# Phylociraptor – A unified computational framework for reproducible phylogenomic inference

**DOI:** 10.1101/2023.09.22.558970

**Authors:** Philipp Resl, Christoph Hahn

**Affiliations:** Institute of Biology, University of Graz, Austria

## Abstract

Multi-stage phylogenomic analyses require complex computational environments. A unified framework for software- and parameter space exploration, data organisation and -deposition is currently lacking. We present phylociraptor (https://github.com/reslp/phylociraptor), a pipeline facilitating fast, reliable, portable (containerized), and reproducible phylogenomics. We demonstrate phylociraptor’s utility and uncover topological discrepancies associated with software and parameter choices in large-scale datasets (31-101 species, 241-2894 genes) addressing enigmatic questions in fungal and vertebrate evolution.

## Main Text

The formal inference of evolutionary relationships based on the state information of homologous characters has come a long way since Willi Hennig founded modern phylogenetic systematics [1]. Since then, DNA and Protein data have become the prime character source, typically drawn from single or a few specifically targeted loci. Now, in the post-genomic era, information from thousands of loci can be utilized while the number of studies under the umbrella of *phylogenomics* increases incessantly.

Phylogenomic inference is a complex multi-stage process principally including orthology inference, multiple sequence alignment (MSA), alignment trimming, and gene tree-, supermatrix- and/or species tree reconstructions. At every stage researchers are confronted with a wide range of software-, parameter-, as well as data filtering choices, each of which may profoundly affect the ultimate result, the phylogenomic tree. Exhaustive evaluation of software- and parameter space is often deemed impractical due to computational limitations (time-, power-, and software dependencies).

In a survey of 50 phylogenomic studies (Suppl. Table on Dryad: https://doi.org/10.5061/dryad.prr4xgxsh) published in leading journals from 2020 onwards all studies employed only a single alignment and trimming strategy (if any), without further justifying the choices or evaluating the effects. Furthermore, the overall data handling and organization was nonuniform, exacerbating problems with reproducibility. While 84% of the investigated studies provided access to at least some crucial supplementary data (single gene alignments, -trees, custom scripts, etc.), the deposited data vary considerably and in some cases it is questionable whether they are sufficient to fully reproduce the published results.

To overcome the above issues we developed a unified computational framework for streamlined, explorative (in terms of software/parameter space), comparative and reproducible phylogenomic inference of large-scale datasets: **phylociraptor** (the rapid phylogenomic tree calculator). Phylociraptor (1) provides a maximum degree of reproducibility, portability and scalability to perform and repeat analyses regardless of computational environment and the underlying dataset, (2) is customizable to account for datasets requiring different parameter settings and software versions and (3) is simple to set up, easy to use and produces comprehensive results.

Phylociraptor performs all steps of typical phylogenomic workflows from raw data download, orthology inference, MSA, trimming, and concatenation, to gene tree-, supermatrix- and species tree reconstructions, complemented with various filtering steps (see details in online-methods). Analyses are steered via an intuitive command-line interface, and internally processes are orchestrated through the workflow management system Snakemake [2]. We provide all third party software tools for our framework as Docker containers [3] (software versions in online-methods), leaving phylociraptor with only two dependencies (Snakemake and Docker or Singularity/Apptainer). This ensures easy installation and portability between systems. Software parameters are specified in a comprehensive YAML configuration file. Upon execution, the particular parameter combinations of an analysis are recorded and encoded as hashes that are stored and incorporated into the output directory structure for full reproducibility. In subsequent analyses with modified parameters phylociraptor automatically determines which parts of the workflow are affected by the changes and executes them.

Phylociraptor is organised into separate modules, which are executed consecutively, adhering to the principial stages of phylogenomic analyses (Figure 1, see also online-methods): `phylociraptor setup` organizes input files, i.e. whole genomes, transcriptomes or proteomes, potentially from different sources, in a uniform directory structure. Input can be provided locally or will be downloaded automatically from NCBI, as specified in a simple comma-separated text file, `phylociraptor orthology` identifies single-copy orthologous genes in all samples using BUSCO v3 [4] or v5 [5]. `phylociraptor filter-orthology` filters the initial set of genes and taxa based on user defined criteria (e.g. amount of missing data for a taxon, or number of taxa in which a given gene is present).

**Figure 1:**
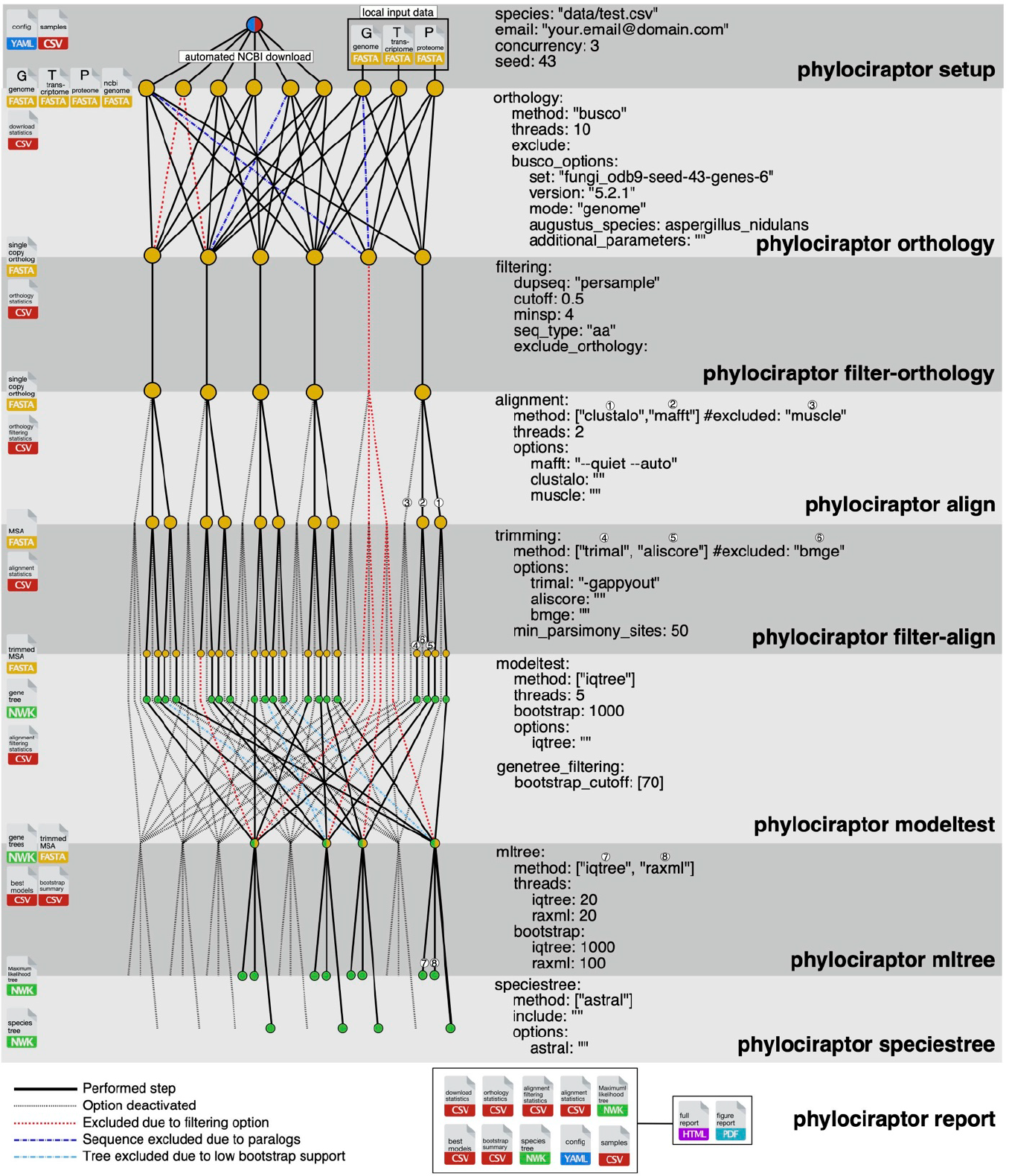
Overview of the phylociraptor framework. Gray boxes delimit individual steps of a phylociraptor analysis. The corresponding phylociraptor commands are provided on the right. The relevant sections of the settings hie for each part are listed in the middle. On the left, side different input and output hies are depicted with different hie symbols. The network graph depicts how analyis steps are connected and which input and output hies are used.

`phylociraptor align` creates MSAs using MAFFT [6], Clustal Omega [7] and MUSCLE [8] for each gene. MS As are automatically trimmed using trimAl [9], AliScore/Alicut [10] and BMGE [11] via `phylociraptor filter-align`, which also filters alignments by length and information content (number of parsimony informative sites and relative composition variability [12]), as specified by the user.

`phylociraptor modeltest` infers the best substitution model for each MSA and calculates gene trees using IQ-Ttee [13]. Phylogenomic inference is performed on concatenated and partitioned MSAs (supermatrix) using maximum-likelihood approaches, via RAxML-NG [14] and/or IQ-Tree (`phylociraptor mltree`), or neighbor-joining, using quicktree [15] (`phylociraptor njtree`), or based on individual gene trees to infer species trees using ASTRAL [16] (`phylociraptor speciestree`). The gene sets considered for these final stages may be automatically filtered by average bootstrap support [17] of the individual gene trees. Throughout, phylociraptor calculates overview statistics of individual steps, which may be summarized in HTML reports (supplementary files) at any point during the analyses via `phylociraptor report` or plotted to publication-ready figures in PDF (see Figure 2). To investigate topological differences, `phylociraptor util` provides several utilities to plot and compare trees. Results, intermediate files and logs of all processes are captured and organised in a coherent directory structure that can be shared with peers or deposited for full reproducibility.

**Figure 2:**
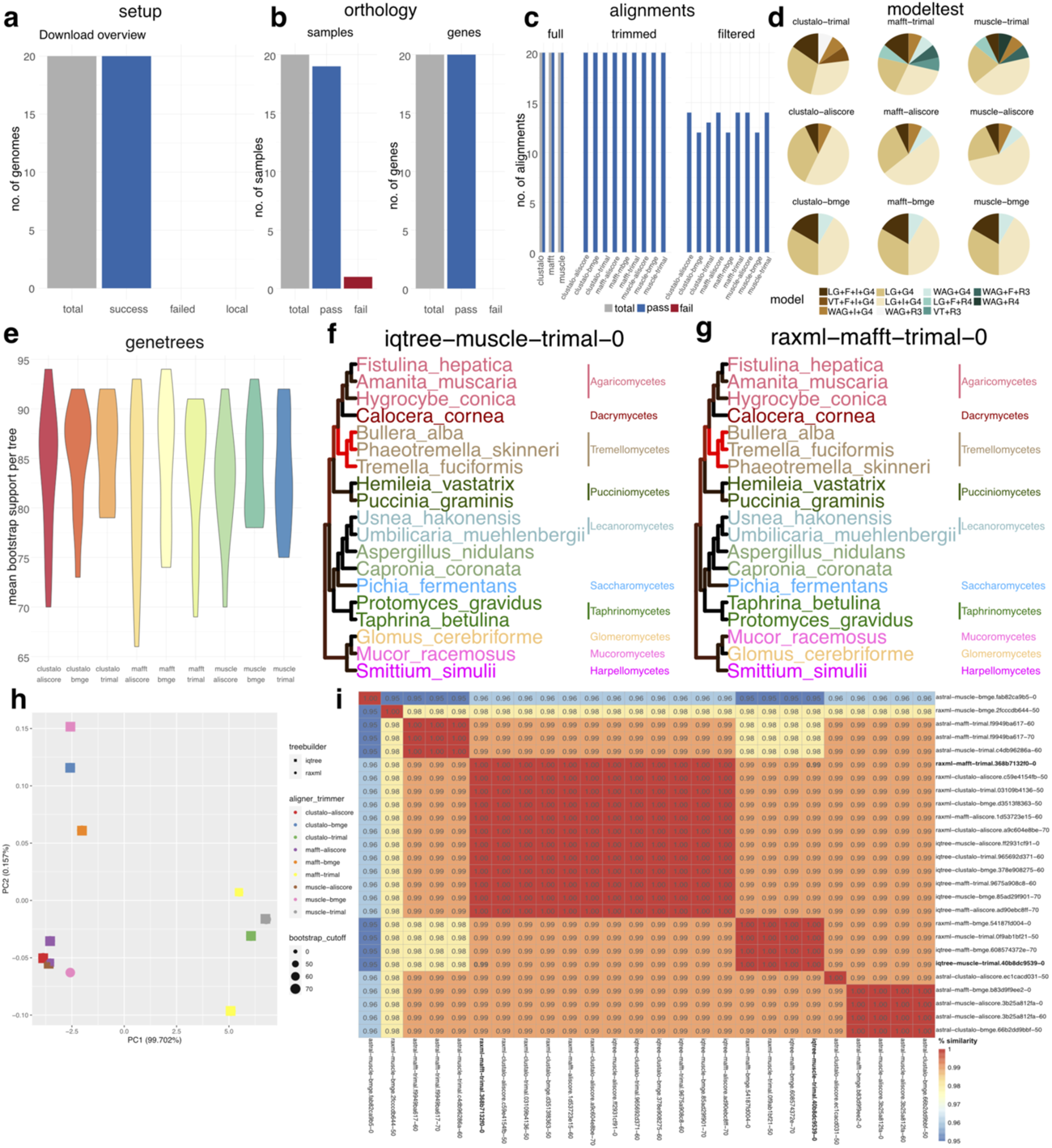
Selection of output created with phylociraptor for a small demo dataset (Drayd: https://doi.org/10.5061/dryad.prr4xgxsh). (a-e) Overview statistics of different steps generated using `phylociraptor report`, (f-g) Two different topologies acquired in a single phylociraptor run. Taxonomic information was retrieved automatically with phylociraptor util get-lineage and monophyletic groupings labelled automatically during plotting. Topological differences between the trees is inferred using quartets of tips with phylociraptor util estimate-conflict and displayed as shades of red along the tree branches. Stronger red color indicates conflicting parts between the two trees, (h) Prinicpal component analyses (PCA) of all tree topologies converted to distance matrices acquired in this run. Different aligners, trimmers, tree-reconstruction methods and average bootstrap cutoff values for gene trees are visualized in different colors and symbols with different sizes. This figure was produced with `phylociraptor util plot-pea`. (i) Heatmap of tree similarities between a selection of trees computed in the example analysis. Red colors indicate more similar trees (with values of 1.00 indicating identical topologies), blue colors more dissimilar trees. This figure was produced with `phylociraptor util plot-heatmap`.

Phylociraptor scales from desktop computers to large High performance computing (HPC) clusters and automatically handles the submission of individual tasks to common HPC queuing systems such as SLURM, SGE and TORQUE to increase parallelization and thus speed.

A complete phylogenomic inference with phylociraptor consists of only seven to eight commands, which, if the full set of alignment and trimming solutions are enabled, would produce nine species trees and/or 18 supermatrix-based maximum-likelihood inferences, each informed by hundreds to thousands of genes (Figure 1 illustrates the process for two plus two aligner/trimmer combinations, yielding four species trees and eight maximum-likelihood trees). If the user had also specified three different average bootstrap support cutoffs in the settings file the same commands would yield 27 species trees and/or 54 supermatrix-based maximum-likelihood trees (Figure 1 illustrates the process with one average bootstrap cutoff of 70).

We tested phylociraptor extensively by analyzing two datasets addressing enigmatic questions in fungal (e.g. the position of the symbiotic Glomeromycota [18, 19, 20, 21, 22]; 101 genomes/transcriptomes) and vertebrate evolution (the closest relative of tetrapods [23, 24, 25, 26, 27]; 31 genomes/proteomes). For each dataset we performed a full phylociraptor analysis using all implemented software combinations and multiple filters, yielding 135 (241-250 genes) and 108 (340-2894 genes) phylogenomic trees, respectively. Our analyses confirm previous results but also uncover topological variation associated with choice of muliple sequence alignment and -trimming algorithms (online-methods). Full analyses, including all intermediate files and logs are deposited with Dryad (https://doi.org/10.5061/dryad.prr4xgxsh).

Phylociraptor is open source (MIT license) and can be obtained from Github (https://github.com/reslp/phylociraptor); version 1.0.0 described here is deposited with Zenodo [https://doi.org/10.5281/zenodo.8365887]). It comes with a minimal test dataset consisting of six fungal genomes. This serves as a tutorial for new users and runs in approximately one hour on a modern desktop computer.

## Supporting information

Phylociraptor report as HTML (zipped) for the small test case presented in the main text (Figure 2).

Short phylociraptor summary report (PDF) for small test case presented in the main text (Figure2).

Phylociraptor report as HTML (zipped) for test-case-1 fungi.

Short phylociraptor summary report (PDF) for test-case-1 fungi.

Phylociraptor report as HTML (zipped) for test-case-2 vertebrates.

Short phylociraptor summary report (PDF) for test-case-2 vertebrates.

## Acknowledgements

We thank Maximilian Wagner, Jonas Lescroart, David Frohlich, Julia Gerasimova and David Díaz Escanden for testing early versions of phylociraptor. We are grateful to Stephan Koblmiiller and Steven Weiss for critical discussions and proofreading.

## Code availability

The full code of phylociraptor is available on GitHub: https://github.com/reslp/phylociraptor. Version 1.0.0 DOI: [https://doi.org/10.5281/zenodo.8365887].

## Data availability

We provide an extensive online-methods document with the main mansucript. Results and additional supplementary data (including all commands to recreate results) are proivided on Dryad: https://doi.org/10.5061/dryad.prr4xgxsh.

## Online Methods

### Introduction

Phylociraptor provides a simple, scalable, customizable and reproducible platform to calculate phylogenomic trees using large-scale genomic, transcriptomic and proteomic datasets. It is based on version-controlled containers of state of the art phylogenetic software connected through a Snakemake workflow and controlled through a single command line interface (CLI) written in Python. Phylociraptor is enhanced by multiple *a-posteriori* tools to compare and visualize analysis results. The only two requirements to run phylociraptor are a container runtime environment (either Singularity/Apptainer or Docker) as well as a Snakemake installation. Phylociraptor is freely available on Github: https://github.com/reslp/phylociraptor. Extensive Online Documentation is available here: https://phylociraptor.readthedocs.io/en/latest/.

### Preparing a phylogenomic analysis

Phylociraptor’s command line interface (CLI) operates based on the settings provided in three config files, of which two are mandatory. The first config file written in the YAML format includes basic settings and parameters to customize different parts of the pipeline, hereafter referred to as settings file (Listing 1, Listing 3, Listing 4, Listing 5, Listing 6, Listing 7). The second mandatory configuration file (hereafter: data file, Listing 2) is a comma-separated text file with at least two mandatory columns: 1) a column labelled species contains the sample name to be used in subsequent analyses, and 2) a column labelled web_local specifies whether NCBI will be automatically queried for a genome sequence or whether a local file is provided. Note that the file was designed to closely resemble the format produced by the NCBI Genome Browser (https://www.ncbi.nlm.nih.gov/genome/browse#!/overview/), and such a file can easily be adjusted to fit the format required by phylociraptor.

**Listing 1:**
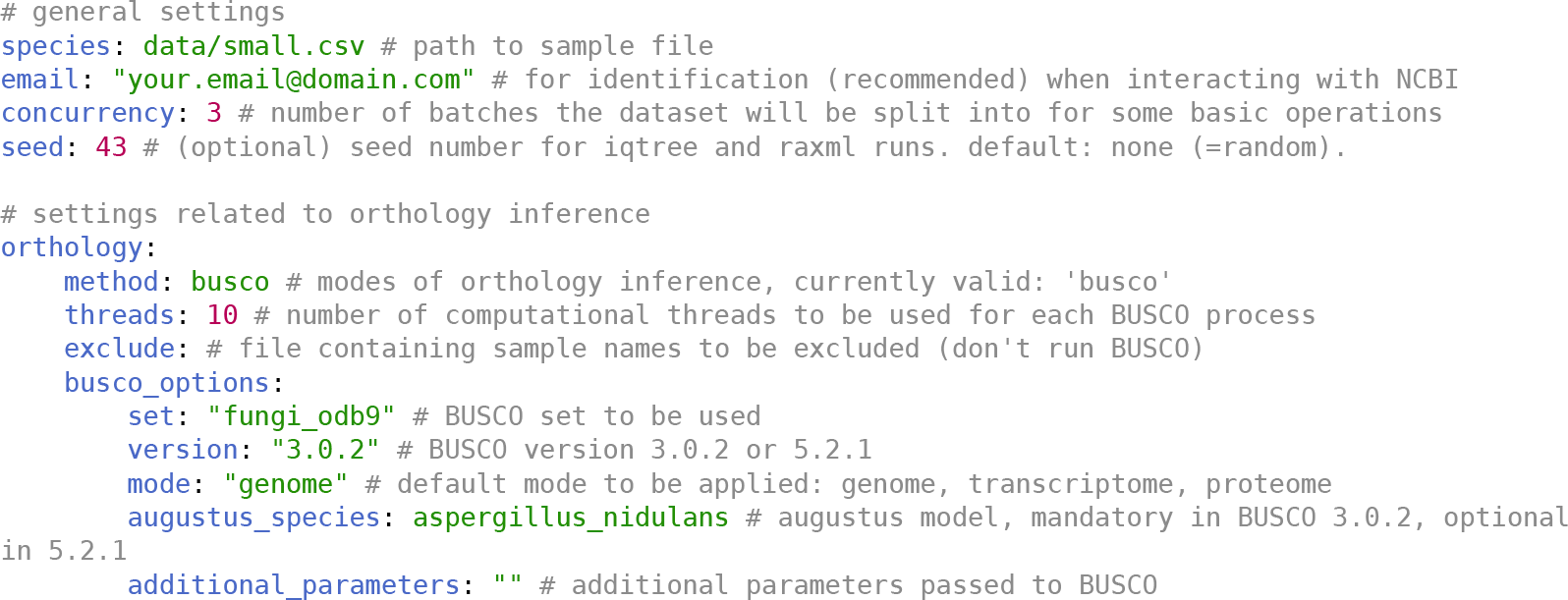
Excerpt of settings file - sections general and orthology

Optionally, the user may also use a cluster configuration file which includes settings and information necessary to interact with common job schedulers and automatically submit individual phylociraptor tasks to a HPC cluster. These settings are specific to the individual computational infrastructure and phylociraptor comes with template files for the common job scheduling systems SLURM, SGE and TORQUE.

**Listing 2:**
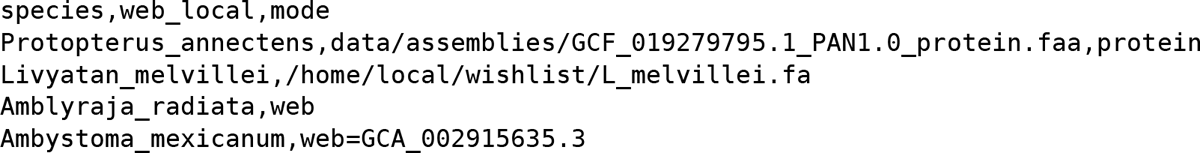
Example data file

A phylociraptor run is separated into distinct stages, corresponding to typical steps familiar from few-gene phylogenetic reconstructions. In the phylociraptor framework this process is expanded automatically to hundreds or thousands of genes and across multiple (exhaustive) combinations of software solutions included in the workflow, and parameters, as specified by the user.

### Setting up a phylociraptor run

The typical initial input of a phylogenomic analyses is genome, transcriptome or protein data, specifically compiled for the current research question. Often, data for key taxa are generated especially for the project in question and combined with hand picked data from reference databases.

`phylociraptor setup` is the entry point for a phylogenomic analysis in phylociraptor and usually the first command a user will run. It performs three main tasks: First, phylociraptor will create the basic directory structure for subsequent analyses. This structure is designed to be self contained, which allows for easy sharing of analyses files. Second, the directory structure will be populated with genome-, transcriptome, and proteome data, as specified by the user in the data file. Column one (`species`) specifies the sample name to be used in subsequent analyses. Column two (`web_local`) specifies whether NCBI will be automatically queried for a genome sequence or whether a local file is provided, in which case this column should contain the path to the local file. Phylociraptor will differentiate between files which are already in (or in a subdirectory of) the current phylocriaptor base directory and files somewhere else in the file system. In the former case (Listing 2, line 2) it will create a symbolic link to the actual file and in the latter case (Listing 2, line 3) it will deposit a copy of the file in the relevant place. If the string in column two starts with the keyword ‘web’ phylociraptor will attempt to automatically download a genome from the NCBI genome database. In this case the species name in the samples file needs to match the binomial under which a genome is deposited at NCBI (Listing 2, lines 4 and 5). If multiple assemblies are available, phylociraptor applies a built-in decision process to select an appropriate assembly to download according to NCBIs nomenclature: First it tries to identify and download a *reference assembly*. If none is available, phylociraptor will download a *representative genome*. For an explanation of these terms see: https://www.ncbi.nlm.nih.gov/assembly/help/#resultsfields. In case there is also no representative genome available, it will download the latest assembly. Alternatively one can also specify a specific assembly accession number (Listing 2, line 5). Genomes will be downloaded in parallel in batches, if specified via the setting concurrency in the general section of the settings file (Listing 1).

Third, phylociraptor downloads the specific BUSCO set, specified in the settings file, the genes of which will serve as basis for subsequent phylogenomic reconstructions.

### Infer single-copy orthologs for all included species

Phylogenetic inference depends on robust establishment of orthologous relationships, `phylociraptor orthology` infers a set of conserved single-copy orthologs in a standardized and well established way for each genome, transcriptome or proteome in the dataset by applying either BUSCO 3.0.2 [1] or 5.2.1 [2]. This step can be highly parallelized: on the one hand, the level of parallelization of a single BUSCO job can be controlled in the settings file (`orthology: busco_settings : threads: 10`), on the other, phylociraptor runs multiple BUSCO jobs in parallel, if the availalbe computational resources allow it (e.g. in HPC environments). After BUSCO has finished successfully for all samples, phylociraptor will extract information on the presence, absence and status of all genes into a table which forms the basis for subsequent filtering of orthogroups. See Listing 1 for the relevant section of the settings file.

### Filter orthogroups

`phylociraptor filter-orthology` gathers data from all samples and generates multi-sequence fasta files for each gene passing the filtering criteria specified in the corresponding section of the settings file (Listing 3). First, it is possible to exclude species for which less than a specified proportion of the reference set of single-copy orthologs could be identified (`filtering: cutoff: 0.5`). The second parameter allows to control the minimum number of species in the dataset which have a specific BUSCO gene so that this gene is included in subsequent analysis steps (`filtering: minsp: 3`). Despite being reported as single-copy, BUSCO occasionally generates multiple sequences for a single BUSCO gene. Since it is unclear which of the sequences should be used for downstream analyses, phylociraptor offers two ways to filter BUSCO gene files for duplicated sequences. Setting `dupseq: “persample”` will instruct phylociraptor to only filter out the duplicated sequences for the affected sample. Alternatively setting `dupseq : “perfile”` will exclude the ortholog from the analysis for all samples. Finally, samples can be excluded all-together from subsequent analyses by specfying a file with a list of sample names (`filtering : exclude_orthology : data/exclude. txt`). See Listing 3 for the relevant section of the settings file.

**Listing 3:**
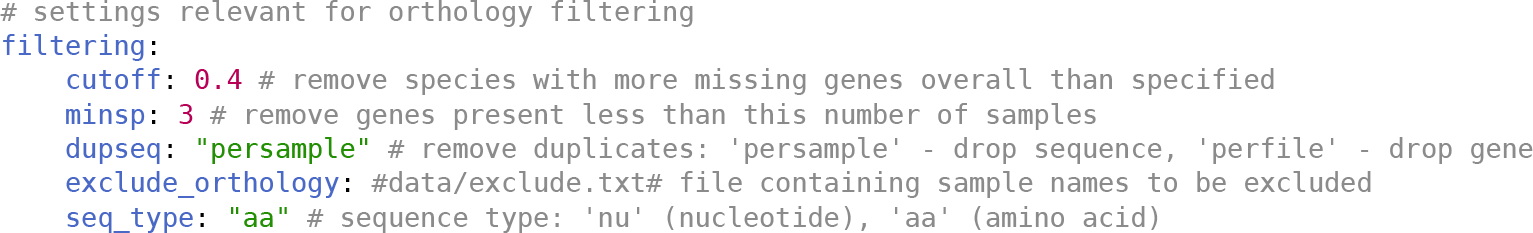
Excerpt of settings file - section orthology filtering

### Multiple Sequence alignment

Alignment is a crucial yet non-trivial step in phylogenetic reconstructions. Poorly aligned regions may impact phylogenetic branch support and could ultimately lead to erroneous tree structures [3]. `phylociraptor align` will apply up to three different multiple sequence alignment algorithms to each gene. Currently supported are MAFFT 7.464 [4], Clustal Omega 1.2.4 [5] and MUSCLE 5.1 [6]. Users can specify the grade of parallelisation as well as any additional aligner specific parameters for each gene alignment in the settings file. Again, phylociraptor will parallelize individual alignment jobs according to the amount of resources available. See Listing 4 for the relevant section of the settings file.

**Listing 4:**
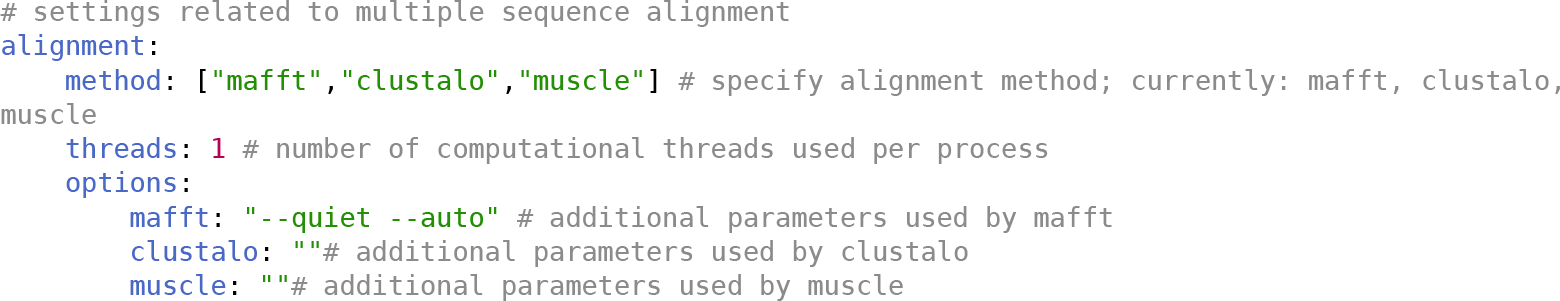
Excerpt of settings file - section alignment

### Alignment trimming and filtering

To remove poorly aligned positions and improve overall alignment quality, alignment trimming is common practice. The alignment algorithms trimAl 1.4.1 [7], AliScore/Alicut 2.31 [8] and BMGE 1.12 [9] will be used to trim poorly aligned positions in each alignment generated by the previous step if the user executes `phylociraptor filter-align`. Individual trimming parameters can be modified in the settings file. Phylociraptor provides two additional alignment filtering options to reduce the amount of missing data and remove uninformative alignments based on the number of parsimony informative sites and relative composition variability (rev; [10]). These filters will be applied to batches of multiple sequence alignments in parallel according to the grade of parallelisation specified via the concurrency: option (Listing 1) in the general part of the settings file. Again, in HPC environments these steps can be highly parallelized. See Listing 5 for the relevant section of the settings file.

**Listing 5:**
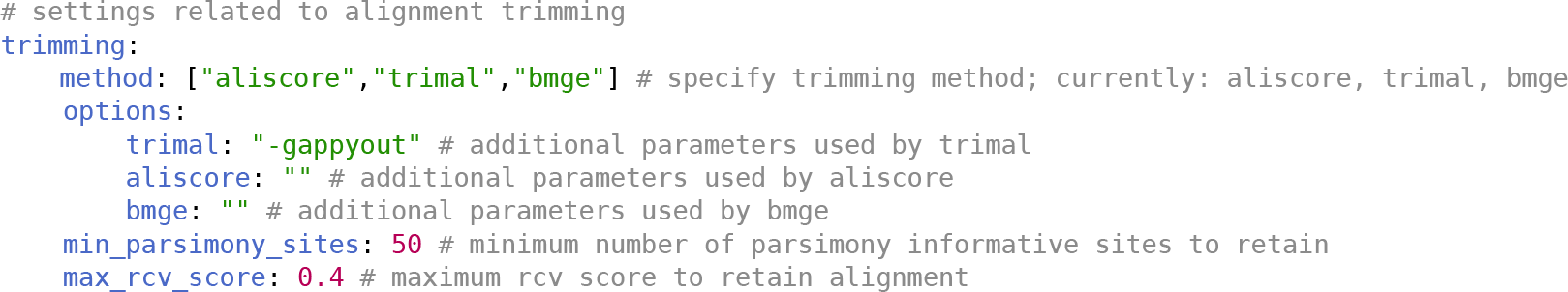
Excerpt of settings file - section alignment trimming

### Substitution model selection and gene tree calculation

Choosing inaccurate substitution models impacts phylogenetic inference [11]. To aid with selecting the best substitution model, phylociraptor uses the modeltest functionality of IQ-Tree [12] to estimate the best fitting substitution model based on the Bayesian information criterion (BIC) for each alignment. During this process, phylociraptor performs a full maximum-likelihood tree search to calculate gene-trees for each alignment. The estimated substitution models and gene trees are saved for subsequent phylogenomic reconstructions (eg. using ASTRAL) and the inferred models will be applied to gene partitions in supermatrix-based maximum-likelihood inference. This stage is triggered by executing `phylociraptor modeltest`. See Listing 6 for the relevant section of the settings file.

**Listing 6:**
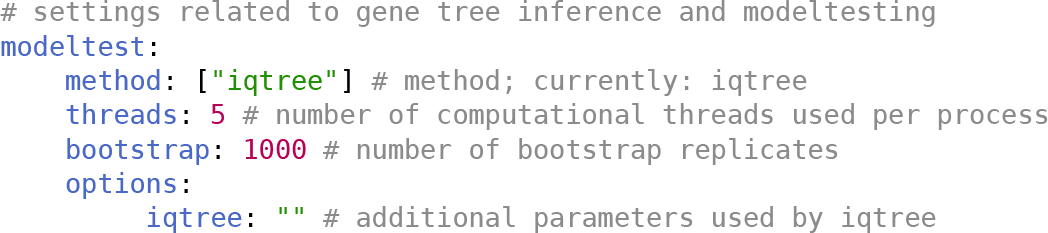
Excerpt of settings file - section modeltest

### Phylogenomic tree reconstruction

Phylociraptor offers several options to calculate phylogenomic trees. Currently supported software includes the maximum-likelihood based software IQ-Ttee 2.0.7 [13], and RAxML-NG 1.1.0 [14], the coalescence based software ASTRAL 5.7.1 [15] as well as the Neighbor-Joining algorithm implemented in QuickTtee 2.5 [16]. Executing `phylociraptor mltree` will run IQ-Tree and/or RAxML-NG based on an alignment supermatrix constructed from the individidual gene alignments incorporating the estimated best models from the previous model testing step. Inferences will be run for each combination of aligners and trimmers specified in the settings file. In the same vein, `phylociraptor speciestree` will gather individual gene trees and infer species trees using ASTRAL for all sets of trees obtained through all enabled aligner/trimmer/filtering options. Finally, `phylociraptor njtree` will gather data and execute QuickTree on each dataset, accordingly.

Each of these programs requires differently formatted input files and different parameters. Phylociraptor handles all of this automatically with extensive options to customize analyses through the settings file. To increase confidence in the final topology, it is possible to filter the genes used in the combined phylogenomic analyses based on the average bootstrap support of the corresponding gene tree as suggested by Salichos & Rokas [17]. See Listing 7 for the relevant section of the settings file.

**Listing 7:**
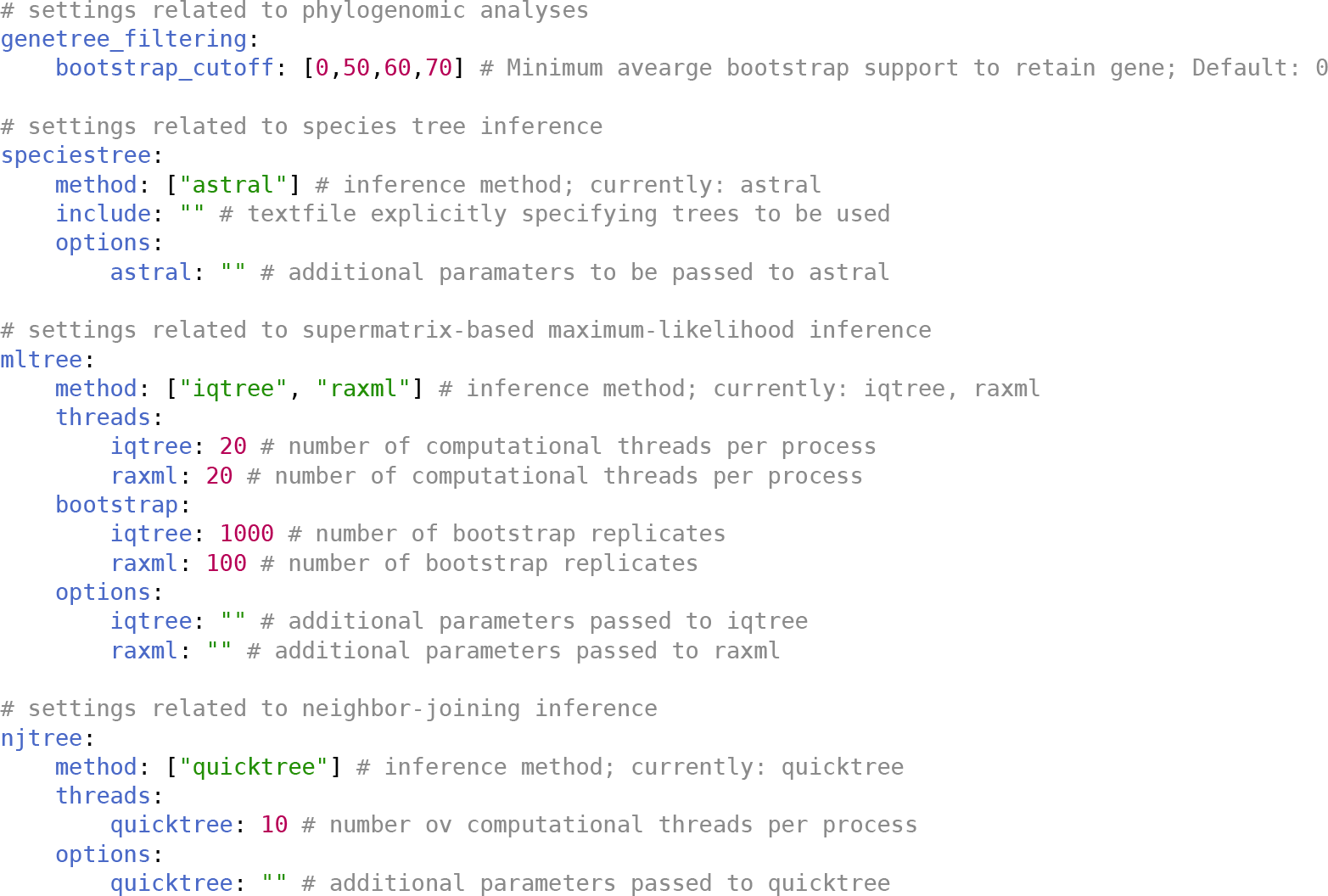
Excerpt of settings file - section phylogenomic inference

### Creating an overview reports

During large phylogenomic analyses a lot of information accumulates. Phylociraptor gathers information from individual steps and creates reports in HTML and PDF formats. The HTML report gives detailed information about the used genomes, alignments, substitution models, gene trees and phylogenomic trees. The report can be created at any time via `phylociraptor report` and helps to inform sensible parameter choices as the analysis progresses. Additionally to the comprehensive HTML report, phylociraptor can produce a publication ready report figure in PDF format (example given in Figure 2 in the main text) summarizing key metrics of the run for quick inspection, via executing `phylociraptor report --figure`.

### Utilities for *a-posteriori* analyses of trees

Due to its comprehensive exploratory approach, phylociraptor can quickly produce large numbers of phylogenomic trees. To facilitate downstream analyses of large phylogenies, phylociraptor offers several utilities (implemented in `phylociraptor util`) to visualize, inspect and compare different trees. Carefully written R and python scripts utilise R packages such as ape [18] and phytools [19] ggplot [20], ggtree [21] and knitr [22] and process phylogenetic data obtained through phylociraptor for visualization. Our scripts solve several challenges encountered when plotting trees with R. By applying a built-in ruleset to automatically decide on sensible plot dimensions based on the number of included taxa phylociraptor’s plotting approach tries to achieve the best possible visualizations for a given dataset. To easily depict taxonomic relationships tips and clades can be annotated with taxonomic lineage information which phylociraptor automatically retrieves from the NCBI taxonomy database.

Phylociraptor can estimate tree similarity in two ways. First, it allows tree comparisons based on the similarity of subtrees containing four taxa, a.k.a. quartet subtrees. This method, originally proposed by George F. Estabrook [23] assumes that two trees are topologically identical if all quartet subtrees are identical. To calculate and compare quartet subtrees, phylociraptor provides a python script interfacing the R package ape through rpy2 (https://rpy2.github.io/) to handle phylogenetic trees. This script compares quartet subtrees between all available pairs of trees created with phylociraptor. Each comparison between two trees will receive a value between 0 and 1 indicating the proportion of identical quartets found. 0 indicates complete dissimilarity (every compared quartet subtree is different) and 1 that the trees are assumed to be topologically identical (every compared quartet subtree is identical). Tree similarity is calculated by running `phylociraptor util estimate-conflict`. This phylociraptor utility is multithreaded to reduce runtime.

With this approach the number of possible quartet subtrees grows quickly as the number of tips increases. This can slow down computation time drastically when comparing large trees. Other implementations using quartet subtree comparisons thus limit the number of tips trees can have. With phylociraptor trees subjected to this approach can have any number of tips. While calculating all possible quartet subtrees could be impractical, to decrease computation time phylociraptor provides multiple stopping criteria such as the maximum number of investigated quartets, the number of identified conflicting quartets or the mean number of times each tip occurs in the sampled quartets. Additionally, the analysis can be limited only to tips belonging to specific taxonomic groups or by providing a list of tips. Estimates of topological conflict can be summarised for all, or a user specified set of, trees as heatmap (`phylociraptor util plot-heatmap`). Pairs of conflicting trees can be visualized via `phylociraptor util plot-conflict` (Figure 2, see also Figure 2 in main text).

A second method to investigate the similarity of trees is based on distances between pairs of tips. This approach assumes that within trees with identical topologies and identical branch lengths, pairs of tips should have the same distance. While there are situations in which this assumption is violated, this approach generally works well in real world scenarios when trees are based on the same underlying alignments. However it should be noted that this comparison is only feasible to identify large scale differences between tree reconstruction methods. Phylociraptor calculates the distance (the sum of the lengths of all branches connecting two tips) for each pair of tips in each tree. This results in a large matrix of tip to tip distances which is visualized through a principal component analysis (phylociraptor plot-pea). This allows us to identify islands of similar and dissimilar trees, which can be investigated further e.g. with phylociraptor’s quartet similarity approach above.

### Case study 1: A phylogenomic tree of the kingdom fungi

To demonstrate how phylociraptor can be used to explore software and parameter combinations with a large phylogenomic datasets, we performed a phylogenomic analysis of the kingdom fungi based on the set of taxa recently published by Strassert and Monaghan (2022; [24]). For 72 taxa we were able to identify the corresponding NCBI Genbank accession IDs based on careful inspections of the data in [24, Suppl. Table 1]. We used these accession numbers to download corresponding genome assemblies automatically from NCBI with phylociraptor. For 24 taxa we downloaded genome and transcriptome assemblies or sets of proteins manually from the corresponding database specified in [24, Suppl. Table 1], and included them as local files in the phylociraptor data file.

For five taxa used by [24] the exact data which would be required for phylociraptor (genome or transcriptome assemblies or sets of protein sequenes) was not deposited in public databases. We therefore downloaded raw sequence reads from NCBI’s Sequence read archive (SRA) according to the information given by [24] and assembled them *de-novo*. Data for *Amaparvus languides, Nuclearia sp*., *Stereomyxa ramosa*, and *Cunea sp*. were assembled with Ttinity 2.10.0 [25] and `-min_kmer_cov 2`. For the latter, it was unclear which specific dataset was originally used, so we used *Cunea sp*. with the SRA accession SRR5396443, which was the only data from *Cunea sp*. available on SRA (last checked Aug. 4, 2022). To deduplicate Trinity assemblies we employed CD-HIT v4.8.1 [26] with -c 0.95. For *Coelornornyces lativittatus* we downloaded trimmed genomic reads from SRA (accession: SRX9369341) and assembled them using SPAdes v3.15.3 [27] with `--k 21,31,51,71 --careful`.

We then performed a full phylogenomic analysis of the 101 taxa from Strassert and Monaghan (2022) using phylociraptor v.0.9.12 (git branch `test-cases`). This analysis (including all config files, settings, intermediate files and commands) is provided in Supplementary Data on Dryad (https://doi.org/10.5061/dryad.prr4xgxsh). An overview of the run is provided in Figure 1. Our analysis entailed BUSCO 5.2.1 searches using the *eukaryota_odbl0* set, multiple sequence alignment using MAFFT, MUSCLE and Clustal Omega, MSA trimming using BMGE, trimAl, AliScore and phylogenomic reconstruction with RAxML-NG, IQ-Ttee and ASTRAL. After trimming, we filtered out alignments with fewer than ten parsimony informative sites and we applied five different boostrap cutoffs (0, 50, 60, 70, 80 and 90) to the filtered alignments and gene trees. This resulted in a total of 136 phylogenomic trees each based on information from 241-250 individual genes. We included the Bayesian consensus trees of three independent Markov-Chains from [24] in our *a-posteriori* analyses of the topologies inferred via phylociraptor because their final consensus tree, as displayed in the article [24,, Figure 1], was not deposited in a textual representation (e.g. Newick) together with the study. Visual comparison with the three deposited Bayesian consensus trees suggests that the tree from Markov Chain 3 has the same branching-order as the tree in Figure 1 of Strassert and Monaghan [24].

**Figure 1:**
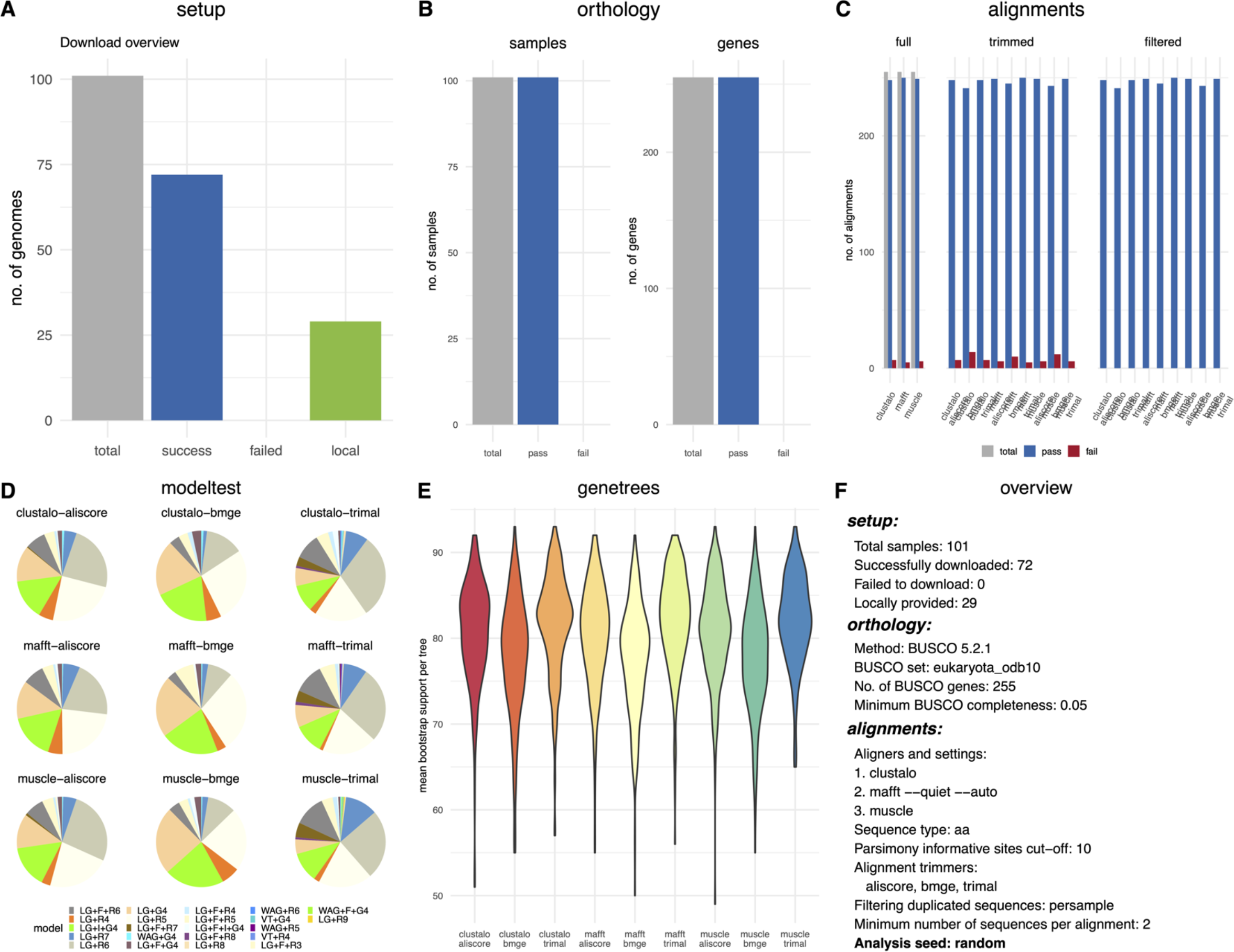
Summary plot of phylociraptor analysis - generated with `phylociraptor report --figure`

Our phylogenomic trees are mostly in accordance with, but also display several differences to, other published phylogenomic trees of fungi, including those by [24]. It is noteworthy, though, that none of the trees recovered by us was topologically identical to the tree presented by [24]. The topologies recovered here differ between 6% and 10% (Figure 4, Figure 2) from the three Baysian consensus trees of different Markov-Chains recovered by [24] based on our analyses of conflicting quartets of tips.

**Figure 2:**
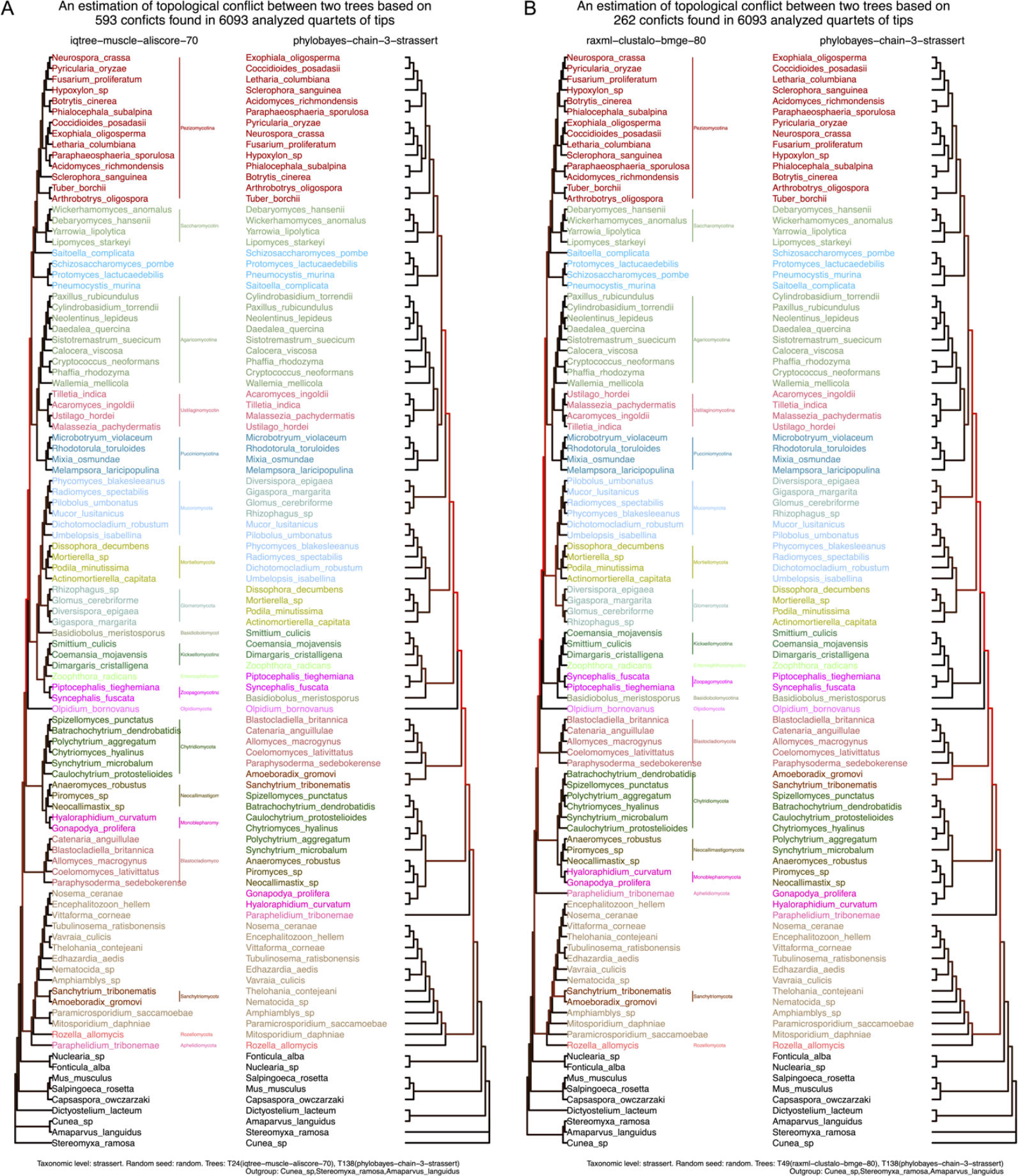
Two examples of differing topologies between the current study and [24]. A) Two trees with approx. 10% conflicting quartets of tips. Left: Topology recovered with phylociraptor using IQ-Tree, MUSCLE and AliScore/Alicut and a bootstrap cutoff of 70. Right: Topology recovered by [24] from the Bayesian Markov-Chain number 3. B) Two trees with approx. 5% conflicting quartets of tips. Left: Topology recovered with phylociraptor using RaxML-NG, Clustal Omega, BMGE and a bootstrap cutoff of 80. Right: Topology recovered by [24] from the Bayesian Markov-Chain number 3. The labeling of the taxonomic groups is based on the groupings suggested by [24]. This plot was generated with `phylociraptor util plot-conflict`.

**Figure 3:**
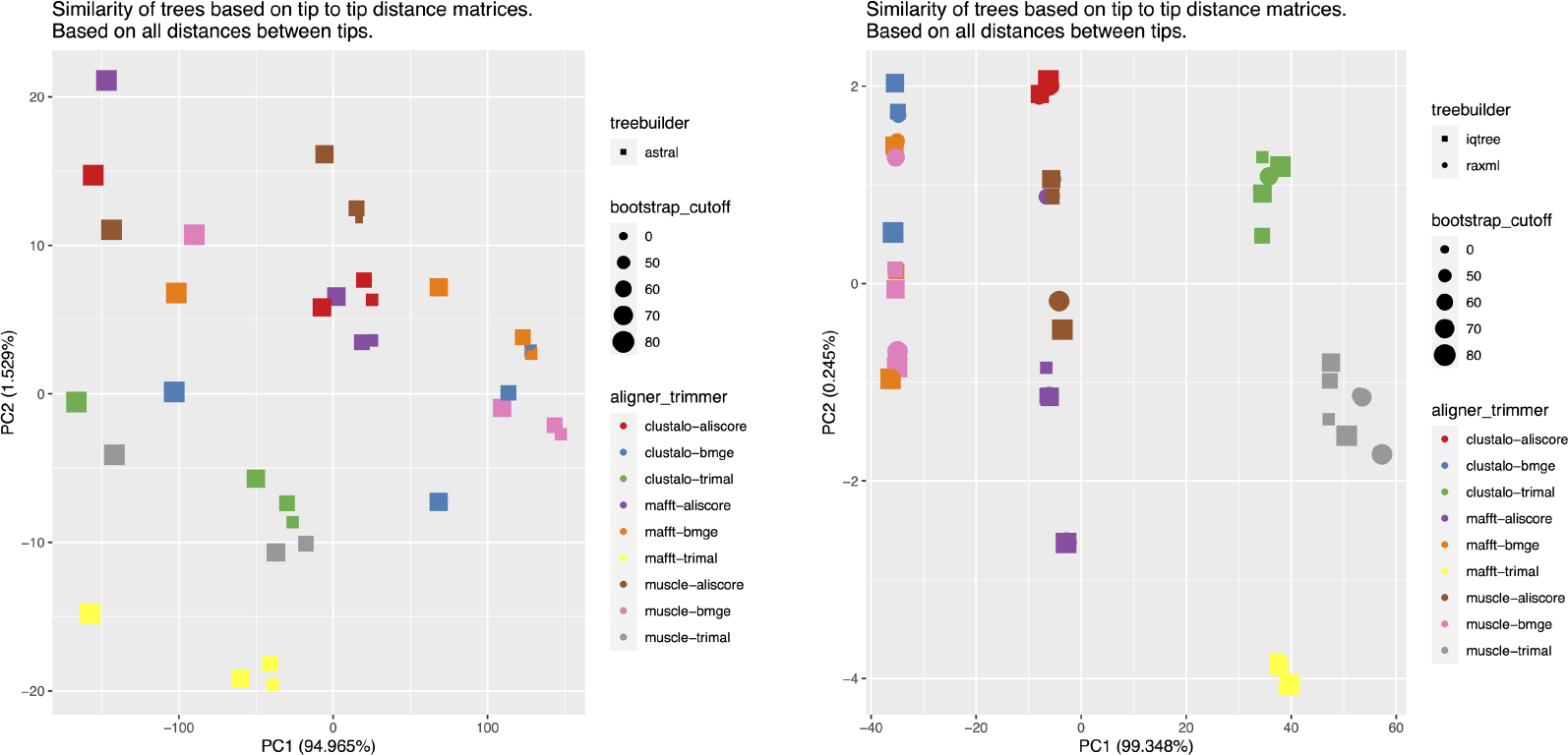
Principal component analyses based on tip2tip distances of phylogenomic trees - generated with `phylociraptor util plot-pca`. Left: All trees from ASTRAL species tree reconstructions. Right: All trees from concatenated Maximum-likelihood reconstructions.

**Figure 4:**
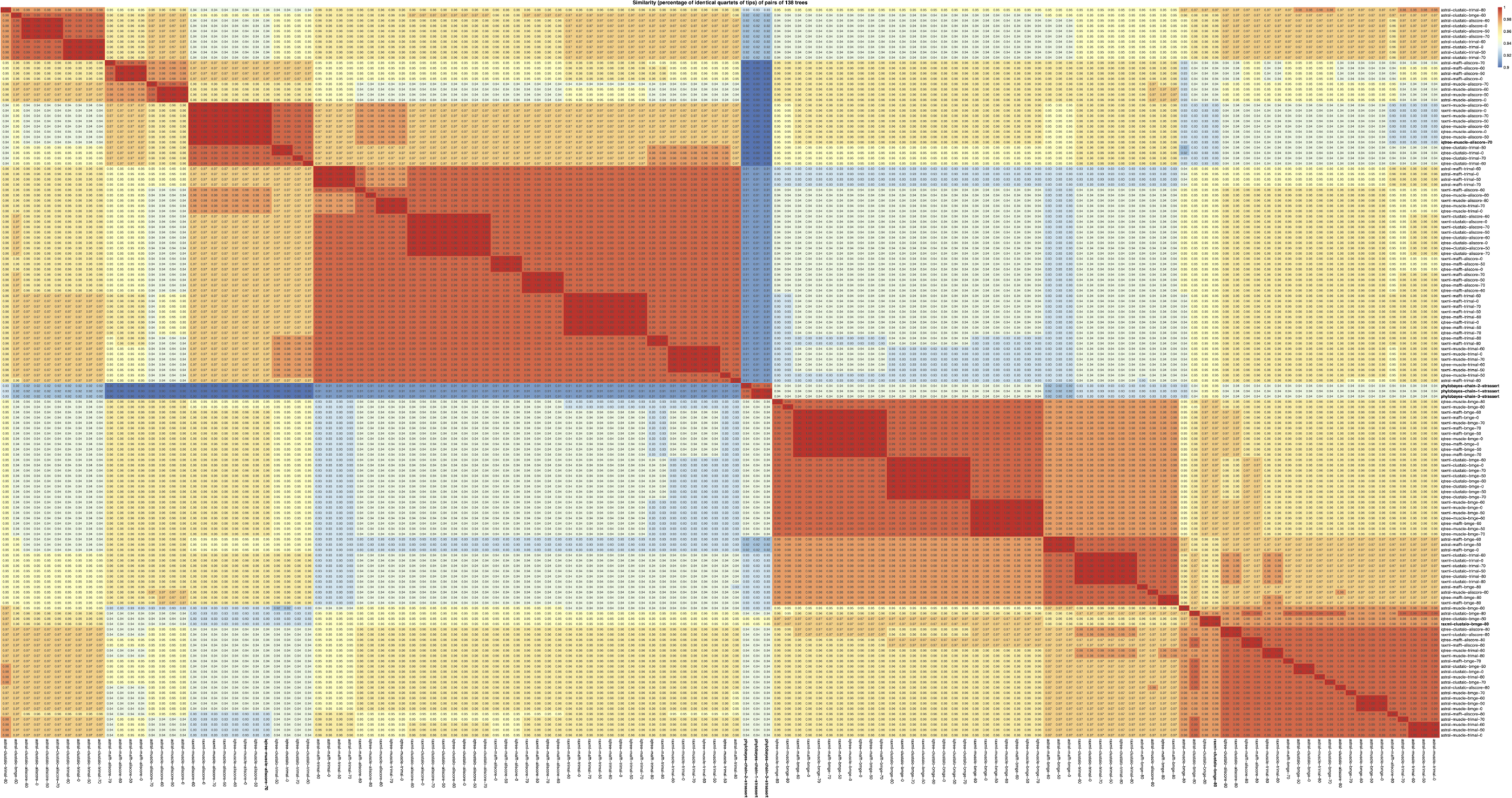
Heatmap of similarities across all phylogenomic trees inferred here including three consensus trees from Bayesian markov chains from [24] - generated with `phylociraptor util plot-heatmap`. In bold are the tree topologies from [24] as well as the trees visualized in Figure 2.

The incongruences were recovered despite using the same set of taxa (to the best of our knowledge) but with different analytical approaches (Figure 3, Figure 4). Strassert & Monaghani (2022) use a set of 320 orthologous genes processed by a consecutive clustering, homology identification and tree building workflow [24]. We utilize the standardized and well curated sets of BUSCO genes. Using this set of genes we tested all combinations of three aligners, alignment trimmers and tree reconstruction methods under five different average bootstrap cutoff values. Combined results obtained by the different analytical approaches taken in the present manuscript and by Strassert and Monaghan (2022) suggests that a considerable amount of phylogenetic uncertainty in this group of organisms remains.

Overall, our 136 phylogenomic analyses resulted in 67 different topologies with up to *7%* different sampled quartets between them (Figure 4), specifically 25, 23, and 19 from ASTRAL, IQTtee and RAxML-NG, respectively. Of 19 topologies that were recovered in at least three different analyses, 16 were consistently inferred from data treated with the same aligner/trimmer combination (supported by multiple average bootstrap cutoffs) and only three were recovered with two aligner/trimmer combinations independently (representatives: iqtree-mafft-bmge-70, iqtree-muscle-trimal-70 and iqtree-muscle-bmge-70). All of these, plus additional seven (total 10), topologies were recovered with both maximum-likelihood software solutions (IQ-Tree and RAxML-NG), and nine with ASTRAL only. Notably, no identical topology was recovered both by maximum-likelihood- and species tree inference methods. The observed topological variability is caused by different branching order within and between fungal groups, occasionally leading to loss of monophyly.

We could confirm the monophyly of several fungal groups over a wide range of software combinations (Table 1). Of the divisions and subdivisions recognized by [24], we consistently recovered Monoblepharomycota, Zoopagomycotina, Mortiellomycota, Mucoromycota, Pucciniomycotina, Glomeromycota, Saccharomycotina, Agaricomycotina, Neocallimastigomycota, Sanchytriomycota, Blastocladiomycota, Pezizomycotina and Ustilaginomycotina as monophyletic, although some groups in some trees had relatively low (<90 BS; <0.95PP) support (https://doi.org/10.5061/dryad.prr4xgxsh). Taphrinomycotina was always recovered as polyphyletic. Microsporidia were also mostly recovered as polyphyletic (because Sanchytriomycota resolved inside it) except for a handful of trees in which it is recovered as monophyletic with low support. The only tree in which Microsporidia forms a well supported monophyletic group is our ASTRAL tree with Clustal Omega and trimAl and a bootstrap cutoff of 80%. Chytridiomycota is recovered as monophyletic in most analyses, however often with poor support. Occasionally it is recovered as being not monophyletic. In topological accordance with [24], 131 of our phylogenies also resolve Chytridiomycota as sister to a clade composed of Neocallimastigomycota and Monoblepharomycota.

**Table 1.**
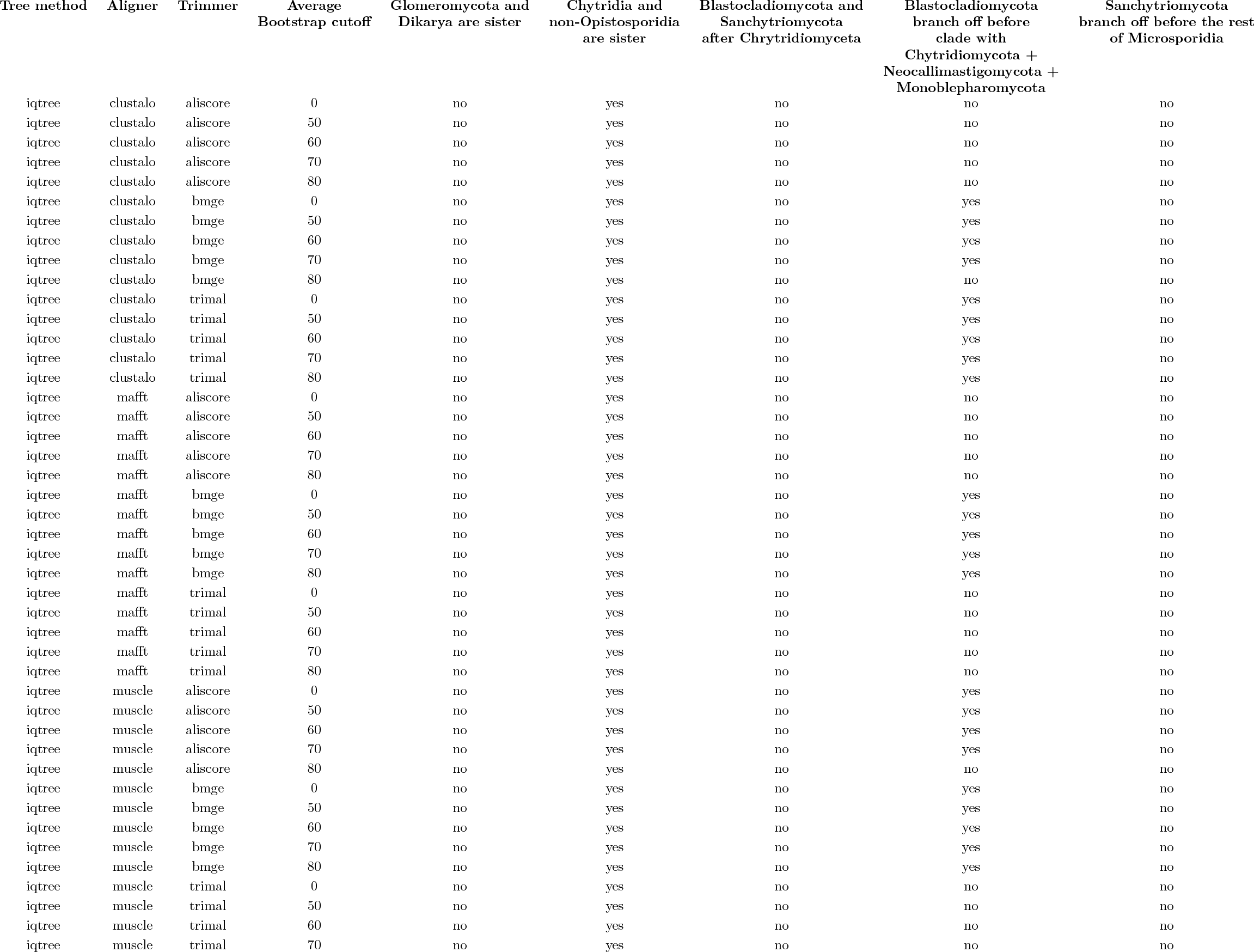
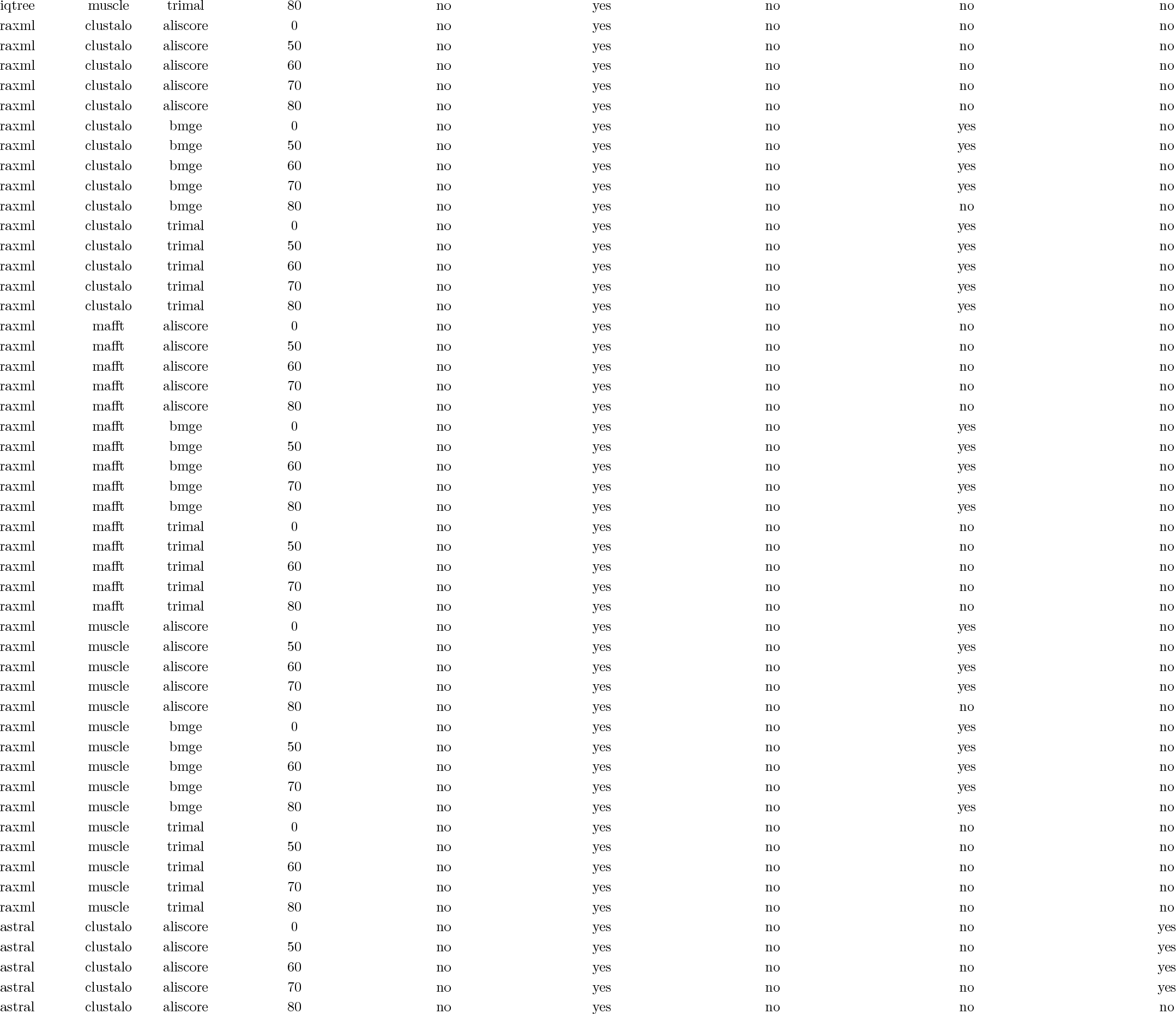
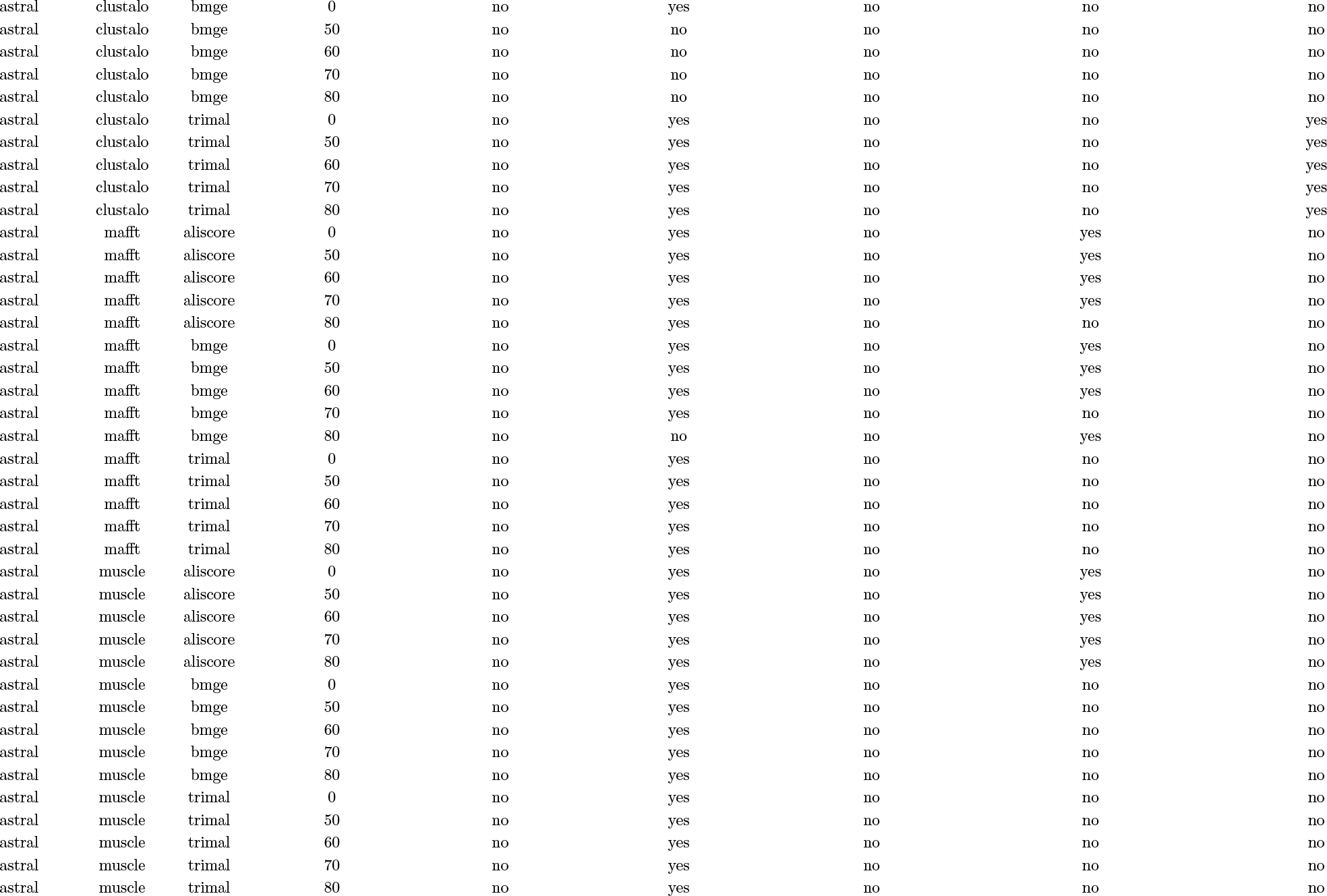
Phylogenetic groups as suggested by [24] and how these groupings are represented in our phylociraptor reanalysis using different aligner, trimmer, bootstrap cutoff and tree inference combinations.

We recover Glomeromycota within a clade together with Mortiellomycota and Mucoromycota and not sister to Dikarya (Table 1). Likewise, none of our trees report the branching order suggested by [24] for the position of a clade composed of Blastocladiomycota and Sanchytriomycota branching off after Chytridiomyceta. (hereafter referred to as clade BSC). Although Blastocladiomycota are consistently recovered as monophyletic, the position of this clade remains unstable. In 77 trees Blastocladiomycota branch off after Chytridiomyceta and Sanchytriomycota, whereas in the remaining 59 trees, Blastocladiomycota branch off before. The two topologies are recovered in roughly equal proportions from data aligned with three different aligners and tree reconstruction methods, however we did notice that the majority (32 of 45) of trees reconstructed from alignments trimmed with BMGE, exhibit a branching order where Blastocladiomycota branch off before the rest of the BSC clade.

Our results suggest that the tree inference method, gene tree filtering and, perhaps surprisingly, alignment and trimming strategies, can have strong influence on node support and branching order of phylogenomic trees. This suggests that on top of other well-established phylogenetic best practices thorough exploration of the software and parameter space is crucial to confidently resolve the evolutionary relationships of fungal groups studied here.

### Case study 2 - the closest living relative of tetrapods

The identity of the closest living relative of tetrapods as part of the evolutionary history of vertebrates has been a long standing question in evolutionary biology. Recent work has for the first time incorporated transcriptome or full genome evidence and consistently recovered lungfishes as the closest living relative of tetrapods ([28, 29, 30, 31, 32]). In this case study we aimed to investigate vertebrate relationships and particularly the position of the closest living relative of tetrapods with a new and extended dataset. The taxon set was chosen to include the same taxa as in the most recent take on the question by Meyer and colleagues [32], with extended taxon sampling to better represent cartilaginous- and slow-evolving ray-finned fishes, as suggested by Takezaki & Nishihara ([31]). We also incorporated jawless fishes (Cyclostomi) into our analyses as an outgroup. Further, our analyses is the first to incorporate both Australian and African lungfish. We used phylociraptor v.0.9.12 (git branch test-cases). The full phylocirpator setup for these analyses (including all config and intermediate files, settings and commands) are provided on Dryad (https://doi.org/10.5061/dryad.prr4xgxsh).

In brief, genome data were automatically downloaded via phylociraptor setup according to NCBI accession numbers as specified in the data file. The proteome of the African lungfish *Protopterus annectens* (Genbank accession: GCF_019279795.1) was downloaded manually and provided as local file. Orthology inference was done using BUSCO 5.2.1, using the *vertebrata_odblO* set, principally with augustus as ab-initio gene predictor with the augustus training set ‘human’. For the large genomes of *Amblyraja radiaba, Arnbystorna mexicanum, Eptatretus burgeri, Neoceratodus forsteri, Rana temporaria, Scyliorhinus canícula*, and *Xenopus tropicalis*, the genefinder metaeuk was used instead of augustus, with additional parameters passed to metaeuk to restrict memory usage (orthology: busco_options : additional_parameters : “--metaeuk_parameters=-- slice-search=1 - -remove-tmp-files=1 --disk-space-limit=3000G --split-mode=0 --split-memory-limit=1500G --metaeuk_rerun_parameters=--slice-search=1 - -remove-tmp-files=1 --disk-space-limit=3000G --split- mode^ - -split-memory-limit=1500G”). Multiple sequence alignment was performed using MAFFT, MUSCLE and Clustal Omega, MSA trimming using BMGE, trimAl, AliScore, excluding any genes with less than 50 parsimony informative sites after trimming, and phylogenomic reconstruction with RAxML-NG, IQ-Tree and ASTRAL, with bootstrap cutoffs 50, 60, 70 (IQ-Tree and ASTRAL only due to extensive runtime of RAxML), 80, and 90.

Our analyses resulted in a total of 108 phylogenomic trees based on 380 to 2,800 individual single copy genes and between 372,000 to 1,459,582 amino acid positions, depending on the bootstrap filtering. For a complete overview of the run including all phylogenomic trees in Newick format see the corresponding phylociraptor report in the Suppl. Data for test case 2 (https://doi.org/10.5061/dryad.prr4xgxsh) and Figure 5 for a summary.

**Figure 5:**
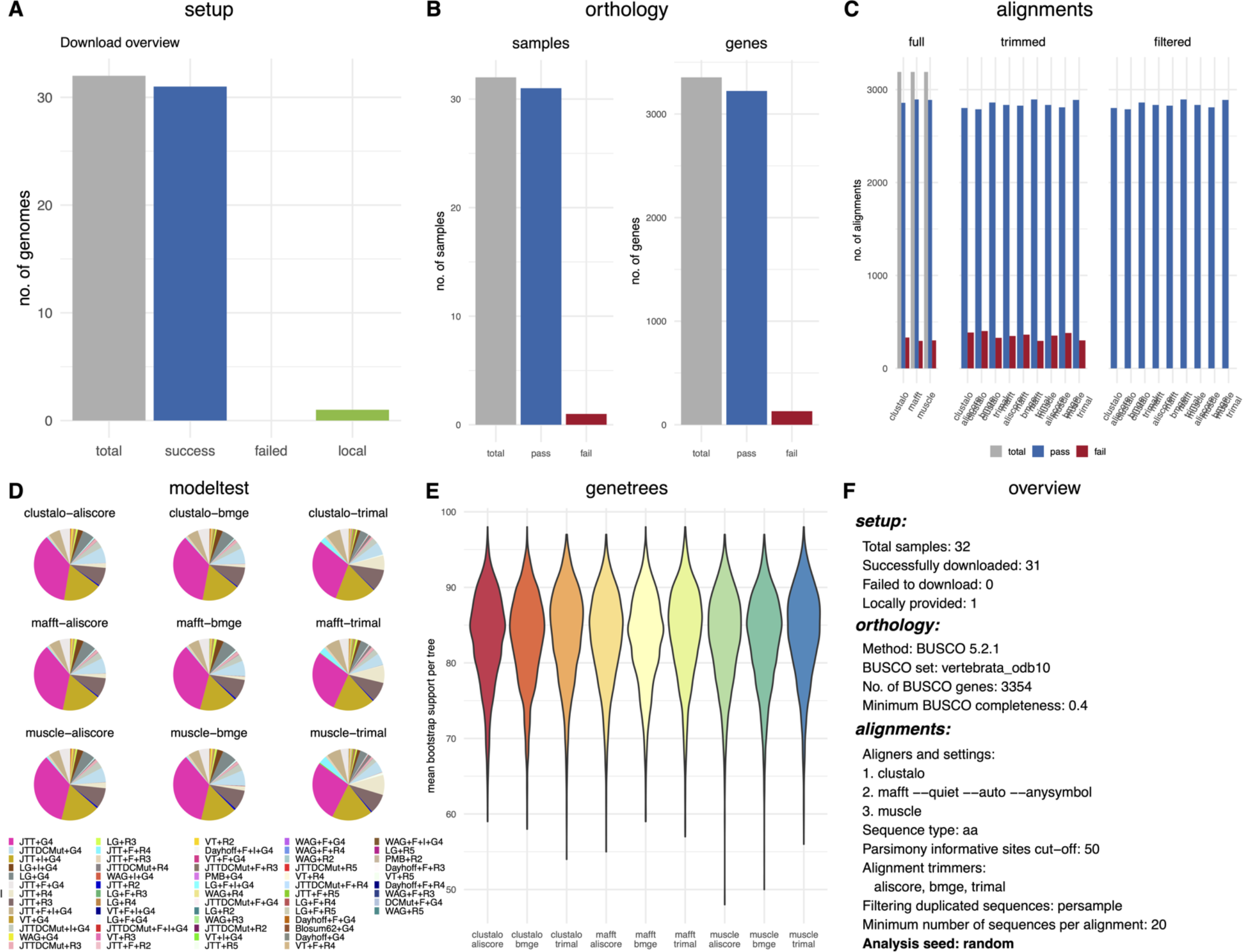
Summary plot of phylociraptor analyst - generated with `phylociraptor report --figure`

The tree topologies thus inferred are highly consistent (Figure 6). Specifically, 105 (97%) of phylogenomic trees, across all software combinations, recovered lungfishes as the closest living relative of tetrapods (Figure 7), in most cases with full statistical support, confirming the most recent results [28, 29, 30, 32]. One alternative topology was recovered by ASTRAL for three datasets processed with Clustal Omega and trimAl and bootstrap cutoffs 50, 60, and 70 (Figure 8). Specifically, in these analyses *L. chalurnnae* groups with lungfishes with reasonable support of 0.85-0.94 PP. However, due to the near unanimous support for the accepted relationships, the relatively low statistical support for the alternative grouping and that it was recovered with only one particular aligner/trimmer combination and based on relatively liberal filtering of trees, we see no reason to challenge the previous conclusions.

**Figure 6:**
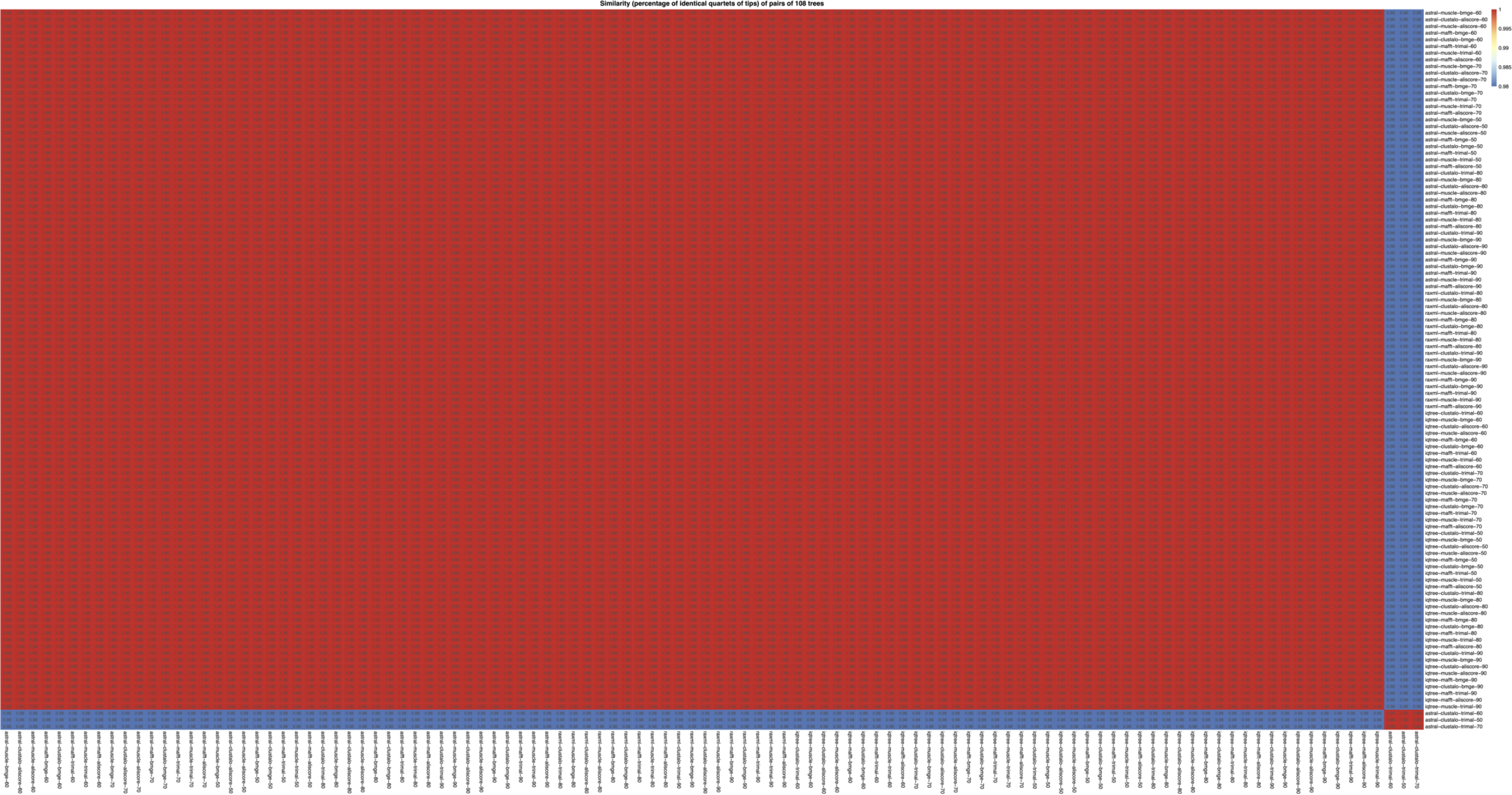
Heatmap of tree similarities across all inferred trees - generated with `phylociraptor util plot-heatmap`

**Figure 7:**
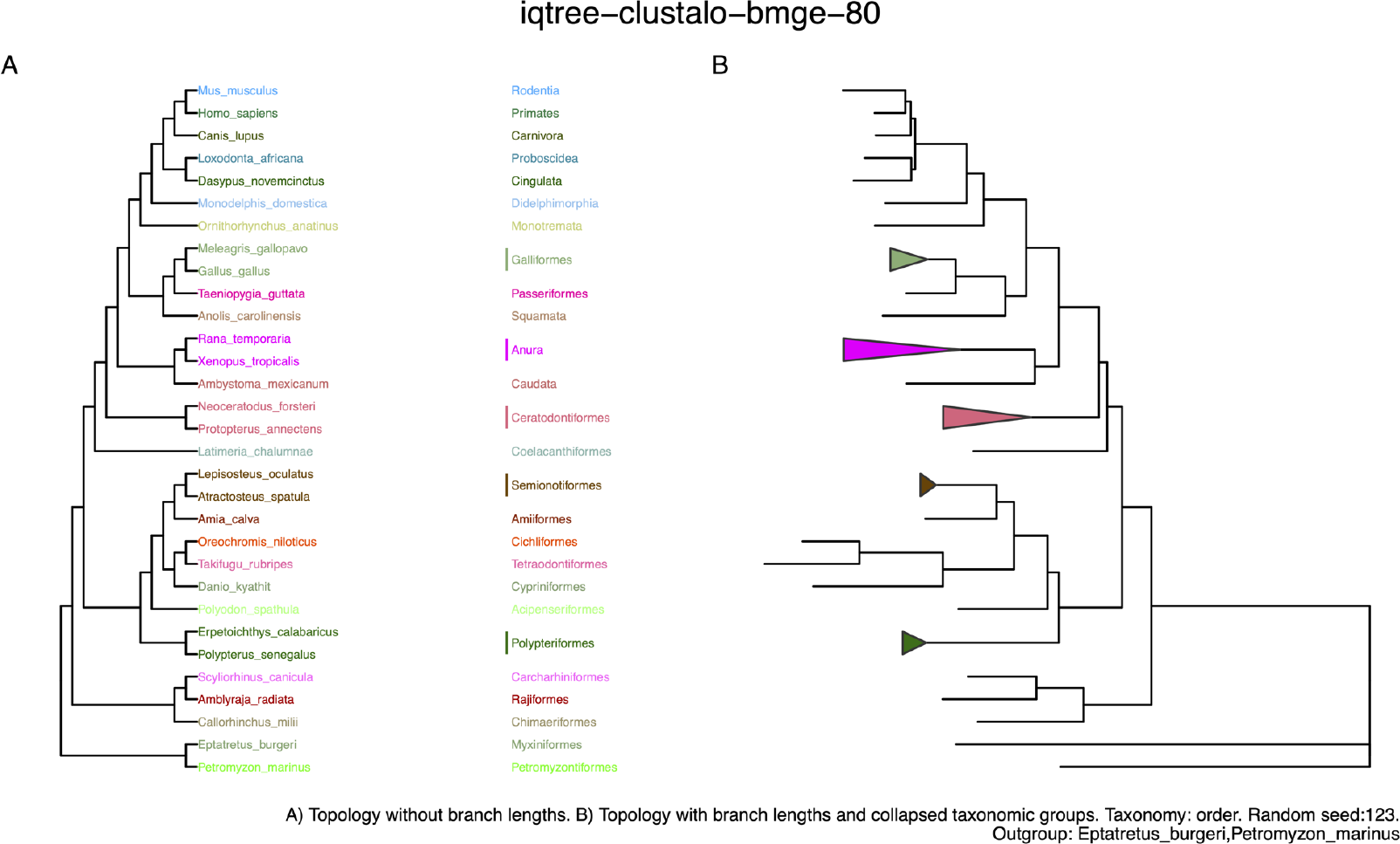
Vertebrate relationships as inferred by the majority of analyses - generated with `phylociraptor util plot-tree`

**Figure 8:**
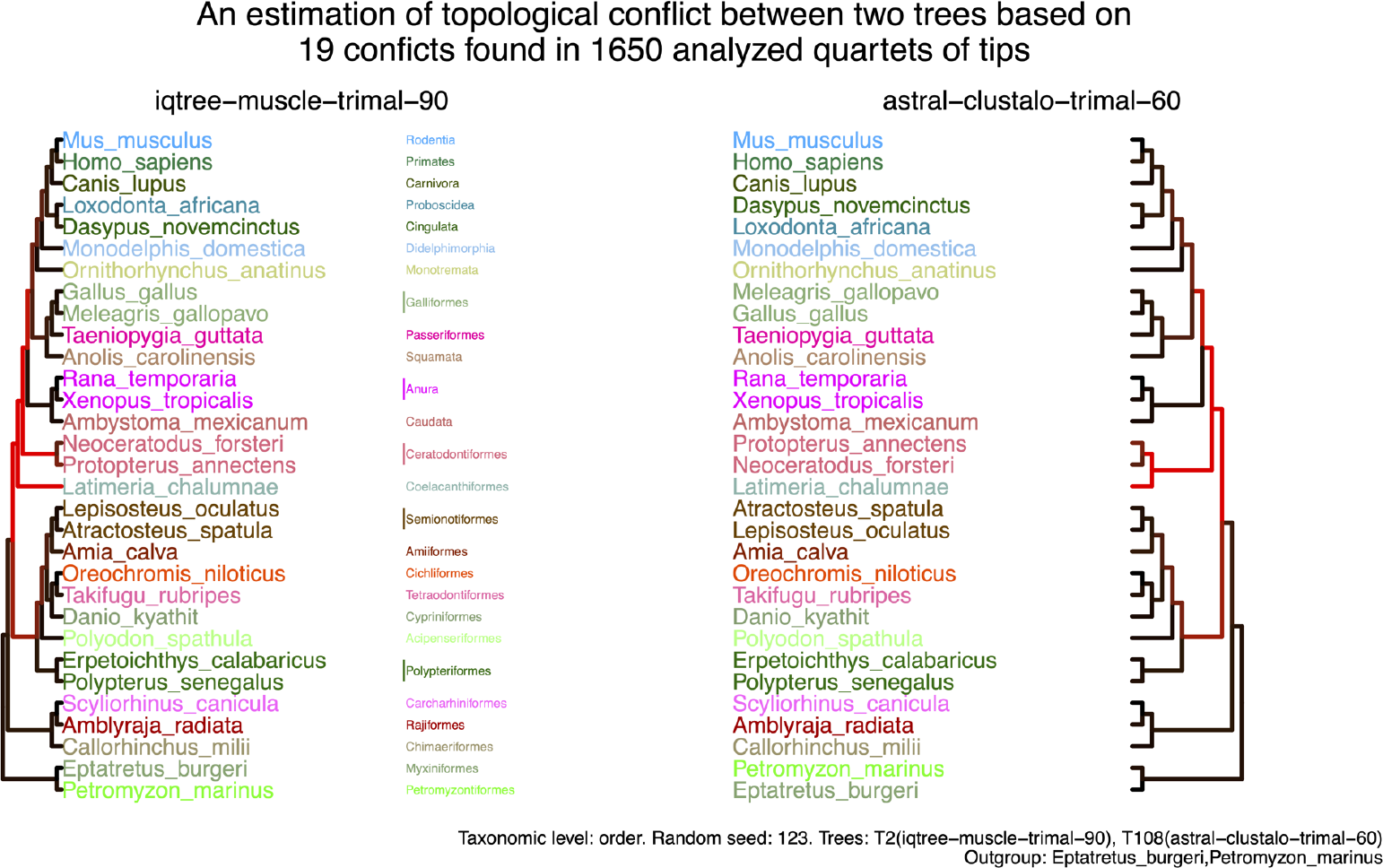
Comparison between the topology inferred by the majority of analyses and an alternative topology - generated with `phylociraptor util plot-conflict`

It does illustrate, however, that high-level choices (in this case likely one software and/or settings) without aggressive filtering can lead to spurious results in phylogenomic studies. While the alternative toplogy is not apparent in the PGA based on tip-to-tip distances of all inferred species trees, it seems that more stringent filtering of gene trees based on average bootstrap support increases the variation of tip distances (Figure 9) between species tree inferences. Whether this is a general phenomenon will need to be investigated further. Interestingly, a similar pattern, albeit less pronounced, was observed in case study 1 (Figure 3). However, datasets processed via trimAl stand out in the PCAs performed for maximum-likelihood based trees with respect to their branch lenghts in both the vertebrate (Figure 9) and the fungi test case (Figure 3). Whether this effect is caused by trimAl generally or the particular trimAl trimming scheme (`-gappyout`) chosen in our analyses remains open, but it appears independent of the multiple sequence alignment software.

**Figure 9:**
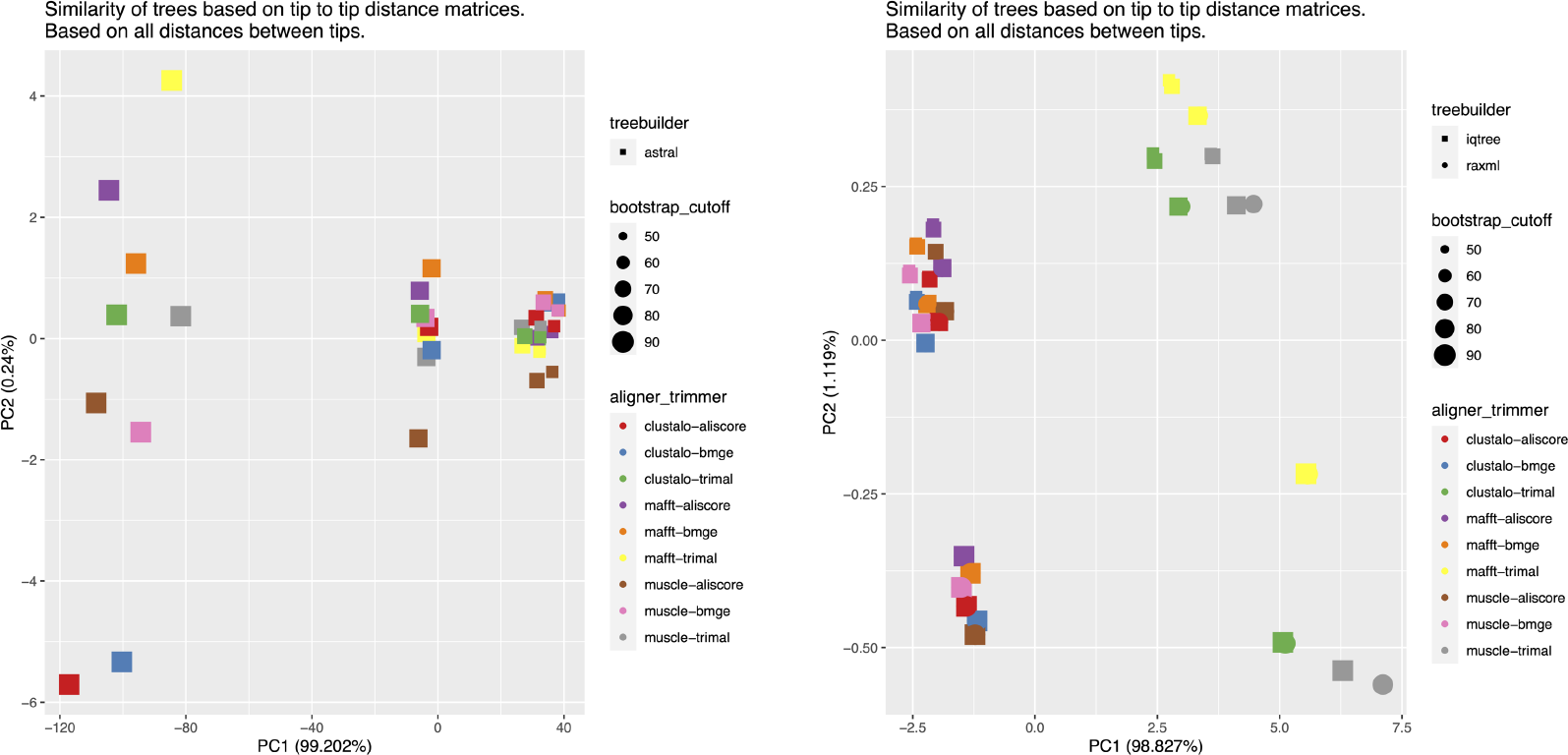
Principal component analysis based on tip2tip distances of species trees - generated with `phylociraptor util plot-pca`. Left: All trees from ASTRAL species tree reconstructions. Right: All trees from concatenated Maximum-likelihood reconstructions.

