## Supplementary material for "Phylociraptor – A unified computational framework for reproducible phylogenomic inference": Phylociraptor report as HTML (zipped) for the small test case presented in the main text (Figure 2).: small-test-report.html


### phylociraptor

**Analysis report - 18 September, 2023 - 10:51 CEST - version: 0.9.13**

### Setup

**Location of genomes files:**

```
results/assemblies
```

**Total number of included genomes:** 20  
**Locally provided genomes:** 0  
**Successfully downloaded genomes:** 20  
**Failed species downloads:** 0  
**Not downloaded:**

**Information on successfully downloaded genomes:**

### Orthology

Analysis hash for step orthology: 9562732d11  
Analysis hash for step filter-orthology: 5039029d55

Orthologous genes were inferred using BUSCO.

**Location of BUSCO results:**

```
results/orthology/busco/busco_runs/fungi_odb9-seed-43-genes-20.9562732d11
```

**Used BUSCO set:** fungi\_odb9-seed-43-genes-20

**Total number of BUSCO genes in BUSCO set:** 20  
**Number of BUSCO genes recovered in too few (3) samples:** 0  
**BUSCO genes below threshold:**  
none

**Number of samples with single-copy BUSCO score below threshold (0.4):** 1  
**Genomes below threshold:**  
Uromyces\_transversalis

**Orthology overview table:**

### Alignments

#### full alignments

**Location of alignment files:**

```
results/alignments/full
```

**Used aligner(s) and settings:**  
**1.** clustalo  
**2.** mafft –quiet –auto  
**3.** muscle  
  
**Parsimony informative sites cutoff (nparsimony):** 50  
**Relative composition variability (rcv) cutoff:** 0.2  
  
**Number of alignments for clustalo.e1f50397f9:** 20   
**Number of alignments for mafft.c40a23812e:** 20   
**Number of alignments for muscle.b39de8829a:** 20   
  
**Overview of individual full alignments:**

#### trimmed alignments

**Location of trimmed alignment files:**

```
results/alignments/trimmed
```

**Parsimony informative sites cutoff:** 50  
**Relative composition variability (rcv) cutoff:** 0.2

**Number of trimmed alignments for clustalo-trimal.77e654e392:** 20   
**Number of trimmed alignments for mafft-trimal.19dd021496:** 20   
**Number of trimmed alignments for muscle-trimal.92654633d8:** 20   
**Number of trimmed alignments for clustalo-aliscore.ff2865ad93:** 20   
**Number of trimmed alignments for mafft-aliscore.43643b3bc4:** 20   
**Number of trimmed alignments for muscle-aliscore.1097e69ca6:** 20   
**Number of trimmed alignments for clustalo-bmge.27f3d280b1:** 20   
**Number of trimmed alignments for mafft-bmge.82198b9885:** 20   
**Number of trimmed alignments for muscle-bmge.c4b681949d:** 20   
  
**Overview of individual trimmed alignments:**

#### filtered alignments

**Location of filtered alignment files:**

```
results/alignments/filtered
```

**Parsimony informative sites cutoff:** 50  
**Relative composition variability (rcv) cutoff:** 0.2

**Number of filtered alignments for clustalo-trimal.49acf35c59:** 13   
**Number of filtered alignments for mafft-trimal.cdca9f0d8d:** 14   
**Number of filtered alignments for muscle-trimal.01f79eaa17:** 14   
**Number of filtered alignments for clustalo-aliscore.adf1309ae8:** 14   
**Number of filtered alignments for mafft-aliscore.114e903deb:** 14   
**Number of filtered alignments for muscle-aliscore.a91abb08a1:** 14   
**Number of filtered alignments for clustalo-bmge.4ebad93401:** 12   
**Number of filtered alignments for mafft-bmge.2528a9aaaf:** 12   
**Number of filtered alignments for muscle-bmge.303f10a856:** 12   
  
**Overview of individual filtered alignments:**

Show additional figures

### Modeltesting

**Location of modeltesting files:**

```
results/modeltest
```

**Overview of individual modeltest results:**

### Genetrees

**Location of gene trees:**

```
results/modeltest
```

**Overview of mean bootstrap values per gene-tree:**   
 Total number of gene trees for clustalo-trimal: 13  
 Total number of gene trees for mafft-trimal: 14  
 Total number of gene trees for muscle-trimal: 14  
 Total number of gene trees for clustalo-aliscore: 14  
 Total number of gene trees for mafft-aliscore: 14  
 Total number of gene trees for muscle-aliscore: 14  
 Total number of gene trees for clustalo-bmge: 12  
 Total number of gene trees for mafft-bmge: 12  
 Total number of gene trees for muscle-bmge: 12

FALSE
FALSE

### Maximum-likelihood trees

**Random seed:** 43

**Location of phylogenetic results:**

```
results/phylogeny/
```

  
**Overview of concatenated Maximum-likelihood phylogenies:**

| software | aligner | trimmer | boostraps | bootstrap-cutoff | ngenes | tree |
| --- | --- | --- | --- | --- | --- | --- |
| iqtree | clustalo | trimal | 1000 | 0 | 13 | Copy tree |
| iqtree | mafft | trimal | 1000 | 0 | 14 | Copy tree |
| iqtree | muscle | trimal | 1000 | 0 | 14 | Copy tree |
| iqtree | clustalo | aliscore | 1000 | 0 | 14 | Copy tree |
| iqtree | mafft | aliscore | 1000 | 0 | 14 | Copy tree |
| iqtree | muscle | aliscore | 1000 | 0 | 14 | Copy tree |
| iqtree | clustalo | bmge | 1000 | 0 | 12 | Copy tree |
| iqtree | mafft | bmge | 1000 | 0 | 12 | Copy tree |
| iqtree | muscle | bmge | 1000 | 0 | 12 | Copy tree |
| raxmlng | clustalo | trimal | 100 | 0 | 13 | Copy tree |
| raxmlng | mafft | trimal | 100 | 0 | 14 | Copy tree |
| raxmlng | muscle | trimal | 100 | 0 | 14 | Copy tree |
| raxmlng | clustalo | aliscore | 100 | 0 | 14 | Copy tree |
| raxmlng | mafft | aliscore | 100 | 0 | 14 | Copy tree |
| raxmlng | muscle | aliscore | 100 | 0 | 14 | Copy tree |
| raxmlng | clustalo | bmge | 100 | 0 | 12 | Copy tree |
| raxmlng | mafft | bmge | 100 | 0 | 12 | Copy tree |
| raxmlng | muscle | bmge | 100 | 0 | 12 | Copy tree |
| iqtree | clustalo | trimal | 1000 | 50 | 13 | Copy tree |
| iqtree | mafft | trimal | 1000 | 50 | 14 | Copy tree |
| iqtree | muscle | trimal | 1000 | 50 | 14 | Copy tree |
| iqtree | clustalo | aliscore | 1000 | 50 | 14 | Copy tree |
| iqtree | mafft | aliscore | 1000 | 50 | 14 | Copy tree |
| iqtree | muscle | aliscore | 1000 | 50 | 14 | Copy tree |
| iqtree | clustalo | bmge | 1000 | 50 | 12 | Copy tree |
| iqtree | mafft | bmge | 1000 | 50 | 12 | Copy tree |
| iqtree | muscle | bmge | 1000 | 50 | 12 | Copy tree |
| raxmlng | clustalo | trimal | 100 | 50 | 13 | Copy tree |
| raxmlng | mafft | trimal | 100 | 50 | 14 | Copy tree |
| raxmlng | muscle | trimal | 100 | 50 | 14 | Copy tree |
| raxmlng | clustalo | aliscore | 100 | 50 | 14 | Copy tree |
| raxmlng | mafft | aliscore | 100 | 50 | 14 | Copy tree |
| raxmlng | muscle | aliscore | 100 | 50 | 14 | Copy tree |
| raxmlng | clustalo | bmge | 100 | 50 | 12 | Copy tree |
| raxmlng | mafft | bmge | 100 | 50 | 12 | Copy tree |
| raxmlng | muscle | bmge | 100 | 50 | 12 | Copy tree |
| iqtree | clustalo | trimal | 1000 | 60 | 13 | Copy tree |
| iqtree | mafft | trimal | 1000 | 60 | 14 | Copy tree |
| iqtree | muscle | trimal | 1000 | 60 | 14 | Copy tree |
| iqtree | clustalo | aliscore | 1000 | 60 | 14 | Copy tree |
| iqtree | mafft | aliscore | 1000 | 60 | 14 | Copy tree |
| iqtree | muscle | aliscore | 1000 | 60 | 14 | Copy tree |
| iqtree | clustalo | bmge | 1000 | 60 | 12 | Copy tree |
| iqtree | mafft | bmge | 1000 | 60 | 12 | Copy tree |
| iqtree | muscle | bmge | 1000 | 60 | 12 | Copy tree |
| raxmlng | clustalo | trimal | 100 | 60 | 13 | Copy tree |
| raxmlng | mafft | trimal | 100 | 60 | 14 | Copy tree |
| raxmlng | muscle | trimal | 100 | 60 | 14 | Copy tree |
| raxmlng | clustalo | aliscore | 100 | 60 | 14 | Copy tree |
| raxmlng | mafft | aliscore | 100 | 60 | 14 | Copy tree |
| raxmlng | muscle | aliscore | 100 | 60 | 14 | Copy tree |
| raxmlng | clustalo | bmge | 100 | 60 | 12 | Copy tree |
| raxmlng | mafft | bmge | 100 | 60 | 12 | Copy tree |
| raxmlng | muscle | bmge | 100 | 60 | 12 | Copy tree |
| iqtree | clustalo | trimal | 1000 | 70 | 13 | Copy tree |
| iqtree | mafft | trimal | 1000 | 70 | 13 | Copy tree |
| iqtree | muscle | trimal | 1000 | 70 | 14 | Copy tree |
| iqtree | clustalo | aliscore | 1000 | 70 | 14 | Copy tree |
| iqtree | mafft | aliscore | 1000 | 70 | 13 | Copy tree |
| iqtree | muscle | aliscore | 1000 | 70 | 14 | Copy tree |
| iqtree | clustalo | bmge | 1000 | 70 | 12 | Copy tree |
| iqtree | mafft | bmge | 1000 | 70 | 12 | Copy tree |
| iqtree | muscle | bmge | 1000 | 70 | 12 | Copy tree |
| raxmlng | clustalo | trimal | 100 | 70 | 13 | Copy tree |
| raxmlng | mafft | trimal | 100 | 70 | 13 | Copy tree |
| raxmlng | muscle | trimal | 100 | 70 | 14 | Copy tree |
| raxmlng | clustalo | aliscore | 100 | 70 | 14 | Copy tree |
| raxmlng | mafft | aliscore | 100 | 70 | 13 | Copy tree |
| raxmlng | muscle | aliscore | 100 | 70 | 14 | Copy tree |
| raxmlng | clustalo | bmge | 100 | 70 | 12 | Copy tree |
| raxmlng | mafft | bmge | 100 | 70 | 12 | Copy tree |
| raxmlng | muscle | bmge | 100 | 70 | 12 | Copy tree |

### Speciestree

**Location of phylogenetic results:**

```
results/phylogeny/
```

  
**Overview of species-tree phylogenies:**

| software | aligner | trimmer | bootstrap-cutoff | ngenes | tree |
| --- | --- | --- | --- | --- | --- |
| astral | clustalo | trimal | 0 | 13 | Copy tree |
| astral | mafft | trimal | 0 | 14 | Copy tree |
| astral | muscle | trimal | 0 | 14 | Copy tree |
| astral | clustalo | aliscore | 0 | 14 | Copy tree |
| astral | mafft | aliscore | 0 | 14 | Copy tree |
| astral | muscle | aliscore | 0 | 14 | Copy tree |
| astral | clustalo | bmge | 0 | 12 | Copy tree |
| astral | mafft | bmge | 0 | 12 | Copy tree |
| astral | muscle | bmge | 0 | 12 | Copy tree |
| astral | clustalo | trimal | 50 | 13 | Copy tree |
| astral | mafft | trimal | 50 | 14 | Copy tree |
| astral | muscle | trimal | 50 | 14 | Copy tree |
| astral | clustalo | aliscore | 50 | 14 | Copy tree |
| astral | mafft | aliscore | 50 | 14 | Copy tree |
| astral | muscle | aliscore | 50 | 14 | Copy tree |
| astral | clustalo | bmge | 50 | 12 | Copy tree |
| astral | mafft | bmge | 50 | 12 | Copy tree |
| astral | muscle | bmge | 50 | 12 | Copy tree |
| astral | clustalo | trimal | 60 | 13 | Copy tree |
| astral | mafft | trimal | 60 | 14 | Copy tree |
| astral | muscle | trimal | 60 | 14 | Copy tree |
| astral | clustalo | aliscore | 60 | 14 | Copy tree |
| astral | mafft | aliscore | 60 | 14 | Copy tree |
| astral | muscle | aliscore | 60 | 14 | Copy tree |
| astral | clustalo | bmge | 60 | 12 | Copy tree |
| astral | mafft | bmge | 60 | 12 | Copy tree |
| astral | muscle | bmge | 60 | 12 | Copy tree |
| astral | clustalo | trimal | 70 | 13 | Copy tree |
| astral | mafft | trimal | 70 | 13 | Copy tree |
| astral | muscle | trimal | 70 | 14 | Copy tree |
| astral | clustalo | aliscore | 70 | 14 | Copy tree |
| astral | mafft | aliscore | 70 | 13 | Copy tree |
| astral | muscle | aliscore | 70 | 14 | Copy tree |
| astral | clustalo | bmge | 70 | 12 | Copy tree |
| astral | mafft | bmge | 70 | 12 | Copy tree |
| astral | muscle | bmge | 70 | 12 | Copy tree |

### Log

```
## Fri Sep  8 14:27:16 CEST 2023 - Setup: Will download species now - batch: 2
## Fri Sep  8 14:28:12 CEST 2023 - Setup: Will download species now - batch: 3
## Fri Sep  8 14:29:01 CEST 2023 - Setup: Will download species now - batch: 1
## Fri Sep  8 14:30:05 CEST 2023 - phylociraptor setup done.
## Fri Sep  8 14:38:46 CEST 2023 Hygrocybe_conica      18 Complete       2 Duplicated       1 Fragmented
## Fri Sep  8 14:39:34 CEST 2023 Usnea_hakonensis      19 Complete       1 Missing
## Fri Sep  8 14:40:24 CEST 2023 Calocera_cornea      19 Complete       1 Missing
## Fri Sep  8 14:40:54 CEST 2023 Protomyces_gravidus      19 Complete       2 Duplicated
## Fri Sep  8 14:41:25 CEST 2023 Taphrina_betulina      19 Complete       1 Missing
## Fri Sep  8 14:43:17 CEST 2023 Uromyces_transversalis       2 Complete       5 Fragmented      13 Missing
## Fri Sep  8 14:44:05 CEST 2023 Aspergillus_nidulans      20 Complete
## Fri Sep  8 14:45:24 CEST 2023 Puccinia_graminis      15 Complete       2 Duplicated       4 Missing
## Fri Sep  8 14:46:22 CEST 2023 Bullera_alba      20 Complete
## Fri Sep  8 14:47:07 CEST 2023 Tremella_fuciformis      18 Complete       2 Fragmented
## Fri Sep  8 14:47:49 CEST 2023 Capronia_coronata      20 Complete
## Fri Sep  8 14:48:42 CEST 2023 Fistulina_hepatica      19 Complete       2 Duplicated
## Fri Sep  8 14:49:22 CEST 2023 Smittium_simulii       9 Complete       2 Duplicated      10 Missing
## Fri Sep  8 14:50:22 CEST 2023 Amanita_muscaria      19 Complete       1 Fragmented
## Fri Sep  8 14:51:59 CEST 2023 Hemileia_vastatrix       9 Complete      11 Missing
## Fri Sep  8 14:52:27 CEST 2023 Pichia_fermentans      16 Complete       1 Fragmented       3 Missing
## Fri Sep  8 14:53:20 CEST 2023 Phaeotremella_skinneri      19 Complete       1 Fragmented
## Fri Sep  8 14:54:21 CEST 2023 Mucor_racemosus      13 Complete      11 Duplicated       1 Fragmented       2 Missing
## Fri Sep  8 14:55:35 CEST 2023 Glomus_cerebriforme      14 Complete       6 Missing
## Fri Sep  8 14:56:35 CEST 2023 Umbilicaria_muehlenbergii      20 Complete
## Fri Sep  8 14:56:37 CEST 2023 - phylociraptor orthology done.
## Fri Sep  8 15:01:59 CEST 2023 - BUSCO files will be filtered on a per-sample basis. This could lower the number of species in the final tree.
## Fri Sep  8 15:02:02 CEST 2023 - Number of BUSCO sequence files: 20
## Fri Sep  8 15:02:02 CEST 2023 - Number of deduplicated BUSCO sequence files: 20
## Fri Sep  8 15:02:03 CEST 2023 - Phylociraptor filter-orthology done.
## Fri Sep  8 15:11:16 CEST 2023 - phylociraptor align done.
## Fri Sep  8 15:11:58 CEST 2023 - The aliscore output file does not exist. Check results for BUSCO: EOG092C3D5H
## Fri Sep  8 15:11:58 CEST 2023 - The aliscore output appears to be empty. Check results for BUSCO: EOG092C3D5H
## Fri Sep  8 15:12:00 CEST 2023 - The aliscore output file does not exist. Check results for BUSCO: EOG092C4JSY
## Fri Sep  8 15:12:00 CEST 2023 - The aliscore output appears to be empty. Check results for BUSCO: EOG092C4JSY
## Fri Sep  8 15:12:00 CEST 2023 - The aliscore output file does not exist. Check results for BUSCO: EOG092C0ZT0
## Fri Sep  8 15:12:00 CEST 2023 - The aliscore output appears to be empty. Check results for BUSCO: EOG092C0ZT0
## Fri Sep  8 15:12:01 CEST 2023 - The aliscore output file does not exist. Check results for BUSCO: EOG092C3WW5
## Fri Sep  8 15:12:01 CEST 2023 - The aliscore output appears to be empty. Check results for BUSCO: EOG092C3WW5
## Fri Sep  8 15:12:06 CEST 2023 - The aliscore output file does not exist. Check results for BUSCO: EOG092C46M0
## Fri Sep  8 15:12:06 CEST 2023 - The aliscore output appears to be empty. Check results for BUSCO: EOG092C46M0
## Fri Sep  8 15:12:08 CEST 2023 - The aliscore output file does not exist. Check results for BUSCO: EOG092C4SHL
## Fri Sep  8 15:12:08 CEST 2023 - The aliscore output appears to be empty. Check results for BUSCO: EOG092C4SHL
## Fri Sep  8 15:12:10 CEST 2023 - The aliscore output file does not exist. Check results for BUSCO: EOG092C0CZK
## Fri Sep  8 15:12:10 CEST 2023 - The aliscore output appears to be empty. Check results for BUSCO: EOG092C0CZK
## Fri Sep  8 15:12:12 CEST 2023 - The aliscore output file does not exist. Check results for BUSCO: EOG092C5E45
## Fri Sep  8 15:12:12 CEST 2023 - The aliscore output appears to be empty. Check results for BUSCO: EOG092C5E45
## Fri Sep  8 15:12:16 CEST 2023 - The aliscore output file does not exist. Check results for BUSCO: EOG092C03KT
## Fri Sep  8 15:12:16 CEST 2023 - The aliscore output appears to be empty. Check results for BUSCO: EOG092C03KT
## Fri Sep  8 15:12:19 CEST 2023 - The aliscore output file does not exist. Check results for BUSCO: EOG092C1PRY
## Fri Sep  8 15:12:19 CEST 2023 - The aliscore output appears to be empty. Check results for BUSCO: EOG092C1PRY
## Fri Sep  8 15:12:31 CEST 2023 - The aliscore output file does not exist. Check results for BUSCO: EOG092C27SX
## Fri Sep  8 15:12:31 CEST 2023 - The aliscore output appears to be empty. Check results for BUSCO: EOG092C27SX
## Fri Sep  8 15:12:34 CEST 2023 - The aliscore output file does not exist. Check results for BUSCO: EOG092C3P8T
## Fri Sep  8 15:12:34 CEST 2023 - The aliscore output appears to be empty. Check results for BUSCO: EOG092C3P8T
## Fri Sep  8 15:12:46 CEST 2023 - The aliscore output file does not exist. Check results for BUSCO: EOG092C48JI
## Fri Sep  8 15:12:46 CEST 2023 - The aliscore output appears to be empty. Check results for BUSCO: EOG092C48JI
## Fri Sep  8 15:12:50 CEST 2023 - The aliscore output file does not exist. Check results for BUSCO: EOG092C5JQL
## Fri Sep  8 15:12:50 CEST 2023 - The aliscore output appears to be empty. Check results for BUSCO: EOG092C5JQL
## Fri Sep  8 15:12:53 CEST 2023 - The aliscore output file does not exist. Check results for BUSCO: EOG092C3D5H
## Fri Sep  8 15:12:53 CEST 2023 - The aliscore output appears to be empty. Check results for BUSCO: EOG092C3D5H
## Fri Sep  8 15:12:54 CEST 2023 - The aliscore output file does not exist. Check results for BUSCO: EOG092C0ZT0
## Fri Sep  8 15:12:54 CEST 2023 - The aliscore output appears to be empty. Check results for BUSCO: EOG092C0ZT0
## Fri Sep  8 15:12:54 CEST 2023 - The aliscore output file does not exist. Check results for BUSCO: EOG092C0CHR
## Fri Sep  8 15:12:54 CEST 2023 - The aliscore output appears to be empty. Check results for BUSCO: EOG092C0CHR
## Fri Sep  8 15:12:56 CEST 2023 - The aliscore output file does not exist. Check results for BUSCO: EOG092C4JSY
## Fri Sep  8 15:12:56 CEST 2023 - The aliscore output appears to be empty. Check results for BUSCO: EOG092C4JSY
## Fri Sep  8 15:13:00 CEST 2023 - The aliscore output file does not exist. Check results for BUSCO: EOG092C5E45
## Fri Sep  8 15:13:00 CEST 2023 - The aliscore output appears to be empty. Check results for BUSCO: EOG092C5E45
## Fri Sep  8 15:13:00 CEST 2023 - The aliscore output file does not exist. Check results for BUSCO: EOG092C48JI
## Fri Sep  8 15:13:00 CEST 2023 - The aliscore output appears to be empty. Check results for BUSCO: EOG092C48JI
## Fri Sep  8 15:13:01 CEST 2023 - The aliscore output file does not exist. Check results for BUSCO: EOG092C08KH
## Fri Sep  8 15:13:02 CEST 2023 - The aliscore output appears to be empty. Check results for BUSCO: EOG092C08KH
## Fri Sep  8 15:13:03 CEST 2023 - The aliscore output file does not exist. Check results for BUSCO: EOG092C46M0
## Fri Sep  8 15:13:03 CEST 2023 - The aliscore output appears to be empty. Check results for BUSCO: EOG092C46M0
## Fri Sep  8 15:13:07 CEST 2023 - The aliscore output file does not exist. Check results for BUSCO: EOG092C3WW5
## Fri Sep  8 15:13:07 CEST 2023 - The aliscore output file does not exist. Check results for BUSCO: EOG092C27SX
## Fri Sep  8 15:13:07 CEST 2023 - The aliscore output appears to be empty. Check results for BUSCO: EOG092C3WW5
## Fri Sep  8 15:13:07 CEST 2023 - The aliscore output appears to be empty. Check results for BUSCO: EOG092C27SX
## Fri Sep  8 15:13:09 CEST 2023 - The aliscore output file does not exist. Check results for BUSCO: EOG092C0CHR
## Fri Sep  8 15:13:09 CEST 2023 - The aliscore output appears to be empty. Check results for BUSCO: EOG092C0CHR
## Fri Sep  8 15:13:13 CEST 2023 - The aliscore output file does not exist. Check results for BUSCO: EOG092C03KT
## Fri Sep  8 15:13:13 CEST 2023 - The aliscore output appears to be empty. Check results for BUSCO: EOG092C03KT
## Fri Sep  8 15:13:14 CEST 2023 - The aliscore output file does not exist. Check results for BUSCO: EOG092C48JI
## Fri Sep  8 15:13:14 CEST 2023 - The aliscore output appears to be empty. Check results for BUSCO: EOG092C48JI
## Fri Sep  8 15:13:14 CEST 2023 - The aliscore output file does not exist. Check results for BUSCO: EOG092C4PJG
## Fri Sep  8 15:13:14 CEST 2023 - The aliscore output appears to be empty. Check results for BUSCO: EOG092C4PJG
## Fri Sep  8 15:13:16 CEST 2023 - The aliscore output file does not exist. Check results for BUSCO: EOG092C5E45
## Fri Sep  8 15:13:16 CEST 2023 - The aliscore output appears to be empty. Check results for BUSCO: EOG092C5E45
## Fri Sep  8 15:13:19 CEST 2023 - The aliscore output file does not exist. Check results for BUSCO: EOG092C0ZT0
## Fri Sep  8 15:13:19 CEST 2023 - The aliscore output appears to be empty. Check results for BUSCO: EOG092C0ZT0
## Fri Sep  8 15:13:21 CEST 2023 - The aliscore output file does not exist. Check results for BUSCO: EOG092C4JSY
## Fri Sep  8 15:13:21 CEST 2023 - The aliscore output appears to be empty. Check results for BUSCO: EOG092C4JSY
## Fri Sep  8 15:13:21 CEST 2023 - The aliscore output file does not exist. Check results for BUSCO: EOG092C0CZK
## Fri Sep  8 15:13:21 CEST 2023 - The aliscore output appears to be empty. Check results for BUSCO: EOG092C0CZK
## Fri Sep  8 15:13:22 CEST 2023 - The aliscore output file does not exist. Check results for BUSCO: EOG092C46M0
## Fri Sep  8 15:13:23 CEST 2023 - The aliscore output appears to be empty. Check results for BUSCO: EOG092C46M0
## Fri Sep  8 15:13:25 CEST 2023 - The aliscore output file does not exist. Check results for BUSCO: EOG092C3D5H
## Fri Sep  8 15:13:25 CEST 2023 - The aliscore output appears to be empty. Check results for BUSCO: EOG092C3D5H
## Fri Sep  8 15:13:27 CEST 2023 - The aliscore output file does not exist. Check results for BUSCO: EOG092C3WW5
## Fri Sep  8 15:13:27 CEST 2023 - The aliscore output appears to be empty. Check results for BUSCO: EOG092C3WW5
## Fri Sep  8 15:13:27 CEST 2023 - The aliscore output file does not exist. Check results for BUSCO: EOG092C27SX
## Fri Sep  8 15:13:27 CEST 2023 - The aliscore output appears to be empty. Check results for BUSCO: EOG092C27SX
## Fri Sep  8 15:13:30 CEST 2023 - The aliscore output file does not exist. Check results for BUSCO: EOG092C08KH
## Fri Sep  8 15:13:30 CEST 2023 - The aliscore output appears to be empty. Check results for BUSCO: EOG092C08KH
## Fri Sep  8 15:13:33 CEST 2023 - The aliscore output file does not exist. Check results for BUSCO: EOG092C5DTV
## Fri Sep  8 15:13:33 CEST 2023 - The aliscore output appears to be empty. Check results for BUSCO: EOG092C5DTV
## Fri Sep  8 15:13:33 CEST 2023 - The aliscore output file does not exist. Check results for BUSCO: EOG092C5G2T
## Fri Sep  8 15:13:33 CEST 2023 - The aliscore output appears to be empty. Check results for BUSCO: EOG092C5G2T
## Fri Sep  8 15:13:34 CEST 2023 - The aliscore output file does not exist. Check results for BUSCO: EOG092C03KT
## Fri Sep  8 15:13:34 CEST 2023 - The aliscore output appears to be empty. Check results for BUSCO: EOG092C03KT
## Fri Sep  8 15:13:36 CEST 2023 - The aliscore output file does not exist. Check results for BUSCO: EOG092C5G2T
## Fri Sep  8 15:13:36 CEST 2023 - The aliscore output appears to be empty. Check results for BUSCO: EOG092C5G2T
## Fri Sep  8 15:13:40 CEST 2023 - The aliscore output file does not exist. Check results for BUSCO: EOG092C4SHL
## Fri Sep  8 15:13:41 CEST 2023 - The aliscore output appears to be empty. Check results for BUSCO: EOG092C4SHL
## Fri Sep  8 15:13:41 CEST 2023 - The aliscore output file does not exist. Check results for BUSCO: EOG092C0CZK
## Fri Sep  8 15:13:41 CEST 2023 - The aliscore output appears to be empty. Check results for BUSCO: EOG092C0CZK
## Fri Sep  8 15:13:43 CEST 2023 - The aliscore output file does not exist. Check results for BUSCO: EOG092C5DTV
## Fri Sep  8 15:13:43 CEST 2023 - The aliscore output appears to be empty. Check results for BUSCO: EOG092C5DTV
## Fri Sep  8 15:13:43 CEST 2023 - The aliscore output file does not exist. Check results for BUSCO: EOG092C1PRY
## Fri Sep  8 15:13:43 CEST 2023 - The aliscore output appears to be empty. Check results for BUSCO: EOG092C1PRY
## Fri Sep  8 15:13:46 CEST 2023 - The aliscore output file does not exist. Check results for BUSCO: EOG092C5DTV
## Fri Sep  8 15:13:46 CEST 2023 - The aliscore output appears to be empty. Check results for BUSCO: EOG092C5DTV
## Fri Sep  8 15:13:48 CEST 2023 - The aliscore output file does not exist. Check results for BUSCO: EOG092C1PRY
## Fri Sep  8 15:13:48 CEST 2023 - The aliscore output appears to be empty. Check results for BUSCO: EOG092C1PRY
## Fri Sep  8 15:13:49 CEST 2023 - The aliscore output file does not exist. Check results for BUSCO: EOG092C3P8T
## Fri Sep  8 15:13:49 CEST 2023 - The aliscore output appears to be empty. Check results for BUSCO: EOG092C3P8T
## Fri Sep  8 15:13:49 CEST 2023 - The aliscore output file does not exist. Check results for BUSCO: EOG092C3P8T
## Fri Sep  8 15:13:49 CEST 2023 - The aliscore output appears to be empty. Check results for BUSCO: EOG092C3P8T
## Fri Sep  8 15:13:54 CEST 2023 - The aliscore output file does not exist. Check results for BUSCO: EOG092C4PJG
## Fri Sep  8 15:13:54 CEST 2023 - The aliscore output appears to be empty. Check results for BUSCO: EOG092C4PJG
## Fri Sep  8 15:13:57 CEST 2023 - The aliscore output file does not exist. Check results for BUSCO: EOG092C0CHR
## Fri Sep  8 15:13:57 CEST 2023 - The aliscore output appears to be empty. Check results for BUSCO: EOG092C0CHR
## Fri Sep  8 15:13:59 CEST 2023 - The aliscore output file does not exist. Check results for BUSCO: EOG092C3VJP
## Fri Sep  8 15:13:59 CEST 2023 - The aliscore output appears to be empty. Check results for BUSCO: EOG092C3VJP
## Fri Sep  8 15:14:05 CEST 2023 - The aliscore output file does not exist. Check results for BUSCO: EOG092C4SHL
## Fri Sep  8 15:14:05 CEST 2023 - The aliscore output appears to be empty. Check results for BUSCO: EOG092C4SHL
## Fri Sep  8 15:14:06 CEST 2023 - The aliscore output file does not exist. Check results for BUSCO: EOG092C5JQL
## Fri Sep  8 15:14:06 CEST 2023 - The aliscore output appears to be empty. Check results for BUSCO: EOG092C5JQL
## Fri Sep  8 15:14:08 CEST 2023 - The aliscore output file does not exist. Check results for BUSCO: EOG092C5JQL
## Fri Sep  8 15:14:08 CEST 2023 - The aliscore output appears to be empty. Check results for BUSCO: EOG092C5JQL
## Fri Sep  8 15:14:09 CEST 2023 - The aliscore output file does not exist. Check results for BUSCO: EOG092C3VJP
## Fri Sep  8 15:14:09 CEST 2023 - The aliscore output appears to be empty. Check results for BUSCO: EOG092C3VJP
## Fri Sep  8 15:14:11 CEST 2023 - The aliscore output file does not exist. Check results for BUSCO: EOG092C3VJP
## Fri Sep  8 15:14:11 CEST 2023 - The aliscore output appears to be empty. Check results for BUSCO: EOG092C3VJP
## Fri Sep  8 15:14:15 CEST 2023 - The aliscore output file does not exist. Check results for BUSCO: EOG092C08KH
## Fri Sep  8 15:14:15 CEST 2023 - The aliscore output appears to be empty. Check results for BUSCO: EOG092C08KH
## Fri Sep  8 15:14:15 CEST 2023 - The aliscore output file does not exist. Check results for BUSCO: EOG092C5G2T
## Fri Sep  8 15:14:15 CEST 2023 - The aliscore output appears to be empty. Check results for BUSCO: EOG092C5G2T
## Fri Sep  8 15:14:35 CEST 2023 - The aliscore output file does not exist. Check results for BUSCO: EOG092C4PJG
## Fri Sep  8 15:14:35 CEST 2023 - The aliscore output appears to be empty. Check results for BUSCO: EOG092C4PJG
## Fri Sep  8 15:14:37 CEST 2023 - Number of alignments (clustalo): 20
## Fri Sep  8 15:14:37 CEST 2023 - Number of trimmed alignments (clustalo - bmge): 0
## Fri Sep  8 15:14:37 CEST 2023 - Number of alignments (clustalo - bmge) after filtering: 20
## Fri Sep  8 15:14:43 CEST 2023 - Number of alignments (mafft): 20
## Fri Sep  8 15:14:43 CEST 2023 - Number of trimmed alignments (mafft - bmge): 0
## Fri Sep  8 15:14:43 CEST 2023 - Number of alignments (mafft - bmge) after filtering: 20
## Fri Sep  8 15:14:45 CEST 2023 - Number of alignments (clustalo): 20
## Fri Sep  8 15:14:45 CEST 2023 - Number of trimmed alignments (clustalo - aliscore): 0
## Fri Sep  8 15:14:45 CEST 2023 - Number of alignments (clustalo - aliscore) after filtering: 20
## Fri Sep  8 15:14:45 CEST 2023 - Number of alignments (mafft): 20
## Fri Sep  8 15:14:45 CEST 2023 - Number of trimmed alignments (mafft - aliscore): 0
## Fri Sep  8 15:14:45 CEST 2023 - Number of alignments (mafft - aliscore) after filtering: 20
## Fri Sep  8 15:14:45 CEST 2023 - Number of alignments (muscle): 20
## Fri Sep  8 15:14:45 CEST 2023 - Number of trimmed alignments (muscle - bmge): 0
## Fri Sep  8 15:14:45 CEST 2023 - Number of alignments (muscle - bmge) after filtering: 20
## Fri Sep  8 15:14:46 CEST 2023 - Number of alignments (clustalo): 20
## Fri Sep  8 15:14:46 CEST 2023 - Number of trimmed alignments (clustalo - trimal): 0
## Fri Sep  8 15:14:46 CEST 2023 - Number of alignments (clustalo - trimal) after filtering: 20
## Fri Sep  8 15:15:03 CEST 2023 - Number of alignments (muscle): 20
## Fri Sep  8 15:15:03 CEST 2023 - Number of trimmed alignments (muscle - aliscore): 0
## Fri Sep  8 15:15:03 CEST 2023 - Number of alignments (muscle - aliscore) after filtering: 20
## Fri Sep  8 15:15:06 CEST 2023 - Number of alignments (mafft): 20
## Fri Sep  8 15:15:06 CEST 2023 - Number of trimmed alignments (mafft - trimal): 0
## Fri Sep  8 15:15:06 CEST 2023 - Number of alignments (mafft - trimal) after filtering: 20
## Fri Sep  8 15:15:15 CEST 2023 - Number of alignments (muscle): 20
## Fri Sep  8 15:15:15 CEST 2023 - Number of trimmed alignments (muscle - trimal): 0
## Fri Sep  8 15:15:15 CEST 2023 - Number of alignments (muscle - trimal) after filtering: 20
## Fri Sep  8 15:15:23 CEST 2023 - phylociraptor filter-align done.
## Fri Sep  8 15:48:15 CEST 2023 - Number of alignments (muscle): 20
## Fri Sep  8 15:48:15 CEST 2023 - Number of trimmed alignments (muscle - trimal): 0
## Fri Sep  8 15:48:15 CEST 2023 - Number of alignments (muscle - trimal) after filtering: 14
## Fri Sep  8 15:48:15 CEST 2023 - Number of alignments (clustalo): 20
## Fri Sep  8 15:48:15 CEST 2023 - Number of trimmed alignments (clustalo - aliscore): 0
## Fri Sep  8 15:48:15 CEST 2023 - Number of alignments (clustalo - aliscore) after filtering: 14
## Fri Sep  8 15:48:15 CEST 2023 - Number of alignments (mafft): 20
## Fri Sep  8 15:48:15 CEST 2023 - Number of alignments (mafft): 20
## Fri Sep  8 15:48:15 CEST 2023 - Number of trimmed alignments (mafft - bmge): 0
## Fri Sep  8 15:48:15 CEST 2023 - Number of trimmed alignments (mafft - trimal): 0
## Fri Sep  8 15:48:15 CEST 2023 - Number of alignments (mafft - bmge) after filtering: 12
## Fri Sep  8 15:48:15 CEST 2023 - Number of alignments (mafft - trimal) after filtering: 14
## Fri Sep  8 15:48:16 CEST 2023 - Number of alignments (muscle): 20
## Fri Sep  8 15:48:16 CEST 2023 - Number of trimmed alignments (muscle - aliscore): 0
## Fri Sep  8 15:48:16 CEST 2023 - Number of alignments (muscle - aliscore) after filtering: 14
## Fri Sep  8 15:48:21 CEST 2023 - Number of alignments (clustalo): 20
## Fri Sep  8 15:48:21 CEST 2023 - Number of trimmed alignments (clustalo - bmge): 0
## Fri Sep  8 15:48:21 CEST 2023 - Number of alignments (clustalo - bmge) after filtering: 12
## Fri Sep  8 15:48:27 CEST 2023 - Number of alignments (mafft): 20
## Fri Sep  8 15:48:27 CEST 2023 - Number of trimmed alignments (mafft - aliscore): 0
## Fri Sep  8 15:48:27 CEST 2023 - Number of alignments (mafft - aliscore) after filtering: 14
## Fri Sep  8 15:48:33 CEST 2023 - Number of alignments (muscle): 20
## Fri Sep  8 15:48:33 CEST 2023 - Number of trimmed alignments (muscle - bmge): 0
## Fri Sep  8 15:48:33 CEST 2023 - Number of alignments (muscle - bmge) after filtering: 12
## Fri Sep  8 15:48:38 CEST 2023 - Number of alignments (clustalo): 20
## Fri Sep  8 15:48:38 CEST 2023 - Number of trimmed alignments (clustalo - trimal): 0
## Fri Sep  8 15:48:38 CEST 2023 - Number of alignments (clustalo - trimal) after filtering: 13
## Fri Sep  8 15:48:47 CEST 2023 - phylociraptor filter-align done.
## Fri Sep  8 16:14:03 CEST 2023 - phylociraptor modeltest done.
## Fri Sep  8 16:18:29 CEST 2023 - phylociraptor will use gene tree filtering based on average bootstrap support value of 70.
## Fri Sep  8 16:18:30 CEST 2023 - phylociraptor will use gene tree filtering based on average bootstrap support value of 0.
## Fri Sep  8 16:18:30 CEST 2023 - phylociraptor will use gene tree filtering based on average bootstrap support value of 50.
## Fri Sep  8 16:18:30 CEST 2023 - phylociraptor will use gene tree filtering based on average bootstrap support value of 60.
## Fri Sep  8 16:18:36 CEST 2023 - phylociraptor will use gene tree filtering based on average bootstrap support value of 60.
## Fri Sep  8 16:18:36 CEST 2023 - phylociraptor will use gene tree filtering based on average bootstrap support value of 60.
## Fri Sep  8 16:18:37 CEST 2023 - phylociraptor will use gene tree filtering based on average bootstrap support value of 50.
## Fri Sep  8 16:18:37 CEST 2023 - phylociraptor will use gene tree filtering based on average bootstrap support value of 70.
## Fri Sep  8 16:18:37 CEST 2023 - phylociraptor will use gene tree filtering based on average bootstrap support value of 70.
## Fri Sep  8 16:18:39 CEST 2023 - phylociraptor will use gene tree filtering based on average bootstrap support value of 50.
## Fri Sep  8 16:18:39 CEST 2023 - phylociraptor will use gene tree filtering based on average bootstrap support value of 0.
## Fri Sep  8 16:18:39 CEST 2023 - phylociraptor will use gene tree filtering based on average bootstrap support value of 50.
## Fri Sep  8 16:18:40 CEST 2023 - phylociraptor will use gene tree filtering based on average bootstrap support value of 0.
## Fri Sep  8 16:18:46 CEST 2023 - phylociraptor will use gene tree filtering based on average bootstrap support value of 0.
## Fri Sep  8 16:18:46 CEST 2023 - phylociraptor will use gene tree filtering based on average bootstrap support value of 70.
## Fri Sep  8 16:18:47 CEST 2023 - phylociraptor will use gene tree filtering based on average bootstrap support value of 70.
## Fri Sep  8 16:18:47 CEST 2023 - phylociraptor will use gene tree filtering based on average bootstrap support value of 0.
## Fri Sep  8 16:18:47 CEST 2023 - phylociraptor will use gene tree filtering based on average bootstrap support value of 60.
## Fri Sep  8 16:18:48 CEST 2023 - phylociraptor will use gene tree filtering based on average bootstrap support value of 50.
## Fri Sep  8 16:18:49 CEST 2023 - phylociraptor will use gene tree filtering based on average bootstrap support value of 50.
## Fri Sep  8 16:18:49 CEST 2023 - phylociraptor will use gene tree filtering based on average bootstrap support value of 60.
## Fri Sep  8 16:18:49 CEST 2023 - phylociraptor will use gene tree filtering based on average bootstrap support value of 70.
## Fri Sep  8 16:18:59 CEST 2023 - phylociraptor will use gene tree filtering based on average bootstrap support value of 70.
## Fri Sep  8 16:19:00 CEST 2023 - phylociraptor will use gene tree filtering based on average bootstrap support value of 60.
## Fri Sep  8 16:19:00 CEST 2023 - phylociraptor will use gene tree filtering based on average bootstrap support value of 0.
## Fri Sep  8 16:19:01 CEST 2023 - phylociraptor will use gene tree filtering based on average bootstrap support value of 50.
## Fri Sep  8 16:19:02 CEST 2023 - phylociraptor will use gene tree filtering based on average bootstrap support value of 70.
## Fri Sep  8 16:19:03 CEST 2023 - phylociraptor will use gene tree filtering based on average bootstrap support value of 0.
## Fri Sep  8 16:19:04 CEST 2023 - phylociraptor will use gene tree filtering based on average bootstrap support value of 0.
## Fri Sep  8 16:19:05 CEST 2023 - phylociraptor will use gene tree filtering based on average bootstrap support value of 50.
## Fri Sep  8 16:19:07 CEST 2023 - phylociraptor will use gene tree filtering based on average bootstrap support value of 60.
## Fri Sep  8 16:19:08 CEST 2023 - phylociraptor will use gene tree filtering based on average bootstrap support value of 0.
## Fri Sep  8 16:19:09 CEST 2023 - phylociraptor will use gene tree filtering based on average bootstrap support value of 70.
## Fri Sep  8 16:19:10 CEST 2023 - phylociraptor will use gene tree filtering based on average bootstrap support value of 50.
## Fri Sep  8 16:19:13 CEST 2023 - phylociraptor will use gene tree filtering based on average bootstrap support value of 60.
## Fri Sep  8 16:19:14 CEST 2023 - phylociraptor will use gene tree filtering based on average bootstrap support value of 60.
## Fri Sep  8 16:19:15 CEST 2023 - phylociraptor speciestree reconstruction done.
## Fri Sep  8 16:23:24 CEST 2023 - prepare_iqtree clustalo-trimal: Will use bootstrap cutoff (50) before creating concatenated alignment
## Fri Sep  8 16:23:24 CEST 2023 - prepare_iqtree muscle-aliscore: Will use bootstrap cutoff (70) before creating concatenated alignment
## Will create NEXUS partition file with model information now.
## Fri Sep  8 16:23:24 CEST 2023 - prepare_iqtree mafft-trimal: Will use bootstrap cutoff (60) before creating concatenated alignment
## Fri Sep  8 16:23:24 CEST 2023 - concatenate muscle-trimal: Will use bootstrap cutoff 60 before creating concatenated alignment
## Fri Sep  8 16:23:24 CEST 2023 - concatenate clustalo-trimal: Will use bootstrap cutoff 0 before creating concatenated alignment
## Fri Sep  8 16:23:24 CEST 2023 - concatenate muscle-bmge: Will use bootstrap cutoff 70 before creating concatenated alignment
## Fri Sep  8 16:23:24 CEST 2023 - concatenate clustalo-bmge: Will use bootstrap cutoff 70 before creating concatenated alignment
## Fri Sep  8 16:23:24 CEST 2023 - concatenate mafft-trimal: Will use bootstrap cutoff 50 before creating concatenated alignment
## Will create NEXUS partition file with model information now.
## Fri Sep  8 16:23:24 CEST 2023 - prepare_iqtree mafft-bmge: Will use bootstrap cutoff (50) before creating concatenated alignment
## Fri Sep  8 16:23:24 CEST 2023 - prepare_iqtree muscle-trimal: Will use bootstrap cutoff (60) before creating concatenated alignment
## Fri Sep  8 16:23:24 CEST 2023 - nexus file for iqtree written.
## Fri Sep  8 16:23:24 CEST 2023 - prepare_iqtree clustalo-bmge: Will use bootstrap cutoff (70) before creating concatenated alignment
## Will create NEXUS partition file with model information now.
## Will create NEXUS partition file with model information now.
## Fri Sep  8 16:23:24 CEST 2023 - concatenate clustalo-trimal: Will use bootstrap cutoff 70 before creating concatenated alignment
## Fri Sep  8 16:23:24 CEST 2023 - concatenate muscle-bmge: Will use bootstrap cutoff 50 before creating concatenated alignment
## Will create NEXUS partition file with model information now.
## Will create NEXUS partition file with model information now.
## Fri Sep  8 16:23:24 CEST 2023 - nexus file for iqtree written.
## Fri Sep  8 16:23:24 CEST 2023 - prepare_iqtree clustalo-trimal: Will use bootstrap cutoff (0) before creating concatenated alignment
## Fri Sep  8 16:23:24 CEST 2023 - concatenate mafft-trimal: Will use bootstrap cutoff 60 before creating concatenated alignment
## Fri Sep  8 16:23:24 CEST 2023 - concatenate muscle-trimal: Will use bootstrap cutoff 70 before creating concatenated alignment
## Fri Sep  8 16:23:24 CEST 2023 - prepare_iqtree clustalo-aliscore: Will use bootstrap cutoff (50) before creating concatenated alignment
## Fri Sep  8 16:23:25 CEST 2023 - nexus file for iqtree written.
## Fri Sep  8 16:23:25 CEST 2023 - prepare_iqtree muscle-aliscore: Will use bootstrap cutoff (60) before creating concatenated alignment
## Fri Sep  8 16:23:25 CEST 2023 - prepare_iqtree mafft-trimal: Will use bootstrap cutoff (70) before creating concatenated alignment
## Fri Sep  8 16:23:25 CEST 2023 - nexus file for iqtree written.
## Fri Sep  8 16:23:25 CEST 2023 - prepare_iqtree muscle-trimal: Will use bootstrap cutoff (50) before creating concatenated alignment
## Will create NEXUS partition file with model information now.
## Fri Sep  8 16:23:25 CEST 2023 - nexus file for iqtree written.
## Fri Sep  8 16:23:25 CEST 2023 - nexus file for iqtree written.
## Will create NEXUS partition file with model information now.
## Will create NEXUS partition file with model information now.
## Will create NEXUS partition file with model information now.
## Will create NEXUS partition file with model information now.
## Fri Sep  8 16:23:25 CEST 2023 - prepare_iqtree mafft-bmge: Will use bootstrap cutoff (70) before creating concatenated alignment
## Fri Sep  8 16:23:25 CEST 2023 - concatenate clustalo-aliscore: Will use bootstrap cutoff 70 before creating concatenated alignment
## Fri Sep  8 16:23:25 CEST 2023 - prepare_iqtree muscle-bmge: Will use bootstrap cutoff (70) before creating concatenated alignment
## Fri Sep  8 16:23:25 CEST 2023 - prepare_iqtree mafft-bmge: Will use bootstrap cutoff (0) before creating concatenated alignment
## Fri Sep  8 16:23:25 CEST 2023 - nexus file for iqtree written.
## Fri Sep  8 16:23:25 CEST 2023 - concatenate mafft-bmge: Will use bootstrap cutoff 0 before creating concatenated alignment
## Fri Sep  8 16:23:25 CEST 2023 - concatenate mafft-aliscore: Will use bootstrap cutoff 60 before creating concatenated alignment
## Will create NEXUS partition file with model information now.
## Will create NEXUS partition file with model information now.
## Fri Sep  8 16:23:25 CEST 2023 - prepare_iqtree muscle-aliscore: Will use bootstrap cutoff (0) before creating concatenated alignment
## Fri Sep  8 16:23:25 CEST 2023 - concatenate muscle-trimal: Will use bootstrap cutoff 0 before creating concatenated alignment
## Fri Sep  8 16:23:25 CEST 2023 - nexus file for iqtree written.
## Fri Sep  8 16:23:25 CEST 2023 - nexus file for iqtree written.
## Will create NEXUS partition file with model information now.
## Fri Sep  8 16:23:25 CEST 2023 - nexus file for iqtree written.
## Fri Sep  8 16:23:25 CEST 2023 - nexus file for iqtree written.
## Will create NEXUS partition file with model information now.
## Fri Sep  8 16:23:25 CEST 2023 - nexus file for iqtree written.
## Fri Sep  8 16:23:25 CEST 2023 - nexus file for iqtree written.
## Fri Sep  8 16:23:25 CEST 2023 - nexus file for iqtree written.
## Fri Sep  8 16:23:26 CEST 2023 - nexus file for iqtree written.
## Fri Sep  8 16:23:26 CEST 2023 - prepare_iqtree mafft-trimal: Will use bootstrap cutoff (50) before creating concatenated alignment
## Fri Sep  8 16:23:26 CEST 2023 - prepare_iqtree clustalo-trimal: Will use bootstrap cutoff (70) before creating concatenated alignment
## Fri Sep  8 16:23:26 CEST 2023 - concatenate mafft-bmge: Will use bootstrap cutoff 60 before creating concatenated alignment
## Fri Sep  8 16:23:26 CEST 2023 - prepare_iqtree clustalo-bmge: Will use bootstrap cutoff (0) before creating concatenated alignment
## Will create NEXUS partition file with model information now.
## Will create NEXUS partition file with model information now.
## Will create NEXUS partition file with model information now.
## Fri Sep  8 16:23:27 CEST 2023 - nexus file for iqtree written.
## Fri Sep  8 16:23:27 CEST 2023 - prepare_iqtree muscle-bmge: Will use bootstrap cutoff (50) before creating concatenated alignment
## Fri Sep  8 16:23:27 CEST 2023 - prepare_iqtree mafft-aliscore: Will use bootstrap cutoff (60) before creating concatenated alignment
## Fri Sep  8 16:23:27 CEST 2023 - nexus file for iqtree written.
## Fri Sep  8 16:23:27 CEST 2023 - nexus file for iqtree written.
## Will create NEXUS partition file with model information now.
## Will create NEXUS partition file with model information now.
## Fri Sep  8 16:23:28 CEST 2023 - nexus file for iqtree written.
## Fri Sep  8 16:23:28 CEST 2023 - nexus file for iqtree written.
## Fri Sep  8 16:23:39 CEST 2023 - concatenate mafft-aliscore: Will use bootstrap cutoff 0 before creating concatenated alignment
## Fri Sep  8 16:23:39 CEST 2023 - concatenate clustalo-trimal: Will use bootstrap cutoff 50 before creating concatenated alignment
## Fri Sep  8 16:23:50 CEST 2023 - 'phylociraptor modeltest' finished successfully before. Will run raxml with best models.
## Fri Sep  8 16:23:50 CEST 2023 - 'phylociraptor modeltest' finished successfully before. Will run raxml with best models.
## Fri Sep  8 16:23:50 CEST 2023 - 'phylociraptor modeltest' finished successfully before. Will run raxml with best models.
## Fri Sep  8 16:23:50 CEST 2023 - 'phylociraptor modeltest' finished successfully before. Will run raxml with best models.
## Fri Sep  8 16:23:50 CEST 2023 - 'phylociraptor modeltest' finished successfully before. Will run raxml with best models.
## Fri Sep  8 16:23:50 CEST 2023 - 'phylociraptor modeltest' finished successfully before. Will run raxml with best models.
## Fri Sep  8 16:23:50 CEST 2023 - 'phylociraptor modeltest' finished successfully before. Will run raxml with best models.
## Fri Sep  8 16:23:50 CEST 2023 - 'phylociraptor modeltest' finished successfully before. Will run raxml with best models.
## Fri Sep  8 16:23:50 CEST 2023 - concatenate mafft-trimal: Will use bootstrap cutoff 0 before creating concatenated alignment
## Fri Sep  8 16:23:50 CEST 2023 - prepare_iqtree clustalo-aliscore: Will use bootstrap cutoff (70) before creating concatenated alignment
## Fri Sep  8 16:23:50 CEST 2023 - prepare_iqtree clustalo-aliscore: Will use bootstrap cutoff (0) before creating concatenated alignment
## Fri Sep  8 16:23:50 CEST 2023 - prepare_iqtree muscle-bmge: Will use bootstrap cutoff (0) before creating concatenated alignment
## Fri Sep  8 16:23:50 CEST 2023 - concatenate mafft-bmge: Will use bootstrap cutoff 50 before creating concatenated alignment
## Fri Sep  8 16:23:50 CEST 2023 - prepare_iqtree muscle-aliscore: Will use bootstrap cutoff (50) before creating concatenated alignment
## Fri Sep  8 16:23:50 CEST 2023 - prepare_iqtree mafft-aliscore: Will use bootstrap cutoff (50) before creating concatenated alignment
## Fri Sep  8 16:23:50 CEST 2023 - concatenate clustalo-bmge: Will use bootstrap cutoff 50 before creating concatenated alignment
## Will create NEXUS partition file with model information now.
## Will create NEXUS partition file with model information now.
## Will create NEXUS partition file with model information now.
## Will create NEXUS partition file with model information now.
## Will create NEXUS partition file with model information now.
## Fri Sep  8 16:23:51 CEST 2023 - prepare_iqtree muscle-bmge: Will use bootstrap cutoff (60) before creating concatenated alignment
## Fri Sep  8 16:23:51 CEST 2023 - nexus file for iqtree written.
## Fri Sep  8 16:23:51 CEST 2023 - nexus file for iqtree written.
## Fri Sep  8 16:23:51 CEST 2023 - concatenate muscle-bmge: Will use bootstrap cutoff 0 before creating concatenated alignment
## Fri Sep  8 16:23:51 CEST 2023 - nexus file for iqtree written.
## Will create NEXUS partition file with model information now.
## Fri Sep  8 16:23:51 CEST 2023 - nexus file for iqtree written.
## Fri Sep  8 16:23:51 CEST 2023 - concatenate clustalo-bmge: Will use bootstrap cutoff 60 before creating concatenated alignment
## Fri Sep  8 16:23:51 CEST 2023 - concatenate clustalo-bmge: Will use bootstrap cutoff 0 before creating concatenated alignment
## Fri Sep  8 16:23:51 CEST 2023 - nexus file for iqtree written.
## Fri Sep  8 16:23:51 CEST 2023 - concatenate muscle-trimal: Will use bootstrap cutoff 50 before creating concatenated alignment
## Fri Sep  8 16:23:51 CEST 2023 - prepare_iqtree mafft-bmge: Will use bootstrap cutoff (60) before creating concatenated alignment
## Fri Sep  8 16:23:51 CEST 2023 - concatenate muscle-aliscore: Will use bootstrap cutoff 70 before creating concatenated alignment
## Fri Sep  8 16:23:51 CEST 2023 - nexus file for iqtree written.
## Fri Sep  8 16:23:51 CEST 2023 - concatenate clustalo-aliscore: Will use bootstrap cutoff 0 before creating concatenated alignment
## Fri Sep  8 16:23:51 CEST 2023 - concatenate clustalo-aliscore: Will use bootstrap cutoff 60 before creating concatenated alignment
## Will create NEXUS partition file with model information now.
## Fri Sep  8 16:23:51 CEST 2023 - concatenate muscle-aliscore: Will use bootstrap cutoff 60 before creating concatenated alignment
## Fri Sep  8 16:23:52 CEST 2023 - prepare_iqtree muscle-trimal: Will use bootstrap cutoff (70) before creating concatenated alignment
## Fri Sep  8 16:23:52 CEST 2023 - concatenate muscle-bmge: Will use bootstrap cutoff 60 before creating concatenated alignment
## Fri Sep  8 16:23:52 CEST 2023 - nexus file for iqtree written.
## Fri Sep  8 16:23:52 CEST 2023 - prepare_iqtree mafft-aliscore: Will use bootstrap cutoff (0) before creating concatenated alignment
## Fri Sep  8 16:23:52 CEST 2023 - concatenate clustalo-aliscore: Will use bootstrap cutoff 50 before creating concatenated alignment
## Will create NEXUS partition file with model information now.
## Will create NEXUS partition file with model information now.
## Fri Sep  8 16:23:52 CEST 2023 - concatenate muscle-aliscore: Will use bootstrap cutoff 50 before creating concatenated alignment
## Fri Sep  8 16:23:52 CEST 2023 - concatenate mafft-bmge: Will use bootstrap cutoff 70 before creating concatenated alignment
## Fri Sep  8 16:23:53 CEST 2023 - nexus file for iqtree written.
## Fri Sep  8 16:23:53 CEST 2023 - nexus file for iqtree written.
## Fri Sep  8 16:23:54 CEST 2023 - prepare_iqtree mafft-trimal: Will use bootstrap cutoff (0) before creating concatenated alignment
## Fri Sep  8 16:23:54 CEST 2023 - prepare_iqtree clustalo-trimal: Will use bootstrap cutoff (60) before creating concatenated alignment
## Will create NEXUS partition file with model information now.
## Will create NEXUS partition file with model information now.
## Fri Sep  8 16:23:54 CEST 2023 - nexus file for iqtree written.
## Fri Sep  8 16:23:54 CEST 2023 - nexus file for iqtree written.
## Fri Sep  8 16:23:55 CEST 2023 - concatenate mafft-aliscore: Will use bootstrap cutoff 70 before creating concatenated alignment
## Fri Sep  8 16:23:55 CEST 2023 - prepare_iqtree mafft-aliscore: Will use bootstrap cutoff (70) before creating concatenated alignment
## Fri Sep  8 16:23:55 CEST 2023 - prepare_iqtree clustalo-bmge: Will use bootstrap cutoff (50) before creating concatenated alignment
## Will create NEXUS partition file with model information now.
## Fri Sep  8 16:23:55 CEST 2023 - prepare_iqtree muscle-trimal: Will use bootstrap cutoff (0) before creating concatenated alignment
## Will create NEXUS partition file with model information now.
## Will create NEXUS partition file with model information now.
## Fri Sep  8 16:23:55 CEST 2023 - nexus file for iqtree written.
## Fri Sep  8 16:23:55 CEST 2023 - nexus file for iqtree written.
## Fri Sep  8 16:23:55 CEST 2023 - nexus file for iqtree written.
## Fri Sep  8 16:24:07 CEST 2023 - prepare_iqtree clustalo-aliscore: Will use bootstrap cutoff (60) before creating concatenated alignment
## Fri Sep  8 16:24:07 CEST 2023 - prepare_iqtree clustalo-bmge: Will use bootstrap cutoff (60) before creating concatenated alignment
## Fri Sep  8 16:24:07 CEST 2023 - concatenate mafft-trimal: Will use bootstrap cutoff 70 before creating concatenated alignment
## Fri Sep  8 16:24:07 CEST 2023 - concatenate muscle-aliscore: Will use bootstrap cutoff 0 before creating concatenated alignment
## Will create NEXUS partition file with model information now.
## Fri Sep  8 16:24:07 CEST 2023 - concatenate mafft-aliscore: Will use bootstrap cutoff 50 before creating concatenated alignment
## Will create NEXUS partition file with model information now.
## Fri Sep  8 16:24:07 CEST 2023 - concatenate clustalo-trimal: Will use bootstrap cutoff 60 before creating concatenated alignment
## Fri Sep  8 16:24:08 CEST 2023 - nexus file for iqtree written.
## Fri Sep  8 16:24:08 CEST 2023 - nexus file for iqtree written.
## Fri Sep  8 16:24:20 CEST 2023 - 'phylociraptor modeltest' finished successfully before. Will run raxml with best models.
## Fri Sep  8 16:24:20 CEST 2023 - 'phylociraptor modeltest' finished successfully before. Will run raxml with best models.
## Fri Sep  8 16:24:20 CEST 2023 - 'phylociraptor modeltest' finished successfully before. Will run raxml with best models.
## Fri Sep  8 16:24:20 CEST 2023 - 'phylociraptor modeltest' finished successfully before. Will run raxml with best models.
## Fri Sep  8 16:24:20 CEST 2023 - 'phylociraptor modeltest' finished successfully before. Will run raxml with best models.
## Fri Sep  8 16:24:20 CEST 2023 - 'phylociraptor modeltest' finished successfully before. Will run raxml with best models.
## Fri Sep  8 16:24:20 CEST 2023 - 'phylociraptor modeltest' finished successfully before. Will run raxml with best models.
## Fri Sep  8 16:24:20 CEST 2023 - 'phylociraptor modeltest' finished successfully before. Will run raxml with best models.
## Fri Sep  8 16:24:20 CEST 2023 - 'phylociraptor modeltest' finished successfully before. Will run raxml with best models.
## Fri Sep  8 16:24:20 CEST 2023 - 'phylociraptor modeltest' finished successfully before. Will run raxml with best models.
## Fri Sep  8 16:24:20 CEST 2023 - 'phylociraptor modeltest' finished successfully before. Will run raxml with best models.
## Fri Sep  8 16:24:20 CEST 2023 - 'phylociraptor modeltest' finished successfully before. Will run raxml with best models.
## Fri Sep  8 16:24:20 CEST 2023 - 'phylociraptor modeltest' finished successfully before. Will run raxml with best models.
## Fri Sep  8 16:24:20 CEST 2023 - 'phylociraptor modeltest' finished successfully before. Will run raxml with best models.
## Fri Sep  8 16:24:24 CEST 2023 - 'phylociraptor modeltest' finished successfully before. Will run raxml with best models.
## Fri Sep  8 16:24:24 CEST 2023 - 'phylociraptor modeltest' finished successfully before. Will run raxml with best models.
## Fri Sep  8 16:24:24 CEST 2023 - 'phylociraptor modeltest' finished successfully before. Will run raxml with best models.
## Fri Sep  8 16:24:24 CEST 2023 - 'phylociraptor modeltest' finished successfully before. Will run raxml with best models.
## Fri Sep  8 16:24:24 CEST 2023 - 'phylociraptor modeltest' finished successfully before. Will run raxml with best models.
## Fri Sep  8 16:24:24 CEST 2023 - 'phylociraptor modeltest' finished successfully before. Will run raxml with best models.
## Fri Sep  8 16:24:24 CEST 2023 - 'phylociraptor modeltest' finished successfully before. Will run raxml with best models.
## Fri Sep  8 16:24:24 CEST 2023 - 'phylociraptor modeltest' finished successfully before. Will run raxml with best models.
## Fri Sep  8 16:24:36 CEST 2023 - 'phylociraptor modeltest' finished successfully before. Will run raxml with best models.
## Fri Sep  8 16:24:36 CEST 2023 - 'phylociraptor modeltest' finished successfully before. Will run raxml with best models.
## Fri Sep  8 16:24:37 CEST 2023 - 'phylociraptor modeltest' finished successfully before. Will run raxml with best models.
## Fri Sep  8 16:24:37 CEST 2023 - 'phylociraptor modeltest' finished successfully before. Will run raxml with best models.
## Fri Sep  8 16:24:37 CEST 2023 - 'phylociraptor modeltest' finished successfully before. Will run raxml with best models.
## Fri Sep  8 16:24:37 CEST 2023 - 'phylociraptor modeltest' finished successfully before. Will run raxml with best models.
## Sat Sep  9 17:14:06 CEST 2023 - phylociraptor mltree (tree) done.
```
