## Supplementary figures and images for "Phylociraptor – A unified computational framework for reproducible phylogenomic inference"

### Short phylociraptor summary report (PDF) for small test case presented in the main text (Figure2).

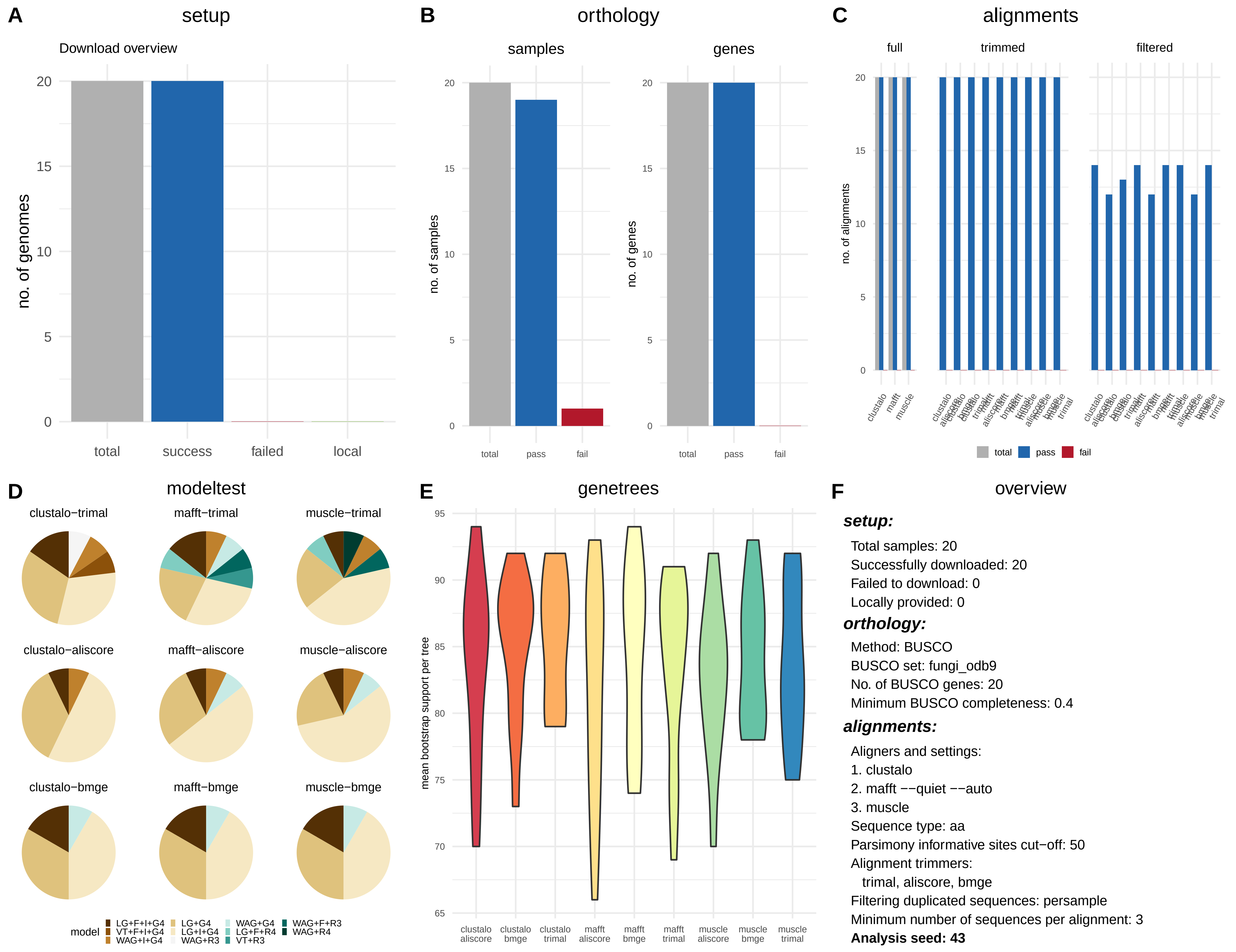

### Short phylociraptor summary report (PDF) for test-case-1 fungi.

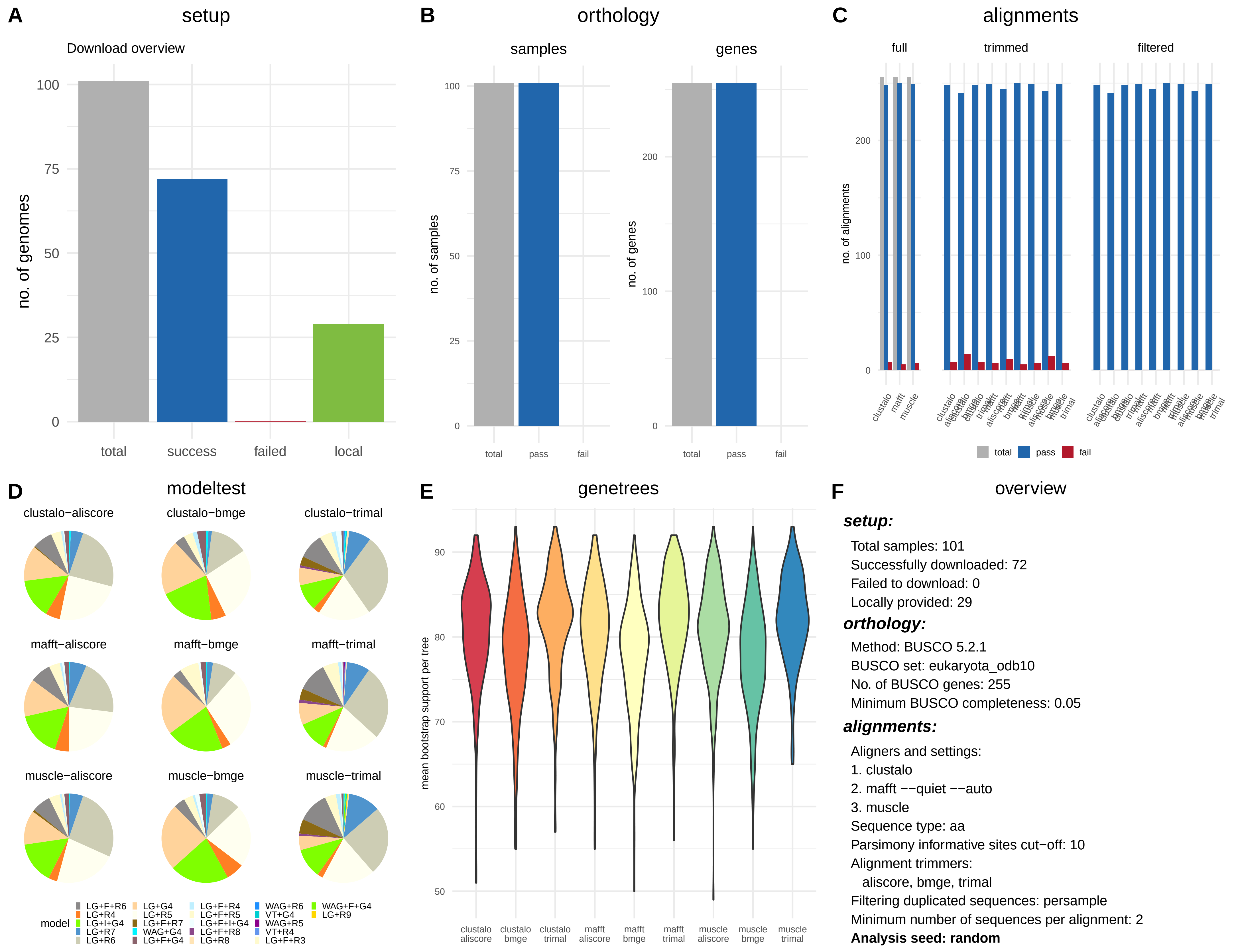

### Short phylociraptor summary report (PDF) for test-case-2 vertebrates.

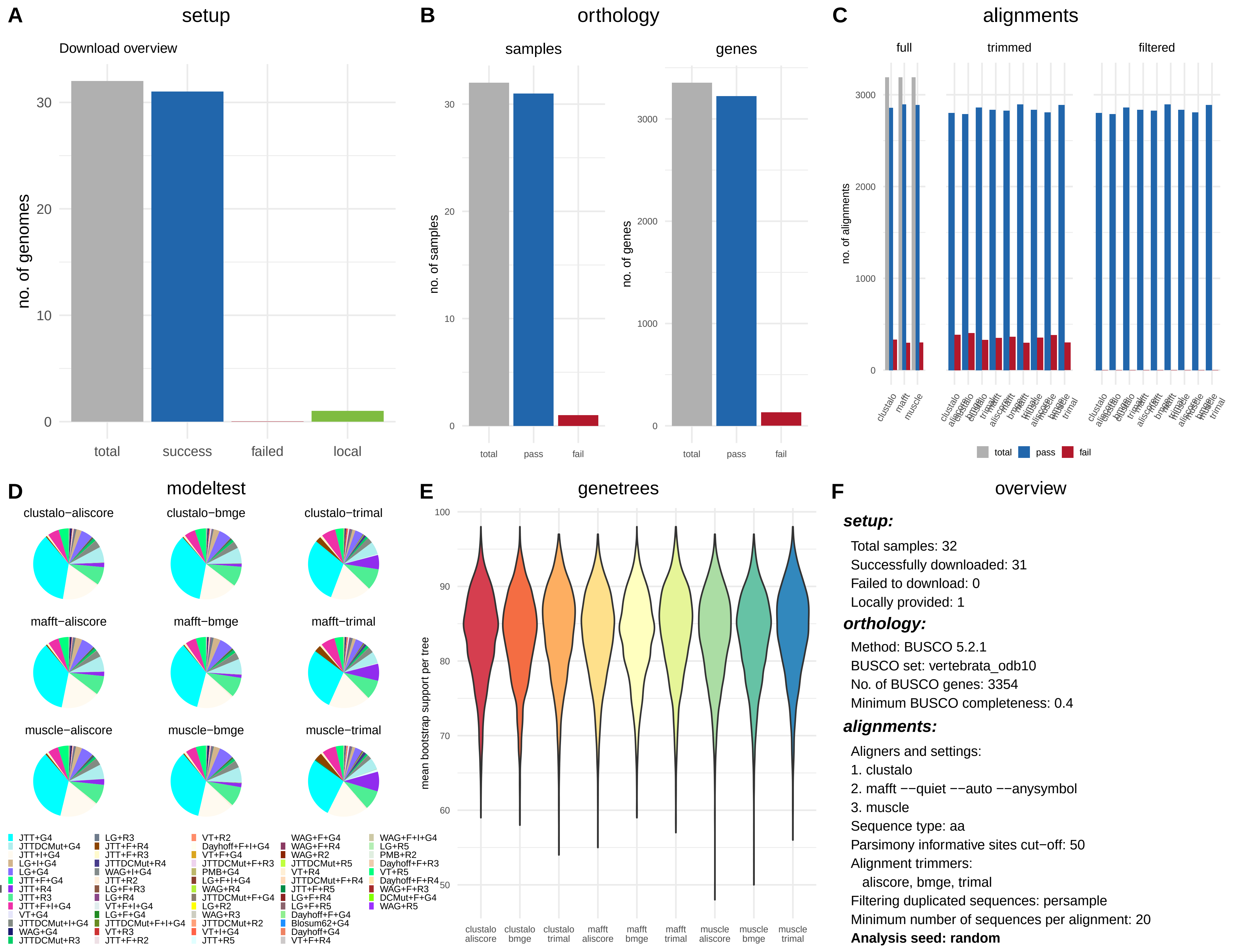
