## Supplementary material for "Phylociraptor – A unified computational framework for reproducible phylogenomic inference": Phylociraptor report as HTML (zipped) for test-case-1 fungi.: test-case1-fungi-report.html


### phylociraptor

**Analysis report - 21 June, 2023 - 11:54 CEST - version: 0.9.12**

### Setup

**Location of genomes files:**

```
results/assemblies
```

**Total number of included genomes:** 101  
**Locally provided genomes:** 29  
**Successfully downloaded genomes:** 72  
**Failed species downloads:** 0  
**Not downloaded:**

**Information on successfully downloaded genomes:**

### Orthology

Orthologous genes were inferred using BUSCO.

**Location of BUSCO results:**

```
results/orthology/busco
```

**Used BUSCO set:** eukaryota\_odb10

**Total number of BUSCO genes in BUSCO set:** 255  
**Number of BUSCO genes recovered in too few (2) samples:** 0  
**BUSCO genes below threshold:**  
none

**Number of samples with single-copy BUSCO score below threshold (0.05):** 0  
**Genomes below threshold:**

**Orthology overview table:**

### Alignments

#### full alignments

**Location of alignment files:**

```
results/alignments/full
```

**Used aligner(s) and settings:**  
**1.** clustalo   
**2.** mafft –quiet –auto  
**3.** muscle  
  
**Parsimony informative sites cutoff:** 10  
  
**Number of alignments for clustalo:** 255   
**Number of alignments for mafft:** 255   
**Number of alignments for muscle:** 255   
  
**Overview of individual full alignments:**

#### trimmed alignments

**Location of trimmed alignment files:**

```
results/alignments/trimmed
```

**Parsimony informative sites cutoff:** 10

**Number of trimmed alignments for clustalo-aliscore:** 255   
**Number of trimmed alignments for clustalo-bmge:** 255   
**Number of trimmed alignments for clustalo-trimal:** 255   
**Number of trimmed alignments for mafft-aliscore:** 255   
**Number of trimmed alignments for mafft-bmge:** 255   
**Number of trimmed alignments for mafft-trimal:** 255   
**Number of trimmed alignments for muscle-aliscore:** 255   
**Number of trimmed alignments for muscle-bmge:** 255   
**Number of trimmed alignments for muscle-trimal:** 255   
  
**Overview of individual trimmed alignments:**

#### filtered alignments

**Location of filtered alignment files:**

```
results/alignments/filtered
```

**Parsimony informative sites cutoff:** 10

**Number of filtered alignments for clustalo-aliscore:** 248   
**Number of filtered alignments for clustalo-bmge:** 241   
**Number of filtered alignments for clustalo-trimal:** 248   
**Number of filtered alignments for mafft-aliscore:** 249   
**Number of filtered alignments for mafft-bmge:** 245   
**Number of filtered alignments for mafft-trimal:** 250   
**Number of filtered alignments for muscle-aliscore:** 249   
**Number of filtered alignments for muscle-bmge:** 243   
**Number of filtered alignments for muscle-trimal:** 249   
  
**Overview of individual filtered alignments:**

### Modeltesting

**Location of modeltesting files:**

```
results/modeltest
```

**Overview of individual modeltest results:**

### Genetrees

**Location of gene trees:**

```
results/modeltest
```

**Overview of mean bootstrap values per gene-tree:**   
 Total number of gene trees for clustalo-aliscore: 248  
 Total number of gene trees for clustalo-bmge: 241  
 Total number of gene trees for clustalo-trimal: 248  
 Total number of gene trees for mafft-aliscore: 249  
 Total number of gene trees for mafft-bmge: 245  
 Total number of gene trees for mafft-trimal: 250  
 Total number of gene trees for muscle-aliscore: 249  
 Total number of gene trees for muscle-bmge: 243  
 Total number of gene trees for muscle-trimal: 249

FALSE
FALSE

### Maximum-likelihood trees

**Random seed:** random

**Location of phylogenetic results:**

```
results/phylogeny-*/
```

  
**Overview of concatenated Maximum-likelihood phylogenies:**

| software | aligner | trimmer | boostraps | bootstrap-cutoff | ngenes | tree |
| --- | --- | --- | --- | --- | --- | --- |
| iqtree | clustalo | aliscore | 1000 | 0 | 248 | Copy tree |
| iqtree | clustalo | aliscore | 1000 | 50 | 248 | Copy tree |
| iqtree | clustalo | aliscore | 1000 | 60 | 246 | Copy tree |
| iqtree | clustalo | aliscore | 1000 | 70 | 238 | Copy tree |
| iqtree | clustalo | aliscore | 1000 | 80 | 153 | Copy tree |
| iqtree | clustalo | aliscore | 1000 | 90 | 8 | Copy tree |
| iqtree | clustalo | bmge | 1000 | 0 | 241 | Copy tree |
| iqtree | clustalo | bmge | 1000 | 50 | 241 | Copy tree |
| iqtree | clustalo | bmge | 1000 | 60 | 236 | Copy tree |
| iqtree | clustalo | bmge | 1000 | 70 | 215 | Copy tree |
| iqtree | clustalo | bmge | 1000 | 80 | 106 | Copy tree |
| iqtree | clustalo | bmge | 1000 | 90 | 3 | Copy tree |
| iqtree | clustalo | trimal | 1000 | 0 | 248 | Copy tree |
| iqtree | clustalo | trimal | 1000 | 50 | 248 | Copy tree |
| iqtree | clustalo | trimal | 1000 | 60 | 246 | Copy tree |
| iqtree | clustalo | trimal | 1000 | 70 | 240 | Copy tree |
| iqtree | clustalo | trimal | 1000 | 80 | 177 | Copy tree |
| iqtree | clustalo | trimal | 1000 | 90 | 15 | Copy tree |
| iqtree | mafft | aliscore | 1000 | 0 | 249 | Copy tree |
| iqtree | mafft | aliscore | 1000 | 50 | 249 | Copy tree |
| iqtree | mafft | aliscore | 1000 | 60 | 247 | Copy tree |
| iqtree | mafft | aliscore | 1000 | 70 | 238 | Copy tree |
| iqtree | mafft | aliscore | 1000 | 80 | 152 | Copy tree |
| iqtree | mafft | aliscore | 1000 | 90 | 9 | Copy tree |
| iqtree | mafft | bmge | 1000 | 0 | 245 | Copy tree |
| iqtree | mafft | bmge | 1000 | 50 | 245 | Copy tree |
| iqtree | mafft | bmge | 1000 | 60 | 242 | Copy tree |
| iqtree | mafft | bmge | 1000 | 70 | 215 | Copy tree |
| iqtree | mafft | bmge | 1000 | 80 | 105 | Copy tree |
| iqtree | mafft | bmge | 1000 | 90 | 2 | Copy tree |
| iqtree | mafft | trimal | 1000 | 0 | 250 | Copy tree |
| iqtree | mafft | trimal | 1000 | 50 | 250 | Copy tree |
| iqtree | mafft | trimal | 1000 | 60 | 249 | Copy tree |
| iqtree | mafft | trimal | 1000 | 70 | 242 | Copy tree |
| iqtree | mafft | trimal | 1000 | 80 | 182 | Copy tree |
| iqtree | mafft | trimal | 1000 | 90 | 17 | Copy tree |
| iqtree | muscle | aliscore | 1000 | 0 | 249 | Copy tree |
| iqtree | muscle | aliscore | 1000 | 50 | 248 | Copy tree |
| iqtree | muscle | aliscore | 1000 | 60 | 248 | Copy tree |
| iqtree | muscle | aliscore | 1000 | 70 | 241 | Copy tree |
| iqtree | muscle | aliscore | 1000 | 80 | 158 | Copy tree |
| iqtree | muscle | aliscore | 1000 | 90 | 7 | Copy tree |
| iqtree | muscle | bmge | 1000 | 0 | 243 | Copy tree |
| iqtree | muscle | bmge | 1000 | 50 | 243 | Copy tree |
| iqtree | muscle | bmge | 1000 | 60 | 241 | Copy tree |
| iqtree | muscle | bmge | 1000 | 70 | 222 | Copy tree |
| iqtree | muscle | bmge | 1000 | 80 | 102 | Copy tree |
| iqtree | muscle | bmge | 1000 | 90 | 3 | Copy tree |
| iqtree | muscle | trimal | 1000 | 0 | 249 | Copy tree |
| iqtree | muscle | trimal | 1000 | 50 | 249 | Copy tree |
| iqtree | muscle | trimal | 1000 | 60 | 249 | Copy tree |
| iqtree | muscle | trimal | 1000 | 70 | 242 | Copy tree |
| iqtree | muscle | trimal | 1000 | 80 | 183 | Copy tree |
| iqtree | muscle | trimal | 1000 | 90 | 12 | Copy tree |
| raxmlng | clustalo | aliscore | 100 | 0 | 248 | Copy tree |
| raxmlng | clustalo | aliscore | 100 | 60 | 246 | Copy tree |
| raxmlng | clustalo | aliscore | 100 | 70 | 238 | Copy tree |
| raxmlng | clustalo | aliscore | 100 | 80 | 153 | Copy tree |
| raxmlng | clustalo | bmge | 100 | 0 | 241 | Copy tree |
| raxmlng | clustalo | bmge | 100 | 50 | 241 | Copy tree |
| raxmlng | clustalo | bmge | 100 | 60 | 236 | Copy tree |
| raxmlng | clustalo | bmge | 100 | 70 | 215 | Copy tree |
| raxmlng | clustalo | bmge | 100 | 80 | 106 | Copy tree |
| raxmlng | clustalo | trimal | 100 | 50 | 248 | Copy tree |
| raxmlng | mafft | aliscore | 100 | 0 | 249 | Copy tree |
| raxmlng | mafft | aliscore | 100 | 60 | 247 | Copy tree |
| raxmlng | mafft | aliscore | 100 | 70 | 238 | Copy tree |
| raxmlng | mafft | aliscore | 100 | 80 | 152 | Copy tree |
| raxmlng | mafft | bmge | 100 | 0 | 245 | Copy tree |
| raxmlng | mafft | bmge | 100 | 50 | 245 | Copy tree |
| raxmlng | mafft | bmge | 100 | 60 | 242 | Copy tree |
| raxmlng | mafft | bmge | 100 | 70 | 215 | Copy tree |
| raxmlng | mafft | bmge | 100 | 80 | 105 | Copy tree |
| raxmlng | mafft | trimal | 100 | 50 | 250 | Copy tree |
| raxmlng | muscle | aliscore | 100 | 0 | 249 | Copy tree |
| raxmlng | muscle | aliscore | 100 | 50 | 248 | Copy tree |
| raxmlng | muscle | aliscore | 100 | 60 | 248 | Copy tree |
| raxmlng | muscle | aliscore | 100 | 70 | 241 | Copy tree |
| raxmlng | muscle | bmge | 100 | 0 | 243 | Copy tree |
| raxmlng | muscle | bmge | 100 | 50 | 243 | Copy tree |
| raxmlng | muscle | bmge | 100 | 60 | 241 | Copy tree |
| raxmlng | muscle | bmge | 100 | 70 | 222 | Copy tree |
| raxmlng | muscle | bmge | 100 | 80 | 102 | Copy tree |
| raxmlng | muscle | trimal | 100 | 0 | 249 | Copy tree |
| raxmlng | muscle | trimal | 100 | 50 | 249 | Copy tree |
| raxmlng | muscle | trimal | 100 | 60 | 249 | Copy tree |

### Speciestree

**Location of phylogenetic results:**

```
results/phylogeny-*/
```

  
**Overview of species-tree phylogenies:**

| software | aligner | trimmer | bootstrap-cutoff | ngenes | tree |
| --- | --- | --- | --- | --- | --- |
| astral | clustalo | aliscore | 0 | 248 | Copy tree |
| astral | clustalo | aliscore | 50 | 248 | Copy tree |
| astral | clustalo | aliscore | 60 | 246 | Copy tree |
| astral | clustalo | aliscore | 70 | 238 | Copy tree |
| astral | clustalo | aliscore | 80 | 153 | Copy tree |
| astral | clustalo | aliscore | 90 | 8 | Copy tree |
| astral | clustalo | bmge | 0 | 241 | Copy tree |
| astral | clustalo | bmge | 50 | 241 | Copy tree |
| astral | clustalo | bmge | 60 | 236 | Copy tree |
| astral | clustalo | bmge | 70 | 215 | Copy tree |
| astral | clustalo | bmge | 80 | 106 | Copy tree |
| astral | clustalo | bmge | 90 | 3 | Copy tree |
| astral | clustalo | trimal | 0 | 248 | Copy tree |
| astral | clustalo | trimal | 50 | 248 | Copy tree |
| astral | clustalo | trimal | 60 | 246 | Copy tree |
| astral | clustalo | trimal | 70 | 240 | Copy tree |
| astral | clustalo | trimal | 80 | 177 | Copy tree |
| astral | clustalo | trimal | 90 | 15 | Copy tree |
| astral | mafft | aliscore | 0 | 249 | Copy tree |
| astral | mafft | aliscore | 50 | 249 | Copy tree |
| astral | mafft | aliscore | 60 | 247 | Copy tree |
| astral | mafft | aliscore | 70 | 238 | Copy tree |
| astral | mafft | aliscore | 80 | 152 | Copy tree |
| astral | mafft | aliscore | 90 | 9 | Copy tree |
| astral | mafft | bmge | 0 | 245 | Copy tree |
| astral | mafft | bmge | 50 | 245 | Copy tree |
| astral | mafft | bmge | 60 | 242 | Copy tree |
| astral | mafft | bmge | 70 | 215 | Copy tree |
| astral | mafft | bmge | 80 | 105 | Copy tree |
| astral | mafft | bmge | 90 | 2 | Copy tree |
| astral | mafft | trimal | 0 | 250 | Copy tree |
| astral | mafft | trimal | 50 | 250 | Copy tree |
| astral | mafft | trimal | 60 | 249 | Copy tree |
| astral | mafft | trimal | 70 | 242 | Copy tree |
| astral | mafft | trimal | 80 | 182 | Copy tree |
| astral | mafft | trimal | 90 | 17 | Copy tree |
| astral | muscle | aliscore | 0 | 249 | Copy tree |
| astral | muscle | aliscore | 50 | 248 | Copy tree |
| astral | muscle | aliscore | 60 | 248 | Copy tree |
| astral | muscle | aliscore | 70 | 241 | Copy tree |
| astral | muscle | aliscore | 80 | 158 | Copy tree |
| astral | muscle | aliscore | 90 | 7 | Copy tree |
| astral | muscle | bmge | 0 | 243 | Copy tree |
| astral | muscle | bmge | 50 | 243 | Copy tree |
| astral | muscle | bmge | 60 | 241 | Copy tree |
| astral | muscle | bmge | 70 | 222 | Copy tree |
| astral | muscle | bmge | 80 | 102 | Copy tree |
| astral | muscle | bmge | 90 | 3 | Copy tree |
| astral | muscle | trimal | 0 | 249 | Copy tree |
| astral | muscle | trimal | 50 | 249 | Copy tree |
| astral | muscle | trimal | 60 | 249 | Copy tree |
| astral | muscle | trimal | 70 | 242 | Copy tree |
| astral | muscle | trimal | 80 | 183 | Copy tree |
| astral | muscle | trimal | 90 | 12 | Copy tree |

### Log

```
## Thu Jul 21 13:48:21 CEST 2022 - Setup: Will download species now - batch: 2
## Thu Jul 21 13:49:39 CEST 2022 - Setup: Will download species now - batch: 1
## Thu Jul 21 13:50:45 CEST 2022 - Setup: Will download species now - batch: 3
## Thu Jul 21 13:51:51 CEST 2022 - Setup: Will download species now - batch: 2
## Thu Jul 21 13:52:18 CEST 2022 - Setup: Will download species now - batch: 1
## Thu Jul 21 13:52:47 CEST 2022 - Setup: Will download species now - batch: 3
## Thu Jul 21 13:54:10 CEST 2022 - Setup: Will download species now - batch: 2
## Thu Jul 21 13:54:37 CEST 2022 - Setup: Will download species now - batch: 1
## Thu Jul 21 13:55:05 CEST 2022 - Setup: Will download species now - batch: 3
## Thu Jul 21 13:56:45 CEST 2022 - phylociraptor setup done.
## Thu Jul 21 15:02:56 CEST 2022 Tubulinosema_ratisbonensis       5 Complete     248 Missing
## Thu Jul 21 15:05:01 CEST 2022 Schizosaccharomyces_pombe     228 Complete       5 Fragmented      20 Missing
## Thu Jul 21 15:05:30 CEST 2022 Amoeboradix_gromovi     118 Complete      14 Fragmented     121 Missing
## Thu Jul 21 15:06:02 CEST 2022 Vavraia_culicis      51 Complete      10 Fragmented     192 Missing
## Thu Jul 21 15:07:22 CEST 2022 Mitosporidium_daphniae      34 Complete       4 Duplicated      14 Fragmented     203 Missing
## Thu Jul 21 15:07:44 CEST 2022 Amphiamblys_sp      35 Complete      11 Fragmented     207 Missing
## Thu Jul 21 15:08:33 CEST 2022 Pneumocystis_murina     199 Complete      18 Fragmented      36 Missing
## Thu Jul 21 15:09:54 CEST 2022 Debaryomyces_hansenii     226 Complete       2 Duplicated       8 Fragmented      18 Missing
## Thu Jul 21 15:10:14 CEST 2022 Protomyces_lactucaedebilis     229 Complete       2 Duplicated       9 Fragmented      14 Missing
## Thu Jul 21 15:11:39 CEST 2022 Spizellomyces_punctatus     227 Complete       2 Duplicated      12 Fragmented      13 Missing
## Thu Jul 21 15:11:44 CEST 2022 Dichotomocladium_robustum     231 Complete       6 Duplicated       8 Fragmented      11 Missing
## Thu Jul 21 15:12:03 CEST 2022 Paramicrosporidium_saccamoebae     134 Complete      32 Fragmented      87 Missing
## Thu Jul 21 15:12:12 CEST 2022 Piptocephalis_tieghemiana     195 Complete       3 Duplicated      16 Fragmented      41 Missing
## Thu Jul 21 15:12:32 CEST 2022 Batrachochytrium_dendrobatidis     235 Complete       6 Duplicated       6 Fragmented       9 Missing
## Thu Jul 21 15:12:46 CEST 2022 Acaromyces_ingoldii     214 Complete      19 Fragmented      20 Missing
## Thu Jul 21 15:14:33 CEST 2022 Capsaspora_owczarzaki     225 Complete       2 Duplicated      13 Fragmented      14 Missing
## Thu Jul 21 15:14:36 CEST 2022 Saitoella_complicata     228 Complete      10 Duplicated      11 Fragmented       9 Missing
## Thu Jul 21 15:14:36 CEST 2022 Coccidioides_posadasii     246 Complete       6 Fragmented       1 Missing
## Thu Jul 21 15:15:11 CEST 2022 Paxillus_rubicundulus     219 Complete      21 Fragmented      13 Missing
## Thu Jul 21 15:16:14 CEST 2022 Dissophora_decumbens     242 Complete       6 Duplicated       4 Fragmented       4 Missing
## Thu Jul 21 15:16:32 CEST 2022 Sanchytrium_tribonematis      68 Complete      21 Fragmented     164 Missing
## Thu Jul 21 15:16:39 CEST 2022 Mixia_osmundae     187 Complete      28 Fragmented      38 Missing
## Thu Jul 21 15:16:50 CEST 2022 Rozella_allomycis      20 Complete      10 Fragmented     223 Missing
## Thu Jul 21 15:16:53 CEST 2022 Mucor_lusitanicus     239 Complete       4 Duplicated       5 Fragmented       7 Missing
## Thu Jul 21 15:18:37 CEST 2022 Gonapodya_prolifera     169 Complete      52 Duplicated      28 Fragmented      30 Missing
## Thu Jul 21 15:19:03 CEST 2022 Hypoxylon_sp     247 Complete       2 Fragmented       4 Missing
## Thu Jul 21 15:20:27 CEST 2022 Nosema_ceranae       6 Complete       3 Fragmented     244 Missing
## Thu Jul 21 15:21:25 CEST 2022 Letharia_columbiana     231 Complete      26 Duplicated       3 Fragmented       6 Missing
## Thu Jul 21 15:21:45 CEST 2022 Nematocida_sp      43 Complete       7 Fragmented     203 Missing
## Thu Jul 21 15:23:12 CEST 2022 Sistotremastrum_suecicum     214 Complete       2 Duplicated      21 Fragmented      17 Missing
## Thu Jul 21 15:23:54 CEST 2022 Thelohania_contejeani      25 Complete      11 Fragmented     217 Missing
## Thu Jul 21 15:24:14 CEST 2022 Piromyces_sp     217 Complete       7 Fragmented      29 Missing
## Thu Jul 21 15:24:44 CEST 2022 Radiomyces_spectabilis     242 Complete       2 Duplicated       6 Fragmented       4 Missing
## Thu Jul 21 15:26:09 CEST 2022 Umbelopsis_isabellina     243 Complete       7 Fragmented       3 Missing
## Thu Jul 21 15:26:13 CEST 2022 Polychytrium_aggregatum     112 Complete     247 Duplicated       8 Fragmented      12 Missing
## Thu Jul 21 15:27:44 CEST 2022 Wickerhamomyces_anomalus     228 Complete       5 Fragmented      20 Missing
## Thu Jul 21 15:28:01 CEST 2022 Edhazardia_aedis      24 Complete      10 Fragmented     219 Missing
## Thu Jul 21 15:28:04 CEST 2022 Dictyostelium_lacteum     223 Complete       4 Duplicated      11 Fragmented      17 Missing
## Thu Jul 21 15:28:09 CEST 2022 Neurospora_crassa     253 Complete
## Thu Jul 21 15:28:36 CEST 2022 Syncephalis_fuscata     193 Complete      40 Duplicated      19 Fragmented      21 Missing
## Thu Jul 21 15:29:31 CEST 2022 Olpidium_bornovanus      86 Complete      36 Duplicated      44 Fragmented     107 Missing
## Thu Jul 21 15:29:32 CEST 2022 Acidomyces_richmondensis     245 Complete       2 Duplicated       5 Fragmented       2 Missing
## Thu Jul 21 15:29:46 CEST 2022 Coemansia_mojavensis     234 Complete      12 Duplicated       3 Fragmented      10 Missing
## Thu Jul 21 15:34:20 CEST 2022 Hyaloraphidium_curvatum     224 Complete      15 Fragmented      14 Missing
## Thu Jul 21 15:34:35 CEST 2022 Paraphysoderma_sedebokerense     205 Complete      26 Duplicated      18 Fragmented      18 Missing
## Thu Jul 21 15:34:39 CEST 2022 Catenaria_anguillulae     115 Complete     137 Duplicated      33 Fragmented      38 Missing
## Thu Jul 21 15:34:53 CEST 2022 Sclerophora_sanguinea     240 Complete       9 Fragmented       4 Missing
## Thu Jul 21 15:36:00 CEST 2022 Exophiala_oligosperma     247 Complete       3 Fragmented       3 Missing
## Thu Jul 21 15:37:44 CEST 2022 Yarrowia_lipolytica     243 Complete       6 Fragmented       4 Missing
## Thu Jul 21 15:37:44 CEST 2022 Cryptococcus_neoformans     221 Complete       2 Duplicated       7 Fragmented      24 Missing
## Thu Jul 21 15:38:04 CEST 2022 Daedalea_quercina     227 Complete      16 Fragmented      10 Missing
## Thu Jul 21 15:38:52 CEST 2022 Encephalitozoon_hellem      59 Complete      12 Fragmented     182 Missing
## Thu Jul 21 15:41:23 CEST 2022 Phaffia_rhodozyma     185 Complete      22 Fragmented      46 Missing
## Thu Jul 21 15:41:49 CEST 2022 Malassezia_pachydermatis     184 Complete      28 Fragmented      41 Missing
## Thu Jul 21 15:42:41 CEST 2022 Caulochytrium_protostelioides      44 Complete      26 Fragmented     183 Missing
## Thu Jul 21 15:42:47 CEST 2022 Vittaforma_corneae      36 Complete       2 Duplicated       8 Fragmented     208 Missing
## Thu Jul 21 15:45:26 CEST 2022 Blastocladiella_britannica     199 Complete      20 Fragmented      34 Missing
## Thu Jul 21 15:45:51 CEST 2022 Cylindrobasidium_torrendii     226 Complete       4 Duplicated      16 Fragmented       9 Missing
## Thu Jul 21 15:46:20 CEST 2022 Tilletia_indica     215 Complete      16 Fragmented      22 Missing
## Thu Jul 21 15:46:37 CEST 2022 Diversispora_epigaea     232 Complete       4 Duplicated       6 Fragmented      13 Missing
## Thu Jul 21 15:46:58 CEST 2022 Phycomyces_blakesleeanus     222 Complete      24 Duplicated      10 Fragmented       9 Missing
## Thu Jul 21 15:47:55 CEST 2022 Chytriomyces_hyalinus     178 Complete      91 Duplicated      12 Fragmented      20 Missing
## Thu Jul 21 15:48:22 CEST 2022 Glomus_cerebriforme     229 Complete       6 Duplicated       6 Fragmented      15 Missing
## Thu Jul 21 15:48:31 CEST 2022 Smittium_culicis     132 Complete     142 Duplicated      18 Fragmented      37 Missing
## Thu Jul 21 15:50:39 CEST 2022 Paraphaeosphaeria_sporulosa     248 Complete       2 Duplicated       4 Fragmented
## Thu Jul 21 15:52:52 CEST 2022 Pilobolus_umbonatus     238 Complete       4 Duplicated       3 Fragmented      10 Missing
## Thu Jul 21 15:53:00 CEST 2022 Neocallimastix_sp     147 Complete     145 Duplicated       3 Fragmented      31 Missing
## Thu Jul 21 15:53:29 CEST 2022 Tuber_borchii     244 Complete       2 Duplicated       4 Fragmented       4 Missing
## Thu Jul 21 15:56:11 CEST 2022 Microbotryum_violaceum     222 Complete       7 Fragmented      24 Missing
## Thu Jul 21 15:56:43 CEST 2022 Dimargaris_cristalligena     217 Complete      13 Fragmented      23 Missing
## Thu Jul 21 15:58:23 CEST 2022 Salpingoeca_rosetta     165 Complete      21 Fragmented      67 Missing
## Thu Jul 21 16:06:19 CEST 2022 Melampsora_laricipopulina     209 Complete       9 Duplicated      12 Fragmented      28 Missing
## Thu Jul 21 16:08:00 CEST 2022 Mortierella_sp     239 Complete      21 Duplicated       3 Fragmented       7 Missing
## Thu Jul 21 16:09:13 CEST 2022 Basidiobolus_meristosporus     208 Complete      67 Duplicated       7 Fragmented       7 Missing
## Thu Jul 21 16:17:43 CEST 2022 Calocera_viscosa     207 Complete      15 Duplicated      20 Fragmented      20 Missing
## Thu Jul 21 16:19:17 CEST 2022 Allomyces_macrogynus      74 Complete     237 Duplicated      24 Fragmented      38 Missing
## Thu Jul 21 17:01:44 CEST 2022 Gigaspora_margarita     226 Complete       6 Duplicated       6 Fragmented      18 Missing
## Thu Jul 21 17:19:23 CEST 2022 Zoophthora_radicans     192 Complete      32 Duplicated      13 Fragmented      32 Missing
## Fri Jul 22 00:36:02 CEST 2022 Mus_musculus     240 Complete      18 Duplicated       3 Fragmented       2 Missing
## Fri Jul 22 09:04:11 CEST 2022 - Setup: Will download species now - batch: 2
## Fri Jul 22 09:04:40 CEST 2022 - Setup: Will download species now - batch: 1
## Fri Jul 22 09:05:09 CEST 2022 - Setup: Will download species now - batch: 3
## Fri 22 Jul 2022 09:05:45 AM CEST - phylociraptor setup done.
## Fri 22 Jul 2022 09:14:08 AM CEST Amaparvus_languidus      40 Complete      82 Duplicated      13 Fragmented     159 Missing
## Fri 22 Jul 2022 09:18:59 AM CEST Nuclearia_sp      84 Complete     282 Duplicated      34 Fragmented       8 Missing
## Fri Jul 22 09:53:02 CEST 2022 - Setup: Will download species now - batch: 2
## Fri Jul 22 09:53:36 CEST 2022 - Setup: Will download species now - batch: 1
## Fri Jul 22 09:54:08 CEST 2022 - Setup: Will download species now - batch: 3
## Fri 22 Jul 2022 09:54:42 AM CEST - phylociraptor setup done.
## Fri 22 Jul 2022 10:14:43 AM CEST Malassezia_pachydermatis     232 Complete       2 Duplicated      14 Fragmented       6 Missing
## Fri 22 Jul 2022 10:16:07 AM CEST Wickerhamomyces_anomalus     238 Complete      12 Duplicated       3 Fragmented       6 Missing
## Fri 22 Jul 2022 10:16:51 AM CEST Mixia_osmundae     219 Complete       2 Duplicated      23 Fragmented      10 Missing
## Fri 22 Jul 2022 10:17:58 AM CEST Saitoella_complicata     237 Complete      14 Duplicated       9 Fragmented
## Fri 22 Jul 2022 10:18:02 AM CEST Nosema_ceranae      67 Complete      15 Fragmented     171 Missing
## Fri 22 Jul 2022 10:19:01 AM CEST Coemansia_mojavensis     234 Complete      18 Duplicated       3 Fragmented       7 Missing
## Fri 22 Jul 2022 10:19:01 AM CEST Sclerophora_sanguinea     251 Complete       2 Fragmented
## Fri 22 Jul 2022 10:19:32 AM CEST Thelohania_contejeani      82 Complete      16 Fragmented     155 Missing
## Fri 22 Jul 2022 10:20:07 AM CEST Dictyostelium_lacteum     231 Complete       2 Duplicated      10 Fragmented      11 Missing
## Fri 22 Jul 2022 10:21:00 AM CEST Amoeboradix_gromovi     150 Complete       4 Duplicated      22 Fragmented      79 Missing
## Fri 22 Jul 2022 10:21:55 AM CEST Vavraia_culicis      56 Complete      15 Fragmented     182 Missing
## Fri 22 Jul 2022 10:22:32 AM CEST Umbelopsis_isabellina     248 Complete       2 Fragmented       3 Missing
## Fri 22 Jul 2022 10:26:04 AM CEST Pilobolus_umbonatus     237 Complete      12 Duplicated       6 Fragmented       4 Missing
## Fri 22 Jul 2022 10:27:58 AM CEST Piptocephalis_tieghemiana     192 Complete      21 Fragmented      40 Missing
## Fri 22 Jul 2022 10:28:15 AM CEST Blastocladiella_britannica     202 Complete      25 Fragmented      26 Missing
## Fri 22 Jul 2022 10:28:21 AM CEST Radiomyces_spectabilis     243 Complete       8 Duplicated       3 Fragmented       3 Missing
## Fri 22 Jul 2022 10:29:40 AM CEST Tilletia_indica     225 Complete       4 Duplicated      12 Fragmented      15 Missing
## Fri 22 Jul 2022 10:30:16 AM CEST Actinomortierella_capitata     245 Complete       5 Fragmented       3 Missing
## Fri 22 Jul 2022 10:30:55 AM CEST Synchytrium_microbalum     223 Complete      19 Fragmented      11 Missing
## Fri 22 Jul 2022 10:31:25 AM CEST Syncephalis_fuscata     177 Complete      41 Duplicated      30 Fragmented      27 Missing
## Fri 22 Jul 2022 10:33:07 AM CEST Amaparvus_languidus      27 Complete     136 Duplicated      58 Fragmented     101 Missing
## Fri Jul 22 10:34:55 CEST 2022 Mucor_lusitanicus     234 Complete      24 Duplicated       4 Fragmented       3 Missing
## Fri 22 Jul 2022 10:34:57 AM CEST Neurospora_crassa     252 Complete       1 Fragmented
## Fri 22 Jul 2022 10:35:30 AM CEST Piromyces_sp     217 Complete      10 Duplicated       4 Fragmented      27 Missing
## Fri 22 Jul 2022 10:36:35 AM CEST Dichotomocladium_robustum     237 Complete       8 Duplicated       7 Fragmented       5 Missing
## Fri 22 Jul 2022 10:36:39 AM CEST Nuclearia_sp      43 Complete     419 Duplicated      35 Fragmented       6 Missing
## Fri 22 Jul 2022 10:37:07 AM CEST Dissophora_decumbens     240 Complete      13 Duplicated       6 Fragmented       1 Missing
## Fri 22 Jul 2022 10:37:50 AM CEST Paraphysoderma_sedebokerense     202 Complete      27 Duplicated      21 Fragmented      18 Missing
## Fri 22 Jul 2022 10:41:28 AM CEST Hypoxylon_sp     246 Complete       2 Duplicated       5 Fragmented       1 Missing
## Fri 22 Jul 2022 10:44:29 AM CEST Phycomyces_blakesleeanus     226 Complete      30 Duplicated       9 Fragmented       3 Missing
## Fri 22 Jul 2022 10:44:31 AM CEST Phialocephala_subalpina     246 Complete      10 Duplicated       1 Fragmented       1 Missing
## Fri 22 Jul 2022 10:50:10 AM CEST Gonapodya_prolifera     188 Complete      74 Duplicated      17 Fragmented      11 Missing
## Fri 22 Jul 2022 10:51:52 AM CEST Stereomyxa_ramosa      39 Complete     211 Duplicated      52 Fragmented      67 Missing
## Fri 22 Jul 2022 11:15:13 AM CEST Diversispora_epigaea     237 Complete       4 Duplicated       9 Fragmented       5 Missing
## Fri 22 Jul 2022 11:29:46 AM CEST Polychytrium_aggregatum     108 Complete     255 Duplicated      16 Fragmented       7 Missing
## Fri 22 Jul 2022 03:03:45 PM CEST Gigaspora_margarita     233 Complete       9 Duplicated       6 Fragmented      10 Missing
## Fri 22 Jul 2022 04:34:23 PM CEST Encephalitozoon_hellem      70 Complete       3 Duplicated      17 Fragmented     165 Missing
## Fri 22 Jul 2022 04:35:37 PM CEST Wallemia_mellicola     231 Complete       2 Duplicated      12 Fragmented       9 Missing
## Fri 22 Jul 2022 04:37:29 PM CEST Debaryomyces_hansenii     233 Complete       2 Duplicated       7 Fragmented      12 Missing
## Fri Jul 22 16:38:01 CEST 2022 Tubulinosema_ratisbonensis      63 Complete       6 Duplicated      16 Fragmented     172 Missing
## Fri 22 Jul 2022 04:39:04 PM CEST Paramicrosporidium_saccamoebae     182 Complete       2 Duplicated      20 Fragmented      50 Missing
## Fri 22 Jul 2022 04:40:19 PM CEST Mitosporidium_daphniae     105 Complete      11 Duplicated      37 Fragmented     106 Missing
## Fri Jul 22 16:43:13 CEST 2022 Ustilago_hordei     248 Complete       4 Fragmented       1 Missing
## Fri 22 Jul 2022 04:43:55 PM CEST Protomyces_lactucaedebilis     238 Complete       7 Fragmented       8 Missing
## Fri 22 Jul 2022 04:46:12 PM CEST Phaffia_rhodozyma     209 Complete       2 Duplicated      28 Fragmented      15 Missing
## Fri Jul 22 16:47:25 CEST 2022 Coccidioides_posadasii     247 Complete       4 Fragmented       2 Missing
## Fri 22 Jul 2022 04:47:29 PM CEST Acidomyces_richmondensis     247 Complete       4 Duplicated       3 Fragmented       1 Missing
## Fri 22 Jul 2022 04:47:31 PM CEST Microbotryum_violaceum     233 Complete       4 Duplicated      13 Fragmented       5 Missing
## Fri 22 Jul 2022 04:49:36 PM CEST Neolentinus_lepideus     233 Complete       9 Duplicated      10 Fragmented       6 Missing
## Fri 22 Jul 2022 04:52:14 PM CEST Exophiala_oligosperma     251 Complete       2 Fragmented
## Fri Jul 22 16:57:11 CEST 2022 Arthrobotrys_oligospora     245 Complete       8 Fragmented
## Fri 22 Jul 2022 04:58:54 PM CEST Sistotremastrum_suecicum     237 Complete       2 Duplicated       9 Fragmented       6 Missing
## Fri 22 Jul 2022 05:03:29 PM CEST Zoophthora_radicans     183 Complete      34 Duplicated      23 Fragmented      30 Missing
## Fri 22 Jul 2022 05:11:19 PM CEST Chytriomyces_hyalinus     177 Complete     100 Duplicated      15 Fragmented      14 Missing
## Fri 22 Jul 2022 05:11:27 PM CEST Edhazardia_aedis      66 Complete      12 Fragmented     175 Missing
## Fri 22 Jul 2022 05:11:56 PM CEST Smittium_culicis     130 Complete     191 Duplicated      17 Fragmented      20 Missing
## Fri 22 Jul 2022 05:12:50 PM CEST Allomyces_macrogynus      77 Complete     306 Duplicated      12 Fragmented      14 Missing
## Fri 22 Jul 2022 05:13:40 PM CEST Glomus_cerebriforme     235 Complete       6 Duplicated       7 Fragmented       8 Missing
## Fri 22 Jul 2022 05:18:50 PM CEST Neocallimastix_sp     128 Complete     193 Duplicated       6 Fragmented      24 Missing
## Fri 22 Jul 2022 05:19:05 PM CEST Olpidium_bornovanus      78 Complete      69 Duplicated      60 Fragmented      87 Missing
## Fri 22 Jul 2022 05:30:16 PM CEST Mortierella_sp     240 Complete      24 Duplicated       4 Fragmented       3 Missing
## Sat 23 Jul 2022 01:18:48 PM CEST Mus_musculus     235 Complete      52 Duplicated       1 Fragmented
## Sun 24 Jul 2022 07:57:40 AM CEST Paraphelidium_tribonemae     124 Complete     225 Duplicated       9 Fragmented      22 Missing
## Sun 24 Jul 2022 08:02:52 AM CEST Nematocida_sp      65 Complete       4 Duplicated       8 Fragmented     178 Missing
## Sun 24 Jul 2022 08:04:00 AM CEST Schizosaccharomyces_pombe     236 Complete       8 Duplicated       5 Fragmented       8 Missing
## Sun 24 Jul 2022 08:09:52 AM CEST Cryptococcus_neoformans     235 Complete       4 Duplicated      10 Fragmented       6 Missing
## Sun Jul 24 08:15:29 CEST 2022 Spizellomyces_punctatus     234 Complete       2 Duplicated       8 Fragmented      10 Missing
## Sun 24 Jul 2022 08:19:19 AM CEST Hyaloraphidium_curvatum     223 Complete      20 Fragmented      10 Missing
## Sun 24 Jul 2022 08:21:10 AM CEST Cylindrobasidium_torrendii     236 Complete      11 Fragmented       6 Missing
## Sun 24 Jul 2022 08:33:10 AM CEST Fonticula_alba     162 Complete       2 Duplicated      30 Fragmented      60 Missing
## Sun Jul 24 08:34:05 CEST 2022 Dimargaris_cristalligena     213 Complete       2 Duplicated      17 Fragmented      22 Missing
## Sun 24 Jul 2022 08:59:53 AM CEST Tuber_borchii     244 Complete       6 Duplicated       4 Fragmented       2 Missing
## Sun 24 Jul 2022 08:35:49 PM CEST Vittaforma_corneae      56 Complete       2 Duplicated      14 Fragmented     182 Missing
## Sun 24 Jul 2022 08:43:03 PM CEST Lipomyces_starkeyi     240 Complete       4 Duplicated       8 Fragmented       3 Missing
## Sun 24 Jul 2022 08:43:43 PM CEST Rozella_allomycis     142 Complete      11 Duplicated      42 Fragmented      64 Missing
## Sun 24 Jul 2022 08:44:54 PM CEST Acaromyces_ingoldii     236 Complete      10 Fragmented       7 Missing
## Sun 24 Jul 2022 08:45:52 PM CEST Sanchytrium_tribonematis     137 Complete      15 Duplicated      30 Fragmented      79 Missing
## Sun 24 Jul 2022 08:47:20 PM CEST Caulochytrium_protostelioides      71 Complete      40 Fragmented     142 Missing
## Sun 24 Jul 2022 08:48:06 PM CEST Calocera_viscosa     232 Complete      14 Duplicated       9 Fragmented       5 Missing
## Sun 24 Jul 2022 08:54:31 PM CEST Paraphaeosphaeria_sporulosa     245 Complete       2 Duplicated       6 Fragmented       1 Missing
## Sun Jul 24 20:58:53 CEST 2022 Pyricularia_oryzae     247 Complete       6 Fragmented
## Sun 24 Jul 2022 08:59:32 PM CEST Capsaspora_owczarzaki     227 Complete      10 Duplicated      11 Fragmented      10 Missing
## Sun Jul 24 21:16:04 CEST 2022 Catenaria_anguillulae     128 Complete     156 Duplicated      27 Fragmented      22 Missing
## Sun 24 Jul 2022 09:16:09 PM CEST Botrytis_cinerea     245 Complete       2 Duplicated       5 Fragmented       2 Missing
## Sun 24 Jul 2022 10:29:45 PM CEST Pneumocystis_murina     239 Complete       2 Duplicated       3 Fragmented      10 Missing
## Sun 24 Jul 2022 10:35:05 PM CEST Yarrowia_lipolytica     242 Complete       4 Duplicated       5 Fragmented       4 Missing
## Sun 24 Jul 2022 10:39:20 PM CEST Amphiamblys_sp      78 Complete      26 Fragmented     149 Missing
## Sun Jul 24 22:40:17 CEST 2022 Anaeromyces_robustus     205 Complete      10 Duplicated       9 Fragmented      34 Missing
## Sun Jul 24 22:41:32 CEST 2022 Daedalea_quercina     244 Complete       6 Fragmented       3 Missing
## Sun 24 Jul 2022 10:50:30 PM CEST Fusarium_proliferatum     248 Complete       4 Duplicated       1 Fragmented       2 Missing
## Sun 24 Jul 2022 10:55:40 PM CEST Paxillus_rubicundulus     235 Complete      10 Fragmented       8 Missing
## Sun 24 Jul 2022 11:01:42 PM CEST Salpingoeca_rosetta     198 Complete       5 Duplicated      15 Fragmented      38 Missing
## Sun 24 Jul 2022 11:19:14 PM CEST Basidiobolus_meristosporus     207 Complete      82 Duplicated       7 Fragmented       1 Missing
## Mon Jul 25 09:17:36 CEST 2022 Batrachochytrium_dendrobatidis     237 Complete       9 Duplicated       7 Fragmented       5 Missing
## Mon Jul 25 09:25:51 CEST 2022 Letharia_columbiana     234 Complete      28 Duplicated       4 Fragmented       1 Missing
## Mon Jul 25 09:51:20 CEST 2022 Rhizophagus_sp     236 Complete       4 Duplicated       8 Fragmented       7 Missing
## Mon Jul 25 10:04:08 CEST 2022 Melampsora_laricipopulina     228 Complete      12 Duplicated      13 Fragmented       6 Missing
## Mon Jul 25 10:33:00 CEST 2022 Rhodotorula_toruloides     224 Complete       2 Duplicated      15 Fragmented      13 Missing
## Mon Jul 25 11:51:20 CEST 2022 Podila_minutissima     243 Complete      12 Duplicated       1 Fragmented       3 Missing
## Mon Jul 25 11:52:26 CEST 2022 - Pipeline part 1 (orthology) done.
## Mon Jul 25 13:10:28 CEST 2022 - BUSCO files will be filtered on a per-sample basis. This could lower the number of species in the final tree.
## Mon Jul 25 13:11:04 CEST 2022 - Number of BUSCO sequence files: 255
## Mon Jul 25 13:11:04 CEST 2022 - Number of deduplicated BUSCO sequence files: 255
## Mon Jul 25 13:11:27 CEST 2022 - Pipeline part filter-orthology done.
## Mon Jul 25 14:12:01 CEST 2022 - Pipeline part 2 (align) done.
## Mon Jul 25 14:20:38 CEST 2022 - The aliscore output file does not exist. Check results for BUSCO: 687505at2759
## Mon Jul 25 14:20:38 CEST 2022 - The aliscore output appears to be empty. Check results for BUSCO: 687505at2759
## Mon Jul 25 14:20:42 CEST 2022 - The aliscore output file does not exist. Check results for BUSCO: 1079130at2759
## Mon Jul 25 14:20:42 CEST 2022 - The aliscore output appears to be empty. Check results for BUSCO: 1079130at2759
## Mon Jul 25 14:20:42 CEST 2022 - The aliscore output file does not exist. Check results for BUSCO: 673132at2759
## Mon Jul 25 14:20:42 CEST 2022 - The aliscore output appears to be empty. Check results for BUSCO: 673132at2759
## Mon Jul 25 14:20:43 CEST 2022 - The aliscore output file does not exist. Check results for BUSCO: 671536at2759
## Mon Jul 25 14:20:43 CEST 2022 - The aliscore output appears to be empty. Check results for BUSCO: 671536at2759
## Mon Jul 25 14:20:45 CEST 2022 - The aliscore output file does not exist. Check results for BUSCO: 324863at2759
## Mon Jul 25 14:20:45 CEST 2022 - The aliscore output appears to be empty. Check results for BUSCO: 324863at2759
## Mon Jul 25 14:20:46 CEST 2022 - The aliscore output file does not exist. Check results for BUSCO: 142542at2759
## Mon Jul 25 14:20:47 CEST 2022 - The aliscore output appears to be empty. Check results for BUSCO: 142542at2759
## Mon Jul 25 14:21:07 CEST 2022 - The aliscore output file does not exist. Check results for BUSCO: 1094121at2759
## Mon Jul 25 14:21:07 CEST 2022 - The aliscore output appears to be empty. Check results for BUSCO: 1094121at2759
## Mon Jul 25 14:21:09 CEST 2022 - The aliscore output file does not exist. Check results for BUSCO: 1053181at2759
## Mon Jul 25 14:21:09 CEST 2022 - The aliscore output file does not exist. Check results for BUSCO: 974865at2759
## Mon Jul 25 14:21:09 CEST 2022 - The aliscore output appears to be empty. Check results for BUSCO: 1053181at2759
## Mon Jul 25 14:21:09 CEST 2022 - The aliscore output appears to be empty. Check results for BUSCO: 974865at2759
## Mon Jul 25 14:21:10 CEST 2022 - The aliscore output file does not exist. Check results for BUSCO: 1222562at2759
## Mon Jul 25 14:21:10 CEST 2022 - The aliscore output appears to be empty. Check results for BUSCO: 1222562at2759
## Mon Jul 25 14:21:10 CEST 2022 - The aliscore output file does not exist. Check results for BUSCO: 1346432at2759
## Mon Jul 25 14:21:10 CEST 2022 - The aliscore output appears to be empty. Check results for BUSCO: 1346432at2759
## Mon Jul 25 14:21:10 CEST 2022 - The aliscore output file does not exist. Check results for BUSCO: 345441at2759
## Mon Jul 25 14:21:10 CEST 2022 - The aliscore output appears to be empty. Check results for BUSCO: 345441at2759
## Mon Jul 25 14:21:12 CEST 2022 - The aliscore output file does not exist. Check results for BUSCO: 975557at2759
## Mon Jul 25 14:21:12 CEST 2022 - The aliscore output file does not exist. Check results for BUSCO: 937275at2759
## Mon Jul 25 14:21:12 CEST 2022 - The aliscore output appears to be empty. Check results for BUSCO: 975557at2759
## Mon Jul 25 14:21:12 CEST 2022 - The aliscore output appears to be empty. Check results for BUSCO: 937275at2759
## Mon Jul 25 14:21:12 CEST 2022 - The aliscore output file does not exist. Check results for BUSCO: 1490022at2759
## Mon Jul 25 14:21:13 CEST 2022 - The aliscore output appears to be empty. Check results for BUSCO: 1490022at2759
## Mon Jul 25 14:21:13 CEST 2022 - The aliscore output file does not exist. Check results for BUSCO: 1552706at2759
## Mon Jul 25 14:21:13 CEST 2022 - The aliscore output appears to be empty. Check results for BUSCO: 1552706at2759
## Mon Jul 25 14:21:13 CEST 2022 - The aliscore output file does not exist. Check results for BUSCO: 388820at2759
## Mon Jul 25 14:21:13 CEST 2022 - The aliscore output appears to be empty. Check results for BUSCO: 388820at2759
## Mon Jul 25 14:21:13 CEST 2022 - The aliscore output file does not exist. Check results for BUSCO: 1395649at2759
## Mon Jul 25 14:21:13 CEST 2022 - The aliscore output appears to be empty. Check results for BUSCO: 1395649at2759
## Mon Jul 25 14:21:13 CEST 2022 - The aliscore output file does not exist. Check results for BUSCO: 453044at2759
## Mon Jul 25 14:21:13 CEST 2022 - The aliscore output file does not exist. Check results for BUSCO: 1593937at2759
## Mon Jul 25 14:21:13 CEST 2022 - The aliscore output appears to be empty. Check results for BUSCO: 453044at2759
## Mon Jul 25 14:21:13 CEST 2022 - The aliscore output appears to be empty. Check results for BUSCO: 1593937at2759
## Mon Jul 25 14:21:13 CEST 2022 - The aliscore output file does not exist. Check results for BUSCO: 1025450at2759
## Mon Jul 25 14:21:13 CEST 2022 - The aliscore output appears to be empty. Check results for BUSCO: 1025450at2759
## Mon Jul 25 14:21:13 CEST 2022 - The aliscore output file does not exist. Check results for BUSCO: 937686at2759
## Mon Jul 25 14:21:14 CEST 2022 - The aliscore output appears to be empty. Check results for BUSCO: 937686at2759
## Mon Jul 25 14:21:14 CEST 2022 - The aliscore output file does not exist. Check results for BUSCO: 923657at2759
## Mon Jul 25 14:21:14 CEST 2022 - The aliscore output appears to be empty. Check results for BUSCO: 923657at2759
## Mon Jul 25 14:21:14 CEST 2022 - The aliscore output file does not exist. Check results for BUSCO: 491869at2759
## Mon Jul 25 14:21:14 CEST 2022 - The aliscore output appears to be empty. Check results for BUSCO: 491869at2759
## Mon Jul 25 14:21:14 CEST 2022 - The aliscore output file does not exist. Check results for BUSCO: 1247641at2759
## Mon Jul 25 14:21:14 CEST 2022 - The aliscore output file does not exist. Check results for BUSCO: 1049599at2759
## Mon Jul 25 14:21:14 CEST 2022 - The aliscore output appears to be empty. Check results for BUSCO: 1247641at2759
## Mon Jul 25 14:21:14 CEST 2022 - The aliscore output file does not exist. Check results for BUSCO: 1032689at2759
## Mon Jul 25 14:21:14 CEST 2022 - The aliscore output file does not exist. Check results for BUSCO: 1442062at2759
## Mon Jul 25 14:21:14 CEST 2022 - The aliscore output appears to be empty. Check results for BUSCO: 1049599at2759
## Mon Jul 25 14:21:14 CEST 2022 - The aliscore output appears to be empty. Check results for BUSCO: 1032689at2759
## Mon Jul 25 14:21:14 CEST 2022 - The aliscore output appears to be empty. Check results for BUSCO: 1442062at2759
## Mon Jul 25 14:21:14 CEST 2022 - The aliscore output file does not exist. Check results for BUSCO: 924753at2759
## Mon Jul 25 14:21:14 CEST 2022 - The aliscore output file does not exist. Check results for BUSCO: 100698at2759
## Mon Jul 25 14:21:14 CEST 2022 - The aliscore output appears to be empty. Check results for BUSCO: 924753at2759
## Mon Jul 25 14:21:14 CEST 2022 - The aliscore output appears to be empty. Check results for BUSCO: 100698at2759
## Mon Jul 25 14:21:15 CEST 2022 - The aliscore output file does not exist. Check results for BUSCO: 1364586at2759
## Mon Jul 25 14:21:15 CEST 2022 - The aliscore output appears to be empty. Check results for BUSCO: 1364586at2759
## Mon Jul 25 14:21:15 CEST 2022 - The aliscore output file does not exist. Check results for BUSCO: 549762at2759
## Mon Jul 25 14:21:15 CEST 2022 - The aliscore output file does not exist. Check results for BUSCO: 331411at2759
## Mon Jul 25 14:21:15 CEST 2022 - The aliscore output appears to be empty. Check results for BUSCO: 549762at2759
## Mon Jul 25 14:21:15 CEST 2022 - The aliscore output appears to be empty. Check results for BUSCO: 331411at2759
## Mon Jul 25 14:21:15 CEST 2022 - The aliscore output file does not exist. Check results for BUSCO: 1645187at2759
## Mon Jul 25 14:21:15 CEST 2022 - The aliscore output appears to be empty. Check results for BUSCO: 1645187at2759
## Mon Jul 25 14:21:15 CEST 2022 - The aliscore output file does not exist. Check results for BUSCO: 1287094at2759
## Mon Jul 25 14:21:15 CEST 2022 - The aliscore output appears to be empty. Check results for BUSCO: 1287094at2759
## Mon Jul 25 14:21:16 CEST 2022 - The aliscore output file does not exist. Check results for BUSCO: 759498at2759
## Mon Jul 25 14:21:16 CEST 2022 - The aliscore output appears to be empty. Check results for BUSCO: 759498at2759
## Mon Jul 25 14:21:16 CEST 2022 - The aliscore output file does not exist. Check results for BUSCO: 844748at2759
## Mon Jul 25 14:21:16 CEST 2022 - The aliscore output appears to be empty. Check results for BUSCO: 844748at2759
## Mon Jul 25 14:21:16 CEST 2022 - The aliscore output file does not exist. Check results for BUSCO: 1057950at2759
## Mon Jul 25 14:21:16 CEST 2022 - The aliscore output appears to be empty. Check results for BUSCO: 1057950at2759
## Mon Jul 25 14:21:17 CEST 2022 - The aliscore output file does not exist. Check results for BUSCO: 1479417at2759
## Mon Jul 25 14:21:17 CEST 2022 - The aliscore output appears to be empty. Check results for BUSCO: 1479417at2759
## Mon Jul 25 14:21:17 CEST 2022 - The aliscore output file does not exist. Check results for BUSCO: 1096688at2759
## Mon Jul 25 14:21:17 CEST 2022 - The aliscore output appears to be empty. Check results for BUSCO: 1096688at2759
## Mon Jul 25 14:21:17 CEST 2022 - The aliscore output file does not exist. Check results for BUSCO: 261419at2759
## Mon Jul 25 14:21:17 CEST 2022 - The aliscore output appears to be empty. Check results for BUSCO: 261419at2759
## Mon Jul 25 14:21:18 CEST 2022 - The aliscore output file does not exist. Check results for BUSCO: 1259741at2759
## Mon Jul 25 14:21:18 CEST 2022 - The aliscore output appears to be empty. Check results for BUSCO: 1259741at2759
## Mon Jul 25 14:21:18 CEST 2022 - The aliscore output file does not exist. Check results for BUSCO: 543764at2759
## Mon Jul 25 14:21:18 CEST 2022 - The aliscore output appears to be empty. Check results for BUSCO: 543764at2759
## Mon Jul 25 14:21:19 CEST 2022 - The aliscore output file does not exist. Check results for BUSCO: 1428854at2759
## Mon Jul 25 14:21:19 CEST 2022 - The aliscore output appears to be empty. Check results for BUSCO: 1428854at2759
## Mon Jul 25 14:21:19 CEST 2022 - The aliscore output file does not exist. Check results for BUSCO: 271586at2759
## Mon Jul 25 14:21:19 CEST 2022 - The aliscore output appears to be empty. Check results for BUSCO: 271586at2759
## Mon Jul 25 14:21:21 CEST 2022 - The aliscore output file does not exist. Check results for BUSCO: 160593at2759
## Mon Jul 25 14:21:21 CEST 2022 - The aliscore output appears to be empty. Check results for BUSCO: 160593at2759
## Mon Jul 25 14:21:22 CEST 2022 - The aliscore output file does not exist. Check results for BUSCO: 166920at2759
## Mon Jul 25 14:21:22 CEST 2022 - The aliscore output appears to be empty. Check results for BUSCO: 166920at2759
## Mon Jul 25 14:21:39 CEST 2022 - The aliscore output file does not exist. Check results for BUSCO: 1338131at2759
## Mon Jul 25 14:21:39 CEST 2022 - The aliscore output appears to be empty. Check results for BUSCO: 1338131at2759
## Mon Jul 25 14:21:40 CEST 2022 - The aliscore output file does not exist. Check results for BUSCO: 865202at2759
## Mon Jul 25 14:21:40 CEST 2022 - The aliscore output appears to be empty. Check results for BUSCO: 865202at2759
## Mon Jul 25 14:21:40 CEST 2022 - The aliscore output file does not exist. Check results for BUSCO: 1138059at2759
## Mon Jul 25 14:21:40 CEST 2022 - The aliscore output appears to be empty. Check results for BUSCO: 1138059at2759
## Mon Jul 25 14:21:40 CEST 2022 - The aliscore output file does not exist. Check results for BUSCO: 1513531at2759
## Mon Jul 25 14:21:40 CEST 2022 - The aliscore output appears to be empty. Check results for BUSCO: 1513531at2759
## Mon Jul 25 14:21:40 CEST 2022 - The aliscore output file does not exist. Check results for BUSCO: 1342242at2759
## Mon Jul 25 14:21:40 CEST 2022 - The aliscore output appears to be empty. Check results for BUSCO: 1342242at2759
## Mon Jul 25 14:21:41 CEST 2022 - The aliscore output file does not exist. Check results for BUSCO: 774318at2759
## Mon Jul 25 14:21:41 CEST 2022 - The aliscore output appears to be empty. Check results for BUSCO: 774318at2759
## Mon Jul 25 14:21:41 CEST 2022 - The aliscore output file does not exist. Check results for BUSCO: 1382842at2759
## Mon Jul 25 14:21:41 CEST 2022 - The aliscore output appears to be empty. Check results for BUSCO: 1382842at2759
## Mon Jul 25 14:21:41 CEST 2022 - The aliscore output file does not exist. Check results for BUSCO: 1223488at2759
## Mon Jul 25 14:21:41 CEST 2022 - The aliscore output file does not exist. Check results for BUSCO: 1200489at2759
## Mon Jul 25 14:21:41 CEST 2022 - The aliscore output appears to be empty. Check results for BUSCO: 1200489at2759
## Mon Jul 25 14:21:41 CEST 2022 - The aliscore output appears to be empty. Check results for BUSCO: 1223488at2759
## Mon Jul 25 14:21:41 CEST 2022 - The aliscore output file does not exist. Check results for BUSCO: 1421503at2759
## Mon Jul 25 14:21:41 CEST 2022 - The aliscore output appears to be empty. Check results for BUSCO: 1421503at2759
## Mon Jul 25 14:21:41 CEST 2022 - The aliscore output file does not exist. Check results for BUSCO: 1370304at2759
## Mon Jul 25 14:21:41 CEST 2022 - The aliscore output appears to be empty. Check results for BUSCO: 1370304at2759
## Mon Jul 25 14:21:42 CEST 2022 - The aliscore output file does not exist. Check results for BUSCO: 751335at2759
## Mon Jul 25 14:21:42 CEST 2022 - The aliscore output appears to be empty. Check results for BUSCO: 751335at2759
## Mon Jul 25 14:21:43 CEST 2022 - The aliscore output file does not exist. Check results for BUSCO: 570797at2759
## Mon Jul 25 14:21:43 CEST 2022 - The aliscore output appears to be empty. Check results for BUSCO: 570797at2759
## Mon Jul 25 14:21:43 CEST 2022 - The aliscore output file does not exist. Check results for BUSCO: 736068at2759
## Mon Jul 25 14:21:43 CEST 2022 - The aliscore output appears to be empty. Check results for BUSCO: 736068at2759
## Mon Jul 25 14:21:43 CEST 2022 - The aliscore output file does not exist. Check results for BUSCO: 355408at2759
## Mon Jul 25 14:21:43 CEST 2022 - The aliscore output appears to be empty. Check results for BUSCO: 355408at2759
## Mon Jul 25 14:21:43 CEST 2022 - The aliscore output file does not exist. Check results for BUSCO: 1538526at2759
## Mon Jul 25 14:21:43 CEST 2022 - The aliscore output appears to be empty. Check results for BUSCO: 1538526at2759
## Mon Jul 25 14:21:44 CEST 2022 - The aliscore output file does not exist. Check results for BUSCO: 1264469at2759
## Mon Jul 25 14:21:44 CEST 2022 - The aliscore output appears to be empty. Check results for BUSCO: 1264469at2759
## Mon Jul 25 14:21:44 CEST 2022 - The aliscore output file does not exist. Check results for BUSCO: 1620056at2759
## Mon Jul 25 14:21:44 CEST 2022 - The aliscore output appears to be empty. Check results for BUSCO: 1620056at2759
## Mon Jul 25 14:21:44 CEST 2022 - The aliscore output file does not exist. Check results for BUSCO: 1377237at2759
## Mon Jul 25 14:21:44 CEST 2022 - The aliscore output file does not exist. Check results for BUSCO: 1432328at2759
## Mon Jul 25 14:21:44 CEST 2022 - The aliscore output appears to be empty. Check results for BUSCO: 1377237at2759
## Mon Jul 25 14:21:44 CEST 2022 - The aliscore output appears to be empty. Check results for BUSCO: 1432328at2759
## Mon Jul 25 14:21:44 CEST 2022 - The aliscore output file does not exist. Check results for BUSCO: 1575179at2759
## Mon Jul 25 14:21:44 CEST 2022 - The aliscore output appears to be empty. Check results for BUSCO: 1575179at2759
## Mon Jul 25 14:21:45 CEST 2022 - The aliscore output file does not exist. Check results for BUSCO: 1233814at2759
## Mon Jul 25 14:21:45 CEST 2022 - The aliscore output appears to be empty. Check results for BUSCO: 1233814at2759
## Mon Jul 25 14:21:45 CEST 2022 - The aliscore output file does not exist. Check results for BUSCO: 931188at2759
## Mon Jul 25 14:21:45 CEST 2022 - The aliscore output file does not exist. Check results for BUSCO: 1339553at2759
## Mon Jul 25 14:21:46 CEST 2022 - The aliscore output appears to be empty. Check results for BUSCO: 1339553at2759
## Mon Jul 25 14:21:46 CEST 2022 - The aliscore output file does not exist. Check results for BUSCO: 1287401at2759
## Mon Jul 25 14:21:46 CEST 2022 - The aliscore output appears to be empty. Check results for BUSCO: 1287401at2759
## Mon Jul 25 14:21:46 CEST 2022 - The aliscore output file does not exist. Check results for BUSCO: 768809at2759
## Mon Jul 25 14:21:46 CEST 2022 - The aliscore output appears to be empty. Check results for BUSCO: 768809at2759
## Mon Jul 25 14:21:46 CEST 2022 - The aliscore output file does not exist. Check results for BUSCO: 1563319at2759
## Mon Jul 25 14:21:46 CEST 2022 - The aliscore output file does not exist. Check results for BUSCO: 919955at2759
## Mon Jul 25 14:21:46 CEST 2022 - The aliscore output appears to be empty. Check results for BUSCO: 1563319at2759
## Mon Jul 25 14:21:46 CEST 2022 - The aliscore output appears to be empty. Check results for BUSCO: 919955at2759
## Mon Jul 25 14:21:47 CEST 2022 - The aliscore output file does not exist. Check results for BUSCO: 1018517at2759
## Mon Jul 25 14:21:47 CEST 2022 - The aliscore output appears to be empty. Check results for BUSCO: 1018517at2759
## Mon Jul 25 14:21:47 CEST 2022 - The aliscore output file does not exist. Check results for BUSCO: 898782at2759
## Mon Jul 25 14:21:48 CEST 2022 - The aliscore output appears to be empty. Check results for BUSCO: 898782at2759
## Mon Jul 25 14:21:48 CEST 2022 - The aliscore output file does not exist. Check results for BUSCO: 836599at2759
## Mon Jul 25 14:21:48 CEST 2022 - The aliscore output appears to be empty. Check results for BUSCO: 836599at2759
## Mon Jul 25 14:21:48 CEST 2022 - The aliscore output file does not exist. Check results for BUSCO: 176625at2759
## Mon Jul 25 14:21:48 CEST 2022 - The aliscore output file does not exist. Check results for BUSCO: 418107at2759
## Mon Jul 25 14:21:48 CEST 2022 - The aliscore output appears to be empty. Check results for BUSCO: 176625at2759
## Mon Jul 25 14:21:48 CEST 2022 - The aliscore output appears to be empty. Check results for BUSCO: 418107at2759
## Mon Jul 25 14:21:48 CEST 2022 - The aliscore output file does not exist. Check results for BUSCO: 1087488at2759
## Mon Jul 25 14:21:48 CEST 2022 - The aliscore output appears to be empty. Check results for BUSCO: 1087488at2759
## Mon Jul 25 14:21:48 CEST 2022 - The aliscore output file does not exist. Check results for BUSCO: 814241at2759
## Mon Jul 25 14:21:48 CEST 2022 - The aliscore output file does not exist. Check results for BUSCO: 1258856at2759
## Mon Jul 25 14:21:48 CEST 2022 - The aliscore output appears to be empty. Check results for BUSCO: 1258856at2759
## Mon Jul 25 14:21:48 CEST 2022 - The aliscore output appears to be empty. Check results for BUSCO: 814241at2759
## Mon Jul 25 14:21:49 CEST 2022 - The aliscore output file does not exist. Check results for BUSCO: 674160at2759
## Mon Jul 25 14:21:49 CEST 2022 - The aliscore output appears to be empty. Check results for BUSCO: 674160at2759
## Mon Jul 25 14:22:06 CEST 2022 - The aliscore output file does not exist. Check results for BUSCO: 1525971at2759
## Mon Jul 25 14:22:07 CEST 2022 - The aliscore output appears to be empty. Check results for BUSCO: 1525971at2759
## Mon Jul 25 14:22:10 CEST 2022 - The aliscore output file does not exist. Check results for BUSCO: 1220881at2759
## Mon Jul 25 14:22:10 CEST 2022 - The aliscore output appears to be empty. Check results for BUSCO: 1220881at2759
## Mon Jul 25 14:22:10 CEST 2022 - The aliscore output file does not exist. Check results for BUSCO: 1284731at2759
## Mon Jul 25 14:22:10 CEST 2022 - The aliscore output appears to be empty. Check results for BUSCO: 1284731at2759
## Mon Jul 25 14:22:11 CEST 2022 - The aliscore output file does not exist. Check results for BUSCO: 1236198at2759
## Mon Jul 25 14:22:11 CEST 2022 - The aliscore output appears to be empty. Check results for BUSCO: 1236198at2759
## Mon Jul 25 14:22:11 CEST 2022 - The aliscore output file does not exist. Check results for BUSCO: 887370at2759
## Mon Jul 25 14:22:11 CEST 2022 - The aliscore output appears to be empty. Check results for BUSCO: 887370at2759
## Mon Jul 25 14:22:13 CEST 2022 - The aliscore output file does not exist. Check results for BUSCO: 1450538at2759
## Mon Jul 25 14:22:13 CEST 2022 - The aliscore output appears to be empty. Check results for BUSCO: 1450538at2759
## Mon Jul 25 14:22:13 CEST 2022 - The aliscore output file does not exist. Check results for BUSCO: 1545004at2759
## Mon Jul 25 14:22:13 CEST 2022 - The aliscore output appears to be empty. Check results for BUSCO: 1545004at2759
## Mon Jul 25 14:22:13 CEST 2022 - The aliscore output file does not exist. Check results for BUSCO: 598949at2759
## Mon Jul 25 14:22:13 CEST 2022 - The aliscore output appears to be empty. Check results for BUSCO: 598949at2759
## Mon Jul 25 14:22:13 CEST 2022 - The aliscore output file does not exist. Check results for BUSCO: 1488235at2759
## Mon Jul 25 14:22:13 CEST 2022 - The aliscore output appears to be empty. Check results for BUSCO: 1488235at2759
## Mon Jul 25 14:22:13 CEST 2022 - The aliscore output file does not exist. Check results for BUSCO: 1488436at2759
## Mon Jul 25 14:22:14 CEST 2022 - The aliscore output appears to be empty. Check results for BUSCO: 1488436at2759
## Mon Jul 25 14:22:14 CEST 2022 - The aliscore output file does not exist. Check results for BUSCO: 1348942at2759
## Mon Jul 25 14:22:14 CEST 2022 - The aliscore output appears to be empty. Check results for BUSCO: 1348942at2759
## Mon Jul 25 14:22:14 CEST 2022 - The aliscore output file does not exist. Check results for BUSCO: 1003258at2759
## Mon Jul 25 14:22:14 CEST 2022 - The aliscore output appears to be empty. Check results for BUSCO: 1003258at2759
## Mon Jul 25 14:22:14 CEST 2022 - The aliscore output file does not exist. Check results for BUSCO: 891751at2759
## Mon Jul 25 14:22:14 CEST 2022 - The aliscore output appears to be empty. Check results for BUSCO: 891751at2759
## Mon Jul 25 14:22:14 CEST 2022 - The aliscore output file does not exist. Check results for BUSCO: 1504863at2759
## Mon Jul 25 14:22:14 CEST 2022 - The aliscore output appears to be empty. Check results for BUSCO: 1504863at2759
## Mon Jul 25 14:22:14 CEST 2022 - The aliscore output file does not exist. Check results for BUSCO: 97116at2759
## Mon Jul 25 14:22:14 CEST 2022 - The aliscore output appears to be empty. Check results for BUSCO: 97116at2759
## Mon Jul 25 14:22:15 CEST 2022 - The aliscore output file does not exist. Check results for BUSCO: 1567796at2759
## Mon Jul 25 14:22:15 CEST 2022 - The aliscore output file does not exist. Check results for BUSCO: 1157302at2759
## Mon Jul 25 14:22:15 CEST 2022 - The aliscore output appears to be empty. Check results for BUSCO: 1567796at2759
## Mon Jul 25 14:22:15 CEST 2022 - The aliscore output file does not exist. Check results for BUSCO: 1576404at2759
## Mon Jul 25 14:22:15 CEST 2022 - The aliscore output file does not exist. Check results for BUSCO: 1322299at2759
## Mon Jul 25 14:22:15 CEST 2022 - The aliscore output appears to be empty. Check results for BUSCO: 1157302at2759
## Mon Jul 25 14:22:15 CEST 2022 - The aliscore output appears to be empty. Check results for BUSCO: 1576404at2759
## Mon Jul 25 14:22:15 CEST 2022 - The aliscore output appears to be empty. Check results for BUSCO: 1322299at2759
## Mon Jul 25 14:22:15 CEST 2022 - The aliscore output file does not exist. Check results for BUSCO: 293315at2759
## Mon Jul 25 14:22:15 CEST 2022 - The aliscore output appears to be empty. Check results for BUSCO: 293315at2759
## Mon Jul 25 14:22:15 CEST 2022 - The aliscore output file does not exist. Check results for BUSCO: 513979at2759
## Mon Jul 25 14:22:15 CEST 2022 - The aliscore output appears to be empty. Check results for BUSCO: 513979at2759
## Mon Jul 25 14:22:15 CEST 2022 - The aliscore output file does not exist. Check results for BUSCO: 976469at2759
## Mon Jul 25 14:22:15 CEST 2022 - The aliscore output appears to be empty. Check results for BUSCO: 976469at2759
## Mon Jul 25 14:22:15 CEST 2022 - The aliscore output file does not exist. Check results for BUSCO: 1626636at2759
## Mon Jul 25 14:22:15 CEST 2022 - The aliscore output appears to be empty. Check results for BUSCO: 1626636at2759
## Mon Jul 25 14:22:15 CEST 2022 - The aliscore output file does not exist. Check results for BUSCO: 1194691at2759
## Mon Jul 25 14:22:15 CEST 2022 - The aliscore output file does not exist. Check results for BUSCO: 779909at2759
## Mon Jul 25 14:22:15 CEST 2022 - The aliscore output appears to be empty. Check results for BUSCO: 1194691at2759
## Mon Jul 25 14:22:15 CEST 2022 - The aliscore output appears to be empty. Check results for BUSCO: 779909at2759
## Mon Jul 25 14:22:15 CEST 2022 - The aliscore output file does not exist. Check results for BUSCO: 655400at2759
## Mon Jul 25 14:22:16 CEST 2022 - The aliscore output appears to be empty. Check results for BUSCO: 655400at2759
## Mon Jul 25 14:22:16 CEST 2022 - The aliscore output file does not exist. Check results for BUSCO: 946128at2759
## Mon Jul 25 14:22:16 CEST 2022 - The aliscore output appears to be empty. Check results for BUSCO: 946128at2759
## Mon Jul 25 14:22:16 CEST 2022 - The aliscore output file does not exist. Check results for BUSCO: 1426075at2759
## Mon Jul 25 14:22:16 CEST 2022 - The aliscore output appears to be empty. Check results for BUSCO: 1426075at2759
## Mon Jul 25 14:22:16 CEST 2022 - The aliscore output file does not exist. Check results for BUSCO: 1193442at2759
## Mon Jul 25 14:22:16 CEST 2022 - The aliscore output appears to be empty. Check results for BUSCO: 1193442at2759
## Mon Jul 25 14:22:17 CEST 2022 - The aliscore output file does not exist. Check results for BUSCO: 1111142at2759
## Mon Jul 25 14:22:17 CEST 2022 - The aliscore output appears to be empty. Check results for BUSCO: 1111142at2759
## Mon Jul 25 14:22:17 CEST 2022 - The aliscore output file does not exist. Check results for BUSCO: 325552at2759
## Mon Jul 25 14:22:17 CEST 2022 - The aliscore output appears to be empty. Check results for BUSCO: 325552at2759
## Mon Jul 25 14:22:17 CEST 2022 - The aliscore output file does not exist. Check results for BUSCO: 1182451at2759
## Mon Jul 25 14:22:17 CEST 2022 - The aliscore output appears to be empty. Check results for BUSCO: 1182451at2759
## Mon Jul 25 14:22:17 CEST 2022 - The aliscore output file does not exist. Check results for BUSCO: 905026at2759
## Mon Jul 25 14:22:17 CEST 2022 - The aliscore output appears to be empty. Check results for BUSCO: 905026at2759
## Mon Jul 25 14:22:19 CEST 2022 - The aliscore output file does not exist. Check results for BUSCO: 296129at2759
## Mon Jul 25 14:22:19 CEST 2022 - The aliscore output appears to be empty. Check results for BUSCO: 296129at2759
## Mon Jul 25 14:22:21 CEST 2022 - The aliscore output file does not exist. Check results for BUSCO: 1314980at2759
## Mon Jul 25 14:22:21 CEST 2022 - The aliscore output appears to be empty. Check results for BUSCO: 1314980at2759
## Mon Jul 25 14:22:22 CEST 2022 - The aliscore output file does not exist. Check results for BUSCO: 1312453at2759
## Mon Jul 25 14:22:22 CEST 2022 - The aliscore output appears to be empty. Check results for BUSCO: 1312453at2759
## Mon Jul 25 14:22:38 CEST 2022 - The aliscore output file does not exist. Check results for BUSCO: 457861at2759
## Mon Jul 25 14:22:38 CEST 2022 - The aliscore output appears to be empty. Check results for BUSCO: 457861at2759
## Mon Jul 25 14:22:38 CEST 2022 - The aliscore output file does not exist. Check results for BUSCO: 1197019at2759
## Mon Jul 25 14:22:38 CEST 2022 - The aliscore output appears to be empty. Check results for BUSCO: 1197019at2759
## Mon Jul 25 14:22:40 CEST 2022 - The aliscore output file does not exist. Check results for BUSCO: 1309031at2759
## Mon Jul 25 14:22:40 CEST 2022 - The aliscore output appears to be empty. Check results for BUSCO: 1309031at2759
## Mon Jul 25 14:22:40 CEST 2022 - The aliscore output file does not exist. Check results for BUSCO: 937686at2759
## Mon Jul 25 14:22:41 CEST 2022 - The aliscore output appears to be empty. Check results for BUSCO: 937686at2759
## Mon Jul 25 14:22:41 CEST 2022 - The aliscore output file does not exist. Check results for BUSCO: 1178688at2759
## Mon Jul 25 14:22:41 CEST 2022 - The aliscore output appears to be empty. Check results for BUSCO: 1178688at2759
## Mon Jul 25 14:22:41 CEST 2022 - The aliscore output file does not exist. Check results for BUSCO: 453044at2759
## Mon Jul 25 14:22:41 CEST 2022 - The aliscore output appears to be empty. Check results for BUSCO: 453044at2759
## Mon Jul 25 14:22:42 CEST 2022 - The aliscore output file does not exist. Check results for BUSCO: 1407446at2759
## Mon Jul 25 14:22:42 CEST 2022 - The aliscore output appears to be empty. Check results for BUSCO: 1407446at2759
## Mon Jul 25 14:22:43 CEST 2022 - The aliscore output file does not exist. Check results for BUSCO: 1076134at2759
## Mon Jul 25 14:22:43 CEST 2022 - The aliscore output appears to be empty. Check results for BUSCO: 1076134at2759
## Mon Jul 25 14:22:43 CEST 2022 - The aliscore output file does not exist. Check results for BUSCO: 464990at2759
## Mon Jul 25 14:22:43 CEST 2022 - The aliscore output appears to be empty. Check results for BUSCO: 464990at2759
## Mon Jul 25 14:22:43 CEST 2022 - The aliscore output file does not exist. Check results for BUSCO: 1442062at2759
## Mon Jul 25 14:22:43 CEST 2022 - The aliscore output appears to be empty. Check results for BUSCO: 1442062at2759
## Mon Jul 25 14:22:43 CEST 2022 - The aliscore output file does not exist. Check results for BUSCO: 1558822at2759
## Mon Jul 25 14:22:43 CEST 2022 - The aliscore output appears to be empty. Check results for BUSCO: 1558822at2759
## Mon Jul 25 14:22:43 CEST 2022 - The aliscore output file does not exist. Check results for BUSCO: 396755at2759
## Mon Jul 25 14:22:43 CEST 2022 - The aliscore output file does not exist. Check results for BUSCO: 844748at2759
## Mon Jul 25 14:22:43 CEST 2022 - The aliscore output appears to be empty. Check results for BUSCO: 396755at2759
## Mon Jul 25 14:22:43 CEST 2022 - The aliscore output appears to be empty. Check results for BUSCO: 844748at2759
## Mon Jul 25 14:22:43 CEST 2022 - The aliscore output file does not exist. Check results for BUSCO: 1041560at2759
## Mon Jul 25 14:22:43 CEST 2022 - The aliscore output appears to be empty. Check results for BUSCO: 1041560at2759
## Mon Jul 25 14:22:44 CEST 2022 - The aliscore output file does not exist. Check results for BUSCO: 1346165at2759
## Mon Jul 25 14:22:44 CEST 2022 - The aliscore output appears to be empty. Check results for BUSCO: 1346165at2759
## Mon Jul 25 14:22:44 CEST 2022 - The aliscore output file does not exist. Check results for BUSCO: 1459797at2759
## Mon Jul 25 14:22:44 CEST 2022 - The aliscore output appears to be empty. Check results for BUSCO: 1459797at2759
## Mon Jul 25 14:22:44 CEST 2022 - The aliscore output file does not exist. Check results for BUSCO: 1014314at2759
## Mon Jul 25 14:22:44 CEST 2022 - The aliscore output appears to be empty. Check results for BUSCO: 1014314at2759
## Mon Jul 25 14:22:44 CEST 2022 - The aliscore output file does not exist. Check results for BUSCO: 1623701at2759
## Mon Jul 25 14:22:44 CEST 2022 - The aliscore output file does not exist. Check results for BUSCO: 1380409at2759
## Mon Jul 25 14:22:44 CEST 2022 - The aliscore output file does not exist. Check results for BUSCO: 1079827at2759
## Mon Jul 25 14:22:44 CEST 2022 - The aliscore output file does not exist. Check results for BUSCO: 1260807at2759
## Mon Jul 25 14:22:44 CEST 2022 - The aliscore output appears to be empty. Check results for BUSCO: 1623701at2759
## Mon Jul 25 14:22:44 CEST 2022 - The aliscore output appears to be empty. Check results for BUSCO: 1380409at2759
## Mon Jul 25 14:22:44 CEST 2022 - The aliscore output appears to be empty. Check results for BUSCO: 1079827at2759
## Mon Jul 25 14:22:45 CEST 2022 - The aliscore output appears to be empty. Check results for BUSCO: 1260807at2759
## Mon Jul 25 14:22:45 CEST 2022 - The aliscore output file does not exist. Check results for BUSCO: 734666at2759
## Mon Jul 25 14:22:45 CEST 2022 - The aliscore output appears to be empty. Check results for BUSCO: 734666at2759
## Mon Jul 25 14:22:45 CEST 2022 - The aliscore output file does not exist. Check results for BUSCO: 1445102at2759
## Mon Jul 25 14:22:45 CEST 2022 - The aliscore output appears to be empty. Check results for BUSCO: 1445102at2759
## Mon Jul 25 14:22:45 CEST 2022 - The aliscore output file does not exist. Check results for BUSCO: 1526152at2759
## Mon Jul 25 14:22:45 CEST 2022 - The aliscore output appears to be empty. Check results for BUSCO: 1526152at2759
## Mon Jul 25 14:22:45 CEST 2022 - The aliscore output file does not exist. Check results for BUSCO: 1645187at2759
## Mon Jul 25 14:22:45 CEST 2022 - The aliscore output appears to be empty. Check results for BUSCO: 1645187at2759
## Mon Jul 25 14:22:45 CEST 2022 - The aliscore output file does not exist. Check results for BUSCO: 1428265at2759
## Mon Jul 25 14:22:45 CEST 2022 - The aliscore output file does not exist. Check results for BUSCO: 1346432at2759
## Mon Jul 25 14:22:45 CEST 2022 - The aliscore output appears to be empty. Check results for BUSCO: 1346432at2759
## Mon Jul 25 14:22:45 CEST 2022 - The aliscore output file does not exist. Check results for BUSCO: 1094121at2759
## Mon Jul 25 14:22:45 CEST 2022 - The aliscore output appears to be empty. Check results for BUSCO: 1428265at2759
## Mon Jul 25 14:22:45 CEST 2022 - The aliscore output appears to be empty. Check results for BUSCO: 1094121at2759
## Mon Jul 25 14:22:45 CEST 2022 - The aliscore output file does not exist. Check results for BUSCO: 975158at2759
## Mon Jul 25 14:22:45 CEST 2022 - The aliscore output appears to be empty. Check results for BUSCO: 975158at2759
## Mon Jul 25 14:22:46 CEST 2022 - The aliscore output file does not exist. Check results for BUSCO: 1257440at2759
## Mon Jul 25 14:22:46 CEST 2022 - The aliscore output appears to be empty. Check results for BUSCO: 1257440at2759
## Mon Jul 25 14:22:46 CEST 2022 - The aliscore output file does not exist. Check results for BUSCO: 1552706at2759
## Mon Jul 25 14:22:46 CEST 2022 - The aliscore output appears to be empty. Check results for BUSCO: 1552706at2759
## Mon Jul 25 14:22:46 CEST 2022 - The aliscore output file does not exist. Check results for BUSCO: 866359at2759
## Mon Jul 25 14:22:46 CEST 2022 - The aliscore output appears to be empty. Check results for BUSCO: 866359at2759
## Mon Jul 25 14:22:46 CEST 2022 - The aliscore output file does not exist. Check results for BUSCO: 388820at2759
## Mon Jul 25 14:22:46 CEST 2022 - The aliscore output file does not exist. Check results for BUSCO: 1454155at2759
## Mon Jul 25 14:22:46 CEST 2022 - The aliscore output appears to be empty. Check results for BUSCO: 388820at2759
## Mon Jul 25 14:22:46 CEST 2022 - The aliscore output appears to be empty. Check results for BUSCO: 1454155at2759
## Mon Jul 25 14:22:46 CEST 2022 - The aliscore output file does not exist. Check results for BUSCO: 1090038at2759
## Mon Jul 25 14:22:46 CEST 2022 - The aliscore output appears to be empty. Check results for BUSCO: 1090038at2759
## Mon Jul 25 14:22:46 CEST 2022 - The aliscore output file does not exist. Check results for BUSCO: 858842at2759
## Mon Jul 25 14:22:46 CEST 2022 - The aliscore output appears to be empty. Check results for BUSCO: 858842at2759
## Mon Jul 25 14:22:46 CEST 2022 - The aliscore output file does not exist. Check results for BUSCO: 1057950at2759
## Mon Jul 25 14:22:46 CEST 2022 - The aliscore output file does not exist. Check results for BUSCO: 708105at2759
## Mon Jul 25 14:22:47 CEST 2022 - The aliscore output appears to be empty. Check results for BUSCO: 1057950at2759
## Mon Jul 25 14:22:47 CEST 2022 - The aliscore output appears to be empty. Check results for BUSCO: 708105at2759
## Mon Jul 25 14:22:47 CEST 2022 - The aliscore output file does not exist. Check results for BUSCO: 1304061at2759
## Mon Jul 25 14:22:47 CEST 2022 - The aliscore output file does not exist. Check results for BUSCO: 1030907at2759
## Mon Jul 25 14:22:47 CEST 2022 - The aliscore output file does not exist. Check results for BUSCO: 923657at2759
## Mon Jul 25 14:22:47 CEST 2022 - The aliscore output appears to be empty. Check results for BUSCO: 1304061at2759
## Mon Jul 25 14:22:47 CEST 2022 - The aliscore output appears to be empty. Check results for BUSCO: 1030907at2759
## Mon Jul 25 14:22:47 CEST 2022 - The aliscore output appears to be empty. Check results for BUSCO: 923657at2759
## Mon Jul 25 14:22:47 CEST 2022 - The aliscore output file does not exist. Check results for BUSCO: 1287094at2759
## Mon Jul 25 14:22:47 CEST 2022 - The aliscore output appears to be empty. Check results for BUSCO: 1287094at2759
## Mon Jul 25 14:22:47 CEST 2022 - The aliscore output file does not exist. Check results for BUSCO: 1441854at2759
## Mon Jul 25 14:22:47 CEST 2022 - The aliscore output file does not exist. Check results for BUSCO: 1364586at2759
## Mon Jul 25 14:22:47 CEST 2022 - The aliscore output appears to be empty. Check results for BUSCO: 1441854at2759
## Mon Jul 25 14:22:47 CEST 2022 - The aliscore output file does not exist. Check results for BUSCO: 1129824at2759
## Mon Jul 25 14:22:47 CEST 2022 - The aliscore output appears to be empty. Check results for BUSCO: 1364586at2759
## Mon Jul 25 14:22:47 CEST 2022 - The aliscore output appears to be empty. Check results for BUSCO: 1129824at2759
## Mon Jul 25 14:22:47 CEST 2022 - The aliscore output file does not exist. Check results for BUSCO: 100698at2759
## Mon Jul 25 14:22:47 CEST 2022 - The aliscore output appears to be empty. Check results for BUSCO: 100698at2759
## Mon Jul 25 14:22:47 CEST 2022 - The aliscore output file does not exist. Check results for BUSCO: 1479417at2759
## Mon Jul 25 14:22:47 CEST 2022 - The aliscore output appears to be empty. Check results for BUSCO: 1479417at2759
## Mon Jul 25 14:22:47 CEST 2022 - The aliscore output file does not exist. Check results for BUSCO: 1173229at2759
## Mon Jul 25 14:22:47 CEST 2022 - The aliscore output file does not exist. Check results for BUSCO: 759498at2759
## Mon Jul 25 14:22:47 CEST 2022 - The aliscore output appears to be empty. Check results for BUSCO: 1173229at2759
## Mon Jul 25 14:22:47 CEST 2022 - The aliscore output appears to be empty. Check results for BUSCO: 759498at2759
## Mon Jul 25 14:22:48 CEST 2022 - The aliscore output file does not exist. Check results for BUSCO: 664730at2759
## Mon Jul 25 14:22:48 CEST 2022 - The aliscore output appears to be empty. Check results for BUSCO: 664730at2759
## Mon Jul 25 14:22:48 CEST 2022 - The aliscore output file does not exist. Check results for BUSCO: 1323575at2759
## Mon Jul 25 14:22:48 CEST 2022 - The aliscore output appears to be empty. Check results for BUSCO: 1323575at2759
## Mon Jul 25 14:22:48 CEST 2022 - The aliscore output file does not exist. Check results for BUSCO: 719531at2759
## Mon Jul 25 14:22:48 CEST 2022 - The aliscore output appears to be empty. Check results for BUSCO: 719531at2759
## Mon Jul 25 14:22:48 CEST 2022 - The aliscore output file does not exist. Check results for BUSCO: 722805at2759
## Mon Jul 25 14:22:48 CEST 2022 - The aliscore output file does not exist. Check results for BUSCO: 1115196at2759
## Mon Jul 25 14:22:48 CEST 2022 - The aliscore output appears to be empty. Check results for BUSCO: 722805at2759
## Mon Jul 25 14:22:48 CEST 2022 - The aliscore output appears to be empty. Check results for BUSCO: 1115196at2759
## Mon Jul 25 14:22:48 CEST 2022 - The aliscore output file does not exist. Check results for BUSCO: 917326at2759
## Mon Jul 25 14:22:48 CEST 2022 - The aliscore output appears to be empty. Check results for BUSCO: 917326at2759
## Mon Jul 25 14:22:49 CEST 2022 - The aliscore output file does not exist. Check results for BUSCO: 720340at2759
## Mon Jul 25 14:22:49 CEST 2022 - The aliscore output appears to be empty. Check results for BUSCO: 720340at2759
## Mon Jul 25 14:22:49 CEST 2022 - The aliscore output file does not exist. Check results for BUSCO: 306227at2759
## Mon Jul 25 14:22:49 CEST 2022 - The aliscore output appears to be empty. Check results for BUSCO: 306227at2759
## Mon Jul 25 14:22:49 CEST 2022 - The aliscore output file does not exist. Check results for BUSCO: 1247641at2759
## Mon Jul 25 14:22:49 CEST 2022 - The aliscore output appears to be empty. Check results for BUSCO: 1247641at2759
## Mon Jul 25 14:22:49 CEST 2022 - The aliscore output file does not exist. Check results for BUSCO: 679187at2759
## Mon Jul 25 14:22:49 CEST 2022 - The aliscore output appears to be empty. Check results for BUSCO: 679187at2759
## Mon Jul 25 14:22:49 CEST 2022 - The aliscore output file does not exist. Check results for BUSCO: 390348at2759
## Mon Jul 25 14:22:49 CEST 2022 - The aliscore output appears to be empty. Check results for BUSCO: 390348at2759
## Mon Jul 25 14:22:49 CEST 2022 - The aliscore output file does not exist. Check results for BUSCO: 1085752at2759
## Mon Jul 25 14:22:49 CEST 2022 - The aliscore output appears to be empty. Check results for BUSCO: 1085752at2759
## Mon Jul 25 14:22:49 CEST 2022 - The aliscore output file does not exist. Check results for BUSCO: 582756at2759
## Mon Jul 25 14:22:49 CEST 2022 - The aliscore output appears to be empty. Check results for BUSCO: 582756at2759
## Mon Jul 25 14:22:50 CEST 2022 - The aliscore output file does not exist. Check results for BUSCO: 426305at2759
## Mon Jul 25 14:22:50 CEST 2022 - The aliscore output file does not exist. Check results for BUSCO: 320059at2759
## Mon Jul 25 14:22:50 CEST 2022 - The aliscore output file does not exist. Check results for BUSCO: 1106766at2759
## Mon Jul 25 14:22:50 CEST 2022 - The aliscore output appears to be empty. Check results for BUSCO: 426305at2759
## Mon Jul 25 14:22:50 CEST 2022 - The aliscore output appears to be empty. Check results for BUSCO: 320059at2759
## Mon Jul 25 14:22:50 CEST 2022 - The aliscore output appears to be empty. Check results for BUSCO: 1106766at2759
## Mon Jul 25 14:22:50 CEST 2022 - The aliscore output file does not exist. Check results for BUSCO: 855834at2759
## Mon Jul 25 14:22:50 CEST 2022 - The aliscore output appears to be empty. Check results for BUSCO: 855834at2759
## Mon Jul 25 14:22:50 CEST 2022 - The aliscore output file does not exist. Check results for BUSCO: 673132at2759
## Mon Jul 25 14:22:50 CEST 2022 - The aliscore output file does not exist. Check results for BUSCO: 450058at2759
## Mon Jul 25 14:22:50 CEST 2022 - The aliscore output file does not exist. Check results for BUSCO: 1324510at2759
## Mon Jul 25 14:22:50 CEST 2022 - The aliscore output appears to be empty. Check results for BUSCO: 673132at2759
## Mon Jul 25 14:22:50 CEST 2022 - The aliscore output appears to be empty. Check results for BUSCO: 450058at2759
## Mon Jul 25 14:22:50 CEST 2022 - The aliscore output appears to be empty. Check results for BUSCO: 1324510at2759
## Mon Jul 25 14:22:50 CEST 2022 - The aliscore output file does not exist. Check results for BUSCO: 692986at2759
## Mon Jul 25 14:22:50 CEST 2022 - The aliscore output appears to be empty. Check results for BUSCO: 692986at2759
## Mon Jul 25 14:22:50 CEST 2022 - The aliscore output file does not exist. Check results for BUSCO: 1194797at2759
## Mon Jul 25 14:22:50 CEST 2022 - The aliscore output appears to be empty. Check results for BUSCO: 1194797at2759
## Mon Jul 25 14:22:50 CEST 2022 - The aliscore output file does not exist. Check results for BUSCO: 261419at2759
## Mon Jul 25 14:22:51 CEST 2022 - The aliscore output appears to be empty. Check results for BUSCO: 261419at2759
## Mon Jul 25 14:22:51 CEST 2022 - The aliscore output file does not exist. Check results for BUSCO: 1379600at2759
## Mon Jul 25 14:22:51 CEST 2022 - The aliscore output appears to be empty. Check results for BUSCO: 1379600at2759
## Mon Jul 25 14:22:51 CEST 2022 - The aliscore output file does not exist. Check results for BUSCO: 1248958at2759
## Mon Jul 25 14:22:51 CEST 2022 - The aliscore output appears to be empty. Check results for BUSCO: 1248958at2759
## Mon Jul 25 14:22:51 CEST 2022 - The aliscore output file does not exist. Check results for BUSCO: 543764at2759
## Mon Jul 25 14:22:51 CEST 2022 - The aliscore output appears to be empty. Check results for BUSCO: 543764at2759
## Mon Jul 25 14:22:51 CEST 2022 - The aliscore output file does not exist. Check results for BUSCO: 87842at2759
## Mon Jul 25 14:22:51 CEST 2022 - The aliscore output appears to be empty. Check results for BUSCO: 87842at2759
## Mon Jul 25 14:22:51 CEST 2022 - The aliscore output file does not exist. Check results for BUSCO: 834694at2759
## Mon Jul 25 14:22:51 CEST 2022 - The aliscore output appears to be empty. Check results for BUSCO: 834694at2759
## Mon Jul 25 14:22:53 CEST 2022 - The aliscore output file does not exist. Check results for BUSCO: 777920at2759
## Mon Jul 25 14:22:53 CEST 2022 - The aliscore output appears to be empty. Check results for BUSCO: 777920at2759
## Mon Jul 25 14:22:53 CEST 2022 - The aliscore output file does not exist. Check results for BUSCO: 271586at2759
## Mon Jul 25 14:22:53 CEST 2022 - The aliscore output appears to be empty. Check results for BUSCO: 271586at2759
## Mon Jul 25 14:22:54 CEST 2022 - The aliscore output file does not exist. Check results for BUSCO: 261328at2759
## Mon Jul 25 14:22:54 CEST 2022 - The aliscore output appears to be empty. Check results for BUSCO: 261328at2759
## Mon Jul 25 14:22:54 CEST 2022 - The aliscore output file does not exist. Check results for BUSCO: 166920at2759
## Mon Jul 25 14:22:54 CEST 2022 - The aliscore output file does not exist. Check results for BUSCO: 324863at2759
## Mon Jul 25 14:22:54 CEST 2022 - The aliscore output appears to be empty. Check results for BUSCO: 166920at2759
## Mon Jul 25 14:22:54 CEST 2022 - The aliscore output appears to be empty. Check results for BUSCO: 324863at2759
## Mon Jul 25 14:22:56 CEST 2022 - The aliscore output file does not exist. Check results for BUSCO: 142542at2759
## Mon Jul 25 14:22:56 CEST 2022 - The aliscore output appears to be empty. Check results for BUSCO: 142542at2759
## Mon Jul 25 14:22:58 CEST 2022 - The aliscore output file does not exist. Check results for BUSCO: 156083at2759
## Mon Jul 25 14:22:58 CEST 2022 - The aliscore output appears to be empty. Check results for BUSCO: 156083at2759
## Mon Jul 25 14:23:08 CEST 2022 - The aliscore output file does not exist. Check results for BUSCO: 1440961at2759
## Mon Jul 25 14:23:08 CEST 2022 - The aliscore output appears to be empty. Check results for BUSCO: 1440961at2759
## Mon Jul 25 14:23:09 CEST 2022 - The aliscore output file does not exist. Check results for BUSCO: 1398309at2759
## Mon Jul 25 14:23:09 CEST 2022 - The aliscore output appears to be empty. Check results for BUSCO: 1398309at2759
## Mon Jul 25 14:23:10 CEST 2022 - The aliscore output file does not exist. Check results for BUSCO: 1454155at2759
## Mon Jul 25 14:23:10 CEST 2022 - The aliscore output appears to be empty. Check results for BUSCO: 1454155at2759
## Mon Jul 25 14:23:10 CEST 2022 - The aliscore output file does not exist. Check results for BUSCO: 1217666at2759
## Mon Jul 25 14:23:10 CEST 2022 - The aliscore output file does not exist. Check results for BUSCO: 1430056at2759
## Mon Jul 25 14:23:10 CEST 2022 - The aliscore output appears to be empty. Check results for BUSCO: 1217666at2759
## Mon Jul 25 14:23:10 CEST 2022 - The aliscore output appears to be empty. Check results for BUSCO: 1430056at2759
## Mon Jul 25 14:23:10 CEST 2022 - The aliscore output file does not exist. Check results for BUSCO: 974865at2759
## Mon Jul 25 14:23:10 CEST 2022 - The aliscore output appears to be empty. Check results for BUSCO: 974865at2759
## Mon Jul 25 14:23:11 CEST 2022 - The aliscore output file does not exist. Check results for BUSCO: 1304061at2759
## Mon Jul 25 14:23:11 CEST 2022 - The aliscore output appears to be empty. Check results for BUSCO: 1304061at2759
## Mon Jul 25 14:23:11 CEST 2022 - The aliscore output file does not exist. Check results for BUSCO: 621827at2759
## Mon Jul 25 14:23:11 CEST 2022 - The aliscore output appears to be empty. Check results for BUSCO: 621827at2759
## Mon Jul 25 14:23:12 CEST 2022 - The aliscore output file does not exist. Check results for BUSCO: 858842at2759
## Mon Jul 25 14:23:12 CEST 2022 - The aliscore output appears to be empty. Check results for BUSCO: 858842at2759
## Mon Jul 25 14:23:12 CEST 2022 - The aliscore output file does not exist. Check results for BUSCO: 1309031at2759
## Mon Jul 25 14:23:12 CEST 2022 - The aliscore output appears to be empty. Check results for BUSCO: 1309031at2759
## Mon Jul 25 14:23:12 CEST 2022 - The aliscore output file does not exist. Check results for BUSCO: 1108845at2759
## Mon Jul 25 14:23:12 CEST 2022 - The aliscore output appears to be empty. Check results for BUSCO: 1108845at2759
## Mon Jul 25 14:23:12 CEST 2022 - The aliscore output file does not exist. Check results for BUSCO: 1530008at2759
## Mon Jul 25 14:23:12 CEST 2022 - The aliscore output appears to be empty. Check results for BUSCO: 1530008at2759
## Mon Jul 25 14:23:13 CEST 2022 - The aliscore output file does not exist. Check results for BUSCO: 426305at2759
## Mon Jul 25 14:23:13 CEST 2022 - The aliscore output appears to be empty. Check results for BUSCO: 426305at2759
## Mon Jul 25 14:23:13 CEST 2022 - The aliscore output file does not exist. Check results for BUSCO: 679187at2759
## Mon Jul 25 14:23:13 CEST 2022 - The aliscore output appears to be empty. Check results for BUSCO: 679187at2759
## Mon Jul 25 14:23:13 CEST 2022 - The aliscore output file does not exist. Check results for BUSCO: 1593937at2759
## Mon Jul 25 14:23:13 CEST 2022 - The aliscore output appears to be empty. Check results for BUSCO: 1593937at2759
## Mon Jul 25 14:23:13 CEST 2022 - The aliscore output file does not exist. Check results for BUSCO: 1324510at2759
## Mon Jul 25 14:23:13 CEST 2022 - The aliscore output appears to be empty. Check results for BUSCO: 1324510at2759
## Mon Jul 25 14:23:13 CEST 2022 - The aliscore output file does not exist. Check results for BUSCO: 891751at2759
## Mon Jul 25 14:23:13 CEST 2022 - The aliscore output appears to be empty. Check results for BUSCO: 891751at2759
## Mon Jul 25 14:23:14 CEST 2022 - The aliscore output file does not exist. Check results for BUSCO: 1405146at2759
## Mon Jul 25 14:23:14 CEST 2022 - The aliscore output appears to be empty. Check results for BUSCO: 1405146at2759
## Mon Jul 25 14:23:14 CEST 2022 - The aliscore output file does not exist. Check results for BUSCO: 1380409at2759
## Mon Jul 25 14:23:14 CEST 2022 - The aliscore output appears to be empty. Check results for BUSCO: 1380409at2759
## Mon Jul 25 14:23:14 CEST 2022 - The aliscore output file does not exist. Check results for BUSCO: 330169at2759
## Mon Jul 25 14:23:14 CEST 2022 - The aliscore output appears to be empty. Check results for BUSCO: 330169at2759
## Mon Jul 25 14:23:14 CEST 2022 - The aliscore output file does not exist. Check results for BUSCO: 1428265at2759
## Mon Jul 25 14:23:14 CEST 2022 - The aliscore output appears to be empty. Check results for BUSCO: 1428265at2759
## Mon Jul 25 14:23:14 CEST 2022 - The aliscore output file does not exist. Check results for BUSCO: 1395649at2759
## Mon Jul 25 14:23:14 CEST 2022 - The aliscore output appears to be empty. Check results for BUSCO: 1395649at2759
## Mon Jul 25 14:23:14 CEST 2022 - The aliscore output file does not exist. Check results for BUSCO: 777920at2759
## Mon Jul 25 14:23:14 CEST 2022 - The aliscore output appears to be empty. Check results for BUSCO: 777920at2759
## Mon Jul 25 14:23:15 CEST 2022 - The aliscore output file does not exist. Check results for BUSCO: 87842at2759
## Mon Jul 25 14:23:15 CEST 2022 - The aliscore output appears to be empty. Check results for BUSCO: 87842at2759
## Mon Jul 25 14:23:15 CEST 2022 - The aliscore output file does not exist. Check results for BUSCO: 1220881at2759
## Mon Jul 25 14:23:15 CEST 2022 - The aliscore output file does not exist. Check results for BUSCO: 671846at2759
## Mon Jul 25 14:23:15 CEST 2022 - The aliscore output file does not exist. Check results for BUSCO: 708105at2759
## Mon Jul 25 14:23:15 CEST 2022 - The aliscore output appears to be empty. Check results for BUSCO: 1220881at2759
## Mon Jul 25 14:23:15 CEST 2022 - The aliscore output appears to be empty. Check results for BUSCO: 671846at2759
## Mon Jul 25 14:23:15 CEST 2022 - The aliscore output appears to be empty. Check results for BUSCO: 708105at2759
## Mon Jul 25 14:23:15 CEST 2022 - The aliscore output file does not exist. Check results for BUSCO: 937275at2759
## Mon Jul 25 14:23:15 CEST 2022 - The aliscore output file does not exist. Check results for BUSCO: 1432328at2759
## Mon Jul 25 14:23:15 CEST 2022 - The aliscore output appears to be empty. Check results for BUSCO: 937275at2759
## Mon Jul 25 14:23:15 CEST 2022 - The aliscore output appears to be empty. Check results for BUSCO: 1432328at2759
## Mon Jul 25 14:23:15 CEST 2022 - The aliscore output file does not exist. Check results for BUSCO: 345441at2759
## Mon Jul 25 14:23:15 CEST 2022 - The aliscore output file does not exist. Check results for BUSCO: 270107at2759
## Mon Jul 25 14:23:15 CEST 2022 - The aliscore output appears to be empty. Check results for BUSCO: 345441at2759
## Mon Jul 25 14:23:15 CEST 2022 - The aliscore output appears to be empty. Check results for BUSCO: 270107at2759
## Mon Jul 25 14:23:15 CEST 2022 - The aliscore output file does not exist. Check results for BUSCO: 1434266at2759
## Mon Jul 25 14:23:15 CEST 2022 - The aliscore output appears to be empty. Check results for BUSCO: 1434266at2759
## Mon Jul 25 14:23:16 CEST 2022 - The aliscore output file does not exist. Check results for BUSCO: 887370at2759
## Mon Jul 25 14:23:16 CEST 2022 - The aliscore output appears to be empty. Check results for BUSCO: 887370at2759
## Mon Jul 25 14:23:16 CEST 2022 - The aliscore output file does not exist. Check results for BUSCO: 1490022at2759
## Mon Jul 25 14:23:16 CEST 2022 - The aliscore output appears to be empty. Check results for BUSCO: 1490022at2759
## Mon Jul 25 14:23:16 CEST 2022 - The aliscore output file does not exist. Check results for BUSCO: 1085752at2759
## Mon Jul 25 14:23:16 CEST 2022 - The aliscore output appears to be empty. Check results for BUSCO: 1085752at2759
## Mon Jul 25 14:23:17 CEST 2022 - The aliscore output file does not exist. Check results for BUSCO: 1001705at2759
## Mon Jul 25 14:23:17 CEST 2022 - The aliscore output file does not exist. Check results for BUSCO: 1370285at2759
## Mon Jul 25 14:23:17 CEST 2022 - The aliscore output appears to be empty. Check results for BUSCO: 1001705at2759
## Mon Jul 25 14:23:17 CEST 2022 - The aliscore output appears to be empty. Check results for BUSCO: 1370285at2759
## Mon Jul 25 14:23:17 CEST 2022 - The aliscore output file does not exist. Check results for BUSCO: 975557at2759
## Mon Jul 25 14:23:17 CEST 2022 - The aliscore output file does not exist. Check results for BUSCO: 1025450at2759
## Mon Jul 25 14:23:17 CEST 2022 - The aliscore output appears to be empty. Check results for BUSCO: 975557at2759
## Mon Jul 25 14:23:17 CEST 2022 - The aliscore output appears to be empty. Check results for BUSCO: 1025450at2759
## Mon Jul 25 14:23:17 CEST 2022 - The aliscore output file does not exist. Check results for BUSCO: 1322642at2759
## Mon Jul 25 14:23:17 CEST 2022 - The aliscore output appears to be empty. Check results for BUSCO: 1322642at2759
## Mon Jul 25 14:23:17 CEST 2022 - The aliscore output file does not exist. Check results for BUSCO: 390348at2759
## Mon Jul 25 14:23:17 CEST 2022 - The aliscore output appears to be empty. Check results for BUSCO: 390348at2759
## Mon Jul 25 14:23:17 CEST 2022 - The aliscore output file does not exist. Check results for BUSCO: 674169at2759
## Mon Jul 25 14:23:18 CEST 2022 - The aliscore output appears to be empty. Check results for BUSCO: 674169at2759
## Mon Jul 25 14:23:18 CEST 2022 - The aliscore output file does not exist. Check results for BUSCO: 918816at2759
## Mon Jul 25 14:23:18 CEST 2022 - The aliscore output appears to be empty. Check results for BUSCO: 918816at2759
## Mon Jul 25 14:23:19 CEST 2022 - The aliscore output file does not exist. Check results for BUSCO: 720340at2759
## Mon Jul 25 14:23:19 CEST 2022 - The aliscore output appears to be empty. Check results for BUSCO: 720340at2759
## Mon Jul 25 14:23:19 CEST 2022 - The aliscore output file does not exist. Check results for BUSCO: 931188at2759
## Mon Jul 25 14:23:19 CEST 2022 - The aliscore output appears to be empty. Check results for BUSCO: 931188at2759
## Mon Jul 25 14:23:20 CEST 2022 - The aliscore output file does not exist. Check results for BUSCO: 1258856at2759
## Mon Jul 25 14:23:20 CEST 2022 - The aliscore output appears to be empty. Check results for BUSCO: 1258856at2759
## Mon Jul 25 14:23:22 CEST 2022 - The aliscore output file does not exist. Check results for BUSCO: 674160at2759
## Mon Jul 25 14:23:22 CEST 2022 - The aliscore output appears to be empty. Check results for BUSCO: 674160at2759
## Mon Jul 25 14:23:39 CEST 2022 - The aliscore output file does not exist. Check results for BUSCO: 1382842at2759
## Mon Jul 25 14:23:39 CEST 2022 - The aliscore output appears to be empty. Check results for BUSCO: 1382842at2759
## Mon Jul 25 14:23:40 CEST 2022 - The aliscore output file does not exist. Check results for BUSCO: 878143at2759
## Mon Jul 25 14:23:40 CEST 2022 - The aliscore output file does not exist. Check results for BUSCO: 1291729at2759
## Mon Jul 25 14:23:40 CEST 2022 - The aliscore output appears to be empty. Check results for BUSCO: 878143at2759
## Mon Jul 25 14:23:40 CEST 2022 - The aliscore output appears to be empty. Check results for BUSCO: 1291729at2759
## Mon Jul 25 14:23:40 CEST 2022 - The aliscore output file does not exist. Check results for BUSCO: 734341at2759
## Mon Jul 25 14:23:40 CEST 2022 - The aliscore output appears to be empty. Check results for BUSCO: 734341at2759
## Mon Jul 25 14:23:40 CEST 2022 - The aliscore output file does not exist. Check results for BUSCO: 1355894at2759
## Mon Jul 25 14:23:40 CEST 2022 - The aliscore output appears to be empty. Check results for BUSCO: 1355894at2759
## Mon Jul 25 14:23:40 CEST 2022 - The aliscore output file does not exist. Check results for BUSCO: 1251252at2759
## Mon Jul 25 14:23:40 CEST 2022 - The aliscore output appears to be empty. Check results for BUSCO: 1251252at2759
## Mon Jul 25 14:23:41 CEST 2022 - The aliscore output file does not exist. Check results for BUSCO: 1051021at2759
## Mon Jul 25 14:23:41 CEST 2022 - The aliscore output file does not exist. Check results for BUSCO: 687505at2759
## Mon Jul 25 14:23:41 CEST 2022 - The aliscore output appears to be empty. Check results for BUSCO: 1051021at2759
## Mon Jul 25 14:23:41 CEST 2022 - The aliscore output appears to be empty. Check results for BUSCO: 687505at2759
## Mon Jul 25 14:23:41 CEST 2022 - The aliscore output file does not exist. Check results for BUSCO: 1228942at2759
## Mon Jul 25 14:23:41 CEST 2022 - The aliscore output file does not exist. Check results for BUSCO: 865202at2759
## Mon Jul 25 14:23:41 CEST 2022 - The aliscore output appears to be empty. Check results for BUSCO: 1228942at2759
## Mon Jul 25 14:23:41 CEST 2022 - The aliscore output appears to be empty. Check results for BUSCO: 865202at2759
## Mon Jul 25 14:23:41 CEST 2022 - The aliscore output file does not exist. Check results for BUSCO: 1588798at2759
## Mon Jul 25 14:23:41 CEST 2022 - The aliscore output appears to be empty. Check results for BUSCO: 1588798at2759
## Mon Jul 25 14:23:41 CEST 2022 - The aliscore output file does not exist. Check results for BUSCO: 1065019at2759
## Mon Jul 25 14:23:41 CEST 2022 - The aliscore output appears to be empty. Check results for BUSCO: 1065019at2759
## Mon Jul 25 14:23:42 CEST 2022 - The aliscore output file does not exist. Check results for BUSCO: 1128607at2759
## Mon Jul 25 14:23:42 CEST 2022 - The aliscore output appears to be empty. Check results for BUSCO: 1128607at2759
## Mon Jul 25 14:23:42 CEST 2022 - The aliscore output file does not exist. Check results for BUSCO: 1346165at2759
## Mon Jul 25 14:23:42 CEST 2022 - The aliscore output appears to be empty. Check results for BUSCO: 1346165at2759
## Mon Jul 25 14:23:42 CEST 2022 - The aliscore output file does not exist. Check results for BUSCO: 341721at2759
## Mon Jul 25 14:23:42 CEST 2022 - The aliscore output appears to be empty. Check results for BUSCO: 341721at2759
## Mon Jul 25 14:23:42 CEST 2022 - The aliscore output file does not exist. Check results for BUSCO: 901894at2759
## Mon Jul 25 14:23:42 CEST 2022 - The aliscore output appears to be empty. Check results for BUSCO: 901894at2759
## Mon Jul 25 14:23:43 CEST 2022 - The aliscore output file does not exist. Check results for BUSCO: 83779at2759
## Mon Jul 25 14:23:43 CEST 2022 - The aliscore output appears to be empty. Check results for BUSCO: 83779at2759
## Mon Jul 25 14:23:43 CEST 2022 - The aliscore output file does not exist. Check results for BUSCO: 1620056at2759
## Mon Jul 25 14:23:43 CEST 2022 - The aliscore output appears to be empty. Check results for BUSCO: 1620056at2759
## Mon Jul 25 14:23:43 CEST 2022 - The aliscore output file does not exist. Check results for BUSCO: 975158at2759
## Mon Jul 25 14:23:43 CEST 2022 - The aliscore output appears to be empty. Check results for BUSCO: 975158at2759
## Mon Jul 25 14:23:44 CEST 2022 - The aliscore output file does not exist. Check results for BUSCO: 1538526at2759
## Mon Jul 25 14:23:44 CEST 2022 - The aliscore output appears to be empty. Check results for BUSCO: 1538526at2759
## Mon Jul 25 14:23:44 CEST 2022 - The aliscore output file does not exist. Check results for BUSCO: 836599at2759
## Mon Jul 25 14:23:44 CEST 2022 - The aliscore output file does not exist. Check results for BUSCO: 1423847at2759
## Mon Jul 25 14:23:44 CEST 2022 - The aliscore output file does not exist. Check results for BUSCO: 973442at2759
## Mon Jul 25 14:23:44 CEST 2022 - The aliscore output appears to be empty. Check results for BUSCO: 973442at2759
## Mon Jul 25 14:23:44 CEST 2022 - The aliscore output appears to be empty. Check results for BUSCO: 1423847at2759
## Mon Jul 25 14:23:44 CEST 2022 - The aliscore output appears to be empty. Check results for BUSCO: 836599at2759
## Mon Jul 25 14:23:44 CEST 2022 - The aliscore output file does not exist. Check results for BUSCO: 1112002at2759
## Mon Jul 25 14:23:44 CEST 2022 - The aliscore output appears to be empty. Check results for BUSCO: 1112002at2759
## Mon Jul 25 14:23:44 CEST 2022 - The aliscore output file does not exist. Check results for BUSCO: 1264469at2759
## Mon Jul 25 14:23:44 CEST 2022 - The aliscore output file does not exist. Check results for BUSCO: 1633672at2759
## Mon Jul 25 14:23:44 CEST 2022 - The aliscore output appears to be empty. Check results for BUSCO: 1264469at2759
## Mon Jul 25 14:23:44 CEST 2022 - The aliscore output appears to be empty. Check results for BUSCO: 1633672at2759
## Mon Jul 25 14:23:44 CEST 2022 - The aliscore output file does not exist. Check results for BUSCO: 369837at2759
## Mon Jul 25 14:23:44 CEST 2022 - The aliscore output appears to be empty. Check results for BUSCO: 369837at2759
## Mon Jul 25 14:23:44 CEST 2022 - The aliscore output file does not exist. Check results for BUSCO: 1575179at2759
## Mon Jul 25 14:23:45 CEST 2022 - The aliscore output appears to be empty. Check results for BUSCO: 1575179at2759
## Mon Jul 25 14:23:45 CEST 2022 - The aliscore output file does not exist. Check results for BUSCO: 981902at2759
## Mon Jul 25 14:23:45 CEST 2022 - The aliscore output appears to be empty. Check results for BUSCO: 981902at2759
## Mon Jul 25 14:23:45 CEST 2022 - The aliscore output file does not exist. Check results for BUSCO: 956854at2759
## Mon Jul 25 14:23:45 CEST 2022 - The aliscore output appears to be empty. Check results for BUSCO: 956854at2759
## Mon Jul 25 14:23:45 CEST 2022 - The aliscore output file does not exist. Check results for BUSCO: 919955at2759
## Mon Jul 25 14:23:45 CEST 2022 - The aliscore output appears to be empty. Check results for BUSCO: 919955at2759
## Mon Jul 25 14:23:45 CEST 2022 - The aliscore output file does not exist. Check results for BUSCO: 721605at2759
## Mon Jul 25 14:23:45 CEST 2022 - The aliscore output file does not exist. Check results for BUSCO: 1370304at2759
## Mon Jul 25 14:23:45 CEST 2022 - The aliscore output appears to be empty. Check results for BUSCO: 721605at2759
## Mon Jul 25 14:23:46 CEST 2022 - The aliscore output appears to be empty. Check results for BUSCO: 1370304at2759
## Mon Jul 25 14:23:46 CEST 2022 - The aliscore output file does not exist. Check results for BUSCO: 1339553at2759
## Mon Jul 25 14:23:46 CEST 2022 - The aliscore output file does not exist. Check results for BUSCO: 1455730at2759
## Mon Jul 25 14:23:46 CEST 2022 - The aliscore output appears to be empty. Check results for BUSCO: 1339553at2759
## Mon Jul 25 14:23:46 CEST 2022 - The aliscore output appears to be empty. Check results for BUSCO: 1455730at2759
## Mon Jul 25 14:23:46 CEST 2022 - The aliscore output file does not exist. Check results for BUSCO: 996662at2759
## Mon Jul 25 14:23:46 CEST 2022 - The aliscore output file does not exist. Check results for BUSCO: 1249159at2759
## Mon Jul 25 14:23:46 CEST 2022 - The aliscore output appears to be empty. Check results for BUSCO: 996662at2759
## Mon Jul 25 14:23:46 CEST 2022 - The aliscore output appears to be empty. Check results for BUSCO: 1249159at2759
## Mon Jul 25 14:23:46 CEST 2022 - The aliscore output file does not exist. Check results for BUSCO: 257318at2759
## Mon Jul 25 14:23:46 CEST 2022 - The aliscore output appears to be empty. Check results for BUSCO: 257318at2759
## Mon Jul 25 14:23:46 CEST 2022 - The aliscore output file does not exist. Check results for BUSCO: 1377237at2759
## Mon Jul 25 14:23:46 CEST 2022 - The aliscore output file does not exist. Check results for BUSCO: 1563319at2759
## Mon Jul 25 14:23:46 CEST 2022 - The aliscore output appears to be empty. Check results for BUSCO: 1377237at2759
## Mon Jul 25 14:23:46 CEST 2022 - The aliscore output file does not exist. Check results for BUSCO: 179362at2759
## Mon Jul 25 14:23:46 CEST 2022 - The aliscore output appears to be empty. Check results for BUSCO: 1563319at2759
## Mon Jul 25 14:23:46 CEST 2022 - The aliscore output appears to be empty. Check results for BUSCO: 179362at2759
## Mon Jul 25 14:23:47 CEST 2022 - The aliscore output file does not exist. Check results for BUSCO: 801857at2759
## Mon Jul 25 14:23:47 CEST 2022 - The aliscore output appears to be empty. Check results for BUSCO: 801857at2759
## Mon Jul 25 14:23:48 CEST 2022 - The aliscore output file does not exist. Check results for BUSCO: 551907at2759
## Mon Jul 25 14:23:48 CEST 2022 - The aliscore output file does not exist. Check results for BUSCO: 375960at2759
## Mon Jul 25 14:23:48 CEST 2022 - The aliscore output appears to be empty. Check results for BUSCO: 551907at2759
## Mon Jul 25 14:23:48 CEST 2022 - The aliscore output file does not exist. Check results for BUSCO: 392369at2759
## Mon Jul 25 14:23:48 CEST 2022 - The aliscore output appears to be empty. Check results for BUSCO: 375960at2759
## Mon Jul 25 14:23:48 CEST 2022 - The aliscore output appears to be empty. Check results for BUSCO: 392369at2759
## Mon Jul 25 14:23:48 CEST 2022 - The aliscore output file does not exist. Check results for BUSCO: 1018517at2759
## Mon Jul 25 14:23:48 CEST 2022 - The aliscore output appears to be empty. Check results for BUSCO: 1018517at2759
## Mon Jul 25 14:23:48 CEST 2022 - The aliscore output file does not exist. Check results for BUSCO: 383503at2759
## Mon Jul 25 14:23:48 CEST 2022 - The aliscore output appears to be empty. Check results for BUSCO: 383503at2759
## Mon Jul 25 14:23:48 CEST 2022 - The aliscore output file does not exist. Check results for BUSCO: 1087488at2759
## Mon Jul 25 14:23:48 CEST 2022 - The aliscore output appears to be empty. Check results for BUSCO: 1087488at2759
## Mon Jul 25 14:23:48 CEST 2022 - The aliscore output file does not exist. Check results for BUSCO: 176625at2759
## Mon Jul 25 14:23:48 CEST 2022 - The aliscore output appears to be empty. Check results for BUSCO: 176625at2759
## Mon Jul 25 14:23:48 CEST 2022 - The aliscore output file does not exist. Check results for BUSCO: 1041560at2759
## Mon Jul 25 14:23:48 CEST 2022 - The aliscore output appears to be empty. Check results for BUSCO: 1041560at2759
## Mon Jul 25 14:23:48 CEST 2022 - The aliscore output file does not exist. Check results for BUSCO: 814241at2759
## Mon Jul 25 14:23:48 CEST 2022 - The aliscore output appears to be empty. Check results for BUSCO: 814241at2759
## Mon Jul 25 14:23:48 CEST 2022 - The aliscore output file does not exist. Check results for BUSCO: 898782at2759
## Mon Jul 25 14:23:48 CEST 2022 - The aliscore output file does not exist. Check results for BUSCO: 418107at2759
## Mon Jul 25 14:23:48 CEST 2022 - The aliscore output appears to be empty. Check results for BUSCO: 898782at2759
## Mon Jul 25 14:23:49 CEST 2022 - The aliscore output appears to be empty. Check results for BUSCO: 418107at2759
## Mon Jul 25 14:23:49 CEST 2022 - The aliscore output file does not exist. Check results for BUSCO: 917326at2759
## Mon Jul 25 14:23:49 CEST 2022 - The aliscore output file does not exist. Check results for BUSCO: 722805at2759
## Mon Jul 25 14:23:49 CEST 2022 - The aliscore output appears to be empty. Check results for BUSCO: 917326at2759
## Mon Jul 25 14:23:49 CEST 2022 - The aliscore output appears to be empty. Check results for BUSCO: 722805at2759
## Mon Jul 25 14:23:49 CEST 2022 - The aliscore output file does not exist. Check results for BUSCO: 245208at2759
## Mon Jul 25 14:23:49 CEST 2022 - The aliscore output appears to be empty. Check results for BUSCO: 245208at2759
## Mon Jul 25 14:23:50 CEST 2022 - The aliscore output file does not exist. Check results for BUSCO: 396755at2759
## Mon Jul 25 14:23:50 CEST 2022 - The aliscore output appears to be empty. Check results for BUSCO: 396755at2759
## Mon Jul 25 14:24:10 CEST 2022 - The aliscore output file does not exist. Check results for BUSCO: 1129824at2759
## Mon Jul 25 14:24:10 CEST 2022 - The aliscore output appears to be empty. Check results for BUSCO: 1129824at2759
## Mon Jul 25 14:24:11 CEST 2022 - The aliscore output file does not exist. Check results for BUSCO: 1178688at2759
## Mon Jul 25 14:24:11 CEST 2022 - The aliscore output appears to be empty. Check results for BUSCO: 1178688at2759
## Mon Jul 25 14:24:12 CEST 2022 - The aliscore output file does not exist. Check results for BUSCO: 761109at2759
## Mon Jul 25 14:24:12 CEST 2022 - The aliscore output appears to be empty. Check results for BUSCO: 761109at2759
## Mon Jul 25 14:24:12 CEST 2022 - The aliscore output file does not exist. Check results for BUSCO: 1432328at2759
## Mon Jul 25 14:24:12 CEST 2022 - The aliscore output appears to be empty. Check results for BUSCO: 1432328at2759
## Mon Jul 25 14:24:12 CEST 2022 - The aliscore output file does not exist. Check results for BUSCO: 290630at2759
## Mon Jul 25 14:24:12 CEST 2022 - The aliscore output appears to be empty. Check results for BUSCO: 290630at2759
## Mon Jul 25 14:24:13 CEST 2022 - The aliscore output file does not exist. Check results for BUSCO: 674169at2759
## Mon Jul 25 14:24:13 CEST 2022 - The aliscore output appears to be empty. Check results for BUSCO: 674169at2759
## Mon Jul 25 14:24:13 CEST 2022 - The aliscore output file does not exist. Check results for BUSCO: 1194797at2759
## Mon Jul 25 14:24:13 CEST 2022 - The aliscore output appears to be empty. Check results for BUSCO: 1194797at2759
## Mon Jul 25 14:24:13 CEST 2022 - The aliscore output file does not exist. Check results for BUSCO: 543764at2759
## Mon Jul 25 14:24:13 CEST 2022 - The aliscore output appears to be empty. Check results for BUSCO: 543764at2759
## Mon Jul 25 14:24:14 CEST 2022 - The aliscore output file does not exist. Check results for BUSCO: 923657at2759
## Mon Jul 25 14:24:14 CEST 2022 - The aliscore output appears to be empty. Check results for BUSCO: 923657at2759
## Mon Jul 25 14:24:14 CEST 2022 - The aliscore output file does not exist. Check results for BUSCO: 1407446at2759
## Mon Jul 25 14:24:14 CEST 2022 - The aliscore output appears to be empty. Check results for BUSCO: 1407446at2759
## Mon Jul 25 14:24:14 CEST 2022 - The aliscore output file does not exist. Check results for BUSCO: 1038775at2759
## Mon Jul 25 14:24:14 CEST 2022 - The aliscore output appears to be empty. Check results for BUSCO: 1038775at2759
## Mon Jul 25 14:24:14 CEST 2022 - The aliscore output file does not exist. Check results for BUSCO: 1530008at2759
## Mon Jul 25 14:24:14 CEST 2022 - The aliscore output appears to be empty. Check results for BUSCO: 1530008at2759
## Mon Jul 25 14:24:14 CEST 2022 - The aliscore output file does not exist. Check results for BUSCO: 1358374at2759
## Mon Jul 25 14:24:15 CEST 2022 - The aliscore output appears to be empty. Check results for BUSCO: 1358374at2759
## Mon Jul 25 14:24:15 CEST 2022 - The aliscore output file does not exist. Check results for BUSCO: 1395649at2759
## Mon Jul 25 14:24:15 CEST 2022 - The aliscore output appears to be empty. Check results for BUSCO: 1395649at2759
## Mon Jul 25 14:24:15 CEST 2022 - The aliscore output file does not exist. Check results for BUSCO: 1434266at2759
## Mon Jul 25 14:24:15 CEST 2022 - The aliscore output appears to be empty. Check results for BUSCO: 1434266at2759
## Mon Jul 25 14:24:15 CEST 2022 - The aliscore output file does not exist. Check results for BUSCO: 1275837at2759
## Mon Jul 25 14:24:15 CEST 2022 - The aliscore output appears to be empty. Check results for BUSCO: 1275837at2759
## Mon Jul 25 14:24:15 CEST 2022 - The aliscore output file does not exist. Check results for BUSCO: 931188at2759
## Mon Jul 25 14:24:15 CEST 2022 - The aliscore output file does not exist. Check results for BUSCO: 1248958at2759
## Mon Jul 25 14:24:15 CEST 2022 - The aliscore output appears to be empty. Check results for BUSCO: 931188at2759
## Mon Jul 25 14:24:15 CEST 2022 - The aliscore output appears to be empty. Check results for BUSCO: 1248958at2759
## Mon Jul 25 14:24:16 CEST 2022 - The aliscore output file does not exist. Check results for BUSCO: 1010730at2759
## Mon Jul 25 14:24:16 CEST 2022 - The aliscore output appears to be empty. Check results for BUSCO: 1010730at2759
## Mon Jul 25 14:24:16 CEST 2022 - The aliscore output file does not exist. Check results for BUSCO: 1593937at2759
## Mon Jul 25 14:24:16 CEST 2022 - The aliscore output appears to be empty. Check results for BUSCO: 1593937at2759
## Mon Jul 25 14:24:16 CEST 2022 - The aliscore output file does not exist. Check results for BUSCO: 944899at2759
## Mon Jul 25 14:24:16 CEST 2022 - The aliscore output appears to be empty. Check results for BUSCO: 944899at2759
## Mon Jul 25 14:24:16 CEST 2022 - The aliscore output file does not exist. Check results for BUSCO: 817008at2759
## Mon Jul 25 14:24:16 CEST 2022 - The aliscore output appears to be empty. Check results for BUSCO: 817008at2759
## Mon Jul 25 14:24:17 CEST 2022 - The aliscore output file does not exist. Check results for BUSCO: 673132at2759
## Mon Jul 25 14:24:17 CEST 2022 - The aliscore output appears to be empty. Check results for BUSCO: 673132at2759
## Mon Jul 25 14:24:17 CEST 2022 - The aliscore output file does not exist. Check results for BUSCO: 261328at2759
## Mon Jul 25 14:24:17 CEST 2022 - The aliscore output appears to be empty. Check results for BUSCO: 261328at2759
## Mon Jul 25 14:24:17 CEST 2022 - The aliscore output file does not exist. Check results for BUSCO: 621827at2759
## Mon Jul 25 14:24:17 CEST 2022 - The aliscore output file does not exist. Check results for BUSCO: 1079130at2759
## Mon Jul 25 14:24:17 CEST 2022 - The aliscore output appears to be empty. Check results for BUSCO: 621827at2759
## Mon Jul 25 14:24:17 CEST 2022 - The aliscore output appears to be empty. Check results for BUSCO: 1079130at2759
## Mon Jul 25 14:24:17 CEST 2022 - The aliscore output file does not exist. Check results for BUSCO: 582756at2759
## Mon Jul 25 14:24:17 CEST 2022 - The aliscore output appears to be empty. Check results for BUSCO: 582756at2759
## Mon Jul 25 14:24:17 CEST 2022 - The aliscore output file does not exist. Check results for BUSCO: 939345at2759
## Mon Jul 25 14:24:17 CEST 2022 - The aliscore output file does not exist. Check results for BUSCO: 1322642at2759
## Mon Jul 25 14:24:17 CEST 2022 - The aliscore output file does not exist. Check results for BUSCO: 671536at2759
## Mon Jul 25 14:24:17 CEST 2022 - The aliscore output appears to be empty. Check results for BUSCO: 939345at2759
## Mon Jul 25 14:24:18 CEST 2022 - The aliscore output appears to be empty. Check results for BUSCO: 1322642at2759
## Mon Jul 25 14:24:18 CEST 2022 - The aliscore output appears to be empty. Check results for BUSCO: 671536at2759
## Mon Jul 25 14:24:18 CEST 2022 - The aliscore output file does not exist. Check results for BUSCO: 604979at2759
## Mon Jul 25 14:24:18 CEST 2022 - The aliscore output appears to be empty. Check results for BUSCO: 604979at2759
## Mon Jul 25 14:24:19 CEST 2022 - The aliscore output file does not exist. Check results for BUSCO: 271586at2759
## Mon Jul 25 14:24:19 CEST 2022 - The aliscore output appears to be empty. Check results for BUSCO: 271586at2759
## Mon Jul 25 14:24:19 CEST 2022 - The aliscore output file does not exist. Check results for BUSCO: 1379600at2759
## Mon Jul 25 14:24:19 CEST 2022 - The aliscore output appears to be empty. Check results for BUSCO: 1379600at2759
## Mon Jul 25 14:24:19 CEST 2022 - The aliscore output file does not exist. Check results for BUSCO: 918816at2759
## Mon Jul 25 14:24:19 CEST 2022 - The aliscore output appears to be empty. Check results for BUSCO: 918816at2759
## Mon Jul 25 14:24:20 CEST 2022 - The aliscore output file does not exist. Check results for BUSCO: 324863at2759
## Mon Jul 25 14:24:20 CEST 2022 - The aliscore output appears to be empty. Check results for BUSCO: 324863at2759
## Mon Jul 25 14:24:20 CEST 2022 - The aliscore output file does not exist. Check results for BUSCO: 39650at2759
## Mon Jul 25 14:24:20 CEST 2022 - The aliscore output appears to be empty. Check results for BUSCO: 39650at2759
## Mon Jul 25 14:24:22 CEST 2022 - The aliscore output file does not exist. Check results for BUSCO: 142542at2759
## Mon Jul 25 14:24:22 CEST 2022 - The aliscore output appears to be empty. Check results for BUSCO: 142542at2759
## Mon Jul 25 14:24:34 CEST 2022 - The aliscore output file does not exist. Check results for BUSCO: 674160at2759
## Mon Jul 25 14:24:34 CEST 2022 - The aliscore output appears to be empty. Check results for BUSCO: 674160at2759
## Mon Jul 25 14:24:39 CEST 2022 - The aliscore output file does not exist. Check results for BUSCO: 1474000at2759
## Mon Jul 25 14:24:39 CEST 2022 - The aliscore output appears to be empty. Check results for BUSCO: 1474000at2759
## Mon Jul 25 14:24:41 CEST 2022 - The aliscore output file does not exist. Check results for BUSCO: 1338131at2759
## Mon Jul 25 14:24:41 CEST 2022 - The aliscore output appears to be empty. Check results for BUSCO: 1338131at2759
## Mon Jul 25 14:24:41 CEST 2022 - The aliscore output file does not exist. Check results for BUSCO: 1107630at2759
## Mon Jul 25 14:24:41 CEST 2022 - The aliscore output appears to be empty. Check results for BUSCO: 1107630at2759
## Mon Jul 25 14:24:42 CEST 2022 - The aliscore output file does not exist. Check results for BUSCO: 1576404at2759
## Mon Jul 25 14:24:42 CEST 2022 - The aliscore output appears to be empty. Check results for BUSCO: 1576404at2759
## Mon Jul 25 14:24:42 CEST 2022 - The aliscore output file does not exist. Check results for BUSCO: 1430056at2759
## Mon Jul 25 14:24:42 CEST 2022 - The aliscore output appears to be empty. Check results for BUSCO: 1430056at2759
## Mon Jul 25 14:24:42 CEST 2022 - The aliscore output file does not exist. Check results for BUSCO: 1348942at2759
## Mon Jul 25 14:24:42 CEST 2022 - The aliscore output appears to be empty. Check results for BUSCO: 1348942at2759
## Mon Jul 25 14:24:42 CEST 2022 - The aliscore output file does not exist. Check results for BUSCO: 937686at2759
## Mon Jul 25 14:24:42 CEST 2022 - The aliscore output file does not exist. Check results for BUSCO: 1266231at2759
## Mon Jul 25 14:24:42 CEST 2022 - The aliscore output file does not exist. Check results for BUSCO: 679771at2759
## Mon Jul 25 14:24:42 CEST 2022 - The aliscore output appears to be empty. Check results for BUSCO: 937686at2759
## Mon Jul 25 14:24:42 CEST 2022 - The aliscore output file does not exist. Check results for BUSCO: 1404162at2759
## Mon Jul 25 14:24:42 CEST 2022 - The aliscore output appears to be empty. Check results for BUSCO: 1266231at2759
## Mon Jul 25 14:24:43 CEST 2022 - The aliscore output appears to be empty. Check results for BUSCO: 679771at2759
## Mon Jul 25 14:24:43 CEST 2022 - The aliscore output file does not exist. Check results for BUSCO: 719531at2759
## Mon Jul 25 14:24:43 CEST 2022 - The aliscore output appears to be empty. Check results for BUSCO: 1404162at2759
## Mon Jul 25 14:24:43 CEST 2022 - The aliscore output appears to be empty. Check results for BUSCO: 719531at2759
## Mon Jul 25 14:24:43 CEST 2022 - The aliscore output file does not exist. Check results for BUSCO: 956854at2759
## Mon Jul 25 14:24:43 CEST 2022 - The aliscore output file does not exist. Check results for BUSCO: 1490022at2759
## Mon Jul 25 14:24:43 CEST 2022 - The aliscore output appears to be empty. Check results for BUSCO: 956854at2759
## Mon Jul 25 14:24:43 CEST 2022 - The aliscore output appears to be empty. Check results for BUSCO: 1490022at2759
## Mon Jul 25 14:24:43 CEST 2022 - The aliscore output file does not exist. Check results for BUSCO: 1405146at2759
## Mon Jul 25 14:24:43 CEST 2022 - The aliscore output appears to be empty. Check results for BUSCO: 1405146at2759
## Mon Jul 25 14:24:43 CEST 2022 - The aliscore output file does not exist. Check results for BUSCO: 1236198at2759
## Mon Jul 25 14:24:43 CEST 2022 - The aliscore output appears to be empty. Check results for BUSCO: 1236198at2759
## Mon Jul 25 14:24:43 CEST 2022 - The aliscore output file does not exist. Check results for BUSCO: 1405073at2759
## Mon Jul 25 14:24:43 CEST 2022 - The aliscore output file does not exist. Check results for BUSCO: 1269244at2759
## Mon Jul 25 14:24:43 CEST 2022 - The aliscore output file does not exist. Check results for BUSCO: 1260807at2759
## Mon Jul 25 14:24:43 CEST 2022 - The aliscore output appears to be empty. Check results for BUSCO: 1405073at2759
## Mon Jul 25 14:24:43 CEST 2022 - The aliscore output appears to be empty. Check results for BUSCO: 1269244at2759
## Mon Jul 25 14:24:43 CEST 2022 - The aliscore output appears to be empty. Check results for BUSCO: 1260807at2759
## Mon Jul 25 14:24:43 CEST 2022 - The aliscore output file does not exist. Check results for BUSCO: 869548at2759
## Mon Jul 25 14:24:44 CEST 2022 - The aliscore output appears to be empty. Check results for BUSCO: 869548at2759
## Mon Jul 25 14:24:44 CEST 2022 - The aliscore output file does not exist. Check results for BUSCO: 293315at2759
## Mon Jul 25 14:24:44 CEST 2022 - The aliscore output file does not exist. Check results for BUSCO: 1106766at2759
## Mon Jul 25 14:24:44 CEST 2022 - The aliscore output appears to be empty. Check results for BUSCO: 293315at2759
## Mon Jul 25 14:24:44 CEST 2022 - The aliscore output appears to be empty. Check results for BUSCO: 1106766at2759
## Mon Jul 25 14:24:44 CEST 2022 - The aliscore output file does not exist. Check results for BUSCO: 1161199at2759
## Mon Jul 25 14:24:44 CEST 2022 - The aliscore output appears to be empty. Check results for BUSCO: 1161199at2759
## Mon Jul 25 14:24:44 CEST 2022 - The aliscore output file does not exist. Check results for BUSCO: 1247641at2759
## Mon Jul 25 14:24:44 CEST 2022 - The aliscore output appears to be empty. Check results for BUSCO: 1247641at2759
## Mon Jul 25 14:24:44 CEST 2022 - The aliscore output file does not exist. Check results for BUSCO: 1370304at2759
## Mon Jul 25 14:24:44 CEST 2022 - The aliscore output appears to be empty. Check results for BUSCO: 1370304at2759
## Mon Jul 25 14:24:44 CEST 2022 - The aliscore output file does not exist. Check results for BUSCO: 1258856at2759
## Mon Jul 25 14:24:44 CEST 2022 - The aliscore output appears to be empty. Check results for BUSCO: 1258856at2759
## Mon Jul 25 14:24:45 CEST 2022 - The aliscore output file does not exist. Check results for BUSCO: 1440961at2759
## Mon Jul 25 14:24:45 CEST 2022 - The aliscore output appears to be empty. Check results for BUSCO: 1440961at2759
## Mon Jul 25 14:24:45 CEST 2022 - The aliscore output file does not exist. Check results for BUSCO: 1269649at2759
## Mon Jul 25 14:24:45 CEST 2022 - The aliscore output appears to be empty. Check results for BUSCO: 1269649at2759
## Mon Jul 25 14:24:45 CEST 2022 - The aliscore output file does not exist. Check results for BUSCO: 1620056at2759
## Mon Jul 25 14:24:45 CEST 2022 - The aliscore output appears to be empty. Check results for BUSCO: 1620056at2759
## Mon Jul 25 14:24:45 CEST 2022 - The aliscore output file does not exist. Check results for BUSCO: 1223488at2759
## Mon Jul 25 14:24:45 CEST 2022 - The aliscore output file does not exist. Check results for BUSCO: 1053181at2759
## Mon Jul 25 14:24:45 CEST 2022 - The aliscore output appears to be empty. Check results for BUSCO: 1223488at2759
## Mon Jul 25 14:24:45 CEST 2022 - The aliscore output file does not exist. Check results for BUSCO: 1617752at2759
## Mon Jul 25 14:24:45 CEST 2022 - The aliscore output appears to be empty. Check results for BUSCO: 1053181at2759
## Mon Jul 25 14:24:45 CEST 2022 - The aliscore output appears to be empty. Check results for BUSCO: 1617752at2759
## Mon Jul 25 14:24:45 CEST 2022 - The aliscore output file does not exist. Check results for BUSCO: 759498at2759
## Mon Jul 25 14:24:45 CEST 2022 - The aliscore output appears to be empty. Check results for BUSCO: 759498at2759
## Mon Jul 25 14:24:45 CEST 2022 - The aliscore output file does not exist. Check results for BUSCO: 97116at2759
## Mon Jul 25 14:24:45 CEST 2022 - The aliscore output appears to be empty. Check results for BUSCO: 97116at2759
## Mon Jul 25 14:24:45 CEST 2022 - The aliscore output file does not exist. Check results for BUSCO: 176625at2759
## Mon Jul 25 14:24:45 CEST 2022 - The aliscore output appears to be empty. Check results for BUSCO: 176625at2759
## Mon Jul 25 14:24:46 CEST 2022 - The aliscore output file does not exist. Check results for BUSCO: 1346432at2759
## Mon Jul 25 14:24:46 CEST 2022 - The aliscore output appears to be empty. Check results for BUSCO: 1346432at2759
## Mon Jul 25 14:24:46 CEST 2022 - The aliscore output file does not exist. Check results for BUSCO: 549762at2759
## Mon Jul 25 14:24:46 CEST 2022 - The aliscore output file does not exist. Check results for BUSCO: 1197019at2759
## Mon Jul 25 14:24:46 CEST 2022 - The aliscore output appears to be empty. Check results for BUSCO: 549762at2759
## Mon Jul 25 14:24:46 CEST 2022 - The aliscore output appears to be empty. Check results for BUSCO: 1197019at2759
## Mon Jul 25 14:24:46 CEST 2022 - The aliscore output file does not exist. Check results for BUSCO: 1421503at2759
## Mon Jul 25 14:24:46 CEST 2022 - The aliscore output appears to be empty. Check results for BUSCO: 1421503at2759
## Mon Jul 25 14:24:46 CEST 2022 - The aliscore output file does not exist. Check results for BUSCO: 974865at2759
## Mon Jul 25 14:24:46 CEST 2022 - The aliscore output file does not exist. Check results for BUSCO: 692986at2759
## Mon Jul 25 14:24:46 CEST 2022 - The aliscore output file does not exist. Check results for BUSCO: 330169at2759
## Mon Jul 25 14:24:46 CEST 2022 - The aliscore output appears to be empty. Check results for BUSCO: 974865at2759
## Mon Jul 25 14:24:46 CEST 2022 - The aliscore output appears to be empty. Check results for BUSCO: 692986at2759
## Mon Jul 25 14:24:46 CEST 2022 - The aliscore output appears to be empty. Check results for BUSCO: 330169at2759
## Mon Jul 25 14:24:46 CEST 2022 - The aliscore output file does not exist. Check results for BUSCO: 1323575at2759
## Mon Jul 25 14:24:46 CEST 2022 - The aliscore output appears to be empty. Check results for BUSCO: 1323575at2759
## Mon Jul 25 14:24:46 CEST 2022 - The aliscore output file does not exist. Check results for BUSCO: 1504863at2759
## Mon Jul 25 14:24:46 CEST 2022 - The aliscore output appears to be empty. Check results for BUSCO: 1504863at2759
## Mon Jul 25 14:24:46 CEST 2022 - The aliscore output file does not exist. Check results for BUSCO: 1019762at2759
## Mon Jul 25 14:24:46 CEST 2022 - The aliscore output file does not exist. Check results for BUSCO: 1538526at2759
## Mon Jul 25 14:24:46 CEST 2022 - The aliscore output appears to be empty. Check results for BUSCO: 1019762at2759
## Mon Jul 25 14:24:46 CEST 2022 - The aliscore output appears to be empty. Check results for BUSCO: 1538526at2759
## Mon Jul 25 14:24:46 CEST 2022 - The aliscore output file does not exist. Check results for BUSCO: 878143at2759
## Mon Jul 25 14:24:46 CEST 2022 - The aliscore output file does not exist. Check results for BUSCO: 1545004at2759
## Mon Jul 25 14:24:46 CEST 2022 - The aliscore output appears to be empty. Check results for BUSCO: 878143at2759
## Mon Jul 25 14:24:46 CEST 2022 - The aliscore output appears to be empty. Check results for BUSCO: 1545004at2759
## Mon Jul 25 14:24:46 CEST 2022 - The aliscore output file does not exist. Check results for BUSCO: 1222562at2759
## Mon Jul 25 14:24:47 CEST 2022 - The aliscore output appears to be empty. Check results for BUSCO: 1222562at2759
## Mon Jul 25 14:24:47 CEST 2022 - The aliscore output file does not exist. Check results for BUSCO: 392369at2759
## Mon Jul 25 14:24:47 CEST 2022 - The aliscore output appears to be empty. Check results for BUSCO: 392369at2759
## Mon Jul 25 14:24:47 CEST 2022 - The aliscore output file does not exist. Check results for BUSCO: 450058at2759
## Mon Jul 25 14:24:47 CEST 2022 - The aliscore output appears to be empty. Check results for BUSCO: 450058at2759
## Mon Jul 25 14:24:47 CEST 2022 - The aliscore output file does not exist. Check results for BUSCO: 911863at2759
## Mon Jul 25 14:24:47 CEST 2022 - The aliscore output appears to be empty. Check results for BUSCO: 911863at2759
## Mon Jul 25 14:24:47 CEST 2022 - The aliscore output file does not exist. Check results for BUSCO: 1370285at2759
## Mon Jul 25 14:24:47 CEST 2022 - The aliscore output file does not exist. Check results for BUSCO: 937275at2759
## Mon Jul 25 14:24:47 CEST 2022 - The aliscore output appears to be empty. Check results for BUSCO: 937275at2759
## Mon Jul 25 14:24:47 CEST 2022 - The aliscore output appears to be empty. Check results for BUSCO: 1370285at2759
## Mon Jul 25 14:24:47 CEST 2022 - The aliscore output file does not exist. Check results for BUSCO: 1450538at2759
## Mon Jul 25 14:24:47 CEST 2022 - The aliscore output appears to be empty. Check results for BUSCO: 1450538at2759
## Mon Jul 25 14:24:47 CEST 2022 - The aliscore output file does not exist. Check results for BUSCO: 1339553at2759
## Mon Jul 25 14:24:47 CEST 2022 - The aliscore output appears to be empty. Check results for BUSCO: 1339553at2759
## Mon Jul 25 14:24:47 CEST 2022 - The aliscore output file does not exist. Check results for BUSCO: 1314980at2759
## Mon Jul 25 14:24:47 CEST 2022 - The aliscore output appears to be empty. Check results for BUSCO: 1314980at2759
## Mon Jul 25 14:24:47 CEST 2022 - The aliscore output file does not exist. Check results for BUSCO: 1096688at2759
## Mon Jul 25 14:24:47 CEST 2022 - The aliscore output appears to be empty. Check results for BUSCO: 1096688at2759
## Mon Jul 25 14:24:48 CEST 2022 - The aliscore output file does not exist. Check results for BUSCO: 320059at2759
## Mon Jul 25 14:24:48 CEST 2022 - The aliscore output appears to be empty. Check results for BUSCO: 320059at2759
## Mon Jul 25 14:24:48 CEST 2022 - The aliscore output file does not exist. Check results for BUSCO: 655400at2759
## Mon Jul 25 14:24:48 CEST 2022 - The aliscore output appears to be empty. Check results for BUSCO: 655400at2759
## Mon Jul 25 14:24:48 CEST 2022 - The aliscore output file does not exist. Check results for BUSCO: 1001705at2759
## Mon Jul 25 14:24:48 CEST 2022 - The aliscore output appears to be empty. Check results for BUSCO: 1001705at2759
## Mon Jul 25 14:24:48 CEST 2022 - The aliscore output file does not exist. Check results for BUSCO: 299766at2759
## Mon Jul 25 14:24:48 CEST 2022 - The aliscore output file does not exist. Check results for BUSCO: 388820at2759
## Mon Jul 25 14:24:48 CEST 2022 - The aliscore output appears to be empty. Check results for BUSCO: 299766at2759
## Mon Jul 25 14:24:48 CEST 2022 - The aliscore output appears to be empty. Check results for BUSCO: 388820at2759
## Mon Jul 25 14:24:48 CEST 2022 - The aliscore output file does not exist. Check results for BUSCO: 345441at2759
## Mon Jul 25 14:24:48 CEST 2022 - The aliscore output appears to be empty. Check results for BUSCO: 345441at2759
## Mon Jul 25 14:24:48 CEST 2022 - The aliscore output file does not exist. Check results for BUSCO: 1423847at2759
## Mon Jul 25 14:24:48 CEST 2022 - The aliscore output appears to be empty. Check results for BUSCO: 1423847at2759
## Mon Jul 25 14:24:48 CEST 2022 - The aliscore output file does not exist. Check results for BUSCO: 855834at2759
## Mon Jul 25 14:24:48 CEST 2022 - The aliscore output file does not exist. Check results for BUSCO: 768809at2759
## Mon Jul 25 14:24:48 CEST 2022 - The aliscore output file does not exist. Check results for BUSCO: 1018517at2759
## Mon Jul 25 14:24:48 CEST 2022 - The aliscore output appears to be empty. Check results for BUSCO: 855834at2759
## Mon Jul 25 14:24:48 CEST 2022 - The aliscore output appears to be empty. Check results for BUSCO: 768809at2759
## Mon Jul 25 14:24:48 CEST 2022 - The aliscore output appears to be empty. Check results for BUSCO: 1018517at2759
## Mon Jul 25 14:24:50 CEST 2022 - The aliscore output file does not exist. Check results for BUSCO: 975557at2759
## Mon Jul 25 14:24:50 CEST 2022 - The aliscore output appears to be empty. Check results for BUSCO: 975557at2759
## Mon Jul 25 14:24:50 CEST 2022 - The aliscore output file does not exist. Check results for BUSCO: 551907at2759
## Mon Jul 25 14:24:50 CEST 2022 - The aliscore output appears to be empty. Check results for BUSCO: 551907at2759
## Mon Jul 25 14:24:50 CEST 2022 - The aliscore output file does not exist. Check results for BUSCO: 671846at2759
## Mon Jul 25 14:24:50 CEST 2022 - The aliscore output appears to be empty. Check results for BUSCO: 671846at2759
## Mon Jul 25 14:24:50 CEST 2022 - The aliscore output file does not exist. Check results for BUSCO: 1025450at2759
## Mon Jul 25 14:24:50 CEST 2022 - The aliscore output file does not exist. Check results for BUSCO: 296129at2759
## Mon Jul 25 14:24:50 CEST 2022 - The aliscore output appears to be empty. Check results for BUSCO: 1025450at2759
## Mon Jul 25 14:24:50 CEST 2022 - The aliscore output appears to be empty. Check results for BUSCO: 296129at2759
## Mon Jul 25 14:24:51 CEST 2022 - The aliscore output file does not exist. Check results for BUSCO: 625387at2759
## Mon Jul 25 14:24:51 CEST 2022 - The aliscore output appears to be empty. Check results for BUSCO: 625387at2759
## Mon Jul 25 14:25:08 CEST 2022 - The aliscore output file does not exist. Check results for BUSCO: 1217666at2759
## Mon Jul 25 14:25:09 CEST 2022 - The aliscore output appears to be empty. Check results for BUSCO: 1217666at2759
## Mon Jul 25 14:25:09 CEST 2022 - The aliscore output file does not exist. Check results for BUSCO: 1382842at2759
## Mon Jul 25 14:25:09 CEST 2022 - The aliscore output appears to be empty. Check results for BUSCO: 1382842at2759
## Mon Jul 25 14:25:11 CEST 2022 - The aliscore output file does not exist. Check results for BUSCO: 1513531at2759
## Mon Jul 25 14:25:11 CEST 2022 - The aliscore output appears to be empty. Check results for BUSCO: 1513531at2759
## Mon Jul 25 14:25:11 CEST 2022 - The aliscore output file does not exist. Check results for BUSCO: 865202at2759
## Mon Jul 25 14:25:11 CEST 2022 - The aliscore output appears to be empty. Check results for BUSCO: 865202at2759
## Mon Jul 25 14:25:12 CEST 2022 - The aliscore output file does not exist. Check results for BUSCO: 1138059at2759
## Mon Jul 25 14:25:12 CEST 2022 - The aliscore output appears to be empty. Check results for BUSCO: 1138059at2759
## Mon Jul 25 14:25:12 CEST 2022 - The aliscore output file does not exist. Check results for BUSCO: 1552706at2759
## Mon Jul 25 14:25:12 CEST 2022 - The aliscore output appears to be empty. Check results for BUSCO: 1552706at2759
## Mon Jul 25 14:25:12 CEST 2022 - The aliscore output file does not exist. Check results for BUSCO: 1079827at2759
## Mon Jul 25 14:25:12 CEST 2022 - The aliscore output appears to be empty. Check results for BUSCO: 1079827at2759
## Mon Jul 25 14:25:12 CEST 2022 - The aliscore output file does not exist. Check results for BUSCO: 1445102at2759
## Mon Jul 25 14:25:12 CEST 2022 - The aliscore output appears to be empty. Check results for BUSCO: 1445102at2759
## Mon Jul 25 14:25:13 CEST 2022 - The aliscore output file does not exist. Check results for BUSCO: 1426075at2759
## Mon Jul 25 14:25:13 CEST 2022 - The aliscore output appears to be empty. Check results for BUSCO: 1426075at2759
## Mon Jul 25 14:25:13 CEST 2022 - The aliscore output file does not exist. Check results for BUSCO: 975158at2759
## Mon Jul 25 14:25:13 CEST 2022 - The aliscore output appears to be empty. Check results for BUSCO: 975158at2759
## Mon Jul 25 14:25:13 CEST 2022 - The aliscore output file does not exist. Check results for BUSCO: 1346165at2759
## Mon Jul 25 14:25:13 CEST 2022 - The aliscore output appears to be empty. Check results for BUSCO: 1346165at2759
## Mon Jul 25 14:25:13 CEST 2022 - The aliscore output file does not exist. Check results for BUSCO: 1128607at2759
## Mon Jul 25 14:25:13 CEST 2022 - The aliscore output appears to be empty. Check results for BUSCO: 1128607at2759
## Mon Jul 25 14:25:14 CEST 2022 - The aliscore output file does not exist. Check results for BUSCO: 1398309at2759
## Mon Jul 25 14:25:14 CEST 2022 - The aliscore output appears to be empty. Check results for BUSCO: 1398309at2759
## Mon Jul 25 14:25:15 CEST 2022 - The aliscore output file does not exist. Check results for BUSCO: 598949at2759
## Mon Jul 25 14:25:15 CEST 2022 - The aliscore output file does not exist. Check results for BUSCO: 331411at2759
## Mon Jul 25 14:25:15 CEST 2022 - The aliscore output appears to be empty. Check results for BUSCO: 331411at2759
## Mon Jul 25 14:25:15 CEST 2022 - The aliscore output appears to be empty. Check results for BUSCO: 598949at2759
## Mon Jul 25 14:25:15 CEST 2022 - The aliscore output file does not exist. Check results for BUSCO: 1626636at2759
## Mon Jul 25 14:25:15 CEST 2022 - The aliscore output appears to be empty. Check results for BUSCO: 1626636at2759
## Mon Jul 25 14:25:15 CEST 2022 - The aliscore output file does not exist. Check results for BUSCO: 1575179at2759
## Mon Jul 25 14:25:15 CEST 2022 - The aliscore output appears to be empty. Check results for BUSCO: 1575179at2759
## Mon Jul 25 14:25:15 CEST 2022 - The aliscore output file does not exist. Check results for BUSCO: 355408at2759
## Mon Jul 25 14:25:15 CEST 2022 - The aliscore output appears to be empty. Check results for BUSCO: 355408at2759
## Mon Jul 25 14:25:15 CEST 2022 - The aliscore output file does not exist. Check results for BUSCO: 919955at2759
## Mon Jul 25 14:25:15 CEST 2022 - The aliscore output appears to be empty. Check results for BUSCO: 919955at2759
## Mon Jul 25 14:25:16 CEST 2022 - The aliscore output file does not exist. Check results for BUSCO: 570797at2759
## Mon Jul 25 14:25:16 CEST 2022 - The aliscore output appears to be empty. Check results for BUSCO: 570797at2759
## Mon Jul 25 14:25:16 CEST 2022 - The aliscore output file does not exist. Check results for BUSCO: 1264469at2759
## Mon Jul 25 14:25:16 CEST 2022 - The aliscore output appears to be empty. Check results for BUSCO: 1264469at2759
## Mon Jul 25 14:25:16 CEST 2022 - The aliscore output file does not exist. Check results for BUSCO: 721605at2759
## Mon Jul 25 14:25:16 CEST 2022 - The aliscore output file does not exist. Check results for BUSCO: 996662at2759
## Mon Jul 25 14:25:16 CEST 2022 - The aliscore output appears to be empty. Check results for BUSCO: 721605at2759
## Mon Jul 25 14:25:16 CEST 2022 - The aliscore output appears to be empty. Check results for BUSCO: 996662at2759
## Mon Jul 25 14:25:16 CEST 2022 - The aliscore output file does not exist. Check results for BUSCO: 1228942at2759
## Mon Jul 25 14:25:16 CEST 2022 - The aliscore output appears to be empty. Check results for BUSCO: 1228942at2759
## Mon Jul 25 14:25:16 CEST 2022 - The aliscore output file does not exist. Check results for BUSCO: 1108845at2759
## Mon Jul 25 14:25:16 CEST 2022 - The aliscore output file does not exist. Check results for BUSCO: 1355894at2759
## Mon Jul 25 14:25:16 CEST 2022 - The aliscore output appears to be empty. Check results for BUSCO: 1108845at2759
## Mon Jul 25 14:25:16 CEST 2022 - The aliscore output appears to be empty. Check results for BUSCO: 1355894at2759
## Mon Jul 25 14:25:16 CEST 2022 - The aliscore output file does not exist. Check results for BUSCO: 1112002at2759
## Mon Jul 25 14:25:16 CEST 2022 - The aliscore output appears to be empty. Check results for BUSCO: 1112002at2759
## Mon Jul 25 14:25:16 CEST 2022 - The aliscore output file does not exist. Check results for BUSCO: 245208at2759
## Mon Jul 25 14:25:16 CEST 2022 - The aliscore output appears to be empty. Check results for BUSCO: 245208at2759
## Mon Jul 25 14:25:16 CEST 2022 - The aliscore output file does not exist. Check results for BUSCO: 736068at2759
## Mon Jul 25 14:25:16 CEST 2022 - The aliscore output appears to be empty. Check results for BUSCO: 736068at2759
## Mon Jul 25 14:25:17 CEST 2022 - The aliscore output file does not exist. Check results for BUSCO: 1563319at2759
## Mon Jul 25 14:25:17 CEST 2022 - The aliscore output appears to be empty. Check results for BUSCO: 1563319at2759
## Mon Jul 25 14:25:17 CEST 2022 - The aliscore output file does not exist. Check results for BUSCO: 1588798at2759
## Mon Jul 25 14:25:17 CEST 2022 - The aliscore output appears to be empty. Check results for BUSCO: 1588798at2759
## Mon Jul 25 14:25:17 CEST 2022 - The aliscore output file does not exist. Check results for BUSCO: 1455730at2759
## Mon Jul 25 14:25:17 CEST 2022 - The aliscore output appears to be empty. Check results for BUSCO: 1455730at2759
## Mon Jul 25 14:25:17 CEST 2022 - The aliscore output file does not exist. Check results for BUSCO: 1111142at2759
## Mon Jul 25 14:25:17 CEST 2022 - The aliscore output appears to be empty. Check results for BUSCO: 1111142at2759
## Mon Jul 25 14:25:18 CEST 2022 - The aliscore output file does not exist. Check results for BUSCO: 418107at2759
## Mon Jul 25 14:25:18 CEST 2022 - The aliscore output appears to be empty. Check results for BUSCO: 418107at2759
## Mon Jul 25 14:25:18 CEST 2022 - The aliscore output file does not exist. Check results for BUSCO: 924753at2759
## Mon Jul 25 14:25:18 CEST 2022 - The aliscore output file does not exist. Check results for BUSCO: 801857at2759
## Mon Jul 25 14:25:18 CEST 2022 - The aliscore output appears to be empty. Check results for BUSCO: 924753at2759
## Mon Jul 25 14:25:18 CEST 2022 - The aliscore output appears to be empty. Check results for BUSCO: 801857at2759
## Mon Jul 25 14:25:18 CEST 2022 - The aliscore output file does not exist. Check results for BUSCO: 1377237at2759
## Mon Jul 25 14:25:18 CEST 2022 - The aliscore output appears to be empty. Check results for BUSCO: 1377237at2759
## Mon Jul 25 14:25:18 CEST 2022 - The aliscore output file does not exist. Check results for BUSCO: 734666at2759
## Mon Jul 25 14:25:18 CEST 2022 - The aliscore output file does not exist. Check results for BUSCO: 1041560at2759
## Mon Jul 25 14:25:18 CEST 2022 - The aliscore output appears to be empty. Check results for BUSCO: 734666at2759
## Mon Jul 25 14:25:18 CEST 2022 - The aliscore output appears to be empty. Check results for BUSCO: 1041560at2759
## Mon Jul 25 14:25:18 CEST 2022 - The aliscore output file does not exist. Check results for BUSCO: 396755at2759
## Mon Jul 25 14:25:18 CEST 2022 - The aliscore output appears to be empty. Check results for BUSCO: 396755at2759
## Mon Jul 25 14:25:19 CEST 2022 - The aliscore output file does not exist. Check results for BUSCO: 341721at2759
## Mon Jul 25 14:25:19 CEST 2022 - The aliscore output appears to be empty. Check results for BUSCO: 341721at2759
## Mon Jul 25 14:25:19 CEST 2022 - The aliscore output file does not exist. Check results for BUSCO: 257318at2759
## Mon Jul 25 14:25:19 CEST 2022 - The aliscore output appears to be empty. Check results for BUSCO: 257318at2759
## Mon Jul 25 14:25:19 CEST 2022 - The aliscore output file does not exist. Check results for BUSCO: 814241at2759
## Mon Jul 25 14:25:19 CEST 2022 - The aliscore output appears to be empty. Check results for BUSCO: 814241at2759
## Mon Jul 25 14:25:20 CEST 2022 - The aliscore output file does not exist. Check results for BUSCO: 917326at2759
## Mon Jul 25 14:25:20 CEST 2022 - The aliscore output appears to be empty. Check results for BUSCO: 917326at2759
## Mon Jul 25 14:25:20 CEST 2022 - The aliscore output file does not exist. Check results for BUSCO: 898782at2759
## Mon Jul 25 14:25:20 CEST 2022 - The aliscore output appears to be empty. Check results for BUSCO: 898782at2759
## Mon Jul 25 14:25:20 CEST 2022 - The aliscore output file does not exist. Check results for BUSCO: 836599at2759
## Mon Jul 25 14:25:20 CEST 2022 - The aliscore output appears to be empty. Check results for BUSCO: 836599at2759
## Mon Jul 25 14:25:20 CEST 2022 - The aliscore output file does not exist. Check results for BUSCO: 722805at2759
## Mon Jul 25 14:25:20 CEST 2022 - The aliscore output appears to be empty. Check results for BUSCO: 722805at2759
## Mon Jul 25 14:25:21 CEST 2022 - The aliscore output file does not exist. Check results for BUSCO: 1087488at2759
## Mon Jul 25 14:25:21 CEST 2022 - The aliscore output appears to be empty. Check results for BUSCO: 1087488at2759
## Mon Jul 25 14:25:22 CEST 2022 - The aliscore output file does not exist. Check results for BUSCO: 973442at2759
## Mon Jul 25 14:25:22 CEST 2022 - The aliscore output appears to be empty. Check results for BUSCO: 973442at2759
## Mon Jul 25 14:25:22 CEST 2022 - The aliscore output file does not exist. Check results for BUSCO: 270107at2759
## Mon Jul 25 14:25:22 CEST 2022 - The aliscore output appears to be empty. Check results for BUSCO: 270107at2759
## Mon Jul 25 14:25:22 CEST 2022 - The aliscore output file does not exist. Check results for BUSCO: 179362at2759
## Mon Jul 25 14:25:23 CEST 2022 - The aliscore output appears to be empty. Check results for BUSCO: 179362at2759
## Mon Jul 25 14:25:27 CEST 2022 - The aliscore output file does not exist. Check results for BUSCO: 156083at2759
## Mon Jul 25 14:25:27 CEST 2022 - The aliscore output appears to be empty. Check results for BUSCO: 156083at2759
## Mon Jul 25 14:25:38 CEST 2022 - The aliscore output file does not exist. Check results for BUSCO: 1223488at2759
## Mon Jul 25 14:25:38 CEST 2022 - The aliscore output appears to be empty. Check results for BUSCO: 1223488at2759
## Mon Jul 25 14:25:38 CEST 2022 - The aliscore output file does not exist. Check results for BUSCO: 1421503at2759
## Mon Jul 25 14:25:39 CEST 2022 - The aliscore output file does not exist. Check results for BUSCO: 1525971at2759
## Mon Jul 25 14:25:39 CEST 2022 - The aliscore output appears to be empty. Check results for BUSCO: 1421503at2759
## Mon Jul 25 14:25:39 CEST 2022 - The aliscore output appears to be empty. Check results for BUSCO: 1525971at2759
## Mon Jul 25 14:25:39 CEST 2022 - The aliscore output file does not exist. Check results for BUSCO: 1049599at2759
## Mon Jul 25 14:25:39 CEST 2022 - The aliscore output appears to be empty. Check results for BUSCO: 1049599at2759
## Mon Jul 25 14:25:40 CEST 2022 - The aliscore output file does not exist. Check results for BUSCO: 1626636at2759
## Mon Jul 25 14:25:40 CEST 2022 - The aliscore output appears to be empty. Check results for BUSCO: 1626636at2759
## Mon Jul 25 14:25:40 CEST 2022 - The aliscore output file does not exist. Check results for BUSCO: 1488436at2759
## Mon Jul 25 14:25:40 CEST 2022 - The aliscore output appears to be empty. Check results for BUSCO: 1488436at2759
## Mon Jul 25 14:25:40 CEST 2022 - The aliscore output file does not exist. Check results for BUSCO: 946128at2759
## Mon Jul 25 14:25:40 CEST 2022 - The aliscore output appears to be empty. Check results for BUSCO: 946128at2759
## Mon Jul 25 14:25:41 CEST 2022 - The aliscore output file does not exist. Check results for BUSCO: 1038775at2759
## Mon Jul 25 14:25:41 CEST 2022 - The aliscore output appears to be empty. Check results for BUSCO: 1038775at2759
## Mon Jul 25 14:25:41 CEST 2022 - The aliscore output file does not exist. Check results for BUSCO: 1358374at2759
## Mon Jul 25 14:25:41 CEST 2022 - The aliscore output appears to be empty. Check results for BUSCO: 1358374at2759
## Mon Jul 25 14:25:42 CEST 2022 - The aliscore output file does not exist. Check results for BUSCO: 1488235at2759
## Mon Jul 25 14:25:42 CEST 2022 - The aliscore output appears to be empty. Check results for BUSCO: 1488235at2759
## Mon Jul 25 14:25:42 CEST 2022 - The aliscore output file does not exist. Check results for BUSCO: 924753at2759
## Mon Jul 25 14:25:42 CEST 2022 - The aliscore output appears to be empty. Check results for BUSCO: 924753at2759
## Mon Jul 25 14:25:42 CEST 2022 - The aliscore output file does not exist. Check results for BUSCO: 290630at2759
## Mon Jul 25 14:25:43 CEST 2022 - The aliscore output appears to be empty. Check results for BUSCO: 290630at2759
## Mon Jul 25 14:25:43 CEST 2022 - The aliscore output file does not exist. Check results for BUSCO: 1200489at2759
## Mon Jul 25 14:25:43 CEST 2022 - The aliscore output appears to be empty. Check results for BUSCO: 1200489at2759
## Mon Jul 25 14:25:43 CEST 2022 - The aliscore output file does not exist. Check results for BUSCO: 1342242at2759
## Mon Jul 25 14:25:43 CEST 2022 - The aliscore output appears to be empty. Check results for BUSCO: 1342242at2759
## Mon Jul 25 14:25:43 CEST 2022 - The aliscore output file does not exist. Check results for BUSCO: 1284731at2759
## Mon Jul 25 14:25:43 CEST 2022 - The aliscore output appears to be empty. Check results for BUSCO: 1284731at2759
## Mon Jul 25 14:25:44 CEST 2022 - The aliscore output file does not exist. Check results for BUSCO: 1249159at2759
## Mon Jul 25 14:25:44 CEST 2022 - The aliscore output appears to be empty. Check results for BUSCO: 1249159at2759
## Mon Jul 25 14:25:44 CEST 2022 - The aliscore output file does not exist. Check results for BUSCO: 976469at2759
## Mon Jul 25 14:25:44 CEST 2022 - The aliscore output appears to be empty. Check results for BUSCO: 976469at2759
## Mon Jul 25 14:25:44 CEST 2022 - The aliscore output file does not exist. Check results for BUSCO: 1287401at2759
## Mon Jul 25 14:25:44 CEST 2022 - The aliscore output file does not exist. Check results for BUSCO: 1251252at2759
## Mon Jul 25 14:25:44 CEST 2022 - The aliscore output appears to be empty. Check results for BUSCO: 1287401at2759
## Mon Jul 25 14:25:44 CEST 2022 - The aliscore output appears to be empty. Check results for BUSCO: 1251252at2759
## Mon Jul 25 14:25:44 CEST 2022 - The aliscore output file does not exist. Check results for BUSCO: 1322299at2759
## Mon Jul 25 14:25:44 CEST 2022 - The aliscore output appears to be empty. Check results for BUSCO: 1322299at2759
## Mon Jul 25 14:25:45 CEST 2022 - The aliscore output file does not exist. Check results for BUSCO: 1233814at2759
## Mon Jul 25 14:25:45 CEST 2022 - The aliscore output appears to be empty. Check results for BUSCO: 1233814at2759
## Mon Jul 25 14:25:45 CEST 2022 - The aliscore output file does not exist. Check results for BUSCO: 570797at2759
## Mon Jul 25 14:25:45 CEST 2022 - The aliscore output appears to be empty. Check results for BUSCO: 570797at2759
## Mon Jul 25 14:25:45 CEST 2022 - The aliscore output file does not exist. Check results for BUSCO: 1138059at2759
## Mon Jul 25 14:25:45 CEST 2022 - The aliscore output appears to be empty. Check results for BUSCO: 1138059at2759
## Mon Jul 25 14:25:45 CEST 2022 - The aliscore output file does not exist. Check results for BUSCO: 1051021at2759
## Mon Jul 25 14:25:45 CEST 2022 - The aliscore output appears to be empty. Check results for BUSCO: 1051021at2759
## Mon Jul 25 14:25:45 CEST 2022 - The aliscore output file does not exist. Check results for BUSCO: 944899at2759
## Mon Jul 25 14:25:45 CEST 2022 - The aliscore output file does not exist. Check results for BUSCO: 1426075at2759
## Mon Jul 25 14:25:45 CEST 2022 - The aliscore output appears to be empty. Check results for BUSCO: 944899at2759
## Mon Jul 25 14:25:46 CEST 2022 - The aliscore output appears to be empty. Check results for BUSCO: 1426075at2759
## Mon Jul 25 14:25:46 CEST 2022 - The aliscore output file does not exist. Check results for BUSCO: 981902at2759
## Mon Jul 25 14:25:46 CEST 2022 - The aliscore output file does not exist. Check results for BUSCO: 160593at2759
## Mon Jul 25 14:25:46 CEST 2022 - The aliscore output appears to be empty. Check results for BUSCO: 981902at2759
## Mon Jul 25 14:25:46 CEST 2022 - The aliscore output appears to be empty. Check results for BUSCO: 160593at2759
## Mon Jul 25 14:25:46 CEST 2022 - The aliscore output file does not exist. Check results for BUSCO: 751335at2759
## Mon Jul 25 14:25:46 CEST 2022 - The aliscore output appears to be empty. Check results for BUSCO: 751335at2759
## Mon Jul 25 14:25:46 CEST 2022 - The aliscore output file does not exist. Check results for BUSCO: 768809at2759
## Mon Jul 25 14:25:46 CEST 2022 - The aliscore output file does not exist. Check results for BUSCO: 383503at2759
## Mon Jul 25 14:25:46 CEST 2022 - The aliscore output appears to be empty. Check results for BUSCO: 768809at2759
## Mon Jul 25 14:25:46 CEST 2022 - The aliscore output appears to be empty. Check results for BUSCO: 383503at2759
## Mon Jul 25 14:25:46 CEST 2022 - The aliscore output file does not exist. Check results for BUSCO: 736068at2759
## Mon Jul 25 14:25:46 CEST 2022 - The aliscore output appears to be empty. Check results for BUSCO: 736068at2759
## Mon Jul 25 14:25:47 CEST 2022 - The aliscore output file does not exist. Check results for BUSCO: 604979at2759
## Mon Jul 25 14:25:47 CEST 2022 - The aliscore output file does not exist. Check results for BUSCO: 774318at2759
## Mon Jul 25 14:25:47 CEST 2022 - The aliscore output appears to be empty. Check results for BUSCO: 604979at2759
## Mon Jul 25 14:25:47 CEST 2022 - The aliscore output appears to be empty. Check results for BUSCO: 774318at2759
## Mon Jul 25 14:25:47 CEST 2022 - The aliscore output file does not exist. Check results for BUSCO: 355408at2759
## Mon Jul 25 14:25:47 CEST 2022 - The aliscore output appears to be empty. Check results for BUSCO: 355408at2759
## Mon Jul 25 14:25:47 CEST 2022 - The aliscore output file does not exist. Check results for BUSCO: 549762at2759
## Mon Jul 25 14:25:47 CEST 2022 - The aliscore output appears to be empty. Check results for BUSCO: 549762at2759
## Mon Jul 25 14:25:48 CEST 2022 - The aliscore output file does not exist. Check results for BUSCO: 83779at2759
## Mon Jul 25 14:25:48 CEST 2022 - The aliscore output appears to be empty. Check results for BUSCO: 83779at2759
## Mon Jul 25 14:25:50 CEST 2022 - The aliscore output file does not exist. Check results for BUSCO: 39650at2759
## Mon Jul 25 14:25:50 CEST 2022 - The aliscore output appears to be empty. Check results for BUSCO: 39650at2759
## Mon Jul 25 14:25:50 CEST 2022 - The aliscore output file does not exist. Check results for BUSCO: 331411at2759
## Mon Jul 25 14:25:50 CEST 2022 - The aliscore output appears to be empty. Check results for BUSCO: 331411at2759
## Mon Jul 25 14:25:51 CEST 2022 - The aliscore output file does not exist. Check results for BUSCO: 1312453at2759
## Mon Jul 25 14:25:51 CEST 2022 - The aliscore output appears to be empty. Check results for BUSCO: 1312453at2759
## Mon Jul 25 14:26:10 CEST 2022 - The aliscore output file does not exist. Check results for BUSCO: 1633672at2759
## Mon Jul 25 14:26:10 CEST 2022 - The aliscore output appears to be empty. Check results for BUSCO: 1633672at2759
## Mon Jul 25 14:26:10 CEST 2022 - The aliscore output file does not exist. Check results for BUSCO: 1459797at2759
## Mon Jul 25 14:26:10 CEST 2022 - The aliscore output appears to be empty. Check results for BUSCO: 1459797at2759
## Mon Jul 25 14:26:10 CEST 2022 - The aliscore output file does not exist. Check results for BUSCO: 1474000at2759
## Mon Jul 25 14:26:10 CEST 2022 - The aliscore output appears to be empty. Check results for BUSCO: 1474000at2759
## Mon Jul 25 14:26:10 CEST 2022 - The aliscore output file does not exist. Check results for BUSCO: 1161199at2759
## Mon Jul 25 14:26:10 CEST 2022 - The aliscore output appears to be empty. Check results for BUSCO: 1161199at2759
## Mon Jul 25 14:26:10 CEST 2022 - The aliscore output file does not exist. Check results for BUSCO: 1430056at2759
## Mon Jul 25 14:26:10 CEST 2022 - The aliscore output appears to be empty. Check results for BUSCO: 1430056at2759
## Mon Jul 25 14:26:11 CEST 2022 - The aliscore output file does not exist. Check results for BUSCO: 1504863at2759
## Mon Jul 25 14:26:11 CEST 2022 - The aliscore output file does not exist. Check results for BUSCO: 1623701at2759
## Mon Jul 25 14:26:11 CEST 2022 - The aliscore output appears to be empty. Check results for BUSCO: 1504863at2759
## Mon Jul 25 14:26:11 CEST 2022 - The aliscore output appears to be empty. Check results for BUSCO: 1623701at2759
## Mon Jul 25 14:26:11 CEST 2022 - The aliscore output file does not exist. Check results for BUSCO: 1558822at2759
## Mon Jul 25 14:26:11 CEST 2022 - The aliscore output appears to be empty. Check results for BUSCO: 1558822at2759
## Mon Jul 25 14:26:11 CEST 2022 - The aliscore output file does not exist. Check results for BUSCO: 1617752at2759
## Mon Jul 25 14:26:11 CEST 2022 - The aliscore output file does not exist. Check results for BUSCO: 1079827at2759
## Mon Jul 25 14:26:11 CEST 2022 - The aliscore output appears to be empty. Check results for BUSCO: 1617752at2759
## Mon Jul 25 14:26:11 CEST 2022 - The aliscore output file does not exist. Check results for BUSCO: 491869at2759
## Mon Jul 25 14:26:11 CEST 2022 - The aliscore output appears to be empty. Check results for BUSCO: 491869at2759
## Mon Jul 25 14:26:11 CEST 2022 - The aliscore output file does not exist. Check results for BUSCO: 1259741at2759
## Mon Jul 25 14:26:11 CEST 2022 - The aliscore output appears to be empty. Check results for BUSCO: 1259741at2759
## Mon Jul 25 14:26:11 CEST 2022 - The aliscore output file does not exist. Check results for BUSCO: 1513531at2759
## Mon Jul 25 14:26:11 CEST 2022 - The aliscore output appears to be empty. Check results for BUSCO: 1513531at2759
## Mon Jul 25 14:26:11 CEST 2022 - The aliscore output file does not exist. Check results for BUSCO: 779909at2759
## Mon Jul 25 14:26:12 CEST 2022 - The aliscore output appears to be empty. Check results for BUSCO: 779909at2759
## Mon Jul 25 14:26:12 CEST 2022 - The aliscore output file does not exist. Check results for BUSCO: 878143at2759
## Mon Jul 25 14:26:12 CEST 2022 - The aliscore output appears to be empty. Check results for BUSCO: 878143at2759
## Mon Jul 25 14:26:12 CEST 2022 - The aliscore output file does not exist. Check results for BUSCO: 1526152at2759
## Mon Jul 25 14:26:12 CEST 2022 - The aliscore output appears to be empty. Check results for BUSCO: 1526152at2759
## Mon Jul 25 14:26:12 CEST 2022 - The aliscore output file does not exist. Check results for BUSCO: 1405073at2759
## Mon Jul 25 14:26:12 CEST 2022 - The aliscore output appears to be empty. Check results for BUSCO: 1405073at2759
## Mon Jul 25 14:26:12 CEST 2022 - The aliscore output file does not exist. Check results for BUSCO: 1445102at2759
## Mon Jul 25 14:26:12 CEST 2022 - The aliscore output appears to be empty. Check results for BUSCO: 1445102at2759
## Mon Jul 25 14:26:12 CEST 2022 - The aliscore output file does not exist. Check results for BUSCO: 869548at2759
## Mon Jul 25 14:26:12 CEST 2022 - The aliscore output appears to be empty. Check results for BUSCO: 869548at2759
## Mon Jul 25 14:26:12 CEST 2022 - The aliscore output file does not exist. Check results for BUSCO: 1370285at2759
## Mon Jul 25 14:26:12 CEST 2022 - The aliscore output file does not exist. Check results for BUSCO: 1001705at2759
## Mon Jul 25 14:26:12 CEST 2022 - The aliscore output appears to be empty. Check results for BUSCO: 1370285at2759
## Mon Jul 25 14:26:12 CEST 2022 - The aliscore output appears to be empty. Check results for BUSCO: 1001705at2759
## Mon Jul 25 14:26:12 CEST 2022 - The aliscore output file does not exist. Check results for BUSCO: 457861at2759
## Mon Jul 25 14:26:12 CEST 2022 - The aliscore output file does not exist. Check results for BUSCO: 1090038at2759
## Mon Jul 25 14:26:12 CEST 2022 - The aliscore output appears to be empty. Check results for BUSCO: 457861at2759
## Mon Jul 25 14:26:12 CEST 2022 - The aliscore output appears to be empty. Check results for BUSCO: 1090038at2759
## Mon Jul 25 14:26:12 CEST 2022 - The aliscore output file does not exist. Check results for BUSCO: 621827at2759
## Mon Jul 25 14:26:13 CEST 2022 - The aliscore output appears to be empty. Check results for BUSCO: 621827at2759
## Mon Jul 25 14:26:13 CEST 2022 - The aliscore output file does not exist. Check results for BUSCO: 664730at2759
## Mon Jul 25 14:26:13 CEST 2022 - The aliscore output appears to be empty. Check results for BUSCO: 664730at2759
## Mon Jul 25 14:26:13 CEST 2022 - The aliscore output file does not exist. Check results for BUSCO: 1107630at2759
## Mon Jul 25 14:26:13 CEST 2022 - The aliscore output appears to be empty. Check results for BUSCO: 1107630at2759
## Mon Jul 25 14:26:13 CEST 2022 - The aliscore output file does not exist. Check results for BUSCO: 1257440at2759
## Mon Jul 25 14:26:13 CEST 2022 - The aliscore output file does not exist. Check results for BUSCO: 866359at2759
## Mon Jul 25 14:26:13 CEST 2022 - The aliscore output appears to be empty. Check results for BUSCO: 1257440at2759
## Mon Jul 25 14:26:13 CEST 2022 - The aliscore output appears to be empty. Check results for BUSCO: 866359at2759
## Mon Jul 25 14:26:13 CEST 2022 - The aliscore output file does not exist. Check results for BUSCO: 1157302at2759
## Mon Jul 25 14:26:13 CEST 2022 - The aliscore output appears to be empty. Check results for BUSCO: 1157302at2759
## Mon Jul 25 14:26:13 CEST 2022 - The aliscore output file does not exist. Check results for BUSCO: 1545004at2759
## Mon Jul 25 14:26:13 CEST 2022 - The aliscore output appears to be empty. Check results for BUSCO: 1545004at2759
## Mon Jul 25 14:26:13 CEST 2022 - The aliscore output file does not exist. Check results for BUSCO: 1014314at2759
## Mon Jul 25 14:26:13 CEST 2022 - The aliscore output file does not exist. Check results for BUSCO: 1182451at2759
## Mon Jul 25 14:26:13 CEST 2022 - The aliscore output file does not exist. Check results for BUSCO: 834694at2759
## Mon Jul 25 14:26:13 CEST 2022 - The aliscore output appears to be empty. Check results for BUSCO: 1014314at2759
## Mon Jul 25 14:26:13 CEST 2022 - The aliscore output appears to be empty. Check results for BUSCO: 1182451at2759
## Mon Jul 25 14:26:13 CEST 2022 - The aliscore output appears to be empty. Check results for BUSCO: 834694at2759
## Mon Jul 25 14:26:13 CEST 2022 - The aliscore output file does not exist. Check results for BUSCO: 1338131at2759
## Mon Jul 25 14:26:13 CEST 2022 - The aliscore output file does not exist. Check results for BUSCO: 679771at2759
## Mon Jul 25 14:26:13 CEST 2022 - The aliscore output appears to be empty. Check results for BUSCO: 1338131at2759
## Mon Jul 25 14:26:13 CEST 2022 - The aliscore output appears to be empty. Check results for BUSCO: 679771at2759
## Mon Jul 25 14:26:14 CEST 2022 - The aliscore output file does not exist. Check results for BUSCO: 464990at2759
## Mon Jul 25 14:26:14 CEST 2022 - The aliscore output appears to be empty. Check results for BUSCO: 464990at2759
## Mon Jul 25 14:26:14 CEST 2022 - The aliscore output file does not exist. Check results for BUSCO: 1111142at2759
## Mon Jul 25 14:26:14 CEST 2022 - The aliscore output appears to be empty. Check results for BUSCO: 1111142at2759
## Mon Jul 25 14:26:14 CEST 2022 - The aliscore output file does not exist. Check results for BUSCO: 1266231at2759
## Mon Jul 25 14:26:14 CEST 2022 - The aliscore output file does not exist. Check results for BUSCO: 341721at2759
## Mon Jul 25 14:26:14 CEST 2022 - The aliscore output appears to be empty. Check results for BUSCO: 1266231at2759
## Mon Jul 25 14:26:14 CEST 2022 - The aliscore output file does not exist. Check results for BUSCO: 1269244at2759
## Mon Jul 25 14:26:14 CEST 2022 - The aliscore output appears to be empty. Check results for BUSCO: 341721at2759
## Mon Jul 25 14:26:14 CEST 2022 - The aliscore output file does not exist. Check results for BUSCO: 97116at2759
## Mon Jul 25 14:26:14 CEST 2022 - The aliscore output appears to be empty. Check results for BUSCO: 1269244at2759
## Mon Jul 25 14:26:14 CEST 2022 - The aliscore output file does not exist. Check results for BUSCO: 375960at2759
## Mon Jul 25 14:26:14 CEST 2022 - The aliscore output appears to be empty. Check results for BUSCO: 97116at2759
## Mon Jul 25 14:26:14 CEST 2022 - The aliscore output file does not exist. Check results for BUSCO: 1269649at2759
## Mon Jul 25 14:26:14 CEST 2022 - The aliscore output file does not exist. Check results for BUSCO: 887370at2759
## Mon Jul 25 14:26:14 CEST 2022 - The aliscore output appears to be empty. Check results for BUSCO: 375960at2759
## Mon Jul 25 14:26:14 CEST 2022 - The aliscore output file does not exist. Check results for BUSCO: 1576404at2759
## Mon Jul 25 14:26:14 CEST 2022 - The aliscore output appears to be empty. Check results for BUSCO: 1269649at2759
## Mon Jul 25 14:26:14 CEST 2022 - The aliscore output appears to be empty. Check results for BUSCO: 1576404at2759
## Mon Jul 25 14:26:14 CEST 2022 - The aliscore output appears to be empty. Check results for BUSCO: 887370at2759
## Mon Jul 25 14:26:14 CEST 2022 - The aliscore output file does not exist. Check results for BUSCO: 918816at2759
## Mon Jul 25 14:26:14 CEST 2022 - The aliscore output file does not exist. Check results for BUSCO: 1115196at2759
## Mon Jul 25 14:26:14 CEST 2022 - The aliscore output appears to be empty. Check results for BUSCO: 918816at2759
## Mon Jul 25 14:26:14 CEST 2022 - The aliscore output appears to be empty. Check results for BUSCO: 1115196at2759
## Mon Jul 25 14:26:14 CEST 2022 - The aliscore output file does not exist. Check results for BUSCO: 1404162at2759
## Mon Jul 25 14:26:14 CEST 2022 - The aliscore output appears to be empty. Check results for BUSCO: 1404162at2759
## Mon Jul 25 14:26:14 CEST 2022 - The aliscore output file does not exist. Check results for BUSCO: 1076134at2759
## Mon Jul 25 14:26:14 CEST 2022 - The aliscore output appears to be empty. Check results for BUSCO: 1076134at2759
## Mon Jul 25 14:26:14 CEST 2022 - The aliscore output file does not exist. Check results for BUSCO: 1173229at2759
## Mon Jul 25 14:26:15 CEST 2022 - The aliscore output appears to be empty. Check results for BUSCO: 1173229at2759
## Mon Jul 25 14:26:15 CEST 2022 - The aliscore output file does not exist. Check results for BUSCO: 1030907at2759
## Mon Jul 25 14:26:15 CEST 2022 - The aliscore output appears to be empty. Check results for BUSCO: 1030907at2759
## Mon Jul 25 14:26:15 CEST 2022 - The aliscore output file does not exist. Check results for BUSCO: 1567796at2759
## Mon Jul 25 14:26:15 CEST 2022 - The aliscore output appears to be empty. Check results for BUSCO: 1567796at2759
## Mon Jul 25 14:26:15 CEST 2022 - The aliscore output file does not exist. Check results for BUSCO: 1220881at2759
## Mon Jul 25 14:26:15 CEST 2022 - The aliscore output file does not exist. Check results for BUSCO: 1032689at2759
## Mon Jul 25 14:26:15 CEST 2022 - The aliscore output appears to be empty. Check results for BUSCO: 1220881at2759
## Mon Jul 25 14:26:15 CEST 2022 - The aliscore output appears to be empty. Check results for BUSCO: 1032689at2759
## Mon Jul 25 14:26:15 CEST 2022 - The aliscore output file does not exist. Check results for BUSCO: 1193442at2759
## Mon Jul 25 14:26:15 CEST 2022 - The aliscore output file does not exist. Check results for BUSCO: 708105at2759
## Mon Jul 25 14:26:15 CEST 2022 - The aliscore output file does not exist. Check results for BUSCO: 270107at2759
## Mon Jul 25 14:26:15 CEST 2022 - The aliscore output appears to be empty. Check results for BUSCO: 1193442at2759
## Mon Jul 25 14:26:15 CEST 2022 - The aliscore output appears to be empty. Check results for BUSCO: 270107at2759
## Mon Jul 25 14:26:15 CEST 2022 - The aliscore output appears to be empty. Check results for BUSCO: 708105at2759
## Mon Jul 25 14:26:15 CEST 2022 - The aliscore output file does not exist. Check results for BUSCO: 1454155at2759
## Mon Jul 25 14:26:15 CEST 2022 - The aliscore output file does not exist. Check results for BUSCO: 858842at2759
## Mon Jul 25 14:26:15 CEST 2022 - The aliscore output file does not exist. Check results for BUSCO: 513979at2759
## Mon Jul 25 14:26:15 CEST 2022 - The aliscore output file does not exist. Check results for BUSCO: 1304061at2759
## Mon Jul 25 14:26:15 CEST 2022 - The aliscore output appears to be empty. Check results for BUSCO: 858842at2759
## Mon Jul 25 14:26:15 CEST 2022 - The aliscore output appears to be empty. Check results for BUSCO: 513979at2759
## Mon Jul 25 14:26:15 CEST 2022 - The aliscore output appears to be empty. Check results for BUSCO: 1304061at2759
## Mon Jul 25 14:26:15 CEST 2022 - The aliscore output appears to be empty. Check results for BUSCO: 1454155at2759
## Mon Jul 25 14:26:15 CEST 2022 - The aliscore output file does not exist. Check results for BUSCO: 390348at2759
## Mon Jul 25 14:26:15 CEST 2022 - The aliscore output appears to be empty. Check results for BUSCO: 390348at2759
## Mon Jul 25 14:26:15 CEST 2022 - The aliscore output file does not exist. Check results for BUSCO: 1194691at2759
## Mon Jul 25 14:26:16 CEST 2022 - The aliscore output appears to be empty. Check results for BUSCO: 1194691at2759
## Mon Jul 25 14:26:16 CEST 2022 - The aliscore output file does not exist. Check results for BUSCO: 1019762at2759
## Mon Jul 25 14:26:16 CEST 2022 - The aliscore output file does not exist. Check results for BUSCO: 598949at2759
## Mon Jul 25 14:26:16 CEST 2022 - The aliscore output file does not exist. Check results for BUSCO: 1441854at2759
## Mon Jul 25 14:26:16 CEST 2022 - The aliscore output appears to be empty. Check results for BUSCO: 1019762at2759
## Mon Jul 25 14:26:16 CEST 2022 - The aliscore output appears to be empty. Check results for BUSCO: 598949at2759
## Mon Jul 25 14:26:16 CEST 2022 - The aliscore output appears to be empty. Check results for BUSCO: 1441854at2759
## Mon Jul 25 14:26:16 CEST 2022 - The aliscore output file does not exist. Check results for BUSCO: 293315at2759
## Mon Jul 25 14:26:16 CEST 2022 - The aliscore output appears to be empty. Check results for BUSCO: 293315at2759
## Mon Jul 25 14:26:16 CEST 2022 - The aliscore output file does not exist. Check results for BUSCO: 1428265at2759
## Mon Jul 25 14:26:16 CEST 2022 - The aliscore output appears to be empty. Check results for BUSCO: 1428265at2759
## Mon Jul 25 14:26:16 CEST 2022 - The aliscore output file does not exist. Check results for BUSCO: 426305at2759
## Mon Jul 25 14:26:16 CEST 2022 - The aliscore output appears to be empty. Check results for BUSCO: 426305at2759
## Mon Jul 25 14:26:16 CEST 2022 - The aliscore output file does not exist. Check results for BUSCO: 720340at2759
## Mon Jul 25 14:26:16 CEST 2022 - The aliscore output appears to be empty. Check results for BUSCO: 720340at2759
## Mon Jul 25 14:26:16 CEST 2022 - The aliscore output file does not exist. Check results for BUSCO: 1003258at2759
## Mon Jul 25 14:26:16 CEST 2022 - The aliscore output appears to be empty. Check results for BUSCO: 1003258at2759
## Mon Jul 25 14:26:16 CEST 2022 - The aliscore output file does not exist. Check results for BUSCO: 1314980at2759
## Mon Jul 25 14:26:16 CEST 2022 - The aliscore output appears to be empty. Check results for BUSCO: 1314980at2759
## Mon Jul 25 14:26:17 CEST 2022 - The aliscore output file does not exist. Check results for BUSCO: 679187at2759
## Mon Jul 25 14:26:17 CEST 2022 - The aliscore output appears to be empty. Check results for BUSCO: 679187at2759
## Mon Jul 25 14:26:17 CEST 2022 - The aliscore output file does not exist. Check results for BUSCO: 306227at2759
## Mon Jul 25 14:26:17 CEST 2022 - The aliscore output appears to be empty. Check results for BUSCO: 306227at2759
## Mon Jul 25 14:26:17 CEST 2022 - The aliscore output file does not exist. Check results for BUSCO: 905026at2759
## Mon Jul 25 14:26:17 CEST 2022 - The aliscore output appears to be empty. Check results for BUSCO: 905026at2759
## Mon Jul 25 14:26:17 CEST 2022 - The aliscore output file does not exist. Check results for BUSCO: 1428854at2759
## Mon Jul 25 14:26:17 CEST 2022 - The aliscore output appears to be empty. Check results for BUSCO: 1428854at2759
## Mon Jul 25 14:26:17 CEST 2022 - The aliscore output file does not exist. Check results for BUSCO: 891751at2759
## Mon Jul 25 14:26:17 CEST 2022 - The aliscore output appears to be empty. Check results for BUSCO: 891751at2759
## Mon Jul 25 14:26:19 CEST 2022 - The aliscore output file does not exist. Check results for BUSCO: 325552at2759
## Mon Jul 25 14:26:19 CEST 2022 - The aliscore output appears to be empty. Check results for BUSCO: 325552at2759
## Mon Jul 25 14:26:29 CEST 2022 - The aliscore output file does not exist. Check results for BUSCO: 156083at2759
## Mon Jul 25 14:26:29 CEST 2022 - The aliscore output appears to be empty. Check results for BUSCO: 156083at2759
## Mon Jul 25 14:26:38 CEST 2022 - The aliscore output file does not exist. Check results for BUSCO: 1236198at2759
## Mon Jul 25 14:26:38 CEST 2022 - The aliscore output appears to be empty. Check results for BUSCO: 1236198at2759
## Mon Jul 25 14:26:39 CEST 2022 - The aliscore output file does not exist. Check results for BUSCO: 1440961at2759
## Mon Jul 25 14:26:39 CEST 2022 - The aliscore output appears to be empty. Check results for BUSCO: 1440961at2759
## Mon Jul 25 14:26:40 CEST 2022 - The aliscore output file does not exist. Check results for BUSCO: 721605at2759
## Mon Jul 25 14:26:40 CEST 2022 - The aliscore output appears to be empty. Check results for BUSCO: 721605at2759
## Mon Jul 25 14:26:40 CEST 2022 - The aliscore output file does not exist. Check results for BUSCO: 1287094at2759
## Mon Jul 25 14:26:40 CEST 2022 - The aliscore output appears to be empty. Check results for BUSCO: 1287094at2759
## Mon Jul 25 14:26:40 CEST 2022 - The aliscore output file does not exist. Check results for BUSCO: 1645187at2759
## Mon Jul 25 14:26:40 CEST 2022 - The aliscore output appears to be empty. Check results for BUSCO: 1645187at2759
## Mon Jul 25 14:26:41 CEST 2022 - The aliscore output file does not exist. Check results for BUSCO: 1049599at2759
## Mon Jul 25 14:26:41 CEST 2022 - The aliscore output appears to be empty. Check results for BUSCO: 1049599at2759
## Mon Jul 25 14:26:41 CEST 2022 - The aliscore output file does not exist. Check results for BUSCO: 1348942at2759
## Mon Jul 25 14:26:41 CEST 2022 - The aliscore output appears to be empty. Check results for BUSCO: 1348942at2759
## Mon Jul 25 14:26:41 CEST 2022 - The aliscore output file does not exist. Check results for BUSCO: 1588798at2759
## Mon Jul 25 14:26:41 CEST 2022 - The aliscore output appears to be empty. Check results for BUSCO: 1588798at2759
## Mon Jul 25 14:26:41 CEST 2022 - The aliscore output file does not exist. Check results for BUSCO: 1233814at2759
## Mon Jul 25 14:26:41 CEST 2022 - The aliscore output file does not exist. Check results for BUSCO: 453044at2759
## Mon Jul 25 14:26:41 CEST 2022 - The aliscore output appears to be empty. Check results for BUSCO: 1233814at2759
## Mon Jul 25 14:26:41 CEST 2022 - The aliscore output appears to be empty. Check results for BUSCO: 453044at2759
## Mon Jul 25 14:26:41 CEST 2022 - The aliscore output file does not exist. Check results for BUSCO: 1405146at2759
## Mon Jul 25 14:26:41 CEST 2022 - The aliscore output appears to be empty. Check results for BUSCO: 1405146at2759
## Mon Jul 25 14:26:42 CEST 2022 - The aliscore output file does not exist. Check results for BUSCO: 674169at2759
## Mon Jul 25 14:26:42 CEST 2022 - The aliscore output appears to be empty. Check results for BUSCO: 674169at2759
## Mon Jul 25 14:26:42 CEST 2022 - The aliscore output file does not exist. Check results for BUSCO: 1434266at2759
## Mon Jul 25 14:26:42 CEST 2022 - The aliscore output appears to be empty. Check results for BUSCO: 1434266at2759
## Mon Jul 25 14:26:42 CEST 2022 - The aliscore output file does not exist. Check results for BUSCO: 1228942at2759
## Mon Jul 25 14:26:42 CEST 2022 - The aliscore output file does not exist. Check results for BUSCO: 1398309at2759
## Mon Jul 25 14:26:42 CEST 2022 - The aliscore output appears to be empty. Check results for BUSCO: 1228942at2759
## Mon Jul 25 14:26:42 CEST 2022 - The aliscore output appears to be empty. Check results for BUSCO: 1398309at2759
## Mon Jul 25 14:26:42 CEST 2022 - The aliscore output file does not exist. Check results for BUSCO: 330169at2759
## Mon Jul 25 14:26:42 CEST 2022 - The aliscore output appears to be empty. Check results for BUSCO: 330169at2759
## Mon Jul 25 14:26:42 CEST 2022 - The aliscore output file does not exist. Check results for BUSCO: 1530008at2759
## Mon Jul 25 14:26:42 CEST 2022 - The aliscore output appears to be empty. Check results for BUSCO: 1530008at2759
## Mon Jul 25 14:26:42 CEST 2022 - The aliscore output file does not exist. Check results for BUSCO: 1342242at2759
## Mon Jul 25 14:26:42 CEST 2022 - The aliscore output appears to be empty. Check results for BUSCO: 1342242at2759
## Mon Jul 25 14:26:42 CEST 2022 - The aliscore output file does not exist. Check results for BUSCO: 1450538at2759
## Mon Jul 25 14:26:42 CEST 2022 - The aliscore output appears to be empty. Check results for BUSCO: 1450538at2759
## Mon Jul 25 14:26:43 CEST 2022 - The aliscore output file does not exist. Check results for BUSCO: 1085752at2759
## Mon Jul 25 14:26:43 CEST 2022 - The aliscore output appears to be empty. Check results for BUSCO: 1085752at2759
## Mon Jul 25 14:26:43 CEST 2022 - The aliscore output file does not exist. Check results for BUSCO: 1442062at2759
## Mon Jul 25 14:26:43 CEST 2022 - The aliscore output appears to be empty. Check results for BUSCO: 1442062at2759
## Mon Jul 25 14:26:43 CEST 2022 - The aliscore output file does not exist. Check results for BUSCO: 1094121at2759
## Mon Jul 25 14:26:43 CEST 2022 - The aliscore output appears to be empty. Check results for BUSCO: 1094121at2759
## Mon Jul 25 14:26:43 CEST 2022 - The aliscore output file does not exist. Check results for BUSCO: 734341at2759
## Mon Jul 25 14:26:43 CEST 2022 - The aliscore output appears to be empty. Check results for BUSCO: 734341at2759
## Mon Jul 25 14:26:43 CEST 2022 - The aliscore output file does not exist. Check results for BUSCO: 100698at2759
## Mon Jul 25 14:26:43 CEST 2022 - The aliscore output appears to be empty. Check results for BUSCO: 100698at2759
## Mon Jul 25 14:26:43 CEST 2022 - The aliscore output file does not exist. Check results for BUSCO: 1309031at2759
## Mon Jul 25 14:26:43 CEST 2022 - The aliscore output appears to be empty. Check results for BUSCO: 1309031at2759
## Mon Jul 25 14:26:44 CEST 2022 - The aliscore output file does not exist. Check results for BUSCO: 956854at2759
## Mon Jul 25 14:26:44 CEST 2022 - The aliscore output appears to be empty. Check results for BUSCO: 956854at2759
## Mon Jul 25 14:26:44 CEST 2022 - The aliscore output file does not exist. Check results for BUSCO: 1128607at2759
## Mon Jul 25 14:26:44 CEST 2022 - The aliscore output file does not exist. Check results for BUSCO: 1057950at2759
## Mon Jul 25 14:26:44 CEST 2022 - The aliscore output appears to be empty. Check results for BUSCO: 1128607at2759
## Mon Jul 25 14:26:44 CEST 2022 - The aliscore output appears to be empty. Check results for BUSCO: 1057950at2759
## Mon Jul 25 14:26:44 CEST 2022 - The aliscore output file does not exist. Check results for BUSCO: 392369at2759
## Mon Jul 25 14:26:44 CEST 2022 - The aliscore output appears to be empty. Check results for BUSCO: 392369at2759
## Mon Jul 25 14:26:44 CEST 2022 - The aliscore output file does not exist. Check results for BUSCO: 1479417at2759
## Mon Jul 25 14:26:44 CEST 2022 - The aliscore output file does not exist. Check results for BUSCO: 774318at2759
## Mon Jul 25 14:26:44 CEST 2022 - The aliscore output appears to be empty. Check results for BUSCO: 1479417at2759
## Mon Jul 25 14:26:44 CEST 2022 - The aliscore output appears to be empty. Check results for BUSCO: 774318at2759
## Mon Jul 25 14:26:44 CEST 2022 - The aliscore output file does not exist. Check results for BUSCO: 1065019at2759
## Mon Jul 25 14:26:44 CEST 2022 - The aliscore output appears to be empty. Check results for BUSCO: 1065019at2759
## Mon Jul 25 14:26:44 CEST 2022 - The aliscore output file does not exist. Check results for BUSCO: 801857at2759
## Mon Jul 25 14:26:44 CEST 2022 - The aliscore output file does not exist. Check results for BUSCO: 655400at2759
## Mon Jul 25 14:26:44 CEST 2022 - The aliscore output appears to be empty. Check results for BUSCO: 801857at2759
## Mon Jul 25 14:26:44 CEST 2022 - The aliscore output appears to be empty. Check results for BUSCO: 655400at2759
## Mon Jul 25 14:26:44 CEST 2022 - The aliscore output file does not exist. Check results for BUSCO: 1364586at2759
## Mon Jul 25 14:26:44 CEST 2022 - The aliscore output appears to be empty. Check results for BUSCO: 1364586at2759
## Mon Jul 25 14:26:45 CEST 2022 - The aliscore output file does not exist. Check results for BUSCO: 1259741at2759
## Mon Jul 25 14:26:45 CEST 2022 - The aliscore output appears to be empty. Check results for BUSCO: 1259741at2759
## Mon Jul 25 14:26:45 CEST 2022 - The aliscore output file does not exist. Check results for BUSCO: 1322642at2759
## Mon Jul 25 14:26:45 CEST 2022 - The aliscore output appears to be empty. Check results for BUSCO: 1322642at2759
## Mon Jul 25 14:26:46 CEST 2022 - The aliscore output file does not exist. Check results for BUSCO: 257318at2759
## Mon Jul 25 14:26:46 CEST 2022 - The aliscore output appears to be empty. Check results for BUSCO: 257318at2759
## Mon Jul 25 14:26:46 CEST 2022 - The aliscore output file does not exist. Check results for BUSCO: 1112002at2759
## Mon Jul 25 14:26:46 CEST 2022 - The aliscore output appears to be empty. Check results for BUSCO: 1112002at2759
## Mon Jul 25 14:26:46 CEST 2022 - The aliscore output file does not exist. Check results for BUSCO: 551907at2759
## Mon Jul 25 14:26:46 CEST 2022 - The aliscore output appears to be empty. Check results for BUSCO: 551907at2759
## Mon Jul 25 14:26:46 CEST 2022 - The aliscore output file does not exist. Check results for BUSCO: 777920at2759
## Mon Jul 25 14:26:46 CEST 2022 - The aliscore output appears to be empty. Check results for BUSCO: 777920at2759
## Mon Jul 25 14:26:47 CEST 2022 - The aliscore output file does not exist. Check results for BUSCO: 901894at2759
## Mon Jul 25 14:26:47 CEST 2022 - The aliscore output appears to be empty. Check results for BUSCO: 901894at2759
## Mon Jul 25 14:26:47 CEST 2022 - The aliscore output file does not exist. Check results for BUSCO: 671846at2759
## Mon Jul 25 14:26:47 CEST 2022 - The aliscore output file does not exist. Check results for BUSCO: 296129at2759
## Mon Jul 25 14:26:47 CEST 2022 - The aliscore output appears to be empty. Check results for BUSCO: 671846at2759
## Mon Jul 25 14:26:47 CEST 2022 - The aliscore output file does not exist. Check results for BUSCO: 1423847at2759
## Mon Jul 25 14:26:47 CEST 2022 - The aliscore output appears to be empty. Check results for BUSCO: 296129at2759
## Mon Jul 25 14:26:47 CEST 2022 - The aliscore output appears to be empty. Check results for BUSCO: 1423847at2759
## Mon Jul 25 14:26:48 CEST 2022 - The aliscore output file does not exist. Check results for BUSCO: 973442at2759
## Mon Jul 25 14:26:48 CEST 2022 - The aliscore output appears to be empty. Check results for BUSCO: 973442at2759
## Mon Jul 25 14:26:48 CEST 2022 - The aliscore output file does not exist. Check results for BUSCO: 369837at2759
## Mon Jul 25 14:26:48 CEST 2022 - The aliscore output appears to be empty. Check results for BUSCO: 369837at2759
## Mon Jul 25 14:26:49 CEST 2022 - The aliscore output file does not exist. Check results for BUSCO: 160593at2759
## Mon Jul 25 14:26:49 CEST 2022 - The aliscore output appears to be empty. Check results for BUSCO: 160593at2759
## Mon Jul 25 14:26:49 CEST 2022 - The aliscore output file does not exist. Check results for BUSCO: 166920at2759
## Mon Jul 25 14:26:49 CEST 2022 - The aliscore output appears to be empty. Check results for BUSCO: 166920at2759
## Mon Jul 25 14:26:50 CEST 2022 - The aliscore output file does not exist. Check results for BUSCO: 179362at2759
## Mon Jul 25 14:26:50 CEST 2022 - The aliscore output appears to be empty. Check results for BUSCO: 179362at2759
## Mon Jul 25 14:27:10 CEST 2022 - The aliscore output file does not exist. Check results for BUSCO: 1380409at2759
## Mon Jul 25 14:27:10 CEST 2022 - The aliscore output appears to be empty. Check results for BUSCO: 1380409at2759
## Mon Jul 25 14:27:11 CEST 2022 - The aliscore output file does not exist. Check results for BUSCO: 1284731at2759
## Mon Jul 25 14:27:11 CEST 2022 - The aliscore output appears to be empty. Check results for BUSCO: 1284731at2759
## Mon Jul 25 14:27:11 CEST 2022 - The aliscore output file does not exist. Check results for BUSCO: 996662at2759
## Mon Jul 25 14:27:11 CEST 2022 - The aliscore output appears to be empty. Check results for BUSCO: 996662at2759
## Mon Jul 25 14:27:11 CEST 2022 - The aliscore output file does not exist. Check results for BUSCO: 1217666at2759
## Mon Jul 25 14:27:11 CEST 2022 - The aliscore output file does not exist. Check results for BUSCO: 1567796at2759
## Mon Jul 25 14:27:11 CEST 2022 - The aliscore output appears to be empty. Check results for BUSCO: 1217666at2759
## Mon Jul 25 14:27:11 CEST 2022 - The aliscore output appears to be empty. Check results for BUSCO: 1567796at2759
## Mon Jul 25 14:27:12 CEST 2022 - The aliscore output file does not exist. Check results for BUSCO: 1488235at2759
## Mon Jul 25 14:27:12 CEST 2022 - The aliscore output appears to be empty. Check results for BUSCO: 1488235at2759
## Mon Jul 25 14:27:12 CEST 2022 - The aliscore output file does not exist. Check results for BUSCO: 1194691at2759
## Mon Jul 25 14:27:12 CEST 2022 - The aliscore output appears to be empty. Check results for BUSCO: 1194691at2759
## Mon Jul 25 14:27:12 CEST 2022 - The aliscore output file does not exist. Check results for BUSCO: 1200489at2759
## Mon Jul 25 14:27:12 CEST 2022 - The aliscore output appears to be empty. Check results for BUSCO: 1200489at2759
## Mon Jul 25 14:27:12 CEST 2022 - The aliscore output file does not exist. Check results for BUSCO: 844748at2759
## Mon Jul 25 14:27:12 CEST 2022 - The aliscore output appears to be empty. Check results for BUSCO: 844748at2759
## Mon Jul 25 14:27:13 CEST 2022 - The aliscore output file does not exist. Check results for BUSCO: 1474000at2759
## Mon Jul 25 14:27:13 CEST 2022 - The aliscore output appears to be empty. Check results for BUSCO: 1474000at2759
## Mon Jul 25 14:27:13 CEST 2022 - The aliscore output file does not exist. Check results for BUSCO: 1161199at2759
## Mon Jul 25 14:27:13 CEST 2022 - The aliscore output appears to be empty. Check results for BUSCO: 1161199at2759
## Mon Jul 25 14:27:13 CEST 2022 - The aliscore output file does not exist. Check results for BUSCO: 1107630at2759
## Mon Jul 25 14:27:13 CEST 2022 - The aliscore output file does not exist. Check results for BUSCO: 325552at2759
## Mon Jul 25 14:27:13 CEST 2022 - The aliscore output appears to be empty. Check results for BUSCO: 1107630at2759
## Mon Jul 25 14:27:13 CEST 2022 - The aliscore output appears to be empty. Check results for BUSCO: 325552at2759
## Mon Jul 25 14:27:14 CEST 2022 - The aliscore output file does not exist. Check results for BUSCO: 1322299at2759
## Mon Jul 25 14:27:14 CEST 2022 - The aliscore output file does not exist. Check results for BUSCO: 1405073at2759
## Mon Jul 25 14:27:14 CEST 2022 - The aliscore output file does not exist. Check results for BUSCO: 1358374at2759
## Mon Jul 25 14:27:14 CEST 2022 - The aliscore output appears to be empty. Check results for BUSCO: 1405073at2759
## Mon Jul 25 14:27:14 CEST 2022 - The aliscore output appears to be empty. Check results for BUSCO: 1322299at2759
## Mon Jul 25 14:27:14 CEST 2022 - The aliscore output appears to be empty. Check results for BUSCO: 1358374at2759
## Mon Jul 25 14:27:14 CEST 2022 - The aliscore output file does not exist. Check results for BUSCO: 1617752at2759
## Mon Jul 25 14:27:14 CEST 2022 - The aliscore output appears to be empty. Check results for BUSCO: 1617752at2759
## Mon Jul 25 14:27:14 CEST 2022 - The aliscore output file does not exist. Check results for BUSCO: 1032689at2759
## Mon Jul 25 14:27:14 CEST 2022 - The aliscore output appears to be empty. Check results for BUSCO: 1032689at2759
## Mon Jul 25 14:27:15 CEST 2022 - The aliscore output file does not exist. Check results for BUSCO: 1355894at2759
## Mon Jul 25 14:27:15 CEST 2022 - The aliscore output file does not exist. Check results for BUSCO: 679771at2759
## Mon Jul 25 14:27:15 CEST 2022 - The aliscore output appears to be empty. Check results for BUSCO: 1355894at2759
## Mon Jul 25 14:27:15 CEST 2022 - The aliscore output appears to be empty. Check results for BUSCO: 679771at2759
## Mon Jul 25 14:27:15 CEST 2022 - The aliscore output file does not exist. Check results for BUSCO: 1157302at2759
## Mon Jul 25 14:27:15 CEST 2022 - The aliscore output appears to be empty. Check results for BUSCO: 1157302at2759
## Mon Jul 25 14:27:15 CEST 2022 - The aliscore output file does not exist. Check results for BUSCO: 290630at2759
## Mon Jul 25 14:27:15 CEST 2022 - The aliscore output appears to be empty. Check results for BUSCO: 290630at2759
## Mon Jul 25 14:27:15 CEST 2022 - The aliscore output file does not exist. Check results for BUSCO: 976469at2759
## Mon Jul 25 14:27:15 CEST 2022 - The aliscore output appears to be empty. Check results for BUSCO: 976469at2759
## Mon Jul 25 14:27:15 CEST 2022 - The aliscore output file does not exist. Check results for BUSCO: 761109at2759
## Mon Jul 25 14:27:15 CEST 2022 - The aliscore output file does not exist. Check results for BUSCO: 1038775at2759
## Mon Jul 25 14:27:15 CEST 2022 - The aliscore output appears to be empty. Check results for BUSCO: 761109at2759
## Mon Jul 25 14:27:15 CEST 2022 - The aliscore output appears to be empty. Check results for BUSCO: 1038775at2759
## Mon Jul 25 14:27:15 CEST 2022 - The aliscore output file does not exist. Check results for BUSCO: 604979at2759
## Mon Jul 25 14:27:15 CEST 2022 - The aliscore output appears to be empty. Check results for BUSCO: 604979at2759
## Mon Jul 25 14:27:16 CEST 2022 - The aliscore output file does not exist. Check results for BUSCO: 817008at2759
## Mon Jul 25 14:27:16 CEST 2022 - The aliscore output appears to be empty. Check results for BUSCO: 817008at2759
## Mon Jul 25 14:27:16 CEST 2022 - The aliscore output file does not exist. Check results for BUSCO: 1287401at2759
## Mon Jul 25 14:27:16 CEST 2022 - The aliscore output file does not exist. Check results for BUSCO: 1108845at2759
## Mon Jul 25 14:27:16 CEST 2022 - The aliscore output appears to be empty. Check results for BUSCO: 1287401at2759
## Mon Jul 25 14:27:16 CEST 2022 - The aliscore output appears to be empty. Check results for BUSCO: 1108845at2759
## Mon Jul 25 14:27:16 CEST 2022 - The aliscore output file does not exist. Check results for BUSCO: 1324510at2759
## Mon Jul 25 14:27:16 CEST 2022 - The aliscore output appears to be empty. Check results for BUSCO: 1324510at2759
## Mon Jul 25 14:27:16 CEST 2022 - The aliscore output file does not exist. Check results for BUSCO: 944899at2759
## Mon Jul 25 14:27:16 CEST 2022 - The aliscore output appears to be empty. Check results for BUSCO: 944899at2759
## Mon Jul 25 14:27:16 CEST 2022 - The aliscore output file does not exist. Check results for BUSCO: 1455730at2759
## Mon Jul 25 14:27:16 CEST 2022 - The aliscore output appears to be empty. Check results for BUSCO: 1455730at2759
## Mon Jul 25 14:27:16 CEST 2022 - The aliscore output file does not exist. Check results for BUSCO: 911863at2759
## Mon Jul 25 14:27:17 CEST 2022 - The aliscore output appears to be empty. Check results for BUSCO: 911863at2759
## Mon Jul 25 14:27:17 CEST 2022 - The aliscore output file does not exist. Check results for BUSCO: 1428854at2759
## Mon Jul 25 14:27:17 CEST 2022 - The aliscore output appears to be empty. Check results for BUSCO: 1428854at2759
## Mon Jul 25 14:27:17 CEST 2022 - The aliscore output file does not exist. Check results for BUSCO: 1182451at2759
## Mon Jul 25 14:27:17 CEST 2022 - The aliscore output appears to be empty. Check results for BUSCO: 1182451at2759
## Mon Jul 25 14:27:17 CEST 2022 - The aliscore output file does not exist. Check results for BUSCO: 751335at2759
## Mon Jul 25 14:27:17 CEST 2022 - The aliscore output appears to be empty. Check results for BUSCO: 751335at2759
## Mon Jul 25 14:27:17 CEST 2022 - The aliscore output file does not exist. Check results for BUSCO: 245208at2759
## Mon Jul 25 14:27:17 CEST 2022 - The aliscore output appears to be empty. Check results for BUSCO: 245208at2759
## Mon Jul 25 14:27:17 CEST 2022 - The aliscore output file does not exist. Check results for BUSCO: 491869at2759
## Mon Jul 25 14:27:17 CEST 2022 - The aliscore output appears to be empty. Check results for BUSCO: 491869at2759
## Mon Jul 25 14:27:18 CEST 2022 - The aliscore output file does not exist. Check results for BUSCO: 87842at2759
## Mon Jul 25 14:27:18 CEST 2022 - The aliscore output appears to be empty. Check results for BUSCO: 87842at2759
## Mon Jul 25 14:27:18 CEST 2022 - The aliscore output file does not exist. Check results for BUSCO: 939345at2759
## Mon Jul 25 14:27:18 CEST 2022 - The aliscore output appears to be empty. Check results for BUSCO: 939345at2759
## Mon Jul 25 14:27:18 CEST 2022 - The aliscore output file does not exist. Check results for BUSCO: 261419at2759
## Mon Jul 25 14:27:18 CEST 2022 - The aliscore output appears to be empty. Check results for BUSCO: 261419at2759
## Mon Jul 25 14:27:19 CEST 2022 - The aliscore output file does not exist. Check results for BUSCO: 39650at2759
## Mon Jul 25 14:27:19 CEST 2022 - The aliscore output appears to be empty. Check results for BUSCO: 39650at2759
## Mon Jul 25 14:27:20 CEST 2022 - The aliscore output file does not exist. Check results for BUSCO: 299766at2759
## Mon Jul 25 14:27:20 CEST 2022 - The aliscore output appears to be empty. Check results for BUSCO: 299766at2759
## Mon Jul 25 14:27:22 CEST 2022 - The aliscore output file does not exist. Check results for BUSCO: 625387at2759
## Mon Jul 25 14:27:23 CEST 2022 - The aliscore output appears to be empty. Check results for BUSCO: 625387at2759
## Mon Jul 25 14:27:25 CEST 2022 - The aliscore output file does not exist. Check results for BUSCO: 1312453at2759
## Mon Jul 25 14:27:25 CEST 2022 - The aliscore output appears to be empty. Check results for BUSCO: 1312453at2759
## Mon Jul 25 14:27:39 CEST 2022 - The aliscore output file does not exist. Check results for BUSCO: 1488436at2759
## Mon Jul 25 14:27:39 CEST 2022 - The aliscore output appears to be empty. Check results for BUSCO: 1488436at2759
## Mon Jul 25 14:27:41 CEST 2022 - The aliscore output file does not exist. Check results for BUSCO: 1633672at2759
## Mon Jul 25 14:27:41 CEST 2022 - The aliscore output file does not exist. Check results for BUSCO: 1526152at2759
## Mon Jul 25 14:27:41 CEST 2022 - The aliscore output appears to be empty. Check results for BUSCO: 1633672at2759
## Mon Jul 25 14:27:41 CEST 2022 - The aliscore output appears to be empty. Check results for BUSCO: 1526152at2759
## Mon Jul 25 14:27:41 CEST 2022 - The aliscore output file does not exist. Check results for BUSCO: 687505at2759
## Mon Jul 25 14:27:41 CEST 2022 - The aliscore output appears to be empty. Check results for BUSCO: 687505at2759
## Mon Jul 25 14:27:41 CEST 2022 - The aliscore output file does not exist. Check results for BUSCO: 779909at2759
## Mon Jul 25 14:27:41 CEST 2022 - The aliscore output appears to be empty. Check results for BUSCO: 779909at2759
## Mon Jul 25 14:27:41 CEST 2022 - The aliscore output file does not exist. Check results for BUSCO: 1197019at2759
## Mon Jul 25 14:27:41 CEST 2022 - The aliscore output appears to be empty. Check results for BUSCO: 1197019at2759
## Mon Jul 25 14:27:41 CEST 2022 - The aliscore output file does not exist. Check results for BUSCO: 1407446at2759
## Mon Jul 25 14:27:41 CEST 2022 - The aliscore output file does not exist. Check results for BUSCO: 1291729at2759
## Mon Jul 25 14:27:41 CEST 2022 - The aliscore output appears to be empty. Check results for BUSCO: 1407446at2759
## Mon Jul 25 14:27:41 CEST 2022 - The aliscore output appears to be empty. Check results for BUSCO: 1291729at2759
## Mon Jul 25 14:27:41 CEST 2022 - The aliscore output file does not exist. Check results for BUSCO: 1257440at2759
## Mon Jul 25 14:27:41 CEST 2022 - The aliscore output appears to be empty. Check results for BUSCO: 1257440at2759
## Mon Jul 25 14:27:42 CEST 2022 - The aliscore output file does not exist. Check results for BUSCO: 1178688at2759
## Mon Jul 25 14:27:42 CEST 2022 - The aliscore output appears to be empty. Check results for BUSCO: 1178688at2759
## Mon Jul 25 14:27:42 CEST 2022 - The aliscore output file does not exist. Check results for BUSCO: 1269244at2759
## Mon Jul 25 14:27:42 CEST 2022 - The aliscore output appears to be empty. Check results for BUSCO: 1269244at2759
## Mon Jul 25 14:27:42 CEST 2022 - The aliscore output file does not exist. Check results for BUSCO: 320059at2759
## Mon Jul 25 14:27:42 CEST 2022 - The aliscore output appears to be empty. Check results for BUSCO: 320059at2759
## Mon Jul 25 14:27:42 CEST 2022 - The aliscore output file does not exist. Check results for BUSCO: 1030907at2759
## Mon Jul 25 14:27:42 CEST 2022 - The aliscore output appears to be empty. Check results for BUSCO: 1030907at2759
## Mon Jul 25 14:27:42 CEST 2022 - The aliscore output file does not exist. Check results for BUSCO: 734341at2759
## Mon Jul 25 14:27:42 CEST 2022 - The aliscore output appears to be empty. Check results for BUSCO: 734341at2759
## Mon Jul 25 14:27:43 CEST 2022 - The aliscore output file does not exist. Check results for BUSCO: 1275837at2759
## Mon Jul 25 14:27:43 CEST 2022 - The aliscore output appears to be empty. Check results for BUSCO: 1275837at2759
## Mon Jul 25 14:27:43 CEST 2022 - The aliscore output file does not exist. Check results for BUSCO: 1173229at2759
## Mon Jul 25 14:27:43 CEST 2022 - The aliscore output file does not exist. Check results for BUSCO: 1051021at2759
## Mon Jul 25 14:27:43 CEST 2022 - The aliscore output appears to be empty. Check results for BUSCO: 1173229at2759
## Mon Jul 25 14:27:43 CEST 2022 - The aliscore output appears to be empty. Check results for BUSCO: 1051021at2759
## Mon Jul 25 14:27:43 CEST 2022 - The aliscore output file does not exist. Check results for BUSCO: 1090038at2759
## Mon Jul 25 14:27:43 CEST 2022 - The aliscore output appears to be empty. Check results for BUSCO: 1090038at2759
## Mon Jul 25 14:27:43 CEST 2022 - The aliscore output file does not exist. Check results for BUSCO: 1010730at2759
## Mon Jul 25 14:27:43 CEST 2022 - The aliscore output appears to be empty. Check results for BUSCO: 1010730at2759
## Mon Jul 25 14:27:43 CEST 2022 - The aliscore output file does not exist. Check results for BUSCO: 1275837at2759
## Mon Jul 25 14:27:43 CEST 2022 - The aliscore output appears to be empty. Check results for BUSCO: 1275837at2759
## Mon Jul 25 14:27:43 CEST 2022 - The aliscore output file does not exist. Check results for BUSCO: 1323575at2759
## Mon Jul 25 14:27:43 CEST 2022 - The aliscore output file does not exist. Check results for BUSCO: 911863at2759
## Mon Jul 25 14:27:43 CEST 2022 - The aliscore output appears to be empty. Check results for BUSCO: 1323575at2759
## Mon Jul 25 14:27:43 CEST 2022 - The aliscore output appears to be empty. Check results for BUSCO: 911863at2759
## Mon Jul 25 14:27:43 CEST 2022 - The aliscore output file does not exist. Check results for BUSCO: 1129824at2759
## Mon Jul 25 14:27:44 CEST 2022 - The aliscore output appears to be empty. Check results for BUSCO: 1129824at2759
## Mon Jul 25 14:27:44 CEST 2022 - The aliscore output file does not exist. Check results for BUSCO: 1260807at2759
## Mon Jul 25 14:27:44 CEST 2022 - The aliscore output file does not exist. Check results for BUSCO: 1525971at2759
## Mon Jul 25 14:27:44 CEST 2022 - The aliscore output appears to be empty. Check results for BUSCO: 1260807at2759
## Mon Jul 25 14:27:44 CEST 2022 - The aliscore output appears to be empty. Check results for BUSCO: 1525971at2759
## Mon Jul 25 14:27:44 CEST 2022 - The aliscore output file does not exist. Check results for BUSCO: 946128at2759
## Mon Jul 25 14:27:44 CEST 2022 - The aliscore output appears to be empty. Check results for BUSCO: 946128at2759
## Mon Jul 25 14:27:44 CEST 2022 - The aliscore output file does not exist. Check results for BUSCO: 1269649at2759
## Mon Jul 25 14:27:44 CEST 2022 - The aliscore output appears to be empty. Check results for BUSCO: 1269649at2759
## Mon Jul 25 14:27:45 CEST 2022 - The aliscore output file does not exist. Check results for BUSCO: 855834at2759
## Mon Jul 25 14:27:45 CEST 2022 - The aliscore output appears to be empty. Check results for BUSCO: 855834at2759
## Mon Jul 25 14:27:45 CEST 2022 - The aliscore output file does not exist. Check results for BUSCO: 981902at2759
## Mon Jul 25 14:27:45 CEST 2022 - The aliscore output appears to be empty. Check results for BUSCO: 981902at2759
## Mon Jul 25 14:27:45 CEST 2022 - The aliscore output file does not exist. Check results for BUSCO: 869548at2759
## Mon Jul 25 14:27:45 CEST 2022 - The aliscore output file does not exist. Check results for BUSCO: 299766at2759
## Mon Jul 25 14:27:45 CEST 2022 - The aliscore output appears to be empty. Check results for BUSCO: 869548at2759
## Mon Jul 25 14:27:45 CEST 2022 - The aliscore output appears to be empty. Check results for BUSCO: 299766at2759
## Mon Jul 25 14:27:45 CEST 2022 - The aliscore output file does not exist. Check results for BUSCO: 719531at2759
## Mon Jul 25 14:27:45 CEST 2022 - The aliscore output appears to be empty. Check results for BUSCO: 719531at2759
## Mon Jul 25 14:27:45 CEST 2022 - The aliscore output file does not exist. Check results for BUSCO: 582756at2759
## Mon Jul 25 14:27:45 CEST 2022 - The aliscore output appears to be empty. Check results for BUSCO: 582756at2759
## Mon Jul 25 14:27:45 CEST 2022 - The aliscore output file does not exist. Check results for BUSCO: 692986at2759
## Mon Jul 25 14:27:45 CEST 2022 - The aliscore output appears to be empty. Check results for BUSCO: 692986at2759
## Mon Jul 25 14:27:45 CEST 2022 - The aliscore output file does not exist. Check results for BUSCO: 1251252at2759
## Mon Jul 25 14:27:45 CEST 2022 - The aliscore output appears to be empty. Check results for BUSCO: 1251252at2759
## Mon Jul 25 14:27:45 CEST 2022 - The aliscore output file does not exist. Check results for BUSCO: 1010730at2759
## Mon Jul 25 14:27:45 CEST 2022 - The aliscore output file does not exist. Check results for BUSCO: 1053181at2759
## Mon Jul 25 14:27:45 CEST 2022 - The aliscore output appears to be empty. Check results for BUSCO: 1010730at2759
## Mon Jul 25 14:27:45 CEST 2022 - The aliscore output file does not exist. Check results for BUSCO: 1291729at2759
## Mon Jul 25 14:27:45 CEST 2022 - The aliscore output appears to be empty. Check results for BUSCO: 1053181at2759
## Mon Jul 25 14:27:45 CEST 2022 - The aliscore output appears to be empty. Check results for BUSCO: 1291729at2759
## Mon Jul 25 14:27:46 CEST 2022 - The aliscore output file does not exist. Check results for BUSCO: 834694at2759
## Mon Jul 25 14:27:46 CEST 2022 - The aliscore output appears to be empty. Check results for BUSCO: 834694at2759
## Mon Jul 25 14:27:46 CEST 2022 - The aliscore output file does not exist. Check results for BUSCO: 450058at2759
## Mon Jul 25 14:27:46 CEST 2022 - The aliscore output appears to be empty. Check results for BUSCO: 450058at2759
## Mon Jul 25 14:27:46 CEST 2022 - The aliscore output file does not exist. Check results for BUSCO: 1249159at2759
## Mon Jul 25 14:27:46 CEST 2022 - The aliscore output appears to be empty. Check results for BUSCO: 1249159at2759
## Mon Jul 25 14:27:46 CEST 2022 - The aliscore output file does not exist. Check results for BUSCO: 817008at2759
## Mon Jul 25 14:27:46 CEST 2022 - The aliscore output appears to be empty. Check results for BUSCO: 817008at2759
## Mon Jul 25 14:27:46 CEST 2022 - The aliscore output file does not exist. Check results for BUSCO: 1222562at2759
## Mon Jul 25 14:27:46 CEST 2022 - The aliscore output appears to be empty. Check results for BUSCO: 1222562at2759
## Mon Jul 25 14:27:47 CEST 2022 - The aliscore output file does not exist. Check results for BUSCO: 1248958at2759
## Mon Jul 25 14:27:47 CEST 2022 - The aliscore output appears to be empty. Check results for BUSCO: 1248958at2759
## Mon Jul 25 14:27:47 CEST 2022 - The aliscore output file does not exist. Check results for BUSCO: 671536at2759
## Mon Jul 25 14:27:47 CEST 2022 - The aliscore output appears to be empty. Check results for BUSCO: 671536at2759
## Mon Jul 25 14:27:47 CEST 2022 - The aliscore output file does not exist. Check results for BUSCO: 905026at2759
## Mon Jul 25 14:27:47 CEST 2022 - The aliscore output appears to be empty. Check results for BUSCO: 905026at2759
## Mon Jul 25 14:27:47 CEST 2022 - The aliscore output file does not exist. Check results for BUSCO: 761109at2759
## Mon Jul 25 14:27:47 CEST 2022 - The aliscore output appears to be empty. Check results for BUSCO: 761109at2759
## Mon Jul 25 14:27:47 CEST 2022 - The aliscore output file does not exist. Check results for BUSCO: 375960at2759
## Mon Jul 25 14:27:47 CEST 2022 - The aliscore output file does not exist. Check results for BUSCO: 1266231at2759
## Mon Jul 25 14:27:47 CEST 2022 - The aliscore output appears to be empty. Check results for BUSCO: 375960at2759
## Mon Jul 25 14:27:47 CEST 2022 - The aliscore output appears to be empty. Check results for BUSCO: 1266231at2759
## Mon Jul 25 14:27:48 CEST 2022 - The aliscore output file does not exist. Check results for BUSCO: 625387at2759
## Mon Jul 25 14:27:48 CEST 2022 - The aliscore output appears to be empty. Check results for BUSCO: 625387at2759
## Mon Jul 25 14:27:48 CEST 2022 - The aliscore output file does not exist. Check results for BUSCO: 1065019at2759
## Mon Jul 25 14:27:48 CEST 2022 - The aliscore output appears to be empty. Check results for BUSCO: 1065019at2759
## Mon Jul 25 14:27:48 CEST 2022 - The aliscore output file does not exist. Check results for BUSCO: 383503at2759
## Mon Jul 25 14:27:48 CEST 2022 - The aliscore output file does not exist. Check results for BUSCO: 939345at2759
## Mon Jul 25 14:27:48 CEST 2022 - The aliscore output appears to be empty. Check results for BUSCO: 383503at2759
## Mon Jul 25 14:27:48 CEST 2022 - The aliscore output appears to be empty. Check results for BUSCO: 939345at2759
## Mon Jul 25 14:27:48 CEST 2022 - The aliscore output file does not exist. Check results for BUSCO: 1193442at2759
## Mon Jul 25 14:27:48 CEST 2022 - The aliscore output appears to be empty. Check results for BUSCO: 1193442at2759
## Mon Jul 25 14:27:48 CEST 2022 - The aliscore output file does not exist. Check results for BUSCO: 1096688at2759
## Mon Jul 25 14:27:48 CEST 2022 - The aliscore output appears to be empty. Check results for BUSCO: 1096688at2759
## Mon Jul 25 14:27:49 CEST 2022 - The aliscore output file does not exist. Check results for BUSCO: 1019762at2759
## Mon Jul 25 14:27:49 CEST 2022 - The aliscore output appears to be empty. Check results for BUSCO: 1019762at2759
## Mon Jul 25 14:27:49 CEST 2022 - The aliscore output file does not exist. Check results for BUSCO: 901894at2759
## Mon Jul 25 14:27:49 CEST 2022 - The aliscore output file does not exist. Check results for BUSCO: 1079130at2759
## Mon Jul 25 14:27:49 CEST 2022 - The aliscore output appears to be empty. Check results for BUSCO: 901894at2759
## Mon Jul 25 14:27:49 CEST 2022 - The aliscore output appears to be empty. Check results for BUSCO: 1079130at2759
## Mon Jul 25 14:27:49 CEST 2022 - The aliscore output file does not exist. Check results for BUSCO: 513979at2759
## Mon Jul 25 14:27:49 CEST 2022 - The aliscore output appears to be empty. Check results for BUSCO: 513979at2759
## Mon Jul 25 14:27:49 CEST 2022 - The aliscore output file does not exist. Check results for BUSCO: 1003258at2759
## Mon Jul 25 14:27:50 CEST 2022 - The aliscore output appears to be empty. Check results for BUSCO: 1003258at2759
## Mon Jul 25 14:27:50 CEST 2022 - The aliscore output file does not exist. Check results for BUSCO: 1194797at2759
## Mon Jul 25 14:27:50 CEST 2022 - The aliscore output appears to be empty. Check results for BUSCO: 1194797at2759
## Mon Jul 25 14:27:51 CEST 2022 - The aliscore output file does not exist. Check results for BUSCO: 83779at2759
## Mon Jul 25 14:27:51 CEST 2022 - The aliscore output appears to be empty. Check results for BUSCO: 83779at2759
## Mon Jul 25 14:27:51 CEST 2022 - The aliscore output file does not exist. Check results for BUSCO: 369837at2759
## Mon Jul 25 14:27:51 CEST 2022 - The aliscore output appears to be empty. Check results for BUSCO: 369837at2759
## Mon Jul 25 14:27:51 CEST 2022 - The aliscore output file does not exist. Check results for BUSCO: 1404162at2759
## Mon Jul 25 14:27:52 CEST 2022 - The aliscore output appears to be empty. Check results for BUSCO: 1404162at2759
## Mon Jul 25 14:28:09 CEST 2022 - The aliscore output file does not exist. Check results for BUSCO: 1558822at2759
## Mon Jul 25 14:28:09 CEST 2022 - The aliscore output appears to be empty. Check results for BUSCO: 1558822at2759
## Mon Jul 25 14:28:09 CEST 2022 - The aliscore output file does not exist. Check results for BUSCO: 1623701at2759
## Mon Jul 25 14:28:09 CEST 2022 - The aliscore output appears to be empty. Check results for BUSCO: 1623701at2759
## Mon Jul 25 14:28:10 CEST 2022 - The aliscore output file does not exist. Check results for BUSCO: 1459797at2759
## Mon Jul 25 14:28:10 CEST 2022 - The aliscore output appears to be empty. Check results for BUSCO: 1459797at2759
## Mon Jul 25 14:28:10 CEST 2022 - The aliscore output file does not exist. Check results for BUSCO: 457861at2759
## Mon Jul 25 14:28:10 CEST 2022 - The aliscore output appears to be empty. Check results for BUSCO: 457861at2759
## Mon Jul 25 14:28:11 CEST 2022 - The aliscore output file does not exist. Check results for BUSCO: 1076134at2759
## Mon Jul 25 14:28:11 CEST 2022 - The aliscore output file does not exist. Check results for BUSCO: 866359at2759
## Mon Jul 25 14:28:11 CEST 2022 - The aliscore output appears to be empty. Check results for BUSCO: 1076134at2759
## Mon Jul 25 14:28:11 CEST 2022 - The aliscore output appears to be empty. Check results for BUSCO: 866359at2759
## Mon Jul 25 14:28:12 CEST 2022 - The aliscore output file does not exist. Check results for BUSCO: 464990at2759
## Mon Jul 25 14:28:12 CEST 2022 - The aliscore output appears to be empty. Check results for BUSCO: 464990at2759
## Mon Jul 25 14:28:12 CEST 2022 - The aliscore output file does not exist. Check results for BUSCO: 734666at2759
## Mon Jul 25 14:28:12 CEST 2022 - The aliscore output appears to be empty. Check results for BUSCO: 734666at2759
## Mon Jul 25 14:28:13 CEST 2022 - The aliscore output file does not exist. Check results for BUSCO: 1115196at2759
## Mon Jul 25 14:28:13 CEST 2022 - The aliscore output appears to be empty. Check results for BUSCO: 1115196at2759
## Mon Jul 25 14:28:13 CEST 2022 - The aliscore output file does not exist. Check results for BUSCO: 1441854at2759
## Mon Jul 25 14:28:13 CEST 2022 - The aliscore output appears to be empty. Check results for BUSCO: 1441854at2759
## Mon Jul 25 14:28:13 CEST 2022 - The aliscore output file does not exist. Check results for BUSCO: 1106766at2759
## Mon Jul 25 14:28:13 CEST 2022 - The aliscore output file does not exist. Check results for BUSCO: 664730at2759
## Mon Jul 25 14:28:13 CEST 2022 - The aliscore output appears to be empty. Check results for BUSCO: 1106766at2759
## Mon Jul 25 14:28:13 CEST 2022 - The aliscore output appears to be empty. Check results for BUSCO: 664730at2759
## Mon Jul 25 14:28:13 CEST 2022 - The aliscore output file does not exist. Check results for BUSCO: 1014314at2759
## Mon Jul 25 14:28:13 CEST 2022 - The aliscore output appears to be empty. Check results for BUSCO: 1014314at2759
## Mon Jul 25 14:28:16 CEST 2022 - The aliscore output file does not exist. Check results for BUSCO: 1379600at2759
## Mon Jul 25 14:28:16 CEST 2022 - The aliscore output appears to be empty. Check results for BUSCO: 1379600at2759
## Mon Jul 25 14:28:16 CEST 2022 - The aliscore output file does not exist. Check results for BUSCO: 306227at2759
## Mon Jul 25 14:28:16 CEST 2022 - The aliscore output appears to be empty. Check results for BUSCO: 306227at2759
## Mon Jul 25 14:28:17 CEST 2022 - The aliscore output file does not exist. Check results for BUSCO: 261328at2759
## Mon Jul 25 14:28:17 CEST 2022 - The aliscore output appears to be empty. Check results for BUSCO: 261328at2759
## Mon Jul 25 14:30:09 CEST 2022 - The aliscore output file does not exist. Check results for BUSCO: 1107630at2759
## Mon Jul 25 14:30:09 CEST 2022 - The aliscore output appears to be empty. Check results for BUSCO: 1107630at2759
## Mon Jul 25 14:30:10 CEST 2022 - The aliscore output file does not exist. Check results for BUSCO: 1398309at2759
## Mon Jul 25 14:30:10 CEST 2022 - The aliscore output file does not exist. Check results for BUSCO: 1157302at2759
## Mon Jul 25 14:30:10 CEST 2022 - The aliscore output appears to be empty. Check results for BUSCO: 1398309at2759
## Mon Jul 25 14:30:10 CEST 2022 - The aliscore output appears to be empty. Check results for BUSCO: 1157302at2759
## Mon Jul 25 14:30:10 CEST 2022 - The aliscore output file does not exist. Check results for BUSCO: 679771at2759
## Mon Jul 25 14:30:10 CEST 2022 - The aliscore output appears to be empty. Check results for BUSCO: 679771at2759
## Mon Jul 25 14:30:10 CEST 2022 - The aliscore output file does not exist. Check results for BUSCO: 1161199at2759
## Mon Jul 25 14:30:10 CEST 2022 - The aliscore output appears to be empty. Check results for BUSCO: 1161199at2759
## Mon Jul 25 14:30:11 CEST 2022 - The aliscore output file does not exist. Check results for BUSCO: 1217666at2759
## Mon Jul 25 14:30:11 CEST 2022 - The aliscore output appears to be empty. Check results for BUSCO: 1217666at2759
## Mon Jul 25 14:30:11 CEST 2022 - The aliscore output file does not exist. Check results for BUSCO: 1129824at2759
## Mon Jul 25 14:30:11 CEST 2022 - The aliscore output appears to be empty. Check results for BUSCO: 1129824at2759
## Mon Jul 25 14:30:11 CEST 2022 - The aliscore output file does not exist. Check results for BUSCO: 923657at2759
## Mon Jul 25 14:30:11 CEST 2022 - The aliscore output file does not exist. Check results for BUSCO: 1182451at2759
## Mon Jul 25 14:30:11 CEST 2022 - The aliscore output appears to be empty. Check results for BUSCO: 923657at2759
## Mon Jul 25 14:30:11 CEST 2022 - The aliscore output appears to be empty. Check results for BUSCO: 1182451at2759
## Mon Jul 25 14:30:12 CEST 2022 - The aliscore output file does not exist. Check results for BUSCO: 1194691at2759
## Mon Jul 25 14:30:12 CEST 2022 - The aliscore output appears to be empty. Check results for BUSCO: 1194691at2759
## Mon Jul 25 14:30:12 CEST 2022 - The aliscore output file does not exist. Check results for BUSCO: 290630at2759
## Mon Jul 25 14:30:12 CEST 2022 - The aliscore output appears to be empty. Check results for BUSCO: 290630at2759
## Mon Jul 25 14:30:13 CEST 2022 - The aliscore output file does not exist. Check results for BUSCO: 97116at2759
## Mon Jul 25 14:30:13 CEST 2022 - The aliscore output appears to be empty. Check results for BUSCO: 97116at2759
## Mon Jul 25 14:30:14 CEST 2022 - The aliscore output file does not exist. Check results for BUSCO: 1322299at2759
## Mon Jul 25 14:30:14 CEST 2022 - The aliscore output appears to be empty. Check results for BUSCO: 1322299at2759
## Mon Jul 25 14:30:14 CEST 2022 - The aliscore output file does not exist. Check results for BUSCO: 1194797at2759
## Mon Jul 25 14:30:14 CEST 2022 - The aliscore output appears to be empty. Check results for BUSCO: 1194797at2759
## Mon Jul 25 14:30:14 CEST 2022 - The aliscore output file does not exist. Check results for BUSCO: 673132at2759
## Mon Jul 25 14:30:14 CEST 2022 - The aliscore output appears to be empty. Check results for BUSCO: 673132at2759
## Mon Jul 25 14:30:14 CEST 2022 - The aliscore output file does not exist. Check results for BUSCO: 1322642at2759
## Mon Jul 25 14:30:14 CEST 2022 - The aliscore output appears to be empty. Check results for BUSCO: 1322642at2759
## Mon Jul 25 14:30:15 CEST 2022 - The aliscore output file does not exist. Check results for BUSCO: 918816at2759
## Mon Jul 25 14:30:15 CEST 2022 - The aliscore output appears to be empty. Check results for BUSCO: 918816at2759
## Mon Jul 25 14:30:15 CEST 2022 - The aliscore output file does not exist. Check results for BUSCO: 325552at2759
## Mon Jul 25 14:30:15 CEST 2022 - The aliscore output appears to be empty. Check results for BUSCO: 325552at2759
## Mon Jul 25 14:30:15 CEST 2022 - The aliscore output file does not exist. Check results for BUSCO: 1423847at2759
## Mon Jul 25 14:30:15 CEST 2022 - The aliscore output appears to be empty. Check results for BUSCO: 1423847at2759
## Mon Jul 25 14:30:15 CEST 2022 - The aliscore output file does not exist. Check results for BUSCO: 1003258at2759
## Mon Jul 25 14:30:15 CEST 2022 - The aliscore output appears to be empty. Check results for BUSCO: 1003258at2759
## Mon Jul 25 14:30:18 CEST 2022 - The aliscore output file does not exist. Check results for BUSCO: 179362at2759
## Mon Jul 25 14:30:18 CEST 2022 - The aliscore output appears to be empty. Check results for BUSCO: 179362at2759
## Mon Jul 25 14:30:18 CEST 2022 - The aliscore output file does not exist. Check results for BUSCO: 973442at2759
## Mon Jul 25 14:30:18 CEST 2022 - The aliscore output appears to be empty. Check results for BUSCO: 973442at2759
## Mon Jul 25 14:30:21 CEST 2022 - The aliscore output file does not exist. Check results for BUSCO: 674160at2759
## Mon Jul 25 14:30:21 CEST 2022 - The aliscore output appears to be empty. Check results for BUSCO: 674160at2759
## Mon Jul 25 14:30:22 CEST 2022 - The aliscore output file does not exist. Check results for BUSCO: 1312453at2759
## Mon Jul 25 14:30:22 CEST 2022 - The aliscore output appears to be empty. Check results for BUSCO: 1312453at2759
## Mon Jul 25 14:30:39 CEST 2022 - The aliscore output file does not exist. Check results for BUSCO: 1474000at2759
## Mon Jul 25 14:30:39 CEST 2022 - The aliscore output appears to be empty. Check results for BUSCO: 1474000at2759
## Mon Jul 25 14:30:40 CEST 2022 - Number of alignments (mafft): 255
## Mon Jul 25 14:30:40 CEST 2022 - Number of alignments (muscle): 255
## Mon Jul 25 14:30:40 CEST 2022 - Number of trimmed alignments (mafft - bmge): 255
## Mon Jul 25 14:30:40 CEST 2022 - The aliscore output file does not exist. Check results for BUSCO: 1459797at2759
## Mon Jul 25 14:30:40 CEST 2022 - Number of trimmed alignments (muscle - bmge): 255
## Mon Jul 25 14:30:40 CEST 2022 - Number of alignments (muscle): 255
## Mon Jul 25 14:30:40 CEST 2022 - The aliscore output appears to be empty. Check results for BUSCO: 1459797at2759
## Mon Jul 25 14:30:40 CEST 2022 - Number of trimmed alignments (muscle - trimal): 255
## Mon Jul 25 14:30:40 CEST 2022 - Number of alignments (mafft - bmge) after filtering: 245
## Mon Jul 25 14:30:40 CEST 2022 - Number of alignments (muscle - bmge) after filtering: 244
## Mon Jul 25 14:30:40 CEST 2022 - Number of alignments (muscle - trimal) after filtering: 249
## Mon Jul 25 14:30:41 CEST 2022 - Number of alignments (mafft): 255
## Mon Jul 25 14:30:41 CEST 2022 - Number of trimmed alignments (mafft - trimal): 255
## Mon Jul 25 14:30:41 CEST 2022 - Number of alignments (mafft - trimal) after filtering: 250
## Mon Jul 25 14:30:41 CEST 2022 - The aliscore output file does not exist. Check results for BUSCO: 1193442at2759
## Mon Jul 25 14:30:41 CEST 2022 - The aliscore output file does not exist. Check results for BUSCO: 1575179at2759
## Mon Jul 25 14:30:41 CEST 2022 - The aliscore output appears to be empty. Check results for BUSCO: 1193442at2759
## Mon Jul 25 14:30:41 CEST 2022 - The aliscore output appears to be empty. Check results for BUSCO: 1575179at2759
## Mon Jul 25 14:30:41 CEST 2022 - The aliscore output file does not exist. Check results for BUSCO: 604979at2759
## Mon Jul 25 14:30:41 CEST 2022 - The aliscore output file does not exist. Check results for BUSCO: 1504863at2759
## Mon Jul 25 14:30:41 CEST 2022 - The aliscore output appears to be empty. Check results for BUSCO: 604979at2759
## Mon Jul 25 14:30:41 CEST 2022 - The aliscore output appears to be empty. Check results for BUSCO: 1504863at2759
## Mon Jul 25 14:30:41 CEST 2022 - The aliscore output file does not exist. Check results for BUSCO: 1200489at2759
## Mon Jul 25 14:30:42 CEST 2022 - The aliscore output appears to be empty. Check results for BUSCO: 1200489at2759
## Mon Jul 25 14:30:42 CEST 2022 - The aliscore output file does not exist. Check results for BUSCO: 768809at2759
## Mon Jul 25 14:30:42 CEST 2022 - The aliscore output appears to be empty. Check results for BUSCO: 768809at2759
## Mon Jul 25 14:30:42 CEST 2022 - The aliscore output file does not exist. Check results for BUSCO: 1233814at2759
## Mon Jul 25 14:30:42 CEST 2022 - The aliscore output appears to be empty. Check results for BUSCO: 1233814at2759
## Mon Jul 25 14:30:42 CEST 2022 - The aliscore output file does not exist. Check results for BUSCO: 1526152at2759
## Mon Jul 25 14:30:42 CEST 2022 - The aliscore output appears to be empty. Check results for BUSCO: 1526152at2759
## Mon Jul 25 14:30:42 CEST 2022 - The aliscore output file does not exist. Check results for BUSCO: 919955at2759
## Mon Jul 25 14:30:42 CEST 2022 - The aliscore output appears to be empty. Check results for BUSCO: 919955at2759
## Mon Jul 25 14:30:42 CEST 2022 - The aliscore output file does not exist. Check results for BUSCO: 1479417at2759
## Mon Jul 25 14:30:42 CEST 2022 - The aliscore output appears to be empty. Check results for BUSCO: 1479417at2759
## Mon Jul 25 14:30:42 CEST 2022 - The aliscore output file does not exist. Check results for BUSCO: 655400at2759
## Mon Jul 25 14:30:42 CEST 2022 - The aliscore output file does not exist. Check results for BUSCO: 1200489at2759
## Mon Jul 25 14:30:42 CEST 2022 - The aliscore output appears to be empty. Check results for BUSCO: 655400at2759
## Mon Jul 25 14:30:42 CEST 2022 - The aliscore output file does not exist. Check results for BUSCO: 1575179at2759
## Mon Jul 25 14:30:42 CEST 2022 - The aliscore output appears to be empty. Check results for BUSCO: 1200489at2759
## Mon Jul 25 14:30:42 CEST 2022 - The aliscore output appears to be empty. Check results for BUSCO: 1575179at2759
## Mon Jul 25 14:30:42 CEST 2022 - The aliscore output file does not exist. Check results for BUSCO: 1526152at2759
## Mon Jul 25 14:30:42 CEST 2022 - The aliscore output appears to be empty. Check results for BUSCO: 1526152at2759
## Mon Jul 25 14:30:42 CEST 2022 - The aliscore output file does not exist. Check results for BUSCO: 1623701at2759
## Mon Jul 25 14:30:43 CEST 2022 - The aliscore output appears to be empty. Check results for BUSCO: 1623701at2759
## Mon Jul 25 14:30:43 CEST 2022 - The aliscore output file does not exist. Check results for BUSCO: 1620056at2759
## Mon Jul 25 14:30:43 CEST 2022 - The aliscore output file does not exist. Check results for BUSCO: 1620056at2759
## Mon Jul 25 14:30:43 CEST 2022 - The aliscore output appears to be empty. Check results for BUSCO: 1620056at2759
## Mon Jul 25 14:30:43 CEST 2022 - The aliscore output appears to be empty. Check results for BUSCO: 1620056at2759
## Mon Jul 25 14:30:43 CEST 2022 - The aliscore output file does not exist. Check results for BUSCO: 1405073at2759
## Mon Jul 25 14:30:43 CEST 2022 - The aliscore output file does not exist. Check results for BUSCO: 919955at2759
## Mon Jul 25 14:30:43 CEST 2022 - The aliscore output appears to be empty. Check results for BUSCO: 919955at2759
## Mon Jul 25 14:30:43 CEST 2022 - The aliscore output appears to be empty. Check results for BUSCO: 1405073at2759
## Mon Jul 25 14:30:43 CEST 2022 - The aliscore output file does not exist. Check results for BUSCO: 464990at2759
## Mon Jul 25 14:30:43 CEST 2022 - The aliscore output file does not exist. Check results for BUSCO: 687505at2759
## Mon Jul 25 14:30:43 CEST 2022 - The aliscore output file does not exist. Check results for BUSCO: 1504863at2759
## Mon Jul 25 14:30:43 CEST 2022 - The aliscore output appears to be empty. Check results for BUSCO: 464990at2759
## Mon Jul 25 14:30:43 CEST 2022 - The aliscore output appears to be empty. Check results for BUSCO: 687505at2759
## Mon Jul 25 14:30:43 CEST 2022 - The aliscore output appears to be empty. Check results for BUSCO: 1504863at2759
## Mon Jul 25 14:30:43 CEST 2022 - The aliscore output file does not exist. Check results for BUSCO: 513979at2759
## Mon Jul 25 14:30:43 CEST 2022 - The aliscore output appears to be empty. Check results for BUSCO: 513979at2759
## Mon Jul 25 14:30:43 CEST 2022 - The aliscore output file does not exist. Check results for BUSCO: 1266231at2759
## Mon Jul 25 14:30:43 CEST 2022 - The aliscore output file does not exist. Check results for BUSCO: 1178688at2759
## Mon Jul 25 14:30:43 CEST 2022 - The aliscore output appears to be empty. Check results for BUSCO: 1266231at2759
## Mon Jul 25 14:30:43 CEST 2022 - The aliscore output appears to be empty. Check results for BUSCO: 1178688at2759
## Mon Jul 25 14:30:43 CEST 2022 - The aliscore output file does not exist. Check results for BUSCO: 1129824at2759
## Mon Jul 25 14:30:43 CEST 2022 - The aliscore output appears to be empty. Check results for BUSCO: 1129824at2759
## Mon Jul 25 14:30:43 CEST 2022 - The aliscore output file does not exist. Check results for BUSCO: 974865at2759
## Mon Jul 25 14:30:43 CEST 2022 - The aliscore output file does not exist. Check results for BUSCO: 1018517at2759
## Mon Jul 25 14:30:43 CEST 2022 - The aliscore output appears to be empty. Check results for BUSCO: 974865at2759
## Mon Jul 25 14:30:43 CEST 2022 - The aliscore output appears to be empty. Check results for BUSCO: 1018517at2759
## Mon Jul 25 14:30:43 CEST 2022 - The aliscore output file does not exist. Check results for BUSCO: 1223488at2759
## Mon Jul 25 14:30:43 CEST 2022 - The aliscore output appears to be empty. Check results for BUSCO: 1223488at2759
## Mon Jul 25 14:30:43 CEST 2022 - The aliscore output file does not exist. Check results for BUSCO: 1645187at2759
## Mon Jul 25 14:30:43 CEST 2022 - The aliscore output appears to be empty. Check results for BUSCO: 1645187at2759
## Mon Jul 25 14:30:43 CEST 2022 - The aliscore output file does not exist. Check results for BUSCO: 1445102at2759
## Mon Jul 25 14:30:43 CEST 2022 - The aliscore output file does not exist. Check results for BUSCO: 1079827at2759
## Mon Jul 25 14:30:43 CEST 2022 - The aliscore output appears to be empty. Check results for BUSCO: 1445102at2759
## Mon Jul 25 14:30:43 CEST 2022 - The aliscore output appears to be empty. Check results for BUSCO: 1079827at2759
## Mon Jul 25 14:30:43 CEST 2022 - The aliscore output file does not exist. Check results for BUSCO: 1287094at2759
## Mon Jul 25 14:30:43 CEST 2022 - The aliscore output appears to be empty. Check results for BUSCO: 1287094at2759
## Mon Jul 25 14:30:43 CEST 2022 - The aliscore output file does not exist. Check results for BUSCO: 1490022at2759
## Mon Jul 25 14:30:43 CEST 2022 - The aliscore output file does not exist. Check results for BUSCO: 1076134at2759
## Mon Jul 25 14:30:44 CEST 2022 - The aliscore output file does not exist. Check results for BUSCO: 1228942at2759
## Mon Jul 25 14:30:44 CEST 2022 - The aliscore output appears to be empty. Check results for BUSCO: 1490022at2759
## Mon Jul 25 14:30:44 CEST 2022 - The aliscore output appears to be empty. Check results for BUSCO: 1076134at2759
## Mon Jul 25 14:30:44 CEST 2022 - The aliscore output appears to be empty. Check results for BUSCO: 1228942at2759
## Mon Jul 25 14:30:44 CEST 2022 - The aliscore output file does not exist. Check results for BUSCO: 1358374at2759
## Mon Jul 25 14:30:44 CEST 2022 - The aliscore output appears to be empty. Check results for BUSCO: 1358374at2759
## Mon Jul 25 14:30:44 CEST 2022 - The aliscore output file does not exist. Check results for BUSCO: 981902at2759
## Mon Jul 25 14:30:44 CEST 2022 - The aliscore output appears to be empty. Check results for BUSCO: 981902at2759
## Mon Jul 25 14:30:44 CEST 2022 - The aliscore output file does not exist. Check results for BUSCO: 814241at2759
## Mon Jul 25 14:30:44 CEST 2022 - The aliscore output file does not exist. Check results for BUSCO: 1266231at2759
## Mon Jul 25 14:30:44 CEST 2022 - The aliscore output appears to be empty. Check results for BUSCO: 814241at2759
## Mon Jul 25 14:30:44 CEST 2022 - The aliscore output appears to be empty. Check results for BUSCO: 1266231at2759
## Mon Jul 25 14:30:44 CEST 2022 - The aliscore output file does not exist. Check results for BUSCO: 1249159at2759
## Mon Jul 25 14:30:44 CEST 2022 - The aliscore output file does not exist. Check results for BUSCO: 1275837at2759
## Mon Jul 25 14:30:44 CEST 2022 - The aliscore output appears to be empty. Check results for BUSCO: 1249159at2759
## Mon Jul 25 14:30:44 CEST 2022 - The aliscore output appears to be empty. Check results for BUSCO: 1275837at2759
## Mon Jul 25 14:30:44 CEST 2022 - The aliscore output file does not exist. Check results for BUSCO: 1030907at2759
## Mon Jul 25 14:30:44 CEST 2022 - The aliscore output file does not exist. Check results for BUSCO: 1018517at2759
## Mon Jul 25 14:30:44 CEST 2022 - The aliscore output appears to be empty. Check results for BUSCO: 1030907at2759
## Mon Jul 25 14:30:44 CEST 2022 - The aliscore output appears to be empty. Check results for BUSCO: 1018517at2759
## Mon Jul 25 14:30:44 CEST 2022 - The aliscore output file does not exist. Check results for BUSCO: 1264469at2759
## Mon Jul 25 14:30:44 CEST 2022 - The aliscore output file does not exist. Check results for BUSCO: 293315at2759
## Mon Jul 25 14:30:44 CEST 2022 - The aliscore output appears to be empty. Check results for BUSCO: 1264469at2759
## Mon Jul 25 14:30:44 CEST 2022 - The aliscore output appears to be empty. Check results for BUSCO: 293315at2759
## Mon Jul 25 14:30:44 CEST 2022 - The aliscore output file does not exist. Check results for BUSCO: 751335at2759
## Mon Jul 25 14:30:44 CEST 2022 - The aliscore output file does not exist. Check results for BUSCO: 1513531at2759
## Mon Jul 25 14:30:44 CEST 2022 - The aliscore output file does not exist. Check results for BUSCO: 1222562at2759
## Mon Jul 25 14:30:44 CEST 2022 - The aliscore output appears to be empty. Check results for BUSCO: 751335at2759
## Mon Jul 25 14:30:44 CEST 2022 - The aliscore output appears to be empty. Check results for BUSCO: 1513531at2759
## Mon Jul 25 14:30:44 CEST 2022 - The aliscore output file does not exist. Check results for BUSCO: 1030907at2759
## Mon Jul 25 14:30:44 CEST 2022 - The aliscore output appears to be empty. Check results for BUSCO: 1222562at2759
## Mon Jul 25 14:30:44 CEST 2022 - The aliscore output file does not exist. Check results for BUSCO: 721605at2759
## Mon Jul 25 14:30:44 CEST 2022 - The aliscore output appears to be empty. Check results for BUSCO: 1030907at2759
## Mon Jul 25 14:30:44 CEST 2022 - The aliscore output appears to be empty. Check results for BUSCO: 721605at2759
## Mon Jul 25 14:30:44 CEST 2022 - The aliscore output file does not exist. Check results for BUSCO: 1233814at2759
## Mon Jul 25 14:30:44 CEST 2022 - The aliscore output file does not exist. Check results for BUSCO: 1249159at2759
## Mon Jul 25 14:30:44 CEST 2022 - The aliscore output appears to be empty. Check results for BUSCO: 1233814at2759
## Mon Jul 25 14:30:44 CEST 2022 - The aliscore output appears to be empty. Check results for BUSCO: 1249159at2759
## Mon Jul 25 14:30:45 CEST 2022 - The aliscore output file does not exist. Check results for BUSCO: 1264469at2759
## Mon Jul 25 14:30:45 CEST 2022 - The aliscore output appears to be empty. Check results for BUSCO: 1264469at2759
## Mon Jul 25 14:30:45 CEST 2022 - The aliscore output file does not exist. Check results for BUSCO: 1479417at2759
## Mon Jul 25 14:30:45 CEST 2022 - The aliscore output appears to be empty. Check results for BUSCO: 1479417at2759
## Mon Jul 25 14:30:45 CEST 2022 - The aliscore output file does not exist. Check results for BUSCO: 817008at2759
## Mon Jul 25 14:30:45 CEST 2022 - The aliscore output appears to be empty. Check results for BUSCO: 817008at2759
## Mon Jul 25 14:30:45 CEST 2022 - The aliscore output file does not exist. Check results for BUSCO: 176625at2759
## Mon Jul 25 14:30:45 CEST 2022 - The aliscore output appears to be empty. Check results for BUSCO: 176625at2759
## Mon Jul 25 14:30:45 CEST 2022 - The aliscore output file does not exist. Check results for BUSCO: 345441at2759
## Mon Jul 25 14:30:45 CEST 2022 - The aliscore output appears to be empty. Check results for BUSCO: 345441at2759
## Mon Jul 25 14:30:45 CEST 2022 - The aliscore output file does not exist. Check results for BUSCO: 664730at2759
## Mon Jul 25 14:30:45 CEST 2022 - The aliscore output appears to be empty. Check results for BUSCO: 664730at2759
## Mon Jul 25 14:30:45 CEST 2022 - The aliscore output file does not exist. Check results for BUSCO: 375960at2759
## Mon Jul 25 14:30:45 CEST 2022 - The aliscore output file does not exist. Check results for BUSCO: 1115196at2759
## Mon Jul 25 14:30:45 CEST 2022 - The aliscore output appears to be empty. Check results for BUSCO: 1115196at2759
## Mon Jul 25 14:30:45 CEST 2022 - The aliscore output appears to be empty. Check results for BUSCO: 375960at2759
## Mon Jul 25 14:30:45 CEST 2022 - The aliscore output file does not exist. Check results for BUSCO: 751335at2759
## Mon Jul 25 14:30:45 CEST 2022 - The aliscore output appears to be empty. Check results for BUSCO: 751335at2759
## Mon Jul 25 14:30:45 CEST 2022 - The aliscore output file does not exist. Check results for BUSCO: 975158at2759
## Mon Jul 25 14:30:45 CEST 2022 - The aliscore output file does not exist. Check results for BUSCO: 1223488at2759
## Mon Jul 25 14:30:45 CEST 2022 - The aliscore output file does not exist. Check results for BUSCO: 722805at2759
## Mon Jul 25 14:30:45 CEST 2022 - The aliscore output appears to be empty. Check results for BUSCO: 975158at2759
## Mon Jul 25 14:30:45 CEST 2022 - The aliscore output file does not exist. Check results for BUSCO: 1220881at2759
## Mon Jul 25 14:30:45 CEST 2022 - The aliscore output appears to be empty. Check results for BUSCO: 1223488at2759
## Mon Jul 25 14:30:45 CEST 2022 - The aliscore output file does not exist. Check results for BUSCO: 1025450at2759
## Mon Jul 25 14:30:45 CEST 2022 - The aliscore output file does not exist. Check results for BUSCO: 1323575at2759
## Mon Jul 25 14:30:45 CEST 2022 - The aliscore output appears to be empty. Check results for BUSCO: 722805at2759
## Mon Jul 25 14:30:45 CEST 2022 - The aliscore output file does not exist. Check results for BUSCO: 418107at2759
## Mon Jul 25 14:30:45 CEST 2022 - The aliscore output appears to be empty. Check results for BUSCO: 1220881at2759
## Mon Jul 25 14:30:45 CEST 2022 - The aliscore output appears to be empty. Check results for BUSCO: 1025450at2759
## Mon Jul 25 14:30:46 CEST 2022 - The aliscore output appears to be empty. Check results for BUSCO: 1323575at2759
## Mon Jul 25 14:30:46 CEST 2022 - The aliscore output appears to be empty. Check results for BUSCO: 418107at2759
## Mon Jul 25 14:30:46 CEST 2022 - The aliscore output file does not exist. Check results for BUSCO: 418107at2759
## Mon Jul 25 14:30:46 CEST 2022 - The aliscore output appears to be empty. Check results for BUSCO: 418107at2759
## Mon Jul 25 14:30:46 CEST 2022 - The aliscore output file does not exist. Check results for BUSCO: 390348at2759
## Mon Jul 25 14:30:46 CEST 2022 - The aliscore output appears to be empty. Check results for BUSCO: 390348at2759
## Mon Jul 25 14:30:46 CEST 2022 - The aliscore output file does not exist. Check results for BUSCO: 1287401at2759
## Mon Jul 25 14:30:46 CEST 2022 - The aliscore output appears to be empty. Check results for BUSCO: 1287401at2759
## Mon Jul 25 14:30:46 CEST 2022 - The aliscore output file does not exist. Check results for BUSCO: 293315at2759
## Mon Jul 25 14:30:46 CEST 2022 - The aliscore output appears to be empty. Check results for BUSCO: 293315at2759
## Mon Jul 25 14:30:46 CEST 2022 - The aliscore output file does not exist. Check results for BUSCO: 1309031at2759
## Mon Jul 25 14:30:46 CEST 2022 - The aliscore output appears to be empty. Check results for BUSCO: 1309031at2759
## Mon Jul 25 14:30:46 CEST 2022 - The aliscore output file does not exist. Check results for BUSCO: 671846at2759
## Mon Jul 25 14:30:46 CEST 2022 - The aliscore output appears to be empty. Check results for BUSCO: 671846at2759
## Mon Jul 25 14:30:46 CEST 2022 - The aliscore output file does not exist. Check results for BUSCO: 383503at2759
## Mon Jul 25 14:30:46 CEST 2022 - The aliscore output appears to be empty. Check results for BUSCO: 383503at2759
## Mon Jul 25 14:30:46 CEST 2022 - The aliscore output file does not exist. Check results for BUSCO: 834694at2759
## Mon Jul 25 14:30:46 CEST 2022 - The aliscore output file does not exist. Check results for BUSCO: 1194797at2759
## Mon Jul 25 14:30:46 CEST 2022 - The aliscore output appears to be empty. Check results for BUSCO: 834694at2759
## Mon Jul 25 14:30:46 CEST 2022 - The aliscore output appears to be empty. Check results for BUSCO: 1194797at2759
## Mon Jul 25 14:30:46 CEST 2022 - The aliscore output file does not exist. Check results for BUSCO: 1248958at2759
## Mon Jul 25 14:30:47 CEST 2022 - The aliscore output file does not exist. Check results for BUSCO: 1087488at2759
## Mon Jul 25 14:30:47 CEST 2022 - The aliscore output file does not exist. Check results for BUSCO: 836599at2759
## Mon Jul 25 14:30:47 CEST 2022 - The aliscore output appears to be empty. Check results for BUSCO: 1248958at2759
## Mon Jul 25 14:30:47 CEST 2022 - The aliscore output file does not exist. Check results for BUSCO: 768809at2759
## Mon Jul 25 14:30:47 CEST 2022 - The aliscore output appears to be empty. Check results for BUSCO: 1087488at2759
## Mon Jul 25 14:30:47 CEST 2022 - The aliscore output appears to be empty. Check results for BUSCO: 836599at2759
## Mon Jul 25 14:30:47 CEST 2022 - The aliscore output appears to be empty. Check results for BUSCO: 768809at2759
## Mon Jul 25 14:30:47 CEST 2022 - The aliscore output file does not exist. Check results for BUSCO: 917326at2759
## Mon Jul 25 14:30:47 CEST 2022 - The aliscore output appears to be empty. Check results for BUSCO: 917326at2759
## Mon Jul 25 14:30:47 CEST 2022 - The aliscore output file does not exist. Check results for BUSCO: 392369at2759
## Mon Jul 25 14:30:47 CEST 2022 - The aliscore output appears to be empty. Check results for BUSCO: 392369at2759
## Mon Jul 25 14:30:47 CEST 2022 - The aliscore output file does not exist. Check results for BUSCO: 759498at2759
## Mon Jul 25 14:30:47 CEST 2022 - The aliscore output file does not exist. Check results for BUSCO: 375960at2759
## Mon Jul 25 14:30:47 CEST 2022 - The aliscore output appears to be empty. Check results for BUSCO: 759498at2759
## Mon Jul 25 14:30:47 CEST 2022 - The aliscore output appears to be empty. Check results for BUSCO: 375960at2759
## Mon Jul 25 14:30:47 CEST 2022 - The aliscore output file does not exist. Check results for BUSCO: 981902at2759
## Mon Jul 25 14:30:47 CEST 2022 - The aliscore output file does not exist. Check results for BUSCO: 176625at2759
## Mon Jul 25 14:30:47 CEST 2022 - The aliscore output appears to be empty. Check results for BUSCO: 981902at2759
## Mon Jul 25 14:30:47 CEST 2022 - The aliscore output appears to be empty. Check results for BUSCO: 176625at2759
## Mon Jul 25 14:30:48 CEST 2022 - The aliscore output file does not exist. Check results for BUSCO: 1258856at2759
## Mon Jul 25 14:30:48 CEST 2022 - The aliscore output file does not exist. Check results for BUSCO: 836599at2759
## Mon Jul 25 14:30:48 CEST 2022 - The aliscore output appears to be empty. Check results for BUSCO: 1258856at2759
## Mon Jul 25 14:30:48 CEST 2022 - The aliscore output appears to be empty. Check results for BUSCO: 836599at2759
## Mon Jul 25 14:30:48 CEST 2022 - The aliscore output file does not exist. Check results for BUSCO: 341721at2759
## Mon Jul 25 14:30:48 CEST 2022 - The aliscore output appears to be empty. Check results for BUSCO: 341721at2759
## Mon Jul 25 14:30:48 CEST 2022 - The aliscore output file does not exist. Check results for BUSCO: 1323575at2759
## Mon Jul 25 14:30:48 CEST 2022 - The aliscore output appears to be empty. Check results for BUSCO: 1323575at2759
## Mon Jul 25 14:30:48 CEST 2022 - The aliscore output file does not exist. Check results for BUSCO: 383503at2759
## Mon Jul 25 14:30:48 CEST 2022 - The aliscore output file does not exist. Check results for BUSCO: 722805at2759
## Mon Jul 25 14:30:48 CEST 2022 - The aliscore output appears to be empty. Check results for BUSCO: 383503at2759
## Mon Jul 25 14:30:48 CEST 2022 - The aliscore output appears to be empty. Check results for BUSCO: 722805at2759
## Mon Jul 25 14:30:48 CEST 2022 - The aliscore output file does not exist. Check results for BUSCO: 1258856at2759
## Mon Jul 25 14:30:48 CEST 2022 - The aliscore output file does not exist. Check results for BUSCO: 777920at2759
## Mon Jul 25 14:30:49 CEST 2022 - The aliscore output file does not exist. Check results for BUSCO: 1314980at2759
## Mon Jul 25 14:30:49 CEST 2022 - The aliscore output appears to be empty. Check results for BUSCO: 1258856at2759
## Mon Jul 25 14:30:49 CEST 2022 - The aliscore output appears to be empty. Check results for BUSCO: 777920at2759
## Mon Jul 25 14:30:49 CEST 2022 - The aliscore output appears to be empty. Check results for BUSCO: 1314980at2759
## Mon Jul 25 14:30:49 CEST 2022 - The aliscore output file does not exist. Check results for BUSCO: 625387at2759
## Mon Jul 25 14:30:49 CEST 2022 - The aliscore output appears to be empty. Check results for BUSCO: 625387at2759
## Mon Jul 25 14:30:49 CEST 2022 - The aliscore output file does not exist. Check results for BUSCO: 834694at2759
## Mon Jul 25 14:30:49 CEST 2022 - The aliscore output appears to be empty. Check results for BUSCO: 834694at2759
## Mon Jul 25 14:30:51 CEST 2022 - The aliscore output file does not exist. Check results for BUSCO: 1314980at2759
## Mon Jul 25 14:30:51 CEST 2022 - The aliscore output appears to be empty. Check results for BUSCO: 1314980at2759
## Mon Jul 25 14:30:52 CEST 2022 - The aliscore output file does not exist. Check results for BUSCO: 160593at2759
## Mon Jul 25 14:30:52 CEST 2022 - The aliscore output appears to be empty. Check results for BUSCO: 160593at2759
## Mon Jul 25 14:30:59 CEST 2022 - The aliscore output file does not exist. Check results for BUSCO: 674160at2759
## Mon Jul 25 14:30:59 CEST 2022 - The aliscore output appears to be empty. Check results for BUSCO: 674160at2759
## Mon Jul 25 14:30:59 CEST 2022 - The aliscore output file does not exist. Check results for BUSCO: 156083at2759
## Mon Jul 25 14:30:59 CEST 2022 - The aliscore output appears to be empty. Check results for BUSCO: 156083at2759
## Mon Jul 25 14:31:09 CEST 2022 - The aliscore output file does not exist. Check results for BUSCO: 1405073at2759
## Mon Jul 25 14:31:10 CEST 2022 - The aliscore output appears to be empty. Check results for BUSCO: 1405073at2759
## Mon Jul 25 14:31:10 CEST 2022 - The aliscore output file does not exist. Check results for BUSCO: 865202at2759
## Mon Jul 25 14:31:10 CEST 2022 - The aliscore output appears to be empty. Check results for BUSCO: 865202at2759
## Mon Jul 25 14:31:10 CEST 2022 - Number of alignments (clustalo): 255
## Mon Jul 25 14:31:10 CEST 2022 - Number of trimmed alignments (clustalo - bmge): 255
## Mon Jul 25 14:31:10 CEST 2022 - Number of alignments (clustalo - bmge) after filtering: 245
## Mon Jul 25 14:31:10 CEST 2022 - The aliscore output file does not exist. Check results for BUSCO: 866359at2759
## Mon Jul 25 14:31:10 CEST 2022 - The aliscore output file does not exist. Check results for BUSCO: 1426075at2759
## Mon Jul 25 14:31:10 CEST 2022 - The aliscore output appears to be empty. Check results for BUSCO: 866359at2759
## Mon Jul 25 14:31:10 CEST 2022 - The aliscore output appears to be empty. Check results for BUSCO: 1426075at2759
## Mon Jul 25 14:31:11 CEST 2022 - The aliscore output file does not exist. Check results for BUSCO: 1398309at2759
## Mon Jul 25 14:31:11 CEST 2022 - The aliscore output appears to be empty. Check results for BUSCO: 1398309at2759
## Mon Jul 25 14:31:11 CEST 2022 - The aliscore output file does not exist. Check results for BUSCO: 1287094at2759
## Mon Jul 25 14:31:11 CEST 2022 - The aliscore output file does not exist. Check results for BUSCO: 1001705at2759
## Mon Jul 25 14:31:11 CEST 2022 - The aliscore output appears to be empty. Check results for BUSCO: 1287094at2759
## Mon Jul 25 14:31:11 CEST 2022 - The aliscore output appears to be empty. Check results for BUSCO: 1001705at2759
## Mon Jul 25 14:31:11 CEST 2022 - The aliscore output file does not exist. Check results for BUSCO: 1157302at2759
## Mon Jul 25 14:31:11 CEST 2022 - The aliscore output appears to be empty. Check results for BUSCO: 1157302at2759
## Mon Jul 25 14:31:11 CEST 2022 - The aliscore output file does not exist. Check results for BUSCO: 917326at2759
## Mon Jul 25 14:31:11 CEST 2022 - The aliscore output appears to be empty. Check results for BUSCO: 917326at2759
## Mon Jul 25 14:31:11 CEST 2022 - The aliscore output file does not exist. Check results for BUSCO: 1459797at2759
## Mon Jul 25 14:31:11 CEST 2022 - The aliscore output appears to be empty. Check results for BUSCO: 1459797at2759
## Mon Jul 25 14:31:11 CEST 2022 - The aliscore output file does not exist. Check results for BUSCO: 865202at2759
## Mon Jul 25 14:31:11 CEST 2022 - The aliscore output appears to be empty. Check results for BUSCO: 865202at2759
## Mon Jul 25 14:31:12 CEST 2022 - The aliscore output file does not exist. Check results for BUSCO: 1217666at2759
## Mon Jul 25 14:31:12 CEST 2022 - The aliscore output file does not exist. Check results for BUSCO: 1178688at2759
## Mon Jul 25 14:31:12 CEST 2022 - The aliscore output appears to be empty. Check results for BUSCO: 1217666at2759
## Mon Jul 25 14:31:12 CEST 2022 - The aliscore output appears to be empty. Check results for BUSCO: 1178688at2759
## Mon Jul 25 14:31:12 CEST 2022 - The aliscore output file does not exist. Check results for BUSCO: 923657at2759
## Mon Jul 25 14:31:12 CEST 2022 - The aliscore output appears to be empty. Check results for BUSCO: 923657at2759
## Mon Jul 25 14:31:12 CEST 2022 - The aliscore output file does not exist. Check results for BUSCO: 1623701at2759
## Mon Jul 25 14:31:12 CEST 2022 - The aliscore output file does not exist. Check results for BUSCO: 1474000at2759
## Mon Jul 25 14:31:12 CEST 2022 - The aliscore output appears to be empty. Check results for BUSCO: 1623701at2759
## Mon Jul 25 14:31:12 CEST 2022 - The aliscore output file does not exist. Check results for BUSCO: 1079827at2759
## Mon Jul 25 14:31:12 CEST 2022 - The aliscore output appears to be empty. Check results for BUSCO: 1079827at2759
## Mon Jul 25 14:31:12 CEST 2022 - The aliscore output file does not exist. Check results for BUSCO: 1222562at2759
## Mon Jul 25 14:31:12 CEST 2022 - The aliscore output appears to be empty. Check results for BUSCO: 1222562at2759
## Mon Jul 25 14:31:12 CEST 2022 - The aliscore output file does not exist. Check results for BUSCO: 1228942at2759
## Mon Jul 25 14:31:12 CEST 2022 - The aliscore output appears to be empty. Check results for BUSCO: 1228942at2759
## Mon Jul 25 14:31:12 CEST 2022 - The aliscore output file does not exist. Check results for BUSCO: 1445102at2759
## Mon Jul 25 14:31:12 CEST 2022 - The aliscore output appears to be empty. Check results for BUSCO: 1445102at2759
## Mon Jul 25 14:31:13 CEST 2022 - The aliscore output file does not exist. Check results for BUSCO: 1645187at2759
## Mon Jul 25 14:31:13 CEST 2022 - The aliscore output appears to be empty. Check results for BUSCO: 1645187at2759
## Mon Jul 25 14:31:13 CEST 2022 - The aliscore output file does not exist. Check results for BUSCO: 1220881at2759
## Mon Jul 25 14:31:13 CEST 2022 - The aliscore output appears to be empty. Check results for BUSCO: 1220881at2759
## Mon Jul 25 14:31:13 CEST 2022 - The aliscore output file does not exist. Check results for BUSCO: 1076134at2759
## Mon Jul 25 14:31:13 CEST 2022 - The aliscore output appears to be empty. Check results for BUSCO: 1076134at2759
## Mon Jul 25 14:31:13 CEST 2022 - The aliscore output file does not exist. Check results for BUSCO: 721605at2759
## Mon Jul 25 14:31:13 CEST 2022 - The aliscore output appears to be empty. Check results for BUSCO: 721605at2759
## Mon Jul 25 14:31:13 CEST 2022 - The aliscore output file does not exist. Check results for BUSCO: 679771at2759
## Mon Jul 25 14:31:13 CEST 2022 - The aliscore output file does not exist. Check results for BUSCO: 1287401at2759
## Mon Jul 25 14:31:13 CEST 2022 - The aliscore output appears to be empty. Check results for BUSCO: 1287401at2759
## Mon Jul 25 14:31:13 CEST 2022 - The aliscore output appears to be empty. Check results for BUSCO: 679771at2759
## Mon Jul 25 14:31:13 CEST 2022 - The aliscore output file does not exist. Check results for BUSCO: 687505at2759
## Mon Jul 25 14:31:13 CEST 2022 - The aliscore output appears to be empty. Check results for BUSCO: 687505at2759
## Mon Jul 25 14:31:13 CEST 2022 - The aliscore output file does not exist. Check results for BUSCO: 1107630at2759
## Mon Jul 25 14:31:13 CEST 2022 - The aliscore output file does not exist. Check results for BUSCO: 1161199at2759
## Mon Jul 25 14:31:13 CEST 2022 - The aliscore output file does not exist. Check results for BUSCO: 974865at2759
## Mon Jul 25 14:31:13 CEST 2022 - The aliscore output appears to be empty. Check results for BUSCO: 1107630at2759
## Mon Jul 25 14:31:13 CEST 2022 - The aliscore output appears to be empty. Check results for BUSCO: 1161199at2759
## Mon Jul 25 14:31:13 CEST 2022 - The aliscore output appears to be empty. Check results for BUSCO: 974865at2759
## Mon Jul 25 14:31:13 CEST 2022 - The aliscore output file does not exist. Check results for BUSCO: 1194691at2759
## Mon Jul 25 14:31:13 CEST 2022 - The aliscore output appears to be empty. Check results for BUSCO: 1194691at2759
## Mon Jul 25 14:31:13 CEST 2022 - The aliscore output file does not exist. Check results for BUSCO: 937686at2759
## Mon Jul 25 14:31:13 CEST 2022 - The aliscore output appears to be empty. Check results for BUSCO: 937686at2759
## Mon Jul 25 14:31:13 CEST 2022 - The aliscore output file does not exist. Check results for BUSCO: 1513531at2759
## Mon Jul 25 14:31:13 CEST 2022 - The aliscore output appears to be empty. Check results for BUSCO: 1513531at2759
## Mon Jul 25 14:31:13 CEST 2022 - The aliscore output file does not exist. Check results for BUSCO: 1087488at2759
## Mon Jul 25 14:31:13 CEST 2022 - The aliscore output appears to be empty. Check results for BUSCO: 1087488at2759
## Mon Jul 25 14:31:14 CEST 2022 - The aliscore output file does not exist. Check results for BUSCO: 866359at2759
## Mon Jul 25 14:31:14 CEST 2022 - The aliscore output file does not exist. Check results for BUSCO: 464990at2759
## Mon Jul 25 14:31:14 CEST 2022 - The aliscore output appears to be empty. Check results for BUSCO: 866359at2759
## Mon Jul 25 14:31:14 CEST 2022 - The aliscore output appears to be empty. Check results for BUSCO: 464990at2759
## Mon Jul 25 14:31:14 CEST 2022 - The aliscore output file does not exist. Check results for BUSCO: 975158at2759
## Mon Jul 25 14:31:14 CEST 2022 - The aliscore output appears to be empty. Check results for BUSCO: 975158at2759
## Mon Jul 25 14:31:14 CEST 2022 - The aliscore output file does not exist. Check results for BUSCO: 1275837at2759
## Mon Jul 25 14:31:14 CEST 2022 - The aliscore output appears to be empty. Check results for BUSCO: 1275837at2759
## Mon Jul 25 14:31:14 CEST 2022 - The aliscore output file does not exist. Check results for BUSCO: 1309031at2759
## Mon Jul 25 14:31:14 CEST 2022 - The aliscore output appears to be empty. Check results for BUSCO: 1309031at2759
## Mon Jul 25 14:31:14 CEST 2022 - The aliscore output file does not exist. Check results for BUSCO: 1182451at2759
## Mon Jul 25 14:31:14 CEST 2022 - The aliscore output appears to be empty. Check results for BUSCO: 1182451at2759
## Mon Jul 25 14:31:14 CEST 2022 - The aliscore output file does not exist. Check results for BUSCO: 1001705at2759
## Mon Jul 25 14:31:14 CEST 2022 - The aliscore output file does not exist. Check results for BUSCO: 1426075at2759
## Mon Jul 25 14:31:14 CEST 2022 - The aliscore output appears to be empty. Check results for BUSCO: 1001705at2759
## Mon Jul 25 14:31:14 CEST 2022 - The aliscore output appears to be empty. Check results for BUSCO: 1426075at2759
## Mon Jul 25 14:31:14 CEST 2022 - The aliscore output file does not exist. Check results for BUSCO: 1322299at2759
## Mon Jul 25 14:31:14 CEST 2022 - The aliscore output file does not exist. Check results for BUSCO: 655400at2759
## Mon Jul 25 14:31:14 CEST 2022 - The aliscore output file does not exist. Check results for BUSCO: 1358374at2759
## Mon Jul 25 14:31:14 CEST 2022 - The aliscore output appears to be empty. Check results for BUSCO: 1322299at2759
## Mon Jul 25 14:31:14 CEST 2022 - The aliscore output appears to be empty. Check results for BUSCO: 655400at2759
## Mon Jul 25 14:31:14 CEST 2022 - The aliscore output file does not exist. Check results for BUSCO: 1193442at2759
## Mon Jul 25 14:31:14 CEST 2022 - The aliscore output appears to be empty. Check results for BUSCO: 1193442at2759
## Mon Jul 25 14:31:15 CEST 2022 - The aliscore output appears to be empty. Check results for BUSCO: 1358374at2759
## Mon Jul 25 14:31:15 CEST 2022 - The aliscore output file does not exist. Check results for BUSCO: 817008at2759
## Mon Jul 25 14:31:15 CEST 2022 - The aliscore output file does not exist. Check results for BUSCO: 937686at2759
## Mon Jul 25 14:31:15 CEST 2022 - The aliscore output appears to be empty. Check results for BUSCO: 817008at2759
## Mon Jul 25 14:31:15 CEST 2022 - The aliscore output appears to be empty. Check results for BUSCO: 937686at2759
## Mon Jul 25 14:31:15 CEST 2022 - The aliscore output file does not exist. Check results for BUSCO: 290630at2759
## Mon Jul 25 14:31:15 CEST 2022 - The aliscore output file does not exist. Check results for BUSCO: 1322642at2759
## Mon Jul 25 14:31:15 CEST 2022 - The aliscore output appears to be empty. Check results for BUSCO: 290630at2759
## Mon Jul 25 14:31:15 CEST 2022 - The aliscore output appears to be empty. Check results for BUSCO: 1322642at2759
## Mon Jul 25 14:31:15 CEST 2022 - The aliscore output file does not exist. Check results for BUSCO: 426305at2759
## Mon Jul 25 14:31:15 CEST 2022 - The aliscore output appears to be empty. Check results for BUSCO: 426305at2759
## Mon Jul 25 14:31:15 CEST 2022 - The aliscore output file does not exist. Check results for BUSCO: 341721at2759
## Mon Jul 25 14:31:15 CEST 2022 - The aliscore output appears to be empty. Check results for BUSCO: 341721at2759
## Mon Jul 25 14:31:15 CEST 2022 - The aliscore output file does not exist. Check results for BUSCO: 1025450at2759
## Mon Jul 25 14:31:15 CEST 2022 - The aliscore output appears to be empty. Check results for BUSCO: 1025450at2759
## Mon Jul 25 14:31:15 CEST 2022 - The aliscore output file does not exist. Check results for BUSCO: 390348at2759
## Mon Jul 25 14:31:15 CEST 2022 - The aliscore output file does not exist. Check results for BUSCO: 673132at2759
## Mon Jul 25 14:31:15 CEST 2022 - The aliscore output file does not exist. Check results for BUSCO: 924753at2759
## Mon Jul 25 14:31:15 CEST 2022 - The aliscore output appears to be empty. Check results for BUSCO: 390348at2759
## Mon Jul 25 14:31:15 CEST 2022 - The aliscore output file does not exist. Check results for BUSCO: 814241at2759
## Mon Jul 25 14:31:15 CEST 2022 - The aliscore output appears to be empty. Check results for BUSCO: 924753at2759
## Mon Jul 25 14:31:15 CEST 2022 - The aliscore output appears to be empty. Check results for BUSCO: 814241at2759
## Mon Jul 25 14:31:15 CEST 2022 - The aliscore output file does not exist. Check results for BUSCO: 513979at2759
## Mon Jul 25 14:31:15 CEST 2022 - The aliscore output file does not exist. Check results for BUSCO: 918816at2759
## Mon Jul 25 14:31:15 CEST 2022 - The aliscore output appears to be empty. Check results for BUSCO: 513979at2759
## Mon Jul 25 14:31:15 CEST 2022 - The aliscore output appears to be empty. Check results for BUSCO: 918816at2759
## Mon Jul 25 14:31:15 CEST 2022 - The aliscore output file does not exist. Check results for BUSCO: 345441at2759
## Mon Jul 25 14:31:16 CEST 2022 - The aliscore output appears to be empty. Check results for BUSCO: 345441at2759
## Mon Jul 25 14:31:16 CEST 2022 - The aliscore output file does not exist. Check results for BUSCO: 331411at2759
## Mon Jul 25 14:31:16 CEST 2022 - The aliscore output file does not exist. Check results for BUSCO: 664730at2759
## Mon Jul 25 14:31:16 CEST 2022 - The aliscore output appears to be empty. Check results for BUSCO: 331411at2759
## Mon Jul 25 14:31:16 CEST 2022 - The aliscore output appears to be empty. Check results for BUSCO: 664730at2759
## Mon Jul 25 14:31:16 CEST 2022 - The aliscore output file does not exist. Check results for BUSCO: 97116at2759
## Mon Jul 25 14:31:16 CEST 2022 - The aliscore output file does not exist. Check results for BUSCO: 570797at2759
## Mon Jul 25 14:31:16 CEST 2022 - The aliscore output file does not exist. Check results for BUSCO: 392369at2759
## Mon Jul 25 14:31:16 CEST 2022 - The aliscore output appears to be empty. Check results for BUSCO: 97116at2759
## Mon Jul 25 14:31:16 CEST 2022 - The aliscore output appears to be empty. Check results for BUSCO: 570797at2759
## Mon Jul 25 14:31:16 CEST 2022 - The aliscore output appears to be empty. Check results for BUSCO: 392369at2759
## Mon Jul 25 14:31:16 CEST 2022 - The aliscore output file does not exist. Check results for BUSCO: 1115196at2759
## Mon Jul 25 14:31:16 CEST 2022 - The aliscore output appears to be empty. Check results for BUSCO: 1115196at2759
## Mon Jul 25 14:31:16 CEST 2022 - The aliscore output file does not exist. Check results for BUSCO: 325552at2759
## Mon Jul 25 14:31:16 CEST 2022 - The aliscore output appears to be empty. Check results for BUSCO: 325552at2759
## Mon Jul 25 14:31:16 CEST 2022 - The aliscore output file does not exist. Check results for BUSCO: 1490022at2759
## Mon Jul 25 14:31:16 CEST 2022 - The aliscore output file does not exist. Check results for BUSCO: 1423847at2759
## Mon Jul 25 14:31:16 CEST 2022 - The aliscore output file does not exist. Check results for BUSCO: 924753at2759
## Mon Jul 25 14:31:16 CEST 2022 - The aliscore output appears to be empty. Check results for BUSCO: 1423847at2759
## Mon Jul 25 14:31:16 CEST 2022 - The aliscore output appears to be empty. Check results for BUSCO: 924753at2759
## Mon Jul 25 14:31:16 CEST 2022 - The aliscore output file does not exist. Check results for BUSCO: 604979at2759
## Mon Jul 25 14:31:16 CEST 2022 - The aliscore output appears to be empty. Check results for BUSCO: 604979at2759
## Mon Jul 25 14:31:16 CEST 2022 - The aliscore output file does not exist. Check results for BUSCO: 759498at2759
## Mon Jul 25 14:31:17 CEST 2022 - The aliscore output appears to be empty. Check results for BUSCO: 759498at2759
## Mon Jul 25 14:31:17 CEST 2022 - The aliscore output appears to be empty. Check results for BUSCO: 1490022at2759
## Mon Jul 25 14:31:17 CEST 2022 - The aliscore output file does not exist. Check results for BUSCO: 973442at2759
## Mon Jul 25 14:31:17 CEST 2022 - The aliscore output file does not exist. Check results for BUSCO: 671846at2759
## Mon Jul 25 14:31:17 CEST 2022 - The aliscore output appears to be empty. Check results for BUSCO: 973442at2759
## Mon Jul 25 14:31:17 CEST 2022 - The aliscore output appears to be empty. Check results for BUSCO: 671846at2759
## Mon Jul 25 14:31:17 CEST 2022 - The aliscore output file does not exist. Check results for BUSCO: 426305at2759
## Mon Jul 25 14:31:17 CEST 2022 - The aliscore output file does not exist. Check results for BUSCO: 1379600at2759
## Mon Jul 25 14:31:17 CEST 2022 - The aliscore output file does not exist. Check results for BUSCO: 1248958at2759
## Mon Jul 25 14:31:17 CEST 2022 - The aliscore output appears to be empty. Check results for BUSCO: 426305at2759
## Mon Jul 25 14:31:17 CEST 2022 - The aliscore output appears to be empty. Check results for BUSCO: 1379600at2759
## Mon Jul 25 14:31:17 CEST 2022 - The aliscore output appears to be empty. Check results for BUSCO: 1248958at2759
## Mon Jul 25 14:31:17 CEST 2022 - The aliscore output file does not exist. Check results for BUSCO: 777920at2759
## Mon Jul 25 14:31:17 CEST 2022 - The aliscore output file does not exist. Check results for BUSCO: 160593at2759
## Mon Jul 25 14:31:17 CEST 2022 - The aliscore output appears to be empty. Check results for BUSCO: 777920at2759
## Mon Jul 25 14:31:17 CEST 2022 - The aliscore output appears to be empty. Check results for BUSCO: 160593at2759
## Mon Jul 25 14:31:18 CEST 2022 - The aliscore output file does not exist. Check results for BUSCO: 331411at2759
## Mon Jul 25 14:31:18 CEST 2022 - The aliscore output appears to be empty. Check results for BUSCO: 331411at2759
## Mon Jul 25 14:31:18 CEST 2022 - The aliscore output file does not exist. Check results for BUSCO: 1003258at2759
## Mon Jul 25 14:31:18 CEST 2022 - The aliscore output appears to be empty. Check results for BUSCO: 1003258at2759
## Mon Jul 25 14:31:18 CEST 2022 - The aliscore output file does not exist. Check results for BUSCO: 570797at2759
## Mon Jul 25 14:31:18 CEST 2022 - The aliscore output appears to be empty. Check results for BUSCO: 570797at2759
## Mon Jul 25 14:31:18 CEST 2022 - The aliscore output file does not exist. Check results for BUSCO: 179362at2759
## Mon Jul 25 14:31:18 CEST 2022 - The aliscore output appears to be empty. Check results for BUSCO: 179362at2759
## Mon Jul 25 14:31:19 CEST 2022 - The aliscore output file does not exist. Check results for BUSCO: 1379600at2759
## Mon Jul 25 14:31:19 CEST 2022 - The aliscore output appears to be empty. Check results for BUSCO: 1379600at2759
## Mon Jul 25 14:31:20 CEST 2022 - The aliscore output file does not exist. Check results for BUSCO: 1312453at2759
## Mon Jul 25 14:31:20 CEST 2022 - The aliscore output appears to be empty. Check results for BUSCO: 1312453at2759
## Mon Jul 25 14:31:21 CEST 2022 - The aliscore output file does not exist. Check results for BUSCO: 625387at2759
## Mon Jul 25 14:31:21 CEST 2022 - The aliscore output appears to be empty. Check results for BUSCO: 625387at2759
## Mon Jul 25 14:31:26 CEST 2022 - The aliscore output file does not exist. Check results for BUSCO: 156083at2759
## Mon Jul 25 14:31:26 CEST 2022 - The aliscore output appears to be empty. Check results for BUSCO: 156083at2759
## Mon Jul 25 14:41:09 CEST 2022 - The aliscore output file does not exist. Check results for BUSCO: 1623701at2759
## Mon Jul 25 14:41:09 CEST 2022 - The aliscore output appears to be empty. Check results for BUSCO: 1623701at2759
## Mon Jul 25 14:41:09 CEST 2022 - The aliscore output file does not exist. Check results for BUSCO: 1504863at2759
## Mon Jul 25 14:41:09 CEST 2022 - The aliscore output appears to be empty. Check results for BUSCO: 1504863at2759
## Mon Jul 25 14:41:10 CEST 2022 - The aliscore output file does not exist. Check results for BUSCO: 1178688at2759
## Mon Jul 25 14:41:10 CEST 2022 - The aliscore output appears to be empty. Check results for BUSCO: 1178688at2759
## Mon Jul 25 14:41:10 CEST 2022 - The aliscore output file does not exist. Check results for BUSCO: 1620056at2759
## Mon Jul 25 14:41:10 CEST 2022 - The aliscore output appears to be empty. Check results for BUSCO: 1620056at2759
## Mon Jul 25 14:41:10 CEST 2022 - The aliscore output file does not exist. Check results for BUSCO: 1030907at2759
## Mon Jul 25 14:41:10 CEST 2022 - The aliscore output appears to be empty. Check results for BUSCO: 1030907at2759
## Mon Jul 25 14:41:11 CEST 2022 - The aliscore output file does not exist. Check results for BUSCO: 974865at2759
## Mon Jul 25 14:41:11 CEST 2022 - The aliscore output file does not exist. Check results for BUSCO: 1233814at2759
## Mon Jul 25 14:41:11 CEST 2022 - The aliscore output appears to be empty. Check results for BUSCO: 974865at2759
## Mon Jul 25 14:41:11 CEST 2022 - The aliscore output appears to be empty. Check results for BUSCO: 1233814at2759
## Mon Jul 25 14:41:11 CEST 2022 - Number of alignments (mafft): 255
## Mon Jul 25 14:41:11 CEST 2022 - Number of trimmed alignments (mafft - aliscore): 255
## Mon Jul 25 14:41:11 CEST 2022 - Number of alignments (mafft - aliscore) after filtering: 248
## Mon Jul 25 14:41:12 CEST 2022 - The aliscore output file does not exist. Check results for BUSCO: 345441at2759
## Mon Jul 25 14:41:12 CEST 2022 - The aliscore output appears to be empty. Check results for BUSCO: 345441at2759
## Mon Jul 25 14:41:12 CEST 2022 - The aliscore output file does not exist. Check results for BUSCO: 1115196at2759
## Mon Jul 25 14:41:12 CEST 2022 - The aliscore output appears to be empty. Check results for BUSCO: 1115196at2759
## Mon Jul 25 14:41:12 CEST 2022 - The aliscore output file does not exist. Check results for BUSCO: 375960at2759
## Mon Jul 25 14:41:12 CEST 2022 - The aliscore output appears to be empty. Check results for BUSCO: 375960at2759
## Mon Jul 25 14:41:14 CEST 2022 - The aliscore output file does not exist. Check results for BUSCO: 383503at2759
## Mon Jul 25 14:41:15 CEST 2022 - The aliscore output appears to be empty. Check results for BUSCO: 383503at2759
## Mon Jul 25 14:41:15 CEST 2022 - The aliscore output file does not exist. Check results for BUSCO: 836599at2759
## Mon Jul 25 14:41:15 CEST 2022 - The aliscore output appears to be empty. Check results for BUSCO: 836599at2759
## Mon Jul 25 14:41:18 CEST 2022 - The aliscore output file does not exist. Check results for BUSCO: 160593at2759
## Mon Jul 25 14:41:18 CEST 2022 - The aliscore output appears to be empty. Check results for BUSCO: 160593at2759
## Mon Jul 25 14:41:39 CEST 2022 - The aliscore output file does not exist. Check results for BUSCO: 1459797at2759
## Mon Jul 25 14:41:39 CEST 2022 - The aliscore output appears to be empty. Check results for BUSCO: 1459797at2759
## Mon Jul 25 14:41:41 CEST 2022 - Number of alignments (clustalo): 255
## Mon Jul 25 14:41:41 CEST 2022 - Number of trimmed alignments (clustalo - trimal): 255
## Mon Jul 25 14:41:41 CEST 2022 - The aliscore output file does not exist. Check results for BUSCO: 1264469at2759
## Mon Jul 25 14:41:41 CEST 2022 - The aliscore output file does not exist. Check results for BUSCO: 1249159at2759
## Mon Jul 25 14:41:41 CEST 2022 - The aliscore output appears to be empty. Check results for BUSCO: 1264469at2759
## Mon Jul 25 14:41:41 CEST 2022 - The aliscore output appears to be empty. Check results for BUSCO: 1249159at2759
## Mon Jul 25 14:41:41 CEST 2022 - Number of alignments (clustalo - trimal) after filtering: 249
## Mon Jul 25 14:41:42 CEST 2022 - The aliscore output file does not exist. Check results for BUSCO: 1575179at2759
## Mon Jul 25 14:41:42 CEST 2022 - The aliscore output appears to be empty. Check results for BUSCO: 1575179at2759
## Mon Jul 25 14:41:42 CEST 2022 - The aliscore output file does not exist. Check results for BUSCO: 1228942at2759
## Mon Jul 25 14:41:42 CEST 2022 - The aliscore output appears to be empty. Check results for BUSCO: 1228942at2759
## Mon Jul 25 14:41:42 CEST 2022 - The aliscore output file does not exist. Check results for BUSCO: 923657at2759
## Mon Jul 25 14:41:42 CEST 2022 - The aliscore output appears to be empty. Check results for BUSCO: 923657at2759
## Mon Jul 25 14:41:42 CEST 2022 - The aliscore output file does not exist. Check results for BUSCO: 919955at2759
## Mon Jul 25 14:41:42 CEST 2022 - The aliscore output appears to be empty. Check results for BUSCO: 919955at2759
## Mon Jul 25 14:41:42 CEST 2022 - The aliscore output file does not exist. Check results for BUSCO: 1526152at2759
## Mon Jul 25 14:41:42 CEST 2022 - The aliscore output appears to be empty. Check results for BUSCO: 1526152at2759
## Mon Jul 25 14:41:42 CEST 2022 - The aliscore output file does not exist. Check results for BUSCO: 1358374at2759
## Mon Jul 25 14:41:42 CEST 2022 - The aliscore output appears to be empty. Check results for BUSCO: 1358374at2759
## Mon Jul 25 14:41:42 CEST 2022 - The aliscore output file does not exist. Check results for BUSCO: 1479417at2759
## Mon Jul 25 14:41:42 CEST 2022 - The aliscore output file does not exist. Check results for BUSCO: 664730at2759
## Mon Jul 25 14:41:42 CEST 2022 - The aliscore output appears to be empty. Check results for BUSCO: 1479417at2759
## Mon Jul 25 14:41:42 CEST 2022 - The aliscore output appears to be empty. Check results for BUSCO: 664730at2759
## Mon Jul 25 14:41:42 CEST 2022 - The aliscore output file does not exist. Check results for BUSCO: 464990at2759
## Mon Jul 25 14:41:42 CEST 2022 - The aliscore output appears to be empty. Check results for BUSCO: 464990at2759
## Mon Jul 25 14:41:42 CEST 2022 - The aliscore output file does not exist. Check results for BUSCO: 1474000at2759
## Mon Jul 25 14:41:42 CEST 2022 - The aliscore output appears to be empty. Check results for BUSCO: 1474000at2759
## Mon Jul 25 14:41:42 CEST 2022 - The aliscore output file does not exist. Check results for BUSCO: 1398309at2759
## Mon Jul 25 14:41:42 CEST 2022 - The aliscore output file does not exist. Check results for BUSCO: 721605at2759
## Mon Jul 25 14:41:43 CEST 2022 - The aliscore output appears to be empty. Check results for BUSCO: 1398309at2759
## Mon Jul 25 14:41:43 CEST 2022 - The aliscore output appears to be empty. Check results for BUSCO: 721605at2759
## Mon Jul 25 14:41:43 CEST 2022 - The aliscore output file does not exist. Check results for BUSCO: 975158at2759
## Mon Jul 25 14:41:43 CEST 2022 - The aliscore output file does not exist. Check results for BUSCO: 1459797at2759
## Mon Jul 25 14:41:43 CEST 2022 - The aliscore output appears to be empty. Check results for BUSCO: 975158at2759
## Mon Jul 25 14:41:43 CEST 2022 - The aliscore output appears to be empty. Check results for BUSCO: 1459797at2759
## Mon Jul 25 14:41:43 CEST 2022 - The aliscore output file does not exist. Check results for BUSCO: 865202at2759
## Mon Jul 25 14:41:43 CEST 2022 - The aliscore output appears to be empty. Check results for BUSCO: 865202at2759
## Mon Jul 25 14:41:43 CEST 2022 - The aliscore output file does not exist. Check results for BUSCO: 721605at2759
## Mon Jul 25 14:41:43 CEST 2022 - The aliscore output appears to be empty. Check results for BUSCO: 721605at2759
## Mon Jul 25 14:41:43 CEST 2022 - The aliscore output file does not exist. Check results for BUSCO: 974865at2759
## Mon Jul 25 14:41:43 CEST 2022 - The aliscore output appears to be empty. Check results for BUSCO: 974865at2759
## Mon Jul 25 14:41:43 CEST 2022 - The aliscore output file does not exist. Check results for BUSCO: 1157302at2759
## Mon Jul 25 14:41:43 CEST 2022 - The aliscore output appears to be empty. Check results for BUSCO: 1157302at2759
## Mon Jul 25 14:41:43 CEST 2022 - The aliscore output file does not exist. Check results for BUSCO: 1194691at2759
## Mon Jul 25 14:41:43 CEST 2022 - The aliscore output file does not exist. Check results for BUSCO: 1217666at2759
## Mon Jul 25 14:41:43 CEST 2022 - The aliscore output appears to be empty. Check results for BUSCO: 1194691at2759
## Mon Jul 25 14:41:43 CEST 2022 - The aliscore output appears to be empty. Check results for BUSCO: 1217666at2759
## Mon Jul 25 14:41:43 CEST 2022 - The aliscore output file does not exist. Check results for BUSCO: 1178688at2759
## Mon Jul 25 14:41:43 CEST 2022 - The aliscore output file does not exist. Check results for BUSCO: 1490022at2759
## Mon Jul 25 14:41:43 CEST 2022 - The aliscore output appears to be empty. Check results for BUSCO: 1178688at2759
## Mon Jul 25 14:41:43 CEST 2022 - The aliscore output file does not exist. Check results for BUSCO: 981902at2759
## Mon Jul 25 14:41:43 CEST 2022 - The aliscore output appears to be empty. Check results for BUSCO: 1490022at2759
## Mon Jul 25 14:41:43 CEST 2022 - The aliscore output file does not exist. Check results for BUSCO: 1287094at2759
## Mon Jul 25 14:41:43 CEST 2022 - The aliscore output appears to be empty. Check results for BUSCO: 981902at2759
## Mon Jul 25 14:41:43 CEST 2022 - The aliscore output file does not exist. Check results for BUSCO: 866359at2759
## Mon Jul 25 14:41:43 CEST 2022 - The aliscore output appears to be empty. Check results for BUSCO: 1287094at2759
## Mon Jul 25 14:41:43 CEST 2022 - The aliscore output appears to be empty. Check results for BUSCO: 866359at2759
## Mon Jul 25 14:41:43 CEST 2022 - The aliscore output file does not exist. Check results for BUSCO: 687505at2759
## Mon Jul 25 14:41:43 CEST 2022 - The aliscore output appears to be empty. Check results for BUSCO: 687505at2759
## Mon Jul 25 14:41:43 CEST 2022 - The aliscore output file does not exist. Check results for BUSCO: 1445102at2759
## Mon Jul 25 14:41:43 CEST 2022 - The aliscore output appears to be empty. Check results for BUSCO: 1445102at2759
## Mon Jul 25 14:41:43 CEST 2022 - The aliscore output file does not exist. Check results for BUSCO: 937686at2759
## Mon Jul 25 14:41:43 CEST 2022 - The aliscore output appears to be empty. Check results for BUSCO: 937686at2759
## Mon Jul 25 14:41:43 CEST 2022 - The aliscore output file does not exist. Check results for BUSCO: 464990at2759
## Mon Jul 25 14:41:44 CEST 2022 - The aliscore output appears to be empty. Check results for BUSCO: 464990at2759
## Mon Jul 25 14:41:44 CEST 2022 - The aliscore output file does not exist. Check results for BUSCO: 1193442at2759
## Mon Jul 25 14:41:44 CEST 2022 - The aliscore output file does not exist. Check results for BUSCO: 1222562at2759
## Mon Jul 25 14:41:44 CEST 2022 - The aliscore output appears to be empty. Check results for BUSCO: 1193442at2759
## Mon Jul 25 14:41:44 CEST 2022 - The aliscore output appears to be empty. Check results for BUSCO: 1222562at2759
## Mon Jul 25 14:41:44 CEST 2022 - The aliscore output file does not exist. Check results for BUSCO: 1513531at2759
## Mon Jul 25 14:41:44 CEST 2022 - The aliscore output file does not exist. Check results for BUSCO: 1322642at2759
## Mon Jul 25 14:41:44 CEST 2022 - The aliscore output appears to be empty. Check results for BUSCO: 1513531at2759
## Mon Jul 25 14:41:44 CEST 2022 - The aliscore output appears to be empty. Check results for BUSCO: 1322642at2759
## Mon Jul 25 14:41:44 CEST 2022 - The aliscore output file does not exist. Check results for BUSCO: 1107630at2759
## Mon Jul 25 14:41:44 CEST 2022 - The aliscore output appears to be empty. Check results for BUSCO: 1107630at2759
## Mon Jul 25 14:41:44 CEST 2022 - The aliscore output file does not exist. Check results for BUSCO: 1490022at2759
## Mon Jul 25 14:41:44 CEST 2022 - The aliscore output file does not exist. Check results for BUSCO: 1076134at2759
## Mon Jul 25 14:41:44 CEST 2022 - The aliscore output file does not exist. Check results for BUSCO: 1266231at2759
## Mon Jul 25 14:41:44 CEST 2022 - The aliscore output appears to be empty. Check results for BUSCO: 1490022at2759
## Mon Jul 25 14:41:44 CEST 2022 - The aliscore output file does not exist. Check results for BUSCO: 1161199at2759
## Mon Jul 25 14:41:44 CEST 2022 - The aliscore output appears to be empty. Check results for BUSCO: 1076134at2759
## Mon Jul 25 14:41:44 CEST 2022 - The aliscore output appears to be empty. Check results for BUSCO: 1266231at2759
## Mon Jul 25 14:41:44 CEST 2022 - The aliscore output appears to be empty. Check results for BUSCO: 1161199at2759
## Mon Jul 25 14:41:44 CEST 2022 - The aliscore output file does not exist. Check results for BUSCO: 1322299at2759
## Mon Jul 25 14:41:44 CEST 2022 - The aliscore output file does not exist. Check results for BUSCO: 1018517at2759
## Mon Jul 25 14:41:44 CEST 2022 - The aliscore output appears to be empty. Check results for BUSCO: 1322299at2759
## Mon Jul 25 14:41:44 CEST 2022 - The aliscore output appears to be empty. Check results for BUSCO: 1018517at2759
## Mon Jul 25 14:41:44 CEST 2022 - The aliscore output file does not exist. Check results for BUSCO: 1275837at2759
## Mon Jul 25 14:41:44 CEST 2022 - The aliscore output appears to be empty. Check results for BUSCO: 1275837at2759
## Mon Jul 25 14:41:45 CEST 2022 - The aliscore output file does not exist. Check results for BUSCO: 513979at2759
## Mon Jul 25 14:41:45 CEST 2022 - The aliscore output appears to be empty. Check results for BUSCO: 513979at2759
## Mon Jul 25 14:41:45 CEST 2022 - The aliscore output file does not exist. Check results for BUSCO: 1287401at2759
## Mon Jul 25 14:41:45 CEST 2022 - The aliscore output appears to be empty. Check results for BUSCO: 1287401at2759
## Mon Jul 25 14:41:45 CEST 2022 - The aliscore output file does not exist. Check results for BUSCO: 1076134at2759
## Mon Jul 25 14:41:45 CEST 2022 - The aliscore output file does not exist. Check results for BUSCO: 1087488at2759
## Mon Jul 25 14:41:45 CEST 2022 - The aliscore output appears to be empty. Check results for BUSCO: 1087488at2759
## Mon Jul 25 14:41:45 CEST 2022 - The aliscore output appears to be empty. Check results for BUSCO: 1076134at2759
## Mon Jul 25 14:41:45 CEST 2022 - The aliscore output file does not exist. Check results for BUSCO: 1003258at2759
## Mon Jul 25 14:41:45 CEST 2022 - The aliscore output file does not exist. Check results for BUSCO: 1426075at2759
## Mon Jul 25 14:41:45 CEST 2022 - The aliscore output appears to be empty. Check results for BUSCO: 1003258at2759
## Mon Jul 25 14:41:45 CEST 2022 - The aliscore output appears to be empty. Check results for BUSCO: 1426075at2759
## Mon Jul 25 14:41:45 CEST 2022 - The aliscore output file does not exist. Check results for BUSCO: 1309031at2759
## Mon Jul 25 14:41:45 CEST 2022 - The aliscore output appears to be empty. Check results for BUSCO: 1309031at2759
## Mon Jul 25 14:41:45 CEST 2022 - The aliscore output file does not exist. Check results for BUSCO: 426305at2759
## Mon Jul 25 14:41:45 CEST 2022 - The aliscore output appears to be empty. Check results for BUSCO: 426305at2759
## Mon Jul 25 14:41:45 CEST 2022 - The aliscore output file does not exist. Check results for BUSCO: 1423847at2759
## Mon Jul 25 14:41:45 CEST 2022 - The aliscore output appears to be empty. Check results for BUSCO: 1423847at2759
## Mon Jul 25 14:41:45 CEST 2022 - The aliscore output file does not exist. Check results for BUSCO: 834694at2759
## Mon Jul 25 14:41:45 CEST 2022 - The aliscore output appears to be empty. Check results for BUSCO: 834694at2759
## Mon Jul 25 14:41:45 CEST 2022 - The aliscore output file does not exist. Check results for BUSCO: 917326at2759
## Mon Jul 25 14:41:45 CEST 2022 - The aliscore output file does not exist. Check results for BUSCO: 866359at2759
## Mon Jul 25 14:41:45 CEST 2022 - The aliscore output appears to be empty. Check results for BUSCO: 917326at2759
## Mon Jul 25 14:41:45 CEST 2022 - The aliscore output appears to be empty. Check results for BUSCO: 866359at2759
## Mon Jul 25 14:41:45 CEST 2022 - The aliscore output file does not exist. Check results for BUSCO: 1001705at2759
## Mon Jul 25 14:41:45 CEST 2022 - The aliscore output appears to be empty. Check results for BUSCO: 1001705at2759
## Mon Jul 25 14:41:45 CEST 2022 - The aliscore output file does not exist. Check results for BUSCO: 418107at2759
## Mon Jul 25 14:41:45 CEST 2022 - The aliscore output appears to be empty. Check results for BUSCO: 418107at2759
## Mon Jul 25 14:41:45 CEST 2022 - The aliscore output file does not exist. Check results for BUSCO: 1182451at2759
## Mon Jul 25 14:41:45 CEST 2022 - The aliscore output appears to be empty. Check results for BUSCO: 1182451at2759
## Mon Jul 25 14:41:45 CEST 2022 - The aliscore output file does not exist. Check results for BUSCO: 937686at2759
## Mon Jul 25 14:41:46 CEST 2022 - The aliscore output appears to be empty. Check results for BUSCO: 937686at2759
## Mon Jul 25 14:41:46 CEST 2022 - The aliscore output file does not exist. Check results for BUSCO: 759498at2759
## Mon Jul 25 14:41:46 CEST 2022 - The aliscore output file does not exist. Check results for BUSCO: 814241at2759
## Mon Jul 25 14:41:46 CEST 2022 - The aliscore output appears to be empty. Check results for BUSCO: 759498at2759
## Mon Jul 25 14:41:46 CEST 2022 - The aliscore output appears to be empty. Check results for BUSCO: 814241at2759
## Mon Jul 25 14:41:46 CEST 2022 - The aliscore output file does not exist. Check results for BUSCO: 325552at2759
## Mon Jul 25 14:41:46 CEST 2022 - The aliscore output file does not exist. Check results for BUSCO: 1025450at2759
## Mon Jul 25 14:41:46 CEST 2022 - The aliscore output appears to be empty. Check results for BUSCO: 325552at2759
## Mon Jul 25 14:41:46 CEST 2022 - The aliscore output file does not exist. Check results for BUSCO: 331411at2759
## Mon Jul 25 14:41:46 CEST 2022 - The aliscore output appears to be empty. Check results for BUSCO: 1025450at2759
## Mon Jul 25 14:41:46 CEST 2022 - The aliscore output appears to be empty. Check results for BUSCO: 331411at2759
## Mon Jul 25 14:41:46 CEST 2022 - The aliscore output file does not exist. Check results for BUSCO: 345441at2759
## Mon Jul 25 14:41:46 CEST 2022 - The aliscore output appears to be empty. Check results for BUSCO: 345441at2759
## Mon Jul 25 14:41:46 CEST 2022 - The aliscore output file does not exist. Check results for BUSCO: 390348at2759
## Mon Jul 25 14:41:46 CEST 2022 - The aliscore output file does not exist. Check results for BUSCO: 673132at2759
## Mon Jul 25 14:41:46 CEST 2022 - The aliscore output file does not exist. Check results for BUSCO: 290630at2759
## Mon Jul 25 14:41:46 CEST 2022 - The aliscore output appears to be empty. Check results for BUSCO: 390348at2759
## Mon Jul 25 14:41:46 CEST 2022 - The aliscore output appears to be empty. Check results for BUSCO: 673132at2759
## Mon Jul 25 14:41:46 CEST 2022 - The aliscore output appears to be empty. Check results for BUSCO: 290630at2759
## Mon Jul 25 14:41:46 CEST 2022 - The aliscore output file does not exist. Check results for BUSCO: 570797at2759
## Mon Jul 25 14:41:46 CEST 2022 - The aliscore output appears to be empty. Check results for BUSCO: 570797at2759
## Mon Jul 25 14:41:46 CEST 2022 - The aliscore output file does not exist. Check results for BUSCO: 671846at2759
## Mon Jul 25 14:41:46 CEST 2022 - The aliscore output file does not exist. Check results for BUSCO: 293315at2759
## Mon Jul 25 14:41:46 CEST 2022 - The aliscore output appears to be empty. Check results for BUSCO: 751335at2759
## Mon Jul 25 14:41:46 CEST 2022 - The aliscore output appears to be empty. Check results for BUSCO: 671846at2759
## Mon Jul 25 14:41:46 CEST 2022 - The aliscore output file does not exist. Check results for BUSCO: 176625at2759
## Mon Jul 25 14:41:46 CEST 2022 - The aliscore output appears to be empty. Check results for BUSCO: 293315at2759
## Mon Jul 25 14:41:46 CEST 2022 - The aliscore output appears to be empty. Check results for BUSCO: 176625at2759
## Mon Jul 25 14:41:46 CEST 2022 - The aliscore output file does not exist. Check results for BUSCO: 341721at2759
## Mon Jul 25 14:41:46 CEST 2022 - The aliscore output file does not exist. Check results for BUSCO: 604979at2759
## Mon Jul 25 14:41:46 CEST 2022 - The aliscore output appears to be empty. Check results for BUSCO: 341721at2759
## Mon Jul 25 14:41:46 CEST 2022 - The aliscore output file does not exist. Check results for BUSCO: 1115196at2759
## Mon Jul 25 14:41:46 CEST 2022 - The aliscore output appears to be empty. Check results for BUSCO: 604979at2759
## Mon Jul 25 14:41:46 CEST 2022 - The aliscore output appears to be empty. Check results for BUSCO: 1115196at2759
## Mon Jul 25 14:41:46 CEST 2022 - The aliscore output file does not exist. Check results for BUSCO: 918816at2759
## Mon Jul 25 14:41:46 CEST 2022 - The aliscore output appears to be empty. Check results for BUSCO: 918816at2759
## Mon Jul 25 14:41:46 CEST 2022 - The aliscore output file does not exist. Check results for BUSCO: 1001705at2759
## Mon Jul 25 14:41:46 CEST 2022 - The aliscore output file does not exist. Check results for BUSCO: 1248958at2759
## Mon Jul 25 14:41:47 CEST 2022 - The aliscore output file does not exist. Check results for BUSCO: 664730at2759
## Mon Jul 25 14:41:47 CEST 2022 - The aliscore output appears to be empty. Check results for BUSCO: 1001705at2759
## Mon Jul 25 14:41:47 CEST 2022 - The aliscore output file does not exist. Check results for BUSCO: 331411at2759
## Mon Jul 25 14:41:47 CEST 2022 - The aliscore output appears to be empty. Check results for BUSCO: 1248958at2759
## Mon Jul 25 14:41:47 CEST 2022 - The aliscore output appears to be empty. Check results for BUSCO: 664730at2759
## Mon Jul 25 14:41:47 CEST 2022 - The aliscore output appears to be empty. Check results for BUSCO: 331411at2759
## Mon Jul 25 14:41:47 CEST 2022 - The aliscore output file does not exist. Check results for BUSCO: 341721at2759
## Mon Jul 25 14:41:47 CEST 2022 - The aliscore output appears to be empty. Check results for BUSCO: 341721at2759
## Mon Jul 25 14:41:47 CEST 2022 - The aliscore output file does not exist. Check results for BUSCO: 1314980at2759
## Mon Jul 25 14:41:47 CEST 2022 - The aliscore output appears to be empty. Check results for BUSCO: 1314980at2759
## Mon Jul 25 14:41:47 CEST 2022 - The aliscore output file does not exist. Check results for BUSCO: 97116at2759
## Mon Jul 25 14:41:47 CEST 2022 - The aliscore output file does not exist. Check results for BUSCO: 1079827at2759
## Mon Jul 25 14:41:47 CEST 2022 - The aliscore output file does not exist. Check results for BUSCO: 722805at2759
## Mon Jul 25 14:41:47 CEST 2022 - The aliscore output appears to be empty. Check results for BUSCO: 97116at2759
## Mon Jul 25 14:41:47 CEST 2022 - The aliscore output appears to be empty. Check results for BUSCO: 722805at2759
## Mon Jul 25 14:41:47 CEST 2022 - The aliscore output appears to be empty. Check results for BUSCO: 1079827at2759
## Mon Jul 25 14:41:47 CEST 2022 - The aliscore output file does not exist. Check results for BUSCO: 777920at2759
## Mon Jul 25 14:41:47 CEST 2022 - The aliscore output file does not exist. Check results for BUSCO: 1220881at2759
## Mon Jul 25 14:41:47 CEST 2022 - The aliscore output appears to be empty. Check results for BUSCO: 777920at2759
## Mon Jul 25 14:41:47 CEST 2022 - The aliscore output appears to be empty. Check results for BUSCO: 1220881at2759
## Mon Jul 25 14:41:47 CEST 2022 - The aliscore output file does not exist. Check results for BUSCO: 814241at2759
## Mon Jul 25 14:41:47 CEST 2022 - The aliscore output appears to be empty. Check results for BUSCO: 814241at2759
## Mon Jul 25 14:41:48 CEST 2022 - The aliscore output file does not exist. Check results for BUSCO: 179362at2759
## Mon Jul 25 14:41:48 CEST 2022 - The aliscore output appears to be empty. Check results for BUSCO: 179362at2759
## Mon Jul 25 14:41:48 CEST 2022 - The aliscore output file does not exist. Check results for BUSCO: 1025450at2759
## Mon Jul 25 14:41:48 CEST 2022 - The aliscore output appears to be empty. Check results for BUSCO: 1025450at2759
## Mon Jul 25 14:41:48 CEST 2022 - The aliscore output file does not exist. Check results for BUSCO: 973442at2759
## Mon Jul 25 14:41:48 CEST 2022 - The aliscore output appears to be empty. Check results for BUSCO: 973442at2759
## Mon Jul 25 14:41:49 CEST 2022 - The aliscore output file does not exist. Check results for BUSCO: 671846at2759
## Mon Jul 25 14:41:49 CEST 2022 - The aliscore output appears to be empty. Check results for BUSCO: 671846at2759
## Mon Jul 25 14:41:49 CEST 2022 - The aliscore output file does not exist. Check results for BUSCO: 1248958at2759
## Mon Jul 25 14:41:49 CEST 2022 - The aliscore output appears to be empty. Check results for BUSCO: 1248958at2759
## Mon Jul 25 14:41:50 CEST 2022 - The aliscore output file does not exist. Check results for BUSCO: 1379600at2759
## Mon Jul 25 14:41:50 CEST 2022 - The aliscore output appears to be empty. Check results for BUSCO: 1379600at2759
## Mon Jul 25 14:41:50 CEST 2022 - The aliscore output file does not exist. Check results for BUSCO: 759498at2759
## Mon Jul 25 14:41:50 CEST 2022 - The aliscore output appears to be empty. Check results for BUSCO: 759498at2759
## Mon Jul 25 14:41:50 CEST 2022 - The aliscore output file does not exist. Check results for BUSCO: 160593at2759
## Mon Jul 25 14:41:50 CEST 2022 - The aliscore output appears to be empty. Check results for BUSCO: 160593at2759
## Mon Jul 25 14:41:51 CEST 2022 - The aliscore output file does not exist. Check results for BUSCO: 1312453at2759
## Mon Jul 25 14:41:51 CEST 2022 - The aliscore output appears to be empty. Check results for BUSCO: 1312453at2759
## Mon Jul 25 14:41:51 CEST 2022 - The aliscore output file does not exist. Check results for BUSCO: 924753at2759
## Mon Jul 25 14:41:51 CEST 2022 - The aliscore output appears to be empty. Check results for BUSCO: 924753at2759
## Mon Jul 25 14:41:54 CEST 2022 - The aliscore output file does not exist. Check results for BUSCO: 1087488at2759
## Mon Jul 25 14:41:54 CEST 2022 - The aliscore output appears to be empty. Check results for BUSCO: 1087488at2759
## Mon Jul 25 14:41:56 CEST 2022 - The aliscore output file does not exist. Check results for BUSCO: 156083at2759
## Mon Jul 25 14:41:56 CEST 2022 - The aliscore output appears to be empty. Check results for BUSCO: 156083at2759
## Mon Jul 25 14:42:11 CEST 2022 - The aliscore output file does not exist. Check results for BUSCO: 1200489at2759
## Mon Jul 25 14:42:11 CEST 2022 - The aliscore output appears to be empty. Check results for BUSCO: 1200489at2759
## Mon Jul 25 14:42:11 CEST 2022 - The aliscore output file does not exist. Check results for BUSCO: 1287094at2759
## Mon Jul 25 14:42:11 CEST 2022 - The aliscore output appears to be empty. Check results for BUSCO: 1287094at2759
## Mon Jul 25 14:42:11 CEST 2022 - The aliscore output file does not exist. Check results for BUSCO: 1079827at2759
## Mon Jul 25 14:42:11 CEST 2022 - The aliscore output appears to be empty. Check results for BUSCO: 1079827at2759
## Mon Jul 25 14:42:11 CEST 2022 - The aliscore output file does not exist. Check results for BUSCO: 679771at2759
## Mon Jul 25 14:42:11 CEST 2022 - The aliscore output appears to be empty. Check results for BUSCO: 679771at2759
## Mon Jul 25 14:42:11 CEST 2022 - The aliscore output file does not exist. Check results for BUSCO: 865202at2759
## Mon Jul 25 14:42:11 CEST 2022 - The aliscore output appears to be empty. Check results for BUSCO: 865202at2759
## Mon Jul 25 14:42:12 CEST 2022 - The aliscore output file does not exist. Check results for BUSCO: 1220881at2759
## Mon Jul 25 14:42:12 CEST 2022 - The aliscore output file does not exist. Check results for BUSCO: 1445102at2759
## Mon Jul 25 14:42:12 CEST 2022 - The aliscore output appears to be empty. Check results for BUSCO: 1220881at2759
## Mon Jul 25 14:42:12 CEST 2022 - The aliscore output file does not exist. Check results for BUSCO: 1623701at2759
## Mon Jul 25 14:42:12 CEST 2022 - The aliscore output appears to be empty. Check results for BUSCO: 1445102at2759
## Mon Jul 25 14:42:12 CEST 2022 - The aliscore output appears to be empty. Check results for BUSCO: 1623701at2759
## Mon Jul 25 14:42:12 CEST 2022 - The aliscore output file does not exist. Check results for BUSCO: 1223488at2759
## Mon Jul 25 14:42:12 CEST 2022 - The aliscore output appears to be empty. Check results for BUSCO: 1223488at2759
## Mon Jul 25 14:42:12 CEST 2022 - The aliscore output file does not exist. Check results for BUSCO: 1222562at2759
## Mon Jul 25 14:42:12 CEST 2022 - The aliscore output appears to be empty. Check results for BUSCO: 1222562at2759
## Mon Jul 25 14:42:12 CEST 2022 - The aliscore output file does not exist. Check results for BUSCO: 687505at2759
## Mon Jul 25 14:42:12 CEST 2022 - The aliscore output appears to be empty. Check results for BUSCO: 687505at2759
## Mon Jul 25 14:42:12 CEST 2022 - The aliscore output file does not exist. Check results for BUSCO: 1526152at2759
## Mon Jul 25 14:42:12 CEST 2022 - The aliscore output appears to be empty. Check results for BUSCO: 1526152at2759
## Mon Jul 25 14:42:12 CEST 2022 - The aliscore output file does not exist. Check results for BUSCO: 1513531at2759
## Mon Jul 25 14:42:12 CEST 2022 - The aliscore output file does not exist. Check results for BUSCO: 1287401at2759
## Mon Jul 25 14:42:12 CEST 2022 - The aliscore output appears to be empty. Check results for BUSCO: 1513531at2759
## Mon Jul 25 14:42:12 CEST 2022 - The aliscore output appears to be empty. Check results for BUSCO: 1287401at2759
## Mon Jul 25 14:42:12 CEST 2022 - The aliscore output file does not exist. Check results for BUSCO: 1575179at2759
## Mon Jul 25 14:42:12 CEST 2022 - The aliscore output appears to be empty. Check results for BUSCO: 1575179at2759
## Mon Jul 25 14:42:13 CEST 2022 - The aliscore output file does not exist. Check results for BUSCO: 1405073at2759
## Mon Jul 25 14:42:13 CEST 2022 - The aliscore output file does not exist. Check results for BUSCO: 1620056at2759
## Mon Jul 25 14:42:13 CEST 2022 - The aliscore output file does not exist. Check results for BUSCO: 1645187at2759
## Mon Jul 25 14:42:13 CEST 2022 - The aliscore output appears to be empty. Check results for BUSCO: 1405073at2759
## Mon Jul 25 14:42:13 CEST 2022 - The aliscore output appears to be empty. Check results for BUSCO: 1620056at2759
## Mon Jul 25 14:42:13 CEST 2022 - The aliscore output file does not exist. Check results for BUSCO: 1275837at2759
## Mon Jul 25 14:42:13 CEST 2022 - The aliscore output appears to be empty. Check results for BUSCO: 1645187at2759
## Mon Jul 25 14:42:13 CEST 2022 - The aliscore output appears to be empty. Check results for BUSCO: 1275837at2759
## Mon Jul 25 14:42:13 CEST 2022 - The aliscore output file does not exist. Check results for BUSCO: 1323575at2759
## Mon Jul 25 14:42:13 CEST 2022 - The aliscore output file does not exist. Check results for BUSCO: 751335at2759
## Mon Jul 25 14:42:13 CEST 2022 - The aliscore output appears to be empty. Check results for BUSCO: 1323575at2759
## Mon Jul 25 14:42:13 CEST 2022 - The aliscore output appears to be empty. Check results for BUSCO: 751335at2759
## Mon Jul 25 14:42:13 CEST 2022 - The aliscore output file does not exist. Check results for BUSCO: 923657at2759
## Mon Jul 25 14:42:13 CEST 2022 - The aliscore output appears to be empty. Check results for BUSCO: 923657at2759
## Mon Jul 25 14:42:13 CEST 2022 - The aliscore output file does not exist. Check results for BUSCO: 1405073at2759
## Mon Jul 25 14:42:13 CEST 2022 - The aliscore output appears to be empty. Check results for BUSCO: 1405073at2759
## Mon Jul 25 14:42:13 CEST 2022 - The aliscore output file does not exist. Check results for BUSCO: 1223488at2759
## Mon Jul 25 14:42:13 CEST 2022 - The aliscore output appears to be empty. Check results for BUSCO: 1223488at2759
## Mon Jul 25 14:42:13 CEST 2022 - The aliscore output file does not exist. Check results for BUSCO: 1249159at2759
## Mon Jul 25 14:42:13 CEST 2022 - The aliscore output appears to be empty. Check results for BUSCO: 1249159at2759
## Mon Jul 25 14:42:13 CEST 2022 - The aliscore output file does not exist. Check results for BUSCO: 1233814at2759
## Mon Jul 25 14:42:13 CEST 2022 - The aliscore output file does not exist. Check results for BUSCO: 1398309at2759
## Mon Jul 25 14:42:13 CEST 2022 - The aliscore output appears to be empty. Check results for BUSCO: 1233814at2759
## Mon Jul 25 14:42:13 CEST 2022 - The aliscore output appears to be empty. Check results for BUSCO: 1398309at2759
## Mon Jul 25 14:42:13 CEST 2022 - The aliscore output file does not exist. Check results for BUSCO: 1358374at2759
## Mon Jul 25 14:42:13 CEST 2022 - The aliscore output appears to be empty. Check results for BUSCO: 1358374at2759
## Mon Jul 25 14:42:13 CEST 2022 - The aliscore output file does not exist. Check results for BUSCO: 679771at2759
## Mon Jul 25 14:42:13 CEST 2022 - The aliscore output appears to be empty. Check results for BUSCO: 679771at2759
## Mon Jul 25 14:42:14 CEST 2022 - The aliscore output file does not exist. Check results for BUSCO: 1264469at2759
## Mon Jul 25 14:42:14 CEST 2022 - The aliscore output file does not exist. Check results for BUSCO: 1228942at2759
## Mon Jul 25 14:42:14 CEST 2022 - The aliscore output file does not exist. Check results for BUSCO: 919955at2759
## Mon Jul 25 14:42:14 CEST 2022 - The aliscore output appears to be empty. Check results for BUSCO: 1264469at2759
## Mon Jul 25 14:42:14 CEST 2022 - The aliscore output file does not exist. Check results for BUSCO: 1474000at2759
## Mon Jul 25 14:42:14 CEST 2022 - The aliscore output file does not exist. Check results for BUSCO: 1645187at2759
## Mon Jul 25 14:42:14 CEST 2022 - The aliscore output appears to be empty. Check results for BUSCO: 1228942at2759
## Mon Jul 25 14:42:14 CEST 2022 - The aliscore output file does not exist. Check results for BUSCO: 1157302at2759
## Mon Jul 25 14:42:14 CEST 2022 - The aliscore output appears to be empty. Check results for BUSCO: 919955at2759
## Mon Jul 25 14:42:14 CEST 2022 - The aliscore output appears to be empty. Check results for BUSCO: 1474000at2759
## Mon Jul 25 14:42:14 CEST 2022 - The aliscore output appears to be empty. Check results for BUSCO: 1645187at2759
## Mon Jul 25 14:42:14 CEST 2022 - The aliscore output appears to be empty. Check results for BUSCO: 1157302at2759
## Mon Jul 25 14:42:14 CEST 2022 - The aliscore output file does not exist. Check results for BUSCO: 1200489at2759
## Mon Jul 25 14:42:14 CEST 2022 - The aliscore output appears to be empty. Check results for BUSCO: 1200489at2759
## Mon Jul 25 14:42:14 CEST 2022 - The aliscore output file does not exist. Check results for BUSCO: 1129824at2759
## Mon Jul 25 14:42:14 CEST 2022 - The aliscore output appears to be empty. Check results for BUSCO: 1129824at2759
## Mon Jul 25 14:42:14 CEST 2022 - The aliscore output appears to be empty. Check results for BUSCO: 1194691at2759
## Mon Jul 25 14:42:14 CEST 2022 - The aliscore output file does not exist. Check results for BUSCO: 390348at2759
## Mon Jul 25 14:42:14 CEST 2022 - The aliscore output file does not exist. Check results for BUSCO: 1107630at2759
## Mon Jul 25 14:42:14 CEST 2022 - The aliscore output appears to be empty. Check results for BUSCO: 390348at2759
## Mon Jul 25 14:42:14 CEST 2022 - The aliscore output appears to be empty. Check results for BUSCO: 1107630at2759
## Mon Jul 25 14:42:14 CEST 2022 - The aliscore output file does not exist. Check results for BUSCO: 1161199at2759
## Mon Jul 25 14:42:14 CEST 2022 - The aliscore output appears to be empty. Check results for BUSCO: 1161199at2759
## Mon Jul 25 14:42:14 CEST 2022 - The aliscore output file does not exist. Check results for BUSCO: 655400at2759
## Mon Jul 25 14:42:14 CEST 2022 - The aliscore output appears to be empty. Check results for BUSCO: 655400at2759
## Mon Jul 25 14:42:14 CEST 2022 - The aliscore output file does not exist. Check results for BUSCO: 975158at2759
## Mon Jul 25 14:42:14 CEST 2022 - The aliscore output appears to be empty. Check results for BUSCO: 975158at2759
## Mon Jul 25 14:42:14 CEST 2022 - The aliscore output file does not exist. Check results for BUSCO: 1217666at2759
## Mon Jul 25 14:42:14 CEST 2022 - The aliscore output appears to be empty. Check results for BUSCO: 1217666at2759
## Mon Jul 25 14:42:14 CEST 2022 - The aliscore output file does not exist. Check results for BUSCO: 1030907at2759
## Mon Jul 25 14:42:14 CEST 2022 - The aliscore output appears to be empty. Check results for BUSCO: 1030907at2759
## Mon Jul 25 14:42:14 CEST 2022 - The aliscore output file does not exist. Check results for BUSCO: 1479417at2759
## Mon Jul 25 14:42:15 CEST 2022 - The aliscore output appears to be empty. Check results for BUSCO: 1479417at2759
## Mon Jul 25 14:42:15 CEST 2022 - The aliscore output file does not exist. Check results for BUSCO: 1018517at2759
## Mon Jul 25 14:42:15 CEST 2022 - The aliscore output appears to be empty. Check results for BUSCO: 1018517at2759
## Mon Jul 25 14:42:15 CEST 2022 - The aliscore output file does not exist. Check results for BUSCO: 768809at2759
## Mon Jul 25 14:42:15 CEST 2022 - The aliscore output file does not exist. Check results for BUSCO: 1266231at2759
## Mon Jul 25 14:42:15 CEST 2022 - The aliscore output appears to be empty. Check results for BUSCO: 1266231at2759
## Mon Jul 25 14:42:15 CEST 2022 - The aliscore output file does not exist. Check results for BUSCO: 817008at2759
## Mon Jul 25 14:42:15 CEST 2022 - The aliscore output file does not exist. Check results for BUSCO: 1129824at2759
## Mon Jul 25 14:42:15 CEST 2022 - The aliscore output appears to be empty. Check results for BUSCO: 817008at2759
## Mon Jul 25 14:42:15 CEST 2022 - The aliscore output appears to be empty. Check results for BUSCO: 1129824at2759
## Mon Jul 25 14:42:15 CEST 2022 - The aliscore output file does not exist. Check results for BUSCO: 290630at2759
## Mon Jul 25 14:42:15 CEST 2022 - The aliscore output appears to be empty. Check results for BUSCO: 290630at2759
## Mon Jul 25 14:42:15 CEST 2022 - The aliscore output file does not exist. Check results for BUSCO: 1323575at2759
## Mon Jul 25 14:42:15 CEST 2022 - The aliscore output appears to be empty. Check results for BUSCO: 418107at2759
## Mon Jul 25 14:42:15 CEST 2022 - The aliscore output file does not exist. Check results for BUSCO: 1322299at2759
## Mon Jul 25 14:42:15 CEST 2022 - The aliscore output file does not exist. Check results for BUSCO: 1309031at2759
## Mon Jul 25 14:42:15 CEST 2022 - The aliscore output appears to be empty. Check results for BUSCO: 1323575at2759
## Mon Jul 25 14:42:15 CEST 2022 - The aliscore output appears to be empty. Check results for BUSCO: 1309031at2759
## Mon Jul 25 14:42:16 CEST 2022 - The aliscore output appears to be empty. Check results for BUSCO: 1322299at2759
## Mon Jul 25 14:42:16 CEST 2022 - The aliscore output file does not exist. Check results for BUSCO: 1258856at2759
## Mon Jul 25 14:42:16 CEST 2022 - The aliscore output appears to be empty. Check results for BUSCO: 1258856at2759
## Mon Jul 25 14:42:16 CEST 2022 - The aliscore output file does not exist. Check results for BUSCO: 655400at2759
## Mon Jul 25 14:42:16 CEST 2022 - The aliscore output appears to be empty. Check results for BUSCO: 655400at2759
## Mon Jul 25 14:42:16 CEST 2022 - The aliscore output file does not exist. Check results for BUSCO: 1426075at2759
## Mon Jul 25 14:42:16 CEST 2022 - The aliscore output appears to be empty. Check results for BUSCO: 1426075at2759
## Mon Jul 25 14:42:16 CEST 2022 - The aliscore output file does not exist. Check results for BUSCO: 1322642at2759
## Mon Jul 25 14:42:16 CEST 2022 - The aliscore output file does not exist. Check results for BUSCO: 1193442at2759
## Mon Jul 25 14:42:16 CEST 2022 - The aliscore output appears to be empty. Check results for BUSCO: 1322642at2759
## Mon Jul 25 14:42:16 CEST 2022 - The aliscore output appears to be empty. Check results for BUSCO: 1193442at2759
## Mon Jul 25 14:42:16 CEST 2022 - The aliscore output file does not exist. Check results for BUSCO: 375960at2759
## Mon Jul 25 14:42:16 CEST 2022 - The aliscore output appears to be empty. Check results for BUSCO: 375960at2759
## Mon Jul 25 14:42:16 CEST 2022 - The aliscore output file does not exist. Check results for BUSCO: 1182451at2759
## Mon Jul 25 14:42:16 CEST 2022 - The aliscore output appears to be empty. Check results for BUSCO: 1182451at2759
## Mon Jul 25 14:42:16 CEST 2022 - The aliscore output file does not exist. Check results for BUSCO: 604979at2759
## Mon Jul 25 14:42:16 CEST 2022 - The aliscore output file does not exist. Check results for BUSCO: 426305at2759
## Mon Jul 25 14:42:16 CEST 2022 - The aliscore output appears to be empty. Check results for BUSCO: 604979at2759
## Mon Jul 25 14:42:16 CEST 2022 - The aliscore output file does not exist. Check results for BUSCO: 981902at2759
## Mon Jul 25 14:42:16 CEST 2022 - The aliscore output appears to be empty. Check results for BUSCO: 426305at2759
## Mon Jul 25 14:42:17 CEST 2022 - The aliscore output appears to be empty. Check results for BUSCO: 981902at2759
## Mon Jul 25 14:42:17 CEST 2022 - The aliscore output file does not exist. Check results for BUSCO: 392369at2759
## Mon Jul 25 14:42:17 CEST 2022 - The aliscore output appears to be empty. Check results for BUSCO: 392369at2759
## Mon Jul 25 14:42:17 CEST 2022 - The aliscore output file does not exist. Check results for BUSCO: 392369at2759
## Mon Jul 25 14:42:17 CEST 2022 - The aliscore output appears to be empty. Check results for BUSCO: 392369at2759
## Mon Jul 25 14:42:17 CEST 2022 - The aliscore output file does not exist. Check results for BUSCO: 1504863at2759
## Mon Jul 25 14:42:17 CEST 2022 - The aliscore output file does not exist. Check results for BUSCO: 777920at2759
## Mon Jul 25 14:42:17 CEST 2022 - The aliscore output appears to be empty. Check results for BUSCO: 1504863at2759
## Mon Jul 25 14:42:17 CEST 2022 - The aliscore output appears to be empty. Check results for BUSCO: 777920at2759
## Mon Jul 25 14:42:17 CEST 2022 - The aliscore output file does not exist. Check results for BUSCO: 1003258at2759
## Mon Jul 25 14:42:17 CEST 2022 - The aliscore output file does not exist. Check results for BUSCO: 924753at2759
## Mon Jul 25 14:42:17 CEST 2022 - The aliscore output appears to be empty. Check results for BUSCO: 924753at2759
## Mon Jul 25 14:42:17 CEST 2022 - The aliscore output appears to be empty. Check results for BUSCO: 1003258at2759
## Mon Jul 25 14:42:17 CEST 2022 - The aliscore output file does not exist. Check results for BUSCO: 1194797at2759
## Mon Jul 25 14:42:17 CEST 2022 - The aliscore output appears to be empty. Check results for BUSCO: 1194797at2759
## Mon Jul 25 14:42:17 CEST 2022 - The aliscore output file does not exist. Check results for BUSCO: 836599at2759
## Mon Jul 25 14:42:17 CEST 2022 - The aliscore output file does not exist. Check results for BUSCO: 673132at2759
## Mon Jul 25 14:42:17 CEST 2022 - The aliscore output appears to be empty. Check results for BUSCO: 836599at2759
## Mon Jul 25 14:42:17 CEST 2022 - The aliscore output appears to be empty. Check results for BUSCO: 673132at2759
## Mon Jul 25 14:42:18 CEST 2022 - The aliscore output file does not exist. Check results for BUSCO: 513979at2759
## Mon Jul 25 14:42:18 CEST 2022 - The aliscore output appears to be empty. Check results for BUSCO: 513979at2759
## Mon Jul 25 14:42:18 CEST 2022 - The aliscore output file does not exist. Check results for BUSCO: 917326at2759
## Mon Jul 25 14:42:18 CEST 2022 - The aliscore output file does not exist. Check results for BUSCO: 1258856at2759
## Mon Jul 25 14:42:18 CEST 2022 - The aliscore output appears to be empty. Check results for BUSCO: 917326at2759
## Mon Jul 25 14:42:18 CEST 2022 - The aliscore output file does not exist. Check results for BUSCO: 176625at2759
## Mon Jul 25 14:42:18 CEST 2022 - The aliscore output file does not exist. Check results for BUSCO: 293315at2759
## Mon Jul 25 14:42:18 CEST 2022 - The aliscore output appears to be empty. Check results for BUSCO: 1258856at2759
## Mon Jul 25 14:42:18 CEST 2022 - The aliscore output file does not exist. Check results for BUSCO: 1379600at2759
## Mon Jul 25 14:42:18 CEST 2022 - The aliscore output appears to be empty. Check results for BUSCO: 176625at2759
## Mon Jul 25 14:42:18 CEST 2022 - The aliscore output appears to be empty. Check results for BUSCO: 293315at2759
## Mon Jul 25 14:42:18 CEST 2022 - The aliscore output appears to be empty. Check results for BUSCO: 1379600at2759
## Mon Jul 25 14:42:18 CEST 2022 - The aliscore output file does not exist. Check results for BUSCO: 570797at2759
## Mon Jul 25 14:42:18 CEST 2022 - The aliscore output appears to be empty. Check results for BUSCO: 570797at2759
## Mon Jul 25 14:42:18 CEST 2022 - The aliscore output file does not exist. Check results for BUSCO: 383503at2759
## Mon Jul 25 14:42:18 CEST 2022 - The aliscore output appears to be empty. Check results for BUSCO: 383503at2759
## Mon Jul 25 14:42:18 CEST 2022 - The aliscore output file does not exist. Check results for BUSCO: 97116at2759
## Mon Jul 25 14:42:18 CEST 2022 - The aliscore output file does not exist. Check results for BUSCO: 918816at2759
## Mon Jul 25 14:42:18 CEST 2022 - The aliscore output appears to be empty. Check results for BUSCO: 97116at2759
## Mon Jul 25 14:42:18 CEST 2022 - The aliscore output appears to be empty. Check results for BUSCO: 918816at2759
## Mon Jul 25 14:42:18 CEST 2022 - The aliscore output file does not exist. Check results for BUSCO: 1423847at2759
## Mon Jul 25 14:42:18 CEST 2022 - The aliscore output appears to be empty. Check results for BUSCO: 1423847at2759
## Mon Jul 25 14:42:19 CEST 2022 - The aliscore output file does not exist. Check results for BUSCO: 1194797at2759
## Mon Jul 25 14:42:19 CEST 2022 - The aliscore output file does not exist. Check results for BUSCO: 834694at2759
## Mon Jul 25 14:42:19 CEST 2022 - The aliscore output appears to be empty. Check results for BUSCO: 1194797at2759
## Mon Jul 25 14:42:19 CEST 2022 - The aliscore output appears to be empty. Check results for BUSCO: 834694at2759
## Mon Jul 25 14:42:19 CEST 2022 - The aliscore output file does not exist. Check results for BUSCO: 325552at2759
## Mon Jul 25 14:42:19 CEST 2022 - The aliscore output appears to be empty. Check results for BUSCO: 325552at2759
## Mon Jul 25 14:42:19 CEST 2022 - The aliscore output file does not exist. Check results for BUSCO: 817008at2759
## Mon Jul 25 14:42:19 CEST 2022 - The aliscore output appears to be empty. Check results for BUSCO: 817008at2759
## Mon Jul 25 14:42:19 CEST 2022 - The aliscore output file does not exist. Check results for BUSCO: 722805at2759
## Mon Jul 25 14:42:19 CEST 2022 - The aliscore output appears to be empty. Check results for BUSCO: 722805at2759
## Mon Jul 25 14:42:20 CEST 2022 - The aliscore output file does not exist. Check results for BUSCO: 768809at2759
## Mon Jul 25 14:42:20 CEST 2022 - The aliscore output appears to be empty. Check results for BUSCO: 768809at2759
## Mon Jul 25 14:42:20 CEST 2022 - The aliscore output file does not exist. Check results for BUSCO: 1314980at2759
## Mon Jul 25 14:42:20 CEST 2022 - The aliscore output appears to be empty. Check results for BUSCO: 1314980at2759
## Mon Jul 25 14:42:20 CEST 2022 - The aliscore output file does not exist. Check results for BUSCO: 179362at2759
## Mon Jul 25 14:42:20 CEST 2022 - The aliscore output appears to be empty. Check results for BUSCO: 179362at2759
## Mon Jul 25 14:42:21 CEST 2022 - The aliscore output file does not exist. Check results for BUSCO: 973442at2759
## Mon Jul 25 14:42:21 CEST 2022 - The aliscore output appears to be empty. Check results for BUSCO: 973442at2759
## Mon Jul 25 14:42:22 CEST 2022 - The aliscore output file does not exist. Check results for BUSCO: 625387at2759
## Mon Jul 25 14:42:22 CEST 2022 - The aliscore output appears to be empty. Check results for BUSCO: 625387at2759
## Mon Jul 25 14:42:23 CEST 2022 - The aliscore output file does not exist. Check results for BUSCO: 625387at2759
## Mon Jul 25 14:42:23 CEST 2022 - The aliscore output appears to be empty. Check results for BUSCO: 625387at2759
## Mon Jul 25 14:42:24 CEST 2022 - The aliscore output file does not exist. Check results for BUSCO: 674160at2759
## Mon Jul 25 14:42:24 CEST 2022 - The aliscore output appears to be empty. Check results for BUSCO: 674160at2759
## Mon Jul 25 14:42:25 CEST 2022 - The aliscore output file does not exist. Check results for BUSCO: 1312453at2759
## Mon Jul 25 14:42:25 CEST 2022 - The aliscore output appears to be empty. Check results for BUSCO: 1312453at2759
## Mon Jul 25 14:42:26 CEST 2022 - The aliscore output file does not exist. Check results for BUSCO: 156083at2759
## Mon Jul 25 14:42:26 CEST 2022 - The aliscore output appears to be empty. Check results for BUSCO: 156083at2759
## Mon Jul 25 14:42:28 CEST 2022 - The aliscore output file does not exist. Check results for BUSCO: 674160at2759
## Mon Jul 25 14:42:28 CEST 2022 - The aliscore output appears to be empty. Check results for BUSCO: 674160at2759
## Mon Jul 25 15:02:11 CEST 2022 - The aliscore output file does not exist. Check results for BUSCO: 1398309at2759
## Mon Jul 25 15:02:11 CEST 2022 - The aliscore output appears to be empty. Check results for BUSCO: 1398309at2759
## Mon Jul 25 15:02:14 CEST 2022 - The aliscore output file does not exist. Check results for BUSCO: 1182451at2759
## Mon Jul 25 15:02:14 CEST 2022 - The aliscore output appears to be empty. Check results for BUSCO: 1182451at2759
## Mon Jul 25 15:02:16 CEST 2022 - The aliscore output file does not exist. Check results for BUSCO: 97116at2759
## Mon Jul 25 15:02:16 CEST 2022 - The aliscore output appears to be empty. Check results for BUSCO: 97116at2759
## Mon Jul 25 15:02:16 CEST 2022 - The aliscore output file does not exist. Check results for BUSCO: 1194797at2759
## Mon Jul 25 15:02:16 CEST 2022 - The aliscore output appears to be empty. Check results for BUSCO: 1194797at2759
## Mon Jul 25 15:02:40 CEST 2022 - The aliscore output file does not exist. Check results for BUSCO: 1157302at2759
## Mon Jul 25 15:02:40 CEST 2022 - The aliscore output appears to be empty. Check results for BUSCO: 1157302at2759
## Mon Jul 25 15:02:42 CEST 2022 - The aliscore output file does not exist. Check results for BUSCO: 1129824at2759
## Mon Jul 25 15:02:42 CEST 2022 - The aliscore output appears to be empty. Check results for BUSCO: 1129824at2759
## Mon Jul 25 15:02:42 CEST 2022 - The aliscore output file does not exist. Check results for BUSCO: 1623701at2759
## Mon Jul 25 15:02:42 CEST 2022 - The aliscore output appears to be empty. Check results for BUSCO: 1623701at2759
## Mon Jul 25 15:02:42 CEST 2022 - The aliscore output file does not exist. Check results for BUSCO: 1459797at2759
## Mon Jul 25 15:02:42 CEST 2022 - The aliscore output appears to be empty. Check results for BUSCO: 1459797at2759
## Mon Jul 25 15:02:43 CEST 2022 - The aliscore output file does not exist. Check results for BUSCO: 1504863at2759
## Mon Jul 25 15:02:43 CEST 2022 - The aliscore output appears to be empty. Check results for BUSCO: 1504863at2759
## Mon Jul 25 15:02:43 CEST 2022 - The aliscore output file does not exist. Check results for BUSCO: 1178688at2759
## Mon Jul 25 15:02:43 CEST 2022 - The aliscore output file does not exist. Check results for BUSCO: 679771at2759
## Mon Jul 25 15:02:43 CEST 2022 - The aliscore output appears to be empty. Check results for BUSCO: 1178688at2759
## Mon Jul 25 15:02:43 CEST 2022 - The aliscore output appears to be empty. Check results for BUSCO: 679771at2759
## Mon Jul 25 15:02:43 CEST 2022 - The aliscore output file does not exist. Check results for BUSCO: 1264469at2759
## Mon Jul 25 15:02:43 CEST 2022 - The aliscore output appears to be empty. Check results for BUSCO: 1264469at2759
## Mon Jul 25 15:02:43 CEST 2022 - The aliscore output file does not exist. Check results for BUSCO: 1030907at2759
## Mon Jul 25 15:02:43 CEST 2022 - The aliscore output appears to be empty. Check results for BUSCO: 1030907at2759
## Mon Jul 25 15:02:43 CEST 2022 - The aliscore output file does not exist. Check results for BUSCO: 1233814at2759
## Mon Jul 25 15:02:43 CEST 2022 - The aliscore output appears to be empty. Check results for BUSCO: 1233814at2759
## Mon Jul 25 15:02:43 CEST 2022 - The aliscore output file does not exist. Check results for BUSCO: 1398309at2759
## Mon Jul 25 15:02:43 CEST 2022 - The aliscore output file does not exist. Check results for BUSCO: 1620056at2759
## Mon Jul 25 15:02:43 CEST 2022 - The aliscore output appears to be empty. Check results for BUSCO: 1398309at2759
## Mon Jul 25 15:02:43 CEST 2022 - The aliscore output file does not exist. Check results for BUSCO: 1161199at2759
## Mon Jul 25 15:02:43 CEST 2022 - The aliscore output appears to be empty. Check results for BUSCO: 1620056at2759
## Mon Jul 25 15:02:43 CEST 2022 - The aliscore output appears to be empty. Check results for BUSCO: 1161199at2759
## Mon Jul 25 15:02:44 CEST 2022 - The aliscore output file does not exist. Check results for BUSCO: 1079827at2759
## Mon Jul 25 15:02:44 CEST 2022 - The aliscore output appears to be empty. Check results for BUSCO: 1079827at2759
## Mon Jul 25 15:02:44 CEST 2022 - The aliscore output file does not exist. Check results for BUSCO: 919955at2759
## Mon Jul 25 15:02:44 CEST 2022 - The aliscore output appears to be empty. Check results for BUSCO: 919955at2759
## Mon Jul 25 15:02:44 CEST 2022 - The aliscore output file does not exist. Check results for BUSCO: 974865at2759
## Mon Jul 25 15:02:44 CEST 2022 - The aliscore output appears to be empty. Check results for BUSCO: 974865at2759
## Mon Jul 25 15:02:44 CEST 2022 - The aliscore output file does not exist. Check results for BUSCO: 375960at2759
## Mon Jul 25 15:02:44 CEST 2022 - The aliscore output appears to be empty. Check results for BUSCO: 375960at2759
## Mon Jul 25 15:02:44 CEST 2022 - The aliscore output file does not exist. Check results for BUSCO: 1107630at2759
## Mon Jul 25 15:02:44 CEST 2022 - The aliscore output file does not exist. Check results for BUSCO: 975158at2759
## Mon Jul 25 15:02:44 CEST 2022 - The aliscore output appears to be empty. Check results for BUSCO: 1107630at2759
## Mon Jul 25 15:02:44 CEST 2022 - The aliscore output appears to be empty. Check results for BUSCO: 975158at2759
## Mon Jul 25 15:02:44 CEST 2022 - The aliscore output file does not exist. Check results for BUSCO: 1233814at2759
## Mon Jul 25 15:02:44 CEST 2022 - The aliscore output file does not exist. Check results for BUSCO: 345441at2759
## Mon Jul 25 15:02:44 CEST 2022 - The aliscore output appears to be empty. Check results for BUSCO: 1233814at2759
## Mon Jul 25 15:02:44 CEST 2022 - The aliscore output appears to be empty. Check results for BUSCO: 345441at2759
## Mon Jul 25 15:02:44 CEST 2022 - The aliscore output file does not exist. Check results for BUSCO: 919955at2759
## Mon Jul 25 15:02:44 CEST 2022 - The aliscore output file does not exist. Check results for BUSCO: 664730at2759
## Mon Jul 25 15:02:44 CEST 2022 - The aliscore output appears to be empty. Check results for BUSCO: 919955at2759
## Mon Jul 25 15:02:44 CEST 2022 - The aliscore output appears to be empty. Check results for BUSCO: 664730at2759
## Mon Jul 25 15:02:45 CEST 2022 - The aliscore output file does not exist. Check results for BUSCO: 1526152at2759
## Mon Jul 25 15:02:45 CEST 2022 - The aliscore output appears to be empty. Check results for BUSCO: 1526152at2759
## Mon Jul 25 15:02:45 CEST 2022 - The aliscore output file does not exist. Check results for BUSCO: 1223488at2759
## Mon Jul 25 15:02:45 CEST 2022 - The aliscore output file does not exist. Check results for BUSCO: 1217666at2759
## Mon Jul 25 15:02:45 CEST 2022 - The aliscore output appears to be empty. Check results for BUSCO: 1223488at2759
## Mon Jul 25 15:02:45 CEST 2022 - The aliscore output appears to be empty. Check results for BUSCO: 1217666at2759
## Mon Jul 25 15:02:45 CEST 2022 - The aliscore output file does not exist. Check results for BUSCO: 1575179at2759
## Mon Jul 25 15:02:45 CEST 2022 - The aliscore output file does not exist. Check results for BUSCO: 1223488at2759
## Mon Jul 25 15:02:45 CEST 2022 - The aliscore output file does not exist. Check results for BUSCO: 1358374at2759
## Mon Jul 25 15:02:45 CEST 2022 - The aliscore output appears to be empty. Check results for BUSCO: 1223488at2759
## Mon Jul 25 15:02:45 CEST 2022 - The aliscore output file does not exist. Check results for BUSCO: 1504863at2759
## Mon Jul 25 15:02:45 CEST 2022 - The aliscore output appears to be empty. Check results for BUSCO: 1358374at2759
## Mon Jul 25 15:02:45 CEST 2022 - The aliscore output file does not exist. Check results for BUSCO: 1129824at2759
## Mon Jul 25 15:02:45 CEST 2022 - The aliscore output file does not exist. Check results for BUSCO: 918816at2759
## Mon Jul 25 15:02:45 CEST 2022 - The aliscore output appears to be empty. Check results for BUSCO: 1504863at2759
## Mon Jul 25 15:02:45 CEST 2022 - The aliscore output appears to be empty. Check results for BUSCO: 918816at2759
## Mon Jul 25 15:02:45 CEST 2022 - The aliscore output appears to be empty. Check results for BUSCO: 1129824at2759
## Mon Jul 25 15:02:45 CEST 2022 - The aliscore output file does not exist. Check results for BUSCO: 1030907at2759
## Mon Jul 25 15:02:45 CEST 2022 - The aliscore output appears to be empty. Check results for BUSCO: 1030907at2759
## Mon Jul 25 15:02:45 CEST 2022 - The aliscore output appears to be empty. Check results for BUSCO: 1575179at2759
## Mon Jul 25 15:02:45 CEST 2022 - The aliscore output file does not exist. Check results for BUSCO: 836599at2759
## Mon Jul 25 15:02:45 CEST 2022 - The aliscore output appears to be empty. Check results for BUSCO: 836599at2759
## Mon Jul 25 15:02:45 CEST 2022 - The aliscore output file does not exist. Check results for BUSCO: 1220881at2759
## Mon Jul 25 15:02:45 CEST 2022 - The aliscore output appears to be empty. Check results for BUSCO: 1220881at2759
## Mon Jul 25 15:02:45 CEST 2022 - The aliscore output file does not exist. Check results for BUSCO: 1200489at2759
## Mon Jul 25 15:02:45 CEST 2022 - The aliscore output file does not exist. Check results for BUSCO: 1479417at2759
## Mon Jul 25 15:02:45 CEST 2022 - The aliscore output appears to be empty. Check results for BUSCO: 1200489at2759
## Mon Jul 25 15:02:45 CEST 2022 - The aliscore output file does not exist. Check results for BUSCO: 1001705at2759
## Mon Jul 25 15:02:45 CEST 2022 - The aliscore output file does not exist. Check results for BUSCO: 1287094at2759
## Mon Jul 25 15:02:45 CEST 2022 - The aliscore output file does not exist. Check results for BUSCO: 1217666at2759
## Mon Jul 25 15:02:45 CEST 2022 - The aliscore output appears to be empty. Check results for BUSCO: 1479417at2759
## Mon Jul 25 15:02:45 CEST 2022 - The aliscore output appears to be empty. Check results for BUSCO: 1001705at2759
## Mon Jul 25 15:02:45 CEST 2022 - The aliscore output appears to be empty. Check results for BUSCO: 1287094at2759
## Mon Jul 25 15:02:45 CEST 2022 - The aliscore output appears to be empty. Check results for BUSCO: 1217666at2759
## Mon Jul 25 15:02:46 CEST 2022 - The aliscore output file does not exist. Check results for BUSCO: 1087488at2759
## Mon Jul 25 15:02:46 CEST 2022 - The aliscore output appears to be empty. Check results for BUSCO: 1087488at2759
## Mon Jul 25 15:02:46 CEST 2022 - The aliscore output file does not exist. Check results for BUSCO: 1479417at2759
## Mon Jul 25 15:02:46 CEST 2022 - The aliscore output file does not exist. Check results for BUSCO: 1264469at2759
## Mon Jul 25 15:02:46 CEST 2022 - The aliscore output appears to be empty. Check results for BUSCO: 1479417at2759
## Mon Jul 25 15:02:46 CEST 2022 - The aliscore output appears to be empty. Check results for BUSCO: 1264469at2759
## Mon Jul 25 15:02:46 CEST 2022 - The aliscore output file does not exist. Check results for BUSCO: 1115196at2759
## Mon Jul 25 15:02:46 CEST 2022 - The aliscore output appears to be empty. Check results for BUSCO: 1115196at2759
## Mon Jul 25 15:02:46 CEST 2022 - The aliscore output file does not exist. Check results for BUSCO: 923657at2759
## Mon Jul 25 15:02:46 CEST 2022 - The aliscore output file does not exist. Check results for BUSCO: 655400at2759
## Mon Jul 25 15:02:46 CEST 2022 - The aliscore output appears to be empty. Check results for BUSCO: 923657at2759
## Mon Jul 25 15:02:46 CEST 2022 - The aliscore output file does not exist. Check results for BUSCO: 1249159at2759
## Mon Jul 25 15:02:46 CEST 2022 - The aliscore output appears to be empty. Check results for BUSCO: 655400at2759
## Mon Jul 25 15:02:46 CEST 2022 - The aliscore output appears to be empty. Check results for BUSCO: 1249159at2759
## Mon Jul 25 15:02:46 CEST 2022 - The aliscore output file does not exist. Check results for BUSCO: 1323575at2759
## Mon Jul 25 15:02:46 CEST 2022 - The aliscore output file does not exist. Check results for BUSCO: 834694at2759
## Mon Jul 25 15:02:46 CEST 2022 - The aliscore output appears to be empty. Check results for BUSCO: 1323575at2759
## Mon Jul 25 15:02:46 CEST 2022 - The aliscore output file does not exist. Check results for BUSCO: 981902at2759
## Mon Jul 25 15:02:46 CEST 2022 - The aliscore output appears to be empty. Check results for BUSCO: 834694at2759
## Mon Jul 25 15:02:46 CEST 2022 - The aliscore output appears to be empty. Check results for BUSCO: 981902at2759
## Mon Jul 25 15:02:46 CEST 2022 - The aliscore output file does not exist. Check results for BUSCO: 1309031at2759
## Mon Jul 25 15:02:46 CEST 2022 - The aliscore output appears to be empty. Check results for BUSCO: 1309031at2759
## Mon Jul 25 15:02:46 CEST 2022 - The aliscore output file does not exist. Check results for BUSCO: 1287094at2759
## Mon Jul 25 15:02:46 CEST 2022 - The aliscore output file does not exist. Check results for BUSCO: 1194691at2759
## Mon Jul 25 15:02:46 CEST 2022 - The aliscore output appears to be empty. Check results for BUSCO: 1287094at2759
## Mon Jul 25 15:02:46 CEST 2022 - The aliscore output appears to be empty. Check results for BUSCO: 1194691at2759
## Mon Jul 25 15:02:46 CEST 2022 - The aliscore output file does not exist. Check results for BUSCO: 1003258at2759
## Mon Jul 25 15:02:47 CEST 2022 - The aliscore output file does not exist. Check results for BUSCO: 1445102at2759
## Mon Jul 25 15:02:46 CEST 2022 - The aliscore output appears to be empty. Check results for BUSCO: 1003258at2759
## Mon Jul 25 15:02:47 CEST 2022 - The aliscore output appears to be empty. Check results for BUSCO: 1445102at2759
## Mon Jul 25 15:02:47 CEST 2022 - The aliscore output file does not exist. Check results for BUSCO: 817008at2759
## Mon Jul 25 15:02:47 CEST 2022 - The aliscore output file does not exist. Check results for BUSCO: 975158at2759
## Mon Jul 25 15:02:47 CEST 2022 - The aliscore output appears to be empty. Check results for BUSCO: 817008at2759
## Mon Jul 25 15:02:47 CEST 2022 - The aliscore output file does not exist. Check results for BUSCO: 1309031at2759
## Mon Jul 25 15:02:47 CEST 2022 - The aliscore output appears to be empty. Check results for BUSCO: 975158at2759
## Mon Jul 25 15:02:47 CEST 2022 - The aliscore output appears to be empty. Check results for BUSCO: 1309031at2759
## Mon Jul 25 15:02:47 CEST 2022 - The aliscore output file does not exist. Check results for BUSCO: 1287401at2759
## Mon Jul 25 15:02:47 CEST 2022 - The aliscore output appears to be empty. Check results for BUSCO: 1287401at2759
## Mon Jul 25 15:02:47 CEST 2022 - The aliscore output file does not exist. Check results for BUSCO: 687505at2759
## Mon Jul 25 15:02:47 CEST 2022 - The aliscore output file does not exist. Check results for BUSCO: 290630at2759
## Mon Jul 25 15:02:47 CEST 2022 - The aliscore output file does not exist. Check results for BUSCO: 1490022at2759
## Mon Jul 25 15:02:47 CEST 2022 - The aliscore output appears to be empty. Check results for BUSCO: 687505at2759
## Mon Jul 25 15:02:47 CEST 2022 - The aliscore output appears to be empty. Check results for BUSCO: 290630at2759
## Mon Jul 25 15:02:47 CEST 2022 - The aliscore output appears to be empty. Check results for BUSCO: 1490022at2759
## Mon Jul 25 15:02:47 CEST 2022 - The aliscore output file does not exist. Check results for BUSCO: 1323575at2759
## Mon Jul 25 15:02:47 CEST 2022 - The aliscore output appears to be empty. Check results for BUSCO: 1323575at2759
## Mon Jul 25 15:02:47 CEST 2022 - The aliscore output file does not exist. Check results for BUSCO: 97116at2759
## Mon Jul 25 15:02:47 CEST 2022 - The aliscore output appears to be empty. Check results for BUSCO: 97116at2759
## Mon Jul 25 15:02:47 CEST 2022 - The aliscore output file does not exist. Check results for BUSCO: 1275837at2759
## Mon Jul 25 15:02:47 CEST 2022 - The aliscore output appears to be empty. Check results for BUSCO: 1275837at2759
## Mon Jul 25 15:02:47 CEST 2022 - The aliscore output file does not exist. Check results for BUSCO: 341721at2759
## Mon Jul 25 15:02:47 CEST 2022 - The aliscore output file does not exist. Check results for BUSCO: 751335at2759
## Mon Jul 25 15:02:47 CEST 2022 - The aliscore output appears to be empty. Check results for BUSCO: 341721at2759
## Mon Jul 25 15:02:47 CEST 2022 - The aliscore output appears to be empty. Check results for BUSCO: 751335at2759
## Mon Jul 25 15:02:47 CEST 2022 - The aliscore output file does not exist. Check results for BUSCO: 418107at2759
## Mon Jul 25 15:02:47 CEST 2022 - The aliscore output file does not exist. Check results for BUSCO: 655400at2759
## Mon Jul 25 15:02:47 CEST 2022 - The aliscore output appears to be empty. Check results for BUSCO: 418107at2759
## Mon Jul 25 15:02:47 CEST 2022 - The aliscore output file does not exist. Check results for BUSCO: 981902at2759
## Mon Jul 25 15:02:47 CEST 2022 - The aliscore output appears to be empty. Check results for BUSCO: 655400at2759
## Mon Jul 25 15:02:47 CEST 2022 - The aliscore output appears to be empty. Check results for BUSCO: 981902at2759
## Mon Jul 25 15:02:48 CEST 2022 - The aliscore output file does not exist. Check results for BUSCO: 768809at2759
## Mon Jul 25 15:02:48 CEST 2022 - The aliscore output file does not exist. Check results for BUSCO: 604979at2759
## Mon Jul 25 15:02:48 CEST 2022 - The aliscore output appears to be empty. Check results for BUSCO: 768809at2759
## Mon Jul 25 15:02:48 CEST 2022 - The aliscore output appears to be empty. Check results for BUSCO: 604979at2759
## Mon Jul 25 15:02:48 CEST 2022 - The aliscore output file does not exist. Check results for BUSCO: 917326at2759
## Mon Jul 25 15:02:48 CEST 2022 - The aliscore output appears to be empty. Check results for BUSCO: 917326at2759
## Mon Jul 25 15:02:48 CEST 2022 - The aliscore output file does not exist. Check results for BUSCO: 1322299at2759
## Mon Jul 25 15:02:48 CEST 2022 - The aliscore output file does not exist. Check results for BUSCO: 375960at2759
## Mon Jul 25 15:02:48 CEST 2022 - The aliscore output appears to be empty. Check results for BUSCO: 1322299at2759
## Mon Jul 25 15:02:48 CEST 2022 - The aliscore output appears to be empty. Check results for BUSCO: 375960at2759
## Mon Jul 25 15:02:48 CEST 2022 - The aliscore output file does not exist. Check results for BUSCO: 1194797at2759
## Mon Jul 25 15:02:48 CEST 2022 - The aliscore output appears to be empty. Check results for BUSCO: 1194797at2759
## Mon Jul 25 15:02:48 CEST 2022 - The aliscore output file does not exist. Check results for BUSCO: 383503at2759
## Mon Jul 25 15:02:48 CEST 2022 - The aliscore output appears to be empty. Check results for BUSCO: 383503at2759
## Mon Jul 25 15:02:48 CEST 2022 - The aliscore output file does not exist. Check results for BUSCO: 1322642at2759
## Mon Jul 25 15:02:48 CEST 2022 - The aliscore output file does not exist. Check results for BUSCO: 1314980at2759
## Mon Jul 25 15:02:48 CEST 2022 - The aliscore output appears to be empty. Check results for BUSCO: 1314980at2759
## Mon Jul 25 15:02:48 CEST 2022 - The aliscore output appears to be empty. Check results for BUSCO: 1322642at2759
## Mon Jul 25 15:02:49 CEST 2022 - The aliscore output file does not exist. Check results for BUSCO: 673132at2759
## Mon Jul 25 15:02:49 CEST 2022 - The aliscore output appears to be empty. Check results for BUSCO: 673132at2759
## Mon Jul 25 15:02:49 CEST 2022 - The aliscore output file does not exist. Check results for BUSCO: 390348at2759
## Mon Jul 25 15:02:49 CEST 2022 - The aliscore output appears to be empty. Check results for BUSCO: 390348at2759
## Mon Jul 25 15:02:50 CEST 2022 - The aliscore output file does not exist. Check results for BUSCO: 1258856at2759
## Mon Jul 25 15:02:50 CEST 2022 - The aliscore output file does not exist. Check results for BUSCO: 325552at2759
## Mon Jul 25 15:02:50 CEST 2022 - The aliscore output appears to be empty. Check results for BUSCO: 1258856at2759
## Mon Jul 25 15:02:50 CEST 2022 - The aliscore output appears to be empty. Check results for BUSCO: 325552at2759
## Mon Jul 25 15:02:50 CEST 2022 - The aliscore output file does not exist. Check results for BUSCO: 768809at2759
## Mon Jul 25 15:02:50 CEST 2022 - The aliscore output file does not exist. Check results for BUSCO: 160593at2759
## Mon Jul 25 15:02:50 CEST 2022 - The aliscore output file does not exist. Check results for BUSCO: 836599at2759
## Mon Jul 25 15:02:50 CEST 2022 - The aliscore output appears to be empty. Check results for BUSCO: 160593at2759
## Mon Jul 25 15:02:50 CEST 2022 - The aliscore output appears to be empty. Check results for BUSCO: 768809at2759
## Mon Jul 25 15:02:50 CEST 2022 - The aliscore output appears to be empty. Check results for BUSCO: 836599at2759
## Mon Jul 25 15:02:50 CEST 2022 - The aliscore output file does not exist. Check results for BUSCO: 383503at2759
## Mon Jul 25 15:02:50 CEST 2022 - The aliscore output appears to be empty. Check results for BUSCO: 383503at2759
## Mon Jul 25 15:02:50 CEST 2022 - The aliscore output file does not exist. Check results for BUSCO: 834694at2759
## Mon Jul 25 15:02:51 CEST 2022 - The aliscore output appears to be empty. Check results for BUSCO: 834694at2759
## Mon Jul 25 15:02:51 CEST 2022 - The aliscore output file does not exist. Check results for BUSCO: 777920at2759
## Mon Jul 25 15:02:51 CEST 2022 - The aliscore output appears to be empty. Check results for BUSCO: 777920at2759
## Mon Jul 25 15:02:51 CEST 2022 - The aliscore output file does not exist. Check results for BUSCO: 973442at2759
## Mon Jul 25 15:02:51 CEST 2022 - The aliscore output appears to be empty. Check results for BUSCO: 973442at2759
## Mon Jul 25 15:02:51 CEST 2022 - The aliscore output file does not exist. Check results for BUSCO: 722805at2759
## Mon Jul 25 15:02:51 CEST 2022 - The aliscore output file does not exist. Check results for BUSCO: 777920at2759
## Mon Jul 25 15:02:51 CEST 2022 - The aliscore output appears to be empty. Check results for BUSCO: 722805at2759
## Mon Jul 25 15:02:51 CEST 2022 - The aliscore output appears to be empty. Check results for BUSCO: 777920at2759
## Mon Jul 25 15:02:51 CEST 2022 - The aliscore output file does not exist. Check results for BUSCO: 293315at2759
## Mon Jul 25 15:02:51 CEST 2022 - The aliscore output appears to be empty. Check results for BUSCO: 293315at2759
## Mon Jul 25 15:02:52 CEST 2022 - The aliscore output file does not exist. Check results for BUSCO: 179362at2759
## Mon Jul 25 15:02:52 CEST 2022 - The aliscore output appears to be empty. Check results for BUSCO: 179362at2759
## Mon Jul 25 15:02:52 CEST 2022 - The aliscore output file does not exist. Check results for BUSCO: 1258856at2759
## Mon Jul 25 15:02:52 CEST 2022 - The aliscore output appears to be empty. Check results for BUSCO: 1258856at2759
## Mon Jul 25 15:02:52 CEST 2022 - The aliscore output file does not exist. Check results for BUSCO: 179362at2759
## Mon Jul 25 15:02:52 CEST 2022 - The aliscore output appears to be empty. Check results for BUSCO: 179362at2759
## Mon Jul 25 15:02:52 CEST 2022 - The aliscore output file does not exist. Check results for BUSCO: 973442at2759
## Mon Jul 25 15:02:52 CEST 2022 - The aliscore output appears to be empty. Check results for BUSCO: 973442at2759
## Mon Jul 25 15:02:53 CEST 2022 - The aliscore output file does not exist. Check results for BUSCO: 1314980at2759
## Mon Jul 25 15:02:53 CEST 2022 - The aliscore output appears to be empty. Check results for BUSCO: 1314980at2759
## Mon Jul 25 15:02:53 CEST 2022 - The aliscore output file does not exist. Check results for BUSCO: 1312453at2759
## Mon Jul 25 15:02:53 CEST 2022 - The aliscore output appears to be empty. Check results for BUSCO: 1312453at2759
## Mon Jul 25 15:02:56 CEST 2022 - The aliscore output file does not exist. Check results for BUSCO: 674160at2759
## Mon Jul 25 15:02:56 CEST 2022 - The aliscore output appears to be empty. Check results for BUSCO: 674160at2759
## Mon Jul 25 15:03:01 CEST 2022 - The aliscore output file does not exist. Check results for BUSCO: 674160at2759
## Mon Jul 25 15:03:01 CEST 2022 - The aliscore output appears to be empty. Check results for BUSCO: 674160at2759
## Mon Jul 25 15:03:02 CEST 2022 - The aliscore output file does not exist. Check results for BUSCO: 156083at2759
## Mon Jul 25 15:03:02 CEST 2022 - The aliscore output appears to be empty. Check results for BUSCO: 156083at2759
## Mon Jul 25 15:03:10 CEST 2022 - The aliscore output file does not exist. Check results for BUSCO: 865202at2759
## Mon Jul 25 15:03:10 CEST 2022 - The aliscore output appears to be empty. Check results for BUSCO: 865202at2759
## Mon Jul 25 15:03:11 CEST 2022 - The aliscore output file does not exist. Check results for BUSCO: 1220881at2759
## Mon Jul 25 15:03:11 CEST 2022 - The aliscore output appears to be empty. Check results for BUSCO: 1220881at2759
## Mon Jul 25 15:03:11 CEST 2022 - The aliscore output file does not exist. Check results for BUSCO: 1645187at2759
## Mon Jul 25 15:03:11 CEST 2022 - The aliscore output appears to be empty. Check results for BUSCO: 1645187at2759
## Mon Jul 25 15:03:12 CEST 2022 - The aliscore output file does not exist. Check results for BUSCO: 464990at2759
## Mon Jul 25 15:03:12 CEST 2022 - The aliscore output appears to be empty. Check results for BUSCO: 464990at2759
## Mon Jul 25 15:03:12 CEST 2022 - The aliscore output file does not exist. Check results for BUSCO: 923657at2759
## Mon Jul 25 15:03:12 CEST 2022 - The aliscore output appears to be empty. Check results for BUSCO: 923657at2759
## Mon Jul 25 15:03:12 CEST 2022 - The aliscore output file does not exist. Check results for BUSCO: 721605at2759
## Mon Jul 25 15:03:12 CEST 2022 - The aliscore output appears to be empty. Check results for BUSCO: 721605at2759
## Mon Jul 25 15:03:12 CEST 2022 - The aliscore output file does not exist. Check results for BUSCO: 1526152at2759
## Mon Jul 25 15:03:12 CEST 2022 - The aliscore output file does not exist. Check results for BUSCO: 1459797at2759
## Mon Jul 25 15:03:12 CEST 2022 - The aliscore output file does not exist. Check results for BUSCO: 1575179at2759
## Mon Jul 25 15:03:13 CEST 2022 - The aliscore output appears to be empty. Check results for BUSCO: 1459797at2759
## Mon Jul 25 15:03:13 CEST 2022 - The aliscore output appears to be empty. Check results for BUSCO: 1575179at2759
## Mon Jul 25 15:03:13 CEST 2022 - The aliscore output file does not exist. Check results for BUSCO: 1018517at2759
## Mon Jul 25 15:03:13 CEST 2022 - The aliscore output appears to be empty. Check results for BUSCO: 1018517at2759
## Mon Jul 25 15:03:13 CEST 2022 - The aliscore output file does not exist. Check results for BUSCO: 866359at2759
## Mon Jul 25 15:03:13 CEST 2022 - The aliscore output appears to be empty. Check results for BUSCO: 866359at2759
## Mon Jul 25 15:03:13 CEST 2022 - The aliscore output file does not exist. Check results for BUSCO: 1358374at2759
## Mon Jul 25 15:03:13 CEST 2022 - The aliscore output appears to be empty. Check results for BUSCO: 1358374at2759
## Mon Jul 25 15:03:13 CEST 2022 - The aliscore output file does not exist. Check results for BUSCO: 1405073at2759
## Mon Jul 25 15:03:13 CEST 2022 - The aliscore output appears to be empty. Check results for BUSCO: 1405073at2759
## Mon Jul 25 15:03:13 CEST 2022 - The aliscore output file does not exist. Check results for BUSCO: 1079827at2759
## Mon Jul 25 15:03:13 CEST 2022 - The aliscore output file does not exist. Check results for BUSCO: 1194691at2759
## Mon Jul 25 15:03:13 CEST 2022 - The aliscore output appears to be empty. Check results for BUSCO: 1079827at2759
## Mon Jul 25 15:03:13 CEST 2022 - The aliscore output file does not exist. Check results for BUSCO: 1405073at2759
## Mon Jul 25 15:03:13 CEST 2022 - The aliscore output appears to be empty. Check results for BUSCO: 1194691at2759
## Mon Jul 25 15:03:13 CEST 2022 - The aliscore output appears to be empty. Check results for BUSCO: 1405073at2759
## Mon Jul 25 15:03:13 CEST 2022 - The aliscore output file does not exist. Check results for BUSCO: 1623701at2759
## Mon Jul 25 15:03:13 CEST 2022 - The aliscore output file does not exist. Check results for BUSCO: 865202at2759
## Mon Jul 25 15:03:13 CEST 2022 - The aliscore output appears to be empty. Check results for BUSCO: 1623701at2759
## Mon Jul 25 15:03:13 CEST 2022 - The aliscore output appears to be empty. Check results for BUSCO: 865202at2759
## Mon Jul 25 15:03:13 CEST 2022 - The aliscore output file does not exist. Check results for BUSCO: 679771at2759
## Mon Jul 25 15:03:13 CEST 2022 - The aliscore output file does not exist. Check results for BUSCO: 1620056at2759
## Mon Jul 25 15:03:14 CEST 2022 - The aliscore output appears to be empty. Check results for BUSCO: 679771at2759
## Mon Jul 25 15:03:14 CEST 2022 - The aliscore output appears to be empty. Check results for BUSCO: 1620056at2759
## Mon Jul 25 15:03:14 CEST 2022 - The aliscore output file does not exist. Check results for BUSCO: 1157302at2759
## Mon Jul 25 15:03:14 CEST 2022 - The aliscore output file does not exist. Check results for BUSCO: 1228942at2759
## Mon Jul 25 15:03:14 CEST 2022 - The aliscore output file does not exist. Check results for BUSCO: 1178688at2759
## Mon Jul 25 15:03:14 CEST 2022 - The aliscore output appears to be empty. Check results for BUSCO: 1157302at2759
## Mon Jul 25 15:03:14 CEST 2022 - The aliscore output appears to be empty. Check results for BUSCO: 1228942at2759
## Mon Jul 25 15:03:14 CEST 2022 - The aliscore output file does not exist. Check results for BUSCO: 1445102at2759
## Mon Jul 25 15:03:14 CEST 2022 - The aliscore output appears to be empty. Check results for BUSCO: 1178688at2759
## Mon Jul 25 15:03:14 CEST 2022 - The aliscore output file does not exist. Check results for BUSCO: 1474000at2759
## Mon Jul 25 15:03:14 CEST 2022 - The aliscore output file does not exist. Check results for BUSCO: 1025450at2759
## Mon Jul 25 15:03:14 CEST 2022 - The aliscore output appears to be empty. Check results for BUSCO: 1445102at2759
## Mon Jul 25 15:03:14 CEST 2022 - The aliscore output appears to be empty. Check results for BUSCO: 1474000at2759
## Mon Jul 25 15:03:14 CEST 2022 - The aliscore output appears to be empty. Check results for BUSCO: 1025450at2759
## Mon Jul 25 15:03:14 CEST 2022 - The aliscore output file does not exist. Check results for BUSCO: 1076134at2759
## Mon Jul 25 15:03:14 CEST 2022 - The aliscore output file does not exist. Check results for BUSCO: 1200489at2759
## Mon Jul 25 15:03:14 CEST 2022 - The aliscore output file does not exist. Check results for BUSCO: 866359at2759
## Mon Jul 25 15:03:14 CEST 2022 - The aliscore output appears to be empty. Check results for BUSCO: 1076134at2759
## Mon Jul 25 15:03:14 CEST 2022 - The aliscore output appears to be empty. Check results for BUSCO: 1200489at2759
## Mon Jul 25 15:03:14 CEST 2022 - The aliscore output appears to be empty. Check results for BUSCO: 866359at2759
## Mon Jul 25 15:03:14 CEST 2022 - The aliscore output file does not exist. Check results for BUSCO: 1161199at2759
## Mon Jul 25 15:03:14 CEST 2022 - The aliscore output appears to be empty. Check results for BUSCO: 1161199at2759
## Mon Jul 25 15:03:14 CEST 2022 - The aliscore output file does not exist. Check results for BUSCO: 1322642at2759
## Mon Jul 25 15:03:14 CEST 2022 - The aliscore output file does not exist. Check results for BUSCO: 1322299at2759
## Mon Jul 25 15:03:14 CEST 2022 - The aliscore output appears to be empty. Check results for BUSCO: 1322642at2759
## Mon Jul 25 15:03:14 CEST 2022 - The aliscore output appears to be empty. Check results for BUSCO: 1322299at2759
## Mon Jul 25 15:03:14 CEST 2022 - The aliscore output file does not exist. Check results for BUSCO: 1645187at2759
## Mon Jul 25 15:03:14 CEST 2022 - The aliscore output appears to be empty. Check results for BUSCO: 1645187at2759
## Mon Jul 25 15:03:14 CEST 2022 - The aliscore output file does not exist. Check results for BUSCO: 1513531at2759
## Mon Jul 25 15:03:14 CEST 2022 - The aliscore output appears to be empty. Check results for BUSCO: 1513531at2759
## Mon Jul 25 15:03:14 CEST 2022 - The aliscore output file does not exist. Check results for BUSCO: 687505at2759
## Mon Jul 25 15:03:14 CEST 2022 - The aliscore output appears to be empty. Check results for BUSCO: 687505at2759
## Mon Jul 25 15:03:14 CEST 2022 - The aliscore output file does not exist. Check results for BUSCO: 1266231at2759
## Mon Jul 25 15:03:14 CEST 2022 - The aliscore output appears to be empty. Check results for BUSCO: 1266231at2759
## Mon Jul 25 15:03:14 CEST 2022 - The aliscore output file does not exist. Check results for BUSCO: 464990at2759
## Mon Jul 25 15:03:14 CEST 2022 - The aliscore output appears to be empty. Check results for BUSCO: 464990at2759
## Mon Jul 25 15:03:14 CEST 2022 - The aliscore output file does not exist. Check results for BUSCO: 817008at2759
## Mon Jul 25 15:03:14 CEST 2022 - The aliscore output file does not exist. Check results for BUSCO: 917326at2759
## Mon Jul 25 15:03:14 CEST 2022 - The aliscore output appears to be empty. Check results for BUSCO: 817008at2759
## Mon Jul 25 15:03:14 CEST 2022 - The aliscore output file does not exist. Check results for BUSCO: 1474000at2759
## Mon Jul 25 15:03:14 CEST 2022 - The aliscore output appears to be empty. Check results for BUSCO: 917326at2759
## Mon Jul 25 15:03:14 CEST 2022 - The aliscore output file does not exist. Check results for BUSCO: 974865at2759
## Mon Jul 25 15:03:14 CEST 2022 - The aliscore output appears to be empty. Check results for BUSCO: 1474000at2759
## Mon Jul 25 15:03:14 CEST 2022 - The aliscore output appears to be empty. Check results for BUSCO: 974865at2759
## Mon Jul 25 15:03:15 CEST 2022 - The aliscore output file does not exist. Check results for BUSCO: 1275837at2759
## Mon Jul 25 15:03:15 CEST 2022 - The aliscore output appears to be empty. Check results for BUSCO: 1275837at2759
## Mon Jul 25 15:03:15 CEST 2022 - The aliscore output file does not exist. Check results for BUSCO: 1076134at2759
## Mon Jul 25 15:03:15 CEST 2022 - The aliscore output appears to be empty. Check results for BUSCO: 1076134at2759
## Mon Jul 25 15:03:15 CEST 2022 - The aliscore output file does not exist. Check results for BUSCO: 1490022at2759
## Mon Jul 25 15:03:15 CEST 2022 - The aliscore output appears to be empty. Check results for BUSCO: 1490022at2759
## Mon Jul 25 15:03:15 CEST 2022 - The aliscore output file does not exist. Check results for BUSCO: 513979at2759
## Mon Jul 25 15:03:15 CEST 2022 - The aliscore output file does not exist. Check results for BUSCO: 1107630at2759
## Mon Jul 25 15:03:15 CEST 2022 - The aliscore output appears to be empty. Check results for BUSCO: 513979at2759
## Mon Jul 25 15:03:15 CEST 2022 - The aliscore output appears to be empty. Check results for BUSCO: 1107630at2759
## Mon Jul 25 15:03:15 CEST 2022 - The aliscore output file does not exist. Check results for BUSCO: 721605at2759
## Mon Jul 25 15:03:15 CEST 2022 - The aliscore output appears to be empty. Check results for BUSCO: 721605at2759
## Mon Jul 25 15:03:15 CEST 2022 - The aliscore output file does not exist. Check results for BUSCO: 751335at2759
## Mon Jul 25 15:03:15 CEST 2022 - The aliscore output file does not exist. Check results for BUSCO: 1287401at2759
## Mon Jul 25 15:03:15 CEST 2022 - The aliscore output appears to be empty. Check results for BUSCO: 751335at2759
## Mon Jul 25 15:03:15 CEST 2022 - The aliscore output appears to be empty. Check results for BUSCO: 1287401at2759
## Mon Jul 25 15:03:15 CEST 2022 - The aliscore output file does not exist. Check results for BUSCO: 1003258at2759
## Mon Jul 25 15:03:15 CEST 2022 - The aliscore output appears to be empty. Check results for BUSCO: 1003258at2759
## Mon Jul 25 15:03:15 CEST 2022 - The aliscore output file does not exist. Check results for BUSCO: 325552at2759
## Mon Jul 25 15:03:16 CEST 2022 - The aliscore output file does not exist. Check results for BUSCO: 1423847at2759
## Mon Jul 25 15:03:16 CEST 2022 - The aliscore output appears to be empty. Check results for BUSCO: 325552at2759
## Mon Jul 25 15:03:16 CEST 2022 - The aliscore output file does not exist. Check results for BUSCO: 1182451at2759
## Mon Jul 25 15:03:16 CEST 2022 - The aliscore output file does not exist. Check results for BUSCO: 1423847at2759
## Mon Jul 25 15:03:16 CEST 2022 - The aliscore output appears to be empty. Check results for BUSCO: 1423847at2759
## Mon Jul 25 15:03:16 CEST 2022 - The aliscore output appears to be empty. Check results for BUSCO: 1423847at2759
## Mon Jul 25 15:03:16 CEST 2022 - The aliscore output file does not exist. Check results for BUSCO: 1266231at2759
## Mon Jul 25 15:03:16 CEST 2022 - The aliscore output file does not exist. Check results for BUSCO: 937686at2759
## Mon Jul 25 15:03:16 CEST 2022 - The aliscore output appears to be empty. Check results for BUSCO: 1182451at2759
## Mon Jul 25 15:03:16 CEST 2022 - The aliscore output appears to be empty. Check results for BUSCO: 1266231at2759
## Mon Jul 25 15:03:16 CEST 2022 - The aliscore output appears to be empty. Check results for BUSCO: 937686at2759
## Mon Jul 25 15:03:16 CEST 2022 - The aliscore output file does not exist. Check results for BUSCO: 290630at2759
## Mon Jul 25 15:03:16 CEST 2022 - The aliscore output appears to be empty. Check results for BUSCO: 290630at2759
## Mon Jul 25 15:03:16 CEST 2022 - The aliscore output file does not exist. Check results for BUSCO: 426305at2759
## Mon Jul 25 15:03:16 CEST 2022 - The aliscore output file does not exist. Check results for BUSCO: 1001705at2759
## Mon Jul 25 15:03:16 CEST 2022 - The aliscore output file does not exist. Check results for BUSCO: 814241at2759
## Mon Jul 25 15:03:16 CEST 2022 - The aliscore output appears to be empty. Check results for BUSCO: 426305at2759
## Mon Jul 25 15:03:16 CEST 2022 - The aliscore output appears to be empty. Check results for BUSCO: 1001705at2759
## Mon Jul 25 15:03:16 CEST 2022 - The aliscore output file does not exist. Check results for BUSCO: 418107at2759
## Mon Jul 25 15:03:16 CEST 2022 - The aliscore output appears to be empty. Check results for BUSCO: 814241at2759
## Mon Jul 25 15:03:16 CEST 2022 - The aliscore output appears to be empty. Check results for BUSCO: 418107at2759
## Mon Jul 25 15:03:16 CEST 2022 - The aliscore output file does not exist. Check results for BUSCO: 1018517at2759
## Mon Jul 25 15:03:16 CEST 2022 - The aliscore output file does not exist. Check results for BUSCO: 176625at2759
## Mon Jul 25 15:03:16 CEST 2022 - The aliscore output appears to be empty. Check results for BUSCO: 1018517at2759
## Mon Jul 25 15:03:16 CEST 2022 - The aliscore output appears to be empty. Check results for BUSCO: 176625at2759
## Mon Jul 25 15:03:16 CEST 2022 - The aliscore output file does not exist. Check results for BUSCO: 759498at2759
## Mon Jul 25 15:03:16 CEST 2022 - The aliscore output appears to be empty. Check results for BUSCO: 759498at2759
## Mon Jul 25 15:03:16 CEST 2022 - The aliscore output file does not exist. Check results for BUSCO: 673132at2759
## Mon Jul 25 15:03:16 CEST 2022 - The aliscore output file does not exist. Check results for BUSCO: 1222562at2759
## Mon Jul 25 15:03:16 CEST 2022 - The aliscore output file does not exist. Check results for BUSCO: 293315at2759
## Mon Jul 25 15:03:16 CEST 2022 - The aliscore output appears to be empty. Check results for BUSCO: 673132at2759
## Mon Jul 25 15:03:16 CEST 2022 - The aliscore output file does not exist. Check results for BUSCO: 664730at2759
## Mon Jul 25 15:03:16 CEST 2022 - The aliscore output appears to be empty. Check results for BUSCO: 1222562at2759
## Mon Jul 25 15:03:16 CEST 2022 - The aliscore output appears to be empty. Check results for BUSCO: 293315at2759
## Mon Jul 25 15:03:16 CEST 2022 - The aliscore output file does not exist. Check results for BUSCO: 671846at2759
## Mon Jul 25 15:03:16 CEST 2022 - The aliscore output appears to be empty. Check results for BUSCO: 664730at2759
## Mon Jul 25 15:03:16 CEST 2022 - The aliscore output appears to be empty. Check results for BUSCO: 671846at2759
## Mon Jul 25 15:03:16 CEST 2022 - The aliscore output file does not exist. Check results for BUSCO: 1426075at2759
## Mon Jul 25 15:03:16 CEST 2022 - The aliscore output file does not exist. Check results for BUSCO: 345441at2759
## Mon Jul 25 15:03:16 CEST 2022 - The aliscore output appears to be empty. Check results for BUSCO: 1426075at2759
## Mon Jul 25 15:03:16 CEST 2022 - The aliscore output appears to be empty. Check results for BUSCO: 345441at2759
## Mon Jul 25 15:03:16 CEST 2022 - The aliscore output file does not exist. Check results for BUSCO: 331411at2759
## Mon Jul 25 15:03:17 CEST 2022 - The aliscore output appears to be empty. Check results for BUSCO: 331411at2759
## Mon Jul 25 15:03:17 CEST 2022 - The aliscore output file does not exist. Check results for BUSCO: 1193442at2759
## Mon Jul 25 15:03:17 CEST 2022 - The aliscore output appears to be empty. Check results for BUSCO: 1193442at2759
## Mon Jul 25 15:03:17 CEST 2022 - The aliscore output file does not exist. Check results for BUSCO: 390348at2759
## Mon Jul 25 15:03:17 CEST 2022 - The aliscore output appears to be empty. Check results for BUSCO: 390348at2759
## Mon Jul 25 15:03:17 CEST 2022 - The aliscore output file does not exist. Check results for BUSCO: 1248958at2759
## Mon Jul 25 15:03:17 CEST 2022 - The aliscore output appears to be empty. Check results for BUSCO: 1248958at2759
## Mon Jul 25 15:03:17 CEST 2022 - The aliscore output file does not exist. Check results for BUSCO: 918816at2759
## Mon Jul 25 15:03:17 CEST 2022 - The aliscore output appears to be empty. Check results for BUSCO: 918816at2759
## Mon Jul 25 15:03:17 CEST 2022 - The aliscore output file does not exist. Check results for BUSCO: 570797at2759
## Mon Jul 25 15:03:17 CEST 2022 - The aliscore output appears to be empty. Check results for BUSCO: 570797at2759
## Mon Jul 25 15:03:17 CEST 2022 - The aliscore output file does not exist. Check results for BUSCO: 341721at2759
## Mon Jul 25 15:03:17 CEST 2022 - The aliscore output appears to be empty. Check results for BUSCO: 341721at2759
## Mon Jul 25 15:03:17 CEST 2022 - The aliscore output file does not exist. Check results for BUSCO: 392369at2759
## Mon Jul 25 15:03:17 CEST 2022 - The aliscore output appears to be empty. Check results for BUSCO: 392369at2759
## Mon Jul 25 15:03:17 CEST 2022 - The aliscore output file does not exist. Check results for BUSCO: 513979at2759
## Mon Jul 25 15:03:17 CEST 2022 - The aliscore output file does not exist. Check results for BUSCO: 722805at2759
## Mon Jul 25 15:03:17 CEST 2022 - The aliscore output appears to be empty. Check results for BUSCO: 513979at2759
## Mon Jul 25 15:03:17 CEST 2022 - The aliscore output appears to be empty. Check results for BUSCO: 722805at2759
## Mon Jul 25 15:03:17 CEST 2022 - The aliscore output file does not exist. Check results for BUSCO: 814241at2759
## Mon Jul 25 15:03:17 CEST 2022 - The aliscore output appears to be empty. Check results for BUSCO: 814241at2759
## Mon Jul 25 15:03:18 CEST 2022 - The aliscore output file does not exist. Check results for BUSCO: 924753at2759
## Mon Jul 25 15:03:18 CEST 2022 - The aliscore output appears to be empty. Check results for BUSCO: 924753at2759
## Mon Jul 25 15:03:18 CEST 2022 - The aliscore output file does not exist. Check results for BUSCO: 1025450at2759
## Mon Jul 25 15:03:18 CEST 2022 - The aliscore output appears to be empty. Check results for BUSCO: 1025450at2759
## Mon Jul 25 15:03:18 CEST 2022 - The aliscore output file does not exist. Check results for BUSCO: 1115196at2759
## Mon Jul 25 15:03:18 CEST 2022 - The aliscore output appears to be empty. Check results for BUSCO: 1115196at2759
## Mon Jul 25 15:03:18 CEST 2022 - The aliscore output file does not exist. Check results for BUSCO: 625387at2759
## Mon Jul 25 15:03:18 CEST 2022 - The aliscore output file does not exist. Check results for BUSCO: 1087488at2759
## Mon Jul 25 15:03:18 CEST 2022 - The aliscore output file does not exist. Check results for BUSCO: 671846at2759
## Mon Jul 25 15:03:19 CEST 2022 - The aliscore output file does not exist. Check results for BUSCO: 759498at2759
## Mon Jul 25 15:03:19 CEST 2022 - The aliscore output appears to be empty. Check results for BUSCO: 1087488at2759
## Mon Jul 25 15:03:19 CEST 2022 - The aliscore output appears to be empty. Check results for BUSCO: 625387at2759
## Mon Jul 25 15:03:19 CEST 2022 - The aliscore output appears to be empty. Check results for BUSCO: 671846at2759
## Mon Jul 25 15:03:19 CEST 2022 - The aliscore output appears to be empty. Check results for BUSCO: 759498at2759
## Mon Jul 25 15:03:19 CEST 2022 - The aliscore output file does not exist. Check results for BUSCO: 604979at2759
## Mon Jul 25 15:03:19 CEST 2022 - The aliscore output appears to be empty. Check results for BUSCO: 604979at2759
## Mon Jul 25 15:03:19 CEST 2022 - The aliscore output file does not exist. Check results for BUSCO: 1248958at2759
## Mon Jul 25 15:03:19 CEST 2022 - The aliscore output appears to be empty. Check results for BUSCO: 1248958at2759
## Mon Jul 25 15:03:20 CEST 2022 - The aliscore output file does not exist. Check results for BUSCO: 160593at2759
## Mon Jul 25 15:03:20 CEST 2022 - The aliscore output appears to be empty. Check results for BUSCO: 160593at2759
## Mon Jul 25 15:03:20 CEST 2022 - The aliscore output file does not exist. Check results for BUSCO: 1379600at2759
## Mon Jul 25 15:03:20 CEST 2022 - The aliscore output appears to be empty. Check results for BUSCO: 1379600at2759
## Mon Jul 25 15:03:22 CEST 2022 - The aliscore output file does not exist. Check results for BUSCO: 1312453at2759
## Mon Jul 25 15:03:22 CEST 2022 - The aliscore output appears to be empty. Check results for BUSCO: 1312453at2759
## Mon Jul 25 15:03:28 CEST 2022 - The aliscore output file does not exist. Check results for BUSCO: 156083at2759
## Mon Jul 25 15:03:28 CEST 2022 - The aliscore output appears to be empty. Check results for BUSCO: 156083at2759
## Mon Jul 25 15:07:38 CEST 2022 - The aliscore output file does not exist. Check results for BUSCO: 1076134at2759
## Mon Jul 25 15:07:38 CEST 2022 - The aliscore output appears to be empty. Check results for BUSCO: 1076134at2759
## Mon Jul 25 15:08:11 CEST 2022 - The aliscore output file does not exist. Check results for BUSCO: 768809at2759
## Mon Jul 25 15:08:11 CEST 2022 - The aliscore output appears to be empty. Check results for BUSCO: 768809at2759
## Mon Jul 25 15:10:42 CEST 2022 - The aliscore output file does not exist. Check results for BUSCO: 974865at2759
## Mon Jul 25 15:10:42 CEST 2022 - The aliscore output appears to be empty. Check results for BUSCO: 974865at2759
## Mon Jul 25 15:10:42 CEST 2022 - The aliscore output file does not exist. Check results for BUSCO: 1623701at2759
## Mon Jul 25 15:10:42 CEST 2022 - The aliscore output appears to be empty. Check results for BUSCO: 1623701at2759
## Mon Jul 25 15:10:42 CEST 2022 - The aliscore output file does not exist. Check results for BUSCO: 1178688at2759
## Mon Jul 25 15:10:42 CEST 2022 - The aliscore output appears to be empty. Check results for BUSCO: 1178688at2759
## Mon Jul 25 15:10:42 CEST 2022 - The aliscore output file does not exist. Check results for BUSCO: 679771at2759
## Mon Jul 25 15:10:42 CEST 2022 - The aliscore output appears to be empty. Check results for BUSCO: 679771at2759
## Mon Jul 25 15:10:42 CEST 2022 - The aliscore output file does not exist. Check results for BUSCO: 1504863at2759
## Mon Jul 25 15:10:42 CEST 2022 - The aliscore output appears to be empty. Check results for BUSCO: 1504863at2759
## Mon Jul 25 15:10:42 CEST 2022 - The aliscore output file does not exist. Check results for BUSCO: 1107630at2759
## Mon Jul 25 15:10:43 CEST 2022 - The aliscore output appears to be empty. Check results for BUSCO: 1107630at2759
## Mon Jul 25 15:10:43 CEST 2022 - The aliscore output file does not exist. Check results for BUSCO: 1645187at2759
## Mon Jul 25 15:10:43 CEST 2022 - The aliscore output appears to be empty. Check results for BUSCO: 1645187at2759
## Mon Jul 25 15:10:43 CEST 2022 - The aliscore output file does not exist. Check results for BUSCO: 1030907at2759
## Mon Jul 25 15:10:43 CEST 2022 - The aliscore output appears to be empty. Check results for BUSCO: 1030907at2759
## Mon Jul 25 15:10:43 CEST 2022 - The aliscore output file does not exist. Check results for BUSCO: 1161199at2759
## Mon Jul 25 15:10:43 CEST 2022 - The aliscore output appears to be empty. Check results for BUSCO: 1161199at2759
## Mon Jul 25 15:10:43 CEST 2022 - The aliscore output file does not exist. Check results for BUSCO: 1264469at2759
## Mon Jul 25 15:10:43 CEST 2022 - The aliscore output appears to be empty. Check results for BUSCO: 1264469at2759
## Mon Jul 25 15:10:43 CEST 2022 - The aliscore output file does not exist. Check results for BUSCO: 1249159at2759
## Mon Jul 25 15:10:43 CEST 2022 - The aliscore output appears to be empty. Check results for BUSCO: 1249159at2759
## Mon Jul 25 15:10:43 CEST 2022 - The aliscore output file does not exist. Check results for BUSCO: 1233814at2759
## Mon Jul 25 15:10:43 CEST 2022 - The aliscore output appears to be empty. Check results for BUSCO: 1233814at2759
## Mon Jul 25 15:10:43 CEST 2022 - The aliscore output file does not exist. Check results for BUSCO: 919955at2759
## Mon Jul 25 15:10:43 CEST 2022 - The aliscore output appears to be empty. Check results for BUSCO: 919955at2759
## Mon Jul 25 15:10:43 CEST 2022 - The aliscore output file does not exist. Check results for BUSCO: 721605at2759
## Mon Jul 25 15:10:43 CEST 2022 - The aliscore output appears to be empty. Check results for BUSCO: 721605at2759
## Mon Jul 25 15:10:44 CEST 2022 - The aliscore output file does not exist. Check results for BUSCO: 1018517at2759
## Mon Jul 25 15:10:44 CEST 2022 - The aliscore output appears to be empty. Check results for BUSCO: 1018517at2759
## Mon Jul 25 15:10:44 CEST 2022 - The aliscore output file does not exist. Check results for BUSCO: 664730at2759
## Mon Jul 25 15:10:44 CEST 2022 - The aliscore output appears to be empty. Check results for BUSCO: 664730at2759
## Mon Jul 25 15:10:44 CEST 2022 - The aliscore output file does not exist. Check results for BUSCO: 814241at2759
## Mon Jul 25 15:10:45 CEST 2022 - The aliscore output file does not exist. Check results for BUSCO: 345441at2759
## Mon Jul 25 15:10:45 CEST 2022 - The aliscore output file does not exist. Check results for BUSCO: 1115196at2759
## Mon Jul 25 15:10:45 CEST 2022 - The aliscore output appears to be empty. Check results for BUSCO: 814241at2759
## Mon Jul 25 15:10:45 CEST 2022 - The aliscore output appears to be empty. Check results for BUSCO: 345441at2759
## Mon Jul 25 15:10:45 CEST 2022 - The aliscore output appears to be empty. Check results for BUSCO: 1115196at2759
## Mon Jul 25 15:10:45 CEST 2022 - The aliscore output file does not exist. Check results for BUSCO: 1025450at2759
## Mon Jul 25 15:10:45 CEST 2022 - The aliscore output appears to be empty. Check results for BUSCO: 1025450at2759
## Mon Jul 25 15:10:45 CEST 2022 - The aliscore output file does not exist. Check results for BUSCO: 375960at2759
## Mon Jul 25 15:10:45 CEST 2022 - The aliscore output appears to be empty. Check results for BUSCO: 375960at2759
## Mon Jul 25 15:10:45 CEST 2022 - The aliscore output file does not exist. Check results for BUSCO: 383503at2759
## Mon Jul 25 15:10:45 CEST 2022 - The aliscore output appears to be empty. Check results for BUSCO: 383503at2759
## Mon Jul 25 15:10:45 CEST 2022 - The aliscore output file does not exist. Check results for BUSCO: 1620056at2759
## Mon Jul 25 15:10:45 CEST 2022 - The aliscore output appears to be empty. Check results for BUSCO: 1620056at2759
## Mon Jul 25 15:10:45 CEST 2022 - The aliscore output file does not exist. Check results for BUSCO: 836599at2759
## Mon Jul 25 15:10:45 CEST 2022 - The aliscore output appears to be empty. Check results for BUSCO: 836599at2759
## Mon Jul 25 15:10:46 CEST 2022 - The aliscore output file does not exist. Check results for BUSCO: 834694at2759
## Mon Jul 25 15:10:46 CEST 2022 - The aliscore output appears to be empty. Check results for BUSCO: 834694at2759
## Mon Jul 25 15:10:46 CEST 2022 - The aliscore output file does not exist. Check results for BUSCO: 981902at2759
## Mon Jul 25 15:10:46 CEST 2022 - The aliscore output file does not exist. Check results for BUSCO: 1087488at2759
## Mon Jul 25 15:10:46 CEST 2022 - The aliscore output appears to be empty. Check results for BUSCO: 981902at2759
## Mon Jul 25 15:10:46 CEST 2022 - The aliscore output appears to be empty. Check results for BUSCO: 1087488at2759
## Mon Jul 25 15:10:46 CEST 2022 - The aliscore output file does not exist. Check results for BUSCO: 917326at2759
## Mon Jul 25 15:10:46 CEST 2022 - The aliscore output appears to be empty. Check results for BUSCO: 917326at2759
## Mon Jul 25 15:10:47 CEST 2022 - The aliscore output file does not exist. Check results for BUSCO: 1003258at2759
## Mon Jul 25 15:10:47 CEST 2022 - The aliscore output appears to be empty. Check results for BUSCO: 1003258at2759
## Mon Jul 25 15:10:47 CEST 2022 - The aliscore output file does not exist. Check results for BUSCO: 1423847at2759
## Mon Jul 25 15:10:47 CEST 2022 - The aliscore output appears to be empty. Check results for BUSCO: 1423847at2759
## Mon Jul 25 15:10:48 CEST 2022 - The aliscore output file does not exist. Check results for BUSCO: 1314980at2759
## Mon Jul 25 15:10:48 CEST 2022 - The aliscore output appears to be empty. Check results for BUSCO: 1314980at2759
## Mon Jul 25 15:10:51 CEST 2022 - The aliscore output file does not exist. Check results for BUSCO: 160593at2759
## Mon Jul 25 15:10:51 CEST 2022 - The aliscore output appears to be empty. Check results for BUSCO: 160593at2759
## Mon Jul 25 16:26:37 CEST 2022 - Pipeline part 2 (align) done.
## Mon Jul 25 16:57:31 CEST 2022 - The aliscore output file does not exist. Check results for BUSCO: 1407446at2759
## Mon Jul 25 16:57:31 CEST 2022 - The aliscore output appears to be empty. Check results for BUSCO: 1407446at2759
## Mon Jul 25 16:57:31 CEST 2022 - The aliscore output file does not exist. Check results for BUSCO: 1197019at2759
## Mon Jul 25 16:57:31 CEST 2022 - The aliscore output appears to be empty. Check results for BUSCO: 1197019at2759
## Mon Jul 25 16:57:31 CEST 2022 - The aliscore output file does not exist. Check results for BUSCO: 1236198at2759
## Mon Jul 25 16:57:32 CEST 2022 - The aliscore output appears to be empty. Check results for BUSCO: 1236198at2759
## Mon Jul 25 16:57:32 CEST 2022 - The aliscore output file does not exist. Check results for BUSCO: 1346432at2759
## Mon Jul 25 16:57:32 CEST 2022 - The aliscore output appears to be empty. Check results for BUSCO: 1346432at2759
## Mon Jul 25 16:57:32 CEST 2022 - The aliscore output file does not exist. Check results for BUSCO: 1178688at2759
## Mon Jul 25 16:57:32 CEST 2022 - The aliscore output appears to be empty. Check results for BUSCO: 1178688at2759
## Mon Jul 25 16:57:32 CEST 2022 - The aliscore output file does not exist. Check results for BUSCO: 1348942at2759
## Mon Jul 25 16:57:32 CEST 2022 - The aliscore output appears to be empty. Check results for BUSCO: 1348942at2759
## Mon Jul 25 16:57:33 CEST 2022 - The aliscore output file does not exist. Check results for BUSCO: 1129824at2759
## Mon Jul 25 16:57:33 CEST 2022 - The aliscore output appears to be empty. Check results for BUSCO: 1129824at2759
## Mon Jul 25 16:57:34 CEST 2022 - The aliscore output file does not exist. Check results for BUSCO: 975158at2759
## Mon Jul 25 16:57:34 CEST 2022 - The aliscore output appears to be empty. Check results for BUSCO: 975158at2759
## Mon Jul 25 16:57:35 CEST 2022 - The aliscore output file does not exist. Check results for BUSCO: 917326at2759
## Mon Jul 25 16:57:35 CEST 2022 - The aliscore output file does not exist. Check results for BUSCO: 1450538at2759
## Mon Jul 25 16:57:35 CEST 2022 - The aliscore output appears to be empty. Check results for BUSCO: 917326at2759
## Mon Jul 25 16:57:35 CEST 2022 - The aliscore output appears to be empty. Check results for BUSCO: 1450538at2759
## Mon Jul 25 16:57:35 CEST 2022 - The aliscore output file does not exist. Check results for BUSCO: 388820at2759
## Mon Jul 25 16:57:35 CEST 2022 - The aliscore output appears to be empty. Check results for BUSCO: 388820at2759
## Mon Jul 25 16:57:35 CEST 2022 - The aliscore output file does not exist. Check results for BUSCO: 719531at2759
## Mon Jul 25 16:57:35 CEST 2022 - The aliscore output appears to be empty. Check results for BUSCO: 719531at2759
## Mon Jul 25 16:57:36 CEST 2022 - The aliscore output file does not exist. Check results for BUSCO: 692986at2759
## Mon Jul 25 16:57:36 CEST 2022 - The aliscore output file does not exist. Check results for BUSCO: 1041560at2759
## Mon Jul 25 16:57:36 CEST 2022 - The aliscore output appears to be empty. Check results for BUSCO: 692986at2759
## Mon Jul 25 16:57:36 CEST 2022 - The aliscore output appears to be empty. Check results for BUSCO: 1041560at2759
## Mon Jul 25 16:57:37 CEST 2022 - The aliscore output file does not exist. Check results for BUSCO: 396755at2759
## Mon Jul 25 16:57:37 CEST 2022 - The aliscore output appears to be empty. Check results for BUSCO: 396755at2759
## Mon Jul 25 16:57:37 CEST 2022 - The aliscore output file does not exist. Check results for BUSCO: 582756at2759
## Mon Jul 25 16:57:37 CEST 2022 - The aliscore output appears to be empty. Check results for BUSCO: 582756at2759
## Mon Jul 25 16:57:37 CEST 2022 - The aliscore output file does not exist. Check results for BUSCO: 97116at2759
## Mon Jul 25 16:57:37 CEST 2022 - The aliscore output appears to be empty. Check results for BUSCO: 97116at2759
## Mon Jul 25 16:57:37 CEST 2022 - The aliscore output file does not exist. Check results for BUSCO: 1247641at2759
## Mon Jul 25 16:57:37 CEST 2022 - The aliscore output appears to be empty. Check results for BUSCO: 1247641at2759
## Mon Jul 25 16:57:38 CEST 2022 - The aliscore output file does not exist. Check results for BUSCO: 855834at2759
## Mon Jul 25 16:57:38 CEST 2022 - The aliscore output appears to be empty. Check results for BUSCO: 855834at2759
## Mon Jul 25 16:57:38 CEST 2022 - The aliscore output file does not exist. Check results for BUSCO: 1194797at2759
## Mon Jul 25 16:57:38 CEST 2022 - The aliscore output appears to be empty. Check results for BUSCO: 1194797at2759
## Mon Jul 25 16:58:01 CEST 2022 - The aliscore output file does not exist. Check results for BUSCO: 1526152at2759
## Mon Jul 25 16:58:01 CEST 2022 - The aliscore output appears to be empty. Check results for BUSCO: 1526152at2759
## Mon Jul 25 16:58:02 CEST 2022 - The aliscore output file does not exist. Check results for BUSCO: 1257440at2759
## Mon Jul 25 16:58:02 CEST 2022 - The aliscore output appears to be empty. Check results for BUSCO: 1257440at2759
## Mon Jul 25 16:58:02 CEST 2022 - The aliscore output file does not exist. Check results for BUSCO: 1545004at2759
## Mon Jul 25 16:58:02 CEST 2022 - The aliscore output appears to be empty. Check results for BUSCO: 1545004at2759
## Mon Jul 25 16:58:02 CEST 2022 - The aliscore output file does not exist. Check results for BUSCO: 1459797at2759
## Mon Jul 25 16:58:02 CEST 2022 - The aliscore output file does not exist. Check results for BUSCO: 1428265at2759
## Mon Jul 25 16:58:02 CEST 2022 - The aliscore output appears to be empty. Check results for BUSCO: 1459797at2759
## Mon Jul 25 16:58:02 CEST 2022 - The aliscore output appears to be empty. Check results for BUSCO: 1428265at2759
## Mon Jul 25 16:58:02 CEST 2022 - The aliscore output file does not exist. Check results for BUSCO: 1079827at2759
## Mon Jul 25 16:58:02 CEST 2022 - The aliscore output appears to be empty. Check results for BUSCO: 1079827at2759
## Mon Jul 25 16:58:02 CEST 2022 - The aliscore output file does not exist. Check results for BUSCO: 1558822at2759
## Mon Jul 25 16:58:03 CEST 2022 - The aliscore output appears to be empty. Check results for BUSCO: 1558822at2759
## Mon Jul 25 16:58:03 CEST 2022 - The aliscore output file does not exist. Check results for BUSCO: 457861at2759
## Mon Jul 25 16:58:03 CEST 2022 - The aliscore output file does not exist. Check results for BUSCO: 1623701at2759
## Mon Jul 25 16:58:03 CEST 2022 - The aliscore output appears to be empty. Check results for BUSCO: 457861at2759
## Mon Jul 25 16:58:03 CEST 2022 - The aliscore output appears to be empty. Check results for BUSCO: 1623701at2759
## Mon Jul 25 16:58:03 CEST 2022 - The aliscore output file does not exist. Check results for BUSCO: 1504863at2759
## Mon Jul 25 16:58:03 CEST 2022 - The aliscore output appears to be empty. Check results for BUSCO: 1504863at2759
## Mon Jul 25 16:58:03 CEST 2022 - The aliscore output file does not exist. Check results for BUSCO: 1076134at2759
## Mon Jul 25 16:58:03 CEST 2022 - The aliscore output appears to be empty. Check results for BUSCO: 1076134at2759
## Mon Jul 25 16:58:03 CEST 2022 - The aliscore output file does not exist. Check results for BUSCO: 1645187at2759
## Mon Jul 25 16:58:03 CEST 2022 - The aliscore output file does not exist. Check results for BUSCO: 1260807at2759
## Mon Jul 25 16:58:03 CEST 2022 - The aliscore output appears to be empty. Check results for BUSCO: 1645187at2759
## Mon Jul 25 16:58:03 CEST 2022 - The aliscore output appears to be empty. Check results for BUSCO: 1260807at2759
## Mon Jul 25 16:58:03 CEST 2022 - The aliscore output file does not exist. Check results for BUSCO: 1445102at2759
## Mon Jul 25 16:58:03 CEST 2022 - The aliscore output appears to be empty. Check results for BUSCO: 1445102at2759
## Mon Jul 25 16:58:04 CEST 2022 - The aliscore output file does not exist. Check results for BUSCO: 866359at2759
## Mon Jul 25 16:58:04 CEST 2022 - The aliscore output appears to be empty. Check results for BUSCO: 866359at2759
## Mon Jul 25 16:58:04 CEST 2022 - The aliscore output file does not exist. Check results for BUSCO: 887370at2759
## Mon Jul 25 16:58:04 CEST 2022 - The aliscore output file does not exist. Check results for BUSCO: 1346165at2759
## Mon Jul 25 16:58:04 CEST 2022 - The aliscore output file does not exist. Check results for BUSCO: 1380409at2759
## Mon Jul 25 16:58:04 CEST 2022 - The aliscore output appears to be empty. Check results for BUSCO: 887370at2759
## Mon Jul 25 16:58:04 CEST 2022 - The aliscore output appears to be empty. Check results for BUSCO: 1346165at2759
## Mon Jul 25 16:58:04 CEST 2022 - The aliscore output appears to be empty. Check results for BUSCO: 1380409at2759
## Mon Jul 25 16:58:04 CEST 2022 - The aliscore output file does not exist. Check results for BUSCO: 453044at2759
## Mon Jul 25 16:58:04 CEST 2022 - The aliscore output file does not exist. Check results for BUSCO: 858842at2759
## Mon Jul 25 16:58:04 CEST 2022 - The aliscore output appears to be empty. Check results for BUSCO: 453044at2759
## Mon Jul 25 16:58:04 CEST 2022 - The aliscore output file does not exist. Check results for BUSCO: 320059at2759
## Mon Jul 25 16:58:04 CEST 2022 - The aliscore output appears to be empty. Check results for BUSCO: 858842at2759
## Mon Jul 25 16:58:04 CEST 2022 - The aliscore output file does not exist. Check results for BUSCO: 598949at2759
## Mon Jul 25 16:58:04 CEST 2022 - The aliscore output appears to be empty. Check results for BUSCO: 320059at2759
## Mon Jul 25 16:58:04 CEST 2022 - The aliscore output file does not exist. Check results for BUSCO: 1552706at2759
## Mon Jul 25 16:58:04 CEST 2022 - The aliscore output appears to be empty. Check results for BUSCO: 598949at2759
## Mon Jul 25 16:58:04 CEST 2022 - The aliscore output file does not exist. Check results for BUSCO: 1576404at2759
## Mon Jul 25 16:58:04 CEST 2022 - The aliscore output appears to be empty. Check results for BUSCO: 1552706at2759
## Mon Jul 25 16:58:04 CEST 2022 - The aliscore output file does not exist. Check results for BUSCO: 1090038at2759
## Mon Jul 25 16:58:04 CEST 2022 - The aliscore output appears to be empty. Check results for BUSCO: 1576404at2759
## Mon Jul 25 16:58:04 CEST 2022 - The aliscore output file does not exist. Check results for BUSCO: 1454155at2759
## Mon Jul 25 16:58:04 CEST 2022 - The aliscore output appears to be empty. Check results for BUSCO: 1090038at2759
## Mon Jul 25 16:58:04 CEST 2022 - The aliscore output file does not exist. Check results for BUSCO: 1030907at2759
## Mon Jul 25 16:58:04 CEST 2022 - The aliscore output file does not exist. Check results for BUSCO: 844748at2759
## Mon Jul 25 16:58:04 CEST 2022 - The aliscore output appears to be empty. Check results for BUSCO: 1030907at2759
## Mon Jul 25 16:58:04 CEST 2022 - The aliscore output file does not exist. Check results for BUSCO: 664730at2759
## Mon Jul 25 16:58:04 CEST 2022 - The aliscore output appears to be empty. Check results for BUSCO: 1454155at2759
## Mon Jul 25 16:58:04 CEST 2022 - The aliscore output appears to be empty. Check results for BUSCO: 844748at2759
## Mon Jul 25 16:58:04 CEST 2022 - The aliscore output appears to be empty. Check results for BUSCO: 664730at2759
## Mon Jul 25 16:58:04 CEST 2022 - The aliscore output file does not exist. Check results for BUSCO: 464990at2759
## Mon Jul 25 16:58:05 CEST 2022 - The aliscore output file does not exist. Check results for BUSCO: 1441854at2759
## Mon Jul 25 16:58:05 CEST 2022 - The aliscore output appears to be empty. Check results for BUSCO: 1441854at2759
## Mon Jul 25 16:58:05 CEST 2022 - The aliscore output file does not exist. Check results for BUSCO: 1220881at2759
## Mon Jul 25 16:58:05 CEST 2022 - The aliscore output appears to be empty. Check results for BUSCO: 464990at2759
## Mon Jul 25 16:58:05 CEST 2022 - The aliscore output appears to be empty. Check results for BUSCO: 1220881at2759
## Mon Jul 25 16:58:05 CEST 2022 - The aliscore output file does not exist. Check results for BUSCO: 1173229at2759
## Mon Jul 25 16:58:05 CEST 2022 - The aliscore output appears to be empty. Check results for BUSCO: 1173229at2759
## Mon Jul 25 16:58:05 CEST 2022 - The aliscore output file does not exist. Check results for BUSCO: 708105at2759
## Mon Jul 25 16:58:05 CEST 2022 - The aliscore output appears to be empty. Check results for BUSCO: 708105at2759
## Mon Jul 25 16:58:05 CEST 2022 - The aliscore output file does not exist. Check results for BUSCO: 1094121at2759
## Mon Jul 25 16:58:05 CEST 2022 - The aliscore output appears to be empty. Check results for BUSCO: 1094121at2759
## Mon Jul 25 16:58:05 CEST 2022 - The aliscore output file does not exist. Check results for BUSCO: 1304061at2759
## Mon Jul 25 16:58:05 CEST 2022 - The aliscore output appears to be empty. Check results for BUSCO: 1304061at2759
## Mon Jul 25 16:58:05 CEST 2022 - The aliscore output file does not exist. Check results for BUSCO: 923657at2759
## Mon Jul 25 16:58:05 CEST 2022 - The aliscore output file does not exist. Check results for BUSCO: 1442062at2759
## Mon Jul 25 16:58:05 CEST 2022 - The aliscore output appears to be empty. Check results for BUSCO: 923657at2759
## Mon Jul 25 16:58:05 CEST 2022 - The aliscore output file does not exist. Check results for BUSCO: 679187at2759
## Mon Jul 25 16:58:06 CEST 2022 - The aliscore output appears to be empty. Check results for BUSCO: 1442062at2759
## Mon Jul 25 16:58:06 CEST 2022 - The aliscore output file does not exist. Check results for BUSCO: 1323575at2759
## Mon Jul 25 16:58:06 CEST 2022 - The aliscore output appears to be empty. Check results for BUSCO: 679187at2759
## Mon Jul 25 16:58:06 CEST 2022 - The aliscore output appears to be empty. Check results for BUSCO: 1323575at2759
## Mon Jul 25 16:58:06 CEST 2022 - The aliscore output file does not exist. Check results for BUSCO: 426305at2759
## Mon Jul 25 16:58:06 CEST 2022 - The aliscore output file does not exist. Check results for BUSCO: 655400at2759
## Mon Jul 25 16:58:06 CEST 2022 - The aliscore output appears to be empty. Check results for BUSCO: 426305at2759
## Mon Jul 25 16:58:06 CEST 2022 - The aliscore output appears to be empty. Check results for BUSCO: 655400at2759
## Mon Jul 25 16:58:06 CEST 2022 - The aliscore output file does not exist. Check results for BUSCO: 937686at2759
## Mon Jul 25 16:58:06 CEST 2022 - The aliscore output file does not exist. Check results for BUSCO: 1324510at2759
## Mon Jul 25 16:58:06 CEST 2022 - The aliscore output appears to be empty. Check results for BUSCO: 937686at2759
## Mon Jul 25 16:58:06 CEST 2022 - The aliscore output appears to be empty. Check results for BUSCO: 1324510at2759
## Mon Jul 25 16:58:06 CEST 2022 - The aliscore output file does not exist. Check results for BUSCO: 1014314at2759
## Mon Jul 25 16:58:06 CEST 2022 - The aliscore output appears to be empty. Check results for BUSCO: 1014314at2759
## Mon Jul 25 16:58:06 CEST 2022 - The aliscore output file does not exist. Check results for BUSCO: 1364586at2759
## Mon Jul 25 16:58:06 CEST 2022 - The aliscore output appears to be empty. Check results for BUSCO: 1364586at2759
## Mon Jul 25 16:58:06 CEST 2022 - The aliscore output file does not exist. Check results for BUSCO: 100698at2759
## Mon Jul 25 16:58:06 CEST 2022 - The aliscore output appears to be empty. Check results for BUSCO: 100698at2759
## Mon Jul 25 16:58:06 CEST 2022 - The aliscore output file does not exist. Check results for BUSCO: 722805at2759
## Mon Jul 25 16:58:07 CEST 2022 - The aliscore output appears to be empty. Check results for BUSCO: 722805at2759
## Mon Jul 25 16:58:07 CEST 2022 - The aliscore output file does not exist. Check results for BUSCO: 734666at2759
## Mon Jul 25 16:58:07 CEST 2022 - The aliscore output appears to be empty. Check results for BUSCO: 734666at2759
## Mon Jul 25 16:58:07 CEST 2022 - The aliscore output file does not exist. Check results for BUSCO: 1287094at2759
## Mon Jul 25 16:58:07 CEST 2022 - The aliscore output appears to be empty. Check results for BUSCO: 1287094at2759
## Mon Jul 25 16:58:07 CEST 2022 - The aliscore output file does not exist. Check results for BUSCO: 1248958at2759
## Mon Jul 25 16:58:07 CEST 2022 - The aliscore output appears to be empty. Check results for BUSCO: 1248958at2759
## Mon Jul 25 16:58:07 CEST 2022 - The aliscore output file does not exist. Check results for BUSCO: 390348at2759
## Mon Jul 25 16:58:07 CEST 2022 - The aliscore output appears to be empty. Check results for BUSCO: 390348at2759
## Mon Jul 25 16:58:07 CEST 2022 - The aliscore output file does not exist. Check results for BUSCO: 1057950at2759
## Mon Jul 25 16:58:07 CEST 2022 - The aliscore output appears to be empty. Check results for BUSCO: 1057950at2759
## Mon Jul 25 16:58:07 CEST 2022 - The aliscore output file does not exist. Check results for BUSCO: 720340at2759
## Mon Jul 25 16:58:07 CEST 2022 - The aliscore output appears to be empty. Check results for BUSCO: 720340at2759
## Mon Jul 25 16:58:07 CEST 2022 - The aliscore output file does not exist. Check results for BUSCO: 293315at2759
## Mon Jul 25 16:58:07 CEST 2022 - The aliscore output file does not exist. Check results for BUSCO: 450058at2759
## Mon Jul 25 16:58:07 CEST 2022 - The aliscore output appears to be empty. Check results for BUSCO: 293315at2759
## Mon Jul 25 16:58:07 CEST 2022 - The aliscore output appears to be empty. Check results for BUSCO: 450058at2759
## Mon Jul 25 16:58:08 CEST 2022 - The aliscore output file does not exist. Check results for BUSCO: 1115196at2759
## Mon Jul 25 16:58:08 CEST 2022 - The aliscore output appears to be empty. Check results for BUSCO: 1115196at2759
## Mon Jul 25 16:58:08 CEST 2022 - The aliscore output file does not exist. Check results for BUSCO: 759498at2759
## Mon Jul 25 16:58:08 CEST 2022 - The aliscore output appears to be empty. Check results for BUSCO: 759498at2759
## Mon Jul 25 16:58:08 CEST 2022 - The aliscore output file does not exist. Check results for BUSCO: 1106766at2759
## Mon Jul 25 16:58:08 CEST 2022 - The aliscore output appears to be empty. Check results for BUSCO: 1106766at2759
## Mon Jul 25 16:58:08 CEST 2022 - The aliscore output file does not exist. Check results for BUSCO: 1479417at2759
## Mon Jul 25 16:58:08 CEST 2022 - The aliscore output file does not exist. Check results for BUSCO: 1309031at2759
## Mon Jul 25 16:58:08 CEST 2022 - The aliscore output appears to be empty. Check results for BUSCO: 1479417at2759
## Mon Jul 25 16:58:08 CEST 2022 - The aliscore output appears to be empty. Check results for BUSCO: 1309031at2759
## Mon Jul 25 16:58:08 CEST 2022 - The aliscore output appears to be empty. Check results for BUSCO: 834694at2759
## Mon Jul 25 16:58:08 CEST 2022 - The aliscore output file does not exist. Check results for BUSCO: 673132at2759
## Mon Jul 25 16:58:08 CEST 2022 - The aliscore output file does not exist. Check results for BUSCO: 299766at2759
## Mon Jul 25 16:58:08 CEST 2022 - The aliscore output appears to be empty. Check results for BUSCO: 673132at2759
## Mon Jul 25 16:58:08 CEST 2022 - The aliscore output appears to be empty. Check results for BUSCO: 299766at2759
## Mon Jul 25 16:58:09 CEST 2022 - The aliscore output file does not exist. Check results for BUSCO: 891751at2759
## Mon Jul 25 16:58:09 CEST 2022 - The aliscore output appears to be empty. Check results for BUSCO: 891751at2759
## Mon Jul 25 16:58:09 CEST 2022 - The aliscore output file does not exist. Check results for BUSCO: 1085752at2759
## Mon Jul 25 16:58:09 CEST 2022 - The aliscore output appears to be empty. Check results for BUSCO: 1085752at2759
## Mon Jul 25 16:58:09 CEST 2022 - The aliscore output file does not exist. Check results for BUSCO: 1379600at2759
## Mon Jul 25 16:58:09 CEST 2022 - The aliscore output appears to be empty. Check results for BUSCO: 1379600at2759
## Mon Jul 25 16:58:10 CEST 2022 - The aliscore output file does not exist. Check results for BUSCO: 296129at2759
## Mon Jul 25 16:58:10 CEST 2022 - The aliscore output appears to be empty. Check results for BUSCO: 296129at2759
## Mon Jul 25 16:58:10 CEST 2022 - The aliscore output file does not exist. Check results for BUSCO: 261419at2759
## Mon Jul 25 16:58:10 CEST 2022 - The aliscore output file does not exist. Check results for BUSCO: 543764at2759
## Mon Jul 25 16:58:10 CEST 2022 - The aliscore output appears to be empty. Check results for BUSCO: 261419at2759
## Mon Jul 25 16:58:10 CEST 2022 - The aliscore output appears to be empty. Check results for BUSCO: 543764at2759
## Mon Jul 25 16:58:11 CEST 2022 - The aliscore output file does not exist. Check results for BUSCO: 87842at2759
## Mon Jul 25 16:58:11 CEST 2022 - The aliscore output file does not exist. Check results for BUSCO: 1314980at2759
## Mon Jul 25 16:58:11 CEST 2022 - The aliscore output appears to be empty. Check results for BUSCO: 87842at2759
## Mon Jul 25 16:58:11 CEST 2022 - The aliscore output appears to be empty. Check results for BUSCO: 1314980at2759
## Mon Jul 25 16:58:11 CEST 2022 - The aliscore output file does not exist. Check results for BUSCO: 306227at2759
## Mon Jul 25 16:58:11 CEST 2022 - The aliscore output appears to be empty. Check results for BUSCO: 306227at2759
## Mon Jul 25 16:58:12 CEST 2022 - The aliscore output file does not exist. Check results for BUSCO: 324863at2759
## Mon Jul 25 16:58:12 CEST 2022 - The aliscore output appears to be empty. Check results for BUSCO: 324863at2759
## Mon Jul 25 16:58:12 CEST 2022 - The aliscore output file does not exist. Check results for BUSCO: 261328at2759
## Mon Jul 25 16:58:12 CEST 2022 - The aliscore output appears to be empty. Check results for BUSCO: 261328at2759
## Mon Jul 25 16:58:12 CEST 2022 - The aliscore output file does not exist. Check results for BUSCO: 777920at2759
## Mon Jul 25 16:58:12 CEST 2022 - The aliscore output appears to be empty. Check results for BUSCO: 777920at2759
## Mon Jul 25 16:58:12 CEST 2022 - The aliscore output file does not exist. Check results for BUSCO: 271586at2759
## Mon Jul 25 16:58:12 CEST 2022 - The aliscore output appears to be empty. Check results for BUSCO: 271586at2759
## Mon Jul 25 16:58:14 CEST 2022 - The aliscore output file does not exist. Check results for BUSCO: 166920at2759
## Mon Jul 25 16:58:14 CEST 2022 - The aliscore output appears to be empty. Check results for BUSCO: 166920at2759
## Mon Jul 25 16:58:15 CEST 2022 - The aliscore output file does not exist. Check results for BUSCO: 142542at2759
## Mon Jul 25 16:58:15 CEST 2022 - The aliscore output appears to be empty. Check results for BUSCO: 142542at2759
## Mon Jul 25 16:58:17 CEST 2022 - The aliscore output file does not exist. Check results for BUSCO: 156083at2759
## Mon Jul 25 16:58:17 CEST 2022 - The aliscore output appears to be empty. Check results for BUSCO: 156083at2759
## Mon Jul 25 16:58:29 CEST 2022 - The aliscore output file does not exist. Check results for BUSCO: 865202at2759
## Mon Jul 25 16:58:29 CEST 2022 - The aliscore output appears to be empty. Check results for BUSCO: 865202at2759
## Mon Jul 25 16:58:30 CEST 2022 - The aliscore output file does not exist. Check results for BUSCO: 1575179at2759
## Mon Jul 25 16:58:30 CEST 2022 - The aliscore output appears to be empty. Check results for BUSCO: 1575179at2759
## Mon Jul 25 16:58:32 CEST 2022 - The aliscore output file does not exist. Check results for BUSCO: 1440961at2759
## Mon Jul 25 16:58:32 CEST 2022 - The aliscore output appears to be empty. Check results for BUSCO: 1440961at2759
## Mon Jul 25 16:58:32 CEST 2022 - The aliscore output file does not exist. Check results for BUSCO: 937275at2759
## Mon Jul 25 16:58:32 CEST 2022 - The aliscore output appears to be empty. Check results for BUSCO: 937275at2759
## Mon Jul 25 16:58:32 CEST 2022 - The aliscore output file does not exist. Check results for BUSCO: 1530008at2759
## Mon Jul 25 16:58:32 CEST 2022 - The aliscore output appears to be empty. Check results for BUSCO: 1530008at2759
## Mon Jul 25 16:58:32 CEST 2022 - The aliscore output file does not exist. Check results for BUSCO: 1112002at2759
## Mon Jul 25 16:58:32 CEST 2022 - The aliscore output file does not exist. Check results for BUSCO: 1620056at2759
## Mon Jul 25 16:58:32 CEST 2022 - The aliscore output file does not exist. Check results for BUSCO: 1398309at2759
## Mon Jul 25 16:58:32 CEST 2022 - The aliscore output appears to be empty. Check results for BUSCO: 1112002at2759
## Mon Jul 25 16:58:32 CEST 2022 - The aliscore output file does not exist. Check results for BUSCO: 708105at2759
## Mon Jul 25 16:58:32 CEST 2022 - The aliscore output appears to be empty. Check results for BUSCO: 1620056at2759
## Mon Jul 25 16:58:32 CEST 2022 - The aliscore output appears to be empty. Check results for BUSCO: 1398309at2759
## Mon Jul 25 16:58:32 CEST 2022 - The aliscore output appears to be empty. Check results for BUSCO: 708105at2759
## Mon Jul 25 16:58:32 CEST 2022 - The aliscore output file does not exist. Check results for BUSCO: 1018517at2759
## Mon Jul 25 16:58:32 CEST 2022 - The aliscore output appears to be empty. Check results for BUSCO: 1018517at2759
## Mon Jul 25 16:58:33 CEST 2022 - The aliscore output file does not exist. Check results for BUSCO: 1395649at2759
## Mon Jul 25 16:58:33 CEST 2022 - The aliscore output appears to be empty. Check results for BUSCO: 1395649at2759
## Mon Jul 25 16:58:33 CEST 2022 - The aliscore output file does not exist. Check results for BUSCO: 974865at2759
## Mon Jul 25 16:58:33 CEST 2022 - The aliscore output file does not exist. Check results for BUSCO: 1432328at2759
## Mon Jul 25 16:58:33 CEST 2022 - The aliscore output appears to be empty. Check results for BUSCO: 974865at2759
## Mon Jul 25 16:58:33 CEST 2022 - The aliscore output appears to be empty. Check results for BUSCO: 1432328at2759
## Mon Jul 25 16:58:33 CEST 2022 - The aliscore output file does not exist. Check results for BUSCO: 1490022at2759
## Mon Jul 25 16:58:33 CEST 2022 - The aliscore output appears to be empty. Check results for BUSCO: 1490022at2759
## Mon Jul 25 16:58:33 CEST 2022 - The aliscore output file does not exist. Check results for BUSCO: 1085752at2759
## Mon Jul 25 16:58:33 CEST 2022 - The aliscore output file does not exist. Check results for BUSCO: 1588798at2759
## Mon Jul 25 16:58:33 CEST 2022 - The aliscore output file does not exist. Check results for BUSCO: 1430056at2759
## Mon Jul 25 16:58:33 CEST 2022 - The aliscore output appears to be empty. Check results for BUSCO: 1085752at2759
## Mon Jul 25 16:58:33 CEST 2022 - The aliscore output appears to be empty. Check results for BUSCO: 1588798at2759
## Mon Jul 25 16:58:33 CEST 2022 - The aliscore output appears to be empty. Check results for BUSCO: 1430056at2759
## Mon Jul 25 16:58:33 CEST 2022 - The aliscore output file does not exist. Check results for BUSCO: 1228942at2759
## Mon Jul 25 16:58:33 CEST 2022 - The aliscore output appears to be empty. Check results for BUSCO: 1228942at2759
## Mon Jul 25 16:58:33 CEST 2022 - The aliscore output file does not exist. Check results for BUSCO: 1405146at2759
## Mon Jul 25 16:58:33 CEST 2022 - The aliscore output appears to be empty. Check results for BUSCO: 1405146at2759
## Mon Jul 25 16:58:33 CEST 2022 - The aliscore output file does not exist. Check results for BUSCO: 1339553at2759
## Mon Jul 25 16:58:33 CEST 2022 - The aliscore output file does not exist. Check results for BUSCO: 1538526at2759
## Mon Jul 25 16:58:33 CEST 2022 - The aliscore output appears to be empty. Check results for BUSCO: 1339553at2759
## Mon Jul 25 16:58:33 CEST 2022 - The aliscore output appears to be empty. Check results for BUSCO: 1538526at2759
## Mon Jul 25 16:58:33 CEST 2022 - The aliscore output file does not exist. Check results for BUSCO: 956854at2759
## 
## Mon Jul 25 16:58:34 CEST 2022 - The aliscore output file does not exist. Check results for BUSCO: 1377237at2759
## Mon Jul 25 16:58:34 CEST 2022 - The aliscore output appears to be empty. Check results for BUSCO: 1454155at2759
## Mon Jul 25 16:58:34 CEST 2022 - The aliscore output appears to be empty. Check results for BUSCO: 956854at2759
## Mon Jul 25 16:58:34 CEST 2022 - The aliscore output appears to be empty. Check results for BUSCO: 1377237at2759
## Mon Jul 25 16:58:34 CEST 2022 - The aliscore output file does not exist. Check results for BUSCO: 898782at2759
## Mon Jul 25 16:58:34 CEST 2022 - The aliscore output file does not exist. Check results for BUSCO: 919955at2759
## Mon Jul 25 16:58:34 CEST 2022 - The aliscore output appears to be empty. Check results for BUSCO: 898782at2759
## Mon Jul 25 16:58:34 CEST 2022 - The aliscore output appears to be empty. Check results for BUSCO: 919955at2759
## Mon Jul 25 16:58:34 CEST 2022 - The aliscore output file does not exist. Check results for BUSCO: 1128607at2759
## Mon Jul 25 16:58:34 CEST 2022 - The aliscore output file does not exist. Check results for BUSCO: 176625at2759
## Mon Jul 25 16:58:34 CEST 2022 - The aliscore output appears to be empty. Check results for BUSCO: 1128607at2759
## Mon Jul 25 16:58:34 CEST 2022 - The aliscore output file does not exist. Check results for BUSCO: 1434266at2759
## Mon Jul 25 16:58:34 CEST 2022 - The aliscore output file does not exist. Check results for BUSCO: 1593937at2759
## Mon Jul 25 16:58:34 CEST 2022 - The aliscore output appears to be empty. Check results for BUSCO: 176625at2759
## Mon Jul 25 16:58:34 CEST 2022 - The aliscore output file does not exist. Check results for BUSCO: 858842at2759
## Mon Jul 25 16:58:34 CEST 2022 - The aliscore output appears to be empty. Check results for BUSCO: 1434266at2759
## Mon Jul 25 16:58:34 CEST 2022 - The aliscore output appears to be empty. Check results for BUSCO: 1593937at2759
## Mon Jul 25 16:58:34 CEST 2022 - The aliscore output appears to be empty. Check results for BUSCO: 858842at2759
## Mon Jul 25 16:58:34 CEST 2022 - The aliscore output file does not exist. Check results for BUSCO: 1370304at2759
## Mon Jul 25 16:58:34 CEST 2022 - The aliscore output appears to be empty. Check results for BUSCO: 1370304at2759
## Mon Jul 25 16:58:34 CEST 2022 - The aliscore output file does not exist. Check results for BUSCO: 426305at2759
## Mon Jul 25 16:58:34 CEST 2022 - The aliscore output file does not exist. Check results for BUSCO: 1304061at2759
## Mon Jul 25 16:58:34 CEST 2022 - The aliscore output file does not exist. Check results for BUSCO: 1428265at2759
## Mon Jul 25 16:58:34 CEST 2022 - The aliscore output appears to be empty. Check results for BUSCO: 426305at2759
## Mon Jul 25 16:58:34 CEST 2022 - The aliscore output appears to be empty. Check results for BUSCO: 1304061at2759
## Mon Jul 25 16:58:34 CEST 2022 - The aliscore output file does not exist. Check results for BUSCO: 1220881at2759
## Mon Jul 25 16:58:34 CEST 2022 - The aliscore output appears to be empty. Check results for BUSCO: 1428265at2759
## Mon Jul 25 16:58:34 CEST 2022 - The aliscore output appears to be empty. Check results for BUSCO: 1220881at2759
## Mon Jul 25 16:58:34 CEST 2022 - The aliscore output file does not exist. Check results for BUSCO: 721605at2759
## Mon Jul 25 16:58:34 CEST 2022 - The aliscore output appears to be empty. Check results for BUSCO: 721605at2759
## Mon Jul 25 16:58:34 CEST 2022 - The aliscore output file does not exist. Check results for BUSCO: 1563319at2759
## Mon Jul 25 16:58:34 CEST 2022 - The aliscore output appears to be empty. Check results for BUSCO: 1563319at2759
## Mon Jul 25 16:58:34 CEST 2022 - The aliscore output file does not exist. Check results for BUSCO: 1264469at2759
## Mon Jul 25 16:58:34 CEST 2022 - The aliscore output appears to be empty. Check results for BUSCO: 1264469at2759
## Mon Jul 25 16:58:34 CEST 2022 - The aliscore output file does not exist. Check results for BUSCO: 330169at2759
## Mon Jul 25 16:58:34 CEST 2022 - The aliscore output appears to be empty. Check results for BUSCO: 330169at2759
## Mon Jul 25 16:58:35 CEST 2022 - The aliscore output file does not exist. Check results for BUSCO: 801857at2759
## Mon Jul 25 16:58:35 CEST 2022 - The aliscore output appears to be empty. Check results for BUSCO: 801857at2759
## Mon Jul 25 16:58:35 CEST 2022 - The aliscore output file does not exist. Check results for BUSCO: 887370at2759
## Mon Jul 25 16:58:35 CEST 2022 - The aliscore output appears to be empty. Check results for BUSCO: 887370at2759
## Mon Jul 25 16:58:35 CEST 2022 - The aliscore output file does not exist. Check results for BUSCO: 878143at2759
## Mon Jul 25 16:58:35 CEST 2022 - The aliscore output appears to be empty. Check results for BUSCO: 878143at2759
## Mon Jul 25 16:58:35 CEST 2022 - The aliscore output file does not exist. Check results for BUSCO: 345441at2759
## Mon Jul 25 16:58:35 CEST 2022 - The aliscore output file does not exist. Check results for BUSCO: 257318at2759
## Mon Jul 25 16:58:35 CEST 2022 - The aliscore output appears to be empty. Check results for BUSCO: 345441at2759
## Mon Jul 25 16:58:35 CEST 2022 - The aliscore output file does not exist. Check results for BUSCO: 1309031at2759
## Mon Jul 25 16:58:35 CEST 2022 - The aliscore output appears to be empty. Check results for BUSCO: 257318at2759
## Mon Jul 25 16:58:35 CEST 2022 - The aliscore output appears to be empty. Check results for BUSCO: 1309031at2759
## Mon Jul 25 16:58:35 CEST 2022 - The aliscore output file does not exist. Check results for BUSCO: 918816at2759
## Mon Jul 25 16:58:36 CEST 2022 - The aliscore output appears to be empty. Check results for BUSCO: 918816at2759
## Mon Jul 25 16:58:36 CEST 2022 - The aliscore output file does not exist. Check results for BUSCO: 1001705at2759
## Mon Jul 25 16:58:36 CEST 2022 - The aliscore output file does not exist. Check results for BUSCO: 621827at2759
## Mon Jul 25 16:58:36 CEST 2022 - The aliscore output file does not exist. Check results for BUSCO: 1370285at2759
## Mon Jul 25 16:58:36 CEST 2022 - The aliscore output appears to be empty. Check results for BUSCO: 1001705at2759
## Mon Jul 25 16:58:36 CEST 2022 - The aliscore output appears to be empty. Check results for BUSCO: 621827at2759
## Mon Jul 25 16:58:36 CEST 2022 - The aliscore output appears to be empty. Check results for BUSCO: 1370285at2759
## Mon Jul 25 16:58:36 CEST 2022 - The aliscore output file does not exist. Check results for BUSCO: 720340at2759
## Mon Jul 25 16:58:36 CEST 2022 - The aliscore output appears to be empty. Check results for BUSCO: 720340at2759
## Mon Jul 25 16:58:36 CEST 2022 - The aliscore output file does not exist. Check results for BUSCO: 814241at2759
## Mon Jul 25 16:58:36 CEST 2022 - The aliscore output appears to be empty. Check results for BUSCO: 814241at2759
## Mon Jul 25 16:58:36 CEST 2022 - The aliscore output file does not exist. Check results for BUSCO: 418107at2759
## Mon Jul 25 16:58:36 CEST 2022 - The aliscore output appears to be empty. Check results for BUSCO: 418107at2759
## Mon Jul 25 16:58:36 CEST 2022 - The aliscore output file does not exist. Check results for BUSCO: 1322642at2759
## Mon Jul 25 16:58:36 CEST 2022 - The aliscore output appears to be empty. Check results for BUSCO: 1322642at2759
## Mon Jul 25 16:58:36 CEST 2022 - The aliscore output file does not exist. Check results for BUSCO: 679187at2759
## Mon Jul 25 16:58:36 CEST 2022 - The aliscore output file does not exist. Check results for BUSCO: 975557at2759
## Mon Jul 25 16:58:36 CEST 2022 - The aliscore output appears to be empty. Check results for BUSCO: 679187at2759
## Mon Jul 25 16:58:36 CEST 2022 - The aliscore output appears to be empty. Check results for BUSCO: 975557at2759
## Mon Jul 25 16:58:36 CEST 2022 - The aliscore output file does not exist. Check results for BUSCO: 674169at2759
## Mon Jul 25 16:58:36 CEST 2022 - The aliscore output appears to be empty. Check results for BUSCO: 674169at2759
## Mon Jul 25 16:58:36 CEST 2022 - The aliscore output file does not exist. Check results for BUSCO: 777920at2759
## Mon Jul 25 16:58:36 CEST 2022 - The aliscore output file does not exist. Check results for BUSCO: 1025450at2759
## Mon Jul 25 16:58:36 CEST 2022 - The aliscore output appears to be empty. Check results for BUSCO: 777920at2759
## Mon Jul 25 16:58:36 CEST 2022 - The aliscore output appears to be empty. Check results for BUSCO: 1025450at2759
## Mon Jul 25 16:58:36 CEST 2022 - The aliscore output file does not exist. Check results for BUSCO: 1258856at2759
## Mon Jul 25 16:58:36 CEST 2022 - The aliscore output appears to be empty. Check results for BUSCO: 1258856at2759
## Mon Jul 25 16:58:36 CEST 2022 - The aliscore output file does not exist. Check results for BUSCO: 341721at2759
## Mon Jul 25 16:58:36 CEST 2022 - The aliscore output file does not exist. Check results for BUSCO: 390348at2759
## Mon Jul 25 16:58:36 CEST 2022 - The aliscore output appears to be empty. Check results for BUSCO: 341721at2759
## Mon Jul 25 16:58:37 CEST 2022 - The aliscore output appears to be empty. Check results for BUSCO: 390348at2759
## Mon Jul 25 16:58:37 CEST 2022 - The aliscore output file does not exist. Check results for BUSCO: 836599at2759
## Mon Jul 25 16:58:37 CEST 2022 - The aliscore output appears to be empty. Check results for BUSCO: 836599at2759
## Mon Jul 25 16:58:37 CEST 2022 - The aliscore output file does not exist. Check results for BUSCO: 392369at2759
## Mon Jul 25 16:58:37 CEST 2022 - The aliscore output appears to be empty. Check results for BUSCO: 392369at2759
## Mon Jul 25 16:58:37 CEST 2022 - The aliscore output file does not exist. Check results for BUSCO: 551907at2759
## Mon Jul 25 16:58:37 CEST 2022 - The aliscore output appears to be empty. Check results for BUSCO: 551907at2759
## Mon Jul 25 16:58:37 CEST 2022 - The aliscore output file does not exist. Check results for BUSCO: 891751at2759
## Mon Jul 25 16:58:37 CEST 2022 - The aliscore output appears to be empty. Check results for BUSCO: 891751at2759
## Mon Jul 25 16:58:37 CEST 2022 - The aliscore output file does not exist. Check results for BUSCO: 1423847at2759
## Mon Jul 25 16:58:37 CEST 2022 - The aliscore output appears to be empty. Check results for BUSCO: 1423847at2759
## Mon Jul 25 16:58:37 CEST 2022 - The aliscore output file does not exist. Check results for BUSCO: 931188at2759
## Mon Jul 25 16:58:37 CEST 2022 - The aliscore output appears to be empty. Check results for BUSCO: 931188at2759
## Mon Jul 25 16:58:37 CEST 2022 - The aliscore output file does not exist. Check results for BUSCO: 671846at2759
## Mon Jul 25 16:58:37 CEST 2022 - The aliscore output appears to be empty. Check results for BUSCO: 671846at2759
## Mon Jul 25 16:58:37 CEST 2022 - The aliscore output file does not exist. Check results for BUSCO: 1087488at2759
## Mon Jul 25 16:58:37 CEST 2022 - The aliscore output appears to be empty. Check results for BUSCO: 1087488at2759
## Mon Jul 25 16:58:38 CEST 2022 - The aliscore output file does not exist. Check results for BUSCO: 973442at2759
## Mon Jul 25 16:58:38 CEST 2022 - The aliscore output file does not exist. Check results for BUSCO: 270107at2759
## Mon Jul 25 16:58:38 CEST 2022 - The aliscore output appears to be empty. Check results for BUSCO: 973442at2759
## Mon Jul 25 16:58:38 CEST 2022 - The aliscore output appears to be empty. Check results for BUSCO: 270107at2759
## Mon Jul 25 16:58:39 CEST 2022 - The aliscore output file does not exist. Check results for BUSCO: 179362at2759
## Mon Jul 25 16:58:39 CEST 2022 - The aliscore output appears to be empty. Check results for BUSCO: 179362at2759
## Mon Jul 25 16:58:44 CEST 2022 - The aliscore output file does not exist. Check results for BUSCO: 674160at2759
## Mon Jul 25 16:58:44 CEST 2022 - The aliscore output appears to be empty. Check results for BUSCO: 674160at2759
## Mon Jul 25 16:59:00 CEST 2022 - The aliscore output file does not exist. Check results for BUSCO: 1382842at2759
## Mon Jul 25 16:59:00 CEST 2022 - The aliscore output appears to be empty. Check results for BUSCO: 1382842at2759
## Mon Jul 25 16:59:01 CEST 2022 - The aliscore output file does not exist. Check results for BUSCO: 944899at2759
## Mon Jul 25 16:59:01 CEST 2022 - The aliscore output appears to be empty. Check results for BUSCO: 944899at2759
## Mon Jul 25 16:59:02 CEST 2022 - The aliscore output file does not exist. Check results for BUSCO: 1178688at2759
## Mon Jul 25 16:59:02 CEST 2022 - The aliscore output appears to be empty. Check results for BUSCO: 1178688at2759
## Mon Jul 25 16:59:02 CEST 2022 - The aliscore output file does not exist. Check results for BUSCO: 1633672at2759
## Mon Jul 25 16:59:02 CEST 2022 - The aliscore output appears to be empty. Check results for BUSCO: 1633672at2759
## Mon Jul 25 16:59:02 CEST 2022 - The aliscore output file does not exist. Check results for BUSCO: 975158at2759
## Mon Jul 25 16:59:02 CEST 2022 - The aliscore output appears to be empty. Check results for BUSCO: 975158at2759
## Mon Jul 25 16:59:02 CEST 2022 - The aliscore output file does not exist. Check results for BUSCO: 383503at2759
## Mon Jul 25 16:59:02 CEST 2022 - The aliscore output appears to be empty. Check results for BUSCO: 383503at2759
## Mon Jul 25 16:59:03 CEST 2022 - The aliscore output file does not exist. Check results for BUSCO: 1407446at2759
## Mon Jul 25 16:59:03 CEST 2022 - The aliscore output appears to be empty. Check results for BUSCO: 1407446at2759
## Mon Jul 25 16:59:03 CEST 2022 - The aliscore output file does not exist. Check results for BUSCO: 1355894at2759
## Mon Jul 25 16:59:03 CEST 2022 - The aliscore output appears to be empty. Check results for BUSCO: 1355894at2759
## Mon Jul 25 16:59:03 CEST 2022 - The aliscore output file does not exist. Check results for BUSCO: 1346165at2759
## Mon Jul 25 16:59:03 CEST 2022 - The aliscore output appears to be empty. Check results for BUSCO: 1346165at2759
## Mon Jul 25 16:59:04 CEST 2022 - The aliscore output file does not exist. Check results for BUSCO: 1291729at2759
## Mon Jul 25 16:59:04 CEST 2022 - The aliscore output appears to be empty. Check results for BUSCO: 1291729at2759
## Mon Jul 25 16:59:04 CEST 2022 - The aliscore output file does not exist. Check results for BUSCO: 734341at2759
## Mon Jul 25 16:59:05 CEST 2022 - The aliscore output appears to be empty. Check results for BUSCO: 734341at2759
## Mon Jul 25 16:59:05 CEST 2022 - The aliscore output file does not exist. Check results for BUSCO: 1010730at2759
## Mon Jul 25 16:59:05 CEST 2022 - The aliscore output appears to be empty. Check results for BUSCO: 1010730at2759
## Mon Jul 25 16:59:05 CEST 2022 - The aliscore output file does not exist. Check results for BUSCO: 1217666at2759
## Mon Jul 25 16:59:05 CEST 2022 - The aliscore output appears to be empty. Check results for BUSCO: 1217666at2759
## Mon Jul 25 16:59:05 CEST 2022 - The aliscore output file does not exist. Check results for BUSCO: 1065019at2759
## Mon Jul 25 16:59:05 CEST 2022 - The aliscore output appears to be empty. Check results for BUSCO: 1065019at2759
## Mon Jul 25 16:59:05 CEST 2022 - The aliscore output file does not exist. Check results for BUSCO: 375960at2759
## Mon Jul 25 16:59:05 CEST 2022 - The aliscore output file does not exist. Check results for BUSCO: 722805at2759
## Mon Jul 25 16:59:05 CEST 2022 - The aliscore output appears to be empty. Check results for BUSCO: 375960at2759
## Mon Jul 25 16:59:05 CEST 2022 - The aliscore output appears to be empty. Check results for BUSCO: 722805at2759
## Mon Jul 25 16:59:06 CEST 2022 - The aliscore output file does not exist. Check results for BUSCO: 996662at2759
## Mon Jul 25 16:59:06 CEST 2022 - The aliscore output appears to be empty. Check results for BUSCO: 996662at2759
## Mon Jul 25 16:59:06 CEST 2022 - The aliscore output appears to be empty. Check results for BUSCO: 582756at2759
## Mon Jul 25 16:59:06 CEST 2022 - The aliscore output file does not exist. Check results for BUSCO: 1275837at2759
## Mon Jul 25 16:59:06 CEST 2022 - The aliscore output appears to be empty. Check results for BUSCO: 1275837at2759
## Mon Jul 25 16:59:06 CEST 2022 - The aliscore output file does not exist. Check results for BUSCO: 1380409at2759
## Mon Jul 25 16:59:06 CEST 2022 - The aliscore output appears to be empty. Check results for BUSCO: 1380409at2759
## Mon Jul 25 16:59:06 CEST 2022 - The aliscore output file does not exist. Check results for BUSCO: 901894at2759
## Mon Jul 25 16:59:06 CEST 2022 - The aliscore output appears to be empty. Check results for BUSCO: 901894at2759
## Mon Jul 25 16:59:06 CEST 2022 - The aliscore output file does not exist. Check results for BUSCO: 1455730at2759
## Mon Jul 25 16:59:06 CEST 2022 - The aliscore output appears to be empty. Check results for BUSCO: 1455730at2759
## Mon Jul 25 16:59:07 CEST 2022 - The aliscore output file does not exist. Check results for BUSCO: 1379600at2759
## Mon Jul 25 16:59:07 CEST 2022 - The aliscore output appears to be empty. Check results for BUSCO: 1379600at2759
## Mon Jul 25 16:59:07 CEST 2022 - The aliscore output file does not exist. Check results for BUSCO: 1248958at2759
## Mon Jul 25 16:59:07 CEST 2022 - The aliscore output appears to be empty. Check results for BUSCO: 1248958at2759
## Mon Jul 25 16:59:07 CEST 2022 - The aliscore output file does not exist. Check results for BUSCO: 396755at2759
## Mon Jul 25 16:59:07 CEST 2022 - The aliscore output appears to be empty. Check results for BUSCO: 396755at2759
## Mon Jul 25 16:59:08 CEST 2022 - The aliscore output file does not exist. Check results for BUSCO: 1108845at2759
## Mon Jul 25 16:59:08 CEST 2022 - The aliscore output appears to be empty. Check results for BUSCO: 1108845at2759
## Mon Jul 25 16:59:08 CEST 2022 - The aliscore output file does not exist. Check results for BUSCO: 1041560at2759
## Mon Jul 25 16:59:08 CEST 2022 - The aliscore output appears to be empty. Check results for BUSCO: 1041560at2759
## Mon Jul 25 16:59:08 CEST 2022 - The aliscore output file does not exist. Check results for BUSCO: 1324510at2759
## Mon Jul 25 16:59:08 CEST 2022 - The aliscore output appears to be empty. Check results for BUSCO: 1324510at2759
## Mon Jul 25 16:59:09 CEST 2022 - The aliscore output file does not exist. Check results for BUSCO: 245208at2759
## Mon Jul 25 16:59:09 CEST 2022 - The aliscore output appears to be empty. Check results for BUSCO: 245208at2759
## Mon Jul 25 16:59:09 CEST 2022 - The aliscore output file does not exist. Check results for BUSCO: 369837at2759
## Mon Jul 25 16:59:09 CEST 2022 - The aliscore output file does not exist. Check results for BUSCO: 917326at2759
## Mon Jul 25 16:59:09 CEST 2022 - The aliscore output appears to be empty. Check results for BUSCO: 369837at2759
## Mon Jul 25 16:59:09 CEST 2022 - The aliscore output file does not exist. Check results for BUSCO: 261328at2759
## Mon Jul 25 16:59:09 CEST 2022 - The aliscore output appears to be empty. Check results for BUSCO: 917326at2759
## Mon Jul 25 16:59:09 CEST 2022 - The aliscore output file does not exist. Check results for BUSCO: 39650at2759
## Mon Jul 25 16:59:09 CEST 2022 - The aliscore output appears to be empty. Check results for BUSCO: 261328at2759
## Mon Jul 25 16:59:09 CEST 2022 - The aliscore output appears to be empty. Check results for BUSCO: 39650at2759
## Mon Jul 25 16:59:10 CEST 2022 - The aliscore output file does not exist. Check results for BUSCO: 87842at2759
## Mon Jul 25 16:59:10 CEST 2022 - The aliscore output appears to be empty. Check results for BUSCO: 87842at2759
## Mon Jul 25 16:59:30 CEST 2022 - The aliscore output file does not exist. Check results for BUSCO: 1249159at2759
## Mon Jul 25 16:59:30 CEST 2022 - The aliscore output appears to be empty. Check results for BUSCO: 1249159at2759
## Mon Jul 25 16:59:31 CEST 2022 - The aliscore output file does not exist. Check results for BUSCO: 1617752at2759
## Mon Jul 25 16:59:31 CEST 2022 - The aliscore output appears to be empty. Check results for BUSCO: 1617752at2759
## Mon Jul 25 16:59:31 CEST 2022 - The aliscore output file does not exist. Check results for BUSCO: 1269244at2759
## Mon Jul 25 16:59:31 CEST 2022 - The aliscore output appears to be empty. Check results for BUSCO: 1269244at2759
## Mon Jul 25 16:59:31 CEST 2022 - The aliscore output file does not exist. Check results for BUSCO: 1474000at2759
## Mon Jul 25 16:59:31 CEST 2022 - The aliscore output appears to be empty. Check results for BUSCO: 1474000at2759
## Mon Jul 25 16:59:31 CEST 2022 - The aliscore output file does not exist. Check results for BUSCO: 1269649at2759
## Mon Jul 25 16:59:31 CEST 2022 - The aliscore output appears to be empty. Check results for BUSCO: 1269649at2759
## Mon Jul 25 16:59:32 CEST 2022 - The aliscore output file does not exist. Check results for BUSCO: 1348942at2759
## Mon Jul 25 16:59:32 CEST 2022 - The aliscore output appears to be empty. Check results for BUSCO: 1348942at2759
## Mon Jul 25 16:59:32 CEST 2022 - The aliscore output file does not exist. Check results for BUSCO: 679771at2759
## Mon Jul 25 16:59:32 CEST 2022 - The aliscore output appears to be empty. Check results for BUSCO: 679771at2759
## Mon Jul 25 16:59:32 CEST 2022 - The aliscore output file does not exist. Check results for BUSCO: 869548at2759
## Mon Jul 25 16:59:32 CEST 2022 - The aliscore output appears to be empty. Check results for BUSCO: 869548at2759
## Mon Jul 25 16:59:32 CEST 2022 - The aliscore output file does not exist. Check results for BUSCO: 1107630at2759
## Mon Jul 25 16:59:32 CEST 2022 - The aliscore output appears to be empty. Check results for BUSCO: 1107630at2759
## Mon Jul 25 16:59:32 CEST 2022 - The aliscore output file does not exist. Check results for BUSCO: 1161199at2759
## Mon Jul 25 16:59:32 CEST 2022 - The aliscore output appears to be empty. Check results for BUSCO: 1161199at2759
## Mon Jul 25 16:59:33 CEST 2022 - The aliscore output file does not exist. Check results for BUSCO: 290630at2759
## Mon Jul 25 16:59:33 CEST 2022 - The aliscore output appears to be empty. Check results for BUSCO: 290630at2759
## Mon Jul 25 16:59:34 CEST 2022 - The aliscore output file does not exist. Check results for BUSCO: 1051021at2759
## Mon Jul 25 16:59:34 CEST 2022 - The aliscore output appears to be empty. Check results for BUSCO: 1051021at2759
## Mon Jul 25 16:59:34 CEST 2022 - The aliscore output file does not exist. Check results for BUSCO: 1430056at2759
## Mon Jul 25 16:59:34 CEST 2022 - The aliscore output appears to be empty. Check results for BUSCO: 1430056at2759
## Mon Jul 25 16:59:34 CEST 2022 - The aliscore output file does not exist. Check results for BUSCO: 1038775at2759
## Mon Jul 25 16:59:34 CEST 2022 - The aliscore output appears to be empty. Check results for BUSCO: 1038775at2759
## Mon Jul 25 16:59:34 CEST 2022 - The aliscore output file does not exist. Check results for BUSCO: 878143at2759
## Mon Jul 25 16:59:34 CEST 2022 - The aliscore output appears to be empty. Check results for BUSCO: 878143at2759
## Mon Jul 25 16:59:34 CEST 2022 - The aliscore output file does not exist. Check results for BUSCO: 1358374at2759
## Mon Jul 25 16:59:34 CEST 2022 - The aliscore output file does not exist. Check results for BUSCO: 923657at2759
## Mon Jul 25 16:59:34 CEST 2022 - The aliscore output appears to be empty. Check results for BUSCO: 923657at2759
## 
## Mon Jul 25 16:59:35 CEST 2022 - The aliscore output file does not exist. Check results for BUSCO: 1129824at2759
## Mon Jul 25 16:59:35 CEST 2022 - The aliscore output appears to be empty. Check results for BUSCO: 1129824at2759
## Mon Jul 25 16:59:35 CEST 2022 - The aliscore output file does not exist. Check results for BUSCO: 97116at2759
## Mon Jul 25 16:59:35 CEST 2022 - The aliscore output appears to be empty. Check results for BUSCO: 97116at2759
## Mon Jul 25 16:59:35 CEST 2022 - The aliscore output file does not exist. Check results for BUSCO: 1251252at2759
## Mon Jul 25 16:59:35 CEST 2022 - The aliscore output appears to be empty. Check results for BUSCO: 1251252at2759
## Mon Jul 25 16:59:36 CEST 2022 - The aliscore output file does not exist. Check results for BUSCO: 621827at2759
## Mon Jul 25 16:59:36 CEST 2022 - The aliscore output appears to be empty. Check results for BUSCO: 621827at2759
## Mon Jul 25 16:59:36 CEST 2022 - The aliscore output file does not exist. Check results for BUSCO: 918816at2759
## Mon Jul 25 16:59:36 CEST 2022 - The aliscore output appears to be empty. Check results for BUSCO: 918816at2759
## Mon Jul 25 16:59:36 CEST 2022 - The aliscore output file does not exist. Check results for BUSCO: 687505at2759
## Mon Jul 25 16:59:36 CEST 2022 - The aliscore output appears to be empty. Check results for BUSCO: 687505at2759
## Mon Jul 25 16:59:36 CEST 2022 - The aliscore output file does not exist. Check results for BUSCO: 673132at2759
## Mon Jul 25 16:59:36 CEST 2022 - The aliscore output appears to be empty. Check results for BUSCO: 673132at2759
## Mon Jul 25 16:59:36 CEST 2022 - The aliscore output file does not exist. Check results for BUSCO: 817008at2759
## Mon Jul 25 16:59:36 CEST 2022 - The aliscore output appears to be empty. Check results for BUSCO: 817008at2759
## Mon Jul 25 16:59:36 CEST 2022 - The aliscore output file does not exist. Check results for BUSCO: 1323575at2759
## Mon Jul 25 16:59:36 CEST 2022 - The aliscore output appears to be empty. Check results for BUSCO: 1323575at2759
## Mon Jul 25 16:59:36 CEST 2022 - The aliscore output file does not exist. Check results for BUSCO: 1404162at2759
## Mon Jul 25 16:59:36 CEST 2022 - The aliscore output file does not exist. Check results for BUSCO: 551907at2759
## Mon Jul 25 16:59:36 CEST 2022 - The aliscore output appears to be empty. Check results for BUSCO: 551907at2759
## Mon Jul 25 16:59:36 CEST 2022 - The aliscore output file does not exist. Check results for BUSCO: 981902at2759
## Mon Jul 25 16:59:36 CEST 2022 - The aliscore output appears to be empty. Check results for BUSCO: 1404162at2759
## Mon Jul 25 16:59:36 CEST 2022 - The aliscore output appears to be empty. Check results for BUSCO: 981902at2759
## Mon Jul 25 16:59:37 CEST 2022 - The aliscore output file does not exist. Check results for BUSCO: 674169at2759
## Mon Jul 25 16:59:37 CEST 2022 - The aliscore output appears to be empty. Check results for BUSCO: 674169at2759
## Mon Jul 25 16:59:37 CEST 2022 - The aliscore output file does not exist. Check results for BUSCO: 1423847at2759
## Mon Jul 25 16:59:37 CEST 2022 - The aliscore output file does not exist. Check results for BUSCO: 692986at2759
## Mon Jul 25 16:59:37 CEST 2022 - The aliscore output file does not exist. Check results for BUSCO: 761109at2759
## Mon Jul 25 16:59:37 CEST 2022 - The aliscore output appears to be empty. Check results for BUSCO: 692986at2759
## Mon Jul 25 16:59:37 CEST 2022 - The aliscore output appears to be empty. Check results for BUSCO: 761109at2759
## Mon Jul 25 16:59:37 CEST 2022 - The aliscore output appears to be empty. Check results for BUSCO: 1423847at2759
## Mon Jul 25 16:59:37 CEST 2022 - The aliscore output file does not exist. Check results for BUSCO: 450058at2759
## Mon Jul 25 16:59:37 CEST 2022 - The aliscore output appears to be empty. Check results for BUSCO: 450058at2759
## Mon Jul 25 16:59:37 CEST 2022 - The aliscore output file does not exist. Check results for BUSCO: 543764at2759
## Mon Jul 25 16:59:37 CEST 2022 - The aliscore output appears to be empty. Check results for BUSCO: 543764at2759
## Mon Jul 25 16:59:37 CEST 2022 - The aliscore output file does not exist. Check results for BUSCO: 83779at2759
## Mon Jul 25 16:59:37 CEST 2022 - The aliscore output file does not exist. Check results for BUSCO: 939345at2759
## Mon Jul 25 16:59:37 CEST 2022 - The aliscore output appears to be empty. Check results for BUSCO: 83779at2759
## Mon Jul 25 16:59:37 CEST 2022 - The aliscore output appears to be empty. Check results for BUSCO: 939345at2759
## Mon Jul 25 16:59:38 CEST 2022 - The aliscore output file does not exist. Check results for BUSCO: 604979at2759
## Mon Jul 25 16:59:38 CEST 2022 - The aliscore output file does not exist. Check results for BUSCO: 320059at2759
## Mon Jul 25 16:59:38 CEST 2022 - The aliscore output appears to be empty. Check results for BUSCO: 604979at2759
## Mon Jul 25 16:59:38 CEST 2022 - The aliscore output appears to be empty. Check results for BUSCO: 320059at2759
## Mon Jul 25 16:59:38 CEST 2022 - The aliscore output file does not exist. Check results for BUSCO: 1079130at2759
## Mon Jul 25 16:59:38 CEST 2022 - The aliscore output appears to be empty. Check results for BUSCO: 1079130at2759
## Mon Jul 25 16:59:39 CEST 2022 - The aliscore output file does not exist. Check results for BUSCO: 671536at2759
## Mon Jul 25 16:59:39 CEST 2022 - The aliscore output appears to be empty. Check results for BUSCO: 671536at2759
## Mon Jul 25 16:59:39 CEST 2022 - The aliscore output file does not exist. Check results for BUSCO: 324863at2759
## Mon Jul 25 16:59:39 CEST 2022 - The aliscore output appears to be empty. Check results for BUSCO: 324863at2759
## Mon Jul 25 16:59:40 CEST 2022 - The aliscore output file does not exist. Check results for BUSCO: 1194797at2759
## Mon Jul 25 16:59:40 CEST 2022 - The aliscore output file does not exist. Check results for BUSCO: 142542at2759
## Mon Jul 25 16:59:40 CEST 2022 - The aliscore output appears to be empty. Check results for BUSCO: 1194797at2759
## Mon Jul 25 16:59:40 CEST 2022 - The aliscore output appears to be empty. Check results for BUSCO: 142542at2759
## Mon Jul 25 16:59:40 CEST 2022 - The aliscore output file does not exist. Check results for BUSCO: 271586at2759
## Mon Jul 25 16:59:40 CEST 2022 - The aliscore output appears to be empty. Check results for BUSCO: 271586at2759
## Mon Jul 25 17:00:02 CEST 2022 - The aliscore output file does not exist. Check results for BUSCO: 1552706at2759
## Mon Jul 25 17:00:02 CEST 2022 - The aliscore output appears to be empty. Check results for BUSCO: 1552706at2759
## Mon Jul 25 17:00:02 CEST 2022 - The aliscore output file does not exist. Check results for BUSCO: 1339553at2759
## Mon Jul 25 17:00:02 CEST 2022 - The aliscore output appears to be empty. Check results for BUSCO: 1339553at2759
## Mon Jul 25 17:00:03 CEST 2022 - The aliscore output file does not exist. Check results for BUSCO: 996662at2759
## Mon Jul 25 17:00:03 CEST 2022 - The aliscore output appears to be empty. Check results for BUSCO: 996662at2759
## Mon Jul 25 17:00:03 CEST 2022 - The aliscore output file does not exist. Check results for BUSCO: 1355894at2759
## Mon Jul 25 17:00:03 CEST 2022 - The aliscore output file does not exist. Check results for BUSCO: 1440961at2759
## Mon Jul 25 17:00:03 CEST 2022 - The aliscore output appears to be empty. Check results for BUSCO: 1355894at2759
## Mon Jul 25 17:00:03 CEST 2022 - The aliscore output appears to be empty. Check results for BUSCO: 1440961at2759
## Mon Jul 25 17:00:03 CEST 2022 - The aliscore output file does not exist. Check results for BUSCO: 1236198at2759
## Mon Jul 25 17:00:03 CEST 2022 - The aliscore output appears to be empty. Check results for BUSCO: 1236198at2759
## Mon Jul 25 17:00:03 CEST 2022 - The aliscore output file does not exist. Check results for BUSCO: 1405073at2759
## Mon Jul 25 17:00:03 CEST 2022 - The aliscore output appears to be empty. Check results for BUSCO: 1405073at2759
## Mon Jul 25 17:00:03 CEST 2022 - The aliscore output file does not exist. Check results for BUSCO: 1217666at2759
## Mon Jul 25 17:00:03 CEST 2022 - The aliscore output file does not exist. Check results for BUSCO: 1545004at2759
## Mon Jul 25 17:00:03 CEST 2022 - The aliscore output file does not exist. Check results for BUSCO: 1620056at2759
## Mon Jul 25 17:00:03 CEST 2022 - The aliscore output appears to be empty. Check results for BUSCO: 1217666at2759
## Mon Jul 25 17:00:03 CEST 2022 - The aliscore output appears to be empty. Check results for BUSCO: 1545004at2759
## Mon Jul 25 17:00:03 CEST 2022 - The aliscore output file does not exist. Check results for BUSCO: 1108845at2759
## Mon Jul 25 17:00:03 CEST 2022 - The aliscore output appears to be empty. Check results for BUSCO: 1620056at2759
## Mon Jul 25 17:00:03 CEST 2022 - The aliscore output file does not exist. Check results for BUSCO: 1395649at2759
## Mon Jul 25 17:00:03 CEST 2022 - The aliscore output appears to be empty. Check results for BUSCO: 1108845at2759
## Mon Jul 25 17:00:03 CEST 2022 - The aliscore output appears to be empty. Check results for BUSCO: 1395649at2759
## Mon Jul 25 17:00:03 CEST 2022 - The aliscore output file does not exist. Check results for BUSCO: 1197019at2759
## Mon Jul 25 17:00:03 CEST 2022 - The aliscore output file does not exist. Check results for BUSCO: 655400at2759
## Mon Jul 25 17:00:03 CEST 2022 - The aliscore output appears to be empty. Check results for BUSCO: 1197019at2759
## Mon Jul 25 17:00:03 CEST 2022 - The aliscore output appears to be empty. Check results for BUSCO: 655400at2759
## Mon Jul 25 17:00:04 CEST 2022 - The aliscore output file does not exist. Check results for BUSCO: 1530008at2759
## Mon Jul 25 17:00:04 CEST 2022 - The aliscore output appears to be empty. Check results for BUSCO: 1530008at2759
## Mon Jul 25 17:00:04 CEST 2022 - The aliscore output file does not exist. Check results for BUSCO: 1247641at2759
## Mon Jul 25 17:00:04 CEST 2022 - The aliscore output appears to be empty. Check results for BUSCO: 1247641at2759
## Mon Jul 25 17:00:04 CEST 2022 - The aliscore output file does not exist. Check results for BUSCO: 1053181at2759
## Mon Jul 25 17:00:04 CEST 2022 - The aliscore output file does not exist. Check results for BUSCO: 1222562at2759
## Mon Jul 25 17:00:04 CEST 2022 - The aliscore output appears to be empty. Check results for BUSCO: 1053181at2759
## Mon Jul 25 17:00:04 CEST 2022 - The aliscore output appears to be empty. Check results for BUSCO: 1222562at2759
## Mon Jul 25 17:00:04 CEST 2022 - The aliscore output file does not exist. Check results for BUSCO: 1455730at2759
## Mon Jul 25 17:00:04 CEST 2022 - The aliscore output appears to be empty. Check results for BUSCO: 1455730at2759
## Mon Jul 25 17:00:04 CEST 2022 - The aliscore output file does not exist. Check results for BUSCO: 1405146at2759
## Mon Jul 25 17:00:04 CEST 2022 - The aliscore output appears to be empty. Check results for BUSCO: 1405146at2759
## Mon Jul 25 17:00:04 CEST 2022 - The aliscore output file does not exist. Check results for BUSCO: 1504863at2759
## Mon Jul 25 17:00:04 CEST 2022 - The aliscore output file does not exist. Check results for BUSCO: 721605at2759
## Mon Jul 25 17:00:04 CEST 2022 - The aliscore output appears to be empty. Check results for BUSCO: 721605at2759
## Mon Jul 25 17:00:04 CEST 2022 - The aliscore output appears to be empty. Check results for BUSCO: 1504863at2759
## Mon Jul 25 17:00:04 CEST 2022 - The aliscore output file does not exist. Check results for BUSCO: 1346432at2759
## Mon Jul 25 17:00:04 CEST 2022 - The aliscore output appears to be empty. Check results for BUSCO: 1346432at2759
## Mon Jul 25 17:00:04 CEST 2022 - The aliscore output file does not exist. Check results for BUSCO: 1338131at2759
## Mon Jul 25 17:00:04 CEST 2022 - The aliscore output appears to be empty. Check results for BUSCO: 1338131at2759
## Mon Jul 25 17:00:04 CEST 2022 - The aliscore output file does not exist. Check results for BUSCO: 1111142at2759
## Mon Jul 25 17:00:04 CEST 2022 - The aliscore output appears to be empty. Check results for BUSCO: 1111142at2759
## Mon Jul 25 17:00:04 CEST 2022 - The aliscore output file does not exist. Check results for BUSCO: 1588798at2759
## Mon Jul 25 17:00:05 CEST 2022 - The aliscore output appears to be empty. Check results for BUSCO: 1588798at2759
## Mon Jul 25 17:00:05 CEST 2022 - The aliscore output file does not exist. Check results for BUSCO: 1228942at2759
## Mon Jul 25 17:00:05 CEST 2022 - The aliscore output file does not exist. Check results for BUSCO: 911863at2759
## Mon Jul 25 17:00:05 CEST 2022 - The aliscore output appears to be empty. Check results for BUSCO: 1228942at2759
## Mon Jul 25 17:00:05 CEST 2022 - The aliscore output appears to be empty. Check results for BUSCO: 911863at2759
## Mon Jul 25 17:00:05 CEST 2022 - The aliscore output file does not exist. Check results for BUSCO: 974865at2759
## Mon Jul 25 17:00:05 CEST 2022 - The aliscore output appears to be empty. Check results for BUSCO: 974865at2759
## Mon Jul 25 17:00:05 CEST 2022 - The aliscore output file does not exist. Check results for BUSCO: 801857at2759
## Mon Jul 25 17:00:05 CEST 2022 - The aliscore output appears to be empty. Check results for BUSCO: 801857at2759
## Mon Jul 25 17:00:05 CEST 2022 - The aliscore output file does not exist. Check results for BUSCO: 1563319at2759
## Mon Jul 25 17:00:05 CEST 2022 - The aliscore output appears to be empty. Check results for BUSCO: 1563319at2759
## Mon Jul 25 17:00:05 CEST 2022 - The aliscore output file does not exist. Check results for BUSCO: 734666at2759
## Mon Jul 25 17:00:05 CEST 2022 - The aliscore output appears to be empty. Check results for BUSCO: 734666at2759
## Mon Jul 25 17:00:05 CEST 2022 - The aliscore output file does not exist. Check results for BUSCO: 1576404at2759
## Mon Jul 25 17:00:05 CEST 2022 - The aliscore output appears to be empty. Check results for BUSCO: 1576404at2759
## Mon Jul 25 17:00:05 CEST 2022 - The aliscore output file does not exist. Check results for BUSCO: 1370285at2759
## Mon Jul 25 17:00:05 CEST 2022 - The aliscore output appears to be empty. Check results for BUSCO: 1370285at2759
## Mon Jul 25 17:00:05 CEST 2022 - The aliscore output file does not exist. Check results for BUSCO: 956854at2759
## Mon Jul 25 17:00:05 CEST 2022 - The aliscore output appears to be empty. Check results for BUSCO: 956854at2759
## Mon Jul 25 17:00:05 CEST 2022 - The aliscore output file does not exist. Check results for BUSCO: 1001705at2759
## Mon Jul 25 17:00:05 CEST 2022 - The aliscore output file does not exist. Check results for BUSCO: 1421503at2759
## Mon Jul 25 17:00:05 CEST 2022 - The aliscore output appears to be empty. Check results for BUSCO: 1001705at2759
## Mon Jul 25 17:00:05 CEST 2022 - The aliscore output file does not exist. Check results for BUSCO: 1377237at2759
## Mon Jul 25 17:00:05 CEST 2022 - The aliscore output appears to be empty. Check results for BUSCO: 1421503at2759
## Mon Jul 25 17:00:05 CEST 2022 - The aliscore output appears to be empty. Check results for BUSCO: 1377237at2759
## Mon Jul 25 17:00:05 CEST 2022 - The aliscore output file does not exist. Check results for BUSCO: 719531at2759
## Mon Jul 25 17:00:06 CEST 2022 - The aliscore output appears to be empty. Check results for BUSCO: 719531at2759
## Mon Jul 25 17:00:06 CEST 2022 - The aliscore output file does not exist. Check results for BUSCO: 1593937at2759
## Mon Jul 25 17:00:06 CEST 2022 - The aliscore output appears to be empty. Check results for BUSCO: 1593937at2759
## Mon Jul 25 17:00:06 CEST 2022 - The aliscore output file does not exist. Check results for BUSCO: 1513531at2759
## Mon Jul 25 17:00:06 CEST 2022 - The aliscore output appears to be empty. Check results for BUSCO: 1513531at2759
## Mon Jul 25 17:00:06 CEST 2022 - The aliscore output file does not exist. Check results for BUSCO: 975158at2759
## Mon Jul 25 17:00:06 CEST 2022 - The aliscore output appears to be empty. Check results for BUSCO: 975158at2759
## Mon Jul 25 17:00:06 CEST 2022 - The aliscore output file does not exist. Check results for BUSCO: 1434266at2759
## Mon Jul 25 17:00:06 CEST 2022 - The aliscore output file does not exist. Check results for BUSCO: 1450538at2759
## Mon Jul 25 17:00:06 CEST 2022 - The aliscore output appears to be empty. Check results for BUSCO: 1434266at2759
## Mon Jul 25 17:00:06 CEST 2022 - The aliscore output appears to be empty. Check results for BUSCO: 1450538at2759
## Mon Jul 25 17:00:06 CEST 2022 - The aliscore output file does not exist. Check results for BUSCO: 1019762at2759
## Mon Jul 25 17:00:06 CEST 2022 - The aliscore output appears to be empty. Check results for BUSCO: 1019762at2759
## Mon Jul 25 17:00:06 CEST 2022 - The aliscore output file does not exist. Check results for BUSCO: 1096688at2759
## Mon Jul 25 17:00:06 CEST 2022 - The aliscore output appears to be empty. Check results for BUSCO: 1096688at2759
## Mon Jul 25 17:00:06 CEST 2022 - The aliscore output file does not exist. Check results for BUSCO: 1260807at2759
## Mon Jul 25 17:00:06 CEST 2022 - The aliscore output appears to be empty. Check results for BUSCO: 1260807at2759
## Mon Jul 25 17:00:06 CEST 2022 - The aliscore output file does not exist. Check results for BUSCO: 330169at2759
## Mon Jul 25 17:00:06 CEST 2022 - The aliscore output appears to be empty. Check results for BUSCO: 330169at2759
## Mon Jul 25 17:00:07 CEST 2022 - The aliscore output file does not exist. Check results for BUSCO: 345441at2759
## Mon Jul 25 17:00:07 CEST 2022 - The aliscore output appears to be empty. Check results for BUSCO: 345441at2759
## Mon Jul 25 17:00:07 CEST 2022 - The aliscore output file does not exist. Check results for BUSCO: 855834at2759
## Mon Jul 25 17:00:07 CEST 2022 - The aliscore output appears to be empty. Check results for BUSCO: 855834at2759
## Mon Jul 25 17:00:07 CEST 2022 - The aliscore output file does not exist. Check results for BUSCO: 299766at2759
## Mon Jul 25 17:00:07 CEST 2022 - The aliscore output appears to be empty. Check results for BUSCO: 299766at2759
## Mon Jul 25 17:00:07 CEST 2022 - The aliscore output file does not exist. Check results for BUSCO: 1432328at2759
## Mon Jul 25 17:00:07 CEST 2022 - The aliscore output appears to be empty. Check results for BUSCO: 1432328at2759
## Mon Jul 25 17:00:07 CEST 2022 - The aliscore output file does not exist. Check results for BUSCO: 975557at2759
## Mon Jul 25 17:00:07 CEST 2022 - The aliscore output file does not exist. Check results for BUSCO: 1370304at2759
## Mon Jul 25 17:00:07 CEST 2022 - The aliscore output appears to be empty. Check results for BUSCO: 975557at2759
## Mon Jul 25 17:00:07 CEST 2022 - The aliscore output appears to be empty. Check results for BUSCO: 1370304at2759
## Mon Jul 25 17:00:07 CEST 2022 - The aliscore output file does not exist. Check results for BUSCO: 293315at2759
## Mon Jul 25 17:00:07 CEST 2022 - The aliscore output appears to be empty. Check results for BUSCO: 293315at2759
## Mon Jul 25 17:00:07 CEST 2022 - The aliscore output file does not exist. Check results for BUSCO: 1106766at2759
## Mon Jul 25 17:00:07 CEST 2022 - The aliscore output appears to be empty. Check results for BUSCO: 1106766at2759
## Mon Jul 25 17:00:07 CEST 2022 - The aliscore output file does not exist. Check results for BUSCO: 598949at2759
## Mon Jul 25 17:00:07 CEST 2022 - The aliscore output file does not exist. Check results for BUSCO: 341721at2759
## Mon Jul 25 17:00:07 CEST 2022 - The aliscore output appears to be empty. Check results for BUSCO: 598949at2759
## Mon Jul 25 17:00:08 CEST 2022 - The aliscore output appears to be empty. Check results for BUSCO: 341721at2759
## Mon Jul 25 17:00:08 CEST 2022 - The aliscore output file does not exist. Check results for BUSCO: 392369at2759
## Mon Jul 25 17:00:08 CEST 2022 - The aliscore output appears to be empty. Check results for BUSCO: 392369at2759
## Mon Jul 25 17:00:08 CEST 2022 - The aliscore output file does not exist. Check results for BUSCO: 671846at2759
## Mon Jul 25 17:00:08 CEST 2022 - The aliscore output appears to be empty. Check results for BUSCO: 671846at2759
## Mon Jul 25 17:00:08 CEST 2022 - The aliscore output file does not exist. Check results for BUSCO: 245208at2759
## Mon Jul 25 17:00:08 CEST 2022 - The aliscore output appears to be empty. Check results for BUSCO: 245208at2759
## Mon Jul 25 17:00:09 CEST 2022 - The aliscore output file does not exist. Check results for BUSCO: 296129at2759
## Mon Jul 25 17:00:09 CEST 2022 - The aliscore output appears to be empty. Check results for BUSCO: 296129at2759
## Mon Jul 25 17:00:09 CEST 2022 - The aliscore output file does not exist. Check results for BUSCO: 1087488at2759
## Mon Jul 25 17:00:09 CEST 2022 - The aliscore output file does not exist. Check results for BUSCO: 1314980at2759
## Mon Jul 25 17:00:09 CEST 2022 - The aliscore output appears to be empty. Check results for BUSCO: 1087488at2759
## Mon Jul 25 17:00:09 CEST 2022 - The aliscore output appears to be empty. Check results for BUSCO: 1314980at2759
## Mon Jul 25 17:00:09 CEST 2022 - The aliscore output file does not exist. Check results for BUSCO: 768809at2759
## Mon Jul 25 17:00:09 CEST 2022 - The aliscore output appears to be empty. Check results for BUSCO: 768809at2759
## Mon Jul 25 17:00:09 CEST 2022 - The aliscore output file does not exist. Check results for BUSCO: 176625at2759
## Mon Jul 25 17:00:09 CEST 2022 - The aliscore output file does not exist. Check results for BUSCO: 270107at2759
## Mon Jul 25 17:00:09 CEST 2022 - The aliscore output appears to be empty. Check results for BUSCO: 176625at2759
## Mon Jul 25 17:00:09 CEST 2022 - The aliscore output appears to be empty. Check results for BUSCO: 270107at2759
## Mon Jul 25 17:00:09 CEST 2022 - The aliscore output file does not exist. Check results for BUSCO: 931188at2759
## Mon Jul 25 17:00:10 CEST 2022 - The aliscore output appears to be empty. Check results for BUSCO: 931188at2759
## Mon Jul 25 17:00:10 CEST 2022 - The aliscore output file does not exist. Check results for BUSCO: 625387at2759
## Mon Jul 25 17:00:10 CEST 2022 - The aliscore output appears to be empty. Check results for BUSCO: 625387at2759
## Mon Jul 25 17:00:10 CEST 2022 - The aliscore output file does not exist. Check results for BUSCO: 396755at2759
## Mon Jul 25 17:00:10 CEST 2022 - The aliscore output appears to be empty. Check results for BUSCO: 396755at2759
## Mon Jul 25 17:00:31 CEST 2022 - The aliscore output file does not exist. Check results for BUSCO: 1200489at2759
## Mon Jul 25 17:00:31 CEST 2022 - The aliscore output file does not exist. Check results for BUSCO: 1079827at2759
## Mon Jul 25 17:00:31 CEST 2022 - The aliscore output appears to be empty. Check results for BUSCO: 1200489at2759
## Mon Jul 25 17:00:31 CEST 2022 - The aliscore output appears to be empty. Check results for BUSCO: 1079827at2759
## Mon Jul 25 17:00:32 CEST 2022 - The aliscore output file does not exist. Check results for BUSCO: 1287401at2759
## Mon Jul 25 17:00:32 CEST 2022 - The aliscore output appears to be empty. Check results for BUSCO: 1287401at2759
## Mon Jul 25 17:00:32 CEST 2022 - The aliscore output file does not exist. Check results for BUSCO: 290630at2759
## Mon Jul 25 17:00:32 CEST 2022 - The aliscore output appears to be empty. Check results for BUSCO: 290630at2759
## Mon Jul 25 17:00:32 CEST 2022 - The aliscore output file does not exist. Check results for BUSCO: 1575179at2759
## Mon Jul 25 17:00:32 CEST 2022 - The aliscore output appears to be empty. Check results for BUSCO: 1575179at2759
## Mon Jul 25 17:00:33 CEST 2022 - The aliscore output file does not exist. Check results for BUSCO: 1426075at2759
## Mon Jul 25 17:00:33 CEST 2022 - The aliscore output appears to be empty. Check results for BUSCO: 1426075at2759
## Mon Jul 25 17:00:33 CEST 2022 - The aliscore output file does not exist. Check results for BUSCO: 919955at2759
## Mon Jul 25 17:00:33 CEST 2022 - The aliscore output appears to be empty. Check results for BUSCO: 919955at2759
## Mon Jul 25 17:00:33 CEST 2022 - The aliscore output file does not exist. Check results for BUSCO: 944899at2759
## Mon Jul 25 17:00:33 CEST 2022 - The aliscore output appears to be empty. Check results for BUSCO: 944899at2759
## Mon Jul 25 17:00:33 CEST 2022 - The aliscore output file does not exist. Check results for BUSCO: 865202at2759
## Mon Jul 25 17:00:33 CEST 2022 - The aliscore output appears to be empty. Check results for BUSCO: 865202at2759
## Mon Jul 25 17:00:33 CEST 2022 - The aliscore output file does not exist. Check results for BUSCO: 1223488at2759
## Mon Jul 25 17:00:33 CEST 2022 - The aliscore output file does not exist. Check results for BUSCO: 1398309at2759
## Mon Jul 25 17:00:33 CEST 2022 - The aliscore output appears to be empty. Check results for BUSCO: 1223488at2759
## Mon Jul 25 17:00:33 CEST 2022 - The aliscore output appears to be empty. Check results for BUSCO: 1398309at2759
## Mon Jul 25 17:00:33 CEST 2022 - The aliscore output file does not exist. Check results for BUSCO: 1538526at2759
## Mon Jul 25 17:00:33 CEST 2022 - The aliscore output appears to be empty. Check results for BUSCO: 1538526at2759
## Mon Jul 25 17:00:34 CEST 2022 - The aliscore output file does not exist. Check results for BUSCO: 1233814at2759
## Mon Jul 25 17:00:34 CEST 2022 - The aliscore output appears to be empty. Check results for BUSCO: 1233814at2759
## Mon Jul 25 17:00:34 CEST 2022 - The aliscore output file does not exist. Check results for BUSCO: 1421503at2759
## Mon Jul 25 17:00:34 CEST 2022 - The aliscore output appears to be empty. Check results for BUSCO: 1421503at2759
## Mon Jul 25 17:00:34 CEST 2022 - The aliscore output file does not exist. Check results for BUSCO: 1128607at2759
## Mon Jul 25 17:00:34 CEST 2022 - The aliscore output appears to be empty. Check results for BUSCO: 1128607at2759
## Mon Jul 25 17:00:35 CEST 2022 - The aliscore output file does not exist. Check results for BUSCO: 1342242at2759
## Mon Jul 25 17:00:35 CEST 2022 - The aliscore output appears to be empty. Check results for BUSCO: 1342242at2759
## Mon Jul 25 17:00:35 CEST 2022 - The aliscore output file does not exist. Check results for BUSCO: 1264469at2759
## Mon Jul 25 17:00:35 CEST 2022 - The aliscore output file does not exist. Check results for BUSCO: 1445102at2759
## Mon Jul 25 17:00:35 CEST 2022 - The aliscore output appears to be empty. Check results for BUSCO: 1445102at2759
## Mon Jul 25 17:00:35 CEST 2022 - The aliscore output appears to be empty. Check results for BUSCO: 1264469at2759
## Mon Jul 25 17:00:35 CEST 2022 - The aliscore output file does not exist. Check results for BUSCO: 1382842at2759
## Mon Jul 25 17:00:35 CEST 2022 - The aliscore output appears to be empty. Check results for BUSCO: 1382842at2759
## Mon Jul 25 17:00:35 CEST 2022 - The aliscore output file does not exist. Check results for BUSCO: 1138059at2759
## Mon Jul 25 17:00:35 CEST 2022 - The aliscore output appears to be empty. Check results for BUSCO: 1138059at2759
## Mon Jul 25 17:00:35 CEST 2022 - The aliscore output file does not exist. Check results for BUSCO: 1018517at2759
## Mon Jul 25 17:00:35 CEST 2022 - The aliscore output appears to be empty. Check results for BUSCO: 1018517at2759
## Mon Jul 25 17:00:35 CEST 2022 - The aliscore output file does not exist. Check results for BUSCO: 751335at2759
## Mon Jul 25 17:00:35 CEST 2022 - The aliscore output appears to be empty. Check results for BUSCO: 751335at2759
## Mon Jul 25 17:00:35 CEST 2022 - The aliscore output file does not exist. Check results for BUSCO: 917326at2759
## Mon Jul 25 17:00:35 CEST 2022 - The aliscore output appears to be empty. Check results for BUSCO: 917326at2759
## Mon Jul 25 17:00:36 CEST 2022 - The aliscore output file does not exist. Check results for BUSCO: 1626636at2759
## Mon Jul 25 17:00:36 CEST 2022 - The aliscore output appears to be empty. Check results for BUSCO: 1626636at2759
## Mon Jul 25 17:00:36 CEST 2022 - The aliscore output file does not exist. Check results for BUSCO: 1112002at2759
## Mon Jul 25 17:00:36 CEST 2022 - The aliscore output appears to be empty. Check results for BUSCO: 1112002at2759
## Mon Jul 25 17:00:36 CEST 2022 - The aliscore output file does not exist. Check results for BUSCO: 937275at2759
## Mon Jul 25 17:00:36 CEST 2022 - The aliscore output appears to be empty. Check results for BUSCO: 937275at2759
## Mon Jul 25 17:00:36 CEST 2022 - The aliscore output file does not exist. Check results for BUSCO: 1490022at2759
## Mon Jul 25 17:00:36 CEST 2022 - The aliscore output appears to be empty. Check results for BUSCO: 1490022at2759
## Mon Jul 25 17:00:36 CEST 2022 - The aliscore output file does not exist. Check results for BUSCO: 1346165at2759
## Mon Jul 25 17:00:36 CEST 2022 - The aliscore output appears to be empty. Check results for BUSCO: 1346165at2759
## Mon Jul 25 17:00:36 CEST 2022 - The aliscore output file does not exist. Check results for BUSCO: 257318at2759
## Mon Jul 25 17:00:36 CEST 2022 - The aliscore output appears to be empty. Check results for BUSCO: 257318at2759
## Mon Jul 25 17:00:37 CEST 2022 - The aliscore output file does not exist. Check results for BUSCO: 774318at2759
## Mon Jul 25 17:00:37 CEST 2022 - The aliscore output appears to be empty. Check results for BUSCO: 774318at2759
## Mon Jul 25 17:00:37 CEST 2022 - The aliscore output file does not exist. Check results for BUSCO: 768809at2759
## Mon Jul 25 17:00:37 CEST 2022 - The aliscore output appears to be empty. Check results for BUSCO: 768809at2759
## Mon Jul 25 17:00:37 CEST 2022 - The aliscore output file does not exist. Check results for BUSCO: 39650at2759
## Mon Jul 25 17:00:37 CEST 2022 - The aliscore output appears to be empty. Check results for BUSCO: 39650at2759
## Mon Jul 25 17:00:37 CEST 2022 - The aliscore output file does not exist. Check results for BUSCO: 924753at2759
## Mon Jul 25 17:00:37 CEST 2022 - The aliscore output appears to be empty. Check results for BUSCO: 924753at2759
## Mon Jul 25 17:00:37 CEST 2022 - The aliscore output file does not exist. Check results for BUSCO: 1025450at2759
## Mon Jul 25 17:00:37 CEST 2022 - The aliscore output appears to be empty. Check results for BUSCO: 1025450at2759
## Mon Jul 25 17:00:38 CEST 2022 - The aliscore output file does not exist. Check results for BUSCO: 814241at2759
## Mon Jul 25 17:00:38 CEST 2022 - The aliscore output appears to be empty. Check results for BUSCO: 814241at2759
## Mon Jul 25 17:00:38 CEST 2022 - The aliscore output file does not exist. Check results for BUSCO: 331411at2759
## Mon Jul 25 17:00:38 CEST 2022 - The aliscore output appears to be empty. Check results for BUSCO: 331411at2759
## Mon Jul 25 17:00:38 CEST 2022 - The aliscore output file does not exist. Check results for BUSCO: 1041560at2759
## Mon Jul 25 17:00:38 CEST 2022 - The aliscore output appears to be empty. Check results for BUSCO: 1041560at2759
## Mon Jul 25 17:00:38 CEST 2022 - The aliscore output file does not exist. Check results for BUSCO: 898782at2759
## Mon Jul 25 17:00:38 CEST 2022 - The aliscore output file does not exist. Check results for BUSCO: 355408at2759
## Mon Jul 25 17:00:38 CEST 2022 - The aliscore output appears to be empty. Check results for BUSCO: 898782at2759
## Mon Jul 25 17:00:38 CEST 2022 - The aliscore output appears to be empty. Check results for BUSCO: 355408at2759
## Mon Jul 25 17:00:38 CEST 2022 - The aliscore output file does not exist. Check results for BUSCO: 418107at2759
## Mon Jul 25 17:00:38 CEST 2022 - The aliscore output appears to be empty. Check results for BUSCO: 418107at2759
## Mon Jul 25 17:00:39 CEST 2022 - The aliscore output file does not exist. Check results for BUSCO: 549762at2759
## Mon Jul 25 17:00:39 CEST 2022 - The aliscore output file does not exist. Check results for BUSCO: 836599at2759
## Mon Jul 25 17:00:39 CEST 2022 - The aliscore output appears to be empty. Check results for BUSCO: 549762at2759
## Mon Jul 25 17:00:39 CEST 2022 - The aliscore output appears to be empty. Check results for BUSCO: 836599at2759
## Mon Jul 25 17:00:39 CEST 2022 - The aliscore output file does not exist. Check results for BUSCO: 570797at2759
## Mon Jul 25 17:00:39 CEST 2022 - The aliscore output appears to be empty. Check results for BUSCO: 570797at2759
## Mon Jul 25 17:00:39 CEST 2022 - The aliscore output file does not exist. Check results for BUSCO: 736068at2759
## Mon Jul 25 17:00:39 CEST 2022 - The aliscore output appears to be empty. Check results for BUSCO: 736068at2759
## Mon Jul 25 17:00:40 CEST 2022 - The aliscore output file does not exist. Check results for BUSCO: 179362at2759
## Mon Jul 25 17:00:40 CEST 2022 - The aliscore output appears to be empty. Check results for BUSCO: 179362at2759
## Mon Jul 25 17:00:40 CEST 2022 - The aliscore output file does not exist. Check results for BUSCO: 722805at2759
## Mon Jul 25 17:00:40 CEST 2022 - The aliscore output appears to be empty. Check results for BUSCO: 722805at2759
## Mon Jul 25 17:00:41 CEST 2022 - The aliscore output file does not exist. Check results for BUSCO: 973442at2759
## Mon Jul 25 17:00:41 CEST 2022 - The aliscore output appears to be empty. Check results for BUSCO: 973442at2759
## Mon Jul 25 17:00:50 CEST 2022 - The aliscore output file does not exist. Check results for BUSCO: 156083at2759
## Mon Jul 25 17:00:50 CEST 2022 - The aliscore output appears to be empty. Check results for BUSCO: 156083at2759
## Mon Jul 25 17:01:01 CEST 2022 - The aliscore output file does not exist. Check results for BUSCO: 1558822at2759
## Mon Jul 25 17:01:01 CEST 2022 - The aliscore output appears to be empty. Check results for BUSCO: 1558822at2759
## Mon Jul 25 17:01:01 CEST 2022 - The aliscore output file does not exist. Check results for BUSCO: 1038775at2759
## Mon Jul 25 17:01:01 CEST 2022 - The aliscore output appears to be empty. Check results for BUSCO: 1038775at2759
## Mon Jul 25 17:01:02 CEST 2022 - The aliscore output file does not exist. Check results for BUSCO: 1049599at2759
## Mon Jul 25 17:01:02 CEST 2022 - The aliscore output file does not exist. Check results for BUSCO: 708105at2759
## Mon Jul 25 17:01:02 CEST 2022 - The aliscore output appears to be empty. Check results for BUSCO: 708105at2759
## Mon Jul 25 17:01:02 CEST 2022 - The aliscore output file does not exist. Check results for BUSCO: 1626636at2759
## Mon Jul 25 17:01:02 CEST 2022 - The aliscore output appears to be empty. Check results for BUSCO: 1626636at2759
## Mon Jul 25 17:01:02 CEST 2022 - The aliscore output appears to be empty. Check results for BUSCO: 1049599at2759
## Mon Jul 25 17:01:03 CEST 2022 - The aliscore output file does not exist. Check results for BUSCO: 1525971at2759
## Mon Jul 25 17:01:03 CEST 2022 - The aliscore output appears to be empty. Check results for BUSCO: 1525971at2759
## Mon Jul 25 17:01:03 CEST 2022 - The aliscore output file does not exist. Check results for BUSCO: 1405073at2759
## Mon Jul 25 17:01:03 CEST 2022 - The aliscore output file does not exist. Check results for BUSCO: 866359at2759
## Mon Jul 25 17:01:03 CEST 2022 - The aliscore output appears to be empty. Check results for BUSCO: 1405073at2759
## Mon Jul 25 17:01:03 CEST 2022 - The aliscore output appears to be empty. Check results for BUSCO: 866359at2759
## Mon Jul 25 17:01:03 CEST 2022 - The aliscore output file does not exist. Check results for BUSCO: 1284731at2759
## Mon Jul 25 17:01:03 CEST 2022 - The aliscore output appears to be empty. Check results for BUSCO: 1284731at2759
## Mon Jul 25 17:01:03 CEST 2022 - The aliscore output file does not exist. Check results for BUSCO: 1269649at2759
## Mon Jul 25 17:01:03 CEST 2022 - The aliscore output file does not exist. Check results for BUSCO: 1474000at2759
## Mon Jul 25 17:01:03 CEST 2022 - The aliscore output appears to be empty. Check results for BUSCO: 1269649at2759
## Mon Jul 25 17:01:03 CEST 2022 - The aliscore output appears to be empty. Check results for BUSCO: 1474000at2759
## Mon Jul 25 17:01:03 CEST 2022 - The aliscore output file does not exist. Check results for BUSCO: 1617752at2759
## Mon Jul 25 17:01:03 CEST 2022 - The aliscore output appears to be empty. Check results for BUSCO: 1617752at2759
## Mon Jul 25 17:01:03 CEST 2022 - The aliscore output file does not exist. Check results for BUSCO: 679771at2759
## Mon Jul 25 17:01:03 CEST 2022 - The aliscore output file does not exist. Check results for BUSCO: 1161199at2759
## Mon Jul 25 17:01:03 CEST 2022 - The aliscore output appears to be empty. Check results for BUSCO: 679771at2759
## Mon Jul 25 17:01:03 CEST 2022 - The aliscore output appears to be empty. Check results for BUSCO: 1161199at2759
## Mon Jul 25 17:01:03 CEST 2022 - The aliscore output file does not exist. Check results for BUSCO: 1545004at2759
## Mon Jul 25 17:01:03 CEST 2022 - The aliscore output file does not exist. Check results for BUSCO: 1488436at2759
## Mon Jul 25 17:01:03 CEST 2022 - The aliscore output appears to be empty. Check results for BUSCO: 1545004at2759
## Mon Jul 25 17:01:03 CEST 2022 - The aliscore output appears to be empty. Check results for BUSCO: 1488436at2759
## Mon Jul 25 17:01:03 CEST 2022 - The aliscore output file does not exist. Check results for BUSCO: 1459797at2759
## Mon Jul 25 17:01:03 CEST 2022 - The aliscore output appears to be empty. Check results for BUSCO: 1459797at2759
## Mon Jul 25 17:01:04 CEST 2022 - The aliscore output file does not exist. Check results for BUSCO: 1107630at2759
## Mon Jul 25 17:01:04 CEST 2022 - The aliscore output file does not exist. Check results for BUSCO: 1249159at2759
## Mon Jul 25 17:01:04 CEST 2022 - The aliscore output appears to be empty. Check results for BUSCO: 1107630at2759
## Mon Jul 25 17:01:04 CEST 2022 - The aliscore output appears to be empty. Check results for BUSCO: 1249159at2759
## Mon Jul 25 17:01:04 CEST 2022 - The aliscore output file does not exist. Check results for BUSCO: 1228942at2759
## Mon Jul 25 17:01:04 CEST 2022 - The aliscore output appears to be empty. Check results for BUSCO: 1228942at2759
## Mon Jul 25 17:01:04 CEST 2022 - The aliscore output file does not exist. Check results for BUSCO: 1304061at2759
## Mon Jul 25 17:01:04 CEST 2022 - The aliscore output appears to be empty. Check results for BUSCO: 1304061at2759
## Mon Jul 25 17:01:04 CEST 2022 - The aliscore output file does not exist. Check results for BUSCO: 976469at2759
## Mon Jul 25 17:01:04 CEST 2022 - The aliscore output file does not exist. Check results for BUSCO: 1269244at2759
## Mon Jul 25 17:01:04 CEST 2022 - The aliscore output appears to be empty. Check results for BUSCO: 976469at2759
## Mon Jul 25 17:01:04 CEST 2022 - The aliscore output appears to be empty. Check results for BUSCO: 1269244at2759
## Mon Jul 25 17:01:04 CEST 2022 - The aliscore output file does not exist. Check results for BUSCO: 721605at2759
## Mon Jul 25 17:01:04 CEST 2022 - The aliscore output appears to be empty. Check results for BUSCO: 721605at2759
## Mon Jul 25 17:01:04 CEST 2022 - The aliscore output file does not exist. Check results for BUSCO: 1138059at2759
## Mon Jul 25 17:01:04 CEST 2022 - The aliscore output file does not exist. Check results for BUSCO: 1032689at2759
## Mon Jul 25 17:01:04 CEST 2022 - The aliscore output file does not exist. Check results for BUSCO: 801857at2759
## Mon Jul 25 17:01:04 CEST 2022 - The aliscore output appears to be empty. Check results for BUSCO: 1138059at2759
## Mon Jul 25 17:01:04 CEST 2022 - The aliscore output appears to be empty. Check results for BUSCO: 1032689at2759
## Mon Jul 25 17:01:04 CEST 2022 - The aliscore output appears to be empty. Check results for BUSCO: 801857at2759
## Mon Jul 25 17:01:04 CEST 2022 - The aliscore output file does not exist. Check results for BUSCO: 1338131at2759
## Mon Jul 25 17:01:04 CEST 2022 - The aliscore output file does not exist. Check results for BUSCO: 491869at2759
## Mon Jul 25 17:01:04 CEST 2022 - The aliscore output file does not exist. Check results for BUSCO: 1588798at2759
## Mon Jul 25 17:01:04 CEST 2022 - The aliscore output appears to be empty. Check results for BUSCO: 1338131at2759
## Mon Jul 25 17:01:04 CEST 2022 - The aliscore output appears to be empty. Check results for BUSCO: 491869at2759
## Mon Jul 25 17:01:04 CEST 2022 - The aliscore output appears to be empty. Check results for BUSCO: 1588798at2759
## Mon Jul 25 17:01:04 CEST 2022 - The aliscore output file does not exist. Check results for BUSCO: 1428854at2759
## Mon Jul 25 17:01:04 CEST 2022 - The aliscore output appears to be empty. Check results for BUSCO: 1428854at2759
## Mon Jul 25 17:01:04 CEST 2022 - The aliscore output file does not exist. Check results for BUSCO: 1358374at2759
## Mon Jul 25 17:01:05 CEST 2022 - The aliscore output file does not exist. Check results for BUSCO: 869548at2759
## Mon Jul 25 17:01:05 CEST 2022 - The aliscore output appears to be empty. Check results for BUSCO: 1358374at2759
## Mon Jul 25 17:01:05 CEST 2022 - The aliscore output appears to be empty. Check results for BUSCO: 869548at2759
## Mon Jul 25 17:01:05 CEST 2022 - The aliscore output file does not exist. Check results for BUSCO: 1014314at2759
## Mon Jul 25 17:01:05 CEST 2022 - The aliscore output appears to be empty. Check results for BUSCO: 1014314at2759
## Mon Jul 25 17:01:05 CEST 2022 - The aliscore output file does not exist. Check results for BUSCO: 604979at2759
## Mon Jul 25 17:01:05 CEST 2022 - The aliscore output appears to be empty. Check results for BUSCO: 604979at2759
## Mon Jul 25 17:01:05 CEST 2022 - The aliscore output file does not exist. Check results for BUSCO: 426305at2759
## Mon Jul 25 17:01:05 CEST 2022 - The aliscore output appears to be empty. Check results for BUSCO: 426305at2759
## Mon Jul 25 17:01:05 CEST 2022 - The aliscore output file does not exist. Check results for BUSCO: 1488235at2759
## Mon Jul 25 17:01:05 CEST 2022 - The aliscore output file does not exist. Check results for BUSCO: 1259741at2759
## Mon Jul 25 17:01:05 CEST 2022 - The aliscore output appears to be empty. Check results for BUSCO: 1488235at2759
## Mon Jul 25 17:01:05 CEST 2022 - The aliscore output appears to be empty. Check results for BUSCO: 1259741at2759
## Mon Jul 25 17:01:06 CEST 2022 - The aliscore output file does not exist. Check results for BUSCO: 1251252at2759
## Mon Jul 25 17:01:06 CEST 2022 - The aliscore output file does not exist. Check results for BUSCO: 946128at2759
## Mon Jul 25 17:01:06 CEST 2022 - The aliscore output appears to be empty. Check results for BUSCO: 1251252at2759
## Mon Jul 25 17:01:06 CEST 2022 - The aliscore output file does not exist. Check results for BUSCO: 1266231at2759
## Mon Jul 25 17:01:06 CEST 2022 - The aliscore output appears to be empty. Check results for BUSCO: 946128at2759
## Mon Jul 25 17:01:06 CEST 2022 - The aliscore output appears to be empty. Check results for BUSCO: 1266231at2759
## Mon Jul 25 17:01:06 CEST 2022 - The aliscore output file does not exist. Check results for BUSCO: 1426075at2759
## Mon Jul 25 17:01:06 CEST 2022 - The aliscore output appears to be empty. Check results for BUSCO: 1426075at2759
## Mon Jul 25 17:01:06 CEST 2022 - The aliscore output file does not exist. Check results for BUSCO: 383503at2759
## Mon Jul 25 17:01:06 CEST 2022 - The aliscore output file does not exist. Check results for BUSCO: 1193442at2759
## Mon Jul 25 17:01:06 CEST 2022 - The aliscore output appears to be empty. Check results for BUSCO: 383503at2759
## Mon Jul 25 17:01:06 CEST 2022 - The aliscore output appears to be empty. Check results for BUSCO: 1193442at2759
## Mon Jul 25 17:01:06 CEST 2022 - The aliscore output file does not exist. Check results for BUSCO: 1322299at2759
## Mon Jul 25 17:01:06 CEST 2022 - The aliscore output file does not exist. Check results for BUSCO: 1051021at2759
## Mon Jul 25 17:01:06 CEST 2022 - The aliscore output appears to be empty. Check results for BUSCO: 1322299at2759
## Mon Jul 25 17:01:06 CEST 2022 - The aliscore output appears to be empty. Check results for BUSCO: 1051021at2759
## Mon Jul 25 17:01:07 CEST 2022 - The aliscore output file does not exist. Check results for BUSCO: 341721at2759
## Mon Jul 25 17:01:07 CEST 2022 - The aliscore output appears to be empty. Check results for BUSCO: 341721at2759
## Mon Jul 25 17:01:07 CEST 2022 - The aliscore output file does not exist. Check results for BUSCO: 905026at2759
## Mon Jul 25 17:01:07 CEST 2022 - The aliscore output appears to be empty. Check results for BUSCO: 905026at2759
## Mon Jul 25 17:01:07 CEST 2022 - The aliscore output file does not exist. Check results for BUSCO: 355408at2759
## Mon Jul 25 17:01:07 CEST 2022 - The aliscore output appears to be empty. Check results for BUSCO: 355408at2759
## Mon Jul 25 17:01:07 CEST 2022 - The aliscore output file does not exist. Check results for BUSCO: 1441854at2759
## Mon Jul 25 17:01:07 CEST 2022 - The aliscore output appears to be empty. Check results for BUSCO: 1441854at2759
## Mon Jul 25 17:01:07 CEST 2022 - The aliscore output file does not exist. Check results for BUSCO: 513979at2759
## Mon Jul 25 17:01:07 CEST 2022 - The aliscore output appears to be empty. Check results for BUSCO: 513979at2759
## Mon Jul 25 17:01:07 CEST 2022 - The aliscore output file does not exist. Check results for BUSCO: 549762at2759
## Mon Jul 25 17:01:07 CEST 2022 - The aliscore output appears to be empty. Check results for BUSCO: 549762at2759
## Mon Jul 25 17:01:07 CEST 2022 - The aliscore output file does not exist. Check results for BUSCO: 1019762at2759
## Mon Jul 25 17:01:07 CEST 2022 - The aliscore output appears to be empty. Check results for BUSCO: 1019762at2759
## Mon Jul 25 17:01:08 CEST 2022 - The aliscore output file does not exist. Check results for BUSCO: 570797at2759
## Mon Jul 25 17:01:08 CEST 2022 - The aliscore output appears to be empty. Check results for BUSCO: 570797at2759
## Mon Jul 25 17:01:08 CEST 2022 - The aliscore output file does not exist. Check results for BUSCO: 924753at2759
## Mon Jul 25 17:01:08 CEST 2022 - The aliscore output appears to be empty. Check results for BUSCO: 924753at2759
## Mon Jul 25 17:01:08 CEST 2022 - The aliscore output file does not exist. Check results for BUSCO: 270107at2759
## Mon Jul 25 17:01:08 CEST 2022 - The aliscore output appears to be empty. Check results for BUSCO: 270107at2759
## Mon Jul 25 17:01:08 CEST 2022 - The aliscore output file does not exist. Check results for BUSCO: 736068at2759
## Mon Jul 25 17:01:08 CEST 2022 - The aliscore output appears to be empty. Check results for BUSCO: 736068at2759
## Mon Jul 25 17:01:08 CEST 2022 - The aliscore output file does not exist. Check results for BUSCO: 981902at2759
## Mon Jul 25 17:01:08 CEST 2022 - The aliscore output appears to be empty. Check results for BUSCO: 981902at2759
## Mon Jul 25 17:01:08 CEST 2022 - The aliscore output file does not exist. Check results for BUSCO: 1003258at2759
## Mon Jul 25 17:01:08 CEST 2022 - The aliscore output appears to be empty. Check results for BUSCO: 1003258at2759
## Mon Jul 25 17:01:08 CEST 2022 - The aliscore output file does not exist. Check results for BUSCO: 331411at2759
## Mon Jul 25 17:01:08 CEST 2022 - The aliscore output appears to be empty. Check results for BUSCO: 331411at2759
## Mon Jul 25 17:01:08 CEST 2022 - The aliscore output file does not exist. Check results for BUSCO: 1404162at2759
## Mon Jul 25 17:01:08 CEST 2022 - The aliscore output appears to be empty. Check results for BUSCO: 1404162at2759
## Mon Jul 25 17:01:09 CEST 2022 - The aliscore output file does not exist. Check results for BUSCO: 83779at2759
## Mon Jul 25 17:01:09 CEST 2022 - The aliscore output appears to be empty. Check results for BUSCO: 83779at2759
## Mon Jul 25 17:01:09 CEST 2022 - The aliscore output file does not exist. Check results for BUSCO: 306227at2759
## Mon Jul 25 17:01:09 CEST 2022 - The aliscore output appears to be empty. Check results for BUSCO: 306227at2759
## Mon Jul 25 17:01:11 CEST 2022 - The aliscore output file does not exist. Check results for BUSCO: 1312453at2759
## Mon Jul 25 17:01:11 CEST 2022 - The aliscore output appears to be empty. Check results for BUSCO: 1312453at2759
## Mon Jul 25 17:01:12 CEST 2022 - The aliscore output file does not exist. Check results for BUSCO: 160593at2759
## Mon Jul 25 17:01:12 CEST 2022 - The aliscore output appears to be empty. Check results for BUSCO: 160593at2759
## Mon Jul 25 17:01:19 CEST 2022 - The aliscore output file does not exist. Check results for BUSCO: 156083at2759
## Mon Jul 25 17:01:19 CEST 2022 - The aliscore output appears to be empty. Check results for BUSCO: 156083at2759
## Mon Jul 25 17:01:31 CEST 2022 - The aliscore output file does not exist. Check results for BUSCO: 1173229at2759
## Mon Jul 25 17:01:31 CEST 2022 - The aliscore output appears to be empty. Check results for BUSCO: 1173229at2759
## Mon Jul 25 17:01:31 CEST 2022 - The aliscore output file does not exist. Check results for BUSCO: 1504863at2759
## Mon Jul 25 17:01:32 CEST 2022 - The aliscore output appears to be empty. Check results for BUSCO: 1504863at2759
## Mon Jul 25 17:01:32 CEST 2022 - The aliscore output file does not exist. Check results for BUSCO: 1623701at2759
## Mon Jul 25 17:01:32 CEST 2022 - The aliscore output appears to be empty. Check results for BUSCO: 1623701at2759
## Mon Jul 25 17:01:32 CEST 2022 - The aliscore output file does not exist. Check results for BUSCO: 1530008at2759
## Mon Jul 25 17:01:32 CEST 2022 - The aliscore output appears to be empty. Check results for BUSCO: 1530008at2759
## Mon Jul 25 17:01:33 CEST 2022 - The aliscore output file does not exist. Check results for BUSCO: 1405146at2759
## Mon Jul 25 17:01:33 CEST 2022 - The aliscore output file does not exist. Check results for BUSCO: 457861at2759
## Mon Jul 25 17:01:33 CEST 2022 - The aliscore output appears to be empty. Check results for BUSCO: 1405146at2759
## Mon Jul 25 17:01:33 CEST 2022 - The aliscore output appears to be empty. Check results for BUSCO: 457861at2759
## Mon Jul 25 17:01:33 CEST 2022 - The aliscore output file does not exist. Check results for BUSCO: 1633672at2759
## Mon Jul 25 17:01:33 CEST 2022 - The aliscore output appears to be empty. Check results for BUSCO: 1633672at2759
## Mon Jul 25 17:01:33 CEST 2022 - The aliscore output file does not exist. Check results for BUSCO: 1434266at2759
## Mon Jul 25 17:01:33 CEST 2022 - The aliscore output appears to be empty. Check results for BUSCO: 1434266at2759
## Mon Jul 25 17:01:33 CEST 2022 - The aliscore output file does not exist. Check results for BUSCO: 1236198at2759
## Mon Jul 25 17:01:34 CEST 2022 - The aliscore output file does not exist. Check results for BUSCO: 734341at2759
## Mon Jul 25 17:01:34 CEST 2022 - The aliscore output appears to be empty. Check results for BUSCO: 1236198at2759
## Mon Jul 25 17:01:34 CEST 2022 - The aliscore output appears to be empty. Check results for BUSCO: 734341at2759
## Mon Jul 25 17:01:34 CEST 2022 - The aliscore output file does not exist. Check results for BUSCO: 655400at2759
## Mon Jul 25 17:01:34 CEST 2022 - The aliscore output appears to be empty. Check results for BUSCO: 655400at2759
## Mon Jul 25 17:01:34 CEST 2022 - The aliscore output file does not exist. Check results for BUSCO: 1445102at2759
## Mon Jul 25 17:01:34 CEST 2022 - The aliscore output appears to be empty. Check results for BUSCO: 1445102at2759
## Mon Jul 25 17:01:34 CEST 2022 - The aliscore output file does not exist. Check results for BUSCO: 330169at2759
## Mon Jul 25 17:01:34 CEST 2022 - The aliscore output file does not exist. Check results for BUSCO: 1076134at2759
## Mon Jul 25 17:01:34 CEST 2022 - The aliscore output appears to be empty. Check results for BUSCO: 1076134at2759
## Mon Jul 25 17:01:34 CEST 2022 - The aliscore output appears to be empty. Check results for BUSCO: 330169at2759
## Mon Jul 25 17:01:34 CEST 2022 - The aliscore output file does not exist. Check results for BUSCO: 1526152at2759
## Mon Jul 25 17:01:34 CEST 2022 - The aliscore output appears to be empty. Check results for BUSCO: 1526152at2759
## Mon Jul 25 17:01:34 CEST 2022 - The aliscore output file does not exist. Check results for BUSCO: 1079827at2759
## Mon Jul 25 17:01:34 CEST 2022 - The aliscore output file does not exist. Check results for BUSCO: 464990at2759
## Mon Jul 25 17:01:34 CEST 2022 - The aliscore output appears to be empty. Check results for BUSCO: 1079827at2759
## Mon Jul 25 17:01:34 CEST 2022 - The aliscore output appears to be empty. Check results for BUSCO: 464990at2759
## Mon Jul 25 17:01:34 CEST 2022 - The aliscore output file does not exist. Check results for BUSCO: 1220881at2759
## Mon Jul 25 17:01:34 CEST 2022 - The aliscore output appears to be empty. Check results for BUSCO: 1220881at2759
## Mon Jul 25 17:01:34 CEST 2022 - The aliscore output file does not exist. Check results for BUSCO: 1115196at2759
## Mon Jul 25 17:01:34 CEST 2022 - The aliscore output file does not exist. Check results for BUSCO: 1440961at2759
## Mon Jul 25 17:01:34 CEST 2022 - The aliscore output appears to be empty. Check results for BUSCO: 1115196at2759
## Mon Jul 25 17:01:34 CEST 2022 - The aliscore output appears to be empty. Check results for BUSCO: 1440961at2759
## Mon Jul 25 17:01:34 CEST 2022 - The aliscore output file does not exist. Check results for BUSCO: 1257440at2759
## Mon Jul 25 17:01:34 CEST 2022 - The aliscore output appears to be empty. Check results for BUSCO: 1257440at2759
## Mon Jul 25 17:01:35 CEST 2022 - The aliscore output file does not exist. Check results for BUSCO: 1428265at2759
## Mon Jul 25 17:01:35 CEST 2022 - The aliscore output file does not exist. Check results for BUSCO: 664730at2759
## Mon Jul 25 17:01:35 CEST 2022 - The aliscore output appears to be empty. Check results for BUSCO: 1428265at2759
## Mon Jul 25 17:01:35 CEST 2022 - The aliscore output appears to be empty. Check results for BUSCO: 664730at2759
## Mon Jul 25 17:01:34 CEST 2022 - The aliscore output file does not exist. Check results for BUSCO: 1450538at2759
## Mon Jul 25 17:01:35 CEST 2022 - The aliscore output appears to be empty. Check results for BUSCO: 1450538at2759
## Mon Jul 25 17:01:35 CEST 2022 - The aliscore output file does not exist. Check results for BUSCO: 1567796at2759
## Mon Jul 25 17:01:35 CEST 2022 - The aliscore output appears to be empty. Check results for BUSCO: 1567796at2759
## Mon Jul 25 17:01:35 CEST 2022 - The aliscore output file does not exist. Check results for BUSCO: 1322642at2759
## Mon Jul 25 17:01:35 CEST 2022 - The aliscore output file does not exist. Check results for BUSCO: 1157302at2759
## Mon Jul 25 17:01:35 CEST 2022 - The aliscore output appears to be empty. Check results for BUSCO: 1322642at2759
## Mon Jul 25 17:01:35 CEST 2022 - The aliscore output appears to be empty. Check results for BUSCO: 1157302at2759
## Mon Jul 25 17:01:35 CEST 2022 - The aliscore output file does not exist. Check results for BUSCO: 887370at2759
## Mon Jul 25 17:01:35 CEST 2022 - The aliscore output file does not exist. Check results for BUSCO: 1348942at2759
## Mon Jul 25 17:01:35 CEST 2022 - The aliscore output appears to be empty. Check results for BUSCO: 887370at2759
## Mon Jul 25 17:01:35 CEST 2022 - The aliscore output file does not exist. Check results for BUSCO: 858842at2759
## Mon Jul 25 17:01:35 CEST 2022 - The aliscore output appears to be empty. Check results for BUSCO: 1348942at2759
## Mon Jul 25 17:01:35 CEST 2022 - The aliscore output file does not exist. Check results for BUSCO: 901894at2759
## Mon Jul 25 17:01:35 CEST 2022 - The aliscore output appears to be empty. Check results for BUSCO: 858842at2759
## Mon Jul 25 17:01:35 CEST 2022 - The aliscore output file does not exist. Check results for BUSCO: 779909at2759
## Mon Jul 25 17:01:35 CEST 2022 - The aliscore output appears to be empty. Check results for BUSCO: 901894at2759
## Mon Jul 25 17:01:35 CEST 2022 - The aliscore output appears to be empty. Check results for BUSCO: 779909at2759
## Mon Jul 25 17:01:35 CEST 2022 - The aliscore output file does not exist. Check results for BUSCO: 1030907at2759
## Mon Jul 25 17:01:35 CEST 2022 - The aliscore output appears to be empty. Check results for BUSCO: 1030907at2759
## Mon Jul 25 17:01:35 CEST 2022 - The aliscore output file does not exist. Check results for BUSCO: 1576404at2759
## Mon Jul 25 17:01:35 CEST 2022 - The aliscore output appears to be empty. Check results for BUSCO: 1576404at2759
## Mon Jul 25 17:01:35 CEST 2022 - The aliscore output file does not exist. Check results for BUSCO: 720340at2759
## Mon Jul 25 17:01:35 CEST 2022 - The aliscore output appears to be empty. Check results for BUSCO: 720340at2759
## Mon Jul 25 17:01:36 CEST 2022 - The aliscore output file does not exist. Check results for BUSCO: 674169at2759
## Mon Jul 25 17:01:36 CEST 2022 - The aliscore output file does not exist. Check results for BUSCO: 1182451at2759
## Mon Jul 25 17:01:36 CEST 2022 - The aliscore output appears to be empty. Check results for BUSCO: 674169at2759
## Mon Jul 25 17:01:36 CEST 2022 - The aliscore output appears to be empty. Check results for BUSCO: 1182451at2759
## Mon Jul 25 17:01:36 CEST 2022 - The aliscore output file does not exist. Check results for BUSCO: 1513531at2759
## Mon Jul 25 17:01:36 CEST 2022 - The aliscore output file does not exist. Check results for BUSCO: 1194691at2759
## Mon Jul 25 17:01:36 CEST 2022 - The aliscore output appears to be empty. Check results for BUSCO: 1513531at2759
## Mon Jul 25 17:01:36 CEST 2022 - The aliscore output appears to be empty. Check results for BUSCO: 1194691at2759
## Mon Jul 25 17:01:36 CEST 2022 - The aliscore output file does not exist. Check results for BUSCO: 621827at2759
## Mon Jul 25 17:01:36 CEST 2022 - The aliscore output file does not exist. Check results for BUSCO: 1090038at2759
## Mon Jul 25 17:01:36 CEST 2022 - The aliscore output appears to be empty. Check results for BUSCO: 621827at2759
## Mon Jul 25 17:01:36 CEST 2022 - The aliscore output appears to be empty. Check results for BUSCO: 1090038at2759
## Mon Jul 25 17:01:36 CEST 2022 - The aliscore output file does not exist. Check results for BUSCO: 296129at2759
## Mon Jul 25 17:01:37 CEST 2022 - The aliscore output appears to be empty. Check results for BUSCO: 296129at2759
## Mon Jul 25 17:01:37 CEST 2022 - The aliscore output file does not exist. Check results for BUSCO: 97116at2759
## Mon Jul 25 17:01:37 CEST 2022 - The aliscore output file does not exist. Check results for BUSCO: 671846at2759
## Mon Jul 25 17:01:37 CEST 2022 - The aliscore output appears to be empty. Check results for BUSCO: 97116at2759
## Mon Jul 25 17:01:37 CEST 2022 - The aliscore output appears to be empty. Check results for BUSCO: 671846at2759
## Mon Jul 25 17:01:37 CEST 2022 - The aliscore output file does not exist. Check results for BUSCO: 1111142at2759
## Mon Jul 25 17:01:37 CEST 2022 - The aliscore output file does not exist. Check results for BUSCO: 1001705at2759
## Mon Jul 25 17:01:37 CEST 2022 - The aliscore output appears to be empty. Check results for BUSCO: 1001705at2759
## Mon Jul 25 17:01:37 CEST 2022 - The aliscore output appears to be empty. Check results for BUSCO: 1111142at2759
## Mon Jul 25 17:01:37 CEST 2022 - The aliscore output file does not exist. Check results for BUSCO: 1065019at2759
## Mon Jul 25 17:01:37 CEST 2022 - The aliscore output file does not exist. Check results for BUSCO: 1370285at2759
## Mon Jul 25 17:01:37 CEST 2022 - The aliscore output appears to be empty. Check results for BUSCO: 1065019at2759
## Mon Jul 25 17:01:37 CEST 2022 - The aliscore output appears to be empty. Check results for BUSCO: 1370285at2759
## Mon Jul 25 17:01:37 CEST 2022 - The aliscore output file does not exist. Check results for BUSCO: 375960at2759
## Mon Jul 25 17:01:37 CEST 2022 - The aliscore output appears to be empty. Check results for BUSCO: 375960at2759
## Mon Jul 25 17:01:37 CEST 2022 - The aliscore output file does not exist. Check results for BUSCO: 834694at2759
## Mon Jul 25 17:01:37 CEST 2022 - The aliscore output appears to be empty. Check results for BUSCO: 834694at2759
## Mon Jul 25 17:01:37 CEST 2022 - The aliscore output file does not exist. Check results for BUSCO: 293315at2759
## Mon Jul 25 17:01:37 CEST 2022 - The aliscore output file does not exist. Check results for BUSCO: 390348at2759
## Mon Jul 25 17:01:38 CEST 2022 - The aliscore output appears to be empty. Check results for BUSCO: 293315at2759
## Mon Jul 25 17:01:38 CEST 2022 - The aliscore output appears to be empty. Check results for BUSCO: 390348at2759
## Mon Jul 25 17:01:38 CEST 2022 - The aliscore output file does not exist. Check results for BUSCO: 918816at2759
## Mon Jul 25 17:01:38 CEST 2022 - The aliscore output appears to be empty. Check results for BUSCO: 918816at2759
## Mon Jul 25 17:01:38 CEST 2022 - The aliscore output file does not exist. Check results for BUSCO: 598949at2759
## Mon Jul 25 17:01:38 CEST 2022 - The aliscore output appears to be empty. Check results for BUSCO: 598949at2759
## Mon Jul 25 17:01:38 CEST 2022 - The aliscore output file does not exist. Check results for BUSCO: 891751at2759
## Mon Jul 25 17:01:38 CEST 2022 - The aliscore output appears to be empty. Check results for BUSCO: 891751at2759
## Mon Jul 25 17:01:38 CEST 2022 - The aliscore output file does not exist. Check results for BUSCO: 369837at2759
## Mon Jul 25 17:01:38 CEST 2022 - The aliscore output appears to be empty. Check results for BUSCO: 369837at2759
## Mon Jul 25 17:01:38 CEST 2022 - The aliscore output file does not exist. Check results for BUSCO: 1322642at2759
## Mon Jul 25 17:01:38 CEST 2022 - The aliscore output appears to be empty. Check results for BUSCO: 1322642at2759
## Mon Jul 25 17:01:39 CEST 2022 - The aliscore output file does not exist. Check results for BUSCO: 245208at2759
## Mon Jul 25 17:01:39 CEST 2022 - The aliscore output appears to be empty. Check results for BUSCO: 245208at2759
## Mon Jul 25 17:01:39 CEST 2022 - The aliscore output file does not exist. Check results for BUSCO: 325552at2759
## Mon Jul 25 17:01:39 CEST 2022 - The aliscore output appears to be empty. Check results for BUSCO: 325552at2759
## Mon Jul 25 17:01:40 CEST 2022 - The aliscore output file does not exist. Check results for BUSCO: 1314980at2759
## Mon Jul 25 17:01:40 CEST 2022 - The aliscore output appears to be empty. Check results for BUSCO: 1314980at2759
## Mon Jul 25 17:02:01 CEST 2022 - The aliscore output file does not exist. Check results for BUSCO: 1405073at2759
## Mon Jul 25 17:02:01 CEST 2022 - The aliscore output appears to be empty. Check results for BUSCO: 1405073at2759
## Mon Jul 25 17:02:01 CEST 2022 - The aliscore output file does not exist. Check results for BUSCO: 996662at2759
## Mon Jul 25 17:02:01 CEST 2022 - The aliscore output appears to be empty. Check results for BUSCO: 996662at2759
## Mon Jul 25 17:02:01 CEST 2022 - The aliscore output file does not exist. Check results for BUSCO: 1217666at2759
## Mon Jul 25 17:02:01 CEST 2022 - The aliscore output appears to be empty. Check results for BUSCO: 1217666at2759
## Mon Jul 25 17:02:01 CEST 2022 - The aliscore output file does not exist. Check results for BUSCO: 1398309at2759
## Mon Jul 25 17:02:01 CEST 2022 - The aliscore output appears to be empty. Check results for BUSCO: 1398309at2759
## Mon Jul 25 17:02:02 CEST 2022 - The aliscore output file does not exist. Check results for BUSCO: 1358374at2759
## Mon Jul 25 17:02:02 CEST 2022 - The aliscore output appears to be empty. Check results for BUSCO: 1358374at2759
## Mon Jul 25 17:02:02 CEST 2022 - The aliscore output file does not exist. Check results for BUSCO: 1617752at2759
## Mon Jul 25 17:02:02 CEST 2022 - The aliscore output appears to be empty. Check results for BUSCO: 1617752at2759
## Mon Jul 25 17:02:02 CEST 2022 - The aliscore output file does not exist. Check results for BUSCO: 1200489at2759
## Mon Jul 25 17:02:02 CEST 2022 - The aliscore output appears to be empty. Check results for BUSCO: 1200489at2759
## Mon Jul 25 17:02:02 CEST 2022 - The aliscore output file does not exist. Check results for BUSCO: 1161199at2759
## Mon Jul 25 17:02:02 CEST 2022 - The aliscore output appears to be empty. Check results for BUSCO: 1161199at2759
## Mon Jul 25 17:02:03 CEST 2022 - The aliscore output file does not exist. Check results for BUSCO: 1342242at2759
## Mon Jul 25 17:02:03 CEST 2022 - The aliscore output appears to be empty. Check results for BUSCO: 1342242at2759
## Mon Jul 25 17:02:03 CEST 2022 - The aliscore output file does not exist. Check results for BUSCO: 1474000at2759
## Mon Jul 25 17:02:03 CEST 2022 - The aliscore output file does not exist. Check results for BUSCO: 679187at2759
## Mon Jul 25 17:02:03 CEST 2022 - The aliscore output appears to be empty. Check results for BUSCO: 679187at2759
## Mon Jul 25 17:02:03 CEST 2022 - The aliscore output appears to be empty. Check results for BUSCO: 1474000at2759
## Mon Jul 25 17:02:03 CEST 2022 - The aliscore output file does not exist. Check results for BUSCO: 257318at2759
## Mon Jul 25 17:02:03 CEST 2022 - The aliscore output appears to be empty. Check results for BUSCO: 257318at2759
## Mon Jul 25 17:02:03 CEST 2022 - The aliscore output file does not exist. Check results for BUSCO: 1038775at2759
## Mon Jul 25 17:02:03 CEST 2022 - The aliscore output appears to be empty. Check results for BUSCO: 1038775at2759
## Mon Jul 25 17:02:03 CEST 2022 - The aliscore output file does not exist. Check results for BUSCO: 1645187at2759
## Mon Jul 25 17:02:03 CEST 2022 - The aliscore output appears to be empty. Check results for BUSCO: 1645187at2759
## Mon Jul 25 17:02:04 CEST 2022 - The aliscore output file does not exist. Check results for BUSCO: 1309031at2759
## Mon Jul 25 17:02:04 CEST 2022 - The aliscore output appears to be empty. Check results for BUSCO: 1309031at2759
## Mon Jul 25 17:02:04 CEST 2022 - The aliscore output file does not exist. Check results for BUSCO: 1094121at2759
## Mon Jul 25 17:02:04 CEST 2022 - The aliscore output appears to be empty. Check results for BUSCO: 1094121at2759
## Mon Jul 25 17:02:04 CEST 2022 - The aliscore output file does not exist. Check results for BUSCO: 817008at2759
## Mon Jul 25 17:02:04 CEST 2022 - The aliscore output appears to be empty. Check results for BUSCO: 817008at2759
## Mon Jul 25 17:02:04 CEST 2022 - The aliscore output file does not exist. Check results for BUSCO: 290630at2759
## Mon Jul 25 17:02:04 CEST 2022 - The aliscore output appears to be empty. Check results for BUSCO: 290630at2759
## Mon Jul 25 17:02:04 CEST 2022 - The aliscore output file does not exist. Check results for BUSCO: 1287094at2759
## Mon Jul 25 17:02:04 CEST 2022 - The aliscore output appears to be empty. Check results for BUSCO: 1287094at2759
## Mon Jul 25 17:02:04 CEST 2022 - The aliscore output file does not exist. Check results for BUSCO: 1108845at2759
## Mon Jul 25 17:02:04 CEST 2022 - The aliscore output appears to be empty. Check results for BUSCO: 1108845at2759
## Mon Jul 25 17:02:05 CEST 2022 - The aliscore output file does not exist. Check results for BUSCO: 1233814at2759
## Mon Jul 25 17:02:05 CEST 2022 - The aliscore output appears to be empty. Check results for BUSCO: 1233814at2759
## Mon Jul 25 17:02:05 CEST 2022 - The aliscore output file does not exist. Check results for BUSCO: 1380409at2759
## Mon Jul 25 17:02:05 CEST 2022 - The aliscore output appears to be empty. Check results for BUSCO: 1380409at2759
## Mon Jul 25 17:02:05 CEST 2022 - The aliscore output file does not exist. Check results for BUSCO: 751335at2759
## Mon Jul 25 17:02:05 CEST 2022 - The aliscore output appears to be empty. Check results for BUSCO: 751335at2759
## Mon Jul 25 17:02:05 CEST 2022 - The aliscore output file does not exist. Check results for BUSCO: 1364586at2759
## Mon Jul 25 17:02:05 CEST 2022 - The aliscore output appears to be empty. Check results for BUSCO: 1364586at2759
## Mon Jul 25 17:02:05 CEST 2022 - The aliscore output file does not exist. Check results for BUSCO: 1049599at2759
## Mon Jul 25 17:02:05 CEST 2022 - The aliscore output appears to be empty. Check results for BUSCO: 1049599at2759
## Mon Jul 25 17:02:05 CEST 2022 - The aliscore output file does not exist. Check results for BUSCO: 956854at2759
## Mon Jul 25 17:02:05 CEST 2022 - The aliscore output appears to be empty. Check results for BUSCO: 956854at2759
## Mon Jul 25 17:02:06 CEST 2022 - The aliscore output file does not exist. Check results for BUSCO: 1442062at2759
## Mon Jul 25 17:02:06 CEST 2022 - The aliscore output appears to be empty. Check results for BUSCO: 1442062at2759
## Mon Jul 25 17:02:06 CEST 2022 - The aliscore output file does not exist. Check results for BUSCO: 844748at2759
## Mon Jul 25 17:02:06 CEST 2022 - The aliscore output appears to be empty. Check results for BUSCO: 844748at2759
## Mon Jul 25 17:02:06 CEST 2022 - The aliscore output file does not exist. Check results for BUSCO: 1057950at2759
## Mon Jul 25 17:02:06 CEST 2022 - The aliscore output appears to be empty. Check results for BUSCO: 1057950at2759
## Mon Jul 25 17:02:06 CEST 2022 - The aliscore output file does not exist. Check results for BUSCO: 1479417at2759
## Mon Jul 25 17:02:06 CEST 2022 - The aliscore output file does not exist. Check results for BUSCO: 1430056at2759
## Mon Jul 25 17:02:06 CEST 2022 - The aliscore output appears to be empty. Check results for BUSCO: 1479417at2759
## Mon Jul 25 17:02:06 CEST 2022 - The aliscore output appears to be empty. Check results for BUSCO: 1430056at2759
## Mon Jul 25 17:02:06 CEST 2022 - The aliscore output file does not exist. Check results for BUSCO: 1454155at2759
## Mon Jul 25 17:02:06 CEST 2022 - The aliscore output appears to be empty. Check results for BUSCO: 1454155at2759
## Mon Jul 25 17:02:06 CEST 2022 - The aliscore output file does not exist. Check results for BUSCO: 453044at2759
## Mon Jul 25 17:02:06 CEST 2022 - The aliscore output appears to be empty. Check results for BUSCO: 453044at2759
## Mon Jul 25 17:02:06 CEST 2022 - The aliscore output file does not exist. Check results for BUSCO: 878143at2759
## Mon Jul 25 17:02:06 CEST 2022 - The aliscore output appears to be empty. Check results for BUSCO: 878143at2759
## Mon Jul 25 17:02:06 CEST 2022 - The aliscore output file does not exist. Check results for BUSCO: 87842at2759
## Mon Jul 25 17:02:06 CEST 2022 - The aliscore output appears to be empty. Check results for BUSCO: 87842at2759
## Mon Jul 25 17:02:06 CEST 2022 - The aliscore output file does not exist. Check results for BUSCO: 100698at2759
## Mon Jul 25 17:02:07 CEST 2022 - The aliscore output appears to be empty. Check results for BUSCO: 100698at2759
## Mon Jul 25 17:02:07 CEST 2022 - The aliscore output file does not exist. Check results for BUSCO: 944899at2759
## Mon Jul 25 17:02:07 CEST 2022 - The aliscore output appears to be empty. Check results for BUSCO: 944899at2759
## Mon Jul 25 17:02:07 CEST 2022 - The aliscore output file does not exist. Check results for BUSCO: 604979at2759
## Mon Jul 25 17:02:07 CEST 2022 - The aliscore output appears to be empty. Check results for BUSCO: 604979at2759
## Mon Jul 25 17:02:07 CEST 2022 - The aliscore output file does not exist. Check results for BUSCO: 1287401at2759
## Mon Jul 25 17:02:07 CEST 2022 - The aliscore output appears to be empty. Check results for BUSCO: 1287401at2759
## Mon Jul 25 17:02:07 CEST 2022 - The aliscore output file does not exist. Check results for BUSCO: 1085752at2759
## Mon Jul 25 17:02:07 CEST 2022 - The aliscore output file does not exist. Check results for BUSCO: 1428854at2759
## Mon Jul 25 17:02:07 CEST 2022 - The aliscore output appears to be empty. Check results for BUSCO: 1085752at2759
## Mon Jul 25 17:02:07 CEST 2022 - The aliscore output appears to be empty. Check results for BUSCO: 1428854at2759
## Mon Jul 25 17:02:07 CEST 2022 - The aliscore output file does not exist. Check results for BUSCO: 392369at2759
## Mon Jul 25 17:02:07 CEST 2022 - The aliscore output appears to be empty. Check results for BUSCO: 392369at2759
## Mon Jul 25 17:02:08 CEST 2022 - The aliscore output file does not exist. Check results for BUSCO: 1032689at2759
## Mon Jul 25 17:02:08 CEST 2022 - The aliscore output appears to be empty. Check results for BUSCO: 1032689at2759
## Mon Jul 25 17:02:08 CEST 2022 - The aliscore output file does not exist. Check results for BUSCO: 1259741at2759
## Mon Jul 25 17:02:08 CEST 2022 - The aliscore output appears to be empty. Check results for BUSCO: 1259741at2759
## Mon Jul 25 17:02:08 CEST 2022 - The aliscore output file does not exist. Check results for BUSCO: 1324510at2759
## Mon Jul 25 17:02:08 CEST 2022 - The aliscore output file does not exist. Check results for BUSCO: 261419at2759
## Mon Jul 25 17:02:08 CEST 2022 - The aliscore output appears to be empty. Check results for BUSCO: 1324510at2759
## Mon Jul 25 17:02:08 CEST 2022 - The aliscore output appears to be empty. Check results for BUSCO: 261419at2759
## Mon Jul 25 17:02:08 CEST 2022 - The aliscore output file does not exist. Check results for BUSCO: 774318at2759
## Mon Jul 25 17:02:08 CEST 2022 - The aliscore output appears to be empty. Check results for BUSCO: 774318at2759
## Mon Jul 25 17:02:09 CEST 2022 - The aliscore output file does not exist. Check results for BUSCO: 551907at2759
## Mon Jul 25 17:02:09 CEST 2022 - The aliscore output appears to be empty. Check results for BUSCO: 551907at2759
## Mon Jul 25 17:02:10 CEST 2022 - The aliscore output file does not exist. Check results for BUSCO: 777920at2759
## Mon Jul 25 17:02:10 CEST 2022 - The aliscore output appears to be empty. Check results for BUSCO: 777920at2759
## Mon Jul 25 17:02:10 CEST 2022 - The aliscore output file does not exist. Check results for BUSCO: 1423847at2759
## Mon Jul 25 17:02:10 CEST 2022 - The aliscore output appears to be empty. Check results for BUSCO: 1423847at2759
## Mon Jul 25 17:02:10 CEST 2022 - The aliscore output file does not exist. Check results for BUSCO: 166920at2759
## Mon Jul 25 17:02:11 CEST 2022 - The aliscore output appears to be empty. Check results for BUSCO: 166920at2759
## Mon Jul 25 17:02:13 CEST 2022 - The aliscore output file does not exist. Check results for BUSCO: 39650at2759
## Mon Jul 25 17:02:13 CEST 2022 - The aliscore output appears to be empty. Check results for BUSCO: 39650at2759
## Mon Jul 25 17:02:19 CEST 2022 - The aliscore output file does not exist. Check results for BUSCO: 674160at2759
## Mon Jul 25 17:02:19 CEST 2022 - The aliscore output appears to be empty. Check results for BUSCO: 674160at2759
## Mon Jul 25 17:02:31 CEST 2022 - The aliscore output file does not exist. Check results for BUSCO: 1488436at2759
## Mon Jul 25 17:02:31 CEST 2022 - The aliscore output appears to be empty. Check results for BUSCO: 1488436at2759
## Mon Jul 25 17:02:31 CEST 2022 - The aliscore output file does not exist. Check results for BUSCO: 869548at2759
## Mon Jul 25 17:02:31 CEST 2022 - The aliscore output appears to be empty. Check results for BUSCO: 869548at2759
## Mon Jul 25 17:02:32 CEST 2022 - The aliscore output file does not exist. Check results for BUSCO: 1269649at2759
## Mon Jul 25 17:02:32 CEST 2022 - The aliscore output appears to be empty. Check results for BUSCO: 1269649at2759
## Mon Jul 25 17:02:32 CEST 2022 - The aliscore output file does not exist. Check results for BUSCO: 1455730at2759
## Mon Jul 25 17:02:32 CEST 2022 - The aliscore output appears to be empty. Check results for BUSCO: 1455730at2759
## Mon Jul 25 17:02:32 CEST 2022 - The aliscore output file does not exist. Check results for BUSCO: 1488235at2759
## Mon Jul 25 17:02:32 CEST 2022 - The aliscore output appears to be empty. Check results for BUSCO: 1488235at2759
## Mon Jul 25 17:02:32 CEST 2022 - The aliscore output file does not exist. Check results for BUSCO: 1525971at2759
## Mon Jul 25 17:02:32 CEST 2022 - The aliscore output appears to be empty. Check results for BUSCO: 1525971at2759
## Mon Jul 25 17:02:33 CEST 2022 - The aliscore output file does not exist. Check results for BUSCO: 976469at2759
## Mon Jul 25 17:02:33 CEST 2022 - The aliscore output appears to be empty. Check results for BUSCO: 976469at2759
## Mon Jul 25 17:02:33 CEST 2022 - The aliscore output file does not exist. Check results for BUSCO: 1633672at2759
## Mon Jul 25 17:02:33 CEST 2022 - The aliscore output appears to be empty. Check results for BUSCO: 1633672at2759
## Mon Jul 25 17:02:33 CEST 2022 - The aliscore output file does not exist. Check results for BUSCO: 1223488at2759
## Mon Jul 25 17:02:34 CEST 2022 - The aliscore output appears to be empty. Check results for BUSCO: 1223488at2759
## Mon Jul 25 17:02:34 CEST 2022 - The aliscore output file does not exist. Check results for BUSCO: 1284731at2759
## Mon Jul 25 17:02:34 CEST 2022 - The aliscore output appears to be empty. Check results for BUSCO: 1284731at2759
## Mon Jul 25 17:02:34 CEST 2022 - The aliscore output file does not exist. Check results for BUSCO: 1193442at2759
## Mon Jul 25 17:02:34 CEST 2022 - The aliscore output appears to be empty. Check results for BUSCO: 1193442at2759
## Mon Jul 25 17:02:34 CEST 2022 - The aliscore output file does not exist. Check results for BUSCO: 1275837at2759
## Mon Jul 25 17:02:34 CEST 2022 - The aliscore output appears to be empty. Check results for BUSCO: 1275837at2759
## Mon Jul 25 17:02:34 CEST 2022 - The aliscore output file does not exist. Check results for BUSCO: 1128607at2759
## Mon Jul 25 17:02:34 CEST 2022 - The aliscore output appears to be empty. Check results for BUSCO: 1128607at2759
## Mon Jul 25 17:02:35 CEST 2022 - The aliscore output file does not exist. Check results for BUSCO: 905026at2759
## Mon Jul 25 17:02:35 CEST 2022 - The aliscore output appears to be empty. Check results for BUSCO: 905026at2759
## Mon Jul 25 17:02:35 CEST 2022 - The aliscore output file does not exist. Check results for BUSCO: 1355894at2759
## Mon Jul 25 17:02:35 CEST 2022 - The aliscore output appears to be empty. Check results for BUSCO: 1355894at2759
## Mon Jul 25 17:02:35 CEST 2022 - The aliscore output file does not exist. Check results for BUSCO: 946128at2759
## Mon Jul 25 17:02:35 CEST 2022 - The aliscore output appears to be empty. Check results for BUSCO: 946128at2759
## Mon Jul 25 17:02:35 CEST 2022 - The aliscore output file does not exist. Check results for BUSCO: 1010730at2759
## Mon Jul 25 17:02:35 CEST 2022 - The aliscore output appears to be empty. Check results for BUSCO: 1010730at2759
## Mon Jul 25 17:02:35 CEST 2022 - The aliscore output file does not exist. Check results for BUSCO: 513979at2759
## Mon Jul 25 17:02:35 CEST 2022 - The aliscore output appears to be empty. Check results for BUSCO: 513979at2759
## Mon Jul 25 17:02:36 CEST 2022 - The aliscore output file does not exist. Check results for BUSCO: 1291729at2759
## Mon Jul 25 17:02:36 CEST 2022 - The aliscore output file does not exist. Check results for BUSCO: 1322299at2759
## Mon Jul 25 17:02:36 CEST 2022 - The aliscore output appears to be empty. Check results for BUSCO: 1322299at2759
## Mon Jul 25 17:02:36 CEST 2022 - The aliscore output appears to be empty. Check results for BUSCO: 1291729at2759
## Mon Jul 25 17:02:36 CEST 2022 - The aliscore output file does not exist. Check results for BUSCO: 383503at2759
## Mon Jul 25 17:02:36 CEST 2022 - The aliscore output appears to be empty. Check results for BUSCO: 383503at2759
## Mon Jul 25 17:02:37 CEST 2022 - The aliscore output file does not exist. Check results for BUSCO: 1157302at2759
## Mon Jul 25 17:02:37 CEST 2022 - The aliscore output appears to be empty. Check results for BUSCO: 1157302at2759
## Mon Jul 25 17:02:37 CEST 2022 - The aliscore output file does not exist. Check results for BUSCO: 779909at2759
## Mon Jul 25 17:02:37 CEST 2022 - The aliscore output appears to be empty. Check results for BUSCO: 779909at2759
## Mon Jul 25 17:02:37 CEST 2022 - The aliscore output file does not exist. Check results for BUSCO: 734341at2759
## Mon Jul 25 17:02:37 CEST 2022 - The aliscore output appears to be empty. Check results for BUSCO: 734341at2759
## Mon Jul 25 17:02:37 CEST 2022 - The aliscore output file does not exist. Check results for BUSCO: 1003258at2759
## Mon Jul 25 17:02:37 CEST 2022 - The aliscore output appears to be empty. Check results for BUSCO: 1003258at2759
## Mon Jul 25 17:02:38 CEST 2022 - The aliscore output file does not exist. Check results for BUSCO: 761109at2759
## Mon Jul 25 17:02:38 CEST 2022 - The aliscore output file does not exist. Check results for BUSCO: 1567796at2759
## Mon Jul 25 17:02:38 CEST 2022 - The aliscore output appears to be empty. Check results for BUSCO: 761109at2759
## Mon Jul 25 17:02:38 CEST 2022 - The aliscore output appears to be empty. Check results for BUSCO: 1567796at2759
## Mon Jul 25 17:02:38 CEST 2022 - The aliscore output file does not exist. Check results for BUSCO: 1182451at2759
## Mon Jul 25 17:02:38 CEST 2022 - The aliscore output appears to be empty. Check results for BUSCO: 1182451at2759
## Mon Jul 25 17:02:38 CEST 2022 - The aliscore output file does not exist. Check results for BUSCO: 1194691at2759
## Mon Jul 25 17:02:38 CEST 2022 - The aliscore output appears to be empty. Check results for BUSCO: 1194691at2759
## Mon Jul 25 17:02:39 CEST 2022 - The aliscore output file does not exist. Check results for BUSCO: 1065019at2759
## Mon Jul 25 17:02:39 CEST 2022 - The aliscore output appears to be empty. Check results for BUSCO: 1065019at2759
## Mon Jul 25 17:02:39 CEST 2022 - The aliscore output file does not exist. Check results for BUSCO: 1112002at2759
## Mon Jul 25 17:02:39 CEST 2022 - The aliscore output file does not exist. Check results for BUSCO: 911863at2759
## Mon Jul 25 17:02:39 CEST 2022 - The aliscore output appears to be empty. Check results for BUSCO: 1112002at2759
## Mon Jul 25 17:02:39 CEST 2022 - The aliscore output appears to be empty. Check results for BUSCO: 911863at2759
## Mon Jul 25 17:02:40 CEST 2022 - The aliscore output file does not exist. Check results for BUSCO: 901894at2759
## Mon Jul 25 17:02:40 CEST 2022 - The aliscore output appears to be empty. Check results for BUSCO: 901894at2759
## Mon Jul 25 17:02:40 CEST 2022 - The aliscore output file does not exist. Check results for BUSCO: 375960at2759
## Mon Jul 25 17:02:40 CEST 2022 - The aliscore output appears to be empty. Check results for BUSCO: 375960at2759
## Mon Jul 25 17:02:40 CEST 2022 - The aliscore output file does not exist. Check results for BUSCO: 939345at2759
## Mon Jul 25 17:02:41 CEST 2022 - The aliscore output appears to be empty. Check results for BUSCO: 939345at2759
## Mon Jul 25 17:02:41 CEST 2022 - The aliscore output file does not exist. Check results for BUSCO: 973442at2759
## Mon Jul 25 17:02:41 CEST 2022 - The aliscore output file does not exist. Check results for BUSCO: 325552at2759
## Mon Jul 25 17:02:41 CEST 2022 - The aliscore output appears to be empty. Check results for BUSCO: 973442at2759
## Mon Jul 25 17:02:41 CEST 2022 - The aliscore output appears to be empty. Check results for BUSCO: 325552at2759
## Mon Jul 25 17:02:41 CEST 2022 - The aliscore output file does not exist. Check results for BUSCO: 179362at2759
## Mon Jul 25 17:02:41 CEST 2022 - The aliscore output appears to be empty. Check results for BUSCO: 179362at2759
## Mon Jul 25 17:02:42 CEST 2022 - The aliscore output file does not exist. Check results for BUSCO: 369837at2759
## Mon Jul 25 17:02:42 CEST 2022 - The aliscore output appears to be empty. Check results for BUSCO: 369837at2759
## Mon Jul 25 17:02:43 CEST 2022 - The aliscore output file does not exist. Check results for BUSCO: 1258856at2759
## Mon Jul 25 17:02:43 CEST 2022 - The aliscore output appears to be empty. Check results for BUSCO: 1258856at2759
## Mon Jul 25 17:02:44 CEST 2022 - The aliscore output file does not exist. Check results for BUSCO: 1312453at2759
## Mon Jul 25 17:02:44 CEST 2022 - The aliscore output appears to be empty. Check results for BUSCO: 1312453at2759
## Mon Jul 25 17:02:45 CEST 2022 - The aliscore output file does not exist. Check results for BUSCO: 625387at2759
## Mon Jul 25 17:02:45 CEST 2022 - The aliscore output appears to be empty. Check results for BUSCO: 625387at2759
## Mon Jul 25 17:03:02 CEST 2022 - The aliscore output file does not exist. Check results for BUSCO: 1459797at2759
## Mon Jul 25 17:03:02 CEST 2022 - The aliscore output appears to be empty. Check results for BUSCO: 1459797at2759
## Mon Jul 25 17:03:02 CEST 2022 - The aliscore output file does not exist. Check results for BUSCO: 1526152at2759
## Mon Jul 25 17:03:02 CEST 2022 - The aliscore output appears to be empty. Check results for BUSCO: 1526152at2759
## Mon Jul 25 17:03:03 CEST 2022 - The aliscore output file does not exist. Check results for BUSCO: 1249159at2759
## Mon Jul 25 17:03:03 CEST 2022 - The aliscore output appears to be empty. Check results for BUSCO: 1249159at2759
## Mon Jul 25 17:03:04 CEST 2022 - The aliscore output file does not exist. Check results for BUSCO: 1257440at2759
## Mon Jul 25 17:03:04 CEST 2022 - The aliscore output appears to be empty. Check results for BUSCO: 1257440at2759
## Mon Jul 25 17:03:04 CEST 2022 - The aliscore output file does not exist. Check results for BUSCO: 1269244at2759
## Mon Jul 25 17:03:04 CEST 2022 - The aliscore output file does not exist. Check results for BUSCO: 1407446at2759
## Mon Jul 25 17:03:04 CEST 2022 - The aliscore output appears to be empty. Check results for BUSCO: 1269244at2759
## Mon Jul 25 17:03:04 CEST 2022 - The aliscore output appears to be empty. Check results for BUSCO: 1407446at2759
## Mon Jul 25 17:03:04 CEST 2022 - The aliscore output file does not exist. Check results for BUSCO: 1030907at2759
## Mon Jul 25 17:03:04 CEST 2022 - The aliscore output file does not exist. Check results for BUSCO: 1197019at2759
## Mon Jul 25 17:03:04 CEST 2022 - The aliscore output appears to be empty. Check results for BUSCO: 1030907at2759
## Mon Jul 25 17:03:04 CEST 2022 - The aliscore output appears to be empty. Check results for BUSCO: 1197019at2759
## Mon Jul 25 17:03:04 CEST 2022 - The aliscore output file does not exist. Check results for BUSCO: 1291729at2759
## Mon Jul 25 17:03:04 CEST 2022 - The aliscore output appears to be empty. Check results for BUSCO: 1291729at2759
## Mon Jul 25 17:03:04 CEST 2022 - The aliscore output file does not exist. Check results for BUSCO: 866359at2759
## Mon Jul 25 17:03:04 CEST 2022 - The aliscore output appears to be empty. Check results for BUSCO: 866359at2759
## Mon Jul 25 17:03:05 CEST 2022 - The aliscore output file does not exist. Check results for BUSCO: 1178688at2759
## Mon Jul 25 17:03:05 CEST 2022 - The aliscore output appears to be empty. Check results for BUSCO: 1178688at2759
## Mon Jul 25 17:03:05 CEST 2022 - The aliscore output file does not exist. Check results for BUSCO: 457861at2759
## Mon Jul 25 17:03:05 CEST 2022 - The aliscore output file does not exist. Check results for BUSCO: 1076134at2759
## Mon Jul 25 17:03:05 CEST 2022 - The aliscore output appears to be empty. Check results for BUSCO: 457861at2759
## Mon Jul 25 17:03:05 CEST 2022 - The aliscore output appears to be empty. Check results for BUSCO: 1076134at2759
## Mon Jul 25 17:03:05 CEST 2022 - The aliscore output file does not exist. Check results for BUSCO: 464990at2759
## Mon Jul 25 17:03:05 CEST 2022 - The aliscore output appears to be empty. Check results for BUSCO: 464990at2759
## Mon Jul 25 17:03:05 CEST 2022 - The aliscore output file does not exist. Check results for BUSCO: 1275837at2759
## Mon Jul 25 17:03:05 CEST 2022 - The aliscore output file does not exist. Check results for BUSCO: 1090038at2759
## Mon Jul 25 17:03:05 CEST 2022 - The aliscore output appears to be empty. Check results for BUSCO: 1275837at2759
## Mon Jul 25 17:03:05 CEST 2022 - The aliscore output appears to be empty. Check results for BUSCO: 1090038at2759
## Mon Jul 25 17:03:06 CEST 2022 - The aliscore output file does not exist. Check results for BUSCO: 1441854at2759
## Mon Jul 25 17:03:06 CEST 2022 - The aliscore output appears to be empty. Check results for BUSCO: 1441854at2759
## Mon Jul 25 17:03:06 CEST 2022 - The aliscore output file does not exist. Check results for BUSCO: 1053181at2759
## Mon Jul 25 17:03:06 CEST 2022 - The aliscore output appears to be empty. Check results for BUSCO: 1053181at2759
## Mon Jul 25 17:03:06 CEST 2022 - The aliscore output file does not exist. Check results for BUSCO: 1623701at2759
## Mon Jul 25 17:03:06 CEST 2022 - The aliscore output appears to be empty. Check results for BUSCO: 1623701at2759
## Mon Jul 25 17:03:06 CEST 2022 - The aliscore output file does not exist. Check results for BUSCO: 1010730at2759
## Mon Jul 25 17:03:06 CEST 2022 - The aliscore output appears to be empty. Check results for BUSCO: 1010730at2759
## Mon Jul 25 17:03:06 CEST 2022 - The aliscore output file does not exist. Check results for BUSCO: 911863at2759
## Mon Jul 25 17:03:06 CEST 2022 - The aliscore output appears to be empty. Check results for BUSCO: 911863at2759
## Mon Jul 25 17:03:06 CEST 2022 - The aliscore output file does not exist. Check results for BUSCO: 664730at2759
## Mon Jul 25 17:03:06 CEST 2022 - The aliscore output file does not exist. Check results for BUSCO: 1014314at2759
## Mon Jul 25 17:03:06 CEST 2022 - The aliscore output appears to be empty. Check results for BUSCO: 664730at2759
## Mon Jul 25 17:03:06 CEST 2022 - The aliscore output file does not exist. Check results for BUSCO: 1129824at2759
## Mon Jul 25 17:03:06 CEST 2022 - The aliscore output appears to be empty. Check results for BUSCO: 1014314at2759
## Mon Jul 25 17:03:06 CEST 2022 - The aliscore output appears to be empty. Check results for BUSCO: 1129824at2759
## Mon Jul 25 17:03:06 CEST 2022 - The aliscore output file does not exist. Check results for BUSCO: 1266231at2759
## Mon Jul 25 17:03:06 CEST 2022 - The aliscore output file does not exist. Check results for BUSCO: 734666at2759
## Mon Jul 25 17:03:06 CEST 2022 - The aliscore output appears to be empty. Check results for BUSCO: 1266231at2759
## Mon Jul 25 17:03:06 CEST 2022 - The aliscore output appears to be empty. Check results for BUSCO: 734666at2759
## Mon Jul 25 17:03:07 CEST 2022 - The aliscore output file does not exist. Check results for BUSCO: 687505at2759
## Mon Jul 25 17:03:07 CEST 2022 - The aliscore output appears to be empty. Check results for BUSCO: 687505at2759
## Mon Jul 25 17:03:07 CEST 2022 - The aliscore output file does not exist. Check results for BUSCO: 1173229at2759
## Mon Jul 25 17:03:07 CEST 2022 - The aliscore output appears to be empty. Check results for BUSCO: 1173229at2759
## Mon Jul 25 17:03:07 CEST 2022 - The aliscore output file does not exist. Check results for BUSCO: 1558822at2759
## Mon Jul 25 17:03:07 CEST 2022 - The aliscore output appears to be empty. Check results for BUSCO: 1558822at2759
## Mon Jul 25 17:03:07 CEST 2022 - The aliscore output file does not exist. Check results for BUSCO: 1222562at2759
## Mon Jul 25 17:03:07 CEST 2022 - The aliscore output appears to be empty. Check results for BUSCO: 1222562at2759
## Mon Jul 25 17:03:07 CEST 2022 - The aliscore output file does not exist. Check results for BUSCO: 1260807at2759
## Mon Jul 25 17:03:07 CEST 2022 - The aliscore output file does not exist. Check results for BUSCO: 299766at2759
## Mon Jul 25 17:03:07 CEST 2022 - The aliscore output appears to be empty. Check results for BUSCO: 1260807at2759
## Mon Jul 25 17:03:07 CEST 2022 - The aliscore output appears to be empty. Check results for BUSCO: 299766at2759
## Mon Jul 25 17:03:07 CEST 2022 - The aliscore output file does not exist. Check results for BUSCO: 1251252at2759
## Mon Jul 25 17:03:07 CEST 2022 - The aliscore output appears to be empty. Check results for BUSCO: 1251252at2759
## Mon Jul 25 17:03:07 CEST 2022 - The aliscore output file does not exist. Check results for BUSCO: 834694at2759
## Mon Jul 25 17:03:07 CEST 2022 - The aliscore output file does not exist. Check results for BUSCO: 1323575at2759
## Mon Jul 25 17:03:07 CEST 2022 - The aliscore output appears to be empty. Check results for BUSCO: 834694at2759
## Mon Jul 25 17:03:07 CEST 2022 - The aliscore output appears to be empty. Check results for BUSCO: 1323575at2759
## Mon Jul 25 17:03:07 CEST 2022 - The aliscore output file does not exist. Check results for BUSCO: 855834at2759
## Mon Jul 25 17:03:08 CEST 2022 - The aliscore output appears to be empty. Check results for BUSCO: 855834at2759
## Mon Jul 25 17:03:08 CEST 2022 - The aliscore output file does not exist. Check results for BUSCO: 761109at2759
## Mon Jul 25 17:03:08 CEST 2022 - The aliscore output appears to be empty. Check results for BUSCO: 761109at2759
## Mon Jul 25 17:03:08 CEST 2022 - The aliscore output file does not exist. Check results for BUSCO: 1096688at2759
## Mon Jul 25 17:03:08 CEST 2022 - The aliscore output appears to be empty. Check results for BUSCO: 1096688at2759
## Mon Jul 25 17:03:08 CEST 2022 - The aliscore output file does not exist. Check results for BUSCO: 817008at2759
## Mon Jul 25 17:03:08 CEST 2022 - The aliscore output file does not exist. Check results for BUSCO: 582756at2759
## Mon Jul 25 17:03:08 CEST 2022 - The aliscore output file does not exist. Check results for BUSCO: 1051021at2759
## Mon Jul 25 17:03:08 CEST 2022 - The aliscore output file does not exist. Check results for BUSCO: 719531at2759
## Mon Jul 25 17:03:08 CEST 2022 - The aliscore output appears to be empty. Check results for BUSCO: 817008at2759
## Mon Jul 25 17:03:08 CEST 2022 - The aliscore output appears to be empty. Check results for BUSCO: 582756at2759
## Mon Jul 25 17:03:08 CEST 2022 - The aliscore output appears to be empty. Check results for BUSCO: 1051021at2759
## Mon Jul 25 17:03:08 CEST 2022 - The aliscore output appears to be empty. Check results for BUSCO: 719531at2759
## Mon Jul 25 17:03:08 CEST 2022 - The aliscore output file does not exist. Check results for BUSCO: 1106766at2759
## Mon Jul 25 17:03:08 CEST 2022 - The aliscore output appears to be empty. Check results for BUSCO: 1106766at2759
## Mon Jul 25 17:03:08 CEST 2022 - The aliscore output file does not exist. Check results for BUSCO: 692986at2759
## Mon Jul 25 17:03:08 CEST 2022 - The aliscore output appears to be empty. Check results for BUSCO: 692986at2759
## Mon Jul 25 17:03:08 CEST 2022 - The aliscore output file does not exist. Check results for BUSCO: 320059at2759
## Mon Jul 25 17:03:08 CEST 2022 - The aliscore output appears to be empty. Check results for BUSCO: 320059at2759
## Mon Jul 25 17:03:08 CEST 2022 - The aliscore output file does not exist. Check results for BUSCO: 450058at2759
## Mon Jul 25 17:03:09 CEST 2022 - The aliscore output appears to be empty. Check results for BUSCO: 450058at2759
## Mon Jul 25 17:03:09 CEST 2022 - The aliscore output file does not exist. Check results for BUSCO: 1019762at2759
## Mon Jul 25 17:03:09 CEST 2022 - The aliscore output appears to be empty. Check results for BUSCO: 1019762at2759
## Mon Jul 25 17:03:09 CEST 2022 - The aliscore output file does not exist. Check results for BUSCO: 1404162at2759
## Mon Jul 25 17:03:09 CEST 2022 - The aliscore output appears to be empty. Check results for BUSCO: 1404162at2759
## Mon Jul 25 17:03:09 CEST 2022 - The aliscore output file does not exist. Check results for BUSCO: 306227at2759
## Mon Jul 25 17:03:09 CEST 2022 - The aliscore output appears to be empty. Check results for BUSCO: 306227at2759
## Mon Jul 25 17:03:09 CEST 2022 - The aliscore output file does not exist. Check results for BUSCO: 939345at2759
## Mon Jul 25 17:03:09 CEST 2022 - The aliscore output appears to be empty. Check results for BUSCO: 939345at2759
## Mon Jul 25 17:03:09 CEST 2022 - The aliscore output file does not exist. Check results for BUSCO: 981902at2759
## Mon Jul 25 17:03:09 CEST 2022 - The aliscore output appears to be empty. Check results for BUSCO: 981902at2759
## Mon Jul 25 17:03:10 CEST 2022 - The aliscore output file does not exist. Check results for BUSCO: 1248958at2759
## Mon Jul 25 17:03:10 CEST 2022 - The aliscore output appears to be empty. Check results for BUSCO: 1248958at2759
## Mon Jul 25 17:03:10 CEST 2022 - The aliscore output file does not exist. Check results for BUSCO: 671536at2759
## Mon Jul 25 17:03:10 CEST 2022 - The aliscore output appears to be empty. Check results for BUSCO: 671536at2759
## Mon Jul 25 17:03:10 CEST 2022 - The aliscore output file does not exist. Check results for BUSCO: 1194797at2759
## Mon Jul 25 17:03:10 CEST 2022 - The aliscore output appears to be empty. Check results for BUSCO: 1194797at2759
## Mon Jul 25 17:03:11 CEST 2022 - The aliscore output file does not exist. Check results for BUSCO: 1115196at2759
## Mon Jul 25 17:03:11 CEST 2022 - The aliscore output appears to be empty. Check results for BUSCO: 1115196at2759
## Mon Jul 25 17:03:11 CEST 2022 - The aliscore output file does not exist. Check results for BUSCO: 625387at2759
## Mon Jul 25 17:03:11 CEST 2022 - The aliscore output appears to be empty. Check results for BUSCO: 625387at2759
## Mon Jul 25 17:03:11 CEST 2022 - The aliscore output file does not exist. Check results for BUSCO: 1079130at2759
## Mon Jul 25 17:03:11 CEST 2022 - The aliscore output appears to be empty. Check results for BUSCO: 1079130at2759
## Mon Jul 25 17:03:11 CEST 2022 - The aliscore output file does not exist. Check results for BUSCO: 83779at2759
## Mon Jul 25 17:03:11 CEST 2022 - The aliscore output appears to be empty. Check results for BUSCO: 83779at2759
## Mon Jul 25 17:03:11 CEST 2022 - The aliscore output file does not exist. Check results for BUSCO: 261328at2759
## Mon Jul 25 17:03:11 CEST 2022 - The aliscore output appears to be empty. Check results for BUSCO: 261328at2759
## Mon Jul 25 17:03:14 CEST 2022 - The aliscore output file does not exist. Check results for BUSCO: 1379600at2759
## Mon Jul 25 17:03:14 CEST 2022 - The aliscore output appears to be empty. Check results for BUSCO: 1379600at2759
## Mon Jul 25 17:03:32 CEST 2022 - The aliscore output file does not exist. Check results for BUSCO: 974865at2759
## Mon Jul 25 17:03:33 CEST 2022 - The aliscore output appears to be empty. Check results for BUSCO: 974865at2759
## Mon Jul 25 17:03:33 CEST 2022 - The aliscore output file does not exist. Check results for BUSCO: 1053181at2759
## Mon Jul 25 17:03:33 CEST 2022 - The aliscore output appears to be empty. Check results for BUSCO: 1053181at2759
## Mon Jul 25 17:03:33 CEST 2022 - The aliscore output file does not exist. Check results for BUSCO: 1395649at2759
## Mon Jul 25 17:03:33 CEST 2022 - The aliscore output appears to be empty. Check results for BUSCO: 1395649at2759
## Mon Jul 25 17:03:33 CEST 2022 - The aliscore output file does not exist. Check results for BUSCO: 937275at2759
## Mon Jul 25 17:03:33 CEST 2022 - The aliscore output appears to be empty. Check results for BUSCO: 937275at2759
## Mon Jul 25 17:03:33 CEST 2022 - The aliscore output file does not exist. Check results for BUSCO: 1346432at2759
## Mon Jul 25 17:03:33 CEST 2022 - The aliscore output appears to be empty. Check results for BUSCO: 1346432at2759
## Mon Jul 25 17:03:33 CEST 2022 - The aliscore output file does not exist. Check results for BUSCO: 1432328at2759
## Mon Jul 25 17:03:33 CEST 2022 - The aliscore output appears to be empty. Check results for BUSCO: 1432328at2759
## Mon Jul 25 17:03:33 CEST 2022 - The aliscore output file does not exist. Check results for BUSCO: 1222562at2759
## Mon Jul 25 17:03:33 CEST 2022 - The aliscore output appears to be empty. Check results for BUSCO: 1222562at2759
## Mon Jul 25 17:03:34 CEST 2022 - The aliscore output file does not exist. Check results for BUSCO: 1266231at2759
## Mon Jul 25 17:03:34 CEST 2022 - The aliscore output appears to be empty. Check results for BUSCO: 1266231at2759
## Mon Jul 25 17:03:34 CEST 2022 - The aliscore output file does not exist. Check results for BUSCO: 1490022at2759
## Mon Jul 25 17:03:34 CEST 2022 - The aliscore output appears to be empty. Check results for BUSCO: 1490022at2759
## Mon Jul 25 17:03:34 CEST 2022 - The aliscore output file does not exist. Check results for BUSCO: 1593937at2759
## Mon Jul 25 17:03:34 CEST 2022 - The aliscore output appears to be empty. Check results for BUSCO: 1593937at2759
## Mon Jul 25 17:03:35 CEST 2022 - The aliscore output file does not exist. Check results for BUSCO: 345441at2759
## Mon Jul 25 17:03:35 CEST 2022 - The aliscore output appears to be empty. Check results for BUSCO: 345441at2759
## Mon Jul 25 17:03:35 CEST 2022 - The aliscore output file does not exist. Check results for BUSCO: 1364586at2759
## Mon Jul 25 17:03:35 CEST 2022 - The aliscore output appears to be empty. Check results for BUSCO: 1364586at2759
## Mon Jul 25 17:03:35 CEST 2022 - The aliscore output file does not exist. Check results for BUSCO: 388820at2759
## Mon Jul 25 17:03:35 CEST 2022 - The aliscore output appears to be empty. Check results for BUSCO: 388820at2759
## Mon Jul 25 17:03:35 CEST 2022 - The aliscore output file does not exist. Check results for BUSCO: 1025450at2759
## Mon Jul 25 17:03:35 CEST 2022 - The aliscore output file does not exist. Check results for BUSCO: 1094121at2759
## Mon Jul 25 17:03:35 CEST 2022 - The aliscore output appears to be empty. Check results for BUSCO: 1025450at2759
## Mon Jul 25 17:03:35 CEST 2022 - The aliscore output appears to be empty. Check results for BUSCO: 1094121at2759
## Mon Jul 25 17:03:35 CEST 2022 - The aliscore output file does not exist. Check results for BUSCO: 1096688at2759
## Mon Jul 25 17:03:35 CEST 2022 - The aliscore output appears to be empty. Check results for BUSCO: 1096688at2759
## Mon Jul 25 17:03:36 CEST 2022 - The aliscore output file does not exist. Check results for BUSCO: 975557at2759
## Mon Jul 25 17:03:36 CEST 2022 - The aliscore output file does not exist. Check results for BUSCO: 931188at2759
## Mon Jul 25 17:03:36 CEST 2022 - The aliscore output appears to be empty. Check results for BUSCO: 975557at2759
## Mon Jul 25 17:03:36 CEST 2022 - The aliscore output appears to be empty. Check results for BUSCO: 931188at2759
## Mon Jul 25 17:03:36 CEST 2022 - The aliscore output file does not exist. Check results for BUSCO: 1287094at2759
## Mon Jul 25 17:03:36 CEST 2022 - The aliscore output file does not exist. Check results for BUSCO: 1057950at2759
## Mon Jul 25 17:03:36 CEST 2022 - The aliscore output appears to be empty. Check results for BUSCO: 1287094at2759
## Mon Jul 25 17:03:36 CEST 2022 - The aliscore output appears to be empty. Check results for BUSCO: 1057950at2759
## Mon Jul 25 17:03:36 CEST 2022 - The aliscore output file does not exist. Check results for BUSCO: 549762at2759
## Mon Jul 25 17:03:36 CEST 2022 - The aliscore output appears to be empty. Check results for BUSCO: 549762at2759
## Mon Jul 25 17:03:36 CEST 2022 - The aliscore output file does not exist. Check results for BUSCO: 1247641at2759
## Mon Jul 25 17:03:36 CEST 2022 - The aliscore output appears to be empty. Check results for BUSCO: 1247641at2759
## Mon Jul 25 17:03:36 CEST 2022 - The aliscore output file does not exist. Check results for BUSCO: 687505at2759
## Mon Jul 25 17:03:36 CEST 2022 - The aliscore output appears to be empty. Check results for BUSCO: 687505at2759
## Mon Jul 25 17:03:37 CEST 2022 - The aliscore output file does not exist. Check results for BUSCO: 543764at2759
## Mon Jul 25 17:03:37 CEST 2022 - The aliscore output appears to be empty. Check results for BUSCO: 543764at2759
## Mon Jul 25 17:03:37 CEST 2022 - The aliscore output file does not exist. Check results for BUSCO: 923657at2759
## Mon Jul 25 17:03:37 CEST 2022 - The aliscore output appears to be empty. Check results for BUSCO: 923657at2759
## Mon Jul 25 17:03:37 CEST 2022 - The aliscore output file does not exist. Check results for BUSCO: 1479417at2759
## Mon Jul 25 17:03:37 CEST 2022 - The aliscore output appears to be empty. Check results for BUSCO: 1479417at2759
## Mon Jul 25 17:03:37 CEST 2022 - The aliscore output file does not exist. Check results for BUSCO: 1258856at2759
## Mon Jul 25 17:03:37 CEST 2022 - The aliscore output appears to be empty. Check results for BUSCO: 1258856at2759
## Mon Jul 25 17:03:37 CEST 2022 - The aliscore output file does not exist. Check results for BUSCO: 271586at2759
## Mon Jul 25 17:03:37 CEST 2022 - The aliscore output appears to be empty. Check results for BUSCO: 271586at2759
## Mon Jul 25 17:03:39 CEST 2022 - The aliscore output file does not exist. Check results for BUSCO: 1079130at2759
## Mon Jul 25 17:03:39 CEST 2022 - The aliscore output appears to be empty. Check results for BUSCO: 1079130at2759
## Mon Jul 25 17:03:40 CEST 2022 - The aliscore output file does not exist. Check results for BUSCO: 673132at2759
## Mon Jul 25 17:03:40 CEST 2022 - The aliscore output appears to be empty. Check results for BUSCO: 673132at2759
## Mon Jul 25 17:03:40 CEST 2022 - The aliscore output file does not exist. Check results for BUSCO: 671536at2759
## Mon Jul 25 17:03:40 CEST 2022 - The aliscore output appears to be empty. Check results for BUSCO: 671536at2759
## Mon Jul 25 17:03:41 CEST 2022 - The aliscore output file does not exist. Check results for BUSCO: 166920at2759
## Mon Jul 25 17:03:41 CEST 2022 - The aliscore output appears to be empty. Check results for BUSCO: 166920at2759
## Mon Jul 25 17:03:43 CEST 2022 - The aliscore output file does not exist. Check results for BUSCO: 324863at2759
## Mon Jul 25 17:03:43 CEST 2022 - The aliscore output appears to be empty. Check results for BUSCO: 324863at2759
## Mon Jul 25 17:03:43 CEST 2022 - The aliscore output file does not exist. Check results for BUSCO: 160593at2759
## Mon Jul 25 17:03:43 CEST 2022 - The aliscore output appears to be empty. Check results for BUSCO: 160593at2759
## Mon Jul 25 17:03:44 CEST 2022 - The aliscore output file does not exist. Check results for BUSCO: 674160at2759
## Mon Jul 25 17:03:44 CEST 2022 - The aliscore output appears to be empty. Check results for BUSCO: 674160at2759
## Mon Jul 25 17:03:44 CEST 2022 - The aliscore output file does not exist. Check results for BUSCO: 142542at2759
## Mon Jul 25 17:03:44 CEST 2022 - The aliscore output appears to be empty. Check results for BUSCO: 142542at2759
## Mon Jul 25 17:04:02 CEST 2022 - The aliscore output file does not exist. Check results for BUSCO: 1138059at2759
## Mon Jul 25 17:04:02 CEST 2022 - The aliscore output appears to be empty. Check results for BUSCO: 1138059at2759
## Mon Jul 25 17:04:02 CEST 2022 - The aliscore output file does not exist. Check results for BUSCO: 1421503at2759
## Mon Jul 25 17:04:02 CEST 2022 - The aliscore output appears to be empty. Check results for BUSCO: 1421503at2759
## Mon Jul 25 17:04:03 CEST 2022 - The aliscore output file does not exist. Check results for BUSCO: 1525971at2759
## Mon Jul 25 17:04:03 CEST 2022 - The aliscore output appears to be empty. Check results for BUSCO: 1525971at2759
## Mon Jul 25 17:04:03 CEST 2022 - The aliscore output file does not exist. Check results for BUSCO: 865202at2759
## Mon Jul 25 17:04:03 CEST 2022 - The aliscore output file does not exist. Check results for BUSCO: 976469at2759
## Mon Jul 25 17:04:03 CEST 2022 - The aliscore output appears to be empty. Check results for BUSCO: 865202at2759
## Mon Jul 25 17:04:03 CEST 2022 - The aliscore output appears to be empty. Check results for BUSCO: 976469at2759
## Mon Jul 25 17:04:04 CEST 2022 - The aliscore output file does not exist. Check results for BUSCO: 946128at2759
## Mon Jul 25 17:04:04 CEST 2022 - The aliscore output appears to be empty. Check results for BUSCO: 946128at2759
## Mon Jul 25 17:04:04 CEST 2022 - The aliscore output file does not exist. Check results for BUSCO: 751335at2759
## Mon Jul 25 17:04:04 CEST 2022 - The aliscore output appears to be empty. Check results for BUSCO: 751335at2759
## Mon Jul 25 17:04:05 CEST 2022 - The aliscore output file does not exist. Check results for BUSCO: 1107630at2759
## Mon Jul 25 17:04:05 CEST 2022 - The aliscore output appears to be empty. Check results for BUSCO: 1107630at2759
## Mon Jul 25 17:04:05 CEST 2022 - The aliscore output file does not exist. Check results for BUSCO: 1626636at2759
## Mon Jul 25 17:04:05 CEST 2022 - The aliscore output appears to be empty. Check results for BUSCO: 1626636at2759
## Mon Jul 25 17:04:05 CEST 2022 - The aliscore output file does not exist. Check results for BUSCO: 1620056at2759
## Mon Jul 25 17:04:05 CEST 2022 - The aliscore output appears to be empty. Check results for BUSCO: 1620056at2759
## Mon Jul 25 17:04:05 CEST 2022 - The aliscore output file does not exist. Check results for BUSCO: 1426075at2759
## Mon Jul 25 17:04:05 CEST 2022 - The aliscore output appears to be empty. Check results for BUSCO: 1426075at2759
## Mon Jul 25 17:04:05 CEST 2022 - The aliscore output file does not exist. Check results for BUSCO: 1338131at2759
## Mon Jul 25 17:04:05 CEST 2022 - The aliscore output file does not exist. Check results for BUSCO: 1182451at2759
## Mon Jul 25 17:04:05 CEST 2022 - The aliscore output appears to be empty. Check results for BUSCO: 1338131at2759
## Mon Jul 25 17:04:05 CEST 2022 - The aliscore output appears to be empty. Check results for BUSCO: 1182451at2759
## Mon Jul 25 17:04:05 CEST 2022 - The aliscore output file does not exist. Check results for BUSCO: 1442062at2759
## Mon Jul 25 17:04:05 CEST 2022 - The aliscore output appears to be empty. Check results for BUSCO: 1442062at2759
## Mon Jul 25 17:04:05 CEST 2022 - The aliscore output file does not exist. Check results for BUSCO: 1513531at2759
## Mon Jul 25 17:04:05 CEST 2022 - The aliscore output appears to be empty. Check results for BUSCO: 1513531at2759
## Mon Jul 25 17:04:05 CEST 2022 - The aliscore output file does not exist. Check results for BUSCO: 1200489at2759
## Mon Jul 25 17:04:05 CEST 2022 - The aliscore output appears to be empty. Check results for BUSCO: 1200489at2759
## Mon Jul 25 17:04:05 CEST 2022 - The aliscore output file does not exist. Check results for BUSCO: 1538526at2759
## Mon Jul 25 17:04:05 CEST 2022 - The aliscore output appears to be empty. Check results for BUSCO: 1538526at2759
## Mon Jul 25 17:04:06 CEST 2022 - The aliscore output file does not exist. Check results for BUSCO: 1049599at2759
## Mon Jul 25 17:04:06 CEST 2022 - The aliscore output appears to be empty. Check results for BUSCO: 1049599at2759
## Mon Jul 25 17:04:06 CEST 2022 - The aliscore output file does not exist. Check results for BUSCO: 1563319at2759
## Mon Jul 25 17:04:06 CEST 2022 - The aliscore output appears to be empty. Check results for BUSCO: 1563319at2759
## Mon Jul 25 17:04:06 CEST 2022 - The aliscore output file does not exist. Check results for BUSCO: 1645187at2759
## Mon Jul 25 17:04:06 CEST 2022 - The aliscore output file does not exist. Check results for BUSCO: 1233814at2759
## Mon Jul 25 17:04:06 CEST 2022 - The aliscore output appears to be empty. Check results for BUSCO: 1233814at2759
## Mon Jul 25 17:04:06 CEST 2022 - The aliscore output appears to be empty. Check results for BUSCO: 1645187at2759
## Mon Jul 25 17:04:06 CEST 2022 - The aliscore output file does not exist. Check results for BUSCO: 1111142at2759
## Mon Jul 25 17:04:06 CEST 2022 - The aliscore output appears to be empty. Check results for BUSCO: 1111142at2759
## Mon Jul 25 17:04:06 CEST 2022 - The aliscore output file does not exist. Check results for BUSCO: 1342242at2759
## Mon Jul 25 17:04:06 CEST 2022 - The aliscore output appears to be empty. Check results for BUSCO: 1342242at2759
## Mon Jul 25 17:04:06 CEST 2022 - The aliscore output file does not exist. Check results for BUSCO: 1223488at2759
## Mon Jul 25 17:04:06 CEST 2022 - The aliscore output appears to be empty. Check results for BUSCO: 1223488at2759
## Mon Jul 25 17:04:06 CEST 2022 - The aliscore output file does not exist. Check results for BUSCO: 937686at2759
## Mon Jul 25 17:04:06 CEST 2022 - The aliscore output appears to be empty. Check results for BUSCO: 937686at2759
## Mon Jul 25 17:04:06 CEST 2022 - The aliscore output file does not exist. Check results for BUSCO: 453044at2759
## Mon Jul 25 17:04:06 CEST 2022 - The aliscore output appears to be empty. Check results for BUSCO: 453044at2759
## Mon Jul 25 17:04:07 CEST 2022 - The aliscore output file does not exist. Check results for BUSCO: 1032689at2759
## Mon Jul 25 17:04:07 CEST 2022 - The aliscore output appears to be empty. Check results for BUSCO: 1032689at2759
## Mon Jul 25 17:04:07 CEST 2022 - The aliscore output file does not exist. Check results for BUSCO: 759498at2759
## Mon Jul 25 17:04:07 CEST 2022 - The aliscore output appears to be empty. Check results for BUSCO: 759498at2759
## Mon Jul 25 17:04:07 CEST 2022 - The aliscore output file does not exist. Check results for BUSCO: 1575179at2759
## Mon Jul 25 17:04:07 CEST 2022 - The aliscore output appears to be empty. Check results for BUSCO: 1575179at2759
## Mon Jul 25 17:04:07 CEST 2022 - The aliscore output file does not exist. Check results for BUSCO: 1377237at2759
## Mon Jul 25 17:04:07 CEST 2022 - The aliscore output appears to be empty. Check results for BUSCO: 1377237at2759
## Mon Jul 25 17:04:07 CEST 2022 - The aliscore output file does not exist. Check results for BUSCO: 1428854at2759
## Mon Jul 25 17:04:07 CEST 2022 - The aliscore output appears to be empty. Check results for BUSCO: 1428854at2759
## Mon Jul 25 17:04:07 CEST 2022 - The aliscore output file does not exist. Check results for BUSCO: 919955at2759
## Mon Jul 25 17:04:07 CEST 2022 - The aliscore output file does not exist. Check results for BUSCO: 1552706at2759
## Mon Jul 25 17:04:07 CEST 2022 - The aliscore output appears to be empty. Check results for BUSCO: 919955at2759
## Mon Jul 25 17:04:07 CEST 2022 - The aliscore output file does not exist. Check results for BUSCO: 844748at2759
## Mon Jul 25 17:04:07 CEST 2022 - The aliscore output appears to be empty. Check results for BUSCO: 1552706at2759
## Mon Jul 25 17:04:07 CEST 2022 - The aliscore output appears to be empty. Check results for BUSCO: 844748at2759
## Mon Jul 25 17:04:07 CEST 2022 - The aliscore output file does not exist. Check results for BUSCO: 937686at2759
## Mon Jul 25 17:04:07 CEST 2022 - The aliscore output appears to be empty. Check results for BUSCO: 937686at2759
## Mon Jul 25 17:04:07 CEST 2022 - The aliscore output file does not exist. Check results for BUSCO: 1264469at2759
## Mon Jul 25 17:04:07 CEST 2022 - The aliscore output file does not exist. Check results for BUSCO: 388820at2759
## Mon Jul 25 17:04:07 CEST 2022 - The aliscore output appears to be empty. Check results for BUSCO: 1264469at2759
## Mon Jul 25 17:04:07 CEST 2022 - The aliscore output appears to be empty. Check results for BUSCO: 388820at2759
## Mon Jul 25 17:04:07 CEST 2022 - The aliscore output file does not exist. Check results for BUSCO: 736068at2759
## Mon Jul 25 17:04:07 CEST 2022 - The aliscore output appears to be empty. Check results for BUSCO: 736068at2759
## Mon Jul 25 17:04:07 CEST 2022 - The aliscore output file does not exist. Check results for BUSCO: 1339553at2759
## Mon Jul 25 17:04:08 CEST 2022 - The aliscore output appears to be empty. Check results for BUSCO: 1339553at2759
## Mon Jul 25 17:04:08 CEST 2022 - The aliscore output file does not exist. Check results for BUSCO: 355408at2759
## Mon Jul 25 17:04:08 CEST 2022 - The aliscore output file does not exist. Check results for BUSCO: 570797at2759
## Mon Jul 25 17:04:08 CEST 2022 - The aliscore output appears to be empty. Check results for BUSCO: 355408at2759
## Mon Jul 25 17:04:08 CEST 2022 - The aliscore output file does not exist. Check results for BUSCO: 1287401at2759
## Mon Jul 25 17:04:08 CEST 2022 - The aliscore output appears to be empty. Check results for BUSCO: 570797at2759
## Mon Jul 25 17:04:08 CEST 2022 - The aliscore output appears to be empty. Check results for BUSCO: 1287401at2759
## Mon Jul 25 17:04:08 CEST 2022 - The aliscore output file does not exist. Check results for BUSCO: 774318at2759
## Mon Jul 25 17:04:08 CEST 2022 - The aliscore output file does not exist. Check results for BUSCO: 331411at2759
## Mon Jul 25 17:04:08 CEST 2022 - The aliscore output appears to be empty. Check results for BUSCO: 774318at2759
## Mon Jul 25 17:04:08 CEST 2022 - The aliscore output file does not exist. Check results for BUSCO: 100698at2759
## Mon Jul 25 17:04:08 CEST 2022 - The aliscore output appears to be empty. Check results for BUSCO: 331411at2759
## Mon Jul 25 17:04:08 CEST 2022 - The aliscore output appears to be empty. Check results for BUSCO: 100698at2759
## Mon Jul 25 17:04:08 CEST 2022 - The aliscore output file does not exist. Check results for BUSCO: 1382842at2759
## Mon Jul 25 17:04:08 CEST 2022 - The aliscore output appears to be empty. Check results for BUSCO: 1382842at2759
## Mon Jul 25 17:04:08 CEST 2022 - The aliscore output file does not exist. Check results for BUSCO: 924753at2759
## Mon Jul 25 17:04:08 CEST 2022 - The aliscore output appears to be empty. Check results for BUSCO: 924753at2759
## Mon Jul 25 17:04:09 CEST 2022 - The aliscore output file does not exist. Check results for BUSCO: 768809at2759
## Mon Jul 25 17:04:09 CEST 2022 - The aliscore output file does not exist. Check results for BUSCO: 1370304at2759
## Mon Jul 25 17:04:09 CEST 2022 - The aliscore output appears to be empty. Check results for BUSCO: 768809at2759
## Mon Jul 25 17:04:09 CEST 2022 - The aliscore output appears to be empty. Check results for BUSCO: 1370304at2759
## Mon Jul 25 17:04:09 CEST 2022 - The aliscore output file does not exist. Check results for BUSCO: 1087488at2759
## Mon Jul 25 17:04:09 CEST 2022 - The aliscore output file does not exist. Check results for BUSCO: 491869at2759
## Mon Jul 25 17:04:09 CEST 2022 - The aliscore output appears to be empty. Check results for BUSCO: 1087488at2759
## Mon Jul 25 17:04:09 CEST 2022 - The aliscore output appears to be empty. Check results for BUSCO: 491869at2759
## Mon Jul 25 17:04:09 CEST 2022 - The aliscore output file does not exist. Check results for BUSCO: 1259741at2759
## Mon Jul 25 17:04:09 CEST 2022 - The aliscore output appears to be empty. Check results for BUSCO: 1259741at2759
## Mon Jul 25 17:04:09 CEST 2022 - The aliscore output file does not exist. Check results for BUSCO: 176625at2759
## Mon Jul 25 17:04:09 CEST 2022 - The aliscore output appears to be empty. Check results for BUSCO: 176625at2759
## Mon Jul 25 17:04:09 CEST 2022 - The aliscore output file does not exist. Check results for BUSCO: 1018517at2759
## Mon Jul 25 17:04:09 CEST 2022 - The aliscore output appears to be empty. Check results for BUSCO: 1018517at2759
## Mon Jul 25 17:04:10 CEST 2022 - The aliscore output file does not exist. Check results for BUSCO: 898782at2759
## Mon Jul 25 17:04:10 CEST 2022 - The aliscore output file does not exist. Check results for BUSCO: 836599at2759
## Mon Jul 25 17:04:10 CEST 2022 - The aliscore output appears to be empty. Check results for BUSCO: 898782at2759
## Mon Jul 25 17:04:10 CEST 2022 - The aliscore output appears to be empty. Check results for BUSCO: 836599at2759
## Mon Jul 25 17:04:10 CEST 2022 - The aliscore output file does not exist. Check results for BUSCO: 814241at2759
## Mon Jul 25 17:04:10 CEST 2022 - The aliscore output appears to be empty. Check results for BUSCO: 814241at2759
## Mon Jul 25 17:04:10 CEST 2022 - The aliscore output file does not exist. Check results for BUSCO: 418107at2759
## Mon Jul 25 17:04:10 CEST 2022 - The aliscore output appears to be empty. Check results for BUSCO: 418107at2759
## Mon Jul 25 17:04:11 CEST 2022 - The aliscore output file does not exist. Check results for BUSCO: 261419at2759
## Mon Jul 25 17:04:11 CEST 2022 - The aliscore output appears to be empty. Check results for BUSCO: 261419at2759
## Mon Jul 25 17:04:11 CEST 2022 - The aliscore output file does not exist. Check results for BUSCO: 759498at2759
## Mon Jul 25 17:04:11 CEST 2022 - The aliscore output appears to be empty. Check results for BUSCO: 759498at2759
## Mon Jul 25 17:04:12 CEST 2022 - The aliscore output file does not exist. Check results for BUSCO: 491869at2759
## Mon Jul 25 17:04:12 CEST 2022 - The aliscore output appears to be empty. Check results for BUSCO: 491869at2759
## Mon Jul 25 17:04:12 CEST 2022 - The aliscore output file does not exist. Check results for BUSCO: 160593at2759
## Mon Jul 25 17:04:12 CEST 2022 - The aliscore output appears to be empty. Check results for BUSCO: 160593at2759
## Mon Jul 25 17:04:31 CEST 2022 - The aliscore output file does not exist. Check results for BUSCO: 1157302at2759
## Mon Jul 25 17:04:31 CEST 2022 - The aliscore output appears to be empty. Check results for BUSCO: 1157302at2759
## Mon Jul 25 17:04:33 CEST 2022 - The aliscore output file does not exist. Check results for BUSCO: 1284731at2759
## Mon Jul 25 17:04:33 CEST 2022 - The aliscore output appears to be empty. Check results for BUSCO: 1284731at2759
## Mon Jul 25 17:04:34 CEST 2022 - The aliscore output file does not exist. Check results for BUSCO: 1488436at2759
## Mon Jul 25 17:04:34 CEST 2022 - The aliscore output appears to be empty. Check results for BUSCO: 1488436at2759
## Mon Jul 25 17:04:34 CEST 2022 - The aliscore output file does not exist. Check results for BUSCO: 679771at2759
## Mon Jul 25 17:04:34 CEST 2022 - The aliscore output appears to be empty. Check results for BUSCO: 679771at2759
## Mon Jul 25 17:04:34 CEST 2022 - The aliscore output file does not exist. Check results for BUSCO: 1567796at2759
## Mon Jul 25 17:04:35 CEST 2022 - The aliscore output appears to be empty. Check results for BUSCO: 1567796at2759
## Mon Jul 25 17:04:35 CEST 2022 - The aliscore output file does not exist. Check results for BUSCO: 1194691at2759
## Mon Jul 25 17:04:35 CEST 2022 - The aliscore output appears to be empty. Check results for BUSCO: 1194691at2759
## Mon Jul 25 17:04:35 CEST 2022 - The aliscore output file does not exist. Check results for BUSCO: 1322299at2759
## Mon Jul 25 17:04:35 CEST 2022 - The aliscore output file does not exist. Check results for BUSCO: 1488235at2759
## Mon Jul 25 17:04:35 CEST 2022 - The aliscore output appears to be empty. Check results for BUSCO: 1322299at2759
## Mon Jul 25 17:04:35 CEST 2022 - The aliscore output appears to be empty. Check results for BUSCO: 1488235at2759
## Mon Jul 25 17:04:36 CEST 2022 - The aliscore output file does not exist. Check results for BUSCO: 779909at2759
## Mon Jul 25 17:04:36 CEST 2022 - The aliscore output appears to be empty. Check results for BUSCO: 779909at2759
## Mon Jul 25 17:04:37 CEST 2022 - The aliscore output file does not exist. Check results for BUSCO: 1193442at2759
## Mon Jul 25 17:04:37 CEST 2022 - The aliscore output appears to be empty. Check results for BUSCO: 1193442at2759
## Mon Jul 25 17:04:37 CEST 2022 - The aliscore output file does not exist. Check results for BUSCO: 1003258at2759
## Mon Jul 25 17:04:37 CEST 2022 - The aliscore output appears to be empty. Check results for BUSCO: 1003258at2759
## Mon Jul 25 17:04:37 CEST 2022 - The aliscore output file does not exist. Check results for BUSCO: 513979at2759
## Mon Jul 25 17:04:37 CEST 2022 - The aliscore output appears to be empty. Check results for BUSCO: 513979at2759
## Mon Jul 25 17:04:38 CEST 2022 - The aliscore output file does not exist. Check results for BUSCO: 325552at2759
## Mon Jul 25 17:04:38 CEST 2022 - The aliscore output appears to be empty. Check results for BUSCO: 325552at2759
## Mon Jul 25 17:04:38 CEST 2022 - The aliscore output file does not exist. Check results for BUSCO: 905026at2759
## Mon Jul 25 17:04:38 CEST 2022 - The aliscore output appears to be empty. Check results for BUSCO: 905026at2759
## Mon Jul 25 17:04:40 CEST 2022 - The aliscore output file does not exist. Check results for BUSCO: 1312453at2759
## Mon Jul 25 17:04:40 CEST 2022 - The aliscore output appears to be empty. Check results for BUSCO: 1312453at2759
## Mon Jul 25 17:05:00 CEST 2022 - Number of alignments (mafft): 255
## Mon Jul 25 17:05:00 CEST 2022 - Number of alignments (clustalo): 255
## Mon Jul 25 17:05:01 CEST 2022 - Number of alignments (mafft): 255
## Mon Jul 25 17:05:01 CEST 2022 - Number of trimmed alignments (mafft - bmge): 255
## Mon Jul 25 17:05:01 CEST 2022 - Number of trimmed alignments (clustalo - trimal): 255
## Mon Jul 25 17:05:01 CEST 2022 - Number of alignments (mafft): 255
## Mon Jul 25 17:05:01 CEST 2022 - Number of trimmed alignments (mafft - trimal): 255
## Mon Jul 25 17:05:01 CEST 2022 - Number of trimmed alignments (mafft - aliscore): 255
## Mon Jul 25 17:05:01 CEST 2022 - Number of alignments (mafft - bmge) after filtering: 245
## Mon Jul 25 17:05:01 CEST 2022 - Number of alignments (clustalo - trimal) after filtering: 249
## Mon Jul 25 17:05:01 CEST 2022 - Number of alignments (mafft - trimal) after filtering: 250
## Mon Jul 25 17:05:01 CEST 2022 - Number of alignments (mafft - aliscore) after filtering: 248
## Mon Jul 25 17:05:30 CEST 2022 - Number of alignments (muscle): 255
## Mon Jul 25 17:05:30 CEST 2022 - Number of alignments (clustalo): 255
## Mon Jul 25 17:05:30 CEST 2022 - Number of alignments (clustalo): 255
## Mon Jul 25 17:05:30 CEST 2022 - Number of trimmed alignments (muscle - trimal): 255
## Mon Jul 25 17:05:30 CEST 2022 - Number of trimmed alignments (clustalo - bmge): 255
## Mon Jul 25 17:05:30 CEST 2022 - Number of trimmed alignments (clustalo - aliscore): 255
## Mon Jul 25 17:05:30 CEST 2022 - Number of alignments (muscle): 255
## Mon Jul 25 17:05:30 CEST 2022 - Number of alignments (muscle): 255
## Mon Jul 25 17:05:30 CEST 2022 - Number of alignments (muscle - trimal) after filtering: 249
## Mon Jul 25 17:05:30 CEST 2022 - Number of trimmed alignments (muscle - aliscore): 255
## Mon Jul 25 17:05:30 CEST 2022 - Number of alignments (clustalo - bmge) after filtering: 245
## Mon Jul 25 17:05:30 CEST 2022 - Number of alignments (clustalo - aliscore) after filtering: 249
## Mon Jul 25 17:05:31 CEST 2022 - Number of trimmed alignments (muscle - bmge): 255
## Mon Jul 25 17:05:31 CEST 2022 - Number of alignments (muscle - aliscore) after filtering: 249
## Mon Jul 25 17:05:31 CEST 2022 - Number of alignments (muscle - bmge) after filtering: 244
## Mon 25 Jul 2022 05:06:26 PM CEST - Pipeline part filter_align done.
## Mon 25 Jul 2022 07:47:41 PM CEST - Pipeline part modeltest (model) done.
## Tue Jul 26 08:02:32 CEST 2022 - concatenate mafft-aliscore: Will use bootstrap cutoff 80 before creating concatenated alignment
## t
## Tue Jul 26 08:02:32 CEST 2022 - prepare_iqtree muscle-aliscore: Will use bootstrap cutoff (80) before creating concatenated alignment
## Tue Jul 26 08:02:32 CEST 2022 - prepare_iqtree clustalo-aliscore: Will use bootstrap cutoff (80) before creating concatenated alignment
## Tue Jul 26 08:02:33 CEST 2022 - prepare_iqtree clustalo-trimal: Will use bootstrap cutoff (90) before creating concatenated alignment
## Tue Jul 26 08:02:33 CEST 2022 - prepare_iqtree mafft-bmge: Will use bootstrap cutoff (0) before creating concatenated alignment
## Tue Jul 26 08:02:33 CEST 2022 - prepare_iqtree clustalo-trimal: Will use bootstrap cutoff (0) before creating concatenated alignment
## Tue Jul 26 08:02:33 CEST 2022 - concatenate clustalo-trimal: Will use bootstrap cutoff 80 before creating concatenated alignment
## Tue Jul 26 08:02:33 CEST 2022 - prepare_iqtree clustalo-aliscore: Will use bootstrap cutoff (0) before creating concatenated alignment
## Tue Jul 26 08:02:33 CEST 2022 - prepare_iqtree clustalo-bmge: Will use bootstrap cutoff (0) before creating concatenated alignment
## Tue Jul 26 08:02:33 CEST 2022 - concatenate mafft-bmge: Will use bootstrap cutoff 80 before creating concatenated alignment
## Tue Jul 26 08:02:33 CEST 2022 - prepare_iqtree muscle-aliscore: Will use bootstrap cutoff (90) before creating concatenated alignment
## Will create NEXUS partition file with model information now.
## Tue Jul 26 08:02:33 CEST 2022 - prepare_iqtree muscle-bmge: Will use bootstrap cutoff (90) before creating concatenated alignment
## Tue Jul 26 08:02:33 CEST 2022 - prepare_iqtree clustalo-aliscore: Will use bootstrap cutoff (90) before creating concatenated alignment
## Tue Jul 26 08:02:33 CEST 2022 - prepare_iqtree mafft-bmge: Will use bootstrap cutoff (50) before creating concatenated alignment
## Will create NEXUS partition file with model information now.
## Tue Jul 26 08:02:33 CEST 2022 - prepare_iqtree muscle-bmge: Will use bootstrap cutoff (0) before creating concatenated alignment
## Tue Jul 26 08:02:33 CEST 2022 - concatenate muscle-bmge: Will use bootstrap cutoff 50 before creating concatenated alignment
## Will create NEXUS partition file with model information now.
## Tue Jul 26 08:02:33 CEST 2022 - concatenate clustalo-aliscore: Will use bootstrap cutoff 50 before creating concatenated alignment
## Will create NEXUS partition file with model information now.
## Tue Jul 26 08:02:33 CEST 2022 - concatenate clustalo-trimal: Will use bootstrap cutoff 50 before creating concatenated alignment
## Tue Jul 26 08:02:33 CEST 2022 - concatenate muscle-aliscore: Will use bootstrap cutoff 50 before creating concatenated alignment
## Tue Jul 26 08:02:34 CEST 2022 - nexus file for iqtree written.
## Tue Jul 26 08:02:34 CEST 2022 - concatenate muscle-bmge: Will use bootstrap cutoff 80 before creating concatenated alignment
## Tue Jul 26 08:02:34 CEST 2022 - nexus file for iqtree written.
## Tue Jul 26 08:02:34 CEST 2022 - concatenate clustalo-trimal: Will use bootstrap cutoff 0 before creating concatenated alignment
## Tue Jul 26 08:02:34 CEST 2022 - concatenate mafft-aliscore: Will use bootstrap cutoff 90 before creating concatenated alignment
## Tue Jul 26 08:02:34 CEST 2022 - concatenate mafft-bmge: Will use bootstrap cutoff 90 before creating concatenated alignment
## Tue Jul 26 08:02:34 CEST 2022 - nexus file for iqtree written.
## Tue Jul 26 08:02:34 CEST 2022 - concatenate clustalo-trimal: Will use bootstrap cutoff 90 before creating concatenated alignment
## Tue Jul 26 08:02:34 CEST 2022 - concatenate clustalo-aliscore: Will use bootstrap cutoff 0 before creating concatenated alignment
## Tue Jul 26 08:02:34 CEST 2022 - nexus file for iqtree written.
## Tue Jul 26 08:02:34 CEST 2022 - concatenate muscle-aliscore: Will use bootstrap cutoff 90 before creating concatenated alignment
## Tue Jul 26 08:02:34 CEST 2022 - concatenate muscle-bmge: Will use bootstrap cutoff 0 before creating concatenated alignment
## Tue Jul 26 08:02:34 CEST 2022 - concatenate muscle-bmge: Will use bootstrap cutoff 90 before creating concatenated alignment
## Tue Jul 26 08:02:34 CEST 2022 - concatenate clustalo-aliscore: Will use bootstrap cutoff 90 before creating concatenated alignment
## Will create NEXUS partition file with model information now.
## Tue Jul 26 08:02:35 CEST 2022 - nexus file for iqtree written.
## Will create NEXUS partition file with model information now.
## Will create NEXUS partition file with model information now.
## Will create NEXUS partition file with model information now.
## Will create NEXUS partition file with model information now.
## Will create NEXUS partition file with model information now.
## Will create NEXUS partition file with model information now.
## Will create NEXUS partition file with model information now.
## Will create NEXUS partition file with model information now.
## Tue Jul 26 08:02:37 CEST 2022 - prepare_iqtree clustalo-trimal: Will use bootstrap cutoff (80) before creating concatenated alignment
## Tue Jul 26 08:02:37 CEST 2022 - concatenate muscle-aliscore: Will use bootstrap cutoff 0 before creating concatenated alignment
## Tue Jul 26 08:02:37 CEST 2022 - concatenate muscle-aliscore: Will use bootstrap cutoff 80 before creating concatenated alignment
## Tue Jul 26 08:02:37 CEST 2022 - concatenate clustalo-aliscore: Will use bootstrap cutoff 80 before creating concatenated alignment
## Tue Jul 26 08:02:38 CEST 2022 - nexus file for iqtree written.
## Tue Jul 26 08:02:38 CEST 2022 - nexus file for iqtree written.
## Tue Jul 26 08:02:38 CEST 2022 - nexus file for iqtree written.
## Tue Jul 26 08:02:38 CEST 2022 - nexus file for iqtree written.
## Tue Jul 26 08:02:38 CEST 2022 - nexus file for iqtree written.
## Tue Jul 26 08:02:39 CEST 2022 - nexus file for iqtree written.
## Will create NEXUS partition file with model information now.
## Tue Jul 26 08:02:39 CEST 2022 - nexus file for iqtree written.
## Tue Jul 26 08:02:39 CEST 2022 - nexus file for iqtree written.
## Tue Jul 26 08:02:40 CEST 2022 - nexus file for iqtree written.
## Tue Jul 26 08:03:01 CEST 2022 - prepare_iqtree mafft-bmge: Will use bootstrap cutoff (60) before creating concatenated alignment
## Tue 26 Jul 2022 08:03:01 AM CEST - 'phylociraptor model' finished successfully before. Will run raxml with best models.
## Tue Jul 26 08:03:01 CEST 2022 - prepare_iqtree clustalo-aliscore: Will use bootstrap cutoff (50) before creating concatenated alignment
## Tue Jul 26 08:03:01 CEST 2022 - prepare_iqtree clustalo-trimal: Will use bootstrap cutoff (50) before creating concatenated alignment
## Tue 26 Jul 2022 08:03:01 AM CEST - 'phylociraptor model' finished successfully before. Will run raxml with best models.
## Tue 26 Jul 2022 08:03:01 AM CEST - 'phylociraptor model' finished successfully before. Will run raxml with best models.
## Tue Jul 26 08:03:01 CEST 2022 - prepare_iqtree clustalo-bmge: Will use bootstrap cutoff (50) before creating concatenated alignment
## Tue 26 Jul 2022 08:03:01 AM CEST - 'phylociraptor model' finished successfully before. Will run raxml with best models.
## Tue 26 Jul 2022 08:03:01 AM CEST - 'phylociraptor model' finished successfully before. Will run raxml with best models.
## Tue 26 Jul 2022 08:03:01 AM CEST - 'phylociraptor model' finished successfully before. Will run raxml with best models.
## Tue 26 Jul 2022 08:03:01 AM CEST - 'phylociraptor model' finished successfully before. Will run raxml with best models.
## Tue 26 Jul 2022 08:03:01 AM CEST - 'phylociraptor model' finished successfully before. Will run raxml with best models.
## Tue 26 Jul 2022 08:03:01 AM CEST - 'phylociraptor model' finished successfully before. Will run raxml with best models.
## Tue 26 Jul 2022 08:03:01 AM CEST - 'phylociraptor model' finished successfully before. Will run raxml with best models.
## Tue 26 Jul 2022 08:03:01 AM CEST - 'phylociraptor model' finished successfully before. Will run raxml with best models.
## Tue 26 Jul 2022 08:03:01 AM CEST - 'phylociraptor model' finished successfully before. Will run raxml with best models.
## Tue 26 Jul 2022 08:03:01 AM CEST - 'phylociraptor model' finished successfully before. Will run raxml with best models.
## Tue 26 Jul 2022 08:03:01 AM CEST - 'phylociraptor model' finished successfully before. Will run raxml with best models.
## Tue 26 Jul 2022 08:03:01 AM CEST - 'phylociraptor model' finished successfully before. Will run raxml with best models.
## Tue Jul 26 08:03:01 CEST 2022 - prepare_iqtree muscle-bmge: Will use bootstrap cutoff (50) before creating concatenated alignment
## Tue Jul 26 08:03:02 CEST 2022 - concatenate muscle-aliscore: Will use bootstrap cutoff 60 before creating concatenated alignment
## Tue Jul 26 08:03:02 CEST 2022 - prepare_iqtree muscle-trimal: Will use bootstrap cutoff (80) before creating concatenated alignment
## Tue Jul 26 08:03:02 CEST 2022 - prepare_iqtree muscle-bmge: Will use bootstrap cutoff (70) before creating concatenated alignment
## Tue Jul 26 08:03:02 CEST 2022 - prepare_iqtree clustalo-aliscore: Will use bootstrap cutoff (70) before creating concatenated alignment
## Tue Jul 26 08:03:02 CEST 2022 - concatenate clustalo-aliscore: Will use bootstrap cutoff 70 before creating concatenated alignment
## Tue Jul 26 08:03:02 CEST 2022 - prepare_iqtree clustalo-trimal: Will use bootstrap cutoff (60) before creating concatenated alignment
## Tue Jul 26 08:03:02 CEST 2022 - prepare_iqtree clustalo-bmge: Will use bootstrap cutoff (90) before creating concatenated alignment
## Tue Jul 26 08:03:02 CEST 2022 - prepare_iqtree clustalo-aliscore: Will use bootstrap cutoff (60) before creating concatenated alignment
## Will create NEXUS partition file with model information now.
## Tue Jul 26 08:03:02 CEST 2022 - prepare_iqtree clustalo-bmge: Will use bootstrap cutoff (80) before creating concatenated alignment
## Tue Jul 26 08:03:02 CEST 2022 - prepare_iqtree clustalo-bmge: Will use bootstrap cutoff (70) before creating concatenated alignment
## Tue Jul 26 08:03:02 CEST 2022 - concatenate muscle-trimal: Will use bootstrap cutoff 90 before creating concatenated alignment
## Tue Jul 26 08:03:02 CEST 2022 - prepare_iqtree muscle-bmge: Will use bootstrap cutoff (60) before creating concatenated alignment
## Tue Jul 26 08:03:02 CEST 2022 - prepare_iqtree mafft-bmge: Will use bootstrap cutoff (70) before creating concatenated alignment
## Tue Jul 26 08:03:02 CEST 2022 - concatenate mafft-bmge: Will use bootstrap cutoff 70 before creating concatenated alignment
## Tue Jul 26 08:03:02 CEST 2022 - concatenate muscle-trimal: Will use bootstrap cutoff 0 before creating concatenated alignment
## Tue Jul 26 08:03:02 CEST 2022 - prepare_iqtree clustalo-bmge: Will use bootstrap cutoff (60) before creating concatenated alignment
## Tue Jul 26 08:03:02 CEST 2022 - nexus file for iqtree written.
## Tue Jul 26 08:03:02 CEST 2022 - concatenate muscle-bmge: Will use bootstrap cutoff 70 before creating concatenated alignment
## Tue Jul 26 08:03:02 CEST 2022 - prepare_iqtree mafft-bmge: Will use bootstrap cutoff (80) before creating concatenated alignment
## Tue Jul 26 08:03:02 CEST 2022 - concatenate muscle-aliscore: Will use bootstrap cutoff 70 before creating concatenated alignment
## Tue Jul 26 08:03:02 CEST 2022 - prepare_iqtree mafft-trimal: Will use bootstrap cutoff (80) before creating concatenated alignment
## Tue Jul 26 08:03:02 CEST 2022 - prepare_iqtree mafft-trimal: Will use bootstrap cutoff (50) before creating concatenated alignment
## Tue Jul 26 08:03:02 CEST 2022 - prepare_iqtree muscle-trimal: Will use bootstrap cutoff (50) before creating concatenated alignment
## Tue Jul 26 08:03:03 CEST 2022 - concatenate clustalo-bmge: Will use bootstrap cutoff 90 before creating concatenated alignment
## Tue Jul 26 08:03:03 CEST 2022 - concatenate clustalo-bmge: Will use bootstrap cutoff 80 before creating concatenated alignment
## Tue Jul 26 08:03:03 CEST 2022 - prepare_iqtree mafft-aliscore: Will use bootstrap cutoff (0) before creating concatenated alignment
## Tue Jul 26 08:03:03 CEST 2022 - prepare_iqtree mafft-aliscore: Will use bootstrap cutoff (80) before creating concatenated alignment
## Tue Jul 26 08:03:03 CEST 2022 - prepare_iqtree mafft-aliscore: Will use bootstrap cutoff (70) before creating concatenated alignment
## Tue Jul 26 08:03:03 CEST 2022 - prepare_iqtree muscle-aliscore: Will use bootstrap cutoff (70) before creating concatenated alignment
## Tue Jul 26 08:03:03 CEST 2022 - prepare_iqtree clustalo-trimal: Will use bootstrap cutoff (70) before creating concatenated alignment
## Tue Jul 26 08:03:03 CEST 2022 - prepare_iqtree muscle-aliscore: Will use bootstrap cutoff (0) before creating concatenated alignment
## Tue Jul 26 08:03:03 CEST 2022 - concatenate mafft-trimal: Will use bootstrap cutoff 80 before creating concatenated alignment
## Tue Jul 26 08:03:03 CEST 2022 - concatenate clustalo-trimal: Will use bootstrap cutoff 70 before creating concatenated alignment
## Tue Jul 26 08:03:03 CEST 2022 - prepare_iqtree mafft-aliscore: Will use bootstrap cutoff (50) before creating concatenated alignment
## Tue Jul 26 08:03:03 CEST 2022 - concatenate muscle-trimal: Will use bootstrap cutoff 80 before creating concatenated alignment
## Tue Jul 26 08:03:03 CEST 2022 - concatenate muscle-bmge: Will use bootstrap cutoff 60 before creating concatenated alignment
## Tue Jul 26 08:03:03 CEST 2022 - concatenate clustalo-trimal: Will use bootstrap cutoff 60 before creating concatenated alignment
## Tue Jul 26 08:03:03 CEST 2022 - concatenate mafft-aliscore: Will use bootstrap cutoff 50 before creating concatenated alignment
## Tue Jul 26 08:03:03 CEST 2022 - prepare_iqtree mafft-trimal: Will use bootstrap cutoff (60) before creating concatenated alignment
## Tue Jul 26 08:03:03 CEST 2022 - concatenate clustalo-bmge: Will use bootstrap cutoff 50 before creating concatenated alignment
## Tue Jul 26 08:03:03 CEST 2022 - concatenate clustalo-aliscore: Will use bootstrap cutoff 60 before creating concatenated alignment
## Tue Jul 26 08:03:03 CEST 2022 - concatenate muscle-trimal: Will use bootstrap cutoff 70 before creating concatenated alignment
## Tue Jul 26 08:03:03 CEST 2022 - concatenate mafft-aliscore: Will use bootstrap cutoff 70 before creating concatenated alignment
## Tue Jul 26 08:03:03 CEST 2022 - prepare_iqtree mafft-trimal: Will use bootstrap cutoff (70) before creating concatenated alignment
## Tue Jul 26 08:03:03 CEST 2022 - concatenate mafft-trimal: Will use bootstrap cutoff 70 before creating concatenated alignment
## Tue Jul 26 08:03:03 CEST 2022 - concatenate mafft-bmge: Will use bootstrap cutoff 60 before creating concatenated alignment
## Tue Jul 26 08:03:03 CEST 2022 - concatenate mafft-aliscore: Will use bootstrap cutoff 0 before creating concatenated alignment
## Will create NEXUS partition file with model information now.
## Will create NEXUS partition file with model information now.
## Will create NEXUS partition file with model information now.
## Tue Jul 26 08:03:04 CEST 2022 - prepare_iqtree mafft-trimal: Will use bootstrap cutoff (0) before creating concatenated alignment
## Tue Jul 26 08:03:04 CEST 2022 - concatenate mafft-trimal: Will use bootstrap cutoff 90 before creating concatenated alignment
## Tue Jul 26 08:03:04 CEST 2022 - prepare_iqtree mafft-trimal: Will use bootstrap cutoff (90) before creating concatenated alignment
## Tue Jul 26 08:03:04 CEST 2022 - concatenate clustalo-bmge: Will use bootstrap cutoff 60 before creating concatenated alignment
## Tue Jul 26 08:03:04 CEST 2022 - concatenate clustalo-bmge: Will use bootstrap cutoff 0 before creating concatenated alignment
## Will create NEXUS partition file with model information now.
## Tue Jul 26 08:03:05 CEST 2022 - prepare_iqtree muscle-trimal: Will use bootstrap cutoff (0) before creating concatenated alignment
## Tue Jul 26 08:03:05 CEST 2022 - prepare_iqtree muscle-trimal: Will use bootstrap cutoff (60) before creating concatenated alignment
## Tue Jul 26 08:03:05 CEST 2022 - prepare_iqtree mafft-aliscore: Will use bootstrap cutoff (90) before creating concatenated alignment
## Tue Jul 26 08:03:05 CEST 2022 - concatenate muscle-trimal: Will use bootstrap cutoff 50 before creating concatenated alignment
## Tue Jul 26 08:03:05 CEST 2022 - concatenate mafft-trimal: Will use bootstrap cutoff 0 before creating concatenated alignment
## Tue Jul 26 08:03:05 CEST 2022 - concatenate mafft-bmge: Will use bootstrap cutoff 50 before creating concatenated alignment
## Tue Jul 26 08:03:05 CEST 2022 - prepare_iqtree muscle-aliscore: Will use bootstrap cutoff (50) before creating concatenated alignment
## Tue Jul 26 08:03:05 CEST 2022 - concatenate mafft-trimal: Will use bootstrap cutoff 60 before creating concatenated alignment
## Will create NEXUS partition file with model information now.
## Will create NEXUS partition file with model information now.
## Tue Jul 26 08:03:05 CEST 2022 - prepare_iqtree muscle-aliscore: Will use bootstrap cutoff (60) before creating concatenated alignment
## Will create NEXUS partition file with model information now.
## Tue Jul 26 08:03:05 CEST 2022 - prepare_iqtree mafft-aliscore: Will use bootstrap cutoff (60) before creating concatenated alignment
## Tue Jul 26 08:03:05 CEST 2022 - concatenate mafft-aliscore: Will use bootstrap cutoff 60 before creating concatenated alignment
## Will create NEXUS partition file with model information now.
## Will create NEXUS partition file with model information now.
## ll use bootstrap cutoff (90) before creating concatenated alignment
## Tue Jul 26 08:03:05 CEST 2022 - concatenate mafft-bmge: Will use bootstrap cutoff 0 before creating concatenated alignment
## Tue Jul 26 08:03:05 CEST 2022 - concatenate mafft-trimal: Will use bootstrap cutoff 50 before creating concatenated alignment
## Will create NEXUS partition file with model information now.
## Tue Jul 26 08:03:05 CEST 2022 - prepare_iqtree muscle-trimal: Will use bootstrap cutoff (70) before creating concatenated alignment
## Tue Jul 26 08:03:05 CEST 2022 - concatenate muscle-trimal: Will use bootstrap cutoff 60 before creating concatenated alignment
## Will create NEXUS partition file with model information now.
## Tue Jul 26 08:03:05 CEST 2022 - concatenate clustalo-bmge: Will use bootstrap cutoff 70 before creating concatenated alignment
## Will create NEXUS partition file with model information now.
## Will create NEXUS partition file with model information now.
## Will create NEXUS partition file with model information now.
## Will create NEXUS partition file with model information now.
## Will create NEXUS partition file with model information now.
## Tue Jul 26 08:03:05 CEST 2022 - nexus file for iqtree written.
## Will create NEXUS partition file with model information now.
## Will create NEXUS partition file with model information now.
## Tue Jul 26 08:03:06 CEST 2022 - nexus file for iqtree written.
## Tue Jul 26 08:03:06 CEST 2022 - nexus file for iqtree written.
## Will create NEXUS partition file with model information now.
## Will create NEXUS partition file with model information now.
## Will create NEXUS partition file with model information now.
## Tue Jul 26 08:03:06 CEST 2022 - nexus file for iqtree written.
## Tue Jul 26 08:03:06 CEST 2022 - nexus file for iqtree written.
## Tue Jul 26 08:03:06 CEST 2022 - nexus file for iqtree written.
## Tue Jul 26 08:03:06 CEST 2022 - nexus file for iqtree written.
## Will create NEXUS partition file with model information now.
## Tue 26 Jul 2022 08:03:07 AM CEST - 'phylociraptor model' finished successfully before. Will run raxml with best models.
## Will create NEXUS partition file with model information now.
## Will create NEXUS partition file with model information now.
## Will create NEXUS partition file with model information now.
## Will create NEXUS partition file with model information now.
## Will create NEXUS partition file with model information now.
## Will create NEXUS partition file with model information now.
## Tue Jul 26 08:03:08 CEST 2022 - nexus file for iqtree written.
## Tue Jul 26 08:03:08 CEST 2022 - nexus file for iqtree written.
## Tue Jul 26 08:03:08 CEST 2022 - nexus file for iqtree written.
## Will create NEXUS partition file with model information now.
## Will create NEXUS partition file with model information now.
## Tue Jul 26 08:03:08 CEST 2022 - nexus file for iqtree written.
## Will create NEXUS partition file with model information now.
## Tue Jul 26 08:03:08 CEST 2022 - nexus file for iqtree written.
## Tue Jul 26 08:03:08 CEST 2022 - nexus file for iqtree written.
## Tue Jul 26 08:03:09 CEST 2022 - nexus file for iqtree written.
## Tue Jul 26 08:03:09 CEST 2022 - nexus file for iqtree written.
## Tue Jul 26 08:03:09 CEST 2022 - nexus file for iqtree written.
## Tue Jul 26 08:03:09 CEST 2022 - nexus file for iqtree written.
## Tue Jul 26 08:03:09 CEST 2022 - nexus file for iqtree written.
## Tue Jul 26 08:03:09 CEST 2022 - nexus file for iqtree written.
## Tue Jul 26 08:03:09 CEST 2022 - nexus file for iqtree written.
## Tue Jul 26 08:03:09 CEST 2022 - nexus file for iqtree written.
## Tue Jul 26 08:03:09 CEST 2022 - nexus file for iqtree written.
## Tue Jul 26 08:03:10 CEST 2022 - nexus file for iqtree written.
## Will create NEXUS partition file with model information now.
## Will create NEXUS partition file with model information now.
## Will create NEXUS partition file with model information now.
## Will create NEXUS partition file with model information now.
## Will create NEXUS partition file with model information now.
## Will create NEXUS partition file with model information now.
## Will create NEXUS partition file with model information now.
## Tue Jul 26 08:03:11 CEST 2022 - nexus file for iqtree written.
## Tue Jul 26 08:03:11 CEST 2022 - nexus file for iqtree written.
## Tue Jul 26 08:03:11 CEST 2022 - nexus file for iqtree written.
## Tue Jul 26 08:03:11 CEST 2022 - nexus file for iqtree written.
## Tue Jul 26 08:03:12 CEST 2022 - nexus file for iqtree written.
## Tue Jul 26 08:03:12 CEST 2022 - nexus file for iqtree written.
## Tue Jul 26 08:03:12 CEST 2022 - nexus file for iqtree written.
## Tue Jul 26 08:03:12 CEST 2022 - nexus file for iqtree written.
## Tue Jul 26 08:03:12 CEST 2022 - nexus file for iqtree written.
## Tue Jul 26 08:03:13 CEST 2022 - nexus file for iqtree written.
## Tue Jul 26 08:03:13 CEST 2022 - nexus file for iqtree written.
## Tue Jul 26 08:03:13 CEST 2022 - nexus file for iqtree written.
## Tue Jul 26 08:03:13 CEST 2022 - nexus file for iqtree written.
## Tue Jul 26 08:03:13 CEST 2022 - nexus file for iqtree written.
## Tue Jul 26 08:03:13 CEST 2022 - nexus file for iqtree written.
## Tue Jul 26 08:03:13 CEST 2022 - nexus file for iqtree written.
## Tue 26 Jul 2022 08:03:30 AM CEST - 'phylociraptor model' finished successfully before. Will run raxml with best models.
## Tue 26 Jul 2022 08:03:30 AM CEST - 'phylociraptor model' finished successfully before. Will run raxml with best models.
## Tue 26 Jul 2022 08:03:30 AM CEST - 'phylociraptor model' finished successfully before. Will run raxml with best models.
## Tue 26 Jul 2022 08:03:31 AM CEST - 'phylociraptor model' finished successfully before. Will run raxml with best models.
## Tue 26 Jul 2022 08:03:31 AM CEST - 'phylociraptor model' finished successfully before. Will run raxml with best models.
## Tue 26 Jul 2022 08:03:31 AM CEST - 'phylociraptor model' finished successfully before. Will run raxml with best models.
## Tue 26 Jul 2022 08:03:31 AM CEST - 'phylociraptor model' finished successfully before. Will run raxml with best models.
## Tue 26 Jul 2022 08:03:31 AM CEST - 'phylociraptor model' finished successfully before. Will run raxml with best models.
## Tue 26 Jul 2022 08:03:31 AM CEST - 'phylociraptor model' finished successfully before. Will run raxml with best models.
## Tue 26 Jul 2022 08:03:31 AM CEST - 'phylociraptor model' finished successfully before. Will run raxml with best models.
## Tue 26 Jul 2022 08:03:31 AM CEST - 'phylociraptor model' finished successfully before. Will run raxml with best models.
## Tue 26 Jul 2022 08:03:31 AM CEST - 'phylociraptor model' finished successfully before. Will run raxml with best models.
## Tue 26 Jul 2022 08:03:31 AM CEST - 'phylociraptor model' finished successfully before. Will run raxml with best models.
## Tue 26 Jul 2022 08:03:31 AM CEST - 'phylociraptor model' finished successfully before. Will run raxml with best models.
## Tue 26 Jul 2022 08:03:31 AM CEST - 'phylociraptor model' finished successfully before. Will run raxml with best models.
## Tue 26 Jul 2022 08:03:31 AM CEST - 'phylociraptor model' finished successfully before. Will run raxml with best models.
## Tue 26 Jul 2022 08:03:31 AM CEST - 'phylociraptor model' finished successfully before. Will run raxml with best models.
## Tue 26 Jul 2022 08:03:31 AM CEST - 'phylociraptor model' finished successfully before. Will run raxml with best models.
## Tue 26 Jul 2022 08:03:31 AM CEST - 'phylociraptor model' finished successfully before. Will run raxml with best models.
## Tue 26 Jul 2022 08:03:31 AM CEST - 'phylociraptor model' finished successfully before. Will run raxml with best models.
## Tue 26 Jul 2022 08:04:01 AM CEST - 'phylociraptor model' finished successfully before. Will run raxml with best models.
## Tue 26 Jul 2022 08:04:01 AM CEST - 'phylociraptor model' finished successfully before. Will run raxml with best models.
## Tue 26 Jul 2022 08:04:01 AM CEST - 'phylociraptor model' finished successfully before. Will run raxml with best models.
## Tue 26 Jul 2022 08:04:01 AM CEST - 'phylociraptor model' finished successfully before. Will run raxml with best models.
## Tue 26 Jul 2022 08:04:01 AM CEST - 'phylociraptor model' finished successfully before. Will run raxml with best models.
## Tue 26 Jul 2022 08:04:01 AM CEST - 'phylociraptor model' finished successfully before. Will run raxml with best models.
## Tue 26 Jul 2022 08:04:01 AM CEST - 'phylociraptor model' finished successfully before. Will run raxml with best models.
## Tue 26 Jul 2022 08:04:01 AM CEST - 'phylociraptor model' finished successfully before. Will run raxml with best models.
## Tue 26 Jul 2022 08:04:01 AM CEST - 'phylociraptor model' finished successfully before. Will run raxml with best models.
## Tue 26 Jul 2022 08:04:01 AM CEST - 'phylociraptor model' finished successfully before. Will run raxml with best models.
## Tue 26 Jul 2022 08:04:01 AM CEST - 'phylociraptor model' finished successfully before. Will run raxml with best models.
## Tue 26 Jul 2022 08:04:01 AM CEST - 'phylociraptor model' finished successfully before. Will run raxml with best models.
## Tue 26 Jul 2022 08:04:01 AM CEST - 'phylociraptor model' finished successfully before. Will run raxml with best models.
## Tue 26 Jul 2022 08:04:01 AM CEST - 'phylociraptor model' finished successfully before. Will run raxml with best models.
## Tue 26 Jul 2022 08:04:01 AM CEST - 'phylociraptor model' finished successfully before. Will run raxml with best models.
## Tue 26 Jul 2022 08:04:01 AM CEST - 'phylociraptor model' finished successfully before. Will run raxml with best models.
## Tue 26 Jul 2022 08:04:01 AM CEST - 'phylociraptor model' finished successfully before. Will run raxml with best models.
## Tue Jul 26 08:53:35 CEST 2022 - concatenate clustalo-trimal: Will use bootstrap cutoff 80 before creating concatenated alignment
## Tue Jul 26 08:53:35 CEST 2022 - concatenate mafft-trimal: Will use bootstrap cutoff 80 before creating concatenated alignment
## Tue Jul 26 08:53:36 CEST 2022 - concatenate mafft-bmge: Will use bootstrap cutoff 80 before creating concatenated alignment
## Tue Jul 26 08:53:36 CEST 2022 - prepare_iqtree mafft-aliscore: Will use bootstrap cutoff (90) before creating concatenated alignment
## Tue Jul 26 08:53:36 CEST 2022 - prepare_iqtree clustalo-trimal: Will use bootstrap cutoff (80) before creating concatenated alignment
## Tue Jul 26 08:53:36 CEST 2022 - prepare_iqtree clustalo-aliscore: Will use bootstrap cutoff (80) before creating concatenated alignment
## Tue Jul 26 08:53:36 CEST 2022 - prepare_iqtree muscle-aliscore: Will use bootstrap cutoff (80) before creating concatenated alignment
## Tue Jul 26 08:53:36 CEST 2022 - prepare_iqtree mafft-aliscore: Will use bootstrap cutoff (80) before creating concatenated alignment
## Tue Jul 26 08:53:36 CEST 2022 - prepare_iqtree clustalo-bmge: Will use bootstrap cutoff (80) before creating concatenated alignment
## Tue Jul 26 08:53:36 CEST 2022 - prepare_iqtree mafft-trimal: Will use bootstrap cutoff (80) before creating concatenated alignment
## Tue Jul 26 08:53:36 CEST 2022 - prepare_iqtree clustalo-bmge: Will use bootstrap cutoff (90) before creating concatenated alignment
## Tue Jul 26 08:53:36 CEST 2022 - prepare_iqtree muscle-bmge: Will use bootstrap cutoff (80) before creating concatenated alignment
## Tue Jul 26 08:53:36 CEST 2022 - prepare_iqtree muscle-bmge: Will use bootstrap cutoff (90) before creating concatenated alignment
## Tue Jul 26 08:53:36 CEST 2022 - prepare_iqtree mafft-bmge: Will use bootstrap cutoff (80) before creating concatenated alignment
## Tue Jul 26 08:53:36 CEST 2022 - concatenate clustalo-bmge: Will use bootstrap cutoff 80 before creating concatenated alignment
## Will create NEXUS partition file with model information now.
## Tue Jul 26 08:53:36 CEST 2022 - prepare_iqtree muscle-trimal: Will use bootstrap cutoff (90) before creating concatenated alignment
## Tue Jul 26 08:53:36 CEST 2022 - prepare_iqtree muscle-trimal: Will use bootstrap cutoff (80) before creating concatenated alignment
## Will create NEXUS partition file with model information now.
## Tue Jul 26 08:53:36 CEST 2022 - concatenate mafft-bmge: Will use bootstrap cutoff 0 before creating concatenated alignment
## Will create NEXUS partition file with model information now.
## Tue Jul 26 08:53:36 CEST 2022 - prepare_iqtree muscle-bmge: Will use bootstrap cutoff (70) before creating concatenated alignment
## Tue Jul 26 08:53:36 CEST 2022 - concatenate muscle-trimal: Will use bootstrap cutoff 0 before creating concatenated alignment
## Tue Jul 26 08:53:36 CEST 2022 - concatenate mafft-aliscore: Will use bootstrap cutoff 0 before creating concatenated alignment
## Will create NEXUS partition file with model information now.
## Tue Jul 26 08:53:36 CEST 2022 - concatenate mafft-trimal: Will use bootstrap cutoff 90 before creating concatenated alignment
## Tue Jul 26 08:53:36 CEST 2022 - prepare_iqtree mafft-trimal: Will use bootstrap cutoff (0) before creating concatenated alignment
## Tue Jul 26 08:53:36 CEST 2022 - concatenate mafft-bmge: Will use bootstrap cutoff 90 before creating concatenated alignment
## Tue Jul 26 08:53:36 CEST 2022 - concatenate clustalo-aliscore: Will use bootstrap cutoff 80 before creating concatenated alignment
## Tue Jul 26 08:53:36 CEST 2022 - nexus file for iqtree written.
## Tue Jul 26 08:53:36 CEST 2022 - nexus file for iqtree written.
## Tue Jul 26 08:53:37 CEST 2022 - nexus file for iqtree written.
## Tue Jul 26 08:53:37 CEST 2022 - prepare_iqtree mafft-trimal: Will use bootstrap cutoff (50) before creating concatenated alignment
## Tue Jul 26 08:53:37 CEST 2022 - prepare_iqtree clustalo-bmge: Will use bootstrap cutoff (0) before creating concatenated alignment
## Tue Jul 26 08:53:37 CEST 2022 - concatenate clustalo-trimal: Will use bootstrap cutoff 60 before creating concatenated alignment
## Tue Jul 26 08:53:37 CEST 2022 - concatenate mafft-bmge: Will use bootstrap cutoff 50 before creating concatenated alignment
## Tue Jul 26 08:53:37 CEST 2022 - prepare_iqtree muscle-bmge: Will use bootstrap cutoff (50) before creating concatenated alignment
## Tue Jul 26 08:53:37 CEST 2022 - concatenate mafft-bmge: Will use bootstrap cutoff 60 before creating concatenated alignment
## Tue Jul 26 08:53:37 CEST 2022 - concatenate muscle-aliscore: Will use bootstrap cutoff 60 before creating concatenated alignment
## Tue Jul 26 08:53:37 CEST 2022 - concatenate clustalo-trimal: Will use bootstrap cutoff 0 before creating concatenated alignment
## Tue Jul 26 08:53:37 CEST 2022 - concatenate mafft-aliscore: Will use bootstrap cutoff 80 before creating concatenated alignment
## Tue Jul 26 08:53:37 CEST 2022 - concatenate muscle-bmge: Will use bootstrap cutoff 50 before creating concatenated alignment
## Tue Jul 26 08:53:37 CEST 2022 - prepare_iqtree clustalo-aliscore: Will use bootstrap cutoff (90) before creating concatenated alignment
## Tue Jul 26 08:53:37 CEST 2022 - nexus file for iqtree written.
## Tue Jul 26 08:53:37 CEST 2022 - concatenate mafft-trimal: Will use bootstrap cutoff 50 before creating concatenated alignment
## Tue Jul 26 08:53:37 CEST 2022 - prepare_iqtree clustalo-trimal: Will use bootstrap cutoff (0) before creating concatenated alignment
## Tue Jul 26 08:53:37 CEST 2022 - prepare_iqtree muscle-aliscore: Will use bootstrap cutoff (50) before creating concatenated alignment
## Tue Jul 26 08:53:37 CEST 2022 - prepare_iqtree clustalo-trimal: Will use bootstrap cutoff (90) before creating concatenated alignment
## Tue Jul 26 08:53:37 CEST 2022 - concatenate mafft-trimal: Will use bootstrap cutoff 0 before creating concatenated alignment
## Tue Jul 26 08:53:37 CEST 2022 - prepare_iqtree mafft-trimal: Will use bootstrap cutoff (90) before creating concatenated alignment
## Tue Jul 26 08:53:37 CEST 2022 - concatenate clustalo-bmge: Will use bootstrap cutoff 50 before creating concatenated alignment
## Tue Jul 26 08:53:37 CEST 2022 - prepare_iqtree clustalo-bmge: Will use bootstrap cutoff (50) before creating concatenated alignment
## Tue Jul 26 08:53:37 CEST 2022 - prepare_iqtree muscle-aliscore: Will use bootstrap cutoff (90) before creating concatenated alignment
## Will create NEXUS partition file with model information now.
## Tue Jul 26 08:53:37 CEST 2022 - prepare_iqtree mafft-bmge: Will use bootstrap cutoff (60) before creating concatenated alignment
## Tue Jul 26 08:53:37 CEST 2022 - prepare_iqtree clustalo-aliscore: Will use bootstrap cutoff (60) before creating concatenated alignment
## Tue Jul 26 08:53:37 CEST 2022 - prepare_iqtree mafft-aliscore: Will use bootstrap cutoff (0) before creating concatenated alignment
## Will create NEXUS partition file with model information now.
## Will create NEXUS partition file with model information now.
## Tue Jul 26 08:53:37 CEST 2022 - concatenate mafft-trimal: Will use bootstrap cutoff 60 before creating concatenated alignment
## Will create NEXUS partition file with model information now.
## Tue Jul 26 08:53:37 CEST 2022 - prepare_iqtree muscle-aliscore: Will use bootstrap cutoff (0) before creating concatenated alignment
## Tue Jul 26 08:53:37 CEST 2022 - concatenate clustalo-aliscore: Will use bootstrap cutoff 60 before creating concatenated alignment
## Tue Jul 26 08:53:37 CEST 2022 - prepare_iqtree mafft-bmge: Will use bootstrap cutoff (0) before creating concatenated alignment
## Tue Jul 26 08:53:37 CEST 2022 - prepare_iqtree clustalo-trimal: Will use bootstrap cutoff (50) before creating concatenated alignment
## Tue Jul 26 08:53:37 CEST 2022 - prepare_iqtree mafft-bmge: Will use bootstrap cutoff (50) before creating concatenated alignment
## Tue Jul 26 08:53:37 CEST 2022 - prepare_iqtree muscle-trimal: Will use bootstrap cutoff (50) before creating concatenated alignment
## Tue Jul 26 08:53:37 CEST 2022 - prepare_iqtree mafft-aliscore: Will use bootstrap cutoff (50) before creating concatenated alignment
## Tue Jul 26 08:53:37 CEST 2022 - prepare_iqtree muscle-trimal: Will use bootstrap cutoff (0) before creating concatenated alignment
## Tue Jul 26 08:53:37 CEST 2022 - prepare_iqtree muscle-bmge: Will use bootstrap cutoff (0) before creating concatenated alignment
## Tue Jul 26 08:53:38 CEST 2022 - concatenate muscle-trimal: Will use bootstrap cutoff 50 before creating concatenated alignment
## Tue Jul 26 08:53:38 CEST 2022 - concatenate mafft-aliscore: Will use bootstrap cutoff 60 before creating concatenated alignment
## Tue Jul 26 08:53:38 CEST 2022 - concatenate clustalo-bmge: Will use bootstrap cutoff 0 before creating concatenated alignment
## Tue Jul 26 08:53:38 CEST 2022 - nexus file for iqtree written.
## Tue Jul 26 08:53:38 CEST 2022 - concatenate mafft-aliscore: Will use bootstrap cutoff 50 before creating concatenated alignment
## Tue Jul 26 08:53:38 CEST 2022 - concatenate clustalo-bmge: Will use bootstrap cutoff 90 before creating concatenated alignment
## Tue Jul 26 08:53:38 CEST 2022 - prepare_iqtree clustalo-aliscore: Will use bootstrap cutoff (70) before creating concatenated alignment
## Tue Jul 26 08:53:38 CEST 2022 - concatenate muscle-trimal: Will use bootstrap cutoff 80 before creating concatenated alignment
## Tue Jul 26 08:53:38 CEST 2022 - concatenate clustalo-aliscore: Will use bootstrap cutoff 90 before creating concatenated alignment
## Tue Jul 26 08:53:38 CEST 2022 - concatenate muscle-bmge: Will use bootstrap cutoff 0 before creating concatenated alignment
## Tue Jul 26 08:53:38 CEST 2022 - prepare_iqtree mafft-bmge: Will use bootstrap cutoff (90) before creating concatenated alignment
## Tue Jul 26 08:53:38 CEST 2022 - prepare_iqtree mafft-bmge: Will use bootstrap cutoff (70) before creating concatenated alignment
## Tue Jul 26 08:53:38 CEST 2022 - prepare_iqtree muscle-aliscore: Will use bootstrap cutoff (70) before creating concatenated alignment
## Tue Jul 26 08:53:38 CEST 2022 - nexus file for iqtree written.
## Will create NEXUS partition file with model information now.
## Tue Jul 26 08:53:38 CEST 2022 - prepare_iqtree muscle-bmge: Will use bootstrap cutoff (60) before creating concatenated alignment
## Tue Jul 26 08:53:38 CEST 2022 - nexus file for iqtree written.
## Tue Jul 26 08:53:38 CEST 2022 - concatenate muscle-bmge: Will use bootstrap cutoff 80 before creating concatenated alignment
## Tue Jul 26 08:53:38 CEST 2022 - prepare_iqtree muscle-aliscore: Will use bootstrap cutoff (60) before creating concatenated alignment
## Will create NEXUS partition file with model information now.
## Tue Jul 26 08:53:38 CEST 2022 - concatenate clustalo-aliscore: Will use bootstrap cutoff 0 before creating concatenated alignment
## Tue Jul 26 08:53:38 CEST 2022 - concatenate muscle-bmge: Will use bootstrap cutoff 90 before creating concatenated alignment
## Tue Jul 26 08:53:38 CEST 2022 - concatenate mafft-aliscore: Will use bootstrap cutoff 90 before creating concatenated alignment
## Tue Jul 26 08:53:38 CEST 2022 - nexus file for iqtree written.
## Tue Jul 26 08:53:38 CEST 2022 - concatenate muscle-trimal: Will use bootstrap cutoff 90 before creating concatenated alignment
## Tue Jul 26 08:53:38 CEST 2022 - concatenate muscle-aliscore: Will use bootstrap cutoff 0 before creating concatenated alignment
## Tue Jul 26 08:53:38 CEST 2022 - concatenate muscle-aliscore: Will use bootstrap cutoff 90 before creating concatenated alignment
## Tue Jul 26 08:53:38 CEST 2022 - concatenate muscle-aliscore: Will use bootstrap cutoff 50 before creating concatenated alignment
## Tue Jul 26 08:53:38 CEST 2022 - concatenate muscle-aliscore: Will use bootstrap cutoff 80 before creating concatenated alignment
## Tue Jul 26 08:53:38 CEST 2022 - concatenate clustalo-aliscore: Will use bootstrap cutoff 50 before creating concatenated alignment
## Tue Jul 26 08:53:38 CEST 2022 - nexus file for iqtree written.
## Will create NEXUS partition file with model information now.
## Tue Jul 26 08:53:38 CEST 2022 - concatenate clustalo-trimal: Will use bootstrap cutoff 50 before creating concatenated alignment
## Tue Jul 26 08:53:38 CEST 2022 - concatenate clustalo-aliscore: Will use bootstrap cutoff 70 before creating concatenated alignment
## Will create NEXUS partition file with model information now.
## Tue Jul 26 08:53:38 CEST 2022 - concatenate clustalo-trimal: Will use bootstrap cutoff 70 before creating concatenated alignment
## Tue Jul 26 08:53:38 CEST 2022 - concatenate muscle-aliscore: Will use bootstrap cutoff 70 before creating concatenated alignment
## Will create NEXUS partition file with model information now.
## Will create NEXUS partition file with model information now.
## Will create NEXUS partition file with model information now.
## Tue Jul 26 08:53:40 CEST 2022 - nexus file for iqtree written.
## Tue Jul 26 08:53:41 CEST 2022 - nexus file for iqtree written.
## Will create NEXUS partition file with model information now.
## Will create NEXUS partition file with model information now.
## Will create NEXUS partition file with model information now.
## Tue Jul 26 08:53:41 CEST 2022 - nexus file for iqtree written.
## Will create NEXUS partition file with model information now.
## Will create NEXUS partition file with model information now.
## Will create NEXUS partition file with model information now.
## Will create NEXUS partition file with model information now.
## Will create NEXUS partition file with model information now.
## Tue Jul 26 08:53:41 CEST 2022 - nexus file for iqtree written.
## Will create NEXUS partition file with model information now.
## Will create NEXUS partition file with model information now.
## Will create NEXUS partition file with model information now.
## Tue Jul 26 08:53:41 CEST 2022 - concatenate clustalo-trimal: Will use bootstrap cutoff 90 before creating concatenated alignment
## Tue Jul 26 08:53:42 CEST 2022 - prepare_iqtree clustalo-aliscore: Will use bootstrap cutoff (0) before creating concatenated alignment
## Tue Jul 26 08:53:42 CEST 2022 - prepare_iqtree clustalo-aliscore: Will use bootstrap cutoff (50) before creating concatenated alignment
## Will create NEXUS partition file with model information now.
## Tue Jul 26 08:53:42 CEST 2022 - nexus file for iqtree written.
## Will create NEXUS partition file with model information now.
## Will create NEXUS partition file with model information now.
## Will create NEXUS partition file with model information now.
## Will create NEXUS partition file with model information now.
## Will create NEXUS partition file with model information now.
## Will create NEXUS partition file with model information now.
## Will create NEXUS partition file with model information now.
## Will create NEXUS partition file with model information now.
## Will create NEXUS partition file with model information now.
## Will create NEXUS partition file with model information now.
## Will create NEXUS partition file with model information now.
## Will create NEXUS partition file with model information now.
## Will create NEXUS partition file with model information now.
## Will create NEXUS partition file with model information now.
## Tue Jul 26 08:53:43 CEST 2022 - nexus file for iqtree written.
## Tue Jul 26 08:53:43 CEST 2022 - nexus file for iqtree written.
## Tue Jul 26 08:53:43 CEST 2022 - nexus file for iqtree written.
## Tue Jul 26 08:53:43 CEST 2022 - nexus file for iqtree written.
## Tue Jul 26 08:53:44 CEST 2022 - nexus file for iqtree written.
## Tue Jul 26 08:53:45 CEST 2022 - nexus file for iqtree written.
## Tue Jul 26 08:53:45 CEST 2022 - nexus file for iqtree written.
## Will create NEXUS partition file with model information now.
## Tue Jul 26 08:53:45 CEST 2022 - nexus file for iqtree written.
## Tue Jul 26 08:53:45 CEST 2022 - nexus file for iqtree written.
## Will create NEXUS partition file with model information now.
## Tue Jul 26 08:53:45 CEST 2022 - nexus file for iqtree written.
## Tue Jul 26 08:53:45 CEST 2022 - nexus file for iqtree written.
## Tue Jul 26 08:53:45 CEST 2022 - nexus file for iqtree written.
## Tue Jul 26 08:53:45 CEST 2022 - nexus file for iqtree written.
## Tue Jul 26 08:53:46 CEST 2022 - nexus file for iqtree written.
## Tue Jul 26 08:53:46 CEST 2022 - nexus file for iqtree written.
## Tue Jul 26 08:53:46 CEST 2022 - nexus file for iqtree written.
## Tue Jul 26 08:53:46 CEST 2022 - nexus file for iqtree written.
## Tue Jul 26 08:53:46 CEST 2022 - nexus file for iqtree written.
## Tue Jul 26 08:53:46 CEST 2022 - nexus file for iqtree written.
## Tue Jul 26 08:53:46 CEST 2022 - nexus file for iqtree written.
## Tue Jul 26 08:53:46 CEST 2022 - nexus file for iqtree written.
## Tue Jul 26 08:53:46 CEST 2022 - nexus file for iqtree written.
## Tue Jul 26 08:53:46 CEST 2022 - nexus file for iqtree written.
## Tue Jul 26 08:53:46 CEST 2022 - nexus file for iqtree written.
## Tue Jul 26 08:53:46 CEST 2022 - nexus file for iqtree written.
## Tue Jul 26 08:53:46 CEST 2022 - nexus file for iqtree written.
## Tue Jul 26 08:53:46 CEST 2022 - nexus file for iqtree written.
## Tue Jul 26 08:53:46 CEST 2022 - nexus file for iqtree written.
## Tue Jul 26 08:53:48 CEST 2022 - nexus file for iqtree written.
## Tue Jul 26 08:53:48 CEST 2022 - nexus file for iqtree written.
## Tue 26 Jul 2022 08:54:04 AM CEST - 'phylociraptor model' finished successfully before. Will run raxml with best models.
## Tue 26 Jul 2022 08:54:04 AM CEST - 'phylociraptor model' finished successfully before. Will run raxml with best models.
## Tue 26 Jul 2022 08:54:04 AM CEST - 'phylociraptor model' finished successfully before. Will run raxml with best models.
## Tue 26 Jul 2022 08:54:04 AM CEST - 'phylociraptor model' finished successfully before. Will run raxml with best models.
## Tue 26 Jul 2022 08:54:04 AM CEST - 'phylociraptor model' finished successfully before. Will run raxml with best models.
## Tue 26 Jul 2022 08:54:04 AM CEST - 'phylociraptor model' finished successfully before. Will run raxml with best models.
## Tue 26 Jul 2022 08:54:04 AM CEST - 'phylociraptor model' finished successfully before. Will run raxml with best models.
## Tue 26 Jul 2022 08:54:04 AM CEST - 'phylociraptor model' finished successfully before. Will run raxml with best models.
## Tue 26 Jul 2022 08:54:04 AM CEST - 'phylociraptor model' finished successfully before. Will run raxml with best models.
## Tue 26 Jul 2022 08:54:04 AM CEST - 'phylociraptor model' finished successfully before. Will run raxml with best models.
## Tue 26 Jul 2022 08:54:04 AM CEST - 'phylociraptor model' finished successfully before. Will run raxml with best models.
## Tue 26 Jul 2022 08:54:04 AM CEST - 'phylociraptor model' finished successfully before. Will run raxml with best models.
## Tue Jul 26 08:54:04 CEST 2022 - prepare_iqtree mafft-trimal: Will use bootstrap cutoff (60) before creating concatenated alignment
## Tue 26 Jul 2022 08:54:04 AM CEST - 'phylociraptor model' finished successfully before. Will run raxml with best models.
## Tue 26 Jul 2022 08:54:04 AM CEST - 'phylociraptor model' finished successfully before. Will run raxml with best models.
## Tue 26 Jul 2022 08:54:04 AM CEST - 'phylociraptor model' finished successfully before. Will run raxml with best models.
## Tue 26 Jul 2022 08:54:04 AM CEST - 'phylociraptor model' finished successfully before. Will run raxml with best models.
## Tue 26 Jul 2022 08:54:04 AM CEST - 'phylociraptor model' finished successfully before. Will run raxml with best models.
## Tue 26 Jul 2022 08:54:04 AM CEST - 'phylociraptor model' finished successfully before. Will run raxml with best models.
## Tue 26 Jul 2022 08:54:04 AM CEST - 'phylociraptor model' finished successfully before. Will run raxml with best models.
## Tue 26 Jul 2022 08:54:04 AM CEST - 'phylociraptor model' finished successfully before. Will run raxml with best models.
## Tue 26 Jul 2022 08:54:04 AM CEST - 'phylociraptor model' finished successfully before. Will run raxml with best models.
## Tue 26 Jul 2022 08:54:04 AM CEST - 'phylociraptor model' finished successfully before. Will run raxml with best models.
## Tue Jul 26 08:54:05 CEST 2022 - concatenate muscle-bmge: Will use bootstrap cutoff 60 before creating concatenated alignment
## Tue Jul 26 08:54:05 CEST 2022 - concatenate muscle-trimal: Will use bootstrap cutoff 60 before creating concatenated alignment
## Tue Jul 26 08:54:05 CEST 2022 - concatenate clustalo-bmge: Will use bootstrap cutoff 60 before creating concatenated alignment
## Tue Jul 26 08:54:05 CEST 2022 - concatenate clustalo-bmge: Will use bootstrap cutoff 70 before creating concatenated alignment
## Tue Jul 26 08:54:05 CEST 2022 - prepare_iqtree clustalo-bmge: Will use bootstrap cutoff (60) before creating concatenated alignment
## Tue Jul 26 08:54:05 CEST 2022 - prepare_iqtree muscle-trimal: Will use bootstrap cutoff (60) before creating concatenated alignment
## Tue Jul 26 08:54:05 CEST 2022 - prepare_iqtree clustalo-trimal: Will use bootstrap cutoff (60) before creating concatenated alignment
## Tue Jul 26 08:54:05 CEST 2022 - concatenate muscle-bmge: Will use bootstrap cutoff 70 before creating concatenated alignment
## Tue Jul 26 08:54:05 CEST 2022 - concatenate muscle-trimal: Will use bootstrap cutoff 70 before creating concatenated alignment
## Tue Jul 26 08:54:05 CEST 2022 - prepare_iqtree mafft-aliscore: Will use bootstrap cutoff (60) before creating concatenated alignment
## Tue Jul 26 08:54:05 CEST 2022 - prepare_iqtree mafft-trimal: Will use bootstrap cutoff (70) before creating concatenated alignment
## Tue Jul 26 08:54:05 CEST 2022 - prepare_iqtree mafft-aliscore: Will use bootstrap cutoff (70) before creating concatenated alignment
## Tue Jul 26 08:54:05 CEST 2022 - concatenate mafft-aliscore: Will use bootstrap cutoff 70 before creating concatenated alignment
## Tue Jul 26 08:54:05 CEST 2022 - concatenate mafft-trimal: Will use bootstrap cutoff 70 before creating concatenated alignment
## Tue Jul 26 08:54:05 CEST 2022 - concatenate mafft-bmge: Will use bootstrap cutoff 70 before creating concatenated alignment
## Tue Jul 26 08:54:05 CEST 2022 - prepare_iqtree clustalo-bmge: Will use bootstrap cutoff (70) before creating concatenated alignment
## Tue Jul 26 08:54:06 CEST 2022 - prepare_iqtree muscle-trimal: Will use bootstrap cutoff (70) before creating concatenated alignment
## Tue Jul 26 08:54:06 CEST 2022 - prepare_iqtree clustalo-trimal: Will use bootstrap cutoff (70) before creating concatenated alignment
## Will create NEXUS partition file with model information now.
## Will create NEXUS partition file with model information now.
## Will create NEXUS partition file with model information now.
## Will create NEXUS partition file with model information now.
## Will create NEXUS partition file with model information now.
## Will create NEXUS partition file with model information now.
## Will create NEXUS partition file with model information now.
## Will create NEXUS partition file with model information now.
## Tue Jul 26 08:54:10 CEST 2022 - nexus file for iqtree written.
## Tue Jul 26 08:54:10 CEST 2022 - nexus file for iqtree written.
## Tue Jul 26 08:54:10 CEST 2022 - nexus file for iqtree written.
## Tue Jul 26 08:54:10 CEST 2022 - nexus file for iqtree written.
## Tue Jul 26 08:54:10 CEST 2022 - nexus file for iqtree written.
## Tue Jul 26 08:54:11 CEST 2022 - nexus file for iqtree written.
## Will create NEXUS partition file with model information now.
## Tue Jul 26 08:54:11 CEST 2022 - nexus file for iqtree written.
## Will create NEXUS partition file with model information now.
## Tue Jul 26 08:54:15 CEST 2022 - nexus file for iqtree written.
## Tue Jul 26 08:54:15 CEST 2022 - nexus file for iqtree written.
## Tue Jul 26 08:54:15 CEST 2022 - nexus file for iqtree written.
## Tue 26 Jul 2022 08:54:34 AM CEST - 'phylociraptor model' finished successfully before. Will run raxml with best models.
## Tue 26 Jul 2022 08:54:34 AM CEST - 'phylociraptor model' finished successfully before. Will run raxml with best models.
## Tue 26 Jul 2022 08:54:34 AM CEST - 'phylociraptor model' finished successfully before. Will run raxml with best models.
## Tue 26 Jul 2022 08:54:34 AM CEST - 'phylociraptor model' finished successfully before. Will run raxml with best models.
## Tue 26 Jul 2022 08:54:34 AM CEST - 'phylociraptor model' finished successfully before. Will run raxml with best models.
## Tue 26 Jul 2022 08:54:34 AM CEST - 'phylociraptor model' finished successfully before. Will run raxml with best models.
## Tue 26 Jul 2022 08:54:34 AM CEST - 'phylociraptor model' finished successfully before. Will run raxml with best models.
## Tue 26 Jul 2022 08:54:34 AM CEST - 'phylociraptor model' finished successfully before. Will run raxml with best models.
## Tue 26 Jul 2022 08:54:34 AM CEST - 'phylociraptor model' finished successfully before. Will run raxml with best models.
## Tue 26 Jul 2022 08:54:34 AM CEST - 'phylociraptor model' finished successfully before. Will run raxml with best models.
## Tue 26 Jul 2022 08:54:34 AM CEST - 'phylociraptor model' finished successfully before. Will run raxml with best models.
## Tue 26 Jul 2022 08:54:34 AM CEST - 'phylociraptor model' finished successfully before. Will run raxml with best models.
## Tue 26 Jul 2022 08:54:34 AM CEST - 'phylociraptor model' finished successfully before. Will run raxml with best models.
## Tue 26 Jul 2022 08:54:34 AM CEST - 'phylociraptor model' finished successfully before. Will run raxml with best models.
## Tue 26 Jul 2022 08:54:34 AM CEST - 'phylociraptor model' finished successfully before. Will run raxml with best models.
## Tue 26 Jul 2022 08:54:34 AM CEST - 'phylociraptor model' finished successfully before. Will run raxml with best models.
## Tue 26 Jul 2022 08:54:34 AM CEST - 'phylociraptor model' finished successfully before. Will run raxml with best models.
## Tue 26 Jul 2022 08:54:34 AM CEST - 'phylociraptor model' finished successfully before. Will run raxml with best models.
## Tue 26 Jul 2022 08:54:34 AM CEST - 'phylociraptor model' finished successfully before. Will run raxml with best models.
## Tue 26 Jul 2022 08:54:34 AM CEST - 'phylociraptor model' finished successfully before. Will run raxml with best models.
## Tue 26 Jul 2022 08:54:34 AM CEST - 'phylociraptor model' finished successfully before. Will run raxml with best models.
## Tue 26 Jul 2022 08:54:34 AM CEST - 'phylociraptor model' finished successfully before. Will run raxml with best models.
## Tue 26 Jul 2022 08:54:34 AM CEST - 'phylociraptor model' finished successfully before. Will run raxml with best models.
## Tue 26 Jul 2022 08:54:34 AM CEST - 'phylociraptor model' finished successfully before. Will run raxml with best models.
## Tue 26 Jul 2022 08:54:34 AM CEST - 'phylociraptor model' finished successfully before. Will run raxml with best models.
## Tue 26 Jul 2022 08:54:34 AM CEST - 'phylociraptor model' finished successfully before. Will run raxml with best models.
## Tue 26 Jul 2022 08:54:34 AM CEST - 'phylociraptor model' finished successfully before. Will run raxml with best models.
## Tue 26 Jul 2022 08:54:34 AM CEST - 'phylociraptor model' finished successfully before. Will run raxml with best models.
## Tue 26 Jul 2022 08:55:04 AM CEST - 'phylociraptor model' finished successfully before. Will run raxml with best models.
## Wed Jul 27 15:34:55 CEST 2022 - Pipeline part 3 (tree) done.
## Wed Jul 27 15:36:24 CEST 2022 - phylociraptor will use gene tree filtering based on average bootstrap support value of 90.
## Wed Jul 27 15:36:24 CEST 2022 - phylociraptor will use gene tree filtering based on average bootstrap support value of 50.
## Wed Jul 27 15:36:25 CEST 2022 - phylociraptor will use gene tree filtering based on average bootstrap support value of 50.
## Wed Jul 27 15:36:25 CEST 2022 - phylociraptor will use gene tree filtering based on average bootstrap support value of 80.
## Wed Jul 27 15:36:25 CEST 2022 - phylociraptor will use gene tree filtering based on average bootstrap support value of 90.
## Wed Jul 27 15:36:25 CEST 2022 - phylociraptor will use gene tree filtering based on average bootstrap support value of 60.
## Wed Jul 27 15:36:25 CEST 2022 - phylociraptor will use gene tree filtering based on average bootstrap support value of 60.
## Wed Jul 27 15:36:25 CEST 2022 - phylociraptor will use gene tree filtering based on average bootstrap support value of 70.
## Wed Jul 27 15:36:25 CEST 2022 - phylociraptor will use gene tree filtering based on average bootstrap support value of 80.
## Wed Jul 27 15:36:25 CEST 2022 - phylociraptor will use gene tree filtering based on average bootstrap support value of 60.
## Wed Jul 27 15:36:25 CEST 2022 - phylociraptor will use gene tree filtering based on average bootstrap support value of 90.
## Wed Jul 27 15:36:25 CEST 2022 - phylociraptor will use gene tree filtering based on average bootstrap support value of 50.
## Wed Jul 27 15:36:25 CEST 2022 - phylociraptor will use gene tree filtering based on average bootstrap support value of 70.
## Wed Jul 27 15:36:25 CEST 2022 - phylociraptor will use gene tree filtering based on average bootstrap support value of 60.
## Wed Jul 27 15:36:25 CEST 2022 - phylociraptor will use gene tree filtering based on average bootstrap support value of 70.
## Wed Jul 27 15:36:25 CEST 2022 - phylociraptor will use gene tree filtering based on average bootstrap support value of 50.
## Wed Jul 27 15:36:25 CEST 2022 - phylociraptor will use gene tree filtering based on average bootstrap support value of 90.
## Wed Jul 27 15:36:25 CEST 2022 - phylociraptor will use gene tree filtering based on average bootstrap support value of 50.
## Wed Jul 27 15:36:25 CEST 2022 - phylociraptor will use gene tree filtering based on average bootstrap support value of 70.
## Wed Jul 27 15:36:25 CEST 2022 - phylociraptor will use gene tree filtering based on average bootstrap support value of 50.
## Wed Jul 27 15:36:25 CEST 2022 - phylociraptor will use gene tree filtering based on average bootstrap support value of 60.
## Wed Jul 27 15:36:25 CEST 2022 - phylociraptor will use gene tree filtering based on average bootstrap support value of 0.
## Wed Jul 27 15:36:25 CEST 2022 - phylociraptor will use gene tree filtering based on average bootstrap support value of 0.
## Wed Jul 27 15:36:25 CEST 2022 - phylociraptor will use gene tree filtering based on average bootstrap support value of 90.
## Wed Jul 27 15:36:25 CEST 2022 - phylociraptor will use gene tree filtering based on average bootstrap support value of 0.
## Wed Jul 27 15:36:25 CEST 2022 - phylociraptor will use gene tree filtering based on average bootstrap support value of 70.
## Wed Jul 27 15:36:25 CEST 2022 - phylociraptor will use gene tree filtering based on average bootstrap support value of 90.
## Wed Jul 27 15:36:25 CEST 2022 - phylociraptor will use gene tree filtering based on average bootstrap support value of 80.
## Wed Jul 27 15:36:25 CEST 2022 - phylociraptor will use gene tree filtering based on average bootstrap support value of 50.
## Wed Jul 27 15:36:25 CEST 2022 - phylociraptor will use gene tree filtering based on average bootstrap support value of 60.
## Wed Jul 27 15:36:25 CEST 2022 - phylociraptor will use gene tree filtering based on average bootstrap support value of 60.
## Wed Jul 27 15:36:25 CEST 2022 - phylociraptor will use gene tree filtering based on average bootstrap support value of 70.
## Wed Jul 27 15:36:25 CEST 2022 - phylociraptor will use gene tree filtering based on average bootstrap support value of 70.
## Wed Jul 27 15:36:25 CEST 2022 - phylociraptor will use gene tree filtering based on average bootstrap support value of 0.
## 
## Wed Jul 27 15:36:25 CEST 2022 - phylociraptor will use gene tree filtering based on average bootstrap support value of 0.
## Wed Jul 27 15:36:25 CEST 2022 - phylociraptor will use gene tree filtering based on average bootstrap support value of 80.
## Wed Jul 27 15:36:25 CEST 2022 - phylociraptor will use gene tree filtering based on average bootstrap support value of 80.
## Wed Jul 27 15:36:25 CEST 2022 - phylociraptor will use gene tree filtering based on average bootstrap support value of 90.
## Wed Jul 27 15:36:25 CEST 2022 - phylociraptor will use gene tree filtering based on average bootstrap support value of 80.
## Wed Jul 27 15:36:25 CEST 2022 - phylociraptor will use gene tree filtering based on average bootstrap support value of 50.
## Wed Jul 27 15:36:25 CEST 2022 - phylociraptor will use gene tree filtering based on average bootstrap support value of 50.
## Wed Jul 27 15:36:25 CEST 2022 - phylociraptor will use gene tree filtering based on average bootstrap support value of 80.
## Wed Jul 27 15:36:25 CEST 2022 - phylociraptor will use gene tree filtering based on average bootstrap support value of 90.
## Wed Jul 27 15:36:25 CEST 2022 - phylociraptor will use gene tree filtering based on average bootstrap support value of 80.
## Wed Jul 27 15:36:25 CEST 2022 - phylociraptor will use gene tree filtering based on average bootstrap support value of 0.
## Wed Jul 27 15:36:26 CEST 2022 - phylociraptor will use gene tree filtering based on average bootstrap support value of 0.
## Wed Jul 27 15:36:26 CEST 2022 - phylociraptor will use gene tree filtering based on average bootstrap support value of 80.
## Wed Jul 27 15:36:30 CEST 2022 - phylociraptor will use gene tree filtering based on average bootstrap support value of 90.
## Wed Jul 27 15:42:24 CEST 2022 - Speciestree reconstruction done.
## Fri Jul 29 11:05:44 CEST 2022 - concatenate mafft-aliscore: Will use bootstrap cutoff 90 before creating concatenated alignment
## Fri Jul 29 11:05:44 CEST 2022 - concatenate clustalo-trimal: Will use bootstrap cutoff 90 before creating concatenated alignment
## Fri Jul 29 11:05:44 CEST 2022 - concatenate clustalo-bmge: Will use bootstrap cutoff 90 before creating concatenated alignment
## Fri Jul 29 11:05:44 CEST 2022 - concatenate mafft-bmge: Will use bootstrap cutoff 90 before creating concatenated alignment
## Fri Jul 29 11:05:44 CEST 2022 - concatenate muscle-aliscore: Will use bootstrap cutoff 90 before creating concatenated alignment
## Fri Jul 29 11:05:44 CEST 2022 - concatenate clustalo-aliscore: Will use bootstrap cutoff 90 before creating concatenated alignment
## Fri Jul 29 11:05:46 CEST 2022 - concatenate muscle-bmge: Will use bootstrap cutoff 90 before creating concatenated alignment
## Fri Jul 29 11:05:47 CEST 2022 - concatenate muscle-trimal: Will use bootstrap cutoff 90 before creating concatenated alignment
## Fri Jul 29 11:05:47 CEST 2022 - concatenate mafft-trimal: Will use bootstrap cutoff 90 before creating concatenated alignment
## Fri Jul 29 11:06:12 CEST 2022 - 'phylociraptor model' finished successfully before. Will run raxml with best models.
## Fri Jul 29 11:06:12 CEST 2022 - 'phylociraptor model' finished successfully before. Will run raxml with best models.
## Fri Jul 29 11:06:12 CEST 2022 - 'phylociraptor model' finished successfully before. Will run raxml with best models.
## Fri Jul 29 11:06:12 CEST 2022 - 'phylociraptor model' finished successfully before. Will run raxml with best models.
## Fri Jul 29 11:06:12 CEST 2022 - 'phylociraptor model' finished successfully before. Will run raxml with best models.
## Fri Jul 29 11:06:13 CEST 2022 - 'phylociraptor model' finished successfully before. Will run raxml with best models.
## Fri Jul 29 11:06:13 CEST 2022 - 'phylociraptor model' finished successfully before. Will run raxml with best models.
## Fri Jul 29 11:06:13 CEST 2022 - 'phylociraptor model' finished successfully before. Will run raxml with best models.
## Fri Jul 29 11:06:13 CEST 2022 - 'phylociraptor model' finished successfully before. Will run raxml with best models.
## Thu Aug  4 19:41:37 CEST 2022 - Setup: Will download species now - batch: 3
## Thu Aug  4 19:41:37 CEST 2022 - Setup: Will download species now - batch: 2
## Thu Aug  4 19:41:37 CEST 2022 - Setup: Will download species now - batch: 1
## Thu 04 Aug 2022 07:42:22 PM CEST - phylociraptor setup done.
## Thu Aug  4 19:47:15 CEST 2022 Paraphelidium_tribonemae     124 Complete     225 Duplicated       9 Fragmented      22 Missing
## Thu Aug  4 19:52:30 CEST 2022 - Setup: Will download species now - batch: 1
## Thu Aug  4 19:52:30 CEST 2022 - Setup: Will download species now - batch: 2
## Thu Aug  4 19:52:30 CEST 2022 - Setup: Will download species now - batch: 3
## Thu 04 Aug 2022 07:53:12 PM CEST - phylociraptor setup done.
## Thu 04 Aug 2022 08:02:13 PM CEST Paraphelidium_tribonemae     124 Complete     225 Duplicated       9 Fragmented      22 Missing
## Thu 04 Aug 2022 08:08:11 PM CEST Encephalitozoon_hellem      70 Complete       3 Duplicated      17 Fragmented     165 Missing
## Thu 04 Aug 2022 08:09:03 PM CEST Nosema_ceranae      67 Complete      15 Fragmented     171 Missing
## Thu 04 Aug 2022 08:09:24 PM CEST Malassezia_pachydermatis     232 Complete       2 Duplicated      14 Fragmented       6 Missing
## Thu 04 Aug 2022 08:10:00 PM CEST Schizosaccharomyces_pombe     236 Complete       8 Duplicated       5 Fragmented       8 Missing
## Thu 04 Aug 2022 08:10:45 PM CEST Wickerhamomyces_anomalus     238 Complete      12 Duplicated       3 Fragmented       6 Missing
## Thu 04 Aug 2022 08:11:22 PM CEST Mixia_osmundae     219 Complete       2 Duplicated      23 Fragmented      10 Missing
## Thu 04 Aug 2022 08:11:53 PM CEST Wallemia_mellicola     231 Complete       2 Duplicated      12 Fragmented       9 Missing
## Thu 04 Aug 2022 08:12:13 PM CEST Thelohania_contejeani      82 Complete      16 Fragmented     155 Missing
## Thu 04 Aug 2022 08:12:20 PM CEST Vavraia_culicis      56 Complete      15 Fragmented     182 Missing
## Thu 04 Aug 2022 08:14:07 PM CEST Cunea_sp      57 Complete       4 Duplicated      49 Fragmented     145 Missing
## Thu 04 Aug 2022 08:15:43 PM CEST Coemansia_mojavensis     234 Complete      18 Duplicated       3 Fragmented       7 Missing
## Thu 04 Aug 2022 08:15:50 PM CEST Sclerophora_sanguinea     251 Complete       2 Fragmented
## Thu 04 Aug 2022 08:15:56 PM CEST Mitosporidium_daphniae     105 Complete      11 Duplicated      37 Fragmented     106 Missing
## Thu 04 Aug 2022 08:16:39 PM CEST Cryptococcus_neoformans     235 Complete       4 Duplicated      10 Fragmented       6 Missing
## Thu 04 Aug 2022 08:18:25 PM CEST Nematocida_sp      65 Complete       4 Duplicated       8 Fragmented     178 Missing
## Thu 04 Aug 2022 08:19:02 PM CEST Rozella_allomycis     142 Complete      11 Duplicated      42 Fragmented      64 Missing
## Thu 04 Aug 2022 08:20:45 PM CEST Dictyostelium_lacteum     231 Complete       2 Duplicated      10 Fragmented      11 Missing
## Thu 04 Aug 2022 08:20:46 PM CEST Debaryomyces_hansenii     233 Complete       2 Duplicated       7 Fragmented      12 Missing
## Thu 04 Aug 2022 08:21:19 PM CEST Calocera_viscosa     232 Complete      14 Duplicated       9 Fragmented       5 Missing
## Thu 04 Aug 2022 08:22:13 PM CEST Anaeromyces_robustus     205 Complete      10 Duplicated       9 Fragmented      34 Missing
## Thu 04 Aug 2022 08:22:26 PM CEST Acidomyces_richmondensis     247 Complete       4 Duplicated       3 Fragmented       1 Missing
## Thu 04 Aug 2022 08:22:50 PM CEST Amaparvus_languidus      91 Complete       7 Duplicated      58 Fragmented     101 Missing
## Thu 04 Aug 2022 08:23:00 PM CEST Yarrowia_lipolytica     242 Complete       4 Duplicated       5 Fragmented       4 Missing
## Thu 04 Aug 2022 08:23:11 PM CEST Daedalea_quercina     244 Complete       6 Fragmented       3 Missing
## Thu 04 Aug 2022 08:24:33 PM CEST Mucor_lusitanicus     234 Complete      24 Duplicated       4 Fragmented       3 Missing
## Thu 04 Aug 2022 08:24:52 PM CEST Pilobolus_umbonatus     237 Complete      12 Duplicated       6 Fragmented       4 Missing
## Thu 04 Aug 2022 08:25:22 PM CEST Lipomyces_starkeyi     240 Complete       4 Duplicated       8 Fragmented       3 Missing
## Thu 04 Aug 2022 08:26:13 PM CEST Tubulinosema_ratisbonensis      63 Complete       6 Duplicated      16 Fragmented     172 Missing
## Thu 04 Aug 2022 08:26:24 PM CEST Ustilago_hordei     248 Complete       4 Fragmented       1 Missing
## Thu 04 Aug 2022 08:27:16 PM CEST Neurospora_crassa     252 Complete       1 Fragmented
## Thu 04 Aug 2022 08:27:27 PM CEST Caulochytrium_protostelioides      71 Complete      40 Fragmented     142 Missing
## Thu 04 Aug 2022 08:27:31 PM CEST Exophiala_oligosperma     251 Complete       2 Fragmented
## Thu 04 Aug 2022 08:27:35 PM CEST Rhodotorula_toruloides     224 Complete       2 Duplicated      15 Fragmented      13 Missing
## Thu 04 Aug 2022 08:27:36 PM CEST Hypoxylon_sp     246 Complete       2 Duplicated       5 Fragmented       1 Missing
## Thu 04 Aug 2022 08:27:53 PM CEST Blastocladiella_britannica     202 Complete      25 Fragmented      26 Missing
## Thu 04 Aug 2022 08:28:30 PM CEST Actinomortierella_capitata     245 Complete       5 Fragmented       3 Missing
## Thu 04 Aug 2022 08:28:35 PM CEST Capsaspora_owczarzaki     227 Complete      10 Duplicated      11 Fragmented      10 Missing
## Thu 04 Aug 2022 08:29:12 PM CEST Letharia_columbiana     234 Complete      28 Duplicated       4 Fragmented       1 Missing
## Thu 04 Aug 2022 08:30:05 PM CEST Cylindrobasidium_torrendii     236 Complete      11 Fragmented       6 Missing
## Thu 04 Aug 2022 08:30:35 PM CEST Synchytrium_microbalum     223 Complete      19 Fragmented      11 Missing
## Thu 04 Aug 2022 08:31:09 PM CEST Nuclearia_sp     163 Complete     109 Duplicated      35 Fragmented       6 Missing
## Thu 04 Aug 2022 08:31:11 PM CEST Syncephalis_fuscata     177 Complete      41 Duplicated      30 Fragmented      27 Missing
## Thu 04 Aug 2022 08:31:54 PM CEST Amphiamblys_sp      78 Complete      26 Fragmented     149 Missing
## Thu 04 Aug 2022 08:32:39 PM CEST Dichotomocladium_robustum     237 Complete       8 Duplicated       7 Fragmented       5 Missing
## Thu 04 Aug 2022 08:33:03 PM CEST Pneumocystis_murina     239 Complete       2 Duplicated       3 Fragmented      10 Missing
## Thu 04 Aug 2022 08:33:14 PM CEST Podila_minutissima     243 Complete      12 Duplicated       1 Fragmented       3 Missing
## Thu 04 Aug 2022 08:33:21 PM CEST Paraphaeosphaeria_sporulosa     245 Complete       2 Duplicated       6 Fragmented       1 Missing
## Thu 04 Aug 2022 08:33:41 PM CEST Spizellomyces_punctatus     234 Complete       2 Duplicated       8 Fragmented      10 Missing
## Thu 04 Aug 2022 08:34:17 PM CEST Acaromyces_ingoldii     236 Complete      10 Fragmented       7 Missing
## Thu 04 Aug 2022 08:34:33 PM CEST Vittaforma_corneae      56 Complete       2 Duplicated      14 Fragmented     182 Missing
## Thu 04 Aug 2022 08:35:45 PM CEST Coelomomyces_lativittatus     163 Complete       4 Duplicated      47 Fragmented      41 Missing
## Thu 04 Aug 2022 08:36:08 PM CEST Sistotremastrum_suecicum     237 Complete       2 Duplicated       9 Fragmented       6 Missing
## Thu 04 Aug 2022 08:36:44 PM CEST Sanchytrium_tribonematis     137 Complete      15 Duplicated      30 Fragmented      79 Missing
## Thu 04 Aug 2022 08:37:54 PM CEST Phaffia_rhodozyma     209 Complete       2 Duplicated      28 Fragmented      15 Missing
## Thu 04 Aug 2022 08:38:05 PM CEST Fonticula_alba     162 Complete       2 Duplicated      30 Fragmented      60 Missing
## Thu 04 Aug 2022 08:38:09 PM CEST Dissophora_decumbens     240 Complete      13 Duplicated       6 Fragmented       1 Missing
## Thu 04 Aug 2022 08:38:48 PM CEST Umbelopsis_isabellina     248 Complete       2 Fragmented       3 Missing
## Thu 04 Aug 2022 08:39:02 PM CEST Paramicrosporidium_saccamoebae     182 Complete       2 Duplicated      20 Fragmented      50 Missing
## Thu 04 Aug 2022 08:39:19 PM CEST Fusarium_proliferatum     248 Complete       4 Duplicated       1 Fragmented       2 Missing
## Thu Aug  4 20:39:27 CEST 2022 Arthrobotrys_oligospora     245 Complete       8 Fragmented
## Thu 04 Aug 2022 08:39:36 PM CEST Saitoella_complicata     237 Complete      14 Duplicated       9 Fragmented
## Thu Aug  4 20:39:43 CEST 2022 Piromyces_sp     217 Complete      10 Duplicated       4 Fragmented      27 Missing
## Thu 04 Aug 2022 08:40:15 PM CEST Amoeboradix_gromovi     150 Complete       4 Duplicated      22 Fragmented      79 Missing
## Thu 04 Aug 2022 08:41:11 PM CEST Microbotryum_violaceum     233 Complete       4 Duplicated      13 Fragmented       5 Missing
## Thu 04 Aug 2022 08:41:25 PM CEST Protomyces_lactucaedebilis     238 Complete       7 Fragmented       8 Missing
## Thu 04 Aug 2022 08:42:05 PM CEST Botrytis_cinerea     245 Complete       2 Duplicated       5 Fragmented       2 Missing
## Thu 04 Aug 2022 08:45:24 PM CEST Tilletia_indica     225 Complete       4 Duplicated      12 Fragmented      15 Missing
## Thu 04 Aug 2022 08:45:25 PM CEST Gonapodya_prolifera     188 Complete      74 Duplicated      17 Fragmented      11 Missing
## Thu 04 Aug 2022 08:46:45 PM CEST Piptocephalis_tieghemiana     192 Complete      21 Fragmented      40 Missing
## Thu 04 Aug 2022 08:46:58 PM CEST Hyaloraphidium_curvatum     223 Complete      20 Fragmented      10 Missing
## Thu 04 Aug 2022 08:48:09 PM CEST Stereomyxa_ramosa     130 Complete      18 Duplicated      51 Fragmented      68 Missing
## Thu 04 Aug 2022 08:49:04 PM CEST Paraphysoderma_sedebokerense     202 Complete      27 Duplicated      21 Fragmented      18 Missing
## Thu 04 Aug 2022 08:49:33 PM CEST Salpingoeca_rosetta     198 Complete       5 Duplicated      15 Fragmented      38 Missing
## Thu 04 Aug 2022 08:49:40 PM CEST Batrachochytrium_dendrobatidis     237 Complete       9 Duplicated       7 Fragmented       5 Missing
## Thu 04 Aug 2022 08:49:42 PM CEST Mortierella_sp     240 Complete      24 Duplicated       4 Fragmented       3 Missing
## Thu 04 Aug 2022 08:50:59 PM CEST Neolentinus_lepideus     233 Complete       9 Duplicated      10 Fragmented       6 Missing
## Thu 04 Aug 2022 08:51:05 PM CEST Radiomyces_spectabilis     243 Complete       8 Duplicated       3 Fragmented       3 Missing
## Thu Aug  4 20:51:36 CEST 2022 Phycomyces_blakesleeanus     226 Complete      30 Duplicated       9 Fragmented       3 Missing
## Thu 04 Aug 2022 08:52:08 PM CEST Pyricularia_oryzae     247 Complete       6 Fragmented
## Thu 04 Aug 2022 08:52:18 PM CEST Phialocephala_subalpina     246 Complete      10 Duplicated       1 Fragmented       1 Missing
## Thu 04 Aug 2022 08:52:19 PM CEST Coccidioides_posadasii     247 Complete       4 Fragmented       2 Missing
## Thu 04 Aug 2022 08:53:28 PM CEST Paxillus_rubicundulus     235 Complete      10 Fragmented       8 Missing
## Thu 04 Aug 2022 08:55:14 PM CEST Olpidium_bornovanus      78 Complete      69 Duplicated      60 Fragmented      87 Missing
## Thu 04 Aug 2022 08:55:58 PM CEST Glomus_cerebriforme     235 Complete       6 Duplicated       7 Fragmented       8 Missing
## Thu 04 Aug 2022 08:56:46 PM CEST Rhizophagus_sp     236 Complete       4 Duplicated       8 Fragmented       7 Missing
## Thu 04 Aug 2022 08:57:22 PM CEST Dimargaris_cristalligena     213 Complete       2 Duplicated      17 Fragmented      22 Missing
## Thu 04 Aug 2022 09:01:13 PM CEST Neocallimastix_sp     128 Complete     193 Duplicated       6 Fragmented      24 Missing
## Thu 04 Aug 2022 09:04:45 PM CEST Melampsora_laricipopulina     228 Complete      12 Duplicated      13 Fragmented       6 Missing
## Thu 04 Aug 2022 09:04:49 PM CEST Allomyces_macrogynus      77 Complete     306 Duplicated      12 Fragmented      14 Missing
## Thu 04 Aug 2022 09:06:30 PM CEST Tuber_borchii     244 Complete       6 Duplicated       4 Fragmented       2 Missing
## Thu 04 Aug 2022 09:10:47 PM CEST Edhazardia_aedis      66 Complete      12 Fragmented     175 Missing
## Thu 04 Aug 2022 09:16:25 PM CEST Chytriomyces_hyalinus     177 Complete     100 Duplicated      15 Fragmented      14 Missing
## Thu 04 Aug 2022 09:19:19 PM CEST Polychytrium_aggregatum     108 Complete     255 Duplicated      16 Fragmented       7 Missing
## Thu 04 Aug 2022 09:19:27 PM CEST Basidiobolus_meristosporus     207 Complete      82 Duplicated       7 Fragmented       1 Missing
## Thu 04 Aug 2022 09:20:37 PM CEST Catenaria_anguillulae     128 Complete     156 Duplicated      27 Fragmented      22 Missing
## Thu 04 Aug 2022 09:23:44 PM CEST Smittium_culicis     130 Complete     191 Duplicated      17 Fragmented      20 Missing
## Thu Aug  4 21:36:24 CEST 2022 Diversispora_epigaea     237 Complete       4 Duplicated       9 Fragmented       5 Missing
## Fri 05 Aug 2022 02:00:49 AM CEST Gigaspora_margarita     233 Complete       9 Duplicated       6 Fragmented      10 Missing
## Fri 05 Aug 2022 03:58:49 AM CEST Zoophthora_radicans     183 Complete      34 Duplicated      23 Fragmented      30 Missing
## Fri 05 Aug 2022 09:06:47 PM CEST Mus_musculus     235 Complete      52 Duplicated       1 Fragmented
## Fri 05 Aug 2022 09:08:11 PM CEST - phylociraptor orthology done.
## Mon Aug  8 09:04:18 CEST 2022 - BUSCO files will be filtered on a per-sample basis. This could lower the number of species in the final tree.
## Mon Aug  8 09:04:53 CEST 2022 - Number of BUSCO sequence files: 255
## Mon Aug  8 09:04:53 CEST 2022 - Number of deduplicated BUSCO sequence files: 255
## Mon Aug  8 09:05:16 CEST 2022 - Phylociraptor filter-orthology done.
## Mon Aug  8 09:46:50 CEST 2022 - phylociraptor align done.
## Mon Aug  8 09:48:28 CEST 2022 - The aliscore output file does not exist. Check results for BUSCO: 1617752at2759
## Mon Aug  8 09:48:28 CEST 2022 - The aliscore output file does not exist. Check results for BUSCO: 1405073at2759
## Mon Aug  8 09:48:28 CEST 2022 - The aliscore output file does not exist. Check results for BUSCO: 1407446at2759
## Mon Aug  8 09:48:28 CEST 2022 - The aliscore output appears to be empty. Check results for BUSCO: 1617752at2759
## Mon Aug  8 09:48:28 CEST 2022 - The aliscore output appears to be empty. Check results for BUSCO: 1405073at2759
## Mon Aug  8 09:48:28 CEST 2022 - The aliscore output appears to be empty. Check results for BUSCO: 1407446at2759
## Mon Aug  8 09:48:29 CEST 2022 - The aliscore output file does not exist. Check results for BUSCO: 1094121at2759
## Mon Aug  8 09:48:29 CEST 2022 - The aliscore output appears to be empty. Check results for BUSCO: 1094121at2759
## Mon Aug  8 09:48:29 CEST 2022 - The aliscore output file does not exist. Check results for BUSCO: 679771at2759
## Mon Aug  8 09:48:29 CEST 2022 - The aliscore output appears to be empty. Check results for BUSCO: 679771at2759
## Mon Aug  8 09:48:29 CEST 2022 - The aliscore output file does not exist. Check results for BUSCO: 1161199at2759
## Mon Aug  8 09:48:29 CEST 2022 - The aliscore output appears to be empty. Check results for BUSCO: 1161199at2759
## Mon Aug  8 09:48:29 CEST 2022 - The aliscore output file does not exist. Check results for BUSCO: 1474000at2759
## Mon Aug  8 09:48:29 CEST 2022 - The aliscore output appears to be empty. Check results for BUSCO: 1474000at2759
## Mon Aug  8 09:48:29 CEST 2022 - The aliscore output file does not exist. Check results for BUSCO: 1178688at2759
## Mon Aug  8 09:48:30 CEST 2022 - The aliscore output appears to be empty. Check results for BUSCO: 1178688at2759
## Mon Aug  8 09:48:30 CEST 2022 - The aliscore output file does not exist. Check results for BUSCO: 1291729at2759
## Mon Aug  8 09:48:30 CEST 2022 - The aliscore output file does not exist. Check results for BUSCO: 1107630at2759
## Mon Aug  8 09:48:30 CEST 2022 - The aliscore output appears to be empty. Check results for BUSCO: 1291729at2759
## Mon Aug  8 09:48:30 CEST 2022 - The aliscore output appears to be empty. Check results for BUSCO: 1107630at2759
## Mon Aug  8 09:48:30 CEST 2022 - The aliscore output file does not exist. Check results for BUSCO: 1287094at2759
## Mon Aug  8 09:48:30 CEST 2022 - The aliscore output appears to be empty. Check results for BUSCO: 1287094at2759
## Mon Aug  8 09:48:30 CEST 2022 - The aliscore output file does not exist. Check results for BUSCO: 1364586at2759
## Mon Aug  8 09:48:30 CEST 2022 - The aliscore output appears to be empty. Check results for BUSCO: 1364586at2759
## Mon Aug  8 09:48:30 CEST 2022 - The aliscore output file does not exist. Check results for BUSCO: 1479417at2759
## Mon Aug  8 09:48:30 CEST 2022 - The aliscore output appears to be empty. Check results for BUSCO: 1479417at2759
## Mon Aug  8 09:48:30 CEST 2022 - The aliscore output file does not exist. Check results for BUSCO: 1057950at2759
## Mon Aug  8 09:48:30 CEST 2022 - The aliscore output appears to be empty. Check results for BUSCO: 1057950at2759
## Mon Aug  8 09:48:31 CEST 2022 - The aliscore output file does not exist. Check results for BUSCO: 1085752at2759
## Mon Aug  8 09:48:31 CEST 2022 - The aliscore output appears to be empty. Check results for BUSCO: 1085752at2759
## Mon Aug  8 09:48:31 CEST 2022 - The aliscore output file does not exist. Check results for BUSCO: 1010730at2759
## Mon Aug  8 09:48:31 CEST 2022 - The aliscore output appears to be empty. Check results for BUSCO: 1010730at2759
## Mon Aug  8 09:48:31 CEST 2022 - The aliscore output file does not exist. Check results for BUSCO: 1275837at2759
## Mon Aug  8 09:48:31 CEST 2022 - The aliscore output appears to be empty. Check results for BUSCO: 1275837at2759
## Mon Aug  8 09:48:32 CEST 2022 - The aliscore output file does not exist. Check results for BUSCO: 582756at2759
## Mon Aug  8 09:48:32 CEST 2022 - The aliscore output appears to be empty. Check results for BUSCO: 582756at2759
## Mon Aug  8 09:48:33 CEST 2022 - The aliscore output file does not exist. Check results for BUSCO: 299766at2759
## Mon Aug  8 09:48:33 CEST 2022 - The aliscore output appears to be empty. Check results for BUSCO: 299766at2759
## Mon Aug  8 09:48:34 CEST 2022 - The aliscore output file does not exist. Check results for BUSCO: 1248958at2759
## Mon Aug  8 09:48:34 CEST 2022 - The aliscore output appears to be empty. Check results for BUSCO: 1248958at2759
## Mon Aug  8 09:48:35 CEST 2022 - The aliscore output file does not exist. Check results for BUSCO: 777920at2759
## Mon Aug  8 09:48:35 CEST 2022 - The aliscore output file does not exist. Check results for BUSCO: 625387at2759
## Mon Aug  8 09:48:35 CEST 2022 - The aliscore output appears to be empty. Check results for BUSCO: 777920at2759
## Mon Aug  8 09:48:35 CEST 2022 - The aliscore output appears to be empty. Check results for BUSCO: 625387at2759
## Mon Aug  8 09:48:35 CEST 2022 - The aliscore output file does not exist. Check results for BUSCO: 166920at2759
## Mon Aug  8 09:48:35 CEST 2022 - The aliscore output appears to be empty. Check results for BUSCO: 166920at2759
## Mon Aug  8 09:48:37 CEST 2022 - The aliscore output file does not exist. Check results for BUSCO: 261328at2759
## Mon Aug  8 09:48:37 CEST 2022 - The aliscore output appears to be empty. Check results for BUSCO: 261328at2759
## Mon Aug  8 09:48:57 CEST 2022 - The aliscore output file does not exist. Check results for BUSCO: 1440961at2759
## Mon Aug  8 09:48:57 CEST 2022 - The aliscore output appears to be empty. Check results for BUSCO: 1440961at2759
## Mon Aug  8 09:48:58 CEST 2022 - The aliscore output file does not exist. Check results for BUSCO: 1558822at2759
## Mon Aug  8 09:48:58 CEST 2022 - The aliscore output appears to be empty. Check results for BUSCO: 1558822at2759
## Mon Aug  8 09:48:59 CEST 2022 - The aliscore output file does not exist. Check results for BUSCO: 1398309at2759
## Mon Aug  8 09:48:59 CEST 2022 - The aliscore output file does not exist. Check results for BUSCO: 1197019at2759
## Mon Aug  8 09:48:59 CEST 2022 - The aliscore output appears to be empty. Check results for BUSCO: 1398309at2759
## Mon Aug  8 09:48:59 CEST 2022 - The aliscore output file does not exist. Check results for BUSCO: 956854at2759
## Mon Aug  8 09:48:59 CEST 2022 - The aliscore output appears to be empty. Check results for BUSCO: 1197019at2759
## Mon Aug  8 09:48:59 CEST 2022 - The aliscore output appears to be empty. Check results for BUSCO: 956854at2759
## Mon Aug  8 09:48:59 CEST 2022 - The aliscore output file does not exist. Check results for BUSCO: 1459797at2759
## Mon Aug  8 09:48:59 CEST 2022 - The aliscore output appears to be empty. Check results for BUSCO: 1459797at2759
## Mon Aug  8 09:48:59 CEST 2022 - The aliscore output file does not exist. Check results for BUSCO: 1380409at2759
## Mon Aug  8 09:49:00 CEST 2022 - The aliscore output appears to be empty. Check results for BUSCO: 1380409at2759
## Mon Aug  8 09:49:00 CEST 2022 - The aliscore output file does not exist. Check results for BUSCO: 1309031at2759
## Mon Aug  8 09:49:01 CEST 2022 - The aliscore output appears to be empty. Check results for BUSCO: 1309031at2759
## Mon Aug  8 09:49:01 CEST 2022 - The aliscore output file does not exist. Check results for BUSCO: 911863at2759
## Mon Aug  8 09:49:01 CEST 2022 - The aliscore output appears to be empty. Check results for BUSCO: 911863at2759
## Mon Aug  8 09:49:01 CEST 2022 - The aliscore output file does not exist. Check results for BUSCO: 866359at2759
## Mon Aug  8 09:49:01 CEST 2022 - The aliscore output appears to be empty. Check results for BUSCO: 866359at2759
## Mon Aug  8 09:49:01 CEST 2022 - The aliscore output file does not exist. Check results for BUSCO: 1526152at2759
## Mon Aug  8 09:49:01 CEST 2022 - The aliscore output appears to be empty. Check results for BUSCO: 1526152at2759
## Mon Aug  8 09:49:01 CEST 2022 - The aliscore output file does not exist. Check results for BUSCO: 87842at2759
## Mon Aug  8 09:49:01 CEST 2022 - The aliscore output file does not exist. Check results for BUSCO: 1128607at2759
## Mon Aug  8 09:49:01 CEST 2022 - The aliscore output file does not exist. Check results for BUSCO: 257318at2759
## Mon Aug  8 09:49:01 CEST 2022 - The aliscore output file does not exist. Check results for BUSCO: 1346165at2759
## Mon Aug  8 09:49:01 CEST 2022 - The aliscore output appears to be empty. Check results for BUSCO: 87842at2759
## Mon Aug  8 09:49:01 CEST 2022 - The aliscore output appears to be empty. Check results for BUSCO: 257318at2759
## Mon Aug  8 09:49:01 CEST 2022 - The aliscore output appears to be empty. Check results for BUSCO: 1346165at2759
## Mon Aug  8 09:49:01 CEST 2022 - The aliscore output appears to be empty. Check results for BUSCO: 1128607at2759
## Mon Aug  8 09:49:01 CEST 2022 - The aliscore output file does not exist. Check results for BUSCO: 1405146at2759
## Mon Aug  8 09:49:01 CEST 2022 - The aliscore output appears to be empty. Check results for BUSCO: 1405146at2759
## Mon Aug  8 09:49:01 CEST 2022 - The aliscore output file does not exist. Check results for BUSCO: 917326at2759
## Mon Aug  8 09:49:01 CEST 2022 - The aliscore output file does not exist. Check results for BUSCO: 996662at2759
## Mon Aug  8 09:49:01 CEST 2022 - The aliscore output appears to be empty. Check results for BUSCO: 917326at2759
## Mon Aug  8 09:49:02 CEST 2022 - The aliscore output appears to be empty. Check results for BUSCO: 996662at2759
## Mon Aug  8 09:49:02 CEST 2022 - The aliscore output file does not exist. Check results for BUSCO: 1001705at2759
## Mon Aug  8 09:49:02 CEST 2022 - The aliscore output file does not exist. Check results for BUSCO: 1441854at2759
## Mon Aug  8 09:49:02 CEST 2022 - The aliscore output appears to be empty. Check results for BUSCO: 1001705at2759
## Mon Aug  8 09:49:02 CEST 2022 - The aliscore output appears to be empty. Check results for BUSCO: 1441854at2759
## Mon Aug  8 09:49:02 CEST 2022 - The aliscore output file does not exist. Check results for BUSCO: 719531at2759
## Mon Aug  8 09:49:02 CEST 2022 - The aliscore output appears to be empty. Check results for BUSCO: 719531at2759
## Mon Aug  8 09:49:02 CEST 2022 - The aliscore output file does not exist. Check results for BUSCO: 1260807at2759
## Mon Aug  8 09:49:02 CEST 2022 - The aliscore output appears to be empty. Check results for BUSCO: 1260807at2759
## Mon Aug  8 09:49:02 CEST 2022 - The aliscore output file does not exist. Check results for BUSCO: 1236198at2759
## Mon Aug  8 09:49:02 CEST 2022 - The aliscore output appears to be empty. Check results for BUSCO: 1236198at2759
## Mon Aug  8 09:49:02 CEST 2022 - The aliscore output file does not exist. Check results for BUSCO: 1445102at2759
## Mon Aug  8 09:49:02 CEST 2022 - The aliscore output appears to be empty. Check results for BUSCO: 1445102at2759
## Mon Aug  8 09:49:02 CEST 2022 - The aliscore output file does not exist. Check results for BUSCO: 734666at2759
## Mon Aug  8 09:49:02 CEST 2022 - The aliscore output appears to be empty. Check results for BUSCO: 734666at2759
## Mon Aug  8 09:49:02 CEST 2022 - The aliscore output file does not exist. Check results for BUSCO: 1079827at2759
## Mon Aug  8 09:49:03 CEST 2022 - The aliscore output file does not exist. Check results for BUSCO: 1090038at2759
## Mon Aug  8 09:49:03 CEST 2022 - The aliscore output file does not exist. Check results for BUSCO: 1257440at2759
## Mon Aug  8 09:49:03 CEST 2022 - The aliscore output appears to be empty. Check results for BUSCO: 1079827at2759
## Mon Aug  8 09:49:03 CEST 2022 - The aliscore output appears to be empty. Check results for BUSCO: 1090038at2759
## Mon Aug  8 09:49:03 CEST 2022 - The aliscore output file does not exist. Check results for BUSCO: 855834at2759
## Mon Aug  8 09:49:03 CEST 2022 - The aliscore output appears to be empty. Check results for BUSCO: 1257440at2759
## Mon Aug  8 09:49:03 CEST 2022 - The aliscore output appears to be empty. Check results for BUSCO: 855834at2759
## Mon Aug  8 09:49:03 CEST 2022 - The aliscore output file does not exist. Check results for BUSCO: 1324510at2759
## Mon Aug  8 09:49:03 CEST 2022 - The aliscore output appears to be empty. Check results for BUSCO: 1324510at2759
## Mon Aug  8 09:49:03 CEST 2022 - The aliscore output file does not exist. Check results for BUSCO: 1014314at2759
## Mon Aug  8 09:49:03 CEST 2022 - The aliscore output file does not exist. Check results for BUSCO: 1030907at2759
## Mon Aug  8 09:49:03 CEST 2022 - The aliscore output file does not exist. Check results for BUSCO: 330169at2759
## Mon Aug  8 09:49:03 CEST 2022 - The aliscore output file does not exist. Check results for BUSCO: 1428265at2759
## Mon Aug  8 09:49:03 CEST 2022 - The aliscore output appears to be empty. Check results for BUSCO: 1030907at2759
## Mon Aug  8 09:49:03 CEST 2022 - The aliscore output appears to be empty. Check results for BUSCO: 330169at2759
## Mon Aug  8 09:49:03 CEST 2022 - The aliscore output appears to be empty. Check results for BUSCO: 1014314at2759
## Mon Aug  8 09:49:03 CEST 2022 - The aliscore output file does not exist. Check results for BUSCO: 1112002at2759
## Mon Aug  8 09:49:03 CEST 2022 - The aliscore output appears to be empty. Check results for BUSCO: 1428265at2759
## Mon Aug  8 09:49:03 CEST 2022 - The aliscore output file does not exist. Check results for BUSCO: 1355894at2759
## Mon Aug  8 09:49:03 CEST 2022 - The aliscore output file does not exist. Check results for BUSCO: 1173229at2759
## Mon Aug  8 09:49:03 CEST 2022 - The aliscore output appears to be empty. Check results for BUSCO: 1112002at2759
## Mon Aug  8 09:49:03 CEST 2022 - The aliscore output appears to be empty. Check results for BUSCO: 1355894at2759
## Mon Aug  8 09:49:03 CEST 2022 - The aliscore output appears to be empty. Check results for BUSCO: 1173229at2759
## Mon Aug  8 09:49:03 CEST 2022 - The aliscore output file does not exist. Check results for BUSCO: 1430056at2759
## Mon Aug  8 09:49:03 CEST 2022 - The aliscore output file does not exist. Check results for BUSCO: 296129at2759
## Mon Aug  8 09:49:03 CEST 2022 - The aliscore output appears to be empty. Check results for BUSCO: 296129at2759
## Mon Aug  8 09:49:03 CEST 2022 - The aliscore output appears to be empty. Check results for BUSCO: 1430056at2759
## Mon Aug  8 09:49:03 CEST 2022 - The aliscore output file does not exist. Check results for BUSCO: 858842at2759
## Mon Aug  8 09:49:03 CEST 2022 - The aliscore output appears to be empty. Check results for BUSCO: 858842at2759
## Mon Aug  8 09:49:03 CEST 2022 - The aliscore output file does not exist. Check results for BUSCO: 975158at2759
## Mon Aug  8 09:49:03 CEST 2022 - The aliscore output file does not exist. Check results for BUSCO: 1348942at2759
## Mon Aug  8 09:49:03 CEST 2022 - The aliscore output appears to be empty. Check results for BUSCO: 975158at2759
## Mon Aug  8 09:49:03 CEST 2022 - The aliscore output appears to be empty. Check results for BUSCO: 1348942at2759
## Mon Aug  8 09:49:03 CEST 2022 - The aliscore output file does not exist. Check results for BUSCO: 1106766at2759
## Mon Aug  8 09:49:03 CEST 2022 - The aliscore output appears to be empty. Check results for BUSCO: 1106766at2759
## Mon Aug  8 09:49:03 CEST 2022 - The aliscore output file does not exist. Check results for BUSCO: 1370285at2759
## Mon Aug  8 09:49:04 CEST 2022 - The aliscore output appears to be empty. Check results for BUSCO: 1370285at2759
## Mon Aug  8 09:49:04 CEST 2022 - The aliscore output file does not exist. Check results for BUSCO: 1217666at2759
## Mon Aug  8 09:49:04 CEST 2022 - The aliscore output appears to be empty. Check results for BUSCO: 1217666at2759
## Mon Aug  8 09:49:04 CEST 2022 - The aliscore output file does not exist. Check results for BUSCO: 1346165at2759
## Mon Aug  8 09:49:04 CEST 2022 - The aliscore output appears to be empty. Check results for BUSCO: 1346165at2759
## Mon Aug  8 09:49:04 CEST 2022 - The aliscore output file does not exist. Check results for BUSCO: 973442at2759
## Mon Aug  8 09:49:04 CEST 2022 - The aliscore output file does not exist. Check results for BUSCO: 1108845at2759
## Mon Aug  8 09:49:04 CEST 2022 - The aliscore output appears to be empty. Check results for BUSCO: 973442at2759
## Mon Aug  8 09:49:04 CEST 2022 - The aliscore output appears to be empty. Check results for BUSCO: 1108845at2759
## Mon Aug  8 09:49:04 CEST 2022 - The aliscore output file does not exist. Check results for BUSCO: 975158at2759
## Mon Aug  8 09:49:04 CEST 2022 - The aliscore output appears to be empty. Check results for BUSCO: 975158at2759
## Mon Aug  8 09:49:05 CEST 2022 - The aliscore output file does not exist. Check results for BUSCO: 1041560at2759
## Mon Aug  8 09:49:05 CEST 2022 - The aliscore output appears to be empty. Check results for BUSCO: 1041560at2759
## Mon Aug  8 09:49:05 CEST 2022 - The aliscore output file does not exist. Check results for BUSCO: 996662at2759
## Mon Aug  8 09:49:05 CEST 2022 - The aliscore output appears to be empty. Check results for BUSCO: 996662at2759
## Mon Aug  8 09:49:05 CEST 2022 - The aliscore output file does not exist. Check results for BUSCO: 1455730at2759
## Mon Aug  8 09:49:05 CEST 2022 - The aliscore output file does not exist. Check results for BUSCO: 1355894at2759
## Mon Aug  8 09:49:05 CEST 2022 - The aliscore output file does not exist. Check results for BUSCO: 671846at2759
## Mon Aug  8 09:49:05 CEST 2022 - The aliscore output file does not exist. Check results for BUSCO: 878143at2759
## Mon Aug  8 09:49:05 CEST 2022 - The aliscore output appears to be empty. Check results for BUSCO: 1355894at2759
## Mon Aug  8 09:49:05 CEST 2022 - The aliscore output appears to be empty. Check results for BUSCO: 671846at2759
## Mon Aug  8 09:49:05 CEST 2022 - The aliscore output file does not exist. Check results for BUSCO: 245208at2759
## Mon Aug  8 09:49:05 CEST 2022 - The aliscore output file does not exist. Check results for BUSCO: 834694at2759
## Mon Aug  8 09:49:05 CEST 2022 - The aliscore output appears to be empty. Check results for BUSCO: 878143at2759
## Mon Aug  8 09:49:05 CEST 2022 - The aliscore output file does not exist. Check results for BUSCO: 1217666at2759
## Mon Aug  8 09:49:05 CEST 2022 - The aliscore output appears to be empty. Check results for BUSCO: 245208at2759
## Mon Aug  8 09:49:05 CEST 2022 - The aliscore output appears to be empty. Check results for BUSCO: 834694at2759
## Mon Aug  8 09:49:05 CEST 2022 - The aliscore output file does not exist. Check results for BUSCO: 1455730at2759
## Mon Aug  8 09:49:05 CEST 2022 - The aliscore output appears to be empty. Check results for BUSCO: 1455730at2759
## Mon Aug  8 09:49:05 CEST 2022 - The aliscore output appears to be empty. Check results for BUSCO: 1455730at2759
## Mon Aug  8 09:49:05 CEST 2022 - The aliscore output file does not exist. Check results for BUSCO: 1041560at2759
## Mon Aug  8 09:49:05 CEST 2022 - The aliscore output file does not exist. Check results for BUSCO: 1379600at2759
## Mon Aug  8 09:49:05 CEST 2022 - The aliscore output appears to be empty. Check results for BUSCO: 1041560at2759
## Mon Aug  8 09:49:05 CEST 2022 - The aliscore output appears to be empty. Check results for BUSCO: 1379600at2759
## Mon Aug  8 09:49:05 CEST 2022 - The aliscore output appears to be empty. Check results for BUSCO: 1217666at2759
## Mon Aug  8 09:49:05 CEST 2022 - The aliscore output file does not exist. Check results for BUSCO: 1450538at2759
## Mon Aug  8 09:49:05 CEST 2022 - The aliscore output file does not exist. Check results for BUSCO: 917326at2759
## Mon Aug  8 09:49:05 CEST 2022 - The aliscore output appears to be empty. Check results for BUSCO: 1450538at2759
## Mon Aug  8 09:49:05 CEST 2022 - The aliscore output appears to be empty. Check results for BUSCO: 917326at2759
## Mon Aug  8 09:49:06 CEST 2022 - The aliscore output file does not exist. Check results for BUSCO: 179362at2759
## Mon Aug  8 09:49:06 CEST 2022 - The aliscore output appears to be empty. Check results for BUSCO: 179362at2759
## Mon Aug  8 09:49:06 CEST 2022 - The aliscore output file does not exist. Check results for BUSCO: 306227at2759
## Mon Aug  8 09:49:06 CEST 2022 - The aliscore output file does not exist. Check results for BUSCO: 722805at2759
## Mon Aug  8 09:49:06 CEST 2022 - The aliscore output appears to be empty. Check results for BUSCO: 306227at2759
## Mon Aug  8 09:49:06 CEST 2022 - The aliscore output appears to be empty. Check results for BUSCO: 722805at2759
## Mon Aug  8 09:49:07 CEST 2022 - The aliscore output file does not exist. Check results for BUSCO: 390348at2759
## Mon Aug  8 09:49:07 CEST 2022 - The aliscore output appears to be empty. Check results for BUSCO: 390348at2759
## Mon Aug  8 09:49:07 CEST 2022 - The aliscore output file does not exist. Check results for BUSCO: 1423847at2759
## Mon Aug  8 09:49:07 CEST 2022 - The aliscore output appears to be empty. Check results for BUSCO: 1423847at2759
## Mon Aug  8 09:49:07 CEST 2022 - The aliscore output file does not exist. Check results for BUSCO: 722805at2759
## Mon Aug  8 09:49:07 CEST 2022 - The aliscore output appears to be empty. Check results for BUSCO: 722805at2759
## Mon Aug  8 09:49:08 CEST 2022 - The aliscore output file does not exist. Check results for BUSCO: 1108845at2759
## Mon Aug  8 09:49:08 CEST 2022 - The aliscore output appears to be empty. Check results for BUSCO: 1108845at2759
## Mon Aug  8 09:49:08 CEST 2022 - The aliscore output file does not exist. Check results for BUSCO: 245208at2759
## Mon Aug  8 09:49:08 CEST 2022 - The aliscore output appears to be empty. Check results for BUSCO: 245208at2759
## Mon Aug  8 09:49:08 CEST 2022 - The aliscore output file does not exist. Check results for BUSCO: 97116at2759
## Mon Aug  8 09:49:08 CEST 2022 - The aliscore output appears to be empty. Check results for BUSCO: 97116at2759
## Mon Aug  8 09:49:08 CEST 2022 - The aliscore output file does not exist. Check results for BUSCO: 551907at2759
## Mon Aug  8 09:49:08 CEST 2022 - The aliscore output appears to be empty. Check results for BUSCO: 551907at2759
## Mon Aug  8 09:49:09 CEST 2022 - The aliscore output file does not exist. Check results for BUSCO: 396755at2759
## Mon Aug  8 09:49:09 CEST 2022 - The aliscore output appears to be empty. Check results for BUSCO: 396755at2759
## Mon Aug  8 09:49:09 CEST 2022 - The aliscore output file does not exist. Check results for BUSCO: 396755at2759
## Mon Aug  8 09:49:09 CEST 2022 - The aliscore output appears to be empty. Check results for BUSCO: 396755at2759
## Mon Aug  8 09:49:22 CEST 2022 - The aliscore output file does not exist. Check results for BUSCO: 156083at2759
## Mon Aug  8 09:49:22 CEST 2022 - The aliscore output appears to be empty. Check results for BUSCO: 156083at2759
## Mon Aug  8 09:49:28 CEST 2022 - The aliscore output file does not exist. Check results for BUSCO: 1620056at2759
## Mon Aug  8 09:49:28 CEST 2022 - The aliscore output appears to be empty. Check results for BUSCO: 1620056at2759
## Mon Aug  8 09:49:30 CEST 2022 - The aliscore output file does not exist. Check results for BUSCO: 1545004at2759
## Mon Aug  8 09:49:30 CEST 2022 - The aliscore output appears to be empty. Check results for BUSCO: 1545004at2759
## Mon Aug  8 09:49:30 CEST 2022 - The aliscore output file does not exist. Check results for BUSCO: 1284731at2759
## Mon Aug  8 09:49:30 CEST 2022 - The aliscore output appears to be empty. Check results for BUSCO: 1284731at2759
## Mon Aug  8 09:49:30 CEST 2022 - The aliscore output file does not exist. Check results for BUSCO: 1488436at2759
## Mon Aug  8 09:49:30 CEST 2022 - The aliscore output appears to be empty. Check results for BUSCO: 1488436at2759
## Mon Aug  8 09:49:30 CEST 2022 - The aliscore output file does not exist. Check results for BUSCO: 176625at2759
## Mon Aug  8 09:49:30 CEST 2022 - The aliscore output appears to be empty. Check results for BUSCO: 176625at2759
## Mon Aug  8 09:49:30 CEST 2022 - The aliscore output file does not exist. Check results for BUSCO: 1525971at2759
## Mon Aug  8 09:49:31 CEST 2022 - The aliscore output appears to be empty. Check results for BUSCO: 1525971at2759
## Mon Aug  8 09:49:31 CEST 2022 - The aliscore output file does not exist. Check results for BUSCO: 293315at2759
## Mon Aug  8 09:49:31 CEST 2022 - The aliscore output appears to be empty. Check results for BUSCO: 293315at2759
## Mon Aug  8 09:49:31 CEST 2022 - The aliscore output file does not exist. Check results for BUSCO: 1504863at2759
## Mon Aug  8 09:49:31 CEST 2022 - The aliscore output file does not exist. Check results for BUSCO: 976469at2759
## Mon Aug  8 09:49:31 CEST 2022 - The aliscore output appears to be empty. Check results for BUSCO: 1504863at2759
## Mon Aug  8 09:49:31 CEST 2022 - The aliscore output appears to be empty. Check results for BUSCO: 976469at2759
## Mon Aug  8 09:49:31 CEST 2022 - The aliscore output file does not exist. Check results for BUSCO: 1488235at2759
## Mon Aug  8 09:49:31 CEST 2022 - The aliscore output appears to be empty. Check results for BUSCO: 1488235at2759
## Mon Aug  8 09:49:31 CEST 2022 - The aliscore output file does not exist. Check results for BUSCO: 1538526at2759
## Mon Aug  8 09:49:31 CEST 2022 - The aliscore output appears to be empty. Check results for BUSCO: 1538526at2759
## Mon Aug  8 09:49:32 CEST 2022 - The aliscore output file does not exist. Check results for BUSCO: 1626636at2759
## Mon Aug  8 09:49:32 CEST 2022 - The aliscore output file does not exist. Check results for BUSCO: 1370304at2759
## Mon Aug  8 09:49:32 CEST 2022 - The aliscore output appears to be empty. Check results for BUSCO: 1626636at2759
## Mon Aug  8 09:49:32 CEST 2022 - The aliscore output appears to be empty. Check results for BUSCO: 1370304at2759
## Mon Aug  8 09:49:32 CEST 2022 - The aliscore output file does not exist. Check results for BUSCO: 946128at2759
## Mon Aug  8 09:49:32 CEST 2022 - The aliscore output appears to be empty. Check results for BUSCO: 946128at2759
## Mon Aug  8 09:49:32 CEST 2022 - The aliscore output file does not exist. Check results for BUSCO: 1426075at2759
## Mon Aug  8 09:49:32 CEST 2022 - The aliscore output file does not exist. Check results for BUSCO: 655400at2759
## Mon Aug  8 09:49:32 CEST 2022 - The aliscore output appears to be empty. Check results for BUSCO: 1426075at2759
## Mon Aug  8 09:49:33 CEST 2022 - The aliscore output appears to be empty. Check results for BUSCO: 655400at2759
## Mon Aug  8 09:49:33 CEST 2022 - The aliscore output file does not exist. Check results for BUSCO: 1322299at2759
## Mon Aug  8 09:49:33 CEST 2022 - The aliscore output appears to be empty. Check results for BUSCO: 1322299at2759
## Mon Aug  8 09:49:33 CEST 2022 - The aliscore output file does not exist. Check results for BUSCO: 598949at2759
## Mon Aug  8 09:49:33 CEST 2022 - The aliscore output appears to be empty. Check results for BUSCO: 598949at2759
## Mon Aug  8 09:49:33 CEST 2022 - The aliscore output file does not exist. Check results for BUSCO: 1338131at2759
## Mon Aug  8 09:49:33 CEST 2022 - The aliscore output appears to be empty. Check results for BUSCO: 1338131at2759
## Mon Aug  8 09:49:33 CEST 2022 - The aliscore output file does not exist. Check results for BUSCO: 865202at2759
## Mon Aug  8 09:49:34 CEST 2022 - The aliscore output appears to be empty. Check results for BUSCO: 865202at2759
## Mon Aug  8 09:49:34 CEST 2022 - The aliscore output file does not exist. Check results for BUSCO: 1377237at2759
## Mon Aug  8 09:49:34 CEST 2022 - The aliscore output appears to be empty. Check results for BUSCO: 1377237at2759
## Mon Aug  8 09:49:34 CEST 2022 - The aliscore output file does not exist. Check results for BUSCO: 1220881at2759
## Mon Aug  8 09:49:34 CEST 2022 - The aliscore output appears to be empty. Check results for BUSCO: 1220881at2759
## Mon Aug  8 09:49:34 CEST 2022 - The aliscore output file does not exist. Check results for BUSCO: 513979at2759
## Mon Aug  8 09:49:34 CEST 2022 - The aliscore output file does not exist. Check results for BUSCO: 1563319at2759
## Mon Aug  8 09:49:34 CEST 2022 - The aliscore output file does not exist. Check results for BUSCO: 355408at2759
## Mon Aug  8 09:49:34 CEST 2022 - The aliscore output appears to be empty. Check results for BUSCO: 513979at2759
## Mon Aug  8 09:49:34 CEST 2022 - The aliscore output file does not exist. Check results for BUSCO: 392369at2759
## Mon Aug  8 09:49:34 CEST 2022 - The aliscore output appears to be empty. Check results for BUSCO: 1563319at2759
## Mon Aug  8 09:49:34 CEST 2022 - The aliscore output appears to be empty. Check results for BUSCO: 392369at2759
## Mon Aug  8 09:49:34 CEST 2022 - The aliscore output appears to be empty. Check results for BUSCO: 355408at2759
## Mon Aug  8 09:49:34 CEST 2022 - The aliscore output file does not exist. Check results for BUSCO: 570797at2759
## Mon Aug  8 09:49:34 CEST 2022 - The aliscore output appears to be empty. Check results for BUSCO: 570797at2759
## Mon Aug  8 09:49:35 CEST 2022 - The aliscore output file does not exist. Check results for BUSCO: 887370at2759
## Mon Aug  8 09:49:35 CEST 2022 - The aliscore output appears to be empty. Check results for BUSCO: 887370at2759
## Mon Aug  8 09:49:35 CEST 2022 - The aliscore output file does not exist. Check results for BUSCO: 1003258at2759
## Mon Aug  8 09:49:35 CEST 2022 - The aliscore output appears to be empty. Check results for BUSCO: 1003258at2759
## Mon Aug  8 09:49:35 CEST 2022 - The aliscore output file does not exist. Check results for BUSCO: 1513531at2759
## Mon Aug  8 09:49:35 CEST 2022 - The aliscore output appears to be empty. Check results for BUSCO: 1513531at2759
## Mon Aug  8 09:49:35 CEST 2022 - The aliscore output file does not exist. Check results for BUSCO: 1138059at2759
## Mon Aug  8 09:49:35 CEST 2022 - The aliscore output appears to be empty. Check results for BUSCO: 1138059at2759
## Mon Aug  8 09:49:35 CEST 2022 - The aliscore output file does not exist. Check results for BUSCO: 1576404at2759
## Mon Aug  8 09:49:35 CEST 2022 - The aliscore output file does not exist. Check results for BUSCO: 736068at2759
## Mon Aug  8 09:49:35 CEST 2022 - The aliscore output appears to be empty. Check results for BUSCO: 736068at2759
## Mon Aug  8 09:49:35 CEST 2022 - The aliscore output appears to be empty. Check results for BUSCO: 1576404at2759
## Mon Aug  8 09:49:36 CEST 2022 - The aliscore output file does not exist. Check results for BUSCO: 1339553at2759
## Mon Aug  8 09:49:36 CEST 2022 - The aliscore output file does not exist. Check results for BUSCO: 814241at2759
## Mon Aug  8 09:49:36 CEST 2022 - The aliscore output appears to be empty. Check results for BUSCO: 814241at2759
## Mon Aug  8 09:49:36 CEST 2022 - The aliscore output appears to be empty. Check results for BUSCO: 1339553at2759
## Mon Aug  8 09:49:36 CEST 2022 - The aliscore output file does not exist. Check results for BUSCO: 1111142at2759
## Mon Aug  8 09:49:36 CEST 2022 - The aliscore output appears to be empty. Check results for BUSCO: 1111142at2759
## Mon Aug  8 09:49:36 CEST 2022 - The aliscore output file does not exist. Check results for BUSCO: 1018517at2759
## Mon Aug  8 09:49:36 CEST 2022 - The aliscore output appears to be empty. Check results for BUSCO: 1018517at2759
## Mon Aug  8 09:49:37 CEST 2022 - The aliscore output file does not exist. Check results for BUSCO: 898782at2759
## Mon Aug  8 09:49:37 CEST 2022 - The aliscore output appears to be empty. Check results for BUSCO: 898782at2759
## Mon Aug  8 09:49:37 CEST 2022 - The aliscore output file does not exist. Check results for BUSCO: 1087488at2759
## Mon Aug  8 09:49:37 CEST 2022 - The aliscore output appears to be empty. Check results for BUSCO: 1087488at2759
## Mon Aug  8 09:49:38 CEST 2022 - The aliscore output file does not exist. Check results for BUSCO: 1314980at2759
## Mon Aug  8 09:49:38 CEST 2022 - The aliscore output file does not exist. Check results for BUSCO: 891751at2759
## Mon Aug  8 09:49:38 CEST 2022 - The aliscore output appears to be empty. Check results for BUSCO: 891751at2759
## Mon Aug  8 09:49:38 CEST 2022 - The aliscore output file does not exist. Check results for BUSCO: 1312453at2759
## Mon Aug  8 09:49:38 CEST 2022 - The aliscore output appears to be empty. Check results for BUSCO: 1314980at2759
## Mon Aug  8 09:49:38 CEST 2022 - The aliscore output appears to be empty. Check results for BUSCO: 1312453at2759
## Mon Aug  8 09:49:59 CEST 2022 - The aliscore output file does not exist. Check results for BUSCO: 1053181at2759
## Mon Aug  8 09:49:59 CEST 2022 - The aliscore output appears to be empty. Check results for BUSCO: 1053181at2759
## Mon Aug  8 09:50:00 CEST 2022 - The aliscore output file does not exist. Check results for BUSCO: 1200489at2759
## Mon Aug  8 09:50:00 CEST 2022 - The aliscore output appears to be empty. Check results for BUSCO: 1200489at2759
## Mon Aug  8 09:50:00 CEST 2022 - The aliscore output file does not exist. Check results for BUSCO: 905026at2759
## Mon Aug  8 09:50:00 CEST 2022 - The aliscore output appears to be empty. Check results for BUSCO: 905026at2759
## Mon Aug  8 09:50:00 CEST 2022 - The aliscore output file does not exist. Check results for BUSCO: 1454155at2759
## Mon Aug  8 09:50:00 CEST 2022 - The aliscore output file does not exist. Check results for BUSCO: 1269649at2759
## Mon Aug  8 09:50:00 CEST 2022 - The aliscore output file does not exist. Check results for BUSCO: 1003258at2759
## Mon Aug  8 09:50:00 CEST 2022 - The aliscore output appears to be empty. Check results for BUSCO: 1269649at2759
## Mon Aug  8 09:50:00 CEST 2022 - The aliscore output appears to be empty. Check results for BUSCO: 1003258at2759
## Mon Aug  8 09:50:00 CEST 2022 - The aliscore output appears to be empty. Check results for BUSCO: 1454155at2759
## Mon Aug  8 09:50:00 CEST 2022 - The aliscore output file does not exist. Check results for BUSCO: 1269244at2759
## Mon Aug  8 09:50:00 CEST 2022 - The aliscore output file does not exist. Check results for BUSCO: 1287401at2759
## Mon Aug  8 09:50:00 CEST 2022 - The aliscore output appears to be empty. Check results for BUSCO: 1269244at2759
## Mon Aug  8 09:50:00 CEST 2022 - The aliscore output file does not exist. Check results for BUSCO: 1395649at2759
## Mon Aug  8 09:50:00 CEST 2022 - The aliscore output appears to be empty. Check results for BUSCO: 1287401at2759
## Mon Aug  8 09:50:00 CEST 2022 - The aliscore output appears to be empty. Check results for BUSCO: 1395649at2759
## Mon Aug  8 09:50:01 CEST 2022 - The aliscore output file does not exist. Check results for BUSCO: 869548at2759
## Mon Aug  8 09:50:01 CEST 2022 - The aliscore output file does not exist. Check results for BUSCO: 1525971at2759
## Mon Aug  8 09:50:01 CEST 2022 - The aliscore output appears to be empty. Check results for BUSCO: 869548at2759
## Mon Aug  8 09:50:01 CEST 2022 - The aliscore output appears to be empty. Check results for BUSCO: 1525971at2759
## Mon Aug  8 09:50:01 CEST 2022 - The aliscore output file does not exist. Check results for BUSCO: 1222562at2759
## Mon Aug  8 09:50:01 CEST 2022 - The aliscore output appears to be empty. Check results for BUSCO: 1222562at2759
## Mon Aug  8 09:50:01 CEST 2022 - The aliscore output file does not exist. Check results for BUSCO: 869548at2759
## Mon Aug  8 09:50:01 CEST 2022 - The aliscore output file does not exist. Check results for BUSCO: 1258856at2759
## Mon Aug  8 09:50:01 CEST 2022 - The aliscore output appears to be empty. Check results for BUSCO: 869548at2759
## Mon Aug  8 09:50:01 CEST 2022 - The aliscore output file does not exist. Check results for BUSCO: 1269649at2759
## Mon Aug  8 09:50:01 CEST 2022 - The aliscore output appears to be empty. Check results for BUSCO: 1258856at2759
## Mon Aug  8 09:50:01 CEST 2022 - The aliscore output appears to be empty. Check results for BUSCO: 1269649at2759
## Mon Aug  8 09:50:02 CEST 2022 - The aliscore output file does not exist. Check results for BUSCO: 1593937at2759
## Mon Aug  8 09:50:02 CEST 2022 - The aliscore output appears to be empty. Check results for BUSCO: 1593937at2759
## Mon Aug  8 09:50:02 CEST 2022 - The aliscore output file does not exist. Check results for BUSCO: 956854at2759
## Mon Aug  8 09:50:02 CEST 2022 - The aliscore output appears to be empty. Check results for BUSCO: 956854at2759
## Mon Aug  8 09:50:03 CEST 2022 - The aliscore output file does not exist. Check results for BUSCO: 768809at2759
## Mon Aug  8 09:50:03 CEST 2022 - The aliscore output appears to be empty. Check results for BUSCO: 768809at2759
## Mon Aug  8 09:50:03 CEST 2022 - The aliscore output file does not exist. Check results for BUSCO: 1266231at2759
## Mon Aug  8 09:50:03 CEST 2022 - The aliscore output file does not exist. Check results for BUSCO: 176625at2759
## Mon Aug  8 09:50:03 CEST 2022 - The aliscore output file does not exist. Check results for BUSCO: 1538526at2759
## Mon Aug  8 09:50:03 CEST 2022 - The aliscore output appears to be empty. Check results for BUSCO: 1266231at2759
## Mon Aug  8 09:50:03 CEST 2022 - The aliscore output appears to be empty. Check results for BUSCO: 176625at2759
## Mon Aug  8 09:50:03 CEST 2022 - The aliscore output appears to be empty. Check results for BUSCO: 1538526at2759
## Mon Aug  8 09:50:03 CEST 2022 - The aliscore output file does not exist. Check results for BUSCO: 1223488at2759
## Mon Aug  8 09:50:03 CEST 2022 - The aliscore output file does not exist. Check results for BUSCO: 946128at2759
## Mon Aug  8 09:50:03 CEST 2022 - The aliscore output appears to be empty. Check results for BUSCO: 1223488at2759
## Mon Aug  8 09:50:03 CEST 2022 - The aliscore output file does not exist. Check results for BUSCO: 1269244at2759
## Mon Aug  8 09:50:03 CEST 2022 - The aliscore output appears to be empty. Check results for BUSCO: 946128at2759
## Mon Aug  8 09:50:03 CEST 2022 - The aliscore output appears to be empty. Check results for BUSCO: 1269244at2759
## Mon Aug  8 09:50:03 CEST 2022 - The aliscore output file does not exist. Check results for BUSCO: 1019762at2759
## Mon Aug  8 09:50:03 CEST 2022 - The aliscore output file does not exist. Check results for BUSCO: 1049599at2759
## Mon Aug  8 09:50:03 CEST 2022 - The aliscore output file does not exist. Check results for BUSCO: 1620056at2759
## Mon Aug  8 09:50:03 CEST 2022 - The aliscore output file does not exist. Check results for BUSCO: 1193442at2759
## Mon Aug  8 09:50:03 CEST 2022 - The aliscore output appears to be empty. Check results for BUSCO: 1019762at2759
## Mon Aug  8 09:50:03 CEST 2022 - The aliscore output appears to be empty. Check results for BUSCO: 1049599at2759
## Mon Aug  8 09:50:03 CEST 2022 - The aliscore output appears to be empty. Check results for BUSCO: 1620056at2759
## Mon Aug  8 09:50:03 CEST 2022 - The aliscore output appears to be empty. Check results for BUSCO: 1193442at2759
## Mon Aug  8 09:50:03 CEST 2022 - The aliscore output file does not exist. Check results for BUSCO: 1421503at2759
## Mon Aug  8 09:50:03 CEST 2022 - The aliscore output appears to be empty. Check results for BUSCO: 1421503at2759
## Mon Aug  8 09:50:04 CEST 2022 - The aliscore output file does not exist. Check results for BUSCO: 905026at2759
## Mon Aug  8 09:50:04 CEST 2022 - The aliscore output appears to be empty. Check results for BUSCO: 905026at2759
## Mon Aug  8 09:50:04 CEST 2022 - The aliscore output file does not exist. Check results for BUSCO: 1107630at2759
## Mon Aug  8 09:50:04 CEST 2022 - The aliscore output file does not exist. Check results for BUSCO: 1342242at2759
## Mon Aug  8 09:50:04 CEST 2022 - The aliscore output appears to be empty. Check results for BUSCO: 1107630at2759
## Mon Aug  8 09:50:04 CEST 2022 - The aliscore output appears to be empty. Check results for BUSCO: 1342242at2759
## Mon Aug  8 09:50:04 CEST 2022 - The aliscore output file does not exist. Check results for BUSCO: 720340at2759
## Mon Aug  8 09:50:04 CEST 2022 - The aliscore output file does not exist. Check results for BUSCO: 1233814at2759
## Mon Aug  8 09:50:04 CEST 2022 - The aliscore output file does not exist. Check results for BUSCO: 679187at2759
## Mon Aug  8 09:50:04 CEST 2022 - The aliscore output appears to be empty. Check results for BUSCO: 720340at2759
## Mon Aug  8 09:50:04 CEST 2022 - The aliscore output appears to be empty. Check results for BUSCO: 1233814at2759
## Mon Aug  8 09:50:04 CEST 2022 - The aliscore output appears to be empty. Check results for BUSCO: 679187at2759
## Mon Aug  8 09:50:04 CEST 2022 - The aliscore output file does not exist. Check results for BUSCO: 1370304at2759
## Mon Aug  8 09:50:04 CEST 2022 - The aliscore output appears to be empty. Check results for BUSCO: 1370304at2759
## Mon Aug  8 09:50:05 CEST 2022 - The aliscore output file does not exist. Check results for BUSCO: 1339553at2759
## Mon Aug  8 09:50:05 CEST 2022 - The aliscore output file does not exist. Check results for BUSCO: 392369at2759
## Mon Aug  8 09:50:05 CEST 2022 - The aliscore output appears to be empty. Check results for BUSCO: 1339553at2759
## Mon Aug  8 09:50:05 CEST 2022 - The aliscore output appears to be empty. Check results for BUSCO: 392369at2759
## Mon Aug  8 09:50:05 CEST 2022 - The aliscore output file does not exist. Check results for BUSCO: 551907at2759
## Mon Aug  8 09:50:05 CEST 2022 - The aliscore output appears to be empty. Check results for BUSCO: 551907at2759
## Mon Aug  8 09:50:05 CEST 2022 - The aliscore output file does not exist. Check results for BUSCO: 1266231at2759
## Mon Aug  8 09:50:05 CEST 2022 - The aliscore output appears to be empty. Check results for BUSCO: 1266231at2759
## Mon Aug  8 09:50:05 CEST 2022 - The aliscore output file does not exist. Check results for BUSCO: 1404162at2759
## Mon Aug  8 09:50:05 CEST 2022 - The aliscore output appears to be empty. Check results for BUSCO: 1404162at2759
## Mon Aug  8 09:50:05 CEST 2022 - The aliscore output file does not exist. Check results for BUSCO: 751335at2759
## Mon Aug  8 09:50:05 CEST 2022 - The aliscore output appears to be empty. Check results for BUSCO: 751335at2759
## Mon Aug  8 09:50:06 CEST 2022 - The aliscore output file does not exist. Check results for BUSCO: 1018517at2759
## Mon Aug  8 09:50:06 CEST 2022 - The aliscore output file does not exist. Check results for BUSCO: 1032689at2759
## Mon Aug  8 09:50:06 CEST 2022 - The aliscore output appears to be empty. Check results for BUSCO: 1018517at2759
## Mon Aug  8 09:50:06 CEST 2022 - The aliscore output appears to be empty. Check results for BUSCO: 1032689at2759
## Mon Aug  8 09:50:06 CEST 2022 - The aliscore output file does not exist. Check results for BUSCO: 491869at2759
## Mon Aug  8 09:50:06 CEST 2022 - The aliscore output appears to be empty. Check results for BUSCO: 491869at2759
## Mon Aug  8 09:50:06 CEST 2022 - The aliscore output file does not exist. Check results for BUSCO: 1096688at2759
## Mon Aug  8 09:50:06 CEST 2022 - The aliscore output appears to be empty. Check results for BUSCO: 1096688at2759
## Mon Aug  8 09:50:07 CEST 2022 - The aliscore output file does not exist. Check results for BUSCO: 774318at2759
## Mon Aug  8 09:50:07 CEST 2022 - The aliscore output appears to be empty. Check results for BUSCO: 774318at2759
## Mon Aug  8 09:50:07 CEST 2022 - The aliscore output file does not exist. Check results for BUSCO: 1430056at2759
## Mon Aug  8 09:50:07 CEST 2022 - The aliscore output appears to be empty. Check results for BUSCO: 1430056at2759
## Mon Aug  8 09:50:07 CEST 2022 - The aliscore output file does not exist. Check results for BUSCO: 1019762at2759
## Mon Aug  8 09:50:07 CEST 2022 - The aliscore output file does not exist. Check results for BUSCO: 1432328at2759
## Mon Aug  8 09:50:07 CEST 2022 - The aliscore output appears to be empty. Check results for BUSCO: 1019762at2759
## Mon Aug  8 09:50:07 CEST 2022 - The aliscore output appears to be empty. Check results for BUSCO: 1432328at2759
## Mon Aug  8 09:50:07 CEST 2022 - The aliscore output file does not exist. Check results for BUSCO: 1259741at2759
## Mon Aug  8 09:50:07 CEST 2022 - The aliscore output appears to be empty. Check results for BUSCO: 1259741at2759
## Mon Aug  8 09:50:09 CEST 2022 - The aliscore output file does not exist. Check results for BUSCO: 878143at2759
## Mon Aug  8 09:50:09 CEST 2022 - The aliscore output file does not exist. Check results for BUSCO: 1193442at2759
## Mon Aug  8 09:50:09 CEST 2022 - The aliscore output appears to be empty. Check results for BUSCO: 1193442at2759
## Mon Aug  8 09:50:09 CEST 2022 - The aliscore output appears to be empty. Check results for BUSCO: 878143at2759
## Mon Aug  8 09:50:10 CEST 2022 - The aliscore output file does not exist. Check results for BUSCO: 1404162at2759
## Mon Aug  8 09:50:10 CEST 2022 - The aliscore output appears to be empty. Check results for BUSCO: 1404162at2759
## Mon Aug  8 09:50:10 CEST 2022 - The aliscore output file does not exist. Check results for BUSCO: 1428854at2759
## Mon Aug  8 09:50:10 CEST 2022 - The aliscore output appears to be empty. Check results for BUSCO: 1428854at2759
## Mon Aug  8 09:50:10 CEST 2022 - The aliscore output file does not exist. Check results for BUSCO: 513979at2759
## Mon Aug  8 09:50:10 CEST 2022 - The aliscore output appears to be empty. Check results for BUSCO: 513979at2759
## Mon Aug  8 09:50:11 CEST 2022 - The aliscore output file does not exist. Check results for BUSCO: 160593at2759
## Mon Aug  8 09:50:11 CEST 2022 - The aliscore output appears to be empty. Check results for BUSCO: 160593at2759
## Mon Aug  8 09:50:11 CEST 2022 - The aliscore output file does not exist. Check results for BUSCO: 931188at2759
## Mon Aug  8 09:50:11 CEST 2022 - The aliscore output appears to be empty. Check results for BUSCO: 931188at2759
## Mon Aug  8 09:50:11 CEST 2022 - The aliscore output file does not exist. Check results for BUSCO: 674160at2759
## Mon Aug  8 09:50:11 CEST 2022 - The aliscore output appears to be empty. Check results for BUSCO: 674160at2759
## Mon Aug  8 09:50:12 CEST 2022 - The aliscore output file does not exist. Check results for BUSCO: 1423847at2759
## Mon Aug  8 09:50:12 CEST 2022 - The aliscore output appears to be empty. Check results for BUSCO: 1423847at2759
## Mon Aug  8 09:50:25 CEST 2022 - The aliscore output file does not exist. Check results for BUSCO: 1633672at2759
## Mon Aug  8 09:50:25 CEST 2022 - The aliscore output appears to be empty. Check results for BUSCO: 1633672at2759
## Mon Aug  8 09:50:26 CEST 2022 - The aliscore output file does not exist. Check results for BUSCO: 1405073at2759
## Mon Aug  8 09:50:26 CEST 2022 - The aliscore output appears to be empty. Check results for BUSCO: 1405073at2759
## Mon Aug  8 09:50:29 CEST 2022 - The aliscore output file does not exist. Check results for BUSCO: 375960at2759
## Mon Aug  8 09:50:29 CEST 2022 - The aliscore output appears to be empty. Check results for BUSCO: 375960at2759
## Mon Aug  8 09:50:31 CEST 2022 - The aliscore output file does not exist. Check results for BUSCO: 679771at2759
## Mon Aug  8 09:50:31 CEST 2022 - The aliscore output appears to be empty. Check results for BUSCO: 679771at2759
## Mon Aug  8 09:50:31 CEST 2022 - The aliscore output file does not exist. Check results for BUSCO: 1617752at2759
## Mon Aug  8 09:50:31 CEST 2022 - The aliscore output file does not exist. Check results for BUSCO: 1161199at2759
## Mon Aug  8 09:50:31 CEST 2022 - The aliscore output appears to be empty. Check results for BUSCO: 1617752at2759
## Mon Aug  8 09:50:31 CEST 2022 - The aliscore output appears to be empty. Check results for BUSCO: 1161199at2759
## Mon Aug  8 09:50:31 CEST 2022 - The aliscore output file does not exist. Check results for BUSCO: 1065019at2759
## Mon Aug  8 09:50:32 CEST 2022 - The aliscore output appears to be empty. Check results for BUSCO: 1065019at2759
## Mon Aug  8 09:50:33 CEST 2022 - The aliscore output file does not exist. Check results for BUSCO: 1474000at2759
## Mon Aug  8 09:50:33 CEST 2022 - The aliscore output appears to be empty. Check results for BUSCO: 1474000at2759
## Mon Aug  8 09:50:34 CEST 2022 - The aliscore output file does not exist. Check results for BUSCO: 1248958at2759
## Mon Aug  8 09:50:34 CEST 2022 - The aliscore output appears to be empty. Check results for BUSCO: 1248958at2759
## Mon Aug  8 09:50:34 CEST 2022 - The aliscore output file does not exist. Check results for BUSCO: 331411at2759
## Mon Aug  8 09:50:34 CEST 2022 - The aliscore output appears to be empty. Check results for BUSCO: 331411at2759
## Mon Aug  8 09:50:35 CEST 2022 - The aliscore output file does not exist. Check results for BUSCO: 734341at2759
## Mon Aug  8 09:50:35 CEST 2022 - The aliscore output appears to be empty. Check results for BUSCO: 734341at2759
## Mon Aug  8 09:50:35 CEST 2022 - The aliscore output file does not exist. Check results for BUSCO: 1178688at2759
## Mon Aug  8 09:50:35 CEST 2022 - The aliscore output appears to be empty. Check results for BUSCO: 1178688at2759
## Mon Aug  8 09:50:35 CEST 2022 - The aliscore output file does not exist. Check results for BUSCO: 1407446at2759
## Mon Aug  8 09:50:35 CEST 2022 - The aliscore output file does not exist. Check results for BUSCO: 1379600at2759
## Mon Aug  8 09:50:35 CEST 2022 - The aliscore output appears to be empty. Check results for BUSCO: 1407446at2759
## Mon Aug  8 09:50:35 CEST 2022 - The aliscore output file does not exist. Check results for BUSCO: 549762at2759
## Mon Aug  8 09:50:35 CEST 2022 - The aliscore output appears to be empty. Check results for BUSCO: 1379600at2759
## Mon Aug  8 09:50:35 CEST 2022 - The aliscore output appears to be empty. Check results for BUSCO: 549762at2759
## Mon Aug  8 09:50:35 CEST 2022 - The aliscore output file does not exist. Check results for BUSCO: 1291729at2759
## Mon Aug  8 09:50:35 CEST 2022 - The aliscore output appears to be empty. Check results for BUSCO: 1291729at2759
## Mon Aug  8 09:50:36 CEST 2022 - The aliscore output file does not exist. Check results for BUSCO: 1275837at2759
## Mon Aug  8 09:50:36 CEST 2022 - The aliscore output appears to be empty. Check results for BUSCO: 1275837at2759
## Mon Aug  8 09:50:36 CEST 2022 - The aliscore output file does not exist. Check results for BUSCO: 261328at2759
## Mon Aug  8 09:50:36 CEST 2022 - The aliscore output appears to be empty. Check results for BUSCO: 261328at2759
## Mon Aug  8 09:50:37 CEST 2022 - The aliscore output file does not exist. Check results for BUSCO: 1010730at2759
## Mon Aug  8 09:50:37 CEST 2022 - The aliscore output appears to be empty. Check results for BUSCO: 1010730at2759
## Mon Aug  8 09:50:37 CEST 2022 - The aliscore output file does not exist. Check results for BUSCO: 901894at2759
## Mon Aug  8 09:50:37 CEST 2022 - The aliscore output appears to be empty. Check results for BUSCO: 901894at2759
## Mon Aug  8 09:50:38 CEST 2022 - The aliscore output file does not exist. Check results for BUSCO: 582756at2759
## Mon Aug  8 09:50:38 CEST 2022 - The aliscore output appears to be empty. Check results for BUSCO: 582756at2759
## Mon Aug  8 09:50:38 CEST 2022 - The aliscore output file does not exist. Check results for BUSCO: 911863at2759
## Mon Aug  8 09:50:38 CEST 2022 - The aliscore output appears to be empty. Check results for BUSCO: 911863at2759
## Mon Aug  8 09:50:40 CEST 2022 - The aliscore output file does not exist. Check results for BUSCO: 369837at2759
## Mon Aug  8 09:50:40 CEST 2022 - The aliscore output appears to be empty. Check results for BUSCO: 369837at2759
## Mon Aug  8 09:50:41 CEST 2022 - The aliscore output file does not exist. Check results for BUSCO: 299766at2759
## Mon Aug  8 09:50:42 CEST 2022 - The aliscore output appears to be empty. Check results for BUSCO: 299766at2759
## Mon Aug  8 09:50:43 CEST 2022 - The aliscore output file does not exist. Check results for BUSCO: 625387at2759
## Mon Aug  8 09:50:43 CEST 2022 - The aliscore output appears to be empty. Check results for BUSCO: 625387at2759
## Mon Aug  8 09:50:55 CEST 2022 - The aliscore output file does not exist. Check results for BUSCO: 1405073at2759
## Mon Aug  8 09:50:55 CEST 2022 - The aliscore output appears to be empty. Check results for BUSCO: 1405073at2759
## Mon Aug  8 09:50:57 CEST 2022 - The aliscore output file does not exist. Check results for BUSCO: 1107630at2759
## Mon Aug  8 09:50:57 CEST 2022 - The aliscore output appears to be empty. Check results for BUSCO: 1107630at2759
## Mon Aug  8 09:50:57 CEST 2022 - The aliscore output file does not exist. Check results for BUSCO: 100698at2759
## Mon Aug  8 09:50:58 CEST 2022 - The aliscore output appears to be empty. Check results for BUSCO: 100698at2759
## Mon Aug  8 09:50:59 CEST 2022 - The aliscore output file does not exist. Check results for BUSCO: 1645187at2759
## Mon Aug  8 09:50:59 CEST 2022 - The aliscore output appears to be empty. Check results for BUSCO: 1645187at2759
## Mon Aug  8 09:50:59 CEST 2022 - The aliscore output file does not exist. Check results for BUSCO: 923657at2759
## Mon Aug  8 09:50:59 CEST 2022 - The aliscore output appears to be empty. Check results for BUSCO: 923657at2759
## Mon Aug  8 09:50:59 CEST 2022 - The aliscore output file does not exist. Check results for BUSCO: 1428265at2759
## Mon Aug  8 09:51:00 CEST 2022 - The aliscore output appears to be empty. Check results for BUSCO: 1428265at2759
## Mon Aug  8 09:51:00 CEST 2022 - The aliscore output file does not exist. Check results for BUSCO: 924753at2759
## Mon Aug  8 09:51:00 CEST 2022 - The aliscore output appears to be empty. Check results for BUSCO: 924753at2759
## Mon Aug  8 09:51:00 CEST 2022 - The aliscore output file does not exist. Check results for BUSCO: 687505at2759
## Mon Aug  8 09:51:00 CEST 2022 - The aliscore output appears to be empty. Check results for BUSCO: 687505at2759
## Mon Aug  8 09:51:01 CEST 2022 - The aliscore output file does not exist. Check results for BUSCO: 1129824at2759
## Mon Aug  8 09:51:01 CEST 2022 - The aliscore output appears to be empty. Check results for BUSCO: 1129824at2759
## Mon Aug  8 09:51:01 CEST 2022 - The aliscore output file does not exist. Check results for BUSCO: 453044at2759
## Mon Aug  8 09:51:01 CEST 2022 - The aliscore output file does not exist. Check results for BUSCO: 1380409at2759
## Mon Aug  8 09:51:01 CEST 2022 - The aliscore output file does not exist. Check results for BUSCO: 1049599at2759
## Mon Aug  8 09:51:01 CEST 2022 - The aliscore output appears to be empty. Check results for BUSCO: 453044at2759
## Mon Aug  8 09:51:01 CEST 2022 - The aliscore output appears to be empty. Check results for BUSCO: 1380409at2759
## Mon Aug  8 09:51:01 CEST 2022 - The aliscore output appears to be empty. Check results for BUSCO: 1049599at2759
## Mon Aug  8 09:51:01 CEST 2022 - The aliscore output file does not exist. Check results for BUSCO: 1309031at2759
## Mon Aug  8 09:51:01 CEST 2022 - The aliscore output appears to be empty. Check results for BUSCO: 1309031at2759
## Mon Aug  8 09:51:01 CEST 2022 - The aliscore output file does not exist. Check results for BUSCO: 923657at2759
## Mon Aug  8 09:51:01 CEST 2022 - The aliscore output appears to be empty. Check results for BUSCO: 923657at2759
## Mon Aug  8 09:51:01 CEST 2022 - The aliscore output file does not exist. Check results for BUSCO: 1526152at2759
## Mon Aug  8 09:51:01 CEST 2022 - The aliscore output file does not exist. Check results for BUSCO: 673132at2759
## Mon Aug  8 09:51:01 CEST 2022 - The aliscore output appears to be empty. Check results for BUSCO: 1526152at2759
## Mon Aug  8 09:51:01 CEST 2022 - The aliscore output appears to be empty. Check results for BUSCO: 673132at2759
## Mon Aug  8 09:51:02 CEST 2022 - The aliscore output file does not exist. Check results for BUSCO: 1364586at2759
## Mon Aug  8 09:51:02 CEST 2022 - The aliscore output appears to be empty. Check results for BUSCO: 1364586at2759
## Mon Aug  8 09:51:02 CEST 2022 - The aliscore output file does not exist. Check results for BUSCO: 1094121at2759
## Mon Aug  8 09:51:02 CEST 2022 - The aliscore output file does not exist. Check results for BUSCO: 844748at2759
## Mon Aug  8 09:51:02 CEST 2022 - The aliscore output file does not exist. Check results for BUSCO: 996662at2759
## Mon Aug  8 09:51:02 CEST 2022 - The aliscore output appears to be empty. Check results for BUSCO: 1094121at2759
## Mon Aug  8 09:51:02 CEST 2022 - The aliscore output appears to be empty. Check results for BUSCO: 844748at2759
## Mon Aug  8 09:51:02 CEST 2022 - The aliscore output appears to be empty. Check results for BUSCO: 996662at2759
## Mon Aug  8 09:51:02 CEST 2022 - The aliscore output file does not exist. Check results for BUSCO: 1257440at2759
## Mon Aug  8 09:51:02 CEST 2022 - The aliscore output appears to be empty. Check results for BUSCO: 1257440at2759
## Mon Aug  8 09:51:02 CEST 2022 - The aliscore output file does not exist. Check results for BUSCO: 1030907at2759
## Mon Aug  8 09:51:02 CEST 2022 - The aliscore output appears to be empty. Check results for BUSCO: 1030907at2759
## Mon Aug  8 09:51:02 CEST 2022 - The aliscore output file does not exist. Check results for BUSCO: 1355894at2759
## Mon Aug  8 09:51:02 CEST 2022 - The aliscore output file does not exist. Check results for BUSCO: 1079130at2759
## Mon Aug  8 09:51:02 CEST 2022 - The aliscore output file does not exist. Check results for BUSCO: 1173229at2759
## Mon Aug  8 09:51:02 CEST 2022 - The aliscore output appears to be empty. Check results for BUSCO: 1355894at2759
## Mon Aug  8 09:51:02 CEST 2022 - The aliscore output file does not exist. Check results for BUSCO: 671536at2759
## Mon Aug  8 09:51:02 CEST 2022 - The aliscore output file does not exist. Check results for BUSCO: 1032689at2759
## Mon Aug  8 09:51:02 CEST 2022 - The aliscore output appears to be empty. Check results for BUSCO: 1079130at2759
## Mon Aug  8 09:51:02 CEST 2022 - The aliscore output appears to be empty. Check results for BUSCO: 1173229at2759
## Mon Aug  8 09:51:03 CEST 2022 - The aliscore output appears to be empty. Check results for BUSCO: 671536at2759
## Mon Aug  8 09:51:03 CEST 2022 - The aliscore output appears to be empty. Check results for BUSCO: 1032689at2759
## Mon Aug  8 09:51:03 CEST 2022 - The aliscore output file does not exist. Check results for BUSCO: 245208at2759
## Mon Aug  8 09:51:03 CEST 2022 - The aliscore output file does not exist. Check results for BUSCO: 1217666at2759
## Mon Aug  8 09:51:03 CEST 2022 - The aliscore output appears to be empty. Check results for BUSCO: 245208at2759
## Mon Aug  8 09:51:03 CEST 2022 - The aliscore output appears to be empty. Check results for BUSCO: 1217666at2759
## Mon Aug  8 09:51:03 CEST 2022 - The aliscore output file does not exist. Check results for BUSCO: 491869at2759
## Mon Aug  8 09:51:03 CEST 2022 - The aliscore output appears to be empty. Check results for BUSCO: 491869at2759
## Mon Aug  8 09:51:03 CEST 2022 - The aliscore output file does not exist. Check results for BUSCO: 1323575at2759
## Mon Aug  8 09:51:03 CEST 2022 - The aliscore output appears to be empty. Check results for BUSCO: 1323575at2759
## Mon Aug  8 09:51:03 CEST 2022 - The aliscore output file does not exist. Check results for BUSCO: 1287094at2759
## Mon Aug  8 09:51:03 CEST 2022 - The aliscore output appears to be empty. Check results for BUSCO: 1287094at2759
## Mon Aug  8 09:51:04 CEST 2022 - The aliscore output file does not exist. Check results for BUSCO: 1617752at2759
## Mon Aug  8 09:51:04 CEST 2022 - The aliscore output file does not exist. Check results for BUSCO: 1090038at2759
## Mon Aug  8 09:51:04 CEST 2022 - The aliscore output appears to be empty. Check results for BUSCO: 1617752at2759
## Mon Aug  8 09:51:04 CEST 2022 - The aliscore output file does not exist. Check results for BUSCO: 1259741at2759
## Mon Aug  8 09:51:04 CEST 2022 - The aliscore output appears to be empty. Check results for BUSCO: 1090038at2759
## Mon Aug  8 09:51:04 CEST 2022 - The aliscore output appears to be empty. Check results for BUSCO: 1259741at2759
## Mon Aug  8 09:51:04 CEST 2022 - The aliscore output file does not exist. Check results for BUSCO: 261419at2759
## Mon Aug  8 09:51:04 CEST 2022 - The aliscore output file does not exist. Check results for BUSCO: 1161199at2759
## Mon Aug  8 09:51:04 CEST 2022 - The aliscore output file does not exist. Check results for BUSCO: 1108845at2759
## Mon Aug  8 09:51:04 CEST 2022 - The aliscore output file does not exist. Check results for BUSCO: 1479417at2759
## Mon Aug  8 09:51:04 CEST 2022 - The aliscore output appears to be empty. Check results for BUSCO: 261419at2759
## Mon Aug  8 09:51:04 CEST 2022 - The aliscore output appears to be empty. Check results for BUSCO: 1161199at2759
## Mon Aug  8 09:51:04 CEST 2022 - The aliscore output appears to be empty. Check results for BUSCO: 1108845at2759
## Mon Aug  8 09:51:04 CEST 2022 - The aliscore output appears to be empty. Check results for BUSCO: 1479417at2759
## Mon Aug  8 09:51:04 CEST 2022 - The aliscore output file does not exist. Check results for BUSCO: 1194797at2759
## Mon Aug  8 09:51:04 CEST 2022 - The aliscore output file does not exist. Check results for BUSCO: 679771at2759
## Mon Aug  8 09:51:04 CEST 2022 - The aliscore output appears to be empty. Check results for BUSCO: 1194797at2759
## Mon Aug  8 09:51:04 CEST 2022 - The aliscore output file does not exist. Check results for BUSCO: 1057950at2759
## Mon Aug  8 09:51:04 CEST 2022 - The aliscore output appears to be empty. Check results for BUSCO: 679771at2759
## Mon Aug  8 09:51:04 CEST 2022 - The aliscore output file does not exist. Check results for BUSCO: 320059at2759
## Mon Aug  8 09:51:04 CEST 2022 - The aliscore output appears to be empty. Check results for BUSCO: 1057950at2759
## Mon Aug  8 09:51:04 CEST 2022 - The aliscore output file does not exist. Check results for BUSCO: 673132at2759
## Mon Aug  8 09:51:04 CEST 2022 - The aliscore output file does not exist. Check results for BUSCO: 1455730at2759
## Mon Aug  8 09:51:04 CEST 2022 - The aliscore output appears to be empty. Check results for BUSCO: 320059at2759
## Mon Aug  8 09:51:04 CEST 2022 - The aliscore output appears to be empty. Check results for BUSCO: 673132at2759
## Mon Aug  8 09:51:04 CEST 2022 - The aliscore output file does not exist. Check results for BUSCO: 324863at2759
## Mon Aug  8 09:51:04 CEST 2022 - The aliscore output appears to be empty. Check results for BUSCO: 1455730at2759
## Mon Aug  8 09:51:04 CEST 2022 - The aliscore output appears to be empty. Check results for BUSCO: 324863at2759
## Mon Aug  8 09:51:04 CEST 2022 - The aliscore output file does not exist. Check results for BUSCO: 692986at2759
## Mon Aug  8 09:51:04 CEST 2022 - The aliscore output appears to be empty. Check results for BUSCO: 692986at2759
## Mon Aug  8 09:51:05 CEST 2022 - The aliscore output file does not exist. Check results for BUSCO: 87842at2759
## Mon Aug  8 09:51:05 CEST 2022 - The aliscore output appears to be empty. Check results for BUSCO: 87842at2759
## Mon Aug  8 09:51:05 CEST 2022 - The aliscore output file does not exist. Check results for BUSCO: 1442062at2759
## Mon Aug  8 09:51:05 CEST 2022 - The aliscore output file does not exist. Check results for BUSCO: 858842at2759
## Mon Aug  8 09:51:05 CEST 2022 - The aliscore output file does not exist. Check results for BUSCO: 1428854at2759
## Mon Aug  8 09:51:05 CEST 2022 - The aliscore output appears to be empty. Check results for BUSCO: 1442062at2759
## Mon Aug  8 09:51:05 CEST 2022 - The aliscore output appears to be empty. Check results for BUSCO: 858842at2759
## Mon Aug  8 09:51:05 CEST 2022 - The aliscore output appears to be empty. Check results for BUSCO: 1428854at2759
## Mon Aug  8 09:51:05 CEST 2022 - The aliscore output file does not exist. Check results for BUSCO: 450058at2759
## Mon Aug  8 09:51:05 CEST 2022 - The aliscore output appears to be empty. Check results for BUSCO: 450058at2759
## Mon Aug  8 09:51:05 CEST 2022 - The aliscore output file does not exist. Check results for BUSCO: 1324510at2759
## Mon Aug  8 09:51:05 CEST 2022 - The aliscore output appears to be empty. Check results for BUSCO: 1324510at2759
## Mon Aug  8 09:51:06 CEST 2022 - The aliscore output file does not exist. Check results for BUSCO: 1474000at2759
## Mon Aug  8 09:51:06 CEST 2022 - The aliscore output file does not exist. Check results for BUSCO: 543764at2759
## Mon Aug  8 09:51:06 CEST 2022 - The aliscore output appears to be empty. Check results for BUSCO: 1474000at2759
## Mon Aug  8 09:51:06 CEST 2022 - The aliscore output appears to be empty. Check results for BUSCO: 543764at2759
## Mon Aug  8 09:51:06 CEST 2022 - The aliscore output file does not exist. Check results for BUSCO: 1085752at2759
## Mon Aug  8 09:51:06 CEST 2022 - The aliscore output appears to be empty. Check results for BUSCO: 1085752at2759
## Mon Aug  8 09:51:06 CEST 2022 - The aliscore output file does not exist. Check results for BUSCO: 142542at2759
## Mon Aug  8 09:51:06 CEST 2022 - The aliscore output appears to be empty. Check results for BUSCO: 142542at2759
## Mon Aug  8 09:51:06 CEST 2022 - The aliscore output file does not exist. Check results for BUSCO: 390348at2759
## Mon Aug  8 09:51:06 CEST 2022 - The aliscore output appears to be empty. Check results for BUSCO: 390348at2759
## Mon Aug  8 09:51:07 CEST 2022 - The aliscore output file does not exist. Check results for BUSCO: 834694at2759
## Mon Aug  8 09:51:07 CEST 2022 - The aliscore output appears to be empty. Check results for BUSCO: 834694at2759
## Mon Aug  8 09:51:08 CEST 2022 - The aliscore output file does not exist. Check results for BUSCO: 777920at2759
## Mon Aug  8 09:51:08 CEST 2022 - The aliscore output appears to be empty. Check results for BUSCO: 777920at2759
## Mon Aug  8 09:51:08 CEST 2022 - The aliscore output file does not exist. Check results for BUSCO: 160593at2759
## Mon Aug  8 09:51:08 CEST 2022 - The aliscore output appears to be empty. Check results for BUSCO: 160593at2759
## Mon Aug  8 09:51:08 CEST 2022 - The aliscore output file does not exist. Check results for BUSCO: 271586at2759
## Mon Aug  8 09:51:08 CEST 2022 - The aliscore output appears to be empty. Check results for BUSCO: 271586at2759
## Mon Aug  8 09:51:09 CEST 2022 - The aliscore output file does not exist. Check results for BUSCO: 324863at2759
## Mon Aug  8 09:51:09 CEST 2022 - The aliscore output appears to be empty. Check results for BUSCO: 324863at2759
## Mon Aug  8 09:51:10 CEST 2022 - The aliscore output file does not exist. Check results for BUSCO: 142542at2759
## Mon Aug  8 09:51:10 CEST 2022 - The aliscore output appears to be empty. Check results for BUSCO: 142542at2759
## Mon Aug  8 09:51:10 CEST 2022 - The aliscore output file does not exist. Check results for BUSCO: 166920at2759
## Mon Aug  8 09:51:10 CEST 2022 - The aliscore output appears to be empty. Check results for BUSCO: 166920at2759
## Mon Aug  8 09:51:26 CEST 2022 - The aliscore output file does not exist. Check results for BUSCO: 457861at2759
## Mon Aug  8 09:51:26 CEST 2022 - The aliscore output appears to be empty. Check results for BUSCO: 457861at2759
## Mon Aug  8 09:51:27 CEST 2022 - The aliscore output file does not exist. Check results for BUSCO: 1076134at2759
## Mon Aug  8 09:51:27 CEST 2022 - The aliscore output appears to be empty. Check results for BUSCO: 1076134at2759
## Mon Aug  8 09:51:28 CEST 2022 - The aliscore output file does not exist. Check results for BUSCO: 1115196at2759
## Mon Aug  8 09:51:28 CEST 2022 - The aliscore output appears to be empty. Check results for BUSCO: 1115196at2759
## Mon Aug  8 09:51:30 CEST 2022 - The aliscore output file does not exist. Check results for BUSCO: 1257440at2759
## Mon Aug  8 09:51:30 CEST 2022 - The aliscore output appears to be empty. Check results for BUSCO: 1257440at2759
## Mon Aug  8 09:51:30 CEST 2022 - The aliscore output file does not exist. Check results for BUSCO: 1094121at2759
## Mon Aug  8 09:51:30 CEST 2022 - The aliscore output appears to be empty. Check results for BUSCO: 1094121at2759
## Mon Aug  8 09:51:30 CEST 2022 - The aliscore output file does not exist. Check results for BUSCO: 464990at2759
## Mon Aug  8 09:51:30 CEST 2022 - The aliscore output file does not exist. Check results for BUSCO: 956854at2759
## Mon Aug  8 09:51:30 CEST 2022 - The aliscore output appears to be empty. Check results for BUSCO: 464990at2759
## Mon Aug  8 09:51:31 CEST 2022 - The aliscore output appears to be empty. Check results for BUSCO: 956854at2759
## Mon Aug  8 09:51:31 CEST 2022 - The aliscore output file does not exist. Check results for BUSCO: 1264469at2759
## Mon Aug  8 09:51:31 CEST 2022 - The aliscore output file does not exist. Check results for BUSCO: 1479417at2759
## Mon Aug  8 09:51:31 CEST 2022 - The aliscore output appears to be empty. Check results for BUSCO: 1479417at2759
## Mon Aug  8 09:51:31 CEST 2022 - The aliscore output appears to be empty. Check results for BUSCO: 1264469at2759
## Mon Aug  8 09:51:31 CEST 2022 - The aliscore output file does not exist. Check results for BUSCO: 919955at2759
## Mon Aug  8 09:51:31 CEST 2022 - The aliscore output file does not exist. Check results for BUSCO: 1057950at2759
## Mon Aug  8 09:51:31 CEST 2022 - The aliscore output appears to be empty. Check results for BUSCO: 919955at2759
## Mon Aug  8 09:51:31 CEST 2022 - The aliscore output appears to be empty. Check results for BUSCO: 1057950at2759
## Mon Aug  8 09:51:31 CEST 2022 - The aliscore output file does not exist. Check results for BUSCO: 1129824at2759
## Mon Aug  8 09:51:31 CEST 2022 - The aliscore output appears to be empty. Check results for BUSCO: 1129824at2759
## Mon Aug  8 09:51:31 CEST 2022 - The aliscore output file does not exist. Check results for BUSCO: 1364586at2759
## Mon Aug  8 09:51:32 CEST 2022 - The aliscore output file does not exist. Check results for BUSCO: 392369at2759
## Mon Aug  8 09:51:32 CEST 2022 - The aliscore output appears to be empty. Check results for BUSCO: 1364586at2759
## Mon Aug  8 09:51:32 CEST 2022 - The aliscore output appears to be empty. Check results for BUSCO: 392369at2759
## Mon Aug  8 09:51:32 CEST 2022 - The aliscore output file does not exist. Check results for BUSCO: 939345at2759
## Mon Aug  8 09:51:32 CEST 2022 - The aliscore output file does not exist. Check results for BUSCO: 721605at2759
## Mon Aug  8 09:51:32 CEST 2022 - The aliscore output appears to be empty. Check results for BUSCO: 721605at2759
## Mon Aug  8 09:51:32 CEST 2022 - The aliscore output file does not exist. Check results for BUSCO: 1287094at2759
## Mon Aug  8 09:51:32 CEST 2022 - The aliscore output appears to be empty. Check results for BUSCO: 939345at2759
## Mon Aug  8 09:51:32 CEST 2022 - The aliscore output appears to be empty. Check results for BUSCO: 1287094at2759
## Mon Aug  8 09:51:32 CEST 2022 - The aliscore output file does not exist. Check results for BUSCO: 836599at2759
## Mon Aug  8 09:51:32 CEST 2022 - The aliscore output appears to be empty. Check results for BUSCO: 836599at2759
## Mon Aug  8 09:51:33 CEST 2022 - The aliscore output file does not exist. Check results for BUSCO: 692986at2759
## Mon Aug  8 09:51:33 CEST 2022 - The aliscore output file does not exist. Check results for BUSCO: 817008at2759
## Mon Aug  8 09:51:33 CEST 2022 - The aliscore output appears to be empty. Check results for BUSCO: 817008at2759
## Mon Aug  8 09:51:33 CEST 2022 - The aliscore output appears to be empty. Check results for BUSCO: 692986at2759
## Mon Aug  8 09:51:33 CEST 2022 - The aliscore output file does not exist. Check results for BUSCO: 418107at2759
## Mon Aug  8 09:51:33 CEST 2022 - The aliscore output appears to be empty. Check results for BUSCO: 418107at2759
## Mon Aug  8 09:51:34 CEST 2022 - The aliscore output file does not exist. Check results for BUSCO: 1454155at2759
## Mon Aug  8 09:51:34 CEST 2022 - The aliscore output appears to be empty. Check results for BUSCO: 1454155at2759
## Mon Aug  8 09:51:34 CEST 2022 - The aliscore output file does not exist. Check results for BUSCO: 1526152at2759
## Mon Aug  8 09:51:34 CEST 2022 - The aliscore output file does not exist. Check results for BUSCO: 543764at2759
## Mon Aug  8 09:51:34 CEST 2022 - The aliscore output appears to be empty. Check results for BUSCO: 1526152at2759
## Mon Aug  8 09:51:34 CEST 2022 - The aliscore output appears to be empty. Check results for BUSCO: 543764at2759
## Mon Aug  8 09:51:35 CEST 2022 - The aliscore output file does not exist. Check results for BUSCO: 1623701at2759
## Mon Aug  8 09:51:35 CEST 2022 - The aliscore output appears to be empty. Check results for BUSCO: 1623701at2759
## Mon Aug  8 09:51:35 CEST 2022 - The aliscore output file does not exist. Check results for BUSCO: 1430056at2759
## Mon Aug  8 09:51:35 CEST 2022 - The aliscore output appears to be empty. Check results for BUSCO: 1430056at2759
## Mon Aug  8 09:51:35 CEST 2022 - The aliscore output file does not exist. Check results for BUSCO: 1090038at2759
## Mon Aug  8 09:51:35 CEST 2022 - The aliscore output appears to be empty. Check results for BUSCO: 1090038at2759
## Mon Aug  8 09:51:35 CEST 2022 - The aliscore output file does not exist. Check results for BUSCO: 341721at2759
## Mon Aug  8 09:51:35 CEST 2022 - The aliscore output appears to be empty. Check results for BUSCO: 341721at2759
## Mon Aug  8 09:51:36 CEST 2022 - The aliscore output file does not exist. Check results for BUSCO: 878143at2759
## Mon Aug  8 09:51:36 CEST 2022 - The aliscore output appears to be empty. Check results for BUSCO: 878143at2759
## Mon Aug  8 09:51:36 CEST 2022 - The aliscore output file does not exist. Check results for BUSCO: 664730at2759
## Mon Aug  8 09:51:36 CEST 2022 - The aliscore output appears to be empty. Check results for BUSCO: 664730at2759
## Mon Aug  8 09:51:36 CEST 2022 - The aliscore output file does not exist. Check results for BUSCO: 1030907at2759
## Mon Aug  8 09:51:36 CEST 2022 - The aliscore output appears to be empty. Check results for BUSCO: 1030907at2759
## Mon Aug  8 09:51:36 CEST 2022 - The aliscore output file does not exist. Check results for BUSCO: 271586at2759
## Mon Aug  8 09:51:36 CEST 2022 - The aliscore output appears to be empty. Check results for BUSCO: 271586at2759
## Mon Aug  8 09:51:36 CEST 2022 - The aliscore output file does not exist. Check results for BUSCO: 1173229at2759
## Mon Aug  8 09:51:36 CEST 2022 - The aliscore output appears to be empty. Check results for BUSCO: 1173229at2759
## Mon Aug  8 09:51:36 CEST 2022 - The aliscore output file does not exist. Check results for BUSCO: 720340at2759
## Mon Aug  8 09:51:36 CEST 2022 - The aliscore output appears to be empty. Check results for BUSCO: 720340at2759
## Mon Aug  8 09:51:36 CEST 2022 - The aliscore output file does not exist. Check results for BUSCO: 679187at2759
## Mon Aug  8 09:51:36 CEST 2022 - The aliscore output appears to be empty. Check results for BUSCO: 679187at2759
## Mon Aug  8 09:51:37 CEST 2022 - The aliscore output file does not exist. Check results for BUSCO: 320059at2759
## Mon Aug  8 09:51:37 CEST 2022 - The aliscore output appears to be empty. Check results for BUSCO: 320059at2759
## Mon Aug  8 09:51:37 CEST 2022 - The aliscore output file does not exist. Check results for BUSCO: 450058at2759
## Mon Aug  8 09:51:37 CEST 2022 - The aliscore output appears to be empty. Check results for BUSCO: 450058at2759
## Mon Aug  8 09:51:37 CEST 2022 - The aliscore output file does not exist. Check results for BUSCO: 761109at2759
## Mon Aug  8 09:51:37 CEST 2022 - The aliscore output file does not exist. Check results for BUSCO: 270107at2759
## Mon Aug  8 09:51:37 CEST 2022 - The aliscore output file does not exist. Check results for BUSCO: 166920at2759
## Mon Aug  8 09:51:37 CEST 2022 - The aliscore output file does not exist. Check results for BUSCO: 1323575at2759
## Mon Aug  8 09:51:37 CEST 2022 - The aliscore output appears to be empty. Check results for BUSCO: 270107at2759
## Mon Aug  8 09:51:37 CEST 2022 - The aliscore output appears to be empty. Check results for BUSCO: 166920at2759
## Mon Aug  8 09:51:37 CEST 2022 - The aliscore output appears to be empty. Check results for BUSCO: 1323575at2759
## Mon Aug  8 09:51:37 CEST 2022 - The aliscore output appears to be empty. Check results for BUSCO: 761109at2759
## Mon Aug  8 09:51:39 CEST 2022 - The aliscore output file does not exist. Check results for BUSCO: 551907at2759
## Mon Aug  8 09:51:39 CEST 2022 - The aliscore output appears to be empty. Check results for BUSCO: 551907at2759
## Mon Aug  8 09:51:39 CEST 2022 - The aliscore output file does not exist. Check results for BUSCO: 1423847at2759
## Mon Aug  8 09:51:39 CEST 2022 - The aliscore output appears to be empty. Check results for BUSCO: 1423847at2759
## Mon Aug  8 09:51:39 CEST 2022 - The aliscore output file does not exist. Check results for BUSCO: 834694at2759
## Mon Aug  8 09:51:39 CEST 2022 - The aliscore output appears to be empty. Check results for BUSCO: 834694at2759
## Mon Aug  8 09:51:40 CEST 2022 - The aliscore output file does not exist. Check results for BUSCO: 1194797at2759
## Mon Aug  8 09:51:40 CEST 2022 - The aliscore output appears to be empty. Check results for BUSCO: 1194797at2759
## Mon Aug  8 09:51:56 CEST 2022 - The aliscore output file does not exist. Check results for BUSCO: 1626636at2759
## Mon Aug  8 09:51:56 CEST 2022 - The aliscore output appears to be empty. Check results for BUSCO: 1626636at2759
## Mon Aug  8 09:51:57 CEST 2022 - The aliscore output file does not exist. Check results for BUSCO: 1525971at2759
## Mon Aug  8 09:51:57 CEST 2022 - The aliscore output appears to be empty. Check results for BUSCO: 1525971at2759
## Mon Aug  8 09:51:59 CEST 2022 - The aliscore output file does not exist. Check results for BUSCO: 865202at2759
## Mon Aug  8 09:51:59 CEST 2022 - The aliscore output appears to be empty. Check results for BUSCO: 865202at2759
## Mon Aug  8 09:51:59 CEST 2022 - The aliscore output file does not exist. Check results for BUSCO: 1003258at2759
## Mon Aug  8 09:52:00 CEST 2022 - The aliscore output appears to be empty. Check results for BUSCO: 1003258at2759
## Mon Aug  8 09:52:00 CEST 2022 - The aliscore output file does not exist. Check results for BUSCO: 1575179at2759
## Mon Aug  8 09:52:00 CEST 2022 - The aliscore output appears to be empty. Check results for BUSCO: 1575179at2759
## Mon Aug  8 09:52:00 CEST 2022 - The aliscore output file does not exist. Check results for BUSCO: 898782at2759
## Mon Aug  8 09:52:00 CEST 2022 - The aliscore output appears to be empty. Check results for BUSCO: 898782at2759
## Mon Aug  8 09:52:01 CEST 2022 - The aliscore output file does not exist. Check results for BUSCO: 1087488at2759
## Mon Aug  8 09:52:01 CEST 2022 - The aliscore output appears to be empty. Check results for BUSCO: 1087488at2759
## Mon Aug  8 09:52:01 CEST 2022 - The aliscore output file does not exist. Check results for BUSCO: 1382842at2759
## Mon Aug  8 09:52:01 CEST 2022 - The aliscore output appears to be empty. Check results for BUSCO: 1382842at2759
## Mon Aug  8 09:52:01 CEST 2022 - The aliscore output file does not exist. Check results for BUSCO: 1228942at2759
## Mon Aug  8 09:52:02 CEST 2022 - The aliscore output appears to be empty. Check results for BUSCO: 1228942at2759
## Mon Aug  8 09:52:02 CEST 2022 - The aliscore output file does not exist. Check results for BUSCO: 1588798at2759
## Mon Aug  8 09:52:02 CEST 2022 - The aliscore output file does not exist. Check results for BUSCO: 1377237at2759
## Mon Aug  8 09:52:02 CEST 2022 - The aliscore output appears to be empty. Check results for BUSCO: 1588798at2759
## Mon Aug  8 09:52:02 CEST 2022 - The aliscore output appears to be empty. Check results for BUSCO: 1377237at2759
## Mon Aug  8 09:52:02 CEST 2022 - The aliscore output file does not exist. Check results for BUSCO: 513979at2759
## Mon Aug  8 09:52:02 CEST 2022 - The aliscore output appears to be empty. Check results for BUSCO: 513979at2759
## Mon Aug  8 09:52:03 CEST 2022 - The aliscore output file does not exist. Check results for BUSCO: 721605at2759
## Mon Aug  8 09:52:03 CEST 2022 - The aliscore output appears to be empty. Check results for BUSCO: 721605at2759
## Mon Aug  8 09:52:03 CEST 2022 - The aliscore output file does not exist. Check results for BUSCO: 708105at2759
## Mon Aug  8 09:52:03 CEST 2022 - The aliscore output file does not exist. Check results for BUSCO: 1223488at2759
## Mon Aug  8 09:52:03 CEST 2022 - The aliscore output file does not exist. Check results for BUSCO: 1421503at2759
## Mon Aug  8 09:52:03 CEST 2022 - The aliscore output appears to be empty. Check results for BUSCO: 708105at2759
## Mon Aug  8 09:52:03 CEST 2022 - The aliscore output appears to be empty. Check results for BUSCO: 1421503at2759
## Mon Aug  8 09:52:03 CEST 2022 - The aliscore output appears to be empty. Check results for BUSCO: 1223488at2759
## Mon Aug  8 09:52:03 CEST 2022 - The aliscore output file does not exist. Check results for BUSCO: 1138059at2759
## Mon Aug  8 09:52:03 CEST 2022 - The aliscore output file does not exist. Check results for BUSCO: 801857at2759
## Mon Aug  8 09:52:03 CEST 2022 - The aliscore output appears to be empty. Check results for BUSCO: 1138059at2759
## Mon Aug  8 09:52:03 CEST 2022 - The aliscore output file does not exist. Check results for BUSCO: 1304061at2759
## Mon Aug  8 09:52:03 CEST 2022 - The aliscore output file does not exist. Check results for BUSCO: 1530008at2759
## Mon Aug  8 09:52:03 CEST 2022 - The aliscore output appears to be empty. Check results for BUSCO: 801857at2759
## Mon Aug  8 09:52:03 CEST 2022 - The aliscore output appears to be empty. Check results for BUSCO: 1304061at2759
## Mon Aug  8 09:52:03 CEST 2022 - The aliscore output appears to be empty. Check results for BUSCO: 1530008at2759
## Mon Aug  8 09:52:04 CEST 2022 - The aliscore output file does not exist. Check results for BUSCO: 1395649at2759
## Mon Aug  8 09:52:04 CEST 2022 - The aliscore output file does not exist. Check results for BUSCO: 801857at2759
## Mon Aug  8 09:52:04 CEST 2022 - The aliscore output appears to be empty. Check results for BUSCO: 1395649at2759
## Mon Aug  8 09:52:04 CEST 2022 - The aliscore output appears to be empty. Check results for BUSCO: 801857at2759
## Mon Aug  8 09:52:04 CEST 2022 - The aliscore output file does not exist. Check results for BUSCO: 355408at2759
## Mon Aug  8 09:52:04 CEST 2022 - The aliscore output file does not exist. Check results for BUSCO: 1193442at2759
## Mon Aug  8 09:52:04 CEST 2022 - The aliscore output appears to be empty. Check results for BUSCO: 355408at2759
## Mon Aug  8 09:52:04 CEST 2022 - The aliscore output appears to be empty. Check results for BUSCO: 1193442at2759
## Mon Aug  8 09:52:05 CEST 2022 - The aliscore output file does not exist. Check results for BUSCO: 341721at2759
## Mon Aug  8 09:52:05 CEST 2022 - The aliscore output appears to be empty. Check results for BUSCO: 341721at2759
## Mon Aug  8 09:52:05 CEST 2022 - The aliscore output file does not exist. Check results for BUSCO: 1434266at2759
## Mon Aug  8 09:52:05 CEST 2022 - The aliscore output file does not exist. Check results for BUSCO: 1432328at2759
## Mon Aug  8 09:52:05 CEST 2022 - The aliscore output file does not exist. Check results for BUSCO: 426305at2759
## Mon Aug  8 09:52:05 CEST 2022 - The aliscore output appears to be empty. Check results for BUSCO: 1434266at2759
## Mon Aug  8 09:52:05 CEST 2022 - The aliscore output appears to be empty. Check results for BUSCO: 1432328at2759
## Mon Aug  8 09:52:05 CEST 2022 - The aliscore output appears to be empty. Check results for BUSCO: 426305at2759
## Mon Aug  8 09:52:05 CEST 2022 - The aliscore output file does not exist. Check results for BUSCO: 736068at2759
## Mon Aug  8 09:52:05 CEST 2022 - The aliscore output appears to be empty. Check results for BUSCO: 736068at2759
## Mon Aug  8 09:52:06 CEST 2022 - The aliscore output file does not exist. Check results for BUSCO: 1426075at2759
## Mon Aug  8 09:52:06 CEST 2022 - The aliscore output file does not exist. Check results for BUSCO: 1593937at2759
## Mon Aug  8 09:52:06 CEST 2022 - The aliscore output appears to be empty. Check results for BUSCO: 1426075at2759
## Mon Aug  8 09:52:06 CEST 2022 - The aliscore output appears to be empty. Check results for BUSCO: 1593937at2759
## Mon Aug  8 09:52:06 CEST 2022 - The aliscore output file does not exist. Check results for BUSCO: 946128at2759
## Mon Aug  8 09:52:06 CEST 2022 - The aliscore output appears to be empty. Check results for BUSCO: 946128at2759
## Mon Aug  8 09:52:06 CEST 2022 - The aliscore output file does not exist. Check results for BUSCO: 621827at2759
## Mon Aug  8 09:52:06 CEST 2022 - The aliscore output appears to be empty. Check results for BUSCO: 621827at2759
## Mon Aug  8 09:52:07 CEST 2022 - The aliscore output file does not exist. Check results for BUSCO: 905026at2759
## Mon Aug  8 09:52:07 CEST 2022 - The aliscore output appears to be empty. Check results for BUSCO: 905026at2759
## Mon Aug  8 09:52:07 CEST 2022 - The aliscore output file does not exist. Check results for BUSCO: 1563319at2759
## Mon Aug  8 09:52:07 CEST 2022 - The aliscore output appears to be empty. Check results for BUSCO: 1563319at2759
## Mon Aug  8 09:52:07 CEST 2022 - The aliscore output file does not exist. Check results for BUSCO: 768809at2759
## Mon Aug  8 09:52:07 CEST 2022 - The aliscore output appears to be empty. Check results for BUSCO: 768809at2759
## Mon Aug  8 09:52:07 CEST 2022 - The aliscore output file does not exist. Check results for BUSCO: 270107at2759
## Mon Aug  8 09:52:07 CEST 2022 - The aliscore output file does not exist. Check results for BUSCO: 1322642at2759
## Mon Aug  8 09:52:08 CEST 2022 - The aliscore output appears to be empty. Check results for BUSCO: 270107at2759
## Mon Aug  8 09:52:08 CEST 2022 - The aliscore output appears to be empty. Check results for BUSCO: 1322642at2759
## Mon Aug  8 09:52:08 CEST 2022 - The aliscore output file does not exist. Check results for BUSCO: 918816at2759
## Mon Aug  8 09:52:08 CEST 2022 - The aliscore output appears to be empty. Check results for BUSCO: 918816at2759
## Mon Aug  8 09:52:08 CEST 2022 - The aliscore output file does not exist. Check results for BUSCO: 1258856at2759
## Mon Aug  8 09:52:08 CEST 2022 - The aliscore output appears to be empty. Check results for BUSCO: 1258856at2759
## Mon Aug  8 09:52:08 CEST 2022 - The aliscore output file does not exist. Check results for BUSCO: 814241at2759
## Mon Aug  8 09:52:08 CEST 2022 - The aliscore output file does not exist. Check results for BUSCO: 674169at2759
## Mon Aug  8 09:52:08 CEST 2022 - The aliscore output appears to be empty. Check results for BUSCO: 814241at2759
## Mon Aug  8 09:52:08 CEST 2022 - The aliscore output file does not exist. Check results for BUSCO: 931188at2759
## Mon Aug  8 09:52:08 CEST 2022 - The aliscore output file does not exist. Check results for BUSCO: 570797at2759
## Mon Aug  8 09:52:08 CEST 2022 - The aliscore output appears to be empty. Check results for BUSCO: 674169at2759
## Mon Aug  8 09:52:08 CEST 2022 - The aliscore output appears to be empty. Check results for BUSCO: 570797at2759
## Mon Aug  8 09:52:08 CEST 2022 - The aliscore output appears to be empty. Check results for BUSCO: 931188at2759
## Mon Aug  8 09:52:19 CEST 2022 - The aliscore output file does not exist. Check results for BUSCO: 674160at2759
## Mon Aug  8 09:52:19 CEST 2022 - The aliscore output appears to be empty. Check results for BUSCO: 674160at2759
## Mon Aug  8 09:52:26 CEST 2022 - The aliscore output file does not exist. Check results for BUSCO: 1079827at2759
## Mon Aug  8 09:52:26 CEST 2022 - The aliscore output appears to be empty. Check results for BUSCO: 1079827at2759
## Mon Aug  8 09:52:31 CEST 2022 - The aliscore output file does not exist. Check results for BUSCO: 891751at2759
## Mon Aug  8 09:52:31 CEST 2022 - The aliscore output appears to be empty. Check results for BUSCO: 891751at2759
## Mon Aug  8 09:52:32 CEST 2022 - The aliscore output file does not exist. Check results for BUSCO: 865202at2759
## Mon Aug  8 09:52:32 CEST 2022 - The aliscore output appears to be empty. Check results for BUSCO: 865202at2759
## Mon Aug  8 09:52:32 CEST 2022 - The aliscore output file does not exist. Check results for BUSCO: 1228942at2759
## Mon Aug  8 09:52:32 CEST 2022 - The aliscore output file does not exist. Check results for BUSCO: 1588798at2759
## Mon Aug  8 09:52:32 CEST 2022 - The aliscore output appears to be empty. Check results for BUSCO: 1228942at2759
## Mon Aug  8 09:52:32 CEST 2022 - The aliscore output file does not exist. Check results for BUSCO: 1194691at2759
## Mon Aug  8 09:52:32 CEST 2022 - The aliscore output appears to be empty. Check results for BUSCO: 1588798at2759
## Mon Aug  8 09:52:32 CEST 2022 - The aliscore output appears to be empty. Check results for BUSCO: 1194691at2759
## Mon Aug  8 09:52:32 CEST 2022 - The aliscore output file does not exist. Check results for BUSCO: 1377237at2759
## Mon Aug  8 09:52:32 CEST 2022 - The aliscore output appears to be empty. Check results for BUSCO: 1377237at2759
## Mon Aug  8 09:52:33 CEST 2022 - The aliscore output file does not exist. Check results for BUSCO: 621827at2759
## Mon Aug  8 09:52:33 CEST 2022 - The aliscore output appears to be empty. Check results for BUSCO: 621827at2759
## Mon Aug  8 09:52:33 CEST 2022 - The aliscore output file does not exist. Check results for BUSCO: 1563319at2759
## Mon Aug  8 09:52:33 CEST 2022 - The aliscore output appears to be empty. Check results for BUSCO: 1563319at2759
## Mon Aug  8 09:52:33 CEST 2022 - The aliscore output file does not exist. Check results for BUSCO: 1291729at2759
## Mon Aug  8 09:52:33 CEST 2022 - The aliscore output appears to be empty. Check results for BUSCO: 1291729at2759
## Mon Aug  8 09:52:34 CEST 2022 - The aliscore output file does not exist. Check results for BUSCO: 1530008at2759
## Mon Aug  8 09:52:34 CEST 2022 - The aliscore output appears to be empty. Check results for BUSCO: 1530008at2759
## Mon Aug  8 09:52:34 CEST 2022 - The aliscore output file does not exist. Check results for BUSCO: 1432328at2759
## Mon Aug  8 09:52:34 CEST 2022 - The aliscore output appears to be empty. Check results for BUSCO: 1432328at2759
## Mon Aug  8 09:52:34 CEST 2022 - The aliscore output file does not exist. Check results for BUSCO: 1275837at2759
## Mon Aug  8 09:52:34 CEST 2022 - The aliscore output appears to be empty. Check results for BUSCO: 1275837at2759
## Mon Aug  8 09:52:34 CEST 2022 - The aliscore output file does not exist. Check results for BUSCO: 1633672at2759
## Mon Aug  8 09:52:35 CEST 2022 - The aliscore output appears to be empty. Check results for BUSCO: 1633672at2759
## Mon Aug  8 09:52:35 CEST 2022 - The aliscore output file does not exist. Check results for BUSCO: 898782at2759
## Mon Aug  8 09:52:35 CEST 2022 - The aliscore output appears to be empty. Check results for BUSCO: 898782at2759
## Mon Aug  8 09:52:35 CEST 2022 - The aliscore output file does not exist. Check results for BUSCO: 1182451at2759
## Mon Aug  8 09:52:35 CEST 2022 - The aliscore output appears to be empty. Check results for BUSCO: 1182451at2759
## Mon Aug  8 09:52:35 CEST 2022 - The aliscore output file does not exist. Check results for BUSCO: 1395649at2759
## Mon Aug  8 09:52:35 CEST 2022 - The aliscore output appears to be empty. Check results for BUSCO: 1395649at2759
## Mon Aug  8 09:52:35 CEST 2022 - The aliscore output file does not exist. Check results for BUSCO: 814241at2759
## Mon Aug  8 09:52:35 CEST 2022 - The aliscore output appears to be empty. Check results for BUSCO: 814241at2759
## Mon Aug  8 09:52:35 CEST 2022 - The aliscore output file does not exist. Check results for BUSCO: 1157302at2759
## Mon Aug  8 09:52:35 CEST 2022 - The aliscore output file does not exist. Check results for BUSCO: 1567796at2759
## Mon Aug  8 09:52:35 CEST 2022 - The aliscore output appears to be empty. Check results for BUSCO: 1567796at2759
## Mon Aug  8 09:52:35 CEST 2022 - The aliscore output appears to be empty. Check results for BUSCO: 1157302at2759
## Mon Aug  8 09:52:36 CEST 2022 - The aliscore output file does not exist. Check results for BUSCO: 1220881at2759
## Mon Aug  8 09:52:36 CEST 2022 - The aliscore output appears to be empty. Check results for BUSCO: 1220881at2759
## Mon Aug  8 09:52:36 CEST 2022 - The aliscore output file does not exist. Check results for BUSCO: 887370at2759
## Mon Aug  8 09:52:36 CEST 2022 - The aliscore output appears to be empty. Check results for BUSCO: 887370at2759
## Mon Aug  8 09:52:36 CEST 2022 - The aliscore output file does not exist. Check results for BUSCO: 1087488at2759
## Mon Aug  8 09:52:36 CEST 2022 - The aliscore output file does not exist. Check results for BUSCO: 1445102at2759
## Mon Aug  8 09:52:36 CEST 2022 - The aliscore output appears to be empty. Check results for BUSCO: 1087488at2759
## Mon Aug  8 09:52:36 CEST 2022 - The aliscore output appears to be empty. Check results for BUSCO: 1445102at2759
## Mon Aug  8 09:52:36 CEST 2022 - The aliscore output file does not exist. Check results for BUSCO: 1434266at2759
## Mon Aug  8 09:52:36 CEST 2022 - The aliscore output appears to be empty. Check results for BUSCO: 1434266at2759
## Mon Aug  8 09:52:37 CEST 2022 - The aliscore output file does not exist. Check results for BUSCO: 918816at2759
## Mon Aug  8 09:52:37 CEST 2022 - The aliscore output appears to be empty. Check results for BUSCO: 918816at2759
## Mon Aug  8 09:52:37 CEST 2022 - The aliscore output file does not exist. Check results for BUSCO: 931188at2759
## Mon Aug  8 09:52:37 CEST 2022 - The aliscore output file does not exist. Check results for BUSCO: 779909at2759
## Mon Aug  8 09:52:37 CEST 2022 - The aliscore output appears to be empty. Check results for BUSCO: 931188at2759
## Mon Aug  8 09:52:37 CEST 2022 - The aliscore output appears to be empty. Check results for BUSCO: 779909at2759
## Mon Aug  8 09:52:37 CEST 2022 - The aliscore output file does not exist. Check results for BUSCO: 1593937at2759
## Mon Aug  8 09:52:37 CEST 2022 - The aliscore output file does not exist. Check results for BUSCO: 734341at2759
## Mon Aug  8 09:52:37 CEST 2022 - The aliscore output appears to be empty. Check results for BUSCO: 1593937at2759
## Mon Aug  8 09:52:37 CEST 2022 - The aliscore output appears to be empty. Check results for BUSCO: 734341at2759
## Mon Aug  8 09:52:37 CEST 2022 - The aliscore output file does not exist. Check results for BUSCO: 674169at2759
## Mon Aug  8 09:52:38 CEST 2022 - The aliscore output appears to be empty. Check results for BUSCO: 674169at2759
## Mon Aug  8 09:52:38 CEST 2022 - The aliscore output file does not exist. Check results for BUSCO: 1322642at2759
## Mon Aug  8 09:52:38 CEST 2022 - The aliscore output appears to be empty. Check results for BUSCO: 1322642at2759
## Mon Aug  8 09:52:38 CEST 2022 - The aliscore output file does not exist. Check results for BUSCO: 1258856at2759
## Mon Aug  8 09:52:38 CEST 2022 - The aliscore output appears to be empty. Check results for BUSCO: 1258856at2759
## Mon Aug  8 09:52:39 CEST 2022 - The aliscore output file does not exist. Check results for BUSCO: 369837at2759
## Mon Aug  8 09:52:39 CEST 2022 - The aliscore output file does not exist. Check results for BUSCO: 1065019at2759
## Mon Aug  8 09:52:39 CEST 2022 - The aliscore output appears to be empty. Check results for BUSCO: 369837at2759
## Mon Aug  8 09:52:39 CEST 2022 - The aliscore output appears to be empty. Check results for BUSCO: 1065019at2759
## Mon Aug  8 09:52:39 CEST 2022 - The aliscore output file does not exist. Check results for BUSCO: 375960at2759
## Mon Aug  8 09:52:39 CEST 2022 - The aliscore output appears to be empty. Check results for BUSCO: 375960at2759
## Mon Aug  8 09:52:40 CEST 2022 - The aliscore output file does not exist. Check results for BUSCO: 901894at2759
## Mon Aug  8 09:52:40 CEST 2022 - The aliscore output file does not exist. Check results for BUSCO: 325552at2759
## Mon Aug  8 09:52:40 CEST 2022 - The aliscore output appears to be empty. Check results for BUSCO: 325552at2759
## Mon Aug  8 09:52:40 CEST 2022 - The aliscore output appears to be empty. Check results for BUSCO: 901894at2759
## Mon Aug  8 09:52:42 CEST 2022 - The aliscore output file does not exist. Check results for BUSCO: 674160at2759
## Mon Aug  8 09:52:42 CEST 2022 - The aliscore output appears to be empty. Check results for BUSCO: 674160at2759
## Mon Aug  8 09:52:46 CEST 2022 - The aliscore output file does not exist. Check results for BUSCO: 156083at2759
## Mon Aug  8 09:52:46 CEST 2022 - The aliscore output appears to be empty. Check results for BUSCO: 156083at2759
## Mon Aug  8 09:52:56 CEST 2022 - The aliscore output file does not exist. Check results for BUSCO: 1053181at2759
## Mon Aug  8 09:52:56 CEST 2022 - The aliscore output appears to be empty. Check results for BUSCO: 1053181at2759
## Mon Aug  8 09:52:58 CEST 2022 - The aliscore output file does not exist. Check results for BUSCO: 1490022at2759
## Mon Aug  8 09:52:58 CEST 2022 - The aliscore output appears to be empty. Check results for BUSCO: 1490022at2759
## Mon Aug  8 09:53:01 CEST 2022 - The aliscore output file does not exist. Check results for BUSCO: 1032689at2759
## Mon Aug  8 09:53:01 CEST 2022 - The aliscore output appears to be empty. Check results for BUSCO: 1032689at2759
## Mon Aug  8 09:53:02 CEST 2022 - The aliscore output file does not exist. Check results for BUSCO: 974865at2759
## Mon Aug  8 09:53:02 CEST 2022 - The aliscore output appears to be empty. Check results for BUSCO: 974865at2759
## Mon Aug  8 09:53:02 CEST 2022 - The aliscore output file does not exist. Check results for BUSCO: 1633672at2759
## Mon Aug  8 09:53:02 CEST 2022 - The aliscore output appears to be empty. Check results for BUSCO: 1633672at2759
## Mon Aug  8 09:53:03 CEST 2022 - The aliscore output file does not exist. Check results for BUSCO: 1259741at2759
## Mon Aug  8 09:53:03 CEST 2022 - The aliscore output appears to be empty. Check results for BUSCO: 1259741at2759
## Mon Aug  8 09:53:03 CEST 2022 - The aliscore output file does not exist. Check results for BUSCO: 1645187at2759
## Mon Aug  8 09:53:03 CEST 2022 - The aliscore output file does not exist. Check results for BUSCO: 1567796at2759
## Mon Aug  8 09:53:03 CEST 2022 - The aliscore output file does not exist. Check results for BUSCO: 937275at2759
## Mon Aug  8 09:53:03 CEST 2022 - The aliscore output file does not exist. Check results for BUSCO: 453044at2759
## Mon Aug  8 09:53:03 CEST 2022 - The aliscore output appears to be empty. Check results for BUSCO: 1645187at2759
## Mon Aug  8 09:53:03 CEST 2022 - The aliscore output appears to be empty. Check results for BUSCO: 1567796at2759
## Mon Aug  8 09:53:03 CEST 2022 - The aliscore output appears to be empty. Check results for BUSCO: 937275at2759
## Mon Aug  8 09:53:03 CEST 2022 - The aliscore output appears to be empty. Check results for BUSCO: 453044at2759
## Mon Aug  8 09:53:03 CEST 2022 - The aliscore output file does not exist. Check results for BUSCO: 1049599at2759
## Mon Aug  8 09:53:03 CEST 2022 - The aliscore output appears to be empty. Check results for BUSCO: 1049599at2759
## Mon Aug  8 09:53:03 CEST 2022 - The aliscore output file does not exist. Check results for BUSCO: 1380409at2759
## Mon Aug  8 09:53:03 CEST 2022 - The aliscore output file does not exist. Check results for BUSCO: 975557at2759
## Mon Aug  8 09:53:03 CEST 2022 - The aliscore output appears to be empty. Check results for BUSCO: 1380409at2759
## Mon Aug  8 09:53:03 CEST 2022 - The aliscore output file does not exist. Check results for BUSCO: 1157302at2759
## Mon Aug  8 09:53:03 CEST 2022 - The aliscore output appears to be empty. Check results for BUSCO: 975557at2759
## Mon Aug  8 09:53:03 CEST 2022 - The aliscore output appears to be empty. Check results for BUSCO: 1157302at2759
## Mon Aug  8 09:53:04 CEST 2022 - The aliscore output file does not exist. Check results for BUSCO: 1025450at2759
## Mon Aug  8 09:53:04 CEST 2022 - The aliscore output file does not exist. Check results for BUSCO: 779909at2759
## Mon Aug  8 09:53:04 CEST 2022 - The aliscore output appears to be empty. Check results for BUSCO: 1025450at2759
## Mon Aug  8 09:53:04 CEST 2022 - The aliscore output file does not exist. Check results for BUSCO: 734341at2759
## Mon Aug  8 09:53:04 CEST 2022 - The aliscore output file does not exist. Check results for BUSCO: 1338131at2759
## Mon Aug  8 09:53:04 CEST 2022 - The aliscore output appears to be empty. Check results for BUSCO: 779909at2759
## Mon Aug  8 09:53:04 CEST 2022 - The aliscore output file does not exist. Check results for BUSCO: 345441at2759
## Mon Aug  8 09:53:04 CEST 2022 - The aliscore output appears to be empty. Check results for BUSCO: 734341at2759
## Mon Aug  8 09:53:04 CEST 2022 - The aliscore output file does not exist. Check results for BUSCO: 549762at2759
## Mon Aug  8 09:53:04 CEST 2022 - The aliscore output appears to be empty. Check results for BUSCO: 1338131at2759
## Mon Aug  8 09:53:04 CEST 2022 - The aliscore output appears to be empty. Check results for BUSCO: 345441at2759
## Mon Aug  8 09:53:04 CEST 2022 - The aliscore output appears to be empty. Check results for BUSCO: 549762at2759
## Mon Aug  8 09:53:04 CEST 2022 - The aliscore output file does not exist. Check results for BUSCO: 1346432at2759
## Mon Aug  8 09:53:04 CEST 2022 - The aliscore output appears to be empty. Check results for BUSCO: 1346432at2759
## Mon Aug  8 09:53:04 CEST 2022 - The aliscore output file does not exist. Check results for BUSCO: 1194691at2759
## Mon Aug  8 09:53:04 CEST 2022 - The aliscore output appears to be empty. Check results for BUSCO: 1194691at2759
## Mon Aug  8 09:53:04 CEST 2022 - The aliscore output file does not exist. Check results for BUSCO: 1010730at2759
## Mon Aug  8 09:53:04 CEST 2022 - The aliscore output appears to be empty. Check results for BUSCO: 1010730at2759
## Mon Aug  8 09:53:04 CEST 2022 - The aliscore output file does not exist. Check results for BUSCO: 924753at2759
## Mon Aug  8 09:53:04 CEST 2022 - The aliscore output appears to be empty. Check results for BUSCO: 924753at2759
## Mon Aug  8 09:53:04 CEST 2022 - The aliscore output file does not exist. Check results for BUSCO: 1513531at2759
## Mon Aug  8 09:53:05 CEST 2022 - The aliscore output appears to be empty. Check results for BUSCO: 1513531at2759
## Mon Aug  8 09:53:05 CEST 2022 - The aliscore output file does not exist. Check results for BUSCO: 491869at2759
## Mon Aug  8 09:53:05 CEST 2022 - The aliscore output file does not exist. Check results for BUSCO: 844748at2759
## Mon Aug  8 09:53:05 CEST 2022 - The aliscore output appears to be empty. Check results for BUSCO: 491869at2759
## Mon Aug  8 09:53:05 CEST 2022 - The aliscore output file does not exist. Check results for BUSCO: 1096688at2759
## Mon Aug  8 09:53:05 CEST 2022 - The aliscore output appears to be empty. Check results for BUSCO: 844748at2759
## Mon Aug  8 09:53:05 CEST 2022 - The aliscore output appears to be empty. Check results for BUSCO: 1096688at2759
## Mon Aug  8 09:53:05 CEST 2022 - The aliscore output file does not exist. Check results for BUSCO: 383503at2759
## Mon Aug  8 09:53:05 CEST 2022 - The aliscore output file does not exist. Check results for BUSCO: 1065019at2759
## Mon Aug  8 09:53:05 CEST 2022 - The aliscore output file does not exist. Check results for BUSCO: 1552706at2759
## Mon Aug  8 09:53:05 CEST 2022 - The aliscore output appears to be empty. Check results for BUSCO: 383503at2759
## Mon Aug  8 09:53:05 CEST 2022 - The aliscore output appears to be empty. Check results for BUSCO: 1552706at2759
## Mon Aug  8 09:53:05 CEST 2022 - The aliscore output appears to be empty. Check results for BUSCO: 1065019at2759
## Mon Aug  8 09:53:06 CEST 2022 - The aliscore output file does not exist. Check results for BUSCO: 1182451at2759
## Mon Aug  8 09:53:06 CEST 2022 - The aliscore output file does not exist. Check results for BUSCO: 1111142at2759
## Mon Aug  8 09:53:06 CEST 2022 - The aliscore output file does not exist. Check results for BUSCO: 375960at2759
## Mon Aug  8 09:53:06 CEST 2022 - The aliscore output appears to be empty. Check results for BUSCO: 1182451at2759
## Mon Aug  8 09:53:06 CEST 2022 - The aliscore output appears to be empty. Check results for BUSCO: 1111142at2759
## Mon Aug  8 09:53:06 CEST 2022 - The aliscore output appears to be empty. Check results for BUSCO: 375960at2759
## Mon Aug  8 09:53:06 CEST 2022 - The aliscore output file does not exist. Check results for BUSCO: 1442062at2759
## Mon Aug  8 09:53:06 CEST 2022 - The aliscore output appears to be empty. Check results for BUSCO: 1442062at2759
## Mon Aug  8 09:53:06 CEST 2022 - The aliscore output file does not exist. Check results for BUSCO: 100698at2759
## Mon Aug  8 09:53:06 CEST 2022 - The aliscore output appears to be empty. Check results for BUSCO: 100698at2759
## Mon Aug  8 09:53:06 CEST 2022 - The aliscore output file does not exist. Check results for BUSCO: 901894at2759
## Mon Aug  8 09:53:06 CEST 2022 - The aliscore output appears to be empty. Check results for BUSCO: 901894at2759
## Mon Aug  8 09:53:06 CEST 2022 - The aliscore output file does not exist. Check results for BUSCO: 937686at2759
## Mon Aug  8 09:53:07 CEST 2022 - The aliscore output appears to be empty. Check results for BUSCO: 937686at2759
## Mon Aug  8 09:53:07 CEST 2022 - The aliscore output file does not exist. Check results for BUSCO: 1428854at2759
## Mon Aug  8 09:53:07 CEST 2022 - The aliscore output appears to be empty. Check results for BUSCO: 1428854at2759
## Mon Aug  8 09:53:07 CEST 2022 - The aliscore output file does not exist. Check results for BUSCO: 388820at2759
## Mon Aug  8 09:53:07 CEST 2022 - The aliscore output appears to be empty. Check results for BUSCO: 388820at2759
## Mon Aug  8 09:53:08 CEST 2022 - The aliscore output file does not exist. Check results for BUSCO: 1324510at2759
## Mon Aug  8 09:53:08 CEST 2022 - The aliscore output appears to be empty. Check results for BUSCO: 1324510at2759
## Mon Aug  8 09:53:08 CEST 2022 - The aliscore output file does not exist. Check results for BUSCO: 369837at2759
## Mon Aug  8 09:53:08 CEST 2022 - The aliscore output appears to be empty. Check results for BUSCO: 369837at2759
## Mon Aug  8 09:53:08 CEST 2022 - The aliscore output file does not exist. Check results for BUSCO: 331411at2759
## Mon Aug  8 09:53:08 CEST 2022 - The aliscore output appears to be empty. Check results for BUSCO: 331411at2759
## Mon Aug  8 09:53:08 CEST 2022 - The aliscore output file does not exist. Check results for BUSCO: 1247641at2759
## Mon Aug  8 09:53:09 CEST 2022 - The aliscore output appears to be empty. Check results for BUSCO: 1247641at2759
## Mon Aug  8 09:53:09 CEST 2022 - The aliscore output file does not exist. Check results for BUSCO: 325552at2759
## Mon Aug  8 09:53:09 CEST 2022 - The aliscore output appears to be empty. Check results for BUSCO: 325552at2759
## Mon Aug  8 09:53:09 CEST 2022 - The aliscore output file does not exist. Check results for BUSCO: 261419at2759
## Mon Aug  8 09:53:09 CEST 2022 - The aliscore output appears to be empty. Check results for BUSCO: 261419at2759
## Mon Aug  8 09:53:09 CEST 2022 - The aliscore output file does not exist. Check results for BUSCO: 87842at2759
## Mon Aug  8 09:53:09 CEST 2022 - The aliscore output appears to be empty. Check results for BUSCO: 87842at2759
## Mon Aug  8 09:53:10 CEST 2022 - The aliscore output file does not exist. Check results for BUSCO: 759498at2759
## Mon Aug  8 09:53:10 CEST 2022 - The aliscore output appears to be empty. Check results for BUSCO: 759498at2759
## Mon Aug  8 09:53:10 CEST 2022 - The aliscore output file does not exist. Check results for BUSCO: 160593at2759
## Mon Aug  8 09:53:10 CEST 2022 - The aliscore output appears to be empty. Check results for BUSCO: 160593at2759
## Mon Aug  8 09:53:26 CEST 2022 - The aliscore output file does not exist. Check results for BUSCO: 1623701at2759
## Mon Aug  8 09:53:26 CEST 2022 - The aliscore output appears to be empty. Check results for BUSCO: 1623701at2759
## Mon Aug  8 09:53:28 CEST 2022 - The aliscore output file does not exist. Check results for BUSCO: 655400at2759
## Mon Aug  8 09:53:28 CEST 2022 - The aliscore output appears to be empty. Check results for BUSCO: 655400at2759
## Mon Aug  8 09:53:28 CEST 2022 - The aliscore output file does not exist. Check results for BUSCO: 453044at2759
## Mon Aug  8 09:53:28 CEST 2022 - The aliscore output appears to be empty. Check results for BUSCO: 453044at2759
## Mon Aug  8 09:53:29 CEST 2022 - The aliscore output file does not exist. Check results for BUSCO: 1504863at2759
## Mon Aug  8 09:53:29 CEST 2022 - The aliscore output appears to be empty. Check results for BUSCO: 1504863at2759
## Mon Aug  8 09:53:30 CEST 2022 - The aliscore output file does not exist. Check results for BUSCO: 1576404at2759
## Mon Aug  8 09:53:30 CEST 2022 - The aliscore output appears to be empty. Check results for BUSCO: 1576404at2759
## Mon Aug  8 09:53:31 CEST 2022 - The aliscore output file does not exist. Check results for BUSCO: 1454155at2759
## Mon Aug  8 09:53:31 CEST 2022 - The aliscore output appears to be empty. Check results for BUSCO: 1454155at2759
## Mon Aug  8 09:53:31 CEST 2022 - The aliscore output file does not exist. Check results for BUSCO: 1236198at2759
## Mon Aug  8 09:53:31 CEST 2022 - The aliscore output file does not exist. Check results for BUSCO: 1251252at2759
## Mon Aug  8 09:53:31 CEST 2022 - The aliscore output appears to be empty. Check results for BUSCO: 1236198at2759
## Mon Aug  8 09:53:31 CEST 2022 - The aliscore output appears to be empty. Check results for BUSCO: 1251252at2759
## Mon Aug  8 09:53:31 CEST 2022 - The aliscore output file does not exist. Check results for BUSCO: 1222562at2759
## Mon Aug  8 09:53:31 CEST 2022 - The aliscore output appears to be empty. Check results for BUSCO: 1222562at2759
## Mon Aug  8 09:53:31 CEST 2022 - The aliscore output file does not exist. Check results for BUSCO: 1346432at2759
## Mon Aug  8 09:53:31 CEST 2022 - The aliscore output file does not exist. Check results for BUSCO: 1249159at2759
## Mon Aug  8 09:53:31 CEST 2022 - The aliscore output appears to be empty. Check results for BUSCO: 1346432at2759
## Mon Aug  8 09:53:32 CEST 2022 - The aliscore output appears to be empty. Check results for BUSCO: 1249159at2759
## Mon Aug  8 09:53:32 CEST 2022 - The aliscore output file does not exist. Check results for BUSCO: 1348942at2759
## Mon Aug  8 09:53:32 CEST 2022 - The aliscore output appears to be empty. Check results for BUSCO: 1348942at2759
## Mon Aug  8 09:53:32 CEST 2022 - The aliscore output file does not exist. Check results for BUSCO: 687505at2759
## Mon Aug  8 09:53:32 CEST 2022 - The aliscore output appears to be empty. Check results for BUSCO: 687505at2759
## Mon Aug  8 09:53:33 CEST 2022 - The aliscore output file does not exist. Check results for BUSCO: 937686at2759
## Mon Aug  8 09:53:33 CEST 2022 - The aliscore output file does not exist. Check results for BUSCO: 1051021at2759
## Mon Aug  8 09:53:33 CEST 2022 - The aliscore output appears to be empty. Check results for BUSCO: 1051021at2759
## Mon Aug  8 09:53:33 CEST 2022 - The aliscore output appears to be empty. Check results for BUSCO: 937686at2759
## Mon Aug  8 09:53:33 CEST 2022 - The aliscore output file does not exist. Check results for BUSCO: 293315at2759
## Mon Aug  8 09:53:33 CEST 2022 - The aliscore output file does not exist. Check results for BUSCO: 923657at2759
## Mon Aug  8 09:53:33 CEST 2022 - The aliscore output appears to be empty. Check results for BUSCO: 293315at2759
## Mon Aug  8 09:53:33 CEST 2022 - The aliscore output appears to be empty. Check results for BUSCO: 923657at2759
## Mon Aug  8 09:53:33 CEST 2022 - The aliscore output file does not exist. Check results for BUSCO: 1450538at2759
## Mon Aug  8 09:53:33 CEST 2022 - The aliscore output file does not exist. Check results for BUSCO: 1197019at2759
## Mon Aug  8 09:53:33 CEST 2022 - The aliscore output appears to be empty. Check results for BUSCO: 1450538at2759
## Mon Aug  8 09:53:33 CEST 2022 - The aliscore output appears to be empty. Check results for BUSCO: 1197019at2759
## Mon Aug  8 09:53:33 CEST 2022 - The aliscore output file does not exist. Check results for BUSCO: 39650at2759
## Mon Aug  8 09:53:34 CEST 2022 - The aliscore output file does not exist. Check results for BUSCO: 324863at2759
## Mon Aug  8 09:53:34 CEST 2022 - The aliscore output appears to be empty. Check results for BUSCO: 39650at2759
## Mon Aug  8 09:53:34 CEST 2022 - The aliscore output appears to be empty. Check results for BUSCO: 324863at2759
## Mon Aug  8 09:53:34 CEST 2022 - The aliscore output file does not exist. Check results for BUSCO: 1197019at2759
## Mon Aug  8 09:53:34 CEST 2022 - The aliscore output file does not exist. Check results for BUSCO: 1545004at2759
## Mon Aug  8 09:53:34 CEST 2022 - The aliscore output appears to be empty. Check results for BUSCO: 1197019at2759
## Mon Aug  8 09:53:34 CEST 2022 - The aliscore output appears to be empty. Check results for BUSCO: 1545004at2759
## Mon Aug  8 09:53:34 CEST 2022 - The aliscore output file does not exist. Check results for BUSCO: 299766at2759
## Mon Aug  8 09:53:34 CEST 2022 - The aliscore output appears to be empty. Check results for BUSCO: 299766at2759
## Mon Aug  8 09:53:34 CEST 2022 - The aliscore output file does not exist. Check results for BUSCO: 671536at2759
## Mon Aug  8 09:53:34 CEST 2022 - The aliscore output appears to be empty. Check results for BUSCO: 671536at2759
## Mon Aug  8 09:53:34 CEST 2022 - The aliscore output file does not exist. Check results for BUSCO: 981902at2759
## Mon Aug  8 09:53:34 CEST 2022 - The aliscore output appears to be empty. Check results for BUSCO: 981902at2759
## Mon Aug  8 09:53:34 CEST 2022 - The aliscore output file does not exist. Check results for BUSCO: 1106766at2759
## Mon Aug  8 09:53:34 CEST 2022 - The aliscore output appears to be empty. Check results for BUSCO: 1106766at2759
## Mon Aug  8 09:53:34 CEST 2022 - The aliscore output file does not exist. Check results for BUSCO: 388820at2759
## Mon Aug  8 09:53:34 CEST 2022 - The aliscore output appears to be empty. Check results for BUSCO: 388820at2759
## Mon Aug  8 09:53:34 CEST 2022 - The aliscore output file does not exist. Check results for BUSCO: 911863at2759
## Mon Aug  8 09:53:34 CEST 2022 - The aliscore output appears to be empty. Check results for BUSCO: 911863at2759
## Mon Aug  8 09:53:34 CEST 2022 - The aliscore output file does not exist. Check results for BUSCO: 457861at2759
## Mon Aug  8 09:53:34 CEST 2022 - The aliscore output file does not exist. Check results for BUSCO: 1247641at2759
## Mon Aug  8 09:53:34 CEST 2022 - The aliscore output appears to be empty. Check results for BUSCO: 457861at2759
## Mon Aug  8 09:53:35 CEST 2022 - The aliscore output appears to be empty. Check results for BUSCO: 1247641at2759
## Mon Aug  8 09:53:35 CEST 2022 - The aliscore output file does not exist. Check results for BUSCO: 679187at2759
## Mon Aug  8 09:53:35 CEST 2022 - The aliscore output appears to be empty. Check results for BUSCO: 679187at2759
## Mon Aug  8 09:53:35 CEST 2022 - The aliscore output file does not exist. Check results for BUSCO: 1358374at2759
## Mon Aug  8 09:53:35 CEST 2022 - The aliscore output file does not exist. Check results for BUSCO: 464990at2759
## Mon Aug  8 09:53:35 CEST 2022 - The aliscore output appears to be empty. Check results for BUSCO: 1358374at2759
## Mon Aug  8 09:53:35 CEST 2022 - The aliscore output file does not exist. Check results for BUSCO: 1076134at2759
## Mon Aug  8 09:53:35 CEST 2022 - The aliscore output appears to be empty. Check results for BUSCO: 464990at2759
## Mon Aug  8 09:53:35 CEST 2022 - The aliscore output appears to be empty. Check results for BUSCO: 1076134at2759
## Mon Aug  8 09:53:35 CEST 2022 - The aliscore output file does not exist. Check results for BUSCO: 944899at2759
## Mon Aug  8 09:53:35 CEST 2022 - The aliscore output appears to be empty. Check results for BUSCO: 944899at2759
## Mon Aug  8 09:53:35 CEST 2022 - The aliscore output file does not exist. Check results for BUSCO: 1645187at2759
## Mon Aug  8 09:53:35 CEST 2022 - The aliscore output appears to be empty. Check results for BUSCO: 1645187at2759
## Mon Aug  8 09:53:35 CEST 2022 - The aliscore output file does not exist. Check results for BUSCO: 83779at2759
## Mon Aug  8 09:53:35 CEST 2022 - The aliscore output file does not exist. Check results for BUSCO: 290630at2759
## Mon Aug  8 09:53:35 CEST 2022 - The aliscore output file does not exist. Check results for BUSCO: 1038775at2759
## Mon Aug  8 09:53:35 CEST 2022 - The aliscore output appears to be empty. Check results for BUSCO: 83779at2759
## Mon Aug  8 09:53:35 CEST 2022 - The aliscore output appears to be empty. Check results for BUSCO: 290630at2759
## Mon Aug  8 09:53:35 CEST 2022 - The aliscore output appears to be empty. Check results for BUSCO: 1038775at2759
## Mon Aug  8 09:53:35 CEST 2022 - The aliscore output file does not exist. Check results for BUSCO: 844748at2759
## Mon Aug  8 09:53:35 CEST 2022 - The aliscore output appears to be empty. Check results for BUSCO: 844748at2759
## Mon Aug  8 09:53:36 CEST 2022 - The aliscore output file does not exist. Check results for BUSCO: 1260807at2759
## Mon Aug  8 09:53:36 CEST 2022 - The aliscore output file does not exist. Check results for BUSCO: 1115196at2759
## Mon Aug  8 09:53:36 CEST 2022 - The aliscore output appears to be empty. Check results for BUSCO: 1260807at2759
## Mon Aug  8 09:53:36 CEST 2022 - The aliscore output appears to be empty. Check results for BUSCO: 1115196at2759
## Mon Aug  8 09:53:36 CEST 2022 - The aliscore output file does not exist. Check results for BUSCO: 734666at2759
## Mon Aug  8 09:53:36 CEST 2022 - The aliscore output file does not exist. Check results for BUSCO: 1442062at2759
## Mon Aug  8 09:53:36 CEST 2022 - The aliscore output appears to be empty. Check results for BUSCO: 734666at2759
## Mon Aug  8 09:53:36 CEST 2022 - The aliscore output appears to be empty. Check results for BUSCO: 1442062at2759
## Mon Aug  8 09:53:36 CEST 2022 - The aliscore output file does not exist. Check results for BUSCO: 97116at2759
## Mon Aug  8 09:53:36 CEST 2022 - The aliscore output appears to be empty. Check results for BUSCO: 97116at2759
## Mon Aug  8 09:53:36 CEST 2022 - The aliscore output file does not exist. Check results for BUSCO: 1260807at2759
## Mon Aug  8 09:53:36 CEST 2022 - The aliscore output file does not exist. Check results for BUSCO: 759498at2759
## Mon Aug  8 09:53:36 CEST 2022 - The aliscore output appears to be empty. Check results for BUSCO: 1260807at2759
## Mon Aug  8 09:53:36 CEST 2022 - The aliscore output appears to be empty. Check results for BUSCO: 759498at2759
## Mon Aug  8 09:53:37 CEST 2022 - The aliscore output file does not exist. Check results for BUSCO: 855834at2759
## Mon Aug  8 09:53:37 CEST 2022 - The aliscore output file does not exist. Check results for BUSCO: 296129at2759
## Mon Aug  8 09:53:37 CEST 2022 - The aliscore output appears to be empty. Check results for BUSCO: 855834at2759
## Mon Aug  8 09:53:37 CEST 2022 - The aliscore output file does not exist. Check results for BUSCO: 543764at2759
## Mon Aug  8 09:53:37 CEST 2022 - The aliscore output appears to be empty. Check results for BUSCO: 296129at2759
## Mon Aug  8 09:53:37 CEST 2022 - The aliscore output file does not exist. Check results for BUSCO: 673132at2759
## Mon Aug  8 09:53:37 CEST 2022 - The aliscore output appears to be empty. Check results for BUSCO: 543764at2759
## Mon Aug  8 09:53:37 CEST 2022 - The aliscore output file does not exist. Check results for BUSCO: 664730at2759
## Mon Aug  8 09:53:37 CEST 2022 - The aliscore output appears to be empty. Check results for BUSCO: 673132at2759
## Mon Aug  8 09:53:37 CEST 2022 - The aliscore output appears to be empty. Check results for BUSCO: 664730at2759
## Mon Aug  8 09:53:37 CEST 2022 - The aliscore output file does not exist. Check results for BUSCO: 598949at2759
## Mon Aug  8 09:53:37 CEST 2022 - The aliscore output appears to be empty. Check results for BUSCO: 598949at2759
## Mon Aug  8 09:53:37 CEST 2022 - The aliscore output file does not exist. Check results for BUSCO: 719531at2759
## Mon Aug  8 09:53:37 CEST 2022 - The aliscore output appears to be empty. Check results for BUSCO: 719531at2759
## Mon Aug  8 09:53:37 CEST 2022 - The aliscore output file does not exist. Check results for BUSCO: 261419at2759
## Mon Aug  8 09:53:37 CEST 2022 - The aliscore output file does not exist. Check results for BUSCO: 604979at2759
## Mon Aug  8 09:53:37 CEST 2022 - The aliscore output file does not exist. Check results for BUSCO: 719531at2759
## Mon Aug  8 09:53:37 CEST 2022 - The aliscore output file does not exist. Check results for BUSCO: 625387at2759
## Mon Aug  8 09:53:37 CEST 2022 - The aliscore output appears to be empty. Check results for BUSCO: 261419at2759
## Mon Aug  8 09:53:37 CEST 2022 - The aliscore output appears to be empty. Check results for BUSCO: 604979at2759
## Mon Aug  8 09:53:38 CEST 2022 - The aliscore output appears to be empty. Check results for BUSCO: 719531at2759
## Mon Aug  8 09:53:38 CEST 2022 - The aliscore output appears to be empty. Check results for BUSCO: 625387at2759
## Mon Aug  8 09:53:38 CEST 2022 - The aliscore output file does not exist. Check results for BUSCO: 100698at2759
## Mon Aug  8 09:53:38 CEST 2022 - The aliscore output file does not exist. Check results for BUSCO: 720340at2759
## Mon Aug  8 09:53:38 CEST 2022 - The aliscore output appears to be empty. Check results for BUSCO: 100698at2759
## Mon Aug  8 09:53:38 CEST 2022 - The aliscore output appears to be empty. Check results for BUSCO: 720340at2759
## Mon Aug  8 09:53:38 CEST 2022 - The aliscore output file does not exist. Check results for BUSCO: 855834at2759
## Mon Aug  8 09:53:38 CEST 2022 - The aliscore output appears to be empty. Check results for BUSCO: 855834at2759
## Mon Aug  8 09:53:38 CEST 2022 - The aliscore output file does not exist. Check results for BUSCO: 271586at2759
## Mon Aug  8 09:53:38 CEST 2022 - The aliscore output appears to be empty. Check results for BUSCO: 271586at2759
## Mon Aug  8 09:53:38 CEST 2022 - The aliscore output file does not exist. Check results for BUSCO: 1552706at2759
## Mon Aug  8 09:53:39 CEST 2022 - The aliscore output appears to be empty. Check results for BUSCO: 1552706at2759
## Mon Aug  8 09:53:39 CEST 2022 - The aliscore output file does not exist. Check results for BUSCO: 1079130at2759
## Mon Aug  8 09:53:39 CEST 2022 - The aliscore output appears to be empty. Check results for BUSCO: 1079130at2759
## Mon Aug  8 09:53:42 CEST 2022 - The aliscore output file does not exist. Check results for BUSCO: 1314980at2759
## Mon Aug  8 09:53:42 CEST 2022 - The aliscore output appears to be empty. Check results for BUSCO: 1314980at2759
## Mon Aug  8 09:53:43 CEST 2022 - The aliscore output file does not exist. Check results for BUSCO: 142542at2759
## Mon Aug  8 09:53:43 CEST 2022 - The aliscore output appears to be empty. Check results for BUSCO: 142542at2759
## Mon Aug  8 09:53:57 CEST 2022 - The aliscore output file does not exist. Check results for BUSCO: 1129824at2759
## Mon Aug  8 09:53:57 CEST 2022 - The aliscore output appears to be empty. Check results for BUSCO: 1129824at2759
## Mon Aug  8 09:53:58 CEST 2022 - The aliscore output file does not exist. Check results for BUSCO: 464990at2759
## Mon Aug  8 09:53:58 CEST 2022 - The aliscore output appears to be empty. Check results for BUSCO: 464990at2759
## Mon Aug  8 09:53:58 CEST 2022 - The aliscore output file does not exist. Check results for BUSCO: 1382842at2759
## Mon Aug  8 09:53:58 CEST 2022 - The aliscore output appears to be empty. Check results for BUSCO: 1382842at2759
## Mon Aug  8 09:53:59 CEST 2022 - The aliscore output file does not exist. Check results for BUSCO: 801857at2759
## Mon Aug  8 09:53:59 CEST 2022 - The aliscore output appears to be empty. Check results for BUSCO: 801857at2759
## Mon Aug  8 09:53:59 CEST 2022 - The aliscore output file does not exist. Check results for BUSCO: 1450538at2759
## Mon Aug  8 09:53:59 CEST 2022 - The aliscore output appears to be empty. Check results for BUSCO: 1450538at2759
## Mon Aug  8 09:53:59 CEST 2022 - The aliscore output file does not exist. Check results for BUSCO: 1459797at2759
## Mon Aug  8 09:53:59 CEST 2022 - The aliscore output appears to be empty. Check results for BUSCO: 1459797at2759
## Mon Aug  8 09:54:00 CEST 2022 - The aliscore output file does not exist. Check results for BUSCO: 866359at2759
## Mon Aug  8 09:54:01 CEST 2022 - The aliscore output appears to be empty. Check results for BUSCO: 866359at2759
## Mon Aug  8 09:54:01 CEST 2022 - The aliscore output file does not exist. Check results for BUSCO: 1575179at2759
## Mon Aug  8 09:54:01 CEST 2022 - The aliscore output appears to be empty. Check results for BUSCO: 1575179at2759
## Mon Aug  8 09:54:01 CEST 2022 - The aliscore output file does not exist. Check results for BUSCO: 1236198at2759
## Mon Aug  8 09:54:01 CEST 2022 - The aliscore output appears to be empty. Check results for BUSCO: 1236198at2759
## Mon Aug  8 09:54:01 CEST 2022 - The aliscore output file does not exist. Check results for BUSCO: 355408at2759
## Mon Aug  8 09:54:01 CEST 2022 - The aliscore output appears to be empty. Check results for BUSCO: 355408at2759
## Mon Aug  8 09:54:01 CEST 2022 - The aliscore output file does not exist. Check results for BUSCO: 1076134at2759
## Mon Aug  8 09:54:01 CEST 2022 - The aliscore output appears to be empty. Check results for BUSCO: 1076134at2759
## Mon Aug  8 09:54:01 CEST 2022 - The aliscore output file does not exist. Check results for BUSCO: 708105at2759
## Mon Aug  8 09:54:01 CEST 2022 - The aliscore output appears to be empty. Check results for BUSCO: 708105at2759
## Mon Aug  8 09:54:01 CEST 2022 - The aliscore output file does not exist. Check results for BUSCO: 1304061at2759
## Mon Aug  8 09:54:01 CEST 2022 - The aliscore output file does not exist. Check results for BUSCO: 1623701at2759
## Mon Aug  8 09:54:02 CEST 2022 - The aliscore output appears to be empty. Check results for BUSCO: 1623701at2759
## Mon Aug  8 09:54:02 CEST 2022 - The aliscore output file does not exist. Check results for BUSCO: 1128607at2759
## Mon Aug  8 09:54:02 CEST 2022 - The aliscore output file does not exist. Check results for BUSCO: 1348942at2759
## Mon Aug  8 09:54:02 CEST 2022 - The aliscore output appears to be empty. Check results for BUSCO: 1128607at2759
## Mon Aug  8 09:54:02 CEST 2022 - The aliscore output file does not exist. Check results for BUSCO: 1014314at2759
## Mon Aug  8 09:54:02 CEST 2022 - The aliscore output appears to be empty. Check results for BUSCO: 1348942at2759
## Mon Aug  8 09:54:02 CEST 2022 - The aliscore output appears to be empty. Check results for BUSCO: 1014314at2759
## Mon Aug  8 09:54:02 CEST 2022 - The aliscore output file does not exist. Check results for BUSCO: 721605at2759
## Mon Aug  8 09:54:02 CEST 2022 - The aliscore output appears to be empty. Check results for BUSCO: 721605at2759
## Mon Aug  8 09:54:02 CEST 2022 - The aliscore output file does not exist. Check results for BUSCO: 1264469at2759
## Mon Aug  8 09:54:02 CEST 2022 - The aliscore output appears to be empty. Check results for BUSCO: 1264469at2759
## Mon Aug  8 09:54:02 CEST 2022 - The aliscore output file does not exist. Check results for BUSCO: 1441854at2759
## Mon Aug  8 09:54:02 CEST 2022 - The aliscore output appears to be empty. Check results for BUSCO: 1441854at2759
## Mon Aug  8 09:54:03 CEST 2022 - The aliscore output file does not exist. Check results for BUSCO: 919955at2759
## Mon Aug  8 09:54:03 CEST 2022 - The aliscore output appears to be empty. Check results for BUSCO: 919955at2759
## Mon Aug  8 09:54:03 CEST 2022 - The aliscore output file does not exist. Check results for BUSCO: 426305at2759
## Mon Aug  8 09:54:03 CEST 2022 - The aliscore output appears to be empty. Check results for BUSCO: 426305at2759
## Mon Aug  8 09:54:03 CEST 2022 - The aliscore output file does not exist. Check results for BUSCO: 761109at2759
## Mon Aug  8 09:54:03 CEST 2022 - The aliscore output appears to be empty. Check results for BUSCO: 761109at2759
## Mon Aug  8 09:54:04 CEST 2022 - The aliscore output file does not exist. Check results for BUSCO: 734666at2759
## Mon Aug  8 09:54:04 CEST 2022 - The aliscore output file does not exist. Check results for BUSCO: 817008at2759
## Mon Aug  8 09:54:04 CEST 2022 - The aliscore output appears to be empty. Check results for BUSCO: 734666at2759
## Mon Aug  8 09:54:04 CEST 2022 - The aliscore output appears to be empty. Check results for BUSCO: 817008at2759
## Mon Aug  8 09:54:04 CEST 2022 - The aliscore output file does not exist. Check results for BUSCO: 1558822at2759
## Mon Aug  8 09:54:04 CEST 2022 - The aliscore output appears to be empty. Check results for BUSCO: 1558822at2759
## Mon Aug  8 09:54:04 CEST 2022 - The aliscore output file does not exist. Check results for BUSCO: 1112002at2759
## Mon Aug  8 09:54:04 CEST 2022 - The aliscore output appears to be empty. Check results for BUSCO: 1112002at2759
## Mon Aug  8 09:54:04 CEST 2022 - The aliscore output file does not exist. Check results for BUSCO: 341721at2759
## Mon Aug  8 09:54:04 CEST 2022 - The aliscore output file does not exist. Check results for BUSCO: 257318at2759
## Mon Aug  8 09:54:04 CEST 2022 - The aliscore output appears to be empty. Check results for BUSCO: 341721at2759
## Mon Aug  8 09:54:04 CEST 2022 - The aliscore output file does not exist. Check results for BUSCO: 97116at2759
## Mon Aug  8 09:54:04 CEST 2022 - The aliscore output appears to be empty. Check results for BUSCO: 257318at2759
## Mon Aug  8 09:54:04 CEST 2022 - The aliscore output appears to be empty. Check results for BUSCO: 97116at2759
## Mon Aug  8 09:54:05 CEST 2022 - The aliscore output file does not exist. Check results for BUSCO: 418107at2759
## Mon Aug  8 09:54:05 CEST 2022 - The aliscore output file does not exist. Check results for BUSCO: 939345at2759
## Mon Aug  8 09:54:05 CEST 2022 - The aliscore output appears to be empty. Check results for BUSCO: 418107at2759
## Mon Aug  8 09:54:05 CEST 2022 - The aliscore output file does not exist. Check results for BUSCO: 1115196at2759
## Mon Aug  8 09:54:05 CEST 2022 - The aliscore output appears to be empty. Check results for BUSCO: 939345at2759
## Mon Aug  8 09:54:05 CEST 2022 - The aliscore output appears to be empty. Check results for BUSCO: 1115196at2759
## Mon Aug  8 09:54:05 CEST 2022 - The aliscore output file does not exist. Check results for BUSCO: 1398309at2759
## Mon Aug  8 09:54:05 CEST 2022 - The aliscore output appears to be empty. Check results for BUSCO: 1398309at2759
## Mon Aug  8 09:54:05 CEST 2022 - The aliscore output file does not exist. Check results for BUSCO: 306227at2759
## Mon Aug  8 09:54:05 CEST 2022 - The aliscore output appears to be empty. Check results for BUSCO: 306227at2759
## Mon Aug  8 09:54:06 CEST 2022 - The aliscore output file does not exist. Check results for BUSCO: 1106766at2759
## Mon Aug  8 09:54:06 CEST 2022 - The aliscore output appears to be empty. Check results for BUSCO: 1106766at2759
## Mon Aug  8 09:54:06 CEST 2022 - The aliscore output file does not exist. Check results for BUSCO: 836599at2759
## Mon Aug  8 09:54:06 CEST 2022 - The aliscore output appears to be empty. Check results for BUSCO: 836599at2759
## Mon Aug  8 09:54:06 CEST 2022 - The aliscore output file does not exist. Check results for BUSCO: 457861at2759
## Mon Aug  8 09:54:06 CEST 2022 - The aliscore output appears to be empty. Check results for BUSCO: 457861at2759
## Mon Aug  8 09:54:06 CEST 2022 - The aliscore output file does not exist. Check results for BUSCO: 320059at2759
## Mon Aug  8 09:54:06 CEST 2022 - The aliscore output file does not exist. Check results for BUSCO: 270107at2759
## Mon Aug  8 09:54:06 CEST 2022 - The aliscore output appears to be empty. Check results for BUSCO: 270107at2759
## Mon Aug  8 09:54:06 CEST 2022 - The aliscore output file does not exist. Check results for BUSCO: 1194797at2759
## Mon Aug  8 09:54:06 CEST 2022 - The aliscore output file does not exist. Check results for BUSCO: 450058at2759
## Mon Aug  8 09:54:06 CEST 2022 - The aliscore output appears to be empty. Check results for BUSCO: 320059at2759
## Mon Aug  8 09:54:06 CEST 2022 - The aliscore output appears to be empty. Check results for BUSCO: 1194797at2759
## Mon Aug  8 09:54:06 CEST 2022 - The aliscore output appears to be empty. Check results for BUSCO: 450058at2759
## Mon Aug  8 09:54:07 CEST 2022 - The aliscore output file does not exist. Check results for BUSCO: 1323575at2759
## Mon Aug  8 09:54:07 CEST 2022 - The aliscore output appears to be empty. Check results for BUSCO: 1323575at2759
## Mon Aug  8 09:54:07 CEST 2022 - The aliscore output file does not exist. Check results for BUSCO: 179362at2759
## Mon Aug  8 09:54:08 CEST 2022 - The aliscore output appears to be empty. Check results for BUSCO: 179362at2759
## Mon Aug  8 09:54:08 CEST 2022 - The aliscore output file does not exist. Check results for BUSCO: 664730at2759
## Mon Aug  8 09:54:08 CEST 2022 - The aliscore output appears to be empty. Check results for BUSCO: 664730at2759
## Mon Aug  8 09:54:08 CEST 2022 - The aliscore output file does not exist. Check results for BUSCO: 973442at2759
## Mon Aug  8 09:54:08 CEST 2022 - The aliscore output appears to be empty. Check results for BUSCO: 973442at2759
## Mon Aug  8 09:54:09 CEST 2022 - The aliscore output file does not exist. Check results for BUSCO: 692986at2759
## Mon Aug  8 09:54:09 CEST 2022 - The aliscore output appears to be empty. Check results for BUSCO: 692986at2759
## Mon Aug  8 09:54:27 CEST 2022 - The aliscore output file does not exist. Check results for BUSCO: 1576404at2759
## Mon Aug  8 09:54:27 CEST 2022 - The aliscore output appears to be empty. Check results for BUSCO: 1576404at2759
## Mon Aug  8 09:54:30 CEST 2022 - The aliscore output file does not exist. Check results for BUSCO: 1112002at2759
## Mon Aug  8 09:54:30 CEST 2022 - The aliscore output appears to be empty. Check results for BUSCO: 1112002at2759
## Mon Aug  8 09:54:30 CEST 2022 - The aliscore output file does not exist. Check results for BUSCO: 919955at2759
## Mon Aug  8 09:54:30 CEST 2022 - The aliscore output appears to be empty. Check results for BUSCO: 919955at2759
## Mon Aug  8 09:54:30 CEST 2022 - The aliscore output file does not exist. Check results for BUSCO: 1220881at2759
## Mon Aug  8 09:54:30 CEST 2022 - The aliscore output appears to be empty. Check results for BUSCO: 1220881at2759
## Mon Aug  8 09:54:30 CEST 2022 - The aliscore output file does not exist. Check results for BUSCO: 1545004at2759
## Mon Aug  8 09:54:31 CEST 2022 - The aliscore output appears to be empty. Check results for BUSCO: 1545004at2759
## Mon Aug  8 09:54:31 CEST 2022 - The aliscore output file does not exist. Check results for BUSCO: 1264469at2759
## Mon Aug  8 09:54:31 CEST 2022 - The aliscore output appears to be empty. Check results for BUSCO: 1264469at2759
## Mon Aug  8 09:54:31 CEST 2022 - The aliscore output file does not exist. Check results for BUSCO: 887370at2759
## Mon Aug  8 09:54:31 CEST 2022 - The aliscore output file does not exist. Check results for BUSCO: 1504863at2759
## Mon Aug  8 09:54:31 CEST 2022 - The aliscore output appears to be empty. Check results for BUSCO: 887370at2759
## Mon Aug  8 09:54:31 CEST 2022 - The aliscore output appears to be empty. Check results for BUSCO: 1504863at2759
## Mon Aug  8 09:54:32 CEST 2022 - The aliscore output file does not exist. Check results for BUSCO: 973442at2759
## Mon Aug  8 09:54:32 CEST 2022 - The aliscore output appears to be empty. Check results for BUSCO: 973442at2759
## Mon Aug  8 09:54:32 CEST 2022 - The aliscore output file does not exist. Check results for BUSCO: 1398309at2759
## Mon Aug  8 09:54:32 CEST 2022 - The aliscore output appears to be empty. Check results for BUSCO: 1398309at2759
## Mon Aug  8 09:54:32 CEST 2022 - The aliscore output file does not exist. Check results for BUSCO: 1228942at2759
## Mon Aug  8 09:54:32 CEST 2022 - The aliscore output file does not exist. Check results for BUSCO: 1588798at2759
## Mon Aug  8 09:54:32 CEST 2022 - The aliscore output appears to be empty. Check results for BUSCO: 1228942at2759
## Mon Aug  8 09:54:32 CEST 2022 - The aliscore output appears to be empty. Check results for BUSCO: 1588798at2759
## Mon Aug  8 09:54:33 CEST 2022 - The aliscore output file does not exist. Check results for BUSCO: 1284731at2759
## Mon Aug  8 09:54:33 CEST 2022 - The aliscore output appears to be empty. Check results for BUSCO: 1284731at2759
## Mon Aug  8 09:54:33 CEST 2022 - The aliscore output file does not exist. Check results for BUSCO: 1626636at2759
## Mon Aug  8 09:54:33 CEST 2022 - The aliscore output appears to be empty. Check results for BUSCO: 1626636at2759
## Mon Aug  8 09:54:33 CEST 2022 - The aliscore output file does not exist. Check results for BUSCO: 836599at2759
## Mon Aug  8 09:54:33 CEST 2022 - The aliscore output appears to be empty. Check results for BUSCO: 836599at2759
## Mon Aug  8 09:54:33 CEST 2022 - The aliscore output file does not exist. Check results for BUSCO: 1346165at2759
## Mon Aug  8 09:54:33 CEST 2022 - The aliscore output appears to be empty. Check results for BUSCO: 1346165at2759
## Mon Aug  8 09:54:33 CEST 2022 - The aliscore output file does not exist. Check results for BUSCO: 975158at2759
## Mon Aug  8 09:54:33 CEST 2022 - The aliscore output appears to be empty. Check results for BUSCO: 975158at2759
## Mon Aug  8 09:54:34 CEST 2022 - The aliscore output file does not exist. Check results for BUSCO: 655400at2759
## Mon Aug  8 09:54:34 CEST 2022 - The aliscore output file does not exist. Check results for BUSCO: 1530008at2759
## Mon Aug  8 09:54:34 CEST 2022 - The aliscore output file does not exist. Check results for BUSCO: 1488235at2759
## Mon Aug  8 09:54:34 CEST 2022 - The aliscore output appears to be empty. Check results for BUSCO: 655400at2759
## Mon Aug  8 09:54:34 CEST 2022 - The aliscore output file does not exist. Check results for BUSCO: 891751at2759
## Mon Aug  8 09:54:34 CEST 2022 - The aliscore output appears to be empty. Check results for BUSCO: 1530008at2759
## Mon Aug  8 09:54:34 CEST 2022 - The aliscore output appears to be empty. Check results for BUSCO: 1488235at2759
## Mon Aug  8 09:54:34 CEST 2022 - The aliscore output appears to be empty. Check results for BUSCO: 891751at2759
## Mon Aug  8 09:54:34 CEST 2022 - The aliscore output file does not exist. Check results for BUSCO: 1575179at2759
## Mon Aug  8 09:54:34 CEST 2022 - The aliscore output file does not exist. Check results for BUSCO: 1309031at2759
## Mon Aug  8 09:54:34 CEST 2022 - The aliscore output appears to be empty. Check results for BUSCO: 1575179at2759
## Mon Aug  8 09:54:34 CEST 2022 - The aliscore output appears to be empty. Check results for BUSCO: 1309031at2759
## Mon Aug  8 09:54:34 CEST 2022 - The aliscore output file does not exist. Check results for BUSCO: 1085752at2759
## Mon Aug  8 09:54:34 CEST 2022 - The aliscore output appears to be empty. Check results for BUSCO: 1085752at2759
## Mon Aug  8 09:54:35 CEST 2022 - The aliscore output file does not exist. Check results for BUSCO: 917326at2759
## Mon Aug  8 09:54:35 CEST 2022 - The aliscore output appears to be empty. Check results for BUSCO: 917326at2759
## Mon Aug  8 09:54:35 CEST 2022 - The aliscore output file does not exist. Check results for BUSCO: 976469at2759
## Mon Aug  8 09:54:35 CEST 2022 - The aliscore output appears to be empty. Check results for BUSCO: 976469at2759
## Mon Aug  8 09:54:35 CEST 2022 - The aliscore output file does not exist. Check results for BUSCO: 1041560at2759
## Mon Aug  8 09:54:35 CEST 2022 - The aliscore output appears to be empty. Check results for BUSCO: 1041560at2759
## Mon Aug  8 09:54:36 CEST 2022 - The aliscore output file does not exist. Check results for BUSCO: 1322299at2759
## Mon Aug  8 09:54:36 CEST 2022 - The aliscore output appears to be empty. Check results for BUSCO: 1322299at2759
## Mon Aug  8 09:54:36 CEST 2022 - The aliscore output file does not exist. Check results for BUSCO: 1434266at2759
## Mon Aug  8 09:54:36 CEST 2022 - The aliscore output file does not exist. Check results for BUSCO: 1128607at2759
## Mon Aug  8 09:54:36 CEST 2022 - The aliscore output appears to be empty. Check results for BUSCO: 1434266at2759
## Mon Aug  8 09:54:36 CEST 2022 - The aliscore output appears to be empty. Check results for BUSCO: 1128607at2759
## Mon Aug  8 09:54:36 CEST 2022 - The aliscore output file does not exist. Check results for BUSCO: 1382842at2759
## Mon Aug  8 09:54:36 CEST 2022 - The aliscore output appears to be empty. Check results for BUSCO: 1382842at2759
## Mon Aug  8 09:54:36 CEST 2022 - The aliscore output file does not exist. Check results for BUSCO: 296129at2759
## Mon Aug  8 09:54:36 CEST 2022 - The aliscore output appears to be empty. Check results for BUSCO: 296129at2759
## Mon Aug  8 09:54:36 CEST 2022 - The aliscore output file does not exist. Check results for BUSCO: 674169at2759
## Mon Aug  8 09:54:36 CEST 2022 - The aliscore output appears to be empty. Check results for BUSCO: 674169at2759
## Mon Aug  8 09:54:36 CEST 2022 - The aliscore output file does not exist. Check results for BUSCO: 722805at2759
## Mon Aug  8 09:54:37 CEST 2022 - The aliscore output appears to be empty. Check results for BUSCO: 722805at2759
## Mon Aug  8 09:54:37 CEST 2022 - The aliscore output file does not exist. Check results for BUSCO: 1322642at2759
## Mon Aug  8 09:54:37 CEST 2022 - The aliscore output file does not exist. Check results for BUSCO: 621827at2759
## Mon Aug  8 09:54:37 CEST 2022 - The aliscore output appears to be empty. Check results for BUSCO: 1322642at2759
## Mon Aug  8 09:54:37 CEST 2022 - The aliscore output appears to be empty. Check results for BUSCO: 621827at2759
## Mon Aug  8 09:54:37 CEST 2022 - The aliscore output file does not exist. Check results for BUSCO: 1314980at2759
## Mon Aug  8 09:54:37 CEST 2022 - The aliscore output appears to be empty. Check results for BUSCO: 1314980at2759
## Mon Aug  8 09:54:37 CEST 2022 - The aliscore output file does not exist. Check results for BUSCO: 418107at2759
## Mon Aug  8 09:54:37 CEST 2022 - The aliscore output file does not exist. Check results for BUSCO: 777920at2759
## Mon Aug  8 09:54:37 CEST 2022 - The aliscore output appears to be empty. Check results for BUSCO: 418107at2759
## Mon Aug  8 09:54:37 CEST 2022 - The aliscore output appears to be empty. Check results for BUSCO: 777920at2759
## Mon Aug  8 09:54:37 CEST 2022 - The aliscore output file does not exist. Check results for BUSCO: 736068at2759
## Mon Aug  8 09:54:37 CEST 2022 - The aliscore output appears to be empty. Check results for BUSCO: 736068at2759
## Mon Aug  8 09:54:37 CEST 2022 - The aliscore output file does not exist. Check results for BUSCO: 918816at2759
## Mon Aug  8 09:54:37 CEST 2022 - The aliscore output appears to be empty. Check results for BUSCO: 918816at2759
## Mon Aug  8 09:54:39 CEST 2022 - The aliscore output file does not exist. Check results for BUSCO: 1312453at2759
## Mon Aug  8 09:54:39 CEST 2022 - The aliscore output appears to be empty. Check results for BUSCO: 1312453at2759
## Mon Aug  8 09:54:40 CEST 2022 - The aliscore output file does not exist. Check results for BUSCO: 396755at2759
## Mon Aug  8 09:54:40 CEST 2022 - The aliscore output file does not exist. Check results for BUSCO: 293315at2759
## Mon Aug  8 09:54:40 CEST 2022 - The aliscore output appears to be empty. Check results for BUSCO: 396755at2759
## Mon Aug  8 09:54:40 CEST 2022 - The aliscore output appears to be empty. Check results for BUSCO: 293315at2759
## Mon Aug  8 09:54:40 CEST 2022 - The aliscore output file does not exist. Check results for BUSCO: 257318at2759
## Mon Aug  8 09:54:40 CEST 2022 - The aliscore output appears to be empty. Check results for BUSCO: 257318at2759
## Mon Aug  8 09:54:40 CEST 2022 - The aliscore output file does not exist. Check results for BUSCO: 598949at2759
## Mon Aug  8 09:54:40 CEST 2022 - The aliscore output appears to be empty. Check results for BUSCO: 598949at2759
## Mon Aug  8 09:54:42 CEST 2022 - The aliscore output file does not exist. Check results for BUSCO: 179362at2759
## Mon Aug  8 09:54:43 CEST 2022 - The aliscore output appears to be empty. Check results for BUSCO: 179362at2759
## Mon Aug  8 09:54:59 CEST 2022 - The aliscore output file does not exist. Check results for BUSCO: 1251252at2759
## Mon Aug  8 09:54:59 CEST 2022 - The aliscore output appears to be empty. Check results for BUSCO: 1251252at2759
## Mon Aug  8 09:55:00 CEST 2022 - The aliscore output file does not exist. Check results for BUSCO: 1269244at2759
## Mon Aug  8 09:55:00 CEST 2022 - The aliscore output appears to be empty. Check results for BUSCO: 1269244at2759
## Mon Aug  8 09:55:01 CEST 2022 - The aliscore output file does not exist. Check results for BUSCO: 976469at2759
## Mon Aug  8 09:55:01 CEST 2022 - The aliscore output appears to be empty. Check results for BUSCO: 976469at2759
## Mon Aug  8 09:55:01 CEST 2022 - The aliscore output file does not exist. Check results for BUSCO: 1266231at2759
## Mon Aug  8 09:55:01 CEST 2022 - The aliscore output appears to be empty. Check results for BUSCO: 1266231at2759
## Mon Aug  8 09:55:02 CEST 2022 - The aliscore output file does not exist. Check results for BUSCO: 981902at2759
## Mon Aug  8 09:55:02 CEST 2022 - The aliscore output appears to be empty. Check results for BUSCO: 981902at2759
## Mon Aug  8 09:55:02 CEST 2022 - The aliscore output file does not exist. Check results for BUSCO: 176625at2759
## Mon Aug  8 09:55:02 CEST 2022 - The aliscore output appears to be empty. Check results for BUSCO: 176625at2759
## Mon Aug  8 09:55:03 CEST 2022 - The aliscore output file does not exist. Check results for BUSCO: 1079827at2759
## Mon Aug  8 09:55:03 CEST 2022 - The aliscore output appears to be empty. Check results for BUSCO: 1079827at2759
## Mon Aug  8 09:55:03 CEST 2022 - The aliscore output file does not exist. Check results for BUSCO: 1620056at2759
## Mon Aug  8 09:55:03 CEST 2022 - The aliscore output appears to be empty. Check results for BUSCO: 1620056at2759
## Mon Aug  8 09:55:03 CEST 2022 - The aliscore output file does not exist. Check results for BUSCO: 1488436at2759
## Mon Aug  8 09:55:03 CEST 2022 - The aliscore output appears to be empty. Check results for BUSCO: 1488436at2759
## Mon Aug  8 09:55:04 CEST 2022 - The aliscore output file does not exist. Check results for BUSCO: 1445102at2759
## Mon Aug  8 09:55:04 CEST 2022 - The aliscore output appears to be empty. Check results for BUSCO: 1445102at2759
## Mon Aug  8 09:55:04 CEST 2022 - The aliscore output file does not exist. Check results for BUSCO: 1538526at2759
## Mon Aug  8 09:55:04 CEST 2022 - The aliscore output appears to be empty. Check results for BUSCO: 1538526at2759
## Mon Aug  8 09:55:04 CEST 2022 - The aliscore output file does not exist. Check results for BUSCO: 1269649at2759
## Mon Aug  8 09:55:05 CEST 2022 - The aliscore output file does not exist. Check results for BUSCO: 1249159at2759
## Mon Aug  8 09:55:05 CEST 2022 - The aliscore output appears to be empty. Check results for BUSCO: 1269649at2759
## Mon Aug  8 09:55:05 CEST 2022 - The aliscore output file does not exist. Check results for BUSCO: 974865at2759
## Mon Aug  8 09:55:05 CEST 2022 - The aliscore output appears to be empty. Check results for BUSCO: 1249159at2759
## Mon Aug  8 09:55:05 CEST 2022 - The aliscore output appears to be empty. Check results for BUSCO: 974865at2759
## Mon Aug  8 09:55:05 CEST 2022 - The aliscore output file does not exist. Check results for BUSCO: 869548at2759
## Mon Aug  8 09:55:05 CEST 2022 - The aliscore output appears to be empty. Check results for BUSCO: 869548at2759
## Mon Aug  8 09:55:05 CEST 2022 - The aliscore output file does not exist. Check results for BUSCO: 1138059at2759
## Mon Aug  8 09:55:05 CEST 2022 - The aliscore output file does not exist. Check results for BUSCO: 937275at2759
## Mon Aug  8 09:55:05 CEST 2022 - The aliscore output appears to be empty. Check results for BUSCO: 937275at2759
## Mon Aug  8 09:55:05 CEST 2022 - The aliscore output appears to be empty. Check results for BUSCO: 1138059at2759
## Mon Aug  8 09:55:05 CEST 2022 - The aliscore output file does not exist. Check results for BUSCO: 1339553at2759
## Mon Aug  8 09:55:05 CEST 2022 - The aliscore output appears to be empty. Check results for BUSCO: 1339553at2759
## Mon Aug  8 09:55:06 CEST 2022 - The aliscore output file does not exist. Check results for BUSCO: 1018517at2759
## Mon Aug  8 09:55:06 CEST 2022 - The aliscore output appears to be empty. Check results for BUSCO: 1018517at2759
## Mon Aug  8 09:55:06 CEST 2022 - The aliscore output file does not exist. Check results for BUSCO: 937275at2759
## Mon Aug  8 09:55:06 CEST 2022 - The aliscore output file does not exist. Check results for BUSCO: 1490022at2759
## Mon Aug  8 09:55:06 CEST 2022 - The aliscore output appears to be empty. Check results for BUSCO: 937275at2759
## Mon Aug  8 09:55:06 CEST 2022 - The aliscore output appears to be empty. Check results for BUSCO: 1490022at2759
## Mon Aug  8 09:55:06 CEST 2022 - The aliscore output file does not exist. Check results for BUSCO: 1426075at2759
## Mon Aug  8 09:55:06 CEST 2022 - The aliscore output appears to be empty. Check results for BUSCO: 1426075at2759
## Mon Aug  8 09:55:06 CEST 2022 - The aliscore output file does not exist. Check results for BUSCO: 1490022at2759
## Mon Aug  8 09:55:06 CEST 2022 - The aliscore output file does not exist. Check results for BUSCO: 1440961at2759
## Mon Aug  8 09:55:06 CEST 2022 - The aliscore output appears to be empty. Check results for BUSCO: 1440961at2759
## Mon Aug  8 09:55:07 CEST 2022 - The aliscore output file does not exist. Check results for BUSCO: 1440961at2759
## Mon Aug  8 09:55:07 CEST 2022 - The aliscore output file does not exist. Check results for BUSCO: 1405146at2759
## Mon Aug  8 09:55:07 CEST 2022 - The aliscore output appears to be empty. Check results for BUSCO: 1440961at2759
## Mon Aug  8 09:55:07 CEST 2022 - The aliscore output appears to be empty. Check results for BUSCO: 1405146at2759
## Mon Aug  8 09:55:07 CEST 2022 - The aliscore output file does not exist. Check results for BUSCO: 345441at2759
## Mon Aug  8 09:55:07 CEST 2022 - The aliscore output appears to be empty. Check results for BUSCO: 345441at2759
## Mon Aug  8 09:55:07 CEST 2022 - The aliscore output file does not exist. Check results for BUSCO: 974865at2759
## Mon Aug  8 09:55:07 CEST 2022 - The aliscore output appears to be empty. Check results for BUSCO: 974865at2759
## Mon Aug  8 09:55:07 CEST 2022 - The aliscore output file does not exist. Check results for BUSCO: 1428265at2759
## Mon Aug  8 09:55:07 CEST 2022 - The aliscore output appears to be empty. Check results for BUSCO: 1428265at2759
## Mon Aug  8 09:55:07 CEST 2022 - The aliscore output file does not exist. Check results for BUSCO: 1405146at2759
## Mon Aug  8 09:55:07 CEST 2022 - The aliscore output appears to be empty. Check results for BUSCO: 1405146at2759
## Mon Aug  8 09:55:07 CEST 2022 - The aliscore output file does not exist. Check results for BUSCO: 83779at2759
## Mon Aug  8 09:55:08 CEST 2022 - The aliscore output appears to be empty. Check results for BUSCO: 83779at2759
## Mon Aug  8 09:55:08 CEST 2022 - The aliscore output file does not exist. Check results for BUSCO: 858842at2759
## Mon Aug  8 09:55:08 CEST 2022 - The aliscore output appears to be empty. Check results for BUSCO: 858842at2759
## Mon Aug  8 09:55:08 CEST 2022 - The aliscore output file does not exist. Check results for BUSCO: 687505at2759
## Mon Aug  8 09:55:08 CEST 2022 - The aliscore output file does not exist. Check results for BUSCO: 975557at2759
## Mon Aug  8 09:55:08 CEST 2022 - The aliscore output appears to be empty. Check results for BUSCO: 687505at2759
## Mon Aug  8 09:55:08 CEST 2022 - The aliscore output file does not exist. Check results for BUSCO: 1312453at2759
## Mon Aug  8 09:55:08 CEST 2022 - The aliscore output appears to be empty. Check results for BUSCO: 1312453at2759
## Mon Aug  8 09:55:08 CEST 2022 - The aliscore output appears to be empty. Check results for BUSCO: 975557at2759
## Mon Aug  8 09:55:08 CEST 2022 - The aliscore output file does not exist. Check results for BUSCO: 924753at2759
## Mon Aug  8 09:55:08 CEST 2022 - The aliscore output appears to be empty. Check results for BUSCO: 924753at2759
## Mon Aug  8 09:55:08 CEST 2022 - The aliscore output file does not exist. Check results for BUSCO: 330169at2759
## Mon Aug  8 09:55:08 CEST 2022 - The aliscore output file does not exist. Check results for BUSCO: 1025450at2759
## Mon Aug  8 09:55:08 CEST 2022 - The aliscore output appears to be empty. Check results for BUSCO: 330169at2759
## Mon Aug  8 09:55:08 CEST 2022 - The aliscore output file does not exist. Check results for BUSCO: 383503at2759
## Mon Aug  8 09:55:08 CEST 2022 - The aliscore output appears to be empty. Check results for BUSCO: 1025450at2759
## Mon Aug  8 09:55:08 CEST 2022 - The aliscore output appears to be empty. Check results for BUSCO: 383503at2759
## Mon Aug  8 09:55:09 CEST 2022 - The aliscore output file does not exist. Check results for BUSCO: 570797at2759
## Mon Aug  8 09:55:09 CEST 2022 - The aliscore output appears to be empty. Check results for BUSCO: 570797at2759
## Mon Aug  8 09:55:09 CEST 2022 - The aliscore output file does not exist. Check results for BUSCO: 1025450at2759
## Mon Aug  8 09:55:09 CEST 2022 - The aliscore output appears to be empty. Check results for BUSCO: 1025450at2759
## Mon Aug  8 09:55:09 CEST 2022 - The aliscore output file does not exist. Check results for BUSCO: 1001705at2759
## Mon Aug  8 09:55:09 CEST 2022 - The aliscore output file does not exist. Check results for BUSCO: 1001705at2759
## Mon Aug  8 09:55:09 CEST 2022 - The aliscore output appears to be empty. Check results for BUSCO: 1001705at2759
## Mon Aug  8 09:55:09 CEST 2022 - The aliscore output appears to be empty. Check results for BUSCO: 1001705at2759
## Mon Aug  8 09:55:09 CEST 2022 - The aliscore output file does not exist. Check results for BUSCO: 1404162at2759
## Mon Aug  8 09:55:09 CEST 2022 - The aliscore output appears to be empty. Check results for BUSCO: 1404162at2759
## Mon Aug  8 09:55:09 CEST 2022 - The aliscore output file does not exist. Check results for BUSCO: 1370285at2759
## Mon Aug  8 09:55:09 CEST 2022 - The aliscore output appears to be empty. Check results for BUSCO: 1370285at2759
## Mon Aug  8 09:55:09 CEST 2022 - The aliscore output file does not exist. Check results for BUSCO: 549762at2759
## Mon Aug  8 09:55:09 CEST 2022 - The aliscore output appears to be empty. Check results for BUSCO: 549762at2759
## Mon Aug  8 09:55:09 CEST 2022 - The aliscore output file does not exist. Check results for BUSCO: 1051021at2759
## Mon Aug  8 09:55:10 CEST 2022 - The aliscore output appears to be empty. Check results for BUSCO: 1051021at2759
## Mon Aug  8 09:55:10 CEST 2022 - The aliscore output file does not exist. Check results for BUSCO: 345441at2759
## Mon Aug  8 09:55:10 CEST 2022 - The aliscore output appears to be empty. Check results for BUSCO: 345441at2759
## Mon Aug  8 09:55:10 CEST 2022 - The aliscore output file does not exist. Check results for BUSCO: 331411at2759
## Mon Aug  8 09:55:10 CEST 2022 - The aliscore output file does not exist. Check results for BUSCO: 1370285at2759
## Mon Aug  8 09:55:10 CEST 2022 - The aliscore output file does not exist. Check results for BUSCO: 330169at2759
## Mon Aug  8 09:55:10 CEST 2022 - The aliscore output appears to be empty. Check results for BUSCO: 331411at2759
## Mon Aug  8 09:55:10 CEST 2022 - The aliscore output appears to be empty. Check results for BUSCO: 1370285at2759
## Mon Aug  8 09:55:10 CEST 2022 - The aliscore output appears to be empty. Check results for BUSCO: 330169at2759
## Mon Aug  8 09:55:10 CEST 2022 - The aliscore output file does not exist. Check results for BUSCO: 390348at2759
## Mon Aug  8 09:55:10 CEST 2022 - The aliscore output appears to be empty. Check results for BUSCO: 390348at2759
## Mon Aug  8 09:55:10 CEST 2022 - The aliscore output file does not exist. Check results for BUSCO: 671536at2759
## Mon Aug  8 09:55:10 CEST 2022 - The aliscore output appears to be empty. Check results for BUSCO: 671536at2759
## Mon Aug  8 09:55:11 CEST 2022 - The aliscore output file does not exist. Check results for BUSCO: 975557at2759
## Mon Aug  8 09:55:11 CEST 2022 - The aliscore output appears to be empty. Check results for BUSCO: 975557at2759
## Mon Aug  8 09:55:11 CEST 2022 - The aliscore output file does not exist. Check results for BUSCO: 671846at2759
## Mon Aug  8 09:55:11 CEST 2022 - The aliscore output appears to be empty. Check results for BUSCO: 671846at2759
## Mon Aug  8 09:55:11 CEST 2022 - The aliscore output file does not exist. Check results for BUSCO: 1079130at2759
## Mon Aug  8 09:55:11 CEST 2022 - The aliscore output appears to be empty. Check results for BUSCO: 1079130at2759
## Mon Aug  8 09:55:12 CEST 2022 - The aliscore output file does not exist. Check results for BUSCO: 671846at2759
## Mon Aug  8 09:55:12 CEST 2022 - The aliscore output appears to be empty. Check results for BUSCO: 671846at2759
## Mon Aug  8 09:55:19 CEST 2022 - The aliscore output file does not exist. Check results for BUSCO: 156083at2759
## Mon Aug  8 09:55:19 CEST 2022 - The aliscore output appears to be empty. Check results for BUSCO: 156083at2759
## Mon Aug  8 09:55:26 CEST 2022 - The aliscore output file does not exist. Check results for BUSCO: 1488436at2759
## Mon Aug  8 09:55:26 CEST 2022 - The aliscore output appears to be empty. Check results for BUSCO: 1488436at2759
## Mon Aug  8 09:55:28 CEST 2022 - The aliscore output file does not exist. Check results for BUSCO: 1051021at2759
## Mon Aug  8 09:55:28 CEST 2022 - The aliscore output file does not exist. Check results for BUSCO: 1157302at2759
## Mon Aug  8 09:55:28 CEST 2022 - The aliscore output appears to be empty. Check results for BUSCO: 1051021at2759
## Mon Aug  8 09:55:28 CEST 2022 - The aliscore output appears to be empty. Check results for BUSCO: 1157302at2759
## Mon Aug  8 09:55:29 CEST 2022 - The aliscore output file does not exist. Check results for BUSCO: 1222562at2759
## Mon Aug  8 09:55:29 CEST 2022 - The aliscore output appears to be empty. Check results for BUSCO: 1222562at2759
## Mon Aug  8 09:55:31 CEST 2022 - The aliscore output file does not exist. Check results for BUSCO: 1284731at2759
## Mon Aug  8 09:55:31 CEST 2022 - The aliscore output appears to be empty. Check results for BUSCO: 1284731at2759
## Mon Aug  8 09:55:31 CEST 2022 - The aliscore output file does not exist. Check results for BUSCO: 1223488at2759
## Mon Aug  8 09:55:31 CEST 2022 - The aliscore output appears to be empty. Check results for BUSCO: 1223488at2759
## Mon Aug  8 09:55:31 CEST 2022 - The aliscore output file does not exist. Check results for BUSCO: 1194691at2759
## Mon Aug  8 09:55:31 CEST 2022 - The aliscore output file does not exist. Check results for BUSCO: 388820at2759
## Mon Aug  8 09:55:31 CEST 2022 - The aliscore output appears to be empty. Check results for BUSCO: 388820at2759
## Mon Aug  8 09:55:31 CEST 2022 - The aliscore output appears to be empty. Check results for BUSCO: 1194691at2759
## Mon Aug  8 09:55:31 CEST 2022 - The aliscore output file does not exist. Check results for BUSCO: 1053181at2759
## Mon Aug  8 09:55:31 CEST 2022 - The aliscore output appears to be empty. Check results for BUSCO: 1053181at2759
## Mon Aug  8 09:55:32 CEST 2022 - The aliscore output file does not exist. Check results for BUSCO: 1233814at2759
## Mon Aug  8 09:55:32 CEST 2022 - The aliscore output appears to be empty. Check results for BUSCO: 1233814at2759
## Mon Aug  8 09:55:32 CEST 2022 - The aliscore output file does not exist. Check results for BUSCO: 1338131at2759
## Mon Aug  8 09:55:32 CEST 2022 - The aliscore output appears to be empty. Check results for BUSCO: 1338131at2759
## Mon Aug  8 09:55:32 CEST 2022 - The aliscore output file does not exist. Check results for BUSCO: 1513531at2759
## Mon Aug  8 09:55:32 CEST 2022 - The aliscore output file does not exist. Check results for BUSCO: 1342242at2759
## Mon Aug  8 09:55:32 CEST 2022 - The aliscore output file does not exist. Check results for BUSCO: 1488235at2759
## Mon Aug  8 09:55:32 CEST 2022 - The aliscore output appears to be empty. Check results for BUSCO: 1513531at2759
## Mon Aug  8 09:55:33 CEST 2022 - The aliscore output appears to be empty. Check results for BUSCO: 1342242at2759
## Mon Aug  8 09:55:33 CEST 2022 - The aliscore output appears to be empty. Check results for BUSCO: 1488235at2759
## Mon Aug  8 09:55:33 CEST 2022 - The aliscore output file does not exist. Check results for BUSCO: 1370304at2759
## Mon Aug  8 09:55:33 CEST 2022 - The aliscore output appears to be empty. Check results for BUSCO: 1370304at2759
## Mon Aug  8 09:55:33 CEST 2022 - The aliscore output file does not exist. Check results for BUSCO: 1019762at2759
## Mon Aug  8 09:55:33 CEST 2022 - The aliscore output appears to be empty. Check results for BUSCO: 1019762at2759
## Mon Aug  8 09:55:33 CEST 2022 - The aliscore output file does not exist. Check results for BUSCO: 1096688at2759
## Mon Aug  8 09:55:33 CEST 2022 - The aliscore output appears to be empty. Check results for BUSCO: 1096688at2759
## Mon Aug  8 09:55:34 CEST 2022 - The aliscore output file does not exist. Check results for BUSCO: 1322299at2759
## Mon Aug  8 09:55:34 CEST 2022 - The aliscore output appears to be empty. Check results for BUSCO: 1322299at2759
## Mon Aug  8 09:55:34 CEST 2022 - The aliscore output file does not exist. Check results for BUSCO: 981902at2759
## Mon Aug  8 09:55:34 CEST 2022 - The aliscore output file does not exist. Check results for BUSCO: 774318at2759
## Mon Aug  8 09:55:34 CEST 2022 - The aliscore output appears to be empty. Check results for BUSCO: 774318at2759
## Mon Aug  8 09:55:34 CEST 2022 - The aliscore output appears to be empty. Check results for BUSCO: 981902at2759
## Mon Aug  8 09:55:35 CEST 2022 - The aliscore output file does not exist. Check results for BUSCO: 1567796at2759
## Mon Aug  8 09:55:35 CEST 2022 - The aliscore output appears to be empty. Check results for BUSCO: 1567796at2759
## Mon Aug  8 09:55:35 CEST 2022 - The aliscore output file does not exist. Check results for BUSCO: 1552706at2759
## Mon Aug  8 09:55:35 CEST 2022 - The aliscore output appears to be empty. Check results for BUSCO: 1552706at2759
## Mon Aug  8 09:55:35 CEST 2022 - The aliscore output file does not exist. Check results for BUSCO: 1251252at2759
## Mon Aug  8 09:55:35 CEST 2022 - The aliscore output appears to be empty. Check results for BUSCO: 1251252at2759
## Mon Aug  8 09:55:35 CEST 2022 - The aliscore output file does not exist. Check results for BUSCO: 1421503at2759
## Mon Aug  8 09:55:35 CEST 2022 - The aliscore output appears to be empty. Check results for BUSCO: 1421503at2759
## Mon Aug  8 09:55:35 CEST 2022 - The aliscore output file does not exist. Check results for BUSCO: 779909at2759
## Mon Aug  8 09:55:35 CEST 2022 - The aliscore output appears to be empty. Check results for BUSCO: 779909at2759
## Mon Aug  8 09:55:36 CEST 2022 - The aliscore output file does not exist. Check results for BUSCO: 1346432at2759
## Mon Aug  8 09:55:36 CEST 2022 - The aliscore output appears to be empty. Check results for BUSCO: 1346432at2759
## Mon Aug  8 09:55:36 CEST 2022 - The aliscore output file does not exist. Check results for BUSCO: 937686at2759
## Mon Aug  8 09:55:36 CEST 2022 - The aliscore output appears to be empty. Check results for BUSCO: 937686at2759
## Mon Aug  8 09:55:37 CEST 2022 - The aliscore output file does not exist. Check results for BUSCO: 1247641at2759
## Mon Aug  8 09:55:37 CEST 2022 - The aliscore output appears to be empty. Check results for BUSCO: 1247641at2759
## Mon Aug  8 09:55:38 CEST 2022 - The aliscore output file does not exist. Check results for BUSCO: 1249159at2759
## Mon Aug  8 09:55:38 CEST 2022 - The aliscore output file does not exist. Check results for BUSCO: 1182451at2759
## Mon Aug  8 09:55:38 CEST 2022 - The aliscore output appears to be empty. Check results for BUSCO: 1249159at2759
## Mon Aug  8 09:55:38 CEST 2022 - The aliscore output appears to be empty. Check results for BUSCO: 1182451at2759
## Mon Aug  8 09:55:38 CEST 2022 - The aliscore output file does not exist. Check results for BUSCO: 83779at2759
## Mon Aug  8 09:55:38 CEST 2022 - The aliscore output appears to be empty. Check results for BUSCO: 83779at2759
## Mon Aug  8 09:55:38 CEST 2022 - The aliscore output file does not exist. Check results for BUSCO: 759498at2759
## Mon Aug  8 09:55:38 CEST 2022 - The aliscore output appears to be empty. Check results for BUSCO: 759498at2759
## Mon Aug  8 09:55:38 CEST 2022 - The aliscore output file does not exist. Check results for BUSCO: 768809at2759
## Mon Aug  8 09:55:38 CEST 2022 - The aliscore output appears to be empty. Check results for BUSCO: 768809at2759
## Mon Aug  8 09:55:39 CEST 2022 - The aliscore output file does not exist. Check results for BUSCO: 325552at2759
## Mon Aug  8 09:55:39 CEST 2022 - The aliscore output appears to be empty. Check results for BUSCO: 325552at2759
## Mon Aug  8 09:55:39 CEST 2022 - The aliscore output file does not exist. Check results for BUSCO: 383503at2759
## Mon Aug  8 09:55:39 CEST 2022 - The aliscore output appears to be empty. Check results for BUSCO: 383503at2759
## Mon Aug  8 09:55:40 CEST 2022 - The aliscore output file does not exist. Check results for BUSCO: 1111142at2759
## Mon Aug  8 09:55:40 CEST 2022 - The aliscore output appears to be empty. Check results for BUSCO: 1111142at2759
## Mon Aug  8 09:55:57 CEST 2022 - The aliscore output file does not exist. Check results for BUSCO: 1358374at2759
## Mon Aug  8 09:55:57 CEST 2022 - The aliscore output appears to be empty. Check results for BUSCO: 1358374at2759
## Mon Aug  8 09:55:59 CEST 2022 - The aliscore output file does not exist. Check results for BUSCO: 944899at2759
## Mon Aug  8 09:55:59 CEST 2022 - The aliscore output appears to be empty. Check results for BUSCO: 944899at2759
## Mon Aug  8 09:56:00 CEST 2022 - The aliscore output file does not exist. Check results for BUSCO: 1038775at2759
## Mon Aug  8 09:56:00 CEST 2022 - The aliscore output appears to be empty. Check results for BUSCO: 1038775at2759
## Mon Aug  8 09:56:01 CEST 2022 - The aliscore output file does not exist. Check results for BUSCO: 1178688at2759
## Mon Aug  8 09:56:01 CEST 2022 - The aliscore output appears to be empty. Check results for BUSCO: 1178688at2759
## Mon Aug  8 09:56:01 CEST 2022 - The aliscore output file does not exist. Check results for BUSCO: 1200489at2759
## Mon Aug  8 09:56:01 CEST 2022 - The aliscore output appears to be empty. Check results for BUSCO: 1200489at2759
## Mon Aug  8 09:56:02 CEST 2022 - The aliscore output file does not exist. Check results for BUSCO: 1459797at2759
## Mon Aug  8 09:56:02 CEST 2022 - The aliscore output appears to be empty. Check results for BUSCO: 1459797at2759
## Mon Aug  8 09:56:03 CEST 2022 - The aliscore output file does not exist. Check results for BUSCO: 1248958at2759
## Mon Aug  8 09:56:03 CEST 2022 - The aliscore output appears to be empty. Check results for BUSCO: 1248958at2759
## Mon Aug  8 09:56:03 CEST 2022 - The aliscore output file does not exist. Check results for BUSCO: 1038775at2759
## Mon Aug  8 09:56:04 CEST 2022 - The aliscore output appears to be empty. Check results for BUSCO: 1038775at2759
## Mon Aug  8 09:56:04 CEST 2022 - The aliscore output file does not exist. Check results for BUSCO: 1407446at2759
## Mon Aug  8 09:56:04 CEST 2022 - The aliscore output appears to be empty. Check results for BUSCO: 1407446at2759
## Mon Aug  8 09:56:04 CEST 2022 - The aliscore output file does not exist. Check results for BUSCO: 1287401at2759
## Mon Aug  8 09:56:04 CEST 2022 - The aliscore output appears to be empty. Check results for BUSCO: 1287401at2759
## Mon Aug  8 09:56:04 CEST 2022 - The aliscore output file does not exist. Check results for BUSCO: 751335at2759
## Mon Aug  8 09:56:04 CEST 2022 - The aliscore output file does not exist. Check results for BUSCO: 1358374at2759
## Mon Aug  8 09:56:05 CEST 2022 - The aliscore output appears to be empty. Check results for BUSCO: 751335at2759
## Mon Aug  8 09:56:05 CEST 2022 - The aliscore output appears to be empty. Check results for BUSCO: 1358374at2759
## Mon Aug  8 09:56:05 CEST 2022 - The aliscore output file does not exist. Check results for BUSCO: 866359at2759
## Mon Aug  8 09:56:05 CEST 2022 - The aliscore output appears to be empty. Check results for BUSCO: 866359at2759
## Mon Aug  8 09:56:05 CEST 2022 - The aliscore output file does not exist. Check results for BUSCO: 708105at2759
## Mon Aug  8 09:56:05 CEST 2022 - The aliscore output file does not exist. Check results for BUSCO: 1304061at2759
## Mon Aug  8 09:56:05 CEST 2022 - The aliscore output appears to be empty. Check results for BUSCO: 708105at2759
## Mon Aug  8 09:56:05 CEST 2022 - The aliscore output appears to be empty. Check results for BUSCO: 1304061at2759
## Mon Aug  8 09:56:05 CEST 2022 - The aliscore output file does not exist. Check results for BUSCO: 1200489at2759
## Mon Aug  8 09:56:05 CEST 2022 - The aliscore output appears to be empty. Check results for BUSCO: 1200489at2759
## Mon Aug  8 09:56:05 CEST 2022 - The aliscore output file does not exist. Check results for BUSCO: 1558822at2759
## Mon Aug  8 09:56:05 CEST 2022 - The aliscore output appears to be empty. Check results for BUSCO: 1558822at2759
## Mon Aug  8 09:56:05 CEST 2022 - The aliscore output file does not exist. Check results for BUSCO: 761109at2759
## Mon Aug  8 09:56:05 CEST 2022 - The aliscore output appears to be empty. Check results for BUSCO: 761109at2759
## Mon Aug  8 09:56:06 CEST 2022 - The aliscore output file does not exist. Check results for BUSCO: 1014314at2759
## Mon Aug  8 09:56:06 CEST 2022 - The aliscore output appears to be empty. Check results for BUSCO: 1014314at2759
## Mon Aug  8 09:56:06 CEST 2022 - The aliscore output file does not exist. Check results for BUSCO: 774318at2759
## Mon Aug  8 09:56:06 CEST 2022 - The aliscore output appears to be empty. Check results for BUSCO: 774318at2759
## Mon Aug  8 09:56:06 CEST 2022 - The aliscore output file does not exist. Check results for BUSCO: 817008at2759
## Mon Aug  8 09:56:06 CEST 2022 - The aliscore output appears to be empty. Check results for BUSCO: 817008at2759
## Mon Aug  8 09:56:07 CEST 2022 - The aliscore output file does not exist. Check results for BUSCO: 1342242at2759
## Mon Aug  8 09:56:07 CEST 2022 - The aliscore output file does not exist. Check results for BUSCO: 1233814at2759
## Mon Aug  8 09:56:07 CEST 2022 - The aliscore output appears to be empty. Check results for BUSCO: 1342242at2759
## Mon Aug  8 09:56:07 CEST 2022 - The aliscore output appears to be empty. Check results for BUSCO: 1233814at2759
## Mon Aug  8 09:56:07 CEST 2022 - The aliscore output file does not exist. Check results for BUSCO: 426305at2759
## Mon Aug  8 09:56:08 CEST 2022 - The aliscore output appears to be empty. Check results for BUSCO: 426305at2759
## Mon Aug  8 09:56:08 CEST 2022 - The aliscore output file does not exist. Check results for BUSCO: 604979at2759
## Mon Aug  8 09:56:08 CEST 2022 - The aliscore output file does not exist. Check results for BUSCO: 939345at2759
## Mon Aug  8 09:56:08 CEST 2022 - The aliscore output appears to be empty. Check results for BUSCO: 939345at2759
## Mon Aug  8 09:56:08 CEST 2022 - The aliscore output appears to be empty. Check results for BUSCO: 604979at2759
## Mon Aug  8 09:56:08 CEST 2022 - The aliscore output file does not exist. Check results for BUSCO: 1287401at2759
## Mon Aug  8 09:56:08 CEST 2022 - The aliscore output file does not exist. Check results for BUSCO: 944899at2759
## Mon Aug  8 09:56:08 CEST 2022 - The aliscore output file does not exist. Check results for BUSCO: 1441854at2759
## Mon Aug  8 09:56:08 CEST 2022 - The aliscore output appears to be empty. Check results for BUSCO: 1287401at2759
## Mon Aug  8 09:56:08 CEST 2022 - The aliscore output appears to be empty. Check results for BUSCO: 944899at2759
## Mon Aug  8 09:56:08 CEST 2022 - The aliscore output appears to be empty. Check results for BUSCO: 1441854at2759
## Mon Aug  8 09:56:08 CEST 2022 - The aliscore output file does not exist. Check results for BUSCO: 290630at2759
## Mon Aug  8 09:56:08 CEST 2022 - The aliscore output appears to be empty. Check results for BUSCO: 290630at2759
## Mon Aug  8 09:56:09 CEST 2022 - The aliscore output file does not exist. Check results for BUSCO: 751335at2759
## Mon Aug  8 09:56:09 CEST 2022 - The aliscore output appears to be empty. Check results for BUSCO: 751335at2759
## Mon Aug  8 09:56:09 CEST 2022 - The aliscore output file does not exist. Check results for BUSCO: 290630at2759
## Mon Aug  8 09:56:09 CEST 2022 - The aliscore output appears to be empty. Check results for BUSCO: 290630at2759
## Mon Aug  8 09:56:09 CEST 2022 - The aliscore output file does not exist. Check results for BUSCO: 39650at2759
## Mon Aug  8 09:56:09 CEST 2022 - The aliscore output appears to be empty. Check results for BUSCO: 39650at2759
## Mon Aug  8 09:56:10 CEST 2022 - The aliscore output file does not exist. Check results for BUSCO: 582756at2759
## Mon Aug  8 09:56:10 CEST 2022 - The aliscore output appears to be empty. Check results for BUSCO: 582756at2759
## Mon Aug  8 09:56:10 CEST 2022 - The aliscore output file does not exist. Check results for BUSCO: 604979at2759
## Mon Aug  8 09:56:10 CEST 2022 - The aliscore output appears to be empty. Check results for BUSCO: 604979at2759
## Mon Aug  8 09:56:11 CEST 2022 - The aliscore output file does not exist. Check results for BUSCO: 306227at2759
## Mon Aug  8 09:56:11 CEST 2022 - The aliscore output appears to be empty. Check results for BUSCO: 306227at2759
## Mon Aug  8 09:56:13 CEST 2022 - The aliscore output file does not exist. Check results for BUSCO: 1379600at2759
## Mon Aug  8 09:56:13 CEST 2022 - The aliscore output appears to be empty. Check results for BUSCO: 1379600at2759
## Mon Aug  8 09:56:13 CEST 2022 - The aliscore output file does not exist. Check results for BUSCO: 39650at2759
## Mon Aug  8 09:56:13 CEST 2022 - The aliscore output appears to be empty. Check results for BUSCO: 39650at2759
## Mon Aug  8 09:56:15 CEST 2022 - The aliscore output file does not exist. Check results for BUSCO: 261328at2759
## Mon Aug  8 09:56:15 CEST 2022 - The aliscore output appears to be empty. Check results for BUSCO: 261328at2759
## Mon Aug  8 09:56:59 CEST 2022 - Number of alignments (clustalo): 255
## Mon Aug  8 09:56:59 CEST 2022 - Number of trimmed alignments (clustalo - bmge): 255
## Mon Aug  8 09:57:00 CEST 2022 - Number of alignments (muscle): 255
## Mon Aug  8 09:57:01 CEST 2022 - Number of trimmed alignments (muscle - aliscore): 255
## Mon Aug  8 09:57:00 CEST 2022 - Number of alignments (clustalo - bmge) after filtering: 241
## Mon Aug  8 09:57:01 CEST 2022 - Number of alignments (mafft): 255
## Mon Aug  8 09:57:01 CEST 2022 - Number of alignments (clustalo): 255
## Mon Aug  8 09:57:01 CEST 2022 - Number of trimmed alignments (mafft - aliscore): 255
## Mon Aug  8 09:57:01 CEST 2022 - Number of trimmed alignments (clustalo - aliscore): 255
## Mon Aug  8 09:57:01 CEST 2022 - Number of alignments (muscle - aliscore) after filtering: 249
## Mon Aug  8 09:57:01 CEST 2022 - Number of alignments (mafft - aliscore) after filtering: 249
## Mon Aug  8 09:57:01 CEST 2022 - Number of alignments (clustalo - aliscore) after filtering: 248
## Mon Aug  8 09:57:29 CEST 2022 - Number of alignments (muscle): 255
## Mon Aug  8 09:57:30 CEST 2022 - Number of trimmed alignments (muscle - bmge): 255
## Mon Aug  8 09:57:30 CEST 2022 - Number of alignments (mafft): 255
## Mon Aug  8 09:57:30 CEST 2022 - Number of alignments (mafft): 255
## Mon Aug  8 09:57:30 CEST 2022 - Number of alignments (muscle): 255
## Mon Aug  8 09:57:30 CEST 2022 - Number of alignments (clustalo): 255
## Mon Aug  8 09:57:30 CEST 2022 - Number of trimmed alignments (mafft - bmge): 255
## Mon Aug  8 09:57:30 CEST 2022 - Number of trimmed alignments (mafft - trimal): 255
## Mon Aug  8 09:57:30 CEST 2022 - Number of trimmed alignments (muscle - trimal): 255
## Mon Aug  8 09:57:30 CEST 2022 - Number of trimmed alignments (clustalo - trimal): 255
## Mon Aug  8 09:57:30 CEST 2022 - Number of alignments (muscle - bmge) after filtering: 243
## Mon Aug  8 09:57:30 CEST 2022 - Number of alignments (mafft - bmge) after filtering: 245
## Mon Aug  8 09:57:30 CEST 2022 - Number of alignments (mafft - trimal) after filtering: 250
## Mon Aug  8 09:57:30 CEST 2022 - Number of alignments (muscle - trimal) after filtering: 249
## Mon Aug  8 09:57:30 CEST 2022 - Number of alignments (clustalo - trimal) after filtering: 248
## Mon Aug  8 09:58:22 CEST 2022 - phylociraptor filter-align done.
## Tue 09 Aug 2022 12:33:18 PM CEST - phylociraptor modeltest done.
## Tue 09 Aug 2022 12:35:34 PM CEST - phylociraptor will use gene tree filtering based on average bootstrap support value of 70.
## Tue 09 Aug 2022 12:35:34 PM CEST - phylociraptor will use gene tree filtering based on average bootstrap support value of 70.
## Tue 09 Aug 2022 12:35:34 PM CEST - phylociraptor will use gene tree filtering based on average bootstrap support value of 80.
## Tue 09 Aug 2022 12:35:34 PM CEST - phylociraptor will use gene tree filtering based on average bootstrap support value of 80.
## Tue 09 Aug 2022 12:35:34 PM CEST - phylociraptor will use gene tree filtering based on average bootstrap support value of 80.
## Tue 09 Aug 2022 12:35:34 PM CEST - phylociraptor will use gene tree filtering based on average bootstrap support value of 70.
## Tue 09 Aug 2022 12:35:34 PM CEST - phylociraptor will use gene tree filtering based on average bootstrap support value of 60.
## Tue 09 Aug 2022 12:35:34 PM CEST - phylociraptor will use gene tree filtering based on average bootstrap support value of 60.
## Tue 09 Aug 2022 12:35:34 PM CEST - phylociraptor will use gene tree filtering based on average bootstrap support value of 50.
## Tue 09 Aug 2022 12:35:34 PM CEST - phylociraptor will use gene tree filtering based on average bootstrap support value of 50.
## Tue 09 Aug 2022 12:35:34 PM CEST - phylociraptor will use gene tree filtering based on average bootstrap support value of 60.
## Tue 09 Aug 2022 12:35:34 PM CEST - phylociraptor will use gene tree filtering based on average bootstrap support value of 50.
## Tue 09 Aug 2022 12:35:34 PM CEST - phylociraptor will use gene tree filtering based on average bootstrap support value of 0.
## Tue 09 Aug 2022 12:35:34 PM CEST - phylociraptor will use gene tree filtering based on average bootstrap support value of 0.
## Tue 09 Aug 2022 12:35:34 PM CEST - phylociraptor will use gene tree filtering based on average bootstrap support value of 80.
## Tue 09 Aug 2022 12:35:34 PM CEST - phylociraptor will use gene tree filtering based on average bootstrap support value of 80.
## Tue 09 Aug 2022 12:35:34 PM CEST - phylociraptor will use gene tree filtering based on average bootstrap support value of 0.
## Tue 09 Aug 2022 12:35:34 PM CEST - phylociraptor will use gene tree filtering based on average bootstrap support value of 80.
## Tue 09 Aug 2022 12:35:34 PM CEST - phylociraptor will use gene tree filtering based on average bootstrap support value of 70.
## Tue 09 Aug 2022 12:35:34 PM CEST - phylociraptor will use gene tree filtering based on average bootstrap support value of 70.
## Tue 09 Aug 2022 12:35:34 PM CEST - phylociraptor will use gene tree filtering based on average bootstrap support value of 60.
## Tue 09 Aug 2022 12:35:34 PM CEST - phylociraptor will use gene tree filtering based on average bootstrap support value of 60.
## Tue 09 Aug 2022 12:35:34 PM CEST - phylociraptor will use gene tree filtering based on average bootstrap support value of 70.
## Tue 09 Aug 2022 12:35:34 PM CEST - phylociraptor will use gene tree filtering based on average bootstrap support value of 60.
## Tue 09 Aug 2022 12:35:34 PM CEST - phylociraptor will use gene tree filtering based on average bootstrap support value of 0.
## Tue 09 Aug 2022 12:35:34 PM CEST - phylociraptor will use gene tree filtering based on average bootstrap support value of 80.
## Tue 09 Aug 2022 12:35:34 PM CEST - phylociraptor will use gene tree filtering based on average bootstrap support value of 80.
## Tue 09 Aug 2022 12:35:34 PM CEST - phylociraptor will use gene tree filtering based on average bootstrap support value of 70.
## Tue 09 Aug 2022 12:35:34 PM CEST - phylociraptor will use gene tree filtering based on average bootstrap support value of 70.
## Tue 09 Aug 2022 12:35:34 PM CEST - phylociraptor will use gene tree filtering based on average bootstrap support value of 80.
## Tue 09 Aug 2022 12:35:35 PM CEST - phylociraptor will use gene tree filtering based on average bootstrap support value of 70.
## Tue 09 Aug 2022 12:35:35 PM CEST - phylociraptor will use gene tree filtering based on average bootstrap support value of 60.
## Tue 09 Aug 2022 12:35:35 PM CEST - phylociraptor will use gene tree filtering based on average bootstrap support value of 60.
## Tue 09 Aug 2022 12:35:35 PM CEST - phylociraptor will use gene tree filtering based on average bootstrap support value of 50.
## Tue 09 Aug 2022 12:35:35 PM CEST - phylociraptor will use gene tree filtering based on average bootstrap support value of 50.
## Tue 09 Aug 2022 12:35:35 PM CEST - phylociraptor will use gene tree filtering based on average bootstrap support value of 60.
## Tue 09 Aug 2022 12:35:35 PM CEST - phylociraptor will use gene tree filtering based on average bootstrap support value of 50.
## Tue 09 Aug 2022 12:35:35 PM CEST - phylociraptor will use gene tree filtering based on average bootstrap support value of 0.
## Tue 09 Aug 2022 12:35:35 PM CEST - phylociraptor will use gene tree filtering based on average bootstrap support value of 0.
## Tue 09 Aug 2022 12:35:35 PM CEST - phylociraptor will use gene tree filtering based on average bootstrap support value of 0.
## Tue 09 Aug 2022 12:35:37 PM CEST - phylociraptor will use gene tree filtering based on average bootstrap support value of 0.
## Tue 09 Aug 2022 12:35:37 PM CEST - phylociraptor will use gene tree filtering based on average bootstrap support value of 50.
## Tue 09 Aug 2022 12:35:37 PM CEST - phylociraptor will use gene tree filtering based on average bootstrap support value of 0.
## Tue 09 Aug 2022 12:35:37 PM CEST - phylociraptor will use gene tree filtering based on average bootstrap support value of 50.
## Tue 09 Aug 2022 12:35:37 PM CEST - phylociraptor will use gene tree filtering based on average bootstrap support value of 50.
## Tue 09 Aug 2022 12:36:36 PM CEST - phylociraptor speciestree reconstruction done.
## Tue Aug  9 12:38:13 CEST 2022 - concatenate muscle-trimal: Will use bootstrap cutoff 0 before creating concatenated alignment
## Tue Aug  9 12:38:13 CEST 2022 - concatenate mafft-aliscore: Will use bootstrap cutoff 60 before creating concatenated alignment
## Tue Aug  9 12:38:13 CEST 2022 - concatenate clustalo-aliscore: Will use bootstrap cutoff 60 before creating concatenated alignment
## Tue Aug  9 12:38:13 CEST 2022 - concatenate mafft-trimal: Will use bootstrap cutoff 60 before creating concatenated alignment
## Tue Aug  9 12:38:13 CEST 2022 - concatenate mafft-trimal: Will use bootstrap cutoff 0 before creating concatenated alignment
## Tue Aug  9 12:38:13 CEST 2022 - concatenate clustalo-trimal: Will use bootstrap cutoff 0 before creating concatenated alignment
## Tue Aug  9 12:38:13 CEST 2022 - concatenate mafft-bmge: Will use bootstrap cutoff 80 before creating concatenated alignment
## Tue Aug  9 12:38:13 CEST 2022 - prepare_iqtree clustalo-aliscore: Will use bootstrap cutoff (0) before creating concatenated alignment
## Tue Aug  9 12:38:13 CEST 2022 - prepare_iqtree clustalo-aliscore: Will use bootstrap cutoff (50) before creating concatenated alignment
## Tue Aug  9 12:38:13 CEST 2022 - prepare_iqtree mafft-bmge: Will use bootstrap cutoff (70) before creating concatenated alignment
## Tue Aug  9 12:38:13 CEST 2022 - prepare_iqtree muscle-aliscore: Will use bootstrap cutoff (0) before creating concatenated alignment
## Tue Aug  9 12:38:13 CEST 2022 - concatenate mafft-trimal: Will use bootstrap cutoff 50 before creating concatenated alignment
## Tue Aug  9 12:38:13 CEST 2022 - prepare_iqtree mafft-bmge: Will use bootstrap cutoff (80) before creating concatenated alignment
## Tue Aug  9 12:38:13 CEST 2022 - prepare_iqtree clustalo-aliscore: Will use bootstrap cutoff (60) before creating concatenated alignment
## Tue Aug  9 12:38:13 CEST 2022 - concatenate clustalo-trimal: Will use bootstrap cutoff 50 before creating concatenated alignment
## Tue Aug  9 12:38:13 CEST 2022 - concatenate muscle-bmge: Will use bootstrap cutoff 80 before creating concatenated alignment
## Tue Aug  9 12:38:13 CEST 2022 - concatenate mafft-aliscore: Will use bootstrap cutoff 70 before creating concatenated alignment
## Tue Aug  9 12:38:13 CEST 2022 - concatenate mafft-bmge: Will use bootstrap cutoff 0 before creating concatenated alignment
## Tue Aug  9 12:38:13 CEST 2022 - concatenate clustalo-aliscore: Will use bootstrap cutoff 70 before creating concatenated alignment
## Tue Aug  9 12:38:13 CEST 2022 - prepare_iqtree clustalo-trimal: Will use bootstrap cutoff (0) before creating concatenated alignment
## Will create NEXUS partition file with model information now.
## Will create NEXUS partition file with model information now.
## Tue Aug  9 12:38:16 CEST 2022 - nexus file for iqtree written.
## Tue Aug  9 12:38:17 CEST 2022 - prepare_iqtree clustalo-aliscore: Will use bootstrap cutoff (80) before creating concatenated alignment
## Will create NEXUS partition file with model information now.
## Tue Aug  9 12:38:17 CEST 2022 - nexus file for iqtree written.
## Tue Aug  9 12:38:18 CEST 2022 - concatenate mafft-bmge: Will use bootstrap cutoff 50 before creating concatenated alignment
## Tue Aug  9 12:38:19 CEST 2022 - nexus file for iqtree written.
## Will create NEXUS partition file with model information now.
## Will create NEXUS partition file with model information now.
## Will create NEXUS partition file with model information now.
## Will create NEXUS partition file with model information now.
## Tue Aug  9 12:38:20 CEST 2022 - prepare_iqtree clustalo-bmge: Will use bootstrap cutoff (70) before creating concatenated alignment
## Will create NEXUS partition file with model information now.
## Tue Aug  9 12:38:21 CEST 2022 - nexus file for iqtree written.
## Tue Aug  9 12:38:21 CEST 2022 - nexus file for iqtree written.
## Tue Aug  9 12:38:21 CEST 2022 - nexus file for iqtree written.
## Tue Aug  9 12:38:21 CEST 2022 - concatenate muscle-aliscore: Will use bootstrap cutoff 60 before creating concatenated alignment
## Tue Aug  9 12:38:21 CEST 2022 - nexus file for iqtree written.
## Tue Aug  9 12:38:22 CEST 2022 - nexus file for iqtree written.
## Tue Aug  9 12:38:22 CEST 2022 - prepare_iqtree muscle-bmge: Will use bootstrap cutoff (70) before creating concatenated alignment
## Tue Aug  9 12:38:22 CEST 2022 - prepare_iqtree mafft-aliscore: Will use bootstrap cutoff (60) before creating concatenated alignment
## Tue Aug  9 12:38:22 CEST 2022 - prepare_iqtree mafft-trimal: Will use bootstrap cutoff (70) before creating concatenated alignment
## Tue Aug  9 12:38:23 CEST 2022 - prepare_iqtree mafft-aliscore: Will use bootstrap cutoff (50) before creating concatenated alignment
## Will create NEXUS partition file with model information now.
## Will create NEXUS partition file with model information now.
## Will create NEXUS partition file with model information now.
## Will create NEXUS partition file with model information now.
## Tue Aug  9 12:38:25 CEST 2022 - nexus file for iqtree written.
## Will create NEXUS partition file with model information now.
## Tue Aug  9 12:38:26 CEST 2022 - concatenate mafft-aliscore: Will use bootstrap cutoff 80 before creating concatenated alignment
## Tue Aug  9 12:38:26 CEST 2022 - nexus file for iqtree written.
## Tue Aug  9 12:38:26 CEST 2022 - nexus file for iqtree written.
## Tue Aug  9 12:38:27 CEST 2022 - nexus file for iqtree written.
## Tue Aug  9 12:38:27 CEST 2022 - nexus file for iqtree written.
## Tue Aug  9 12:38:27 CEST 2022 - prepare_iqtree mafft-trimal: Will use bootstrap cutoff (60) before creating concatenated alignment
## Tue Aug  9 12:38:27 CEST 2022 - prepare_iqtree clustalo-trimal: Will use bootstrap cutoff (50) before creating concatenated alignment
## Tue Aug  9 12:38:28 CEST 2022 - concatenate muscle-bmge: Will use bootstrap cutoff 50 before creating concatenated alignment
## Tue Aug  9 12:38:28 CEST 2022 - concatenate muscle-aliscore: Will use bootstrap cutoff 70 before creating concatenated alignment
## Tue Aug  9 12:38:28 CEST 2022 - prepare_iqtree muscle-bmge: Will use bootstrap cutoff (80) before creating concatenated alignment
## Tue Aug  9 12:38:28 CEST 2022 - concatenate mafft-trimal: Will use bootstrap cutoff 70 before creating concatenated alignment
## Will create NEXUS partition file with model information now.
## Will create NEXUS partition file with model information now.
## Will create NEXUS partition file with model information now.
## Tue Aug  9 12:38:29 CEST 2022 - nexus file for iqtree written.
## Tue Aug  9 12:38:30 CEST 2022 - prepare_iqtree clustalo-trimal: Will use bootstrap cutoff (60) before creating concatenated alignment
## Tue Aug  9 12:38:31 CEST 2022 - nexus file for iqtree written.
## Tue Aug  9 12:38:31 CEST 2022 - nexus file for iqtree written.
## Tue Aug  9 12:38:32 CEST 2022 - concatenate muscle-trimal: Will use bootstrap cutoff 70 before creating concatenated alignment
## Will create NEXUS partition file with model information now.
## Tue Aug  9 12:38:32 CEST 2022 - prepare_iqtree clustalo-trimal: Will use bootstrap cutoff (70) before creating concatenated alignment
## Will create NEXUS partition file with model information now.
## Tue Aug  9 12:38:34 CEST 2022 - nexus file for iqtree written.
## Tue Aug  9 12:38:35 CEST 2022 - prepare_iqtree muscle-trimal: Will use bootstrap cutoff (80) before creating concatenated alignment
## Tue Aug  9 12:38:35 CEST 2022 - nexus file for iqtree written.
## Will create NEXUS partition file with model information now.
## Tue Aug  9 12:38:36 CEST 2022 - prepare_iqtree muscle-trimal: Will use bootstrap cutoff (60) before creating concatenated alignment
## Tue Aug  9 12:38:37 CEST 2022 - prepare_iqtree muscle-aliscore: Will use bootstrap cutoff (70) before creating concatenated alignment
## Tue Aug  9 12:38:37 CEST 2022 - nexus file for iqtree written.
## Will create NEXUS partition file with model information now.
## Tue Aug  9 12:38:38 CEST 2022 - prepare_iqtree clustalo-bmge: Will use bootstrap cutoff (50) before creating concatenated alignment
## Will create NEXUS partition file with model information now.
## Tue Aug  9 12:38:40 CEST 2022 - nexus file for iqtree written.
## Will create NEXUS partition file with model information now.
## Tue Aug  9 12:38:40 CEST 2022 - nexus file for iqtree written.
## Tue Aug  9 12:38:41 CEST 2022 - concatenate mafft-bmge: Will use bootstrap cutoff 60 before creating concatenated alignment
## Tue Aug  9 12:38:41 CEST 2022 - prepare_iqtree mafft-trimal: Will use bootstrap cutoff (0) before creating concatenated alignment
## Tue Aug  9 12:38:41 CEST 2022 - concatenate clustalo-bmge: Will use bootstrap cutoff 80 before creating concatenated alignment
## Tue Aug  9 12:38:42 CEST 2022 - nexus file for iqtree written.
## Tue 09 Aug 2022 12:38:42 PM CEST - 'phylociraptor model' finished successfully before. Will run raxml with best models.
## Tue Aug  9 12:38:42 CEST 2022 - concatenate clustalo-trimal: Will use bootstrap cutoff 80 before creating concatenated alignment
## Tue 09 Aug 2022 12:38:42 PM CEST - 'phylociraptor model' finished successfully before. Will run raxml with best models.
## Tue Aug  9 12:38:42 CEST 2022 - concatenate clustalo-trimal: Will use bootstrap cutoff 70 before creating concatenated alignment
## Will create NEXUS partition file with model information now.
## Tue Aug  9 12:38:43 CEST 2022 - concatenate muscle-aliscore: Will use bootstrap cutoff 50 before creating concatenated alignment
## Tue 09 Aug 2022 12:38:43 PM CEST - 'phylociraptor model' finished successfully before. Will run raxml with best models.
## Tue Aug  9 12:38:43 CEST 2022 - prepare_iqtree muscle-bmge: Will use bootstrap cutoff (60) before creating concatenated alignment
## Tue Aug  9 12:38:44 CEST 2022 - concatenate mafft-aliscore: Will use bootstrap cutoff 50 before creating concatenated alignment
## Tue Aug  9 12:38:44 CEST 2022 - prepare_iqtree clustalo-trimal: Will use bootstrap cutoff (80) before creating concatenated alignment
## Tue Aug  9 12:38:44 CEST 2022 - concatenate mafft-aliscore: Will use bootstrap cutoff 0 before creating concatenated alignment
## Tue Aug  9 12:38:44 CEST 2022 - concatenate muscle-aliscore: Will use bootstrap cutoff 0 before creating concatenated alignment
## Tue Aug  9 12:38:44 CEST 2022 - prepare_iqtree mafft-aliscore: Will use bootstrap cutoff (70) before creating concatenated alignment
## Tue 09 Aug 2022 12:38:44 PM CEST - 'phylociraptor model' finished successfully before. Will run raxml with best models.
## Tue Aug  9 12:38:45 CEST 2022 - nexus file for iqtree written.
## Tue Aug  9 12:38:45 CEST 2022 - concatenate muscle-trimal: Will use bootstrap cutoff 60 before creating concatenated alignment
## Will create NEXUS partition file with model information now.
## Will create NEXUS partition file with model information now.
## Tue Aug  9 12:38:45 CEST 2022 - concatenate clustalo-aliscore: Will use bootstrap cutoff 50 before creating concatenated alignment
## Tue Aug  9 12:38:46 CEST 2022 - prepare_iqtree mafft-trimal: Will use bootstrap cutoff (80) before creating concatenated alignment
## Will create NEXUS partition file with model information now.
## Tue Aug  9 12:38:46 CEST 2022 - concatenate clustalo-trimal: Will use bootstrap cutoff 60 before creating concatenated alignment
## Tue Aug  9 12:38:47 CEST 2022 - nexus file for iqtree written.
## Will create NEXUS partition file with model information now.
## Tue Aug  9 12:38:47 CEST 2022 - nexus file for iqtree written.
## Tue Aug  9 12:38:48 CEST 2022 - prepare_iqtree muscle-bmge: Will use bootstrap cutoff (0) before creating concatenated alignment
## Tue Aug  9 12:38:48 CEST 2022 - nexus file for iqtree written.
## Tue Aug  9 12:38:48 CEST 2022 - concatenate muscle-aliscore: Will use bootstrap cutoff 80 before creating concatenated alignment
## Tue Aug  9 12:38:49 CEST 2022 - nexus file for iqtree written.
## Tue Aug  9 12:38:49 CEST 2022 - prepare_iqtree muscle-aliscore: Will use bootstrap cutoff (50) before creating concatenated alignment
## Tue 09 Aug 2022 12:38:49 PM CEST - 'phylociraptor model' finished successfully before. Will run raxml with best models.
## Will create NEXUS partition file with model information now.
## Tue Aug  9 12:38:49 CEST 2022 - prepare_iqtree mafft-aliscore: Will use bootstrap cutoff (0) before creating concatenated alignment
## Tue Aug  9 12:38:50 CEST 2022 - prepare_iqtree mafft-bmge: Will use bootstrap cutoff (50) before creating concatenated alignment
## Tue Aug  9 12:38:50 CEST 2022 - prepare_iqtree mafft-aliscore: Will use bootstrap cutoff (80) before creating concatenated alignment
## Will create NEXUS partition file with model information now.
## Will create NEXUS partition file with model information now.
## Will create NEXUS partition file with model information now.
## Tue Aug  9 12:38:51 CEST 2022 - nexus file for iqtree written.
## Tue 09 Aug 2022 12:38:52 PM CEST - 'phylociraptor model' finished successfully before. Will run raxml with best models.
## Will create NEXUS partition file with model information now.
## Tue Aug  9 12:38:52 CEST 2022 - concatenate clustalo-bmge: Will use bootstrap cutoff 50 before creating concatenated alignment
## Tue Aug  9 12:38:52 CEST 2022 - nexus file for iqtree written.
## Tue Aug  9 12:38:53 CEST 2022 - nexus file for iqtree written.
## Tue 09 Aug 2022 12:38:53 PM CEST - 'phylociraptor model' finished successfully before. Will run raxml with best models.
## Tue 09 Aug 2022 12:38:53 PM CEST - 'phylociraptor model' finished successfully before. Will run raxml with best models.
## Tue 09 Aug 2022 12:38:53 PM CEST - 'phylociraptor model' finished successfully before. Will run raxml with best models.
## Tue Aug  9 12:38:53 CEST 2022 - prepare_iqtree clustalo-aliscore: Will use bootstrap cutoff (70) before creating concatenated alignment
## Tue Aug  9 12:38:53 CEST 2022 - nexus file for iqtree written.
## Tue Aug  9 12:38:54 CEST 2022 - concatenate muscle-bmge: Will use bootstrap cutoff 70 before creating concatenated alignment
## Tue Aug  9 12:38:54 CEST 2022 - nexus file for iqtree written.
## Tue Aug  9 12:38:54 CEST 2022 - concatenate mafft-bmge: Will use bootstrap cutoff 70 before creating concatenated alignment
## Tue Aug  9 12:38:54 CEST 2022 - prepare_iqtree mafft-bmge: Will use bootstrap cutoff (60) before creating concatenated alignment
## Tue Aug  9 12:38:55 CEST 2022 - prepare_iqtree muscle-aliscore: Will use bootstrap cutoff (80) before creating concatenated alignment
## Will create NEXUS partition file with model information now.
## Tue 09 Aug 2022 12:38:55 PM CEST - 'phylociraptor model' finished successfully before. Will run raxml with best models.
## Tue 09 Aug 2022 12:38:55 PM CEST - 'phylociraptor model' finished successfully before. Will run raxml with best models.
## Tue Aug  9 12:38:56 CEST 2022 - prepare_iqtree muscle-trimal: Will use bootstrap cutoff (50) before creating concatenated alignment
## Will create NEXUS partition file with model information now.
## Will create NEXUS partition file with model information now.
## Tue Aug  9 12:38:56 CEST 2022 - concatenate mafft-trimal: Will use bootstrap cutoff 80 before creating concatenated alignment
## Tue Aug  9 12:38:57 CEST 2022 - nexus file for iqtree written.
## Tue Aug  9 12:38:57 CEST 2022 - nexus file for iqtree written.
## Tue 09 Aug 2022 12:38:57 PM CEST - 'phylociraptor model' finished successfully before. Will run raxml with best models.
## Will create NEXUS partition file with model information now.
## Tue Aug  9 12:38:58 CEST 2022 - concatenate clustalo-aliscore: Will use bootstrap cutoff 80 before creating concatenated alignment
## Tue Aug  9 12:38:58 CEST 2022 - prepare_iqtree mafft-trimal: Will use bootstrap cutoff (50) before creating concatenated alignment
## Tue Aug  9 12:38:58 CEST 2022 - nexus file for iqtree written.
## Tue Aug  9 12:38:59 CEST 2022 - concatenate muscle-bmge: Will use bootstrap cutoff 60 before creating concatenated alignment
## Tue Aug  9 12:38:59 CEST 2022 - prepare_iqtree muscle-trimal: Will use bootstrap cutoff (70) before creating concatenated alignment
## Tue Aug  9 12:38:59 CEST 2022 - nexus file for iqtree written.
## Will create NEXUS partition file with model information now.
## Tue Aug  9 12:39:00 CEST 2022 - prepare_iqtree clustalo-bmge: Will use bootstrap cutoff (80) before creating concatenated alignment
## Will create NEXUS partition file with model information now.
## Will create NEXUS partition file with model information now.
## Tue Aug  9 12:39:02 CEST 2022 - nexus file for iqtree written.
## Tue Aug  9 12:39:02 CEST 2022 - nexus file for iqtree written.
## Tue 09 Aug 2022 12:39:02 PM CEST - 'phylociraptor model' finished successfully before. Will run raxml with best models.
## Tue 09 Aug 2022 12:39:02 PM CEST - 'phylociraptor model' finished successfully before. Will run raxml with best models.
## Tue Aug  9 12:39:02 CEST 2022 - concatenate clustalo-bmge: Will use bootstrap cutoff 70 before creating concatenated alignment
## Tue 09 Aug 2022 12:39:03 PM CEST - 'phylociraptor model' finished successfully before. Will run raxml with best models.
## Tue Aug  9 12:39:03 CEST 2022 - prepare_iqtree clustalo-bmge: Will use bootstrap cutoff (60) before creating concatenated alignment
## Tue Aug  9 12:39:03 CEST 2022 - nexus file for iqtree written.
## Tue Aug  9 12:39:03 CEST 2022 - concatenate clustalo-aliscore: Will use bootstrap cutoff 0 before creating concatenated alignment
## Tue Aug  9 12:39:04 CEST 2022 - prepare_iqtree clustalo-bmge: Will use bootstrap cutoff (0) before creating concatenated alignment
## Tue Aug  9 12:39:04 CEST 2022 - prepare_iqtree mafft-bmge: Will use bootstrap cutoff (0) before creating concatenated alignment
## Will create NEXUS partition file with model information now.
## Will create NEXUS partition file with model information now.
## Will create NEXUS partition file with model information now.
## Tue Aug  9 12:39:06 CEST 2022 - nexus file for iqtree written.
## Tue 09 Aug 2022 12:39:06 PM CEST - 'phylociraptor model' finished successfully before. Will run raxml with best models.
## Tue 09 Aug 2022 12:39:06 PM CEST - 'phylociraptor model' finished successfully before. Will run raxml with best models.
## Tue 09 Aug 2022 12:39:07 PM CEST - 'phylociraptor model' finished successfully before. Will run raxml with best models.
## Tue Aug  9 12:39:07 CEST 2022 - nexus file for iqtree written.
## Tue 09 Aug 2022 12:39:07 PM CEST - 'phylociraptor model' finished successfully before. Will run raxml with best models.
## Tue Aug  9 12:39:07 CEST 2022 - concatenate muscle-trimal: Will use bootstrap cutoff 50 before creating concatenated alignment
## Tue 09 Aug 2022 12:39:07 PM CEST - 'phylociraptor model' finished successfully before. Will run raxml with best models.
## Tue Aug  9 12:39:07 CEST 2022 - nexus file for iqtree written.
## Tue 09 Aug 2022 12:39:08 PM CEST - 'phylociraptor model' finished successfully before. Will run raxml with best models.
## Tue 09 Aug 2022 12:39:08 PM CEST - 'phylociraptor model' finished successfully before. Will run raxml with best models.
## Tue 09 Aug 2022 12:39:08 PM CEST - 'phylociraptor model' finished successfully before. Will run raxml with best models.
## Tue Aug  9 12:39:08 CEST 2022 - prepare_iqtree muscle-aliscore: Will use bootstrap cutoff (60) before creating concatenated alignment
## Tue Aug  9 12:39:08 CEST 2022 - concatenate clustalo-bmge: Will use bootstrap cutoff 60 before creating concatenated alignment
## Tue Aug  9 12:39:09 CEST 2022 - prepare_iqtree muscle-trimal: Will use bootstrap cutoff (0) before creating concatenated alignment
## Tue 09 Aug 2022 12:39:09 PM CEST - 'phylociraptor model' finished successfully before. Will run raxml with best models.
## Tue Aug  9 12:39:10 CEST 2022 - concatenate clustalo-bmge: Will use bootstrap cutoff 0 before creating concatenated alignment
## Will create NEXUS partition file with model information now.
## Will create NEXUS partition file with model information now.
## Tue Aug  9 12:39:11 CEST 2022 - concatenate muscle-bmge: Will use bootstrap cutoff 0 before creating concatenated alignment
## Tue 09 Aug 2022 12:39:11 PM CEST - 'phylociraptor model' finished successfully before. Will run raxml with best models.
## Tue 09 Aug 2022 12:39:12 PM CEST - 'phylociraptor model' finished successfully before. Will run raxml with best models.
## Tue 09 Aug 2022 12:39:12 PM CEST - 'phylociraptor model' finished successfully before. Will run raxml with best models.
## Tue 09 Aug 2022 12:39:12 PM CEST - 'phylociraptor model' finished successfully before. Will run raxml with best models.
## Tue 09 Aug 2022 12:39:12 PM CEST - 'phylociraptor model' finished successfully before. Will run raxml with best models.
## Tue Aug  9 12:39:12 CEST 2022 - nexus file for iqtree written.
## Tue Aug  9 12:39:12 CEST 2022 - prepare_iqtree muscle-bmge: Will use bootstrap cutoff (50) before creating concatenated alignment
## Tue Aug  9 12:39:12 CEST 2022 - nexus file for iqtree written.
## Tue Aug  9 12:39:12 CEST 2022 - concatenate muscle-trimal: Will use bootstrap cutoff 80 before creating concatenated alignment
## Tue 09 Aug 2022 12:39:13 PM CEST - 'phylociraptor model' finished successfully before. Will run raxml with best models.
## Tue 09 Aug 2022 12:39:13 PM CEST - 'phylociraptor model' finished successfully before. Will run raxml with best models.
## Tue 09 Aug 2022 12:39:13 PM CEST - 'phylociraptor model' finished successfully before. Will run raxml with best models.
## Tue 09 Aug 2022 12:39:13 PM CEST - 'phylociraptor model' finished successfully before. Will run raxml with best models.
## Tue 09 Aug 2022 12:39:14 PM CEST - 'phylociraptor model' finished successfully before. Will run raxml with best models.
## Will create NEXUS partition file with model information now.
## Tue 09 Aug 2022 12:39:14 PM CEST - 'phylociraptor model' finished successfully before. Will run raxml with best models.
## Tue Aug  9 12:39:15 CEST 2022 - nexus file for iqtree written.
## Tue 09 Aug 2022 12:39:16 PM CEST - 'phylociraptor model' finished successfully before. Will run raxml with best models.
## Tue 09 Aug 2022 12:39:17 PM CEST - 'phylociraptor model' finished successfully before. Will run raxml with best models.
## Tue 09 Aug 2022 12:39:20 PM CEST - 'phylociraptor model' finished successfully before. Will run raxml with best models.
## Tue 09 Aug 2022 12:39:22 PM CEST - 'phylociraptor model' finished successfully before. Will run raxml with best models.
## Tue 09 Aug 2022 12:39:27 PM CEST - 'phylociraptor model' finished successfully before. Will run raxml with best models.
## Tue 09 Aug 2022 12:39:28 PM CEST - 'phylociraptor model' finished successfully before. Will run raxml with best models.
## Tue 09 Aug 2022 12:39:29 PM CEST - 'phylociraptor model' finished successfully before. Will run raxml with best models.
## Tue 09 Aug 2022 12:39:30 PM CEST - 'phylociraptor model' finished successfully before. Will run raxml with best models.
## Tue 09 Aug 2022 12:39:31 PM CEST - 'phylociraptor model' finished successfully before. Will run raxml with best models.
## Wed Aug 10 10:16:24 CEST 2022 - 'phylociraptor model' finished successfully before. Will run raxml with best models.
```
