## Supplementary material for "Phylociraptor – A unified computational framework for reproducible phylogenomic inference": Phylociraptor report as HTML (zipped) for test-case-2 vertebrates.: test-case2-vertebrates-report.html


### phylociraptor

**Analysis report - 06 September, 2023 - 21:08 CEST - version: 0.9.12**

### Setup

**Location of genomes files:**

```
results/assemblies
```

**Total number of included genomes:** 32  
**Locally provided genomes:** 1  
**Successfully downloaded genomes:** 31  
**Failed species downloads:** 0  
**Not downloaded:**

**Information on successfully downloaded genomes:**

### Orthology

Orthologous genes were inferred using BUSCO.

**Location of BUSCO results:**

```
results/orthology/busco
```

**Used BUSCO set:** vertebrata\_odb10

**Total number of BUSCO genes in BUSCO set:** 3354  
**Number of BUSCO genes recovered in too few (20) samples:** 131  
**BUSCO genes below threshold:**  
54574at7742, 232256at7742, 24468at7742, 263280at7742, 339849at7742, 284429at7742, 245463at7742, 398107at7742, 329837at7742, 157016at7742, 259004at7742, 110339at7742, 82717at7742, 169126at7742, 120673at7742, 310175at7742, 299674at7742, 47273at7742, 65834at7742, 66884at7742, 429826at7742, 278895at7742, 102515at7742, 173113at7742, 328736at7742, 269163at7742, 202675at7742, 55268at7742, 27280at7742, 181216at7742, 393128at7742, 275603at7742, 222334at7742, 154432at7742, 111501at7742, 58069at7742, 77215at7742, 12004at7742, 388825at7742, 73984at7742, 181872at7742, 220441at7742, 41125at7742, 62592at7742, 166540at7742, 200053at7742, 64479at7742, 128901at7742, 275050at7742, 384835at7742, 354526at7742, 68501at7742, 175798at7742, 130943at7742, 129046at7742, 221780at7742, 266944at7742, 287310at7742, 319460at7742, 396414at7742, 81595at7742, 142850at7742, 72937at7742, 127445at7742, 129939at7742, 15083at7742, 113409at7742, 229692at7742, 62307at7742, 78292at7742, 248600at7742, 228730at7742, 105242at7742, 230622at7742, 123936at7742, 163855at7742, 309843at7742, 19085at7742, 393381at7742, 131366at7742, 438295at7742, 307170at7742, 12613at7742, 383831at7742, 140367at7742, 144812at7742, 212720at7742, 265085at7742, 233465at7742, 402679at7742, 346365at7742, 150696at7742, 262337at7742, 152820at7742, 134368at7742, 204865at7742, 302828at7742, 360502at7742, 164538at7742, 333671at7742, 5520at7742, 126705at7742, 208665at7742, 205835at7742, 436864at7742, 377136at7742, 120857at7742, 211124at7742, 171462at7742, 269700at7742, 394294at7742, 9484at7742, 88436at7742, 201616at7742, 42975at7742, 419694at7742, 239772at7742, 10163at7742, 188041at7742, 210433at7742, 65838at7742, 53835at7742, 107971at7742, 336001at7742, 57359at7742, 400457at7742, 69330at7742, 379901at7742, 124775at7742, 320238at7742, 381399at7742

**Number of samples with single-copy BUSCO score below threshold (0.4):** 1  
**Genomes below threshold:**  
Notamacropus\_eugenii

**Orthology overview table:**

### Alignments

#### full alignments

**Location of alignment files:**

```
results/alignments/full
```

**Used aligner(s) and settings:**  
**1.** clustalo   
**2.** mafft –quiet –auto –anysymbol  
**3.** muscle  
  
**Parsimony informative sites cutoff:** 50  
  
**Number of alignments for clustalo:** 3190   
**Number of alignments for mafft:** 3190   
**Number of alignments for muscle:** 3190   
  
**Overview of individual full alignments:**

#### trimmed alignments

**Location of trimmed alignment files:**

```
results/alignments/trimmed
```

**Parsimony informative sites cutoff:** 50

**Number of trimmed alignments for clustalo-aliscore:** 3188   
**Number of trimmed alignments for clustalo-bmge:** 3190   
**Number of trimmed alignments for clustalo-trimal:** 3190   
**Number of trimmed alignments for mafft-aliscore:** 3184   
**Number of trimmed alignments for mafft-bmge:** 3190   
**Number of trimmed alignments for mafft-trimal:** 3190   
**Number of trimmed alignments for muscle-aliscore:** 3188   
**Number of trimmed alignments for muscle-bmge:** 3190   
**Number of trimmed alignments for muscle-trimal:** 3190   
  
**Overview of individual trimmed alignments:**

#### filtered alignments

**Location of filtered alignment files:**

```
results/alignments/filtered
```

**Parsimony informative sites cutoff:** 50

**Number of filtered alignments for clustalo-aliscore:** 2802   
**Number of filtered alignments for clustalo-bmge:** 2788   
**Number of filtered alignments for clustalo-trimal:** 2861   
**Number of filtered alignments for mafft-aliscore:** 2835   
**Number of filtered alignments for mafft-bmge:** 2827   
**Number of filtered alignments for mafft-trimal:** 2894   
**Number of filtered alignments for muscle-aliscore:** 2835   
**Number of filtered alignments for muscle-bmge:** 2809   
**Number of filtered alignments for muscle-trimal:** 2889   
  
**Overview of individual filtered alignments:**

### Modeltesting

**Location of modeltesting files:**

```
results/modeltest
```

**Overview of individual modeltest results:**

### Genetrees

**Location of gene trees:**

```
results/modeltest
```

**Overview of mean bootstrap values per gene-tree:**   
 Total number of gene trees for clustalo-aliscore: 2802  
 Total number of gene trees for clustalo-bmge: 2788  
 Total number of gene trees for clustalo-trimal: 2861  
 Total number of gene trees for mafft-aliscore: 2835  
 Total number of gene trees for mafft-bmge: 2827  
 Total number of gene trees for mafft-trimal: 2894  
 Total number of gene trees for muscle-aliscore: 2835  
 Total number of gene trees for muscle-bmge: 2809  
 Total number of gene trees for muscle-trimal: 2889

FALSE
FALSE

### Maximum-likelihood trees

**Random seed:** random

**Location of phylogenetic results:**

```
results/phylogeny-*/
```

  
**Overview of concatenated Maximum-likelihood phylogenies:**

| software | aligner | trimmer | boostraps | bootstrap-cutoff | ngenes | tree |
| --- | --- | --- | --- | --- | --- | --- |
| iqtree | clustalo | aliscore | 1000 | 50 | 2802 | Copy tree |
| iqtree | clustalo | aliscore | 1000 | 60 | 2801 | Copy tree |
| iqtree | clustalo | aliscore | 1000 | 70 | 2730 | Copy tree |
| iqtree | clustalo | aliscore | 1000 | 80 | 2141 | Copy tree |
| iqtree | clustalo | aliscore | 1000 | 90 | 386 | Copy tree |
| iqtree | clustalo | bmge | 1000 | 50 | 2788 | Copy tree |
| iqtree | clustalo | bmge | 1000 | 60 | 2785 | Copy tree |
| iqtree | clustalo | bmge | 1000 | 70 | 2722 | Copy tree |
| iqtree | clustalo | bmge | 1000 | 80 | 2108 | Copy tree |
| iqtree | clustalo | bmge | 1000 | 90 | 369 | Copy tree |
| iqtree | clustalo | trimal | 1000 | 50 | 2861 | Copy tree |
| iqtree | clustalo | trimal | 1000 | 60 | 2857 | Copy tree |
| iqtree | clustalo | trimal | 1000 | 70 | 2804 | Copy tree |
| iqtree | clustalo | trimal | 1000 | 80 | 2213 | Copy tree |
| iqtree | clustalo | trimal | 1000 | 90 | 487 | Copy tree |
| iqtree | mafft | aliscore | 1000 | 50 | 2835 | Copy tree |
| iqtree | mafft | aliscore | 1000 | 60 | 2831 | Copy tree |
| iqtree | mafft | aliscore | 1000 | 70 | 2770 | Copy tree |
| iqtree | mafft | aliscore | 1000 | 80 | 2158 | Copy tree |
| iqtree | mafft | aliscore | 1000 | 90 | 389 | Copy tree |
| iqtree | mafft | bmge | 1000 | 50 | 2827 | Copy tree |
| iqtree | mafft | bmge | 1000 | 60 | 2826 | Copy tree |
| iqtree | mafft | bmge | 1000 | 70 | 2769 | Copy tree |
| iqtree | mafft | bmge | 1000 | 80 | 2142 | Copy tree |
| iqtree | mafft | bmge | 1000 | 90 | 371 | Copy tree |
| iqtree | mafft | trimal | 1000 | 50 | 2894 | Copy tree |
| iqtree | mafft | trimal | 1000 | 60 | 2888 | Copy tree |
| iqtree | mafft | trimal | 1000 | 70 | 2828 | Copy tree |
| iqtree | mafft | trimal | 1000 | 80 | 2281 | Copy tree |
| iqtree | mafft | trimal | 1000 | 90 | 496 | Copy tree |
| iqtree | muscle | aliscore | 1000 | 50 | 2833 | Copy tree |
| iqtree | muscle | aliscore | 1000 | 60 | 2831 | Copy tree |
| iqtree | muscle | aliscore | 1000 | 70 | 2773 | Copy tree |
| iqtree | muscle | aliscore | 1000 | 80 | 2137 | Copy tree |
| iqtree | muscle | aliscore | 1000 | 90 | 393 | Copy tree |
| iqtree | muscle | bmge | 1000 | 50 | 2809 | Copy tree |
| iqtree | muscle | bmge | 1000 | 60 | 2806 | Copy tree |
| iqtree | muscle | bmge | 1000 | 70 | 2745 | Copy tree |
| iqtree | muscle | bmge | 1000 | 80 | 2113 | Copy tree |
| iqtree | muscle | bmge | 1000 | 90 | 340 | Copy tree |
| iqtree | muscle | trimal | 1000 | 50 | 2889 | Copy tree |
| iqtree | muscle | trimal | 1000 | 60 | 2886 | Copy tree |
| iqtree | muscle | trimal | 1000 | 70 | 2823 | Copy tree |
| iqtree | muscle | trimal | 1000 | 80 | 2273 | Copy tree |
| iqtree | muscle | trimal | 1000 | 90 | 493 | Copy tree |
| raxmlng | clustalo | aliscore | 100 | 80 | 2141 | Copy tree |
| raxmlng | clustalo | aliscore | 100 | 90 | 386 | Copy tree |
| raxmlng | clustalo | bmge | 100 | 80 | 2108 | Copy tree |
| raxmlng | clustalo | bmge | 100 | 90 | 369 | Copy tree |
| raxmlng | clustalo | trimal | 100 | 80 | 2213 | Copy tree |
| raxmlng | clustalo | trimal | 100 | 90 | 487 | Copy tree |
| raxmlng | mafft | aliscore | 100 | 80 | 2158 | Copy tree |
| raxmlng | mafft | aliscore | 100 | 90 | 389 | Copy tree |
| raxmlng | mafft | bmge | 100 | 80 | 2142 | Copy tree |
| raxmlng | mafft | bmge | 100 | 90 | 371 | Copy tree |
| raxmlng | mafft | trimal | 100 | 80 | 2281 | Copy tree |
| raxmlng | mafft | trimal | 100 | 90 | 496 | Copy tree |
| raxmlng | muscle | aliscore | 100 | 80 | 2137 | Copy tree |
| raxmlng | muscle | aliscore | 100 | 90 | 393 | Copy tree |
| raxmlng | muscle | bmge | 100 | 80 | 2113 | Copy tree |
| raxmlng | muscle | bmge | 100 | 90 | 340 | Copy tree |
| raxmlng | muscle | trimal | 100 | 80 | 2273 | Copy tree |
| raxmlng | muscle | trimal | 100 | 90 | 493 | Copy tree |

### Speciestree

**Location of phylogenetic results:**

```
results/phylogeny-*/
```

  
**Overview of species-tree phylogenies:**

| software | aligner | trimmer | bootstrap-cutoff | ngenes | tree |
| --- | --- | --- | --- | --- | --- |
| astral | clustalo | aliscore | 50 | 2802 | Copy tree |
| astral | clustalo | aliscore | 60 | 2801 | Copy tree |
| astral | clustalo | aliscore | 70 | 2730 | Copy tree |
| astral | clustalo | aliscore | 80 | 2141 | Copy tree |
| astral | clustalo | aliscore | 90 | 386 | Copy tree |
| astral | clustalo | bmge | 50 | 2788 | Copy tree |
| astral | clustalo | bmge | 60 | 2785 | Copy tree |
| astral | clustalo | bmge | 70 | 2722 | Copy tree |
| astral | clustalo | bmge | 80 | 2108 | Copy tree |
| astral | clustalo | bmge | 90 | 369 | Copy tree |
| astral | clustalo | trimal | 50 | 2861 | Copy tree |
| astral | clustalo | trimal | 60 | 2857 | Copy tree |
| astral | clustalo | trimal | 70 | 2804 | Copy tree |
| astral | clustalo | trimal | 80 | 2213 | Copy tree |
| astral | clustalo | trimal | 90 | 487 | Copy tree |
| astral | mafft | aliscore | 50 | 2835 | Copy tree |
| astral | mafft | aliscore | 60 | 2831 | Copy tree |
| astral | mafft | aliscore | 70 | 2770 | Copy tree |
| astral | mafft | aliscore | 80 | 2158 | Copy tree |
| astral | mafft | aliscore | 90 | 389 | Copy tree |
| astral | mafft | bmge | 50 | 2827 | Copy tree |
| astral | mafft | bmge | 60 | 2826 | Copy tree |
| astral | mafft | bmge | 70 | 2769 | Copy tree |
| astral | mafft | bmge | 80 | 2142 | Copy tree |
| astral | mafft | bmge | 90 | 371 | Copy tree |
| astral | mafft | trimal | 50 | 2894 | Copy tree |
| astral | mafft | trimal | 60 | 2888 | Copy tree |
| astral | mafft | trimal | 70 | 2828 | Copy tree |
| astral | mafft | trimal | 80 | 2281 | Copy tree |
| astral | mafft | trimal | 90 | 496 | Copy tree |
| astral | muscle | aliscore | 50 | 2833 | Copy tree |
| astral | muscle | aliscore | 60 | 2831 | Copy tree |
| astral | muscle | aliscore | 70 | 2773 | Copy tree |
| astral | muscle | aliscore | 80 | 2137 | Copy tree |
| astral | muscle | aliscore | 90 | 393 | Copy tree |
| astral | muscle | bmge | 50 | 2809 | Copy tree |
| astral | muscle | bmge | 60 | 2806 | Copy tree |
| astral | muscle | bmge | 70 | 2745 | Copy tree |
| astral | muscle | bmge | 80 | 2113 | Copy tree |
| astral | muscle | bmge | 90 | 340 | Copy tree |
| astral | muscle | trimal | 50 | 2889 | Copy tree |
| astral | muscle | trimal | 60 | 2886 | Copy tree |
| astral | muscle | trimal | 70 | 2823 | Copy tree |
| astral | muscle | trimal | 80 | 2273 | Copy tree |
| astral | muscle | trimal | 90 | 493 | Copy tree |

### Log

```
## Tue Jul 12 00:55:56 CEST 2022 - Setup: Will download species now - batch: 4
## Tue Jul 12 00:55:56 CEST 2022 - Setup: Will download species now - batch: 3
## Tue Jul 12 00:55:56 CEST 2022 - Setup: Will download species now - batch: 5
## Tue Jul 12 00:55:56 CEST 2022 - Setup: Will download species now - batch: 2
## Tue Jul 12 00:55:57 CEST 2022 - Setup: Will download species now - batch: 1
## Tue Jul 12 00:58:05 CEST 2022 - Setup: Will download species now - batch: 4
## Tue Jul 12 00:58:05 CEST 2022 - Setup: Will download species now - batch: 3
## Tue Jul 12 00:58:05 CEST 2022 - Setup: Will download species now - batch: 5
## Tue Jul 12 00:58:05 CEST 2022 - Setup: Will download species now - batch: 2
## Tue Jul 12 00:58:05 CEST 2022 - Setup: Will download species now - batch: 1
## Tue Jul 12 01:04:23 CEST 2022 - Setup: Will download species now - batch: 1
## Tue Jul 12 01:06:06 CEST 2022 - phylociraptor setup done.
## Tue Jul 12 01:18:52 CEST 2022 - Pipeline part 1 (orthology) done.
## Tue Jul 12 02:58:29 CEST 2022 - BUSCO files will be filtered on a per-sample basis. This could lower the number of species in the final tree.
## Tue Jul 12 03:08:20 CEST 2022 - Number of BUSCO sequence files: 3201
## Tue Jul 12 03:08:21 CEST 2022 - Number of deduplicated BUSCO sequence files: 3168
## Tue Jul 12 03:08:25 CEST 2022 - Pipeline part filter-orthology done.
## Wed Jul 13 04:33:39 CEST 2022 - Pipeline part 2 (align) done.
## Wed Jul 13 23:48:18 CEST 2022 - The aliscore output file does not exist. Check results for BUSCO: 3764at7742
## Wed Jul 13 23:48:18 CEST 2022 - The aliscore output appears to be empty. Check results for BUSCO: 3764at7742
## Wed Jul 13 23:48:19 CEST 2022 - The aliscore output file does not exist. Check results for BUSCO: 107720at7742
## Wed Jul 13 23:48:19 CEST 2022 - The aliscore output appears to be empty. Check results for BUSCO: 107720at7742
## Wed Jul 13 23:48:19 CEST 2022 - The aliscore output file does not exist. Check results for BUSCO: 17781at7742
## Wed Jul 13 23:48:19 CEST 2022 - The aliscore output appears to be empty. Check results for BUSCO: 17781at7742
## Wed Jul 13 23:48:20 CEST 2022 - The aliscore output file does not exist. Check results for BUSCO: 12616at7742
## Wed Jul 13 23:48:20 CEST 2022 - The aliscore output appears to be empty. Check results for BUSCO: 12616at7742
## Wed Jul 13 23:48:20 CEST 2022 - The aliscore output file does not exist. Check results for BUSCO: 12616at7742
## Wed Jul 13 23:48:20 CEST 2022 - The aliscore output appears to be empty. Check results for BUSCO: 12616at7742
## Wed Jul 13 23:48:21 CEST 2022 - The aliscore output file does not exist. Check results for BUSCO: 12616at7742
## Wed Jul 13 23:48:21 CEST 2022 - The aliscore output appears to be empty. Check results for BUSCO: 12616at7742
## Wed Jul 13 23:48:21 CEST 2022 - The aliscore output file does not exist. Check results for BUSCO: 3764at7742
## Wed Jul 13 23:48:21 CEST 2022 - The aliscore output appears to be empty. Check results for BUSCO: 3764at7742
## Wed Jul 13 23:48:25 CEST 2022 - The aliscore output file does not exist. Check results for BUSCO: 17781at7742
## Wed Jul 13 23:48:25 CEST 2022 - The aliscore output appears to be empty. Check results for BUSCO: 17781at7742
## Wed Jul 13 23:48:26 CEST 2022 - The aliscore output file does not exist. Check results for BUSCO: 107720at7742
## Wed Jul 13 23:48:26 CEST 2022 - The aliscore output appears to be empty. Check results for BUSCO: 107720at7742
## Wed Jul 13 23:48:26 CEST 2022 - The aliscore output file does not exist. Check results for BUSCO: 1034at7742
## Wed Jul 13 23:48:26 CEST 2022 - The aliscore output appears to be empty. Check results for BUSCO: 1034at7742
## Wed Jul 13 23:48:26 CEST 2022 - The aliscore output file does not exist. Check results for BUSCO: 44193at7742
## Wed Jul 13 23:48:26 CEST 2022 - The aliscore output appears to be empty. Check results for BUSCO: 44193at7742
## Wed Jul 13 23:50:01 CEST 2022 - The aliscore output file does not exist. Check results for BUSCO: 2178at7742
## Wed Jul 13 23:50:01 CEST 2022 - The aliscore output appears to be empty. Check results for BUSCO: 2178at7742
## Wed Jul 13 23:50:02 CEST 2022 - The aliscore output file does not exist. Check results for BUSCO: 2178at7742
## Wed Jul 13 23:50:02 CEST 2022 - The aliscore output appears to be empty. Check results for BUSCO: 2178at7742
## Wed Jul 13 23:50:02 CEST 2022 - The aliscore output file does not exist. Check results for BUSCO: 107720at7742
## Wed Jul 13 23:50:02 CEST 2022 - The aliscore output appears to be empty. Check results for BUSCO: 107720at7742
## Wed Jul 13 23:50:03 CEST 2022 - The aliscore output file does not exist. Check results for BUSCO: 17781at7742
## Wed Jul 13 23:50:03 CEST 2022 - The aliscore output appears to be empty. Check results for BUSCO: 17781at7742
## Wed Jul 13 23:50:03 CEST 2022 - The aliscore output file does not exist. Check results for BUSCO: 1034at7742
## Wed Jul 13 23:50:03 CEST 2022 - The aliscore output appears to be empty. Check results for BUSCO: 1034at7742
## Wed Jul 13 23:50:04 CEST 2022 - The aliscore output file does not exist. Check results for BUSCO: 1034at7742
## Wed Jul 13 23:50:04 CEST 2022 - The aliscore output appears to be empty. Check results for BUSCO: 1034at7742
## Wed Jul 13 23:50:04 CEST 2022 - The aliscore output file does not exist. Check results for BUSCO: 107720at7742
## Wed Jul 13 23:50:04 CEST 2022 - The aliscore output appears to be empty. Check results for BUSCO: 107720at7742
## Wed Jul 13 23:50:04 CEST 2022 - The aliscore output file does not exist. Check results for BUSCO: 1034at7742
## Wed Jul 13 23:50:04 CEST 2022 - The aliscore output appears to be empty. Check results for BUSCO: 1034at7742
## Wed Jul 13 23:50:04 CEST 2022 - The aliscore output file does not exist. Check results for BUSCO: 2845at7742
## Wed Jul 13 23:50:04 CEST 2022 - The aliscore output appears to be empty. Check results for BUSCO: 2845at7742
## Wed Jul 13 23:50:05 CEST 2022 - The aliscore output file does not exist. Check results for BUSCO: 2178at7742
## Wed Jul 13 23:50:05 CEST 2022 - The aliscore output appears to be empty. Check results for BUSCO: 2178at7742
## Wed Jul 13 23:50:05 CEST 2022 - The aliscore output file does not exist. Check results for BUSCO: 44193at7742
## Wed Jul 13 23:50:05 CEST 2022 - The aliscore output appears to be empty. Check results for BUSCO: 44193at7742
## Wed Jul 13 23:50:07 CEST 2022 - The aliscore output file does not exist. Check results for BUSCO: 2066at7742
## Wed Jul 13 23:50:07 CEST 2022 - The aliscore output appears to be empty. Check results for BUSCO: 2066at7742
## Wed Jul 13 23:50:08 CEST 2022 - The aliscore output file does not exist. Check results for BUSCO: 3764at7742
## Wed Jul 13 23:50:08 CEST 2022 - The aliscore output appears to be empty. Check results for BUSCO: 3764at7742
## Wed Jul 13 23:50:10 CEST 2022 - The aliscore output file does not exist. Check results for BUSCO: 2066at7742
## Wed Jul 13 23:50:11 CEST 2022 - The aliscore output appears to be empty. Check results for BUSCO: 2066at7742
## Wed Jul 13 23:50:13 CEST 2022 - The aliscore output file does not exist. Check results for BUSCO: 29959at7742
## Wed Jul 13 23:50:13 CEST 2022 - The aliscore output appears to be empty. Check results for BUSCO: 29959at7742
## Wed Jul 13 23:50:13 CEST 2022 - The aliscore output file does not exist. Check results for BUSCO: 2845at7742
## Wed Jul 13 23:50:13 CEST 2022 - The aliscore output appears to be empty. Check results for BUSCO: 2845at7742
## Wed Jul 13 23:51:31 CEST 2022 - The aliscore output file does not exist. Check results for BUSCO: 3022at7742
## Wed Jul 13 23:51:31 CEST 2022 - The aliscore output appears to be empty. Check results for BUSCO: 3022at7742
## Wed Jul 13 23:51:31 CEST 2022 - The aliscore output file does not exist. Check results for BUSCO: 1662at7742
## Wed Jul 13 23:51:31 CEST 2022 - The aliscore output appears to be empty. Check results for BUSCO: 1662at7742
## Wed Jul 13 23:51:32 CEST 2022 - The aliscore output file does not exist. Check results for BUSCO: 29959at7742
## Wed Jul 13 23:51:32 CEST 2022 - The aliscore output appears to be empty. Check results for BUSCO: 29959at7742
## Wed Jul 13 23:51:32 CEST 2022 - The aliscore output file does not exist. Check results for BUSCO: 44193at7742
## Wed Jul 13 23:51:32 CEST 2022 - The aliscore output appears to be empty. Check results for BUSCO: 44193at7742
## Wed Jul 13 23:51:32 CEST 2022 - The aliscore output file does not exist. Check results for BUSCO: 2066at7742
## Wed Jul 13 23:51:32 CEST 2022 - The aliscore output appears to be empty. Check results for BUSCO: 2066at7742
## Wed Jul 13 23:51:33 CEST 2022 - The aliscore output file does not exist. Check results for BUSCO: 1662at7742
## Wed Jul 13 23:51:33 CEST 2022 - The aliscore output appears to be empty. Check results for BUSCO: 1662at7742
## Wed Jul 13 23:51:33 CEST 2022 - The aliscore output file does not exist. Check results for BUSCO: 243571at7742
## Wed Jul 13 23:51:34 CEST 2022 - The aliscore output appears to be empty. Check results for BUSCO: 243571at7742
## Wed Jul 13 23:51:34 CEST 2022 - The aliscore output file does not exist. Check results for BUSCO: 1662at7742
## Wed Jul 13 23:51:34 CEST 2022 - The aliscore output file does not exist. Check results for BUSCO: 2845at7742
## Wed Jul 13 23:51:34 CEST 2022 - The aliscore output appears to be empty. Check results for BUSCO: 1662at7742
## Wed Jul 13 23:51:34 CEST 2022 - The aliscore output appears to be empty. Check results for BUSCO: 2845at7742
## Wed Jul 13 23:51:35 CEST 2022 - The aliscore output file does not exist. Check results for BUSCO: 44193at7742
## Wed Jul 13 23:51:35 CEST 2022 - The aliscore output appears to be empty. Check results for BUSCO: 44193at7742
## Wed Jul 13 23:51:38 CEST 2022 - The aliscore output file does not exist. Check results for BUSCO: 243571at7742
## Wed Jul 13 23:51:38 CEST 2022 - The aliscore output appears to be empty. Check results for BUSCO: 243571at7742
## Wed Jul 13 23:51:38 CEST 2022 - The aliscore output file does not exist. Check results for BUSCO: 29959at7742
## Wed Jul 13 23:51:38 CEST 2022 - The aliscore output appears to be empty. Check results for BUSCO: 29959at7742
## Wed Jul 13 23:51:41 CEST 2022 - The aliscore output file does not exist. Check results for BUSCO: 243571at7742
## Wed Jul 13 23:51:41 CEST 2022 - The aliscore output appears to be empty. Check results for BUSCO: 243571at7742
## Wed Jul 13 23:51:42 CEST 2022 - The aliscore output file does not exist. Check results for BUSCO: 62216at7742
## Wed Jul 13 23:51:42 CEST 2022 - The aliscore output appears to be empty. Check results for BUSCO: 62216at7742
## Wed Jul 13 23:51:43 CEST 2022 - The aliscore output file does not exist. Check results for BUSCO: 3022at7742
## Wed Jul 13 23:51:43 CEST 2022 - The aliscore output appears to be empty. Check results for BUSCO: 3022at7742
## Wed Jul 13 23:51:43 CEST 2022 - The aliscore output file does not exist. Check results for BUSCO: 62216at7742
## Wed Jul 13 23:51:43 CEST 2022 - The aliscore output appears to be empty. Check results for BUSCO: 62216at7742
## Wed Jul 13 23:51:43 CEST 2022 - The aliscore output file does not exist. Check results for BUSCO: 18037at7742
## Wed Jul 13 23:51:43 CEST 2022 - The aliscore output appears to be empty. Check results for BUSCO: 18037at7742
## Wed Jul 13 23:51:48 CEST 2022 - The aliscore output file does not exist. Check results for BUSCO: 173114at7742
## Wed Jul 13 23:51:48 CEST 2022 - The aliscore output appears to be empty. Check results for BUSCO: 173114at7742
## Wed Jul 13 23:51:50 CEST 2022 - The aliscore output file does not exist. Check results for BUSCO: 62216at7742
## Wed Jul 13 23:51:50 CEST 2022 - The aliscore output appears to be empty. Check results for BUSCO: 62216at7742
## Wed Jul 13 23:51:53 CEST 2022 - The aliscore output file does not exist. Check results for BUSCO: 7262at7742
## Wed Jul 13 23:51:53 CEST 2022 - The aliscore output appears to be empty. Check results for BUSCO: 7262at7742
## Wed Jul 13 23:51:55 CEST 2022 - The aliscore output file does not exist. Check results for BUSCO: 3022at7742
## Wed Jul 13 23:51:55 CEST 2022 - The aliscore output appears to be empty. Check results for BUSCO: 3022at7742
## Wed Jul 13 23:51:55 CEST 2022 - The aliscore output file does not exist. Check results for BUSCO: 21841at7742
## Wed Jul 13 23:51:55 CEST 2022 - The aliscore output appears to be empty. Check results for BUSCO: 21841at7742
## Wed Jul 13 23:51:56 CEST 2022 - The aliscore output file does not exist. Check results for BUSCO: 7262at7742
## Wed Jul 13 23:51:56 CEST 2022 - The aliscore output appears to be empty. Check results for BUSCO: 7262at7742
## Wed Jul 13 23:51:56 CEST 2022 - The aliscore output file does not exist. Check results for BUSCO: 19351at7742
## Wed Jul 13 23:51:56 CEST 2022 - The aliscore output appears to be empty. Check results for BUSCO: 19351at7742
## Wed Jul 13 23:52:00 CEST 2022 - The aliscore output file does not exist. Check results for BUSCO: 2503at7742
## Wed Jul 13 23:52:00 CEST 2022 - The aliscore output appears to be empty. Check results for BUSCO: 2503at7742
## Wed Jul 13 23:52:01 CEST 2022 - The aliscore output file does not exist. Check results for BUSCO: 2503at7742
## Wed Jul 13 23:52:01 CEST 2022 - The aliscore output appears to be empty. Check results for BUSCO: 2503at7742
## Wed Jul 13 23:52:03 CEST 2022 - The aliscore output file does not exist. Check results for BUSCO: 45247at7742
## Wed Jul 13 23:52:03 CEST 2022 - The aliscore output appears to be empty. Check results for BUSCO: 45247at7742
## Wed Jul 13 23:52:05 CEST 2022 - The aliscore output file does not exist. Check results for BUSCO: 23457at7742
## Wed Jul 13 23:52:05 CEST 2022 - The aliscore output appears to be empty. Check results for BUSCO: 23457at7742
## Wed Jul 13 23:52:07 CEST 2022 - The aliscore output file does not exist. Check results for BUSCO: 10485at7742
## Wed Jul 13 23:52:07 CEST 2022 - The aliscore output appears to be empty. Check results for BUSCO: 10485at7742
## Wed Jul 13 23:52:08 CEST 2022 - The aliscore output file does not exist. Check results for BUSCO: 7262at7742
## Wed Jul 13 23:52:08 CEST 2022 - The aliscore output appears to be empty. Check results for BUSCO: 7262at7742
## Wed Jul 13 23:52:09 CEST 2022 - The aliscore output file does not exist. Check results for BUSCO: 37552at7742
## Wed Jul 13 23:52:09 CEST 2022 - The aliscore output appears to be empty. Check results for BUSCO: 37552at7742
## Wed Jul 13 23:52:10 CEST 2022 - The aliscore output file does not exist. Check results for BUSCO: 5018at7742
## Wed Jul 13 23:52:10 CEST 2022 - The aliscore output appears to be empty. Check results for BUSCO: 5018at7742
## Wed Jul 13 23:52:14 CEST 2022 - The aliscore output file does not exist. Check results for BUSCO: 2503at7742
## Wed Jul 13 23:52:14 CEST 2022 - The aliscore output appears to be empty. Check results for BUSCO: 2503at7742
## Wed Jul 13 23:52:15 CEST 2022 - The aliscore output file does not exist. Check results for BUSCO: 21841at7742
## Wed Jul 13 23:52:15 CEST 2022 - The aliscore output appears to be empty. Check results for BUSCO: 21841at7742
## Wed Jul 13 23:52:15 CEST 2022 - The aliscore output file does not exist. Check results for BUSCO: 173114at7742
## Wed Jul 13 23:52:16 CEST 2022 - The aliscore output appears to be empty. Check results for BUSCO: 173114at7742
## Wed Jul 13 23:52:18 CEST 2022 - The aliscore output file does not exist. Check results for BUSCO: 11550at7742
## Wed Jul 13 23:52:18 CEST 2022 - The aliscore output appears to be empty. Check results for BUSCO: 11550at7742
## Wed Jul 13 23:52:18 CEST 2022 - The aliscore output file does not exist. Check results for BUSCO: 6253at7742
## Wed Jul 13 23:52:18 CEST 2022 - The aliscore output appears to be empty. Check results for BUSCO: 6253at7742
## Wed Jul 13 23:52:19 CEST 2022 - The aliscore output file does not exist. Check results for BUSCO: 48106at7742
## Wed Jul 13 23:52:19 CEST 2022 - The aliscore output appears to be empty. Check results for BUSCO: 48106at7742
## Wed Jul 13 23:52:22 CEST 2022 - The aliscore output file does not exist. Check results for BUSCO: 1270at7742
## Wed Jul 13 23:52:22 CEST 2022 - The aliscore output appears to be empty. Check results for BUSCO: 1270at7742
## Wed Jul 13 23:52:23 CEST 2022 - The aliscore output file does not exist. Check results for BUSCO: 64048at7742
## Wed Jul 13 23:52:23 CEST 2022 - The aliscore output appears to be empty. Check results for BUSCO: 64048at7742
## Wed Jul 13 23:52:24 CEST 2022 - The aliscore output file does not exist. Check results for BUSCO: 15711at7742
## Wed Jul 13 23:52:24 CEST 2022 - The aliscore output appears to be empty. Check results for BUSCO: 15711at7742
## Wed Jul 13 23:52:25 CEST 2022 - The aliscore output file does not exist. Check results for BUSCO: 1270at7742
## Wed Jul 13 23:52:25 CEST 2022 - The aliscore output appears to be empty. Check results for BUSCO: 1270at7742
## Wed Jul 13 23:52:27 CEST 2022 - The aliscore output file does not exist. Check results for BUSCO: 2461at7742
## Wed Jul 13 23:52:28 CEST 2022 - The aliscore output appears to be empty. Check results for BUSCO: 2461at7742
## Wed Jul 13 23:52:28 CEST 2022 - The aliscore output file does not exist. Check results for BUSCO: 1270at7742
## Wed Jul 13 23:52:28 CEST 2022 - The aliscore output appears to be empty. Check results for BUSCO: 1270at7742
## Wed Jul 13 23:52:28 CEST 2022 - The aliscore output file does not exist. Check results for BUSCO: 4577at7742
## Wed Jul 13 23:52:28 CEST 2022 - The aliscore output appears to be empty. Check results for BUSCO: 4577at7742
## Wed Jul 13 23:52:31 CEST 2022 - The aliscore output file does not exist. Check results for BUSCO: 21841at7742
## Wed Jul 13 23:52:31 CEST 2022 - The aliscore output appears to be empty. Check results for BUSCO: 21841at7742
## Wed Jul 13 23:52:34 CEST 2022 - The aliscore output file does not exist. Check results for BUSCO: 16236at7742
## Wed Jul 13 23:52:34 CEST 2022 - The aliscore output appears to be empty. Check results for BUSCO: 16236at7742
## Wed Jul 13 23:52:35 CEST 2022 - The aliscore output file does not exist. Check results for BUSCO: 2461at7742
## Wed Jul 13 23:52:35 CEST 2022 - The aliscore output appears to be empty. Check results for BUSCO: 2461at7742
## Wed Jul 13 23:52:37 CEST 2022 - The aliscore output file does not exist. Check results for BUSCO: 19351at7742
## Wed Jul 13 23:52:37 CEST 2022 - The aliscore output appears to be empty. Check results for BUSCO: 19351at7742
## Wed Jul 13 23:52:37 CEST 2022 - The aliscore output file does not exist. Check results for BUSCO: 3190at7742
## Wed Jul 13 23:52:37 CEST 2022 - The aliscore output appears to be empty. Check results for BUSCO: 3190at7742
## Wed Jul 13 23:52:41 CEST 2022 - The aliscore output file does not exist. Check results for BUSCO: 10003at7742
## Wed Jul 13 23:52:41 CEST 2022 - The aliscore output appears to be empty. Check results for BUSCO: 10003at7742
## Wed Jul 13 23:52:42 CEST 2022 - The aliscore output file does not exist. Check results for BUSCO: 11550at7742
## Wed Jul 13 23:52:42 CEST 2022 - The aliscore output appears to be empty. Check results for BUSCO: 11550at7742
## Wed Jul 13 23:52:42 CEST 2022 - The aliscore output file does not exist. Check results for BUSCO: 4577at7742
## Wed Jul 13 23:52:42 CEST 2022 - The aliscore output appears to be empty. Check results for BUSCO: 4577at7742
## Wed Jul 13 23:52:44 CEST 2022 - The aliscore output file does not exist. Check results for BUSCO: 3190at7742
## Wed Jul 13 23:52:44 CEST 2022 - The aliscore output appears to be empty. Check results for BUSCO: 3190at7742
## Wed Jul 13 23:52:46 CEST 2022 - The aliscore output file does not exist. Check results for BUSCO: 10485at7742
## Wed Jul 13 23:52:46 CEST 2022 - The aliscore output appears to be empty. Check results for BUSCO: 10485at7742
## Wed Jul 13 23:52:46 CEST 2022 - The aliscore output file does not exist. Check results for BUSCO: 44104at7742
## Wed Jul 13 23:52:46 CEST 2022 - The aliscore output appears to be empty. Check results for BUSCO: 44104at7742
## Wed Jul 13 23:52:48 CEST 2022 - The aliscore output file does not exist. Check results for BUSCO: 5018at7742
## Wed Jul 13 23:52:48 CEST 2022 - The aliscore output appears to be empty. Check results for BUSCO: 5018at7742
## Wed Jul 13 23:52:50 CEST 2022 - The aliscore output file does not exist. Check results for BUSCO: 2461at7742
## Wed Jul 13 23:52:50 CEST 2022 - The aliscore output appears to be empty. Check results for BUSCO: 2461at7742
## Wed Jul 13 23:52:50 CEST 2022 - The aliscore output file does not exist. Check results for BUSCO: 4577at7742
## Wed Jul 13 23:52:50 CEST 2022 - The aliscore output appears to be empty. Check results for BUSCO: 4577at7742
## Wed Jul 13 23:52:54 CEST 2022 - The aliscore output file does not exist. Check results for BUSCO: 17848at7742
## Wed Jul 13 23:52:54 CEST 2022 - The aliscore output appears to be empty. Check results for BUSCO: 17848at7742
## Wed Jul 13 23:52:55 CEST 2022 - The aliscore output file does not exist. Check results for BUSCO: 10003at7742
## Wed Jul 13 23:52:55 CEST 2022 - The aliscore output appears to be empty. Check results for BUSCO: 10003at7742
## Wed Jul 13 23:52:55 CEST 2022 - The aliscore output file does not exist. Check results for BUSCO: 19351at7742
## Wed Jul 13 23:52:55 CEST 2022 - The aliscore output appears to be empty. Check results for BUSCO: 19351at7742
## Wed Jul 13 23:52:57 CEST 2022 - The aliscore output file does not exist. Check results for BUSCO: 115534at7742
## Wed Jul 13 23:52:57 CEST 2022 - The aliscore output appears to be empty. Check results for BUSCO: 115534at7742
## Wed Jul 13 23:52:58 CEST 2022 - The aliscore output file does not exist. Check results for BUSCO: 390830at7742
## Wed Jul 13 23:52:58 CEST 2022 - The aliscore output appears to be empty. Check results for BUSCO: 390830at7742
## Wed Jul 13 23:52:59 CEST 2022 - The aliscore output file does not exist. Check results for BUSCO: 48106at7742
## Wed Jul 13 23:52:59 CEST 2022 - The aliscore output appears to be empty. Check results for BUSCO: 48106at7742
## Wed Jul 13 23:53:02 CEST 2022 - The aliscore output file does not exist. Check results for BUSCO: 44104at7742
## Wed Jul 13 23:53:02 CEST 2022 - The aliscore output appears to be empty. Check results for BUSCO: 44104at7742
## Wed Jul 13 23:53:02 CEST 2022 - The aliscore output file does not exist. Check results for BUSCO: 39061at7742
## Wed Jul 13 23:53:02 CEST 2022 - The aliscore output appears to be empty. Check results for BUSCO: 39061at7742
## Wed Jul 13 23:53:06 CEST 2022 - The aliscore output file does not exist. Check results for BUSCO: 64048at7742
## Wed Jul 13 23:53:06 CEST 2022 - The aliscore output appears to be empty. Check results for BUSCO: 64048at7742
## Wed Jul 13 23:53:09 CEST 2022 - The aliscore output file does not exist. Check results for BUSCO: 3190at7742
## Wed Jul 13 23:53:09 CEST 2022 - The aliscore output appears to be empty. Check results for BUSCO: 3190at7742
## Wed Jul 13 23:53:10 CEST 2022 - The aliscore output file does not exist. Check results for BUSCO: 64048at7742
## Wed Jul 13 23:53:10 CEST 2022 - The aliscore output appears to be empty. Check results for BUSCO: 64048at7742
## Wed Jul 13 23:53:11 CEST 2022 - The aliscore output file does not exist. Check results for BUSCO: 37552at7742
## Wed Jul 13 23:53:11 CEST 2022 - The aliscore output appears to be empty. Check results for BUSCO: 37552at7742
## Wed Jul 13 23:53:12 CEST 2022 - The aliscore output file does not exist. Check results for BUSCO: 6305at7742
## Wed Jul 13 23:53:12 CEST 2022 - The aliscore output appears to be empty. Check results for BUSCO: 6305at7742
## Wed Jul 13 23:53:13 CEST 2022 - The aliscore output file does not exist. Check results for BUSCO: 8757at7742
## Wed Jul 13 23:53:13 CEST 2022 - The aliscore output appears to be empty. Check results for BUSCO: 8757at7742
## Wed Jul 13 23:53:13 CEST 2022 - The aliscore output file does not exist. Check results for BUSCO: 5018at7742
## Wed Jul 13 23:53:13 CEST 2022 - The aliscore output appears to be empty. Check results for BUSCO: 5018at7742
## Wed Jul 13 23:53:16 CEST 2022 - The aliscore output file does not exist. Check results for BUSCO: 50582at7742
## Wed Jul 13 23:53:16 CEST 2022 - The aliscore output appears to be empty. Check results for BUSCO: 50582at7742
## Wed Jul 13 23:53:17 CEST 2022 - Alignment of BUSCO: 50582at7742 contains empty sequence after aliscore/alicut. This sequence will be removed.
## Wed Jul 13 23:53:18 CEST 2022 - The aliscore output file does not exist. Check results for BUSCO: 173114at7742
## Wed Jul 13 23:53:18 CEST 2022 - The aliscore output appears to be empty. Check results for BUSCO: 173114at7742
## Wed Jul 13 23:53:19 CEST 2022 - The aliscore output file does not exist. Check results for BUSCO: 11550at7742
## Wed Jul 13 23:53:19 CEST 2022 - The aliscore output appears to be empty. Check results for BUSCO: 11550at7742
## Wed Jul 13 23:53:21 CEST 2022 - The aliscore output file does not exist. Check results for BUSCO: 17848at7742
## Wed Jul 13 23:53:21 CEST 2022 - The aliscore output appears to be empty. Check results for BUSCO: 17848at7742
## Wed Jul 13 23:53:23 CEST 2022 - The aliscore output file does not exist. Check results for BUSCO: 10003at7742
## Wed Jul 13 23:53:23 CEST 2022 - The aliscore output appears to be empty. Check results for BUSCO: 10003at7742
## Wed Jul 13 23:53:24 CEST 2022 - The aliscore output file does not exist. Check results for BUSCO: 17789at7742
## Wed Jul 13 23:53:24 CEST 2022 - The aliscore output appears to be empty. Check results for BUSCO: 17789at7742
## Wed Jul 13 23:53:24 CEST 2022 - The aliscore output file does not exist. Check results for BUSCO: 45247at7742
## Wed Jul 13 23:53:24 CEST 2022 - The aliscore output appears to be empty. Check results for BUSCO: 45247at7742
## Wed Jul 13 23:53:27 CEST 2022 - The aliscore output file does not exist. Check results for BUSCO: 10485at7742
## Wed Jul 13 23:53:27 CEST 2022 - The aliscore output appears to be empty. Check results for BUSCO: 10485at7742
## Wed Jul 13 23:53:28 CEST 2022 - The aliscore output file does not exist. Check results for BUSCO: 37552at7742
## Wed Jul 13 23:53:28 CEST 2022 - The aliscore output appears to be empty. Check results for BUSCO: 37552at7742
## Wed Jul 13 23:53:29 CEST 2022 - The aliscore output file does not exist. Check results for BUSCO: 48106at7742
## Wed Jul 13 23:53:29 CEST 2022 - The aliscore output appears to be empty. Check results for BUSCO: 48106at7742
## Wed Jul 13 23:53:30 CEST 2022 - The aliscore output file does not exist. Check results for BUSCO: 18037at7742
## Wed Jul 13 23:53:30 CEST 2022 - The aliscore output appears to be empty. Check results for BUSCO: 18037at7742
## Wed Jul 13 23:53:33 CEST 2022 - The aliscore output file does not exist. Check results for BUSCO: 39061at7742
## Wed Jul 13 23:53:33 CEST 2022 - The aliscore output appears to be empty. Check results for BUSCO: 39061at7742
## Wed Jul 13 23:53:35 CEST 2022 - The aliscore output file does not exist. Check results for BUSCO: 23457at7742
## Wed Jul 13 23:53:35 CEST 2022 - The aliscore output appears to be empty. Check results for BUSCO: 23457at7742
## Wed Jul 13 23:53:35 CEST 2022 - The aliscore output file does not exist. Check results for BUSCO: 8258at7742
## Wed Jul 13 23:53:35 CEST 2022 - The aliscore output appears to be empty. Check results for BUSCO: 8258at7742
## Wed Jul 13 23:53:38 CEST 2022 - The aliscore output file does not exist. Check results for BUSCO: 15604at7742
## Wed Jul 13 23:53:38 CEST 2022 - The aliscore output appears to be empty. Check results for BUSCO: 15604at7742
## Wed Jul 13 23:53:38 CEST 2022 - The aliscore output file does not exist. Check results for BUSCO: 15711at7742
## Wed Jul 13 23:53:38 CEST 2022 - The aliscore output appears to be empty. Check results for BUSCO: 15711at7742
## Wed Jul 13 23:53:40 CEST 2022 - The aliscore output file does not exist. Check results for BUSCO: 20246at7742
## Wed Jul 13 23:53:40 CEST 2022 - The aliscore output appears to be empty. Check results for BUSCO: 20246at7742
## Wed Jul 13 23:53:40 CEST 2022 - The aliscore output file does not exist. Check results for BUSCO: 1653at7742
## Wed Jul 13 23:53:40 CEST 2022 - The aliscore output appears to be empty. Check results for BUSCO: 1653at7742
## Wed Jul 13 23:53:42 CEST 2022 - The aliscore output file does not exist. Check results for BUSCO: 1653at7742
## Wed Jul 13 23:53:42 CEST 2022 - The aliscore output appears to be empty. Check results for BUSCO: 1653at7742
## Wed Jul 13 23:53:45 CEST 2022 - The aliscore output file does not exist. Check results for BUSCO: 13963at7742
## Wed Jul 13 23:53:45 CEST 2022 - The aliscore output appears to be empty. Check results for BUSCO: 13963at7742
## Wed Jul 13 23:53:46 CEST 2022 - The aliscore output file does not exist. Check results for BUSCO: 7732at7742
## Wed Jul 13 23:53:46 CEST 2022 - The aliscore output appears to be empty. Check results for BUSCO: 7732at7742
## Wed Jul 13 23:53:47 CEST 2022 - The aliscore output file does not exist. Check results for BUSCO: 50582at7742
## Wed Jul 13 23:53:47 CEST 2022 - The aliscore output appears to be empty. Check results for BUSCO: 50582at7742
## Wed Jul 13 23:53:48 CEST 2022 - The aliscore output file does not exist. Check results for BUSCO: 8757at7742
## Wed Jul 13 23:53:48 CEST 2022 - The aliscore output appears to be empty. Check results for BUSCO: 8757at7742
## Wed Jul 13 23:53:50 CEST 2022 - The aliscore output file does not exist. Check results for BUSCO: 16347at7742
## Wed Jul 13 23:53:50 CEST 2022 - The aliscore output appears to be empty. Check results for BUSCO: 16347at7742
## Wed Jul 13 23:53:51 CEST 2022 - The aliscore output file does not exist. Check results for BUSCO: 14102at7742
## Wed Jul 13 23:53:51 CEST 2022 - The aliscore output appears to be empty. Check results for BUSCO: 14102at7742
## Wed Jul 13 23:53:52 CEST 2022 - The aliscore output file does not exist. Check results for BUSCO: 45247at7742
## Wed Jul 13 23:53:52 CEST 2022 - The aliscore output appears to be empty. Check results for BUSCO: 45247at7742
## Wed Jul 13 23:53:53 CEST 2022 - The aliscore output file does not exist. Check results for BUSCO: 1653at7742
## Wed Jul 13 23:53:53 CEST 2022 - The aliscore output appears to be empty. Check results for BUSCO: 1653at7742
## Wed Jul 13 23:53:55 CEST 2022 - The aliscore output file does not exist. Check results for BUSCO: 6305at7742
## Wed Jul 13 23:53:55 CEST 2022 - The aliscore output appears to be empty. Check results for BUSCO: 6305at7742
## Wed Jul 13 23:53:56 CEST 2022 - The aliscore output file does not exist. Check results for BUSCO: 32307at7742
## Wed Jul 13 23:53:56 CEST 2022 - The aliscore output appears to be empty. Check results for BUSCO: 32307at7742
## Wed Jul 13 23:54:00 CEST 2022 - The aliscore output file does not exist. Check results for BUSCO: 6253at7742
## Wed Jul 13 23:54:00 CEST 2022 - The aliscore output appears to be empty. Check results for BUSCO: 6253at7742
## Wed Jul 13 23:54:00 CEST 2022 - The aliscore output file does not exist. Check results for BUSCO: 15711at7742
## Wed Jul 13 23:54:00 CEST 2022 - The aliscore output appears to be empty. Check results for BUSCO: 15711at7742
## Wed Jul 13 23:54:02 CEST 2022 - The aliscore output file does not exist. Check results for BUSCO: 8874at7742
## Wed Jul 13 23:54:02 CEST 2022 - The aliscore output appears to be empty. Check results for BUSCO: 8874at7742
## Wed Jul 13 23:54:04 CEST 2022 - The aliscore output file does not exist. Check results for BUSCO: 7732at7742
## Wed Jul 13 23:54:04 CEST 2022 - The aliscore output appears to be empty. Check results for BUSCO: 7732at7742
## Wed Jul 13 23:54:05 CEST 2022 - The aliscore output file does not exist. Check results for BUSCO: 14102at7742
## Wed Jul 13 23:54:05 CEST 2022 - The aliscore output appears to be empty. Check results for BUSCO: 14102at7742
## Wed Jul 13 23:54:06 CEST 2022 - The aliscore output file does not exist. Check results for BUSCO: 16236at7742
## Wed Jul 13 23:54:06 CEST 2022 - The aliscore output appears to be empty. Check results for BUSCO: 16236at7742
## Wed Jul 13 23:54:08 CEST 2022 - The aliscore output file does not exist. Check results for BUSCO: 17848at7742
## Wed Jul 13 23:54:08 CEST 2022 - The aliscore output appears to be empty. Check results for BUSCO: 17848at7742
## Wed Jul 13 23:54:09 CEST 2022 - The aliscore output file does not exist. Check results for BUSCO: 23457at7742
## Wed Jul 13 23:54:09 CEST 2022 - The aliscore output appears to be empty. Check results for BUSCO: 23457at7742
## Wed Jul 13 23:54:11 CEST 2022 - The aliscore output file does not exist. Check results for BUSCO: 6305at7742
## Wed Jul 13 23:54:11 CEST 2022 - The aliscore output appears to be empty. Check results for BUSCO: 6305at7742
## Wed Jul 13 23:54:12 CEST 2022 - The aliscore output file does not exist. Check results for BUSCO: 44104at7742
## Wed Jul 13 23:54:12 CEST 2022 - The aliscore output appears to be empty. Check results for BUSCO: 44104at7742
## Wed Jul 13 23:54:15 CEST 2022 - The aliscore output file does not exist. Check results for BUSCO: 14102at7742
## Wed Jul 13 23:54:15 CEST 2022 - The aliscore output appears to be empty. Check results for BUSCO: 14102at7742
## Wed Jul 13 23:54:16 CEST 2022 - The aliscore output file does not exist. Check results for BUSCO: 18037at7742
## Wed Jul 13 23:54:16 CEST 2022 - The aliscore output appears to be empty. Check results for BUSCO: 18037at7742
## Wed Jul 13 23:54:16 CEST 2022 - The aliscore output file does not exist. Check results for BUSCO: 115534at7742
## Wed Jul 13 23:54:16 CEST 2022 - The aliscore output appears to be empty. Check results for BUSCO: 115534at7742
## Wed Jul 13 23:54:17 CEST 2022 - The aliscore output file does not exist. Check results for BUSCO: 7732at7742
## Wed Jul 13 23:54:17 CEST 2022 - The aliscore output appears to be empty. Check results for BUSCO: 7732at7742
## Wed Jul 13 23:54:21 CEST 2022 - The aliscore output file does not exist. Check results for BUSCO: 8757at7742
## Wed Jul 13 23:54:21 CEST 2022 - The aliscore output appears to be empty. Check results for BUSCO: 8757at7742
## Wed Jul 13 23:54:21 CEST 2022 - The aliscore output file does not exist. Check results for BUSCO: 20518at7742
## Wed Jul 13 23:54:21 CEST 2022 - The aliscore output appears to be empty. Check results for BUSCO: 20518at7742
## Wed Jul 13 23:54:23 CEST 2022 - The aliscore output file does not exist. Check results for BUSCO: 15604at7742
## Wed Jul 13 23:54:23 CEST 2022 - The aliscore output appears to be empty. Check results for BUSCO: 15604at7742
## Wed Jul 13 23:54:25 CEST 2022 - The aliscore output file does not exist. Check results for BUSCO: 5095at7742
## Wed Jul 13 23:54:25 CEST 2022 - The aliscore output appears to be empty. Check results for BUSCO: 5095at7742
## Wed Jul 13 23:54:28 CEST 2022 - The aliscore output file does not exist. Check results for BUSCO: 202098at7742
## Wed Jul 13 23:54:28 CEST 2022 - The aliscore output file does not exist. Check results for BUSCO: 52390at7742
## Wed Jul 13 23:54:28 CEST 2022 - The aliscore output appears to be empty. Check results for BUSCO: 202098at7742
## Wed Jul 13 23:54:28 CEST 2022 - The aliscore output appears to be empty. Check results for BUSCO: 52390at7742
## Wed Jul 13 23:54:31 CEST 2022 - The aliscore output file does not exist. Check results for BUSCO: 8409at7742
## Wed Jul 13 23:54:31 CEST 2022 - The aliscore output appears to be empty. Check results for BUSCO: 8409at7742
## Wed Jul 13 23:54:32 CEST 2022 - The aliscore output file does not exist. Check results for BUSCO: 199410at7742
## Wed Jul 13 23:54:32 CEST 2022 - The aliscore output appears to be empty. Check results for BUSCO: 199410at7742
## Wed Jul 13 23:54:32 CEST 2022 - The aliscore output file does not exist. Check results for BUSCO: 365702at7742
## Wed Jul 13 23:54:32 CEST 2022 - The aliscore output appears to be empty. Check results for BUSCO: 365702at7742
## Wed Jul 13 23:54:33 CEST 2022 - The aliscore output file does not exist. Check results for BUSCO: 5465at7742
## Wed Jul 13 23:54:33 CEST 2022 - The aliscore output appears to be empty. Check results for BUSCO: 5465at7742
## Wed Jul 13 23:54:36 CEST 2022 - The aliscore output file does not exist. Check results for BUSCO: 32732at7742
## Wed Jul 13 23:54:36 CEST 2022 - The aliscore output appears to be empty. Check results for BUSCO: 32732at7742
## Wed Jul 13 23:54:38 CEST 2022 - The aliscore output file does not exist. Check results for BUSCO: 17720at7742
## Wed Jul 13 23:54:38 CEST 2022 - The aliscore output appears to be empty. Check results for BUSCO: 17720at7742
## Wed Jul 13 23:54:38 CEST 2022 - The aliscore output file does not exist. Check results for BUSCO: 15257at7742
## Wed Jul 13 23:54:38 CEST 2022 - The aliscore output appears to be empty. Check results for BUSCO: 15257at7742
## Wed Jul 13 23:54:39 CEST 2022 - The aliscore output file does not exist. Check results for BUSCO: 39865at7742
## Wed Jul 13 23:54:39 CEST 2022 - The aliscore output appears to be empty. Check results for BUSCO: 39865at7742
## Wed Jul 13 23:54:41 CEST 2022 - The aliscore output file does not exist. Check results for BUSCO: 16236at7742
## Wed Jul 13 23:54:41 CEST 2022 - The aliscore output appears to be empty. Check results for BUSCO: 16236at7742
## Wed Jul 13 23:54:42 CEST 2022 - The aliscore output file does not exist. Check results for BUSCO: 5465at7742
## Wed Jul 13 23:54:42 CEST 2022 - The aliscore output appears to be empty. Check results for BUSCO: 5465at7742
## Wed Jul 13 23:54:44 CEST 2022 - The aliscore output file does not exist. Check results for BUSCO: 390830at7742
## Wed Jul 13 23:54:44 CEST 2022 - The aliscore output appears to be empty. Check results for BUSCO: 390830at7742
## Wed Jul 13 23:54:47 CEST 2022 - The aliscore output file does not exist. Check results for BUSCO: 6253at7742
## Wed Jul 13 23:54:47 CEST 2022 - The aliscore output appears to be empty. Check results for BUSCO: 6253at7742
## Wed Jul 13 23:54:48 CEST 2022 - The aliscore output file does not exist. Check results for BUSCO: 52059at7742
## Wed Jul 13 23:54:48 CEST 2022 - The aliscore output appears to be empty. Check results for BUSCO: 52059at7742
## Wed Jul 13 23:54:49 CEST 2022 - The aliscore output file does not exist. Check results for BUSCO: 4372at7742
## Wed Jul 13 23:54:49 CEST 2022 - The aliscore output appears to be empty. Check results for BUSCO: 4372at7742
## Wed Jul 13 23:54:50 CEST 2022 - The aliscore output file does not exist. Check results for BUSCO: 20246at7742
## Wed Jul 13 23:54:50 CEST 2022 - The aliscore output appears to be empty. Check results for BUSCO: 20246at7742
## Wed Jul 13 23:54:52 CEST 2022 - The aliscore output file does not exist. Check results for BUSCO: 9579at7742
## Wed Jul 13 23:54:52 CEST 2022 - The aliscore output appears to be empty. Check results for BUSCO: 9579at7742
## Wed Jul 13 23:54:53 CEST 2022 - The aliscore output file does not exist. Check results for BUSCO: 39061at7742
## Wed Jul 13 23:54:53 CEST 2022 - The aliscore output appears to be empty. Check results for BUSCO: 39061at7742
## Wed Jul 13 23:54:55 CEST 2022 - The aliscore output file does not exist. Check results for BUSCO: 15257at7742
## Wed Jul 13 23:54:55 CEST 2022 - The aliscore output appears to be empty. Check results for BUSCO: 15257at7742
## Wed Jul 13 23:54:55 CEST 2022 - The aliscore output file does not exist. Check results for BUSCO: 115534at7742
## Wed Jul 13 23:54:55 CEST 2022 - The aliscore output appears to be empty. Check results for BUSCO: 115534at7742
## Wed Jul 13 23:54:59 CEST 2022 - The aliscore output file does not exist. Check results for BUSCO: 17720at7742
## Wed Jul 13 23:54:59 CEST 2022 - The aliscore output appears to be empty. Check results for BUSCO: 17720at7742
## Wed Jul 13 23:54:59 CEST 2022 - The aliscore output file does not exist. Check results for BUSCO: 20518at7742
## Wed Jul 13 23:54:59 CEST 2022 - The aliscore output appears to be empty. Check results for BUSCO: 20518at7742
## Wed Jul 13 23:55:00 CEST 2022 - The aliscore output file does not exist. Check results for BUSCO: 4372at7742
## Wed Jul 13 23:55:00 CEST 2022 - The aliscore output appears to be empty. Check results for BUSCO: 4372at7742
## Wed Jul 13 23:55:01 CEST 2022 - The aliscore output file does not exist. Check results for BUSCO: 17789at7742
## Wed Jul 13 23:55:01 CEST 2022 - The aliscore output appears to be empty. Check results for BUSCO: 17789at7742
## Wed Jul 13 23:55:02 CEST 2022 - The aliscore output file does not exist. Check results for BUSCO: 5465at7742
## Wed Jul 13 23:55:02 CEST 2022 - The aliscore output appears to be empty. Check results for BUSCO: 5465at7742
## Wed Jul 13 23:55:06 CEST 2022 - The aliscore output file does not exist. Check results for BUSCO: 8258at7742
## Wed Jul 13 23:55:06 CEST 2022 - The aliscore output appears to be empty. Check results for BUSCO: 8258at7742
## Wed Jul 13 23:55:07 CEST 2022 - The aliscore output file does not exist. Check results for BUSCO: 20518at7742
## Wed Jul 13 23:55:07 CEST 2022 - The aliscore output appears to be empty. Check results for BUSCO: 20518at7742
## Wed Jul 13 23:55:12 CEST 2022 - The aliscore output file does not exist. Check results for BUSCO: 4372at7742
## Wed Jul 13 23:55:12 CEST 2022 - The aliscore output file does not exist. Check results for BUSCO: 388101at7742
## Wed Jul 13 23:55:12 CEST 2022 - The aliscore output appears to be empty. Check results for BUSCO: 4372at7742
## Wed Jul 13 23:55:12 CEST 2022 - The aliscore output appears to be empty. Check results for BUSCO: 388101at7742
## Wed Jul 13 23:55:12 CEST 2022 - The aliscore output file does not exist. Check results for BUSCO: 75557at7742
## Wed Jul 13 23:55:12 CEST 2022 - The aliscore output appears to be empty. Check results for BUSCO: 75557at7742
## Wed Jul 13 23:55:13 CEST 2022 - The aliscore output file does not exist. Check results for BUSCO: 388101at7742
## Wed Jul 13 23:55:13 CEST 2022 - The aliscore output appears to be empty. Check results for BUSCO: 388101at7742
## Wed Jul 13 23:55:13 CEST 2022 - The aliscore output file does not exist. Check results for BUSCO: 8258at7742
## Wed Jul 13 23:55:13 CEST 2022 - The aliscore output appears to be empty. Check results for BUSCO: 8258at7742
## Wed Jul 13 23:55:15 CEST 2022 - The aliscore output file does not exist. Check results for BUSCO: 34549at7742
## Wed Jul 13 23:55:15 CEST 2022 - The aliscore output appears to be empty. Check results for BUSCO: 34549at7742
## Wed Jul 13 23:55:18 CEST 2022 - The aliscore output file does not exist. Check results for BUSCO: 52390at7742
## Wed Jul 13 23:55:18 CEST 2022 - The aliscore output appears to be empty. Check results for BUSCO: 52390at7742
## Wed Jul 13 23:55:21 CEST 2022 - The aliscore output file does not exist. Check results for BUSCO: 23531at7742
## Wed Jul 13 23:55:21 CEST 2022 - The aliscore output appears to be empty. Check results for BUSCO: 23531at7742
## Wed Jul 13 23:55:22 CEST 2022 - The aliscore output file does not exist. Check results for BUSCO: 5095at7742
## Wed Jul 13 23:55:22 CEST 2022 - The aliscore output appears to be empty. Check results for BUSCO: 5095at7742
## Wed Jul 13 23:55:23 CEST 2022 - The aliscore output file does not exist. Check results for BUSCO: 388101at7742
## Wed Jul 13 23:55:23 CEST 2022 - The aliscore output appears to be empty. Check results for BUSCO: 388101at7742
## Wed Jul 13 23:55:23 CEST 2022 - The aliscore output file does not exist. Check results for BUSCO: 20246at7742
## Wed Jul 13 23:55:23 CEST 2022 - The aliscore output appears to be empty. Check results for BUSCO: 20246at7742
## Wed Jul 13 23:55:24 CEST 2022 - The aliscore output file does not exist. Check results for BUSCO: 17720at7742
## Wed Jul 13 23:55:24 CEST 2022 - The aliscore output appears to be empty. Check results for BUSCO: 17720at7742
## Wed Jul 13 23:55:26 CEST 2022 - The aliscore output file does not exist. Check results for BUSCO: 15604at7742
## Wed Jul 13 23:55:26 CEST 2022 - The aliscore output appears to be empty. Check results for BUSCO: 15604at7742
## Wed Jul 13 23:55:28 CEST 2022 - The aliscore output file does not exist. Check results for BUSCO: 15257at7742
## Wed Jul 13 23:55:28 CEST 2022 - The aliscore output appears to be empty. Check results for BUSCO: 15257at7742
## Wed Jul 13 23:55:30 CEST 2022 - The aliscore output file does not exist. Check results for BUSCO: 45908at7742
## Wed Jul 13 23:55:30 CEST 2022 - The aliscore output appears to be empty. Check results for BUSCO: 45908at7742
## Wed Jul 13 23:55:32 CEST 2022 - The aliscore output file does not exist. Check results for BUSCO: 33527at7742
## Wed Jul 13 23:55:32 CEST 2022 - The aliscore output appears to be empty. Check results for BUSCO: 33527at7742
## Wed Jul 13 23:55:32 CEST 2022 - The aliscore output file does not exist. Check results for BUSCO: 8874at7742
## Wed Jul 13 23:55:33 CEST 2022 - The aliscore output appears to be empty. Check results for BUSCO: 8874at7742
## Wed Jul 13 23:55:33 CEST 2022 - The aliscore output file does not exist. Check results for BUSCO: 52390at7742
## Wed Jul 13 23:55:33 CEST 2022 - The aliscore output appears to be empty. Check results for BUSCO: 52390at7742
## Wed Jul 13 23:55:36 CEST 2022 - The aliscore output file does not exist. Check results for BUSCO: 5422at7742
## Wed Jul 13 23:55:36 CEST 2022 - The aliscore output appears to be empty. Check results for BUSCO: 5422at7742
## Wed Jul 13 23:55:38 CEST 2022 - The aliscore output file does not exist. Check results for BUSCO: 32307at7742
## Wed Jul 13 23:55:38 CEST 2022 - The aliscore output appears to be empty. Check results for BUSCO: 32307at7742
## Wed Jul 13 23:55:38 CEST 2022 - The aliscore output file does not exist. Check results for BUSCO: 5422at7742
## Wed Jul 13 23:55:38 CEST 2022 - The aliscore output appears to be empty. Check results for BUSCO: 5422at7742
## Wed Jul 13 23:55:40 CEST 2022 - The aliscore output file does not exist. Check results for BUSCO: 365702at7742
## Wed Jul 13 23:55:40 CEST 2022 - The aliscore output appears to be empty. Check results for BUSCO: 365702at7742
## Wed Jul 13 23:55:42 CEST 2022 - The aliscore output file does not exist. Check results for BUSCO: 44744at7742
## Wed Jul 13 23:55:42 CEST 2022 - The aliscore output appears to be empty. Check results for BUSCO: 44744at7742
## Wed Jul 13 23:55:43 CEST 2022 - The aliscore output file does not exist. Check results for BUSCO: 8874at7742
## Wed Jul 13 23:55:43 CEST 2022 - The aliscore output appears to be empty. Check results for BUSCO: 8874at7742
## Wed Jul 13 23:55:46 CEST 2022 - The aliscore output file does not exist. Check results for BUSCO: 5095at7742
## Wed Jul 13 23:55:46 CEST 2022 - The aliscore output appears to be empty. Check results for BUSCO: 5095at7742
## Wed Jul 13 23:55:46 CEST 2022 - The aliscore output file does not exist. Check results for BUSCO: 32983at7742
## Wed Jul 13 23:55:47 CEST 2022 - The aliscore output appears to be empty. Check results for BUSCO: 32983at7742
## Wed Jul 13 23:55:47 CEST 2022 - The aliscore output file does not exist. Check results for BUSCO: 17789at7742
## Wed Jul 13 23:55:47 CEST 2022 - The aliscore output appears to be empty. Check results for BUSCO: 17789at7742
## Wed Jul 13 23:55:48 CEST 2022 - The aliscore output file does not exist. Check results for BUSCO: 36859at7742
## Wed Jul 13 23:55:48 CEST 2022 - The aliscore output appears to be empty. Check results for BUSCO: 36859at7742
## Wed Jul 13 23:55:51 CEST 2022 - The aliscore output file does not exist. Check results for BUSCO: 65757at7742
## Wed Jul 13 23:55:51 CEST 2022 - The aliscore output appears to be empty. Check results for BUSCO: 65757at7742
## Wed Jul 13 23:55:53 CEST 2022 - The aliscore output file does not exist. Check results for BUSCO: 390830at7742
## Wed Jul 13 23:55:53 CEST 2022 - The aliscore output appears to be empty. Check results for BUSCO: 390830at7742
## Wed Jul 13 23:55:54 CEST 2022 - The aliscore output file does not exist. Check results for BUSCO: 11472at7742
## Wed Jul 13 23:55:54 CEST 2022 - The aliscore output appears to be empty. Check results for BUSCO: 11472at7742
## Wed Jul 13 23:55:56 CEST 2022 - The aliscore output file does not exist. Check results for BUSCO: 16061at7742
## Wed Jul 13 23:55:56 CEST 2022 - The aliscore output appears to be empty. Check results for BUSCO: 16061at7742
## Wed Jul 13 23:55:58 CEST 2022 - The aliscore output file does not exist. Check results for BUSCO: 202098at7742
## Wed Jul 13 23:55:58 CEST 2022 - The aliscore output appears to be empty. Check results for BUSCO: 202098at7742
## Wed Jul 13 23:56:00 CEST 2022 - The aliscore output file does not exist. Check results for BUSCO: 16347at7742
## Wed Jul 13 23:56:00 CEST 2022 - The aliscore output appears to be empty. Check results for BUSCO: 16347at7742
## Wed Jul 13 23:56:01 CEST 2022 - The aliscore output file does not exist. Check results for BUSCO: 202098at7742
## Wed Jul 13 23:56:01 CEST 2022 - The aliscore output appears to be empty. Check results for BUSCO: 202098at7742
## Wed Jul 13 23:56:02 CEST 2022 - The aliscore output file does not exist. Check results for BUSCO: 5422at7742
## Wed Jul 13 23:56:02 CEST 2022 - The aliscore output appears to be empty. Check results for BUSCO: 5422at7742
## Wed Jul 13 23:56:04 CEST 2022 - The aliscore output file does not exist. Check results for BUSCO: 73826at7742
## Wed Jul 13 23:56:04 CEST 2022 - The aliscore output appears to be empty. Check results for BUSCO: 73826at7742
## Wed Jul 13 23:56:04 CEST 2022 - The aliscore output file does not exist. Check results for BUSCO: 365702at7742
## Wed Jul 13 23:56:04 CEST 2022 - The aliscore output appears to be empty. Check results for BUSCO: 365702at7742
## Wed Jul 13 23:56:07 CEST 2022 - The aliscore output file does not exist. Check results for BUSCO: 27385at7742
## Wed Jul 13 23:56:07 CEST 2022 - The aliscore output appears to be empty. Check results for BUSCO: 27385at7742
## Wed Jul 13 23:56:08 CEST 2022 - The aliscore output file does not exist. Check results for BUSCO: 8995at7742
## Wed Jul 13 23:56:08 CEST 2022 - The aliscore output appears to be empty. Check results for BUSCO: 8995at7742
## Wed Jul 13 23:56:08 CEST 2022 - The aliscore output file does not exist. Check results for BUSCO: 52059at7742
## Wed Jul 13 23:56:08 CEST 2022 - The aliscore output appears to be empty. Check results for BUSCO: 52059at7742
## Wed Jul 13 23:56:11 CEST 2022 - The aliscore output file does not exist. Check results for BUSCO: 199410at7742
## Wed Jul 13 23:56:11 CEST 2022 - The aliscore output appears to be empty. Check results for BUSCO: 199410at7742
## Wed Jul 13 23:56:12 CEST 2022 - The aliscore output file does not exist. Check results for BUSCO: 13963at7742
## Wed Jul 13 23:56:12 CEST 2022 - The aliscore output appears to be empty. Check results for BUSCO: 13963at7742
## Wed Jul 13 23:56:13 CEST 2022 - The aliscore output file does not exist. Check results for BUSCO: 16347at7742
## Wed Jul 13 23:56:14 CEST 2022 - The aliscore output appears to be empty. Check results for BUSCO: 16347at7742
## Wed Jul 13 23:56:17 CEST 2022 - The aliscore output file does not exist. Check results for BUSCO: 32732at7742
## Wed Jul 13 23:56:17 CEST 2022 - The aliscore output appears to be empty. Check results for BUSCO: 32732at7742
## Wed Jul 13 23:56:18 CEST 2022 - The aliscore output file does not exist. Check results for BUSCO: 150509at7742
## Wed Jul 13 23:56:18 CEST 2022 - The aliscore output appears to be empty. Check results for BUSCO: 150509at7742
## Wed Jul 13 23:56:19 CEST 2022 - The aliscore output file does not exist. Check results for BUSCO: 150509at7742
## Wed Jul 13 23:56:19 CEST 2022 - The aliscore output appears to be empty. Check results for BUSCO: 150509at7742
## Wed Jul 13 23:56:20 CEST 2022 - The aliscore output file does not exist. Check results for BUSCO: 143232at7742
## Wed Jul 13 23:56:21 CEST 2022 - The aliscore output appears to be empty. Check results for BUSCO: 143232at7742
## Wed Jul 13 23:56:22 CEST 2022 - The aliscore output file does not exist. Check results for BUSCO: 64985at7742
## Wed Jul 13 23:56:22 CEST 2022 - The aliscore output appears to be empty. Check results for BUSCO: 64985at7742
## Wed Jul 13 23:56:23 CEST 2022 - The aliscore output file does not exist. Check results for BUSCO: 4900at7742
## Wed Jul 13 23:56:23 CEST 2022 - The aliscore output appears to be empty. Check results for BUSCO: 4900at7742
## Wed Jul 13 23:56:24 CEST 2022 - The aliscore output file does not exist. Check results for BUSCO: 150509at7742
## Wed Jul 13 23:56:24 CEST 2022 - The aliscore output appears to be empty. Check results for BUSCO: 150509at7742
## Wed Jul 13 23:56:26 CEST 2022 - The aliscore output file does not exist. Check results for BUSCO: 33527at7742
## Wed Jul 13 23:56:26 CEST 2022 - The aliscore output appears to be empty. Check results for BUSCO: 33527at7742
## Wed Jul 13 23:56:29 CEST 2022 - The aliscore output file does not exist. Check results for BUSCO: 24985at7742
## Wed Jul 13 23:56:29 CEST 2022 - The aliscore output appears to be empty. Check results for BUSCO: 24985at7742
## Wed Jul 13 23:56:29 CEST 2022 - The aliscore output file does not exist. Check results for BUSCO: 21252at7742
## Wed Jul 13 23:56:29 CEST 2022 - The aliscore output appears to be empty. Check results for BUSCO: 21252at7742
## Wed Jul 13 23:56:31 CEST 2022 - The aliscore output file does not exist. Check results for BUSCO: 50582at7742
## Wed Jul 13 23:56:31 CEST 2022 - The aliscore output appears to be empty. Check results for BUSCO: 50582at7742
## Wed Jul 13 23:56:31 CEST 2022 - The aliscore output file does not exist. Check results for BUSCO: 32732at7742
## Wed Jul 13 23:56:31 CEST 2022 - The aliscore output appears to be empty. Check results for BUSCO: 32732at7742
## Wed Jul 13 23:56:35 CEST 2022 - The aliscore output file does not exist. Check results for BUSCO: 8995at7742
## Wed Jul 13 23:56:35 CEST 2022 - The aliscore output appears to be empty. Check results for BUSCO: 8995at7742
## Wed Jul 13 23:56:35 CEST 2022 - The aliscore output file does not exist. Check results for BUSCO: 9579at7742
## Wed Jul 13 23:56:35 CEST 2022 - The aliscore output appears to be empty. Check results for BUSCO: 9579at7742
## Wed Jul 13 23:56:37 CEST 2022 - The aliscore output file does not exist. Check results for BUSCO: 129936at7742
## Wed Jul 13 23:56:37 CEST 2022 - The aliscore output appears to be empty. Check results for BUSCO: 129936at7742
## Wed Jul 13 23:56:38 CEST 2022 - The aliscore output file does not exist. Check results for BUSCO: 32307at7742
## Wed Jul 13 23:56:38 CEST 2022 - The aliscore output appears to be empty. Check results for BUSCO: 32307at7742
## Wed Jul 13 23:56:40 CEST 2022 - The aliscore output file does not exist. Check results for BUSCO: 74593at7742
## Wed Jul 13 23:56:40 CEST 2022 - The aliscore output appears to be empty. Check results for BUSCO: 74593at7742
## Wed Jul 13 23:56:42 CEST 2022 - The aliscore output file does not exist. Check results for BUSCO: 4900at7742
## Wed Jul 13 23:56:43 CEST 2022 - The aliscore output appears to be empty. Check results for BUSCO: 4900at7742
## Wed Jul 13 23:56:43 CEST 2022 - The aliscore output file does not exist. Check results for BUSCO: 75557at7742
## Wed Jul 13 23:56:43 CEST 2022 - The aliscore output appears to be empty. Check results for BUSCO: 75557at7742
## Wed Jul 13 23:56:45 CEST 2022 - The aliscore output file does not exist. Check results for BUSCO: 13963at7742
## Wed Jul 13 23:56:45 CEST 2022 - The aliscore output appears to be empty. Check results for BUSCO: 13963at7742
## Wed Jul 13 23:56:46 CEST 2022 - The aliscore output file does not exist. Check results for BUSCO: 4900at7742
## Wed Jul 13 23:56:46 CEST 2022 - The aliscore output appears to be empty. Check results for BUSCO: 4900at7742
## Wed Jul 13 23:56:49 CEST 2022 - The aliscore output file does not exist. Check results for BUSCO: 52059at7742
## Wed Jul 13 23:56:49 CEST 2022 - The aliscore output appears to be empty. Check results for BUSCO: 52059at7742
## Wed Jul 13 23:56:50 CEST 2022 - The aliscore output file does not exist. Check results for BUSCO: 33527at7742
## Wed Jul 13 23:56:50 CEST 2022 - The aliscore output appears to be empty. Check results for BUSCO: 33527at7742
## Wed Jul 13 23:56:51 CEST 2022 - The aliscore output file does not exist. Check results for BUSCO: 9579at7742
## Wed Jul 13 23:56:51 CEST 2022 - The aliscore output appears to be empty. Check results for BUSCO: 9579at7742
## Wed Jul 13 23:56:53 CEST 2022 - The aliscore output file does not exist. Check results for BUSCO: 33364at7742
## Wed Jul 13 23:56:53 CEST 2022 - The aliscore output appears to be empty. Check results for BUSCO: 33364at7742
## Wed Jul 13 23:56:55 CEST 2022 - The aliscore output file does not exist. Check results for BUSCO: 34549at7742
## Wed Jul 13 23:56:55 CEST 2022 - The aliscore output appears to be empty. Check results for BUSCO: 34549at7742
## Wed Jul 13 23:56:56 CEST 2022 - The aliscore output file does not exist. Check results for BUSCO: 15408at7742
## Wed Jul 13 23:56:56 CEST 2022 - The aliscore output appears to be empty. Check results for BUSCO: 15408at7742
## Wed Jul 13 23:56:57 CEST 2022 - The aliscore output file does not exist. Check results for BUSCO: 8995at7742
## Wed Jul 13 23:56:57 CEST 2022 - The aliscore output appears to be empty. Check results for BUSCO: 8995at7742
## Wed Jul 13 23:56:59 CEST 2022 - The aliscore output file does not exist. Check results for BUSCO: 15408at7742
## Wed Jul 13 23:57:00 CEST 2022 - The aliscore output appears to be empty. Check results for BUSCO: 15408at7742
## Wed Jul 13 23:57:01 CEST 2022 - The aliscore output file does not exist. Check results for BUSCO: 11472at7742
## Wed Jul 13 23:57:01 CEST 2022 - The aliscore output appears to be empty. Check results for BUSCO: 11472at7742
## Wed Jul 13 23:57:03 CEST 2022 - The aliscore output file does not exist. Check results for BUSCO: 58421at7742
## Wed Jul 13 23:57:03 CEST 2022 - The aliscore output appears to be empty. Check results for BUSCO: 58421at7742
## Wed Jul 13 23:57:03 CEST 2022 - The aliscore output file does not exist. Check results for BUSCO: 23531at7742
## Wed Jul 13 23:57:03 CEST 2022 - The aliscore output appears to be empty. Check results for BUSCO: 23531at7742
## Wed Jul 13 23:57:06 CEST 2022 - The aliscore output file does not exist. Check results for BUSCO: 103093at7742
## Wed Jul 13 23:57:06 CEST 2022 - The aliscore output appears to be empty. Check results for BUSCO: 103093at7742
## Wed Jul 13 23:57:07 CEST 2022 - The aliscore output file does not exist. Check results for BUSCO: 15408at7742
## Wed Jul 13 23:57:07 CEST 2022 - The aliscore output appears to be empty. Check results for BUSCO: 15408at7742
## Wed Jul 13 23:57:10 CEST 2022 - The aliscore output file does not exist. Check results for BUSCO: 32983at7742
## Wed Jul 13 23:57:10 CEST 2022 - The aliscore output appears to be empty. Check results for BUSCO: 32983at7742
## Wed Jul 13 23:57:10 CEST 2022 - The aliscore output file does not exist. Check results for BUSCO: 18541at7742
## Wed Jul 13 23:57:10 CEST 2022 - The aliscore output appears to be empty. Check results for BUSCO: 18541at7742
## Wed Jul 13 23:57:13 CEST 2022 - The aliscore output file does not exist. Check results for BUSCO: 34549at7742
## Wed Jul 13 23:57:13 CEST 2022 - The aliscore output appears to be empty. Check results for BUSCO: 34549at7742
## Wed Jul 13 23:57:14 CEST 2022 - The aliscore output file does not exist. Check results for BUSCO: 13987at7742
## Wed Jul 13 23:57:14 CEST 2022 - The aliscore output appears to be empty. Check results for BUSCO: 13987at7742
## Wed Jul 13 23:57:14 CEST 2022 - The aliscore output file does not exist. Check results for BUSCO: 11472at7742
## Wed Jul 13 23:57:14 CEST 2022 - The aliscore output appears to be empty. Check results for BUSCO: 11472at7742
## Wed Jul 13 23:57:17 CEST 2022 - The aliscore output file does not exist. Check results for BUSCO: 27364at7742
## Wed Jul 13 23:57:17 CEST 2022 - The aliscore output appears to be empty. Check results for BUSCO: 27364at7742
## Wed Jul 13 23:57:19 CEST 2022 - The aliscore output file does not exist. Check results for BUSCO: 75557at7742
## Wed Jul 13 23:57:19 CEST 2022 - The aliscore output appears to be empty. Check results for BUSCO: 75557at7742
## Wed Jul 13 23:57:20 CEST 2022 - The aliscore output file does not exist. Check results for BUSCO: 45908at7742
## Wed Jul 13 23:57:20 CEST 2022 - The aliscore output appears to be empty. Check results for BUSCO: 45908at7742
## Wed Jul 13 23:57:21 CEST 2022 - The aliscore output file does not exist. Check results for BUSCO: 8409at7742
## Wed Jul 13 23:57:21 CEST 2022 - The aliscore output appears to be empty. Check results for BUSCO: 8409at7742
## Wed Jul 13 23:57:22 CEST 2022 - The aliscore output file does not exist. Check results for BUSCO: 50917at7742
## Wed Jul 13 23:57:22 CEST 2022 - The aliscore output appears to be empty. Check results for BUSCO: 50917at7742
## Wed Jul 13 23:57:22 CEST 2022 - The aliscore output file does not exist. Check results for BUSCO: 143232at7742
## Wed Jul 13 23:57:22 CEST 2022 - The aliscore output appears to be empty. Check results for BUSCO: 143232at7742
## Wed Jul 13 23:57:24 CEST 2022 - The aliscore output file does not exist. Check results for BUSCO: 23531at7742
## Wed Jul 13 23:57:24 CEST 2022 - The aliscore output appears to be empty. Check results for BUSCO: 23531at7742
## Wed Jul 13 23:57:26 CEST 2022 - The aliscore output file does not exist. Check results for BUSCO: 6108at7742
## Wed Jul 13 23:57:26 CEST 2022 - The aliscore output appears to be empty. Check results for BUSCO: 6108at7742
## Wed Jul 13 23:57:28 CEST 2022 - The aliscore output file does not exist. Check results for BUSCO: 6701at7742
## Wed Jul 13 23:57:28 CEST 2022 - The aliscore output appears to be empty. Check results for BUSCO: 6701at7742
## Wed Jul 13 23:57:29 CEST 2022 - The aliscore output file does not exist. Check results for BUSCO: 39865at7742
## Wed Jul 13 23:57:29 CEST 2022 - The aliscore output appears to be empty. Check results for BUSCO: 39865at7742
## Wed Jul 13 23:57:30 CEST 2022 - The aliscore output file does not exist. Check results for BUSCO: 129936at7742
## Wed Jul 13 23:57:30 CEST 2022 - The aliscore output appears to be empty. Check results for BUSCO: 129936at7742
## Wed Jul 13 23:57:33 CEST 2022 - The aliscore output file does not exist. Check results for BUSCO: 64333at7742
## Wed Jul 13 23:57:33 CEST 2022 - The aliscore output appears to be empty. Check results for BUSCO: 64333at7742
## Wed Jul 13 23:57:34 CEST 2022 - The aliscore output file does not exist. Check results for BUSCO: 6108at7742
## Wed Jul 13 23:57:34 CEST 2022 - The aliscore output appears to be empty. Check results for BUSCO: 6108at7742
## Wed Jul 13 23:57:35 CEST 2022 - The aliscore output file does not exist. Check results for BUSCO: 24985at7742
## Wed Jul 13 23:57:35 CEST 2022 - The aliscore output appears to be empty. Check results for BUSCO: 24985at7742
## Wed Jul 13 23:57:37 CEST 2022 - The aliscore output file does not exist. Check results for BUSCO: 32983at7742
## Wed Jul 13 23:57:37 CEST 2022 - The aliscore output appears to be empty. Check results for BUSCO: 32983at7742
## Wed Jul 13 23:57:38 CEST 2022 - The aliscore output file does not exist. Check results for BUSCO: 16061at7742
## Wed Jul 13 23:57:38 CEST 2022 - The aliscore output appears to be empty. Check results for BUSCO: 16061at7742
## Wed Jul 13 23:57:40 CEST 2022 - The aliscore output file does not exist. Check results for BUSCO: 18541at7742
## Wed Jul 13 23:57:40 CEST 2022 - The aliscore output appears to be empty. Check results for BUSCO: 18541at7742
## Wed Jul 13 23:57:41 CEST 2022 - The aliscore output file does not exist. Check results for BUSCO: 129936at7742
## Wed Jul 13 23:57:41 CEST 2022 - The aliscore output appears to be empty. Check results for BUSCO: 129936at7742
## Wed Jul 13 23:57:42 CEST 2022 - The aliscore output file does not exist. Check results for BUSCO: 6108at7742
## Wed Jul 13 23:57:42 CEST 2022 - The aliscore output appears to be empty. Check results for BUSCO: 6108at7742
## Wed Jul 13 23:57:44 CEST 2022 - The aliscore output file does not exist. Check results for BUSCO: 84691at7742
## Wed Jul 13 23:57:44 CEST 2022 - The aliscore output appears to be empty. Check results for BUSCO: 84691at7742
## Wed Jul 13 23:57:46 CEST 2022 - The aliscore output file does not exist. Check results for BUSCO: 84691at7742
## Wed Jul 13 23:57:46 CEST 2022 - The aliscore output appears to be empty. Check results for BUSCO: 84691at7742
## Wed Jul 13 23:57:47 CEST 2022 - The aliscore output file does not exist. Check results for BUSCO: 199410at7742
## Wed Jul 13 23:57:47 CEST 2022 - The aliscore output appears to be empty. Check results for BUSCO: 199410at7742
## Wed Jul 13 23:57:48 CEST 2022 - The aliscore output file does not exist. Check results for BUSCO: 16601at7742
## Wed Jul 13 23:57:48 CEST 2022 - The aliscore output appears to be empty. Check results for BUSCO: 16601at7742
## Wed Jul 13 23:57:50 CEST 2022 - The aliscore output file does not exist. Check results for BUSCO: 146754at7742
## Wed Jul 13 23:57:50 CEST 2022 - The aliscore output appears to be empty. Check results for BUSCO: 146754at7742
## Wed Jul 13 23:57:50 CEST 2022 - The aliscore output file does not exist. Check results for BUSCO: 21252at7742
## Wed Jul 13 23:57:50 CEST 2022 - The aliscore output appears to be empty. Check results for BUSCO: 21252at7742
## Wed Jul 13 23:57:52 CEST 2022 - The aliscore output file does not exist. Check results for BUSCO: 13881at7742
## Wed Jul 13 23:57:52 CEST 2022 - The aliscore output appears to be empty. Check results for BUSCO: 13881at7742
## Wed Jul 13 23:57:53 CEST 2022 - The aliscore output file does not exist. Check results for BUSCO: 143232at7742
## Wed Jul 13 23:57:53 CEST 2022 - The aliscore output appears to be empty. Check results for BUSCO: 143232at7742
## Wed Jul 13 23:57:53 CEST 2022 - The aliscore output file does not exist. Check results for BUSCO: 36859at7742
## Wed Jul 13 23:57:53 CEST 2022 - The aliscore output appears to be empty. Check results for BUSCO: 36859at7742
## Wed Jul 13 23:57:57 CEST 2022 - The aliscore output file does not exist. Check results for BUSCO: 6701at7742
## Wed Jul 13 23:57:57 CEST 2022 - The aliscore output appears to be empty. Check results for BUSCO: 6701at7742
## Wed Jul 13 23:57:59 CEST 2022 - The aliscore output file does not exist. Check results for BUSCO: 50917at7742
## Wed Jul 13 23:57:59 CEST 2022 - The aliscore output appears to be empty. Check results for BUSCO: 50917at7742
## Wed Jul 13 23:58:01 CEST 2022 - The aliscore output file does not exist. Check results for BUSCO: 8409at7742
## Wed Jul 13 23:58:01 CEST 2022 - The aliscore output appears to be empty. Check results for BUSCO: 8409at7742
## Wed Jul 13 23:58:03 CEST 2022 - The aliscore output file does not exist. Check results for BUSCO: 39865at7742
## Wed Jul 13 23:58:03 CEST 2022 - The aliscore output appears to be empty. Check results for BUSCO: 39865at7742
## Wed Jul 13 23:58:03 CEST 2022 - The aliscore output file does not exist. Check results for BUSCO: 45908at7742
## Wed Jul 13 23:58:03 CEST 2022 - The aliscore output appears to be empty. Check results for BUSCO: 45908at7742
## Wed Jul 13 23:58:03 CEST 2022 - The aliscore output file does not exist. Check results for BUSCO: 11505at7742
## Wed Jul 13 23:58:03 CEST 2022 - The aliscore output appears to be empty. Check results for BUSCO: 11505at7742
## Wed Jul 13 23:58:03 CEST 2022 - The aliscore output file does not exist. Check results for BUSCO: 6701at7742
## Wed Jul 13 23:58:03 CEST 2022 - The aliscore output appears to be empty. Check results for BUSCO: 6701at7742
## Wed Jul 13 23:58:09 CEST 2022 - The aliscore output file does not exist. Check results for BUSCO: 21252at7742
## Wed Jul 13 23:58:09 CEST 2022 - The aliscore output appears to be empty. Check results for BUSCO: 21252at7742
## Wed Jul 13 23:58:12 CEST 2022 - The aliscore output file does not exist. Check results for BUSCO: 22187at7742
## Wed Jul 13 23:58:12 CEST 2022 - The aliscore output appears to be empty. Check results for BUSCO: 22187at7742
## Wed Jul 13 23:58:12 CEST 2022 - The aliscore output file does not exist. Check results for BUSCO: 27364at7742
## Wed Jul 13 23:58:12 CEST 2022 - The aliscore output appears to be empty. Check results for BUSCO: 27364at7742
## Wed Jul 13 23:58:13 CEST 2022 - The aliscore output file does not exist. Check results for BUSCO: 27364at7742
## Wed Jul 13 23:58:13 CEST 2022 - The aliscore output appears to be empty. Check results for BUSCO: 27364at7742
## Wed Jul 13 23:58:13 CEST 2022 - The aliscore output file does not exist. Check results for BUSCO: 65634at7742
## Wed Jul 13 23:58:13 CEST 2022 - The aliscore output appears to be empty. Check results for BUSCO: 65634at7742
## Wed Jul 13 23:58:13 CEST 2022 - The aliscore output file does not exist. Check results for BUSCO: 146754at7742
## Wed Jul 13 23:58:13 CEST 2022 - The aliscore output appears to be empty. Check results for BUSCO: 146754at7742
## Wed Jul 13 23:58:17 CEST 2022 - The aliscore output file does not exist. Check results for BUSCO: 18946at7742
## Wed Jul 13 23:58:17 CEST 2022 - The aliscore output appears to be empty. Check results for BUSCO: 18946at7742
## Wed Jul 13 23:58:19 CEST 2022 - The aliscore output file does not exist. Check results for BUSCO: 74593at7742
## Wed Jul 13 23:58:19 CEST 2022 - The aliscore output appears to be empty. Check results for BUSCO: 74593at7742
## Wed Jul 13 23:58:19 CEST 2022 - The aliscore output file does not exist. Check results for BUSCO: 142339at7742
## Wed Jul 13 23:58:20 CEST 2022 - The aliscore output appears to be empty. Check results for BUSCO: 142339at7742
## Wed Jul 13 23:58:22 CEST 2022 - The aliscore output file does not exist. Check results for BUSCO: 156140at7742
## Wed Jul 13 23:58:22 CEST 2022 - The aliscore output appears to be empty. Check results for BUSCO: 156140at7742
## Wed Jul 13 23:58:23 CEST 2022 - The aliscore output file does not exist. Check results for BUSCO: 27385at7742
## Wed Jul 13 23:58:23 CEST 2022 - The aliscore output appears to be empty. Check results for BUSCO: 27385at7742
## Wed Jul 13 23:58:24 CEST 2022 - The aliscore output file does not exist. Check results for BUSCO: 41902at7742
## Wed Jul 13 23:58:24 CEST 2022 - The aliscore output appears to be empty. Check results for BUSCO: 41902at7742
## Wed Jul 13 23:58:26 CEST 2022 - The aliscore output file does not exist. Check results for BUSCO: 74593at7742
## Wed Jul 13 23:58:26 CEST 2022 - The aliscore output appears to be empty. Check results for BUSCO: 74593at7742
## Wed Jul 13 23:58:27 CEST 2022 - The aliscore output file does not exist. Check results for BUSCO: 33525at7742
## Wed Jul 13 23:58:27 CEST 2022 - The aliscore output appears to be empty. Check results for BUSCO: 33525at7742
## Wed Jul 13 23:58:29 CEST 2022 - The aliscore output file does not exist. Check results for BUSCO: 146754at7742
## Wed Jul 13 23:58:29 CEST 2022 - The aliscore output appears to be empty. Check results for BUSCO: 146754at7742
## Wed Jul 13 23:58:31 CEST 2022 - The aliscore output file does not exist. Check results for BUSCO: 16061at7742
## Wed Jul 13 23:58:31 CEST 2022 - The aliscore output appears to be empty. Check results for BUSCO: 16061at7742
## Wed Jul 13 23:58:32 CEST 2022 - The aliscore output file does not exist. Check results for BUSCO: 36859at7742
## Wed Jul 13 23:58:32 CEST 2022 - The aliscore output appears to be empty. Check results for BUSCO: 36859at7742
## Wed Jul 13 23:58:33 CEST 2022 - The aliscore output file does not exist. Check results for BUSCO: 27385at7742
## Wed Jul 13 23:58:33 CEST 2022 - The aliscore output appears to be empty. Check results for BUSCO: 27385at7742
## Wed Jul 13 23:58:34 CEST 2022 - The aliscore output file does not exist. Check results for BUSCO: 66047at7742
## Wed Jul 13 23:58:34 CEST 2022 - The aliscore output appears to be empty. Check results for BUSCO: 66047at7742
## Wed Jul 13 23:58:35 CEST 2022 - The aliscore output file does not exist. Check results for BUSCO: 12491at7742
## Wed Jul 13 23:58:35 CEST 2022 - The aliscore output appears to be empty. Check results for BUSCO: 12491at7742
## Wed Jul 13 23:58:38 CEST 2022 - The aliscore output file does not exist. Check results for BUSCO: 12491at7742
## Wed Jul 13 23:58:38 CEST 2022 - The aliscore output appears to be empty. Check results for BUSCO: 12491at7742
## Wed Jul 13 23:58:38 CEST 2022 - The aliscore output file does not exist. Check results for BUSCO: 321789at7742
## Wed Jul 13 23:58:39 CEST 2022 - The aliscore output appears to be empty. Check results for BUSCO: 321789at7742
## Wed Jul 13 23:58:42 CEST 2022 - The aliscore output file does not exist. Check results for BUSCO: 24985at7742
## Wed Jul 13 23:58:42 CEST 2022 - The aliscore output appears to be empty. Check results for BUSCO: 24985at7742
## Wed Jul 13 23:58:42 CEST 2022 - The aliscore output file does not exist. Check results for BUSCO: 108312at7742
## Wed Jul 13 23:58:42 CEST 2022 - The aliscore output appears to be empty. Check results for BUSCO: 108312at7742
## Wed Jul 13 23:58:44 CEST 2022 - The aliscore output file does not exist. Check results for BUSCO: 18541at7742
## Wed Jul 13 23:58:44 CEST 2022 - The aliscore output appears to be empty. Check results for BUSCO: 18541at7742
## Wed Jul 13 23:58:45 CEST 2022 - The aliscore output file does not exist. Check results for BUSCO: 73826at7742
## Wed Jul 13 23:58:45 CEST 2022 - The aliscore output appears to be empty. Check results for BUSCO: 73826at7742
## Wed Jul 13 23:58:45 CEST 2022 - The aliscore output file does not exist. Check results for BUSCO: 43051at7742
## Wed Jul 13 23:58:45 CEST 2022 - The aliscore output appears to be empty. Check results for BUSCO: 43051at7742
## Wed Jul 13 23:58:47 CEST 2022 - The aliscore output file does not exist. Check results for BUSCO: 38686at7742
## Wed Jul 13 23:58:47 CEST 2022 - The aliscore output appears to be empty. Check results for BUSCO: 38686at7742
## Wed Jul 13 23:58:50 CEST 2022 - The aliscore output file does not exist. Check results for BUSCO: 22187at7742
## Wed Jul 13 23:58:50 CEST 2022 - The aliscore output appears to be empty. Check results for BUSCO: 22187at7742
## Wed Jul 13 23:58:51 CEST 2022 - The aliscore output file does not exist. Check results for BUSCO: 33364at7742
## Wed Jul 13 23:58:51 CEST 2022 - The aliscore output appears to be empty. Check results for BUSCO: 33364at7742
## Wed Jul 13 23:58:52 CEST 2022 - The aliscore output file does not exist. Check results for BUSCO: 104785at7742
## Wed Jul 13 23:58:52 CEST 2022 - The aliscore output appears to be empty. Check results for BUSCO: 104785at7742
## Wed Jul 13 23:58:55 CEST 2022 - The aliscore output file does not exist. Check results for BUSCO: 108312at7742
## Wed Jul 13 23:58:56 CEST 2022 - The aliscore output appears to be empty. Check results for BUSCO: 108312at7742
## Wed Jul 13 23:58:56 CEST 2022 - The aliscore output file does not exist. Check results for BUSCO: 73826at7742
## Wed Jul 13 23:58:56 CEST 2022 - The aliscore output appears to be empty. Check results for BUSCO: 73826at7742
## Wed Jul 13 23:58:56 CEST 2022 - The aliscore output file does not exist. Check results for BUSCO: 9744at7742
## Wed Jul 13 23:58:56 CEST 2022 - The aliscore output appears to be empty. Check results for BUSCO: 9744at7742
## Wed Jul 13 23:58:58 CEST 2022 - The aliscore output file does not exist. Check results for BUSCO: 108312at7742
## Wed Jul 13 23:58:58 CEST 2022 - The aliscore output appears to be empty. Check results for BUSCO: 108312at7742
## Wed Jul 13 23:59:00 CEST 2022 - The aliscore output file does not exist. Check results for BUSCO: 13480at7742
## Wed Jul 13 23:59:00 CEST 2022 - The aliscore output appears to be empty. Check results for BUSCO: 13480at7742
## Wed Jul 13 23:59:01 CEST 2022 - The aliscore output file does not exist. Check results for BUSCO: 33364at7742
## Wed Jul 13 23:59:01 CEST 2022 - The aliscore output appears to be empty. Check results for BUSCO: 33364at7742
## Wed Jul 13 23:59:03 CEST 2022 - The aliscore output file does not exist. Check results for BUSCO: 22187at7742
## Wed Jul 13 23:59:03 CEST 2022 - The aliscore output appears to be empty. Check results for BUSCO: 22187at7742
## Wed Jul 13 23:59:05 CEST 2022 - The aliscore output file does not exist. Check results for BUSCO: 33525at7742
## Wed Jul 13 23:59:05 CEST 2022 - The aliscore output appears to be empty. Check results for BUSCO: 33525at7742
## Wed Jul 13 23:59:07 CEST 2022 - The aliscore output file does not exist. Check results for BUSCO: 103093at7742
## Wed Jul 13 23:59:07 CEST 2022 - The aliscore output file does not exist. Check results for BUSCO: 13881at7742
## Wed Jul 13 23:59:07 CEST 2022 - The aliscore output appears to be empty. Check results for BUSCO: 103093at7742
## Wed Jul 13 23:59:07 CEST 2022 - The aliscore output appears to be empty. Check results for BUSCO: 13881at7742
## Wed Jul 13 23:59:07 CEST 2022 - The aliscore output file does not exist. Check results for BUSCO: 11505at7742
## Wed Jul 13 23:59:07 CEST 2022 - The aliscore output appears to be empty. Check results for BUSCO: 11505at7742
## Wed Jul 13 23:59:12 CEST 2022 - The aliscore output file does not exist. Check results for BUSCO: 185277at7742
## Wed Jul 13 23:59:12 CEST 2022 - The aliscore output appears to be empty. Check results for BUSCO: 185277at7742
## Wed Jul 13 23:59:12 CEST 2022 - The aliscore output file does not exist. Check results for BUSCO: 33525at7742
## Wed Jul 13 23:59:13 CEST 2022 - The aliscore output appears to be empty. Check results for BUSCO: 33525at7742
## Wed Jul 13 23:59:13 CEST 2022 - The aliscore output file does not exist. Check results for BUSCO: 11505at7742
## Wed Jul 13 23:59:13 CEST 2022 - The aliscore output appears to be empty. Check results for BUSCO: 11505at7742
## Wed Jul 13 23:59:16 CEST 2022 - The aliscore output file does not exist. Check results for BUSCO: 103093at7742
## Wed Jul 13 23:59:16 CEST 2022 - The aliscore output appears to be empty. Check results for BUSCO: 103093at7742
## Wed Jul 13 23:59:17 CEST 2022 - The aliscore output file does not exist. Check results for BUSCO: 18946at7742
## Wed Jul 13 23:59:17 CEST 2022 - The aliscore output appears to be empty. Check results for BUSCO: 18946at7742
## Wed Jul 13 23:59:17 CEST 2022 - The aliscore output file does not exist. Check results for BUSCO: 225400at7742
## Wed Jul 13 23:59:17 CEST 2022 - The aliscore output appears to be empty. Check results for BUSCO: 225400at7742
## Wed Jul 13 23:59:20 CEST 2022 - The aliscore output file does not exist. Check results for BUSCO: 156140at7742
## Wed Jul 13 23:59:20 CEST 2022 - The aliscore output appears to be empty. Check results for BUSCO: 156140at7742
## Wed Jul 13 23:59:20 CEST 2022 - The aliscore output file does not exist. Check results for BUSCO: 30973at7742
## Wed Jul 13 23:59:20 CEST 2022 - The aliscore output appears to be empty. Check results for BUSCO: 30973at7742
## Wed Jul 13 23:59:22 CEST 2022 - The aliscore output file does not exist. Check results for BUSCO: 231143at7742
## Wed Jul 13 23:59:22 CEST 2022 - The aliscore output appears to be empty. Check results for BUSCO: 231143at7742
## Wed Jul 13 23:59:22 CEST 2022 - The aliscore output file does not exist. Check results for BUSCO: 153465at7742
## Wed Jul 13 23:59:22 CEST 2022 - The aliscore output appears to be empty. Check results for BUSCO: 153465at7742
## Wed Jul 13 23:59:27 CEST 2022 - The aliscore output file does not exist. Check results for BUSCO: 38686at7742
## Wed Jul 13 23:59:27 CEST 2022 - The aliscore output appears to be empty. Check results for BUSCO: 38686at7742
## Wed Jul 13 23:59:27 CEST 2022 - The aliscore output file does not exist. Check results for BUSCO: 159513at7742
## Wed Jul 13 23:59:27 CEST 2022 - The aliscore output appears to be empty. Check results for BUSCO: 159513at7742
## Wed Jul 13 23:59:28 CEST 2022 - The aliscore output file does not exist. Check results for BUSCO: 79715at7742
## Wed Jul 13 23:59:28 CEST 2022 - The aliscore output appears to be empty. Check results for BUSCO: 79715at7742
## Wed Jul 13 23:59:28 CEST 2022 - The aliscore output file does not exist. Check results for BUSCO: 6126at7742
## Wed Jul 13 23:59:28 CEST 2022 - The aliscore output appears to be empty. Check results for BUSCO: 6126at7742
## Wed Jul 13 23:59:29 CEST 2022 - The aliscore output file does not exist. Check results for BUSCO: 10136at7742
## Wed Jul 13 23:59:29 CEST 2022 - The aliscore output appears to be empty. Check results for BUSCO: 10136at7742
## Wed Jul 13 23:59:32 CEST 2022 - The aliscore output file does not exist. Check results for BUSCO: 36449at7742
## Wed Jul 13 23:59:32 CEST 2022 - The aliscore output appears to be empty. Check results for BUSCO: 36449at7742
## Wed Jul 13 23:59:33 CEST 2022 - The aliscore output file does not exist. Check results for BUSCO: 102945at7742
## Wed Jul 13 23:59:33 CEST 2022 - The aliscore output appears to be empty. Check results for BUSCO: 102945at7742
## Wed Jul 13 23:59:35 CEST 2022 - The aliscore output file does not exist. Check results for BUSCO: 46117at7742
## Wed Jul 13 23:59:35 CEST 2022 - The aliscore output appears to be empty. Check results for BUSCO: 46117at7742
## Wed Jul 13 23:59:36 CEST 2022 - The aliscore output file does not exist. Check results for BUSCO: 97499at7742
## Wed Jul 13 23:59:37 CEST 2022 - The aliscore output appears to be empty. Check results for BUSCO: 97499at7742
## Wed Jul 13 23:59:39 CEST 2022 - The aliscore output file does not exist. Check results for BUSCO: 41902at7742
## Wed Jul 13 23:59:39 CEST 2022 - The aliscore output appears to be empty. Check results for BUSCO: 41902at7742
## Wed Jul 13 23:59:40 CEST 2022 - The aliscore output file does not exist. Check results for BUSCO: 62617at7742
## Wed Jul 13 23:59:40 CEST 2022 - The aliscore output appears to be empty. Check results for BUSCO: 62617at7742
## Wed Jul 13 23:59:42 CEST 2022 - The aliscore output file does not exist. Check results for BUSCO: 189431at7742
## Wed Jul 13 23:59:42 CEST 2022 - The aliscore output appears to be empty. Check results for BUSCO: 189431at7742
## Wed Jul 13 23:59:42 CEST 2022 - The aliscore output file does not exist. Check results for BUSCO: 6126at7742
## Wed Jul 13 23:59:42 CEST 2022 - The aliscore output appears to be empty. Check results for BUSCO: 6126at7742
## Wed Jul 13 23:59:45 CEST 2022 - The aliscore output file does not exist. Check results for BUSCO: 156140at7742
## Wed Jul 13 23:59:45 CEST 2022 - The aliscore output appears to be empty. Check results for BUSCO: 156140at7742
## Wed Jul 13 23:59:47 CEST 2022 - The aliscore output file does not exist. Check results for BUSCO: 41902at7742
## Wed Jul 13 23:59:47 CEST 2022 - The aliscore output appears to be empty. Check results for BUSCO: 41902at7742
## Wed Jul 13 23:59:49 CEST 2022 - The aliscore output file does not exist. Check results for BUSCO: 38686at7742
## Wed Jul 13 23:59:49 CEST 2022 - The aliscore output appears to be empty. Check results for BUSCO: 38686at7742
## Wed Jul 13 23:59:49 CEST 2022 - The aliscore output file does not exist. Check results for BUSCO: 62617at7742
## Wed Jul 13 23:59:50 CEST 2022 - The aliscore output appears to be empty. Check results for BUSCO: 62617at7742
## Wed Jul 13 23:59:50 CEST 2022 - The aliscore output file does not exist. Check results for BUSCO: 13987at7742
## Wed Jul 13 23:59:50 CEST 2022 - The aliscore output appears to be empty. Check results for BUSCO: 13987at7742
## Wed Jul 13 23:59:52 CEST 2022 - The aliscore output file does not exist. Check results for BUSCO: 16601at7742
## Wed Jul 13 23:59:52 CEST 2022 - The aliscore output appears to be empty. Check results for BUSCO: 16601at7742
## Wed Jul 13 23:59:54 CEST 2022 - The aliscore output file does not exist. Check results for BUSCO: 62617at7742
## Wed Jul 13 23:59:54 CEST 2022 - The aliscore output appears to be empty. Check results for BUSCO: 62617at7742
## Wed Jul 13 23:59:56 CEST 2022 - The aliscore output file does not exist. Check results for BUSCO: 6126at7742
## Wed Jul 13 23:59:56 CEST 2022 - The aliscore output appears to be empty. Check results for BUSCO: 6126at7742
## Wed Jul 13 23:59:57 CEST 2022 - The aliscore output file does not exist. Check results for BUSCO: 14442at7742
## Wed Jul 13 23:59:57 CEST 2022 - The aliscore output appears to be empty. Check results for BUSCO: 14442at7742
## Wed Jul 13 23:59:58 CEST 2022 - The aliscore output file does not exist. Check results for BUSCO: 185277at7742
## Wed Jul 13 23:59:58 CEST 2022 - The aliscore output appears to be empty. Check results for BUSCO: 185277at7742
## Wed Jul 13 23:59:59 CEST 2022 - The aliscore output file does not exist. Check results for BUSCO: 16601at7742
## Wed Jul 13 23:59:59 CEST 2022 - The aliscore output appears to be empty. Check results for BUSCO: 16601at7742
## Thu Jul 14 00:00:01 CEST 2022 - The aliscore output file does not exist. Check results for BUSCO: 25109at7742
## Thu Jul 14 00:00:01 CEST 2022 - The aliscore output appears to be empty. Check results for BUSCO: 25109at7742
## Thu Jul 14 00:00:02 CEST 2022 - The aliscore output file does not exist. Check results for BUSCO: 185277at7742
## Thu Jul 14 00:00:02 CEST 2022 - The aliscore output appears to be empty. Check results for BUSCO: 185277at7742
## Thu Jul 14 00:00:04 CEST 2022 - The aliscore output file does not exist. Check results for BUSCO: 12300at7742
## Thu Jul 14 00:00:04 CEST 2022 - The aliscore output appears to be empty. Check results for BUSCO: 12300at7742
## Thu Jul 14 00:00:06 CEST 2022 - The aliscore output file does not exist. Check results for BUSCO: 44744at7742
## Thu Jul 14 00:00:06 CEST 2022 - The aliscore output appears to be empty. Check results for BUSCO: 44744at7742
## Thu Jul 14 00:00:06 CEST 2022 - The aliscore output file does not exist. Check results for BUSCO: 83333at7742
## Thu Jul 14 00:00:06 CEST 2022 - The aliscore output appears to be empty. Check results for BUSCO: 83333at7742
## Thu Jul 14 00:00:09 CEST 2022 - The aliscore output file does not exist. Check results for BUSCO: 64985at7742
## Thu Jul 14 00:00:09 CEST 2022 - The aliscore output appears to be empty. Check results for BUSCO: 64985at7742
## Thu Jul 14 00:00:09 CEST 2022 - The aliscore output file does not exist. Check results for BUSCO: 9744at7742
## Thu Jul 14 00:00:09 CEST 2022 - The aliscore output appears to be empty. Check results for BUSCO: 9744at7742
## Thu Jul 14 00:00:10 CEST 2022 - The aliscore output file does not exist. Check results for BUSCO: 13881at7742
## Thu Jul 14 00:00:10 CEST 2022 - The aliscore output appears to be empty. Check results for BUSCO: 13881at7742
## Thu Jul 14 00:00:10 CEST 2022 - The aliscore output file does not exist. Check results for BUSCO: 104785at7742
## Thu Jul 14 00:00:10 CEST 2022 - The aliscore output appears to be empty. Check results for BUSCO: 104785at7742
## Thu Jul 14 00:00:13 CEST 2022 - The aliscore output file does not exist. Check results for BUSCO: 64985at7742
## Thu Jul 14 00:00:14 CEST 2022 - The aliscore output appears to be empty. Check results for BUSCO: 64985at7742
## Thu Jul 14 00:00:16 CEST 2022 - The aliscore output file does not exist. Check results for BUSCO: 159513at7742
## Thu Jul 14 00:00:16 CEST 2022 - The aliscore output appears to be empty. Check results for BUSCO: 159513at7742
## Thu Jul 14 00:00:18 CEST 2022 - The aliscore output file does not exist. Check results for BUSCO: 47172at7742
## Thu Jul 14 00:00:18 CEST 2022 - The aliscore output appears to be empty. Check results for BUSCO: 47172at7742
## Thu Jul 14 00:00:19 CEST 2022 - The aliscore output file does not exist. Check results for BUSCO: 42353at7742
## Thu Jul 14 00:00:19 CEST 2022 - The aliscore output appears to be empty. Check results for BUSCO: 42353at7742
## Thu Jul 14 00:00:19 CEST 2022 - The aliscore output file does not exist. Check results for BUSCO: 58421at7742
## Thu Jul 14 00:00:19 CEST 2022 - The aliscore output appears to be empty. Check results for BUSCO: 58421at7742
## Thu Jul 14 00:00:19 CEST 2022 - The aliscore output file does not exist. Check results for BUSCO: 66047at7742
## Thu Jul 14 00:00:19 CEST 2022 - The aliscore output appears to be empty. Check results for BUSCO: 66047at7742
## Thu Jul 14 00:00:21 CEST 2022 - The aliscore output file does not exist. Check results for BUSCO: 66047at7742
## Thu Jul 14 00:00:21 CEST 2022 - The aliscore output appears to be empty. Check results for BUSCO: 66047at7742
## Thu Jul 14 00:00:24 CEST 2022 - The aliscore output file does not exist. Check results for BUSCO: 29855at7742
## Thu Jul 14 00:00:24 CEST 2022 - The aliscore output appears to be empty. Check results for BUSCO: 29855at7742
## Thu Jul 14 00:00:24 CEST 2022 - The aliscore output file does not exist. Check results for BUSCO: 142339at7742
## Thu Jul 14 00:00:24 CEST 2022 - The aliscore output appears to be empty. Check results for BUSCO: 142339at7742
## Thu Jul 14 00:00:26 CEST 2022 - The aliscore output file does not exist. Check results for BUSCO: 16405at7742
## Thu Jul 14 00:00:26 CEST 2022 - The aliscore output appears to be empty. Check results for BUSCO: 16405at7742
## Thu Jul 14 00:00:30 CEST 2022 - The aliscore output file does not exist. Check results for BUSCO: 11895at7742
## Thu Jul 14 00:00:30 CEST 2022 - The aliscore output appears to be empty. Check results for BUSCO: 11895at7742
## Thu Jul 14 00:00:30 CEST 2022 - The aliscore output file does not exist. Check results for BUSCO: 5981at7742
## Thu Jul 14 00:00:30 CEST 2022 - The aliscore output file does not exist. Check results for BUSCO: 30479at7742
## Thu Jul 14 00:00:30 CEST 2022 - The aliscore output appears to be empty. Check results for BUSCO: 5981at7742
## Thu Jul 14 00:00:30 CEST 2022 - The aliscore output appears to be empty. Check results for BUSCO: 30479at7742
## Thu Jul 14 00:00:31 CEST 2022 - The aliscore output file does not exist. Check results for BUSCO: 50917at7742
## Thu Jul 14 00:00:31 CEST 2022 - The aliscore output appears to be empty. Check results for BUSCO: 50917at7742
## Thu Jul 14 00:00:33 CEST 2022 - The aliscore output file does not exist. Check results for BUSCO: 58421at7742
## Thu Jul 14 00:00:33 CEST 2022 - The aliscore output appears to be empty. Check results for BUSCO: 58421at7742
## Thu Jul 14 00:00:34 CEST 2022 - The aliscore output file does not exist. Check results for BUSCO: 14442at7742
## Thu Jul 14 00:00:34 CEST 2022 - The aliscore output appears to be empty. Check results for BUSCO: 14442at7742
## Thu Jul 14 00:00:37 CEST 2022 - The aliscore output file does not exist. Check results for BUSCO: 11895at7742
## Thu Jul 14 00:00:37 CEST 2022 - The aliscore output appears to be empty. Check results for BUSCO: 11895at7742
## Thu Jul 14 00:00:39 CEST 2022 - The aliscore output file does not exist. Check results for BUSCO: 142339at7742
## Thu Jul 14 00:00:39 CEST 2022 - The aliscore output appears to be empty. Check results for BUSCO: 142339at7742
## Thu Jul 14 00:00:40 CEST 2022 - The aliscore output file does not exist. Check results for BUSCO: 9744at7742
## Thu Jul 14 00:00:40 CEST 2022 - The aliscore output file does not exist. Check results for BUSCO: 5981at7742
## Thu Jul 14 00:00:40 CEST 2022 - The aliscore output appears to be empty. Check results for BUSCO: 9744at7742
## Thu Jul 14 00:00:40 CEST 2022 - The aliscore output appears to be empty. Check results for BUSCO: 5981at7742
## Thu Jul 14 00:00:40 CEST 2022 - The aliscore output file does not exist. Check results for BUSCO: 73684at7742
## Thu Jul 14 00:00:40 CEST 2022 - The aliscore output appears to be empty. Check results for BUSCO: 73684at7742
## Thu Jul 14 00:00:44 CEST 2022 - The aliscore output file does not exist. Check results for BUSCO: 53999at7742
## Thu Jul 14 00:00:44 CEST 2022 - The aliscore output appears to be empty. Check results for BUSCO: 53999at7742
## Thu Jul 14 00:00:44 CEST 2022 - The aliscore output file does not exist. Check results for BUSCO: 91290at7742
## Thu Jul 14 00:00:44 CEST 2022 - The aliscore output appears to be empty. Check results for BUSCO: 91290at7742
## Thu Jul 14 00:00:50 CEST 2022 - The aliscore output file does not exist. Check results for BUSCO: 44744at7742
## Thu Jul 14 00:00:50 CEST 2022 - The aliscore output appears to be empty. Check results for BUSCO: 44744at7742
## Thu Jul 14 00:00:51 CEST 2022 - The aliscore output file does not exist. Check results for BUSCO: 159513at7742
## Thu Jul 14 00:00:51 CEST 2022 - The aliscore output appears to be empty. Check results for BUSCO: 159513at7742
## Thu Jul 14 00:00:51 CEST 2022 - The aliscore output file does not exist. Check results for BUSCO: 136464at7742
## Thu Jul 14 00:00:51 CEST 2022 - The aliscore output appears to be empty. Check results for BUSCO: 136464at7742
## Thu Jul 14 00:00:51 CEST 2022 - The aliscore output file does not exist. Check results for BUSCO: 10136at7742
## Thu Jul 14 00:00:51 CEST 2022 - The aliscore output appears to be empty. Check results for BUSCO: 10136at7742
## Thu Jul 14 00:00:52 CEST 2022 - The aliscore output file does not exist. Check results for BUSCO: 11895at7742
## Thu Jul 14 00:00:52 CEST 2022 - The aliscore output appears to be empty. Check results for BUSCO: 11895at7742
## Thu Jul 14 00:00:53 CEST 2022 - The aliscore output file does not exist. Check results for BUSCO: 25109at7742
## Thu Jul 14 00:00:53 CEST 2022 - The aliscore output appears to be empty. Check results for BUSCO: 25109at7742
## Thu Jul 14 00:00:53 CEST 2022 - The aliscore output file does not exist. Check results for BUSCO: 136464at7742
## Thu Jul 14 00:00:53 CEST 2022 - The aliscore output appears to be empty. Check results for BUSCO: 136464at7742
## Thu Jul 14 00:00:54 CEST 2022 - The aliscore output file does not exist. Check results for BUSCO: 16405at7742
## Thu Jul 14 00:00:54 CEST 2022 - The aliscore output appears to be empty. Check results for BUSCO: 16405at7742
## Thu Jul 14 00:00:58 CEST 2022 - The aliscore output file does not exist. Check results for BUSCO: 81824at7742
## Thu Jul 14 00:00:59 CEST 2022 - The aliscore output appears to be empty. Check results for BUSCO: 81824at7742
## Thu Jul 14 00:01:01 CEST 2022 - The aliscore output file does not exist. Check results for BUSCO: 28532at7742
## Thu Jul 14 00:01:01 CEST 2022 - The aliscore output appears to be empty. Check results for BUSCO: 28532at7742
## Thu Jul 14 00:01:01 CEST 2022 - The aliscore output file does not exist. Check results for BUSCO: 73684at7742
## Thu Jul 14 00:01:01 CEST 2022 - The aliscore output appears to be empty. Check results for BUSCO: 73684at7742
## Thu Jul 14 00:01:03 CEST 2022 - The aliscore output file does not exist. Check results for BUSCO: 136464at7742
## Thu Jul 14 00:01:03 CEST 2022 - The aliscore output appears to be empty. Check results for BUSCO: 136464at7742
## Thu Jul 14 00:01:05 CEST 2022 - The aliscore output file does not exist. Check results for BUSCO: 29855at7742
## Thu Jul 14 00:01:05 CEST 2022 - The aliscore output appears to be empty. Check results for BUSCO: 29855at7742
## Thu Jul 14 00:01:06 CEST 2022 - The aliscore output file does not exist. Check results for BUSCO: 104785at7742
## Thu Jul 14 00:01:06 CEST 2022 - The aliscore output appears to be empty. Check results for BUSCO: 104785at7742
## Thu Jul 14 00:01:09 CEST 2022 - The aliscore output file does not exist. Check results for BUSCO: 30973at7742
## Thu Jul 14 00:01:09 CEST 2022 - The aliscore output appears to be empty. Check results for BUSCO: 30973at7742
## Thu Jul 14 00:01:10 CEST 2022 - The aliscore output file does not exist. Check results for BUSCO: 13987at7742
## Thu Jul 14 00:01:10 CEST 2022 - The aliscore output appears to be empty. Check results for BUSCO: 13987at7742
## Thu Jul 14 00:01:10 CEST 2022 - The aliscore output file does not exist. Check results for BUSCO: 5981at7742
## Thu Jul 14 00:01:10 CEST 2022 - The aliscore output appears to be empty. Check results for BUSCO: 5981at7742
## Thu Jul 14 00:01:11 CEST 2022 - The aliscore output file does not exist. Check results for BUSCO: 14442at7742
## Thu Jul 14 00:01:12 CEST 2022 - The aliscore output appears to be empty. Check results for BUSCO: 14442at7742
## Thu Jul 14 00:01:12 CEST 2022 - The aliscore output file does not exist. Check results for BUSCO: 29855at7742
## Thu Jul 14 00:01:12 CEST 2022 - The aliscore output appears to be empty. Check results for BUSCO: 29855at7742
## Thu Jul 14 00:01:17 CEST 2022 - The aliscore output file does not exist. Check results for BUSCO: 73684at7742
## Thu Jul 14 00:01:17 CEST 2022 - The aliscore output appears to be empty. Check results for BUSCO: 73684at7742
## Thu Jul 14 00:01:18 CEST 2022 - The aliscore output file does not exist. Check results for BUSCO: 26808at7742
## Thu Jul 14 00:01:18 CEST 2022 - The aliscore output appears to be empty. Check results for BUSCO: 26808at7742
## Thu Jul 14 00:01:19 CEST 2022 - The aliscore output file does not exist. Check results for BUSCO: 13670at7742
## Thu Jul 14 00:01:19 CEST 2022 - The aliscore output appears to be empty. Check results for BUSCO: 13670at7742
## Thu Jul 14 00:01:20 CEST 2022 - The aliscore output file does not exist. Check results for BUSCO: 13670at7742
## Thu Jul 14 00:01:20 CEST 2022 - The aliscore output appears to be empty. Check results for BUSCO: 13670at7742
## Thu Jul 14 00:01:20 CEST 2022 - The aliscore output file does not exist. Check results for BUSCO: 16405at7742
## Thu Jul 14 00:01:20 CEST 2022 - The aliscore output appears to be empty. Check results for BUSCO: 16405at7742
## Thu Jul 14 00:01:24 CEST 2022 - The aliscore output file does not exist. Check results for BUSCO: 25109at7742
## Thu Jul 14 00:01:24 CEST 2022 - The aliscore output appears to be empty. Check results for BUSCO: 25109at7742
## Thu Jul 14 00:01:25 CEST 2022 - The aliscore output file does not exist. Check results for BUSCO: 172364at7742
## Thu Jul 14 00:01:25 CEST 2022 - The aliscore output appears to be empty. Check results for BUSCO: 172364at7742
## Thu Jul 14 00:01:26 CEST 2022 - The aliscore output file does not exist. Check results for BUSCO: 36449at7742
## Thu Jul 14 00:01:26 CEST 2022 - The aliscore output appears to be empty. Check results for BUSCO: 36449at7742
## Thu Jul 14 00:01:26 CEST 2022 - The aliscore output file does not exist. Check results for BUSCO: 359556at7742
## Thu Jul 14 00:01:26 CEST 2022 - The aliscore output appears to be empty. Check results for BUSCO: 359556at7742
## Thu Jul 14 00:01:30 CEST 2022 - The aliscore output file does not exist. Check results for BUSCO: 30973at7742
## Thu Jul 14 00:01:30 CEST 2022 - The aliscore output appears to be empty. Check results for BUSCO: 30973at7742
## Thu Jul 14 00:01:31 CEST 2022 - The aliscore output file does not exist. Check results for BUSCO: 12300at7742
## Thu Jul 14 00:01:31 CEST 2022 - The aliscore output appears to be empty. Check results for BUSCO: 12300at7742
## Thu Jul 14 00:01:32 CEST 2022 - The aliscore output file does not exist. Check results for BUSCO: 65634at7742
## Thu Jul 14 00:01:32 CEST 2022 - The aliscore output appears to be empty. Check results for BUSCO: 65634at7742
## Thu Jul 14 00:01:33 CEST 2022 - The aliscore output file does not exist. Check results for BUSCO: 23675at7742
## Thu Jul 14 00:01:33 CEST 2022 - The aliscore output appears to be empty. Check results for BUSCO: 23675at7742
## Thu Jul 14 00:01:35 CEST 2022 - The aliscore output file does not exist. Check results for BUSCO: 18946at7742
## Thu Jul 14 00:01:35 CEST 2022 - The aliscore output appears to be empty. Check results for BUSCO: 18946at7742
## Thu Jul 14 00:01:36 CEST 2022 - The aliscore output file does not exist. Check results for BUSCO: 13480at7742
## Thu Jul 14 00:01:36 CEST 2022 - The aliscore output appears to be empty. Check results for BUSCO: 13480at7742
## Thu Jul 14 00:01:38 CEST 2022 - The aliscore output file does not exist. Check results for BUSCO: 215138at7742
## Thu Jul 14 00:01:38 CEST 2022 - The aliscore output appears to be empty. Check results for BUSCO: 215138at7742
## Thu Jul 14 00:01:40 CEST 2022 - The aliscore output file does not exist. Check results for BUSCO: 12491at7742
## Thu Jul 14 00:01:40 CEST 2022 - The aliscore output appears to be empty. Check results for BUSCO: 12491at7742
## Thu Jul 14 00:01:42 CEST 2022 - The aliscore output file does not exist. Check results for BUSCO: 13670at7742
## Thu Jul 14 00:01:42 CEST 2022 - The aliscore output appears to be empty. Check results for BUSCO: 13670at7742
## Thu Jul 14 00:01:42 CEST 2022 - The aliscore output file does not exist. Check results for BUSCO: 13480at7742
## Thu Jul 14 00:01:42 CEST 2022 - The aliscore output file does not exist. Check results for BUSCO: 9110at7742
## Thu Jul 14 00:01:42 CEST 2022 - The aliscore output appears to be empty. Check results for BUSCO: 13480at7742
## Thu Jul 14 00:01:42 CEST 2022 - The aliscore output appears to be empty. Check results for BUSCO: 9110at7742
## Thu Jul 14 00:01:45 CEST 2022 - The aliscore output file does not exist. Check results for BUSCO: 9110at7742
## Thu Jul 14 00:01:45 CEST 2022 - The aliscore output appears to be empty. Check results for BUSCO: 9110at7742
## Thu Jul 14 00:01:45 CEST 2022 - The aliscore output file does not exist. Check results for BUSCO: 92414at7742
## Thu Jul 14 00:01:45 CEST 2022 - The aliscore output appears to be empty. Check results for BUSCO: 92414at7742
## Thu Jul 14 00:01:48 CEST 2022 - The aliscore output file does not exist. Check results for BUSCO: 81824at7742
## Thu Jul 14 00:01:48 CEST 2022 - The aliscore output appears to be empty. Check results for BUSCO: 81824at7742
## Thu Jul 14 00:01:49 CEST 2022 - The aliscore output file does not exist. Check results for BUSCO: 36449at7742
## Thu Jul 14 00:01:49 CEST 2022 - The aliscore output appears to be empty. Check results for BUSCO: 36449at7742
## Thu Jul 14 00:01:49 CEST 2022 - The aliscore output file does not exist. Check results for BUSCO: 40997at7742
## Thu Jul 14 00:01:49 CEST 2022 - The aliscore output appears to be empty. Check results for BUSCO: 40997at7742
## Thu Jul 14 00:01:51 CEST 2022 - The aliscore output file does not exist. Check results for BUSCO: 285264at7742
## Thu Jul 14 00:01:51 CEST 2022 - The aliscore output appears to be empty. Check results for BUSCO: 285264at7742
## Thu Jul 14 00:01:51 CEST 2022 - The aliscore output file does not exist. Check results for BUSCO: 211831at7742
## Thu Jul 14 00:01:51 CEST 2022 - The aliscore output appears to be empty. Check results for BUSCO: 211831at7742
## Thu Jul 14 00:01:53 CEST 2022 - The aliscore output file does not exist. Check results for BUSCO: 10136at7742
## Thu Jul 14 00:01:53 CEST 2022 - The aliscore output appears to be empty. Check results for BUSCO: 10136at7742
## Thu Jul 14 00:01:54 CEST 2022 - The aliscore output file does not exist. Check results for BUSCO: 175198at7742
## Thu Jul 14 00:01:54 CEST 2022 - The aliscore output appears to be empty. Check results for BUSCO: 175198at7742
## Thu Jul 14 00:01:58 CEST 2022 - The aliscore output file does not exist. Check results for BUSCO: 81824at7742
## Thu Jul 14 00:01:58 CEST 2022 - The aliscore output appears to be empty. Check results for BUSCO: 81824at7742
## Thu Jul 14 00:01:58 CEST 2022 - The aliscore output file does not exist. Check results for BUSCO: 122318at7742
## Thu Jul 14 00:01:58 CEST 2022 - The aliscore output appears to be empty. Check results for BUSCO: 122318at7742
## Thu Jul 14 00:02:02 CEST 2022 - The aliscore output file does not exist. Check results for BUSCO: 9110at7742
## Thu Jul 14 00:02:02 CEST 2022 - The aliscore output appears to be empty. Check results for BUSCO: 9110at7742
## Thu Jul 14 00:02:03 CEST 2022 - The aliscore output file does not exist. Check results for BUSCO: 12300at7742
## Thu Jul 14 00:02:03 CEST 2022 - The aliscore output appears to be empty. Check results for BUSCO: 12300at7742
## Thu Jul 14 00:02:03 CEST 2022 - The aliscore output file does not exist. Check results for BUSCO: 26808at7742
## Thu Jul 14 00:02:03 CEST 2022 - The aliscore output appears to be empty. Check results for BUSCO: 26808at7742
## Thu Jul 14 00:02:04 CEST 2022 - The aliscore output file does not exist. Check results for BUSCO: 46117at7742
## Thu Jul 14 00:02:04 CEST 2022 - The aliscore output file does not exist. Check results for BUSCO: 65634at7742
## Thu Jul 14 00:02:04 CEST 2022 - The aliscore output appears to be empty. Check results for BUSCO: 46117at7742
## Thu Jul 14 00:02:04 CEST 2022 - The aliscore output appears to be empty. Check results for BUSCO: 65634at7742
## Thu Jul 14 00:02:04 CEST 2022 - The aliscore output file does not exist. Check results for BUSCO: 321789at7742
## Thu Jul 14 00:02:04 CEST 2022 - The aliscore output appears to be empty. Check results for BUSCO: 321789at7742
## Thu Jul 14 00:02:09 CEST 2022 - The aliscore output file does not exist. Check results for BUSCO: 30479at7742
## Thu Jul 14 00:02:09 CEST 2022 - The aliscore output appears to be empty. Check results for BUSCO: 30479at7742
## Thu Jul 14 00:02:10 CEST 2022 - The aliscore output file does not exist. Check results for BUSCO: 141334at7742
## Thu Jul 14 00:02:10 CEST 2022 - The aliscore output appears to be empty. Check results for BUSCO: 141334at7742
## Thu Jul 14 00:02:12 CEST 2022 - The aliscore output file does not exist. Check results for BUSCO: 28532at7742
## Thu Jul 14 00:02:12 CEST 2022 - The aliscore output file does not exist. Check results for BUSCO: 189431at7742
## Thu Jul 14 00:02:12 CEST 2022 - The aliscore output appears to be empty. Check results for BUSCO: 28532at7742
## Thu Jul 14 00:02:12 CEST 2022 - The aliscore output appears to be empty. Check results for BUSCO: 189431at7742
## Thu Jul 14 00:02:13 CEST 2022 - The aliscore output file does not exist. Check results for BUSCO: 102945at7742
## Thu Jul 14 00:02:13 CEST 2022 - The aliscore output appears to be empty. Check results for BUSCO: 102945at7742
## Thu Jul 14 00:02:14 CEST 2022 - The aliscore output file does not exist. Check results for BUSCO: 99006at7742
## Thu Jul 14 00:02:14 CEST 2022 - The aliscore output appears to be empty. Check results for BUSCO: 99006at7742
## Thu Jul 14 00:02:17 CEST 2022 - The aliscore output file does not exist. Check results for BUSCO: 49885at7742
## Thu Jul 14 00:02:17 CEST 2022 - The aliscore output appears to be empty. Check results for BUSCO: 49885at7742
## Thu Jul 14 00:02:19 CEST 2022 - The aliscore output file does not exist. Check results for BUSCO: 99006at7742
## Thu Jul 14 00:02:19 CEST 2022 - The aliscore output appears to be empty. Check results for BUSCO: 99006at7742
## Thu Jul 14 00:02:20 CEST 2022 - The aliscore output file does not exist. Check results for BUSCO: 83333at7742
## Thu Jul 14 00:02:20 CEST 2022 - The aliscore output appears to be empty. Check results for BUSCO: 83333at7742
## Thu Jul 14 00:02:22 CEST 2022 - The aliscore output file does not exist. Check results for BUSCO: 27786at7742
## Thu Jul 14 00:02:22 CEST 2022 - The aliscore output appears to be empty. Check results for BUSCO: 27786at7742
## Thu Jul 14 00:02:23 CEST 2022 - The aliscore output file does not exist. Check results for BUSCO: 99006at7742
## Thu Jul 14 00:02:23 CEST 2022 - The aliscore output appears to be empty. Check results for BUSCO: 99006at7742
## Thu Jul 14 00:02:24 CEST 2022 - The aliscore output file does not exist. Check results for BUSCO: 23675at7742
## Thu Jul 14 00:02:24 CEST 2022 - The aliscore output appears to be empty. Check results for BUSCO: 23675at7742
## Thu Jul 14 00:02:26 CEST 2022 - The aliscore output file does not exist. Check results for BUSCO: 215138at7742
## Thu Jul 14 00:02:26 CEST 2022 - The aliscore output appears to be empty. Check results for BUSCO: 215138at7742
## Thu Jul 14 00:02:27 CEST 2022 - The aliscore output file does not exist. Check results for BUSCO: 141334at7742
## Thu Jul 14 00:02:27 CEST 2022 - The aliscore output appears to be empty. Check results for BUSCO: 141334at7742
## Thu Jul 14 00:02:29 CEST 2022 - The aliscore output file does not exist. Check results for BUSCO: 27463at7742
## Thu Jul 14 00:02:29 CEST 2022 - The aliscore output appears to be empty. Check results for BUSCO: 27463at7742
## Thu Jul 14 00:02:31 CEST 2022 - The aliscore output file does not exist. Check results for BUSCO: 23692at7742
## Thu Jul 14 00:02:31 CEST 2022 - The aliscore output appears to be empty. Check results for BUSCO: 23692at7742
## Thu Jul 14 00:02:32 CEST 2022 - The aliscore output file does not exist. Check results for BUSCO: 26808at7742
## Thu Jul 14 00:02:32 CEST 2022 - The aliscore output appears to be empty. Check results for BUSCO: 26808at7742
## Thu Jul 14 00:02:33 CEST 2022 - The aliscore output file does not exist. Check results for BUSCO: 153465at7742
## Thu Jul 14 00:02:33 CEST 2022 - The aliscore output appears to be empty. Check results for BUSCO: 153465at7742
## Thu Jul 14 00:02:33 CEST 2022 - The aliscore output file does not exist. Check results for BUSCO: 53999at7742
## Thu Jul 14 00:02:33 CEST 2022 - The aliscore output appears to be empty. Check results for BUSCO: 53999at7742
## Thu Jul 14 00:02:36 CEST 2022 - The aliscore output file does not exist. Check results for BUSCO: 177448at7742
## Thu Jul 14 00:02:36 CEST 2022 - The aliscore output appears to be empty. Check results for BUSCO: 177448at7742
## Thu Jul 14 00:02:37 CEST 2022 - The aliscore output file does not exist. Check results for BUSCO: 215138at7742
## Thu Jul 14 00:02:37 CEST 2022 - The aliscore output appears to be empty. Check results for BUSCO: 215138at7742
## Thu Jul 14 00:02:39 CEST 2022 - The aliscore output file does not exist. Check results for BUSCO: 172364at7742
## Thu Jul 14 00:02:39 CEST 2022 - The aliscore output appears to be empty. Check results for BUSCO: 172364at7742
## Thu Jul 14 00:02:40 CEST 2022 - The aliscore output file does not exist. Check results for BUSCO: 193224at7742
## Thu Jul 14 00:02:40 CEST 2022 - The aliscore output appears to be empty. Check results for BUSCO: 193224at7742
## Thu Jul 14 00:02:41 CEST 2022 - The aliscore output file does not exist. Check results for BUSCO: 225400at7742
## Thu Jul 14 00:02:41 CEST 2022 - The aliscore output appears to be empty. Check results for BUSCO: 225400at7742
## Thu Jul 14 00:02:42 CEST 2022 - The aliscore output file does not exist. Check results for BUSCO: 231143at7742
## Thu Jul 14 00:02:42 CEST 2022 - The aliscore output appears to be empty. Check results for BUSCO: 231143at7742
## Thu Jul 14 00:02:42 CEST 2022 - The aliscore output file does not exist. Check results for BUSCO: 4106at7742
## Thu Jul 14 00:02:42 CEST 2022 - The aliscore output appears to be empty. Check results for BUSCO: 4106at7742
## Thu Jul 14 00:02:45 CEST 2022 - The aliscore output file does not exist. Check results for BUSCO: 90587at7742
## Thu Jul 14 00:02:45 CEST 2022 - The aliscore output appears to be empty. Check results for BUSCO: 90587at7742
## Thu Jul 14 00:02:46 CEST 2022 - The aliscore output file does not exist. Check results for BUSCO: 23996at7742
## Thu Jul 14 00:02:46 CEST 2022 - The aliscore output appears to be empty. Check results for BUSCO: 23996at7742
## Thu Jul 14 00:02:50 CEST 2022 - The aliscore output file does not exist. Check results for BUSCO: 11696at7742
## Thu Jul 14 00:02:50 CEST 2022 - The aliscore output appears to be empty. Check results for BUSCO: 11696at7742
## Thu Jul 14 00:02:50 CEST 2022 - The aliscore output file does not exist. Check results for BUSCO: 17577at7742
## Thu Jul 14 00:02:50 CEST 2022 - The aliscore output appears to be empty. Check results for BUSCO: 17577at7742
## Thu Jul 14 00:02:52 CEST 2022 - The aliscore output file does not exist. Check results for BUSCO: 97499at7742
## Thu Jul 14 00:02:52 CEST 2022 - The aliscore output appears to be empty. Check results for BUSCO: 97499at7742
## Thu Jul 14 00:02:52 CEST 2022 - The aliscore output file does not exist. Check results for BUSCO: 42353at7742
## Thu Jul 14 00:02:52 CEST 2022 - The aliscore output appears to be empty. Check results for BUSCO: 42353at7742
## Thu Jul 14 00:02:55 CEST 2022 - The aliscore output file does not exist. Check results for BUSCO: 92414at7742
## Thu Jul 14 00:02:55 CEST 2022 - The aliscore output appears to be empty. Check results for BUSCO: 92414at7742
## Thu Jul 14 00:02:57 CEST 2022 - The aliscore output file does not exist. Check results for BUSCO: 84691at7742
## Thu Jul 14 00:02:57 CEST 2022 - The aliscore output appears to be empty. Check results for BUSCO: 84691at7742
## Thu Jul 14 00:02:57 CEST 2022 - The aliscore output file does not exist. Check results for BUSCO: 4106at7742
## Thu Jul 14 00:02:57 CEST 2022 - The aliscore output appears to be empty. Check results for BUSCO: 4106at7742
## Thu Jul 14 00:02:58 CEST 2022 - The aliscore output file does not exist. Check results for BUSCO: 321789at7742
## Thu Jul 14 00:02:58 CEST 2022 - The aliscore output appears to be empty. Check results for BUSCO: 321789at7742
## Thu Jul 14 00:03:02 CEST 2022 - The aliscore output file does not exist. Check results for BUSCO: 10899at7742
## Thu Jul 14 00:03:02 CEST 2022 - The aliscore output appears to be empty. Check results for BUSCO: 10899at7742
## Thu Jul 14 00:03:02 CEST 2022 - The aliscore output file does not exist. Check results for BUSCO: 62115at7742
## Thu Jul 14 00:03:02 CEST 2022 - The aliscore output appears to be empty. Check results for BUSCO: 62115at7742
## Thu Jul 14 00:03:02 CEST 2022 - The aliscore output file does not exist. Check results for BUSCO: 83333at7742
## Thu Jul 14 00:03:02 CEST 2022 - The aliscore output appears to be empty. Check results for BUSCO: 83333at7742
## Thu Jul 14 00:03:02 CEST 2022 - The aliscore output file does not exist. Check results for BUSCO: 102945at7742
## Thu Jul 14 00:03:02 CEST 2022 - The aliscore output appears to be empty. Check results for BUSCO: 102945at7742
## Thu Jul 14 00:03:06 CEST 2022 - The aliscore output file does not exist. Check results for BUSCO: 47172at7742
## Thu Jul 14 00:03:06 CEST 2022 - The aliscore output appears to be empty. Check results for BUSCO: 47172at7742
## Thu Jul 14 00:03:07 CEST 2022 - The aliscore output file does not exist. Check results for BUSCO: 30479at7742
## Thu Jul 14 00:03:07 CEST 2022 - The aliscore output appears to be empty. Check results for BUSCO: 30479at7742
## Thu Jul 14 00:03:08 CEST 2022 - The aliscore output file does not exist. Check results for BUSCO: 55030at7742
## Thu Jul 14 00:03:08 CEST 2022 - The aliscore output appears to be empty. Check results for BUSCO: 55030at7742
## Thu Jul 14 00:03:12 CEST 2022 - The aliscore output file does not exist. Check results for BUSCO: 231143at7742
## Thu Jul 14 00:03:12 CEST 2022 - The aliscore output appears to be empty. Check results for BUSCO: 231143at7742
## Thu Jul 14 00:03:12 CEST 2022 - The aliscore output file does not exist. Check results for BUSCO: 194852at7742
## Thu Jul 14 00:03:12 CEST 2022 - The aliscore output appears to be empty. Check results for BUSCO: 194852at7742
## Thu Jul 14 00:03:13 CEST 2022 - The aliscore output file does not exist. Check results for BUSCO: 211831at7742
## Thu Jul 14 00:03:13 CEST 2022 - The aliscore output file does not exist. Check results for BUSCO: 141334at7742
## Thu Jul 14 00:03:13 CEST 2022 - The aliscore output appears to be empty. Check results for BUSCO: 211831at7742
## Thu Jul 14 00:03:13 CEST 2022 - The aliscore output appears to be empty. Check results for BUSCO: 141334at7742
## Thu Jul 14 00:03:14 CEST 2022 - The aliscore output file does not exist. Check results for BUSCO: 92414at7742
## Thu Jul 14 00:03:14 CEST 2022 - The aliscore output appears to be empty. Check results for BUSCO: 92414at7742
## Thu Jul 14 00:03:18 CEST 2022 - The aliscore output file does not exist. Check results for BUSCO: 106853at7742
## Thu Jul 14 00:03:18 CEST 2022 - The aliscore output appears to be empty. Check results for BUSCO: 106853at7742
## Thu Jul 14 00:03:18 CEST 2022 - The aliscore output file does not exist. Check results for BUSCO: 211831at7742
## Thu Jul 14 00:03:18 CEST 2022 - The aliscore output appears to be empty. Check results for BUSCO: 211831at7742
## Thu Jul 14 00:03:22 CEST 2022 - The aliscore output file does not exist. Check results for BUSCO: 24868at7742
## Thu Jul 14 00:03:22 CEST 2022 - The aliscore output appears to be empty. Check results for BUSCO: 24868at7742
## Thu Jul 14 00:03:22 CEST 2022 - The aliscore output file does not exist. Check results for BUSCO: 102393at7742
## Thu Jul 14 00:03:22 CEST 2022 - The aliscore output appears to be empty. Check results for BUSCO: 102393at7742
## Thu Jul 14 00:03:24 CEST 2022 - The aliscore output file does not exist. Check results for BUSCO: 42353at7742
## Thu Jul 14 00:03:24 CEST 2022 - The aliscore output file does not exist. Check results for BUSCO: 4106at7742
## Thu Jul 14 00:03:24 CEST 2022 - The aliscore output appears to be empty. Check results for BUSCO: 42353at7742
## Thu Jul 14 00:03:24 CEST 2022 - The aliscore output appears to be empty. Check results for BUSCO: 4106at7742
## Thu Jul 14 00:03:27 CEST 2022 - The aliscore output file does not exist. Check results for BUSCO: 189431at7742
## Thu Jul 14 00:03:27 CEST 2022 - The aliscore output appears to be empty. Check results for BUSCO: 189431at7742
## Thu Jul 14 00:03:27 CEST 2022 - The aliscore output file does not exist. Check results for BUSCO: 36236at7742
## Thu Jul 14 00:03:27 CEST 2022 - The aliscore output appears to be empty. Check results for BUSCO: 36236at7742
## Thu Jul 14 00:03:28 CEST 2022 - The aliscore output file does not exist. Check results for BUSCO: 23675at7742
## Thu Jul 14 00:03:28 CEST 2022 - The aliscore output appears to be empty. Check results for BUSCO: 23675at7742
## Thu Jul 14 00:03:31 CEST 2022 - The aliscore output file does not exist. Check results for BUSCO: 42342at7742
## Thu Jul 14 00:03:31 CEST 2022 - The aliscore output appears to be empty. Check results for BUSCO: 42342at7742
## Thu Jul 14 00:03:34 CEST 2022 - The aliscore output file does not exist. Check results for BUSCO: 359556at7742
## Thu Jul 14 00:03:34 CEST 2022 - The aliscore output appears to be empty. Check results for BUSCO: 359556at7742
## Thu Jul 14 00:03:34 CEST 2022 - The aliscore output file does not exist. Check results for BUSCO: 16779at7742
## Thu Jul 14 00:03:34 CEST 2022 - The aliscore output appears to be empty. Check results for BUSCO: 16779at7742
## Thu Jul 14 00:03:34 CEST 2022 - The aliscore output file does not exist. Check results for BUSCO: 56852at7742
## Thu Jul 14 00:03:34 CEST 2022 - The aliscore output appears to be empty. Check results for BUSCO: 56852at7742
## Thu Jul 14 00:03:35 CEST 2022 - The aliscore output file does not exist. Check results for BUSCO: 102393at7742
## Thu Jul 14 00:03:35 CEST 2022 - The aliscore output appears to be empty. Check results for BUSCO: 102393at7742
## Thu Jul 14 00:03:37 CEST 2022 - The aliscore output file does not exist. Check results for BUSCO: 14788at7742
## Thu Jul 14 00:03:37 CEST 2022 - The aliscore output appears to be empty. Check results for BUSCO: 14788at7742
## Thu Jul 14 00:03:38 CEST 2022 - The aliscore output file does not exist. Check results for BUSCO: 33771at7742
## Thu Jul 14 00:03:38 CEST 2022 - The aliscore output appears to be empty. Check results for BUSCO: 33771at7742
## Thu Jul 14 00:03:40 CEST 2022 - The aliscore output file does not exist. Check results for BUSCO: 8503at7742
## Thu Jul 14 00:03:40 CEST 2022 - The aliscore output appears to be empty. Check results for BUSCO: 8503at7742
## Thu Jul 14 00:03:42 CEST 2022 - The aliscore output file does not exist. Check results for BUSCO: 11696at7742
## Thu Jul 14 00:03:42 CEST 2022 - The aliscore output appears to be empty. Check results for BUSCO: 11696at7742
## Thu Jul 14 00:03:44 CEST 2022 - The aliscore output file does not exist. Check results for BUSCO: 46117at7742
## Thu Jul 14 00:03:44 CEST 2022 - The aliscore output appears to be empty. Check results for BUSCO: 46117at7742
## Thu Jul 14 00:03:45 CEST 2022 - The aliscore output file does not exist. Check results for BUSCO: 122318at7742
## Thu Jul 14 00:03:45 CEST 2022 - The aliscore output appears to be empty. Check results for BUSCO: 122318at7742
## Thu Jul 14 00:03:45 CEST 2022 - The aliscore output file does not exist. Check results for BUSCO: 64333at7742
## Thu Jul 14 00:03:45 CEST 2022 - The aliscore output appears to be empty. Check results for BUSCO: 64333at7742
## Thu Jul 14 00:03:46 CEST 2022 - The aliscore output file does not exist. Check results for BUSCO: 153465at7742
## Thu Jul 14 00:03:46 CEST 2022 - The aliscore output appears to be empty. Check results for BUSCO: 153465at7742
## Thu Jul 14 00:03:49 CEST 2022 - The aliscore output file does not exist. Check results for BUSCO: 102393at7742
## Thu Jul 14 00:03:50 CEST 2022 - The aliscore output appears to be empty. Check results for BUSCO: 102393at7742
## Thu Jul 14 00:03:50 CEST 2022 - The aliscore output file does not exist. Check results for BUSCO: 177448at7742
## Thu Jul 14 00:03:50 CEST 2022 - The aliscore output appears to be empty. Check results for BUSCO: 177448at7742
## Thu Jul 14 00:03:52 CEST 2022 - The aliscore output file does not exist. Check results for BUSCO: 133758at7742
## Thu Jul 14 00:03:52 CEST 2022 - The aliscore output appears to be empty. Check results for BUSCO: 133758at7742
## Thu Jul 14 00:03:54 CEST 2022 - The aliscore output file does not exist. Check results for BUSCO: 96327at7742
## Thu Jul 14 00:03:54 CEST 2022 - The aliscore output appears to be empty. Check results for BUSCO: 96327at7742
## Thu Jul 14 00:03:54 CEST 2022 - The aliscore output file does not exist. Check results for BUSCO: 41154at7742
## Thu Jul 14 00:03:54 CEST 2022 - The aliscore output appears to be empty. Check results for BUSCO: 41154at7742
## Thu Jul 14 00:03:55 CEST 2022 - The aliscore output file does not exist. Check results for BUSCO: 11696at7742
## Thu Jul 14 00:03:55 CEST 2022 - The aliscore output appears to be empty. Check results for BUSCO: 11696at7742
## Thu Jul 14 00:03:57 CEST 2022 - The aliscore output file does not exist. Check results for BUSCO: 11195at7742
## Thu Jul 14 00:03:57 CEST 2022 - The aliscore output appears to be empty. Check results for BUSCO: 11195at7742
## Thu Jul 14 00:03:59 CEST 2022 - The aliscore output file does not exist. Check results for BUSCO: 53999at7742
## Thu Jul 14 00:03:59 CEST 2022 - The aliscore output appears to be empty. Check results for BUSCO: 53999at7742
## Thu Jul 14 00:04:00 CEST 2022 - The aliscore output file does not exist. Check results for BUSCO: 11195at7742
## Thu Jul 14 00:04:00 CEST 2022 - The aliscore output appears to be empty. Check results for BUSCO: 11195at7742
## Thu Jul 14 00:04:02 CEST 2022 - The aliscore output file does not exist. Check results for BUSCO: 90690at7742
## Thu Jul 14 00:04:02 CEST 2022 - The aliscore output appears to be empty. Check results for BUSCO: 90690at7742
## Thu Jul 14 00:04:03 CEST 2022 - The aliscore output file does not exist. Check results for BUSCO: 79715at7742
## Thu Jul 14 00:04:03 CEST 2022 - The aliscore output appears to be empty. Check results for BUSCO: 79715at7742
## Thu Jul 14 00:04:04 CEST 2022 - The aliscore output file does not exist. Check results for BUSCO: 91290at7742
## Thu Jul 14 00:04:04 CEST 2022 - The aliscore output appears to be empty. Check results for BUSCO: 91290at7742
## Thu Jul 14 00:04:04 CEST 2022 - The aliscore output file does not exist. Check results for BUSCO: 28267at7742
## Thu Jul 14 00:04:04 CEST 2022 - The aliscore output appears to be empty. Check results for BUSCO: 28267at7742
## Thu Jul 14 00:04:07 CEST 2022 - The aliscore output file does not exist. Check results for BUSCO: 27786at7742
## Thu Jul 14 00:04:07 CEST 2022 - The aliscore output appears to be empty. Check results for BUSCO: 27786at7742
## Thu Jul 14 00:04:08 CEST 2022 - The aliscore output file does not exist. Check results for BUSCO: 42342at7742
## Thu Jul 14 00:04:08 CEST 2022 - The aliscore output appears to be empty. Check results for BUSCO: 42342at7742
## Thu Jul 14 00:04:10 CEST 2022 - The aliscore output file does not exist. Check results for BUSCO: 172364at7742
## Thu Jul 14 00:04:10 CEST 2022 - The aliscore output appears to be empty. Check results for BUSCO: 172364at7742
## Thu Jul 14 00:04:11 CEST 2022 - The aliscore output file does not exist. Check results for BUSCO: 65671at7742
## Thu Jul 14 00:04:11 CEST 2022 - The aliscore output appears to be empty. Check results for BUSCO: 65671at7742
## Thu Jul 14 00:04:12 CEST 2022 - The aliscore output file does not exist. Check results for BUSCO: 23173at7742
## Thu Jul 14 00:04:12 CEST 2022 - The aliscore output appears to be empty. Check results for BUSCO: 23173at7742
## Thu Jul 14 00:04:13 CEST 2022 - The aliscore output file does not exist. Check results for BUSCO: 12659at7742
## Thu Jul 14 00:04:13 CEST 2022 - The aliscore output appears to be empty. Check results for BUSCO: 12659at7742
## Thu Jul 14 00:04:13 CEST 2022 - The aliscore output file does not exist. Check results for BUSCO: 211943at7742
## Thu Jul 14 00:04:13 CEST 2022 - The aliscore output appears to be empty. Check results for BUSCO: 211943at7742
## Thu Jul 14 00:04:15 CEST 2022 - The aliscore output file does not exist. Check results for BUSCO: 164002at7742
## Thu Jul 14 00:04:15 CEST 2022 - The aliscore output appears to be empty. Check results for BUSCO: 164002at7742
## Thu Jul 14 00:04:18 CEST 2022 - The aliscore output file does not exist. Check results for BUSCO: 48764at7742
## Thu Jul 14 00:04:18 CEST 2022 - The aliscore output appears to be empty. Check results for BUSCO: 48764at7742
## Thu Jul 14 00:04:20 CEST 2022 - The aliscore output file does not exist. Check results for BUSCO: 55030at7742
## Thu Jul 14 00:04:20 CEST 2022 - The aliscore output appears to be empty. Check results for BUSCO: 55030at7742
## Thu Jul 14 00:04:20 CEST 2022 - The aliscore output file does not exist. Check results for BUSCO: 122318at7742
## Thu Jul 14 00:04:20 CEST 2022 - The aliscore output appears to be empty. Check results for BUSCO: 122318at7742
## Thu Jul 14 00:04:23 CEST 2022 - The aliscore output file does not exist. Check results for BUSCO: 42342at7742
## Thu Jul 14 00:04:23 CEST 2022 - The aliscore output appears to be empty. Check results for BUSCO: 42342at7742
## Thu Jul 14 00:04:24 CEST 2022 - The aliscore output file does not exist. Check results for BUSCO: 11195at7742
## Thu Jul 14 00:04:24 CEST 2022 - The aliscore output appears to be empty. Check results for BUSCO: 11195at7742
## Thu Jul 14 00:04:25 CEST 2022 - The aliscore output file does not exist. Check results for BUSCO: 90587at7742
## Thu Jul 14 00:04:25 CEST 2022 - The aliscore output appears to be empty. Check results for BUSCO: 90587at7742
## Thu Jul 14 00:04:27 CEST 2022 - The aliscore output file does not exist. Check results for BUSCO: 79715at7742
## Thu Jul 14 00:04:27 CEST 2022 - The aliscore output appears to be empty. Check results for BUSCO: 79715at7742
## Thu Jul 14 00:04:27 CEST 2022 - The aliscore output file does not exist. Check results for BUSCO: 12659at7742
## Thu Jul 14 00:04:28 CEST 2022 - The aliscore output appears to be empty. Check results for BUSCO: 12659at7742
## Thu Jul 14 00:04:30 CEST 2022 - The aliscore output file does not exist. Check results for BUSCO: 8503at7742
## Thu Jul 14 00:04:30 CEST 2022 - The aliscore output appears to be empty. Check results for BUSCO: 8503at7742
## Thu Jul 14 00:04:31 CEST 2022 - The aliscore output file does not exist. Check results for BUSCO: 17577at7742
## Thu Jul 14 00:04:31 CEST 2022 - The aliscore output appears to be empty. Check results for BUSCO: 17577at7742
## Thu Jul 14 00:04:33 CEST 2022 - The aliscore output file does not exist. Check results for BUSCO: 10757at7742
## Thu Jul 14 00:04:33 CEST 2022 - The aliscore output appears to be empty. Check results for BUSCO: 10757at7742
## Thu Jul 14 00:04:33 CEST 2022 - The aliscore output file does not exist. Check results for BUSCO: 10899at7742
## Thu Jul 14 00:04:33 CEST 2022 - The aliscore output appears to be empty. Check results for BUSCO: 10899at7742
## Thu Jul 14 00:04:34 CEST 2022 - The aliscore output file does not exist. Check results for BUSCO: 55709at7742
## Thu Jul 14 00:04:34 CEST 2022 - The aliscore output appears to be empty. Check results for BUSCO: 55709at7742
## Thu Jul 14 00:04:37 CEST 2022 - The aliscore output file does not exist. Check results for BUSCO: 23692at7742
## Thu Jul 14 00:04:37 CEST 2022 - The aliscore output appears to be empty. Check results for BUSCO: 23692at7742
## Thu Jul 14 00:04:38 CEST 2022 - The aliscore output file does not exist. Check results for BUSCO: 10757at7742
## Thu Jul 14 00:04:38 CEST 2022 - The aliscore output appears to be empty. Check results for BUSCO: 10757at7742
## Thu Jul 14 00:04:41 CEST 2022 - The aliscore output file does not exist. Check results for BUSCO: 23692at7742
## Thu Jul 14 00:04:41 CEST 2022 - The aliscore output appears to be empty. Check results for BUSCO: 23692at7742
## Thu Jul 14 00:04:42 CEST 2022 - The aliscore output file does not exist. Check results for BUSCO: 16779at7742
## Thu Jul 14 00:04:42 CEST 2022 - The aliscore output appears to be empty. Check results for BUSCO: 16779at7742
## Thu Jul 14 00:04:43 CEST 2022 - The aliscore output file does not exist. Check results for BUSCO: 8503at7742
## Thu Jul 14 00:04:43 CEST 2022 - The aliscore output appears to be empty. Check results for BUSCO: 8503at7742
## Thu Jul 14 00:04:44 CEST 2022 - The aliscore output file does not exist. Check results for BUSCO: 38882at7742
## Thu Jul 14 00:04:44 CEST 2022 - The aliscore output appears to be empty. Check results for BUSCO: 38882at7742
## Thu Jul 14 00:04:44 CEST 2022 - The aliscore output file does not exist. Check results for BUSCO: 12659at7742
## Thu Jul 14 00:04:44 CEST 2022 - The aliscore output appears to be empty. Check results for BUSCO: 12659at7742
## Thu Jul 14 00:04:47 CEST 2022 - The aliscore output file does not exist. Check results for BUSCO: 88880at7742
## Thu Jul 14 00:04:47 CEST 2022 - The aliscore output appears to be empty. Check results for BUSCO: 88880at7742
## Thu Jul 14 00:04:49 CEST 2022 - The aliscore output file does not exist. Check results for BUSCO: 91290at7742
## Thu Jul 14 00:04:49 CEST 2022 - The aliscore output appears to be empty. Check results for BUSCO: 91290at7742
## Thu Jul 14 00:04:51 CEST 2022 - The aliscore output file does not exist. Check results for BUSCO: 26210at7742
## Thu Jul 14 00:04:51 CEST 2022 - The aliscore output appears to be empty. Check results for BUSCO: 26210at7742
## Thu Jul 14 00:04:51 CEST 2022 - The aliscore output file does not exist. Check results for BUSCO: 28267at7742
## Thu Jul 14 00:04:51 CEST 2022 - The aliscore output appears to be empty. Check results for BUSCO: 28267at7742
## Thu Jul 14 00:04:54 CEST 2022 - The aliscore output file does not exist. Check results for BUSCO: 140317at7742
## Thu Jul 14 00:04:54 CEST 2022 - The aliscore output appears to be empty. Check results for BUSCO: 140317at7742
## Thu Jul 14 00:04:54 CEST 2022 - The aliscore output file does not exist. Check results for BUSCO: 43051at7742
## Thu Jul 14 00:04:54 CEST 2022 - The aliscore output appears to be empty. Check results for BUSCO: 43051at7742
## Thu Jul 14 00:04:56 CEST 2022 - The aliscore output file does not exist. Check results for BUSCO: 27786at7742
## Thu Jul 14 00:04:56 CEST 2022 - The aliscore output appears to be empty. Check results for BUSCO: 27786at7742
## Thu Jul 14 00:04:57 CEST 2022 - The aliscore output file does not exist. Check results for BUSCO: 141686at7742
## Thu Jul 14 00:04:57 CEST 2022 - The aliscore output appears to be empty. Check results for BUSCO: 141686at7742
## Thu Jul 14 00:05:00 CEST 2022 - The aliscore output file does not exist. Check results for BUSCO: 16779at7742
## Thu Jul 14 00:05:00 CEST 2022 - The aliscore output appears to be empty. Check results for BUSCO: 16779at7742
## Thu Jul 14 00:05:01 CEST 2022 - The aliscore output file does not exist. Check results for BUSCO: 64333at7742
## Thu Jul 14 00:05:01 CEST 2022 - The aliscore output appears to be empty. Check results for BUSCO: 64333at7742
## Thu Jul 14 00:05:03 CEST 2022 - The aliscore output file does not exist. Check results for BUSCO: 27463at7742
## Thu Jul 14 00:05:03 CEST 2022 - The aliscore output appears to be empty. Check results for BUSCO: 27463at7742
## Thu Jul 14 00:05:05 CEST 2022 - The aliscore output file does not exist. Check results for BUSCO: 16923at7742
## Thu Jul 14 00:05:05 CEST 2022 - The aliscore output appears to be empty. Check results for BUSCO: 16923at7742
## Thu Jul 14 00:05:05 CEST 2022 - The aliscore output file does not exist. Check results for BUSCO: 38882at7742
## Thu Jul 14 00:05:05 CEST 2022 - The aliscore output appears to be empty. Check results for BUSCO: 38882at7742
## Thu Jul 14 00:05:08 CEST 2022 - The aliscore output file does not exist. Check results for BUSCO: 177448at7742
## Thu Jul 14 00:05:08 CEST 2022 - The aliscore output appears to be empty. Check results for BUSCO: 177448at7742
## Thu Jul 14 00:05:09 CEST 2022 - The aliscore output file does not exist. Check results for BUSCO: 24868at7742
## Thu Jul 14 00:05:09 CEST 2022 - The aliscore output appears to be empty. Check results for BUSCO: 24868at7742
## Thu Jul 14 00:05:10 CEST 2022 - The aliscore output file does not exist. Check results for BUSCO: 32511at7742
## Thu Jul 14 00:05:11 CEST 2022 - The aliscore output appears to be empty. Check results for BUSCO: 32511at7742
## Thu Jul 14 00:05:11 CEST 2022 - The aliscore output file does not exist. Check results for BUSCO: 41154at7742
## Thu Jul 14 00:05:11 CEST 2022 - The aliscore output appears to be empty. Check results for BUSCO: 41154at7742
## Thu Jul 14 00:05:13 CEST 2022 - The aliscore output file does not exist. Check results for BUSCO: 106344at7742
## Thu Jul 14 00:05:13 CEST 2022 - The aliscore output appears to be empty. Check results for BUSCO: 106344at7742
## Thu Jul 14 00:05:15 CEST 2022 - The aliscore output file does not exist. Check results for BUSCO: 56852at7742
## Thu Jul 14 00:05:15 CEST 2022 - The aliscore output appears to be empty. Check results for BUSCO: 56852at7742
## Thu Jul 14 00:05:16 CEST 2022 - The aliscore output file does not exist. Check results for BUSCO: 152085at7742
## Thu Jul 14 00:05:16 CEST 2022 - The aliscore output appears to be empty. Check results for BUSCO: 152085at7742
## Thu Jul 14 00:05:17 CEST 2022 - The aliscore output file does not exist. Check results for BUSCO: 16923at7742
## Thu Jul 14 00:05:17 CEST 2022 - The aliscore output appears to be empty. Check results for BUSCO: 16923at7742
## Thu Jul 14 00:05:19 CEST 2022 - The aliscore output file does not exist. Check results for BUSCO: 23996at7742
## Thu Jul 14 00:05:19 CEST 2022 - The aliscore output appears to be empty. Check results for BUSCO: 23996at7742
## Thu Jul 14 00:05:19 CEST 2022 - The aliscore output file does not exist. Check results for BUSCO: 106853at7742
## Thu Jul 14 00:05:19 CEST 2022 - The aliscore output appears to be empty. Check results for BUSCO: 106853at7742
## Thu Jul 14 00:05:23 CEST 2022 - The aliscore output file does not exist. Check results for BUSCO: 65671at7742
## Thu Jul 14 00:05:23 CEST 2022 - The aliscore output appears to be empty. Check results for BUSCO: 65671at7742
## Thu Jul 14 00:05:23 CEST 2022 - The aliscore output file does not exist. Check results for BUSCO: 97499at7742
## Thu Jul 14 00:05:23 CEST 2022 - The aliscore output appears to be empty. Check results for BUSCO: 97499at7742
## Thu Jul 14 00:05:25 CEST 2022 - The aliscore output file does not exist. Check results for BUSCO: 276779at7742
## Thu Jul 14 00:05:26 CEST 2022 - The aliscore output appears to be empty. Check results for BUSCO: 276779at7742
## Thu Jul 14 00:05:27 CEST 2022 - The aliscore output file does not exist. Check results for BUSCO: 55709at7742
## Thu Jul 14 00:05:27 CEST 2022 - The aliscore output appears to be empty. Check results for BUSCO: 55709at7742
## Thu Jul 14 00:05:28 CEST 2022 - The aliscore output file does not exist. Check results for BUSCO: 43051at7742
## Thu Jul 14 00:05:28 CEST 2022 - The aliscore output appears to be empty. Check results for BUSCO: 43051at7742
## Thu Jul 14 00:05:30 CEST 2022 - The aliscore output file does not exist. Check results for BUSCO: 28532at7742
## Thu Jul 14 00:05:30 CEST 2022 - The aliscore output appears to be empty. Check results for BUSCO: 28532at7742
## Thu Jul 14 00:05:31 CEST 2022 - The aliscore output file does not exist. Check results for BUSCO: 33771at7742
## Thu Jul 14 00:05:31 CEST 2022 - The aliscore output appears to be empty. Check results for BUSCO: 33771at7742
## Thu Jul 14 00:05:33 CEST 2022 - The aliscore output file does not exist. Check results for BUSCO: 133758at7742
## Thu Jul 14 00:05:33 CEST 2022 - The aliscore output appears to be empty. Check results for BUSCO: 133758at7742
## Thu Jul 14 00:05:34 CEST 2022 - The aliscore output file does not exist. Check results for BUSCO: 10757at7742
## Thu Jul 14 00:05:34 CEST 2022 - The aliscore output appears to be empty. Check results for BUSCO: 10757at7742
## Thu Jul 14 00:05:35 CEST 2022 - The aliscore output file does not exist. Check results for BUSCO: 11104at7742
## Thu Jul 14 00:05:35 CEST 2022 - The aliscore output appears to be empty. Check results for BUSCO: 11104at7742
## Thu Jul 14 00:05:36 CEST 2022 - The aliscore output file does not exist. Check results for BUSCO: 11104at7742
## Thu Jul 14 00:05:36 CEST 2022 - The aliscore output appears to be empty. Check results for BUSCO: 11104at7742
## Thu Jul 14 00:05:37 CEST 2022 - The aliscore output file does not exist. Check results for BUSCO: 27595at7742
## Thu Jul 14 00:05:37 CEST 2022 - The aliscore output appears to be empty. Check results for BUSCO: 27595at7742
## Thu Jul 14 00:05:39 CEST 2022 - The aliscore output file does not exist. Check results for BUSCO: 48764at7742
## Thu Jul 14 00:05:39 CEST 2022 - The aliscore output appears to be empty. Check results for BUSCO: 48764at7742
## Thu Jul 14 00:05:40 CEST 2022 - The aliscore output file does not exist. Check results for BUSCO: 11104at7742
## Thu Jul 14 00:05:40 CEST 2022 - The aliscore output appears to be empty. Check results for BUSCO: 11104at7742
## Thu Jul 14 00:05:41 CEST 2022 - The aliscore output file does not exist. Check results for BUSCO: 31951at7742
## Thu Jul 14 00:05:41 CEST 2022 - The aliscore output appears to be empty. Check results for BUSCO: 31951at7742
## Thu Jul 14 00:05:44 CEST 2022 - The aliscore output file does not exist. Check results for BUSCO: 46737at7742
## Thu Jul 14 00:05:44 CEST 2022 - The aliscore output appears to be empty. Check results for BUSCO: 46737at7742
## Thu Jul 14 00:05:47 CEST 2022 - The aliscore output file does not exist. Check results for BUSCO: 16923at7742
## Thu Jul 14 00:05:47 CEST 2022 - The aliscore output appears to be empty. Check results for BUSCO: 16923at7742
## Thu Jul 14 00:05:48 CEST 2022 - The aliscore output file does not exist. Check results for BUSCO: 38882at7742
## Thu Jul 14 00:05:48 CEST 2022 - The aliscore output appears to be empty. Check results for BUSCO: 38882at7742
## Thu Jul 14 00:05:48 CEST 2022 - The aliscore output file does not exist. Check results for BUSCO: 47172at7742
## Thu Jul 14 00:05:48 CEST 2022 - The aliscore output appears to be empty. Check results for BUSCO: 47172at7742
## Thu Jul 14 00:05:50 CEST 2022 - The aliscore output file does not exist. Check results for BUSCO: 211943at7742
## Thu Jul 14 00:05:50 CEST 2022 - The aliscore output appears to be empty. Check results for BUSCO: 211943at7742
## Thu Jul 14 00:05:51 CEST 2022 - The aliscore output file does not exist. Check results for BUSCO: 23996at7742
## Thu Jul 14 00:05:51 CEST 2022 - The aliscore output appears to be empty. Check results for BUSCO: 23996at7742
## Thu Jul 14 00:05:52 CEST 2022 - The aliscore output file does not exist. Check results for BUSCO: 55709at7742
## Thu Jul 14 00:05:52 CEST 2022 - The aliscore output appears to be empty. Check results for BUSCO: 55709at7742
## Thu Jul 14 00:05:53 CEST 2022 - The aliscore output file does not exist. Check results for BUSCO: 22554at7742
## Thu Jul 14 00:05:53 CEST 2022 - The aliscore output appears to be empty. Check results for BUSCO: 22554at7742
## Thu Jul 14 00:05:56 CEST 2022 - The aliscore output file does not exist. Check results for BUSCO: 50444at7742
## Thu Jul 14 00:05:56 CEST 2022 - The aliscore output appears to be empty. Check results for BUSCO: 50444at7742
## Thu Jul 14 00:05:56 CEST 2022 - The aliscore output file does not exist. Check results for BUSCO: 58913at7742
## Thu Jul 14 00:05:56 CEST 2022 - The aliscore output appears to be empty. Check results for BUSCO: 58913at7742
## Thu Jul 14 00:05:57 CEST 2022 - The aliscore output file does not exist. Check results for BUSCO: 28267at7742
## Thu Jul 14 00:05:57 CEST 2022 - The aliscore output appears to be empty. Check results for BUSCO: 28267at7742
## Thu Jul 14 00:06:00 CEST 2022 - The aliscore output file does not exist. Check results for BUSCO: 406184at7742
## Thu Jul 14 00:06:00 CEST 2022 - The aliscore output file does not exist. Check results for BUSCO: 10899at7742
## Thu Jul 14 00:06:00 CEST 2022 - The aliscore output appears to be empty. Check results for BUSCO: 10899at7742
## Thu Jul 14 00:06:00 CEST 2022 - The aliscore output appears to be empty. Check results for BUSCO: 406184at7742
## Thu Jul 14 00:06:02 CEST 2022 - The aliscore output file does not exist. Check results for BUSCO: 26210at7742
## Thu Jul 14 00:06:02 CEST 2022 - The aliscore output appears to be empty. Check results for BUSCO: 26210at7742
## Thu Jul 14 00:06:04 CEST 2022 - The aliscore output file does not exist. Check results for BUSCO: 90690at7742
## Thu Jul 14 00:06:04 CEST 2022 - The aliscore output appears to be empty. Check results for BUSCO: 90690at7742
## Thu Jul 14 00:06:05 CEST 2022 - The aliscore output file does not exist. Check results for BUSCO: 179099at7742
## Thu Jul 14 00:06:05 CEST 2022 - The aliscore output appears to be empty. Check results for BUSCO: 179099at7742
## Thu Jul 14 00:06:07 CEST 2022 - The aliscore output file does not exist. Check results for BUSCO: 74327at7742
## Thu Jul 14 00:06:07 CEST 2022 - The aliscore output appears to be empty. Check results for BUSCO: 74327at7742
## Thu Jul 14 00:06:07 CEST 2022 - The aliscore output file does not exist. Check results for BUSCO: 175198at7742
## Thu Jul 14 00:06:07 CEST 2022 - The aliscore output appears to be empty. Check results for BUSCO: 175198at7742
## Thu Jul 14 00:06:08 CEST 2022 - The aliscore output file does not exist. Check results for BUSCO: 56852at7742
## Thu Jul 14 00:06:09 CEST 2022 - The aliscore output appears to be empty. Check results for BUSCO: 56852at7742
## Thu Jul 14 00:06:09 CEST 2022 - The aliscore output file does not exist. Check results for BUSCO: 61914at7742
## Thu Jul 14 00:06:09 CEST 2022 - The aliscore output appears to be empty. Check results for BUSCO: 61914at7742
## Thu Jul 14 00:06:12 CEST 2022 - The aliscore output file does not exist. Check results for BUSCO: 58913at7742
## Thu Jul 14 00:06:12 CEST 2022 - The aliscore output appears to be empty. Check results for BUSCO: 58913at7742
## Thu Jul 14 00:06:14 CEST 2022 - The aliscore output file does not exist. Check results for BUSCO: 17577at7742
## Thu Jul 14 00:06:14 CEST 2022 - The aliscore output appears to be empty. Check results for BUSCO: 17577at7742
## Thu Jul 14 00:06:16 CEST 2022 - The aliscore output file does not exist. Check results for BUSCO: 99281at7742
## Thu Jul 14 00:06:16 CEST 2022 - The aliscore output appears to be empty. Check results for BUSCO: 99281at7742
## Thu Jul 14 00:06:16 CEST 2022 - The aliscore output file does not exist. Check results for BUSCO: 41154at7742
## Thu Jul 14 00:06:16 CEST 2022 - The aliscore output appears to be empty. Check results for BUSCO: 41154at7742
## Thu Jul 14 00:06:19 CEST 2022 - The aliscore output file does not exist. Check results for BUSCO: 124120at7742
## Thu Jul 14 00:06:19 CEST 2022 - The aliscore output appears to be empty. Check results for BUSCO: 124120at7742
## Thu Jul 14 00:06:19 CEST 2022 - The aliscore output file does not exist. Check results for BUSCO: 74327at7742
## Thu Jul 14 00:06:19 CEST 2022 - The aliscore output appears to be empty. Check results for BUSCO: 74327at7742
## Thu Jul 14 00:06:19 CEST 2022 - The aliscore output file does not exist. Check results for BUSCO: 104791at7742
## Thu Jul 14 00:06:19 CEST 2022 - The aliscore output appears to be empty. Check results for BUSCO: 104791at7742
## Thu Jul 14 00:06:21 CEST 2022 - The aliscore output file does not exist. Check results for BUSCO: 24868at7742
## Thu Jul 14 00:06:21 CEST 2022 - The aliscore output appears to be empty. Check results for BUSCO: 24868at7742
## Thu Jul 14 00:06:22 CEST 2022 - The aliscore output file does not exist. Check results for BUSCO: 193224at7742
## Thu Jul 14 00:06:22 CEST 2022 - The aliscore output appears to be empty. Check results for BUSCO: 193224at7742
## Thu Jul 14 00:06:25 CEST 2022 - The aliscore output file does not exist. Check results for BUSCO: 331581at7742
## Thu Jul 14 00:06:25 CEST 2022 - The aliscore output appears to be empty. Check results for BUSCO: 331581at7742
## Thu Jul 14 00:06:25 CEST 2022 - The aliscore output file does not exist. Check results for BUSCO: 48764at7742
## Thu Jul 14 00:06:26 CEST 2022 - The aliscore output appears to be empty. Check results for BUSCO: 48764at7742
## Thu Jul 14 00:06:28 CEST 2022 - The aliscore output file does not exist. Check results for BUSCO: 50444at7742
## Thu Jul 14 00:06:28 CEST 2022 - The aliscore output appears to be empty. Check results for BUSCO: 50444at7742
## Thu Jul 14 00:06:28 CEST 2022 - The aliscore output file does not exist. Check results for BUSCO: 58913at7742
## Thu Jul 14 00:06:28 CEST 2022 - The aliscore output appears to be empty. Check results for BUSCO: 58913at7742
## Thu Jul 14 00:06:30 CEST 2022 - The aliscore output file does not exist. Check results for BUSCO: 138901at7742
## Thu Jul 14 00:06:30 CEST 2022 - The aliscore output appears to be empty. Check results for BUSCO: 138901at7742
## Thu Jul 14 00:06:30 CEST 2022 - The aliscore output file does not exist. Check results for BUSCO: 28708at7742
## Thu Jul 14 00:06:30 CEST 2022 - The aliscore output appears to be empty. Check results for BUSCO: 28708at7742
## Thu Jul 14 00:06:33 CEST 2022 - The aliscore output file does not exist. Check results for BUSCO: 98298at7742
## Thu Jul 14 00:06:33 CEST 2022 - The aliscore output appears to be empty. Check results for BUSCO: 98298at7742
## Thu Jul 14 00:06:34 CEST 2022 - The aliscore output file does not exist. Check results for BUSCO: 41150at7742
## Thu Jul 14 00:06:34 CEST 2022 - The aliscore output appears to be empty. Check results for BUSCO: 41150at7742
## Thu Jul 14 00:06:36 CEST 2022 - The aliscore output file does not exist. Check results for BUSCO: 96327at7742
## Thu Jul 14 00:06:36 CEST 2022 - The aliscore output appears to be empty. Check results for BUSCO: 96327at7742
## Thu Jul 14 00:06:39 CEST 2022 - The aliscore output file does not exist. Check results for BUSCO: 331581at7742
## Thu Jul 14 00:06:39 CEST 2022 - The aliscore output file does not exist. Check results for BUSCO: 28386at7742
## Thu Jul 14 00:06:39 CEST 2022 - The aliscore output appears to be empty. Check results for BUSCO: 331581at7742
## Thu Jul 14 00:06:39 CEST 2022 - The aliscore output appears to be empty. Check results for BUSCO: 28386at7742
## Thu Jul 14 00:06:40 CEST 2022 - The aliscore output file does not exist. Check results for BUSCO: 28708at7742
## Thu Jul 14 00:06:40 CEST 2022 - The aliscore output appears to be empty. Check results for BUSCO: 28708at7742
## Thu Jul 14 00:06:40 CEST 2022 - The aliscore output file does not exist. Check results for BUSCO: 26485at7742
## Thu Jul 14 00:06:40 CEST 2022 - The aliscore output appears to be empty. Check results for BUSCO: 26485at7742
## Thu Jul 14 00:06:42 CEST 2022 - The aliscore output file does not exist. Check results for BUSCO: 44800at7742
## Thu Jul 14 00:06:42 CEST 2022 - The aliscore output appears to be empty. Check results for BUSCO: 44800at7742
## Thu Jul 14 00:06:45 CEST 2022 - The aliscore output file does not exist. Check results for BUSCO: 186292at7742
## Thu Jul 14 00:06:45 CEST 2022 - The aliscore output appears to be empty. Check results for BUSCO: 186292at7742
## Thu Jul 14 00:06:45 CEST 2022 - The aliscore output file does not exist. Check results for BUSCO: 26210at7742
## Thu Jul 14 00:06:45 CEST 2022 - The aliscore output appears to be empty. Check results for BUSCO: 26210at7742
## Thu Jul 14 00:06:49 CEST 2022 - The aliscore output file does not exist. Check results for BUSCO: 126446at7742
## Thu Jul 14 00:06:49 CEST 2022 - The aliscore output appears to be empty. Check results for BUSCO: 126446at7742
## Thu Jul 14 00:06:49 CEST 2022 - The aliscore output file does not exist. Check results for BUSCO: 276779at7742
## Thu Jul 14 00:06:49 CEST 2022 - The aliscore output appears to be empty. Check results for BUSCO: 276779at7742
## Thu Jul 14 00:06:50 CEST 2022 - The aliscore output file does not exist. Check results for BUSCO: 79862at7742
## Thu Jul 14 00:06:50 CEST 2022 - The aliscore output appears to be empty. Check results for BUSCO: 79862at7742
## Thu Jul 14 00:06:50 CEST 2022 - The aliscore output file does not exist. Check results for BUSCO: 32152at7742
## Thu Jul 14 00:06:51 CEST 2022 - The aliscore output appears to be empty. Check results for BUSCO: 32152at7742
## Thu Jul 14 00:06:53 CEST 2022 - The aliscore output file does not exist. Check results for BUSCO: 28708at7742
## Thu Jul 14 00:06:53 CEST 2022 - The aliscore output appears to be empty. Check results for BUSCO: 28708at7742
## Thu Jul 14 00:06:54 CEST 2022 - The aliscore output file does not exist. Check results for BUSCO: 65671at7742
## Thu Jul 14 00:06:54 CEST 2022 - The aliscore output appears to be empty. Check results for BUSCO: 65671at7742
## Thu Jul 14 00:06:56 CEST 2022 - The aliscore output file does not exist. Check results for BUSCO: 27463at7742
## Thu Jul 14 00:06:56 CEST 2022 - The aliscore output appears to be empty. Check results for BUSCO: 27463at7742
## Thu Jul 14 00:06:59 CEST 2022 - The aliscore output file does not exist. Check results for BUSCO: 30224at7742
## Thu Jul 14 00:06:59 CEST 2022 - The aliscore output file does not exist. Check results for BUSCO: 406184at7742
## Thu Jul 14 00:06:59 CEST 2022 - The aliscore output appears to be empty. Check results for BUSCO: 30224at7742
## Thu Jul 14 00:06:59 CEST 2022 - The aliscore output appears to be empty. Check results for BUSCO: 406184at7742
## Thu Jul 14 00:07:00 CEST 2022 - The aliscore output file does not exist. Check results for BUSCO: 36236at7742
## Thu Jul 14 00:07:00 CEST 2022 - The aliscore output appears to be empty. Check results for BUSCO: 36236at7742
## Thu Jul 14 00:07:00 CEST 2022 - The aliscore output file does not exist. Check results for BUSCO: 23173at7742
## Thu Jul 14 00:07:00 CEST 2022 - The aliscore output appears to be empty. Check results for BUSCO: 23173at7742
## Thu Jul 14 00:07:01 CEST 2022 - The aliscore output file does not exist. Check results for BUSCO: 79862at7742
## Thu Jul 14 00:07:01 CEST 2022 - The aliscore output appears to be empty. Check results for BUSCO: 79862at7742
## Thu Jul 14 00:07:04 CEST 2022 - The aliscore output file does not exist. Check results for BUSCO: 130025at7742
## Thu Jul 14 00:07:04 CEST 2022 - The aliscore output appears to be empty. Check results for BUSCO: 130025at7742
## Thu Jul 14 00:07:07 CEST 2022 - The aliscore output file does not exist. Check results for BUSCO: 331581at7742
## Thu Jul 14 00:07:07 CEST 2022 - The aliscore output appears to be empty. Check results for BUSCO: 331581at7742
## Thu Jul 14 00:07:08 CEST 2022 - The aliscore output file does not exist. Check results for BUSCO: 26485at7742
## Thu Jul 14 00:07:08 CEST 2022 - The aliscore output appears to be empty. Check results for BUSCO: 26485at7742
## Thu Jul 14 00:07:10 CEST 2022 - The aliscore output file does not exist. Check results for BUSCO: 41890at7742
## Thu Jul 14 00:07:10 CEST 2022 - The aliscore output appears to be empty. Check results for BUSCO: 41890at7742
## Thu Jul 14 00:07:10 CEST 2022 - The aliscore output file does not exist. Check results for BUSCO: 135024at7742
## Thu Jul 14 00:07:10 CEST 2022 - The aliscore output appears to be empty. Check results for BUSCO: 135024at7742
## Thu Jul 14 00:07:10 CEST 2022 - The aliscore output file does not exist. Check results for BUSCO: 62115at7742
## Thu Jul 14 00:07:10 CEST 2022 - The aliscore output appears to be empty. Check results for BUSCO: 62115at7742
## Thu Jul 14 00:07:11 CEST 2022 - The aliscore output file does not exist. Check results for BUSCO: 41890at7742
## Thu Jul 14 00:07:11 CEST 2022 - The aliscore output appears to be empty. Check results for BUSCO: 41890at7742
## Thu Jul 14 00:07:15 CEST 2022 - The aliscore output file does not exist. Check results for BUSCO: 46737at7742
## Thu Jul 14 00:07:15 CEST 2022 - The aliscore output appears to be empty. Check results for BUSCO: 46737at7742
## Thu Jul 14 00:07:17 CEST 2022 - The aliscore output file does not exist. Check results for BUSCO: 406184at7742
## Thu Jul 14 00:07:17 CEST 2022 - The aliscore output appears to be empty. Check results for BUSCO: 406184at7742
## Thu Jul 14 00:07:19 CEST 2022 - The aliscore output file does not exist. Check results for BUSCO: 13296at7742
## Thu Jul 14 00:07:19 CEST 2022 - The aliscore output appears to be empty. Check results for BUSCO: 13296at7742
## Thu Jul 14 00:07:21 CEST 2022 - The aliscore output file does not exist. Check results for BUSCO: 106853at7742
## Thu Jul 14 00:07:21 CEST 2022 - The aliscore output appears to be empty. Check results for BUSCO: 106853at7742
## Thu Jul 14 00:07:21 CEST 2022 - The aliscore output file does not exist. Check results for BUSCO: 14788at7742
## Thu Jul 14 00:07:21 CEST 2022 - The aliscore output appears to be empty. Check results for BUSCO: 14788at7742
## Thu Jul 14 00:07:22 CEST 2022 - The aliscore output file does not exist. Check results for BUSCO: 113103at7742
## Thu Jul 14 00:07:22 CEST 2022 - The aliscore output appears to be empty. Check results for BUSCO: 113103at7742
## Thu Jul 14 00:07:22 CEST 2022 - The aliscore output file does not exist. Check results for BUSCO: 151895at7742
## Thu Jul 14 00:07:22 CEST 2022 - The aliscore output appears to be empty. Check results for BUSCO: 151895at7742
## Thu Jul 14 00:07:22 CEST 2022 - The aliscore output file does not exist. Check results for BUSCO: 22554at7742
## Thu Jul 14 00:07:22 CEST 2022 - The aliscore output appears to be empty. Check results for BUSCO: 22554at7742
## Thu Jul 14 00:07:27 CEST 2022 - The aliscore output file does not exist. Check results for BUSCO: 138901at7742
## Thu Jul 14 00:07:27 CEST 2022 - The aliscore output appears to be empty. Check results for BUSCO: 138901at7742
## Thu Jul 14 00:07:27 CEST 2022 - The aliscore output file does not exist. Check results for BUSCO: 23173at7742
## Thu Jul 14 00:07:27 CEST 2022 - The aliscore output appears to be empty. Check results for BUSCO: 23173at7742
## Thu Jul 14 00:07:29 CEST 2022 - The aliscore output file does not exist. Check results for BUSCO: 40843at7742
## Thu Jul 14 00:07:30 CEST 2022 - The aliscore output appears to be empty. Check results for BUSCO: 40843at7742
## Thu Jul 14 00:07:31 CEST 2022 - The aliscore output file does not exist. Check results for BUSCO: 26485at7742
## Thu Jul 14 00:07:31 CEST 2022 - The aliscore output appears to be empty. Check results for BUSCO: 26485at7742
## Thu Jul 14 00:07:32 CEST 2022 - The aliscore output file does not exist. Check results for BUSCO: 74327at7742
## Thu Jul 14 00:07:32 CEST 2022 - The aliscore output appears to be empty. Check results for BUSCO: 74327at7742
## Thu Jul 14 00:07:32 CEST 2022 - The aliscore output file does not exist. Check results for BUSCO: 66052at7742
## Thu Jul 14 00:07:32 CEST 2022 - The aliscore output appears to be empty. Check results for BUSCO: 66052at7742
## Thu Jul 14 00:07:33 CEST 2022 - The aliscore output file does not exist. Check results for BUSCO: 68853at7742
## Thu Jul 14 00:07:33 CEST 2022 - The aliscore output appears to be empty. Check results for BUSCO: 68853at7742
## Thu Jul 14 00:07:37 CEST 2022 - The aliscore output file does not exist. Check results for BUSCO: 19072at7742
## Thu Jul 14 00:07:37 CEST 2022 - The aliscore output appears to be empty. Check results for BUSCO: 19072at7742
## Thu Jul 14 00:07:39 CEST 2022 - The aliscore output file does not exist. Check results for BUSCO: 170437at7742
## Thu Jul 14 00:07:40 CEST 2022 - The aliscore output appears to be empty. Check results for BUSCO: 170437at7742
## Thu Jul 14 00:07:40 CEST 2022 - The aliscore output file does not exist. Check results for BUSCO: 197548at7742
## Thu Jul 14 00:07:40 CEST 2022 - The aliscore output appears to be empty. Check results for BUSCO: 197548at7742
## Thu Jul 14 00:07:42 CEST 2022 - The aliscore output file does not exist. Check results for BUSCO: 40997at7742
## Thu Jul 14 00:07:42 CEST 2022 - The aliscore output appears to be empty. Check results for BUSCO: 40997at7742
## Thu Jul 14 00:07:43 CEST 2022 - The aliscore output file does not exist. Check results for BUSCO: 37715at7742
## Thu Jul 14 00:07:43 CEST 2022 - The aliscore output appears to be empty. Check results for BUSCO: 37715at7742
## Thu Jul 14 00:07:43 CEST 2022 - The aliscore output file does not exist. Check results for BUSCO: 13296at7742
## Thu Jul 14 00:07:43 CEST 2022 - The aliscore output appears to be empty. Check results for BUSCO: 13296at7742
## Thu Jul 14 00:07:43 CEST 2022 - The aliscore output file does not exist. Check results for BUSCO: 40997at7742
## Thu Jul 14 00:07:43 CEST 2022 - The aliscore output appears to be empty. Check results for BUSCO: 40997at7742
## Thu Jul 14 00:07:44 CEST 2022 - The aliscore output file does not exist. Check results for BUSCO: 23673at7742
## Thu Jul 14 00:07:44 CEST 2022 - The aliscore output appears to be empty. Check results for BUSCO: 23673at7742
## Thu Jul 14 00:07:49 CEST 2022 - The aliscore output file does not exist. Check results for BUSCO: 22554at7742
## Thu Jul 14 00:07:49 CEST 2022 - The aliscore output appears to be empty. Check results for BUSCO: 22554at7742
## Thu Jul 14 00:07:49 CEST 2022 - The aliscore output file does not exist. Check results for BUSCO: 32511at7742
## Thu Jul 14 00:07:49 CEST 2022 - The aliscore output appears to be empty. Check results for BUSCO: 32511at7742
## Thu Jul 14 00:07:52 CEST 2022 - The aliscore output file does not exist. Check results for BUSCO: 28386at7742
## Thu Jul 14 00:07:52 CEST 2022 - The aliscore output appears to be empty. Check results for BUSCO: 28386at7742
## Thu Jul 14 00:07:52 CEST 2022 - The aliscore output file does not exist. Check results for BUSCO: 55030at7742
## Thu Jul 14 00:07:52 CEST 2022 - The aliscore output appears to be empty. Check results for BUSCO: 55030at7742
## Thu Jul 14 00:07:53 CEST 2022 - The aliscore output file does not exist. Check results for BUSCO: 14788at7742
## Thu Jul 14 00:07:53 CEST 2022 - The aliscore output appears to be empty. Check results for BUSCO: 14788at7742
## Thu Jul 14 00:07:54 CEST 2022 - The aliscore output file does not exist. Check results for BUSCO: 32152at7742
## Thu Jul 14 00:07:54 CEST 2022 - The aliscore output appears to be empty. Check results for BUSCO: 32152at7742
## Thu Jul 14 00:07:57 CEST 2022 - The aliscore output file does not exist. Check results for BUSCO: 174426at7742
## Thu Jul 14 00:07:58 CEST 2022 - The aliscore output appears to be empty. Check results for BUSCO: 174426at7742
## Thu Jul 14 00:07:58 CEST 2022 - The aliscore output file does not exist. Check results for BUSCO: 24566at7742
## Thu Jul 14 00:07:58 CEST 2022 - The aliscore output appears to be empty. Check results for BUSCO: 24566at7742
## Thu Jul 14 00:07:58 CEST 2022 - The aliscore output file does not exist. Check results for BUSCO: 36236at7742
## Thu Jul 14 00:07:58 CEST 2022 - The aliscore output appears to be empty. Check results for BUSCO: 36236at7742
## Thu Jul 14 00:08:01 CEST 2022 - The aliscore output file does not exist. Check results for BUSCO: 138901at7742
## Thu Jul 14 00:08:01 CEST 2022 - The aliscore output appears to be empty. Check results for BUSCO: 138901at7742
## Thu Jul 14 00:08:03 CEST 2022 - The aliscore output file does not exist. Check results for BUSCO: 90587at7742
## Thu Jul 14 00:08:03 CEST 2022 - The aliscore output file does not exist. Check results for BUSCO: 174426at7742
## Thu Jul 14 00:08:03 CEST 2022 - The aliscore output appears to be empty. Check results for BUSCO: 90587at7742
## Thu Jul 14 00:08:03 CEST 2022 - The aliscore output appears to be empty. Check results for BUSCO: 174426at7742
## Thu Jul 14 00:08:04 CEST 2022 - The aliscore output file does not exist. Check results for BUSCO: 174426at7742
## Thu Jul 14 00:08:04 CEST 2022 - The aliscore output appears to be empty. Check results for BUSCO: 174426at7742
## Thu Jul 14 00:08:06 CEST 2022 - The aliscore output file does not exist. Check results for BUSCO: 82622at7742
## Thu Jul 14 00:08:06 CEST 2022 - The aliscore output appears to be empty. Check results for BUSCO: 82622at7742
## Thu Jul 14 00:08:08 CEST 2022 - The aliscore output file does not exist. Check results for BUSCO: 98298at7742
## Thu Jul 14 00:08:08 CEST 2022 - The aliscore output appears to be empty. Check results for BUSCO: 98298at7742
## Thu Jul 14 00:08:09 CEST 2022 - The aliscore output file does not exist. Check results for BUSCO: 65757at7742
## Thu Jul 14 00:08:09 CEST 2022 - The aliscore output appears to be empty. Check results for BUSCO: 65757at7742
## Thu Jul 14 00:08:10 CEST 2022 - The aliscore output file does not exist. Check results for BUSCO: 65757at7742
## Thu Jul 14 00:08:10 CEST 2022 - The aliscore output appears to be empty. Check results for BUSCO: 65757at7742
## Thu Jul 14 00:08:13 CEST 2022 - The aliscore output file does not exist. Check results for BUSCO: 186464at7742
## Thu Jul 14 00:08:13 CEST 2022 - The aliscore output appears to be empty. Check results for BUSCO: 186464at7742
## Thu Jul 14 00:08:14 CEST 2022 - The aliscore output file does not exist. Check results for BUSCO: 30224at7742
## Thu Jul 14 00:08:14 CEST 2022 - The aliscore output appears to be empty. Check results for BUSCO: 30224at7742
## Thu Jul 14 00:08:14 CEST 2022 - The aliscore output file does not exist. Check results for BUSCO: 83097at7742
## Thu Jul 14 00:08:14 CEST 2022 - The aliscore output appears to be empty. Check results for BUSCO: 83097at7742
## Thu Jul 14 00:08:16 CEST 2022 - The aliscore output file does not exist. Check results for BUSCO: 171211at7742
## Thu Jul 14 00:08:16 CEST 2022 - The aliscore output appears to be empty. Check results for BUSCO: 171211at7742
## Thu Jul 14 00:08:17 CEST 2022 - The aliscore output file does not exist. Check results for BUSCO: 98298at7742
## Thu Jul 14 00:08:17 CEST 2022 - The aliscore output appears to be empty. Check results for BUSCO: 98298at7742
## Thu Jul 14 00:08:19 CEST 2022 - The aliscore output file does not exist. Check results for BUSCO: 62115at7742
## Thu Jul 14 00:08:19 CEST 2022 - The aliscore output appears to be empty. Check results for BUSCO: 62115at7742
## Thu Jul 14 00:08:19 CEST 2022 - The aliscore output file does not exist. Check results for BUSCO: 87643at7742
## Thu Jul 14 00:08:20 CEST 2022 - The aliscore output appears to be empty. Check results for BUSCO: 87643at7742
## Thu Jul 14 00:08:23 CEST 2022 - The aliscore output file does not exist. Check results for BUSCO: 212520at7742
## Thu Jul 14 00:08:23 CEST 2022 - The aliscore output file does not exist. Check results for BUSCO: 276779at7742
## Thu Jul 14 00:08:23 CEST 2022 - The aliscore output appears to be empty. Check results for BUSCO: 276779at7742
## Thu Jul 14 00:08:23 CEST 2022 - The aliscore output appears to be empty. Check results for BUSCO: 212520at7742
## Thu Jul 14 00:08:23 CEST 2022 - The aliscore output file does not exist. Check results for BUSCO: 33771at7742
## Thu Jul 14 00:08:23 CEST 2022 - The aliscore output appears to be empty. Check results for BUSCO: 33771at7742
## Thu Jul 14 00:08:26 CEST 2022 - The aliscore output file does not exist. Check results for BUSCO: 46737at7742
## Thu Jul 14 00:08:26 CEST 2022 - The aliscore output appears to be empty. Check results for BUSCO: 46737at7742
## Thu Jul 14 00:08:27 CEST 2022 - The aliscore output file does not exist. Check results for BUSCO: 106344at7742
## Thu Jul 14 00:08:27 CEST 2022 - The aliscore output appears to be empty. Check results for BUSCO: 106344at7742
## Thu Jul 14 00:08:28 CEST 2022 - The aliscore output file does not exist. Check results for BUSCO: 106943at7742
## Thu Jul 14 00:08:28 CEST 2022 - The aliscore output appears to be empty. Check results for BUSCO: 106943at7742
## Thu Jul 14 00:08:30 CEST 2022 - The aliscore output file does not exist. Check results for BUSCO: 130025at7742
## Thu Jul 14 00:08:30 CEST 2022 - The aliscore output appears to be empty. Check results for BUSCO: 130025at7742
## Thu Jul 14 00:08:34 CEST 2022 - The aliscore output file does not exist. Check results for BUSCO: 114216at7742
## Thu Jul 14 00:08:34 CEST 2022 - The aliscore output appears to be empty. Check results for BUSCO: 114216at7742
## Thu Jul 14 00:08:34 CEST 2022 - The aliscore output file does not exist. Check results for BUSCO: 45658at7742
## Thu Jul 14 00:08:34 CEST 2022 - The aliscore output appears to be empty. Check results for BUSCO: 45658at7742
## Thu Jul 14 00:08:35 CEST 2022 - The aliscore output file does not exist. Check results for BUSCO: 41150at7742
## Thu Jul 14 00:08:35 CEST 2022 - The aliscore output appears to be empty. Check results for BUSCO: 41150at7742
## Thu Jul 14 00:08:35 CEST 2022 - The aliscore output file does not exist. Check results for BUSCO: 90690at7742
## Thu Jul 14 00:08:35 CEST 2022 - The aliscore output appears to be empty. Check results for BUSCO: 90690at7742
## Thu Jul 14 00:08:37 CEST 2022 - The aliscore output file does not exist. Check results for BUSCO: 40843at7742
## Thu Jul 14 00:08:37 CEST 2022 - The aliscore output appears to be empty. Check results for BUSCO: 40843at7742
## Thu Jul 14 00:08:38 CEST 2022 - The aliscore output file does not exist. Check results for BUSCO: 83097at7742
## Thu Jul 14 00:08:38 CEST 2022 - The aliscore output appears to be empty. Check results for BUSCO: 83097at7742
## Thu Jul 14 00:08:39 CEST 2022 - The aliscore output file does not exist. Check results for BUSCO: 82448at7742
## Thu Jul 14 00:08:39 CEST 2022 - The aliscore output appears to be empty. Check results for BUSCO: 82448at7742
## Thu Jul 14 00:08:42 CEST 2022 - The aliscore output file does not exist. Check results for BUSCO: 61914at7742
## Thu Jul 14 00:08:42 CEST 2022 - The aliscore output appears to be empty. Check results for BUSCO: 61914at7742
## Thu Jul 14 00:08:42 CEST 2022 - The aliscore output file does not exist. Check results for BUSCO: 113103at7742
## Thu Jul 14 00:08:42 CEST 2022 - The aliscore output appears to be empty. Check results for BUSCO: 113103at7742
## Thu Jul 14 00:08:43 CEST 2022 - The aliscore output file does not exist. Check results for BUSCO: 186292at7742
## Thu Jul 14 00:08:43 CEST 2022 - The aliscore output appears to be empty. Check results for BUSCO: 186292at7742
## Thu Jul 14 00:08:46 CEST 2022 - The aliscore output file does not exist. Check results for BUSCO: 30224at7742
## Thu Jul 14 00:08:46 CEST 2022 - The aliscore output appears to be empty. Check results for BUSCO: 30224at7742
## Thu Jul 14 00:08:46 CEST 2022 - The aliscore output file does not exist. Check results for BUSCO: 31951at7742
## Thu Jul 14 00:08:46 CEST 2022 - The aliscore output appears to be empty. Check results for BUSCO: 31951at7742
## Thu Jul 14 00:08:48 CEST 2022 - The aliscore output file does not exist. Check results for BUSCO: 41890at7742
## Thu Jul 14 00:08:48 CEST 2022 - The aliscore output appears to be empty. Check results for BUSCO: 41890at7742
## Thu Jul 14 00:08:50 CEST 2022 - The aliscore output file does not exist. Check results for BUSCO: 179099at7742
## Thu Jul 14 00:08:50 CEST 2022 - The aliscore output appears to be empty. Check results for BUSCO: 179099at7742
## Thu Jul 14 00:08:52 CEST 2022 - The aliscore output file does not exist. Check results for BUSCO: 152085at7742
## Thu Jul 14 00:08:52 CEST 2022 - The aliscore output appears to be empty. Check results for BUSCO: 152085at7742
## Thu Jul 14 00:08:52 CEST 2022 - The aliscore output file does not exist. Check results for BUSCO: 83097at7742
## Thu Jul 14 00:08:52 CEST 2022 - The aliscore output appears to be empty. Check results for BUSCO: 83097at7742
## Thu Jul 14 00:08:54 CEST 2022 - The aliscore output file does not exist. Check results for BUSCO: 28386at7742
## Thu Jul 14 00:08:54 CEST 2022 - The aliscore output appears to be empty. Check results for BUSCO: 28386at7742
## Thu Jul 14 00:08:55 CEST 2022 - The aliscore output file does not exist. Check results for BUSCO: 32152at7742
## Thu Jul 14 00:08:55 CEST 2022 - The aliscore output appears to be empty. Check results for BUSCO: 32152at7742
## Thu Jul 14 00:08:56 CEST 2022 - The aliscore output file does not exist. Check results for BUSCO: 50444at7742
## Thu Jul 14 00:08:56 CEST 2022 - The aliscore output appears to be empty. Check results for BUSCO: 50444at7742
## Thu Jul 14 00:08:58 CEST 2022 - The aliscore output file does not exist. Check results for BUSCO: 82704at7742
## Thu Jul 14 00:08:58 CEST 2022 - The aliscore output appears to be empty. Check results for BUSCO: 82704at7742
## Thu Jul 14 00:08:59 CEST 2022 - The aliscore output file does not exist. Check results for BUSCO: 49885at7742
## Thu Jul 14 00:08:59 CEST 2022 - The aliscore output appears to be empty. Check results for BUSCO: 49885at7742
## Thu Jul 14 00:09:01 CEST 2022 - The aliscore output file does not exist. Check results for BUSCO: 253119at7742
## Thu Jul 14 00:09:01 CEST 2022 - The aliscore output appears to be empty. Check results for BUSCO: 253119at7742
## Thu Jul 14 00:09:03 CEST 2022 - The aliscore output file does not exist. Check results for BUSCO: 101309at7742
## Thu Jul 14 00:09:03 CEST 2022 - The aliscore output appears to be empty. Check results for BUSCO: 101309at7742
## Thu Jul 14 00:09:03 CEST 2022 - The aliscore output file does not exist. Check results for BUSCO: 162634at7742
## Thu Jul 14 00:09:03 CEST 2022 - The aliscore output appears to be empty. Check results for BUSCO: 162634at7742
## Thu Jul 14 00:09:04 CEST 2022 - The aliscore output file does not exist. Check results for BUSCO: 109869at7742
## Thu Jul 14 00:09:04 CEST 2022 - The aliscore output appears to be empty. Check results for BUSCO: 109869at7742
## Thu Jul 14 00:09:05 CEST 2022 - The aliscore output file does not exist. Check results for BUSCO: 185168at7742
## Thu Jul 14 00:09:05 CEST 2022 - The aliscore output appears to be empty. Check results for BUSCO: 185168at7742
## Thu Jul 14 00:09:07 CEST 2022 - The aliscore output file does not exist. Check results for BUSCO: 82448at7742
## Thu Jul 14 00:09:07 CEST 2022 - The aliscore output appears to be empty. Check results for BUSCO: 82448at7742
## Thu Jul 14 00:09:09 CEST 2022 - The aliscore output file does not exist. Check results for BUSCO: 26047at7742
## Thu Jul 14 00:09:09 CEST 2022 - The aliscore output appears to be empty. Check results for BUSCO: 26047at7742
## Thu Jul 14 00:09:10 CEST 2022 - The aliscore output file does not exist. Check results for BUSCO: 82448at7742
## Thu Jul 14 00:09:10 CEST 2022 - The aliscore output appears to be empty. Check results for BUSCO: 82448at7742
## Thu Jul 14 00:09:12 CEST 2022 - The aliscore output file does not exist. Check results for BUSCO: 197548at7742
## Thu Jul 14 00:09:12 CEST 2022 - The aliscore output appears to be empty. Check results for BUSCO: 197548at7742
## Thu Jul 14 00:09:13 CEST 2022 - The aliscore output file does not exist. Check results for BUSCO: 13296at7742
## Thu Jul 14 00:09:13 CEST 2022 - The aliscore output appears to be empty. Check results for BUSCO: 13296at7742
## Thu Jul 14 00:09:15 CEST 2022 - The aliscore output file does not exist. Check results for BUSCO: 162634at7742
## Thu Jul 14 00:09:15 CEST 2022 - The aliscore output appears to be empty. Check results for BUSCO: 162634at7742
## Thu Jul 14 00:09:15 CEST 2022 - The aliscore output file does not exist. Check results for BUSCO: 63488at7742
## Thu Jul 14 00:09:15 CEST 2022 - The aliscore output appears to be empty. Check results for BUSCO: 63488at7742
## Thu Jul 14 00:09:18 CEST 2022 - The aliscore output file does not exist. Check results for BUSCO: 130025at7742
## Thu Jul 14 00:09:18 CEST 2022 - The aliscore output appears to be empty. Check results for BUSCO: 130025at7742
## Thu Jul 14 00:09:18 CEST 2022 - The aliscore output file does not exist. Check results for BUSCO: 175198at7742
## Thu Jul 14 00:09:18 CEST 2022 - The aliscore output appears to be empty. Check results for BUSCO: 175198at7742
## Thu Jul 14 00:09:19 CEST 2022 - The aliscore output file does not exist. Check results for BUSCO: 185168at7742
## Thu Jul 14 00:09:19 CEST 2022 - The aliscore output appears to be empty. Check results for BUSCO: 185168at7742
## Thu Jul 14 00:09:22 CEST 2022 - The aliscore output file does not exist. Check results for BUSCO: 104791at7742
## Thu Jul 14 00:09:22 CEST 2022 - The aliscore output appears to be empty. Check results for BUSCO: 104791at7742
## Thu Jul 14 00:09:23 CEST 2022 - The aliscore output file does not exist. Check results for BUSCO: 104791at7742
## Thu Jul 14 00:09:23 CEST 2022 - The aliscore output appears to be empty. Check results for BUSCO: 104791at7742
## Thu Jul 14 00:09:24 CEST 2022 - The aliscore output file does not exist. Check results for BUSCO: 96327at7742
## Thu Jul 14 00:09:24 CEST 2022 - The aliscore output appears to be empty. Check results for BUSCO: 96327at7742
## Thu Jul 14 00:09:25 CEST 2022 - The aliscore output file does not exist. Check results for BUSCO: 162634at7742
## Thu Jul 14 00:09:25 CEST 2022 - The aliscore output appears to be empty. Check results for BUSCO: 162634at7742
## Thu Jul 14 00:09:26 CEST 2022 - The aliscore output file does not exist. Check results for BUSCO: 73217at7742
## Thu Jul 14 00:09:26 CEST 2022 - The aliscore output appears to be empty. Check results for BUSCO: 73217at7742
## Thu Jul 14 00:09:28 CEST 2022 - The aliscore output file does not exist. Check results for BUSCO: 106301at7742
## Thu Jul 14 00:09:28 CEST 2022 - The aliscore output appears to be empty. Check results for BUSCO: 106301at7742
## Thu Jul 14 00:09:30 CEST 2022 - The aliscore output file does not exist. Check results for BUSCO: 155652at7742
## Thu Jul 14 00:09:30 CEST 2022 - The aliscore output appears to be empty. Check results for BUSCO: 155652at7742
## Thu Jul 14 00:09:30 CEST 2022 - The aliscore output file does not exist. Check results for BUSCO: 55500at7742
## Thu Jul 14 00:09:30 CEST 2022 - The aliscore output appears to be empty. Check results for BUSCO: 55500at7742
## Thu Jul 14 00:09:31 CEST 2022 - The aliscore output file does not exist. Check results for BUSCO: 193224at7742
## Thu Jul 14 00:09:31 CEST 2022 - The aliscore output appears to be empty. Check results for BUSCO: 193224at7742
## Thu Jul 14 00:09:34 CEST 2022 - The aliscore output file does not exist. Check results for BUSCO: 43568at7742
## Thu Jul 14 00:09:34 CEST 2022 - The aliscore output appears to be empty. Check results for BUSCO: 43568at7742
## Thu Jul 14 00:09:35 CEST 2022 - The aliscore output file does not exist. Check results for BUSCO: 49993at7742
## Thu Jul 14 00:09:35 CEST 2022 - The aliscore output appears to be empty. Check results for BUSCO: 49993at7742
## Thu Jul 14 00:09:36 CEST 2022 - The aliscore output file does not exist. Check results for BUSCO: 285264at7742
## Thu Jul 14 00:09:36 CEST 2022 - The aliscore output appears to be empty. Check results for BUSCO: 285264at7742
## Thu Jul 14 00:09:37 CEST 2022 - The aliscore output file does not exist. Check results for BUSCO: 26047at7742
## Thu Jul 14 00:09:37 CEST 2022 - The aliscore output appears to be empty. Check results for BUSCO: 26047at7742
## Thu Jul 14 00:09:39 CEST 2022 - The aliscore output file does not exist. Check results for BUSCO: 158865at7742
## Thu Jul 14 00:09:39 CEST 2022 - The aliscore output appears to be empty. Check results for BUSCO: 158865at7742
## Thu Jul 14 00:09:41 CEST 2022 - The aliscore output file does not exist. Check results for BUSCO: 44800at7742
## Thu Jul 14 00:09:41 CEST 2022 - The aliscore output appears to be empty. Check results for BUSCO: 44800at7742
## Thu Jul 14 00:09:42 CEST 2022 - The aliscore output file does not exist. Check results for BUSCO: 359556at7742
## Thu Jul 14 00:09:42 CEST 2022 - The aliscore output appears to be empty. Check results for BUSCO: 359556at7742
## Thu Jul 14 00:09:44 CEST 2022 - The aliscore output file does not exist. Check results for BUSCO: 194852at7742
## Thu Jul 14 00:09:44 CEST 2022 - The aliscore output appears to be empty. Check results for BUSCO: 194852at7742
## Thu Jul 14 00:09:46 CEST 2022 - The aliscore output file does not exist. Check results for BUSCO: 24566at7742
## Thu Jul 14 00:09:46 CEST 2022 - The aliscore output appears to be empty. Check results for BUSCO: 24566at7742
## Thu Jul 14 00:09:46 CEST 2022 - The aliscore output file does not exist. Check results for BUSCO: 211943at7742
## Thu Jul 14 00:09:46 CEST 2022 - The aliscore output appears to be empty. Check results for BUSCO: 211943at7742
## Thu Jul 14 00:09:48 CEST 2022 - The aliscore output file does not exist. Check results for BUSCO: 140317at7742
## Thu Jul 14 00:09:48 CEST 2022 - The aliscore output appears to be empty. Check results for BUSCO: 140317at7742
## Thu Jul 14 00:09:50 CEST 2022 - The aliscore output file does not exist. Check results for BUSCO: 42469at7742
## Thu Jul 14 00:09:50 CEST 2022 - The aliscore output appears to be empty. Check results for BUSCO: 42469at7742
## Thu Jul 14 00:09:50 CEST 2022 - The aliscore output file does not exist. Check results for BUSCO: 170024at7742
## Thu Jul 14 00:09:50 CEST 2022 - The aliscore output appears to be empty. Check results for BUSCO: 170024at7742
## Thu Jul 14 00:09:53 CEST 2022 - The aliscore output file does not exist. Check results for BUSCO: 25752at7742
## Thu Jul 14 00:09:53 CEST 2022 - The aliscore output appears to be empty. Check results for BUSCO: 25752at7742
## Thu Jul 14 00:09:55 CEST 2022 - The aliscore output file does not exist. Check results for BUSCO: 57077at7742
## Thu Jul 14 00:09:55 CEST 2022 - The aliscore output appears to be empty. Check results for BUSCO: 57077at7742
## Thu Jul 14 00:09:56 CEST 2022 - The aliscore output file does not exist. Check results for BUSCO: 26047at7742
## Thu Jul 14 00:09:56 CEST 2022 - The aliscore output appears to be empty. Check results for BUSCO: 26047at7742
## Thu Jul 14 00:09:57 CEST 2022 - The aliscore output file does not exist. Check results for BUSCO: 43750at7742
## Thu Jul 14 00:09:57 CEST 2022 - The aliscore output appears to be empty. Check results for BUSCO: 43750at7742
## Thu Jul 14 00:09:57 CEST 2022 - The aliscore output file does not exist. Check results for BUSCO: 113103at7742
## Thu Jul 14 00:09:57 CEST 2022 - The aliscore output appears to be empty. Check results for BUSCO: 113103at7742
## Thu Jul 14 00:09:59 CEST 2022 - The aliscore output file does not exist. Check results for BUSCO: 170658at7742
## Thu Jul 14 00:09:59 CEST 2022 - The aliscore output appears to be empty. Check results for BUSCO: 170658at7742
## Thu Jul 14 00:10:00 CEST 2022 - The aliscore output file does not exist. Check results for BUSCO: 120920at7742
## Thu Jul 14 00:10:00 CEST 2022 - The aliscore output appears to be empty. Check results for BUSCO: 120920at7742
## Thu Jul 14 00:10:02 CEST 2022 - The aliscore output file does not exist. Check results for BUSCO: 30774at7742
## Thu Jul 14 00:10:02 CEST 2022 - The aliscore output appears to be empty. Check results for BUSCO: 30774at7742
## Thu Jul 14 00:10:02 CEST 2022 - The aliscore output file does not exist. Check results for BUSCO: 311450at7742
## Thu Jul 14 00:10:02 CEST 2022 - The aliscore output appears to be empty. Check results for BUSCO: 311450at7742
## Thu Jul 14 00:10:05 CEST 2022 - The aliscore output file does not exist. Check results for BUSCO: 35547at7742
## Thu Jul 14 00:10:05 CEST 2022 - The aliscore output file does not exist. Check results for BUSCO: 125886at7742
## Thu Jul 14 00:10:05 CEST 2022 - The aliscore output appears to be empty. Check results for BUSCO: 35547at7742
## Thu Jul 14 00:10:05 CEST 2022 - The aliscore output appears to be empty. Check results for BUSCO: 125886at7742
## Thu Jul 14 00:10:09 CEST 2022 - The aliscore output file does not exist. Check results for BUSCO: 61914at7742
## Thu Jul 14 00:10:09 CEST 2022 - The aliscore output appears to be empty. Check results for BUSCO: 61914at7742
## Thu Jul 14 00:10:09 CEST 2022 - The aliscore output file does not exist. Check results for BUSCO: 186464at7742
## Thu Jul 14 00:10:09 CEST 2022 - The aliscore output appears to be empty. Check results for BUSCO: 186464at7742
## Thu Jul 14 00:10:09 CEST 2022 - The aliscore output file does not exist. Check results for BUSCO: 41150at7742
## Thu Jul 14 00:10:09 CEST 2022 - The aliscore output appears to be empty. Check results for BUSCO: 41150at7742
## Thu Jul 14 00:10:11 CEST 2022 - The aliscore output file does not exist. Check results for BUSCO: 31951at7742
## Thu Jul 14 00:10:11 CEST 2022 - The aliscore output appears to be empty. Check results for BUSCO: 31951at7742
## Thu Jul 14 00:10:13 CEST 2022 - The aliscore output file does not exist. Check results for BUSCO: 44800at7742
## Thu Jul 14 00:10:13 CEST 2022 - The aliscore output appears to be empty. Check results for BUSCO: 44800at7742
## Thu Jul 14 00:10:16 CEST 2022 - The aliscore output file does not exist. Check results for BUSCO: 106943at7742
## Thu Jul 14 00:10:16 CEST 2022 - The aliscore output appears to be empty. Check results for BUSCO: 106943at7742
## Thu Jul 14 00:10:16 CEST 2022 - The aliscore output file does not exist. Check results for BUSCO: 49318at7742
## Thu Jul 14 00:10:16 CEST 2022 - The aliscore output appears to be empty. Check results for BUSCO: 49318at7742
## Thu Jul 14 00:10:21 CEST 2022 - The aliscore output file does not exist. Check results for BUSCO: 146445at7742
## Thu Jul 14 00:10:21 CEST 2022 - The aliscore output appears to be empty. Check results for BUSCO: 146445at7742
## Thu Jul 14 00:10:22 CEST 2022 - The aliscore output file does not exist. Check results for BUSCO: 25752at7742
## Thu Jul 14 00:10:22 CEST 2022 - The aliscore output appears to be empty. Check results for BUSCO: 25752at7742
## Thu Jul 14 00:10:22 CEST 2022 - The aliscore output file does not exist. Check results for BUSCO: 23673at7742
## Thu Jul 14 00:10:22 CEST 2022 - The aliscore output appears to be empty. Check results for BUSCO: 23673at7742
## Thu Jul 14 00:10:23 CEST 2022 - The aliscore output file does not exist. Check results for BUSCO: 185168at7742
## Thu Jul 14 00:10:23 CEST 2022 - The aliscore output appears to be empty. Check results for BUSCO: 185168at7742
## Thu Jul 14 00:10:24 CEST 2022 - The aliscore output file does not exist. Check results for BUSCO: 19367at7742
## Thu Jul 14 00:10:24 CEST 2022 - The aliscore output appears to be empty. Check results for BUSCO: 19367at7742
## Thu Jul 14 00:10:24 CEST 2022 - The aliscore output file does not exist. Check results for BUSCO: 45658at7742
## Thu Jul 14 00:10:24 CEST 2022 - The aliscore output appears to be empty. Check results for BUSCO: 45658at7742
## Thu Jul 14 00:10:25 CEST 2022 - The aliscore output file does not exist. Check results for BUSCO: 285264at7742
## Thu Jul 14 00:10:25 CEST 2022 - The aliscore output appears to be empty. Check results for BUSCO: 285264at7742
## Thu Jul 14 00:10:28 CEST 2022 - The aliscore output file does not exist. Check results for BUSCO: 106943at7742
## Thu Jul 14 00:10:28 CEST 2022 - The aliscore output appears to be empty. Check results for BUSCO: 106943at7742
## Thu Jul 14 00:10:29 CEST 2022 - The aliscore output file does not exist. Check results for BUSCO: 35041at7742
## Thu Jul 14 00:10:29 CEST 2022 - The aliscore output appears to be empty. Check results for BUSCO: 35041at7742
## Thu Jul 14 00:10:30 CEST 2022 - The aliscore output file does not exist. Check results for BUSCO: 155652at7742
## Thu Jul 14 00:10:30 CEST 2022 - The aliscore output appears to be empty. Check results for BUSCO: 155652at7742
## Thu Jul 14 00:10:32 CEST 2022 - The aliscore output file does not exist. Check results for BUSCO: 57077at7742
## Thu Jul 14 00:10:32 CEST 2022 - The aliscore output appears to be empty. Check results for BUSCO: 57077at7742
## Thu Jul 14 00:10:33 CEST 2022 - The aliscore output file does not exist. Check results for BUSCO: 30774at7742
## Thu Jul 14 00:10:33 CEST 2022 - The aliscore output appears to be empty. Check results for BUSCO: 30774at7742
## Thu Jul 14 00:10:35 CEST 2022 - The aliscore output file does not exist. Check results for BUSCO: 25752at7742
## Thu Jul 14 00:10:35 CEST 2022 - The aliscore output appears to be empty. Check results for BUSCO: 25752at7742
## Thu Jul 14 00:10:37 CEST 2022 - The aliscore output file does not exist. Check results for BUSCO: 226989at7742
## Thu Jul 14 00:10:37 CEST 2022 - The aliscore output appears to be empty. Check results for BUSCO: 226989at7742
## Thu Jul 14 00:10:38 CEST 2022 - The aliscore output file does not exist. Check results for BUSCO: 55500at7742
## Thu Jul 14 00:10:38 CEST 2022 - The aliscore output appears to be empty. Check results for BUSCO: 55500at7742
## Thu Jul 14 00:10:38 CEST 2022 - The aliscore output file does not exist. Check results for BUSCO: 197548at7742
## Thu Jul 14 00:10:38 CEST 2022 - The aliscore output appears to be empty. Check results for BUSCO: 197548at7742
## Thu Jul 14 00:10:40 CEST 2022 - The aliscore output file does not exist. Check results for BUSCO: 106344at7742
## Thu Jul 14 00:10:40 CEST 2022 - The aliscore output appears to be empty. Check results for BUSCO: 106344at7742
## Thu Jul 14 00:10:42 CEST 2022 - The aliscore output file does not exist. Check results for BUSCO: 19072at7742
## Thu Jul 14 00:10:42 CEST 2022 - The aliscore output appears to be empty. Check results for BUSCO: 19072at7742
## Thu Jul 14 00:10:43 CEST 2022 - The aliscore output file does not exist. Check results for BUSCO: 170024at7742
## Thu Jul 14 00:10:43 CEST 2022 - The aliscore output appears to be empty. Check results for BUSCO: 170024at7742
## Thu Jul 14 00:10:44 CEST 2022 - The aliscore output file does not exist. Check results for BUSCO: 80808at7742
## Thu Jul 14 00:10:44 CEST 2022 - The aliscore output appears to be empty. Check results for BUSCO: 80808at7742
## Thu Jul 14 00:10:46 CEST 2022 - The aliscore output file does not exist. Check results for BUSCO: 40843at7742
## Thu Jul 14 00:10:46 CEST 2022 - The aliscore output appears to be empty. Check results for BUSCO: 40843at7742
## Thu Jul 14 00:10:48 CEST 2022 - The aliscore output file does not exist. Check results for BUSCO: 114216at7742
## Thu Jul 14 00:10:48 CEST 2022 - The aliscore output appears to be empty. Check results for BUSCO: 114216at7742
## Thu Jul 14 00:10:48 CEST 2022 - The aliscore output file does not exist. Check results for BUSCO: 46928at7742
## Thu Jul 14 00:10:48 CEST 2022 - The aliscore output appears to be empty. Check results for BUSCO: 46928at7742
## Thu Jul 14 00:10:49 CEST 2022 - The aliscore output file does not exist. Check results for BUSCO: 122255at7742
## Thu Jul 14 00:10:49 CEST 2022 - The aliscore output appears to be empty. Check results for BUSCO: 122255at7742
## Thu Jul 14 00:10:50 CEST 2022 - The aliscore output file does not exist. Check results for BUSCO: 102786at7742
## Thu Jul 14 00:10:50 CEST 2022 - The aliscore output appears to be empty. Check results for BUSCO: 102786at7742
## Thu Jul 14 00:10:53 CEST 2022 - The aliscore output file does not exist. Check results for BUSCO: 111254at7742
## Thu Jul 14 00:10:53 CEST 2022 - The aliscore output appears to be empty. Check results for BUSCO: 111254at7742
## Thu Jul 14 00:10:54 CEST 2022 - The aliscore output file does not exist. Check results for BUSCO: 73217at7742
## Thu Jul 14 00:10:54 CEST 2022 - The aliscore output appears to be empty. Check results for BUSCO: 73217at7742
## Thu Jul 14 00:10:55 CEST 2022 - The aliscore output file does not exist. Check results for BUSCO: 23673at7742
## Thu Jul 14 00:10:55 CEST 2022 - The aliscore output appears to be empty. Check results for BUSCO: 23673at7742
## Thu Jul 14 00:10:58 CEST 2022 - The aliscore output file does not exist. Check results for BUSCO: 37715at7742
## Thu Jul 14 00:10:58 CEST 2022 - The aliscore output appears to be empty. Check results for BUSCO: 37715at7742
## Thu Jul 14 00:10:58 CEST 2022 - The aliscore output file does not exist. Check results for BUSCO: 170024at7742
## Thu Jul 14 00:10:58 CEST 2022 - The aliscore output file does not exist. Check results for BUSCO: 164002at7742
## Thu Jul 14 00:10:58 CEST 2022 - The aliscore output appears to be empty. Check results for BUSCO: 170024at7742
## Thu Jul 14 00:10:58 CEST 2022 - The aliscore output appears to be empty. Check results for BUSCO: 164002at7742
## Thu Jul 14 00:11:00 CEST 2022 - The aliscore output file does not exist. Check results for BUSCO: 43568at7742
## Thu Jul 14 00:11:00 CEST 2022 - The aliscore output appears to be empty. Check results for BUSCO: 43568at7742
## Thu Jul 14 00:11:02 CEST 2022 - The aliscore output file does not exist. Check results for BUSCO: 87643at7742
## Thu Jul 14 00:11:02 CEST 2022 - The aliscore output appears to be empty. Check results for BUSCO: 87643at7742
## Thu Jul 14 00:11:02 CEST 2022 - The aliscore output file does not exist. Check results for BUSCO: 179099at7742
## Thu Jul 14 00:11:02 CEST 2022 - The aliscore output appears to be empty. Check results for BUSCO: 179099at7742
## Thu Jul 14 00:11:06 CEST 2022 - The aliscore output file does not exist. Check results for BUSCO: 56293at7742
## Thu Jul 14 00:11:06 CEST 2022 - The aliscore output appears to be empty. Check results for BUSCO: 56293at7742
## Thu Jul 14 00:11:07 CEST 2022 - The aliscore output file does not exist. Check results for BUSCO: 43750at7742
## Thu Jul 14 00:11:07 CEST 2022 - The aliscore output appears to be empty. Check results for BUSCO: 43750at7742
## Thu Jul 14 00:11:10 CEST 2022 - The aliscore output file does not exist. Check results for BUSCO: 68853at7742
## Thu Jul 14 00:11:10 CEST 2022 - The aliscore output appears to be empty. Check results for BUSCO: 68853at7742
## Thu Jul 14 00:11:11 CEST 2022 - The aliscore output file does not exist. Check results for BUSCO: 32511at7742
## Thu Jul 14 00:11:11 CEST 2022 - The aliscore output appears to be empty. Check results for BUSCO: 32511at7742
## Thu Jul 14 00:11:11 CEST 2022 - The aliscore output file does not exist. Check results for BUSCO: 152085at7742
## Thu Jul 14 00:11:11 CEST 2022 - The aliscore output appears to be empty. Check results for BUSCO: 152085at7742
## Thu Jul 14 00:11:13 CEST 2022 - The aliscore output file does not exist. Check results for BUSCO: 135024at7742
## Thu Jul 14 00:11:13 CEST 2022 - The aliscore output appears to be empty. Check results for BUSCO: 135024at7742
## Thu Jul 14 00:11:14 CEST 2022 - The aliscore output file does not exist. Check results for BUSCO: 24566at7742
## Thu Jul 14 00:11:14 CEST 2022 - The aliscore output appears to be empty. Check results for BUSCO: 24566at7742
## Thu Jul 14 00:11:15 CEST 2022 - The aliscore output file does not exist. Check results for BUSCO: 164002at7742
## Thu Jul 14 00:11:15 CEST 2022 - The aliscore output appears to be empty. Check results for BUSCO: 164002at7742
## Thu Jul 14 00:11:16 CEST 2022 - The aliscore output file does not exist. Check results for BUSCO: 46538at7742
## Thu Jul 14 00:11:16 CEST 2022 - The aliscore output appears to be empty. Check results for BUSCO: 46538at7742
## Thu Jul 14 00:11:19 CEST 2022 - The aliscore output file does not exist. Check results for BUSCO: 216073at7742
## Thu Jul 14 00:11:19 CEST 2022 - The aliscore output appears to be empty. Check results for BUSCO: 216073at7742
## Thu Jul 14 00:11:20 CEST 2022 - The aliscore output file does not exist. Check results for BUSCO: 75522at7742
## Thu Jul 14 00:11:20 CEST 2022 - The aliscore output appears to be empty. Check results for BUSCO: 75522at7742
## Thu Jul 14 00:11:20 CEST 2022 - The aliscore output file does not exist. Check results for BUSCO: 55500at7742
## Thu Jul 14 00:11:20 CEST 2022 - The aliscore output appears to be empty. Check results for BUSCO: 55500at7742
## Thu Jul 14 00:11:21 CEST 2022 - The aliscore output file does not exist. Check results for BUSCO: 76024at7742
## Thu Jul 14 00:11:21 CEST 2022 - The aliscore output appears to be empty. Check results for BUSCO: 76024at7742
## Thu Jul 14 00:11:24 CEST 2022 - The aliscore output file does not exist. Check results for BUSCO: 19072at7742
## Thu Jul 14 00:11:24 CEST 2022 - The aliscore output appears to be empty. Check results for BUSCO: 19072at7742
## Thu Jul 14 00:11:25 CEST 2022 - The aliscore output file does not exist. Check results for BUSCO: 124120at7742
## Thu Jul 14 00:11:25 CEST 2022 - The aliscore output appears to be empty. Check results for BUSCO: 124120at7742
## Thu Jul 14 00:11:28 CEST 2022 - The aliscore output file does not exist. Check results for BUSCO: 53792at7742
## Thu Jul 14 00:11:28 CEST 2022 - The aliscore output appears to be empty. Check results for BUSCO: 53792at7742
## Thu Jul 14 00:11:29 CEST 2022 - The aliscore output file does not exist. Check results for BUSCO: 42346at7742
## Thu Jul 14 00:11:29 CEST 2022 - The aliscore output appears to be empty. Check results for BUSCO: 42346at7742
## Thu Jul 14 00:11:30 CEST 2022 - The aliscore output file does not exist. Check results for BUSCO: 151895at7742
## Thu Jul 14 00:11:30 CEST 2022 - The aliscore output appears to be empty. Check results for BUSCO: 151895at7742
## Thu Jul 14 00:11:30 CEST 2022 - The aliscore output file does not exist. Check results for BUSCO: 74629at7742
## Thu Jul 14 00:11:30 CEST 2022 - The aliscore output file does not exist. Check results for BUSCO: 199826at7742
## Thu Jul 14 00:11:30 CEST 2022 - The aliscore output appears to be empty. Check results for BUSCO: 74629at7742
## Thu Jul 14 00:11:30 CEST 2022 - The aliscore output appears to be empty. Check results for BUSCO: 199826at7742
## Thu Jul 14 00:11:33 CEST 2022 - The aliscore output file does not exist. Check results for BUSCO: 104369at7742
## Thu Jul 14 00:11:33 CEST 2022 - The aliscore output appears to be empty. Check results for BUSCO: 104369at7742
## Thu Jul 14 00:11:34 CEST 2022 - The aliscore output file does not exist. Check results for BUSCO: 78893at7742
## Thu Jul 14 00:11:34 CEST 2022 - The aliscore output appears to be empty. Check results for BUSCO: 78893at7742
## Thu Jul 14 00:11:35 CEST 2022 - The aliscore output file does not exist. Check results for BUSCO: 124120at7742
## Thu Jul 14 00:11:35 CEST 2022 - The aliscore output appears to be empty. Check results for BUSCO: 124120at7742
## Thu Jul 14 00:11:37 CEST 2022 - The aliscore output file does not exist. Check results for BUSCO: 76947at7742
## Thu Jul 14 00:11:37 CEST 2022 - The aliscore output appears to be empty. Check results for BUSCO: 76947at7742
## Thu Jul 14 00:11:39 CEST 2022 - The aliscore output file does not exist. Check results for BUSCO: 106301at7742
## Thu Jul 14 00:11:39 CEST 2022 - The aliscore output appears to be empty. Check results for BUSCO: 106301at7742
## Thu Jul 14 00:11:40 CEST 2022 - The aliscore output file does not exist. Check results for BUSCO: 139811at7742
## Thu Jul 14 00:11:40 CEST 2022 - The aliscore output appears to be empty. Check results for BUSCO: 139811at7742
## Thu Jul 14 00:11:41 CEST 2022 - The aliscore output file does not exist. Check results for BUSCO: 75780at7742
## Thu Jul 14 00:11:41 CEST 2022 - The aliscore output appears to be empty. Check results for BUSCO: 75780at7742
## Thu Jul 14 00:11:44 CEST 2022 - The aliscore output file does not exist. Check results for BUSCO: 109869at7742
## Thu Jul 14 00:11:44 CEST 2022 - The aliscore output appears to be empty. Check results for BUSCO: 109869at7742
## Thu Jul 14 00:11:45 CEST 2022 - The aliscore output file does not exist. Check results for BUSCO: 27308at7742
## Thu Jul 14 00:11:45 CEST 2022 - The aliscore output appears to be empty. Check results for BUSCO: 27308at7742
## Thu Jul 14 00:11:46 CEST 2022 - The aliscore output file does not exist. Check results for BUSCO: 27308at7742
## Thu Jul 14 00:11:46 CEST 2022 - The aliscore output appears to be empty. Check results for BUSCO: 27308at7742
## Thu Jul 14 00:11:47 CEST 2022 - The aliscore output file does not exist. Check results for BUSCO: 74629at7742
## Thu Jul 14 00:11:47 CEST 2022 - The aliscore output appears to be empty. Check results for BUSCO: 74629at7742
## Thu Jul 14 00:11:49 CEST 2022 - The aliscore output file does not exist. Check results for BUSCO: 75522at7742
## Thu Jul 14 00:11:49 CEST 2022 - The aliscore output appears to be empty. Check results for BUSCO: 75522at7742
## Thu Jul 14 00:11:50 CEST 2022 - The aliscore output file does not exist. Check results for BUSCO: 186292at7742
## Thu Jul 14 00:11:50 CEST 2022 - The aliscore output appears to be empty. Check results for BUSCO: 186292at7742
## Thu Jul 14 00:11:52 CEST 2022 - The aliscore output file does not exist. Check results for BUSCO: 188180at7742
## Thu Jul 14 00:11:52 CEST 2022 - The aliscore output appears to be empty. Check results for BUSCO: 188180at7742
## Thu Jul 14 00:11:54 CEST 2022 - The aliscore output file does not exist. Check results for BUSCO: 56293at7742
## Thu Jul 14 00:11:54 CEST 2022 - The aliscore output appears to be empty. Check results for BUSCO: 56293at7742
## Thu Jul 14 00:11:54 CEST 2022 - The aliscore output file does not exist. Check results for BUSCO: 101309at7742
## Thu Jul 14 00:11:54 CEST 2022 - The aliscore output appears to be empty. Check results for BUSCO: 101309at7742
## Thu Jul 14 00:11:57 CEST 2022 - The aliscore output file does not exist. Check results for BUSCO: 79862at7742
## Thu Jul 14 00:11:57 CEST 2022 - The aliscore output appears to be empty. Check results for BUSCO: 79862at7742
## Thu Jul 14 00:11:58 CEST 2022 - The aliscore output file does not exist. Check results for BUSCO: 49885at7742
## Thu Jul 14 00:11:58 CEST 2022 - The aliscore output appears to be empty. Check results for BUSCO: 49885at7742
## Thu Jul 14 00:12:01 CEST 2022 - The aliscore output file does not exist. Check results for BUSCO: 60277at7742
## Thu Jul 14 00:12:01 CEST 2022 - The aliscore output appears to be empty. Check results for BUSCO: 60277at7742
## Thu Jul 14 00:12:01 CEST 2022 - The aliscore output file does not exist. Check results for BUSCO: 34542at7742
## Thu Jul 14 00:12:01 CEST 2022 - The aliscore output appears to be empty. Check results for BUSCO: 34542at7742
## Thu Jul 14 00:12:02 CEST 2022 - The aliscore output file does not exist. Check results for BUSCO: 29623at7742
## Thu Jul 14 00:12:02 CEST 2022 - The aliscore output appears to be empty. Check results for BUSCO: 29623at7742
## Thu Jul 14 00:12:04 CEST 2022 - The aliscore output file does not exist. Check results for BUSCO: 27595at7742
## Thu Jul 14 00:12:04 CEST 2022 - The aliscore output appears to be empty. Check results for BUSCO: 27595at7742
## Thu Jul 14 00:12:05 CEST 2022 - The aliscore output file does not exist. Check results for BUSCO: 163696at7742
## Thu Jul 14 00:12:05 CEST 2022 - The aliscore output appears to be empty. Check results for BUSCO: 163696at7742
## Thu Jul 14 00:12:06 CEST 2022 - The aliscore output file does not exist. Check results for BUSCO: 23568at7742
## Thu Jul 14 00:12:06 CEST 2022 - The aliscore output appears to be empty. Check results for BUSCO: 23568at7742
## Thu Jul 14 00:12:09 CEST 2022 - The aliscore output file does not exist. Check results for BUSCO: 35041at7742
## Thu Jul 14 00:12:09 CEST 2022 - The aliscore output appears to be empty. Check results for BUSCO: 35041at7742
## Thu Jul 14 00:12:10 CEST 2022 - The aliscore output file does not exist. Check results for BUSCO: 22898at7742
## Thu Jul 14 00:12:10 CEST 2022 - The aliscore output appears to be empty. Check results for BUSCO: 22898at7742
## Thu Jul 14 00:12:11 CEST 2022 - The aliscore output file does not exist. Check results for BUSCO: 69398at7742
## Thu Jul 14 00:12:11 CEST 2022 - The aliscore output appears to be empty. Check results for BUSCO: 69398at7742
## Thu Jul 14 00:12:11 CEST 2022 - The aliscore output file does not exist. Check results for BUSCO: 141686at7742
## Thu Jul 14 00:12:11 CEST 2022 - The aliscore output appears to be empty. Check results for BUSCO: 141686at7742
## Thu Jul 14 00:12:14 CEST 2022 - The aliscore output file does not exist. Check results for BUSCO: 53792at7742
## Thu Jul 14 00:12:14 CEST 2022 - The aliscore output appears to be empty. Check results for BUSCO: 53792at7742
## Thu Jul 14 00:12:14 CEST 2022 - The aliscore output file does not exist. Check results for BUSCO: 186464at7742
## Thu Jul 14 00:12:14 CEST 2022 - The aliscore output file does not exist. Check results for BUSCO: 158865at7742
## Thu Jul 14 00:12:14 CEST 2022 - The aliscore output appears to be empty. Check results for BUSCO: 186464at7742
## Thu Jul 14 00:12:14 CEST 2022 - The aliscore output appears to be empty. Check results for BUSCO: 158865at7742
## Thu Jul 14 00:12:18 CEST 2022 - The aliscore output file does not exist. Check results for BUSCO: 155652at7742
## Thu Jul 14 00:12:18 CEST 2022 - The aliscore output appears to be empty. Check results for BUSCO: 155652at7742
## Thu Jul 14 00:12:20 CEST 2022 - The aliscore output file does not exist. Check results for BUSCO: 43750at7742
## Thu Jul 14 00:12:20 CEST 2022 - The aliscore output appears to be empty. Check results for BUSCO: 43750at7742
## Thu Jul 14 00:12:21 CEST 2022 - The aliscore output file does not exist. Check results for BUSCO: 42346at7742
## Thu Jul 14 00:12:21 CEST 2022 - The aliscore output appears to be empty. Check results for BUSCO: 42346at7742
## Thu Jul 14 00:12:23 CEST 2022 - The aliscore output file does not exist. Check results for BUSCO: 34542at7742
## Thu Jul 14 00:12:23 CEST 2022 - The aliscore output appears to be empty. Check results for BUSCO: 34542at7742
## Thu Jul 14 00:12:24 CEST 2022 - The aliscore output file does not exist. Check results for BUSCO: 106855at7742
## Thu Jul 14 00:12:24 CEST 2022 - The aliscore output appears to be empty. Check results for BUSCO: 106855at7742
## Thu Jul 14 00:12:24 CEST 2022 - The aliscore output file does not exist. Check results for BUSCO: 37715at7742
## Thu Jul 14 00:12:24 CEST 2022 - The aliscore output appears to be empty. Check results for BUSCO: 37715at7742
## Thu Jul 14 00:12:24 CEST 2022 - The aliscore output file does not exist. Check results for BUSCO: 105918at7742
## Thu Jul 14 00:12:24 CEST 2022 - The aliscore output appears to be empty. Check results for BUSCO: 105918at7742
## Thu Jul 14 00:12:28 CEST 2022 - The aliscore output file does not exist. Check results for BUSCO: 57077at7742
## Thu Jul 14 00:12:28 CEST 2022 - The aliscore output appears to be empty. Check results for BUSCO: 57077at7742
## Thu Jul 14 00:12:28 CEST 2022 - The aliscore output file does not exist. Check results for BUSCO: 56293at7742
## Thu Jul 14 00:12:28 CEST 2022 - The aliscore output appears to be empty. Check results for BUSCO: 56293at7742
## Thu Jul 14 00:12:30 CEST 2022 - The aliscore output file does not exist. Check results for BUSCO: 22108at7742
## Thu Jul 14 00:12:30 CEST 2022 - The aliscore output appears to be empty. Check results for BUSCO: 22108at7742
## Thu Jul 14 00:12:31 CEST 2022 - The aliscore output file does not exist. Check results for BUSCO: 35547at7742
## Thu Jul 14 00:12:31 CEST 2022 - The aliscore output appears to be empty. Check results for BUSCO: 35547at7742
## Thu Jul 14 00:12:34 CEST 2022 - The aliscore output file does not exist. Check results for BUSCO: 138498at7742
## Thu Jul 14 00:12:34 CEST 2022 - The aliscore output appears to be empty. Check results for BUSCO: 138498at7742
## Thu Jul 14 00:12:34 CEST 2022 - The aliscore output file does not exist. Check results for BUSCO: 104369at7742
## Thu Jul 14 00:12:34 CEST 2022 - The aliscore output appears to be empty. Check results for BUSCO: 104369at7742
## Thu Jul 14 00:12:35 CEST 2022 - The aliscore output file does not exist. Check results for BUSCO: 74629at7742
## Thu Jul 14 00:12:35 CEST 2022 - The aliscore output appears to be empty. Check results for BUSCO: 74629at7742
## Thu Jul 14 00:12:37 CEST 2022 - The aliscore output file does not exist. Check results for BUSCO: 101309at7742
## Thu Jul 14 00:12:37 CEST 2022 - The aliscore output appears to be empty. Check results for BUSCO: 101309at7742
## Thu Jul 14 00:12:38 CEST 2022 - The aliscore output file does not exist. Check results for BUSCO: 27595at7742
## Thu Jul 14 00:12:39 CEST 2022 - The aliscore output appears to be empty. Check results for BUSCO: 27595at7742
## Thu Jul 14 00:12:41 CEST 2022 - The aliscore output file does not exist. Check results for BUSCO: 30774at7742
## Thu Jul 14 00:12:41 CEST 2022 - The aliscore output appears to be empty. Check results for BUSCO: 30774at7742
## Thu Jul 14 00:12:42 CEST 2022 - The aliscore output file does not exist. Check results for BUSCO: 68853at7742
## Thu Jul 14 00:12:42 CEST 2022 - The aliscore output appears to be empty. Check results for BUSCO: 68853at7742
## Thu Jul 14 00:12:45 CEST 2022 - The aliscore output file does not exist. Check results for BUSCO: 22898at7742
## Thu Jul 14 00:12:45 CEST 2022 - The aliscore output appears to be empty. Check results for BUSCO: 22898at7742
## Thu Jul 14 00:12:46 CEST 2022 - The aliscore output file does not exist. Check results for BUSCO: 34542at7742
## Thu Jul 14 00:12:46 CEST 2022 - The aliscore output appears to be empty. Check results for BUSCO: 34542at7742
## Thu Jul 14 00:12:47 CEST 2022 - The aliscore output file does not exist. Check results for BUSCO: 21546at7742
## Thu Jul 14 00:12:47 CEST 2022 - The aliscore output appears to be empty. Check results for BUSCO: 21546at7742
## Thu Jul 14 00:12:49 CEST 2022 - The aliscore output file does not exist. Check results for BUSCO: 76138at7742
## Thu Jul 14 00:12:49 CEST 2022 - The aliscore output appears to be empty. Check results for BUSCO: 76138at7742
## Thu Jul 14 00:12:50 CEST 2022 - The aliscore output file does not exist. Check results for BUSCO: 53792at7742
## Thu Jul 14 00:12:50 CEST 2022 - The aliscore output appears to be empty. Check results for BUSCO: 53792at7742
## Thu Jul 14 00:12:52 CEST 2022 - The aliscore output file does not exist. Check results for BUSCO: 49935at7742
## Thu Jul 14 00:12:52 CEST 2022 - The aliscore output appears to be empty. Check results for BUSCO: 49935at7742
## Thu Jul 14 00:12:52 CEST 2022 - The aliscore output file does not exist. Check results for BUSCO: 33320at7742
## Thu Jul 14 00:12:52 CEST 2022 - The aliscore output appears to be empty. Check results for BUSCO: 33320at7742
## Thu Jul 14 00:12:55 CEST 2022 - The aliscore output file does not exist. Check results for BUSCO: 36011at7742
## Thu Jul 14 00:12:55 CEST 2022 - The aliscore output appears to be empty. Check results for BUSCO: 36011at7742
## Thu Jul 14 00:12:55 CEST 2022 - The aliscore output file does not exist. Check results for BUSCO: 135024at7742
## Thu Jul 14 00:12:55 CEST 2022 - The aliscore output appears to be empty. Check results for BUSCO: 135024at7742
## Thu Jul 14 00:12:56 CEST 2022 - The aliscore output file does not exist. Check results for BUSCO: 106301at7742
## Thu Jul 14 00:12:56 CEST 2022 - The aliscore output appears to be empty. Check results for BUSCO: 106301at7742
## Thu Jul 14 00:12:57 CEST 2022 - The aliscore output file does not exist. Check results for BUSCO: 75522at7742
## Thu Jul 14 00:12:57 CEST 2022 - The aliscore output appears to be empty. Check results for BUSCO: 75522at7742
## Thu Jul 14 00:12:59 CEST 2022 - The aliscore output file does not exist. Check results for BUSCO: 17657at7742
## Thu Jul 14 00:12:59 CEST 2022 - The aliscore output appears to be empty. Check results for BUSCO: 17657at7742
## Thu Jul 14 00:13:00 CEST 2022 - The aliscore output file does not exist. Check results for BUSCO: 158865at7742
## Thu Jul 14 00:13:00 CEST 2022 - The aliscore output appears to be empty. Check results for BUSCO: 158865at7742
## Thu Jul 14 00:13:03 CEST 2022 - The aliscore output file does not exist. Check results for BUSCO: 223702at7742
## Thu Jul 14 00:13:03 CEST 2022 - The aliscore output appears to be empty. Check results for BUSCO: 223702at7742
## Thu Jul 14 00:13:05 CEST 2022 - The aliscore output file does not exist. Check results for BUSCO: 81797at7742
## Thu Jul 14 00:13:05 CEST 2022 - The aliscore output appears to be empty. Check results for BUSCO: 81797at7742
## Thu Jul 14 00:13:05 CEST 2022 - The aliscore output file does not exist. Check results for BUSCO: 36011at7742
## Thu Jul 14 00:13:05 CEST 2022 - The aliscore output appears to be empty. Check results for BUSCO: 36011at7742
## Thu Jul 14 00:13:06 CEST 2022 - The aliscore output file does not exist. Check results for BUSCO: 126446at7742
## Thu Jul 14 00:13:06 CEST 2022 - The aliscore output appears to be empty. Check results for BUSCO: 126446at7742
## Thu Jul 14 00:13:08 CEST 2022 - The aliscore output file does not exist. Check results for BUSCO: 200035at7742
## Thu Jul 14 00:13:08 CEST 2022 - The aliscore output appears to be empty. Check results for BUSCO: 200035at7742
## Thu Jul 14 00:13:09 CEST 2022 - The aliscore output file does not exist. Check results for BUSCO: 137061at7742
## Thu Jul 14 00:13:09 CEST 2022 - The aliscore output appears to be empty. Check results for BUSCO: 137061at7742
## Thu Jul 14 00:13:10 CEST 2022 - The aliscore output file does not exist. Check results for BUSCO: 27308at7742
## Thu Jul 14 00:13:10 CEST 2022 - The aliscore output appears to be empty. Check results for BUSCO: 27308at7742
## Thu Jul 14 00:13:13 CEST 2022 - The aliscore output file does not exist. Check results for BUSCO: 226989at7742
## Thu Jul 14 00:13:13 CEST 2022 - The aliscore output appears to be empty. Check results for BUSCO: 226989at7742
## Thu Jul 14 00:13:14 CEST 2022 - The aliscore output file does not exist. Check results for BUSCO: 24621at7742
## Thu Jul 14 00:13:15 CEST 2022 - The aliscore output appears to be empty. Check results for BUSCO: 24621at7742
## Thu Jul 14 00:13:15 CEST 2022 - The aliscore output file does not exist. Check results for BUSCO: 42346at7742
## Thu Jul 14 00:13:15 CEST 2022 - The aliscore output appears to be empty. Check results for BUSCO: 42346at7742
## Thu Jul 14 00:13:15 CEST 2022 - The aliscore output file does not exist. Check results for BUSCO: 76947at7742
## Thu Jul 14 00:13:15 CEST 2022 - The aliscore output appears to be empty. Check results for BUSCO: 76947at7742
## Thu Jul 14 00:13:18 CEST 2022 - The aliscore output file does not exist. Check results for BUSCO: 42469at7742
## Thu Jul 14 00:13:18 CEST 2022 - The aliscore output appears to be empty. Check results for BUSCO: 42469at7742
## Thu Jul 14 00:13:20 CEST 2022 - The aliscore output file does not exist. Check results for BUSCO: 226989at7742
## Thu Jul 14 00:13:20 CEST 2022 - The aliscore output appears to be empty. Check results for BUSCO: 226989at7742
## Thu Jul 14 00:13:21 CEST 2022 - The aliscore output file does not exist. Check results for BUSCO: 32396at7742
## Thu Jul 14 00:13:21 CEST 2022 - The aliscore output appears to be empty. Check results for BUSCO: 32396at7742
## Thu Jul 14 00:13:21 CEST 2022 - The aliscore output file does not exist. Check results for BUSCO: 102786at7742
## Thu Jul 14 00:13:21 CEST 2022 - The aliscore output appears to be empty. Check results for BUSCO: 102786at7742
## Thu Jul 14 00:13:24 CEST 2022 - The aliscore output file does not exist. Check results for BUSCO: 42469at7742
## Thu Jul 14 00:13:24 CEST 2022 - The aliscore output appears to be empty. Check results for BUSCO: 42469at7742
## Thu Jul 14 00:13:24 CEST 2022 - The aliscore output file does not exist. Check results for BUSCO: 76138at7742
## Thu Jul 14 00:13:24 CEST 2022 - The aliscore output appears to be empty. Check results for BUSCO: 76138at7742
## Thu Jul 14 00:13:25 CEST 2022 - The aliscore output file does not exist. Check results for BUSCO: 98098at7742
## Thu Jul 14 00:13:25 CEST 2022 - The aliscore output appears to be empty. Check results for BUSCO: 98098at7742
## Thu Jul 14 00:13:28 CEST 2022 - The aliscore output file does not exist. Check results for BUSCO: 21546at7742
## Thu Jul 14 00:13:28 CEST 2022 - The aliscore output appears to be empty. Check results for BUSCO: 21546at7742
## Thu Jul 14 00:13:28 CEST 2022 - The aliscore output file does not exist. Check results for BUSCO: 141686at7742
## Thu Jul 14 00:13:28 CEST 2022 - The aliscore output appears to be empty. Check results for BUSCO: 141686at7742
## Thu Jul 14 00:13:29 CEST 2022 - The aliscore output file does not exist. Check results for BUSCO: 141264at7742
## Thu Jul 14 00:13:30 CEST 2022 - The aliscore output appears to be empty. Check results for BUSCO: 141264at7742
## Thu Jul 14 00:13:30 CEST 2022 - The aliscore output file does not exist. Check results for BUSCO: 87643at7742
## Thu Jul 14 00:13:30 CEST 2022 - The aliscore output appears to be empty. Check results for BUSCO: 87643at7742
## Thu Jul 14 00:13:34 CEST 2022 - The aliscore output file does not exist. Check results for BUSCO: 71417at7742
## Thu Jul 14 00:13:34 CEST 2022 - The aliscore output appears to be empty. Check results for BUSCO: 71417at7742
## Thu Jul 14 00:13:34 CEST 2022 - The aliscore output file does not exist. Check results for BUSCO: 32840at7742
## Thu Jul 14 00:13:34 CEST 2022 - The aliscore output appears to be empty. Check results for BUSCO: 32840at7742
## Thu Jul 14 00:13:35 CEST 2022 - The aliscore output file does not exist. Check results for BUSCO: 35041at7742
## Thu Jul 14 00:13:35 CEST 2022 - The aliscore output appears to be empty. Check results for BUSCO: 35041at7742
## Thu Jul 14 00:13:35 CEST 2022 - The aliscore output file does not exist. Check results for BUSCO: 137301at7742
## Thu Jul 14 00:13:35 CEST 2022 - The aliscore output appears to be empty. Check results for BUSCO: 137301at7742
## Thu Jul 14 00:13:38 CEST 2022 - The aliscore output file does not exist. Check results for BUSCO: 29623at7742
## Thu Jul 14 00:13:38 CEST 2022 - The aliscore output appears to be empty. Check results for BUSCO: 29623at7742
## Thu Jul 14 00:13:39 CEST 2022 - The aliscore output file does not exist. Check results for BUSCO: 73217at7742
## Thu Jul 14 00:13:39 CEST 2022 - The aliscore output appears to be empty. Check results for BUSCO: 73217at7742
## Thu Jul 14 00:13:40 CEST 2022 - The aliscore output file does not exist. Check results for BUSCO: 311450at7742
## Thu Jul 14 00:13:40 CEST 2022 - The aliscore output appears to be empty. Check results for BUSCO: 311450at7742
## Thu Jul 14 00:13:41 CEST 2022 - The aliscore output file does not exist. Check results for BUSCO: 76947at7742
## Thu Jul 14 00:13:41 CEST 2022 - The aliscore output appears to be empty. Check results for BUSCO: 76947at7742
## Thu Jul 14 00:13:44 CEST 2022 - The aliscore output file does not exist. Check results for BUSCO: 35547at7742
## Thu Jul 14 00:13:44 CEST 2022 - The aliscore output appears to be empty. Check results for BUSCO: 35547at7742
## Thu Jul 14 00:13:44 CEST 2022 - The aliscore output file does not exist. Check results for BUSCO: 178561at7742
## Thu Jul 14 00:13:44 CEST 2022 - The aliscore output appears to be empty. Check results for BUSCO: 178561at7742
## Thu Jul 14 00:13:46 CEST 2022 - The aliscore output file does not exist. Check results for BUSCO: 109869at7742
## Thu Jul 14 00:13:46 CEST 2022 - The aliscore output appears to be empty. Check results for BUSCO: 109869at7742
## Thu Jul 14 00:13:47 CEST 2022 - The aliscore output file does not exist. Check results for BUSCO: 82704at7742
## Thu Jul 14 00:13:47 CEST 2022 - The aliscore output appears to be empty. Check results for BUSCO: 82704at7742
## Thu Jul 14 00:13:49 CEST 2022 - The aliscore output file does not exist. Check results for BUSCO: 21546at7742
## Thu Jul 14 00:13:49 CEST 2022 - The aliscore output appears to be empty. Check results for BUSCO: 21546at7742
## Thu Jul 14 00:13:52 CEST 2022 - The aliscore output file does not exist. Check results for BUSCO: 55780at7742
## Thu Jul 14 00:13:52 CEST 2022 - The aliscore output appears to be empty. Check results for BUSCO: 55780at7742
## Thu Jul 14 00:13:52 CEST 2022 - The aliscore output file does not exist. Check results for BUSCO: 88880at7742
## Thu Jul 14 00:13:53 CEST 2022 - The aliscore output appears to be empty. Check results for BUSCO: 88880at7742
## Thu Jul 14 00:13:53 CEST 2022 - The aliscore output file does not exist. Check results for BUSCO: 36011at7742
## Thu Jul 14 00:13:53 CEST 2022 - The aliscore output appears to be empty. Check results for BUSCO: 36011at7742
## Thu Jul 14 00:13:54 CEST 2022 - The aliscore output file does not exist. Check results for BUSCO: 104081at7742
## Thu Jul 14 00:13:54 CEST 2022 - The aliscore output appears to be empty. Check results for BUSCO: 104081at7742
## Thu Jul 14 00:13:55 CEST 2022 - The aliscore output file does not exist. Check results for BUSCO: 53273at7742
## Thu Jul 14 00:13:55 CEST 2022 - The aliscore output appears to be empty. Check results for BUSCO: 53273at7742
## Thu Jul 14 00:13:57 CEST 2022 - The aliscore output file does not exist. Check results for BUSCO: 171211at7742
## Thu Jul 14 00:13:58 CEST 2022 - The aliscore output appears to be empty. Check results for BUSCO: 171211at7742
## Thu Jul 14 00:13:58 CEST 2022 - The aliscore output file does not exist. Check results for BUSCO: 43568at7742
## Thu Jul 14 00:13:58 CEST 2022 - The aliscore output appears to be empty. Check results for BUSCO: 43568at7742
## Thu Jul 14 00:14:00 CEST 2022 - The aliscore output file does not exist. Check results for BUSCO: 200035at7742
## Thu Jul 14 00:14:00 CEST 2022 - The aliscore output appears to be empty. Check results for BUSCO: 200035at7742
## Thu Jul 14 00:14:02 CEST 2022 - The aliscore output file does not exist. Check results for BUSCO: 45658at7742
## Thu Jul 14 00:14:02 CEST 2022 - The aliscore output appears to be empty. Check results for BUSCO: 45658at7742
## Thu Jul 14 00:14:03 CEST 2022 - The aliscore output file does not exist. Check results for BUSCO: 253119at7742
## Thu Jul 14 00:14:03 CEST 2022 - The aliscore output appears to be empty. Check results for BUSCO: 253119at7742
## Thu Jul 14 00:14:07 CEST 2022 - The aliscore output file does not exist. Check results for BUSCO: 46839at7742
## Thu Jul 14 00:14:07 CEST 2022 - The aliscore output appears to be empty. Check results for BUSCO: 46839at7742
## Thu Jul 14 00:14:07 CEST 2022 - The aliscore output file does not exist. Check results for BUSCO: 47899at7742
## Thu Jul 14 00:14:08 CEST 2022 - The aliscore output appears to be empty. Check results for BUSCO: 47899at7742
## Thu Jul 14 00:14:08 CEST 2022 - The aliscore output file does not exist. Check results for BUSCO: 23568at7742
## Thu Jul 14 00:14:08 CEST 2022 - The aliscore output appears to be empty. Check results for BUSCO: 23568at7742
## Thu Jul 14 00:14:09 CEST 2022 - The aliscore output file does not exist. Check results for BUSCO: 156102at7742
## Thu Jul 14 00:14:09 CEST 2022 - The aliscore output appears to be empty. Check results for BUSCO: 156102at7742
## Thu Jul 14 00:14:10 CEST 2022 - The aliscore output file does not exist. Check results for BUSCO: 94920at7742
## Thu Jul 14 00:14:10 CEST 2022 - The aliscore output appears to be empty. Check results for BUSCO: 94920at7742
## Thu Jul 14 00:14:11 CEST 2022 - The aliscore output file does not exist. Check results for BUSCO: 64168at7742
## Thu Jul 14 00:14:11 CEST 2022 - The aliscore output appears to be empty. Check results for BUSCO: 64168at7742
## Thu Jul 14 00:14:13 CEST 2022 - The aliscore output file does not exist. Check results for BUSCO: 137061at7742
## Thu Jul 14 00:14:13 CEST 2022 - The aliscore output appears to be empty. Check results for BUSCO: 137061at7742
## Thu Jul 14 00:14:15 CEST 2022 - The aliscore output file does not exist. Check results for BUSCO: 23568at7742
## Thu Jul 14 00:14:15 CEST 2022 - The aliscore output appears to be empty. Check results for BUSCO: 23568at7742
## Thu Jul 14 00:14:16 CEST 2022 - The aliscore output file does not exist. Check results for BUSCO: 32840at7742
## Thu Jul 14 00:14:16 CEST 2022 - The aliscore output appears to be empty. Check results for BUSCO: 32840at7742
## Thu Jul 14 00:14:16 CEST 2022 - The aliscore output file does not exist. Check results for BUSCO: 96827at7742
## Thu Jul 14 00:14:16 CEST 2022 - The aliscore output appears to be empty. Check results for BUSCO: 96827at7742
## Thu Jul 14 00:14:19 CEST 2022 - The aliscore output file does not exist. Check results for BUSCO: 24621at7742
## Thu Jul 14 00:14:19 CEST 2022 - The aliscore output appears to be empty. Check results for BUSCO: 24621at7742
## Thu Jul 14 00:14:19 CEST 2022 - The aliscore output file does not exist. Check results for BUSCO: 46928at7742
## Thu Jul 14 00:14:19 CEST 2022 - The aliscore output appears to be empty. Check results for BUSCO: 46928at7742
## Thu Jul 14 00:14:21 CEST 2022 - The aliscore output file does not exist. Check results for BUSCO: 96827at7742
## Thu Jul 14 00:14:21 CEST 2022 - The aliscore output appears to be empty. Check results for BUSCO: 96827at7742
## Thu Jul 14 00:14:22 CEST 2022 - The aliscore output file does not exist. Check results for BUSCO: 249530at7742
## Thu Jul 14 00:14:22 CEST 2022 - The aliscore output appears to be empty. Check results for BUSCO: 249530at7742
## Thu Jul 14 00:14:23 CEST 2022 - The aliscore output file does not exist. Check results for BUSCO: 249530at7742
## Thu Jul 14 00:14:23 CEST 2022 - The aliscore output appears to be empty. Check results for BUSCO: 249530at7742
## Thu Jul 14 00:14:25 CEST 2022 - The aliscore output file does not exist. Check results for BUSCO: 105918at7742
## Thu Jul 14 00:14:25 CEST 2022 - The aliscore output appears to be empty. Check results for BUSCO: 105918at7742
## Thu Jul 14 00:14:26 CEST 2022 - The aliscore output file does not exist. Check results for BUSCO: 35830at7742
## Thu Jul 14 00:14:26 CEST 2022 - The aliscore output appears to be empty. Check results for BUSCO: 35830at7742
## Thu Jul 14 00:14:27 CEST 2022 - The aliscore output file does not exist. Check results for BUSCO: 99281at7742
## Thu Jul 14 00:14:27 CEST 2022 - The aliscore output appears to be empty. Check results for BUSCO: 99281at7742
## Thu Jul 14 00:14:29 CEST 2022 - The aliscore output file does not exist. Check results for BUSCO: 33320at7742
## Thu Jul 14 00:14:29 CEST 2022 - The aliscore output appears to be empty. Check results for BUSCO: 33320at7742
## Thu Jul 14 00:14:31 CEST 2022 - The aliscore output file does not exist. Check results for BUSCO: 200035at7742
## Thu Jul 14 00:14:31 CEST 2022 - The aliscore output appears to be empty. Check results for BUSCO: 200035at7742
## Thu Jul 14 00:14:32 CEST 2022 - The aliscore output file does not exist. Check results for BUSCO: 19367at7742
## Thu Jul 14 00:14:32 CEST 2022 - The aliscore output appears to be empty. Check results for BUSCO: 19367at7742
## Thu Jul 14 00:14:33 CEST 2022 - The aliscore output file does not exist. Check results for BUSCO: 106855at7742
## Thu Jul 14 00:14:33 CEST 2022 - The aliscore output appears to be empty. Check results for BUSCO: 106855at7742
## Thu Jul 14 00:14:36 CEST 2022 - The aliscore output file does not exist. Check results for BUSCO: 59233at7742
## Thu Jul 14 00:14:36 CEST 2022 - The aliscore output appears to be empty. Check results for BUSCO: 59233at7742
## Thu Jul 14 00:14:36 CEST 2022 - The aliscore output file does not exist. Check results for BUSCO: 32396at7742
## Thu Jul 14 00:14:36 CEST 2022 - The aliscore output appears to be empty. Check results for BUSCO: 32396at7742
## Thu Jul 14 00:14:37 CEST 2022 - The aliscore output file does not exist. Check results for BUSCO: 46928at7742
## Thu Jul 14 00:14:37 CEST 2022 - The aliscore output appears to be empty. Check results for BUSCO: 46928at7742
## Thu Jul 14 00:14:40 CEST 2022 - The aliscore output file does not exist. Check results for BUSCO: 66052at7742
## Thu Jul 14 00:14:40 CEST 2022 - The aliscore output appears to be empty. Check results for BUSCO: 66052at7742
## Thu Jul 14 00:14:40 CEST 2022 - The aliscore output file does not exist. Check results for BUSCO: 19367at7742
## Thu Jul 14 00:14:40 CEST 2022 - The aliscore output appears to be empty. Check results for BUSCO: 19367at7742
## Thu Jul 14 00:14:42 CEST 2022 - The aliscore output file does not exist. Check results for BUSCO: 57179at7742
## Thu Jul 14 00:14:42 CEST 2022 - The aliscore output appears to be empty. Check results for BUSCO: 57179at7742
## Thu Jul 14 00:14:43 CEST 2022 - The aliscore output file does not exist. Check results for BUSCO: 249530at7742
## Thu Jul 14 00:14:43 CEST 2022 - The aliscore output appears to be empty. Check results for BUSCO: 249530at7742
## Thu Jul 14 00:14:44 CEST 2022 - The aliscore output file does not exist. Check results for BUSCO: 143698at7742
## Thu Jul 14 00:14:44 CEST 2022 - The aliscore output appears to be empty. Check results for BUSCO: 143698at7742
## Thu Jul 14 00:14:46 CEST 2022 - The aliscore output file does not exist. Check results for BUSCO: 55780at7742
## Thu Jul 14 00:14:46 CEST 2022 - The aliscore output appears to be empty. Check results for BUSCO: 55780at7742
## Thu Jul 14 00:14:48 CEST 2022 - The aliscore output file does not exist. Check results for BUSCO: 32840at7742
## Thu Jul 14 00:14:48 CEST 2022 - The aliscore output appears to be empty. Check results for BUSCO: 32840at7742
## Thu Jul 14 00:14:49 CEST 2022 - The aliscore output file does not exist. Check results for BUSCO: 76138at7742
## Thu Jul 14 00:14:49 CEST 2022 - The aliscore output appears to be empty. Check results for BUSCO: 76138at7742
## Thu Jul 14 00:14:50 CEST 2022 - The aliscore output file does not exist. Check results for BUSCO: 22898at7742
## Thu Jul 14 00:14:50 CEST 2022 - The aliscore output appears to be empty. Check results for BUSCO: 22898at7742
## Thu Jul 14 00:14:51 CEST 2022 - The aliscore output file does not exist. Check results for BUSCO: 140317at7742
## Thu Jul 14 00:14:52 CEST 2022 - The aliscore output appears to be empty. Check results for BUSCO: 140317at7742
## Thu Jul 14 00:14:52 CEST 2022 - The aliscore output file does not exist. Check results for BUSCO: 91440at7742
## Thu Jul 14 00:14:52 CEST 2022 - The aliscore output appears to be empty. Check results for BUSCO: 91440at7742
## Thu Jul 14 00:14:55 CEST 2022 - The aliscore output file does not exist. Check results for BUSCO: 341080at7742
## Thu Jul 14 00:14:55 CEST 2022 - The aliscore output appears to be empty. Check results for BUSCO: 341080at7742
## Thu Jul 14 00:14:55 CEST 2022 - The aliscore output file does not exist. Check results for BUSCO: 47899at7742
## Thu Jul 14 00:14:55 CEST 2022 - The aliscore output appears to be empty. Check results for BUSCO: 47899at7742
## Thu Jul 14 00:14:57 CEST 2022 - The aliscore output file does not exist. Check results for BUSCO: 29623at7742
## Thu Jul 14 00:14:57 CEST 2022 - The aliscore output appears to be empty. Check results for BUSCO: 29623at7742
## Thu Jul 14 00:14:58 CEST 2022 - The aliscore output file does not exist. Check results for BUSCO: 96827at7742
## Thu Jul 14 00:14:58 CEST 2022 - The aliscore output appears to be empty. Check results for BUSCO: 96827at7742
## Thu Jul 14 00:14:58 CEST 2022 - The aliscore output file does not exist. Check results for BUSCO: 44485at7742
## Thu Jul 14 00:14:59 CEST 2022 - The aliscore output appears to be empty. Check results for BUSCO: 44485at7742
## Thu Jul 14 00:15:01 CEST 2022 - The aliscore output file does not exist. Check results for BUSCO: 34300at7742
## Thu Jul 14 00:15:01 CEST 2022 - The aliscore output appears to be empty. Check results for BUSCO: 34300at7742
## Thu Jul 14 00:15:02 CEST 2022 - The aliscore output file does not exist. Check results for BUSCO: 102786at7742
## Thu Jul 14 00:15:02 CEST 2022 - The aliscore output appears to be empty. Check results for BUSCO: 102786at7742
## Thu Jul 14 00:15:04 CEST 2022 - The aliscore output file does not exist. Check results for BUSCO: 64168at7742
## Thu Jul 14 00:15:04 CEST 2022 - The aliscore output appears to be empty. Check results for BUSCO: 64168at7742
## Thu Jul 14 00:15:05 CEST 2022 - The aliscore output file does not exist. Check results for BUSCO: 125886at7742
## Thu Jul 14 00:15:06 CEST 2022 - The aliscore output appears to be empty. Check results for BUSCO: 125886at7742
## Thu Jul 14 00:15:06 CEST 2022 - The aliscore output file does not exist. Check results for BUSCO: 44485at7742
## Thu Jul 14 00:15:06 CEST 2022 - The aliscore output appears to be empty. Check results for BUSCO: 44485at7742
## Thu Jul 14 00:15:08 CEST 2022 - The aliscore output file does not exist. Check results for BUSCO: 166801at7742
## Thu Jul 14 00:15:08 CEST 2022 - The aliscore output appears to be empty. Check results for BUSCO: 166801at7742
## Thu Jul 14 00:15:10 CEST 2022 - The aliscore output file does not exist. Check results for BUSCO: 49993at7742
## Thu Jul 14 00:15:10 CEST 2022 - The aliscore output appears to be empty. Check results for BUSCO: 49993at7742
## Thu Jul 14 00:15:11 CEST 2022 - The aliscore output file does not exist. Check results for BUSCO: 80808at7742
## Thu Jul 14 00:15:11 CEST 2022 - The aliscore output appears to be empty. Check results for BUSCO: 80808at7742
## Thu Jul 14 00:15:13 CEST 2022 - The aliscore output file does not exist. Check results for BUSCO: 41524at7742
## Thu Jul 14 00:15:13 CEST 2022 - The aliscore output appears to be empty. Check results for BUSCO: 41524at7742
## Thu Jul 14 00:15:14 CEST 2022 - The aliscore output file does not exist. Check results for BUSCO: 35830at7742
## Thu Jul 14 00:15:14 CEST 2022 - The aliscore output appears to be empty. Check results for BUSCO: 35830at7742
## Thu Jul 14 00:15:14 CEST 2022 - The aliscore output file does not exist. Check results for BUSCO: 54892at7742
## Thu Jul 14 00:15:14 CEST 2022 - The aliscore output appears to be empty. Check results for BUSCO: 54892at7742
## Thu Jul 14 00:15:17 CEST 2022 - The aliscore output file does not exist. Check results for BUSCO: 49935at7742
## Thu Jul 14 00:15:18 CEST 2022 - The aliscore output appears to be empty. Check results for BUSCO: 49935at7742
## Thu Jul 14 00:15:18 CEST 2022 - The aliscore output file does not exist. Check results for BUSCO: 34233at7742
## Thu Jul 14 00:15:18 CEST 2022 - The aliscore output appears to be empty. Check results for BUSCO: 34233at7742
## Thu Jul 14 00:15:18 CEST 2022 - The aliscore output file does not exist. Check results for BUSCO: 260562at7742
## Thu Jul 14 00:15:18 CEST 2022 - The aliscore output appears to be empty. Check results for BUSCO: 260562at7742
## Thu Jul 14 00:15:20 CEST 2022 - The aliscore output file does not exist. Check results for BUSCO: 247622at7742
## Thu Jul 14 00:15:20 CEST 2022 - The aliscore output appears to be empty. Check results for BUSCO: 247622at7742
## Thu Jul 14 00:15:21 CEST 2022 - The aliscore output file does not exist. Check results for BUSCO: 51434at7742
## Thu Jul 14 00:15:21 CEST 2022 - The aliscore output appears to be empty. Check results for BUSCO: 51434at7742
## Thu Jul 14 00:15:23 CEST 2022 - The aliscore output file does not exist. Check results for BUSCO: 82622at7742
## Thu Jul 14 00:15:23 CEST 2022 - The aliscore output appears to be empty. Check results for BUSCO: 82622at7742
## Thu Jul 14 00:15:25 CEST 2022 - The aliscore output file does not exist. Check results for BUSCO: 116202at7742
## Thu Jul 14 00:15:25 CEST 2022 - The aliscore output appears to be empty. Check results for BUSCO: 116202at7742
## Thu Jul 14 00:15:27 CEST 2022 - The aliscore output file does not exist. Check results for BUSCO: 125886at7742
## Thu Jul 14 00:15:27 CEST 2022 - The aliscore output appears to be empty. Check results for BUSCO: 125886at7742
## Thu Jul 14 00:15:28 CEST 2022 - The aliscore output file does not exist. Check results for BUSCO: 315842at7742
## Thu Jul 14 00:15:29 CEST 2022 - The aliscore output appears to be empty. Check results for BUSCO: 315842at7742
## Thu Jul 14 00:15:29 CEST 2022 - The aliscore output file does not exist. Check results for BUSCO: 22108at7742
## Thu Jul 14 00:15:29 CEST 2022 - The aliscore output appears to be empty. Check results for BUSCO: 22108at7742
## Thu Jul 14 00:15:31 CEST 2022 - The aliscore output file does not exist. Check results for BUSCO: 35830at7742
## Thu Jul 14 00:15:31 CEST 2022 - The aliscore output appears to be empty. Check results for BUSCO: 35830at7742
## Thu Jul 14 00:15:31 CEST 2022 - The aliscore output file does not exist. Check results for BUSCO: 104369at7742
## Thu Jul 14 00:15:32 CEST 2022 - The aliscore output appears to be empty. Check results for BUSCO: 104369at7742
## Thu Jul 14 00:15:34 CEST 2022 - The aliscore output file does not exist. Check results for BUSCO: 34300at7742
## Thu Jul 14 00:15:34 CEST 2022 - The aliscore output appears to be empty. Check results for BUSCO: 34300at7742
## Thu Jul 14 00:15:35 CEST 2022 - The aliscore output file does not exist. Check results for BUSCO: 98098at7742
## Thu Jul 14 00:15:35 CEST 2022 - The aliscore output appears to be empty. Check results for BUSCO: 98098at7742
## Thu Jul 14 00:15:35 CEST 2022 - The aliscore output file does not exist. Check results for BUSCO: 46839at7742
## Thu Jul 14 00:15:35 CEST 2022 - The aliscore output appears to be empty. Check results for BUSCO: 46839at7742
## Thu Jul 14 00:15:38 CEST 2022 - The aliscore output file does not exist. Check results for BUSCO: 80808at7742
## Thu Jul 14 00:15:38 CEST 2022 - The aliscore output appears to be empty. Check results for BUSCO: 80808at7742
## Thu Jul 14 00:15:38 CEST 2022 - The aliscore output file does not exist. Check results for BUSCO: 78893at7742
## Thu Jul 14 00:15:38 CEST 2022 - The aliscore output appears to be empty. Check results for BUSCO: 78893at7742
## Thu Jul 14 00:15:42 CEST 2022 - The aliscore output file does not exist. Check results for BUSCO: 133758at7742
## Thu Jul 14 00:15:42 CEST 2022 - The aliscore output appears to be empty. Check results for BUSCO: 133758at7742
## Thu Jul 14 00:15:43 CEST 2022 - The aliscore output file does not exist. Check results for BUSCO: 199826at7742
## Thu Jul 14 00:15:43 CEST 2022 - The aliscore output appears to be empty. Check results for BUSCO: 199826at7742
## Thu Jul 14 00:15:44 CEST 2022 - The aliscore output file does not exist. Check results for BUSCO: 225400at7742
## Thu Jul 14 00:15:44 CEST 2022 - The aliscore output appears to be empty. Check results for BUSCO: 225400at7742
## Thu Jul 14 00:15:45 CEST 2022 - The aliscore output file does not exist. Check results for BUSCO: 341080at7742
## Thu Jul 14 00:15:45 CEST 2022 - The aliscore output appears to be empty. Check results for BUSCO: 341080at7742
## Thu Jul 14 00:15:48 CEST 2022 - The aliscore output file does not exist. Check results for BUSCO: 24621at7742
## Thu Jul 14 00:15:48 CEST 2022 - The aliscore output appears to be empty. Check results for BUSCO: 24621at7742
## Thu Jul 14 00:15:48 CEST 2022 - The aliscore output file does not exist. Check results for BUSCO: 105918at7742
## Thu Jul 14 00:15:48 CEST 2022 - The aliscore output appears to be empty. Check results for BUSCO: 105918at7742
## Thu Jul 14 00:15:50 CEST 2022 - The aliscore output file does not exist. Check results for BUSCO: 260562at7742
## Thu Jul 14 00:15:50 CEST 2022 - The aliscore output appears to be empty. Check results for BUSCO: 260562at7742
## Thu Jul 14 00:15:51 CEST 2022 - The aliscore output file does not exist. Check results for BUSCO: 106855at7742
## Thu Jul 14 00:15:51 CEST 2022 - The aliscore output appears to be empty. Check results for BUSCO: 106855at7742
## Thu Jul 14 00:15:53 CEST 2022 - The aliscore output file does not exist. Check results for BUSCO: 36575at7742
## Thu Jul 14 00:15:53 CEST 2022 - The aliscore output appears to be empty. Check results for BUSCO: 36575at7742
## Thu Jul 14 00:15:53 CEST 2022 - The aliscore output file does not exist. Check results for BUSCO: 286974at7742
## Thu Jul 14 00:15:53 CEST 2022 - The aliscore output appears to be empty. Check results for BUSCO: 286974at7742
## Thu Jul 14 00:15:54 CEST 2022 - The aliscore output file does not exist. Check results for BUSCO: 164399at7742
## Thu Jul 14 00:15:55 CEST 2022 - The aliscore output appears to be empty. Check results for BUSCO: 164399at7742
## Thu Jul 14 00:15:57 CEST 2022 - The aliscore output file does not exist. Check results for BUSCO: 41524at7742
## Thu Jul 14 00:15:57 CEST 2022 - The aliscore output appears to be empty. Check results for BUSCO: 41524at7742
## Thu Jul 14 00:15:59 CEST 2022 - The aliscore output file does not exist. Check results for BUSCO: 91440at7742
## Thu Jul 14 00:15:59 CEST 2022 - The aliscore output appears to be empty. Check results for BUSCO: 91440at7742
## Thu Jul 14 00:16:01 CEST 2022 - The aliscore output file does not exist. Check results for BUSCO: 71963at7742
## Thu Jul 14 00:16:01 CEST 2022 - The aliscore output appears to be empty. Check results for BUSCO: 71963at7742
## Thu Jul 14 00:16:02 CEST 2022 - The aliscore output file does not exist. Check results for BUSCO: 98098at7742
## Thu Jul 14 00:16:02 CEST 2022 - The aliscore output appears to be empty. Check results for BUSCO: 98098at7742
## Thu Jul 14 00:16:02 CEST 2022 - The aliscore output file does not exist. Check results for BUSCO: 120920at7742
## Thu Jul 14 00:16:02 CEST 2022 - The aliscore output appears to be empty. Check results for BUSCO: 120920at7742
## Thu Jul 14 00:16:04 CEST 2022 - The aliscore output file does not exist. Check results for BUSCO: 55780at7742
## Thu Jul 14 00:16:04 CEST 2022 - The aliscore output appears to be empty. Check results for BUSCO: 55780at7742
## Thu Jul 14 00:16:06 CEST 2022 - The aliscore output file does not exist. Check results for BUSCO: 49993at7742
## Thu Jul 14 00:16:07 CEST 2022 - The aliscore output appears to be empty. Check results for BUSCO: 49993at7742
## Thu Jul 14 00:16:07 CEST 2022 - The aliscore output file does not exist. Check results for BUSCO: 164399at7742
## Thu Jul 14 00:16:07 CEST 2022 - The aliscore output appears to be empty. Check results for BUSCO: 164399at7742
## Thu Jul 14 00:16:08 CEST 2022 - The aliscore output file does not exist. Check results for BUSCO: 60277at7742
## Thu Jul 14 00:16:08 CEST 2022 - The aliscore output appears to be empty. Check results for BUSCO: 60277at7742
## Thu Jul 14 00:16:08 CEST 2022 - The aliscore output file does not exist. Check results for BUSCO: 44485at7742
## Thu Jul 14 00:16:08 CEST 2022 - The aliscore output appears to be empty. Check results for BUSCO: 44485at7742
## Thu Jul 14 00:16:10 CEST 2022 - The aliscore output file does not exist. Check results for BUSCO: 57824at7742
## Thu Jul 14 00:16:10 CEST 2022 - The aliscore output appears to be empty. Check results for BUSCO: 57824at7742
## Thu Jul 14 00:16:14 CEST 2022 - The aliscore output file does not exist. Check results for BUSCO: 164399at7742
## Thu Jul 14 00:16:14 CEST 2022 - The aliscore output appears to be empty. Check results for BUSCO: 164399at7742
## Thu Jul 14 00:16:14 CEST 2022 - The aliscore output file does not exist. Check results for BUSCO: 33222at7742
## Thu Jul 14 00:16:14 CEST 2022 - The aliscore output appears to be empty. Check results for BUSCO: 33222at7742
## Thu Jul 14 00:16:15 CEST 2022 - The aliscore output file does not exist. Check results for BUSCO: 152755at7742
## Thu Jul 14 00:16:15 CEST 2022 - The aliscore output appears to be empty. Check results for BUSCO: 152755at7742
## Thu Jul 14 00:16:18 CEST 2022 - The aliscore output file does not exist. Check results for BUSCO: 33320at7742
## Thu Jul 14 00:16:18 CEST 2022 - The aliscore output appears to be empty. Check results for BUSCO: 33320at7742
## Thu Jul 14 00:16:19 CEST 2022 - The aliscore output file does not exist. Check results for BUSCO: 20356at7742
## Thu Jul 14 00:16:19 CEST 2022 - The aliscore output appears to be empty. Check results for BUSCO: 20356at7742
## Thu Jul 14 00:16:21 CEST 2022 - The aliscore output file does not exist. Check results for BUSCO: 156689at7742
## Thu Jul 14 00:16:21 CEST 2022 - The aliscore output appears to be empty. Check results for BUSCO: 156689at7742
## Thu Jul 14 00:16:22 CEST 2022 - The aliscore output file does not exist. Check results for BUSCO: 26714at7742
## Thu Jul 14 00:16:22 CEST 2022 - The aliscore output appears to be empty. Check results for BUSCO: 26714at7742
## Thu Jul 14 00:16:24 CEST 2022 - The aliscore output file does not exist. Check results for BUSCO: 41524at7742
## Thu Jul 14 00:16:24 CEST 2022 - The aliscore output appears to be empty. Check results for BUSCO: 41524at7742
## Thu Jul 14 00:16:24 CEST 2022 - The aliscore output file does not exist. Check results for BUSCO: 104081at7742
## Thu Jul 14 00:16:24 CEST 2022 - The aliscore output appears to be empty. Check results for BUSCO: 104081at7742
## Thu Jul 14 00:16:25 CEST 2022 - The aliscore output file does not exist. Check results for BUSCO: 120920at7742
## Thu Jul 14 00:16:25 CEST 2022 - The aliscore output appears to be empty. Check results for BUSCO: 120920at7742
## Thu Jul 14 00:16:27 CEST 2022 - The aliscore output file does not exist. Check results for BUSCO: 162368at7742
## Thu Jul 14 00:16:27 CEST 2022 - The aliscore output appears to be empty. Check results for BUSCO: 162368at7742
## Thu Jul 14 00:16:28 CEST 2022 - The aliscore output file does not exist. Check results for BUSCO: 74297at7742
## Thu Jul 14 00:16:28 CEST 2022 - The aliscore output appears to be empty. Check results for BUSCO: 74297at7742
## Thu Jul 14 00:16:30 CEST 2022 - The aliscore output file does not exist. Check results for BUSCO: 260562at7742
## Thu Jul 14 00:16:30 CEST 2022 - The aliscore output appears to be empty. Check results for BUSCO: 260562at7742
## Thu Jul 14 00:16:31 CEST 2022 - The aliscore output file does not exist. Check results for BUSCO: 162368at7742
## Thu Jul 14 00:16:31 CEST 2022 - The aliscore output appears to be empty. Check results for BUSCO: 162368at7742
## Thu Jul 14 00:16:33 CEST 2022 - The aliscore output file does not exist. Check results for BUSCO: 341080at7742
## Thu Jul 14 00:16:33 CEST 2022 - The aliscore output appears to be empty. Check results for BUSCO: 341080at7742
## Thu Jul 14 00:16:34 CEST 2022 - The aliscore output file does not exist. Check results for BUSCO: 140098at7742
## Thu Jul 14 00:16:34 CEST 2022 - The aliscore output appears to be empty. Check results for BUSCO: 140098at7742
## Thu Jul 14 00:16:34 CEST 2022 - The aliscore output file does not exist. Check results for BUSCO: 104484at7742
## Thu Jul 14 00:16:34 CEST 2022 - The aliscore output appears to be empty. Check results for BUSCO: 104484at7742
## Thu Jul 14 00:16:38 CEST 2022 - The aliscore output file does not exist. Check results for BUSCO: 315842at7742
## Thu Jul 14 00:16:38 CEST 2022 - The aliscore output appears to be empty. Check results for BUSCO: 315842at7742
## Thu Jul 14 00:16:38 CEST 2022 - The aliscore output file does not exist. Check results for BUSCO: 151895at7742
## Thu Jul 14 00:16:38 CEST 2022 - The aliscore output appears to be empty. Check results for BUSCO: 151895at7742
## Thu Jul 14 00:16:41 CEST 2022 - The aliscore output file does not exist. Check results for BUSCO: 127587at7742
## Thu Jul 14 00:16:41 CEST 2022 - The aliscore output appears to be empty. Check results for BUSCO: 127587at7742
## Thu Jul 14 00:16:41 CEST 2022 - The aliscore output file does not exist. Check results for BUSCO: 20356at7742
## Thu Jul 14 00:16:41 CEST 2022 - The aliscore output appears to be empty. Check results for BUSCO: 20356at7742
## Thu Jul 14 00:16:41 CEST 2022 - The aliscore output file does not exist. Check results for BUSCO: 34117at7742
## Thu Jul 14 00:16:42 CEST 2022 - The aliscore output appears to be empty. Check results for BUSCO: 34117at7742
## Thu Jul 14 00:16:43 CEST 2022 - The aliscore output file does not exist. Check results for BUSCO: 25377at7742
## Thu Jul 14 00:16:43 CEST 2022 - The aliscore output appears to be empty. Check results for BUSCO: 25377at7742
## Thu Jul 14 00:16:43 CEST 2022 - The aliscore output file does not exist. Check results for BUSCO: 152755at7742
## Thu Jul 14 00:16:43 CEST 2022 - The aliscore output appears to be empty. Check results for BUSCO: 152755at7742
## Thu Jul 14 00:16:47 CEST 2022 - The aliscore output file does not exist. Check results for BUSCO: 63488at7742
## Thu Jul 14 00:16:47 CEST 2022 - The aliscore output appears to be empty. Check results for BUSCO: 63488at7742
## Thu Jul 14 00:16:47 CEST 2022 - The aliscore output file does not exist. Check results for BUSCO: 22108at7742
## Thu Jul 14 00:16:47 CEST 2022 - The aliscore output appears to be empty. Check results for BUSCO: 22108at7742
## Thu Jul 14 00:16:49 CEST 2022 - The aliscore output file does not exist. Check results for BUSCO: 137301at7742
## Thu Jul 14 00:16:49 CEST 2022 - The aliscore output appears to be empty. Check results for BUSCO: 137301at7742
## Thu Jul 14 00:16:50 CEST 2022 - The aliscore output file does not exist. Check results for BUSCO: 145564at7742
## Thu Jul 14 00:16:50 CEST 2022 - The aliscore output appears to be empty. Check results for BUSCO: 145564at7742
## Thu Jul 14 00:16:52 CEST 2022 - The aliscore output file does not exist. Check results for BUSCO: 103478at7742
## Thu Jul 14 00:16:52 CEST 2022 - The aliscore output appears to be empty. Check results for BUSCO: 103478at7742
## Thu Jul 14 00:16:53 CEST 2022 - The aliscore output file does not exist. Check results for BUSCO: 100129at7742
## Thu Jul 14 00:16:53 CEST 2022 - The aliscore output appears to be empty. Check results for BUSCO: 100129at7742
## Thu Jul 14 00:16:53 CEST 2022 - The aliscore output file does not exist. Check results for BUSCO: 17657at7742
## Thu Jul 14 00:16:53 CEST 2022 - The aliscore output appears to be empty. Check results for BUSCO: 17657at7742
## Thu Jul 14 00:16:56 CEST 2022 - The aliscore output file does not exist. Check results for BUSCO: 32396at7742
## Thu Jul 14 00:16:56 CEST 2022 - The aliscore output appears to be empty. Check results for BUSCO: 32396at7742
## Thu Jul 14 00:16:57 CEST 2022 - The aliscore output file does not exist. Check results for BUSCO: 253119at7742
## Thu Jul 14 00:16:57 CEST 2022 - The aliscore output appears to be empty. Check results for BUSCO: 253119at7742
## Thu Jul 14 00:16:58 CEST 2022 - The aliscore output file does not exist. Check results for BUSCO: 34233at7742
## Thu Jul 14 00:16:58 CEST 2022 - The aliscore output appears to be empty. Check results for BUSCO: 34233at7742
## Thu Jul 14 00:16:58 CEST 2022 - The aliscore output file does not exist. Check results for BUSCO: 218806at7742
## Thu Jul 14 00:16:58 CEST 2022 - The aliscore output appears to be empty. Check results for BUSCO: 218806at7742
## Thu Jul 14 00:17:01 CEST 2022 - The aliscore output file does not exist. Check results for BUSCO: 25377at7742
## Thu Jul 14 00:17:01 CEST 2022 - The aliscore output appears to be empty. Check results for BUSCO: 25377at7742
## Thu Jul 14 00:17:03 CEST 2022 - The aliscore output file does not exist. Check results for BUSCO: 66052at7742
## Thu Jul 14 00:17:03 CEST 2022 - The aliscore output appears to be empty. Check results for BUSCO: 66052at7742
## Thu Jul 14 00:17:04 CEST 2022 - The aliscore output file does not exist. Check results for BUSCO: 20356at7742
## Thu Jul 14 00:17:04 CEST 2022 - The aliscore output appears to be empty. Check results for BUSCO: 20356at7742
## Thu Jul 14 00:17:06 CEST 2022 - The aliscore output file does not exist. Check results for BUSCO: 194852at7742
## Thu Jul 14 00:17:06 CEST 2022 - The aliscore output appears to be empty. Check results for BUSCO: 194852at7742
## Thu Jul 14 00:17:07 CEST 2022 - The aliscore output file does not exist. Check results for BUSCO: 104081at7742
## Thu Jul 14 00:17:07 CEST 2022 - The aliscore output appears to be empty. Check results for BUSCO: 104081at7742
## Thu Jul 14 00:17:08 CEST 2022 - The aliscore output file does not exist. Check results for BUSCO: 91440at7742
## Thu Jul 14 00:17:08 CEST 2022 - The aliscore output appears to be empty. Check results for BUSCO: 91440at7742
## Thu Jul 14 00:17:10 CEST 2022 - The aliscore output file does not exist. Check results for BUSCO: 82704at7742
## Thu Jul 14 00:17:10 CEST 2022 - The aliscore output appears to be empty. Check results for BUSCO: 82704at7742
## Thu Jul 14 00:17:11 CEST 2022 - The aliscore output file does not exist. Check results for BUSCO: 43285at7742
## Thu Jul 14 00:17:11 CEST 2022 - The aliscore output appears to be empty. Check results for BUSCO: 43285at7742
## Thu Jul 14 00:17:12 CEST 2022 - The aliscore output file does not exist. Check results for BUSCO: 88880at7742
## Thu Jul 14 00:17:12 CEST 2022 - The aliscore output appears to be empty. Check results for BUSCO: 88880at7742
## Thu Jul 14 00:17:13 CEST 2022 - The aliscore output file does not exist. Check results for BUSCO: 25377at7742
## Thu Jul 14 00:17:13 CEST 2022 - The aliscore output appears to be empty. Check results for BUSCO: 25377at7742
## Thu Jul 14 00:17:14 CEST 2022 - The aliscore output file does not exist. Check results for BUSCO: 82622at7742
## Thu Jul 14 00:17:14 CEST 2022 - The aliscore output file does not exist. Check results for BUSCO: 34117at7742
## Thu Jul 14 00:17:14 CEST 2022 - The aliscore output appears to be empty. Check results for BUSCO: 82622at7742
## Thu Jul 14 00:17:14 CEST 2022 - The aliscore output appears to be empty. Check results for BUSCO: 34117at7742
## Thu Jul 14 00:17:17 CEST 2022 - The aliscore output file does not exist. Check results for BUSCO: 57824at7742
## Thu Jul 14 00:17:17 CEST 2022 - The aliscore output appears to be empty. Check results for BUSCO: 57824at7742
## Thu Jul 14 00:17:18 CEST 2022 - The aliscore output file does not exist. Check results for BUSCO: 45427at7742
## Thu Jul 14 00:17:18 CEST 2022 - The aliscore output appears to be empty. Check results for BUSCO: 45427at7742
## Thu Jul 14 00:17:20 CEST 2022 - The aliscore output file does not exist. Check results for BUSCO: 177338at7742
## Thu Jul 14 00:17:20 CEST 2022 - The aliscore output appears to be empty. Check results for BUSCO: 177338at7742
## Thu Jul 14 00:17:21 CEST 2022 - The aliscore output file does not exist. Check results for BUSCO: 61105at7742
## Thu Jul 14 00:17:21 CEST 2022 - The aliscore output appears to be empty. Check results for BUSCO: 61105at7742
## Thu Jul 14 00:17:23 CEST 2022 - The aliscore output file does not exist. Check results for BUSCO: 26714at7742
## Thu Jul 14 00:17:23 CEST 2022 - The aliscore output appears to be empty. Check results for BUSCO: 26714at7742
## Thu Jul 14 00:17:23 CEST 2022 - The aliscore output file does not exist. Check results for BUSCO: 162368at7742
## Thu Jul 14 00:17:23 CEST 2022 - The aliscore output appears to be empty. Check results for BUSCO: 162368at7742
## Thu Jul 14 00:17:24 CEST 2022 - The aliscore output file does not exist. Check results for BUSCO: 311450at7742
## Thu Jul 14 00:17:24 CEST 2022 - The aliscore output appears to be empty. Check results for BUSCO: 311450at7742
## Thu Jul 14 00:17:27 CEST 2022 - The aliscore output file does not exist. Check results for BUSCO: 212520at7742
## Thu Jul 14 00:17:27 CEST 2022 - The aliscore output appears to be empty. Check results for BUSCO: 212520at7742
## Thu Jul 14 00:17:29 CEST 2022 - The aliscore output file does not exist. Check results for BUSCO: 153497at7742
## Thu Jul 14 00:17:29 CEST 2022 - The aliscore output appears to be empty. Check results for BUSCO: 153497at7742
## Thu Jul 14 00:17:30 CEST 2022 - The aliscore output file does not exist. Check results for BUSCO: 34117at7742
## Thu Jul 14 00:17:30 CEST 2022 - The aliscore output appears to be empty. Check results for BUSCO: 34117at7742
## Thu Jul 14 00:17:31 CEST 2022 - The aliscore output file does not exist. Check results for BUSCO: 140683at7742
## Thu Jul 14 00:17:31 CEST 2022 - The aliscore output appears to be empty. Check results for BUSCO: 140683at7742
## Thu Jul 14 00:17:33 CEST 2022 - The aliscore output file does not exist. Check results for BUSCO: 32286at7742
## Thu Jul 14 00:17:33 CEST 2022 - The aliscore output appears to be empty. Check results for BUSCO: 32286at7742
## Thu Jul 14 00:17:34 CEST 2022 - The aliscore output file does not exist. Check results for BUSCO: 117682at7742
## Thu Jul 14 00:17:34 CEST 2022 - The aliscore output appears to be empty. Check results for BUSCO: 117682at7742
## Thu Jul 14 00:17:35 CEST 2022 - The aliscore output file does not exist. Check results for BUSCO: 77231at7742
## Thu Jul 14 00:17:35 CEST 2022 - The aliscore output appears to be empty. Check results for BUSCO: 77231at7742
## Thu Jul 14 00:17:37 CEST 2022 - The aliscore output file does not exist. Check results for BUSCO: 137061at7742
## Thu Jul 14 00:17:37 CEST 2022 - The aliscore output appears to be empty. Check results for BUSCO: 137061at7742
## Thu Jul 14 00:17:37 CEST 2022 - The aliscore output file does not exist. Check results for BUSCO: 26890at7742
## Thu Jul 14 00:17:38 CEST 2022 - The aliscore output appears to be empty. Check results for BUSCO: 26890at7742
## Thu Jul 14 00:17:40 CEST 2022 - The aliscore output file does not exist. Check results for BUSCO: 48275at7742
## Thu Jul 14 00:17:41 CEST 2022 - The aliscore output appears to be empty. Check results for BUSCO: 48275at7742
## Thu Jul 14 00:17:41 CEST 2022 - The aliscore output file does not exist. Check results for BUSCO: 139811at7742
## Thu Jul 14 00:17:41 CEST 2022 - The aliscore output appears to be empty. Check results for BUSCO: 139811at7742
## Thu Jul 14 00:17:42 CEST 2022 - The aliscore output file does not exist. Check results for BUSCO: 223702at7742
## Thu Jul 14 00:17:42 CEST 2022 - The aliscore output appears to be empty. Check results for BUSCO: 223702at7742
## Thu Jul 14 00:17:43 CEST 2022 - The aliscore output file does not exist. Check results for BUSCO: 47838at7742
## Thu Jul 14 00:17:43 CEST 2022 - The aliscore output appears to be empty. Check results for BUSCO: 47838at7742
## Thu Jul 14 00:17:45 CEST 2022 - The aliscore output file does not exist. Check results for BUSCO: 163696at7742
## Thu Jul 14 00:17:45 CEST 2022 - The aliscore output appears to be empty. Check results for BUSCO: 163696at7742
## Thu Jul 14 00:17:47 CEST 2022 - The aliscore output file does not exist. Check results for BUSCO: 34233at7742
## Thu Jul 14 00:17:47 CEST 2022 - The aliscore output appears to be empty. Check results for BUSCO: 34233at7742
## Thu Jul 14 00:17:47 CEST 2022 - The aliscore output file does not exist. Check results for BUSCO: 150697at7742
## Thu Jul 14 00:17:48 CEST 2022 - The aliscore output appears to be empty. Check results for BUSCO: 150697at7742
## Thu Jul 14 00:17:50 CEST 2022 - The aliscore output file does not exist. Check results for BUSCO: 76024at7742
## Thu Jul 14 00:17:50 CEST 2022 - The aliscore output appears to be empty. Check results for BUSCO: 76024at7742
## Thu Jul 14 00:17:51 CEST 2022 - The aliscore output file does not exist. Check results for BUSCO: 315842at7742
## Thu Jul 14 00:17:51 CEST 2022 - The aliscore output appears to be empty. Check results for BUSCO: 315842at7742
## Thu Jul 14 00:17:52 CEST 2022 - The aliscore output file does not exist. Check results for BUSCO: 152755at7742
## Thu Jul 14 00:17:52 CEST 2022 - The aliscore output appears to be empty. Check results for BUSCO: 152755at7742
## Thu Jul 14 00:17:53 CEST 2022 - The aliscore output file does not exist. Check results for BUSCO: 404163at7742
## Thu Jul 14 00:17:53 CEST 2022 - The aliscore output appears to be empty. Check results for BUSCO: 404163at7742
## Thu Jul 14 00:17:55 CEST 2022 - The aliscore output file does not exist. Check results for BUSCO: 20951at7742
## Thu Jul 14 00:17:55 CEST 2022 - The aliscore output appears to be empty. Check results for BUSCO: 20951at7742
## Thu Jul 14 00:17:57 CEST 2022 - The aliscore output file does not exist. Check results for BUSCO: 36575at7742
## Thu Jul 14 00:17:57 CEST 2022 - The aliscore output appears to be empty. Check results for BUSCO: 36575at7742
## Thu Jul 14 00:17:58 CEST 2022 - The aliscore output file does not exist. Check results for BUSCO: 111509at7742
## Thu Jul 14 00:17:58 CEST 2022 - The aliscore output appears to be empty. Check results for BUSCO: 111509at7742
## Thu Jul 14 00:18:00 CEST 2022 - The aliscore output file does not exist. Check results for BUSCO: 35211at7742
## Thu Jul 14 00:18:00 CEST 2022 - The aliscore output appears to be empty. Check results for BUSCO: 35211at7742
## Thu Jul 14 00:18:00 CEST 2022 - The aliscore output file does not exist. Check results for BUSCO: 47899at7742
## Thu Jul 14 00:18:00 CEST 2022 - The aliscore output appears to be empty. Check results for BUSCO: 47899at7742
## Thu Jul 14 00:18:01 CEST 2022 - The aliscore output file does not exist. Check results for BUSCO: 137301at7742
## Thu Jul 14 00:18:01 CEST 2022 - The aliscore output appears to be empty. Check results for BUSCO: 137301at7742
## Thu Jul 14 00:18:03 CEST 2022 - The aliscore output file does not exist. Check results for BUSCO: 47838at7742
## Thu Jul 14 00:18:03 CEST 2022 - The aliscore output appears to be empty. Check results for BUSCO: 47838at7742
## Thu Jul 14 00:18:03 CEST 2022 - The aliscore output file does not exist. Check results for BUSCO: 44698at7742
## Thu Jul 14 00:18:03 CEST 2022 - The aliscore output appears to be empty. Check results for BUSCO: 44698at7742
## Thu Jul 14 00:18:07 CEST 2022 - The aliscore output file does not exist. Check results for BUSCO: 47838at7742
## Thu Jul 14 00:18:07 CEST 2022 - The aliscore output appears to be empty. Check results for BUSCO: 47838at7742
## Thu Jul 14 00:18:07 CEST 2022 - The aliscore output file does not exist. Check results for BUSCO: 56414at7742
## Thu Jul 14 00:18:07 CEST 2022 - The aliscore output appears to be empty. Check results for BUSCO: 56414at7742
## Thu Jul 14 00:18:09 CEST 2022 - The aliscore output file does not exist. Check results for BUSCO: 49104at7742
## Thu Jul 14 00:18:09 CEST 2022 - The aliscore output appears to be empty. Check results for BUSCO: 49104at7742
## Thu Jul 14 00:18:09 CEST 2022 - The aliscore output file does not exist. Check results for BUSCO: 169538at7742
## Thu Jul 14 00:18:10 CEST 2022 - The aliscore output appears to be empty. Check results for BUSCO: 169538at7742
## Thu Jul 14 00:18:11 CEST 2022 - The aliscore output file does not exist. Check results for BUSCO: 35338at7742
## Thu Jul 14 00:18:11 CEST 2022 - The aliscore output appears to be empty. Check results for BUSCO: 35338at7742
## Thu Jul 14 00:18:13 CEST 2022 - The aliscore output file does not exist. Check results for BUSCO: 106972at7742
## Thu Jul 14 00:18:13 CEST 2022 - The aliscore output appears to be empty. Check results for BUSCO: 106972at7742
## Thu Jul 14 00:18:15 CEST 2022 - The aliscore output file does not exist. Check results for BUSCO: 122255at7742
## Thu Jul 14 00:18:15 CEST 2022 - The aliscore output appears to be empty. Check results for BUSCO: 122255at7742
## Thu Jul 14 00:18:15 CEST 2022 - The aliscore output file does not exist. Check results for BUSCO: 111684at7742
## Thu Jul 14 00:18:15 CEST 2022 - The aliscore output appears to be empty. Check results for BUSCO: 111684at7742
## Thu Jul 14 00:18:16 CEST 2022 - The aliscore output file does not exist. Check results for BUSCO: 35338at7742
## Thu Jul 14 00:18:16 CEST 2022 - The aliscore output appears to be empty. Check results for BUSCO: 35338at7742
## Thu Jul 14 00:18:19 CEST 2022 - The aliscore output file does not exist. Check results for BUSCO: 20951at7742
## Thu Jul 14 00:18:19 CEST 2022 - The aliscore output appears to be empty. Check results for BUSCO: 20951at7742
## Thu Jul 14 00:18:20 CEST 2022 - The aliscore output file does not exist. Check results for BUSCO: 26890at7742
## Thu Jul 14 00:18:20 CEST 2022 - The aliscore output appears to be empty. Check results for BUSCO: 26890at7742
## Thu Jul 14 00:18:21 CEST 2022 - The aliscore output file does not exist. Check results for BUSCO: 99281at7742
## Thu Jul 14 00:18:21 CEST 2022 - The aliscore output appears to be empty. Check results for BUSCO: 99281at7742
## Thu Jul 14 00:18:21 CEST 2022 - The aliscore output file does not exist. Check results for BUSCO: 59233at7742
## Thu Jul 14 00:18:21 CEST 2022 - The aliscore output appears to be empty. Check results for BUSCO: 59233at7742
## Thu Jul 14 00:18:25 CEST 2022 - The aliscore output file does not exist. Check results for BUSCO: 43798at7742
## Thu Jul 14 00:18:25 CEST 2022 - The aliscore output appears to be empty. Check results for BUSCO: 43798at7742
## Thu Jul 14 00:18:26 CEST 2022 - The aliscore output file does not exist. Check results for BUSCO: 46538at7742
## Thu Jul 14 00:18:26 CEST 2022 - The aliscore output appears to be empty. Check results for BUSCO: 46538at7742
## Thu Jul 14 00:18:27 CEST 2022 - The aliscore output file does not exist. Check results for BUSCO: 404163at7742
## Thu Jul 14 00:18:27 CEST 2022 - The aliscore output appears to be empty. Check results for BUSCO: 404163at7742
## Thu Jul 14 00:18:29 CEST 2022 - The aliscore output file does not exist. Check results for BUSCO: 114216at7742
## Thu Jul 14 00:18:29 CEST 2022 - The aliscore output appears to be empty. Check results for BUSCO: 114216at7742
## Thu Jul 14 00:18:29 CEST 2022 - The aliscore output file does not exist. Check results for BUSCO: 17657at7742
## Thu Jul 14 00:18:29 CEST 2022 - The aliscore output appears to be empty. Check results for BUSCO: 17657at7742
## Thu Jul 14 00:18:31 CEST 2022 - The aliscore output file does not exist. Check results for BUSCO: 80917at7742
## Thu Jul 14 00:18:31 CEST 2022 - The aliscore output appears to be empty. Check results for BUSCO: 80917at7742
## Thu Jul 14 00:18:33 CEST 2022 - The aliscore output file does not exist. Check results for BUSCO: 218806at7742
## Thu Jul 14 00:18:33 CEST 2022 - The aliscore output appears to be empty. Check results for BUSCO: 218806at7742
## Thu Jul 14 00:18:34 CEST 2022 - The aliscore output file does not exist. Check results for BUSCO: 26970at7742
## Thu Jul 14 00:18:34 CEST 2022 - The aliscore output appears to be empty. Check results for BUSCO: 26970at7742
## Thu Jul 14 00:18:36 CEST 2022 - The aliscore output file does not exist. Check results for BUSCO: 81797at7742
## Thu Jul 14 00:18:36 CEST 2022 - The aliscore output appears to be empty. Check results for BUSCO: 81797at7742
## Thu Jul 14 00:18:37 CEST 2022 - The aliscore output file does not exist. Check results for BUSCO: 44698at7742
## Thu Jul 14 00:18:37 CEST 2022 - The aliscore output appears to be empty. Check results for BUSCO: 44698at7742
## Thu Jul 14 00:18:39 CEST 2022 - The aliscore output file does not exist. Check results for BUSCO: 49318at7742
## Thu Jul 14 00:18:39 CEST 2022 - The aliscore output file does not exist. Check results for BUSCO: 26714at7742
## Thu Jul 14 00:18:39 CEST 2022 - The aliscore output appears to be empty. Check results for BUSCO: 49318at7742
## Thu Jul 14 00:18:39 CEST 2022 - The aliscore output appears to be empty. Check results for BUSCO: 26714at7742
## Thu Jul 14 00:18:41 CEST 2022 - The aliscore output file does not exist. Check results for BUSCO: 29952at7742
## Thu Jul 14 00:18:41 CEST 2022 - The aliscore output appears to be empty. Check results for BUSCO: 29952at7742
## Thu Jul 14 00:18:42 CEST 2022 - The aliscore output file does not exist. Check results for BUSCO: 77231at7742
## Thu Jul 14 00:18:42 CEST 2022 - The aliscore output appears to be empty. Check results for BUSCO: 77231at7742
## Thu Jul 14 00:18:43 CEST 2022 - The aliscore output file does not exist. Check results for BUSCO: 111509at7742
## Thu Jul 14 00:18:43 CEST 2022 - The aliscore output appears to be empty. Check results for BUSCO: 111509at7742
## Thu Jul 14 00:18:47 CEST 2022 - The aliscore output file does not exist. Check results for BUSCO: 171211at7742
## Thu Jul 14 00:18:47 CEST 2022 - The aliscore output appears to be empty. Check results for BUSCO: 171211at7742
## Thu Jul 14 00:18:48 CEST 2022 - The aliscore output file does not exist. Check results for BUSCO: 38426at7742
## Thu Jul 14 00:18:48 CEST 2022 - The aliscore output appears to be empty. Check results for BUSCO: 38426at7742
## Thu Jul 14 00:18:49 CEST 2022 - The aliscore output file does not exist. Check results for BUSCO: 126446at7742
## Thu Jul 14 00:18:49 CEST 2022 - The aliscore output appears to be empty. Check results for BUSCO: 126446at7742
## Thu Jul 14 00:18:51 CEST 2022 - The aliscore output file does not exist. Check results for BUSCO: 138116at7742
## Thu Jul 14 00:18:51 CEST 2022 - The aliscore output appears to be empty. Check results for BUSCO: 138116at7742
## Thu Jul 14 00:18:52 CEST 2022 - The aliscore output file does not exist. Check results for BUSCO: 48275at7742
## Thu Jul 14 00:18:52 CEST 2022 - The aliscore output appears to be empty. Check results for BUSCO: 48275at7742
## Thu Jul 14 00:18:54 CEST 2022 - The aliscore output file does not exist. Check results for BUSCO: 404163at7742
## Thu Jul 14 00:18:54 CEST 2022 - The aliscore output appears to be empty. Check results for BUSCO: 404163at7742
## Thu Jul 14 00:18:56 CEST 2022 - The aliscore output file does not exist. Check results for BUSCO: 105636at7742
## Thu Jul 14 00:18:56 CEST 2022 - The aliscore output appears to be empty. Check results for BUSCO: 105636at7742
## Thu Jul 14 00:18:56 CEST 2022 - The aliscore output file does not exist. Check results for BUSCO: 26970at7742
## Thu Jul 14 00:18:56 CEST 2022 - The aliscore output appears to be empty. Check results for BUSCO: 26970at7742
## Thu Jul 14 00:18:58 CEST 2022 - The aliscore output file does not exist. Check results for BUSCO: 104484at7742
## Thu Jul 14 00:18:58 CEST 2022 - The aliscore output appears to be empty. Check results for BUSCO: 104484at7742
## Thu Jul 14 00:18:59 CEST 2022 - The aliscore output file does not exist. Check results for BUSCO: 58866at7742
## Thu Jul 14 00:18:59 CEST 2022 - The aliscore output appears to be empty. Check results for BUSCO: 58866at7742
## Thu Jul 14 00:19:00 CEST 2022 - The aliscore output file does not exist. Check results for BUSCO: 191412at7742
## Thu Jul 14 00:19:00 CEST 2022 - The aliscore output appears to be empty. Check results for BUSCO: 191412at7742
## Thu Jul 14 00:19:02 CEST 2022 - The aliscore output file does not exist. Check results for BUSCO: 23496at7742
## Thu Jul 14 00:19:02 CEST 2022 - The aliscore output appears to be empty. Check results for BUSCO: 23496at7742
## Thu Jul 14 00:19:04 CEST 2022 - The aliscore output file does not exist. Check results for BUSCO: 170437at7742
## Thu Jul 14 00:19:04 CEST 2022 - The aliscore output appears to be empty. Check results for BUSCO: 170437at7742
## Thu Jul 14 00:19:04 CEST 2022 - The aliscore output file does not exist. Check results for BUSCO: 104992at7742
## Thu Jul 14 00:19:04 CEST 2022 - The aliscore output appears to be empty. Check results for BUSCO: 104992at7742
## Thu Jul 14 00:19:07 CEST 2022 - The aliscore output file does not exist. Check results for BUSCO: 94920at7742
## Thu Jul 14 00:19:07 CEST 2022 - The aliscore output appears to be empty. Check results for BUSCO: 94920at7742
## Thu Jul 14 00:19:09 CEST 2022 - The aliscore output file does not exist. Check results for BUSCO: 74281at7742
## Thu Jul 14 00:19:09 CEST 2022 - The aliscore output appears to be empty. Check results for BUSCO: 74281at7742
## Thu Jul 14 00:19:09 CEST 2022 - The aliscore output file does not exist. Check results for BUSCO: 46839at7742
## Thu Jul 14 00:19:09 CEST 2022 - The aliscore output appears to be empty. Check results for BUSCO: 46839at7742
## Thu Jul 14 00:19:11 CEST 2022 - The aliscore output file does not exist. Check results for BUSCO: 218806at7742
## Thu Jul 14 00:19:11 CEST 2022 - The aliscore output appears to be empty. Check results for BUSCO: 218806at7742
## Thu Jul 14 00:19:13 CEST 2022 - The aliscore output file does not exist. Check results for BUSCO: 23496at7742
## Thu Jul 14 00:19:13 CEST 2022 - The aliscore output appears to be empty. Check results for BUSCO: 23496at7742
## Thu Jul 14 00:19:13 CEST 2022 - The aliscore output file does not exist. Check results for BUSCO: 170437at7742
## Thu Jul 14 00:19:13 CEST 2022 - The aliscore output appears to be empty. Check results for BUSCO: 170437at7742
## Thu Jul 14 00:19:15 CEST 2022 - The aliscore output file does not exist. Check results for BUSCO: 160770at7742
## Thu Jul 14 00:19:15 CEST 2022 - The aliscore output appears to be empty. Check results for BUSCO: 160770at7742
## Thu Jul 14 00:19:17 CEST 2022 - The aliscore output file does not exist. Check results for BUSCO: 361964at7742
## Thu Jul 14 00:19:17 CEST 2022 - The aliscore output appears to be empty. Check results for BUSCO: 361964at7742
## Thu Jul 14 00:19:18 CEST 2022 - The aliscore output file does not exist. Check results for BUSCO: 71417at7742
## Thu Jul 14 00:19:18 CEST 2022 - The aliscore output appears to be empty. Check results for BUSCO: 71417at7742
## Thu Jul 14 00:19:19 CEST 2022 - The aliscore output file does not exist. Check results for BUSCO: 75780at7742
## Thu Jul 14 00:19:19 CEST 2022 - The aliscore output appears to be empty. Check results for BUSCO: 75780at7742
## Thu Jul 14 00:19:22 CEST 2022 - The aliscore output file does not exist. Check results for BUSCO: 150497at7742
## Thu Jul 14 00:19:22 CEST 2022 - The aliscore output file does not exist. Check results for BUSCO: 38426at7742
## Thu Jul 14 00:19:22 CEST 2022 - The aliscore output appears to be empty. Check results for BUSCO: 150497at7742
## Thu Jul 14 00:19:22 CEST 2022 - The aliscore output appears to be empty. Check results for BUSCO: 38426at7742
## Thu Jul 14 00:19:24 CEST 2022 - The aliscore output file does not exist. Check results for BUSCO: 128918at7742
## Thu Jul 14 00:19:25 CEST 2022 - The aliscore output appears to be empty. Check results for BUSCO: 128918at7742
## Thu Jul 14 00:19:25 CEST 2022 - The aliscore output file does not exist. Check results for BUSCO: 98179at7742
## Thu Jul 14 00:19:25 CEST 2022 - The aliscore output appears to be empty. Check results for BUSCO: 98179at7742
## Thu Jul 14 00:19:27 CEST 2022 - The aliscore output file does not exist. Check results for BUSCO: 52093at7742
## Thu Jul 14 00:19:27 CEST 2022 - The aliscore output appears to be empty. Check results for BUSCO: 52093at7742
## Thu Jul 14 00:19:27 CEST 2022 - The aliscore output file does not exist. Check results for BUSCO: 111667at7742
## Thu Jul 14 00:19:27 CEST 2022 - The aliscore output appears to be empty. Check results for BUSCO: 111667at7742
## Thu Jul 14 00:19:30 CEST 2022 - The aliscore output file does not exist. Check results for BUSCO: 51434at7742
## Thu Jul 14 00:19:30 CEST 2022 - The aliscore output appears to be empty. Check results for BUSCO: 51434at7742
## Thu Jul 14 00:19:30 CEST 2022 - The aliscore output file does not exist. Check results for BUSCO: 94920at7742
## Thu Jul 14 00:19:30 CEST 2022 - The aliscore output appears to be empty. Check results for BUSCO: 94920at7742
## Thu Jul 14 00:19:32 CEST 2022 - The aliscore output file does not exist. Check results for BUSCO: 57824at7742
## Thu Jul 14 00:19:32 CEST 2022 - The aliscore output appears to be empty. Check results for BUSCO: 57824at7742
## Thu Jul 14 00:19:33 CEST 2022 - The aliscore output file does not exist. Check results for BUSCO: 64860at7742
## Thu Jul 14 00:19:33 CEST 2022 - The aliscore output appears to be empty. Check results for BUSCO: 64860at7742
## Thu Jul 14 00:19:35 CEST 2022 - The aliscore output file does not exist. Check results for BUSCO: 188180at7742
## Thu Jul 14 00:19:35 CEST 2022 - The aliscore output appears to be empty. Check results for BUSCO: 188180at7742
## Thu Jul 14 00:19:37 CEST 2022 - The aliscore output file does not exist. Check results for BUSCO: 48275at7742
## Thu Jul 14 00:19:37 CEST 2022 - The aliscore output appears to be empty. Check results for BUSCO: 48275at7742
## Thu Jul 14 00:19:37 CEST 2022 - The aliscore output file does not exist. Check results for BUSCO: 45427at7742
## Thu Jul 14 00:19:37 CEST 2022 - The aliscore output appears to be empty. Check results for BUSCO: 45427at7742
## Thu Jul 14 00:19:39 CEST 2022 - The aliscore output file does not exist. Check results for BUSCO: 49776at7742
## Thu Jul 14 00:19:39 CEST 2022 - The aliscore output appears to be empty. Check results for BUSCO: 49776at7742
## Thu Jul 14 00:19:40 CEST 2022 - The aliscore output file does not exist. Check results for BUSCO: 199826at7742
## Thu Jul 14 00:19:40 CEST 2022 - The aliscore output appears to be empty. Check results for BUSCO: 199826at7742
## Thu Jul 14 00:19:42 CEST 2022 - The aliscore output file does not exist. Check results for BUSCO: 34300at7742
## Thu Jul 14 00:19:42 CEST 2022 - The aliscore output appears to be empty. Check results for BUSCO: 34300at7742
## Thu Jul 14 00:19:44 CEST 2022 - The aliscore output file does not exist. Check results for BUSCO: 176410at7742
## Thu Jul 14 00:19:45 CEST 2022 - The aliscore output appears to be empty. Check results for BUSCO: 176410at7742
## Thu Jul 14 00:19:45 CEST 2022 - The aliscore output file does not exist. Check results for BUSCO: 71417at7742
## Thu Jul 14 00:19:45 CEST 2022 - The aliscore output appears to be empty. Check results for BUSCO: 71417at7742
## Thu Jul 14 00:19:47 CEST 2022 - The aliscore output file does not exist. Check results for BUSCO: 26536at7742
## Thu Jul 14 00:19:47 CEST 2022 - The aliscore output appears to be empty. Check results for BUSCO: 26536at7742
## Thu Jul 14 00:19:48 CEST 2022 - The aliscore output file does not exist. Check results for BUSCO: 170658at7742
## Thu Jul 14 00:19:48 CEST 2022 - The aliscore output appears to be empty. Check results for BUSCO: 170658at7742
## Thu Jul 14 00:19:49 CEST 2022 - The aliscore output file does not exist. Check results for BUSCO: 54892at7742
## Thu Jul 14 00:19:49 CEST 2022 - The aliscore output appears to be empty. Check results for BUSCO: 54892at7742
## Thu Jul 14 00:19:52 CEST 2022 - The aliscore output file does not exist. Check results for BUSCO: 114807at7742
## Thu Jul 14 00:19:52 CEST 2022 - The aliscore output appears to be empty. Check results for BUSCO: 114807at7742
## Thu Jul 14 00:19:52 CEST 2022 - The aliscore output file does not exist. Check results for BUSCO: 29952at7742
## Thu Jul 14 00:19:52 CEST 2022 - The aliscore output appears to be empty. Check results for BUSCO: 29952at7742
## Thu Jul 14 00:19:54 CEST 2022 - The aliscore output file does not exist. Check results for BUSCO: 20951at7742
## Thu Jul 14 00:19:54 CEST 2022 - The aliscore output appears to be empty. Check results for BUSCO: 20951at7742
## Thu Jul 14 00:19:57 CEST 2022 - The aliscore output file does not exist. Check results for BUSCO: 212520at7742
## Thu Jul 14 00:19:57 CEST 2022 - The aliscore output appears to be empty. Check results for BUSCO: 212520at7742
## Thu Jul 14 00:19:57 CEST 2022 - The aliscore output file does not exist. Check results for BUSCO: 78822at7742
## Thu Jul 14 00:19:57 CEST 2022 - The aliscore output appears to be empty. Check results for BUSCO: 78822at7742
## Thu Jul 14 00:19:58 CEST 2022 - The aliscore output file does not exist. Check results for BUSCO: 114807at7742
## Thu Jul 14 00:19:58 CEST 2022 - The aliscore output appears to be empty. Check results for BUSCO: 114807at7742
## Thu Jul 14 00:20:01 CEST 2022 - The aliscore output file does not exist. Check results for BUSCO: 93412at7742
## Thu Jul 14 00:20:01 CEST 2022 - The aliscore output appears to be empty. Check results for BUSCO: 93412at7742
## Thu Jul 14 00:20:01 CEST 2022 - The aliscore output file does not exist. Check results for BUSCO: 141264at7742
## Thu Jul 14 00:20:01 CEST 2022 - The aliscore output appears to be empty. Check results for BUSCO: 141264at7742
## Thu Jul 14 00:20:02 CEST 2022 - The aliscore output file does not exist. Check results for BUSCO: 49776at7742
## Thu Jul 14 00:20:02 CEST 2022 - The aliscore output appears to be empty. Check results for BUSCO: 49776at7742
## Thu Jul 14 00:20:04 CEST 2022 - The aliscore output file does not exist. Check results for BUSCO: 46538at7742
## Thu Jul 14 00:20:04 CEST 2022 - The aliscore output appears to be empty. Check results for BUSCO: 46538at7742
## Thu Jul 14 00:20:05 CEST 2022 - The aliscore output file does not exist. Check results for BUSCO: 52093at7742
## Thu Jul 14 00:20:05 CEST 2022 - The aliscore output appears to be empty. Check results for BUSCO: 52093at7742
## Thu Jul 14 00:20:07 CEST 2022 - The aliscore output file does not exist. Check results for BUSCO: 36575at7742
## Thu Jul 14 00:20:07 CEST 2022 - The aliscore output appears to be empty. Check results for BUSCO: 36575at7742
## Thu Jul 14 00:20:08 CEST 2022 - The aliscore output file does not exist. Check results for BUSCO: 38426at7742
## Thu Jul 14 00:20:08 CEST 2022 - The aliscore output appears to be empty. Check results for BUSCO: 38426at7742
## Thu Jul 14 00:20:09 CEST 2022 - The aliscore output file does not exist. Check results for BUSCO: 64168at7742
## Thu Jul 14 00:20:09 CEST 2022 - The aliscore output appears to be empty. Check results for BUSCO: 64168at7742
## Thu Jul 14 00:20:11 CEST 2022 - The aliscore output file does not exist. Check results for BUSCO: 26970at7742
## Thu Jul 14 00:20:11 CEST 2022 - The aliscore output appears to be empty. Check results for BUSCO: 26970at7742
## Thu Jul 14 00:20:13 CEST 2022 - The aliscore output file does not exist. Check results for BUSCO: 111509at7742
## Thu Jul 14 00:20:13 CEST 2022 - The aliscore output appears to be empty. Check results for BUSCO: 111509at7742
## Thu Jul 14 00:20:14 CEST 2022 - The aliscore output file does not exist. Check results for BUSCO: 49318at7742
## Thu Jul 14 00:20:14 CEST 2022 - The aliscore output appears to be empty. Check results for BUSCO: 49318at7742
## Thu Jul 14 00:20:15 CEST 2022 - The aliscore output file does not exist. Check results for BUSCO: 247622at7742
## Thu Jul 14 00:20:15 CEST 2022 - The aliscore output appears to be empty. Check results for BUSCO: 247622at7742
## Thu Jul 14 00:20:17 CEST 2022 - The aliscore output file does not exist. Check results for BUSCO: 347249at7742
## Thu Jul 14 00:20:17 CEST 2022 - The aliscore output appears to be empty. Check results for BUSCO: 347249at7742
## Thu Jul 14 00:20:18 CEST 2022 - The aliscore output file does not exist. Check results for BUSCO: 54892at7742
## Thu Jul 14 00:20:18 CEST 2022 - The aliscore output appears to be empty. Check results for BUSCO: 54892at7742
## Thu Jul 14 00:20:19 CEST 2022 - The aliscore output file does not exist. Check results for BUSCO: 49776at7742
## Thu Jul 14 00:20:19 CEST 2022 - The aliscore output appears to be empty. Check results for BUSCO: 49776at7742
## Thu Jul 14 00:20:20 CEST 2022 - The aliscore output file does not exist. Check results for BUSCO: 45427at7742
## Thu Jul 14 00:20:20 CEST 2022 - The aliscore output appears to be empty. Check results for BUSCO: 45427at7742
## Thu Jul 14 00:20:23 CEST 2022 - The aliscore output file does not exist. Check results for BUSCO: 31418at7742
## Thu Jul 14 00:20:23 CEST 2022 - The aliscore output appears to be empty. Check results for BUSCO: 31418at7742
## Thu Jul 14 00:20:24 CEST 2022 - The aliscore output file does not exist. Check results for BUSCO: 76024at7742
## Thu Jul 14 00:20:24 CEST 2022 - The aliscore output appears to be empty. Check results for BUSCO: 76024at7742
## Thu Jul 14 00:20:29 CEST 2022 - The aliscore output file does not exist. Check results for BUSCO: 57179at7742
## Thu Jul 14 00:20:29 CEST 2022 - The aliscore output appears to be empty. Check results for BUSCO: 57179at7742
## Thu Jul 14 00:20:30 CEST 2022 - The aliscore output file does not exist. Check results for BUSCO: 37335at7742
## Thu Jul 14 00:20:30 CEST 2022 - The aliscore output appears to be empty. Check results for BUSCO: 37335at7742
## Thu Jul 14 00:20:30 CEST 2022 - The aliscore output file does not exist. Check results for BUSCO: 160770at7742
## Thu Jul 14 00:20:30 CEST 2022 - The aliscore output appears to be empty. Check results for BUSCO: 160770at7742
## Thu Jul 14 00:20:31 CEST 2022 - The aliscore output file does not exist. Check results for BUSCO: 78893at7742
## Thu Jul 14 00:20:31 CEST 2022 - The aliscore output appears to be empty. Check results for BUSCO: 78893at7742
## Thu Jul 14 00:20:33 CEST 2022 - The aliscore output file does not exist. Check results for BUSCO: 63939at7742
## Thu Jul 14 00:20:33 CEST 2022 - The aliscore output appears to be empty. Check results for BUSCO: 63939at7742
## Thu Jul 14 00:20:34 CEST 2022 - The aliscore output file does not exist. Check results for BUSCO: 50070at7742
## Thu Jul 14 00:20:34 CEST 2022 - The aliscore output appears to be empty. Check results for BUSCO: 50070at7742
## Thu Jul 14 00:20:34 CEST 2022 - The aliscore output file does not exist. Check results for BUSCO: 108510at7742
## Thu Jul 14 00:20:34 CEST 2022 - The aliscore output appears to be empty. Check results for BUSCO: 108510at7742
## Thu Jul 14 00:20:39 CEST 2022 - The aliscore output file does not exist. Check results for BUSCO: 48975at7742
## Thu Jul 14 00:20:39 CEST 2022 - The aliscore output appears to be empty. Check results for BUSCO: 48975at7742
## Thu Jul 14 00:20:39 CEST 2022 - The aliscore output file does not exist. Check results for BUSCO: 111684at7742
## Thu Jul 14 00:20:39 CEST 2022 - The aliscore output file does not exist. Check results for BUSCO: 102147at7742
## Thu Jul 14 00:20:39 CEST 2022 - The aliscore output appears to be empty. Check results for BUSCO: 111684at7742
## Thu Jul 14 00:20:39 CEST 2022 - The aliscore output appears to be empty. Check results for BUSCO: 102147at7742
## Thu Jul 14 00:20:40 CEST 2022 - The aliscore output file does not exist. Check results for BUSCO: 128918at7742
## Thu Jul 14 00:20:40 CEST 2022 - The aliscore output appears to be empty. Check results for BUSCO: 128918at7742
## Thu Jul 14 00:20:42 CEST 2022 - The aliscore output file does not exist. Check results for BUSCO: 60277at7742
## Thu Jul 14 00:20:43 CEST 2022 - The aliscore output appears to be empty. Check results for BUSCO: 60277at7742
## Thu Jul 14 00:20:44 CEST 2022 - The aliscore output file does not exist. Check results for BUSCO: 176414at7742
## Thu Jul 14 00:20:44 CEST 2022 - The aliscore output appears to be empty. Check results for BUSCO: 176414at7742
## Thu Jul 14 00:20:44 CEST 2022 - The aliscore output file does not exist. Check results for BUSCO: 23496at7742
## Thu Jul 14 00:20:44 CEST 2022 - The aliscore output appears to be empty. Check results for BUSCO: 23496at7742
## Thu Jul 14 00:20:47 CEST 2022 - The aliscore output file does not exist. Check results for BUSCO: 170658at7742
## Thu Jul 14 00:20:47 CEST 2022 - The aliscore output appears to be empty. Check results for BUSCO: 170658at7742
## Thu Jul 14 00:20:47 CEST 2022 - The aliscore output file does not exist. Check results for BUSCO: 114807at7742
## Thu Jul 14 00:20:47 CEST 2022 - The aliscore output appears to be empty. Check results for BUSCO: 114807at7742
## Thu Jul 14 00:20:49 CEST 2022 - The aliscore output file does not exist. Check results for BUSCO: 32286at7742
## Thu Jul 14 00:20:49 CEST 2022 - The aliscore output appears to be empty. Check results for BUSCO: 32286at7742
## Thu Jul 14 00:20:50 CEST 2022 - The aliscore output file does not exist. Check results for BUSCO: 82217at7742
## Thu Jul 14 00:20:50 CEST 2022 - The aliscore output appears to be empty. Check results for BUSCO: 82217at7742
## Thu Jul 14 00:20:54 CEST 2022 - The aliscore output file does not exist. Check results for BUSCO: 50838at7742
## Thu Jul 14 00:20:54 CEST 2022 - The aliscore output appears to be empty. Check results for BUSCO: 50838at7742
## Thu Jul 14 00:20:54 CEST 2022 - The aliscore output file does not exist. Check results for BUSCO: 223702at7742
## Thu Jul 14 00:20:54 CEST 2022 - The aliscore output appears to be empty. Check results for BUSCO: 223702at7742
## Thu Jul 14 00:20:54 CEST 2022 - The aliscore output file does not exist. Check results for BUSCO: 97645at7742
## Thu Jul 14 00:20:54 CEST 2022 - The aliscore output appears to be empty. Check results for BUSCO: 97645at7742
## Thu Jul 14 00:20:56 CEST 2022 - The aliscore output file does not exist. Check results for BUSCO: 111254at7742
## Thu Jul 14 00:20:56 CEST 2022 - The aliscore output appears to be empty. Check results for BUSCO: 111254at7742
## Thu Jul 14 00:20:57 CEST 2022 - The aliscore output file does not exist. Check results for BUSCO: 223348at7742
## Thu Jul 14 00:20:57 CEST 2022 - The aliscore output appears to be empty. Check results for BUSCO: 223348at7742
## Thu Jul 14 00:20:59 CEST 2022 - The aliscore output file does not exist. Check results for BUSCO: 49935at7742
## Thu Jul 14 00:20:59 CEST 2022 - The aliscore output appears to be empty. Check results for BUSCO: 49935at7742
## Thu Jul 14 00:21:01 CEST 2022 - The aliscore output file does not exist. Check results for BUSCO: 26890at7742
## Thu Jul 14 00:21:01 CEST 2022 - The aliscore output appears to be empty. Check results for BUSCO: 26890at7742
## Thu Jul 14 00:21:03 CEST 2022 - The aliscore output file does not exist. Check results for BUSCO: 42483at7742
## Thu Jul 14 00:21:03 CEST 2022 - The aliscore output appears to be empty. Check results for BUSCO: 42483at7742
## Thu Jul 14 00:21:03 CEST 2022 - The aliscore output file does not exist. Check results for BUSCO: 286974at7742
## Thu Jul 14 00:21:04 CEST 2022 - The aliscore output appears to be empty. Check results for BUSCO: 286974at7742
## Thu Jul 14 00:21:04 CEST 2022 - The aliscore output file does not exist. Check results for BUSCO: 216073at7742
## Thu Jul 14 00:21:04 CEST 2022 - The aliscore output appears to be empty. Check results for BUSCO: 216073at7742
## Thu Jul 14 00:21:04 CEST 2022 - The aliscore output file does not exist. Check results for BUSCO: 26536at7742
## Thu Jul 14 00:21:04 CEST 2022 - The aliscore output appears to be empty. Check results for BUSCO: 26536at7742
## Thu Jul 14 00:21:05 CEST 2022 - The aliscore output file does not exist. Check results for BUSCO: 50284at7742
## Thu Jul 14 00:21:05 CEST 2022 - The aliscore output appears to be empty. Check results for BUSCO: 50284at7742
## Thu Jul 14 00:21:08 CEST 2022 - The aliscore output file does not exist. Check results for BUSCO: 74297at7742
## Thu Jul 14 00:21:08 CEST 2022 - The aliscore output appears to be empty. Check results for BUSCO: 74297at7742
## Thu Jul 14 00:21:10 CEST 2022 - The aliscore output file does not exist. Check results for BUSCO: 177338at7742
## Thu Jul 14 00:21:10 CEST 2022 - The aliscore output appears to be empty. Check results for BUSCO: 177338at7742
## Thu Jul 14 00:21:11 CEST 2022 - The aliscore output file does not exist. Check results for BUSCO: 28500at7742
## Thu Jul 14 00:21:11 CEST 2022 - The aliscore output appears to be empty. Check results for BUSCO: 28500at7742
## Thu Jul 14 00:21:13 CEST 2022 - The aliscore output file does not exist. Check results for BUSCO: 50284at7742
## Thu Jul 14 00:21:13 CEST 2022 - The aliscore output appears to be empty. Check results for BUSCO: 50284at7742
## Thu Jul 14 00:21:13 CEST 2022 - The aliscore output file does not exist. Check results for BUSCO: 156102at7742
## Thu Jul 14 00:21:13 CEST 2022 - The aliscore output appears to be empty. Check results for BUSCO: 156102at7742
## Thu Jul 14 00:21:14 CEST 2022 - The aliscore output file does not exist. Check results for BUSCO: 235319at7742
## Thu Jul 14 00:21:14 CEST 2022 - The aliscore output appears to be empty. Check results for BUSCO: 235319at7742
## Thu Jul 14 00:21:16 CEST 2022 - The aliscore output file does not exist. Check results for BUSCO: 160945at7742
## Thu Jul 14 00:21:16 CEST 2022 - The aliscore output appears to be empty. Check results for BUSCO: 160945at7742
## Thu Jul 14 00:21:18 CEST 2022 - The aliscore output file does not exist. Check results for BUSCO: 403293at7742
## Thu Jul 14 00:21:18 CEST 2022 - The aliscore output appears to be empty. Check results for BUSCO: 403293at7742
## Thu Jul 14 00:21:18 CEST 2022 - The aliscore output file does not exist. Check results for BUSCO: 111684at7742
## Thu Jul 14 00:21:18 CEST 2022 - The aliscore output appears to be empty. Check results for BUSCO: 111684at7742
## Thu Jul 14 00:21:20 CEST 2022 - The aliscore output file does not exist. Check results for BUSCO: 141264at7742
## Thu Jul 14 00:21:20 CEST 2022 - The aliscore output appears to be empty. Check results for BUSCO: 141264at7742
## Thu Jul 14 00:21:23 CEST 2022 - The aliscore output file does not exist. Check results for BUSCO: 268813at7742
## Thu Jul 14 00:21:23 CEST 2022 - The aliscore output appears to be empty. Check results for BUSCO: 268813at7742
## Thu Jul 14 00:21:25 CEST 2022 - The aliscore output file does not exist. Check results for BUSCO: 287179at7742
## Thu Jul 14 00:21:25 CEST 2022 - The aliscore output file does not exist. Check results for BUSCO: 59233at7742
## Thu Jul 14 00:21:25 CEST 2022 - The aliscore output appears to be empty. Check results for BUSCO: 287179at7742
## Thu Jul 14 00:21:25 CEST 2022 - The aliscore output appears to be empty. Check results for BUSCO: 59233at7742
## Thu Jul 14 00:21:27 CEST 2022 - The aliscore output file does not exist. Check results for BUSCO: 52093at7742
## Thu Jul 14 00:21:27 CEST 2022 - The aliscore output appears to be empty. Check results for BUSCO: 52093at7742
## Thu Jul 14 00:21:28 CEST 2022 - The aliscore output file does not exist. Check results for BUSCO: 163696at7742
## Thu Jul 14 00:21:28 CEST 2022 - The aliscore output appears to be empty. Check results for BUSCO: 163696at7742
## Thu Jul 14 00:21:31 CEST 2022 - The aliscore output file does not exist. Check results for BUSCO: 129880at7742
## Thu Jul 14 00:21:31 CEST 2022 - The aliscore output appears to be empty. Check results for BUSCO: 129880at7742
## Thu Jul 14 00:21:31 CEST 2022 - The aliscore output file does not exist. Check results for BUSCO: 81797at7742
## Thu Jul 14 00:21:31 CEST 2022 - The aliscore output appears to be empty. Check results for BUSCO: 81797at7742
## Thu Jul 14 00:21:35 CEST 2022 - The aliscore output file does not exist. Check results for BUSCO: 42483at7742
## Thu Jul 14 00:21:35 CEST 2022 - The aliscore output appears to be empty. Check results for BUSCO: 42483at7742
## Thu Jul 14 00:21:35 CEST 2022 - The aliscore output file does not exist. Check results for BUSCO: 32286at7742
## Thu Jul 14 00:21:35 CEST 2022 - The aliscore output appears to be empty. Check results for BUSCO: 32286at7742
## Thu Jul 14 00:21:36 CEST 2022 - The aliscore output file does not exist. Check results for BUSCO: 29952at7742
## Thu Jul 14 00:21:36 CEST 2022 - The aliscore output appears to be empty. Check results for BUSCO: 29952at7742
## Thu Jul 14 00:21:38 CEST 2022 - The aliscore output file does not exist. Check results for BUSCO: 111667at7742
## Thu Jul 14 00:21:38 CEST 2022 - The aliscore output appears to be empty. Check results for BUSCO: 111667at7742
## Thu Jul 14 00:21:39 CEST 2022 - The aliscore output file does not exist. Check results for BUSCO: 156102at7742
## Thu Jul 14 00:21:39 CEST 2022 - The aliscore output appears to be empty. Check results for BUSCO: 156102at7742
## Thu Jul 14 00:21:40 CEST 2022 - The aliscore output file does not exist. Check results for BUSCO: 73527at7742
## Thu Jul 14 00:21:40 CEST 2022 - The aliscore output appears to be empty. Check results for BUSCO: 73527at7742
## Thu Jul 14 00:21:42 CEST 2022 - The aliscore output file does not exist. Check results for BUSCO: 25839at7742
## Thu Jul 14 00:21:42 CEST 2022 - The aliscore output appears to be empty. Check results for BUSCO: 25839at7742
## Thu Jul 14 00:21:42 CEST 2022 - The aliscore output file does not exist. Check results for BUSCO: 188180at7742
## Thu Jul 14 00:21:42 CEST 2022 - The aliscore output appears to be empty. Check results for BUSCO: 188180at7742
## Thu Jul 14 00:21:45 CEST 2022 - The aliscore output file does not exist. Check results for BUSCO: 39313at7742
## Thu Jul 14 00:21:45 CEST 2022 - The aliscore output appears to be empty. Check results for BUSCO: 39313at7742
## Thu Jul 14 00:21:45 CEST 2022 - The aliscore output file does not exist. Check results for BUSCO: 35338at7742
## Thu Jul 14 00:21:45 CEST 2022 - The aliscore output appears to be empty. Check results for BUSCO: 35338at7742
## Thu Jul 14 00:21:45 CEST 2022 - The aliscore output file does not exist. Check results for BUSCO: 128918at7742
## Thu Jul 14 00:21:46 CEST 2022 - The aliscore output appears to be empty. Check results for BUSCO: 128918at7742
## Thu Jul 14 00:21:48 CEST 2022 - The aliscore output file does not exist. Check results for BUSCO: 361964at7742
## Thu Jul 14 00:21:48 CEST 2022 - The aliscore output appears to be empty. Check results for BUSCO: 361964at7742
## Thu Jul 14 00:21:49 CEST 2022 - The aliscore output file does not exist. Check results for BUSCO: 75305at7742
## Thu Jul 14 00:21:49 CEST 2022 - The aliscore output appears to be empty. Check results for BUSCO: 75305at7742
## Thu Jul 14 00:21:50 CEST 2022 - The aliscore output file does not exist. Check results for BUSCO: 105636at7742
## Thu Jul 14 00:21:50 CEST 2022 - The aliscore output appears to be empty. Check results for BUSCO: 105636at7742
## Thu Jul 14 00:21:52 CEST 2022 - The aliscore output file does not exist. Check results for BUSCO: 403293at7742
## Thu Jul 14 00:21:52 CEST 2022 - The aliscore output appears to be empty. Check results for BUSCO: 403293at7742
## Thu Jul 14 00:21:55 CEST 2022 - The aliscore output file does not exist. Check results for BUSCO: 82833at7742
## Thu Jul 14 00:21:55 CEST 2022 - The aliscore output file does not exist. Check results for BUSCO: 16096at7742
## Thu Jul 14 00:21:55 CEST 2022 - The aliscore output appears to be empty. Check results for BUSCO: 82833at7742
## Thu Jul 14 00:21:55 CEST 2022 - The aliscore output appears to be empty. Check results for BUSCO: 16096at7742
## Thu Jul 14 00:21:55 CEST 2022 - The aliscore output file does not exist. Check results for BUSCO: 277286at7742
## Thu Jul 14 00:21:55 CEST 2022 - The aliscore output appears to be empty. Check results for BUSCO: 277286at7742
## Thu Jul 14 00:21:56 CEST 2022 - The aliscore output file does not exist. Check results for BUSCO: 191412at7742
## Thu Jul 14 00:21:56 CEST 2022 - The aliscore output appears to be empty. Check results for BUSCO: 191412at7742
## Thu Jul 14 00:22:01 CEST 2022 - The aliscore output file does not exist. Check results for BUSCO: 55967at7742
## Thu Jul 14 00:22:01 CEST 2022 - The aliscore output appears to be empty. Check results for BUSCO: 55967at7742
## Thu Jul 14 00:22:03 CEST 2022 - The aliscore output file does not exist. Check results for BUSCO: 26536at7742
## Thu Jul 14 00:22:03 CEST 2022 - The aliscore output appears to be empty. Check results for BUSCO: 26536at7742
## Thu Jul 14 00:22:03 CEST 2022 - The aliscore output file does not exist. Check results for BUSCO: 55967at7742
## Thu Jul 14 00:22:03 CEST 2022 - The aliscore output appears to be empty. Check results for BUSCO: 55967at7742
## Thu Jul 14 00:22:05 CEST 2022 - The aliscore output file does not exist. Check results for BUSCO: 82833at7742
## Thu Jul 14 00:22:05 CEST 2022 - The aliscore output appears to be empty. Check results for BUSCO: 82833at7742
## Thu Jul 14 00:22:05 CEST 2022 - The aliscore output file does not exist. Check results for BUSCO: 85263at7742
## Thu Jul 14 00:22:05 CEST 2022 - The aliscore output appears to be empty. Check results for BUSCO: 85263at7742
## Thu Jul 14 00:22:06 CEST 2022 - The aliscore output file does not exist. Check results for BUSCO: 63488at7742
## Thu Jul 14 00:22:06 CEST 2022 - The aliscore output file does not exist. Check results for BUSCO: 82833at7742
## Thu Jul 14 00:22:06 CEST 2022 - The aliscore output appears to be empty. Check results for BUSCO: 63488at7742
## Thu Jul 14 00:22:06 CEST 2022 - The aliscore output appears to be empty. Check results for BUSCO: 82833at7742
## Thu Jul 14 00:22:09 CEST 2022 - The aliscore output file does not exist. Check results for BUSCO: 31418at7742
## Thu Jul 14 00:22:09 CEST 2022 - The aliscore output appears to be empty. Check results for BUSCO: 31418at7742
## Thu Jul 14 00:22:10 CEST 2022 - The aliscore output file does not exist. Check results for BUSCO: 56414at7742
## Thu Jul 14 00:22:10 CEST 2022 - The aliscore output appears to be empty. Check results for BUSCO: 56414at7742
## Thu Jul 14 00:22:12 CEST 2022 - The aliscore output file does not exist. Check results for BUSCO: 287179at7742
## Thu Jul 14 00:22:12 CEST 2022 - The aliscore output appears to be empty. Check results for BUSCO: 287179at7742
## Thu Jul 14 00:22:13 CEST 2022 - The aliscore output file does not exist. Check results for BUSCO: 103284at7742
## Thu Jul 14 00:22:14 CEST 2022 - The aliscore output appears to be empty. Check results for BUSCO: 103284at7742
## Thu Jul 14 00:22:15 CEST 2022 - The aliscore output file does not exist. Check results for BUSCO: 75780at7742
## Thu Jul 14 00:22:16 CEST 2022 - The aliscore output appears to be empty. Check results for BUSCO: 75780at7742
## Thu Jul 14 00:22:16 CEST 2022 - The aliscore output file does not exist. Check results for BUSCO: 51434at7742
## Thu Jul 14 00:22:16 CEST 2022 - The aliscore output appears to be empty. Check results for BUSCO: 51434at7742
## Thu Jul 14 00:22:18 CEST 2022 - The aliscore output file does not exist. Check results for BUSCO: 24404at7742
## Thu Jul 14 00:22:18 CEST 2022 - The aliscore output appears to be empty. Check results for BUSCO: 24404at7742
## Thu Jul 14 00:22:18 CEST 2022 - The aliscore output file does not exist. Check results for BUSCO: 146445at7742
## Thu Jul 14 00:22:18 CEST 2022 - The aliscore output appears to be empty. Check results for BUSCO: 146445at7742
## Thu Jul 14 00:22:20 CEST 2022 - The aliscore output file does not exist. Check results for BUSCO: 111667at7742
## Thu Jul 14 00:22:20 CEST 2022 - The aliscore output appears to be empty. Check results for BUSCO: 111667at7742
## Thu Jul 14 00:22:22 CEST 2022 - The aliscore output file does not exist. Check results for BUSCO: 50284at7742
## Thu Jul 14 00:22:22 CEST 2022 - The aliscore output appears to be empty. Check results for BUSCO: 50284at7742
## Thu Jul 14 00:22:24 CEST 2022 - The aliscore output file does not exist. Check results for BUSCO: 49104at7742
## Thu Jul 14 00:22:24 CEST 2022 - The aliscore output appears to be empty. Check results for BUSCO: 49104at7742
## Thu Jul 14 00:22:26 CEST 2022 - The aliscore output file does not exist. Check results for BUSCO: 102917at7742
## Thu Jul 14 00:22:26 CEST 2022 - The aliscore output appears to be empty. Check results for BUSCO: 102917at7742
## Thu Jul 14 00:22:26 CEST 2022 - The aliscore output file does not exist. Check results for BUSCO: 286974at7742
## Thu Jul 14 00:22:26 CEST 2022 - The aliscore output appears to be empty. Check results for BUSCO: 286974at7742
## Thu Jul 14 00:22:30 CEST 2022 - The aliscore output file does not exist. Check results for BUSCO: 37335at7742
## Thu Jul 14 00:22:30 CEST 2022 - The aliscore output appears to be empty. Check results for BUSCO: 37335at7742
## Thu Jul 14 00:22:30 CEST 2022 - The aliscore output file does not exist. Check results for BUSCO: 31574at7742
## Thu Jul 14 00:22:30 CEST 2022 - The aliscore output appears to be empty. Check results for BUSCO: 31574at7742
## Thu Jul 14 00:22:32 CEST 2022 - The aliscore output file does not exist. Check results for BUSCO: 403293at7742
## Thu Jul 14 00:22:32 CEST 2022 - The aliscore output appears to be empty. Check results for BUSCO: 403293at7742
## Thu Jul 14 00:22:33 CEST 2022 - The aliscore output file does not exist. Check results for BUSCO: 103689at7742
## Thu Jul 14 00:22:33 CEST 2022 - The aliscore output appears to be empty. Check results for BUSCO: 103689at7742
## Thu Jul 14 00:22:34 CEST 2022 - The aliscore output file does not exist. Check results for BUSCO: 139811at7742
## Thu Jul 14 00:22:34 CEST 2022 - The aliscore output appears to be empty. Check results for BUSCO: 139811at7742
## Thu Jul 14 00:22:36 CEST 2022 - The aliscore output file does not exist. Check results for BUSCO: 72648at7742
## Thu Jul 14 00:22:37 CEST 2022 - The aliscore output appears to be empty. Check results for BUSCO: 72648at7742
## Thu Jul 14 00:22:37 CEST 2022 - The aliscore output file does not exist. Check results for BUSCO: 132279at7742
## Thu Jul 14 00:22:37 CEST 2022 - The aliscore output appears to be empty. Check results for BUSCO: 132279at7742
## Thu Jul 14 00:22:37 CEST 2022 - The aliscore output file does not exist. Check results for BUSCO: 72648at7742
## Thu Jul 14 00:22:37 CEST 2022 - The aliscore output appears to be empty. Check results for BUSCO: 72648at7742
## Thu Jul 14 00:22:40 CEST 2022 - The aliscore output file does not exist. Check results for BUSCO: 287179at7742
## Thu Jul 14 00:22:40 CEST 2022 - The aliscore output appears to be empty. Check results for BUSCO: 287179at7742
## Thu Jul 14 00:22:42 CEST 2022 - The aliscore output file does not exist. Check results for BUSCO: 172713at7742
## Thu Jul 14 00:22:42 CEST 2022 - The aliscore output appears to be empty. Check results for BUSCO: 172713at7742
## Thu Jul 14 00:22:43 CEST 2022 - The aliscore output file does not exist. Check results for BUSCO: 277134at7742
## Thu Jul 14 00:22:43 CEST 2022 - The aliscore output appears to be empty. Check results for BUSCO: 277134at7742
## Thu Jul 14 00:22:45 CEST 2022 - The aliscore output file does not exist. Check results for BUSCO: 334890at7742
## Thu Jul 14 00:22:45 CEST 2022 - The aliscore output appears to be empty. Check results for BUSCO: 334890at7742
## Thu Jul 14 00:22:46 CEST 2022 - The aliscore output file does not exist. Check results for BUSCO: 247622at7742
## Thu Jul 14 00:22:46 CEST 2022 - The aliscore output file does not exist. Check results for BUSCO: 25839at7742
## Thu Jul 14 00:22:46 CEST 2022 - The aliscore output appears to be empty. Check results for BUSCO: 247622at7742
## Thu Jul 14 00:22:46 CEST 2022 - The aliscore output appears to be empty. Check results for BUSCO: 25839at7742
## Thu Jul 14 00:22:48 CEST 2022 - The aliscore output file does not exist. Check results for BUSCO: 95435at7742
## Thu Jul 14 00:22:48 CEST 2022 - The aliscore output appears to be empty. Check results for BUSCO: 95435at7742
## Thu Jul 14 00:22:50 CEST 2022 - The aliscore output file does not exist. Check results for BUSCO: 156689at7742
## Thu Jul 14 00:22:50 CEST 2022 - The aliscore output appears to be empty. Check results for BUSCO: 156689at7742
## Thu Jul 14 00:22:51 CEST 2022 - The aliscore output file does not exist. Check results for BUSCO: 129062at7742
## Thu Jul 14 00:22:51 CEST 2022 - The aliscore output appears to be empty. Check results for BUSCO: 129062at7742
## Thu Jul 14 00:22:52 CEST 2022 - The aliscore output file does not exist. Check results for BUSCO: 49005at7742
## Thu Jul 14 00:22:52 CEST 2022 - The aliscore output appears to be empty. Check results for BUSCO: 49005at7742
## Thu Jul 14 00:22:55 CEST 2022 - The aliscore output file does not exist. Check results for BUSCO: 334890at7742
## Thu Jul 14 00:22:55 CEST 2022 - The aliscore output appears to be empty. Check results for BUSCO: 334890at7742
## Thu Jul 14 00:22:57 CEST 2022 - The aliscore output file does not exist. Check results for BUSCO: 149121at7742
## Thu Jul 14 00:22:57 CEST 2022 - The aliscore output appears to be empty. Check results for BUSCO: 149121at7742
## Thu Jul 14 00:22:58 CEST 2022 - The aliscore output file does not exist. Check results for BUSCO: 28500at7742
## Thu Jul 14 00:22:58 CEST 2022 - The aliscore output appears to be empty. Check results for BUSCO: 28500at7742
## Thu Jul 14 00:23:00 CEST 2022 - The aliscore output file does not exist. Check results for BUSCO: 39313at7742
## Thu Jul 14 00:23:00 CEST 2022 - The aliscore output appears to be empty. Check results for BUSCO: 39313at7742
## Thu Jul 14 00:23:00 CEST 2022 - The aliscore output file does not exist. Check results for BUSCO: 73313at7742
## Thu Jul 14 00:23:01 CEST 2022 - The aliscore output appears to be empty. Check results for BUSCO: 73313at7742
## Thu Jul 14 00:23:02 CEST 2022 - The aliscore output file does not exist. Check results for BUSCO: 111254at7742
## Thu Jul 14 00:23:03 CEST 2022 - The aliscore output appears to be empty. Check results for BUSCO: 111254at7742
## Thu Jul 14 00:23:03 CEST 2022 - The aliscore output file does not exist. Check results for BUSCO: 58866at7742
## Thu Jul 14 00:23:03 CEST 2022 - The aliscore output appears to be empty. Check results for BUSCO: 58866at7742
## Thu Jul 14 00:23:06 CEST 2022 - The aliscore output file does not exist. Check results for BUSCO: 95643at7742
## Thu Jul 14 00:23:06 CEST 2022 - The aliscore output appears to be empty. Check results for BUSCO: 95643at7742
## Thu Jul 14 00:23:08 CEST 2022 - The aliscore output file does not exist. Check results for BUSCO: 50838at7742
## Thu Jul 14 00:23:08 CEST 2022 - The aliscore output appears to be empty. Check results for BUSCO: 50838at7742
## Thu Jul 14 00:23:08 CEST 2022 - The aliscore output file does not exist. Check results for BUSCO: 117682at7742
## Thu Jul 14 00:23:08 CEST 2022 - The aliscore output appears to be empty. Check results for BUSCO: 117682at7742
## Thu Jul 14 00:23:09 CEST 2022 - The aliscore output file does not exist. Check results for BUSCO: 49104at7742
## Thu Jul 14 00:23:09 CEST 2022 - The aliscore output appears to be empty. Check results for BUSCO: 49104at7742
## Thu Jul 14 00:23:11 CEST 2022 - The aliscore output file does not exist. Check results for BUSCO: 73527at7742
## Thu Jul 14 00:23:11 CEST 2022 - The aliscore output appears to be empty. Check results for BUSCO: 73527at7742
## Thu Jul 14 00:23:12 CEST 2022 - The aliscore output file does not exist. Check results for BUSCO: 77231at7742
## Thu Jul 14 00:23:12 CEST 2022 - The aliscore output appears to be empty. Check results for BUSCO: 77231at7742
## Thu Jul 14 00:23:13 CEST 2022 - The aliscore output file does not exist. Check results for BUSCO: 37782at7742
## Thu Jul 14 00:23:13 CEST 2022 - The aliscore output appears to be empty. Check results for BUSCO: 37782at7742
## Thu Jul 14 00:23:14 CEST 2022 - The aliscore output file does not exist. Check results for BUSCO: 37782at7742
## Thu Jul 14 00:23:14 CEST 2022 - The aliscore output appears to be empty. Check results for BUSCO: 37782at7742
## Thu Jul 14 00:23:17 CEST 2022 - The aliscore output file does not exist. Check results for BUSCO: 31574at7742
## Thu Jul 14 00:23:17 CEST 2022 - The aliscore output appears to be empty. Check results for BUSCO: 31574at7742
## Thu Jul 14 00:23:17 CEST 2022 - The aliscore output file does not exist. Check results for BUSCO: 177338at7742
## Thu Jul 14 00:23:17 CEST 2022 - The aliscore output appears to be empty. Check results for BUSCO: 177338at7742
## Thu Jul 14 00:23:18 CEST 2022 - The aliscore output file does not exist. Check results for BUSCO: 46657at7742
## Thu Jul 14 00:23:18 CEST 2022 - The aliscore output appears to be empty. Check results for BUSCO: 46657at7742
## Thu Jul 14 00:23:18 CEST 2022 - The aliscore output file does not exist. Check results for BUSCO: 98179at7742
## Thu Jul 14 00:23:19 CEST 2022 - The aliscore output appears to be empty. Check results for BUSCO: 98179at7742
## Thu Jul 14 00:23:21 CEST 2022 - The aliscore output file does not exist. Check results for BUSCO: 17900at7742
## Thu Jul 14 00:23:21 CEST 2022 - The aliscore output appears to be empty. Check results for BUSCO: 17900at7742
## Thu Jul 14 00:23:23 CEST 2022 - The aliscore output file does not exist. Check results for BUSCO: 48975at7742
## Thu Jul 14 00:23:23 CEST 2022 - The aliscore output appears to be empty. Check results for BUSCO: 48975at7742
## Thu Jul 14 00:23:24 CEST 2022 - The aliscore output file does not exist. Check results for BUSCO: 98179at7742
## Thu Jul 14 00:23:24 CEST 2022 - The aliscore output appears to be empty. Check results for BUSCO: 98179at7742
## Thu Jul 14 00:23:26 CEST 2022 - The aliscore output file does not exist. Check results for BUSCO: 76178at7742
## Thu Jul 14 00:23:26 CEST 2022 - The aliscore output appears to be empty. Check results for BUSCO: 76178at7742
## Thu Jul 14 00:23:26 CEST 2022 - The aliscore output file does not exist. Check results for BUSCO: 142324at7742
## Thu Jul 14 00:23:26 CEST 2022 - The aliscore output appears to be empty. Check results for BUSCO: 142324at7742
## Thu Jul 14 00:23:28 CEST 2022 - The aliscore output file does not exist. Check results for BUSCO: 156439at7742
## Thu Jul 14 00:23:28 CEST 2022 - The aliscore output appears to be empty. Check results for BUSCO: 156439at7742
## Thu Jul 14 00:23:29 CEST 2022 - The aliscore output file does not exist. Check results for BUSCO: 105636at7742
## Thu Jul 14 00:23:29 CEST 2022 - The aliscore output appears to be empty. Check results for BUSCO: 105636at7742
## Thu Jul 14 00:23:32 CEST 2022 - The aliscore output file does not exist. Check results for BUSCO: 69398at7742
## Thu Jul 14 00:23:32 CEST 2022 - The aliscore output appears to be empty. Check results for BUSCO: 69398at7742
## Thu Jul 14 00:23:33 CEST 2022 - The aliscore output file does not exist. Check results for BUSCO: 68016at7742
## Thu Jul 14 00:23:33 CEST 2022 - The aliscore output appears to be empty. Check results for BUSCO: 68016at7742
## Thu Jul 14 00:23:34 CEST 2022 - The aliscore output file does not exist. Check results for BUSCO: 61105at7742
## Thu Jul 14 00:23:34 CEST 2022 - The aliscore output appears to be empty. Check results for BUSCO: 61105at7742
## Thu Jul 14 00:23:36 CEST 2022 - The aliscore output file does not exist. Check results for BUSCO: 33222at7742
## Thu Jul 14 00:23:36 CEST 2022 - The aliscore output appears to be empty. Check results for BUSCO: 33222at7742
## Thu Jul 14 00:23:38 CEST 2022 - The aliscore output file does not exist. Check results for BUSCO: 31574at7742
## Thu Jul 14 00:23:38 CEST 2022 - The aliscore output appears to be empty. Check results for BUSCO: 31574at7742
## Thu Jul 14 00:23:39 CEST 2022 - The aliscore output file does not exist. Check results for BUSCO: 43285at7742
## Thu Jul 14 00:23:39 CEST 2022 - The aliscore output appears to be empty. Check results for BUSCO: 43285at7742
## Thu Jul 14 00:23:40 CEST 2022 - The aliscore output file does not exist. Check results for BUSCO: 19307at7742
## Thu Jul 14 00:23:40 CEST 2022 - The aliscore output appears to be empty. Check results for BUSCO: 19307at7742
## Thu Jul 14 00:23:42 CEST 2022 - The aliscore output file does not exist. Check results for BUSCO: 104484at7742
## Thu Jul 14 00:23:42 CEST 2022 - The aliscore output appears to be empty. Check results for BUSCO: 104484at7742
## Thu Jul 14 00:23:42 CEST 2022 - The aliscore output file does not exist. Check results for BUSCO: 122017at7742
## Thu Jul 14 00:23:42 CEST 2022 - The aliscore output appears to be empty. Check results for BUSCO: 122017at7742
## Thu Jul 14 00:23:44 CEST 2022 - The aliscore output file does not exist. Check results for BUSCO: 160770at7742
## Thu Jul 14 00:23:44 CEST 2022 - The aliscore output appears to be empty. Check results for BUSCO: 160770at7742
## Thu Jul 14 00:23:47 CEST 2022 - The aliscore output file does not exist. Check results for BUSCO: 172713at7742
## Thu Jul 14 00:23:47 CEST 2022 - The aliscore output appears to be empty. Check results for BUSCO: 172713at7742
## Thu Jul 14 00:23:48 CEST 2022 - The aliscore output file does not exist. Check results for BUSCO: 166801at7742
## Thu Jul 14 00:23:48 CEST 2022 - The aliscore output appears to be empty. Check results for BUSCO: 166801at7742
## Thu Jul 14 00:23:49 CEST 2022 - The aliscore output file does not exist. Check results for BUSCO: 55967at7742
## Thu Jul 14 00:23:49 CEST 2022 - The aliscore output appears to be empty. Check results for BUSCO: 55967at7742
## Thu Jul 14 00:23:49 CEST 2022 - The aliscore output file does not exist. Check results for BUSCO: 235319at7742
## Thu Jul 14 00:23:49 CEST 2022 - The aliscore output appears to be empty. Check results for BUSCO: 235319at7742
## Thu Jul 14 00:23:52 CEST 2022 - The aliscore output file does not exist. Check results for BUSCO: 102147at7742
## Thu Jul 14 00:23:52 CEST 2022 - The aliscore output appears to be empty. Check results for BUSCO: 102147at7742
## Thu Jul 14 00:23:54 CEST 2022 - The aliscore output file does not exist. Check results for BUSCO: 92591at7742
## Thu Jul 14 00:23:54 CEST 2022 - The aliscore output appears to be empty. Check results for BUSCO: 92591at7742
## Thu Jul 14 00:23:54 CEST 2022 - The aliscore output file does not exist. Check results for BUSCO: 79920at7742
## Thu Jul 14 00:23:55 CEST 2022 - The aliscore output appears to be empty. Check results for BUSCO: 79920at7742
## Thu Jul 14 00:23:58 CEST 2022 - The aliscore output file does not exist. Check results for BUSCO: 19307at7742
## Thu Jul 14 00:23:58 CEST 2022 - The aliscore output appears to be empty. Check results for BUSCO: 19307at7742
## Thu Jul 14 00:23:59 CEST 2022 - The aliscore output file does not exist. Check results for BUSCO: 79920at7742
## Thu Jul 14 00:23:59 CEST 2022 - The aliscore output appears to be empty. Check results for BUSCO: 79920at7742
## Thu Jul 14 00:24:00 CEST 2022 - The aliscore output file does not exist. Check results for BUSCO: 273631at7742
## Thu Jul 14 00:24:00 CEST 2022 - The aliscore output appears to be empty. Check results for BUSCO: 273631at7742
## Thu Jul 14 00:24:00 CEST 2022 - The aliscore output file does not exist. Check results for BUSCO: 47023at7742
## Thu Jul 14 00:24:00 CEST 2022 - The aliscore output appears to be empty. Check results for BUSCO: 47023at7742
## Thu Jul 14 00:24:01 CEST 2022 - The aliscore output file does not exist. Check results for BUSCO: 37782at7742
## Thu Jul 14 00:24:01 CEST 2022 - The aliscore output appears to be empty. Check results for BUSCO: 37782at7742
## Thu Jul 14 00:24:04 CEST 2022 - The aliscore output file does not exist. Check results for BUSCO: 74297at7742
## Thu Jul 14 00:24:04 CEST 2022 - The aliscore output appears to be empty. Check results for BUSCO: 74297at7742
## Thu Jul 14 00:24:04 CEST 2022 - The aliscore output file does not exist. Check results for BUSCO: 137258at7742
## Thu Jul 14 00:24:04 CEST 2022 - The aliscore output appears to be empty. Check results for BUSCO: 137258at7742
## Thu Jul 14 00:24:06 CEST 2022 - The aliscore output file does not exist. Check results for BUSCO: 155749at7742
## Thu Jul 14 00:24:06 CEST 2022 - The aliscore output appears to be empty. Check results for BUSCO: 155749at7742
## Thu Jul 14 00:24:08 CEST 2022 - The aliscore output file does not exist. Check results for BUSCO: 166801at7742
## Thu Jul 14 00:24:08 CEST 2022 - The aliscore output appears to be empty. Check results for BUSCO: 166801at7742
## Thu Jul 14 00:24:10 CEST 2022 - The aliscore output file does not exist. Check results for BUSCO: 28500at7742
## Thu Jul 14 00:24:10 CEST 2022 - The aliscore output appears to be empty. Check results for BUSCO: 28500at7742
## Thu Jul 14 00:24:10 CEST 2022 - The aliscore output file does not exist. Check results for BUSCO: 63207at7742
## Thu Jul 14 00:24:10 CEST 2022 - The aliscore output appears to be empty. Check results for BUSCO: 63207at7742
## Thu Jul 14 00:24:12 CEST 2022 - The aliscore output file does not exist. Check results for BUSCO: 268813at7742
## Thu Jul 14 00:24:12 CEST 2022 - The aliscore output appears to be empty. Check results for BUSCO: 268813at7742
## Thu Jul 14 00:24:13 CEST 2022 - The aliscore output file does not exist. Check results for BUSCO: 17900at7742
## Thu Jul 14 00:24:13 CEST 2022 - The aliscore output appears to be empty. Check results for BUSCO: 17900at7742
## Thu Jul 14 00:24:15 CEST 2022 - The aliscore output file does not exist. Check results for BUSCO: 102147at7742
## Thu Jul 14 00:24:15 CEST 2022 - The aliscore output appears to be empty. Check results for BUSCO: 102147at7742
## Thu Jul 14 00:24:15 CEST 2022 - The aliscore output file does not exist. Check results for BUSCO: 17900at7742
## Thu Jul 14 00:24:16 CEST 2022 - The aliscore output appears to be empty. Check results for BUSCO: 17900at7742
## Thu Jul 14 00:24:18 CEST 2022 - The aliscore output file does not exist. Check results for BUSCO: 104992at7742
## Thu Jul 14 00:24:18 CEST 2022 - The aliscore output appears to be empty. Check results for BUSCO: 104992at7742
## Thu Jul 14 00:24:19 CEST 2022 - The aliscore output file does not exist. Check results for BUSCO: 95166at7742
## Thu Jul 14 00:24:19 CEST 2022 - The aliscore output appears to be empty. Check results for BUSCO: 95166at7742
## Thu Jul 14 00:24:20 CEST 2022 - The aliscore output file does not exist. Check results for BUSCO: 31418at7742
## Thu Jul 14 00:24:20 CEST 2022 - The aliscore output appears to be empty. Check results for BUSCO: 31418at7742
## Thu Jul 14 00:24:22 CEST 2022 - The aliscore output file does not exist. Check results for BUSCO: 103478at7742
## Thu Jul 14 00:24:22 CEST 2022 - The aliscore output appears to be empty. Check results for BUSCO: 103478at7742
## Thu Jul 14 00:24:24 CEST 2022 - The aliscore output file does not exist. Check results for BUSCO: 69398at7742
## Thu Jul 14 00:24:24 CEST 2022 - The aliscore output appears to be empty. Check results for BUSCO: 69398at7742
## Thu Jul 14 00:24:25 CEST 2022 - The aliscore output file does not exist. Check results for BUSCO: 87762at7742
## Thu Jul 14 00:24:25 CEST 2022 - The aliscore output appears to be empty. Check results for BUSCO: 87762at7742
## Thu Jul 14 00:24:27 CEST 2022 - The aliscore output file does not exist. Check results for BUSCO: 139572at7742
## Thu Jul 14 00:24:27 CEST 2022 - The aliscore output appears to be empty. Check results for BUSCO: 139572at7742
## Thu Jul 14 00:24:27 CEST 2022 - The aliscore output file does not exist. Check results for BUSCO: 234786at7742
## Thu Jul 14 00:24:27 CEST 2022 - The aliscore output appears to be empty. Check results for BUSCO: 234786at7742
## Thu Jul 14 00:24:31 CEST 2022 - The aliscore output file does not exist. Check results for BUSCO: 36274at7742
## Thu Jul 14 00:24:31 CEST 2022 - The aliscore output appears to be empty. Check results for BUSCO: 36274at7742
## Thu Jul 14 00:24:31 CEST 2022 - The aliscore output file does not exist. Check results for BUSCO: 119886at7742
## Thu Jul 14 00:24:31 CEST 2022 - The aliscore output appears to be empty. Check results for BUSCO: 119886at7742
## Thu Jul 14 00:24:31 CEST 2022 - The aliscore output file does not exist. Check results for BUSCO: 108510at7742
## Thu Jul 14 00:24:31 CEST 2022 - The aliscore output appears to be empty. Check results for BUSCO: 108510at7742
## Thu Jul 14 00:24:33 CEST 2022 - The aliscore output file does not exist. Check results for BUSCO: 273629at7742
## Thu Jul 14 00:24:33 CEST 2022 - The aliscore output appears to be empty. Check results for BUSCO: 273629at7742
## Thu Jul 14 00:24:35 CEST 2022 - The aliscore output file does not exist. Check results for BUSCO: 175472at7742
## Thu Jul 14 00:24:35 CEST 2022 - The aliscore output appears to be empty. Check results for BUSCO: 175472at7742
## Thu Jul 14 00:24:37 CEST 2022 - The aliscore output file does not exist. Check results for BUSCO: 277134at7742
## Thu Jul 14 00:24:37 CEST 2022 - The aliscore output appears to be empty. Check results for BUSCO: 277134at7742
## Thu Jul 14 00:24:38 CEST 2022 - The aliscore output file does not exist. Check results for BUSCO: 156112at7742
## Thu Jul 14 00:24:38 CEST 2022 - The aliscore output appears to be empty. Check results for BUSCO: 156112at7742
## Thu Jul 14 00:24:39 CEST 2022 - The aliscore output file does not exist. Check results for BUSCO: 126913at7742
## Thu Jul 14 00:24:39 CEST 2022 - The aliscore output appears to be empty. Check results for BUSCO: 126913at7742
## Thu Jul 14 00:24:40 CEST 2022 - The aliscore output file does not exist. Check results for BUSCO: 43285at7742
## Thu Jul 14 00:24:41 CEST 2022 - The aliscore output appears to be empty. Check results for BUSCO: 43285at7742
## Thu Jul 14 00:24:43 CEST 2022 - The aliscore output file does not exist. Check results for BUSCO: 137258at7742
## Thu Jul 14 00:24:43 CEST 2022 - The aliscore output appears to be empty. Check results for BUSCO: 137258at7742
## Thu Jul 14 00:24:43 CEST 2022 - The aliscore output file does not exist. Check results for BUSCO: 50502at7742
## Thu Jul 14 00:24:43 CEST 2022 - The aliscore output appears to be empty. Check results for BUSCO: 50502at7742
## Thu Jul 14 00:24:44 CEST 2022 - The aliscore output file does not exist. Check results for BUSCO: 37335at7742
## Thu Jul 14 00:24:44 CEST 2022 - The aliscore output appears to be empty. Check results for BUSCO: 37335at7742
## Thu Jul 14 00:24:48 CEST 2022 - The aliscore output file does not exist. Check results for BUSCO: 82217at7742
## Thu Jul 14 00:24:48 CEST 2022 - The aliscore output appears to be empty. Check results for BUSCO: 82217at7742
## Thu Jul 14 00:24:48 CEST 2022 - The aliscore output file does not exist. Check results for BUSCO: 44698at7742
## Thu Jul 14 00:24:48 CEST 2022 - The aliscore output appears to be empty. Check results for BUSCO: 44698at7742
## Thu Jul 14 00:24:50 CEST 2022 - The aliscore output file does not exist. Check results for BUSCO: 106972at7742
## Thu Jul 14 00:24:50 CEST 2022 - The aliscore output appears to be empty. Check results for BUSCO: 106972at7742
## Thu Jul 14 00:24:50 CEST 2022 - The aliscore output file does not exist. Check results for BUSCO: 29096at7742
## Thu Jul 14 00:24:50 CEST 2022 - The aliscore output appears to be empty. Check results for BUSCO: 29096at7742
## Thu Jul 14 00:24:50 CEST 2022 - The aliscore output file does not exist. Check results for BUSCO: 122017at7742
## Thu Jul 14 00:24:50 CEST 2022 - The aliscore output appears to be empty. Check results for BUSCO: 122017at7742
## Thu Jul 14 00:24:54 CEST 2022 - The aliscore output file does not exist. Check results for BUSCO: 29096at7742
## Thu Jul 14 00:24:54 CEST 2022 - The aliscore output appears to be empty. Check results for BUSCO: 29096at7742
## Thu Jul 14 00:24:56 CEST 2022 - The aliscore output file does not exist. Check results for BUSCO: 139572at7742
## Thu Jul 14 00:24:56 CEST 2022 - The aliscore output appears to be empty. Check results for BUSCO: 139572at7742
## Thu Jul 14 00:24:56 CEST 2022 - The aliscore output file does not exist. Check results for BUSCO: 172963at7742
## Thu Jul 14 00:24:56 CEST 2022 - The aliscore output appears to be empty. Check results for BUSCO: 172963at7742
## Thu Jul 14 00:24:58 CEST 2022 - The aliscore output file does not exist. Check results for BUSCO: 95166at7742
## Thu Jul 14 00:24:58 CEST 2022 - The aliscore output appears to be empty. Check results for BUSCO: 95166at7742
## Thu Jul 14 00:24:59 CEST 2022 - The aliscore output file does not exist. Check results for BUSCO: 268813at7742
## Thu Jul 14 00:24:59 CEST 2022 - The aliscore output appears to be empty. Check results for BUSCO: 268813at7742
## Thu Jul 14 00:25:01 CEST 2022 - The aliscore output file does not exist. Check results for BUSCO: 29096at7742
## Thu Jul 14 00:25:01 CEST 2022 - The aliscore output appears to be empty. Check results for BUSCO: 29096at7742
## Thu Jul 14 00:25:01 CEST 2022 - The aliscore output file does not exist. Check results for BUSCO: 60899at7742
## Thu Jul 14 00:25:01 CEST 2022 - The aliscore output appears to be empty. Check results for BUSCO: 60899at7742
## Thu Jul 14 00:25:03 CEST 2022 - The aliscore output file does not exist. Check results for BUSCO: 28859at7742
## Thu Jul 14 00:25:03 CEST 2022 - The aliscore output appears to be empty. Check results for BUSCO: 28859at7742
## Thu Jul 14 00:25:05 CEST 2022 - The aliscore output file does not exist. Check results for BUSCO: 79920at7742
## Thu Jul 14 00:25:05 CEST 2022 - The aliscore output appears to be empty. Check results for BUSCO: 79920at7742
## Thu Jul 14 00:25:09 CEST 2022 - The aliscore output file does not exist. Check results for BUSCO: 42483at7742
## Thu Jul 14 00:25:09 CEST 2022 - The aliscore output appears to be empty. Check results for BUSCO: 42483at7742
## Thu Jul 14 00:25:09 CEST 2022 - The aliscore output file does not exist. Check results for BUSCO: 105968at7742
## Thu Jul 14 00:25:09 CEST 2022 - The aliscore output appears to be empty. Check results for BUSCO: 105968at7742
## Thu Jul 14 00:25:09 CEST 2022 - The aliscore output file does not exist. Check results for BUSCO: 25839at7742
## Thu Jul 14 00:25:09 CEST 2022 - The aliscore output appears to be empty. Check results for BUSCO: 25839at7742
## Thu Jul 14 00:25:09 CEST 2022 - The aliscore output file does not exist. Check results for BUSCO: 134632at7742
## Thu Jul 14 00:25:09 CEST 2022 - The aliscore output appears to be empty. Check results for BUSCO: 134632at7742
## Thu Jul 14 00:25:10 CEST 2022 - The aliscore output file does not exist. Check results for BUSCO: 50838at7742
## Thu Jul 14 00:25:10 CEST 2022 - The aliscore output appears to be empty. Check results for BUSCO: 50838at7742
## Thu Jul 14 00:25:14 CEST 2022 - The aliscore output file does not exist. Check results for BUSCO: 31066at7742
## Thu Jul 14 00:25:14 CEST 2022 - The aliscore output appears to be empty. Check results for BUSCO: 31066at7742
## Thu Jul 14 00:25:15 CEST 2022 - The aliscore output file does not exist. Check results for BUSCO: 57179at7742
## Thu Jul 14 00:25:15 CEST 2022 - The aliscore output appears to be empty. Check results for BUSCO: 57179at7742
## Thu Jul 14 00:25:15 CEST 2022 - The aliscore output file does not exist. Check results for BUSCO: 156439at7742
## Thu Jul 14 00:25:15 CEST 2022 - The aliscore output appears to be empty. Check results for BUSCO: 156439at7742
## Thu Jul 14 00:25:18 CEST 2022 - The aliscore output file does not exist. Check results for BUSCO: 24404at7742
## Thu Jul 14 00:25:18 CEST 2022 - The aliscore output appears to be empty. Check results for BUSCO: 24404at7742
## Thu Jul 14 00:25:19 CEST 2022 - The aliscore output file does not exist. Check results for BUSCO: 28859at7742
## Thu Jul 14 00:25:19 CEST 2022 - The aliscore output appears to be empty. Check results for BUSCO: 28859at7742
## Thu Jul 14 00:25:19 CEST 2022 - The aliscore output file does not exist. Check results for BUSCO: 191412at7742
## Thu Jul 14 00:25:19 CEST 2022 - The aliscore output appears to be empty. Check results for BUSCO: 191412at7742
## Thu Jul 14 00:25:20 CEST 2022 - The aliscore output file does not exist. Check results for BUSCO: 53816at7742
## Thu Jul 14 00:25:20 CEST 2022 - The aliscore output appears to be empty. Check results for BUSCO: 53816at7742
## Thu Jul 14 00:25:20 CEST 2022 - The aliscore output file does not exist. Check results for BUSCO: 122017at7742
## Thu Jul 14 00:25:20 CEST 2022 - The aliscore output appears to be empty. Check results for BUSCO: 122017at7742
## Thu Jul 14 00:25:23 CEST 2022 - The aliscore output file does not exist. Check results for BUSCO: 169538at7742
## Thu Jul 14 00:25:23 CEST 2022 - The aliscore output appears to be empty. Check results for BUSCO: 169538at7742
## Thu Jul 14 00:25:25 CEST 2022 - The aliscore output file does not exist. Check results for BUSCO: 46657at7742
## Thu Jul 14 00:25:26 CEST 2022 - The aliscore output appears to be empty. Check results for BUSCO: 46657at7742
## Thu Jul 14 00:25:27 CEST 2022 - The aliscore output file does not exist. Check results for BUSCO: 261794at7742
## Thu Jul 14 00:25:27 CEST 2022 - The aliscore output appears to be empty. Check results for BUSCO: 261794at7742
## Thu Jul 14 00:25:29 CEST 2022 - The aliscore output file does not exist. Check results for BUSCO: 150697at7742
## Thu Jul 14 00:25:29 CEST 2022 - The aliscore output appears to be empty. Check results for BUSCO: 150697at7742
## Thu Jul 14 00:25:31 CEST 2022 - The aliscore output file does not exist. Check results for BUSCO: 156112at7742
## Thu Jul 14 00:25:31 CEST 2022 - The aliscore output appears to be empty. Check results for BUSCO: 156112at7742
## Thu Jul 14 00:25:31 CEST 2022 - The aliscore output file does not exist. Check results for BUSCO: 122255at7742
## Thu Jul 14 00:25:31 CEST 2022 - The aliscore output appears to be empty. Check results for BUSCO: 122255at7742
## Thu Jul 14 00:25:32 CEST 2022 - The aliscore output file does not exist. Check results for BUSCO: 207707at7742
## Thu Jul 14 00:25:32 CEST 2022 - The aliscore output appears to be empty. Check results for BUSCO: 207707at7742
## Thu Jul 14 00:25:34 CEST 2022 - The aliscore output file does not exist. Check results for BUSCO: 33222at7742
## Thu Jul 14 00:25:34 CEST 2022 - The aliscore output appears to be empty. Check results for BUSCO: 33222at7742
## Thu Jul 14 00:25:36 CEST 2022 - The aliscore output file does not exist. Check results for BUSCO: 216073at7742
## Thu Jul 14 00:25:36 CEST 2022 - The aliscore output appears to be empty. Check results for BUSCO: 216073at7742
## Thu Jul 14 00:25:38 CEST 2022 - The aliscore output file does not exist. Check results for BUSCO: 24404at7742
## Thu Jul 14 00:25:38 CEST 2022 - The aliscore output appears to be empty. Check results for BUSCO: 24404at7742
## Thu Jul 14 00:25:39 CEST 2022 - The aliscore output file does not exist. Check results for BUSCO: 43798at7742
## Thu Jul 14 00:25:39 CEST 2022 - The aliscore output appears to be empty. Check results for BUSCO: 43798at7742
## Thu Jul 14 00:25:41 CEST 2022 - The aliscore output file does not exist. Check results for BUSCO: 80917at7742
## Thu Jul 14 00:25:41 CEST 2022 - The aliscore output appears to be empty. Check results for BUSCO: 80917at7742
## Thu Jul 14 00:25:41 CEST 2022 - The aliscore output file does not exist. Check results for BUSCO: 143698at7742
## Thu Jul 14 00:25:41 CEST 2022 - The aliscore output appears to be empty. Check results for BUSCO: 143698at7742
## Thu Jul 14 00:25:42 CEST 2022 - The aliscore output file does not exist. Check results for BUSCO: 53273at7742
## Thu Jul 14 00:25:42 CEST 2022 - The aliscore output appears to be empty. Check results for BUSCO: 53273at7742
## Thu Jul 14 00:25:46 CEST 2022 - The aliscore output file does not exist. Check results for BUSCO: 153497at7742
## Thu Jul 14 00:25:46 CEST 2022 - The aliscore output appears to be empty. Check results for BUSCO: 153497at7742
## Thu Jul 14 00:25:48 CEST 2022 - The aliscore output file does not exist. Check results for BUSCO: 155749at7742
## Thu Jul 14 00:25:48 CEST 2022 - The aliscore output appears to be empty. Check results for BUSCO: 155749at7742
## Thu Jul 14 00:25:48 CEST 2022 - The aliscore output file does not exist. Check results for BUSCO: 19307at7742
## Thu Jul 14 00:25:49 CEST 2022 - The aliscore output appears to be empty. Check results for BUSCO: 19307at7742
## Thu Jul 14 00:25:49 CEST 2022 - The aliscore output file does not exist. Check results for BUSCO: 35861at7742
## Thu Jul 14 00:25:49 CEST 2022 - The aliscore output appears to be empty. Check results for BUSCO: 35861at7742
## Thu Jul 14 00:25:51 CEST 2022 - The aliscore output file does not exist. Check results for BUSCO: 95643at7742
## Thu Jul 14 00:25:51 CEST 2022 - The aliscore output appears to be empty. Check results for BUSCO: 95643at7742
## Thu Jul 14 00:25:51 CEST 2022 - The aliscore output file does not exist. Check results for BUSCO: 169538at7742
## Thu Jul 14 00:25:51 CEST 2022 - The aliscore output appears to be empty. Check results for BUSCO: 169538at7742
## Thu Jul 14 00:25:51 CEST 2022 - The aliscore output file does not exist. Check results for BUSCO: 64860at7742
## Thu Jul 14 00:25:52 CEST 2022 - The aliscore output appears to be empty. Check results for BUSCO: 64860at7742
## Thu Jul 14 00:25:52 CEST 2022 - The aliscore output file does not exist. Check results for BUSCO: 28859at7742
## Thu Jul 14 00:25:52 CEST 2022 - The aliscore output appears to be empty. Check results for BUSCO: 28859at7742
## Thu Jul 14 00:25:55 CEST 2022 - The aliscore output file does not exist. Check results for BUSCO: 33940at7742
## Thu Jul 14 00:25:55 CEST 2022 - The aliscore output appears to be empty. Check results for BUSCO: 33940at7742
## Thu Jul 14 00:25:58 CEST 2022 - The aliscore output file does not exist. Check results for BUSCO: 31066at7742
## Thu Jul 14 00:25:58 CEST 2022 - The aliscore output appears to be empty. Check results for BUSCO: 31066at7742
## Thu Jul 14 00:25:59 CEST 2022 - The aliscore output file does not exist. Check results for BUSCO: 137258at7742
## Thu Jul 14 00:25:59 CEST 2022 - The aliscore output appears to be empty. Check results for BUSCO: 137258at7742
## Thu Jul 14 00:26:00 CEST 2022 - The aliscore output file does not exist. Check results for BUSCO: 344416at7742
## Thu Jul 14 00:26:00 CEST 2022 - The aliscore output appears to be empty. Check results for BUSCO: 344416at7742
## Thu Jul 14 00:26:00 CEST 2022 - The aliscore output file does not exist. Check results for BUSCO: 155749at7742
## Thu Jul 14 00:26:00 CEST 2022 - The aliscore output appears to be empty. Check results for BUSCO: 155749at7742
## Thu Jul 14 00:26:02 CEST 2022 - The aliscore output file does not exist. Check results for BUSCO: 39313at7742
## Thu Jul 14 00:26:02 CEST 2022 - The aliscore output appears to be empty. Check results for BUSCO: 39313at7742
## Thu Jul 14 00:26:04 CEST 2022 - The aliscore output file does not exist. Check results for BUSCO: 33940at7742
## Thu Jul 14 00:26:04 CEST 2022 - The aliscore output appears to be empty. Check results for BUSCO: 33940at7742
## Thu Jul 14 00:26:06 CEST 2022 - The aliscore output file does not exist. Check results for BUSCO: 138116at7742
## Thu Jul 14 00:26:06 CEST 2022 - The aliscore output appears to be empty. Check results for BUSCO: 138116at7742
## Thu Jul 14 00:26:07 CEST 2022 - The aliscore output file does not exist. Check results for BUSCO: 129039at7742
## Thu Jul 14 00:26:07 CEST 2022 - The aliscore output appears to be empty. Check results for BUSCO: 129039at7742
## Thu Jul 14 00:26:10 CEST 2022 - The aliscore output file does not exist. Check results for BUSCO: 52065at7742
## Thu Jul 14 00:26:10 CEST 2022 - The aliscore output appears to be empty. Check results for BUSCO: 52065at7742
## Thu Jul 14 00:26:10 CEST 2022 - The aliscore output file does not exist. Check results for BUSCO: 139572at7742
## Thu Jul 14 00:26:10 CEST 2022 - The aliscore output appears to be empty. Check results for BUSCO: 139572at7742
## Thu Jul 14 00:26:11 CEST 2022 - The aliscore output file does not exist. Check results for BUSCO: 108510at7742
## Thu Jul 14 00:26:11 CEST 2022 - The aliscore output appears to be empty. Check results for BUSCO: 108510at7742
## Thu Jul 14 00:26:12 CEST 2022 - The aliscore output file does not exist. Check results for BUSCO: 63939at7742
## Thu Jul 14 00:26:13 CEST 2022 - The aliscore output appears to be empty. Check results for BUSCO: 63939at7742
## Thu Jul 14 00:26:17 CEST 2022 - The aliscore output file does not exist. Check results for BUSCO: 35211at7742
## Thu Jul 14 00:26:17 CEST 2022 - The aliscore output appears to be empty. Check results for BUSCO: 35211at7742
## Thu Jul 14 00:26:17 CEST 2022 - The aliscore output file does not exist. Check results for BUSCO: 261794at7742
## Thu Jul 14 00:26:17 CEST 2022 - The aliscore output appears to be empty. Check results for BUSCO: 261794at7742
## Thu Jul 14 00:26:18 CEST 2022 - The aliscore output file does not exist. Check results for BUSCO: 53816at7742
## Thu Jul 14 00:26:18 CEST 2022 - The aliscore output appears to be empty. Check results for BUSCO: 53816at7742
## Thu Jul 14 00:26:21 CEST 2022 - The aliscore output file does not exist. Check results for BUSCO: 39393at7742
## Thu Jul 14 00:26:21 CEST 2022 - The aliscore output appears to be empty. Check results for BUSCO: 39393at7742
## Thu Jul 14 00:26:22 CEST 2022 - The aliscore output file does not exist. Check results for BUSCO: 361964at7742
## Thu Jul 14 00:26:22 CEST 2022 - The aliscore output file does not exist. Check results for BUSCO: 46657at7742
## Thu Jul 14 00:26:22 CEST 2022 - The aliscore output appears to be empty. Check results for BUSCO: 361964at7742
## Thu Jul 14 00:26:22 CEST 2022 - The aliscore output appears to be empty. Check results for BUSCO: 46657at7742
## Thu Jul 14 00:26:24 CEST 2022 - The aliscore output file does not exist. Check results for BUSCO: 150497at7742
## Thu Jul 14 00:26:24 CEST 2022 - The aliscore output appears to be empty. Check results for BUSCO: 150497at7742
## Thu Jul 14 00:26:26 CEST 2022 - The aliscore output file does not exist. Check results for BUSCO: 92481at7742
## Thu Jul 14 00:26:26 CEST 2022 - The aliscore output appears to be empty. Check results for BUSCO: 92481at7742
## Thu Jul 14 00:26:28 CEST 2022 - The aliscore output file does not exist. Check results for BUSCO: 47023at7742
## Thu Jul 14 00:26:28 CEST 2022 - The aliscore output appears to be empty. Check results for BUSCO: 47023at7742
## Thu Jul 14 00:26:28 CEST 2022 - The aliscore output file does not exist. Check results for BUSCO: 87117at7742
## Thu Jul 14 00:26:28 CEST 2022 - The aliscore output appears to be empty. Check results for BUSCO: 87117at7742
## Thu Jul 14 00:26:32 CEST 2022 - The aliscore output file does not exist. Check results for BUSCO: 299948at7742
## Thu Jul 14 00:26:32 CEST 2022 - The aliscore output appears to be empty. Check results for BUSCO: 299948at7742
## Thu Jul 14 00:26:32 CEST 2022 - The aliscore output file does not exist. Check results for BUSCO: 52094at7742
## Thu Jul 14 00:26:32 CEST 2022 - The aliscore output appears to be empty. Check results for BUSCO: 52094at7742
## Thu Jul 14 00:26:32 CEST 2022 - The aliscore output file does not exist. Check results for BUSCO: 95435at7742
## Thu Jul 14 00:26:32 CEST 2022 - The aliscore output appears to be empty. Check results for BUSCO: 95435at7742
## Thu Jul 14 00:26:32 CEST 2022 - The aliscore output file does not exist. Check results for BUSCO: 174474at7742
## Thu Jul 14 00:26:32 CEST 2022 - The aliscore output appears to be empty. Check results for BUSCO: 174474at7742
## Thu Jul 14 00:26:35 CEST 2022 - The aliscore output file does not exist. Check results for BUSCO: 319198at7742
## Thu Jul 14 00:26:35 CEST 2022 - The aliscore output appears to be empty. Check results for BUSCO: 319198at7742
## Thu Jul 14 00:26:38 CEST 2022 - The aliscore output file does not exist. Check results for BUSCO: 253823at7742
## Thu Jul 14 00:26:38 CEST 2022 - The aliscore output appears to be empty. Check results for BUSCO: 253823at7742
## Thu Jul 14 00:26:39 CEST 2022 - The aliscore output file does not exist. Check results for BUSCO: 53816at7742
## Thu Jul 14 00:26:39 CEST 2022 - The aliscore output appears to be empty. Check results for BUSCO: 53816at7742
## Thu Jul 14 00:26:41 CEST 2022 - The aliscore output file does not exist. Check results for BUSCO: 140683at7742
## Thu Jul 14 00:26:41 CEST 2022 - The aliscore output appears to be empty. Check results for BUSCO: 140683at7742
## Thu Jul 14 00:26:42 CEST 2022 - The aliscore output file does not exist. Check results for BUSCO: 82217at7742
## Thu Jul 14 00:26:42 CEST 2022 - The aliscore output appears to be empty. Check results for BUSCO: 82217at7742
## Thu Jul 14 00:26:43 CEST 2022 - The aliscore output file does not exist. Check results for BUSCO: 143864at7742
## Thu Jul 14 00:26:43 CEST 2022 - The aliscore output appears to be empty. Check results for BUSCO: 143864at7742
## Thu Jul 14 00:26:44 CEST 2022 - The aliscore output file does not exist. Check results for BUSCO: 63939at7742
## Thu Jul 14 00:26:44 CEST 2022 - The aliscore output appears to be empty. Check results for BUSCO: 63939at7742
## Thu Jul 14 00:26:45 CEST 2022 - The aliscore output file does not exist. Check results for BUSCO: 95166at7742
## Thu Jul 14 00:26:45 CEST 2022 - The aliscore output appears to be empty. Check results for BUSCO: 95166at7742
## Thu Jul 14 00:26:47 CEST 2022 - The aliscore output file does not exist. Check results for BUSCO: 334890at7742
## Thu Jul 14 00:26:47 CEST 2022 - The aliscore output appears to be empty. Check results for BUSCO: 334890at7742
## Thu Jul 14 00:26:48 CEST 2022 - The aliscore output file does not exist. Check results for BUSCO: 36274at7742
## Thu Jul 14 00:26:48 CEST 2022 - The aliscore output appears to be empty. Check results for BUSCO: 36274at7742
## Thu Jul 14 00:26:51 CEST 2022 - The aliscore output file does not exist. Check results for BUSCO: 52094at7742
## Thu Jul 14 00:26:51 CEST 2022 - The aliscore output appears to be empty. Check results for BUSCO: 52094at7742
## Thu Jul 14 00:26:52 CEST 2022 - The aliscore output file does not exist. Check results for BUSCO: 235319at7742
## Thu Jul 14 00:26:52 CEST 2022 - The aliscore output file does not exist. Check results for BUSCO: 138498at7742
## Thu Jul 14 00:26:52 CEST 2022 - The aliscore output appears to be empty. Check results for BUSCO: 235319at7742
## Thu Jul 14 00:26:52 CEST 2022 - The aliscore output appears to be empty. Check results for BUSCO: 138498at7742
## Thu Jul 14 00:26:55 CEST 2022 - The aliscore output file does not exist. Check results for BUSCO: 49005at7742
## Thu Jul 14 00:26:55 CEST 2022 - The aliscore output appears to be empty. Check results for BUSCO: 49005at7742
## Thu Jul 14 00:26:56 CEST 2022 - The aliscore output file does not exist. Check results for BUSCO: 39393at7742
## Thu Jul 14 00:26:56 CEST 2022 - The aliscore output appears to be empty. Check results for BUSCO: 39393at7742
## Thu Jul 14 00:26:57 CEST 2022 - The aliscore output file does not exist. Check results for BUSCO: 35861at7742
## Thu Jul 14 00:26:57 CEST 2022 - The aliscore output appears to be empty. Check results for BUSCO: 35861at7742
## Thu Jul 14 00:26:57 CEST 2022 - The aliscore output file does not exist. Check results for BUSCO: 42971at7742
## Thu Jul 14 00:26:57 CEST 2022 - The aliscore output appears to be empty. Check results for BUSCO: 42971at7742
## Thu Jul 14 00:27:00 CEST 2022 - The aliscore output file does not exist. Check results for BUSCO: 74922at7742
## Thu Jul 14 00:27:00 CEST 2022 - The aliscore output appears to be empty. Check results for BUSCO: 74922at7742
## Thu Jul 14 00:27:00 CEST 2022 - The aliscore output file does not exist. Check results for BUSCO: 39393at7742
## Thu Jul 14 00:27:00 CEST 2022 - The aliscore output appears to be empty. Check results for BUSCO: 39393at7742
## Thu Jul 14 00:27:02 CEST 2022 - The aliscore output file does not exist. Check results for BUSCO: 68589at7742
## Thu Jul 14 00:27:02 CEST 2022 - The aliscore output appears to be empty. Check results for BUSCO: 68589at7742
## Thu Jul 14 00:27:04 CEST 2022 - The aliscore output file does not exist. Check results for BUSCO: 137788at7742
## Thu Jul 14 00:27:04 CEST 2022 - The aliscore output appears to be empty. Check results for BUSCO: 137788at7742
## Thu Jul 14 00:27:06 CEST 2022 - The aliscore output file does not exist. Check results for BUSCO: 52094at7742
## Thu Jul 14 00:27:06 CEST 2022 - The aliscore output appears to be empty. Check results for BUSCO: 52094at7742
## Thu Jul 14 00:27:07 CEST 2022 - The aliscore output file does not exist. Check results for BUSCO: 100129at7742
## Thu Jul 14 00:27:07 CEST 2022 - The aliscore output file does not exist. Check results for BUSCO: 87762at7742
## Thu Jul 14 00:27:07 CEST 2022 - The aliscore output appears to be empty. Check results for BUSCO: 100129at7742
## Thu Jul 14 00:27:07 CEST 2022 - The aliscore output appears to be empty. Check results for BUSCO: 87762at7742
## Thu Jul 14 00:27:10 CEST 2022 - The aliscore output file does not exist. Check results for BUSCO: 224567at7742
## Thu Jul 14 00:27:10 CEST 2022 - The aliscore output appears to be empty. Check results for BUSCO: 224567at7742
## Thu Jul 14 00:27:12 CEST 2022 - The aliscore output file does not exist. Check results for BUSCO: 108633at7742
## Thu Jul 14 00:27:12 CEST 2022 - The aliscore output appears to be empty. Check results for BUSCO: 108633at7742
## Thu Jul 14 00:27:14 CEST 2022 - The aliscore output file does not exist. Check results for BUSCO: 299948at7742
## Thu Jul 14 00:27:14 CEST 2022 - The aliscore output appears to be empty. Check results for BUSCO: 299948at7742
## Thu Jul 14 00:27:15 CEST 2022 - The aliscore output file does not exist. Check results for BUSCO: 299948at7742
## Thu Jul 14 00:27:15 CEST 2022 - The aliscore output appears to be empty. Check results for BUSCO: 299948at7742
## Thu Jul 14 00:27:17 CEST 2022 - The aliscore output file does not exist. Check results for BUSCO: 224567at7742
## Thu Jul 14 00:27:17 CEST 2022 - The aliscore output appears to be empty. Check results for BUSCO: 224567at7742
## Thu Jul 14 00:27:17 CEST 2022 - The aliscore output file does not exist. Check results for BUSCO: 224692at7742
## Thu Jul 14 00:27:17 CEST 2022 - The aliscore output appears to be empty. Check results for BUSCO: 224692at7742
## Thu Jul 14 00:27:17 CEST 2022 - The aliscore output file does not exist. Check results for BUSCO: 61105at7742
## Thu Jul 14 00:27:17 CEST 2022 - The aliscore output appears to be empty. Check results for BUSCO: 61105at7742
## Thu Jul 14 00:27:21 CEST 2022 - The aliscore output file does not exist. Check results for BUSCO: 72648at7742
## Thu Jul 14 00:27:21 CEST 2022 - The aliscore output appears to be empty. Check results for BUSCO: 72648at7742
## Thu Jul 14 00:27:21 CEST 2022 - The aliscore output file does not exist. Check results for BUSCO: 31066at7742
## Thu Jul 14 00:27:22 CEST 2022 - The aliscore output appears to be empty. Check results for BUSCO: 31066at7742
## Thu Jul 14 00:27:23 CEST 2022 - The aliscore output file does not exist. Check results for BUSCO: 105968at7742
## Thu Jul 14 00:27:23 CEST 2022 - The aliscore output appears to be empty. Check results for BUSCO: 105968at7742
## Thu Jul 14 00:27:26 CEST 2022 - The aliscore output file does not exist. Check results for BUSCO: 43798at7742
## Thu Jul 14 00:27:26 CEST 2022 - The aliscore output appears to be empty. Check results for BUSCO: 43798at7742
## Thu Jul 14 00:27:26 CEST 2022 - The aliscore output file does not exist. Check results for BUSCO: 35861at7742
## Thu Jul 14 00:27:26 CEST 2022 - The aliscore output appears to be empty. Check results for BUSCO: 35861at7742
## Thu Jul 14 00:27:27 CEST 2022 - The aliscore output file does not exist. Check results for BUSCO: 344416at7742
## Thu Jul 14 00:27:27 CEST 2022 - The aliscore output appears to be empty. Check results for BUSCO: 344416at7742
## Thu Jul 14 00:27:29 CEST 2022 - The aliscore output file does not exist. Check results for BUSCO: 123570at7742
## Thu Jul 14 00:27:29 CEST 2022 - The aliscore output appears to be empty. Check results for BUSCO: 123570at7742
## Thu Jul 14 00:27:30 CEST 2022 - The aliscore output file does not exist. Check results for BUSCO: 138116at7742
## Thu Jul 14 00:27:30 CEST 2022 - The aliscore output appears to be empty. Check results for BUSCO: 138116at7742
## Thu Jul 14 00:27:32 CEST 2022 - The aliscore output file does not exist. Check results for BUSCO: 46728at7742
## Thu Jul 14 00:27:32 CEST 2022 - The aliscore output file does not exist. Check results for BUSCO: 246595at7742
## Thu Jul 14 00:27:32 CEST 2022 - The aliscore output appears to be empty. Check results for BUSCO: 46728at7742
## Thu Jul 14 00:27:33 CEST 2022 - The aliscore output appears to be empty. Check results for BUSCO: 246595at7742
## Thu Jul 14 00:27:36 CEST 2022 - The aliscore output file does not exist. Check results for BUSCO: 103284at7742
## Thu Jul 14 00:27:36 CEST 2022 - The aliscore output appears to be empty. Check results for BUSCO: 103284at7742
## Thu Jul 14 00:27:37 CEST 2022 - The aliscore output file does not exist. Check results for BUSCO: 156689at7742
## Thu Jul 14 00:27:37 CEST 2022 - The aliscore output appears to be empty. Check results for BUSCO: 156689at7742
## Thu Jul 14 00:27:37 CEST 2022 - The aliscore output file does not exist. Check results for BUSCO: 229239at7742
## Thu Jul 14 00:27:37 CEST 2022 - The aliscore output appears to be empty. Check results for BUSCO: 229239at7742
## Thu Jul 14 00:27:39 CEST 2022 - The aliscore output file does not exist. Check results for BUSCO: 224567at7742
## Thu Jul 14 00:27:39 CEST 2022 - The aliscore output appears to be empty. Check results for BUSCO: 224567at7742
## Thu Jul 14 00:27:39 CEST 2022 - The aliscore output file does not exist. Check results for BUSCO: 126913at7742
## Thu Jul 14 00:27:39 CEST 2022 - The aliscore output appears to be empty. Check results for BUSCO: 126913at7742
## Thu Jul 14 00:27:40 CEST 2022 - The aliscore output file does not exist. Check results for BUSCO: 121053at7742
## Thu Jul 14 00:27:40 CEST 2022 - The aliscore output appears to be empty. Check results for BUSCO: 121053at7742
## Thu Jul 14 00:27:42 CEST 2022 - The aliscore output file does not exist. Check results for BUSCO: 121375at7742
## Thu Jul 14 00:27:42 CEST 2022 - The aliscore output appears to be empty. Check results for BUSCO: 121375at7742
## Thu Jul 14 00:27:45 CEST 2022 - The aliscore output file does not exist. Check results for BUSCO: 41344at7742
## Thu Jul 14 00:27:45 CEST 2022 - The aliscore output appears to be empty. Check results for BUSCO: 41344at7742
## Thu Jul 14 00:27:45 CEST 2022 - The aliscore output file does not exist. Check results for BUSCO: 319198at7742
## Thu Jul 14 00:27:45 CEST 2022 - The aliscore output appears to be empty. Check results for BUSCO: 319198at7742
## Thu Jul 14 00:27:48 CEST 2022 - The aliscore output file does not exist. Check results for BUSCO: 35211at7742
## Thu Jul 14 00:27:48 CEST 2022 - The aliscore output appears to be empty. Check results for BUSCO: 35211at7742
## Thu Jul 14 00:27:48 CEST 2022 - The aliscore output file does not exist. Check results for BUSCO: 95963at7742
## Thu Jul 14 00:27:48 CEST 2022 - The aliscore output appears to be empty. Check results for BUSCO: 95963at7742
## Thu Jul 14 00:27:49 CEST 2022 - The aliscore output file does not exist. Check results for BUSCO: 53273at7742
## Thu Jul 14 00:27:49 CEST 2022 - The aliscore output appears to be empty. Check results for BUSCO: 53273at7742
## Thu Jul 14 00:27:49 CEST 2022 - The aliscore output file does not exist. Check results for BUSCO: 153497at7742
## Thu Jul 14 00:27:49 CEST 2022 - The aliscore output appears to be empty. Check results for BUSCO: 153497at7742
## Thu Jul 14 00:27:52 CEST 2022 - The aliscore output file does not exist. Check results for BUSCO: 116202at7742
## Thu Jul 14 00:27:52 CEST 2022 - The aliscore output appears to be empty. Check results for BUSCO: 116202at7742
## Thu Jul 14 00:27:54 CEST 2022 - The aliscore output file does not exist. Check results for BUSCO: 48975at7742
## Thu Jul 14 00:27:54 CEST 2022 - The aliscore output appears to be empty. Check results for BUSCO: 48975at7742
## Thu Jul 14 00:27:55 CEST 2022 - The aliscore output file does not exist. Check results for BUSCO: 58275at7742
## Thu Jul 14 00:27:55 CEST 2022 - The aliscore output appears to be empty. Check results for BUSCO: 58275at7742
## Thu Jul 14 00:27:55 CEST 2022 - The aliscore output file does not exist. Check results for BUSCO: 91371at7742
## Thu Jul 14 00:27:56 CEST 2022 - The aliscore output appears to be empty. Check results for BUSCO: 91371at7742
## Thu Jul 14 00:27:58 CEST 2022 - The aliscore output file does not exist. Check results for BUSCO: 108633at7742
## Thu Jul 14 00:27:58 CEST 2022 - The aliscore output appears to be empty. Check results for BUSCO: 108633at7742
## Thu Jul 14 00:28:00 CEST 2022 - The aliscore output file does not exist. Check results for BUSCO: 103478at7742
## Thu Jul 14 00:28:00 CEST 2022 - The aliscore output appears to be empty. Check results for BUSCO: 103478at7742
## Thu Jul 14 00:28:01 CEST 2022 - The aliscore output file does not exist. Check results for BUSCO: 198404at7742
## Thu Jul 14 00:28:01 CEST 2022 - The aliscore output appears to be empty. Check results for BUSCO: 198404at7742
## Thu Jul 14 00:28:01 CEST 2022 - The aliscore output file does not exist. Check results for BUSCO: 238925at7742
## Thu Jul 14 00:28:02 CEST 2022 - The aliscore output appears to be empty. Check results for BUSCO: 238925at7742
## Thu Jul 14 00:28:04 CEST 2022 - The aliscore output file does not exist. Check results for BUSCO: 60899at7742
## Thu Jul 14 00:28:04 CEST 2022 - The aliscore output appears to be empty. Check results for BUSCO: 60899at7742
## Thu Jul 14 00:28:06 CEST 2022 - The aliscore output file does not exist. Check results for BUSCO: 127587at7742
## Thu Jul 14 00:28:06 CEST 2022 - The aliscore output appears to be empty. Check results for BUSCO: 127587at7742
## Thu Jul 14 00:28:07 CEST 2022 - The aliscore output file does not exist. Check results for BUSCO: 93412at7742
## Thu Jul 14 00:28:07 CEST 2022 - The aliscore output appears to be empty. Check results for BUSCO: 93412at7742
## Thu Jul 14 00:28:08 CEST 2022 - The aliscore output file does not exist. Check results for BUSCO: 95310at7742
## Thu Jul 14 00:28:08 CEST 2022 - The aliscore output appears to be empty. Check results for BUSCO: 95310at7742
## Thu Jul 14 00:28:10 CEST 2022 - The aliscore output file does not exist. Check results for BUSCO: 103284at7742
## Thu Jul 14 00:28:10 CEST 2022 - The aliscore output appears to be empty. Check results for BUSCO: 103284at7742
## Thu Jul 14 00:28:11 CEST 2022 - The aliscore output file does not exist. Check results for BUSCO: 246595at7742
## Thu Jul 14 00:28:11 CEST 2022 - The aliscore output appears to be empty. Check results for BUSCO: 246595at7742
## Thu Jul 14 00:28:12 CEST 2022 - The aliscore output file does not exist. Check results for BUSCO: 33940at7742
## Thu Jul 14 00:28:12 CEST 2022 - The aliscore output appears to be empty. Check results for BUSCO: 33940at7742
## Thu Jul 14 00:28:14 CEST 2022 - The aliscore output file does not exist. Check results for BUSCO: 160945at7742
## Thu Jul 14 00:28:14 CEST 2022 - The aliscore output appears to be empty. Check results for BUSCO: 160945at7742
## Thu Jul 14 00:28:15 CEST 2022 - The aliscore output file does not exist. Check results for BUSCO: 58275at7742
## Thu Jul 14 00:28:15 CEST 2022 - The aliscore output appears to be empty. Check results for BUSCO: 58275at7742
## Thu Jul 14 00:28:17 CEST 2022 - The aliscore output file does not exist. Check results for BUSCO: 104501at7742
## Thu Jul 14 00:28:17 CEST 2022 - The aliscore output appears to be empty. Check results for BUSCO: 104501at7742
## Thu Jul 14 00:28:17 CEST 2022 - The aliscore output file does not exist. Check results for BUSCO: 117682at7742
## Thu Jul 14 00:28:17 CEST 2022 - The aliscore output appears to be empty. Check results for BUSCO: 117682at7742
## Thu Jul 14 00:28:19 CEST 2022 - The aliscore output file does not exist. Check results for BUSCO: 172963at7742
## Thu Jul 14 00:28:19 CEST 2022 - The aliscore output appears to be empty. Check results for BUSCO: 172963at7742
## Thu Jul 14 00:28:20 CEST 2022 - The aliscore output file does not exist. Check results for BUSCO: 140683at7742
## Thu Jul 14 00:28:20 CEST 2022 - The aliscore output appears to be empty. Check results for BUSCO: 140683at7742
## Thu Jul 14 00:28:23 CEST 2022 - The aliscore output file does not exist. Check results for BUSCO: 34483at7742
## Thu Jul 14 00:28:23 CEST 2022 - The aliscore output appears to be empty. Check results for BUSCO: 34483at7742
## Thu Jul 14 00:28:23 CEST 2022 - The aliscore output file does not exist. Check results for BUSCO: 114695at7742
## Thu Jul 14 00:28:23 CEST 2022 - The aliscore output appears to be empty. Check results for BUSCO: 114695at7742
## Thu Jul 14 00:28:25 CEST 2022 - The aliscore output file does not exist. Check results for BUSCO: 87279at7742
## Thu Jul 14 00:28:25 CEST 2022 - The aliscore output appears to be empty. Check results for BUSCO: 87279at7742
## Thu Jul 14 00:28:25 CEST 2022 - The aliscore output file does not exist. Check results for BUSCO: 277134at7742
## Thu Jul 14 00:28:25 CEST 2022 - The aliscore output appears to be empty. Check results for BUSCO: 277134at7742
## Thu Jul 14 00:28:28 CEST 2022 - The aliscore output file does not exist. Check results for BUSCO: 49005at7742
## Thu Jul 14 00:28:28 CEST 2022 - The aliscore output appears to be empty. Check results for BUSCO: 49005at7742
## Thu Jul 14 00:28:30 CEST 2022 - The aliscore output file does not exist. Check results for BUSCO: 76822at7742
## Thu Jul 14 00:28:30 CEST 2022 - The aliscore output appears to be empty. Check results for BUSCO: 76822at7742
## Thu Jul 14 00:28:31 CEST 2022 - The aliscore output file does not exist. Check results for BUSCO: 106972at7742
## Thu Jul 14 00:28:31 CEST 2022 - The aliscore output appears to be empty. Check results for BUSCO: 106972at7742
## Thu Jul 14 00:28:33 CEST 2022 - The aliscore output file does not exist. Check results for BUSCO: 131825at7742
## Thu Jul 14 00:28:33 CEST 2022 - The aliscore output appears to be empty. Check results for BUSCO: 131825at7742
## Thu Jul 14 00:28:35 CEST 2022 - The aliscore output file does not exist. Check results for BUSCO: 116202at7742
## Thu Jul 14 00:28:35 CEST 2022 - The aliscore output appears to be empty. Check results for BUSCO: 116202at7742
## Thu Jul 14 00:28:35 CEST 2022 - The aliscore output file does not exist. Check results for BUSCO: 34483at7742
## Thu Jul 14 00:28:35 CEST 2022 - The aliscore output appears to be empty. Check results for BUSCO: 34483at7742
## Thu Jul 14 00:28:36 CEST 2022 - The aliscore output file does not exist. Check results for BUSCO: 77923at7742
## Thu Jul 14 00:28:36 CEST 2022 - The aliscore output appears to be empty. Check results for BUSCO: 77923at7742
## Thu Jul 14 00:28:38 CEST 2022 - The aliscore output file does not exist. Check results for BUSCO: 129880at7742
## Thu Jul 14 00:28:38 CEST 2022 - The aliscore output appears to be empty. Check results for BUSCO: 129880at7742
## Thu Jul 14 00:28:40 CEST 2022 - The aliscore output file does not exist. Check results for BUSCO: 127587at7742
## Thu Jul 14 00:28:40 CEST 2022 - The aliscore output appears to be empty. Check results for BUSCO: 127587at7742
## Thu Jul 14 00:28:40 CEST 2022 - The aliscore output file does not exist. Check results for BUSCO: 84802at7742
## Thu Jul 14 00:28:40 CEST 2022 - The aliscore output appears to be empty. Check results for BUSCO: 84802at7742
## Thu Jul 14 00:28:43 CEST 2022 - The aliscore output file does not exist. Check results for BUSCO: 92591at7742
## Thu Jul 14 00:28:43 CEST 2022 - The aliscore output appears to be empty. Check results for BUSCO: 92591at7742
## Thu Jul 14 00:28:43 CEST 2022 - The aliscore output file does not exist. Check results for BUSCO: 64359at7742
## Thu Jul 14 00:28:43 CEST 2022 - The aliscore output appears to be empty. Check results for BUSCO: 64359at7742
## Thu Jul 14 00:28:46 CEST 2022 - The aliscore output file does not exist. Check results for BUSCO: 153924at7742
## Thu Jul 14 00:28:46 CEST 2022 - The aliscore output appears to be empty. Check results for BUSCO: 153924at7742
## Thu Jul 14 00:28:46 CEST 2022 - The aliscore output file does not exist. Check results for BUSCO: 94269at7742
## Thu Jul 14 00:28:46 CEST 2022 - The aliscore output appears to be empty. Check results for BUSCO: 94269at7742
## Thu Jul 14 00:28:47 CEST 2022 - The aliscore output file does not exist. Check results for BUSCO: 64860at7742
## Thu Jul 14 00:28:47 CEST 2022 - The aliscore output appears to be empty. Check results for BUSCO: 64860at7742
## Thu Jul 14 00:28:48 CEST 2022 - The aliscore output file does not exist. Check results for BUSCO: 103690at7742
## Thu Jul 14 00:28:48 CEST 2022 - The aliscore output appears to be empty. Check results for BUSCO: 103690at7742
## Thu Jul 14 00:28:50 CEST 2022 - The aliscore output file does not exist. Check results for BUSCO: 172713at7742
## Thu Jul 14 00:28:50 CEST 2022 - The aliscore output appears to be empty. Check results for BUSCO: 172713at7742
## Thu Jul 14 00:28:52 CEST 2022 - The aliscore output file does not exist. Check results for BUSCO: 108633at7742
## Thu Jul 14 00:28:53 CEST 2022 - The aliscore output appears to be empty. Check results for BUSCO: 108633at7742
## Thu Jul 14 00:28:54 CEST 2022 - The aliscore output file does not exist. Check results for BUSCO: 52065at7742
## Thu Jul 14 00:28:54 CEST 2022 - The aliscore output file does not exist. Check results for BUSCO: 97645at7742
## Thu Jul 14 00:28:54 CEST 2022 - The aliscore output appears to be empty. Check results for BUSCO: 52065at7742
## Thu Jul 14 00:28:54 CEST 2022 - The aliscore output appears to be empty. Check results for BUSCO: 97645at7742
## Thu Jul 14 00:28:56 CEST 2022 - The aliscore output file does not exist. Check results for BUSCO: 100129at7742
## Thu Jul 14 00:28:56 CEST 2022 - The aliscore output appears to be empty. Check results for BUSCO: 100129at7742
## Thu Jul 14 00:28:57 CEST 2022 - The aliscore output file does not exist. Check results for BUSCO: 104501at7742
## Thu Jul 14 00:28:57 CEST 2022 - The aliscore output appears to be empty. Check results for BUSCO: 104501at7742
## Thu Jul 14 00:28:57 CEST 2022 - The aliscore output file does not exist. Check results for BUSCO: 102917at7742
## Thu Jul 14 00:28:57 CEST 2022 - The aliscore output appears to be empty. Check results for BUSCO: 102917at7742
## Thu Jul 14 00:28:59 CEST 2022 - The aliscore output file does not exist. Check results for BUSCO: 156439at7742
## Thu Jul 14 00:28:59 CEST 2022 - The aliscore output appears to be empty. Check results for BUSCO: 156439at7742
## Thu Jul 14 00:29:02 CEST 2022 - The aliscore output file does not exist. Check results for BUSCO: 112232at7742
## Thu Jul 14 00:29:02 CEST 2022 - The aliscore output appears to be empty. Check results for BUSCO: 112232at7742
## Thu Jul 14 00:29:04 CEST 2022 - The aliscore output file does not exist. Check results for BUSCO: 261794at7742
## Thu Jul 14 00:29:04 CEST 2022 - The aliscore output appears to be empty. Check results for BUSCO: 261794at7742
## Thu Jul 14 00:29:06 CEST 2022 - The aliscore output file does not exist. Check results for BUSCO: 47023at7742
## Thu Jul 14 00:29:06 CEST 2022 - The aliscore output appears to be empty. Check results for BUSCO: 47023at7742
## Thu Jul 14 00:29:07 CEST 2022 - The aliscore output file does not exist. Check results for BUSCO: 37796at7742
## Thu Jul 14 00:29:07 CEST 2022 - The aliscore output appears to be empty. Check results for BUSCO: 37796at7742
## Thu Jul 14 00:29:07 CEST 2022 - The aliscore output file does not exist. Check results for BUSCO: 347249at7742
## Thu Jul 14 00:29:08 CEST 2022 - The aliscore output appears to be empty. Check results for BUSCO: 347249at7742
## Thu Jul 14 00:29:08 CEST 2022 - The aliscore output file does not exist. Check results for BUSCO: 163788at7742
## Thu Jul 14 00:29:08 CEST 2022 - The aliscore output appears to be empty. Check results for BUSCO: 163788at7742
## Thu Jul 14 00:29:10 CEST 2022 - The aliscore output file does not exist. Check results for BUSCO: 163788at7742
## Thu Jul 14 00:29:10 CEST 2022 - The aliscore output appears to be empty. Check results for BUSCO: 163788at7742
## Thu Jul 14 00:29:12 CEST 2022 - The aliscore output file does not exist. Check results for BUSCO: 85263at7742
## Thu Jul 14 00:29:12 CEST 2022 - The aliscore output appears to be empty. Check results for BUSCO: 85263at7742
## Thu Jul 14 00:29:13 CEST 2022 - The aliscore output file does not exist. Check results for BUSCO: 401756at7742
## Thu Jul 14 00:29:14 CEST 2022 - The aliscore output appears to be empty. Check results for BUSCO: 401756at7742
## Thu Jul 14 00:29:14 CEST 2022 - The aliscore output file does not exist. Check results for BUSCO: 157863at7742
## Thu Jul 14 00:29:14 CEST 2022 - The aliscore output appears to be empty. Check results for BUSCO: 157863at7742
## Thu Jul 14 00:29:16 CEST 2022 - The aliscore output file does not exist. Check results for BUSCO: 87276at7742
## Thu Jul 14 00:29:16 CEST 2022 - The aliscore output appears to be empty. Check results for BUSCO: 87276at7742
## Thu Jul 14 00:29:18 CEST 2022 - The aliscore output file does not exist. Check results for BUSCO: 58866at7742
## Thu Jul 14 00:29:18 CEST 2022 - The aliscore output appears to be empty. Check results for BUSCO: 58866at7742
## Thu Jul 14 00:29:20 CEST 2022 - The aliscore output file does not exist. Check results for BUSCO: 76822at7742
## Thu Jul 14 00:29:20 CEST 2022 - The aliscore output appears to be empty. Check results for BUSCO: 76822at7742
## Thu Jul 14 00:29:20 CEST 2022 - The aliscore output file does not exist. Check results for BUSCO: 100655at7742
## Thu Jul 14 00:29:20 CEST 2022 - The aliscore output appears to be empty. Check results for BUSCO: 100655at7742
## Thu Jul 14 00:29:22 CEST 2022 - The aliscore output file does not exist. Check results for BUSCO: 40409at7742
## Thu Jul 14 00:29:22 CEST 2022 - The aliscore output appears to be empty. Check results for BUSCO: 40409at7742
## Thu Jul 14 00:29:23 CEST 2022 - The aliscore output file does not exist. Check results for BUSCO: 157863at7742
## Thu Jul 14 00:29:23 CEST 2022 - The aliscore output appears to be empty. Check results for BUSCO: 157863at7742
## Thu Jul 14 00:29:24 CEST 2022 - The aliscore output file does not exist. Check results for BUSCO: 77680at7742
## Thu Jul 14 00:29:24 CEST 2022 - The aliscore output appears to be empty. Check results for BUSCO: 77680at7742
## Thu Jul 14 00:29:26 CEST 2022 - The aliscore output file does not exist. Check results for BUSCO: 156112at7742
## Thu Jul 14 00:29:26 CEST 2022 - The aliscore output appears to be empty. Check results for BUSCO: 156112at7742
## Thu Jul 14 00:29:27 CEST 2022 - The aliscore output file does not exist. Check results for BUSCO: 80917at7742
## Thu Jul 14 00:29:27 CEST 2022 - The aliscore output appears to be empty. Check results for BUSCO: 80917at7742
## Thu Jul 14 00:29:27 CEST 2022 - The aliscore output file does not exist. Check results for BUSCO: 163788at7742
## Thu Jul 14 00:29:27 CEST 2022 - The aliscore output appears to be empty. Check results for BUSCO: 163788at7742
## Thu Jul 14 00:29:31 CEST 2022 - The aliscore output file does not exist. Check results for BUSCO: 112232at7742
## Thu Jul 14 00:29:31 CEST 2022 - The aliscore output appears to be empty. Check results for BUSCO: 112232at7742
## Thu Jul 14 00:29:32 CEST 2022 - The aliscore output file does not exist. Check results for BUSCO: 63207at7742
## Thu Jul 14 00:29:32 CEST 2022 - The aliscore output appears to be empty. Check results for BUSCO: 63207at7742
## Thu Jul 14 00:29:33 CEST 2022 - The aliscore output file does not exist. Check results for BUSCO: 223348at7742
## Thu Jul 14 00:29:33 CEST 2022 - The aliscore output appears to be empty. Check results for BUSCO: 223348at7742
## Thu Jul 14 00:29:34 CEST 2022 - The aliscore output file does not exist. Check results for BUSCO: 93412at7742
## Thu Jul 14 00:29:34 CEST 2022 - The aliscore output appears to be empty. Check results for BUSCO: 93412at7742
## Thu Jul 14 00:29:34 CEST 2022 - The aliscore output file does not exist. Check results for BUSCO: 146661at7742
## Thu Jul 14 00:29:34 CEST 2022 - The aliscore output appears to be empty. Check results for BUSCO: 146661at7742
## Thu Jul 14 00:29:37 CEST 2022 - The aliscore output file does not exist. Check results for BUSCO: 195010at7742
## Thu Jul 14 00:29:37 CEST 2022 - The aliscore output appears to be empty. Check results for BUSCO: 195010at7742
## Thu Jul 14 00:29:37 CEST 2022 - The aliscore output file does not exist. Check results for BUSCO: 50070at7742
## Thu Jul 14 00:29:38 CEST 2022 - The aliscore output appears to be empty. Check results for BUSCO: 50070at7742
## Thu Jul 14 00:29:39 CEST 2022 - The aliscore output file does not exist. Check results for BUSCO: 78822at7742
## Thu Jul 14 00:29:39 CEST 2022 - The aliscore output appears to be empty. Check results for BUSCO: 78822at7742
## Thu Jul 14 00:29:41 CEST 2022 - The aliscore output file does not exist. Check results for BUSCO: 160945at7742
## Thu Jul 14 00:29:41 CEST 2022 - The aliscore output appears to be empty. Check results for BUSCO: 160945at7742
## Thu Jul 14 00:29:43 CEST 2022 - The aliscore output file does not exist. Check results for BUSCO: 47069at7742
## Thu Jul 14 00:29:43 CEST 2022 - The aliscore output appears to be empty. Check results for BUSCO: 47069at7742
## Thu Jul 14 00:29:44 CEST 2022 - The aliscore output file does not exist. Check results for BUSCO: 102917at7742
## Thu Jul 14 00:29:44 CEST 2022 - The aliscore output appears to be empty. Check results for BUSCO: 102917at7742
## Thu Jul 14 00:29:47 CEST 2022 - The aliscore output file does not exist. Check results for BUSCO: 97302at7742
## Thu Jul 14 00:29:47 CEST 2022 - The aliscore output appears to be empty. Check results for BUSCO: 97302at7742
## Thu Jul 14 00:29:47 CEST 2022 - The aliscore output file does not exist. Check results for BUSCO: 90360at7742
## Thu Jul 14 00:29:47 CEST 2022 - The aliscore output appears to be empty. Check results for BUSCO: 90360at7742
## Thu Jul 14 00:29:47 CEST 2022 - The aliscore output file does not exist. Check results for BUSCO: 97302at7742
## Thu Jul 14 00:29:47 CEST 2022 - The aliscore output appears to be empty. Check results for BUSCO: 97302at7742
## Thu Jul 14 00:29:50 CEST 2022 - The aliscore output file does not exist. Check results for BUSCO: 75305at7742
## Thu Jul 14 00:29:50 CEST 2022 - The aliscore output appears to be empty. Check results for BUSCO: 75305at7742
## Thu Jul 14 00:29:51 CEST 2022 - The aliscore output file does not exist. Check results for BUSCO: 129062at7742
## Thu Jul 14 00:29:51 CEST 2022 - The aliscore output appears to be empty. Check results for BUSCO: 129062at7742
## Thu Jul 14 00:29:53 CEST 2022 - The aliscore output file does not exist. Check results for BUSCO: 60899at7742
## Thu Jul 14 00:29:53 CEST 2022 - The aliscore output file does not exist. Check results for BUSCO: 80119at7742
## Thu Jul 14 00:29:53 CEST 2022 - The aliscore output appears to be empty. Check results for BUSCO: 60899at7742
## Thu Jul 14 00:29:53 CEST 2022 - The aliscore output appears to be empty. Check results for BUSCO: 80119at7742
## Thu Jul 14 00:29:56 CEST 2022 - The aliscore output file does not exist. Check results for BUSCO: 155361at7742
## Thu Jul 14 00:29:56 CEST 2022 - The aliscore output appears to be empty. Check results for BUSCO: 155361at7742
## Thu Jul 14 00:29:57 CEST 2022 - The aliscore output file does not exist. Check results for BUSCO: 114695at7742
## Thu Jul 14 00:29:57 CEST 2022 - The aliscore output appears to be empty. Check results for BUSCO: 114695at7742
## Thu Jul 14 00:29:57 CEST 2022 - The aliscore output file does not exist. Check results for BUSCO: 249120at7742
## Thu Jul 14 00:29:57 CEST 2022 - The aliscore output appears to be empty. Check results for BUSCO: 249120at7742
## Thu Jul 14 00:29:59 CEST 2022 - The aliscore output file does not exist. Check results for BUSCO: 266951at7742
## Thu Jul 14 00:29:59 CEST 2022 - The aliscore output appears to be empty. Check results for BUSCO: 266951at7742
## Thu Jul 14 00:30:01 CEST 2022 - The aliscore output file does not exist. Check results for BUSCO: 101303at7742
## Thu Jul 14 00:30:01 CEST 2022 - The aliscore output appears to be empty. Check results for BUSCO: 101303at7742
## Thu Jul 14 00:30:03 CEST 2022 - The aliscore output file does not exist. Check results for BUSCO: 90853at7742
## Thu Jul 14 00:30:03 CEST 2022 - The aliscore output appears to be empty. Check results for BUSCO: 90853at7742
## Thu Jul 14 00:30:03 CEST 2022 - The aliscore output file does not exist. Check results for BUSCO: 123570at7742
## Thu Jul 14 00:30:03 CEST 2022 - The aliscore output appears to be empty. Check results for BUSCO: 123570at7742
## Thu Jul 14 00:30:04 CEST 2022 - The aliscore output file does not exist. Check results for BUSCO: 157863at7742
## Thu Jul 14 00:30:04 CEST 2022 - The aliscore output appears to be empty. Check results for BUSCO: 157863at7742
## Thu Jul 14 00:30:06 CEST 2022 - The aliscore output file does not exist. Check results for BUSCO: 46728at7742
## Thu Jul 14 00:30:06 CEST 2022 - The aliscore output appears to be empty. Check results for BUSCO: 46728at7742
## Thu Jul 14 00:30:08 CEST 2022 - The aliscore output file does not exist. Check results for BUSCO: 71963at7742
## Thu Jul 14 00:30:08 CEST 2022 - The aliscore output appears to be empty. Check results for BUSCO: 71963at7742
## Thu Jul 14 00:30:10 CEST 2022 - The aliscore output file does not exist. Check results for BUSCO: 178561at7742
## Thu Jul 14 00:30:10 CEST 2022 - The aliscore output appears to be empty. Check results for BUSCO: 178561at7742
## Thu Jul 14 00:30:13 CEST 2022 - The aliscore output file does not exist. Check results for BUSCO: 16096at7742
## Thu Jul 14 00:30:13 CEST 2022 - The aliscore output appears to be empty. Check results for BUSCO: 16096at7742
## Thu Jul 14 00:30:13 CEST 2022 - The aliscore output file does not exist. Check results for BUSCO: 176414at7742
## Thu Jul 14 00:30:13 CEST 2022 - The aliscore output appears to be empty. Check results for BUSCO: 176414at7742
## Thu Jul 14 00:30:13 CEST 2022 - The aliscore output file does not exist. Check results for BUSCO: 42033at7742
## Thu Jul 14 00:30:13 CEST 2022 - The aliscore output appears to be empty. Check results for BUSCO: 42033at7742
## Thu Jul 14 00:30:14 CEST 2022 - The aliscore output file does not exist. Check results for BUSCO: 111204at7742
## Thu Jul 14 00:30:14 CEST 2022 - The aliscore output appears to be empty. Check results for BUSCO: 111204at7742
## Thu Jul 14 00:30:14 CEST 2022 - The aliscore output file does not exist. Check results for BUSCO: 315794at7742
## Thu Jul 14 00:30:14 CEST 2022 - The aliscore output appears to be empty. Check results for BUSCO: 315794at7742
## Thu Jul 14 00:30:15 CEST 2022 - The aliscore output file does not exist. Check results for BUSCO: 92591at7742
## Thu Jul 14 00:30:15 CEST 2022 - The aliscore output appears to be empty. Check results for BUSCO: 92591at7742
## Thu Jul 14 00:30:18 CEST 2022 - The aliscore output file does not exist. Check results for BUSCO: 114819at7742
## Thu Jul 14 00:30:18 CEST 2022 - The aliscore output appears to be empty. Check results for BUSCO: 114819at7742
## Thu Jul 14 00:30:21 CEST 2022 - The aliscore output file does not exist. Check results for BUSCO: 28425at7742
## Thu Jul 14 00:30:21 CEST 2022 - The aliscore output appears to be empty. Check results for BUSCO: 28425at7742
## Thu Jul 14 00:30:23 CEST 2022 - The aliscore output file does not exist. Check results for BUSCO: 68589at7742
## Thu Jul 14 00:30:23 CEST 2022 - The aliscore output appears to be empty. Check results for BUSCO: 68589at7742
## Thu Jul 14 00:30:24 CEST 2022 - The aliscore output file does not exist. Check results for BUSCO: 90360at7742
## Thu Jul 14 00:30:25 CEST 2022 - The aliscore output appears to be empty. Check results for BUSCO: 90360at7742
## Thu Jul 14 00:30:25 CEST 2022 - The aliscore output file does not exist. Check results for BUSCO: 188889at7742
## Thu Jul 14 00:30:25 CEST 2022 - The aliscore output file does not exist. Check results for BUSCO: 146445at7742
## Thu Jul 14 00:30:25 CEST 2022 - The aliscore output appears to be empty. Check results for BUSCO: 188889at7742
## Thu Jul 14 00:30:25 CEST 2022 - The aliscore output appears to be empty. Check results for BUSCO: 146445at7742
## Thu Jul 14 00:30:25 CEST 2022 - The aliscore output file does not exist. Check results for BUSCO: 175033at7742
## Thu Jul 14 00:30:25 CEST 2022 - The aliscore output appears to be empty. Check results for BUSCO: 175033at7742
## Thu Jul 14 00:30:28 CEST 2022 - The aliscore output file does not exist. Check results for BUSCO: 319198at7742
## Thu Jul 14 00:30:28 CEST 2022 - The aliscore output appears to be empty. Check results for BUSCO: 319198at7742
## Thu Jul 14 00:30:32 CEST 2022 - The aliscore output file does not exist. Check results for BUSCO: 76178at7742
## Thu Jul 14 00:30:32 CEST 2022 - The aliscore output appears to be empty. Check results for BUSCO: 76178at7742
## Thu Jul 14 00:30:32 CEST 2022 - The aliscore output file does not exist. Check results for BUSCO: 111204at7742
## Thu Jul 14 00:30:32 CEST 2022 - The aliscore output appears to be empty. Check results for BUSCO: 111204at7742
## Thu Jul 14 00:30:32 CEST 2022 - The aliscore output file does not exist. Check results for BUSCO: 134632at7742
## Thu Jul 14 00:30:32 CEST 2022 - The aliscore output appears to be empty. Check results for BUSCO: 134632at7742
## Thu Jul 14 00:30:34 CEST 2022 - The aliscore output file does not exist. Check results for BUSCO: 121375at7742
## Thu Jul 14 00:30:34 CEST 2022 - The aliscore output appears to be empty. Check results for BUSCO: 121375at7742
## Thu Jul 14 00:30:34 CEST 2022 - The aliscore output file does not exist. Check results for BUSCO: 129062at7742
## Thu Jul 14 00:30:34 CEST 2022 - The aliscore output appears to be empty. Check results for BUSCO: 129062at7742
## Thu Jul 14 00:30:35 CEST 2022 - The aliscore output file does not exist. Check results for BUSCO: 41344at7742
## Thu Jul 14 00:30:35 CEST 2022 - The aliscore output appears to be empty. Check results for BUSCO: 41344at7742
## Thu Jul 14 00:30:38 CEST 2022 - The aliscore output file does not exist. Check results for BUSCO: 28425at7742
## Thu Jul 14 00:30:38 CEST 2022 - The aliscore output appears to be empty. Check results for BUSCO: 28425at7742
## Thu Jul 14 00:30:41 CEST 2022 - The aliscore output file does not exist. Check results for BUSCO: 112232at7742
## Thu Jul 14 00:30:41 CEST 2022 - The aliscore output appears to be empty. Check results for BUSCO: 112232at7742
## Thu Jul 14 00:30:42 CEST 2022 - The aliscore output file does not exist. Check results for BUSCO: 70459at7742
## Thu Jul 14 00:30:42 CEST 2022 - The aliscore output appears to be empty. Check results for BUSCO: 70459at7742
## Thu Jul 14 00:30:43 CEST 2022 - The aliscore output file does not exist. Check results for BUSCO: 36274at7742
## Thu Jul 14 00:30:43 CEST 2022 - The aliscore output appears to be empty. Check results for BUSCO: 36274at7742
## Thu Jul 14 00:30:45 CEST 2022 - The aliscore output file does not exist. Check results for BUSCO: 107610at7742
## Thu Jul 14 00:30:45 CEST 2022 - The aliscore output appears to be empty. Check results for BUSCO: 107610at7742
## Thu Jul 14 00:30:46 CEST 2022 - The aliscore output file does not exist. Check results for BUSCO: 142324at7742
## Thu Jul 14 00:30:46 CEST 2022 - The aliscore output appears to be empty. Check results for BUSCO: 142324at7742
## Thu Jul 14 00:30:46 CEST 2022 - The aliscore output file does not exist. Check results for BUSCO: 75305at7742
## Thu Jul 14 00:30:46 CEST 2022 - The aliscore output appears to be empty. Check results for BUSCO: 75305at7742
## Thu Jul 14 00:30:47 CEST 2022 - The aliscore output file does not exist. Check results for BUSCO: 87276at7742
## Thu Jul 14 00:30:47 CEST 2022 - The aliscore output appears to be empty. Check results for BUSCO: 87276at7742
## Thu Jul 14 00:30:50 CEST 2022 - The aliscore output file does not exist. Check results for BUSCO: 87279at7742
## Thu Jul 14 00:30:50 CEST 2022 - The aliscore output appears to be empty. Check results for BUSCO: 87279at7742
## Thu Jul 14 00:30:51 CEST 2022 - The aliscore output file does not exist. Check results for BUSCO: 315794at7742
## Thu Jul 14 00:30:51 CEST 2022 - The aliscore output appears to be empty. Check results for BUSCO: 315794at7742
## Thu Jul 14 00:30:54 CEST 2022 - The aliscore output file does not exist. Check results for BUSCO: 133969at7742
## Thu Jul 14 00:30:54 CEST 2022 - The aliscore output appears to be empty. Check results for BUSCO: 133969at7742
## Thu Jul 14 00:30:55 CEST 2022 - The aliscore output file does not exist. Check results for BUSCO: 138885at7742
## Thu Jul 14 00:30:55 CEST 2022 - The aliscore output appears to be empty. Check results for BUSCO: 138885at7742
## Thu Jul 14 00:30:55 CEST 2022 - The aliscore output file does not exist. Check results for BUSCO: 85263at7742
## Thu Jul 14 00:30:55 CEST 2022 - The aliscore output appears to be empty. Check results for BUSCO: 85263at7742
## Thu Jul 14 00:30:56 CEST 2022 - The aliscore output file does not exist. Check results for BUSCO: 76822at7742
## Thu Jul 14 00:30:57 CEST 2022 - The aliscore output appears to be empty. Check results for BUSCO: 76822at7742
## Thu Jul 14 00:30:59 CEST 2022 - The aliscore output file does not exist. Check results for BUSCO: 123570at7742
## Thu Jul 14 00:30:59 CEST 2022 - The aliscore output file does not exist. Check results for BUSCO: 142324at7742
## Thu Jul 14 00:30:59 CEST 2022 - The aliscore output appears to be empty. Check results for BUSCO: 142324at7742
## Thu Jul 14 00:30:59 CEST 2022 - The aliscore output appears to be empty. Check results for BUSCO: 123570at7742
## Thu Jul 14 00:31:01 CEST 2022 - The aliscore output file does not exist. Check results for BUSCO: 58275at7742
## Thu Jul 14 00:31:01 CEST 2022 - The aliscore output appears to be empty. Check results for BUSCO: 58275at7742
## Thu Jul 14 00:31:02 CEST 2022 - The aliscore output file does not exist. Check results for BUSCO: 37904at7742
## Thu Jul 14 00:31:02 CEST 2022 - The aliscore output appears to be empty. Check results for BUSCO: 37904at7742
## Thu Jul 14 00:31:04 CEST 2022 - The aliscore output file does not exist. Check results for BUSCO: 111204at7742
## Thu Jul 14 00:31:04 CEST 2022 - The aliscore output appears to be empty. Check results for BUSCO: 111204at7742
## Thu Jul 14 00:31:05 CEST 2022 - The aliscore output file does not exist. Check results for BUSCO: 56414at7742
## Thu Jul 14 00:31:05 CEST 2022 - The aliscore output appears to be empty. Check results for BUSCO: 56414at7742
## Thu Jul 14 00:31:05 CEST 2022 - The aliscore output file does not exist. Check results for BUSCO: 437512at7742
## Thu Jul 14 00:31:05 CEST 2022 - The aliscore output appears to be empty. Check results for BUSCO: 437512at7742
## Thu Jul 14 00:31:07 CEST 2022 - The aliscore output file does not exist. Check results for BUSCO: 40409at7742
## Thu Jul 14 00:31:07 CEST 2022 - The aliscore output appears to be empty. Check results for BUSCO: 40409at7742
## Thu Jul 14 00:31:11 CEST 2022 - The aliscore output file does not exist. Check results for BUSCO: 80119at7742
## Thu Jul 14 00:31:11 CEST 2022 - The aliscore output file does not exist. Check results for BUSCO: 21970at7742
## Thu Jul 14 00:31:11 CEST 2022 - The aliscore output appears to be empty. Check results for BUSCO: 80119at7742
## Thu Jul 14 00:31:11 CEST 2022 - The aliscore output appears to be empty. Check results for BUSCO: 21970at7742
## Thu Jul 14 00:31:14 CEST 2022 - The aliscore output file does not exist. Check results for BUSCO: 234786at7742
## Thu Jul 14 00:31:14 CEST 2022 - The aliscore output appears to be empty. Check results for BUSCO: 234786at7742
## Thu Jul 14 00:31:15 CEST 2022 - The aliscore output file does not exist. Check results for BUSCO: 131825at7742
## Thu Jul 14 00:31:15 CEST 2022 - The aliscore output appears to be empty. Check results for BUSCO: 131825at7742
## Thu Jul 14 00:31:15 CEST 2022 - The aliscore output file does not exist. Check results for BUSCO: 40897at7742
## Thu Jul 14 00:31:15 CEST 2022 - The aliscore output file does not exist. Check results for BUSCO: 21970at7742
## Thu Jul 14 00:31:15 CEST 2022 - The aliscore output appears to be empty. Check results for BUSCO: 40897at7742
## Thu Jul 14 00:31:15 CEST 2022 - The aliscore output appears to be empty. Check results for BUSCO: 21970at7742
## Thu Jul 14 00:31:16 CEST 2022 - The aliscore output file does not exist. Check results for BUSCO: 21970at7742
## Thu Jul 14 00:31:16 CEST 2022 - The aliscore output appears to be empty. Check results for BUSCO: 21970at7742
## Thu Jul 14 00:31:19 CEST 2022 - The aliscore output file does not exist. Check results for BUSCO: 114695at7742
## Thu Jul 14 00:31:19 CEST 2022 - The aliscore output appears to be empty. Check results for BUSCO: 114695at7742
## Thu Jul 14 00:31:21 CEST 2022 - The aliscore output file does not exist. Check results for BUSCO: 91878at7742
## Thu Jul 14 00:31:21 CEST 2022 - The aliscore output appears to be empty. Check results for BUSCO: 91878at7742
## Thu Jul 14 00:31:23 CEST 2022 - The aliscore output file does not exist. Check results for BUSCO: 176414at7742
## Thu Jul 14 00:31:23 CEST 2022 - The aliscore output appears to be empty. Check results for BUSCO: 176414at7742
## Thu Jul 14 00:31:23 CEST 2022 - The aliscore output file does not exist. Check results for BUSCO: 321778at7742
## Thu Jul 14 00:31:23 CEST 2022 - The aliscore output appears to be empty. Check results for BUSCO: 321778at7742
## Thu Jul 14 00:31:25 CEST 2022 - The aliscore output file does not exist. Check results for BUSCO: 101303at7742
## Thu Jul 14 00:31:25 CEST 2022 - The aliscore output appears to be empty. Check results for BUSCO: 101303at7742
## Thu Jul 14 00:31:25 CEST 2022 - The aliscore output file does not exist. Check results for BUSCO: 80119at7742
## Thu Jul 14 00:31:25 CEST 2022 - The aliscore output appears to be empty. Check results for BUSCO: 80119at7742
## Thu Jul 14 00:31:27 CEST 2022 - The aliscore output file does not exist. Check results for BUSCO: 42033at7742
## Thu Jul 14 00:31:27 CEST 2022 - The aliscore output appears to be empty. Check results for BUSCO: 42033at7742
## Thu Jul 14 00:31:28 CEST 2022 - The aliscore output file does not exist. Check results for BUSCO: 97302at7742
## Thu Jul 14 00:31:29 CEST 2022 - The aliscore output appears to be empty. Check results for BUSCO: 97302at7742
## Thu Jul 14 00:31:30 CEST 2022 - The aliscore output file does not exist. Check results for BUSCO: 28425at7742
## Thu Jul 14 00:31:30 CEST 2022 - The aliscore output appears to be empty. Check results for BUSCO: 28425at7742
## Thu Jul 14 00:31:32 CEST 2022 - The aliscore output file does not exist. Check results for BUSCO: 68016at7742
## Thu Jul 14 00:31:32 CEST 2022 - The aliscore output appears to be empty. Check results for BUSCO: 68016at7742
## Thu Jul 14 00:31:33 CEST 2022 - The aliscore output file does not exist. Check results for BUSCO: 78812at7742
## Thu Jul 14 00:31:33 CEST 2022 - The aliscore output appears to be empty. Check results for BUSCO: 78812at7742
## Thu Jul 14 00:31:34 CEST 2022 - The aliscore output file does not exist. Check results for BUSCO: 87276at7742
## Thu Jul 14 00:31:34 CEST 2022 - The aliscore output appears to be empty. Check results for BUSCO: 87276at7742
## Thu Jul 14 00:31:35 CEST 2022 - The aliscore output file does not exist. Check results for BUSCO: 88390at7742
## Thu Jul 14 00:31:35 CEST 2022 - The aliscore output appears to be empty. Check results for BUSCO: 88390at7742
## Thu Jul 14 00:31:37 CEST 2022 - The aliscore output file does not exist. Check results for BUSCO: 61069at7742
## Thu Jul 14 00:31:37 CEST 2022 - The aliscore output appears to be empty. Check results for BUSCO: 61069at7742
## Thu Jul 14 00:31:40 CEST 2022 - The aliscore output file does not exist. Check results for BUSCO: 74281at7742
## Thu Jul 14 00:31:40 CEST 2022 - The aliscore output appears to be empty. Check results for BUSCO: 74281at7742
## Thu Jul 14 00:31:41 CEST 2022 - The aliscore output file does not exist. Check results for BUSCO: 249120at7742
## Thu Jul 14 00:31:41 CEST 2022 - The aliscore output appears to be empty. Check results for BUSCO: 249120at7742
## Thu Jul 14 00:31:41 CEST 2022 - The aliscore output file does not exist. Check results for BUSCO: 126913at7742
## Thu Jul 14 00:31:41 CEST 2022 - The aliscore output appears to be empty. Check results for BUSCO: 126913at7742
## Thu Jul 14 00:31:42 CEST 2022 - The aliscore output file does not exist. Check results for BUSCO: 77680at7742
## Thu Jul 14 00:31:42 CEST 2022 - The aliscore output appears to be empty. Check results for BUSCO: 77680at7742
## Thu Jul 14 00:31:45 CEST 2022 - The aliscore output file does not exist. Check results for BUSCO: 90360at7742
## Thu Jul 14 00:31:45 CEST 2022 - The aliscore output appears to be empty. Check results for BUSCO: 90360at7742
## Thu Jul 14 00:31:46 CEST 2022 - The aliscore output file does not exist. Check results for BUSCO: 37904at7742
## Thu Jul 14 00:31:46 CEST 2022 - The aliscore output appears to be empty. Check results for BUSCO: 37904at7742
## Thu Jul 14 00:31:47 CEST 2022 - The aliscore output file does not exist. Check results for BUSCO: 69813at7742
## Thu Jul 14 00:31:47 CEST 2022 - The aliscore output appears to be empty. Check results for BUSCO: 69813at7742
## Thu Jul 14 00:31:49 CEST 2022 - The aliscore output file does not exist. Check results for BUSCO: 335206at7742
## Thu Jul 14 00:31:49 CEST 2022 - The aliscore output appears to be empty. Check results for BUSCO: 335206at7742
## Thu Jul 14 00:31:50 CEST 2022 - The aliscore output file does not exist. Check results for BUSCO: 52065at7742
## Thu Jul 14 00:31:50 CEST 2022 - The aliscore output appears to be empty. Check results for BUSCO: 52065at7742
## Thu Jul 14 00:31:51 CEST 2022 - The aliscore output file does not exist. Check results for BUSCO: 286498at7742
## Thu Jul 14 00:31:51 CEST 2022 - The aliscore output appears to be empty. Check results for BUSCO: 286498at7742
## Thu Jul 14 00:31:52 CEST 2022 - The aliscore output file does not exist. Check results for BUSCO: 123337at7742
## Thu Jul 14 00:31:52 CEST 2022 - The aliscore output appears to be empty. Check results for BUSCO: 123337at7742
## Thu Jul 14 00:31:54 CEST 2022 - The aliscore output file does not exist. Check results for BUSCO: 234786at7742
## Thu Jul 14 00:31:54 CEST 2022 - The aliscore output appears to be empty. Check results for BUSCO: 234786at7742
## Thu Jul 14 00:31:56 CEST 2022 - The aliscore output file does not exist. Check results for BUSCO: 132279at7742
## Thu Jul 14 00:31:56 CEST 2022 - The aliscore output appears to be empty. Check results for BUSCO: 132279at7742
## Thu Jul 14 00:31:57 CEST 2022 - The aliscore output file does not exist. Check results for BUSCO: 29337at7742
## Thu Jul 14 00:31:57 CEST 2022 - The aliscore output appears to be empty. Check results for BUSCO: 29337at7742
## Thu Jul 14 00:31:58 CEST 2022 - The aliscore output file does not exist. Check results for BUSCO: 46317at7742
## Thu Jul 14 00:31:58 CEST 2022 - The aliscore output appears to be empty. Check results for BUSCO: 46317at7742
## Thu Jul 14 00:32:00 CEST 2022 - The aliscore output file does not exist. Check results for BUSCO: 71963at7742
## Thu Jul 14 00:32:00 CEST 2022 - The aliscore output appears to be empty. Check results for BUSCO: 71963at7742
## Thu Jul 14 00:32:01 CEST 2022 - The aliscore output file does not exist. Check results for BUSCO: 92481at7742
## Thu Jul 14 00:32:02 CEST 2022 - The aliscore output appears to be empty. Check results for BUSCO: 92481at7742
## Thu Jul 14 00:32:04 CEST 2022 - The aliscore output file does not exist. Check results for BUSCO: 58506at7742
## Thu Jul 14 00:32:04 CEST 2022 - The aliscore output appears to be empty. Check results for BUSCO: 58506at7742
## Thu Jul 14 00:32:04 CEST 2022 - The aliscore output file does not exist. Check results for BUSCO: 50070at7742
## Thu Jul 14 00:32:04 CEST 2022 - The aliscore output appears to be empty. Check results for BUSCO: 50070at7742
## Thu Jul 14 00:32:05 CEST 2022 - The aliscore output file does not exist. Check results for BUSCO: 129880at7742
## Thu Jul 14 00:32:05 CEST 2022 - The aliscore output appears to be empty. Check results for BUSCO: 129880at7742
## Thu Jul 14 00:32:07 CEST 2022 - The aliscore output file does not exist. Check results for BUSCO: 29337at7742
## Thu Jul 14 00:32:08 CEST 2022 - The aliscore output appears to be empty. Check results for BUSCO: 29337at7742
## Thu Jul 14 00:32:08 CEST 2022 - The aliscore output file does not exist. Check results for BUSCO: 209493at7742
## Thu Jul 14 00:32:08 CEST 2022 - The aliscore output appears to be empty. Check results for BUSCO: 209493at7742
## Thu Jul 14 00:32:08 CEST 2022 - The aliscore output file does not exist. Check results for BUSCO: 143698at7742
## Thu Jul 14 00:32:08 CEST 2022 - The aliscore output appears to be empty. Check results for BUSCO: 143698at7742
## Thu Jul 14 00:32:11 CEST 2022 - The aliscore output file does not exist. Check results for BUSCO: 60783at7742
## Thu Jul 14 00:32:11 CEST 2022 - The aliscore output appears to be empty. Check results for BUSCO: 60783at7742
## Thu Jul 14 00:32:13 CEST 2022 - The aliscore output file does not exist. Check results for BUSCO: 184785at7742
## Thu Jul 14 00:32:13 CEST 2022 - The aliscore output appears to be empty. Check results for BUSCO: 184785at7742
## Thu Jul 14 00:32:13 CEST 2022 - The aliscore output file does not exist. Check results for BUSCO: 87279at7742
## Thu Jul 14 00:32:13 CEST 2022 - The aliscore output appears to be empty. Check results for BUSCO: 87279at7742
## Thu Jul 14 00:32:15 CEST 2022 - The aliscore output file does not exist. Check results for BUSCO: 42971at7742
## Thu Jul 14 00:32:15 CEST 2022 - The aliscore output appears to be empty. Check results for BUSCO: 42971at7742
## Thu Jul 14 00:32:17 CEST 2022 - The aliscore output file does not exist. Check results for BUSCO: 141754at7742
## Thu Jul 14 00:32:17 CEST 2022 - The aliscore output appears to be empty. Check results for BUSCO: 141754at7742
## Thu Jul 14 00:32:18 CEST 2022 - The aliscore output file does not exist. Check results for BUSCO: 128709at7742
## Thu Jul 14 00:32:18 CEST 2022 - The aliscore output appears to be empty. Check results for BUSCO: 128709at7742
## Thu Jul 14 00:32:19 CEST 2022 - The aliscore output file does not exist. Check results for BUSCO: 91371at7742
## Thu Jul 14 00:32:19 CEST 2022 - The aliscore output appears to be empty. Check results for BUSCO: 91371at7742
## Thu Jul 14 00:32:22 CEST 2022 - The aliscore output file does not exist. Check results for BUSCO: 212876at7742
## Thu Jul 14 00:32:22 CEST 2022 - The aliscore output appears to be empty. Check results for BUSCO: 212876at7742
## Thu Jul 14 00:32:22 CEST 2022 - The aliscore output file does not exist. Check results for BUSCO: 132279at7742
## Thu Jul 14 00:32:22 CEST 2022 - The aliscore output file does not exist. Check results for BUSCO: 103731at7742
## Thu Jul 14 00:32:22 CEST 2022 - The aliscore output appears to be empty. Check results for BUSCO: 132279at7742
## Thu Jul 14 00:32:22 CEST 2022 - The aliscore output appears to be empty. Check results for BUSCO: 103731at7742
## Thu Jul 14 00:32:24 CEST 2022 - The aliscore output file does not exist. Check results for BUSCO: 286498at7742
## Thu Jul 14 00:32:24 CEST 2022 - The aliscore output appears to be empty. Check results for BUSCO: 286498at7742
## Thu Jul 14 00:32:26 CEST 2022 - The aliscore output file does not exist. Check results for BUSCO: 42033at7742
## Thu Jul 14 00:32:26 CEST 2022 - The aliscore output appears to be empty. Check results for BUSCO: 42033at7742
## Thu Jul 14 00:32:28 CEST 2022 - The aliscore output file does not exist. Check results for BUSCO: 95435at7742
## Thu Jul 14 00:32:28 CEST 2022 - The aliscore output appears to be empty. Check results for BUSCO: 95435at7742
## Thu Jul 14 00:32:28 CEST 2022 - The aliscore output file does not exist. Check results for BUSCO: 42971at7742
## Thu Jul 14 00:32:29 CEST 2022 - The aliscore output appears to be empty. Check results for BUSCO: 42971at7742
## Thu Jul 14 00:32:30 CEST 2022 - The aliscore output file does not exist. Check results for BUSCO: 175841at7742
## Thu Jul 14 00:32:30 CEST 2022 - The aliscore output appears to be empty. Check results for BUSCO: 175841at7742
## Thu Jul 14 00:32:31 CEST 2022 - The aliscore output file does not exist. Check results for BUSCO: 105968at7742
## Thu Jul 14 00:32:31 CEST 2022 - The aliscore output appears to be empty. Check results for BUSCO: 105968at7742
## Thu Jul 14 00:32:32 CEST 2022 - The aliscore output file does not exist. Check results for BUSCO: 121375at7742
## Thu Jul 14 00:32:32 CEST 2022 - The aliscore output appears to be empty. Check results for BUSCO: 121375at7742
## Thu Jul 14 00:32:35 CEST 2022 - The aliscore output file does not exist. Check results for BUSCO: 59009at7742
## Thu Jul 14 00:32:35 CEST 2022 - The aliscore output appears to be empty. Check results for BUSCO: 59009at7742
## Thu Jul 14 00:32:37 CEST 2022 - The aliscore output file does not exist. Check results for BUSCO: 103689at7742
## Thu Jul 14 00:32:37 CEST 2022 - The aliscore output appears to be empty. Check results for BUSCO: 103689at7742
## Thu Jul 14 00:32:38 CEST 2022 - The aliscore output file does not exist. Check results for BUSCO: 119886at7742
## Thu Jul 14 00:32:38 CEST 2022 - The aliscore output appears to be empty. Check results for BUSCO: 119886at7742
## Thu Jul 14 00:32:39 CEST 2022 - The aliscore output file does not exist. Check results for BUSCO: 41344at7742
## Thu Jul 14 00:32:39 CEST 2022 - The aliscore output appears to be empty. Check results for BUSCO: 41344at7742
## Thu Jul 14 00:32:42 CEST 2022 - The aliscore output file does not exist. Check results for BUSCO: 16096at7742
## Thu Jul 14 00:32:42 CEST 2022 - The aliscore output file does not exist. Check results for BUSCO: 37796at7742
## Thu Jul 14 00:32:42 CEST 2022 - The aliscore output appears to be empty. Check results for BUSCO: 16096at7742
## Thu Jul 14 00:32:42 CEST 2022 - The aliscore output appears to be empty. Check results for BUSCO: 37796at7742
## Thu Jul 14 00:32:43 CEST 2022 - The aliscore output file does not exist. Check results for BUSCO: 78822at7742
## Thu Jul 14 00:32:43 CEST 2022 - The aliscore output appears to be empty. Check results for BUSCO: 78822at7742
## Thu Jul 14 00:32:44 CEST 2022 - The aliscore output file does not exist. Check results for BUSCO: 37904at7742
## Thu Jul 14 00:32:44 CEST 2022 - The aliscore output appears to be empty. Check results for BUSCO: 37904at7742
## Thu Jul 14 00:32:47 CEST 2022 - The aliscore output file does not exist. Check results for BUSCO: 128709at7742
## Thu Jul 14 00:32:47 CEST 2022 - The aliscore output file does not exist. Check results for BUSCO: 103731at7742
## Thu Jul 14 00:32:47 CEST 2022 - The aliscore output appears to be empty. Check results for BUSCO: 128709at7742
## Thu Jul 14 00:32:47 CEST 2022 - The aliscore output appears to be empty. Check results for BUSCO: 103731at7742
## Thu Jul 14 00:32:47 CEST 2022 - The aliscore output file does not exist. Check results for BUSCO: 35271at7742
## Thu Jul 14 00:32:47 CEST 2022 - The aliscore output appears to be empty. Check results for BUSCO: 35271at7742
## Thu Jul 14 00:32:48 CEST 2022 - The aliscore output file does not exist. Check results for BUSCO: 35271at7742
## Thu Jul 14 00:32:48 CEST 2022 - The aliscore output appears to be empty. Check results for BUSCO: 35271at7742
## Thu Jul 14 00:32:52 CEST 2022 - The aliscore output file does not exist. Check results for BUSCO: 64591at7742
## Thu Jul 14 00:32:52 CEST 2022 - The aliscore output appears to be empty. Check results for BUSCO: 64591at7742
## Thu Jul 14 00:32:53 CEST 2022 - The aliscore output file does not exist. Check results for BUSCO: 170533at7742
## Thu Jul 14 00:32:53 CEST 2022 - The aliscore output appears to be empty. Check results for BUSCO: 170533at7742
## Thu Jul 14 00:32:55 CEST 2022 - The aliscore output file does not exist. Check results for BUSCO: 88390at7742
## Thu Jul 14 00:32:55 CEST 2022 - The aliscore output appears to be empty. Check results for BUSCO: 88390at7742
## Thu Jul 14 00:32:55 CEST 2022 - The aliscore output file does not exist. Check results for BUSCO: 41189at7742
## Thu Jul 14 00:32:55 CEST 2022 - The aliscore output appears to be empty. Check results for BUSCO: 41189at7742
## Thu Jul 14 00:32:57 CEST 2022 - The aliscore output file does not exist. Check results for BUSCO: 315794at7742
## Thu Jul 14 00:32:57 CEST 2022 - The aliscore output appears to be empty. Check results for BUSCO: 315794at7742
## Thu Jul 14 00:32:59 CEST 2022 - The aliscore output file does not exist. Check results for BUSCO: 114459at7742
## Thu Jul 14 00:32:59 CEST 2022 - The aliscore output appears to be empty. Check results for BUSCO: 114459at7742
## Thu Jul 14 00:33:01 CEST 2022 - The aliscore output file does not exist. Check results for BUSCO: 87762at7742
## Thu Jul 14 00:33:01 CEST 2022 - The aliscore output appears to be empty. Check results for BUSCO: 87762at7742
## Thu Jul 14 00:33:02 CEST 2022 - The aliscore output file does not exist. Check results for BUSCO: 46317at7742
## Thu Jul 14 00:33:02 CEST 2022 - The aliscore output appears to be empty. Check results for BUSCO: 46317at7742
## Thu Jul 14 00:33:02 CEST 2022 - The aliscore output file does not exist. Check results for BUSCO: 115220at7742
## Thu Jul 14 00:33:02 CEST 2022 - The aliscore output appears to be empty. Check results for BUSCO: 115220at7742
## Thu Jul 14 00:33:03 CEST 2022 - The aliscore output file does not exist. Check results for BUSCO: 61069at7742
## Thu Jul 14 00:33:03 CEST 2022 - The aliscore output appears to be empty. Check results for BUSCO: 61069at7742
## Thu Jul 14 00:33:07 CEST 2022 - The aliscore output file does not exist. Check results for BUSCO: 128709at7742
## Thu Jul 14 00:33:07 CEST 2022 - The aliscore output appears to be empty. Check results for BUSCO: 128709at7742
## Thu Jul 14 00:33:07 CEST 2022 - The aliscore output file does not exist. Check results for BUSCO: 131825at7742
## Thu Jul 14 00:33:07 CEST 2022 - The aliscore output appears to be empty. Check results for BUSCO: 131825at7742
## Thu Jul 14 00:33:08 CEST 2022 - The aliscore output file does not exist. Check results for BUSCO: 66726at7742
## Thu Jul 14 00:33:08 CEST 2022 - The aliscore output appears to be empty. Check results for BUSCO: 66726at7742
## Thu Jul 14 00:33:11 CEST 2022 - The aliscore output file does not exist. Check results for BUSCO: 150697at7742
## Thu Jul 14 00:33:11 CEST 2022 - The aliscore output appears to be empty. Check results for BUSCO: 150697at7742
## Thu Jul 14 00:33:12 CEST 2022 - The aliscore output file does not exist. Check results for BUSCO: 156420at7742
## Thu Jul 14 00:33:12 CEST 2022 - The aliscore output appears to be empty. Check results for BUSCO: 156420at7742
## Thu Jul 14 00:33:13 CEST 2022 - The aliscore output file does not exist. Check results for BUSCO: 154706at7742
## Thu Jul 14 00:33:13 CEST 2022 - The aliscore output appears to be empty. Check results for BUSCO: 154706at7742
## Thu Jul 14 00:33:13 CEST 2022 - The aliscore output file does not exist. Check results for BUSCO: 121053at7742
## Thu Jul 14 00:33:13 CEST 2022 - The aliscore output appears to be empty. Check results for BUSCO: 121053at7742
## Thu Jul 14 00:33:16 CEST 2022 - The aliscore output file does not exist. Check results for BUSCO: 143864at7742
## Thu Jul 14 00:33:16 CEST 2022 - The aliscore output appears to be empty. Check results for BUSCO: 143864at7742
## Thu Jul 14 00:33:16 CEST 2022 - The aliscore output file does not exist. Check results for BUSCO: 51094at7742
## Thu Jul 14 00:33:16 CEST 2022 - The aliscore output appears to be empty. Check results for BUSCO: 51094at7742
## Thu Jul 14 00:33:17 CEST 2022 - The aliscore output file does not exist. Check results for BUSCO: 235274at7742
## Thu Jul 14 00:33:17 CEST 2022 - The aliscore output appears to be empty. Check results for BUSCO: 235274at7742
## Thu Jul 14 00:33:21 CEST 2022 - The aliscore output file does not exist. Check results for BUSCO: 46728at7742
## Thu Jul 14 00:33:21 CEST 2022 - The aliscore output appears to be empty. Check results for BUSCO: 46728at7742
## Thu Jul 14 00:33:22 CEST 2022 - The aliscore output file does not exist. Check results for BUSCO: 251511at7742
## Thu Jul 14 00:33:22 CEST 2022 - The aliscore output file does not exist. Check results for BUSCO: 132656at7742
## Thu Jul 14 00:33:22 CEST 2022 - The aliscore output appears to be empty. Check results for BUSCO: 251511at7742
## Thu Jul 14 00:33:22 CEST 2022 - The aliscore output appears to be empty. Check results for BUSCO: 132656at7742
## Thu Jul 14 00:33:23 CEST 2022 - The aliscore output file does not exist. Check results for BUSCO: 319486at7742
## Thu Jul 14 00:33:23 CEST 2022 - The aliscore output appears to be empty. Check results for BUSCO: 319486at7742
## Thu Jul 14 00:33:23 CEST 2022 - The aliscore output file does not exist. Check results for BUSCO: 138498at7742
## Thu Jul 14 00:33:23 CEST 2022 - The aliscore output appears to be empty. Check results for BUSCO: 138498at7742
## Thu Jul 14 00:33:26 CEST 2022 - The aliscore output file does not exist. Check results for BUSCO: 134048at7742
## Thu Jul 14 00:33:26 CEST 2022 - The aliscore output appears to be empty. Check results for BUSCO: 134048at7742
## Thu Jul 14 00:33:30 CEST 2022 - The aliscore output file does not exist. Check results for BUSCO: 103731at7742
## Thu Jul 14 00:33:30 CEST 2022 - The aliscore output appears to be empty. Check results for BUSCO: 103731at7742
## Thu Jul 14 00:33:30 CEST 2022 - The aliscore output file does not exist. Check results for BUSCO: 76178at7742
## Thu Jul 14 00:33:30 CEST 2022 - The aliscore output appears to be empty. Check results for BUSCO: 76178at7742
## Thu Jul 14 00:33:32 CEST 2022 - The aliscore output file does not exist. Check results for BUSCO: 122963at7742
## Thu Jul 14 00:33:32 CEST 2022 - The aliscore output appears to be empty. Check results for BUSCO: 122963at7742
## Thu Jul 14 00:33:34 CEST 2022 - The aliscore output file does not exist. Check results for BUSCO: 35271at7742
## Thu Jul 14 00:33:34 CEST 2022 - The aliscore output appears to be empty. Check results for BUSCO: 35271at7742
## Thu Jul 14 00:33:34 CEST 2022 - The aliscore output file does not exist. Check results for BUSCO: 56477at7742
## Thu Jul 14 00:33:34 CEST 2022 - The aliscore output appears to be empty. Check results for BUSCO: 56477at7742
## Thu Jul 14 00:33:35 CEST 2022 - The aliscore output file does not exist. Check results for BUSCO: 167134at7742
## Thu Jul 14 00:33:35 CEST 2022 - The aliscore output appears to be empty. Check results for BUSCO: 167134at7742
## Thu Jul 14 00:33:36 CEST 2022 - The aliscore output file does not exist. Check results for BUSCO: 286498at7742
## Thu Jul 14 00:33:36 CEST 2022 - The aliscore output appears to be empty. Check results for BUSCO: 286498at7742
## Thu Jul 14 00:33:37 CEST 2022 - The aliscore output file does not exist. Check results for BUSCO: 66726at7742
## Thu Jul 14 00:33:37 CEST 2022 - The aliscore output appears to be empty. Check results for BUSCO: 66726at7742
## Thu Jul 14 00:33:39 CEST 2022 - The aliscore output file does not exist. Check results for BUSCO: 107610at7742
## Thu Jul 14 00:33:39 CEST 2022 - The aliscore output appears to be empty. Check results for BUSCO: 107610at7742
## Thu Jul 14 00:33:41 CEST 2022 - The aliscore output file does not exist. Check results for BUSCO: 133006at7742
## Thu Jul 14 00:33:41 CEST 2022 - The aliscore output appears to be empty. Check results for BUSCO: 133006at7742
## Thu Jul 14 00:33:41 CEST 2022 - The aliscore output file does not exist. Check results for BUSCO: 176410at7742
## Thu Jul 14 00:33:41 CEST 2022 - The aliscore output appears to be empty. Check results for BUSCO: 176410at7742
## Thu Jul 14 00:33:42 CEST 2022 - The aliscore output file does not exist. Check results for BUSCO: 232468at7742
## Thu Jul 14 00:33:42 CEST 2022 - The aliscore output appears to be empty. Check results for BUSCO: 232468at7742
## Thu Jul 14 00:33:44 CEST 2022 - The aliscore output file does not exist. Check results for BUSCO: 437512at7742
## Thu Jul 14 00:33:44 CEST 2022 - The aliscore output appears to be empty. Check results for BUSCO: 437512at7742
## Thu Jul 14 00:33:47 CEST 2022 - The aliscore output file does not exist. Check results for BUSCO: 107107at7742
## Thu Jul 14 00:33:47 CEST 2022 - The aliscore output appears to be empty. Check results for BUSCO: 107107at7742
## Thu Jul 14 00:33:47 CEST 2022 - The aliscore output file does not exist. Check results for BUSCO: 266951at7742
## Thu Jul 14 00:33:47 CEST 2022 - The aliscore output appears to be empty. Check results for BUSCO: 266951at7742
## Thu Jul 14 00:33:51 CEST 2022 - The aliscore output file does not exist. Check results for BUSCO: 159524at7742
## Thu Jul 14 00:33:51 CEST 2022 - The aliscore output appears to be empty. Check results for BUSCO: 159524at7742
## Thu Jul 14 00:33:51 CEST 2022 - The aliscore output file does not exist. Check results for BUSCO: 69813at7742
## Thu Jul 14 00:33:51 CEST 2022 - The aliscore output appears to be empty. Check results for BUSCO: 69813at7742
## Thu Jul 14 00:33:51 CEST 2022 - The aliscore output file does not exist. Check results for BUSCO: 125236at7742
## Thu Jul 14 00:33:51 CEST 2022 - The aliscore output appears to be empty. Check results for BUSCO: 125236at7742
## Thu Jul 14 00:33:52 CEST 2022 - The aliscore output file does not exist. Check results for BUSCO: 124962at7742
## Thu Jul 14 00:33:52 CEST 2022 - The aliscore output appears to be empty. Check results for BUSCO: 124962at7742
## Thu Jul 14 00:33:54 CEST 2022 - The aliscore output file does not exist. Check results for BUSCO: 290517at7742
## Thu Jul 14 00:33:54 CEST 2022 - The aliscore output appears to be empty. Check results for BUSCO: 290517at7742
## Thu Jul 14 00:33:56 CEST 2022 - The aliscore output file does not exist. Check results for BUSCO: 46317at7742
## Thu Jul 14 00:33:56 CEST 2022 - The aliscore output appears to be empty. Check results for BUSCO: 46317at7742
## Thu Jul 14 00:33:56 CEST 2022 - The aliscore output file does not exist. Check results for BUSCO: 437512at7742
## Thu Jul 14 00:33:56 CEST 2022 - The aliscore output appears to be empty. Check results for BUSCO: 437512at7742
## Thu Jul 14 00:33:58 CEST 2022 - The aliscore output file does not exist. Check results for BUSCO: 227556at7742
## Thu Jul 14 00:33:58 CEST 2022 - The aliscore output appears to be empty. Check results for BUSCO: 227556at7742
## Thu Jul 14 00:34:00 CEST 2022 - The aliscore output file does not exist. Check results for BUSCO: 61069at7742
## Thu Jul 14 00:34:00 CEST 2022 - The aliscore output appears to be empty. Check results for BUSCO: 61069at7742
## Thu Jul 14 00:34:02 CEST 2022 - The aliscore output file does not exist. Check results for BUSCO: 19730at7742
## Thu Jul 14 00:34:02 CEST 2022 - The aliscore output file does not exist. Check results for BUSCO: 103689at7742
## Thu Jul 14 00:34:02 CEST 2022 - The aliscore output appears to be empty. Check results for BUSCO: 19730at7742
## Thu Jul 14 00:34:02 CEST 2022 - The aliscore output appears to be empty. Check results for BUSCO: 103689at7742
## Thu Jul 14 00:34:02 CEST 2022 - The aliscore output file does not exist. Check results for BUSCO: 52541at7742
## Thu Jul 14 00:34:02 CEST 2022 - The aliscore output appears to be empty. Check results for BUSCO: 52541at7742
## Thu Jul 14 00:34:05 CEST 2022 - The aliscore output file does not exist. Check results for BUSCO: 141015at7742
## Thu Jul 14 00:34:05 CEST 2022 - The aliscore output appears to be empty. Check results for BUSCO: 141015at7742
## Thu Jul 14 00:34:07 CEST 2022 - The aliscore output file does not exist. Check results for BUSCO: 34483at7742
## Thu Jul 14 00:34:07 CEST 2022 - The aliscore output appears to be empty. Check results for BUSCO: 34483at7742
## Thu Jul 14 00:34:10 CEST 2022 - The aliscore output file does not exist. Check results for BUSCO: 68016at7742
## Thu Jul 14 00:34:10 CEST 2022 - The aliscore output appears to be empty. Check results for BUSCO: 68016at7742
## Thu Jul 14 00:34:11 CEST 2022 - The aliscore output file does not exist. Check results for BUSCO: 170032at7742
## Thu Jul 14 00:34:11 CEST 2022 - The aliscore output appears to be empty. Check results for BUSCO: 170032at7742
## Thu Jul 14 00:34:11 CEST 2022 - The aliscore output file does not exist. Check results for BUSCO: 177227at7742
## Thu Jul 14 00:34:11 CEST 2022 - The aliscore output appears to be empty. Check results for BUSCO: 177227at7742
## Thu Jul 14 00:34:12 CEST 2022 - The aliscore output file does not exist. Check results for BUSCO: 134048at7742
## Thu Jul 14 00:34:12 CEST 2022 - The aliscore output appears to be empty. Check results for BUSCO: 134048at7742
## Thu Jul 14 00:34:13 CEST 2022 - The aliscore output file does not exist. Check results for BUSCO: 68589at7742
## Thu Jul 14 00:34:13 CEST 2022 - The aliscore output appears to be empty. Check results for BUSCO: 68589at7742
## Thu Jul 14 00:34:16 CEST 2022 - The aliscore output file does not exist. Check results for BUSCO: 178561at7742
## Thu Jul 14 00:34:16 CEST 2022 - The aliscore output appears to be empty. Check results for BUSCO: 178561at7742
## Thu Jul 14 00:34:16 CEST 2022 - The aliscore output file does not exist. Check results for BUSCO: 39294at7742
## Thu Jul 14 00:34:16 CEST 2022 - The aliscore output appears to be empty. Check results for BUSCO: 39294at7742
## Thu Jul 14 00:34:17 CEST 2022 - The aliscore output file does not exist. Check results for BUSCO: 77680at7742
## Thu Jul 14 00:34:17 CEST 2022 - The aliscore output appears to be empty. Check results for BUSCO: 77680at7742
## Thu Jul 14 00:34:18 CEST 2022 - The aliscore output file does not exist. Check results for BUSCO: 66726at7742
## Thu Jul 14 00:34:18 CEST 2022 - The aliscore output appears to be empty. Check results for BUSCO: 66726at7742
## Thu Jul 14 00:34:22 CEST 2022 - The aliscore output file does not exist. Check results for BUSCO: 172963at7742
## Thu Jul 14 00:34:22 CEST 2022 - The aliscore output appears to be empty. Check results for BUSCO: 172963at7742
## Thu Jul 14 00:34:22 CEST 2022 - The aliscore output file does not exist. Check results for BUSCO: 40409at7742
## Thu Jul 14 00:34:22 CEST 2022 - The aliscore output appears to be empty. Check results for BUSCO: 40409at7742
## Thu Jul 14 00:34:23 CEST 2022 - The aliscore output file does not exist. Check results for BUSCO: 169833at7742
## Thu Jul 14 00:34:23 CEST 2022 - The aliscore output appears to be empty. Check results for BUSCO: 169833at7742
## Thu Jul 14 00:34:25 CEST 2022 - The aliscore output file does not exist. Check results for BUSCO: 319486at7742
## Thu Jul 14 00:34:25 CEST 2022 - The aliscore output appears to be empty. Check results for BUSCO: 319486at7742
## Thu Jul 14 00:34:26 CEST 2022 - The aliscore output file does not exist. Check results for BUSCO: 319486at7742
## Thu Jul 14 00:34:26 CEST 2022 - The aliscore output appears to be empty. Check results for BUSCO: 319486at7742
## Thu Jul 14 00:34:27 CEST 2022 - The aliscore output file does not exist. Check results for BUSCO: 47275at7742
## Thu Jul 14 00:34:27 CEST 2022 - The aliscore output appears to be empty. Check results for BUSCO: 47275at7742
## Thu Jul 14 00:34:30 CEST 2022 - The aliscore output file does not exist. Check results for BUSCO: 296592at7742
## Thu Jul 14 00:34:30 CEST 2022 - The aliscore output appears to be empty. Check results for BUSCO: 296592at7742
## Thu Jul 14 00:34:31 CEST 2022 - The aliscore output file does not exist. Check results for BUSCO: 160468at7742
## Thu Jul 14 00:34:31 CEST 2022 - The aliscore output appears to be empty. Check results for BUSCO: 160468at7742
## Thu Jul 14 00:34:32 CEST 2022 - The aliscore output file does not exist. Check results for BUSCO: 249120at7742
## Thu Jul 14 00:34:32 CEST 2022 - The aliscore output appears to be empty. Check results for BUSCO: 249120at7742
## Thu Jul 14 00:34:32 CEST 2022 - The aliscore output file does not exist. Check results for BUSCO: 134048at7742
## Thu Jul 14 00:34:32 CEST 2022 - The aliscore output appears to be empty. Check results for BUSCO: 134048at7742
## Thu Jul 14 00:34:35 CEST 2022 - The aliscore output file does not exist. Check results for BUSCO: 238925at7742
## Thu Jul 14 00:34:35 CEST 2022 - The aliscore output appears to be empty. Check results for BUSCO: 238925at7742
## Thu Jul 14 00:34:35 CEST 2022 - The aliscore output file does not exist. Check results for BUSCO: 169833at7742
## Thu Jul 14 00:34:36 CEST 2022 - The aliscore output appears to be empty. Check results for BUSCO: 169833at7742
## Thu Jul 14 00:34:38 CEST 2022 - The aliscore output file does not exist. Check results for BUSCO: 176410at7742
## Thu Jul 14 00:34:38 CEST 2022 - The aliscore output appears to be empty. Check results for BUSCO: 176410at7742
## Thu Jul 14 00:34:40 CEST 2022 - The aliscore output file does not exist. Check results for BUSCO: 40897at7742
## Thu Jul 14 00:34:40 CEST 2022 - The aliscore output appears to be empty. Check results for BUSCO: 40897at7742
## Thu Jul 14 00:34:42 CEST 2022 - The aliscore output file does not exist. Check results for BUSCO: 253823at7742
## Thu Jul 14 00:34:42 CEST 2022 - The aliscore output appears to be empty. Check results for BUSCO: 253823at7742
## Thu Jul 14 00:34:42 CEST 2022 - The aliscore output file does not exist. Check results for BUSCO: 100256at7742
## Thu Jul 14 00:34:42 CEST 2022 - The aliscore output file does not exist. Check results for BUSCO: 109081at7742
## Thu Jul 14 00:34:42 CEST 2022 - The aliscore output appears to be empty. Check results for BUSCO: 100256at7742
## Thu Jul 14 00:34:42 CEST 2022 - The aliscore output appears to be empty. Check results for BUSCO: 109081at7742
## Thu Jul 14 00:34:45 CEST 2022 - The aliscore output file does not exist. Check results for BUSCO: 73527at7742
## Thu Jul 14 00:34:45 CEST 2022 - The aliscore output appears to be empty. Check results for BUSCO: 73527at7742
## Thu Jul 14 00:34:46 CEST 2022 - The aliscore output file does not exist. Check results for BUSCO: 327742at7742
## Thu Jul 14 00:34:46 CEST 2022 - The aliscore output appears to be empty. Check results for BUSCO: 327742at7742
## Thu Jul 14 00:34:47 CEST 2022 - The aliscore output file does not exist. Check results for BUSCO: 70039at7742
## Thu Jul 14 00:34:47 CEST 2022 - The aliscore output appears to be empty. Check results for BUSCO: 70039at7742
## Thu Jul 14 00:34:49 CEST 2022 - The aliscore output file does not exist. Check results for BUSCO: 146661at7742
## Thu Jul 14 00:34:49 CEST 2022 - The aliscore output appears to be empty. Check results for BUSCO: 146661at7742
## Thu Jul 14 00:34:51 CEST 2022 - The aliscore output file does not exist. Check results for BUSCO: 157162at7742
## Thu Jul 14 00:34:51 CEST 2022 - The aliscore output appears to be empty. Check results for BUSCO: 157162at7742
## Thu Jul 14 00:34:53 CEST 2022 - The aliscore output file does not exist. Check results for BUSCO: 73313at7742
## Thu Jul 14 00:34:53 CEST 2022 - The aliscore output file does not exist. Check results for BUSCO: 117939at7742
## Thu Jul 14 00:34:53 CEST 2022 - The aliscore output appears to be empty. Check results for BUSCO: 73313at7742
## Thu Jul 14 00:34:53 CEST 2022 - The aliscore output appears to be empty. Check results for BUSCO: 117939at7742
## Thu Jul 14 00:34:54 CEST 2022 - The aliscore output file does not exist. Check results for BUSCO: 146661at7742
## Thu Jul 14 00:34:54 CEST 2022 - The aliscore output appears to be empty. Check results for BUSCO: 146661at7742
## Thu Jul 14 00:34:57 CEST 2022 - The aliscore output file does not exist. Check results for BUSCO: 59333at7742
## Thu Jul 14 00:34:57 CEST 2022 - The aliscore output appears to be empty. Check results for BUSCO: 59333at7742
## Thu Jul 14 00:34:58 CEST 2022 - The aliscore output file does not exist. Check results for BUSCO: 140098at7742
## Thu Jul 14 00:34:58 CEST 2022 - The aliscore output appears to be empty. Check results for BUSCO: 140098at7742
## Thu Jul 14 00:34:59 CEST 2022 - The aliscore output file does not exist. Check results for BUSCO: 164340at7742
## Thu Jul 14 00:34:59 CEST 2022 - The aliscore output appears to be empty. Check results for BUSCO: 164340at7742
## Thu Jul 14 00:35:01 CEST 2022 - The aliscore output file does not exist. Check results for BUSCO: 37796at7742
## Thu Jul 14 00:35:01 CEST 2022 - The aliscore output appears to be empty. Check results for BUSCO: 37796at7742
## Thu Jul 14 00:35:02 CEST 2022 - The aliscore output file does not exist. Check results for BUSCO: 175472at7742
## Thu Jul 14 00:35:02 CEST 2022 - The aliscore output appears to be empty. Check results for BUSCO: 175472at7742
## Thu Jul 14 00:35:04 CEST 2022 - The aliscore output file does not exist. Check results for BUSCO: 103690at7742
## Thu Jul 14 00:35:04 CEST 2022 - The aliscore output appears to be empty. Check results for BUSCO: 103690at7742
## Thu Jul 14 00:35:05 CEST 2022 - The aliscore output file does not exist. Check results for BUSCO: 207140at7742
## Thu Jul 14 00:35:05 CEST 2022 - The aliscore output appears to be empty. Check results for BUSCO: 207140at7742
## Thu Jul 14 00:35:05 CEST 2022 - The aliscore output file does not exist. Check results for BUSCO: 87266at7742
## Thu Jul 14 00:35:06 CEST 2022 - The aliscore output appears to be empty. Check results for BUSCO: 87266at7742
## Thu Jul 14 00:35:08 CEST 2022 - The aliscore output file does not exist. Check results for BUSCO: 108576at7742
## Thu Jul 14 00:35:08 CEST 2022 - The aliscore output appears to be empty. Check results for BUSCO: 108576at7742
## Thu Jul 14 00:35:08 CEST 2022 - The aliscore output file does not exist. Check results for BUSCO: 29337at7742
## Thu Jul 14 00:35:08 CEST 2022 - The aliscore output appears to be empty. Check results for BUSCO: 29337at7742
## Thu Jul 14 00:35:11 CEST 2022 - The aliscore output file does not exist. Check results for BUSCO: 171639at7742
## Thu Jul 14 00:35:11 CEST 2022 - The aliscore output appears to be empty. Check results for BUSCO: 171639at7742
## Thu Jul 14 00:35:11 CEST 2022 - The aliscore output file does not exist. Check results for BUSCO: 107107at7742
## Thu Jul 14 00:35:11 CEST 2022 - The aliscore output appears to be empty. Check results for BUSCO: 107107at7742
## Thu Jul 14 00:35:13 CEST 2022 - The aliscore output file does not exist. Check results for BUSCO: 95643at7742
## Thu Jul 14 00:35:13 CEST 2022 - The aliscore output appears to be empty. Check results for BUSCO: 95643at7742
## Thu Jul 14 00:35:13 CEST 2022 - The aliscore output file does not exist. Check results for BUSCO: 41189at7742
## Thu Jul 14 00:35:13 CEST 2022 - The aliscore output appears to be empty. Check results for BUSCO: 41189at7742
## Thu Jul 14 00:35:16 CEST 2022 - The aliscore output file does not exist. Check results for BUSCO: 69813at7742
## Thu Jul 14 00:35:17 CEST 2022 - The aliscore output appears to be empty. Check results for BUSCO: 69813at7742
## Thu Jul 14 00:35:18 CEST 2022 - The aliscore output file does not exist. Check results for BUSCO: 80163at7742
## Thu Jul 14 00:35:18 CEST 2022 - The aliscore output appears to be empty. Check results for BUSCO: 80163at7742
## Thu Jul 14 00:35:21 CEST 2022 - The aliscore output file does not exist. Check results for BUSCO: 159524at7742
## Thu Jul 14 00:35:21 CEST 2022 - The aliscore output appears to be empty. Check results for BUSCO: 159524at7742
## Thu Jul 14 00:35:21 CEST 2022 - The aliscore output file does not exist. Check results for BUSCO: 124962at7742
## Thu Jul 14 00:35:21 CEST 2022 - The aliscore output appears to be empty. Check results for BUSCO: 124962at7742
## Thu Jul 14 00:35:22 CEST 2022 - The aliscore output file does not exist. Check results for BUSCO: 101303at7742
## Thu Jul 14 00:35:22 CEST 2022 - The aliscore output appears to be empty. Check results for BUSCO: 101303at7742
## Thu Jul 14 00:35:22 CEST 2022 - The aliscore output file does not exist. Check results for BUSCO: 161769at7742
## Thu Jul 14 00:35:22 CEST 2022 - The aliscore output appears to be empty. Check results for BUSCO: 161769at7742
## Thu Jul 14 00:35:25 CEST 2022 - The aliscore output file does not exist. Check results for BUSCO: 189983at7742
## Thu Jul 14 00:35:25 CEST 2022 - The aliscore output appears to be empty. Check results for BUSCO: 189983at7742
## Thu Jul 14 00:35:25 CEST 2022 - The aliscore output file does not exist. Check results for BUSCO: 97645at7742
## Thu Jul 14 00:35:25 CEST 2022 - The aliscore output appears to be empty. Check results for BUSCO: 97645at7742
## Thu Jul 14 00:35:28 CEST 2022 - The aliscore output file does not exist. Check results for BUSCO: 131585at7742
## Thu Jul 14 00:35:28 CEST 2022 - The aliscore output appears to be empty. Check results for BUSCO: 131585at7742
## Thu Jul 14 00:35:28 CEST 2022 - The aliscore output file does not exist. Check results for BUSCO: 303797at7742
## Thu Jul 14 00:35:28 CEST 2022 - The aliscore output appears to be empty. Check results for BUSCO: 303797at7742
## Thu Jul 14 00:35:30 CEST 2022 - The aliscore output file does not exist. Check results for BUSCO: 53548at7742
## Thu Jul 14 00:35:30 CEST 2022 - The aliscore output appears to be empty. Check results for BUSCO: 53548at7742
## Thu Jul 14 00:35:30 CEST 2022 - The aliscore output file does not exist. Check results for BUSCO: 51094at7742
## Thu Jul 14 00:35:30 CEST 2022 - The aliscore output appears to be empty. Check results for BUSCO: 51094at7742
## Thu Jul 14 00:35:31 CEST 2022 - The aliscore output file does not exist. Check results for BUSCO: 141015at7742
## Thu Jul 14 00:35:31 CEST 2022 - The aliscore output appears to be empty. Check results for BUSCO: 141015at7742
## Thu Jul 14 00:35:33 CEST 2022 - The aliscore output file does not exist. Check results for BUSCO: 84802at7742
## Thu Jul 14 00:35:33 CEST 2022 - The aliscore output appears to be empty. Check results for BUSCO: 84802at7742
## Thu Jul 14 00:35:34 CEST 2022 - The aliscore output file does not exist. Check results for BUSCO: 235274at7742
## Thu Jul 14 00:35:34 CEST 2022 - The aliscore output appears to be empty. Check results for BUSCO: 235274at7742
## Thu Jul 14 00:35:37 CEST 2022 - The aliscore output file does not exist. Check results for BUSCO: 165703at7742
## Thu Jul 14 00:35:37 CEST 2022 - The aliscore output appears to be empty. Check results for BUSCO: 165703at7742
## Thu Jul 14 00:35:40 CEST 2022 - The aliscore output file does not exist. Check results for BUSCO: 39294at7742
## Thu Jul 14 00:35:40 CEST 2022 - The aliscore output appears to be empty. Check results for BUSCO: 39294at7742
## Thu Jul 14 00:35:41 CEST 2022 - The aliscore output file does not exist. Check results for BUSCO: 82221at7742
## Thu Jul 14 00:35:41 CEST 2022 - The aliscore output file does not exist. Check results for BUSCO: 88390at7742
## Thu Jul 14 00:35:41 CEST 2022 - The aliscore output appears to be empty. Check results for BUSCO: 82221at7742
## Thu Jul 14 00:35:41 CEST 2022 - The aliscore output appears to be empty. Check results for BUSCO: 88390at7742
## Thu Jul 14 00:35:41 CEST 2022 - The aliscore output file does not exist. Check results for BUSCO: 127789at7742
## Thu Jul 14 00:35:42 CEST 2022 - The aliscore output appears to be empty. Check results for BUSCO: 127789at7742
## Thu Jul 14 00:35:44 CEST 2022 - The aliscore output file does not exist. Check results for BUSCO: 344416at7742
## Thu Jul 14 00:35:44 CEST 2022 - The aliscore output appears to be empty. Check results for BUSCO: 344416at7742
## Thu Jul 14 00:35:45 CEST 2022 - The aliscore output file does not exist. Check results for BUSCO: 133969at7742
## Thu Jul 14 00:35:45 CEST 2022 - The aliscore output appears to be empty. Check results for BUSCO: 133969at7742
## Thu Jul 14 00:35:47 CEST 2022 - The aliscore output file does not exist. Check results for BUSCO: 77923at7742
## Thu Jul 14 00:35:47 CEST 2022 - The aliscore output appears to be empty. Check results for BUSCO: 77923at7742
## Thu Jul 14 00:35:49 CEST 2022 - The aliscore output file does not exist. Check results for BUSCO: 278954at7742
## Thu Jul 14 00:35:49 CEST 2022 - The aliscore output appears to be empty. Check results for BUSCO: 278954at7742
## Thu Jul 14 00:35:50 CEST 2022 - The aliscore output file does not exist. Check results for BUSCO: 104992at7742
## Thu Jul 14 00:35:50 CEST 2022 - The aliscore output appears to be empty. Check results for BUSCO: 104992at7742
## Thu Jul 14 00:35:51 CEST 2022 - The aliscore output file does not exist. Check results for BUSCO: 270327at7742
## Thu Jul 14 00:35:51 CEST 2022 - The aliscore output appears to be empty. Check results for BUSCO: 270327at7742
## Thu Jul 14 00:35:51 CEST 2022 - The aliscore output file does not exist. Check results for BUSCO: 114623at7742
## Thu Jul 14 00:35:51 CEST 2022 - The aliscore output appears to be empty. Check results for BUSCO: 114623at7742
## Thu Jul 14 00:35:53 CEST 2022 - The aliscore output file does not exist. Check results for BUSCO: 59333at7742
## Thu Jul 14 00:35:53 CEST 2022 - The aliscore output appears to be empty. Check results for BUSCO: 59333at7742
## Thu Jul 14 00:35:55 CEST 2022 - The aliscore output file does not exist. Check results for BUSCO: 137788at7742
## Thu Jul 14 00:35:55 CEST 2022 - The aliscore output appears to be empty. Check results for BUSCO: 137788at7742
## Thu Jul 14 00:35:58 CEST 2022 - The aliscore output file does not exist. Check results for BUSCO: 22496at7742
## Thu Jul 14 00:35:58 CEST 2022 - The aliscore output appears to be empty. Check results for BUSCO: 22496at7742
## Thu Jul 14 00:35:59 CEST 2022 - The aliscore output file does not exist. Check results for BUSCO: 120085at7742
## Thu Jul 14 00:35:59 CEST 2022 - The aliscore output appears to be empty. Check results for BUSCO: 120085at7742
## Thu Jul 14 00:35:59 CEST 2022 - The aliscore output file does not exist. Check results for BUSCO: 73313at7742
## Thu Jul 14 00:35:59 CEST 2022 - The aliscore output appears to be empty. Check results for BUSCO: 73313at7742
## Thu Jul 14 00:36:01 CEST 2022 - The aliscore output file does not exist. Check results for BUSCO: 74922at7742
## Thu Jul 14 00:36:01 CEST 2022 - The aliscore output appears to be empty. Check results for BUSCO: 74922at7742
## Thu Jul 14 00:36:01 CEST 2022 - The aliscore output file does not exist. Check results for BUSCO: 290517at7742
## Thu Jul 14 00:36:01 CEST 2022 - The aliscore output appears to be empty. Check results for BUSCO: 290517at7742
## Thu Jul 14 00:36:02 CEST 2022 - The aliscore output file does not exist. Check results for BUSCO: 86277at7742
## Thu Jul 14 00:36:02 CEST 2022 - The aliscore output appears to be empty. Check results for BUSCO: 86277at7742
## Thu Jul 14 00:36:04 CEST 2022 - The aliscore output file does not exist. Check results for BUSCO: 60672at7742
## Thu Jul 14 00:36:04 CEST 2022 - The aliscore output appears to be empty. Check results for BUSCO: 60672at7742
## Thu Jul 14 00:36:05 CEST 2022 - The aliscore output file does not exist. Check results for BUSCO: 87266at7742
## Thu Jul 14 00:36:05 CEST 2022 - The aliscore output appears to be empty. Check results for BUSCO: 87266at7742
## Thu Jul 14 00:36:07 CEST 2022 - The aliscore output file does not exist. Check results for BUSCO: 175033at7742
## Thu Jul 14 00:36:07 CEST 2022 - The aliscore output appears to be empty. Check results for BUSCO: 175033at7742
## Thu Jul 14 00:36:09 CEST 2022 - The aliscore output file does not exist. Check results for BUSCO: 39294at7742
## Thu Jul 14 00:36:09 CEST 2022 - The aliscore output appears to be empty. Check results for BUSCO: 39294at7742
## Thu Jul 14 00:36:10 CEST 2022 - The aliscore output file does not exist. Check results for BUSCO: 294047at7742
## Thu Jul 14 00:36:10 CEST 2022 - The aliscore output file does not exist. Check results for BUSCO: 19730at7742
## Thu Jul 14 00:36:10 CEST 2022 - The aliscore output appears to be empty. Check results for BUSCO: 294047at7742
## Thu Jul 14 00:36:10 CEST 2022 - The aliscore output appears to be empty. Check results for BUSCO: 19730at7742
## Thu Jul 14 00:36:12 CEST 2022 - The aliscore output file does not exist. Check results for BUSCO: 212876at7742
## Thu Jul 14 00:36:12 CEST 2022 - The aliscore output appears to be empty. Check results for BUSCO: 212876at7742
## Thu Jul 14 00:36:15 CEST 2022 - The aliscore output file does not exist. Check results for BUSCO: 175472at7742
## Thu Jul 14 00:36:15 CEST 2022 - The aliscore output appears to be empty. Check results for BUSCO: 175472at7742
## Thu Jul 14 00:36:16 CEST 2022 - The aliscore output file does not exist. Check results for BUSCO: 114459at7742
## Thu Jul 14 00:36:16 CEST 2022 - The aliscore output appears to be empty. Check results for BUSCO: 114459at7742
## Thu Jul 14 00:36:17 CEST 2022 - The aliscore output file does not exist. Check results for BUSCO: 235284at7742
## Thu Jul 14 00:36:17 CEST 2022 - The aliscore output appears to be empty. Check results for BUSCO: 235284at7742
## Thu Jul 14 00:36:17 CEST 2022 - The aliscore output file does not exist. Check results for BUSCO: 59663at7742
## Thu Jul 14 00:36:17 CEST 2022 - The aliscore output appears to be empty. Check results for BUSCO: 59663at7742
## Thu Jul 14 00:36:19 CEST 2022 - The aliscore output file does not exist. Check results for BUSCO: 223032at7742
## Thu Jul 14 00:36:19 CEST 2022 - The aliscore output appears to be empty. Check results for BUSCO: 223032at7742
## Thu Jul 14 00:36:20 CEST 2022 - The aliscore output file does not exist. Check results for BUSCO: 253823at7742
## Thu Jul 14 00:36:20 CEST 2022 - The aliscore output appears to be empty. Check results for BUSCO: 253823at7742
## Thu Jul 14 00:36:22 CEST 2022 - The aliscore output file does not exist. Check results for BUSCO: 143864at7742
## Thu Jul 14 00:36:22 CEST 2022 - The aliscore output appears to be empty. Check results for BUSCO: 143864at7742
## Thu Jul 14 00:36:24 CEST 2022 - The aliscore output file does not exist. Check results for BUSCO: 105115at7742
## Thu Jul 14 00:36:24 CEST 2022 - The aliscore output appears to be empty. Check results for BUSCO: 105115at7742
## Thu Jul 14 00:36:26 CEST 2022 - The aliscore output file does not exist. Check results for BUSCO: 169833at7742
## Thu Jul 14 00:36:26 CEST 2022 - The aliscore output appears to be empty. Check results for BUSCO: 169833at7742
## Thu Jul 14 00:36:27 CEST 2022 - The aliscore output file does not exist. Check results for BUSCO: 84802at7742
## Thu Jul 14 00:36:27 CEST 2022 - The aliscore output appears to be empty. Check results for BUSCO: 84802at7742
## Thu Jul 14 00:36:27 CEST 2022 - The aliscore output file does not exist. Check results for BUSCO: 146804at7742
## Thu Jul 14 00:36:27 CEST 2022 - The aliscore output appears to be empty. Check results for BUSCO: 146804at7742
## Thu Jul 14 00:36:29 CEST 2022 - The aliscore output file does not exist. Check results for BUSCO: 143004at7742
## Thu Jul 14 00:36:29 CEST 2022 - The aliscore output appears to be empty. Check results for BUSCO: 143004at7742
## Thu Jul 14 00:36:29 CEST 2022 - The aliscore output file does not exist. Check results for BUSCO: 42710at7742
## Thu Jul 14 00:36:29 CEST 2022 - The aliscore output appears to be empty. Check results for BUSCO: 42710at7742
## Thu Jul 14 00:36:33 CEST 2022 - The aliscore output file does not exist. Check results for BUSCO: 133969at7742
## Thu Jul 14 00:36:33 CEST 2022 - The aliscore output appears to be empty. Check results for BUSCO: 133969at7742
## Thu Jul 14 00:36:33 CEST 2022 - The aliscore output file does not exist. Check results for BUSCO: 114819at7742
## Thu Jul 14 00:36:33 CEST 2022 - The aliscore output appears to be empty. Check results for BUSCO: 114819at7742
## Thu Jul 14 00:36:35 CEST 2022 - The aliscore output file does not exist. Check results for BUSCO: 241590at7742
## Thu Jul 14 00:36:35 CEST 2022 - The aliscore output appears to be empty. Check results for BUSCO: 241590at7742
## Thu Jul 14 00:36:37 CEST 2022 - The aliscore output file does not exist. Check results for BUSCO: 87117at7742
## Thu Jul 14 00:36:37 CEST 2022 - The aliscore output appears to be empty. Check results for BUSCO: 87117at7742
## Thu Jul 14 00:36:38 CEST 2022 - The aliscore output file does not exist. Check results for BUSCO: 238925at7742
## Thu Jul 14 00:36:38 CEST 2022 - The aliscore output appears to be empty. Check results for BUSCO: 238925at7742
## Thu Jul 14 00:36:39 CEST 2022 - The aliscore output file does not exist. Check results for BUSCO: 74281at7742
## Thu Jul 14 00:36:39 CEST 2022 - The aliscore output appears to be empty. Check results for BUSCO: 74281at7742
## Thu Jul 14 00:36:42 CEST 2022 - The aliscore output file does not exist. Check results for BUSCO: 161395at7742
## Thu Jul 14 00:36:42 CEST 2022 - The aliscore output appears to be empty. Check results for BUSCO: 161395at7742
## Thu Jul 14 00:36:43 CEST 2022 - The aliscore output file does not exist. Check results for BUSCO: 80527at7742
## Thu Jul 14 00:36:43 CEST 2022 - The aliscore output appears to be empty. Check results for BUSCO: 80527at7742
## Thu Jul 14 00:36:46 CEST 2022 - The aliscore output file does not exist. Check results for BUSCO: 104501at7742
## Thu Jul 14 00:36:46 CEST 2022 - The aliscore output appears to be empty. Check results for BUSCO: 104501at7742
## Thu Jul 14 00:36:47 CEST 2022 - The aliscore output file does not exist. Check results for BUSCO: 127789at7742
## Thu Jul 14 00:36:47 CEST 2022 - The aliscore output appears to be empty. Check results for BUSCO: 127789at7742
## Thu Jul 14 00:36:48 CEST 2022 - The aliscore output file does not exist. Check results for BUSCO: 235274at7742
## Thu Jul 14 00:36:48 CEST 2022 - The aliscore output appears to be empty. Check results for BUSCO: 235274at7742
## Thu Jul 14 00:36:49 CEST 2022 - The aliscore output file does not exist. Check results for BUSCO: 160468at7742
## Thu Jul 14 00:36:49 CEST 2022 - The aliscore output appears to be empty. Check results for BUSCO: 160468at7742
## Thu Jul 14 00:36:50 CEST 2022 - The aliscore output file does not exist. Check results for BUSCO: 194260at7742
## Thu Jul 14 00:36:50 CEST 2022 - The aliscore output appears to be empty. Check results for BUSCO: 194260at7742
## Thu Jul 14 00:36:50 CEST 2022 - The aliscore output file does not exist. Check results for BUSCO: 60672at7742
## Thu Jul 14 00:36:50 CEST 2022 - The aliscore output appears to be empty. Check results for BUSCO: 60672at7742
## Thu Jul 14 00:36:53 CEST 2022 - The aliscore output file does not exist. Check results for BUSCO: 56477at7742
## Thu Jul 14 00:36:53 CEST 2022 - The aliscore output appears to be empty. Check results for BUSCO: 56477at7742
## Thu Jul 14 00:36:54 CEST 2022 - The aliscore output file does not exist. Check results for BUSCO: 58506at7742
## Thu Jul 14 00:36:54 CEST 2022 - The aliscore output appears to be empty. Check results for BUSCO: 58506at7742
## Thu Jul 14 00:36:56 CEST 2022 - The aliscore output file does not exist. Check results for BUSCO: 176579at7742
## Thu Jul 14 00:36:56 CEST 2022 - The aliscore output appears to be empty. Check results for BUSCO: 176579at7742
## Thu Jul 14 00:36:57 CEST 2022 - The aliscore output file does not exist. Check results for BUSCO: 203180at7742
## Thu Jul 14 00:36:57 CEST 2022 - The aliscore output appears to be empty. Check results for BUSCO: 203180at7742
## Thu Jul 14 00:36:57 CEST 2022 - The aliscore output file does not exist. Check results for BUSCO: 124764at7742
## Thu Jul 14 00:36:57 CEST 2022 - The aliscore output appears to be empty. Check results for BUSCO: 124764at7742
## Thu Jul 14 00:37:01 CEST 2022 - The aliscore output file does not exist. Check results for BUSCO: 60672at7742
## Thu Jul 14 00:37:01 CEST 2022 - The aliscore output appears to be empty. Check results for BUSCO: 60672at7742
## Thu Jul 14 00:37:01 CEST 2022 - The aliscore output file does not exist. Check results for BUSCO: 63207at7742
## Thu Jul 14 00:37:01 CEST 2022 - The aliscore output appears to be empty. Check results for BUSCO: 63207at7742
## Thu Jul 14 00:37:03 CEST 2022 - The aliscore output file does not exist. Check results for BUSCO: 327742at7742
## Thu Jul 14 00:37:03 CEST 2022 - The aliscore output file does not exist. Check results for BUSCO: 159524at7742
## Thu Jul 14 00:37:03 CEST 2022 - The aliscore output appears to be empty. Check results for BUSCO: 327742at7742
## Thu Jul 14 00:37:03 CEST 2022 - The aliscore output appears to be empty. Check results for BUSCO: 159524at7742
## Thu Jul 14 00:37:05 CEST 2022 - The aliscore output file does not exist. Check results for BUSCO: 47069at7742
## Thu Jul 14 00:37:05 CEST 2022 - The aliscore output appears to be empty. Check results for BUSCO: 47069at7742
## Thu Jul 14 00:37:06 CEST 2022 - The aliscore output file does not exist. Check results for BUSCO: 58896at7742
## Thu Jul 14 00:37:06 CEST 2022 - The aliscore output appears to be empty. Check results for BUSCO: 58896at7742
## Thu Jul 14 00:37:09 CEST 2022 - The aliscore output file does not exist. Check results for BUSCO: 64359at7742
## Thu Jul 14 00:37:09 CEST 2022 - The aliscore output appears to be empty. Check results for BUSCO: 64359at7742
## Thu Jul 14 00:37:09 CEST 2022 - The aliscore output file does not exist. Check results for BUSCO: 59241at7742
## Thu Jul 14 00:37:09 CEST 2022 - The aliscore output appears to be empty. Check results for BUSCO: 59241at7742
## Thu Jul 14 00:37:10 CEST 2022 - The aliscore output file does not exist. Check results for BUSCO: 50502at7742
## Thu Jul 14 00:37:10 CEST 2022 - The aliscore output appears to be empty. Check results for BUSCO: 50502at7742
## Thu Jul 14 00:37:12 CEST 2022 - The aliscore output file does not exist. Check results for BUSCO: 133596at7742
## Thu Jul 14 00:37:12 CEST 2022 - The aliscore output appears to be empty. Check results for BUSCO: 133596at7742
## Thu Jul 14 00:37:13 CEST 2022 - The aliscore output file does not exist. Check results for BUSCO: 241590at7742
## Thu Jul 14 00:37:13 CEST 2022 - The aliscore output appears to be empty. Check results for BUSCO: 241590at7742
## Thu Jul 14 00:37:14 CEST 2022 - The aliscore output file does not exist. Check results for BUSCO: 207993at7742
## Thu Jul 14 00:37:14 CEST 2022 - The aliscore output appears to be empty. Check results for BUSCO: 207993at7742
## Thu Jul 14 00:37:16 CEST 2022 - The aliscore output file does not exist. Check results for BUSCO: 207993at7742
## Thu Jul 14 00:37:16 CEST 2022 - The aliscore output appears to be empty. Check results for BUSCO: 207993at7742
## Thu Jul 14 00:37:17 CEST 2022 - The aliscore output file does not exist. Check results for BUSCO: 48716at7742
## Thu Jul 14 00:37:17 CEST 2022 - The aliscore output appears to be empty. Check results for BUSCO: 48716at7742
## Thu Jul 14 00:37:20 CEST 2022 - The aliscore output file does not exist. Check results for BUSCO: 51066at7742
## Thu Jul 14 00:37:20 CEST 2022 - The aliscore output appears to be empty. Check results for BUSCO: 51066at7742
## Thu Jul 14 00:37:21 CEST 2022 - The aliscore output file does not exist. Check results for BUSCO: 277286at7742
## Thu Jul 14 00:37:21 CEST 2022 - The aliscore output appears to be empty. Check results for BUSCO: 277286at7742
## Thu Jul 14 00:37:21 CEST 2022 - The aliscore output file does not exist. Check results for BUSCO: 50502at7742
## Thu Jul 14 00:37:21 CEST 2022 - The aliscore output appears to be empty. Check results for BUSCO: 50502at7742
## Thu Jul 14 00:37:23 CEST 2022 - The aliscore output file does not exist. Check results for BUSCO: 41189at7742
## Thu Jul 14 00:37:23 CEST 2022 - The aliscore output appears to be empty. Check results for BUSCO: 41189at7742
## Thu Jul 14 00:37:24 CEST 2022 - The aliscore output file does not exist. Check results for BUSCO: 396665at7742
## Thu Jul 14 00:37:24 CEST 2022 - The aliscore output appears to be empty. Check results for BUSCO: 396665at7742
## Thu Jul 14 00:37:26 CEST 2022 - The aliscore output file does not exist. Check results for BUSCO: 401756at7742
## Thu Jul 14 00:37:26 CEST 2022 - The aliscore output appears to be empty. Check results for BUSCO: 401756at7742
## Thu Jul 14 00:37:28 CEST 2022 - The aliscore output file does not exist. Check results for BUSCO: 123742at7742
## Thu Jul 14 00:37:28 CEST 2022 - The aliscore output appears to be empty. Check results for BUSCO: 123742at7742
## Thu Jul 14 00:37:29 CEST 2022 - The aliscore output file does not exist. Check results for BUSCO: 207993at7742
## Thu Jul 14 00:37:29 CEST 2022 - The aliscore output appears to be empty. Check results for BUSCO: 207993at7742
## Thu Jul 14 00:37:30 CEST 2022 - The aliscore output file does not exist. Check results for BUSCO: 54353at7742
## Thu Jul 14 00:37:30 CEST 2022 - The aliscore output appears to be empty. Check results for BUSCO: 54353at7742
## Thu Jul 14 00:37:31 CEST 2022 - The aliscore output file does not exist. Check results for BUSCO: 162375at7742
## Thu Jul 14 00:37:31 CEST 2022 - The aliscore output appears to be empty. Check results for BUSCO: 162375at7742
## Thu Jul 14 00:37:33 CEST 2022 - The aliscore output file does not exist. Check results for BUSCO: 47275at7742
## Thu Jul 14 00:37:33 CEST 2022 - The aliscore output appears to be empty. Check results for BUSCO: 47275at7742
## Thu Jul 14 00:37:35 CEST 2022 - The aliscore output file does not exist. Check results for BUSCO: 109081at7742
## Thu Jul 14 00:37:35 CEST 2022 - The aliscore output appears to be empty. Check results for BUSCO: 109081at7742
## Thu Jul 14 00:37:35 CEST 2022 - The aliscore output file does not exist. Check results for BUSCO: 175036at7742
## Thu Jul 14 00:37:35 CEST 2022 - The aliscore output appears to be empty. Check results for BUSCO: 175036at7742
## Thu Jul 14 00:37:37 CEST 2022 - The aliscore output file does not exist. Check results for BUSCO: 175033at7742
## Thu Jul 14 00:37:38 CEST 2022 - The aliscore output appears to be empty. Check results for BUSCO: 175033at7742
## Thu Jul 14 00:37:39 CEST 2022 - The aliscore output file does not exist. Check results for BUSCO: 207707at7742
## Thu Jul 14 00:37:39 CEST 2022 - The aliscore output appears to be empty. Check results for BUSCO: 207707at7742
## Thu Jul 14 00:37:39 CEST 2022 - The aliscore output file does not exist. Check results for BUSCO: 396665at7742
## Thu Jul 14 00:37:39 CEST 2022 - The aliscore output appears to be empty. Check results for BUSCO: 396665at7742
## Thu Jul 14 00:37:39 CEST 2022 - The aliscore output file does not exist. Check results for BUSCO: 64591at7742
## Thu Jul 14 00:37:39 CEST 2022 - The aliscore output appears to be empty. Check results for BUSCO: 64591at7742
## Thu Jul 14 00:37:43 CEST 2022 - The aliscore output file does not exist. Check results for BUSCO: 158068at7742
## Thu Jul 14 00:37:43 CEST 2022 - The aliscore output appears to be empty. Check results for BUSCO: 158068at7742
## Thu Jul 14 00:37:44 CEST 2022 - The aliscore output file does not exist. Check results for BUSCO: 87266at7742
## Thu Jul 14 00:37:44 CEST 2022 - The aliscore output appears to be empty. Check results for BUSCO: 87266at7742
## Thu Jul 14 00:37:46 CEST 2022 - The aliscore output file does not exist. Check results for BUSCO: 170533at7742
## Thu Jul 14 00:37:46 CEST 2022 - The aliscore output appears to be empty. Check results for BUSCO: 170533at7742
## Thu Jul 14 00:37:49 CEST 2022 - The aliscore output file does not exist. Check results for BUSCO: 225592at7742
## Thu Jul 14 00:37:49 CEST 2022 - The aliscore output appears to be empty. Check results for BUSCO: 225592at7742
## Thu Jul 14 00:37:49 CEST 2022 - The aliscore output file does not exist. Check results for BUSCO: 189983at7742
## Thu Jul 14 00:37:49 CEST 2022 - The aliscore output appears to be empty. Check results for BUSCO: 189983at7742
## Thu Jul 14 00:37:50 CEST 2022 - The aliscore output file does not exist. Check results for BUSCO: 303814at7742
## Thu Jul 14 00:37:50 CEST 2022 - The aliscore output appears to be empty. Check results for BUSCO: 303814at7742
## Thu Jul 14 00:37:51 CEST 2022 - The aliscore output file does not exist. Check results for BUSCO: 225592at7742
## Thu Jul 14 00:37:52 CEST 2022 - The aliscore output appears to be empty. Check results for BUSCO: 225592at7742
## Thu Jul 14 00:37:52 CEST 2022 - The aliscore output file does not exist. Check results for BUSCO: 124764at7742
## Thu Jul 14 00:37:52 CEST 2022 - The aliscore output appears to be empty. Check results for BUSCO: 124764at7742
## Thu Jul 14 00:37:53 CEST 2022 - The aliscore output file does not exist. Check results for BUSCO: 41071at7742
## Thu Jul 14 00:37:53 CEST 2022 - The aliscore output appears to be empty. Check results for BUSCO: 41071at7742
## Thu Jul 14 00:37:55 CEST 2022 - The aliscore output file does not exist. Check results for BUSCO: 189983at7742
## Thu Jul 14 00:37:55 CEST 2022 - The aliscore output appears to be empty. Check results for BUSCO: 189983at7742
## Thu Jul 14 00:37:56 CEST 2022 - The aliscore output file does not exist. Check results for BUSCO: 325663at7742
## Thu Jul 14 00:37:56 CEST 2022 - The aliscore output appears to be empty. Check results for BUSCO: 325663at7742
## Thu Jul 14 00:37:58 CEST 2022 - The aliscore output file does not exist. Check results for BUSCO: 119886at7742
## Thu Jul 14 00:37:58 CEST 2022 - The aliscore output appears to be empty. Check results for BUSCO: 119886at7742
## Thu Jul 14 00:37:58 CEST 2022 - The aliscore output file does not exist. Check results for BUSCO: 203180at7742
## Thu Jul 14 00:37:58 CEST 2022 - The aliscore output appears to be empty. Check results for BUSCO: 203180at7742
## Thu Jul 14 00:37:58 CEST 2022 - The aliscore output file does not exist. Check results for BUSCO: 396665at7742
## Thu Jul 14 00:37:59 CEST 2022 - The aliscore output appears to be empty. Check results for BUSCO: 396665at7742
## Thu Jul 14 00:38:02 CEST 2022 - The aliscore output file does not exist. Check results for BUSCO: 153051at7742
## Thu Jul 14 00:38:02 CEST 2022 - The aliscore output appears to be empty. Check results for BUSCO: 153051at7742
## Thu Jul 14 00:38:03 CEST 2022 - The aliscore output file does not exist. Check results for BUSCO: 163295at7742
## Thu Jul 14 00:38:04 CEST 2022 - The aliscore output appears to be empty. Check results for BUSCO: 163295at7742
## Thu Jul 14 00:38:05 CEST 2022 - The aliscore output file does not exist. Check results for BUSCO: 155361at7742
## Thu Jul 14 00:38:05 CEST 2022 - The aliscore output appears to be empty. Check results for BUSCO: 155361at7742
## Thu Jul 14 00:38:07 CEST 2022 - The aliscore output file does not exist. Check results for BUSCO: 103690at7742
## Thu Jul 14 00:38:07 CEST 2022 - The aliscore output appears to be empty. Check results for BUSCO: 103690at7742
## Thu Jul 14 00:38:07 CEST 2022 - The aliscore output file does not exist. Check results for BUSCO: 290517at7742
## Thu Jul 14 00:38:07 CEST 2022 - The aliscore output appears to be empty. Check results for BUSCO: 290517at7742
## Thu Jul 14 00:38:08 CEST 2022 - The aliscore output file does not exist. Check results for BUSCO: 45067at7742
## Thu Jul 14 00:38:08 CEST 2022 - The aliscore output file does not exist. Check results for BUSCO: 140098at7742
## Thu Jul 14 00:38:08 CEST 2022 - The aliscore output appears to be empty. Check results for BUSCO: 45067at7742
## Thu Jul 14 00:38:08 CEST 2022 - The aliscore output appears to be empty. Check results for BUSCO: 140098at7742
## Thu Jul 14 00:38:11 CEST 2022 - The aliscore output file does not exist. Check results for BUSCO: 207707at7742
## Thu Jul 14 00:38:11 CEST 2022 - The aliscore output appears to be empty. Check results for BUSCO: 207707at7742
## Thu Jul 14 00:38:13 CEST 2022 - The aliscore output file does not exist. Check results for BUSCO: 40567at7742
## Thu Jul 14 00:38:13 CEST 2022 - The aliscore output appears to be empty. Check results for BUSCO: 40567at7742
## Thu Jul 14 00:38:13 CEST 2022 - The aliscore output file does not exist. Check results for BUSCO: 294047at7742
## Thu Jul 14 00:38:13 CEST 2022 - The aliscore output appears to be empty. Check results for BUSCO: 294047at7742
## Thu Jul 14 00:38:15 CEST 2022 - The aliscore output file does not exist. Check results for BUSCO: 142117at7742
## Thu Jul 14 00:38:16 CEST 2022 - The aliscore output appears to be empty. Check results for BUSCO: 142117at7742
## Thu Jul 14 00:38:16 CEST 2022 - The aliscore output file does not exist. Check results for BUSCO: 137681at7742
## Thu Jul 14 00:38:16 CEST 2022 - The aliscore output appears to be empty. Check results for BUSCO: 137681at7742
## Thu Jul 14 00:38:17 CEST 2022 - The aliscore output file does not exist. Check results for BUSCO: 64807at7742
## Thu Jul 14 00:38:17 CEST 2022 - The aliscore output appears to be empty. Check results for BUSCO: 64807at7742
## Thu Jul 14 00:38:17 CEST 2022 - The aliscore output file does not exist. Check results for BUSCO: 87513at7742
## Thu Jul 14 00:38:17 CEST 2022 - The aliscore output appears to be empty. Check results for BUSCO: 87513at7742
## Thu Jul 14 00:38:20 CEST 2022 - The aliscore output file does not exist. Check results for BUSCO: 129039at7742
## Thu Jul 14 00:38:20 CEST 2022 - The aliscore output appears to be empty. Check results for BUSCO: 129039at7742
## Thu Jul 14 00:38:21 CEST 2022 - The aliscore output file does not exist. Check results for BUSCO: 129039at7742
## Thu Jul 14 00:38:21 CEST 2022 - The aliscore output appears to be empty. Check results for BUSCO: 129039at7742
## Thu Jul 14 00:38:25 CEST 2022 - The aliscore output file does not exist. Check results for BUSCO: 246595at7742
## Thu Jul 14 00:38:25 CEST 2022 - The aliscore output appears to be empty. Check results for BUSCO: 246595at7742
## Thu Jul 14 00:38:25 CEST 2022 - The aliscore output file does not exist. Check results for BUSCO: 80163at7742
## Thu Jul 14 00:38:25 CEST 2022 - The aliscore output appears to be empty. Check results for BUSCO: 80163at7742
## Thu Jul 14 00:38:26 CEST 2022 - The aliscore output file does not exist. Check results for BUSCO: 58506at7742
## Thu Jul 14 00:38:26 CEST 2022 - The aliscore output appears to be empty. Check results for BUSCO: 58506at7742
## Thu Jul 14 00:38:27 CEST 2022 - The aliscore output file does not exist. Check results for BUSCO: 185735at7742
## Thu Jul 14 00:38:27 CEST 2022 - The aliscore output appears to be empty. Check results for BUSCO: 185735at7742
## Thu Jul 14 00:38:27 CEST 2022 - The aliscore output file does not exist. Check results for BUSCO: 156420at7742
## Thu Jul 14 00:38:27 CEST 2022 - The aliscore output appears to be empty. Check results for BUSCO: 156420at7742
## Thu Jul 14 00:38:28 CEST 2022 - The aliscore output file does not exist. Check results for BUSCO: 124962at7742
## Thu Jul 14 00:38:28 CEST 2022 - The aliscore output appears to be empty. Check results for BUSCO: 124962at7742
## Thu Jul 14 00:38:31 CEST 2022 - The aliscore output file does not exist. Check results for BUSCO: 134632at7742
## Thu Jul 14 00:38:32 CEST 2022 - The aliscore output appears to be empty. Check results for BUSCO: 134632at7742
## Thu Jul 14 00:38:34 CEST 2022 - The aliscore output file does not exist. Check results for BUSCO: 95511at7742
## Thu Jul 14 00:38:34 CEST 2022 - The aliscore output appears to be empty. Check results for BUSCO: 95511at7742
## Thu Jul 14 00:38:34 CEST 2022 - The aliscore output file does not exist. Check results for BUSCO: 170533at7742
## Thu Jul 14 00:38:34 CEST 2022 - The aliscore output appears to be empty. Check results for BUSCO: 170533at7742
## Thu Jul 14 00:38:36 CEST 2022 - The aliscore output file does not exist. Check results for BUSCO: 19730at7742
## Thu Jul 14 00:38:36 CEST 2022 - The aliscore output appears to be empty. Check results for BUSCO: 19730at7742
## Thu Jul 14 00:38:37 CEST 2022 - The aliscore output file does not exist. Check results for BUSCO: 70459at7742
## Thu Jul 14 00:38:37 CEST 2022 - The aliscore output appears to be empty. Check results for BUSCO: 70459at7742
## Thu Jul 14 00:38:37 CEST 2022 - The aliscore output file does not exist. Check results for BUSCO: 153924at7742
## Thu Jul 14 00:38:37 CEST 2022 - The aliscore output appears to be empty. Check results for BUSCO: 153924at7742
## Thu Jul 14 00:38:38 CEST 2022 - The aliscore output file does not exist. Check results for BUSCO: 60142at7742
## Thu Jul 14 00:38:38 CEST 2022 - The aliscore output appears to be empty. Check results for BUSCO: 60142at7742
## Thu Jul 14 00:38:41 CEST 2022 - The aliscore output file does not exist. Check results for BUSCO: 155361at7742
## Thu Jul 14 00:38:41 CEST 2022 - The aliscore output appears to be empty. Check results for BUSCO: 155361at7742
## Thu Jul 14 00:38:42 CEST 2022 - The aliscore output file does not exist. Check results for BUSCO: 212876at7742
## Thu Jul 14 00:38:42 CEST 2022 - The aliscore output appears to be empty. Check results for BUSCO: 212876at7742
## Thu Jul 14 00:38:45 CEST 2022 - The aliscore output file does not exist. Check results for BUSCO: 92481at7742
## Thu Jul 14 00:38:45 CEST 2022 - The aliscore output appears to be empty. Check results for BUSCO: 92481at7742
## Thu Jul 14 00:38:47 CEST 2022 - The aliscore output file does not exist. Check results for BUSCO: 157162at7742
## Thu Jul 14 00:38:47 CEST 2022 - The aliscore output appears to be empty. Check results for BUSCO: 157162at7742
## Thu Jul 14 00:38:47 CEST 2022 - The aliscore output file does not exist. Check results for BUSCO: 154706at7742
## Thu Jul 14 00:38:47 CEST 2022 - The aliscore output appears to be empty. Check results for BUSCO: 154706at7742
## Thu Jul 14 00:38:48 CEST 2022 - The aliscore output file does not exist. Check results for BUSCO: 77322at7742
## Thu Jul 14 00:38:48 CEST 2022 - The aliscore output appears to be empty. Check results for BUSCO: 77322at7742
## Thu Jul 14 00:38:49 CEST 2022 - The aliscore output file does not exist. Check results for BUSCO: 122527at7742
## Thu Jul 14 00:38:49 CEST 2022 - The aliscore output appears to be empty. Check results for BUSCO: 122527at7742
## Thu Jul 14 00:38:49 CEST 2022 - The aliscore output file does not exist. Check results for BUSCO: 40897at7742
## Thu Jul 14 00:38:49 CEST 2022 - The aliscore output appears to be empty. Check results for BUSCO: 40897at7742
## Thu Jul 14 00:38:51 CEST 2022 - The aliscore output file does not exist. Check results for BUSCO: 77473at7742
## Thu Jul 14 00:38:51 CEST 2022 - The aliscore output appears to be empty. Check results for BUSCO: 77473at7742
## Thu Jul 14 00:38:53 CEST 2022 - The aliscore output file does not exist. Check results for BUSCO: 195017at7742
## Thu Jul 14 00:38:53 CEST 2022 - The aliscore output appears to be empty. Check results for BUSCO: 195017at7742
## Thu Jul 14 00:38:53 CEST 2022 - The aliscore output file does not exist. Check results for BUSCO: 168949at7742
## Thu Jul 14 00:38:53 CEST 2022 - The aliscore output appears to be empty. Check results for BUSCO: 168949at7742
## Thu Jul 14 00:38:55 CEST 2022 - The aliscore output file does not exist. Check results for BUSCO: 111583at7742
## Thu Jul 14 00:38:55 CEST 2022 - The aliscore output appears to be empty. Check results for BUSCO: 111583at7742
## Thu Jul 14 00:38:56 CEST 2022 - The aliscore output file does not exist. Check results for BUSCO: 59009at7742
## Thu Jul 14 00:38:56 CEST 2022 - The aliscore output appears to be empty. Check results for BUSCO: 59009at7742
## Thu Jul 14 00:38:56 CEST 2022 - The aliscore output file does not exist. Check results for BUSCO: 268563at7742
## Thu Jul 14 00:38:56 CEST 2022 - The aliscore output appears to be empty. Check results for BUSCO: 268563at7742
## Thu Jul 14 00:38:58 CEST 2022 - The aliscore output file does not exist. Check results for BUSCO: 51066at7742
## Thu Jul 14 00:38:58 CEST 2022 - The aliscore output appears to be empty. Check results for BUSCO: 51066at7742
## Thu Jul 14 00:38:59 CEST 2022 - The aliscore output file does not exist. Check results for BUSCO: 51094at7742
## Thu Jul 14 00:39:00 CEST 2022 - The aliscore output appears to be empty. Check results for BUSCO: 51094at7742
## Thu Jul 14 00:39:01 CEST 2022 - The aliscore output file does not exist. Check results for BUSCO: 160674at7742
## Thu Jul 14 00:39:01 CEST 2022 - The aliscore output appears to be empty. Check results for BUSCO: 160674at7742
## Thu Jul 14 00:39:04 CEST 2022 - The aliscore output file does not exist. Check results for BUSCO: 117939at7742
## Thu Jul 14 00:39:04 CEST 2022 - The aliscore output appears to be empty. Check results for BUSCO: 117939at7742
## Thu Jul 14 00:39:06 CEST 2022 - The aliscore output file does not exist. Check results for BUSCO: 40567at7742
## Thu Jul 14 00:39:06 CEST 2022 - The aliscore output appears to be empty. Check results for BUSCO: 40567at7742
## Thu Jul 14 00:39:06 CEST 2022 - The aliscore output file does not exist. Check results for BUSCO: 145564at7742
## Thu Jul 14 00:39:06 CEST 2022 - The aliscore output appears to be empty. Check results for BUSCO: 145564at7742
## Thu Jul 14 00:39:07 CEST 2022 - The aliscore output file does not exist. Check results for BUSCO: 77322at7742
## Thu Jul 14 00:39:07 CEST 2022 - The aliscore output appears to be empty. Check results for BUSCO: 77322at7742
## Thu Jul 14 00:39:08 CEST 2022 - The aliscore output file does not exist. Check results for BUSCO: 214140at7742
## Thu Jul 14 00:39:08 CEST 2022 - The aliscore output appears to be empty. Check results for BUSCO: 214140at7742
## Thu Jul 14 00:39:10 CEST 2022 - The aliscore output file does not exist. Check results for BUSCO: 223874at7742
## Thu Jul 14 00:39:10 CEST 2022 - The aliscore output appears to be empty. Check results for BUSCO: 223874at7742
## Thu Jul 14 00:39:10 CEST 2022 - The aliscore output file does not exist. Check results for BUSCO: 41071at7742
## Thu Jul 14 00:39:10 CEST 2022 - The aliscore output appears to be empty. Check results for BUSCO: 41071at7742
## Thu Jul 14 00:39:12 CEST 2022 - The aliscore output file does not exist. Check results for BUSCO: 232468at7742
## Thu Jul 14 00:39:12 CEST 2022 - The aliscore output appears to be empty. Check results for BUSCO: 232468at7742
## Thu Jul 14 00:39:14 CEST 2022 - The aliscore output file does not exist. Check results for BUSCO: 224692at7742
## Thu Jul 14 00:39:14 CEST 2022 - The aliscore output appears to be empty. Check results for BUSCO: 224692at7742
## Thu Jul 14 00:39:14 CEST 2022 - The aliscore output file does not exist. Check results for BUSCO: 137727at7742
## Thu Jul 14 00:39:14 CEST 2022 - The aliscore output appears to be empty. Check results for BUSCO: 137727at7742
## Thu Jul 14 00:39:16 CEST 2022 - The aliscore output file does not exist. Check results for BUSCO: 39255at7742
## Thu Jul 14 00:39:17 CEST 2022 - The aliscore output appears to be empty. Check results for BUSCO: 39255at7742
## Thu Jul 14 00:39:18 CEST 2022 - The aliscore output file does not exist. Check results for BUSCO: 138885at7742
## Thu Jul 14 00:39:18 CEST 2022 - The aliscore output appears to be empty. Check results for BUSCO: 138885at7742
## Thu Jul 14 00:39:20 CEST 2022 - The aliscore output file does not exist. Check results for BUSCO: 277286at7742
## Thu Jul 14 00:39:20 CEST 2022 - The aliscore output appears to be empty. Check results for BUSCO: 277286at7742
## Thu Jul 14 00:39:20 CEST 2022 - The aliscore output file does not exist. Check results for BUSCO: 153924at7742
## Thu Jul 14 00:39:20 CEST 2022 - The aliscore output appears to be empty. Check results for BUSCO: 153924at7742
## Thu Jul 14 00:39:22 CEST 2022 - The aliscore output file does not exist. Check results for BUSCO: 78812at7742
## Thu Jul 14 00:39:22 CEST 2022 - The aliscore output appears to be empty. Check results for BUSCO: 78812at7742
## Thu Jul 14 00:39:22 CEST 2022 - The aliscore output file does not exist. Check results for BUSCO: 100256at7742
## Thu Jul 14 00:39:22 CEST 2022 - The aliscore output file does not exist. Check results for BUSCO: 77322at7742
## Thu Jul 14 00:39:22 CEST 2022 - The aliscore output appears to be empty. Check results for BUSCO: 100256at7742
## Thu Jul 14 00:39:22 CEST 2022 - The aliscore output appears to be empty. Check results for BUSCO: 77322at7742
## Thu Jul 14 00:39:24 CEST 2022 - The aliscore output file does not exist. Check results for BUSCO: 114819at7742
## Thu Jul 14 00:39:24 CEST 2022 - The aliscore output appears to be empty. Check results for BUSCO: 114819at7742
## Thu Jul 14 00:39:27 CEST 2022 - The aliscore output file does not exist. Check results for BUSCO: 107610at7742
## Thu Jul 14 00:39:27 CEST 2022 - The aliscore output appears to be empty. Check results for BUSCO: 107610at7742
## Thu Jul 14 00:39:29 CEST 2022 - The aliscore output file does not exist. Check results for BUSCO: 162155at7742
## Thu Jul 14 00:39:29 CEST 2022 - The aliscore output appears to be empty. Check results for BUSCO: 162155at7742
## Thu Jul 14 00:39:29 CEST 2022 - The aliscore output file does not exist. Check results for BUSCO: 150497at7742
## Thu Jul 14 00:39:29 CEST 2022 - The aliscore output appears to be empty. Check results for BUSCO: 150497at7742
## Thu Jul 14 00:39:31 CEST 2022 - The aliscore output file does not exist. Check results for BUSCO: 42945at7742
## Thu Jul 14 00:39:31 CEST 2022 - The aliscore output appears to be empty. Check results for BUSCO: 42945at7742
## Thu Jul 14 00:39:31 CEST 2022 - The aliscore output file does not exist. Check results for BUSCO: 335206at7742
## Thu Jul 14 00:39:31 CEST 2022 - The aliscore output appears to be empty. Check results for BUSCO: 335206at7742
## Thu Jul 14 00:39:32 CEST 2022 - The aliscore output file does not exist. Check results for BUSCO: 60783at7742
## Thu Jul 14 00:39:32 CEST 2022 - The aliscore output appears to be empty. Check results for BUSCO: 60783at7742
## Thu Jul 14 00:39:34 CEST 2022 - The aliscore output file does not exist. Check results for BUSCO: 121053at7742
## Thu Jul 14 00:39:34 CEST 2022 - The aliscore output appears to be empty. Check results for BUSCO: 121053at7742
## Thu Jul 14 00:39:37 CEST 2022 - The aliscore output file does not exist. Check results for BUSCO: 22496at7742
## Thu Jul 14 00:39:37 CEST 2022 - The aliscore output file does not exist. Check results for BUSCO: 60870at7742
## Thu Jul 14 00:39:37 CEST 2022 - The aliscore output appears to be empty. Check results for BUSCO: 22496at7742
## Thu Jul 14 00:39:37 CEST 2022 - The aliscore output appears to be empty. Check results for BUSCO: 60870at7742
## Thu Jul 14 00:39:38 CEST 2022 - The aliscore output file does not exist. Check results for BUSCO: 105115at7742
## Thu Jul 14 00:39:38 CEST 2022 - The aliscore output appears to be empty. Check results for BUSCO: 105115at7742
## Thu Jul 14 00:39:40 CEST 2022 - The aliscore output file does not exist. Check results for BUSCO: 41071at7742
## Thu Jul 14 00:39:40 CEST 2022 - The aliscore output appears to be empty. Check results for BUSCO: 41071at7742
## Thu Jul 14 00:39:41 CEST 2022 - The aliscore output file does not exist. Check results for BUSCO: 161425at7742
## Thu Jul 14 00:39:41 CEST 2022 - The aliscore output appears to be empty. Check results for BUSCO: 161425at7742
## Thu Jul 14 00:39:41 CEST 2022 - The aliscore output file does not exist. Check results for BUSCO: 347249at7742
## Thu Jul 14 00:39:41 CEST 2022 - The aliscore output appears to be empty. Check results for BUSCO: 347249at7742
## Thu Jul 14 00:39:43 CEST 2022 - The aliscore output file does not exist. Check results for BUSCO: 149121at7742
## Thu Jul 14 00:39:43 CEST 2022 - The aliscore output appears to be empty. Check results for BUSCO: 149121at7742
## Thu Jul 14 00:39:46 CEST 2022 - The aliscore output file does not exist. Check results for BUSCO: 49394at7742
## Thu Jul 14 00:39:46 CEST 2022 - The aliscore output appears to be empty. Check results for BUSCO: 49394at7742
## Thu Jul 14 00:39:47 CEST 2022 - The aliscore output file does not exist. Check results for BUSCO: 56477at7742
## Thu Jul 14 00:39:47 CEST 2022 - The aliscore output appears to be empty. Check results for BUSCO: 56477at7742
## Thu Jul 14 00:39:47 CEST 2022 - The aliscore output file does not exist. Check results for BUSCO: 58896at7742
## Thu Jul 14 00:39:47 CEST 2022 - The aliscore output appears to be empty. Check results for BUSCO: 58896at7742
## Thu Jul 14 00:39:49 CEST 2022 - The aliscore output file does not exist. Check results for BUSCO: 110832at7742
## Thu Jul 14 00:39:49 CEST 2022 - The aliscore output appears to be empty. Check results for BUSCO: 110832at7742
## Thu Jul 14 00:39:49 CEST 2022 - The aliscore output file does not exist. Check results for BUSCO: 45067at7742
## Thu Jul 14 00:39:49 CEST 2022 - The aliscore output appears to be empty. Check results for BUSCO: 45067at7742
## Thu Jul 14 00:39:50 CEST 2022 - The aliscore output file does not exist. Check results for BUSCO: 108576at7742
## Thu Jul 14 00:39:50 CEST 2022 - The aliscore output appears to be empty. Check results for BUSCO: 108576at7742
## Thu Jul 14 00:39:53 CEST 2022 - The aliscore output file does not exist. Check results for BUSCO: 42945at7742
## Thu Jul 14 00:39:53 CEST 2022 - The aliscore output appears to be empty. Check results for BUSCO: 42945at7742
## Thu Jul 14 00:39:56 CEST 2022 - The aliscore output file does not exist. Check results for BUSCO: 162703at7742
## Thu Jul 14 00:39:56 CEST 2022 - The aliscore output appears to be empty. Check results for BUSCO: 162703at7742
## Thu Jul 14 00:39:56 CEST 2022 - The aliscore output file does not exist. Check results for BUSCO: 64807at7742
## Thu Jul 14 00:39:56 CEST 2022 - The aliscore output appears to be empty. Check results for BUSCO: 64807at7742
## Thu Jul 14 00:39:57 CEST 2022 - The aliscore output file does not exist. Check results for BUSCO: 79344at7742
## Thu Jul 14 00:39:57 CEST 2022 - The aliscore output appears to be empty. Check results for BUSCO: 79344at7742
## Thu Jul 14 00:39:57 CEST 2022 - The aliscore output file does not exist. Check results for BUSCO: 203180at7742
## Thu Jul 14 00:39:58 CEST 2022 - The aliscore output appears to be empty. Check results for BUSCO: 203180at7742
## Thu Jul 14 00:39:58 CEST 2022 - The aliscore output file does not exist. Check results for BUSCO: 347981at7742
## Thu Jul 14 00:39:58 CEST 2022 - The aliscore output appears to be empty. Check results for BUSCO: 347981at7742
## Thu Jul 14 00:39:58 CEST 2022 - The aliscore output file does not exist. Check results for BUSCO: 165703at7742
## Thu Jul 14 00:39:58 CEST 2022 - The aliscore output file does not exist. Check results for BUSCO: 60870at7742
## Thu Jul 14 00:39:58 CEST 2022 - The aliscore output appears to be empty. Check results for BUSCO: 165703at7742
## Thu Jul 14 00:39:58 CEST 2022 - The aliscore output appears to be empty. Check results for BUSCO: 60870at7742
## Thu Jul 14 00:40:02 CEST 2022 - The aliscore output file does not exist. Check results for BUSCO: 111583at7742
## Thu Jul 14 00:40:02 CEST 2022 - The aliscore output appears to be empty. Check results for BUSCO: 111583at7742
## Thu Jul 14 00:40:02 CEST 2022 - The aliscore output file does not exist. Check results for BUSCO: 168901at7742
## Thu Jul 14 00:40:02 CEST 2022 - The aliscore output appears to be empty. Check results for BUSCO: 168901at7742
## Thu Jul 14 00:40:05 CEST 2022 - The aliscore output file does not exist. Check results for BUSCO: 60142at7742
## Thu Jul 14 00:40:05 CEST 2022 - The aliscore output appears to be empty. Check results for BUSCO: 60142at7742
## Thu Jul 14 00:40:06 CEST 2022 - The aliscore output file does not exist. Check results for BUSCO: 74887at7742
## Thu Jul 14 00:40:06 CEST 2022 - The aliscore output appears to be empty. Check results for BUSCO: 74887at7742
## Thu Jul 14 00:40:08 CEST 2022 - The aliscore output file does not exist. Check results for BUSCO: 76254at7742
## Thu Jul 14 00:40:08 CEST 2022 - The aliscore output appears to be empty. Check results for BUSCO: 76254at7742
## Thu Jul 14 00:40:09 CEST 2022 - The aliscore output file does not exist. Check results for BUSCO: 171708at7742
## Thu Jul 14 00:40:09 CEST 2022 - The aliscore output appears to be empty. Check results for BUSCO: 171708at7742
## Thu Jul 14 00:40:10 CEST 2022 - The aliscore output file does not exist. Check results for BUSCO: 91371at7742
## Thu Jul 14 00:40:10 CEST 2022 - The aliscore output appears to be empty. Check results for BUSCO: 91371at7742
## Thu Jul 14 00:40:11 CEST 2022 - The aliscore output file does not exist. Check results for BUSCO: 64807at7742
## Thu Jul 14 00:40:11 CEST 2022 - The aliscore output appears to be empty. Check results for BUSCO: 64807at7742
## Thu Jul 14 00:40:12 CEST 2022 - The aliscore output file does not exist. Check results for BUSCO: 138885at7742
## Thu Jul 14 00:40:12 CEST 2022 - The aliscore output appears to be empty. Check results for BUSCO: 138885at7742
## Thu Jul 14 00:40:13 CEST 2022 - The aliscore output file does not exist. Check results for BUSCO: 257726at7742
## Thu Jul 14 00:40:13 CEST 2022 - The aliscore output appears to be empty. Check results for BUSCO: 257726at7742
## Thu Jul 14 00:40:15 CEST 2022 - The aliscore output file does not exist. Check results for BUSCO: 99383at7742
## Thu Jul 14 00:40:15 CEST 2022 - The aliscore output appears to be empty. Check results for BUSCO: 99383at7742
## Thu Jul 14 00:40:16 CEST 2022 - The aliscore output file does not exist. Check results for BUSCO: 209493at7742
## Thu Jul 14 00:40:16 CEST 2022 - The aliscore output appears to be empty. Check results for BUSCO: 209493at7742
## Thu Jul 14 00:40:16 CEST 2022 - The aliscore output file does not exist. Check results for BUSCO: 117939at7742
## Thu Jul 14 00:40:17 CEST 2022 - The aliscore output appears to be empty. Check results for BUSCO: 117939at7742
## Thu Jul 14 00:40:20 CEST 2022 - The aliscore output file does not exist. Check results for BUSCO: 84543at7742
## Thu Jul 14 00:40:20 CEST 2022 - The aliscore output appears to be empty. Check results for BUSCO: 84543at7742
## Thu Jul 14 00:40:20 CEST 2022 - The aliscore output file does not exist. Check results for BUSCO: 77473at7742
## Thu Jul 14 00:40:20 CEST 2022 - The aliscore output appears to be empty. Check results for BUSCO: 77473at7742
## Thu Jul 14 00:40:21 CEST 2022 - The aliscore output file does not exist. Check results for BUSCO: 42945at7742
## Thu Jul 14 00:40:21 CEST 2022 - The aliscore output appears to be empty. Check results for BUSCO: 42945at7742
## Thu Jul 14 00:40:24 CEST 2022 - The aliscore output file does not exist. Check results for BUSCO: 167017at7742
## Thu Jul 14 00:40:24 CEST 2022 - The aliscore output appears to be empty. Check results for BUSCO: 167017at7742
## Thu Jul 14 00:40:24 CEST 2022 - The aliscore output file does not exist. Check results for BUSCO: 92370at7742
## Thu Jul 14 00:40:24 CEST 2022 - The aliscore output appears to be empty. Check results for BUSCO: 92370at7742
## Thu Jul 14 00:40:26 CEST 2022 - The aliscore output file does not exist. Check results for BUSCO: 100655at7742
## Thu Jul 14 00:40:26 CEST 2022 - The aliscore output appears to be empty. Check results for BUSCO: 100655at7742
## Thu Jul 14 00:40:26 CEST 2022 - The aliscore output file does not exist. Check results for BUSCO: 123337at7742
## Thu Jul 14 00:40:26 CEST 2022 - The aliscore output appears to be empty. Check results for BUSCO: 123337at7742
## Thu Jul 14 00:40:29 CEST 2022 - The aliscore output file does not exist. Check results for BUSCO: 204919at7742
## Thu Jul 14 00:40:29 CEST 2022 - The aliscore output appears to be empty. Check results for BUSCO: 204919at7742
## Thu Jul 14 00:40:29 CEST 2022 - The aliscore output file does not exist. Check results for BUSCO: 94269at7742
## Thu Jul 14 00:40:29 CEST 2022 - The aliscore output appears to be empty. Check results for BUSCO: 94269at7742
## Thu Jul 14 00:40:29 CEST 2022 - The aliscore output file does not exist. Check results for BUSCO: 64359at7742
## Thu Jul 14 00:40:29 CEST 2022 - The aliscore output appears to be empty. Check results for BUSCO: 64359at7742
## Thu Jul 14 00:40:33 CEST 2022 - The aliscore output file does not exist. Check results for BUSCO: 96933at7742
## Thu Jul 14 00:40:33 CEST 2022 - The aliscore output appears to be empty. Check results for BUSCO: 96933at7742
## Thu Jul 14 00:40:34 CEST 2022 - The aliscore output file does not exist. Check results for BUSCO: 401756at7742
## Thu Jul 14 00:40:34 CEST 2022 - The aliscore output file does not exist. Check results for BUSCO: 161769at7742
## Thu Jul 14 00:40:34 CEST 2022 - The aliscore output appears to be empty. Check results for BUSCO: 401756at7742
## Thu Jul 14 00:40:34 CEST 2022 - The aliscore output appears to be empty. Check results for BUSCO: 161769at7742
## Thu Jul 14 00:40:35 CEST 2022 - The aliscore output file does not exist. Check results for BUSCO: 60870at7742
## Thu Jul 14 00:40:35 CEST 2022 - The aliscore output appears to be empty. Check results for BUSCO: 60870at7742
## Thu Jul 14 00:40:36 CEST 2022 - The aliscore output file does not exist. Check results for BUSCO: 72074at7742
## Thu Jul 14 00:40:36 CEST 2022 - The aliscore output appears to be empty. Check results for BUSCO: 72074at7742
## Thu Jul 14 00:40:37 CEST 2022 - The aliscore output file does not exist. Check results for BUSCO: 52541at7742
## Thu Jul 14 00:40:37 CEST 2022 - The aliscore output appears to be empty. Check results for BUSCO: 52541at7742
## Thu Jul 14 00:40:40 CEST 2022 - The aliscore output file does not exist. Check results for BUSCO: 99383at7742
## Thu Jul 14 00:40:40 CEST 2022 - The aliscore output appears to be empty. Check results for BUSCO: 99383at7742
## Thu Jul 14 00:40:40 CEST 2022 - The aliscore output file does not exist. Check results for BUSCO: 161769at7742
## Thu Jul 14 00:40:40 CEST 2022 - The aliscore output appears to be empty. Check results for BUSCO: 161769at7742
## Thu Jul 14 00:40:43 CEST 2022 - The aliscore output file does not exist. Check results for BUSCO: 160468at7742
## Thu Jul 14 00:40:43 CEST 2022 - The aliscore output appears to be empty. Check results for BUSCO: 160468at7742
## Thu Jul 14 00:40:43 CEST 2022 - The aliscore output file does not exist. Check results for BUSCO: 39255at7742
## Thu Jul 14 00:40:43 CEST 2022 - The aliscore output appears to be empty. Check results for BUSCO: 39255at7742
## Thu Jul 14 00:40:46 CEST 2022 - The aliscore output file does not exist. Check results for BUSCO: 44653at7742
## Thu Jul 14 00:40:46 CEST 2022 - The aliscore output appears to be empty. Check results for BUSCO: 44653at7742
## Thu Jul 14 00:40:46 CEST 2022 - The aliscore output file does not exist. Check results for BUSCO: 251511at7742
## Thu Jul 14 00:40:46 CEST 2022 - The aliscore output appears to be empty. Check results for BUSCO: 251511at7742
## Thu Jul 14 00:40:47 CEST 2022 - The aliscore output file does not exist. Check results for BUSCO: 70459at7742
## Thu Jul 14 00:40:47 CEST 2022 - The aliscore output appears to be empty. Check results for BUSCO: 70459at7742
## Thu Jul 14 00:40:48 CEST 2022 - The aliscore output file does not exist. Check results for BUSCO: 123337at7742
## Thu Jul 14 00:40:48 CEST 2022 - The aliscore output appears to be empty. Check results for BUSCO: 123337at7742
## Thu Jul 14 00:40:51 CEST 2022 - The aliscore output file does not exist. Check results for BUSCO: 58896at7742
## Thu Jul 14 00:40:51 CEST 2022 - The aliscore output appears to be empty. Check results for BUSCO: 58896at7742
## Thu Jul 14 00:40:52 CEST 2022 - The aliscore output file does not exist. Check results for BUSCO: 64591at7742
## Thu Jul 14 00:40:52 CEST 2022 - The aliscore output appears to be empty. Check results for BUSCO: 64591at7742
## Thu Jul 14 00:40:52 CEST 2022 - The aliscore output file does not exist. Check results for BUSCO: 109426at7742
## Thu Jul 14 00:40:52 CEST 2022 - The aliscore output appears to be empty. Check results for BUSCO: 109426at7742
## Thu Jul 14 00:40:54 CEST 2022 - The aliscore output file does not exist. Check results for BUSCO: 227556at7742
## Thu Jul 14 00:40:54 CEST 2022 - The aliscore output appears to be empty. Check results for BUSCO: 227556at7742
## Thu Jul 14 00:40:54 CEST 2022 - The aliscore output file does not exist. Check results for BUSCO: 40567at7742
## Thu Jul 14 00:40:55 CEST 2022 - The aliscore output appears to be empty. Check results for BUSCO: 40567at7742
## Thu Jul 14 00:40:56 CEST 2022 - The aliscore output file does not exist. Check results for BUSCO: 97786at7742
## Thu Jul 14 00:40:56 CEST 2022 - The aliscore output appears to be empty. Check results for BUSCO: 97786at7742
## Thu Jul 14 00:40:57 CEST 2022 - The aliscore output file does not exist. Check results for BUSCO: 108576at7742
## Thu Jul 14 00:40:57 CEST 2022 - The aliscore output appears to be empty. Check results for BUSCO: 108576at7742
## Thu Jul 14 00:41:00 CEST 2022 - The aliscore output file does not exist. Check results for BUSCO: 51066at7742
## Thu Jul 14 00:41:00 CEST 2022 - The aliscore output appears to be empty. Check results for BUSCO: 51066at7742
## Thu Jul 14 00:41:01 CEST 2022 - The aliscore output file does not exist. Check results for BUSCO: 154706at7742
## Thu Jul 14 00:41:01 CEST 2022 - The aliscore output appears to be empty. Check results for BUSCO: 154706at7742
## Thu Jul 14 00:41:01 CEST 2022 - The aliscore output file does not exist. Check results for BUSCO: 82221at7742
## Thu Jul 14 00:41:01 CEST 2022 - The aliscore output appears to be empty. Check results for BUSCO: 82221at7742
## Thu Jul 14 00:41:02 CEST 2022 - The aliscore output file does not exist. Check results for BUSCO: 157162at7742
## Thu Jul 14 00:41:02 CEST 2022 - The aliscore output appears to be empty. Check results for BUSCO: 157162at7742
## Thu Jul 14 00:41:05 CEST 2022 - The aliscore output file does not exist. Check results for BUSCO: 47069at7742
## Thu Jul 14 00:41:05 CEST 2022 - The aliscore output appears to be empty. Check results for BUSCO: 47069at7742
## Thu Jul 14 00:41:06 CEST 2022 - The aliscore output file does not exist. Check results for BUSCO: 74887at7742
## Thu Jul 14 00:41:06 CEST 2022 - The aliscore output appears to be empty. Check results for BUSCO: 74887at7742
## Thu Jul 14 00:41:08 CEST 2022 - The aliscore output file does not exist. Check results for BUSCO: 49394at7742
## Thu Jul 14 00:41:08 CEST 2022 - The aliscore output appears to be empty. Check results for BUSCO: 49394at7742
## Thu Jul 14 00:41:08 CEST 2022 - The aliscore output file does not exist. Check results for BUSCO: 108458at7742
## Thu Jul 14 00:41:09 CEST 2022 - The aliscore output appears to be empty. Check results for BUSCO: 108458at7742
## Thu Jul 14 00:41:10 CEST 2022 - The aliscore output file does not exist. Check results for BUSCO: 63198at7742
## Thu Jul 14 00:41:10 CEST 2022 - The aliscore output appears to be empty. Check results for BUSCO: 63198at7742
## Thu Jul 14 00:41:11 CEST 2022 - The aliscore output file does not exist. Check results for BUSCO: 111583at7742
## Thu Jul 14 00:41:11 CEST 2022 - The aliscore output appears to be empty. Check results for BUSCO: 111583at7742
## Thu Jul 14 00:41:12 CEST 2022 - The aliscore output file does not exist. Check results for BUSCO: 54353at7742
## Thu Jul 14 00:41:12 CEST 2022 - The aliscore output appears to be empty. Check results for BUSCO: 54353at7742
## Thu Jul 14 00:41:12 CEST 2022 - The aliscore output file does not exist. Check results for BUSCO: 63922at7742
## Thu Jul 14 00:41:12 CEST 2022 - The aliscore output appears to be empty. Check results for BUSCO: 63922at7742
## Thu Jul 14 00:41:16 CEST 2022 - The aliscore output file does not exist. Check results for BUSCO: 99383at7742
## Thu Jul 14 00:41:16 CEST 2022 - The aliscore output appears to be empty. Check results for BUSCO: 99383at7742
## Thu Jul 14 00:41:16 CEST 2022 - The aliscore output file does not exist. Check results for BUSCO: 162426at7742
## Thu Jul 14 00:41:16 CEST 2022 - The aliscore output appears to be empty. Check results for BUSCO: 162426at7742
## Thu Jul 14 00:41:20 CEST 2022 - The aliscore output file does not exist. Check results for BUSCO: 110839at7742
## Thu Jul 14 00:41:20 CEST 2022 - The aliscore output appears to be empty. Check results for BUSCO: 110839at7742
## Thu Jul 14 00:41:20 CEST 2022 - The aliscore output file does not exist. Check results for BUSCO: 109426at7742
## Thu Jul 14 00:41:20 CEST 2022 - The aliscore output appears to be empty. Check results for BUSCO: 109426at7742
## Thu Jul 14 00:41:21 CEST 2022 - The aliscore output file does not exist. Check results for BUSCO: 59663at7742
## Thu Jul 14 00:41:21 CEST 2022 - The aliscore output appears to be empty. Check results for BUSCO: 59663at7742
## Thu Jul 14 00:41:21 CEST 2022 - The aliscore output file does not exist. Check results for BUSCO: 223348at7742
## Thu Jul 14 00:41:21 CEST 2022 - The aliscore output appears to be empty. Check results for BUSCO: 223348at7742
## Thu Jul 14 00:41:21 CEST 2022 - The aliscore output file does not exist. Check results for BUSCO: 59241at7742
## Thu Jul 14 00:41:21 CEST 2022 - The aliscore output appears to be empty. Check results for BUSCO: 59241at7742
## Thu Jul 14 00:41:22 CEST 2022 - The aliscore output file does not exist. Check results for BUSCO: 209493at7742
## Thu Jul 14 00:41:22 CEST 2022 - The aliscore output appears to be empty. Check results for BUSCO: 209493at7742
## Thu Jul 14 00:41:26 CEST 2022 - The aliscore output file does not exist. Check results for BUSCO: 195017at7742
## Thu Jul 14 00:41:26 CEST 2022 - The aliscore output appears to be empty. Check results for BUSCO: 195017at7742
## Thu Jul 14 00:41:26 CEST 2022 - The aliscore output file does not exist. Check results for BUSCO: 69988at7742
## Thu Jul 14 00:41:26 CEST 2022 - The aliscore output appears to be empty. Check results for BUSCO: 69988at7742
## Thu Jul 14 00:41:29 CEST 2022 - The aliscore output file does not exist. Check results for BUSCO: 110839at7742
## Thu Jul 14 00:41:29 CEST 2022 - The aliscore output appears to be empty. Check results for BUSCO: 110839at7742
## Thu Jul 14 00:41:31 CEST 2022 - The aliscore output file does not exist. Check results for BUSCO: 135517at7742
## Thu Jul 14 00:41:31 CEST 2022 - The aliscore output appears to be empty. Check results for BUSCO: 135517at7742
## Thu Jul 14 00:41:31 CEST 2022 - The aliscore output file does not exist. Check results for BUSCO: 123387at7742
## Thu Jul 14 00:41:31 CEST 2022 - The aliscore output appears to be empty. Check results for BUSCO: 123387at7742
## Thu Jul 14 00:41:31 CEST 2022 - The aliscore output file does not exist. Check results for BUSCO: 44653at7742
## Thu Jul 14 00:41:31 CEST 2022 - The aliscore output appears to be empty. Check results for BUSCO: 44653at7742
## Thu Jul 14 00:41:33 CEST 2022 - The aliscore output file does not exist. Check results for BUSCO: 225592at7742
## Thu Jul 14 00:41:33 CEST 2022 - The aliscore output appears to be empty. Check results for BUSCO: 225592at7742
## Thu Jul 14 00:41:35 CEST 2022 - The aliscore output file does not exist. Check results for BUSCO: 123291at7742
## Thu Jul 14 00:41:35 CEST 2022 - The aliscore output appears to be empty. Check results for BUSCO: 123291at7742
## Thu Jul 14 00:41:36 CEST 2022 - The aliscore output file does not exist. Check results for BUSCO: 59241at7742
## Thu Jul 14 00:41:36 CEST 2022 - The aliscore output appears to be empty. Check results for BUSCO: 59241at7742
## Thu Jul 14 00:41:37 CEST 2022 - The aliscore output file does not exist. Check results for BUSCO: 48716at7742
## Thu Jul 14 00:41:37 CEST 2022 - The aliscore output appears to be empty. Check results for BUSCO: 48716at7742
## Thu Jul 14 00:41:38 CEST 2022 - The aliscore output file does not exist. Check results for BUSCO: 78812at7742
## Thu Jul 14 00:41:38 CEST 2022 - The aliscore output appears to be empty. Check results for BUSCO: 78812at7742
## Thu Jul 14 00:41:40 CEST 2022 - The aliscore output file does not exist. Check results for BUSCO: 195017at7742
## Thu Jul 14 00:41:40 CEST 2022 - The aliscore output appears to be empty. Check results for BUSCO: 195017at7742
## Thu Jul 14 00:41:40 CEST 2022 - The aliscore output file does not exist. Check results for BUSCO: 54075at7742
## Thu Jul 14 00:41:40 CEST 2022 - The aliscore output appears to be empty. Check results for BUSCO: 54075at7742
## Thu Jul 14 00:41:42 CEST 2022 - The aliscore output file does not exist. Check results for BUSCO: 236610at7742
## Thu Jul 14 00:41:42 CEST 2022 - The aliscore output appears to be empty. Check results for BUSCO: 236610at7742
## Thu Jul 14 00:41:45 CEST 2022 - The aliscore output file does not exist. Check results for BUSCO: 92370at7742
## Thu Jul 14 00:41:45 CEST 2022 - The aliscore output appears to be empty. Check results for BUSCO: 92370at7742
## Thu Jul 14 00:41:46 CEST 2022 - The aliscore output file does not exist. Check results for BUSCO: 224692at7742
## Thu Jul 14 00:41:46 CEST 2022 - The aliscore output appears to be empty. Check results for BUSCO: 224692at7742
## Thu Jul 14 00:41:46 CEST 2022 - The aliscore output file does not exist. Check results for BUSCO: 122918at7742
## Thu Jul 14 00:41:46 CEST 2022 - The aliscore output appears to be empty. Check results for BUSCO: 122918at7742
## Thu Jul 14 00:41:48 CEST 2022 - The aliscore output file does not exist. Check results for BUSCO: 54353at7742
## Thu Jul 14 00:41:48 CEST 2022 - The aliscore output appears to be empty. Check results for BUSCO: 54353at7742
## Thu Jul 14 00:41:49 CEST 2022 - The aliscore output file does not exist. Check results for BUSCO: 188889at7742
## Thu Jul 14 00:41:49 CEST 2022 - The aliscore output appears to be empty. Check results for BUSCO: 188889at7742
## Thu Jul 14 00:41:50 CEST 2022 - The aliscore output file does not exist. Check results for BUSCO: 210369at7742
## Thu Jul 14 00:41:50 CEST 2022 - The aliscore output appears to be empty. Check results for BUSCO: 210369at7742
## Thu Jul 14 00:41:50 CEST 2022 - The aliscore output file does not exist. Check results for BUSCO: 124764at7742
## Thu Jul 14 00:41:50 CEST 2022 - The aliscore output appears to be empty. Check results for BUSCO: 124764at7742
## Thu Jul 14 00:41:53 CEST 2022 - The aliscore output file does not exist. Check results for BUSCO: 133006at7742
## Thu Jul 14 00:41:53 CEST 2022 - The aliscore output appears to be empty. Check results for BUSCO: 133006at7742
## Thu Jul 14 00:41:54 CEST 2022 - The aliscore output file does not exist. Check results for BUSCO: 141754at7742
## Thu Jul 14 00:41:54 CEST 2022 - The aliscore output appears to be empty. Check results for BUSCO: 141754at7742
## Thu Jul 14 00:41:55 CEST 2022 - The aliscore output file does not exist. Check results for BUSCO: 162375at7742
## Thu Jul 14 00:41:55 CEST 2022 - The aliscore output appears to be empty. Check results for BUSCO: 162375at7742
## Thu Jul 14 00:41:56 CEST 2022 - The aliscore output file does not exist. Check results for BUSCO: 162375at7742
## Thu Jul 14 00:41:56 CEST 2022 - The aliscore output appears to be empty. Check results for BUSCO: 162375at7742
## Thu Jul 14 00:41:58 CEST 2022 - The aliscore output file does not exist. Check results for BUSCO: 108967at7742
## Thu Jul 14 00:41:58 CEST 2022 - The aliscore output appears to be empty. Check results for BUSCO: 108967at7742
## Thu Jul 14 00:41:59 CEST 2022 - The aliscore output file does not exist. Check results for BUSCO: 214140at7742
## Thu Jul 14 00:41:59 CEST 2022 - The aliscore output appears to be empty. Check results for BUSCO: 214140at7742
## Thu Jul 14 00:42:01 CEST 2022 - The aliscore output file does not exist. Check results for BUSCO: 52541at7742
## Thu Jul 14 00:42:01 CEST 2022 - The aliscore output appears to be empty. Check results for BUSCO: 52541at7742
## Thu Jul 14 00:42:02 CEST 2022 - The aliscore output file does not exist. Check results for BUSCO: 330757at7742
## Thu Jul 14 00:42:02 CEST 2022 - The aliscore output appears to be empty. Check results for BUSCO: 330757at7742
## Thu Jul 14 00:42:04 CEST 2022 - The aliscore output file does not exist. Check results for BUSCO: 149121at7742
## Thu Jul 14 00:42:04 CEST 2022 - The aliscore output appears to be empty. Check results for BUSCO: 149121at7742
## Thu Jul 14 00:42:05 CEST 2022 - The aliscore output file does not exist. Check results for BUSCO: 123387at7742
## Thu Jul 14 00:42:05 CEST 2022 - The aliscore output appears to be empty. Check results for BUSCO: 123387at7742
## Thu Jul 14 00:42:06 CEST 2022 - The aliscore output file does not exist. Check results for BUSCO: 109426at7742
## Thu Jul 14 00:42:06 CEST 2022 - The aliscore output appears to be empty. Check results for BUSCO: 109426at7742
## Thu Jul 14 00:42:06 CEST 2022 - The aliscore output file does not exist. Check results for BUSCO: 173159at7742
## Thu Jul 14 00:42:06 CEST 2022 - The aliscore output appears to be empty. Check results for BUSCO: 173159at7742
## Thu Jul 14 00:42:09 CEST 2022 - The aliscore output file does not exist. Check results for BUSCO: 149083at7742
## Thu Jul 14 00:42:09 CEST 2022 - The aliscore output appears to be empty. Check results for BUSCO: 149083at7742
## Thu Jul 14 00:42:10 CEST 2022 - The aliscore output file does not exist. Check results for BUSCO: 74922at7742
## Thu Jul 14 00:42:10 CEST 2022 - The aliscore output appears to be empty. Check results for BUSCO: 74922at7742
## Thu Jul 14 00:42:11 CEST 2022 - The aliscore output file does not exist. Check results for BUSCO: 223032at7742
## Thu Jul 14 00:42:11 CEST 2022 - The aliscore output appears to be empty. Check results for BUSCO: 223032at7742
## Thu Jul 14 00:42:12 CEST 2022 - The aliscore output file does not exist. Check results for BUSCO: 128400at7742
## Thu Jul 14 00:42:12 CEST 2022 - The aliscore output appears to be empty. Check results for BUSCO: 128400at7742
## Thu Jul 14 00:42:13 CEST 2022 - The aliscore output file does not exist. Check results for BUSCO: 47275at7742
## Thu Jul 14 00:42:13 CEST 2022 - The aliscore output appears to be empty. Check results for BUSCO: 47275at7742
## Thu Jul 14 00:42:14 CEST 2022 - The aliscore output file does not exist. Check results for BUSCO: 177227at7742
## Thu Jul 14 00:42:14 CEST 2022 - The aliscore output appears to be empty. Check results for BUSCO: 177227at7742
## Thu Jul 14 00:42:18 CEST 2022 - The aliscore output file does not exist. Check results for BUSCO: 172631at7742
## Thu Jul 14 00:42:18 CEST 2022 - The aliscore output appears to be empty. Check results for BUSCO: 172631at7742
## Thu Jul 14 00:42:19 CEST 2022 - The aliscore output file does not exist. Check results for BUSCO: 151961at7742
## Thu Jul 14 00:42:20 CEST 2022 - The aliscore output appears to be empty. Check results for BUSCO: 151961at7742
## Thu Jul 14 00:42:20 CEST 2022 - The aliscore output file does not exist. Check results for BUSCO: 172631at7742
## Thu Jul 14 00:42:20 CEST 2022 - The aliscore output appears to be empty. Check results for BUSCO: 172631at7742
## Thu Jul 14 00:42:23 CEST 2022 - The aliscore output file does not exist. Check results for BUSCO: 168901at7742
## Thu Jul 14 00:42:23 CEST 2022 - The aliscore output appears to be empty. Check results for BUSCO: 168901at7742
## Thu Jul 14 00:42:26 CEST 2022 - The aliscore output file does not exist. Check results for BUSCO: 104543at7742
## Thu Jul 14 00:42:26 CEST 2022 - The aliscore output file does not exist. Check results for BUSCO: 112360at7742
## Thu Jul 14 00:42:26 CEST 2022 - The aliscore output appears to be empty. Check results for BUSCO: 104543at7742
## Thu Jul 14 00:42:26 CEST 2022 - The aliscore output appears to be empty. Check results for BUSCO: 112360at7742
## Thu Jul 14 00:42:26 CEST 2022 - The aliscore output file does not exist. Check results for BUSCO: 22496at7742
## Thu Jul 14 00:42:26 CEST 2022 - The aliscore output appears to be empty. Check results for BUSCO: 22496at7742
## Thu Jul 14 00:42:27 CEST 2022 - The aliscore output file does not exist. Check results for BUSCO: 49394at7742
## Thu Jul 14 00:42:27 CEST 2022 - The aliscore output appears to be empty. Check results for BUSCO: 49394at7742
## Thu Jul 14 00:42:29 CEST 2022 - The aliscore output file does not exist. Check results for BUSCO: 63922at7742
## Thu Jul 14 00:42:29 CEST 2022 - The aliscore output appears to be empty. Check results for BUSCO: 63922at7742
## Thu Jul 14 00:42:29 CEST 2022 - The aliscore output file does not exist. Check results for BUSCO: 273629at7742
## Thu Jul 14 00:42:29 CEST 2022 - The aliscore output appears to be empty. Check results for BUSCO: 273629at7742
## Thu Jul 14 00:42:31 CEST 2022 - The aliscore output file does not exist. Check results for BUSCO: 165703at7742
## Thu Jul 14 00:42:31 CEST 2022 - The aliscore output appears to be empty. Check results for BUSCO: 165703at7742
## Thu Jul 14 00:42:32 CEST 2022 - The aliscore output file does not exist. Check results for BUSCO: 53548at7742
## Thu Jul 14 00:42:32 CEST 2022 - The aliscore output appears to be empty. Check results for BUSCO: 53548at7742
## Thu Jul 14 00:42:33 CEST 2022 - The aliscore output file does not exist. Check results for BUSCO: 114626at7742
## Thu Jul 14 00:42:33 CEST 2022 - The aliscore output appears to be empty. Check results for BUSCO: 114626at7742
## Thu Jul 14 00:42:34 CEST 2022 - The aliscore output file does not exist. Check results for BUSCO: 59333at7742
## Thu Jul 14 00:42:34 CEST 2022 - The aliscore output appears to be empty. Check results for BUSCO: 59333at7742
## Thu Jul 14 00:42:38 CEST 2022 - The aliscore output file does not exist. Check results for BUSCO: 127789at7742
## Thu Jul 14 00:42:38 CEST 2022 - The aliscore output appears to be empty. Check results for BUSCO: 127789at7742
## Thu Jul 14 00:42:38 CEST 2022 - The aliscore output file does not exist. Check results for BUSCO: 278954at7742
## Thu Jul 14 00:42:38 CEST 2022 - The aliscore output appears to be empty. Check results for BUSCO: 278954at7742
## Thu Jul 14 00:42:39 CEST 2022 - The aliscore output file does not exist. Check results for BUSCO: 93430at7742
## Thu Jul 14 00:42:40 CEST 2022 - The aliscore output appears to be empty. Check results for BUSCO: 93430at7742
## Thu Jul 14 00:42:41 CEST 2022 - The aliscore output file does not exist. Check results for BUSCO: 84817at7742
## Thu Jul 14 00:42:41 CEST 2022 - The aliscore output appears to be empty. Check results for BUSCO: 84817at7742
## Thu Jul 14 00:42:42 CEST 2022 - The aliscore output file does not exist. Check results for BUSCO: 205786at7742
## Thu Jul 14 00:42:42 CEST 2022 - The aliscore output appears to be empty. Check results for BUSCO: 205786at7742
## Thu Jul 14 00:42:43 CEST 2022 - The aliscore output file does not exist. Check results for BUSCO: 210369at7742
## Thu Jul 14 00:42:43 CEST 2022 - The aliscore output appears to be empty. Check results for BUSCO: 210369at7742
## Thu Jul 14 00:42:43 CEST 2022 - The aliscore output file does not exist. Check results for BUSCO: 44653at7742
## Thu Jul 14 00:42:43 CEST 2022 - The aliscore output appears to be empty. Check results for BUSCO: 44653at7742
## Thu Jul 14 00:42:45 CEST 2022 - The aliscore output file does not exist. Check results for BUSCO: 214140at7742
## Thu Jul 14 00:42:45 CEST 2022 - The aliscore output appears to be empty. Check results for BUSCO: 214140at7742
## Thu Jul 14 00:42:46 CEST 2022 - The aliscore output file does not exist. Check results for BUSCO: 95963at7742
## Thu Jul 14 00:42:46 CEST 2022 - The aliscore output appears to be empty. Check results for BUSCO: 95963at7742
## Thu Jul 14 00:42:49 CEST 2022 - The aliscore output file does not exist. Check results for BUSCO: 351213at7742
## Thu Jul 14 00:42:49 CEST 2022 - The aliscore output appears to be empty. Check results for BUSCO: 351213at7742
## Thu Jul 14 00:42:49 CEST 2022 - The aliscore output file does not exist. Check results for BUSCO: 303797at7742
## Thu Jul 14 00:42:49 CEST 2022 - The aliscore output appears to be empty. Check results for BUSCO: 303797at7742
## Thu Jul 14 00:42:50 CEST 2022 - The aliscore output file does not exist. Check results for BUSCO: 133006at7742
## Thu Jul 14 00:42:50 CEST 2022 - The aliscore output appears to be empty. Check results for BUSCO: 133006at7742
## Thu Jul 14 00:42:52 CEST 2022 - The aliscore output file does not exist. Check results for BUSCO: 172631at7742
## Thu Jul 14 00:42:52 CEST 2022 - The aliscore output appears to be empty. Check results for BUSCO: 172631at7742
## Thu Jul 14 00:42:52 CEST 2022 - The aliscore output file does not exist. Check results for BUSCO: 109650at7742
## Thu Jul 14 00:42:52 CEST 2022 - The aliscore output appears to be empty. Check results for BUSCO: 109650at7742
## Thu Jul 14 00:42:54 CEST 2022 - The aliscore output file does not exist. Check results for BUSCO: 321778at7742
## Thu Jul 14 00:42:54 CEST 2022 - The aliscore output appears to be empty. Check results for BUSCO: 321778at7742
## Thu Jul 14 00:42:55 CEST 2022 - The aliscore output file does not exist. Check results for BUSCO: 87513at7742
## Thu Jul 14 00:42:55 CEST 2022 - The aliscore output appears to be empty. Check results for BUSCO: 87513at7742
## Thu Jul 14 00:42:56 CEST 2022 - The aliscore output file does not exist. Check results for BUSCO: 142117at7742
## Thu Jul 14 00:42:56 CEST 2022 - The aliscore output appears to be empty. Check results for BUSCO: 142117at7742
## Thu Jul 14 00:42:57 CEST 2022 - The aliscore output file does not exist. Check results for BUSCO: 45067at7742
## Thu Jul 14 00:42:57 CEST 2022 - The aliscore output appears to be empty. Check results for BUSCO: 45067at7742
## Thu Jul 14 00:43:00 CEST 2022 - The aliscore output file does not exist. Check results for BUSCO: 59009at7742
## Thu Jul 14 00:43:01 CEST 2022 - The aliscore output appears to be empty. Check results for BUSCO: 59009at7742
## Thu Jul 14 00:43:02 CEST 2022 - The aliscore output file does not exist. Check results for BUSCO: 170032at7742
## Thu Jul 14 00:43:02 CEST 2022 - The aliscore output appears to be empty. Check results for BUSCO: 170032at7742
## Thu Jul 14 00:43:02 CEST 2022 - The aliscore output file does not exist. Check results for BUSCO: 119040at7742
## Thu Jul 14 00:43:02 CEST 2022 - The aliscore output appears to be empty. Check results for BUSCO: 119040at7742
## Thu Jul 14 00:43:04 CEST 2022 - The aliscore output file does not exist. Check results for BUSCO: 39465at7742
## Thu Jul 14 00:43:04 CEST 2022 - The aliscore output appears to be empty. Check results for BUSCO: 39465at7742
## Thu Jul 14 00:43:04 CEST 2022 - The aliscore output file does not exist. Check results for BUSCO: 115220at7742
## Thu Jul 14 00:43:05 CEST 2022 - The aliscore output appears to be empty. Check results for BUSCO: 115220at7742
## Thu Jul 14 00:43:06 CEST 2022 - The aliscore output file does not exist. Check results for BUSCO: 191130at7742
## Thu Jul 14 00:43:06 CEST 2022 - The aliscore output appears to be empty. Check results for BUSCO: 191130at7742
## Thu Jul 14 00:43:09 CEST 2022 - The aliscore output file does not exist. Check results for BUSCO: 92115at7742
## Thu Jul 14 00:43:09 CEST 2022 - The aliscore output appears to be empty. Check results for BUSCO: 92115at7742
## Thu Jul 14 00:43:10 CEST 2022 - The aliscore output file does not exist. Check results for BUSCO: 156420at7742
## Thu Jul 14 00:43:10 CEST 2022 - The aliscore output appears to be empty. Check results for BUSCO: 156420at7742
## Thu Jul 14 00:43:11 CEST 2022 - The aliscore output file does not exist. Check results for BUSCO: 210369at7742
## Thu Jul 14 00:43:11 CEST 2022 - The aliscore output appears to be empty. Check results for BUSCO: 210369at7742
## Thu Jul 14 00:43:11 CEST 2022 - The aliscore output file does not exist. Check results for BUSCO: 39465at7742
## Thu Jul 14 00:43:11 CEST 2022 - The aliscore output appears to be empty. Check results for BUSCO: 39465at7742
## Thu Jul 14 00:43:12 CEST 2022 - The aliscore output file does not exist. Check results for BUSCO: 273631at7742
## Thu Jul 14 00:43:12 CEST 2022 - The aliscore output appears to be empty. Check results for BUSCO: 273631at7742
## Thu Jul 14 00:43:14 CEST 2022 - The aliscore output file does not exist. Check results for BUSCO: 232468at7742
## Thu Jul 14 00:43:14 CEST 2022 - The aliscore output appears to be empty. Check results for BUSCO: 232468at7742
## Thu Jul 14 00:43:16 CEST 2022 - The aliscore output file does not exist. Check results for BUSCO: 141754at7742
## Thu Jul 14 00:43:16 CEST 2022 - The aliscore output appears to be empty. Check results for BUSCO: 141754at7742
## Thu Jul 14 00:43:17 CEST 2022 - The aliscore output file does not exist. Check results for BUSCO: 90853at7742
## Thu Jul 14 00:43:17 CEST 2022 - The aliscore output appears to be empty. Check results for BUSCO: 90853at7742
## Thu Jul 14 00:43:19 CEST 2022 - The aliscore output file does not exist. Check results for BUSCO: 46694at7742
## Thu Jul 14 00:43:19 CEST 2022 - The aliscore output appears to be empty. Check results for BUSCO: 46694at7742
## Thu Jul 14 00:43:21 CEST 2022 - The aliscore output file does not exist. Check results for BUSCO: 205786at7742
## Thu Jul 14 00:43:21 CEST 2022 - The aliscore output appears to be empty. Check results for BUSCO: 205786at7742
## Thu Jul 14 00:43:22 CEST 2022 - The aliscore output file does not exist. Check results for BUSCO: 60142at7742
## Thu Jul 14 00:43:22 CEST 2022 - The aliscore output appears to be empty. Check results for BUSCO: 60142at7742
## Thu Jul 14 00:43:22 CEST 2022 - The aliscore output file does not exist. Check results for BUSCO: 123291at7742
## Thu Jul 14 00:43:22 CEST 2022 - The aliscore output appears to be empty. Check results for BUSCO: 123291at7742
## Thu Jul 14 00:43:23 CEST 2022 - The aliscore output file does not exist. Check results for BUSCO: 227556at7742
## Thu Jul 14 00:43:23 CEST 2022 - The aliscore output appears to be empty. Check results for BUSCO: 227556at7742
## Thu Jul 14 00:43:26 CEST 2022 - The aliscore output file does not exist. Check results for BUSCO: 131585at7742
## Thu Jul 14 00:43:26 CEST 2022 - The aliscore output appears to be empty. Check results for BUSCO: 131585at7742
## Thu Jul 14 00:43:28 CEST 2022 - The aliscore output file does not exist. Check results for BUSCO: 107107at7742
## Thu Jul 14 00:43:28 CEST 2022 - The aliscore output appears to be empty. Check results for BUSCO: 107107at7742
## Thu Jul 14 00:43:28 CEST 2022 - The aliscore output file does not exist. Check results for BUSCO: 163295at7742
## Thu Jul 14 00:43:28 CEST 2022 - The aliscore output appears to be empty. Check results for BUSCO: 163295at7742
## Thu Jul 14 00:43:30 CEST 2022 - The aliscore output file does not exist. Check results for BUSCO: 115220at7742
## Thu Jul 14 00:43:30 CEST 2022 - The aliscore output appears to be empty. Check results for BUSCO: 115220at7742
## Thu Jul 14 00:43:31 CEST 2022 - The aliscore output file does not exist. Check results for BUSCO: 39255at7742
## Thu Jul 14 00:43:31 CEST 2022 - The aliscore output appears to be empty. Check results for BUSCO: 39255at7742
## Thu Jul 14 00:43:33 CEST 2022 - The aliscore output file does not exist. Check results for BUSCO: 58280at7742
## Thu Jul 14 00:43:33 CEST 2022 - The aliscore output appears to be empty. Check results for BUSCO: 58280at7742
## Thu Jul 14 00:43:33 CEST 2022 - The aliscore output file does not exist. Check results for BUSCO: 84543at7742
## Thu Jul 14 00:43:33 CEST 2022 - The aliscore output appears to be empty. Check results for BUSCO: 84543at7742
## Thu Jul 14 00:43:35 CEST 2022 - The aliscore output file does not exist. Check results for BUSCO: 294047at7742
## Thu Jul 14 00:43:35 CEST 2022 - The aliscore output appears to be empty. Check results for BUSCO: 294047at7742
## Thu Jul 14 00:43:35 CEST 2022 - The aliscore output file does not exist. Check results for BUSCO: 124028at7742
## Thu Jul 14 00:43:35 CEST 2022 - The aliscore output appears to be empty. Check results for BUSCO: 124028at7742
## Thu Jul 14 00:43:38 CEST 2022 - The aliscore output file does not exist. Check results for BUSCO: 126277at7742
## Thu Jul 14 00:43:38 CEST 2022 - The aliscore output appears to be empty. Check results for BUSCO: 126277at7742
## Thu Jul 14 00:43:39 CEST 2022 - The aliscore output file does not exist. Check results for BUSCO: 124028at7742
## Thu Jul 14 00:43:39 CEST 2022 - The aliscore output appears to be empty. Check results for BUSCO: 124028at7742
## Thu Jul 14 00:43:39 CEST 2022 - The aliscore output file does not exist. Check results for BUSCO: 198404at7742
## Thu Jul 14 00:43:39 CEST 2022 - The aliscore output appears to be empty. Check results for BUSCO: 198404at7742
## Thu Jul 14 00:43:40 CEST 2022 - The aliscore output file does not exist. Check results for BUSCO: 100655at7742
## Thu Jul 14 00:43:40 CEST 2022 - The aliscore output appears to be empty. Check results for BUSCO: 100655at7742
## Thu Jul 14 00:43:44 CEST 2022 - The aliscore output file does not exist. Check results for BUSCO: 50786at7742
## Thu Jul 14 00:43:44 CEST 2022 - The aliscore output appears to be empty. Check results for BUSCO: 50786at7742
## Thu Jul 14 00:43:45 CEST 2022 - The aliscore output file does not exist. Check results for BUSCO: 73229at7742
## Thu Jul 14 00:43:45 CEST 2022 - The aliscore output appears to be empty. Check results for BUSCO: 73229at7742
## Thu Jul 14 00:43:46 CEST 2022 - The aliscore output file does not exist. Check results for BUSCO: 92115at7742
## Thu Jul 14 00:43:46 CEST 2022 - The aliscore output appears to be empty. Check results for BUSCO: 92115at7742
## Thu Jul 14 00:43:47 CEST 2022 - The aliscore output file does not exist. Check results for BUSCO: 126277at7742
## Thu Jul 14 00:43:47 CEST 2022 - The aliscore output file does not exist. Check results for BUSCO: 251511at7742
## Thu Jul 14 00:43:47 CEST 2022 - The aliscore output appears to be empty. Check results for BUSCO: 126277at7742
## Thu Jul 14 00:43:47 CEST 2022 - The aliscore output appears to be empty. Check results for BUSCO: 251511at7742
## Thu Jul 14 00:43:48 CEST 2022 - The aliscore output file does not exist. Check results for BUSCO: 124028at7742
## Thu Jul 14 00:43:48 CEST 2022 - The aliscore output appears to be empty. Check results for BUSCO: 124028at7742
## Thu Jul 14 00:43:49 CEST 2022 - The aliscore output file does not exist. Check results for BUSCO: 73229at7742
## Thu Jul 14 00:43:49 CEST 2022 - The aliscore output appears to be empty. Check results for BUSCO: 73229at7742
## Thu Jul 14 00:43:52 CEST 2022 - The aliscore output file does not exist. Check results for BUSCO: 132224at7742
## Thu Jul 14 00:43:52 CEST 2022 - The aliscore output appears to be empty. Check results for BUSCO: 132224at7742
## Thu Jul 14 00:43:54 CEST 2022 - The aliscore output file does not exist. Check results for BUSCO: 120085at7742
## Thu Jul 14 00:43:54 CEST 2022 - The aliscore output appears to be empty. Check results for BUSCO: 120085at7742
## Thu Jul 14 00:43:56 CEST 2022 - The aliscore output file does not exist. Check results for BUSCO: 127830at7742
## Thu Jul 14 00:43:56 CEST 2022 - The aliscore output appears to be empty. Check results for BUSCO: 127830at7742
## Thu Jul 14 00:43:57 CEST 2022 - The aliscore output file does not exist. Check results for BUSCO: 229239at7742
## Thu Jul 14 00:43:57 CEST 2022 - The aliscore output appears to be empty. Check results for BUSCO: 229239at7742
## Thu Jul 14 00:43:57 CEST 2022 - The aliscore output file does not exist. Check results for BUSCO: 79344at7742
## Thu Jul 14 00:43:57 CEST 2022 - The aliscore output appears to be empty. Check results for BUSCO: 79344at7742
## Thu Jul 14 00:43:58 CEST 2022 - The aliscore output file does not exist. Check results for BUSCO: 404307at7742
## Thu Jul 14 00:43:58 CEST 2022 - The aliscore output appears to be empty. Check results for BUSCO: 404307at7742
## Thu Jul 14 00:44:01 CEST 2022 - The aliscore output file does not exist. Check results for BUSCO: 235284at7742
## Thu Jul 14 00:44:01 CEST 2022 - The aliscore output appears to be empty. Check results for BUSCO: 235284at7742
## Thu Jul 14 00:44:01 CEST 2022 - The aliscore output file does not exist. Check results for BUSCO: 158068at7742
## Thu Jul 14 00:44:01 CEST 2022 - The aliscore output appears to be empty. Check results for BUSCO: 158068at7742
## Thu Jul 14 00:44:02 CEST 2022 - The aliscore output file does not exist. Check results for BUSCO: 69988at7742
## Thu Jul 14 00:44:02 CEST 2022 - The aliscore output appears to be empty. Check results for BUSCO: 69988at7742
## Thu Jul 14 00:44:04 CEST 2022 - The aliscore output file does not exist. Check results for BUSCO: 130375at7742
## Thu Jul 14 00:44:04 CEST 2022 - The aliscore output appears to be empty. Check results for BUSCO: 130375at7742
## Thu Jul 14 00:44:06 CEST 2022 - The aliscore output file does not exist. Check results for BUSCO: 137788at7742
## Thu Jul 14 00:44:06 CEST 2022 - The aliscore output appears to be empty. Check results for BUSCO: 137788at7742
## Thu Jul 14 00:44:06 CEST 2022 - The aliscore output file does not exist. Check results for BUSCO: 46227at7742
## Thu Jul 14 00:44:06 CEST 2022 - The aliscore output appears to be empty. Check results for BUSCO: 46227at7742
## Thu Jul 14 00:44:07 CEST 2022 - The aliscore output file does not exist. Check results for BUSCO: 131585at7742
## Thu Jul 14 00:44:07 CEST 2022 - The aliscore output appears to be empty. Check results for BUSCO: 131585at7742
## Thu Jul 14 00:44:10 CEST 2022 - The aliscore output file does not exist. Check results for BUSCO: 188889at7742
## Thu Jul 14 00:44:10 CEST 2022 - The aliscore output appears to be empty. Check results for BUSCO: 188889at7742
## Thu Jul 14 00:44:11 CEST 2022 - The aliscore output file does not exist. Check results for BUSCO: 76254at7742
## Thu Jul 14 00:44:11 CEST 2022 - The aliscore output appears to be empty. Check results for BUSCO: 76254at7742
## Thu Jul 14 00:44:11 CEST 2022 - The aliscore output file does not exist. Check results for BUSCO: 123404at7742
## Thu Jul 14 00:44:11 CEST 2022 - The aliscore output appears to be empty. Check results for BUSCO: 123404at7742
## Thu Jul 14 00:44:13 CEST 2022 - The aliscore output file does not exist. Check results for BUSCO: 39465at7742
## Thu Jul 14 00:44:13 CEST 2022 - The aliscore output appears to be empty. Check results for BUSCO: 39465at7742
## Thu Jul 14 00:44:14 CEST 2022 - The aliscore output file does not exist. Check results for BUSCO: 273631at7742
## Thu Jul 14 00:44:14 CEST 2022 - The aliscore output appears to be empty. Check results for BUSCO: 273631at7742
## Thu Jul 14 00:44:16 CEST 2022 - The aliscore output file does not exist. Check results for BUSCO: 39501at7742
## Thu Jul 14 00:44:16 CEST 2022 - The aliscore output appears to be empty. Check results for BUSCO: 39501at7742
## Thu Jul 14 00:44:16 CEST 2022 - The aliscore output file does not exist. Check results for BUSCO: 143004at7742
## Thu Jul 14 00:44:16 CEST 2022 - The aliscore output appears to be empty. Check results for BUSCO: 143004at7742
## Thu Jul 14 00:44:19 CEST 2022 - The aliscore output file does not exist. Check results for BUSCO: 81484at7742
## Thu Jul 14 00:44:19 CEST 2022 - The aliscore output appears to be empty. Check results for BUSCO: 81484at7742
## Thu Jul 14 00:44:19 CEST 2022 - The aliscore output file does not exist. Check results for BUSCO: 168901at7742
## Thu Jul 14 00:44:19 CEST 2022 - The aliscore output appears to be empty. Check results for BUSCO: 168901at7742
## Thu Jul 14 00:44:20 CEST 2022 - The aliscore output file does not exist. Check results for BUSCO: 59663at7742
## Thu Jul 14 00:44:20 CEST 2022 - The aliscore output appears to be empty. Check results for BUSCO: 59663at7742
## Thu Jul 14 00:44:23 CEST 2022 - The aliscore output file does not exist. Check results for BUSCO: 226528at7742
## Thu Jul 14 00:44:23 CEST 2022 - The aliscore output appears to be empty. Check results for BUSCO: 226528at7742
## Thu Jul 14 00:44:23 CEST 2022 - The aliscore output file does not exist. Check results for BUSCO: 72074at7742
## Thu Jul 14 00:44:23 CEST 2022 - The aliscore output appears to be empty. Check results for BUSCO: 72074at7742
## Thu Jul 14 00:44:25 CEST 2022 - The aliscore output file does not exist. Check results for BUSCO: 170032at7742
## Thu Jul 14 00:44:25 CEST 2022 - The aliscore output appears to be empty. Check results for BUSCO: 170032at7742
## Thu Jul 14 00:44:29 CEST 2022 - The aliscore output file does not exist. Check results for BUSCO: 185735at7742
## Thu Jul 14 00:44:29 CEST 2022 - The aliscore output appears to be empty. Check results for BUSCO: 185735at7742
## Thu Jul 14 00:44:29 CEST 2022 - The aliscore output file does not exist. Check results for BUSCO: 212288at7742
## Thu Jul 14 00:44:29 CEST 2022 - The aliscore output appears to be empty. Check results for BUSCO: 212288at7742
## Thu Jul 14 00:44:29 CEST 2022 - The aliscore output file does not exist. Check results for BUSCO: 114459at7742
## Thu Jul 14 00:44:29 CEST 2022 - The aliscore output appears to be empty. Check results for BUSCO: 114459at7742
## Thu Jul 14 00:44:31 CEST 2022 - The aliscore output file does not exist. Check results for BUSCO: 162703at7742
## Thu Jul 14 00:44:31 CEST 2022 - The aliscore output appears to be empty. Check results for BUSCO: 162703at7742
## Thu Jul 14 00:44:31 CEST 2022 - The aliscore output file does not exist. Check results for BUSCO: 229787at7742
## Thu Jul 14 00:44:31 CEST 2022 - The aliscore output appears to be empty. Check results for BUSCO: 229787at7742
## Thu Jul 14 00:44:32 CEST 2022 - The aliscore output file does not exist. Check results for BUSCO: 95310at7742
## Thu Jul 14 00:44:32 CEST 2022 - The aliscore output appears to be empty. Check results for BUSCO: 95310at7742
## Thu Jul 14 00:44:34 CEST 2022 - The aliscore output file does not exist. Check results for BUSCO: 194260at7742
## Thu Jul 14 00:44:34 CEST 2022 - The aliscore output appears to be empty. Check results for BUSCO: 194260at7742
## Thu Jul 14 00:44:34 CEST 2022 - The aliscore output file does not exist. Check results for BUSCO: 73229at7742
## Thu Jul 14 00:44:34 CEST 2022 - The aliscore output appears to be empty. Check results for BUSCO: 73229at7742
## Thu Jul 14 00:44:37 CEST 2022 - The aliscore output file does not exist. Check results for BUSCO: 110839at7742
## Thu Jul 14 00:44:37 CEST 2022 - The aliscore output appears to be empty. Check results for BUSCO: 110839at7742
## Thu Jul 14 00:44:39 CEST 2022 - The aliscore output file does not exist. Check results for BUSCO: 63198at7742
## Thu Jul 14 00:44:39 CEST 2022 - The aliscore output appears to be empty. Check results for BUSCO: 63198at7742
## Thu Jul 14 00:44:39 CEST 2022 - The aliscore output file does not exist. Check results for BUSCO: 39501at7742
## Thu Jul 14 00:44:39 CEST 2022 - The aliscore output appears to be empty. Check results for BUSCO: 39501at7742
## Thu Jul 14 00:44:39 CEST 2022 - The aliscore output file does not exist. Check results for BUSCO: 114623at7742
## Thu Jul 14 00:44:39 CEST 2022 - The aliscore output appears to be empty. Check results for BUSCO: 114623at7742
## Thu Jul 14 00:44:42 CEST 2022 - The aliscore output file does not exist. Check results for BUSCO: 100256at7742
## Thu Jul 14 00:44:42 CEST 2022 - The aliscore output appears to be empty. Check results for BUSCO: 100256at7742
## Thu Jul 14 00:44:42 CEST 2022 - The aliscore output file does not exist. Check results for BUSCO: 122527at7742
## Thu Jul 14 00:44:42 CEST 2022 - The aliscore output appears to be empty. Check results for BUSCO: 122527at7742
## Thu Jul 14 00:44:47 CEST 2022 - The aliscore output file does not exist. Check results for BUSCO: 226571at7742
## Thu Jul 14 00:44:47 CEST 2022 - The aliscore output appears to be empty. Check results for BUSCO: 226571at7742
## Thu Jul 14 00:44:48 CEST 2022 - The aliscore output file does not exist. Check results for BUSCO: 236610at7742
## Thu Jul 14 00:44:48 CEST 2022 - The aliscore output appears to be empty. Check results for BUSCO: 236610at7742
## Thu Jul 14 00:44:48 CEST 2022 - The aliscore output file does not exist. Check results for BUSCO: 80163at7742
## Thu Jul 14 00:44:48 CEST 2022 - The aliscore output appears to be empty. Check results for BUSCO: 80163at7742
## Thu Jul 14 00:44:48 CEST 2022 - The aliscore output file does not exist. Check results for BUSCO: 42710at7742
## Thu Jul 14 00:44:48 CEST 2022 - The aliscore output appears to be empty. Check results for BUSCO: 42710at7742
## Thu Jul 14 00:44:49 CEST 2022 - The aliscore output file does not exist. Check results for BUSCO: 239731at7742
## Thu Jul 14 00:44:49 CEST 2022 - The aliscore output appears to be empty. Check results for BUSCO: 239731at7742
## Thu Jul 14 00:44:49 CEST 2022 - The aliscore output file does not exist. Check results for BUSCO: 95310at7742
## Thu Jul 14 00:44:49 CEST 2022 - The aliscore output appears to be empty. Check results for BUSCO: 95310at7742
## Thu Jul 14 00:44:53 CEST 2022 - The aliscore output file does not exist. Check results for BUSCO: 207140at7742
## Thu Jul 14 00:44:53 CEST 2022 - The aliscore output appears to be empty. Check results for BUSCO: 207140at7742
## Thu Jul 14 00:44:54 CEST 2022 - The aliscore output file does not exist. Check results for BUSCO: 137681at7742
## Thu Jul 14 00:44:54 CEST 2022 - The aliscore output appears to be empty. Check results for BUSCO: 137681at7742
## Thu Jul 14 00:44:56 CEST 2022 - The aliscore output file does not exist. Check results for BUSCO: 100816at7742
## Thu Jul 14 00:44:56 CEST 2022 - The aliscore output appears to be empty. Check results for BUSCO: 100816at7742
## Thu Jul 14 00:44:57 CEST 2022 - The aliscore output file does not exist. Check results for BUSCO: 84817at7742
## Thu Jul 14 00:44:57 CEST 2022 - The aliscore output appears to be empty. Check results for BUSCO: 84817at7742
## Thu Jul 14 00:44:58 CEST 2022 - The aliscore output file does not exist. Check results for BUSCO: 337981at7742
## Thu Jul 14 00:44:58 CEST 2022 - The aliscore output appears to be empty. Check results for BUSCO: 337981at7742
## Thu Jul 14 00:44:58 CEST 2022 - The aliscore output file does not exist. Check results for BUSCO: 123291at7742
## Thu Jul 14 00:44:58 CEST 2022 - The aliscore output appears to be empty. Check results for BUSCO: 123291at7742
## Thu Jul 14 00:44:59 CEST 2022 - The aliscore output file does not exist. Check results for BUSCO: 92115at7742
## Thu Jul 14 00:44:59 CEST 2022 - The aliscore output appears to be empty. Check results for BUSCO: 92115at7742
## Thu Jul 14 00:45:02 CEST 2022 - The aliscore output file does not exist. Check results for BUSCO: 123387at7742
## Thu Jul 14 00:45:02 CEST 2022 - The aliscore output appears to be empty. Check results for BUSCO: 123387at7742
## Thu Jul 14 00:45:03 CEST 2022 - The aliscore output file does not exist. Check results for BUSCO: 280447at7742
## Thu Jul 14 00:45:03 CEST 2022 - The aliscore output appears to be empty. Check results for BUSCO: 280447at7742
## Thu Jul 14 00:45:04 CEST 2022 - The aliscore output file does not exist. Check results for BUSCO: 175841at7742
## Thu Jul 14 00:45:04 CEST 2022 - The aliscore output appears to be empty. Check results for BUSCO: 175841at7742
## Thu Jul 14 00:45:04 CEST 2022 - The aliscore output file does not exist. Check results for BUSCO: 48716at7742
## Thu Jul 14 00:45:04 CEST 2022 - The aliscore output appears to be empty. Check results for BUSCO: 48716at7742
## Thu Jul 14 00:45:09 CEST 2022 - The aliscore output file does not exist. Check results for BUSCO: 77923at7742
## Thu Jul 14 00:45:09 CEST 2022 - The aliscore output appears to be empty. Check results for BUSCO: 77923at7742
## Thu Jul 14 00:45:09 CEST 2022 - The aliscore output file does not exist. Check results for BUSCO: 145946at7742
## Thu Jul 14 00:45:09 CEST 2022 - The aliscore output appears to be empty. Check results for BUSCO: 145946at7742
## Thu Jul 14 00:45:09 CEST 2022 - The aliscore output file does not exist. Check results for BUSCO: 278954at7742
## Thu Jul 14 00:45:09 CEST 2022 - The aliscore output appears to be empty. Check results for BUSCO: 278954at7742
## Thu Jul 14 00:45:11 CEST 2022 - The aliscore output file does not exist. Check results for BUSCO: 125236at7742
## Thu Jul 14 00:45:11 CEST 2022 - The aliscore output appears to be empty. Check results for BUSCO: 125236at7742
## Thu Jul 14 00:45:13 CEST 2022 - The aliscore output file does not exist. Check results for BUSCO: 330367at7742
## Thu Jul 14 00:45:13 CEST 2022 - The aliscore output appears to be empty. Check results for BUSCO: 330367at7742
## Thu Jul 14 00:45:14 CEST 2022 - The aliscore output file does not exist. Check results for BUSCO: 141015at7742
## Thu Jul 14 00:45:14 CEST 2022 - The aliscore output appears to be empty. Check results for BUSCO: 141015at7742
## Thu Jul 14 00:45:16 CEST 2022 - The aliscore output file does not exist. Check results for BUSCO: 236610at7742
## Thu Jul 14 00:45:16 CEST 2022 - The aliscore output file does not exist. Check results for BUSCO: 212765at7742
## Thu Jul 14 00:45:16 CEST 2022 - The aliscore output appears to be empty. Check results for BUSCO: 236610at7742
## Thu Jul 14 00:45:16 CEST 2022 - The aliscore output appears to be empty. Check results for BUSCO: 212765at7742
## Thu Jul 14 00:45:18 CEST 2022 - The aliscore output file does not exist. Check results for BUSCO: 190480at7742
## Thu Jul 14 00:45:18 CEST 2022 - The aliscore output appears to be empty. Check results for BUSCO: 190480at7742
## Thu Jul 14 00:45:19 CEST 2022 - The aliscore output file does not exist. Check results for BUSCO: 63922at7742
## Thu Jul 14 00:45:19 CEST 2022 - The aliscore output appears to be empty. Check results for BUSCO: 63922at7742
## Thu Jul 14 00:45:20 CEST 2022 - The aliscore output file does not exist. Check results for BUSCO: 196659at7742
## Thu Jul 14 00:45:21 CEST 2022 - The aliscore output appears to be empty. Check results for BUSCO: 196659at7742
## Thu Jul 14 00:45:21 CEST 2022 - The aliscore output file does not exist. Check results for BUSCO: 203744at7742
## Thu Jul 14 00:45:21 CEST 2022 - The aliscore output appears to be empty. Check results for BUSCO: 203744at7742
## Thu Jul 14 00:45:23 CEST 2022 - The aliscore output file does not exist. Check results for BUSCO: 74887at7742
## Thu Jul 14 00:45:23 CEST 2022 - The aliscore output appears to be empty. Check results for BUSCO: 74887at7742
## Thu Jul 14 00:45:23 CEST 2022 - The aliscore output file does not exist. Check results for BUSCO: 212765at7742
## Thu Jul 14 00:45:23 CEST 2022 - The aliscore output appears to be empty. Check results for BUSCO: 212765at7742
## Thu Jul 14 00:45:26 CEST 2022 - The aliscore output file does not exist. Check results for BUSCO: 120085at7742
## Thu Jul 14 00:45:26 CEST 2022 - The aliscore output appears to be empty. Check results for BUSCO: 120085at7742
## Thu Jul 14 00:45:26 CEST 2022 - The aliscore output file does not exist. Check results for BUSCO: 150068at7742
## Thu Jul 14 00:45:26 CEST 2022 - The aliscore output file does not exist. Check results for BUSCO: 66866at7742
## Thu Jul 14 00:45:26 CEST 2022 - The aliscore output appears to be empty. Check results for BUSCO: 150068at7742
## Thu Jul 14 00:45:26 CEST 2022 - The aliscore output appears to be empty. Check results for BUSCO: 66866at7742
## Thu Jul 14 00:45:29 CEST 2022 - The aliscore output file does not exist. Check results for BUSCO: 229239at7742
## Thu Jul 14 00:45:29 CEST 2022 - The aliscore output appears to be empty. Check results for BUSCO: 229239at7742
## Thu Jul 14 00:45:30 CEST 2022 - The aliscore output file does not exist. Check results for BUSCO: 266951at7742
## Thu Jul 14 00:45:31 CEST 2022 - The aliscore output appears to be empty. Check results for BUSCO: 266951at7742
## Thu Jul 14 00:45:32 CEST 2022 - The aliscore output file does not exist. Check results for BUSCO: 69988at7742
## Thu Jul 14 00:45:32 CEST 2022 - The aliscore output appears to be empty. Check results for BUSCO: 69988at7742
## Thu Jul 14 00:45:34 CEST 2022 - The aliscore output file does not exist. Check results for BUSCO: 131862at7742
## Thu Jul 14 00:45:34 CEST 2022 - The aliscore output appears to be empty. Check results for BUSCO: 131862at7742
## Thu Jul 14 00:45:35 CEST 2022 - The aliscore output file does not exist. Check results for BUSCO: 163295at7742
## Thu Jul 14 00:45:35 CEST 2022 - The aliscore output appears to be empty. Check results for BUSCO: 163295at7742
## Thu Jul 14 00:45:36 CEST 2022 - The aliscore output file does not exist. Check results for BUSCO: 177142at7742
## Thu Jul 14 00:45:36 CEST 2022 - The aliscore output appears to be empty. Check results for BUSCO: 177142at7742
## Thu Jul 14 00:45:36 CEST 2022 - The aliscore output file does not exist. Check results for BUSCO: 150068at7742
## Thu Jul 14 00:45:36 CEST 2022 - The aliscore output appears to be empty. Check results for BUSCO: 150068at7742
## Thu Jul 14 00:45:39 CEST 2022 - The aliscore output file does not exist. Check results for BUSCO: 233490at7742
## Thu Jul 14 00:45:39 CEST 2022 - The aliscore output appears to be empty. Check results for BUSCO: 233490at7742
## Thu Jul 14 00:45:41 CEST 2022 - The aliscore output file does not exist. Check results for BUSCO: 126277at7742
## Thu Jul 14 00:45:41 CEST 2022 - The aliscore output appears to be empty. Check results for BUSCO: 126277at7742
## Thu Jul 14 00:45:41 CEST 2022 - The aliscore output file does not exist. Check results for BUSCO: 76254at7742
## Thu Jul 14 00:45:41 CEST 2022 - The aliscore output appears to be empty. Check results for BUSCO: 76254at7742
## Thu Jul 14 00:45:41 CEST 2022 - The aliscore output file does not exist. Check results for BUSCO: 46694at7742
## Thu Jul 14 00:45:41 CEST 2022 - The aliscore output appears to be empty. Check results for BUSCO: 46694at7742
## Thu Jul 14 00:45:44 CEST 2022 - The aliscore output file does not exist. Check results for BUSCO: 143015at7742
## Thu Jul 14 00:45:44 CEST 2022 - The aliscore output appears to be empty. Check results for BUSCO: 143015at7742
## Thu Jul 14 00:45:46 CEST 2022 - The aliscore output file does not exist. Check results for BUSCO: 87513at7742
## Thu Jul 14 00:45:46 CEST 2022 - The aliscore output appears to be empty. Check results for BUSCO: 87513at7742
## Thu Jul 14 00:45:46 CEST 2022 - The aliscore output file does not exist. Check results for BUSCO: 165094at7742
## Thu Jul 14 00:45:46 CEST 2022 - The aliscore output appears to be empty. Check results for BUSCO: 165094at7742
## Thu Jul 14 00:45:47 CEST 2022 - The aliscore output file does not exist. Check results for BUSCO: 76998at7742
## Thu Jul 14 00:45:47 CEST 2022 - The aliscore output appears to be empty. Check results for BUSCO: 76998at7742
## Thu Jul 14 00:45:48 CEST 2022 - The aliscore output file does not exist. Check results for BUSCO: 367008at7742
## Thu Jul 14 00:45:48 CEST 2022 - The aliscore output appears to be empty. Check results for BUSCO: 367008at7742
## Thu Jul 14 00:45:49 CEST 2022 - The aliscore output file does not exist. Check results for BUSCO: 105115at7742
## Thu Jul 14 00:45:49 CEST 2022 - The aliscore output appears to be empty. Check results for BUSCO: 105115at7742
## Thu Jul 14 00:45:50 CEST 2022 - The aliscore output file does not exist. Check results for BUSCO: 63198at7742
## Thu Jul 14 00:45:50 CEST 2022 - The aliscore output appears to be empty. Check results for BUSCO: 63198at7742
## Thu Jul 14 00:45:52 CEST 2022 - The aliscore output file does not exist. Check results for BUSCO: 113847at7742
## Thu Jul 14 00:45:52 CEST 2022 - The aliscore output file does not exist. Check results for BUSCO: 151961at7742
## Thu Jul 14 00:45:52 CEST 2022 - The aliscore output appears to be empty. Check results for BUSCO: 113847at7742
## Thu Jul 14 00:45:52 CEST 2022 - The aliscore output appears to be empty. Check results for BUSCO: 151961at7742
## Thu Jul 14 00:45:54 CEST 2022 - The aliscore output file does not exist. Check results for BUSCO: 108967at7742
## Thu Jul 14 00:45:54 CEST 2022 - The aliscore output appears to be empty. Check results for BUSCO: 108967at7742
## Thu Jul 14 00:45:57 CEST 2022 - The aliscore output file does not exist. Check results for BUSCO: 54075at7742
## Thu Jul 14 00:45:57 CEST 2022 - The aliscore output file does not exist. Check results for BUSCO: 129943at7742
## Thu Jul 14 00:45:57 CEST 2022 - The aliscore output appears to be empty. Check results for BUSCO: 54075at7742
## Thu Jul 14 00:45:57 CEST 2022 - The aliscore output appears to be empty. Check results for BUSCO: 129943at7742
## Thu Jul 14 00:45:59 CEST 2022 - The aliscore output file does not exist. Check results for BUSCO: 122918at7742
## Thu Jul 14 00:45:59 CEST 2022 - The aliscore output appears to be empty. Check results for BUSCO: 122918at7742
## Thu Jul 14 00:46:00 CEST 2022 - The aliscore output file does not exist. Check results for BUSCO: 57918at7742
## Thu Jul 14 00:46:00 CEST 2022 - The aliscore output appears to be empty. Check results for BUSCO: 57918at7742
## Thu Jul 14 00:46:02 CEST 2022 - The aliscore output file does not exist. Check results for BUSCO: 127291at7742
## Thu Jul 14 00:46:02 CEST 2022 - The aliscore output appears to be empty. Check results for BUSCO: 127291at7742
## Thu Jul 14 00:46:02 CEST 2022 - The aliscore output file does not exist. Check results for BUSCO: 125127at7742
## Thu Jul 14 00:46:02 CEST 2022 - The aliscore output file does not exist. Check results for BUSCO: 330367at7742
## Thu Jul 14 00:46:02 CEST 2022 - The aliscore output appears to be empty. Check results for BUSCO: 125127at7742
## Thu Jul 14 00:46:02 CEST 2022 - The aliscore output appears to be empty. Check results for BUSCO: 330367at7742
## Thu Jul 14 00:46:05 CEST 2022 - The aliscore output file does not exist. Check results for BUSCO: 97786at7742
## Thu Jul 14 00:46:05 CEST 2022 - The aliscore output appears to be empty. Check results for BUSCO: 97786at7742
## Thu Jul 14 00:46:06 CEST 2022 - The aliscore output file does not exist. Check results for BUSCO: 162703at7742
## Thu Jul 14 00:46:06 CEST 2022 - The aliscore output appears to be empty. Check results for BUSCO: 162703at7742
## Thu Jul 14 00:46:08 CEST 2022 - The aliscore output file does not exist. Check results for BUSCO: 91878at7742
## Thu Jul 14 00:46:08 CEST 2022 - The aliscore output appears to be empty. Check results for BUSCO: 91878at7742
## Thu Jul 14 00:46:09 CEST 2022 - The aliscore output file does not exist. Check results for BUSCO: 229787at7742
## Thu Jul 14 00:46:09 CEST 2022 - The aliscore output appears to be empty. Check results for BUSCO: 229787at7742
## Thu Jul 14 00:46:10 CEST 2022 - The aliscore output file does not exist. Check results for BUSCO: 53548at7742
## Thu Jul 14 00:46:11 CEST 2022 - The aliscore output appears to be empty. Check results for BUSCO: 53548at7742
## Thu Jul 14 00:46:12 CEST 2022 - The aliscore output file does not exist. Check results for BUSCO: 180758at7742
## Thu Jul 14 00:46:12 CEST 2022 - The aliscore output file does not exist. Check results for BUSCO: 87193at7742
## Thu Jul 14 00:46:12 CEST 2022 - The aliscore output appears to be empty. Check results for BUSCO: 180758at7742
## Thu Jul 14 00:46:12 CEST 2022 - The aliscore output appears to be empty. Check results for BUSCO: 87193at7742
## Thu Jul 14 00:46:13 CEST 2022 - The aliscore output file does not exist. Check results for BUSCO: 115767at7742
## Thu Jul 14 00:46:13 CEST 2022 - The aliscore output appears to be empty. Check results for BUSCO: 115767at7742
## Thu Jul 14 00:46:15 CEST 2022 - The aliscore output file does not exist. Check results for BUSCO: 198404at7742
## Thu Jul 14 00:46:15 CEST 2022 - The aliscore output file does not exist. Check results for BUSCO: 330367at7742
## Thu Jul 14 00:46:15 CEST 2022 - The aliscore output appears to be empty. Check results for BUSCO: 198404at7742
## Thu Jul 14 00:46:15 CEST 2022 - The aliscore output appears to be empty. Check results for BUSCO: 330367at7742
## Thu Jul 14 00:46:18 CEST 2022 - The aliscore output file does not exist. Check results for BUSCO: 367008at7742
## Thu Jul 14 00:46:18 CEST 2022 - The aliscore output appears to be empty. Check results for BUSCO: 367008at7742
## Thu Jul 14 00:46:20 CEST 2022 - The aliscore output file does not exist. Check results for BUSCO: 46227at7742
## Thu Jul 14 00:46:20 CEST 2022 - The aliscore output appears to be empty. Check results for BUSCO: 46227at7742
## Thu Jul 14 00:46:21 CEST 2022 - The aliscore output file does not exist. Check results for BUSCO: 158236at7742
## Thu Jul 14 00:46:21 CEST 2022 - The aliscore output appears to be empty. Check results for BUSCO: 158236at7742
## Thu Jul 14 00:46:21 CEST 2022 - The aliscore output file does not exist. Check results for BUSCO: 87236at7742
## Thu Jul 14 00:46:22 CEST 2022 - The aliscore output appears to be empty. Check results for BUSCO: 87236at7742
## Thu Jul 14 00:46:22 CEST 2022 - The aliscore output file does not exist. Check results for BUSCO: 100658at7742
## Thu Jul 14 00:46:22 CEST 2022 - The aliscore output appears to be empty. Check results for BUSCO: 100658at7742
## Thu Jul 14 00:46:24 CEST 2022 - The aliscore output file does not exist. Check results for BUSCO: 57918at7742
## Thu Jul 14 00:46:24 CEST 2022 - The aliscore output appears to be empty. Check results for BUSCO: 57918at7742
## Thu Jul 14 00:46:26 CEST 2022 - The aliscore output file does not exist. Check results for BUSCO: 350970at7742
## Thu Jul 14 00:46:27 CEST 2022 - The aliscore output appears to be empty. Check results for BUSCO: 350970at7742
## Thu Jul 14 00:46:28 CEST 2022 - The aliscore output file does not exist. Check results for BUSCO: 94269at7742
## Thu Jul 14 00:46:28 CEST 2022 - The aliscore output appears to be empty. Check results for BUSCO: 94269at7742
## Thu Jul 14 00:46:28 CEST 2022 - The aliscore output file does not exist. Check results for BUSCO: 213614at7742
## Thu Jul 14 00:46:28 CEST 2022 - The aliscore output appears to be empty. Check results for BUSCO: 213614at7742
## Thu Jul 14 00:46:29 CEST 2022 - The aliscore output file does not exist. Check results for BUSCO: 126566at7742
## Thu Jul 14 00:46:29 CEST 2022 - The aliscore output appears to be empty. Check results for BUSCO: 126566at7742
## Thu Jul 14 00:46:31 CEST 2022 - The aliscore output file does not exist. Check results for BUSCO: 178398at7742
## Thu Jul 14 00:46:31 CEST 2022 - The aliscore output appears to be empty. Check results for BUSCO: 178398at7742
## Thu Jul 14 00:46:32 CEST 2022 - The aliscore output file does not exist. Check results for BUSCO: 111099at7742
## Thu Jul 14 00:46:32 CEST 2022 - The aliscore output appears to be empty. Check results for BUSCO: 111099at7742
## Thu Jul 14 00:46:34 CEST 2022 - The aliscore output file does not exist. Check results for BUSCO: 305045at7742
## Thu Jul 14 00:46:34 CEST 2022 - The aliscore output appears to be empty. Check results for BUSCO: 305045at7742
## Thu Jul 14 00:46:36 CEST 2022 - The aliscore output file does not exist. Check results for BUSCO: 46740at7742
## Thu Jul 14 00:46:36 CEST 2022 - The aliscore output appears to be empty. Check results for BUSCO: 46740at7742
## Thu Jul 14 00:46:37 CEST 2022 - The aliscore output file does not exist. Check results for BUSCO: 93430at7742
## Thu Jul 14 00:46:37 CEST 2022 - The aliscore output appears to be empty. Check results for BUSCO: 93430at7742
## Thu Jul 14 00:46:37 CEST 2022 - The aliscore output file does not exist. Check results for BUSCO: 101457at7742
## Thu Jul 14 00:46:37 CEST 2022 - The aliscore output appears to be empty. Check results for BUSCO: 101457at7742
## Thu Jul 14 00:46:39 CEST 2022 - The aliscore output file does not exist. Check results for BUSCO: 97786at7742
## Thu Jul 14 00:46:39 CEST 2022 - The aliscore output appears to be empty. Check results for BUSCO: 97786at7742
## Thu Jul 14 00:46:40 CEST 2022 - The aliscore output file does not exist. Check results for BUSCO: 59584at7742
## Thu Jul 14 00:46:40 CEST 2022 - The aliscore output appears to be empty. Check results for BUSCO: 59584at7742
## Thu Jul 14 00:46:42 CEST 2022 - The aliscore output file does not exist. Check results for BUSCO: 142495at7742
## Thu Jul 14 00:46:42 CEST 2022 - The aliscore output appears to be empty. Check results for BUSCO: 142495at7742
## Thu Jul 14 00:46:43 CEST 2022 - The aliscore output file does not exist. Check results for BUSCO: 145962at7742
## Thu Jul 14 00:46:43 CEST 2022 - The aliscore output appears to be empty. Check results for BUSCO: 145962at7742
## Thu Jul 14 00:46:44 CEST 2022 - The aliscore output file does not exist. Check results for BUSCO: 90853at7742
## Thu Jul 14 00:46:44 CEST 2022 - The aliscore output appears to be empty. Check results for BUSCO: 90853at7742
## Thu Jul 14 00:46:45 CEST 2022 - The aliscore output file does not exist. Check results for BUSCO: 182992at7742
## Thu Jul 14 00:46:45 CEST 2022 - The aliscore output appears to be empty. Check results for BUSCO: 182992at7742
## Thu Jul 14 00:46:49 CEST 2022 - The aliscore output file does not exist. Check results for BUSCO: 46227at7742
## Thu Jul 14 00:46:49 CEST 2022 - The aliscore output appears to be empty. Check results for BUSCO: 46227at7742
## Thu Jul 14 00:46:49 CEST 2022 - The aliscore output file does not exist. Check results for BUSCO: 95963at7742
## Thu Jul 14 00:46:49 CEST 2022 - The aliscore output file does not exist. Check results for BUSCO: 143004at7742
## Thu Jul 14 00:46:49 CEST 2022 - The aliscore output appears to be empty. Check results for BUSCO: 95963at7742
## Thu Jul 14 00:46:49 CEST 2022 - The aliscore output appears to be empty. Check results for BUSCO: 143004at7742
## Thu Jul 14 00:46:51 CEST 2022 - The aliscore output file does not exist. Check results for BUSCO: 174858at7742
## Thu Jul 14 00:46:51 CEST 2022 - The aliscore output appears to be empty. Check results for BUSCO: 174858at7742
## Thu Jul 14 00:46:53 CEST 2022 - The aliscore output file does not exist. Check results for BUSCO: 258445at7742
## Thu Jul 14 00:46:53 CEST 2022 - The aliscore output appears to be empty. Check results for BUSCO: 258445at7742
## Thu Jul 14 00:46:54 CEST 2022 - The aliscore output file does not exist. Check results for BUSCO: 264645at7742
## Thu Jul 14 00:46:54 CEST 2022 - The aliscore output appears to be empty. Check results for BUSCO: 264645at7742
## Thu Jul 14 00:46:55 CEST 2022 - The aliscore output file does not exist. Check results for BUSCO: 145104at7742
## Thu Jul 14 00:46:55 CEST 2022 - The aliscore output appears to be empty. Check results for BUSCO: 145104at7742
## Thu Jul 14 00:47:00 CEST 2022 - The aliscore output file does not exist. Check results for BUSCO: 122963at7742
## Thu Jul 14 00:47:00 CEST 2022 - The aliscore output appears to be empty. Check results for BUSCO: 122963at7742
## Thu Jul 14 00:47:00 CEST 2022 - The aliscore output file does not exist. Check results for BUSCO: 93430at7742
## Thu Jul 14 00:47:00 CEST 2022 - The aliscore output appears to be empty. Check results for BUSCO: 93430at7742
## Thu Jul 14 00:47:00 CEST 2022 - The aliscore output file does not exist. Check results for BUSCO: 145104at7742
## Thu Jul 14 00:47:00 CEST 2022 - The aliscore output appears to be empty. Check results for BUSCO: 145104at7742
## Thu Jul 14 00:47:00 CEST 2022 - The aliscore output file does not exist. Check results for BUSCO: 189794at7742
## Thu Jul 14 00:47:00 CEST 2022 - The aliscore output appears to be empty. Check results for BUSCO: 189794at7742
## Thu Jul 14 00:47:01 CEST 2022 - The aliscore output file does not exist. Check results for BUSCO: 152476at7742
## Thu Jul 14 00:47:01 CEST 2022 - The aliscore output appears to be empty. Check results for BUSCO: 152476at7742
## Thu Jul 14 00:47:03 CEST 2022 - The aliscore output file does not exist. Check results for BUSCO: 241590at7742
## Thu Jul 14 00:47:04 CEST 2022 - The aliscore output appears to be empty. Check results for BUSCO: 241590at7742
## Thu Jul 14 00:47:06 CEST 2022 - The aliscore output file does not exist. Check results for BUSCO: 68766at7742
## Thu Jul 14 00:47:06 CEST 2022 - The aliscore output appears to be empty. Check results for BUSCO: 68766at7742
## Thu Jul 14 00:47:06 CEST 2022 - The aliscore output file does not exist. Check results for BUSCO: 123847at7742
## Thu Jul 14 00:47:06 CEST 2022 - The aliscore output appears to be empty. Check results for BUSCO: 123847at7742
## Thu Jul 14 00:47:09 CEST 2022 - The aliscore output file does not exist. Check results for BUSCO: 59584at7742
## Thu Jul 14 00:47:09 CEST 2022 - The aliscore output file does not exist. Check results for BUSCO: 95511at7742
## Thu Jul 14 00:47:09 CEST 2022 - The aliscore output appears to be empty. Check results for BUSCO: 59584at7742
## Thu Jul 14 00:47:09 CEST 2022 - The aliscore output appears to be empty. Check results for BUSCO: 95511at7742
## Thu Jul 14 00:47:09 CEST 2022 - The aliscore output file does not exist. Check results for BUSCO: 141724at7742
## Thu Jul 14 00:47:09 CEST 2022 - The aliscore output appears to be empty. Check results for BUSCO: 141724at7742
## Thu Jul 14 00:47:12 CEST 2022 - The aliscore output file does not exist. Check results for BUSCO: 171708at7742
## Thu Jul 14 00:47:12 CEST 2022 - The aliscore output appears to be empty. Check results for BUSCO: 171708at7742
## Thu Jul 14 00:47:12 CEST 2022 - The aliscore output file does not exist. Check results for BUSCO: 54075at7742
## Thu Jul 14 00:47:12 CEST 2022 - The aliscore output appears to be empty. Check results for BUSCO: 54075at7742
## Thu Jul 14 00:47:14 CEST 2022 - The aliscore output file does not exist. Check results for BUSCO: 165094at7742
## Thu Jul 14 00:47:14 CEST 2022 - The aliscore output appears to be empty. Check results for BUSCO: 165094at7742
## Thu Jul 14 00:47:16 CEST 2022 - The aliscore output file does not exist. Check results for BUSCO: 376015at7742
## Thu Jul 14 00:47:16 CEST 2022 - The aliscore output appears to be empty. Check results for BUSCO: 376015at7742
## Thu Jul 14 00:47:17 CEST 2022 - The aliscore output file does not exist. Check results for BUSCO: 165094at7742
## Thu Jul 14 00:47:17 CEST 2022 - The aliscore output appears to be empty. Check results for BUSCO: 165094at7742
## Thu Jul 14 00:47:18 CEST 2022 - The aliscore output file does not exist. Check results for BUSCO: 100658at7742
## Thu Jul 14 00:47:18 CEST 2022 - The aliscore output appears to be empty. Check results for BUSCO: 100658at7742
## Thu Jul 14 00:47:19 CEST 2022 - The aliscore output file does not exist. Check results for BUSCO: 190480at7742
## Thu Jul 14 00:47:19 CEST 2022 - The aliscore output appears to be empty. Check results for BUSCO: 190480at7742
## Thu Jul 14 00:47:19 CEST 2022 - The aliscore output file does not exist. Check results for BUSCO: 275273at7742
## Thu Jul 14 00:47:19 CEST 2022 - The aliscore output appears to be empty. Check results for BUSCO: 275273at7742
## Thu Jul 14 00:47:21 CEST 2022 - The aliscore output file does not exist. Check results for BUSCO: 222975at7742
## Thu Jul 14 00:47:21 CEST 2022 - The aliscore output appears to be empty. Check results for BUSCO: 222975at7742
## Thu Jul 14 00:47:22 CEST 2022 - The aliscore output file does not exist. Check results for BUSCO: 143015at7742
## Thu Jul 14 00:47:22 CEST 2022 - The aliscore output appears to be empty. Check results for BUSCO: 143015at7742
## Thu Jul 14 00:47:24 CEST 2022 - The aliscore output file does not exist. Check results for BUSCO: 254316at7742
## Thu Jul 14 00:47:24 CEST 2022 - The aliscore output appears to be empty. Check results for BUSCO: 254316at7742
## Thu Jul 14 00:47:26 CEST 2022 - The aliscore output file does not exist. Check results for BUSCO: 90735at7742
## Thu Jul 14 00:47:26 CEST 2022 - The aliscore output appears to be empty. Check results for BUSCO: 90735at7742
## Thu Jul 14 00:47:27 CEST 2022 - The aliscore output file does not exist. Check results for BUSCO: 368834at7742
## Thu Jul 14 00:47:27 CEST 2022 - The aliscore output appears to be empty. Check results for BUSCO: 368834at7742
## Thu Jul 14 00:47:27 CEST 2022 - The aliscore output file does not exist. Check results for BUSCO: 133596at7742
## Thu Jul 14 00:47:27 CEST 2022 - The aliscore output appears to be empty. Check results for BUSCO: 133596at7742
## Thu Jul 14 00:47:28 CEST 2022 - The aliscore output file does not exist. Check results for BUSCO: 157513at7742
## Thu Jul 14 00:47:28 CEST 2022 - The aliscore output appears to be empty. Check results for BUSCO: 157513at7742
## Thu Jul 14 00:47:29 CEST 2022 - The aliscore output file does not exist. Check results for BUSCO: 142117at7742
## Thu Jul 14 00:47:29 CEST 2022 - The aliscore output appears to be empty. Check results for BUSCO: 142117at7742
## Thu Jul 14 00:47:31 CEST 2022 - The aliscore output file does not exist. Check results for BUSCO: 67119at7742
## Thu Jul 14 00:47:31 CEST 2022 - The aliscore output appears to be empty. Check results for BUSCO: 67119at7742
## Thu Jul 14 00:47:33 CEST 2022 - The aliscore output file does not exist. Check results for BUSCO: 42538at7742
## Thu Jul 14 00:47:33 CEST 2022 - The aliscore output appears to be empty. Check results for BUSCO: 42538at7742
## Thu Jul 14 00:47:34 CEST 2022 - The aliscore output file does not exist. Check results for BUSCO: 82221at7742
## Thu Jul 14 00:47:34 CEST 2022 - The aliscore output appears to be empty. Check results for BUSCO: 82221at7742
## Thu Jul 14 00:47:36 CEST 2022 - The aliscore output file does not exist. Check results for BUSCO: 89000at7742
## Thu Jul 14 00:47:36 CEST 2022 - The aliscore output appears to be empty. Check results for BUSCO: 89000at7742
## Thu Jul 14 00:47:37 CEST 2022 - The aliscore output file does not exist. Check results for BUSCO: 87117at7742
## Thu Jul 14 00:47:37 CEST 2022 - The aliscore output appears to be empty. Check results for BUSCO: 87117at7742
## Thu Jul 14 00:47:37 CEST 2022 - The aliscore output file does not exist. Check results for BUSCO: 137681at7742
## Thu Jul 14 00:47:38 CEST 2022 - The aliscore output appears to be empty. Check results for BUSCO: 137681at7742
## Thu Jul 14 00:47:38 CEST 2022 - The aliscore output file does not exist. Check results for BUSCO: 180758at7742
## Thu Jul 14 00:47:38 CEST 2022 - The aliscore output appears to be empty. Check results for BUSCO: 180758at7742
## Thu Jul 14 00:47:40 CEST 2022 - The aliscore output file does not exist. Check results for BUSCO: 317124at7742
## Thu Jul 14 00:47:40 CEST 2022 - The aliscore output appears to be empty. Check results for BUSCO: 317124at7742
## Thu Jul 14 00:47:41 CEST 2022 - The aliscore output file does not exist. Check results for BUSCO: 83474at7742
## Thu Jul 14 00:47:41 CEST 2022 - The aliscore output appears to be empty. Check results for BUSCO: 83474at7742
## Thu Jul 14 00:47:42 CEST 2022 - The aliscore output file does not exist. Check results for BUSCO: 60783at7742
## Thu Jul 14 00:47:42 CEST 2022 - The aliscore output appears to be empty. Check results for BUSCO: 60783at7742
## Thu Jul 14 00:47:45 CEST 2022 - The aliscore output file does not exist. Check results for BUSCO: 171639at7742
## Thu Jul 14 00:47:45 CEST 2022 - The aliscore output appears to be empty. Check results for BUSCO: 171639at7742
## Thu Jul 14 00:47:46 CEST 2022 - The aliscore output file does not exist. Check results for BUSCO: 174858at7742
## Thu Jul 14 00:47:46 CEST 2022 - The aliscore output appears to be empty. Check results for BUSCO: 174858at7742
## Thu Jul 14 00:47:47 CEST 2022 - The aliscore output file does not exist. Check results for BUSCO: 195010at7742
## Thu Jul 14 00:47:47 CEST 2022 - The aliscore output appears to be empty. Check results for BUSCO: 195010at7742
## Thu Jul 14 00:47:47 CEST 2022 - The aliscore output file does not exist. Check results for BUSCO: 145564at7742
## Thu Jul 14 00:47:47 CEST 2022 - The aliscore output appears to be empty. Check results for BUSCO: 145564at7742
## Thu Jul 14 00:47:50 CEST 2022 - The aliscore output file does not exist. Check results for BUSCO: 132224at7742
## Thu Jul 14 00:47:50 CEST 2022 - The aliscore output appears to be empty. Check results for BUSCO: 132224at7742
## Thu Jul 14 00:47:51 CEST 2022 - The aliscore output file does not exist. Check results for BUSCO: 79086at7742
## Thu Jul 14 00:47:51 CEST 2022 - The aliscore output appears to be empty. Check results for BUSCO: 79086at7742
## Thu Jul 14 00:47:53 CEST 2022 - The aliscore output file does not exist. Check results for BUSCO: 194260at7742
## Thu Jul 14 00:47:53 CEST 2022 - The aliscore output appears to be empty. Check results for BUSCO: 194260at7742
## Thu Jul 14 00:47:53 CEST 2022 - The aliscore output file does not exist. Check results for BUSCO: 134433at7742
## Thu Jul 14 00:47:53 CEST 2022 - The aliscore output appears to be empty. Check results for BUSCO: 134433at7742
## Thu Jul 14 00:47:54 CEST 2022 - The aliscore output file does not exist. Check results for BUSCO: 208314at7742
## Thu Jul 14 00:47:55 CEST 2022 - The aliscore output appears to be empty. Check results for BUSCO: 208314at7742
## Thu Jul 14 00:47:56 CEST 2022 - The aliscore output file does not exist. Check results for BUSCO: 84543at7742
## Thu Jul 14 00:47:56 CEST 2022 - The aliscore output appears to be empty. Check results for BUSCO: 84543at7742
## Thu Jul 14 00:47:56 CEST 2022 - The aliscore output file does not exist. Check results for BUSCO: 223032at7742
## Thu Jul 14 00:47:56 CEST 2022 - The aliscore output appears to be empty. Check results for BUSCO: 223032at7742
## Thu Jul 14 00:47:56 CEST 2022 - The aliscore output file does not exist. Check results for BUSCO: 151961at7742
## Thu Jul 14 00:47:56 CEST 2022 - The aliscore output appears to be empty. Check results for BUSCO: 151961at7742
## Thu Jul 14 00:48:01 CEST 2022 - The aliscore output file does not exist. Check results for BUSCO: 174858at7742
## Thu Jul 14 00:48:01 CEST 2022 - The aliscore output appears to be empty. Check results for BUSCO: 174858at7742
## Thu Jul 14 00:48:01 CEST 2022 - The aliscore output file does not exist. Check results for BUSCO: 108458at7742
## Thu Jul 14 00:48:01 CEST 2022 - The aliscore output appears to be empty. Check results for BUSCO: 108458at7742
## Thu Jul 14 00:48:02 CEST 2022 - The aliscore output file does not exist. Check results for BUSCO: 125236at7742
## Thu Jul 14 00:48:02 CEST 2022 - The aliscore output appears to be empty. Check results for BUSCO: 125236at7742
## Thu Jul 14 00:48:05 CEST 2022 - The aliscore output file does not exist. Check results for BUSCO: 81484at7742
## Thu Jul 14 00:48:05 CEST 2022 - The aliscore output appears to be empty. Check results for BUSCO: 81484at7742
## Thu Jul 14 00:48:06 CEST 2022 - The aliscore output file does not exist. Check results for BUSCO: 58280at7742
## Thu Jul 14 00:48:06 CEST 2022 - The aliscore output appears to be empty. Check results for BUSCO: 58280at7742
## Thu Jul 14 00:48:06 CEST 2022 - The aliscore output file does not exist. Check results for BUSCO: 70359at7742
## Thu Jul 14 00:48:06 CEST 2022 - The aliscore output appears to be empty. Check results for BUSCO: 70359at7742
## Thu Jul 14 00:48:08 CEST 2022 - The aliscore output file does not exist. Check results for BUSCO: 53153at7742
## Thu Jul 14 00:48:08 CEST 2022 - The aliscore output appears to be empty. Check results for BUSCO: 53153at7742
## Thu Jul 14 00:48:10 CEST 2022 - The aliscore output file does not exist. Check results for BUSCO: 299207at7742
## Thu Jul 14 00:48:10 CEST 2022 - The aliscore output appears to be empty. Check results for BUSCO: 299207at7742
## Thu Jul 14 00:48:12 CEST 2022 - The aliscore output file does not exist. Check results for BUSCO: 79344at7742
## Thu Jul 14 00:48:12 CEST 2022 - The aliscore output appears to be empty. Check results for BUSCO: 79344at7742
## Thu Jul 14 00:48:12 CEST 2022 - The aliscore output file does not exist. Check results for BUSCO: 160674at7742
## Thu Jul 14 00:48:12 CEST 2022 - The aliscore output appears to be empty. Check results for BUSCO: 160674at7742
## Thu Jul 14 00:48:13 CEST 2022 - The aliscore output file does not exist. Check results for BUSCO: 109081at7742
## Thu Jul 14 00:48:13 CEST 2022 - The aliscore output appears to be empty. Check results for BUSCO: 109081at7742
## Thu Jul 14 00:48:15 CEST 2022 - The aliscore output file does not exist. Check results for BUSCO: 100494at7742
## Thu Jul 14 00:48:15 CEST 2022 - The aliscore output appears to be empty. Check results for BUSCO: 100494at7742
## Thu Jul 14 00:48:17 CEST 2022 - The aliscore output file does not exist. Check results for BUSCO: 264645at7742
## Thu Jul 14 00:48:17 CEST 2022 - The aliscore output appears to be empty. Check results for BUSCO: 264645at7742
## Thu Jul 14 00:48:17 CEST 2022 - The aliscore output file does not exist. Check results for BUSCO: 110832at7742
## Thu Jul 14 00:48:17 CEST 2022 - The aliscore output appears to be empty. Check results for BUSCO: 110832at7742
## Thu Jul 14 00:48:18 CEST 2022 - The aliscore output file does not exist. Check results for BUSCO: 128400at7742
## Thu Jul 14 00:48:18 CEST 2022 - The aliscore output appears to be empty. Check results for BUSCO: 128400at7742
## Thu Jul 14 00:48:21 CEST 2022 - The aliscore output file does not exist. Check results for BUSCO: 64461at7742
## Thu Jul 14 00:48:21 CEST 2022 - The aliscore output appears to be empty. Check results for BUSCO: 64461at7742
## Thu Jul 14 00:48:21 CEST 2022 - The aliscore output file does not exist. Check results for BUSCO: 212765at7742
## Thu Jul 14 00:48:21 CEST 2022 - The aliscore output appears to be empty. Check results for BUSCO: 212765at7742
## Thu Jul 14 00:48:21 CEST 2022 - The aliscore output file does not exist. Check results for BUSCO: 132656at7742
## Thu Jul 14 00:48:21 CEST 2022 - The aliscore output appears to be empty. Check results for BUSCO: 132656at7742
## Thu Jul 14 00:48:24 CEST 2022 - The aliscore output file does not exist. Check results for BUSCO: 64473at7742
## Thu Jul 14 00:48:24 CEST 2022 - The aliscore output appears to be empty. Check results for BUSCO: 64473at7742
## Thu Jul 14 00:48:24 CEST 2022 - The aliscore output file does not exist. Check results for BUSCO: 59584at7742
## Thu Jul 14 00:48:24 CEST 2022 - The aliscore output appears to be empty. Check results for BUSCO: 59584at7742
## Thu Jul 14 00:48:27 CEST 2022 - The aliscore output file does not exist. Check results for BUSCO: 57918at7742
## Thu Jul 14 00:48:27 CEST 2022 - The aliscore output appears to be empty. Check results for BUSCO: 57918at7742
## Thu Jul 14 00:48:31 CEST 2022 - The aliscore output file does not exist. Check results for BUSCO: 131862at7742
## Thu Jul 14 00:48:31 CEST 2022 - The aliscore output appears to be empty. Check results for BUSCO: 131862at7742
## Thu Jul 14 00:48:31 CEST 2022 - The aliscore output file does not exist. Check results for BUSCO: 134433at7742
## Thu Jul 14 00:48:31 CEST 2022 - The aliscore output appears to be empty. Check results for BUSCO: 134433at7742
## Thu Jul 14 00:48:31 CEST 2022 - The aliscore output file does not exist. Check results for BUSCO: 46740at7742
## Thu Jul 14 00:48:32 CEST 2022 - The aliscore output appears to be empty. Check results for BUSCO: 46740at7742
## Thu Jul 14 00:48:32 CEST 2022 - The aliscore output file does not exist. Check results for BUSCO: 88808at7742
## Thu Jul 14 00:48:32 CEST 2022 - The aliscore output appears to be empty. Check results for BUSCO: 88808at7742
## Thu Jul 14 00:48:32 CEST 2022 - The aliscore output file does not exist. Check results for BUSCO: 258916at7742
## Thu Jul 14 00:48:32 CEST 2022 - The aliscore output appears to be empty. Check results for BUSCO: 258916at7742
## Thu Jul 14 00:48:33 CEST 2022 - The aliscore output file does not exist. Check results for BUSCO: 67465at7742
## Thu Jul 14 00:48:33 CEST 2022 - The aliscore output appears to be empty. Check results for BUSCO: 67465at7742
## Thu Jul 14 00:48:37 CEST 2022 - The aliscore output file does not exist. Check results for BUSCO: 146804at7742
## Thu Jul 14 00:48:37 CEST 2022 - The aliscore output appears to be empty. Check results for BUSCO: 146804at7742
## Thu Jul 14 00:48:38 CEST 2022 - The aliscore output file does not exist. Check results for BUSCO: 50786at7742
## Thu Jul 14 00:48:38 CEST 2022 - The aliscore output appears to be empty. Check results for BUSCO: 50786at7742
## Thu Jul 14 00:48:39 CEST 2022 - The aliscore output file does not exist. Check results for BUSCO: 167134at7742
## Thu Jul 14 00:48:40 CEST 2022 - The aliscore output appears to be empty. Check results for BUSCO: 167134at7742
## Thu Jul 14 00:48:41 CEST 2022 - The aliscore output file does not exist. Check results for BUSCO: 127929at7742
## Thu Jul 14 00:48:41 CEST 2022 - The aliscore output appears to be empty. Check results for BUSCO: 127929at7742
## Thu Jul 14 00:48:41 CEST 2022 - The aliscore output file does not exist. Check results for BUSCO: 182992at7742
## Thu Jul 14 00:48:41 CEST 2022 - The aliscore output appears to be empty. Check results for BUSCO: 182992at7742
## Thu Jul 14 00:48:41 CEST 2022 - The aliscore output file does not exist. Check results for BUSCO: 204311at7742
## Thu Jul 14 00:48:41 CEST 2022 - The aliscore output file does not exist. Check results for BUSCO: 264645at7742
## Thu Jul 14 00:48:41 CEST 2022 - The aliscore output appears to be empty. Check results for BUSCO: 204311at7742
## Thu Jul 14 00:48:41 CEST 2022 - The aliscore output appears to be empty. Check results for BUSCO: 264645at7742
## Thu Jul 14 00:48:42 CEST 2022 - The aliscore output file does not exist. Check results for BUSCO: 76998at7742
## Thu Jul 14 00:48:42 CEST 2022 - The aliscore output appears to be empty. Check results for BUSCO: 76998at7742
## Thu Jul 14 00:48:47 CEST 2022 - The aliscore output file does not exist. Check results for BUSCO: 80527at7742
## Thu Jul 14 00:48:47 CEST 2022 - The aliscore output appears to be empty. Check results for BUSCO: 80527at7742
## Thu Jul 14 00:48:48 CEST 2022 - The aliscore output file does not exist. Check results for BUSCO: 350970at7742
## Thu Jul 14 00:48:48 CEST 2022 - The aliscore output appears to be empty. Check results for BUSCO: 350970at7742
## Thu Jul 14 00:48:51 CEST 2022 - The aliscore output file does not exist. Check results for BUSCO: 90735at7742
## Thu Jul 14 00:48:51 CEST 2022 - The aliscore output file does not exist. Check results for BUSCO: 72074at7742
## Thu Jul 14 00:48:51 CEST 2022 - The aliscore output appears to be empty. Check results for BUSCO: 90735at7742
## Thu Jul 14 00:48:51 CEST 2022 - The aliscore output appears to be empty. Check results for BUSCO: 72074at7742
## Thu Jul 14 00:48:51 CEST 2022 - The aliscore output file does not exist. Check results for BUSCO: 123872at7742
## Thu Jul 14 00:48:51 CEST 2022 - The aliscore output appears to be empty. Check results for BUSCO: 123872at7742
## Thu Jul 14 00:48:52 CEST 2022 - The aliscore output file does not exist. Check results for BUSCO: 149377at7742
## Thu Jul 14 00:48:52 CEST 2022 - The aliscore output appears to be empty. Check results for BUSCO: 149377at7742
## Thu Jul 14 00:48:53 CEST 2022 - The aliscore output file does not exist. Check results for BUSCO: 53153at7742
## Thu Jul 14 00:48:54 CEST 2022 - The aliscore output appears to be empty. Check results for BUSCO: 53153at7742
## Thu Jul 14 00:48:55 CEST 2022 - The aliscore output file does not exist. Check results for BUSCO: 350970at7742
## Thu Jul 14 00:48:55 CEST 2022 - The aliscore output appears to be empty. Check results for BUSCO: 350970at7742
## Thu Jul 14 00:48:59 CEST 2022 - The aliscore output file does not exist. Check results for BUSCO: 277807at7742
## Thu Jul 14 00:48:59 CEST 2022 - The aliscore output appears to be empty. Check results for BUSCO: 277807at7742
## Thu Jul 14 00:48:59 CEST 2022 - The aliscore output file does not exist. Check results for BUSCO: 100623at7742
## Thu Jul 14 00:48:59 CEST 2022 - The aliscore output file does not exist. Check results for BUSCO: 89000at7742
## Thu Jul 14 00:48:59 CEST 2022 - The aliscore output appears to be empty. Check results for BUSCO: 100623at7742
## Thu Jul 14 00:48:59 CEST 2022 - The aliscore output appears to be empty. Check results for BUSCO: 89000at7742
## Thu Jul 14 00:48:59 CEST 2022 - The aliscore output file does not exist. Check results for BUSCO: 176119at7742
## Thu Jul 14 00:48:59 CEST 2022 - The aliscore output appears to be empty. Check results for BUSCO: 176119at7742
## Thu Jul 14 00:49:00 CEST 2022 - The aliscore output file does not exist. Check results for BUSCO: 258445at7742
## Thu Jul 14 00:49:00 CEST 2022 - The aliscore output appears to be empty. Check results for BUSCO: 258445at7742
## Thu Jul 14 00:49:01 CEST 2022 - The aliscore output file does not exist. Check results for BUSCO: 95511at7742
## Thu Jul 14 00:49:01 CEST 2022 - The aliscore output appears to be empty. Check results for BUSCO: 95511at7742
## Thu Jul 14 00:49:04 CEST 2022 - The aliscore output file does not exist. Check results for BUSCO: 108458at7742
## Thu Jul 14 00:49:05 CEST 2022 - The aliscore output appears to be empty. Check results for BUSCO: 108458at7742
## Thu Jul 14 00:49:06 CEST 2022 - The aliscore output file does not exist. Check results for BUSCO: 367008at7742
## Thu Jul 14 00:49:06 CEST 2022 - The aliscore output appears to be empty. Check results for BUSCO: 367008at7742
## Thu Jul 14 00:49:07 CEST 2022 - The aliscore output file does not exist. Check results for BUSCO: 46694at7742
## Thu Jul 14 00:49:07 CEST 2022 - The aliscore output appears to be empty. Check results for BUSCO: 46694at7742
## Thu Jul 14 00:49:08 CEST 2022 - The aliscore output file does not exist. Check results for BUSCO: 79542at7742
## Thu Jul 14 00:49:08 CEST 2022 - The aliscore output appears to be empty. Check results for BUSCO: 79542at7742
## Thu Jul 14 00:49:10 CEST 2022 - The aliscore output file does not exist. Check results for BUSCO: 88808at7742
## Thu Jul 14 00:49:10 CEST 2022 - The aliscore output appears to be empty. Check results for BUSCO: 88808at7742
## Thu Jul 14 00:49:10 CEST 2022 - The aliscore output file does not exist. Check results for BUSCO: 122527at7742
## Thu Jul 14 00:49:10 CEST 2022 - The aliscore output appears to be empty. Check results for BUSCO: 122527at7742
## Thu Jul 14 00:49:13 CEST 2022 - The aliscore output file does not exist. Check results for BUSCO: 42538at7742
## Thu Jul 14 00:49:13 CEST 2022 - The aliscore output appears to be empty. Check results for BUSCO: 42538at7742
## Thu Jul 14 00:49:14 CEST 2022 - The aliscore output file does not exist. Check results for BUSCO: 130375at7742
## Thu Jul 14 00:49:14 CEST 2022 - The aliscore output appears to be empty. Check results for BUSCO: 130375at7742
## Thu Jul 14 00:49:16 CEST 2022 - The aliscore output file does not exist. Check results for BUSCO: 92370at7742
## Thu Jul 14 00:49:16 CEST 2022 - The aliscore output appears to be empty. Check results for BUSCO: 92370at7742
## Thu Jul 14 00:49:17 CEST 2022 - The aliscore output file does not exist. Check results for BUSCO: 153711at7742
## Thu Jul 14 00:49:17 CEST 2022 - The aliscore output appears to be empty. Check results for BUSCO: 153711at7742
## Thu Jul 14 00:49:18 CEST 2022 - The aliscore output file does not exist. Check results for BUSCO: 108033at7742
## Thu Jul 14 00:49:18 CEST 2022 - The aliscore output appears to be empty. Check results for BUSCO: 108033at7742
## Thu Jul 14 00:49:18 CEST 2022 - The aliscore output file does not exist. Check results for BUSCO: 330757at7742
## Thu Jul 14 00:49:18 CEST 2022 - The aliscore output appears to be empty. Check results for BUSCO: 330757at7742
## Thu Jul 14 00:49:19 CEST 2022 - The aliscore output file does not exist. Check results for BUSCO: 111099at7742
## Thu Jul 14 00:49:19 CEST 2022 - The aliscore output appears to be empty. Check results for BUSCO: 111099at7742
## Thu Jul 14 00:49:21 CEST 2022 - The aliscore output file does not exist. Check results for BUSCO: 205786at7742
## Thu Jul 14 00:49:21 CEST 2022 - The aliscore output appears to be empty. Check results for BUSCO: 205786at7742
## Thu Jul 14 00:49:22 CEST 2022 - The aliscore output file does not exist. Check results for BUSCO: 42710at7742
## Thu Jul 14 00:49:22 CEST 2022 - The aliscore output appears to be empty. Check results for BUSCO: 42710at7742
## Thu Jul 14 00:49:23 CEST 2022 - The aliscore output file does not exist. Check results for BUSCO: 229787at7742
## Thu Jul 14 00:49:23 CEST 2022 - The aliscore output appears to be empty. Check results for BUSCO: 229787at7742
## Thu Jul 14 00:49:24 CEST 2022 - The aliscore output file does not exist. Check results for BUSCO: 168949at7742
## Thu Jul 14 00:49:24 CEST 2022 - The aliscore output appears to be empty. Check results for BUSCO: 168949at7742
## Thu Jul 14 00:49:25 CEST 2022 - The aliscore output file does not exist. Check results for BUSCO: 277807at7742
## Thu Jul 14 00:49:25 CEST 2022 - The aliscore output appears to be empty. Check results for BUSCO: 277807at7742
## Thu Jul 14 00:49:28 CEST 2022 - The aliscore output file does not exist. Check results for BUSCO: 277807at7742
## Thu Jul 14 00:49:28 CEST 2022 - The aliscore output appears to be empty. Check results for BUSCO: 277807at7742
## Thu Jul 14 00:49:29 CEST 2022 - The aliscore output file does not exist. Check results for BUSCO: 114623at7742
## Thu Jul 14 00:49:29 CEST 2022 - The aliscore output appears to be empty. Check results for BUSCO: 114623at7742
## Thu Jul 14 00:49:29 CEST 2022 - The aliscore output file does not exist. Check results for BUSCO: 39501at7742
## Thu Jul 14 00:49:29 CEST 2022 - The aliscore output appears to be empty. Check results for BUSCO: 39501at7742
## Thu Jul 14 00:49:31 CEST 2022 - The aliscore output file does not exist. Check results for BUSCO: 136018at7742
## Thu Jul 14 00:49:31 CEST 2022 - The aliscore output appears to be empty. Check results for BUSCO: 136018at7742
## Thu Jul 14 00:49:32 CEST 2022 - The aliscore output file does not exist. Check results for BUSCO: 153051at7742
## Thu Jul 14 00:49:32 CEST 2022 - The aliscore output appears to be empty. Check results for BUSCO: 153051at7742
## Thu Jul 14 00:49:34 CEST 2022 - The aliscore output file does not exist. Check results for BUSCO: 152476at7742
## Thu Jul 14 00:49:34 CEST 2022 - The aliscore output appears to be empty. Check results for BUSCO: 152476at7742
## Thu Jul 14 00:49:35 CEST 2022 - The aliscore output file does not exist. Check results for BUSCO: 91759at7742
## Thu Jul 14 00:49:35 CEST 2022 - The aliscore output appears to be empty. Check results for BUSCO: 91759at7742
## Thu Jul 14 00:49:37 CEST 2022 - The aliscore output file does not exist. Check results for BUSCO: 46740at7742
## Thu Jul 14 00:49:37 CEST 2022 - The aliscore output appears to be empty. Check results for BUSCO: 46740at7742
## Thu Jul 14 00:49:38 CEST 2022 - The aliscore output file does not exist. Check results for BUSCO: 28954at7742
## Thu Jul 14 00:49:38 CEST 2022 - The aliscore output appears to be empty. Check results for BUSCO: 28954at7742
## Thu Jul 14 00:49:40 CEST 2022 - The aliscore output file does not exist. Check results for BUSCO: 219710at7742
## Thu Jul 14 00:49:40 CEST 2022 - The aliscore output appears to be empty. Check results for BUSCO: 219710at7742
## Thu Jul 14 00:49:40 CEST 2022 - The aliscore output file does not exist. Check results for BUSCO: 53153at7742
## Thu Jul 14 00:49:40 CEST 2022 - The aliscore output appears to be empty. Check results for BUSCO: 53153at7742
## Thu Jul 14 00:49:43 CEST 2022 - The aliscore output file does not exist. Check results for BUSCO: 90735at7742
## Thu Jul 14 00:49:43 CEST 2022 - The aliscore output appears to be empty. Check results for BUSCO: 90735at7742
## Thu Jul 14 00:49:43 CEST 2022 - The aliscore output file does not exist. Check results for BUSCO: 91759at7742
## Thu Jul 14 00:49:43 CEST 2022 - The aliscore output appears to be empty. Check results for BUSCO: 91759at7742
## Thu Jul 14 00:49:45 CEST 2022 - The aliscore output file does not exist. Check results for BUSCO: 91638at7742
## Thu Jul 14 00:49:45 CEST 2022 - The aliscore output appears to be empty. Check results for BUSCO: 91638at7742
## Thu Jul 14 00:49:46 CEST 2022 - The aliscore output file does not exist. Check results for BUSCO: 178398at7742
## Thu Jul 14 00:49:46 CEST 2022 - The aliscore output appears to be empty. Check results for BUSCO: 178398at7742
## Thu Jul 14 00:49:48 CEST 2022 - The aliscore output file does not exist. Check results for BUSCO: 96933at7742
## Thu Jul 14 00:49:48 CEST 2022 - The aliscore output appears to be empty. Check results for BUSCO: 96933at7742
## Thu Jul 14 00:49:48 CEST 2022 - The aliscore output file does not exist. Check results for BUSCO: 91759at7742
## Thu Jul 14 00:49:48 CEST 2022 - The aliscore output appears to be empty. Check results for BUSCO: 91759at7742
## Thu Jul 14 00:49:50 CEST 2022 - The aliscore output file does not exist. Check results for BUSCO: 235284at7742
## Thu Jul 14 00:49:50 CEST 2022 - The aliscore output appears to be empty. Check results for BUSCO: 235284at7742
## Thu Jul 14 00:49:51 CEST 2022 - The aliscore output file does not exist. Check results for BUSCO: 134433at7742
## Thu Jul 14 00:49:51 CEST 2022 - The aliscore output appears to be empty. Check results for BUSCO: 134433at7742
## Thu Jul 14 00:49:51 CEST 2022 - The aliscore output file does not exist. Check results for BUSCO: 110832at7742
## Thu Jul 14 00:49:51 CEST 2022 - The aliscore output appears to be empty. Check results for BUSCO: 110832at7742
## Thu Jul 14 00:49:54 CEST 2022 - The aliscore output file does not exist. Check results for BUSCO: 88808at7742
## Thu Jul 14 00:49:54 CEST 2022 - The aliscore output appears to be empty. Check results for BUSCO: 88808at7742
## Thu Jul 14 00:49:55 CEST 2022 - The aliscore output file does not exist. Check results for BUSCO: 121505at7742
## Thu Jul 14 00:49:55 CEST 2022 - The aliscore output appears to be empty. Check results for BUSCO: 121505at7742
## Thu Jul 14 00:49:56 CEST 2022 - The aliscore output file does not exist. Check results for BUSCO: 373324at7742
## Thu Jul 14 00:49:56 CEST 2022 - The aliscore output appears to be empty. Check results for BUSCO: 373324at7742
## Thu Jul 14 00:49:56 CEST 2022 - The aliscore output file does not exist. Check results for BUSCO: 178398at7742
## Thu Jul 14 00:49:56 CEST 2022 - The aliscore output appears to be empty. Check results for BUSCO: 178398at7742
## Thu Jul 14 00:49:59 CEST 2022 - The aliscore output file does not exist. Check results for BUSCO: 76998at7742
## Thu Jul 14 00:49:59 CEST 2022 - The aliscore output appears to be empty. Check results for BUSCO: 76998at7742
## Thu Jul 14 00:50:00 CEST 2022 - The aliscore output file does not exist. Check results for BUSCO: 60688at7742
## Thu Jul 14 00:50:00 CEST 2022 - The aliscore output appears to be empty. Check results for BUSCO: 60688at7742
## Thu Jul 14 00:50:01 CEST 2022 - The aliscore output file does not exist. Check results for BUSCO: 96764at7742
## Thu Jul 14 00:50:01 CEST 2022 - The aliscore output appears to be empty. Check results for BUSCO: 96764at7742
## Thu Jul 14 00:50:03 CEST 2022 - The aliscore output file does not exist. Check results for BUSCO: 111099at7742
## Thu Jul 14 00:50:03 CEST 2022 - The aliscore output appears to be empty. Check results for BUSCO: 111099at7742
## Thu Jul 14 00:50:03 CEST 2022 - The aliscore output file does not exist. Check results for BUSCO: 182992at7742
## Thu Jul 14 00:50:03 CEST 2022 - The aliscore output appears to be empty. Check results for BUSCO: 182992at7742
## Thu Jul 14 00:50:06 CEST 2022 - The aliscore output file does not exist. Check results for BUSCO: 60688at7742
## Thu Jul 14 00:50:06 CEST 2022 - The aliscore output appears to be empty. Check results for BUSCO: 60688at7742
## Thu Jul 14 00:50:06 CEST 2022 - The aliscore output file does not exist. Check results for BUSCO: 321778at7742
## Thu Jul 14 00:50:06 CEST 2022 - The aliscore output appears to be empty. Check results for BUSCO: 321778at7742
## Thu Jul 14 00:50:07 CEST 2022 - The aliscore output file does not exist. Check results for BUSCO: 185735at7742
## Thu Jul 14 00:50:07 CEST 2022 - The aliscore output appears to be empty. Check results for BUSCO: 185735at7742
## Thu Jul 14 00:50:09 CEST 2022 - The aliscore output file does not exist. Check results for BUSCO: 145104at7742
## Thu Jul 14 00:50:09 CEST 2022 - The aliscore output appears to be empty. Check results for BUSCO: 145104at7742
## Thu Jul 14 00:50:11 CEST 2022 - The aliscore output file does not exist. Check results for BUSCO: 106009at7742
## Thu Jul 14 00:50:11 CEST 2022 - The aliscore output appears to be empty. Check results for BUSCO: 106009at7742
## Thu Jul 14 00:50:13 CEST 2022 - The aliscore output file does not exist. Check results for BUSCO: 328263at7742
## Thu Jul 14 00:50:13 CEST 2022 - The aliscore output appears to be empty. Check results for BUSCO: 328263at7742
## Thu Jul 14 00:50:13 CEST 2022 - The aliscore output file does not exist. Check results for BUSCO: 158068at7742
## Thu Jul 14 00:50:13 CEST 2022 - The aliscore output appears to be empty. Check results for BUSCO: 158068at7742
## Thu Jul 14 00:50:15 CEST 2022 - The aliscore output file does not exist. Check results for BUSCO: 80914at7742
## Thu Jul 14 00:50:15 CEST 2022 - The aliscore output appears to be empty. Check results for BUSCO: 80914at7742
## Thu Jul 14 00:50:16 CEST 2022 - The aliscore output file does not exist. Check results for BUSCO: 56183at7742
## Thu Jul 14 00:50:16 CEST 2022 - The aliscore output appears to be empty. Check results for BUSCO: 56183at7742
## Thu Jul 14 00:50:17 CEST 2022 - The aliscore output file does not exist. Check results for BUSCO: 303797at7742
## Thu Jul 14 00:50:17 CEST 2022 - The aliscore output appears to be empty. Check results for BUSCO: 303797at7742
## Thu Jul 14 00:50:19 CEST 2022 - The aliscore output file does not exist. Check results for BUSCO: 280447at7742
## Thu Jul 14 00:50:19 CEST 2022 - The aliscore output appears to be empty. Check results for BUSCO: 280447at7742
## Thu Jul 14 00:50:19 CEST 2022 - The aliscore output file does not exist. Check results for BUSCO: 84817at7742
## Thu Jul 14 00:50:20 CEST 2022 - The aliscore output appears to be empty. Check results for BUSCO: 84817at7742
## Thu Jul 14 00:50:22 CEST 2022 - The aliscore output file does not exist. Check results for BUSCO: 177142at7742
## Thu Jul 14 00:50:22 CEST 2022 - The aliscore output appears to be empty. Check results for BUSCO: 177142at7742
## Thu Jul 14 00:50:22 CEST 2022 - The aliscore output file does not exist. Check results for BUSCO: 160275at7742
## Thu Jul 14 00:50:22 CEST 2022 - The aliscore output appears to be empty. Check results for BUSCO: 160275at7742
## Thu Jul 14 00:50:24 CEST 2022 - The aliscore output file does not exist. Check results for BUSCO: 99049at7742
## Thu Jul 14 00:50:24 CEST 2022 - The aliscore output appears to be empty. Check results for BUSCO: 99049at7742
## Thu Jul 14 00:50:25 CEST 2022 - The aliscore output file does not exist. Check results for BUSCO: 335206at7742
## Thu Jul 14 00:50:25 CEST 2022 - The aliscore output appears to be empty. Check results for BUSCO: 335206at7742
## Thu Jul 14 00:50:27 CEST 2022 - The aliscore output file does not exist. Check results for BUSCO: 199449at7742
## Thu Jul 14 00:50:27 CEST 2022 - The aliscore output appears to be empty. Check results for BUSCO: 199449at7742
## Thu Jul 14 00:50:28 CEST 2022 - The aliscore output file does not exist. Check results for BUSCO: 42538at7742
## Thu Jul 14 00:50:28 CEST 2022 - The aliscore output appears to be empty. Check results for BUSCO: 42538at7742
## Thu Jul 14 00:50:29 CEST 2022 - The aliscore output file does not exist. Check results for BUSCO: 143015at7742
## Thu Jul 14 00:50:29 CEST 2022 - The aliscore output appears to be empty. Check results for BUSCO: 143015at7742
## Thu Jul 14 00:50:31 CEST 2022 - The aliscore output file does not exist. Check results for BUSCO: 180758at7742
## Thu Jul 14 00:50:31 CEST 2022 - The aliscore output appears to be empty. Check results for BUSCO: 180758at7742
## Thu Jul 14 00:50:32 CEST 2022 - The aliscore output file does not exist. Check results for BUSCO: 337981at7742
## Thu Jul 14 00:50:32 CEST 2022 - The aliscore output appears to be empty. Check results for BUSCO: 337981at7742
## Thu Jul 14 00:50:33 CEST 2022 - The aliscore output file does not exist. Check results for BUSCO: 56183at7742
## Thu Jul 14 00:50:33 CEST 2022 - The aliscore output appears to be empty. Check results for BUSCO: 56183at7742
## Thu Jul 14 00:50:36 CEST 2022 - The aliscore output file does not exist. Check results for BUSCO: 233490at7742
## Thu Jul 14 00:50:36 CEST 2022 - The aliscore output appears to be empty. Check results for BUSCO: 233490at7742
## Thu Jul 14 00:50:36 CEST 2022 - The aliscore output file does not exist. Check results for BUSCO: 91878at7742
## Thu Jul 14 00:50:36 CEST 2022 - The aliscore output appears to be empty. Check results for BUSCO: 91878at7742
## Thu Jul 14 00:50:37 CEST 2022 - The aliscore output file does not exist. Check results for BUSCO: 143627at7742
## Thu Jul 14 00:50:37 CEST 2022 - The aliscore output appears to be empty. Check results for BUSCO: 143627at7742
## Thu Jul 14 00:50:40 CEST 2022 - The aliscore output file does not exist. Check results for BUSCO: 226528at7742
## Thu Jul 14 00:50:40 CEST 2022 - The aliscore output appears to be empty. Check results for BUSCO: 226528at7742
## Thu Jul 14 00:50:40 CEST 2022 - The aliscore output file does not exist. Check results for BUSCO: 99049at7742
## Thu Jul 14 00:50:40 CEST 2022 - The aliscore output appears to be empty. Check results for BUSCO: 99049at7742
## Thu Jul 14 00:50:41 CEST 2022 - The aliscore output file does not exist. Check results for BUSCO: 305955at7742
## Thu Jul 14 00:50:41 CEST 2022 - The aliscore output appears to be empty. Check results for BUSCO: 305955at7742
## Thu Jul 14 00:50:41 CEST 2022 - The aliscore output file does not exist. Check results for BUSCO: 80527at7742
## Thu Jul 14 00:50:41 CEST 2022 - The aliscore output appears to be empty. Check results for BUSCO: 80527at7742
## Thu Jul 14 00:50:43 CEST 2022 - The aliscore output file does not exist. Check results for BUSCO: 87193at7742
## Thu Jul 14 00:50:43 CEST 2022 - The aliscore output appears to be empty. Check results for BUSCO: 87193at7742
## Thu Jul 14 00:50:44 CEST 2022 - The aliscore output file does not exist. Check results for BUSCO: 100658at7742
## Thu Jul 14 00:50:44 CEST 2022 - The aliscore output appears to be empty. Check results for BUSCO: 100658at7742
## Thu Jul 14 00:50:46 CEST 2022 - The aliscore output file does not exist. Check results for BUSCO: 136018at7742
## Thu Jul 14 00:50:46 CEST 2022 - The aliscore output appears to be empty. Check results for BUSCO: 136018at7742
## Thu Jul 14 00:50:47 CEST 2022 - The aliscore output file does not exist. Check results for BUSCO: 133596at7742
## Thu Jul 14 00:50:47 CEST 2022 - The aliscore output appears to be empty. Check results for BUSCO: 133596at7742
## Thu Jul 14 00:50:49 CEST 2022 - The aliscore output file does not exist. Check results for BUSCO: 337981at7742
## Thu Jul 14 00:50:49 CEST 2022 - The aliscore output appears to be empty. Check results for BUSCO: 337981at7742
## Thu Jul 14 00:50:50 CEST 2022 - The aliscore output file does not exist. Check results for BUSCO: 280261at7742
## Thu Jul 14 00:50:50 CEST 2022 - The aliscore output appears to be empty. Check results for BUSCO: 280261at7742
## Thu Jul 14 00:50:50 CEST 2022 - The aliscore output file does not exist. Check results for BUSCO: 328263at7742
## Thu Jul 14 00:50:50 CEST 2022 - The aliscore output appears to be empty. Check results for BUSCO: 328263at7742
## Thu Jul 14 00:50:53 CEST 2022 - The aliscore output file does not exist. Check results for BUSCO: 190480at7742
## Thu Jul 14 00:50:53 CEST 2022 - The aliscore output appears to be empty. Check results for BUSCO: 190480at7742
## Thu Jul 14 00:50:53 CEST 2022 - The aliscore output file does not exist. Check results for BUSCO: 152020at7742
## Thu Jul 14 00:50:53 CEST 2022 - The aliscore output appears to be empty. Check results for BUSCO: 152020at7742
## Thu Jul 14 00:50:55 CEST 2022 - The aliscore output file does not exist. Check results for BUSCO: 28693at7742
## Thu Jul 14 00:50:55 CEST 2022 - The aliscore output appears to be empty. Check results for BUSCO: 28693at7742
## Thu Jul 14 00:50:58 CEST 2022 - The aliscore output file does not exist. Check results for BUSCO: 149128at7742
## Thu Jul 14 00:50:58 CEST 2022 - The aliscore output appears to be empty. Check results for BUSCO: 149128at7742
## Thu Jul 14 00:50:58 CEST 2022 - The aliscore output file does not exist. Check results for BUSCO: 81484at7742
## Thu Jul 14 00:50:58 CEST 2022 - The aliscore output appears to be empty. Check results for BUSCO: 81484at7742
## Thu Jul 14 00:51:00 CEST 2022 - The aliscore output file does not exist. Check results for BUSCO: 347981at7742
## Thu Jul 14 00:51:00 CEST 2022 - The aliscore output appears to be empty. Check results for BUSCO: 347981at7742
## Thu Jul 14 00:51:02 CEST 2022 - The aliscore output file does not exist. Check results for BUSCO: 79542at7742
## Thu Jul 14 00:51:02 CEST 2022 - The aliscore output appears to be empty. Check results for BUSCO: 79542at7742
## Thu Jul 14 00:51:02 CEST 2022 - The aliscore output file does not exist. Check results for BUSCO: 80791at7742
## Thu Jul 14 00:51:02 CEST 2022 - The aliscore output appears to be empty. Check results for BUSCO: 80791at7742
## Thu Jul 14 00:51:03 CEST 2022 - The aliscore output file does not exist. Check results for BUSCO: 303814at7742
## Thu Jul 14 00:51:03 CEST 2022 - The aliscore output appears to be empty. Check results for BUSCO: 303814at7742
## Thu Jul 14 00:51:05 CEST 2022 - The aliscore output file does not exist. Check results for BUSCO: 103919at7742
## Thu Jul 14 00:51:05 CEST 2022 - The aliscore output appears to be empty. Check results for BUSCO: 103919at7742
## Thu Jul 14 00:51:06 CEST 2022 - The aliscore output file does not exist. Check results for BUSCO: 67465at7742
## Thu Jul 14 00:51:06 CEST 2022 - The aliscore output appears to be empty. Check results for BUSCO: 67465at7742
## Thu Jul 14 00:51:08 CEST 2022 - The aliscore output file does not exist. Check results for BUSCO: 99037at7742
## Thu Jul 14 00:51:08 CEST 2022 - The aliscore output appears to be empty. Check results for BUSCO: 99037at7742
## Thu Jul 14 00:51:09 CEST 2022 - The aliscore output file does not exist. Check results for BUSCO: 59168at7742
## Thu Jul 14 00:51:09 CEST 2022 - The aliscore output appears to be empty. Check results for BUSCO: 59168at7742
## Thu Jul 14 00:51:09 CEST 2022 - The aliscore output file does not exist. Check results for BUSCO: 64473at7742
## Thu Jul 14 00:51:09 CEST 2022 - The aliscore output appears to be empty. Check results for BUSCO: 64473at7742
## Thu Jul 14 00:51:12 CEST 2022 - The aliscore output file does not exist. Check results for BUSCO: 68766at7742
## Thu Jul 14 00:51:12 CEST 2022 - The aliscore output appears to be empty. Check results for BUSCO: 68766at7742
## Thu Jul 14 00:51:13 CEST 2022 - The aliscore output file does not exist. Check results for BUSCO: 79542at7742
## Thu Jul 14 00:51:13 CEST 2022 - The aliscore output appears to be empty. Check results for BUSCO: 79542at7742
## Thu Jul 14 00:51:13 CEST 2022 - The aliscore output file does not exist. Check results for BUSCO: 264650at7742
## Thu Jul 14 00:51:13 CEST 2022 - The aliscore output appears to be empty. Check results for BUSCO: 264650at7742
## Thu Jul 14 00:51:16 CEST 2022 - The aliscore output file does not exist. Check results for BUSCO: 167134at7742
## Thu Jul 14 00:51:17 CEST 2022 - The aliscore output appears to be empty. Check results for BUSCO: 167134at7742
## Thu Jul 14 00:51:18 CEST 2022 - The aliscore output file does not exist. Check results for BUSCO: 146804at7742
## Thu Jul 14 00:51:18 CEST 2022 - The aliscore output appears to be empty. Check results for BUSCO: 146804at7742
## Thu Jul 14 00:51:18 CEST 2022 - The aliscore output file does not exist. Check results for BUSCO: 152476at7742
## Thu Jul 14 00:51:18 CEST 2022 - The aliscore output appears to be empty. Check results for BUSCO: 152476at7742
## Thu Jul 14 00:51:19 CEST 2022 - The aliscore output file does not exist. Check results for BUSCO: 161164at7742
## Thu Jul 14 00:51:19 CEST 2022 - The aliscore output appears to be empty. Check results for BUSCO: 161164at7742
## Thu Jul 14 00:51:21 CEST 2022 - The aliscore output file does not exist. Check results for BUSCO: 175841at7742
## Thu Jul 14 00:51:21 CEST 2022 - The aliscore output appears to be empty. Check results for BUSCO: 175841at7742
## Thu Jul 14 00:51:22 CEST 2022 - The aliscore output file does not exist. Check results for BUSCO: 153051at7742
## Thu Jul 14 00:51:22 CEST 2022 - The aliscore output appears to be empty. Check results for BUSCO: 153051at7742
## Thu Jul 14 00:51:23 CEST 2022 - The aliscore output file does not exist. Check results for BUSCO: 258916at7742
## Thu Jul 14 00:51:23 CEST 2022 - The aliscore output appears to be empty. Check results for BUSCO: 258916at7742
## Thu Jul 14 00:51:26 CEST 2022 - The aliscore output file does not exist. Check results for BUSCO: 128400at7742
## Thu Jul 14 00:51:26 CEST 2022 - The aliscore output appears to be empty. Check results for BUSCO: 128400at7742
## Thu Jul 14 00:51:28 CEST 2022 - The aliscore output file does not exist. Check results for BUSCO: 289598at7742
## Thu Jul 14 00:51:28 CEST 2022 - The aliscore output appears to be empty. Check results for BUSCO: 289598at7742
## Thu Jul 14 00:51:28 CEST 2022 - The aliscore output file does not exist. Check results for BUSCO: 184910at7742
## Thu Jul 14 00:51:28 CEST 2022 - The aliscore output appears to be empty. Check results for BUSCO: 184910at7742
## Thu Jul 14 00:51:28 CEST 2022 - The aliscore output file does not exist. Check results for BUSCO: 328263at7742
## Thu Jul 14 00:51:28 CEST 2022 - The aliscore output appears to be empty. Check results for BUSCO: 328263at7742
## Thu Jul 14 00:51:31 CEST 2022 - The aliscore output file does not exist. Check results for BUSCO: 161395at7742
## Thu Jul 14 00:51:31 CEST 2022 - The aliscore output appears to be empty. Check results for BUSCO: 161395at7742
## Thu Jul 14 00:51:31 CEST 2022 - The aliscore output file does not exist. Check results for BUSCO: 28693at7742
## Thu Jul 14 00:51:32 CEST 2022 - The aliscore output appears to be empty. Check results for BUSCO: 28693at7742
## Thu Jul 14 00:51:35 CEST 2022 - The aliscore output file does not exist. Check results for BUSCO: 171708at7742
## Thu Jul 14 00:51:35 CEST 2022 - The aliscore output appears to be empty. Check results for BUSCO: 171708at7742
## Thu Jul 14 00:51:36 CEST 2022 - The aliscore output file does not exist. Check results for BUSCO: 139083at7742
## Thu Jul 14 00:51:36 CEST 2022 - The aliscore output appears to be empty. Check results for BUSCO: 139083at7742
## Thu Jul 14 00:51:38 CEST 2022 - The aliscore output file does not exist. Check results for BUSCO: 258916at7742
## Thu Jul 14 00:51:38 CEST 2022 - The aliscore output appears to be empty. Check results for BUSCO: 258916at7742
## Thu Jul 14 00:51:39 CEST 2022 - The aliscore output file does not exist. Check results for BUSCO: 117635at7742
## Thu Jul 14 00:51:39 CEST 2022 - The aliscore output appears to be empty. Check results for BUSCO: 117635at7742
## Thu Jul 14 00:51:39 CEST 2022 - The aliscore output file does not exist. Check results for BUSCO: 87236at7742
## Thu Jul 14 00:51:39 CEST 2022 - The aliscore output appears to be empty. Check results for BUSCO: 87236at7742
## Thu Jul 14 00:51:39 CEST 2022 - The aliscore output file does not exist. Check results for BUSCO: 116065at7742
## Thu Jul 14 00:51:39 CEST 2022 - The aliscore output appears to be empty. Check results for BUSCO: 116065at7742
## Thu Jul 14 00:51:42 CEST 2022 - The aliscore output file does not exist. Check results for BUSCO: 351213at7742
## Thu Jul 14 00:51:42 CEST 2022 - The aliscore output appears to be empty. Check results for BUSCO: 351213at7742
## Thu Jul 14 00:51:44 CEST 2022 - The aliscore output file does not exist. Check results for BUSCO: 91638at7742
## Thu Jul 14 00:51:44 CEST 2022 - The aliscore output appears to be empty. Check results for BUSCO: 91638at7742
## Thu Jul 14 00:51:44 CEST 2022 - The aliscore output file does not exist. Check results for BUSCO: 311908at7742
## Thu Jul 14 00:51:44 CEST 2022 - The aliscore output appears to be empty. Check results for BUSCO: 311908at7742
## Thu Jul 14 00:51:46 CEST 2022 - The aliscore output file does not exist. Check results for BUSCO: 177142at7742
## Thu Jul 14 00:51:46 CEST 2022 - The aliscore output appears to be empty. Check results for BUSCO: 177142at7742
## Thu Jul 14 00:51:48 CEST 2022 - The aliscore output file does not exist. Check results for BUSCO: 132656at7742
## Thu Jul 14 00:51:48 CEST 2022 - The aliscore output appears to be empty. Check results for BUSCO: 132656at7742
## Thu Jul 14 00:51:48 CEST 2022 - The aliscore output file does not exist. Check results for BUSCO: 85369at7742
## Thu Jul 14 00:51:48 CEST 2022 - The aliscore output appears to be empty. Check results for BUSCO: 85369at7742
## Thu Jul 14 00:51:48 CEST 2022 - The aliscore output file does not exist. Check results for BUSCO: 125127at7742
## Thu Jul 14 00:51:48 CEST 2022 - The aliscore output appears to be empty. Check results for BUSCO: 125127at7742
## Thu Jul 14 00:51:50 CEST 2022 - The aliscore output file does not exist. Check results for BUSCO: 114626at7742
## Thu Jul 14 00:51:50 CEST 2022 - The aliscore output appears to be empty. Check results for BUSCO: 114626at7742
## Thu Jul 14 00:51:53 CEST 2022 - The aliscore output file does not exist. Check results for BUSCO: 158042at7742
## Thu Jul 14 00:51:53 CEST 2022 - The aliscore output appears to be empty. Check results for BUSCO: 158042at7742
## Thu Jul 14 00:51:54 CEST 2022 - The aliscore output file does not exist. Check results for BUSCO: 204919at7742
## Thu Jul 14 00:51:55 CEST 2022 - The aliscore output appears to be empty. Check results for BUSCO: 204919at7742
## Thu Jul 14 00:51:57 CEST 2022 - The aliscore output file does not exist. Check results for BUSCO: 28693at7742
## Thu Jul 14 00:51:57 CEST 2022 - The aliscore output appears to be empty. Check results for BUSCO: 28693at7742
## Thu Jul 14 00:51:57 CEST 2022 - The aliscore output file does not exist. Check results for BUSCO: 139946at7742
## Thu Jul 14 00:51:57 CEST 2022 - The aliscore output appears to be empty. Check results for BUSCO: 139946at7742
## Thu Jul 14 00:51:57 CEST 2022 - The aliscore output file does not exist. Check results for BUSCO: 195010at7742
## Thu Jul 14 00:51:57 CEST 2022 - The aliscore output appears to be empty. Check results for BUSCO: 195010at7742
## Thu Jul 14 00:51:57 CEST 2022 - The aliscore output file does not exist. Check results for BUSCO: 149377at7742
## Thu Jul 14 00:51:57 CEST 2022 - The aliscore output appears to be empty. Check results for BUSCO: 149377at7742
## Thu Jul 14 00:52:00 CEST 2022 - The aliscore output file does not exist. Check results for BUSCO: 176369at7742
## Thu Jul 14 00:52:00 CEST 2022 - The aliscore output appears to be empty. Check results for BUSCO: 176369at7742
## Thu Jul 14 00:52:01 CEST 2022 - The aliscore output file does not exist. Check results for BUSCO: 123404at7742
## Thu Jul 14 00:52:01 CEST 2022 - The aliscore output appears to be empty. Check results for BUSCO: 123404at7742
## Thu Jul 14 00:52:02 CEST 2022 - The aliscore output file does not exist. Check results for BUSCO: 127392at7742
## Thu Jul 14 00:52:02 CEST 2022 - The aliscore output appears to be empty. Check results for BUSCO: 127392at7742
## Thu Jul 14 00:52:05 CEST 2022 - The aliscore output file does not exist. Check results for BUSCO: 211495at7742
## Thu Jul 14 00:52:05 CEST 2022 - The aliscore output appears to be empty. Check results for BUSCO: 211495at7742
## Thu Jul 14 00:52:06 CEST 2022 - The aliscore output file does not exist. Check results for BUSCO: 236895at7742
## Thu Jul 14 00:52:06 CEST 2022 - The aliscore output appears to be empty. Check results for BUSCO: 236895at7742
## Thu Jul 14 00:52:07 CEST 2022 - The aliscore output file does not exist. Check results for BUSCO: 175036at7742
## Thu Jul 14 00:52:07 CEST 2022 - The aliscore output appears to be empty. Check results for BUSCO: 175036at7742
## Thu Jul 14 00:52:08 CEST 2022 - The aliscore output file does not exist. Check results for BUSCO: 64473at7742
## Thu Jul 14 00:52:08 CEST 2022 - The aliscore output appears to be empty. Check results for BUSCO: 64473at7742
## Thu Jul 14 00:52:09 CEST 2022 - The aliscore output file does not exist. Check results for BUSCO: 191192at7742
## Thu Jul 14 00:52:09 CEST 2022 - The aliscore output appears to be empty. Check results for BUSCO: 191192at7742
## Thu Jul 14 00:52:09 CEST 2022 - The aliscore output file does not exist. Check results for BUSCO: 126566at7742
## Thu Jul 14 00:52:10 CEST 2022 - The aliscore output appears to be empty. Check results for BUSCO: 126566at7742
## Thu Jul 14 00:52:12 CEST 2022 - The aliscore output file does not exist. Check results for BUSCO: 213614at7742
## Thu Jul 14 00:52:12 CEST 2022 - The aliscore output appears to be empty. Check results for BUSCO: 213614at7742
## Thu Jul 14 00:52:12 CEST 2022 - The aliscore output file does not exist. Check results for BUSCO: 117499at7742
## Thu Jul 14 00:52:12 CEST 2022 - The aliscore output appears to be empty. Check results for BUSCO: 117499at7742
## Thu Jul 14 00:52:16 CEST 2022 - The aliscore output file does not exist. Check results for BUSCO: 152020at7742
## Thu Jul 14 00:52:16 CEST 2022 - The aliscore output appears to be empty. Check results for BUSCO: 152020at7742
## Thu Jul 14 00:52:16 CEST 2022 - The aliscore output file does not exist. Check results for BUSCO: 58280at7742
## Thu Jul 14 00:52:16 CEST 2022 - The aliscore output appears to be empty. Check results for BUSCO: 58280at7742
## Thu Jul 14 00:52:17 CEST 2022 - The aliscore output file does not exist. Check results for BUSCO: 213614at7742
## Thu Jul 14 00:52:17 CEST 2022 - The aliscore output appears to be empty. Check results for BUSCO: 213614at7742
## Thu Jul 14 00:52:18 CEST 2022 - The aliscore output file does not exist. Check results for BUSCO: 202578at7742
## Thu Jul 14 00:52:18 CEST 2022 - The aliscore output appears to be empty. Check results for BUSCO: 202578at7742
## Thu Jul 14 00:52:19 CEST 2022 - The aliscore output file does not exist. Check results for BUSCO: 123847at7742
## Thu Jul 14 00:52:19 CEST 2022 - The aliscore output appears to be empty. Check results for BUSCO: 123847at7742
## Thu Jul 14 00:52:20 CEST 2022 - The aliscore output file does not exist. Check results for BUSCO: 194866at7742
## Thu Jul 14 00:52:20 CEST 2022 - The aliscore output appears to be empty. Check results for BUSCO: 194866at7742
## Thu Jul 14 00:52:22 CEST 2022 - The aliscore output file does not exist. Check results for BUSCO: 60688at7742
## Thu Jul 14 00:52:22 CEST 2022 - The aliscore output appears to be empty. Check results for BUSCO: 60688at7742
## Thu Jul 14 00:52:22 CEST 2022 - The aliscore output file does not exist. Check results for BUSCO: 135517at7742
## Thu Jul 14 00:52:22 CEST 2022 - The aliscore output appears to be empty. Check results for BUSCO: 135517at7742
## Thu Jul 14 00:52:24 CEST 2022 - The aliscore output file does not exist. Check results for BUSCO: 283522at7742
## Thu Jul 14 00:52:25 CEST 2022 - The aliscore output appears to be empty. Check results for BUSCO: 283522at7742
## Thu Jul 14 00:52:26 CEST 2022 - The aliscore output file does not exist. Check results for BUSCO: 160674at7742
## Thu Jul 14 00:52:26 CEST 2022 - The aliscore output appears to be empty. Check results for BUSCO: 160674at7742
## Thu Jul 14 00:52:27 CEST 2022 - The aliscore output file does not exist. Check results for BUSCO: 82034at7742
## Thu Jul 14 00:52:27 CEST 2022 - The aliscore output appears to be empty. Check results for BUSCO: 82034at7742
## Thu Jul 14 00:52:28 CEST 2022 - The aliscore output file does not exist. Check results for BUSCO: 203744at7742
## Thu Jul 14 00:52:28 CEST 2022 - The aliscore output appears to be empty. Check results for BUSCO: 203744at7742
## Thu Jul 14 00:52:31 CEST 2022 - The aliscore output file does not exist. Check results for BUSCO: 168479at7742
## Thu Jul 14 00:52:31 CEST 2022 - The aliscore output appears to be empty. Check results for BUSCO: 168479at7742
## Thu Jul 14 00:52:31 CEST 2022 - The aliscore output file does not exist. Check results for BUSCO: 219710at7742
## Thu Jul 14 00:52:31 CEST 2022 - The aliscore output appears to be empty. Check results for BUSCO: 219710at7742
## Thu Jul 14 00:52:32 CEST 2022 - The aliscore output file does not exist. Check results for BUSCO: 70359at7742
## Thu Jul 14 00:52:32 CEST 2022 - The aliscore output appears to be empty. Check results for BUSCO: 70359at7742
## Thu Jul 14 00:52:34 CEST 2022 - The aliscore output file does not exist. Check results for BUSCO: 95757at7742
## Thu Jul 14 00:52:34 CEST 2022 - The aliscore output appears to be empty. Check results for BUSCO: 95757at7742
## Thu Jul 14 00:52:36 CEST 2022 - The aliscore output file does not exist. Check results for BUSCO: 112360at7742
## Thu Jul 14 00:52:36 CEST 2022 - The aliscore output appears to be empty. Check results for BUSCO: 112360at7742
## Thu Jul 14 00:52:36 CEST 2022 - The aliscore output file does not exist. Check results for BUSCO: 174474at7742
## Thu Jul 14 00:52:36 CEST 2022 - The aliscore output appears to be empty. Check results for BUSCO: 174474at7742
## Thu Jul 14 00:52:37 CEST 2022 - The aliscore output file does not exist. Check results for BUSCO: 116359at7742
## Thu Jul 14 00:52:37 CEST 2022 - The aliscore output appears to be empty. Check results for BUSCO: 116359at7742
## Thu Jul 14 00:52:38 CEST 2022 - The aliscore output file does not exist. Check results for BUSCO: 83474at7742
## Thu Jul 14 00:52:39 CEST 2022 - The aliscore output appears to be empty. Check results for BUSCO: 83474at7742
## Thu Jul 14 00:52:41 CEST 2022 - The aliscore output file does not exist. Check results for BUSCO: 366345at7742
## Thu Jul 14 00:52:41 CEST 2022 - The aliscore output appears to be empty. Check results for BUSCO: 366345at7742
## Thu Jul 14 00:52:42 CEST 2022 - The aliscore output file does not exist. Check results for BUSCO: 139083at7742
## Thu Jul 14 00:52:42 CEST 2022 - The aliscore output appears to be empty. Check results for BUSCO: 139083at7742
## Thu Jul 14 00:52:43 CEST 2022 - The aliscore output file does not exist. Check results for BUSCO: 270327at7742
## Thu Jul 14 00:52:43 CEST 2022 - The aliscore output appears to be empty. Check results for BUSCO: 270327at7742
## Thu Jul 14 00:52:45 CEST 2022 - The aliscore output file does not exist. Check results for BUSCO: 293720at7742
## Thu Jul 14 00:52:45 CEST 2022 - The aliscore output appears to be empty. Check results for BUSCO: 293720at7742
## Thu Jul 14 00:52:45 CEST 2022 - The aliscore output file does not exist. Check results for BUSCO: 65316at7742
## Thu Jul 14 00:52:45 CEST 2022 - The aliscore output appears to be empty. Check results for BUSCO: 65316at7742
## Thu Jul 14 00:52:45 CEST 2022 - The aliscore output file does not exist. Check results for BUSCO: 210019at7742
## Thu Jul 14 00:52:45 CEST 2022 - The aliscore output appears to be empty. Check results for BUSCO: 210019at7742
## Thu Jul 14 00:52:48 CEST 2022 - The aliscore output file does not exist. Check results for BUSCO: 187791at7742
## Thu Jul 14 00:52:48 CEST 2022 - The aliscore output appears to be empty. Check results for BUSCO: 187791at7742
## Thu Jul 14 00:52:50 CEST 2022 - The aliscore output file does not exist. Check results for BUSCO: 171639at7742
## Thu Jul 14 00:52:50 CEST 2022 - The aliscore output appears to be empty. Check results for BUSCO: 171639at7742
## Thu Jul 14 00:52:51 CEST 2022 - The aliscore output file does not exist. Check results for BUSCO: 263251at7742
## Thu Jul 14 00:52:51 CEST 2022 - The aliscore output appears to be empty. Check results for BUSCO: 263251at7742
## Thu Jul 14 00:52:52 CEST 2022 - The aliscore output file does not exist. Check results for BUSCO: 293720at7742
## Thu Jul 14 00:52:52 CEST 2022 - The aliscore output appears to be empty. Check results for BUSCO: 293720at7742
## Thu Jul 14 00:52:53 CEST 2022 - The aliscore output file does not exist. Check results for BUSCO: 268563at7742
## Thu Jul 14 00:52:53 CEST 2022 - The aliscore output appears to be empty. Check results for BUSCO: 268563at7742
## Thu Jul 14 00:52:54 CEST 2022 - The aliscore output file does not exist. Check results for BUSCO: 122918at7742
## Thu Jul 14 00:52:54 CEST 2022 - The aliscore output appears to be empty. Check results for BUSCO: 122918at7742
## Thu Jul 14 00:52:54 CEST 2022 - The aliscore output file does not exist. Check results for BUSCO: 162426at7742
## Thu Jul 14 00:52:54 CEST 2022 - The aliscore output appears to be empty. Check results for BUSCO: 162426at7742
## Thu Jul 14 00:52:59 CEST 2022 - The aliscore output file does not exist. Check results for BUSCO: 228455at7742
## Thu Jul 14 00:52:59 CEST 2022 - The aliscore output appears to be empty. Check results for BUSCO: 228455at7742
## Thu Jul 14 00:52:59 CEST 2022 - The aliscore output file does not exist. Check results for BUSCO: 59168at7742
## Thu Jul 14 00:52:59 CEST 2022 - The aliscore output appears to be empty. Check results for BUSCO: 59168at7742
## Thu Jul 14 00:53:00 CEST 2022 - The aliscore output file does not exist. Check results for BUSCO: 219710at7742
## Thu Jul 14 00:53:00 CEST 2022 - The aliscore output appears to be empty. Check results for BUSCO: 219710at7742
## Thu Jul 14 00:53:02 CEST 2022 - The aliscore output file does not exist. Check results for BUSCO: 109054at7742
## Thu Jul 14 00:53:02 CEST 2022 - The aliscore output appears to be empty. Check results for BUSCO: 109054at7742
## Thu Jul 14 00:53:05 CEST 2022 - The aliscore output file does not exist. Check results for BUSCO: 152020at7742
## Thu Jul 14 00:53:05 CEST 2022 - The aliscore output appears to be empty. Check results for BUSCO: 152020at7742
## Thu Jul 14 00:53:05 CEST 2022 - The aliscore output file does not exist. Check results for BUSCO: 77473at7742
## Thu Jul 14 00:53:05 CEST 2022 - The aliscore output appears to be empty. Check results for BUSCO: 77473at7742
## Thu Jul 14 00:53:06 CEST 2022 - The aliscore output file does not exist. Check results for BUSCO: 208931at7742
## Thu Jul 14 00:53:06 CEST 2022 - The aliscore output appears to be empty. Check results for BUSCO: 208931at7742
## Thu Jul 14 00:53:08 CEST 2022 - The aliscore output file does not exist. Check results for BUSCO: 89615at7742
## Thu Jul 14 00:53:08 CEST 2022 - The aliscore output appears to be empty. Check results for BUSCO: 89615at7742
## Thu Jul 14 00:53:10 CEST 2022 - The aliscore output file does not exist. Check results for BUSCO: 146203at7742
## Thu Jul 14 00:53:10 CEST 2022 - The aliscore output appears to be empty. Check results for BUSCO: 146203at7742
## Thu Jul 14 00:53:11 CEST 2022 - The aliscore output file does not exist. Check results for BUSCO: 92625at7742
## Thu Jul 14 00:53:11 CEST 2022 - The aliscore output appears to be empty. Check results for BUSCO: 92625at7742
## Thu Jul 14 00:53:12 CEST 2022 - The aliscore output file does not exist. Check results for BUSCO: 402344at7742
## Thu Jul 14 00:53:13 CEST 2022 - The aliscore output appears to be empty. Check results for BUSCO: 402344at7742
## Thu Jul 14 00:53:14 CEST 2022 - The aliscore output file does not exist. Check results for BUSCO: 211495at7742
## Thu Jul 14 00:53:14 CEST 2022 - The aliscore output appears to be empty. Check results for BUSCO: 211495at7742
## Thu Jul 14 00:53:14 CEST 2022 - The aliscore output file does not exist. Check results for BUSCO: 53077at7742
## Thu Jul 14 00:53:14 CEST 2022 - The aliscore output appears to be empty. Check results for BUSCO: 53077at7742
## Thu Jul 14 00:53:16 CEST 2022 - The aliscore output file does not exist. Check results for BUSCO: 123404at7742
## Thu Jul 14 00:53:16 CEST 2022 - The aliscore output appears to be empty. Check results for BUSCO: 123404at7742
## Thu Jul 14 00:53:17 CEST 2022 - The aliscore output file does not exist. Check results for BUSCO: 68766at7742
## Thu Jul 14 00:53:17 CEST 2022 - The aliscore output appears to be empty. Check results for BUSCO: 68766at7742
## Thu Jul 14 00:53:18 CEST 2022 - The aliscore output file does not exist. Check results for BUSCO: 161425at7742
## Thu Jul 14 00:53:18 CEST 2022 - The aliscore output appears to be empty. Check results for BUSCO: 161425at7742
## Thu Jul 14 00:53:19 CEST 2022 - The aliscore output file does not exist. Check results for BUSCO: 172204at7742
## Thu Jul 14 00:53:19 CEST 2022 - The aliscore output appears to be empty. Check results for BUSCO: 172204at7742
## Thu Jul 14 00:53:21 CEST 2022 - The aliscore output file does not exist. Check results for BUSCO: 139083at7742
## Thu Jul 14 00:53:21 CEST 2022 - The aliscore output appears to be empty. Check results for BUSCO: 139083at7742
## Thu Jul 14 00:53:21 CEST 2022 - The aliscore output file does not exist. Check results for BUSCO: 141724at7742
## Thu Jul 14 00:53:21 CEST 2022 - The aliscore output appears to be empty. Check results for BUSCO: 141724at7742
## Thu Jul 14 00:53:23 CEST 2022 - The aliscore output file does not exist. Check results for BUSCO: 86277at7742
## Thu Jul 14 00:53:24 CEST 2022 - The aliscore output appears to be empty. Check results for BUSCO: 86277at7742
## Thu Jul 14 00:53:24 CEST 2022 - The aliscore output file does not exist. Check results for BUSCO: 80791at7742
## Thu Jul 14 00:53:24 CEST 2022 - The aliscore output appears to be empty. Check results for BUSCO: 80791at7742
## Thu Jul 14 00:53:24 CEST 2022 - The aliscore output file does not exist. Check results for BUSCO: 298175at7742
## Thu Jul 14 00:53:24 CEST 2022 - The aliscore output appears to be empty. Check results for BUSCO: 298175at7742
## Thu Jul 14 00:53:27 CEST 2022 - The aliscore output file does not exist. Check results for BUSCO: 174966at7742
## Thu Jul 14 00:53:27 CEST 2022 - The aliscore output appears to be empty. Check results for BUSCO: 174966at7742
## Thu Jul 14 00:53:28 CEST 2022 - The aliscore output file does not exist. Check results for BUSCO: 214498at7742
## Thu Jul 14 00:53:28 CEST 2022 - The aliscore output appears to be empty. Check results for BUSCO: 214498at7742
## Thu Jul 14 00:53:30 CEST 2022 - The aliscore output file does not exist. Check results for BUSCO: 177227at7742
## Thu Jul 14 00:53:30 CEST 2022 - The aliscore output appears to be empty. Check results for BUSCO: 177227at7742
## Thu Jul 14 00:53:31 CEST 2022 - The aliscore output file does not exist. Check results for BUSCO: 127929at7742
## Thu Jul 14 00:53:31 CEST 2022 - The aliscore output appears to be empty. Check results for BUSCO: 127929at7742
## Thu Jul 14 00:53:31 CEST 2022 - The aliscore output file does not exist. Check results for BUSCO: 174474at7742
## Thu Jul 14 00:53:32 CEST 2022 - The aliscore output appears to be empty. Check results for BUSCO: 174474at7742
## Thu Jul 14 00:53:33 CEST 2022 - The aliscore output file does not exist. Check results for BUSCO: 149083at7742
## Thu Jul 14 00:53:33 CEST 2022 - The aliscore output appears to be empty. Check results for BUSCO: 149083at7742
## Thu Jul 14 00:53:36 CEST 2022 - The aliscore output file does not exist. Check results for BUSCO: 221197at7742
## Thu Jul 14 00:53:36 CEST 2022 - The aliscore output appears to be empty. Check results for BUSCO: 221197at7742
## Thu Jul 14 00:53:36 CEST 2022 - The aliscore output file does not exist. Check results for BUSCO: 213511at7742
## Thu Jul 14 00:53:36 CEST 2022 - The aliscore output appears to be empty. Check results for BUSCO: 213511at7742
## Thu Jul 14 00:53:38 CEST 2022 - The aliscore output file does not exist. Check results for BUSCO: 226528at7742
## Thu Jul 14 00:53:38 CEST 2022 - The aliscore output appears to be empty. Check results for BUSCO: 226528at7742
## Thu Jul 14 00:53:39 CEST 2022 - The aliscore output file does not exist. Check results for BUSCO: 303814at7742
## Thu Jul 14 00:53:39 CEST 2022 - The aliscore output appears to be empty. Check results for BUSCO: 303814at7742
## Thu Jul 14 00:53:40 CEST 2022 - The aliscore output file does not exist. Check results for BUSCO: 173159at7742
## Thu Jul 14 00:53:40 CEST 2022 - The aliscore output appears to be empty. Check results for BUSCO: 173159at7742
## Thu Jul 14 00:53:40 CEST 2022 - The aliscore output file does not exist. Check results for BUSCO: 424377at7742
## Thu Jul 14 00:53:40 CEST 2022 - The aliscore output appears to be empty. Check results for BUSCO: 424377at7742
## Thu Jul 14 00:53:44 CEST 2022 - The aliscore output file does not exist. Check results for BUSCO: 174966at7742
## Thu Jul 14 00:53:44 CEST 2022 - The aliscore output appears to be empty. Check results for BUSCO: 174966at7742
## Thu Jul 14 00:53:44 CEST 2022 - The aliscore output file does not exist. Check results for BUSCO: 174966at7742
## Thu Jul 14 00:53:44 CEST 2022 - The aliscore output appears to be empty. Check results for BUSCO: 174966at7742
## Thu Jul 14 00:53:46 CEST 2022 - The aliscore output file does not exist. Check results for BUSCO: 128396at7742
## Thu Jul 14 00:53:46 CEST 2022 - The aliscore output appears to be empty. Check results for BUSCO: 128396at7742
## Thu Jul 14 00:53:48 CEST 2022 - The aliscore output file does not exist. Check results for BUSCO: 75881at7742
## Thu Jul 14 00:53:48 CEST 2022 - The aliscore output appears to be empty. Check results for BUSCO: 75881at7742
## Thu Jul 14 00:53:48 CEST 2022 - The aliscore output file does not exist. Check results for BUSCO: 73001at7742
## Thu Jul 14 00:53:48 CEST 2022 - The aliscore output appears to be empty. Check results for BUSCO: 73001at7742
## Thu Jul 14 00:53:48 CEST 2022 - The aliscore output file does not exist. Check results for BUSCO: 127830at7742
## Thu Jul 14 00:53:49 CEST 2022 - The aliscore output appears to be empty. Check results for BUSCO: 127830at7742
## Thu Jul 14 00:53:51 CEST 2022 - The aliscore output file does not exist. Check results for BUSCO: 53843at7742
## Thu Jul 14 00:53:51 CEST 2022 - The aliscore output appears to be empty. Check results for BUSCO: 53843at7742
## Thu Jul 14 00:53:51 CEST 2022 - The aliscore output file does not exist. Check results for BUSCO: 67465at7742
## Thu Jul 14 00:53:51 CEST 2022 - The aliscore output appears to be empty. Check results for BUSCO: 67465at7742
## Thu Jul 14 00:53:52 CEST 2022 - The aliscore output file does not exist. Check results for BUSCO: 228455at7742
## Thu Jul 14 00:53:52 CEST 2022 - The aliscore output appears to be empty. Check results for BUSCO: 228455at7742
## Thu Jul 14 00:53:54 CEST 2022 - The aliscore output file does not exist. Check results for BUSCO: 59168at7742
## Thu Jul 14 00:53:54 CEST 2022 - The aliscore output appears to be empty. Check results for BUSCO: 59168at7742
## Thu Jul 14 00:53:55 CEST 2022 - The aliscore output file does not exist. Check results for BUSCO: 210019at7742
## Thu Jul 14 00:53:55 CEST 2022 - The aliscore output appears to be empty. Check results for BUSCO: 210019at7742
## Thu Jul 14 00:53:57 CEST 2022 - The aliscore output file does not exist. Check results for BUSCO: 107664at7742
## Thu Jul 14 00:53:57 CEST 2022 - The aliscore output appears to be empty. Check results for BUSCO: 107664at7742
## Thu Jul 14 00:53:58 CEST 2022 - The aliscore output file does not exist. Check results for BUSCO: 329621at7742
## Thu Jul 14 00:53:58 CEST 2022 - The aliscore output appears to be empty. Check results for BUSCO: 329621at7742
## Thu Jul 14 00:53:58 CEST 2022 - The aliscore output file does not exist. Check results for BUSCO: 91638at7742
## Thu Jul 14 00:53:58 CEST 2022 - The aliscore output appears to be empty. Check results for BUSCO: 91638at7742
## Thu Jul 14 00:54:01 CEST 2022 - The aliscore output file does not exist. Check results for BUSCO: 245053at7742
## Thu Jul 14 00:54:01 CEST 2022 - The aliscore output appears to be empty. Check results for BUSCO: 245053at7742
## Thu Jul 14 00:54:02 CEST 2022 - The aliscore output file does not exist. Check results for BUSCO: 53077at7742
## Thu Jul 14 00:54:02 CEST 2022 - The aliscore output appears to be empty. Check results for BUSCO: 53077at7742
## Thu Jul 14 00:54:04 CEST 2022 - The aliscore output file does not exist. Check results for BUSCO: 104543at7742
## Thu Jul 14 00:54:05 CEST 2022 - The aliscore output appears to be empty. Check results for BUSCO: 104543at7742
## Thu Jul 14 00:54:05 CEST 2022 - The aliscore output file does not exist. Check results for BUSCO: 75874at7742
## Thu Jul 14 00:54:05 CEST 2022 - The aliscore output appears to be empty. Check results for BUSCO: 75874at7742
## Thu Jul 14 00:54:06 CEST 2022 - The aliscore output file does not exist. Check results for BUSCO: 97039at7742
## Thu Jul 14 00:54:06 CEST 2022 - The aliscore output appears to be empty. Check results for BUSCO: 97039at7742
## Thu Jul 14 00:54:08 CEST 2022 - The aliscore output file does not exist. Check results for BUSCO: 87783at7742
## Thu Jul 14 00:54:08 CEST 2022 - The aliscore output appears to be empty. Check results for BUSCO: 87783at7742
## Thu Jul 14 00:54:09 CEST 2022 - The aliscore output file does not exist. Check results for BUSCO: 87236at7742
## Thu Jul 14 00:54:09 CEST 2022 - The aliscore output appears to be empty. Check results for BUSCO: 87236at7742
## Thu Jul 14 00:54:10 CEST 2022 - The aliscore output file does not exist. Check results for BUSCO: 245445at7742
## Thu Jul 14 00:54:10 CEST 2022 - The aliscore output appears to be empty. Check results for BUSCO: 245445at7742
## Thu Jul 14 00:54:11 CEST 2022 - The aliscore output file does not exist. Check results for BUSCO: 96764at7742
## Thu Jul 14 00:54:12 CEST 2022 - The aliscore output appears to be empty. Check results for BUSCO: 96764at7742
## Thu Jul 14 00:54:13 CEST 2022 - The aliscore output file does not exist. Check results for BUSCO: 208931at7742
## Thu Jul 14 00:54:13 CEST 2022 - The aliscore output appears to be empty. Check results for BUSCO: 208931at7742
## Thu Jul 14 00:54:13 CEST 2022 - The aliscore output file does not exist. Check results for BUSCO: 42045at7742
## Thu Jul 14 00:54:13 CEST 2022 - The aliscore output appears to be empty. Check results for BUSCO: 42045at7742
## Thu Jul 14 00:54:16 CEST 2022 - The aliscore output file does not exist. Check results for BUSCO: 64461at7742
## Thu Jul 14 00:54:16 CEST 2022 - The aliscore output appears to be empty. Check results for BUSCO: 64461at7742
## Thu Jul 14 00:54:16 CEST 2022 - The aliscore output file does not exist. Check results for BUSCO: 202578at7742
## Thu Jul 14 00:54:16 CEST 2022 - The aliscore output appears to be empty. Check results for BUSCO: 202578at7742
## Thu Jul 14 00:54:18 CEST 2022 - The aliscore output file does not exist. Check results for BUSCO: 100494at7742
## Thu Jul 14 00:54:18 CEST 2022 - The aliscore output appears to be empty. Check results for BUSCO: 100494at7742
## Thu Jul 14 00:54:19 CEST 2022 - The aliscore output file does not exist. Check results for BUSCO: 104543at7742
## Thu Jul 14 00:54:19 CEST 2022 - The aliscore output appears to be empty. Check results for BUSCO: 104543at7742
## Thu Jul 14 00:54:20 CEST 2022 - The aliscore output file does not exist. Check results for BUSCO: 92625at7742
## Thu Jul 14 00:54:20 CEST 2022 - The aliscore output appears to be empty. Check results for BUSCO: 92625at7742
## Thu Jul 14 00:54:22 CEST 2022 - The aliscore output file does not exist. Check results for BUSCO: 53077at7742
## Thu Jul 14 00:54:22 CEST 2022 - The aliscore output appears to be empty. Check results for BUSCO: 53077at7742
## Thu Jul 14 00:54:24 CEST 2022 - The aliscore output file does not exist. Check results for BUSCO: 128396at7742
## Thu Jul 14 00:54:24 CEST 2022 - The aliscore output appears to be empty. Check results for BUSCO: 128396at7742
## Thu Jul 14 00:54:25 CEST 2022 - The aliscore output file does not exist. Check results for BUSCO: 51955at7742
## Thu Jul 14 00:54:25 CEST 2022 - The aliscore output appears to be empty. Check results for BUSCO: 51955at7742
## Thu Jul 14 00:54:25 CEST 2022 - The aliscore output file does not exist. Check results for BUSCO: 143938at7742
## Thu Jul 14 00:54:25 CEST 2022 - The aliscore output appears to be empty. Check results for BUSCO: 143938at7742
## Thu Jul 14 00:54:25 CEST 2022 - The aliscore output file does not exist. Check results for BUSCO: 56183at7742
## Thu Jul 14 00:54:25 CEST 2022 - The aliscore output appears to be empty. Check results for BUSCO: 56183at7742
## Thu Jul 14 00:54:27 CEST 2022 - The aliscore output file does not exist. Check results for BUSCO: 168949at7742
## Thu Jul 14 00:54:27 CEST 2022 - The aliscore output appears to be empty. Check results for BUSCO: 168949at7742
## Thu Jul 14 00:54:30 CEST 2022 - The aliscore output file does not exist. Check results for BUSCO: 131862at7742
## Thu Jul 14 00:54:30 CEST 2022 - The aliscore output appears to be empty. Check results for BUSCO: 131862at7742
## Thu Jul 14 00:54:30 CEST 2022 - The aliscore output file does not exist. Check results for BUSCO: 214498at7742
## Thu Jul 14 00:54:30 CEST 2022 - The aliscore output appears to be empty. Check results for BUSCO: 214498at7742
## Thu Jul 14 00:54:31 CEST 2022 - The aliscore output file does not exist. Check results for BUSCO: 75881at7742
## Thu Jul 14 00:54:32 CEST 2022 - The aliscore output appears to be empty. Check results for BUSCO: 75881at7742
## Thu Jul 14 00:54:33 CEST 2022 - The aliscore output file does not exist. Check results for BUSCO: 175036at7742
## Thu Jul 14 00:54:33 CEST 2022 - The aliscore output appears to be empty. Check results for BUSCO: 175036at7742
## Thu Jul 14 00:54:35 CEST 2022 - The aliscore output file does not exist. Check results for BUSCO: 70359at7742
## Thu Jul 14 00:54:35 CEST 2022 - The aliscore output appears to be empty. Check results for BUSCO: 70359at7742
## Thu Jul 14 00:54:36 CEST 2022 - The aliscore output file does not exist. Check results for BUSCO: 115767at7742
## Thu Jul 14 00:54:36 CEST 2022 - The aliscore output appears to be empty. Check results for BUSCO: 115767at7742
## Thu Jul 14 00:54:36 CEST 2022 - The aliscore output file does not exist. Check results for BUSCO: 78565at7742
## Thu Jul 14 00:54:36 CEST 2022 - The aliscore output appears to be empty. Check results for BUSCO: 78565at7742
## Thu Jul 14 00:54:38 CEST 2022 - The aliscore output file does not exist. Check results for BUSCO: 50786at7742
## Thu Jul 14 00:54:38 CEST 2022 - The aliscore output appears to be empty. Check results for BUSCO: 50786at7742
## Thu Jul 14 00:54:39 CEST 2022 - The aliscore output file does not exist. Check results for BUSCO: 367334at7742
## Thu Jul 14 00:54:39 CEST 2022 - The aliscore output appears to be empty. Check results for BUSCO: 367334at7742
## Thu Jul 14 00:54:40 CEST 2022 - The aliscore output file does not exist. Check results for BUSCO: 80914at7742
## Thu Jul 14 00:54:40 CEST 2022 - The aliscore output appears to be empty. Check results for BUSCO: 80914at7742
## Thu Jul 14 00:54:42 CEST 2022 - The aliscore output file does not exist. Check results for BUSCO: 127291at7742
## Thu Jul 14 00:54:42 CEST 2022 - The aliscore output appears to be empty. Check results for BUSCO: 127291at7742
## Thu Jul 14 00:54:42 CEST 2022 - The aliscore output file does not exist. Check results for BUSCO: 311908at7742
## Thu Jul 14 00:54:42 CEST 2022 - The aliscore output appears to be empty. Check results for BUSCO: 311908at7742
## Thu Jul 14 00:54:43 CEST 2022 - The aliscore output file does not exist. Check results for BUSCO: 123446at7742
## Thu Jul 14 00:54:43 CEST 2022 - The aliscore output appears to be empty. Check results for BUSCO: 123446at7742
## Thu Jul 14 00:54:45 CEST 2022 - The aliscore output file does not exist. Check results for BUSCO: 135770at7742
## Thu Jul 14 00:54:45 CEST 2022 - The aliscore output appears to be empty. Check results for BUSCO: 135770at7742
## Thu Jul 14 00:54:45 CEST 2022 - The aliscore output file does not exist. Check results for BUSCO: 249853at7742
## Thu Jul 14 00:54:45 CEST 2022 - The aliscore output appears to be empty. Check results for BUSCO: 249853at7742
## Thu Jul 14 00:54:49 CEST 2022 - The aliscore output file does not exist. Check results for BUSCO: 236607at7742
## Thu Jul 14 00:54:49 CEST 2022 - The aliscore output appears to be empty. Check results for BUSCO: 236607at7742
## Thu Jul 14 00:54:49 CEST 2022 - The aliscore output file does not exist. Check results for BUSCO: 100494at7742
## Thu Jul 14 00:54:49 CEST 2022 - The aliscore output appears to be empty. Check results for BUSCO: 100494at7742
## Thu Jul 14 00:54:50 CEST 2022 - The aliscore output file does not exist. Check results for BUSCO: 70039at7742
## Thu Jul 14 00:54:50 CEST 2022 - The aliscore output appears to be empty. Check results for BUSCO: 70039at7742
## Thu Jul 14 00:54:53 CEST 2022 - The aliscore output file does not exist. Check results for BUSCO: 28954at7742
## Thu Jul 14 00:54:53 CEST 2022 - The aliscore output appears to be empty. Check results for BUSCO: 28954at7742
## Thu Jul 14 00:54:53 CEST 2022 - The aliscore output file does not exist. Check results for BUSCO: 145962at7742
## Thu Jul 14 00:54:53 CEST 2022 - The aliscore output appears to be empty. Check results for BUSCO: 145962at7742
## Thu Jul 14 00:54:54 CEST 2022 - The aliscore output file does not exist. Check results for BUSCO: 221391at7742
## Thu Jul 14 00:54:54 CEST 2022 - The aliscore output appears to be empty. Check results for BUSCO: 221391at7742
## Thu Jul 14 00:54:55 CEST 2022 - The aliscore output file does not exist. Check results for BUSCO: 228455at7742
## Thu Jul 14 00:54:55 CEST 2022 - The aliscore output appears to be empty. Check results for BUSCO: 228455at7742
## Thu Jul 14 00:54:57 CEST 2022 - The aliscore output file does not exist. Check results for BUSCO: 116065at7742
## Thu Jul 14 00:54:57 CEST 2022 - The aliscore output appears to be empty. Check results for BUSCO: 116065at7742
## Thu Jul 14 00:54:59 CEST 2022 - The aliscore output file does not exist. Check results for BUSCO: 123847at7742
## Thu Jul 14 00:54:59 CEST 2022 - The aliscore output appears to be empty. Check results for BUSCO: 123847at7742
## Thu Jul 14 00:55:00 CEST 2022 - The aliscore output file does not exist. Check results for BUSCO: 189794at7742
## Thu Jul 14 00:55:00 CEST 2022 - The aliscore output appears to be empty. Check results for BUSCO: 189794at7742
## Thu Jul 14 00:55:00 CEST 2022 - The aliscore output file does not exist. Check results for BUSCO: 155845at7742
## Thu Jul 14 00:55:00 CEST 2022 - The aliscore output appears to be empty. Check results for BUSCO: 155845at7742
## Thu Jul 14 00:55:01 CEST 2022 - The aliscore output file does not exist. Check results for BUSCO: 115767at7742
## Thu Jul 14 00:55:01 CEST 2022 - The aliscore output appears to be empty. Check results for BUSCO: 115767at7742
## Thu Jul 14 00:55:02 CEST 2022 - The aliscore output file does not exist. Check results for BUSCO: 168479at7742
## Thu Jul 14 00:55:02 CEST 2022 - The aliscore output appears to be empty. Check results for BUSCO: 168479at7742
## Thu Jul 14 00:55:05 CEST 2022 - The aliscore output file does not exist. Check results for BUSCO: 153711at7742
## Thu Jul 14 00:55:05 CEST 2022 - The aliscore output appears to be empty. Check results for BUSCO: 153711at7742
## Thu Jul 14 00:55:06 CEST 2022 - The aliscore output file does not exist. Check results for BUSCO: 100952at7742
## Thu Jul 14 00:55:06 CEST 2022 - The aliscore output appears to be empty. Check results for BUSCO: 100952at7742
## Thu Jul 14 00:55:07 CEST 2022 - The aliscore output file does not exist. Check results for BUSCO: 150068at7742
## Thu Jul 14 00:55:07 CEST 2022 - The aliscore output appears to be empty. Check results for BUSCO: 150068at7742
## Thu Jul 14 00:55:07 CEST 2022 - The aliscore output file does not exist. Check results for BUSCO: 214498at7742
## Thu Jul 14 00:55:07 CEST 2022 - The aliscore output appears to be empty. Check results for BUSCO: 214498at7742
## Thu Jul 14 00:55:09 CEST 2022 - The aliscore output file does not exist. Check results for BUSCO: 295275at7742
## Thu Jul 14 00:55:09 CEST 2022 - The aliscore output appears to be empty. Check results for BUSCO: 295275at7742
## Thu Jul 14 00:55:11 CEST 2022 - The aliscore output file does not exist. Check results for BUSCO: 237620at7742
## Thu Jul 14 00:55:11 CEST 2022 - The aliscore output appears to be empty. Check results for BUSCO: 237620at7742
## Thu Jul 14 00:55:11 CEST 2022 - The aliscore output file does not exist. Check results for BUSCO: 117635at7742
## Thu Jul 14 00:55:11 CEST 2022 - The aliscore output appears to be empty. Check results for BUSCO: 117635at7742
## Thu Jul 14 00:55:14 CEST 2022 - The aliscore output file does not exist. Check results for BUSCO: 70039at7742
## Thu Jul 14 00:55:14 CEST 2022 - The aliscore output appears to be empty. Check results for BUSCO: 70039at7742
## Thu Jul 14 00:55:14 CEST 2022 - The aliscore output file does not exist. Check results for BUSCO: 95757at7742
## Thu Jul 14 00:55:14 CEST 2022 - The aliscore output appears to be empty. Check results for BUSCO: 95757at7742
## Thu Jul 14 00:55:16 CEST 2022 - The aliscore output file does not exist. Check results for BUSCO: 221197at7742
## Thu Jul 14 00:55:16 CEST 2022 - The aliscore output appears to be empty. Check results for BUSCO: 221197at7742
## Thu Jul 14 00:55:18 CEST 2022 - The aliscore output file does not exist. Check results for BUSCO: 83474at7742
## Thu Jul 14 00:55:18 CEST 2022 - The aliscore output appears to be empty. Check results for BUSCO: 83474at7742
## Thu Jul 14 00:55:19 CEST 2022 - The aliscore output file does not exist. Check results for BUSCO: 100952at7742
## Thu Jul 14 00:55:19 CEST 2022 - The aliscore output appears to be empty. Check results for BUSCO: 100952at7742
## Thu Jul 14 00:55:20 CEST 2022 - The aliscore output file does not exist. Check results for BUSCO: 99049at7742
## Thu Jul 14 00:55:20 CEST 2022 - The aliscore output appears to be empty. Check results for BUSCO: 99049at7742
## Thu Jul 14 00:55:21 CEST 2022 - The aliscore output file does not exist. Check results for BUSCO: 106009at7742
## Thu Jul 14 00:55:21 CEST 2022 - The aliscore output appears to be empty. Check results for BUSCO: 106009at7742
## Thu Jul 14 00:55:23 CEST 2022 - The aliscore output file does not exist. Check results for BUSCO: 129214at7742
## Thu Jul 14 00:55:23 CEST 2022 - The aliscore output appears to be empty. Check results for BUSCO: 129214at7742
## Thu Jul 14 00:55:24 CEST 2022 - The aliscore output file does not exist. Check results for BUSCO: 161467at7742
## Thu Jul 14 00:55:24 CEST 2022 - The aliscore output appears to be empty. Check results for BUSCO: 161467at7742
## Thu Jul 14 00:55:25 CEST 2022 - The aliscore output file does not exist. Check results for BUSCO: 42045at7742
## Thu Jul 14 00:55:25 CEST 2022 - The aliscore output appears to be empty. Check results for BUSCO: 42045at7742
## Thu Jul 14 00:55:28 CEST 2022 - The aliscore output file does not exist. Check results for BUSCO: 182480at7742
## Thu Jul 14 00:55:28 CEST 2022 - The aliscore output appears to be empty. Check results for BUSCO: 182480at7742
## Thu Jul 14 00:55:28 CEST 2022 - The aliscore output file does not exist. Check results for BUSCO: 145962at7742
## Thu Jul 14 00:55:28 CEST 2022 - The aliscore output appears to be empty. Check results for BUSCO: 145962at7742
## Thu Jul 14 00:55:29 CEST 2022 - The aliscore output file does not exist. Check results for BUSCO: 70131at7742
## Thu Jul 14 00:55:29 CEST 2022 - The aliscore output appears to be empty. Check results for BUSCO: 70131at7742
## Thu Jul 14 00:55:30 CEST 2022 - The aliscore output file does not exist. Check results for BUSCO: 86856at7742
## Thu Jul 14 00:55:30 CEST 2022 - The aliscore output appears to be empty. Check results for BUSCO: 86856at7742
## Thu Jul 14 00:55:31 CEST 2022 - The aliscore output file does not exist. Check results for BUSCO: 42045at7742
## Thu Jul 14 00:55:31 CEST 2022 - The aliscore output appears to be empty. Check results for BUSCO: 42045at7742
## Thu Jul 14 00:55:34 CEST 2022 - The aliscore output file does not exist. Check results for BUSCO: 210019at7742
## Thu Jul 14 00:55:34 CEST 2022 - The aliscore output appears to be empty. Check results for BUSCO: 210019at7742
## Thu Jul 14 00:55:34 CEST 2022 - The aliscore output file does not exist. Check results for BUSCO: 245053at7742
## Thu Jul 14 00:55:34 CEST 2022 - The aliscore output appears to be empty. Check results for BUSCO: 245053at7742
## Thu Jul 14 00:55:36 CEST 2022 - The aliscore output file does not exist. Check results for BUSCO: 122030at7742
## Thu Jul 14 00:55:36 CEST 2022 - The aliscore output appears to be empty. Check results for BUSCO: 122030at7742
## Thu Jul 14 00:55:38 CEST 2022 - The aliscore output file does not exist. Check results for BUSCO: 299207at7742
## Thu Jul 14 00:55:38 CEST 2022 - The aliscore output appears to be empty. Check results for BUSCO: 299207at7742
## Thu Jul 14 00:55:38 CEST 2022 - The aliscore output file does not exist. Check results for BUSCO: 78565at7742
## Thu Jul 14 00:55:38 CEST 2022 - The aliscore output appears to be empty. Check results for BUSCO: 78565at7742
## Thu Jul 14 00:55:39 CEST 2022 - The aliscore output file does not exist. Check results for BUSCO: 89615at7742
## Thu Jul 14 00:55:39 CEST 2022 - The aliscore output appears to be empty. Check results for BUSCO: 89615at7742
## Thu Jul 14 00:55:40 CEST 2022 - The aliscore output file does not exist. Check results for BUSCO: 184785at7742
## Thu Jul 14 00:55:40 CEST 2022 - The aliscore output appears to be empty. Check results for BUSCO: 184785at7742
## Thu Jul 14 00:55:42 CEST 2022 - The aliscore output file does not exist. Check results for BUSCO: 97039at7742
## Thu Jul 14 00:55:42 CEST 2022 - The aliscore output appears to be empty. Check results for BUSCO: 97039at7742
## Thu Jul 14 00:55:44 CEST 2022 - The aliscore output file does not exist. Check results for BUSCO: 300767at7742
## Thu Jul 14 00:55:44 CEST 2022 - The aliscore output appears to be empty. Check results for BUSCO: 300767at7742
## Thu Jul 14 00:55:46 CEST 2022 - The aliscore output file does not exist. Check results for BUSCO: 117499at7742
## Thu Jul 14 00:55:46 CEST 2022 - The aliscore output appears to be empty. Check results for BUSCO: 117499at7742
## Thu Jul 14 00:55:46 CEST 2022 - The aliscore output file does not exist. Check results for BUSCO: 351213at7742
## Thu Jul 14 00:55:46 CEST 2022 - The aliscore output appears to be empty. Check results for BUSCO: 351213at7742
## Thu Jul 14 00:55:48 CEST 2022 - The aliscore output file does not exist. Check results for BUSCO: 203744at7742
## Thu Jul 14 00:55:48 CEST 2022 - The aliscore output appears to be empty. Check results for BUSCO: 203744at7742
## Thu Jul 14 00:55:48 CEST 2022 - The aliscore output file does not exist. Check results for BUSCO: 125127at7742
## Thu Jul 14 00:55:48 CEST 2022 - The aliscore output appears to be empty. Check results for BUSCO: 125127at7742
## Thu Jul 14 00:55:50 CEST 2022 - The aliscore output file does not exist. Check results for BUSCO: 162155at7742
## Thu Jul 14 00:55:50 CEST 2022 - The aliscore output appears to be empty. Check results for BUSCO: 162155at7742
## Thu Jul 14 00:55:51 CEST 2022 - The aliscore output file does not exist. Check results for BUSCO: 75881at7742
## Thu Jul 14 00:55:51 CEST 2022 - The aliscore output appears to be empty. Check results for BUSCO: 75881at7742
## Thu Jul 14 00:55:53 CEST 2022 - The aliscore output file does not exist. Check results for BUSCO: 117635at7742
## Thu Jul 14 00:55:53 CEST 2022 - The aliscore output appears to be empty. Check results for BUSCO: 117635at7742
## Thu Jul 14 00:55:55 CEST 2022 - The aliscore output file does not exist. Check results for BUSCO: 73001at7742
## Thu Jul 14 00:55:55 CEST 2022 - The aliscore output appears to be empty. Check results for BUSCO: 73001at7742
## Thu Jul 14 00:55:55 CEST 2022 - The aliscore output file does not exist. Check results for BUSCO: 248515at7742
## Thu Jul 14 00:55:55 CEST 2022 - The aliscore output appears to be empty. Check results for BUSCO: 248515at7742
## Thu Jul 14 00:55:56 CEST 2022 - The aliscore output file does not exist. Check results for BUSCO: 155845at7742
## Thu Jul 14 00:55:56 CEST 2022 - The aliscore output appears to be empty. Check results for BUSCO: 155845at7742
## Thu Jul 14 00:55:56 CEST 2022 - The aliscore output file does not exist. Check results for BUSCO: 135517at7742
## Thu Jul 14 00:55:56 CEST 2022 - The aliscore output appears to be empty. Check results for BUSCO: 135517at7742
## Thu Jul 14 00:55:59 CEST 2022 - The aliscore output file does not exist. Check results for BUSCO: 329870at7742
## Thu Jul 14 00:55:59 CEST 2022 - The aliscore output appears to be empty. Check results for BUSCO: 329870at7742
## Thu Jul 14 00:56:00 CEST 2022 - The aliscore output file does not exist. Check results for BUSCO: 51955at7742
## Thu Jul 14 00:56:00 CEST 2022 - The aliscore output appears to be empty. Check results for BUSCO: 51955at7742
## Thu Jul 14 00:56:01 CEST 2022 - The aliscore output file does not exist. Check results for BUSCO: 129943at7742
## Thu Jul 14 00:56:01 CEST 2022 - The aliscore output appears to be empty. Check results for BUSCO: 129943at7742
## Thu Jul 14 00:56:03 CEST 2022 - The aliscore output file does not exist. Check results for BUSCO: 128396at7742
## Thu Jul 14 00:56:03 CEST 2022 - The aliscore output appears to be empty. Check results for BUSCO: 128396at7742
## Thu Jul 14 00:56:03 CEST 2022 - The aliscore output file does not exist. Check results for BUSCO: 405703at7742
## Thu Jul 14 00:56:03 CEST 2022 - The aliscore output appears to be empty. Check results for BUSCO: 405703at7742
## Thu Jul 14 00:56:04 CEST 2022 - The aliscore output file does not exist. Check results for BUSCO: 100623at7742
## Thu Jul 14 00:56:04 CEST 2022 - The aliscore output appears to be empty. Check results for BUSCO: 100623at7742
## Thu Jul 14 00:56:06 CEST 2022 - The aliscore output file does not exist. Check results for BUSCO: 100623at7742
## Thu Jul 14 00:56:06 CEST 2022 - The aliscore output appears to be empty. Check results for BUSCO: 100623at7742
## Thu Jul 14 00:56:06 CEST 2022 - The aliscore output file does not exist. Check results for BUSCO: 122963at7742
## Thu Jul 14 00:56:06 CEST 2022 - The aliscore output appears to be empty. Check results for BUSCO: 122963at7742
## Thu Jul 14 00:56:08 CEST 2022 - The aliscore output file does not exist. Check results for BUSCO: 95927at7742
## Thu Jul 14 00:56:08 CEST 2022 - The aliscore output appears to be empty. Check results for BUSCO: 95927at7742
## Thu Jul 14 00:56:09 CEST 2022 - The aliscore output file does not exist. Check results for BUSCO: 91624at7742
## Thu Jul 14 00:56:09 CEST 2022 - The aliscore output appears to be empty. Check results for BUSCO: 91624at7742
## Thu Jul 14 00:56:10 CEST 2022 - The aliscore output file does not exist. Check results for BUSCO: 249853at7742
## Thu Jul 14 00:56:10 CEST 2022 - The aliscore output appears to be empty. Check results for BUSCO: 249853at7742
## Thu Jul 14 00:56:13 CEST 2022 - The aliscore output file does not exist. Check results for BUSCO: 149377at7742
## Thu Jul 14 00:56:13 CEST 2022 - The aliscore output appears to be empty. Check results for BUSCO: 149377at7742
## Thu Jul 14 00:56:14 CEST 2022 - The aliscore output file does not exist. Check results for BUSCO: 223874at7742
## Thu Jul 14 00:56:14 CEST 2022 - The aliscore output appears to be empty. Check results for BUSCO: 223874at7742
## Thu Jul 14 00:56:14 CEST 2022 - The aliscore output file does not exist. Check results for BUSCO: 126306at7742
## Thu Jul 14 00:56:14 CEST 2022 - The aliscore output appears to be empty. Check results for BUSCO: 126306at7742
## Thu Jul 14 00:56:16 CEST 2022 - The aliscore output file does not exist. Check results for BUSCO: 373324at7742
## Thu Jul 14 00:56:16 CEST 2022 - The aliscore output appears to be empty. Check results for BUSCO: 373324at7742
## Thu Jul 14 00:56:17 CEST 2022 - The aliscore output file does not exist. Check results for BUSCO: 281891at7742
## Thu Jul 14 00:56:17 CEST 2022 - The aliscore output appears to be empty. Check results for BUSCO: 281891at7742
## Thu Jul 14 00:56:18 CEST 2022 - The aliscore output file does not exist. Check results for BUSCO: 206434at7742
## Thu Jul 14 00:56:18 CEST 2022 - The aliscore output appears to be empty. Check results for BUSCO: 206434at7742
## Thu Jul 14 00:56:20 CEST 2022 - The aliscore output file does not exist. Check results for BUSCO: 192234at7742
## Thu Jul 14 00:56:20 CEST 2022 - The aliscore output appears to be empty. Check results for BUSCO: 192234at7742
## Thu Jul 14 00:56:20 CEST 2022 - The aliscore output file does not exist. Check results for BUSCO: 28954at7742
## Thu Jul 14 00:56:20 CEST 2022 - The aliscore output appears to be empty. Check results for BUSCO: 28954at7742
## Thu Jul 14 00:56:23 CEST 2022 - The aliscore output file does not exist. Check results for BUSCO: 358181at7742
## Thu Jul 14 00:56:23 CEST 2022 - The aliscore output appears to be empty. Check results for BUSCO: 358181at7742
## Thu Jul 14 00:56:25 CEST 2022 - The aliscore output file does not exist. Check results for BUSCO: 127929at7742
## Thu Jul 14 00:56:25 CEST 2022 - The aliscore output appears to be empty. Check results for BUSCO: 127929at7742
## Thu Jul 14 00:56:25 CEST 2022 - The aliscore output file does not exist. Check results for BUSCO: 79086at7742
## Thu Jul 14 00:56:25 CEST 2022 - The aliscore output appears to be empty. Check results for BUSCO: 79086at7742
## Thu Jul 14 00:56:26 CEST 2022 - The aliscore output file does not exist. Check results for BUSCO: 75797at7742
## Thu Jul 14 00:56:26 CEST 2022 - The aliscore output appears to be empty. Check results for BUSCO: 75797at7742
## Thu Jul 14 00:56:28 CEST 2022 - The aliscore output file does not exist. Check results for BUSCO: 347981at7742
## Thu Jul 14 00:56:28 CEST 2022 - The aliscore output appears to be empty. Check results for BUSCO: 347981at7742
## Thu Jul 14 00:56:28 CEST 2022 - The aliscore output file does not exist. Check results for BUSCO: 123872at7742
## Thu Jul 14 00:56:28 CEST 2022 - The aliscore output appears to be empty. Check results for BUSCO: 123872at7742
## Thu Jul 14 00:56:30 CEST 2022 - The aliscore output file does not exist. Check results for BUSCO: 96933at7742
## Thu Jul 14 00:56:30 CEST 2022 - The aliscore output appears to be empty. Check results for BUSCO: 96933at7742
## Thu Jul 14 00:56:31 CEST 2022 - The aliscore output file does not exist. Check results for BUSCO: 327742at7742
## Thu Jul 14 00:56:31 CEST 2022 - The aliscore output appears to be empty. Check results for BUSCO: 327742at7742
## Thu Jul 14 00:56:32 CEST 2022 - The aliscore output file does not exist. Check results for BUSCO: 119040at7742
## Thu Jul 14 00:56:32 CEST 2022 - The aliscore output appears to be empty. Check results for BUSCO: 119040at7742
## Thu Jul 14 00:56:33 CEST 2022 - The aliscore output file does not exist. Check results for BUSCO: 87193at7742
## Thu Jul 14 00:56:33 CEST 2022 - The aliscore output appears to be empty. Check results for BUSCO: 87193at7742
## Thu Jul 14 00:56:34 CEST 2022 - The aliscore output file does not exist. Check results for BUSCO: 319001at7742
## Thu Jul 14 00:56:34 CEST 2022 - The aliscore output appears to be empty. Check results for BUSCO: 319001at7742
## Thu Jul 14 00:56:37 CEST 2022 - The aliscore output file does not exist. Check results for BUSCO: 424377at7742
## Thu Jul 14 00:56:37 CEST 2022 - The aliscore output appears to be empty. Check results for BUSCO: 424377at7742
## Thu Jul 14 00:56:39 CEST 2022 - The aliscore output file does not exist. Check results for BUSCO: 184785at7742
## Thu Jul 14 00:56:39 CEST 2022 - The aliscore output appears to be empty. Check results for BUSCO: 184785at7742
## Thu Jul 14 00:56:39 CEST 2022 - The aliscore output file does not exist. Check results for BUSCO: 108967at7742
## Thu Jul 14 00:56:39 CEST 2022 - The aliscore output appears to be empty. Check results for BUSCO: 108967at7742
## Thu Jul 14 00:56:40 CEST 2022 - The aliscore output file does not exist. Check results for BUSCO: 53843at7742
## Thu Jul 14 00:56:40 CEST 2022 - The aliscore output appears to be empty. Check results for BUSCO: 53843at7742
## Thu Jul 14 00:56:41 CEST 2022 - The aliscore output file does not exist. Check results for BUSCO: 117499at7742
## Thu Jul 14 00:56:41 CEST 2022 - The aliscore output appears to be empty. Check results for BUSCO: 117499at7742
## Thu Jul 14 00:56:42 CEST 2022 - The aliscore output file does not exist. Check results for BUSCO: 151777at7742
## Thu Jul 14 00:56:43 CEST 2022 - The aliscore output appears to be empty. Check results for BUSCO: 151777at7742
## Thu Jul 14 00:56:43 CEST 2022 - The aliscore output file does not exist. Check results for BUSCO: 51573at7742
## Thu Jul 14 00:56:43 CEST 2022 - The aliscore output appears to be empty. Check results for BUSCO: 51573at7742
## Thu Jul 14 00:56:46 CEST 2022 - The aliscore output file does not exist. Check results for BUSCO: 225564at7742
## Thu Jul 14 00:56:46 CEST 2022 - The aliscore output appears to be empty. Check results for BUSCO: 225564at7742
## Thu Jul 14 00:56:47 CEST 2022 - The aliscore output file does not exist. Check results for BUSCO: 424377at7742
## Thu Jul 14 00:56:47 CEST 2022 - The aliscore output appears to be empty. Check results for BUSCO: 424377at7742
## Thu Jul 14 00:56:47 CEST 2022 - The aliscore output file does not exist. Check results for BUSCO: 211156at7742
## Thu Jul 14 00:56:47 CEST 2022 - The aliscore output appears to be empty. Check results for BUSCO: 211156at7742
## Thu Jul 14 00:56:50 CEST 2022 - The aliscore output file does not exist. Check results for BUSCO: 127830at7742
## Thu Jul 14 00:56:50 CEST 2022 - The aliscore output appears to be empty. Check results for BUSCO: 127830at7742
## Thu Jul 14 00:56:51 CEST 2022 - The aliscore output file does not exist. Check results for BUSCO: 149552at7742
## Thu Jul 14 00:56:51 CEST 2022 - The aliscore output appears to be empty. Check results for BUSCO: 149552at7742
## Thu Jul 14 00:56:52 CEST 2022 - The aliscore output file does not exist. Check results for BUSCO: 189794at7742
## Thu Jul 14 00:56:52 CEST 2022 - The aliscore output appears to be empty. Check results for BUSCO: 189794at7742
## Thu Jul 14 00:56:53 CEST 2022 - The aliscore output file does not exist. Check results for BUSCO: 367334at7742
## Thu Jul 14 00:56:53 CEST 2022 - The aliscore output appears to be empty. Check results for BUSCO: 367334at7742
## Thu Jul 14 00:56:54 CEST 2022 - The aliscore output file does not exist. Check results for BUSCO: 103919at7742
## Thu Jul 14 00:56:54 CEST 2022 - The aliscore output appears to be empty. Check results for BUSCO: 103919at7742
## Thu Jul 14 00:56:55 CEST 2022 - The aliscore output file does not exist. Check results for BUSCO: 91624at7742
## Thu Jul 14 00:56:55 CEST 2022 - The aliscore output appears to be empty. Check results for BUSCO: 91624at7742
## Thu Jul 14 00:56:55 CEST 2022 - The aliscore output file does not exist. Check results for BUSCO: 146337at7742
## Thu Jul 14 00:56:56 CEST 2022 - The aliscore output appears to be empty. Check results for BUSCO: 146337at7742
## Thu Jul 14 00:56:58 CEST 2022 - The aliscore output file does not exist. Check results for BUSCO: 226571at7742
## Thu Jul 14 00:56:58 CEST 2022 - The aliscore output appears to be empty. Check results for BUSCO: 226571at7742
## Thu Jul 14 00:57:00 CEST 2022 - The aliscore output file does not exist. Check results for BUSCO: 87423at7742
## Thu Jul 14 00:57:00 CEST 2022 - The aliscore output appears to be empty. Check results for BUSCO: 87423at7742
## Thu Jul 14 00:57:00 CEST 2022 - The aliscore output file does not exist. Check results for BUSCO: 162426at7742
## Thu Jul 14 00:57:00 CEST 2022 - The aliscore output appears to be empty. Check results for BUSCO: 162426at7742
## Thu Jul 14 00:57:01 CEST 2022 - The aliscore output file does not exist. Check results for BUSCO: 75464at7742
## Thu Jul 14 00:57:01 CEST 2022 - The aliscore output appears to be empty. Check results for BUSCO: 75464at7742
## Thu Jul 14 00:57:02 CEST 2022 - The aliscore output file does not exist. Check results for BUSCO: 254316at7742
## Thu Jul 14 00:57:02 CEST 2022 - The aliscore output appears to be empty. Check results for BUSCO: 254316at7742
## Thu Jul 14 00:57:04 CEST 2022 - The aliscore output file does not exist. Check results for BUSCO: 67119at7742
## Thu Jul 14 00:57:04 CEST 2022 - The aliscore output appears to be empty. Check results for BUSCO: 67119at7742
## Thu Jul 14 00:57:05 CEST 2022 - The aliscore output file does not exist. Check results for BUSCO: 127291at7742
## Thu Jul 14 00:57:05 CEST 2022 - The aliscore output appears to be empty. Check results for BUSCO: 127291at7742
## Thu Jul 14 00:57:08 CEST 2022 - The aliscore output file does not exist. Check results for BUSCO: 233490at7742
## Thu Jul 14 00:57:09 CEST 2022 - The aliscore output appears to be empty. Check results for BUSCO: 233490at7742
## Thu Jul 14 00:57:09 CEST 2022 - The aliscore output file does not exist. Check results for BUSCO: 143938at7742
## Thu Jul 14 00:57:09 CEST 2022 - The aliscore output appears to be empty. Check results for BUSCO: 143938at7742
## Thu Jul 14 00:57:10 CEST 2022 - The aliscore output file does not exist. Check results for BUSCO: 204919at7742
## Thu Jul 14 00:57:10 CEST 2022 - The aliscore output appears to be empty. Check results for BUSCO: 204919at7742
## Thu Jul 14 00:57:10 CEST 2022 - The aliscore output file does not exist. Check results for BUSCO: 95927at7742
## Thu Jul 14 00:57:11 CEST 2022 - The aliscore output appears to be empty. Check results for BUSCO: 95927at7742
## Thu Jul 14 00:57:13 CEST 2022 - The aliscore output file does not exist. Check results for BUSCO: 94359at7742
## Thu Jul 14 00:57:13 CEST 2022 - The aliscore output appears to be empty. Check results for BUSCO: 94359at7742
## Thu Jul 14 00:57:14 CEST 2022 - The aliscore output file does not exist. Check results for BUSCO: 161467at7742
## Thu Jul 14 00:57:14 CEST 2022 - The aliscore output appears to be empty. Check results for BUSCO: 161467at7742
## Thu Jul 14 00:57:16 CEST 2022 - The aliscore output file does not exist. Check results for BUSCO: 176579at7742
## Thu Jul 14 00:57:16 CEST 2022 - The aliscore output appears to be empty. Check results for BUSCO: 176579at7742
## Thu Jul 14 00:57:17 CEST 2022 - The aliscore output file does not exist. Check results for BUSCO: 299207at7742
## Thu Jul 14 00:57:17 CEST 2022 - The aliscore output appears to be empty. Check results for BUSCO: 299207at7742
## Thu Jul 14 00:57:18 CEST 2022 - The aliscore output file does not exist. Check results for BUSCO: 173102at7742
## Thu Jul 14 00:57:18 CEST 2022 - The aliscore output appears to be empty. Check results for BUSCO: 173102at7742
## Thu Jul 14 00:57:19 CEST 2022 - The aliscore output file does not exist. Check results for BUSCO: 155845at7742
## Thu Jul 14 00:57:19 CEST 2022 - The aliscore output appears to be empty. Check results for BUSCO: 155845at7742
## Thu Jul 14 00:57:19 CEST 2022 - The aliscore output file does not exist. Check results for BUSCO: 275273at7742
## Thu Jul 14 00:57:19 CEST 2022 - The aliscore output appears to be empty. Check results for BUSCO: 275273at7742
## Thu Jul 14 00:57:21 CEST 2022 - The aliscore output file does not exist. Check results for BUSCO: 65316at7742
## Thu Jul 14 00:57:21 CEST 2022 - The aliscore output appears to be empty. Check results for BUSCO: 65316at7742
## Thu Jul 14 00:57:23 CEST 2022 - The aliscore output file does not exist. Check results for BUSCO: 137727at7742
## Thu Jul 14 00:57:23 CEST 2022 - The aliscore output appears to be empty. Check results for BUSCO: 137727at7742
## Thu Jul 14 00:57:24 CEST 2022 - The aliscore output file does not exist. Check results for BUSCO: 192114at7742
## Thu Jul 14 00:57:25 CEST 2022 - The aliscore output appears to be empty. Check results for BUSCO: 192114at7742
## Thu Jul 14 00:57:26 CEST 2022 - The aliscore output file does not exist. Check results for BUSCO: 145946at7742
## Thu Jul 14 00:57:26 CEST 2022 - The aliscore output appears to be empty. Check results for BUSCO: 145946at7742
## Thu Jul 14 00:57:30 CEST 2022 - The aliscore output file does not exist. Check results for BUSCO: 177660at7742
## Thu Jul 14 00:57:30 CEST 2022 - The aliscore output appears to be empty. Check results for BUSCO: 177660at7742
## Thu Jul 14 00:57:30 CEST 2022 - The aliscore output file does not exist. Check results for BUSCO: 122841at7742
## Thu Jul 14 00:57:30 CEST 2022 - The aliscore output appears to be empty. Check results for BUSCO: 122841at7742
## Thu Jul 14 00:57:30 CEST 2022 - The aliscore output file does not exist. Check results for BUSCO: 132224at7742
## Thu Jul 14 00:57:30 CEST 2022 - The aliscore output appears to be empty. Check results for BUSCO: 132224at7742
## Thu Jul 14 00:57:31 CEST 2022 - The aliscore output file does not exist. Check results for BUSCO: 191192at7742
## Thu Jul 14 00:57:31 CEST 2022 - The aliscore output appears to be empty. Check results for BUSCO: 191192at7742
## Thu Jul 14 00:57:33 CEST 2022 - The aliscore output file does not exist. Check results for BUSCO: 73001at7742
## Thu Jul 14 00:57:33 CEST 2022 - The aliscore output appears to be empty. Check results for BUSCO: 73001at7742
## Thu Jul 14 00:57:33 CEST 2022 - The aliscore output file does not exist. Check results for BUSCO: 367334at7742
## Thu Jul 14 00:57:33 CEST 2022 - The aliscore output appears to be empty. Check results for BUSCO: 367334at7742
## Thu Jul 14 00:57:34 CEST 2022 - The aliscore output file does not exist. Check results for BUSCO: 149552at7742
## Thu Jul 14 00:57:34 CEST 2022 - The aliscore output appears to be empty. Check results for BUSCO: 149552at7742
## Thu Jul 14 00:57:38 CEST 2022 - The aliscore output file does not exist. Check results for BUSCO: 82295at7742
## Thu Jul 14 00:57:38 CEST 2022 - The aliscore output appears to be empty. Check results for BUSCO: 82295at7742
## Thu Jul 14 00:57:38 CEST 2022 - The aliscore output file does not exist. Check results for BUSCO: 149083at7742
## Thu Jul 14 00:57:38 CEST 2022 - The aliscore output appears to be empty. Check results for BUSCO: 149083at7742
## Thu Jul 14 00:57:38 CEST 2022 - The aliscore output file does not exist. Check results for BUSCO: 194866at7742
## Thu Jul 14 00:57:38 CEST 2022 - The aliscore output appears to be empty. Check results for BUSCO: 194866at7742
## Thu Jul 14 00:57:39 CEST 2022 - The aliscore output file does not exist. Check results for BUSCO: 129943at7742
## Thu Jul 14 00:57:39 CEST 2022 - The aliscore output appears to be empty. Check results for BUSCO: 129943at7742
## Thu Jul 14 00:57:39 CEST 2022 - The aliscore output file does not exist. Check results for BUSCO: 107664at7742
## Thu Jul 14 00:57:39 CEST 2022 - The aliscore output appears to be empty. Check results for BUSCO: 107664at7742
## Thu Jul 14 00:57:41 CEST 2022 - The aliscore output file does not exist. Check results for BUSCO: 293720at7742
## Thu Jul 14 00:57:41 CEST 2022 - The aliscore output appears to be empty. Check results for BUSCO: 293720at7742
## Thu Jul 14 00:57:43 CEST 2022 - The aliscore output file does not exist. Check results for BUSCO: 161467at7742
## Thu Jul 14 00:57:43 CEST 2022 - The aliscore output appears to be empty. Check results for BUSCO: 161467at7742
## Thu Jul 14 00:57:45 CEST 2022 - The aliscore output file does not exist. Check results for BUSCO: 113847at7742
## Thu Jul 14 00:57:45 CEST 2022 - The aliscore output appears to be empty. Check results for BUSCO: 113847at7742
## Thu Jul 14 00:57:47 CEST 2022 - The aliscore output file does not exist. Check results for BUSCO: 65316at7742
## Thu Jul 14 00:57:47 CEST 2022 - The aliscore output appears to be empty. Check results for BUSCO: 65316at7742
## Thu Jul 14 00:57:47 CEST 2022 - The aliscore output file does not exist. Check results for BUSCO: 112360at7742
## Thu Jul 14 00:57:47 CEST 2022 - The aliscore output appears to be empty. Check results for BUSCO: 112360at7742
## Thu Jul 14 00:57:48 CEST 2022 - The aliscore output file does not exist. Check results for BUSCO: 184910at7742
## Thu Jul 14 00:57:48 CEST 2022 - The aliscore output appears to be empty. Check results for BUSCO: 184910at7742
## Thu Jul 14 00:57:48 CEST 2022 - The aliscore output file does not exist. Check results for BUSCO: 404307at7742
## Thu Jul 14 00:57:48 CEST 2022 - The aliscore output appears to be empty. Check results for BUSCO: 404307at7742
## Thu Jul 14 00:57:51 CEST 2022 - The aliscore output file does not exist. Check results for BUSCO: 75797at7742
## Thu Jul 14 00:57:51 CEST 2022 - The aliscore output appears to be empty. Check results for BUSCO: 75797at7742
## Thu Jul 14 00:57:53 CEST 2022 - The aliscore output file does not exist. Check results for BUSCO: 75797at7742
## Thu Jul 14 00:57:53 CEST 2022 - The aliscore output appears to be empty. Check results for BUSCO: 75797at7742
## Thu Jul 14 00:57:54 CEST 2022 - The aliscore output file does not exist. Check results for BUSCO: 160275at7742
## Thu Jul 14 00:57:54 CEST 2022 - The aliscore output appears to be empty. Check results for BUSCO: 160275at7742
## Thu Jul 14 00:57:54 CEST 2022 - The aliscore output file does not exist. Check results for BUSCO: 143938at7742
## Thu Jul 14 00:57:54 CEST 2022 - The aliscore output appears to be empty. Check results for BUSCO: 143938at7742
## Thu Jul 14 00:57:56 CEST 2022 - The aliscore output file does not exist. Check results for BUSCO: 212288at7742
## Thu Jul 14 00:57:57 CEST 2022 - The aliscore output appears to be empty. Check results for BUSCO: 212288at7742
## Thu Jul 14 00:57:57 CEST 2022 - The aliscore output file does not exist. Check results for BUSCO: 116065at7742
## Thu Jul 14 00:57:57 CEST 2022 - The aliscore output appears to be empty. Check results for BUSCO: 116065at7742
## Thu Jul 14 00:57:57 CEST 2022 - The aliscore output file does not exist. Check results for BUSCO: 123872at7742
## Thu Jul 14 00:57:57 CEST 2022 - The aliscore output appears to be empty. Check results for BUSCO: 123872at7742
## Thu Jul 14 00:57:58 CEST 2022 - The aliscore output file does not exist. Check results for BUSCO: 283449at7742
## Thu Jul 14 00:57:58 CEST 2022 - The aliscore output appears to be empty. Check results for BUSCO: 283449at7742
## Thu Jul 14 00:58:01 CEST 2022 - The aliscore output file does not exist. Check results for BUSCO: 94359at7742
## Thu Jul 14 00:58:01 CEST 2022 - The aliscore output appears to be empty. Check results for BUSCO: 94359at7742
## Thu Jul 14 00:58:02 CEST 2022 - The aliscore output file does not exist. Check results for BUSCO: 108033at7742
## Thu Jul 14 00:58:02 CEST 2022 - The aliscore output appears to be empty. Check results for BUSCO: 108033at7742
## Thu Jul 14 00:58:05 CEST 2022 - The aliscore output file does not exist. Check results for BUSCO: 268237at7742
## Thu Jul 14 00:58:05 CEST 2022 - The aliscore output appears to be empty. Check results for BUSCO: 268237at7742
## Thu Jul 14 00:58:06 CEST 2022 - The aliscore output file does not exist. Check results for BUSCO: 130375at7742
## Thu Jul 14 00:58:06 CEST 2022 - The aliscore output appears to be empty. Check results for BUSCO: 130375at7742
## Thu Jul 14 00:58:06 CEST 2022 - The aliscore output file does not exist. Check results for BUSCO: 116245at7742
## Thu Jul 14 00:58:06 CEST 2022 - The aliscore output appears to be empty. Check results for BUSCO: 116245at7742
## Thu Jul 14 00:58:06 CEST 2022 - The aliscore output file does not exist. Check results for BUSCO: 130161at7742
## Thu Jul 14 00:58:06 CEST 2022 - The aliscore output appears to be empty. Check results for BUSCO: 130161at7742
## Thu Jul 14 00:58:07 CEST 2022 - The aliscore output file does not exist. Check results for BUSCO: 82295at7742
## Thu Jul 14 00:58:07 CEST 2022 - The aliscore output appears to be empty. Check results for BUSCO: 82295at7742
## Thu Jul 14 00:58:09 CEST 2022 - The aliscore output file does not exist. Check results for BUSCO: 134679at7742
## Thu Jul 14 00:58:09 CEST 2022 - The aliscore output appears to be empty. Check results for BUSCO: 134679at7742
## Thu Jul 14 00:58:10 CEST 2022 - The aliscore output file does not exist. Check results for BUSCO: 132931at7742
## Thu Jul 14 00:58:10 CEST 2022 - The aliscore output appears to be empty. Check results for BUSCO: 132931at7742
## Thu Jul 14 00:58:12 CEST 2022 - The aliscore output file does not exist. Check results for BUSCO: 192234at7742
## Thu Jul 14 00:58:12 CEST 2022 - The aliscore output appears to be empty. Check results for BUSCO: 192234at7742
## Thu Jul 14 00:58:13 CEST 2022 - The aliscore output file does not exist. Check results for BUSCO: 213224at7742
## Thu Jul 14 00:58:13 CEST 2022 - The aliscore output appears to be empty. Check results for BUSCO: 213224at7742
## Thu Jul 14 00:58:16 CEST 2022 - The aliscore output file does not exist. Check results for BUSCO: 254316at7742
## Thu Jul 14 00:58:16 CEST 2022 - The aliscore output appears to be empty. Check results for BUSCO: 254316at7742
## Thu Jul 14 00:58:16 CEST 2022 - The aliscore output file does not exist. Check results for BUSCO: 63057at7742
## Thu Jul 14 00:58:16 CEST 2022 - The aliscore output appears to be empty. Check results for BUSCO: 63057at7742
## Thu Jul 14 00:58:16 CEST 2022 - The aliscore output file does not exist. Check results for BUSCO: 87114at7742
## Thu Jul 14 00:58:16 CEST 2022 - The aliscore output appears to be empty. Check results for BUSCO: 87114at7742
## Thu Jul 14 00:58:19 CEST 2022 - The aliscore output file does not exist. Check results for BUSCO: 94359at7742
## Thu Jul 14 00:58:19 CEST 2022 - The aliscore output appears to be empty. Check results for BUSCO: 94359at7742
## Thu Jul 14 00:58:20 CEST 2022 - The aliscore output file does not exist. Check results for BUSCO: 325335at7742
## Thu Jul 14 00:58:20 CEST 2022 - The aliscore output appears to be empty. Check results for BUSCO: 325335at7742
## Thu Jul 14 00:58:21 CEST 2022 - The aliscore output file does not exist. Check results for BUSCO: 313611at7742
## Thu Jul 14 00:58:21 CEST 2022 - The aliscore output appears to be empty. Check results for BUSCO: 313611at7742
## Thu Jul 14 00:58:24 CEST 2022 - The aliscore output file does not exist. Check results for BUSCO: 319001at7742
## Thu Jul 14 00:58:24 CEST 2022 - The aliscore output appears to be empty. Check results for BUSCO: 319001at7742
## Thu Jul 14 00:58:25 CEST 2022 - The aliscore output file does not exist. Check results for BUSCO: 190917at7742
## Thu Jul 14 00:58:25 CEST 2022 - The aliscore output appears to be empty. Check results for BUSCO: 190917at7742
## Thu Jul 14 00:58:25 CEST 2022 - The aliscore output file does not exist. Check results for BUSCO: 160275at7742
## Thu Jul 14 00:58:25 CEST 2022 - The aliscore output appears to be empty. Check results for BUSCO: 160275at7742
## Thu Jul 14 00:58:25 CEST 2022 - The aliscore output file does not exist. Check results for BUSCO: 202578at7742
## Thu Jul 14 00:58:25 CEST 2022 - The aliscore output appears to be empty. Check results for BUSCO: 202578at7742
## Thu Jul 14 00:58:27 CEST 2022 - The aliscore output file does not exist. Check results for BUSCO: 319001at7742
## Thu Jul 14 00:58:27 CEST 2022 - The aliscore output appears to be empty. Check results for BUSCO: 319001at7742
## Thu Jul 14 00:58:29 CEST 2022 - The aliscore output file does not exist. Check results for BUSCO: 358171at7742
## Thu Jul 14 00:58:29 CEST 2022 - The aliscore output appears to be empty. Check results for BUSCO: 358171at7742
## Thu Jul 14 00:58:31 CEST 2022 - The aliscore output file does not exist. Check results for BUSCO: 153711at7742
## Thu Jul 14 00:58:31 CEST 2022 - The aliscore output appears to be empty. Check results for BUSCO: 153711at7742
## Thu Jul 14 00:58:34 CEST 2022 - The aliscore output file does not exist. Check results for BUSCO: 135349at7742
## Thu Jul 14 00:58:34 CEST 2022 - The aliscore output file does not exist. Check results for BUSCO: 149552at7742
## Thu Jul 14 00:58:34 CEST 2022 - The aliscore output appears to be empty. Check results for BUSCO: 135349at7742
## Thu Jul 14 00:58:34 CEST 2022 - The aliscore output appears to be empty. Check results for BUSCO: 149552at7742
## Thu Jul 14 00:58:35 CEST 2022 - The aliscore output file does not exist. Check results for BUSCO: 63057at7742
## Thu Jul 14 00:58:35 CEST 2022 - The aliscore output appears to be empty. Check results for BUSCO: 63057at7742
## Thu Jul 14 00:58:35 CEST 2022 - The aliscore output file does not exist. Check results for BUSCO: 109650at7742
## Thu Jul 14 00:58:35 CEST 2022 - The aliscore output appears to be empty. Check results for BUSCO: 109650at7742
## Thu Jul 14 00:58:37 CEST 2022 - The aliscore output file does not exist. Check results for BUSCO: 358171at7742
## Thu Jul 14 00:58:37 CEST 2022 - The aliscore output appears to be empty. Check results for BUSCO: 358171at7742
## Thu Jul 14 00:58:37 CEST 2022 - The aliscore output file does not exist. Check results for BUSCO: 221391at7742
## Thu Jul 14 00:58:37 CEST 2022 - The aliscore output appears to be empty. Check results for BUSCO: 221391at7742
## Thu Jul 14 00:58:39 CEST 2022 - The aliscore output file does not exist. Check results for BUSCO: 192234at7742
## Thu Jul 14 00:58:39 CEST 2022 - The aliscore output appears to be empty. Check results for BUSCO: 192234at7742
## Thu Jul 14 00:58:40 CEST 2022 - The aliscore output file does not exist. Check results for BUSCO: 106313at7742
## Thu Jul 14 00:58:41 CEST 2022 - The aliscore output appears to be empty. Check results for BUSCO: 106313at7742
## Thu Jul 14 00:58:44 CEST 2022 - The aliscore output file does not exist. Check results for BUSCO: 176119at7742
## Thu Jul 14 00:58:44 CEST 2022 - The aliscore output appears to be empty. Check results for BUSCO: 176119at7742
## Thu Jul 14 00:58:44 CEST 2022 - The aliscore output file does not exist. Check results for BUSCO: 140310at7742
## Thu Jul 14 00:58:44 CEST 2022 - The aliscore output appears to be empty. Check results for BUSCO: 140310at7742
## Thu Jul 14 00:58:44 CEST 2022 - The aliscore output file does not exist. Check results for BUSCO: 168479at7742
## Thu Jul 14 00:58:44 CEST 2022 - The aliscore output appears to be empty. Check results for BUSCO: 168479at7742
## Thu Jul 14 00:58:45 CEST 2022 - The aliscore output file does not exist. Check results for BUSCO: 226571at7742
## Thu Jul 14 00:58:45 CEST 2022 - The aliscore output appears to be empty. Check results for BUSCO: 226571at7742
## Thu Jul 14 00:58:46 CEST 2022 - The aliscore output file does not exist. Check results for BUSCO: 173159at7742
## Thu Jul 14 00:58:47 CEST 2022 - The aliscore output appears to be empty. Check results for BUSCO: 173159at7742
## Thu Jul 14 00:58:49 CEST 2022 - The aliscore output file does not exist. Check results for BUSCO: 183360at7742
## Thu Jul 14 00:58:49 CEST 2022 - The aliscore output appears to be empty. Check results for BUSCO: 183360at7742
## Thu Jul 14 00:58:50 CEST 2022 - The aliscore output file does not exist. Check results for BUSCO: 272309at7742
## Thu Jul 14 00:58:50 CEST 2022 - The aliscore output appears to be empty. Check results for BUSCO: 272309at7742
## Thu Jul 14 00:58:52 CEST 2022 - The aliscore output file does not exist. Check results for BUSCO: 96764at7742
## Thu Jul 14 00:58:52 CEST 2022 - The aliscore output appears to be empty. Check results for BUSCO: 96764at7742
## Thu Jul 14 00:58:53 CEST 2022 - The aliscore output file does not exist. Check results for BUSCO: 221197at7742
## Thu Jul 14 00:58:53 CEST 2022 - The aliscore output file does not exist. Check results for BUSCO: 80914at7742
## Thu Jul 14 00:58:53 CEST 2022 - The aliscore output appears to be empty. Check results for BUSCO: 221197at7742
## Thu Jul 14 00:58:53 CEST 2022 - The aliscore output appears to be empty. Check results for BUSCO: 80914at7742
## Thu Jul 14 00:58:56 CEST 2022 - The aliscore output file does not exist. Check results for BUSCO: 311908at7742
## Thu Jul 14 00:58:56 CEST 2022 - The aliscore output appears to be empty. Check results for BUSCO: 311908at7742
## Thu Jul 14 00:58:57 CEST 2022 - The aliscore output file does not exist. Check results for BUSCO: 80791at7742
## Thu Jul 14 00:58:57 CEST 2022 - The aliscore output appears to be empty. Check results for BUSCO: 80791at7742
## Thu Jul 14 00:58:59 CEST 2022 - The aliscore output file does not exist. Check results for BUSCO: 75464at7742
## Thu Jul 14 00:58:59 CEST 2022 - The aliscore output appears to be empty. Check results for BUSCO: 75464at7742
## Thu Jul 14 00:58:59 CEST 2022 - The aliscore output file does not exist. Check results for BUSCO: 280447at7742
## Thu Jul 14 00:58:59 CEST 2022 - The aliscore output appears to be empty. Check results for BUSCO: 280447at7742
## Thu Jul 14 00:59:01 CEST 2022 - The aliscore output file does not exist. Check results for BUSCO: 126566at7742
## Thu Jul 14 00:59:01 CEST 2022 - The aliscore output appears to be empty. Check results for BUSCO: 126566at7742
## Thu Jul 14 00:59:01 CEST 2022 - The aliscore output file does not exist. Check results for BUSCO: 132972at7742
## Thu Jul 14 00:59:01 CEST 2022 - The aliscore output appears to be empty. Check results for BUSCO: 132972at7742
## Thu Jul 14 00:59:02 CEST 2022 - The aliscore output file does not exist. Check results for BUSCO: 245053at7742
## Thu Jul 14 00:59:02 CEST 2022 - The aliscore output appears to be empty. Check results for BUSCO: 245053at7742
## Thu Jul 14 00:59:04 CEST 2022 - The aliscore output file does not exist. Check results for BUSCO: 66866at7742
## Thu Jul 14 00:59:04 CEST 2022 - The aliscore output appears to be empty. Check results for BUSCO: 66866at7742
## Thu Jul 14 00:59:05 CEST 2022 - The aliscore output file does not exist. Check results for BUSCO: 89615at7742
## Thu Jul 14 00:59:05 CEST 2022 - The aliscore output appears to be empty. Check results for BUSCO: 89615at7742
## Thu Jul 14 00:59:07 CEST 2022 - The aliscore output file does not exist. Check results for BUSCO: 95261at7742
## Thu Jul 14 00:59:07 CEST 2022 - The aliscore output appears to be empty. Check results for BUSCO: 95261at7742
## Thu Jul 14 00:59:08 CEST 2022 - The aliscore output file does not exist. Check results for BUSCO: 187791at7742
## Thu Jul 14 00:59:08 CEST 2022 - The aliscore output appears to be empty. Check results for BUSCO: 187791at7742
## Thu Jul 14 00:59:10 CEST 2022 - The aliscore output file does not exist. Check results for BUSCO: 51955at7742
## Thu Jul 14 00:59:10 CEST 2022 - The aliscore output appears to be empty. Check results for BUSCO: 51955at7742
## Thu Jul 14 00:59:11 CEST 2022 - The aliscore output file does not exist. Check results for BUSCO: 105001at7742
## Thu Jul 14 00:59:11 CEST 2022 - The aliscore output appears to be empty. Check results for BUSCO: 105001at7742
## Thu Jul 14 00:59:11 CEST 2022 - The aliscore output file does not exist. Check results for BUSCO: 105001at7742
## Thu Jul 14 00:59:11 CEST 2022 - The aliscore output appears to be empty. Check results for BUSCO: 105001at7742
## Thu Jul 14 00:59:11 CEST 2022 - The aliscore output file does not exist. Check results for BUSCO: 290897at7742
## Thu Jul 14 00:59:11 CEST 2022 - The aliscore output appears to be empty. Check results for BUSCO: 290897at7742
## Thu Jul 14 00:59:14 CEST 2022 - The aliscore output file does not exist. Check results for BUSCO: 137296at7742
## Thu Jul 14 00:59:14 CEST 2022 - The aliscore output appears to be empty. Check results for BUSCO: 137296at7742
## Thu Jul 14 00:59:14 CEST 2022 - The aliscore output file does not exist. Check results for BUSCO: 51573at7742
## Thu Jul 14 00:59:14 CEST 2022 - The aliscore output appears to be empty. Check results for BUSCO: 51573at7742
## Thu Jul 14 00:59:19 CEST 2022 - The aliscore output file does not exist. Check results for BUSCO: 89000at7742
## Thu Jul 14 00:59:19 CEST 2022 - The aliscore output appears to be empty. Check results for BUSCO: 89000at7742
## Thu Jul 14 00:59:19 CEST 2022 - The aliscore output file does not exist. Check results for BUSCO: 163089at7742
## Thu Jul 14 00:59:19 CEST 2022 - The aliscore output appears to be empty. Check results for BUSCO: 163089at7742
## Thu Jul 14 00:59:20 CEST 2022 - The aliscore output file does not exist. Check results for BUSCO: 81669at7742
## Thu Jul 14 00:59:20 CEST 2022 - The aliscore output appears to be empty. Check results for BUSCO: 81669at7742
## Thu Jul 14 00:59:20 CEST 2022 - The aliscore output file does not exist. Check results for BUSCO: 51573at7742
## Thu Jul 14 00:59:20 CEST 2022 - The aliscore output appears to be empty. Check results for BUSCO: 51573at7742
## Thu Jul 14 00:59:20 CEST 2022 - The aliscore output file does not exist. Check results for BUSCO: 86856at7742
## Thu Jul 14 00:59:20 CEST 2022 - The aliscore output appears to be empty. Check results for BUSCO: 86856at7742
## Thu Jul 14 00:59:20 CEST 2022 - The aliscore output file does not exist. Check results for BUSCO: 223874at7742
## Thu Jul 14 00:59:20 CEST 2022 - The aliscore output appears to be empty. Check results for BUSCO: 223874at7742
## Thu Jul 14 00:59:24 CEST 2022 - The aliscore output file does not exist. Check results for BUSCO: 176579at7742
## Thu Jul 14 00:59:24 CEST 2022 - The aliscore output appears to be empty. Check results for BUSCO: 176579at7742
## Thu Jul 14 00:59:26 CEST 2022 - The aliscore output file does not exist. Check results for BUSCO: 142495at7742
## Thu Jul 14 00:59:26 CEST 2022 - The aliscore output appears to be empty. Check results for BUSCO: 142495at7742
## Thu Jul 14 00:59:27 CEST 2022 - The aliscore output file does not exist. Check results for BUSCO: 146337at7742
## Thu Jul 14 00:59:27 CEST 2022 - The aliscore output appears to be empty. Check results for BUSCO: 146337at7742
## Thu Jul 14 00:59:28 CEST 2022 - The aliscore output file does not exist. Check results for BUSCO: 99037at7742
## Thu Jul 14 00:59:28 CEST 2022 - The aliscore output appears to be empty. Check results for BUSCO: 99037at7742
## Thu Jul 14 00:59:29 CEST 2022 - The aliscore output file does not exist. Check results for BUSCO: 163089at7742
## Thu Jul 14 00:59:29 CEST 2022 - The aliscore output appears to be empty. Check results for BUSCO: 163089at7742
## Thu Jul 14 00:59:30 CEST 2022 - The aliscore output file does not exist. Check results for BUSCO: 384977at7742
## Thu Jul 14 00:59:30 CEST 2022 - The aliscore output appears to be empty. Check results for BUSCO: 384977at7742
## Thu Jul 14 00:59:30 CEST 2022 - The aliscore output file does not exist. Check results for BUSCO: 78565at7742
## Thu Jul 14 00:59:30 CEST 2022 - The aliscore output appears to be empty. Check results for BUSCO: 78565at7742
## Thu Jul 14 00:59:32 CEST 2022 - The aliscore output file does not exist. Check results for BUSCO: 64603at7742
## Thu Jul 14 00:59:32 CEST 2022 - The aliscore output appears to be empty. Check results for BUSCO: 64603at7742
## Thu Jul 14 00:59:33 CEST 2022 - The aliscore output file does not exist. Check results for BUSCO: 325663at7742
## Thu Jul 14 00:59:33 CEST 2022 - The aliscore output appears to be empty. Check results for BUSCO: 325663at7742
## Thu Jul 14 00:59:35 CEST 2022 - The aliscore output file does not exist. Check results for BUSCO: 116245at7742
## Thu Jul 14 00:59:35 CEST 2022 - The aliscore output appears to be empty. Check results for BUSCO: 116245at7742
## Thu Jul 14 00:59:38 CEST 2022 - The aliscore output file does not exist. Check results for BUSCO: 81669at7742
## Thu Jul 14 00:59:38 CEST 2022 - The aliscore output file does not exist. Check results for BUSCO: 264816at7742
## Thu Jul 14 00:59:38 CEST 2022 - The aliscore output appears to be empty. Check results for BUSCO: 81669at7742
## Thu Jul 14 00:59:38 CEST 2022 - The aliscore output appears to be empty. Check results for BUSCO: 264816at7742
## Thu Jul 14 00:59:38 CEST 2022 - The aliscore output file does not exist. Check results for BUSCO: 305045at7742
## Thu Jul 14 00:59:38 CEST 2022 - The aliscore output file does not exist. Check results for BUSCO: 114626at7742
## Thu Jul 14 00:59:38 CEST 2022 - The aliscore output appears to be empty. Check results for BUSCO: 305045at7742
## Thu Jul 14 00:59:38 CEST 2022 - The aliscore output appears to be empty. Check results for BUSCO: 114626at7742
## Thu Jul 14 00:59:41 CEST 2022 - The aliscore output file does not exist. Check results for BUSCO: 67119at7742
## Thu Jul 14 00:59:41 CEST 2022 - The aliscore output appears to be empty. Check results for BUSCO: 67119at7742
## Thu Jul 14 00:59:43 CEST 2022 - The aliscore output file does not exist. Check results for BUSCO: 281413at7742
## Thu Jul 14 00:59:43 CEST 2022 - The aliscore output appears to be empty. Check results for BUSCO: 281413at7742
## Thu Jul 14 00:59:43 CEST 2022 - The aliscore output file does not exist. Check results for BUSCO: 155999at7742
## Thu Jul 14 00:59:43 CEST 2022 - The aliscore output appears to be empty. Check results for BUSCO: 155999at7742
## Thu Jul 14 00:59:44 CEST 2022 - The aliscore output file does not exist. Check results for BUSCO: 113535at7742
## Thu Jul 14 00:59:44 CEST 2022 - The aliscore output appears to be empty. Check results for BUSCO: 113535at7742
## Thu Jul 14 00:59:47 CEST 2022 - The aliscore output file does not exist. Check results for BUSCO: 405703at7742
## Thu Jul 14 00:59:47 CEST 2022 - The aliscore output appears to be empty. Check results for BUSCO: 405703at7742
## Thu Jul 14 00:59:47 CEST 2022 - The aliscore output file does not exist. Check results for BUSCO: 96693at7742
## Thu Jul 14 00:59:47 CEST 2022 - The aliscore output appears to be empty. Check results for BUSCO: 96693at7742
## Thu Jul 14 00:59:47 CEST 2022 - The aliscore output file does not exist. Check results for BUSCO: 97039at7742
## Thu Jul 14 00:59:47 CEST 2022 - The aliscore output appears to be empty. Check results for BUSCO: 97039at7742
## Thu Jul 14 00:59:49 CEST 2022 - The aliscore output file does not exist. Check results for BUSCO: 64603at7742
## Thu Jul 14 00:59:49 CEST 2022 - The aliscore output appears to be empty. Check results for BUSCO: 64603at7742
## Thu Jul 14 00:59:50 CEST 2022 - The aliscore output file does not exist. Check results for BUSCO: 79086at7742
## Thu Jul 14 00:59:50 CEST 2022 - The aliscore output appears to be empty. Check results for BUSCO: 79086at7742
## Thu Jul 14 00:59:52 CEST 2022 - The aliscore output file does not exist. Check results for BUSCO: 109264at7742
## Thu Jul 14 00:59:52 CEST 2022 - The aliscore output appears to be empty. Check results for BUSCO: 109264at7742
## Thu Jul 14 00:59:55 CEST 2022 - The aliscore output file does not exist. Check results for BUSCO: 184873at7742
## Thu Jul 14 00:59:55 CEST 2022 - The aliscore output appears to be empty. Check results for BUSCO: 184873at7742
## Thu Jul 14 00:59:55 CEST 2022 - The aliscore output file does not exist. Check results for BUSCO: 163089at7742
## Thu Jul 14 00:59:55 CEST 2022 - The aliscore output appears to be empty. Check results for BUSCO: 163089at7742
## Thu Jul 14 00:59:55 CEST 2022 - The aliscore output file does not exist. Check results for BUSCO: 142495at7742
## Thu Jul 14 00:59:55 CEST 2022 - The aliscore output appears to be empty. Check results for BUSCO: 142495at7742
## Thu Jul 14 00:59:56 CEST 2022 - The aliscore output file does not exist. Check results for BUSCO: 330757at7742
## Thu Jul 14 00:59:56 CEST 2022 - The aliscore output appears to be empty. Check results for BUSCO: 330757at7742
## Thu Jul 14 00:59:58 CEST 2022 - The aliscore output file does not exist. Check results for BUSCO: 82295at7742
## Thu Jul 14 00:59:58 CEST 2022 - The aliscore output appears to be empty. Check results for BUSCO: 82295at7742
## Thu Jul 14 01:00:00 CEST 2022 - The aliscore output file does not exist. Check results for BUSCO: 323828at7742
## Thu Jul 14 01:00:00 CEST 2022 - The aliscore output appears to be empty. Check results for BUSCO: 323828at7742
## Thu Jul 14 01:00:00 CEST 2022 - The aliscore output file does not exist. Check results for BUSCO: 62345at7742
## Thu Jul 14 01:00:00 CEST 2022 - The aliscore output appears to be empty. Check results for BUSCO: 62345at7742
## Thu Jul 14 01:00:02 CEST 2022 - The aliscore output file does not exist. Check results for BUSCO: 106060at7742
## Thu Jul 14 01:00:02 CEST 2022 - The aliscore output appears to be empty. Check results for BUSCO: 106060at7742
## Thu Jul 14 01:00:05 CEST 2022 - The aliscore output file does not exist. Check results for BUSCO: 64461at7742
## Thu Jul 14 01:00:05 CEST 2022 - The aliscore output appears to be empty. Check results for BUSCO: 64461at7742
## Thu Jul 14 01:00:07 CEST 2022 - The aliscore output file does not exist. Check results for BUSCO: 345012at7742
## Thu Jul 14 01:00:07 CEST 2022 - The aliscore output appears to be empty. Check results for BUSCO: 345012at7742
## Thu Jul 14 01:00:07 CEST 2022 - The aliscore output file does not exist. Check results for BUSCO: 106009at7742
## Thu Jul 14 01:00:07 CEST 2022 - The aliscore output appears to be empty. Check results for BUSCO: 106009at7742
## Thu Jul 14 01:00:07 CEST 2022 - The aliscore output file does not exist. Check results for BUSCO: 136018at7742
## Thu Jul 14 01:00:07 CEST 2022 - The aliscore output appears to be empty. Check results for BUSCO: 136018at7742
## Thu Jul 14 01:00:07 CEST 2022 - The aliscore output file does not exist. Check results for BUSCO: 129053at7742
## Thu Jul 14 01:00:08 CEST 2022 - The aliscore output appears to be empty. Check results for BUSCO: 129053at7742
## Thu Jul 14 01:00:09 CEST 2022 - The aliscore output file does not exist. Check results for BUSCO: 169724at7742
## Thu Jul 14 01:00:09 CEST 2022 - The aliscore output appears to be empty. Check results for BUSCO: 169724at7742
## Thu Jul 14 01:00:11 CEST 2022 - The aliscore output file does not exist. Check results for BUSCO: 211156at7742
## Thu Jul 14 01:00:11 CEST 2022 - The aliscore output appears to be empty. Check results for BUSCO: 211156at7742
## Thu Jul 14 01:00:13 CEST 2022 - The aliscore output file does not exist. Check results for BUSCO: 289598at7742
## Thu Jul 14 01:00:13 CEST 2022 - The aliscore output appears to be empty. Check results for BUSCO: 289598at7742
## Thu Jul 14 01:00:14 CEST 2022 - The aliscore output file does not exist. Check results for BUSCO: 147455at7742
## Thu Jul 14 01:00:14 CEST 2022 - The aliscore output appears to be empty. Check results for BUSCO: 147455at7742
## Thu Jul 14 01:00:14 CEST 2022 - The aliscore output file does not exist. Check results for BUSCO: 54203at7742
## Thu Jul 14 01:00:14 CEST 2022 - The aliscore output appears to be empty. Check results for BUSCO: 54203at7742
## Thu Jul 14 01:00:15 CEST 2022 - The aliscore output file does not exist. Check results for BUSCO: 263251at7742
## Thu Jul 14 01:00:15 CEST 2022 - The aliscore output appears to be empty. Check results for BUSCO: 263251at7742
## Thu Jul 14 01:00:16 CEST 2022 - The aliscore output file does not exist. Check results for BUSCO: 91624at7742
## Thu Jul 14 01:00:16 CEST 2022 - The aliscore output appears to be empty. Check results for BUSCO: 91624at7742
## Thu Jul 14 01:00:19 CEST 2022 - The aliscore output file does not exist. Check results for BUSCO: 95757at7742
## Thu Jul 14 01:00:19 CEST 2022 - The aliscore output appears to be empty. Check results for BUSCO: 95757at7742
## Thu Jul 14 01:00:19 CEST 2022 - The aliscore output file does not exist. Check results for BUSCO: 132931at7742
## Thu Jul 14 01:00:19 CEST 2022 - The aliscore output appears to be empty. Check results for BUSCO: 132931at7742
## Thu Jul 14 01:00:21 CEST 2022 - The aliscore output file does not exist. Check results for BUSCO: 89391at7742
## Thu Jul 14 01:00:21 CEST 2022 - The aliscore output appears to be empty. Check results for BUSCO: 89391at7742
## Thu Jul 14 01:00:22 CEST 2022 - The aliscore output file does not exist. Check results for BUSCO: 99440at7742
## Thu Jul 14 01:00:22 CEST 2022 - The aliscore output appears to be empty. Check results for BUSCO: 99440at7742
## Thu Jul 14 01:00:23 CEST 2022 - The aliscore output file does not exist. Check results for BUSCO: 129979at7742
## Thu Jul 14 01:00:23 CEST 2022 - The aliscore output appears to be empty. Check results for BUSCO: 129979at7742
## Thu Jul 14 01:00:24 CEST 2022 - The aliscore output file does not exist. Check results for BUSCO: 236895at7742
## Thu Jul 14 01:00:24 CEST 2022 - The aliscore output appears to be empty. Check results for BUSCO: 236895at7742
## Thu Jul 14 01:00:25 CEST 2022 - The aliscore output file does not exist. Check results for BUSCO: 54203at7742
## Thu Jul 14 01:00:25 CEST 2022 - The aliscore output appears to be empty. Check results for BUSCO: 54203at7742
## Thu Jul 14 01:00:28 CEST 2022 - The aliscore output file does not exist. Check results for BUSCO: 221391at7742
## Thu Jul 14 01:00:28 CEST 2022 - The aliscore output appears to be empty. Check results for BUSCO: 221391at7742
## Thu Jul 14 01:00:29 CEST 2022 - The aliscore output file does not exist. Check results for BUSCO: 63057at7742
## Thu Jul 14 01:00:29 CEST 2022 - The aliscore output appears to be empty. Check results for BUSCO: 63057at7742
## Thu Jul 14 01:00:29 CEST 2022 - The aliscore output file does not exist. Check results for BUSCO: 146337at7742
## Thu Jul 14 01:00:29 CEST 2022 - The aliscore output appears to be empty. Check results for BUSCO: 146337at7742
## Thu Jul 14 01:00:31 CEST 2022 - The aliscore output file does not exist. Check results for BUSCO: 81669at7742
## Thu Jul 14 01:00:31 CEST 2022 - The aliscore output appears to be empty. Check results for BUSCO: 81669at7742
## Thu Jul 14 01:00:32 CEST 2022 - The aliscore output file does not exist. Check results for BUSCO: 248515at7742
## Thu Jul 14 01:00:32 CEST 2022 - The aliscore output appears to be empty. Check results for BUSCO: 248515at7742
## Thu Jul 14 01:00:32 CEST 2022 - The aliscore output file does not exist. Check results for BUSCO: 77080at7742
## Thu Jul 14 01:00:32 CEST 2022 - The aliscore output appears to be empty. Check results for BUSCO: 77080at7742
## Thu Jul 14 01:00:34 CEST 2022 - The aliscore output file does not exist. Check results for BUSCO: 126473at7742
## Thu Jul 14 01:00:34 CEST 2022 - The aliscore output appears to be empty. Check results for BUSCO: 126473at7742
## Thu Jul 14 01:00:36 CEST 2022 - The aliscore output file does not exist. Check results for BUSCO: 257487at7742
## Thu Jul 14 01:00:36 CEST 2022 - The aliscore output appears to be empty. Check results for BUSCO: 257487at7742
## Thu Jul 14 01:00:36 CEST 2022 - The aliscore output file does not exist. Check results for BUSCO: 171487at7742
## Thu Jul 14 01:00:36 CEST 2022 - The aliscore output appears to be empty. Check results for BUSCO: 171487at7742
## Thu Jul 14 01:00:39 CEST 2022 - The aliscore output file does not exist. Check results for BUSCO: 202773at7742
## Thu Jul 14 01:00:39 CEST 2022 - The aliscore output appears to be empty. Check results for BUSCO: 202773at7742
## Thu Jul 14 01:00:40 CEST 2022 - The aliscore output file does not exist. Check results for BUSCO: 110226at7742
## Thu Jul 14 01:00:40 CEST 2022 - The aliscore output appears to be empty. Check results for BUSCO: 110226at7742
## Thu Jul 14 01:00:41 CEST 2022 - The aliscore output file does not exist. Check results for BUSCO: 99037at7742
## Thu Jul 14 01:00:41 CEST 2022 - The aliscore output appears to be empty. Check results for BUSCO: 99037at7742
## Thu Jul 14 01:00:41 CEST 2022 - The aliscore output file does not exist. Check results for BUSCO: 164526at7742
## Thu Jul 14 01:00:41 CEST 2022 - The aliscore output appears to be empty. Check results for BUSCO: 164526at7742
## Thu Jul 14 01:00:43 CEST 2022 - The aliscore output file does not exist. Check results for BUSCO: 244702at7742
## Thu Jul 14 01:00:43 CEST 2022 - The aliscore output appears to be empty. Check results for BUSCO: 244702at7742
## Thu Jul 14 01:00:45 CEST 2022 - The aliscore output file does not exist. Check results for BUSCO: 172376at7742
## Thu Jul 14 01:00:45 CEST 2022 - The aliscore output appears to be empty. Check results for BUSCO: 172376at7742
## Thu Jul 14 01:00:47 CEST 2022 - The aliscore output file does not exist. Check results for BUSCO: 176119at7742
## Thu Jul 14 01:00:47 CEST 2022 - The aliscore output appears to be empty. Check results for BUSCO: 176119at7742
## Thu Jul 14 01:00:47 CEST 2022 - The aliscore output file does not exist. Check results for BUSCO: 62525at7742
## Thu Jul 14 01:00:47 CEST 2022 - The aliscore output appears to be empty. Check results for BUSCO: 62525at7742
## Thu Jul 14 01:00:49 CEST 2022 - The aliscore output file does not exist. Check results for BUSCO: 141724at7742
## Thu Jul 14 01:00:49 CEST 2022 - The aliscore output appears to be empty. Check results for BUSCO: 141724at7742
## Thu Jul 14 01:00:50 CEST 2022 - The aliscore output file does not exist. Check results for BUSCO: 113805at7742
## Thu Jul 14 01:00:50 CEST 2022 - The aliscore output appears to be empty. Check results for BUSCO: 113805at7742
## Thu Jul 14 01:00:50 CEST 2022 - The aliscore output file does not exist. Check results for BUSCO: 289598at7742
## Thu Jul 14 01:00:50 CEST 2022 - The aliscore output appears to be empty. Check results for BUSCO: 289598at7742
## Thu Jul 14 01:00:51 CEST 2022 - The aliscore output file does not exist. Check results for BUSCO: 121505at7742
## Thu Jul 14 01:00:52 CEST 2022 - The aliscore output appears to be empty. Check results for BUSCO: 121505at7742
## Thu Jul 14 01:00:54 CEST 2022 - The aliscore output file does not exist. Check results for BUSCO: 69819at7742
## Thu Jul 14 01:00:54 CEST 2022 - The aliscore output appears to be empty. Check results for BUSCO: 69819at7742
## Thu Jul 14 01:00:56 CEST 2022 - The aliscore output file does not exist. Check results for BUSCO: 95927at7742
## Thu Jul 14 01:00:56 CEST 2022 - The aliscore output file does not exist. Check results for BUSCO: 170750at7742
## Thu Jul 14 01:00:56 CEST 2022 - The aliscore output appears to be empty. Check results for BUSCO: 95927at7742
## Thu Jul 14 01:00:56 CEST 2022 - The aliscore output appears to be empty. Check results for BUSCO: 170750at7742
## Thu Jul 14 01:00:56 CEST 2022 - The aliscore output file does not exist. Check results for BUSCO: 381001at7742
## Thu Jul 14 01:00:56 CEST 2022 - The aliscore output appears to be empty. Check results for BUSCO: 381001at7742
## Thu Jul 14 01:00:59 CEST 2022 - The aliscore output file does not exist. Check results for BUSCO: 62345at7742
## Thu Jul 14 01:00:59 CEST 2022 - The aliscore output appears to be empty. Check results for BUSCO: 62345at7742
## Thu Jul 14 01:01:00 CEST 2022 - The aliscore output file does not exist. Check results for BUSCO: 143627at7742
## Thu Jul 14 01:01:00 CEST 2022 - The aliscore output appears to be empty. Check results for BUSCO: 143627at7742
## Thu Jul 14 01:01:03 CEST 2022 - The aliscore output file does not exist. Check results for BUSCO: 106423at7742
## Thu Jul 14 01:01:03 CEST 2022 - The aliscore output appears to be empty. Check results for BUSCO: 106423at7742
## Thu Jul 14 01:01:04 CEST 2022 - The aliscore output file does not exist. Check results for BUSCO: 122030at7742
## Thu Jul 14 01:01:04 CEST 2022 - The aliscore output appears to be empty. Check results for BUSCO: 122030at7742
## Thu Jul 14 01:01:06 CEST 2022 - The aliscore output file does not exist. Check results for BUSCO: 174574at7742
## Thu Jul 14 01:01:06 CEST 2022 - The aliscore output appears to be empty. Check results for BUSCO: 174574at7742
## Thu Jul 14 01:01:06 CEST 2022 - The aliscore output file does not exist. Check results for BUSCO: 144645at7742
## Thu Jul 14 01:01:06 CEST 2022 - The aliscore output appears to be empty. Check results for BUSCO: 144645at7742
## Thu Jul 14 01:01:07 CEST 2022 - The aliscore output file does not exist. Check results for BUSCO: 126473at7742
## Thu Jul 14 01:01:07 CEST 2022 - The aliscore output appears to be empty. Check results for BUSCO: 126473at7742
## Thu Jul 14 01:01:08 CEST 2022 - The aliscore output file does not exist. Check results for BUSCO: 174574at7742
## Thu Jul 14 01:01:08 CEST 2022 - The aliscore output appears to be empty. Check results for BUSCO: 174574at7742
## Thu Jul 14 01:01:10 CEST 2022 - The aliscore output file does not exist. Check results for BUSCO: 100816at7742
## Thu Jul 14 01:01:10 CEST 2022 - The aliscore output appears to be empty. Check results for BUSCO: 100816at7742
## Thu Jul 14 01:01:11 CEST 2022 - The aliscore output file does not exist. Check results for BUSCO: 258445at7742
## Thu Jul 14 01:01:11 CEST 2022 - The aliscore output appears to be empty. Check results for BUSCO: 258445at7742
## Thu Jul 14 01:01:14 CEST 2022 - The aliscore output file does not exist. Check results for BUSCO: 64603at7742
## Thu Jul 14 01:01:14 CEST 2022 - The aliscore output appears to be empty. Check results for BUSCO: 64603at7742
## Thu Jul 14 01:01:14 CEST 2022 - The aliscore output file does not exist. Check results for BUSCO: 109264at7742
## Thu Jul 14 01:01:14 CEST 2022 - The aliscore output appears to be empty. Check results for BUSCO: 109264at7742
## Thu Jul 14 01:01:14 CEST 2022 - The aliscore output file does not exist. Check results for BUSCO: 212288at7742
## Thu Jul 14 01:01:14 CEST 2022 - The aliscore output appears to be empty. Check results for BUSCO: 212288at7742
## Thu Jul 14 01:01:15 CEST 2022 - The aliscore output file does not exist. Check results for BUSCO: 86277at7742
## Thu Jul 14 01:01:15 CEST 2022 - The aliscore output appears to be empty. Check results for BUSCO: 86277at7742
## Thu Jul 14 01:01:18 CEST 2022 - The aliscore output file does not exist. Check results for BUSCO: 130161at7742
## Thu Jul 14 01:01:18 CEST 2022 - The aliscore output appears to be empty. Check results for BUSCO: 130161at7742
## Thu Jul 14 01:01:19 CEST 2022 - The aliscore output file does not exist. Check results for BUSCO: 200403at7742
## Thu Jul 14 01:01:19 CEST 2022 - The aliscore output appears to be empty. Check results for BUSCO: 200403at7742
## Thu Jul 14 01:01:20 CEST 2022 - The aliscore output file does not exist. Check results for BUSCO: 184910at7742
## Thu Jul 14 01:01:20 CEST 2022 - The aliscore output file does not exist. Check results for BUSCO: 60330at7742
## Thu Jul 14 01:01:21 CEST 2022 - The aliscore output appears to be empty. Check results for BUSCO: 184910at7742
## Thu Jul 14 01:01:21 CEST 2022 - The aliscore output appears to be empty. Check results for BUSCO: 60330at7742
## Thu Jul 14 01:01:21 CEST 2022 - The aliscore output file does not exist. Check results for BUSCO: 85369at7742
## Thu Jul 14 01:01:21 CEST 2022 - The aliscore output appears to be empty. Check results for BUSCO: 85369at7742
## Thu Jul 14 01:01:24 CEST 2022 - The aliscore output file does not exist. Check results for BUSCO: 275273at7742
## Thu Jul 14 01:01:24 CEST 2022 - The aliscore output appears to be empty. Check results for BUSCO: 275273at7742
## Thu Jul 14 01:01:24 CEST 2022 - The aliscore output file does not exist. Check results for BUSCO: 82034at7742
## Thu Jul 14 01:01:24 CEST 2022 - The aliscore output file does not exist. Check results for BUSCO: 86001at7742
## Thu Jul 14 01:01:24 CEST 2022 - The aliscore output appears to be empty. Check results for BUSCO: 82034at7742
## Thu Jul 14 01:01:24 CEST 2022 - The aliscore output appears to be empty. Check results for BUSCO: 86001at7742
## Thu Jul 14 01:01:27 CEST 2022 - The aliscore output file does not exist. Check results for BUSCO: 325335at7742
## Thu Jul 14 01:01:27 CEST 2022 - The aliscore output appears to be empty. Check results for BUSCO: 325335at7742
## Thu Jul 14 01:01:27 CEST 2022 - The aliscore output file does not exist. Check results for BUSCO: 89391at7742
## Thu Jul 14 01:01:27 CEST 2022 - The aliscore output appears to be empty. Check results for BUSCO: 89391at7742
## Thu Jul 14 01:01:30 CEST 2022 - The aliscore output file does not exist. Check results for BUSCO: 75464at7742
## Thu Jul 14 01:01:30 CEST 2022 - The aliscore output appears to be empty. Check results for BUSCO: 75464at7742
## Thu Jul 14 01:01:31 CEST 2022 - The aliscore output file does not exist. Check results for BUSCO: 89229at7742
## Thu Jul 14 01:01:31 CEST 2022 - The aliscore output appears to be empty. Check results for BUSCO: 89229at7742
## Thu Jul 14 01:01:32 CEST 2022 - The aliscore output file does not exist. Check results for BUSCO: 116359at7742
## Thu Jul 14 01:01:32 CEST 2022 - The aliscore output appears to be empty. Check results for BUSCO: 116359at7742
## Thu Jul 14 01:01:33 CEST 2022 - The aliscore output file does not exist. Check results for BUSCO: 158236at7742
## Thu Jul 14 01:01:33 CEST 2022 - The aliscore output appears to be empty. Check results for BUSCO: 158236at7742
## Thu Jul 14 01:01:35 CEST 2022 - The aliscore output file does not exist. Check results for BUSCO: 162155at7742
## Thu Jul 14 01:01:35 CEST 2022 - The aliscore output appears to be empty. Check results for BUSCO: 162155at7742
## Thu Jul 14 01:01:35 CEST 2022 - The aliscore output file does not exist. Check results for BUSCO: 62525at7742
## Thu Jul 14 01:01:35 CEST 2022 - The aliscore output appears to be empty. Check results for BUSCO: 62525at7742
## Thu Jul 14 01:01:36 CEST 2022 - The aliscore output file does not exist. Check results for BUSCO: 126473at7742
## Thu Jul 14 01:01:36 CEST 2022 - The aliscore output appears to be empty. Check results for BUSCO: 126473at7742
## Thu Jul 14 01:01:39 CEST 2022 - The aliscore output file does not exist. Check results for BUSCO: 86001at7742
## Thu Jul 14 01:01:39 CEST 2022 - The aliscore output appears to be empty. Check results for BUSCO: 86001at7742
## Thu Jul 14 01:01:39 CEST 2022 - The aliscore output file does not exist. Check results for BUSCO: 155999at7742
## Thu Jul 14 01:01:39 CEST 2022 - The aliscore output appears to be empty. Check results for BUSCO: 155999at7742
## Thu Jul 14 01:01:42 CEST 2022 - The aliscore output file does not exist. Check results for BUSCO: 376015at7742
## Thu Jul 14 01:01:42 CEST 2022 - The aliscore output appears to be empty. Check results for BUSCO: 376015at7742
## Thu Jul 14 01:01:42 CEST 2022 - The aliscore output file does not exist. Check results for BUSCO: 202773at7742
## Thu Jul 14 01:01:42 CEST 2022 - The aliscore output appears to be empty. Check results for BUSCO: 202773at7742
## Thu Jul 14 01:01:42 CEST 2022 - The aliscore output file does not exist. Check results for BUSCO: 100816at7742
## Thu Jul 14 01:01:42 CEST 2022 - The aliscore output appears to be empty. Check results for BUSCO: 100816at7742
## Thu Jul 14 01:01:43 CEST 2022 - The aliscore output file does not exist. Check results for BUSCO: 236910at7742
## Thu Jul 14 01:01:43 CEST 2022 - The aliscore output appears to be empty. Check results for BUSCO: 236910at7742
## Thu Jul 14 01:01:44 CEST 2022 - The aliscore output file does not exist. Check results for BUSCO: 317124at7742
## Thu Jul 14 01:01:44 CEST 2022 - The aliscore output appears to be empty. Check results for BUSCO: 317124at7742
## Thu Jul 14 01:01:48 CEST 2022 - The aliscore output file does not exist. Check results for BUSCO: 194866at7742
## Thu Jul 14 01:01:48 CEST 2022 - The aliscore output appears to be empty. Check results for BUSCO: 194866at7742
## Thu Jul 14 01:01:48 CEST 2022 - The aliscore output file does not exist. Check results for BUSCO: 368834at7742
## Thu Jul 14 01:01:48 CEST 2022 - The aliscore output appears to be empty. Check results for BUSCO: 368834at7742
## Thu Jul 14 01:01:51 CEST 2022 - The aliscore output file does not exist. Check results for BUSCO: 220642at7742
## Thu Jul 14 01:01:51 CEST 2022 - The aliscore output appears to be empty. Check results for BUSCO: 220642at7742
## Thu Jul 14 01:01:51 CEST 2022 - The aliscore output file does not exist. Check results for BUSCO: 150433at7742
## Thu Jul 14 01:01:51 CEST 2022 - The aliscore output appears to be empty. Check results for BUSCO: 150433at7742
## Thu Jul 14 01:01:52 CEST 2022 - The aliscore output file does not exist. Check results for BUSCO: 140588at7742
## Thu Jul 14 01:01:52 CEST 2022 - The aliscore output appears to be empty. Check results for BUSCO: 140588at7742
## Thu Jul 14 01:01:52 CEST 2022 - The aliscore output file does not exist. Check results for BUSCO: 181384at7742
## Thu Jul 14 01:01:52 CEST 2022 - The aliscore output appears to be empty. Check results for BUSCO: 181384at7742
## Thu Jul 14 01:01:54 CEST 2022 - The aliscore output file does not exist. Check results for BUSCO: 329870at7742
## Thu Jul 14 01:01:54 CEST 2022 - The aliscore output appears to be empty. Check results for BUSCO: 329870at7742
## Thu Jul 14 01:01:56 CEST 2022 - The aliscore output file does not exist. Check results for BUSCO: 211495at7742
## Thu Jul 14 01:01:56 CEST 2022 - The aliscore output appears to be empty. Check results for BUSCO: 211495at7742
## Thu Jul 14 01:01:57 CEST 2022 - The aliscore output file does not exist. Check results for BUSCO: 54203at7742
## Thu Jul 14 01:01:57 CEST 2022 - The aliscore output appears to be empty. Check results for BUSCO: 54203at7742
## Thu Jul 14 01:01:59 CEST 2022 - The aliscore output file does not exist. Check results for BUSCO: 69819at7742
## Thu Jul 14 01:01:59 CEST 2022 - The aliscore output appears to be empty. Check results for BUSCO: 69819at7742
## Thu Jul 14 01:02:00 CEST 2022 - The aliscore output file does not exist. Check results for BUSCO: 172376at7742
## Thu Jul 14 01:02:00 CEST 2022 - The aliscore output appears to be empty. Check results for BUSCO: 172376at7742
## Thu Jul 14 01:02:01 CEST 2022 - The aliscore output file does not exist. Check results for BUSCO: 225564at7742
## Thu Jul 14 01:02:01 CEST 2022 - The aliscore output appears to be empty. Check results for BUSCO: 225564at7742
## Thu Jul 14 01:02:01 CEST 2022 - The aliscore output file does not exist. Check results for BUSCO: 117494at7742
## Thu Jul 14 01:02:01 CEST 2022 - The aliscore output appears to be empty. Check results for BUSCO: 117494at7742
## Thu Jul 14 01:02:03 CEST 2022 - The aliscore output file does not exist. Check results for BUSCO: 114303at7742
## Thu Jul 14 01:02:03 CEST 2022 - The aliscore output appears to be empty. Check results for BUSCO: 114303at7742
## Thu Jul 14 01:02:04 CEST 2022 - The aliscore output file does not exist. Check results for BUSCO: 313611at7742
## Thu Jul 14 01:02:04 CEST 2022 - The aliscore output appears to be empty. Check results for BUSCO: 313611at7742
## Thu Jul 14 01:02:06 CEST 2022 - The aliscore output file does not exist. Check results for BUSCO: 204311at7742
## Thu Jul 14 01:02:06 CEST 2022 - The aliscore output appears to be empty. Check results for BUSCO: 204311at7742
## Thu Jul 14 01:02:07 CEST 2022 - The aliscore output file does not exist. Check results for BUSCO: 236895at7742
## Thu Jul 14 01:02:07 CEST 2022 - The aliscore output appears to be empty. Check results for BUSCO: 236895at7742
## Thu Jul 14 01:02:10 CEST 2022 - The aliscore output file does not exist. Check results for BUSCO: 122030at7742
## Thu Jul 14 01:02:10 CEST 2022 - The aliscore output appears to be empty. Check results for BUSCO: 122030at7742
## Thu Jul 14 01:02:10 CEST 2022 - The aliscore output file does not exist. Check results for BUSCO: 158042at7742
## Thu Jul 14 01:02:10 CEST 2022 - The aliscore output appears to be empty. Check results for BUSCO: 158042at7742
## Thu Jul 14 01:02:10 CEST 2022 - The aliscore output file does not exist. Check results for BUSCO: 53843at7742
## Thu Jul 14 01:02:10 CEST 2022 - The aliscore output file does not exist. Check results for BUSCO: 324714at7742
## Thu Jul 14 01:02:10 CEST 2022 - The aliscore output appears to be empty. Check results for BUSCO: 53843at7742
## Thu Jul 14 01:02:10 CEST 2022 - The aliscore output appears to be empty. Check results for BUSCO: 324714at7742
## Thu Jul 14 01:02:12 CEST 2022 - The aliscore output file does not exist. Check results for BUSCO: 77080at7742
## Thu Jul 14 01:02:12 CEST 2022 - The aliscore output appears to be empty. Check results for BUSCO: 77080at7742
## Thu Jul 14 01:02:14 CEST 2022 - The aliscore output file does not exist. Check results for BUSCO: 70131at7742
## Thu Jul 14 01:02:14 CEST 2022 - The aliscore output appears to be empty. Check results for BUSCO: 70131at7742
## Thu Jul 14 01:02:15 CEST 2022 - The aliscore output file does not exist. Check results for BUSCO: 109054at7742
## Thu Jul 14 01:02:15 CEST 2022 - The aliscore output appears to be empty. Check results for BUSCO: 109054at7742
## Thu Jul 14 01:02:17 CEST 2022 - The aliscore output file does not exist. Check results for BUSCO: 207140at7742
## Thu Jul 14 01:02:17 CEST 2022 - The aliscore output appears to be empty. Check results for BUSCO: 207140at7742
## Thu Jul 14 01:02:19 CEST 2022 - The aliscore output file does not exist. Check results for BUSCO: 109650at7742
## Thu Jul 14 01:02:19 CEST 2022 - The aliscore output appears to be empty. Check results for BUSCO: 109650at7742
## Thu Jul 14 01:02:19 CEST 2022 - The aliscore output file does not exist. Check results for BUSCO: 116359at7742
## Thu Jul 14 01:02:19 CEST 2022 - The aliscore output appears to be empty. Check results for BUSCO: 116359at7742
## Thu Jul 14 01:02:20 CEST 2022 - The aliscore output file does not exist. Check results for BUSCO: 384977at7742
## Thu Jul 14 01:02:20 CEST 2022 - The aliscore output appears to be empty. Check results for BUSCO: 384977at7742
## Thu Jul 14 01:02:21 CEST 2022 - The aliscore output file does not exist. Check results for BUSCO: 89391at7742
## Thu Jul 14 01:02:21 CEST 2022 - The aliscore output appears to be empty. Check results for BUSCO: 89391at7742
## Thu Jul 14 01:02:23 CEST 2022 - The aliscore output file does not exist. Check results for BUSCO: 249853at7742
## Thu Jul 14 01:02:23 CEST 2022 - The aliscore output appears to be empty. Check results for BUSCO: 249853at7742
## Thu Jul 14 01:02:24 CEST 2022 - The aliscore output file does not exist. Check results for BUSCO: 123742at7742
## Thu Jul 14 01:02:24 CEST 2022 - The aliscore output appears to be empty. Check results for BUSCO: 123742at7742
## Thu Jul 14 01:02:25 CEST 2022 - The aliscore output file does not exist. Check results for BUSCO: 105001at7742
## Thu Jul 14 01:02:25 CEST 2022 - The aliscore output appears to be empty. Check results for BUSCO: 105001at7742
## Thu Jul 14 01:02:26 CEST 2022 - The aliscore output file does not exist. Check results for BUSCO: 352256at7742
## Thu Jul 14 01:02:26 CEST 2022 - The aliscore output appears to be empty. Check results for BUSCO: 352256at7742
## Thu Jul 14 01:02:28 CEST 2022 - The aliscore output file does not exist. Check results for BUSCO: 101457at7742
## Thu Jul 14 01:02:28 CEST 2022 - The aliscore output appears to be empty. Check results for BUSCO: 101457at7742
## Thu Jul 14 01:02:30 CEST 2022 - The aliscore output file does not exist. Check results for BUSCO: 113805at7742
## Thu Jul 14 01:02:30 CEST 2022 - The aliscore output appears to be empty. Check results for BUSCO: 113805at7742
## Thu Jul 14 01:02:32 CEST 2022 - The aliscore output file does not exist. Check results for BUSCO: 126306at7742
## Thu Jul 14 01:02:32 CEST 2022 - The aliscore output appears to be empty. Check results for BUSCO: 126306at7742
## Thu Jul 14 01:02:33 CEST 2022 - The aliscore output file does not exist. Check results for BUSCO: 109264at7742
## Thu Jul 14 01:02:33 CEST 2022 - The aliscore output appears to be empty. Check results for BUSCO: 109264at7742
## Thu Jul 14 01:02:33 CEST 2022 - The aliscore output file does not exist. Check results for BUSCO: 158445at7742
## Thu Jul 14 01:02:33 CEST 2022 - The aliscore output appears to be empty. Check results for BUSCO: 158445at7742
## Thu Jul 14 01:02:36 CEST 2022 - The aliscore output file does not exist. Check results for BUSCO: 126329at7742
## Thu Jul 14 01:02:36 CEST 2022 - The aliscore output appears to be empty. Check results for BUSCO: 126329at7742
## Thu Jul 14 01:02:36 CEST 2022 - The aliscore output file does not exist. Check results for BUSCO: 376554at7742
## Thu Jul 14 01:02:36 CEST 2022 - The aliscore output appears to be empty. Check results for BUSCO: 376554at7742
## Thu Jul 14 01:02:38 CEST 2022 - The aliscore output file does not exist. Check results for BUSCO: 116835at7742
## Thu Jul 14 01:02:38 CEST 2022 - The aliscore output appears to be empty. Check results for BUSCO: 116835at7742
## Thu Jul 14 01:02:38 CEST 2022 - The aliscore output file does not exist. Check results for BUSCO: 116245at7742
## Thu Jul 14 01:02:38 CEST 2022 - The aliscore output appears to be empty. Check results for BUSCO: 116245at7742
## Thu Jul 14 01:02:40 CEST 2022 - The aliscore output file does not exist. Check results for BUSCO: 106858at7742
## Thu Jul 14 01:02:40 CEST 2022 - The aliscore output appears to be empty. Check results for BUSCO: 106858at7742
## Thu Jul 14 01:02:41 CEST 2022 - The aliscore output file does not exist. Check results for BUSCO: 87012at7742
## Thu Jul 14 01:02:41 CEST 2022 - The aliscore output appears to be empty. Check results for BUSCO: 87012at7742
## Thu Jul 14 01:02:43 CEST 2022 - The aliscore output file does not exist. Check results for BUSCO: 227299at7742
## Thu Jul 14 01:02:43 CEST 2022 - The aliscore output appears to be empty. Check results for BUSCO: 227299at7742
## Thu Jul 14 01:02:43 CEST 2022 - The aliscore output file does not exist. Check results for BUSCO: 120131at7742
## Thu Jul 14 01:02:43 CEST 2022 - The aliscore output appears to be empty. Check results for BUSCO: 120131at7742
## Thu Jul 14 01:02:45 CEST 2022 - The aliscore output file does not exist. Check results for BUSCO: 130549at7742
## Thu Jul 14 01:02:45 CEST 2022 - The aliscore output appears to be empty. Check results for BUSCO: 130549at7742
## Thu Jul 14 01:02:45 CEST 2022 - The aliscore output file does not exist. Check results for BUSCO: 381001at7742
## Thu Jul 14 01:02:45 CEST 2022 - The aliscore output appears to be empty. Check results for BUSCO: 381001at7742
## Thu Jul 14 01:02:47 CEST 2022 - The aliscore output file does not exist. Check results for BUSCO: 143627at7742
## Thu Jul 14 01:02:47 CEST 2022 - The aliscore output appears to be empty. Check results for BUSCO: 143627at7742
## Thu Jul 14 01:02:50 CEST 2022 - The aliscore output file does not exist. Check results for BUSCO: 257726at7742
## Thu Jul 14 01:02:50 CEST 2022 - The aliscore output appears to be empty. Check results for BUSCO: 257726at7742
## Thu Jul 14 01:02:51 CEST 2022 - The aliscore output file does not exist. Check results for BUSCO: 177660at7742
## Thu Jul 14 01:02:51 CEST 2022 - The aliscore output file does not exist. Check results for BUSCO: 222975at7742
## Thu Jul 14 01:02:51 CEST 2022 - The aliscore output appears to be empty. Check results for BUSCO: 177660at7742
## Thu Jul 14 01:02:51 CEST 2022 - The aliscore output appears to be empty. Check results for BUSCO: 222975at7742
## Thu Jul 14 01:02:52 CEST 2022 - The aliscore output file does not exist. Check results for BUSCO: 151923at7742
## Thu Jul 14 01:02:52 CEST 2022 - The aliscore output appears to be empty. Check results for BUSCO: 151923at7742
## Thu Jul 14 01:02:53 CEST 2022 - The aliscore output file does not exist. Check results for BUSCO: 92625at7742
## Thu Jul 14 01:02:53 CEST 2022 - The aliscore output appears to be empty. Check results for BUSCO: 92625at7742
## Thu Jul 14 01:02:54 CEST 2022 - The aliscore output file does not exist. Check results for BUSCO: 183360at7742
## Thu Jul 14 01:02:54 CEST 2022 - The aliscore output appears to be empty. Check results for BUSCO: 183360at7742
## Thu Jul 14 01:02:55 CEST 2022 - The aliscore output file does not exist. Check results for BUSCO: 204311at7742
## Thu Jul 14 01:02:55 CEST 2022 - The aliscore output appears to be empty. Check results for BUSCO: 204311at7742
## Thu Jul 14 01:02:56 CEST 2022 - The aliscore output file does not exist. Check results for BUSCO: 376015at7742
## Thu Jul 14 01:02:57 CEST 2022 - The aliscore output appears to be empty. Check results for BUSCO: 376015at7742
## Thu Jul 14 01:02:58 CEST 2022 - The aliscore output file does not exist. Check results for BUSCO: 62345at7742
## Thu Jul 14 01:02:58 CEST 2022 - The aliscore output appears to be empty. Check results for BUSCO: 62345at7742
## Thu Jul 14 01:02:59 CEST 2022 - The aliscore output file does not exist. Check results for BUSCO: 107188at7742
## Thu Jul 14 01:02:59 CEST 2022 - The aliscore output appears to be empty. Check results for BUSCO: 107188at7742
## Thu Jul 14 01:03:01 CEST 2022 - The aliscore output file does not exist. Check results for BUSCO: 237620at7742
## Thu Jul 14 01:03:01 CEST 2022 - The aliscore output appears to be empty. Check results for BUSCO: 237620at7742
## Thu Jul 14 01:03:02 CEST 2022 - The aliscore output file does not exist. Check results for BUSCO: 87114at7742
## Thu Jul 14 01:03:02 CEST 2022 - The aliscore output appears to be empty. Check results for BUSCO: 87114at7742
## Thu Jul 14 01:03:03 CEST 2022 - The aliscore output file does not exist. Check results for BUSCO: 202773at7742
## Thu Jul 14 01:03:03 CEST 2022 - The aliscore output appears to be empty. Check results for BUSCO: 202773at7742
## Thu Jul 14 01:03:04 CEST 2022 - The aliscore output file does not exist. Check results for BUSCO: 68557at7742
## Thu Jul 14 01:03:04 CEST 2022 - The aliscore output appears to be empty. Check results for BUSCO: 68557at7742
## Thu Jul 14 01:03:06 CEST 2022 - The aliscore output file does not exist. Check results for BUSCO: 273629at7742
## Thu Jul 14 01:03:06 CEST 2022 - The aliscore output appears to be empty. Check results for BUSCO: 273629at7742
## Thu Jul 14 01:03:06 CEST 2022 - The aliscore output file does not exist. Check results for BUSCO: 404307at7742
## Thu Jul 14 01:03:06 CEST 2022 - The aliscore output appears to be empty. Check results for BUSCO: 404307at7742
## Thu Jul 14 01:03:07 CEST 2022 - The aliscore output file does not exist. Check results for BUSCO: 87423at7742
## Thu Jul 14 01:03:07 CEST 2022 - The aliscore output appears to be empty. Check results for BUSCO: 87423at7742
## Thu Jul 14 01:03:11 CEST 2022 - The aliscore output file does not exist. Check results for BUSCO: 87114at7742
## Thu Jul 14 01:03:11 CEST 2022 - The aliscore output appears to be empty. Check results for BUSCO: 87114at7742
## Thu Jul 14 01:03:13 CEST 2022 - The aliscore output file does not exist. Check results for BUSCO: 183058at7742
## Thu Jul 14 01:03:13 CEST 2022 - The aliscore output appears to be empty. Check results for BUSCO: 183058at7742
## Thu Jul 14 01:03:13 CEST 2022 - The aliscore output file does not exist. Check results for BUSCO: 121505at7742
## Thu Jul 14 01:03:13 CEST 2022 - The aliscore output appears to be empty. Check results for BUSCO: 121505at7742
## Thu Jul 14 01:03:13 CEST 2022 - The aliscore output file does not exist. Check results for BUSCO: 208314at7742
## Thu Jul 14 01:03:14 CEST 2022 - The aliscore output appears to be empty. Check results for BUSCO: 208314at7742
## Thu Jul 14 01:03:16 CEST 2022 - The aliscore output file does not exist. Check results for BUSCO: 308660at7742
## Thu Jul 14 01:03:16 CEST 2022 - The aliscore output appears to be empty. Check results for BUSCO: 308660at7742
## Thu Jul 14 01:03:17 CEST 2022 - The aliscore output file does not exist. Check results for BUSCO: 127392at7742
## Thu Jul 14 01:03:17 CEST 2022 - The aliscore output appears to be empty. Check results for BUSCO: 127392at7742
## Thu Jul 14 01:03:17 CEST 2022 - The aliscore output file does not exist. Check results for BUSCO: 126306at7742
## Thu Jul 14 01:03:17 CEST 2022 - The aliscore output appears to be empty. Check results for BUSCO: 126306at7742
## Thu Jul 14 01:03:20 CEST 2022 - The aliscore output file does not exist. Check results for BUSCO: 298175at7742
## Thu Jul 14 01:03:20 CEST 2022 - The aliscore output appears to be empty. Check results for BUSCO: 298175at7742
## Thu Jul 14 01:03:20 CEST 2022 - The aliscore output file does not exist. Check results for BUSCO: 192715at7742
## Thu Jul 14 01:03:20 CEST 2022 - The aliscore output appears to be empty. Check results for BUSCO: 192715at7742
## Thu Jul 14 01:03:21 CEST 2022 - The aliscore output file does not exist. Check results for BUSCO: 268563at7742
## Thu Jul 14 01:03:21 CEST 2022 - The aliscore output appears to be empty. Check results for BUSCO: 268563at7742
## Thu Jul 14 01:03:22 CEST 2022 - The aliscore output file does not exist. Check results for BUSCO: 268947at7742
## Thu Jul 14 01:03:22 CEST 2022 - The aliscore output appears to be empty. Check results for BUSCO: 268947at7742
## Thu Jul 14 01:03:24 CEST 2022 - The aliscore output file does not exist. Check results for BUSCO: 131983at7742
## Thu Jul 14 01:03:24 CEST 2022 - The aliscore output appears to be empty. Check results for BUSCO: 131983at7742
## Thu Jul 14 01:03:25 CEST 2022 - The aliscore output file does not exist. Check results for BUSCO: 113535at7742
## Thu Jul 14 01:03:25 CEST 2022 - The aliscore output appears to be empty. Check results for BUSCO: 113535at7742
## Thu Jul 14 01:03:25 CEST 2022 - The aliscore output file does not exist. Check results for BUSCO: 129053at7742
## Thu Jul 14 01:03:25 CEST 2022 - The aliscore output appears to be empty. Check results for BUSCO: 129053at7742
## Thu Jul 14 01:03:27 CEST 2022 - The aliscore output file does not exist. Check results for BUSCO: 108033at7742
## Thu Jul 14 01:03:27 CEST 2022 - The aliscore output appears to be empty. Check results for BUSCO: 108033at7742
## Thu Jul 14 01:03:27 CEST 2022 - The aliscore output file does not exist. Check results for BUSCO: 127392at7742
## Thu Jul 14 01:03:27 CEST 2022 - The aliscore output appears to be empty. Check results for BUSCO: 127392at7742
## Thu Jul 14 01:03:28 CEST 2022 - The aliscore output file does not exist. Check results for BUSCO: 106060at7742
## Thu Jul 14 01:03:28 CEST 2022 - The aliscore output appears to be empty. Check results for BUSCO: 106060at7742
## Thu Jul 14 01:03:31 CEST 2022 - The aliscore output file does not exist. Check results for BUSCO: 245445at7742
## Thu Jul 14 01:03:31 CEST 2022 - The aliscore output appears to be empty. Check results for BUSCO: 245445at7742
## Thu Jul 14 01:03:33 CEST 2022 - The aliscore output file does not exist. Check results for BUSCO: 257726at7742
## Thu Jul 14 01:03:33 CEST 2022 - The aliscore output appears to be empty. Check results for BUSCO: 257726at7742
## Thu Jul 14 01:03:33 CEST 2022 - The aliscore output file does not exist. Check results for BUSCO: 137296at7742
## Thu Jul 14 01:03:33 CEST 2022 - The aliscore output appears to be empty. Check results for BUSCO: 137296at7742
## Thu Jul 14 01:03:35 CEST 2022 - The aliscore output file does not exist. Check results for BUSCO: 161395at7742
## Thu Jul 14 01:03:35 CEST 2022 - The aliscore output appears to be empty. Check results for BUSCO: 161395at7742
## Thu Jul 14 01:03:36 CEST 2022 - The aliscore output file does not exist. Check results for BUSCO: 87783at7742
## Thu Jul 14 01:03:36 CEST 2022 - The aliscore output appears to be empty. Check results for BUSCO: 87783at7742
## Thu Jul 14 01:03:36 CEST 2022 - The aliscore output file does not exist. Check results for BUSCO: 384977at7742
## Thu Jul 14 01:03:36 CEST 2022 - The aliscore output appears to be empty. Check results for BUSCO: 384977at7742
## Thu Jul 14 01:03:39 CEST 2022 - The aliscore output file does not exist. Check results for BUSCO: 210721at7742
## Thu Jul 14 01:03:39 CEST 2022 - The aliscore output appears to be empty. Check results for BUSCO: 210721at7742
## Thu Jul 14 01:03:40 CEST 2022 - The aliscore output file does not exist. Check results for BUSCO: 200403at7742
## Thu Jul 14 01:03:40 CEST 2022 - The aliscore output appears to be empty. Check results for BUSCO: 200403at7742
## Thu Jul 14 01:03:41 CEST 2022 - The aliscore output file does not exist. Check results for BUSCO: 113847at7742
## Thu Jul 14 01:03:41 CEST 2022 - The aliscore output appears to be empty. Check results for BUSCO: 113847at7742
## Thu Jul 14 01:03:42 CEST 2022 - The aliscore output file does not exist. Check results for BUSCO: 329870at7742
## Thu Jul 14 01:03:42 CEST 2022 - The aliscore output appears to be empty. Check results for BUSCO: 329870at7742
## Thu Jul 14 01:03:43 CEST 2022 - The aliscore output file does not exist. Check results for BUSCO: 211156at7742
## Thu Jul 14 01:03:43 CEST 2022 - The aliscore output appears to be empty. Check results for BUSCO: 211156at7742
## Thu Jul 14 01:03:45 CEST 2022 - The aliscore output file does not exist. Check results for BUSCO: 69819at7742
## Thu Jul 14 01:03:46 CEST 2022 - The aliscore output appears to be empty. Check results for BUSCO: 69819at7742
## Thu Jul 14 01:03:46 CEST 2022 - The aliscore output file does not exist. Check results for BUSCO: 107664at7742
## Thu Jul 14 01:03:46 CEST 2022 - The aliscore output appears to be empty. Check results for BUSCO: 107664at7742
## Thu Jul 14 01:03:46 CEST 2022 - The aliscore output file does not exist. Check results for BUSCO: 213224at7742
## Thu Jul 14 01:03:46 CEST 2022 - The aliscore output appears to be empty. Check results for BUSCO: 213224at7742
## Thu Jul 14 01:03:48 CEST 2022 - The aliscore output file does not exist. Check results for BUSCO: 196659at7742
## Thu Jul 14 01:03:48 CEST 2022 - The aliscore output appears to be empty. Check results for BUSCO: 196659at7742
## Thu Jul 14 01:03:49 CEST 2022 - The aliscore output file does not exist. Check results for BUSCO: 106858at7742
## Thu Jul 14 01:03:49 CEST 2022 - The aliscore output appears to be empty. Check results for BUSCO: 106858at7742
## Thu Jul 14 01:03:51 CEST 2022 - The aliscore output file does not exist. Check results for BUSCO: 87783at7742
## Thu Jul 14 01:03:51 CEST 2022 - The aliscore output appears to be empty. Check results for BUSCO: 87783at7742
## Thu Jul 14 01:03:52 CEST 2022 - The aliscore output file does not exist. Check results for BUSCO: 191192at7742
## Thu Jul 14 01:03:53 CEST 2022 - The aliscore output appears to be empty. Check results for BUSCO: 191192at7742
## Thu Jul 14 01:03:53 CEST 2022 - The aliscore output file does not exist. Check results for BUSCO: 115748at7742
## Thu Jul 14 01:03:53 CEST 2022 - The aliscore output appears to be empty. Check results for BUSCO: 115748at7742
## Thu Jul 14 01:03:54 CEST 2022 - The aliscore output file does not exist. Check results for BUSCO: 257487at7742
## Thu Jul 14 01:03:54 CEST 2022 - The aliscore output appears to be empty. Check results for BUSCO: 257487at7742
## Thu Jul 14 01:03:56 CEST 2022 - The aliscore output file does not exist. Check results for BUSCO: 86001at7742
## Thu Jul 14 01:03:56 CEST 2022 - The aliscore output appears to be empty. Check results for BUSCO: 86001at7742
## Thu Jul 14 01:03:57 CEST 2022 - The aliscore output file does not exist. Check results for BUSCO: 264650at7742
## Thu Jul 14 01:03:57 CEST 2022 - The aliscore output appears to be empty. Check results for BUSCO: 264650at7742
## Thu Jul 14 01:03:57 CEST 2022 - The aliscore output file does not exist. Check results for BUSCO: 368834at7742
## Thu Jul 14 01:03:57 CEST 2022 - The aliscore output appears to be empty. Check results for BUSCO: 368834at7742
## Thu Jul 14 01:03:58 CEST 2022 - The aliscore output file does not exist. Check results for BUSCO: 386844at7742
## Thu Jul 14 01:03:59 CEST 2022 - The aliscore output appears to be empty. Check results for BUSCO: 386844at7742
## Thu Jul 14 01:04:01 CEST 2022 - The aliscore output file does not exist. Check results for BUSCO: 244028at7742
## Thu Jul 14 01:04:01 CEST 2022 - The aliscore output appears to be empty. Check results for BUSCO: 244028at7742
## Thu Jul 14 01:04:03 CEST 2022 - The aliscore output file does not exist. Check results for BUSCO: 57538at7742
## Thu Jul 14 01:04:03 CEST 2022 - The aliscore output appears to be empty. Check results for BUSCO: 57538at7742
## Thu Jul 14 01:04:03 CEST 2022 - The aliscore output file does not exist. Check results for BUSCO: 280261at7742
## Thu Jul 14 01:04:03 CEST 2022 - The aliscore output appears to be empty. Check results for BUSCO: 280261at7742
## Thu Jul 14 01:04:04 CEST 2022 - The aliscore output file does not exist. Check results for BUSCO: 195929at7742
## Thu Jul 14 01:04:04 CEST 2022 - The aliscore output appears to be empty. Check results for BUSCO: 195929at7742
## Thu Jul 14 01:04:07 CEST 2022 - The aliscore output file does not exist. Check results for BUSCO: 283522at7742
## Thu Jul 14 01:04:07 CEST 2022 - The aliscore output appears to be empty. Check results for BUSCO: 283522at7742
## Thu Jul 14 01:04:07 CEST 2022 - The aliscore output file does not exist. Check results for BUSCO: 185742at7742
## Thu Jul 14 01:04:07 CEST 2022 - The aliscore output appears to be empty. Check results for BUSCO: 185742at7742
## Thu Jul 14 01:04:09 CEST 2022 - The aliscore output file does not exist. Check results for BUSCO: 181591at7742
## Thu Jul 14 01:04:09 CEST 2022 - The aliscore output appears to be empty. Check results for BUSCO: 181591at7742
## Thu Jul 14 01:04:10 CEST 2022 - The aliscore output file does not exist. Check results for BUSCO: 161425at7742
## Thu Jul 14 01:04:10 CEST 2022 - The aliscore output appears to be empty. Check results for BUSCO: 161425at7742
## Thu Jul 14 01:04:10 CEST 2022 - The aliscore output file does not exist. Check results for BUSCO: 257487at7742
## Thu Jul 14 01:04:10 CEST 2022 - The aliscore output appears to be empty. Check results for BUSCO: 257487at7742
## Thu Jul 14 01:04:11 CEST 2022 - The aliscore output file does not exist. Check results for BUSCO: 268947at7742
## Thu Jul 14 01:04:11 CEST 2022 - The aliscore output appears to be empty. Check results for BUSCO: 268947at7742
## Thu Jul 14 01:04:14 CEST 2022 - The aliscore output file does not exist. Check results for BUSCO: 378656at7742
## Thu Jul 14 01:04:14 CEST 2022 - The aliscore output appears to be empty. Check results for BUSCO: 378656at7742
## Thu Jul 14 01:04:15 CEST 2022 - The aliscore output file does not exist. Check results for BUSCO: 384788at7742
## Thu Jul 14 01:04:15 CEST 2022 - The aliscore output appears to be empty. Check results for BUSCO: 384788at7742
## Thu Jul 14 01:04:16 CEST 2022 - The aliscore output file does not exist. Check results for BUSCO: 95871at7742
## Thu Jul 14 01:04:17 CEST 2022 - The aliscore output appears to be empty. Check results for BUSCO: 95871at7742
## Thu Jul 14 01:04:20 CEST 2022 - The aliscore output file does not exist. Check results for BUSCO: 85986at7742
## Thu Jul 14 01:04:20 CEST 2022 - The aliscore output appears to be empty. Check results for BUSCO: 85986at7742
## Thu Jul 14 01:04:20 CEST 2022 - The aliscore output file does not exist. Check results for BUSCO: 256267at7742
## Thu Jul 14 01:04:20 CEST 2022 - The aliscore output appears to be empty. Check results for BUSCO: 256267at7742
## Thu Jul 14 01:04:20 CEST 2022 - The aliscore output file does not exist. Check results for BUSCO: 248515at7742
## Thu Jul 14 01:04:20 CEST 2022 - The aliscore output appears to be empty. Check results for BUSCO: 248515at7742
## Thu Jul 14 01:04:21 CEST 2022 - The aliscore output file does not exist. Check results for BUSCO: 145946at7742
## Thu Jul 14 01:04:21 CEST 2022 - The aliscore output appears to be empty. Check results for BUSCO: 145946at7742
## Thu Jul 14 01:04:23 CEST 2022 - The aliscore output file does not exist. Check results for BUSCO: 276337at7742
## Thu Jul 14 01:04:23 CEST 2022 - The aliscore output appears to be empty. Check results for BUSCO: 276337at7742
## Thu Jul 14 01:04:24 CEST 2022 - The aliscore output file does not exist. Check results for BUSCO: 233794at7742
## Thu Jul 14 01:04:24 CEST 2022 - The aliscore output appears to be empty. Check results for BUSCO: 233794at7742
## Thu Jul 14 01:04:25 CEST 2022 - The aliscore output file does not exist. Check results for BUSCO: 115007at7742
## Thu Jul 14 01:04:25 CEST 2022 - The aliscore output appears to be empty. Check results for BUSCO: 115007at7742
## Thu Jul 14 01:04:26 CEST 2022 - The aliscore output file does not exist. Check results for BUSCO: 236607at7742
## Thu Jul 14 01:04:26 CEST 2022 - The aliscore output appears to be empty. Check results for BUSCO: 236607at7742
## Thu Jul 14 01:04:27 CEST 2022 - The aliscore output file does not exist. Check results for BUSCO: 198831at7742
## Thu Jul 14 01:04:27 CEST 2022 - The aliscore output appears to be empty. Check results for BUSCO: 198831at7742
## Thu Jul 14 01:04:28 CEST 2022 - The aliscore output file does not exist. Check results for BUSCO: 270426at7742
## Thu Jul 14 01:04:28 CEST 2022 - The aliscore output appears to be empty. Check results for BUSCO: 270426at7742
## Thu Jul 14 01:04:29 CEST 2022 - The aliscore output file does not exist. Check results for BUSCO: 123446at7742
## Thu Jul 14 01:04:29 CEST 2022 - The aliscore output appears to be empty. Check results for BUSCO: 123446at7742
## Thu Jul 14 01:04:33 CEST 2022 - The aliscore output file does not exist. Check results for BUSCO: 102691at7742
## Thu Jul 14 01:04:33 CEST 2022 - The aliscore output appears to be empty. Check results for BUSCO: 102691at7742
## Thu Jul 14 01:04:33 CEST 2022 - The aliscore output file does not exist. Check results for BUSCO: 151923at7742
## Thu Jul 14 01:04:33 CEST 2022 - The aliscore output appears to be empty. Check results for BUSCO: 151923at7742
## Thu Jul 14 01:04:34 CEST 2022 - The aliscore output file does not exist. Check results for BUSCO: 138249at7742
## Thu Jul 14 01:04:34 CEST 2022 - The aliscore output appears to be empty. Check results for BUSCO: 138249at7742
## Thu Jul 14 01:04:34 CEST 2022 - The aliscore output file does not exist. Check results for BUSCO: 172922at7742
## Thu Jul 14 01:04:34 CEST 2022 - The aliscore output appears to be empty. Check results for BUSCO: 172922at7742
## Thu Jul 14 01:04:35 CEST 2022 - The aliscore output file does not exist. Check results for BUSCO: 60330at7742
## Thu Jul 14 01:04:35 CEST 2022 - The aliscore output appears to be empty. Check results for BUSCO: 60330at7742
## Thu Jul 14 01:04:37 CEST 2022 - The aliscore output file does not exist. Check results for BUSCO: 381001at7742
## Thu Jul 14 01:04:37 CEST 2022 - The aliscore output appears to be empty. Check results for BUSCO: 381001at7742
## Thu Jul 14 01:04:38 CEST 2022 - The aliscore output file does not exist. Check results for BUSCO: 301755at7742
## Thu Jul 14 01:04:38 CEST 2022 - The aliscore output appears to be empty. Check results for BUSCO: 301755at7742
## Thu Jul 14 01:04:41 CEST 2022 - The aliscore output file does not exist. Check results for BUSCO: 174574at7742
## Thu Jul 14 01:04:41 CEST 2022 - The aliscore output appears to be empty. Check results for BUSCO: 174574at7742
## Thu Jul 14 01:04:41 CEST 2022 - The aliscore output file does not exist. Check results for BUSCO: 300767at7742
## Thu Jul 14 01:04:41 CEST 2022 - The aliscore output appears to be empty. Check results for BUSCO: 300767at7742
## Thu Jul 14 01:04:42 CEST 2022 - The aliscore output file does not exist. Check results for BUSCO: 87012at7742
## Thu Jul 14 01:04:42 CEST 2022 - The aliscore output appears to be empty. Check results for BUSCO: 87012at7742
## Thu Jul 14 01:04:45 CEST 2022 - The aliscore output file does not exist. Check results for BUSCO: 290897at7742
## Thu Jul 14 01:04:45 CEST 2022 - The aliscore output appears to be empty. Check results for BUSCO: 290897at7742
## Thu Jul 14 01:04:46 CEST 2022 - The aliscore output file does not exist. Check results for BUSCO: 173102at7742
## Thu Jul 14 01:04:46 CEST 2022 - The aliscore output appears to be empty. Check results for BUSCO: 173102at7742
## Thu Jul 14 01:04:46 CEST 2022 - The aliscore output file does not exist. Check results for BUSCO: 270426at7742
## Thu Jul 14 01:04:46 CEST 2022 - The aliscore output appears to be empty. Check results for BUSCO: 270426at7742
## Thu Jul 14 01:04:47 CEST 2022 - The aliscore output file does not exist. Check results for BUSCO: 137727at7742
## Thu Jul 14 01:04:47 CEST 2022 - The aliscore output appears to be empty. Check results for BUSCO: 137727at7742
## Thu Jul 14 01:04:48 CEST 2022 - The aliscore output file does not exist. Check results for BUSCO: 68557at7742
## Thu Jul 14 01:04:48 CEST 2022 - The aliscore output appears to be empty. Check results for BUSCO: 68557at7742
## Thu Jul 14 01:04:49 CEST 2022 - The aliscore output file does not exist. Check results for BUSCO: 241923at7742
## Thu Jul 14 01:04:49 CEST 2022 - The aliscore output appears to be empty. Check results for BUSCO: 241923at7742
## Thu Jul 14 01:04:50 CEST 2022 - The aliscore output file does not exist. Check results for BUSCO: 125985at7742
## Thu Jul 14 01:04:50 CEST 2022 - The aliscore output appears to be empty. Check results for BUSCO: 125985at7742
## Thu Jul 14 01:04:52 CEST 2022 - The aliscore output file does not exist. Check results for BUSCO: 147455at7742
## Thu Jul 14 01:04:52 CEST 2022 - The aliscore output appears to be empty. Check results for BUSCO: 147455at7742
## Thu Jul 14 01:04:53 CEST 2022 - The aliscore output file does not exist. Check results for BUSCO: 114924at7742
## Thu Jul 14 01:04:53 CEST 2022 - The aliscore output appears to be empty. Check results for BUSCO: 114924at7742
## Thu Jul 14 01:04:54 CEST 2022 - The aliscore output file does not exist. Check results for BUSCO: 192114at7742
## Thu Jul 14 01:04:54 CEST 2022 - The aliscore output appears to be empty. Check results for BUSCO: 192114at7742
## Thu Jul 14 01:04:56 CEST 2022 - The aliscore output file does not exist. Check results for BUSCO: 87599at7742
## Thu Jul 14 01:04:56 CEST 2022 - The aliscore output appears to be empty. Check results for BUSCO: 87599at7742
## Thu Jul 14 01:04:56 CEST 2022 - The aliscore output file does not exist. Check results for BUSCO: 104126at7742
## Thu Jul 14 01:04:57 CEST 2022 - The aliscore output appears to be empty. Check results for BUSCO: 104126at7742
## Thu Jul 14 01:04:58 CEST 2022 - The aliscore output file does not exist. Check results for BUSCO: 113805at7742
## Thu Jul 14 01:04:58 CEST 2022 - The aliscore output appears to be empty. Check results for BUSCO: 113805at7742
## Thu Jul 14 01:04:59 CEST 2022 - The aliscore output file does not exist. Check results for BUSCO: 100952at7742
## Thu Jul 14 01:04:59 CEST 2022 - The aliscore output appears to be empty. Check results for BUSCO: 100952at7742
## Thu Jul 14 01:05:01 CEST 2022 - The aliscore output file does not exist. Check results for BUSCO: 190917at7742
## Thu Jul 14 01:05:01 CEST 2022 - The aliscore output appears to be empty. Check results for BUSCO: 190917at7742
## Thu Jul 14 01:05:03 CEST 2022 - The aliscore output file does not exist. Check results for BUSCO: 135349at7742
## Thu Jul 14 01:05:03 CEST 2022 - The aliscore output appears to be empty. Check results for BUSCO: 135349at7742
## Thu Jul 14 01:05:03 CEST 2022 - The aliscore output file does not exist. Check results for BUSCO: 208314at7742
## Thu Jul 14 01:05:03 CEST 2022 - The aliscore output appears to be empty. Check results for BUSCO: 208314at7742
## Thu Jul 14 01:05:05 CEST 2022 - The aliscore output file does not exist. Check results for BUSCO: 305045at7742
## Thu Jul 14 01:05:05 CEST 2022 - The aliscore output appears to be empty. Check results for BUSCO: 305045at7742
## Thu Jul 14 01:05:06 CEST 2022 - The aliscore output file does not exist. Check results for BUSCO: 268237at7742
## Thu Jul 14 01:05:06 CEST 2022 - The aliscore output appears to be empty. Check results for BUSCO: 268237at7742
## Thu Jul 14 01:05:06 CEST 2022 - The aliscore output file does not exist. Check results for BUSCO: 115007at7742
## Thu Jul 14 01:05:06 CEST 2022 - The aliscore output appears to be empty. Check results for BUSCO: 115007at7742
## Thu Jul 14 01:05:07 CEST 2022 - The aliscore output file does not exist. Check results for BUSCO: 123446at7742
## Thu Jul 14 01:05:07 CEST 2022 - The aliscore output appears to be empty. Check results for BUSCO: 123446at7742
## Thu Jul 14 01:05:09 CEST 2022 - The aliscore output file does not exist. Check results for BUSCO: 132931at7742
## Thu Jul 14 01:05:09 CEST 2022 - The aliscore output file does not exist. Check results for BUSCO: 308660at7742
## Thu Jul 14 01:05:10 CEST 2022 - The aliscore output appears to be empty. Check results for BUSCO: 132931at7742
## Thu Jul 14 01:05:10 CEST 2022 - The aliscore output appears to be empty. Check results for BUSCO: 308660at7742
## Thu Jul 14 01:05:11 CEST 2022 - The aliscore output file does not exist. Check results for BUSCO: 172922at7742
## Thu Jul 14 01:05:11 CEST 2022 - The aliscore output appears to be empty. Check results for BUSCO: 172922at7742
## Thu Jul 14 01:05:14 CEST 2022 - The aliscore output file does not exist. Check results for BUSCO: 87012at7742
## Thu Jul 14 01:05:14 CEST 2022 - The aliscore output appears to be empty. Check results for BUSCO: 87012at7742
## Thu Jul 14 01:05:15 CEST 2022 - The aliscore output file does not exist. Check results for BUSCO: 119040at7742
## Thu Jul 14 01:05:15 CEST 2022 - The aliscore output file does not exist. Check results for BUSCO: 161164at7742
## Thu Jul 14 01:05:15 CEST 2022 - The aliscore output appears to be empty. Check results for BUSCO: 119040at7742
## Thu Jul 14 01:05:15 CEST 2022 - The aliscore output appears to be empty. Check results for BUSCO: 161164at7742
## Thu Jul 14 01:05:16 CEST 2022 - The aliscore output file does not exist. Check results for BUSCO: 203050at7742
## Thu Jul 14 01:05:16 CEST 2022 - The aliscore output appears to be empty. Check results for BUSCO: 203050at7742
## Thu Jul 14 01:05:19 CEST 2022 - The aliscore output file does not exist. Check results for BUSCO: 405703at7742
## Thu Jul 14 01:05:19 CEST 2022 - The aliscore output appears to be empty. Check results for BUSCO: 405703at7742
## Thu Jul 14 01:05:19 CEST 2022 - The aliscore output file does not exist. Check results for BUSCO: 198831at7742
## Thu Jul 14 01:05:19 CEST 2022 - The aliscore output appears to be empty. Check results for BUSCO: 198831at7742
## Thu Jul 14 01:05:21 CEST 2022 - The aliscore output file does not exist. Check results for BUSCO: 106313at7742
## Thu Jul 14 01:05:21 CEST 2022 - The aliscore output appears to be empty. Check results for BUSCO: 106313at7742
## Thu Jul 14 01:05:21 CEST 2022 - The aliscore output file does not exist. Check results for BUSCO: 60330at7742
## Thu Jul 14 01:05:21 CEST 2022 - The aliscore output appears to be empty. Check results for BUSCO: 60330at7742
## Thu Jul 14 01:05:24 CEST 2022 - The aliscore output file does not exist. Check results for BUSCO: 241923at7742
## Thu Jul 14 01:05:24 CEST 2022 - The aliscore output appears to be empty. Check results for BUSCO: 241923at7742
## Thu Jul 14 01:05:25 CEST 2022 - The aliscore output file does not exist. Check results for BUSCO: 187791at7742
## Thu Jul 14 01:05:25 CEST 2022 - The aliscore output appears to be empty. Check results for BUSCO: 187791at7742
## Thu Jul 14 01:05:26 CEST 2022 - The aliscore output file does not exist. Check results for BUSCO: 57538at7742
## Thu Jul 14 01:05:26 CEST 2022 - The aliscore output appears to be empty. Check results for BUSCO: 57538at7742
## Thu Jul 14 01:05:30 CEST 2022 - The aliscore output file does not exist. Check results for BUSCO: 161164at7742
## Thu Jul 14 01:05:30 CEST 2022 - The aliscore output appears to be empty. Check results for BUSCO: 161164at7742
## Thu Jul 14 01:05:30 CEST 2022 - The aliscore output file does not exist. Check results for BUSCO: 158236at7742
## Thu Jul 14 01:05:30 CEST 2022 - The aliscore output appears to be empty. Check results for BUSCO: 158236at7742
## Thu Jul 14 01:05:32 CEST 2022 - The aliscore output file does not exist. Check results for BUSCO: 296592at7742
## Thu Jul 14 01:05:32 CEST 2022 - The aliscore output appears to be empty. Check results for BUSCO: 296592at7742
## Thu Jul 14 01:05:32 CEST 2022 - The aliscore output file does not exist. Check results for BUSCO: 102222at7742
## Thu Jul 14 01:05:32 CEST 2022 - The aliscore output appears to be empty. Check results for BUSCO: 102222at7742
## Thu Jul 14 01:05:34 CEST 2022 - The aliscore output file does not exist. Check results for BUSCO: 241923at7742
## Thu Jul 14 01:05:34 CEST 2022 - The aliscore output appears to be empty. Check results for BUSCO: 241923at7742
## Thu Jul 14 01:05:35 CEST 2022 - The aliscore output file does not exist. Check results for BUSCO: 129053at7742
## Thu Jul 14 01:05:35 CEST 2022 - The aliscore output appears to be empty. Check results for BUSCO: 129053at7742
## Thu Jul 14 01:05:35 CEST 2022 - The aliscore output file does not exist. Check results for BUSCO: 140310at7742
## Thu Jul 14 01:05:35 CEST 2022 - The aliscore output appears to be empty. Check results for BUSCO: 140310at7742
## Thu Jul 14 01:05:37 CEST 2022 - The aliscore output file does not exist. Check results for BUSCO: 281785at7742
## Thu Jul 14 01:05:37 CEST 2022 - The aliscore output appears to be empty. Check results for BUSCO: 281785at7742
## Thu Jul 14 01:05:37 CEST 2022 - The aliscore output file does not exist. Check results for BUSCO: 103640at7742
## Thu Jul 14 01:05:37 CEST 2022 - The aliscore output appears to be empty. Check results for BUSCO: 103640at7742
## Thu Jul 14 01:05:39 CEST 2022 - The aliscore output file does not exist. Check results for BUSCO: 106858at7742
## Thu Jul 14 01:05:39 CEST 2022 - The aliscore output appears to be empty. Check results for BUSCO: 106858at7742
## Thu Jul 14 01:05:41 CEST 2022 - The aliscore output file does not exist. Check results for BUSCO: 134188at7742
## Thu Jul 14 01:05:41 CEST 2022 - The aliscore output appears to be empty. Check results for BUSCO: 134188at7742
## Thu Jul 14 01:05:44 CEST 2022 - The aliscore output file does not exist. Check results for BUSCO: 60569at7742
## Thu Jul 14 01:05:44 CEST 2022 - The aliscore output appears to be empty. Check results for BUSCO: 60569at7742
## Thu Jul 14 01:05:44 CEST 2022 - The aliscore output file does not exist. Check results for BUSCO: 157513at7742
## Thu Jul 14 01:05:44 CEST 2022 - The aliscore output appears to be empty. Check results for BUSCO: 157513at7742
## Thu Jul 14 01:05:44 CEST 2022 - The aliscore output file does not exist. Check results for BUSCO: 183360at7742
## Thu Jul 14 01:05:44 CEST 2022 - The aliscore output appears to be empty. Check results for BUSCO: 183360at7742
## Thu Jul 14 01:05:46 CEST 2022 - The aliscore output file does not exist. Check results for BUSCO: 102691at7742
## Thu Jul 14 01:05:46 CEST 2022 - The aliscore output appears to be empty. Check results for BUSCO: 102691at7742
## Thu Jul 14 01:05:47 CEST 2022 - The aliscore output file does not exist. Check results for BUSCO: 77080at7742
## Thu Jul 14 01:05:47 CEST 2022 - The aliscore output appears to be empty. Check results for BUSCO: 77080at7742
## Thu Jul 14 01:05:48 CEST 2022 - The aliscore output file does not exist. Check results for BUSCO: 87599at7742
## Thu Jul 14 01:05:48 CEST 2022 - The aliscore output appears to be empty. Check results for BUSCO: 87599at7742
## Thu Jul 14 01:05:49 CEST 2022 - The aliscore output file does not exist. Check results for BUSCO: 60569at7742
## Thu Jul 14 01:05:49 CEST 2022 - The aliscore output appears to be empty. Check results for BUSCO: 60569at7742
## Thu Jul 14 01:05:52 CEST 2022 - The aliscore output file does not exist. Check results for BUSCO: 103640at7742
## Thu Jul 14 01:05:52 CEST 2022 - The aliscore output appears to be empty. Check results for BUSCO: 103640at7742
## Thu Jul 14 01:05:52 CEST 2022 - The aliscore output file does not exist. Check results for BUSCO: 281413at7742
## Thu Jul 14 01:05:52 CEST 2022 - The aliscore output file does not exist. Check results for BUSCO: 87599at7742
## Thu Jul 14 01:05:52 CEST 2022 - The aliscore output appears to be empty. Check results for BUSCO: 281413at7742
## Thu Jul 14 01:05:52 CEST 2022 - The aliscore output appears to be empty. Check results for BUSCO: 87599at7742
## Thu Jul 14 01:05:55 CEST 2022 - The aliscore output file does not exist. Check results for BUSCO: 135770at7742
## Thu Jul 14 01:05:55 CEST 2022 - The aliscore output appears to be empty. Check results for BUSCO: 135770at7742
## Thu Jul 14 01:05:55 CEST 2022 - The aliscore output file does not exist. Check results for BUSCO: 236910at7742
## Thu Jul 14 01:05:55 CEST 2022 - The aliscore output appears to be empty. Check results for BUSCO: 236910at7742
## Thu Jul 14 01:05:56 CEST 2022 - The aliscore output file does not exist. Check results for BUSCO: 123742at7742
## Thu Jul 14 01:05:56 CEST 2022 - The aliscore output appears to be empty. Check results for BUSCO: 123742at7742
## Thu Jul 14 01:05:58 CEST 2022 - The aliscore output file does not exist. Check results for BUSCO: 245445at7742
## Thu Jul 14 01:05:58 CEST 2022 - The aliscore output appears to be empty. Check results for BUSCO: 245445at7742
## Thu Jul 14 01:05:58 CEST 2022 - The aliscore output file does not exist. Check results for BUSCO: 165456at7742
## Thu Jul 14 01:05:58 CEST 2022 - The aliscore output appears to be empty. Check results for BUSCO: 165456at7742
## Thu Jul 14 01:06:01 CEST 2022 - The aliscore output file does not exist. Check results for BUSCO: 95261at7742
## Thu Jul 14 01:06:01 CEST 2022 - The aliscore output appears to be empty. Check results for BUSCO: 95261at7742
## Thu Jul 14 01:06:02 CEST 2022 - The aliscore output file does not exist. Check results for BUSCO: 136089at7742
## Thu Jul 14 01:06:02 CEST 2022 - The aliscore output appears to be empty. Check results for BUSCO: 136089at7742
## Thu Jul 14 01:06:04 CEST 2022 - The aliscore output file does not exist. Check results for BUSCO: 135349at7742
## Thu Jul 14 01:06:04 CEST 2022 - The aliscore output appears to be empty. Check results for BUSCO: 135349at7742
## Thu Jul 14 01:06:05 CEST 2022 - The aliscore output file does not exist. Check results for BUSCO: 208931at7742
## Thu Jul 14 01:06:05 CEST 2022 - The aliscore output appears to be empty. Check results for BUSCO: 208931at7742
## Thu Jul 14 01:06:07 CEST 2022 - The aliscore output file does not exist. Check results for BUSCO: 158445at7742
## Thu Jul 14 01:06:07 CEST 2022 - The aliscore output appears to be empty. Check results for BUSCO: 158445at7742
## Thu Jul 14 01:06:07 CEST 2022 - The aliscore output file does not exist. Check results for BUSCO: 172922at7742
## Thu Jul 14 01:06:07 CEST 2022 - The aliscore output appears to be empty. Check results for BUSCO: 172922at7742
## Thu Jul 14 01:06:08 CEST 2022 - The aliscore output file does not exist. Check results for BUSCO: 129214at7742
## Thu Jul 14 01:06:08 CEST 2022 - The aliscore output appears to be empty. Check results for BUSCO: 129214at7742
## Thu Jul 14 01:06:10 CEST 2022 - The aliscore output file does not exist. Check results for BUSCO: 173398at7742
## Thu Jul 14 01:06:10 CEST 2022 - The aliscore output file does not exist. Check results for BUSCO: 131898at7742
## Thu Jul 14 01:06:10 CEST 2022 - The aliscore output appears to be empty. Check results for BUSCO: 173398at7742
## Thu Jul 14 01:06:10 CEST 2022 - The aliscore output appears to be empty. Check results for BUSCO: 131898at7742
## Thu Jul 14 01:06:13 CEST 2022 - The aliscore output file does not exist. Check results for BUSCO: 267345at7742
## Thu Jul 14 01:06:13 CEST 2022 - The aliscore output appears to be empty. Check results for BUSCO: 267345at7742
## Thu Jul 14 01:06:13 CEST 2022 - The aliscore output file does not exist. Check results for BUSCO: 202558at7742
## Thu Jul 14 01:06:13 CEST 2022 - The aliscore output appears to be empty. Check results for BUSCO: 202558at7742
## Thu Jul 14 01:06:16 CEST 2022 - The aliscore output file does not exist. Check results for BUSCO: 181591at7742
## Thu Jul 14 01:06:16 CEST 2022 - The aliscore output appears to be empty. Check results for BUSCO: 181591at7742
## Thu Jul 14 01:06:17 CEST 2022 - The aliscore output file does not exist. Check results for BUSCO: 149128at7742
## Thu Jul 14 01:06:17 CEST 2022 - The aliscore output appears to be empty. Check results for BUSCO: 149128at7742
## Thu Jul 14 01:06:18 CEST 2022 - The aliscore output file does not exist. Check results for BUSCO: 170750at7742
## Thu Jul 14 01:06:18 CEST 2022 - The aliscore output appears to be empty. Check results for BUSCO: 170750at7742
## Thu Jul 14 01:06:19 CEST 2022 - The aliscore output file does not exist. Check results for BUSCO: 82034at7742
## Thu Jul 14 01:06:19 CEST 2022 - The aliscore output appears to be empty. Check results for BUSCO: 82034at7742
## Thu Jul 14 01:06:20 CEST 2022 - The aliscore output file does not exist. Check results for BUSCO: 126329at7742
## Thu Jul 14 01:06:20 CEST 2022 - The aliscore output appears to be empty. Check results for BUSCO: 126329at7742
## Thu Jul 14 01:06:21 CEST 2022 - The aliscore output file does not exist. Check results for BUSCO: 290897at7742
## Thu Jul 14 01:06:21 CEST 2022 - The aliscore output appears to be empty. Check results for BUSCO: 290897at7742
## Thu Jul 14 01:06:22 CEST 2022 - The aliscore output file does not exist. Check results for BUSCO: 130161at7742
## Thu Jul 14 01:06:22 CEST 2022 - The aliscore output appears to be empty. Check results for BUSCO: 130161at7742
## Thu Jul 14 01:06:23 CEST 2022 - The aliscore output file does not exist. Check results for BUSCO: 166600at7742
## Thu Jul 14 01:06:23 CEST 2022 - The aliscore output appears to be empty. Check results for BUSCO: 166600at7742
## Thu Jul 14 01:06:25 CEST 2022 - The aliscore output file does not exist. Check results for BUSCO: 222821at7742
## Thu Jul 14 01:06:25 CEST 2022 - The aliscore output appears to be empty. Check results for BUSCO: 222821at7742
## Thu Jul 14 01:06:26 CEST 2022 - The aliscore output file does not exist. Check results for BUSCO: 303765at7742
## Thu Jul 14 01:06:26 CEST 2022 - The aliscore output appears to be empty. Check results for BUSCO: 303765at7742
## Thu Jul 14 01:06:27 CEST 2022 - The aliscore output file does not exist. Check results for BUSCO: 270426at7742
## Thu Jul 14 01:06:27 CEST 2022 - The aliscore output appears to be empty. Check results for BUSCO: 270426at7742
## Thu Jul 14 01:06:29 CEST 2022 - The aliscore output file does not exist. Check results for BUSCO: 379829at7742
## Thu Jul 14 01:06:29 CEST 2022 - The aliscore output appears to be empty. Check results for BUSCO: 379829at7742
## Thu Jul 14 01:06:29 CEST 2022 - The aliscore output file does not exist. Check results for BUSCO: 198831at7742
## Thu Jul 14 01:06:29 CEST 2022 - The aliscore output appears to be empty. Check results for BUSCO: 198831at7742
## Thu Jul 14 01:06:32 CEST 2022 - The aliscore output file does not exist. Check results for BUSCO: 131898at7742
## Thu Jul 14 01:06:32 CEST 2022 - The aliscore output appears to be empty. Check results for BUSCO: 131898at7742
## Thu Jul 14 01:06:32 CEST 2022 - The aliscore output file does not exist. Check results for BUSCO: 115007at7742
## Thu Jul 14 01:06:32 CEST 2022 - The aliscore output appears to be empty. Check results for BUSCO: 115007at7742
## Thu Jul 14 01:06:33 CEST 2022 - The aliscore output file does not exist. Check results for BUSCO: 93727at7742
## Thu Jul 14 01:06:33 CEST 2022 - The aliscore output appears to be empty. Check results for BUSCO: 93727at7742
## Thu Jul 14 01:06:34 CEST 2022 - The aliscore output file does not exist. Check results for BUSCO: 131983at7742
## Thu Jul 14 01:06:34 CEST 2022 - The aliscore output appears to be empty. Check results for BUSCO: 131983at7742
## Thu Jul 14 01:06:35 CEST 2022 - The aliscore output file does not exist. Check results for BUSCO: 95871at7742
## Thu Jul 14 01:06:35 CEST 2022 - The aliscore output appears to be empty. Check results for BUSCO: 95871at7742
## Thu Jul 14 01:06:38 CEST 2022 - The aliscore output file does not exist. Check results for BUSCO: 210721at7742
## Thu Jul 14 01:06:38 CEST 2022 - The aliscore output appears to be empty. Check results for BUSCO: 210721at7742
## Thu Jul 14 01:06:39 CEST 2022 - The aliscore output file does not exist. Check results for BUSCO: 117494at7742
## Thu Jul 14 01:06:39 CEST 2022 - The aliscore output appears to be empty. Check results for BUSCO: 117494at7742
## Thu Jul 14 01:06:40 CEST 2022 - The aliscore output file does not exist. Check results for BUSCO: 101457at7742
## Thu Jul 14 01:06:40 CEST 2022 - The aliscore output appears to be empty. Check results for BUSCO: 101457at7742
## Thu Jul 14 01:06:41 CEST 2022 - The aliscore output file does not exist. Check results for BUSCO: 68557at7742
## Thu Jul 14 01:06:41 CEST 2022 - The aliscore output appears to be empty. Check results for BUSCO: 68557at7742
## Thu Jul 14 01:06:42 CEST 2022 - The aliscore output file does not exist. Check results for BUSCO: 270327at7742
## Thu Jul 14 01:06:42 CEST 2022 - The aliscore output appears to be empty. Check results for BUSCO: 270327at7742
## Thu Jul 14 01:06:43 CEST 2022 - The aliscore output file does not exist. Check results for BUSCO: 191130at7742
## Thu Jul 14 01:06:43 CEST 2022 - The aliscore output appears to be empty. Check results for BUSCO: 191130at7742
## Thu Jul 14 01:06:45 CEST 2022 - The aliscore output file does not exist. Check results for BUSCO: 96693at7742
## Thu Jul 14 01:06:45 CEST 2022 - The aliscore output appears to be empty. Check results for BUSCO: 96693at7742
## Thu Jul 14 01:06:47 CEST 2022 - The aliscore output file does not exist. Check results for BUSCO: 106423at7742
## Thu Jul 14 01:06:47 CEST 2022 - The aliscore output appears to be empty. Check results for BUSCO: 106423at7742
## Thu Jul 14 01:06:47 CEST 2022 - The aliscore output file does not exist. Check results for BUSCO: 66866at7742
## Thu Jul 14 01:06:47 CEST 2022 - The aliscore output appears to be empty. Check results for BUSCO: 66866at7742
## Thu Jul 14 01:06:48 CEST 2022 - The aliscore output file does not exist. Check results for BUSCO: 288361at7742
## Thu Jul 14 01:06:48 CEST 2022 - The aliscore output appears to be empty. Check results for BUSCO: 288361at7742
## Thu Jul 14 01:06:50 CEST 2022 - The aliscore output file does not exist. Check results for BUSCO: 122841at7742
## Thu Jul 14 01:06:50 CEST 2022 - The aliscore output appears to be empty. Check results for BUSCO: 122841at7742
## Thu Jul 14 01:06:51 CEST 2022 - The aliscore output file does not exist. Check results for BUSCO: 104126at7742
## Thu Jul 14 01:06:51 CEST 2022 - The aliscore output appears to be empty. Check results for BUSCO: 104126at7742
## Thu Jul 14 01:06:52 CEST 2022 - The aliscore output file does not exist. Check results for BUSCO: 166600at7742
## Thu Jul 14 01:06:52 CEST 2022 - The aliscore output appears to be empty. Check results for BUSCO: 166600at7742
## Thu Jul 14 01:06:54 CEST 2022 - The aliscore output file does not exist. Check results for BUSCO: 162265at7742
## Thu Jul 14 01:06:54 CEST 2022 - The aliscore output appears to be empty. Check results for BUSCO: 162265at7742
## Thu Jul 14 01:06:55 CEST 2022 - The aliscore output file does not exist. Check results for BUSCO: 252557at7742
## Thu Jul 14 01:06:55 CEST 2022 - The aliscore output appears to be empty. Check results for BUSCO: 252557at7742
## Thu Jul 14 01:06:55 CEST 2022 - The aliscore output file does not exist. Check results for BUSCO: 181384at7742
## Thu Jul 14 01:06:55 CEST 2022 - The aliscore output appears to be empty. Check results for BUSCO: 181384at7742
## Thu Jul 14 01:06:57 CEST 2022 - The aliscore output file does not exist. Check results for BUSCO: 137296at7742
## Thu Jul 14 01:06:57 CEST 2022 - The aliscore output appears to be empty. Check results for BUSCO: 137296at7742
## Thu Jul 14 01:06:59 CEST 2022 - The aliscore output file does not exist. Check results for BUSCO: 300767at7742
## Thu Jul 14 01:06:59 CEST 2022 - The aliscore output appears to be empty. Check results for BUSCO: 300767at7742
## Thu Jul 14 01:06:59 CEST 2022 - The aliscore output file does not exist. Check results for BUSCO: 264650at7742
## Thu Jul 14 01:06:59 CEST 2022 - The aliscore output appears to be empty. Check results for BUSCO: 264650at7742
## Thu Jul 14 01:07:02 CEST 2022 - The aliscore output file does not exist. Check results for BUSCO: 120131at7742
## Thu Jul 14 01:07:02 CEST 2022 - The aliscore output appears to be empty. Check results for BUSCO: 120131at7742
## Thu Jul 14 01:07:03 CEST 2022 - The aliscore output file does not exist. Check results for BUSCO: 115070at7742
## Thu Jul 14 01:07:03 CEST 2022 - The aliscore output appears to be empty. Check results for BUSCO: 115070at7742
## Thu Jul 14 01:07:03 CEST 2022 - The aliscore output file does not exist. Check results for BUSCO: 112966at7742
## Thu Jul 14 01:07:03 CEST 2022 - The aliscore output appears to be empty. Check results for BUSCO: 112966at7742
## Thu Jul 14 01:07:04 CEST 2022 - The aliscore output file does not exist. Check results for BUSCO: 155999at7742
## Thu Jul 14 01:07:04 CEST 2022 - The aliscore output appears to be empty. Check results for BUSCO: 155999at7742
## Thu Jul 14 01:07:06 CEST 2022 - The aliscore output file does not exist. Check results for BUSCO: 308333at7742
## Thu Jul 14 01:07:06 CEST 2022 - The aliscore output appears to be empty. Check results for BUSCO: 308333at7742
## Thu Jul 14 01:07:08 CEST 2022 - The aliscore output file does not exist. Check results for BUSCO: 281785at7742
## Thu Jul 14 01:07:08 CEST 2022 - The aliscore output appears to be empty. Check results for BUSCO: 281785at7742
## Thu Jul 14 01:07:09 CEST 2022 - The aliscore output file does not exist. Check results for BUSCO: 75874at7742
## Thu Jul 14 01:07:09 CEST 2022 - The aliscore output appears to be empty. Check results for BUSCO: 75874at7742
## Thu Jul 14 01:07:10 CEST 2022 - The aliscore output file does not exist. Check results for BUSCO: 223515at7742
## Thu Jul 14 01:07:11 CEST 2022 - The aliscore output appears to be empty. Check results for BUSCO: 223515at7742
## Thu Jul 14 01:07:11 CEST 2022 - The aliscore output file does not exist. Check results for BUSCO: 70131at7742
## Thu Jul 14 01:07:11 CEST 2022 - The aliscore output appears to be empty. Check results for BUSCO: 70131at7742
## Thu Jul 14 01:07:13 CEST 2022 - The aliscore output file does not exist. Check results for BUSCO: 113535at7742
## Thu Jul 14 01:07:13 CEST 2022 - The aliscore output appears to be empty. Check results for BUSCO: 113535at7742
## Thu Jul 14 01:07:13 CEST 2022 - The aliscore output file does not exist. Check results for BUSCO: 194479at7742
## Thu Jul 14 01:07:13 CEST 2022 - The aliscore output appears to be empty. Check results for BUSCO: 194479at7742
## Thu Jul 14 01:07:16 CEST 2022 - The aliscore output file does not exist. Check results for BUSCO: 150433at7742
## Thu Jul 14 01:07:16 CEST 2022 - The aliscore output appears to be empty. Check results for BUSCO: 150433at7742
## Thu Jul 14 01:07:17 CEST 2022 - The aliscore output file does not exist. Check results for BUSCO: 102691at7742
## Thu Jul 14 01:07:17 CEST 2022 - The aliscore output appears to be empty. Check results for BUSCO: 102691at7742
## Thu Jul 14 01:07:19 CEST 2022 - The aliscore output file does not exist. Check results for BUSCO: 171487at7742
## Thu Jul 14 01:07:19 CEST 2022 - The aliscore output appears to be empty. Check results for BUSCO: 171487at7742
## Thu Jul 14 01:07:20 CEST 2022 - The aliscore output file does not exist. Check results for BUSCO: 414653at7742
## Thu Jul 14 01:07:20 CEST 2022 - The aliscore output appears to be empty. Check results for BUSCO: 414653at7742
## Thu Jul 14 01:07:20 CEST 2022 - The aliscore output file does not exist. Check results for BUSCO: 134453at7742
## Thu Jul 14 01:07:20 CEST 2022 - The aliscore output appears to be empty. Check results for BUSCO: 134453at7742
## Thu Jul 14 01:07:20 CEST 2022 - The aliscore output file does not exist. Check results for BUSCO: 240065at7742
## Thu Jul 14 01:07:20 CEST 2022 - The aliscore output appears to be empty. Check results for BUSCO: 240065at7742
## Thu Jul 14 01:07:22 CEST 2022 - The aliscore output file does not exist. Check results for BUSCO: 78859at7742
## Thu Jul 14 01:07:22 CEST 2022 - The aliscore output appears to be empty. Check results for BUSCO: 78859at7742
## Thu Jul 14 01:07:23 CEST 2022 - The aliscore output file does not exist. Check results for BUSCO: 84209at7742
## Thu Jul 14 01:07:23 CEST 2022 - The aliscore output appears to be empty. Check results for BUSCO: 84209at7742
## Thu Jul 14 01:07:26 CEST 2022 - The aliscore output file does not exist. Check results for BUSCO: 119066at7742
## Thu Jul 14 01:07:26 CEST 2022 - The aliscore output appears to be empty. Check results for BUSCO: 119066at7742
## Thu Jul 14 01:07:26 CEST 2022 - The aliscore output file does not exist. Check results for BUSCO: 123814at7742
## Thu Jul 14 01:07:26 CEST 2022 - The aliscore output appears to be empty. Check results for BUSCO: 123814at7742
## Thu Jul 14 01:07:27 CEST 2022 - The aliscore output file does not exist. Check results for BUSCO: 95871at7742
## Thu Jul 14 01:07:27 CEST 2022 - The aliscore output appears to be empty. Check results for BUSCO: 95871at7742
## Thu Jul 14 01:07:28 CEST 2022 - The aliscore output file does not exist. Check results for BUSCO: 158042at7742
## Thu Jul 14 01:07:28 CEST 2022 - The aliscore output appears to be empty. Check results for BUSCO: 158042at7742
## Thu Jul 14 01:07:28 CEST 2022 - The aliscore output file does not exist. Check results for BUSCO: 140310at7742
## Thu Jul 14 01:07:28 CEST 2022 - The aliscore output appears to be empty. Check results for BUSCO: 140310at7742
## Thu Jul 14 01:07:29 CEST 2022 - The aliscore output file does not exist. Check results for BUSCO: 325663at7742
## Thu Jul 14 01:07:30 CEST 2022 - The aliscore output appears to be empty. Check results for BUSCO: 325663at7742
## Thu Jul 14 01:07:32 CEST 2022 - The aliscore output file does not exist. Check results for BUSCO: 170750at7742
## Thu Jul 14 01:07:32 CEST 2022 - The aliscore output appears to be empty. Check results for BUSCO: 170750at7742
## Thu Jul 14 01:07:35 CEST 2022 - The aliscore output file does not exist. Check results for BUSCO: 303207at7742
## Thu Jul 14 01:07:35 CEST 2022 - The aliscore output appears to be empty. Check results for BUSCO: 303207at7742
## Thu Jul 14 01:07:37 CEST 2022 - The aliscore output file does not exist. Check results for BUSCO: 132847at7742
## Thu Jul 14 01:07:37 CEST 2022 - The aliscore output appears to be empty. Check results for BUSCO: 132847at7742
## Thu Jul 14 01:07:37 CEST 2022 - The aliscore output file does not exist. Check results for BUSCO: 252557at7742
## Thu Jul 14 01:07:37 CEST 2022 - The aliscore output appears to be empty. Check results for BUSCO: 252557at7742
## Thu Jul 14 01:07:37 CEST 2022 - The aliscore output file does not exist. Check results for BUSCO: 102408at7742
## Thu Jul 14 01:07:37 CEST 2022 - The aliscore output appears to be empty. Check results for BUSCO: 102408at7742
## Thu Jul 14 01:07:38 CEST 2022 - The aliscore output file does not exist. Check results for BUSCO: 81711at7742
## Thu Jul 14 01:07:38 CEST 2022 - The aliscore output appears to be empty. Check results for BUSCO: 81711at7742
## Thu Jul 14 01:07:38 CEST 2022 - The aliscore output file does not exist. Check results for BUSCO: 84209at7742
## Thu Jul 14 01:07:38 CEST 2022 - The aliscore output appears to be empty. Check results for BUSCO: 84209at7742
## Thu Jul 14 01:07:42 CEST 2022 - The aliscore output file does not exist. Check results for BUSCO: 122955at7742
## Thu Jul 14 01:07:42 CEST 2022 - The aliscore output appears to be empty. Check results for BUSCO: 122955at7742
## Thu Jul 14 01:07:42 CEST 2022 - The aliscore output file does not exist. Check results for BUSCO: 103640at7742
## Thu Jul 14 01:07:42 CEST 2022 - The aliscore output appears to be empty. Check results for BUSCO: 103640at7742
## Thu Jul 14 01:07:43 CEST 2022 - The aliscore output file does not exist. Check results for BUSCO: 263251at7742
## Thu Jul 14 01:07:43 CEST 2022 - The aliscore output appears to be empty. Check results for BUSCO: 263251at7742
## Thu Jul 14 01:07:44 CEST 2022 - The aliscore output file does not exist. Check results for BUSCO: 132010at7742
## Thu Jul 14 01:07:44 CEST 2022 - The aliscore output appears to be empty. Check results for BUSCO: 132010at7742
## Thu Jul 14 01:07:45 CEST 2022 - The aliscore output file does not exist. Check results for BUSCO: 303765at7742
## Thu Jul 14 01:07:45 CEST 2022 - The aliscore output appears to be empty. Check results for BUSCO: 303765at7742
## Thu Jul 14 01:07:46 CEST 2022 - The aliscore output file does not exist. Check results for BUSCO: 294082at7742
## Thu Jul 14 01:07:46 CEST 2022 - The aliscore output appears to be empty. Check results for BUSCO: 294082at7742
## Thu Jul 14 01:07:47 CEST 2022 - The aliscore output file does not exist. Check results for BUSCO: 139307at7742
## Thu Jul 14 01:07:47 CEST 2022 - The aliscore output file does not exist. Check results for BUSCO: 402344at7742
## Thu Jul 14 01:07:47 CEST 2022 - The aliscore output appears to be empty. Check results for BUSCO: 139307at7742
## Thu Jul 14 01:07:47 CEST 2022 - The aliscore output appears to be empty. Check results for BUSCO: 402344at7742
## Thu Jul 14 01:07:51 CEST 2022 - The aliscore output file does not exist. Check results for BUSCO: 236439at7742
## Thu Jul 14 01:07:51 CEST 2022 - The aliscore output appears to be empty. Check results for BUSCO: 236439at7742
## Thu Jul 14 01:07:52 CEST 2022 - The aliscore output file does not exist. Check results for BUSCO: 192715at7742
## Thu Jul 14 01:07:52 CEST 2022 - The aliscore output appears to be empty. Check results for BUSCO: 192715at7742
## Thu Jul 14 01:07:54 CEST 2022 - The aliscore output file does not exist. Check results for BUSCO: 277665at7742
## Thu Jul 14 01:07:54 CEST 2022 - The aliscore output appears to be empty. Check results for BUSCO: 277665at7742
## Thu Jul 14 01:07:55 CEST 2022 - The aliscore output file does not exist. Check results for BUSCO: 203861at7742
## Thu Jul 14 01:07:55 CEST 2022 - The aliscore output appears to be empty. Check results for BUSCO: 203861at7742
## Thu Jul 14 01:07:55 CEST 2022 - The aliscore output file does not exist. Check results for BUSCO: 62525at7742
## Thu Jul 14 01:07:55 CEST 2022 - The aliscore output appears to be empty. Check results for BUSCO: 62525at7742
## Thu Jul 14 01:07:55 CEST 2022 - The aliscore output file does not exist. Check results for BUSCO: 327776at7742
## Thu Jul 14 01:07:55 CEST 2022 - The aliscore output appears to be empty. Check results for BUSCO: 327776at7742
## Thu Jul 14 01:07:58 CEST 2022 - The aliscore output file does not exist. Check results for BUSCO: 285589at7742
## Thu Jul 14 01:07:58 CEST 2022 - The aliscore output appears to be empty. Check results for BUSCO: 285589at7742
## Thu Jul 14 01:07:59 CEST 2022 - The aliscore output file does not exist. Check results for BUSCO: 303765at7742
## Thu Jul 14 01:07:59 CEST 2022 - The aliscore output appears to be empty. Check results for BUSCO: 303765at7742
## Thu Jul 14 01:08:00 CEST 2022 - The aliscore output file does not exist. Check results for BUSCO: 206434at7742
## Thu Jul 14 01:08:00 CEST 2022 - The aliscore output appears to be empty. Check results for BUSCO: 206434at7742
## Thu Jul 14 01:08:02 CEST 2022 - The aliscore output file does not exist. Check results for BUSCO: 104126at7742
## Thu Jul 14 01:08:02 CEST 2022 - The aliscore output appears to be empty. Check results for BUSCO: 104126at7742
## Thu Jul 14 01:08:03 CEST 2022 - The aliscore output file does not exist. Check results for BUSCO: 202558at7742
## Thu Jul 14 01:08:03 CEST 2022 - The aliscore output appears to be empty. Check results for BUSCO: 202558at7742
## Thu Jul 14 01:08:04 CEST 2022 - The aliscore output file does not exist. Check results for BUSCO: 414653at7742
## Thu Jul 14 01:08:04 CEST 2022 - The aliscore output appears to be empty. Check results for BUSCO: 414653at7742
## Thu Jul 14 01:08:05 CEST 2022 - The aliscore output file does not exist. Check results for BUSCO: 115042at7742
## Thu Jul 14 01:08:05 CEST 2022 - The aliscore output appears to be empty. Check results for BUSCO: 115042at7742
## Thu Jul 14 01:08:07 CEST 2022 - The aliscore output file does not exist. Check results for BUSCO: 232857at7742
## Thu Jul 14 01:08:07 CEST 2022 - The aliscore output appears to be empty. Check results for BUSCO: 232857at7742
## Thu Jul 14 01:08:08 CEST 2022 - The aliscore output file does not exist. Check results for BUSCO: 135770at7742
## Thu Jul 14 01:08:08 CEST 2022 - The aliscore output appears to be empty. Check results for BUSCO: 135770at7742
## Thu Jul 14 01:08:10 CEST 2022 - The aliscore output file does not exist. Check results for BUSCO: 366345at7742
## Thu Jul 14 01:08:10 CEST 2022 - The aliscore output appears to be empty. Check results for BUSCO: 366345at7742
## Thu Jul 14 01:08:11 CEST 2022 - The aliscore output file does not exist. Check results for BUSCO: 194479at7742
## Thu Jul 14 01:08:11 CEST 2022 - The aliscore output appears to be empty. Check results for BUSCO: 194479at7742
## Thu Jul 14 01:08:12 CEST 2022 - The aliscore output file does not exist. Check results for BUSCO: 81711at7742
## Thu Jul 14 01:08:12 CEST 2022 - The aliscore output appears to be empty. Check results for BUSCO: 81711at7742
## Thu Jul 14 01:08:13 CEST 2022 - The aliscore output file does not exist. Check results for BUSCO: 224267at7742
## Thu Jul 14 01:08:13 CEST 2022 - The aliscore output appears to be empty. Check results for BUSCO: 224267at7742
## Thu Jul 14 01:08:13 CEST 2022 - The aliscore output file does not exist. Check results for BUSCO: 283469at7742
## Thu Jul 14 01:08:13 CEST 2022 - The aliscore output appears to be empty. Check results for BUSCO: 283469at7742
## Thu Jul 14 01:08:14 CEST 2022 - The aliscore output file does not exist. Check results for BUSCO: 239108at7742
## Thu Jul 14 01:08:14 CEST 2022 - The aliscore output appears to be empty. Check results for BUSCO: 239108at7742
## Thu Jul 14 01:08:15 CEST 2022 - The aliscore output file does not exist. Check results for BUSCO: 236607at7742
## Thu Jul 14 01:08:15 CEST 2022 - The aliscore output appears to be empty. Check results for BUSCO: 236607at7742
## Thu Jul 14 01:08:18 CEST 2022 - The aliscore output file does not exist. Check results for BUSCO: 366345at7742
## Thu Jul 14 01:08:18 CEST 2022 - The aliscore output appears to be empty. Check results for BUSCO: 366345at7742
## Thu Jul 14 01:08:19 CEST 2022 - The aliscore output file does not exist. Check results for BUSCO: 181384at7742
## Thu Jul 14 01:08:19 CEST 2022 - The aliscore output appears to be empty. Check results for BUSCO: 181384at7742
## Thu Jul 14 01:08:19 CEST 2022 - The aliscore output file does not exist. Check results for BUSCO: 129214at7742
## Thu Jul 14 01:08:19 CEST 2022 - The aliscore output appears to be empty. Check results for BUSCO: 129214at7742
## Thu Jul 14 01:08:20 CEST 2022 - The aliscore output file does not exist. Check results for BUSCO: 253625at7742
## Thu Jul 14 01:08:20 CEST 2022 - The aliscore output appears to be empty. Check results for BUSCO: 253625at7742
## Thu Jul 14 01:08:21 CEST 2022 - The aliscore output file does not exist. Check results for BUSCO: 200616at7742
## Thu Jul 14 01:08:21 CEST 2022 - The aliscore output appears to be empty. Check results for BUSCO: 200616at7742
## Thu Jul 14 01:08:22 CEST 2022 - The aliscore output file does not exist. Check results for BUSCO: 57538at7742
## Thu Jul 14 01:08:22 CEST 2022 - The aliscore output appears to be empty. Check results for BUSCO: 57538at7742
## Thu Jul 14 01:08:22 CEST 2022 - The aliscore output file does not exist. Check results for BUSCO: 384788at7742
## Thu Jul 14 01:08:23 CEST 2022 - The aliscore output appears to be empty. Check results for BUSCO: 384788at7742
## Thu Jul 14 01:08:24 CEST 2022 - The aliscore output file does not exist. Check results for BUSCO: 109054at7742
## Thu Jul 14 01:08:24 CEST 2022 - The aliscore output appears to be empty. Check results for BUSCO: 109054at7742
## Thu Jul 14 01:08:26 CEST 2022 - The aliscore output file does not exist. Check results for BUSCO: 191130at7742
## Thu Jul 14 01:08:26 CEST 2022 - The aliscore output appears to be empty. Check results for BUSCO: 191130at7742
## Thu Jul 14 01:08:27 CEST 2022 - The aliscore output file does not exist. Check results for BUSCO: 308333at7742
## Thu Jul 14 01:08:27 CEST 2022 - The aliscore output appears to be empty. Check results for BUSCO: 308333at7742
## Thu Jul 14 01:08:28 CEST 2022 - The aliscore output file does not exist. Check results for BUSCO: 172339at7742
## Thu Jul 14 01:08:28 CEST 2022 - The aliscore output appears to be empty. Check results for BUSCO: 172339at7742
## Thu Jul 14 01:08:29 CEST 2022 - The aliscore output file does not exist. Check results for BUSCO: 81711at7742
## Thu Jul 14 01:08:29 CEST 2022 - The aliscore output appears to be empty. Check results for BUSCO: 81711at7742
## Thu Jul 14 01:08:31 CEST 2022 - The aliscore output file does not exist. Check results for BUSCO: 56885at7742
## Thu Jul 14 01:08:31 CEST 2022 - The aliscore output appears to be empty. Check results for BUSCO: 56885at7742
## Thu Jul 14 01:08:31 CEST 2022 - The aliscore output file does not exist. Check results for BUSCO: 280261at7742
## Thu Jul 14 01:08:31 CEST 2022 - The aliscore output appears to be empty. Check results for BUSCO: 280261at7742
## Thu Jul 14 01:08:33 CEST 2022 - The aliscore output file does not exist. Check results for BUSCO: 199449at7742
## Thu Jul 14 01:08:33 CEST 2022 - The aliscore output appears to be empty. Check results for BUSCO: 199449at7742
## Thu Jul 14 01:08:35 CEST 2022 - The aliscore output file does not exist. Check results for BUSCO: 180447at7742
## Thu Jul 14 01:08:35 CEST 2022 - The aliscore output appears to be empty. Check results for BUSCO: 180447at7742
## Thu Jul 14 01:08:35 CEST 2022 - The aliscore output file does not exist. Check results for BUSCO: 78859at7742
## Thu Jul 14 01:08:35 CEST 2022 - The aliscore output appears to be empty. Check results for BUSCO: 78859at7742
## Thu Jul 14 01:08:38 CEST 2022 - The aliscore output file does not exist. Check results for BUSCO: 281785at7742
## Thu Jul 14 01:08:38 CEST 2022 - The aliscore output appears to be empty. Check results for BUSCO: 281785at7742
## Thu Jul 14 01:08:38 CEST 2022 - The aliscore output file does not exist. Check results for BUSCO: 384788at7742
## Thu Jul 14 01:08:38 CEST 2022 - The aliscore output appears to be empty. Check results for BUSCO: 384788at7742
## Thu Jul 14 01:08:40 CEST 2022 - The aliscore output file does not exist. Check results for BUSCO: 93269at7742
## Thu Jul 14 01:08:40 CEST 2022 - The aliscore output appears to be empty. Check results for BUSCO: 93269at7742
## Thu Jul 14 01:08:41 CEST 2022 - The aliscore output file does not exist. Check results for BUSCO: 180416at7742
## Thu Jul 14 01:08:41 CEST 2022 - The aliscore output appears to be empty. Check results for BUSCO: 180416at7742
## Thu Jul 14 01:08:42 CEST 2022 - The aliscore output file does not exist. Check results for BUSCO: 164678at7742
## Thu Jul 14 01:08:42 CEST 2022 - The aliscore output appears to be empty. Check results for BUSCO: 164678at7742
## Thu Jul 14 01:08:43 CEST 2022 - The aliscore output file does not exist. Check results for BUSCO: 146203at7742
## Thu Jul 14 01:08:43 CEST 2022 - The aliscore output appears to be empty. Check results for BUSCO: 146203at7742
## Thu Jul 14 01:08:45 CEST 2022 - The aliscore output file does not exist. Check results for BUSCO: 384476at7742
## Thu Jul 14 01:08:45 CEST 2022 - The aliscore output appears to be empty. Check results for BUSCO: 384476at7742
## Thu Jul 14 01:08:47 CEST 2022 - The aliscore output file does not exist. Check results for BUSCO: 103919at7742
## Thu Jul 14 01:08:47 CEST 2022 - The aliscore output appears to be empty. Check results for BUSCO: 103919at7742
## Thu Jul 14 01:08:47 CEST 2022 - The aliscore output file does not exist. Check results for BUSCO: 114303at7742
## Thu Jul 14 01:08:47 CEST 2022 - The aliscore output appears to be empty. Check results for BUSCO: 114303at7742
## Thu Jul 14 01:08:48 CEST 2022 - The aliscore output file does not exist. Check results for BUSCO: 114924at7742
## Thu Jul 14 01:08:48 CEST 2022 - The aliscore output appears to be empty. Check results for BUSCO: 114924at7742
## Thu Jul 14 01:08:50 CEST 2022 - The aliscore output file does not exist. Check results for BUSCO: 373324at7742
## Thu Jul 14 01:08:50 CEST 2022 - The aliscore output appears to be empty. Check results for BUSCO: 373324at7742
## Thu Jul 14 01:08:50 CEST 2022 - The aliscore output file does not exist. Check results for BUSCO: 310707at7742
## Thu Jul 14 01:08:50 CEST 2022 - The aliscore output appears to be empty. Check results for BUSCO: 310707at7742
## Thu Jul 14 01:08:52 CEST 2022 - The aliscore output file does not exist. Check results for BUSCO: 227299at7742
## Thu Jul 14 01:08:52 CEST 2022 - The aliscore output appears to be empty. Check results for BUSCO: 227299at7742
## Thu Jul 14 01:08:53 CEST 2022 - The aliscore output file does not exist. Check results for BUSCO: 177660at7742
## Thu Jul 14 01:08:53 CEST 2022 - The aliscore output appears to be empty. Check results for BUSCO: 177660at7742
## Thu Jul 14 01:08:55 CEST 2022 - The aliscore output file does not exist. Check results for BUSCO: 72241at7742
## Thu Jul 14 01:08:55 CEST 2022 - The aliscore output appears to be empty. Check results for BUSCO: 72241at7742
## Thu Jul 14 01:08:56 CEST 2022 - The aliscore output file does not exist. Check results for BUSCO: 190917at7742
## Thu Jul 14 01:08:56 CEST 2022 - The aliscore output appears to be empty. Check results for BUSCO: 190917at7742
## Thu Jul 14 01:08:58 CEST 2022 - The aliscore output file does not exist. Check results for BUSCO: 281413at7742
## Thu Jul 14 01:08:58 CEST 2022 - The aliscore output appears to be empty. Check results for BUSCO: 281413at7742
## Thu Jul 14 01:08:59 CEST 2022 - The aliscore output file does not exist. Check results for BUSCO: 85986at7742
## Thu Jul 14 01:08:59 CEST 2022 - The aliscore output appears to be empty. Check results for BUSCO: 85986at7742
## Thu Jul 14 01:09:00 CEST 2022 - The aliscore output file does not exist. Check results for BUSCO: 213224at7742
## Thu Jul 14 01:09:00 CEST 2022 - The aliscore output appears to be empty. Check results for BUSCO: 213224at7742
## Thu Jul 14 01:09:00 CEST 2022 - The aliscore output file does not exist. Check results for BUSCO: 400635at7742
## Thu Jul 14 01:09:00 CEST 2022 - The aliscore output appears to be empty. Check results for BUSCO: 400635at7742
## Thu Jul 14 01:09:03 CEST 2022 - The aliscore output file does not exist. Check results for BUSCO: 85986at7742
## Thu Jul 14 01:09:03 CEST 2022 - The aliscore output appears to be empty. Check results for BUSCO: 85986at7742
## Thu Jul 14 01:09:03 CEST 2022 - The aliscore output file does not exist. Check results for BUSCO: 170526at7742
## Thu Jul 14 01:09:03 CEST 2022 - The aliscore output appears to be empty. Check results for BUSCO: 170526at7742
## Thu Jul 14 01:09:04 CEST 2022 - The aliscore output file does not exist. Check results for BUSCO: 158445at7742
## Thu Jul 14 01:09:04 CEST 2022 - The aliscore output appears to be empty. Check results for BUSCO: 158445at7742
## Thu Jul 14 01:09:06 CEST 2022 - The aliscore output file does not exist. Check results for BUSCO: 206434at7742
## Thu Jul 14 01:09:06 CEST 2022 - The aliscore output appears to be empty. Check results for BUSCO: 206434at7742
## Thu Jul 14 01:09:07 CEST 2022 - The aliscore output file does not exist. Check results for BUSCO: 154679at7742
## Thu Jul 14 01:09:08 CEST 2022 - The aliscore output appears to be empty. Check results for BUSCO: 154679at7742
## Thu Jul 14 01:09:09 CEST 2022 - The aliscore output file does not exist. Check results for BUSCO: 114924at7742
## Thu Jul 14 01:09:09 CEST 2022 - The aliscore output appears to be empty. Check results for BUSCO: 114924at7742
## Thu Jul 14 01:09:09 CEST 2022 - The aliscore output file does not exist. Check results for BUSCO: 115748at7742
## Thu Jul 14 01:09:09 CEST 2022 - The aliscore output appears to be empty. Check results for BUSCO: 115748at7742
## Thu Jul 14 01:09:12 CEST 2022 - The aliscore output file does not exist. Check results for BUSCO: 95261at7742
## Thu Jul 14 01:09:12 CEST 2022 - The aliscore output appears to be empty. Check results for BUSCO: 95261at7742
## Thu Jul 14 01:09:13 CEST 2022 - The aliscore output file does not exist. Check results for BUSCO: 252557at7742
## Thu Jul 14 01:09:13 CEST 2022 - The aliscore output appears to be empty. Check results for BUSCO: 252557at7742
## Thu Jul 14 01:09:15 CEST 2022 - The aliscore output file does not exist. Check results for BUSCO: 139946at7742
## Thu Jul 14 01:09:15 CEST 2022 - The aliscore output appears to be empty. Check results for BUSCO: 139946at7742
## Thu Jul 14 01:09:16 CEST 2022 - The aliscore output file does not exist. Check results for BUSCO: 169034at7742
## Thu Jul 14 01:09:16 CEST 2022 - The aliscore output appears to be empty. Check results for BUSCO: 169034at7742
## Thu Jul 14 01:09:17 CEST 2022 - The aliscore output file does not exist. Check results for BUSCO: 202558at7742
## Thu Jul 14 01:09:17 CEST 2022 - The aliscore output appears to be empty. Check results for BUSCO: 202558at7742
## Thu Jul 14 01:09:18 CEST 2022 - The aliscore output file does not exist. Check results for BUSCO: 115070at7742
## Thu Jul 14 01:09:18 CEST 2022 - The aliscore output appears to be empty. Check results for BUSCO: 115070at7742
## Thu Jul 14 01:09:19 CEST 2022 - The aliscore output file does not exist. Check results for BUSCO: 329621at7742
## Thu Jul 14 01:09:19 CEST 2022 - The aliscore output appears to be empty. Check results for BUSCO: 329621at7742
## Thu Jul 14 01:09:20 CEST 2022 - The aliscore output file does not exist. Check results for BUSCO: 400635at7742
## Thu Jul 14 01:09:20 CEST 2022 - The aliscore output appears to be empty. Check results for BUSCO: 400635at7742
## Thu Jul 14 01:09:21 CEST 2022 - The aliscore output file does not exist. Check results for BUSCO: 239731at7742
## Thu Jul 14 01:09:21 CEST 2022 - The aliscore output appears to be empty. Check results for BUSCO: 239731at7742
## Thu Jul 14 01:09:21 CEST 2022 - The aliscore output file does not exist. Check results for BUSCO: 85369at7742
## Thu Jul 14 01:09:22 CEST 2022 - The aliscore output appears to be empty. Check results for BUSCO: 85369at7742
## Thu Jul 14 01:09:24 CEST 2022 - The aliscore output file does not exist. Check results for BUSCO: 324714at7742
## Thu Jul 14 01:09:24 CEST 2022 - The aliscore output appears to be empty. Check results for BUSCO: 324714at7742
## Thu Jul 14 01:09:26 CEST 2022 - The aliscore output file does not exist. Check results for BUSCO: 139974at7742
## Thu Jul 14 01:09:26 CEST 2022 - The aliscore output appears to be empty. Check results for BUSCO: 139974at7742
## Thu Jul 14 01:09:27 CEST 2022 - The aliscore output file does not exist. Check results for BUSCO: 222975at7742
## Thu Jul 14 01:09:27 CEST 2022 - The aliscore output appears to be empty. Check results for BUSCO: 222975at7742
## Thu Jul 14 01:09:28 CEST 2022 - The aliscore output file does not exist. Check results for BUSCO: 126329at7742
## Thu Jul 14 01:09:28 CEST 2022 - The aliscore output appears to be empty. Check results for BUSCO: 126329at7742
## Thu Jul 14 01:09:28 CEST 2022 - The aliscore output file does not exist. Check results for BUSCO: 268947at7742
## Thu Jul 14 01:09:28 CEST 2022 - The aliscore output appears to be empty. Check results for BUSCO: 268947at7742
## Thu Jul 14 01:09:30 CEST 2022 - The aliscore output file does not exist. Check results for BUSCO: 181591at7742
## Thu Jul 14 01:09:30 CEST 2022 - The aliscore output appears to be empty. Check results for BUSCO: 181591at7742
## Thu Jul 14 01:09:32 CEST 2022 - The aliscore output file does not exist. Check results for BUSCO: 174607at7742
## Thu Jul 14 01:09:32 CEST 2022 - The aliscore output appears to be empty. Check results for BUSCO: 174607at7742
## Thu Jul 14 01:09:32 CEST 2022 - The aliscore output file does not exist. Check results for BUSCO: 196659at7742
## Thu Jul 14 01:09:32 CEST 2022 - The aliscore output appears to be empty. Check results for BUSCO: 196659at7742
## Thu Jul 14 01:09:33 CEST 2022 - The aliscore output file does not exist. Check results for BUSCO: 198300at7742
## Thu Jul 14 01:09:33 CEST 2022 - The aliscore output appears to be empty. Check results for BUSCO: 198300at7742
## Thu Jul 14 01:09:36 CEST 2022 - The aliscore output file does not exist. Check results for BUSCO: 106423at7742
## Thu Jul 14 01:09:36 CEST 2022 - The aliscore output appears to be empty. Check results for BUSCO: 106423at7742
## Thu Jul 14 01:09:36 CEST 2022 - The aliscore output file does not exist. Check results for BUSCO: 280159at7742
## Thu Jul 14 01:09:36 CEST 2022 - The aliscore output appears to be empty. Check results for BUSCO: 280159at7742
## Thu Jul 14 01:09:39 CEST 2022 - The aliscore output file does not exist. Check results for BUSCO: 93020at7742
## Thu Jul 14 01:09:39 CEST 2022 - The aliscore output appears to be empty. Check results for BUSCO: 93020at7742
## Thu Jul 14 01:09:40 CEST 2022 - The aliscore output file does not exist. Check results for BUSCO: 161077at7742
## Thu Jul 14 01:09:40 CEST 2022 - The aliscore output appears to be empty. Check results for BUSCO: 161077at7742
## Thu Jul 14 01:09:40 CEST 2022 - The aliscore output file does not exist. Check results for BUSCO: 98464at7742
## Thu Jul 14 01:09:41 CEST 2022 - The aliscore output appears to be empty. Check results for BUSCO: 98464at7742
## Thu Jul 14 01:09:41 CEST 2022 - The aliscore output file does not exist. Check results for BUSCO: 96693at7742
## Thu Jul 14 01:09:41 CEST 2022 - The aliscore output appears to be empty. Check results for BUSCO: 96693at7742
## Thu Jul 14 01:09:43 CEST 2022 - The aliscore output file does not exist. Check results for BUSCO: 324312at7742
## Thu Jul 14 01:09:43 CEST 2022 - The aliscore output appears to be empty. Check results for BUSCO: 324312at7742
## Thu Jul 14 01:09:44 CEST 2022 - The aliscore output file does not exist. Check results for BUSCO: 233794at7742
## Thu Jul 14 01:09:44 CEST 2022 - The aliscore output appears to be empty. Check results for BUSCO: 233794at7742
## Thu Jul 14 01:09:46 CEST 2022 - The aliscore output file does not exist. Check results for BUSCO: 192715at7742
## Thu Jul 14 01:09:46 CEST 2022 - The aliscore output appears to be empty. Check results for BUSCO: 192715at7742
## Thu Jul 14 01:09:46 CEST 2022 - The aliscore output file does not exist. Check results for BUSCO: 173398at7742
## Thu Jul 14 01:09:46 CEST 2022 - The aliscore output appears to be empty. Check results for BUSCO: 173398at7742
## Thu Jul 14 01:09:49 CEST 2022 - The aliscore output file does not exist. Check results for BUSCO: 59090at7742
## Thu Jul 14 01:09:49 CEST 2022 - The aliscore output appears to be empty. Check results for BUSCO: 59090at7742
## Thu Jul 14 01:09:49 CEST 2022 - The aliscore output file does not exist. Check results for BUSCO: 136142at7742
## Thu Jul 14 01:09:50 CEST 2022 - The aliscore output appears to be empty. Check results for BUSCO: 136142at7742
## Thu Jul 14 01:09:50 CEST 2022 - The aliscore output file does not exist. Check results for BUSCO: 122841at7742
## Thu Jul 14 01:09:51 CEST 2022 - The aliscore output appears to be empty. Check results for BUSCO: 122841at7742
## Thu Jul 14 01:09:52 CEST 2022 - The aliscore output file does not exist. Check results for BUSCO: 130549at7742
## Thu Jul 14 01:09:52 CEST 2022 - The aliscore output appears to be empty. Check results for BUSCO: 130549at7742
## Thu Jul 14 01:09:53 CEST 2022 - The aliscore output file does not exist. Check results for BUSCO: 151923at7742
## Thu Jul 14 01:09:53 CEST 2022 - The aliscore output appears to be empty. Check results for BUSCO: 151923at7742
## Thu Jul 14 01:09:55 CEST 2022 - The aliscore output file does not exist. Check results for BUSCO: 131898at7742
## Thu Jul 14 01:09:55 CEST 2022 - The aliscore output appears to be empty. Check results for BUSCO: 131898at7742
## Thu Jul 14 01:09:55 CEST 2022 - The aliscore output file does not exist. Check results for BUSCO: 370236at7742
## Thu Jul 14 01:09:55 CEST 2022 - The aliscore output appears to be empty. Check results for BUSCO: 370236at7742
## Thu Jul 14 01:09:57 CEST 2022 - The aliscore output file does not exist. Check results for BUSCO: 173398at7742
## Thu Jul 14 01:09:57 CEST 2022 - The aliscore output appears to be empty. Check results for BUSCO: 173398at7742
## Thu Jul 14 01:09:57 CEST 2022 - The aliscore output file does not exist. Check results for BUSCO: 139974at7742
## Thu Jul 14 01:09:57 CEST 2022 - The aliscore output appears to be empty. Check results for BUSCO: 139974at7742
## Thu Jul 14 01:09:59 CEST 2022 - The aliscore output file does not exist. Check results for BUSCO: 86856at7742
## Thu Jul 14 01:09:59 CEST 2022 - The aliscore output appears to be empty. Check results for BUSCO: 86856at7742
## Thu Jul 14 01:10:00 CEST 2022 - The aliscore output file does not exist. Check results for BUSCO: 256457at7742
## Thu Jul 14 01:10:00 CEST 2022 - The aliscore output appears to be empty. Check results for BUSCO: 256457at7742
## Thu Jul 14 01:10:01 CEST 2022 - The aliscore output file does not exist. Check results for BUSCO: 272794at7742
## Thu Jul 14 01:10:01 CEST 2022 - The aliscore output appears to be empty. Check results for BUSCO: 272794at7742
## Thu Jul 14 01:10:02 CEST 2022 - The aliscore output file does not exist. Check results for BUSCO: 203050at7742
## Thu Jul 14 01:10:02 CEST 2022 - The aliscore output appears to be empty. Check results for BUSCO: 203050at7742
## Thu Jul 14 01:10:03 CEST 2022 - The aliscore output file does not exist. Check results for BUSCO: 291807at7742
## Thu Jul 14 01:10:03 CEST 2022 - The aliscore output appears to be empty. Check results for BUSCO: 291807at7742
## Thu Jul 14 01:10:05 CEST 2022 - The aliscore output file does not exist. Check results for BUSCO: 151777at7742
## Thu Jul 14 01:10:05 CEST 2022 - The aliscore output appears to be empty. Check results for BUSCO: 151777at7742
## Thu Jul 14 01:10:07 CEST 2022 - The aliscore output file does not exist. Check results for BUSCO: 175755at7742
## Thu Jul 14 01:10:07 CEST 2022 - The aliscore output file does not exist. Check results for BUSCO: 367625at7742
## Thu Jul 14 01:10:07 CEST 2022 - The aliscore output appears to be empty. Check results for BUSCO: 175755at7742
## Thu Jul 14 01:10:07 CEST 2022 - The aliscore output appears to be empty. Check results for BUSCO: 367625at7742
## Thu Jul 14 01:10:09 CEST 2022 - The aliscore output file does not exist. Check results for BUSCO: 102222at7742
## Thu Jul 14 01:10:09 CEST 2022 - The aliscore output appears to be empty. Check results for BUSCO: 102222at7742
## Thu Jul 14 01:10:11 CEST 2022 - The aliscore output file does not exist. Check results for BUSCO: 102594at7742
## Thu Jul 14 01:10:11 CEST 2022 - The aliscore output appears to be empty. Check results for BUSCO: 102594at7742
## Thu Jul 14 01:10:12 CEST 2022 - The aliscore output file does not exist. Check results for BUSCO: 172737at7742
## Thu Jul 14 01:10:12 CEST 2022 - The aliscore output appears to be empty. Check results for BUSCO: 172737at7742
## Thu Jul 14 01:10:12 CEST 2022 - The aliscore output file does not exist. Check results for BUSCO: 171487at7742
## Thu Jul 14 01:10:12 CEST 2022 - The aliscore output appears to be empty. Check results for BUSCO: 171487at7742
## Thu Jul 14 01:10:15 CEST 2022 - The aliscore output file does not exist. Check results for BUSCO: 325335at7742
## Thu Jul 14 01:10:15 CEST 2022 - The aliscore output appears to be empty. Check results for BUSCO: 325335at7742
## Thu Jul 14 01:10:17 CEST 2022 - The aliscore output file does not exist. Check results for BUSCO: 66867at7742
## Thu Jul 14 01:10:17 CEST 2022 - The aliscore output appears to be empty. Check results for BUSCO: 66867at7742
## Thu Jul 14 01:10:18 CEST 2022 - The aliscore output file does not exist. Check results for BUSCO: 140588at7742
## Thu Jul 14 01:10:18 CEST 2022 - The aliscore output appears to be empty. Check results for BUSCO: 140588at7742
## Thu Jul 14 01:10:18 CEST 2022 - The aliscore output file does not exist. Check results for BUSCO: 163908at7742
## Thu Jul 14 01:10:18 CEST 2022 - The aliscore output appears to be empty. Check results for BUSCO: 163908at7742
## Thu Jul 14 01:10:19 CEST 2022 - The aliscore output file does not exist. Check results for BUSCO: 183058at7742
## Thu Jul 14 01:10:19 CEST 2022 - The aliscore output appears to be empty. Check results for BUSCO: 183058at7742
## Thu Jul 14 01:10:20 CEST 2022 - The aliscore output file does not exist. Check results for BUSCO: 299549at7742
## Thu Jul 14 01:10:20 CEST 2022 - The aliscore output appears to be empty. Check results for BUSCO: 299549at7742
## Thu Jul 14 01:10:23 CEST 2022 - The aliscore output file does not exist. Check results for BUSCO: 116667at7742
## Thu Jul 14 01:10:23 CEST 2022 - The aliscore output appears to be empty. Check results for BUSCO: 116667at7742
## Thu Jul 14 01:10:23 CEST 2022 - The aliscore output file does not exist. Check results for BUSCO: 223515at7742
## Thu Jul 14 01:10:23 CEST 2022 - The aliscore output appears to be empty. Check results for BUSCO: 223515at7742
## Thu Jul 14 01:10:24 CEST 2022 - The aliscore output file does not exist. Check results for BUSCO: 117494at7742
## Thu Jul 14 01:10:24 CEST 2022 - The aliscore output appears to be empty. Check results for BUSCO: 117494at7742
## Thu Jul 14 01:10:24 CEST 2022 - The aliscore output file does not exist. Check results for BUSCO: 143678at7742
## Thu Jul 14 01:10:24 CEST 2022 - The aliscore output appears to be empty. Check results for BUSCO: 143678at7742
## Thu Jul 14 01:10:25 CEST 2022 - The aliscore output file does not exist. Check results for BUSCO: 343527at7742
## Thu Jul 14 01:10:25 CEST 2022 - The aliscore output appears to be empty. Check results for BUSCO: 343527at7742
## Thu Jul 14 01:10:28 CEST 2022 - The aliscore output file does not exist. Check results for BUSCO: 209187at7742
## Thu Jul 14 01:10:28 CEST 2022 - The aliscore output appears to be empty. Check results for BUSCO: 209187at7742
## Thu Jul 14 01:10:28 CEST 2022 - The aliscore output file does not exist. Check results for BUSCO: 119066at7742
## Thu Jul 14 01:10:28 CEST 2022 - The aliscore output appears to be empty. Check results for BUSCO: 119066at7742
## Thu Jul 14 01:10:31 CEST 2022 - The aliscore output file does not exist. Check results for BUSCO: 237620at7742
## Thu Jul 14 01:10:31 CEST 2022 - The aliscore output appears to be empty. Check results for BUSCO: 237620at7742
## Thu Jul 14 01:10:31 CEST 2022 - The aliscore output file does not exist. Check results for BUSCO: 340519at7742
## Thu Jul 14 01:10:31 CEST 2022 - The aliscore output appears to be empty. Check results for BUSCO: 340519at7742
## Thu Jul 14 01:10:32 CEST 2022 - The aliscore output file does not exist. Check results for BUSCO: 164526at7742
## Thu Jul 14 01:10:32 CEST 2022 - The aliscore output appears to be empty. Check results for BUSCO: 164526at7742
## Thu Jul 14 01:10:33 CEST 2022 - The aliscore output file does not exist. Check results for BUSCO: 220642at7742
## Thu Jul 14 01:10:33 CEST 2022 - The aliscore output file does not exist. Check results for BUSCO: 120131at7742
## Thu Jul 14 01:10:33 CEST 2022 - The aliscore output appears to be empty. Check results for BUSCO: 220642at7742
## Thu Jul 14 01:10:33 CEST 2022 - The aliscore output appears to be empty. Check results for BUSCO: 120131at7742
## Thu Jul 14 01:10:35 CEST 2022 - The aliscore output file does not exist. Check results for BUSCO: 131983at7742
## Thu Jul 14 01:10:35 CEST 2022 - The aliscore output appears to be empty. Check results for BUSCO: 131983at7742
## Thu Jul 14 01:10:36 CEST 2022 - The aliscore output file does not exist. Check results for BUSCO: 123156at7742
## Thu Jul 14 01:10:36 CEST 2022 - The aliscore output appears to be empty. Check results for BUSCO: 123156at7742
## Thu Jul 14 01:10:37 CEST 2022 - The aliscore output file does not exist. Check results for BUSCO: 313611at7742
## Thu Jul 14 01:10:37 CEST 2022 - The aliscore output appears to be empty. Check results for BUSCO: 313611at7742
## Thu Jul 14 01:10:41 CEST 2022 - The aliscore output file does not exist. Check results for BUSCO: 137629at7742
## Thu Jul 14 01:10:41 CEST 2022 - The aliscore output appears to be empty. Check results for BUSCO: 137629at7742
## Thu Jul 14 01:10:41 CEST 2022 - The aliscore output file does not exist. Check results for BUSCO: 162840at7742
## Thu Jul 14 01:10:41 CEST 2022 - The aliscore output appears to be empty. Check results for BUSCO: 162840at7742
## Thu Jul 14 01:10:41 CEST 2022 - The aliscore output file does not exist. Check results for BUSCO: 223515at7742
## Thu Jul 14 01:10:41 CEST 2022 - The aliscore output appears to be empty. Check results for BUSCO: 223515at7742
## Thu Jul 14 01:10:42 CEST 2022 - The aliscore output file does not exist. Check results for BUSCO: 325029at7742
## Thu Jul 14 01:10:42 CEST 2022 - The aliscore output appears to be empty. Check results for BUSCO: 325029at7742
## Thu Jul 14 01:10:43 CEST 2022 - The aliscore output file does not exist. Check results for BUSCO: 194479at7742
## Thu Jul 14 01:10:43 CEST 2022 - The aliscore output appears to be empty. Check results for BUSCO: 194479at7742
## Thu Jul 14 01:10:45 CEST 2022 - The aliscore output file does not exist. Check results for BUSCO: 130549at7742
## Thu Jul 14 01:10:45 CEST 2022 - The aliscore output appears to be empty. Check results for BUSCO: 130549at7742
## Thu Jul 14 01:10:47 CEST 2022 - The aliscore output file does not exist. Check results for BUSCO: 133877at7742
## Thu Jul 14 01:10:47 CEST 2022 - The aliscore output appears to be empty. Check results for BUSCO: 133877at7742
## Thu Jul 14 01:10:47 CEST 2022 - The aliscore output file does not exist. Check results for BUSCO: 239108at7742
## Thu Jul 14 01:10:47 CEST 2022 - The aliscore output appears to be empty. Check results for BUSCO: 239108at7742
## Thu Jul 14 01:10:49 CEST 2022 - The aliscore output file does not exist. Check results for BUSCO: 151777at7742
## Thu Jul 14 01:10:49 CEST 2022 - The aliscore output appears to be empty. Check results for BUSCO: 151777at7742
## Thu Jul 14 01:10:50 CEST 2022 - The aliscore output file does not exist. Check results for BUSCO: 90151at7742
## Thu Jul 14 01:10:50 CEST 2022 - The aliscore output appears to be empty. Check results for BUSCO: 90151at7742
## Thu Jul 14 01:10:50 CEST 2022 - The aliscore output file does not exist. Check results for BUSCO: 114214at7742
## Thu Jul 14 01:10:50 CEST 2022 - The aliscore output appears to be empty. Check results for BUSCO: 114214at7742
## Thu Jul 14 01:10:51 CEST 2022 - The aliscore output file does not exist. Check results for BUSCO: 59090at7742
## Thu Jul 14 01:10:51 CEST 2022 - The aliscore output appears to be empty. Check results for BUSCO: 59090at7742
## Thu Jul 14 01:10:54 CEST 2022 - The aliscore output file does not exist. Check results for BUSCO: 134453at7742
## Thu Jul 14 01:10:54 CEST 2022 - The aliscore output file does not exist. Check results for BUSCO: 220786at7742
## Thu Jul 14 01:10:54 CEST 2022 - The aliscore output appears to be empty. Check results for BUSCO: 134453at7742
## Thu Jul 14 01:10:54 CEST 2022 - The aliscore output appears to be empty. Check results for BUSCO: 220786at7742
## Thu Jul 14 01:10:56 CEST 2022 - The aliscore output file does not exist. Check results for BUSCO: 253625at7742
## Thu Jul 14 01:10:56 CEST 2022 - The aliscore output appears to be empty. Check results for BUSCO: 253625at7742
## Thu Jul 14 01:10:57 CEST 2022 - The aliscore output file does not exist. Check results for BUSCO: 101858at7742
## Thu Jul 14 01:10:57 CEST 2022 - The aliscore output appears to be empty. Check results for BUSCO: 101858at7742
## Thu Jul 14 01:10:57 CEST 2022 - The aliscore output file does not exist. Check results for BUSCO: 236910at7742
## Thu Jul 14 01:10:57 CEST 2022 - The aliscore output appears to be empty. Check results for BUSCO: 236910at7742
## Thu Jul 14 01:10:59 CEST 2022 - The aliscore output file does not exist. Check results for BUSCO: 164526at7742
## Thu Jul 14 01:10:59 CEST 2022 - The aliscore output appears to be empty. Check results for BUSCO: 164526at7742
## Thu Jul 14 01:11:00 CEST 2022 - The aliscore output file does not exist. Check results for BUSCO: 157513at7742
## Thu Jul 14 01:11:00 CEST 2022 - The aliscore output appears to be empty. Check results for BUSCO: 157513at7742
## Thu Jul 14 01:11:02 CEST 2022 - The aliscore output file does not exist. Check results for BUSCO: 239731at7742
## Thu Jul 14 01:11:02 CEST 2022 - The aliscore output appears to be empty. Check results for BUSCO: 239731at7742
## Thu Jul 14 01:11:02 CEST 2022 - The aliscore output file does not exist. Check results for BUSCO: 272309at7742
## Thu Jul 14 01:11:03 CEST 2022 - The aliscore output appears to be empty. Check results for BUSCO: 272309at7742
## Thu Jul 14 01:11:06 CEST 2022 - The aliscore output file does not exist. Check results for BUSCO: 337175at7742
## Thu Jul 14 01:11:06 CEST 2022 - The aliscore output appears to be empty. Check results for BUSCO: 337175at7742
## Thu Jul 14 01:11:06 CEST 2022 - The aliscore output file does not exist. Check results for BUSCO: 358181at7742
## Thu Jul 14 01:11:06 CEST 2022 - The aliscore output appears to be empty. Check results for BUSCO: 358181at7742
## Thu Jul 14 01:11:06 CEST 2022 - The aliscore output file does not exist. Check results for BUSCO: 139974at7742
## Thu Jul 14 01:11:06 CEST 2022 - The aliscore output appears to be empty. Check results for BUSCO: 139974at7742
## Thu Jul 14 01:11:07 CEST 2022 - The aliscore output file does not exist. Check results for BUSCO: 203050at7742
## Thu Jul 14 01:11:07 CEST 2022 - The aliscore output appears to be empty. Check results for BUSCO: 203050at7742
## Thu Jul 14 01:11:07 CEST 2022 - The aliscore output file does not exist. Check results for BUSCO: 78859at7742
## Thu Jul 14 01:11:07 CEST 2022 - The aliscore output appears to be empty. Check results for BUSCO: 78859at7742
## Thu Jul 14 01:11:10 CEST 2022 - The aliscore output file does not exist. Check results for BUSCO: 170526at7742
## Thu Jul 14 01:11:10 CEST 2022 - The aliscore output appears to be empty. Check results for BUSCO: 170526at7742
## Thu Jul 14 01:11:11 CEST 2022 - The aliscore output file does not exist. Check results for BUSCO: 60569at7742
## Thu Jul 14 01:11:11 CEST 2022 - The aliscore output appears to be empty. Check results for BUSCO: 60569at7742
## Thu Jul 14 01:11:13 CEST 2022 - The aliscore output file does not exist. Check results for BUSCO: 223283at7742
## Thu Jul 14 01:11:13 CEST 2022 - The aliscore output appears to be empty. Check results for BUSCO: 223283at7742
## Thu Jul 14 01:11:15 CEST 2022 - The aliscore output file does not exist. Check results for BUSCO: 102408at7742
## Thu Jul 14 01:11:15 CEST 2022 - The aliscore output appears to be empty. Check results for BUSCO: 102408at7742
## Thu Jul 14 01:11:15 CEST 2022 - The aliscore output file does not exist. Check results for BUSCO: 183058at7742
## Thu Jul 14 01:11:15 CEST 2022 - The aliscore output appears to be empty. Check results for BUSCO: 183058at7742
## Thu Jul 14 01:11:16 CEST 2022 - The aliscore output file does not exist. Check results for BUSCO: 141965at7742
## Thu Jul 14 01:11:16 CEST 2022 - The aliscore output appears to be empty. Check results for BUSCO: 141965at7742
## Thu Jul 14 01:11:16 CEST 2022 - The aliscore output file does not exist. Check results for BUSCO: 164340at7742
## Thu Jul 14 01:11:16 CEST 2022 - The aliscore output appears to be empty. Check results for BUSCO: 164340at7742
## Thu Jul 14 01:11:20 CEST 2022 - The aliscore output file does not exist. Check results for BUSCO: 270710at7742
## Thu Jul 14 01:11:20 CEST 2022 - The aliscore output appears to be empty. Check results for BUSCO: 270710at7742
## Thu Jul 14 01:11:20 CEST 2022 - The aliscore output file does not exist. Check results for BUSCO: 236439at7742
## Thu Jul 14 01:11:20 CEST 2022 - The aliscore output appears to be empty. Check results for BUSCO: 236439at7742
## Thu Jul 14 01:11:21 CEST 2022 - The aliscore output file does not exist. Check results for BUSCO: 376554at7742
## Thu Jul 14 01:11:21 CEST 2022 - The aliscore output file does not exist. Check results for BUSCO: 172376at7742
## Thu Jul 14 01:11:21 CEST 2022 - The aliscore output appears to be empty. Check results for BUSCO: 376554at7742
## Thu Jul 14 01:11:21 CEST 2022 - The aliscore output appears to be empty. Check results for BUSCO: 172376at7742
## Thu Jul 14 01:11:22 CEST 2022 - The aliscore output file does not exist. Check results for BUSCO: 334896at7742
## Thu Jul 14 01:11:22 CEST 2022 - The aliscore output appears to be empty. Check results for BUSCO: 334896at7742
## Thu Jul 14 01:11:25 CEST 2022 - The aliscore output file does not exist. Check results for BUSCO: 130415at7742
## Thu Jul 14 01:11:25 CEST 2022 - The aliscore output appears to be empty. Check results for BUSCO: 130415at7742
## Thu Jul 14 01:11:26 CEST 2022 - The aliscore output file does not exist. Check results for BUSCO: 172339at7742
## Thu Jul 14 01:11:26 CEST 2022 - The aliscore output appears to be empty. Check results for BUSCO: 172339at7742
## Thu Jul 14 01:11:26 CEST 2022 - The aliscore output file does not exist. Check results for BUSCO: 93727at7742
## Thu Jul 14 01:11:26 CEST 2022 - The aliscore output appears to be empty. Check results for BUSCO: 93727at7742
## Thu Jul 14 01:11:29 CEST 2022 - The aliscore output file does not exist. Check results for BUSCO: 115070at7742
## Thu Jul 14 01:11:29 CEST 2022 - The aliscore output appears to be empty. Check results for BUSCO: 115070at7742
## Thu Jul 14 01:11:30 CEST 2022 - The aliscore output file does not exist. Check results for BUSCO: 123814at7742
## Thu Jul 14 01:11:30 CEST 2022 - The aliscore output appears to be empty. Check results for BUSCO: 123814at7742
## Thu Jul 14 01:11:30 CEST 2022 - The aliscore output file does not exist. Check results for BUSCO: 162701at7742
## Thu Jul 14 01:11:30 CEST 2022 - The aliscore output appears to be empty. Check results for BUSCO: 162701at7742
## Thu Jul 14 01:11:32 CEST 2022 - The aliscore output file does not exist. Check results for BUSCO: 291147at7742
## Thu Jul 14 01:11:32 CEST 2022 - The aliscore output appears to be empty. Check results for BUSCO: 291147at7742
## Thu Jul 14 01:11:33 CEST 2022 - The aliscore output file does not exist. Check results for BUSCO: 132847at7742
## Thu Jul 14 01:11:33 CEST 2022 - The aliscore output appears to be empty. Check results for BUSCO: 132847at7742
## Thu Jul 14 01:11:34 CEST 2022 - The aliscore output file does not exist. Check results for BUSCO: 220642at7742
## Thu Jul 14 01:11:34 CEST 2022 - The aliscore output appears to be empty. Check results for BUSCO: 220642at7742
## Thu Jul 14 01:11:36 CEST 2022 - The aliscore output file does not exist. Check results for BUSCO: 83010at7742
## Thu Jul 14 01:11:36 CEST 2022 - The aliscore output appears to be empty. Check results for BUSCO: 83010at7742
## Thu Jul 14 01:11:36 CEST 2022 - The aliscore output file does not exist. Check results for BUSCO: 224105at7742
## Thu Jul 14 01:11:36 CEST 2022 - The aliscore output appears to be empty. Check results for BUSCO: 224105at7742
## Thu Jul 14 01:11:38 CEST 2022 - The aliscore output file does not exist. Check results for BUSCO: 162701at7742
## Thu Jul 14 01:11:38 CEST 2022 - The aliscore output appears to be empty. Check results for BUSCO: 162701at7742
## Thu Jul 14 01:11:39 CEST 2022 - The aliscore output file does not exist. Check results for BUSCO: 250310at7742
## Thu Jul 14 01:11:39 CEST 2022 - The aliscore output appears to be empty. Check results for BUSCO: 250310at7742
## Thu Jul 14 01:11:40 CEST 2022 - The aliscore output file does not exist. Check results for BUSCO: 99014at7742
## Thu Jul 14 01:11:40 CEST 2022 - The aliscore output appears to be empty. Check results for BUSCO: 99014at7742
## Thu Jul 14 01:11:42 CEST 2022 - The aliscore output file does not exist. Check results for BUSCO: 376554at7742
## Thu Jul 14 01:11:42 CEST 2022 - The aliscore output appears to be empty. Check results for BUSCO: 376554at7742
## Thu Jul 14 01:11:43 CEST 2022 - The aliscore output file does not exist. Check results for BUSCO: 88636at7742
## Thu Jul 14 01:11:43 CEST 2022 - The aliscore output appears to be empty. Check results for BUSCO: 88636at7742
## Thu Jul 14 01:11:44 CEST 2022 - The aliscore output file does not exist. Check results for BUSCO: 134188at7742
## Thu Jul 14 01:11:44 CEST 2022 - The aliscore output appears to be empty. Check results for BUSCO: 134188at7742
## Thu Jul 14 01:11:46 CEST 2022 - The aliscore output file does not exist. Check results for BUSCO: 180447at7742
## Thu Jul 14 01:11:46 CEST 2022 - The aliscore output appears to be empty. Check results for BUSCO: 180447at7742
## Thu Jul 14 01:11:47 CEST 2022 - The aliscore output file does not exist. Check results for BUSCO: 239108at7742
## Thu Jul 14 01:11:47 CEST 2022 - The aliscore output appears to be empty. Check results for BUSCO: 239108at7742
## Thu Jul 14 01:11:47 CEST 2022 - The aliscore output file does not exist. Check results for BUSCO: 123814at7742
## Thu Jul 14 01:11:47 CEST 2022 - The aliscore output appears to be empty. Check results for BUSCO: 123814at7742
## Thu Jul 14 01:11:49 CEST 2022 - The aliscore output file does not exist. Check results for BUSCO: 111128at7742
## Thu Jul 14 01:11:49 CEST 2022 - The aliscore output appears to be empty. Check results for BUSCO: 111128at7742
## Thu Jul 14 01:11:50 CEST 2022 - The aliscore output file does not exist. Check results for BUSCO: 296592at7742
## Thu Jul 14 01:11:50 CEST 2022 - The aliscore output appears to be empty. Check results for BUSCO: 296592at7742
## Thu Jul 14 01:11:51 CEST 2022 - The aliscore output file does not exist. Check results for BUSCO: 106060at7742
## Thu Jul 14 01:11:51 CEST 2022 - The aliscore output appears to be empty. Check results for BUSCO: 106060at7742
## Thu Jul 14 01:11:52 CEST 2022 - The aliscore output file does not exist. Check results for BUSCO: 167017at7742
## Thu Jul 14 01:11:52 CEST 2022 - The aliscore output appears to be empty. Check results for BUSCO: 167017at7742
## Thu Jul 14 01:11:53 CEST 2022 - The aliscore output file does not exist. Check results for BUSCO: 119066at7742
## Thu Jul 14 01:11:53 CEST 2022 - The aliscore output appears to be empty. Check results for BUSCO: 119066at7742
## Thu Jul 14 01:11:55 CEST 2022 - The aliscore output file does not exist. Check results for BUSCO: 227299at7742
## Thu Jul 14 01:11:55 CEST 2022 - The aliscore output appears to be empty. Check results for BUSCO: 227299at7742
## Thu Jul 14 01:11:55 CEST 2022 - The aliscore output file does not exist. Check results for BUSCO: 199120at7742
## Thu Jul 14 01:11:55 CEST 2022 - The aliscore output appears to be empty. Check results for BUSCO: 199120at7742
## Thu Jul 14 01:11:57 CEST 2022 - The aliscore output file does not exist. Check results for BUSCO: 159169at7742
## Thu Jul 14 01:11:57 CEST 2022 - The aliscore output appears to be empty. Check results for BUSCO: 159169at7742
## Thu Jul 14 01:11:58 CEST 2022 - The aliscore output file does not exist. Check results for BUSCO: 184498at7742
## Thu Jul 14 01:11:58 CEST 2022 - The aliscore output appears to be empty. Check results for BUSCO: 184498at7742
## Thu Jul 14 01:12:00 CEST 2022 - The aliscore output file does not exist. Check results for BUSCO: 103208at7742
## Thu Jul 14 01:12:00 CEST 2022 - The aliscore output appears to be empty. Check results for BUSCO: 103208at7742
## Thu Jul 14 01:12:01 CEST 2022 - The aliscore output file does not exist. Check results for BUSCO: 197420at7742
## Thu Jul 14 01:12:01 CEST 2022 - The aliscore output appears to be empty. Check results for BUSCO: 197420at7742
## Thu Jul 14 01:12:01 CEST 2022 - The aliscore output file does not exist. Check results for BUSCO: 208949at7742
## Thu Jul 14 01:12:01 CEST 2022 - The aliscore output appears to be empty. Check results for BUSCO: 208949at7742
## Thu Jul 14 01:12:03 CEST 2022 - The aliscore output file does not exist. Check results for BUSCO: 102408at7742
## Thu Jul 14 01:12:03 CEST 2022 - The aliscore output appears to be empty. Check results for BUSCO: 102408at7742
## Thu Jul 14 01:12:04 CEST 2022 - The aliscore output file does not exist. Check results for BUSCO: 384476at7742
## Thu Jul 14 01:12:04 CEST 2022 - The aliscore output appears to be empty. Check results for BUSCO: 384476at7742
## Thu Jul 14 01:12:04 CEST 2022 - The aliscore output file does not exist. Check results for BUSCO: 253625at7742
## Thu Jul 14 01:12:04 CEST 2022 - The aliscore output appears to be empty. Check results for BUSCO: 253625at7742
## Thu Jul 14 01:12:07 CEST 2022 - The aliscore output file does not exist. Check results for BUSCO: 139946at7742
## Thu Jul 14 01:12:07 CEST 2022 - The aliscore output appears to be empty. Check results for BUSCO: 139946at7742
## Thu Jul 14 01:12:08 CEST 2022 - The aliscore output file does not exist. Check results for BUSCO: 98464at7742
## Thu Jul 14 01:12:08 CEST 2022 - The aliscore output appears to be empty. Check results for BUSCO: 98464at7742
## Thu Jul 14 01:12:09 CEST 2022 - The aliscore output file does not exist. Check results for BUSCO: 116835at7742
## Thu Jul 14 01:12:09 CEST 2022 - The aliscore output appears to be empty. Check results for BUSCO: 116835at7742
## Thu Jul 14 01:12:11 CEST 2022 - The aliscore output file does not exist. Check results for BUSCO: 250310at7742
## Thu Jul 14 01:12:11 CEST 2022 - The aliscore output appears to be empty. Check results for BUSCO: 250310at7742
## Thu Jul 14 01:12:11 CEST 2022 - The aliscore output file does not exist. Check results for BUSCO: 119796at7742
## Thu Jul 14 01:12:11 CEST 2022 - The aliscore output appears to be empty. Check results for BUSCO: 119796at7742
## Thu Jul 14 01:12:12 CEST 2022 - The aliscore output file does not exist. Check results for BUSCO: 56885at7742
## Thu Jul 14 01:12:12 CEST 2022 - The aliscore output appears to be empty. Check results for BUSCO: 56885at7742
## Thu Jul 14 01:12:13 CEST 2022 - The aliscore output file does not exist. Check results for BUSCO: 56885at7742
## Thu Jul 14 01:12:13 CEST 2022 - The aliscore output appears to be empty. Check results for BUSCO: 56885at7742
## Thu Jul 14 01:12:14 CEST 2022 - The aliscore output file does not exist. Check results for BUSCO: 166600at7742
## Thu Jul 14 01:12:14 CEST 2022 - The aliscore output appears to be empty. Check results for BUSCO: 166600at7742
## Thu Jul 14 01:12:15 CEST 2022 - The aliscore output file does not exist. Check results for BUSCO: 172339at7742
## Thu Jul 14 01:12:15 CEST 2022 - The aliscore output appears to be empty. Check results for BUSCO: 172339at7742
## Thu Jul 14 01:12:16 CEST 2022 - The aliscore output file does not exist. Check results for BUSCO: 172737at7742
## Thu Jul 14 01:12:16 CEST 2022 - The aliscore output appears to be empty. Check results for BUSCO: 172737at7742
## Thu Jul 14 01:12:19 CEST 2022 - The aliscore output file does not exist. Check results for BUSCO: 402344at7742
## Thu Jul 14 01:12:19 CEST 2022 - The aliscore output appears to be empty. Check results for BUSCO: 402344at7742
## Thu Jul 14 01:12:20 CEST 2022 - The aliscore output file does not exist. Check results for BUSCO: 59090at7742
## Thu Jul 14 01:12:20 CEST 2022 - The aliscore output appears to be empty. Check results for BUSCO: 59090at7742
## Thu Jul 14 01:12:20 CEST 2022 - The aliscore output file does not exist. Check results for BUSCO: 168697at7742
## Thu Jul 14 01:12:20 CEST 2022 - The aliscore output appears to be empty. Check results for BUSCO: 168697at7742
## Thu Jul 14 01:12:22 CEST 2022 - The aliscore output file does not exist. Check results for BUSCO: 99014at7742
## Thu Jul 14 01:12:22 CEST 2022 - The aliscore output appears to be empty. Check results for BUSCO: 99014at7742
## Thu Jul 14 01:12:23 CEST 2022 - The aliscore output file does not exist. Check results for BUSCO: 269302at7742
## Thu Jul 14 01:12:24 CEST 2022 - The aliscore output appears to be empty. Check results for BUSCO: 269302at7742
## Thu Jul 14 01:12:24 CEST 2022 - The aliscore output file does not exist. Check results for BUSCO: 141965at7742
## Thu Jul 14 01:12:24 CEST 2022 - The aliscore output appears to be empty. Check results for BUSCO: 141965at7742
## Thu Jul 14 01:12:25 CEST 2022 - The aliscore output file does not exist. Check results for BUSCO: 109623at7742
## Thu Jul 14 01:12:25 CEST 2022 - The aliscore output appears to be empty. Check results for BUSCO: 109623at7742
## Thu Jul 14 01:12:26 CEST 2022 - The aliscore output file does not exist. Check results for BUSCO: 90151at7742
## Thu Jul 14 01:12:26 CEST 2022 - The aliscore output appears to be empty. Check results for BUSCO: 90151at7742
## Thu Jul 14 01:12:28 CEST 2022 - The aliscore output file does not exist. Check results for BUSCO: 169034at7742
## Thu Jul 14 01:12:28 CEST 2022 - The aliscore output appears to be empty. Check results for BUSCO: 169034at7742
## Thu Jul 14 01:12:29 CEST 2022 - The aliscore output file does not exist. Check results for BUSCO: 147455at7742
## Thu Jul 14 01:12:29 CEST 2022 - The aliscore output appears to be empty. Check results for BUSCO: 147455at7742
## Thu Jul 14 01:12:31 CEST 2022 - The aliscore output file does not exist. Check results for BUSCO: 141965at7742
## Thu Jul 14 01:12:31 CEST 2022 - The aliscore output appears to be empty. Check results for BUSCO: 141965at7742
## Thu Jul 14 01:12:32 CEST 2022 - The aliscore output file does not exist. Check results for BUSCO: 173102at7742
## Thu Jul 14 01:12:32 CEST 2022 - The aliscore output appears to be empty. Check results for BUSCO: 173102at7742
## Thu Jul 14 01:12:32 CEST 2022 - The aliscore output file does not exist. Check results for BUSCO: 101858at7742
## Thu Jul 14 01:12:32 CEST 2022 - The aliscore output appears to be empty. Check results for BUSCO: 101858at7742
## Thu Jul 14 01:12:34 CEST 2022 - The aliscore output file does not exist. Check results for BUSCO: 283522at7742
## Thu Jul 14 01:12:34 CEST 2022 - The aliscore output appears to be empty. Check results for BUSCO: 283522at7742
## Thu Jul 14 01:12:36 CEST 2022 - The aliscore output file does not exist. Check results for BUSCO: 90151at7742
## Thu Jul 14 01:12:36 CEST 2022 - The aliscore output appears to be empty. Check results for BUSCO: 90151at7742
## Thu Jul 14 01:12:37 CEST 2022 - The aliscore output file does not exist. Check results for BUSCO: 414653at7742
## Thu Jul 14 01:12:37 CEST 2022 - The aliscore output appears to be empty. Check results for BUSCO: 414653at7742
## Thu Jul 14 01:12:37 CEST 2022 - The aliscore output file does not exist. Check results for BUSCO: 291289at7742
## Thu Jul 14 01:12:37 CEST 2022 - The aliscore output appears to be empty. Check results for BUSCO: 291289at7742
## Thu Jul 14 01:12:38 CEST 2022 - The aliscore output file does not exist. Check results for BUSCO: 250310at7742
## Thu Jul 14 01:12:39 CEST 2022 - The aliscore output appears to be empty. Check results for BUSCO: 250310at7742
## Thu Jul 14 01:12:39 CEST 2022 - The aliscore output file does not exist. Check results for BUSCO: 160923at7742
## Thu Jul 14 01:12:39 CEST 2022 - The aliscore output appears to be empty. Check results for BUSCO: 160923at7742
## Thu Jul 14 01:12:40 CEST 2022 - The aliscore output file does not exist. Check results for BUSCO: 110226at7742
## Thu Jul 14 01:12:40 CEST 2022 - The aliscore output appears to be empty. Check results for BUSCO: 110226at7742
## Thu Jul 14 01:12:44 CEST 2022 - The aliscore output file does not exist. Check results for BUSCO: 210721at7742
## Thu Jul 14 01:12:44 CEST 2022 - The aliscore output appears to be empty. Check results for BUSCO: 210721at7742
## Thu Jul 14 01:12:44 CEST 2022 - The aliscore output file does not exist. Check results for BUSCO: 93269at7742
## Thu Jul 14 01:12:45 CEST 2022 - The aliscore output appears to be empty. Check results for BUSCO: 93269at7742
## Thu Jul 14 01:12:45 CEST 2022 - The aliscore output file does not exist. Check results for BUSCO: 213511at7742
## Thu Jul 14 01:12:45 CEST 2022 - The aliscore output appears to be empty. Check results for BUSCO: 213511at7742
## Thu Jul 14 01:12:46 CEST 2022 - The aliscore output file does not exist. Check results for BUSCO: 93727at7742
## Thu Jul 14 01:12:46 CEST 2022 - The aliscore output file does not exist. Check results for BUSCO: 261612at7742
## Thu Jul 14 01:12:46 CEST 2022 - The aliscore output appears to be empty. Check results for BUSCO: 93727at7742
## Thu Jul 14 01:12:46 CEST 2022 - The aliscore output appears to be empty. Check results for BUSCO: 261612at7742
## Thu Jul 14 01:12:46 CEST 2022 - The aliscore output file does not exist. Check results for BUSCO: 229422at7742
## Thu Jul 14 01:12:46 CEST 2022 - The aliscore output appears to be empty. Check results for BUSCO: 229422at7742
## Thu Jul 14 01:12:49 CEST 2022 - The aliscore output file does not exist. Check results for BUSCO: 102135at7742
## Thu Jul 14 01:12:49 CEST 2022 - The aliscore output appears to be empty. Check results for BUSCO: 102135at7742
## Thu Jul 14 01:12:50 CEST 2022 - The aliscore output file does not exist. Check results for BUSCO: 305955at7742
## Thu Jul 14 01:12:50 CEST 2022 - The aliscore output appears to be empty. Check results for BUSCO: 305955at7742
## Thu Jul 14 01:12:53 CEST 2022 - The aliscore output file does not exist. Check results for BUSCO: 136089at7742
## Thu Jul 14 01:12:53 CEST 2022 - The aliscore output appears to be empty. Check results for BUSCO: 136089at7742
## Thu Jul 14 01:12:54 CEST 2022 - The aliscore output file does not exist. Check results for BUSCO: 317124at7742
## Thu Jul 14 01:12:54 CEST 2022 - The aliscore output file does not exist. Check results for BUSCO: 130062at7742
## Thu Jul 14 01:12:54 CEST 2022 - The aliscore output appears to be empty. Check results for BUSCO: 317124at7742
## Thu Jul 14 01:12:54 CEST 2022 - The aliscore output appears to be empty. Check results for BUSCO: 130062at7742
## Thu Jul 14 01:12:54 CEST 2022 - The aliscore output file does not exist. Check results for BUSCO: 373540at7742
## Thu Jul 14 01:12:54 CEST 2022 - The aliscore output appears to be empty. Check results for BUSCO: 373540at7742
## Thu Jul 14 01:12:54 CEST 2022 - The aliscore output file does not exist. Check results for BUSCO: 75874at7742
## Thu Jul 14 01:12:54 CEST 2022 - The aliscore output appears to be empty. Check results for BUSCO: 75874at7742
## Thu Jul 14 01:12:56 CEST 2022 - The aliscore output file does not exist. Check results for BUSCO: 244028at7742
## Thu Jul 14 01:12:56 CEST 2022 - The aliscore output appears to be empty. Check results for BUSCO: 244028at7742
## Thu Jul 14 01:12:59 CEST 2022 - The aliscore output file does not exist. Check results for BUSCO: 190356at7742
## Thu Jul 14 01:12:59 CEST 2022 - The aliscore output appears to be empty. Check results for BUSCO: 190356at7742
## Thu Jul 14 01:13:00 CEST 2022 - The aliscore output file does not exist. Check results for BUSCO: 248762at7742
## Thu Jul 14 01:13:00 CEST 2022 - The aliscore output appears to be empty. Check results for BUSCO: 248762at7742
## Thu Jul 14 01:13:03 CEST 2022 - The aliscore output file does not exist. Check results for BUSCO: 214501at7742
## Thu Jul 14 01:13:03 CEST 2022 - The aliscore output appears to be empty. Check results for BUSCO: 214501at7742
## Thu Jul 14 01:13:03 CEST 2022 - The aliscore output file does not exist. Check results for BUSCO: 139307at7742
## Thu Jul 14 01:13:03 CEST 2022 - The aliscore output appears to be empty. Check results for BUSCO: 139307at7742
## Thu Jul 14 01:13:04 CEST 2022 - The aliscore output file does not exist. Check results for BUSCO: 200403at7742
## Thu Jul 14 01:13:04 CEST 2022 - The aliscore output appears to be empty. Check results for BUSCO: 200403at7742
## Thu Jul 14 01:13:04 CEST 2022 - The aliscore output file does not exist. Check results for BUSCO: 214501at7742
## Thu Jul 14 01:13:04 CEST 2022 - The aliscore output appears to be empty. Check results for BUSCO: 214501at7742
## Thu Jul 14 01:13:06 CEST 2022 - The aliscore output file does not exist. Check results for BUSCO: 188851at7742
## Thu Jul 14 01:13:06 CEST 2022 - The aliscore output appears to be empty. Check results for BUSCO: 188851at7742
## Thu Jul 14 01:13:07 CEST 2022 - The aliscore output file does not exist. Check results for BUSCO: 84209at7742
## Thu Jul 14 01:13:07 CEST 2022 - The aliscore output appears to be empty. Check results for BUSCO: 84209at7742
## Thu Jul 14 01:13:09 CEST 2022 - The aliscore output file does not exist. Check results for BUSCO: 87423at7742
## Thu Jul 14 01:13:09 CEST 2022 - The aliscore output appears to be empty. Check results for BUSCO: 87423at7742
## Thu Jul 14 01:13:09 CEST 2022 - The aliscore output file does not exist. Check results for BUSCO: 358181at7742
## Thu Jul 14 01:13:09 CEST 2022 - The aliscore output appears to be empty. Check results for BUSCO: 358181at7742
## Thu Jul 14 01:13:11 CEST 2022 - The aliscore output file does not exist. Check results for BUSCO: 233794at7742
## Thu Jul 14 01:13:11 CEST 2022 - The aliscore output appears to be empty. Check results for BUSCO: 233794at7742
## Thu Jul 14 01:13:12 CEST 2022 - The aliscore output file does not exist. Check results for BUSCO: 283449at7742
## Thu Jul 14 01:13:12 CEST 2022 - The aliscore output appears to be empty. Check results for BUSCO: 283449at7742
## Thu Jul 14 01:13:12 CEST 2022 - The aliscore output file does not exist. Check results for BUSCO: 102594at7742
## Thu Jul 14 01:13:13 CEST 2022 - The aliscore output appears to be empty. Check results for BUSCO: 102594at7742
## Thu Jul 14 01:13:14 CEST 2022 - The aliscore output file does not exist. Check results for BUSCO: 193294at7742
## Thu Jul 14 01:13:14 CEST 2022 - The aliscore output appears to be empty. Check results for BUSCO: 193294at7742
## Thu Jul 14 01:13:16 CEST 2022 - The aliscore output file does not exist. Check results for BUSCO: 140588at7742
## Thu Jul 14 01:13:16 CEST 2022 - The aliscore output appears to be empty. Check results for BUSCO: 140588at7742
## Thu Jul 14 01:13:17 CEST 2022 - The aliscore output file does not exist. Check results for BUSCO: 161484at7742
## Thu Jul 14 01:13:17 CEST 2022 - The aliscore output appears to be empty. Check results for BUSCO: 161484at7742
## Thu Jul 14 01:13:18 CEST 2022 - The aliscore output file does not exist. Check results for BUSCO: 66867at7742
## Thu Jul 14 01:13:18 CEST 2022 - The aliscore output appears to be empty. Check results for BUSCO: 66867at7742
## Thu Jul 14 01:13:19 CEST 2022 - The aliscore output file does not exist. Check results for BUSCO: 93566at7742
## Thu Jul 14 01:13:19 CEST 2022 - The aliscore output appears to be empty. Check results for BUSCO: 93566at7742
## Thu Jul 14 01:13:20 CEST 2022 - The aliscore output file does not exist. Check results for BUSCO: 129979at7742
## Thu Jul 14 01:13:20 CEST 2022 - The aliscore output appears to be empty. Check results for BUSCO: 129979at7742
## Thu Jul 14 01:13:21 CEST 2022 - The aliscore output file does not exist. Check results for BUSCO: 162174at7742
## Thu Jul 14 01:13:21 CEST 2022 - The aliscore output appears to be empty. Check results for BUSCO: 162174at7742
## Thu Jul 14 01:13:22 CEST 2022 - The aliscore output file does not exist. Check results for BUSCO: 132847at7742
## Thu Jul 14 01:13:22 CEST 2022 - The aliscore output appears to be empty. Check results for BUSCO: 132847at7742
## Thu Jul 14 01:13:23 CEST 2022 - The aliscore output file does not exist. Check results for BUSCO: 102135at7742
## Thu Jul 14 01:13:23 CEST 2022 - The aliscore output appears to be empty. Check results for BUSCO: 102135at7742
## Thu Jul 14 01:13:24 CEST 2022 - The aliscore output file does not exist. Check results for BUSCO: 365677at7742
## Thu Jul 14 01:13:24 CEST 2022 - The aliscore output appears to be empty. Check results for BUSCO: 365677at7742
## Thu Jul 14 01:13:25 CEST 2022 - The aliscore output file does not exist. Check results for BUSCO: 136089at7742
## Thu Jul 14 01:13:25 CEST 2022 - The aliscore output appears to be empty. Check results for BUSCO: 136089at7742
## Thu Jul 14 01:13:26 CEST 2022 - The aliscore output file does not exist. Check results for BUSCO: 229422at7742
## Thu Jul 14 01:13:26 CEST 2022 - The aliscore output appears to be empty. Check results for BUSCO: 229422at7742
## Thu Jul 14 01:13:29 CEST 2022 - The aliscore output file does not exist. Check results for BUSCO: 150433at7742
## Thu Jul 14 01:13:29 CEST 2022 - The aliscore output appears to be empty. Check results for BUSCO: 150433at7742
## Thu Jul 14 01:13:29 CEST 2022 - The aliscore output file does not exist. Check results for BUSCO: 103208at7742
## Thu Jul 14 01:13:30 CEST 2022 - The aliscore output appears to be empty. Check results for BUSCO: 103208at7742
## Thu Jul 14 01:13:32 CEST 2022 - The aliscore output file does not exist. Check results for BUSCO: 180447at7742
## Thu Jul 14 01:13:32 CEST 2022 - The aliscore output appears to be empty. Check results for BUSCO: 180447at7742
## Thu Jul 14 01:13:33 CEST 2022 - The aliscore output file does not exist. Check results for BUSCO: 283910at7742
## Thu Jul 14 01:13:33 CEST 2022 - The aliscore output appears to be empty. Check results for BUSCO: 283910at7742
## Thu Jul 14 01:13:35 CEST 2022 - The aliscore output file does not exist. Check results for BUSCO: 154679at7742
## Thu Jul 14 01:13:35 CEST 2022 - The aliscore output appears to be empty. Check results for BUSCO: 154679at7742
## Thu Jul 14 01:13:35 CEST 2022 - The aliscore output file does not exist. Check results for BUSCO: 161077at7742
## Thu Jul 14 01:13:35 CEST 2022 - The aliscore output appears to be empty. Check results for BUSCO: 161077at7742
## Thu Jul 14 01:13:36 CEST 2022 - The aliscore output file does not exist. Check results for BUSCO: 224537at7742
## Thu Jul 14 01:13:36 CEST 2022 - The aliscore output appears to be empty. Check results for BUSCO: 224537at7742
## Thu Jul 14 01:13:38 CEST 2022 - The aliscore output file does not exist. Check results for BUSCO: 116667at7742
## Thu Jul 14 01:13:38 CEST 2022 - The aliscore output appears to be empty. Check results for BUSCO: 116667at7742
## Thu Jul 14 01:13:39 CEST 2022 - The aliscore output file does not exist. Check results for BUSCO: 88636at7742
## Thu Jul 14 01:13:40 CEST 2022 - The aliscore output appears to be empty. Check results for BUSCO: 88636at7742
## Thu Jul 14 01:13:40 CEST 2022 - The aliscore output file does not exist. Check results for BUSCO: 216380at7742
## Thu Jul 14 01:13:41 CEST 2022 - The aliscore output appears to be empty. Check results for BUSCO: 216380at7742
## Thu Jul 14 01:13:41 CEST 2022 - The aliscore output file does not exist. Check results for BUSCO: 365677at7742
## Thu Jul 14 01:13:41 CEST 2022 - The aliscore output appears to be empty. Check results for BUSCO: 365677at7742
## Thu Jul 14 01:13:43 CEST 2022 - The aliscore output file does not exist. Check results for BUSCO: 99440at7742
## Thu Jul 14 01:13:43 CEST 2022 - The aliscore output appears to be empty. Check results for BUSCO: 99440at7742
## Thu Jul 14 01:13:44 CEST 2022 - The aliscore output file does not exist. Check results for BUSCO: 116667at7742
## Thu Jul 14 01:13:44 CEST 2022 - The aliscore output appears to be empty. Check results for BUSCO: 116667at7742
## Thu Jul 14 01:13:45 CEST 2022 - The aliscore output file does not exist. Check results for BUSCO: 88636at7742
## Thu Jul 14 01:13:46 CEST 2022 - The aliscore output appears to be empty. Check results for BUSCO: 88636at7742
## Thu Jul 14 01:13:46 CEST 2022 - The aliscore output file does not exist. Check results for BUSCO: 72241at7742
## Thu Jul 14 01:13:46 CEST 2022 - The aliscore output appears to be empty. Check results for BUSCO: 72241at7742
## Thu Jul 14 01:13:47 CEST 2022 - The aliscore output file does not exist. Check results for BUSCO: 81278at7742
## Thu Jul 14 01:13:47 CEST 2022 - The aliscore output appears to be empty. Check results for BUSCO: 81278at7742
## Thu Jul 14 01:13:49 CEST 2022 - The aliscore output file does not exist. Check results for BUSCO: 164340at7742
## Thu Jul 14 01:13:49 CEST 2022 - The aliscore output file does not exist. Check results for BUSCO: 269540at7742
## Thu Jul 14 01:13:49 CEST 2022 - The aliscore output appears to be empty. Check results for BUSCO: 164340at7742
## Thu Jul 14 01:13:49 CEST 2022 - The aliscore output appears to be empty. Check results for BUSCO: 269540at7742
## Thu Jul 14 01:13:51 CEST 2022 - The aliscore output file does not exist. Check results for BUSCO: 192114at7742
## Thu Jul 14 01:13:51 CEST 2022 - The aliscore output appears to be empty. Check results for BUSCO: 192114at7742
## Thu Jul 14 01:13:52 CEST 2022 - The aliscore output file does not exist. Check results for BUSCO: 115748at7742
## Thu Jul 14 01:13:52 CEST 2022 - The aliscore output appears to be empty. Check results for BUSCO: 115748at7742
## Thu Jul 14 01:13:54 CEST 2022 - The aliscore output file does not exist. Check results for BUSCO: 288361at7742
## Thu Jul 14 01:13:54 CEST 2022 - The aliscore output appears to be empty. Check results for BUSCO: 288361at7742
## Thu Jul 14 01:13:54 CEST 2022 - The aliscore output file does not exist. Check results for BUSCO: 215159at7742
## Thu Jul 14 01:13:54 CEST 2022 - The aliscore output appears to be empty. Check results for BUSCO: 215159at7742
## Thu Jul 14 01:13:56 CEST 2022 - The aliscore output file does not exist. Check results for BUSCO: 122990at7742
## Thu Jul 14 01:13:56 CEST 2022 - The aliscore output appears to be empty. Check results for BUSCO: 122990at7742
## Thu Jul 14 01:13:57 CEST 2022 - The aliscore output file does not exist. Check results for BUSCO: 222821at7742
## Thu Jul 14 01:13:57 CEST 2022 - The aliscore output appears to be empty. Check results for BUSCO: 222821at7742
## Thu Jul 14 01:13:58 CEST 2022 - The aliscore output file does not exist. Check results for BUSCO: 198300at7742
## Thu Jul 14 01:13:58 CEST 2022 - The aliscore output appears to be empty. Check results for BUSCO: 198300at7742
## Thu Jul 14 01:14:00 CEST 2022 - The aliscore output file does not exist. Check results for BUSCO: 232857at7742
## Thu Jul 14 01:14:00 CEST 2022 - The aliscore output appears to be empty. Check results for BUSCO: 232857at7742
## Thu Jul 14 01:14:00 CEST 2022 - The aliscore output file does not exist. Check results for BUSCO: 303207at7742
## Thu Jul 14 01:14:00 CEST 2022 - The aliscore output appears to be empty. Check results for BUSCO: 303207at7742
## Thu Jul 14 01:14:00 CEST 2022 - The aliscore output file does not exist. Check results for BUSCO: 192313at7742
## Thu Jul 14 01:14:00 CEST 2022 - The aliscore output appears to be empty. Check results for BUSCO: 192313at7742
## Thu Jul 14 01:14:02 CEST 2022 - The aliscore output file does not exist. Check results for BUSCO: 345012at7742
## Thu Jul 14 01:14:02 CEST 2022 - The aliscore output appears to be empty. Check results for BUSCO: 345012at7742
## Thu Jul 14 01:14:03 CEST 2022 - The aliscore output file does not exist. Check results for BUSCO: 215427at7742
## Thu Jul 14 01:14:03 CEST 2022 - The aliscore output appears to be empty. Check results for BUSCO: 215427at7742
## Thu Jul 14 01:14:05 CEST 2022 - The aliscore output file does not exist. Check results for BUSCO: 170526at7742
## Thu Jul 14 01:14:05 CEST 2022 - The aliscore output appears to be empty. Check results for BUSCO: 170526at7742
## Thu Jul 14 01:14:06 CEST 2022 - The aliscore output file does not exist. Check results for BUSCO: 199553at7742
## Thu Jul 14 01:14:06 CEST 2022 - The aliscore output appears to be empty. Check results for BUSCO: 199553at7742
## Thu Jul 14 01:14:06 CEST 2022 - The aliscore output file does not exist. Check results for BUSCO: 140071at7742
## Thu Jul 14 01:14:06 CEST 2022 - The aliscore output appears to be empty. Check results for BUSCO: 140071at7742
## Thu Jul 14 01:14:09 CEST 2022 - The aliscore output file does not exist. Check results for BUSCO: 358171at7742
## Thu Jul 14 01:14:09 CEST 2022 - The aliscore output appears to be empty. Check results for BUSCO: 358171at7742
## Thu Jul 14 01:14:10 CEST 2022 - The aliscore output file does not exist. Check results for BUSCO: 134453at7742
## Thu Jul 14 01:14:10 CEST 2022 - The aliscore output appears to be empty. Check results for BUSCO: 134453at7742
## Thu Jul 14 01:14:12 CEST 2022 - The aliscore output file does not exist. Check results for BUSCO: 163908at7742
## Thu Jul 14 01:14:12 CEST 2022 - The aliscore output appears to be empty. Check results for BUSCO: 163908at7742
## Thu Jul 14 01:14:12 CEST 2022 - The aliscore output file does not exist. Check results for BUSCO: 149128at7742
## Thu Jul 14 01:14:12 CEST 2022 - The aliscore output appears to be empty. Check results for BUSCO: 149128at7742
## Thu Jul 14 01:14:14 CEST 2022 - The aliscore output file does not exist. Check results for BUSCO: 114115at7742
## Thu Jul 14 01:14:14 CEST 2022 - The aliscore output appears to be empty. Check results for BUSCO: 114115at7742
## Thu Jul 14 01:14:15 CEST 2022 - The aliscore output file does not exist. Check results for BUSCO: 66867at7742
## Thu Jul 14 01:14:15 CEST 2022 - The aliscore output appears to be empty. Check results for BUSCO: 66867at7742
## Thu Jul 14 01:14:16 CEST 2022 - The aliscore output file does not exist. Check results for BUSCO: 99440at7742
## Thu Jul 14 01:14:16 CEST 2022 - The aliscore output appears to be empty. Check results for BUSCO: 99440at7742
## Thu Jul 14 01:14:17 CEST 2022 - The aliscore output file does not exist. Check results for BUSCO: 166688at7742
## Thu Jul 14 01:14:17 CEST 2022 - The aliscore output appears to be empty. Check results for BUSCO: 166688at7742
## Thu Jul 14 01:14:19 CEST 2022 - The aliscore output file does not exist. Check results for BUSCO: 93566at7742
## Thu Jul 14 01:14:19 CEST 2022 - The aliscore output appears to be empty. Check results for BUSCO: 93566at7742
## Thu Jul 14 01:14:20 CEST 2022 - The aliscore output file does not exist. Check results for BUSCO: 199553at7742
## Thu Jul 14 01:14:20 CEST 2022 - The aliscore output appears to be empty. Check results for BUSCO: 199553at7742
## Thu Jul 14 01:14:22 CEST 2022 - The aliscore output file does not exist. Check results for BUSCO: 217728at7742
## Thu Jul 14 01:14:22 CEST 2022 - The aliscore output appears to be empty. Check results for BUSCO: 217728at7742
## Thu Jul 14 01:14:23 CEST 2022 - The aliscore output file does not exist. Check results for BUSCO: 209187at7742
## Thu Jul 14 01:14:23 CEST 2022 - The aliscore output appears to be empty. Check results for BUSCO: 209187at7742
## Thu Jul 14 01:14:24 CEST 2022 - The aliscore output file does not exist. Check results for BUSCO: 416929at7742
## Thu Jul 14 01:14:24 CEST 2022 - The aliscore output appears to be empty. Check results for BUSCO: 416929at7742
## Thu Jul 14 01:14:26 CEST 2022 - The aliscore output file does not exist. Check results for BUSCO: 144445at7742
## Thu Jul 14 01:14:26 CEST 2022 - The aliscore output appears to be empty. Check results for BUSCO: 144445at7742
## Thu Jul 14 01:14:27 CEST 2022 - The aliscore output file does not exist. Check results for BUSCO: 199553at7742
## Thu Jul 14 01:14:27 CEST 2022 - The aliscore output appears to be empty. Check results for BUSCO: 199553at7742
## Thu Jul 14 01:14:28 CEST 2022 - The aliscore output file does not exist. Check results for BUSCO: 203861at7742
## Thu Jul 14 01:14:28 CEST 2022 - The aliscore output appears to be empty. Check results for BUSCO: 203861at7742
## Thu Jul 14 01:14:29 CEST 2022 - The aliscore output file does not exist. Check results for BUSCO: 116812at7742
## Thu Jul 14 01:14:29 CEST 2022 - The aliscore output appears to be empty. Check results for BUSCO: 116812at7742
## Thu Jul 14 01:14:31 CEST 2022 - The aliscore output file does not exist. Check results for BUSCO: 197420at7742
## Thu Jul 14 01:14:31 CEST 2022 - The aliscore output appears to be empty. Check results for BUSCO: 197420at7742
## Thu Jul 14 01:14:32 CEST 2022 - The aliscore output file does not exist. Check results for BUSCO: 81278at7742
## Thu Jul 14 01:14:32 CEST 2022 - The aliscore output appears to be empty. Check results for BUSCO: 81278at7742
## Thu Jul 14 01:14:33 CEST 2022 - The aliscore output file does not exist. Check results for BUSCO: 229422at7742
## Thu Jul 14 01:14:33 CEST 2022 - The aliscore output appears to be empty. Check results for BUSCO: 229422at7742
## Thu Jul 14 01:14:36 CEST 2022 - The aliscore output file does not exist. Check results for BUSCO: 384476at7742
## Thu Jul 14 01:14:36 CEST 2022 - The aliscore output appears to be empty. Check results for BUSCO: 384476at7742
## Thu Jul 14 01:14:36 CEST 2022 - The aliscore output file does not exist. Check results for BUSCO: 268237at7742
## Thu Jul 14 01:14:36 CEST 2022 - The aliscore output appears to be empty. Check results for BUSCO: 268237at7742
## Thu Jul 14 01:14:37 CEST 2022 - The aliscore output file does not exist. Check results for BUSCO: 159169at7742
## Thu Jul 14 01:14:37 CEST 2022 - The aliscore output appears to be empty. Check results for BUSCO: 159169at7742
## Thu Jul 14 01:14:39 CEST 2022 - The aliscore output file does not exist. Check results for BUSCO: 162701at7742
## Thu Jul 14 01:14:39 CEST 2022 - The aliscore output appears to be empty. Check results for BUSCO: 162701at7742
## Thu Jul 14 01:14:39 CEST 2022 - The aliscore output file does not exist. Check results for BUSCO: 197883at7742
## Thu Jul 14 01:14:39 CEST 2022 - The aliscore output appears to be empty. Check results for BUSCO: 197883at7742
## Thu Jul 14 01:14:39 CEST 2022 - The aliscore output file does not exist. Check results for BUSCO: 93229at7742
## Thu Jul 14 01:14:40 CEST 2022 - The aliscore output appears to be empty. Check results for BUSCO: 93229at7742
## Thu Jul 14 01:14:42 CEST 2022 - The aliscore output file does not exist. Check results for BUSCO: 416929at7742
## Thu Jul 14 01:14:42 CEST 2022 - The aliscore output appears to be empty. Check results for BUSCO: 416929at7742
## Thu Jul 14 01:14:42 CEST 2022 - The aliscore output file does not exist. Check results for BUSCO: 164950at7742
## Thu Jul 14 01:14:42 CEST 2022 - The aliscore output appears to be empty. Check results for BUSCO: 164950at7742
## Thu Jul 14 01:14:45 CEST 2022 - The aliscore output file does not exist. Check results for BUSCO: 199449at7742
## Thu Jul 14 01:14:45 CEST 2022 - The aliscore output appears to be empty. Check results for BUSCO: 199449at7742
## Thu Jul 14 01:14:45 CEST 2022 - The aliscore output file does not exist. Check results for BUSCO: 139050at7742
## Thu Jul 14 01:14:45 CEST 2022 - The aliscore output appears to be empty. Check results for BUSCO: 139050at7742
## Thu Jul 14 01:14:46 CEST 2022 - The aliscore output file does not exist. Check results for BUSCO: 103422at7742
## Thu Jul 14 01:14:46 CEST 2022 - The aliscore output appears to be empty. Check results for BUSCO: 103422at7742
## Thu Jul 14 01:14:49 CEST 2022 - The aliscore output file does not exist. Check results for BUSCO: 225564at7742
## Thu Jul 14 01:14:49 CEST 2022 - The aliscore output file does not exist. Check results for BUSCO: 139307at7742
## Thu Jul 14 01:14:49 CEST 2022 - The aliscore output appears to be empty. Check results for BUSCO: 225564at7742
## Thu Jul 14 01:14:49 CEST 2022 - The aliscore output appears to be empty. Check results for BUSCO: 139307at7742
## Thu Jul 14 01:14:49 CEST 2022 - The aliscore output file does not exist. Check results for BUSCO: 83010at7742
## Thu Jul 14 01:14:49 CEST 2022 - The aliscore output appears to be empty. Check results for BUSCO: 83010at7742
## Thu Jul 14 01:14:51 CEST 2022 - The aliscore output file does not exist. Check results for BUSCO: 238721at7742
## Thu Jul 14 01:14:51 CEST 2022 - The aliscore output appears to be empty. Check results for BUSCO: 238721at7742
## Thu Jul 14 01:14:52 CEST 2022 - The aliscore output file does not exist. Check results for BUSCO: 324714at7742
## Thu Jul 14 01:14:52 CEST 2022 - The aliscore output appears to be empty. Check results for BUSCO: 324714at7742
## Thu Jul 14 01:14:53 CEST 2022 - The aliscore output file does not exist. Check results for BUSCO: 132010at7742
## Thu Jul 14 01:14:53 CEST 2022 - The aliscore output appears to be empty. Check results for BUSCO: 132010at7742
## Thu Jul 14 01:14:56 CEST 2022 - The aliscore output file does not exist. Check results for BUSCO: 167017at7742
## Thu Jul 14 01:14:56 CEST 2022 - The aliscore output appears to be empty. Check results for BUSCO: 167017at7742
## Thu Jul 14 01:14:57 CEST 2022 - The aliscore output file does not exist. Check results for BUSCO: 236439at7742
## Thu Jul 14 01:14:57 CEST 2022 - The aliscore output appears to be empty. Check results for BUSCO: 236439at7742
## Thu Jul 14 01:14:57 CEST 2022 - The aliscore output file does not exist. Check results for BUSCO: 277665at7742
## Thu Jul 14 01:14:57 CEST 2022 - The aliscore output appears to be empty. Check results for BUSCO: 277665at7742
## Thu Jul 14 01:14:59 CEST 2022 - The aliscore output file does not exist. Check results for BUSCO: 237959at7742
## Thu Jul 14 01:14:59 CEST 2022 - The aliscore output appears to be empty. Check results for BUSCO: 237959at7742
## Thu Jul 14 01:14:59 CEST 2022 - The aliscore output file does not exist. Check results for BUSCO: 81278at7742
## Thu Jul 14 01:14:59 CEST 2022 - The aliscore output appears to be empty. Check results for BUSCO: 81278at7742
## Thu Jul 14 01:15:00 CEST 2022 - The aliscore output file does not exist. Check results for BUSCO: 98464at7742
## Thu Jul 14 01:15:00 CEST 2022 - The aliscore output appears to be empty. Check results for BUSCO: 98464at7742
## Thu Jul 14 01:15:02 CEST 2022 - The aliscore output file does not exist. Check results for BUSCO: 162840at7742
## Thu Jul 14 01:15:02 CEST 2022 - The aliscore output appears to be empty. Check results for BUSCO: 162840at7742
## Thu Jul 14 01:15:04 CEST 2022 - The aliscore output file does not exist. Check results for BUSCO: 129979at7742
## Thu Jul 14 01:15:04 CEST 2022 - The aliscore output appears to be empty. Check results for BUSCO: 129979at7742
## Thu Jul 14 01:15:04 CEST 2022 - The aliscore output file does not exist. Check results for BUSCO: 164678at7742
## Thu Jul 14 01:15:05 CEST 2022 - The aliscore output appears to be empty. Check results for BUSCO: 164678at7742
## Thu Jul 14 01:15:05 CEST 2022 - The aliscore output file does not exist. Check results for BUSCO: 110226at7742
## Thu Jul 14 01:15:05 CEST 2022 - The aliscore output appears to be empty. Check results for BUSCO: 110226at7742
## Thu Jul 14 01:15:05 CEST 2022 - The aliscore output file does not exist. Check results for BUSCO: 218283at7742
## Thu Jul 14 01:15:05 CEST 2022 - The aliscore output appears to be empty. Check results for BUSCO: 218283at7742
## Thu Jul 14 01:15:08 CEST 2022 - The aliscore output file does not exist. Check results for BUSCO: 93229at7742
## Thu Jul 14 01:15:08 CEST 2022 - The aliscore output appears to be empty. Check results for BUSCO: 93229at7742
## Thu Jul 14 01:15:10 CEST 2022 - The aliscore output file does not exist. Check results for BUSCO: 154679at7742
## Thu Jul 14 01:15:10 CEST 2022 - The aliscore output appears to be empty. Check results for BUSCO: 154679at7742
## Thu Jul 14 01:15:11 CEST 2022 - The aliscore output file does not exist. Check results for BUSCO: 303207at7742
## Thu Jul 14 01:15:11 CEST 2022 - The aliscore output appears to be empty. Check results for BUSCO: 303207at7742
## Thu Jul 14 01:15:13 CEST 2022 - The aliscore output file does not exist. Check results for BUSCO: 174607at7742
## Thu Jul 14 01:15:13 CEST 2022 - The aliscore output appears to be empty. Check results for BUSCO: 174607at7742
## Thu Jul 14 01:15:13 CEST 2022 - The aliscore output file does not exist. Check results for BUSCO: 102594at7742
## Thu Jul 14 01:15:13 CEST 2022 - The aliscore output appears to be empty. Check results for BUSCO: 102594at7742
## Thu Jul 14 01:15:14 CEST 2022 - The aliscore output file does not exist. Check results for BUSCO: 203861at7742
## Thu Jul 14 01:15:14 CEST 2022 - The aliscore output appears to be empty. Check results for BUSCO: 203861at7742
## Thu Jul 14 01:15:16 CEST 2022 - The aliscore output file does not exist. Check results for BUSCO: 123758at7742
## Thu Jul 14 01:15:16 CEST 2022 - The aliscore output appears to be empty. Check results for BUSCO: 123758at7742
## Thu Jul 14 01:15:17 CEST 2022 - The aliscore output file does not exist. Check results for BUSCO: 159169at7742
## Thu Jul 14 01:15:17 CEST 2022 - The aliscore output appears to be empty. Check results for BUSCO: 159169at7742
## Thu Jul 14 01:15:18 CEST 2022 - The aliscore output file does not exist. Check results for BUSCO: 367625at7742
## Thu Jul 14 01:15:18 CEST 2022 - The aliscore output appears to be empty. Check results for BUSCO: 367625at7742
## Thu Jul 14 01:15:20 CEST 2022 - The aliscore output file does not exist. Check results for BUSCO: 83010at7742
## Thu Jul 14 01:15:20 CEST 2022 - The aliscore output appears to be empty. Check results for BUSCO: 83010at7742
## Thu Jul 14 01:15:21 CEST 2022 - The aliscore output file does not exist. Check results for BUSCO: 131564at7742
## Thu Jul 14 01:15:21 CEST 2022 - The aliscore output appears to be empty. Check results for BUSCO: 131564at7742
## Thu Jul 14 01:15:23 CEST 2022 - The aliscore output file does not exist. Check results for BUSCO: 362170at7742
## Thu Jul 14 01:15:23 CEST 2022 - The aliscore output appears to be empty. Check results for BUSCO: 362170at7742
## Thu Jul 14 01:15:23 CEST 2022 - The aliscore output file does not exist. Check results for BUSCO: 53269at7742
## Thu Jul 14 01:15:23 CEST 2022 - The aliscore output appears to be empty. Check results for BUSCO: 53269at7742
## Thu Jul 14 01:15:24 CEST 2022 - The aliscore output file does not exist. Check results for BUSCO: 205393at7742
## Thu Jul 14 01:15:25 CEST 2022 - The aliscore output appears to be empty. Check results for BUSCO: 205393at7742
## Thu Jul 14 01:15:27 CEST 2022 - The aliscore output file does not exist. Check results for BUSCO: 161077at7742
## Thu Jul 14 01:15:27 CEST 2022 - The aliscore output appears to be empty. Check results for BUSCO: 161077at7742
## Thu Jul 14 01:15:27 CEST 2022 - The aliscore output file does not exist. Check results for BUSCO: 138249at7742
## Thu Jul 14 01:15:27 CEST 2022 - The aliscore output appears to be empty. Check results for BUSCO: 138249at7742
## Thu Jul 14 01:15:28 CEST 2022 - The aliscore output file does not exist. Check results for BUSCO: 116835at7742
## Thu Jul 14 01:15:28 CEST 2022 - The aliscore output appears to be empty. Check results for BUSCO: 116835at7742
## Thu Jul 14 01:15:30 CEST 2022 - The aliscore output file does not exist. Check results for BUSCO: 144645at7742
## Thu Jul 14 01:15:30 CEST 2022 - The aliscore output appears to be empty. Check results for BUSCO: 144645at7742
## Thu Jul 14 01:15:30 CEST 2022 - The aliscore output file does not exist. Check results for BUSCO: 238721at7742
## Thu Jul 14 01:15:30 CEST 2022 - The aliscore output appears to be empty. Check results for BUSCO: 238721at7742
## Thu Jul 14 01:15:31 CEST 2022 - The aliscore output file does not exist. Check results for BUSCO: 198300at7742
## Thu Jul 14 01:15:31 CEST 2022 - The aliscore output appears to be empty. Check results for BUSCO: 198300at7742
## Thu Jul 14 01:15:33 CEST 2022 - The aliscore output file does not exist. Check results for BUSCO: 206370at7742
## Thu Jul 14 01:15:33 CEST 2022 - The aliscore output appears to be empty. Check results for BUSCO: 206370at7742
## Thu Jul 14 01:15:34 CEST 2022 - The aliscore output file does not exist. Check results for BUSCO: 99014at7742
## Thu Jul 14 01:15:34 CEST 2022 - The aliscore output appears to be empty. Check results for BUSCO: 99014at7742
## Thu Jul 14 01:15:35 CEST 2022 - The aliscore output file does not exist. Check results for BUSCO: 162840at7742
## Thu Jul 14 01:15:35 CEST 2022 - The aliscore output appears to be empty. Check results for BUSCO: 162840at7742
## Thu Jul 14 01:15:37 CEST 2022 - The aliscore output file does not exist. Check results for BUSCO: 298175at7742
## Thu Jul 14 01:15:37 CEST 2022 - The aliscore output appears to be empty. Check results for BUSCO: 298175at7742
## Thu Jul 14 01:15:38 CEST 2022 - The aliscore output file does not exist. Check results for BUSCO: 367625at7742
## Thu Jul 14 01:15:38 CEST 2022 - The aliscore output appears to be empty. Check results for BUSCO: 367625at7742
## Thu Jul 14 01:15:39 CEST 2022 - The aliscore output file does not exist. Check results for BUSCO: 169034at7742
## Thu Jul 14 01:15:39 CEST 2022 - The aliscore output appears to be empty. Check results for BUSCO: 169034at7742
## Thu Jul 14 01:15:42 CEST 2022 - The aliscore output file does not exist. Check results for BUSCO: 193294at7742
## Thu Jul 14 01:15:42 CEST 2022 - The aliscore output appears to be empty. Check results for BUSCO: 193294at7742
## Thu Jul 14 01:15:42 CEST 2022 - The aliscore output file does not exist. Check results for BUSCO: 204261at7742
## Thu Jul 14 01:15:42 CEST 2022 - The aliscore output appears to be empty. Check results for BUSCO: 204261at7742
## Thu Jul 14 01:15:43 CEST 2022 - The aliscore output file does not exist. Check results for BUSCO: 308333at7742
## Thu Jul 14 01:15:43 CEST 2022 - The aliscore output appears to be empty. Check results for BUSCO: 308333at7742
## Thu Jul 14 01:15:44 CEST 2022 - The aliscore output file does not exist. Check results for BUSCO: 168697at7742
## Thu Jul 14 01:15:44 CEST 2022 - The aliscore output appears to be empty. Check results for BUSCO: 168697at7742
## Thu Jul 14 01:15:45 CEST 2022 - The aliscore output file does not exist. Check results for BUSCO: 192142at7742
## Thu Jul 14 01:15:46 CEST 2022 - The aliscore output appears to be empty. Check results for BUSCO: 192142at7742
## Thu Jul 14 01:15:46 CEST 2022 - The aliscore output file does not exist. Check results for BUSCO: 197420at7742
## Thu Jul 14 01:15:46 CEST 2022 - The aliscore output appears to be empty. Check results for BUSCO: 197420at7742
## Thu Jul 14 01:15:48 CEST 2022 - The aliscore output file does not exist. Check results for BUSCO: 311134at7742
## Thu Jul 14 01:15:48 CEST 2022 - The aliscore output appears to be empty. Check results for BUSCO: 311134at7742
## Thu Jul 14 01:15:48 CEST 2022 - The aliscore output file does not exist. Check results for BUSCO: 190356at7742
## Thu Jul 14 01:15:48 CEST 2022 - The aliscore output appears to be empty. Check results for BUSCO: 190356at7742
## Thu Jul 14 01:15:51 CEST 2022 - The aliscore output file does not exist. Check results for BUSCO: 137629at7742
## Thu Jul 14 01:15:51 CEST 2022 - The aliscore output appears to be empty. Check results for BUSCO: 137629at7742
## Thu Jul 14 01:15:51 CEST 2022 - The aliscore output file does not exist. Check results for BUSCO: 72241at7742
## Thu Jul 14 01:15:51 CEST 2022 - The aliscore output appears to be empty. Check results for BUSCO: 72241at7742
## Thu Jul 14 01:15:51 CEST 2022 - The aliscore output file does not exist. Check results for BUSCO: 106271at7742
## Thu Jul 14 01:15:51 CEST 2022 - The aliscore output appears to be empty. Check results for BUSCO: 106271at7742
## Thu Jul 14 01:15:53 CEST 2022 - The aliscore output file does not exist. Check results for BUSCO: 139653at7742
## Thu Jul 14 01:15:53 CEST 2022 - The aliscore output appears to be empty. Check results for BUSCO: 139653at7742
## Thu Jul 14 01:15:55 CEST 2022 - The aliscore output file does not exist. Check results for BUSCO: 244028at7742
## Thu Jul 14 01:15:56 CEST 2022 - The aliscore output appears to be empty. Check results for BUSCO: 244028at7742
## Thu Jul 14 01:15:56 CEST 2022 - The aliscore output file does not exist. Check results for BUSCO: 266521at7742
## Thu Jul 14 01:15:56 CEST 2022 - The aliscore output appears to be empty. Check results for BUSCO: 266521at7742
## Thu Jul 14 01:15:58 CEST 2022 - The aliscore output file does not exist. Check results for BUSCO: 197883at7742
## Thu Jul 14 01:15:58 CEST 2022 - The aliscore output file does not exist. Check results for BUSCO: 340971at7742
## Thu Jul 14 01:15:58 CEST 2022 - The aliscore output appears to be empty. Check results for BUSCO: 197883at7742
## Thu Jul 14 01:15:58 CEST 2022 - The aliscore output appears to be empty. Check results for BUSCO: 340971at7742
## Thu Jul 14 01:15:59 CEST 2022 - The aliscore output file does not exist. Check results for BUSCO: 172204at7742
## Thu Jul 14 01:15:59 CEST 2022 - The aliscore output appears to be empty. Check results for BUSCO: 172204at7742
## Thu Jul 14 01:16:01 CEST 2022 - The aliscore output file does not exist. Check results for BUSCO: 416929at7742
## Thu Jul 14 01:16:01 CEST 2022 - The aliscore output appears to be empty. Check results for BUSCO: 416929at7742
## Thu Jul 14 01:16:03 CEST 2022 - The aliscore output file does not exist. Check results for BUSCO: 116812at7742
## Thu Jul 14 01:16:03 CEST 2022 - The aliscore output appears to be empty. Check results for BUSCO: 116812at7742
## Thu Jul 14 01:16:03 CEST 2022 - The aliscore output file does not exist. Check results for BUSCO: 125985at7742
## Thu Jul 14 01:16:03 CEST 2022 - The aliscore output appears to be empty. Check results for BUSCO: 125985at7742
## Thu Jul 14 01:16:04 CEST 2022 - The aliscore output file does not exist. Check results for BUSCO: 238721at7742
## Thu Jul 14 01:16:04 CEST 2022 - The aliscore output appears to be empty. Check results for BUSCO: 238721at7742
## Thu Jul 14 01:16:05 CEST 2022 - The aliscore output file does not exist. Check results for BUSCO: 140962at7742
## Thu Jul 14 01:16:05 CEST 2022 - The aliscore output appears to be empty. Check results for BUSCO: 140962at7742
## Thu Jul 14 01:16:06 CEST 2022 - The aliscore output file does not exist. Check results for BUSCO: 344884at7742
## Thu Jul 14 01:16:06 CEST 2022 - The aliscore output appears to be empty. Check results for BUSCO: 344884at7742
## Thu Jul 14 01:16:08 CEST 2022 - The aliscore output file does not exist. Check results for BUSCO: 258449at7742
## Thu Jul 14 01:16:08 CEST 2022 - The aliscore output appears to be empty. Check results for BUSCO: 258449at7742
## Thu Jul 14 01:16:09 CEST 2022 - The aliscore output file does not exist. Check results for BUSCO: 224078at7742
## Thu Jul 14 01:16:09 CEST 2022 - The aliscore output appears to be empty. Check results for BUSCO: 224078at7742
## Thu Jul 14 01:16:10 CEST 2022 - The aliscore output file does not exist. Check results for BUSCO: 232857at7742
## Thu Jul 14 01:16:10 CEST 2022 - The aliscore output appears to be empty. Check results for BUSCO: 232857at7742
## Thu Jul 14 01:16:11 CEST 2022 - The aliscore output file does not exist. Check results for BUSCO: 142074at7742
## Thu Jul 14 01:16:11 CEST 2022 - The aliscore output appears to be empty. Check results for BUSCO: 142074at7742
## Thu Jul 14 01:16:12 CEST 2022 - The aliscore output file does not exist. Check results for BUSCO: 174607at7742
## Thu Jul 14 01:16:12 CEST 2022 - The aliscore output appears to be empty. Check results for BUSCO: 174607at7742
## Thu Jul 14 01:16:14 CEST 2022 - The aliscore output file does not exist. Check results for BUSCO: 288361at7742
## Thu Jul 14 01:16:14 CEST 2022 - The aliscore output appears to be empty. Check results for BUSCO: 288361at7742
## Thu Jul 14 01:16:16 CEST 2022 - The aliscore output file does not exist. Check results for BUSCO: 107391at7742
## Thu Jul 14 01:16:16 CEST 2022 - The aliscore output appears to be empty. Check results for BUSCO: 107391at7742
## Thu Jul 14 01:16:16 CEST 2022 - The aliscore output file does not exist. Check results for BUSCO: 54598at7742
## Thu Jul 14 01:16:16 CEST 2022 - The aliscore output appears to be empty. Check results for BUSCO: 54598at7742
## Thu Jul 14 01:16:16 CEST 2022 - The aliscore output file does not exist. Check results for BUSCO: 88086at7742
## Thu Jul 14 01:16:16 CEST 2022 - The aliscore output appears to be empty. Check results for BUSCO: 88086at7742
## Thu Jul 14 01:16:18 CEST 2022 - The aliscore output file does not exist. Check results for BUSCO: 370224at7742
## Thu Jul 14 01:16:18 CEST 2022 - The aliscore output appears to be empty. Check results for BUSCO: 370224at7742
## Thu Jul 14 01:16:21 CEST 2022 - The aliscore output file does not exist. Check results for BUSCO: 132972at7742
## Thu Jul 14 01:16:21 CEST 2022 - The aliscore output appears to be empty. Check results for BUSCO: 132972at7742
## Thu Jul 14 01:16:21 CEST 2022 - The aliscore output file does not exist. Check results for BUSCO: 93269at7742
## Thu Jul 14 01:16:21 CEST 2022 - The aliscore output appears to be empty. Check results for BUSCO: 93269at7742
## Thu Jul 14 01:16:23 CEST 2022 - The aliscore output file does not exist. Check results for BUSCO: 293554at7742
## Thu Jul 14 01:16:23 CEST 2022 - The aliscore output appears to be empty. Check results for BUSCO: 293554at7742
## Thu Jul 14 01:16:25 CEST 2022 - The aliscore output file does not exist. Check results for BUSCO: 209187at7742
## Thu Jul 14 01:16:25 CEST 2022 - The aliscore output appears to be empty. Check results for BUSCO: 209187at7742
## Thu Jul 14 01:16:25 CEST 2022 - The aliscore output file does not exist. Check results for BUSCO: 176601at7742
## Thu Jul 14 01:16:25 CEST 2022 - The aliscore output appears to be empty. Check results for BUSCO: 176601at7742
## Thu Jul 14 01:16:25 CEST 2022 - The aliscore output file does not exist. Check results for BUSCO: 75578at7742
## Thu Jul 14 01:16:25 CEST 2022 - The aliscore output appears to be empty. Check results for BUSCO: 75578at7742
## Thu Jul 14 01:16:28 CEST 2022 - The aliscore output file does not exist. Check results for BUSCO: 213511at7742
## Thu Jul 14 01:16:28 CEST 2022 - The aliscore output appears to be empty. Check results for BUSCO: 213511at7742
## Thu Jul 14 01:16:28 CEST 2022 - The aliscore output file does not exist. Check results for BUSCO: 169724at7742
## Thu Jul 14 01:16:28 CEST 2022 - The aliscore output appears to be empty. Check results for BUSCO: 169724at7742
## Thu Jul 14 01:16:29 CEST 2022 - The aliscore output file does not exist. Check results for BUSCO: 119120at7742
## Thu Jul 14 01:16:29 CEST 2022 - The aliscore output appears to be empty. Check results for BUSCO: 119120at7742
## Thu Jul 14 01:16:31 CEST 2022 - The aliscore output file does not exist. Check results for BUSCO: 144445at7742
## Thu Jul 14 01:16:31 CEST 2022 - The aliscore output appears to be empty. Check results for BUSCO: 144445at7742
## Thu Jul 14 01:16:33 CEST 2022 - The aliscore output file does not exist. Check results for BUSCO: 134679at7742
## Thu Jul 14 01:16:33 CEST 2022 - The aliscore output appears to be empty. Check results for BUSCO: 134679at7742
## Thu Jul 14 01:16:33 CEST 2022 - The aliscore output file does not exist. Check results for BUSCO: 81756at7742
## Thu Jul 14 01:16:34 CEST 2022 - The aliscore output appears to be empty. Check results for BUSCO: 81756at7742
## Thu Jul 14 01:16:34 CEST 2022 - The aliscore output file does not exist. Check results for BUSCO: 161167at7742
## Thu Jul 14 01:16:34 CEST 2022 - The aliscore output appears to be empty. Check results for BUSCO: 161167at7742
## Thu Jul 14 01:16:35 CEST 2022 - The aliscore output file does not exist. Check results for BUSCO: 81689at7742
## Thu Jul 14 01:16:35 CEST 2022 - The aliscore output appears to be empty. Check results for BUSCO: 81689at7742
## Thu Jul 14 01:16:39 CEST 2022 - The aliscore output file does not exist. Check results for BUSCO: 97043at7742
## Thu Jul 14 01:16:39 CEST 2022 - The aliscore output appears to be empty. Check results for BUSCO: 97043at7742
## Thu Jul 14 01:16:39 CEST 2022 - The aliscore output file does not exist. Check results for BUSCO: 114303at7742
## Thu Jul 14 01:16:39 CEST 2022 - The aliscore output appears to be empty. Check results for BUSCO: 114303at7742
## Thu Jul 14 01:16:41 CEST 2022 - The aliscore output file does not exist. Check results for BUSCO: 93231at7742
## Thu Jul 14 01:16:41 CEST 2022 - The aliscore output appears to be empty. Check results for BUSCO: 93231at7742
## Thu Jul 14 01:16:41 CEST 2022 - The aliscore output file does not exist. Check results for BUSCO: 272309at7742
## Thu Jul 14 01:16:41 CEST 2022 - The aliscore output appears to be empty. Check results for BUSCO: 272309at7742
## Thu Jul 14 01:16:42 CEST 2022 - The aliscore output file does not exist. Check results for BUSCO: 106271at7742
## Thu Jul 14 01:16:42 CEST 2022 - The aliscore output appears to be empty. Check results for BUSCO: 106271at7742
## Thu Jul 14 01:16:44 CEST 2022 - The aliscore output file does not exist. Check results for BUSCO: 219513at7742
## Thu Jul 14 01:16:44 CEST 2022 - The aliscore output appears to be empty. Check results for BUSCO: 219513at7742
## Thu Jul 14 01:16:45 CEST 2022 - The aliscore output file does not exist. Check results for BUSCO: 139653at7742
## Thu Jul 14 01:16:45 CEST 2022 - The aliscore output appears to be empty. Check results for BUSCO: 139653at7742
## Thu Jul 14 01:16:47 CEST 2022 - The aliscore output file does not exist. Check results for BUSCO: 240065at7742
## Thu Jul 14 01:16:47 CEST 2022 - The aliscore output appears to be empty. Check results for BUSCO: 240065at7742
## Thu Jul 14 01:16:48 CEST 2022 - The aliscore output file does not exist. Check results for BUSCO: 256267at7742
## Thu Jul 14 01:16:48 CEST 2022 - The aliscore output appears to be empty. Check results for BUSCO: 256267at7742
## Thu Jul 14 01:16:50 CEST 2022 - The aliscore output file does not exist. Check results for BUSCO: 197883at7742
## Thu Jul 14 01:16:50 CEST 2022 - The aliscore output appears to be empty. Check results for BUSCO: 197883at7742
## Thu Jul 14 01:16:51 CEST 2022 - The aliscore output file does not exist. Check results for BUSCO: 301755at7742
## Thu Jul 14 01:16:51 CEST 2022 - The aliscore output appears to be empty. Check results for BUSCO: 301755at7742
## Thu Jul 14 01:16:52 CEST 2022 - The aliscore output file does not exist. Check results for BUSCO: 327776at7742
## Thu Jul 14 01:16:52 CEST 2022 - The aliscore output appears to be empty. Check results for BUSCO: 327776at7742
## Thu Jul 14 01:16:53 CEST 2022 - The aliscore output file does not exist. Check results for BUSCO: 132010at7742
## Thu Jul 14 01:16:53 CEST 2022 - The aliscore output appears to be empty. Check results for BUSCO: 132010at7742
## Thu Jul 14 01:16:55 CEST 2022 - The aliscore output file does not exist. Check results for BUSCO: 299549at7742
## Thu Jul 14 01:16:55 CEST 2022 - The aliscore output appears to be empty. Check results for BUSCO: 299549at7742
## Thu Jul 14 01:16:56 CEST 2022 - The aliscore output file does not exist. Check results for BUSCO: 80688at7742
## Thu Jul 14 01:16:56 CEST 2022 - The aliscore output appears to be empty. Check results for BUSCO: 80688at7742
## Thu Jul 14 01:16:56 CEST 2022 - The aliscore output file does not exist. Check results for BUSCO: 80688at7742
## Thu Jul 14 01:16:56 CEST 2022 - The aliscore output appears to be empty. Check results for BUSCO: 80688at7742
## Thu Jul 14 01:16:58 CEST 2022 - The aliscore output file does not exist. Check results for BUSCO: 378656at7742
## Thu Jul 14 01:16:58 CEST 2022 - The aliscore output appears to be empty. Check results for BUSCO: 378656at7742
## Thu Jul 14 01:16:59 CEST 2022 - The aliscore output file does not exist. Check results for BUSCO: 193456at7742
## Thu Jul 14 01:16:59 CEST 2022 - The aliscore output appears to be empty. Check results for BUSCO: 193456at7742
## Thu Jul 14 01:16:59 CEST 2022 - The aliscore output file does not exist. Check results for BUSCO: 88086at7742
## Thu Jul 14 01:16:59 CEST 2022 - The aliscore output appears to be empty. Check results for BUSCO: 88086at7742
## Thu Jul 14 01:17:01 CEST 2022 - The aliscore output file does not exist. Check results for BUSCO: 107188at7742
## Thu Jul 14 01:17:01 CEST 2022 - The aliscore output appears to be empty. Check results for BUSCO: 107188at7742
## Thu Jul 14 01:17:02 CEST 2022 - The aliscore output file does not exist. Check results for BUSCO: 340971at7742
## Thu Jul 14 01:17:02 CEST 2022 - The aliscore output appears to be empty. Check results for BUSCO: 340971at7742
## Thu Jul 14 01:17:05 CEST 2022 - The aliscore output file does not exist. Check results for BUSCO: 138249at7742
## Thu Jul 14 01:17:05 CEST 2022 - The aliscore output appears to be empty. Check results for BUSCO: 138249at7742
## Thu Jul 14 01:17:05 CEST 2022 - The aliscore output file does not exist. Check results for BUSCO: 108553at7742
## Thu Jul 14 01:17:06 CEST 2022 - The aliscore output appears to be empty. Check results for BUSCO: 108553at7742
## Thu Jul 14 01:17:07 CEST 2022 - The aliscore output file does not exist. Check results for BUSCO: 93229at7742
## Thu Jul 14 01:17:07 CEST 2022 - The aliscore output appears to be empty. Check results for BUSCO: 93229at7742
## Thu Jul 14 01:17:07 CEST 2022 - The aliscore output file does not exist. Check results for BUSCO: 112966at7742
## Thu Jul 14 01:17:07 CEST 2022 - The aliscore output appears to be empty. Check results for BUSCO: 112966at7742
## Thu Jul 14 01:17:09 CEST 2022 - The aliscore output file does not exist. Check results for BUSCO: 308660at7742
## Thu Jul 14 01:17:09 CEST 2022 - The aliscore output appears to be empty. Check results for BUSCO: 308660at7742
## Thu Jul 14 01:17:10 CEST 2022 - The aliscore output file does not exist. Check results for BUSCO: 291147at7742
## Thu Jul 14 01:17:10 CEST 2022 - The aliscore output appears to be empty. Check results for BUSCO: 291147at7742
## Thu Jul 14 01:17:11 CEST 2022 - The aliscore output file does not exist. Check results for BUSCO: 170012at7742
## Thu Jul 14 01:17:11 CEST 2022 - The aliscore output appears to be empty. Check results for BUSCO: 170012at7742
## Thu Jul 14 01:17:13 CEST 2022 - The aliscore output file does not exist. Check results for BUSCO: 109623at7742
## Thu Jul 14 01:17:13 CEST 2022 - The aliscore output file does not exist. Check results for BUSCO: 102135at7742
## Thu Jul 14 01:17:13 CEST 2022 - The aliscore output appears to be empty. Check results for BUSCO: 109623at7742
## Thu Jul 14 01:17:13 CEST 2022 - The aliscore output appears to be empty. Check results for BUSCO: 102135at7742
## Thu Jul 14 01:17:16 CEST 2022 - The aliscore output file does not exist. Check results for BUSCO: 75578at7742
## Thu Jul 14 01:17:16 CEST 2022 - The aliscore output appears to be empty. Check results for BUSCO: 75578at7742
## Thu Jul 14 01:17:17 CEST 2022 - The aliscore output file does not exist. Check results for BUSCO: 131817at7742
## Thu Jul 14 01:17:17 CEST 2022 - The aliscore output appears to be empty. Check results for BUSCO: 131817at7742
## Thu Jul 14 01:17:20 CEST 2022 - The aliscore output file does not exist. Check results for BUSCO: 123758at7742
## Thu Jul 14 01:17:20 CEST 2022 - The aliscore output file does not exist. Check results for BUSCO: 161484at7742
## Thu Jul 14 01:17:20 CEST 2022 - The aliscore output appears to be empty. Check results for BUSCO: 123758at7742
## Thu Jul 14 01:17:20 CEST 2022 - The aliscore output appears to be empty. Check results for BUSCO: 161484at7742
## Thu Jul 14 01:17:21 CEST 2022 - The aliscore output file does not exist. Check results for BUSCO: 81756at7742
## Thu Jul 14 01:17:21 CEST 2022 - The aliscore output appears to be empty. Check results for BUSCO: 81756at7742
## Thu Jul 14 01:17:22 CEST 2022 - The aliscore output file does not exist. Check results for BUSCO: 106313at7742
## Thu Jul 14 01:17:22 CEST 2022 - The aliscore output appears to be empty. Check results for BUSCO: 106313at7742
## Thu Jul 14 01:17:23 CEST 2022 - The aliscore output file does not exist. Check results for BUSCO: 159349at7742
## Thu Jul 14 01:17:23 CEST 2022 - The aliscore output appears to be empty. Check results for BUSCO: 159349at7742
## Thu Jul 14 01:17:24 CEST 2022 - The aliscore output file does not exist. Check results for BUSCO: 360487at7742
## Thu Jul 14 01:17:24 CEST 2022 - The aliscore output appears to be empty. Check results for BUSCO: 360487at7742
## Thu Jul 14 01:17:25 CEST 2022 - The aliscore output file does not exist. Check results for BUSCO: 329621at7742
## Thu Jul 14 01:17:25 CEST 2022 - The aliscore output appears to be empty. Check results for BUSCO: 329621at7742
## Thu Jul 14 01:17:25 CEST 2022 - The aliscore output file does not exist. Check results for BUSCO: 209425at7742
## Thu Jul 14 01:17:25 CEST 2022 - The aliscore output appears to be empty. Check results for BUSCO: 209425at7742
## Thu Jul 14 01:17:27 CEST 2022 - The aliscore output file does not exist. Check results for BUSCO: 245662at7742
## Thu Jul 14 01:17:27 CEST 2022 - The aliscore output appears to be empty. Check results for BUSCO: 245662at7742
## Thu Jul 14 01:17:29 CEST 2022 - The aliscore output file does not exist. Check results for BUSCO: 140071at7742
## Thu Jul 14 01:17:29 CEST 2022 - The aliscore output appears to be empty. Check results for BUSCO: 140071at7742
## Thu Jul 14 01:17:31 CEST 2022 - The aliscore output file does not exist. Check results for BUSCO: 351544at7742
## Thu Jul 14 01:17:31 CEST 2022 - The aliscore output appears to be empty. Check results for BUSCO: 351544at7742
## Thu Jul 14 01:17:31 CEST 2022 - The aliscore output file does not exist. Check results for BUSCO: 88086at7742
## Thu Jul 14 01:17:31 CEST 2022 - The aliscore output appears to be empty. Check results for BUSCO: 88086at7742
## Thu Jul 14 01:17:33 CEST 2022 - The aliscore output file does not exist. Check results for BUSCO: 333967at7742
## Thu Jul 14 01:17:33 CEST 2022 - The aliscore output appears to be empty. Check results for BUSCO: 333967at7742
## Thu Jul 14 01:17:34 CEST 2022 - The aliscore output file does not exist. Check results for BUSCO: 93020at7742
## Thu Jul 14 01:17:34 CEST 2022 - The aliscore output appears to be empty. Check results for BUSCO: 93020at7742
## Thu Jul 14 01:17:34 CEST 2022 - The aliscore output file does not exist. Check results for BUSCO: 82541at7742
## Thu Jul 14 01:17:34 CEST 2022 - The aliscore output appears to be empty. Check results for BUSCO: 82541at7742
## Thu Jul 14 01:17:37 CEST 2022 - The aliscore output file does not exist. Check results for BUSCO: 172737at7742
## Thu Jul 14 01:17:37 CEST 2022 - The aliscore output appears to be empty. Check results for BUSCO: 172737at7742
## Thu Jul 14 01:17:38 CEST 2022 - The aliscore output file does not exist. Check results for BUSCO: 284061at7742
## Thu Jul 14 01:17:38 CEST 2022 - The aliscore output appears to be empty. Check results for BUSCO: 284061at7742
## Thu Jul 14 01:17:38 CEST 2022 - The aliscore output file does not exist. Check results for BUSCO: 103422at7742
## Thu Jul 14 01:17:38 CEST 2022 - The aliscore output appears to be empty. Check results for BUSCO: 103422at7742
## Thu Jul 14 01:17:40 CEST 2022 - The aliscore output file does not exist. Check results for BUSCO: 334896at7742
## Thu Jul 14 01:17:40 CEST 2022 - The aliscore output appears to be empty. Check results for BUSCO: 334896at7742
## Thu Jul 14 01:17:42 CEST 2022 - The aliscore output file does not exist. Check results for BUSCO: 325029at7742
## Thu Jul 14 01:17:42 CEST 2022 - The aliscore output appears to be empty. Check results for BUSCO: 325029at7742
## Thu Jul 14 01:17:43 CEST 2022 - The aliscore output file does not exist. Check results for BUSCO: 276337at7742
## Thu Jul 14 01:17:43 CEST 2022 - The aliscore output appears to be empty. Check results for BUSCO: 276337at7742
## Thu Jul 14 01:17:44 CEST 2022 - The aliscore output file does not exist. Check results for BUSCO: 195437at7742
## Thu Jul 14 01:17:44 CEST 2022 - The aliscore output appears to be empty. Check results for BUSCO: 195437at7742
## Thu Jul 14 01:17:45 CEST 2022 - The aliscore output file does not exist. Check results for BUSCO: 101858at7742
## Thu Jul 14 01:17:45 CEST 2022 - The aliscore output appears to be empty. Check results for BUSCO: 101858at7742
## Thu Jul 14 01:17:46 CEST 2022 - The aliscore output file does not exist. Check results for BUSCO: 62905at7742
## Thu Jul 14 01:17:46 CEST 2022 - The aliscore output appears to be empty. Check results for BUSCO: 62905at7742
## Thu Jul 14 01:17:48 CEST 2022 - The aliscore output file does not exist. Check results for BUSCO: 143678at7742
## Thu Jul 14 01:17:48 CEST 2022 - The aliscore output appears to be empty. Check results for BUSCO: 143678at7742
## Thu Jul 14 01:17:48 CEST 2022 - The aliscore output file does not exist. Check results for BUSCO: 337175at7742
## Thu Jul 14 01:17:48 CEST 2022 - The aliscore output appears to be empty. Check results for BUSCO: 337175at7742
## Thu Jul 14 01:17:50 CEST 2022 - The aliscore output file does not exist. Check results for BUSCO: 169724at7742
## Thu Jul 14 01:17:50 CEST 2022 - The aliscore output appears to be empty. Check results for BUSCO: 169724at7742
## Thu Jul 14 01:17:50 CEST 2022 - The aliscore output file does not exist. Check results for BUSCO: 176317at7742
## Thu Jul 14 01:17:50 CEST 2022 - The aliscore output appears to be empty. Check results for BUSCO: 176317at7742
## Thu Jul 14 01:17:51 CEST 2022 - The aliscore output file does not exist. Check results for BUSCO: 79012at7742
## Thu Jul 14 01:17:51 CEST 2022 - The aliscore output appears to be empty. Check results for BUSCO: 79012at7742
## Thu Jul 14 01:17:52 CEST 2022 - The aliscore output file does not exist. Check results for BUSCO: 81756at7742
## Thu Jul 14 01:17:52 CEST 2022 - The aliscore output appears to be empty. Check results for BUSCO: 81756at7742
## Thu Jul 14 01:17:53 CEST 2022 - The aliscore output file does not exist. Check results for BUSCO: 75578at7742
## Thu Jul 14 01:17:53 CEST 2022 - The aliscore output appears to be empty. Check results for BUSCO: 75578at7742
## Thu Jul 14 01:17:54 CEST 2022 - The aliscore output file does not exist. Check results for BUSCO: 89229at7742
## Thu Jul 14 01:17:54 CEST 2022 - The aliscore output appears to be empty. Check results for BUSCO: 89229at7742
## Thu Jul 14 01:17:58 CEST 2022 - The aliscore output file does not exist. Check results for BUSCO: 256267at7742
## Thu Jul 14 01:17:58 CEST 2022 - The aliscore output appears to be empty. Check results for BUSCO: 256267at7742
## Thu Jul 14 01:17:58 CEST 2022 - The aliscore output file does not exist. Check results for BUSCO: 176369at7742
## Thu Jul 14 01:17:58 CEST 2022 - The aliscore output file does not exist. Check results for BUSCO: 277665at7742
## Thu Jul 14 01:17:58 CEST 2022 - The aliscore output appears to be empty. Check results for BUSCO: 176369at7742
## Thu Jul 14 01:17:58 CEST 2022 - The aliscore output appears to be empty. Check results for BUSCO: 277665at7742
## Thu Jul 14 01:17:59 CEST 2022 - The aliscore output file does not exist. Check results for BUSCO: 95886at7742
## Thu Jul 14 01:17:59 CEST 2022 - The aliscore output appears to be empty. Check results for BUSCO: 95886at7742
## Thu Jul 14 01:18:01 CEST 2022 - The aliscore output file does not exist. Check results for BUSCO: 209714at7742
## Thu Jul 14 01:18:01 CEST 2022 - The aliscore output appears to be empty. Check results for BUSCO: 209714at7742
## Thu Jul 14 01:18:01 CEST 2022 - The aliscore output file does not exist. Check results for BUSCO: 81689at7742
## Thu Jul 14 01:18:01 CEST 2022 - The aliscore output appears to be empty. Check results for BUSCO: 81689at7742
## Thu Jul 14 01:18:03 CEST 2022 - The aliscore output file does not exist. Check results for BUSCO: 156795at7742
## Thu Jul 14 01:18:03 CEST 2022 - The aliscore output appears to be empty. Check results for BUSCO: 156795at7742
## Thu Jul 14 01:18:03 CEST 2022 - The aliscore output file does not exist. Check results for BUSCO: 195929at7742
## Thu Jul 14 01:18:03 CEST 2022 - The aliscore output appears to be empty. Check results for BUSCO: 195929at7742
## Thu Jul 14 01:18:04 CEST 2022 - The aliscore output file does not exist. Check results for BUSCO: 164950at7742
## Thu Jul 14 01:18:05 CEST 2022 - The aliscore output appears to be empty. Check results for BUSCO: 164950at7742
## Thu Jul 14 01:18:07 CEST 2022 - The aliscore output file does not exist. Check results for BUSCO: 101941at7742
## Thu Jul 14 01:18:07 CEST 2022 - The aliscore output appears to be empty. Check results for BUSCO: 101941at7742
## Thu Jul 14 01:18:08 CEST 2022 - The aliscore output file does not exist. Check results for BUSCO: 233977at7742
## Thu Jul 14 01:18:08 CEST 2022 - The aliscore output appears to be empty. Check results for BUSCO: 233977at7742
## Thu Jul 14 01:18:10 CEST 2022 - The aliscore output file does not exist. Check results for BUSCO: 135511at7742
## Thu Jul 14 01:18:10 CEST 2022 - The aliscore output appears to be empty. Check results for BUSCO: 135511at7742
## Thu Jul 14 01:18:11 CEST 2022 - The aliscore output file does not exist. Check results for BUSCO: 240075at7742
## Thu Jul 14 01:18:11 CEST 2022 - The aliscore output appears to be empty. Check results for BUSCO: 240075at7742
## Thu Jul 14 01:18:11 CEST 2022 - The aliscore output file does not exist. Check results for BUSCO: 170944at7742
## Thu Jul 14 01:18:11 CEST 2022 - The aliscore output appears to be empty. Check results for BUSCO: 170944at7742
## Thu Jul 14 01:18:13 CEST 2022 - The aliscore output file does not exist. Check results for BUSCO: 114115at7742
## Thu Jul 14 01:18:13 CEST 2022 - The aliscore output appears to be empty. Check results for BUSCO: 114115at7742
## Thu Jul 14 01:18:15 CEST 2022 - The aliscore output file does not exist. Check results for BUSCO: 133642at7742
## Thu Jul 14 01:18:15 CEST 2022 - The aliscore output appears to be empty. Check results for BUSCO: 133642at7742
## Thu Jul 14 01:18:16 CEST 2022 - The aliscore output file does not exist. Check results for BUSCO: 248762at7742
## Thu Jul 14 01:18:16 CEST 2022 - The aliscore output appears to be empty. Check results for BUSCO: 248762at7742
## Thu Jul 14 01:18:16 CEST 2022 - The aliscore output file does not exist. Check results for BUSCO: 62905at7742
## Thu Jul 14 01:18:16 CEST 2022 - The aliscore output appears to be empty. Check results for BUSCO: 62905at7742
## Thu Jul 14 01:18:18 CEST 2022 - The aliscore output file does not exist. Check results for BUSCO: 104856at7742
## Thu Jul 14 01:18:18 CEST 2022 - The aliscore output appears to be empty. Check results for BUSCO: 104856at7742
## Thu Jul 14 01:18:20 CEST 2022 - The aliscore output file does not exist. Check results for BUSCO: 81689at7742
## Thu Jul 14 01:18:20 CEST 2022 - The aliscore output appears to be empty. Check results for BUSCO: 81689at7742
## Thu Jul 14 01:18:20 CEST 2022 - The aliscore output file does not exist. Check results for BUSCO: 216380at7742
## Thu Jul 14 01:18:21 CEST 2022 - The aliscore output appears to be empty. Check results for BUSCO: 216380at7742
## Thu Jul 14 01:18:22 CEST 2022 - The aliscore output file does not exist. Check results for BUSCO: 323828at7742
## Thu Jul 14 01:18:22 CEST 2022 - The aliscore output appears to be empty. Check results for BUSCO: 323828at7742
## Thu Jul 14 01:18:23 CEST 2022 - The aliscore output file does not exist. Check results for BUSCO: 239989at7742
## Thu Jul 14 01:18:23 CEST 2022 - The aliscore output appears to be empty. Check results for BUSCO: 239989at7742
## Thu Jul 14 01:18:24 CEST 2022 - The aliscore output file does not exist. Check results for BUSCO: 168697at7742
## Thu Jul 14 01:18:24 CEST 2022 - The aliscore output appears to be empty. Check results for BUSCO: 168697at7742
## Thu Jul 14 01:18:24 CEST 2022 - The aliscore output file does not exist. Check results for BUSCO: 89229at7742
## Thu Jul 14 01:18:24 CEST 2022 - The aliscore output appears to be empty. Check results for BUSCO: 89229at7742
## Thu Jul 14 01:18:27 CEST 2022 - The aliscore output file does not exist. Check results for BUSCO: 102222at7742
## Thu Jul 14 01:18:27 CEST 2022 - The aliscore output appears to be empty. Check results for BUSCO: 102222at7742
## Thu Jul 14 01:18:29 CEST 2022 - The aliscore output file does not exist. Check results for BUSCO: 220786at7742
## Thu Jul 14 01:18:29 CEST 2022 - The aliscore output appears to be empty. Check results for BUSCO: 220786at7742
## Thu Jul 14 01:18:30 CEST 2022 - The aliscore output file does not exist. Check results for BUSCO: 163908at7742
## Thu Jul 14 01:18:30 CEST 2022 - The aliscore output appears to be empty. Check results for BUSCO: 163908at7742
## Thu Jul 14 01:18:31 CEST 2022 - The aliscore output file does not exist. Check results for BUSCO: 300687at7742
## Thu Jul 14 01:18:31 CEST 2022 - The aliscore output appears to be empty. Check results for BUSCO: 300687at7742
## Thu Jul 14 01:18:32 CEST 2022 - The aliscore output file does not exist. Check results for BUSCO: 116812at7742
## Thu Jul 14 01:18:32 CEST 2022 - The aliscore output appears to be empty. Check results for BUSCO: 116812at7742
## Thu Jul 14 01:18:33 CEST 2022 - The aliscore output file does not exist. Check results for BUSCO: 115042at7742
## Thu Jul 14 01:18:33 CEST 2022 - The aliscore output appears to be empty. Check results for BUSCO: 115042at7742
## Thu Jul 14 01:18:33 CEST 2022 - The aliscore output file does not exist. Check results for BUSCO: 80688at7742
## Thu Jul 14 01:18:33 CEST 2022 - The aliscore output appears to be empty. Check results for BUSCO: 80688at7742
## Thu Jul 14 01:18:36 CEST 2022 - The aliscore output file does not exist. Check results for BUSCO: 139050at7742
## Thu Jul 14 01:18:36 CEST 2022 - The aliscore output appears to be empty. Check results for BUSCO: 139050at7742
## Thu Jul 14 01:18:36 CEST 2022 - The aliscore output file does not exist. Check results for BUSCO: 239989at7742
## Thu Jul 14 01:18:36 CEST 2022 - The aliscore output appears to be empty. Check results for BUSCO: 239989at7742
## Thu Jul 14 01:18:38 CEST 2022 - The aliscore output file does not exist. Check results for BUSCO: 109623at7742
## Thu Jul 14 01:18:38 CEST 2022 - The aliscore output appears to be empty. Check results for BUSCO: 109623at7742
## Thu Jul 14 01:18:39 CEST 2022 - The aliscore output file does not exist. Check results for BUSCO: 125985at7742
## Thu Jul 14 01:18:39 CEST 2022 - The aliscore output appears to be empty. Check results for BUSCO: 125985at7742
## Thu Jul 14 01:18:39 CEST 2022 - The aliscore output file does not exist. Check results for BUSCO: 112966at7742
## Thu Jul 14 01:18:39 CEST 2022 - The aliscore output appears to be empty. Check results for BUSCO: 112966at7742
## Thu Jul 14 01:18:40 CEST 2022 - The aliscore output file does not exist. Check results for BUSCO: 182481at7742
## Thu Jul 14 01:18:40 CEST 2022 - The aliscore output appears to be empty. Check results for BUSCO: 182481at7742
## Thu Jul 14 01:18:41 CEST 2022 - The aliscore output file does not exist. Check results for BUSCO: 79012at7742
## Thu Jul 14 01:18:41 CEST 2022 - The aliscore output appears to be empty. Check results for BUSCO: 79012at7742
## Thu Jul 14 01:18:43 CEST 2022 - The aliscore output file does not exist. Check results for BUSCO: 136946at7742
## Thu Jul 14 01:18:43 CEST 2022 - The aliscore output appears to be empty. Check results for BUSCO: 136946at7742
## Thu Jul 14 01:18:46 CEST 2022 - The aliscore output file does not exist. Check results for BUSCO: 248762at7742
## Thu Jul 14 01:18:46 CEST 2022 - The aliscore output appears to be empty. Check results for BUSCO: 248762at7742
## Thu Jul 14 01:18:46 CEST 2022 - The aliscore output file does not exist. Check results for BUSCO: 145949at7742
## Thu Jul 14 01:18:46 CEST 2022 - The aliscore output appears to be empty. Check results for BUSCO: 145949at7742
## Thu Jul 14 01:18:47 CEST 2022 - The aliscore output file does not exist. Check results for BUSCO: 133877at7742
## Thu Jul 14 01:18:47 CEST 2022 - The aliscore output appears to be empty. Check results for BUSCO: 133877at7742
## Thu Jul 14 01:18:49 CEST 2022 - The aliscore output file does not exist. Check results for BUSCO: 131817at7742
## Thu Jul 14 01:18:49 CEST 2022 - The aliscore output appears to be empty. Check results for BUSCO: 131817at7742
## Thu Jul 14 01:18:49 CEST 2022 - The aliscore output file does not exist. Check results for BUSCO: 284061at7742
## Thu Jul 14 01:18:49 CEST 2022 - The aliscore output appears to be empty. Check results for BUSCO: 284061at7742
## Thu Jul 14 01:18:50 CEST 2022 - The aliscore output file does not exist. Check results for BUSCO: 54598at7742
## Thu Jul 14 01:18:50 CEST 2022 - The aliscore output appears to be empty. Check results for BUSCO: 54598at7742
## Thu Jul 14 01:18:51 CEST 2022 - The aliscore output file does not exist. Check results for BUSCO: 301755at7742
## Thu Jul 14 01:18:51 CEST 2022 - The aliscore output appears to be empty. Check results for BUSCO: 301755at7742
## Thu Jul 14 01:18:52 CEST 2022 - The aliscore output file does not exist. Check results for BUSCO: 132972at7742
## Thu Jul 14 01:18:52 CEST 2022 - The aliscore output appears to be empty. Check results for BUSCO: 132972at7742
## Thu Jul 14 01:18:54 CEST 2022 - The aliscore output file does not exist. Check results for BUSCO: 114214at7742
## Thu Jul 14 01:18:54 CEST 2022 - The aliscore output appears to be empty. Check results for BUSCO: 114214at7742
## Thu Jul 14 01:18:54 CEST 2022 - The aliscore output file does not exist. Check results for BUSCO: 332125at7742
## Thu Jul 14 01:18:54 CEST 2022 - The aliscore output appears to be empty. Check results for BUSCO: 332125at7742
## Thu Jul 14 01:18:57 CEST 2022 - The aliscore output file does not exist. Check results for BUSCO: 107391at7742
## Thu Jul 14 01:18:57 CEST 2022 - The aliscore output appears to be empty. Check results for BUSCO: 107391at7742
## Thu Jul 14 01:18:57 CEST 2022 - The aliscore output file does not exist. Check results for BUSCO: 283449at7742
## Thu Jul 14 01:18:58 CEST 2022 - The aliscore output appears to be empty. Check results for BUSCO: 283449at7742
## Thu Jul 14 01:18:59 CEST 2022 - The aliscore output file does not exist. Check results for BUSCO: 183332at7742
## Thu Jul 14 01:18:59 CEST 2022 - The aliscore output appears to be empty. Check results for BUSCO: 183332at7742
## Thu Jul 14 01:19:00 CEST 2022 - The aliscore output file does not exist. Check results for BUSCO: 296707at7742
## Thu Jul 14 01:19:00 CEST 2022 - The aliscore output appears to be empty. Check results for BUSCO: 296707at7742
## Thu Jul 14 01:19:02 CEST 2022 - The aliscore output file does not exist. Check results for BUSCO: 145949at7742
## Thu Jul 14 01:19:02 CEST 2022 - The aliscore output appears to be empty. Check results for BUSCO: 145949at7742
## Thu Jul 14 01:19:02 CEST 2022 - The aliscore output file does not exist. Check results for BUSCO: 79012at7742
## Thu Jul 14 01:19:02 CEST 2022 - The aliscore output appears to be empty. Check results for BUSCO: 79012at7742
## Thu Jul 14 01:19:05 CEST 2022 - The aliscore output file does not exist. Check results for BUSCO: 200616at7742
## Thu Jul 14 01:19:05 CEST 2022 - The aliscore output file does not exist. Check results for BUSCO: 176369at7742
## Thu Jul 14 01:19:05 CEST 2022 - The aliscore output appears to be empty. Check results for BUSCO: 176369at7742
## Thu Jul 14 01:19:05 CEST 2022 - The aliscore output appears to be empty. Check results for BUSCO: 200616at7742
## Thu Jul 14 01:19:05 CEST 2022 - The aliscore output file does not exist. Check results for BUSCO: 184701at7742
## Thu Jul 14 01:19:05 CEST 2022 - The aliscore output appears to be empty. Check results for BUSCO: 184701at7742
## Thu Jul 14 01:19:07 CEST 2022 - The aliscore output file does not exist. Check results for BUSCO: 130415at7742
## Thu Jul 14 01:19:07 CEST 2022 - The aliscore output appears to be empty. Check results for BUSCO: 130415at7742
## Thu Jul 14 01:19:09 CEST 2022 - The aliscore output file does not exist. Check results for BUSCO: 284061at7742
## Thu Jul 14 01:19:09 CEST 2022 - The aliscore output appears to be empty. Check results for BUSCO: 284061at7742
## Thu Jul 14 01:19:10 CEST 2022 - The aliscore output file does not exist. Check results for BUSCO: 283910at7742
## Thu Jul 14 01:19:10 CEST 2022 - The aliscore output appears to be empty. Check results for BUSCO: 283910at7742
## Thu Jul 14 01:19:12 CEST 2022 - The aliscore output file does not exist. Check results for BUSCO: 344884at7742
## Thu Jul 14 01:19:12 CEST 2022 - The aliscore output appears to be empty. Check results for BUSCO: 344884at7742
## Thu Jul 14 01:19:13 CEST 2022 - The aliscore output file does not exist. Check results for BUSCO: 139053at7742
## Thu Jul 14 01:19:13 CEST 2022 - The aliscore output appears to be empty. Check results for BUSCO: 139053at7742
## Thu Jul 14 01:19:14 CEST 2022 - The aliscore output file does not exist. Check results for BUSCO: 108730at7742
## Thu Jul 14 01:19:14 CEST 2022 - The aliscore output appears to be empty. Check results for BUSCO: 108730at7742
## Thu Jul 14 01:19:14 CEST 2022 - The aliscore output file does not exist. Check results for BUSCO: 310707at7742
## Thu Jul 14 01:19:14 CEST 2022 - The aliscore output appears to be empty. Check results for BUSCO: 310707at7742
## Thu Jul 14 01:19:16 CEST 2022 - The aliscore output file does not exist. Check results for BUSCO: 193294at7742
## Thu Jul 14 01:19:16 CEST 2022 - The aliscore output appears to be empty. Check results for BUSCO: 193294at7742
## Thu Jul 14 01:19:19 CEST 2022 - The aliscore output file does not exist. Check results for BUSCO: 232849at7742
## Thu Jul 14 01:19:19 CEST 2022 - The aliscore output appears to be empty. Check results for BUSCO: 232849at7742
## Thu Jul 14 01:19:20 CEST 2022 - The aliscore output file does not exist. Check results for BUSCO: 107391at7742
## Thu Jul 14 01:19:20 CEST 2022 - The aliscore output appears to be empty. Check results for BUSCO: 107391at7742
## Thu Jul 14 01:19:21 CEST 2022 - The aliscore output file does not exist. Check results for BUSCO: 233977at7742
## Thu Jul 14 01:19:21 CEST 2022 - The aliscore output appears to be empty. Check results for BUSCO: 233977at7742
## Thu Jul 14 01:19:21 CEST 2022 - The aliscore output file does not exist. Check results for BUSCO: 122990at7742
## Thu Jul 14 01:19:21 CEST 2022 - The aliscore output appears to be empty. Check results for BUSCO: 122990at7742
## Thu Jul 14 01:19:22 CEST 2022 - The aliscore output file does not exist. Check results for BUSCO: 345012at7742
## Thu Jul 14 01:19:22 CEST 2022 - The aliscore output file does not exist. Check results for BUSCO: 327776at7742
## Thu Jul 14 01:19:22 CEST 2022 - The aliscore output appears to be empty. Check results for BUSCO: 345012at7742
## Thu Jul 14 01:19:22 CEST 2022 - The aliscore output appears to be empty. Check results for BUSCO: 327776at7742
## Thu Jul 14 01:19:23 CEST 2022 - The aliscore output file does not exist. Check results for BUSCO: 378656at7742
## Thu Jul 14 01:19:23 CEST 2022 - The aliscore output appears to be empty. Check results for BUSCO: 378656at7742
## Thu Jul 14 01:19:25 CEST 2022 - The aliscore output file does not exist. Check results for BUSCO: 218284at7742
## Thu Jul 14 01:19:25 CEST 2022 - The aliscore output appears to be empty. Check results for BUSCO: 218284at7742
## Thu Jul 14 01:19:27 CEST 2022 - The aliscore output file does not exist. Check results for BUSCO: 244702at7742
## Thu Jul 14 01:19:27 CEST 2022 - The aliscore output appears to be empty. Check results for BUSCO: 244702at7742
## Thu Jul 14 01:19:27 CEST 2022 - The aliscore output file does not exist. Check results for BUSCO: 133877at7742
## Thu Jul 14 01:19:27 CEST 2022 - The aliscore output appears to be empty. Check results for BUSCO: 133877at7742
## Thu Jul 14 01:19:31 CEST 2022 - The aliscore output file does not exist. Check results for BUSCO: 107188at7742
## Thu Jul 14 01:19:31 CEST 2022 - The aliscore output appears to be empty. Check results for BUSCO: 107188at7742
## Thu Jul 14 01:19:31 CEST 2022 - The aliscore output file does not exist. Check results for BUSCO: 97043at7742
## Thu Jul 14 01:19:31 CEST 2022 - The aliscore output appears to be empty. Check results for BUSCO: 97043at7742
## Thu Jul 14 01:19:31 CEST 2022 - The aliscore output file does not exist. Check results for BUSCO: 108730at7742
## Thu Jul 14 01:19:31 CEST 2022 - The aliscore output appears to be empty. Check results for BUSCO: 108730at7742
## Thu Jul 14 01:19:32 CEST 2022 - The aliscore output file does not exist. Check results for BUSCO: 82541at7742
## Thu Jul 14 01:19:32 CEST 2022 - The aliscore output appears to be empty. Check results for BUSCO: 82541at7742
## Thu Jul 14 01:19:32 CEST 2022 - The aliscore output file does not exist. Check results for BUSCO: 224105at7742
## Thu Jul 14 01:19:32 CEST 2022 - The aliscore output appears to be empty. Check results for BUSCO: 224105at7742
## Thu Jul 14 01:19:34 CEST 2022 - The aliscore output file does not exist. Check results for BUSCO: 218283at7742
## Thu Jul 14 01:19:34 CEST 2022 - The aliscore output appears to be empty. Check results for BUSCO: 218283at7742
## Thu Jul 14 01:19:36 CEST 2022 - The aliscore output file does not exist. Check results for BUSCO: 114115at7742
## Thu Jul 14 01:19:36 CEST 2022 - The aliscore output appears to be empty. Check results for BUSCO: 114115at7742
## Thu Jul 14 01:19:37 CEST 2022 - The aliscore output file does not exist. Check results for BUSCO: 147736at7742
## Thu Jul 14 01:19:37 CEST 2022 - The aliscore output appears to be empty. Check results for BUSCO: 147736at7742
## Thu Jul 14 01:19:39 CEST 2022 - The aliscore output file does not exist. Check results for BUSCO: 197023at7742
## Thu Jul 14 01:19:39 CEST 2022 - The aliscore output appears to be empty. Check results for BUSCO: 197023at7742
## Thu Jul 14 01:19:39 CEST 2022 - The aliscore output file does not exist. Check results for BUSCO: 108730at7742
## Thu Jul 14 01:19:39 CEST 2022 - The aliscore output appears to be empty. Check results for BUSCO: 108730at7742
## Thu Jul 14 01:19:42 CEST 2022 - The aliscore output file does not exist. Check results for BUSCO: 97043at7742
## Thu Jul 14 01:19:42 CEST 2022 - The aliscore output appears to be empty. Check results for BUSCO: 97043at7742
## Thu Jul 14 01:19:43 CEST 2022 - The aliscore output file does not exist. Check results for BUSCO: 103208at7742
## Thu Jul 14 01:19:43 CEST 2022 - The aliscore output appears to be empty. Check results for BUSCO: 103208at7742
## Thu Jul 14 01:19:44 CEST 2022 - The aliscore output file does not exist. Check results for BUSCO: 352256at7742
## Thu Jul 14 01:19:45 CEST 2022 - The aliscore output appears to be empty. Check results for BUSCO: 352256at7742
## Thu Jul 14 01:19:45 CEST 2022 - The aliscore output file does not exist. Check results for BUSCO: 206370at7742
## Thu Jul 14 01:19:45 CEST 2022 - The aliscore output appears to be empty. Check results for BUSCO: 206370at7742
## Thu Jul 14 01:19:47 CEST 2022 - The aliscore output file does not exist. Check results for BUSCO: 157460at7742
## Thu Jul 14 01:19:47 CEST 2022 - The aliscore output appears to be empty. Check results for BUSCO: 157460at7742
## Thu Jul 14 01:19:48 CEST 2022 - The aliscore output file does not exist. Check results for BUSCO: 134188at7742
## Thu Jul 14 01:19:48 CEST 2022 - The aliscore output appears to be empty. Check results for BUSCO: 134188at7742
## Thu Jul 14 01:19:48 CEST 2022 - The aliscore output file does not exist. Check results for BUSCO: 122955at7742
## Thu Jul 14 01:19:48 CEST 2022 - The aliscore output appears to be empty. Check results for BUSCO: 122955at7742
## Thu Jul 14 01:19:50 CEST 2022 - The aliscore output file does not exist. Check results for BUSCO: 272794at7742
## Thu Jul 14 01:19:50 CEST 2022 - The aliscore output appears to be empty. Check results for BUSCO: 272794at7742
## Thu Jul 14 01:19:50 CEST 2022 - The aliscore output file does not exist. Check results for BUSCO: 239989at7742
## Thu Jul 14 01:19:50 CEST 2022 - The aliscore output appears to be empty. Check results for BUSCO: 239989at7742
## Thu Jul 14 01:19:52 CEST 2022 - The aliscore output file does not exist. Check results for BUSCO: 155482at7742
## Thu Jul 14 01:19:52 CEST 2022 - The aliscore output appears to be empty. Check results for BUSCO: 155482at7742
## Thu Jul 14 01:19:53 CEST 2022 - The aliscore output file does not exist. Check results for BUSCO: 115042at7742
## Thu Jul 14 01:19:53 CEST 2022 - The aliscore output appears to be empty. Check results for BUSCO: 115042at7742
## Thu Jul 14 01:19:54 CEST 2022 - The aliscore output file does not exist. Check results for BUSCO: 239783at7742
## Thu Jul 14 01:19:54 CEST 2022 - The aliscore output appears to be empty. Check results for BUSCO: 239783at7742
## Thu Jul 14 01:19:56 CEST 2022 - The aliscore output file does not exist. Check results for BUSCO: 210385at7742
## Thu Jul 14 01:19:56 CEST 2022 - The aliscore output appears to be empty. Check results for BUSCO: 210385at7742
## Thu Jul 14 01:19:56 CEST 2022 - The aliscore output file does not exist. Check results for BUSCO: 93231at7742
## Thu Jul 14 01:19:56 CEST 2022 - The aliscore output appears to be empty. Check results for BUSCO: 93231at7742
## Thu Jul 14 01:19:57 CEST 2022 - The aliscore output file does not exist. Check results for BUSCO: 119120at7742
## Thu Jul 14 01:19:57 CEST 2022 - The aliscore output appears to be empty. Check results for BUSCO: 119120at7742
## Thu Jul 14 01:19:59 CEST 2022 - The aliscore output file does not exist. Check results for BUSCO: 122793at7742
## Thu Jul 14 01:19:59 CEST 2022 - The aliscore output appears to be empty. Check results for BUSCO: 122793at7742
## Thu Jul 14 01:20:00 CEST 2022 - The aliscore output file does not exist. Check results for BUSCO: 205393at7742
## Thu Jul 14 01:20:01 CEST 2022 - The aliscore output appears to be empty. Check results for BUSCO: 205393at7742
## Thu Jul 14 01:20:01 CEST 2022 - The aliscore output file does not exist. Check results for BUSCO: 166688at7742
## Thu Jul 14 01:20:01 CEST 2022 - The aliscore output appears to be empty. Check results for BUSCO: 166688at7742
## Thu Jul 14 01:20:03 CEST 2022 - The aliscore output file does not exist. Check results for BUSCO: 280833at7742
## Thu Jul 14 01:20:03 CEST 2022 - The aliscore output appears to be empty. Check results for BUSCO: 280833at7742
## Thu Jul 14 01:20:04 CEST 2022 - The aliscore output file does not exist. Check results for BUSCO: 116710at7742
## Thu Jul 14 01:20:04 CEST 2022 - The aliscore output appears to be empty. Check results for BUSCO: 116710at7742
## Thu Jul 14 01:20:05 CEST 2022 - The aliscore output file does not exist. Check results for BUSCO: 296707at7742
## Thu Jul 14 01:20:05 CEST 2022 - The aliscore output appears to be empty. Check results for BUSCO: 296707at7742
## Thu Jul 14 01:20:07 CEST 2022 - The aliscore output file does not exist. Check results for BUSCO: 222631at7742
## Thu Jul 14 01:20:07 CEST 2022 - The aliscore output appears to be empty. Check results for BUSCO: 222631at7742
## Thu Jul 14 01:20:08 CEST 2022 - The aliscore output file does not exist. Check results for BUSCO: 162265at7742
## Thu Jul 14 01:20:08 CEST 2022 - The aliscore output appears to be empty. Check results for BUSCO: 162265at7742
## Thu Jul 14 01:20:10 CEST 2022 - The aliscore output file does not exist. Check results for BUSCO: 150456at7742
## Thu Jul 14 01:20:10 CEST 2022 - The aliscore output appears to be empty. Check results for BUSCO: 150456at7742
## Thu Jul 14 01:20:11 CEST 2022 - The aliscore output file does not exist. Check results for BUSCO: 175755at7742
## Thu Jul 14 01:20:11 CEST 2022 - The aliscore output appears to be empty. Check results for BUSCO: 175755at7742
## Thu Jul 14 01:20:12 CEST 2022 - The aliscore output file does not exist. Check results for BUSCO: 332125at7742
## Thu Jul 14 01:20:12 CEST 2022 - The aliscore output appears to be empty. Check results for BUSCO: 332125at7742
## Thu Jul 14 01:20:13 CEST 2022 - The aliscore output file does not exist. Check results for BUSCO: 239747at7742
## Thu Jul 14 01:20:13 CEST 2022 - The aliscore output appears to be empty. Check results for BUSCO: 239747at7742
## Thu Jul 14 01:20:13 CEST 2022 - The aliscore output file does not exist. Check results for BUSCO: 171737at7742
## Thu Jul 14 01:20:13 CEST 2022 - The aliscore output appears to be empty. Check results for BUSCO: 171737at7742
## Thu Jul 14 01:20:17 CEST 2022 - The aliscore output file does not exist. Check results for BUSCO: 82541at7742
## Thu Jul 14 01:20:17 CEST 2022 - The aliscore output appears to be empty. Check results for BUSCO: 82541at7742
## Thu Jul 14 01:20:17 CEST 2022 - The aliscore output file does not exist. Check results for BUSCO: 155526at7742
## Thu Jul 14 01:20:17 CEST 2022 - The aliscore output appears to be empty. Check results for BUSCO: 155526at7742
## Thu Jul 14 01:20:17 CEST 2022 - The aliscore output file does not exist. Check results for BUSCO: 233977at7742
## Thu Jul 14 01:20:17 CEST 2022 - The aliscore output appears to be empty. Check results for BUSCO: 233977at7742
## Thu Jul 14 01:20:20 CEST 2022 - The aliscore output file does not exist. Check results for BUSCO: 53269at7742
## Thu Jul 14 01:20:20 CEST 2022 - The aliscore output appears to be empty. Check results for BUSCO: 53269at7742
## Thu Jul 14 01:20:21 CEST 2022 - The aliscore output file does not exist. Check results for BUSCO: 53269at7742
## Thu Jul 14 01:20:21 CEST 2022 - The aliscore output appears to be empty. Check results for BUSCO: 53269at7742
## Thu Jul 14 01:20:22 CEST 2022 - The aliscore output file does not exist. Check results for BUSCO: 294082at7742
## Thu Jul 14 01:20:22 CEST 2022 - The aliscore output appears to be empty. Check results for BUSCO: 294082at7742
## Thu Jul 14 01:20:22 CEST 2022 - The aliscore output file does not exist. Check results for BUSCO: 98637at7742
## Thu Jul 14 01:20:22 CEST 2022 - The aliscore output appears to be empty. Check results for BUSCO: 98637at7742
## Thu Jul 14 01:20:24 CEST 2022 - The aliscore output file does not exist. Check results for BUSCO: 119120at7742
## Thu Jul 14 01:20:24 CEST 2022 - The aliscore output appears to be empty. Check results for BUSCO: 119120at7742
## Thu Jul 14 01:20:24 CEST 2022 - The aliscore output file does not exist. Check results for BUSCO: 156795at7742
## Thu Jul 14 01:20:24 CEST 2022 - The aliscore output appears to be empty. Check results for BUSCO: 156795at7742
## Thu Jul 14 01:20:27 CEST 2022 - The aliscore output file does not exist. Check results for BUSCO: 93778at7742
## Thu Jul 14 01:20:27 CEST 2022 - The aliscore output appears to be empty. Check results for BUSCO: 93778at7742
## Thu Jul 14 01:20:28 CEST 2022 - The aliscore output file does not exist. Check results for BUSCO: 191875at7742
## Thu Jul 14 01:20:28 CEST 2022 - The aliscore output appears to be empty. Check results for BUSCO: 191875at7742
## Thu Jul 14 01:20:29 CEST 2022 - The aliscore output file does not exist. Check results for BUSCO: 140071at7742
## Thu Jul 14 01:20:29 CEST 2022 - The aliscore output appears to be empty. Check results for BUSCO: 140071at7742
## Thu Jul 14 01:20:29 CEST 2022 - The aliscore output file does not exist. Check results for BUSCO: 291147at7742
## Thu Jul 14 01:20:29 CEST 2022 - The aliscore output appears to be empty. Check results for BUSCO: 291147at7742
## Thu Jul 14 01:20:32 CEST 2022 - The aliscore output file does not exist. Check results for BUSCO: 93778at7742
## Thu Jul 14 01:20:32 CEST 2022 - The aliscore output appears to be empty. Check results for BUSCO: 93778at7742
## Thu Jul 14 01:20:33 CEST 2022 - The aliscore output file does not exist. Check results for BUSCO: 240075at7742
## Thu Jul 14 01:20:33 CEST 2022 - The aliscore output appears to be empty. Check results for BUSCO: 240075at7742
## Thu Jul 14 01:20:33 CEST 2022 - The aliscore output file does not exist. Check results for BUSCO: 139653at7742
## Thu Jul 14 01:20:33 CEST 2022 - The aliscore output appears to be empty. Check results for BUSCO: 139653at7742
## Thu Jul 14 01:20:36 CEST 2022 - The aliscore output file does not exist. Check results for BUSCO: 64971at7742
## Thu Jul 14 01:20:36 CEST 2022 - The aliscore output appears to be empty. Check results for BUSCO: 64971at7742
## Thu Jul 14 01:20:36 CEST 2022 - The aliscore output file does not exist. Check results for BUSCO: 301301at7742
## Thu Jul 14 01:20:36 CEST 2022 - The aliscore output appears to be empty. Check results for BUSCO: 301301at7742
## Thu Jul 14 01:20:37 CEST 2022 - The aliscore output file does not exist. Check results for BUSCO: 261612at7742
## Thu Jul 14 01:20:37 CEST 2022 - The aliscore output file does not exist. Check results for BUSCO: 291464at7742
## Thu Jul 14 01:20:37 CEST 2022 - The aliscore output appears to be empty. Check results for BUSCO: 261612at7742
## Thu Jul 14 01:20:37 CEST 2022 - The aliscore output appears to be empty. Check results for BUSCO: 291464at7742
## Thu Jul 14 01:20:40 CEST 2022 - The aliscore output file does not exist. Check results for BUSCO: 215440at7742
## Thu Jul 14 01:20:40 CEST 2022 - The aliscore output appears to be empty. Check results for BUSCO: 215440at7742
## Thu Jul 14 01:20:40 CEST 2022 - The aliscore output file does not exist. Check results for BUSCO: 344884at7742
## Thu Jul 14 01:20:40 CEST 2022 - The aliscore output appears to be empty. Check results for BUSCO: 344884at7742
## Thu Jul 14 01:20:41 CEST 2022 - The aliscore output file does not exist. Check results for BUSCO: 240075at7742
## Thu Jul 14 01:20:41 CEST 2022 - The aliscore output appears to be empty. Check results for BUSCO: 240075at7742
## Thu Jul 14 01:20:44 CEST 2022 - The aliscore output file does not exist. Check results for BUSCO: 266927at7742
## Thu Jul 14 01:20:44 CEST 2022 - The aliscore output appears to be empty. Check results for BUSCO: 266927at7742
## Thu Jul 14 01:20:44 CEST 2022 - The aliscore output file does not exist. Check results for BUSCO: 260905at7742
## Thu Jul 14 01:20:44 CEST 2022 - The aliscore output appears to be empty. Check results for BUSCO: 260905at7742
## Thu Jul 14 01:20:44 CEST 2022 - The aliscore output file does not exist. Check results for BUSCO: 352256at7742
## Thu Jul 14 01:20:45 CEST 2022 - The aliscore output appears to be empty. Check results for BUSCO: 352256at7742
## Thu Jul 14 01:20:46 CEST 2022 - The aliscore output file does not exist. Check results for BUSCO: 146203at7742
## Thu Jul 14 01:20:46 CEST 2022 - The aliscore output appears to be empty. Check results for BUSCO: 146203at7742
## Thu Jul 14 01:20:48 CEST 2022 - The aliscore output file does not exist. Check results for BUSCO: 294332at7742
## Thu Jul 14 01:20:48 CEST 2022 - The aliscore output appears to be empty. Check results for BUSCO: 294332at7742
## Thu Jul 14 01:20:49 CEST 2022 - The aliscore output file does not exist. Check results for BUSCO: 144645at7742
## Thu Jul 14 01:20:49 CEST 2022 - The aliscore output file does not exist. Check results for BUSCO: 221393at7742
## Thu Jul 14 01:20:49 CEST 2022 - The aliscore output appears to be empty. Check results for BUSCO: 144645at7742
## Thu Jul 14 01:20:49 CEST 2022 - The aliscore output appears to be empty. Check results for BUSCO: 221393at7742
## Thu Jul 14 01:20:50 CEST 2022 - The aliscore output file does not exist. Check results for BUSCO: 195929at7742
## Thu Jul 14 01:20:50 CEST 2022 - The aliscore output appears to be empty. Check results for BUSCO: 195929at7742
## Thu Jul 14 01:20:52 CEST 2022 - The aliscore output file does not exist. Check results for BUSCO: 153401at7742
## Thu Jul 14 01:20:52 CEST 2022 - The aliscore output appears to be empty. Check results for BUSCO: 153401at7742
## Thu Jul 14 01:20:52 CEST 2022 - The aliscore output file does not exist. Check results for BUSCO: 408649at7742
## Thu Jul 14 01:20:52 CEST 2022 - The aliscore output appears to be empty. Check results for BUSCO: 408649at7742
## Thu Jul 14 01:20:52 CEST 2022 - The aliscore output file does not exist. Check results for BUSCO: 161520at7742
## Thu Jul 14 01:20:52 CEST 2022 - The aliscore output appears to be empty. Check results for BUSCO: 161520at7742
## Thu Jul 14 01:20:55 CEST 2022 - The aliscore output file does not exist. Check results for BUSCO: 155795at7742
## Thu Jul 14 01:20:55 CEST 2022 - The aliscore output appears to be empty. Check results for BUSCO: 155795at7742
## Thu Jul 14 01:20:56 CEST 2022 - The aliscore output file does not exist. Check results for BUSCO: 196730at7742
## Thu Jul 14 01:20:56 CEST 2022 - The aliscore output appears to be empty. Check results for BUSCO: 196730at7742
## Thu Jul 14 01:20:57 CEST 2022 - The aliscore output file does not exist. Check results for BUSCO: 111128at7742
## Thu Jul 14 01:20:57 CEST 2022 - The aliscore output appears to be empty. Check results for BUSCO: 111128at7742
## Thu Jul 14 01:20:58 CEST 2022 - The aliscore output file does not exist. Check results for BUSCO: 224105at7742
## Thu Jul 14 01:20:58 CEST 2022 - The aliscore output appears to be empty. Check results for BUSCO: 224105at7742
## Thu Jul 14 01:21:00 CEST 2022 - The aliscore output file does not exist. Check results for BUSCO: 96292at7742
## Thu Jul 14 01:21:00 CEST 2022 - The aliscore output appears to be empty. Check results for BUSCO: 96292at7742
## Thu Jul 14 01:21:00 CEST 2022 - The aliscore output file does not exist. Check results for BUSCO: 93566at7742
## Thu Jul 14 01:21:00 CEST 2022 - The aliscore output appears to be empty. Check results for BUSCO: 93566at7742
## Thu Jul 14 01:21:02 CEST 2022 - The aliscore output file does not exist. Check results for BUSCO: 182480at7742
## Thu Jul 14 01:21:02 CEST 2022 - The aliscore output appears to be empty. Check results for BUSCO: 182480at7742
## Thu Jul 14 01:21:04 CEST 2022 - The aliscore output file does not exist. Check results for BUSCO: 123156at7742
## Thu Jul 14 01:21:04 CEST 2022 - The aliscore output appears to be empty. Check results for BUSCO: 123156at7742
## Thu Jul 14 01:21:04 CEST 2022 - The aliscore output file does not exist. Check results for BUSCO: 256473at7742
## Thu Jul 14 01:21:04 CEST 2022 - The aliscore output appears to be empty. Check results for BUSCO: 256473at7742
## Thu Jul 14 01:21:07 CEST 2022 - The aliscore output file does not exist. Check results for BUSCO: 170944at7742
## Thu Jul 14 01:21:07 CEST 2022 - The aliscore output appears to be empty. Check results for BUSCO: 170944at7742
## Thu Jul 14 01:21:07 CEST 2022 - The aliscore output file does not exist. Check results for BUSCO: 273954at7742
## Thu Jul 14 01:21:07 CEST 2022 - The aliscore output appears to be empty. Check results for BUSCO: 273954at7742
## Thu Jul 14 01:21:09 CEST 2022 - The aliscore output file does not exist. Check results for BUSCO: 93020at7742
## Thu Jul 14 01:21:09 CEST 2022 - The aliscore output appears to be empty. Check results for BUSCO: 93020at7742
## Thu Jul 14 01:21:09 CEST 2022 - The aliscore output file does not exist. Check results for BUSCO: 192313at7742
## Thu Jul 14 01:21:09 CEST 2022 - The aliscore output appears to be empty. Check results for BUSCO: 192313at7742
## Thu Jul 14 01:21:12 CEST 2022 - The aliscore output file does not exist. Check results for BUSCO: 128562at7742
## Thu Jul 14 01:21:12 CEST 2022 - The aliscore output appears to be empty. Check results for BUSCO: 128562at7742
## Thu Jul 14 01:21:12 CEST 2022 - The aliscore output file does not exist. Check results for BUSCO: 400635at7742
## Thu Jul 14 01:21:12 CEST 2022 - The aliscore output appears to be empty. Check results for BUSCO: 400635at7742
## Thu Jul 14 01:21:12 CEST 2022 - The aliscore output file does not exist. Check results for BUSCO: 128562at7742
## Thu Jul 14 01:21:12 CEST 2022 - The aliscore output appears to be empty. Check results for BUSCO: 128562at7742
## Thu Jul 14 01:21:14 CEST 2022 - The aliscore output file does not exist. Check results for BUSCO: 288402at7742
## Thu Jul 14 01:21:14 CEST 2022 - The aliscore output appears to be empty. Check results for BUSCO: 288402at7742
## Thu Jul 14 01:21:16 CEST 2022 - The aliscore output file does not exist. Check results for BUSCO: 64971at7742
## Thu Jul 14 01:21:16 CEST 2022 - The aliscore output appears to be empty. Check results for BUSCO: 64971at7742
## Thu Jul 14 01:21:17 CEST 2022 - The aliscore output file does not exist. Check results for BUSCO: 190356at7742
## Thu Jul 14 01:21:17 CEST 2022 - The aliscore output appears to be empty. Check results for BUSCO: 190356at7742
## Thu Jul 14 01:21:17 CEST 2022 - The aliscore output file does not exist. Check results for BUSCO: 197023at7742
## Thu Jul 14 01:21:17 CEST 2022 - The aliscore output appears to be empty. Check results for BUSCO: 197023at7742
## Thu Jul 14 01:21:18 CEST 2022 - The aliscore output file does not exist. Check results for BUSCO: 197023at7742
## Thu Jul 14 01:21:18 CEST 2022 - The aliscore output appears to be empty. Check results for BUSCO: 197023at7742
## Thu Jul 14 01:21:20 CEST 2022 - The aliscore output file does not exist. Check results for BUSCO: 104856at7742
## Thu Jul 14 01:21:20 CEST 2022 - The aliscore output appears to be empty. Check results for BUSCO: 104856at7742
## Thu Jul 14 01:21:22 CEST 2022 - The aliscore output file does not exist. Check results for BUSCO: 325029at7742
## Thu Jul 14 01:21:22 CEST 2022 - The aliscore output appears to be empty. Check results for BUSCO: 325029at7742
## Thu Jul 14 01:21:23 CEST 2022 - The aliscore output file does not exist. Check results for BUSCO: 137629at7742
## Thu Jul 14 01:21:23 CEST 2022 - The aliscore output appears to be empty. Check results for BUSCO: 137629at7742
## Thu Jul 14 01:21:24 CEST 2022 - The aliscore output file does not exist. Check results for BUSCO: 122955at7742
## Thu Jul 14 01:21:24 CEST 2022 - The aliscore output appears to be empty. Check results for BUSCO: 122955at7742
## Thu Jul 14 01:21:25 CEST 2022 - The aliscore output file does not exist. Check results for BUSCO: 97588at7742
## Thu Jul 14 01:21:25 CEST 2022 - The aliscore output appears to be empty. Check results for BUSCO: 97588at7742
## Thu Jul 14 01:21:27 CEST 2022 - The aliscore output file does not exist. Check results for BUSCO: 316777at7742
## Thu Jul 14 01:21:27 CEST 2022 - The aliscore output appears to be empty. Check results for BUSCO: 316777at7742
## Thu Jul 14 01:21:27 CEST 2022 - The aliscore output file does not exist. Check results for BUSCO: 54598at7742
## Thu Jul 14 01:21:27 CEST 2022 - The aliscore output appears to be empty. Check results for BUSCO: 54598at7742
## Thu Jul 14 01:21:29 CEST 2022 - The aliscore output file does not exist. Check results for BUSCO: 98753at7742
## Thu Jul 14 01:21:29 CEST 2022 - The aliscore output appears to be empty. Check results for BUSCO: 98753at7742
## Thu Jul 14 01:21:30 CEST 2022 - The aliscore output file does not exist. Check results for BUSCO: 131564at7742
## Thu Jul 14 01:21:30 CEST 2022 - The aliscore output appears to be empty. Check results for BUSCO: 131564at7742
## Thu Jul 14 01:21:30 CEST 2022 - The aliscore output file does not exist. Check results for BUSCO: 103422at7742
## Thu Jul 14 01:21:30 CEST 2022 - The aliscore output appears to be empty. Check results for BUSCO: 103422at7742
## Thu Jul 14 01:21:31 CEST 2022 - The aliscore output file does not exist. Check results for BUSCO: 337175at7742
## Thu Jul 14 01:21:31 CEST 2022 - The aliscore output appears to be empty. Check results for BUSCO: 337175at7742
## Thu Jul 14 01:21:33 CEST 2022 - The aliscore output file does not exist. Check results for BUSCO: 176317at7742
## Thu Jul 14 01:21:33 CEST 2022 - The aliscore output appears to be empty. Check results for BUSCO: 176317at7742
## Thu Jul 14 01:21:35 CEST 2022 - The aliscore output file does not exist. Check results for BUSCO: 224267at7742
## Thu Jul 14 01:21:35 CEST 2022 - The aliscore output appears to be empty. Check results for BUSCO: 224267at7742
## Thu Jul 14 01:21:37 CEST 2022 - The aliscore output file does not exist. Check results for BUSCO: 156795at7742
## Thu Jul 14 01:21:37 CEST 2022 - The aliscore output appears to be empty. Check results for BUSCO: 156795at7742
## Thu Jul 14 01:21:37 CEST 2022 - The aliscore output file does not exist. Check results for BUSCO: 373540at7742
## Thu Jul 14 01:21:37 CEST 2022 - The aliscore output appears to be empty. Check results for BUSCO: 373540at7742
## Thu Jul 14 01:21:37 CEST 2022 - The aliscore output file does not exist. Check results for BUSCO: 296707at7742
## Thu Jul 14 01:21:37 CEST 2022 - The aliscore output appears to be empty. Check results for BUSCO: 296707at7742
## Thu Jul 14 01:21:39 CEST 2022 - The aliscore output file does not exist. Check results for BUSCO: 340508at7742
## Thu Jul 14 01:21:39 CEST 2022 - The aliscore output appears to be empty. Check results for BUSCO: 340508at7742
## Thu Jul 14 01:21:42 CEST 2022 - The aliscore output file does not exist. Check results for BUSCO: 143678at7742
## Thu Jul 14 01:21:42 CEST 2022 - The aliscore output appears to be empty. Check results for BUSCO: 143678at7742
## Thu Jul 14 01:21:42 CEST 2022 - The aliscore output file does not exist. Check results for BUSCO: 288766at7742
## Thu Jul 14 01:21:42 CEST 2022 - The aliscore output appears to be empty. Check results for BUSCO: 288766at7742
## Thu Jul 14 01:21:43 CEST 2022 - The aliscore output file does not exist. Check results for BUSCO: 172204at7742
## Thu Jul 14 01:21:43 CEST 2022 - The aliscore output appears to be empty. Check results for BUSCO: 172204at7742
## Thu Jul 14 01:21:46 CEST 2022 - The aliscore output file does not exist. Check results for BUSCO: 130062at7742
## Thu Jul 14 01:21:46 CEST 2022 - The aliscore output appears to be empty. Check results for BUSCO: 130062at7742
## Thu Jul 14 01:21:46 CEST 2022 - The aliscore output file does not exist. Check results for BUSCO: 108553at7742
## Thu Jul 14 01:21:46 CEST 2022 - The aliscore output appears to be empty. Check results for BUSCO: 108553at7742
## Thu Jul 14 01:21:47 CEST 2022 - The aliscore output file does not exist. Check results for BUSCO: 144445at7742
## Thu Jul 14 01:21:47 CEST 2022 - The aliscore output appears to be empty. Check results for BUSCO: 144445at7742
## Thu Jul 14 01:21:48 CEST 2022 - The aliscore output file does not exist. Check results for BUSCO: 123156at7742
## Thu Jul 14 01:21:48 CEST 2022 - The aliscore output appears to be empty. Check results for BUSCO: 123156at7742
## Thu Jul 14 01:21:50 CEST 2022 - The aliscore output file does not exist. Check results for BUSCO: 359184at7742
## Thu Jul 14 01:21:50 CEST 2022 - The aliscore output appears to be empty. Check results for BUSCO: 359184at7742
## Thu Jul 14 01:21:51 CEST 2022 - The aliscore output file does not exist. Check results for BUSCO: 218673at7742
## Thu Jul 14 01:21:51 CEST 2022 - The aliscore output file does not exist. Check results for BUSCO: 291807at7742
## Thu Jul 14 01:21:51 CEST 2022 - The aliscore output appears to be empty. Check results for BUSCO: 218673at7742
## Thu Jul 14 01:21:51 CEST 2022 - The aliscore output appears to be empty. Check results for BUSCO: 291807at7742
## Thu Jul 14 01:21:54 CEST 2022 - The aliscore output file does not exist. Check results for BUSCO: 199227at7742
## Thu Jul 14 01:21:54 CEST 2022 - The aliscore output appears to be empty. Check results for BUSCO: 199227at7742
## Thu Jul 14 01:21:54 CEST 2022 - The aliscore output file does not exist. Check results for BUSCO: 174048at7742
## Thu Jul 14 01:21:54 CEST 2022 - The aliscore output appears to be empty. Check results for BUSCO: 174048at7742
## Thu Jul 14 01:21:55 CEST 2022 - The aliscore output file does not exist. Check results for BUSCO: 62905at7742
## Thu Jul 14 01:21:55 CEST 2022 - The aliscore output appears to be empty. Check results for BUSCO: 62905at7742
## Thu Jul 14 01:21:58 CEST 2022 - The aliscore output file does not exist. Check results for BUSCO: 151349at7742
## Thu Jul 14 01:21:58 CEST 2022 - The aliscore output appears to be empty. Check results for BUSCO: 151349at7742
## Thu Jul 14 01:21:58 CEST 2022 - The aliscore output file does not exist. Check results for BUSCO: 95886at7742
## Thu Jul 14 01:21:58 CEST 2022 - The aliscore output appears to be empty. Check results for BUSCO: 95886at7742
## Thu Jul 14 01:22:01 CEST 2022 - The aliscore output file does not exist. Check results for BUSCO: 240065at7742
## Thu Jul 14 01:22:01 CEST 2022 - The aliscore output appears to be empty. Check results for BUSCO: 240065at7742
## Thu Jul 14 01:22:01 CEST 2022 - The aliscore output file does not exist. Check results for BUSCO: 155435at7742
## Thu Jul 14 01:22:01 CEST 2022 - The aliscore output appears to be empty. Check results for BUSCO: 155435at7742
## Thu Jul 14 01:22:01 CEST 2022 - The aliscore output file does not exist. Check results for BUSCO: 283910at7742
## Thu Jul 14 01:22:02 CEST 2022 - The aliscore output appears to be empty. Check results for BUSCO: 283910at7742
## Thu Jul 14 01:22:02 CEST 2022 - The aliscore output file does not exist. Check results for BUSCO: 269540at7742
## Thu Jul 14 01:22:02 CEST 2022 - The aliscore output appears to be empty. Check results for BUSCO: 269540at7742
## Thu Jul 14 01:22:04 CEST 2022 - The aliscore output file does not exist. Check results for BUSCO: 313142at7742
## Thu Jul 14 01:22:04 CEST 2022 - The aliscore output appears to be empty. Check results for BUSCO: 313142at7742
## Thu Jul 14 01:22:05 CEST 2022 - The aliscore output file does not exist. Check results for BUSCO: 162265at7742
## Thu Jul 14 01:22:05 CEST 2022 - The aliscore output appears to be empty. Check results for BUSCO: 162265at7742
## Thu Jul 14 01:22:06 CEST 2022 - The aliscore output file does not exist. Check results for BUSCO: 160923at7742
## Thu Jul 14 01:22:06 CEST 2022 - The aliscore output appears to be empty. Check results for BUSCO: 160923at7742
## Thu Jul 14 01:22:08 CEST 2022 - The aliscore output file does not exist. Check results for BUSCO: 280159at7742
## Thu Jul 14 01:22:08 CEST 2022 - The aliscore output appears to be empty. Check results for BUSCO: 280159at7742
## Thu Jul 14 01:22:08 CEST 2022 - The aliscore output file does not exist. Check results for BUSCO: 200616at7742
## Thu Jul 14 01:22:08 CEST 2022 - The aliscore output appears to be empty. Check results for BUSCO: 200616at7742
## Thu Jul 14 01:22:11 CEST 2022 - The aliscore output file does not exist. Check results for BUSCO: 155435at7742
## Thu Jul 14 01:22:11 CEST 2022 - The aliscore output appears to be empty. Check results for BUSCO: 155435at7742
## Thu Jul 14 01:22:11 CEST 2022 - The aliscore output file does not exist. Check results for BUSCO: 136142at7742
## Thu Jul 14 01:22:11 CEST 2022 - The aliscore output appears to be empty. Check results for BUSCO: 136142at7742
## Thu Jul 14 01:22:11 CEST 2022 - The aliscore output file does not exist. Check results for BUSCO: 182489at7742
## Thu Jul 14 01:22:11 CEST 2022 - The aliscore output appears to be empty. Check results for BUSCO: 182489at7742
## Thu Jul 14 01:22:13 CEST 2022 - The aliscore output file does not exist. Check results for BUSCO: 294082at7742
## Thu Jul 14 01:22:13 CEST 2022 - The aliscore output appears to be empty. Check results for BUSCO: 294082at7742
## Thu Jul 14 01:22:14 CEST 2022 - The aliscore output file does not exist. Check results for BUSCO: 106271at7742
## Thu Jul 14 01:22:14 CEST 2022 - The aliscore output appears to be empty. Check results for BUSCO: 106271at7742
## Thu Jul 14 01:22:16 CEST 2022 - The aliscore output file does not exist. Check results for BUSCO: 164950at7742
## Thu Jul 14 01:22:16 CEST 2022 - The aliscore output appears to be empty. Check results for BUSCO: 164950at7742
## Thu Jul 14 01:22:17 CEST 2022 - The aliscore output file does not exist. Check results for BUSCO: 318895at7742
## Thu Jul 14 01:22:17 CEST 2022 - The aliscore output appears to be empty. Check results for BUSCO: 318895at7742
## Thu Jul 14 01:22:18 CEST 2022 - The aliscore output file does not exist. Check results for BUSCO: 155795at7742
## Thu Jul 14 01:22:19 CEST 2022 - The aliscore output appears to be empty. Check results for BUSCO: 155795at7742
## Thu Jul 14 01:22:19 CEST 2022 - The aliscore output file does not exist. Check results for BUSCO: 184498at7742
## Thu Jul 14 01:22:19 CEST 2022 - The aliscore output appears to be empty. Check results for BUSCO: 184498at7742
## Thu Jul 14 01:22:20 CEST 2022 - The aliscore output file does not exist. Check results for BUSCO: 187969at7742
## Thu Jul 14 01:22:20 CEST 2022 - The aliscore output appears to be empty. Check results for BUSCO: 187969at7742
## Thu Jul 14 01:22:22 CEST 2022 - The aliscore output file does not exist. Check results for BUSCO: 134679at7742
## Thu Jul 14 01:22:22 CEST 2022 - The aliscore output appears to be empty. Check results for BUSCO: 134679at7742
## Thu Jul 14 01:22:22 CEST 2022 - The aliscore output file does not exist. Check results for BUSCO: 224267at7742
## Thu Jul 14 01:22:22 CEST 2022 - The aliscore output appears to be empty. Check results for BUSCO: 224267at7742
## Thu Jul 14 01:22:23 CEST 2022 - The aliscore output file does not exist. Check results for BUSCO: 184498at7742
## Thu Jul 14 01:22:23 CEST 2022 - The aliscore output appears to be empty. Check results for BUSCO: 184498at7742
## Thu Jul 14 01:22:26 CEST 2022 - The aliscore output file does not exist. Check results for BUSCO: 318965at7742
## Thu Jul 14 01:22:26 CEST 2022 - The aliscore output appears to be empty. Check results for BUSCO: 318965at7742
## Thu Jul 14 01:22:27 CEST 2022 - The aliscore output file does not exist. Check results for BUSCO: 109231at7742
## Thu Jul 14 01:22:27 CEST 2022 - The aliscore output appears to be empty. Check results for BUSCO: 109231at7742
## Thu Jul 14 01:22:28 CEST 2022 - The aliscore output file does not exist. Check results for BUSCO: 206370at7742
## Thu Jul 14 01:22:28 CEST 2022 - The aliscore output appears to be empty. Check results for BUSCO: 206370at7742
## Thu Jul 14 01:22:29 CEST 2022 - The aliscore output file does not exist. Check results for BUSCO: 155526at7742
## Thu Jul 14 01:22:29 CEST 2022 - The aliscore output appears to be empty. Check results for BUSCO: 155526at7742
## Thu Jul 14 01:22:30 CEST 2022 - The aliscore output file does not exist. Check results for BUSCO: 373540at7742
## Thu Jul 14 01:22:30 CEST 2022 - The aliscore output appears to be empty. Check results for BUSCO: 373540at7742
## Thu Jul 14 01:22:32 CEST 2022 - The aliscore output file does not exist. Check results for BUSCO: 260905at7742
## Thu Jul 14 01:22:32 CEST 2022 - The aliscore output appears to be empty. Check results for BUSCO: 260905at7742
## Thu Jul 14 01:22:33 CEST 2022 - The aliscore output file does not exist. Check results for BUSCO: 139050at7742
## Thu Jul 14 01:22:33 CEST 2022 - The aliscore output appears to be empty. Check results for BUSCO: 139050at7742
## Thu Jul 14 01:22:34 CEST 2022 - The aliscore output file does not exist. Check results for BUSCO: 122990at7742
## Thu Jul 14 01:22:34 CEST 2022 - The aliscore output appears to be empty. Check results for BUSCO: 122990at7742
## Thu Jul 14 01:22:34 CEST 2022 - The aliscore output file does not exist. Check results for BUSCO: 130062at7742
## Thu Jul 14 01:22:34 CEST 2022 - The aliscore output appears to be empty. Check results for BUSCO: 130062at7742
## Thu Jul 14 01:22:36 CEST 2022 - The aliscore output file does not exist. Check results for BUSCO: 220786at7742
## Thu Jul 14 01:22:36 CEST 2022 - The aliscore output appears to be empty. Check results for BUSCO: 220786at7742
## Thu Jul 14 01:22:38 CEST 2022 - The aliscore output file does not exist. Check results for BUSCO: 111128at7742
## Thu Jul 14 01:22:38 CEST 2022 - The aliscore output appears to be empty. Check results for BUSCO: 111128at7742
## Thu Jul 14 01:22:38 CEST 2022 - The aliscore output file does not exist. Check results for BUSCO: 369435at7742
## Thu Jul 14 01:22:38 CEST 2022 - The aliscore output appears to be empty. Check results for BUSCO: 369435at7742
## Thu Jul 14 01:22:40 CEST 2022 - The aliscore output file does not exist. Check results for BUSCO: 323828at7742
## Thu Jul 14 01:22:40 CEST 2022 - The aliscore output appears to be empty. Check results for BUSCO: 323828at7742
## Thu Jul 14 01:22:42 CEST 2022 - The aliscore output file does not exist. Check results for BUSCO: 155526at7742
## Thu Jul 14 01:22:42 CEST 2022 - The aliscore output appears to be empty. Check results for BUSCO: 155526at7742
## Thu Jul 14 01:22:43 CEST 2022 - The aliscore output file does not exist. Check results for BUSCO: 183332at7742
## Thu Jul 14 01:22:43 CEST 2022 - The aliscore output appears to be empty. Check results for BUSCO: 183332at7742
## Thu Jul 14 01:22:44 CEST 2022 - The aliscore output file does not exist. Check results for BUSCO: 305417at7742
## Thu Jul 14 01:22:44 CEST 2022 - The aliscore output appears to be empty. Check results for BUSCO: 305417at7742
## Thu Jul 14 01:22:45 CEST 2022 - The aliscore output file does not exist. Check results for BUSCO: 147736at7742
## Thu Jul 14 01:22:45 CEST 2022 - The aliscore output appears to be empty. Check results for BUSCO: 147736at7742
## Thu Jul 14 01:22:46 CEST 2022 - The aliscore output file does not exist. Check results for BUSCO: 123758at7742
## Thu Jul 14 01:22:46 CEST 2022 - The aliscore output appears to be empty. Check results for BUSCO: 123758at7742
## Thu Jul 14 01:22:46 CEST 2022 - The aliscore output file does not exist. Check results for BUSCO: 218524at7742
## Thu Jul 14 01:22:46 CEST 2022 - The aliscore output appears to be empty. Check results for BUSCO: 218524at7742
## Thu Jul 14 01:22:47 CEST 2022 - The aliscore output file does not exist. Check results for BUSCO: 119371at7742
## Thu Jul 14 01:22:47 CEST 2022 - The aliscore output appears to be empty. Check results for BUSCO: 119371at7742
## Thu Jul 14 01:22:50 CEST 2022 - The aliscore output file does not exist. Check results for BUSCO: 281891at7742
## Thu Jul 14 01:22:50 CEST 2022 - The aliscore output appears to be empty. Check results for BUSCO: 281891at7742
## Thu Jul 14 01:22:52 CEST 2022 - The aliscore output file does not exist. Check results for BUSCO: 199120at7742
## Thu Jul 14 01:22:52 CEST 2022 - The aliscore output appears to be empty. Check results for BUSCO: 199120at7742
## Thu Jul 14 01:22:52 CEST 2022 - The aliscore output file does not exist. Check results for BUSCO: 340971at7742
## Thu Jul 14 01:22:52 CEST 2022 - The aliscore output appears to be empty. Check results for BUSCO: 340971at7742
## Thu Jul 14 01:22:53 CEST 2022 - The aliscore output file does not exist. Check results for BUSCO: 263171at7742
## Thu Jul 14 01:22:53 CEST 2022 - The aliscore output appears to be empty. Check results for BUSCO: 263171at7742
## Thu Jul 14 01:22:54 CEST 2022 - The aliscore output file does not exist. Check results for BUSCO: 64971at7742
## Thu Jul 14 01:22:54 CEST 2022 - The aliscore output appears to be empty. Check results for BUSCO: 64971at7742
## Thu Jul 14 01:22:56 CEST 2022 - The aliscore output file does not exist. Check results for BUSCO: 379829at7742
## Thu Jul 14 01:22:56 CEST 2022 - The aliscore output appears to be empty. Check results for BUSCO: 379829at7742
## Thu Jul 14 01:22:56 CEST 2022 - The aliscore output file does not exist. Check results for BUSCO: 367197at7742
## Thu Jul 14 01:22:56 CEST 2022 - The aliscore output appears to be empty. Check results for BUSCO: 367197at7742
## Thu Jul 14 01:22:57 CEST 2022 - The aliscore output file does not exist. Check results for BUSCO: 145949at7742
## Thu Jul 14 01:22:57 CEST 2022 - The aliscore output appears to be empty. Check results for BUSCO: 145949at7742
## Thu Jul 14 01:22:59 CEST 2022 - The aliscore output file does not exist. Check results for BUSCO: 219513at7742
## Thu Jul 14 01:22:59 CEST 2022 - The aliscore output appears to be empty. Check results for BUSCO: 219513at7742
## Thu Jul 14 01:23:00 CEST 2022 - The aliscore output file does not exist. Check results for BUSCO: 93231at7742
## Thu Jul 14 01:23:00 CEST 2022 - The aliscore output appears to be empty. Check results for BUSCO: 93231at7742
## Thu Jul 14 01:23:01 CEST 2022 - The aliscore output file does not exist. Check results for BUSCO: 369435at7742
## Thu Jul 14 01:23:01 CEST 2022 - The aliscore output appears to be empty. Check results for BUSCO: 369435at7742
## Thu Jul 14 01:23:02 CEST 2022 - The aliscore output file does not exist. Check results for BUSCO: 218283at7742
## Thu Jul 14 01:23:02 CEST 2022 - The aliscore output appears to be empty. Check results for BUSCO: 218283at7742
## Thu Jul 14 01:23:04 CEST 2022 - The aliscore output file does not exist. Check results for BUSCO: 142902at7742
## Thu Jul 14 01:23:04 CEST 2022 - The aliscore output appears to be empty. Check results for BUSCO: 142902at7742
## Thu Jul 14 01:23:05 CEST 2022 - The aliscore output file does not exist. Check results for BUSCO: 326687at7742
## Thu Jul 14 01:23:05 CEST 2022 - The aliscore output appears to be empty. Check results for BUSCO: 326687at7742
## Thu Jul 14 01:23:06 CEST 2022 - The aliscore output file does not exist. Check results for BUSCO: 185742at7742
## Thu Jul 14 01:23:06 CEST 2022 - The aliscore output appears to be empty. Check results for BUSCO: 185742at7742
## Thu Jul 14 01:23:07 CEST 2022 - The aliscore output file does not exist. Check results for BUSCO: 263171at7742
## Thu Jul 14 01:23:07 CEST 2022 - The aliscore output appears to be empty. Check results for BUSCO: 263171at7742
## Thu Jul 14 01:23:07 CEST 2022 - The aliscore output file does not exist. Check results for BUSCO: 68376at7742
## Thu Jul 14 01:23:07 CEST 2022 - The aliscore output appears to be empty. Check results for BUSCO: 68376at7742
## Thu Jul 14 01:23:11 CEST 2022 - The aliscore output file does not exist. Check results for BUSCO: 112051at7742
## Thu Jul 14 01:23:11 CEST 2022 - The aliscore output file does not exist. Check results for BUSCO: 221061at7742
## Thu Jul 14 01:23:11 CEST 2022 - The aliscore output appears to be empty. Check results for BUSCO: 112051at7742
## Thu Jul 14 01:23:11 CEST 2022 - The aliscore output appears to be empty. Check results for BUSCO: 221061at7742
## Thu Jul 14 01:23:11 CEST 2022 - The aliscore output file does not exist. Check results for BUSCO: 256457at7742
## Thu Jul 14 01:23:11 CEST 2022 - The aliscore output appears to be empty. Check results for BUSCO: 256457at7742
## Thu Jul 14 01:23:13 CEST 2022 - The aliscore output file does not exist. Check results for BUSCO: 97588at7742
## Thu Jul 14 01:23:13 CEST 2022 - The aliscore output appears to be empty. Check results for BUSCO: 97588at7742
## Thu Jul 14 01:23:15 CEST 2022 - The aliscore output file does not exist. Check results for BUSCO: 258449at7742
## Thu Jul 14 01:23:15 CEST 2022 - The aliscore output appears to be empty. Check results for BUSCO: 258449at7742
## Thu Jul 14 01:23:15 CEST 2022 - The aliscore output file does not exist. Check results for BUSCO: 101941at7742
## Thu Jul 14 01:23:15 CEST 2022 - The aliscore output appears to be empty. Check results for BUSCO: 101941at7742
## Thu Jul 14 01:23:17 CEST 2022 - The aliscore output file does not exist. Check results for BUSCO: 104856at7742
## Thu Jul 14 01:23:17 CEST 2022 - The aliscore output appears to be empty. Check results for BUSCO: 104856at7742
## Thu Jul 14 01:23:18 CEST 2022 - The aliscore output file does not exist. Check results for BUSCO: 388724at7742
## Thu Jul 14 01:23:18 CEST 2022 - The aliscore output appears to be empty. Check results for BUSCO: 388724at7742
## Thu Jul 14 01:23:18 CEST 2022 - The aliscore output file does not exist. Check results for BUSCO: 369435at7742
## Thu Jul 14 01:23:18 CEST 2022 - The aliscore output appears to be empty. Check results for BUSCO: 369435at7742
## Thu Jul 14 01:23:21 CEST 2022 - The aliscore output file does not exist. Check results for BUSCO: 92374at7742
## Thu Jul 14 01:23:21 CEST 2022 - The aliscore output appears to be empty. Check results for BUSCO: 92374at7742
## Thu Jul 14 01:23:24 CEST 2022 - The aliscore output file does not exist. Check results for BUSCO: 128562at7742
## Thu Jul 14 01:23:24 CEST 2022 - The aliscore output appears to be empty. Check results for BUSCO: 128562at7742
## Thu Jul 14 01:23:24 CEST 2022 - The aliscore output file does not exist. Check results for BUSCO: 125500at7742
## Thu Jul 14 01:23:24 CEST 2022 - The aliscore output appears to be empty. Check results for BUSCO: 125500at7742
## Thu Jul 14 01:23:24 CEST 2022 - The aliscore output file does not exist. Check results for BUSCO: 174048at7742
## Thu Jul 14 01:23:24 CEST 2022 - The aliscore output appears to be empty. Check results for BUSCO: 174048at7742
## Thu Jul 14 01:23:25 CEST 2022 - The aliscore output file does not exist. Check results for BUSCO: 351544at7742
## Thu Jul 14 01:23:25 CEST 2022 - The aliscore output appears to be empty. Check results for BUSCO: 351544at7742
## Thu Jul 14 01:23:26 CEST 2022 - The aliscore output file does not exist. Check results for BUSCO: 365677at7742
## Thu Jul 14 01:23:26 CEST 2022 - The aliscore output appears to be empty. Check results for BUSCO: 365677at7742
## Thu Jul 14 01:23:28 CEST 2022 - The aliscore output file does not exist. Check results for BUSCO: 260905at7742
## Thu Jul 14 01:23:28 CEST 2022 - The aliscore output appears to be empty. Check results for BUSCO: 260905at7742
## Thu Jul 14 01:23:28 CEST 2022 - The aliscore output file does not exist. Check results for BUSCO: 321918at7742
## Thu Jul 14 01:23:28 CEST 2022 - The aliscore output appears to be empty. Check results for BUSCO: 321918at7742
## Thu Jul 14 01:23:31 CEST 2022 - The aliscore output file does not exist. Check results for BUSCO: 256348at7742
## Thu Jul 14 01:23:31 CEST 2022 - The aliscore output appears to be empty. Check results for BUSCO: 256348at7742
## Thu Jul 14 01:23:31 CEST 2022 - The aliscore output file does not exist. Check results for BUSCO: 96459at7742
## Thu Jul 14 01:23:31 CEST 2022 - The aliscore output file does not exist. Check results for BUSCO: 119532at7742
## Thu Jul 14 01:23:31 CEST 2022 - The aliscore output appears to be empty. Check results for BUSCO: 96459at7742
## Thu Jul 14 01:23:31 CEST 2022 - The aliscore output appears to be empty. Check results for BUSCO: 119532at7742
## Thu Jul 14 01:23:33 CEST 2022 - The aliscore output file does not exist. Check results for BUSCO: 219513at7742
## Thu Jul 14 01:23:33 CEST 2022 - The aliscore output appears to be empty. Check results for BUSCO: 219513at7742
## Thu Jul 14 01:23:34 CEST 2022 - The aliscore output file does not exist. Check results for BUSCO: 205318at7742
## Thu Jul 14 01:23:34 CEST 2022 - The aliscore output appears to be empty. Check results for BUSCO: 205318at7742
## Thu Jul 14 01:23:35 CEST 2022 - The aliscore output file does not exist. Check results for BUSCO: 144578at7742
## Thu Jul 14 01:23:35 CEST 2022 - The aliscore output appears to be empty. Check results for BUSCO: 144578at7742
## Thu Jul 14 01:23:37 CEST 2022 - The aliscore output file does not exist. Check results for BUSCO: 224101at7742
## Thu Jul 14 01:23:37 CEST 2022 - The aliscore output appears to be empty. Check results for BUSCO: 224101at7742
## Thu Jul 14 01:23:39 CEST 2022 - The aliscore output file does not exist. Check results for BUSCO: 128068at7742
## Thu Jul 14 01:23:39 CEST 2022 - The aliscore output appears to be empty. Check results for BUSCO: 128068at7742
## Thu Jul 14 01:23:40 CEST 2022 - The aliscore output file does not exist. Check results for BUSCO: 336679at7742
## Thu Jul 14 01:23:40 CEST 2022 - The aliscore output appears to be empty. Check results for BUSCO: 336679at7742
## Thu Jul 14 01:23:40 CEST 2022 - The aliscore output file does not exist. Check results for BUSCO: 199227at7742
## Thu Jul 14 01:23:40 CEST 2022 - The aliscore output appears to be empty. Check results for BUSCO: 199227at7742
## Thu Jul 14 01:23:41 CEST 2022 - The aliscore output file does not exist. Check results for BUSCO: 256457at7742
## Thu Jul 14 01:23:41 CEST 2022 - The aliscore output appears to be empty. Check results for BUSCO: 256457at7742
## Thu Jul 14 01:23:41 CEST 2022 - The aliscore output file does not exist. Check results for BUSCO: 210385at7742
## Thu Jul 14 01:23:41 CEST 2022 - The aliscore output appears to be empty. Check results for BUSCO: 210385at7742
## Thu Jul 14 01:23:44 CEST 2022 - The aliscore output file does not exist. Check results for BUSCO: 96459at7742
## Thu Jul 14 01:23:44 CEST 2022 - The aliscore output appears to be empty. Check results for BUSCO: 96459at7742
## Thu Jul 14 01:23:45 CEST 2022 - The aliscore output file does not exist. Check results for BUSCO: 154457at7742
## Thu Jul 14 01:23:45 CEST 2022 - The aliscore output appears to be empty. Check results for BUSCO: 154457at7742
## Thu Jul 14 01:23:48 CEST 2022 - The aliscore output file does not exist. Check results for BUSCO: 108553at7742
## Thu Jul 14 01:23:48 CEST 2022 - The aliscore output file does not exist. Check results for BUSCO: 207031at7742
## Thu Jul 14 01:23:48 CEST 2022 - The aliscore output appears to be empty. Check results for BUSCO: 108553at7742
## Thu Jul 14 01:23:48 CEST 2022 - The aliscore output appears to be empty. Check results for BUSCO: 207031at7742
## Thu Jul 14 01:23:48 CEST 2022 - The aliscore output file does not exist. Check results for BUSCO: 109231at7742
## Thu Jul 14 01:23:48 CEST 2022 - The aliscore output appears to be empty. Check results for BUSCO: 109231at7742
## Thu Jul 14 01:23:48 CEST 2022 - The aliscore output file does not exist. Check results for BUSCO: 147736at7742
## Thu Jul 14 01:23:48 CEST 2022 - The aliscore output appears to be empty. Check results for BUSCO: 147736at7742
## Thu Jul 14 01:23:48 CEST 2022 - The aliscore output file does not exist. Check results for BUSCO: 142902at7742
## Thu Jul 14 01:23:49 CEST 2022 - The aliscore output appears to be empty. Check results for BUSCO: 142902at7742
## Thu Jul 14 01:23:49 CEST 2022 - The aliscore output file does not exist. Check results for BUSCO: 184873at7742
## Thu Jul 14 01:23:49 CEST 2022 - The aliscore output appears to be empty. Check results for BUSCO: 184873at7742
## Thu Jul 14 01:23:52 CEST 2022 - The aliscore output file does not exist. Check results for BUSCO: 119796at7742
## Thu Jul 14 01:23:52 CEST 2022 - The aliscore output appears to be empty. Check results for BUSCO: 119796at7742
## Thu Jul 14 01:23:57 CEST 2022 - The aliscore output file does not exist. Check results for BUSCO: 97588at7742
## Thu Jul 14 01:23:57 CEST 2022 - The aliscore output appears to be empty. Check results for BUSCO: 97588at7742
## Thu Jul 14 01:23:57 CEST 2022 - The aliscore output file does not exist. Check results for BUSCO: 150306at7742
## Thu Jul 14 01:23:57 CEST 2022 - The aliscore output appears to be empty. Check results for BUSCO: 150306at7742
## Thu Jul 14 01:23:57 CEST 2022 - The aliscore output file does not exist. Check results for BUSCO: 318895at7742
## Thu Jul 14 01:23:57 CEST 2022 - The aliscore output appears to be empty. Check results for BUSCO: 318895at7742
## Thu Jul 14 01:23:58 CEST 2022 - The aliscore output file does not exist. Check results for BUSCO: 325307at7742
## Thu Jul 14 01:23:58 CEST 2022 - The aliscore output appears to be empty. Check results for BUSCO: 325307at7742
## Thu Jul 14 01:23:58 CEST 2022 - The aliscore output file does not exist. Check results for BUSCO: 320126at7742
## Thu Jul 14 01:23:58 CEST 2022 - The aliscore output appears to be empty. Check results for BUSCO: 320126at7742
## Thu Jul 14 01:23:58 CEST 2022 - The aliscore output file does not exist. Check results for BUSCO: 195437at7742
## Thu Jul 14 01:23:58 CEST 2022 - The aliscore output appears to be empty. Check results for BUSCO: 195437at7742
## Thu Jul 14 01:23:59 CEST 2022 - The aliscore output file does not exist. Check results for BUSCO: 119532at7742
## Thu Jul 14 01:23:59 CEST 2022 - The aliscore output appears to be empty. Check results for BUSCO: 119532at7742
## Thu Jul 14 01:24:02 CEST 2022 - The aliscore output file does not exist. Check results for BUSCO: 386844at7742
## Thu Jul 14 01:24:02 CEST 2022 - The aliscore output appears to be empty. Check results for BUSCO: 386844at7742
## Thu Jul 14 01:24:03 CEST 2022 - The aliscore output file does not exist. Check results for BUSCO: 324312at7742
## Thu Jul 14 01:24:03 CEST 2022 - The aliscore output appears to be empty. Check results for BUSCO: 324312at7742
## Thu Jul 14 01:24:05 CEST 2022 - The aliscore output file does not exist. Check results for BUSCO: 112051at7742
## Thu Jul 14 01:24:05 CEST 2022 - The aliscore output appears to be empty. Check results for BUSCO: 112051at7742
## Thu Jul 14 01:24:06 CEST 2022 - The aliscore output file does not exist. Check results for BUSCO: 109231at7742
## Thu Jul 14 01:24:06 CEST 2022 - The aliscore output appears to be empty. Check results for BUSCO: 109231at7742
## Thu Jul 14 01:24:06 CEST 2022 - The aliscore output file does not exist. Check results for BUSCO: 285589at7742
## Thu Jul 14 01:24:06 CEST 2022 - The aliscore output appears to be empty. Check results for BUSCO: 285589at7742
## Thu Jul 14 01:24:09 CEST 2022 - The aliscore output file does not exist. Check results for BUSCO: 180416at7742
## Thu Jul 14 01:24:09 CEST 2022 - The aliscore output appears to be empty. Check results for BUSCO: 180416at7742
## Thu Jul 14 01:24:10 CEST 2022 - The aliscore output file does not exist. Check results for BUSCO: 259790at7742
## Thu Jul 14 01:24:10 CEST 2022 - The aliscore output appears to be empty. Check results for BUSCO: 259790at7742
## Thu Jul 14 01:24:11 CEST 2022 - The aliscore output file does not exist. Check results for BUSCO: 386844at7742
## Thu Jul 14 01:24:11 CEST 2022 - The aliscore output appears to be empty. Check results for BUSCO: 386844at7742
## Thu Jul 14 01:24:12 CEST 2022 - The aliscore output file does not exist. Check results for BUSCO: 362121at7742
## Thu Jul 14 01:24:12 CEST 2022 - The aliscore output appears to be empty. Check results for BUSCO: 362121at7742
## Thu Jul 14 01:24:14 CEST 2022 - The aliscore output file does not exist. Check results for BUSCO: 125500at7742
## Thu Jul 14 01:24:14 CEST 2022 - The aliscore output file does not exist. Check results for BUSCO: 174048at7742
## Thu Jul 14 01:24:14 CEST 2022 - The aliscore output appears to be empty. Check results for BUSCO: 125500at7742
## Thu Jul 14 01:24:14 CEST 2022 - The aliscore output appears to be empty. Check results for BUSCO: 174048at7742
## Thu Jul 14 01:24:15 CEST 2022 - The aliscore output file does not exist. Check results for BUSCO: 114214at7742
## Thu Jul 14 01:24:15 CEST 2022 - The aliscore output appears to be empty. Check results for BUSCO: 114214at7742
## Thu Jul 14 01:24:15 CEST 2022 - The aliscore output file does not exist. Check results for BUSCO: 116482at7742
## Thu Jul 14 01:24:15 CEST 2022 - The aliscore output appears to be empty. Check results for BUSCO: 116482at7742
## Thu Jul 14 01:24:18 CEST 2022 - The aliscore output file does not exist. Check results for BUSCO: 93522at7742
## Thu Jul 14 01:24:18 CEST 2022 - The aliscore output appears to be empty. Check results for BUSCO: 93522at7742
## Thu Jul 14 01:24:19 CEST 2022 - The aliscore output file does not exist. Check results for BUSCO: 93778at7742
## Thu Jul 14 01:24:19 CEST 2022 - The aliscore output appears to be empty. Check results for BUSCO: 93778at7742
## Thu Jul 14 01:24:20 CEST 2022 - The aliscore output file does not exist. Check results for BUSCO: 259790at7742
## Thu Jul 14 01:24:20 CEST 2022 - The aliscore output appears to be empty. Check results for BUSCO: 259790at7742
## Thu Jul 14 01:24:21 CEST 2022 - The aliscore output file does not exist. Check results for BUSCO: 168022at7742
## Thu Jul 14 01:24:21 CEST 2022 - The aliscore output appears to be empty. Check results for BUSCO: 168022at7742
## Thu Jul 14 01:24:22 CEST 2022 - The aliscore output file does not exist. Check results for BUSCO: 327391at7742
## Thu Jul 14 01:24:22 CEST 2022 - The aliscore output appears to be empty. Check results for BUSCO: 327391at7742
## Thu Jul 14 01:24:24 CEST 2022 - The aliscore output file does not exist. Check results for BUSCO: 142902at7742
## Thu Jul 14 01:24:24 CEST 2022 - The aliscore output appears to be empty. Check results for BUSCO: 142902at7742
## Thu Jul 14 01:24:24 CEST 2022 - The aliscore output file does not exist. Check results for BUSCO: 263171at7742
## Thu Jul 14 01:24:24 CEST 2022 - The aliscore output appears to be empty. Check results for BUSCO: 263171at7742
## Thu Jul 14 01:24:27 CEST 2022 - The aliscore output file does not exist. Check results for BUSCO: 153242at7742
## Thu Jul 14 01:24:27 CEST 2022 - The aliscore output appears to be empty. Check results for BUSCO: 153242at7742
## Thu Jul 14 01:24:28 CEST 2022 - The aliscore output file does not exist. Check results for BUSCO: 283469at7742
## Thu Jul 14 01:24:28 CEST 2022 - The aliscore output appears to be empty. Check results for BUSCO: 283469at7742
## Thu Jul 14 01:24:28 CEST 2022 - The aliscore output file does not exist. Check results for BUSCO: 215427at7742
## Thu Jul 14 01:24:28 CEST 2022 - The aliscore output appears to be empty. Check results for BUSCO: 215427at7742
## Thu Jul 14 01:24:31 CEST 2022 - The aliscore output file does not exist. Check results for BUSCO: 334616at7742
## Thu Jul 14 01:24:31 CEST 2022 - The aliscore output appears to be empty. Check results for BUSCO: 334616at7742
## Thu Jul 14 01:24:32 CEST 2022 - The aliscore output file does not exist. Check results for BUSCO: 241953at7742
## Thu Jul 14 01:24:32 CEST 2022 - The aliscore output appears to be empty. Check results for BUSCO: 241953at7742
## Thu Jul 14 01:24:32 CEST 2022 - The aliscore output file does not exist. Check results for BUSCO: 324312at7742
## Thu Jul 14 01:24:32 CEST 2022 - The aliscore output appears to be empty. Check results for BUSCO: 324312at7742
## Thu Jul 14 01:24:34 CEST 2022 - The aliscore output file does not exist. Check results for BUSCO: 332125at7742
## Thu Jul 14 01:24:34 CEST 2022 - The aliscore output file does not exist. Check results for BUSCO: 161315at7742
## Thu Jul 14 01:24:34 CEST 2022 - The aliscore output appears to be empty. Check results for BUSCO: 332125at7742
## Thu Jul 14 01:24:34 CEST 2022 - The aliscore output appears to be empty. Check results for BUSCO: 161315at7742
## Thu Jul 14 01:24:36 CEST 2022 - The aliscore output file does not exist. Check results for BUSCO: 119796at7742
## Thu Jul 14 01:24:36 CEST 2022 - The aliscore output appears to be empty. Check results for BUSCO: 119796at7742
## Thu Jul 14 01:24:36 CEST 2022 - The aliscore output file does not exist. Check results for BUSCO: 256473at7742
## Thu Jul 14 01:24:36 CEST 2022 - The aliscore output appears to be empty. Check results for BUSCO: 256473at7742
## Thu Jul 14 01:24:39 CEST 2022 - The aliscore output file does not exist. Check results for BUSCO: 150456at7742
## Thu Jul 14 01:24:39 CEST 2022 - The aliscore output appears to be empty. Check results for BUSCO: 150456at7742
## Thu Jul 14 01:24:40 CEST 2022 - The aliscore output file does not exist. Check results for BUSCO: 272794at7742
## Thu Jul 14 01:24:40 CEST 2022 - The aliscore output appears to be empty. Check results for BUSCO: 272794at7742
## Thu Jul 14 01:24:42 CEST 2022 - The aliscore output file does not exist. Check results for BUSCO: 137318at7742
## Thu Jul 14 01:24:42 CEST 2022 - The aliscore output appears to be empty. Check results for BUSCO: 137318at7742
## Thu Jul 14 01:24:43 CEST 2022 - The aliscore output file does not exist. Check results for BUSCO: 161167at7742
## Thu Jul 14 01:24:43 CEST 2022 - The aliscore output appears to be empty. Check results for BUSCO: 161167at7742
## Thu Jul 14 01:24:44 CEST 2022 - The aliscore output file does not exist. Check results for BUSCO: 168022at7742
## Thu Jul 14 01:24:44 CEST 2022 - The aliscore output appears to be empty. Check results for BUSCO: 168022at7742
## Thu Jul 14 01:24:44 CEST 2022 - The aliscore output file does not exist. Check results for BUSCO: 164678at7742
## Thu Jul 14 01:24:44 CEST 2022 - The aliscore output appears to be empty. Check results for BUSCO: 164678at7742
## Thu Jul 14 01:24:46 CEST 2022 - The aliscore output file does not exist. Check results for BUSCO: 103124at7742
## Thu Jul 14 01:24:46 CEST 2022 - The aliscore output appears to be empty. Check results for BUSCO: 103124at7742
## Thu Jul 14 01:24:46 CEST 2022 - The aliscore output file does not exist. Check results for BUSCO: 208122at7742
## Thu Jul 14 01:24:46 CEST 2022 - The aliscore output appears to be empty. Check results for BUSCO: 208122at7742
## Thu Jul 14 01:24:47 CEST 2022 - The aliscore output file does not exist. Check results for BUSCO: 362170at7742
## Thu Jul 14 01:24:47 CEST 2022 - The aliscore output appears to be empty. Check results for BUSCO: 362170at7742
## Thu Jul 14 01:24:48 CEST 2022 - The aliscore output file does not exist. Check results for BUSCO: 293554at7742
## Thu Jul 14 01:24:48 CEST 2022 - The aliscore output appears to be empty. Check results for BUSCO: 293554at7742
## Thu Jul 14 01:24:49 CEST 2022 - The aliscore output file does not exist. Check results for BUSCO: 188851at7742
## Thu Jul 14 01:24:49 CEST 2022 - The aliscore output appears to be empty. Check results for BUSCO: 188851at7742
## Thu Jul 14 01:24:51 CEST 2022 - The aliscore output file does not exist. Check results for BUSCO: 215159at7742
## Thu Jul 14 01:24:51 CEST 2022 - The aliscore output appears to be empty. Check results for BUSCO: 215159at7742
## Thu Jul 14 01:24:52 CEST 2022 - The aliscore output file does not exist. Check results for BUSCO: 259790at7742
## Thu Jul 14 01:24:52 CEST 2022 - The aliscore output appears to be empty. Check results for BUSCO: 259790at7742
## Thu Jul 14 01:24:52 CEST 2022 - The aliscore output file does not exist. Check results for BUSCO: 137318at7742
## Thu Jul 14 01:24:52 CEST 2022 - The aliscore output appears to be empty. Check results for BUSCO: 137318at7742
## Thu Jul 14 01:24:54 CEST 2022 - The aliscore output file does not exist. Check results for BUSCO: 96292at7742
## Thu Jul 14 01:24:54 CEST 2022 - The aliscore output appears to be empty. Check results for BUSCO: 96292at7742
## Thu Jul 14 01:24:56 CEST 2022 - The aliscore output file does not exist. Check results for BUSCO: 168022at7742
## Thu Jul 14 01:24:56 CEST 2022 - The aliscore output appears to be empty. Check results for BUSCO: 168022at7742
## Thu Jul 14 01:24:56 CEST 2022 - The aliscore output file does not exist. Check results for BUSCO: 72731at7742
## Thu Jul 14 01:24:56 CEST 2022 - The aliscore output appears to be empty. Check results for BUSCO: 72731at7742
## Thu Jul 14 01:24:57 CEST 2022 - The aliscore output file does not exist. Check results for BUSCO: 95886at7742
## Thu Jul 14 01:24:57 CEST 2022 - The aliscore output appears to be empty. Check results for BUSCO: 95886at7742
## Thu Jul 14 01:25:01 CEST 2022 - The aliscore output file does not exist. Check results for BUSCO: 112051at7742
## Thu Jul 14 01:25:01 CEST 2022 - The aliscore output appears to be empty. Check results for BUSCO: 112051at7742
## Thu Jul 14 01:25:02 CEST 2022 - The aliscore output file does not exist. Check results for BUSCO: 300687at7742
## Thu Jul 14 01:25:02 CEST 2022 - The aliscore output appears to be empty. Check results for BUSCO: 300687at7742
## Thu Jul 14 01:25:02 CEST 2022 - The aliscore output file does not exist. Check results for BUSCO: 374963at7742
## Thu Jul 14 01:25:03 CEST 2022 - The aliscore output appears to be empty. Check results for BUSCO: 374963at7742
## Thu Jul 14 01:25:03 CEST 2022 - The aliscore output file does not exist. Check results for BUSCO: 311134at7742
## Thu Jul 14 01:25:03 CEST 2022 - The aliscore output appears to be empty. Check results for BUSCO: 311134at7742
## Thu Jul 14 01:25:04 CEST 2022 - The aliscore output file does not exist. Check results for BUSCO: 184873at7742
## Thu Jul 14 01:25:04 CEST 2022 - The aliscore output appears to be empty. Check results for BUSCO: 184873at7742
## Thu Jul 14 01:25:04 CEST 2022 - The aliscore output file does not exist. Check results for BUSCO: 291702at7742
## Thu Jul 14 01:25:04 CEST 2022 - The aliscore output appears to be empty. Check results for BUSCO: 291702at7742
## Thu Jul 14 01:25:05 CEST 2022 - The aliscore output file does not exist. Check results for BUSCO: 98753at7742
## Thu Jul 14 01:25:05 CEST 2022 - The aliscore output appears to be empty. Check results for BUSCO: 98753at7742
## Thu Jul 14 01:25:07 CEST 2022 - The aliscore output file does not exist. Check results for BUSCO: 125500at7742
## Thu Jul 14 01:25:07 CEST 2022 - The aliscore output appears to be empty. Check results for BUSCO: 125500at7742
## Thu Jul 14 01:25:08 CEST 2022 - The aliscore output file does not exist. Check results for BUSCO: 155435at7742
## Thu Jul 14 01:25:08 CEST 2022 - The aliscore output appears to be empty. Check results for BUSCO: 155435at7742
## Thu Jul 14 01:25:09 CEST 2022 - The aliscore output file does not exist. Check results for BUSCO: 130415at7742
## Thu Jul 14 01:25:09 CEST 2022 - The aliscore output appears to be empty. Check results for BUSCO: 130415at7742
## Thu Jul 14 01:25:10 CEST 2022 - The aliscore output file does not exist. Check results for BUSCO: 311134at7742
## Thu Jul 14 01:25:10 CEST 2022 - The aliscore output appears to be empty. Check results for BUSCO: 311134at7742
## Thu Jul 14 01:25:13 CEST 2022 - The aliscore output file does not exist. Check results for BUSCO: 140962at7742
## Thu Jul 14 01:25:13 CEST 2022 - The aliscore output appears to be empty. Check results for BUSCO: 140962at7742
## Thu Jul 14 01:25:13 CEST 2022 - The aliscore output file does not exist. Check results for BUSCO: 68376at7742
## Thu Jul 14 01:25:13 CEST 2022 - The aliscore output appears to be empty. Check results for BUSCO: 68376at7742
## Thu Jul 14 01:25:13 CEST 2022 - The aliscore output file does not exist. Check results for BUSCO: 118406at7742
## Thu Jul 14 01:25:13 CEST 2022 - The aliscore output appears to be empty. Check results for BUSCO: 118406at7742
## Thu Jul 14 01:25:15 CEST 2022 - The aliscore output file does not exist. Check results for BUSCO: 249629at7742
## Thu Jul 14 01:25:15 CEST 2022 - The aliscore output appears to be empty. Check results for BUSCO: 249629at7742
## Thu Jul 14 01:25:16 CEST 2022 - The aliscore output file does not exist. Check results for BUSCO: 201876at7742
## Thu Jul 14 01:25:16 CEST 2022 - The aliscore output appears to be empty. Check results for BUSCO: 201876at7742
## Thu Jul 14 01:25:19 CEST 2022 - The aliscore output file does not exist. Check results for BUSCO: 108881at7742
## Thu Jul 14 01:25:19 CEST 2022 - The aliscore output appears to be empty. Check results for BUSCO: 108881at7742
## Thu Jul 14 01:25:21 CEST 2022 - The aliscore output file does not exist. Check results for BUSCO: 310011at7742
## Thu Jul 14 01:25:21 CEST 2022 - The aliscore output appears to be empty. Check results for BUSCO: 310011at7742
## Thu Jul 14 01:25:22 CEST 2022 - The aliscore output file does not exist. Check results for BUSCO: 108881at7742
## Thu Jul 14 01:25:22 CEST 2022 - The aliscore output appears to be empty. Check results for BUSCO: 108881at7742
## Thu Jul 14 01:25:22 CEST 2022 - The aliscore output file does not exist. Check results for BUSCO: 192462at7742
## Thu Jul 14 01:25:22 CEST 2022 - The aliscore output appears to be empty. Check results for BUSCO: 192462at7742
## Thu Jul 14 01:25:23 CEST 2022 - The aliscore output file does not exist. Check results for BUSCO: 68376at7742
## Thu Jul 14 01:25:23 CEST 2022 - The aliscore output appears to be empty. Check results for BUSCO: 68376at7742
## Thu Jul 14 01:25:23 CEST 2022 - The aliscore output file does not exist. Check results for BUSCO: 192462at7742
## Thu Jul 14 01:25:23 CEST 2022 - The aliscore output appears to be empty. Check results for BUSCO: 192462at7742
## Thu Jul 14 01:25:25 CEST 2022 - The aliscore output file does not exist. Check results for BUSCO: 246371at7742
## Thu Jul 14 01:25:25 CEST 2022 - The aliscore output appears to be empty. Check results for BUSCO: 246371at7742
## Thu Jul 14 01:25:26 CEST 2022 - The aliscore output file does not exist. Check results for BUSCO: 237959at7742
## Thu Jul 14 01:25:26 CEST 2022 - The aliscore output appears to be empty. Check results for BUSCO: 237959at7742
## Thu Jul 14 01:25:28 CEST 2022 - The aliscore output file does not exist. Check results for BUSCO: 98753at7742
## Thu Jul 14 01:25:28 CEST 2022 - The aliscore output appears to be empty. Check results for BUSCO: 98753at7742
## Thu Jul 14 01:25:28 CEST 2022 - The aliscore output file does not exist. Check results for BUSCO: 182480at7742
## Thu Jul 14 01:25:28 CEST 2022 - The aliscore output appears to be empty. Check results for BUSCO: 182480at7742
## Thu Jul 14 01:25:29 CEST 2022 - The aliscore output file does not exist. Check results for BUSCO: 135511at7742
## Thu Jul 14 01:25:30 CEST 2022 - The aliscore output appears to be empty. Check results for BUSCO: 135511at7742
## Thu Jul 14 01:25:30 CEST 2022 - The aliscore output file does not exist. Check results for BUSCO: 139567at7742
## Thu Jul 14 01:25:30 CEST 2022 - The aliscore output appears to be empty. Check results for BUSCO: 139567at7742
## Thu Jul 14 01:25:30 CEST 2022 - The aliscore output file does not exist. Check results for BUSCO: 182481at7742
## Thu Jul 14 01:25:30 CEST 2022 - The aliscore output appears to be empty. Check results for BUSCO: 182481at7742
## Thu Jul 14 01:25:32 CEST 2022 - The aliscore output file does not exist. Check results for BUSCO: 257616at7742
## Thu Jul 14 01:25:32 CEST 2022 - The aliscore output appears to be empty. Check results for BUSCO: 257616at7742
## Thu Jul 14 01:25:34 CEST 2022 - The aliscore output file does not exist. Check results for BUSCO: 208291at7742
## Thu Jul 14 01:25:34 CEST 2022 - The aliscore output appears to be empty. Check results for BUSCO: 208291at7742
## Thu Jul 14 01:25:36 CEST 2022 - The aliscore output file does not exist. Check results for BUSCO: 72731at7742
## Thu Jul 14 01:25:36 CEST 2022 - The aliscore output appears to be empty. Check results for BUSCO: 72731at7742
## Thu Jul 14 01:25:36 CEST 2022 - The aliscore output file does not exist. Check results for BUSCO: 322442at7742
## Thu Jul 14 01:25:36 CEST 2022 - The aliscore output appears to be empty. Check results for BUSCO: 322442at7742
## Thu Jul 14 01:25:37 CEST 2022 - The aliscore output file does not exist. Check results for BUSCO: 256473at7742
## Thu Jul 14 01:25:37 CEST 2022 - The aliscore output appears to be empty. Check results for BUSCO: 256473at7742
## Thu Jul 14 01:25:40 CEST 2022 - The aliscore output file does not exist. Check results for BUSCO: 101941at7742
## Thu Jul 14 01:25:40 CEST 2022 - The aliscore output appears to be empty. Check results for BUSCO: 101941at7742
## Thu Jul 14 01:25:40 CEST 2022 - The aliscore output file does not exist. Check results for BUSCO: 122793at7742
## Thu Jul 14 01:25:40 CEST 2022 - The aliscore output appears to be empty. Check results for BUSCO: 122793at7742
## Thu Jul 14 01:25:42 CEST 2022 - The aliscore output file does not exist. Check results for BUSCO: 294332at7742
## Thu Jul 14 01:25:42 CEST 2022 - The aliscore output appears to be empty. Check results for BUSCO: 294332at7742
## Thu Jul 14 01:25:43 CEST 2022 - The aliscore output file does not exist. Check results for BUSCO: 276337at7742
## Thu Jul 14 01:25:43 CEST 2022 - The aliscore output appears to be empty. Check results for BUSCO: 276337at7742
## Thu Jul 14 01:25:45 CEST 2022 - The aliscore output file does not exist. Check results for BUSCO: 392098at7742
## Thu Jul 14 01:25:45 CEST 2022 - The aliscore output appears to be empty. Check results for BUSCO: 392098at7742
## Thu Jul 14 01:25:45 CEST 2022 - The aliscore output file does not exist. Check results for BUSCO: 137318at7742
## Thu Jul 14 01:25:45 CEST 2022 - The aliscore output appears to be empty. Check results for BUSCO: 137318at7742
## Thu Jul 14 01:25:46 CEST 2022 - The aliscore output file does not exist. Check results for BUSCO: 223283at7742
## Thu Jul 14 01:25:46 CEST 2022 - The aliscore output appears to be empty. Check results for BUSCO: 223283at7742
## Thu Jul 14 01:25:47 CEST 2022 - The aliscore output file does not exist. Check results for BUSCO: 256348at7742
## Thu Jul 14 01:25:47 CEST 2022 - The aliscore output appears to be empty. Check results for BUSCO: 256348at7742
## Thu Jul 14 01:25:48 CEST 2022 - The aliscore output file does not exist. Check results for BUSCO: 307789at7742
## Thu Jul 14 01:25:48 CEST 2022 - The aliscore output appears to be empty. Check results for BUSCO: 307789at7742
## Thu Jul 14 01:25:48 CEST 2022 - The aliscore output file does not exist. Check results for BUSCO: 113992at7742
## Thu Jul 14 01:25:48 CEST 2022 - The aliscore output appears to be empty. Check results for BUSCO: 113992at7742
## Thu Jul 14 01:25:52 CEST 2022 - The aliscore output file does not exist. Check results for BUSCO: 179036at7742
## Thu Jul 14 01:25:52 CEST 2022 - The aliscore output appears to be empty. Check results for BUSCO: 179036at7742
## Thu Jul 14 01:25:54 CEST 2022 - The aliscore output file does not exist. Check results for BUSCO: 103124at7742
## Thu Jul 14 01:25:54 CEST 2022 - The aliscore output appears to be empty. Check results for BUSCO: 103124at7742
## Thu Jul 14 01:25:54 CEST 2022 - The aliscore output file does not exist. Check results for BUSCO: 312837at7742
## Thu Jul 14 01:25:54 CEST 2022 - The aliscore output appears to be empty. Check results for BUSCO: 312837at7742
## Thu Jul 14 01:25:54 CEST 2022 - The aliscore output file does not exist. Check results for BUSCO: 266521at7742
## Thu Jul 14 01:25:54 CEST 2022 - The aliscore output appears to be empty. Check results for BUSCO: 266521at7742
## Thu Jul 14 01:25:55 CEST 2022 - The aliscore output file does not exist. Check results for BUSCO: 293554at7742
## Thu Jul 14 01:25:55 CEST 2022 - The aliscore output appears to be empty. Check results for BUSCO: 293554at7742
## Thu Jul 14 01:25:55 CEST 2022 - The aliscore output file does not exist. Check results for BUSCO: 135786at7742
## Thu Jul 14 01:25:55 CEST 2022 - The aliscore output appears to be empty. Check results for BUSCO: 135786at7742
## Thu Jul 14 01:25:57 CEST 2022 - The aliscore output file does not exist. Check results for BUSCO: 256510at7742
## Thu Jul 14 01:25:57 CEST 2022 - The aliscore output appears to be empty. Check results for BUSCO: 256510at7742
## Thu Jul 14 01:25:58 CEST 2022 - The aliscore output file does not exist. Check results for BUSCO: 225242at7742
## Thu Jul 14 01:25:58 CEST 2022 - The aliscore output appears to be empty. Check results for BUSCO: 225242at7742
## Thu Jul 14 01:26:01 CEST 2022 - The aliscore output file does not exist. Check results for BUSCO: 135786at7742
## Thu Jul 14 01:26:01 CEST 2022 - The aliscore output file does not exist. Check results for BUSCO: 139250at7742
## Thu Jul 14 01:26:01 CEST 2022 - The aliscore output appears to be empty. Check results for BUSCO: 135786at7742
## Thu Jul 14 01:26:01 CEST 2022 - The aliscore output appears to be empty. Check results for BUSCO: 139250at7742
## Thu Jul 14 01:26:03 CEST 2022 - The aliscore output file does not exist. Check results for BUSCO: 291702at7742
## Thu Jul 14 01:26:03 CEST 2022 - The aliscore output appears to be empty. Check results for BUSCO: 291702at7742
## Thu Jul 14 01:26:03 CEST 2022 - The aliscore output file does not exist. Check results for BUSCO: 175755at7742
## Thu Jul 14 01:26:03 CEST 2022 - The aliscore output appears to be empty. Check results for BUSCO: 175755at7742
## Thu Jul 14 01:26:04 CEST 2022 - The aliscore output file does not exist. Check results for BUSCO: 136457at7742
## Thu Jul 14 01:26:04 CEST 2022 - The aliscore output appears to be empty. Check results for BUSCO: 136457at7742
## Thu Jul 14 01:26:06 CEST 2022 - The aliscore output file does not exist. Check results for BUSCO: 239747at7742
## Thu Jul 14 01:26:06 CEST 2022 - The aliscore output appears to be empty. Check results for BUSCO: 239747at7742
## Thu Jul 14 01:26:07 CEST 2022 - The aliscore output file does not exist. Check results for BUSCO: 122793at7742
## Thu Jul 14 01:26:08 CEST 2022 - The aliscore output appears to be empty. Check results for BUSCO: 122793at7742
## Thu Jul 14 01:26:08 CEST 2022 - The aliscore output file does not exist. Check results for BUSCO: 136457at7742
## Thu Jul 14 01:26:08 CEST 2022 - The aliscore output appears to be empty. Check results for BUSCO: 136457at7742
## Thu Jul 14 01:26:08 CEST 2022 - The aliscore output file does not exist. Check results for BUSCO: 96292at7742
## Thu Jul 14 01:26:08 CEST 2022 - The aliscore output appears to be empty. Check results for BUSCO: 96292at7742
## Thu Jul 14 01:26:10 CEST 2022 - The aliscore output file does not exist. Check results for BUSCO: 343527at7742
## Thu Jul 14 01:26:10 CEST 2022 - The aliscore output appears to be empty. Check results for BUSCO: 343527at7742
## Thu Jul 14 01:26:11 CEST 2022 - The aliscore output file does not exist. Check results for BUSCO: 108888at7742
## Thu Jul 14 01:26:11 CEST 2022 - The aliscore output appears to be empty. Check results for BUSCO: 108888at7742
## Thu Jul 14 01:26:12 CEST 2022 - The aliscore output file does not exist. Check results for BUSCO: 142074at7742
## Thu Jul 14 01:26:12 CEST 2022 - The aliscore output appears to be empty. Check results for BUSCO: 142074at7742
## Thu Jul 14 01:26:14 CEST 2022 - The aliscore output file does not exist. Check results for BUSCO: 107743at7742
## Thu Jul 14 01:26:14 CEST 2022 - The aliscore output appears to be empty. Check results for BUSCO: 107743at7742
## Thu Jul 14 01:26:14 CEST 2022 - The aliscore output file does not exist. Check results for BUSCO: 170012at7742
## Thu Jul 14 01:26:14 CEST 2022 - The aliscore output appears to be empty. Check results for BUSCO: 170012at7742
## Thu Jul 14 01:26:15 CEST 2022 - The aliscore output file does not exist. Check results for BUSCO: 130130at7742
## Thu Jul 14 01:26:15 CEST 2022 - The aliscore output appears to be empty. Check results for BUSCO: 130130at7742
## Thu Jul 14 01:26:16 CEST 2022 - The aliscore output file does not exist. Check results for BUSCO: 254818at7742
## Thu Jul 14 01:26:16 CEST 2022 - The aliscore output appears to be empty. Check results for BUSCO: 254818at7742
## Thu Jul 14 01:26:18 CEST 2022 - The aliscore output file does not exist. Check results for BUSCO: 136457at7742
## Thu Jul 14 01:26:18 CEST 2022 - The aliscore output appears to be empty. Check results for BUSCO: 136457at7742
## Thu Jul 14 01:26:19 CEST 2022 - The aliscore output file does not exist. Check results for BUSCO: 334616at7742
## Thu Jul 14 01:26:19 CEST 2022 - The aliscore output appears to be empty. Check results for BUSCO: 334616at7742
## Thu Jul 14 01:26:20 CEST 2022 - The aliscore output file does not exist. Check results for BUSCO: 162174at7742
## Thu Jul 14 01:26:20 CEST 2022 - The aliscore output appears to be empty. Check results for BUSCO: 162174at7742
## Thu Jul 14 01:26:21 CEST 2022 - The aliscore output file does not exist. Check results for BUSCO: 119532at7742
## Thu Jul 14 01:26:21 CEST 2022 - The aliscore output appears to be empty. Check results for BUSCO: 119532at7742
## Thu Jul 14 01:26:23 CEST 2022 - The aliscore output file does not exist. Check results for BUSCO: 310011at7742
## Thu Jul 14 01:26:23 CEST 2022 - The aliscore output appears to be empty. Check results for BUSCO: 310011at7742
## Thu Jul 14 01:26:24 CEST 2022 - The aliscore output file does not exist. Check results for BUSCO: 93522at7742
## Thu Jul 14 01:26:24 CEST 2022 - The aliscore output appears to be empty. Check results for BUSCO: 93522at7742
## Thu Jul 14 01:26:24 CEST 2022 - The aliscore output file does not exist. Check results for BUSCO: 408649at7742
## Thu Jul 14 01:26:24 CEST 2022 - The aliscore output appears to be empty. Check results for BUSCO: 408649at7742
## Thu Jul 14 01:26:26 CEST 2022 - The aliscore output file does not exist. Check results for BUSCO: 103124at7742
## Thu Jul 14 01:26:26 CEST 2022 - The aliscore output appears to be empty. Check results for BUSCO: 103124at7742
## Thu Jul 14 01:26:27 CEST 2022 - The aliscore output file does not exist. Check results for BUSCO: 108881at7742
## Thu Jul 14 01:26:27 CEST 2022 - The aliscore output appears to be empty. Check results for BUSCO: 108881at7742
## Thu Jul 14 01:26:28 CEST 2022 - The aliscore output file does not exist. Check results for BUSCO: 273954at7742
## Thu Jul 14 01:26:28 CEST 2022 - The aliscore output appears to be empty. Check results for BUSCO: 273954at7742
## Thu Jul 14 01:26:29 CEST 2022 - The aliscore output file does not exist. Check results for BUSCO: 295275at7742
## Thu Jul 14 01:26:29 CEST 2022 - The aliscore output appears to be empty. Check results for BUSCO: 295275at7742
## Thu Jul 14 01:26:30 CEST 2022 - The aliscore output file does not exist. Check results for BUSCO: 222821at7742
## Thu Jul 14 01:26:31 CEST 2022 - The aliscore output appears to be empty. Check results for BUSCO: 222821at7742
## Thu Jul 14 01:26:33 CEST 2022 - The aliscore output file does not exist. Check results for BUSCO: 305417at7742
## Thu Jul 14 01:26:33 CEST 2022 - The aliscore output appears to be empty. Check results for BUSCO: 305417at7742
## Thu Jul 14 01:26:33 CEST 2022 - The aliscore output file does not exist. Check results for BUSCO: 380097at7742
## Thu Jul 14 01:26:33 CEST 2022 - The aliscore output appears to be empty. Check results for BUSCO: 380097at7742
## Thu Jul 14 01:26:34 CEST 2022 - The aliscore output file does not exist. Check results for BUSCO: 72731at7742
## Thu Jul 14 01:26:34 CEST 2022 - The aliscore output appears to be empty. Check results for BUSCO: 72731at7742
## Thu Jul 14 01:26:35 CEST 2022 - The aliscore output file does not exist. Check results for BUSCO: 222631at7742
## Thu Jul 14 01:26:35 CEST 2022 - The aliscore output appears to be empty. Check results for BUSCO: 222631at7742
## Thu Jul 14 01:26:37 CEST 2022 - The aliscore output file does not exist. Check results for BUSCO: 281891at7742
## Thu Jul 14 01:26:37 CEST 2022 - The aliscore output appears to be empty. Check results for BUSCO: 281891at7742
## Thu Jul 14 01:26:38 CEST 2022 - The aliscore output file does not exist. Check results for BUSCO: 249629at7742
## Thu Jul 14 01:26:38 CEST 2022 - The aliscore output appears to be empty. Check results for BUSCO: 249629at7742
## Thu Jul 14 01:26:38 CEST 2022 - The aliscore output file does not exist. Check results for BUSCO: 232849at7742
## Thu Jul 14 01:26:38 CEST 2022 - The aliscore output appears to be empty. Check results for BUSCO: 232849at7742
## Thu Jul 14 01:26:40 CEST 2022 - The aliscore output file does not exist. Check results for BUSCO: 318965at7742
## Thu Jul 14 01:26:40 CEST 2022 - The aliscore output appears to be empty. Check results for BUSCO: 318965at7742
## Thu Jul 14 01:26:42 CEST 2022 - The aliscore output file does not exist. Check results for BUSCO: 224537at7742
## Thu Jul 14 01:26:42 CEST 2022 - The aliscore output appears to be empty. Check results for BUSCO: 224537at7742
## Thu Jul 14 01:26:42 CEST 2022 - The aliscore output file does not exist. Check results for BUSCO: 218524at7742
## Thu Jul 14 01:26:42 CEST 2022 - The aliscore output appears to be empty. Check results for BUSCO: 218524at7742
## Thu Jul 14 01:26:44 CEST 2022 - The aliscore output file does not exist. Check results for BUSCO: 275256at7742
## Thu Jul 14 01:26:44 CEST 2022 - The aliscore output appears to be empty. Check results for BUSCO: 275256at7742
## Thu Jul 14 01:26:44 CEST 2022 - The aliscore output file does not exist. Check results for BUSCO: 98637at7742
## Thu Jul 14 01:26:44 CEST 2022 - The aliscore output appears to be empty. Check results for BUSCO: 98637at7742
## Thu Jul 14 01:26:47 CEST 2022 - The aliscore output file does not exist. Check results for BUSCO: 136946at7742
## Thu Jul 14 01:26:47 CEST 2022 - The aliscore output appears to be empty. Check results for BUSCO: 136946at7742
## Thu Jul 14 01:26:48 CEST 2022 - The aliscore output file does not exist. Check results for BUSCO: 269540at7742
## Thu Jul 14 01:26:48 CEST 2022 - The aliscore output appears to be empty. Check results for BUSCO: 269540at7742
## Thu Jul 14 01:26:48 CEST 2022 - The aliscore output file does not exist. Check results for BUSCO: 362121at7742
## Thu Jul 14 01:26:48 CEST 2022 - The aliscore output appears to be empty. Check results for BUSCO: 362121at7742
## Thu Jul 14 01:26:49 CEST 2022 - The aliscore output file does not exist. Check results for BUSCO: 215427at7742
## Thu Jul 14 01:26:49 CEST 2022 - The aliscore output appears to be empty. Check results for BUSCO: 215427at7742
## Thu Jul 14 01:26:50 CEST 2022 - The aliscore output file does not exist. Check results for BUSCO: 157460at7742
## Thu Jul 14 01:26:50 CEST 2022 - The aliscore output appears to be empty. Check results for BUSCO: 157460at7742
## Thu Jul 14 01:26:52 CEST 2022 - The aliscore output file does not exist. Check results for BUSCO: 161520at7742
## Thu Jul 14 01:26:52 CEST 2022 - The aliscore output appears to be empty. Check results for BUSCO: 161520at7742
## Thu Jul 14 01:26:54 CEST 2022 - The aliscore output file does not exist. Check results for BUSCO: 199341at7742
## Thu Jul 14 01:26:54 CEST 2022 - The aliscore output appears to be empty. Check results for BUSCO: 199341at7742
## Thu Jul 14 01:26:55 CEST 2022 - The aliscore output file does not exist. Check results for BUSCO: 117384at7742
## Thu Jul 14 01:26:55 CEST 2022 - The aliscore output appears to be empty. Check results for BUSCO: 117384at7742
## Thu Jul 14 01:26:58 CEST 2022 - The aliscore output file does not exist. Check results for BUSCO: 266521at7742
## Thu Jul 14 01:26:58 CEST 2022 - The aliscore output appears to be empty. Check results for BUSCO: 266521at7742
## Thu Jul 14 01:26:58 CEST 2022 - The aliscore output file does not exist. Check results for BUSCO: 183332at7742
## Thu Jul 14 01:26:58 CEST 2022 - The aliscore output appears to be empty. Check results for BUSCO: 183332at7742
## Thu Jul 14 01:26:59 CEST 2022 - The aliscore output file does not exist. Check results for BUSCO: 325307at7742
## Thu Jul 14 01:26:59 CEST 2022 - The aliscore output appears to be empty. Check results for BUSCO: 325307at7742
## Thu Jul 14 01:26:59 CEST 2022 - The aliscore output file does not exist. Check results for BUSCO: 161167at7742
## Thu Jul 14 01:26:59 CEST 2022 - The aliscore output appears to be empty. Check results for BUSCO: 161167at7742
## Thu Jul 14 01:27:00 CEST 2022 - The aliscore output file does not exist. Check results for BUSCO: 291289at7742
## Thu Jul 14 01:27:00 CEST 2022 - The aliscore output appears to be empty. Check results for BUSCO: 291289at7742
## Thu Jul 14 01:27:01 CEST 2022 - The aliscore output file does not exist. Check results for BUSCO: 178680at7742
## Thu Jul 14 01:27:01 CEST 2022 - The aliscore output appears to be empty. Check results for BUSCO: 178680at7742
## Thu Jul 14 01:27:02 CEST 2022 - The aliscore output file does not exist. Check results for BUSCO: 208291at7742
## Thu Jul 14 01:27:02 CEST 2022 - The aliscore output appears to be empty. Check results for BUSCO: 208291at7742
## Thu Jul 14 01:27:03 CEST 2022 - The aliscore output file does not exist. Check results for BUSCO: 191875at7742
## Thu Jul 14 01:27:03 CEST 2022 - The aliscore output appears to be empty. Check results for BUSCO: 191875at7742
## Thu Jul 14 01:27:04 CEST 2022 - The aliscore output file does not exist. Check results for BUSCO: 139567at7742
## Thu Jul 14 01:27:04 CEST 2022 - The aliscore output appears to be empty. Check results for BUSCO: 139567at7742
## Thu Jul 14 01:27:05 CEST 2022 - The aliscore output file does not exist. Check results for BUSCO: 280833at7742
## Thu Jul 14 01:27:06 CEST 2022 - The aliscore output appears to be empty. Check results for BUSCO: 280833at7742
## Thu Jul 14 01:27:06 CEST 2022 - The aliscore output file does not exist. Check results for BUSCO: 192331at7742
## Thu Jul 14 01:27:06 CEST 2022 - The aliscore output appears to be empty. Check results for BUSCO: 192331at7742
## Thu Jul 14 01:27:08 CEST 2022 - The aliscore output file does not exist. Check results for BUSCO: 131564at7742
## Thu Jul 14 01:27:08 CEST 2022 - The aliscore output appears to be empty. Check results for BUSCO: 131564at7742
## Thu Jul 14 01:27:09 CEST 2022 - The aliscore output file does not exist. Check results for BUSCO: 343527at7742
## Thu Jul 14 01:27:09 CEST 2022 - The aliscore output appears to be empty. Check results for BUSCO: 343527at7742
## Thu Jul 14 01:27:10 CEST 2022 - The aliscore output file does not exist. Check results for BUSCO: 239747at7742
## Thu Jul 14 01:27:10 CEST 2022 - The aliscore output appears to be empty. Check results for BUSCO: 239747at7742
## Thu Jul 14 01:27:12 CEST 2022 - The aliscore output file does not exist. Check results for BUSCO: 221061at7742
## Thu Jul 14 01:27:12 CEST 2022 - The aliscore output appears to be empty. Check results for BUSCO: 221061at7742
## Thu Jul 14 01:27:12 CEST 2022 - The aliscore output file does not exist. Check results for BUSCO: 370224at7742
## Thu Jul 14 01:27:13 CEST 2022 - The aliscore output appears to be empty. Check results for BUSCO: 370224at7742
## Thu Jul 14 01:27:14 CEST 2022 - The aliscore output file does not exist. Check results for BUSCO: 318895at7742
## Thu Jul 14 01:27:14 CEST 2022 - The aliscore output appears to be empty. Check results for BUSCO: 318895at7742
## Thu Jul 14 01:27:14 CEST 2022 - The aliscore output file does not exist. Check results for BUSCO: 326687at7742
## Thu Jul 14 01:27:14 CEST 2022 - The aliscore output appears to be empty. Check results for BUSCO: 326687at7742
## Thu Jul 14 01:27:16 CEST 2022 - The aliscore output file does not exist. Check results for BUSCO: 204010at7742
## Thu Jul 14 01:27:16 CEST 2022 - The aliscore output appears to be empty. Check results for BUSCO: 204010at7742
## Thu Jul 14 01:27:17 CEST 2022 - The aliscore output file does not exist. Check results for BUSCO: 210107at7742
## Thu Jul 14 01:27:17 CEST 2022 - The aliscore output appears to be empty. Check results for BUSCO: 210107at7742
## Thu Jul 14 01:27:19 CEST 2022 - The aliscore output file does not exist. Check results for BUSCO: 166657at7742
## Thu Jul 14 01:27:19 CEST 2022 - The aliscore output appears to be empty. Check results for BUSCO: 166657at7742
## Thu Jul 14 01:27:21 CEST 2022 - The aliscore output file does not exist. Check results for BUSCO: 354251at7742
## Thu Jul 14 01:27:21 CEST 2022 - The aliscore output appears to be empty. Check results for BUSCO: 354251at7742
## Thu Jul 14 01:27:22 CEST 2022 - The aliscore output file does not exist. Check results for BUSCO: 221592at7742
## Thu Jul 14 01:27:22 CEST 2022 - The aliscore output appears to be empty. Check results for BUSCO: 221592at7742
## Thu Jul 14 01:27:22 CEST 2022 - The aliscore output file does not exist. Check results for BUSCO: 178259at7742
## Thu Jul 14 01:27:22 CEST 2022 - The aliscore output appears to be empty. Check results for BUSCO: 178259at7742
## Thu Jul 14 01:27:23 CEST 2022 - The aliscore output file does not exist. Check results for BUSCO: 159349at7742
## Thu Jul 14 01:27:23 CEST 2022 - The aliscore output appears to be empty. Check results for BUSCO: 159349at7742
## Thu Jul 14 01:27:24 CEST 2022 - The aliscore output file does not exist. Check results for BUSCO: 115533at7742
## Thu Jul 14 01:27:24 CEST 2022 - The aliscore output appears to be empty. Check results for BUSCO: 115533at7742
## Thu Jul 14 01:27:26 CEST 2022 - The aliscore output file does not exist. Check results for BUSCO: 362121at7742
## Thu Jul 14 01:27:26 CEST 2022 - The aliscore output appears to be empty. Check results for BUSCO: 362121at7742
## Thu Jul 14 01:27:28 CEST 2022 - The aliscore output file does not exist. Check results for BUSCO: 159463at7742
## Thu Jul 14 01:27:28 CEST 2022 - The aliscore output appears to be empty. Check results for BUSCO: 159463at7742
## Thu Jul 14 01:27:29 CEST 2022 - The aliscore output file does not exist. Check results for BUSCO: 291702at7742
## Thu Jul 14 01:27:29 CEST 2022 - The aliscore output appears to be empty. Check results for BUSCO: 291702at7742
## Thu Jul 14 01:27:31 CEST 2022 - The aliscore output file does not exist. Check results for BUSCO: 203960at7742
## Thu Jul 14 01:27:31 CEST 2022 - The aliscore output appears to be empty. Check results for BUSCO: 203960at7742
## Thu Jul 14 01:27:31 CEST 2022 - The aliscore output file does not exist. Check results for BUSCO: 143888at7742
## Thu Jul 14 01:27:31 CEST 2022 - The aliscore output appears to be empty. Check results for BUSCO: 143888at7742
## Thu Jul 14 01:27:33 CEST 2022 - The aliscore output file does not exist. Check results for BUSCO: 370236at7742
## Thu Jul 14 01:27:33 CEST 2022 - The aliscore output appears to be empty. Check results for BUSCO: 370236at7742
## Thu Jul 14 01:27:33 CEST 2022 - The aliscore output file does not exist. Check results for BUSCO: 299549at7742
## Thu Jul 14 01:27:33 CEST 2022 - The aliscore output appears to be empty. Check results for BUSCO: 299549at7742
## Thu Jul 14 01:27:35 CEST 2022 - The aliscore output file does not exist. Check results for BUSCO: 129835at7742
## Thu Jul 14 01:27:35 CEST 2022 - The aliscore output appears to be empty. Check results for BUSCO: 129835at7742
## Thu Jul 14 01:27:35 CEST 2022 - The aliscore output file does not exist. Check results for BUSCO: 266927at7742
## Thu Jul 14 01:27:35 CEST 2022 - The aliscore output appears to be empty. Check results for BUSCO: 266927at7742
## Thu Jul 14 01:27:36 CEST 2022 - The aliscore output file does not exist. Check results for BUSCO: 241002at7742
## Thu Jul 14 01:27:36 CEST 2022 - The aliscore output appears to be empty. Check results for BUSCO: 241002at7742
## Thu Jul 14 01:27:37 CEST 2022 - The aliscore output file does not exist. Check results for BUSCO: 192313at7742
## Thu Jul 14 01:27:37 CEST 2022 - The aliscore output appears to be empty. Check results for BUSCO: 192313at7742
## Thu Jul 14 01:27:39 CEST 2022 - The aliscore output file does not exist. Check results for BUSCO: 131817at7742
## Thu Jul 14 01:27:39 CEST 2022 - The aliscore output appears to be empty. Check results for BUSCO: 131817at7742
## Thu Jul 14 01:27:39 CEST 2022 - The aliscore output file does not exist. Check results for BUSCO: 188869at7742
## Thu Jul 14 01:27:39 CEST 2022 - The aliscore output appears to be empty. Check results for BUSCO: 188869at7742
## Thu Jul 14 01:27:40 CEST 2022 - The aliscore output file does not exist. Check results for BUSCO: 140962at7742
## Thu Jul 14 01:27:40 CEST 2022 - The aliscore output appears to be empty. Check results for BUSCO: 140962at7742
## Thu Jul 14 01:27:43 CEST 2022 - The aliscore output file does not exist. Check results for BUSCO: 142074at7742
## Thu Jul 14 01:27:43 CEST 2022 - The aliscore output appears to be empty. Check results for BUSCO: 142074at7742
## Thu Jul 14 01:27:43 CEST 2022 - The aliscore output file does not exist. Check results for BUSCO: 96459at7742
## Thu Jul 14 01:27:44 CEST 2022 - The aliscore output appears to be empty. Check results for BUSCO: 96459at7742
## Thu Jul 14 01:27:44 CEST 2022 - The aliscore output file does not exist. Check results for BUSCO: 171737at7742
## Thu Jul 14 01:27:44 CEST 2022 - The aliscore output appears to be empty. Check results for BUSCO: 171737at7742
## Thu Jul 14 01:27:45 CEST 2022 - The aliscore output file does not exist. Check results for BUSCO: 176497at7742
## Thu Jul 14 01:27:45 CEST 2022 - The aliscore output appears to be empty. Check results for BUSCO: 176497at7742
## Thu Jul 14 01:27:47 CEST 2022 - The aliscore output file does not exist. Check results for BUSCO: 305955at7742
## Thu Jul 14 01:27:47 CEST 2022 - The aliscore output appears to be empty. Check results for BUSCO: 305955at7742
## Thu Jul 14 01:27:47 CEST 2022 - The aliscore output file does not exist. Check results for BUSCO: 139567at7742
## Thu Jul 14 01:27:47 CEST 2022 - The aliscore output appears to be empty. Check results for BUSCO: 139567at7742
## Thu Jul 14 01:27:50 CEST 2022 - The aliscore output file does not exist. Check results for BUSCO: 248693at7742
## Thu Jul 14 01:27:50 CEST 2022 - The aliscore output appears to be empty. Check results for BUSCO: 248693at7742
## Thu Jul 14 01:27:52 CEST 2022 - The aliscore output file does not exist. Check results for BUSCO: 270710at7742
## Thu Jul 14 01:27:52 CEST 2022 - The aliscore output appears to be empty. Check results for BUSCO: 270710at7742
## Thu Jul 14 01:27:52 CEST 2022 - The aliscore output file does not exist. Check results for BUSCO: 438106at7742
## Thu Jul 14 01:27:52 CEST 2022 - The aliscore output appears to be empty. Check results for BUSCO: 438106at7742
## Thu Jul 14 01:27:54 CEST 2022 - The aliscore output file does not exist. Check results for BUSCO: 159463at7742
## Thu Jul 14 01:27:54 CEST 2022 - The aliscore output appears to be empty. Check results for BUSCO: 159463at7742
## Thu Jul 14 01:27:54 CEST 2022 - The aliscore output file does not exist. Check results for BUSCO: 93522at7742
## Thu Jul 14 01:27:54 CEST 2022 - The aliscore output appears to be empty. Check results for BUSCO: 93522at7742
## Thu Jul 14 01:27:55 CEST 2022 - The aliscore output file does not exist. Check results for BUSCO: 176497at7742
## Thu Jul 14 01:27:55 CEST 2022 - The aliscore output appears to be empty. Check results for BUSCO: 176497at7742
## Thu Jul 14 01:27:57 CEST 2022 - The aliscore output file does not exist. Check results for BUSCO: 367197at7742
## Thu Jul 14 01:27:57 CEST 2022 - The aliscore output appears to be empty. Check results for BUSCO: 367197at7742
## Thu Jul 14 01:27:58 CEST 2022 - The aliscore output file does not exist. Check results for BUSCO: 136946at7742
## Thu Jul 14 01:27:58 CEST 2022 - The aliscore output appears to be empty. Check results for BUSCO: 136946at7742
## Thu Jul 14 01:28:00 CEST 2022 - The aliscore output file does not exist. Check results for BUSCO: 199227at7742
## Thu Jul 14 01:28:00 CEST 2022 - The aliscore output appears to be empty. Check results for BUSCO: 199227at7742
## Thu Jul 14 01:28:03 CEST 2022 - The aliscore output file does not exist. Check results for BUSCO: 139053at7742
## Thu Jul 14 01:28:03 CEST 2022 - The aliscore output appears to be empty. Check results for BUSCO: 139053at7742
## Thu Jul 14 01:28:04 CEST 2022 - The aliscore output file does not exist. Check results for BUSCO: 227436at7742
## Thu Jul 14 01:28:04 CEST 2022 - The aliscore output appears to be empty. Check results for BUSCO: 227436at7742
## Thu Jul 14 01:28:05 CEST 2022 - The aliscore output file does not exist. Check results for BUSCO: 155235at7742
## Thu Jul 14 01:28:05 CEST 2022 - The aliscore output appears to be empty. Check results for BUSCO: 155235at7742
## Thu Jul 14 01:28:05 CEST 2022 - The aliscore output file does not exist. Check results for BUSCO: 354251at7742
## Thu Jul 14 01:28:05 CEST 2022 - The aliscore output appears to be empty. Check results for BUSCO: 354251at7742
## Thu Jul 14 01:28:06 CEST 2022 - The aliscore output file does not exist. Check results for BUSCO: 109315at7742
## Thu Jul 14 01:28:06 CEST 2022 - The aliscore output appears to be empty. Check results for BUSCO: 109315at7742
## Thu Jul 14 01:28:06 CEST 2022 - The aliscore output file does not exist. Check results for BUSCO: 246371at7742
## Thu Jul 14 01:28:06 CEST 2022 - The aliscore output appears to be empty. Check results for BUSCO: 246371at7742
## Thu Jul 14 01:28:07 CEST 2022 - The aliscore output file does not exist. Check results for BUSCO: 182489at7742
## Thu Jul 14 01:28:07 CEST 2022 - The aliscore output appears to be empty. Check results for BUSCO: 182489at7742
## Thu Jul 14 01:28:08 CEST 2022 - The aliscore output file does not exist. Check results for BUSCO: 257782at7742
## Thu Jul 14 01:28:08 CEST 2022 - The aliscore output appears to be empty. Check results for BUSCO: 257782at7742
## Thu Jul 14 01:28:10 CEST 2022 - The aliscore output file does not exist. Check results for BUSCO: 185742at7742
## Thu Jul 14 01:28:11 CEST 2022 - The aliscore output appears to be empty. Check results for BUSCO: 185742at7742
## Thu Jul 14 01:28:11 CEST 2022 - The aliscore output file does not exist. Check results for BUSCO: 276660at7742
## Thu Jul 14 01:28:11 CEST 2022 - The aliscore output appears to be empty. Check results for BUSCO: 276660at7742
## Thu Jul 14 01:28:13 CEST 2022 - The aliscore output file does not exist. Check results for BUSCO: 158142at7742
## Thu Jul 14 01:28:13 CEST 2022 - The aliscore output appears to be empty. Check results for BUSCO: 158142at7742
## Thu Jul 14 01:28:13 CEST 2022 - The aliscore output file does not exist. Check results for BUSCO: 340519at7742
## Thu Jul 14 01:28:14 CEST 2022 - The aliscore output appears to be empty. Check results for BUSCO: 340519at7742
## Thu Jul 14 01:28:14 CEST 2022 - The aliscore output file does not exist. Check results for BUSCO: 161520at7742
## Thu Jul 14 01:28:14 CEST 2022 - The aliscore output appears to be empty. Check results for BUSCO: 161520at7742
## Thu Jul 14 01:28:16 CEST 2022 - The aliscore output file does not exist. Check results for BUSCO: 162174at7742
## Thu Jul 14 01:28:16 CEST 2022 - The aliscore output appears to be empty. Check results for BUSCO: 162174at7742
## Thu Jul 14 01:28:17 CEST 2022 - The aliscore output file does not exist. Check results for BUSCO: 108888at7742
## Thu Jul 14 01:28:17 CEST 2022 - The aliscore output appears to be empty. Check results for BUSCO: 108888at7742
## Thu Jul 14 01:28:19 CEST 2022 - The aliscore output file does not exist. Check results for BUSCO: 281807at7742
## Thu Jul 14 01:28:19 CEST 2022 - The aliscore output file does not exist. Check results for BUSCO: 117384at7742
## Thu Jul 14 01:28:19 CEST 2022 - The aliscore output appears to be empty. Check results for BUSCO: 281807at7742
## Thu Jul 14 01:28:19 CEST 2022 - The aliscore output appears to be empty. Check results for BUSCO: 117384at7742
## Thu Jul 14 01:28:19 CEST 2022 - The aliscore output file does not exist. Check results for BUSCO: 220601at7742
## Thu Jul 14 01:28:19 CEST 2022 - The aliscore output appears to be empty. Check results for BUSCO: 220601at7742
## Thu Jul 14 01:28:22 CEST 2022 - The aliscore output file does not exist. Check results for BUSCO: 275720at7742
## Thu Jul 14 01:28:22 CEST 2022 - The aliscore output appears to be empty. Check results for BUSCO: 275720at7742
## Thu Jul 14 01:28:23 CEST 2022 - The aliscore output file does not exist. Check results for BUSCO: 93035at7742
## Thu Jul 14 01:28:23 CEST 2022 - The aliscore output file does not exist. Check results for BUSCO: 208949at7742
## Thu Jul 14 01:28:23 CEST 2022 - The aliscore output appears to be empty. Check results for BUSCO: 93035at7742
## Thu Jul 14 01:28:23 CEST 2022 - The aliscore output appears to be empty. Check results for BUSCO: 208949at7742
## Thu Jul 14 01:28:25 CEST 2022 - The aliscore output file does not exist. Check results for BUSCO: 98637at7742
## Thu Jul 14 01:28:25 CEST 2022 - The aliscore output appears to be empty. Check results for BUSCO: 98637at7742
## Thu Jul 14 01:28:27 CEST 2022 - The aliscore output file does not exist. Check results for BUSCO: 143888at7742
## Thu Jul 14 01:28:27 CEST 2022 - The aliscore output appears to be empty. Check results for BUSCO: 143888at7742
## Thu Jul 14 01:28:28 CEST 2022 - The aliscore output file does not exist. Check results for BUSCO: 170012at7742
## Thu Jul 14 01:28:28 CEST 2022 - The aliscore output appears to be empty. Check results for BUSCO: 170012at7742
## Thu Jul 14 01:28:28 CEST 2022 - The aliscore output file does not exist. Check results for BUSCO: 225148at7742
## Thu Jul 14 01:28:28 CEST 2022 - The aliscore output appears to be empty. Check results for BUSCO: 225148at7742
## Thu Jul 14 01:28:28 CEST 2022 - The aliscore output file does not exist. Check results for BUSCO: 205393at7742
## Thu Jul 14 01:28:28 CEST 2022 - The aliscore output appears to be empty. Check results for BUSCO: 205393at7742
## Thu Jul 14 01:28:30 CEST 2022 - The aliscore output file does not exist. Check results for BUSCO: 187969at7742
## Thu Jul 14 01:28:30 CEST 2022 - The aliscore output appears to be empty. Check results for BUSCO: 187969at7742
## Thu Jul 14 01:28:31 CEST 2022 - The aliscore output file does not exist. Check results for BUSCO: 294332at7742
## Thu Jul 14 01:28:31 CEST 2022 - The aliscore output appears to be empty. Check results for BUSCO: 294332at7742
## Thu Jul 14 01:28:33 CEST 2022 - The aliscore output file does not exist. Check results for BUSCO: 279373at7742
## Thu Jul 14 01:28:33 CEST 2022 - The aliscore output appears to be empty. Check results for BUSCO: 279373at7742
## Thu Jul 14 01:28:34 CEST 2022 - The aliscore output file does not exist. Check results for BUSCO: 108802at7742
## Thu Jul 14 01:28:34 CEST 2022 - The aliscore output appears to be empty. Check results for BUSCO: 108802at7742
## Thu Jul 14 01:28:35 CEST 2022 - The aliscore output file does not exist. Check results for BUSCO: 160923at7742
## Thu Jul 14 01:28:35 CEST 2022 - The aliscore output appears to be empty. Check results for BUSCO: 160923at7742
## Thu Jul 14 01:28:35 CEST 2022 - The aliscore output file does not exist. Check results for BUSCO: 205318at7742
## Thu Jul 14 01:28:35 CEST 2022 - The aliscore output appears to be empty. Check results for BUSCO: 205318at7742
## Thu Jul 14 01:28:38 CEST 2022 - The aliscore output file does not exist. Check results for BUSCO: 150306at7742
## Thu Jul 14 01:28:38 CEST 2022 - The aliscore output appears to be empty. Check results for BUSCO: 150306at7742
## Thu Jul 14 01:28:38 CEST 2022 - The aliscore output file does not exist. Check results for BUSCO: 269302at7742
## Thu Jul 14 01:28:38 CEST 2022 - The aliscore output appears to be empty. Check results for BUSCO: 269302at7742
## Thu Jul 14 01:28:40 CEST 2022 - The aliscore output file does not exist. Check results for BUSCO: 316777at7742
## Thu Jul 14 01:28:40 CEST 2022 - The aliscore output appears to be empty. Check results for BUSCO: 316777at7742
## Thu Jul 14 01:28:41 CEST 2022 - The aliscore output file does not exist. Check results for BUSCO: 143888at7742
## Thu Jul 14 01:28:41 CEST 2022 - The aliscore output appears to be empty. Check results for BUSCO: 143888at7742
## Thu Jul 14 01:28:43 CEST 2022 - The aliscore output file does not exist. Check results for BUSCO: 313142at7742
## Thu Jul 14 01:28:43 CEST 2022 - The aliscore output appears to be empty. Check results for BUSCO: 313142at7742
## Thu Jul 14 01:28:43 CEST 2022 - The aliscore output file does not exist. Check results for BUSCO: 221031at7742
## Thu Jul 14 01:28:43 CEST 2022 - The aliscore output appears to be empty. Check results for BUSCO: 221031at7742
## Thu Jul 14 01:28:44 CEST 2022 - The aliscore output file does not exist. Check results for BUSCO: 163177at7742
## Thu Jul 14 01:28:44 CEST 2022 - The aliscore output appears to be empty. Check results for BUSCO: 163177at7742
## Thu Jul 14 01:28:45 CEST 2022 - The aliscore output file does not exist. Check results for BUSCO: 166688at7742
## Thu Jul 14 01:28:45 CEST 2022 - The aliscore output appears to be empty. Check results for BUSCO: 166688at7742
## Thu Jul 14 01:28:47 CEST 2022 - The aliscore output file does not exist. Check results for BUSCO: 155235at7742
## Thu Jul 14 01:28:47 CEST 2022 - The aliscore output appears to be empty. Check results for BUSCO: 155235at7742
## Thu Jul 14 01:28:48 CEST 2022 - The aliscore output file does not exist. Check results for BUSCO: 178680at7742
## Thu Jul 14 01:28:48 CEST 2022 - The aliscore output appears to be empty. Check results for BUSCO: 178680at7742
## Thu Jul 14 01:28:49 CEST 2022 - The aliscore output file does not exist. Check results for BUSCO: 224101at7742
## Thu Jul 14 01:28:49 CEST 2022 - The aliscore output appears to be empty. Check results for BUSCO: 224101at7742
## Thu Jul 14 01:28:51 CEST 2022 - The aliscore output file does not exist. Check results for BUSCO: 310707at7742
## Thu Jul 14 01:28:51 CEST 2022 - The aliscore output appears to be empty. Check results for BUSCO: 310707at7742
## Thu Jul 14 01:28:52 CEST 2022 - The aliscore output file does not exist. Check results for BUSCO: 145938at7742
## Thu Jul 14 01:28:52 CEST 2022 - The aliscore output appears to be empty. Check results for BUSCO: 145938at7742
## Thu Jul 14 01:28:53 CEST 2022 - The aliscore output file does not exist. Check results for BUSCO: 264239at7742
## Thu Jul 14 01:28:53 CEST 2022 - The aliscore output appears to be empty. Check results for BUSCO: 264239at7742
## Thu Jul 14 01:28:53 CEST 2022 - The aliscore output file does not exist. Check results for BUSCO: 128255at7742
## Thu Jul 14 01:28:53 CEST 2022 - The aliscore output appears to be empty. Check results for BUSCO: 128255at7742
## Thu Jul 14 01:28:55 CEST 2022 - The aliscore output file does not exist. Check results for BUSCO: 269302at7742
## Thu Jul 14 01:28:55 CEST 2022 - The aliscore output appears to be empty. Check results for BUSCO: 269302at7742
## Thu Jul 14 01:28:57 CEST 2022 - The aliscore output file does not exist. Check results for BUSCO: 255610at7742
## Thu Jul 14 01:28:57 CEST 2022 - The aliscore output appears to be empty. Check results for BUSCO: 255610at7742
## Thu Jul 14 01:28:59 CEST 2022 - The aliscore output file does not exist. Check results for BUSCO: 316777at7742
## Thu Jul 14 01:28:59 CEST 2022 - The aliscore output appears to be empty. Check results for BUSCO: 316777at7742
## Thu Jul 14 01:29:00 CEST 2022 - The aliscore output file does not exist. Check results for BUSCO: 214501at7742
## Thu Jul 14 01:29:00 CEST 2022 - The aliscore output appears to be empty. Check results for BUSCO: 214501at7742
## Thu Jul 14 01:29:02 CEST 2022 - The aliscore output file does not exist. Check results for BUSCO: 124354at7742
## Thu Jul 14 01:29:02 CEST 2022 - The aliscore output appears to be empty. Check results for BUSCO: 124354at7742
## Thu Jul 14 01:29:02 CEST 2022 - The aliscore output file does not exist. Check results for BUSCO: 295275at7742
## Thu Jul 14 01:29:02 CEST 2022 - The aliscore output appears to be empty. Check results for BUSCO: 295275at7742
## Thu Jul 14 01:29:04 CEST 2022 - The aliscore output file does not exist. Check results for BUSCO: 178259at7742
## Thu Jul 14 01:29:04 CEST 2022 - The aliscore output appears to be empty. Check results for BUSCO: 178259at7742
## Thu Jul 14 01:29:04 CEST 2022 - The aliscore output file does not exist. Check results for BUSCO: 118353at7742
## Thu Jul 14 01:29:04 CEST 2022 - The aliscore output appears to be empty. Check results for BUSCO: 118353at7742
## Thu Jul 14 01:29:04 CEST 2022 - The aliscore output file does not exist. Check results for BUSCO: 204261at7742
## Thu Jul 14 01:29:04 CEST 2022 - The aliscore output appears to be empty. Check results for BUSCO: 204261at7742
## Thu Jul 14 01:29:06 CEST 2022 - The aliscore output file does not exist. Check results for BUSCO: 218284at7742
## Thu Jul 14 01:29:06 CEST 2022 - The aliscore output appears to be empty. Check results for BUSCO: 218284at7742
## Thu Jul 14 01:29:08 CEST 2022 - The aliscore output file does not exist. Check results for BUSCO: 116710at7742
## Thu Jul 14 01:29:08 CEST 2022 - The aliscore output appears to be empty. Check results for BUSCO: 116710at7742
## Thu Jul 14 01:29:09 CEST 2022 - The aliscore output file does not exist. Check results for BUSCO: 174872at7742
## Thu Jul 14 01:29:09 CEST 2022 - The aliscore output appears to be empty. Check results for BUSCO: 174872at7742
## Thu Jul 14 01:29:09 CEST 2022 - The aliscore output file does not exist. Check results for BUSCO: 225241at7742
## Thu Jul 14 01:29:09 CEST 2022 - The aliscore output appears to be empty. Check results for BUSCO: 225241at7742
## Thu Jul 14 01:29:11 CEST 2022 - The aliscore output file does not exist. Check results for BUSCO: 153401at7742
## Thu Jul 14 01:29:11 CEST 2022 - The aliscore output appears to be empty. Check results for BUSCO: 153401at7742
## Thu Jul 14 01:29:12 CEST 2022 - The aliscore output file does not exist. Check results for BUSCO: 199120at7742
## Thu Jul 14 01:29:12 CEST 2022 - The aliscore output appears to be empty. Check results for BUSCO: 199120at7742
## Thu Jul 14 01:29:13 CEST 2022 - The aliscore output file does not exist. Check results for BUSCO: 188869at7742
## Thu Jul 14 01:29:13 CEST 2022 - The aliscore output appears to be empty. Check results for BUSCO: 188869at7742
## Thu Jul 14 01:29:15 CEST 2022 - The aliscore output file does not exist. Check results for BUSCO: 344933at7742
## Thu Jul 14 01:29:15 CEST 2022 - The aliscore output appears to be empty. Check results for BUSCO: 344933at7742
## Thu Jul 14 01:29:16 CEST 2022 - The aliscore output file does not exist. Check results for BUSCO: 136142at7742
## Thu Jul 14 01:29:16 CEST 2022 - The aliscore output appears to be empty. Check results for BUSCO: 136142at7742
## Thu Jul 14 01:29:17 CEST 2022 - The aliscore output file does not exist. Check results for BUSCO: 261612at7742
## Thu Jul 14 01:29:17 CEST 2022 - The aliscore output appears to be empty. Check results for BUSCO: 261612at7742
## Thu Jul 14 01:29:18 CEST 2022 - The aliscore output file does not exist. Check results for BUSCO: 283821at7742
## Thu Jul 14 01:29:19 CEST 2022 - The aliscore output appears to be empty. Check results for BUSCO: 283821at7742
## Thu Jul 14 01:29:21 CEST 2022 - The aliscore output file does not exist. Check results for BUSCO: 307789at7742
## Thu Jul 14 01:29:21 CEST 2022 - The aliscore output appears to be empty. Check results for BUSCO: 307789at7742
## Thu Jul 14 01:29:22 CEST 2022 - The aliscore output file does not exist. Check results for BUSCO: 202155at7742
## Thu Jul 14 01:29:22 CEST 2022 - The aliscore output appears to be empty. Check results for BUSCO: 202155at7742
## Thu Jul 14 01:29:22 CEST 2022 - The aliscore output file does not exist. Check results for BUSCO: 116482at7742
## Thu Jul 14 01:29:22 CEST 2022 - The aliscore output appears to be empty. Check results for BUSCO: 116482at7742
## Thu Jul 14 01:29:23 CEST 2022 - The aliscore output file does not exist. Check results for BUSCO: 370236at7742
## Thu Jul 14 01:29:23 CEST 2022 - The aliscore output appears to be empty. Check results for BUSCO: 370236at7742
## Thu Jul 14 01:29:25 CEST 2022 - The aliscore output file does not exist. Check results for BUSCO: 253042at7742
## Thu Jul 14 01:29:25 CEST 2022 - The aliscore output appears to be empty. Check results for BUSCO: 253042at7742
## Thu Jul 14 01:29:26 CEST 2022 - The aliscore output file does not exist. Check results for BUSCO: 283469at7742
## Thu Jul 14 01:29:26 CEST 2022 - The aliscore output appears to be empty. Check results for BUSCO: 283469at7742
## Thu Jul 14 01:29:27 CEST 2022 - The aliscore output file does not exist. Check results for BUSCO: 245932at7742
## Thu Jul 14 01:29:27 CEST 2022 - The aliscore output appears to be empty. Check results for BUSCO: 245932at7742
## Thu Jul 14 01:29:29 CEST 2022 - The aliscore output file does not exist. Check results for BUSCO: 205394at7742
## Thu Jul 14 01:29:29 CEST 2022 - The aliscore output appears to be empty. Check results for BUSCO: 205394at7742
## Thu Jul 14 01:29:30 CEST 2022 - The aliscore output file does not exist. Check results for BUSCO: 342319at7742
## Thu Jul 14 01:29:30 CEST 2022 - The aliscore output appears to be empty. Check results for BUSCO: 342319at7742
## Thu Jul 14 01:29:30 CEST 2022 - The aliscore output file does not exist. Check results for BUSCO: 88027at7742
## Thu Jul 14 01:29:30 CEST 2022 - The aliscore output appears to be empty. Check results for BUSCO: 88027at7742
## Thu Jul 14 01:29:33 CEST 2022 - The aliscore output file does not exist. Check results for BUSCO: 291807at7742
## Thu Jul 14 01:29:33 CEST 2022 - The aliscore output appears to be empty. Check results for BUSCO: 291807at7742
## Thu Jul 14 01:29:33 CEST 2022 - The aliscore output file does not exist. Check results for BUSCO: 209714at7742
## Thu Jul 14 01:29:33 CEST 2022 - The aliscore output appears to be empty. Check results for BUSCO: 209714at7742
## Thu Jul 14 01:29:34 CEST 2022 - The aliscore output file does not exist. Check results for BUSCO: 108888at7742
## Thu Jul 14 01:29:34 CEST 2022 - The aliscore output appears to be empty. Check results for BUSCO: 108888at7742
## Thu Jul 14 01:29:35 CEST 2022 - The aliscore output file does not exist. Check results for BUSCO: 187969at7742
## Thu Jul 14 01:29:35 CEST 2022 - The aliscore output appears to be empty. Check results for BUSCO: 187969at7742
## Thu Jul 14 01:29:37 CEST 2022 - The aliscore output file does not exist. Check results for BUSCO: 179036at7742
## Thu Jul 14 01:29:37 CEST 2022 - The aliscore output appears to be empty. Check results for BUSCO: 179036at7742
## Thu Jul 14 01:29:37 CEST 2022 - The aliscore output file does not exist. Check results for BUSCO: 199341at7742
## Thu Jul 14 01:29:37 CEST 2022 - The aliscore output appears to be empty. Check results for BUSCO: 199341at7742
## Thu Jul 14 01:29:38 CEST 2022 - The aliscore output file does not exist. Check results for BUSCO: 300687at7742
## Thu Jul 14 01:29:38 CEST 2022 - The aliscore output appears to be empty. Check results for BUSCO: 300687at7742
## Thu Jul 14 01:29:40 CEST 2022 - The aliscore output file does not exist. Check results for BUSCO: 122495at7742
## Thu Jul 14 01:29:40 CEST 2022 - The aliscore output appears to be empty. Check results for BUSCO: 122495at7742
## Thu Jul 14 01:29:40 CEST 2022 - The aliscore output file does not exist. Check results for BUSCO: 241002at7742
## Thu Jul 14 01:29:40 CEST 2022 - The aliscore output appears to be empty. Check results for BUSCO: 241002at7742
## Thu Jul 14 01:29:41 CEST 2022 - The aliscore output file does not exist. Check results for BUSCO: 122495at7742
## Thu Jul 14 01:29:41 CEST 2022 - The aliscore output appears to be empty. Check results for BUSCO: 122495at7742
## Thu Jul 14 01:29:44 CEST 2022 - The aliscore output file does not exist. Check results for BUSCO: 261602at7742
## Thu Jul 14 01:29:44 CEST 2022 - The aliscore output appears to be empty. Check results for BUSCO: 261602at7742
## Thu Jul 14 01:29:44 CEST 2022 - The aliscore output file does not exist. Check results for BUSCO: 118460at7742
## Thu Jul 14 01:29:44 CEST 2022 - The aliscore output appears to be empty. Check results for BUSCO: 118460at7742
## Thu Jul 14 01:29:46 CEST 2022 - The aliscore output file does not exist. Check results for BUSCO: 218524at7742
## Thu Jul 14 01:29:46 CEST 2022 - The aliscore output appears to be empty. Check results for BUSCO: 218524at7742
## Thu Jul 14 01:29:47 CEST 2022 - The aliscore output file does not exist. Check results for BUSCO: 227436at7742
## Thu Jul 14 01:29:47 CEST 2022 - The aliscore output appears to be empty. Check results for BUSCO: 227436at7742
## Thu Jul 14 01:29:47 CEST 2022 - The aliscore output file does not exist. Check results for BUSCO: 193249at7742
## Thu Jul 14 01:29:47 CEST 2022 - The aliscore output appears to be empty. Check results for BUSCO: 193249at7742
## Thu Jul 14 01:29:49 CEST 2022 - The aliscore output file does not exist. Check results for BUSCO: 275875at7742
## Thu Jul 14 01:29:49 CEST 2022 - The aliscore output appears to be empty. Check results for BUSCO: 275875at7742
## Thu Jul 14 01:29:52 CEST 2022 - The aliscore output file does not exist. Check results for BUSCO: 130130at7742
## Thu Jul 14 01:29:52 CEST 2022 - The aliscore output appears to be empty. Check results for BUSCO: 130130at7742
## Thu Jul 14 01:29:52 CEST 2022 - The aliscore output file does not exist. Check results for BUSCO: 291289at7742
## Thu Jul 14 01:29:52 CEST 2022 - The aliscore output appears to be empty. Check results for BUSCO: 291289at7742
## Thu Jul 14 01:29:53 CEST 2022 - The aliscore output file does not exist. Check results for BUSCO: 210107at7742
## Thu Jul 14 01:29:53 CEST 2022 - The aliscore output appears to be empty. Check results for BUSCO: 210107at7742
## Thu Jul 14 01:29:55 CEST 2022 - The aliscore output file does not exist. Check results for BUSCO: 182489at7742
## Thu Jul 14 01:29:55 CEST 2022 - The aliscore output appears to be empty. Check results for BUSCO: 182489at7742
## Thu Jul 14 01:29:56 CEST 2022 - The aliscore output file does not exist. Check results for BUSCO: 139241at7742
## Thu Jul 14 01:29:56 CEST 2022 - The aliscore output appears to be empty. Check results for BUSCO: 139241at7742
## Thu Jul 14 01:29:56 CEST 2022 - The aliscore output file does not exist. Check results for BUSCO: 216982at7742
## Thu Jul 14 01:29:56 CEST 2022 - The aliscore output appears to be empty. Check results for BUSCO: 216982at7742
## Thu Jul 14 01:29:57 CEST 2022 - The aliscore output file does not exist. Check results for BUSCO: 329437at7742
## Thu Jul 14 01:29:57 CEST 2022 - The aliscore output appears to be empty. Check results for BUSCO: 329437at7742
## Thu Jul 14 01:29:58 CEST 2022 - The aliscore output file does not exist. Check results for BUSCO: 198411at7742
## Thu Jul 14 01:29:58 CEST 2022 - The aliscore output appears to be empty. Check results for BUSCO: 198411at7742
## Thu Jul 14 01:29:58 CEST 2022 - The aliscore output file does not exist. Check results for BUSCO: 199341at7742
## Thu Jul 14 01:29:58 CEST 2022 - The aliscore output appears to be empty. Check results for BUSCO: 199341at7742
## Thu Jul 14 01:30:00 CEST 2022 - The aliscore output file does not exist. Check results for BUSCO: 264816at7742
## Thu Jul 14 01:30:00 CEST 2022 - The aliscore output appears to be empty. Check results for BUSCO: 264816at7742
## Thu Jul 14 01:30:01 CEST 2022 - The aliscore output file does not exist. Check results for BUSCO: 192142at7742
## Thu Jul 14 01:30:01 CEST 2022 - The aliscore output appears to be empty. Check results for BUSCO: 192142at7742
## Thu Jul 14 01:30:03 CEST 2022 - The aliscore output file does not exist. Check results for BUSCO: 180416at7742
## Thu Jul 14 01:30:03 CEST 2022 - The aliscore output appears to be empty. Check results for BUSCO: 180416at7742
## Thu Jul 14 01:30:04 CEST 2022 - The aliscore output file does not exist. Check results for BUSCO: 287723at7742
## Thu Jul 14 01:30:04 CEST 2022 - The aliscore output appears to be empty. Check results for BUSCO: 287723at7742
## Thu Jul 14 01:30:05 CEST 2022 - The aliscore output file does not exist. Check results for BUSCO: 301301at7742
## Thu Jul 14 01:30:05 CEST 2022 - The aliscore output appears to be empty. Check results for BUSCO: 301301at7742
## Thu Jul 14 01:30:05 CEST 2022 - The aliscore output file does not exist. Check results for BUSCO: 389352at7742
## Thu Jul 14 01:30:05 CEST 2022 - The aliscore output appears to be empty. Check results for BUSCO: 389352at7742
## Thu Jul 14 01:30:06 CEST 2022 - The aliscore output file does not exist. Check results for BUSCO: 113992at7742
## Thu Jul 14 01:30:06 CEST 2022 - The aliscore output appears to be empty. Check results for BUSCO: 113992at7742
## Thu Jul 14 01:30:08 CEST 2022 - The aliscore output file does not exist. Check results for BUSCO: 164982at7742
## Thu Jul 14 01:30:08 CEST 2022 - The aliscore output appears to be empty. Check results for BUSCO: 164982at7742
## Thu Jul 14 01:30:09 CEST 2022 - The aliscore output file does not exist. Check results for BUSCO: 116482at7742
## Thu Jul 14 01:30:09 CEST 2022 - The aliscore output appears to be empty. Check results for BUSCO: 116482at7742
## Thu Jul 14 01:30:10 CEST 2022 - The aliscore output file does not exist. Check results for BUSCO: 264816at7742
## Thu Jul 14 01:30:10 CEST 2022 - The aliscore output appears to be empty. Check results for BUSCO: 264816at7742
## Thu Jul 14 01:30:12 CEST 2022 - The aliscore output file does not exist. Check results for BUSCO: 246371at7742
## Thu Jul 14 01:30:12 CEST 2022 - The aliscore output appears to be empty. Check results for BUSCO: 246371at7742
## Thu Jul 14 01:30:12 CEST 2022 - The aliscore output file does not exist. Check results for BUSCO: 105032at7742
## Thu Jul 14 01:30:13 CEST 2022 - The aliscore output appears to be empty. Check results for BUSCO: 105032at7742
## Thu Jul 14 01:30:13 CEST 2022 - The aliscore output file does not exist. Check results for BUSCO: 221031at7742
## Thu Jul 14 01:30:13 CEST 2022 - The aliscore output appears to be empty. Check results for BUSCO: 221031at7742
## Thu Jul 14 01:30:13 CEST 2022 - The aliscore output file does not exist. Check results for BUSCO: 116710at7742
## Thu Jul 14 01:30:13 CEST 2022 - The aliscore output appears to be empty. Check results for BUSCO: 116710at7742
## Thu Jul 14 01:30:15 CEST 2022 - The aliscore output file does not exist. Check results for BUSCO: 160371at7742
## Thu Jul 14 01:30:15 CEST 2022 - The aliscore output appears to be empty. Check results for BUSCO: 160371at7742
## Thu Jul 14 01:30:18 CEST 2022 - The aliscore output file does not exist. Check results for BUSCO: 201359at7742
## Thu Jul 14 01:30:18 CEST 2022 - The aliscore output appears to be empty. Check results for BUSCO: 201359at7742
## Thu Jul 14 01:30:18 CEST 2022 - The aliscore output file does not exist. Check results for BUSCO: 312347at7742
## Thu Jul 14 01:30:18 CEST 2022 - The aliscore output appears to be empty. Check results for BUSCO: 312347at7742
## Thu Jul 14 01:30:18 CEST 2022 - The aliscore output file does not exist. Check results for BUSCO: 223283at7742
## Thu Jul 14 01:30:18 CEST 2022 - The aliscore output appears to be empty. Check results for BUSCO: 223283at7742
## Thu Jul 14 01:30:20 CEST 2022 - The aliscore output file does not exist. Check results for BUSCO: 231455at7742
## Thu Jul 14 01:30:20 CEST 2022 - The aliscore output appears to be empty. Check results for BUSCO: 231455at7742
## Thu Jul 14 01:30:21 CEST 2022 - The aliscore output file does not exist. Check results for BUSCO: 215159at7742
## Thu Jul 14 01:30:21 CEST 2022 - The aliscore output appears to be empty. Check results for BUSCO: 215159at7742
## Thu Jul 14 01:30:22 CEST 2022 - The aliscore output file does not exist. Check results for BUSCO: 334616at7742
## Thu Jul 14 01:30:23 CEST 2022 - The aliscore output appears to be empty. Check results for BUSCO: 334616at7742
## Thu Jul 14 01:30:23 CEST 2022 - The aliscore output file does not exist. Check results for BUSCO: 327369at7742
## Thu Jul 14 01:30:23 CEST 2022 - The aliscore output appears to be empty. Check results for BUSCO: 327369at7742
## Thu Jul 14 01:30:24 CEST 2022 - The aliscore output file does not exist. Check results for BUSCO: 241002at7742
## Thu Jul 14 01:30:24 CEST 2022 - The aliscore output appears to be empty. Check results for BUSCO: 241002at7742
## Thu Jul 14 01:30:27 CEST 2022 - The aliscore output file does not exist. Check results for BUSCO: 171530at7742
## Thu Jul 14 01:30:27 CEST 2022 - The aliscore output file does not exist. Check results for BUSCO: 118460at7742
## Thu Jul 14 01:30:27 CEST 2022 - The aliscore output appears to be empty. Check results for BUSCO: 171530at7742
## Thu Jul 14 01:30:27 CEST 2022 - The aliscore output appears to be empty. Check results for BUSCO: 118460at7742
## Thu Jul 14 01:30:30 CEST 2022 - The aliscore output file does not exist. Check results for BUSCO: 221061at7742
## Thu Jul 14 01:30:30 CEST 2022 - The aliscore output appears to be empty. Check results for BUSCO: 221061at7742
## Thu Jul 14 01:30:30 CEST 2022 - The aliscore output file does not exist. Check results for BUSCO: 88027at7742
## Thu Jul 14 01:30:30 CEST 2022 - The aliscore output appears to be empty. Check results for BUSCO: 88027at7742
## Thu Jul 14 01:30:31 CEST 2022 - The aliscore output file does not exist. Check results for BUSCO: 130130at7742
## Thu Jul 14 01:30:31 CEST 2022 - The aliscore output appears to be empty. Check results for BUSCO: 130130at7742
## Thu Jul 14 01:30:32 CEST 2022 - The aliscore output file does not exist. Check results for BUSCO: 150306at7742
## Thu Jul 14 01:30:32 CEST 2022 - The aliscore output appears to be empty. Check results for BUSCO: 150306at7742
## Thu Jul 14 01:30:33 CEST 2022 - The aliscore output file does not exist. Check results for BUSCO: 169728at7742
## Thu Jul 14 01:30:33 CEST 2022 - The aliscore output appears to be empty. Check results for BUSCO: 169728at7742
## Thu Jul 14 01:30:34 CEST 2022 - The aliscore output file does not exist. Check results for BUSCO: 334896at7742
## Thu Jul 14 01:30:34 CEST 2022 - The aliscore output appears to be empty. Check results for BUSCO: 334896at7742
## Thu Jul 14 01:30:35 CEST 2022 - The aliscore output file does not exist. Check results for BUSCO: 438106at7742
## Thu Jul 14 01:30:35 CEST 2022 - The aliscore output appears to be empty. Check results for BUSCO: 438106at7742
## Thu Jul 14 01:30:37 CEST 2022 - The aliscore output file does not exist. Check results for BUSCO: 88027at7742
## Thu Jul 14 01:30:37 CEST 2022 - The aliscore output appears to be empty. Check results for BUSCO: 88027at7742
## Thu Jul 14 01:30:38 CEST 2022 - The aliscore output file does not exist. Check results for BUSCO: 176601at7742
## Thu Jul 14 01:30:38 CEST 2022 - The aliscore output appears to be empty. Check results for BUSCO: 176601at7742
## Thu Jul 14 01:30:39 CEST 2022 - The aliscore output file does not exist. Check results for BUSCO: 153242at7742
## Thu Jul 14 01:30:39 CEST 2022 - The aliscore output appears to be empty. Check results for BUSCO: 153242at7742
## Thu Jul 14 01:30:41 CEST 2022 - The aliscore output file does not exist. Check results for BUSCO: 128712at7742
## Thu Jul 14 01:30:41 CEST 2022 - The aliscore output appears to be empty. Check results for BUSCO: 128712at7742
## Thu Jul 14 01:30:43 CEST 2022 - The aliscore output file does not exist. Check results for BUSCO: 138339at7742
## Thu Jul 14 01:30:43 CEST 2022 - The aliscore output appears to be empty. Check results for BUSCO: 138339at7742
## Thu Jul 14 01:30:43 CEST 2022 - The aliscore output file does not exist. Check results for BUSCO: 163878at7742
## Thu Jul 14 01:30:43 CEST 2022 - The aliscore output appears to be empty. Check results for BUSCO: 163878at7742
## Thu Jul 14 01:30:44 CEST 2022 - The aliscore output file does not exist. Check results for BUSCO: 171530at7742
## Thu Jul 14 01:30:44 CEST 2022 - The aliscore output appears to be empty. Check results for BUSCO: 171530at7742
## Thu Jul 14 01:30:44 CEST 2022 - The aliscore output file does not exist. Check results for BUSCO: 388724at7742
## Thu Jul 14 01:30:44 CEST 2022 - The aliscore output appears to be empty. Check results for BUSCO: 388724at7742
## Thu Jul 14 01:30:46 CEST 2022 - The aliscore output file does not exist. Check results for BUSCO: 119371at7742
## Thu Jul 14 01:30:46 CEST 2022 - The aliscore output appears to be empty. Check results for BUSCO: 119371at7742
## Thu Jul 14 01:30:48 CEST 2022 - The aliscore output file does not exist. Check results for BUSCO: 137723at7742
## Thu Jul 14 01:30:48 CEST 2022 - The aliscore output appears to be empty. Check results for BUSCO: 137723at7742
## Thu Jul 14 01:30:49 CEST 2022 - The aliscore output file does not exist. Check results for BUSCO: 196730at7742
## Thu Jul 14 01:30:50 CEST 2022 - The aliscore output appears to be empty. Check results for BUSCO: 196730at7742
## Thu Jul 14 01:30:50 CEST 2022 - The aliscore output file does not exist. Check results for BUSCO: 176497at7742
## Thu Jul 14 01:30:50 CEST 2022 - The aliscore output appears to be empty. Check results for BUSCO: 176497at7742
## Thu Jul 14 01:30:53 CEST 2022 - The aliscore output file does not exist. Check results for BUSCO: 153242at7742
## Thu Jul 14 01:30:53 CEST 2022 - The aliscore output appears to be empty. Check results for BUSCO: 153242at7742
## Thu Jul 14 01:30:53 CEST 2022 - The aliscore output file does not exist. Check results for BUSCO: 105032at7742
## Thu Jul 14 01:30:53 CEST 2022 - The aliscore output appears to be empty. Check results for BUSCO: 105032at7742
## Thu Jul 14 01:30:54 CEST 2022 - The aliscore output file does not exist. Check results for BUSCO: 164982at7742
## Thu Jul 14 01:30:55 CEST 2022 - The aliscore output appears to be empty. Check results for BUSCO: 164982at7742
## Thu Jul 14 01:30:56 CEST 2022 - The aliscore output file does not exist. Check results for BUSCO: 354251at7742
## Thu Jul 14 01:30:56 CEST 2022 - The aliscore output appears to be empty. Check results for BUSCO: 354251at7742
## Thu Jul 14 01:30:57 CEST 2022 - The aliscore output file does not exist. Check results for BUSCO: 410850at7742
## Thu Jul 14 01:30:57 CEST 2022 - The aliscore output appears to be empty. Check results for BUSCO: 410850at7742
## Thu Jul 14 01:30:57 CEST 2022 - The aliscore output file does not exist. Check results for BUSCO: 285297at7742
## Thu Jul 14 01:30:57 CEST 2022 - The aliscore output appears to be empty. Check results for BUSCO: 285297at7742
## Thu Jul 14 01:30:59 CEST 2022 - The aliscore output file does not exist. Check results for BUSCO: 275720at7742
## Thu Jul 14 01:30:59 CEST 2022 - The aliscore output appears to be empty. Check results for BUSCO: 275720at7742
## Thu Jul 14 01:31:01 CEST 2022 - The aliscore output file does not exist. Check results for BUSCO: 402515at7742
## Thu Jul 14 01:31:01 CEST 2022 - The aliscore output appears to be empty. Check results for BUSCO: 402515at7742
## Thu Jul 14 01:31:02 CEST 2022 - The aliscore output file does not exist. Check results for BUSCO: 210385at7742
## Thu Jul 14 01:31:02 CEST 2022 - The aliscore output appears to be empty. Check results for BUSCO: 210385at7742
## Thu Jul 14 01:31:03 CEST 2022 - The aliscore output file does not exist. Check results for BUSCO: 135511at7742
## Thu Jul 14 01:31:03 CEST 2022 - The aliscore output appears to be empty. Check results for BUSCO: 135511at7742
## Thu Jul 14 01:31:03 CEST 2022 - The aliscore output file does not exist. Check results for BUSCO: 291464at7742
## Thu Jul 14 01:31:03 CEST 2022 - The aliscore output appears to be empty. Check results for BUSCO: 291464at7742
## Thu Jul 14 01:31:05 CEST 2022 - The aliscore output file does not exist. Check results for BUSCO: 117384at7742
## Thu Jul 14 01:31:05 CEST 2022 - The aliscore output appears to be empty. Check results for BUSCO: 117384at7742
## Thu Jul 14 01:31:06 CEST 2022 - The aliscore output file does not exist. Check results for BUSCO: 92374at7742
## Thu Jul 14 01:31:06 CEST 2022 - The aliscore output appears to be empty. Check results for BUSCO: 92374at7742
## Thu Jul 14 01:31:07 CEST 2022 - The aliscore output file does not exist. Check results for BUSCO: 351544at7742
## Thu Jul 14 01:31:07 CEST 2022 - The aliscore output file does not exist. Check results for BUSCO: 260711at7742
## Thu Jul 14 01:31:07 CEST 2022 - The aliscore output appears to be empty. Check results for BUSCO: 351544at7742
## Thu Jul 14 01:31:07 CEST 2022 - The aliscore output appears to be empty. Check results for BUSCO: 260711at7742
## Thu Jul 14 01:31:08 CEST 2022 - The aliscore output file does not exist. Check results for BUSCO: 370224at7742
## Thu Jul 14 01:31:08 CEST 2022 - The aliscore output appears to be empty. Check results for BUSCO: 370224at7742
## Thu Jul 14 01:31:10 CEST 2022 - The aliscore output file does not exist. Check results for BUSCO: 223898at7742
## Thu Jul 14 01:31:11 CEST 2022 - The aliscore output appears to be empty. Check results for BUSCO: 223898at7742
## Thu Jul 14 01:31:13 CEST 2022 - The aliscore output file does not exist. Check results for BUSCO: 151349at7742
## Thu Jul 14 01:31:13 CEST 2022 - The aliscore output appears to be empty. Check results for BUSCO: 151349at7742
## Thu Jul 14 01:31:13 CEST 2022 - The aliscore output file does not exist. Check results for BUSCO: 129684at7742
## Thu Jul 14 01:31:13 CEST 2022 - The aliscore output appears to be empty. Check results for BUSCO: 129684at7742
## Thu Jul 14 01:31:13 CEST 2022 - The aliscore output file does not exist. Check results for BUSCO: 273954at7742
## Thu Jul 14 01:31:14 CEST 2022 - The aliscore output appears to be empty. Check results for BUSCO: 273954at7742
## Thu Jul 14 01:31:14 CEST 2022 - The aliscore output file does not exist. Check results for BUSCO: 320126at7742
## Thu Jul 14 01:31:14 CEST 2022 - The aliscore output appears to be empty. Check results for BUSCO: 320126at7742
## Thu Jul 14 01:31:16 CEST 2022 - The aliscore output file does not exist. Check results for BUSCO: 336679at7742
## Thu Jul 14 01:31:16 CEST 2022 - The aliscore output appears to be empty. Check results for BUSCO: 336679at7742
## Thu Jul 14 01:31:18 CEST 2022 - The aliscore output file does not exist. Check results for BUSCO: 227544at7742
## Thu Jul 14 01:31:18 CEST 2022 - The aliscore output appears to be empty. Check results for BUSCO: 227544at7742
## Thu Jul 14 01:31:18 CEST 2022 - The aliscore output file does not exist. Check results for BUSCO: 318965at7742
## Thu Jul 14 01:31:18 CEST 2022 - The aliscore output appears to be empty. Check results for BUSCO: 318965at7742
## Thu Jul 14 01:31:20 CEST 2022 - The aliscore output file does not exist. Check results for BUSCO: 291801at7742
## Thu Jul 14 01:31:20 CEST 2022 - The aliscore output appears to be empty. Check results for BUSCO: 291801at7742
## Thu Jul 14 01:31:21 CEST 2022 - The aliscore output file does not exist. Check results for BUSCO: 310011at7742
## Thu Jul 14 01:31:21 CEST 2022 - The aliscore output appears to be empty. Check results for BUSCO: 310011at7742
## Thu Jul 14 01:31:22 CEST 2022 - The aliscore output file does not exist. Check results for BUSCO: 275875at7742
## Thu Jul 14 01:31:22 CEST 2022 - The aliscore output appears to be empty. Check results for BUSCO: 275875at7742
## Thu Jul 14 01:31:22 CEST 2022 - The aliscore output file does not exist. Check results for BUSCO: 176601at7742
## Thu Jul 14 01:31:23 CEST 2022 - The aliscore output appears to be empty. Check results for BUSCO: 176601at7742
## Thu Jul 14 01:31:25 CEST 2022 - The aliscore output file does not exist. Check results for BUSCO: 388724at7742
## Thu Jul 14 01:31:25 CEST 2022 - The aliscore output appears to be empty. Check results for BUSCO: 388724at7742
## Thu Jul 14 01:31:26 CEST 2022 - The aliscore output file does not exist. Check results for BUSCO: 93035at7742
## Thu Jul 14 01:31:26 CEST 2022 - The aliscore output appears to be empty. Check results for BUSCO: 93035at7742
## Thu Jul 14 01:31:27 CEST 2022 - The aliscore output file does not exist. Check results for BUSCO: 105032at7742
## Thu Jul 14 01:31:27 CEST 2022 - The aliscore output appears to be empty. Check results for BUSCO: 105032at7742
## Thu Jul 14 01:31:28 CEST 2022 - The aliscore output file does not exist. Check results for BUSCO: 192462at7742
## Thu Jul 14 01:31:28 CEST 2022 - The aliscore output appears to be empty. Check results for BUSCO: 192462at7742
## Thu Jul 14 01:31:29 CEST 2022 - The aliscore output file does not exist. Check results for BUSCO: 328913at7742
## Thu Jul 14 01:31:29 CEST 2022 - The aliscore output appears to be empty. Check results for BUSCO: 328913at7742
## Thu Jul 14 01:31:31 CEST 2022 - The aliscore output file does not exist. Check results for BUSCO: 305085at7742
## Thu Jul 14 01:31:31 CEST 2022 - The aliscore output appears to be empty. Check results for BUSCO: 305085at7742
## Thu Jul 14 01:31:31 CEST 2022 - The aliscore output file does not exist. Check results for BUSCO: 242072at7742
## Thu Jul 14 01:31:31 CEST 2022 - The aliscore output appears to be empty. Check results for BUSCO: 242072at7742
## Thu Jul 14 01:31:33 CEST 2022 - The aliscore output file does not exist. Check results for BUSCO: 214745at7742
## Thu Jul 14 01:31:33 CEST 2022 - The aliscore output appears to be empty. Check results for BUSCO: 214745at7742
## Thu Jul 14 01:31:35 CEST 2022 - The aliscore output file does not exist. Check results for BUSCO: 159463at7742
## Thu Jul 14 01:31:36 CEST 2022 - The aliscore output appears to be empty. Check results for BUSCO: 159463at7742
## Thu Jul 14 01:31:36 CEST 2022 - The aliscore output file does not exist. Check results for BUSCO: 133255at7742
## Thu Jul 14 01:31:36 CEST 2022 - The aliscore output appears to be empty. Check results for BUSCO: 133255at7742
## Thu Jul 14 01:31:37 CEST 2022 - The aliscore output file does not exist. Check results for BUSCO: 357871at7742
## Thu Jul 14 01:31:37 CEST 2022 - The aliscore output file does not exist. Check results for BUSCO: 215440at7742
## Thu Jul 14 01:31:37 CEST 2022 - The aliscore output appears to be empty. Check results for BUSCO: 357871at7742
## Thu Jul 14 01:31:37 CEST 2022 - The aliscore output appears to be empty. Check results for BUSCO: 215440at7742
## Thu Jul 14 01:31:38 CEST 2022 - The aliscore output file does not exist. Check results for BUSCO: 357871at7742
## Thu Jul 14 01:31:38 CEST 2022 - The aliscore output appears to be empty. Check results for BUSCO: 357871at7742
## Thu Jul 14 01:31:39 CEST 2022 - The aliscore output file does not exist. Check results for BUSCO: 363574at7742
## Thu Jul 14 01:31:39 CEST 2022 - The aliscore output appears to be empty. Check results for BUSCO: 363574at7742
## Thu Jul 14 01:31:42 CEST 2022 - The aliscore output file does not exist. Check results for BUSCO: 221393at7742
## Thu Jul 14 01:31:42 CEST 2022 - The aliscore output appears to be empty. Check results for BUSCO: 221393at7742
## Thu Jul 14 01:31:43 CEST 2022 - The aliscore output file does not exist. Check results for BUSCO: 325307at7742
## Thu Jul 14 01:31:43 CEST 2022 - The aliscore output appears to be empty. Check results for BUSCO: 325307at7742
## Thu Jul 14 01:31:43 CEST 2022 - The aliscore output file does not exist. Check results for BUSCO: 174872at7742
## Thu Jul 14 01:31:43 CEST 2022 - The aliscore output appears to be empty. Check results for BUSCO: 174872at7742
## Thu Jul 14 01:31:45 CEST 2022 - The aliscore output file does not exist. Check results for BUSCO: 215440at7742
## Thu Jul 14 01:31:45 CEST 2022 - The aliscore output appears to be empty. Check results for BUSCO: 215440at7742
## Thu Jul 14 01:31:45 CEST 2022 - The aliscore output file does not exist. Check results for BUSCO: 239682at7742
## Thu Jul 14 01:31:45 CEST 2022 - The aliscore output appears to be empty. Check results for BUSCO: 239682at7742
## Thu Jul 14 01:31:47 CEST 2022 - The aliscore output file does not exist. Check results for BUSCO: 285297at7742
## Thu Jul 14 01:31:47 CEST 2022 - The aliscore output appears to be empty. Check results for BUSCO: 285297at7742
## Thu Jul 14 01:31:47 CEST 2022 - The aliscore output file does not exist. Check results for BUSCO: 208122at7742
## Thu Jul 14 01:31:47 CEST 2022 - The aliscore output appears to be empty. Check results for BUSCO: 208122at7742
## Thu Jul 14 01:31:49 CEST 2022 - The aliscore output file does not exist. Check results for BUSCO: 224078at7742
## Thu Jul 14 01:31:49 CEST 2022 - The aliscore output appears to be empty. Check results for BUSCO: 224078at7742
## Thu Jul 14 01:31:51 CEST 2022 - The aliscore output file does not exist. Check results for BUSCO: 165456at7742
## Thu Jul 14 01:31:51 CEST 2022 - The aliscore output appears to be empty. Check results for BUSCO: 165456at7742
## Thu Jul 14 01:31:51 CEST 2022 - The aliscore output file does not exist. Check results for BUSCO: 163878at7742
## Thu Jul 14 01:31:51 CEST 2022 - The aliscore output appears to be empty. Check results for BUSCO: 163878at7742
## Thu Jul 14 01:31:52 CEST 2022 - The aliscore output file does not exist. Check results for BUSCO: 208122at7742
## Thu Jul 14 01:31:52 CEST 2022 - The aliscore output appears to be empty. Check results for BUSCO: 208122at7742
## Thu Jul 14 01:31:53 CEST 2022 - The aliscore output file does not exist. Check results for BUSCO: 208291at7742
## Thu Jul 14 01:31:53 CEST 2022 - The aliscore output appears to be empty. Check results for BUSCO: 208291at7742
## Thu Jul 14 01:31:56 CEST 2022 - The aliscore output file does not exist. Check results for BUSCO: 256510at7742
## Thu Jul 14 01:31:56 CEST 2022 - The aliscore output appears to be empty. Check results for BUSCO: 256510at7742
## Thu Jul 14 01:31:57 CEST 2022 - The aliscore output file does not exist. Check results for BUSCO: 118353at7742
## Thu Jul 14 01:31:57 CEST 2022 - The aliscore output appears to be empty. Check results for BUSCO: 118353at7742
## Thu Jul 14 01:31:58 CEST 2022 - The aliscore output file does not exist. Check results for BUSCO: 195437at7742
## Thu Jul 14 01:31:58 CEST 2022 - The aliscore output appears to be empty. Check results for BUSCO: 195437at7742
## Thu Jul 14 01:32:00 CEST 2022 - The aliscore output file does not exist. Check results for BUSCO: 346924at7742
## Thu Jul 14 01:32:00 CEST 2022 - The aliscore output appears to be empty. Check results for BUSCO: 346924at7742
## Thu Jul 14 01:32:00 CEST 2022 - The aliscore output file does not exist. Check results for BUSCO: 174148at7742
## Thu Jul 14 01:32:00 CEST 2022 - The aliscore output appears to be empty. Check results for BUSCO: 174148at7742
## Thu Jul 14 01:32:01 CEST 2022 - The aliscore output file does not exist. Check results for BUSCO: 155235at7742
## Thu Jul 14 01:32:01 CEST 2022 - The aliscore output appears to be empty. Check results for BUSCO: 155235at7742
## Thu Jul 14 01:32:02 CEST 2022 - The aliscore output file does not exist. Check results for BUSCO: 128255at7742
## Thu Jul 14 01:32:02 CEST 2022 - The aliscore output appears to be empty. Check results for BUSCO: 128255at7742
## Thu Jul 14 01:32:02 CEST 2022 - The aliscore output file does not exist. Check results for BUSCO: 201876at7742
## Thu Jul 14 01:32:02 CEST 2022 - The aliscore output appears to be empty. Check results for BUSCO: 201876at7742
## Thu Jul 14 01:32:05 CEST 2022 - The aliscore output file does not exist. Check results for BUSCO: 288402at7742
## Thu Jul 14 01:32:05 CEST 2022 - The aliscore output appears to be empty. Check results for BUSCO: 288402at7742
## Thu Jul 14 01:32:06 CEST 2022 - The aliscore output file does not exist. Check results for BUSCO: 138339at7742
## Thu Jul 14 01:32:06 CEST 2022 - The aliscore output appears to be empty. Check results for BUSCO: 138339at7742
## Thu Jul 14 01:32:07 CEST 2022 - The aliscore output file does not exist. Check results for BUSCO: 204010at7742
## Thu Jul 14 01:32:07 CEST 2022 - The aliscore output appears to be empty. Check results for BUSCO: 204010at7742
## Thu Jul 14 01:32:08 CEST 2022 - The aliscore output file does not exist. Check results for BUSCO: 200462at7742
## Thu Jul 14 01:32:08 CEST 2022 - The aliscore output appears to be empty. Check results for BUSCO: 200462at7742
## Thu Jul 14 01:32:08 CEST 2022 - The aliscore output file does not exist. Check results for BUSCO: 93035at7742
## Thu Jul 14 01:32:08 CEST 2022 - The aliscore output appears to be empty. Check results for BUSCO: 93035at7742
## Thu Jul 14 01:32:11 CEST 2022 - The aliscore output file does not exist. Check results for BUSCO: 139053at7742
## Thu Jul 14 01:32:11 CEST 2022 - The aliscore output appears to be empty. Check results for BUSCO: 139053at7742
## Thu Jul 14 01:32:11 CEST 2022 - The aliscore output file does not exist. Check results for BUSCO: 122495at7742
## Thu Jul 14 01:32:11 CEST 2022 - The aliscore output appears to be empty. Check results for BUSCO: 122495at7742
## Thu Jul 14 01:32:12 CEST 2022 - The aliscore output file does not exist. Check results for BUSCO: 174148at7742
## Thu Jul 14 01:32:12 CEST 2022 - The aliscore output appears to be empty. Check results for BUSCO: 174148at7742
## Thu Jul 14 01:32:14 CEST 2022 - The aliscore output file does not exist. Check results for BUSCO: 326687at7742
## Thu Jul 14 01:32:14 CEST 2022 - The aliscore output appears to be empty. Check results for BUSCO: 326687at7742
## Thu Jul 14 01:32:14 CEST 2022 - The aliscore output file does not exist. Check results for BUSCO: 188851at7742
## Thu Jul 14 01:32:14 CEST 2022 - The aliscore output appears to be empty. Check results for BUSCO: 188851at7742
## Thu Jul 14 01:32:15 CEST 2022 - The aliscore output file does not exist. Check results for BUSCO: 203699at7742
## Thu Jul 14 01:32:16 CEST 2022 - The aliscore output appears to be empty. Check results for BUSCO: 203699at7742
## Thu Jul 14 01:32:17 CEST 2022 - The aliscore output file does not exist. Check results for BUSCO: 113992at7742
## Thu Jul 14 01:32:17 CEST 2022 - The aliscore output appears to be empty. Check results for BUSCO: 113992at7742
## Thu Jul 14 01:32:18 CEST 2022 - The aliscore output file does not exist. Check results for BUSCO: 260711at7742
## Thu Jul 14 01:32:18 CEST 2022 - The aliscore output appears to be empty. Check results for BUSCO: 260711at7742
## Thu Jul 14 01:32:19 CEST 2022 - The aliscore output file does not exist. Check results for BUSCO: 313142at7742
## Thu Jul 14 01:32:19 CEST 2022 - The aliscore output appears to be empty. Check results for BUSCO: 313142at7742
## Thu Jul 14 01:32:20 CEST 2022 - The aliscore output file does not exist. Check results for BUSCO: 357871at7742
## Thu Jul 14 01:32:20 CEST 2022 - The aliscore output appears to be empty. Check results for BUSCO: 357871at7742
## Thu Jul 14 01:32:22 CEST 2022 - The aliscore output file does not exist. Check results for BUSCO: 163878at7742
## Thu Jul 14 01:32:22 CEST 2022 - The aliscore output appears to be empty. Check results for BUSCO: 163878at7742
## Thu Jul 14 01:32:23 CEST 2022 - The aliscore output file does not exist. Check results for BUSCO: 285297at7742
## Thu Jul 14 01:32:23 CEST 2022 - The aliscore output appears to be empty. Check results for BUSCO: 285297at7742
## Thu Jul 14 01:32:23 CEST 2022 - The aliscore output file does not exist. Check results for BUSCO: 133255at7742
## Thu Jul 14 01:32:23 CEST 2022 - The aliscore output appears to be empty. Check results for BUSCO: 133255at7742
## Thu Jul 14 01:32:26 CEST 2022 - The aliscore output file does not exist. Check results for BUSCO: 216380at7742
## Thu Jul 14 01:32:26 CEST 2022 - The aliscore output appears to be empty. Check results for BUSCO: 216380at7742
## Thu Jul 14 01:32:27 CEST 2022 - The aliscore output file does not exist. Check results for BUSCO: 161315at7742
## Thu Jul 14 01:32:27 CEST 2022 - The aliscore output appears to be empty. Check results for BUSCO: 161315at7742
## Thu Jul 14 01:32:27 CEST 2022 - The aliscore output file does not exist. Check results for BUSCO: 159349at7742
## Thu Jul 14 01:32:27 CEST 2022 - The aliscore output appears to be empty. Check results for BUSCO: 159349at7742
## Thu Jul 14 01:32:28 CEST 2022 - The aliscore output file does not exist. Check results for BUSCO: 307789at7742
## Thu Jul 14 01:32:28 CEST 2022 - The aliscore output appears to be empty. Check results for BUSCO: 307789at7742
## Thu Jul 14 01:32:29 CEST 2022 - The aliscore output file does not exist. Check results for BUSCO: 108802at7742
## Thu Jul 14 01:32:29 CEST 2022 - The aliscore output appears to be empty. Check results for BUSCO: 108802at7742
## Thu Jul 14 01:32:31 CEST 2022 - The aliscore output file does not exist. Check results for BUSCO: 227436at7742
## Thu Jul 14 01:32:31 CEST 2022 - The aliscore output appears to be empty. Check results for BUSCO: 227436at7742
## Thu Jul 14 01:32:32 CEST 2022 - The aliscore output file does not exist. Check results for BUSCO: 239783at7742
## Thu Jul 14 01:32:32 CEST 2022 - The aliscore output appears to be empty. Check results for BUSCO: 239783at7742
## Thu Jul 14 01:32:32 CEST 2022 - The aliscore output file does not exist. Check results for BUSCO: 353522at7742
## Thu Jul 14 01:32:32 CEST 2022 - The aliscore output appears to be empty. Check results for BUSCO: 353522at7742
## Thu Jul 14 01:32:33 CEST 2022 - The aliscore output file does not exist. Check results for BUSCO: 209714at7742
## Thu Jul 14 01:32:33 CEST 2022 - The aliscore output appears to be empty. Check results for BUSCO: 209714at7742
## Thu Jul 14 01:32:35 CEST 2022 - The aliscore output file does not exist. Check results for BUSCO: 275875at7742
## Thu Jul 14 01:32:35 CEST 2022 - The aliscore output appears to be empty. Check results for BUSCO: 275875at7742
## Thu Jul 14 01:32:37 CEST 2022 - The aliscore output file does not exist. Check results for BUSCO: 171530at7742
## Thu Jul 14 01:32:37 CEST 2022 - The aliscore output appears to be empty. Check results for BUSCO: 171530at7742
## Thu Jul 14 01:32:37 CEST 2022 - The aliscore output file does not exist. Check results for BUSCO: 239783at7742
## Thu Jul 14 01:32:37 CEST 2022 - The aliscore output appears to be empty. Check results for BUSCO: 239783at7742
## Thu Jul 14 01:32:38 CEST 2022 - The aliscore output file does not exist. Check results for BUSCO: 188869at7742
## Thu Jul 14 01:32:38 CEST 2022 - The aliscore output appears to be empty. Check results for BUSCO: 188869at7742
## Thu Jul 14 01:32:40 CEST 2022 - The aliscore output file does not exist. Check results for BUSCO: 204261at7742
## Thu Jul 14 01:32:40 CEST 2022 - The aliscore output appears to be empty. Check results for BUSCO: 204261at7742
## Thu Jul 14 01:32:42 CEST 2022 - The aliscore output file does not exist. Check results for BUSCO: 118406at7742
## Thu Jul 14 01:32:42 CEST 2022 - The aliscore output appears to be empty. Check results for BUSCO: 118406at7742
## Thu Jul 14 01:32:43 CEST 2022 - The aliscore output file does not exist. Check results for BUSCO: 174872at7742
## Thu Jul 14 01:32:43 CEST 2022 - The aliscore output appears to be empty. Check results for BUSCO: 174872at7742
## Thu Jul 14 01:32:43 CEST 2022 - The aliscore output file does not exist. Check results for BUSCO: 336679at7742
## Thu Jul 14 01:32:43 CEST 2022 - The aliscore output appears to be empty. Check results for BUSCO: 336679at7742
## Thu Jul 14 01:32:45 CEST 2022 - The aliscore output file does not exist. Check results for BUSCO: 133255at7742
## Thu Jul 14 01:32:45 CEST 2022 - The aliscore output appears to be empty. Check results for BUSCO: 133255at7742
## Thu Jul 14 01:32:45 CEST 2022 - The aliscore output file does not exist. Check results for BUSCO: 254818at7742
## Thu Jul 14 01:32:46 CEST 2022 - The aliscore output appears to be empty. Check results for BUSCO: 254818at7742
## Thu Jul 14 01:32:47 CEST 2022 - The aliscore output file does not exist. Check results for BUSCO: 184701at7742
## Thu Jul 14 01:32:47 CEST 2022 - The aliscore output appears to be empty. Check results for BUSCO: 184701at7742
## Thu Jul 14 01:32:49 CEST 2022 - The aliscore output file does not exist. Check results for BUSCO: 158142at7742
## Thu Jul 14 01:32:49 CEST 2022 - The aliscore output appears to be empty. Check results for BUSCO: 158142at7742
## Thu Jul 14 01:32:49 CEST 2022 - The aliscore output file does not exist. Check results for BUSCO: 129835at7742
## Thu Jul 14 01:32:49 CEST 2022 - The aliscore output appears to be empty. Check results for BUSCO: 129835at7742
## Thu Jul 14 01:32:51 CEST 2022 - The aliscore output file does not exist. Check results for BUSCO: 209425at7742
## Thu Jul 14 01:32:51 CEST 2022 - The aliscore output appears to be empty. Check results for BUSCO: 209425at7742
## Thu Jul 14 01:32:52 CEST 2022 - The aliscore output file does not exist. Check results for BUSCO: 150456at7742
## Thu Jul 14 01:32:52 CEST 2022 - The aliscore output appears to be empty. Check results for BUSCO: 150456at7742
## Thu Jul 14 01:32:53 CEST 2022 - The aliscore output file does not exist. Check results for BUSCO: 78340at7742
## Thu Jul 14 01:32:54 CEST 2022 - The aliscore output appears to be empty. Check results for BUSCO: 78340at7742
## Thu Jul 14 01:32:54 CEST 2022 - The aliscore output file does not exist. Check results for BUSCO: 205318at7742
## Thu Jul 14 01:32:54 CEST 2022 - The aliscore output file does not exist. Check results for BUSCO: 78340at7742
## Thu Jul 14 01:32:54 CEST 2022 - The aliscore output appears to be empty. Check results for BUSCO: 205318at7742
## Thu Jul 14 01:32:54 CEST 2022 - The aliscore output appears to be empty. Check results for BUSCO: 78340at7742
## Thu Jul 14 01:32:56 CEST 2022 - The aliscore output file does not exist. Check results for BUSCO: 124354at7742
## Thu Jul 14 01:32:57 CEST 2022 - The aliscore output appears to be empty. Check results for BUSCO: 124354at7742
## Thu Jul 14 01:32:57 CEST 2022 - The aliscore output file does not exist. Check results for BUSCO: 244702at7742
## Thu Jul 14 01:32:57 CEST 2022 - The aliscore output appears to be empty. Check results for BUSCO: 244702at7742
## Thu Jul 14 01:32:59 CEST 2022 - The aliscore output file does not exist. Check results for BUSCO: 287723at7742
## Thu Jul 14 01:32:59 CEST 2022 - The aliscore output appears to be empty. Check results for BUSCO: 287723at7742
## Thu Jul 14 01:33:00 CEST 2022 - The aliscore output file does not exist. Check results for BUSCO: 360487at7742
## Thu Jul 14 01:33:00 CEST 2022 - The aliscore output appears to be empty. Check results for BUSCO: 360487at7742
## Thu Jul 14 01:33:00 CEST 2022 - The aliscore output file does not exist. Check results for BUSCO: 329437at7742
## Thu Jul 14 01:33:00 CEST 2022 - The aliscore output appears to be empty. Check results for BUSCO: 329437at7742
## Thu Jul 14 01:33:03 CEST 2022 - The aliscore output file does not exist. Check results for BUSCO: 131994at7742
## Thu Jul 14 01:33:03 CEST 2022 - The aliscore output appears to be empty. Check results for BUSCO: 131994at7742
## Thu Jul 14 01:33:03 CEST 2022 - The aliscore output file does not exist. Check results for BUSCO: 139250at7742
## Thu Jul 14 01:33:04 CEST 2022 - The aliscore output appears to be empty. Check results for BUSCO: 139250at7742
## Thu Jul 14 01:33:06 CEST 2022 - The aliscore output file does not exist. Check results for BUSCO: 139250at7742
## Thu Jul 14 01:33:06 CEST 2022 - The aliscore output appears to be empty. Check results for BUSCO: 139250at7742
## Thu Jul 14 01:33:07 CEST 2022 - The aliscore output file does not exist. Check results for BUSCO: 163142at7742
## Thu Jul 14 01:33:07 CEST 2022 - The aliscore output appears to be empty. Check results for BUSCO: 163142at7742
## Thu Jul 14 01:33:08 CEST 2022 - The aliscore output file does not exist. Check results for BUSCO: 176317at7742
## Thu Jul 14 01:33:08 CEST 2022 - The aliscore output appears to be empty. Check results for BUSCO: 176317at7742
## Thu Jul 14 01:33:09 CEST 2022 - The aliscore output file does not exist. Check results for BUSCO: 175700at7742
## Thu Jul 14 01:33:09 CEST 2022 - The aliscore output appears to be empty. Check results for BUSCO: 175700at7742
## Thu Jul 14 01:33:10 CEST 2022 - The aliscore output file does not exist. Check results for BUSCO: 196730at7742
## Thu Jul 14 01:33:10 CEST 2022 - The aliscore output appears to be empty. Check results for BUSCO: 196730at7742
## Thu Jul 14 01:33:12 CEST 2022 - The aliscore output file does not exist. Check results for BUSCO: 288402at7742
## Thu Jul 14 01:33:12 CEST 2022 - The aliscore output appears to be empty. Check results for BUSCO: 288402at7742
## Thu Jul 14 01:33:12 CEST 2022 - The aliscore output file does not exist. Check results for BUSCO: 205394at7742
## Thu Jul 14 01:33:12 CEST 2022 - The aliscore output appears to be empty. Check results for BUSCO: 205394at7742
## Thu Jul 14 01:33:12 CEST 2022 - The aliscore output file does not exist. Check results for BUSCO: 78340at7742
## Thu Jul 14 01:33:12 CEST 2022 - The aliscore output appears to be empty. Check results for BUSCO: 78340at7742
## Thu Jul 14 01:33:14 CEST 2022 - The aliscore output file does not exist. Check results for BUSCO: 340519at7742
## Thu Jul 14 01:33:14 CEST 2022 - The aliscore output appears to be empty. Check results for BUSCO: 340519at7742
## Thu Jul 14 01:33:15 CEST 2022 - The aliscore output file does not exist. Check results for BUSCO: 330064at7742
## Thu Jul 14 01:33:15 CEST 2022 - The aliscore output appears to be empty. Check results for BUSCO: 330064at7742
## Thu Jul 14 01:33:16 CEST 2022 - The aliscore output file does not exist. Check results for BUSCO: 174148at7742
## Thu Jul 14 01:33:16 CEST 2022 - The aliscore output appears to be empty. Check results for BUSCO: 174148at7742
## Thu Jul 14 01:33:17 CEST 2022 - The aliscore output file does not exist. Check results for BUSCO: 205394at7742
## Thu Jul 14 01:33:17 CEST 2022 - The aliscore output appears to be empty. Check results for BUSCO: 205394at7742
## Thu Jul 14 01:33:18 CEST 2022 - The aliscore output file does not exist. Check results for BUSCO: 163177at7742
## Thu Jul 14 01:33:18 CEST 2022 - The aliscore output appears to be empty. Check results for BUSCO: 163177at7742
## Thu Jul 14 01:33:19 CEST 2022 - The aliscore output file does not exist. Check results for BUSCO: 182481at7742
## Thu Jul 14 01:33:19 CEST 2022 - The aliscore output appears to be empty. Check results for BUSCO: 182481at7742
## Thu Jul 14 01:33:21 CEST 2022 - The aliscore output file does not exist. Check results for BUSCO: 434390at7742
## Thu Jul 14 01:33:21 CEST 2022 - The aliscore output file does not exist. Check results for BUSCO: 142719at7742
## Thu Jul 14 01:33:21 CEST 2022 - The aliscore output appears to be empty. Check results for BUSCO: 434390at7742
## Thu Jul 14 01:33:21 CEST 2022 - The aliscore output appears to be empty. Check results for BUSCO: 142719at7742
## Thu Jul 14 01:33:23 CEST 2022 - The aliscore output file does not exist. Check results for BUSCO: 329437at7742
## Thu Jul 14 01:33:23 CEST 2022 - The aliscore output appears to be empty. Check results for BUSCO: 329437at7742
## Thu Jul 14 01:33:24 CEST 2022 - The aliscore output file does not exist. Check results for BUSCO: 241953at7742
## Thu Jul 14 01:33:24 CEST 2022 - The aliscore output appears to be empty. Check results for BUSCO: 241953at7742
## Thu Jul 14 01:33:24 CEST 2022 - The aliscore output file does not exist. Check results for BUSCO: 243851at7742
## Thu Jul 14 01:33:24 CEST 2022 - The aliscore output appears to be empty. Check results for BUSCO: 243851at7742
## Thu Jul 14 01:33:25 CEST 2022 - The aliscore output file does not exist. Check results for BUSCO: 243851at7742
## Thu Jul 14 01:33:25 CEST 2022 - The aliscore output appears to be empty. Check results for BUSCO: 243851at7742
## Thu Jul 14 01:33:28 CEST 2022 - The aliscore output file does not exist. Check results for BUSCO: 224537at7742
## Thu Jul 14 01:33:28 CEST 2022 - The aliscore output appears to be empty. Check results for BUSCO: 224537at7742
## Thu Jul 14 01:33:29 CEST 2022 - The aliscore output file does not exist. Check results for BUSCO: 192665at7742
## Thu Jul 14 01:33:29 CEST 2022 - The aliscore output appears to be empty. Check results for BUSCO: 192665at7742
## Thu Jul 14 01:33:30 CEST 2022 - The aliscore output file does not exist. Check results for BUSCO: 118460at7742
## Thu Jul 14 01:33:30 CEST 2022 - The aliscore output appears to be empty. Check results for BUSCO: 118460at7742
## Thu Jul 14 01:33:32 CEST 2022 - The aliscore output file does not exist. Check results for BUSCO: 155795at7742
## Thu Jul 14 01:33:32 CEST 2022 - The aliscore output file does not exist. Check results for BUSCO: 128255at7742
## Thu Jul 14 01:33:32 CEST 2022 - The aliscore output appears to be empty. Check results for BUSCO: 155795at7742
## Thu Jul 14 01:33:32 CEST 2022 - The aliscore output appears to be empty. Check results for BUSCO: 128255at7742
## Thu Jul 14 01:33:34 CEST 2022 - The aliscore output file does not exist. Check results for BUSCO: 115533at7742
## Thu Jul 14 01:33:34 CEST 2022 - The aliscore output appears to be empty. Check results for BUSCO: 115533at7742
## Thu Jul 14 01:33:34 CEST 2022 - The aliscore output file does not exist. Check results for BUSCO: 170944at7742
## Thu Jul 14 01:33:34 CEST 2022 - The aliscore output appears to be empty. Check results for BUSCO: 170944at7742
## Thu Jul 14 01:33:35 CEST 2022 - The aliscore output file does not exist. Check results for BUSCO: 129835at7742
## Thu Jul 14 01:33:35 CEST 2022 - The aliscore output appears to be empty. Check results for BUSCO: 129835at7742
## Thu Jul 14 01:33:36 CEST 2022 - The aliscore output file does not exist. Check results for BUSCO: 248693at7742
## Thu Jul 14 01:33:36 CEST 2022 - The aliscore output appears to be empty. Check results for BUSCO: 248693at7742
## Thu Jul 14 01:33:38 CEST 2022 - The aliscore output file does not exist. Check results for BUSCO: 227816at7742
## Thu Jul 14 01:33:38 CEST 2022 - The aliscore output appears to be empty. Check results for BUSCO: 227816at7742
## Thu Jul 14 01:33:40 CEST 2022 - The aliscore output file does not exist. Check results for BUSCO: 208949at7742
## Thu Jul 14 01:33:40 CEST 2022 - The aliscore output appears to be empty. Check results for BUSCO: 208949at7742
## Thu Jul 14 01:33:40 CEST 2022 - The aliscore output file does not exist. Check results for BUSCO: 133642at7742
## Thu Jul 14 01:33:41 CEST 2022 - The aliscore output appears to be empty. Check results for BUSCO: 133642at7742
## Thu Jul 14 01:33:42 CEST 2022 - The aliscore output file does not exist. Check results for BUSCO: 280833at7742
## Thu Jul 14 01:33:42 CEST 2022 - The aliscore output appears to be empty. Check results for BUSCO: 280833at7742
## Thu Jul 14 01:33:43 CEST 2022 - The aliscore output file does not exist. Check results for BUSCO: 92374at7742
## Thu Jul 14 01:33:43 CEST 2022 - The aliscore output appears to be empty. Check results for BUSCO: 92374at7742
## Thu Jul 14 01:33:44 CEST 2022 - The aliscore output file does not exist. Check results for BUSCO: 206547at7742
## Thu Jul 14 01:33:44 CEST 2022 - The aliscore output appears to be empty. Check results for BUSCO: 206547at7742
## Thu Jul 14 01:33:44 CEST 2022 - The aliscore output file does not exist. Check results for BUSCO: 175700at7742
## Thu Jul 14 01:33:44 CEST 2022 - The aliscore output appears to be empty. Check results for BUSCO: 175700at7742
## Thu Jul 14 01:33:46 CEST 2022 - The aliscore output file does not exist. Check results for BUSCO: 218284at7742
## Thu Jul 14 01:33:46 CEST 2022 - The aliscore output appears to be empty. Check results for BUSCO: 218284at7742
## Thu Jul 14 01:33:47 CEST 2022 - The aliscore output file does not exist. Check results for BUSCO: 124354at7742
## Thu Jul 14 01:33:47 CEST 2022 - The aliscore output appears to be empty. Check results for BUSCO: 124354at7742
## Thu Jul 14 01:33:47 CEST 2022 - The aliscore output file does not exist. Check results for BUSCO: 161484at7742
## Thu Jul 14 01:33:48 CEST 2022 - The aliscore output appears to be empty. Check results for BUSCO: 161484at7742
## Thu Jul 14 01:33:49 CEST 2022 - The aliscore output file does not exist. Check results for BUSCO: 187916at7742
## Thu Jul 14 01:33:49 CEST 2022 - The aliscore output appears to be empty. Check results for BUSCO: 187916at7742
## Thu Jul 14 01:33:50 CEST 2022 - The aliscore output file does not exist. Check results for BUSCO: 249399at7742
## Thu Jul 14 01:33:50 CEST 2022 - The aliscore output appears to be empty. Check results for BUSCO: 249399at7742
## Thu Jul 14 01:33:53 CEST 2022 - The aliscore output file does not exist. Check results for BUSCO: 260711at7742
## Thu Jul 14 01:33:53 CEST 2022 - The aliscore output appears to be empty. Check results for BUSCO: 260711at7742
## Thu Jul 14 01:33:53 CEST 2022 - The aliscore output file does not exist. Check results for BUSCO: 321010at7742
## Thu Jul 14 01:33:53 CEST 2022 - The aliscore output appears to be empty. Check results for BUSCO: 321010at7742
## Thu Jul 14 01:33:55 CEST 2022 - The aliscore output file does not exist. Check results for BUSCO: 248693at7742
## Thu Jul 14 01:33:55 CEST 2022 - The aliscore output appears to be empty. Check results for BUSCO: 248693at7742
## Thu Jul 14 01:33:55 CEST 2022 - The aliscore output file does not exist. Check results for BUSCO: 223898at7742
## Thu Jul 14 01:33:55 CEST 2022 - The aliscore output appears to be empty. Check results for BUSCO: 223898at7742
## Thu Jul 14 01:33:56 CEST 2022 - The aliscore output file does not exist. Check results for BUSCO: 193525at7742
## Thu Jul 14 01:33:56 CEST 2022 - The aliscore output appears to be empty. Check results for BUSCO: 193525at7742
## Thu Jul 14 01:33:59 CEST 2022 - The aliscore output file does not exist. Check results for BUSCO: 241953at7742
## Thu Jul 14 01:33:59 CEST 2022 - The aliscore output appears to be empty. Check results for BUSCO: 241953at7742
## Thu Jul 14 01:33:59 CEST 2022 - The aliscore output file does not exist. Check results for BUSCO: 210107at7742
## Thu Jul 14 01:33:59 CEST 2022 - The aliscore output appears to be empty. Check results for BUSCO: 210107at7742
## Thu Jul 14 01:34:00 CEST 2022 - The aliscore output file does not exist. Check results for BUSCO: 118365at7742
## Thu Jul 14 01:34:00 CEST 2022 - The aliscore output appears to be empty. Check results for BUSCO: 118365at7742
## Thu Jul 14 01:34:01 CEST 2022 - The aliscore output file does not exist. Check results for BUSCO: 166657at7742
## Thu Jul 14 01:34:02 CEST 2022 - The aliscore output appears to be empty. Check results for BUSCO: 166657at7742
## Thu Jul 14 01:34:02 CEST 2022 - The aliscore output file does not exist. Check results for BUSCO: 255610at7742
## Thu Jul 14 01:34:02 CEST 2022 - The aliscore output appears to be empty. Check results for BUSCO: 255610at7742
## Thu Jul 14 01:34:04 CEST 2022 - The aliscore output file does not exist. Check results for BUSCO: 305417at7742
## Thu Jul 14 01:34:04 CEST 2022 - The aliscore output appears to be empty. Check results for BUSCO: 305417at7742
## Thu Jul 14 01:34:05 CEST 2022 - The aliscore output file does not exist. Check results for BUSCO: 161315at7742
## Thu Jul 14 01:34:05 CEST 2022 - The aliscore output appears to be empty. Check results for BUSCO: 161315at7742
## Thu Jul 14 01:34:06 CEST 2022 - The aliscore output file does not exist. Check results for BUSCO: 338075at7742
## Thu Jul 14 01:34:06 CEST 2022 - The aliscore output appears to be empty. Check results for BUSCO: 338075at7742
## Thu Jul 14 01:34:06 CEST 2022 - The aliscore output file does not exist. Check results for BUSCO: 171737at7742
## Thu Jul 14 01:34:06 CEST 2022 - The aliscore output appears to be empty. Check results for BUSCO: 171737at7742
## Thu Jul 14 01:34:08 CEST 2022 - The aliscore output file does not exist. Check results for BUSCO: 338746at7742
## Thu Jul 14 01:34:08 CEST 2022 - The aliscore output appears to be empty. Check results for BUSCO: 338746at7742
## Thu Jul 14 01:34:08 CEST 2022 - The aliscore output file does not exist. Check results for BUSCO: 201876at7742
## Thu Jul 14 01:34:08 CEST 2022 - The aliscore output appears to be empty. Check results for BUSCO: 201876at7742
## Thu Jul 14 01:34:09 CEST 2022 - The aliscore output file does not exist. Check results for BUSCO: 144578at7742
## Thu Jul 14 01:34:09 CEST 2022 - The aliscore output appears to be empty. Check results for BUSCO: 144578at7742
## Thu Jul 14 01:34:10 CEST 2022 - The aliscore output file does not exist. Check results for BUSCO: 333967at7742
## Thu Jul 14 01:34:10 CEST 2022 - The aliscore output appears to be empty. Check results for BUSCO: 333967at7742
## Thu Jul 14 01:34:12 CEST 2022 - The aliscore output file does not exist. Check results for BUSCO: 224472at7742
## Thu Jul 14 01:34:12 CEST 2022 - The aliscore output appears to be empty. Check results for BUSCO: 224472at7742
## Thu Jul 14 01:34:13 CEST 2022 - The aliscore output file does not exist. Check results for BUSCO: 255610at7742
## Thu Jul 14 01:34:13 CEST 2022 - The aliscore output appears to be empty. Check results for BUSCO: 255610at7742
## Thu Jul 14 01:34:14 CEST 2022 - The aliscore output file does not exist. Check results for BUSCO: 367197at7742
## Thu Jul 14 01:34:14 CEST 2022 - The aliscore output appears to be empty. Check results for BUSCO: 367197at7742
## Thu Jul 14 01:34:16 CEST 2022 - The aliscore output file does not exist. Check results for BUSCO: 228275at7742
## Thu Jul 14 01:34:16 CEST 2022 - The aliscore output appears to be empty. Check results for BUSCO: 228275at7742
## Thu Jul 14 01:34:17 CEST 2022 - The aliscore output file does not exist. Check results for BUSCO: 165456at7742
## Thu Jul 14 01:34:17 CEST 2022 - The aliscore output appears to be empty. Check results for BUSCO: 165456at7742
## Thu Jul 14 01:34:18 CEST 2022 - The aliscore output file does not exist. Check results for BUSCO: 192350at7742
## Thu Jul 14 01:34:18 CEST 2022 - The aliscore output appears to be empty. Check results for BUSCO: 192350at7742
## Thu Jul 14 01:34:18 CEST 2022 - The aliscore output file does not exist. Check results for BUSCO: 225242at7742
## Thu Jul 14 01:34:18 CEST 2022 - The aliscore output appears to be empty. Check results for BUSCO: 225242at7742
## Thu Jul 14 01:34:21 CEST 2022 - The aliscore output file does not exist. Check results for BUSCO: 338746at7742
## Thu Jul 14 01:34:21 CEST 2022 - The aliscore output appears to be empty. Check results for BUSCO: 338746at7742
## Thu Jul 14 01:34:22 CEST 2022 - The aliscore output file does not exist. Check results for BUSCO: 408649at7742
## Thu Jul 14 01:34:22 CEST 2022 - The aliscore output appears to be empty. Check results for BUSCO: 408649at7742
## Thu Jul 14 01:34:23 CEST 2022 - The aliscore output file does not exist. Check results for BUSCO: 208300at7742
## Thu Jul 14 01:34:23 CEST 2022 - The aliscore output appears to be empty. Check results for BUSCO: 208300at7742
## Thu Jul 14 01:34:23 CEST 2022 - The aliscore output file does not exist. Check results for BUSCO: 275720at7742
## Thu Jul 14 01:34:23 CEST 2022 - The aliscore output appears to be empty. Check results for BUSCO: 275720at7742
## Thu Jul 14 01:34:24 CEST 2022 - The aliscore output file does not exist. Check results for BUSCO: 209951at7742
## Thu Jul 14 01:34:24 CEST 2022 - The aliscore output appears to be empty. Check results for BUSCO: 209951at7742
## Thu Jul 14 01:34:25 CEST 2022 - The aliscore output file does not exist. Check results for BUSCO: 239682at7742
## Thu Jul 14 01:34:25 CEST 2022 - The aliscore output appears to be empty. Check results for BUSCO: 239682at7742
## Thu Jul 14 01:34:26 CEST 2022 - The aliscore output file does not exist. Check results for BUSCO: 225148at7742
## Thu Jul 14 01:34:26 CEST 2022 - The aliscore output appears to be empty. Check results for BUSCO: 225148at7742
## Thu Jul 14 01:34:29 CEST 2022 - The aliscore output file does not exist. Check results for BUSCO: 193657at7742
## Thu Jul 14 01:34:29 CEST 2022 - The aliscore output appears to be empty. Check results for BUSCO: 193657at7742
## Thu Jul 14 01:34:29 CEST 2022 - The aliscore output file does not exist. Check results for BUSCO: 338075at7742
## Thu Jul 14 01:34:29 CEST 2022 - The aliscore output appears to be empty. Check results for BUSCO: 338075at7742
## Thu Jul 14 01:34:31 CEST 2022 - The aliscore output file does not exist. Check results for BUSCO: 264510at7742
## Thu Jul 14 01:34:31 CEST 2022 - The aliscore output file does not exist. Check results for BUSCO: 178680at7742
## Thu Jul 14 01:34:31 CEST 2022 - The aliscore output appears to be empty. Check results for BUSCO: 264510at7742
## Thu Jul 14 01:34:31 CEST 2022 - The aliscore output appears to be empty. Check results for BUSCO: 178680at7742
## Thu Jul 14 01:34:32 CEST 2022 - The aliscore output file does not exist. Check results for BUSCO: 139241at7742
## Thu Jul 14 01:34:32 CEST 2022 - The aliscore output appears to be empty. Check results for BUSCO: 139241at7742
## Thu Jul 14 01:34:33 CEST 2022 - The aliscore output file does not exist. Check results for BUSCO: 221393at7742
## Thu Jul 14 01:34:34 CEST 2022 - The aliscore output appears to be empty. Check results for BUSCO: 221393at7742
## Thu Jul 14 01:34:34 CEST 2022 - The aliscore output file does not exist. Check results for BUSCO: 209951at7742
## Thu Jul 14 01:34:34 CEST 2022 - The aliscore output appears to be empty. Check results for BUSCO: 209951at7742
## Thu Jul 14 01:34:36 CEST 2022 - The aliscore output file does not exist. Check results for BUSCO: 270710at7742
## Thu Jul 14 01:34:36 CEST 2022 - The aliscore output appears to be empty. Check results for BUSCO: 270710at7742
## Thu Jul 14 01:34:37 CEST 2022 - The aliscore output file does not exist. Check results for BUSCO: 272002at7742
## Thu Jul 14 01:34:37 CEST 2022 - The aliscore output appears to be empty. Check results for BUSCO: 272002at7742
## Thu Jul 14 01:34:38 CEST 2022 - The aliscore output file does not exist. Check results for BUSCO: 118406at7742
## Thu Jul 14 01:34:38 CEST 2022 - The aliscore output appears to be empty. Check results for BUSCO: 118406at7742
## Thu Jul 14 01:34:40 CEST 2022 - The aliscore output file does not exist. Check results for BUSCO: 256348at7742
## Thu Jul 14 01:34:40 CEST 2022 - The aliscore output appears to be empty. Check results for BUSCO: 256348at7742
## Thu Jul 14 01:34:40 CEST 2022 - The aliscore output file does not exist. Check results for BUSCO: 312347at7742
## Thu Jul 14 01:34:40 CEST 2022 - The aliscore output appears to be empty. Check results for BUSCO: 312347at7742
## Thu Jul 14 01:34:40 CEST 2022 - The aliscore output file does not exist. Check results for BUSCO: 251667at7742
## Thu Jul 14 01:34:41 CEST 2022 - The aliscore output appears to be empty. Check results for BUSCO: 251667at7742
## Thu Jul 14 01:34:41 CEST 2022 - The aliscore output file does not exist. Check results for BUSCO: 338746at7742
## Thu Jul 14 01:34:42 CEST 2022 - The aliscore output appears to be empty. Check results for BUSCO: 338746at7742
## Thu Jul 14 01:34:44 CEST 2022 - The aliscore output file does not exist. Check results for BUSCO: 193657at7742
## Thu Jul 14 01:34:44 CEST 2022 - The aliscore output appears to be empty. Check results for BUSCO: 193657at7742
## Thu Jul 14 01:34:46 CEST 2022 - The aliscore output file does not exist. Check results for BUSCO: 434390at7742
## Thu Jul 14 01:34:46 CEST 2022 - The aliscore output appears to be empty. Check results for BUSCO: 434390at7742
## Thu Jul 14 01:34:46 CEST 2022 - The aliscore output file does not exist. Check results for BUSCO: 257616at7742
## Thu Jul 14 01:34:46 CEST 2022 - The aliscore output appears to be empty. Check results for BUSCO: 257616at7742
## Thu Jul 14 01:34:47 CEST 2022 - The aliscore output file does not exist. Check results for BUSCO: 142719at7742
## Thu Jul 14 01:34:47 CEST 2022 - The aliscore output appears to be empty. Check results for BUSCO: 142719at7742
## Thu Jul 14 01:34:48 CEST 2022 - The aliscore output file does not exist. Check results for BUSCO: 207031at7742
## Thu Jul 14 01:34:48 CEST 2022 - The aliscore output appears to be empty. Check results for BUSCO: 207031at7742
## Thu Jul 14 01:34:51 CEST 2022 - The aliscore output file does not exist. Check results for BUSCO: 203699at7742
## Thu Jul 14 01:34:51 CEST 2022 - The aliscore output file does not exist. Check results for BUSCO: 94127at7742
## Thu Jul 14 01:34:51 CEST 2022 - The aliscore output appears to be empty. Check results for BUSCO: 203699at7742
## Thu Jul 14 01:34:51 CEST 2022 - The aliscore output appears to be empty. Check results for BUSCO: 94127at7742
## Thu Jul 14 01:34:51 CEST 2022 - The aliscore output file does not exist. Check results for BUSCO: 327369at7742
## Thu Jul 14 01:34:51 CEST 2022 - The aliscore output appears to be empty. Check results for BUSCO: 327369at7742
## Thu Jul 14 01:34:53 CEST 2022 - The aliscore output file does not exist. Check results for BUSCO: 138339at7742
## Thu Jul 14 01:34:53 CEST 2022 - The aliscore output appears to be empty. Check results for BUSCO: 138339at7742
## Thu Jul 14 01:34:55 CEST 2022 - The aliscore output file does not exist. Check results for BUSCO: 113550at7742
## Thu Jul 14 01:34:55 CEST 2022 - The aliscore output appears to be empty. Check results for BUSCO: 113550at7742
## Thu Jul 14 01:34:56 CEST 2022 - The aliscore output file does not exist. Check results for BUSCO: 224078at7742
## Thu Jul 14 01:34:56 CEST 2022 - The aliscore output appears to be empty. Check results for BUSCO: 224078at7742
## Thu Jul 14 01:34:56 CEST 2022 - The aliscore output file does not exist. Check results for BUSCO: 128068at7742
## Thu Jul 14 01:34:56 CEST 2022 - The aliscore output appears to be empty. Check results for BUSCO: 128068at7742
## Thu Jul 14 01:34:57 CEST 2022 - The aliscore output file does not exist. Check results for BUSCO: 224472at7742
## Thu Jul 14 01:34:57 CEST 2022 - The aliscore output appears to be empty. Check results for BUSCO: 224472at7742
## Thu Jul 14 01:34:59 CEST 2022 - The aliscore output file does not exist. Check results for BUSCO: 136144at7742
## Thu Jul 14 01:34:59 CEST 2022 - The aliscore output appears to be empty. Check results for BUSCO: 136144at7742
## Thu Jul 14 01:35:00 CEST 2022 - The aliscore output file does not exist. Check results for BUSCO: 241220at7742
## Thu Jul 14 01:35:00 CEST 2022 - The aliscore output appears to be empty. Check results for BUSCO: 241220at7742
## Thu Jul 14 01:35:01 CEST 2022 - The aliscore output file does not exist. Check results for BUSCO: 164982at7742
## Thu Jul 14 01:35:01 CEST 2022 - The aliscore output appears to be empty. Check results for BUSCO: 164982at7742
## Thu Jul 14 01:35:02 CEST 2022 - The aliscore output file does not exist. Check results for BUSCO: 193657at7742
## Thu Jul 14 01:35:02 CEST 2022 - The aliscore output appears to be empty. Check results for BUSCO: 193657at7742
## Thu Jul 14 01:35:03 CEST 2022 - The aliscore output file does not exist. Check results for BUSCO: 318035at7742
## Thu Jul 14 01:35:03 CEST 2022 - The aliscore output appears to be empty. Check results for BUSCO: 318035at7742
## Thu Jul 14 01:35:04 CEST 2022 - The aliscore output file does not exist. Check results for BUSCO: 136144at7742
## Thu Jul 14 01:35:04 CEST 2022 - The aliscore output appears to be empty. Check results for BUSCO: 136144at7742
## Thu Jul 14 01:35:05 CEST 2022 - The aliscore output file does not exist. Check results for BUSCO: 135786at7742
## Thu Jul 14 01:35:05 CEST 2022 - The aliscore output appears to be empty. Check results for BUSCO: 135786at7742
## Thu Jul 14 01:35:06 CEST 2022 - The aliscore output file does not exist. Check results for BUSCO: 160371at7742
## Thu Jul 14 01:35:06 CEST 2022 - The aliscore output appears to be empty. Check results for BUSCO: 160371at7742
## Thu Jul 14 01:35:06 CEST 2022 - The aliscore output file does not exist. Check results for BUSCO: 165997at7742
## Thu Jul 14 01:35:06 CEST 2022 - The aliscore output appears to be empty. Check results for BUSCO: 165997at7742
## Thu Jul 14 01:35:09 CEST 2022 - The aliscore output file does not exist. Check results for BUSCO: 213005at7742
## Thu Jul 14 01:35:09 CEST 2022 - The aliscore output appears to be empty. Check results for BUSCO: 213005at7742
## Thu Jul 14 01:35:11 CEST 2022 - The aliscore output file does not exist. Check results for BUSCO: 80137at7742
## Thu Jul 14 01:35:11 CEST 2022 - The aliscore output appears to be empty. Check results for BUSCO: 80137at7742
## Thu Jul 14 01:35:12 CEST 2022 - The aliscore output file does not exist. Check results for BUSCO: 182742at7742
## Thu Jul 14 01:35:12 CEST 2022 - The aliscore output appears to be empty. Check results for BUSCO: 182742at7742
## Thu Jul 14 01:35:12 CEST 2022 - The aliscore output file does not exist. Check results for BUSCO: 154457at7742
## Thu Jul 14 01:35:12 CEST 2022 - The aliscore output appears to be empty. Check results for BUSCO: 154457at7742
## Thu Jul 14 01:35:12 CEST 2022 - The aliscore output file does not exist. Check results for BUSCO: 192518at7742
## Thu Jul 14 01:35:12 CEST 2022 - The aliscore output appears to be empty. Check results for BUSCO: 192518at7742
## Thu Jul 14 01:35:14 CEST 2022 - The aliscore output file does not exist. Check results for BUSCO: 281807at7742
## Thu Jul 14 01:35:15 CEST 2022 - The aliscore output appears to be empty. Check results for BUSCO: 281807at7742
## Thu Jul 14 01:35:15 CEST 2022 - The aliscore output file does not exist. Check results for BUSCO: 143665at7742
## Thu Jul 14 01:35:15 CEST 2022 - The aliscore output appears to be empty. Check results for BUSCO: 143665at7742
## Thu Jul 14 01:35:16 CEST 2022 - The aliscore output file does not exist. Check results for BUSCO: 217728at7742
## Thu Jul 14 01:35:16 CEST 2022 - The aliscore output appears to be empty. Check results for BUSCO: 217728at7742
## Thu Jul 14 01:35:19 CEST 2022 - The aliscore output file does not exist. Check results for BUSCO: 389352at7742
## Thu Jul 14 01:35:19 CEST 2022 - The aliscore output appears to be empty. Check results for BUSCO: 389352at7742
## Thu Jul 14 01:35:19 CEST 2022 - The aliscore output file does not exist. Check results for BUSCO: 389667at7742
## Thu Jul 14 01:35:19 CEST 2022 - The aliscore output appears to be empty. Check results for BUSCO: 389667at7742
## Thu Jul 14 01:35:19 CEST 2022 - The aliscore output file does not exist. Check results for BUSCO: 189447at7742
## Thu Jul 14 01:35:19 CEST 2022 - The aliscore output appears to be empty. Check results for BUSCO: 189447at7742
## Thu Jul 14 01:35:21 CEST 2022 - The aliscore output file does not exist. Check results for BUSCO: 214728at7742
## Thu Jul 14 01:35:22 CEST 2022 - The aliscore output appears to be empty. Check results for BUSCO: 214728at7742
## Thu Jul 14 01:35:22 CEST 2022 - The aliscore output file does not exist. Check results for BUSCO: 296646at7742
## Thu Jul 14 01:35:22 CEST 2022 - The aliscore output appears to be empty. Check results for BUSCO: 296646at7742
## Thu Jul 14 01:35:23 CEST 2022 - The aliscore output file does not exist. Check results for BUSCO: 190526at7742
## Thu Jul 14 01:35:23 CEST 2022 - The aliscore output appears to be empty. Check results for BUSCO: 190526at7742
## Thu Jul 14 01:35:24 CEST 2022 - The aliscore output file does not exist. Check results for BUSCO: 118365at7742
## Thu Jul 14 01:35:24 CEST 2022 - The aliscore output appears to be empty. Check results for BUSCO: 118365at7742
## Thu Jul 14 01:35:25 CEST 2022 - The aliscore output file does not exist. Check results for BUSCO: 203699at7742
## Thu Jul 14 01:35:25 CEST 2022 - The aliscore output appears to be empty. Check results for BUSCO: 203699at7742
## Thu Jul 14 01:35:28 CEST 2022 - The aliscore output file does not exist. Check results for BUSCO: 273630at7742
## Thu Jul 14 01:35:28 CEST 2022 - The aliscore output file does not exist. Check results for BUSCO: 157460at7742
## Thu Jul 14 01:35:28 CEST 2022 - The aliscore output appears to be empty. Check results for BUSCO: 273630at7742
## Thu Jul 14 01:35:28 CEST 2022 - The aliscore output appears to be empty. Check results for BUSCO: 157460at7742
## Thu Jul 14 01:35:28 CEST 2022 - The aliscore output file does not exist. Check results for BUSCO: 145875at7742
## Thu Jul 14 01:35:28 CEST 2022 - The aliscore output appears to be empty. Check results for BUSCO: 145875at7742
## Thu Jul 14 01:35:28 CEST 2022 - The aliscore output file does not exist. Check results for BUSCO: 138669at7742
## Thu Jul 14 01:35:28 CEST 2022 - The aliscore output appears to be empty. Check results for BUSCO: 138669at7742
## Thu Jul 14 01:35:30 CEST 2022 - The aliscore output file does not exist. Check results for BUSCO: 206547at7742
## Thu Jul 14 01:35:30 CEST 2022 - The aliscore output appears to be empty. Check results for BUSCO: 206547at7742
## Thu Jul 14 01:35:33 CEST 2022 - The aliscore output file does not exist. Check results for BUSCO: 321359at7742
## Thu Jul 14 01:35:33 CEST 2022 - The aliscore output appears to be empty. Check results for BUSCO: 321359at7742
## Thu Jul 14 01:35:33 CEST 2022 - The aliscore output file does not exist. Check results for BUSCO: 257699at7742
## Thu Jul 14 01:35:33 CEST 2022 - The aliscore output appears to be empty. Check results for BUSCO: 257699at7742
## Thu Jul 14 01:35:35 CEST 2022 - The aliscore output file does not exist. Check results for BUSCO: 178259at7742
## Thu Jul 14 01:35:35 CEST 2022 - The aliscore output appears to be empty. Check results for BUSCO: 178259at7742
## Thu Jul 14 01:35:36 CEST 2022 - The aliscore output file does not exist. Check results for BUSCO: 145938at7742
## Thu Jul 14 01:35:36 CEST 2022 - The aliscore output appears to be empty. Check results for BUSCO: 145938at7742
## Thu Jul 14 01:35:37 CEST 2022 - The aliscore output file does not exist. Check results for BUSCO: 192142at7742
## Thu Jul 14 01:35:37 CEST 2022 - The aliscore output appears to be empty. Check results for BUSCO: 192142at7742
## Thu Jul 14 01:35:37 CEST 2022 - The aliscore output file does not exist. Check results for BUSCO: 256936at7742
## Thu Jul 14 01:35:37 CEST 2022 - The aliscore output appears to be empty. Check results for BUSCO: 256936at7742
## Thu Jul 14 01:35:40 CEST 2022 - The aliscore output file does not exist. Check results for BUSCO: 230477at7742
## Thu Jul 14 01:35:40 CEST 2022 - The aliscore output appears to be empty. Check results for BUSCO: 230477at7742
## Thu Jul 14 01:35:41 CEST 2022 - The aliscore output file does not exist. Check results for BUSCO: 118353at7742
## Thu Jul 14 01:35:41 CEST 2022 - The aliscore output appears to be empty. Check results for BUSCO: 118353at7742
## Thu Jul 14 01:35:41 CEST 2022 - The aliscore output file does not exist. Check results for BUSCO: 330064at7742
## Thu Jul 14 01:35:41 CEST 2022 - The aliscore output appears to be empty. Check results for BUSCO: 330064at7742
## Thu Jul 14 01:35:44 CEST 2022 - The aliscore output file does not exist. Check results for BUSCO: 239682at7742
## Thu Jul 14 01:35:44 CEST 2022 - The aliscore output appears to be empty. Check results for BUSCO: 239682at7742
## Thu Jul 14 01:35:46 CEST 2022 - The aliscore output file does not exist. Check results for BUSCO: 155482at7742
## Thu Jul 14 01:35:46 CEST 2022 - The aliscore output appears to be empty. Check results for BUSCO: 155482at7742
## Thu Jul 14 01:35:46 CEST 2022 - The aliscore output file does not exist. Check results for BUSCO: 266927at7742
## Thu Jul 14 01:35:46 CEST 2022 - The aliscore output appears to be empty. Check results for BUSCO: 266927at7742
## Thu Jul 14 01:35:46 CEST 2022 - The aliscore output file does not exist. Check results for BUSCO: 128068at7742
## Thu Jul 14 01:35:46 CEST 2022 - The aliscore output appears to be empty. Check results for BUSCO: 128068at7742
## Thu Jul 14 01:35:47 CEST 2022 - The aliscore output file does not exist. Check results for BUSCO: 115533at7742
## Thu Jul 14 01:35:47 CEST 2022 - The aliscore output appears to be empty. Check results for BUSCO: 115533at7742
## Thu Jul 14 01:35:47 CEST 2022 - The aliscore output file does not exist. Check results for BUSCO: 163142at7742
## Thu Jul 14 01:35:47 CEST 2022 - The aliscore output appears to be empty. Check results for BUSCO: 163142at7742
## Thu Jul 14 01:35:51 CEST 2022 - The aliscore output file does not exist. Check results for BUSCO: 230477at7742
## Thu Jul 14 01:35:51 CEST 2022 - The aliscore output appears to be empty. Check results for BUSCO: 230477at7742
## Thu Jul 14 01:35:51 CEST 2022 - The aliscore output file does not exist. Check results for BUSCO: 320126at7742
## Thu Jul 14 01:35:51 CEST 2022 - The aliscore output appears to be empty. Check results for BUSCO: 320126at7742
## Thu Jul 14 01:35:52 CEST 2022 - The aliscore output file does not exist. Check results for BUSCO: 327391at7742
## Thu Jul 14 01:35:52 CEST 2022 - The aliscore output appears to be empty. Check results for BUSCO: 327391at7742
## Thu Jul 14 01:35:53 CEST 2022 - The aliscore output file does not exist. Check results for BUSCO: 355364at7742
## Thu Jul 14 01:35:53 CEST 2022 - The aliscore output appears to be empty. Check results for BUSCO: 355364at7742
## Thu Jul 14 01:35:54 CEST 2022 - The aliscore output file does not exist. Check results for BUSCO: 142214at7742
## Thu Jul 14 01:35:54 CEST 2022 - The aliscore output appears to be empty. Check results for BUSCO: 142214at7742
## Thu Jul 14 01:35:56 CEST 2022 - The aliscore output file does not exist. Check results for BUSCO: 80137at7742
## Thu Jul 14 01:35:56 CEST 2022 - The aliscore output appears to be empty. Check results for BUSCO: 80137at7742
## Thu Jul 14 01:35:56 CEST 2022 - The aliscore output file does not exist. Check results for BUSCO: 207031at7742
## Thu Jul 14 01:35:56 CEST 2022 - The aliscore output appears to be empty. Check results for BUSCO: 207031at7742
## Thu Jul 14 01:35:59 CEST 2022 - The aliscore output file does not exist. Check results for BUSCO: 128712at7742
## Thu Jul 14 01:35:59 CEST 2022 - The aliscore output appears to be empty. Check results for BUSCO: 128712at7742
## Thu Jul 14 01:36:00 CEST 2022 - The aliscore output file does not exist. Check results for BUSCO: 163142at7742
## Thu Jul 14 01:36:00 CEST 2022 - The aliscore output appears to be empty. Check results for BUSCO: 163142at7742
## Thu Jul 14 01:36:01 CEST 2022 - The aliscore output file does not exist. Check results for BUSCO: 330054at7742
## Thu Jul 14 01:36:01 CEST 2022 - The aliscore output appears to be empty. Check results for BUSCO: 330054at7742
## Thu Jul 14 01:36:01 CEST 2022 - The aliscore output file does not exist. Check results for BUSCO: 131994at7742
## Thu Jul 14 01:36:01 CEST 2022 - The aliscore output appears to be empty. Check results for BUSCO: 131994at7742
## Thu Jul 14 01:36:02 CEST 2022 - The aliscore output file does not exist. Check results for BUSCO: 129684at7742
## Thu Jul 14 01:36:02 CEST 2022 - The aliscore output appears to be empty. Check results for BUSCO: 129684at7742
## Thu Jul 14 01:36:03 CEST 2022 - The aliscore output file does not exist. Check results for BUSCO: 267345at7742
## Thu Jul 14 01:36:03 CEST 2022 - The aliscore output appears to be empty. Check results for BUSCO: 267345at7742
## Thu Jul 14 01:36:04 CEST 2022 - The aliscore output file does not exist. Check results for BUSCO: 143665at7742
## Thu Jul 14 01:36:04 CEST 2022 - The aliscore output appears to be empty. Check results for BUSCO: 143665at7742
## Thu Jul 14 01:36:07 CEST 2022 - The aliscore output file does not exist. Check results for BUSCO: 279170at7742
## Thu Jul 14 01:36:07 CEST 2022 - The aliscore output appears to be empty. Check results for BUSCO: 279170at7742
## Thu Jul 14 01:36:10 CEST 2022 - The aliscore output file does not exist. Check results for BUSCO: 228390at7742
## Thu Jul 14 01:36:10 CEST 2022 - The aliscore output appears to be empty. Check results for BUSCO: 228390at7742
## Thu Jul 14 01:36:11 CEST 2022 - The aliscore output file does not exist. Check results for BUSCO: 200462at7742
## Thu Jul 14 01:36:11 CEST 2022 - The aliscore output appears to be empty. Check results for BUSCO: 200462at7742
## Thu Jul 14 01:36:11 CEST 2022 - The aliscore output file does not exist. Check results for BUSCO: 344933at7742
## Thu Jul 14 01:36:12 CEST 2022 - The aliscore output appears to be empty. Check results for BUSCO: 344933at7742
## Thu Jul 14 01:36:12 CEST 2022 - The aliscore output file does not exist. Check results for BUSCO: 133642at7742
## Thu Jul 14 01:36:12 CEST 2022 - The aliscore output appears to be empty. Check results for BUSCO: 133642at7742
## Thu Jul 14 01:36:12 CEST 2022 - The aliscore output file does not exist. Check results for BUSCO: 139241at7742
## Thu Jul 14 01:36:13 CEST 2022 - The aliscore output appears to be empty. Check results for BUSCO: 139241at7742
## Thu Jul 14 01:36:13 CEST 2022 - The aliscore output file does not exist. Check results for BUSCO: 98908at7742
## Thu Jul 14 01:36:13 CEST 2022 - The aliscore output appears to be empty. Check results for BUSCO: 98908at7742
## Thu Jul 14 01:36:14 CEST 2022 - The aliscore output file does not exist. Check results for BUSCO: 353522at7742
## Thu Jul 14 01:36:14 CEST 2022 - The aliscore output appears to be empty. Check results for BUSCO: 353522at7742
## Thu Jul 14 01:36:15 CEST 2022 - The aliscore output file does not exist. Check results for BUSCO: 267345at7742
## Thu Jul 14 01:36:15 CEST 2022 - The aliscore output appears to be empty. Check results for BUSCO: 267345at7742
## Thu Jul 14 01:36:17 CEST 2022 - The aliscore output file does not exist. Check results for BUSCO: 80137at7742
## Thu Jul 14 01:36:17 CEST 2022 - The aliscore output appears to be empty. Check results for BUSCO: 80137at7742
## Thu Jul 14 01:36:19 CEST 2022 - The aliscore output file does not exist. Check results for BUSCO: 281807at7742
## Thu Jul 14 01:36:19 CEST 2022 - The aliscore output file does not exist. Check results for BUSCO: 94127at7742
## Thu Jul 14 01:36:19 CEST 2022 - The aliscore output appears to be empty. Check results for BUSCO: 281807at7742
## Thu Jul 14 01:36:19 CEST 2022 - The aliscore output appears to be empty. Check results for BUSCO: 94127at7742
## Thu Jul 14 01:36:21 CEST 2022 - The aliscore output file does not exist. Check results for BUSCO: 102567at7742
## Thu Jul 14 01:36:21 CEST 2022 - The aliscore output appears to be empty. Check results for BUSCO: 102567at7742
## Thu Jul 14 01:36:21 CEST 2022 - The aliscore output file does not exist. Check results for BUSCO: 222631at7742
## Thu Jul 14 01:36:22 CEST 2022 - The aliscore output appears to be empty. Check results for BUSCO: 222631at7742
## Thu Jul 14 01:36:23 CEST 2022 - The aliscore output file does not exist. Check results for BUSCO: 206547at7742
## Thu Jul 14 01:36:23 CEST 2022 - The aliscore output appears to be empty. Check results for BUSCO: 206547at7742
## Thu Jul 14 01:36:25 CEST 2022 - The aliscore output file does not exist. Check results for BUSCO: 291801at7742
## Thu Jul 14 01:36:25 CEST 2022 - The aliscore output appears to be empty. Check results for BUSCO: 291801at7742
## Thu Jul 14 01:36:26 CEST 2022 - The aliscore output file does not exist. Check results for BUSCO: 318035at7742
## Thu Jul 14 01:36:26 CEST 2022 - The aliscore output appears to be empty. Check results for BUSCO: 318035at7742
## Thu Jul 14 01:36:26 CEST 2022 - The aliscore output file does not exist. Check results for BUSCO: 286595at7742
## Thu Jul 14 01:36:26 CEST 2022 - The aliscore output appears to be empty. Check results for BUSCO: 286595at7742
## Thu Jul 14 01:36:28 CEST 2022 - The aliscore output file does not exist. Check results for BUSCO: 249629at7742
## Thu Jul 14 01:36:28 CEST 2022 - The aliscore output file does not exist. Check results for BUSCO: 94127at7742
## Thu Jul 14 01:36:28 CEST 2022 - The aliscore output appears to be empty. Check results for BUSCO: 249629at7742
## Thu Jul 14 01:36:28 CEST 2022 - The aliscore output appears to be empty. Check results for BUSCO: 94127at7742
## Thu Jul 14 01:36:31 CEST 2022 - The aliscore output file does not exist. Check results for BUSCO: 138669at7742
## Thu Jul 14 01:36:31 CEST 2022 - The aliscore output appears to be empty. Check results for BUSCO: 138669at7742
## Thu Jul 14 01:36:32 CEST 2022 - The aliscore output file does not exist. Check results for BUSCO: 118365at7742
## Thu Jul 14 01:36:32 CEST 2022 - The aliscore output appears to be empty. Check results for BUSCO: 118365at7742
## Thu Jul 14 01:36:33 CEST 2022 - The aliscore output file does not exist. Check results for BUSCO: 113550at7742
## Thu Jul 14 01:36:33 CEST 2022 - The aliscore output appears to be empty. Check results for BUSCO: 113550at7742
## Thu Jul 14 01:36:34 CEST 2022 - The aliscore output file does not exist. Check results for BUSCO: 301608at7742
## Thu Jul 14 01:36:34 CEST 2022 - The aliscore output appears to be empty. Check results for BUSCO: 301608at7742
## Thu Jul 14 01:36:34 CEST 2022 - The aliscore output file does not exist. Check results for BUSCO: 118164at7742
## Thu Jul 14 01:36:34 CEST 2022 - The aliscore output appears to be empty. Check results for BUSCO: 118164at7742
## Thu Jul 14 01:36:35 CEST 2022 - The aliscore output file does not exist. Check results for BUSCO: 136144at7742
## Thu Jul 14 01:36:35 CEST 2022 - The aliscore output appears to be empty. Check results for BUSCO: 136144at7742
## Thu Jul 14 01:36:35 CEST 2022 - The aliscore output file does not exist. Check results for BUSCO: 192518at7742
## Thu Jul 14 01:36:35 CEST 2022 - The aliscore output appears to be empty. Check results for BUSCO: 192518at7742
## Thu Jul 14 01:36:38 CEST 2022 - The aliscore output file does not exist. Check results for BUSCO: 153401at7742
## Thu Jul 14 01:36:38 CEST 2022 - The aliscore output appears to be empty. Check results for BUSCO: 153401at7742
## Thu Jul 14 01:36:40 CEST 2022 - The aliscore output file does not exist. Check results for BUSCO: 128712at7742
## Thu Jul 14 01:36:40 CEST 2022 - The aliscore output appears to be empty. Check results for BUSCO: 128712at7742
## Thu Jul 14 01:36:40 CEST 2022 - The aliscore output file does not exist. Check results for BUSCO: 107743at7742
## Thu Jul 14 01:36:40 CEST 2022 - The aliscore output appears to be empty. Check results for BUSCO: 107743at7742
## Thu Jul 14 01:36:41 CEST 2022 - The aliscore output file does not exist. Check results for BUSCO: 254818at7742
## Thu Jul 14 01:36:41 CEST 2022 - The aliscore output appears to be empty. Check results for BUSCO: 254818at7742
## Thu Jul 14 01:36:44 CEST 2022 - The aliscore output file does not exist. Check results for BUSCO: 249399at7742
## Thu Jul 14 01:36:44 CEST 2022 - The aliscore output appears to be empty. Check results for BUSCO: 249399at7742
## Thu Jul 14 01:36:44 CEST 2022 - The aliscore output file does not exist. Check results for BUSCO: 113537at7742
## Thu Jul 14 01:36:44 CEST 2022 - The aliscore output appears to be empty. Check results for BUSCO: 113537at7742
## Thu Jul 14 01:36:47 CEST 2022 - The aliscore output file does not exist. Check results for BUSCO: 283821at7742
## Thu Jul 14 01:36:47 CEST 2022 - The aliscore output appears to be empty. Check results for BUSCO: 283821at7742
## Thu Jul 14 01:36:48 CEST 2022 - The aliscore output file does not exist. Check results for BUSCO: 128299at7742
## Thu Jul 14 01:36:48 CEST 2022 - The aliscore output appears to be empty. Check results for BUSCO: 128299at7742
## Thu Jul 14 01:36:48 CEST 2022 - The aliscore output file does not exist. Check results for BUSCO: 183927at7742
## Thu Jul 14 01:36:48 CEST 2022 - The aliscore output appears to be empty. Check results for BUSCO: 183927at7742
## Thu Jul 14 01:36:49 CEST 2022 - The aliscore output file does not exist. Check results for BUSCO: 145875at7742
## Thu Jul 14 01:36:49 CEST 2022 - The aliscore output appears to be empty. Check results for BUSCO: 145875at7742
## Thu Jul 14 01:36:50 CEST 2022 - The aliscore output file does not exist. Check results for BUSCO: 256936at7742
## Thu Jul 14 01:36:50 CEST 2022 - The aliscore output appears to be empty. Check results for BUSCO: 256936at7742
## Thu Jul 14 01:36:51 CEST 2022 - The aliscore output file does not exist. Check results for BUSCO: 265814at7742
## Thu Jul 14 01:36:51 CEST 2022 - The aliscore output appears to be empty. Check results for BUSCO: 265814at7742
## Thu Jul 14 01:36:52 CEST 2022 - The aliscore output file does not exist. Check results for BUSCO: 177658at7742
## Thu Jul 14 01:36:52 CEST 2022 - The aliscore output appears to be empty. Check results for BUSCO: 177658at7742
## Thu Jul 14 01:36:53 CEST 2022 - The aliscore output file does not exist. Check results for BUSCO: 130337at7742
## Thu Jul 14 01:36:53 CEST 2022 - The aliscore output appears to be empty. Check results for BUSCO: 130337at7742
## Thu Jul 14 01:36:54 CEST 2022 - The aliscore output file does not exist. Check results for BUSCO: 193525at7742
## Thu Jul 14 01:36:55 CEST 2022 - The aliscore output appears to be empty. Check results for BUSCO: 193525at7742
## Thu Jul 14 01:36:56 CEST 2022 - The aliscore output file does not exist. Check results for BUSCO: 128299at7742
## Thu Jul 14 01:36:57 CEST 2022 - The aliscore output appears to be empty. Check results for BUSCO: 128299at7742
## Thu Jul 14 01:36:58 CEST 2022 - The aliscore output file does not exist. Check results for BUSCO: 229564at7742
## Thu Jul 14 01:36:58 CEST 2022 - The aliscore output appears to be empty. Check results for BUSCO: 229564at7742
## Thu Jul 14 01:36:58 CEST 2022 - The aliscore output file does not exist. Check results for BUSCO: 239324at7742
## Thu Jul 14 01:36:58 CEST 2022 - The aliscore output appears to be empty. Check results for BUSCO: 239324at7742
## Thu Jul 14 01:36:59 CEST 2022 - The aliscore output file does not exist. Check results for BUSCO: 107743at7742
## Thu Jul 14 01:36:59 CEST 2022 - The aliscore output appears to be empty. Check results for BUSCO: 107743at7742
## Thu Jul 14 01:37:00 CEST 2022 - The aliscore output file does not exist. Check results for BUSCO: 239338at7742
## Thu Jul 14 01:37:00 CEST 2022 - The aliscore output appears to be empty. Check results for BUSCO: 239338at7742
## Thu Jul 14 01:37:01 CEST 2022 - The aliscore output file does not exist. Check results for BUSCO: 243851at7742
## Thu Jul 14 01:37:01 CEST 2022 - The aliscore output appears to be empty. Check results for BUSCO: 243851at7742
## Thu Jul 14 01:37:02 CEST 2022 - The aliscore output file does not exist. Check results for BUSCO: 189447at7742
## Thu Jul 14 01:37:02 CEST 2022 - The aliscore output appears to be empty. Check results for BUSCO: 189447at7742
## Thu Jul 14 01:37:04 CEST 2022 - The aliscore output file does not exist. Check results for BUSCO: 155482at7742
## Thu Jul 14 01:37:04 CEST 2022 - The aliscore output appears to be empty. Check results for BUSCO: 155482at7742
## Thu Jul 14 01:37:04 CEST 2022 - The aliscore output file does not exist. Check results for BUSCO: 98908at7742
## Thu Jul 14 01:37:04 CEST 2022 - The aliscore output appears to be empty. Check results for BUSCO: 98908at7742
## Thu Jul 14 01:37:06 CEST 2022 - The aliscore output file does not exist. Check results for BUSCO: 248682at7742
## Thu Jul 14 01:37:06 CEST 2022 - The aliscore output appears to be empty. Check results for BUSCO: 248682at7742
## Thu Jul 14 01:37:08 CEST 2022 - The aliscore output file does not exist. Check results for BUSCO: 292028at7742
## Thu Jul 14 01:37:08 CEST 2022 - The aliscore output appears to be empty. Check results for BUSCO: 292028at7742
## Thu Jul 14 01:37:08 CEST 2022 - The aliscore output file does not exist. Check results for BUSCO: 308334at7742
## Thu Jul 14 01:37:08 CEST 2022 - The aliscore output appears to be empty. Check results for BUSCO: 308334at7742
## Thu Jul 14 01:37:10 CEST 2022 - The aliscore output file does not exist. Check results for BUSCO: 131994at7742
## Thu Jul 14 01:37:10 CEST 2022 - The aliscore output appears to be empty. Check results for BUSCO: 131994at7742
## Thu Jul 14 01:37:12 CEST 2022 - The aliscore output file does not exist. Check results for BUSCO: 113550at7742
## Thu Jul 14 01:37:12 CEST 2022 - The aliscore output appears to be empty. Check results for BUSCO: 113550at7742
## Thu Jul 14 01:37:12 CEST 2022 - The aliscore output file does not exist. Check results for BUSCO: 301301at7742
## Thu Jul 14 01:37:12 CEST 2022 - The aliscore output appears to be empty. Check results for BUSCO: 301301at7742
## Thu Jul 14 01:37:13 CEST 2022 - The aliscore output file does not exist. Check results for BUSCO: 308334at7742
## Thu Jul 14 01:37:13 CEST 2022 - The aliscore output appears to be empty. Check results for BUSCO: 308334at7742
## Thu Jul 14 01:37:14 CEST 2022 - The aliscore output file does not exist. Check results for BUSCO: 327369at7742
## Thu Jul 14 01:37:14 CEST 2022 - The aliscore output appears to be empty. Check results for BUSCO: 327369at7742
## Thu Jul 14 01:37:15 CEST 2022 - The aliscore output file does not exist. Check results for BUSCO: 402515at7742
## Thu Jul 14 01:37:15 CEST 2022 - The aliscore output appears to be empty. Check results for BUSCO: 402515at7742
## Thu Jul 14 01:37:15 CEST 2022 - The aliscore output file does not exist. Check results for BUSCO: 239338at7742
## Thu Jul 14 01:37:15 CEST 2022 - The aliscore output appears to be empty. Check results for BUSCO: 239338at7742
## Thu Jul 14 01:37:18 CEST 2022 - The aliscore output file does not exist. Check results for BUSCO: 143611at7742
## Thu Jul 14 01:37:18 CEST 2022 - The aliscore output appears to be empty. Check results for BUSCO: 143611at7742
## Thu Jul 14 01:37:19 CEST 2022 - The aliscore output file does not exist. Check results for BUSCO: 218673at7742
## Thu Jul 14 01:37:19 CEST 2022 - The aliscore output appears to be empty. Check results for BUSCO: 218673at7742
## Thu Jul 14 01:37:21 CEST 2022 - The aliscore output file does not exist. Check results for BUSCO: 192518at7742
## Thu Jul 14 01:37:21 CEST 2022 - The aliscore output appears to be empty. Check results for BUSCO: 192518at7742
## Thu Jul 14 01:37:21 CEST 2022 - The aliscore output file does not exist. Check results for BUSCO: 154457at7742
## Thu Jul 14 01:37:21 CEST 2022 - The aliscore output appears to be empty. Check results for BUSCO: 154457at7742
## Thu Jul 14 01:37:22 CEST 2022 - The aliscore output file does not exist. Check results for BUSCO: 254424at7742
## Thu Jul 14 01:37:22 CEST 2022 - The aliscore output appears to be empty. Check results for BUSCO: 254424at7742
## Thu Jul 14 01:37:22 CEST 2022 - The aliscore output file does not exist. Check results for BUSCO: 322442at7742
## Thu Jul 14 01:37:22 CEST 2022 - The aliscore output appears to be empty. Check results for BUSCO: 322442at7742
## Thu Jul 14 01:37:24 CEST 2022 - The aliscore output file does not exist. Check results for BUSCO: 291464at7742
## Thu Jul 14 01:37:24 CEST 2022 - The aliscore output appears to be empty. Check results for BUSCO: 291464at7742
## Thu Jul 14 01:37:26 CEST 2022 - The aliscore output file does not exist. Check results for BUSCO: 184701at7742
## Thu Jul 14 01:37:26 CEST 2022 - The aliscore output appears to be empty. Check results for BUSCO: 184701at7742
## Thu Jul 14 01:37:27 CEST 2022 - The aliscore output file does not exist. Check results for BUSCO: 401448at7742
## Thu Jul 14 01:37:27 CEST 2022 - The aliscore output appears to be empty. Check results for BUSCO: 401448at7742
## Thu Jul 14 01:37:28 CEST 2022 - The aliscore output file does not exist. Check results for BUSCO: 237959at7742
## Thu Jul 14 01:37:28 CEST 2022 - The aliscore output appears to be empty. Check results for BUSCO: 237959at7742
## Thu Jul 14 01:37:29 CEST 2022 - The aliscore output file does not exist. Check results for BUSCO: 102567at7742
## Thu Jul 14 01:37:29 CEST 2022 - The aliscore output appears to be empty. Check results for BUSCO: 102567at7742
## Thu Jul 14 01:37:29 CEST 2022 - The aliscore output file does not exist. Check results for BUSCO: 154851at7742
## Thu Jul 14 01:37:29 CEST 2022 - The aliscore output appears to be empty. Check results for BUSCO: 154851at7742
## Thu Jul 14 01:37:32 CEST 2022 - The aliscore output file does not exist. Check results for BUSCO: 328571at7742
## Thu Jul 14 01:37:32 CEST 2022 - The aliscore output appears to be empty. Check results for BUSCO: 328571at7742
## Thu Jul 14 01:37:33 CEST 2022 - The aliscore output file does not exist. Check results for BUSCO: 308334at7742
## Thu Jul 14 01:37:33 CEST 2022 - The aliscore output appears to be empty. Check results for BUSCO: 308334at7742
## Thu Jul 14 01:37:33 CEST 2022 - The aliscore output file does not exist. Check results for BUSCO: 295741at7742
## Thu Jul 14 01:37:33 CEST 2022 - The aliscore output appears to be empty. Check results for BUSCO: 295741at7742
## Thu Jul 14 01:37:35 CEST 2022 - The aliscore output file does not exist. Check results for BUSCO: 160371at7742
## Thu Jul 14 01:37:35 CEST 2022 - The aliscore output appears to be empty. Check results for BUSCO: 160371at7742
## Thu Jul 14 01:37:38 CEST 2022 - The aliscore output file does not exist. Check results for BUSCO: 245932at7742
## Thu Jul 14 01:37:38 CEST 2022 - The aliscore output appears to be empty. Check results for BUSCO: 245932at7742
## Thu Jul 14 01:37:38 CEST 2022 - The aliscore output file does not exist. Check results for BUSCO: 434390at7742
## Thu Jul 14 01:37:38 CEST 2022 - The aliscore output file does not exist. Check results for BUSCO: 109315at7742
## Thu Jul 14 01:37:38 CEST 2022 - The aliscore output file does not exist. Check results for BUSCO: 199024at7742
## Thu Jul 14 01:37:38 CEST 2022 - The aliscore output appears to be empty. Check results for BUSCO: 434390at7742
## Thu Jul 14 01:37:38 CEST 2022 - The aliscore output appears to be empty. Check results for BUSCO: 109315at7742
## Thu Jul 14 01:37:38 CEST 2022 - The aliscore output appears to be empty. Check results for BUSCO: 199024at7742
## Thu Jul 14 01:37:40 CEST 2022 - The aliscore output file does not exist. Check results for BUSCO: 348500at7742
## Thu Jul 14 01:37:40 CEST 2022 - The aliscore output appears to be empty. Check results for BUSCO: 348500at7742
## Thu Jul 14 01:37:42 CEST 2022 - The aliscore output file does not exist. Check results for BUSCO: 333967at7742
## Thu Jul 14 01:37:42 CEST 2022 - The aliscore output appears to be empty. Check results for BUSCO: 333967at7742
## Thu Jul 14 01:37:42 CEST 2022 - The aliscore output file does not exist. Check results for BUSCO: 137723at7742
## Thu Jul 14 01:37:42 CEST 2022 - The aliscore output appears to be empty. Check results for BUSCO: 137723at7742
## Thu Jul 14 01:37:45 CEST 2022 - The aliscore output file does not exist. Check results for BUSCO: 178102at7742
## Thu Jul 14 01:37:45 CEST 2022 - The aliscore output appears to be empty. Check results for BUSCO: 178102at7742
## Thu Jul 14 01:37:46 CEST 2022 - The aliscore output file does not exist. Check results for BUSCO: 318035at7742
## Thu Jul 14 01:37:46 CEST 2022 - The aliscore output appears to be empty. Check results for BUSCO: 318035at7742
## Thu Jul 14 01:37:46 CEST 2022 - The aliscore output file does not exist. Check results for BUSCO: 232849at7742
## Thu Jul 14 01:37:46 CEST 2022 - The aliscore output appears to be empty. Check results for BUSCO: 232849at7742
## Thu Jul 14 01:37:48 CEST 2022 - The aliscore output file does not exist. Check results for BUSCO: 417555at7742
## Thu Jul 14 01:37:48 CEST 2022 - The aliscore output appears to be empty. Check results for BUSCO: 417555at7742
## Thu Jul 14 01:37:48 CEST 2022 - The aliscore output file does not exist. Check results for BUSCO: 321918at7742
## Thu Jul 14 01:37:48 CEST 2022 - The aliscore output appears to be empty. Check results for BUSCO: 321918at7742
## Thu Jul 14 01:37:50 CEST 2022 - The aliscore output file does not exist. Check results for BUSCO: 288407at7742
## Thu Jul 14 01:37:50 CEST 2022 - The aliscore output appears to be empty. Check results for BUSCO: 288407at7742
## Thu Jul 14 01:37:51 CEST 2022 - The aliscore output file does not exist. Check results for BUSCO: 113537at7742
## Thu Jul 14 01:37:51 CEST 2022 - The aliscore output appears to be empty. Check results for BUSCO: 113537at7742
## Thu Jul 14 01:37:51 CEST 2022 - The aliscore output file does not exist. Check results for BUSCO: 193525at7742
## Thu Jul 14 01:37:52 CEST 2022 - The aliscore output appears to be empty. Check results for BUSCO: 193525at7742
## Thu Jul 14 01:37:53 CEST 2022 - The aliscore output file does not exist. Check results for BUSCO: 245932at7742
## Thu Jul 14 01:37:53 CEST 2022 - The aliscore output file does not exist. Check results for BUSCO: 225125at7742
## Thu Jul 14 01:37:53 CEST 2022 - The aliscore output appears to be empty. Check results for BUSCO: 245932at7742
## Thu Jul 14 01:37:53 CEST 2022 - The aliscore output file does not exist. Check results for BUSCO: 316762at7742
## Thu Jul 14 01:37:53 CEST 2022 - The aliscore output appears to be empty. Check results for BUSCO: 225125at7742
## Thu Jul 14 01:37:53 CEST 2022 - The aliscore output appears to be empty. Check results for BUSCO: 316762at7742
## Thu Jul 14 01:37:56 CEST 2022 - The aliscore output file does not exist. Check results for BUSCO: 366274at7742
## Thu Jul 14 01:37:56 CEST 2022 - The aliscore output appears to be empty. Check results for BUSCO: 366274at7742
## Thu Jul 14 01:37:58 CEST 2022 - The aliscore output file does not exist. Check results for BUSCO: 154520at7742
## Thu Jul 14 01:37:58 CEST 2022 - The aliscore output appears to be empty. Check results for BUSCO: 154520at7742
## Thu Jul 14 01:37:58 CEST 2022 - The aliscore output file does not exist. Check results for BUSCO: 143665at7742
## Thu Jul 14 01:37:58 CEST 2022 - The aliscore output appears to be empty. Check results for BUSCO: 143665at7742
## Thu Jul 14 01:37:59 CEST 2022 - The aliscore output file does not exist. Check results for BUSCO: 144578at7742
## Thu Jul 14 01:37:59 CEST 2022 - The aliscore output appears to be empty. Check results for BUSCO: 144578at7742
## Thu Jul 14 01:38:01 CEST 2022 - The aliscore output file does not exist. Check results for BUSCO: 207618at7742
## Thu Jul 14 01:38:01 CEST 2022 - The aliscore output appears to be empty. Check results for BUSCO: 207618at7742
## Thu Jul 14 01:38:02 CEST 2022 - The aliscore output file does not exist. Check results for BUSCO: 143611at7742
## Thu Jul 14 01:38:02 CEST 2022 - The aliscore output appears to be empty. Check results for BUSCO: 143611at7742
## Thu Jul 14 01:38:03 CEST 2022 - The aliscore output file does not exist. Check results for BUSCO: 217951at7742
## Thu Jul 14 01:38:03 CEST 2022 - The aliscore output appears to be empty. Check results for BUSCO: 217951at7742
## Thu Jul 14 01:38:04 CEST 2022 - The aliscore output file does not exist. Check results for BUSCO: 287723at7742
## Thu Jul 14 01:38:04 CEST 2022 - The aliscore output appears to be empty. Check results for BUSCO: 287723at7742
## Thu Jul 14 01:38:05 CEST 2022 - The aliscore output file does not exist. Check results for BUSCO: 121516at7742
## Thu Jul 14 01:38:05 CEST 2022 - The aliscore output appears to be empty. Check results for BUSCO: 121516at7742
## Thu Jul 14 01:38:08 CEST 2022 - The aliscore output file does not exist. Check results for BUSCO: 113537at7742
## Thu Jul 14 01:38:08 CEST 2022 - The aliscore output appears to be empty. Check results for BUSCO: 113537at7742
## Thu Jul 14 01:38:09 CEST 2022 - The aliscore output file does not exist. Check results for BUSCO: 239308at7742
## Thu Jul 14 01:38:09 CEST 2022 - The aliscore output appears to be empty. Check results for BUSCO: 239308at7742
## Thu Jul 14 01:38:09 CEST 2022 - The aliscore output file does not exist. Check results for BUSCO: 291239at7742
## Thu Jul 14 01:38:09 CEST 2022 - The aliscore output appears to be empty. Check results for BUSCO: 291239at7742
## Thu Jul 14 01:38:10 CEST 2022 - The aliscore output file does not exist. Check results for BUSCO: 325908at7742
## Thu Jul 14 01:38:10 CEST 2022 - The aliscore output appears to be empty. Check results for BUSCO: 325908at7742
## Thu Jul 14 01:38:11 CEST 2022 - The aliscore output file does not exist. Check results for BUSCO: 349414at7742
## Thu Jul 14 01:38:12 CEST 2022 - The aliscore output appears to be empty. Check results for BUSCO: 349414at7742
## Thu Jul 14 01:38:12 CEST 2022 - The aliscore output file does not exist. Check results for BUSCO: 151349at7742
## Thu Jul 14 01:38:12 CEST 2022 - The aliscore output appears to be empty. Check results for BUSCO: 151349at7742
## Thu Jul 14 01:38:14 CEST 2022 - The aliscore output file does not exist. Check results for BUSCO: 128299at7742
## Thu Jul 14 01:38:14 CEST 2022 - The aliscore output appears to be empty. Check results for BUSCO: 128299at7742
## Thu Jul 14 01:38:15 CEST 2022 - The aliscore output file does not exist. Check results for BUSCO: 224472at7742
## Thu Jul 14 01:38:15 CEST 2022 - The aliscore output appears to be empty. Check results for BUSCO: 224472at7742
## Thu Jul 14 01:38:15 CEST 2022 - The aliscore output file does not exist. Check results for BUSCO: 245662at7742
## Thu Jul 14 01:38:16 CEST 2022 - The aliscore output appears to be empty. Check results for BUSCO: 245662at7742
## Thu Jul 14 01:38:18 CEST 2022 - The aliscore output file does not exist. Check results for BUSCO: 217951at7742
## Thu Jul 14 01:38:18 CEST 2022 - The aliscore output appears to be empty. Check results for BUSCO: 217951at7742
## Thu Jul 14 01:38:18 CEST 2022 - The aliscore output file does not exist. Check results for BUSCO: 360487at7742
## Thu Jul 14 01:38:18 CEST 2022 - The aliscore output appears to be empty. Check results for BUSCO: 360487at7742
## Thu Jul 14 01:38:18 CEST 2022 - The aliscore output file does not exist. Check results for BUSCO: 328913at7742
## Thu Jul 14 01:38:18 CEST 2022 - The aliscore output appears to be empty. Check results for BUSCO: 328913at7742
## Thu Jul 14 01:38:20 CEST 2022 - The aliscore output file does not exist. Check results for BUSCO: 145938at7742
## Thu Jul 14 01:38:20 CEST 2022 - The aliscore output appears to be empty. Check results for BUSCO: 145938at7742
## Thu Jul 14 01:38:22 CEST 2022 - The aliscore output file does not exist. Check results for BUSCO: 294443at7742
## Thu Jul 14 01:38:22 CEST 2022 - The aliscore output appears to be empty. Check results for BUSCO: 294443at7742
## Thu Jul 14 01:38:24 CEST 2022 - The aliscore output file does not exist. Check results for BUSCO: 360372at7742
## Thu Jul 14 01:38:24 CEST 2022 - The aliscore output appears to be empty. Check results for BUSCO: 360372at7742
## Thu Jul 14 01:38:26 CEST 2022 - The aliscore output file does not exist. Check results for BUSCO: 173047at7742
## Thu Jul 14 01:38:26 CEST 2022 - The aliscore output appears to be empty. Check results for BUSCO: 173047at7742
## Thu Jul 14 01:38:26 CEST 2022 - The aliscore output file does not exist. Check results for BUSCO: 250198at7742
## Thu Jul 14 01:38:26 CEST 2022 - The aliscore output appears to be empty. Check results for BUSCO: 250198at7742
## Thu Jul 14 01:38:26 CEST 2022 - The aliscore output file does not exist. Check results for BUSCO: 166657at7742
## Thu Jul 14 01:38:26 CEST 2022 - The aliscore output appears to be empty. Check results for BUSCO: 166657at7742
## Thu Jul 14 01:38:26 CEST 2022 - The aliscore output file does not exist. Check results for BUSCO: 224101at7742
## Thu Jul 14 01:38:27 CEST 2022 - The aliscore output appears to be empty. Check results for BUSCO: 224101at7742
## Thu Jul 14 01:38:29 CEST 2022 - The aliscore output file does not exist. Check results for BUSCO: 231455at7742
## Thu Jul 14 01:38:29 CEST 2022 - The aliscore output appears to be empty. Check results for BUSCO: 231455at7742
## Thu Jul 14 01:38:29 CEST 2022 - The aliscore output file does not exist. Check results for BUSCO: 182049at7742
## Thu Jul 14 01:38:29 CEST 2022 - The aliscore output appears to be empty. Check results for BUSCO: 182049at7742
## Thu Jul 14 01:38:32 CEST 2022 - The aliscore output file does not exist. Check results for BUSCO: 312347at7742
## Thu Jul 14 01:38:32 CEST 2022 - The aliscore output appears to be empty. Check results for BUSCO: 312347at7742
## Thu Jul 14 01:38:33 CEST 2022 - The aliscore output file does not exist. Check results for BUSCO: 386739at7742
## Thu Jul 14 01:38:33 CEST 2022 - The aliscore output appears to be empty. Check results for BUSCO: 386739at7742
## Thu Jul 14 01:38:34 CEST 2022 - The aliscore output file does not exist. Check results for BUSCO: 203960at7742
## Thu Jul 14 01:38:34 CEST 2022 - The aliscore output appears to be empty. Check results for BUSCO: 203960at7742
## Thu Jul 14 01:38:34 CEST 2022 - The aliscore output file does not exist. Check results for BUSCO: 438106at7742
## Thu Jul 14 01:38:34 CEST 2022 - The aliscore output appears to be empty. Check results for BUSCO: 438106at7742
## Thu Jul 14 01:38:35 CEST 2022 - The aliscore output file does not exist. Check results for BUSCO: 102567at7742
## Thu Jul 14 01:38:35 CEST 2022 - The aliscore output appears to be empty. Check results for BUSCO: 102567at7742
## Thu Jul 14 01:38:35 CEST 2022 - The aliscore output file does not exist. Check results for BUSCO: 80213at7742
## Thu Jul 14 01:38:35 CEST 2022 - The aliscore output appears to be empty. Check results for BUSCO: 80213at7742
## Thu Jul 14 01:38:37 CEST 2022 - The aliscore output file does not exist. Check results for BUSCO: 412581at7742
## Thu Jul 14 01:38:37 CEST 2022 - The aliscore output appears to be empty. Check results for BUSCO: 412581at7742
## Thu Jul 14 01:38:38 CEST 2022 - The aliscore output file does not exist. Check results for BUSCO: 145875at7742
## Thu Jul 14 01:38:38 CEST 2022 - The aliscore output appears to be empty. Check results for BUSCO: 145875at7742
## Thu Jul 14 01:38:39 CEST 2022 - The aliscore output file does not exist. Check results for BUSCO: 143555at7742
## Thu Jul 14 01:38:39 CEST 2022 - The aliscore output appears to be empty. Check results for BUSCO: 143555at7742
## Thu Jul 14 01:38:40 CEST 2022 - The aliscore output file does not exist. Check results for BUSCO: 143555at7742
## Thu Jul 14 01:38:40 CEST 2022 - The aliscore output appears to be empty. Check results for BUSCO: 143555at7742
## Thu Jul 14 01:38:41 CEST 2022 - The aliscore output file does not exist. Check results for BUSCO: 173172at7742
## Thu Jul 14 01:38:41 CEST 2022 - The aliscore output appears to be empty. Check results for BUSCO: 173172at7742
## Thu Jul 14 01:38:43 CEST 2022 - The aliscore output file does not exist. Check results for BUSCO: 183995at7742
## Thu Jul 14 01:38:43 CEST 2022 - The aliscore output appears to be empty. Check results for BUSCO: 183995at7742
## Thu Jul 14 01:38:44 CEST 2022 - The aliscore output file does not exist. Check results for BUSCO: 349414at7742
## Thu Jul 14 01:38:44 CEST 2022 - The aliscore output appears to be empty. Check results for BUSCO: 349414at7742
## Thu Jul 14 01:38:45 CEST 2022 - The aliscore output file does not exist. Check results for BUSCO: 191875at7742
## Thu Jul 14 01:38:45 CEST 2022 - The aliscore output appears to be empty. Check results for BUSCO: 191875at7742
## Thu Jul 14 01:38:47 CEST 2022 - The aliscore output file does not exist. Check results for BUSCO: 177658at7742
## Thu Jul 14 01:38:47 CEST 2022 - The aliscore output appears to be empty. Check results for BUSCO: 177658at7742
## Thu Jul 14 01:38:48 CEST 2022 - The aliscore output file does not exist. Check results for BUSCO: 193456at7742
## Thu Jul 14 01:38:48 CEST 2022 - The aliscore output appears to be empty. Check results for BUSCO: 193456at7742
## Thu Jul 14 01:38:49 CEST 2022 - The aliscore output file does not exist. Check results for BUSCO: 285172at7742
## Thu Jul 14 01:38:49 CEST 2022 - The aliscore output appears to be empty. Check results for BUSCO: 285172at7742
## Thu Jul 14 01:38:50 CEST 2022 - The aliscore output file does not exist. Check results for BUSCO: 237617at7742
## Thu Jul 14 01:38:50 CEST 2022 - The aliscore output appears to be empty. Check results for BUSCO: 237617at7742
## Thu Jul 14 01:38:51 CEST 2022 - The aliscore output file does not exist. Check results for BUSCO: 321918at7742
## Thu Jul 14 01:38:51 CEST 2022 - The aliscore output appears to be empty. Check results for BUSCO: 321918at7742
## Thu Jul 14 01:38:52 CEST 2022 - The aliscore output file does not exist. Check results for BUSCO: 83843at7742
## Thu Jul 14 01:38:52 CEST 2022 - The aliscore output appears to be empty. Check results for BUSCO: 83843at7742
## Thu Jul 14 01:38:54 CEST 2022 - The aliscore output file does not exist. Check results for BUSCO: 256936at7742
## Thu Jul 14 01:38:54 CEST 2022 - The aliscore output appears to be empty. Check results for BUSCO: 256936at7742
## Thu Jul 14 01:38:54 CEST 2022 - The aliscore output file does not exist. Check results for BUSCO: 80213at7742
## Thu Jul 14 01:38:54 CEST 2022 - The aliscore output appears to be empty. Check results for BUSCO: 80213at7742
## Thu Jul 14 01:38:56 CEST 2022 - The aliscore output file does not exist. Check results for BUSCO: 198411at7742
## Thu Jul 14 01:38:56 CEST 2022 - The aliscore output appears to be empty. Check results for BUSCO: 198411at7742
## Thu Jul 14 01:38:56 CEST 2022 - The aliscore output file does not exist. Check results for BUSCO: 120479at7742
## Thu Jul 14 01:38:56 CEST 2022 - The aliscore output appears to be empty. Check results for BUSCO: 120479at7742
## Thu Jul 14 01:38:58 CEST 2022 - The aliscore output file does not exist. Check results for BUSCO: 177658at7742
## Thu Jul 14 01:38:58 CEST 2022 - The aliscore output appears to be empty. Check results for BUSCO: 177658at7742
## Thu Jul 14 01:38:59 CEST 2022 - The aliscore output file does not exist. Check results for BUSCO: 160035at7742
## Thu Jul 14 01:38:59 CEST 2022 - The aliscore output appears to be empty. Check results for BUSCO: 160035at7742
## Thu Jul 14 01:39:00 CEST 2022 - The aliscore output file does not exist. Check results for BUSCO: 138435at7742
## Thu Jul 14 01:39:00 CEST 2022 - The aliscore output appears to be empty. Check results for BUSCO: 138435at7742
## Thu Jul 14 01:39:01 CEST 2022 - The aliscore output file does not exist. Check results for BUSCO: 209425at7742
## Thu Jul 14 01:39:01 CEST 2022 - The aliscore output appears to be empty. Check results for BUSCO: 209425at7742
## Thu Jul 14 01:39:03 CEST 2022 - The aliscore output file does not exist. Check results for BUSCO: 154851at7742
## Thu Jul 14 01:39:03 CEST 2022 - The aliscore output appears to be empty. Check results for BUSCO: 154851at7742
## Thu Jul 14 01:39:04 CEST 2022 - The aliscore output file does not exist. Check results for BUSCO: 208840at7742
## Thu Jul 14 01:39:04 CEST 2022 - The aliscore output appears to be empty. Check results for BUSCO: 208840at7742
## Thu Jul 14 01:39:05 CEST 2022 - The aliscore output file does not exist. Check results for BUSCO: 217951at7742
## Thu Jul 14 01:39:05 CEST 2022 - The aliscore output appears to be empty. Check results for BUSCO: 217951at7742
## Thu Jul 14 01:39:07 CEST 2022 - The aliscore output file does not exist. Check results for BUSCO: 232798at7742
## Thu Jul 14 01:39:07 CEST 2022 - The aliscore output appears to be empty. Check results for BUSCO: 232798at7742
## Thu Jul 14 01:39:07 CEST 2022 - The aliscore output file does not exist. Check results for BUSCO: 80213at7742
## Thu Jul 14 01:39:07 CEST 2022 - The aliscore output appears to be empty. Check results for BUSCO: 80213at7742
## Thu Jul 14 01:39:08 CEST 2022 - The aliscore output file does not exist. Check results for BUSCO: 368813at7742
## Thu Jul 14 01:39:08 CEST 2022 - The aliscore output appears to be empty. Check results for BUSCO: 368813at7742
## Thu Jul 14 01:39:10 CEST 2022 - The aliscore output file does not exist. Check results for BUSCO: 101854at7742
## Thu Jul 14 01:39:10 CEST 2022 - The aliscore output appears to be empty. Check results for BUSCO: 101854at7742
## Thu Jul 14 01:39:12 CEST 2022 - The aliscore output file does not exist. Check results for BUSCO: 123145at7742
## Thu Jul 14 01:39:12 CEST 2022 - The aliscore output appears to be empty. Check results for BUSCO: 123145at7742
## Thu Jul 14 01:39:12 CEST 2022 - The aliscore output file does not exist. Check results for BUSCO: 105171at7742
## Thu Jul 14 01:39:12 CEST 2022 - The aliscore output appears to be empty. Check results for BUSCO: 105171at7742
## Thu Jul 14 01:39:14 CEST 2022 - The aliscore output file does not exist. Check results for BUSCO: 280159at7742
## Thu Jul 14 01:39:14 CEST 2022 - The aliscore output appears to be empty. Check results for BUSCO: 280159at7742
## Thu Jul 14 01:39:15 CEST 2022 - The aliscore output file does not exist. Check results for BUSCO: 291801at7742
## Thu Jul 14 01:39:15 CEST 2022 - The aliscore output appears to be empty. Check results for BUSCO: 291801at7742
## Thu Jul 14 01:39:17 CEST 2022 - The aliscore output file does not exist. Check results for BUSCO: 202155at7742
## Thu Jul 14 01:39:17 CEST 2022 - The aliscore output appears to be empty. Check results for BUSCO: 202155at7742
## Thu Jul 14 01:39:17 CEST 2022 - The aliscore output file does not exist. Check results for BUSCO: 207648at7742
## Thu Jul 14 01:39:17 CEST 2022 - The aliscore output appears to be empty. Check results for BUSCO: 207648at7742
## Thu Jul 14 01:39:18 CEST 2022 - The aliscore output file does not exist. Check results for BUSCO: 105171at7742
## Thu Jul 14 01:39:18 CEST 2022 - The aliscore output appears to be empty. Check results for BUSCO: 105171at7742
## Thu Jul 14 01:39:20 CEST 2022 - The aliscore output file does not exist. Check results for BUSCO: 192350at7742
## Thu Jul 14 01:39:20 CEST 2022 - The aliscore output appears to be empty. Check results for BUSCO: 192350at7742
## Thu Jul 14 01:39:21 CEST 2022 - The aliscore output file does not exist. Check results for BUSCO: 189447at7742
## Thu Jul 14 01:39:22 CEST 2022 - The aliscore output appears to be empty. Check results for BUSCO: 189447at7742
## Thu Jul 14 01:39:22 CEST 2022 - The aliscore output file does not exist. Check results for BUSCO: 165997at7742
## Thu Jul 14 01:39:22 CEST 2022 - The aliscore output appears to be empty. Check results for BUSCO: 165997at7742
## Thu Jul 14 01:39:22 CEST 2022 - The aliscore output file does not exist. Check results for BUSCO: 202970at7742
## Thu Jul 14 01:39:22 CEST 2022 - The aliscore output appears to be empty. Check results for BUSCO: 202970at7742
## Thu Jul 14 01:39:23 CEST 2022 - The aliscore output file does not exist. Check results for BUSCO: 302596at7742
## Thu Jul 14 01:39:24 CEST 2022 - The aliscore output appears to be empty. Check results for BUSCO: 302596at7742
## Thu Jul 14 01:39:26 CEST 2022 - The aliscore output file does not exist. Check results for BUSCO: 143611at7742
## Thu Jul 14 01:39:26 CEST 2022 - The aliscore output appears to be empty. Check results for BUSCO: 143611at7742
## Thu Jul 14 01:39:27 CEST 2022 - The aliscore output file does not exist. Check results for BUSCO: 108802at7742
## Thu Jul 14 01:39:27 CEST 2022 - The aliscore output appears to be empty. Check results for BUSCO: 108802at7742
## Thu Jul 14 01:39:29 CEST 2022 - The aliscore output file does not exist. Check results for BUSCO: 121830at7742
## Thu Jul 14 01:39:29 CEST 2022 - The aliscore output appears to be empty. Check results for BUSCO: 121830at7742
## Thu Jul 14 01:39:29 CEST 2022 - The aliscore output file does not exist. Check results for BUSCO: 105171at7742
## Thu Jul 14 01:39:29 CEST 2022 - The aliscore output appears to be empty. Check results for BUSCO: 105171at7742
## Thu Jul 14 01:39:30 CEST 2022 - The aliscore output file does not exist. Check results for BUSCO: 402515at7742
## Thu Jul 14 01:39:30 CEST 2022 - The aliscore output appears to be empty. Check results for BUSCO: 402515at7742
## Thu Jul 14 01:39:32 CEST 2022 - The aliscore output file does not exist. Check results for BUSCO: 160383at7742
## Thu Jul 14 01:39:32 CEST 2022 - The aliscore output appears to be empty. Check results for BUSCO: 160383at7742
## Thu Jul 14 01:39:34 CEST 2022 - The aliscore output file does not exist. Check results for BUSCO: 377100at7742
## Thu Jul 14 01:39:34 CEST 2022 - The aliscore output appears to be empty. Check results for BUSCO: 377100at7742
## Thu Jul 14 01:39:35 CEST 2022 - The aliscore output file does not exist. Check results for BUSCO: 179036at7742
## Thu Jul 14 01:39:35 CEST 2022 - The aliscore output appears to be empty. Check results for BUSCO: 179036at7742
## Thu Jul 14 01:39:36 CEST 2022 - The aliscore output file does not exist. Check results for BUSCO: 312837at7742
## Thu Jul 14 01:39:36 CEST 2022 - The aliscore output appears to be empty. Check results for BUSCO: 312837at7742
## Thu Jul 14 01:39:37 CEST 2022 - The aliscore output file does not exist. Check results for BUSCO: 417555at7742
## Thu Jul 14 01:39:37 CEST 2022 - The aliscore output appears to be empty. Check results for BUSCO: 417555at7742
## Thu Jul 14 01:39:38 CEST 2022 - The aliscore output file does not exist. Check results for BUSCO: 417555at7742
## Thu Jul 14 01:39:38 CEST 2022 - The aliscore output appears to be empty. Check results for BUSCO: 417555at7742
## Thu Jul 14 01:39:39 CEST 2022 - The aliscore output file does not exist. Check results for BUSCO: 133713at7742
## Thu Jul 14 01:39:39 CEST 2022 - The aliscore output appears to be empty. Check results for BUSCO: 133713at7742
## Thu Jul 14 01:39:40 CEST 2022 - The aliscore output file does not exist. Check results for BUSCO: 160035at7742
## Thu Jul 14 01:39:40 CEST 2022 - The aliscore output appears to be empty. Check results for BUSCO: 160035at7742
## Thu Jul 14 01:39:42 CEST 2022 - The aliscore output file does not exist. Check results for BUSCO: 253042at7742
## Thu Jul 14 01:39:42 CEST 2022 - The aliscore output appears to be empty. Check results for BUSCO: 253042at7742
## Thu Jul 14 01:39:43 CEST 2022 - The aliscore output file does not exist. Check results for BUSCO: 138669at7742
## Thu Jul 14 01:39:43 CEST 2022 - The aliscore output appears to be empty. Check results for BUSCO: 138669at7742
## Thu Jul 14 01:39:44 CEST 2022 - The aliscore output file does not exist. Check results for BUSCO: 360372at7742
## Thu Jul 14 01:39:44 CEST 2022 - The aliscore output appears to be empty. Check results for BUSCO: 360372at7742
## Thu Jul 14 01:39:45 CEST 2022 - The aliscore output file does not exist. Check results for BUSCO: 362170at7742
## Thu Jul 14 01:39:45 CEST 2022 - The aliscore output appears to be empty. Check results for BUSCO: 362170at7742
## Thu Jul 14 01:39:46 CEST 2022 - The aliscore output file does not exist. Check results for BUSCO: 121516at7742
## Thu Jul 14 01:39:46 CEST 2022 - The aliscore output appears to be empty. Check results for BUSCO: 121516at7742
## Thu Jul 14 01:39:47 CEST 2022 - The aliscore output file does not exist. Check results for BUSCO: 264239at7742
## Thu Jul 14 01:39:47 CEST 2022 - The aliscore output appears to be empty. Check results for BUSCO: 264239at7742
## Thu Jul 14 01:39:48 CEST 2022 - The aliscore output file does not exist. Check results for BUSCO: 123145at7742
## Thu Jul 14 01:39:48 CEST 2022 - The aliscore output appears to be empty. Check results for BUSCO: 123145at7742
## Thu Jul 14 01:39:50 CEST 2022 - The aliscore output file does not exist. Check results for BUSCO: 145137at7742
## Thu Jul 14 01:39:50 CEST 2022 - The aliscore output appears to be empty. Check results for BUSCO: 145137at7742
## Thu Jul 14 01:39:50 CEST 2022 - The aliscore output file does not exist. Check results for BUSCO: 389352at7742
## Thu Jul 14 01:39:50 CEST 2022 - The aliscore output appears to be empty. Check results for BUSCO: 389352at7742
## Thu Jul 14 01:39:53 CEST 2022 - The aliscore output file does not exist. Check results for BUSCO: 160859at7742
## Thu Jul 14 01:39:53 CEST 2022 - The aliscore output appears to be empty. Check results for BUSCO: 160859at7742
## Thu Jul 14 01:39:53 CEST 2022 - The aliscore output file does not exist. Check results for BUSCO: 307639at7742
## Thu Jul 14 01:39:53 CEST 2022 - The aliscore output appears to be empty. Check results for BUSCO: 307639at7742
## Thu Jul 14 01:39:54 CEST 2022 - The aliscore output file does not exist. Check results for BUSCO: 242072at7742
## Thu Jul 14 01:39:54 CEST 2022 - The aliscore output appears to be empty. Check results for BUSCO: 242072at7742
## Thu Jul 14 01:39:55 CEST 2022 - The aliscore output file does not exist. Check results for BUSCO: 316762at7742
## Thu Jul 14 01:39:55 CEST 2022 - The aliscore output appears to be empty. Check results for BUSCO: 316762at7742
## Thu Jul 14 01:39:57 CEST 2022 - The aliscore output file does not exist. Check results for BUSCO: 185697at7742
## Thu Jul 14 01:39:57 CEST 2022 - The aliscore output file does not exist. Check results for BUSCO: 154851at7742
## Thu Jul 14 01:39:57 CEST 2022 - The aliscore output appears to be empty. Check results for BUSCO: 185697at7742
## Thu Jul 14 01:39:57 CEST 2022 - The aliscore output appears to be empty. Check results for BUSCO: 154851at7742
## Thu Jul 14 01:39:59 CEST 2022 - The aliscore output file does not exist. Check results for BUSCO: 193456at7742
## Thu Jul 14 01:39:59 CEST 2022 - The aliscore output appears to be empty. Check results for BUSCO: 193456at7742
## Thu Jul 14 01:40:00 CEST 2022 - The aliscore output file does not exist. Check results for BUSCO: 220062at7742
## Thu Jul 14 01:40:00 CEST 2022 - The aliscore output appears to be empty. Check results for BUSCO: 220062at7742
## Thu Jul 14 01:40:01 CEST 2022 - The aliscore output file does not exist. Check results for BUSCO: 290261at7742
## Thu Jul 14 01:40:01 CEST 2022 - The aliscore output appears to be empty. Check results for BUSCO: 290261at7742
## Thu Jul 14 01:40:02 CEST 2022 - The aliscore output file does not exist. Check results for BUSCO: 199723at7742
## Thu Jul 14 01:40:02 CEST 2022 - The aliscore output appears to be empty. Check results for BUSCO: 199723at7742
## Thu Jul 14 01:40:03 CEST 2022 - The aliscore output file does not exist. Check results for BUSCO: 123145at7742
## Thu Jul 14 01:40:03 CEST 2022 - The aliscore output appears to be empty. Check results for BUSCO: 123145at7742
## Thu Jul 14 01:40:04 CEST 2022 - The aliscore output file does not exist. Check results for BUSCO: 274707at7742
## Thu Jul 14 01:40:04 CEST 2022 - The aliscore output appears to be empty. Check results for BUSCO: 274707at7742
## Thu Jul 14 01:40:06 CEST 2022 - The aliscore output file does not exist. Check results for BUSCO: 242641at7742
## Thu Jul 14 01:40:06 CEST 2022 - The aliscore output appears to be empty. Check results for BUSCO: 242641at7742
## Thu Jul 14 01:40:07 CEST 2022 - The aliscore output file does not exist. Check results for BUSCO: 340508at7742
## Thu Jul 14 01:40:07 CEST 2022 - The aliscore output appears to be empty. Check results for BUSCO: 340508at7742
## Thu Jul 14 01:40:08 CEST 2022 - The aliscore output file does not exist. Check results for BUSCO: 298884at7742
## Thu Jul 14 01:40:08 CEST 2022 - The aliscore output appears to be empty. Check results for BUSCO: 298884at7742
## Thu Jul 14 01:40:09 CEST 2022 - The aliscore output file does not exist. Check results for BUSCO: 389667at7742
## Thu Jul 14 01:40:09 CEST 2022 - The aliscore output appears to be empty. Check results for BUSCO: 389667at7742
## Thu Jul 14 01:40:11 CEST 2022 - The aliscore output file does not exist. Check results for BUSCO: 267268at7742
## Thu Jul 14 01:40:11 CEST 2022 - The aliscore output appears to be empty. Check results for BUSCO: 267268at7742
## Thu Jul 14 01:40:11 CEST 2022 - The aliscore output file does not exist. Check results for BUSCO: 252806at7742
## Thu Jul 14 01:40:11 CEST 2022 - The aliscore output appears to be empty. Check results for BUSCO: 252806at7742
## Thu Jul 14 01:40:13 CEST 2022 - The aliscore output file does not exist. Check results for BUSCO: 208300at7742
## Thu Jul 14 01:40:13 CEST 2022 - The aliscore output appears to be empty. Check results for BUSCO: 208300at7742
## Thu Jul 14 01:40:13 CEST 2022 - The aliscore output file does not exist. Check results for BUSCO: 249877at7742
## Thu Jul 14 01:40:13 CEST 2022 - The aliscore output appears to be empty. Check results for BUSCO: 249877at7742
## Thu Jul 14 01:40:14 CEST 2022 - The aliscore output file does not exist. Check results for BUSCO: 385740at7742
## Thu Jul 14 01:40:14 CEST 2022 - The aliscore output appears to be empty. Check results for BUSCO: 385740at7742
## Thu Jul 14 01:40:17 CEST 2022 - The aliscore output file does not exist. Check results for BUSCO: 162212at7742
## Thu Jul 14 01:40:17 CEST 2022 - The aliscore output appears to be empty. Check results for BUSCO: 162212at7742
## Thu Jul 14 01:40:18 CEST 2022 - The aliscore output file does not exist. Check results for BUSCO: 160859at7742
## Thu Jul 14 01:40:18 CEST 2022 - The aliscore output appears to be empty. Check results for BUSCO: 160859at7742
## Thu Jul 14 01:40:19 CEST 2022 - The aliscore output file does not exist. Check results for BUSCO: 173172at7742
## Thu Jul 14 01:40:19 CEST 2022 - The aliscore output appears to be empty. Check results for BUSCO: 173172at7742
## Thu Jul 14 01:40:20 CEST 2022 - The aliscore output file does not exist. Check results for BUSCO: 228390at7742
## Thu Jul 14 01:40:20 CEST 2022 - The aliscore output appears to be empty. Check results for BUSCO: 228390at7742
## Thu Jul 14 01:40:21 CEST 2022 - The aliscore output file does not exist. Check results for BUSCO: 248682at7742
## Thu Jul 14 01:40:21 CEST 2022 - The aliscore output appears to be empty. Check results for BUSCO: 248682at7742
## Thu Jul 14 01:40:23 CEST 2022 - The aliscore output file does not exist. Check results for BUSCO: 223898at7742
## Thu Jul 14 01:40:23 CEST 2022 - The aliscore output appears to be empty. Check results for BUSCO: 223898at7742
## Thu Jul 14 01:40:24 CEST 2022 - The aliscore output file does not exist. Check results for BUSCO: 221592at7742
## Thu Jul 14 01:40:24 CEST 2022 - The aliscore output appears to be empty. Check results for BUSCO: 221592at7742
## Thu Jul 14 01:40:25 CEST 2022 - The aliscore output file does not exist. Check results for BUSCO: 130337at7742
## Thu Jul 14 01:40:25 CEST 2022 - The aliscore output appears to be empty. Check results for BUSCO: 130337at7742
## Thu Jul 14 01:40:26 CEST 2022 - The aliscore output file does not exist. Check results for BUSCO: 208840at7742
## Thu Jul 14 01:40:26 CEST 2022 - The aliscore output appears to be empty. Check results for BUSCO: 208840at7742
## Thu Jul 14 01:40:28 CEST 2022 - The aliscore output file does not exist. Check results for BUSCO: 189840at7742
## Thu Jul 14 01:40:28 CEST 2022 - The aliscore output appears to be empty. Check results for BUSCO: 189840at7742
## Thu Jul 14 01:40:30 CEST 2022 - The aliscore output file does not exist. Check results for BUSCO: 349414at7742
## Thu Jul 14 01:40:30 CEST 2022 - The aliscore output appears to be empty. Check results for BUSCO: 349414at7742
## Thu Jul 14 01:40:30 CEST 2022 - The aliscore output file does not exist. Check results for BUSCO: 326191at7742
## Thu Jul 14 01:40:30 CEST 2022 - The aliscore output appears to be empty. Check results for BUSCO: 326191at7742
## Thu Jul 14 01:40:30 CEST 2022 - The aliscore output file does not exist. Check results for BUSCO: 312837at7742
## Thu Jul 14 01:40:30 CEST 2022 - The aliscore output appears to be empty. Check results for BUSCO: 312837at7742
## Thu Jul 14 01:40:31 CEST 2022 - The aliscore output file does not exist. Check results for BUSCO: 256510at7742
## Thu Jul 14 01:40:31 CEST 2022 - The aliscore output appears to be empty. Check results for BUSCO: 256510at7742
## Thu Jul 14 01:40:32 CEST 2022 - The aliscore output file does not exist. Check results for BUSCO: 261602at7742
## Thu Jul 14 01:40:32 CEST 2022 - The aliscore output appears to be empty. Check results for BUSCO: 261602at7742
## Thu Jul 14 01:40:34 CEST 2022 - The aliscore output file does not exist. Check results for BUSCO: 266219at7742
## Thu Jul 14 01:40:34 CEST 2022 - The aliscore output appears to be empty. Check results for BUSCO: 266219at7742
## Thu Jul 14 01:40:36 CEST 2022 - The aliscore output file does not exist. Check results for BUSCO: 193249at7742
## Thu Jul 14 01:40:36 CEST 2022 - The aliscore output appears to be empty. Check results for BUSCO: 193249at7742
## Thu Jul 14 01:40:37 CEST 2022 - The aliscore output file does not exist. Check results for BUSCO: 385740at7742
## Thu Jul 14 01:40:38 CEST 2022 - The aliscore output appears to be empty. Check results for BUSCO: 385740at7742
## Thu Jul 14 01:40:39 CEST 2022 - The aliscore output file does not exist. Check results for BUSCO: 192331at7742
## Thu Jul 14 01:40:39 CEST 2022 - The aliscore output appears to be empty. Check results for BUSCO: 192331at7742
## Thu Jul 14 01:40:39 CEST 2022 - The aliscore output file does not exist. Check results for BUSCO: 245482at7742
## Thu Jul 14 01:40:40 CEST 2022 - The aliscore output appears to be empty. Check results for BUSCO: 245482at7742
## Thu Jul 14 01:40:41 CEST 2022 - The aliscore output file does not exist. Check results for BUSCO: 221592at7742
## Thu Jul 14 01:40:41 CEST 2022 - The aliscore output appears to be empty. Check results for BUSCO: 221592at7742
## Thu Jul 14 01:40:41 CEST 2022 - The aliscore output file does not exist. Check results for BUSCO: 188059at7742
## Thu Jul 14 01:40:41 CEST 2022 - The aliscore output appears to be empty. Check results for BUSCO: 188059at7742
## Thu Jul 14 01:40:43 CEST 2022 - The aliscore output file does not exist. Check results for BUSCO: 114119at7742
## Thu Jul 14 01:40:43 CEST 2022 - The aliscore output appears to be empty. Check results for BUSCO: 114119at7742
## Thu Jul 14 01:40:44 CEST 2022 - The aliscore output file does not exist. Check results for BUSCO: 227544at7742
## Thu Jul 14 01:40:44 CEST 2022 - The aliscore output appears to be empty. Check results for BUSCO: 227544at7742
## Thu Jul 14 01:40:45 CEST 2022 - The aliscore output file does not exist. Check results for BUSCO: 203253at7742
## Thu Jul 14 01:40:46 CEST 2022 - The aliscore output appears to be empty. Check results for BUSCO: 203253at7742
## Thu Jul 14 01:40:46 CEST 2022 - The aliscore output file does not exist. Check results for BUSCO: 118164at7742
## Thu Jul 14 01:40:46 CEST 2022 - The aliscore output appears to be empty. Check results for BUSCO: 118164at7742
## Thu Jul 14 01:40:47 CEST 2022 - The aliscore output file does not exist. Check results for BUSCO: 251667at7742
## Thu Jul 14 01:40:47 CEST 2022 - The aliscore output appears to be empty. Check results for BUSCO: 251667at7742
## Thu Jul 14 01:40:49 CEST 2022 - The aliscore output file does not exist. Check results for BUSCO: 144764at7742
## Thu Jul 14 01:40:49 CEST 2022 - The aliscore output appears to be empty. Check results for BUSCO: 144764at7742
## Thu Jul 14 01:40:51 CEST 2022 - The aliscore output file does not exist. Check results for BUSCO: 251465at7742
## Thu Jul 14 01:40:51 CEST 2022 - The aliscore output appears to be empty. Check results for BUSCO: 251465at7742
## Thu Jul 14 01:40:52 CEST 2022 - The aliscore output file does not exist. Check results for BUSCO: 264772at7742
## Thu Jul 14 01:40:52 CEST 2022 - The aliscore output file does not exist. Check results for BUSCO: 205403at7742
## Thu Jul 14 01:40:52 CEST 2022 - The aliscore output appears to be empty. Check results for BUSCO: 264772at7742
## Thu Jul 14 01:40:52 CEST 2022 - The aliscore output appears to be empty. Check results for BUSCO: 205403at7742
## Thu Jul 14 01:40:53 CEST 2022 - The aliscore output file does not exist. Check results for BUSCO: 245056at7742
## Thu Jul 14 01:40:54 CEST 2022 - The aliscore output appears to be empty. Check results for BUSCO: 245056at7742
## Thu Jul 14 01:40:54 CEST 2022 - The aliscore output file does not exist. Check results for BUSCO: 175700at7742
## Thu Jul 14 01:40:55 CEST 2022 - The aliscore output appears to be empty. Check results for BUSCO: 175700at7742
## Thu Jul 14 01:40:57 CEST 2022 - The aliscore output file does not exist. Check results for BUSCO: 114119at7742
## Thu Jul 14 01:40:57 CEST 2022 - The aliscore output appears to be empty. Check results for BUSCO: 114119at7742
## Thu Jul 14 01:40:58 CEST 2022 - The aliscore output file does not exist. Check results for BUSCO: 368433at7742
## Thu Jul 14 01:40:58 CEST 2022 - The aliscore output appears to be empty. Check results for BUSCO: 368433at7742
## Thu Jul 14 01:40:58 CEST 2022 - The aliscore output file does not exist. Check results for BUSCO: 158142at7742
## Thu Jul 14 01:40:58 CEST 2022 - The aliscore output appears to be empty. Check results for BUSCO: 158142at7742
## Thu Jul 14 01:40:59 CEST 2022 - The aliscore output file does not exist. Check results for BUSCO: 202970at7742
## Thu Jul 14 01:40:59 CEST 2022 - The aliscore output appears to be empty. Check results for BUSCO: 202970at7742
## Thu Jul 14 01:41:00 CEST 2022 - The aliscore output file does not exist. Check results for BUSCO: 114119at7742
## Thu Jul 14 01:41:00 CEST 2022 - The aliscore output appears to be empty. Check results for BUSCO: 114119at7742
## Thu Jul 14 01:41:01 CEST 2022 - The aliscore output file does not exist. Check results for BUSCO: 410850at7742
## Thu Jul 14 01:41:01 CEST 2022 - The aliscore output appears to be empty. Check results for BUSCO: 410850at7742
## Thu Jul 14 01:41:02 CEST 2022 - The aliscore output file does not exist. Check results for BUSCO: 386739at7742
## Thu Jul 14 01:41:02 CEST 2022 - The aliscore output appears to be empty. Check results for BUSCO: 386739at7742
## Thu Jul 14 01:41:05 CEST 2022 - The aliscore output file does not exist. Check results for BUSCO: 208627at7742
## Thu Jul 14 01:41:05 CEST 2022 - The aliscore output appears to be empty. Check results for BUSCO: 208627at7742
## Thu Jul 14 01:41:05 CEST 2022 - The aliscore output file does not exist. Check results for BUSCO: 160859at7742
## Thu Jul 14 01:41:05 CEST 2022 - The aliscore output appears to be empty. Check results for BUSCO: 160859at7742
## Thu Jul 14 01:41:06 CEST 2022 - The aliscore output file does not exist. Check results for BUSCO: 120479at7742
## Thu Jul 14 01:41:06 CEST 2022 - The aliscore output appears to be empty. Check results for BUSCO: 120479at7742
## Thu Jul 14 01:41:07 CEST 2022 - The aliscore output file does not exist. Check results for BUSCO: 182982at7742
## Thu Jul 14 01:41:07 CEST 2022 - The aliscore output appears to be empty. Check results for BUSCO: 182982at7742
## Thu Jul 14 01:41:09 CEST 2022 - The aliscore output file does not exist. Check results for BUSCO: 230477at7742
## Thu Jul 14 01:41:09 CEST 2022 - The aliscore output appears to be empty. Check results for BUSCO: 230477at7742
## Thu Jul 14 01:41:09 CEST 2022 - The aliscore output file does not exist. Check results for BUSCO: 184999at7742
## Thu Jul 14 01:41:10 CEST 2022 - The aliscore output appears to be empty. Check results for BUSCO: 184999at7742
## Thu Jul 14 01:41:10 CEST 2022 - The aliscore output file does not exist. Check results for BUSCO: 216982at7742
## Thu Jul 14 01:41:10 CEST 2022 - The aliscore output appears to be empty. Check results for BUSCO: 216982at7742
## Thu Jul 14 01:41:12 CEST 2022 - The aliscore output file does not exist. Check results for BUSCO: 173172at7742
## Thu Jul 14 01:41:12 CEST 2022 - The aliscore output appears to be empty. Check results for BUSCO: 173172at7742
## Thu Jul 14 01:41:12 CEST 2022 - The aliscore output file does not exist. Check results for BUSCO: 129684at7742
## Thu Jul 14 01:41:12 CEST 2022 - The aliscore output appears to be empty. Check results for BUSCO: 129684at7742
## Thu Jul 14 01:41:15 CEST 2022 - The aliscore output file does not exist. Check results for BUSCO: 208300at7742
## Thu Jul 14 01:41:15 CEST 2022 - The aliscore output appears to be empty. Check results for BUSCO: 208300at7742
## Thu Jul 14 01:41:15 CEST 2022 - The aliscore output file does not exist. Check results for BUSCO: 258449at7742
## Thu Jul 14 01:41:15 CEST 2022 - The aliscore output appears to be empty. Check results for BUSCO: 258449at7742
## Thu Jul 14 01:41:17 CEST 2022 - The aliscore output file does not exist. Check results for BUSCO: 232798at7742
## Thu Jul 14 01:41:17 CEST 2022 - The aliscore output appears to be empty. Check results for BUSCO: 232798at7742
## Thu Jul 14 01:41:18 CEST 2022 - The aliscore output file does not exist. Check results for BUSCO: 145137at7742
## Thu Jul 14 01:41:18 CEST 2022 - The aliscore output appears to be empty. Check results for BUSCO: 145137at7742
## Thu Jul 14 01:41:18 CEST 2022 - The aliscore output file does not exist. Check results for BUSCO: 340508at7742
## Thu Jul 14 01:41:18 CEST 2022 - The aliscore output appears to be empty. Check results for BUSCO: 340508at7742
## Thu Jul 14 01:41:19 CEST 2022 - The aliscore output file does not exist. Check results for BUSCO: 363574at7742
## Thu Jul 14 01:41:19 CEST 2022 - The aliscore output appears to be empty. Check results for BUSCO: 363574at7742
## Thu Jul 14 01:41:22 CEST 2022 - The aliscore output file does not exist. Check results for BUSCO: 133713at7742
## Thu Jul 14 01:41:22 CEST 2022 - The aliscore output file does not exist. Check results for BUSCO: 260359at7742
## Thu Jul 14 01:41:22 CEST 2022 - The aliscore output appears to be empty. Check results for BUSCO: 133713at7742
## Thu Jul 14 01:41:22 CEST 2022 - The aliscore output appears to be empty. Check results for BUSCO: 260359at7742
## Thu Jul 14 01:41:22 CEST 2022 - The aliscore output file does not exist. Check results for BUSCO: 337325at7742
## Thu Jul 14 01:41:22 CEST 2022 - The aliscore output appears to be empty. Check results for BUSCO: 337325at7742
## Thu Jul 14 01:41:24 CEST 2022 - The aliscore output file does not exist. Check results for BUSCO: 338075at7742
## Thu Jul 14 01:41:24 CEST 2022 - The aliscore output appears to be empty. Check results for BUSCO: 338075at7742
## Thu Jul 14 01:41:25 CEST 2022 - The aliscore output file does not exist. Check results for BUSCO: 218673at7742
## Thu Jul 14 01:41:25 CEST 2022 - The aliscore output appears to be empty. Check results for BUSCO: 218673at7742
## Thu Jul 14 01:41:27 CEST 2022 - The aliscore output file does not exist. Check results for BUSCO: 250198at7742
## Thu Jul 14 01:41:27 CEST 2022 - The aliscore output appears to be empty. Check results for BUSCO: 250198at7742
## Thu Jul 14 01:41:28 CEST 2022 - The aliscore output file does not exist. Check results for BUSCO: 143555at7742
## Thu Jul 14 01:41:28 CEST 2022 - The aliscore output appears to be empty. Check results for BUSCO: 143555at7742
## Thu Jul 14 01:41:28 CEST 2022 - The aliscore output file does not exist. Check results for BUSCO: 182742at7742
## Thu Jul 14 01:41:28 CEST 2022 - The aliscore output appears to be empty. Check results for BUSCO: 182742at7742
## Thu Jul 14 01:41:30 CEST 2022 - The aliscore output file does not exist. Check results for BUSCO: 182982at7742
## Thu Jul 14 01:41:30 CEST 2022 - The aliscore output appears to be empty. Check results for BUSCO: 182982at7742
## Thu Jul 14 01:41:31 CEST 2022 - The aliscore output file does not exist. Check results for BUSCO: 229564at7742
## Thu Jul 14 01:41:31 CEST 2022 - The aliscore output appears to be empty. Check results for BUSCO: 229564at7742
## Thu Jul 14 01:41:33 CEST 2022 - The aliscore output file does not exist. Check results for BUSCO: 308423at7742
## Thu Jul 14 01:41:33 CEST 2022 - The aliscore output appears to be empty. Check results for BUSCO: 308423at7742
## Thu Jul 14 01:41:33 CEST 2022 - The aliscore output file does not exist. Check results for BUSCO: 161775at7742
## Thu Jul 14 01:41:33 CEST 2022 - The aliscore output appears to be empty. Check results for BUSCO: 161775at7742
## Thu Jul 14 01:41:33 CEST 2022 - The aliscore output file does not exist. Check results for BUSCO: 379193at7742
## Thu Jul 14 01:41:33 CEST 2022 - The aliscore output appears to be empty. Check results for BUSCO: 379193at7742
## Thu Jul 14 01:41:35 CEST 2022 - The aliscore output file does not exist. Check results for BUSCO: 208238at7742
## Thu Jul 14 01:41:35 CEST 2022 - The aliscore output appears to be empty. Check results for BUSCO: 208238at7742
## Thu Jul 14 01:41:35 CEST 2022 - The aliscore output file does not exist. Check results for BUSCO: 169728at7742
## Thu Jul 14 01:41:35 CEST 2022 - The aliscore output appears to be empty. Check results for BUSCO: 169728at7742
## Thu Jul 14 01:41:37 CEST 2022 - The aliscore output file does not exist. Check results for BUSCO: 162212at7742
## Thu Jul 14 01:41:37 CEST 2022 - The aliscore output appears to be empty. Check results for BUSCO: 162212at7742
## Thu Jul 14 01:41:39 CEST 2022 - The aliscore output file does not exist. Check results for BUSCO: 106667at7742
## Thu Jul 14 01:41:39 CEST 2022 - The aliscore output appears to be empty. Check results for BUSCO: 106667at7742
## Thu Jul 14 01:41:41 CEST 2022 - The aliscore output file does not exist. Check results for BUSCO: 192350at7742
## Thu Jul 14 01:41:41 CEST 2022 - The aliscore output appears to be empty. Check results for BUSCO: 192350at7742
## Thu Jul 14 01:41:41 CEST 2022 - The aliscore output file does not exist. Check results for BUSCO: 190526at7742
## Thu Jul 14 01:41:41 CEST 2022 - The aliscore output appears to be empty. Check results for BUSCO: 190526at7742
## Thu Jul 14 01:41:43 CEST 2022 - The aliscore output file does not exist. Check results for BUSCO: 158802at7742
## Thu Jul 14 01:41:43 CEST 2022 - The aliscore output appears to be empty. Check results for BUSCO: 158802at7742
## Thu Jul 14 01:41:43 CEST 2022 - The aliscore output file does not exist. Check results for BUSCO: 296646at7742
## Thu Jul 14 01:41:43 CEST 2022 - The aliscore output appears to be empty. Check results for BUSCO: 296646at7742
## Thu Jul 14 01:41:44 CEST 2022 - The aliscore output file does not exist. Check results for BUSCO: 225242at7742
## Thu Jul 14 01:41:44 CEST 2022 - The aliscore output appears to be empty. Check results for BUSCO: 225242at7742
## Thu Jul 14 01:41:46 CEST 2022 - The aliscore output file does not exist. Check results for BUSCO: 337000at7742
## Thu Jul 14 01:41:46 CEST 2022 - The aliscore output appears to be empty. Check results for BUSCO: 337000at7742
## Thu Jul 14 01:41:47 CEST 2022 - The aliscore output file does not exist. Check results for BUSCO: 208238at7742
## Thu Jul 14 01:41:47 CEST 2022 - The aliscore output appears to be empty. Check results for BUSCO: 208238at7742
## Thu Jul 14 01:41:48 CEST 2022 - The aliscore output file does not exist. Check results for BUSCO: 296646at7742
## Thu Jul 14 01:41:48 CEST 2022 - The aliscore output appears to be empty. Check results for BUSCO: 296646at7742
## Thu Jul 14 01:41:53 CEST 2022 - The aliscore output file does not exist. Check results for BUSCO: 159866at7742
## Thu Jul 14 01:41:53 CEST 2022 - The aliscore output appears to be empty. Check results for BUSCO: 159866at7742
## Thu Jul 14 01:41:54 CEST 2022 - The aliscore output file does not exist. Check results for BUSCO: 308423at7742
## Thu Jul 14 01:41:54 CEST 2022 - The aliscore output appears to be empty. Check results for BUSCO: 308423at7742
## Thu Jul 14 01:41:54 CEST 2022 - The aliscore output file does not exist. Check results for BUSCO: 239308at7742
## Thu Jul 14 01:41:54 CEST 2022 - The aliscore output appears to be empty. Check results for BUSCO: 239308at7742
## Thu Jul 14 01:41:54 CEST 2022 - The aliscore output file does not exist. Check results for BUSCO: 188059at7742
## Thu Jul 14 01:41:54 CEST 2022 - The aliscore output appears to be empty. Check results for BUSCO: 188059at7742
## Thu Jul 14 01:41:55 CEST 2022 - The aliscore output file does not exist. Check results for BUSCO: 328913at7742
## Thu Jul 14 01:41:55 CEST 2022 - The aliscore output appears to be empty. Check results for BUSCO: 328913at7742
## Thu Jul 14 01:41:55 CEST 2022 - The aliscore output file does not exist. Check results for BUSCO: 239324at7742
## Thu Jul 14 01:41:55 CEST 2022 - The aliscore output appears to be empty. Check results for BUSCO: 239324at7742
## Thu Jul 14 01:41:57 CEST 2022 - The aliscore output file does not exist. Check results for BUSCO: 379829at7742
## Thu Jul 14 01:41:57 CEST 2022 - The aliscore output appears to be empty. Check results for BUSCO: 379829at7742
## Thu Jul 14 01:41:59 CEST 2022 - The aliscore output file does not exist. Check results for BUSCO: 278370at7742
## Thu Jul 14 01:41:59 CEST 2022 - The aliscore output appears to be empty. Check results for BUSCO: 278370at7742
## Thu Jul 14 01:41:59 CEST 2022 - The aliscore output file does not exist. Check results for BUSCO: 173047at7742
## Thu Jul 14 01:41:59 CEST 2022 - The aliscore output appears to be empty. Check results for BUSCO: 173047at7742
## Thu Jul 14 01:42:00 CEST 2022 - The aliscore output file does not exist. Check results for BUSCO: 98908at7742
## Thu Jul 14 01:42:00 CEST 2022 - The aliscore output appears to be empty. Check results for BUSCO: 98908at7742
## Thu Jul 14 01:42:01 CEST 2022 - The aliscore output file does not exist. Check results for BUSCO: 241220at7742
## Thu Jul 14 01:42:01 CEST 2022 - The aliscore output appears to be empty. Check results for BUSCO: 241220at7742
## Thu Jul 14 01:42:02 CEST 2022 - The aliscore output file does not exist. Check results for BUSCO: 257782at7742
## Thu Jul 14 01:42:02 CEST 2022 - The aliscore output appears to be empty. Check results for BUSCO: 257782at7742
## Thu Jul 14 01:42:04 CEST 2022 - The aliscore output file does not exist. Check results for BUSCO: 101854at7742
## Thu Jul 14 01:42:04 CEST 2022 - The aliscore output appears to be empty. Check results for BUSCO: 101854at7742
## Thu Jul 14 01:42:06 CEST 2022 - The aliscore output file does not exist. Check results for BUSCO: 106667at7742
## Thu Jul 14 01:42:06 CEST 2022 - The aliscore output appears to be empty. Check results for BUSCO: 106667at7742
## Thu Jul 14 01:42:08 CEST 2022 - The aliscore output file does not exist. Check results for BUSCO: 159866at7742
## Thu Jul 14 01:42:08 CEST 2022 - The aliscore output appears to be empty. Check results for BUSCO: 159866at7742
## Thu Jul 14 01:42:08 CEST 2022 - The aliscore output file does not exist. Check results for BUSCO: 208627at7742
## Thu Jul 14 01:42:09 CEST 2022 - The aliscore output appears to be empty. Check results for BUSCO: 208627at7742
## Thu Jul 14 01:42:10 CEST 2022 - The aliscore output file does not exist. Check results for BUSCO: 130337at7742
## Thu Jul 14 01:42:10 CEST 2022 - The aliscore output appears to be empty. Check results for BUSCO: 130337at7742
## Thu Jul 14 01:42:11 CEST 2022 - The aliscore output file does not exist. Check results for BUSCO: 288766at7742
## Thu Jul 14 01:42:12 CEST 2022 - The aliscore output appears to be empty. Check results for BUSCO: 288766at7742
## Thu Jul 14 01:42:13 CEST 2022 - The aliscore output file does not exist. Check results for BUSCO: 182742at7742
## Thu Jul 14 01:42:13 CEST 2022 - The aliscore output file does not exist. Check results for BUSCO: 385085at7742
## Thu Jul 14 01:42:13 CEST 2022 - The aliscore output appears to be empty. Check results for BUSCO: 182742at7742
## Thu Jul 14 01:42:13 CEST 2022 - The aliscore output appears to be empty. Check results for BUSCO: 385085at7742
## Thu Jul 14 01:42:14 CEST 2022 - The aliscore output file does not exist. Check results for BUSCO: 359184at7742
## Thu Jul 14 01:42:14 CEST 2022 - The aliscore output appears to be empty. Check results for BUSCO: 359184at7742
## Thu Jul 14 01:42:15 CEST 2022 - The aliscore output file does not exist. Check results for BUSCO: 337000at7742
## Thu Jul 14 01:42:15 CEST 2022 - The aliscore output appears to be empty. Check results for BUSCO: 337000at7742
## Thu Jul 14 01:42:16 CEST 2022 - The aliscore output file does not exist. Check results for BUSCO: 220062at7742
## Thu Jul 14 01:42:16 CEST 2022 - The aliscore output appears to be empty. Check results for BUSCO: 220062at7742
## Thu Jul 14 01:42:17 CEST 2022 - The aliscore output file does not exist. Check results for BUSCO: 319402at7742
## Thu Jul 14 01:42:17 CEST 2022 - The aliscore output appears to be empty. Check results for BUSCO: 319402at7742
## Thu Jul 14 01:42:20 CEST 2022 - The aliscore output file does not exist. Check results for BUSCO: 238259at7742
## Thu Jul 14 01:42:20 CEST 2022 - The aliscore output appears to be empty. Check results for BUSCO: 238259at7742
## Thu Jul 14 01:42:21 CEST 2022 - The aliscore output file does not exist. Check results for BUSCO: 227080at7742
## Thu Jul 14 01:42:21 CEST 2022 - The aliscore output appears to be empty. Check results for BUSCO: 227080at7742
## Thu Jul 14 01:42:22 CEST 2022 - The aliscore output file does not exist. Check results for BUSCO: 192058at7742
## Thu Jul 14 01:42:22 CEST 2022 - The aliscore output appears to be empty. Check results for BUSCO: 192058at7742
## Thu Jul 14 01:42:22 CEST 2022 - The aliscore output file does not exist. Check results for BUSCO: 188059at7742
## Thu Jul 14 01:42:22 CEST 2022 - The aliscore output appears to be empty. Check results for BUSCO: 188059at7742
## Thu Jul 14 01:42:24 CEST 2022 - The aliscore output file does not exist. Check results for BUSCO: 203960at7742
## Thu Jul 14 01:42:24 CEST 2022 - The aliscore output appears to be empty. Check results for BUSCO: 203960at7742
## Thu Jul 14 01:42:25 CEST 2022 - The aliscore output file does not exist. Check results for BUSCO: 106667at7742
## Thu Jul 14 01:42:25 CEST 2022 - The aliscore output appears to be empty. Check results for BUSCO: 106667at7742
## Thu Jul 14 01:42:26 CEST 2022 - The aliscore output file does not exist. Check results for BUSCO: 148672at7742
## Thu Jul 14 01:42:27 CEST 2022 - The aliscore output appears to be empty. Check results for BUSCO: 148672at7742
## Thu Jul 14 01:42:27 CEST 2022 - The aliscore output file does not exist. Check results for BUSCO: 158801at7742
## Thu Jul 14 01:42:27 CEST 2022 - The aliscore output appears to be empty. Check results for BUSCO: 158801at7742
## Thu Jul 14 01:42:28 CEST 2022 - The aliscore output file does not exist. Check results for BUSCO: 109315at7742
## Thu Jul 14 01:42:28 CEST 2022 - The aliscore output appears to be empty. Check results for BUSCO: 109315at7742
## Thu Jul 14 01:42:29 CEST 2022 - The aliscore output file does not exist. Check results for BUSCO: 276660at7742
## Thu Jul 14 01:42:29 CEST 2022 - The aliscore output appears to be empty. Check results for BUSCO: 276660at7742
## Thu Jul 14 01:42:30 CEST 2022 - The aliscore output file does not exist. Check results for BUSCO: 202302at7742
## Thu Jul 14 01:42:30 CEST 2022 - The aliscore output appears to be empty. Check results for BUSCO: 202302at7742
## Thu Jul 14 01:42:32 CEST 2022 - The aliscore output file does not exist. Check results for BUSCO: 330054at7742
## Thu Jul 14 01:42:32 CEST 2022 - The aliscore output appears to be empty. Check results for BUSCO: 330054at7742
## Thu Jul 14 01:42:34 CEST 2022 - The aliscore output file does not exist. Check results for BUSCO: 242641at7742
## Thu Jul 14 01:42:34 CEST 2022 - The aliscore output appears to be empty. Check results for BUSCO: 242641at7742
## Thu Jul 14 01:42:34 CEST 2022 - The aliscore output file does not exist. Check results for BUSCO: 162212at7742
## Thu Jul 14 01:42:34 CEST 2022 - The aliscore output appears to be empty. Check results for BUSCO: 162212at7742
## Thu Jul 14 01:42:37 CEST 2022 - The aliscore output file does not exist. Check results for BUSCO: 274707at7742
## Thu Jul 14 01:42:37 CEST 2022 - The aliscore output appears to be empty. Check results for BUSCO: 274707at7742
## Thu Jul 14 01:42:37 CEST 2022 - The aliscore output file does not exist. Check results for BUSCO: 192331at7742
## Thu Jul 14 01:42:37 CEST 2022 - The aliscore output appears to be empty. Check results for BUSCO: 192331at7742
## Thu Jul 14 01:42:37 CEST 2022 - The aliscore output file does not exist. Check results for BUSCO: 231455at7742
## Thu Jul 14 01:42:37 CEST 2022 - The aliscore output appears to be empty. Check results for BUSCO: 231455at7742
## Thu Jul 14 01:42:39 CEST 2022 - The aliscore output file does not exist. Check results for BUSCO: 142719at7742
## Thu Jul 14 01:42:39 CEST 2022 - The aliscore output appears to be empty. Check results for BUSCO: 142719at7742
## Thu Jul 14 01:42:41 CEST 2022 - The aliscore output file does not exist. Check results for BUSCO: 220601at7742
## Thu Jul 14 01:42:41 CEST 2022 - The aliscore output appears to be empty. Check results for BUSCO: 220601at7742
## Thu Jul 14 01:42:41 CEST 2022 - The aliscore output file does not exist. Check results for BUSCO: 160383at7742
## Thu Jul 14 01:42:41 CEST 2022 - The aliscore output appears to be empty. Check results for BUSCO: 160383at7742
## Thu Jul 14 01:42:42 CEST 2022 - The aliscore output file does not exist. Check results for BUSCO: 220601at7742
## Thu Jul 14 01:42:42 CEST 2022 - The aliscore output appears to be empty. Check results for BUSCO: 220601at7742
## Thu Jul 14 01:42:45 CEST 2022 - The aliscore output file does not exist. Check results for BUSCO: 191044at7742
## Thu Jul 14 01:42:45 CEST 2022 - The aliscore output appears to be empty. Check results for BUSCO: 191044at7742
## Thu Jul 14 01:42:45 CEST 2022 - The aliscore output file does not exist. Check results for BUSCO: 311983at7742
## Thu Jul 14 01:42:45 CEST 2022 - The aliscore output appears to be empty. Check results for BUSCO: 311983at7742
## Thu Jul 14 01:42:46 CEST 2022 - The aliscore output file does not exist. Check results for BUSCO: 363574at7742
## Thu Jul 14 01:42:46 CEST 2022 - The aliscore output appears to be empty. Check results for BUSCO: 363574at7742
## Thu Jul 14 01:42:47 CEST 2022 - The aliscore output file does not exist. Check results for BUSCO: 228390at7742
## Thu Jul 14 01:42:47 CEST 2022 - The aliscore output appears to be empty. Check results for BUSCO: 228390at7742
## Thu Jul 14 01:42:48 CEST 2022 - The aliscore output file does not exist. Check results for BUSCO: 251465at7742
## Thu Jul 14 01:42:48 CEST 2022 - The aliscore output appears to be empty. Check results for BUSCO: 251465at7742
## Thu Jul 14 01:42:51 CEST 2022 - The aliscore output file does not exist. Check results for BUSCO: 361935at7742
## Thu Jul 14 01:42:51 CEST 2022 - The aliscore output appears to be empty. Check results for BUSCO: 361935at7742
## Thu Jul 14 01:42:51 CEST 2022 - The aliscore output file does not exist. Check results for BUSCO: 361935at7742
## Thu Jul 14 01:42:52 CEST 2022 - The aliscore output appears to be empty. Check results for BUSCO: 361935at7742
## Thu Jul 14 01:42:52 CEST 2022 - The aliscore output file does not exist. Check results for BUSCO: 160732at7742
## Thu Jul 14 01:42:52 CEST 2022 - The aliscore output appears to be empty. Check results for BUSCO: 160732at7742
## Thu Jul 14 01:42:53 CEST 2022 - The aliscore output file does not exist. Check results for BUSCO: 161775at7742
## Thu Jul 14 01:42:53 CEST 2022 - The aliscore output appears to be empty. Check results for BUSCO: 161775at7742
## Thu Jul 14 01:42:55 CEST 2022 - The aliscore output file does not exist. Check results for BUSCO: 260359at7742
## Thu Jul 14 01:42:55 CEST 2022 - The aliscore output appears to be empty. Check results for BUSCO: 260359at7742
## Thu Jul 14 01:42:55 CEST 2022 - The aliscore output file does not exist. Check results for BUSCO: 169338at7742
## Thu Jul 14 01:42:55 CEST 2022 - The aliscore output appears to be empty. Check results for BUSCO: 169338at7742
## Thu Jul 14 01:42:58 CEST 2022 - The aliscore output file does not exist. Check results for BUSCO: 237617at7742
## Thu Jul 14 01:42:58 CEST 2022 - The aliscore output appears to be empty. Check results for BUSCO: 237617at7742
## Thu Jul 14 01:42:59 CEST 2022 - The aliscore output file does not exist. Check results for BUSCO: 386739at7742
## Thu Jul 14 01:42:59 CEST 2022 - The aliscore output appears to be empty. Check results for BUSCO: 386739at7742
## Thu Jul 14 01:43:00 CEST 2022 - The aliscore output file does not exist. Check results for BUSCO: 245117at7742
## Thu Jul 14 01:43:00 CEST 2022 - The aliscore output appears to be empty. Check results for BUSCO: 245117at7742
## Thu Jul 14 01:43:01 CEST 2022 - The aliscore output file does not exist. Check results for BUSCO: 202155at7742
## Thu Jul 14 01:43:01 CEST 2022 - The aliscore output appears to be empty. Check results for BUSCO: 202155at7742
## Thu Jul 14 01:43:03 CEST 2022 - The aliscore output file does not exist. Check results for BUSCO: 165642at7742
## Thu Jul 14 01:43:03 CEST 2022 - The aliscore output appears to be empty. Check results for BUSCO: 165642at7742
## Thu Jul 14 01:43:03 CEST 2022 - The aliscore output file does not exist. Check results for BUSCO: 199024at7742
## Thu Jul 14 01:43:03 CEST 2022 - The aliscore output appears to be empty. Check results for BUSCO: 199024at7742
## Thu Jul 14 01:43:06 CEST 2022 - The aliscore output file does not exist. Check results for BUSCO: 227544at7742
## Thu Jul 14 01:43:06 CEST 2022 - The aliscore output appears to be empty. Check results for BUSCO: 227544at7742
## Thu Jul 14 01:43:06 CEST 2022 - The aliscore output file does not exist. Check results for BUSCO: 260359at7742
## Thu Jul 14 01:43:06 CEST 2022 - The aliscore output appears to be empty. Check results for BUSCO: 260359at7742
## Thu Jul 14 01:43:08 CEST 2022 - The aliscore output file does not exist. Check results for BUSCO: 182049at7742
## Thu Jul 14 01:43:08 CEST 2022 - The aliscore output appears to be empty. Check results for BUSCO: 182049at7742
## Thu Jul 14 01:43:08 CEST 2022 - The aliscore output file does not exist. Check results for BUSCO: 137723at7742
## Thu Jul 14 01:43:08 CEST 2022 - The aliscore output appears to be empty. Check results for BUSCO: 137723at7742
## Thu Jul 14 01:43:10 CEST 2022 - The aliscore output file does not exist. Check results for BUSCO: 242895at7742
## Thu Jul 14 01:43:10 CEST 2022 - The aliscore output appears to be empty. Check results for BUSCO: 242895at7742
## Thu Jul 14 01:43:11 CEST 2022 - The aliscore output file does not exist. Check results for BUSCO: 189753at7742
## Thu Jul 14 01:43:12 CEST 2022 - The aliscore output appears to be empty. Check results for BUSCO: 189753at7742
## Thu Jul 14 01:43:13 CEST 2022 - The aliscore output file does not exist. Check results for BUSCO: 160732at7742
## Thu Jul 14 01:43:13 CEST 2022 - The aliscore output appears to be empty. Check results for BUSCO: 160732at7742
## Thu Jul 14 01:43:14 CEST 2022 - The aliscore output file does not exist. Check results for BUSCO: 212696at7742
## Thu Jul 14 01:43:14 CEST 2022 - The aliscore output appears to be empty. Check results for BUSCO: 212696at7742
## Thu Jul 14 01:43:16 CEST 2022 - The aliscore output file does not exist. Check results for BUSCO: 208627at7742
## Thu Jul 14 01:43:16 CEST 2022 - The aliscore output appears to be empty. Check results for BUSCO: 208627at7742
## Thu Jul 14 01:43:16 CEST 2022 - The aliscore output file does not exist. Check results for BUSCO: 159866at7742
## Thu Jul 14 01:43:16 CEST 2022 - The aliscore output appears to be empty. Check results for BUSCO: 159866at7742
## Thu Jul 14 01:43:17 CEST 2022 - The aliscore output file does not exist. Check results for BUSCO: 298884at7742
## Thu Jul 14 01:43:17 CEST 2022 - The aliscore output appears to be empty. Check results for BUSCO: 298884at7742
## Thu Jul 14 01:43:18 CEST 2022 - The aliscore output file does not exist. Check results for BUSCO: 265814at7742
## Thu Jul 14 01:43:18 CEST 2022 - The aliscore output appears to be empty. Check results for BUSCO: 265814at7742
## Thu Jul 14 01:43:20 CEST 2022 - The aliscore output file does not exist. Check results for BUSCO: 83843at7742
## Thu Jul 14 01:43:20 CEST 2022 - The aliscore output appears to be empty. Check results for BUSCO: 83843at7742
## Thu Jul 14 01:43:21 CEST 2022 - The aliscore output file does not exist. Check results for BUSCO: 225125at7742
## Thu Jul 14 01:43:21 CEST 2022 - The aliscore output appears to be empty. Check results for BUSCO: 225125at7742
## Thu Jul 14 01:43:21 CEST 2022 - The aliscore output file does not exist. Check results for BUSCO: 232798at7742
## Thu Jul 14 01:43:21 CEST 2022 - The aliscore output appears to be empty. Check results for BUSCO: 232798at7742
## Thu Jul 14 01:43:24 CEST 2022 - The aliscore output file does not exist. Check results for BUSCO: 119371at7742
## Thu Jul 14 01:43:24 CEST 2022 - The aliscore output appears to be empty. Check results for BUSCO: 119371at7742
## Thu Jul 14 01:43:24 CEST 2022 - The aliscore output file does not exist. Check results for BUSCO: 229564at7742
## Thu Jul 14 01:43:24 CEST 2022 - The aliscore output appears to be empty. Check results for BUSCO: 229564at7742
## Thu Jul 14 01:43:25 CEST 2022 - The aliscore output file does not exist. Check results for BUSCO: 191044at7742
## Thu Jul 14 01:43:25 CEST 2022 - The aliscore output appears to be empty. Check results for BUSCO: 191044at7742
## Thu Jul 14 01:43:27 CEST 2022 - The aliscore output file does not exist. Check results for BUSCO: 273630at7742
## Thu Jul 14 01:43:27 CEST 2022 - The aliscore output appears to be empty. Check results for BUSCO: 273630at7742
## Thu Jul 14 01:43:29 CEST 2022 - The aliscore output file does not exist. Check results for BUSCO: 83843at7742
## Thu Jul 14 01:43:29 CEST 2022 - The aliscore output appears to be empty. Check results for BUSCO: 83843at7742
## Thu Jul 14 01:43:29 CEST 2022 - The aliscore output file does not exist. Check results for BUSCO: 359184at7742
## Thu Jul 14 01:43:29 CEST 2022 - The aliscore output appears to be empty. Check results for BUSCO: 359184at7742
## Thu Jul 14 01:43:29 CEST 2022 - The aliscore output file does not exist. Check results for BUSCO: 327391at7742
## Thu Jul 14 01:43:29 CEST 2022 - The aliscore output appears to be empty. Check results for BUSCO: 327391at7742
## Thu Jul 14 01:43:30 CEST 2022 - The aliscore output file does not exist. Check results for BUSCO: 296930at7742
## Thu Jul 14 01:43:30 CEST 2022 - The aliscore output appears to be empty. Check results for BUSCO: 296930at7742
## Thu Jul 14 01:43:31 CEST 2022 - The aliscore output file does not exist. Check results for BUSCO: 254424at7742
## Thu Jul 14 01:43:31 CEST 2022 - The aliscore output appears to be empty. Check results for BUSCO: 254424at7742
## Thu Jul 14 01:43:33 CEST 2022 - The aliscore output file does not exist. Check results for BUSCO: 290906at7742
## Thu Jul 14 01:43:33 CEST 2022 - The aliscore output appears to be empty. Check results for BUSCO: 290906at7742
## Thu Jul 14 01:43:35 CEST 2022 - The aliscore output file does not exist. Check results for BUSCO: 145137at7742
## Thu Jul 14 01:43:35 CEST 2022 - The aliscore output appears to be empty. Check results for BUSCO: 145137at7742
## Thu Jul 14 01:43:36 CEST 2022 - The aliscore output file does not exist. Check results for BUSCO: 198321at7742
## Thu Jul 14 01:43:36 CEST 2022 - The aliscore output appears to be empty. Check results for BUSCO: 198321at7742
## Thu Jul 14 01:43:37 CEST 2022 - The aliscore output file does not exist. Check results for BUSCO: 220484at7742
## Thu Jul 14 01:43:37 CEST 2022 - The aliscore output appears to be empty. Check results for BUSCO: 220484at7742
## Thu Jul 14 01:43:38 CEST 2022 - The aliscore output file does not exist. Check results for BUSCO: 356998at7742
## Thu Jul 14 01:43:38 CEST 2022 - The aliscore output appears to be empty. Check results for BUSCO: 356998at7742
## Thu Jul 14 01:43:39 CEST 2022 - The aliscore output file does not exist. Check results for BUSCO: 161775at7742
## Thu Jul 14 01:43:39 CEST 2022 - The aliscore output appears to be empty. Check results for BUSCO: 161775at7742
## Thu Jul 14 01:43:41 CEST 2022 - The aliscore output file does not exist. Check results for BUSCO: 242072at7742
## Thu Jul 14 01:43:41 CEST 2022 - The aliscore output file does not exist. Check results for BUSCO: 169728at7742
## Thu Jul 14 01:43:41 CEST 2022 - The aliscore output appears to be empty. Check results for BUSCO: 242072at7742
## Thu Jul 14 01:43:41 CEST 2022 - The aliscore output appears to be empty. Check results for BUSCO: 169728at7742
## Thu Jul 14 01:43:42 CEST 2022 - The aliscore output file does not exist. Check results for BUSCO: 163177at7742
## Thu Jul 14 01:43:42 CEST 2022 - The aliscore output appears to be empty. Check results for BUSCO: 163177at7742
## Thu Jul 14 01:43:42 CEST 2022 - The aliscore output file does not exist. Check results for BUSCO: 275256at7742
## Thu Jul 14 01:43:42 CEST 2022 - The aliscore output appears to be empty. Check results for BUSCO: 275256at7742
## Thu Jul 14 01:43:44 CEST 2022 - The aliscore output file does not exist. Check results for BUSCO: 185697at7742
## Thu Jul 14 01:43:44 CEST 2022 - The aliscore output appears to be empty. Check results for BUSCO: 185697at7742
## Thu Jul 14 01:43:46 CEST 2022 - The aliscore output file does not exist. Check results for BUSCO: 257616at7742
## Thu Jul 14 01:43:46 CEST 2022 - The aliscore output appears to be empty. Check results for BUSCO: 257616at7742
## Thu Jul 14 01:43:47 CEST 2022 - The aliscore output file does not exist. Check results for BUSCO: 227816at7742
## Thu Jul 14 01:43:47 CEST 2022 - The aliscore output appears to be empty. Check results for BUSCO: 227816at7742
## Thu Jul 14 01:43:47 CEST 2022 - The aliscore output file does not exist. Check results for BUSCO: 272002at7742
## Thu Jul 14 01:43:47 CEST 2022 - The aliscore output appears to be empty. Check results for BUSCO: 272002at7742
## Thu Jul 14 01:43:49 CEST 2022 - The aliscore output file does not exist. Check results for BUSCO: 202970at7742
## Thu Jul 14 01:43:49 CEST 2022 - The aliscore output appears to be empty. Check results for BUSCO: 202970at7742
## Thu Jul 14 01:43:50 CEST 2022 - The aliscore output file does not exist. Check results for BUSCO: 353601at7742
## Thu Jul 14 01:43:50 CEST 2022 - The aliscore output appears to be empty. Check results for BUSCO: 353601at7742
## Thu Jul 14 01:43:50 CEST 2022 - The aliscore output file does not exist. Check results for BUSCO: 117231at7742
## Thu Jul 14 01:43:50 CEST 2022 - The aliscore output appears to be empty. Check results for BUSCO: 117231at7742
## Thu Jul 14 01:43:52 CEST 2022 - The aliscore output file does not exist. Check results for BUSCO: 355364at7742
## Thu Jul 14 01:43:52 CEST 2022 - The aliscore output appears to be empty. Check results for BUSCO: 355364at7742
## Thu Jul 14 01:43:53 CEST 2022 - The aliscore output file does not exist. Check results for BUSCO: 274707at7742
## Thu Jul 14 01:43:53 CEST 2022 - The aliscore output appears to be empty. Check results for BUSCO: 274707at7742
## Thu Jul 14 01:43:54 CEST 2022 - The aliscore output file does not exist. Check results for BUSCO: 245056at7742
## Thu Jul 14 01:43:54 CEST 2022 - The aliscore output appears to be empty. Check results for BUSCO: 245056at7742
## Thu Jul 14 01:43:56 CEST 2022 - The aliscore output file does not exist. Check results for BUSCO: 245056at7742
## Thu Jul 14 01:43:56 CEST 2022 - The aliscore output appears to be empty. Check results for BUSCO: 245056at7742
## Thu Jul 14 01:43:57 CEST 2022 - The aliscore output file does not exist. Check results for BUSCO: 198321at7742
## Thu Jul 14 01:43:57 CEST 2022 - The aliscore output appears to be empty. Check results for BUSCO: 198321at7742
## Thu Jul 14 01:43:57 CEST 2022 - The aliscore output file does not exist. Check results for BUSCO: 201359at7742
## Thu Jul 14 01:43:57 CEST 2022 - The aliscore output appears to be empty. Check results for BUSCO: 201359at7742
## Thu Jul 14 01:43:58 CEST 2022 - The aliscore output file does not exist. Check results for BUSCO: 286595at7742
## Thu Jul 14 01:43:58 CEST 2022 - The aliscore output appears to be empty. Check results for BUSCO: 286595at7742
## Thu Jul 14 01:44:00 CEST 2022 - The aliscore output file does not exist. Check results for BUSCO: 189224at7742
## Thu Jul 14 01:44:00 CEST 2022 - The aliscore output appears to be empty. Check results for BUSCO: 189224at7742
## Thu Jul 14 01:44:02 CEST 2022 - The aliscore output file does not exist. Check results for BUSCO: 121516at7742
## Thu Jul 14 01:44:02 CEST 2022 - The aliscore output appears to be empty. Check results for BUSCO: 121516at7742
## Thu Jul 14 01:44:02 CEST 2022 - The aliscore output file does not exist. Check results for BUSCO: 380097at7742
## Thu Jul 14 01:44:02 CEST 2022 - The aliscore output appears to be empty. Check results for BUSCO: 380097at7742
## Thu Jul 14 01:44:04 CEST 2022 - The aliscore output file does not exist. Check results for BUSCO: 190766at7742
## Thu Jul 14 01:44:04 CEST 2022 - The aliscore output appears to be empty. Check results for BUSCO: 190766at7742
## Thu Jul 14 01:44:06 CEST 2022 - The aliscore output file does not exist. Check results for BUSCO: 261602at7742
## Thu Jul 14 01:44:06 CEST 2022 - The aliscore output appears to be empty. Check results for BUSCO: 261602at7742
## Thu Jul 14 01:44:06 CEST 2022 - The aliscore output file does not exist. Check results for BUSCO: 235765at7742
## Thu Jul 14 01:44:06 CEST 2022 - The aliscore output appears to be empty. Check results for BUSCO: 235765at7742
## Thu Jul 14 01:44:08 CEST 2022 - The aliscore output file does not exist. Check results for BUSCO: 310260at7742
## Thu Jul 14 01:44:08 CEST 2022 - The aliscore output appears to be empty. Check results for BUSCO: 310260at7742
## Thu Jul 14 01:44:08 CEST 2022 - The aliscore output file does not exist. Check results for BUSCO: 160732at7742
## Thu Jul 14 01:44:08 CEST 2022 - The aliscore output file does not exist. Check results for BUSCO: 154520at7742
## Thu Jul 14 01:44:08 CEST 2022 - The aliscore output appears to be empty. Check results for BUSCO: 160732at7742
## Thu Jul 14 01:44:08 CEST 2022 - The aliscore output appears to be empty. Check results for BUSCO: 154520at7742
## Thu Jul 14 01:44:10 CEST 2022 - The aliscore output file does not exist. Check results for BUSCO: 277067at7742
## Thu Jul 14 01:44:10 CEST 2022 - The aliscore output appears to be empty. Check results for BUSCO: 277067at7742
## Thu Jul 14 01:44:11 CEST 2022 - The aliscore output file does not exist. Check results for BUSCO: 101854at7742
## Thu Jul 14 01:44:11 CEST 2022 - The aliscore output appears to be empty. Check results for BUSCO: 101854at7742
## Thu Jul 14 01:44:13 CEST 2022 - The aliscore output file does not exist. Check results for BUSCO: 182463at7742
## Thu Jul 14 01:44:13 CEST 2022 - The aliscore output appears to be empty. Check results for BUSCO: 182463at7742
## Thu Jul 14 01:44:13 CEST 2022 - The aliscore output file does not exist. Check results for BUSCO: 178490at7742
## Thu Jul 14 01:44:13 CEST 2022 - The aliscore output appears to be empty. Check results for BUSCO: 178490at7742
## Thu Jul 14 01:44:13 CEST 2022 - The aliscore output file does not exist. Check results for BUSCO: 117231at7742
## Thu Jul 14 01:44:13 CEST 2022 - The aliscore output appears to be empty. Check results for BUSCO: 117231at7742
## Thu Jul 14 01:44:16 CEST 2022 - The aliscore output file does not exist. Check results for BUSCO: 165642at7742
## Thu Jul 14 01:44:16 CEST 2022 - The aliscore output appears to be empty. Check results for BUSCO: 165642at7742
## Thu Jul 14 01:44:17 CEST 2022 - The aliscore output file does not exist. Check results for BUSCO: 327404at7742
## Thu Jul 14 01:44:17 CEST 2022 - The aliscore output appears to be empty. Check results for BUSCO: 327404at7742
## Thu Jul 14 01:44:19 CEST 2022 - The aliscore output file does not exist. Check results for BUSCO: 219805at7742
## Thu Jul 14 01:44:19 CEST 2022 - The aliscore output appears to be empty. Check results for BUSCO: 219805at7742
## Thu Jul 14 01:44:19 CEST 2022 - The aliscore output file does not exist. Check results for BUSCO: 262798at7742
## Thu Jul 14 01:44:20 CEST 2022 - The aliscore output appears to be empty. Check results for BUSCO: 262798at7742
## Thu Jul 14 01:44:21 CEST 2022 - The aliscore output file does not exist. Check results for BUSCO: 225148at7742
## Thu Jul 14 01:44:21 CEST 2022 - The aliscore output appears to be empty. Check results for BUSCO: 225148at7742
## Thu Jul 14 01:44:21 CEST 2022 - The aliscore output file does not exist. Check results for BUSCO: 285172at7742
## Thu Jul 14 01:44:22 CEST 2022 - The aliscore output appears to be empty. Check results for BUSCO: 285172at7742
## Thu Jul 14 01:44:23 CEST 2022 - The aliscore output file does not exist. Check results for BUSCO: 325908at7742
## Thu Jul 14 01:44:23 CEST 2022 - The aliscore output appears to be empty. Check results for BUSCO: 325908at7742
## Thu Jul 14 01:44:24 CEST 2022 - The aliscore output file does not exist. Check results for BUSCO: 238259at7742
## Thu Jul 14 01:44:24 CEST 2022 - The aliscore output appears to be empty. Check results for BUSCO: 238259at7742
## Thu Jul 14 01:44:25 CEST 2022 - The aliscore output file does not exist. Check results for BUSCO: 394021at7742
## Thu Jul 14 01:44:25 CEST 2022 - The aliscore output appears to be empty. Check results for BUSCO: 394021at7742
## Thu Jul 14 01:44:27 CEST 2022 - The aliscore output file does not exist. Check results for BUSCO: 165642at7742
## Thu Jul 14 01:44:27 CEST 2022 - The aliscore output appears to be empty. Check results for BUSCO: 165642at7742
## Thu Jul 14 01:44:28 CEST 2022 - The aliscore output file does not exist. Check results for BUSCO: 266219at7742
## Thu Jul 14 01:44:28 CEST 2022 - The aliscore output appears to be empty. Check results for BUSCO: 266219at7742
## Thu Jul 14 01:44:29 CEST 2022 - The aliscore output file does not exist. Check results for BUSCO: 133153at7742
## Thu Jul 14 01:44:29 CEST 2022 - The aliscore output appears to be empty. Check results for BUSCO: 133153at7742
## Thu Jul 14 01:44:29 CEST 2022 - The aliscore output file does not exist. Check results for BUSCO: 276691at7742
## Thu Jul 14 01:44:29 CEST 2022 - The aliscore output appears to be empty. Check results for BUSCO: 276691at7742
## Thu Jul 14 01:44:31 CEST 2022 - The aliscore output file does not exist. Check results for BUSCO: 234228at7742
## Thu Jul 14 01:44:31 CEST 2022 - The aliscore output appears to be empty. Check results for BUSCO: 234228at7742
## Thu Jul 14 01:44:32 CEST 2022 - The aliscore output file does not exist. Check results for BUSCO: 200462at7742
## Thu Jul 14 01:44:32 CEST 2022 - The aliscore output appears to be empty. Check results for BUSCO: 200462at7742
## Thu Jul 14 01:44:33 CEST 2022 - The aliscore output file does not exist. Check results for BUSCO: 275256at7742
## Thu Jul 14 01:44:33 CEST 2022 - The aliscore output appears to be empty. Check results for BUSCO: 275256at7742
## Thu Jul 14 01:44:34 CEST 2022 - The aliscore output file does not exist. Check results for BUSCO: 265814at7742
## Thu Jul 14 01:44:34 CEST 2022 - The aliscore output appears to be empty. Check results for BUSCO: 265814at7742
## Thu Jul 14 01:44:36 CEST 2022 - The aliscore output file does not exist. Check results for BUSCO: 204708at7742
## Thu Jul 14 01:44:36 CEST 2022 - The aliscore output appears to be empty. Check results for BUSCO: 204708at7742
## Thu Jul 14 01:44:36 CEST 2022 - The aliscore output file does not exist. Check results for BUSCO: 241220at7742
## Thu Jul 14 01:44:36 CEST 2022 - The aliscore output appears to be empty. Check results for BUSCO: 241220at7742
## Thu Jul 14 01:44:37 CEST 2022 - The aliscore output file does not exist. Check results for BUSCO: 118164at7742
## Thu Jul 14 01:44:37 CEST 2022 - The aliscore output appears to be empty. Check results for BUSCO: 118164at7742
## Thu Jul 14 01:44:38 CEST 2022 - The aliscore output file does not exist. Check results for BUSCO: 330064at7742
## Thu Jul 14 01:44:38 CEST 2022 - The aliscore output appears to be empty. Check results for BUSCO: 330064at7742
## Thu Jul 14 01:44:39 CEST 2022 - The aliscore output file does not exist. Check results for BUSCO: 305085at7742
## Thu Jul 14 01:44:39 CEST 2022 - The aliscore output appears to be empty. Check results for BUSCO: 305085at7742
## Thu Jul 14 01:44:41 CEST 2022 - The aliscore output file does not exist. Check results for BUSCO: 207618at7742
## Thu Jul 14 01:44:41 CEST 2022 - The aliscore output appears to be empty. Check results for BUSCO: 207618at7742
## Thu Jul 14 01:44:42 CEST 2022 - The aliscore output file does not exist. Check results for BUSCO: 144352at7742
## Thu Jul 14 01:44:42 CEST 2022 - The aliscore output appears to be empty. Check results for BUSCO: 144352at7742
## Thu Jul 14 01:44:44 CEST 2022 - The aliscore output file does not exist. Check results for BUSCO: 199780at7742
## Thu Jul 14 01:44:44 CEST 2022 - The aliscore output appears to be empty. Check results for BUSCO: 199780at7742
## Thu Jul 14 01:44:45 CEST 2022 - The aliscore output file does not exist. Check results for BUSCO: 223443at7742
## Thu Jul 14 01:44:45 CEST 2022 - The aliscore output appears to be empty. Check results for BUSCO: 223443at7742
## Thu Jul 14 01:44:45 CEST 2022 - The aliscore output file does not exist. Check results for BUSCO: 361935at7742
## Thu Jul 14 01:44:45 CEST 2022 - The aliscore output appears to be empty. Check results for BUSCO: 361935at7742
## Thu Jul 14 01:44:47 CEST 2022 - The aliscore output file does not exist. Check results for BUSCO: 322442at7742
## Thu Jul 14 01:44:47 CEST 2022 - The aliscore output appears to be empty. Check results for BUSCO: 322442at7742
## Thu Jul 14 01:44:47 CEST 2022 - The aliscore output file does not exist. Check results for BUSCO: 262798at7742
## Thu Jul 14 01:44:47 CEST 2022 - The aliscore output appears to be empty. Check results for BUSCO: 262798at7742
## Thu Jul 14 01:44:50 CEST 2022 - The aliscore output file does not exist. Check results for BUSCO: 148672at7742
## Thu Jul 14 01:44:50 CEST 2022 - The aliscore output appears to be empty. Check results for BUSCO: 148672at7742
## Thu Jul 14 01:44:51 CEST 2022 - The aliscore output file does not exist. Check results for BUSCO: 213005at7742
## Thu Jul 14 01:44:51 CEST 2022 - The aliscore output appears to be empty. Check results for BUSCO: 213005at7742
## Thu Jul 14 01:44:51 CEST 2022 - The aliscore output file does not exist. Check results for BUSCO: 243611at7742
## Thu Jul 14 01:44:51 CEST 2022 - The aliscore output appears to be empty. Check results for BUSCO: 243611at7742
## Thu Jul 14 01:44:53 CEST 2022 - The aliscore output file does not exist. Check results for BUSCO: 234228at7742
## Thu Jul 14 01:44:53 CEST 2022 - The aliscore output appears to be empty. Check results for BUSCO: 234228at7742
## Thu Jul 14 01:44:54 CEST 2022 - The aliscore output file does not exist. Check results for BUSCO: 337000at7742
## Thu Jul 14 01:44:54 CEST 2022 - The aliscore output appears to be empty. Check results for BUSCO: 337000at7742
## Thu Jul 14 01:44:55 CEST 2022 - The aliscore output file does not exist. Check results for BUSCO: 199723at7742
## Thu Jul 14 01:44:55 CEST 2022 - The aliscore output appears to be empty. Check results for BUSCO: 199723at7742
## Thu Jul 14 01:44:56 CEST 2022 - The aliscore output file does not exist. Check results for BUSCO: 154520at7742
## Thu Jul 14 01:44:56 CEST 2022 - The aliscore output appears to be empty. Check results for BUSCO: 154520at7742
## Thu Jul 14 01:44:56 CEST 2022 - The aliscore output file does not exist. Check results for BUSCO: 208840at7742
## Thu Jul 14 01:44:56 CEST 2022 - The aliscore output appears to be empty. Check results for BUSCO: 208840at7742
## Thu Jul 14 01:44:59 CEST 2022 - The aliscore output file does not exist. Check results for BUSCO: 344933at7742
## Thu Jul 14 01:45:00 CEST 2022 - The aliscore output appears to be empty. Check results for BUSCO: 344933at7742
## Thu Jul 14 01:45:00 CEST 2022 - The aliscore output file does not exist. Check results for BUSCO: 192665at7742
## Thu Jul 14 01:45:00 CEST 2022 - The aliscore output appears to be empty. Check results for BUSCO: 192665at7742
## Thu Jul 14 01:45:01 CEST 2022 - The aliscore output file does not exist. Check results for BUSCO: 410850at7742
## Thu Jul 14 01:45:01 CEST 2022 - The aliscore output appears to be empty. Check results for BUSCO: 410850at7742
## Thu Jul 14 01:45:02 CEST 2022 - The aliscore output file does not exist. Check results for BUSCO: 223443at7742
## Thu Jul 14 01:45:02 CEST 2022 - The aliscore output appears to be empty. Check results for BUSCO: 223443at7742
## Thu Jul 14 01:45:03 CEST 2022 - The aliscore output file does not exist. Check results for BUSCO: 169313at7742
## Thu Jul 14 01:45:03 CEST 2022 - The aliscore output appears to be empty. Check results for BUSCO: 169313at7742
## Thu Jul 14 01:45:04 CEST 2022 - The aliscore output file does not exist. Check results for BUSCO: 229590at7742
## Thu Jul 14 01:45:04 CEST 2022 - The aliscore output appears to be empty. Check results for BUSCO: 229590at7742
## Thu Jul 14 01:45:06 CEST 2022 - The aliscore output file does not exist. Check results for BUSCO: 182982at7742
## Thu Jul 14 01:45:06 CEST 2022 - The aliscore output appears to be empty. Check results for BUSCO: 182982at7742
## Thu Jul 14 01:45:07 CEST 2022 - The aliscore output file does not exist. Check results for BUSCO: 386982at7742
## Thu Jul 14 01:45:07 CEST 2022 - The aliscore output appears to be empty. Check results for BUSCO: 386982at7742
## Thu Jul 14 01:45:08 CEST 2022 - The aliscore output file does not exist. Check results for BUSCO: 319641at7742
## Thu Jul 14 01:45:08 CEST 2022 - The aliscore output appears to be empty. Check results for BUSCO: 319641at7742
## Thu Jul 14 01:45:09 CEST 2022 - The aliscore output file does not exist. Check results for BUSCO: 191426at7742
## Thu Jul 14 01:45:09 CEST 2022 - The aliscore output appears to be empty. Check results for BUSCO: 191426at7742
## Thu Jul 14 01:45:10 CEST 2022 - The aliscore output file does not exist. Check results for BUSCO: 238259at7742
## Thu Jul 14 01:45:10 CEST 2022 - The aliscore output appears to be empty. Check results for BUSCO: 238259at7742
## Thu Jul 14 01:45:13 CEST 2022 - The aliscore output file does not exist. Check results for BUSCO: 251465at7742
## Thu Jul 14 01:45:13 CEST 2022 - The aliscore output appears to be empty. Check results for BUSCO: 251465at7742
## Thu Jul 14 01:45:13 CEST 2022 - The aliscore output file does not exist. Check results for BUSCO: 253042at7742
## Thu Jul 14 01:45:13 CEST 2022 - The aliscore output appears to be empty. Check results for BUSCO: 253042at7742
## Thu Jul 14 01:45:14 CEST 2022 - The aliscore output file does not exist. Check results for BUSCO: 275566at7742
## Thu Jul 14 01:45:14 CEST 2022 - The aliscore output appears to be empty. Check results for BUSCO: 275566at7742
## Thu Jul 14 01:45:16 CEST 2022 - The aliscore output file does not exist. Check results for BUSCO: 198411at7742
## Thu Jul 14 01:45:16 CEST 2022 - The aliscore output appears to be empty. Check results for BUSCO: 198411at7742
## Thu Jul 14 01:45:16 CEST 2022 - The aliscore output file does not exist. Check results for BUSCO: 239338at7742
## Thu Jul 14 01:45:16 CEST 2022 - The aliscore output appears to be empty. Check results for BUSCO: 239338at7742
## Thu Jul 14 01:45:16 CEST 2022 - The aliscore output file does not exist. Check results for BUSCO: 165997at7742
## Thu Jul 14 01:45:16 CEST 2022 - The aliscore output appears to be empty. Check results for BUSCO: 165997at7742
## Thu Jul 14 01:45:18 CEST 2022 - The aliscore output file does not exist. Check results for BUSCO: 239308at7742
## Thu Jul 14 01:45:18 CEST 2022 - The aliscore output appears to be empty. Check results for BUSCO: 239308at7742
## Thu Jul 14 01:45:20 CEST 2022 - The aliscore output file does not exist. Check results for BUSCO: 140754at7742
## Thu Jul 14 01:45:20 CEST 2022 - The aliscore output file does not exist. Check results for BUSCO: 293668at7742
## Thu Jul 14 01:45:20 CEST 2022 - The aliscore output appears to be empty. Check results for BUSCO: 140754at7742
## Thu Jul 14 01:45:20 CEST 2022 - The aliscore output appears to be empty. Check results for BUSCO: 293668at7742
## Thu Jul 14 01:45:22 CEST 2022 - The aliscore output file does not exist. Check results for BUSCO: 158802at7742
## Thu Jul 14 01:45:23 CEST 2022 - The aliscore output appears to be empty. Check results for BUSCO: 158802at7742
## Thu Jul 14 01:45:23 CEST 2022 - The aliscore output file does not exist. Check results for BUSCO: 220484at7742
## Thu Jul 14 01:45:23 CEST 2022 - The aliscore output appears to be empty. Check results for BUSCO: 220484at7742
## Thu Jul 14 01:45:24 CEST 2022 - The aliscore output file does not exist. Check results for BUSCO: 131719at7742
## Thu Jul 14 01:45:24 CEST 2022 - The aliscore output appears to be empty. Check results for BUSCO: 131719at7742
## Thu Jul 14 01:45:24 CEST 2022 - The aliscore output file does not exist. Check results for BUSCO: 117231at7742
## Thu Jul 14 01:45:24 CEST 2022 - The aliscore output appears to be empty. Check results for BUSCO: 117231at7742
## Thu Jul 14 01:45:25 CEST 2022 - The aliscore output file does not exist. Check results for BUSCO: 394021at7742
## Thu Jul 14 01:45:25 CEST 2022 - The aliscore output appears to be empty. Check results for BUSCO: 394021at7742
## Thu Jul 14 01:45:27 CEST 2022 - The aliscore output file does not exist. Check results for BUSCO: 360372at7742
## Thu Jul 14 01:45:27 CEST 2022 - The aliscore output appears to be empty. Check results for BUSCO: 360372at7742
## Thu Jul 14 01:45:28 CEST 2022 - The aliscore output file does not exist. Check results for BUSCO: 276660at7742
## Thu Jul 14 01:45:28 CEST 2022 - The aliscore output appears to be empty. Check results for BUSCO: 276660at7742
## Thu Jul 14 01:45:28 CEST 2022 - The aliscore output file does not exist. Check results for BUSCO: 243575at7742
## Thu Jul 14 01:45:28 CEST 2022 - The aliscore output appears to be empty. Check results for BUSCO: 243575at7742
## Thu Jul 14 01:45:30 CEST 2022 - The aliscore output file does not exist. Check results for BUSCO: 288766at7742
## Thu Jul 14 01:45:30 CEST 2022 - The aliscore output appears to be empty. Check results for BUSCO: 288766at7742
## Thu Jul 14 01:45:30 CEST 2022 - The aliscore output file does not exist. Check results for BUSCO: 404952at7742
## Thu Jul 14 01:45:30 CEST 2022 - The aliscore output appears to be empty. Check results for BUSCO: 404952at7742
## Thu Jul 14 01:45:32 CEST 2022 - The aliscore output file does not exist. Check results for BUSCO: 131719at7742
## Thu Jul 14 01:45:32 CEST 2022 - The aliscore output appears to be empty. Check results for BUSCO: 131719at7742
## Thu Jul 14 01:45:33 CEST 2022 - The aliscore output file does not exist. Check results for BUSCO: 160035at7742
## Thu Jul 14 01:45:33 CEST 2022 - The aliscore output appears to be empty. Check results for BUSCO: 160035at7742
## Thu Jul 14 01:45:35 CEST 2022 - The aliscore output file does not exist. Check results for BUSCO: 290906at7742
## Thu Jul 14 01:45:35 CEST 2022 - The aliscore output appears to be empty. Check results for BUSCO: 290906at7742
## Thu Jul 14 01:45:35 CEST 2022 - The aliscore output file does not exist. Check results for BUSCO: 214745at7742
## Thu Jul 14 01:45:35 CEST 2022 - The aliscore output appears to be empty. Check results for BUSCO: 214745at7742
## Thu Jul 14 01:45:36 CEST 2022 - The aliscore output file does not exist. Check results for BUSCO: 236098at7742
## Thu Jul 14 01:45:36 CEST 2022 - The aliscore output appears to be empty. Check results for BUSCO: 236098at7742
## Thu Jul 14 01:45:37 CEST 2022 - The aliscore output file does not exist. Check results for BUSCO: 178490at7742
## Thu Jul 14 01:45:38 CEST 2022 - The aliscore output appears to be empty. Check results for BUSCO: 178490at7742
## Thu Jul 14 01:45:40 CEST 2022 - The aliscore output file does not exist. Check results for BUSCO: 220062at7742
## Thu Jul 14 01:45:40 CEST 2022 - The aliscore output appears to be empty. Check results for BUSCO: 220062at7742
## Thu Jul 14 01:45:40 CEST 2022 - The aliscore output file does not exist. Check results for BUSCO: 193249at7742
## Thu Jul 14 01:45:40 CEST 2022 - The aliscore output appears to be empty. Check results for BUSCO: 193249at7742
## Thu Jul 14 01:45:40 CEST 2022 - The aliscore output file does not exist. Check results for BUSCO: 140754at7742
## Thu Jul 14 01:45:40 CEST 2022 - The aliscore output appears to be empty. Check results for BUSCO: 140754at7742
## Thu Jul 14 01:45:40 CEST 2022 - The aliscore output file does not exist. Check results for BUSCO: 346924at7742
## Thu Jul 14 01:45:40 CEST 2022 - The aliscore output appears to be empty. Check results for BUSCO: 346924at7742
## Thu Jul 14 01:45:43 CEST 2022 - The aliscore output file does not exist. Check results for BUSCO: 208238at7742
## Thu Jul 14 01:45:43 CEST 2022 - The aliscore output appears to be empty. Check results for BUSCO: 208238at7742
## Thu Jul 14 01:45:45 CEST 2022 - The aliscore output file does not exist. Check results for BUSCO: 385085at7742
## Thu Jul 14 01:45:45 CEST 2022 - The aliscore output appears to be empty. Check results for BUSCO: 385085at7742
## Thu Jul 14 01:45:46 CEST 2022 - The aliscore output file does not exist. Check results for BUSCO: 305085at7742
## Thu Jul 14 01:45:46 CEST 2022 - The aliscore output appears to be empty. Check results for BUSCO: 305085at7742
## Thu Jul 14 01:45:46 CEST 2022 - The aliscore output file does not exist. Check results for BUSCO: 173047at7742
## Thu Jul 14 01:45:46 CEST 2022 - The aliscore output appears to be empty. Check results for BUSCO: 173047at7742
## Thu Jul 14 01:45:48 CEST 2022 - The aliscore output file does not exist. Check results for BUSCO: 115171at7742
## Thu Jul 14 01:45:48 CEST 2022 - The aliscore output appears to be empty. Check results for BUSCO: 115171at7742
## Thu Jul 14 01:45:49 CEST 2022 - The aliscore output file does not exist. Check results for BUSCO: 156291at7742
## Thu Jul 14 01:45:49 CEST 2022 - The aliscore output appears to be empty. Check results for BUSCO: 156291at7742
## Thu Jul 14 01:45:49 CEST 2022 - The aliscore output file does not exist. Check results for BUSCO: 368433at7742
## Thu Jul 14 01:45:49 CEST 2022 - The aliscore output appears to be empty. Check results for BUSCO: 368433at7742
## Thu Jul 14 01:45:51 CEST 2022 - The aliscore output file does not exist. Check results for BUSCO: 213005at7742
## Thu Jul 14 01:45:51 CEST 2022 - The aliscore output appears to be empty. Check results for BUSCO: 213005at7742
## Thu Jul 14 01:45:53 CEST 2022 - The aliscore output file does not exist. Check results for BUSCO: 115171at7742
## Thu Jul 14 01:45:53 CEST 2022 - The aliscore output appears to be empty. Check results for BUSCO: 115171at7742
## Thu Jul 14 01:45:53 CEST 2022 - The aliscore output file does not exist. Check results for BUSCO: 185467at7742
## Thu Jul 14 01:45:53 CEST 2022 - The aliscore output appears to be empty. Check results for BUSCO: 185467at7742
## Thu Jul 14 01:45:54 CEST 2022 - The aliscore output file does not exist. Check results for BUSCO: 301292at7742
## Thu Jul 14 01:45:54 CEST 2022 - The aliscore output appears to be empty. Check results for BUSCO: 301292at7742
## Thu Jul 14 01:45:55 CEST 2022 - The aliscore output file does not exist. Check results for BUSCO: 389667at7742
## Thu Jul 14 01:45:55 CEST 2022 - The aliscore output appears to be empty. Check results for BUSCO: 389667at7742
## Thu Jul 14 01:45:56 CEST 2022 - The aliscore output file does not exist. Check results for BUSCO: 283821at7742
## Thu Jul 14 01:45:56 CEST 2022 - The aliscore output appears to be empty. Check results for BUSCO: 283821at7742
## Thu Jul 14 01:45:57 CEST 2022 - The aliscore output file does not exist. Check results for BUSCO: 120479at7742
## Thu Jul 14 01:45:57 CEST 2022 - The aliscore output appears to be empty. Check results for BUSCO: 120479at7742
## Thu Jul 14 01:45:58 CEST 2022 - The aliscore output file does not exist. Check results for BUSCO: 262798at7742
## Thu Jul 14 01:45:58 CEST 2022 - The aliscore output appears to be empty. Check results for BUSCO: 262798at7742
## Thu Jul 14 01:46:01 CEST 2022 - The aliscore output file does not exist. Check results for BUSCO: 216982at7742
## Thu Jul 14 01:46:01 CEST 2022 - The aliscore output appears to be empty. Check results for BUSCO: 216982at7742
## Thu Jul 14 01:46:02 CEST 2022 - The aliscore output file does not exist. Check results for BUSCO: 227080at7742
## Thu Jul 14 01:46:02 CEST 2022 - The aliscore output appears to be empty. Check results for BUSCO: 227080at7742
## Thu Jul 14 01:46:03 CEST 2022 - The aliscore output file does not exist. Check results for BUSCO: 412581at7742
## Thu Jul 14 01:46:03 CEST 2022 - The aliscore output appears to be empty. Check results for BUSCO: 412581at7742
## Thu Jul 14 01:46:03 CEST 2022 - The aliscore output file does not exist. Check results for BUSCO: 121830at7742
## Thu Jul 14 01:46:03 CEST 2022 - The aliscore output appears to be empty. Check results for BUSCO: 121830at7742
## Thu Jul 14 01:46:04 CEST 2022 - The aliscore output file does not exist. Check results for BUSCO: 262221at7742
## Thu Jul 14 01:46:04 CEST 2022 - The aliscore output appears to be empty. Check results for BUSCO: 262221at7742
## Thu Jul 14 01:46:04 CEST 2022 - The aliscore output file does not exist. Check results for BUSCO: 131719at7742
## Thu Jul 14 01:46:04 CEST 2022 - The aliscore output appears to be empty. Check results for BUSCO: 131719at7742
## Thu Jul 14 01:46:06 CEST 2022 - The aliscore output file does not exist. Check results for BUSCO: 252806at7742
## Thu Jul 14 01:46:06 CEST 2022 - The aliscore output appears to be empty. Check results for BUSCO: 252806at7742
## Thu Jul 14 01:46:06 CEST 2022 - The aliscore output file does not exist. Check results for BUSCO: 198321at7742
## Thu Jul 14 01:46:06 CEST 2022 - The aliscore output appears to be empty. Check results for BUSCO: 198321at7742
## Thu Jul 14 01:46:08 CEST 2022 - The aliscore output file does not exist. Check results for BUSCO: 294443at7742
## Thu Jul 14 01:46:08 CEST 2022 - The aliscore output appears to be empty. Check results for BUSCO: 294443at7742
## Thu Jul 14 01:46:10 CEST 2022 - The aliscore output file does not exist. Check results for BUSCO: 184999at7742
## Thu Jul 14 01:46:10 CEST 2022 - The aliscore output appears to be empty. Check results for BUSCO: 184999at7742
## Thu Jul 14 01:46:11 CEST 2022 - The aliscore output file does not exist. Check results for BUSCO: 219805at7742
## Thu Jul 14 01:46:11 CEST 2022 - The aliscore output appears to be empty. Check results for BUSCO: 219805at7742
## Thu Jul 14 01:46:12 CEST 2022 - The aliscore output file does not exist. Check results for BUSCO: 205403at7742
## Thu Jul 14 01:46:12 CEST 2022 - The aliscore output appears to be empty. Check results for BUSCO: 205403at7742
## Thu Jul 14 01:46:13 CEST 2022 - The aliscore output file does not exist. Check results for BUSCO: 191426at7742
## Thu Jul 14 01:46:13 CEST 2022 - The aliscore output appears to be empty. Check results for BUSCO: 191426at7742
## Thu Jul 14 01:46:14 CEST 2022 - The aliscore output file does not exist. Check results for BUSCO: 197724at7742
## Thu Jul 14 01:46:14 CEST 2022 - The aliscore output file does not exist. Check results for BUSCO: 249877at7742
## Thu Jul 14 01:46:14 CEST 2022 - The aliscore output appears to be empty. Check results for BUSCO: 197724at7742
## Thu Jul 14 01:46:14 CEST 2022 - The aliscore output appears to be empty. Check results for BUSCO: 249877at7742
## Thu Jul 14 01:46:16 CEST 2022 - The aliscore output file does not exist. Check results for BUSCO: 327404at7742
## Thu Jul 14 01:46:16 CEST 2022 - The aliscore output appears to be empty. Check results for BUSCO: 327404at7742
## Thu Jul 14 01:46:17 CEST 2022 - The aliscore output file does not exist. Check results for BUSCO: 162664at7742
## Thu Jul 14 01:46:17 CEST 2022 - The aliscore output appears to be empty. Check results for BUSCO: 162664at7742
## Thu Jul 14 01:46:19 CEST 2022 - The aliscore output file does not exist. Check results for BUSCO: 252398at7742
## Thu Jul 14 01:46:19 CEST 2022 - The aliscore output appears to be empty. Check results for BUSCO: 252398at7742
## Thu Jul 14 01:46:19 CEST 2022 - The aliscore output file does not exist. Check results for BUSCO: 185467at7742
## Thu Jul 14 01:46:19 CEST 2022 - The aliscore output appears to be empty. Check results for BUSCO: 185467at7742
## Thu Jul 14 01:46:19 CEST 2022 - The aliscore output file does not exist. Check results for BUSCO: 133713at7742
## Thu Jul 14 01:46:19 CEST 2022 - The aliscore output appears to be empty. Check results for BUSCO: 133713at7742
## Thu Jul 14 01:46:22 CEST 2022 - The aliscore output file does not exist. Check results for BUSCO: 285589at7742
## Thu Jul 14 01:46:22 CEST 2022 - The aliscore output appears to be empty. Check results for BUSCO: 285589at7742
## Thu Jul 14 01:46:22 CEST 2022 - The aliscore output file does not exist. Check results for BUSCO: 330054at7742
## Thu Jul 14 01:46:22 CEST 2022 - The aliscore output appears to be empty. Check results for BUSCO: 330054at7742
## Thu Jul 14 01:46:24 CEST 2022 - The aliscore output file does not exist. Check results for BUSCO: 176266at7742
## Thu Jul 14 01:46:24 CEST 2022 - The aliscore output appears to be empty. Check results for BUSCO: 176266at7742
## Thu Jul 14 01:46:26 CEST 2022 - The aliscore output file does not exist. Check results for BUSCO: 177602at7742
## Thu Jul 14 01:46:26 CEST 2022 - The aliscore output appears to be empty. Check results for BUSCO: 177602at7742
## Thu Jul 14 01:46:27 CEST 2022 - The aliscore output file does not exist. Check results for BUSCO: 158802at7742
## Thu Jul 14 01:46:27 CEST 2022 - The aliscore output appears to be empty. Check results for BUSCO: 158802at7742
## Thu Jul 14 01:46:27 CEST 2022 - The aliscore output file does not exist. Check results for BUSCO: 290261at7742
## Thu Jul 14 01:46:27 CEST 2022 - The aliscore output appears to be empty. Check results for BUSCO: 290261at7742
## Thu Jul 14 01:46:29 CEST 2022 - The aliscore output file does not exist. Check results for BUSCO: 115171at7742
## Thu Jul 14 01:46:29 CEST 2022 - The aliscore output appears to be empty. Check results for BUSCO: 115171at7742
## Thu Jul 14 01:46:29 CEST 2022 - The aliscore output file does not exist. Check results for BUSCO: 236417at7742
## Thu Jul 14 01:46:29 CEST 2022 - The aliscore output appears to be empty. Check results for BUSCO: 236417at7742
## Thu Jul 14 01:46:31 CEST 2022 - The aliscore output file does not exist. Check results for BUSCO: 243575at7742
## Thu Jul 14 01:46:31 CEST 2022 - The aliscore output appears to be empty. Check results for BUSCO: 243575at7742
## Thu Jul 14 01:46:31 CEST 2022 - The aliscore output file does not exist. Check results for BUSCO: 375400at7742
## Thu Jul 14 01:46:31 CEST 2022 - The aliscore output appears to be empty. Check results for BUSCO: 375400at7742
## Thu Jul 14 01:46:35 CEST 2022 - The aliscore output file does not exist. Check results for BUSCO: 183927at7742
## Thu Jul 14 01:46:35 CEST 2022 - The aliscore output appears to be empty. Check results for BUSCO: 183927at7742
## Thu Jul 14 01:46:36 CEST 2022 - The aliscore output file does not exist. Check results for BUSCO: 192058at7742
## Thu Jul 14 01:46:36 CEST 2022 - The aliscore output appears to be empty. Check results for BUSCO: 192058at7742
## Thu Jul 14 01:46:36 CEST 2022 - The aliscore output file does not exist. Check results for BUSCO: 199780at7742
## Thu Jul 14 01:46:37 CEST 2022 - The aliscore output appears to be empty. Check results for BUSCO: 199780at7742
## Thu Jul 14 01:46:37 CEST 2022 - The aliscore output file does not exist. Check results for BUSCO: 185648at7742
## Thu Jul 14 01:46:37 CEST 2022 - The aliscore output appears to be empty. Check results for BUSCO: 185648at7742
## Thu Jul 14 01:46:37 CEST 2022 - The aliscore output file does not exist. Check results for BUSCO: 178490at7742
## Thu Jul 14 01:46:37 CEST 2022 - The aliscore output appears to be empty. Check results for BUSCO: 178490at7742
## Thu Jul 14 01:46:39 CEST 2022 - The aliscore output file does not exist. Check results for BUSCO: 263794at7742
## Thu Jul 14 01:46:39 CEST 2022 - The aliscore output appears to be empty. Check results for BUSCO: 263794at7742
## Thu Jul 14 01:46:39 CEST 2022 - The aliscore output file does not exist. Check results for BUSCO: 206342at7742
## Thu Jul 14 01:46:39 CEST 2022 - The aliscore output appears to be empty. Check results for BUSCO: 206342at7742
## Thu Jul 14 01:46:40 CEST 2022 - The aliscore output file does not exist. Check results for BUSCO: 205403at7742
## Thu Jul 14 01:46:40 CEST 2022 - The aliscore output appears to be empty. Check results for BUSCO: 205403at7742
## Thu Jul 14 01:46:42 CEST 2022 - The aliscore output file does not exist. Check results for BUSCO: 264239at7742
## Thu Jul 14 01:46:42 CEST 2022 - The aliscore output appears to be empty. Check results for BUSCO: 264239at7742
## Thu Jul 14 01:46:42 CEST 2022 - The aliscore output file does not exist. Check results for BUSCO: 261523at7742
## Thu Jul 14 01:46:42 CEST 2022 - The aliscore output appears to be empty. Check results for BUSCO: 261523at7742
## Thu Jul 14 01:46:44 CEST 2022 - The aliscore output file does not exist. Check results for BUSCO: 252398at7742
## Thu Jul 14 01:46:44 CEST 2022 - The aliscore output appears to be empty. Check results for BUSCO: 252398at7742
## Thu Jul 14 01:46:45 CEST 2022 - The aliscore output file does not exist. Check results for BUSCO: 376114at7742
## Thu Jul 14 01:46:45 CEST 2022 - The aliscore output appears to be empty. Check results for BUSCO: 376114at7742
## Thu Jul 14 01:46:47 CEST 2022 - The aliscore output file does not exist. Check results for BUSCO: 191426at7742
## Thu Jul 14 01:46:47 CEST 2022 - The aliscore output appears to be empty. Check results for BUSCO: 191426at7742
## Thu Jul 14 01:46:48 CEST 2022 - The aliscore output file does not exist. Check results for BUSCO: 401448at7742
## Thu Jul 14 01:46:48 CEST 2022 - The aliscore output appears to be empty. Check results for BUSCO: 401448at7742
## Thu Jul 14 01:46:50 CEST 2022 - The aliscore output file does not exist. Check results for BUSCO: 360850at7742
## Thu Jul 14 01:46:50 CEST 2022 - The aliscore output appears to be empty. Check results for BUSCO: 360850at7742
## Thu Jul 14 01:46:50 CEST 2022 - The aliscore output file does not exist. Check results for BUSCO: 169465at7742
## Thu Jul 14 01:46:50 CEST 2022 - The aliscore output appears to be empty. Check results for BUSCO: 169465at7742
## Thu Jul 14 01:46:52 CEST 2022 - The aliscore output file does not exist. Check results for BUSCO: 171308at7742
## Thu Jul 14 01:46:52 CEST 2022 - The aliscore output appears to be empty. Check results for BUSCO: 171308at7742
## Thu Jul 14 01:46:53 CEST 2022 - The aliscore output file does not exist. Check results for BUSCO: 326191at7742
## Thu Jul 14 01:46:53 CEST 2022 - The aliscore output appears to be empty. Check results for BUSCO: 326191at7742
## Thu Jul 14 01:46:53 CEST 2022 - The aliscore output file does not exist. Check results for BUSCO: 288407at7742
## Thu Jul 14 01:46:53 CEST 2022 - The aliscore output appears to be empty. Check results for BUSCO: 288407at7742
## Thu Jul 14 01:46:55 CEST 2022 - The aliscore output file does not exist. Check results for BUSCO: 267268at7742
## Thu Jul 14 01:46:55 CEST 2022 - The aliscore output appears to be empty. Check results for BUSCO: 267268at7742
## Thu Jul 14 01:46:56 CEST 2022 - The aliscore output file does not exist. Check results for BUSCO: 369343at7742
## Thu Jul 14 01:46:56 CEST 2022 - The aliscore output appears to be empty. Check results for BUSCO: 369343at7742
## Thu Jul 14 01:46:57 CEST 2022 - The aliscore output file does not exist. Check results for BUSCO: 133326at7742
## Thu Jul 14 01:46:57 CEST 2022 - The aliscore output appears to be empty. Check results for BUSCO: 133326at7742
## Thu Jul 14 01:46:58 CEST 2022 - The aliscore output file does not exist. Check results for BUSCO: 207618at7742
## Thu Jul 14 01:46:58 CEST 2022 - The aliscore output appears to be empty. Check results for BUSCO: 207618at7742
## Thu Jul 14 01:46:59 CEST 2022 - The aliscore output file does not exist. Check results for BUSCO: 187916at7742
## Thu Jul 14 01:46:59 CEST 2022 - The aliscore output appears to be empty. Check results for BUSCO: 187916at7742
## Thu Jul 14 01:47:00 CEST 2022 - The aliscore output file does not exist. Check results for BUSCO: 159334at7742
## Thu Jul 14 01:47:00 CEST 2022 - The aliscore output appears to be empty. Check results for BUSCO: 159334at7742
## Thu Jul 14 01:47:01 CEST 2022 - The aliscore output file does not exist. Check results for BUSCO: 259058at7742
## Thu Jul 14 01:47:01 CEST 2022 - The aliscore output appears to be empty. Check results for BUSCO: 259058at7742
## Thu Jul 14 01:47:03 CEST 2022 - The aliscore output file does not exist. Check results for BUSCO: 149216at7742
## Thu Jul 14 01:47:03 CEST 2022 - The aliscore output appears to be empty. Check results for BUSCO: 149216at7742
## Thu Jul 14 01:47:04 CEST 2022 - The aliscore output file does not exist. Check results for BUSCO: 177602at7742
## Thu Jul 14 01:47:04 CEST 2022 - The aliscore output appears to be empty. Check results for BUSCO: 177602at7742
## Thu Jul 14 01:47:05 CEST 2022 - The aliscore output file does not exist. Check results for BUSCO: 412581at7742
## Thu Jul 14 01:47:05 CEST 2022 - The aliscore output appears to be empty. Check results for BUSCO: 412581at7742
## Thu Jul 14 01:47:06 CEST 2022 - The aliscore output file does not exist. Check results for BUSCO: 192058at7742
## Thu Jul 14 01:47:06 CEST 2022 - The aliscore output appears to be empty. Check results for BUSCO: 192058at7742
## Thu Jul 14 01:47:06 CEST 2022 - The aliscore output file does not exist. Check results for BUSCO: 239324at7742
## Thu Jul 14 01:47:06 CEST 2022 - The aliscore output appears to be empty. Check results for BUSCO: 239324at7742
## Thu Jul 14 01:47:07 CEST 2022 - The aliscore output file does not exist. Check results for BUSCO: 142214at7742
## Thu Jul 14 01:47:07 CEST 2022 - The aliscore output appears to be empty. Check results for BUSCO: 142214at7742
## Thu Jul 14 01:47:09 CEST 2022 - The aliscore output file does not exist. Check results for BUSCO: 292028at7742
## Thu Jul 14 01:47:09 CEST 2022 - The aliscore output appears to be empty. Check results for BUSCO: 292028at7742
## Thu Jul 14 01:47:09 CEST 2022 - The aliscore output file does not exist. Check results for BUSCO: 245662at7742
## Thu Jul 14 01:47:09 CEST 2022 - The aliscore output appears to be empty. Check results for BUSCO: 245662at7742
## Thu Jul 14 01:47:11 CEST 2022 - The aliscore output file does not exist. Check results for BUSCO: 220484at7742
## Thu Jul 14 01:47:11 CEST 2022 - The aliscore output appears to be empty. Check results for BUSCO: 220484at7742
## Thu Jul 14 01:47:12 CEST 2022 - The aliscore output file does not exist. Check results for BUSCO: 257699at7742
## Thu Jul 14 01:47:12 CEST 2022 - The aliscore output appears to be empty. Check results for BUSCO: 257699at7742
## Thu Jul 14 01:47:13 CEST 2022 - The aliscore output file does not exist. Check results for BUSCO: 227816at7742
## Thu Jul 14 01:47:13 CEST 2022 - The aliscore output appears to be empty. Check results for BUSCO: 227816at7742
## Thu Jul 14 01:47:13 CEST 2022 - The aliscore output file does not exist. Check results for BUSCO: 250198at7742
## Thu Jul 14 01:47:13 CEST 2022 - The aliscore output appears to be empty. Check results for BUSCO: 250198at7742
## Thu Jul 14 01:47:15 CEST 2022 - The aliscore output file does not exist. Check results for BUSCO: 144764at7742
## Thu Jul 14 01:47:15 CEST 2022 - The aliscore output appears to be empty. Check results for BUSCO: 144764at7742
## Thu Jul 14 01:47:16 CEST 2022 - The aliscore output file does not exist. Check results for BUSCO: 144764at7742
## Thu Jul 14 01:47:16 CEST 2022 - The aliscore output appears to be empty. Check results for BUSCO: 144764at7742
## Thu Jul 14 01:47:17 CEST 2022 - The aliscore output file does not exist. Check results for BUSCO: 230223at7742
## Thu Jul 14 01:47:17 CEST 2022 - The aliscore output appears to be empty. Check results for BUSCO: 230223at7742
## Thu Jul 14 01:47:18 CEST 2022 - The aliscore output file does not exist. Check results for BUSCO: 221031at7742
## Thu Jul 14 01:47:18 CEST 2022 - The aliscore output appears to be empty. Check results for BUSCO: 221031at7742
## Thu Jul 14 01:47:20 CEST 2022 - The aliscore output file does not exist. Check results for BUSCO: 162664at7742
## Thu Jul 14 01:47:20 CEST 2022 - The aliscore output appears to be empty. Check results for BUSCO: 162664at7742
## Thu Jul 14 01:47:20 CEST 2022 - The aliscore output file does not exist. Check results for BUSCO: 244439at7742
## Thu Jul 14 01:47:20 CEST 2022 - The aliscore output appears to be empty. Check results for BUSCO: 244439at7742
## Thu Jul 14 01:47:21 CEST 2022 - The aliscore output file does not exist. Check results for BUSCO: 321010at7742
## Thu Jul 14 01:47:21 CEST 2022 - The aliscore output appears to be empty. Check results for BUSCO: 321010at7742
## Thu Jul 14 01:47:21 CEST 2022 - The aliscore output file does not exist. Check results for BUSCO: 379193at7742
## Thu Jul 14 01:47:21 CEST 2022 - The aliscore output appears to be empty. Check results for BUSCO: 379193at7742
## Thu Jul 14 01:47:24 CEST 2022 - The aliscore output file does not exist. Check results for BUSCO: 392098at7742
## Thu Jul 14 01:47:24 CEST 2022 - The aliscore output appears to be empty. Check results for BUSCO: 392098at7742
## Thu Jul 14 01:47:25 CEST 2022 - The aliscore output file does not exist. Check results for BUSCO: 133326at7742
## Thu Jul 14 01:47:25 CEST 2022 - The aliscore output appears to be empty. Check results for BUSCO: 133326at7742
## Thu Jul 14 01:47:25 CEST 2022 - The aliscore output file does not exist. Check results for BUSCO: 300443at7742
## Thu Jul 14 01:47:25 CEST 2022 - The aliscore output appears to be empty. Check results for BUSCO: 300443at7742
## Thu Jul 14 01:47:28 CEST 2022 - The aliscore output file does not exist. Check results for BUSCO: 133326at7742
## Thu Jul 14 01:47:28 CEST 2022 - The aliscore output appears to be empty. Check results for BUSCO: 133326at7742
## Thu Jul 14 01:47:29 CEST 2022 - The aliscore output file does not exist. Check results for BUSCO: 185467at7742
## Thu Jul 14 01:47:29 CEST 2022 - The aliscore output appears to be empty. Check results for BUSCO: 185467at7742
## Thu Jul 14 01:47:29 CEST 2022 - The aliscore output file does not exist. Check results for BUSCO: 169313at7742
## Thu Jul 14 01:47:29 CEST 2022 - The aliscore output appears to be empty. Check results for BUSCO: 169313at7742
## Thu Jul 14 01:47:29 CEST 2022 - The aliscore output file does not exist. Check results for BUSCO: 207782at7742
## Thu Jul 14 01:47:29 CEST 2022 - The aliscore output appears to be empty. Check results for BUSCO: 207782at7742
## Thu Jul 14 01:47:31 CEST 2022 - The aliscore output file does not exist. Check results for BUSCO: 353522at7742
## Thu Jul 14 01:47:31 CEST 2022 - The aliscore output appears to be empty. Check results for BUSCO: 353522at7742
## Thu Jul 14 01:47:32 CEST 2022 - The aliscore output file does not exist. Check results for BUSCO: 377100at7742
## Thu Jul 14 01:47:32 CEST 2022 - The aliscore output appears to be empty. Check results for BUSCO: 377100at7742
## Thu Jul 14 01:47:33 CEST 2022 - The aliscore output file does not exist. Check results for BUSCO: 321295at7742
## Thu Jul 14 01:47:33 CEST 2022 - The aliscore output appears to be empty. Check results for BUSCO: 321295at7742
## Thu Jul 14 01:47:36 CEST 2022 - The aliscore output file does not exist. Check results for BUSCO: 161007at7742
## Thu Jul 14 01:47:36 CEST 2022 - The aliscore output appears to be empty. Check results for BUSCO: 161007at7742
## Thu Jul 14 01:47:36 CEST 2022 - The aliscore output file does not exist. Check results for BUSCO: 243575at7742
## Thu Jul 14 01:47:36 CEST 2022 - The aliscore output appears to be empty. Check results for BUSCO: 243575at7742
## Thu Jul 14 01:47:36 CEST 2022 - The aliscore output file does not exist. Check results for BUSCO: 288407at7742
## Thu Jul 14 01:47:36 CEST 2022 - The aliscore output appears to be empty. Check results for BUSCO: 288407at7742
## Thu Jul 14 01:47:36 CEST 2022 - The aliscore output file does not exist. Check results for BUSCO: 225241at7742
## Thu Jul 14 01:47:36 CEST 2022 - The aliscore output appears to be empty. Check results for BUSCO: 225241at7742
## Thu Jul 14 01:47:38 CEST 2022 - The aliscore output file does not exist. Check results for BUSCO: 121830at7742
## Thu Jul 14 01:47:38 CEST 2022 - The aliscore output appears to be empty. Check results for BUSCO: 121830at7742
## Thu Jul 14 01:47:39 CEST 2022 - The aliscore output file does not exist. Check results for BUSCO: 219805at7742
## Thu Jul 14 01:47:39 CEST 2022 - The aliscore output appears to be empty. Check results for BUSCO: 219805at7742
## Thu Jul 14 01:47:39 CEST 2022 - The aliscore output file does not exist. Check results for BUSCO: 199987at7742
## Thu Jul 14 01:47:39 CEST 2022 - The aliscore output appears to be empty. Check results for BUSCO: 199987at7742
## Thu Jul 14 01:47:43 CEST 2022 - The aliscore output file does not exist. Check results for BUSCO: 229590at7742
## Thu Jul 14 01:47:43 CEST 2022 - The aliscore output appears to be empty. Check results for BUSCO: 229590at7742
## Thu Jul 14 01:47:43 CEST 2022 - The aliscore output file does not exist. Check results for BUSCO: 221887at7742
## Thu Jul 14 01:47:43 CEST 2022 - The aliscore output appears to be empty. Check results for BUSCO: 221887at7742
## Thu Jul 14 01:47:44 CEST 2022 - The aliscore output file does not exist. Check results for BUSCO: 245433at7742
## Thu Jul 14 01:47:44 CEST 2022 - The aliscore output appears to be empty. Check results for BUSCO: 245433at7742
## Thu Jul 14 01:47:45 CEST 2022 - The aliscore output file does not exist. Check results for BUSCO: 227080at7742
## Thu Jul 14 01:47:45 CEST 2022 - The aliscore output appears to be empty. Check results for BUSCO: 227080at7742
## Thu Jul 14 01:47:45 CEST 2022 - The aliscore output file does not exist. Check results for BUSCO: 310872at7742
## Thu Jul 14 01:47:45 CEST 2022 - The aliscore output appears to be empty. Check results for BUSCO: 310872at7742
## Thu Jul 14 01:47:47 CEST 2022 - The aliscore output file does not exist. Check results for BUSCO: 228275at7742
## Thu Jul 14 01:47:47 CEST 2022 - The aliscore output appears to be empty. Check results for BUSCO: 228275at7742
## Thu Jul 14 01:47:49 CEST 2022 - The aliscore output file does not exist. Check results for BUSCO: 156291at7742
## Thu Jul 14 01:47:49 CEST 2022 - The aliscore output appears to be empty. Check results for BUSCO: 156291at7742
## Thu Jul 14 01:47:49 CEST 2022 - The aliscore output file does not exist. Check results for BUSCO: 189753at7742
## Thu Jul 14 01:47:50 CEST 2022 - The aliscore output appears to be empty. Check results for BUSCO: 189753at7742
## Thu Jul 14 01:47:50 CEST 2022 - The aliscore output file does not exist. Check results for BUSCO: 327404at7742
## Thu Jul 14 01:47:50 CEST 2022 - The aliscore output appears to be empty. Check results for BUSCO: 327404at7742
## Thu Jul 14 01:47:51 CEST 2022 - The aliscore output file does not exist. Check results for BUSCO: 277067at7742
## Thu Jul 14 01:47:51 CEST 2022 - The aliscore output appears to be empty. Check results for BUSCO: 277067at7742
## Thu Jul 14 01:47:52 CEST 2022 - The aliscore output file does not exist. Check results for BUSCO: 308423at7742
## Thu Jul 14 01:47:52 CEST 2022 - The aliscore output appears to be empty. Check results for BUSCO: 308423at7742
## Thu Jul 14 01:47:52 CEST 2022 - The aliscore output file does not exist. Check results for BUSCO: 189840at7742
## Thu Jul 14 01:47:52 CEST 2022 - The aliscore output appears to be empty. Check results for BUSCO: 189840at7742
## Thu Jul 14 01:47:57 CEST 2022 - The aliscore output file does not exist. Check results for BUSCO: 207648at7742
## Thu Jul 14 01:47:57 CEST 2022 - The aliscore output appears to be empty. Check results for BUSCO: 207648at7742
## Thu Jul 14 01:47:57 CEST 2022 - The aliscore output file does not exist. Check results for BUSCO: 140754at7742
## Thu Jul 14 01:47:57 CEST 2022 - The aliscore output appears to be empty. Check results for BUSCO: 140754at7742
## Thu Jul 14 01:47:57 CEST 2022 - The aliscore output file does not exist. Check results for BUSCO: 190526at7742
## Thu Jul 14 01:47:58 CEST 2022 - The aliscore output appears to be empty. Check results for BUSCO: 190526at7742
## Thu Jul 14 01:47:58 CEST 2022 - The aliscore output file does not exist. Check results for BUSCO: 238523at7742
## Thu Jul 14 01:47:58 CEST 2022 - The aliscore output appears to be empty. Check results for BUSCO: 238523at7742
## Thu Jul 14 01:47:59 CEST 2022 - The aliscore output file does not exist. Check results for BUSCO: 142214at7742
## Thu Jul 14 01:47:59 CEST 2022 - The aliscore output appears to be empty. Check results for BUSCO: 142214at7742
## Thu Jul 14 01:47:59 CEST 2022 - The aliscore output file does not exist. Check results for BUSCO: 185697at7742
## Thu Jul 14 01:47:59 CEST 2022 - The aliscore output appears to be empty. Check results for BUSCO: 185697at7742
## Thu Jul 14 01:48:01 CEST 2022 - The aliscore output file does not exist. Check results for BUSCO: 169313at7742
## Thu Jul 14 01:48:01 CEST 2022 - The aliscore output appears to be empty. Check results for BUSCO: 169313at7742
## Thu Jul 14 01:48:01 CEST 2022 - The aliscore output file does not exist. Check results for BUSCO: 248682at7742
## Thu Jul 14 01:48:02 CEST 2022 - The aliscore output appears to be empty. Check results for BUSCO: 248682at7742
## Thu Jul 14 01:48:04 CEST 2022 - The aliscore output file does not exist. Check results for BUSCO: 259863at7742
## Thu Jul 14 01:48:04 CEST 2022 - The aliscore output appears to be empty. Check results for BUSCO: 259863at7742
## Thu Jul 14 01:48:06 CEST 2022 - The aliscore output file does not exist. Check results for BUSCO: 133153at7742
## Thu Jul 14 01:48:06 CEST 2022 - The aliscore output appears to be empty. Check results for BUSCO: 133153at7742
## Thu Jul 14 01:48:07 CEST 2022 - The aliscore output file does not exist. Check results for BUSCO: 197724at7742
## Thu Jul 14 01:48:07 CEST 2022 - The aliscore output appears to be empty. Check results for BUSCO: 197724at7742
## Thu Jul 14 01:48:07 CEST 2022 - The aliscore output file does not exist. Check results for BUSCO: 235765at7742
## Thu Jul 14 01:48:07 CEST 2022 - The aliscore output appears to be empty. Check results for BUSCO: 235765at7742
## Thu Jul 14 01:48:08 CEST 2022 - The aliscore output file does not exist. Check results for BUSCO: 238523at7742
## Thu Jul 14 01:48:08 CEST 2022 - The aliscore output appears to be empty. Check results for BUSCO: 238523at7742
## Thu Jul 14 01:48:08 CEST 2022 - The aliscore output file does not exist. Check results for BUSCO: 133965at7742
## Thu Jul 14 01:48:08 CEST 2022 - The aliscore output appears to be empty. Check results for BUSCO: 133965at7742
## Thu Jul 14 01:48:11 CEST 2022 - The aliscore output file does not exist. Check results for BUSCO: 235827at7742
## Thu Jul 14 01:48:11 CEST 2022 - The aliscore output appears to be empty. Check results for BUSCO: 235827at7742
## Thu Jul 14 01:48:11 CEST 2022 - The aliscore output file does not exist. Check results for BUSCO: 343795at7742
## Thu Jul 14 01:48:11 CEST 2022 - The aliscore output appears to be empty. Check results for BUSCO: 343795at7742
## Thu Jul 14 01:48:12 CEST 2022 - The aliscore output file does not exist. Check results for BUSCO: 379193at7742
## Thu Jul 14 01:48:12 CEST 2022 - The aliscore output appears to be empty. Check results for BUSCO: 379193at7742
## Thu Jul 14 01:48:12 CEST 2022 - The aliscore output file does not exist. Check results for BUSCO: 273630at7742
## Thu Jul 14 01:48:12 CEST 2022 - The aliscore output appears to be empty. Check results for BUSCO: 273630at7742
## Thu Jul 14 01:48:15 CEST 2022 - The aliscore output file does not exist. Check results for BUSCO: 279373at7742
## Thu Jul 14 01:48:15 CEST 2022 - The aliscore output appears to be empty. Check results for BUSCO: 279373at7742
## Thu Jul 14 01:48:15 CEST 2022 - The aliscore output file does not exist. Check results for BUSCO: 293668at7742
## Thu Jul 14 01:48:15 CEST 2022 - The aliscore output appears to be empty. Check results for BUSCO: 293668at7742
## Thu Jul 14 01:48:16 CEST 2022 - The aliscore output file does not exist. Check results for BUSCO: 194403at7742
## Thu Jul 14 01:48:16 CEST 2022 - The aliscore output appears to be empty. Check results for BUSCO: 194403at7742
## Thu Jul 14 01:48:19 CEST 2022 - The aliscore output file does not exist. Check results for BUSCO: 243611at7742
## Thu Jul 14 01:48:19 CEST 2022 - The aliscore output appears to be empty. Check results for BUSCO: 243611at7742
## Thu Jul 14 01:48:19 CEST 2022 - The aliscore output file does not exist. Check results for BUSCO: 381258at7742
## Thu Jul 14 01:48:19 CEST 2022 - The aliscore output appears to be empty. Check results for BUSCO: 381258at7742
## Thu Jul 14 01:48:20 CEST 2022 - The aliscore output file does not exist. Check results for BUSCO: 135248at7742
## Thu Jul 14 01:48:20 CEST 2022 - The aliscore output appears to be empty. Check results for BUSCO: 135248at7742
## Thu Jul 14 01:48:21 CEST 2022 - The aliscore output file does not exist. Check results for BUSCO: 230223at7742
## Thu Jul 14 01:48:21 CEST 2022 - The aliscore output appears to be empty. Check results for BUSCO: 230223at7742
## Thu Jul 14 01:48:22 CEST 2022 - The aliscore output file does not exist. Check results for BUSCO: 144352at7742
## Thu Jul 14 01:48:22 CEST 2022 - The aliscore output appears to be empty. Check results for BUSCO: 144352at7742
## Thu Jul 14 01:48:24 CEST 2022 - The aliscore output file does not exist. Check results for BUSCO: 192665at7742
## Thu Jul 14 01:48:24 CEST 2022 - The aliscore output appears to be empty. Check results for BUSCO: 192665at7742
## Thu Jul 14 01:48:25 CEST 2022 - The aliscore output file does not exist. Check results for BUSCO: 318959at7742
## Thu Jul 14 01:48:25 CEST 2022 - The aliscore output appears to be empty. Check results for BUSCO: 318959at7742
## Thu Jul 14 01:48:25 CEST 2022 - The aliscore output file does not exist. Check results for BUSCO: 292857at7742
## Thu Jul 14 01:48:25 CEST 2022 - The aliscore output appears to be empty. Check results for BUSCO: 292857at7742
## Thu Jul 14 01:48:27 CEST 2022 - The aliscore output file does not exist. Check results for BUSCO: 199024at7742
## Thu Jul 14 01:48:27 CEST 2022 - The aliscore output appears to be empty. Check results for BUSCO: 199024at7742
## Thu Jul 14 01:48:27 CEST 2022 - The aliscore output file does not exist. Check results for BUSCO: 252398at7742
## Thu Jul 14 01:48:27 CEST 2022 - The aliscore output appears to be empty. Check results for BUSCO: 252398at7742
## Thu Jul 14 01:48:28 CEST 2022 - The aliscore output file does not exist. Check results for BUSCO: 242951at7742
## Thu Jul 14 01:48:28 CEST 2022 - The aliscore output appears to be empty. Check results for BUSCO: 242951at7742
## Thu Jul 14 01:48:29 CEST 2022 - The aliscore output file does not exist. Check results for BUSCO: 224305at7742
## Thu Jul 14 01:48:29 CEST 2022 - The aliscore output appears to be empty. Check results for BUSCO: 224305at7742
## Thu Jul 14 01:48:29 CEST 2022 - The aliscore output file does not exist. Check results for BUSCO: 252546at7742
## Thu Jul 14 01:48:29 CEST 2022 - The aliscore output appears to be empty. Check results for BUSCO: 252546at7742
## Thu Jul 14 01:48:31 CEST 2022 - The aliscore output file does not exist. Check results for BUSCO: 238694at7742
## Thu Jul 14 01:48:31 CEST 2022 - The aliscore output appears to be empty. Check results for BUSCO: 238694at7742
## Thu Jul 14 01:48:33 CEST 2022 - The aliscore output file does not exist. Check results for BUSCO: 169338at7742
## Thu Jul 14 01:48:33 CEST 2022 - The aliscore output appears to be empty. Check results for BUSCO: 169338at7742
## Thu Jul 14 01:48:34 CEST 2022 - The aliscore output file does not exist. Check results for BUSCO: 199780at7742
## Thu Jul 14 01:48:34 CEST 2022 - The aliscore output appears to be empty. Check results for BUSCO: 199780at7742
## Thu Jul 14 01:48:35 CEST 2022 - The aliscore output file does not exist. Check results for BUSCO: 229590at7742
## Thu Jul 14 01:48:35 CEST 2022 - The aliscore output appears to be empty. Check results for BUSCO: 229590at7742
## Thu Jul 14 01:48:37 CEST 2022 - The aliscore output file does not exist. Check results for BUSCO: 192799at7742
## Thu Jul 14 01:48:37 CEST 2022 - The aliscore output appears to be empty. Check results for BUSCO: 192799at7742
## Thu Jul 14 01:48:37 CEST 2022 - The aliscore output file does not exist. Check results for BUSCO: 421821at7742
## Thu Jul 14 01:48:37 CEST 2022 - The aliscore output appears to be empty. Check results for BUSCO: 421821at7742
## Thu Jul 14 01:48:37 CEST 2022 - The aliscore output file does not exist. Check results for BUSCO: 374963at7742
## Thu Jul 14 01:48:37 CEST 2022 - The aliscore output appears to be empty. Check results for BUSCO: 374963at7742
## Thu Jul 14 01:48:38 CEST 2022 - The aliscore output file does not exist. Check results for BUSCO: 206342at7742
## Thu Jul 14 01:48:38 CEST 2022 - The aliscore output appears to be empty. Check results for BUSCO: 206342at7742
## Thu Jul 14 01:48:40 CEST 2022 - The aliscore output file does not exist. Check results for BUSCO: 169338at7742
## Thu Jul 14 01:48:40 CEST 2022 - The aliscore output appears to be empty. Check results for BUSCO: 169338at7742
## Thu Jul 14 01:48:41 CEST 2022 - The aliscore output file does not exist. Check results for BUSCO: 257212at7742
## Thu Jul 14 01:48:42 CEST 2022 - The aliscore output appears to be empty. Check results for BUSCO: 257212at7742
## Thu Jul 14 01:48:43 CEST 2022 - The aliscore output file does not exist. Check results for BUSCO: 178102at7742
## Thu Jul 14 01:48:43 CEST 2022 - The aliscore output appears to be empty. Check results for BUSCO: 178102at7742
## Thu Jul 14 01:48:44 CEST 2022 - The aliscore output file does not exist. Check results for BUSCO: 270024at7742
## Thu Jul 14 01:48:44 CEST 2022 - The aliscore output appears to be empty. Check results for BUSCO: 270024at7742
## Thu Jul 14 01:48:45 CEST 2022 - The aliscore output file does not exist. Check results for BUSCO: 239289at7742
## Thu Jul 14 01:48:45 CEST 2022 - The aliscore output appears to be empty. Check results for BUSCO: 239289at7742
## Thu Jul 14 01:48:45 CEST 2022 - The aliscore output file does not exist. Check results for BUSCO: 221671at7742
## Thu Jul 14 01:48:45 CEST 2022 - The aliscore output appears to be empty. Check results for BUSCO: 221671at7742
## Thu Jul 14 01:48:47 CEST 2022 - The aliscore output file does not exist. Check results for BUSCO: 368798at7742
## Thu Jul 14 01:48:47 CEST 2022 - The aliscore output appears to be empty. Check results for BUSCO: 368798at7742
## Thu Jul 14 01:48:49 CEST 2022 - The aliscore output file does not exist. Check results for BUSCO: 201359at7742
## Thu Jul 14 01:48:49 CEST 2022 - The aliscore output appears to be empty. Check results for BUSCO: 201359at7742
## Thu Jul 14 01:48:49 CEST 2022 - The aliscore output file does not exist. Check results for BUSCO: 204010at7742
## Thu Jul 14 01:48:49 CEST 2022 - The aliscore output appears to be empty. Check results for BUSCO: 204010at7742
## Thu Jul 14 01:48:50 CEST 2022 - The aliscore output file does not exist. Check results for BUSCO: 310260at7742
## Thu Jul 14 01:48:50 CEST 2022 - The aliscore output appears to be empty. Check results for BUSCO: 310260at7742
## Thu Jul 14 01:48:51 CEST 2022 - The aliscore output file does not exist. Check results for BUSCO: 183927at7742
## Thu Jul 14 01:48:51 CEST 2022 - The aliscore output appears to be empty. Check results for BUSCO: 183927at7742
## Thu Jul 14 01:48:51 CEST 2022 - The aliscore output file does not exist. Check results for BUSCO: 249877at7742
## Thu Jul 14 01:48:51 CEST 2022 - The aliscore output appears to be empty. Check results for BUSCO: 249877at7742
## Thu Jul 14 01:48:52 CEST 2022 - The aliscore output file does not exist. Check results for BUSCO: 230223at7742
## Thu Jul 14 01:48:52 CEST 2022 - The aliscore output appears to be empty. Check results for BUSCO: 230223at7742
## Thu Jul 14 01:48:55 CEST 2022 - The aliscore output file does not exist. Check results for BUSCO: 374963at7742
## Thu Jul 14 01:48:55 CEST 2022 - The aliscore output appears to be empty. Check results for BUSCO: 374963at7742
## Thu Jul 14 01:48:57 CEST 2022 - The aliscore output file does not exist. Check results for BUSCO: 318212at7742
## Thu Jul 14 01:48:57 CEST 2022 - The aliscore output appears to be empty. Check results for BUSCO: 318212at7742
## Thu Jul 14 01:48:57 CEST 2022 - The aliscore output file does not exist. Check results for BUSCO: 245117at7742
## Thu Jul 14 01:48:57 CEST 2022 - The aliscore output appears to be empty. Check results for BUSCO: 245117at7742
## Thu Jul 14 01:48:58 CEST 2022 - The aliscore output file does not exist. Check results for BUSCO: 217728at7742
## Thu Jul 14 01:48:58 CEST 2022 - The aliscore output appears to be empty. Check results for BUSCO: 217728at7742
## Thu Jul 14 01:48:59 CEST 2022 - The aliscore output file does not exist. Check results for BUSCO: 307639at7742
## Thu Jul 14 01:48:59 CEST 2022 - The aliscore output appears to be empty. Check results for BUSCO: 307639at7742
## Thu Jul 14 01:49:00 CEST 2022 - The aliscore output file does not exist. Check results for BUSCO: 149216at7742
## Thu Jul 14 01:49:00 CEST 2022 - The aliscore output appears to be empty. Check results for BUSCO: 149216at7742
## Thu Jul 14 01:49:02 CEST 2022 - The aliscore output file does not exist. Check results for BUSCO: 156291at7742
## Thu Jul 14 01:49:02 CEST 2022 - The aliscore output appears to be empty. Check results for BUSCO: 156291at7742
## Thu Jul 14 01:49:04 CEST 2022 - The aliscore output file does not exist. Check results for BUSCO: 319641at7742
## Thu Jul 14 01:49:04 CEST 2022 - The aliscore output appears to be empty. Check results for BUSCO: 319641at7742
## Thu Jul 14 01:49:04 CEST 2022 - The aliscore output file does not exist. Check results for BUSCO: 97370at7742
## Thu Jul 14 01:49:04 CEST 2022 - The aliscore output appears to be empty. Check results for BUSCO: 97370at7742
## Thu Jul 14 01:49:05 CEST 2022 - The aliscore output file does not exist. Check results for BUSCO: 261523at7742
## Thu Jul 14 01:49:05 CEST 2022 - The aliscore output file does not exist. Check results for BUSCO: 144352at7742
## Thu Jul 14 01:49:05 CEST 2022 - The aliscore output appears to be empty. Check results for BUSCO: 261523at7742
## Thu Jul 14 01:49:05 CEST 2022 - The aliscore output appears to be empty. Check results for BUSCO: 144352at7742
## Thu Jul 14 01:49:05 CEST 2022 - The aliscore output file does not exist. Check results for BUSCO: 205949at7742
## Thu Jul 14 01:49:06 CEST 2022 - The aliscore output appears to be empty. Check results for BUSCO: 205949at7742
## Thu Jul 14 01:49:07 CEST 2022 - The aliscore output file does not exist. Check results for BUSCO: 302555at7742
## Thu Jul 14 01:49:07 CEST 2022 - The aliscore output appears to be empty. Check results for BUSCO: 302555at7742
## Thu Jul 14 01:49:10 CEST 2022 - The aliscore output file does not exist. Check results for BUSCO: 160383at7742
## Thu Jul 14 01:49:10 CEST 2022 - The aliscore output appears to be empty. Check results for BUSCO: 160383at7742
## Thu Jul 14 01:49:10 CEST 2022 - The aliscore output file does not exist. Check results for BUSCO: 182463at7742
## Thu Jul 14 01:49:10 CEST 2022 - The aliscore output appears to be empty. Check results for BUSCO: 182463at7742
## Thu Jul 14 01:49:11 CEST 2022 - The aliscore output file does not exist. Check results for BUSCO: 245482at7742
## Thu Jul 14 01:49:11 CEST 2022 - The aliscore output appears to be empty. Check results for BUSCO: 245482at7742
## Thu Jul 14 01:49:13 CEST 2022 - The aliscore output file does not exist. Check results for BUSCO: 330809at7742
## Thu Jul 14 01:49:13 CEST 2022 - The aliscore output appears to be empty. Check results for BUSCO: 330809at7742
## Thu Jul 14 01:49:13 CEST 2022 - The aliscore output file does not exist. Check results for BUSCO: 237617at7742
## Thu Jul 14 01:49:13 CEST 2022 - The aliscore output appears to be empty. Check results for BUSCO: 237617at7742
## Thu Jul 14 01:49:16 CEST 2022 - The aliscore output file does not exist. Check results for BUSCO: 353601at7742
## Thu Jul 14 01:49:16 CEST 2022 - The aliscore output appears to be empty. Check results for BUSCO: 353601at7742
## Thu Jul 14 01:49:17 CEST 2022 - The aliscore output file does not exist. Check results for BUSCO: 323966at7742
## Thu Jul 14 01:49:17 CEST 2022 - The aliscore output appears to be empty. Check results for BUSCO: 323966at7742
## Thu Jul 14 01:49:17 CEST 2022 - The aliscore output file does not exist. Check results for BUSCO: 248773at7742
## Thu Jul 14 01:49:17 CEST 2022 - The aliscore output appears to be empty. Check results for BUSCO: 248773at7742
## Thu Jul 14 01:49:18 CEST 2022 - The aliscore output file does not exist. Check results for BUSCO: 202302at7742
## Thu Jul 14 01:49:18 CEST 2022 - The aliscore output appears to be empty. Check results for BUSCO: 202302at7742
## Thu Jul 14 01:49:18 CEST 2022 - The aliscore output file does not exist. Check results for BUSCO: 323966at7742
## Thu Jul 14 01:49:18 CEST 2022 - The aliscore output appears to be empty. Check results for BUSCO: 323966at7742
## Thu Jul 14 01:49:19 CEST 2022 - The aliscore output file does not exist. Check results for BUSCO: 204708at7742
## Thu Jul 14 01:49:19 CEST 2022 - The aliscore output appears to be empty. Check results for BUSCO: 204708at7742
## Thu Jul 14 01:49:21 CEST 2022 - The aliscore output file does not exist. Check results for BUSCO: 241473at7742
## Thu Jul 14 01:49:21 CEST 2022 - The aliscore output appears to be empty. Check results for BUSCO: 241473at7742
## Thu Jul 14 01:49:21 CEST 2022 - The aliscore output file does not exist. Check results for BUSCO: 204708at7742
## Thu Jul 14 01:49:21 CEST 2022 - The aliscore output appears to be empty. Check results for BUSCO: 204708at7742
## Thu Jul 14 01:49:23 CEST 2022 - The aliscore output file does not exist. Check results for BUSCO: 148672at7742
## Thu Jul 14 01:49:23 CEST 2022 - The aliscore output appears to be empty. Check results for BUSCO: 148672at7742
## Thu Jul 14 01:49:25 CEST 2022 - The aliscore output file does not exist. Check results for BUSCO: 257699at7742
## Thu Jul 14 01:49:25 CEST 2022 - The aliscore output appears to be empty. Check results for BUSCO: 257699at7742
## Thu Jul 14 01:49:26 CEST 2022 - The aliscore output file does not exist. Check results for BUSCO: 97370at7742
## Thu Jul 14 01:49:26 CEST 2022 - The aliscore output appears to be empty. Check results for BUSCO: 97370at7742
## Thu Jul 14 01:49:27 CEST 2022 - The aliscore output file does not exist. Check results for BUSCO: 185648at7742
## Thu Jul 14 01:49:27 CEST 2022 - The aliscore output appears to be empty. Check results for BUSCO: 185648at7742
## Thu Jul 14 01:49:27 CEST 2022 - The aliscore output file does not exist. Check results for BUSCO: 342319at7742
## Thu Jul 14 01:49:27 CEST 2022 - The aliscore output appears to be empty. Check results for BUSCO: 342319at7742
## Thu Jul 14 01:49:28 CEST 2022 - The aliscore output file does not exist. Check results for BUSCO: 385740at7742
## Thu Jul 14 01:49:28 CEST 2022 - The aliscore output appears to be empty. Check results for BUSCO: 385740at7742
## Thu Jul 14 01:49:28 CEST 2022 - The aliscore output file does not exist. Check results for BUSCO: 169465at7742
## Thu Jul 14 01:49:28 CEST 2022 - The aliscore output appears to be empty. Check results for BUSCO: 169465at7742
## Thu Jul 14 01:49:30 CEST 2022 - The aliscore output file does not exist. Check results for BUSCO: 206342at7742
## Thu Jul 14 01:49:30 CEST 2022 - The aliscore output appears to be empty. Check results for BUSCO: 206342at7742
## Thu Jul 14 01:49:32 CEST 2022 - The aliscore output file does not exist. Check results for BUSCO: 298884at7742
## Thu Jul 14 01:49:32 CEST 2022 - The aliscore output appears to be empty. Check results for BUSCO: 298884at7742
## Thu Jul 14 01:49:33 CEST 2022 - The aliscore output file does not exist. Check results for BUSCO: 130125at7742
## Thu Jul 14 01:49:33 CEST 2022 - The aliscore output appears to be empty. Check results for BUSCO: 130125at7742
## Thu Jul 14 01:49:34 CEST 2022 - The aliscore output file does not exist. Check results for BUSCO: 195717at7742
## Thu Jul 14 01:49:34 CEST 2022 - The aliscore output appears to be empty. Check results for BUSCO: 195717at7742
## Thu Jul 14 01:49:34 CEST 2022 - The aliscore output file does not exist. Check results for BUSCO: 376721at7742
## Thu Jul 14 01:49:34 CEST 2022 - The aliscore output appears to be empty. Check results for BUSCO: 376721at7742
## Thu Jul 14 01:49:34 CEST 2022 - The aliscore output file does not exist. Check results for BUSCO: 243611at7742
## Thu Jul 14 01:49:34 CEST 2022 - The aliscore output appears to be empty. Check results for BUSCO: 243611at7742
## Thu Jul 14 01:49:37 CEST 2022 - The aliscore output file does not exist. Check results for BUSCO: 355364at7742
## Thu Jul 14 01:49:37 CEST 2022 - The aliscore output appears to be empty. Check results for BUSCO: 355364at7742
## Thu Jul 14 01:49:37 CEST 2022 - The aliscore output file does not exist. Check results for BUSCO: 310872at7742
## Thu Jul 14 01:49:37 CEST 2022 - The aliscore output appears to be empty. Check results for BUSCO: 310872at7742
## Thu Jul 14 01:49:40 CEST 2022 - The aliscore output file does not exist. Check results for BUSCO: 205949at7742
## Thu Jul 14 01:49:40 CEST 2022 - The aliscore output appears to be empty. Check results for BUSCO: 205949at7742
## Thu Jul 14 01:49:40 CEST 2022 - The aliscore output file does not exist. Check results for BUSCO: 318212at7742
## Thu Jul 14 01:49:40 CEST 2022 - The aliscore output appears to be empty. Check results for BUSCO: 318212at7742
## Thu Jul 14 01:49:40 CEST 2022 - The aliscore output file does not exist. Check results for BUSCO: 130125at7742
## Thu Jul 14 01:49:40 CEST 2022 - The aliscore output appears to be empty. Check results for BUSCO: 130125at7742
## Thu Jul 14 01:49:42 CEST 2022 - The aliscore output file does not exist. Check results for BUSCO: 135248at7742
## Thu Jul 14 01:49:42 CEST 2022 - The aliscore output appears to be empty. Check results for BUSCO: 135248at7742
## Thu Jul 14 01:49:43 CEST 2022 - The aliscore output file does not exist. Check results for BUSCO: 229738at7742
## Thu Jul 14 01:49:43 CEST 2022 - The aliscore output appears to be empty. Check results for BUSCO: 229738at7742
## Thu Jul 14 01:49:43 CEST 2022 - The aliscore output file does not exist. Check results for BUSCO: 220049at7742
## Thu Jul 14 01:49:43 CEST 2022 - The aliscore output appears to be empty. Check results for BUSCO: 220049at7742
## Thu Jul 14 01:49:45 CEST 2022 - The aliscore output file does not exist. Check results for BUSCO: 249399at7742
## Thu Jul 14 01:49:45 CEST 2022 - The aliscore output appears to be empty. Check results for BUSCO: 249399at7742
## Thu Jul 14 01:49:48 CEST 2022 - The aliscore output file does not exist. Check results for BUSCO: 310260at7742
## Thu Jul 14 01:49:48 CEST 2022 - The aliscore output appears to be empty. Check results for BUSCO: 310260at7742
## Thu Jul 14 01:49:48 CEST 2022 - The aliscore output file does not exist. Check results for BUSCO: 273897at7742
## Thu Jul 14 01:49:48 CEST 2022 - The aliscore output appears to be empty. Check results for BUSCO: 273897at7742
## Thu Jul 14 01:49:48 CEST 2022 - The aliscore output file does not exist. Check results for BUSCO: 178102at7742
## Thu Jul 14 01:49:48 CEST 2022 - The aliscore output appears to be empty. Check results for BUSCO: 178102at7742
## Thu Jul 14 01:49:49 CEST 2022 - The aliscore output file does not exist. Check results for BUSCO: 223512at7742
## Thu Jul 14 01:49:49 CEST 2022 - The aliscore output appears to be empty. Check results for BUSCO: 223512at7742
## Thu Jul 14 01:49:51 CEST 2022 - The aliscore output file does not exist. Check results for BUSCO: 380483at7742
## Thu Jul 14 01:49:52 CEST 2022 - The aliscore output appears to be empty. Check results for BUSCO: 380483at7742
## Thu Jul 14 01:49:52 CEST 2022 - The aliscore output file does not exist. Check results for BUSCO: 191044at7742
## Thu Jul 14 01:49:52 CEST 2022 - The aliscore output appears to be empty. Check results for BUSCO: 191044at7742
## Thu Jul 14 01:49:54 CEST 2022 - The aliscore output file does not exist. Check results for BUSCO: 404952at7742
## Thu Jul 14 01:49:54 CEST 2022 - The aliscore output appears to be empty. Check results for BUSCO: 404952at7742
## Thu Jul 14 01:49:54 CEST 2022 - The aliscore output file does not exist. Check results for BUSCO: 318212at7742
## Thu Jul 14 01:49:54 CEST 2022 - The aliscore output appears to be empty. Check results for BUSCO: 318212at7742
## Thu Jul 14 01:49:55 CEST 2022 - The aliscore output file does not exist. Check results for BUSCO: 195717at7742
## Thu Jul 14 01:49:55 CEST 2022 - The aliscore output appears to be empty. Check results for BUSCO: 195717at7742
## Thu Jul 14 01:49:56 CEST 2022 - The aliscore output file does not exist. Check results for BUSCO: 149216at7742
## Thu Jul 14 01:49:56 CEST 2022 - The aliscore output appears to be empty. Check results for BUSCO: 149216at7742
## Thu Jul 14 01:49:56 CEST 2022 - The aliscore output file does not exist. Check results for BUSCO: 375400at7742
## Thu Jul 14 01:49:56 CEST 2022 - The aliscore output appears to be empty. Check results for BUSCO: 375400at7742
## Thu Jul 14 01:49:57 CEST 2022 - The aliscore output file does not exist. Check results for BUSCO: 195717at7742
## Thu Jul 14 01:49:57 CEST 2022 - The aliscore output appears to be empty. Check results for BUSCO: 195717at7742
## Thu Jul 14 01:49:59 CEST 2022 - The aliscore output file does not exist. Check results for BUSCO: 97370at7742
## Thu Jul 14 01:49:59 CEST 2022 - The aliscore output appears to be empty. Check results for BUSCO: 97370at7742
## Thu Jul 14 01:50:01 CEST 2022 - The aliscore output file does not exist. Check results for BUSCO: 385085at7742
## Thu Jul 14 01:50:01 CEST 2022 - The aliscore output appears to be empty. Check results for BUSCO: 385085at7742
## Thu Jul 14 01:50:03 CEST 2022 - The aliscore output file does not exist. Check results for BUSCO: 300443at7742
## Thu Jul 14 01:50:03 CEST 2022 - The aliscore output appears to be empty. Check results for BUSCO: 300443at7742
## Thu Jul 14 01:50:03 CEST 2022 - The aliscore output file does not exist. Check results for BUSCO: 130125at7742
## Thu Jul 14 01:50:03 CEST 2022 - The aliscore output appears to be empty. Check results for BUSCO: 130125at7742
## Thu Jul 14 01:50:03 CEST 2022 - The aliscore output file does not exist. Check results for BUSCO: 254455at7742
## Thu Jul 14 01:50:03 CEST 2022 - The aliscore output appears to be empty. Check results for BUSCO: 254455at7742
## Thu Jul 14 01:50:05 CEST 2022 - The aliscore output file does not exist. Check results for BUSCO: 135248at7742
## Thu Jul 14 01:50:05 CEST 2022 - The aliscore output appears to be empty. Check results for BUSCO: 135248at7742
## Thu Jul 14 01:50:05 CEST 2022 - The aliscore output file does not exist. Check results for BUSCO: 264510at7742
## Thu Jul 14 01:50:05 CEST 2022 - The aliscore output appears to be empty. Check results for BUSCO: 264510at7742
## Thu Jul 14 01:50:07 CEST 2022 - The aliscore output file does not exist. Check results for BUSCO: 321359at7742
## Thu Jul 14 01:50:07 CEST 2022 - The aliscore output appears to be empty. Check results for BUSCO: 321359at7742
## Thu Jul 14 01:50:08 CEST 2022 - The aliscore output file does not exist. Check results for BUSCO: 171308at7742
## Thu Jul 14 01:50:08 CEST 2022 - The aliscore output appears to be empty. Check results for BUSCO: 171308at7742
## Thu Jul 14 01:50:09 CEST 2022 - The aliscore output file does not exist. Check results for BUSCO: 172079at7742
## Thu Jul 14 01:50:09 CEST 2022 - The aliscore output appears to be empty. Check results for BUSCO: 172079at7742
## Thu Jul 14 01:50:11 CEST 2022 - The aliscore output file does not exist. Check results for BUSCO: 189753at7742
## Thu Jul 14 01:50:11 CEST 2022 - The aliscore output appears to be empty. Check results for BUSCO: 189753at7742
## Thu Jul 14 01:50:11 CEST 2022 - The aliscore output file does not exist. Check results for BUSCO: 221671at7742
## Thu Jul 14 01:50:12 CEST 2022 - The aliscore output appears to be empty. Check results for BUSCO: 221671at7742
## Thu Jul 14 01:50:12 CEST 2022 - The aliscore output file does not exist. Check results for BUSCO: 267268at7742
## Thu Jul 14 01:50:12 CEST 2022 - The aliscore output file does not exist. Check results for BUSCO: 328109at7742
## Thu Jul 14 01:50:12 CEST 2022 - The aliscore output appears to be empty. Check results for BUSCO: 267268at7742
## Thu Jul 14 01:50:12 CEST 2022 - The aliscore output appears to be empty. Check results for BUSCO: 328109at7742
## Thu Jul 14 01:50:12 CEST 2022 - The aliscore output file does not exist. Check results for BUSCO: 328571at7742
## Thu Jul 14 01:50:12 CEST 2022 - The aliscore output appears to be empty. Check results for BUSCO: 328571at7742
## Thu Jul 14 01:50:15 CEST 2022 - The aliscore output file does not exist. Check results for BUSCO: 202302at7742
## Thu Jul 14 01:50:15 CEST 2022 - The aliscore output appears to be empty. Check results for BUSCO: 202302at7742
## Thu Jul 14 01:50:18 CEST 2022 - The aliscore output file does not exist. Check results for BUSCO: 330809at7742
## Thu Jul 14 01:50:18 CEST 2022 - The aliscore output appears to be empty. Check results for BUSCO: 330809at7742
## Thu Jul 14 01:50:18 CEST 2022 - The aliscore output file does not exist. Check results for BUSCO: 192799at7742
## Thu Jul 14 01:50:18 CEST 2022 - The aliscore output appears to be empty. Check results for BUSCO: 192799at7742
## Thu Jul 14 01:50:19 CEST 2022 - The aliscore output file does not exist. Check results for BUSCO: 238523at7742
## Thu Jul 14 01:50:19 CEST 2022 - The aliscore output appears to be empty. Check results for BUSCO: 238523at7742
## Thu Jul 14 01:50:19 CEST 2022 - The aliscore output file does not exist. Check results for BUSCO: 213068at7742
## Thu Jul 14 01:50:20 CEST 2022 - The aliscore output appears to be empty. Check results for BUSCO: 213068at7742
## Thu Jul 14 01:50:20 CEST 2022 - The aliscore output file does not exist. Check results for BUSCO: 284666at7742
## Thu Jul 14 01:50:20 CEST 2022 - The aliscore output appears to be empty. Check results for BUSCO: 284666at7742
## Thu Jul 14 01:50:22 CEST 2022 - The aliscore output file does not exist. Check results for BUSCO: 376114at7742
## Thu Jul 14 01:50:22 CEST 2022 - The aliscore output appears to be empty. Check results for BUSCO: 376114at7742
## Thu Jul 14 01:50:23 CEST 2022 - The aliscore output file does not exist. Check results for BUSCO: 337325at7742
## Thu Jul 14 01:50:23 CEST 2022 - The aliscore output appears to be empty. Check results for BUSCO: 337325at7742
## Thu Jul 14 01:50:24 CEST 2022 - The aliscore output file does not exist. Check results for BUSCO: 404952at7742
## Thu Jul 14 01:50:24 CEST 2022 - The aliscore output appears to be empty. Check results for BUSCO: 404952at7742
## Thu Jul 14 01:50:25 CEST 2022 - The aliscore output file does not exist. Check results for BUSCO: 376114at7742
## Thu Jul 14 01:50:25 CEST 2022 - The aliscore output appears to be empty. Check results for BUSCO: 376114at7742
## Thu Jul 14 01:50:27 CEST 2022 - The aliscore output file does not exist. Check results for BUSCO: 187916at7742
## Thu Jul 14 01:50:27 CEST 2022 - The aliscore output appears to be empty. Check results for BUSCO: 187916at7742
## Thu Jul 14 01:50:28 CEST 2022 - The aliscore output file does not exist. Check results for BUSCO: 190766at7742
## Thu Jul 14 01:50:28 CEST 2022 - The aliscore output appears to be empty. Check results for BUSCO: 190766at7742
## Thu Jul 14 01:50:29 CEST 2022 - The aliscore output file does not exist. Check results for BUSCO: 242895at7742
## Thu Jul 14 01:50:29 CEST 2022 - The aliscore output appears to be empty. Check results for BUSCO: 242895at7742
## Thu Jul 14 01:50:29 CEST 2022 - The aliscore output file does not exist. Check results for BUSCO: 203253at7742
## Thu Jul 14 01:50:29 CEST 2022 - The aliscore output appears to be empty. Check results for BUSCO: 203253at7742
## Thu Jul 14 01:50:31 CEST 2022 - The aliscore output file does not exist. Check results for BUSCO: 235827at7742
## Thu Jul 14 01:50:31 CEST 2022 - The aliscore output appears to be empty. Check results for BUSCO: 235827at7742
## Thu Jul 14 01:50:33 CEST 2022 - The aliscore output file does not exist. Check results for BUSCO: 211328at7742
## Thu Jul 14 01:50:33 CEST 2022 - The aliscore output appears to be empty. Check results for BUSCO: 211328at7742
## Thu Jul 14 01:50:33 CEST 2022 - The aliscore output file does not exist. Check results for BUSCO: 185648at7742
## Thu Jul 14 01:50:34 CEST 2022 - The aliscore output appears to be empty. Check results for BUSCO: 185648at7742
## Thu Jul 14 01:50:34 CEST 2022 - The aliscore output file does not exist. Check results for BUSCO: 278370at7742
## Thu Jul 14 01:50:34 CEST 2022 - The aliscore output appears to be empty. Check results for BUSCO: 278370at7742
## Thu Jul 14 01:50:35 CEST 2022 - The aliscore output file does not exist. Check results for BUSCO: 353601at7742
## Thu Jul 14 01:50:35 CEST 2022 - The aliscore output appears to be empty. Check results for BUSCO: 353601at7742
## Thu Jul 14 01:50:36 CEST 2022 - The aliscore output file does not exist. Check results for BUSCO: 309596at7742
## Thu Jul 14 01:50:36 CEST 2022 - The aliscore output appears to be empty. Check results for BUSCO: 309596at7742
## Thu Jul 14 01:50:37 CEST 2022 - The aliscore output file does not exist. Check results for BUSCO: 283441at7742
## Thu Jul 14 01:50:37 CEST 2022 - The aliscore output appears to be empty. Check results for BUSCO: 283441at7742
## Thu Jul 14 01:50:39 CEST 2022 - The aliscore output file does not exist. Check results for BUSCO: 276401at7742
## Thu Jul 14 01:50:39 CEST 2022 - The aliscore output appears to be empty. Check results for BUSCO: 276401at7742
## Thu Jul 14 01:50:40 CEST 2022 - The aliscore output file does not exist. Check results for BUSCO: 224305at7742
## Thu Jul 14 01:50:40 CEST 2022 - The aliscore output appears to be empty. Check results for BUSCO: 224305at7742
## Thu Jul 14 01:50:41 CEST 2022 - The aliscore output file does not exist. Check results for BUSCO: 375400at7742
## Thu Jul 14 01:50:41 CEST 2022 - The aliscore output appears to be empty. Check results for BUSCO: 375400at7742
## Thu Jul 14 01:50:42 CEST 2022 - The aliscore output file does not exist. Check results for BUSCO: 234228at7742
## Thu Jul 14 01:50:42 CEST 2022 - The aliscore output appears to be empty. Check results for BUSCO: 234228at7742
## Thu Jul 14 01:50:42 CEST 2022 - The aliscore output file does not exist. Check results for BUSCO: 369343at7742
## Thu Jul 14 01:50:42 CEST 2022 - The aliscore output appears to be empty. Check results for BUSCO: 369343at7742
## Thu Jul 14 01:50:45 CEST 2022 - The aliscore output file does not exist. Check results for BUSCO: 376721at7742
## Thu Jul 14 01:50:45 CEST 2022 - The aliscore output appears to be empty. Check results for BUSCO: 376721at7742
## Thu Jul 14 01:50:45 CEST 2022 - The aliscore output file does not exist. Check results for BUSCO: 277524at7742
## Thu Jul 14 01:50:45 CEST 2022 - The aliscore output appears to be empty. Check results for BUSCO: 277524at7742
## Thu Jul 14 01:50:47 CEST 2022 - The aliscore output file does not exist. Check results for BUSCO: 180713at7742
## Thu Jul 14 01:50:47 CEST 2022 - The aliscore output appears to be empty. Check results for BUSCO: 180713at7742
## Thu Jul 14 01:50:48 CEST 2022 - The aliscore output file does not exist. Check results for BUSCO: 277067at7742
## Thu Jul 14 01:50:48 CEST 2022 - The aliscore output appears to be empty. Check results for BUSCO: 277067at7742
## Thu Jul 14 01:50:48 CEST 2022 - The aliscore output file does not exist. Check results for BUSCO: 225241at7742
## Thu Jul 14 01:50:48 CEST 2022 - The aliscore output appears to be empty. Check results for BUSCO: 225241at7742
## Thu Jul 14 01:50:48 CEST 2022 - The aliscore output file does not exist. Check results for BUSCO: 376010at7742
## Thu Jul 14 01:50:48 CEST 2022 - The aliscore output appears to be empty. Check results for BUSCO: 376010at7742
## Thu Jul 14 01:50:50 CEST 2022 - The aliscore output file does not exist. Check results for BUSCO: 172079at7742
## Thu Jul 14 01:50:50 CEST 2022 - The aliscore output appears to be empty. Check results for BUSCO: 172079at7742
## Thu Jul 14 01:50:52 CEST 2022 - The aliscore output file does not exist. Check results for BUSCO: 239289at7742
## Thu Jul 14 01:50:52 CEST 2022 - The aliscore output appears to be empty. Check results for BUSCO: 239289at7742
## Thu Jul 14 01:50:53 CEST 2022 - The aliscore output file does not exist. Check results for BUSCO: 264772at7742
## Thu Jul 14 01:50:54 CEST 2022 - The aliscore output appears to be empty. Check results for BUSCO: 264772at7742
## Thu Jul 14 01:50:54 CEST 2022 - The aliscore output file does not exist. Check results for BUSCO: 368798at7742
## Thu Jul 14 01:50:54 CEST 2022 - The aliscore output appears to be empty. Check results for BUSCO: 368798at7742
## Thu Jul 14 01:50:54 CEST 2022 - The aliscore output file does not exist. Check results for BUSCO: 254455at7742
## Thu Jul 14 01:50:54 CEST 2022 - The aliscore output appears to be empty. Check results for BUSCO: 254455at7742
## Thu Jul 14 01:50:56 CEST 2022 - The aliscore output file does not exist. Check results for BUSCO: 264056at7742
## Thu Jul 14 01:50:56 CEST 2022 - The aliscore output appears to be empty. Check results for BUSCO: 264056at7742
## Thu Jul 14 01:50:57 CEST 2022 - The aliscore output file does not exist. Check results for BUSCO: 337325at7742
## Thu Jul 14 01:50:57 CEST 2022 - The aliscore output appears to be empty. Check results for BUSCO: 337325at7742
## Thu Jul 14 01:50:58 CEST 2022 - The aliscore output file does not exist. Check results for BUSCO: 277226at7742
## Thu Jul 14 01:50:58 CEST 2022 - The aliscore output appears to be empty. Check results for BUSCO: 277226at7742
## Thu Jul 14 01:50:59 CEST 2022 - The aliscore output file does not exist. Check results for BUSCO: 189224at7742
## Thu Jul 14 01:50:59 CEST 2022 - The aliscore output appears to be empty. Check results for BUSCO: 189224at7742
## Thu Jul 14 01:50:59 CEST 2022 - The aliscore output file does not exist. Check results for BUSCO: 366274at7742
## Thu Jul 14 01:50:59 CEST 2022 - The aliscore output appears to be empty. Check results for BUSCO: 366274at7742
## Thu Jul 14 01:51:02 CEST 2022 - The aliscore output file does not exist. Check results for BUSCO: 242641at7742
## Thu Jul 14 01:51:02 CEST 2022 - The aliscore output appears to be empty. Check results for BUSCO: 242641at7742
## Thu Jul 14 01:51:02 CEST 2022 - The aliscore output file does not exist. Check results for BUSCO: 276691at7742
## Thu Jul 14 01:51:02 CEST 2022 - The aliscore output appears to be empty. Check results for BUSCO: 276691at7742
## Thu Jul 14 01:51:03 CEST 2022 - The aliscore output file does not exist. Check results for BUSCO: 279373at7742
## Thu Jul 14 01:51:03 CEST 2022 - The aliscore output appears to be empty. Check results for BUSCO: 279373at7742
## Thu Jul 14 01:51:05 CEST 2022 - The aliscore output file does not exist. Check results for BUSCO: 275566at7742
## Thu Jul 14 01:51:05 CEST 2022 - The aliscore output appears to be empty. Check results for BUSCO: 275566at7742
## Thu Jul 14 01:51:05 CEST 2022 - The aliscore output file does not exist. Check results for BUSCO: 174424at7742
## Thu Jul 14 01:51:05 CEST 2022 - The aliscore output appears to be empty. Check results for BUSCO: 174424at7742
## Thu Jul 14 01:51:06 CEST 2022 - The aliscore output file does not exist. Check results for BUSCO: 345950at7742
## Thu Jul 14 01:51:06 CEST 2022 - The aliscore output appears to be empty. Check results for BUSCO: 345950at7742
## Thu Jul 14 01:51:09 CEST 2022 - The aliscore output file does not exist. Check results for BUSCO: 220049at7742
## Thu Jul 14 01:51:09 CEST 2022 - The aliscore output appears to be empty. Check results for BUSCO: 220049at7742
## Thu Jul 14 01:51:10 CEST 2022 - The aliscore output file does not exist. Check results for BUSCO: 207782at7742
## Thu Jul 14 01:51:10 CEST 2022 - The aliscore output appears to be empty. Check results for BUSCO: 207782at7742
## Thu Jul 14 01:51:11 CEST 2022 - The aliscore output file does not exist. Check results for BUSCO: 199723at7742
## Thu Jul 14 01:51:11 CEST 2022 - The aliscore output appears to be empty. Check results for BUSCO: 199723at7742
## Thu Jul 14 01:51:11 CEST 2022 - The aliscore output file does not exist. Check results for BUSCO: 287466at7742
## Thu Jul 14 01:51:11 CEST 2022 - The aliscore output appears to be empty. Check results for BUSCO: 287466at7742
## Thu Jul 14 01:51:12 CEST 2022 - The aliscore output file does not exist. Check results for BUSCO: 254455at7742
## Thu Jul 14 01:51:12 CEST 2022 - The aliscore output appears to be empty. Check results for BUSCO: 254455at7742
## Thu Jul 14 01:51:12 CEST 2022 - The aliscore output file does not exist. Check results for BUSCO: 182463at7742
## Thu Jul 14 01:51:12 CEST 2022 - The aliscore output appears to be empty. Check results for BUSCO: 182463at7742
## Thu Jul 14 01:51:14 CEST 2022 - The aliscore output file does not exist. Check results for BUSCO: 309596at7742
## Thu Jul 14 01:51:14 CEST 2022 - The aliscore output appears to be empty. Check results for BUSCO: 309596at7742
## Thu Jul 14 01:51:16 CEST 2022 - The aliscore output file does not exist. Check results for BUSCO: 180713at7742
## Thu Jul 14 01:51:16 CEST 2022 - The aliscore output appears to be empty. Check results for BUSCO: 180713at7742
## Thu Jul 14 01:51:17 CEST 2022 - The aliscore output file does not exist. Check results for BUSCO: 172079at7742
## Thu Jul 14 01:51:17 CEST 2022 - The aliscore output appears to be empty. Check results for BUSCO: 172079at7742
## Thu Jul 14 01:51:18 CEST 2022 - The aliscore output file does not exist. Check results for BUSCO: 297362at7742
## Thu Jul 14 01:51:18 CEST 2022 - The aliscore output appears to be empty. Check results for BUSCO: 297362at7742
## Thu Jul 14 01:51:18 CEST 2022 - The aliscore output file does not exist. Check results for BUSCO: 328109at7742
## Thu Jul 14 01:51:18 CEST 2022 - The aliscore output appears to be empty. Check results for BUSCO: 328109at7742
## Thu Jul 14 01:51:20 CEST 2022 - The aliscore output file does not exist. Check results for BUSCO: 325935at7742
## Thu Jul 14 01:51:20 CEST 2022 - The aliscore output appears to be empty. Check results for BUSCO: 325935at7742
## Thu Jul 14 01:51:21 CEST 2022 - The aliscore output file does not exist. Check results for BUSCO: 179120at7742
## Thu Jul 14 01:51:21 CEST 2022 - The aliscore output appears to be empty. Check results for BUSCO: 179120at7742
## Thu Jul 14 01:51:21 CEST 2022 - The aliscore output file does not exist. Check results for BUSCO: 209951at7742
## Thu Jul 14 01:51:22 CEST 2022 - The aliscore output appears to be empty. Check results for BUSCO: 209951at7742
## Thu Jul 14 01:51:24 CEST 2022 - The aliscore output file does not exist. Check results for BUSCO: 133153at7742
## Thu Jul 14 01:51:24 CEST 2022 - The aliscore output appears to be empty. Check results for BUSCO: 133153at7742
## Thu Jul 14 01:51:24 CEST 2022 - The aliscore output file does not exist. Check results for BUSCO: 342319at7742
## Thu Jul 14 01:51:24 CEST 2022 - The aliscore output appears to be empty. Check results for BUSCO: 342319at7742
## Thu Jul 14 01:51:26 CEST 2022 - The aliscore output file does not exist. Check results for BUSCO: 363898at7742
## Thu Jul 14 01:51:26 CEST 2022 - The aliscore output appears to be empty. Check results for BUSCO: 363898at7742
## Thu Jul 14 01:51:26 CEST 2022 - The aliscore output file does not exist. Check results for BUSCO: 366847at7742
## Thu Jul 14 01:51:26 CEST 2022 - The aliscore output appears to be empty. Check results for BUSCO: 366847at7742
## Thu Jul 14 01:51:26 CEST 2022 - The aliscore output file does not exist. Check results for BUSCO: 257782at7742
## Thu Jul 14 01:51:26 CEST 2022 - The aliscore output appears to be empty. Check results for BUSCO: 257782at7742
## Thu Jul 14 01:51:28 CEST 2022 - The aliscore output file does not exist. Check results for BUSCO: 202013at7742
## Thu Jul 14 01:51:28 CEST 2022 - The aliscore output appears to be empty. Check results for BUSCO: 202013at7742
## Thu Jul 14 01:51:30 CEST 2022 - The aliscore output file does not exist. Check results for BUSCO: 264510at7742
## Thu Jul 14 01:51:30 CEST 2022 - The aliscore output appears to be empty. Check results for BUSCO: 264510at7742
## Thu Jul 14 01:51:32 CEST 2022 - The aliscore output file does not exist. Check results for BUSCO: 292437at7742
## Thu Jul 14 01:51:32 CEST 2022 - The aliscore output appears to be empty. Check results for BUSCO: 292437at7742
## Thu Jul 14 01:51:32 CEST 2022 - The aliscore output file does not exist. Check results for BUSCO: 311983at7742
## Thu Jul 14 01:51:32 CEST 2022 - The aliscore output appears to be empty. Check results for BUSCO: 311983at7742
## Thu Jul 14 01:51:33 CEST 2022 - The aliscore output file does not exist. Check results for BUSCO: 169465at7742
## Thu Jul 14 01:51:33 CEST 2022 - The aliscore output appears to be empty. Check results for BUSCO: 169465at7742
## Thu Jul 14 01:51:34 CEST 2022 - The aliscore output file does not exist. Check results for BUSCO: 182049at7742
## Thu Jul 14 01:51:34 CEST 2022 - The aliscore output appears to be empty. Check results for BUSCO: 182049at7742
## Thu Jul 14 01:51:34 CEST 2022 - The aliscore output file does not exist. Check results for BUSCO: 214728at7742
## Thu Jul 14 01:51:35 CEST 2022 - The aliscore output appears to be empty. Check results for BUSCO: 214728at7742
## Thu Jul 14 01:51:36 CEST 2022 - The aliscore output file does not exist. Check results for BUSCO: 348500at7742
## Thu Jul 14 01:51:36 CEST 2022 - The aliscore output appears to be empty. Check results for BUSCO: 348500at7742
## Thu Jul 14 01:51:37 CEST 2022 - The aliscore output file does not exist. Check results for BUSCO: 270024at7742
## Thu Jul 14 01:51:37 CEST 2022 - The aliscore output appears to be empty. Check results for BUSCO: 270024at7742
## Thu Jul 14 01:51:39 CEST 2022 - The aliscore output file does not exist. Check results for BUSCO: 319402at7742
## Thu Jul 14 01:51:39 CEST 2022 - The aliscore output appears to be empty. Check results for BUSCO: 319402at7742
## Thu Jul 14 01:51:39 CEST 2022 - The aliscore output file does not exist. Check results for BUSCO: 236098at7742
## Thu Jul 14 01:51:39 CEST 2022 - The aliscore output appears to be empty. Check results for BUSCO: 236098at7742
## Thu Jul 14 01:51:41 CEST 2022 - The aliscore output file does not exist. Check results for BUSCO: 407881at7742
## Thu Jul 14 01:51:41 CEST 2022 - The aliscore output appears to be empty. Check results for BUSCO: 407881at7742
## Thu Jul 14 01:51:42 CEST 2022 - The aliscore output file does not exist. Check results for BUSCO: 174424at7742
## Thu Jul 14 01:51:42 CEST 2022 - The aliscore output file does not exist. Check results for BUSCO: 219315at7742
## Thu Jul 14 01:51:42 CEST 2022 - The aliscore output appears to be empty. Check results for BUSCO: 174424at7742
## Thu Jul 14 01:51:42 CEST 2022 - The aliscore output appears to be empty. Check results for BUSCO: 219315at7742
## Thu Jul 14 01:51:42 CEST 2022 - The aliscore output file does not exist. Check results for BUSCO: 345950at7742
## Thu Jul 14 01:51:42 CEST 2022 - The aliscore output appears to be empty. Check results for BUSCO: 345950at7742
## Thu Jul 14 01:51:45 CEST 2022 - The aliscore output file does not exist. Check results for BUSCO: 318959at7742
## Thu Jul 14 01:51:45 CEST 2022 - The aliscore output appears to be empty. Check results for BUSCO: 318959at7742
## Thu Jul 14 01:51:46 CEST 2022 - The aliscore output file does not exist. Check results for BUSCO: 211150at7742
## Thu Jul 14 01:51:46 CEST 2022 - The aliscore output appears to be empty. Check results for BUSCO: 211150at7742
## Thu Jul 14 01:51:47 CEST 2022 - The aliscore output file does not exist. Check results for BUSCO: 319641at7742
## Thu Jul 14 01:51:47 CEST 2022 - The aliscore output appears to be empty. Check results for BUSCO: 319641at7742
## Thu Jul 14 01:51:48 CEST 2022 - The aliscore output file does not exist. Check results for BUSCO: 288248at7742
## Thu Jul 14 01:51:48 CEST 2022 - The aliscore output appears to be empty. Check results for BUSCO: 288248at7742
## Thu Jul 14 01:51:50 CEST 2022 - The aliscore output file does not exist. Check results for BUSCO: 159879at7742
## Thu Jul 14 01:51:50 CEST 2022 - The aliscore output appears to be empty. Check results for BUSCO: 159879at7742
## Thu Jul 14 01:51:50 CEST 2022 - The aliscore output file does not exist. Check results for BUSCO: 214745at7742
## Thu Jul 14 01:51:50 CEST 2022 - The aliscore output appears to be empty. Check results for BUSCO: 214745at7742
## Thu Jul 14 01:51:52 CEST 2022 - The aliscore output file does not exist. Check results for BUSCO: 291239at7742
## Thu Jul 14 01:51:52 CEST 2022 - The aliscore output appears to be empty. Check results for BUSCO: 291239at7742
## Thu Jul 14 01:51:53 CEST 2022 - The aliscore output file does not exist. Check results for BUSCO: 328582at7742
## Thu Jul 14 01:51:53 CEST 2022 - The aliscore output appears to be empty. Check results for BUSCO: 328582at7742
## Thu Jul 14 01:51:53 CEST 2022 - The aliscore output file does not exist. Check results for BUSCO: 194403at7742
## Thu Jul 14 01:51:53 CEST 2022 - The aliscore output appears to be empty. Check results for BUSCO: 194403at7742
## Thu Jul 14 01:51:54 CEST 2022 - The aliscore output file does not exist. Check results for BUSCO: 366847at7742
## Thu Jul 14 01:51:54 CEST 2022 - The aliscore output appears to be empty. Check results for BUSCO: 366847at7742
## Thu Jul 14 01:51:57 CEST 2022 - The aliscore output file does not exist. Check results for BUSCO: 133604at7742
## Thu Jul 14 01:51:57 CEST 2022 - The aliscore output appears to be empty. Check results for BUSCO: 133604at7742
## Thu Jul 14 01:51:58 CEST 2022 - The aliscore output file does not exist. Check results for BUSCO: 345950at7742
## Thu Jul 14 01:51:58 CEST 2022 - The aliscore output appears to be empty. Check results for BUSCO: 345950at7742
## Thu Jul 14 01:51:59 CEST 2022 - The aliscore output file does not exist. Check results for BUSCO: 207648at7742
## Thu Jul 14 01:51:59 CEST 2022 - The aliscore output appears to be empty. Check results for BUSCO: 207648at7742
## Thu Jul 14 01:52:00 CEST 2022 - The aliscore output file does not exist. Check results for BUSCO: 190562at7742
## Thu Jul 14 01:52:00 CEST 2022 - The aliscore output appears to be empty. Check results for BUSCO: 190562at7742
## Thu Jul 14 01:52:01 CEST 2022 - The aliscore output file does not exist. Check results for BUSCO: 235765at7742
## Thu Jul 14 01:52:01 CEST 2022 - The aliscore output appears to be empty. Check results for BUSCO: 235765at7742
## Thu Jul 14 01:52:01 CEST 2022 - The aliscore output file does not exist. Check results for BUSCO: 242487at7742
## Thu Jul 14 01:52:01 CEST 2022 - The aliscore output appears to be empty. Check results for BUSCO: 242487at7742
## Thu Jul 14 01:52:02 CEST 2022 - The aliscore output file does not exist. Check results for BUSCO: 236476at7742
## Thu Jul 14 01:52:02 CEST 2022 - The aliscore output appears to be empty. Check results for BUSCO: 236476at7742
## Thu Jul 14 01:52:03 CEST 2022 - The aliscore output file does not exist. Check results for BUSCO: 199987at7742
## Thu Jul 14 01:52:03 CEST 2022 - The aliscore output appears to be empty. Check results for BUSCO: 199987at7742
## Thu Jul 14 01:52:05 CEST 2022 - The aliscore output file does not exist. Check results for BUSCO: 373266at7742
## Thu Jul 14 01:52:05 CEST 2022 - The aliscore output appears to be empty. Check results for BUSCO: 373266at7742
## Thu Jul 14 01:52:05 CEST 2022 - The aliscore output file does not exist. Check results for BUSCO: 325908at7742
## Thu Jul 14 01:52:05 CEST 2022 - The aliscore output appears to be empty. Check results for BUSCO: 325908at7742
## Thu Jul 14 01:52:06 CEST 2022 - The aliscore output file does not exist. Check results for BUSCO: 369343at7742
## Thu Jul 14 01:52:07 CEST 2022 - The aliscore output appears to be empty. Check results for BUSCO: 369343at7742
## Thu Jul 14 01:52:07 CEST 2022 - The aliscore output file does not exist. Check results for BUSCO: 138435at7742
## Thu Jul 14 01:52:07 CEST 2022 - The aliscore output appears to be empty. Check results for BUSCO: 138435at7742
## Thu Jul 14 01:52:08 CEST 2022 - The aliscore output file does not exist. Check results for BUSCO: 376010at7742
## Thu Jul 14 01:52:08 CEST 2022 - The aliscore output appears to be empty. Check results for BUSCO: 376010at7742
## Thu Jul 14 01:52:10 CEST 2022 - The aliscore output file does not exist. Check results for BUSCO: 236098at7742
## Thu Jul 14 01:52:10 CEST 2022 - The aliscore output appears to be empty. Check results for BUSCO: 236098at7742
## Thu Jul 14 01:52:10 CEST 2022 - The aliscore output file does not exist. Check results for BUSCO: 186913at7742
## Thu Jul 14 01:52:10 CEST 2022 - The aliscore output appears to be empty. Check results for BUSCO: 186913at7742
## Thu Jul 14 01:52:12 CEST 2022 - The aliscore output file does not exist. Check results for BUSCO: 149872at7742
## Thu Jul 14 01:52:12 CEST 2022 - The aliscore output appears to be empty. Check results for BUSCO: 149872at7742
## Thu Jul 14 01:52:12 CEST 2022 - The aliscore output file does not exist. Check results for BUSCO: 291774at7742
## Thu Jul 14 01:52:12 CEST 2022 - The aliscore output appears to be empty. Check results for BUSCO: 291774at7742
## Thu Jul 14 01:52:14 CEST 2022 - The aliscore output file does not exist. Check results for BUSCO: 162664at7742
## Thu Jul 14 01:52:14 CEST 2022 - The aliscore output appears to be empty. Check results for BUSCO: 162664at7742
## Thu Jul 14 01:52:16 CEST 2022 - The aliscore output file does not exist. Check results for BUSCO: 214728at7742
## Thu Jul 14 01:52:17 CEST 2022 - The aliscore output appears to be empty. Check results for BUSCO: 214728at7742
## Thu Jul 14 01:52:17 CEST 2022 - The aliscore output file does not exist. Check results for BUSCO: 205949at7742
## Thu Jul 14 01:52:17 CEST 2022 - The aliscore output appears to be empty. Check results for BUSCO: 205949at7742
## Thu Jul 14 01:52:19 CEST 2022 - The aliscore output file does not exist. Check results for BUSCO: 183251at7742
## Thu Jul 14 01:52:19 CEST 2022 - The aliscore output appears to be empty. Check results for BUSCO: 183251at7742
## Thu Jul 14 01:52:20 CEST 2022 - The aliscore output file does not exist. Check results for BUSCO: 242951at7742
## Thu Jul 14 01:52:20 CEST 2022 - The aliscore output appears to be empty. Check results for BUSCO: 242951at7742
## Thu Jul 14 01:52:20 CEST 2022 - The aliscore output file does not exist. Check results for BUSCO: 174424at7742
## Thu Jul 14 01:52:20 CEST 2022 - The aliscore output appears to be empty. Check results for BUSCO: 174424at7742
## Thu Jul 14 01:52:20 CEST 2022 - The aliscore output file does not exist. Check results for BUSCO: 252806at7742
## Thu Jul 14 01:52:20 CEST 2022 - The aliscore output appears to be empty. Check results for BUSCO: 252806at7742
## Thu Jul 14 01:52:21 CEST 2022 - The aliscore output file does not exist. Check results for BUSCO: 166002at7742
## Thu Jul 14 01:52:21 CEST 2022 - The aliscore output appears to be empty. Check results for BUSCO: 166002at7742
## Thu Jul 14 01:52:23 CEST 2022 - The aliscore output file does not exist. Check results for BUSCO: 279455at7742
## Thu Jul 14 01:52:23 CEST 2022 - The aliscore output appears to be empty. Check results for BUSCO: 279455at7742
## Thu Jul 14 01:52:24 CEST 2022 - The aliscore output file does not exist. Check results for BUSCO: 296065at7742
## Thu Jul 14 01:52:24 CEST 2022 - The aliscore output appears to be empty. Check results for BUSCO: 296065at7742
## Thu Jul 14 01:52:24 CEST 2022 - The aliscore output file does not exist. Check results for BUSCO: 213068at7742
## Thu Jul 14 01:52:24 CEST 2022 - The aliscore output appears to be empty. Check results for BUSCO: 213068at7742
## Thu Jul 14 01:52:26 CEST 2022 - The aliscore output file does not exist. Check results for BUSCO: 240403at7742
## Thu Jul 14 01:52:26 CEST 2022 - The aliscore output appears to be empty. Check results for BUSCO: 240403at7742
## Thu Jul 14 01:52:28 CEST 2022 - The aliscore output file does not exist. Check results for BUSCO: 318662at7742
## Thu Jul 14 01:52:28 CEST 2022 - The aliscore output appears to be empty. Check results for BUSCO: 318662at7742
## Thu Jul 14 01:52:28 CEST 2022 - The aliscore output file does not exist. Check results for BUSCO: 254424at7742
## Thu Jul 14 01:52:28 CEST 2022 - The aliscore output appears to be empty. Check results for BUSCO: 254424at7742
## Thu Jul 14 01:52:28 CEST 2022 - The aliscore output file does not exist. Check results for BUSCO: 272002at7742
## Thu Jul 14 01:52:28 CEST 2022 - The aliscore output appears to be empty. Check results for BUSCO: 272002at7742
## Thu Jul 14 01:52:30 CEST 2022 - The aliscore output file does not exist. Check results for BUSCO: 266056at7742
## Thu Jul 14 01:52:30 CEST 2022 - The aliscore output appears to be empty. Check results for BUSCO: 266056at7742
## Thu Jul 14 01:52:31 CEST 2022 - The aliscore output file does not exist. Check results for BUSCO: 300584at7742
## Thu Jul 14 01:52:31 CEST 2022 - The aliscore output appears to be empty. Check results for BUSCO: 300584at7742
## Thu Jul 14 01:52:31 CEST 2022 - The aliscore output file does not exist. Check results for BUSCO: 146782at7742
## Thu Jul 14 01:52:31 CEST 2022 - The aliscore output appears to be empty. Check results for BUSCO: 146782at7742
## Thu Jul 14 01:52:34 CEST 2022 - The aliscore output file does not exist. Check results for BUSCO: 229738at7742
## Thu Jul 14 01:52:34 CEST 2022 - The aliscore output appears to be empty. Check results for BUSCO: 229738at7742
## Thu Jul 14 01:52:34 CEST 2022 - The aliscore output file does not exist. Check results for BUSCO: 165972at7742
## Thu Jul 14 01:52:34 CEST 2022 - The aliscore output appears to be empty. Check results for BUSCO: 165972at7742
## Thu Jul 14 01:52:35 CEST 2022 - The aliscore output file does not exist. Check results for BUSCO: 165972at7742
## Thu Jul 14 01:52:35 CEST 2022 - The aliscore output appears to be empty. Check results for BUSCO: 165972at7742
## Thu Jul 14 01:52:36 CEST 2022 - The aliscore output file does not exist. Check results for BUSCO: 350211at7742
## Thu Jul 14 01:52:37 CEST 2022 - The aliscore output appears to be empty. Check results for BUSCO: 350211at7742
## Thu Jul 14 01:52:37 CEST 2022 - The aliscore output file does not exist. Check results for BUSCO: 307638at7742
## Thu Jul 14 01:52:37 CEST 2022 - The aliscore output appears to be empty. Check results for BUSCO: 307638at7742
## Thu Jul 14 01:52:37 CEST 2022 - The aliscore output file does not exist. Check results for BUSCO: 350211at7742
## Thu Jul 14 01:52:37 CEST 2022 - The aliscore output appears to be empty. Check results for BUSCO: 350211at7742
## Thu Jul 14 01:52:41 CEST 2022 - The aliscore output file does not exist. Check results for BUSCO: 376448at7742
## Thu Jul 14 01:52:41 CEST 2022 - The aliscore output appears to be empty. Check results for BUSCO: 376448at7742
## Thu Jul 14 01:52:41 CEST 2022 - The aliscore output file does not exist. Check results for BUSCO: 285172at7742
## Thu Jul 14 01:52:42 CEST 2022 - The aliscore output appears to be empty. Check results for BUSCO: 285172at7742
## Thu Jul 14 01:52:42 CEST 2022 - The aliscore output file does not exist. Check results for BUSCO: 270024at7742
## Thu Jul 14 01:52:42 CEST 2022 - The aliscore output appears to be empty. Check results for BUSCO: 270024at7742
## Thu Jul 14 01:52:43 CEST 2022 - The aliscore output file does not exist. Check results for BUSCO: 292857at7742
## Thu Jul 14 01:52:43 CEST 2022 - The aliscore output appears to be empty. Check results for BUSCO: 292857at7742
## Thu Jul 14 01:52:43 CEST 2022 - The aliscore output file does not exist. Check results for BUSCO: 266228at7742
## Thu Jul 14 01:52:43 CEST 2022 - The aliscore output appears to be empty. Check results for BUSCO: 266228at7742
## Thu Jul 14 01:52:44 CEST 2022 - The aliscore output file does not exist. Check results for BUSCO: 192284at7742
## Thu Jul 14 01:52:44 CEST 2022 - The aliscore output appears to be empty. Check results for BUSCO: 192284at7742
## Thu Jul 14 01:52:46 CEST 2022 - The aliscore output file does not exist. Check results for BUSCO: 221887at7742
## Thu Jul 14 01:52:46 CEST 2022 - The aliscore output appears to be empty. Check results for BUSCO: 221887at7742
## Thu Jul 14 01:52:48 CEST 2022 - The aliscore output file does not exist. Check results for BUSCO: 224305at7742
## Thu Jul 14 01:52:48 CEST 2022 - The aliscore output appears to be empty. Check results for BUSCO: 224305at7742
## Thu Jul 14 01:52:49 CEST 2022 - The aliscore output file does not exist. Check results for BUSCO: 159334at7742
## Thu Jul 14 01:52:49 CEST 2022 - The aliscore output appears to be empty. Check results for BUSCO: 159334at7742
## Thu Jul 14 01:52:50 CEST 2022 - The aliscore output file does not exist. Check results for BUSCO: 363812at7742
## Thu Jul 14 01:52:50 CEST 2022 - The aliscore output appears to be empty. Check results for BUSCO: 363812at7742
## Thu Jul 14 01:52:51 CEST 2022 - The aliscore output file does not exist. Check results for BUSCO: 292028at7742
## Thu Jul 14 01:52:51 CEST 2022 - The aliscore output appears to be empty. Check results for BUSCO: 292028at7742
## Thu Jul 14 01:52:51 CEST 2022 - The aliscore output file does not exist. Check results for BUSCO: 264772at7742
## Thu Jul 14 01:52:51 CEST 2022 - The aliscore output appears to be empty. Check results for BUSCO: 264772at7742
## Thu Jul 14 01:52:51 CEST 2022 - The aliscore output file does not exist. Check results for BUSCO: 294443at7742
## Thu Jul 14 01:52:51 CEST 2022 - The aliscore output appears to be empty. Check results for BUSCO: 294443at7742
## Thu Jul 14 01:52:53 CEST 2022 - The aliscore output file does not exist. Check results for BUSCO: 283441at7742
## Thu Jul 14 01:52:53 CEST 2022 - The aliscore output appears to be empty. Check results for BUSCO: 283441at7742
## Thu Jul 14 01:52:56 CEST 2022 - The aliscore output file does not exist. Check results for BUSCO: 177602at7742
## Thu Jul 14 01:52:56 CEST 2022 - The aliscore output appears to be empty. Check results for BUSCO: 177602at7742
## Thu Jul 14 01:52:58 CEST 2022 - The aliscore output file does not exist. Check results for BUSCO: 330809at7742
## Thu Jul 14 01:52:58 CEST 2022 - The aliscore output appears to be empty. Check results for BUSCO: 330809at7742
## Thu Jul 14 01:52:58 CEST 2022 - The aliscore output file does not exist. Check results for BUSCO: 307639at7742
## Thu Jul 14 01:52:58 CEST 2022 - The aliscore output appears to be empty. Check results for BUSCO: 307639at7742
## Thu Jul 14 01:52:59 CEST 2022 - The aliscore output file does not exist. Check results for BUSCO: 281159at7742
## Thu Jul 14 01:52:59 CEST 2022 - The aliscore output appears to be empty. Check results for BUSCO: 281159at7742
## Thu Jul 14 01:53:00 CEST 2022 - The aliscore output file does not exist. Check results for BUSCO: 249694at7742
## Thu Jul 14 01:53:00 CEST 2022 - The aliscore output appears to be empty. Check results for BUSCO: 249694at7742
## Thu Jul 14 01:53:01 CEST 2022 - The aliscore output file does not exist. Check results for BUSCO: 159879at7742
## Thu Jul 14 01:53:01 CEST 2022 - The aliscore output appears to be empty. Check results for BUSCO: 159879at7742
## Thu Jul 14 01:53:01 CEST 2022 - The aliscore output file does not exist. Check results for BUSCO: 267724at7742
## Thu Jul 14 01:53:01 CEST 2022 - The aliscore output appears to be empty. Check results for BUSCO: 267724at7742
## Thu Jul 14 01:53:03 CEST 2022 - The aliscore output file does not exist. Check results for BUSCO: 146782at7742
## Thu Jul 14 01:53:03 CEST 2022 - The aliscore output appears to be empty. Check results for BUSCO: 146782at7742
## Thu Jul 14 01:53:03 CEST 2022 - The aliscore output file does not exist. Check results for BUSCO: 215706at7742
## Thu Jul 14 01:53:03 CEST 2022 - The aliscore output appears to be empty. Check results for BUSCO: 215706at7742
## Thu Jul 14 01:53:05 CEST 2022 - The aliscore output file does not exist. Check results for BUSCO: 245482at7742
## Thu Jul 14 01:53:05 CEST 2022 - The aliscore output appears to be empty. Check results for BUSCO: 245482at7742
## Thu Jul 14 01:53:07 CEST 2022 - The aliscore output file does not exist. Check results for BUSCO: 309596at7742
## Thu Jul 14 01:53:07 CEST 2022 - The aliscore output appears to be empty. Check results for BUSCO: 309596at7742
## Thu Jul 14 01:53:07 CEST 2022 - The aliscore output file does not exist. Check results for BUSCO: 158698at7742
## Thu Jul 14 01:53:07 CEST 2022 - The aliscore output appears to be empty. Check results for BUSCO: 158698at7742
## Thu Jul 14 01:53:07 CEST 2022 - The aliscore output file does not exist. Check results for BUSCO: 350211at7742
## Thu Jul 14 01:53:07 CEST 2022 - The aliscore output appears to be empty. Check results for BUSCO: 350211at7742
## Thu Jul 14 01:53:07 CEST 2022 - The aliscore output file does not exist. Check results for BUSCO: 236476at7742
## Thu Jul 14 01:53:07 CEST 2022 - The aliscore output appears to be empty. Check results for BUSCO: 236476at7742
## Thu Jul 14 01:53:10 CEST 2022 - The aliscore output file does not exist. Check results for BUSCO: 190562at7742
## Thu Jul 14 01:53:10 CEST 2022 - The aliscore output appears to be empty. Check results for BUSCO: 190562at7742
## Thu Jul 14 01:53:12 CEST 2022 - The aliscore output file does not exist. Check results for BUSCO: 159879at7742
## Thu Jul 14 01:53:12 CEST 2022 - The aliscore output appears to be empty. Check results for BUSCO: 159879at7742
## Thu Jul 14 01:53:13 CEST 2022 - The aliscore output file does not exist. Check results for BUSCO: 274480at7742
## Thu Jul 14 01:53:13 CEST 2022 - The aliscore output appears to be empty. Check results for BUSCO: 274480at7742
## Thu Jul 14 01:53:14 CEST 2022 - The aliscore output file does not exist. Check results for BUSCO: 236476at7742
## Thu Jul 14 01:53:14 CEST 2022 - The aliscore output appears to be empty. Check results for BUSCO: 236476at7742
## Thu Jul 14 01:53:14 CEST 2022 - The aliscore output file does not exist. Check results for BUSCO: 368433at7742
## Thu Jul 14 01:53:14 CEST 2022 - The aliscore output appears to be empty. Check results for BUSCO: 368433at7742
## Thu Jul 14 01:53:15 CEST 2022 - The aliscore output file does not exist. Check results for BUSCO: 278370at7742
## Thu Jul 14 01:53:15 CEST 2022 - The aliscore output appears to be empty. Check results for BUSCO: 278370at7742
## Thu Jul 14 01:53:15 CEST 2022 - The aliscore output file does not exist. Check results for BUSCO: 311983at7742
## Thu Jul 14 01:53:15 CEST 2022 - The aliscore output appears to be empty. Check results for BUSCO: 311983at7742
## Thu Jul 14 01:53:17 CEST 2022 - The aliscore output file does not exist. Check results for BUSCO: 244439at7742
## Thu Jul 14 01:53:17 CEST 2022 - The aliscore output appears to be empty. Check results for BUSCO: 244439at7742
## Thu Jul 14 01:53:17 CEST 2022 - The aliscore output file does not exist. Check results for BUSCO: 292857at7742
## Thu Jul 14 01:53:17 CEST 2022 - The aliscore output appears to be empty. Check results for BUSCO: 292857at7742
## Thu Jul 14 01:53:20 CEST 2022 - The aliscore output file does not exist. Check results for BUSCO: 321010at7742
## Thu Jul 14 01:53:20 CEST 2022 - The aliscore output appears to be empty. Check results for BUSCO: 321010at7742
## Thu Jul 14 01:53:21 CEST 2022 - The aliscore output file does not exist. Check results for BUSCO: 223426at7742
## Thu Jul 14 01:53:21 CEST 2022 - The aliscore output appears to be empty. Check results for BUSCO: 223426at7742
## Thu Jul 14 01:53:21 CEST 2022 - The aliscore output file does not exist. Check results for BUSCO: 207782at7742
## Thu Jul 14 01:53:21 CEST 2022 - The aliscore output appears to be empty. Check results for BUSCO: 207782at7742
## Thu Jul 14 01:53:22 CEST 2022 - The aliscore output file does not exist. Check results for BUSCO: 229738at7742
## Thu Jul 14 01:53:22 CEST 2022 - The aliscore output appears to be empty. Check results for BUSCO: 229738at7742
## Thu Jul 14 01:53:23 CEST 2022 - The aliscore output file does not exist. Check results for BUSCO: 248773at7742
## Thu Jul 14 01:53:23 CEST 2022 - The aliscore output appears to be empty. Check results for BUSCO: 248773at7742
## Thu Jul 14 01:53:24 CEST 2022 - The aliscore output file does not exist. Check results for BUSCO: 275566at7742
## Thu Jul 14 01:53:24 CEST 2022 - The aliscore output appears to be empty. Check results for BUSCO: 275566at7742
## Thu Jul 14 01:53:25 CEST 2022 - The aliscore output file does not exist. Check results for BUSCO: 133965at7742
## Thu Jul 14 01:53:25 CEST 2022 - The aliscore output appears to be empty. Check results for BUSCO: 133965at7742
## Thu Jul 14 01:53:26 CEST 2022 - The aliscore output file does not exist. Check results for BUSCO: 394021at7742
## Thu Jul 14 01:53:26 CEST 2022 - The aliscore output appears to be empty. Check results for BUSCO: 394021at7742
## Thu Jul 14 01:53:28 CEST 2022 - The aliscore output file does not exist. Check results for BUSCO: 189224at7742
## Thu Jul 14 01:53:28 CEST 2022 - The aliscore output appears to be empty. Check results for BUSCO: 189224at7742
## Thu Jul 14 01:53:28 CEST 2022 - The aliscore output file does not exist. Check results for BUSCO: 196409at7742
## Thu Jul 14 01:53:28 CEST 2022 - The aliscore output appears to be empty. Check results for BUSCO: 196409at7742
## Thu Jul 14 01:53:29 CEST 2022 - The aliscore output file does not exist. Check results for BUSCO: 197724at7742
## Thu Jul 14 01:53:29 CEST 2022 - The aliscore output appears to be empty. Check results for BUSCO: 197724at7742
## Thu Jul 14 01:53:30 CEST 2022 - The aliscore output file does not exist. Check results for BUSCO: 175714at7742
## Thu Jul 14 01:53:30 CEST 2022 - The aliscore output appears to be empty. Check results for BUSCO: 175714at7742
## Thu Jul 14 01:53:31 CEST 2022 - The aliscore output file does not exist. Check results for BUSCO: 251667at7742
## Thu Jul 14 01:53:31 CEST 2022 - The aliscore output appears to be empty. Check results for BUSCO: 251667at7742
## Thu Jul 14 01:53:34 CEST 2022 - The aliscore output file does not exist. Check results for BUSCO: 213618at7742
## Thu Jul 14 01:53:34 CEST 2022 - The aliscore output appears to be empty. Check results for BUSCO: 213618at7742
## Thu Jul 14 01:53:34 CEST 2022 - The aliscore output file does not exist. Check results for BUSCO: 133604at7742
## Thu Jul 14 01:53:34 CEST 2022 - The aliscore output appears to be empty. Check results for BUSCO: 133604at7742
## Thu Jul 14 01:53:36 CEST 2022 - The aliscore output file does not exist. Check results for BUSCO: 220049at7742
## Thu Jul 14 01:53:36 CEST 2022 - The aliscore output appears to be empty. Check results for BUSCO: 220049at7742
## Thu Jul 14 01:53:37 CEST 2022 - The aliscore output file does not exist. Check results for BUSCO: 198672at7742
## Thu Jul 14 01:53:37 CEST 2022 - The aliscore output appears to be empty. Check results for BUSCO: 198672at7742
## Thu Jul 14 01:53:37 CEST 2022 - The aliscore output file does not exist. Check results for BUSCO: 159334at7742
## Thu Jul 14 01:53:37 CEST 2022 - The aliscore output appears to be empty. Check results for BUSCO: 159334at7742
## Thu Jul 14 01:53:38 CEST 2022 - The aliscore output file does not exist. Check results for BUSCO: 225125at7742
## Thu Jul 14 01:53:39 CEST 2022 - The aliscore output appears to be empty. Check results for BUSCO: 225125at7742
## Thu Jul 14 01:53:39 CEST 2022 - The aliscore output file does not exist. Check results for BUSCO: 165972at7742
## Thu Jul 14 01:53:39 CEST 2022 - The aliscore output appears to be empty. Check results for BUSCO: 165972at7742
## Thu Jul 14 01:53:40 CEST 2022 - The aliscore output file does not exist. Check results for BUSCO: 262221at7742
## Thu Jul 14 01:53:40 CEST 2022 - The aliscore output appears to be empty. Check results for BUSCO: 262221at7742
## Thu Jul 14 01:53:40 CEST 2022 - The aliscore output file does not exist. Check results for BUSCO: 138435at7742
## Thu Jul 14 01:53:40 CEST 2022 - The aliscore output appears to be empty. Check results for BUSCO: 138435at7742
## Thu Jul 14 01:53:42 CEST 2022 - The aliscore output file does not exist. Check results for BUSCO: 187431at7742
## Thu Jul 14 01:53:42 CEST 2022 - The aliscore output appears to be empty. Check results for BUSCO: 187431at7742
## Thu Jul 14 01:53:43 CEST 2022 - The aliscore output file does not exist. Check results for BUSCO: 381258at7742
## Thu Jul 14 01:53:43 CEST 2022 - The aliscore output appears to be empty. Check results for BUSCO: 381258at7742
## Thu Jul 14 01:53:43 CEST 2022 - The aliscore output file does not exist. Check results for BUSCO: 362410at7742
## Thu Jul 14 01:53:43 CEST 2022 - The aliscore output appears to be empty. Check results for BUSCO: 362410at7742
## Thu Jul 14 01:53:45 CEST 2022 - The aliscore output file does not exist. Check results for BUSCO: 184999at7742
## Thu Jul 14 01:53:45 CEST 2022 - The aliscore output appears to be empty. Check results for BUSCO: 184999at7742
## Thu Jul 14 01:53:46 CEST 2022 - The aliscore output file does not exist. Check results for BUSCO: 158801at7742
## Thu Jul 14 01:53:46 CEST 2022 - The aliscore output appears to be empty. Check results for BUSCO: 158801at7742
## Thu Jul 14 01:53:47 CEST 2022 - The aliscore output file does not exist. Check results for BUSCO: 176266at7742
## Thu Jul 14 01:53:48 CEST 2022 - The aliscore output appears to be empty. Check results for BUSCO: 176266at7742
## Thu Jul 14 01:53:50 CEST 2022 - The aliscore output file does not exist. Check results for BUSCO: 178275at7742
## Thu Jul 14 01:53:50 CEST 2022 - The aliscore output appears to be empty. Check results for BUSCO: 178275at7742
## Thu Jul 14 01:53:50 CEST 2022 - The aliscore output file does not exist. Check results for BUSCO: 178275at7742
## Thu Jul 14 01:53:50 CEST 2022 - The aliscore output appears to be empty. Check results for BUSCO: 178275at7742
## Thu Jul 14 01:53:51 CEST 2022 - The aliscore output file does not exist. Check results for BUSCO: 361680at7742
## Thu Jul 14 01:53:51 CEST 2022 - The aliscore output appears to be empty. Check results for BUSCO: 361680at7742
## Thu Jul 14 01:53:51 CEST 2022 - The aliscore output file does not exist. Check results for BUSCO: 236145at7742
## Thu Jul 14 01:53:51 CEST 2022 - The aliscore output appears to be empty. Check results for BUSCO: 236145at7742
## Thu Jul 14 01:53:53 CEST 2022 - The aliscore output file does not exist. Check results for BUSCO: 291153at7742
## Thu Jul 14 01:53:53 CEST 2022 - The aliscore output appears to be empty. Check results for BUSCO: 291153at7742
## Thu Jul 14 01:53:55 CEST 2022 - The aliscore output file does not exist. Check results for BUSCO: 314547at7742
## Thu Jul 14 01:53:55 CEST 2022 - The aliscore output appears to be empty. Check results for BUSCO: 314547at7742
## Thu Jul 14 01:53:55 CEST 2022 - The aliscore output file does not exist. Check results for BUSCO: 291774at7742
## Thu Jul 14 01:53:55 CEST 2022 - The aliscore output appears to be empty. Check results for BUSCO: 291774at7742
## Thu Jul 14 01:53:57 CEST 2022 - The aliscore output file does not exist. Check results for BUSCO: 124571at7742
## Thu Jul 14 01:53:57 CEST 2022 - The aliscore output appears to be empty. Check results for BUSCO: 124571at7742
## Thu Jul 14 01:53:58 CEST 2022 - The aliscore output file does not exist. Check results for BUSCO: 380483at7742
## Thu Jul 14 01:53:58 CEST 2022 - The aliscore output appears to be empty. Check results for BUSCO: 380483at7742
## Thu Jul 14 01:53:58 CEST 2022 - The aliscore output file does not exist. Check results for BUSCO: 248773at7742
## Thu Jul 14 01:53:58 CEST 2022 - The aliscore output appears to be empty. Check results for BUSCO: 248773at7742
## Thu Jul 14 01:54:00 CEST 2022 - The aliscore output file does not exist. Check results for BUSCO: 173248at7742
## Thu Jul 14 01:54:00 CEST 2022 - The aliscore output appears to be empty. Check results for BUSCO: 173248at7742
## Thu Jul 14 01:54:01 CEST 2022 - The aliscore output file does not exist. Check results for BUSCO: 302596at7742
## Thu Jul 14 01:54:01 CEST 2022 - The aliscore output appears to be empty. Check results for BUSCO: 302596at7742
## Thu Jul 14 01:54:01 CEST 2022 - The aliscore output file does not exist. Check results for BUSCO: 194403at7742
## Thu Jul 14 01:54:01 CEST 2022 - The aliscore output appears to be empty. Check results for BUSCO: 194403at7742
## Thu Jul 14 01:54:03 CEST 2022 - The aliscore output file does not exist. Check results for BUSCO: 366847at7742
## Thu Jul 14 01:54:03 CEST 2022 - The aliscore output appears to be empty. Check results for BUSCO: 366847at7742
## Thu Jul 14 01:54:05 CEST 2022 - The aliscore output file does not exist. Check results for BUSCO: 321359at7742
## Thu Jul 14 01:54:05 CEST 2022 - The aliscore output appears to be empty. Check results for BUSCO: 321359at7742
## Thu Jul 14 01:54:06 CEST 2022 - The aliscore output file does not exist. Check results for BUSCO: 158698at7742
## Thu Jul 14 01:54:06 CEST 2022 - The aliscore output appears to be empty. Check results for BUSCO: 158698at7742
## Thu Jul 14 01:54:06 CEST 2022 - The aliscore output file does not exist. Check results for BUSCO: 133604at7742
## Thu Jul 14 01:54:06 CEST 2022 - The aliscore output appears to be empty. Check results for BUSCO: 133604at7742
## Thu Jul 14 01:54:07 CEST 2022 - The aliscore output file does not exist. Check results for BUSCO: 332304at7742
## Thu Jul 14 01:54:07 CEST 2022 - The aliscore output appears to be empty. Check results for BUSCO: 332304at7742
## Thu Jul 14 01:54:09 CEST 2022 - The aliscore output file does not exist. Check results for BUSCO: 268274at7742
## Thu Jul 14 01:54:09 CEST 2022 - The aliscore output appears to be empty. Check results for BUSCO: 268274at7742
## Thu Jul 14 01:54:09 CEST 2022 - The aliscore output file does not exist. Check results for BUSCO: 242487at7742
## Thu Jul 14 01:54:09 CEST 2022 - The aliscore output appears to be empty. Check results for BUSCO: 242487at7742
## Thu Jul 14 01:54:10 CEST 2022 - The aliscore output file does not exist. Check results for BUSCO: 175714at7742
## Thu Jul 14 01:54:10 CEST 2022 - The aliscore output appears to be empty. Check results for BUSCO: 175714at7742
## Thu Jul 14 01:54:12 CEST 2022 - The aliscore output file does not exist. Check results for BUSCO: 290906at7742
## Thu Jul 14 01:54:12 CEST 2022 - The aliscore output appears to be empty. Check results for BUSCO: 290906at7742
## Thu Jul 14 01:54:13 CEST 2022 - The aliscore output file does not exist. Check results for BUSCO: 192284at7742
## Thu Jul 14 01:54:13 CEST 2022 - The aliscore output appears to be empty. Check results for BUSCO: 192284at7742
## Thu Jul 14 01:54:14 CEST 2022 - The aliscore output file does not exist. Check results for BUSCO: 160476at7742
## Thu Jul 14 01:54:14 CEST 2022 - The aliscore output appears to be empty. Check results for BUSCO: 160476at7742
## Thu Jul 14 01:54:16 CEST 2022 - The aliscore output file does not exist. Check results for BUSCO: 178275at7742
## Thu Jul 14 01:54:16 CEST 2022 - The aliscore output appears to be empty. Check results for BUSCO: 178275at7742
## Thu Jul 14 01:54:16 CEST 2022 - The aliscore output file does not exist. Check results for BUSCO: 226375at7742
## Thu Jul 14 01:54:16 CEST 2022 - The aliscore output appears to be empty. Check results for BUSCO: 226375at7742
## Thu Jul 14 01:54:16 CEST 2022 - The aliscore output file does not exist. Check results for BUSCO: 266219at7742
## Thu Jul 14 01:54:16 CEST 2022 - The aliscore output appears to be empty. Check results for BUSCO: 266219at7742
## Thu Jul 14 01:54:18 CEST 2022 - The aliscore output file does not exist. Check results for BUSCO: 264056at7742
## Thu Jul 14 01:54:18 CEST 2022 - The aliscore output appears to be empty. Check results for BUSCO: 264056at7742
## Thu Jul 14 01:54:20 CEST 2022 - The aliscore output file does not exist. Check results for BUSCO: 291774at7742
## Thu Jul 14 01:54:20 CEST 2022 - The aliscore output appears to be empty. Check results for BUSCO: 291774at7742
## Thu Jul 14 01:54:20 CEST 2022 - The aliscore output file does not exist. Check results for BUSCO: 187431at7742
## Thu Jul 14 01:54:20 CEST 2022 - The aliscore output appears to be empty. Check results for BUSCO: 187431at7742
## Thu Jul 14 01:54:22 CEST 2022 - The aliscore output file does not exist. Check results for BUSCO: 151964at7742
## Thu Jul 14 01:54:22 CEST 2022 - The aliscore output appears to be empty. Check results for BUSCO: 151964at7742
## Thu Jul 14 01:54:22 CEST 2022 - The aliscore output file does not exist. Check results for BUSCO: 300584at7742
## Thu Jul 14 01:54:22 CEST 2022 - The aliscore output appears to be empty. Check results for BUSCO: 300584at7742
## Thu Jul 14 01:54:23 CEST 2022 - The aliscore output file does not exist. Check results for BUSCO: 242895at7742
## Thu Jul 14 01:54:23 CEST 2022 - The aliscore output appears to be empty. Check results for BUSCO: 242895at7742
## Thu Jul 14 01:54:24 CEST 2022 - The aliscore output file does not exist. Check results for BUSCO: 287042at7742
## Thu Jul 14 01:54:24 CEST 2022 - The aliscore output appears to be empty. Check results for BUSCO: 287042at7742
## Thu Jul 14 01:54:26 CEST 2022 - The aliscore output file does not exist. Check results for BUSCO: 161007at7742
## Thu Jul 14 01:54:26 CEST 2022 - The aliscore output appears to be empty. Check results for BUSCO: 161007at7742
## Thu Jul 14 01:54:28 CEST 2022 - The aliscore output file does not exist. Check results for BUSCO: 288248at7742
## Thu Jul 14 01:54:28 CEST 2022 - The aliscore output appears to be empty. Check results for BUSCO: 288248at7742
## Thu Jul 14 01:54:28 CEST 2022 - The aliscore output file does not exist. Check results for BUSCO: 286595at7742
## Thu Jul 14 01:54:28 CEST 2022 - The aliscore output appears to be empty. Check results for BUSCO: 286595at7742
## Thu Jul 14 01:54:29 CEST 2022 - The aliscore output file does not exist. Check results for BUSCO: 244439at7742
## Thu Jul 14 01:54:29 CEST 2022 - The aliscore output appears to be empty. Check results for BUSCO: 244439at7742
## Thu Jul 14 01:54:30 CEST 2022 - The aliscore output file does not exist. Check results for BUSCO: 260107at7742
## Thu Jul 14 01:54:30 CEST 2022 - The aliscore output appears to be empty. Check results for BUSCO: 260107at7742
## Thu Jul 14 01:54:31 CEST 2022 - The aliscore output file does not exist. Check results for BUSCO: 261976at7742
## Thu Jul 14 01:54:31 CEST 2022 - The aliscore output appears to be empty. Check results for BUSCO: 261976at7742
## Thu Jul 14 01:54:32 CEST 2022 - The aliscore output file does not exist. Check results for BUSCO: 284666at7742
## Thu Jul 14 01:54:32 CEST 2022 - The aliscore output file does not exist. Check results for BUSCO: 362410at7742
## Thu Jul 14 01:54:32 CEST 2022 - The aliscore output appears to be empty. Check results for BUSCO: 284666at7742
## Thu Jul 14 01:54:32 CEST 2022 - The aliscore output appears to be empty. Check results for BUSCO: 362410at7742
## Thu Jul 14 01:54:34 CEST 2022 - The aliscore output file does not exist. Check results for BUSCO: 123660at7742
## Thu Jul 14 01:54:34 CEST 2022 - The aliscore output appears to be empty. Check results for BUSCO: 123660at7742
## Thu Jul 14 01:54:35 CEST 2022 - The aliscore output file does not exist. Check results for BUSCO: 237266at7742
## Thu Jul 14 01:54:35 CEST 2022 - The aliscore output appears to be empty. Check results for BUSCO: 237266at7742
## Thu Jul 14 01:54:36 CEST 2022 - The aliscore output file does not exist. Check results for BUSCO: 219315at7742
## Thu Jul 14 01:54:36 CEST 2022 - The aliscore output appears to be empty. Check results for BUSCO: 219315at7742
## Thu Jul 14 01:54:37 CEST 2022 - The aliscore output file does not exist. Check results for BUSCO: 321295at7742
## Thu Jul 14 01:54:37 CEST 2022 - The aliscore output appears to be empty. Check results for BUSCO: 321295at7742
## Thu Jul 14 01:54:37 CEST 2022 - The aliscore output file does not exist. Check results for BUSCO: 173248at7742
## Thu Jul 14 01:54:37 CEST 2022 - The aliscore output appears to be empty. Check results for BUSCO: 173248at7742
## Thu Jul 14 01:54:38 CEST 2022 - The aliscore output file does not exist. Check results for BUSCO: 158801at7742
## Thu Jul 14 01:54:38 CEST 2022 - The aliscore output appears to be empty. Check results for BUSCO: 158801at7742
## Thu Jul 14 01:54:40 CEST 2022 - The aliscore output file does not exist. Check results for BUSCO: 282590at7742
## Thu Jul 14 01:54:40 CEST 2022 - The aliscore output appears to be empty. Check results for BUSCO: 282590at7742
## Thu Jul 14 01:54:42 CEST 2022 - The aliscore output file does not exist. Check results for BUSCO: 161959at7742
## Thu Jul 14 01:54:42 CEST 2022 - The aliscore output appears to be empty. Check results for BUSCO: 161959at7742
## Thu Jul 14 01:54:43 CEST 2022 - The aliscore output file does not exist. Check results for BUSCO: 296065at7742
## Thu Jul 14 01:54:43 CEST 2022 - The aliscore output appears to be empty. Check results for BUSCO: 296065at7742
## Thu Jul 14 01:54:43 CEST 2022 - The aliscore output file does not exist. Check results for BUSCO: 310872at7742
## Thu Jul 14 01:54:43 CEST 2022 - The aliscore output appears to be empty. Check results for BUSCO: 310872at7742
## Thu Jul 14 01:54:43 CEST 2022 - The aliscore output file does not exist. Check results for BUSCO: 366274at7742
## Thu Jul 14 01:54:43 CEST 2022 - The aliscore output appears to be empty. Check results for BUSCO: 366274at7742
## Thu Jul 14 01:54:45 CEST 2022 - The aliscore output file does not exist. Check results for BUSCO: 137854at7742
## Thu Jul 14 01:54:45 CEST 2022 - The aliscore output appears to be empty. Check results for BUSCO: 137854at7742
## Thu Jul 14 01:54:47 CEST 2022 - The aliscore output file does not exist. Check results for BUSCO: 343795at7742
## Thu Jul 14 01:54:47 CEST 2022 - The aliscore output appears to be empty. Check results for BUSCO: 343795at7742
## Thu Jul 14 01:54:48 CEST 2022 - The aliscore output file does not exist. Check results for BUSCO: 223426at7742
## Thu Jul 14 01:54:48 CEST 2022 - The aliscore output appears to be empty. Check results for BUSCO: 223426at7742
## Thu Jul 14 01:54:50 CEST 2022 - The aliscore output file does not exist. Check results for BUSCO: 281858at7742
## Thu Jul 14 01:54:50 CEST 2022 - The aliscore output appears to be empty. Check results for BUSCO: 281858at7742
## Thu Jul 14 01:54:51 CEST 2022 - The aliscore output file does not exist. Check results for BUSCO: 294328at7742
## Thu Jul 14 01:54:51 CEST 2022 - The aliscore output appears to be empty. Check results for BUSCO: 294328at7742
## Thu Jul 14 01:54:51 CEST 2022 - The aliscore output file does not exist. Check results for BUSCO: 261523at7742
## Thu Jul 14 01:54:51 CEST 2022 - The aliscore output appears to be empty. Check results for BUSCO: 261523at7742
## Thu Jul 14 01:54:51 CEST 2022 - The aliscore output file does not exist. Check results for BUSCO: 302596at7742
## Thu Jul 14 01:54:52 CEST 2022 - The aliscore output appears to be empty. Check results for BUSCO: 302596at7742
## Thu Jul 14 01:54:52 CEST 2022 - The aliscore output file does not exist. Check results for BUSCO: 189840at7742
## Thu Jul 14 01:54:52 CEST 2022 - The aliscore output appears to be empty. Check results for BUSCO: 189840at7742
## Thu Jul 14 01:54:55 CEST 2022 - The aliscore output file does not exist. Check results for BUSCO: 221887at7742
## Thu Jul 14 01:54:55 CEST 2022 - The aliscore output appears to be empty. Check results for BUSCO: 221887at7742
## Thu Jul 14 01:54:56 CEST 2022 - The aliscore output file does not exist. Check results for BUSCO: 407881at7742
## Thu Jul 14 01:54:56 CEST 2022 - The aliscore output appears to be empty. Check results for BUSCO: 407881at7742
## Thu Jul 14 01:54:57 CEST 2022 - The aliscore output file does not exist. Check results for BUSCO: 300443at7742
## Thu Jul 14 01:54:57 CEST 2022 - The aliscore output appears to be empty. Check results for BUSCO: 300443at7742
## Thu Jul 14 01:54:57 CEST 2022 - The aliscore output file does not exist. Check results for BUSCO: 202512at7742
## Thu Jul 14 01:54:57 CEST 2022 - The aliscore output appears to be empty. Check results for BUSCO: 202512at7742
## Thu Jul 14 01:54:58 CEST 2022 - The aliscore output file does not exist. Check results for BUSCO: 183995at7742
## Thu Jul 14 01:54:58 CEST 2022 - The aliscore output appears to be empty. Check results for BUSCO: 183995at7742
## Thu Jul 14 01:54:58 CEST 2022 - The aliscore output file does not exist. Check results for BUSCO: 214858at7742
## Thu Jul 14 01:54:58 CEST 2022 - The aliscore output appears to be empty. Check results for BUSCO: 214858at7742
## Thu Jul 14 01:54:59 CEST 2022 - The aliscore output file does not exist. Check results for BUSCO: 268484at7742
## Thu Jul 14 01:54:59 CEST 2022 - The aliscore output appears to be empty. Check results for BUSCO: 268484at7742
## Thu Jul 14 01:55:00 CEST 2022 - The aliscore output file does not exist. Check results for BUSCO: 396312at7742
## Thu Jul 14 01:55:00 CEST 2022 - The aliscore output appears to be empty. Check results for BUSCO: 396312at7742
## Thu Jul 14 01:55:02 CEST 2022 - The aliscore output file does not exist. Check results for BUSCO: 237266at7742
## Thu Jul 14 01:55:02 CEST 2022 - The aliscore output appears to be empty. Check results for BUSCO: 237266at7742
## Thu Jul 14 01:55:03 CEST 2022 - The aliscore output file does not exist. Check results for BUSCO: 213618at7742
## Thu Jul 14 01:55:03 CEST 2022 - The aliscore output appears to be empty. Check results for BUSCO: 213618at7742
## Thu Jul 14 01:55:05 CEST 2022 - The aliscore output file does not exist. Check results for BUSCO: 407881at7742
## Thu Jul 14 01:55:05 CEST 2022 - The aliscore output appears to be empty. Check results for BUSCO: 407881at7742
## Thu Jul 14 01:55:05 CEST 2022 - The aliscore output file does not exist. Check results for BUSCO: 328514at7742
## Thu Jul 14 01:55:05 CEST 2022 - The aliscore output appears to be empty. Check results for BUSCO: 328514at7742
## Thu Jul 14 01:55:08 CEST 2022 - The aliscore output file does not exist. Check results for BUSCO: 171308at7742
## Thu Jul 14 01:55:08 CEST 2022 - The aliscore output appears to be empty. Check results for BUSCO: 171308at7742
## Thu Jul 14 01:55:08 CEST 2022 - The aliscore output file does not exist. Check results for BUSCO: 232734at7742
## Thu Jul 14 01:55:08 CEST 2022 - The aliscore output appears to be empty. Check results for BUSCO: 232734at7742
## Thu Jul 14 01:55:09 CEST 2022 - The aliscore output file does not exist. Check results for BUSCO: 215706at7742
## Thu Jul 14 01:55:09 CEST 2022 - The aliscore output appears to be empty. Check results for BUSCO: 215706at7742
## Thu Jul 14 01:55:10 CEST 2022 - The aliscore output file does not exist. Check results for BUSCO: 316762at7742
## Thu Jul 14 01:55:10 CEST 2022 - The aliscore output appears to be empty. Check results for BUSCO: 316762at7742
## Thu Jul 14 01:55:10 CEST 2022 - The aliscore output file does not exist. Check results for BUSCO: 151964at7742
## Thu Jul 14 01:55:10 CEST 2022 - The aliscore output appears to be empty. Check results for BUSCO: 151964at7742
## Thu Jul 14 01:55:12 CEST 2022 - The aliscore output file does not exist. Check results for BUSCO: 239289at7742
## Thu Jul 14 01:55:12 CEST 2022 - The aliscore output appears to be empty. Check results for BUSCO: 239289at7742
## Thu Jul 14 01:55:13 CEST 2022 - The aliscore output file does not exist. Check results for BUSCO: 123660at7742
## Thu Jul 14 01:55:13 CEST 2022 - The aliscore output appears to be empty. Check results for BUSCO: 123660at7742
## Thu Jul 14 01:55:13 CEST 2022 - The aliscore output file does not exist. Check results for BUSCO: 267724at7742
## Thu Jul 14 01:55:13 CEST 2022 - The aliscore output appears to be empty. Check results for BUSCO: 267724at7742
## Thu Jul 14 01:55:14 CEST 2022 - The aliscore output file does not exist. Check results for BUSCO: 166002at7742
## Thu Jul 14 01:55:14 CEST 2022 - The aliscore output appears to be empty. Check results for BUSCO: 166002at7742
## Thu Jul 14 01:55:16 CEST 2022 - The aliscore output file does not exist. Check results for BUSCO: 377100at7742
## Thu Jul 14 01:55:16 CEST 2022 - The aliscore output appears to be empty. Check results for BUSCO: 377100at7742
## Thu Jul 14 01:55:16 CEST 2022 - The aliscore output file does not exist. Check results for BUSCO: 214100at7742
## Thu Jul 14 01:55:16 CEST 2022 - The aliscore output appears to be empty. Check results for BUSCO: 214100at7742
## Thu Jul 14 01:55:17 CEST 2022 - The aliscore output file does not exist. Check results for BUSCO: 226375at7742
## Thu Jul 14 01:55:17 CEST 2022 - The aliscore output appears to be empty. Check results for BUSCO: 226375at7742
## Thu Jul 14 01:55:19 CEST 2022 - The aliscore output file does not exist. Check results for BUSCO: 211328at7742
## Thu Jul 14 01:55:19 CEST 2022 - The aliscore output appears to be empty. Check results for BUSCO: 211328at7742
## Thu Jul 14 01:55:20 CEST 2022 - The aliscore output file does not exist. Check results for BUSCO: 180713at7742
## Thu Jul 14 01:55:20 CEST 2022 - The aliscore output appears to be empty. Check results for BUSCO: 180713at7742
## Thu Jul 14 01:55:20 CEST 2022 - The aliscore output file does not exist. Check results for BUSCO: 319288at7742
## Thu Jul 14 01:55:20 CEST 2022 - The aliscore output appears to be empty. Check results for BUSCO: 319288at7742
## Thu Jul 14 01:55:22 CEST 2022 - The aliscore output file does not exist. Check results for BUSCO: 312288at7742
## Thu Jul 14 01:55:22 CEST 2022 - The aliscore output appears to be empty. Check results for BUSCO: 312288at7742
## Thu Jul 14 01:55:22 CEST 2022 - The aliscore output file does not exist. Check results for BUSCO: 301292at7742
## Thu Jul 14 01:55:23 CEST 2022 - The aliscore output appears to be empty. Check results for BUSCO: 301292at7742
## Thu Jul 14 01:55:23 CEST 2022 - The aliscore output file does not exist. Check results for BUSCO: 350246at7742
## Thu Jul 14 01:55:23 CEST 2022 - The aliscore output appears to be empty. Check results for BUSCO: 350246at7742
## Thu Jul 14 01:55:24 CEST 2022 - The aliscore output file does not exist. Check results for BUSCO: 366844at7742
## Thu Jul 14 01:55:24 CEST 2022 - The aliscore output appears to be empty. Check results for BUSCO: 366844at7742
## Thu Jul 14 01:55:25 CEST 2022 - The aliscore output file does not exist. Check results for BUSCO: 173248at7742
## Thu Jul 14 01:55:25 CEST 2022 - The aliscore output appears to be empty. Check results for BUSCO: 173248at7742
## Thu Jul 14 01:55:27 CEST 2022 - The aliscore output file does not exist. Check results for BUSCO: 314547at7742
## Thu Jul 14 01:55:27 CEST 2022 - The aliscore output appears to be empty. Check results for BUSCO: 314547at7742
## Thu Jul 14 01:55:27 CEST 2022 - The aliscore output file does not exist. Check results for BUSCO: 229857at7742
## Thu Jul 14 01:55:27 CEST 2022 - The aliscore output appears to be empty. Check results for BUSCO: 229857at7742
## Thu Jul 14 01:55:29 CEST 2022 - The aliscore output file does not exist. Check results for BUSCO: 266056at7742
## Thu Jul 14 01:55:29 CEST 2022 - The aliscore output appears to be empty. Check results for BUSCO: 266056at7742
## Thu Jul 14 01:55:30 CEST 2022 - The aliscore output file does not exist. Check results for BUSCO: 160476at7742
## Thu Jul 14 01:55:30 CEST 2022 - The aliscore output appears to be empty. Check results for BUSCO: 160476at7742
## Thu Jul 14 01:55:32 CEST 2022 - The aliscore output file does not exist. Check results for BUSCO: 223152at7742
## Thu Jul 14 01:55:32 CEST 2022 - The aliscore output appears to be empty. Check results for BUSCO: 223152at7742
## Thu Jul 14 01:55:32 CEST 2022 - The aliscore output file does not exist. Check results for BUSCO: 176266at7742
## Thu Jul 14 01:55:32 CEST 2022 - The aliscore output appears to be empty. Check results for BUSCO: 176266at7742
## Thu Jul 14 01:55:33 CEST 2022 - The aliscore output file does not exist. Check results for BUSCO: 381258at7742
## Thu Jul 14 01:55:33 CEST 2022 - The aliscore output appears to be empty. Check results for BUSCO: 381258at7742
## Thu Jul 14 01:55:33 CEST 2022 - The aliscore output file does not exist. Check results for BUSCO: 242951at7742
## Thu Jul 14 01:55:33 CEST 2022 - The aliscore output appears to be empty. Check results for BUSCO: 242951at7742
## Thu Jul 14 01:55:34 CEST 2022 - The aliscore output file does not exist. Check results for BUSCO: 215706at7742
## Thu Jul 14 01:55:34 CEST 2022 - The aliscore output appears to be empty. Check results for BUSCO: 215706at7742
## Thu Jul 14 01:55:37 CEST 2022 - The aliscore output file does not exist. Check results for BUSCO: 166429at7742
## Thu Jul 14 01:55:37 CEST 2022 - The aliscore output appears to be empty. Check results for BUSCO: 166429at7742
## Thu Jul 14 01:55:37 CEST 2022 - The aliscore output file does not exist. Check results for BUSCO: 370499at7742
## Thu Jul 14 01:55:37 CEST 2022 - The aliscore output appears to be empty. Check results for BUSCO: 370499at7742
## Thu Jul 14 01:55:39 CEST 2022 - The aliscore output file does not exist. Check results for BUSCO: 236145at7742
## Thu Jul 14 01:55:39 CEST 2022 - The aliscore output appears to be empty. Check results for BUSCO: 236145at7742
## Thu Jul 14 01:55:39 CEST 2022 - The aliscore output file does not exist. Check results for BUSCO: 241208at7742
## Thu Jul 14 01:55:39 CEST 2022 - The aliscore output appears to be empty. Check results for BUSCO: 241208at7742
## Thu Jul 14 01:55:40 CEST 2022 - The aliscore output file does not exist. Check results for BUSCO: 183251at7742
## Thu Jul 14 01:55:40 CEST 2022 - The aliscore output appears to be empty. Check results for BUSCO: 183251at7742
## Thu Jul 14 01:55:41 CEST 2022 - The aliscore output file does not exist. Check results for BUSCO: 290261at7742
## Thu Jul 14 01:55:41 CEST 2022 - The aliscore output appears to be empty. Check results for BUSCO: 290261at7742
## Thu Jul 14 01:55:42 CEST 2022 - The aliscore output file does not exist. Check results for BUSCO: 314547at7742
## Thu Jul 14 01:55:42 CEST 2022 - The aliscore output appears to be empty. Check results for BUSCO: 314547at7742
## Thu Jul 14 01:55:43 CEST 2022 - The aliscore output file does not exist. Check results for BUSCO: 350246at7742
## Thu Jul 14 01:55:43 CEST 2022 - The aliscore output appears to be empty. Check results for BUSCO: 350246at7742
## Thu Jul 14 01:55:44 CEST 2022 - The aliscore output file does not exist. Check results for BUSCO: 282590at7742
## Thu Jul 14 01:55:44 CEST 2022 - The aliscore output appears to be empty. Check results for BUSCO: 282590at7742
## Thu Jul 14 01:55:46 CEST 2022 - The aliscore output file does not exist. Check results for BUSCO: 228275at7742
## Thu Jul 14 01:55:46 CEST 2022 - The aliscore output appears to be empty. Check results for BUSCO: 228275at7742
## Thu Jul 14 01:55:47 CEST 2022 - The aliscore output file does not exist. Check results for BUSCO: 360850at7742
## Thu Jul 14 01:55:47 CEST 2022 - The aliscore output file does not exist. Check results for BUSCO: 196409at7742
## Thu Jul 14 01:55:47 CEST 2022 - The aliscore output appears to be empty. Check results for BUSCO: 360850at7742
## Thu Jul 14 01:55:47 CEST 2022 - The aliscore output appears to be empty. Check results for BUSCO: 196409at7742
## Thu Jul 14 01:55:48 CEST 2022 - The aliscore output file does not exist. Check results for BUSCO: 158698at7742
## Thu Jul 14 01:55:48 CEST 2022 - The aliscore output appears to be empty. Check results for BUSCO: 158698at7742
## Thu Jul 14 01:55:49 CEST 2022 - The aliscore output file does not exist. Check results for BUSCO: 239469at7742
## Thu Jul 14 01:55:49 CEST 2022 - The aliscore output appears to be empty. Check results for BUSCO: 239469at7742
## Thu Jul 14 01:55:51 CEST 2022 - The aliscore output file does not exist. Check results for BUSCO: 300584at7742
## Thu Jul 14 01:55:51 CEST 2022 - The aliscore output appears to be empty. Check results for BUSCO: 300584at7742
## Thu Jul 14 01:55:53 CEST 2022 - The aliscore output file does not exist. Check results for BUSCO: 148962at7742
## Thu Jul 14 01:55:53 CEST 2022 - The aliscore output appears to be empty. Check results for BUSCO: 148962at7742
## Thu Jul 14 01:55:53 CEST 2022 - The aliscore output file does not exist. Check results for BUSCO: 187431at7742
## Thu Jul 14 01:55:53 CEST 2022 - The aliscore output appears to be empty. Check results for BUSCO: 187431at7742
## Thu Jul 14 01:55:54 CEST 2022 - The aliscore output file does not exist. Check results for BUSCO: 202512at7742
## Thu Jul 14 01:55:54 CEST 2022 - The aliscore output appears to be empty. Check results for BUSCO: 202512at7742
## Thu Jul 14 01:55:54 CEST 2022 - The aliscore output file does not exist. Check results for BUSCO: 320621at7742
## Thu Jul 14 01:55:54 CEST 2022 - The aliscore output appears to be empty. Check results for BUSCO: 320621at7742
## Thu Jul 14 01:55:56 CEST 2022 - The aliscore output file does not exist. Check results for BUSCO: 223426at7742
## Thu Jul 14 01:55:56 CEST 2022 - The aliscore output appears to be empty. Check results for BUSCO: 223426at7742
## Thu Jul 14 01:55:56 CEST 2022 - The aliscore output file does not exist. Check results for BUSCO: 229857at7742
## Thu Jul 14 01:55:56 CEST 2022 - The aliscore output appears to be empty. Check results for BUSCO: 229857at7742
## Thu Jul 14 01:55:58 CEST 2022 - The aliscore output file does not exist. Check results for BUSCO: 261091at7742
## Thu Jul 14 01:55:58 CEST 2022 - The aliscore output appears to be empty. Check results for BUSCO: 261091at7742
## Thu Jul 14 01:55:59 CEST 2022 - The aliscore output file does not exist. Check results for BUSCO: 359693at7742
## Thu Jul 14 01:55:59 CEST 2022 - The aliscore output appears to be empty. Check results for BUSCO: 359693at7742
## Thu Jul 14 01:55:59 CEST 2022 - The aliscore output file does not exist. Check results for BUSCO: 252307at7742
## Thu Jul 14 01:55:59 CEST 2022 - The aliscore output appears to be empty. Check results for BUSCO: 252307at7742
## Thu Jul 14 01:56:00 CEST 2022 - The aliscore output file does not exist. Check results for BUSCO: 133965at7742
## Thu Jul 14 01:56:00 CEST 2022 - The aliscore output appears to be empty. Check results for BUSCO: 133965at7742
## Thu Jul 14 01:56:01 CEST 2022 - The aliscore output file does not exist. Check results for BUSCO: 292556at7742
## Thu Jul 14 01:56:01 CEST 2022 - The aliscore output appears to be empty. Check results for BUSCO: 292556at7742
## Thu Jul 14 01:56:03 CEST 2022 - The aliscore output file does not exist. Check results for BUSCO: 333162at7742
## Thu Jul 14 01:56:03 CEST 2022 - The aliscore output appears to be empty. Check results for BUSCO: 333162at7742
## Thu Jul 14 01:56:03 CEST 2022 - The aliscore output file does not exist. Check results for BUSCO: 186913at7742
## Thu Jul 14 01:56:03 CEST 2022 - The aliscore output appears to be empty. Check results for BUSCO: 186913at7742
## Thu Jul 14 01:56:04 CEST 2022 - The aliscore output file does not exist. Check results for BUSCO: 229857at7742
## Thu Jul 14 01:56:04 CEST 2022 - The aliscore output appears to be empty. Check results for BUSCO: 229857at7742
## Thu Jul 14 01:56:06 CEST 2022 - The aliscore output file does not exist. Check results for BUSCO: 202512at7742
## Thu Jul 14 01:56:06 CEST 2022 - The aliscore output appears to be empty. Check results for BUSCO: 202512at7742
## Thu Jul 14 01:56:07 CEST 2022 - The aliscore output file does not exist. Check results for BUSCO: 366862at7742
## Thu Jul 14 01:56:07 CEST 2022 - The aliscore output appears to be empty. Check results for BUSCO: 366862at7742
## Thu Jul 14 01:56:08 CEST 2022 - The aliscore output file does not exist. Check results for BUSCO: 245433at7742
## Thu Jul 14 01:56:08 CEST 2022 - The aliscore output appears to be empty. Check results for BUSCO: 245433at7742
## Thu Jul 14 01:56:09 CEST 2022 - The aliscore output file does not exist. Check results for BUSCO: 209386at7742
## Thu Jul 14 01:56:09 CEST 2022 - The aliscore output appears to be empty. Check results for BUSCO: 209386at7742
## Thu Jul 14 01:56:09 CEST 2022 - The aliscore output file does not exist. Check results for BUSCO: 340002at7742
## Thu Jul 14 01:56:09 CEST 2022 - The aliscore output appears to be empty. Check results for BUSCO: 340002at7742
## Thu Jul 14 01:56:11 CEST 2022 - The aliscore output file does not exist. Check results for BUSCO: 151964at7742
## Thu Jul 14 01:56:11 CEST 2022 - The aliscore output appears to be empty. Check results for BUSCO: 151964at7742
## Thu Jul 14 01:56:11 CEST 2022 - The aliscore output file does not exist. Check results for BUSCO: 346924at7742
## Thu Jul 14 01:56:11 CEST 2022 - The aliscore output appears to be empty. Check results for BUSCO: 346924at7742
## Thu Jul 14 01:56:13 CEST 2022 - The aliscore output file does not exist. Check results for BUSCO: 291589at7742
## Thu Jul 14 01:56:13 CEST 2022 - The aliscore output appears to be empty. Check results for BUSCO: 291589at7742
## Thu Jul 14 01:56:16 CEST 2022 - The aliscore output file does not exist. Check results for BUSCO: 223443at7742
## Thu Jul 14 01:56:16 CEST 2022 - The aliscore output appears to be empty. Check results for BUSCO: 223443at7742
## Thu Jul 14 01:56:17 CEST 2022 - The aliscore output file does not exist. Check results for BUSCO: 218109at7742
## Thu Jul 14 01:56:17 CEST 2022 - The aliscore output appears to be empty. Check results for BUSCO: 218109at7742
## Thu Jul 14 01:56:17 CEST 2022 - The aliscore output file does not exist. Check results for BUSCO: 299177at7742
## Thu Jul 14 01:56:17 CEST 2022 - The aliscore output appears to be empty. Check results for BUSCO: 299177at7742
## Thu Jul 14 01:56:19 CEST 2022 - The aliscore output file does not exist. Check results for BUSCO: 149872at7742
## Thu Jul 14 01:56:19 CEST 2022 - The aliscore output appears to be empty. Check results for BUSCO: 149872at7742
## Thu Jul 14 01:56:19 CEST 2022 - The aliscore output file does not exist. Check results for BUSCO: 154805at7742
## Thu Jul 14 01:56:19 CEST 2022 - The aliscore output appears to be empty. Check results for BUSCO: 154805at7742
## Thu Jul 14 01:56:21 CEST 2022 - The aliscore output file does not exist. Check results for BUSCO: 301608at7742
## Thu Jul 14 01:56:21 CEST 2022 - The aliscore output appears to be empty. Check results for BUSCO: 301608at7742
## Thu Jul 14 01:56:22 CEST 2022 - The aliscore output file does not exist. Check results for BUSCO: 179120at7742
## Thu Jul 14 01:56:22 CEST 2022 - The aliscore output appears to be empty. Check results for BUSCO: 179120at7742
## Thu Jul 14 01:56:22 CEST 2022 - The aliscore output file does not exist. Check results for BUSCO: 221671at7742
## Thu Jul 14 01:56:22 CEST 2022 - The aliscore output appears to be empty. Check results for BUSCO: 221671at7742
## Thu Jul 14 01:56:23 CEST 2022 - The aliscore output file does not exist. Check results for BUSCO: 356132at7742
## Thu Jul 14 01:56:23 CEST 2022 - The aliscore output appears to be empty. Check results for BUSCO: 356132at7742
## Thu Jul 14 01:56:24 CEST 2022 - The aliscore output file does not exist. Check results for BUSCO: 166002at7742
## Thu Jul 14 01:56:25 CEST 2022 - The aliscore output file does not exist. Check results for BUSCO: 325448at7742
## Thu Jul 14 01:56:25 CEST 2022 - The aliscore output appears to be empty. Check results for BUSCO: 166002at7742
## Thu Jul 14 01:56:25 CEST 2022 - The aliscore output appears to be empty. Check results for BUSCO: 325448at7742
## Thu Jul 14 01:56:26 CEST 2022 - The aliscore output file does not exist. Check results for BUSCO: 250326at7742
## Thu Jul 14 01:56:26 CEST 2022 - The aliscore output appears to be empty. Check results for BUSCO: 250326at7742
## Thu Jul 14 01:56:28 CEST 2022 - The aliscore output file does not exist. Check results for BUSCO: 155537at7742
## Thu Jul 14 01:56:28 CEST 2022 - The aliscore output appears to be empty. Check results for BUSCO: 155537at7742
## Thu Jul 14 01:56:28 CEST 2022 - The aliscore output file does not exist. Check results for BUSCO: 376010at7742
## Thu Jul 14 01:56:28 CEST 2022 - The aliscore output appears to be empty. Check results for BUSCO: 376010at7742
## Thu Jul 14 01:56:30 CEST 2022 - The aliscore output file does not exist. Check results for BUSCO: 229063at7742
## Thu Jul 14 01:56:30 CEST 2022 - The aliscore output appears to be empty. Check results for BUSCO: 229063at7742
## Thu Jul 14 01:56:30 CEST 2022 - The aliscore output file does not exist. Check results for BUSCO: 282590at7742
## Thu Jul 14 01:56:30 CEST 2022 - The aliscore output appears to be empty. Check results for BUSCO: 282590at7742
## Thu Jul 14 01:56:32 CEST 2022 - The aliscore output file does not exist. Check results for BUSCO: 197208at7742
## Thu Jul 14 01:56:32 CEST 2022 - The aliscore output appears to be empty. Check results for BUSCO: 197208at7742
## Thu Jul 14 01:56:32 CEST 2022 - The aliscore output file does not exist. Check results for BUSCO: 197208at7742
## Thu Jul 14 01:56:33 CEST 2022 - The aliscore output appears to be empty. Check results for BUSCO: 197208at7742
## Thu Jul 14 01:56:33 CEST 2022 - The aliscore output file does not exist. Check results for BUSCO: 291768at7742
## Thu Jul 14 01:56:33 CEST 2022 - The aliscore output appears to be empty. Check results for BUSCO: 291768at7742
## Thu Jul 14 01:56:36 CEST 2022 - The aliscore output file does not exist. Check results for BUSCO: 291239at7742
## Thu Jul 14 01:56:36 CEST 2022 - The aliscore output appears to be empty. Check results for BUSCO: 291239at7742
## Thu Jul 14 01:56:36 CEST 2022 - The aliscore output file does not exist. Check results for BUSCO: 427282at7742
## Thu Jul 14 01:56:36 CEST 2022 - The aliscore output appears to be empty. Check results for BUSCO: 427282at7742
## Thu Jul 14 01:56:37 CEST 2022 - The aliscore output file does not exist. Check results for BUSCO: 202160at7742
## Thu Jul 14 01:56:37 CEST 2022 - The aliscore output appears to be empty. Check results for BUSCO: 202160at7742
## Thu Jul 14 01:56:38 CEST 2022 - The aliscore output file does not exist. Check results for BUSCO: 175714at7742
## Thu Jul 14 01:56:38 CEST 2022 - The aliscore output appears to be empty. Check results for BUSCO: 175714at7742
## Thu Jul 14 01:56:38 CEST 2022 - The aliscore output file does not exist. Check results for BUSCO: 267724at7742
## Thu Jul 14 01:56:38 CEST 2022 - The aliscore output appears to be empty. Check results for BUSCO: 267724at7742
## Thu Jul 14 01:56:40 CEST 2022 - The aliscore output file does not exist. Check results for BUSCO: 293668at7742
## Thu Jul 14 01:56:40 CEST 2022 - The aliscore output appears to be empty. Check results for BUSCO: 293668at7742
## Thu Jul 14 01:56:40 CEST 2022 - The aliscore output file does not exist. Check results for BUSCO: 286230at7742
## Thu Jul 14 01:56:40 CEST 2022 - The aliscore output appears to be empty. Check results for BUSCO: 286230at7742
## Thu Jul 14 01:56:42 CEST 2022 - The aliscore output file does not exist. Check results for BUSCO: 349261at7742
## Thu Jul 14 01:56:42 CEST 2022 - The aliscore output appears to be empty. Check results for BUSCO: 349261at7742
## Thu Jul 14 01:56:43 CEST 2022 - The aliscore output file does not exist. Check results for BUSCO: 180612at7742
## Thu Jul 14 01:56:43 CEST 2022 - The aliscore output appears to be empty. Check results for BUSCO: 180612at7742
## Thu Jul 14 01:56:45 CEST 2022 - The aliscore output file does not exist. Check results for BUSCO: 192284at7742
## Thu Jul 14 01:56:45 CEST 2022 - The aliscore output appears to be empty. Check results for BUSCO: 192284at7742
## Thu Jul 14 01:56:45 CEST 2022 - The aliscore output file does not exist. Check results for BUSCO: 146782at7742
## Thu Jul 14 01:56:45 CEST 2022 - The aliscore output appears to be empty. Check results for BUSCO: 146782at7742
## Thu Jul 14 01:56:47 CEST 2022 - The aliscore output file does not exist. Check results for BUSCO: 309592at7742
## Thu Jul 14 01:56:47 CEST 2022 - The aliscore output appears to be empty. Check results for BUSCO: 309592at7742
## Thu Jul 14 01:56:47 CEST 2022 - The aliscore output file does not exist. Check results for BUSCO: 318320at7742
## Thu Jul 14 01:56:47 CEST 2022 - The aliscore output appears to be empty. Check results for BUSCO: 318320at7742
## Thu Jul 14 01:56:48 CEST 2022 - The aliscore output file does not exist. Check results for BUSCO: 319402at7742
## Thu Jul 14 01:56:48 CEST 2022 - The aliscore output appears to be empty. Check results for BUSCO: 319402at7742
## Thu Jul 14 01:56:50 CEST 2022 - The aliscore output file does not exist. Check results for BUSCO: 295741at7742
## Thu Jul 14 01:56:50 CEST 2022 - The aliscore output appears to be empty. Check results for BUSCO: 295741at7742
## Thu Jul 14 01:56:51 CEST 2022 - The aliscore output file does not exist. Check results for BUSCO: 213618at7742
## Thu Jul 14 01:56:51 CEST 2022 - The aliscore output appears to be empty. Check results for BUSCO: 213618at7742
## Thu Jul 14 01:56:53 CEST 2022 - The aliscore output file does not exist. Check results for BUSCO: 325935at7742
## Thu Jul 14 01:56:53 CEST 2022 - The aliscore output appears to be empty. Check results for BUSCO: 325935at7742
## Thu Jul 14 01:56:53 CEST 2022 - The aliscore output file does not exist. Check results for BUSCO: 197208at7742
## Thu Jul 14 01:56:53 CEST 2022 - The aliscore output appears to be empty. Check results for BUSCO: 197208at7742
## Thu Jul 14 01:56:53 CEST 2022 - The aliscore output file does not exist. Check results for BUSCO: 229063at7742
## Thu Jul 14 01:56:53 CEST 2022 - The aliscore output appears to be empty. Check results for BUSCO: 229063at7742
## Thu Jul 14 01:56:55 CEST 2022 - The aliscore output file does not exist. Check results for BUSCO: 123660at7742
## Thu Jul 14 01:56:55 CEST 2022 - The aliscore output file does not exist. Check results for BUSCO: 368798at7742
## Thu Jul 14 01:56:55 CEST 2022 - The aliscore output appears to be empty. Check results for BUSCO: 123660at7742
## Thu Jul 14 01:56:55 CEST 2022 - The aliscore output appears to be empty. Check results for BUSCO: 368798at7742
## Thu Jul 14 01:56:58 CEST 2022 - The aliscore output file does not exist. Check results for BUSCO: 169787at7742
## Thu Jul 14 01:56:58 CEST 2022 - The aliscore output appears to be empty. Check results for BUSCO: 169787at7742
## Thu Jul 14 01:57:00 CEST 2022 - The aliscore output file does not exist. Check results for BUSCO: 223512at7742
## Thu Jul 14 01:57:00 CEST 2022 - The aliscore output appears to be empty. Check results for BUSCO: 223512at7742
## Thu Jul 14 01:57:00 CEST 2022 - The aliscore output file does not exist. Check results for BUSCO: 183251at7742
## Thu Jul 14 01:57:00 CEST 2022 - The aliscore output appears to be empty. Check results for BUSCO: 183251at7742
## Thu Jul 14 01:57:01 CEST 2022 - The aliscore output file does not exist. Check results for BUSCO: 350246at7742
## Thu Jul 14 01:57:01 CEST 2022 - The aliscore output appears to be empty. Check results for BUSCO: 350246at7742
## Thu Jul 14 01:57:02 CEST 2022 - The aliscore output file does not exist. Check results for BUSCO: 401448at7742
## Thu Jul 14 01:57:02 CEST 2022 - The aliscore output appears to be empty. Check results for BUSCO: 401448at7742
## Thu Jul 14 01:57:04 CEST 2022 - The aliscore output file does not exist. Check results for BUSCO: 376721at7742
## Thu Jul 14 01:57:04 CEST 2022 - The aliscore output appears to be empty. Check results for BUSCO: 376721at7742
## Thu Jul 14 01:57:04 CEST 2022 - The aliscore output file does not exist. Check results for BUSCO: 196409at7742
## Thu Jul 14 01:57:04 CEST 2022 - The aliscore output appears to be empty. Check results for BUSCO: 196409at7742
## Thu Jul 14 01:57:04 CEST 2022 - The aliscore output file does not exist. Check results for BUSCO: 155537at7742
## Thu Jul 14 01:57:04 CEST 2022 - The aliscore output appears to be empty. Check results for BUSCO: 155537at7742
## Thu Jul 14 01:57:07 CEST 2022 - The aliscore output file does not exist. Check results for BUSCO: 281858at7742
## Thu Jul 14 01:57:07 CEST 2022 - The aliscore output appears to be empty. Check results for BUSCO: 281858at7742
## Thu Jul 14 01:57:07 CEST 2022 - The aliscore output file does not exist. Check results for BUSCO: 242487at7742
## Thu Jul 14 01:57:07 CEST 2022 - The aliscore output appears to be empty. Check results for BUSCO: 242487at7742
## Thu Jul 14 01:57:08 CEST 2022 - The aliscore output file does not exist. Check results for BUSCO: 214100at7742
## Thu Jul 14 01:57:08 CEST 2022 - The aliscore output appears to be empty. Check results for BUSCO: 214100at7742
## Thu Jul 14 01:57:08 CEST 2022 - The aliscore output file does not exist. Check results for BUSCO: 195766at7742
## Thu Jul 14 01:57:08 CEST 2022 - The aliscore output appears to be empty. Check results for BUSCO: 195766at7742
## Thu Jul 14 01:57:10 CEST 2022 - The aliscore output file does not exist. Check results for BUSCO: 392098at7742
## Thu Jul 14 01:57:10 CEST 2022 - The aliscore output appears to be empty. Check results for BUSCO: 392098at7742
## Thu Jul 14 01:57:11 CEST 2022 - The aliscore output file does not exist. Check results for BUSCO: 161007at7742
## Thu Jul 14 01:57:12 CEST 2022 - The aliscore output appears to be empty. Check results for BUSCO: 161007at7742
## Thu Jul 14 01:57:12 CEST 2022 - The aliscore output file does not exist. Check results for BUSCO: 190562at7742
## Thu Jul 14 01:57:12 CEST 2022 - The aliscore output appears to be empty. Check results for BUSCO: 190562at7742
## Thu Jul 14 01:57:13 CEST 2022 - The aliscore output file does not exist. Check results for BUSCO: 302110at7742
## Thu Jul 14 01:57:13 CEST 2022 - The aliscore output appears to be empty. Check results for BUSCO: 302110at7742
## Thu Jul 14 01:57:14 CEST 2022 - The aliscore output file does not exist. Check results for BUSCO: 223512at7742
## Thu Jul 14 01:57:14 CEST 2022 - The aliscore output appears to be empty. Check results for BUSCO: 223512at7742
## Thu Jul 14 01:57:16 CEST 2022 - The aliscore output file does not exist. Check results for BUSCO: 299177at7742
## Thu Jul 14 01:57:16 CEST 2022 - The aliscore output appears to be empty. Check results for BUSCO: 299177at7742
## Thu Jul 14 01:57:18 CEST 2022 - The aliscore output file does not exist. Check results for BUSCO: 236145at7742
## Thu Jul 14 01:57:18 CEST 2022 - The aliscore output appears to be empty. Check results for BUSCO: 236145at7742
## Thu Jul 14 01:57:18 CEST 2022 - The aliscore output file does not exist. Check results for BUSCO: 345746at7742
## Thu Jul 14 01:57:18 CEST 2022 - The aliscore output appears to be empty. Check results for BUSCO: 345746at7742
## Thu Jul 14 01:57:18 CEST 2022 - The aliscore output file does not exist. Check results for BUSCO: 124571at7742
## Thu Jul 14 01:57:18 CEST 2022 - The aliscore output appears to be empty. Check results for BUSCO: 124571at7742
## Thu Jul 14 01:57:19 CEST 2022 - The aliscore output file does not exist. Check results for BUSCO: 279455at7742
## Thu Jul 14 01:57:19 CEST 2022 - The aliscore output appears to be empty. Check results for BUSCO: 279455at7742
## Thu Jul 14 01:57:20 CEST 2022 - The aliscore output file does not exist. Check results for BUSCO: 154805at7742
## Thu Jul 14 01:57:20 CEST 2022 - The aliscore output appears to be empty. Check results for BUSCO: 154805at7742
## Thu Jul 14 01:57:21 CEST 2022 - The aliscore output file does not exist. Check results for BUSCO: 239469at7742
## Thu Jul 14 01:57:21 CEST 2022 - The aliscore output appears to be empty. Check results for BUSCO: 239469at7742
## Thu Jul 14 01:57:23 CEST 2022 - The aliscore output file does not exist. Check results for BUSCO: 238694at7742
## Thu Jul 14 01:57:23 CEST 2022 - The aliscore output appears to be empty. Check results for BUSCO: 238694at7742
## Thu Jul 14 01:57:24 CEST 2022 - The aliscore output file does not exist. Check results for BUSCO: 202160at7742
## Thu Jul 14 01:57:24 CEST 2022 - The aliscore output appears to be empty. Check results for BUSCO: 202160at7742
## Thu Jul 14 01:57:26 CEST 2022 - The aliscore output file does not exist. Check results for BUSCO: 254847at7742
## Thu Jul 14 01:57:26 CEST 2022 - The aliscore output appears to be empty. Check results for BUSCO: 254847at7742
## Thu Jul 14 01:57:27 CEST 2022 - The aliscore output file does not exist. Check results for BUSCO: 245117at7742
## Thu Jul 14 01:57:27 CEST 2022 - The aliscore output appears to be empty. Check results for BUSCO: 245117at7742
## Thu Jul 14 01:57:27 CEST 2022 - The aliscore output file does not exist. Check results for BUSCO: 178109at7742
## Thu Jul 14 01:57:27 CEST 2022 - The aliscore output appears to be empty. Check results for BUSCO: 178109at7742
## Thu Jul 14 01:57:28 CEST 2022 - The aliscore output file does not exist. Check results for BUSCO: 274480at7742
## Thu Jul 14 01:57:28 CEST 2022 - The aliscore output appears to be empty. Check results for BUSCO: 274480at7742
## Thu Jul 14 01:57:30 CEST 2022 - The aliscore output file does not exist. Check results for BUSCO: 281858at7742
## Thu Jul 14 01:57:30 CEST 2022 - The aliscore output appears to be empty. Check results for BUSCO: 281858at7742
## Thu Jul 14 01:57:30 CEST 2022 - The aliscore output file does not exist. Check results for BUSCO: 279455at7742
## Thu Jul 14 01:57:30 CEST 2022 - The aliscore output appears to be empty. Check results for BUSCO: 279455at7742
## Thu Jul 14 01:57:30 CEST 2022 - The aliscore output file does not exist. Check results for BUSCO: 180612at7742
## Thu Jul 14 01:57:31 CEST 2022 - The aliscore output appears to be empty. Check results for BUSCO: 180612at7742
## Thu Jul 14 01:57:32 CEST 2022 - The aliscore output file does not exist. Check results for BUSCO: 305792at7742
## Thu Jul 14 01:57:32 CEST 2022 - The aliscore output appears to be empty. Check results for BUSCO: 305792at7742
## Thu Jul 14 01:57:33 CEST 2022 - The aliscore output file does not exist. Check results for BUSCO: 287466at7742
## Thu Jul 14 01:57:33 CEST 2022 - The aliscore output appears to be empty. Check results for BUSCO: 287466at7742
## Thu Jul 14 01:57:34 CEST 2022 - The aliscore output file does not exist. Check results for BUSCO: 203253at7742
## Thu Jul 14 01:57:34 CEST 2022 - The aliscore output appears to be empty. Check results for BUSCO: 203253at7742
## Thu Jul 14 01:57:35 CEST 2022 - The aliscore output file does not exist. Check results for BUSCO: 218430at7742
## Thu Jul 14 01:57:35 CEST 2022 - The aliscore output appears to be empty. Check results for BUSCO: 218430at7742
## Thu Jul 14 01:57:37 CEST 2022 - The aliscore output file does not exist. Check results for BUSCO: 259863at7742
## Thu Jul 14 01:57:37 CEST 2022 - The aliscore output appears to be empty. Check results for BUSCO: 259863at7742
## Thu Jul 14 01:57:37 CEST 2022 - The aliscore output file does not exist. Check results for BUSCO: 257212at7742
## Thu Jul 14 01:57:37 CEST 2022 - The aliscore output appears to be empty. Check results for BUSCO: 257212at7742
## Thu Jul 14 01:57:37 CEST 2022 - The aliscore output file does not exist. Check results for BUSCO: 211150at7742
## Thu Jul 14 01:57:37 CEST 2022 - The aliscore output appears to be empty. Check results for BUSCO: 211150at7742
## Thu Jul 14 01:57:39 CEST 2022 - The aliscore output file does not exist. Check results for BUSCO: 229063at7742
## Thu Jul 14 01:57:40 CEST 2022 - The aliscore output file does not exist. Check results for BUSCO: 137854at7742
## Thu Jul 14 01:57:40 CEST 2022 - The aliscore output appears to be empty. Check results for BUSCO: 229063at7742
## Thu Jul 14 01:57:40 CEST 2022 - The aliscore output appears to be empty. Check results for BUSCO: 137854at7742
## Thu Jul 14 01:57:41 CEST 2022 - The aliscore output file does not exist. Check results for BUSCO: 286230at7742
## Thu Jul 14 01:57:41 CEST 2022 - The aliscore output appears to be empty. Check results for BUSCO: 286230at7742
## Thu Jul 14 01:57:42 CEST 2022 - The aliscore output file does not exist. Check results for BUSCO: 148962at7742
## Thu Jul 14 01:57:42 CEST 2022 - The aliscore output appears to be empty. Check results for BUSCO: 148962at7742
## Thu Jul 14 01:57:43 CEST 2022 - The aliscore output file does not exist. Check results for BUSCO: 326191at7742
## Thu Jul 14 01:57:43 CEST 2022 - The aliscore output appears to be empty. Check results for BUSCO: 326191at7742
## Thu Jul 14 01:57:45 CEST 2022 - The aliscore output file does not exist. Check results for BUSCO: 239538at7742
## Thu Jul 14 01:57:45 CEST 2022 - The aliscore output appears to be empty. Check results for BUSCO: 239538at7742
## Thu Jul 14 01:57:45 CEST 2022 - The aliscore output file does not exist. Check results for BUSCO: 178109at7742
## Thu Jul 14 01:57:45 CEST 2022 - The aliscore output appears to be empty. Check results for BUSCO: 178109at7742
## Thu Jul 14 01:57:46 CEST 2022 - The aliscore output file does not exist. Check results for BUSCO: 364320at7742
## Thu Jul 14 01:57:46 CEST 2022 - The aliscore output appears to be empty. Check results for BUSCO: 364320at7742
## Thu Jul 14 01:57:47 CEST 2022 - The aliscore output file does not exist. Check results for BUSCO: 128421at7742
## Thu Jul 14 01:57:47 CEST 2022 - The aliscore output appears to be empty. Check results for BUSCO: 128421at7742
## Thu Jul 14 01:57:48 CEST 2022 - The aliscore output file does not exist. Check results for BUSCO: 213068at7742
## Thu Jul 14 01:57:48 CEST 2022 - The aliscore output appears to be empty. Check results for BUSCO: 213068at7742
## Thu Jul 14 01:57:48 CEST 2022 - The aliscore output file does not exist. Check results for BUSCO: 259058at7742
## Thu Jul 14 01:57:48 CEST 2022 - The aliscore output appears to be empty. Check results for BUSCO: 259058at7742
## Thu Jul 14 01:57:50 CEST 2022 - The aliscore output file does not exist. Check results for BUSCO: 293487at7742
## Thu Jul 14 01:57:50 CEST 2022 - The aliscore output appears to be empty. Check results for BUSCO: 293487at7742
## Thu Jul 14 01:57:51 CEST 2022 - The aliscore output file does not exist. Check results for BUSCO: 281159at7742
## Thu Jul 14 01:57:52 CEST 2022 - The aliscore output appears to be empty. Check results for BUSCO: 281159at7742
## Thu Jul 14 01:57:53 CEST 2022 - The aliscore output file does not exist. Check results for BUSCO: 183995at7742
## Thu Jul 14 01:57:53 CEST 2022 - The aliscore output appears to be empty. Check results for BUSCO: 183995at7742
## Thu Jul 14 01:57:54 CEST 2022 - The aliscore output file does not exist. Check results for BUSCO: 320621at7742
## Thu Jul 14 01:57:54 CEST 2022 - The aliscore output appears to be empty. Check results for BUSCO: 320621at7742
## Thu Jul 14 01:57:54 CEST 2022 - The aliscore output file does not exist. Check results for BUSCO: 151078at7742
## Thu Jul 14 01:57:54 CEST 2022 - The aliscore output appears to be empty. Check results for BUSCO: 151078at7742
## Thu Jul 14 01:57:56 CEST 2022 - The aliscore output file does not exist. Check results for BUSCO: 317891at7742
## Thu Jul 14 01:57:56 CEST 2022 - The aliscore output appears to be empty. Check results for BUSCO: 317891at7742
## Thu Jul 14 01:57:56 CEST 2022 - The aliscore output file does not exist. Check results for BUSCO: 167517at7742
## Thu Jul 14 01:57:56 CEST 2022 - The aliscore output appears to be empty. Check results for BUSCO: 167517at7742
## Thu Jul 14 01:57:57 CEST 2022 - The aliscore output file does not exist. Check results for BUSCO: 279170at7742
## Thu Jul 14 01:57:57 CEST 2022 - The aliscore output appears to be empty. Check results for BUSCO: 279170at7742
## Thu Jul 14 01:57:58 CEST 2022 - The aliscore output file does not exist. Check results for BUSCO: 295741at7742
## Thu Jul 14 01:57:58 CEST 2022 - The aliscore output appears to be empty. Check results for BUSCO: 295741at7742
## Thu Jul 14 01:57:59 CEST 2022 - The aliscore output file does not exist. Check results for BUSCO: 178109at7742
## Thu Jul 14 01:57:59 CEST 2022 - The aliscore output appears to be empty. Check results for BUSCO: 178109at7742
## Thu Jul 14 01:58:00 CEST 2022 - The aliscore output file does not exist. Check results for BUSCO: 151078at7742
## Thu Jul 14 01:58:00 CEST 2022 - The aliscore output appears to be empty. Check results for BUSCO: 151078at7742
## Thu Jul 14 01:58:01 CEST 2022 - The aliscore output file does not exist. Check results for BUSCO: 211150at7742
## Thu Jul 14 01:58:01 CEST 2022 - The aliscore output appears to be empty. Check results for BUSCO: 211150at7742
## Thu Jul 14 01:58:03 CEST 2022 - The aliscore output file does not exist. Check results for BUSCO: 207153at7742
## Thu Jul 14 01:58:03 CEST 2022 - The aliscore output appears to be empty. Check results for BUSCO: 207153at7742
## Thu Jul 14 01:58:03 CEST 2022 - The aliscore output file does not exist. Check results for BUSCO: 356998at7742
## Thu Jul 14 01:58:03 CEST 2022 - The aliscore output appears to be empty. Check results for BUSCO: 356998at7742
## Thu Jul 14 01:58:04 CEST 2022 - The aliscore output file does not exist. Check results for BUSCO: 264056at7742
## Thu Jul 14 01:58:04 CEST 2022 - The aliscore output appears to be empty. Check results for BUSCO: 264056at7742
## Thu Jul 14 01:58:04 CEST 2022 - The aliscore output file does not exist. Check results for BUSCO: 317891at7742
## Thu Jul 14 01:58:04 CEST 2022 - The aliscore output appears to be empty. Check results for BUSCO: 317891at7742
## Thu Jul 14 01:58:05 CEST 2022 - The aliscore output file does not exist. Check results for BUSCO: 213350at7742
## Thu Jul 14 01:58:05 CEST 2022 - The aliscore output appears to be empty. Check results for BUSCO: 213350at7742
## Thu Jul 14 01:58:07 CEST 2022 - The aliscore output file does not exist. Check results for BUSCO: 328571at7742
## Thu Jul 14 01:58:07 CEST 2022 - The aliscore output appears to be empty. Check results for BUSCO: 328571at7742
## Thu Jul 14 01:58:08 CEST 2022 - The aliscore output file does not exist. Check results for BUSCO: 209386at7742
## Thu Jul 14 01:58:08 CEST 2022 - The aliscore output appears to be empty. Check results for BUSCO: 209386at7742
## Thu Jul 14 01:58:09 CEST 2022 - The aliscore output file does not exist. Check results for BUSCO: 201177at7742
## Thu Jul 14 01:58:09 CEST 2022 - The aliscore output appears to be empty. Check results for BUSCO: 201177at7742
## Thu Jul 14 01:58:10 CEST 2022 - The aliscore output file does not exist. Check results for BUSCO: 250442at7742
## Thu Jul 14 01:58:10 CEST 2022 - The aliscore output appears to be empty. Check results for BUSCO: 250442at7742
## Thu Jul 14 01:58:11 CEST 2022 - The aliscore output file does not exist. Check results for BUSCO: 286230at7742
## Thu Jul 14 01:58:11 CEST 2022 - The aliscore output appears to be empty. Check results for BUSCO: 286230at7742
## Thu Jul 14 01:58:12 CEST 2022 - The aliscore output file does not exist. Check results for BUSCO: 368813at7742
## Thu Jul 14 01:58:12 CEST 2022 - The aliscore output appears to be empty. Check results for BUSCO: 368813at7742
## Thu Jul 14 01:58:12 CEST 2022 - The aliscore output file does not exist. Check results for BUSCO: 377163at7742
## Thu Jul 14 01:58:12 CEST 2022 - The aliscore output appears to be empty. Check results for BUSCO: 377163at7742
## Thu Jul 14 01:58:14 CEST 2022 - The aliscore output file does not exist. Check results for BUSCO: 128421at7742
## Thu Jul 14 01:58:14 CEST 2022 - The aliscore output appears to be empty. Check results for BUSCO: 128421at7742
## Thu Jul 14 01:58:14 CEST 2022 - The aliscore output file does not exist. Check results for BUSCO: 160476at7742
## Thu Jul 14 01:58:14 CEST 2022 - The aliscore output appears to be empty. Check results for BUSCO: 160476at7742
## Thu Jul 14 01:58:17 CEST 2022 - The aliscore output file does not exist. Check results for BUSCO: 137854at7742
## Thu Jul 14 01:58:17 CEST 2022 - The aliscore output appears to be empty. Check results for BUSCO: 137854at7742
## Thu Jul 14 01:58:17 CEST 2022 - The aliscore output file does not exist. Check results for BUSCO: 315274at7742
## Thu Jul 14 01:58:17 CEST 2022 - The aliscore output appears to be empty. Check results for BUSCO: 315274at7742
## Thu Jul 14 01:58:18 CEST 2022 - The aliscore output file does not exist. Check results for BUSCO: 287339at7742
## Thu Jul 14 01:58:18 CEST 2022 - The aliscore output appears to be empty. Check results for BUSCO: 287339at7742
## Thu Jul 14 01:58:19 CEST 2022 - The aliscore output file does not exist. Check results for BUSCO: 249694at7742
## Thu Jul 14 01:58:19 CEST 2022 - The aliscore output appears to be empty. Check results for BUSCO: 249694at7742
## Thu Jul 14 01:58:20 CEST 2022 - The aliscore output file does not exist. Check results for BUSCO: 166722at7742
## Thu Jul 14 01:58:20 CEST 2022 - The aliscore output appears to be empty. Check results for BUSCO: 166722at7742
## Thu Jul 14 01:58:20 CEST 2022 - The aliscore output file does not exist. Check results for BUSCO: 180612at7742
## Thu Jul 14 01:58:20 CEST 2022 - The aliscore output appears to be empty. Check results for BUSCO: 180612at7742
## Thu Jul 14 01:58:22 CEST 2022 - The aliscore output file does not exist. Check results for BUSCO: 264251at7742
## Thu Jul 14 01:58:22 CEST 2022 - The aliscore output appears to be empty. Check results for BUSCO: 264251at7742
## Thu Jul 14 01:58:23 CEST 2022 - The aliscore output file does not exist. Check results for BUSCO: 218430at7742
## Thu Jul 14 01:58:23 CEST 2022 - The aliscore output appears to be empty. Check results for BUSCO: 218430at7742
## Thu Jul 14 01:58:23 CEST 2022 - The aliscore output file does not exist. Check results for BUSCO: 235827at7742
## Thu Jul 14 01:58:23 CEST 2022 - The aliscore output appears to be empty. Check results for BUSCO: 235827at7742
## Thu Jul 14 01:58:24 CEST 2022 - The aliscore output file does not exist. Check results for BUSCO: 274480at7742
## Thu Jul 14 01:58:24 CEST 2022 - The aliscore output appears to be empty. Check results for BUSCO: 274480at7742
## Thu Jul 14 01:58:26 CEST 2022 - The aliscore output file does not exist. Check results for BUSCO: 325935at7742
## Thu Jul 14 01:58:26 CEST 2022 - The aliscore output appears to be empty. Check results for BUSCO: 325935at7742
## Thu Jul 14 01:58:28 CEST 2022 - The aliscore output file does not exist. Check results for BUSCO: 276229at7742
## Thu Jul 14 01:58:28 CEST 2022 - The aliscore output appears to be empty. Check results for BUSCO: 276229at7742
## Thu Jul 14 01:58:28 CEST 2022 - The aliscore output file does not exist. Check results for BUSCO: 276229at7742
## Thu Jul 14 01:58:28 CEST 2022 - The aliscore output appears to be empty. Check results for BUSCO: 276229at7742
## Thu Jul 14 01:58:29 CEST 2022 - The aliscore output file does not exist. Check results for BUSCO: 377163at7742
## Thu Jul 14 01:58:29 CEST 2022 - The aliscore output file does not exist. Check results for BUSCO: 211328at7742
## Thu Jul 14 01:58:29 CEST 2022 - The aliscore output appears to be empty. Check results for BUSCO: 377163at7742
## Thu Jul 14 01:58:29 CEST 2022 - The aliscore output appears to be empty. Check results for BUSCO: 211328at7742
## Thu Jul 14 01:58:30 CEST 2022 - The aliscore output file does not exist. Check results for BUSCO: 315274at7742
## Thu Jul 14 01:58:30 CEST 2022 - The aliscore output appears to be empty. Check results for BUSCO: 315274at7742
## Thu Jul 14 01:58:31 CEST 2022 - The aliscore output file does not exist. Check results for BUSCO: 177937at7742
## Thu Jul 14 01:58:31 CEST 2022 - The aliscore output appears to be empty. Check results for BUSCO: 177937at7742
## Thu Jul 14 01:58:32 CEST 2022 - The aliscore output file does not exist. Check results for BUSCO: 209086at7742
## Thu Jul 14 01:58:32 CEST 2022 - The aliscore output appears to be empty. Check results for BUSCO: 209086at7742
## Thu Jul 14 01:58:33 CEST 2022 - The aliscore output file does not exist. Check results for BUSCO: 223152at7742
## Thu Jul 14 01:58:33 CEST 2022 - The aliscore output appears to be empty. Check results for BUSCO: 223152at7742
## Thu Jul 14 01:58:35 CEST 2022 - The aliscore output file does not exist. Check results for BUSCO: 223152at7742
## Thu Jul 14 01:58:35 CEST 2022 - The aliscore output appears to be empty. Check results for BUSCO: 223152at7742
## Thu Jul 14 01:58:35 CEST 2022 - The aliscore output file does not exist. Check results for BUSCO: 209931at7742
## Thu Jul 14 01:58:35 CEST 2022 - The aliscore output appears to be empty. Check results for BUSCO: 209931at7742
## Thu Jul 14 01:58:36 CEST 2022 - The aliscore output file does not exist. Check results for BUSCO: 340020at7742
## Thu Jul 14 01:58:36 CEST 2022 - The aliscore output appears to be empty. Check results for BUSCO: 340020at7742
## Thu Jul 14 01:58:37 CEST 2022 - The aliscore output file does not exist. Check results for BUSCO: 291153at7742
## Thu Jul 14 01:58:37 CEST 2022 - The aliscore output appears to be empty. Check results for BUSCO: 291153at7742
## Thu Jul 14 01:58:39 CEST 2022 - The aliscore output file does not exist. Check results for BUSCO: 226375at7742
## Thu Jul 14 01:58:39 CEST 2022 - The aliscore output appears to be empty. Check results for BUSCO: 226375at7742
## Thu Jul 14 01:58:39 CEST 2022 - The aliscore output file does not exist. Check results for BUSCO: 380097at7742
## Thu Jul 14 01:58:39 CEST 2022 - The aliscore output appears to be empty. Check results for BUSCO: 380097at7742
## Thu Jul 14 01:58:39 CEST 2022 - The aliscore output file does not exist. Check results for BUSCO: 328514at7742
## Thu Jul 14 01:58:39 CEST 2022 - The aliscore output appears to be empty. Check results for BUSCO: 328514at7742
## Thu Jul 14 01:58:41 CEST 2022 - The aliscore output file does not exist. Check results for BUSCO: 328514at7742
## Thu Jul 14 01:58:41 CEST 2022 - The aliscore output appears to be empty. Check results for BUSCO: 328514at7742
## Thu Jul 14 01:58:41 CEST 2022 - The aliscore output file does not exist. Check results for BUSCO: 263652at7742
## Thu Jul 14 01:58:41 CEST 2022 - The aliscore output appears to be empty. Check results for BUSCO: 263652at7742
## Thu Jul 14 01:58:42 CEST 2022 - The aliscore output file does not exist. Check results for BUSCO: 253661at7742
## Thu Jul 14 01:58:42 CEST 2022 - The aliscore output appears to be empty. Check results for BUSCO: 253661at7742
## Thu Jul 14 01:58:43 CEST 2022 - The aliscore output file does not exist. Check results for BUSCO: 239469at7742
## Thu Jul 14 01:58:43 CEST 2022 - The aliscore output appears to be empty. Check results for BUSCO: 239469at7742
## Thu Jul 14 01:58:45 CEST 2022 - The aliscore output file does not exist. Check results for BUSCO: 155537at7742
## Thu Jul 14 01:58:45 CEST 2022 - The aliscore output appears to be empty. Check results for BUSCO: 155537at7742
## Thu Jul 14 01:58:45 CEST 2022 - The aliscore output file does not exist. Check results for BUSCO: 214100at7742
## Thu Jul 14 01:58:45 CEST 2022 - The aliscore output appears to be empty. Check results for BUSCO: 214100at7742
## Thu Jul 14 01:58:48 CEST 2022 - The aliscore output file does not exist. Check results for BUSCO: 186469at7742
## Thu Jul 14 01:58:48 CEST 2022 - The aliscore output file does not exist. Check results for BUSCO: 381534at7742
## Thu Jul 14 01:58:48 CEST 2022 - The aliscore output appears to be empty. Check results for BUSCO: 186469at7742
## Thu Jul 14 01:58:48 CEST 2022 - The aliscore output appears to be empty. Check results for BUSCO: 381534at7742
## Thu Jul 14 01:58:49 CEST 2022 - The aliscore output file does not exist. Check results for BUSCO: 301608at7742
## Thu Jul 14 01:58:49 CEST 2022 - The aliscore output appears to be empty. Check results for BUSCO: 301608at7742
## Thu Jul 14 01:58:50 CEST 2022 - The aliscore output file does not exist. Check results for BUSCO: 283441at7742
## Thu Jul 14 01:58:50 CEST 2022 - The aliscore output appears to be empty. Check results for BUSCO: 283441at7742
## Thu Jul 14 01:58:51 CEST 2022 - The aliscore output file does not exist. Check results for BUSCO: 202160at7742
## Thu Jul 14 01:58:51 CEST 2022 - The aliscore output appears to be empty. Check results for BUSCO: 202160at7742
## Thu Jul 14 01:58:51 CEST 2022 - The aliscore output file does not exist. Check results for BUSCO: 257212at7742
## Thu Jul 14 01:58:51 CEST 2022 - The aliscore output appears to be empty. Check results for BUSCO: 257212at7742
## Thu Jul 14 01:58:53 CEST 2022 - The aliscore output file does not exist. Check results for BUSCO: 241208at7742
## Thu Jul 14 01:58:53 CEST 2022 - The aliscore output appears to be empty. Check results for BUSCO: 241208at7742
## Thu Jul 14 01:58:53 CEST 2022 - The aliscore output file does not exist. Check results for BUSCO: 238694at7742
## Thu Jul 14 01:58:53 CEST 2022 - The aliscore output appears to be empty. Check results for BUSCO: 238694at7742
## Thu Jul 14 01:58:55 CEST 2022 - The aliscore output file does not exist. Check results for BUSCO: 339768at7742
## Thu Jul 14 01:58:55 CEST 2022 - The aliscore output appears to be empty. Check results for BUSCO: 339768at7742
## Thu Jul 14 01:58:56 CEST 2022 - The aliscore output file does not exist. Check results for BUSCO: 297771at7742
## Thu Jul 14 01:58:56 CEST 2022 - The aliscore output appears to be empty. Check results for BUSCO: 297771at7742
## Thu Jul 14 01:58:58 CEST 2022 - The aliscore output file does not exist. Check results for BUSCO: 328558at7742
## Thu Jul 14 01:58:58 CEST 2022 - The aliscore output appears to be empty. Check results for BUSCO: 328558at7742
## Thu Jul 14 01:58:58 CEST 2022 - The aliscore output file does not exist. Check results for BUSCO: 241208at7742
## Thu Jul 14 01:58:58 CEST 2022 - The aliscore output appears to be empty. Check results for BUSCO: 241208at7742
## Thu Jul 14 01:58:59 CEST 2022 - The aliscore output file does not exist. Check results for BUSCO: 347924at7742
## Thu Jul 14 01:58:59 CEST 2022 - The aliscore output appears to be empty. Check results for BUSCO: 347924at7742
## Thu Jul 14 01:59:00 CEST 2022 - The aliscore output file does not exist. Check results for BUSCO: 168866at7742
## Thu Jul 14 01:59:00 CEST 2022 - The aliscore output appears to be empty. Check results for BUSCO: 168866at7742
## Thu Jul 14 01:59:01 CEST 2022 - The aliscore output file does not exist. Check results for BUSCO: 124571at7742
## Thu Jul 14 01:59:01 CEST 2022 - The aliscore output appears to be empty. Check results for BUSCO: 124571at7742
## Thu Jul 14 01:59:01 CEST 2022 - The aliscore output file does not exist. Check results for BUSCO: 199987at7742
## Thu Jul 14 01:59:01 CEST 2022 - The aliscore output appears to be empty. Check results for BUSCO: 199987at7742
## Thu Jul 14 01:59:02 CEST 2022 - The aliscore output file does not exist. Check results for BUSCO: 373266at7742
## Thu Jul 14 01:59:02 CEST 2022 - The aliscore output appears to be empty. Check results for BUSCO: 373266at7742
## Thu Jul 14 01:59:03 CEST 2022 - The aliscore output file does not exist. Check results for BUSCO: 186469at7742
## Thu Jul 14 01:59:03 CEST 2022 - The aliscore output appears to be empty. Check results for BUSCO: 186469at7742
## Thu Jul 14 01:59:03 CEST 2022 - The aliscore output file does not exist. Check results for BUSCO: 209086at7742
## Thu Jul 14 01:59:04 CEST 2022 - The aliscore output appears to be empty. Check results for BUSCO: 209086at7742
## Thu Jul 14 01:59:04 CEST 2022 - The aliscore output file does not exist. Check results for BUSCO: 363898at7742
## Thu Jul 14 01:59:04 CEST 2022 - The aliscore output appears to be empty. Check results for BUSCO: 363898at7742
## Thu Jul 14 01:59:06 CEST 2022 - The aliscore output file does not exist. Check results for BUSCO: 244272at7742
## Thu Jul 14 01:59:06 CEST 2022 - The aliscore output appears to be empty. Check results for BUSCO: 244272at7742
## Thu Jul 14 01:59:08 CEST 2022 - The aliscore output file does not exist. Check results for BUSCO: 296930at7742
## Thu Jul 14 01:59:08 CEST 2022 - The aliscore output appears to be empty. Check results for BUSCO: 296930at7742
## Thu Jul 14 01:59:08 CEST 2022 - The aliscore output file does not exist. Check results for BUSCO: 349840at7742
## Thu Jul 14 01:59:08 CEST 2022 - The aliscore output appears to be empty. Check results for BUSCO: 349840at7742
## Thu Jul 14 01:59:09 CEST 2022 - The aliscore output file does not exist. Check results for BUSCO: 249694at7742
## Thu Jul 14 01:59:09 CEST 2022 - The aliscore output appears to be empty. Check results for BUSCO: 249694at7742
## Thu Jul 14 01:59:10 CEST 2022 - The aliscore output file does not exist. Check results for BUSCO: 239538at7742
## Thu Jul 14 01:59:10 CEST 2022 - The aliscore output appears to be empty. Check results for BUSCO: 239538at7742
## Thu Jul 14 01:59:11 CEST 2022 - The aliscore output file does not exist. Check results for BUSCO: 281159at7742
## Thu Jul 14 01:59:11 CEST 2022 - The aliscore output appears to be empty. Check results for BUSCO: 281159at7742
## Thu Jul 14 01:59:14 CEST 2022 - The aliscore output file does not exist. Check results for BUSCO: 167517at7742
## Thu Jul 14 01:59:14 CEST 2022 - The aliscore output appears to be empty. Check results for BUSCO: 167517at7742
## Thu Jul 14 01:59:14 CEST 2022 - The aliscore output file does not exist. Check results for BUSCO: 202013at7742
## Thu Jul 14 01:59:14 CEST 2022 - The aliscore output appears to be empty. Check results for BUSCO: 202013at7742
## Thu Jul 14 01:59:15 CEST 2022 - The aliscore output file does not exist. Check results for BUSCO: 297362at7742
## Thu Jul 14 01:59:15 CEST 2022 - The aliscore output appears to be empty. Check results for BUSCO: 297362at7742
## Thu Jul 14 01:59:16 CEST 2022 - The aliscore output file does not exist. Check results for BUSCO: 214136at7742
## Thu Jul 14 01:59:16 CEST 2022 - The aliscore output appears to be empty. Check results for BUSCO: 214136at7742
## Thu Jul 14 01:59:17 CEST 2022 - The aliscore output file does not exist. Check results for BUSCO: 192368at7742
## Thu Jul 14 01:59:17 CEST 2022 - The aliscore output appears to be empty. Check results for BUSCO: 192368at7742
## Thu Jul 14 01:59:17 CEST 2022 - The aliscore output file does not exist. Check results for BUSCO: 277524at7742
## Thu Jul 14 01:59:17 CEST 2022 - The aliscore output appears to be empty. Check results for BUSCO: 277524at7742
## Thu Jul 14 01:59:19 CEST 2022 - The aliscore output file does not exist. Check results for BUSCO: 185249at7742
## Thu Jul 14 01:59:19 CEST 2022 - The aliscore output file does not exist. Check results for BUSCO: 166722at7742
## Thu Jul 14 01:59:19 CEST 2022 - The aliscore output appears to be empty. Check results for BUSCO: 185249at7742
## Thu Jul 14 01:59:19 CEST 2022 - The aliscore output appears to be empty. Check results for BUSCO: 166722at7742
## Thu Jul 14 01:59:20 CEST 2022 - The aliscore output file does not exist. Check results for BUSCO: 279227at7742
## Thu Jul 14 01:59:20 CEST 2022 - The aliscore output appears to be empty. Check results for BUSCO: 279227at7742
## Thu Jul 14 01:59:21 CEST 2022 - The aliscore output file does not exist. Check results for BUSCO: 381011at7742
## Thu Jul 14 01:59:21 CEST 2022 - The aliscore output appears to be empty. Check results for BUSCO: 381011at7742
## Thu Jul 14 01:59:23 CEST 2022 - The aliscore output file does not exist. Check results for BUSCO: 262221at7742
## Thu Jul 14 01:59:23 CEST 2022 - The aliscore output appears to be empty. Check results for BUSCO: 262221at7742
## Thu Jul 14 01:59:23 CEST 2022 - The aliscore output file does not exist. Check results for BUSCO: 184707at7742
## Thu Jul 14 01:59:23 CEST 2022 - The aliscore output appears to be empty. Check results for BUSCO: 184707at7742
## Thu Jul 14 01:59:24 CEST 2022 - The aliscore output file does not exist. Check results for BUSCO: 264251at7742
## Thu Jul 14 01:59:24 CEST 2022 - The aliscore output appears to be empty. Check results for BUSCO: 264251at7742
## Thu Jul 14 01:59:26 CEST 2022 - The aliscore output file does not exist. Check results for BUSCO: 232734at7742
## Thu Jul 14 01:59:26 CEST 2022 - The aliscore output appears to be empty. Check results for BUSCO: 232734at7742
## Thu Jul 14 01:59:27 CEST 2022 - The aliscore output file does not exist. Check results for BUSCO: 232734at7742
## Thu Jul 14 01:59:27 CEST 2022 - The aliscore output appears to be empty. Check results for BUSCO: 232734at7742
## Thu Jul 14 01:59:28 CEST 2022 - The aliscore output file does not exist. Check results for BUSCO: 312288at7742
## Thu Jul 14 01:59:28 CEST 2022 - The aliscore output appears to be empty. Check results for BUSCO: 312288at7742
## Thu Jul 14 01:59:30 CEST 2022 - The aliscore output file does not exist. Check results for BUSCO: 240011at7742
## Thu Jul 14 01:59:30 CEST 2022 - The aliscore output appears to be empty. Check results for BUSCO: 240011at7742
## Thu Jul 14 01:59:30 CEST 2022 - The aliscore output file does not exist. Check results for BUSCO: 330348at7742
## Thu Jul 14 01:59:30 CEST 2022 - The aliscore output appears to be empty. Check results for BUSCO: 330348at7742
## Thu Jul 14 01:59:31 CEST 2022 - The aliscore output file does not exist. Check results for BUSCO: 205317at7742
## Thu Jul 14 01:59:31 CEST 2022 - The aliscore output appears to be empty. Check results for BUSCO: 205317at7742
## Thu Jul 14 01:59:31 CEST 2022 - The aliscore output file does not exist. Check results for BUSCO: 186913at7742
## Thu Jul 14 01:59:31 CEST 2022 - The aliscore output appears to be empty. Check results for BUSCO: 186913at7742
## Thu Jul 14 01:59:33 CEST 2022 - The aliscore output file does not exist. Check results for BUSCO: 252546at7742
## Thu Jul 14 01:59:33 CEST 2022 - The aliscore output appears to be empty. Check results for BUSCO: 252546at7742
## Thu Jul 14 01:59:33 CEST 2022 - The aliscore output file does not exist. Check results for BUSCO: 281421at7742
## Thu Jul 14 01:59:33 CEST 2022 - The aliscore output appears to be empty. Check results for BUSCO: 281421at7742
## Thu Jul 14 01:59:35 CEST 2022 - The aliscore output file does not exist. Check results for BUSCO: 264251at7742
## Thu Jul 14 01:59:35 CEST 2022 - The aliscore output appears to be empty. Check results for BUSCO: 264251at7742
## Thu Jul 14 01:59:37 CEST 2022 - The aliscore output file does not exist. Check results for BUSCO: 286081at7742
## Thu Jul 14 01:59:37 CEST 2022 - The aliscore output appears to be empty. Check results for BUSCO: 286081at7742
## Thu Jul 14 01:59:37 CEST 2022 - The aliscore output file does not exist. Check results for BUSCO: 258381at7742
## Thu Jul 14 01:59:37 CEST 2022 - The aliscore output appears to be empty. Check results for BUSCO: 258381at7742
## Thu Jul 14 01:59:38 CEST 2022 - The aliscore output file does not exist. Check results for BUSCO: 260439at7742
## Thu Jul 14 01:59:38 CEST 2022 - The aliscore output appears to be empty. Check results for BUSCO: 260439at7742
## Thu Jul 14 01:59:39 CEST 2022 - The aliscore output file does not exist. Check results for BUSCO: 243425at7742
## Thu Jul 14 01:59:39 CEST 2022 - The aliscore output appears to be empty. Check results for BUSCO: 243425at7742
## Thu Jul 14 01:59:39 CEST 2022 - The aliscore output file does not exist. Check results for BUSCO: 307158at7742
## Thu Jul 14 01:59:39 CEST 2022 - The aliscore output appears to be empty. Check results for BUSCO: 307158at7742
## Thu Jul 14 01:59:40 CEST 2022 - The aliscore output file does not exist. Check results for BUSCO: 377163at7742
## Thu Jul 14 01:59:41 CEST 2022 - The aliscore output appears to be empty. Check results for BUSCO: 377163at7742
## Thu Jul 14 01:59:42 CEST 2022 - The aliscore output file does not exist. Check results for BUSCO: 244272at7742
## Thu Jul 14 01:59:42 CEST 2022 - The aliscore output appears to be empty. Check results for BUSCO: 244272at7742
## Thu Jul 14 01:59:44 CEST 2022 - The aliscore output file does not exist. Check results for BUSCO: 161959at7742
## Thu Jul 14 01:59:44 CEST 2022 - The aliscore output appears to be empty. Check results for BUSCO: 161959at7742
## Thu Jul 14 01:59:45 CEST 2022 - The aliscore output file does not exist. Check results for BUSCO: 281421at7742
## Thu Jul 14 01:59:45 CEST 2022 - The aliscore output appears to be empty. Check results for BUSCO: 281421at7742
## Thu Jul 14 01:59:45 CEST 2022 - The aliscore output file does not exist. Check results for BUSCO: 407084at7742
## Thu Jul 14 01:59:45 CEST 2022 - The aliscore output appears to be empty. Check results for BUSCO: 407084at7742
## Thu Jul 14 01:59:47 CEST 2022 - The aliscore output file does not exist. Check results for BUSCO: 244272at7742
## Thu Jul 14 01:59:47 CEST 2022 - The aliscore output appears to be empty. Check results for BUSCO: 244272at7742
## Thu Jul 14 01:59:47 CEST 2022 - The aliscore output file does not exist. Check results for BUSCO: 209931at7742
## Thu Jul 14 01:59:47 CEST 2022 - The aliscore output appears to be empty. Check results for BUSCO: 209931at7742
## Thu Jul 14 01:59:49 CEST 2022 - The aliscore output file does not exist. Check results for BUSCO: 403876at7742
## Thu Jul 14 01:59:49 CEST 2022 - The aliscore output appears to be empty. Check results for BUSCO: 403876at7742
## Thu Jul 14 01:59:49 CEST 2022 - The aliscore output file does not exist. Check results for BUSCO: 183663at7742
## Thu Jul 14 01:59:49 CEST 2022 - The aliscore output appears to be empty. Check results for BUSCO: 183663at7742
## Thu Jul 14 01:59:50 CEST 2022 - The aliscore output file does not exist. Check results for BUSCO: 322275at7742
## Thu Jul 14 01:59:50 CEST 2022 - The aliscore output appears to be empty. Check results for BUSCO: 322275at7742
## Thu Jul 14 01:59:51 CEST 2022 - The aliscore output file does not exist. Check results for BUSCO: 157219at7742
## Thu Jul 14 01:59:52 CEST 2022 - The aliscore output appears to be empty. Check results for BUSCO: 157219at7742
## Thu Jul 14 01:59:52 CEST 2022 - The aliscore output file does not exist. Check results for BUSCO: 309958at7742
## Thu Jul 14 01:59:52 CEST 2022 - The aliscore output appears to be empty. Check results for BUSCO: 309958at7742
## Thu Jul 14 01:59:53 CEST 2022 - The aliscore output file does not exist. Check results for BUSCO: 409719at7742
## Thu Jul 14 01:59:53 CEST 2022 - The aliscore output appears to be empty. Check results for BUSCO: 409719at7742
## Thu Jul 14 01:59:53 CEST 2022 - The aliscore output file does not exist. Check results for BUSCO: 173682at7742
## Thu Jul 14 01:59:53 CEST 2022 - The aliscore output appears to be empty. Check results for BUSCO: 173682at7742
## Thu Jul 14 01:59:56 CEST 2022 - The aliscore output file does not exist. Check results for BUSCO: 291229at7742
## Thu Jul 14 01:59:56 CEST 2022 - The aliscore output appears to be empty. Check results for BUSCO: 291229at7742
## Thu Jul 14 01:59:56 CEST 2022 - The aliscore output file does not exist. Check results for BUSCO: 119219at7742
## Thu Jul 14 01:59:57 CEST 2022 - The aliscore output appears to be empty. Check results for BUSCO: 119219at7742
## Thu Jul 14 01:59:58 CEST 2022 - The aliscore output file does not exist. Check results for BUSCO: 307554at7742
## Thu Jul 14 01:59:58 CEST 2022 - The aliscore output appears to be empty. Check results for BUSCO: 307554at7742
## Thu Jul 14 01:59:58 CEST 2022 - The aliscore output file does not exist. Check results for BUSCO: 345746at7742
## Thu Jul 14 01:59:58 CEST 2022 - The aliscore output appears to be empty. Check results for BUSCO: 345746at7742
## Thu Jul 14 01:59:59 CEST 2022 - The aliscore output file does not exist. Check results for BUSCO: 177937at7742
## Thu Jul 14 01:59:59 CEST 2022 - The aliscore output appears to be empty. Check results for BUSCO: 177937at7742
## Thu Jul 14 02:00:00 CEST 2022 - The aliscore output file does not exist. Check results for BUSCO: 148962at7742
## Thu Jul 14 02:00:00 CEST 2022 - The aliscore output appears to be empty. Check results for BUSCO: 148962at7742
## Thu Jul 14 02:00:02 CEST 2022 - The aliscore output file does not exist. Check results for BUSCO: 296065at7742
## Thu Jul 14 02:00:02 CEST 2022 - The aliscore output file does not exist. Check results for BUSCO: 239538at7742
## Thu Jul 14 02:00:02 CEST 2022 - The aliscore output appears to be empty. Check results for BUSCO: 296065at7742
## Thu Jul 14 02:00:02 CEST 2022 - The aliscore output appears to be empty. Check results for BUSCO: 239538at7742
## Thu Jul 14 02:00:03 CEST 2022 - The aliscore output file does not exist. Check results for BUSCO: 349840at7742
## Thu Jul 14 02:00:03 CEST 2022 - The aliscore output appears to be empty. Check results for BUSCO: 349840at7742
## Thu Jul 14 02:00:03 CEST 2022 - The aliscore output file does not exist. Check results for BUSCO: 317891at7742
## Thu Jul 14 02:00:04 CEST 2022 - The aliscore output appears to be empty. Check results for BUSCO: 317891at7742
## Thu Jul 14 02:00:05 CEST 2022 - The aliscore output file does not exist. Check results for BUSCO: 211818at7742
## Thu Jul 14 02:00:05 CEST 2022 - The aliscore output appears to be empty. Check results for BUSCO: 211818at7742
## Thu Jul 14 02:00:06 CEST 2022 - The aliscore output file does not exist. Check results for BUSCO: 328582at7742
## Thu Jul 14 02:00:06 CEST 2022 - The aliscore output appears to be empty. Check results for BUSCO: 328582at7742
## Thu Jul 14 02:00:07 CEST 2022 - The aliscore output file does not exist. Check results for BUSCO: 323076at7742
## Thu Jul 14 02:00:07 CEST 2022 - The aliscore output appears to be empty. Check results for BUSCO: 323076at7742
## Thu Jul 14 02:00:07 CEST 2022 - The aliscore output file does not exist. Check results for BUSCO: 177937at7742
## Thu Jul 14 02:00:07 CEST 2022 - The aliscore output appears to be empty. Check results for BUSCO: 177937at7742
## Thu Jul 14 02:00:10 CEST 2022 - The aliscore output file does not exist. Check results for BUSCO: 273485at7742
## Thu Jul 14 02:00:10 CEST 2022 - The aliscore output file does not exist. Check results for BUSCO: 167517at7742
## Thu Jul 14 02:00:10 CEST 2022 - The aliscore output appears to be empty. Check results for BUSCO: 273485at7742
## Thu Jul 14 02:00:10 CEST 2022 - The aliscore output appears to be empty. Check results for BUSCO: 167517at7742
## Thu Jul 14 02:00:12 CEST 2022 - The aliscore output file does not exist. Check results for BUSCO: 398926at7742
## Thu Jul 14 02:00:12 CEST 2022 - The aliscore output appears to be empty. Check results for BUSCO: 398926at7742
## Thu Jul 14 02:00:12 CEST 2022 - The aliscore output file does not exist. Check results for BUSCO: 309592at7742
## Thu Jul 14 02:00:12 CEST 2022 - The aliscore output appears to be empty. Check results for BUSCO: 309592at7742
## Thu Jul 14 02:00:13 CEST 2022 - The aliscore output file does not exist. Check results for BUSCO: 258381at7742
## Thu Jul 14 02:00:13 CEST 2022 - The aliscore output file does not exist. Check results for BUSCO: 332227at7742
## Thu Jul 14 02:00:13 CEST 2022 - The aliscore output appears to be empty. Check results for BUSCO: 258381at7742
## Thu Jul 14 02:00:13 CEST 2022 - The aliscore output appears to be empty. Check results for BUSCO: 332227at7742
## Thu Jul 14 02:00:14 CEST 2022 - The aliscore output file does not exist. Check results for BUSCO: 328109at7742
## Thu Jul 14 02:00:14 CEST 2022 - The aliscore output appears to be empty. Check results for BUSCO: 328109at7742
## Thu Jul 14 02:00:17 CEST 2022 - The aliscore output file does not exist. Check results for BUSCO: 248163at7742
## Thu Jul 14 02:00:17 CEST 2022 - The aliscore output appears to be empty. Check results for BUSCO: 248163at7742
## Thu Jul 14 02:00:18 CEST 2022 - The aliscore output file does not exist. Check results for BUSCO: 173797at7742
## Thu Jul 14 02:00:18 CEST 2022 - The aliscore output appears to be empty. Check results for BUSCO: 173797at7742
## Thu Jul 14 02:00:19 CEST 2022 - The aliscore output file does not exist. Check results for BUSCO: 198672at7742
## Thu Jul 14 02:00:20 CEST 2022 - The aliscore output appears to be empty. Check results for BUSCO: 198672at7742
## Thu Jul 14 02:00:20 CEST 2022 - The aliscore output file does not exist. Check results for BUSCO: 240011at7742
## Thu Jul 14 02:00:20 CEST 2022 - The aliscore output appears to be empty. Check results for BUSCO: 240011at7742
## Thu Jul 14 02:00:20 CEST 2022 - The aliscore output file does not exist. Check results for BUSCO: 228525at7742
## Thu Jul 14 02:00:20 CEST 2022 - The aliscore output appears to be empty. Check results for BUSCO: 228525at7742
## Thu Jul 14 02:00:21 CEST 2022 - The aliscore output file does not exist. Check results for BUSCO: 266228at7742
## Thu Jul 14 02:00:21 CEST 2022 - The aliscore output appears to be empty. Check results for BUSCO: 266228at7742
## Thu Jul 14 02:00:23 CEST 2022 - The aliscore output file does not exist. Check results for BUSCO: 421394at7742
## Thu Jul 14 02:00:23 CEST 2022 - The aliscore output appears to be empty. Check results for BUSCO: 421394at7742
## Thu Jul 14 02:00:24 CEST 2022 - The aliscore output file does not exist. Check results for BUSCO: 328582at7742
## Thu Jul 14 02:00:24 CEST 2022 - The aliscore output appears to be empty. Check results for BUSCO: 328582at7742
## Thu Jul 14 02:00:25 CEST 2022 - The aliscore output file does not exist. Check results for BUSCO: 198672at7742
## Thu Jul 14 02:00:25 CEST 2022 - The aliscore output appears to be empty. Check results for BUSCO: 198672at7742
## Thu Jul 14 02:00:26 CEST 2022 - The aliscore output file does not exist. Check results for BUSCO: 287042at7742
## Thu Jul 14 02:00:26 CEST 2022 - The aliscore output appears to be empty. Check results for BUSCO: 287042at7742
## Thu Jul 14 02:00:26 CEST 2022 - The aliscore output file does not exist. Check results for BUSCO: 266228at7742
## Thu Jul 14 02:00:26 CEST 2022 - The aliscore output appears to be empty. Check results for BUSCO: 266228at7742
## Thu Jul 14 02:00:27 CEST 2022 - The aliscore output file does not exist. Check results for BUSCO: 370499at7742
## Thu Jul 14 02:00:27 CEST 2022 - The aliscore output appears to be empty. Check results for BUSCO: 370499at7742
## Thu Jul 14 02:00:28 CEST 2022 - The aliscore output file does not exist. Check results for BUSCO: 169787at7742
## Thu Jul 14 02:00:28 CEST 2022 - The aliscore output appears to be empty. Check results for BUSCO: 169787at7742
## Thu Jul 14 02:00:30 CEST 2022 - The aliscore output file does not exist. Check results for BUSCO: 318959at7742
## Thu Jul 14 02:00:30 CEST 2022 - The aliscore output appears to be empty. Check results for BUSCO: 318959at7742
## Thu Jul 14 02:00:31 CEST 2022 - The aliscore output file does not exist. Check results for BUSCO: 157219at7742
## Thu Jul 14 02:00:31 CEST 2022 - The aliscore output appears to be empty. Check results for BUSCO: 157219at7742
## Thu Jul 14 02:00:32 CEST 2022 - The aliscore output file does not exist. Check results for BUSCO: 301292at7742
## Thu Jul 14 02:00:32 CEST 2022 - The aliscore output appears to be empty. Check results for BUSCO: 301292at7742
## Thu Jul 14 02:00:33 CEST 2022 - The aliscore output file does not exist. Check results for BUSCO: 254847at7742
## Thu Jul 14 02:00:33 CEST 2022 - The aliscore output appears to be empty. Check results for BUSCO: 254847at7742
## Thu Jul 14 02:00:33 CEST 2022 - The aliscore output file does not exist. Check results for BUSCO: 192368at7742
## Thu Jul 14 02:00:33 CEST 2022 - The aliscore output appears to be empty. Check results for BUSCO: 192368at7742
## Thu Jul 14 02:00:33 CEST 2022 - The aliscore output file does not exist. Check results for BUSCO: 128421at7742
## Thu Jul 14 02:00:33 CEST 2022 - The aliscore output appears to be empty. Check results for BUSCO: 128421at7742
## Thu Jul 14 02:00:35 CEST 2022 - The aliscore output file does not exist. Check results for BUSCO: 248028at7742
## Thu Jul 14 02:00:35 CEST 2022 - The aliscore output appears to be empty. Check results for BUSCO: 248028at7742
## Thu Jul 14 02:00:37 CEST 2022 - The aliscore output file does not exist. Check results for BUSCO: 219315at7742
## Thu Jul 14 02:00:37 CEST 2022 - The aliscore output appears to be empty. Check results for BUSCO: 219315at7742
## Thu Jul 14 02:00:37 CEST 2022 - The aliscore output file does not exist. Check results for BUSCO: 374996at7742
## Thu Jul 14 02:00:37 CEST 2022 - The aliscore output appears to be empty. Check results for BUSCO: 374996at7742
## Thu Jul 14 02:00:39 CEST 2022 - The aliscore output file does not exist. Check results for BUSCO: 207153at7742
## Thu Jul 14 02:00:39 CEST 2022 - The aliscore output appears to be empty. Check results for BUSCO: 207153at7742
## Thu Jul 14 02:00:40 CEST 2022 - The aliscore output file does not exist. Check results for BUSCO: 213286at7742
## Thu Jul 14 02:00:40 CEST 2022 - The aliscore output appears to be empty. Check results for BUSCO: 213286at7742
## Thu Jul 14 02:00:41 CEST 2022 - The aliscore output file does not exist. Check results for BUSCO: 248163at7742
## Thu Jul 14 02:00:41 CEST 2022 - The aliscore output appears to be empty. Check results for BUSCO: 248163at7742
## Thu Jul 14 02:00:42 CEST 2022 - The aliscore output file does not exist. Check results for BUSCO: 277217at7742
## Thu Jul 14 02:00:42 CEST 2022 - The aliscore output appears to be empty. Check results for BUSCO: 277217at7742
## Thu Jul 14 02:00:42 CEST 2022 - The aliscore output file does not exist. Check results for BUSCO: 218430at7742
## Thu Jul 14 02:00:42 CEST 2022 - The aliscore output appears to be empty. Check results for BUSCO: 218430at7742
## Thu Jul 14 02:00:44 CEST 2022 - The aliscore output file does not exist. Check results for BUSCO: 351419at7742
## Thu Jul 14 02:00:44 CEST 2022 - The aliscore output appears to be empty. Check results for BUSCO: 351419at7742
## Thu Jul 14 02:00:44 CEST 2022 - The aliscore output file does not exist. Check results for BUSCO: 214858at7742
## Thu Jul 14 02:00:44 CEST 2022 - The aliscore output appears to be empty. Check results for BUSCO: 214858at7742
## Thu Jul 14 02:00:46 CEST 2022 - The aliscore output file does not exist. Check results for BUSCO: 214858at7742
## Thu Jul 14 02:00:46 CEST 2022 - The aliscore output appears to be empty. Check results for BUSCO: 214858at7742
## Thu Jul 14 02:00:46 CEST 2022 - The aliscore output file does not exist. Check results for BUSCO: 268484at7742
## Thu Jul 14 02:00:46 CEST 2022 - The aliscore output appears to be empty. Check results for BUSCO: 268484at7742
## Thu Jul 14 02:00:47 CEST 2022 - The aliscore output file does not exist. Check results for BUSCO: 209931at7742
## Thu Jul 14 02:00:47 CEST 2022 - The aliscore output appears to be empty. Check results for BUSCO: 209931at7742
## Thu Jul 14 02:00:48 CEST 2022 - The aliscore output file does not exist. Check results for BUSCO: 252546at7742
## Thu Jul 14 02:00:48 CEST 2022 - The aliscore output appears to be empty. Check results for BUSCO: 252546at7742
## Thu Jul 14 02:00:49 CEST 2022 - The aliscore output file does not exist. Check results for BUSCO: 365226at7742
## Thu Jul 14 02:00:49 CEST 2022 - The aliscore output appears to be empty. Check results for BUSCO: 365226at7742
## Thu Jul 14 02:00:51 CEST 2022 - The aliscore output file does not exist. Check results for BUSCO: 241288at7742
## Thu Jul 14 02:00:51 CEST 2022 - The aliscore output appears to be empty. Check results for BUSCO: 241288at7742
## Thu Jul 14 02:00:52 CEST 2022 - The aliscore output file does not exist. Check results for BUSCO: 192451at7742
## Thu Jul 14 02:00:52 CEST 2022 - The aliscore output appears to be empty. Check results for BUSCO: 192451at7742
## Thu Jul 14 02:00:52 CEST 2022 - The aliscore output file does not exist. Check results for BUSCO: 399601at7742
## Thu Jul 14 02:00:52 CEST 2022 - The aliscore output appears to be empty. Check results for BUSCO: 399601at7742
## Thu Jul 14 02:00:53 CEST 2022 - The aliscore output file does not exist. Check results for BUSCO: 288248at7742
## Thu Jul 14 02:00:53 CEST 2022 - The aliscore output appears to be empty. Check results for BUSCO: 288248at7742
## Thu Jul 14 02:00:54 CEST 2022 - The aliscore output file does not exist. Check results for BUSCO: 241887at7742
## Thu Jul 14 02:00:54 CEST 2022 - The aliscore output appears to be empty. Check results for BUSCO: 241887at7742
## Thu Jul 14 02:00:55 CEST 2022 - The aliscore output file does not exist. Check results for BUSCO: 267296at7742
## Thu Jul 14 02:00:55 CEST 2022 - The aliscore output appears to be empty. Check results for BUSCO: 267296at7742
## Thu Jul 14 02:00:57 CEST 2022 - The aliscore output file does not exist. Check results for BUSCO: 206784at7742
## Thu Jul 14 02:00:57 CEST 2022 - The aliscore output appears to be empty. Check results for BUSCO: 206784at7742
## Thu Jul 14 02:00:57 CEST 2022 - The aliscore output file does not exist. Check results for BUSCO: 276691at7742
## Thu Jul 14 02:00:57 CEST 2022 - The aliscore output appears to be empty. Check results for BUSCO: 276691at7742
## Thu Jul 14 02:00:59 CEST 2022 - The aliscore output file does not exist. Check results for BUSCO: 332036at7742
## Thu Jul 14 02:00:59 CEST 2022 - The aliscore output appears to be empty. Check results for BUSCO: 332036at7742
## Thu Jul 14 02:00:59 CEST 2022 - The aliscore output file does not exist. Check results for BUSCO: 211012at7742
## Thu Jul 14 02:00:59 CEST 2022 - The aliscore output appears to be empty. Check results for BUSCO: 211012at7742
## Thu Jul 14 02:01:00 CEST 2022 - The aliscore output file does not exist. Check results for BUSCO: 237266at7742
## Thu Jul 14 02:01:00 CEST 2022 - The aliscore output appears to be empty. Check results for BUSCO: 237266at7742
## Thu Jul 14 02:01:01 CEST 2022 - The aliscore output file does not exist. Check results for BUSCO: 173797at7742
## Thu Jul 14 02:01:01 CEST 2022 - The aliscore output appears to be empty. Check results for BUSCO: 173797at7742
## Thu Jul 14 02:01:02 CEST 2022 - The aliscore output file does not exist. Check results for BUSCO: 268484at7742
## Thu Jul 14 02:01:02 CEST 2022 - The aliscore output appears to be empty. Check results for BUSCO: 268484at7742
## Thu Jul 14 02:01:03 CEST 2022 - The aliscore output file does not exist. Check results for BUSCO: 381011at7742
## Thu Jul 14 02:01:03 CEST 2022 - The aliscore output appears to be empty. Check results for BUSCO: 381011at7742
## Thu Jul 14 02:01:03 CEST 2022 - The aliscore output file does not exist. Check results for BUSCO: 268926at7742
## Thu Jul 14 02:01:03 CEST 2022 - The aliscore output appears to be empty. Check results for BUSCO: 268926at7742
## Thu Jul 14 02:01:06 CEST 2022 - The aliscore output file does not exist. Check results for BUSCO: 209386at7742
## Thu Jul 14 02:01:06 CEST 2022 - The aliscore output appears to be empty. Check results for BUSCO: 209386at7742
## Thu Jul 14 02:01:06 CEST 2022 - The aliscore output file does not exist. Check results for BUSCO: 317161at7742
## Thu Jul 14 02:01:06 CEST 2022 - The aliscore output appears to be empty. Check results for BUSCO: 317161at7742
## Thu Jul 14 02:01:08 CEST 2022 - The aliscore output file does not exist. Check results for BUSCO: 211818at7742
## Thu Jul 14 02:01:08 CEST 2022 - The aliscore output appears to be empty. Check results for BUSCO: 211818at7742
## Thu Jul 14 02:01:09 CEST 2022 - The aliscore output file does not exist. Check results for BUSCO: 303510at7742
## Thu Jul 14 02:01:09 CEST 2022 - The aliscore output appears to be empty. Check results for BUSCO: 303510at7742
## Thu Jul 14 02:01:10 CEST 2022 - The aliscore output file does not exist. Check results for BUSCO: 151078at7742
## Thu Jul 14 02:01:10 CEST 2022 - The aliscore output appears to be empty. Check results for BUSCO: 151078at7742
## Thu Jul 14 02:01:10 CEST 2022 - The aliscore output file does not exist. Check results for BUSCO: 345549at7742
## Thu Jul 14 02:01:11 CEST 2022 - The aliscore output appears to be empty. Check results for BUSCO: 345549at7742
## Thu Jul 14 02:01:12 CEST 2022 - The aliscore output file does not exist. Check results for BUSCO: 348500at7742
## Thu Jul 14 02:01:12 CEST 2022 - The aliscore output appears to be empty. Check results for BUSCO: 348500at7742
## Thu Jul 14 02:01:13 CEST 2022 - The aliscore output file does not exist. Check results for BUSCO: 166429at7742
## Thu Jul 14 02:01:13 CEST 2022 - The aliscore output appears to be empty. Check results for BUSCO: 166429at7742
## Thu Jul 14 02:01:13 CEST 2022 - The aliscore output file does not exist. Check results for BUSCO: 377177at7742
## Thu Jul 14 02:01:13 CEST 2022 - The aliscore output appears to be empty. Check results for BUSCO: 377177at7742
## Thu Jul 14 02:01:14 CEST 2022 - The aliscore output file does not exist. Check results for BUSCO: 236417at7742
## Thu Jul 14 02:01:14 CEST 2022 - The aliscore output appears to be empty. Check results for BUSCO: 236417at7742
## Thu Jul 14 02:01:16 CEST 2022 - The aliscore output file does not exist. Check results for BUSCO: 214136at7742
## Thu Jul 14 02:01:16 CEST 2022 - The aliscore output appears to be empty. Check results for BUSCO: 214136at7742
## Thu Jul 14 02:01:16 CEST 2022 - The aliscore output file does not exist. Check results for BUSCO: 380303at7742
## Thu Jul 14 02:01:16 CEST 2022 - The aliscore output appears to be empty. Check results for BUSCO: 380303at7742
## Thu Jul 14 02:01:18 CEST 2022 - The aliscore output file does not exist. Check results for BUSCO: 173682at7742
## Thu Jul 14 02:01:18 CEST 2022 - The aliscore output appears to be empty. Check results for BUSCO: 173682at7742
## Thu Jul 14 02:01:19 CEST 2022 - The aliscore output file does not exist. Check results for BUSCO: 202013at7742
## Thu Jul 14 02:01:19 CEST 2022 - The aliscore output appears to be empty. Check results for BUSCO: 202013at7742
## Thu Jul 14 02:01:20 CEST 2022 - The aliscore output file does not exist. Check results for BUSCO: 318704at7742
## Thu Jul 14 02:01:20 CEST 2022 - The aliscore output appears to be empty. Check results for BUSCO: 318704at7742
## Thu Jul 14 02:01:22 CEST 2022 - The aliscore output file does not exist. Check results for BUSCO: 263794at7742
## Thu Jul 14 02:01:22 CEST 2022 - The aliscore output appears to be empty. Check results for BUSCO: 263794at7742
## Thu Jul 14 02:01:23 CEST 2022 - The aliscore output file does not exist. Check results for BUSCO: 333162at7742
## Thu Jul 14 02:01:23 CEST 2022 - The aliscore output appears to be empty. Check results for BUSCO: 333162at7742
## Thu Jul 14 02:01:24 CEST 2022 - The aliscore output file does not exist. Check results for BUSCO: 261976at7742
## Thu Jul 14 02:01:24 CEST 2022 - The aliscore output appears to be empty. Check results for BUSCO: 261976at7742
## Thu Jul 14 02:01:25 CEST 2022 - The aliscore output file does not exist. Check results for BUSCO: 149872at7742
## Thu Jul 14 02:01:25 CEST 2022 - The aliscore output appears to be empty. Check results for BUSCO: 149872at7742
## Thu Jul 14 02:01:26 CEST 2022 - The aliscore output file does not exist. Check results for BUSCO: 186469at7742
## Thu Jul 14 02:01:26 CEST 2022 - The aliscore output appears to be empty. Check results for BUSCO: 186469at7742
## Thu Jul 14 02:01:27 CEST 2022 - The aliscore output file does not exist. Check results for BUSCO: 266056at7742
## Thu Jul 14 02:01:27 CEST 2022 - The aliscore output appears to be empty. Check results for BUSCO: 266056at7742
## Thu Jul 14 02:01:27 CEST 2022 - The aliscore output file does not exist. Check results for BUSCO: 154805at7742
## Thu Jul 14 02:01:27 CEST 2022 - The aliscore output appears to be empty. Check results for BUSCO: 154805at7742
## Thu Jul 14 02:01:29 CEST 2022 - The aliscore output file does not exist. Check results for BUSCO: 268274at7742
## Thu Jul 14 02:01:29 CEST 2022 - The aliscore output appears to be empty. Check results for BUSCO: 268274at7742
## Thu Jul 14 02:01:29 CEST 2022 - The aliscore output file does not exist. Check results for BUSCO: 378093at7742
## Thu Jul 14 02:01:29 CEST 2022 - The aliscore output appears to be empty. Check results for BUSCO: 378093at7742
## Thu Jul 14 02:01:30 CEST 2022 - The aliscore output file does not exist. Check results for BUSCO: 284666at7742
## Thu Jul 14 02:01:30 CEST 2022 - The aliscore output appears to be empty. Check results for BUSCO: 284666at7742
## Thu Jul 14 02:01:31 CEST 2022 - The aliscore output file does not exist. Check results for BUSCO: 253661at7742
## Thu Jul 14 02:01:31 CEST 2022 - The aliscore output appears to be empty. Check results for BUSCO: 253661at7742
## Thu Jul 14 02:01:32 CEST 2022 - The aliscore output file does not exist. Check results for BUSCO: 185249at7742
## Thu Jul 14 02:01:32 CEST 2022 - The aliscore output appears to be empty. Check results for BUSCO: 185249at7742
## Thu Jul 14 02:01:32 CEST 2022 - The aliscore output file does not exist. Check results for BUSCO: 307158at7742
## Thu Jul 14 02:01:32 CEST 2022 - The aliscore output appears to be empty. Check results for BUSCO: 307158at7742
## Thu Jul 14 02:01:34 CEST 2022 - The aliscore output file does not exist. Check results for BUSCO: 309592at7742
## Thu Jul 14 02:01:34 CEST 2022 - The aliscore output appears to be empty. Check results for BUSCO: 309592at7742
## Thu Jul 14 02:01:34 CEST 2022 - The aliscore output file does not exist. Check results for BUSCO: 277226at7742
## Thu Jul 14 02:01:34 CEST 2022 - The aliscore output appears to be empty. Check results for BUSCO: 277226at7742
## Thu Jul 14 02:01:36 CEST 2022 - The aliscore output file does not exist. Check results for BUSCO: 287466at7742
## Thu Jul 14 02:01:36 CEST 2022 - The aliscore output appears to be empty. Check results for BUSCO: 287466at7742
## Thu Jul 14 02:01:36 CEST 2022 - The aliscore output file does not exist. Check results for BUSCO: 281421at7742
## Thu Jul 14 02:01:36 CEST 2022 - The aliscore output appears to be empty. Check results for BUSCO: 281421at7742
## Thu Jul 14 02:01:38 CEST 2022 - The aliscore output file does not exist. Check results for BUSCO: 226799at7742
## Thu Jul 14 02:01:38 CEST 2022 - The aliscore output appears to be empty. Check results for BUSCO: 226799at7742
## Thu Jul 14 02:01:38 CEST 2022 - The aliscore output file does not exist. Check results for BUSCO: 168866at7742
## Thu Jul 14 02:01:38 CEST 2022 - The aliscore output appears to be empty. Check results for BUSCO: 168866at7742
## Thu Jul 14 02:01:40 CEST 2022 - The aliscore output file does not exist. Check results for BUSCO: 314402at7742
## Thu Jul 14 02:01:40 CEST 2022 - The aliscore output appears to be empty. Check results for BUSCO: 314402at7742
## Thu Jul 14 02:01:41 CEST 2022 - The aliscore output file does not exist. Check results for BUSCO: 294328at7742
## Thu Jul 14 02:01:41 CEST 2022 - The aliscore output appears to be empty. Check results for BUSCO: 294328at7742
## Thu Jul 14 02:01:41 CEST 2022 - The aliscore output file does not exist. Check results for BUSCO: 261091at7742
## Thu Jul 14 02:01:42 CEST 2022 - The aliscore output appears to be empty. Check results for BUSCO: 261091at7742
## Thu Jul 14 02:01:42 CEST 2022 - The aliscore output file does not exist. Check results for BUSCO: 294328at7742
## Thu Jul 14 02:01:42 CEST 2022 - The aliscore output appears to be empty. Check results for BUSCO: 294328at7742
## Thu Jul 14 02:01:43 CEST 2022 - The aliscore output file does not exist. Check results for BUSCO: 262915at7742
## Thu Jul 14 02:01:43 CEST 2022 - The aliscore output appears to be empty. Check results for BUSCO: 262915at7742
## Thu Jul 14 02:01:45 CEST 2022 - The aliscore output file does not exist. Check results for BUSCO: 307415at7742
## Thu Jul 14 02:01:45 CEST 2022 - The aliscore output appears to be empty. Check results for BUSCO: 307415at7742
## Thu Jul 14 02:01:46 CEST 2022 - The aliscore output file does not exist. Check results for BUSCO: 252307at7742
## Thu Jul 14 02:01:46 CEST 2022 - The aliscore output appears to be empty. Check results for BUSCO: 252307at7742
## Thu Jul 14 02:01:47 CEST 2022 - The aliscore output file does not exist. Check results for BUSCO: 277217at7742
## Thu Jul 14 02:01:47 CEST 2022 - The aliscore output appears to be empty. Check results for BUSCO: 277217at7742
## Thu Jul 14 02:01:48 CEST 2022 - The aliscore output file does not exist. Check results for BUSCO: 248163at7742
## Thu Jul 14 02:01:48 CEST 2022 - The aliscore output appears to be empty. Check results for BUSCO: 248163at7742
## Thu Jul 14 02:01:49 CEST 2022 - The aliscore output file does not exist. Check results for BUSCO: 209086at7742
## Thu Jul 14 02:01:49 CEST 2022 - The aliscore output appears to be empty. Check results for BUSCO: 209086at7742
## Thu Jul 14 02:01:49 CEST 2022 - The aliscore output file does not exist. Check results for BUSCO: 391055at7742
## Thu Jul 14 02:01:49 CEST 2022 - The aliscore output appears to be empty. Check results for BUSCO: 391055at7742
## Thu Jul 14 02:01:50 CEST 2022 - The aliscore output file does not exist. Check results for BUSCO: 293860at7742
## Thu Jul 14 02:01:50 CEST 2022 - The aliscore output appears to be empty. Check results for BUSCO: 293860at7742
## Thu Jul 14 02:01:52 CEST 2022 - The aliscore output file does not exist. Check results for BUSCO: 363898at7742
## Thu Jul 14 02:01:52 CEST 2022 - The aliscore output appears to be empty. Check results for BUSCO: 363898at7742
## Thu Jul 14 02:01:53 CEST 2022 - The aliscore output file does not exist. Check results for BUSCO: 322871at7742
## Thu Jul 14 02:01:53 CEST 2022 - The aliscore output appears to be empty. Check results for BUSCO: 322871at7742
## Thu Jul 14 02:01:53 CEST 2022 - The aliscore output file does not exist. Check results for BUSCO: 133692at7742
## Thu Jul 14 02:01:53 CEST 2022 - The aliscore output appears to be empty. Check results for BUSCO: 133692at7742
## Thu Jul 14 02:01:54 CEST 2022 - The aliscore output file does not exist. Check results for BUSCO: 240403at7742
## Thu Jul 14 02:01:54 CEST 2022 - The aliscore output appears to be empty. Check results for BUSCO: 240403at7742
## Thu Jul 14 02:01:56 CEST 2022 - The aliscore output file does not exist. Check results for BUSCO: 205317at7742
## Thu Jul 14 02:01:56 CEST 2022 - The aliscore output appears to be empty. Check results for BUSCO: 205317at7742
## Thu Jul 14 02:01:57 CEST 2022 - The aliscore output file does not exist. Check results for BUSCO: 261091at7742
## Thu Jul 14 02:01:57 CEST 2022 - The aliscore output appears to be empty. Check results for BUSCO: 261091at7742
## Thu Jul 14 02:01:58 CEST 2022 - The aliscore output file does not exist. Check results for BUSCO: 261684at7742
## Thu Jul 14 02:01:58 CEST 2022 - The aliscore output appears to be empty. Check results for BUSCO: 261684at7742
## Thu Jul 14 02:01:59 CEST 2022 - The aliscore output file does not exist. Check results for BUSCO: 214005at7742
## Thu Jul 14 02:01:59 CEST 2022 - The aliscore output appears to be empty. Check results for BUSCO: 214005at7742
## Thu Jul 14 02:01:59 CEST 2022 - The aliscore output file does not exist. Check results for BUSCO: 339203at7742
## Thu Jul 14 02:02:00 CEST 2022 - The aliscore output appears to be empty. Check results for BUSCO: 339203at7742
## Thu Jul 14 02:02:01 CEST 2022 - The aliscore output file does not exist. Check results for BUSCO: 376448at7742
## Thu Jul 14 02:02:01 CEST 2022 - The aliscore output appears to be empty. Check results for BUSCO: 376448at7742
## Thu Jul 14 02:02:02 CEST 2022 - The aliscore output file does not exist. Check results for BUSCO: 380483at7742
## Thu Jul 14 02:02:02 CEST 2022 - The aliscore output appears to be empty. Check results for BUSCO: 380483at7742
## Thu Jul 14 02:02:03 CEST 2022 - The aliscore output file does not exist. Check results for BUSCO: 195300at7742
## Thu Jul 14 02:02:03 CEST 2022 - The aliscore output appears to be empty. Check results for BUSCO: 195300at7742
## Thu Jul 14 02:02:04 CEST 2022 - The aliscore output file does not exist. Check results for BUSCO: 321295at7742
## Thu Jul 14 02:02:04 CEST 2022 - The aliscore output appears to be empty. Check results for BUSCO: 321295at7742
## Thu Jul 14 02:02:05 CEST 2022 - The aliscore output file does not exist. Check results for BUSCO: 307554at7742
## Thu Jul 14 02:02:05 CEST 2022 - The aliscore output appears to be empty. Check results for BUSCO: 307554at7742
## Thu Jul 14 02:02:06 CEST 2022 - The aliscore output file does not exist. Check results for BUSCO: 320621at7742
## Thu Jul 14 02:02:06 CEST 2022 - The aliscore output appears to be empty. Check results for BUSCO: 320621at7742
## Thu Jul 14 02:02:07 CEST 2022 - The aliscore output file does not exist. Check results for BUSCO: 241473at7742
## Thu Jul 14 02:02:07 CEST 2022 - The aliscore output appears to be empty. Check results for BUSCO: 241473at7742
## Thu Jul 14 02:02:08 CEST 2022 - The aliscore output file does not exist. Check results for BUSCO: 258381at7742
## Thu Jul 14 02:02:08 CEST 2022 - The aliscore output appears to be empty. Check results for BUSCO: 258381at7742
## Thu Jul 14 02:02:09 CEST 2022 - The aliscore output file does not exist. Check results for BUSCO: 166301at7742
## Thu Jul 14 02:02:09 CEST 2022 - The aliscore output appears to be empty. Check results for BUSCO: 166301at7742
## Thu Jul 14 02:02:12 CEST 2022 - The aliscore output file does not exist. Check results for BUSCO: 250442at7742
## Thu Jul 14 02:02:12 CEST 2022 - The aliscore output appears to be empty. Check results for BUSCO: 250442at7742
## Thu Jul 14 02:02:12 CEST 2022 - The aliscore output file does not exist. Check results for BUSCO: 250326at7742
## Thu Jul 14 02:02:12 CEST 2022 - The aliscore output appears to be empty. Check results for BUSCO: 250326at7742
## Thu Jul 14 02:02:12 CEST 2022 - The aliscore output file does not exist. Check results for BUSCO: 213372at7742
## Thu Jul 14 02:02:13 CEST 2022 - The aliscore output appears to be empty. Check results for BUSCO: 213372at7742
## Thu Jul 14 02:02:14 CEST 2022 - The aliscore output file does not exist. Check results for BUSCO: 192368at7742
## Thu Jul 14 02:02:14 CEST 2022 - The aliscore output appears to be empty. Check results for BUSCO: 192368at7742
## Thu Jul 14 02:02:14 CEST 2022 - The aliscore output file does not exist. Check results for BUSCO: 268926at7742
## Thu Jul 14 02:02:14 CEST 2022 - The aliscore output appears to be empty. Check results for BUSCO: 268926at7742
## Thu Jul 14 02:02:15 CEST 2022 - The aliscore output file does not exist. Check results for BUSCO: 238829at7742
## Thu Jul 14 02:02:15 CEST 2022 - The aliscore output appears to be empty. Check results for BUSCO: 238829at7742
## Thu Jul 14 02:02:16 CEST 2022 - The aliscore output file does not exist. Check results for BUSCO: 196649at7742
## Thu Jul 14 02:02:16 CEST 2022 - The aliscore output appears to be empty. Check results for BUSCO: 196649at7742
## Thu Jul 14 02:02:17 CEST 2022 - The aliscore output file does not exist. Check results for BUSCO: 293860at7742
## Thu Jul 14 02:02:17 CEST 2022 - The aliscore output appears to be empty. Check results for BUSCO: 293860at7742
## Thu Jul 14 02:02:18 CEST 2022 - The aliscore output file does not exist. Check results for BUSCO: 234511at7742
## Thu Jul 14 02:02:18 CEST 2022 - The aliscore output appears to be empty. Check results for BUSCO: 234511at7742
## Thu Jul 14 02:02:19 CEST 2022 - The aliscore output file does not exist. Check results for BUSCO: 234511at7742
## Thu Jul 14 02:02:19 CEST 2022 - The aliscore output appears to be empty. Check results for BUSCO: 234511at7742
## Thu Jul 14 02:02:20 CEST 2022 - The aliscore output file does not exist. Check results for BUSCO: 213350at7742
## Thu Jul 14 02:02:20 CEST 2022 - The aliscore output appears to be empty. Check results for BUSCO: 213350at7742
## Thu Jul 14 02:02:22 CEST 2022 - The aliscore output file does not exist. Check results for BUSCO: 218109at7742
## Thu Jul 14 02:02:22 CEST 2022 - The aliscore output appears to be empty. Check results for BUSCO: 218109at7742
## Thu Jul 14 02:02:22 CEST 2022 - The aliscore output file does not exist. Check results for BUSCO: 292437at7742
## Thu Jul 14 02:02:22 CEST 2022 - The aliscore output appears to be empty. Check results for BUSCO: 292437at7742
## Thu Jul 14 02:02:24 CEST 2022 - The aliscore output file does not exist. Check results for BUSCO: 228525at7742
## Thu Jul 14 02:02:24 CEST 2022 - The aliscore output appears to be empty. Check results for BUSCO: 228525at7742
## Thu Jul 14 02:02:24 CEST 2022 - The aliscore output file does not exist. Check results for BUSCO: 241887at7742
## Thu Jul 14 02:02:24 CEST 2022 - The aliscore output appears to be empty. Check results for BUSCO: 241887at7742
## Thu Jul 14 02:02:25 CEST 2022 - The aliscore output file does not exist. Check results for BUSCO: 173682at7742
## Thu Jul 14 02:02:25 CEST 2022 - The aliscore output appears to be empty. Check results for BUSCO: 173682at7742
## Thu Jul 14 02:02:26 CEST 2022 - The aliscore output file does not exist. Check results for BUSCO: 241473at7742
## Thu Jul 14 02:02:26 CEST 2022 - The aliscore output appears to be empty. Check results for BUSCO: 241473at7742
## Thu Jul 14 02:02:29 CEST 2022 - The aliscore output file does not exist. Check results for BUSCO: 248028at7742
## Thu Jul 14 02:02:29 CEST 2022 - The aliscore output file does not exist. Check results for BUSCO: 332227at7742
## Thu Jul 14 02:02:29 CEST 2022 - The aliscore output appears to be empty. Check results for BUSCO: 248028at7742
## Thu Jul 14 02:02:29 CEST 2022 - The aliscore output appears to be empty. Check results for BUSCO: 332227at7742
## Thu Jul 14 02:02:30 CEST 2022 - The aliscore output file does not exist. Check results for BUSCO: 234511at7742
## Thu Jul 14 02:02:30 CEST 2022 - The aliscore output appears to be empty. Check results for BUSCO: 234511at7742
## Thu Jul 14 02:02:31 CEST 2022 - The aliscore output file does not exist. Check results for BUSCO: 356998at7742
## Thu Jul 14 02:02:31 CEST 2022 - The aliscore output appears to be empty. Check results for BUSCO: 356998at7742
## Thu Jul 14 02:02:32 CEST 2022 - The aliscore output file does not exist. Check results for BUSCO: 277217at7742
## Thu Jul 14 02:02:32 CEST 2022 - The aliscore output appears to be empty. Check results for BUSCO: 277217at7742
## Thu Jul 14 02:02:33 CEST 2022 - The aliscore output file does not exist. Check results for BUSCO: 205317at7742
## Thu Jul 14 02:02:33 CEST 2022 - The aliscore output appears to be empty. Check results for BUSCO: 205317at7742
## Thu Jul 14 02:02:35 CEST 2022 - The aliscore output file does not exist. Check results for BUSCO: 243425at7742
## Thu Jul 14 02:02:35 CEST 2022 - The aliscore output appears to be empty. Check results for BUSCO: 243425at7742
## Thu Jul 14 02:02:35 CEST 2022 - The aliscore output file does not exist. Check results for BUSCO: 228525at7742
## Thu Jul 14 02:02:35 CEST 2022 - The aliscore output appears to be empty. Check results for BUSCO: 228525at7742
## Thu Jul 14 02:02:37 CEST 2022 - The aliscore output file does not exist. Check results for BUSCO: 297409at7742
## Thu Jul 14 02:02:37 CEST 2022 - The aliscore output appears to be empty. Check results for BUSCO: 297409at7742
## Thu Jul 14 02:02:38 CEST 2022 - The aliscore output file does not exist. Check results for BUSCO: 273485at7742
## Thu Jul 14 02:02:38 CEST 2022 - The aliscore output appears to be empty. Check results for BUSCO: 273485at7742
## Thu Jul 14 02:02:38 CEST 2022 - The aliscore output file does not exist. Check results for BUSCO: 297362at7742
## Thu Jul 14 02:02:38 CEST 2022 - The aliscore output appears to be empty. Check results for BUSCO: 297362at7742
## Thu Jul 14 02:02:40 CEST 2022 - The aliscore output file does not exist. Check results for BUSCO: 364320at7742
## Thu Jul 14 02:02:40 CEST 2022 - The aliscore output appears to be empty. Check results for BUSCO: 364320at7742
## Thu Jul 14 02:02:41 CEST 2022 - The aliscore output file does not exist. Check results for BUSCO: 353474at7742
## Thu Jul 14 02:02:41 CEST 2022 - The aliscore output appears to be empty. Check results for BUSCO: 353474at7742
## Thu Jul 14 02:02:42 CEST 2022 - The aliscore output file does not exist. Check results for BUSCO: 192799at7742
## Thu Jul 14 02:02:42 CEST 2022 - The aliscore output appears to be empty. Check results for BUSCO: 192799at7742
## Thu Jul 14 02:02:43 CEST 2022 - The aliscore output file does not exist. Check results for BUSCO: 270938at7742
## Thu Jul 14 02:02:43 CEST 2022 - The aliscore output appears to be empty. Check results for BUSCO: 270938at7742
## Thu Jul 14 02:02:44 CEST 2022 - The aliscore output file does not exist. Check results for BUSCO: 259863at7742
## Thu Jul 14 02:02:44 CEST 2022 - The aliscore output appears to be empty. Check results for BUSCO: 259863at7742
## Thu Jul 14 02:02:46 CEST 2022 - The aliscore output file does not exist. Check results for BUSCO: 299177at7742
## Thu Jul 14 02:02:46 CEST 2022 - The aliscore output appears to be empty. Check results for BUSCO: 299177at7742
## Thu Jul 14 02:02:46 CEST 2022 - The aliscore output file does not exist. Check results for BUSCO: 413952at7742
## Thu Jul 14 02:02:46 CEST 2022 - The aliscore output appears to be empty. Check results for BUSCO: 413952at7742
## Thu Jul 14 02:02:46 CEST 2022 - The aliscore output file does not exist. Check results for BUSCO: 274906at7742
## Thu Jul 14 02:02:46 CEST 2022 - The aliscore output appears to be empty. Check results for BUSCO: 274906at7742
## Thu Jul 14 02:02:48 CEST 2022 - The aliscore output file does not exist. Check results for BUSCO: 168866at7742
## Thu Jul 14 02:02:48 CEST 2022 - The aliscore output appears to be empty. Check results for BUSCO: 168866at7742
## Thu Jul 14 02:02:49 CEST 2022 - The aliscore output file does not exist. Check results for BUSCO: 325448at7742
## Thu Jul 14 02:02:49 CEST 2022 - The aliscore output appears to be empty. Check results for BUSCO: 325448at7742
## Thu Jul 14 02:02:50 CEST 2022 - The aliscore output file does not exist. Check results for BUSCO: 179120at7742
## Thu Jul 14 02:02:50 CEST 2022 - The aliscore output appears to be empty. Check results for BUSCO: 179120at7742
## Thu Jul 14 02:02:51 CEST 2022 - The aliscore output file does not exist. Check results for BUSCO: 296930at7742
## Thu Jul 14 02:02:51 CEST 2022 - The aliscore output appears to be empty. Check results for BUSCO: 296930at7742
## Thu Jul 14 02:02:52 CEST 2022 - The aliscore output file does not exist. Check results for BUSCO: 192451at7742
## Thu Jul 14 02:02:52 CEST 2022 - The aliscore output appears to be empty. Check results for BUSCO: 192451at7742
## Thu Jul 14 02:02:53 CEST 2022 - The aliscore output file does not exist. Check results for BUSCO: 380303at7742
## Thu Jul 14 02:02:53 CEST 2022 - The aliscore output appears to be empty. Check results for BUSCO: 380303at7742
## Thu Jul 14 02:02:55 CEST 2022 - The aliscore output file does not exist. Check results for BUSCO: 290048at7742
## Thu Jul 14 02:02:55 CEST 2022 - The aliscore output appears to be empty. Check results for BUSCO: 290048at7742
## Thu Jul 14 02:02:56 CEST 2022 - The aliscore output file does not exist. Check results for BUSCO: 339203at7742
## Thu Jul 14 02:02:56 CEST 2022 - The aliscore output appears to be empty. Check results for BUSCO: 339203at7742
## Thu Jul 14 02:02:56 CEST 2022 - The aliscore output file does not exist. Check results for BUSCO: 359289at7742
## Thu Jul 14 02:02:56 CEST 2022 - The aliscore output appears to be empty. Check results for BUSCO: 359289at7742
## Thu Jul 14 02:02:57 CEST 2022 - The aliscore output file does not exist. Check results for BUSCO: 366844at7742
## Thu Jul 14 02:02:57 CEST 2022 - The aliscore output appears to be empty. Check results for BUSCO: 366844at7742
## Thu Jul 14 02:02:59 CEST 2022 - The aliscore output file does not exist. Check results for BUSCO: 207153at7742
## Thu Jul 14 02:02:59 CEST 2022 - The aliscore output file does not exist. Check results for BUSCO: 363812at7742
## Thu Jul 14 02:02:59 CEST 2022 - The aliscore output appears to be empty. Check results for BUSCO: 207153at7742
## Thu Jul 14 02:02:59 CEST 2022 - The aliscore output appears to be empty. Check results for BUSCO: 363812at7742
## Thu Jul 14 02:03:01 CEST 2022 - The aliscore output file does not exist. Check results for BUSCO: 260107at7742
## Thu Jul 14 02:03:01 CEST 2022 - The aliscore output appears to be empty. Check results for BUSCO: 260107at7742
## Thu Jul 14 02:03:01 CEST 2022 - The aliscore output file does not exist. Check results for BUSCO: 362410at7742
## Thu Jul 14 02:03:01 CEST 2022 - The aliscore output appears to be empty. Check results for BUSCO: 362410at7742
## Thu Jul 14 02:03:03 CEST 2022 - The aliscore output file does not exist. Check results for BUSCO: 311241at7742
## Thu Jul 14 02:03:03 CEST 2022 - The aliscore output appears to be empty. Check results for BUSCO: 311241at7742
## Thu Jul 14 02:03:03 CEST 2022 - The aliscore output file does not exist. Check results for BUSCO: 360846at7742
## Thu Jul 14 02:03:03 CEST 2022 - The aliscore output appears to be empty. Check results for BUSCO: 360846at7742
## Thu Jul 14 02:03:04 CEST 2022 - The aliscore output file does not exist. Check results for BUSCO: 258712at7742
## Thu Jul 14 02:03:04 CEST 2022 - The aliscore output appears to be empty. Check results for BUSCO: 258712at7742
## Thu Jul 14 02:03:05 CEST 2022 - The aliscore output file does not exist. Check results for BUSCO: 291589at7742
## Thu Jul 14 02:03:05 CEST 2022 - The aliscore output appears to be empty. Check results for BUSCO: 291589at7742
## Thu Jul 14 02:03:06 CEST 2022 - The aliscore output file does not exist. Check results for BUSCO: 270566at7742
## Thu Jul 14 02:03:06 CEST 2022 - The aliscore output appears to be empty. Check results for BUSCO: 270566at7742
## Thu Jul 14 02:03:07 CEST 2022 - The aliscore output file does not exist. Check results for BUSCO: 254847at7742
## Thu Jul 14 02:03:07 CEST 2022 - The aliscore output appears to be empty. Check results for BUSCO: 254847at7742
## Thu Jul 14 02:03:09 CEST 2022 - The aliscore output file does not exist. Check results for BUSCO: 399601at7742
## Thu Jul 14 02:03:09 CEST 2022 - The aliscore output appears to be empty. Check results for BUSCO: 399601at7742
## Thu Jul 14 02:03:10 CEST 2022 - The aliscore output file does not exist. Check results for BUSCO: 223079at7742
## Thu Jul 14 02:03:10 CEST 2022 - The aliscore output appears to be empty. Check results for BUSCO: 223079at7742
## Thu Jul 14 02:03:11 CEST 2022 - The aliscore output file does not exist. Check results for BUSCO: 161959at7742
## Thu Jul 14 02:03:11 CEST 2022 - The aliscore output file does not exist. Check results for BUSCO: 433490at7742
## Thu Jul 14 02:03:11 CEST 2022 - The aliscore output appears to be empty. Check results for BUSCO: 161959at7742
## Thu Jul 14 02:03:11 CEST 2022 - The aliscore output appears to be empty. Check results for BUSCO: 433490at7742
## Thu Jul 14 02:03:13 CEST 2022 - The aliscore output file does not exist. Check results for BUSCO: 195766at7742
## Thu Jul 14 02:03:13 CEST 2022 - The aliscore output appears to be empty. Check results for BUSCO: 195766at7742
## Thu Jul 14 02:03:14 CEST 2022 - The aliscore output file does not exist. Check results for BUSCO: 240011at7742
## Thu Jul 14 02:03:14 CEST 2022 - The aliscore output appears to be empty. Check results for BUSCO: 240011at7742
## Thu Jul 14 02:03:15 CEST 2022 - The aliscore output file does not exist. Check results for BUSCO: 212696at7742
## Thu Jul 14 02:03:15 CEST 2022 - The aliscore output appears to be empty. Check results for BUSCO: 212696at7742
## Thu Jul 14 02:03:15 CEST 2022 - The aliscore output file does not exist. Check results for BUSCO: 349261at7742
## Thu Jul 14 02:03:15 CEST 2022 - The aliscore output appears to be empty. Check results for BUSCO: 349261at7742
## Thu Jul 14 02:03:16 CEST 2022 - The aliscore output file does not exist. Check results for BUSCO: 343795at7742
## Thu Jul 14 02:03:16 CEST 2022 - The aliscore output appears to be empty. Check results for BUSCO: 343795at7742
## Thu Jul 14 02:03:17 CEST 2022 - The aliscore output file does not exist. Check results for BUSCO: 359289at7742
## Thu Jul 14 02:03:17 CEST 2022 - The aliscore output appears to be empty. Check results for BUSCO: 359289at7742
## Thu Jul 14 02:03:18 CEST 2022 - The aliscore output file does not exist. Check results for BUSCO: 184707at7742
## Thu Jul 14 02:03:18 CEST 2022 - The aliscore output appears to be empty. Check results for BUSCO: 184707at7742
## Thu Jul 14 02:03:19 CEST 2022 - The aliscore output file does not exist. Check results for BUSCO: 174124at7742
## Thu Jul 14 02:03:19 CEST 2022 - The aliscore output file does not exist. Check results for BUSCO: 238829at7742
## Thu Jul 14 02:03:19 CEST 2022 - The aliscore output appears to be empty. Check results for BUSCO: 174124at7742
## Thu Jul 14 02:03:19 CEST 2022 - The aliscore output appears to be empty. Check results for BUSCO: 238829at7742
## Thu Jul 14 02:03:21 CEST 2022 - The aliscore output file does not exist. Check results for BUSCO: 276401at7742
## Thu Jul 14 02:03:21 CEST 2022 - The aliscore output appears to be empty. Check results for BUSCO: 276401at7742
## Thu Jul 14 02:03:22 CEST 2022 - The aliscore output file does not exist. Check results for BUSCO: 417286at7742
## Thu Jul 14 02:03:22 CEST 2022 - The aliscore output appears to be empty. Check results for BUSCO: 417286at7742
## Thu Jul 14 02:03:23 CEST 2022 - The aliscore output file does not exist. Check results for BUSCO: 119219at7742
## Thu Jul 14 02:03:23 CEST 2022 - The aliscore output appears to be empty. Check results for BUSCO: 119219at7742
## Thu Jul 14 02:03:24 CEST 2022 - The aliscore output file does not exist. Check results for BUSCO: 320817at7742
## Thu Jul 14 02:03:24 CEST 2022 - The aliscore output appears to be empty. Check results for BUSCO: 320817at7742
## Thu Jul 14 02:03:24 CEST 2022 - The aliscore output file does not exist. Check results for BUSCO: 332227at7742
## Thu Jul 14 02:03:24 CEST 2022 - The aliscore output appears to be empty. Check results for BUSCO: 332227at7742
## Thu Jul 14 02:03:25 CEST 2022 - The aliscore output file does not exist. Check results for BUSCO: 166301at7742
## Thu Jul 14 02:03:25 CEST 2022 - The aliscore output appears to be empty. Check results for BUSCO: 166301at7742
## Thu Jul 14 02:03:26 CEST 2022 - The aliscore output file does not exist. Check results for BUSCO: 278070at7742
## Thu Jul 14 02:03:26 CEST 2022 - The aliscore output appears to be empty. Check results for BUSCO: 278070at7742
## Thu Jul 14 02:03:27 CEST 2022 - The aliscore output file does not exist. Check results for BUSCO: 133692at7742
## Thu Jul 14 02:03:27 CEST 2022 - The aliscore output appears to be empty. Check results for BUSCO: 133692at7742
## Thu Jul 14 02:03:28 CEST 2022 - The aliscore output file does not exist. Check results for BUSCO: 214136at7742
## Thu Jul 14 02:03:29 CEST 2022 - The aliscore output appears to be empty. Check results for BUSCO: 214136at7742
## Thu Jul 14 02:03:29 CEST 2022 - The aliscore output file does not exist. Check results for BUSCO: 218088at7742
## Thu Jul 14 02:03:29 CEST 2022 - The aliscore output appears to be empty. Check results for BUSCO: 218088at7742
## Thu Jul 14 02:03:30 CEST 2022 - The aliscore output file does not exist. Check results for BUSCO: 260439at7742
## Thu Jul 14 02:03:30 CEST 2022 - The aliscore output appears to be empty. Check results for BUSCO: 260439at7742
## Thu Jul 14 02:03:30 CEST 2022 - The aliscore output file does not exist. Check results for BUSCO: 340002at7742
## Thu Jul 14 02:03:30 CEST 2022 - The aliscore output appears to be empty. Check results for BUSCO: 340002at7742
## Thu Jul 14 02:03:31 CEST 2022 - The aliscore output file does not exist. Check results for BUSCO: 259058at7742
## Thu Jul 14 02:03:31 CEST 2022 - The aliscore output appears to be empty. Check results for BUSCO: 259058at7742
## Thu Jul 14 02:03:31 CEST 2022 - The aliscore output file does not exist. Check results for BUSCO: 380303at7742
## Thu Jul 14 02:03:31 CEST 2022 - The aliscore output appears to be empty. Check results for BUSCO: 380303at7742
## Thu Jul 14 02:03:33 CEST 2022 - The aliscore output file does not exist. Check results for BUSCO: 293313at7742
## Thu Jul 14 02:03:33 CEST 2022 - The aliscore output appears to be empty. Check results for BUSCO: 293313at7742
## Thu Jul 14 02:03:35 CEST 2022 - The aliscore output file does not exist. Check results for BUSCO: 318320at7742
## Thu Jul 14 02:03:35 CEST 2022 - The aliscore output appears to be empty. Check results for BUSCO: 318320at7742
## Thu Jul 14 02:03:36 CEST 2022 - The aliscore output file does not exist. Check results for BUSCO: 277109at7742
## Thu Jul 14 02:03:36 CEST 2022 - The aliscore output appears to be empty. Check results for BUSCO: 277109at7742
## Thu Jul 14 02:03:37 CEST 2022 - The aliscore output file does not exist. Check results for BUSCO: 190766at7742
## Thu Jul 14 02:03:37 CEST 2022 - The aliscore output appears to be empty. Check results for BUSCO: 190766at7742
## Thu Jul 14 02:03:38 CEST 2022 - The aliscore output file does not exist. Check results for BUSCO: 214005at7742
## Thu Jul 14 02:03:38 CEST 2022 - The aliscore output file does not exist. Check results for BUSCO: 240403at7742
## Thu Jul 14 02:03:38 CEST 2022 - The aliscore output appears to be empty. Check results for BUSCO: 214005at7742
## Thu Jul 14 02:03:38 CEST 2022 - The aliscore output appears to be empty. Check results for BUSCO: 240403at7742
## Thu Jul 14 02:03:39 CEST 2022 - The aliscore output file does not exist. Check results for BUSCO: 270566at7742
## Thu Jul 14 02:03:39 CEST 2022 - The aliscore output appears to be empty. Check results for BUSCO: 270566at7742
## Thu Jul 14 02:03:41 CEST 2022 - The aliscore output file does not exist. Check results for BUSCO: 122064at7742
## Thu Jul 14 02:03:41 CEST 2022 - The aliscore output appears to be empty. Check results for BUSCO: 122064at7742
## Thu Jul 14 02:03:42 CEST 2022 - The aliscore output file does not exist. Check results for BUSCO: 273897at7742
## Thu Jul 14 02:03:42 CEST 2022 - The aliscore output appears to be empty. Check results for BUSCO: 273897at7742
## Thu Jul 14 02:03:43 CEST 2022 - The aliscore output file does not exist. Check results for BUSCO: 359693at7742
## Thu Jul 14 02:03:43 CEST 2022 - The aliscore output appears to be empty. Check results for BUSCO: 359693at7742
## Thu Jul 14 02:03:44 CEST 2022 - The aliscore output file does not exist. Check results for BUSCO: 335788at7742
## Thu Jul 14 02:03:44 CEST 2022 - The aliscore output appears to be empty. Check results for BUSCO: 335788at7742
## Thu Jul 14 02:03:45 CEST 2022 - The aliscore output file does not exist. Check results for BUSCO: 412231at7742
## Thu Jul 14 02:03:45 CEST 2022 - The aliscore output appears to be empty. Check results for BUSCO: 412231at7742
## Thu Jul 14 02:03:46 CEST 2022 - The aliscore output file does not exist. Check results for BUSCO: 245921at7742
## Thu Jul 14 02:03:46 CEST 2022 - The aliscore output appears to be empty. Check results for BUSCO: 245921at7742
## Thu Jul 14 02:03:47 CEST 2022 - The aliscore output file does not exist. Check results for BUSCO: 152925at7742
## Thu Jul 14 02:03:47 CEST 2022 - The aliscore output appears to be empty. Check results for BUSCO: 152925at7742
## Thu Jul 14 02:03:48 CEST 2022 - The aliscore output file does not exist. Check results for BUSCO: 254518at7742
## Thu Jul 14 02:03:48 CEST 2022 - The aliscore output appears to be empty. Check results for BUSCO: 254518at7742
## Thu Jul 14 02:03:49 CEST 2022 - The aliscore output file does not exist. Check results for BUSCO: 192276at7742
## Thu Jul 14 02:03:49 CEST 2022 - The aliscore output appears to be empty. Check results for BUSCO: 192276at7742
## Thu Jul 14 02:03:51 CEST 2022 - The aliscore output file does not exist. Check results for BUSCO: 122064at7742
## Thu Jul 14 02:03:51 CEST 2022 - The aliscore output appears to be empty. Check results for BUSCO: 122064at7742
## Thu Jul 14 02:03:52 CEST 2022 - The aliscore output file does not exist. Check results for BUSCO: 260439at7742
## Thu Jul 14 02:03:52 CEST 2022 - The aliscore output appears to be empty. Check results for BUSCO: 260439at7742
## Thu Jul 14 02:03:53 CEST 2022 - The aliscore output file does not exist. Check results for BUSCO: 301932at7742
## Thu Jul 14 02:03:53 CEST 2022 - The aliscore output appears to be empty. Check results for BUSCO: 301932at7742
## Thu Jul 14 02:03:54 CEST 2022 - The aliscore output file does not exist. Check results for BUSCO: 273897at7742
## Thu Jul 14 02:03:54 CEST 2022 - The aliscore output appears to be empty. Check results for BUSCO: 273897at7742
## Thu Jul 14 02:03:55 CEST 2022 - The aliscore output file does not exist. Check results for BUSCO: 213286at7742
## Thu Jul 14 02:03:55 CEST 2022 - The aliscore output appears to be empty. Check results for BUSCO: 213286at7742
## Thu Jul 14 02:03:56 CEST 2022 - The aliscore output file does not exist. Check results for BUSCO: 434093at7742
## Thu Jul 14 02:03:56 CEST 2022 - The aliscore output appears to be empty. Check results for BUSCO: 434093at7742
## Thu Jul 14 02:03:57 CEST 2022 - The aliscore output file does not exist. Check results for BUSCO: 236417at7742
## Thu Jul 14 02:03:57 CEST 2022 - The aliscore output appears to be empty. Check results for BUSCO: 236417at7742
## Thu Jul 14 02:03:57 CEST 2022 - The aliscore output file does not exist. Check results for BUSCO: 291153at7742
## Thu Jul 14 02:03:57 CEST 2022 - The aliscore output appears to be empty. Check results for BUSCO: 291153at7742
## Thu Jul 14 02:03:59 CEST 2022 - The aliscore output file does not exist. Check results for BUSCO: 360850at7742
## Thu Jul 14 02:03:59 CEST 2022 - The aliscore output appears to be empty. Check results for BUSCO: 360850at7742
## Thu Jul 14 02:04:00 CEST 2022 - The aliscore output file does not exist. Check results for BUSCO: 293313at7742
## Thu Jul 14 02:04:00 CEST 2022 - The aliscore output appears to be empty. Check results for BUSCO: 293313at7742
## Thu Jul 14 02:04:00 CEST 2022 - The aliscore output file does not exist. Check results for BUSCO: 196963at7742
## Thu Jul 14 02:04:00 CEST 2022 - The aliscore output appears to be empty. Check results for BUSCO: 196963at7742
## Thu Jul 14 02:04:02 CEST 2022 - The aliscore output file does not exist. Check results for BUSCO: 152925at7742
## Thu Jul 14 02:04:02 CEST 2022 - The aliscore output appears to be empty. Check results for BUSCO: 152925at7742
## Thu Jul 14 02:04:04 CEST 2022 - The aliscore output file does not exist. Check results for BUSCO: 291403at7742
## Thu Jul 14 02:04:04 CEST 2022 - The aliscore output appears to be empty. Check results for BUSCO: 291403at7742
## Thu Jul 14 02:04:04 CEST 2022 - The aliscore output file does not exist. Check results for BUSCO: 268430at7742
## Thu Jul 14 02:04:04 CEST 2022 - The aliscore output file does not exist. Check results for BUSCO: 173797at7742
## Thu Jul 14 02:04:04 CEST 2022 - The aliscore output appears to be empty. Check results for BUSCO: 268430at7742
## Thu Jul 14 02:04:04 CEST 2022 - The aliscore output appears to be empty. Check results for BUSCO: 173797at7742
## Thu Jul 14 02:04:08 CEST 2022 - The aliscore output file does not exist. Check results for BUSCO: 183663at7742
## Thu Jul 14 02:04:08 CEST 2022 - The aliscore output appears to be empty. Check results for BUSCO: 183663at7742
## Thu Jul 14 02:04:08 CEST 2022 - The aliscore output file does not exist. Check results for BUSCO: 226477at7742
## Thu Jul 14 02:04:08 CEST 2022 - The aliscore output appears to be empty. Check results for BUSCO: 226477at7742
## Thu Jul 14 02:04:08 CEST 2022 - The aliscore output file does not exist. Check results for BUSCO: 226477at7742
## Thu Jul 14 02:04:09 CEST 2022 - The aliscore output appears to be empty. Check results for BUSCO: 226477at7742
## Thu Jul 14 02:04:09 CEST 2022 - The aliscore output file does not exist. Check results for BUSCO: 207751at7742
## Thu Jul 14 02:04:09 CEST 2022 - The aliscore output appears to be empty. Check results for BUSCO: 207751at7742
## Thu Jul 14 02:04:11 CEST 2022 - The aliscore output file does not exist. Check results for BUSCO: 241352at7742
## Thu Jul 14 02:04:11 CEST 2022 - The aliscore output appears to be empty. Check results for BUSCO: 241352at7742
## Thu Jul 14 02:04:11 CEST 2022 - The aliscore output file does not exist. Check results for BUSCO: 317167at7742
## Thu Jul 14 02:04:11 CEST 2022 - The aliscore output appears to be empty. Check results for BUSCO: 317167at7742
## Thu Jul 14 02:04:12 CEST 2022 - The aliscore output file does not exist. Check results for BUSCO: 366862at7742
## Thu Jul 14 02:04:12 CEST 2022 - The aliscore output appears to be empty. Check results for BUSCO: 366862at7742
## Thu Jul 14 02:04:12 CEST 2022 - The aliscore output file does not exist. Check results for BUSCO: 245433at7742
## Thu Jul 14 02:04:12 CEST 2022 - The aliscore output appears to be empty. Check results for BUSCO: 245433at7742
## Thu Jul 14 02:04:14 CEST 2022 - The aliscore output file does not exist. Check results for BUSCO: 291768at7742
## Thu Jul 14 02:04:14 CEST 2022 - The aliscore output appears to be empty. Check results for BUSCO: 291768at7742
## Thu Jul 14 02:04:15 CEST 2022 - The aliscore output file does not exist. Check results for BUSCO: 196554at7742
## Thu Jul 14 02:04:15 CEST 2022 - The aliscore output appears to be empty. Check results for BUSCO: 196554at7742
## Thu Jul 14 02:04:16 CEST 2022 - The aliscore output file does not exist. Check results for BUSCO: 248028at7742
## Thu Jul 14 02:04:16 CEST 2022 - The aliscore output appears to be empty. Check results for BUSCO: 248028at7742
## Thu Jul 14 02:04:17 CEST 2022 - The aliscore output file does not exist. Check results for BUSCO: 267121at7742
## Thu Jul 14 02:04:17 CEST 2022 - The aliscore output appears to be empty. Check results for BUSCO: 267121at7742
## Thu Jul 14 02:04:18 CEST 2022 - The aliscore output file does not exist. Check results for BUSCO: 152925at7742
## Thu Jul 14 02:04:18 CEST 2022 - The aliscore output appears to be empty. Check results for BUSCO: 152925at7742
## Thu Jul 14 02:04:18 CEST 2022 - The aliscore output file does not exist. Check results for BUSCO: 314402at7742
## Thu Jul 14 02:04:18 CEST 2022 - The aliscore output appears to be empty. Check results for BUSCO: 314402at7742
## Thu Jul 14 02:04:21 CEST 2022 - The aliscore output file does not exist. Check results for BUSCO: 307638at7742
## Thu Jul 14 02:04:21 CEST 2022 - The aliscore output appears to be empty. Check results for BUSCO: 307638at7742
## Thu Jul 14 02:04:22 CEST 2022 - The aliscore output file does not exist. Check results for BUSCO: 201177at7742
## Thu Jul 14 02:04:22 CEST 2022 - The aliscore output appears to be empty. Check results for BUSCO: 201177at7742
## Thu Jul 14 02:04:22 CEST 2022 - The aliscore output file does not exist. Check results for BUSCO: 399601at7742
## Thu Jul 14 02:04:22 CEST 2022 - The aliscore output appears to be empty. Check results for BUSCO: 399601at7742
## Thu Jul 14 02:04:23 CEST 2022 - The aliscore output file does not exist. Check results for BUSCO: 203819at7742
## Thu Jul 14 02:04:23 CEST 2022 - The aliscore output appears to be empty. Check results for BUSCO: 203819at7742
## Thu Jul 14 02:04:24 CEST 2022 - The aliscore output file does not exist. Check results for BUSCO: 160097at7742
## Thu Jul 14 02:04:24 CEST 2022 - The aliscore output appears to be empty. Check results for BUSCO: 160097at7742
## Thu Jul 14 02:04:26 CEST 2022 - The aliscore output file does not exist. Check results for BUSCO: 196649at7742
## Thu Jul 14 02:04:27 CEST 2022 - The aliscore output appears to be empty. Check results for BUSCO: 196649at7742
## Thu Jul 14 02:04:28 CEST 2022 - The aliscore output file does not exist. Check results for BUSCO: 359289at7742
## Thu Jul 14 02:04:28 CEST 2022 - The aliscore output appears to be empty. Check results for BUSCO: 359289at7742
## Thu Jul 14 02:04:28 CEST 2022 - The aliscore output file does not exist. Check results for BUSCO: 318320at7742
## Thu Jul 14 02:04:28 CEST 2022 - The aliscore output appears to be empty. Check results for BUSCO: 318320at7742
## Thu Jul 14 02:04:29 CEST 2022 - The aliscore output file does not exist. Check results for BUSCO: 162268at7742
## Thu Jul 14 02:04:29 CEST 2022 - The aliscore output appears to be empty. Check results for BUSCO: 162268at7742
## Thu Jul 14 02:04:29 CEST 2022 - The aliscore output file does not exist. Check results for BUSCO: 257196at7742
## Thu Jul 14 02:04:29 CEST 2022 - The aliscore output appears to be empty. Check results for BUSCO: 257196at7742
## Thu Jul 14 02:04:30 CEST 2022 - The aliscore output file does not exist. Check results for BUSCO: 219276at7742
## Thu Jul 14 02:04:30 CEST 2022 - The aliscore output appears to be empty. Check results for BUSCO: 219276at7742
## Thu Jul 14 02:04:31 CEST 2022 - The aliscore output file does not exist. Check results for BUSCO: 166722at7742
## Thu Jul 14 02:04:31 CEST 2022 - The aliscore output appears to be empty. Check results for BUSCO: 166722at7742
## Thu Jul 14 02:04:32 CEST 2022 - The aliscore output file does not exist. Check results for BUSCO: 323737at7742
## Thu Jul 14 02:04:32 CEST 2022 - The aliscore output appears to be empty. Check results for BUSCO: 323737at7742
## Thu Jul 14 02:04:36 CEST 2022 - The aliscore output file does not exist. Check results for BUSCO: 307554at7742
## Thu Jul 14 02:04:36 CEST 2022 - The aliscore output appears to be empty. Check results for BUSCO: 307554at7742
## Thu Jul 14 02:04:36 CEST 2022 - The aliscore output file does not exist. Check results for BUSCO: 339203at7742
## Thu Jul 14 02:04:36 CEST 2022 - The aliscore output appears to be empty. Check results for BUSCO: 339203at7742
## Thu Jul 14 02:04:36 CEST 2022 - The aliscore output file does not exist. Check results for BUSCO: 119219at7742
## Thu Jul 14 02:04:36 CEST 2022 - The aliscore output appears to be empty. Check results for BUSCO: 119219at7742
## Thu Jul 14 02:04:36 CEST 2022 - The aliscore output file does not exist. Check results for BUSCO: 263794at7742
## Thu Jul 14 02:04:36 CEST 2022 - The aliscore output appears to be empty. Check results for BUSCO: 263794at7742
## Thu Jul 14 02:04:38 CEST 2022 - The aliscore output file does not exist. Check results for BUSCO: 205632at7742
## Thu Jul 14 02:04:38 CEST 2022 - The aliscore output appears to be empty. Check results for BUSCO: 205632at7742
## Thu Jul 14 02:04:39 CEST 2022 - The aliscore output file does not exist. Check results for BUSCO: 281526at7742
## Thu Jul 14 02:04:39 CEST 2022 - The aliscore output appears to be empty. Check results for BUSCO: 281526at7742
## Thu Jul 14 02:04:39 CEST 2022 - The aliscore output file does not exist. Check results for BUSCO: 196649at7742
## Thu Jul 14 02:04:39 CEST 2022 - The aliscore output appears to be empty. Check results for BUSCO: 196649at7742
## Thu Jul 14 02:04:41 CEST 2022 - The aliscore output file does not exist. Check results for BUSCO: 302110at7742
## Thu Jul 14 02:04:41 CEST 2022 - The aliscore output appears to be empty. Check results for BUSCO: 302110at7742
## Thu Jul 14 02:04:42 CEST 2022 - The aliscore output file does not exist. Check results for BUSCO: 378093at7742
## Thu Jul 14 02:04:42 CEST 2022 - The aliscore output appears to be empty. Check results for BUSCO: 378093at7742
## Thu Jul 14 02:04:43 CEST 2022 - The aliscore output file does not exist. Check results for BUSCO: 211818at7742
## Thu Jul 14 02:04:43 CEST 2022 - The aliscore output appears to be empty. Check results for BUSCO: 211818at7742
## Thu Jul 14 02:04:45 CEST 2022 - The aliscore output file does not exist. Check results for BUSCO: 268926at7742
## Thu Jul 14 02:04:45 CEST 2022 - The aliscore output appears to be empty. Check results for BUSCO: 268926at7742
## Thu Jul 14 02:04:45 CEST 2022 - The aliscore output file does not exist. Check results for BUSCO: 418700at7742
## Thu Jul 14 02:04:45 CEST 2022 - The aliscore output appears to be empty. Check results for BUSCO: 418700at7742
## Thu Jul 14 02:04:46 CEST 2022 - The aliscore output file does not exist. Check results for BUSCO: 133692at7742
## Thu Jul 14 02:04:46 CEST 2022 - The aliscore output appears to be empty. Check results for BUSCO: 133692at7742
## Thu Jul 14 02:04:47 CEST 2022 - The aliscore output file does not exist. Check results for BUSCO: 322345at7742
## Thu Jul 14 02:04:47 CEST 2022 - The aliscore output appears to be empty. Check results for BUSCO: 322345at7742
## Thu Jul 14 02:04:48 CEST 2022 - The aliscore output file does not exist. Check results for BUSCO: 211012at7742
## Thu Jul 14 02:04:48 CEST 2022 - The aliscore output appears to be empty. Check results for BUSCO: 211012at7742
## Thu Jul 14 02:04:49 CEST 2022 - The aliscore output file does not exist. Check results for BUSCO: 293860at7742
## Thu Jul 14 02:04:49 CEST 2022 - The aliscore output appears to be empty. Check results for BUSCO: 293860at7742
## Thu Jul 14 02:04:51 CEST 2022 - The aliscore output file does not exist. Check results for BUSCO: 219276at7742
## Thu Jul 14 02:04:51 CEST 2022 - The aliscore output appears to be empty. Check results for BUSCO: 219276at7742
## Thu Jul 14 02:04:51 CEST 2022 - The aliscore output file does not exist. Check results for BUSCO: 365226at7742
## Thu Jul 14 02:04:52 CEST 2022 - The aliscore output appears to be empty. Check results for BUSCO: 365226at7742
## Thu Jul 14 02:04:54 CEST 2022 - The aliscore output file does not exist. Check results for BUSCO: 157219at7742
## Thu Jul 14 02:04:54 CEST 2022 - The aliscore output appears to be empty. Check results for BUSCO: 157219at7742
## Thu Jul 14 02:04:54 CEST 2022 - The aliscore output file does not exist. Check results for BUSCO: 267121at7742
## Thu Jul 14 02:04:54 CEST 2022 - The aliscore output appears to be empty. Check results for BUSCO: 267121at7742
## Thu Jul 14 02:04:56 CEST 2022 - The aliscore output file does not exist. Check results for BUSCO: 166301at7742
## Thu Jul 14 02:04:56 CEST 2022 - The aliscore output appears to be empty. Check results for BUSCO: 166301at7742
## Thu Jul 14 02:04:56 CEST 2022 - The aliscore output file does not exist. Check results for BUSCO: 237455at7742
## Thu Jul 14 02:04:56 CEST 2022 - The aliscore output appears to be empty. Check results for BUSCO: 237455at7742
## Thu Jul 14 02:04:58 CEST 2022 - The aliscore output file does not exist. Check results for BUSCO: 281526at7742
## Thu Jul 14 02:04:58 CEST 2022 - The aliscore output appears to be empty. Check results for BUSCO: 281526at7742
## Thu Jul 14 02:04:58 CEST 2022 - The aliscore output file does not exist. Check results for BUSCO: 211012at7742
## Thu Jul 14 02:04:58 CEST 2022 - The aliscore output appears to be empty. Check results for BUSCO: 211012at7742
## Thu Jul 14 02:04:58 CEST 2022 - The aliscore output file does not exist. Check results for BUSCO: 340073at7742
## Thu Jul 14 02:04:58 CEST 2022 - The aliscore output appears to be empty. Check results for BUSCO: 340073at7742
## Thu Jul 14 02:05:00 CEST 2022 - The aliscore output file does not exist. Check results for BUSCO: 328558at7742
## Thu Jul 14 02:05:00 CEST 2022 - The aliscore output appears to be empty. Check results for BUSCO: 328558at7742
## Thu Jul 14 02:05:01 CEST 2022 - The aliscore output file does not exist. Check results for BUSCO: 238829at7742
## Thu Jul 14 02:05:01 CEST 2022 - The aliscore output appears to be empty. Check results for BUSCO: 238829at7742
## Thu Jul 14 02:05:02 CEST 2022 - The aliscore output file does not exist. Check results for BUSCO: 337736at7742
## Thu Jul 14 02:05:02 CEST 2022 - The aliscore output appears to be empty. Check results for BUSCO: 337736at7742
## Thu Jul 14 02:05:03 CEST 2022 - The aliscore output file does not exist. Check results for BUSCO: 291589at7742
## Thu Jul 14 02:05:03 CEST 2022 - The aliscore output appears to be empty. Check results for BUSCO: 291589at7742
## Thu Jul 14 02:05:03 CEST 2022 - The aliscore output file does not exist. Check results for BUSCO: 212065at7742
## Thu Jul 14 02:05:03 CEST 2022 - The aliscore output appears to be empty. Check results for BUSCO: 212065at7742
## Thu Jul 14 02:05:05 CEST 2022 - The aliscore output file does not exist. Check results for BUSCO: 213372at7742
## Thu Jul 14 02:05:05 CEST 2022 - The aliscore output appears to be empty. Check results for BUSCO: 213372at7742
## Thu Jul 14 02:05:06 CEST 2022 - The aliscore output file does not exist. Check results for BUSCO: 218088at7742
## Thu Jul 14 02:05:06 CEST 2022 - The aliscore output appears to be empty. Check results for BUSCO: 218088at7742
## Thu Jul 14 02:05:07 CEST 2022 - The aliscore output file does not exist. Check results for BUSCO: 267121at7742
## Thu Jul 14 02:05:07 CEST 2022 - The aliscore output appears to be empty. Check results for BUSCO: 267121at7742
## Thu Jul 14 02:05:07 CEST 2022 - The aliscore output file does not exist. Check results for BUSCO: 315274at7742
## Thu Jul 14 02:05:07 CEST 2022 - The aliscore output appears to be empty. Check results for BUSCO: 315274at7742
## Thu Jul 14 02:05:09 CEST 2022 - The aliscore output file does not exist. Check results for BUSCO: 196416at7742
## Thu Jul 14 02:05:09 CEST 2022 - The aliscore output appears to be empty. Check results for BUSCO: 196416at7742
## Thu Jul 14 02:05:10 CEST 2022 - The aliscore output file does not exist. Check results for BUSCO: 373266at7742
## Thu Jul 14 02:05:10 CEST 2022 - The aliscore output appears to be empty. Check results for BUSCO: 373266at7742
## Thu Jul 14 02:05:11 CEST 2022 - The aliscore output file does not exist. Check results for BUSCO: 317375at7742
## Thu Jul 14 02:05:11 CEST 2022 - The aliscore output appears to be empty. Check results for BUSCO: 317375at7742
## Thu Jul 14 02:05:12 CEST 2022 - The aliscore output file does not exist. Check results for BUSCO: 263652at7742
## Thu Jul 14 02:05:12 CEST 2022 - The aliscore output appears to be empty. Check results for BUSCO: 263652at7742
## Thu Jul 14 02:05:13 CEST 2022 - The aliscore output file does not exist. Check results for BUSCO: 169787at7742
## Thu Jul 14 02:05:13 CEST 2022 - The aliscore output appears to be empty. Check results for BUSCO: 169787at7742
## Thu Jul 14 02:05:13 CEST 2022 - The aliscore output file does not exist. Check results for BUSCO: 307158at7742
## Thu Jul 14 02:05:13 CEST 2022 - The aliscore output appears to be empty. Check results for BUSCO: 307158at7742
## Thu Jul 14 02:05:16 CEST 2022 - The aliscore output file does not exist. Check results for BUSCO: 287339at7742
## Thu Jul 14 02:05:16 CEST 2022 - The aliscore output appears to be empty. Check results for BUSCO: 287339at7742
## Thu Jul 14 02:05:17 CEST 2022 - The aliscore output file does not exist. Check results for BUSCO: 174124at7742
## Thu Jul 14 02:05:17 CEST 2022 - The aliscore output appears to be empty. Check results for BUSCO: 174124at7742
## Thu Jul 14 02:05:17 CEST 2022 - The aliscore output file does not exist. Check results for BUSCO: 277616at7742
## Thu Jul 14 02:05:17 CEST 2022 - The aliscore output appears to be empty. Check results for BUSCO: 277616at7742
## Thu Jul 14 02:05:20 CEST 2022 - The aliscore output file does not exist. Check results for BUSCO: 290543at7742
## Thu Jul 14 02:05:20 CEST 2022 - The aliscore output appears to be empty. Check results for BUSCO: 290543at7742
## Thu Jul 14 02:05:20 CEST 2022 - The aliscore output file does not exist. Check results for BUSCO: 421394at7742
## Thu Jul 14 02:05:20 CEST 2022 - The aliscore output appears to be empty. Check results for BUSCO: 421394at7742
## Thu Jul 14 02:05:21 CEST 2022 - The aliscore output file does not exist. Check results for BUSCO: 213286at7742
## Thu Jul 14 02:05:21 CEST 2022 - The aliscore output appears to be empty. Check results for BUSCO: 213286at7742
## Thu Jul 14 02:05:21 CEST 2022 - The aliscore output file does not exist. Check results for BUSCO: 363812at7742
## Thu Jul 14 02:05:21 CEST 2022 - The aliscore output appears to be empty. Check results for BUSCO: 363812at7742
## Thu Jul 14 02:05:21 CEST 2022 - The aliscore output file does not exist. Check results for BUSCO: 195300at7742
## Thu Jul 14 02:05:21 CEST 2022 - The aliscore output appears to be empty. Check results for BUSCO: 195300at7742
## Thu Jul 14 02:05:23 CEST 2022 - The aliscore output file does not exist. Check results for BUSCO: 417286at7742
## Thu Jul 14 02:05:23 CEST 2022 - The aliscore output appears to be empty. Check results for BUSCO: 417286at7742
## Thu Jul 14 02:05:24 CEST 2022 - The aliscore output file does not exist. Check results for BUSCO: 304804at7742
## Thu Jul 14 02:05:24 CEST 2022 - The aliscore output appears to be empty. Check results for BUSCO: 304804at7742
## Thu Jul 14 02:05:27 CEST 2022 - The aliscore output file does not exist. Check results for BUSCO: 427282at7742
## Thu Jul 14 02:05:27 CEST 2022 - The aliscore output appears to be empty. Check results for BUSCO: 427282at7742
## Thu Jul 14 02:05:27 CEST 2022 - The aliscore output file does not exist. Check results for BUSCO: 252307at7742
## Thu Jul 14 02:05:27 CEST 2022 - The aliscore output appears to be empty. Check results for BUSCO: 252307at7742
## Thu Jul 14 02:05:28 CEST 2022 - The aliscore output file does not exist. Check results for BUSCO: 203819at7742
## Thu Jul 14 02:05:28 CEST 2022 - The aliscore output appears to be empty. Check results for BUSCO: 203819at7742
## Thu Jul 14 02:05:28 CEST 2022 - The aliscore output file does not exist. Check results for BUSCO: 205632at7742
## Thu Jul 14 02:05:28 CEST 2022 - The aliscore output appears to be empty. Check results for BUSCO: 205632at7742
## Thu Jul 14 02:05:30 CEST 2022 - The aliscore output file does not exist. Check results for BUSCO: 180473at7742
## Thu Jul 14 02:05:30 CEST 2022 - The ALICUT output appears to be empty. Will touch an empty file so the pipeline will continue. Check results for BUSCO: 180473at7742
## Thu Jul 14 02:05:30 CEST 2022 - The aliscore output file does not exist. Check results for BUSCO: 180473at7742
## Thu Jul 14 02:05:30 CEST 2022 - The ALICUT output appears to be empty. Will touch an empty file so the pipeline will continue. Check results for BUSCO: 180473at7742
## Thu Jul 14 02:05:31 CEST 2022 - The aliscore output file does not exist. Check results for BUSCO: 192451at7742
## Thu Jul 14 02:05:31 CEST 2022 - The aliscore output appears to be empty. Check results for BUSCO: 192451at7742
## Thu Jul 14 02:05:33 CEST 2022 - The aliscore output file does not exist. Check results for BUSCO: 434093at7742
## Thu Jul 14 02:05:33 CEST 2022 - The aliscore output appears to be empty. Check results for BUSCO: 434093at7742
## Thu Jul 14 02:05:35 CEST 2022 - The aliscore output file does not exist. Check results for BUSCO: 218088at7742
## Thu Jul 14 02:05:35 CEST 2022 - The aliscore output appears to be empty. Check results for BUSCO: 218088at7742
## Thu Jul 14 02:05:35 CEST 2022 - The aliscore output file does not exist. Check results for BUSCO: 293487at7742
## Thu Jul 14 02:05:35 CEST 2022 - The aliscore output appears to be empty. Check results for BUSCO: 293487at7742
## Thu Jul 14 02:05:35 CEST 2022 - The aliscore output file does not exist. Check results for BUSCO: 180473at7742
## Thu Jul 14 02:05:35 CEST 2022 - The ALICUT output appears to be empty. Will touch an empty file so the pipeline will continue. Check results for BUSCO: 180473at7742
## Thu Jul 14 02:05:37 CEST 2022 - The aliscore output file does not exist. Check results for BUSCO: 195766at7742
## Thu Jul 14 02:05:37 CEST 2022 - The aliscore output appears to be empty. Check results for BUSCO: 195766at7742
## Thu Jul 14 02:05:37 CEST 2022 - The aliscore output file does not exist. Check results for BUSCO: 250326at7742
## Thu Jul 14 02:05:37 CEST 2022 - The aliscore output appears to be empty. Check results for BUSCO: 250326at7742
## Thu Jul 14 02:05:40 CEST 2022 - The aliscore output file does not exist. Check results for BUSCO: 319229at7742
## Thu Jul 14 02:05:40 CEST 2022 - The aliscore output appears to be empty. Check results for BUSCO: 319229at7742
## Thu Jul 14 02:05:40 CEST 2022 - The aliscore output file does not exist. Check results for BUSCO: 122064at7742
## Thu Jul 14 02:05:40 CEST 2022 - The aliscore output appears to be empty. Check results for BUSCO: 122064at7742
## Thu Jul 14 02:05:42 CEST 2022 - The aliscore output file does not exist. Check results for BUSCO: 260062at7742
## Thu Jul 14 02:05:42 CEST 2022 - The aliscore output appears to be empty. Check results for BUSCO: 260062at7742
## Thu Jul 14 02:05:42 CEST 2022 - The aliscore output file does not exist. Check results for BUSCO: 260062at7742
## Thu Jul 14 02:05:42 CEST 2022 - The aliscore output appears to be empty. Check results for BUSCO: 260062at7742
## Thu Jul 14 02:05:42 CEST 2022 - The aliscore output file does not exist. Check results for BUSCO: 324341at7742
## Thu Jul 14 02:05:42 CEST 2022 - The aliscore output appears to be empty. Check results for BUSCO: 324341at7742
## Thu Jul 14 02:05:44 CEST 2022 - The aliscore output file does not exist. Check results for BUSCO: 234804at7742
## Thu Jul 14 02:05:44 CEST 2022 - The aliscore output appears to be empty. Check results for BUSCO: 234804at7742
## Thu Jul 14 02:05:45 CEST 2022 - The aliscore output file does not exist. Check results for BUSCO: 335129at7742
## Thu Jul 14 02:05:45 CEST 2022 - The aliscore output appears to be empty. Check results for BUSCO: 335129at7742
## Thu Jul 14 02:05:46 CEST 2022 - The aliscore output file does not exist. Check results for BUSCO: 261976at7742
## Thu Jul 14 02:05:46 CEST 2022 - The aliscore output appears to be empty. Check results for BUSCO: 261976at7742
## Thu Jul 14 02:05:47 CEST 2022 - The aliscore output file does not exist. Check results for BUSCO: 421394at7742
## Thu Jul 14 02:05:47 CEST 2022 - The aliscore output appears to be empty. Check results for BUSCO: 421394at7742
## Thu Jul 14 02:05:49 CEST 2022 - The aliscore output file does not exist. Check results for BUSCO: 226799at7742
## Thu Jul 14 02:05:49 CEST 2022 - The aliscore output appears to be empty. Check results for BUSCO: 226799at7742
## Thu Jul 14 02:05:49 CEST 2022 - The aliscore output file does not exist. Check results for BUSCO: 303129at7742
## Thu Jul 14 02:05:49 CEST 2022 - The aliscore output appears to be empty. Check results for BUSCO: 303129at7742
## Thu Jul 14 02:05:50 CEST 2022 - The aliscore output file does not exist. Check results for BUSCO: 243222at7742
## Thu Jul 14 02:05:50 CEST 2022 - The aliscore output appears to be empty. Check results for BUSCO: 243222at7742
## Thu Jul 14 02:05:50 CEST 2022 - The aliscore output file does not exist. Check results for BUSCO: 349261at7742
## Thu Jul 14 02:05:50 CEST 2022 - The aliscore output appears to be empty. Check results for BUSCO: 349261at7742
## Thu Jul 14 02:05:51 CEST 2022 - The aliscore output file does not exist. Check results for BUSCO: 182326at7742
## Thu Jul 14 02:05:51 CEST 2022 - The aliscore output appears to be empty. Check results for BUSCO: 182326at7742
## Thu Jul 14 02:05:52 CEST 2022 - The aliscore output file does not exist. Check results for BUSCO: 305792at7742
## Thu Jul 14 02:05:52 CEST 2022 - The aliscore output appears to be empty. Check results for BUSCO: 305792at7742
## Thu Jul 14 02:05:54 CEST 2022 - The aliscore output file does not exist. Check results for BUSCO: 219276at7742
## Thu Jul 14 02:05:54 CEST 2022 - The aliscore output appears to be empty. Check results for BUSCO: 219276at7742
## Thu Jul 14 02:05:55 CEST 2022 - The aliscore output file does not exist. Check results for BUSCO: 207751at7742
## Thu Jul 14 02:05:55 CEST 2022 - The aliscore output appears to be empty. Check results for BUSCO: 207751at7742
## Thu Jul 14 02:05:55 CEST 2022 - The aliscore output file does not exist. Check results for BUSCO: 312288at7742
## Thu Jul 14 02:05:55 CEST 2022 - The aliscore output appears to be empty. Check results for BUSCO: 312288at7742
## Thu Jul 14 02:05:56 CEST 2022 - The aliscore output file does not exist. Check results for BUSCO: 253322at7742
## Thu Jul 14 02:05:56 CEST 2022 - The aliscore output appears to be empty. Check results for BUSCO: 253322at7742
## Thu Jul 14 02:05:57 CEST 2022 - The aliscore output file does not exist. Check results for BUSCO: 278076at7742
## Thu Jul 14 02:05:57 CEST 2022 - The aliscore output appears to be empty. Check results for BUSCO: 278076at7742
## Thu Jul 14 02:05:58 CEST 2022 - The aliscore output file does not exist. Check results for BUSCO: 407084at7742
## Thu Jul 14 02:05:58 CEST 2022 - The aliscore output appears to be empty. Check results for BUSCO: 407084at7742
## Thu Jul 14 02:06:00 CEST 2022 - The aliscore output file does not exist. Check results for BUSCO: 302110at7742
## Thu Jul 14 02:06:00 CEST 2022 - The aliscore output appears to be empty. Check results for BUSCO: 302110at7742
## Thu Jul 14 02:06:00 CEST 2022 - The aliscore output file does not exist. Check results for BUSCO: 260062at7742
## Thu Jul 14 02:06:00 CEST 2022 - The aliscore output appears to be empty. Check results for BUSCO: 260062at7742
## Thu Jul 14 02:06:02 CEST 2022 - The aliscore output file does not exist. Check results for BUSCO: 311889at7742
## Thu Jul 14 02:06:02 CEST 2022 - The aliscore output appears to be empty. Check results for BUSCO: 311889at7742
## Thu Jul 14 02:06:03 CEST 2022 - The aliscore output file does not exist. Check results for BUSCO: 226949at7742
## Thu Jul 14 02:06:03 CEST 2022 - The aliscore output appears to be empty. Check results for BUSCO: 226949at7742
## Thu Jul 14 02:06:04 CEST 2022 - The aliscore output file does not exist. Check results for BUSCO: 225169at7742
## Thu Jul 14 02:06:04 CEST 2022 - The aliscore output appears to be empty. Check results for BUSCO: 225169at7742
## Thu Jul 14 02:06:04 CEST 2022 - The aliscore output file does not exist. Check results for BUSCO: 323966at7742
## Thu Jul 14 02:06:04 CEST 2022 - The aliscore output appears to be empty. Check results for BUSCO: 323966at7742
## Thu Jul 14 02:06:07 CEST 2022 - The aliscore output file does not exist. Check results for BUSCO: 247398at7742
## Thu Jul 14 02:06:07 CEST 2022 - The aliscore output appears to be empty. Check results for BUSCO: 247398at7742
## Thu Jul 14 02:06:07 CEST 2022 - The aliscore output file does not exist. Check results for BUSCO: 280835at7742
## Thu Jul 14 02:06:07 CEST 2022 - The aliscore output appears to be empty. Check results for BUSCO: 280835at7742
## Thu Jul 14 02:06:07 CEST 2022 - The aliscore output file does not exist. Check results for BUSCO: 361974at7742
## Thu Jul 14 02:06:07 CEST 2022 - The aliscore output appears to be empty. Check results for BUSCO: 361974at7742
## Thu Jul 14 02:06:09 CEST 2022 - The aliscore output file does not exist. Check results for BUSCO: 251278at7742
## Thu Jul 14 02:06:09 CEST 2022 - The aliscore output appears to be empty. Check results for BUSCO: 251278at7742
## Thu Jul 14 02:06:11 CEST 2022 - The aliscore output file does not exist. Check results for BUSCO: 162268at7742
## Thu Jul 14 02:06:11 CEST 2022 - The aliscore output appears to be empty. Check results for BUSCO: 162268at7742
## Thu Jul 14 02:06:12 CEST 2022 - The aliscore output file does not exist. Check results for BUSCO: 365136at7742
## Thu Jul 14 02:06:12 CEST 2022 - The aliscore output appears to be empty. Check results for BUSCO: 365136at7742
## Thu Jul 14 02:06:13 CEST 2022 - The aliscore output file does not exist. Check results for BUSCO: 277226at7742
## Thu Jul 14 02:06:13 CEST 2022 - The aliscore output appears to be empty. Check results for BUSCO: 277226at7742
## Thu Jul 14 02:06:13 CEST 2022 - The aliscore output file does not exist. Check results for BUSCO: 291768at7742
## Thu Jul 14 02:06:13 CEST 2022 - The aliscore output appears to be empty. Check results for BUSCO: 291768at7742
## Thu Jul 14 02:06:15 CEST 2022 - The aliscore output file does not exist. Check results for BUSCO: 207751at7742
## Thu Jul 14 02:06:15 CEST 2022 - The aliscore output appears to be empty. Check results for BUSCO: 207751at7742
## Thu Jul 14 02:06:16 CEST 2022 - The aliscore output file does not exist. Check results for BUSCO: 277109at7742
## Thu Jul 14 02:06:16 CEST 2022 - The aliscore output appears to be empty. Check results for BUSCO: 277109at7742
## Thu Jul 14 02:06:17 CEST 2022 - The aliscore output file does not exist. Check results for BUSCO: 162268at7742
## Thu Jul 14 02:06:17 CEST 2022 - The aliscore output appears to be empty. Check results for BUSCO: 162268at7742
## Thu Jul 14 02:06:18 CEST 2022 - The aliscore output file does not exist. Check results for BUSCO: 234637at7742
## Thu Jul 14 02:06:18 CEST 2022 - The aliscore output appears to be empty. Check results for BUSCO: 234637at7742
## Thu Jul 14 02:06:18 CEST 2022 - The aliscore output file does not exist. Check results for BUSCO: 319288at7742
## Thu Jul 14 02:06:18 CEST 2022 - The aliscore output appears to be empty. Check results for BUSCO: 319288at7742
## Thu Jul 14 02:06:19 CEST 2022 - The aliscore output file does not exist. Check results for BUSCO: 316568at7742
## Thu Jul 14 02:06:19 CEST 2022 - The aliscore output appears to be empty. Check results for BUSCO: 316568at7742
## Thu Jul 14 02:06:20 CEST 2022 - The aliscore output file does not exist. Check results for BUSCO: 263991at7742
## Thu Jul 14 02:06:20 CEST 2022 - The aliscore output appears to be empty. Check results for BUSCO: 263991at7742
## Thu Jul 14 02:06:21 CEST 2022 - The aliscore output file does not exist. Check results for BUSCO: 242394at7742
## Thu Jul 14 02:06:21 CEST 2022 - The aliscore output appears to be empty. Check results for BUSCO: 242394at7742
## Thu Jul 14 02:06:24 CEST 2022 - The aliscore output file does not exist. Check results for BUSCO: 313364at7742
## Thu Jul 14 02:06:24 CEST 2022 - The aliscore output appears to be empty. Check results for BUSCO: 313364at7742
## Thu Jul 14 02:06:24 CEST 2022 - The aliscore output file does not exist. Check results for BUSCO: 280835at7742
## Thu Jul 14 02:06:24 CEST 2022 - The aliscore output appears to be empty. Check results for BUSCO: 280835at7742
## Thu Jul 14 02:06:26 CEST 2022 - The aliscore output file does not exist. Check results for BUSCO: 243222at7742
## Thu Jul 14 02:06:26 CEST 2022 - The aliscore output appears to be empty. Check results for BUSCO: 243222at7742
## Thu Jul 14 02:06:26 CEST 2022 - The aliscore output file does not exist. Check results for BUSCO: 229159at7742
## Thu Jul 14 02:06:26 CEST 2022 - The aliscore output appears to be empty. Check results for BUSCO: 229159at7742
## Thu Jul 14 02:06:26 CEST 2022 - The aliscore output file does not exist. Check results for BUSCO: 303510at7742
## Thu Jul 14 02:06:26 CEST 2022 - The aliscore output appears to be empty. Check results for BUSCO: 303510at7742
## Thu Jul 14 02:06:28 CEST 2022 - The aliscore output file does not exist. Check results for BUSCO: 250963at7742
## Thu Jul 14 02:06:28 CEST 2022 - The aliscore output appears to be empty. Check results for BUSCO: 250963at7742
## Thu Jul 14 02:06:29 CEST 2022 - The aliscore output file does not exist. Check results for BUSCO: 159802at7742
## Thu Jul 14 02:06:29 CEST 2022 - The aliscore output appears to be empty. Check results for BUSCO: 159802at7742
## Thu Jul 14 02:06:29 CEST 2022 - The aliscore output file does not exist. Check results for BUSCO: 274732at7742
## Thu Jul 14 02:06:29 CEST 2022 - The aliscore output appears to be empty. Check results for BUSCO: 274732at7742
## Thu Jul 14 02:06:31 CEST 2022 - The aliscore output file does not exist. Check results for BUSCO: 196416at7742
## Thu Jul 14 02:06:31 CEST 2022 - The aliscore output appears to be empty. Check results for BUSCO: 196416at7742
## Thu Jul 14 02:06:32 CEST 2022 - The aliscore output file does not exist. Check results for BUSCO: 292437at7742
## Thu Jul 14 02:06:32 CEST 2022 - The aliscore output appears to be empty. Check results for BUSCO: 292437at7742
## Thu Jul 14 02:06:33 CEST 2022 - The aliscore output file does not exist. Check results for BUSCO: 185249at7742
## Thu Jul 14 02:06:33 CEST 2022 - The aliscore output appears to be empty. Check results for BUSCO: 185249at7742
## Thu Jul 14 02:06:37 CEST 2022 - The aliscore output file does not exist. Check results for BUSCO: 366844at7742
## Thu Jul 14 02:06:37 CEST 2022 - The aliscore output appears to be empty. Check results for BUSCO: 366844at7742
## Thu Jul 14 02:06:37 CEST 2022 - The aliscore output file does not exist. Check results for BUSCO: 213372at7742
## Thu Jul 14 02:06:37 CEST 2022 - The aliscore output appears to be empty. Check results for BUSCO: 213372at7742
## Thu Jul 14 02:06:37 CEST 2022 - The aliscore output file does not exist. Check results for BUSCO: 309974at7742
## Thu Jul 14 02:06:38 CEST 2022 - The aliscore output appears to be empty. Check results for BUSCO: 309974at7742
## Thu Jul 14 02:06:38 CEST 2022 - The aliscore output file does not exist. Check results for BUSCO: 205632at7742
## Thu Jul 14 02:06:38 CEST 2022 - The aliscore output appears to be empty. Check results for BUSCO: 205632at7742
## Thu Jul 14 02:06:38 CEST 2022 - The aliscore output file does not exist. Check results for BUSCO: 241288at7742
## Thu Jul 14 02:06:38 CEST 2022 - The aliscore output appears to be empty. Check results for BUSCO: 241288at7742
## Thu Jul 14 02:06:39 CEST 2022 - The aliscore output file does not exist. Check results for BUSCO: 286081at7742
## Thu Jul 14 02:06:39 CEST 2022 - The aliscore output appears to be empty. Check results for BUSCO: 286081at7742
## Thu Jul 14 02:06:40 CEST 2022 - The aliscore output file does not exist. Check results for BUSCO: 434093at7742
## Thu Jul 14 02:06:40 CEST 2022 - The aliscore output appears to be empty. Check results for BUSCO: 434093at7742
## Thu Jul 14 02:06:42 CEST 2022 - The aliscore output file does not exist. Check results for BUSCO: 329248at7742
## Thu Jul 14 02:06:42 CEST 2022 - The aliscore output appears to be empty. Check results for BUSCO: 329248at7742
## Thu Jul 14 02:06:43 CEST 2022 - The aliscore output file does not exist. Check results for BUSCO: 203767at7742
## Thu Jul 14 02:06:43 CEST 2022 - The aliscore output appears to be empty. Check results for BUSCO: 203767at7742
## Thu Jul 14 02:06:44 CEST 2022 - The aliscore output file does not exist. Check results for BUSCO: 320379at7742
## Thu Jul 14 02:06:44 CEST 2022 - The aliscore output appears to be empty. Check results for BUSCO: 320379at7742
## Thu Jul 14 02:06:45 CEST 2022 - The aliscore output file does not exist. Check results for BUSCO: 212696at7742
## Thu Jul 14 02:06:45 CEST 2022 - The aliscore output file does not exist. Check results for BUSCO: 369081at7742
## Thu Jul 14 02:06:45 CEST 2022 - The aliscore output appears to be empty. Check results for BUSCO: 212696at7742
## Thu Jul 14 02:06:45 CEST 2022 - The aliscore output appears to be empty. Check results for BUSCO: 369081at7742
## Thu Jul 14 02:06:46 CEST 2022 - The aliscore output file does not exist. Check results for BUSCO: 366264at7742
## Thu Jul 14 02:06:46 CEST 2022 - The aliscore output appears to be empty. Check results for BUSCO: 366264at7742
## Thu Jul 14 02:06:47 CEST 2022 - The aliscore output file does not exist. Check results for BUSCO: 195300at7742
## Thu Jul 14 02:06:47 CEST 2022 - The aliscore output appears to be empty. Check results for BUSCO: 195300at7742
## Thu Jul 14 02:06:49 CEST 2022 - The aliscore output file does not exist. Check results for BUSCO: 279227at7742
## Thu Jul 14 02:06:49 CEST 2022 - The aliscore output appears to be empty. Check results for BUSCO: 279227at7742
## Thu Jul 14 02:06:50 CEST 2022 - The aliscore output file does not exist. Check results for BUSCO: 173977at7742
## Thu Jul 14 02:06:50 CEST 2022 - The aliscore output appears to be empty. Check results for BUSCO: 173977at7742
## Thu Jul 14 02:06:52 CEST 2022 - The aliscore output file does not exist. Check results for BUSCO: 237258at7742
## Thu Jul 14 02:06:52 CEST 2022 - The aliscore output appears to be empty. Check results for BUSCO: 237258at7742
## Thu Jul 14 02:06:52 CEST 2022 - The aliscore output file does not exist. Check results for BUSCO: 221575at7742
## Thu Jul 14 02:06:52 CEST 2022 - The aliscore output appears to be empty. Check results for BUSCO: 221575at7742
## Thu Jul 14 02:06:53 CEST 2022 - The aliscore output file does not exist. Check results for BUSCO: 342620at7742
## Thu Jul 14 02:06:53 CEST 2022 - The aliscore output appears to be empty. Check results for BUSCO: 342620at7742
## Thu Jul 14 02:06:54 CEST 2022 - The aliscore output file does not exist. Check results for BUSCO: 184707at7742
## Thu Jul 14 02:06:54 CEST 2022 - The aliscore output appears to be empty. Check results for BUSCO: 184707at7742
## Thu Jul 14 02:06:55 CEST 2022 - The aliscore output file does not exist. Check results for BUSCO: 427282at7742
## Thu Jul 14 02:06:55 CEST 2022 - The aliscore output appears to be empty. Check results for BUSCO: 427282at7742
## Thu Jul 14 02:06:56 CEST 2022 - The aliscore output file does not exist. Check results for BUSCO: 318662at7742
## Thu Jul 14 02:06:56 CEST 2022 - The aliscore output appears to be empty. Check results for BUSCO: 318662at7742
## Thu Jul 14 02:06:58 CEST 2022 - The aliscore output file does not exist. Check results for BUSCO: 245921at7742
## Thu Jul 14 02:06:58 CEST 2022 - The aliscore output appears to be empty. Check results for BUSCO: 245921at7742
## Thu Jul 14 02:06:58 CEST 2022 - The aliscore output file does not exist. Check results for BUSCO: 379932at7742
## Thu Jul 14 02:06:58 CEST 2022 - The aliscore output appears to be empty. Check results for BUSCO: 379932at7742
## Thu Jul 14 02:06:59 CEST 2022 - The aliscore output file does not exist. Check results for BUSCO: 276229at7742
## Thu Jul 14 02:06:59 CEST 2022 - The aliscore output appears to be empty. Check results for BUSCO: 276229at7742
## Thu Jul 14 02:06:59 CEST 2022 - The aliscore output file does not exist. Check results for BUSCO: 166429at7742
## Thu Jul 14 02:06:59 CEST 2022 - The aliscore output appears to be empty. Check results for BUSCO: 166429at7742
## Thu Jul 14 02:07:00 CEST 2022 - The aliscore output file does not exist. Check results for BUSCO: 210234at7742
## Thu Jul 14 02:07:00 CEST 2022 - The aliscore output appears to be empty. Check results for BUSCO: 210234at7742
## Thu Jul 14 02:07:01 CEST 2022 - The aliscore output file does not exist. Check results for BUSCO: 277616at7742
## Thu Jul 14 02:07:01 CEST 2022 - The aliscore output appears to be empty. Check results for BUSCO: 277616at7742
## Thu Jul 14 02:07:03 CEST 2022 - The aliscore output file does not exist. Check results for BUSCO: 221575at7742
## Thu Jul 14 02:07:03 CEST 2022 - The aliscore output appears to be empty. Check results for BUSCO: 221575at7742
## Thu Jul 14 02:07:04 CEST 2022 - The aliscore output file does not exist. Check results for BUSCO: 339768at7742
## Thu Jul 14 02:07:05 CEST 2022 - The aliscore output appears to be empty. Check results for BUSCO: 339768at7742
## Thu Jul 14 02:07:05 CEST 2022 - The aliscore output file does not exist. Check results for BUSCO: 305943at7742
## Thu Jul 14 02:07:05 CEST 2022 - The aliscore output appears to be empty. Check results for BUSCO: 305943at7742
## Thu Jul 14 02:07:06 CEST 2022 - The aliscore output file does not exist. Check results for BUSCO: 339768at7742
## Thu Jul 14 02:07:06 CEST 2022 - The aliscore output appears to be empty. Check results for BUSCO: 339768at7742
## Thu Jul 14 02:07:07 CEST 2022 - The aliscore output file does not exist. Check results for BUSCO: 326776at7742
## Thu Jul 14 02:07:07 CEST 2022 - The aliscore output appears to be empty. Check results for BUSCO: 326776at7742
## Thu Jul 14 02:07:07 CEST 2022 - The aliscore output file does not exist. Check results for BUSCO: 145683at7742
## Thu Jul 14 02:07:07 CEST 2022 - The aliscore output appears to be empty. Check results for BUSCO: 145683at7742
## Thu Jul 14 02:07:09 CEST 2022 - The aliscore output file does not exist. Check results for BUSCO: 280430at7742
## Thu Jul 14 02:07:09 CEST 2022 - The aliscore output appears to be empty. Check results for BUSCO: 280430at7742
## Thu Jul 14 02:07:10 CEST 2022 - The aliscore output file does not exist. Check results for BUSCO: 276401at7742
## Thu Jul 14 02:07:10 CEST 2022 - The aliscore output appears to be empty. Check results for BUSCO: 276401at7742
## Thu Jul 14 02:07:11 CEST 2022 - The aliscore output file does not exist. Check results for BUSCO: 229159at7742
## Thu Jul 14 02:07:11 CEST 2022 - The aliscore output appears to be empty. Check results for BUSCO: 229159at7742
## Thu Jul 14 02:07:12 CEST 2022 - The aliscore output file does not exist. Check results for BUSCO: 212065at7742
## Thu Jul 14 02:07:12 CEST 2022 - The aliscore output appears to be empty. Check results for BUSCO: 212065at7742
## Thu Jul 14 02:07:13 CEST 2022 - The aliscore output file does not exist. Check results for BUSCO: 318704at7742
## Thu Jul 14 02:07:13 CEST 2022 - The aliscore output appears to be empty. Check results for BUSCO: 318704at7742
## Thu Jul 14 02:07:14 CEST 2022 - The aliscore output file does not exist. Check results for BUSCO: 324341at7742
## Thu Jul 14 02:07:14 CEST 2022 - The aliscore output appears to be empty. Check results for BUSCO: 324341at7742
## Thu Jul 14 02:07:14 CEST 2022 - The aliscore output file does not exist. Check results for BUSCO: 182599at7742
## Thu Jul 14 02:07:14 CEST 2022 - The aliscore output appears to be empty. Check results for BUSCO: 182599at7742
## Thu Jul 14 02:07:16 CEST 2022 - The aliscore output file does not exist. Check results for BUSCO: 433490at7742
## Thu Jul 14 02:07:16 CEST 2022 - The aliscore output appears to be empty. Check results for BUSCO: 433490at7742
## Thu Jul 14 02:07:17 CEST 2022 - The aliscore output file does not exist. Check results for BUSCO: 340499at7742
## Thu Jul 14 02:07:17 CEST 2022 - The aliscore output appears to be empty. Check results for BUSCO: 340499at7742
## Thu Jul 14 02:07:18 CEST 2022 - The aliscore output file does not exist. Check results for BUSCO: 320379at7742
## Thu Jul 14 02:07:18 CEST 2022 - The aliscore output appears to be empty. Check results for BUSCO: 320379at7742
## Thu Jul 14 02:07:18 CEST 2022 - The aliscore output file does not exist. Check results for BUSCO: 234804at7742
## Thu Jul 14 02:07:18 CEST 2022 - The aliscore output appears to be empty. Check results for BUSCO: 234804at7742
## Thu Jul 14 02:07:20 CEST 2022 - The aliscore output file does not exist. Check results for BUSCO: 370499at7742
## Thu Jul 14 02:07:20 CEST 2022 - The aliscore output appears to be empty. Check results for BUSCO: 370499at7742
## Thu Jul 14 02:07:21 CEST 2022 - The aliscore output file does not exist. Check results for BUSCO: 214005at7742
## Thu Jul 14 02:07:21 CEST 2022 - The aliscore output appears to be empty. Check results for BUSCO: 214005at7742
## Thu Jul 14 02:07:23 CEST 2022 - The aliscore output file does not exist. Check results for BUSCO: 296225at7742
## Thu Jul 14 02:07:23 CEST 2022 - The aliscore output appears to be empty. Check results for BUSCO: 296225at7742
## Thu Jul 14 02:07:23 CEST 2022 - The aliscore output file does not exist. Check results for BUSCO: 296225at7742
## Thu Jul 14 02:07:23 CEST 2022 - The aliscore output appears to be empty. Check results for BUSCO: 296225at7742
## Thu Jul 14 02:07:23 CEST 2022 - The aliscore output file does not exist. Check results for BUSCO: 318704at7742
## Thu Jul 14 02:07:24 CEST 2022 - The aliscore output appears to be empty. Check results for BUSCO: 318704at7742
## Thu Jul 14 02:07:25 CEST 2022 - The aliscore output file does not exist. Check results for BUSCO: 323416at7742
## Thu Jul 14 02:07:26 CEST 2022 - The aliscore output appears to be empty. Check results for BUSCO: 323416at7742
## Thu Jul 14 02:07:26 CEST 2022 - The aliscore output file does not exist. Check results for BUSCO: 222957at7742
## Thu Jul 14 02:07:26 CEST 2022 - The aliscore output appears to be empty. Check results for BUSCO: 222957at7742
## Thu Jul 14 02:07:26 CEST 2022 - The aliscore output file does not exist. Check results for BUSCO: 258712at7742
## Thu Jul 14 02:07:27 CEST 2022 - The aliscore output appears to be empty. Check results for BUSCO: 258712at7742
## Thu Jul 14 02:07:27 CEST 2022 - The aliscore output file does not exist. Check results for BUSCO: 262790at7742
## Thu Jul 14 02:07:27 CEST 2022 - The aliscore output appears to be empty. Check results for BUSCO: 262790at7742
## Thu Jul 14 02:07:29 CEST 2022 - The aliscore output file does not exist. Check results for BUSCO: 274906at7742
## Thu Jul 14 02:07:29 CEST 2022 - The aliscore output appears to be empty. Check results for BUSCO: 274906at7742
## Thu Jul 14 02:07:30 CEST 2022 - The aliscore output file does not exist. Check results for BUSCO: 160097at7742
## Thu Jul 14 02:07:30 CEST 2022 - The aliscore output appears to be empty. Check results for BUSCO: 160097at7742
## Thu Jul 14 02:07:33 CEST 2022 - The aliscore output file does not exist. Check results for BUSCO: 251278at7742
## Thu Jul 14 02:07:33 CEST 2022 - The aliscore output appears to be empty. Check results for BUSCO: 251278at7742
## Thu Jul 14 02:07:34 CEST 2022 - The aliscore output file does not exist. Check results for BUSCO: 194806at7742
## Thu Jul 14 02:07:34 CEST 2022 - The aliscore output appears to be empty. Check results for BUSCO: 194806at7742
## Thu Jul 14 02:07:34 CEST 2022 - The aliscore output file does not exist. Check results for BUSCO: 338146at7742
## Thu Jul 14 02:07:34 CEST 2022 - The aliscore output appears to be empty. Check results for BUSCO: 338146at7742
## Thu Jul 14 02:07:34 CEST 2022 - The aliscore output file does not exist. Check results for BUSCO: 196416at7742
## Thu Jul 14 02:07:34 CEST 2022 - The aliscore output appears to be empty. Check results for BUSCO: 196416at7742
## Thu Jul 14 02:07:34 CEST 2022 - The aliscore output file does not exist. Check results for BUSCO: 330348at7742
## Thu Jul 14 02:07:34 CEST 2022 - The aliscore output appears to be empty. Check results for BUSCO: 330348at7742
## Thu Jul 14 02:07:34 CEST 2022 - The aliscore output file does not exist. Check results for BUSCO: 345746at7742
## Thu Jul 14 02:07:34 CEST 2022 - The aliscore output appears to be empty. Check results for BUSCO: 345746at7742
## Thu Jul 14 02:07:37 CEST 2022 - The aliscore output file does not exist. Check results for BUSCO: 173977at7742
## Thu Jul 14 02:07:37 CEST 2022 - The aliscore output appears to be empty. Check results for BUSCO: 173977at7742
## Thu Jul 14 02:07:38 CEST 2022 - The aliscore output file does not exist. Check results for BUSCO: 278076at7742
## Thu Jul 14 02:07:38 CEST 2022 - The aliscore output appears to be empty. Check results for BUSCO: 278076at7742
## Thu Jul 14 02:07:40 CEST 2022 - The aliscore output file does not exist. Check results for BUSCO: 257976at7742
## Thu Jul 14 02:07:40 CEST 2022 - The aliscore output appears to be empty. Check results for BUSCO: 257976at7742
## Thu Jul 14 02:07:40 CEST 2022 - The aliscore output file does not exist. Check results for BUSCO: 237455at7742
## Thu Jul 14 02:07:40 CEST 2022 - The aliscore output appears to be empty. Check results for BUSCO: 237455at7742
## Thu Jul 14 02:07:41 CEST 2022 - The aliscore output file does not exist. Check results for BUSCO: 360846at7742
## Thu Jul 14 02:07:41 CEST 2022 - The aliscore output appears to be empty. Check results for BUSCO: 360846at7742
## Thu Jul 14 02:07:42 CEST 2022 - The aliscore output file does not exist. Check results for BUSCO: 239559at7742
## Thu Jul 14 02:07:42 CEST 2022 - The aliscore output appears to be empty. Check results for BUSCO: 239559at7742
## Thu Jul 14 02:07:43 CEST 2022 - The aliscore output file does not exist. Check results for BUSCO: 239559at7742
## Thu Jul 14 02:07:43 CEST 2022 - The aliscore output appears to be empty. Check results for BUSCO: 239559at7742
## Thu Jul 14 02:07:44 CEST 2022 - The aliscore output file does not exist. Check results for BUSCO: 309962at7742
## Thu Jul 14 02:07:44 CEST 2022 - The aliscore output appears to be empty. Check results for BUSCO: 309962at7742
## Thu Jul 14 02:07:45 CEST 2022 - The aliscore output file does not exist. Check results for BUSCO: 381011at7742
## Thu Jul 14 02:07:45 CEST 2022 - The aliscore output appears to be empty. Check results for BUSCO: 381011at7742
## Thu Jul 14 02:07:47 CEST 2022 - The aliscore output file does not exist. Check results for BUSCO: 287339at7742
## Thu Jul 14 02:07:47 CEST 2022 - The aliscore output appears to be empty. Check results for BUSCO: 287339at7742
## Thu Jul 14 02:07:47 CEST 2022 - The aliscore output file does not exist. Check results for BUSCO: 274732at7742
## Thu Jul 14 02:07:47 CEST 2022 - The aliscore output appears to be empty. Check results for BUSCO: 274732at7742
## Thu Jul 14 02:07:48 CEST 2022 - The aliscore output file does not exist. Check results for BUSCO: 311519at7742
## Thu Jul 14 02:07:48 CEST 2022 - The aliscore output appears to be empty. Check results for BUSCO: 311519at7742
## Thu Jul 14 02:07:49 CEST 2022 - The aliscore output file does not exist. Check results for BUSCO: 340020at7742
## Thu Jul 14 02:07:49 CEST 2022 - The aliscore output appears to be empty. Check results for BUSCO: 340020at7742
## Thu Jul 14 02:07:50 CEST 2022 - The aliscore output file does not exist. Check results for BUSCO: 317375at7742
## Thu Jul 14 02:07:50 CEST 2022 - The aliscore output appears to be empty. Check results for BUSCO: 317375at7742
## Thu Jul 14 02:07:52 CEST 2022 - The aliscore output file does not exist. Check results for BUSCO: 243836at7742
## Thu Jul 14 02:07:52 CEST 2022 - The aliscore output appears to be empty. Check results for BUSCO: 243836at7742
## Thu Jul 14 02:07:54 CEST 2022 - The aliscore output file does not exist. Check results for BUSCO: 307638at7742
## Thu Jul 14 02:07:54 CEST 2022 - The aliscore output appears to be empty. Check results for BUSCO: 307638at7742
## Thu Jul 14 02:07:54 CEST 2022 - The aliscore output file does not exist. Check results for BUSCO: 234804at7742
## Thu Jul 14 02:07:54 CEST 2022 - The aliscore output appears to be empty. Check results for BUSCO: 234804at7742
## Thu Jul 14 02:07:55 CEST 2022 - The aliscore output file does not exist. Check results for BUSCO: 270566at7742
## Thu Jul 14 02:07:55 CEST 2022 - The aliscore output appears to be empty. Check results for BUSCO: 270566at7742
## Thu Jul 14 02:07:56 CEST 2022 - The aliscore output file does not exist. Check results for BUSCO: 253661at7742
## Thu Jul 14 02:07:56 CEST 2022 - The aliscore output appears to be empty. Check results for BUSCO: 253661at7742
## Thu Jul 14 02:07:56 CEST 2022 - The aliscore output file does not exist. Check results for BUSCO: 145683at7742
## Thu Jul 14 02:07:56 CEST 2022 - The aliscore output appears to be empty. Check results for BUSCO: 145683at7742
## Thu Jul 14 02:07:57 CEST 2022 - The aliscore output file does not exist. Check results for BUSCO: 221029at7742
## Thu Jul 14 02:07:57 CEST 2022 - The aliscore output appears to be empty. Check results for BUSCO: 221029at7742
## Thu Jul 14 02:07:59 CEST 2022 - The aliscore output file does not exist. Check results for BUSCO: 338146at7742
## Thu Jul 14 02:07:59 CEST 2022 - The aliscore output appears to be empty. Check results for BUSCO: 338146at7742
## Thu Jul 14 02:08:01 CEST 2022 - The aliscore output file does not exist. Check results for BUSCO: 160097at7742
## Thu Jul 14 02:08:01 CEST 2022 - The aliscore output appears to be empty. Check results for BUSCO: 160097at7742
## Thu Jul 14 02:08:01 CEST 2022 - The aliscore output file does not exist. Check results for BUSCO: 223079at7742
## Thu Jul 14 02:08:01 CEST 2022 - The aliscore output file does not exist. Check results for BUSCO: 332304at7742
## Thu Jul 14 02:08:01 CEST 2022 - The aliscore output appears to be empty. Check results for BUSCO: 223079at7742
## Thu Jul 14 02:08:01 CEST 2022 - The aliscore output appears to be empty. Check results for BUSCO: 332304at7742
## Thu Jul 14 02:08:03 CEST 2022 - The aliscore output file does not exist. Check results for BUSCO: 161763at7742
## Thu Jul 14 02:08:03 CEST 2022 - The aliscore output appears to be empty. Check results for BUSCO: 161763at7742
## Thu Jul 14 02:08:04 CEST 2022 - The aliscore output file does not exist. Check results for BUSCO: 330751at7742
## Thu Jul 14 02:08:04 CEST 2022 - The aliscore output appears to be empty. Check results for BUSCO: 330751at7742
## Thu Jul 14 02:08:05 CEST 2022 - The aliscore output file does not exist. Check results for BUSCO: 335129at7742
## Thu Jul 14 02:08:06 CEST 2022 - The aliscore output appears to be empty. Check results for BUSCO: 335129at7742
## Thu Jul 14 02:08:06 CEST 2022 - The aliscore output file does not exist. Check results for BUSCO: 145683at7742
## Thu Jul 14 02:08:06 CEST 2022 - The aliscore output appears to be empty. Check results for BUSCO: 145683at7742
## Thu Jul 14 02:08:07 CEST 2022 - The aliscore output file does not exist. Check results for BUSCO: 309958at7742
## Thu Jul 14 02:08:07 CEST 2022 - The aliscore output appears to be empty. Check results for BUSCO: 309958at7742
## Thu Jul 14 02:08:08 CEST 2022 - The aliscore output file does not exist. Check results for BUSCO: 221029at7742
## Thu Jul 14 02:08:08 CEST 2022 - The aliscore output appears to be empty. Check results for BUSCO: 221029at7742
## Thu Jul 14 02:08:08 CEST 2022 - The aliscore output file does not exist. Check results for BUSCO: 311889at7742
## Thu Jul 14 02:08:08 CEST 2022 - The aliscore output appears to be empty. Check results for BUSCO: 311889at7742
## Thu Jul 14 02:08:10 CEST 2022 - The aliscore output file does not exist. Check results for BUSCO: 357094at7742
## Thu Jul 14 02:08:10 CEST 2022 - The aliscore output file does not exist. Check results for BUSCO: 221575at7742
## Thu Jul 14 02:08:10 CEST 2022 - The aliscore output appears to be empty. Check results for BUSCO: 357094at7742
## Thu Jul 14 02:08:10 CEST 2022 - The aliscore output appears to be empty. Check results for BUSCO: 221575at7742
## Thu Jul 14 02:08:11 CEST 2022 - The aliscore output file does not exist. Check results for BUSCO: 339693at7742
## Thu Jul 14 02:08:11 CEST 2022 - The aliscore output appears to be empty. Check results for BUSCO: 339693at7742
## Thu Jul 14 02:08:13 CEST 2022 - The aliscore output file does not exist. Check results for BUSCO: 206164at7742
## Thu Jul 14 02:08:13 CEST 2022 - The aliscore output appears to be empty. Check results for BUSCO: 206164at7742
## Thu Jul 14 02:08:14 CEST 2022 - The aliscore output file does not exist. Check results for BUSCO: 329248at7742
## Thu Jul 14 02:08:14 CEST 2022 - The aliscore output appears to be empty. Check results for BUSCO: 329248at7742
## Thu Jul 14 02:08:15 CEST 2022 - The aliscore output file does not exist. Check results for BUSCO: 190989at7742
## Thu Jul 14 02:08:15 CEST 2022 - The aliscore output appears to be empty. Check results for BUSCO: 190989at7742
## Thu Jul 14 02:08:16 CEST 2022 - The aliscore output file does not exist. Check results for BUSCO: 318662at7742
## Thu Jul 14 02:08:16 CEST 2022 - The aliscore output appears to be empty. Check results for BUSCO: 318662at7742
## Thu Jul 14 02:08:16 CEST 2022 - The aliscore output file does not exist. Check results for BUSCO: 322275at7742
## Thu Jul 14 02:08:16 CEST 2022 - The aliscore output appears to be empty. Check results for BUSCO: 322275at7742
## Thu Jul 14 02:08:16 CEST 2022 - The aliscore output file does not exist. Check results for BUSCO: 226477at7742
## Thu Jul 14 02:08:16 CEST 2022 - The aliscore output appears to be empty. Check results for BUSCO: 226477at7742
## Thu Jul 14 02:08:19 CEST 2022 - The aliscore output file does not exist. Check results for BUSCO: 261684at7742
## Thu Jul 14 02:08:19 CEST 2022 - The aliscore output appears to be empty. Check results for BUSCO: 261684at7742
## Thu Jul 14 02:08:20 CEST 2022 - The aliscore output file does not exist. Check results for BUSCO: 331411at7742
## Thu Jul 14 02:08:20 CEST 2022 - The aliscore output appears to be empty. Check results for BUSCO: 331411at7742
## Thu Jul 14 02:08:20 CEST 2022 - The aliscore output file does not exist. Check results for BUSCO: 225169at7742
## Thu Jul 14 02:08:20 CEST 2022 - The aliscore output appears to be empty. Check results for BUSCO: 225169at7742
## Thu Jul 14 02:08:21 CEST 2022 - The aliscore output file does not exist. Check results for BUSCO: 262915at7742
## Thu Jul 14 02:08:21 CEST 2022 - The aliscore output appears to be empty. Check results for BUSCO: 262915at7742
## Thu Jul 14 02:08:22 CEST 2022 - The aliscore output file does not exist. Check results for BUSCO: 270435at7742
## Thu Jul 14 02:08:22 CEST 2022 - The aliscore output appears to be empty. Check results for BUSCO: 270435at7742
## Thu Jul 14 02:08:24 CEST 2022 - The aliscore output file does not exist. Check results for BUSCO: 301932at7742
## Thu Jul 14 02:08:24 CEST 2022 - The aliscore output appears to be empty. Check results for BUSCO: 301932at7742
## Thu Jul 14 02:08:25 CEST 2022 - The aliscore output file does not exist. Check results for BUSCO: 237455at7742
## Thu Jul 14 02:08:25 CEST 2022 - The aliscore output appears to be empty. Check results for BUSCO: 237455at7742
## Thu Jul 14 02:08:25 CEST 2022 - The aliscore output file does not exist. Check results for BUSCO: 240294at7742
## Thu Jul 14 02:08:26 CEST 2022 - The aliscore output appears to be empty. Check results for BUSCO: 240294at7742
## Thu Jul 14 02:08:27 CEST 2022 - The aliscore output file does not exist. Check results for BUSCO: 190989at7742
## Thu Jul 14 02:08:27 CEST 2022 - The aliscore output appears to be empty. Check results for BUSCO: 190989at7742
## Thu Jul 14 02:08:28 CEST 2022 - The aliscore output file does not exist. Check results for BUSCO: 223079at7742
## Thu Jul 14 02:08:28 CEST 2022 - The aliscore output appears to be empty. Check results for BUSCO: 223079at7742
## Thu Jul 14 02:08:29 CEST 2022 - The aliscore output file does not exist. Check results for BUSCO: 263068at7742
## Thu Jul 14 02:08:29 CEST 2022 - The aliscore output appears to be empty. Check results for BUSCO: 263068at7742
## Thu Jul 14 02:08:29 CEST 2022 - The aliscore output file does not exist. Check results for BUSCO: 263991at7742
## Thu Jul 14 02:08:30 CEST 2022 - The aliscore output appears to be empty. Check results for BUSCO: 263991at7742
## Thu Jul 14 02:08:31 CEST 2022 - The aliscore output file does not exist. Check results for BUSCO: 368813at7742
## Thu Jul 14 02:08:31 CEST 2022 - The aliscore output appears to be empty. Check results for BUSCO: 368813at7742
## Thu Jul 14 02:08:31 CEST 2022 - The aliscore output file does not exist. Check results for BUSCO: 192276at7742
## Thu Jul 14 02:08:31 CEST 2022 - The aliscore output appears to be empty. Check results for BUSCO: 192276at7742
## Thu Jul 14 02:08:32 CEST 2022 - The aliscore output file does not exist. Check results for BUSCO: 333162at7742
## Thu Jul 14 02:08:32 CEST 2022 - The aliscore output appears to be empty. Check results for BUSCO: 333162at7742
## Thu Jul 14 02:08:33 CEST 2022 - The aliscore output file does not exist. Check results for BUSCO: 262790at7742
## Thu Jul 14 02:08:33 CEST 2022 - The aliscore output appears to be empty. Check results for BUSCO: 262790at7742
## Thu Jul 14 02:08:35 CEST 2022 - The aliscore output file does not exist. Check results for BUSCO: 262790at7742
## Thu Jul 14 02:08:35 CEST 2022 - The aliscore output appears to be empty. Check results for BUSCO: 262790at7742
## Thu Jul 14 02:08:35 CEST 2022 - The aliscore output file does not exist. Check results for BUSCO: 240294at7742
## Thu Jul 14 02:08:35 CEST 2022 - The aliscore output appears to be empty. Check results for BUSCO: 240294at7742
## Thu Jul 14 02:08:36 CEST 2022 - The aliscore output file does not exist. Check results for BUSCO: 293313at7742
## Thu Jul 14 02:08:36 CEST 2022 - The aliscore output appears to be empty. Check results for BUSCO: 293313at7742
## Thu Jul 14 02:08:38 CEST 2022 - The aliscore output file does not exist. Check results for BUSCO: 305943at7742
## Thu Jul 14 02:08:38 CEST 2022 - The aliscore output appears to be empty. Check results for BUSCO: 305943at7742
## Thu Jul 14 02:08:39 CEST 2022 - The aliscore output file does not exist. Check results for BUSCO: 205135at7742
## Thu Jul 14 02:08:39 CEST 2022 - The aliscore output appears to be empty. Check results for BUSCO: 205135at7742
## Thu Jul 14 02:08:39 CEST 2022 - The aliscore output file does not exist. Check results for BUSCO: 380513at7742
## Thu Jul 14 02:08:39 CEST 2022 - The aliscore output appears to be empty. Check results for BUSCO: 380513at7742
## Thu Jul 14 02:08:42 CEST 2022 - The aliscore output file does not exist. Check results for BUSCO: 398926at7742
## Thu Jul 14 02:08:42 CEST 2022 - The aliscore output appears to be empty. Check results for BUSCO: 398926at7742
## Thu Jul 14 02:08:43 CEST 2022 - The aliscore output file does not exist. Check results for BUSCO: 226949at7742
## Thu Jul 14 02:08:43 CEST 2022 - The aliscore output appears to be empty. Check results for BUSCO: 226949at7742
## Thu Jul 14 02:08:43 CEST 2022 - The aliscore output file does not exist. Check results for BUSCO: 221029at7742
## Thu Jul 14 02:08:43 CEST 2022 - The aliscore output appears to be empty. Check results for BUSCO: 221029at7742
## Thu Jul 14 02:08:45 CEST 2022 - The aliscore output file does not exist. Check results for BUSCO: 270432at7742
## Thu Jul 14 02:08:45 CEST 2022 - The aliscore output appears to be empty. Check results for BUSCO: 270432at7742
## Thu Jul 14 02:08:46 CEST 2022 - The aliscore output file does not exist. Check results for BUSCO: 280649at7742
## Thu Jul 14 02:08:46 CEST 2022 - The aliscore output appears to be empty. Check results for BUSCO: 280649at7742
## Thu Jul 14 02:08:46 CEST 2022 - The aliscore output file does not exist. Check results for BUSCO: 223375at7742
## Thu Jul 14 02:08:46 CEST 2022 - The aliscore output appears to be empty. Check results for BUSCO: 223375at7742
## Thu Jul 14 02:08:47 CEST 2022 - The aliscore output file does not exist. Check results for BUSCO: 286081at7742
## Thu Jul 14 02:08:47 CEST 2022 - The aliscore output appears to be empty. Check results for BUSCO: 286081at7742
## Thu Jul 14 02:08:49 CEST 2022 - The aliscore output file does not exist. Check results for BUSCO: 229159at7742
## Thu Jul 14 02:08:49 CEST 2022 - The aliscore output appears to be empty. Check results for BUSCO: 229159at7742
## Thu Jul 14 02:08:49 CEST 2022 - The aliscore output file does not exist. Check results for BUSCO: 240294at7742
## Thu Jul 14 02:08:50 CEST 2022 - The aliscore output appears to be empty. Check results for BUSCO: 240294at7742
## Thu Jul 14 02:08:50 CEST 2022 - The aliscore output file does not exist. Check results for BUSCO: 268430at7742
## Thu Jul 14 02:08:50 CEST 2022 - The aliscore output appears to be empty. Check results for BUSCO: 268430at7742
## Thu Jul 14 02:08:51 CEST 2022 - The aliscore output file does not exist. Check results for BUSCO: 273485at7742
## Thu Jul 14 02:08:51 CEST 2022 - The aliscore output appears to be empty. Check results for BUSCO: 273485at7742
## Thu Jul 14 02:08:52 CEST 2022 - The aliscore output file does not exist. Check results for BUSCO: 334363at7742
## Thu Jul 14 02:08:52 CEST 2022 - The aliscore output appears to be empty. Check results for BUSCO: 334363at7742
## Thu Jul 14 02:08:54 CEST 2022 - The aliscore output file does not exist. Check results for BUSCO: 161763at7742
## Thu Jul 14 02:08:54 CEST 2022 - The aliscore output appears to be empty. Check results for BUSCO: 161763at7742
## Thu Jul 14 02:08:55 CEST 2022 - The aliscore output file does not exist. Check results for BUSCO: 161763at7742
## Thu Jul 14 02:08:55 CEST 2022 - The aliscore output appears to be empty. Check results for BUSCO: 161763at7742
## Thu Jul 14 02:08:56 CEST 2022 - The aliscore output file does not exist. Check results for BUSCO: 330348at7742
## Thu Jul 14 02:08:56 CEST 2022 - The aliscore output appears to be empty. Check results for BUSCO: 330348at7742
## Thu Jul 14 02:08:57 CEST 2022 - The aliscore output file does not exist. Check results for BUSCO: 274906at7742
## Thu Jul 14 02:08:57 CEST 2022 - The aliscore output appears to be empty. Check results for BUSCO: 274906at7742
## Thu Jul 14 02:08:58 CEST 2022 - The aliscore output file does not exist. Check results for BUSCO: 203819at7742
## Thu Jul 14 02:08:58 CEST 2022 - The aliscore output appears to be empty. Check results for BUSCO: 203819at7742
## Thu Jul 14 02:08:58 CEST 2022 - The aliscore output file does not exist. Check results for BUSCO: 190989at7742
## Thu Jul 14 02:08:58 CEST 2022 - The aliscore output appears to be empty. Check results for BUSCO: 190989at7742
## Thu Jul 14 02:09:00 CEST 2022 - The aliscore output file does not exist. Check results for BUSCO: 296225at7742
## Thu Jul 14 02:09:00 CEST 2022 - The aliscore output appears to be empty. Check results for BUSCO: 296225at7742
## Thu Jul 14 02:09:01 CEST 2022 - The aliscore output file does not exist. Check results for BUSCO: 278076at7742
## Thu Jul 14 02:09:01 CEST 2022 - The aliscore output appears to be empty. Check results for BUSCO: 278076at7742
## Thu Jul 14 02:09:01 CEST 2022 - The aliscore output file does not exist. Check results for BUSCO: 190981at7742
## Thu Jul 14 02:09:01 CEST 2022 - The aliscore output appears to be empty. Check results for BUSCO: 190981at7742
## Thu Jul 14 02:09:03 CEST 2022 - The aliscore output file does not exist. Check results for BUSCO: 335788at7742
## Thu Jul 14 02:09:03 CEST 2022 - The aliscore output appears to be empty. Check results for BUSCO: 335788at7742
## Thu Jul 14 02:09:03 CEST 2022 - The aliscore output file does not exist. Check results for BUSCO: 335788at7742
## Thu Jul 14 02:09:03 CEST 2022 - The aliscore output appears to be empty. Check results for BUSCO: 335788at7742
## Thu Jul 14 02:09:05 CEST 2022 - The aliscore output file does not exist. Check results for BUSCO: 159802at7742
## Thu Jul 14 02:09:05 CEST 2022 - The aliscore output appears to be empty. Check results for BUSCO: 159802at7742
## Thu Jul 14 02:09:06 CEST 2022 - The aliscore output file does not exist. Check results for BUSCO: 245921at7742
## Thu Jul 14 02:09:06 CEST 2022 - The aliscore output appears to be empty. Check results for BUSCO: 245921at7742
## Thu Jul 14 02:09:07 CEST 2022 - The aliscore output file does not exist. Check results for BUSCO: 205135at7742
## Thu Jul 14 02:09:07 CEST 2022 - The aliscore output appears to be empty. Check results for BUSCO: 205135at7742
## Thu Jul 14 02:09:07 CEST 2022 - The aliscore output file does not exist. Check results for BUSCO: 290523at7742
## Thu Jul 14 02:09:07 CEST 2022 - The aliscore output appears to be empty. Check results for BUSCO: 290523at7742
## Thu Jul 14 02:09:09 CEST 2022 - The aliscore output file does not exist. Check results for BUSCO: 261684at7742
## Thu Jul 14 02:09:10 CEST 2022 - The aliscore output appears to be empty. Check results for BUSCO: 261684at7742
## Thu Jul 14 02:09:10 CEST 2022 - The aliscore output file does not exist. Check results for BUSCO: 317170at7742
## Thu Jul 14 02:09:10 CEST 2022 - The aliscore output appears to be empty. Check results for BUSCO: 317170at7742
## Thu Jul 14 02:09:12 CEST 2022 - The aliscore output file does not exist. Check results for BUSCO: 309974at7742
## Thu Jul 14 02:09:12 CEST 2022 - The aliscore output file does not exist. Check results for BUSCO: 284651at7742
## Thu Jul 14 02:09:12 CEST 2022 - The aliscore output appears to be empty. Check results for BUSCO: 309974at7742
## Thu Jul 14 02:09:12 CEST 2022 - The aliscore output appears to be empty. Check results for BUSCO: 284651at7742
## Thu Jul 14 02:09:12 CEST 2022 - The aliscore output file does not exist. Check results for BUSCO: 190981at7742
## Thu Jul 14 02:09:12 CEST 2022 - The aliscore output appears to be empty. Check results for BUSCO: 190981at7742
## Thu Jul 14 02:09:13 CEST 2022 - The aliscore output file does not exist. Check results for BUSCO: 237258at7742
## Thu Jul 14 02:09:13 CEST 2022 - The aliscore output appears to be empty. Check results for BUSCO: 237258at7742
## Thu Jul 14 02:09:15 CEST 2022 - The aliscore output file does not exist. Check results for BUSCO: 182326at7742
## Thu Jul 14 02:09:15 CEST 2022 - The aliscore output appears to be empty. Check results for BUSCO: 182326at7742
## Thu Jul 14 02:09:15 CEST 2022 - The aliscore output file does not exist. Check results for BUSCO: 242394at7742
## Thu Jul 14 02:09:15 CEST 2022 - The aliscore output appears to be empty. Check results for BUSCO: 242394at7742
## Thu Jul 14 02:09:16 CEST 2022 - The aliscore output file does not exist. Check results for BUSCO: 376448at7742
## Thu Jul 14 02:09:16 CEST 2022 - The aliscore output appears to be empty. Check results for BUSCO: 376448at7742
## Thu Jul 14 02:09:18 CEST 2022 - The aliscore output file does not exist. Check results for BUSCO: 281245at7742
## Thu Jul 14 02:09:18 CEST 2022 - The aliscore output appears to be empty. Check results for BUSCO: 281245at7742
## Thu Jul 14 02:09:19 CEST 2022 - The aliscore output file does not exist. Check results for BUSCO: 386982at7742
## Thu Jul 14 02:09:19 CEST 2022 - The aliscore output appears to be empty. Check results for BUSCO: 386982at7742
## Thu Jul 14 02:09:19 CEST 2022 - The aliscore output file does not exist. Check results for BUSCO: 212065at7742
## Thu Jul 14 02:09:19 CEST 2022 - The aliscore output appears to be empty. Check results for BUSCO: 212065at7742
## Thu Jul 14 02:09:20 CEST 2022 - The aliscore output file does not exist. Check results for BUSCO: 320379at7742
## Thu Jul 14 02:09:20 CEST 2022 - The aliscore output appears to be empty. Check results for BUSCO: 320379at7742
## Thu Jul 14 02:09:21 CEST 2022 - The aliscore output file does not exist. Check results for BUSCO: 213350at7742
## Thu Jul 14 02:09:21 CEST 2022 - The aliscore output appears to be empty. Check results for BUSCO: 213350at7742
## Thu Jul 14 02:09:23 CEST 2022 - The aliscore output file does not exist. Check results for BUSCO: 147171at7742
## Thu Jul 14 02:09:23 CEST 2022 - The aliscore output appears to be empty. Check results for BUSCO: 147171at7742
## Thu Jul 14 02:09:24 CEST 2022 - The aliscore output file does not exist. Check results for BUSCO: 293073at7742
## Thu Jul 14 02:09:24 CEST 2022 - The aliscore output appears to be empty. Check results for BUSCO: 293073at7742
## Thu Jul 14 02:09:24 CEST 2022 - The aliscore output file does not exist. Check results for BUSCO: 349840at7742
## Thu Jul 14 02:09:24 CEST 2022 - The aliscore output appears to be empty. Check results for BUSCO: 349840at7742
## Thu Jul 14 02:09:25 CEST 2022 - The aliscore output file does not exist. Check results for BUSCO: 325448at7742
## Thu Jul 14 02:09:26 CEST 2022 - The aliscore output appears to be empty. Check results for BUSCO: 325448at7742
## Thu Jul 14 02:09:26 CEST 2022 - The aliscore output file does not exist. Check results for BUSCO: 226949at7742
## Thu Jul 14 02:09:26 CEST 2022 - The aliscore output appears to be empty. Check results for BUSCO: 226949at7742
## Thu Jul 14 02:09:27 CEST 2022 - The aliscore output file does not exist. Check results for BUSCO: 250963at7742
## Thu Jul 14 02:09:27 CEST 2022 - The aliscore output appears to be empty. Check results for BUSCO: 250963at7742
## Thu Jul 14 02:09:29 CEST 2022 - The aliscore output file does not exist. Check results for BUSCO: 250963at7742
## Thu Jul 14 02:09:29 CEST 2022 - The aliscore output appears to be empty. Check results for BUSCO: 250963at7742
## Thu Jul 14 02:09:30 CEST 2022 - The aliscore output file does not exist. Check results for BUSCO: 210234at7742
## Thu Jul 14 02:09:30 CEST 2022 - The aliscore output appears to be empty. Check results for BUSCO: 210234at7742
## Thu Jul 14 02:09:30 CEST 2022 - The aliscore output file does not exist. Check results for BUSCO: 174124at7742
## Thu Jul 14 02:09:30 CEST 2022 - The aliscore output appears to be empty. Check results for BUSCO: 174124at7742
## Thu Jul 14 02:09:31 CEST 2022 - The aliscore output file does not exist. Check results for BUSCO: 192276at7742
## Thu Jul 14 02:09:31 CEST 2022 - The aliscore output appears to be empty. Check results for BUSCO: 192276at7742
## Thu Jul 14 02:09:32 CEST 2022 - The aliscore output file does not exist. Check results for BUSCO: 356688at7742
## Thu Jul 14 02:09:32 CEST 2022 - The aliscore output appears to be empty. Check results for BUSCO: 356688at7742
## Thu Jul 14 02:09:33 CEST 2022 - The aliscore output file does not exist. Check results for BUSCO: 356688at7742
## Thu Jul 14 02:09:33 CEST 2022 - The aliscore output appears to be empty. Check results for BUSCO: 356688at7742
## Thu Jul 14 02:09:35 CEST 2022 - The aliscore output file does not exist. Check results for BUSCO: 159802at7742
## Thu Jul 14 02:09:35 CEST 2022 - The aliscore output appears to be empty. Check results for BUSCO: 159802at7742
## Thu Jul 14 02:09:36 CEST 2022 - The aliscore output file does not exist. Check results for BUSCO: 290048at7742
## Thu Jul 14 02:09:36 CEST 2022 - The aliscore output appears to be empty. Check results for BUSCO: 290048at7742
## Thu Jul 14 02:09:38 CEST 2022 - The aliscore output file does not exist. Check results for BUSCO: 328558at7742
## Thu Jul 14 02:09:38 CEST 2022 - The aliscore output appears to be empty. Check results for BUSCO: 328558at7742
## Thu Jul 14 02:09:38 CEST 2022 - The aliscore output file does not exist. Check results for BUSCO: 182326at7742
## Thu Jul 14 02:09:39 CEST 2022 - The aliscore output appears to be empty. Check results for BUSCO: 182326at7742
## Thu Jul 14 02:09:39 CEST 2022 - The aliscore output file does not exist. Check results for BUSCO: 257196at7742
## Thu Jul 14 02:09:39 CEST 2022 - The aliscore output appears to be empty. Check results for BUSCO: 257196at7742
## Thu Jul 14 02:09:40 CEST 2022 - The aliscore output file does not exist. Check results for BUSCO: 239695at7742
## Thu Jul 14 02:09:41 CEST 2022 - The aliscore output appears to be empty. Check results for BUSCO: 239695at7742
## Thu Jul 14 02:09:41 CEST 2022 - The aliscore output file does not exist. Check results for BUSCO: 241887at7742
## Thu Jul 14 02:09:41 CEST 2022 - The aliscore output appears to be empty. Check results for BUSCO: 241887at7742
## Thu Jul 14 02:09:42 CEST 2022 - The aliscore output file does not exist. Check results for BUSCO: 194806at7742
## Thu Jul 14 02:09:42 CEST 2022 - The aliscore output appears to be empty. Check results for BUSCO: 194806at7742
## Thu Jul 14 02:09:44 CEST 2022 - The aliscore output file does not exist. Check results for BUSCO: 277109at7742
## Thu Jul 14 02:09:44 CEST 2022 - The aliscore output appears to be empty. Check results for BUSCO: 277109at7742
## Thu Jul 14 02:09:45 CEST 2022 - The aliscore output file does not exist. Check results for BUSCO: 206784at7742
## Thu Jul 14 02:09:46 CEST 2022 - The aliscore output appears to be empty. Check results for BUSCO: 206784at7742
## Thu Jul 14 02:09:46 CEST 2022 - The aliscore output file does not exist. Check results for BUSCO: 433490at7742
## Thu Jul 14 02:09:46 CEST 2022 - The aliscore output appears to be empty. Check results for BUSCO: 433490at7742
## Thu Jul 14 02:09:47 CEST 2022 - The aliscore output file does not exist. Check results for BUSCO: 268274at7742
## Thu Jul 14 02:09:47 CEST 2022 - The aliscore output appears to be empty. Check results for BUSCO: 268274at7742
## Thu Jul 14 02:09:48 CEST 2022 - The aliscore output file does not exist. Check results for BUSCO: 196554at7742
## Thu Jul 14 02:09:48 CEST 2022 - The aliscore output appears to be empty. Check results for BUSCO: 196554at7742
## Thu Jul 14 02:09:48 CEST 2022 - The aliscore output file does not exist. Check results for BUSCO: 287042at7742
## Thu Jul 14 02:09:48 CEST 2022 - The aliscore output appears to be empty. Check results for BUSCO: 287042at7742
## Thu Jul 14 02:09:50 CEST 2022 - The aliscore output file does not exist. Check results for BUSCO: 291229at7742
## Thu Jul 14 02:09:50 CEST 2022 - The aliscore output appears to be empty. Check results for BUSCO: 291229at7742
## Thu Jul 14 02:09:50 CEST 2022 - The aliscore output file does not exist. Check results for BUSCO: 178357at7742
## Thu Jul 14 02:09:50 CEST 2022 - The aliscore output appears to be empty. Check results for BUSCO: 178357at7742
## Thu Jul 14 02:09:51 CEST 2022 - The aliscore output file does not exist. Check results for BUSCO: 313364at7742
## Thu Jul 14 02:09:51 CEST 2022 - The aliscore output appears to be empty. Check results for BUSCO: 313364at7742
## Thu Jul 14 02:09:53 CEST 2022 - The aliscore output file does not exist. Check results for BUSCO: 357094at7742
## Thu Jul 14 02:09:53 CEST 2022 - The aliscore output appears to be empty. Check results for BUSCO: 357094at7742
## Thu Jul 14 02:09:54 CEST 2022 - The aliscore output file does not exist. Check results for BUSCO: 196554at7742
## Thu Jul 14 02:09:54 CEST 2022 - The aliscore output appears to be empty. Check results for BUSCO: 196554at7742
## Thu Jul 14 02:09:55 CEST 2022 - The aliscore output file does not exist. Check results for BUSCO: 263068at7742
## Thu Jul 14 02:09:55 CEST 2022 - The aliscore output file does not exist. Check results for BUSCO: 236514at7742
## Thu Jul 14 02:09:55 CEST 2022 - The aliscore output appears to be empty. Check results for BUSCO: 263068at7742
## Thu Jul 14 02:09:55 CEST 2022 - The aliscore output appears to be empty. Check results for BUSCO: 236514at7742
## Thu Jul 14 02:09:56 CEST 2022 - The aliscore output file does not exist. Check results for BUSCO: 331411at7742
## Thu Jul 14 02:09:56 CEST 2022 - The aliscore output appears to be empty. Check results for BUSCO: 331411at7742
## Thu Jul 14 02:09:57 CEST 2022 - The aliscore output file does not exist. Check results for BUSCO: 356688at7742
## Thu Jul 14 02:09:57 CEST 2022 - The aliscore output appears to be empty. Check results for BUSCO: 356688at7742
## Thu Jul 14 02:09:59 CEST 2022 - The aliscore output file does not exist. Check results for BUSCO: 317021at7742
## Thu Jul 14 02:09:59 CEST 2022 - The aliscore output appears to be empty. Check results for BUSCO: 317021at7742
## Thu Jul 14 02:10:01 CEST 2022 - The aliscore output file does not exist. Check results for BUSCO: 209555at7742
## Thu Jul 14 02:10:01 CEST 2022 - The aliscore output appears to be empty. Check results for BUSCO: 209555at7742
## Thu Jul 14 02:10:01 CEST 2022 - The aliscore output file does not exist. Check results for BUSCO: 360846at7742
## Thu Jul 14 02:10:01 CEST 2022 - The aliscore output appears to be empty. Check results for BUSCO: 360846at7742
## Thu Jul 14 02:10:01 CEST 2022 - The aliscore output file does not exist. Check results for BUSCO: 380513at7742
## Thu Jul 14 02:10:01 CEST 2022 - The aliscore output appears to be empty. Check results for BUSCO: 380513at7742
## Thu Jul 14 02:10:02 CEST 2022 - The aliscore output file does not exist. Check results for BUSCO: 305943at7742
## Thu Jul 14 02:10:02 CEST 2022 - The aliscore output appears to be empty. Check results for BUSCO: 305943at7742
## Thu Jul 14 02:10:04 CEST 2022 - The aliscore output file does not exist. Check results for BUSCO: 222957at7742
## Thu Jul 14 02:10:04 CEST 2022 - The aliscore output appears to be empty. Check results for BUSCO: 222957at7742
## Thu Jul 14 02:10:05 CEST 2022 - The aliscore output file does not exist. Check results for BUSCO: 311241at7742
## Thu Jul 14 02:10:05 CEST 2022 - The aliscore output appears to be empty. Check results for BUSCO: 311241at7742
## Thu Jul 14 02:10:05 CEST 2022 - The aliscore output file does not exist. Check results for BUSCO: 317167at7742
## Thu Jul 14 02:10:05 CEST 2022 - The aliscore output appears to be empty. Check results for BUSCO: 317167at7742
## Thu Jul 14 02:10:07 CEST 2022 - The aliscore output file does not exist. Check results for BUSCO: 167825at7742
## Thu Jul 14 02:10:07 CEST 2022 - The aliscore output appears to be empty. Check results for BUSCO: 167825at7742
## Thu Jul 14 02:10:07 CEST 2022 - The aliscore output file does not exist. Check results for BUSCO: 206164at7742
## Thu Jul 14 02:10:08 CEST 2022 - The aliscore output appears to be empty. Check results for BUSCO: 206164at7742
## Thu Jul 14 02:10:08 CEST 2022 - The aliscore output file does not exist. Check results for BUSCO: 412231at7742
## Thu Jul 14 02:10:08 CEST 2022 - The aliscore output appears to be empty. Check results for BUSCO: 412231at7742
## Thu Jul 14 02:10:10 CEST 2022 - The aliscore output file does not exist. Check results for BUSCO: 183663at7742
## Thu Jul 14 02:10:10 CEST 2022 - The aliscore output appears to be empty. Check results for BUSCO: 183663at7742
## Thu Jul 14 02:10:11 CEST 2022 - The aliscore output file does not exist. Check results for BUSCO: 178357at7742
## Thu Jul 14 02:10:11 CEST 2022 - The aliscore output appears to be empty. Check results for BUSCO: 178357at7742
## Thu Jul 14 02:10:13 CEST 2022 - The aliscore output file does not exist. Check results for BUSCO: 332036at7742
## Thu Jul 14 02:10:13 CEST 2022 - The aliscore output appears to be empty. Check results for BUSCO: 332036at7742
## Thu Jul 14 02:10:13 CEST 2022 - The aliscore output file does not exist. Check results for BUSCO: 380379at7742
## Thu Jul 14 02:10:13 CEST 2022 - The aliscore output appears to be empty. Check results for BUSCO: 380379at7742
## Thu Jul 14 02:10:14 CEST 2022 - The aliscore output file does not exist. Check results for BUSCO: 268430at7742
## Thu Jul 14 02:10:14 CEST 2022 - The aliscore output appears to be empty. Check results for BUSCO: 268430at7742
## Thu Jul 14 02:10:15 CEST 2022 - The aliscore output file does not exist. Check results for BUSCO: 201177at7742
## Thu Jul 14 02:10:15 CEST 2022 - The aliscore output appears to be empty. Check results for BUSCO: 201177at7742
## Thu Jul 14 02:10:15 CEST 2022 - The aliscore output file does not exist. Check results for BUSCO: 167825at7742
## Thu Jul 14 02:10:15 CEST 2022 - The aliscore output appears to be empty. Check results for BUSCO: 167825at7742
## Thu Jul 14 02:10:18 CEST 2022 - The aliscore output file does not exist. Check results for BUSCO: 178357at7742
## Thu Jul 14 02:10:18 CEST 2022 - The aliscore output appears to be empty. Check results for BUSCO: 178357at7742
## Thu Jul 14 02:10:19 CEST 2022 - The aliscore output file does not exist. Check results for BUSCO: 332304at7742
## Thu Jul 14 02:10:19 CEST 2022 - The aliscore output appears to be empty. Check results for BUSCO: 332304at7742
## Thu Jul 14 02:10:19 CEST 2022 - The aliscore output file does not exist. Check results for BUSCO: 339693at7742
## Thu Jul 14 02:10:19 CEST 2022 - The aliscore output appears to be empty. Check results for BUSCO: 339693at7742
## Thu Jul 14 02:10:20 CEST 2022 - The aliscore output file does not exist. Check results for BUSCO: 322871at7742
## Thu Jul 14 02:10:20 CEST 2022 - The aliscore output appears to be empty. Check results for BUSCO: 322871at7742
## Thu Jul 14 02:10:21 CEST 2022 - The aliscore output file does not exist. Check results for BUSCO: 274732at7742
## Thu Jul 14 02:10:21 CEST 2022 - The aliscore output appears to be empty. Check results for BUSCO: 274732at7742
## Thu Jul 14 02:10:22 CEST 2022 - The aliscore output file does not exist. Check results for BUSCO: 356132at7742
## Thu Jul 14 02:10:22 CEST 2022 - The aliscore output appears to be empty. Check results for BUSCO: 356132at7742
## Thu Jul 14 02:10:23 CEST 2022 - The aliscore output file does not exist. Check results for BUSCO: 324664at7742
## Thu Jul 14 02:10:23 CEST 2022 - The aliscore output appears to be empty. Check results for BUSCO: 324664at7742
## Thu Jul 14 02:10:24 CEST 2022 - The aliscore output file does not exist. Check results for BUSCO: 205135at7742
## Thu Jul 14 02:10:24 CEST 2022 - The aliscore output appears to be empty. Check results for BUSCO: 205135at7742
## Thu Jul 14 02:10:25 CEST 2022 - The aliscore output file does not exist. Check results for BUSCO: 437044at7742
## Thu Jul 14 02:10:25 CEST 2022 - The aliscore output appears to be empty. Check results for BUSCO: 437044at7742
## Thu Jul 14 02:10:26 CEST 2022 - The aliscore output file does not exist. Check results for BUSCO: 360873at7742
## Thu Jul 14 02:10:26 CEST 2022 - The aliscore output appears to be empty. Check results for BUSCO: 360873at7742
## Thu Jul 14 02:10:27 CEST 2022 - The aliscore output file does not exist. Check results for BUSCO: 332036at7742
## Thu Jul 14 02:10:27 CEST 2022 - The aliscore output appears to be empty. Check results for BUSCO: 332036at7742
## Thu Jul 14 02:10:28 CEST 2022 - The aliscore output file does not exist. Check results for BUSCO: 299624at7742
## Thu Jul 14 02:10:28 CEST 2022 - The aliscore output appears to be empty. Check results for BUSCO: 299624at7742
## Thu Jul 14 02:10:29 CEST 2022 - The aliscore output file does not exist. Check results for BUSCO: 247398at7742
## Thu Jul 14 02:10:29 CEST 2022 - The aliscore output appears to be empty. Check results for BUSCO: 247398at7742
## Thu Jul 14 02:10:30 CEST 2022 - The aliscore output file does not exist. Check results for BUSCO: 359738at7742
## Thu Jul 14 02:10:30 CEST 2022 - The aliscore output appears to be empty. Check results for BUSCO: 359738at7742
## Thu Jul 14 02:10:31 CEST 2022 - The aliscore output file does not exist. Check results for BUSCO: 366264at7742
## Thu Jul 14 02:10:31 CEST 2022 - The aliscore output appears to be empty. Check results for BUSCO: 366264at7742
## Thu Jul 14 02:10:32 CEST 2022 - The aliscore output file does not exist. Check results for BUSCO: 309958at7742
## Thu Jul 14 02:10:32 CEST 2022 - The aliscore output appears to be empty. Check results for BUSCO: 309958at7742
## Thu Jul 14 02:10:34 CEST 2022 - The aliscore output file does not exist. Check results for BUSCO: 333576at7742
## Thu Jul 14 02:10:34 CEST 2022 - The aliscore output appears to be empty. Check results for BUSCO: 333576at7742
## Thu Jul 14 02:10:34 CEST 2022 - The aliscore output file does not exist. Check results for BUSCO: 297409at7742
## Thu Jul 14 02:10:34 CEST 2022 - The aliscore output appears to be empty. Check results for BUSCO: 297409at7742
## Thu Jul 14 02:10:35 CEST 2022 - The aliscore output file does not exist. Check results for BUSCO: 359693at7742
## Thu Jul 14 02:10:35 CEST 2022 - The aliscore output appears to be empty. Check results for BUSCO: 359693at7742
## Thu Jul 14 02:10:36 CEST 2022 - The aliscore output file does not exist. Check results for BUSCO: 279400at7742
## Thu Jul 14 02:10:36 CEST 2022 - The aliscore output appears to be empty. Check results for BUSCO: 279400at7742
## Thu Jul 14 02:10:38 CEST 2022 - The aliscore output file does not exist. Check results for BUSCO: 333716at7742
## Thu Jul 14 02:10:38 CEST 2022 - The aliscore output appears to be empty. Check results for BUSCO: 333716at7742
## Thu Jul 14 02:10:38 CEST 2022 - The aliscore output file does not exist. Check results for BUSCO: 147171at7742
## Thu Jul 14 02:10:38 CEST 2022 - The aliscore output appears to be empty. Check results for BUSCO: 147171at7742
## Thu Jul 14 02:10:39 CEST 2022 - The aliscore output file does not exist. Check results for BUSCO: 260107at7742
## Thu Jul 14 02:10:40 CEST 2022 - The aliscore output appears to be empty. Check results for BUSCO: 260107at7742
## Thu Jul 14 02:10:40 CEST 2022 - The aliscore output file does not exist. Check results for BUSCO: 262915at7742
## Thu Jul 14 02:10:40 CEST 2022 - The aliscore output appears to be empty. Check results for BUSCO: 262915at7742
## Thu Jul 14 02:10:41 CEST 2022 - The aliscore output file does not exist. Check results for BUSCO: 229681at7742
## Thu Jul 14 02:10:41 CEST 2022 - The aliscore output appears to be empty. Check results for BUSCO: 229681at7742
## Thu Jul 14 02:10:42 CEST 2022 - The aliscore output file does not exist. Check results for BUSCO: 297771at7742
## Thu Jul 14 02:10:42 CEST 2022 - The aliscore output appears to be empty. Check results for BUSCO: 297771at7742
## Thu Jul 14 02:10:44 CEST 2022 - The aliscore output file does not exist. Check results for BUSCO: 313364at7742
## Thu Jul 14 02:10:44 CEST 2022 - The aliscore output appears to be empty. Check results for BUSCO: 313364at7742
## Thu Jul 14 02:10:44 CEST 2022 - The aliscore output file does not exist. Check results for BUSCO: 182599at7742
## Thu Jul 14 02:10:44 CEST 2022 - The aliscore output appears to be empty. Check results for BUSCO: 182599at7742
## Thu Jul 14 02:10:46 CEST 2022 - The aliscore output file does not exist. Check results for BUSCO: 361974at7742
## Thu Jul 14 02:10:46 CEST 2022 - The aliscore output appears to be empty. Check results for BUSCO: 361974at7742
## Thu Jul 14 02:10:46 CEST 2022 - The aliscore output file does not exist. Check results for BUSCO: 340002at7742
## Thu Jul 14 02:10:46 CEST 2022 - The aliscore output appears to be empty. Check results for BUSCO: 340002at7742
## Thu Jul 14 02:10:47 CEST 2022 - The aliscore output file does not exist. Check results for BUSCO: 297409at7742
## Thu Jul 14 02:10:47 CEST 2022 - The aliscore output appears to be empty. Check results for BUSCO: 297409at7742
## Thu Jul 14 02:10:48 CEST 2022 - The aliscore output file does not exist. Check results for BUSCO: 279400at7742
## Thu Jul 14 02:10:48 CEST 2022 - The aliscore output appears to be empty. Check results for BUSCO: 279400at7742
## Thu Jul 14 02:10:49 CEST 2022 - The aliscore output file does not exist. Check results for BUSCO: 184721at7742
## Thu Jul 14 02:10:49 CEST 2022 - The aliscore output appears to be empty. Check results for BUSCO: 184721at7742
## Thu Jul 14 02:10:50 CEST 2022 - The aliscore output file does not exist. Check results for BUSCO: 239559at7742
## Thu Jul 14 02:10:50 CEST 2022 - The aliscore output appears to be empty. Check results for BUSCO: 239559at7742
## Thu Jul 14 02:10:51 CEST 2022 - The aliscore output file does not exist. Check results for BUSCO: 218109at7742
## Thu Jul 14 02:10:51 CEST 2022 - The aliscore output appears to be empty. Check results for BUSCO: 218109at7742
## Thu Jul 14 02:10:52 CEST 2022 - The aliscore output file does not exist. Check results for BUSCO: 319288at7742
## Thu Jul 14 02:10:52 CEST 2022 - The aliscore output appears to be empty. Check results for BUSCO: 319288at7742
## Thu Jul 14 02:10:54 CEST 2022 - The aliscore output file does not exist. Check results for BUSCO: 229681at7742
## Thu Jul 14 02:10:54 CEST 2022 - The aliscore output appears to be empty. Check results for BUSCO: 229681at7742
## Thu Jul 14 02:10:55 CEST 2022 - The aliscore output file does not exist. Check results for BUSCO: 279170at7742
## Thu Jul 14 02:10:55 CEST 2022 - The aliscore output appears to be empty. Check results for BUSCO: 279170at7742
## Thu Jul 14 02:10:55 CEST 2022 - The aliscore output file does not exist. Check results for BUSCO: 209555at7742
## Thu Jul 14 02:10:55 CEST 2022 - The aliscore output appears to be empty. Check results for BUSCO: 209555at7742
## Thu Jul 14 02:10:57 CEST 2022 - The aliscore output file does not exist. Check results for BUSCO: 257976at7742
## Thu Jul 14 02:10:57 CEST 2022 - The aliscore output appears to be empty. Check results for BUSCO: 257976at7742
## Thu Jul 14 02:10:57 CEST 2022 - The aliscore output file does not exist. Check results for BUSCO: 258712at7742
## Thu Jul 14 02:10:57 CEST 2022 - The aliscore output appears to be empty. Check results for BUSCO: 258712at7742
## Thu Jul 14 02:10:59 CEST 2022 - The aliscore output file does not exist. Check results for BUSCO: 225169at7742
## Thu Jul 14 02:10:59 CEST 2022 - The aliscore output appears to be empty. Check results for BUSCO: 225169at7742
## Thu Jul 14 02:11:00 CEST 2022 - The aliscore output file does not exist. Check results for BUSCO: 290048at7742
## Thu Jul 14 02:11:00 CEST 2022 - The aliscore output appears to be empty. Check results for BUSCO: 290048at7742
## Thu Jul 14 02:11:00 CEST 2022 - The aliscore output file does not exist. Check results for BUSCO: 426882at7742
## Thu Jul 14 02:11:00 CEST 2022 - The aliscore output appears to be empty. Check results for BUSCO: 426882at7742
## Thu Jul 14 02:11:01 CEST 2022 - The aliscore output file does not exist. Check results for BUSCO: 303129at7742
## Thu Jul 14 02:11:01 CEST 2022 - The aliscore output appears to be empty. Check results for BUSCO: 303129at7742
## Thu Jul 14 02:11:02 CEST 2022 - The aliscore output file does not exist. Check results for BUSCO: 184721at7742
## Thu Jul 14 02:11:02 CEST 2022 - The aliscore output appears to be empty. Check results for BUSCO: 184721at7742
## Thu Jul 14 02:11:03 CEST 2022 - The aliscore output file does not exist. Check results for BUSCO: 380340at7742
## Thu Jul 14 02:11:03 CEST 2022 - The aliscore output appears to be empty. Check results for BUSCO: 380340at7742
## Thu Jul 14 02:11:04 CEST 2022 - The aliscore output file does not exist. Check results for BUSCO: 380513at7742
## Thu Jul 14 02:11:04 CEST 2022 - The aliscore output appears to be empty. Check results for BUSCO: 380513at7742
## Thu Jul 14 02:11:05 CEST 2022 - The aliscore output file does not exist. Check results for BUSCO: 236514at7742
## Thu Jul 14 02:11:05 CEST 2022 - The aliscore output appears to be empty. Check results for BUSCO: 236514at7742
## Thu Jul 14 02:11:06 CEST 2022 - The aliscore output file does not exist. Check results for BUSCO: 147171at7742
## Thu Jul 14 02:11:06 CEST 2022 - The aliscore output appears to be empty. Check results for BUSCO: 147171at7742
## Thu Jul 14 02:11:07 CEST 2022 - The aliscore output file does not exist. Check results for BUSCO: 263991at7742
## Thu Jul 14 02:11:07 CEST 2022 - The aliscore output appears to be empty. Check results for BUSCO: 263991at7742
## Thu Jul 14 02:11:08 CEST 2022 - The aliscore output file does not exist. Check results for BUSCO: 159771at7742
## Thu Jul 14 02:11:08 CEST 2022 - The aliscore output appears to be empty. Check results for BUSCO: 159771at7742
## Thu Jul 14 02:11:11 CEST 2022 - The aliscore output file does not exist. Check results for BUSCO: 277616at7742
## Thu Jul 14 02:11:11 CEST 2022 - The aliscore output appears to be empty. Check results for BUSCO: 277616at7742
## Thu Jul 14 02:11:11 CEST 2022 - The aliscore output file does not exist. Check results for BUSCO: 391492at7742
## Thu Jul 14 02:11:11 CEST 2022 - The aliscore output appears to be empty. Check results for BUSCO: 391492at7742
## Thu Jul 14 02:11:12 CEST 2022 - The aliscore output file does not exist. Check results for BUSCO: 257976at7742
## Thu Jul 14 02:11:12 CEST 2022 - The aliscore output appears to be empty. Check results for BUSCO: 257976at7742
## Thu Jul 14 02:11:12 CEST 2022 - The aliscore output file does not exist. Check results for BUSCO: 305792at7742
## Thu Jul 14 02:11:12 CEST 2022 - The aliscore output appears to be empty. Check results for BUSCO: 305792at7742
## Thu Jul 14 02:11:14 CEST 2022 - The aliscore output file does not exist. Check results for BUSCO: 239695at7742
## Thu Jul 14 02:11:14 CEST 2022 - The aliscore output appears to be empty. Check results for BUSCO: 239695at7742
## Thu Jul 14 02:11:15 CEST 2022 - The aliscore output file does not exist. Check results for BUSCO: 226799at7742
## Thu Jul 14 02:11:15 CEST 2022 - The aliscore output appears to be empty. Check results for BUSCO: 226799at7742
## Thu Jul 14 02:11:17 CEST 2022 - The aliscore output file does not exist. Check results for BUSCO: 386982at7742
## Thu Jul 14 02:11:17 CEST 2022 - The aliscore output appears to be empty. Check results for BUSCO: 386982at7742
## Thu Jul 14 02:11:18 CEST 2022 - The aliscore output file does not exist. Check results for BUSCO: 286295at7742
## Thu Jul 14 02:11:18 CEST 2022 - The aliscore output appears to be empty. Check results for BUSCO: 286295at7742
## Thu Jul 14 02:11:19 CEST 2022 - The aliscore output file does not exist. Check results for BUSCO: 324341at7742
## Thu Jul 14 02:11:19 CEST 2022 - The aliscore output appears to be empty. Check results for BUSCO: 324341at7742
## Thu Jul 14 02:11:19 CEST 2022 - The aliscore output file does not exist. Check results for BUSCO: 247398at7742
## Thu Jul 14 02:11:19 CEST 2022 - The aliscore output appears to be empty. Check results for BUSCO: 247398at7742
## Thu Jul 14 02:11:19 CEST 2022 - The aliscore output file does not exist. Check results for BUSCO: 323416at7742
## Thu Jul 14 02:11:19 CEST 2022 - The aliscore output appears to be empty. Check results for BUSCO: 323416at7742
## Thu Jul 14 02:11:21 CEST 2022 - The aliscore output file does not exist. Check results for BUSCO: 253322at7742
## Thu Jul 14 02:11:21 CEST 2022 - The aliscore output appears to be empty. Check results for BUSCO: 253322at7742
## Thu Jul 14 02:11:23 CEST 2022 - The aliscore output file does not exist. Check results for BUSCO: 321845at7742
## Thu Jul 14 02:11:23 CEST 2022 - The aliscore output appears to be empty. Check results for BUSCO: 321845at7742
## Thu Jul 14 02:11:23 CEST 2022 - The aliscore output file does not exist. Check results for BUSCO: 184721at7742
## Thu Jul 14 02:11:23 CEST 2022 - The aliscore output appears to be empty. Check results for BUSCO: 184721at7742
## Thu Jul 14 02:11:24 CEST 2022 - The aliscore output file does not exist. Check results for BUSCO: 328352at7742
## Thu Jul 14 02:11:24 CEST 2022 - The aliscore output appears to be empty. Check results for BUSCO: 328352at7742
## Thu Jul 14 02:11:25 CEST 2022 - The aliscore output file does not exist. Check results for BUSCO: 201147at7742
## Thu Jul 14 02:11:25 CEST 2022 - The aliscore output appears to be empty. Check results for BUSCO: 201147at7742
## Thu Jul 14 02:11:26 CEST 2022 - The aliscore output file does not exist. Check results for BUSCO: 335620at7742
## Thu Jul 14 02:11:26 CEST 2022 - The aliscore output appears to be empty. Check results for BUSCO: 335620at7742
## Thu Jul 14 02:11:26 CEST 2022 - The aliscore output file does not exist. Check results for BUSCO: 214392at7742
## Thu Jul 14 02:11:26 CEST 2022 - The aliscore output appears to be empty. Check results for BUSCO: 214392at7742
## Thu Jul 14 02:11:28 CEST 2022 - The aliscore output file does not exist. Check results for BUSCO: 159771at7742
## Thu Jul 14 02:11:28 CEST 2022 - The aliscore output appears to be empty. Check results for BUSCO: 159771at7742
## Thu Jul 14 02:11:29 CEST 2022 - The aliscore output file does not exist. Check results for BUSCO: 237258at7742
## Thu Jul 14 02:11:29 CEST 2022 - The aliscore output appears to be empty. Check results for BUSCO: 237258at7742
## Thu Jul 14 02:11:31 CEST 2022 - The aliscore output file does not exist. Check results for BUSCO: 271425at7742
## Thu Jul 14 02:11:31 CEST 2022 - The aliscore output appears to be empty. Check results for BUSCO: 271425at7742
## Thu Jul 14 02:11:31 CEST 2022 - The aliscore output file does not exist. Check results for BUSCO: 250442at7742
## Thu Jul 14 02:11:31 CEST 2022 - The aliscore output appears to be empty. Check results for BUSCO: 250442at7742
## Thu Jul 14 02:11:32 CEST 2022 - The aliscore output file does not exist. Check results for BUSCO: 276761at7742
## Thu Jul 14 02:11:32 CEST 2022 - The aliscore output appears to be empty. Check results for BUSCO: 276761at7742
## Thu Jul 14 02:11:32 CEST 2022 - The aliscore output file does not exist. Check results for BUSCO: 279997at7742
## Thu Jul 14 02:11:32 CEST 2022 - The aliscore output appears to be empty. Check results for BUSCO: 279997at7742
## Thu Jul 14 02:11:33 CEST 2022 - The aliscore output file does not exist. Check results for BUSCO: 377238at7742
## Thu Jul 14 02:11:33 CEST 2022 - The aliscore output appears to be empty. Check results for BUSCO: 377238at7742
## Thu Jul 14 02:11:35 CEST 2022 - The aliscore output file does not exist. Check results for BUSCO: 173977at7742
## Thu Jul 14 02:11:35 CEST 2022 - The aliscore output appears to be empty. Check results for BUSCO: 173977at7742
## Thu Jul 14 02:11:35 CEST 2022 - The aliscore output file does not exist. Check results for BUSCO: 243836at7742
## Thu Jul 14 02:11:35 CEST 2022 - The aliscore output appears to be empty. Check results for BUSCO: 243836at7742
## Thu Jul 14 02:11:36 CEST 2022 - The aliscore output file does not exist. Check results for BUSCO: 167825at7742
## Thu Jul 14 02:11:36 CEST 2022 - The aliscore output appears to be empty. Check results for BUSCO: 167825at7742
## Thu Jul 14 02:11:38 CEST 2022 - The aliscore output file does not exist. Check results for BUSCO: 323416at7742
## Thu Jul 14 02:11:38 CEST 2022 - The aliscore output appears to be empty. Check results for BUSCO: 323416at7742
## Thu Jul 14 02:11:38 CEST 2022 - The aliscore output file does not exist. Check results for BUSCO: 423159at7742
## Thu Jul 14 02:11:39 CEST 2022 - The aliscore output appears to be empty. Check results for BUSCO: 423159at7742
## Thu Jul 14 02:11:40 CEST 2022 - The aliscore output file does not exist. Check results for BUSCO: 258747at7742
## Thu Jul 14 02:11:40 CEST 2022 - The aliscore output appears to be empty. Check results for BUSCO: 258747at7742
## Thu Jul 14 02:11:41 CEST 2022 - The aliscore output file does not exist. Check results for BUSCO: 329979at7742
## Thu Jul 14 02:11:41 CEST 2022 - The aliscore output appears to be empty. Check results for BUSCO: 329979at7742
## Thu Jul 14 02:11:42 CEST 2022 - The aliscore output file does not exist. Check results for BUSCO: 378093at7742
## Thu Jul 14 02:11:42 CEST 2022 - The aliscore output appears to be empty. Check results for BUSCO: 378093at7742
## Thu Jul 14 02:11:44 CEST 2022 - The aliscore output file does not exist. Check results for BUSCO: 239444at7742
## Thu Jul 14 02:11:44 CEST 2022 - The aliscore output appears to be empty. Check results for BUSCO: 239444at7742
## Thu Jul 14 02:11:46 CEST 2022 - The aliscore output file does not exist. Check results for BUSCO: 265369at7742
## Thu Jul 14 02:11:46 CEST 2022 - The aliscore output appears to be empty. Check results for BUSCO: 265369at7742
## Thu Jul 14 02:11:46 CEST 2022 - The aliscore output file does not exist. Check results for BUSCO: 239444at7742
## Thu Jul 14 02:11:46 CEST 2022 - The aliscore output appears to be empty. Check results for BUSCO: 239444at7742
## Thu Jul 14 02:11:46 CEST 2022 - The aliscore output file does not exist. Check results for BUSCO: 263652at7742
## Thu Jul 14 02:11:46 CEST 2022 - The aliscore output appears to be empty. Check results for BUSCO: 263652at7742
## Thu Jul 14 02:11:48 CEST 2022 - The aliscore output file does not exist. Check results for BUSCO: 159771at7742
## Thu Jul 14 02:11:48 CEST 2022 - The aliscore output appears to be empty. Check results for BUSCO: 159771at7742
## Thu Jul 14 02:11:49 CEST 2022 - The aliscore output file does not exist. Check results for BUSCO: 214392at7742
## Thu Jul 14 02:11:49 CEST 2022 - The aliscore output appears to be empty. Check results for BUSCO: 214392at7742
## Thu Jul 14 02:11:50 CEST 2022 - The aliscore output file does not exist. Check results for BUSCO: 386558at7742
## Thu Jul 14 02:11:50 CEST 2022 - The aliscore output appears to be empty. Check results for BUSCO: 386558at7742
## Thu Jul 14 02:11:51 CEST 2022 - The aliscore output file does not exist. Check results for BUSCO: 317401at7742
## Thu Jul 14 02:11:51 CEST 2022 - The aliscore output appears to be empty. Check results for BUSCO: 317401at7742
## Thu Jul 14 02:11:53 CEST 2022 - The aliscore output file does not exist. Check results for BUSCO: 322871at7742
## Thu Jul 14 02:11:53 CEST 2022 - The aliscore output appears to be empty. Check results for BUSCO: 322871at7742
## Thu Jul 14 02:11:53 CEST 2022 - The aliscore output file does not exist. Check results for BUSCO: 380340at7742
## Thu Jul 14 02:11:53 CEST 2022 - The aliscore output file does not exist. Check results for BUSCO: 196963at7742
## Thu Jul 14 02:11:53 CEST 2022 - The aliscore output appears to be empty. Check results for BUSCO: 380340at7742
## Thu Jul 14 02:11:53 CEST 2022 - The aliscore output appears to be empty. Check results for BUSCO: 196963at7742
## Thu Jul 14 02:11:54 CEST 2022 - The aliscore output file does not exist. Check results for BUSCO: 290543at7742
## Thu Jul 14 02:11:54 CEST 2022 - The aliscore output appears to be empty. Check results for BUSCO: 290543at7742
## Thu Jul 14 02:11:56 CEST 2022 - The aliscore output file does not exist. Check results for BUSCO: 290523at7742
## Thu Jul 14 02:11:56 CEST 2022 - The aliscore output file does not exist. Check results for BUSCO: 341195at7742
## Thu Jul 14 02:11:56 CEST 2022 - The aliscore output appears to be empty. Check results for BUSCO: 290523at7742
## Thu Jul 14 02:11:56 CEST 2022 - The aliscore output appears to be empty. Check results for BUSCO: 341195at7742
## Thu Jul 14 02:11:57 CEST 2022 - The aliscore output file does not exist. Check results for BUSCO: 333672at7742
## Thu Jul 14 02:11:57 CEST 2022 - The aliscore output appears to be empty. Check results for BUSCO: 333672at7742
## Thu Jul 14 02:11:59 CEST 2022 - The aliscore output file does not exist. Check results for BUSCO: 299624at7742
## Thu Jul 14 02:11:59 CEST 2022 - The aliscore output appears to be empty. Check results for BUSCO: 299624at7742
## Thu Jul 14 02:12:01 CEST 2022 - The aliscore output file does not exist. Check results for BUSCO: 299624at7742
## Thu Jul 14 02:12:01 CEST 2022 - The aliscore output appears to be empty. Check results for BUSCO: 299624at7742
## Thu Jul 14 02:12:01 CEST 2022 - The aliscore output file does not exist. Check results for BUSCO: 325125at7742
## Thu Jul 14 02:12:01 CEST 2022 - The aliscore output appears to be empty. Check results for BUSCO: 325125at7742
## Thu Jul 14 02:12:03 CEST 2022 - The aliscore output file does not exist. Check results for BUSCO: 293487at7742
## Thu Jul 14 02:12:03 CEST 2022 - The aliscore output appears to be empty. Check results for BUSCO: 293487at7742
## Thu Jul 14 02:12:03 CEST 2022 - The aliscore output file does not exist. Check results for BUSCO: 205146at7742
## Thu Jul 14 02:12:03 CEST 2022 - The aliscore output appears to be empty. Check results for BUSCO: 205146at7742
## Thu Jul 14 02:12:03 CEST 2022 - The aliscore output file does not exist. Check results for BUSCO: 239444at7742
## Thu Jul 14 02:12:03 CEST 2022 - The aliscore output appears to be empty. Check results for BUSCO: 239444at7742
## Thu Jul 14 02:12:04 CEST 2022 - The aliscore output file does not exist. Check results for BUSCO: 317167at7742
## Thu Jul 14 02:12:04 CEST 2022 - The aliscore output appears to be empty. Check results for BUSCO: 317167at7742
## Thu Jul 14 02:12:06 CEST 2022 - The aliscore output file does not exist. Check results for BUSCO: 252714at7742
## Thu Jul 14 02:12:06 CEST 2022 - The aliscore output appears to be empty. Check results for BUSCO: 252714at7742
## Thu Jul 14 02:12:07 CEST 2022 - The aliscore output file does not exist. Check results for BUSCO: 298587at7742
## Thu Jul 14 02:12:07 CEST 2022 - The aliscore output appears to be empty. Check results for BUSCO: 298587at7742
## Thu Jul 14 02:12:07 CEST 2022 - The aliscore output file does not exist. Check results for BUSCO: 252714at7742
## Thu Jul 14 02:12:07 CEST 2022 - The aliscore output appears to be empty. Check results for BUSCO: 252714at7742
## Thu Jul 14 02:12:09 CEST 2022 - The aliscore output file does not exist. Check results for BUSCO: 396303at7742
## Thu Jul 14 02:12:09 CEST 2022 - The aliscore output appears to be empty. Check results for BUSCO: 396303at7742
## Thu Jul 14 02:12:10 CEST 2022 - The aliscore output file does not exist. Check results for BUSCO: 293073at7742
## Thu Jul 14 02:12:10 CEST 2022 - The aliscore output appears to be empty. Check results for BUSCO: 293073at7742
## Thu Jul 14 02:12:10 CEST 2022 - The aliscore output file does not exist. Check results for BUSCO: 398926at7742
## Thu Jul 14 02:12:10 CEST 2022 - The aliscore output appears to be empty. Check results for BUSCO: 398926at7742
## Thu Jul 14 02:12:11 CEST 2022 - The aliscore output file does not exist. Check results for BUSCO: 324664at7742
## Thu Jul 14 02:12:11 CEST 2022 - The aliscore output appears to be empty. Check results for BUSCO: 324664at7742
## Thu Jul 14 02:12:12 CEST 2022 - The aliscore output file does not exist. Check results for BUSCO: 302555at7742
## Thu Jul 14 02:12:12 CEST 2022 - The aliscore output appears to be empty. Check results for BUSCO: 302555at7742
## Thu Jul 14 02:12:13 CEST 2022 - The aliscore output file does not exist. Check results for BUSCO: 267296at7742
## Thu Jul 14 02:12:13 CEST 2022 - The aliscore output appears to be empty. Check results for BUSCO: 267296at7742
## Thu Jul 14 02:12:13 CEST 2022 - The aliscore output file does not exist. Check results for BUSCO: 329248at7742
## Thu Jul 14 02:12:13 CEST 2022 - The aliscore output appears to be empty. Check results for BUSCO: 329248at7742
## Thu Jul 14 02:12:15 CEST 2022 - The aliscore output file does not exist. Check results for BUSCO: 398201at7742
## Thu Jul 14 02:12:15 CEST 2022 - The aliscore output appears to be empty. Check results for BUSCO: 398201at7742
## Thu Jul 14 02:12:16 CEST 2022 - The aliscore output file does not exist. Check results for BUSCO: 190981at7742
## Thu Jul 14 02:12:16 CEST 2022 - The aliscore output appears to be empty. Check results for BUSCO: 190981at7742
## Thu Jul 14 02:12:18 CEST 2022 - The aliscore output file does not exist. Check results for BUSCO: 286295at7742
## Thu Jul 14 02:12:18 CEST 2022 - The aliscore output appears to be empty. Check results for BUSCO: 286295at7742
## Thu Jul 14 02:12:19 CEST 2022 - The aliscore output file does not exist. Check results for BUSCO: 423159at7742
## Thu Jul 14 02:12:19 CEST 2022 - The aliscore output appears to be empty. Check results for BUSCO: 423159at7742
## Thu Jul 14 02:12:20 CEST 2022 - The aliscore output file does not exist. Check results for BUSCO: 380340at7742
## Thu Jul 14 02:12:20 CEST 2022 - The aliscore output appears to be empty. Check results for BUSCO: 380340at7742
## Thu Jul 14 02:12:20 CEST 2022 - The aliscore output file does not exist. Check results for BUSCO: 338146at7742
## Thu Jul 14 02:12:20 CEST 2022 - The aliscore output appears to be empty. Check results for BUSCO: 338146at7742
## Thu Jul 14 02:12:22 CEST 2022 - The aliscore output file does not exist. Check results for BUSCO: 328134at7742
## Thu Jul 14 02:12:22 CEST 2022 - The aliscore output appears to be empty. Check results for BUSCO: 328134at7742
## Thu Jul 14 02:12:22 CEST 2022 - The aliscore output file does not exist. Check results for BUSCO: 366862at7742
## Thu Jul 14 02:12:22 CEST 2022 - The aliscore output appears to be empty. Check results for BUSCO: 366862at7742
## Thu Jul 14 02:12:24 CEST 2022 - The aliscore output file does not exist. Check results for BUSCO: 263068at7742
## Thu Jul 14 02:12:24 CEST 2022 - The aliscore output appears to be empty. Check results for BUSCO: 263068at7742
## Thu Jul 14 02:12:25 CEST 2022 - The aliscore output file does not exist. Check results for BUSCO: 197667at7742
## Thu Jul 14 02:12:25 CEST 2022 - The aliscore output appears to be empty. Check results for BUSCO: 197667at7742
## Thu Jul 14 02:12:25 CEST 2022 - The aliscore output file does not exist. Check results for BUSCO: 406718at7742
## Thu Jul 14 02:12:25 CEST 2022 - The aliscore output appears to be empty. Check results for BUSCO: 406718at7742
## Thu Jul 14 02:12:27 CEST 2022 - The aliscore output file does not exist. Check results for BUSCO: 251278at7742
## Thu Jul 14 02:12:27 CEST 2022 - The aliscore output file does not exist. Check results for BUSCO: 266826at7742
## Thu Jul 14 02:12:27 CEST 2022 - The aliscore output appears to be empty. Check results for BUSCO: 251278at7742
## Thu Jul 14 02:12:27 CEST 2022 - The aliscore output appears to be empty. Check results for BUSCO: 266826at7742
## Thu Jul 14 02:12:29 CEST 2022 - The aliscore output file does not exist. Check results for BUSCO: 196963at7742
## Thu Jul 14 02:12:29 CEST 2022 - The aliscore output appears to be empty. Check results for BUSCO: 196963at7742
## Thu Jul 14 02:12:30 CEST 2022 - The aliscore output file does not exist. Check results for BUSCO: 333576at7742
## Thu Jul 14 02:12:30 CEST 2022 - The aliscore output appears to be empty. Check results for BUSCO: 333576at7742
## Thu Jul 14 02:12:30 CEST 2022 - The aliscore output file does not exist. Check results for BUSCO: 333576at7742
## Thu Jul 14 02:12:30 CEST 2022 - The aliscore output appears to be empty. Check results for BUSCO: 333576at7742
## Thu Jul 14 02:12:31 CEST 2022 - The aliscore output file does not exist. Check results for BUSCO: 399664at7742
## Thu Jul 14 02:12:31 CEST 2022 - The aliscore output appears to be empty. Check results for BUSCO: 399664at7742
## Thu Jul 14 02:12:33 CEST 2022 - The aliscore output file does not exist. Check results for BUSCO: 345549at7742
## Thu Jul 14 02:12:33 CEST 2022 - The aliscore output appears to be empty. Check results for BUSCO: 345549at7742
## Thu Jul 14 02:12:33 CEST 2022 - The aliscore output file does not exist. Check results for BUSCO: 270868at7742
## Thu Jul 14 02:12:33 CEST 2022 - The aliscore output appears to be empty. Check results for BUSCO: 270868at7742
## Thu Jul 14 02:12:34 CEST 2022 - The aliscore output file does not exist. Check results for BUSCO: 222957at7742
## Thu Jul 14 02:12:34 CEST 2022 - The aliscore output appears to be empty. Check results for BUSCO: 222957at7742
## Thu Jul 14 02:12:35 CEST 2022 - The aliscore output file does not exist. Check results for BUSCO: 291403at7742
## Thu Jul 14 02:12:35 CEST 2022 - The aliscore output appears to be empty. Check results for BUSCO: 291403at7742
## Thu Jul 14 02:12:36 CEST 2022 - The aliscore output file does not exist. Check results for BUSCO: 275715at7742
## Thu Jul 14 02:12:36 CEST 2022 - The aliscore output appears to be empty. Check results for BUSCO: 275715at7742
## Thu Jul 14 02:12:38 CEST 2022 - The aliscore output file does not exist. Check results for BUSCO: 335517at7742
## Thu Jul 14 02:12:38 CEST 2022 - The aliscore output appears to be empty. Check results for BUSCO: 335517at7742
## Thu Jul 14 02:12:39 CEST 2022 - The aliscore output file does not exist. Check results for BUSCO: 198118at7742
## Thu Jul 14 02:12:39 CEST 2022 - The aliscore output appears to be empty. Check results for BUSCO: 198118at7742
## Thu Jul 14 02:12:40 CEST 2022 - The aliscore output file does not exist. Check results for BUSCO: 307415at7742
## Thu Jul 14 02:12:40 CEST 2022 - The aliscore output appears to be empty. Check results for BUSCO: 307415at7742
## Thu Jul 14 02:12:40 CEST 2022 - The aliscore output file does not exist. Check results for BUSCO: 279997at7742
## Thu Jul 14 02:12:40 CEST 2022 - The aliscore output appears to be empty. Check results for BUSCO: 279997at7742
## Thu Jul 14 02:12:41 CEST 2022 - The aliscore output file does not exist. Check results for BUSCO: 194806at7742
## Thu Jul 14 02:12:41 CEST 2022 - The aliscore output appears to be empty. Check results for BUSCO: 194806at7742
## Thu Jul 14 02:12:42 CEST 2022 - The aliscore output file does not exist. Check results for BUSCO: 214392at7742
## Thu Jul 14 02:12:42 CEST 2022 - The aliscore output appears to be empty. Check results for BUSCO: 214392at7742
## Thu Jul 14 02:12:43 CEST 2022 - The aliscore output file does not exist. Check results for BUSCO: 324120at7742
## Thu Jul 14 02:12:43 CEST 2022 - The aliscore output appears to be empty. Check results for BUSCO: 324120at7742
## Thu Jul 14 02:12:45 CEST 2022 - The aliscore output file does not exist. Check results for BUSCO: 305344at7742
## Thu Jul 14 02:12:45 CEST 2022 - The aliscore output appears to be empty. Check results for BUSCO: 305344at7742
## Thu Jul 14 02:12:45 CEST 2022 - The aliscore output file does not exist. Check results for BUSCO: 265369at7742
## Thu Jul 14 02:12:45 CEST 2022 - The aliscore output appears to be empty. Check results for BUSCO: 265369at7742
## Thu Jul 14 02:12:46 CEST 2022 - The aliscore output file does not exist. Check results for BUSCO: 223813at7742
## Thu Jul 14 02:12:46 CEST 2022 - The aliscore output appears to be empty. Check results for BUSCO: 223813at7742
## Thu Jul 14 02:12:47 CEST 2022 - The aliscore output file does not exist. Check results for BUSCO: 205603at7742
## Thu Jul 14 02:12:47 CEST 2022 - The aliscore output appears to be empty. Check results for BUSCO: 205603at7742
## Thu Jul 14 02:12:47 CEST 2022 - The aliscore output file does not exist. Check results for BUSCO: 314402at7742
## Thu Jul 14 02:12:47 CEST 2022 - The aliscore output appears to be empty. Check results for BUSCO: 314402at7742
## Thu Jul 14 02:12:49 CEST 2022 - The aliscore output file does not exist. Check results for BUSCO: 286295at7742
## Thu Jul 14 02:12:49 CEST 2022 - The aliscore output appears to be empty. Check results for BUSCO: 286295at7742
## Thu Jul 14 02:12:50 CEST 2022 - The aliscore output file does not exist. Check results for BUSCO: 381534at7742
## Thu Jul 14 02:12:50 CEST 2022 - The aliscore output appears to be empty. Check results for BUSCO: 381534at7742
## Thu Jul 14 02:12:51 CEST 2022 - The aliscore output file does not exist. Check results for BUSCO: 298043at7742
## Thu Jul 14 02:12:51 CEST 2022 - The aliscore output appears to be empty. Check results for BUSCO: 298043at7742
## Thu Jul 14 02:12:52 CEST 2022 - The aliscore output file does not exist. Check results for BUSCO: 279997at7742
## Thu Jul 14 02:12:52 CEST 2022 - The aliscore output appears to be empty. Check results for BUSCO: 279997at7742
## Thu Jul 14 02:12:53 CEST 2022 - The aliscore output file does not exist. Check results for BUSCO: 281526at7742
## Thu Jul 14 02:12:53 CEST 2022 - The aliscore output appears to be empty. Check results for BUSCO: 281526at7742
## Thu Jul 14 02:12:53 CEST 2022 - The aliscore output file does not exist. Check results for BUSCO: 166380at7742
## Thu Jul 14 02:12:53 CEST 2022 - The aliscore output appears to be empty. Check results for BUSCO: 166380at7742
## Thu Jul 14 02:12:54 CEST 2022 - The aliscore output file does not exist. Check results for BUSCO: 166380at7742
## Thu Jul 14 02:12:54 CEST 2022 - The aliscore output appears to be empty. Check results for BUSCO: 166380at7742
## Thu Jul 14 02:12:57 CEST 2022 - The aliscore output file does not exist. Check results for BUSCO: 203767at7742
## Thu Jul 14 02:12:57 CEST 2022 - The aliscore output file does not exist. Check results for BUSCO: 336178at7742
## Thu Jul 14 02:12:57 CEST 2022 - The aliscore output appears to be empty. Check results for BUSCO: 203767at7742
## Thu Jul 14 02:12:57 CEST 2022 - The aliscore output appears to be empty. Check results for BUSCO: 336178at7742
## Thu Jul 14 02:12:57 CEST 2022 - The aliscore output file does not exist. Check results for BUSCO: 288347at7742
## Thu Jul 14 02:12:57 CEST 2022 - The aliscore output appears to be empty. Check results for BUSCO: 288347at7742
## Thu Jul 14 02:12:59 CEST 2022 - The aliscore output file does not exist. Check results for BUSCO: 197276at7742
## Thu Jul 14 02:12:59 CEST 2022 - The aliscore output appears to be empty. Check results for BUSCO: 197276at7742
## Thu Jul 14 02:12:59 CEST 2022 - The aliscore output file does not exist. Check results for BUSCO: 286089at7742
## Thu Jul 14 02:12:59 CEST 2022 - The aliscore output appears to be empty. Check results for BUSCO: 286089at7742
## Thu Jul 14 02:13:01 CEST 2022 - The aliscore output file does not exist. Check results for BUSCO: 413350at7742
## Thu Jul 14 02:13:01 CEST 2022 - The aliscore output appears to be empty. Check results for BUSCO: 413350at7742
## Thu Jul 14 02:13:01 CEST 2022 - The aliscore output file does not exist. Check results for BUSCO: 280642at7742
## Thu Jul 14 02:13:01 CEST 2022 - The aliscore output appears to be empty. Check results for BUSCO: 280642at7742
## Thu Jul 14 02:13:03 CEST 2022 - The aliscore output file does not exist. Check results for BUSCO: 267296at7742
## Thu Jul 14 02:13:03 CEST 2022 - The aliscore output appears to be empty. Check results for BUSCO: 267296at7742
## Thu Jul 14 02:13:04 CEST 2022 - The aliscore output file does not exist. Check results for BUSCO: 166380at7742
## Thu Jul 14 02:13:04 CEST 2022 - The aliscore output appears to be empty. Check results for BUSCO: 166380at7742
## Thu Jul 14 02:13:05 CEST 2022 - The aliscore output file does not exist. Check results for BUSCO: 317170at7742
## Thu Jul 14 02:13:05 CEST 2022 - The aliscore output appears to be empty. Check results for BUSCO: 317170at7742
## Thu Jul 14 02:13:06 CEST 2022 - The aliscore output file does not exist. Check results for BUSCO: 251896at7742
## Thu Jul 14 02:13:06 CEST 2022 - The aliscore output appears to be empty. Check results for BUSCO: 251896at7742
## Thu Jul 14 02:13:07 CEST 2022 - The aliscore output file does not exist. Check results for BUSCO: 210234at7742
## Thu Jul 14 02:13:07 CEST 2022 - The aliscore output appears to be empty. Check results for BUSCO: 210234at7742
## Thu Jul 14 02:13:07 CEST 2022 - The aliscore output file does not exist. Check results for BUSCO: 223813at7742
## Thu Jul 14 02:13:07 CEST 2022 - The aliscore output appears to be empty. Check results for BUSCO: 223813at7742
## Thu Jul 14 02:13:09 CEST 2022 - The aliscore output file does not exist. Check results for BUSCO: 324664at7742
## Thu Jul 14 02:13:09 CEST 2022 - The aliscore output appears to be empty. Check results for BUSCO: 324664at7742
## Thu Jul 14 02:13:09 CEST 2022 - The aliscore output file does not exist. Check results for BUSCO: 246297at7742
## Thu Jul 14 02:13:09 CEST 2022 - The aliscore output appears to be empty. Check results for BUSCO: 246297at7742
## Thu Jul 14 02:13:10 CEST 2022 - The aliscore output file does not exist. Check results for BUSCO: 254518at7742
## Thu Jul 14 02:13:10 CEST 2022 - The aliscore output appears to be empty. Check results for BUSCO: 254518at7742
## Thu Jul 14 02:13:12 CEST 2022 - The aliscore output file does not exist. Check results for BUSCO: 239695at7742
## Thu Jul 14 02:13:12 CEST 2022 - The aliscore output appears to be empty. Check results for BUSCO: 239695at7742
## Thu Jul 14 02:13:14 CEST 2022 - The aliscore output file does not exist. Check results for BUSCO: 182599at7742
## Thu Jul 14 02:13:14 CEST 2022 - The aliscore output appears to be empty. Check results for BUSCO: 182599at7742
## Thu Jul 14 02:13:16 CEST 2022 - The aliscore output file does not exist. Check results for BUSCO: 223375at7742
## Thu Jul 14 02:13:16 CEST 2022 - The aliscore output appears to be empty. Check results for BUSCO: 223375at7742
## Thu Jul 14 02:13:16 CEST 2022 - The aliscore output file does not exist. Check results for BUSCO: 321845at7742
## Thu Jul 14 02:13:16 CEST 2022 - The aliscore output appears to be empty. Check results for BUSCO: 321845at7742
## Thu Jul 14 02:13:16 CEST 2022 - The aliscore output file does not exist. Check results for BUSCO: 356132at7742
## Thu Jul 14 02:13:16 CEST 2022 - The aliscore output appears to be empty. Check results for BUSCO: 356132at7742
## Thu Jul 14 02:13:18 CEST 2022 - The aliscore output file does not exist. Check results for BUSCO: 229308at7742
## Thu Jul 14 02:13:18 CEST 2022 - The aliscore output appears to be empty. Check results for BUSCO: 229308at7742
## Thu Jul 14 02:13:18 CEST 2022 - The aliscore output file does not exist. Check results for BUSCO: 288347at7742
## Thu Jul 14 02:13:18 CEST 2022 - The aliscore output appears to be empty. Check results for BUSCO: 288347at7742
## Thu Jul 14 02:13:18 CEST 2022 - The aliscore output file does not exist. Check results for BUSCO: 271228at7742
## Thu Jul 14 02:13:18 CEST 2022 - The aliscore output appears to be empty. Check results for BUSCO: 271228at7742
## Thu Jul 14 02:13:18 CEST 2022 - The aliscore output file does not exist. Check results for BUSCO: 412231at7742
## Thu Jul 14 02:13:19 CEST 2022 - The aliscore output appears to be empty. Check results for BUSCO: 412231at7742
## Thu Jul 14 02:13:20 CEST 2022 - The aliscore output file does not exist. Check results for BUSCO: 361680at7742
## Thu Jul 14 02:13:20 CEST 2022 - The aliscore output appears to be empty. Check results for BUSCO: 361680at7742
## Thu Jul 14 02:13:21 CEST 2022 - The aliscore output file does not exist. Check results for BUSCO: 279400at7742
## Thu Jul 14 02:13:21 CEST 2022 - The aliscore output appears to be empty. Check results for BUSCO: 279400at7742
## Thu Jul 14 02:13:23 CEST 2022 - The aliscore output file does not exist. Check results for BUSCO: 275715at7742
## Thu Jul 14 02:13:23 CEST 2022 - The aliscore output appears to be empty. Check results for BUSCO: 275715at7742
## Thu Jul 14 02:13:23 CEST 2022 - The aliscore output file does not exist. Check results for BUSCO: 229308at7742
## Thu Jul 14 02:13:23 CEST 2022 - The aliscore output appears to be empty. Check results for BUSCO: 229308at7742
## Thu Jul 14 02:13:24 CEST 2022 - The aliscore output file does not exist. Check results for BUSCO: 275715at7742
## Thu Jul 14 02:13:24 CEST 2022 - The aliscore output appears to be empty. Check results for BUSCO: 275715at7742
## Thu Jul 14 02:13:25 CEST 2022 - The aliscore output file does not exist. Check results for BUSCO: 317401at7742
## Thu Jul 14 02:13:26 CEST 2022 - The aliscore output appears to be empty. Check results for BUSCO: 317401at7742
## Thu Jul 14 02:13:27 CEST 2022 - The aliscore output file does not exist. Check results for BUSCO: 298043at7742
## Thu Jul 14 02:13:27 CEST 2022 - The aliscore output appears to be empty. Check results for BUSCO: 298043at7742
## Thu Jul 14 02:13:28 CEST 2022 - The aliscore output file does not exist. Check results for BUSCO: 197667at7742
## Thu Jul 14 02:13:28 CEST 2022 - The aliscore output appears to be empty. Check results for BUSCO: 197667at7742
## Thu Jul 14 02:13:29 CEST 2022 - The aliscore output file does not exist. Check results for BUSCO: 325125at7742
## Thu Jul 14 02:13:29 CEST 2022 - The aliscore output appears to be empty. Check results for BUSCO: 325125at7742
## Thu Jul 14 02:13:30 CEST 2022 - The aliscore output file does not exist. Check results for BUSCO: 280642at7742
## Thu Jul 14 02:13:30 CEST 2022 - The aliscore output appears to be empty. Check results for BUSCO: 280642at7742
## Thu Jul 14 02:13:31 CEST 2022 - The aliscore output file does not exist. Check results for BUSCO: 280649at7742
## Thu Jul 14 02:13:31 CEST 2022 - The aliscore output appears to be empty. Check results for BUSCO: 280649at7742
## Thu Jul 14 02:13:32 CEST 2022 - The aliscore output file does not exist. Check results for BUSCO: 357094at7742
## Thu Jul 14 02:13:32 CEST 2022 - The aliscore output appears to be empty. Check results for BUSCO: 357094at7742
## Thu Jul 14 02:13:33 CEST 2022 - The aliscore output file does not exist. Check results for BUSCO: 331959at7742
## Thu Jul 14 02:13:33 CEST 2022 - The aliscore output appears to be empty. Check results for BUSCO: 331959at7742
## Thu Jul 14 02:13:34 CEST 2022 - The aliscore output file does not exist. Check results for BUSCO: 271060at7742
## Thu Jul 14 02:13:34 CEST 2022 - The aliscore output appears to be empty. Check results for BUSCO: 271060at7742
## Thu Jul 14 02:13:35 CEST 2022 - The aliscore output file does not exist. Check results for BUSCO: 205146at7742
## Thu Jul 14 02:13:35 CEST 2022 - The aliscore output appears to be empty. Check results for BUSCO: 205146at7742
## Thu Jul 14 02:13:36 CEST 2022 - The aliscore output file does not exist. Check results for BUSCO: 255497at7742
## Thu Jul 14 02:13:36 CEST 2022 - The aliscore output appears to be empty. Check results for BUSCO: 255497at7742
## Thu Jul 14 02:13:37 CEST 2022 - The aliscore output file does not exist. Check results for BUSCO: 364320at7742
## Thu Jul 14 02:13:37 CEST 2022 - The aliscore output appears to be empty. Check results for BUSCO: 364320at7742
## Thu Jul 14 02:13:38 CEST 2022 - The aliscore output file does not exist. Check results for BUSCO: 270432at7742
## Thu Jul 14 02:13:38 CEST 2022 - The aliscore output appears to be empty. Check results for BUSCO: 270432at7742
## Thu Jul 14 02:13:39 CEST 2022 - The aliscore output file does not exist. Check results for BUSCO: 321845at7742
## Thu Jul 14 02:13:39 CEST 2022 - The aliscore output appears to be empty. Check results for BUSCO: 321845at7742
## Thu Jul 14 02:13:39 CEST 2022 - The aliscore output file does not exist. Check results for BUSCO: 303047at7742
## Thu Jul 14 02:13:40 CEST 2022 - The aliscore output appears to be empty. Check results for BUSCO: 303047at7742
## Thu Jul 14 02:13:40 CEST 2022 - The aliscore output file does not exist. Check results for BUSCO: 336615at7742
## Thu Jul 14 02:13:40 CEST 2022 - The aliscore output appears to be empty. Check results for BUSCO: 336615at7742
## Thu Jul 14 02:13:41 CEST 2022 - The aliscore output file does not exist. Check results for BUSCO: 423159at7742
## Thu Jul 14 02:13:41 CEST 2022 - The aliscore output appears to be empty. Check results for BUSCO: 423159at7742
## Thu Jul 14 02:13:42 CEST 2022 - The aliscore output file does not exist. Check results for BUSCO: 227342at7742
## Thu Jul 14 02:13:42 CEST 2022 - The aliscore output appears to be empty. Check results for BUSCO: 227342at7742
## Thu Jul 14 02:13:43 CEST 2022 - The aliscore output file does not exist. Check results for BUSCO: 266826at7742
## Thu Jul 14 02:13:43 CEST 2022 - The aliscore output appears to be empty. Check results for BUSCO: 266826at7742
## Thu Jul 14 02:13:44 CEST 2022 - The aliscore output file does not exist. Check results for BUSCO: 236514at7742
## Thu Jul 14 02:13:44 CEST 2022 - The aliscore output appears to be empty. Check results for BUSCO: 236514at7742
## Thu Jul 14 02:13:45 CEST 2022 - The aliscore output file does not exist. Check results for BUSCO: 209555at7742
## Thu Jul 14 02:13:45 CEST 2022 - The aliscore output appears to be empty. Check results for BUSCO: 209555at7742
## Thu Jul 14 02:13:46 CEST 2022 - The aliscore output file does not exist. Check results for BUSCO: 300333at7742
## Thu Jul 14 02:13:46 CEST 2022 - The aliscore output appears to be empty. Check results for BUSCO: 300333at7742
## Thu Jul 14 02:13:47 CEST 2022 - The aliscore output file does not exist. Check results for BUSCO: 249392at7742
## Thu Jul 14 02:13:47 CEST 2022 - The aliscore output appears to be empty. Check results for BUSCO: 249392at7742
## Thu Jul 14 02:13:47 CEST 2022 - The aliscore output file does not exist. Check results for BUSCO: 340073at7742
## Thu Jul 14 02:13:47 CEST 2022 - The aliscore output appears to be empty. Check results for BUSCO: 340073at7742
## Thu Jul 14 02:13:49 CEST 2022 - The aliscore output file does not exist. Check results for BUSCO: 243977at7742
## Thu Jul 14 02:13:49 CEST 2022 - The aliscore output appears to be empty. Check results for BUSCO: 243977at7742
## Thu Jul 14 02:13:50 CEST 2022 - The aliscore output file does not exist. Check results for BUSCO: 369081at7742
## Thu Jul 14 02:13:50 CEST 2022 - The aliscore output appears to be empty. Check results for BUSCO: 369081at7742
## Thu Jul 14 02:13:52 CEST 2022 - The aliscore output file does not exist. Check results for BUSCO: 229681at7742
## Thu Jul 14 02:13:52 CEST 2022 - The aliscore output appears to be empty. Check results for BUSCO: 229681at7742
## Thu Jul 14 02:13:52 CEST 2022 - The aliscore output file does not exist. Check results for BUSCO: 407084at7742
## Thu Jul 14 02:13:52 CEST 2022 - The aliscore output appears to be empty. Check results for BUSCO: 407084at7742
## Thu Jul 14 02:13:53 CEST 2022 - The aliscore output file does not exist. Check results for BUSCO: 270435at7742
## Thu Jul 14 02:13:53 CEST 2022 - The aliscore output appears to be empty. Check results for BUSCO: 270435at7742
## Thu Jul 14 02:13:54 CEST 2022 - The aliscore output file does not exist. Check results for BUSCO: 270868at7742
## Thu Jul 14 02:13:54 CEST 2022 - The aliscore output file does not exist. Check results for BUSCO: 227342at7742
## Thu Jul 14 02:13:54 CEST 2022 - The aliscore output appears to be empty. Check results for BUSCO: 270868at7742
## Thu Jul 14 02:13:55 CEST 2022 - The aliscore output appears to be empty. Check results for BUSCO: 227342at7742
## Thu Jul 14 02:13:56 CEST 2022 - The aliscore output file does not exist. Check results for BUSCO: 241288at7742
## Thu Jul 14 02:13:56 CEST 2022 - The aliscore output appears to be empty. Check results for BUSCO: 241288at7742
## Thu Jul 14 02:13:56 CEST 2022 - The aliscore output file does not exist. Check results for BUSCO: 336615at7742
## Thu Jul 14 02:13:56 CEST 2022 - The aliscore output appears to be empty. Check results for BUSCO: 336615at7742
## Thu Jul 14 02:13:58 CEST 2022 - The aliscore output file does not exist. Check results for BUSCO: 417286at7742
## Thu Jul 14 02:13:58 CEST 2022 - The aliscore output appears to be empty. Check results for BUSCO: 417286at7742
## Thu Jul 14 02:14:00 CEST 2022 - The aliscore output file does not exist. Check results for BUSCO: 223813at7742
## Thu Jul 14 02:14:00 CEST 2022 - The aliscore output appears to be empty. Check results for BUSCO: 223813at7742
## Thu Jul 14 02:14:00 CEST 2022 - The aliscore output file does not exist. Check results for BUSCO: 330751at7742
## Thu Jul 14 02:14:00 CEST 2022 - The aliscore output appears to be empty. Check results for BUSCO: 330751at7742
## Thu Jul 14 02:14:00 CEST 2022 - The aliscore output file does not exist. Check results for BUSCO: 257196at7742
## Thu Jul 14 02:14:00 CEST 2022 - The aliscore output appears to be empty. Check results for BUSCO: 257196at7742
## Thu Jul 14 02:14:02 CEST 2022 - The aliscore output file does not exist. Check results for BUSCO: 317375at7742
## Thu Jul 14 02:14:02 CEST 2022 - The aliscore output appears to be empty. Check results for BUSCO: 317375at7742
## Thu Jul 14 02:14:02 CEST 2022 - The aliscore output file does not exist. Check results for BUSCO: 240465at7742
## Thu Jul 14 02:14:02 CEST 2022 - The aliscore output appears to be empty. Check results for BUSCO: 240465at7742
## Thu Jul 14 02:14:04 CEST 2022 - The aliscore output file does not exist. Check results for BUSCO: 325125at7742
## Thu Jul 14 02:14:04 CEST 2022 - The aliscore output appears to be empty. Check results for BUSCO: 325125at7742
## Thu Jul 14 02:14:05 CEST 2022 - The aliscore output file does not exist. Check results for BUSCO: 359330at7742
## Thu Jul 14 02:14:05 CEST 2022 - The aliscore output appears to be empty. Check results for BUSCO: 359330at7742
## Thu Jul 14 02:14:05 CEST 2022 - The aliscore output file does not exist. Check results for BUSCO: 253322at7742
## Thu Jul 14 02:14:05 CEST 2022 - The aliscore output appears to be empty. Check results for BUSCO: 253322at7742
## Thu Jul 14 02:14:07 CEST 2022 - The aliscore output file does not exist. Check results for BUSCO: 216474at7742
## Thu Jul 14 02:14:07 CEST 2022 - The aliscore output appears to be empty. Check results for BUSCO: 216474at7742
## Thu Jul 14 02:14:08 CEST 2022 - The aliscore output file does not exist. Check results for BUSCO: 291229at7742
## Thu Jul 14 02:14:08 CEST 2022 - The aliscore output appears to be empty. Check results for BUSCO: 291229at7742
## Thu Jul 14 02:14:08 CEST 2022 - The aliscore output file does not exist. Check results for BUSCO: 201147at7742
## Thu Jul 14 02:14:08 CEST 2022 - The aliscore output appears to be empty. Check results for BUSCO: 201147at7742
## Thu Jul 14 02:14:10 CEST 2022 - The aliscore output file does not exist. Check results for BUSCO: 357819at7742
## Thu Jul 14 02:14:10 CEST 2022 - The aliscore output appears to be empty. Check results for BUSCO: 357819at7742
## Thu Jul 14 02:14:12 CEST 2022 - The aliscore output file does not exist. Check results for BUSCO: 262189at7742
## Thu Jul 14 02:14:12 CEST 2022 - The aliscore output appears to be empty. Check results for BUSCO: 262189at7742
## Thu Jul 14 02:14:12 CEST 2022 - The aliscore output file does not exist. Check results for BUSCO: 391492at7742
## Thu Jul 14 02:14:12 CEST 2022 - The aliscore output appears to be empty. Check results for BUSCO: 391492at7742
## Thu Jul 14 02:14:13 CEST 2022 - The aliscore output file does not exist. Check results for BUSCO: 243836at7742
## Thu Jul 14 02:14:13 CEST 2022 - The aliscore output appears to be empty. Check results for BUSCO: 243836at7742
## Thu Jul 14 02:14:14 CEST 2022 - The aliscore output file does not exist. Check results for BUSCO: 197667at7742
## Thu Jul 14 02:14:14 CEST 2022 - The aliscore output appears to be empty. Check results for BUSCO: 197667at7742
## Thu Jul 14 02:14:16 CEST 2022 - The aliscore output file does not exist. Check results for BUSCO: 252714at7742
## Thu Jul 14 02:14:16 CEST 2022 - The aliscore output appears to be empty. Check results for BUSCO: 252714at7742
## Thu Jul 14 02:14:16 CEST 2022 - The aliscore output file does not exist. Check results for BUSCO: 293076at7742
## Thu Jul 14 02:14:16 CEST 2022 - The aliscore output appears to be empty. Check results for BUSCO: 293076at7742
## Thu Jul 14 02:14:17 CEST 2022 - The aliscore output file does not exist. Check results for BUSCO: 203767at7742
## Thu Jul 14 02:14:17 CEST 2022 - The aliscore output appears to be empty. Check results for BUSCO: 203767at7742
## Thu Jul 14 02:14:19 CEST 2022 - The aliscore output file does not exist. Check results for BUSCO: 216474at7742
## Thu Jul 14 02:14:19 CEST 2022 - The aliscore output appears to be empty. Check results for BUSCO: 216474at7742
## Thu Jul 14 02:14:19 CEST 2022 - The aliscore output file does not exist. Check results for BUSCO: 403632at7742
## Thu Jul 14 02:14:19 CEST 2022 - The aliscore output appears to be empty. Check results for BUSCO: 403632at7742
## Thu Jul 14 02:14:20 CEST 2022 - The aliscore output file does not exist. Check results for BUSCO: 317161at7742
## Thu Jul 14 02:14:20 CEST 2022 - The aliscore output appears to be empty. Check results for BUSCO: 317161at7742
## Thu Jul 14 02:14:21 CEST 2022 - The aliscore output file does not exist. Check results for BUSCO: 358920at7742
## Thu Jul 14 02:14:21 CEST 2022 - The aliscore output appears to be empty. Check results for BUSCO: 358920at7742
## Thu Jul 14 02:14:21 CEST 2022 - The aliscore output file does not exist. Check results for BUSCO: 340020at7742
## Thu Jul 14 02:14:21 CEST 2022 - The aliscore output appears to be empty. Check results for BUSCO: 340020at7742
## Thu Jul 14 02:14:22 CEST 2022 - The aliscore output file does not exist. Check results for BUSCO: 326776at7742
## Thu Jul 14 02:14:22 CEST 2022 - The aliscore output appears to be empty. Check results for BUSCO: 326776at7742
## Thu Jul 14 02:14:24 CEST 2022 - The aliscore output file does not exist. Check results for BUSCO: 305356at7742
## Thu Jul 14 02:14:24 CEST 2022 - The aliscore output appears to be empty. Check results for BUSCO: 305356at7742
## Thu Jul 14 02:14:25 CEST 2022 - The aliscore output file does not exist. Check results for BUSCO: 303047at7742
## Thu Jul 14 02:14:25 CEST 2022 - The aliscore output appears to be empty. Check results for BUSCO: 303047at7742
## Thu Jul 14 02:14:25 CEST 2022 - The aliscore output file does not exist. Check results for BUSCO: 240465at7742
## Thu Jul 14 02:14:26 CEST 2022 - The aliscore output appears to be empty. Check results for BUSCO: 240465at7742
## Thu Jul 14 02:14:27 CEST 2022 - The aliscore output file does not exist. Check results for BUSCO: 289044at7742
## Thu Jul 14 02:14:27 CEST 2022 - The aliscore output appears to be empty. Check results for BUSCO: 289044at7742
## Thu Jul 14 02:14:28 CEST 2022 - The aliscore output file does not exist. Check results for BUSCO: 351419at7742
## Thu Jul 14 02:14:28 CEST 2022 - The aliscore output appears to be empty. Check results for BUSCO: 351419at7742
## Thu Jul 14 02:14:29 CEST 2022 - The aliscore output file does not exist. Check results for BUSCO: 359032at7742
## Thu Jul 14 02:14:29 CEST 2022 - The aliscore output appears to be empty. Check results for BUSCO: 359032at7742
## Thu Jul 14 02:14:30 CEST 2022 - The aliscore output file does not exist. Check results for BUSCO: 302536at7742
## Thu Jul 14 02:14:30 CEST 2022 - The aliscore output appears to be empty. Check results for BUSCO: 302536at7742
## Thu Jul 14 02:14:31 CEST 2022 - The aliscore output file does not exist. Check results for BUSCO: 280649at7742
## Thu Jul 14 02:14:31 CEST 2022 - The aliscore output appears to be empty. Check results for BUSCO: 280649at7742
## Thu Jul 14 02:14:31 CEST 2022 - The aliscore output file does not exist. Check results for BUSCO: 359330at7742
## Thu Jul 14 02:14:31 CEST 2022 - The aliscore output appears to be empty. Check results for BUSCO: 359330at7742
## Thu Jul 14 02:14:34 CEST 2022 - The aliscore output file does not exist. Check results for BUSCO: 307415at7742
## Thu Jul 14 02:14:34 CEST 2022 - The aliscore output appears to be empty. Check results for BUSCO: 307415at7742
## Thu Jul 14 02:14:35 CEST 2022 - The aliscore output file does not exist. Check results for BUSCO: 405299at7742
## Thu Jul 14 02:14:35 CEST 2022 - The aliscore output appears to be empty. Check results for BUSCO: 405299at7742
## Thu Jul 14 02:14:36 CEST 2022 - The aliscore output file does not exist. Check results for BUSCO: 386558at7742
## Thu Jul 14 02:14:36 CEST 2022 - The aliscore output appears to be empty. Check results for BUSCO: 386558at7742
## Thu Jul 14 02:14:36 CEST 2022 - The aliscore output file does not exist. Check results for BUSCO: 365226at7742
## Thu Jul 14 02:14:36 CEST 2022 - The aliscore output appears to be empty. Check results for BUSCO: 365226at7742
## Thu Jul 14 02:14:37 CEST 2022 - The aliscore output file does not exist. Check results for BUSCO: 252200at7742
## Thu Jul 14 02:14:37 CEST 2022 - The aliscore output appears to be empty. Check results for BUSCO: 252200at7742
## Thu Jul 14 02:14:38 CEST 2022 - The aliscore output file does not exist. Check results for BUSCO: 195562at7742
## Thu Jul 14 02:14:38 CEST 2022 - The aliscore output appears to be empty. Check results for BUSCO: 195562at7742
## Thu Jul 14 02:14:39 CEST 2022 - The aliscore output file does not exist. Check results for BUSCO: 197276at7742
## Thu Jul 14 02:14:39 CEST 2022 - The aliscore output appears to be empty. Check results for BUSCO: 197276at7742
## Thu Jul 14 02:14:40 CEST 2022 - The aliscore output file does not exist. Check results for BUSCO: 268420at7742
## Thu Jul 14 02:14:40 CEST 2022 - The aliscore output appears to be empty. Check results for BUSCO: 268420at7742
## Thu Jul 14 02:14:41 CEST 2022 - The aliscore output file does not exist. Check results for BUSCO: 361680at7742
## Thu Jul 14 02:14:41 CEST 2022 - The aliscore output appears to be empty. Check results for BUSCO: 361680at7742
## Thu Jul 14 02:14:42 CEST 2022 - The aliscore output file does not exist. Check results for BUSCO: 336615at7742
## Thu Jul 14 02:14:42 CEST 2022 - The aliscore output appears to be empty. Check results for BUSCO: 336615at7742
## Thu Jul 14 02:14:44 CEST 2022 - The aliscore output file does not exist. Check results for BUSCO: 317401at7742
## Thu Jul 14 02:14:44 CEST 2022 - The aliscore output appears to be empty. Check results for BUSCO: 317401at7742
## Thu Jul 14 02:14:44 CEST 2022 - The aliscore output file does not exist. Check results for BUSCO: 371168at7742
## Thu Jul 14 02:14:44 CEST 2022 - The aliscore output appears to be empty. Check results for BUSCO: 371168at7742
## Thu Jul 14 02:14:46 CEST 2022 - The aliscore output file does not exist. Check results for BUSCO: 365136at7742
## Thu Jul 14 02:14:46 CEST 2022 - The aliscore output appears to be empty. Check results for BUSCO: 365136at7742
## Thu Jul 14 02:14:47 CEST 2022 - The aliscore output file does not exist. Check results for BUSCO: 386558at7742
## Thu Jul 14 02:14:47 CEST 2022 - The aliscore output appears to be empty. Check results for BUSCO: 386558at7742
## Thu Jul 14 02:14:48 CEST 2022 - The aliscore output file does not exist. Check results for BUSCO: 261158at7742
## Thu Jul 14 02:14:48 CEST 2022 - The aliscore output appears to be empty. Check results for BUSCO: 261158at7742
## Thu Jul 14 02:14:48 CEST 2022 - The aliscore output file does not exist. Check results for BUSCO: 259157at7742
## Thu Jul 14 02:14:48 CEST 2022 - The aliscore output appears to be empty. Check results for BUSCO: 259157at7742
## Thu Jul 14 02:14:50 CEST 2022 - The aliscore output file does not exist. Check results for BUSCO: 316568at7742
## Thu Jul 14 02:14:50 CEST 2022 - The aliscore output appears to be empty. Check results for BUSCO: 316568at7742
## Thu Jul 14 02:14:51 CEST 2022 - The aliscore output file does not exist. Check results for BUSCO: 374996at7742
## Thu Jul 14 02:14:51 CEST 2022 - The aliscore output appears to be empty. Check results for BUSCO: 374996at7742
## Thu Jul 14 02:14:51 CEST 2022 - The aliscore output file does not exist. Check results for BUSCO: 220264at7742
## Thu Jul 14 02:14:51 CEST 2022 - The aliscore output appears to be empty. Check results for BUSCO: 220264at7742
## Thu Jul 14 02:14:54 CEST 2022 - The aliscore output file does not exist. Check results for BUSCO: 353474at7742
## Thu Jul 14 02:14:54 CEST 2022 - The aliscore output appears to be empty. Check results for BUSCO: 353474at7742
## Thu Jul 14 02:14:54 CEST 2022 - The aliscore output file does not exist. Check results for BUSCO: 376105at7742
## Thu Jul 14 02:14:54 CEST 2022 - The aliscore output appears to be empty. Check results for BUSCO: 376105at7742
## Thu Jul 14 02:14:55 CEST 2022 - The aliscore output file does not exist. Check results for BUSCO: 276761at7742
## Thu Jul 14 02:14:55 CEST 2022 - The aliscore output appears to be empty. Check results for BUSCO: 276761at7742
## Thu Jul 14 02:14:56 CEST 2022 - The aliscore output file does not exist. Check results for BUSCO: 266826at7742
## Thu Jul 14 02:14:56 CEST 2022 - The aliscore output appears to be empty. Check results for BUSCO: 266826at7742
## Thu Jul 14 02:14:56 CEST 2022 - The aliscore output file does not exist. Check results for BUSCO: 229308at7742
## Thu Jul 14 02:14:57 CEST 2022 - The aliscore output appears to be empty. Check results for BUSCO: 229308at7742
## Thu Jul 14 02:14:58 CEST 2022 - The aliscore output file does not exist. Check results for BUSCO: 245299at7742
## Thu Jul 14 02:14:58 CEST 2022 - The aliscore output appears to be empty. Check results for BUSCO: 245299at7742
## Thu Jul 14 02:14:58 CEST 2022 - The aliscore output file does not exist. Check results for BUSCO: 265605at7742
## Thu Jul 14 02:14:58 CEST 2022 - The aliscore output appears to be empty. Check results for BUSCO: 265605at7742
## Thu Jul 14 02:14:59 CEST 2022 - The aliscore output file does not exist. Check results for BUSCO: 409719at7742
## Thu Jul 14 02:14:59 CEST 2022 - The aliscore output appears to be empty. Check results for BUSCO: 409719at7742
## Thu Jul 14 02:15:00 CEST 2022 - The aliscore output file does not exist. Check results for BUSCO: 377177at7742
## Thu Jul 14 02:15:00 CEST 2022 - The aliscore output appears to be empty. Check results for BUSCO: 377177at7742
## Thu Jul 14 02:15:02 CEST 2022 - The aliscore output file does not exist. Check results for BUSCO: 243425at7742
## Thu Jul 14 02:15:02 CEST 2022 - The aliscore output file does not exist. Check results for BUSCO: 237440at7742
## Thu Jul 14 02:15:02 CEST 2022 - The aliscore output appears to be empty. Check results for BUSCO: 243425at7742
## Thu Jul 14 02:15:02 CEST 2022 - The aliscore output appears to be empty. Check results for BUSCO: 237440at7742
## Thu Jul 14 02:15:04 CEST 2022 - The aliscore output file does not exist. Check results for BUSCO: 270868at7742
## Thu Jul 14 02:15:04 CEST 2022 - The aliscore output appears to be empty. Check results for BUSCO: 270868at7742
## Thu Jul 14 02:15:05 CEST 2022 - The aliscore output file does not exist. Check results for BUSCO: 303510at7742
## Thu Jul 14 02:15:05 CEST 2022 - The aliscore output appears to be empty. Check results for BUSCO: 303510at7742
## Thu Jul 14 02:15:05 CEST 2022 - The aliscore output file does not exist. Check results for BUSCO: 323737at7742
## Thu Jul 14 02:15:05 CEST 2022 - The aliscore output appears to be empty. Check results for BUSCO: 323737at7742
## Thu Jul 14 02:15:06 CEST 2022 - The aliscore output file does not exist. Check results for BUSCO: 309935at7742
## Thu Jul 14 02:15:07 CEST 2022 - The aliscore output appears to be empty. Check results for BUSCO: 309935at7742
## Thu Jul 14 02:15:08 CEST 2022 - The aliscore output file does not exist. Check results for BUSCO: 335129at7742
## Thu Jul 14 02:15:08 CEST 2022 - The aliscore output appears to be empty. Check results for BUSCO: 335129at7742
## Thu Jul 14 02:15:08 CEST 2022 - The aliscore output file does not exist. Check results for BUSCO: 360873at7742
## Thu Jul 14 02:15:08 CEST 2022 - The aliscore output appears to be empty. Check results for BUSCO: 360873at7742
## Thu Jul 14 02:15:10 CEST 2022 - The aliscore output file does not exist. Check results for BUSCO: 277524at7742
## Thu Jul 14 02:15:10 CEST 2022 - The aliscore output appears to be empty. Check results for BUSCO: 277524at7742
## Thu Jul 14 02:15:11 CEST 2022 - The aliscore output file does not exist. Check results for BUSCO: 326524at7742
## Thu Jul 14 02:15:11 CEST 2022 - The aliscore output appears to be empty. Check results for BUSCO: 326524at7742
## Thu Jul 14 02:15:11 CEST 2022 - The aliscore output file does not exist. Check results for BUSCO: 206164at7742
## Thu Jul 14 02:15:11 CEST 2022 - The aliscore output appears to be empty. Check results for BUSCO: 206164at7742
## Thu Jul 14 02:15:13 CEST 2022 - The aliscore output file does not exist. Check results for BUSCO: 371168at7742
## Thu Jul 14 02:15:13 CEST 2022 - The aliscore output appears to be empty. Check results for BUSCO: 371168at7742
## Thu Jul 14 02:15:13 CEST 2022 - The aliscore output file does not exist. Check results for BUSCO: 298137at7742
## Thu Jul 14 02:15:13 CEST 2022 - The aliscore output appears to be empty. Check results for BUSCO: 298137at7742
## Thu Jul 14 02:15:14 CEST 2022 - The aliscore output file does not exist. Check results for BUSCO: 195562at7742
## Thu Jul 14 02:15:14 CEST 2022 - The aliscore output appears to be empty. Check results for BUSCO: 195562at7742
## Thu Jul 14 02:15:15 CEST 2022 - The aliscore output file does not exist. Check results for BUSCO: 375593at7742
## Thu Jul 14 02:15:15 CEST 2022 - The aliscore output appears to be empty. Check results for BUSCO: 375593at7742
## Thu Jul 14 02:15:16 CEST 2022 - The aliscore output file does not exist. Check results for BUSCO: 320985at7742
## Thu Jul 14 02:15:16 CEST 2022 - The aliscore output appears to be empty. Check results for BUSCO: 320985at7742
## Thu Jul 14 02:15:16 CEST 2022 - The aliscore output file does not exist. Check results for BUSCO: 322345at7742
## Thu Jul 14 02:15:17 CEST 2022 - The aliscore output appears to be empty. Check results for BUSCO: 322345at7742
## Thu Jul 14 02:15:17 CEST 2022 - The aliscore output file does not exist. Check results for BUSCO: 293076at7742
## Thu Jul 14 02:15:17 CEST 2022 - The aliscore output appears to be empty. Check results for BUSCO: 293076at7742
## Thu Jul 14 02:15:19 CEST 2022 - The aliscore output file does not exist. Check results for BUSCO: 414201at7742
## Thu Jul 14 02:15:19 CEST 2022 - The aliscore output appears to be empty. Check results for BUSCO: 414201at7742
## Thu Jul 14 02:15:19 CEST 2022 - The aliscore output file does not exist. Check results for BUSCO: 375859at7742
## Thu Jul 14 02:15:19 CEST 2022 - The aliscore output appears to be empty. Check results for BUSCO: 375859at7742
## Thu Jul 14 02:15:20 CEST 2022 - The aliscore output file does not exist. Check results for BUSCO: 268420at7742
## Thu Jul 14 02:15:21 CEST 2022 - The aliscore output appears to be empty. Check results for BUSCO: 268420at7742
## Thu Jul 14 02:15:21 CEST 2022 - The aliscore output file does not exist. Check results for BUSCO: 254518at7742
## Thu Jul 14 02:15:21 CEST 2022 - The aliscore output appears to be empty. Check results for BUSCO: 254518at7742
## Thu Jul 14 02:15:23 CEST 2022 - The aliscore output file does not exist. Check results for BUSCO: 286089at7742
## Thu Jul 14 02:15:23 CEST 2022 - The aliscore output appears to be empty. Check results for BUSCO: 286089at7742
## Thu Jul 14 02:15:24 CEST 2022 - The aliscore output file does not exist. Check results for BUSCO: 197276at7742
## Thu Jul 14 02:15:24 CEST 2022 - The aliscore output appears to be empty. Check results for BUSCO: 197276at7742
## Thu Jul 14 02:15:25 CEST 2022 - The aliscore output file does not exist. Check results for BUSCO: 250138at7742
## Thu Jul 14 02:15:25 CEST 2022 - The aliscore output appears to be empty. Check results for BUSCO: 250138at7742
## Thu Jul 14 02:15:25 CEST 2022 - The aliscore output file does not exist. Check results for BUSCO: 317161at7742
## Thu Jul 14 02:15:25 CEST 2022 - The aliscore output appears to be empty. Check results for BUSCO: 317161at7742
## Thu Jul 14 02:15:26 CEST 2022 - The aliscore output file does not exist. Check results for BUSCO: 245299at7742
## Thu Jul 14 02:15:26 CEST 2022 - The aliscore output appears to be empty. Check results for BUSCO: 245299at7742
## Thu Jul 14 02:15:28 CEST 2022 - The aliscore output file does not exist. Check results for BUSCO: 298137at7742
## Thu Jul 14 02:15:28 CEST 2022 - The aliscore output appears to be empty. Check results for BUSCO: 298137at7742
## Thu Jul 14 02:15:29 CEST 2022 - The aliscore output file does not exist. Check results for BUSCO: 190657at7742
## Thu Jul 14 02:15:29 CEST 2022 - The aliscore output appears to be empty. Check results for BUSCO: 190657at7742
## Thu Jul 14 02:15:30 CEST 2022 - The aliscore output file does not exist. Check results for BUSCO: 249392at7742
## Thu Jul 14 02:15:30 CEST 2022 - The aliscore output appears to be empty. Check results for BUSCO: 249392at7742
## Thu Jul 14 02:15:31 CEST 2022 - The aliscore output file does not exist. Check results for BUSCO: 201147at7742
## Thu Jul 14 02:15:31 CEST 2022 - The aliscore output appears to be empty. Check results for BUSCO: 201147at7742
## Thu Jul 14 02:15:31 CEST 2022 - The aliscore output file does not exist. Check results for BUSCO: 292556at7742
## Thu Jul 14 02:15:31 CEST 2022 - The aliscore output appears to be empty. Check results for BUSCO: 292556at7742
## Thu Jul 14 02:15:34 CEST 2022 - The aliscore output file does not exist. Check results for BUSCO: 319229at7742
## Thu Jul 14 02:15:34 CEST 2022 - The aliscore output appears to be empty. Check results for BUSCO: 319229at7742
## Thu Jul 14 02:15:34 CEST 2022 - The aliscore output file does not exist. Check results for BUSCO: 302536at7742
## Thu Jul 14 02:15:34 CEST 2022 - The aliscore output appears to be empty. Check results for BUSCO: 302536at7742
## Thu Jul 14 02:15:35 CEST 2022 - The aliscore output file does not exist. Check results for BUSCO: 221230at7742
## Thu Jul 14 02:15:35 CEST 2022 - The aliscore output appears to be empty. Check results for BUSCO: 221230at7742
## Thu Jul 14 02:15:36 CEST 2022 - The aliscore output file does not exist. Check results for BUSCO: 216474at7742
## Thu Jul 14 02:15:36 CEST 2022 - The aliscore output appears to be empty. Check results for BUSCO: 216474at7742
## Thu Jul 14 02:15:37 CEST 2022 - The aliscore output file does not exist. Check results for BUSCO: 331959at7742
## Thu Jul 14 02:15:37 CEST 2022 - The aliscore output appears to be empty. Check results for BUSCO: 331959at7742
## Thu Jul 14 02:15:37 CEST 2022 - The aliscore output file does not exist. Check results for BUSCO: 326524at7742
## Thu Jul 14 02:15:37 CEST 2022 - The aliscore output appears to be empty. Check results for BUSCO: 326524at7742
## Thu Jul 14 02:15:38 CEST 2022 - The aliscore output file does not exist. Check results for BUSCO: 293704at7742
## Thu Jul 14 02:15:38 CEST 2022 - The aliscore output appears to be empty. Check results for BUSCO: 293704at7742
## Thu Jul 14 02:15:39 CEST 2022 - The aliscore output file does not exist. Check results for BUSCO: 371168at7742
## Thu Jul 14 02:15:39 CEST 2022 - The aliscore output appears to be empty. Check results for BUSCO: 371168at7742
## Thu Jul 14 02:15:41 CEST 2022 - The aliscore output file does not exist. Check results for BUSCO: 297599at7742
## Thu Jul 14 02:15:41 CEST 2022 - The aliscore output appears to be empty. Check results for BUSCO: 297599at7742
## Thu Jul 14 02:15:41 CEST 2022 - The aliscore output file does not exist. Check results for BUSCO: 195562at7742
## Thu Jul 14 02:15:41 CEST 2022 - The aliscore output appears to be empty. Check results for BUSCO: 195562at7742
## Thu Jul 14 02:15:43 CEST 2022 - The aliscore output file does not exist. Check results for BUSCO: 298137at7742
## Thu Jul 14 02:15:43 CEST 2022 - The aliscore output appears to be empty. Check results for BUSCO: 298137at7742
## Thu Jul 14 02:15:43 CEST 2022 - The aliscore output file does not exist. Check results for BUSCO: 405299at7742
## Thu Jul 14 02:15:43 CEST 2022 - The aliscore output appears to be empty. Check results for BUSCO: 405299at7742
## Thu Jul 14 02:15:45 CEST 2022 - The aliscore output file does not exist. Check results for BUSCO: 293704at7742
## Thu Jul 14 02:15:45 CEST 2022 - The aliscore output appears to be empty. Check results for BUSCO: 293704at7742
## Thu Jul 14 02:15:45 CEST 2022 - The aliscore output file does not exist. Check results for BUSCO: 261266at7742
## Thu Jul 14 02:15:45 CEST 2022 - The aliscore output appears to be empty. Check results for BUSCO: 261266at7742
## Thu Jul 14 02:15:45 CEST 2022 - The aliscore output file does not exist. Check results for BUSCO: 284527at7742
## Thu Jul 14 02:15:45 CEST 2022 - The aliscore output appears to be empty. Check results for BUSCO: 284527at7742
## Thu Jul 14 02:15:47 CEST 2022 - The aliscore output file does not exist. Check results for BUSCO: 265369at7742
## Thu Jul 14 02:15:47 CEST 2022 - The aliscore output appears to be empty. Check results for BUSCO: 265369at7742
## Thu Jul 14 02:15:48 CEST 2022 - The aliscore output file does not exist. Check results for BUSCO: 345549at7742
## Thu Jul 14 02:15:48 CEST 2022 - The aliscore output appears to be empty. Check results for BUSCO: 345549at7742
## Thu Jul 14 02:15:49 CEST 2022 - The aliscore output file does not exist. Check results for BUSCO: 256968at7742
## Thu Jul 14 02:15:49 CEST 2022 - The aliscore output appears to be empty. Check results for BUSCO: 256968at7742
## Thu Jul 14 02:15:50 CEST 2022 - The aliscore output file does not exist. Check results for BUSCO: 325689at7742
## Thu Jul 14 02:15:50 CEST 2022 - The aliscore output file does not exist. Check results for BUSCO: 375859at7742
## Thu Jul 14 02:15:50 CEST 2022 - The aliscore output appears to be empty. Check results for BUSCO: 325689at7742
## Thu Jul 14 02:15:50 CEST 2022 - The aliscore output appears to be empty. Check results for BUSCO: 375859at7742
## Thu Jul 14 02:15:52 CEST 2022 - The aliscore output file does not exist. Check results for BUSCO: 265605at7742
## Thu Jul 14 02:15:52 CEST 2022 - The aliscore output appears to be empty. Check results for BUSCO: 265605at7742
## Thu Jul 14 02:15:53 CEST 2022 - The aliscore output file does not exist. Check results for BUSCO: 258747at7742
## Thu Jul 14 02:15:53 CEST 2022 - The aliscore output appears to be empty. Check results for BUSCO: 258747at7742
## Thu Jul 14 02:15:53 CEST 2022 - The aliscore output file does not exist. Check results for BUSCO: 211842at7742
## Thu Jul 14 02:15:53 CEST 2022 - The aliscore output appears to be empty. Check results for BUSCO: 211842at7742
## Thu Jul 14 02:15:55 CEST 2022 - The aliscore output file does not exist. Check results for BUSCO: 316568at7742
## Thu Jul 14 02:15:55 CEST 2022 - The aliscore output appears to be empty. Check results for BUSCO: 316568at7742
## Thu Jul 14 02:15:55 CEST 2022 - The aliscore output file does not exist. Check results for BUSCO: 353318at7742
## Thu Jul 14 02:15:55 CEST 2022 - The aliscore output appears to be empty. Check results for BUSCO: 353318at7742
## Thu Jul 14 02:15:56 CEST 2022 - The aliscore output file does not exist. Check results for BUSCO: 376736at7742
## Thu Jul 14 02:15:56 CEST 2022 - The aliscore output appears to be empty. Check results for BUSCO: 376736at7742
## Thu Jul 14 02:15:58 CEST 2022 - The aliscore output file does not exist. Check results for BUSCO: 284527at7742
## Thu Jul 14 02:15:58 CEST 2022 - The aliscore output appears to be empty. Check results for BUSCO: 284527at7742
## Thu Jul 14 02:15:58 CEST 2022 - The aliscore output file does not exist. Check results for BUSCO: 240465at7742
## Thu Jul 14 02:15:58 CEST 2022 - The aliscore output appears to be empty. Check results for BUSCO: 240465at7742
## Thu Jul 14 02:15:59 CEST 2022 - The aliscore output file does not exist. Check results for BUSCO: 269102at7742
## Thu Jul 14 02:15:59 CEST 2022 - The aliscore output appears to be empty. Check results for BUSCO: 269102at7742
## Thu Jul 14 02:16:00 CEST 2022 - The aliscore output file does not exist. Check results for BUSCO: 269102at7742
## Thu Jul 14 02:16:00 CEST 2022 - The aliscore output appears to be empty. Check results for BUSCO: 269102at7742
## Thu Jul 14 02:16:02 CEST 2022 - The aliscore output file does not exist. Check results for BUSCO: 243222at7742
## Thu Jul 14 02:16:02 CEST 2022 - The aliscore output appears to be empty. Check results for BUSCO: 243222at7742
## Thu Jul 14 02:16:02 CEST 2022 - The aliscore output file does not exist. Check results for BUSCO: 326776at7742
## Thu Jul 14 02:16:02 CEST 2022 - The aliscore output appears to be empty. Check results for BUSCO: 326776at7742
## Thu Jul 14 02:16:04 CEST 2022 - The aliscore output file does not exist. Check results for BUSCO: 305356at7742
## Thu Jul 14 02:16:04 CEST 2022 - The aliscore output appears to be empty. Check results for BUSCO: 305356at7742
## Thu Jul 14 02:16:04 CEST 2022 - The aliscore output file does not exist. Check results for BUSCO: 272432at7742
## Thu Jul 14 02:16:04 CEST 2022 - The aliscore output appears to be empty. Check results for BUSCO: 272432at7742
## Thu Jul 14 02:16:06 CEST 2022 - The aliscore output file does not exist. Check results for BUSCO: 301932at7742
## Thu Jul 14 02:16:06 CEST 2022 - The aliscore output appears to be empty. Check results for BUSCO: 301932at7742
## Thu Jul 14 02:16:06 CEST 2022 - The aliscore output file does not exist. Check results for BUSCO: 284527at7742
## Thu Jul 14 02:16:06 CEST 2022 - The aliscore output appears to be empty. Check results for BUSCO: 284527at7742
## Thu Jul 14 02:16:07 CEST 2022 - The aliscore output file does not exist. Check results for BUSCO: 280835at7742
## Thu Jul 14 02:16:08 CEST 2022 - The aliscore output appears to be empty. Check results for BUSCO: 280835at7742
## Thu Jul 14 02:16:08 CEST 2022 - The aliscore output file does not exist. Check results for BUSCO: 259089at7742
## Thu Jul 14 02:16:08 CEST 2022 - The aliscore output appears to be empty. Check results for BUSCO: 259089at7742
## Thu Jul 14 02:16:09 CEST 2022 - The aliscore output file does not exist. Check results for BUSCO: 206784at7742
## Thu Jul 14 02:16:09 CEST 2022 - The aliscore output appears to be empty. Check results for BUSCO: 206784at7742
## Thu Jul 14 02:16:10 CEST 2022 - The aliscore output file does not exist. Check results for BUSCO: 374996at7742
## Thu Jul 14 02:16:10 CEST 2022 - The aliscore output appears to be empty. Check results for BUSCO: 374996at7742
## Thu Jul 14 02:16:11 CEST 2022 - The aliscore output file does not exist. Check results for BUSCO: 251896at7742
## Thu Jul 14 02:16:11 CEST 2022 - The aliscore output appears to be empty. Check results for BUSCO: 251896at7742
## Thu Jul 14 02:16:11 CEST 2022 - The aliscore output file does not exist. Check results for BUSCO: 317021at7742
## Thu Jul 14 02:16:11 CEST 2022 - The aliscore output appears to be empty. Check results for BUSCO: 317021at7742
## Thu Jul 14 02:16:13 CEST 2022 - The aliscore output file does not exist. Check results for BUSCO: 377238at7742
## Thu Jul 14 02:16:13 CEST 2022 - The aliscore output appears to be empty. Check results for BUSCO: 377238at7742
## Thu Jul 14 02:16:16 CEST 2022 - The aliscore output file does not exist. Check results for BUSCO: 382669at7742
## Thu Jul 14 02:16:16 CEST 2022 - The aliscore output appears to be empty. Check results for BUSCO: 382669at7742
## Thu Jul 14 02:16:16 CEST 2022 - The aliscore output file does not exist. Check results for BUSCO: 286089at7742
## Thu Jul 14 02:16:16 CEST 2022 - The aliscore output appears to be empty. Check results for BUSCO: 286089at7742
## Thu Jul 14 02:16:17 CEST 2022 - The aliscore output file does not exist. Check results for BUSCO: 205603at7742
## Thu Jul 14 02:16:17 CEST 2022 - The aliscore output appears to be empty. Check results for BUSCO: 205603at7742
## Thu Jul 14 02:16:17 CEST 2022 - The aliscore output file does not exist. Check results for BUSCO: 382456at7742
## Thu Jul 14 02:16:17 CEST 2022 - The aliscore output appears to be empty. Check results for BUSCO: 382456at7742
## Thu Jul 14 02:16:18 CEST 2022 - The aliscore output file does not exist. Check results for BUSCO: 262189at7742
## Thu Jul 14 02:16:18 CEST 2022 - The aliscore output appears to be empty. Check results for BUSCO: 262189at7742
## Thu Jul 14 02:16:19 CEST 2022 - The aliscore output file does not exist. Check results for BUSCO: 261658at7742
## Thu Jul 14 02:16:19 CEST 2022 - The aliscore output appears to be empty. Check results for BUSCO: 261658at7742
## Thu Jul 14 02:16:20 CEST 2022 - The aliscore output file does not exist. Check results for BUSCO: 336331at7742
## Thu Jul 14 02:16:20 CEST 2022 - The aliscore output appears to be empty. Check results for BUSCO: 336331at7742
## Thu Jul 14 02:16:21 CEST 2022 - The aliscore output file does not exist. Check results for BUSCO: 304804at7742
## Thu Jul 14 02:16:21 CEST 2022 - The aliscore output appears to be empty. Check results for BUSCO: 304804at7742
## Thu Jul 14 02:16:22 CEST 2022 - The aliscore output file does not exist. Check results for BUSCO: 265605at7742
## Thu Jul 14 02:16:22 CEST 2022 - The aliscore output appears to be empty. Check results for BUSCO: 265605at7742
## Thu Jul 14 02:16:23 CEST 2022 - The aliscore output file does not exist. Check results for BUSCO: 339693at7742
## Thu Jul 14 02:16:23 CEST 2022 - The aliscore output appears to be empty. Check results for BUSCO: 339693at7742
## Thu Jul 14 02:16:24 CEST 2022 - The aliscore output file does not exist. Check results for BUSCO: 252200at7742
## Thu Jul 14 02:16:24 CEST 2022 - The aliscore output appears to be empty. Check results for BUSCO: 252200at7742
## Thu Jul 14 02:16:24 CEST 2022 - The aliscore output file does not exist. Check results for BUSCO: 198118at7742
## Thu Jul 14 02:16:24 CEST 2022 - The aliscore output appears to be empty. Check results for BUSCO: 198118at7742
## Thu Jul 14 02:16:25 CEST 2022 - The aliscore output file does not exist. Check results for BUSCO: 198118at7742
## Thu Jul 14 02:16:25 CEST 2022 - The aliscore output appears to be empty. Check results for BUSCO: 198118at7742
## Thu Jul 14 02:16:26 CEST 2022 - The aliscore output file does not exist. Check results for BUSCO: 309962at7742
## Thu Jul 14 02:16:26 CEST 2022 - The aliscore output appears to be empty. Check results for BUSCO: 309962at7742
## Thu Jul 14 02:16:28 CEST 2022 - The aliscore output file does not exist. Check results for BUSCO: 320985at7742
## Thu Jul 14 02:16:28 CEST 2022 - The aliscore output appears to be empty. Check results for BUSCO: 320985at7742
## Thu Jul 14 02:16:29 CEST 2022 - The aliscore output file does not exist. Check results for BUSCO: 320985at7742
## Thu Jul 14 02:16:29 CEST 2022 - The aliscore output appears to be empty. Check results for BUSCO: 320985at7742
## Thu Jul 14 02:16:30 CEST 2022 - The aliscore output file does not exist. Check results for BUSCO: 305356at7742
## Thu Jul 14 02:16:30 CEST 2022 - The aliscore output appears to be empty. Check results for BUSCO: 305356at7742
## Thu Jul 14 02:16:31 CEST 2022 - The aliscore output file does not exist. Check results for BUSCO: 261658at7742
## Thu Jul 14 02:16:31 CEST 2022 - The aliscore output appears to be empty. Check results for BUSCO: 261658at7742
## Thu Jul 14 02:16:32 CEST 2022 - The aliscore output file does not exist. Check results for BUSCO: 327840at7742
## Thu Jul 14 02:16:32 CEST 2022 - The aliscore output appears to be empty. Check results for BUSCO: 327840at7742
## Thu Jul 14 02:16:32 CEST 2022 - The aliscore output file does not exist. Check results for BUSCO: 328134at7742
## Thu Jul 14 02:16:32 CEST 2022 - The aliscore output appears to be empty. Check results for BUSCO: 328134at7742
## Thu Jul 14 02:16:34 CEST 2022 - The aliscore output file does not exist. Check results for BUSCO: 324120at7742
## Thu Jul 14 02:16:34 CEST 2022 - The aliscore output appears to be empty. Check results for BUSCO: 324120at7742
## Thu Jul 14 02:16:34 CEST 2022 - The aliscore output file does not exist. Check results for BUSCO: 407723at7742
## Thu Jul 14 02:16:34 CEST 2022 - The aliscore output appears to be empty. Check results for BUSCO: 407723at7742
## Thu Jul 14 02:16:35 CEST 2022 - The aliscore output file does not exist. Check results for BUSCO: 437044at7742
## Thu Jul 14 02:16:35 CEST 2022 - The aliscore output appears to be empty. Check results for BUSCO: 437044at7742
## Thu Jul 14 02:16:36 CEST 2022 - The aliscore output file does not exist. Check results for BUSCO: 336564at7742
## Thu Jul 14 02:16:37 CEST 2022 - The aliscore output appears to be empty. Check results for BUSCO: 336564at7742
## Thu Jul 14 02:16:38 CEST 2022 - The aliscore output file does not exist. Check results for BUSCO: 396303at7742
## Thu Jul 14 02:16:38 CEST 2022 - The aliscore output appears to be empty. Check results for BUSCO: 396303at7742
## Thu Jul 14 02:16:39 CEST 2022 - The aliscore output file does not exist. Check results for BUSCO: 334381at7742
## Thu Jul 14 02:16:39 CEST 2022 - The aliscore output appears to be empty. Check results for BUSCO: 334381at7742
## Thu Jul 14 02:16:40 CEST 2022 - The aliscore output file does not exist. Check results for BUSCO: 276761at7742
## Thu Jul 14 02:16:40 CEST 2022 - The aliscore output appears to be empty. Check results for BUSCO: 276761at7742
## Thu Jul 14 02:16:41 CEST 2022 - The aliscore output file does not exist. Check results for BUSCO: 340499at7742
## Thu Jul 14 02:16:41 CEST 2022 - The aliscore output appears to be empty. Check results for BUSCO: 340499at7742
## Thu Jul 14 02:16:41 CEST 2022 - The aliscore output file does not exist. Check results for BUSCO: 252200at7742
## Thu Jul 14 02:16:41 CEST 2022 - The aliscore output appears to be empty. Check results for BUSCO: 252200at7742
## Thu Jul 14 02:16:42 CEST 2022 - The aliscore output file does not exist. Check results for BUSCO: 290213at7742
## Thu Jul 14 02:16:42 CEST 2022 - The aliscore output appears to be empty. Check results for BUSCO: 290213at7742
## Thu Jul 14 02:16:44 CEST 2022 - The aliscore output file does not exist. Check results for BUSCO: 284498at7742
## Thu Jul 14 02:16:44 CEST 2022 - The aliscore output appears to be empty. Check results for BUSCO: 284498at7742
## Thu Jul 14 02:16:44 CEST 2022 - The aliscore output file does not exist. Check results for BUSCO: 284498at7742
## Thu Jul 14 02:16:44 CEST 2022 - The aliscore output appears to be empty. Check results for BUSCO: 284498at7742
## Thu Jul 14 02:16:44 CEST 2022 - The aliscore output file does not exist. Check results for BUSCO: 190657at7742
## Thu Jul 14 02:16:45 CEST 2022 - The aliscore output appears to be empty. Check results for BUSCO: 190657at7742
## Thu Jul 14 02:16:47 CEST 2022 - The aliscore output file does not exist. Check results for BUSCO: 382669at7742
## Thu Jul 14 02:16:47 CEST 2022 - The aliscore output appears to be empty. Check results for BUSCO: 382669at7742
## Thu Jul 14 02:16:47 CEST 2022 - The aliscore output file does not exist. Check results for BUSCO: 336564at7742
## Thu Jul 14 02:16:47 CEST 2022 - The aliscore output appears to be empty. Check results for BUSCO: 336564at7742
## Thu Jul 14 02:16:48 CEST 2022 - The aliscore output file does not exist. Check results for BUSCO: 205603at7742
## Thu Jul 14 02:16:48 CEST 2022 - The aliscore output appears to be empty. Check results for BUSCO: 205603at7742
## Thu Jul 14 02:16:49 CEST 2022 - The aliscore output file does not exist. Check results for BUSCO: 321199at7742
## Thu Jul 14 02:16:49 CEST 2022 - The aliscore output appears to be empty. Check results for BUSCO: 321199at7742
## Thu Jul 14 02:16:50 CEST 2022 - The aliscore output file does not exist. Check results for BUSCO: 335620at7742
## Thu Jul 14 02:16:50 CEST 2022 - The aliscore output appears to be empty. Check results for BUSCO: 335620at7742
## Thu Jul 14 02:16:51 CEST 2022 - The aliscore output file does not exist. Check results for BUSCO: 320817at7742
## Thu Jul 14 02:16:51 CEST 2022 - The aliscore output appears to be empty. Check results for BUSCO: 320817at7742
## Thu Jul 14 02:16:52 CEST 2022 - The aliscore output file does not exist. Check results for BUSCO: 271228at7742
## Thu Jul 14 02:16:52 CEST 2022 - The aliscore output appears to be empty. Check results for BUSCO: 271228at7742
## Thu Jul 14 02:16:52 CEST 2022 - The aliscore output file does not exist. Check results for BUSCO: 309962at7742
## Thu Jul 14 02:16:52 CEST 2022 - The aliscore output appears to be empty. Check results for BUSCO: 309962at7742
## Thu Jul 14 02:16:53 CEST 2022 - The aliscore output file does not exist. Check results for BUSCO: 391055at7742
## Thu Jul 14 02:16:54 CEST 2022 - The aliscore output appears to be empty. Check results for BUSCO: 391055at7742
## Thu Jul 14 02:16:56 CEST 2022 - The aliscore output file does not exist. Check results for BUSCO: 365136at7742
## Thu Jul 14 02:16:56 CEST 2022 - The aliscore output appears to be empty. Check results for BUSCO: 365136at7742
## Thu Jul 14 02:16:57 CEST 2022 - The aliscore output file does not exist. Check results for BUSCO: 191806at7742
## Thu Jul 14 02:16:57 CEST 2022 - The aliscore output appears to be empty. Check results for BUSCO: 191806at7742
## Thu Jul 14 02:16:58 CEST 2022 - The aliscore output file does not exist. Check results for BUSCO: 311241at7742
## Thu Jul 14 02:16:58 CEST 2022 - The aliscore output appears to be empty. Check results for BUSCO: 311241at7742
## Thu Jul 14 02:16:58 CEST 2022 - The aliscore output file does not exist. Check results for BUSCO: 382669at7742
## Thu Jul 14 02:16:58 CEST 2022 - The aliscore output appears to be empty. Check results for BUSCO: 382669at7742
## Thu Jul 14 02:16:59 CEST 2022 - The aliscore output file does not exist. Check results for BUSCO: 246297at7742
## Thu Jul 14 02:16:59 CEST 2022 - The aliscore output appears to be empty. Check results for BUSCO: 246297at7742
## Thu Jul 14 02:17:00 CEST 2022 - The aliscore output file does not exist. Check results for BUSCO: 337045at7742
## Thu Jul 14 02:17:00 CEST 2022 - The aliscore output appears to be empty. Check results for BUSCO: 337045at7742
## Thu Jul 14 02:17:00 CEST 2022 - The aliscore output file does not exist. Check results for BUSCO: 333672at7742
## Thu Jul 14 02:17:00 CEST 2022 - The aliscore output appears to be empty. Check results for BUSCO: 333672at7742
## Thu Jul 14 02:17:03 CEST 2022 - The aliscore output file does not exist. Check results for BUSCO: 323076at7742
## Thu Jul 14 02:17:03 CEST 2022 - The aliscore output appears to be empty. Check results for BUSCO: 323076at7742
## Thu Jul 14 02:17:03 CEST 2022 - The aliscore output file does not exist. Check results for BUSCO: 317054at7742
## Thu Jul 14 02:17:03 CEST 2022 - The aliscore output appears to be empty. Check results for BUSCO: 317054at7742
## Thu Jul 14 02:17:03 CEST 2022 - The aliscore output file does not exist. Check results for BUSCO: 257379at7742
## Thu Jul 14 02:17:03 CEST 2022 - The aliscore output appears to be empty. Check results for BUSCO: 257379at7742
## Thu Jul 14 02:17:06 CEST 2022 - The aliscore output file does not exist. Check results for BUSCO: 205146at7742
## Thu Jul 14 02:17:06 CEST 2022 - The aliscore output appears to be empty. Check results for BUSCO: 205146at7742
## Thu Jul 14 02:17:06 CEST 2022 - The aliscore output file does not exist. Check results for BUSCO: 381534at7742
## Thu Jul 14 02:17:07 CEST 2022 - The aliscore output appears to be empty. Check results for BUSCO: 381534at7742
## Thu Jul 14 02:17:07 CEST 2022 - The aliscore output file does not exist. Check results for BUSCO: 227342at7742
## Thu Jul 14 02:17:07 CEST 2022 - The aliscore output appears to be empty. Check results for BUSCO: 227342at7742
## Thu Jul 14 02:17:07 CEST 2022 - The aliscore output file does not exist. Check results for BUSCO: 284651at7742
## Thu Jul 14 02:17:07 CEST 2022 - The aliscore output appears to be empty. Check results for BUSCO: 284651at7742
## Thu Jul 14 02:17:09 CEST 2022 - The aliscore output file does not exist. Check results for BUSCO: 379564at7742
## Thu Jul 14 02:17:09 CEST 2022 - The aliscore output appears to be empty. Check results for BUSCO: 379564at7742
## Thu Jul 14 02:17:11 CEST 2022 - The aliscore output file does not exist. Check results for BUSCO: 309948at7742
## Thu Jul 14 02:17:11 CEST 2022 - The aliscore output appears to be empty. Check results for BUSCO: 309948at7742
## Thu Jul 14 02:17:12 CEST 2022 - The aliscore output file does not exist. Check results for BUSCO: 315823at7742
## Thu Jul 14 02:17:12 CEST 2022 - The aliscore output appears to be empty. Check results for BUSCO: 315823at7742
## Thu Jul 14 02:17:12 CEST 2022 - The aliscore output file does not exist. Check results for BUSCO: 309948at7742
## Thu Jul 14 02:17:12 CEST 2022 - The aliscore output appears to be empty. Check results for BUSCO: 309948at7742
## Thu Jul 14 02:17:13 CEST 2022 - The aliscore output file does not exist. Check results for BUSCO: 284498at7742
## Thu Jul 14 02:17:13 CEST 2022 - The aliscore output appears to be empty. Check results for BUSCO: 284498at7742
## Thu Jul 14 02:17:15 CEST 2022 - The aliscore output file does not exist. Check results for BUSCO: 374832at7742
## Thu Jul 14 02:17:15 CEST 2022 - The aliscore output appears to be empty. Check results for BUSCO: 374832at7742
## Thu Jul 14 02:17:16 CEST 2022 - The aliscore output file does not exist. Check results for BUSCO: 384842at7742
## Thu Jul 14 02:17:16 CEST 2022 - The aliscore output appears to be empty. Check results for BUSCO: 384842at7742
## Thu Jul 14 02:17:16 CEST 2022 - The aliscore output file does not exist. Check results for BUSCO: 379932at7742
## Thu Jul 14 02:17:16 CEST 2022 - The aliscore output appears to be empty. Check results for BUSCO: 379932at7742
## Thu Jul 14 02:17:18 CEST 2022 - The aliscore output file does not exist. Check results for BUSCO: 256968at7742
## Thu Jul 14 02:17:19 CEST 2022 - The aliscore output appears to be empty. Check results for BUSCO: 256968at7742
## Thu Jul 14 02:17:19 CEST 2022 - The aliscore output file does not exist. Check results for BUSCO: 231838at7742
## Thu Jul 14 02:17:19 CEST 2022 - The aliscore output appears to be empty. Check results for BUSCO: 231838at7742
## Thu Jul 14 02:17:20 CEST 2022 - The aliscore output file does not exist. Check results for BUSCO: 309935at7742
## Thu Jul 14 02:17:20 CEST 2022 - The aliscore output appears to be empty. Check results for BUSCO: 309935at7742
## Thu Jul 14 02:17:21 CEST 2022 - The aliscore output file does not exist. Check results for BUSCO: 279092at7742
## Thu Jul 14 02:17:21 CEST 2022 - The aliscore output appears to be empty. Check results for BUSCO: 279092at7742
## Thu Jul 14 02:17:22 CEST 2022 - The aliscore output file does not exist. Check results for BUSCO: 342620at7742
## Thu Jul 14 02:17:22 CEST 2022 - The aliscore output appears to be empty. Check results for BUSCO: 342620at7742
## Thu Jul 14 02:17:22 CEST 2022 - The aliscore output file does not exist. Check results for BUSCO: 278070at7742
## Thu Jul 14 02:17:22 CEST 2022 - The aliscore output appears to be empty. Check results for BUSCO: 278070at7742
## Thu Jul 14 02:17:24 CEST 2022 - The aliscore output file does not exist. Check results for BUSCO: 270432at7742
## Thu Jul 14 02:17:24 CEST 2022 - The aliscore output appears to be empty. Check results for BUSCO: 270432at7742
## Thu Jul 14 02:17:25 CEST 2022 - The aliscore output file does not exist. Check results for BUSCO: 379932at7742
## Thu Jul 14 02:17:25 CEST 2022 - The aliscore output appears to be empty. Check results for BUSCO: 379932at7742
## Thu Jul 14 02:17:25 CEST 2022 - The aliscore output file does not exist. Check results for BUSCO: 331959at7742
## Thu Jul 14 02:17:25 CEST 2022 - The aliscore output appears to be empty. Check results for BUSCO: 331959at7742
## Thu Jul 14 02:17:27 CEST 2022 - The aliscore output file does not exist. Check results for BUSCO: 309974at7742
## Thu Jul 14 02:17:27 CEST 2022 - The aliscore output appears to be empty. Check results for BUSCO: 309974at7742
## Thu Jul 14 02:17:28 CEST 2022 - The aliscore output file does not exist. Check results for BUSCO: 278070at7742
## Thu Jul 14 02:17:28 CEST 2022 - The aliscore output appears to be empty. Check results for BUSCO: 278070at7742
## Thu Jul 14 02:17:28 CEST 2022 - The aliscore output file does not exist. Check results for BUSCO: 308917at7742
## Thu Jul 14 02:17:28 CEST 2022 - The aliscore output appears to be empty. Check results for BUSCO: 308917at7742
## Thu Jul 14 02:17:28 CEST 2022 - The aliscore output file does not exist. Check results for BUSCO: 325689at7742
## Thu Jul 14 02:17:29 CEST 2022 - The aliscore output appears to be empty. Check results for BUSCO: 325689at7742
## Thu Jul 14 02:17:30 CEST 2022 - The aliscore output file does not exist. Check results for BUSCO: 403632at7742
## Thu Jul 14 02:17:30 CEST 2022 - The aliscore output appears to be empty. Check results for BUSCO: 403632at7742
## Thu Jul 14 02:17:32 CEST 2022 - The aliscore output file does not exist. Check results for BUSCO: 361974at7742
## Thu Jul 14 02:17:32 CEST 2022 - The aliscore output appears to be empty. Check results for BUSCO: 361974at7742
## Thu Jul 14 02:17:32 CEST 2022 - The aliscore output file does not exist. Check results for BUSCO: 270938at7742
## Thu Jul 14 02:17:32 CEST 2022 - The aliscore output appears to be empty. Check results for BUSCO: 270938at7742
## Thu Jul 14 02:17:33 CEST 2022 - The aliscore output file does not exist. Check results for BUSCO: 280642at7742
## Thu Jul 14 02:17:33 CEST 2022 - The aliscore output appears to be empty. Check results for BUSCO: 280642at7742
## Thu Jul 14 02:17:34 CEST 2022 - The aliscore output file does not exist. Check results for BUSCO: 376105at7742
## Thu Jul 14 02:17:34 CEST 2022 - The aliscore output appears to be empty. Check results for BUSCO: 376105at7742
## Thu Jul 14 02:17:34 CEST 2022 - The aliscore output file does not exist. Check results for BUSCO: 242369at7742
## Thu Jul 14 02:17:34 CEST 2022 - The aliscore output appears to be empty. Check results for BUSCO: 242369at7742
## Thu Jul 14 02:17:34 CEST 2022 - The aliscore output file does not exist. Check results for BUSCO: 231838at7742
## Thu Jul 14 02:17:35 CEST 2022 - The aliscore output appears to be empty. Check results for BUSCO: 231838at7742
## Thu Jul 14 02:17:38 CEST 2022 - The aliscore output file does not exist. Check results for BUSCO: 298043at7742
## Thu Jul 14 02:17:38 CEST 2022 - The aliscore output appears to be empty. Check results for BUSCO: 298043at7742
## Thu Jul 14 02:17:39 CEST 2022 - The aliscore output file does not exist. Check results for BUSCO: 333716at7742
## Thu Jul 14 02:17:39 CEST 2022 - The aliscore output appears to be empty. Check results for BUSCO: 333716at7742
## Thu Jul 14 02:17:39 CEST 2022 - The aliscore output file does not exist. Check results for BUSCO: 359738at7742
## Thu Jul 14 02:17:39 CEST 2022 - The aliscore output file does not exist. Check results for BUSCO: 205589at7742
## Thu Jul 14 02:17:39 CEST 2022 - The aliscore output appears to be empty. Check results for BUSCO: 359738at7742
## Thu Jul 14 02:17:39 CEST 2022 - The aliscore output appears to be empty. Check results for BUSCO: 205589at7742
## Thu Jul 14 02:17:41 CEST 2022 - The aliscore output file does not exist. Check results for BUSCO: 262189at7742
## Thu Jul 14 02:17:41 CEST 2022 - The aliscore output appears to be empty. Check results for BUSCO: 262189at7742
## Thu Jul 14 02:17:41 CEST 2022 - The aliscore output file does not exist. Check results for BUSCO: 255497at7742
## Thu Jul 14 02:17:41 CEST 2022 - The aliscore output appears to be empty. Check results for BUSCO: 255497at7742
## Thu Jul 14 02:17:43 CEST 2022 - The aliscore output file does not exist. Check results for BUSCO: 335517at7742
## Thu Jul 14 02:17:43 CEST 2022 - The aliscore output appears to be empty. Check results for BUSCO: 335517at7742
## Thu Jul 14 02:17:43 CEST 2022 - The aliscore output file does not exist. Check results for BUSCO: 323970at7742
## Thu Jul 14 02:17:43 CEST 2022 - The aliscore output appears to be empty. Check results for BUSCO: 323970at7742
## Thu Jul 14 02:17:45 CEST 2022 - The aliscore output file does not exist. Check results for BUSCO: 245299at7742
## Thu Jul 14 02:17:45 CEST 2022 - The aliscore output appears to be empty. Check results for BUSCO: 245299at7742
## Thu Jul 14 02:17:45 CEST 2022 - The aliscore output file does not exist. Check results for BUSCO: 225646at7742
## Thu Jul 14 02:17:45 CEST 2022 - The aliscore output appears to be empty. Check results for BUSCO: 225646at7742
## Thu Jul 14 02:17:45 CEST 2022 - The aliscore output file does not exist. Check results for BUSCO: 245611at7742
## Thu Jul 14 02:17:45 CEST 2022 - The aliscore output appears to be empty. Check results for BUSCO: 245611at7742
## Thu Jul 14 02:17:46 CEST 2022 - The aliscore output file does not exist. Check results for BUSCO: 330614at7742
## Thu Jul 14 02:17:46 CEST 2022 - The aliscore output appears to be empty. Check results for BUSCO: 330614at7742
## Thu Jul 14 02:17:47 CEST 2022 - The aliscore output file does not exist. Check results for BUSCO: 237440at7742
## Thu Jul 14 02:17:47 CEST 2022 - The aliscore output appears to be empty. Check results for BUSCO: 237440at7742
## Thu Jul 14 02:17:48 CEST 2022 - The aliscore output file does not exist. Check results for BUSCO: 293076at7742
## Thu Jul 14 02:17:48 CEST 2022 - The aliscore output appears to be empty. Check results for BUSCO: 293076at7742
## Thu Jul 14 02:17:50 CEST 2022 - The aliscore output file does not exist. Check results for BUSCO: 251896at7742
## Thu Jul 14 02:17:50 CEST 2022 - The aliscore output appears to be empty. Check results for BUSCO: 251896at7742
## Thu Jul 14 02:17:50 CEST 2022 - The aliscore output file does not exist. Check results for BUSCO: 432107at7742
## Thu Jul 14 02:17:50 CEST 2022 - The aliscore output appears to be empty. Check results for BUSCO: 432107at7742
## Thu Jul 14 02:17:51 CEST 2022 - The aliscore output file does not exist. Check results for BUSCO: 357819at7742
## Thu Jul 14 02:17:51 CEST 2022 - The aliscore output appears to be empty. Check results for BUSCO: 357819at7742
## Thu Jul 14 02:17:52 CEST 2022 - The aliscore output file does not exist. Check results for BUSCO: 323076at7742
## Thu Jul 14 02:17:52 CEST 2022 - The aliscore output appears to be empty. Check results for BUSCO: 323076at7742
## Thu Jul 14 02:17:53 CEST 2022 - The aliscore output file does not exist. Check results for BUSCO: 370111at7742
## Thu Jul 14 02:17:53 CEST 2022 - The aliscore output appears to be empty. Check results for BUSCO: 370111at7742
## Thu Jul 14 02:17:54 CEST 2022 - The aliscore output file does not exist. Check results for BUSCO: 290543at7742
## Thu Jul 14 02:17:54 CEST 2022 - The aliscore output appears to be empty. Check results for BUSCO: 290543at7742
## Thu Jul 14 02:17:56 CEST 2022 - The aliscore output file does not exist. Check results for BUSCO: 234247at7742
## Thu Jul 14 02:17:56 CEST 2022 - The aliscore output appears to be empty. Check results for BUSCO: 234247at7742
## Thu Jul 14 02:17:56 CEST 2022 - The aliscore output file does not exist. Check results for BUSCO: 241352at7742
## Thu Jul 14 02:17:56 CEST 2022 - The aliscore output appears to be empty. Check results for BUSCO: 241352at7742
## Thu Jul 14 02:17:57 CEST 2022 - The aliscore output file does not exist. Check results for BUSCO: 340073at7742
## Thu Jul 14 02:17:57 CEST 2022 - The aliscore output appears to be empty. Check results for BUSCO: 340073at7742
## Thu Jul 14 02:17:58 CEST 2022 - The aliscore output file does not exist. Check results for BUSCO: 192574at7742
## Thu Jul 14 02:17:58 CEST 2022 - The aliscore output appears to be empty. Check results for BUSCO: 192574at7742
## Thu Jul 14 02:17:59 CEST 2022 - The aliscore output file does not exist. Check results for BUSCO: 237440at7742
## Thu Jul 14 02:17:59 CEST 2022 - The aliscore output appears to be empty. Check results for BUSCO: 237440at7742
## Thu Jul 14 02:18:00 CEST 2022 - The aliscore output file does not exist. Check results for BUSCO: 268420at7742
## Thu Jul 14 02:18:00 CEST 2022 - The aliscore output appears to be empty. Check results for BUSCO: 268420at7742
## Thu Jul 14 02:18:01 CEST 2022 - The aliscore output file does not exist. Check results for BUSCO: 121142at7742
## Thu Jul 14 02:18:01 CEST 2022 - The aliscore output appears to be empty. Check results for BUSCO: 121142at7742
## Thu Jul 14 02:18:02 CEST 2022 - The aliscore output file does not exist. Check results for BUSCO: 288347at7742
## Thu Jul 14 02:18:02 CEST 2022 - The aliscore output appears to be empty. Check results for BUSCO: 288347at7742
## Thu Jul 14 02:18:03 CEST 2022 - The aliscore output file does not exist. Check results for BUSCO: 290567at7742
## Thu Jul 14 02:18:03 CEST 2022 - The aliscore output appears to be empty. Check results for BUSCO: 290567at7742
## Thu Jul 14 02:18:04 CEST 2022 - The aliscore output file does not exist. Check results for BUSCO: 192574at7742
## Thu Jul 14 02:18:04 CEST 2022 - The aliscore output appears to be empty. Check results for BUSCO: 192574at7742
## Thu Jul 14 02:18:04 CEST 2022 - The aliscore output file does not exist. Check results for BUSCO: 225646at7742
## Thu Jul 14 02:18:04 CEST 2022 - The aliscore output appears to be empty. Check results for BUSCO: 225646at7742
## Thu Jul 14 02:18:06 CEST 2022 - The aliscore output file does not exist. Check results for BUSCO: 231838at7742
## Thu Jul 14 02:18:06 CEST 2022 - The aliscore output appears to be empty. Check results for BUSCO: 231838at7742
## Thu Jul 14 02:18:07 CEST 2022 - The aliscore output file does not exist. Check results for BUSCO: 403876at7742
## Thu Jul 14 02:18:07 CEST 2022 - The aliscore output appears to be empty. Check results for BUSCO: 403876at7742
## Thu Jul 14 02:18:07 CEST 2022 - The aliscore output file does not exist. Check results for BUSCO: 191806at7742
## Thu Jul 14 02:18:07 CEST 2022 - The aliscore output appears to be empty. Check results for BUSCO: 191806at7742
## Thu Jul 14 02:18:09 CEST 2022 - The aliscore output file does not exist. Check results for BUSCO: 297771at7742
## Thu Jul 14 02:18:09 CEST 2022 - The aliscore output appears to be empty. Check results for BUSCO: 297771at7742
## Thu Jul 14 02:18:09 CEST 2022 - The aliscore output file does not exist. Check results for BUSCO: 337736at7742
## Thu Jul 14 02:18:09 CEST 2022 - The aliscore output appears to be empty. Check results for BUSCO: 337736at7742
## Thu Jul 14 02:18:10 CEST 2022 - The aliscore output file does not exist. Check results for BUSCO: 261266at7742
## Thu Jul 14 02:18:10 CEST 2022 - The aliscore output appears to be empty. Check results for BUSCO: 261266at7742
## Thu Jul 14 02:18:11 CEST 2022 - The aliscore output file does not exist. Check results for BUSCO: 292271at7742
## Thu Jul 14 02:18:11 CEST 2022 - The aliscore output appears to be empty. Check results for BUSCO: 292271at7742
## Thu Jul 14 02:18:12 CEST 2022 - The aliscore output file does not exist. Check results for BUSCO: 376384at7742
## Thu Jul 14 02:18:12 CEST 2022 - The aliscore output appears to be empty. Check results for BUSCO: 376384at7742
## Thu Jul 14 02:18:13 CEST 2022 - The aliscore output file does not exist. Check results for BUSCO: 264124at7742
## Thu Jul 14 02:18:13 CEST 2022 - The aliscore output appears to be empty. Check results for BUSCO: 264124at7742
## Thu Jul 14 02:18:14 CEST 2022 - The aliscore output file does not exist. Check results for BUSCO: 347924at7742
## Thu Jul 14 02:18:15 CEST 2022 - The aliscore output appears to be empty. Check results for BUSCO: 347924at7742
## Thu Jul 14 02:18:15 CEST 2022 - The aliscore output file does not exist. Check results for BUSCO: 347924at7742
## Thu Jul 14 02:18:15 CEST 2022 - The aliscore output appears to be empty. Check results for BUSCO: 347924at7742
## Thu Jul 14 02:18:16 CEST 2022 - The aliscore output file does not exist. Check results for BUSCO: 234247at7742
## Thu Jul 14 02:18:16 CEST 2022 - The aliscore output appears to be empty. Check results for BUSCO: 234247at7742
## Thu Jul 14 02:18:16 CEST 2022 - The aliscore output file does not exist. Check results for BUSCO: 261158at7742
## Thu Jul 14 02:18:16 CEST 2022 - The aliscore output appears to be empty. Check results for BUSCO: 261158at7742
## Thu Jul 14 02:18:17 CEST 2022 - The aliscore output file does not exist. Check results for BUSCO: 280430at7742
## Thu Jul 14 02:18:17 CEST 2022 - The aliscore output appears to be empty. Check results for BUSCO: 280430at7742
## Thu Jul 14 02:18:20 CEST 2022 - The aliscore output file does not exist. Check results for BUSCO: 402327at7742
## Thu Jul 14 02:18:20 CEST 2022 - The aliscore output appears to be empty. Check results for BUSCO: 402327at7742
## Thu Jul 14 02:18:20 CEST 2022 - The aliscore output file does not exist. Check results for BUSCO: 281245at7742
## Thu Jul 14 02:18:20 CEST 2022 - The aliscore output appears to be empty. Check results for BUSCO: 281245at7742
## Thu Jul 14 02:18:21 CEST 2022 - The aliscore output file does not exist. Check results for BUSCO: 290523at7742
## Thu Jul 14 02:18:21 CEST 2022 - The aliscore output appears to be empty. Check results for BUSCO: 290523at7742
## Thu Jul 14 02:18:22 CEST 2022 - The aliscore output file does not exist. Check results for BUSCO: 121142at7742
## Thu Jul 14 02:18:22 CEST 2022 - The aliscore output appears to be empty. Check results for BUSCO: 121142at7742
## Thu Jul 14 02:18:23 CEST 2022 - The aliscore output file does not exist. Check results for BUSCO: 311889at7742
## Thu Jul 14 02:18:23 CEST 2022 - The aliscore output file does not exist. Check results for BUSCO: 297599at7742
## Thu Jul 14 02:18:23 CEST 2022 - The aliscore output appears to be empty. Check results for BUSCO: 311889at7742
## Thu Jul 14 02:18:23 CEST 2022 - The aliscore output appears to be empty. Check results for BUSCO: 297599at7742
## Thu Jul 14 02:18:25 CEST 2022 - The aliscore output file does not exist. Check results for BUSCO: 432107at7742
## Thu Jul 14 02:18:25 CEST 2022 - The aliscore output appears to be empty. Check results for BUSCO: 432107at7742
## Thu Jul 14 02:18:27 CEST 2022 - The aliscore output file does not exist. Check results for BUSCO: 301630at7742
## Thu Jul 14 02:18:27 CEST 2022 - The aliscore output appears to be empty. Check results for BUSCO: 301630at7742
## Thu Jul 14 02:18:27 CEST 2022 - The aliscore output file does not exist. Check results for BUSCO: 333716at7742
## Thu Jul 14 02:18:27 CEST 2022 - The aliscore output appears to be empty. Check results for BUSCO: 333716at7742
## Thu Jul 14 02:18:27 CEST 2022 - The aliscore output file does not exist. Check results for BUSCO: 331411at7742
## Thu Jul 14 02:18:27 CEST 2022 - The aliscore output appears to be empty. Check results for BUSCO: 331411at7742
## Thu Jul 14 02:18:28 CEST 2022 - The aliscore output file does not exist. Check results for BUSCO: 409719at7742
## Thu Jul 14 02:18:28 CEST 2022 - The aliscore output appears to be empty. Check results for BUSCO: 409719at7742
## Thu Jul 14 02:18:30 CEST 2022 - The aliscore output file does not exist. Check results for BUSCO: 336619at7742
## Thu Jul 14 02:18:30 CEST 2022 - The aliscore output appears to be empty. Check results for BUSCO: 336619at7742
## Thu Jul 14 02:18:31 CEST 2022 - The aliscore output file does not exist. Check results for BUSCO: 297599at7742
## Thu Jul 14 02:18:31 CEST 2022 - The aliscore output appears to be empty. Check results for BUSCO: 297599at7742
## Thu Jul 14 02:18:32 CEST 2022 - The aliscore output file does not exist. Check results for BUSCO: 305378at7742
## Thu Jul 14 02:18:32 CEST 2022 - The aliscore output appears to be empty. Check results for BUSCO: 305378at7742
## Thu Jul 14 02:18:33 CEST 2022 - The aliscore output file does not exist. Check results for BUSCO: 242394at7742
## Thu Jul 14 02:18:33 CEST 2022 - The aliscore output appears to be empty. Check results for BUSCO: 242394at7742
## Thu Jul 14 02:18:33 CEST 2022 - The aliscore output file does not exist. Check results for BUSCO: 271425at7742
## Thu Jul 14 02:18:34 CEST 2022 - The aliscore output appears to be empty. Check results for BUSCO: 271425at7742
## Thu Jul 14 02:18:34 CEST 2022 - The aliscore output file does not exist. Check results for BUSCO: 334381at7742
## Thu Jul 14 02:18:34 CEST 2022 - The aliscore output appears to be empty. Check results for BUSCO: 334381at7742
## Thu Jul 14 02:18:34 CEST 2022 - The aliscore output file does not exist. Check results for BUSCO: 333672at7742
## Thu Jul 14 02:18:34 CEST 2022 - The aliscore output appears to be empty. Check results for BUSCO: 333672at7742
## Thu Jul 14 02:18:36 CEST 2022 - The aliscore output file does not exist. Check results for BUSCO: 407723at7742
## Thu Jul 14 02:18:37 CEST 2022 - The aliscore output appears to be empty. Check results for BUSCO: 407723at7742
## Thu Jul 14 02:18:38 CEST 2022 - The aliscore output file does not exist. Check results for BUSCO: 216850at7742
## Thu Jul 14 02:18:38 CEST 2022 - The aliscore output appears to be empty. Check results for BUSCO: 216850at7742
## Thu Jul 14 02:18:38 CEST 2022 - The aliscore output file does not exist. Check results for BUSCO: 261658at7742
## Thu Jul 14 02:18:38 CEST 2022 - The aliscore output appears to be empty. Check results for BUSCO: 261658at7742
## Thu Jul 14 02:18:40 CEST 2022 - The aliscore output file does not exist. Check results for BUSCO: 302555at7742
## Thu Jul 14 02:18:40 CEST 2022 - The aliscore output appears to be empty. Check results for BUSCO: 302555at7742
## Thu Jul 14 02:18:42 CEST 2022 - The aliscore output file does not exist. Check results for BUSCO: 243977at7742
## Thu Jul 14 02:18:42 CEST 2022 - The aliscore output file does not exist. Check results for BUSCO: 409736at7742
## Thu Jul 14 02:18:42 CEST 2022 - The aliscore output appears to be empty. Check results for BUSCO: 243977at7742
## Thu Jul 14 02:18:42 CEST 2022 - The aliscore output appears to be empty. Check results for BUSCO: 409736at7742
## Thu Jul 14 02:18:43 CEST 2022 - The aliscore output file does not exist. Check results for BUSCO: 192574at7742
## Thu Jul 14 02:18:43 CEST 2022 - The aliscore output appears to be empty. Check results for BUSCO: 192574at7742
## Thu Jul 14 02:18:43 CEST 2022 - The aliscore output file does not exist. Check results for BUSCO: 271060at7742
## Thu Jul 14 02:18:43 CEST 2022 - The aliscore output appears to be empty. Check results for BUSCO: 271060at7742
## Thu Jul 14 02:18:45 CEST 2022 - The aliscore output file does not exist. Check results for BUSCO: 311206at7742
## Thu Jul 14 02:18:46 CEST 2022 - The aliscore output appears to be empty. Check results for BUSCO: 311206at7742
## Thu Jul 14 02:18:46 CEST 2022 - The aliscore output file does not exist. Check results for BUSCO: 322345at7742
## Thu Jul 14 02:18:46 CEST 2022 - The aliscore output appears to be empty. Check results for BUSCO: 322345at7742
## Thu Jul 14 02:18:47 CEST 2022 - The aliscore output file does not exist. Check results for BUSCO: 264221at7742
## Thu Jul 14 02:18:47 CEST 2022 - The aliscore output appears to be empty. Check results for BUSCO: 264221at7742
## Thu Jul 14 02:18:48 CEST 2022 - The aliscore output file does not exist. Check results for BUSCO: 333698at7742
## Thu Jul 14 02:18:48 CEST 2022 - The aliscore output appears to be empty. Check results for BUSCO: 333698at7742
## Thu Jul 14 02:18:48 CEST 2022 - The aliscore output file does not exist. Check results for BUSCO: 301630at7742
## Thu Jul 14 02:18:48 CEST 2022 - The aliscore output appears to be empty. Check results for BUSCO: 301630at7742
## Thu Jul 14 02:18:49 CEST 2022 - The aliscore output file does not exist. Check results for BUSCO: 406718at7742
## Thu Jul 14 02:18:49 CEST 2022 - The aliscore output appears to be empty. Check results for BUSCO: 406718at7742
## Thu Jul 14 02:18:49 CEST 2022 - The aliscore output file does not exist. Check results for BUSCO: 398201at7742
## Thu Jul 14 02:18:49 CEST 2022 - The aliscore output appears to be empty. Check results for BUSCO: 398201at7742
## Thu Jul 14 02:18:52 CEST 2022 - The aliscore output file does not exist. Check results for BUSCO: 303047at7742
## Thu Jul 14 02:18:52 CEST 2022 - The aliscore output appears to be empty. Check results for BUSCO: 303047at7742
## Thu Jul 14 02:18:52 CEST 2022 - The aliscore output file does not exist. Check results for BUSCO: 319229at7742
## Thu Jul 14 02:18:52 CEST 2022 - The aliscore output appears to be empty. Check results for BUSCO: 319229at7742
## Thu Jul 14 02:18:53 CEST 2022 - The aliscore output file does not exist. Check results for BUSCO: 376105at7742
## Thu Jul 14 02:18:53 CEST 2022 - The aliscore output appears to be empty. Check results for BUSCO: 376105at7742
## Thu Jul 14 02:18:53 CEST 2022 - The aliscore output file does not exist. Check results for BUSCO: 220264at7742
## Thu Jul 14 02:18:53 CEST 2022 - The aliscore output appears to be empty. Check results for BUSCO: 220264at7742
## Thu Jul 14 02:18:54 CEST 2022 - The aliscore output file does not exist. Check results for BUSCO: 220264at7742
## Thu Jul 14 02:18:54 CEST 2022 - The aliscore output appears to be empty. Check results for BUSCO: 220264at7742
## Thu Jul 14 02:18:56 CEST 2022 - The aliscore output file does not exist. Check results for BUSCO: 398201at7742
## Thu Jul 14 02:18:56 CEST 2022 - The aliscore output appears to be empty. Check results for BUSCO: 398201at7742
## Thu Jul 14 02:18:57 CEST 2022 - The aliscore output file does not exist. Check results for BUSCO: 234247at7742
## Thu Jul 14 02:18:57 CEST 2022 - The aliscore output appears to be empty. Check results for BUSCO: 234247at7742
## Thu Jul 14 02:18:58 CEST 2022 - The aliscore output file does not exist. Check results for BUSCO: 187843at7742
## Thu Jul 14 02:18:58 CEST 2022 - The aliscore output appears to be empty. Check results for BUSCO: 187843at7742
## Thu Jul 14 02:19:00 CEST 2022 - The aliscore output file does not exist. Check results for BUSCO: 258997at7742
## Thu Jul 14 02:19:00 CEST 2022 - The aliscore output appears to be empty. Check results for BUSCO: 258997at7742
## Thu Jul 14 02:19:01 CEST 2022 - The aliscore output file does not exist. Check results for BUSCO: 205589at7742
## Thu Jul 14 02:19:01 CEST 2022 - The aliscore output appears to be empty. Check results for BUSCO: 205589at7742
## Thu Jul 14 02:19:01 CEST 2022 - The aliscore output file does not exist. Check results for BUSCO: 336178at7742
## Thu Jul 14 02:19:01 CEST 2022 - The aliscore output appears to be empty. Check results for BUSCO: 336178at7742
## Thu Jul 14 02:19:02 CEST 2022 - The aliscore output file does not exist. Check results for BUSCO: 250138at7742
## Thu Jul 14 02:19:02 CEST 2022 - The aliscore output appears to be empty. Check results for BUSCO: 250138at7742
## Thu Jul 14 02:19:02 CEST 2022 - The aliscore output file does not exist. Check results for BUSCO: 249392at7742
## Thu Jul 14 02:19:02 CEST 2022 - The aliscore output appears to be empty. Check results for BUSCO: 249392at7742
## Thu Jul 14 02:19:04 CEST 2022 - The aliscore output file does not exist. Check results for BUSCO: 241352at7742
## Thu Jul 14 02:19:04 CEST 2022 - The aliscore output appears to be empty. Check results for BUSCO: 241352at7742
## Thu Jul 14 02:19:05 CEST 2022 - The aliscore output file does not exist. Check results for BUSCO: 432107at7742
## Thu Jul 14 02:19:05 CEST 2022 - The aliscore output appears to be empty. Check results for BUSCO: 432107at7742
## Thu Jul 14 02:19:06 CEST 2022 - The aliscore output file does not exist. Check results for BUSCO: 399664at7742
## Thu Jul 14 02:19:06 CEST 2022 - The aliscore output appears to be empty. Check results for BUSCO: 399664at7742
## Thu Jul 14 02:19:07 CEST 2022 - The aliscore output file does not exist. Check results for BUSCO: 397671at7742
## Thu Jul 14 02:19:07 CEST 2022 - The aliscore output appears to be empty. Check results for BUSCO: 397671at7742
## Thu Jul 14 02:19:08 CEST 2022 - The aliscore output file does not exist. Check results for BUSCO: 325689at7742
## Thu Jul 14 02:19:08 CEST 2022 - The aliscore output appears to be empty. Check results for BUSCO: 325689at7742
## Thu Jul 14 02:19:09 CEST 2022 - The aliscore output file does not exist. Check results for BUSCO: 259157at7742
## Thu Jul 14 02:19:09 CEST 2022 - The aliscore output appears to be empty. Check results for BUSCO: 259157at7742
## Thu Jul 14 02:19:10 CEST 2022 - The aliscore output file does not exist. Check results for BUSCO: 271228at7742
## Thu Jul 14 02:19:10 CEST 2022 - The aliscore output appears to be empty. Check results for BUSCO: 271228at7742
## Thu Jul 14 02:19:11 CEST 2022 - The aliscore output file does not exist. Check results for BUSCO: 368306at7742
## Thu Jul 14 02:19:11 CEST 2022 - The aliscore output appears to be empty. Check results for BUSCO: 368306at7742
## Thu Jul 14 02:19:12 CEST 2022 - The aliscore output file does not exist. Check results for BUSCO: 360873at7742
## Thu Jul 14 02:19:12 CEST 2022 - The aliscore output appears to be empty. Check results for BUSCO: 360873at7742
## Thu Jul 14 02:19:12 CEST 2022 - The aliscore output file does not exist. Check results for BUSCO: 259089at7742
## Thu Jul 14 02:19:12 CEST 2022 - The aliscore output appears to be empty. Check results for BUSCO: 259089at7742
## Thu Jul 14 02:19:13 CEST 2022 - The aliscore output file does not exist. Check results for BUSCO: 234637at7742
## Thu Jul 14 02:19:13 CEST 2022 - The aliscore output appears to be empty. Check results for BUSCO: 234637at7742
## Thu Jul 14 02:19:15 CEST 2022 - The aliscore output file does not exist. Check results for BUSCO: 413350at7742
## Thu Jul 14 02:19:15 CEST 2022 - The aliscore output appears to be empty. Check results for BUSCO: 413350at7742
## Thu Jul 14 02:19:15 CEST 2022 - The aliscore output file does not exist. Check results for BUSCO: 318720at7742
## Thu Jul 14 02:19:15 CEST 2022 - The aliscore output appears to be empty. Check results for BUSCO: 318720at7742
## Thu Jul 14 02:19:15 CEST 2022 - The aliscore output file does not exist. Check results for BUSCO: 191806at7742
## Thu Jul 14 02:19:15 CEST 2022 - The aliscore output appears to be empty. Check results for BUSCO: 191806at7742
## Thu Jul 14 02:19:17 CEST 2022 - The aliscore output file does not exist. Check results for BUSCO: 269168at7742
## Thu Jul 14 02:19:17 CEST 2022 - The aliscore output appears to be empty. Check results for BUSCO: 269168at7742
## Thu Jul 14 02:19:19 CEST 2022 - The aliscore output file does not exist. Check results for BUSCO: 370111at7742
## Thu Jul 14 02:19:19 CEST 2022 - The aliscore output appears to be empty. Check results for BUSCO: 370111at7742
## Thu Jul 14 02:19:20 CEST 2022 - The aliscore output file does not exist. Check results for BUSCO: 304827at7742
## Thu Jul 14 02:19:20 CEST 2022 - The aliscore output appears to be empty. Check results for BUSCO: 304827at7742
## Thu Jul 14 02:19:21 CEST 2022 - The aliscore output file does not exist. Check results for BUSCO: 368306at7742
## Thu Jul 14 02:19:21 CEST 2022 - The aliscore output appears to be empty. Check results for BUSCO: 368306at7742
## Thu Jul 14 02:19:22 CEST 2022 - The aliscore output file does not exist. Check results for BUSCO: 396521at7742
## Thu Jul 14 02:19:22 CEST 2022 - The aliscore output appears to be empty. Check results for BUSCO: 396521at7742
## Thu Jul 14 02:19:22 CEST 2022 - The aliscore output file does not exist. Check results for BUSCO: 205589at7742
## Thu Jul 14 02:19:22 CEST 2022 - The aliscore output appears to be empty. Check results for BUSCO: 205589at7742
## Thu Jul 14 02:19:23 CEST 2022 - The aliscore output file does not exist. Check results for BUSCO: 304588at7742
## Thu Jul 14 02:19:23 CEST 2022 - The aliscore output appears to be empty. Check results for BUSCO: 304588at7742
## Thu Jul 14 02:19:24 CEST 2022 - The aliscore output file does not exist. Check results for BUSCO: 323737at7742
## Thu Jul 14 02:19:24 CEST 2022 - The aliscore output appears to be empty. Check results for BUSCO: 323737at7742
## Thu Jul 14 02:19:25 CEST 2022 - The aliscore output file does not exist. Check results for BUSCO: 264221at7742
## Thu Jul 14 02:19:25 CEST 2022 - The aliscore output appears to be empty. Check results for BUSCO: 264221at7742
## Thu Jul 14 02:19:26 CEST 2022 - The aliscore output file does not exist. Check results for BUSCO: 216850at7742
## Thu Jul 14 02:19:26 CEST 2022 - The aliscore output appears to be empty. Check results for BUSCO: 216850at7742
## Thu Jul 14 02:19:27 CEST 2022 - The aliscore output file does not exist. Check results for BUSCO: 279227at7742
## Thu Jul 14 02:19:27 CEST 2022 - The aliscore output appears to be empty. Check results for BUSCO: 279227at7742
## Thu Jul 14 02:19:29 CEST 2022 - The aliscore output file does not exist. Check results for BUSCO: 234637at7742
## Thu Jul 14 02:19:29 CEST 2022 - The aliscore output appears to be empty. Check results for BUSCO: 234637at7742
## Thu Jul 14 02:19:29 CEST 2022 - The aliscore output file does not exist. Check results for BUSCO: 261266at7742
## Thu Jul 14 02:19:29 CEST 2022 - The aliscore output appears to be empty. Check results for BUSCO: 261266at7742
## Thu Jul 14 02:19:30 CEST 2022 - The aliscore output file does not exist. Check results for BUSCO: 377238at7742
## Thu Jul 14 02:19:30 CEST 2022 - The aliscore output appears to be empty. Check results for BUSCO: 377238at7742
## Thu Jul 14 02:19:32 CEST 2022 - The aliscore output file does not exist. Check results for BUSCO: 121142at7742
## Thu Jul 14 02:19:32 CEST 2022 - The aliscore output appears to be empty. Check results for BUSCO: 121142at7742
## Thu Jul 14 02:19:32 CEST 2022 - The aliscore output file does not exist. Check results for BUSCO: 270938at7742
## Thu Jul 14 02:19:32 CEST 2022 - The aliscore output appears to be empty. Check results for BUSCO: 270938at7742
## Thu Jul 14 02:19:34 CEST 2022 - The aliscore output file does not exist. Check results for BUSCO: 378519at7742
## Thu Jul 14 02:19:34 CEST 2022 - The aliscore output appears to be empty. Check results for BUSCO: 378519at7742
## Thu Jul 14 02:19:35 CEST 2022 - The aliscore output file does not exist. Check results for BUSCO: 272582at7742
## Thu Jul 14 02:19:35 CEST 2022 - The aliscore output appears to be empty. Check results for BUSCO: 272582at7742
## Thu Jul 14 02:19:36 CEST 2022 - The aliscore output file does not exist. Check results for BUSCO: 298587at7742
## Thu Jul 14 02:19:36 CEST 2022 - The aliscore output appears to be empty. Check results for BUSCO: 298587at7742
## Thu Jul 14 02:19:36 CEST 2022 - The aliscore output file does not exist. Check results for BUSCO: 384842at7742
## Thu Jul 14 02:19:36 CEST 2022 - The aliscore output file does not exist. Check results for BUSCO: 271060at7742
## Thu Jul 14 02:19:36 CEST 2022 - The aliscore output appears to be empty. Check results for BUSCO: 384842at7742
## Thu Jul 14 02:19:36 CEST 2022 - The aliscore output appears to be empty. Check results for BUSCO: 271060at7742
## Thu Jul 14 02:19:39 CEST 2022 - The aliscore output file does not exist. Check results for BUSCO: 329979at7742
## Thu Jul 14 02:19:39 CEST 2022 - The aliscore output appears to be empty. Check results for BUSCO: 329979at7742
## Thu Jul 14 02:19:40 CEST 2022 - The aliscore output file does not exist. Check results for BUSCO: 264320at7742
## Thu Jul 14 02:19:40 CEST 2022 - The aliscore output appears to be empty. Check results for BUSCO: 264320at7742
## Thu Jul 14 02:19:40 CEST 2022 - The aliscore output file does not exist. Check results for BUSCO: 235735at7742
## Thu Jul 14 02:19:40 CEST 2022 - The aliscore output appears to be empty. Check results for BUSCO: 235735at7742
## Thu Jul 14 02:19:42 CEST 2022 - The aliscore output file does not exist. Check results for BUSCO: 234920at7742
## Thu Jul 14 02:19:42 CEST 2022 - The aliscore output appears to be empty. Check results for BUSCO: 234920at7742
## Thu Jul 14 02:19:43 CEST 2022 - The aliscore output file does not exist. Check results for BUSCO: 320817at7742
## Thu Jul 14 02:19:43 CEST 2022 - The aliscore output appears to be empty. Check results for BUSCO: 320817at7742
## Thu Jul 14 02:19:43 CEST 2022 - The aliscore output file does not exist. Check results for BUSCO: 342301at7742
## Thu Jul 14 02:19:43 CEST 2022 - The aliscore output appears to be empty. Check results for BUSCO: 342301at7742
## Thu Jul 14 02:19:45 CEST 2022 - The aliscore output file does not exist. Check results for BUSCO: 366264at7742
## Thu Jul 14 02:19:45 CEST 2022 - The aliscore output appears to be empty. Check results for BUSCO: 366264at7742
## Thu Jul 14 02:19:46 CEST 2022 - The aliscore output file does not exist. Check results for BUSCO: 301630at7742
## Thu Jul 14 02:19:46 CEST 2022 - The aliscore output appears to be empty. Check results for BUSCO: 301630at7742
## Thu Jul 14 02:19:46 CEST 2022 - The aliscore output file does not exist. Check results for BUSCO: 336178at7742
## Thu Jul 14 02:19:46 CEST 2022 - The aliscore output appears to be empty. Check results for BUSCO: 336178at7742
## Thu Jul 14 02:19:48 CEST 2022 - The aliscore output file does not exist. Check results for BUSCO: 292556at7742
## Thu Jul 14 02:19:48 CEST 2022 - The aliscore output appears to be empty. Check results for BUSCO: 292556at7742
## Thu Jul 14 02:19:48 CEST 2022 - The aliscore output file does not exist. Check results for BUSCO: 304804at7742
## Thu Jul 14 02:19:48 CEST 2022 - The aliscore output appears to be empty. Check results for BUSCO: 304804at7742
## Thu Jul 14 02:19:50 CEST 2022 - The aliscore output file does not exist. Check results for BUSCO: 272432at7742
## Thu Jul 14 02:19:50 CEST 2022 - The aliscore output appears to be empty. Check results for BUSCO: 272432at7742
## Thu Jul 14 02:19:51 CEST 2022 - The aliscore output file does not exist. Check results for BUSCO: 340499at7742
## Thu Jul 14 02:19:51 CEST 2022 - The aliscore output appears to be empty. Check results for BUSCO: 340499at7742
## Thu Jul 14 02:19:52 CEST 2022 - The aliscore output file does not exist. Check results for BUSCO: 293073at7742
## Thu Jul 14 02:19:52 CEST 2022 - The aliscore output appears to be empty. Check results for BUSCO: 293073at7742
## Thu Jul 14 02:19:53 CEST 2022 - The aliscore output file does not exist. Check results for BUSCO: 264320at7742
## Thu Jul 14 02:19:53 CEST 2022 - The aliscore output appears to be empty. Check results for BUSCO: 264320at7742
## Thu Jul 14 02:19:54 CEST 2022 - The aliscore output file does not exist. Check results for BUSCO: 405299at7742
## Thu Jul 14 02:19:54 CEST 2022 - The aliscore output appears to be empty. Check results for BUSCO: 405299at7742
## Thu Jul 14 02:19:54 CEST 2022 - The aliscore output file does not exist. Check results for BUSCO: 336564at7742
## Thu Jul 14 02:19:54 CEST 2022 - The aliscore output appears to be empty. Check results for BUSCO: 336564at7742
## Thu Jul 14 02:19:56 CEST 2022 - The aliscore output file does not exist. Check results for BUSCO: 326524at7742
## Thu Jul 14 02:19:56 CEST 2022 - The aliscore output appears to be empty. Check results for BUSCO: 326524at7742
## Thu Jul 14 02:19:56 CEST 2022 - The aliscore output file does not exist. Check results for BUSCO: 216850at7742
## Thu Jul 14 02:19:57 CEST 2022 - The aliscore output appears to be empty. Check results for BUSCO: 216850at7742
## Thu Jul 14 02:19:58 CEST 2022 - The aliscore output file does not exist. Check results for BUSCO: 303129at7742
## Thu Jul 14 02:19:58 CEST 2022 - The aliscore output appears to be empty. Check results for BUSCO: 303129at7742
## Thu Jul 14 02:19:59 CEST 2022 - The aliscore output file does not exist. Check results for BUSCO: 349595at7742
## Thu Jul 14 02:19:59 CEST 2022 - The aliscore output appears to be empty. Check results for BUSCO: 349595at7742
## Thu Jul 14 02:19:59 CEST 2022 - The aliscore output file does not exist. Check results for BUSCO: 221230at7742
## Thu Jul 14 02:20:00 CEST 2022 - The aliscore output appears to be empty. Check results for BUSCO: 221230at7742
## Thu Jul 14 02:20:00 CEST 2022 - The aliscore output file does not exist. Check results for BUSCO: 403632at7742
## Thu Jul 14 02:20:00 CEST 2022 - The aliscore output appears to be empty. Check results for BUSCO: 403632at7742
## Thu Jul 14 02:20:02 CEST 2022 - The aliscore output file does not exist. Check results for BUSCO: 380379at7742
## Thu Jul 14 02:20:02 CEST 2022 - The aliscore output appears to be empty. Check results for BUSCO: 380379at7742
## Thu Jul 14 02:20:04 CEST 2022 - The aliscore output file does not exist. Check results for BUSCO: 273941at7742
## Thu Jul 14 02:20:04 CEST 2022 - The aliscore output appears to be empty. Check results for BUSCO: 273941at7742
## Thu Jul 14 02:20:05 CEST 2022 - The aliscore output file does not exist. Check results for BUSCO: 302536at7742
## Thu Jul 14 02:20:05 CEST 2022 - The aliscore output appears to be empty. Check results for BUSCO: 302536at7742
## Thu Jul 14 02:20:05 CEST 2022 - The aliscore output file does not exist. Check results for BUSCO: 373281at7742
## Thu Jul 14 02:20:05 CEST 2022 - The aliscore output appears to be empty. Check results for BUSCO: 373281at7742
## Thu Jul 14 02:20:06 CEST 2022 - The aliscore output file does not exist. Check results for BUSCO: 341195at7742
## Thu Jul 14 02:20:06 CEST 2022 - The aliscore output appears to be empty. Check results for BUSCO: 341195at7742
## Thu Jul 14 02:20:06 CEST 2022 - The aliscore output file does not exist. Check results for BUSCO: 396312at7742
## Thu Jul 14 02:20:06 CEST 2022 - The aliscore output appears to be empty. Check results for BUSCO: 396312at7742
## Thu Jul 14 02:20:07 CEST 2022 - The aliscore output file does not exist. Check results for BUSCO: 270381at7742
## Thu Jul 14 02:20:07 CEST 2022 - The aliscore output appears to be empty. Check results for BUSCO: 270381at7742
## Thu Jul 14 02:20:08 CEST 2022 - The aliscore output file does not exist. Check results for BUSCO: 418700at7742
## Thu Jul 14 02:20:08 CEST 2022 - The aliscore output appears to be empty. Check results for BUSCO: 418700at7742
## Thu Jul 14 02:20:10 CEST 2022 - The aliscore output file does not exist. Check results for BUSCO: 261158at7742
## Thu Jul 14 02:20:10 CEST 2022 - The aliscore output appears to be empty. Check results for BUSCO: 261158at7742
## Thu Jul 14 02:20:10 CEST 2022 - The aliscore output file does not exist. Check results for BUSCO: 357819at7742
## Thu Jul 14 02:20:10 CEST 2022 - The aliscore output appears to be empty. Check results for BUSCO: 357819at7742
## Thu Jul 14 02:20:12 CEST 2022 - The aliscore output file does not exist. Check results for BUSCO: 384842at7742
## Thu Jul 14 02:20:12 CEST 2022 - The aliscore output appears to be empty. Check results for BUSCO: 384842at7742
## Thu Jul 14 02:20:12 CEST 2022 - The aliscore output file does not exist. Check results for BUSCO: 328352at7742
## Thu Jul 14 02:20:12 CEST 2022 - The aliscore output appears to be empty. Check results for BUSCO: 328352at7742
## Thu Jul 14 02:20:14 CEST 2022 - The aliscore output file does not exist. Check results for BUSCO: 335620at7742
## Thu Jul 14 02:20:14 CEST 2022 - The aliscore output appears to be empty. Check results for BUSCO: 335620at7742
## Thu Jul 14 02:20:15 CEST 2022 - The aliscore output file does not exist. Check results for BUSCO: 250138at7742
## Thu Jul 14 02:20:15 CEST 2022 - The aliscore output appears to be empty. Check results for BUSCO: 250138at7742
## Thu Jul 14 02:20:15 CEST 2022 - The aliscore output file does not exist. Check results for BUSCO: 421821at7742
## Thu Jul 14 02:20:15 CEST 2022 - The aliscore output appears to be empty. Check results for BUSCO: 421821at7742
## Thu Jul 14 02:20:17 CEST 2022 - The aliscore output file does not exist. Check results for BUSCO: 294706at7742
## Thu Jul 14 02:20:17 CEST 2022 - The aliscore output appears to be empty. Check results for BUSCO: 294706at7742
## Thu Jul 14 02:20:18 CEST 2022 - The aliscore output file does not exist. Check results for BUSCO: 223375at7742
## Thu Jul 14 02:20:19 CEST 2022 - The aliscore output appears to be empty. Check results for BUSCO: 223375at7742
## Thu Jul 14 02:20:19 CEST 2022 - The aliscore output file does not exist. Check results for BUSCO: 317170at7742
## Thu Jul 14 02:20:19 CEST 2022 - The aliscore output appears to be empty. Check results for BUSCO: 317170at7742
## Thu Jul 14 02:20:20 CEST 2022 - The aliscore output file does not exist. Check results for BUSCO: 343725at7742
## Thu Jul 14 02:20:20 CEST 2022 - The aliscore output appears to be empty. Check results for BUSCO: 343725at7742
## Thu Jul 14 02:20:21 CEST 2022 - The aliscore output file does not exist. Check results for BUSCO: 311519at7742
## Thu Jul 14 02:20:21 CEST 2022 - The aliscore output appears to be empty. Check results for BUSCO: 311519at7742
## Thu Jul 14 02:20:21 CEST 2022 - The aliscore output file does not exist. Check results for BUSCO: 321199at7742
## Thu Jul 14 02:20:21 CEST 2022 - The aliscore output appears to be empty. Check results for BUSCO: 321199at7742
## Thu Jul 14 02:20:21 CEST 2022 - The aliscore output file does not exist. Check results for BUSCO: 396312at7742
## Thu Jul 14 02:20:22 CEST 2022 - The aliscore output appears to be empty. Check results for BUSCO: 396312at7742
## Thu Jul 14 02:20:23 CEST 2022 - The aliscore output file does not exist. Check results for BUSCO: 390186at7742
## Thu Jul 14 02:20:23 CEST 2022 - The aliscore output appears to be empty. Check results for BUSCO: 390186at7742
## Thu Jul 14 02:20:24 CEST 2022 - The aliscore output file does not exist. Check results for BUSCO: 256968at7742
## Thu Jul 14 02:20:24 CEST 2022 - The aliscore output appears to be empty. Check results for BUSCO: 256968at7742
## Thu Jul 14 02:20:25 CEST 2022 - The aliscore output file does not exist. Check results for BUSCO: 295973at7742
## Thu Jul 14 02:20:25 CEST 2022 - The aliscore output appears to be empty. Check results for BUSCO: 295973at7742
## Thu Jul 14 02:20:25 CEST 2022 - The aliscore output file does not exist. Check results for BUSCO: 317021at7742
## Thu Jul 14 02:20:25 CEST 2022 - The aliscore output appears to be empty. Check results for BUSCO: 317021at7742
## Thu Jul 14 02:20:27 CEST 2022 - The aliscore output file does not exist. Check results for BUSCO: 388102at7742
## Thu Jul 14 02:20:27 CEST 2022 - The aliscore output appears to be empty. Check results for BUSCO: 388102at7742
## Thu Jul 14 02:20:28 CEST 2022 - The aliscore output file does not exist. Check results for BUSCO: 309935at7742
## Thu Jul 14 02:20:29 CEST 2022 - The aliscore output appears to be empty. Check results for BUSCO: 309935at7742
## Thu Jul 14 02:20:30 CEST 2022 - The aliscore output file does not exist. Check results for BUSCO: 318720at7742
## Thu Jul 14 02:20:30 CEST 2022 - The aliscore output appears to be empty. Check results for BUSCO: 318720at7742
## Thu Jul 14 02:20:30 CEST 2022 - The aliscore output file does not exist. Check results for BUSCO: 272582at7742
## Thu Jul 14 02:20:30 CEST 2022 - The aliscore output appears to be empty. Check results for BUSCO: 272582at7742
## Thu Jul 14 02:20:32 CEST 2022 - The aliscore output file does not exist. Check results for BUSCO: 414238at7742
## Thu Jul 14 02:20:32 CEST 2022 - The aliscore output appears to be empty. Check results for BUSCO: 414238at7742
## Thu Jul 14 02:20:32 CEST 2022 - The aliscore output file does not exist. Check results for BUSCO: 255497at7742
## Thu Jul 14 02:20:32 CEST 2022 - The aliscore output appears to be empty. Check results for BUSCO: 255497at7742
## Thu Jul 14 02:20:32 CEST 2022 - The aliscore output file does not exist. Check results for BUSCO: 271425at7742
## Thu Jul 14 02:20:32 CEST 2022 - The aliscore output appears to be empty. Check results for BUSCO: 271425at7742
## Thu Jul 14 02:20:34 CEST 2022 - The aliscore output file does not exist. Check results for BUSCO: 407384at7742
## Thu Jul 14 02:20:34 CEST 2022 - The aliscore output appears to be empty. Check results for BUSCO: 407384at7742
## Thu Jul 14 02:20:35 CEST 2022 - The aliscore output file does not exist. Check results for BUSCO: 258747at7742
## Thu Jul 14 02:20:35 CEST 2022 - The aliscore output appears to be empty. Check results for BUSCO: 258747at7742
## Thu Jul 14 02:20:36 CEST 2022 - The aliscore output file does not exist. Check results for BUSCO: 284651at7742
## Thu Jul 14 02:20:36 CEST 2022 - The aliscore output appears to be empty. Check results for BUSCO: 284651at7742
## Thu Jul 14 02:20:36 CEST 2022 - The aliscore output file does not exist. Check results for BUSCO: 374832at7742
## Thu Jul 14 02:20:36 CEST 2022 - The aliscore output appears to be empty. Check results for BUSCO: 374832at7742
## Thu Jul 14 02:20:37 CEST 2022 - The aliscore output file does not exist. Check results for BUSCO: 305344at7742
## Thu Jul 14 02:20:37 CEST 2022 - The aliscore output appears to be empty. Check results for BUSCO: 305344at7742
## Thu Jul 14 02:20:39 CEST 2022 - The aliscore output file does not exist. Check results for BUSCO: 330751at7742
## Thu Jul 14 02:20:39 CEST 2022 - The aliscore output appears to be empty. Check results for BUSCO: 330751at7742
## Thu Jul 14 02:20:39 CEST 2022 - The aliscore output file does not exist. Check results for BUSCO: 260364at7742
## Thu Jul 14 02:20:39 CEST 2022 - The aliscore output appears to be empty. Check results for BUSCO: 260364at7742
## Thu Jul 14 02:20:41 CEST 2022 - The aliscore output file does not exist. Check results for BUSCO: 262895at7742
## Thu Jul 14 02:20:41 CEST 2022 - The aliscore output appears to be empty. Check results for BUSCO: 262895at7742
## Thu Jul 14 02:20:42 CEST 2022 - The aliscore output file does not exist. Check results for BUSCO: 372062at7742
## Thu Jul 14 02:20:42 CEST 2022 - The aliscore output appears to be empty. Check results for BUSCO: 372062at7742
## Thu Jul 14 02:20:42 CEST 2022 - The aliscore output file does not exist. Check results for BUSCO: 273993at7742
## Thu Jul 14 02:20:42 CEST 2022 - The aliscore output appears to be empty. Check results for BUSCO: 273993at7742
## Thu Jul 14 02:20:45 CEST 2022 - The aliscore output file does not exist. Check results for BUSCO: 270435at7742
## Thu Jul 14 02:20:45 CEST 2022 - The aliscore output appears to be empty. Check results for BUSCO: 270435at7742
## Thu Jul 14 02:20:45 CEST 2022 - The aliscore output file does not exist. Check results for BUSCO: 258997at7742
## Thu Jul 14 02:20:46 CEST 2022 - The aliscore output appears to be empty. Check results for BUSCO: 258997at7742
## Thu Jul 14 02:20:46 CEST 2022 - The aliscore output file does not exist. Check results for BUSCO: 326316at7742
## Thu Jul 14 02:20:46 CEST 2022 - The aliscore output appears to be empty. Check results for BUSCO: 326316at7742
## Thu Jul 14 02:20:47 CEST 2022 - The aliscore output file does not exist. Check results for BUSCO: 272432at7742
## Thu Jul 14 02:20:48 CEST 2022 - The aliscore output appears to be empty. Check results for BUSCO: 272432at7742
## Thu Jul 14 02:20:48 CEST 2022 - The aliscore output file does not exist. Check results for BUSCO: 400935at7742
## Thu Jul 14 02:20:48 CEST 2022 - The aliscore output appears to be empty. Check results for BUSCO: 400935at7742
## Thu Jul 14 02:20:49 CEST 2022 - The aliscore output file does not exist. Check results for BUSCO: 281245at7742
## Thu Jul 14 02:20:49 CEST 2022 - The aliscore output appears to be empty. Check results for BUSCO: 281245at7742
## Thu Jul 14 02:20:50 CEST 2022 - The aliscore output file does not exist. Check results for BUSCO: 257379at7742
## Thu Jul 14 02:20:50 CEST 2022 - The aliscore output appears to be empty. Check results for BUSCO: 257379at7742
## Thu Jul 14 02:20:51 CEST 2022 - The aliscore output file does not exist. Check results for BUSCO: 413350at7742
## Thu Jul 14 02:20:51 CEST 2022 - The aliscore output appears to be empty. Check results for BUSCO: 413350at7742
## Thu Jul 14 02:20:52 CEST 2022 - The aliscore output file does not exist. Check results for BUSCO: 341195at7742
## Thu Jul 14 02:20:52 CEST 2022 - The aliscore output appears to be empty. Check results for BUSCO: 341195at7742
## Thu Jul 14 02:20:53 CEST 2022 - The aliscore output file does not exist. Check results for BUSCO: 294990at7742
## Thu Jul 14 02:20:53 CEST 2022 - The aliscore output appears to be empty. Check results for BUSCO: 294990at7742
## Thu Jul 14 02:20:54 CEST 2022 - The aliscore output file does not exist. Check results for BUSCO: 391492at7742
## Thu Jul 14 02:20:54 CEST 2022 - The aliscore output appears to be empty. Check results for BUSCO: 391492at7742
## Thu Jul 14 02:20:56 CEST 2022 - The aliscore output file does not exist. Check results for BUSCO: 234920at7742
## Thu Jul 14 02:20:56 CEST 2022 - The aliscore output appears to be empty. Check results for BUSCO: 234920at7742
## Thu Jul 14 02:20:56 CEST 2022 - The aliscore output file does not exist. Check results for BUSCO: 375859at7742
## Thu Jul 14 02:20:56 CEST 2022 - The aliscore output appears to be empty. Check results for BUSCO: 375859at7742
## Thu Jul 14 02:20:57 CEST 2022 - The aliscore output file does not exist. Check results for BUSCO: 315823at7742
## Thu Jul 14 02:20:57 CEST 2022 - The aliscore output appears to be empty. Check results for BUSCO: 315823at7742
## Thu Jul 14 02:20:58 CEST 2022 - The aliscore output file does not exist. Check results for BUSCO: 190657at7742
## Thu Jul 14 02:20:58 CEST 2022 - The aliscore output appears to be empty. Check results for BUSCO: 190657at7742
## Thu Jul 14 02:20:59 CEST 2022 - The aliscore output file does not exist. Check results for BUSCO: 376384at7742
## Thu Jul 14 02:20:59 CEST 2022 - The aliscore output appears to be empty. Check results for BUSCO: 376384at7742
## Thu Jul 14 02:21:00 CEST 2022 - The aliscore output file does not exist. Check results for BUSCO: 311206at7742
## Thu Jul 14 02:21:00 CEST 2022 - The aliscore output appears to be empty. Check results for BUSCO: 311206at7742
## Thu Jul 14 02:21:00 CEST 2022 - The aliscore output file does not exist. Check results for BUSCO: 292271at7742
## Thu Jul 14 02:21:00 CEST 2022 - The aliscore output appears to be empty. Check results for BUSCO: 292271at7742
## Thu Jul 14 02:21:02 CEST 2022 - The aliscore output file does not exist. Check results for BUSCO: 259157at7742
## Thu Jul 14 02:21:02 CEST 2022 - The aliscore output appears to be empty. Check results for BUSCO: 259157at7742
## Thu Jul 14 02:21:03 CEST 2022 - The aliscore output file does not exist. Check results for BUSCO: 242979at7742
## Thu Jul 14 02:21:03 CEST 2022 - The aliscore output appears to be empty. Check results for BUSCO: 242979at7742
## Thu Jul 14 02:21:03 CEST 2022 - The aliscore output file does not exist. Check results for BUSCO: 323970at7742
## Thu Jul 14 02:21:03 CEST 2022 - The aliscore output appears to be empty. Check results for BUSCO: 323970at7742
## Thu Jul 14 02:21:06 CEST 2022 - The aliscore output file does not exist. Check results for BUSCO: 349204at7742
## Thu Jul 14 02:21:06 CEST 2022 - The aliscore output file does not exist. Check results for BUSCO: 414238at7742
## Thu Jul 14 02:21:06 CEST 2022 - The aliscore output appears to be empty. Check results for BUSCO: 349204at7742
## Thu Jul 14 02:21:06 CEST 2022 - The aliscore output appears to be empty. Check results for BUSCO: 414238at7742
## Thu Jul 14 02:21:07 CEST 2022 - The aliscore output file does not exist. Check results for BUSCO: 403876at7742
## Thu Jul 14 02:21:07 CEST 2022 - The aliscore output appears to be empty. Check results for BUSCO: 403876at7742
## Thu Jul 14 02:21:08 CEST 2022 - The aliscore output file does not exist. Check results for BUSCO: 211842at7742
## Thu Jul 14 02:21:09 CEST 2022 - The aliscore output appears to be empty. Check results for BUSCO: 211842at7742
## Thu Jul 14 02:21:09 CEST 2022 - The aliscore output file does not exist. Check results for BUSCO: 359330at7742
## Thu Jul 14 02:21:09 CEST 2022 - The aliscore output appears to be empty. Check results for BUSCO: 359330at7742
## Thu Jul 14 02:21:11 CEST 2022 - The aliscore output file does not exist. Check results for BUSCO: 426882at7742
## Thu Jul 14 02:21:11 CEST 2022 - The aliscore output appears to be empty. Check results for BUSCO: 426882at7742
## Thu Jul 14 02:21:11 CEST 2022 - The aliscore output file does not exist. Check results for BUSCO: 399664at7742
## Thu Jul 14 02:21:11 CEST 2022 - The aliscore output appears to be empty. Check results for BUSCO: 399664at7742
## Thu Jul 14 02:21:12 CEST 2022 - The aliscore output file does not exist. Check results for BUSCO: 271121at7742
## Thu Jul 14 02:21:12 CEST 2022 - The aliscore output appears to be empty. Check results for BUSCO: 271121at7742
## Thu Jul 14 02:21:13 CEST 2022 - The aliscore output file does not exist. Check results for BUSCO: 318720at7742
## Thu Jul 14 02:21:13 CEST 2022 - The aliscore output appears to be empty. Check results for BUSCO: 318720at7742
## Thu Jul 14 02:21:14 CEST 2022 - The aliscore output file does not exist. Check results for BUSCO: 317722at7742
## Thu Jul 14 02:21:15 CEST 2022 - The aliscore output appears to be empty. Check results for BUSCO: 317722at7742
## Thu Jul 14 02:21:15 CEST 2022 - The aliscore output file does not exist. Check results for BUSCO: 257379at7742
## Thu Jul 14 02:21:15 CEST 2022 - The aliscore output appears to be empty. Check results for BUSCO: 257379at7742
## Thu Jul 14 02:21:15 CEST 2022 - The aliscore output file does not exist. Check results for BUSCO: 354226at7742
## Thu Jul 14 02:21:15 CEST 2022 - The aliscore output appears to be empty. Check results for BUSCO: 354226at7742
## Thu Jul 14 02:21:17 CEST 2022 - The aliscore output file does not exist. Check results for BUSCO: 264221at7742
## Thu Jul 14 02:21:17 CEST 2022 - The aliscore output appears to be empty. Check results for BUSCO: 264221at7742
## Thu Jul 14 02:21:18 CEST 2022 - The aliscore output file does not exist. Check results for BUSCO: 217762at7742
## Thu Jul 14 02:21:18 CEST 2022 - The aliscore output appears to be empty. Check results for BUSCO: 217762at7742
## Thu Jul 14 02:21:19 CEST 2022 - The aliscore output file does not exist. Check results for BUSCO: 373281at7742
## Thu Jul 14 02:21:19 CEST 2022 - The aliscore output appears to be empty. Check results for BUSCO: 373281at7742
## Thu Jul 14 02:21:20 CEST 2022 - The aliscore output file does not exist. Check results for BUSCO: 328949at7742
## Thu Jul 14 02:21:20 CEST 2022 - The aliscore output appears to be empty. Check results for BUSCO: 328949at7742
## Thu Jul 14 02:21:22 CEST 2022 - The aliscore output file does not exist. Check results for BUSCO: 400935at7742
## Thu Jul 14 02:21:22 CEST 2022 - The aliscore output appears to be empty. Check results for BUSCO: 400935at7742
## Thu Jul 14 02:21:22 CEST 2022 - The aliscore output file does not exist. Check results for BUSCO: 328352at7742
## Thu Jul 14 02:21:22 CEST 2022 - The aliscore output appears to be empty. Check results for BUSCO: 328352at7742
## Thu Jul 14 02:21:22 CEST 2022 - The aliscore output file does not exist. Check results for BUSCO: 246301at7742
## Thu Jul 14 02:21:22 CEST 2022 - The aliscore output appears to be empty. Check results for BUSCO: 246301at7742
## Thu Jul 14 02:21:24 CEST 2022 - The aliscore output file does not exist. Check results for BUSCO: 328134at7742
## Thu Jul 14 02:21:24 CEST 2022 - The aliscore output appears to be empty. Check results for BUSCO: 328134at7742
## Thu Jul 14 02:21:25 CEST 2022 - The aliscore output file does not exist. Check results for BUSCO: 269168at7742
## Thu Jul 14 02:21:25 CEST 2022 - The aliscore output appears to be empty. Check results for BUSCO: 269168at7742
## Thu Jul 14 02:21:26 CEST 2022 - The aliscore output file does not exist. Check results for BUSCO: 317722at7742
## Thu Jul 14 02:21:26 CEST 2022 - The aliscore output appears to be empty. Check results for BUSCO: 317722at7742
## Thu Jul 14 02:21:26 CEST 2022 - The aliscore output file does not exist. Check results for BUSCO: 388183at7742
## Thu Jul 14 02:21:26 CEST 2022 - The aliscore output appears to be empty. Check results for BUSCO: 388183at7742
## Thu Jul 14 02:21:27 CEST 2022 - The aliscore output file does not exist. Check results for BUSCO: 333698at7742
## Thu Jul 14 02:21:27 CEST 2022 - The aliscore output appears to be empty. Check results for BUSCO: 333698at7742
## Thu Jul 14 02:21:28 CEST 2022 - The aliscore output file does not exist. Check results for BUSCO: 303740at7742
## Thu Jul 14 02:21:28 CEST 2022 - The aliscore output appears to be empty. Check results for BUSCO: 303740at7742
## Thu Jul 14 02:21:30 CEST 2022 - The aliscore output file does not exist. Check results for BUSCO: 187843at7742
## Thu Jul 14 02:21:30 CEST 2022 - The aliscore output appears to be empty. Check results for BUSCO: 187843at7742
## Thu Jul 14 02:21:30 CEST 2022 - The aliscore output file does not exist. Check results for BUSCO: 336346at7742
## Thu Jul 14 02:21:30 CEST 2022 - The aliscore output appears to be empty. Check results for BUSCO: 336346at7742
## Thu Jul 14 02:21:32 CEST 2022 - The aliscore output file does not exist. Check results for BUSCO: 258997at7742
## Thu Jul 14 02:21:32 CEST 2022 - The aliscore output appears to be empty. Check results for BUSCO: 258997at7742
## Thu Jul 14 02:21:32 CEST 2022 - The aliscore output file does not exist. Check results for BUSCO: 334381at7742
## Thu Jul 14 02:21:32 CEST 2022 - The aliscore output appears to be empty. Check results for BUSCO: 334381at7742
## Thu Jul 14 02:21:34 CEST 2022 - The aliscore output file does not exist. Check results for BUSCO: 304827at7742
## Thu Jul 14 02:21:34 CEST 2022 - The aliscore output file does not exist. Check results for BUSCO: 336764at7742
## Thu Jul 14 02:21:34 CEST 2022 - The aliscore output appears to be empty. Check results for BUSCO: 304827at7742
## Thu Jul 14 02:21:34 CEST 2022 - The aliscore output appears to be empty. Check results for BUSCO: 336764at7742
## Thu Jul 14 02:21:35 CEST 2022 - The aliscore output file does not exist. Check results for BUSCO: 291403at7742
## Thu Jul 14 02:21:35 CEST 2022 - The aliscore output appears to be empty. Check results for BUSCO: 291403at7742
## Thu Jul 14 02:21:37 CEST 2022 - The aliscore output file does not exist. Check results for BUSCO: 373281at7742
## Thu Jul 14 02:21:37 CEST 2022 - The aliscore output appears to be empty. Check results for BUSCO: 373281at7742
## Thu Jul 14 02:21:38 CEST 2022 - The aliscore output file does not exist. Check results for BUSCO: 305378at7742
## Thu Jul 14 02:21:38 CEST 2022 - The aliscore output appears to be empty. Check results for BUSCO: 305378at7742
## Thu Jul 14 02:21:39 CEST 2022 - The aliscore output file does not exist. Check results for BUSCO: 304588at7742
## Thu Jul 14 02:21:39 CEST 2022 - The aliscore output appears to be empty. Check results for BUSCO: 304588at7742
## Thu Jul 14 02:21:39 CEST 2022 - The aliscore output file does not exist. Check results for BUSCO: 311519at7742
## Thu Jul 14 02:21:39 CEST 2022 - The aliscore output appears to be empty. Check results for BUSCO: 311519at7742
## Thu Jul 14 02:21:42 CEST 2022 - The aliscore output file does not exist. Check results for BUSCO: 396303at7742
## Thu Jul 14 02:21:42 CEST 2022 - The aliscore output appears to be empty. Check results for BUSCO: 396303at7742
## Thu Jul 14 02:21:42 CEST 2022 - The aliscore output file does not exist. Check results for BUSCO: 368306at7742
## Thu Jul 14 02:21:42 CEST 2022 - The aliscore output appears to be empty. Check results for BUSCO: 368306at7742
## Thu Jul 14 02:21:42 CEST 2022 - The aliscore output file does not exist. Check results for BUSCO: 310457at7742
## Thu Jul 14 02:21:42 CEST 2022 - The aliscore output appears to be empty. Check results for BUSCO: 310457at7742
## Thu Jul 14 02:21:43 CEST 2022 - The aliscore output file does not exist. Check results for BUSCO: 315823at7742
## Thu Jul 14 02:21:43 CEST 2022 - The aliscore output appears to be empty. Check results for BUSCO: 315823at7742
## Thu Jul 14 02:21:44 CEST 2022 - The aliscore output file does not exist. Check results for BUSCO: 293704at7742
## Thu Jul 14 02:21:44 CEST 2022 - The aliscore output appears to be empty. Check results for BUSCO: 293704at7742
## Thu Jul 14 02:21:45 CEST 2022 - The aliscore output file does not exist. Check results for BUSCO: 317722at7742
## Thu Jul 14 02:21:45 CEST 2022 - The aliscore output appears to be empty. Check results for BUSCO: 317722at7742
## Thu Jul 14 02:21:46 CEST 2022 - The aliscore output file does not exist. Check results for BUSCO: 305344at7742
## Thu Jul 14 02:21:46 CEST 2022 - The aliscore output appears to be empty. Check results for BUSCO: 305344at7742
## Thu Jul 14 02:21:47 CEST 2022 - The aliscore output file does not exist. Check results for BUSCO: 337045at7742
## Thu Jul 14 02:21:47 CEST 2022 - The aliscore output appears to be empty. Check results for BUSCO: 337045at7742
## Thu Jul 14 02:21:49 CEST 2022 - The aliscore output file does not exist. Check results for BUSCO: 280430at7742
## Thu Jul 14 02:21:49 CEST 2022 - The aliscore output appears to be empty. Check results for BUSCO: 280430at7742
## Thu Jul 14 02:21:49 CEST 2022 - The aliscore output file does not exist. Check results for BUSCO: 290213at7742
## Thu Jul 14 02:21:49 CEST 2022 - The aliscore output appears to be empty. Check results for BUSCO: 290213at7742
## Thu Jul 14 02:21:50 CEST 2022 - The aliscore output file does not exist. Check results for BUSCO: 426882at7742
## Thu Jul 14 02:21:50 CEST 2022 - The aliscore output appears to be empty. Check results for BUSCO: 426882at7742
## Thu Jul 14 02:21:52 CEST 2022 - The aliscore output file does not exist. Check results for BUSCO: 288175at7742
## Thu Jul 14 02:21:52 CEST 2022 - The aliscore output appears to be empty. Check results for BUSCO: 288175at7742
## Thu Jul 14 02:21:52 CEST 2022 - The aliscore output file does not exist. Check results for BUSCO: 277361at7742
## Thu Jul 14 02:21:52 CEST 2022 - The aliscore output appears to be empty. Check results for BUSCO: 277361at7742
## Thu Jul 14 02:21:53 CEST 2022 - The aliscore output file does not exist. Check results for BUSCO: 365149at7742
## Thu Jul 14 02:21:53 CEST 2022 - The aliscore output appears to be empty. Check results for BUSCO: 365149at7742
## Thu Jul 14 02:21:53 CEST 2022 - The aliscore output file does not exist. Check results for BUSCO: 271121at7742
## Thu Jul 14 02:21:53 CEST 2022 - The aliscore output appears to be empty. Check results for BUSCO: 271121at7742
## Thu Jul 14 02:21:56 CEST 2022 - The aliscore output file does not exist. Check results for BUSCO: 292271at7742
## Thu Jul 14 02:21:56 CEST 2022 - The aliscore output appears to be empty. Check results for BUSCO: 292271at7742
## Thu Jul 14 02:21:56 CEST 2022 - The aliscore output file does not exist. Check results for BUSCO: 359032at7742
## Thu Jul 14 02:21:56 CEST 2022 - The aliscore output appears to be empty. Check results for BUSCO: 359032at7742
## Thu Jul 14 02:21:56 CEST 2022 - The aliscore output file does not exist. Check results for BUSCO: 246297at7742
## Thu Jul 14 02:21:56 CEST 2022 - The aliscore output appears to be empty. Check results for BUSCO: 246297at7742
## Thu Jul 14 02:21:57 CEST 2022 - The aliscore output file does not exist. Check results for BUSCO: 249652at7742
## Thu Jul 14 02:21:57 CEST 2022 - The aliscore output appears to be empty. Check results for BUSCO: 249652at7742
## Thu Jul 14 02:21:58 CEST 2022 - The aliscore output file does not exist. Check results for BUSCO: 271121at7742
## Thu Jul 14 02:21:58 CEST 2022 - The aliscore output appears to be empty. Check results for BUSCO: 271121at7742
## Thu Jul 14 02:21:58 CEST 2022 - The aliscore output file does not exist. Check results for BUSCO: 349858at7742
## Thu Jul 14 02:21:58 CEST 2022 - The aliscore output appears to be empty. Check results for BUSCO: 349858at7742
## Thu Jul 14 02:21:59 CEST 2022 - The aliscore output file does not exist. Check results for BUSCO: 187843at7742
## Thu Jul 14 02:21:59 CEST 2022 - The aliscore output appears to be empty. Check results for BUSCO: 187843at7742
## Thu Jul 14 02:22:03 CEST 2022 - The aliscore output file does not exist. Check results for BUSCO: 327840at7742
## Thu Jul 14 02:22:03 CEST 2022 - The aliscore output appears to be empty. Check results for BUSCO: 327840at7742
## Thu Jul 14 02:22:03 CEST 2022 - The aliscore output file does not exist. Check results for BUSCO: 372431at7742
## Thu Jul 14 02:22:03 CEST 2022 - The aliscore output appears to be empty. Check results for BUSCO: 372431at7742
## Thu Jul 14 02:22:03 CEST 2022 - The aliscore output file does not exist. Check results for BUSCO: 241207at7742
## Thu Jul 14 02:22:03 CEST 2022 - The aliscore output file does not exist. Check results for BUSCO: 324120at7742
## Thu Jul 14 02:22:03 CEST 2022 - The aliscore output appears to be empty. Check results for BUSCO: 241207at7742
## Thu Jul 14 02:22:03 CEST 2022 - The aliscore output appears to be empty. Check results for BUSCO: 324120at7742
## Thu Jul 14 02:22:05 CEST 2022 - The aliscore output file does not exist. Check results for BUSCO: 298587at7742
## Thu Jul 14 02:22:05 CEST 2022 - The aliscore output appears to be empty. Check results for BUSCO: 298587at7742
## Thu Jul 14 02:22:06 CEST 2022 - The aliscore output file does not exist. Check results for BUSCO: 407384at7742
## Thu Jul 14 02:22:06 CEST 2022 - The aliscore output appears to be empty. Check results for BUSCO: 407384at7742
## Thu Jul 14 02:22:09 CEST 2022 - The aliscore output file does not exist. Check results for BUSCO: 409736at7742
## Thu Jul 14 02:22:10 CEST 2022 - The aliscore output appears to be empty. Check results for BUSCO: 409736at7742
## Thu Jul 14 02:22:10 CEST 2022 - The aliscore output file does not exist. Check results for BUSCO: 323574at7742
## Thu Jul 14 02:22:10 CEST 2022 - The aliscore output appears to be empty. Check results for BUSCO: 323574at7742
## Thu Jul 14 02:22:10 CEST 2022 - The aliscore output file does not exist. Check results for BUSCO: 273993at7742
## Thu Jul 14 02:22:10 CEST 2022 - The aliscore output appears to be empty. Check results for BUSCO: 273993at7742
## Thu Jul 14 02:22:11 CEST 2022 - The aliscore output file does not exist. Check results for BUSCO: 415130at7742
## Thu Jul 14 02:22:11 CEST 2022 - The aliscore output appears to be empty. Check results for BUSCO: 415130at7742
## Thu Jul 14 02:22:11 CEST 2022 - The aliscore output file does not exist. Check results for BUSCO: 323970at7742
## Thu Jul 14 02:22:11 CEST 2022 - The aliscore output appears to be empty. Check results for BUSCO: 323970at7742
## Thu Jul 14 02:22:11 CEST 2022 - The aliscore output file does not exist. Check results for BUSCO: 336346at7742
## Thu Jul 14 02:22:11 CEST 2022 - The aliscore output appears to be empty. Check results for BUSCO: 336346at7742
## Thu Jul 14 02:22:11 CEST 2022 - The aliscore output file does not exist. Check results for BUSCO: 418700at7742
## Thu Jul 14 02:22:11 CEST 2022 - The aliscore output appears to be empty. Check results for BUSCO: 418700at7742
## Thu Jul 14 02:22:17 CEST 2022 - The aliscore output file does not exist. Check results for BUSCO: 344436at7742
## Thu Jul 14 02:22:17 CEST 2022 - The aliscore output file does not exist. Check results for BUSCO: 391948at7742
## Thu Jul 14 02:22:17 CEST 2022 - The aliscore output appears to be empty. Check results for BUSCO: 344436at7742
## Thu Jul 14 02:22:17 CEST 2022 - The aliscore output appears to be empty. Check results for BUSCO: 391948at7742
## Thu Jul 14 02:22:17 CEST 2022 - The aliscore output file does not exist. Check results for BUSCO: 351419at7742
## Thu Jul 14 02:22:17 CEST 2022 - The aliscore output appears to be empty. Check results for BUSCO: 351419at7742
## Thu Jul 14 02:22:17 CEST 2022 - The aliscore output file does not exist. Check results for BUSCO: 321199at7742
## Thu Jul 14 02:22:17 CEST 2022 - The aliscore output appears to be empty. Check results for BUSCO: 321199at7742
## Thu Jul 14 02:22:17 CEST 2022 - The aliscore output file does not exist. Check results for BUSCO: 329979at7742
## Thu Jul 14 02:22:17 CEST 2022 - The aliscore output file does not exist. Check results for BUSCO: 397455at7742
## Thu Jul 14 02:22:17 CEST 2022 - The aliscore output appears to be empty. Check results for BUSCO: 329979at7742
## Thu Jul 14 02:22:18 CEST 2022 - The aliscore output appears to be empty. Check results for BUSCO: 397455at7742
## Thu Jul 14 02:22:21 CEST 2022 - The aliscore output file does not exist. Check results for BUSCO: 420085at7742
## Thu Jul 14 02:22:21 CEST 2022 - The aliscore output appears to be empty. Check results for BUSCO: 420085at7742
## Thu Jul 14 02:22:21 CEST 2022 - The aliscore output file does not exist. Check results for BUSCO: 235335at7742
## Thu Jul 14 02:22:21 CEST 2022 - The aliscore output appears to be empty. Check results for BUSCO: 235335at7742
## Thu Jul 14 02:22:22 CEST 2022 - The aliscore output file does not exist. Check results for BUSCO: 376384at7742
## Thu Jul 14 02:22:22 CEST 2022 - The aliscore output appears to be empty. Check results for BUSCO: 376384at7742
## Thu Jul 14 02:22:23 CEST 2022 - The aliscore output file does not exist. Check results for BUSCO: 343085at7742
## Thu Jul 14 02:22:23 CEST 2022 - The aliscore output appears to be empty. Check results for BUSCO: 343085at7742
## Thu Jul 14 02:22:23 CEST 2022 - The aliscore output file does not exist. Check results for BUSCO: 407723at7742
## Thu Jul 14 02:22:23 CEST 2022 - The aliscore output appears to be empty. Check results for BUSCO: 407723at7742
## Thu Jul 14 02:22:24 CEST 2022 - The aliscore output file does not exist. Check results for BUSCO: 334363at7742
## Thu Jul 14 02:22:24 CEST 2022 - The aliscore output appears to be empty. Check results for BUSCO: 334363at7742
## Thu Jul 14 02:22:25 CEST 2022 - The aliscore output file does not exist. Check results for BUSCO: 259089at7742
## Thu Jul 14 02:22:25 CEST 2022 - The aliscore output appears to be empty. Check results for BUSCO: 259089at7742
## Thu Jul 14 02:22:27 CEST 2022 - The aliscore output file does not exist. Check results for BUSCO: 309948at7742
## Thu Jul 14 02:22:27 CEST 2022 - The aliscore output appears to be empty. Check results for BUSCO: 309948at7742
## Thu Jul 14 02:22:27 CEST 2022 - The aliscore output file does not exist. Check results for BUSCO: 279092at7742
## Thu Jul 14 02:22:27 CEST 2022 - The aliscore output appears to be empty. Check results for BUSCO: 279092at7742
## Thu Jul 14 02:22:31 CEST 2022 - The aliscore output file does not exist. Check results for BUSCO: 376479at7742
## Thu Jul 14 02:22:31 CEST 2022 - The aliscore output appears to be empty. Check results for BUSCO: 376479at7742
## Thu Jul 14 02:22:31 CEST 2022 - The aliscore output file does not exist. Check results for BUSCO: 273941at7742
## Thu Jul 14 02:22:31 CEST 2022 - The aliscore output file does not exist. Check results for BUSCO: 376479at7742
## Thu Jul 14 02:22:31 CEST 2022 - The aliscore output appears to be empty. Check results for BUSCO: 273941at7742
## Thu Jul 14 02:22:31 CEST 2022 - The aliscore output appears to be empty. Check results for BUSCO: 376479at7742
## Thu Jul 14 02:22:32 CEST 2022 - The aliscore output file does not exist. Check results for BUSCO: 243977at7742
## Thu Jul 14 02:22:32 CEST 2022 - The aliscore output appears to be empty. Check results for BUSCO: 243977at7742
## Thu Jul 14 02:22:33 CEST 2022 - The aliscore output file does not exist. Check results for BUSCO: 271622at7742
## Thu Jul 14 02:22:33 CEST 2022 - The aliscore output file does not exist. Check results for BUSCO: 326316at7742
## Thu Jul 14 02:22:33 CEST 2022 - The aliscore output appears to be empty. Check results for BUSCO: 271622at7742
## Thu Jul 14 02:22:33 CEST 2022 - The aliscore output appears to be empty. Check results for BUSCO: 326316at7742
## Thu Jul 14 02:22:35 CEST 2022 - The aliscore output file does not exist. Check results for BUSCO: 241207at7742
## Thu Jul 14 02:22:35 CEST 2022 - The aliscore output appears to be empty. Check results for BUSCO: 241207at7742
## Thu Jul 14 02:22:37 CEST 2022 - The aliscore output file does not exist. Check results for BUSCO: 336331at7742
## Thu Jul 14 02:22:37 CEST 2022 - The aliscore output appears to be empty. Check results for BUSCO: 336331at7742
## Thu Jul 14 02:22:37 CEST 2022 - The aliscore output file does not exist. Check results for BUSCO: 245611at7742
## Thu Jul 14 02:22:37 CEST 2022 - The aliscore output appears to be empty. Check results for BUSCO: 245611at7742
## Thu Jul 14 02:22:38 CEST 2022 - The aliscore output file does not exist. Check results for BUSCO: 363264at7742
## Thu Jul 14 02:22:38 CEST 2022 - The aliscore output appears to be empty. Check results for BUSCO: 363264at7742
## Thu Jul 14 02:22:38 CEST 2022 - The aliscore output file does not exist. Check results for BUSCO: 264124at7742
## Thu Jul 14 02:22:38 CEST 2022 - The aliscore output appears to be empty. Check results for BUSCO: 264124at7742
## Thu Jul 14 02:22:38 CEST 2022 - The aliscore output file does not exist. Check results for BUSCO: 246301at7742
## Thu Jul 14 02:22:38 CEST 2022 - The aliscore output appears to be empty. Check results for BUSCO: 246301at7742
## Thu Jul 14 02:22:40 CEST 2022 - The aliscore output file does not exist. Check results for BUSCO: 337042at7742
## Thu Jul 14 02:22:40 CEST 2022 - The aliscore output appears to be empty. Check results for BUSCO: 337042at7742
## Thu Jul 14 02:22:42 CEST 2022 - The aliscore output file does not exist. Check results for BUSCO: 370608at7742
## Thu Jul 14 02:22:42 CEST 2022 - The aliscore output file does not exist. Check results for BUSCO: 337045at7742
## Thu Jul 14 02:22:42 CEST 2022 - The aliscore output appears to be empty. Check results for BUSCO: 370608at7742
## Thu Jul 14 02:22:42 CEST 2022 - The aliscore output appears to be empty. Check results for BUSCO: 337045at7742
## Thu Jul 14 02:22:43 CEST 2022 - The aliscore output file does not exist. Check results for BUSCO: 298740at7742
## Thu Jul 14 02:22:43 CEST 2022 - The aliscore output appears to be empty. Check results for BUSCO: 298740at7742
## Thu Jul 14 02:22:45 CEST 2022 - The aliscore output file does not exist. Check results for BUSCO: 270381at7742
## Thu Jul 14 02:22:45 CEST 2022 - The aliscore output appears to be empty. Check results for BUSCO: 270381at7742
## Thu Jul 14 02:22:45 CEST 2022 - The aliscore output file does not exist. Check results for BUSCO: 235335at7742
## Thu Jul 14 02:22:45 CEST 2022 - The aliscore output appears to be empty. Check results for BUSCO: 235335at7742
## Thu Jul 14 02:22:45 CEST 2022 - The aliscore output file does not exist. Check results for BUSCO: 270381at7742
## Thu Jul 14 02:22:45 CEST 2022 - The aliscore output appears to be empty. Check results for BUSCO: 270381at7742
## Thu Jul 14 02:22:46 CEST 2022 - The aliscore output file does not exist. Check results for BUSCO: 388183at7742
## Thu Jul 14 02:22:46 CEST 2022 - The aliscore output appears to be empty. Check results for BUSCO: 388183at7742
## Thu Jul 14 02:22:47 CEST 2022 - The aliscore output file does not exist. Check results for BUSCO: 295973at7742
## Thu Jul 14 02:22:47 CEST 2022 - The aliscore output appears to be empty. Check results for BUSCO: 295973at7742
## Thu Jul 14 02:22:48 CEST 2022 - The aliscore output file does not exist. Check results for BUSCO: 221230at7742
## Thu Jul 14 02:22:48 CEST 2022 - The aliscore output appears to be empty. Check results for BUSCO: 221230at7742
## Thu Jul 14 02:22:50 CEST 2022 - The aliscore output file does not exist. Check results for BUSCO: 336346at7742
## Thu Jul 14 02:22:50 CEST 2022 - The aliscore output appears to be empty. Check results for BUSCO: 336346at7742
## Thu Jul 14 02:22:51 CEST 2022 - The aliscore output file does not exist. Check results for BUSCO: 377177at7742
## Thu Jul 14 02:22:52 CEST 2022 - The aliscore output appears to be empty. Check results for BUSCO: 377177at7742
## Thu Jul 14 02:22:52 CEST 2022 - The aliscore output file does not exist. Check results for BUSCO: 262895at7742
## Thu Jul 14 02:22:52 CEST 2022 - The aliscore output appears to be empty. Check results for BUSCO: 262895at7742
## Thu Jul 14 02:22:52 CEST 2022 - The aliscore output file does not exist. Check results for BUSCO: 357356at7742
## Thu Jul 14 02:22:52 CEST 2022 - The aliscore output appears to be empty. Check results for BUSCO: 357356at7742
## Thu Jul 14 02:22:54 CEST 2022 - The aliscore output file does not exist. Check results for BUSCO: 217762at7742
## Thu Jul 14 02:22:54 CEST 2022 - The aliscore output appears to be empty. Check results for BUSCO: 217762at7742
## Thu Jul 14 02:22:55 CEST 2022 - The aliscore output file does not exist. Check results for BUSCO: 329385at7742
## Thu Jul 14 02:22:55 CEST 2022 - The aliscore output appears to be empty. Check results for BUSCO: 329385at7742
## Thu Jul 14 02:22:55 CEST 2022 - The aliscore output file does not exist. Check results for BUSCO: 271622at7742
## Thu Jul 14 02:22:55 CEST 2022 - The aliscore output file does not exist. Check results for BUSCO: 279092at7742
## Thu Jul 14 02:22:56 CEST 2022 - The aliscore output appears to be empty. Check results for BUSCO: 271622at7742
## Thu Jul 14 02:22:56 CEST 2022 - The aliscore output appears to be empty. Check results for BUSCO: 279092at7742
## Thu Jul 14 02:22:57 CEST 2022 - The aliscore output file does not exist. Check results for BUSCO: 277361at7742
## Thu Jul 14 02:22:57 CEST 2022 - The aliscore output appears to be empty. Check results for BUSCO: 277361at7742
## Thu Jul 14 02:22:58 CEST 2022 - The aliscore output file does not exist. Check results for BUSCO: 316190at7742
## Thu Jul 14 02:22:59 CEST 2022 - The aliscore output appears to be empty. Check results for BUSCO: 316190at7742
## Thu Jul 14 02:23:00 CEST 2022 - The aliscore output file does not exist. Check results for BUSCO: 338050at7742
## Thu Jul 14 02:23:00 CEST 2022 - The aliscore output appears to be empty. Check results for BUSCO: 338050at7742
## Thu Jul 14 02:23:00 CEST 2022 - The aliscore output file does not exist. Check results for BUSCO: 289878at7742
## Thu Jul 14 02:23:00 CEST 2022 - The aliscore output appears to be empty. Check results for BUSCO: 289878at7742
## Thu Jul 14 02:23:02 CEST 2022 - The aliscore output file does not exist. Check results for BUSCO: 225646at7742
## Thu Jul 14 02:23:02 CEST 2022 - The aliscore output appears to be empty. Check results for BUSCO: 225646at7742
## Thu Jul 14 02:23:03 CEST 2022 - The aliscore output file does not exist. Check results for BUSCO: 273993at7742
## Thu Jul 14 02:23:03 CEST 2022 - The aliscore output appears to be empty. Check results for BUSCO: 273993at7742
## Thu Jul 14 02:23:03 CEST 2022 - The aliscore output file does not exist. Check results for BUSCO: 349204at7742
## Thu Jul 14 02:23:03 CEST 2022 - The aliscore output appears to be empty. Check results for BUSCO: 349204at7742
## Thu Jul 14 02:23:04 CEST 2022 - The aliscore output file does not exist. Check results for BUSCO: 242369at7742
## Thu Jul 14 02:23:04 CEST 2022 - The aliscore output appears to be empty. Check results for BUSCO: 242369at7742
## Thu Jul 14 02:23:05 CEST 2022 - The aliscore output file does not exist. Check results for BUSCO: 369081at7742
## Thu Jul 14 02:23:05 CEST 2022 - The aliscore output appears to be empty. Check results for BUSCO: 369081at7742
## Thu Jul 14 02:23:05 CEST 2022 - The aliscore output file does not exist. Check results for BUSCO: 304827at7742
## Thu Jul 14 02:23:05 CEST 2022 - The aliscore output file does not exist. Check results for BUSCO: 370914at7742
## Thu Jul 14 02:23:05 CEST 2022 - The aliscore output appears to be empty. Check results for BUSCO: 304827at7742
## Thu Jul 14 02:23:05 CEST 2022 - The aliscore output appears to be empty. Check results for BUSCO: 370914at7742
## Thu Jul 14 02:23:06 CEST 2022 - The aliscore output file does not exist. Check results for BUSCO: 388719at7742
## Thu Jul 14 02:23:06 CEST 2022 - The aliscore output appears to be empty. Check results for BUSCO: 388719at7742
## Thu Jul 14 02:23:09 CEST 2022 - The aliscore output file does not exist. Check results for BUSCO: 271622at7742
## Thu Jul 14 02:23:09 CEST 2022 - The aliscore output appears to be empty. Check results for BUSCO: 271622at7742
## Thu Jul 14 02:23:10 CEST 2022 - The aliscore output file does not exist. Check results for BUSCO: 344151at7742
## Thu Jul 14 02:23:10 CEST 2022 - The aliscore output appears to be empty. Check results for BUSCO: 344151at7742
## Thu Jul 14 02:23:10 CEST 2022 - The aliscore output file does not exist. Check results for BUSCO: 384054at7742
## Thu Jul 14 02:23:10 CEST 2022 - The aliscore output appears to be empty. Check results for BUSCO: 384054at7742
## Thu Jul 14 02:23:12 CEST 2022 - The aliscore output file does not exist. Check results for BUSCO: 245611at7742
## Thu Jul 14 02:23:12 CEST 2022 - The aliscore output appears to be empty. Check results for BUSCO: 245611at7742
## Thu Jul 14 02:23:12 CEST 2022 - The aliscore output file does not exist. Check results for BUSCO: 289044at7742
## Thu Jul 14 02:23:12 CEST 2022 - The aliscore output appears to be empty. Check results for BUSCO: 289044at7742
## Thu Jul 14 02:23:13 CEST 2022 - The aliscore output file does not exist. Check results for BUSCO: 376038at7742
## Thu Jul 14 02:23:13 CEST 2022 - The aliscore output appears to be empty. Check results for BUSCO: 376038at7742
## Thu Jul 14 02:23:15 CEST 2022 - The aliscore output file does not exist. Check results for BUSCO: 357059at7742
## Thu Jul 14 02:23:15 CEST 2022 - The aliscore output appears to be empty. Check results for BUSCO: 357059at7742
## Thu Jul 14 02:23:16 CEST 2022 - The aliscore output file does not exist. Check results for BUSCO: 342301at7742
## Thu Jul 14 02:23:16 CEST 2022 - The aliscore output appears to be empty. Check results for BUSCO: 342301at7742
## Thu Jul 14 02:23:17 CEST 2022 - The aliscore output file does not exist. Check results for BUSCO: 277361at7742
## Thu Jul 14 02:23:17 CEST 2022 - The aliscore output appears to be empty. Check results for BUSCO: 277361at7742
## Thu Jul 14 02:23:17 CEST 2022 - The aliscore output file does not exist. Check results for BUSCO: 365186at7742
## Thu Jul 14 02:23:17 CEST 2022 - The aliscore output appears to be empty. Check results for BUSCO: 365186at7742
## Thu Jul 14 02:23:19 CEST 2022 - The aliscore output file does not exist. Check results for BUSCO: 264320at7742
## Thu Jul 14 02:23:19 CEST 2022 - The aliscore output appears to be empty. Check results for BUSCO: 264320at7742
## Thu Jul 14 02:23:21 CEST 2022 - The aliscore output file does not exist. Check results for BUSCO: 241207at7742
## Thu Jul 14 02:23:21 CEST 2022 - The aliscore output appears to be empty. Check results for BUSCO: 241207at7742
## Thu Jul 14 02:23:21 CEST 2022 - The aliscore output file does not exist. Check results for BUSCO: 320707at7742
## Thu Jul 14 02:23:21 CEST 2022 - The aliscore output appears to be empty. Check results for BUSCO: 320707at7742
## Thu Jul 14 02:23:23 CEST 2022 - The aliscore output file does not exist. Check results for BUSCO: 373808at7742
## Thu Jul 14 02:23:23 CEST 2022 - The aliscore output appears to be empty. Check results for BUSCO: 373808at7742
## Thu Jul 14 02:23:23 CEST 2022 - The aliscore output file does not exist. Check results for BUSCO: 298740at7742
## Thu Jul 14 02:23:23 CEST 2022 - The aliscore output appears to be empty. Check results for BUSCO: 298740at7742
## Thu Jul 14 02:23:23 CEST 2022 - The aliscore output file does not exist. Check results for BUSCO: 289878at7742
## Thu Jul 14 02:23:23 CEST 2022 - The aliscore output file does not exist. Check results for BUSCO: 326316at7742
## Thu Jul 14 02:23:23 CEST 2022 - The aliscore output appears to be empty. Check results for BUSCO: 289878at7742
## Thu Jul 14 02:23:23 CEST 2022 - The aliscore output appears to be empty. Check results for BUSCO: 326316at7742
## Thu Jul 14 02:23:25 CEST 2022 - The aliscore output file does not exist. Check results for BUSCO: 342301at7742
## Thu Jul 14 02:23:25 CEST 2022 - The aliscore output appears to be empty. Check results for BUSCO: 342301at7742
## Thu Jul 14 02:23:26 CEST 2022 - The aliscore output file does not exist. Check results for BUSCO: 234920at7742
## Thu Jul 14 02:23:26 CEST 2022 - The aliscore output appears to be empty. Check results for BUSCO: 234920at7742
## Thu Jul 14 02:23:27 CEST 2022 - The aliscore output file does not exist. Check results for BUSCO: 343085at7742
## Thu Jul 14 02:23:27 CEST 2022 - The aliscore output appears to be empty. Check results for BUSCO: 343085at7742
## Thu Jul 14 02:23:29 CEST 2022 - The aliscore output file does not exist. Check results for BUSCO: 357356at7742
## Thu Jul 14 02:23:29 CEST 2022 - The aliscore output appears to be empty. Check results for BUSCO: 357356at7742
## Thu Jul 14 02:23:30 CEST 2022 - The aliscore output file does not exist. Check results for BUSCO: 353474at7742
## Thu Jul 14 02:23:30 CEST 2022 - The aliscore output appears to be empty. Check results for BUSCO: 353474at7742
## Thu Jul 14 02:23:30 CEST 2022 - The aliscore output file does not exist. Check results for BUSCO: 342923at7742
## Thu Jul 14 02:23:30 CEST 2022 - The aliscore output appears to be empty. Check results for BUSCO: 342923at7742
## Thu Jul 14 02:23:30 CEST 2022 - The aliscore output file does not exist. Check results for BUSCO: 338050at7742
## Thu Jul 14 02:23:31 CEST 2022 - The aliscore output appears to be empty. Check results for BUSCO: 338050at7742
## Thu Jul 14 02:23:32 CEST 2022 - The aliscore output file does not exist. Check results for BUSCO: 384878at7742
## Thu Jul 14 02:23:32 CEST 2022 - The aliscore output appears to be empty. Check results for BUSCO: 384878at7742
## Thu Jul 14 02:23:32 CEST 2022 - The aliscore output file does not exist. Check results for BUSCO: 269168at7742
## Thu Jul 14 02:23:32 CEST 2022 - The aliscore output appears to be empty. Check results for BUSCO: 269168at7742
## Thu Jul 14 02:23:33 CEST 2022 - The aliscore output file does not exist. Check results for BUSCO: 320707at7742
## Thu Jul 14 02:23:33 CEST 2022 - The aliscore output appears to be empty. Check results for BUSCO: 320707at7742
## Thu Jul 14 02:23:35 CEST 2022 - The aliscore output file does not exist. Check results for BUSCO: 262895at7742
## Thu Jul 14 02:23:35 CEST 2022 - The aliscore output appears to be empty. Check results for BUSCO: 262895at7742
## Thu Jul 14 02:23:37 CEST 2022 - The aliscore output file does not exist. Check results for BUSCO: 349858at7742
## Thu Jul 14 02:23:37 CEST 2022 - The aliscore output appears to be empty. Check results for BUSCO: 349858at7742
## Thu Jul 14 02:23:37 CEST 2022 - The aliscore output file does not exist. Check results for BUSCO: 272582at7742
## Thu Jul 14 02:23:37 CEST 2022 - The aliscore output appears to be empty. Check results for BUSCO: 272582at7742
## Thu Jul 14 02:23:37 CEST 2022 - The aliscore output file does not exist. Check results for BUSCO: 217762at7742
## Thu Jul 14 02:23:37 CEST 2022 - The aliscore output appears to be empty. Check results for BUSCO: 217762at7742
## Thu Jul 14 02:23:40 CEST 2022 - The aliscore output file does not exist. Check results for BUSCO: 385022at7742
## Thu Jul 14 02:23:40 CEST 2022 - The aliscore output appears to be empty. Check results for BUSCO: 385022at7742
## Thu Jul 14 02:23:41 CEST 2022 - The aliscore output file does not exist. Check results for BUSCO: 294706at7742
## Thu Jul 14 02:23:41 CEST 2022 - The aliscore output appears to be empty. Check results for BUSCO: 294706at7742
## Thu Jul 14 02:23:41 CEST 2022 - The aliscore output file does not exist. Check results for BUSCO: 294706at7742
## Thu Jul 14 02:23:42 CEST 2022 - The aliscore output appears to be empty. Check results for BUSCO: 294706at7742
## Thu Jul 14 02:23:42 CEST 2022 - The aliscore output file does not exist. Check results for BUSCO: 377323at7742
## Thu Jul 14 02:23:42 CEST 2022 - The aliscore output appears to be empty. Check results for BUSCO: 377323at7742
## Thu Jul 14 02:23:43 CEST 2022 - The aliscore output file does not exist. Check results for BUSCO: 413952at7742
## Thu Jul 14 02:23:43 CEST 2022 - The aliscore output appears to be empty. Check results for BUSCO: 413952at7742
## Thu Jul 14 02:23:44 CEST 2022 - The aliscore output file does not exist. Check results for BUSCO: 323574at7742
## Thu Jul 14 02:23:44 CEST 2022 - The aliscore output appears to be empty. Check results for BUSCO: 323574at7742
## Thu Jul 14 02:23:44 CEST 2022 - The aliscore output file does not exist. Check results for BUSCO: 344151at7742
## Thu Jul 14 02:23:45 CEST 2022 - The aliscore output appears to be empty. Check results for BUSCO: 344151at7742
## Thu Jul 14 02:23:45 CEST 2022 - The aliscore output file does not exist. Check results for BUSCO: 322275at7742
## Thu Jul 14 02:23:45 CEST 2022 - The aliscore output appears to be empty. Check results for BUSCO: 322275at7742
## Thu Jul 14 02:23:47 CEST 2022 - The aliscore output file does not exist. Check results for BUSCO: 320550at7742
## Thu Jul 14 02:23:47 CEST 2022 - The aliscore output appears to be empty. Check results for BUSCO: 320550at7742
## Thu Jul 14 02:23:47 CEST 2022 - The aliscore output file does not exist. Check results for BUSCO: 392383at7742
## Thu Jul 14 02:23:47 CEST 2022 - The aliscore output appears to be empty. Check results for BUSCO: 392383at7742
## Thu Jul 14 02:23:49 CEST 2022 - The aliscore output file does not exist. Check results for BUSCO: 289878at7742
## Thu Jul 14 02:23:49 CEST 2022 - The aliscore output appears to be empty. Check results for BUSCO: 289878at7742
## Thu Jul 14 02:23:50 CEST 2022 - The aliscore output file does not exist. Check results for BUSCO: 211842at7742
## Thu Jul 14 02:23:50 CEST 2022 - The aliscore output appears to be empty. Check results for BUSCO: 211842at7742
## Thu Jul 14 02:23:52 CEST 2022 - The aliscore output file does not exist. Check results for BUSCO: 384878at7742
## Thu Jul 14 02:23:52 CEST 2022 - The aliscore output appears to be empty. Check results for BUSCO: 384878at7742
## Thu Jul 14 02:23:52 CEST 2022 - The aliscore output file does not exist. Check results for BUSCO: 397671at7742
## Thu Jul 14 02:23:52 CEST 2022 - The aliscore output appears to be empty. Check results for BUSCO: 397671at7742
## Thu Jul 14 02:23:53 CEST 2022 - The aliscore output file does not exist. Check results for BUSCO: 235335at7742
## Thu Jul 14 02:23:53 CEST 2022 - The aliscore output appears to be empty. Check results for BUSCO: 235335at7742
## Thu Jul 14 02:23:54 CEST 2022 - The aliscore output file does not exist. Check results for BUSCO: 304593at7742
## Thu Jul 14 02:23:54 CEST 2022 - The aliscore output appears to be empty. Check results for BUSCO: 304593at7742
## Thu Jul 14 02:23:54 CEST 2022 - The aliscore output file does not exist. Check results for BUSCO: 304593at7742
## Thu Jul 14 02:23:54 CEST 2022 - The aliscore output appears to be empty. Check results for BUSCO: 304593at7742
## Thu Jul 14 02:23:56 CEST 2022 - The aliscore output file does not exist. Check results for BUSCO: 395686at7742
## Thu Jul 14 02:23:56 CEST 2022 - The aliscore output appears to be empty. Check results for BUSCO: 395686at7742
## Thu Jul 14 02:23:59 CEST 2022 - The aliscore output file does not exist. Check results for BUSCO: 357059at7742
## Thu Jul 14 02:23:59 CEST 2022 - The aliscore output appears to be empty. Check results for BUSCO: 357059at7742
## Thu Jul 14 02:23:59 CEST 2022 - The aliscore output file does not exist. Check results for BUSCO: 329385at7742
## Thu Jul 14 02:23:59 CEST 2022 - The aliscore output appears to be empty. Check results for BUSCO: 329385at7742
## Thu Jul 14 02:23:59 CEST 2022 - The aliscore output file does not exist. Check results for BUSCO: 407384at7742
## Thu Jul 14 02:23:59 CEST 2022 - The aliscore output appears to be empty. Check results for BUSCO: 407384at7742
## Thu Jul 14 02:23:59 CEST 2022 - The aliscore output file does not exist. Check results for BUSCO: 327840at7742
## Thu Jul 14 02:23:59 CEST 2022 - The aliscore output appears to be empty. Check results for BUSCO: 327840at7742
## Thu Jul 14 02:24:01 CEST 2022 - The aliscore output file does not exist. Check results for BUSCO: 390186at7742
## Thu Jul 14 02:24:01 CEST 2022 - The aliscore output appears to be empty. Check results for BUSCO: 390186at7742
## Thu Jul 14 02:24:02 CEST 2022 - The aliscore output file does not exist. Check results for BUSCO: 370111at7742
## Thu Jul 14 02:24:02 CEST 2022 - The aliscore output appears to be empty. Check results for BUSCO: 370111at7742
## Thu Jul 14 02:24:03 CEST 2022 - The aliscore output file does not exist. Check results for BUSCO: 246301at7742
## Thu Jul 14 02:24:03 CEST 2022 - The aliscore output appears to be empty. Check results for BUSCO: 246301at7742
## Thu Jul 14 02:24:04 CEST 2022 - The aliscore output file does not exist. Check results for BUSCO: 303740at7742
## Thu Jul 14 02:24:04 CEST 2022 - The aliscore output appears to be empty. Check results for BUSCO: 303740at7742
## Thu Jul 14 02:24:05 CEST 2022 - The aliscore output file does not exist. Check results for BUSCO: 235735at7742
## Thu Jul 14 02:24:05 CEST 2022 - The aliscore output appears to be empty. Check results for BUSCO: 235735at7742
## Thu Jul 14 02:24:06 CEST 2022 - The aliscore output file does not exist. Check results for BUSCO: 242979at7742
## Thu Jul 14 02:24:06 CEST 2022 - The aliscore output appears to be empty. Check results for BUSCO: 242979at7742
## Thu Jul 14 02:24:07 CEST 2022 - The aliscore output file does not exist. Check results for BUSCO: 344436at7742
## Thu Jul 14 02:24:07 CEST 2022 - The aliscore output appears to be empty. Check results for BUSCO: 344436at7742
## Thu Jul 14 02:24:07 CEST 2022 - The aliscore output file does not exist. Check results for BUSCO: 349858at7742
## Thu Jul 14 02:24:07 CEST 2022 - The aliscore output appears to be empty. Check results for BUSCO: 349858at7742
## Thu Jul 14 02:24:09 CEST 2022 - The aliscore output file does not exist. Check results for BUSCO: 316190at7742
## Thu Jul 14 02:24:09 CEST 2022 - The aliscore output appears to be empty. Check results for BUSCO: 316190at7742
## Thu Jul 14 02:24:10 CEST 2022 - The aliscore output file does not exist. Check results for BUSCO: 316190at7742
## Thu Jul 14 02:24:10 CEST 2022 - The aliscore output appears to be empty. Check results for BUSCO: 316190at7742
## Thu Jul 14 02:24:10 CEST 2022 - The aliscore output file does not exist. Check results for BUSCO: 361842at7742
## Thu Jul 14 02:24:10 CEST 2022 - The aliscore output appears to be empty. Check results for BUSCO: 361842at7742
## Thu Jul 14 02:24:12 CEST 2022 - The aliscore output file does not exist. Check results for BUSCO: 278787at7742
## Thu Jul 14 02:24:12 CEST 2022 - The aliscore output appears to be empty. Check results for BUSCO: 278787at7742
## Thu Jul 14 02:24:13 CEST 2022 - The aliscore output file does not exist. Check results for BUSCO: 373808at7742
## Thu Jul 14 02:24:13 CEST 2022 - The aliscore output appears to be empty. Check results for BUSCO: 373808at7742
## Thu Jul 14 02:24:13 CEST 2022 - The aliscore output file does not exist. Check results for BUSCO: 294646at7742
## Thu Jul 14 02:24:13 CEST 2022 - The aliscore output appears to be empty. Check results for BUSCO: 294646at7742
## Thu Jul 14 02:24:14 CEST 2022 - The aliscore output file does not exist. Check results for BUSCO: 388719at7742
## Thu Jul 14 02:24:14 CEST 2022 - The aliscore output appears to be empty. Check results for BUSCO: 388719at7742
## Thu Jul 14 02:24:16 CEST 2022 - The aliscore output file does not exist. Check results for BUSCO: 349204at7742
## Thu Jul 14 02:24:16 CEST 2022 - The aliscore output appears to be empty. Check results for BUSCO: 349204at7742
## Thu Jul 14 02:24:16 CEST 2022 - The aliscore output file does not exist. Check results for BUSCO: 289044at7742
## Thu Jul 14 02:24:17 CEST 2022 - The aliscore output appears to be empty. Check results for BUSCO: 289044at7742
## Thu Jul 14 02:24:18 CEST 2022 - The aliscore output file does not exist. Check results for BUSCO: 354226at7742
## Thu Jul 14 02:24:18 CEST 2022 - The aliscore output appears to be empty. Check results for BUSCO: 354226at7742
## Thu Jul 14 02:24:19 CEST 2022 - The aliscore output file does not exist. Check results for BUSCO: 359541at7742
## Thu Jul 14 02:24:19 CEST 2022 - The aliscore output appears to be empty. Check results for BUSCO: 359541at7742
## Thu Jul 14 02:24:19 CEST 2022 - The aliscore output file does not exist. Check results for BUSCO: 373808at7742
## Thu Jul 14 02:24:19 CEST 2022 - The aliscore output appears to be empty. Check results for BUSCO: 373808at7742
## Thu Jul 14 02:24:21 CEST 2022 - The aliscore output file does not exist. Check results for BUSCO: 308357at7742
## Thu Jul 14 02:24:21 CEST 2022 - The aliscore output appears to be empty. Check results for BUSCO: 308357at7742
## Thu Jul 14 02:24:21 CEST 2022 - The aliscore output file does not exist. Check results for BUSCO: 326788at7742
## Thu Jul 14 02:24:21 CEST 2022 - The aliscore output appears to be empty. Check results for BUSCO: 326788at7742
## Thu Jul 14 02:24:22 CEST 2022 - The aliscore output file does not exist. Check results for BUSCO: 329385at7742
## Thu Jul 14 02:24:22 CEST 2022 - The aliscore output appears to be empty. Check results for BUSCO: 329385at7742
## Thu Jul 14 02:24:23 CEST 2022 - The aliscore output file does not exist. Check results for BUSCO: 374832at7742
## Thu Jul 14 02:24:23 CEST 2022 - The aliscore output appears to be empty. Check results for BUSCO: 374832at7742
## Thu Jul 14 02:24:25 CEST 2022 - The aliscore output file does not exist. Check results for BUSCO: 381867at7742
## Thu Jul 14 02:24:25 CEST 2022 - The aliscore output appears to be empty. Check results for BUSCO: 381867at7742
## Thu Jul 14 02:24:25 CEST 2022 - The aliscore output file does not exist. Check results for BUSCO: 342620at7742
## Thu Jul 14 02:24:25 CEST 2022 - The aliscore output appears to be empty. Check results for BUSCO: 342620at7742
## Thu Jul 14 02:24:25 CEST 2022 - The aliscore output file does not exist. Check results for BUSCO: 284145at7742
## Thu Jul 14 02:24:26 CEST 2022 - The aliscore output appears to be empty. Check results for BUSCO: 284145at7742
## Thu Jul 14 02:24:26 CEST 2022 - The aliscore output file does not exist. Check results for BUSCO: 359032at7742
## Thu Jul 14 02:24:26 CEST 2022 - The aliscore output appears to be empty. Check results for BUSCO: 359032at7742
## Thu Jul 14 02:24:28 CEST 2022 - The aliscore output file does not exist. Check results for BUSCO: 311206at7742
## Thu Jul 14 02:24:28 CEST 2022 - The aliscore output appears to be empty. Check results for BUSCO: 311206at7742
## Thu Jul 14 02:24:30 CEST 2022 - The aliscore output file does not exist. Check results for BUSCO: 373482at7742
## Thu Jul 14 02:24:30 CEST 2022 - The aliscore output appears to be empty. Check results for BUSCO: 373482at7742
## Thu Jul 14 02:24:32 CEST 2022 - The aliscore output file does not exist. Check results for BUSCO: 339398at7742
## Thu Jul 14 02:24:32 CEST 2022 - The aliscore output appears to be empty. Check results for BUSCO: 339398at7742
## Thu Jul 14 02:24:32 CEST 2022 - The aliscore output file does not exist. Check results for BUSCO: 388719at7742
## Thu Jul 14 02:24:32 CEST 2022 - The aliscore output appears to be empty. Check results for BUSCO: 388719at7742
## Thu Jul 14 02:24:32 CEST 2022 - The aliscore output file does not exist. Check results for BUSCO: 378519at7742
## Thu Jul 14 02:24:32 CEST 2022 - The aliscore output appears to be empty. Check results for BUSCO: 378519at7742
## Thu Jul 14 02:24:33 CEST 2022 - The aliscore output file does not exist. Check results for BUSCO: 305378at7742
## Thu Jul 14 02:24:33 CEST 2022 - The aliscore output appears to be empty. Check results for BUSCO: 305378at7742
## Thu Jul 14 02:24:34 CEST 2022 - The aliscore output file does not exist. Check results for BUSCO: 380542at7742
## Thu Jul 14 02:24:34 CEST 2022 - The aliscore output appears to be empty. Check results for BUSCO: 380542at7742
## Thu Jul 14 02:24:36 CEST 2022 - The aliscore output file does not exist. Check results for BUSCO: 320550at7742
## Thu Jul 14 02:24:36 CEST 2022 - The aliscore output appears to be empty. Check results for BUSCO: 320550at7742
## Thu Jul 14 02:24:36 CEST 2022 - The aliscore output file does not exist. Check results for BUSCO: 317054at7742
## Thu Jul 14 02:24:36 CEST 2022 - The aliscore output appears to be empty. Check results for BUSCO: 317054at7742
## Thu Jul 14 02:24:37 CEST 2022 - The aliscore output file does not exist. Check results for BUSCO: 415822at7742
## Thu Jul 14 02:24:37 CEST 2022 - The aliscore output appears to be empty. Check results for BUSCO: 415822at7742
## Thu Jul 14 02:24:38 CEST 2022 - The aliscore output file does not exist. Check results for BUSCO: 384558at7742
## Thu Jul 14 02:24:38 CEST 2022 - The aliscore output appears to be empty. Check results for BUSCO: 384558at7742
## Thu Jul 14 02:24:38 CEST 2022 - The aliscore output file does not exist. Check results for BUSCO: 346443at7742
## Thu Jul 14 02:24:38 CEST 2022 - The aliscore output appears to be empty. Check results for BUSCO: 346443at7742
## Thu Jul 14 02:24:38 CEST 2022 - The aliscore output file does not exist. Check results for BUSCO: 320550at7742
## Thu Jul 14 02:24:38 CEST 2022 - The aliscore output appears to be empty. Check results for BUSCO: 320550at7742
## Thu Jul 14 02:24:39 CEST 2022 - The aliscore output file does not exist. Check results for BUSCO: 376736at7742
## Thu Jul 14 02:24:39 CEST 2022 - The aliscore output appears to be empty. Check results for BUSCO: 376736at7742
## Thu Jul 14 02:24:40 CEST 2022 - The aliscore output file does not exist. Check results for BUSCO: 354226at7742
## Thu Jul 14 02:24:40 CEST 2022 - The aliscore output appears to be empty. Check results for BUSCO: 354226at7742
## Thu Jul 14 02:24:42 CEST 2022 - The aliscore output file does not exist. Check results for BUSCO: 249652at7742
## Thu Jul 14 02:24:42 CEST 2022 - The aliscore output appears to be empty. Check results for BUSCO: 249652at7742
## Thu Jul 14 02:24:43 CEST 2022 - The aliscore output file does not exist. Check results for BUSCO: 395686at7742
## Thu Jul 14 02:24:43 CEST 2022 - The aliscore output appears to be empty. Check results for BUSCO: 395686at7742
## Thu Jul 14 02:24:43 CEST 2022 - The aliscore output file does not exist. Check results for BUSCO: 301464at7742
## Thu Jul 14 02:24:43 CEST 2022 - The aliscore output appears to be empty. Check results for BUSCO: 301464at7742
## Thu Jul 14 02:24:45 CEST 2022 - The aliscore output file does not exist. Check results for BUSCO: 361842at7742
## Thu Jul 14 02:24:45 CEST 2022 - The aliscore output appears to be empty. Check results for BUSCO: 361842at7742
## Thu Jul 14 02:24:46 CEST 2022 - The aliscore output file does not exist. Check results for BUSCO: 272130at7742
## Thu Jul 14 02:24:46 CEST 2022 - The aliscore output appears to be empty. Check results for BUSCO: 272130at7742
## Thu Jul 14 02:24:46 CEST 2022 - The aliscore output file does not exist. Check results for BUSCO: 336619at7742
## Thu Jul 14 02:24:46 CEST 2022 - The aliscore output appears to be empty. Check results for BUSCO: 336619at7742
## Thu Jul 14 02:24:47 CEST 2022 - The aliscore output file does not exist. Check results for BUSCO: 334363at7742
## Thu Jul 14 02:24:47 CEST 2022 - The aliscore output appears to be empty. Check results for BUSCO: 334363at7742
## Thu Jul 14 02:24:48 CEST 2022 - The aliscore output file does not exist. Check results for BUSCO: 295973at7742
## Thu Jul 14 02:24:48 CEST 2022 - The aliscore output appears to be empty. Check results for BUSCO: 295973at7742
## Thu Jul 14 02:24:50 CEST 2022 - The aliscore output file does not exist. Check results for BUSCO: 420085at7742
## Thu Jul 14 02:24:50 CEST 2022 - The aliscore output appears to be empty. Check results for BUSCO: 420085at7742
## Thu Jul 14 02:24:52 CEST 2022 - The aliscore output file does not exist. Check results for BUSCO: 343489at7742
## Thu Jul 14 02:24:52 CEST 2022 - The aliscore output appears to be empty. Check results for BUSCO: 343489at7742
## Thu Jul 14 02:24:52 CEST 2022 - The aliscore output file does not exist. Check results for BUSCO: 296937at7742
## Thu Jul 14 02:24:52 CEST 2022 - The aliscore output appears to be empty. Check results for BUSCO: 296937at7742
## Thu Jul 14 02:24:52 CEST 2022 - The aliscore output file does not exist. Check results for BUSCO: 342923at7742
## Thu Jul 14 02:24:52 CEST 2022 - The aliscore output appears to be empty. Check results for BUSCO: 342923at7742
## Thu Jul 14 02:24:54 CEST 2022 - The aliscore output file does not exist. Check results for BUSCO: 335473at7742
## Thu Jul 14 02:24:54 CEST 2022 - The aliscore output appears to be empty. Check results for BUSCO: 335473at7742
## Thu Jul 14 02:24:55 CEST 2022 - The aliscore output file does not exist. Check results for BUSCO: 333698at7742
## Thu Jul 14 02:24:55 CEST 2022 - The aliscore output appears to be empty. Check results for BUSCO: 333698at7742
## Thu Jul 14 02:24:56 CEST 2022 - The aliscore output file does not exist. Check results for BUSCO: 249652at7742
## Thu Jul 14 02:24:56 CEST 2022 - The aliscore output appears to be empty. Check results for BUSCO: 249652at7742
## Thu Jul 14 02:24:57 CEST 2022 - The aliscore output file does not exist. Check results for BUSCO: 414201at7742
## Thu Jul 14 02:24:57 CEST 2022 - The aliscore output file does not exist. Check results for BUSCO: 376479at7742
## Thu Jul 14 02:24:57 CEST 2022 - The aliscore output appears to be empty. Check results for BUSCO: 414201at7742
## Thu Jul 14 02:24:57 CEST 2022 - The aliscore output appears to be empty. Check results for BUSCO: 376479at7742
## Thu Jul 14 02:24:59 CEST 2022 - The aliscore output file does not exist. Check results for BUSCO: 342641at7742
## Thu Jul 14 02:24:59 CEST 2022 - The aliscore output appears to be empty. Check results for BUSCO: 342641at7742
## Thu Jul 14 02:24:59 CEST 2022 - The aliscore output file does not exist. Check results for BUSCO: 391948at7742
## Thu Jul 14 02:25:00 CEST 2022 - The aliscore output appears to be empty. Check results for BUSCO: 391948at7742
## Thu Jul 14 02:25:00 CEST 2022 - The aliscore output file does not exist. Check results for BUSCO: 375593at7742
## Thu Jul 14 02:25:00 CEST 2022 - The aliscore output appears to be empty. Check results for BUSCO: 375593at7742
## Thu Jul 14 02:25:01 CEST 2022 - The aliscore output file does not exist. Check results for BUSCO: 391055at7742
## Thu Jul 14 02:25:01 CEST 2022 - The aliscore output appears to be empty. Check results for BUSCO: 391055at7742
## Thu Jul 14 02:25:02 CEST 2022 - The aliscore output file does not exist. Check results for BUSCO: 308357at7742
## Thu Jul 14 02:25:02 CEST 2022 - The aliscore output appears to be empty. Check results for BUSCO: 308357at7742
## Thu Jul 14 02:25:06 CEST 2022 - The aliscore output file does not exist. Check results for BUSCO: 344151at7742
## Thu Jul 14 02:25:06 CEST 2022 - The aliscore output appears to be empty. Check results for BUSCO: 344151at7742
## Thu Jul 14 02:25:06 CEST 2022 - The aliscore output file does not exist. Check results for BUSCO: 284145at7742
## Thu Jul 14 02:25:06 CEST 2022 - The aliscore output appears to be empty. Check results for BUSCO: 284145at7742
## Thu Jul 14 02:25:06 CEST 2022 - The aliscore output file does not exist. Check results for BUSCO: 273271at7742
## Thu Jul 14 02:25:06 CEST 2022 - The aliscore output appears to be empty. Check results for BUSCO: 273271at7742
## Thu Jul 14 02:25:06 CEST 2022 - The aliscore output file does not exist. Check results for BUSCO: 242369at7742
## Thu Jul 14 02:25:06 CEST 2022 - The aliscore output appears to be empty. Check results for BUSCO: 242369at7742
## Thu Jul 14 02:25:08 CEST 2022 - The aliscore output file does not exist. Check results for BUSCO: 342923at7742
## Thu Jul 14 02:25:08 CEST 2022 - The aliscore output appears to be empty. Check results for BUSCO: 342923at7742
## Thu Jul 14 02:25:08 CEST 2022 - The aliscore output file does not exist. Check results for BUSCO: 343725at7742
## Thu Jul 14 02:25:08 CEST 2022 - The aliscore output appears to be empty. Check results for BUSCO: 343725at7742
## Thu Jul 14 02:25:08 CEST 2022 - The aliscore output file does not exist. Check results for BUSCO: 350839at7742
## Thu Jul 14 02:25:08 CEST 2022 - The aliscore output appears to be empty. Check results for BUSCO: 350839at7742
## Thu Jul 14 02:25:10 CEST 2022 - The aliscore output file does not exist. Check results for BUSCO: 320707at7742
## Thu Jul 14 02:25:10 CEST 2022 - The aliscore output appears to be empty. Check results for BUSCO: 320707at7742
## Thu Jul 14 02:25:11 CEST 2022 - The aliscore output file does not exist. Check results for BUSCO: 375593at7742
## Thu Jul 14 02:25:11 CEST 2022 - The aliscore output appears to be empty. Check results for BUSCO: 375593at7742
## Thu Jul 14 02:25:12 CEST 2022 - The aliscore output file does not exist. Check results for BUSCO: 388102at7742
## Thu Jul 14 02:25:12 CEST 2022 - The aliscore output appears to be empty. Check results for BUSCO: 388102at7742
## Thu Jul 14 02:25:12 CEST 2022 - The aliscore output file does not exist. Check results for BUSCO: 372431at7742
## Thu Jul 14 02:25:12 CEST 2022 - The aliscore output appears to be empty. Check results for BUSCO: 372431at7742
## Thu Jul 14 02:25:14 CEST 2022 - The aliscore output file does not exist. Check results for BUSCO: 300333at7742
## Thu Jul 14 02:25:14 CEST 2022 - The aliscore output appears to be empty. Check results for BUSCO: 300333at7742
## Thu Jul 14 02:25:14 CEST 2022 - The aliscore output file does not exist. Check results for BUSCO: 273941at7742
## Thu Jul 14 02:25:14 CEST 2022 - The aliscore output appears to be empty. Check results for BUSCO: 273941at7742
## Thu Jul 14 02:25:16 CEST 2022 - The aliscore output file does not exist. Check results for BUSCO: 343489at7742
## Thu Jul 14 02:25:16 CEST 2022 - The aliscore output appears to be empty. Check results for BUSCO: 343489at7742
## Thu Jul 14 02:25:16 CEST 2022 - The aliscore output file does not exist. Check results for BUSCO: 331452at7742
## Thu Jul 14 02:25:16 CEST 2022 - The aliscore output appears to be empty. Check results for BUSCO: 331452at7742
## Thu Jul 14 02:25:19 CEST 2022 - The aliscore output file does not exist. Check results for BUSCO: 264124at7742
## Thu Jul 14 02:25:19 CEST 2022 - The aliscore output appears to be empty. Check results for BUSCO: 264124at7742
## Thu Jul 14 02:25:20 CEST 2022 - The aliscore output file does not exist. Check results for BUSCO: 380542at7742
## Thu Jul 14 02:25:20 CEST 2022 - The aliscore output appears to be empty. Check results for BUSCO: 380542at7742
## Thu Jul 14 02:25:21 CEST 2022 - The aliscore output file does not exist. Check results for BUSCO: 304593at7742
## Thu Jul 14 02:25:21 CEST 2022 - The aliscore output appears to be empty. Check results for BUSCO: 304593at7742
## Thu Jul 14 02:25:21 CEST 2022 - The aliscore output file does not exist. Check results for BUSCO: 290567at7742
## Thu Jul 14 02:25:21 CEST 2022 - The aliscore output appears to be empty. Check results for BUSCO: 290567at7742
## Thu Jul 14 02:25:21 CEST 2022 - The aliscore output file does not exist. Check results for BUSCO: 269102at7742
## Thu Jul 14 02:25:22 CEST 2022 - The aliscore output appears to be empty. Check results for BUSCO: 269102at7742
## Thu Jul 14 02:25:23 CEST 2022 - The aliscore output file does not exist. Check results for BUSCO: 415822at7742
## Thu Jul 14 02:25:23 CEST 2022 - The aliscore output appears to be empty. Check results for BUSCO: 415822at7742
## Thu Jul 14 02:25:24 CEST 2022 - The aliscore output file does not exist. Check results for BUSCO: 385398at7742
## Thu Jul 14 02:25:24 CEST 2022 - The aliscore output appears to be empty. Check results for BUSCO: 385398at7742
## Thu Jul 14 02:25:25 CEST 2022 - The aliscore output file does not exist. Check results for BUSCO: 345264at7742
## Thu Jul 14 02:25:25 CEST 2022 - The aliscore output appears to be empty. Check results for BUSCO: 345264at7742
## Thu Jul 14 02:25:26 CEST 2022 - The aliscore output file does not exist. Check results for BUSCO: 345556at7742
## Thu Jul 14 02:25:26 CEST 2022 - The aliscore output appears to be empty. Check results for BUSCO: 345556at7742
## Thu Jul 14 02:25:26 CEST 2022 - The aliscore output file does not exist. Check results for BUSCO: 389786at7742
## Thu Jul 14 02:25:26 CEST 2022 - The aliscore output appears to be empty. Check results for BUSCO: 389786at7742
## Thu Jul 14 02:25:26 CEST 2022 - The aliscore output file does not exist. Check results for BUSCO: 349595at7742
## Thu Jul 14 02:25:26 CEST 2022 - The aliscore output appears to be empty. Check results for BUSCO: 349595at7742
## Thu Jul 14 02:25:28 CEST 2022 - The aliscore output file does not exist. Check results for BUSCO: 315366at7742
## Thu Jul 14 02:25:28 CEST 2022 - The aliscore output appears to be empty. Check results for BUSCO: 315366at7742
## Thu Jul 14 02:25:30 CEST 2022 - The aliscore output file does not exist. Check results for BUSCO: 339398at7742
## Thu Jul 14 02:25:30 CEST 2022 - The aliscore output file does not exist. Check results for BUSCO: 335517at7742
## Thu Jul 14 02:25:30 CEST 2022 - The aliscore output appears to be empty. Check results for BUSCO: 339398at7742
## Thu Jul 14 02:25:30 CEST 2022 - The aliscore output appears to be empty. Check results for BUSCO: 335517at7742
## Thu Jul 14 02:25:31 CEST 2022 - The aliscore output file does not exist. Check results for BUSCO: 381528at7742
## Thu Jul 14 02:25:31 CEST 2022 - The aliscore output appears to be empty. Check results for BUSCO: 381528at7742
## Thu Jul 14 02:25:32 CEST 2022 - The aliscore output file does not exist. Check results for BUSCO: 300333at7742
## Thu Jul 14 02:25:32 CEST 2022 - The aliscore output appears to be empty. Check results for BUSCO: 300333at7742
## Thu Jul 14 02:25:33 CEST 2022 - The aliscore output file does not exist. Check results for BUSCO: 326788at7742
## Thu Jul 14 02:25:33 CEST 2022 - The aliscore output appears to be empty. Check results for BUSCO: 326788at7742
## Thu Jul 14 02:25:33 CEST 2022 - The aliscore output file does not exist. Check results for BUSCO: 308357at7742
## Thu Jul 14 02:25:33 CEST 2022 - The aliscore output appears to be empty. Check results for BUSCO: 308357at7742
## Thu Jul 14 02:25:36 CEST 2022 - The aliscore output file does not exist. Check results for BUSCO: 372062at7742
## Thu Jul 14 02:25:36 CEST 2022 - The aliscore output appears to be empty. Check results for BUSCO: 372062at7742
## Thu Jul 14 02:25:36 CEST 2022 - The aliscore output file does not exist. Check results for BUSCO: 299314at7742
## Thu Jul 14 02:25:36 CEST 2022 - The aliscore output appears to be empty. Check results for BUSCO: 299314at7742
## Thu Jul 14 02:25:37 CEST 2022 - The aliscore output file does not exist. Check results for BUSCO: 378519at7742
## Thu Jul 14 02:25:37 CEST 2022 - The aliscore output appears to be empty. Check results for BUSCO: 378519at7742
## Thu Jul 14 02:25:39 CEST 2022 - The aliscore output file does not exist. Check results for BUSCO: 342641at7742
## Thu Jul 14 02:25:39 CEST 2022 - The aliscore output appears to be empty. Check results for BUSCO: 342641at7742
## Thu Jul 14 02:25:40 CEST 2022 - The aliscore output file does not exist. Check results for BUSCO: 376607at7742
## Thu Jul 14 02:25:40 CEST 2022 - The aliscore output appears to be empty. Check results for BUSCO: 376607at7742
## Thu Jul 14 02:25:41 CEST 2022 - The aliscore output file does not exist. Check results for BUSCO: 295101at7742
## Thu Jul 14 02:25:41 CEST 2022 - The aliscore output appears to be empty. Check results for BUSCO: 295101at7742
## Thu Jul 14 02:25:41 CEST 2022 - The aliscore output file does not exist. Check results for BUSCO: 338050at7742
## Thu Jul 14 02:25:41 CEST 2022 - The aliscore output appears to be empty. Check results for BUSCO: 338050at7742
## Thu Jul 14 02:25:41 CEST 2022 - The aliscore output file does not exist. Check results for BUSCO: 376736at7742
## Thu Jul 14 02:25:41 CEST 2022 - The aliscore output appears to be empty. Check results for BUSCO: 376736at7742
## Thu Jul 14 02:25:43 CEST 2022 - The aliscore output file does not exist. Check results for BUSCO: 349595at7742
## Thu Jul 14 02:25:43 CEST 2022 - The aliscore output appears to be empty. Check results for BUSCO: 349595at7742
## Thu Jul 14 02:25:45 CEST 2022 - The aliscore output file does not exist. Check results for BUSCO: 331452at7742
## Thu Jul 14 02:25:45 CEST 2022 - The aliscore output appears to be empty. Check results for BUSCO: 331452at7742
## Thu Jul 14 02:25:47 CEST 2022 - The aliscore output file does not exist. Check results for BUSCO: 315366at7742
## Thu Jul 14 02:25:47 CEST 2022 - The aliscore output appears to be empty. Check results for BUSCO: 315366at7742
## Thu Jul 14 02:25:47 CEST 2022 - The aliscore output file does not exist. Check results for BUSCO: 373482at7742
## Thu Jul 14 02:25:47 CEST 2022 - The aliscore output file does not exist. Check results for BUSCO: 339398at7742
## Thu Jul 14 02:25:47 CEST 2022 - The aliscore output appears to be empty. Check results for BUSCO: 373482at7742
## Thu Jul 14 02:25:47 CEST 2022 - The aliscore output appears to be empty. Check results for BUSCO: 339398at7742
## Thu Jul 14 02:25:48 CEST 2022 - The aliscore output file does not exist. Check results for BUSCO: 290567at7742
## Thu Jul 14 02:25:48 CEST 2022 - The aliscore output appears to be empty. Check results for BUSCO: 290567at7742
## Thu Jul 14 02:25:49 CEST 2022 - The aliscore output file does not exist. Check results for BUSCO: 322470at7742
## Thu Jul 14 02:25:49 CEST 2022 - The aliscore output appears to be empty. Check results for BUSCO: 322470at7742
## Thu Jul 14 02:25:50 CEST 2022 - The aliscore output file does not exist. Check results for BUSCO: 362453at7742
## Thu Jul 14 02:25:50 CEST 2022 - The aliscore output appears to be empty. Check results for BUSCO: 362453at7742
## Thu Jul 14 02:25:51 CEST 2022 - The aliscore output file does not exist. Check results for BUSCO: 346443at7742
## Thu Jul 14 02:25:51 CEST 2022 - The aliscore output appears to be empty. Check results for BUSCO: 346443at7742
## Thu Jul 14 02:25:52 CEST 2022 - The aliscore output file does not exist. Check results for BUSCO: 385022at7742
## Thu Jul 14 02:25:52 CEST 2022 - The aliscore output appears to be empty. Check results for BUSCO: 385022at7742
## Thu Jul 14 02:25:53 CEST 2022 - The aliscore output file does not exist. Check results for BUSCO: 398565at7742
## Thu Jul 14 02:25:53 CEST 2022 - The aliscore output appears to be empty. Check results for BUSCO: 398565at7742
## Thu Jul 14 02:25:54 CEST 2022 - The aliscore output file does not exist. Check results for BUSCO: 277748at7742
## Thu Jul 14 02:25:54 CEST 2022 - The aliscore output appears to be empty. Check results for BUSCO: 277748at7742
## Thu Jul 14 02:25:54 CEST 2022 - The aliscore output file does not exist. Check results for BUSCO: 421415at7742
## Thu Jul 14 02:25:54 CEST 2022 - The aliscore output appears to be empty. Check results for BUSCO: 421415at7742
## Thu Jul 14 02:25:55 CEST 2022 - The aliscore output file does not exist. Check results for BUSCO: 346443at7742
## Thu Jul 14 02:25:55 CEST 2022 - The aliscore output file does not exist. Check results for BUSCO: 361842at7742
## Thu Jul 14 02:25:55 CEST 2022 - The aliscore output appears to be empty. Check results for BUSCO: 346443at7742
## Thu Jul 14 02:25:55 CEST 2022 - The aliscore output appears to be empty. Check results for BUSCO: 361842at7742
## Thu Jul 14 02:25:59 CEST 2022 - The aliscore output file does not exist. Check results for BUSCO: 257768at7742
## Thu Jul 14 02:25:59 CEST 2022 - The aliscore output appears to be empty. Check results for BUSCO: 257768at7742
## Thu Jul 14 02:25:59 CEST 2022 - The aliscore output file does not exist. Check results for BUSCO: 396521at7742
## Thu Jul 14 02:25:59 CEST 2022 - The aliscore output appears to be empty. Check results for BUSCO: 396521at7742
## Thu Jul 14 02:26:00 CEST 2022 - The aliscore output file does not exist. Check results for BUSCO: 379564at7742
## Thu Jul 14 02:26:00 CEST 2022 - The aliscore output file does not exist. Check results for BUSCO: 255423at7742
## Thu Jul 14 02:26:00 CEST 2022 - The aliscore output appears to be empty. Check results for BUSCO: 379564at7742
## Thu Jul 14 02:26:00 CEST 2022 - The aliscore output appears to be empty. Check results for BUSCO: 255423at7742
## Thu Jul 14 02:26:01 CEST 2022 - The aliscore output file does not exist. Check results for BUSCO: 373954at7742
## Thu Jul 14 02:26:01 CEST 2022 - The aliscore output appears to be empty. Check results for BUSCO: 373954at7742
## Thu Jul 14 02:26:01 CEST 2022 - The aliscore output file does not exist. Check results for BUSCO: 242979at7742
## Thu Jul 14 02:26:01 CEST 2022 - The aliscore output appears to be empty. Check results for BUSCO: 242979at7742
## Thu Jul 14 02:26:04 CEST 2022 - The aliscore output file does not exist. Check results for BUSCO: 344436at7742
## Thu Jul 14 02:26:04 CEST 2022 - The aliscore output appears to be empty. Check results for BUSCO: 344436at7742
## Thu Jul 14 02:26:04 CEST 2022 - The aliscore output file does not exist. Check results for BUSCO: 384054at7742
## Thu Jul 14 02:26:04 CEST 2022 - The aliscore output appears to be empty. Check results for BUSCO: 384054at7742
## Thu Jul 14 02:26:05 CEST 2022 - The aliscore output file does not exist. Check results for BUSCO: 298740at7742
## Thu Jul 14 02:26:05 CEST 2022 - The aliscore output appears to be empty. Check results for BUSCO: 298740at7742
## Thu Jul 14 02:26:07 CEST 2022 - The aliscore output file does not exist. Check results for BUSCO: 279816at7742
## Thu Jul 14 02:26:07 CEST 2022 - The aliscore output appears to be empty. Check results for BUSCO: 279816at7742
## Thu Jul 14 02:26:08 CEST 2022 - The aliscore output file does not exist. Check results for BUSCO: 337736at7742
## Thu Jul 14 02:26:08 CEST 2022 - The aliscore output appears to be empty. Check results for BUSCO: 337736at7742
## Thu Jul 14 02:26:08 CEST 2022 - The aliscore output file does not exist. Check results for BUSCO: 343013at7742
## Thu Jul 14 02:26:08 CEST 2022 - The aliscore output appears to be empty. Check results for BUSCO: 343013at7742
## Thu Jul 14 02:26:09 CEST 2022 - The aliscore output file does not exist. Check results for BUSCO: 273271at7742
## Thu Jul 14 02:26:09 CEST 2022 - The aliscore output appears to be empty. Check results for BUSCO: 273271at7742
## Thu Jul 14 02:26:10 CEST 2022 - The aliscore output file does not exist. Check results for BUSCO: 313967at7742
## Thu Jul 14 02:26:10 CEST 2022 - The aliscore output appears to be empty. Check results for BUSCO: 313967at7742
## Thu Jul 14 02:26:10 CEST 2022 - The aliscore output file does not exist. Check results for BUSCO: 261072at7742
## Thu Jul 14 02:26:10 CEST 2022 - The aliscore output appears to be empty. Check results for BUSCO: 261072at7742
## Thu Jul 14 02:26:12 CEST 2022 - The aliscore output file does not exist. Check results for BUSCO: 303740at7742
## Thu Jul 14 02:26:12 CEST 2022 - The aliscore output appears to be empty. Check results for BUSCO: 303740at7742
## Thu Jul 14 02:26:12 CEST 2022 - The aliscore output file does not exist. Check results for BUSCO: 328147at7742
## Thu Jul 14 02:26:12 CEST 2022 - The aliscore output appears to be empty. Check results for BUSCO: 328147at7742
## Thu Jul 14 02:26:14 CEST 2022 - The aliscore output file does not exist. Check results for BUSCO: 365186at7742
## Thu Jul 14 02:26:15 CEST 2022 - The aliscore output appears to be empty. Check results for BUSCO: 365186at7742
## Thu Jul 14 02:26:15 CEST 2022 - The aliscore output file does not exist. Check results for BUSCO: 310903at7742
## Thu Jul 14 02:26:15 CEST 2022 - The aliscore output appears to be empty. Check results for BUSCO: 310903at7742
## Thu Jul 14 02:26:15 CEST 2022 - The aliscore output file does not exist. Check results for BUSCO: 315832at7742
## Thu Jul 14 02:26:15 CEST 2022 - The aliscore output appears to be empty. Check results for BUSCO: 315832at7742
## Thu Jul 14 02:26:17 CEST 2022 - The aliscore output file does not exist. Check results for BUSCO: 235046at7742
## Thu Jul 14 02:26:17 CEST 2022 - The aliscore output appears to be empty. Check results for BUSCO: 235046at7742
## Thu Jul 14 02:26:18 CEST 2022 - The aliscore output file does not exist. Check results for BUSCO: 342641at7742
## Thu Jul 14 02:26:18 CEST 2022 - The aliscore output appears to be empty. Check results for BUSCO: 342641at7742
## Thu Jul 14 02:26:19 CEST 2022 - The aliscore output file does not exist. Check results for BUSCO: 389786at7742
## Thu Jul 14 02:26:19 CEST 2022 - The aliscore output appears to be empty. Check results for BUSCO: 389786at7742
## Thu Jul 14 02:26:21 CEST 2022 - The aliscore output file does not exist. Check results for BUSCO: 392383at7742
## Thu Jul 14 02:26:21 CEST 2022 - The aliscore output appears to be empty. Check results for BUSCO: 392383at7742
## Thu Jul 14 02:26:21 CEST 2022 - The aliscore output file does not exist. Check results for BUSCO: 400534at7742
## Thu Jul 14 02:26:21 CEST 2022 - The aliscore output appears to be empty. Check results for BUSCO: 400534at7742
## Thu Jul 14 02:26:22 CEST 2022 - The aliscore output file does not exist. Check results for BUSCO: 343725at7742
## Thu Jul 14 02:26:22 CEST 2022 - The aliscore output appears to be empty. Check results for BUSCO: 343725at7742
## Thu Jul 14 02:26:23 CEST 2022 - The aliscore output file does not exist. Check results for BUSCO: 325026at7742
## Thu Jul 14 02:26:23 CEST 2022 - The aliscore output appears to be empty. Check results for BUSCO: 325026at7742
## Thu Jul 14 02:26:23 CEST 2022 - The aliscore output file does not exist. Check results for BUSCO: 363264at7742
## Thu Jul 14 02:26:23 CEST 2022 - The aliscore output appears to be empty. Check results for BUSCO: 363264at7742
## Thu Jul 14 02:26:24 CEST 2022 - The aliscore output file does not exist. Check results for BUSCO: 358777at7742
## Thu Jul 14 02:26:24 CEST 2022 - The aliscore output appears to be empty. Check results for BUSCO: 358777at7742
## Thu Jul 14 02:26:25 CEST 2022 - The aliscore output file does not exist. Check results for BUSCO: 290213at7742
## Thu Jul 14 02:26:25 CEST 2022 - The aliscore output appears to be empty. Check results for BUSCO: 290213at7742
## Thu Jul 14 02:26:28 CEST 2022 - The aliscore output file does not exist. Check results for BUSCO: 390186at7742
## Thu Jul 14 02:26:28 CEST 2022 - The aliscore output appears to be empty. Check results for BUSCO: 390186at7742
## Thu Jul 14 02:26:29 CEST 2022 - The aliscore output file does not exist. Check results for BUSCO: 278787at7742
## Thu Jul 14 02:26:29 CEST 2022 - The aliscore output appears to be empty. Check results for BUSCO: 278787at7742
## Thu Jul 14 02:26:29 CEST 2022 - The aliscore output file does not exist. Check results for BUSCO: 337686at7742
## Thu Jul 14 02:26:29 CEST 2022 - The aliscore output appears to be empty. Check results for BUSCO: 337686at7742
## Thu Jul 14 02:26:29 CEST 2022 - The aliscore output file does not exist. Check results for BUSCO: 414238at7742
## Thu Jul 14 02:26:29 CEST 2022 - The aliscore output appears to be empty. Check results for BUSCO: 414238at7742
## Thu Jul 14 02:26:30 CEST 2022 - The aliscore output file does not exist. Check results for BUSCO: 384558at7742
## Thu Jul 14 02:26:30 CEST 2022 - The aliscore output appears to be empty. Check results for BUSCO: 384558at7742
## Thu Jul 14 02:26:31 CEST 2022 - The aliscore output file does not exist. Check results for BUSCO: 353318at7742
## Thu Jul 14 02:26:31 CEST 2022 - The aliscore output appears to be empty. Check results for BUSCO: 353318at7742
## Thu Jul 14 02:26:32 CEST 2022 - The aliscore output file does not exist. Check results for BUSCO: 235046at7742
## Thu Jul 14 02:26:32 CEST 2022 - The aliscore output appears to be empty. Check results for BUSCO: 235046at7742
## Thu Jul 14 02:26:33 CEST 2022 - The aliscore output file does not exist. Check results for BUSCO: 301464at7742
## Thu Jul 14 02:26:33 CEST 2022 - The aliscore output appears to be empty. Check results for BUSCO: 301464at7742
## Thu Jul 14 02:26:34 CEST 2022 - The aliscore output file does not exist. Check results for BUSCO: 285180at7742
## Thu Jul 14 02:26:34 CEST 2022 - The aliscore output appears to be empty. Check results for BUSCO: 285180at7742
## Thu Jul 14 02:26:35 CEST 2022 - The aliscore output file does not exist. Check results for BUSCO: 317054at7742
## Thu Jul 14 02:26:35 CEST 2022 - The aliscore output appears to be empty. Check results for BUSCO: 317054at7742
## Thu Jul 14 02:26:37 CEST 2022 - The aliscore output file does not exist. Check results for BUSCO: 288175at7742
## Thu Jul 14 02:26:37 CEST 2022 - The aliscore output appears to be empty. Check results for BUSCO: 288175at7742
## Thu Jul 14 02:26:37 CEST 2022 - The aliscore output file does not exist. Check results for BUSCO: 272130at7742
## Thu Jul 14 02:26:37 CEST 2022 - The aliscore output appears to be empty. Check results for BUSCO: 272130at7742
## Thu Jul 14 02:26:38 CEST 2022 - The aliscore output file does not exist. Check results for BUSCO: 273483at7742
## Thu Jul 14 02:26:38 CEST 2022 - The aliscore output appears to be empty. Check results for BUSCO: 273483at7742
## Thu Jul 14 02:26:39 CEST 2022 - The aliscore output file does not exist. Check results for BUSCO: 273271at7742
## Thu Jul 14 02:26:39 CEST 2022 - The aliscore output appears to be empty. Check results for BUSCO: 273271at7742
## Thu Jul 14 02:26:41 CEST 2022 - The aliscore output file does not exist. Check results for BUSCO: 265210at7742
## Thu Jul 14 02:26:41 CEST 2022 - The aliscore output appears to be empty. Check results for BUSCO: 265210at7742
## Thu Jul 14 02:26:41 CEST 2022 - The aliscore output file does not exist. Check results for BUSCO: 365149at7742
## Thu Jul 14 02:26:41 CEST 2022 - The aliscore output appears to be empty. Check results for BUSCO: 365149at7742
## Thu Jul 14 02:26:43 CEST 2022 - The aliscore output file does not exist. Check results for BUSCO: 386039at7742
## Thu Jul 14 02:26:43 CEST 2022 - The aliscore output appears to be empty. Check results for BUSCO: 386039at7742
## Thu Jul 14 02:26:44 CEST 2022 - The aliscore output file does not exist. Check results for BUSCO: 255469at7742
## Thu Jul 14 02:26:44 CEST 2022 - The aliscore output appears to be empty. Check results for BUSCO: 255469at7742
## Thu Jul 14 02:26:44 CEST 2022 - The aliscore output file does not exist. Check results for BUSCO: 371581at7742
## Thu Jul 14 02:26:44 CEST 2022 - The aliscore output appears to be empty. Check results for BUSCO: 371581at7742
## Thu Jul 14 02:26:45 CEST 2022 - The aliscore output file does not exist. Check results for BUSCO: 352334at7742
## Thu Jul 14 02:26:45 CEST 2022 - The aliscore output appears to be empty. Check results for BUSCO: 352334at7742
## Thu Jul 14 02:26:47 CEST 2022 - The aliscore output file does not exist. Check results for BUSCO: 257768at7742
## Thu Jul 14 02:26:47 CEST 2022 - The aliscore output appears to be empty. Check results for BUSCO: 257768at7742
## Thu Jul 14 02:26:47 CEST 2022 - The aliscore output file does not exist. Check results for BUSCO: 376038at7742
## Thu Jul 14 02:26:47 CEST 2022 - The aliscore output appears to be empty. Check results for BUSCO: 376038at7742
## Thu Jul 14 02:26:48 CEST 2022 - The aliscore output file does not exist. Check results for BUSCO: 315366at7742
## Thu Jul 14 02:26:48 CEST 2022 - The aliscore output appears to be empty. Check results for BUSCO: 315366at7742
## Thu Jul 14 02:26:49 CEST 2022 - The aliscore output file does not exist. Check results for BUSCO: 326283at7742
## Thu Jul 14 02:26:49 CEST 2022 - The aliscore output appears to be empty. Check results for BUSCO: 326283at7742
## Thu Jul 14 02:26:50 CEST 2022 - The aliscore output file does not exist. Check results for BUSCO: 365186at7742
## Thu Jul 14 02:26:50 CEST 2022 - The aliscore output appears to be empty. Check results for BUSCO: 365186at7742
## Thu Jul 14 02:26:50 CEST 2022 - The aliscore output file does not exist. Check results for BUSCO: 255423at7742
## Thu Jul 14 02:26:50 CEST 2022 - The aliscore output appears to be empty. Check results for BUSCO: 255423at7742
## Thu Jul 14 02:26:51 CEST 2022 - The aliscore output file does not exist. Check results for BUSCO: 388403at7742
## Thu Jul 14 02:26:51 CEST 2022 - The aliscore output appears to be empty. Check results for BUSCO: 388403at7742
## Thu Jul 14 02:26:53 CEST 2022 - The aliscore output file does not exist. Check results for BUSCO: 325026at7742
## Thu Jul 14 02:26:53 CEST 2022 - The aliscore output appears to be empty. Check results for BUSCO: 325026at7742
## Thu Jul 14 02:26:55 CEST 2022 - The aliscore output file does not exist. Check results for BUSCO: 315832at7742
## Thu Jul 14 02:26:55 CEST 2022 - The aliscore output appears to be empty. Check results for BUSCO: 315832at7742
## Thu Jul 14 02:26:55 CEST 2022 - The aliscore output file does not exist. Check results for BUSCO: 362141at7742
## Thu Jul 14 02:26:55 CEST 2022 - The aliscore output appears to be empty. Check results for BUSCO: 362141at7742
## Thu Jul 14 02:26:55 CEST 2022 - The aliscore output file does not exist. Check results for BUSCO: 380542at7742
## Thu Jul 14 02:26:55 CEST 2022 - The aliscore output appears to be empty. Check results for BUSCO: 380542at7742
## Thu Jul 14 02:26:56 CEST 2022 - The aliscore output file does not exist. Check results for BUSCO: 306544at7742
## Thu Jul 14 02:26:56 CEST 2022 - The aliscore output appears to be empty. Check results for BUSCO: 306544at7742
## Thu Jul 14 02:26:56 CEST 2022 - The aliscore output file does not exist. Check results for BUSCO: 265210at7742
## Thu Jul 14 02:26:56 CEST 2022 - The aliscore output appears to be empty. Check results for BUSCO: 265210at7742
## Thu Jul 14 02:26:57 CEST 2022 - The aliscore output file does not exist. Check results for BUSCO: 343085at7742
## Thu Jul 14 02:26:57 CEST 2022 - The aliscore output appears to be empty. Check results for BUSCO: 343085at7742
## Thu Jul 14 02:26:59 CEST 2022 - The aliscore output file does not exist. Check results for BUSCO: 345264at7742
## Thu Jul 14 02:26:59 CEST 2022 - The aliscore output appears to be empty. Check results for BUSCO: 345264at7742
## Thu Jul 14 02:27:01 CEST 2022 - The aliscore output file does not exist. Check results for BUSCO: 352334at7742
## Thu Jul 14 02:27:01 CEST 2022 - The aliscore output appears to be empty. Check results for BUSCO: 352334at7742
## Thu Jul 14 02:27:02 CEST 2022 - The aliscore output file does not exist. Check results for BUSCO: 384878at7742
## Thu Jul 14 02:27:02 CEST 2022 - The aliscore output appears to be empty. Check results for BUSCO: 384878at7742
## Thu Jul 14 02:27:02 CEST 2022 - The aliscore output file does not exist. Check results for BUSCO: 235046at7742
## Thu Jul 14 02:27:02 CEST 2022 - The aliscore output appears to be empty. Check results for BUSCO: 235046at7742
## Thu Jul 14 02:27:03 CEST 2022 - The aliscore output file does not exist. Check results for BUSCO: 353318at7742
## Thu Jul 14 02:27:03 CEST 2022 - The aliscore output appears to be empty. Check results for BUSCO: 353318at7742
## Thu Jul 14 02:27:04 CEST 2022 - The aliscore output file does not exist. Check results for BUSCO: 385594at7742
## Thu Jul 14 02:27:04 CEST 2022 - The aliscore output appears to be empty. Check results for BUSCO: 385594at7742
## Thu Jul 14 02:27:05 CEST 2022 - The aliscore output file does not exist. Check results for BUSCO: 373954at7742
## Thu Jul 14 02:27:05 CEST 2022 - The aliscore output appears to be empty. Check results for BUSCO: 373954at7742
## Thu Jul 14 02:27:07 CEST 2022 - The aliscore output file does not exist. Check results for BUSCO: 387408at7742
## Thu Jul 14 02:27:07 CEST 2022 - The aliscore output appears to be empty. Check results for BUSCO: 387408at7742
## Thu Jul 14 02:27:08 CEST 2022 - The aliscore output file does not exist. Check results for BUSCO: 294646at7742
## Thu Jul 14 02:27:08 CEST 2022 - The aliscore output appears to be empty. Check results for BUSCO: 294646at7742
## Thu Jul 14 02:27:09 CEST 2022 - The aliscore output file does not exist. Check results for BUSCO: 393010at7742
## Thu Jul 14 02:27:09 CEST 2022 - The aliscore output file does not exist. Check results for BUSCO: 357059at7742
## Thu Jul 14 02:27:09 CEST 2022 - The aliscore output appears to be empty. Check results for BUSCO: 393010at7742
## Thu Jul 14 02:27:09 CEST 2022 - The aliscore output appears to be empty. Check results for BUSCO: 357059at7742
## Thu Jul 14 02:27:10 CEST 2022 - The aliscore output file does not exist. Check results for BUSCO: 328949at7742
## Thu Jul 14 02:27:10 CEST 2022 - The aliscore output appears to be empty. Check results for BUSCO: 328949at7742
## Thu Jul 14 02:27:12 CEST 2022 - The aliscore output file does not exist. Check results for BUSCO: 422403at7742
## Thu Jul 14 02:27:12 CEST 2022 - The aliscore output file does not exist. Check results for BUSCO: 296937at7742
## Thu Jul 14 02:27:12 CEST 2022 - The aliscore output appears to be empty. Check results for BUSCO: 422403at7742
## Thu Jul 14 02:27:12 CEST 2022 - The aliscore output appears to be empty. Check results for BUSCO: 296937at7742
## Thu Jul 14 02:27:12 CEST 2022 - The aliscore output file does not exist. Check results for BUSCO: 382456at7742
## Thu Jul 14 02:27:13 CEST 2022 - The aliscore output appears to be empty. Check results for BUSCO: 382456at7742
## Thu Jul 14 02:27:15 CEST 2022 - The aliscore output file does not exist. Check results for BUSCO: 295101at7742
## Thu Jul 14 02:27:15 CEST 2022 - The aliscore output appears to be empty. Check results for BUSCO: 295101at7742
## Thu Jul 14 02:27:16 CEST 2022 - The aliscore output file does not exist. Check results for BUSCO: 299314at7742
## Thu Jul 14 02:27:16 CEST 2022 - The aliscore output appears to be empty. Check results for BUSCO: 299314at7742
## Thu Jul 14 02:27:16 CEST 2022 - The aliscore output file does not exist. Check results for BUSCO: 278647at7742
## Thu Jul 14 02:27:16 CEST 2022 - The aliscore output appears to be empty. Check results for BUSCO: 278647at7742
## Thu Jul 14 02:27:16 CEST 2022 - The aliscore output file does not exist. Check results for BUSCO: 272130at7742
## Thu Jul 14 02:27:16 CEST 2022 - The aliscore output appears to be empty. Check results for BUSCO: 272130at7742
## Thu Jul 14 02:27:17 CEST 2022 - The aliscore output file does not exist. Check results for BUSCO: 328949at7742
## Thu Jul 14 02:27:17 CEST 2022 - The aliscore output appears to be empty. Check results for BUSCO: 328949at7742
## Thu Jul 14 02:27:19 CEST 2022 - The aliscore output file does not exist. Check results for BUSCO: 261213at7742
## Thu Jul 14 02:27:19 CEST 2022 - The aliscore output file does not exist. Check results for BUSCO: 358777at7742
## Thu Jul 14 02:27:19 CEST 2022 - The aliscore output appears to be empty. Check results for BUSCO: 261213at7742
## Thu Jul 14 02:27:19 CEST 2022 - The aliscore output appears to be empty. Check results for BUSCO: 358777at7742
## Thu Jul 14 02:27:21 CEST 2022 - The aliscore output file does not exist. Check results for BUSCO: 387526at7742
## Thu Jul 14 02:27:21 CEST 2022 - The aliscore output appears to be empty. Check results for BUSCO: 387526at7742
## Thu Jul 14 02:27:21 CEST 2022 - The aliscore output file does not exist. Check results for BUSCO: 352334at7742
## Thu Jul 14 02:27:21 CEST 2022 - The aliscore output appears to be empty. Check results for BUSCO: 352334at7742
## Thu Jul 14 02:27:22 CEST 2022 - The aliscore output file does not exist. Check results for BUSCO: 255469at7742
## Thu Jul 14 02:27:22 CEST 2022 - The aliscore output appears to be empty. Check results for BUSCO: 255469at7742
## Thu Jul 14 02:27:22 CEST 2022 - The aliscore output file does not exist. Check results for BUSCO: 361537at7742
## Thu Jul 14 02:27:22 CEST 2022 - The aliscore output appears to be empty. Check results for BUSCO: 361537at7742
## Thu Jul 14 02:27:24 CEST 2022 - The aliscore output file does not exist. Check results for BUSCO: 326141at7742
## Thu Jul 14 02:27:24 CEST 2022 - The aliscore output appears to be empty. Check results for BUSCO: 326141at7742
## Thu Jul 14 02:27:24 CEST 2022 - The aliscore output file does not exist. Check results for BUSCO: 421821at7742
## Thu Jul 14 02:27:24 CEST 2022 - The aliscore output appears to be empty. Check results for BUSCO: 421821at7742
## Thu Jul 14 02:27:26 CEST 2022 - The aliscore output file does not exist. Check results for BUSCO: 338803at7742
## Thu Jul 14 02:27:26 CEST 2022 - The aliscore output appears to be empty. Check results for BUSCO: 338803at7742
## Thu Jul 14 02:27:26 CEST 2022 - The aliscore output file does not exist. Check results for BUSCO: 386039at7742
## Thu Jul 14 02:27:26 CEST 2022 - The aliscore output appears to be empty. Check results for BUSCO: 386039at7742
## Thu Jul 14 02:27:27 CEST 2022 - The aliscore output file does not exist. Check results for BUSCO: 324567at7742
## Thu Jul 14 02:27:27 CEST 2022 - The aliscore output appears to be empty. Check results for BUSCO: 324567at7742
## Thu Jul 14 02:27:29 CEST 2022 - The aliscore output file does not exist. Check results for BUSCO: 406396at7742
## Thu Jul 14 02:27:29 CEST 2022 - The aliscore output appears to be empty. Check results for BUSCO: 406396at7742
## Thu Jul 14 02:27:30 CEST 2022 - The aliscore output file does not exist. Check results for BUSCO: 373954at7742
## Thu Jul 14 02:27:30 CEST 2022 - The aliscore output file does not exist. Check results for BUSCO: 406718at7742
## Thu Jul 14 02:27:30 CEST 2022 - The aliscore output appears to be empty. Check results for BUSCO: 373954at7742
## Thu Jul 14 02:27:30 CEST 2022 - The aliscore output appears to be empty. Check results for BUSCO: 406718at7742
## Thu Jul 14 02:27:32 CEST 2022 - The aliscore output file does not exist. Check results for BUSCO: 308917at7742
## Thu Jul 14 02:27:32 CEST 2022 - The aliscore output appears to be empty. Check results for BUSCO: 308917at7742
## Thu Jul 14 02:27:33 CEST 2022 - The aliscore output file does not exist. Check results for BUSCO: 370914at7742
## Thu Jul 14 02:27:33 CEST 2022 - The aliscore output appears to be empty. Check results for BUSCO: 370914at7742
## Thu Jul 14 02:27:33 CEST 2022 - The aliscore output file does not exist. Check results for BUSCO: 366454at7742
## Thu Jul 14 02:27:33 CEST 2022 - The aliscore output appears to be empty. Check results for BUSCO: 366454at7742
## Thu Jul 14 02:27:35 CEST 2022 - The aliscore output file does not exist. Check results for BUSCO: 336619at7742
## Thu Jul 14 02:27:35 CEST 2022 - The aliscore output appears to be empty. Check results for BUSCO: 336619at7742
## Thu Jul 14 02:27:36 CEST 2022 - The aliscore output file does not exist. Check results for BUSCO: 372764at7742
## Thu Jul 14 02:27:36 CEST 2022 - The aliscore output appears to be empty. Check results for BUSCO: 372764at7742
## Thu Jul 14 02:27:36 CEST 2022 - The aliscore output file does not exist. Check results for BUSCO: 265210at7742
## Thu Jul 14 02:27:36 CEST 2022 - The aliscore output appears to be empty. Check results for BUSCO: 265210at7742
## Thu Jul 14 02:27:36 CEST 2022 - The aliscore output file does not exist. Check results for BUSCO: 273483at7742
## Thu Jul 14 02:27:36 CEST 2022 - The aliscore output appears to be empty. Check results for BUSCO: 273483at7742
## Thu Jul 14 02:27:38 CEST 2022 - The aliscore output file does not exist. Check results for BUSCO: 279816at7742
## Thu Jul 14 02:27:38 CEST 2022 - The aliscore output appears to be empty. Check results for BUSCO: 279816at7742
## Thu Jul 14 02:27:39 CEST 2022 - The aliscore output file does not exist. Check results for BUSCO: 336331at7742
## Thu Jul 14 02:27:39 CEST 2022 - The aliscore output appears to be empty. Check results for BUSCO: 336331at7742
## Thu Jul 14 02:27:41 CEST 2022 - The aliscore output file does not exist. Check results for BUSCO: 384054at7742
## Thu Jul 14 02:27:41 CEST 2022 - The aliscore output file does not exist. Check results for BUSCO: 285180at7742
## Thu Jul 14 02:27:41 CEST 2022 - The aliscore output appears to be empty. Check results for BUSCO: 384054at7742
## Thu Jul 14 02:27:41 CEST 2022 - The aliscore output appears to be empty. Check results for BUSCO: 285180at7742
## Thu Jul 14 02:27:42 CEST 2022 - The aliscore output file does not exist. Check results for BUSCO: 400935at7742
## Thu Jul 14 02:27:42 CEST 2022 - The aliscore output appears to be empty. Check results for BUSCO: 400935at7742
## Thu Jul 14 02:27:43 CEST 2022 - The aliscore output file does not exist. Check results for BUSCO: 352195at7742
## Thu Jul 14 02:27:43 CEST 2022 - The aliscore output appears to be empty. Check results for BUSCO: 352195at7742
## Thu Jul 14 02:27:44 CEST 2022 - The aliscore output file does not exist. Check results for BUSCO: 389786at7742
## Thu Jul 14 02:27:44 CEST 2022 - The aliscore output appears to be empty. Check results for BUSCO: 389786at7742
## Thu Jul 14 02:27:45 CEST 2022 - The aliscore output file does not exist. Check results for BUSCO: 280287at7742
## Thu Jul 14 02:27:45 CEST 2022 - The aliscore output appears to be empty. Check results for BUSCO: 280287at7742
## Thu Jul 14 02:27:46 CEST 2022 - The aliscore output file does not exist. Check results for BUSCO: 296937at7742
## Thu Jul 14 02:27:46 CEST 2022 - The aliscore output appears to be empty. Check results for BUSCO: 296937at7742
## Thu Jul 14 02:27:46 CEST 2022 - The aliscore output file does not exist. Check results for BUSCO: 370914at7742
## Thu Jul 14 02:27:46 CEST 2022 - The aliscore output appears to be empty. Check results for BUSCO: 370914at7742
## Thu Jul 14 02:27:46 CEST 2022 - The aliscore output file does not exist. Check results for BUSCO: 261072at7742
## Thu Jul 14 02:27:46 CEST 2022 - The aliscore output appears to be empty. Check results for BUSCO: 261072at7742
## Thu Jul 14 02:27:47 CEST 2022 - The aliscore output file does not exist. Check results for BUSCO: 398565at7742
## Thu Jul 14 02:27:47 CEST 2022 - The aliscore output appears to be empty. Check results for BUSCO: 398565at7742
## Thu Jul 14 02:27:50 CEST 2022 - The aliscore output file does not exist. Check results for BUSCO: 358777at7742
## Thu Jul 14 02:27:50 CEST 2022 - The aliscore output appears to be empty. Check results for BUSCO: 358777at7742
## Thu Jul 14 02:27:50 CEST 2022 - The aliscore output file does not exist. Check results for BUSCO: 365149at7742
## Thu Jul 14 02:27:50 CEST 2022 - The aliscore output appears to be empty. Check results for BUSCO: 365149at7742
## Thu Jul 14 02:27:51 CEST 2022 - The aliscore output file does not exist. Check results for BUSCO: 363264at7742
## Thu Jul 14 02:27:51 CEST 2022 - The aliscore output appears to be empty. Check results for BUSCO: 363264at7742
## Thu Jul 14 02:27:51 CEST 2022 - The aliscore output file does not exist. Check results for BUSCO: 324567at7742
## Thu Jul 14 02:27:51 CEST 2022 - The aliscore output appears to be empty. Check results for BUSCO: 324567at7742
## Thu Jul 14 02:27:53 CEST 2022 - The aliscore output file does not exist. Check results for BUSCO: 362141at7742
## Thu Jul 14 02:27:53 CEST 2022 - The aliscore output appears to be empty. Check results for BUSCO: 362141at7742
## Thu Jul 14 02:27:53 CEST 2022 - The aliscore output file does not exist. Check results for BUSCO: 306544at7742
## Thu Jul 14 02:27:53 CEST 2022 - The aliscore output appears to be empty. Check results for BUSCO: 306544at7742
## Thu Jul 14 02:27:54 CEST 2022 - The aliscore output file does not exist. Check results for BUSCO: 260364at7742
## Thu Jul 14 02:27:54 CEST 2022 - The aliscore output appears to be empty. Check results for BUSCO: 260364at7742
## Thu Jul 14 02:27:54 CEST 2022 - The aliscore output file does not exist. Check results for BUSCO: 255469at7742
## Thu Jul 14 02:27:54 CEST 2022 - The aliscore output appears to be empty. Check results for BUSCO: 255469at7742
## Thu Jul 14 02:27:57 CEST 2022 - The aliscore output file does not exist. Check results for BUSCO: 400534at7742
## Thu Jul 14 02:27:57 CEST 2022 - The aliscore output appears to be empty. Check results for BUSCO: 400534at7742
## Thu Jul 14 02:27:57 CEST 2022 - The aliscore output file does not exist. Check results for BUSCO: 323574at7742
## Thu Jul 14 02:27:57 CEST 2022 - The aliscore output appears to be empty. Check results for BUSCO: 323574at7742
## Thu Jul 14 02:27:58 CEST 2022 - The aliscore output file does not exist. Check results for BUSCO: 310457at7742
## Thu Jul 14 02:27:58 CEST 2022 - The aliscore output appears to be empty. Check results for BUSCO: 310457at7742
## Thu Jul 14 02:27:59 CEST 2022 - The aliscore output file does not exist. Check results for BUSCO: 255423at7742
## Thu Jul 14 02:27:59 CEST 2022 - The aliscore output appears to be empty. Check results for BUSCO: 255423at7742
## Thu Jul 14 02:27:59 CEST 2022 - The aliscore output file does not exist. Check results for BUSCO: 345264at7742
## Thu Jul 14 02:28:00 CEST 2022 - The aliscore output appears to be empty. Check results for BUSCO: 345264at7742
## Thu Jul 14 02:28:01 CEST 2022 - The aliscore output file does not exist. Check results for BUSCO: 414201at7742
## Thu Jul 14 02:28:01 CEST 2022 - The aliscore output appears to be empty. Check results for BUSCO: 414201at7742
## Thu Jul 14 02:28:03 CEST 2022 - The aliscore output file does not exist. Check results for BUSCO: 397671at7742
## Thu Jul 14 02:28:03 CEST 2022 - The aliscore output appears to be empty. Check results for BUSCO: 397671at7742
## Thu Jul 14 02:28:04 CEST 2022 - The aliscore output file does not exist. Check results for BUSCO: 388403at7742
## Thu Jul 14 02:28:04 CEST 2022 - The aliscore output appears to be empty. Check results for BUSCO: 388403at7742
## Thu Jul 14 02:28:05 CEST 2022 - The aliscore output file does not exist. Check results for BUSCO: 235735at7742
## Thu Jul 14 02:28:05 CEST 2022 - The aliscore output appears to be empty. Check results for BUSCO: 235735at7742
## Thu Jul 14 02:28:06 CEST 2022 - The aliscore output file does not exist. Check results for BUSCO: 391948at7742
## Thu Jul 14 02:28:06 CEST 2022 - The aliscore output appears to be empty. Check results for BUSCO: 391948at7742
## Thu Jul 14 02:28:06 CEST 2022 - The aliscore output file does not exist. Check results for BUSCO: 301464at7742
## Thu Jul 14 02:28:06 CEST 2022 - The aliscore output appears to be empty. Check results for BUSCO: 301464at7742
## Thu Jul 14 02:28:06 CEST 2022 - The aliscore output file does not exist. Check results for BUSCO: 257925at7742
## Thu Jul 14 02:28:06 CEST 2022 - The aliscore output appears to be empty. Check results for BUSCO: 257925at7742
## Thu Jul 14 02:28:08 CEST 2022 - The aliscore output file does not exist. Check results for BUSCO: 299314at7742
## Thu Jul 14 02:28:08 CEST 2022 - The aliscore output appears to be empty. Check results for BUSCO: 299314at7742
## Thu Jul 14 02:28:09 CEST 2022 - The aliscore output file does not exist. Check results for BUSCO: 406306at7742
## Thu Jul 14 02:28:10 CEST 2022 - The aliscore output appears to be empty. Check results for BUSCO: 406306at7742
## Thu Jul 14 02:28:10 CEST 2022 - The aliscore output file does not exist. Check results for BUSCO: 294990at7742
## Thu Jul 14 02:28:10 CEST 2022 - The aliscore output appears to be empty. Check results for BUSCO: 294990at7742
## Thu Jul 14 02:28:10 CEST 2022 - The aliscore output file does not exist. Check results for BUSCO: 406396at7742
## Thu Jul 14 02:28:10 CEST 2022 - The aliscore output appears to be empty. Check results for BUSCO: 406396at7742
## Thu Jul 14 02:28:12 CEST 2022 - The aliscore output file does not exist. Check results for BUSCO: 338803at7742
## Thu Jul 14 02:28:12 CEST 2022 - The aliscore output appears to be empty. Check results for BUSCO: 338803at7742
## Thu Jul 14 02:28:13 CEST 2022 - The aliscore output file does not exist. Check results for BUSCO: 398565at7742
## Thu Jul 14 02:28:13 CEST 2022 - The aliscore output appears to be empty. Check results for BUSCO: 398565at7742
## Thu Jul 14 02:28:14 CEST 2022 - The aliscore output file does not exist. Check results for BUSCO: 352195at7742
## Thu Jul 14 02:28:14 CEST 2022 - The aliscore output appears to be empty. Check results for BUSCO: 352195at7742
## Thu Jul 14 02:28:15 CEST 2022 - The aliscore output file does not exist. Check results for BUSCO: 337042at7742
## Thu Jul 14 02:28:15 CEST 2022 - The aliscore output appears to be empty. Check results for BUSCO: 337042at7742
## Thu Jul 14 02:28:16 CEST 2022 - The aliscore output file does not exist. Check results for BUSCO: 310903at7742
## Thu Jul 14 02:28:16 CEST 2022 - The aliscore output appears to be empty. Check results for BUSCO: 310903at7742
## Thu Jul 14 02:28:17 CEST 2022 - The aliscore output file does not exist. Check results for BUSCO: 387408at7742
## Thu Jul 14 02:28:17 CEST 2022 - The aliscore output appears to be empty. Check results for BUSCO: 387408at7742
## Thu Jul 14 02:28:17 CEST 2022 - The aliscore output file does not exist. Check results for BUSCO: 413352at7742
## Thu Jul 14 02:28:17 CEST 2022 - The aliscore output appears to be empty. Check results for BUSCO: 413352at7742
## Thu Jul 14 02:28:18 CEST 2022 - The aliscore output file does not exist. Check results for BUSCO: 304588at7742
## Thu Jul 14 02:28:19 CEST 2022 - The aliscore output appears to be empty. Check results for BUSCO: 304588at7742
## Thu Jul 14 02:28:19 CEST 2022 - The aliscore output file does not exist. Check results for BUSCO: 306544at7742
## Thu Jul 14 02:28:19 CEST 2022 - The aliscore output appears to be empty. Check results for BUSCO: 306544at7742
## Thu Jul 14 02:28:20 CEST 2022 - The aliscore output file does not exist. Check results for BUSCO: 347954at7742
## Thu Jul 14 02:28:21 CEST 2022 - The aliscore output appears to be empty. Check results for BUSCO: 347954at7742
## Thu Jul 14 02:28:21 CEST 2022 - The aliscore output file does not exist. Check results for BUSCO: 338803at7742
## Thu Jul 14 02:28:21 CEST 2022 - The aliscore output appears to be empty. Check results for BUSCO: 338803at7742
## Thu Jul 14 02:28:22 CEST 2022 - The aliscore output file does not exist. Check results for BUSCO: 421415at7742
## Thu Jul 14 02:28:22 CEST 2022 - The aliscore output appears to be empty. Check results for BUSCO: 421415at7742
## Thu Jul 14 02:28:23 CEST 2022 - The aliscore output file does not exist. Check results for BUSCO: 437044at7742
## Thu Jul 14 02:28:23 CEST 2022 - The aliscore output appears to be empty. Check results for BUSCO: 437044at7742
## Thu Jul 14 02:28:24 CEST 2022 - The aliscore output file does not exist. Check results for BUSCO: 404882at7742
## Thu Jul 14 02:28:24 CEST 2022 - The aliscore output appears to be empty. Check results for BUSCO: 404882at7742
## Thu Jul 14 02:28:25 CEST 2022 - The aliscore output file does not exist. Check results for BUSCO: 406396at7742
## Thu Jul 14 02:28:25 CEST 2022 - The aliscore output appears to be empty. Check results for BUSCO: 406396at7742
## Thu Jul 14 02:28:26 CEST 2022 - The aliscore output file does not exist. Check results for BUSCO: 325026at7742
## Thu Jul 14 02:28:26 CEST 2022 - The aliscore output appears to be empty. Check results for BUSCO: 325026at7742
## Thu Jul 14 02:28:27 CEST 2022 - The aliscore output file does not exist. Check results for BUSCO: 398379at7742
## Thu Jul 14 02:28:27 CEST 2022 - The aliscore output appears to be empty. Check results for BUSCO: 398379at7742
## Thu Jul 14 02:28:28 CEST 2022 - The aliscore output file does not exist. Check results for BUSCO: 257925at7742
## Thu Jul 14 02:28:28 CEST 2022 - The aliscore output appears to be empty. Check results for BUSCO: 257925at7742
## Thu Jul 14 02:28:29 CEST 2022 - The aliscore output file does not exist. Check results for BUSCO: 352195at7742
## Thu Jul 14 02:28:29 CEST 2022 - The aliscore output appears to be empty. Check results for BUSCO: 352195at7742
## Thu Jul 14 02:28:30 CEST 2022 - The aliscore output file does not exist. Check results for BUSCO: 343013at7742
## Thu Jul 14 02:28:30 CEST 2022 - The aliscore output appears to be empty. Check results for BUSCO: 343013at7742
## Thu Jul 14 02:28:30 CEST 2022 - The aliscore output file does not exist. Check results for BUSCO: 294646at7742
## Thu Jul 14 02:28:30 CEST 2022 - The aliscore output appears to be empty. Check results for BUSCO: 294646at7742
## Thu Jul 14 02:28:31 CEST 2022 - The aliscore output file does not exist. Check results for BUSCO: 278647at7742
## Thu Jul 14 02:28:31 CEST 2022 - The aliscore output appears to be empty. Check results for BUSCO: 278647at7742
## Thu Jul 14 02:28:33 CEST 2022 - The aliscore output file does not exist. Check results for BUSCO: 337686at7742
## Thu Jul 14 02:28:33 CEST 2022 - The aliscore output appears to be empty. Check results for BUSCO: 337686at7742
## Thu Jul 14 02:28:34 CEST 2022 - The aliscore output file does not exist. Check results for BUSCO: 277508at7742
## Thu Jul 14 02:28:34 CEST 2022 - The aliscore output appears to be empty. Check results for BUSCO: 277508at7742
## Thu Jul 14 02:28:35 CEST 2022 - The aliscore output file does not exist. Check results for BUSCO: 300899at7742
## Thu Jul 14 02:28:35 CEST 2022 - The aliscore output appears to be empty. Check results for BUSCO: 300899at7742
## Thu Jul 14 02:28:35 CEST 2022 - The aliscore output file does not exist. Check results for BUSCO: 303403at7742
## Thu Jul 14 02:28:35 CEST 2022 - The aliscore output appears to be empty. Check results for BUSCO: 303403at7742
## Thu Jul 14 02:28:36 CEST 2022 - The aliscore output file does not exist. Check results for BUSCO: 388183at7742
## Thu Jul 14 02:28:36 CEST 2022 - The aliscore output appears to be empty. Check results for BUSCO: 388183at7742
## Thu Jul 14 02:28:36 CEST 2022 - The aliscore output file does not exist. Check results for BUSCO: 280287at7742
## Thu Jul 14 02:28:36 CEST 2022 - The aliscore output appears to be empty. Check results for BUSCO: 280287at7742
## Thu Jul 14 02:28:37 CEST 2022 - The aliscore output file does not exist. Check results for BUSCO: 320027at7742
## Thu Jul 14 02:28:37 CEST 2022 - The aliscore output appears to be empty. Check results for BUSCO: 320027at7742
## Thu Jul 14 02:28:39 CEST 2022 - The aliscore output file does not exist. Check results for BUSCO: 358920at7742
## Thu Jul 14 02:28:40 CEST 2022 - The aliscore output appears to be empty. Check results for BUSCO: 358920at7742
## Thu Jul 14 02:28:40 CEST 2022 - The aliscore output file does not exist. Check results for BUSCO: 313967at7742
## Thu Jul 14 02:28:40 CEST 2022 - The aliscore output appears to be empty. Check results for BUSCO: 313967at7742
## Thu Jul 14 02:28:41 CEST 2022 - The aliscore output file does not exist. Check results for BUSCO: 348744at7742
## Thu Jul 14 02:28:41 CEST 2022 - The aliscore output appears to be empty. Check results for BUSCO: 348744at7742
## Thu Jul 14 02:28:42 CEST 2022 - The aliscore output file does not exist. Check results for BUSCO: 278787at7742
## Thu Jul 14 02:28:42 CEST 2022 - The aliscore output appears to be empty. Check results for BUSCO: 278787at7742
## Thu Jul 14 02:28:43 CEST 2022 - The aliscore output file does not exist. Check results for BUSCO: 284145at7742
## Thu Jul 14 02:28:43 CEST 2022 - The aliscore output appears to be empty. Check results for BUSCO: 284145at7742
## Thu Jul 14 02:28:43 CEST 2022 - The aliscore output file does not exist. Check results for BUSCO: 345556at7742
## Thu Jul 14 02:28:43 CEST 2022 - The aliscore output appears to be empty. Check results for BUSCO: 345556at7742
## Thu Jul 14 02:28:45 CEST 2022 - The aliscore output file does not exist. Check results for BUSCO: 257925at7742
## Thu Jul 14 02:28:45 CEST 2022 - The aliscore output appears to be empty. Check results for BUSCO: 257925at7742
## Thu Jul 14 02:28:46 CEST 2022 - The aliscore output file does not exist. Check results for BUSCO: 331986at7742
## Thu Jul 14 02:28:46 CEST 2022 - The aliscore output appears to be empty. Check results for BUSCO: 331986at7742
## Thu Jul 14 02:28:47 CEST 2022 - The aliscore output file does not exist. Check results for BUSCO: 328506at7742
## Thu Jul 14 02:28:47 CEST 2022 - The aliscore output appears to be empty. Check results for BUSCO: 328506at7742
## Thu Jul 14 02:28:47 CEST 2022 - The aliscore output file does not exist. Check results for BUSCO: 368105at7742
## Thu Jul 14 02:28:47 CEST 2022 - The aliscore output appears to be empty. Check results for BUSCO: 368105at7742
## Thu Jul 14 02:28:49 CEST 2022 - The aliscore output file does not exist. Check results for BUSCO: 308917at7742
## Thu Jul 14 02:28:49 CEST 2022 - The aliscore output appears to be empty. Check results for BUSCO: 308917at7742
## Thu Jul 14 02:28:50 CEST 2022 - The aliscore output file does not exist. Check results for BUSCO: 379564at7742
## Thu Jul 14 02:28:50 CEST 2022 - The aliscore output appears to be empty. Check results for BUSCO: 379564at7742
## Thu Jul 14 02:28:51 CEST 2022 - The aliscore output file does not exist. Check results for BUSCO: 343013at7742
## Thu Jul 14 02:28:51 CEST 2022 - The aliscore output appears to be empty. Check results for BUSCO: 343013at7742
## Thu Jul 14 02:28:51 CEST 2022 - The aliscore output file does not exist. Check results for BUSCO: 326788at7742
## Thu Jul 14 02:28:51 CEST 2022 - The aliscore output appears to be empty. Check results for BUSCO: 326788at7742
## Thu Jul 14 02:28:52 CEST 2022 - The aliscore output file does not exist. Check results for BUSCO: 273483at7742
## Thu Jul 14 02:28:52 CEST 2022 - The aliscore output appears to be empty. Check results for BUSCO: 273483at7742
## Thu Jul 14 02:28:53 CEST 2022 - The aliscore output file does not exist. Check results for BUSCO: 385594at7742
## Thu Jul 14 02:28:53 CEST 2022 - The aliscore output appears to be empty. Check results for BUSCO: 385594at7742
## Thu Jul 14 02:28:54 CEST 2022 - The aliscore output file does not exist. Check results for BUSCO: 372764at7742
## Thu Jul 14 02:28:54 CEST 2022 - The aliscore output appears to be empty. Check results for BUSCO: 372764at7742
## Thu Jul 14 02:28:55 CEST 2022 - The aliscore output file does not exist. Check results for BUSCO: 331986at7742
## Thu Jul 14 02:28:56 CEST 2022 - The aliscore output appears to be empty. Check results for BUSCO: 331986at7742
## Thu Jul 14 02:28:56 CEST 2022 - The aliscore output file does not exist. Check results for BUSCO: 391345at7742
## Thu Jul 14 02:28:56 CEST 2022 - The aliscore output appears to be empty. Check results for BUSCO: 391345at7742
## Thu Jul 14 02:28:56 CEST 2022 - The aliscore output file does not exist. Check results for BUSCO: 388102at7742
## Thu Jul 14 02:28:56 CEST 2022 - The aliscore output appears to be empty. Check results for BUSCO: 388102at7742
## Thu Jul 14 02:28:57 CEST 2022 - The aliscore output file does not exist. Check results for BUSCO: 299325at7742
## Thu Jul 14 02:28:57 CEST 2022 - The aliscore output appears to be empty. Check results for BUSCO: 299325at7742
## Thu Jul 14 02:28:58 CEST 2022 - The aliscore output file does not exist. Check results for BUSCO: 277748at7742
## Thu Jul 14 02:28:58 CEST 2022 - The aliscore output appears to be empty. Check results for BUSCO: 277748at7742
## Thu Jul 14 02:29:00 CEST 2022 - The aliscore output file does not exist. Check results for BUSCO: 320027at7742
## Thu Jul 14 02:29:00 CEST 2022 - The aliscore output appears to be empty. Check results for BUSCO: 320027at7742
## Thu Jul 14 02:29:01 CEST 2022 - The aliscore output file does not exist. Check results for BUSCO: 336764at7742
## Thu Jul 14 02:29:01 CEST 2022 - The aliscore output appears to be empty. Check results for BUSCO: 336764at7742
## Thu Jul 14 02:29:01 CEST 2022 - The aliscore output file does not exist. Check results for BUSCO: 372431at7742
## Thu Jul 14 02:29:01 CEST 2022 - The aliscore output appears to be empty. Check results for BUSCO: 372431at7742
## Thu Jul 14 02:29:02 CEST 2022 - The aliscore output file does not exist. Check results for BUSCO: 370608at7742
## Thu Jul 14 02:29:02 CEST 2022 - The aliscore output appears to be empty. Check results for BUSCO: 370608at7742
## Thu Jul 14 02:29:03 CEST 2022 - The aliscore output file does not exist. Check results for BUSCO: 310903at7742
## Thu Jul 14 02:29:03 CEST 2022 - The aliscore output appears to be empty. Check results for BUSCO: 310903at7742
## Thu Jul 14 02:29:05 CEST 2022 - The aliscore output file does not exist. Check results for BUSCO: 273166at7742
## Thu Jul 14 02:29:05 CEST 2022 - The aliscore output appears to be empty. Check results for BUSCO: 273166at7742
## Thu Jul 14 02:29:05 CEST 2022 - The aliscore output file does not exist. Check results for BUSCO: 277748at7742
## Thu Jul 14 02:29:05 CEST 2022 - The aliscore output appears to be empty. Check results for BUSCO: 277748at7742
## Thu Jul 14 02:29:07 CEST 2022 - The aliscore output file does not exist. Check results for BUSCO: 310457at7742
## Thu Jul 14 02:29:07 CEST 2022 - The aliscore output appears to be empty. Check results for BUSCO: 310457at7742
## Thu Jul 14 02:29:07 CEST 2022 - The aliscore output file does not exist. Check results for BUSCO: 331452at7742
## Thu Jul 14 02:29:07 CEST 2022 - The aliscore output appears to be empty. Check results for BUSCO: 331452at7742
## Thu Jul 14 02:29:09 CEST 2022 - The aliscore output file does not exist. Check results for BUSCO: 326283at7742
## Thu Jul 14 02:29:09 CEST 2022 - The aliscore output file does not exist. Check results for BUSCO: 373482at7742
## Thu Jul 14 02:29:09 CEST 2022 - The aliscore output appears to be empty. Check results for BUSCO: 326283at7742
## Thu Jul 14 02:29:09 CEST 2022 - The aliscore output appears to be empty. Check results for BUSCO: 373482at7742
## Thu Jul 14 02:29:11 CEST 2022 - The aliscore output file does not exist. Check results for BUSCO: 313967at7742
## Thu Jul 14 02:29:11 CEST 2022 - The aliscore output file does not exist. Check results for BUSCO: 307391at7742
## Thu Jul 14 02:29:11 CEST 2022 - The aliscore output appears to be empty. Check results for BUSCO: 313967at7742
## Thu Jul 14 02:29:11 CEST 2022 - The aliscore output appears to be empty. Check results for BUSCO: 307391at7742
## Thu Jul 14 02:29:12 CEST 2022 - The aliscore output file does not exist. Check results for BUSCO: 402327at7742
## Thu Jul 14 02:29:12 CEST 2022 - The aliscore output appears to be empty. Check results for BUSCO: 402327at7742
## Thu Jul 14 02:29:13 CEST 2022 - The aliscore output file does not exist. Check results for BUSCO: 261213at7742
## Thu Jul 14 02:29:13 CEST 2022 - The aliscore output appears to be empty. Check results for BUSCO: 261213at7742
## Thu Jul 14 02:29:14 CEST 2022 - The aliscore output file does not exist. Check results for BUSCO: 257768at7742
## Thu Jul 14 02:29:14 CEST 2022 - The aliscore output appears to be empty. Check results for BUSCO: 257768at7742
## Thu Jul 14 02:29:15 CEST 2022 - The aliscore output file does not exist. Check results for BUSCO: 381528at7742
## Thu Jul 14 02:29:15 CEST 2022 - The aliscore output appears to be empty. Check results for BUSCO: 381528at7742
## Thu Jul 14 02:29:17 CEST 2022 - The aliscore output file does not exist. Check results for BUSCO: 312646at7742
## Thu Jul 14 02:29:17 CEST 2022 - The aliscore output file does not exist. Check results for BUSCO: 368948at7742
## Thu Jul 14 02:29:17 CEST 2022 - The aliscore output appears to be empty. Check results for BUSCO: 312646at7742
## Thu Jul 14 02:29:17 CEST 2022 - The aliscore output appears to be empty. Check results for BUSCO: 368948at7742
## Thu Jul 14 02:29:18 CEST 2022 - The aliscore output file does not exist. Check results for BUSCO: 261072at7742
## Thu Jul 14 02:29:19 CEST 2022 - The aliscore output appears to be empty. Check results for BUSCO: 261072at7742
## Thu Jul 14 02:29:19 CEST 2022 - The aliscore output file does not exist. Check results for BUSCO: 238049at7742
## Thu Jul 14 02:29:19 CEST 2022 - The aliscore output appears to be empty. Check results for BUSCO: 238049at7742
## Thu Jul 14 02:29:19 CEST 2022 - The aliscore output file does not exist. Check results for BUSCO: 294990at7742
## Thu Jul 14 02:29:20 CEST 2022 - The aliscore output appears to be empty. Check results for BUSCO: 294990at7742
## Thu Jul 14 02:29:21 CEST 2022 - The aliscore output file does not exist. Check results for BUSCO: 274709at7742
## Thu Jul 14 02:29:21 CEST 2022 - The aliscore output appears to be empty. Check results for BUSCO: 274709at7742
## Thu Jul 14 02:29:22 CEST 2022 - The aliscore output file does not exist. Check results for BUSCO: 395686at7742
## Thu Jul 14 02:29:22 CEST 2022 - The aliscore output appears to be empty. Check results for BUSCO: 395686at7742
## Thu Jul 14 02:29:24 CEST 2022 - The aliscore output file does not exist. Check results for BUSCO: 368105at7742
## Thu Jul 14 02:29:24 CEST 2022 - The aliscore output appears to be empty. Check results for BUSCO: 368105at7742
## Thu Jul 14 02:29:24 CEST 2022 - The aliscore output file does not exist. Check results for BUSCO: 408438at7742
## Thu Jul 14 02:29:24 CEST 2022 - The aliscore output appears to be empty. Check results for BUSCO: 408438at7742
## Thu Jul 14 02:29:25 CEST 2022 - The aliscore output file does not exist. Check results for BUSCO: 387526at7742
## Thu Jul 14 02:29:25 CEST 2022 - The aliscore output appears to be empty. Check results for BUSCO: 387526at7742
## Thu Jul 14 02:29:26 CEST 2022 - The aliscore output file does not exist. Check results for BUSCO: 397031at7742
## Thu Jul 14 02:29:26 CEST 2022 - The aliscore output file does not exist. Check results for BUSCO: 368948at7742
## Thu Jul 14 02:29:26 CEST 2022 - The aliscore output appears to be empty. Check results for BUSCO: 397031at7742
## Thu Jul 14 02:29:26 CEST 2022 - The aliscore output appears to be empty. Check results for BUSCO: 368948at7742
## Thu Jul 14 02:29:27 CEST 2022 - The aliscore output file does not exist. Check results for BUSCO: 328506at7742
## Thu Jul 14 02:29:27 CEST 2022 - The aliscore output appears to be empty. Check results for BUSCO: 328506at7742
## Thu Jul 14 02:29:29 CEST 2022 - The aliscore output file does not exist. Check results for BUSCO: 362453at7742
## Thu Jul 14 02:29:29 CEST 2022 - The aliscore output appears to be empty. Check results for BUSCO: 362453at7742
## Thu Jul 14 02:29:29 CEST 2022 - The aliscore output file does not exist. Check results for BUSCO: 343489at7742
## Thu Jul 14 02:29:29 CEST 2022 - The aliscore output appears to be empty. Check results for BUSCO: 343489at7742
## Thu Jul 14 02:29:32 CEST 2022 - The aliscore output file does not exist. Check results for BUSCO: 361537at7742
## Thu Jul 14 02:29:32 CEST 2022 - The aliscore output appears to be empty. Check results for BUSCO: 361537at7742
## Thu Jul 14 02:29:33 CEST 2022 - The aliscore output file does not exist. Check results for BUSCO: 328147at7742
## Thu Jul 14 02:29:33 CEST 2022 - The aliscore output file does not exist. Check results for BUSCO: 361537at7742
## Thu Jul 14 02:29:33 CEST 2022 - The aliscore output appears to be empty. Check results for BUSCO: 328147at7742
## Thu Jul 14 02:29:33 CEST 2022 - The aliscore output appears to be empty. Check results for BUSCO: 361537at7742
## Thu Jul 14 02:29:34 CEST 2022 - The aliscore output file does not exist. Check results for BUSCO: 315832at7742
## Thu Jul 14 02:29:34 CEST 2022 - The aliscore output appears to be empty. Check results for BUSCO: 315832at7742
## Thu Jul 14 02:29:35 CEST 2022 - The aliscore output file does not exist. Check results for BUSCO: 386508at7742
## Thu Jul 14 02:29:35 CEST 2022 - The aliscore output appears to be empty. Check results for BUSCO: 386508at7742
## Thu Jul 14 02:29:35 CEST 2022 - The aliscore output file does not exist. Check results for BUSCO: 274709at7742
## Thu Jul 14 02:29:35 CEST 2022 - The aliscore output appears to be empty. Check results for BUSCO: 274709at7742
## Thu Jul 14 02:29:36 CEST 2022 - The aliscore output file does not exist. Check results for BUSCO: 274709at7742
## Thu Jul 14 02:29:36 CEST 2022 - The aliscore output appears to be empty. Check results for BUSCO: 274709at7742
## Thu Jul 14 02:29:37 CEST 2022 - The aliscore output file does not exist. Check results for BUSCO: 359738at7742
## Thu Jul 14 02:29:37 CEST 2022 - The aliscore output appears to be empty. Check results for BUSCO: 359738at7742
## Thu Jul 14 02:29:38 CEST 2022 - The aliscore output file does not exist. Check results for BUSCO: 279816at7742
## Thu Jul 14 02:29:38 CEST 2022 - The aliscore output appears to be empty. Check results for BUSCO: 279816at7742
## Thu Jul 14 02:29:39 CEST 2022 - The aliscore output file does not exist. Check results for BUSCO: 413952at7742
## Thu Jul 14 02:29:39 CEST 2022 - The aliscore output appears to be empty. Check results for BUSCO: 413952at7742
## Thu Jul 14 02:29:40 CEST 2022 - The aliscore output file does not exist. Check results for BUSCO: 347416at7742
## Thu Jul 14 02:29:40 CEST 2022 - The aliscore output appears to be empty. Check results for BUSCO: 347416at7742
## Thu Jul 14 02:29:40 CEST 2022 - The aliscore output file does not exist. Check results for BUSCO: 316119at7742
## Thu Jul 14 02:29:41 CEST 2022 - The aliscore output appears to be empty. Check results for BUSCO: 316119at7742
## Thu Jul 14 02:29:42 CEST 2022 - The aliscore output file does not exist. Check results for BUSCO: 384558at7742
## Thu Jul 14 02:29:42 CEST 2022 - The aliscore output appears to be empty. Check results for BUSCO: 384558at7742
## Thu Jul 14 02:29:42 CEST 2022 - The aliscore output file does not exist. Check results for BUSCO: 413352at7742
## Thu Jul 14 02:29:42 CEST 2022 - The aliscore output appears to be empty. Check results for BUSCO: 413352at7742
## Thu Jul 14 02:29:43 CEST 2022 - The aliscore output file does not exist. Check results for BUSCO: 307391at7742
## Thu Jul 14 02:29:44 CEST 2022 - The aliscore output appears to be empty. Check results for BUSCO: 307391at7742
## Thu Jul 14 02:29:44 CEST 2022 - The aliscore output file does not exist. Check results for BUSCO: 317712at7742
## Thu Jul 14 02:29:44 CEST 2022 - The aliscore output appears to be empty. Check results for BUSCO: 317712at7742
## Thu Jul 14 02:29:45 CEST 2022 - The aliscore output file does not exist. Check results for BUSCO: 208372at7742
## Thu Jul 14 02:29:45 CEST 2022 - The aliscore output appears to be empty. Check results for BUSCO: 208372at7742
## Thu Jul 14 02:29:46 CEST 2022 - The aliscore output file does not exist. Check results for BUSCO: 400874at7742
## Thu Jul 14 02:29:46 CEST 2022 - The aliscore output appears to be empty. Check results for BUSCO: 400874at7742
## Thu Jul 14 02:29:48 CEST 2022 - The aliscore output file does not exist. Check results for BUSCO: 415928at7742
## Thu Jul 14 02:29:48 CEST 2022 - The aliscore output appears to be empty. Check results for BUSCO: 415928at7742
## Thu Jul 14 02:29:48 CEST 2022 - The aliscore output file does not exist. Check results for BUSCO: 368105at7742
## Thu Jul 14 02:29:48 CEST 2022 - The aliscore output appears to be empty. Check results for BUSCO: 368105at7742
## Thu Jul 14 02:29:49 CEST 2022 - The aliscore output file does not exist. Check results for BUSCO: 381867at7742
## Thu Jul 14 02:29:49 CEST 2022 - The aliscore output appears to be empty. Check results for BUSCO: 381867at7742
## Thu Jul 14 02:29:50 CEST 2022 - The aliscore output file does not exist. Check results for BUSCO: 330996at7742
## Thu Jul 14 02:29:50 CEST 2022 - The aliscore output file does not exist. Check results for BUSCO: 319099at7742
## Thu Jul 14 02:29:50 CEST 2022 - The aliscore output appears to be empty. Check results for BUSCO: 330996at7742
## Thu Jul 14 02:29:50 CEST 2022 - The aliscore output appears to be empty. Check results for BUSCO: 319099at7742
## Thu Jul 14 02:29:52 CEST 2022 - The aliscore output file does not exist. Check results for BUSCO: 386039at7742
## Thu Jul 14 02:29:52 CEST 2022 - The aliscore output appears to be empty. Check results for BUSCO: 386039at7742
## Thu Jul 14 02:29:52 CEST 2022 - The aliscore output file does not exist. Check results for BUSCO: 300899at7742
## Thu Jul 14 02:29:52 CEST 2022 - The aliscore output appears to be empty. Check results for BUSCO: 300899at7742
## Thu Jul 14 02:29:53 CEST 2022 - The aliscore output file does not exist. Check results for BUSCO: 320027at7742
## Thu Jul 14 02:29:53 CEST 2022 - The aliscore output appears to be empty. Check results for BUSCO: 320027at7742
## Thu Jul 14 02:29:54 CEST 2022 - The aliscore output file does not exist. Check results for BUSCO: 293276at7742
## Thu Jul 14 02:29:54 CEST 2022 - The aliscore output appears to be empty. Check results for BUSCO: 293276at7742
## Thu Jul 14 02:29:56 CEST 2022 - The aliscore output file does not exist. Check results for BUSCO: 288175at7742
## Thu Jul 14 02:29:56 CEST 2022 - The aliscore output appears to be empty. Check results for BUSCO: 288175at7742
## Thu Jul 14 02:29:56 CEST 2022 - The aliscore output file does not exist. Check results for BUSCO: 330996at7742
## Thu Jul 14 02:29:56 CEST 2022 - The aliscore output appears to be empty. Check results for BUSCO: 330996at7742
## Thu Jul 14 02:29:58 CEST 2022 - The aliscore output file does not exist. Check results for BUSCO: 295613at7742
## Thu Jul 14 02:29:58 CEST 2022 - The aliscore output appears to be empty. Check results for BUSCO: 295613at7742
## Thu Jul 14 02:29:58 CEST 2022 - The aliscore output file does not exist. Check results for BUSCO: 346436at7742
## Thu Jul 14 02:29:58 CEST 2022 - The aliscore output appears to be empty. Check results for BUSCO: 346436at7742
## Thu Jul 14 02:30:00 CEST 2022 - The aliscore output file does not exist. Check results for BUSCO: 261213at7742
## Thu Jul 14 02:30:00 CEST 2022 - The aliscore output appears to be empty. Check results for BUSCO: 261213at7742
## Thu Jul 14 02:30:00 CEST 2022 - The aliscore output file does not exist. Check results for BUSCO: 359541at7742
## Thu Jul 14 02:30:00 CEST 2022 - The aliscore output appears to be empty. Check results for BUSCO: 359541at7742
## Thu Jul 14 02:30:01 CEST 2022 - The aliscore output file does not exist. Check results for BUSCO: 326141at7742
## Thu Jul 14 02:30:01 CEST 2022 - The aliscore output appears to be empty. Check results for BUSCO: 326141at7742
## Thu Jul 14 02:30:02 CEST 2022 - The aliscore output file does not exist. Check results for BUSCO: 306239at7742
## Thu Jul 14 02:30:02 CEST 2022 - The aliscore output appears to be empty. Check results for BUSCO: 306239at7742
## Thu Jul 14 02:30:03 CEST 2022 - The aliscore output file does not exist. Check results for BUSCO: 322470at7742
## Thu Jul 14 02:30:03 CEST 2022 - The aliscore output appears to be empty. Check results for BUSCO: 322470at7742
## Thu Jul 14 02:30:05 CEST 2022 - The aliscore output file does not exist. Check results for BUSCO: 240576at7742
## Thu Jul 14 02:30:05 CEST 2022 - The aliscore output appears to be empty. Check results for BUSCO: 240576at7742
## Thu Jul 14 02:30:05 CEST 2022 - The aliscore output file does not exist. Check results for BUSCO: 404882at7742
## Thu Jul 14 02:30:05 CEST 2022 - The aliscore output appears to be empty. Check results for BUSCO: 404882at7742
## Thu Jul 14 02:30:07 CEST 2022 - The aliscore output file does not exist. Check results for BUSCO: 373225at7742
## Thu Jul 14 02:30:07 CEST 2022 - The aliscore output appears to be empty. Check results for BUSCO: 373225at7742
## Thu Jul 14 02:30:07 CEST 2022 - The aliscore output file does not exist. Check results for BUSCO: 255237at7742
## Thu Jul 14 02:30:07 CEST 2022 - The aliscore output appears to be empty. Check results for BUSCO: 255237at7742
## Thu Jul 14 02:30:08 CEST 2022 - The aliscore output file does not exist. Check results for BUSCO: 319137at7742
## Thu Jul 14 02:30:08 CEST 2022 - The aliscore output appears to be empty. Check results for BUSCO: 319137at7742
## Thu Jul 14 02:30:08 CEST 2022 - The aliscore output file does not exist. Check results for BUSCO: 328147at7742
## Thu Jul 14 02:30:08 CEST 2022 - The aliscore output appears to be empty. Check results for BUSCO: 328147at7742
## Thu Jul 14 02:30:09 CEST 2022 - The aliscore output file does not exist. Check results for BUSCO: 298518at7742
## Thu Jul 14 02:30:09 CEST 2022 - The aliscore output appears to be empty. Check results for BUSCO: 298518at7742
## Thu Jul 14 02:30:11 CEST 2022 - The aliscore output file does not exist. Check results for BUSCO: 326141at7742
## Thu Jul 14 02:30:11 CEST 2022 - The aliscore output appears to be empty. Check results for BUSCO: 326141at7742
## Thu Jul 14 02:30:12 CEST 2022 - The aliscore output file does not exist. Check results for BUSCO: 331986at7742
## Thu Jul 14 02:30:12 CEST 2022 - The aliscore output appears to be empty. Check results for BUSCO: 331986at7742
## Thu Jul 14 02:30:12 CEST 2022 - The aliscore output file does not exist. Check results for BUSCO: 364898at7742
## Thu Jul 14 02:30:12 CEST 2022 - The aliscore output appears to be empty. Check results for BUSCO: 364898at7742
## Thu Jul 14 02:30:12 CEST 2022 - The aliscore output file does not exist. Check results for BUSCO: 260364at7742
## Thu Jul 14 02:30:12 CEST 2022 - The aliscore output appears to be empty. Check results for BUSCO: 260364at7742
## Thu Jul 14 02:30:14 CEST 2022 - The aliscore output file does not exist. Check results for BUSCO: 346605at7742
## Thu Jul 14 02:30:14 CEST 2022 - The aliscore output appears to be empty. Check results for BUSCO: 346605at7742
## Thu Jul 14 02:30:15 CEST 2022 - The aliscore output file does not exist. Check results for BUSCO: 415822at7742
## Thu Jul 14 02:30:15 CEST 2022 - The aliscore output appears to be empty. Check results for BUSCO: 415822at7742
## Thu Jul 14 02:30:16 CEST 2022 - The aliscore output file does not exist. Check results for BUSCO: 271347at7742
## Thu Jul 14 02:30:16 CEST 2022 - The aliscore output appears to be empty. Check results for BUSCO: 271347at7742
## Thu Jul 14 02:30:17 CEST 2022 - The aliscore output file does not exist. Check results for BUSCO: 208372at7742
## Thu Jul 14 02:30:17 CEST 2022 - The aliscore output appears to be empty. Check results for BUSCO: 208372at7742
## Thu Jul 14 02:30:18 CEST 2022 - The aliscore output file does not exist. Check results for BUSCO: 300899at7742
## Thu Jul 14 02:30:18 CEST 2022 - The aliscore output appears to be empty. Check results for BUSCO: 300899at7742
## Thu Jul 14 02:30:18 CEST 2022 - The aliscore output file does not exist. Check results for BUSCO: 255237at7742
## Thu Jul 14 02:30:18 CEST 2022 - The aliscore output appears to be empty. Check results for BUSCO: 255237at7742
## Thu Jul 14 02:30:19 CEST 2022 - The aliscore output file does not exist. Check results for BUSCO: 280287at7742
## Thu Jul 14 02:30:19 CEST 2022 - The aliscore output appears to be empty. Check results for BUSCO: 280287at7742
## Thu Jul 14 02:30:20 CEST 2022 - The aliscore output file does not exist. Check results for BUSCO: 297797at7742
## Thu Jul 14 02:30:20 CEST 2022 - The aliscore output appears to be empty. Check results for BUSCO: 297797at7742
## Thu Jul 14 02:30:20 CEST 2022 - The aliscore output file does not exist. Check results for BUSCO: 327515at7742
## Thu Jul 14 02:30:20 CEST 2022 - The aliscore output appears to be empty. Check results for BUSCO: 327515at7742
## Thu Jul 14 02:30:22 CEST 2022 - The aliscore output file does not exist. Check results for BUSCO: 337042at7742
## Thu Jul 14 02:30:22 CEST 2022 - The aliscore output appears to be empty. Check results for BUSCO: 337042at7742
## Thu Jul 14 02:30:23 CEST 2022 - The aliscore output file does not exist. Check results for BUSCO: 208372at7742
## Thu Jul 14 02:30:23 CEST 2022 - The aliscore output appears to be empty. Check results for BUSCO: 208372at7742
## Thu Jul 14 02:30:24 CEST 2022 - The aliscore output file does not exist. Check results for BUSCO: 307391at7742
## Thu Jul 14 02:30:24 CEST 2022 - The aliscore output appears to be empty. Check results for BUSCO: 307391at7742
## Thu Jul 14 02:30:24 CEST 2022 - The aliscore output file does not exist. Check results for BUSCO: 299325at7742
## Thu Jul 14 02:30:24 CEST 2022 - The aliscore output appears to be empty. Check results for BUSCO: 299325at7742
## Thu Jul 14 02:30:26 CEST 2022 - The aliscore output file does not exist. Check results for BUSCO: 266654at7742
## Thu Jul 14 02:30:26 CEST 2022 - The aliscore output appears to be empty. Check results for BUSCO: 266654at7742
## Thu Jul 14 02:30:26 CEST 2022 - The aliscore output file does not exist. Check results for BUSCO: 285180at7742
## Thu Jul 14 02:30:26 CEST 2022 - The aliscore output appears to be empty. Check results for BUSCO: 285180at7742
## Thu Jul 14 02:30:27 CEST 2022 - The aliscore output file does not exist. Check results for BUSCO: 345556at7742
## Thu Jul 14 02:30:28 CEST 2022 - The aliscore output appears to be empty. Check results for BUSCO: 345556at7742
## Thu Jul 14 02:30:29 CEST 2022 - The aliscore output file does not exist. Check results for BUSCO: 333591at7742
## Thu Jul 14 02:30:29 CEST 2022 - The aliscore output appears to be empty. Check results for BUSCO: 333591at7742
## Thu Jul 14 02:30:29 CEST 2022 - The aliscore output file does not exist. Check results for BUSCO: 414793at7742
## Thu Jul 14 02:30:30 CEST 2022 - The aliscore output appears to be empty. Check results for BUSCO: 414793at7742
## Thu Jul 14 02:30:31 CEST 2022 - The aliscore output file does not exist. Check results for BUSCO: 325223at7742
## Thu Jul 14 02:30:31 CEST 2022 - The aliscore output appears to be empty. Check results for BUSCO: 325223at7742
## Thu Jul 14 02:30:31 CEST 2022 - The aliscore output file does not exist. Check results for BUSCO: 358562at7742
## Thu Jul 14 02:30:31 CEST 2022 - The aliscore output appears to be empty. Check results for BUSCO: 358562at7742
## Thu Jul 14 02:30:32 CEST 2022 - The aliscore output file does not exist. Check results for BUSCO: 291607at7742
## Thu Jul 14 02:30:32 CEST 2022 - The aliscore output appears to be empty. Check results for BUSCO: 291607at7742
## Thu Jul 14 02:30:34 CEST 2022 - The aliscore output file does not exist. Check results for BUSCO: 358920at7742
## Thu Jul 14 02:30:34 CEST 2022 - The aliscore output file does not exist. Check results for BUSCO: 306239at7742
## Thu Jul 14 02:30:34 CEST 2022 - The aliscore output appears to be empty. Check results for BUSCO: 358920at7742
## Thu Jul 14 02:30:34 CEST 2022 - The aliscore output appears to be empty. Check results for BUSCO: 306239at7742
## Thu Jul 14 02:30:36 CEST 2022 - The aliscore output file does not exist. Check results for BUSCO: 255237at7742
## Thu Jul 14 02:30:36 CEST 2022 - The aliscore output appears to be empty. Check results for BUSCO: 255237at7742
## Thu Jul 14 02:30:36 CEST 2022 - The aliscore output file does not exist. Check results for BUSCO: 278384at7742
## Thu Jul 14 02:30:36 CEST 2022 - The aliscore output appears to be empty. Check results for BUSCO: 278384at7742
## Thu Jul 14 02:30:37 CEST 2022 - The aliscore output file does not exist. Check results for BUSCO: 342192at7742
## Thu Jul 14 02:30:37 CEST 2022 - The aliscore output appears to be empty. Check results for BUSCO: 342192at7742
## Thu Jul 14 02:30:38 CEST 2022 - The aliscore output file does not exist. Check results for BUSCO: 240576at7742
## Thu Jul 14 02:30:38 CEST 2022 - The aliscore output appears to be empty. Check results for BUSCO: 240576at7742
## Thu Jul 14 02:30:39 CEST 2022 - The aliscore output file does not exist. Check results for BUSCO: 315746at7742
## Thu Jul 14 02:30:39 CEST 2022 - The aliscore output appears to be empty. Check results for BUSCO: 315746at7742
## Thu Jul 14 02:30:39 CEST 2022 - The aliscore output file does not exist. Check results for BUSCO: 315746at7742
## Thu Jul 14 02:30:39 CEST 2022 - The aliscore output appears to be empty. Check results for BUSCO: 315746at7742
## Thu Jul 14 02:30:40 CEST 2022 - The aliscore output file does not exist. Check results for BUSCO: 413352at7742
## Thu Jul 14 02:30:40 CEST 2022 - The aliscore output appears to be empty. Check results for BUSCO: 413352at7742
## Thu Jul 14 02:30:42 CEST 2022 - The aliscore output file does not exist. Check results for BUSCO: 238049at7742
## Thu Jul 14 02:30:42 CEST 2022 - The aliscore output appears to be empty. Check results for BUSCO: 238049at7742
## Thu Jul 14 02:30:43 CEST 2022 - The aliscore output file does not exist. Check results for BUSCO: 406306at7742
## Thu Jul 14 02:30:43 CEST 2022 - The aliscore output appears to be empty. Check results for BUSCO: 406306at7742
## Thu Jul 14 02:30:44 CEST 2022 - The aliscore output file does not exist. Check results for BUSCO: 401745at7742
## Thu Jul 14 02:30:44 CEST 2022 - The aliscore output appears to be empty. Check results for BUSCO: 401745at7742
## Thu Jul 14 02:30:45 CEST 2022 - The aliscore output file does not exist. Check results for BUSCO: 297797at7742
## Thu Jul 14 02:30:45 CEST 2022 - The aliscore output appears to be empty. Check results for BUSCO: 297797at7742
## Thu Jul 14 02:30:45 CEST 2022 - The aliscore output file does not exist. Check results for BUSCO: 325223at7742
## Thu Jul 14 02:30:45 CEST 2022 - The aliscore output appears to be empty. Check results for BUSCO: 325223at7742
## Thu Jul 14 02:30:46 CEST 2022 - The aliscore output file does not exist. Check results for BUSCO: 306239at7742
## Thu Jul 14 02:30:46 CEST 2022 - The aliscore output appears to be empty. Check results for BUSCO: 306239at7742
## Thu Jul 14 02:30:47 CEST 2022 - The aliscore output file does not exist. Check results for BUSCO: 330614at7742
## Thu Jul 14 02:30:47 CEST 2022 - The aliscore output appears to be empty. Check results for BUSCO: 330614at7742
## Thu Jul 14 02:30:48 CEST 2022 - The aliscore output file does not exist. Check results for BUSCO: 393010at7742
## Thu Jul 14 02:30:48 CEST 2022 - The aliscore output appears to be empty. Check results for BUSCO: 393010at7742
## Thu Jul 14 02:30:49 CEST 2022 - The aliscore output file does not exist. Check results for BUSCO: 335473at7742
## Thu Jul 14 02:30:49 CEST 2022 - The aliscore output appears to be empty. Check results for BUSCO: 335473at7742
## Thu Jul 14 02:30:50 CEST 2022 - The aliscore output file does not exist. Check results for BUSCO: 336764at7742
## Thu Jul 14 02:30:50 CEST 2022 - The aliscore output appears to be empty. Check results for BUSCO: 336764at7742
## Thu Jul 14 02:30:50 CEST 2022 - The aliscore output file does not exist. Check results for BUSCO: 327515at7742
## Thu Jul 14 02:30:50 CEST 2022 - The aliscore output appears to be empty. Check results for BUSCO: 327515at7742
## Thu Jul 14 02:30:52 CEST 2022 - The aliscore output file does not exist. Check results for BUSCO: 265531at7742
## Thu Jul 14 02:30:52 CEST 2022 - The aliscore output appears to be empty. Check results for BUSCO: 265531at7742
## Thu Jul 14 02:30:53 CEST 2022 - The aliscore output file does not exist. Check results for BUSCO: 373225at7742
## Thu Jul 14 02:30:53 CEST 2022 - The aliscore output appears to be empty. Check results for BUSCO: 373225at7742
## Thu Jul 14 02:30:53 CEST 2022 - The aliscore output file does not exist. Check results for BUSCO: 295101at7742
## Thu Jul 14 02:30:53 CEST 2022 - The aliscore output appears to be empty. Check results for BUSCO: 295101at7742
## Thu Jul 14 02:30:55 CEST 2022 - The aliscore output file does not exist. Check results for BUSCO: 387408at7742
## Thu Jul 14 02:30:55 CEST 2022 - The aliscore output appears to be empty. Check results for BUSCO: 387408at7742
## Thu Jul 14 02:30:55 CEST 2022 - The aliscore output file does not exist. Check results for BUSCO: 325223at7742
## Thu Jul 14 02:30:55 CEST 2022 - The aliscore output appears to be empty. Check results for BUSCO: 325223at7742
## Thu Jul 14 02:30:56 CEST 2022 - The aliscore output file does not exist. Check results for BUSCO: 278647at7742
## Thu Jul 14 02:30:56 CEST 2022 - The aliscore output appears to be empty. Check results for BUSCO: 278647at7742
## Thu Jul 14 02:30:57 CEST 2022 - The aliscore output file does not exist. Check results for BUSCO: 388403at7742
## Thu Jul 14 02:30:57 CEST 2022 - The aliscore output appears to be empty. Check results for BUSCO: 388403at7742
## Thu Jul 14 02:30:58 CEST 2022 - The aliscore output file does not exist. Check results for BUSCO: 285488at7742
## Thu Jul 14 02:30:58 CEST 2022 - The aliscore output appears to be empty. Check results for BUSCO: 285488at7742
## Thu Jul 14 02:31:00 CEST 2022 - The aliscore output file does not exist. Check results for BUSCO: 355430at7742
## Thu Jul 14 02:31:00 CEST 2022 - The aliscore output appears to be empty. Check results for BUSCO: 355430at7742
## Thu Jul 14 02:31:01 CEST 2022 - The aliscore output file does not exist. Check results for BUSCO: 273713at7742
## Thu Jul 14 02:31:01 CEST 2022 - The aliscore output appears to be empty. Check results for BUSCO: 273713at7742
## Thu Jul 14 02:31:02 CEST 2022 - The aliscore output file does not exist. Check results for BUSCO: 271347at7742
## Thu Jul 14 02:31:02 CEST 2022 - The aliscore output appears to be empty. Check results for BUSCO: 271347at7742
## Thu Jul 14 02:31:03 CEST 2022 - The aliscore output file does not exist. Check results for BUSCO: 312987at7742
## Thu Jul 14 02:31:03 CEST 2022 - The aliscore output appears to be empty. Check results for BUSCO: 312987at7742
## Thu Jul 14 02:31:03 CEST 2022 - The aliscore output file does not exist. Check results for BUSCO: 365662at7742
## Thu Jul 14 02:31:03 CEST 2022 - The aliscore output appears to be empty. Check results for BUSCO: 365662at7742
## Thu Jul 14 02:31:04 CEST 2022 - The aliscore output file does not exist. Check results for BUSCO: 309129at7742
## Thu Jul 14 02:31:04 CEST 2022 - The aliscore output appears to be empty. Check results for BUSCO: 309129at7742
## Thu Jul 14 02:31:04 CEST 2022 - The aliscore output file does not exist. Check results for BUSCO: 414793at7742
## Thu Jul 14 02:31:04 CEST 2022 - The aliscore output appears to be empty. Check results for BUSCO: 414793at7742
## Thu Jul 14 02:31:05 CEST 2022 - The aliscore output file does not exist. Check results for BUSCO: 376607at7742
## Thu Jul 14 02:31:05 CEST 2022 - The aliscore output appears to be empty. Check results for BUSCO: 376607at7742
## Thu Jul 14 02:31:06 CEST 2022 - The aliscore output file does not exist. Check results for BUSCO: 342192at7742
## Thu Jul 14 02:31:06 CEST 2022 - The aliscore output appears to be empty. Check results for BUSCO: 342192at7742
## Thu Jul 14 02:31:08 CEST 2022 - The aliscore output file does not exist. Check results for BUSCO: 291607at7742
## Thu Jul 14 02:31:08 CEST 2022 - The aliscore output file does not exist. Check results for BUSCO: 283060at7742
## Thu Jul 14 02:31:08 CEST 2022 - The aliscore output appears to be empty. Check results for BUSCO: 291607at7742
## Thu Jul 14 02:31:08 CEST 2022 - The aliscore output appears to be empty. Check results for BUSCO: 283060at7742
## Thu Jul 14 02:31:09 CEST 2022 - The aliscore output file does not exist. Check results for BUSCO: 408438at7742
## Thu Jul 14 02:31:09 CEST 2022 - The aliscore output appears to be empty. Check results for BUSCO: 408438at7742
## Thu Jul 14 02:31:10 CEST 2022 - The aliscore output file does not exist. Check results for BUSCO: 357356at7742
## Thu Jul 14 02:31:10 CEST 2022 - The aliscore output appears to be empty. Check results for BUSCO: 357356at7742
## Thu Jul 14 02:31:11 CEST 2022 - The aliscore output file does not exist. Check results for BUSCO: 389517at7742
## Thu Jul 14 02:31:11 CEST 2022 - The aliscore output appears to be empty. Check results for BUSCO: 389517at7742
## Thu Jul 14 02:31:13 CEST 2022 - The aliscore output file does not exist. Check results for BUSCO: 321832at7742
## Thu Jul 14 02:31:13 CEST 2022 - The aliscore output appears to be empty. Check results for BUSCO: 321832at7742
## Thu Jul 14 02:31:13 CEST 2022 - The aliscore output file does not exist. Check results for BUSCO: 356030at7742
## Thu Jul 14 02:31:13 CEST 2022 - The aliscore output appears to be empty. Check results for BUSCO: 356030at7742
## Thu Jul 14 02:31:15 CEST 2022 - The aliscore output file does not exist. Check results for BUSCO: 286842at7742
## Thu Jul 14 02:31:15 CEST 2022 - The aliscore output appears to be empty. Check results for BUSCO: 286842at7742
## Thu Jul 14 02:31:15 CEST 2022 - The aliscore output file does not exist. Check results for BUSCO: 352153at7742
## Thu Jul 14 02:31:15 CEST 2022 - The aliscore output appears to be empty. Check results for BUSCO: 352153at7742
## Thu Jul 14 02:31:16 CEST 2022 - The aliscore output file does not exist. Check results for BUSCO: 286842at7742
## Thu Jul 14 02:31:16 CEST 2022 - The aliscore output appears to be empty. Check results for BUSCO: 286842at7742
## Thu Jul 14 02:31:16 CEST 2022 - The aliscore output file does not exist. Check results for BUSCO: 295613at7742
## Thu Jul 14 02:31:16 CEST 2022 - The aliscore output appears to be empty. Check results for BUSCO: 295613at7742
## Thu Jul 14 02:31:17 CEST 2022 - The aliscore output file does not exist. Check results for BUSCO: 286842at7742
## Thu Jul 14 02:31:18 CEST 2022 - The aliscore output appears to be empty. Check results for BUSCO: 286842at7742
## Thu Jul 14 02:31:20 CEST 2022 - The aliscore output file does not exist. Check results for BUSCO: 403644at7742
## Thu Jul 14 02:31:20 CEST 2022 - The aliscore output appears to be empty. Check results for BUSCO: 403644at7742
## Thu Jul 14 02:31:20 CEST 2022 - The aliscore output file does not exist. Check results for BUSCO: 403644at7742
## Thu Jul 14 02:31:20 CEST 2022 - The aliscore output appears to be empty. Check results for BUSCO: 403644at7742
## Thu Jul 14 02:31:21 CEST 2022 - The aliscore output file does not exist. Check results for BUSCO: 316119at7742
## Thu Jul 14 02:31:21 CEST 2022 - The aliscore output appears to be empty. Check results for BUSCO: 316119at7742
## Thu Jul 14 02:31:21 CEST 2022 - The aliscore output file does not exist. Check results for BUSCO: 299325at7742
## Thu Jul 14 02:31:21 CEST 2022 - The aliscore output appears to be empty. Check results for BUSCO: 299325at7742
## Thu Jul 14 02:31:22 CEST 2022 - The aliscore output file does not exist. Check results for BUSCO: 277508at7742
## Thu Jul 14 02:31:22 CEST 2022 - The aliscore output appears to be empty. Check results for BUSCO: 277508at7742
## Thu Jul 14 02:31:23 CEST 2022 - The aliscore output file does not exist. Check results for BUSCO: 280016at7742
## Thu Jul 14 02:31:24 CEST 2022 - The aliscore output appears to be empty. Check results for BUSCO: 280016at7742
## Thu Jul 14 02:31:24 CEST 2022 - The aliscore output file does not exist. Check results for BUSCO: 316119at7742
## Thu Jul 14 02:31:24 CEST 2022 - The aliscore output appears to be empty. Check results for BUSCO: 316119at7742
## Thu Jul 14 02:31:25 CEST 2022 - The aliscore output file does not exist. Check results for BUSCO: 355430at7742
## Thu Jul 14 02:31:25 CEST 2022 - The aliscore output appears to be empty. Check results for BUSCO: 355430at7742
## Thu Jul 14 02:31:27 CEST 2022 - The aliscore output file does not exist. Check results for BUSCO: 397031at7742
## Thu Jul 14 02:31:27 CEST 2022 - The aliscore output appears to be empty. Check results for BUSCO: 397031at7742
## Thu Jul 14 02:31:28 CEST 2022 - The aliscore output file does not exist. Check results for BUSCO: 414793at7742
## Thu Jul 14 02:31:28 CEST 2022 - The aliscore output appears to be empty. Check results for BUSCO: 414793at7742
## Thu Jul 14 02:31:29 CEST 2022 - The aliscore output file does not exist. Check results for BUSCO: 271347at7742
## Thu Jul 14 02:31:29 CEST 2022 - The aliscore output appears to be empty. Check results for BUSCO: 271347at7742
## Thu Jul 14 02:31:29 CEST 2022 - The aliscore output file does not exist. Check results for BUSCO: 323530at7742
## Thu Jul 14 02:31:29 CEST 2022 - The aliscore output appears to be empty. Check results for BUSCO: 323530at7742
## Thu Jul 14 02:31:31 CEST 2022 - The aliscore output file does not exist. Check results for BUSCO: 297797at7742
## Thu Jul 14 02:31:31 CEST 2022 - The aliscore output appears to be empty. Check results for BUSCO: 297797at7742
## Thu Jul 14 02:31:31 CEST 2022 - The aliscore output file does not exist. Check results for BUSCO: 323530at7742
## Thu Jul 14 02:31:31 CEST 2022 - The aliscore output appears to be empty. Check results for BUSCO: 323530at7742
## Thu Jul 14 02:31:31 CEST 2022 - The aliscore output file does not exist. Check results for BUSCO: 265531at7742
## Thu Jul 14 02:31:32 CEST 2022 - The aliscore output appears to be empty. Check results for BUSCO: 265531at7742
## Thu Jul 14 02:31:33 CEST 2022 - The aliscore output file does not exist. Check results for BUSCO: 291607at7742
## Thu Jul 14 02:31:33 CEST 2022 - The aliscore output appears to be empty. Check results for BUSCO: 291607at7742
## Thu Jul 14 02:31:33 CEST 2022 - The aliscore output file does not exist. Check results for BUSCO: 240576at7742
## Thu Jul 14 02:31:33 CEST 2022 - The aliscore output appears to be empty. Check results for BUSCO: 240576at7742
## Thu Jul 14 02:31:34 CEST 2022 - The aliscore output file does not exist. Check results for BUSCO: 344798at7742
## Thu Jul 14 02:31:34 CEST 2022 - The aliscore output appears to be empty. Check results for BUSCO: 344798at7742
## Thu Jul 14 02:31:36 CEST 2022 - The aliscore output file does not exist. Check results for BUSCO: 384470at7742
## Thu Jul 14 02:31:36 CEST 2022 - The aliscore output file does not exist. Check results for BUSCO: 333591at7742
## Thu Jul 14 02:31:36 CEST 2022 - The aliscore output appears to be empty. Check results for BUSCO: 384470at7742
## Thu Jul 14 02:31:36 CEST 2022 - The aliscore output appears to be empty. Check results for BUSCO: 333591at7742
## Thu Jul 14 02:31:37 CEST 2022 - The aliscore output file does not exist. Check results for BUSCO: 342192at7742
## Thu Jul 14 02:31:37 CEST 2022 - The aliscore output appears to be empty. Check results for BUSCO: 342192at7742
## Thu Jul 14 02:31:39 CEST 2022 - The aliscore output file does not exist. Check results for BUSCO: 397807at7742
## Thu Jul 14 02:31:39 CEST 2022 - The aliscore output appears to be empty. Check results for BUSCO: 397807at7742
## Thu Jul 14 02:31:39 CEST 2022 - The aliscore output file does not exist. Check results for BUSCO: 393341at7742
## Thu Jul 14 02:31:39 CEST 2022 - The aliscore output appears to be empty. Check results for BUSCO: 393341at7742
## Thu Jul 14 02:31:41 CEST 2022 - The aliscore output file does not exist. Check results for BUSCO: 316607at7742
## Thu Jul 14 02:31:41 CEST 2022 - The aliscore output appears to be empty. Check results for BUSCO: 316607at7742
## Thu Jul 14 02:31:41 CEST 2022 - The aliscore output file does not exist. Check results for BUSCO: 316607at7742
## Thu Jul 14 02:31:41 CEST 2022 - The aliscore output appears to be empty. Check results for BUSCO: 316607at7742
## Thu Jul 14 02:31:43 CEST 2022 - The aliscore output file does not exist. Check results for BUSCO: 335473at7742
## Thu Jul 14 02:31:43 CEST 2022 - The aliscore output appears to be empty. Check results for BUSCO: 335473at7742
## Thu Jul 14 02:31:43 CEST 2022 - The aliscore output file does not exist. Check results for BUSCO: 389517at7742
## Thu Jul 14 02:31:43 CEST 2022 - The aliscore output appears to be empty. Check results for BUSCO: 389517at7742
## Thu Jul 14 02:31:44 CEST 2022 - The aliscore output file does not exist. Check results for BUSCO: 315750at7742
## Thu Jul 14 02:31:44 CEST 2022 - The aliscore output appears to be empty. Check results for BUSCO: 315750at7742
## Thu Jul 14 02:31:45 CEST 2022 - The aliscore output file does not exist. Check results for BUSCO: 365662at7742
## Thu Jul 14 02:31:45 CEST 2022 - The aliscore output appears to be empty. Check results for BUSCO: 365662at7742
## Thu Jul 14 02:31:46 CEST 2022 - The aliscore output file does not exist. Check results for BUSCO: 315746at7742
## Thu Jul 14 02:31:46 CEST 2022 - The aliscore output appears to be empty. Check results for BUSCO: 315746at7742
## Thu Jul 14 02:31:48 CEST 2022 - The aliscore output file does not exist. Check results for BUSCO: 362453at7742
## Thu Jul 14 02:31:48 CEST 2022 - The aliscore output appears to be empty. Check results for BUSCO: 362453at7742
## Thu Jul 14 02:31:48 CEST 2022 - The aliscore output file does not exist. Check results for BUSCO: 280016at7742
## Thu Jul 14 02:31:49 CEST 2022 - The aliscore output appears to be empty. Check results for BUSCO: 280016at7742
## Thu Jul 14 02:31:49 CEST 2022 - The aliscore output file does not exist. Check results for BUSCO: 280016at7742
## Thu Jul 14 02:31:49 CEST 2022 - The aliscore output file does not exist. Check results for BUSCO: 363627at7742
## Thu Jul 14 02:31:49 CEST 2022 - The aliscore output appears to be empty. Check results for BUSCO: 280016at7742
## Thu Jul 14 02:31:49 CEST 2022 - The aliscore output appears to be empty. Check results for BUSCO: 363627at7742
## Thu Jul 14 02:31:50 CEST 2022 - The aliscore output file does not exist. Check results for BUSCO: 265531at7742
## Thu Jul 14 02:31:50 CEST 2022 - The ALICUT output appears to be empty. Will touch an empty file so the pipeline will continue. Check results for BUSCO: 265531at7742
## Thu Jul 14 02:31:53 CEST 2022 - The aliscore output file does not exist. Check results for BUSCO: 390017at7742
## Thu Jul 14 02:31:53 CEST 2022 - The aliscore output appears to be empty. Check results for BUSCO: 390017at7742
## Thu Jul 14 02:31:53 CEST 2022 - The aliscore output file does not exist. Check results for BUSCO: 399109at7742
## Thu Jul 14 02:31:53 CEST 2022 - The aliscore output appears to be empty. Check results for BUSCO: 399109at7742
## Thu Jul 14 02:31:54 CEST 2022 - The aliscore output file does not exist. Check results for BUSCO: 280704at7742
## Thu Jul 14 02:31:54 CEST 2022 - The aliscore output appears to be empty. Check results for BUSCO: 280704at7742
## Thu Jul 14 02:31:54 CEST 2022 - The aliscore output file does not exist. Check results for BUSCO: 376038at7742
## Thu Jul 14 02:31:54 CEST 2022 - The aliscore output appears to be empty. Check results for BUSCO: 376038at7742
## Thu Jul 14 02:31:55 CEST 2022 - The aliscore output file does not exist. Check results for BUSCO: 295613at7742
## Thu Jul 14 02:31:55 CEST 2022 - The aliscore output appears to be empty. Check results for BUSCO: 295613at7742
## Thu Jul 14 02:31:58 CEST 2022 - The aliscore output file does not exist. Check results for BUSCO: 359273at7742
## Thu Jul 14 02:31:58 CEST 2022 - The aliscore output appears to be empty. Check results for BUSCO: 359273at7742
## Thu Jul 14 02:31:58 CEST 2022 - The aliscore output file does not exist. Check results for BUSCO: 359541at7742
## Thu Jul 14 02:31:58 CEST 2022 - The aliscore output appears to be empty. Check results for BUSCO: 359541at7742
## Thu Jul 14 02:31:58 CEST 2022 - The aliscore output file does not exist. Check results for BUSCO: 387095at7742
## Thu Jul 14 02:31:58 CEST 2022 - The aliscore output appears to be empty. Check results for BUSCO: 387095at7742
## Thu Jul 14 02:32:00 CEST 2022 - The aliscore output file does not exist. Check results for BUSCO: 360409at7742
## Thu Jul 14 02:32:00 CEST 2022 - The aliscore output appears to be empty. Check results for BUSCO: 360409at7742
## Thu Jul 14 02:32:01 CEST 2022 - The aliscore output file does not exist. Check results for BUSCO: 400325at7742
## Thu Jul 14 02:32:01 CEST 2022 - The aliscore output appears to be empty. Check results for BUSCO: 400325at7742
## Thu Jul 14 02:32:01 CEST 2022 - The aliscore output file does not exist. Check results for BUSCO: 323530at7742
## Thu Jul 14 02:32:01 CEST 2022 - The aliscore output appears to be empty. Check results for BUSCO: 323530at7742
## Thu Jul 14 02:32:03 CEST 2022 - The aliscore output file does not exist. Check results for BUSCO: 407088at7742
## Thu Jul 14 02:32:03 CEST 2022 - The aliscore output appears to be empty. Check results for BUSCO: 407088at7742
## Thu Jul 14 02:32:04 CEST 2022 - The aliscore output file does not exist. Check results for BUSCO: 393599at7742
## Thu Jul 14 02:32:04 CEST 2022 - The aliscore output appears to be empty. Check results for BUSCO: 393599at7742
## Thu Jul 14 02:32:04 CEST 2022 - The aliscore output file does not exist. Check results for BUSCO: 287059at7742
## Thu Jul 14 02:32:04 CEST 2022 - The aliscore output appears to be empty. Check results for BUSCO: 287059at7742
## Thu Jul 14 02:32:04 CEST 2022 - The aliscore output file does not exist. Check results for BUSCO: 365662at7742
## Thu Jul 14 02:32:05 CEST 2022 - The aliscore output appears to be empty. Check results for BUSCO: 365662at7742
## Thu Jul 14 02:32:06 CEST 2022 - The aliscore output file does not exist. Check results for BUSCO: 403644at7742
## Thu Jul 14 02:32:06 CEST 2022 - The aliscore output appears to be empty. Check results for BUSCO: 403644at7742
## Thu Jul 14 02:32:07 CEST 2022 - The aliscore output file does not exist. Check results for BUSCO: 277508at7742
## Thu Jul 14 02:32:07 CEST 2022 - The aliscore output appears to be empty. Check results for BUSCO: 277508at7742
## Thu Jul 14 02:32:08 CEST 2022 - The aliscore output file does not exist. Check results for BUSCO: 353461at7742
## Thu Jul 14 02:32:08 CEST 2022 - The aliscore output appears to be empty. Check results for BUSCO: 353461at7742
## Thu Jul 14 02:32:09 CEST 2022 - The aliscore output file does not exist. Check results for BUSCO: 393010at7742
## Thu Jul 14 02:32:09 CEST 2022 - The aliscore output appears to be empty. Check results for BUSCO: 393010at7742
## Thu Jul 14 02:32:10 CEST 2022 - The aliscore output file does not exist. Check results for BUSCO: 404882at7742
## Thu Jul 14 02:32:10 CEST 2022 - The aliscore output appears to be empty. Check results for BUSCO: 404882at7742
## Thu Jul 14 02:32:11 CEST 2022 - The aliscore output file does not exist. Check results for BUSCO: 371207at7742
## Thu Jul 14 02:32:11 CEST 2022 - The aliscore output appears to be empty. Check results for BUSCO: 371207at7742
## Thu Jul 14 02:32:11 CEST 2022 - The aliscore output file does not exist. Check results for BUSCO: 387095at7742
## Thu Jul 14 02:32:12 CEST 2022 - The aliscore output appears to be empty. Check results for BUSCO: 387095at7742
## Thu Jul 14 02:32:12 CEST 2022 - The aliscore output file does not exist. Check results for BUSCO: 395764at7742
## Thu Jul 14 02:32:12 CEST 2022 - The aliscore output appears to be empty. Check results for BUSCO: 395764at7742
## Thu Jul 14 02:32:12 CEST 2022 - The aliscore output file does not exist. Check results for BUSCO: 382456at7742
## Thu Jul 14 02:32:12 CEST 2022 - The aliscore output appears to be empty. Check results for BUSCO: 382456at7742
## Thu Jul 14 02:32:13 CEST 2022 - The aliscore output file does not exist. Check results for BUSCO: 321832at7742
## Thu Jul 14 02:32:13 CEST 2022 - The aliscore output appears to be empty. Check results for BUSCO: 321832at7742
## Thu Jul 14 02:32:14 CEST 2022 - The aliscore output file does not exist. Check results for BUSCO: 380379at7742
## Thu Jul 14 02:32:14 CEST 2022 - The aliscore output appears to be empty. Check results for BUSCO: 380379at7742
## Thu Jul 14 02:32:16 CEST 2022 - The aliscore output file does not exist. Check results for BUSCO: 386595at7742
## Thu Jul 14 02:32:16 CEST 2022 - The aliscore output appears to be empty. Check results for BUSCO: 386595at7742
## Thu Jul 14 02:32:17 CEST 2022 - The aliscore output file does not exist. Check results for BUSCO: 372062at7742
## Thu Jul 14 02:32:17 CEST 2022 - The aliscore output appears to be empty. Check results for BUSCO: 372062at7742
## Thu Jul 14 02:32:17 CEST 2022 - The aliscore output file does not exist. Check results for BUSCO: 317712at7742
## Thu Jul 14 02:32:17 CEST 2022 - The aliscore output appears to be empty. Check results for BUSCO: 317712at7742
## Thu Jul 14 02:32:19 CEST 2022 - The aliscore output file does not exist. Check results for BUSCO: 239094at7742
## Thu Jul 14 02:32:19 CEST 2022 - The aliscore output appears to be empty. Check results for BUSCO: 239094at7742
## Thu Jul 14 02:32:19 CEST 2022 - The aliscore output file does not exist. Check results for BUSCO: 404437at7742
## Thu Jul 14 02:32:19 CEST 2022 - The aliscore output appears to be empty. Check results for BUSCO: 404437at7742
## Thu Jul 14 02:32:19 CEST 2022 - The aliscore output file does not exist. Check results for BUSCO: 312987at7742
## Thu Jul 14 02:32:19 CEST 2022 - The aliscore output appears to be empty. Check results for BUSCO: 312987at7742
## Thu Jul 14 02:32:21 CEST 2022 - The aliscore output file does not exist. Check results for BUSCO: 341187at7742
## Thu Jul 14 02:32:21 CEST 2022 - The aliscore output appears to be empty. Check results for BUSCO: 341187at7742
## Thu Jul 14 02:32:22 CEST 2022 - The aliscore output file does not exist. Check results for BUSCO: 368948at7742
## Thu Jul 14 02:32:22 CEST 2022 - The aliscore output appears to be empty. Check results for BUSCO: 368948at7742
## Thu Jul 14 02:32:23 CEST 2022 - The aliscore output file does not exist. Check results for BUSCO: 363627at7742
## Thu Jul 14 02:32:23 CEST 2022 - The aliscore output appears to be empty. Check results for BUSCO: 363627at7742
## Thu Jul 14 02:32:23 CEST 2022 - The aliscore output file does not exist. Check results for BUSCO: 402327at7742
## Thu Jul 14 02:32:24 CEST 2022 - The aliscore output appears to be empty. Check results for BUSCO: 402327at7742
## Thu Jul 14 02:32:24 CEST 2022 - The aliscore output file does not exist. Check results for BUSCO: 404437at7742
## Thu Jul 14 02:32:24 CEST 2022 - The aliscore output appears to be empty. Check results for BUSCO: 404437at7742
## Thu Jul 14 02:32:26 CEST 2022 - The aliscore output file does not exist. Check results for BUSCO: 238417at7742
## Thu Jul 14 02:32:26 CEST 2022 - The aliscore output appears to be empty. Check results for BUSCO: 238417at7742
## Thu Jul 14 02:32:26 CEST 2022 - The aliscore output file does not exist. Check results for BUSCO: 238417at7742
## Thu Jul 14 02:32:26 CEST 2022 - The aliscore output appears to be empty. Check results for BUSCO: 238417at7742
## Thu Jul 14 02:32:27 CEST 2022 - The aliscore output file does not exist. Check results for BUSCO: 346436at7742
## Thu Jul 14 02:32:27 CEST 2022 - The aliscore output appears to be empty. Check results for BUSCO: 346436at7742
## Thu Jul 14 02:32:29 CEST 2022 - The aliscore output file does not exist. Check results for BUSCO: 386595at7742
## Thu Jul 14 02:32:29 CEST 2022 - The aliscore output appears to be empty. Check results for BUSCO: 386595at7742
## Thu Jul 14 02:32:30 CEST 2022 - The aliscore output file does not exist. Check results for BUSCO: 422403at7742
## Thu Jul 14 02:32:30 CEST 2022 - The aliscore output appears to be empty. Check results for BUSCO: 422403at7742
## Thu Jul 14 02:32:30 CEST 2022 - The aliscore output file does not exist. Check results for BUSCO: 312987at7742
## Thu Jul 14 02:32:30 CEST 2022 - The aliscore output appears to be empty. Check results for BUSCO: 312987at7742
## Thu Jul 14 02:32:30 CEST 2022 - The aliscore output file does not exist. Check results for BUSCO: 348744at7742
## Thu Jul 14 02:32:31 CEST 2022 - The aliscore output appears to be empty. Check results for BUSCO: 348744at7742
## Thu Jul 14 02:32:32 CEST 2022 - The aliscore output file does not exist. Check results for BUSCO: 422403at7742
## Thu Jul 14 02:32:32 CEST 2022 - The aliscore output appears to be empty. Check results for BUSCO: 422403at7742
## Thu Jul 14 02:32:34 CEST 2022 - The aliscore output file does not exist. Check results for BUSCO: 238417at7742
## Thu Jul 14 02:32:34 CEST 2022 - The aliscore output appears to be empty. Check results for BUSCO: 238417at7742
## Thu Jul 14 02:32:35 CEST 2022 - The aliscore output file does not exist. Check results for BUSCO: 371581at7742
## Thu Jul 14 02:32:35 CEST 2022 - The aliscore output appears to be empty. Check results for BUSCO: 371581at7742
## Thu Jul 14 02:32:36 CEST 2022 - The aliscore output file does not exist. Check results for BUSCO: 305009at7742
## Thu Jul 14 02:32:36 CEST 2022 - The aliscore output appears to be empty. Check results for BUSCO: 305009at7742
## Thu Jul 14 02:32:37 CEST 2022 - The aliscore output file does not exist. Check results for BUSCO: 396521at7742
## Thu Jul 14 02:32:37 CEST 2022 - The aliscore output file does not exist. Check results for BUSCO: 317712at7742
## Thu Jul 14 02:32:37 CEST 2022 - The aliscore output appears to be empty. Check results for BUSCO: 396521at7742
## Thu Jul 14 02:32:37 CEST 2022 - The aliscore output appears to be empty. Check results for BUSCO: 317712at7742
## Thu Jul 14 02:32:38 CEST 2022 - The aliscore output file does not exist. Check results for BUSCO: 358276at7742
## Thu Jul 14 02:32:38 CEST 2022 - The aliscore output appears to be empty. Check results for BUSCO: 358276at7742
## Thu Jul 14 02:32:38 CEST 2022 - The aliscore output file does not exist. Check results for BUSCO: 273713at7742
## Thu Jul 14 02:32:38 CEST 2022 - The aliscore output appears to be empty. Check results for BUSCO: 273713at7742
## Thu Jul 14 02:32:40 CEST 2022 - The aliscore output file does not exist. Check results for BUSCO: 273713at7742
## Thu Jul 14 02:32:40 CEST 2022 - The aliscore output appears to be empty. Check results for BUSCO: 273713at7742
## Thu Jul 14 02:32:41 CEST 2022 - The aliscore output file does not exist. Check results for BUSCO: 386595at7742
## Thu Jul 14 02:32:41 CEST 2022 - The aliscore output appears to be empty. Check results for BUSCO: 386595at7742
## Thu Jul 14 02:32:41 CEST 2022 - The aliscore output file does not exist. Check results for BUSCO: 337686at7742
## Thu Jul 14 02:32:41 CEST 2022 - The aliscore output appears to be empty. Check results for BUSCO: 337686at7742
## Thu Jul 14 02:32:42 CEST 2022 - The aliscore output file does not exist. Check results for BUSCO: 421415at7742
## Thu Jul 14 02:32:42 CEST 2022 - The aliscore output appears to be empty. Check results for BUSCO: 421415at7742
## Thu Jul 14 02:32:44 CEST 2022 - The aliscore output file does not exist. Check results for BUSCO: 239094at7742
## Thu Jul 14 02:32:44 CEST 2022 - The aliscore output appears to be empty. Check results for BUSCO: 239094at7742
## Thu Jul 14 02:32:44 CEST 2022 - The aliscore output file does not exist. Check results for BUSCO: 344798at7742
## Thu Jul 14 02:32:44 CEST 2022 - The aliscore output appears to be empty. Check results for BUSCO: 344798at7742
## Thu Jul 14 02:32:45 CEST 2022 - The aliscore output file does not exist. Check results for BUSCO: 362141at7742
## Thu Jul 14 02:32:45 CEST 2022 - The aliscore output appears to be empty. Check results for BUSCO: 362141at7742
## Thu Jul 14 02:32:45 CEST 2022 - The aliscore output file does not exist. Check results for BUSCO: 359273at7742
## Thu Jul 14 02:32:45 CEST 2022 - The aliscore output appears to be empty. Check results for BUSCO: 359273at7742
## Thu Jul 14 02:32:48 CEST 2022 - The aliscore output file does not exist. Check results for BUSCO: 285301at7742
## Thu Jul 14 02:32:48 CEST 2022 - The aliscore output file does not exist. Check results for BUSCO: 354349at7742
## Thu Jul 14 02:32:48 CEST 2022 - The aliscore output appears to be empty. Check results for BUSCO: 285301at7742
## Thu Jul 14 02:32:48 CEST 2022 - The aliscore output appears to be empty. Check results for BUSCO: 354349at7742
## Thu Jul 14 02:32:48 CEST 2022 - The aliscore output file does not exist. Check results for BUSCO: 324567at7742
## Thu Jul 14 02:32:48 CEST 2022 - The aliscore output appears to be empty. Check results for BUSCO: 324567at7742
## Thu Jul 14 02:32:50 CEST 2022 - The aliscore output file does not exist. Check results for BUSCO: 319137at7742
## Thu Jul 14 02:32:50 CEST 2022 - The aliscore output appears to be empty. Check results for BUSCO: 319137at7742
## Thu Jul 14 02:32:50 CEST 2022 - The aliscore output file does not exist. Check results for BUSCO: 405236at7742
## Thu Jul 14 02:32:51 CEST 2022 - The aliscore output appears to be empty. Check results for BUSCO: 405236at7742
## Thu Jul 14 02:32:51 CEST 2022 - The aliscore output file does not exist. Check results for BUSCO: 385594at7742
## Thu Jul 14 02:32:51 CEST 2022 - The aliscore output appears to be empty. Check results for BUSCO: 385594at7742
## Thu Jul 14 02:32:52 CEST 2022 - The aliscore output file does not exist. Check results for BUSCO: 370608at7742
## Thu Jul 14 02:32:52 CEST 2022 - The aliscore output appears to be empty. Check results for BUSCO: 370608at7742
## Thu Jul 14 02:32:54 CEST 2022 - The aliscore output file does not exist. Check results for BUSCO: 387526at7742
## Thu Jul 14 02:32:54 CEST 2022 - The aliscore output appears to be empty. Check results for BUSCO: 387526at7742
## Thu Jul 14 02:32:55 CEST 2022 - The aliscore output file does not exist. Check results for BUSCO: 389517at7742
## Thu Jul 14 02:32:55 CEST 2022 - The aliscore output file does not exist. Check results for BUSCO: 386508at7742
## Thu Jul 14 02:32:55 CEST 2022 - The aliscore output file does not exist. Check results for BUSCO: 381431at7742
## Thu Jul 14 02:32:55 CEST 2022 - The aliscore output appears to be empty. Check results for BUSCO: 389517at7742
## Thu Jul 14 02:32:55 CEST 2022 - The aliscore output appears to be empty. Check results for BUSCO: 386508at7742
## Thu Jul 14 02:32:55 CEST 2022 - The aliscore output appears to be empty. Check results for BUSCO: 381431at7742
## Thu Jul 14 02:32:57 CEST 2022 - The aliscore output file does not exist. Check results for BUSCO: 397455at7742
## Thu Jul 14 02:32:57 CEST 2022 - The aliscore output appears to be empty. Check results for BUSCO: 397455at7742
## Thu Jul 14 02:32:58 CEST 2022 - The aliscore output file does not exist. Check results for BUSCO: 239094at7742
## Thu Jul 14 02:32:58 CEST 2022 - The aliscore output appears to be empty. Check results for BUSCO: 239094at7742
## Thu Jul 14 02:32:59 CEST 2022 - The aliscore output file does not exist. Check results for BUSCO: 400325at7742
## Thu Jul 14 02:32:59 CEST 2022 - The aliscore output appears to be empty. Check results for BUSCO: 400325at7742
## Thu Jul 14 02:33:00 CEST 2022 - The aliscore output file does not exist. Check results for BUSCO: 361358at7742
## Thu Jul 14 02:33:00 CEST 2022 - The aliscore output appears to be empty. Check results for BUSCO: 361358at7742
## Thu Jul 14 02:33:00 CEST 2022 - The aliscore output file does not exist. Check results for BUSCO: 280704at7742
## Thu Jul 14 02:33:00 CEST 2022 - The aliscore output appears to be empty. Check results for BUSCO: 280704at7742
## Thu Jul 14 02:33:01 CEST 2022 - The aliscore output file does not exist. Check results for BUSCO: 385758at7742
## Thu Jul 14 02:33:01 CEST 2022 - The aliscore output appears to be empty. Check results for BUSCO: 385758at7742
## Thu Jul 14 02:33:02 CEST 2022 - The aliscore output file does not exist. Check results for BUSCO: 401723at7742
## Thu Jul 14 02:33:02 CEST 2022 - The aliscore output appears to be empty. Check results for BUSCO: 401723at7742
## Thu Jul 14 02:33:03 CEST 2022 - The aliscore output file does not exist. Check results for BUSCO: 422259at7742
## Thu Jul 14 02:33:03 CEST 2022 - The aliscore output appears to be empty. Check results for BUSCO: 422259at7742
## Thu Jul 14 02:33:03 CEST 2022 - The aliscore output file does not exist. Check results for BUSCO: 354349at7742
## Thu Jul 14 02:33:03 CEST 2022 - The aliscore output appears to be empty. Check results for BUSCO: 354349at7742
## Thu Jul 14 02:33:04 CEST 2022 - The aliscore output file does not exist. Check results for BUSCO: 333591at7742
## Thu Jul 14 02:33:04 CEST 2022 - The aliscore output appears to be empty. Check results for BUSCO: 333591at7742
## Thu Jul 14 02:33:06 CEST 2022 - The aliscore output file does not exist. Check results for BUSCO: 362133at7742
## Thu Jul 14 02:33:06 CEST 2022 - The aliscore output appears to be empty. Check results for BUSCO: 362133at7742
## Thu Jul 14 02:33:06 CEST 2022 - The aliscore output file does not exist. Check results for BUSCO: 280704at7742
## Thu Jul 14 02:33:06 CEST 2022 - The aliscore output appears to be empty. Check results for BUSCO: 280704at7742
## Thu Jul 14 02:33:07 CEST 2022 - The aliscore output file does not exist. Check results for BUSCO: 372764at7742
## Thu Jul 14 02:33:07 CEST 2022 - The aliscore output appears to be empty. Check results for BUSCO: 372764at7742
## Thu Jul 14 02:33:09 CEST 2022 - The aliscore output file does not exist. Check results for BUSCO: 293643at7742
## Thu Jul 14 02:33:09 CEST 2022 - The aliscore output appears to be empty. Check results for BUSCO: 293643at7742
## Thu Jul 14 02:33:09 CEST 2022 - The aliscore output file does not exist. Check results for BUSCO: 411659at7742
## Thu Jul 14 02:33:09 CEST 2022 - The aliscore output appears to be empty. Check results for BUSCO: 411659at7742
## Thu Jul 14 02:33:10 CEST 2022 - The aliscore output file does not exist. Check results for BUSCO: 293643at7742
## Thu Jul 14 02:33:10 CEST 2022 - The aliscore output appears to be empty. Check results for BUSCO: 293643at7742
## Thu Jul 14 02:33:11 CEST 2022 - The aliscore output file does not exist. Check results for BUSCO: 219560at7742
## Thu Jul 14 02:33:11 CEST 2022 - The ALICUT output appears to be empty. Will touch an empty file so the pipeline will continue. Check results for BUSCO: 219560at7742
## Thu Jul 14 02:33:12 CEST 2022 - The aliscore output file does not exist. Check results for BUSCO: 305009at7742
## Thu Jul 14 02:33:12 CEST 2022 - The aliscore output appears to be empty. Check results for BUSCO: 305009at7742
## Thu Jul 14 02:33:13 CEST 2022 - The aliscore output file does not exist. Check results for BUSCO: 309129at7742
## Thu Jul 14 02:33:13 CEST 2022 - The aliscore output appears to be empty. Check results for BUSCO: 309129at7742
## Thu Jul 14 02:33:13 CEST 2022 - The aliscore output file does not exist. Check results for BUSCO: 339885at7742
## Thu Jul 14 02:33:14 CEST 2022 - The aliscore output appears to be empty. Check results for BUSCO: 339885at7742
## Thu Jul 14 02:33:15 CEST 2022 - The aliscore output file does not exist. Check results for BUSCO: 329799at7742
## Thu Jul 14 02:33:15 CEST 2022 - The aliscore output appears to be empty. Check results for BUSCO: 329799at7742
## Thu Jul 14 02:33:15 CEST 2022 - The aliscore output file does not exist. Check results for BUSCO: 339885at7742
## Thu Jul 14 02:33:15 CEST 2022 - The aliscore output appears to be empty. Check results for BUSCO: 339885at7742
## Thu Jul 14 02:33:17 CEST 2022 - The aliscore output file does not exist. Check results for BUSCO: 400534at7742
## Thu Jul 14 02:33:17 CEST 2022 - The aliscore output appears to be empty. Check results for BUSCO: 400534at7742
## Thu Jul 14 02:33:17 CEST 2022 - The aliscore output file does not exist. Check results for BUSCO: 219560at7742
## Thu Jul 14 02:33:17 CEST 2022 - The aliscore output appears to be empty. Check results for BUSCO: 219560at7742
## Thu Jul 14 02:33:19 CEST 2022 - The aliscore output file does not exist. Check results for BUSCO: 395765at7742
## Thu Jul 14 02:33:19 CEST 2022 - The aliscore output file does not exist. Check results for BUSCO: 390810at7742
## Thu Jul 14 02:33:19 CEST 2022 - The aliscore output appears to be empty. Check results for BUSCO: 390810at7742
## Thu Jul 14 02:33:19 CEST 2022 - The aliscore output appears to be empty. Check results for BUSCO: 395765at7742
## Thu Jul 14 02:33:20 CEST 2022 - The aliscore output file does not exist. Check results for BUSCO: 376607at7742
## Thu Jul 14 02:33:20 CEST 2022 - The aliscore output appears to be empty. Check results for BUSCO: 376607at7742
## Thu Jul 14 02:33:20 CEST 2022 - The aliscore output file does not exist. Check results for BUSCO: 330996at7742
## Thu Jul 14 02:33:20 CEST 2022 - The aliscore output appears to be empty. Check results for BUSCO: 330996at7742
## Thu Jul 14 02:33:22 CEST 2022 - The aliscore output file does not exist. Check results for BUSCO: 355430at7742
## Thu Jul 14 02:33:22 CEST 2022 - The aliscore output file does not exist. Check results for BUSCO: 319137at7742
## Thu Jul 14 02:33:22 CEST 2022 - The aliscore output appears to be empty. Check results for BUSCO: 355430at7742
## Thu Jul 14 02:33:22 CEST 2022 - The aliscore output appears to be empty. Check results for BUSCO: 319137at7742
## Thu Jul 14 02:33:23 CEST 2022 - The aliscore output file does not exist. Check results for BUSCO: 329799at7742
## Thu Jul 14 02:33:23 CEST 2022 - The aliscore output appears to be empty. Check results for BUSCO: 329799at7742
## Thu Jul 14 02:33:25 CEST 2022 - The aliscore output file does not exist. Check results for BUSCO: 235347at7742
## Thu Jul 14 02:33:25 CEST 2022 - The aliscore output appears to be empty. Check results for BUSCO: 235347at7742
## Thu Jul 14 02:33:26 CEST 2022 - The aliscore output file does not exist. Check results for BUSCO: 238049at7742
## Thu Jul 14 02:33:26 CEST 2022 - The aliscore output appears to be empty. Check results for BUSCO: 238049at7742
## Thu Jul 14 02:33:26 CEST 2022 - The aliscore output file does not exist. Check results for BUSCO: 398403at7742
## Thu Jul 14 02:33:26 CEST 2022 - The aliscore output appears to be empty. Check results for BUSCO: 398403at7742
## Thu Jul 14 02:33:27 CEST 2022 - The aliscore output file does not exist. Check results for BUSCO: 330614at7742
## Thu Jul 14 02:33:27 CEST 2022 - The aliscore output appears to be empty. Check results for BUSCO: 330614at7742
## Thu Jul 14 02:33:28 CEST 2022 - The aliscore output file does not exist. Check results for BUSCO: 323567at7742
## Thu Jul 14 02:33:28 CEST 2022 - The aliscore output appears to be empty. Check results for BUSCO: 323567at7742
## Thu Jul 14 02:33:28 CEST 2022 - The aliscore output file does not exist. Check results for BUSCO: 339885at7742
## Thu Jul 14 02:33:28 CEST 2022 - The aliscore output appears to be empty. Check results for BUSCO: 339885at7742
## Thu Jul 14 02:33:29 CEST 2022 - The aliscore output file does not exist. Check results for BUSCO: 309129at7742
## Thu Jul 14 02:33:30 CEST 2022 - The aliscore output appears to be empty. Check results for BUSCO: 309129at7742
## Thu Jul 14 02:33:31 CEST 2022 - The aliscore output file does not exist. Check results for BUSCO: 381867at7742
## Thu Jul 14 02:33:31 CEST 2022 - The aliscore output appears to be empty. Check results for BUSCO: 381867at7742
## Thu Jul 14 02:33:31 CEST 2022 - The aliscore output file does not exist. Check results for BUSCO: 362133at7742
## Thu Jul 14 02:33:31 CEST 2022 - The aliscore output appears to be empty. Check results for BUSCO: 362133at7742
## Thu Jul 14 02:33:33 CEST 2022 - The aliscore output file does not exist. Check results for BUSCO: 358276at7742
## Thu Jul 14 02:33:33 CEST 2022 - The aliscore output appears to be empty. Check results for BUSCO: 358276at7742
## Thu Jul 14 02:33:33 CEST 2022 - The aliscore output file does not exist. Check results for BUSCO: 298518at7742
## Thu Jul 14 02:33:33 CEST 2022 - The aliscore output appears to be empty. Check results for BUSCO: 298518at7742
## Thu Jul 14 02:33:34 CEST 2022 - The aliscore output file does not exist. Check results for BUSCO: 341187at7742
## Thu Jul 14 02:33:34 CEST 2022 - The aliscore output appears to be empty. Check results for BUSCO: 341187at7742
## Thu Jul 14 02:33:36 CEST 2022 - The aliscore output file does not exist. Check results for BUSCO: 219560at7742
## Thu Jul 14 02:33:36 CEST 2022 - The ALICUT output appears to be empty. Will touch an empty file so the pipeline will continue. Check results for BUSCO: 219560at7742
## Thu Jul 14 02:33:36 CEST 2022 - The aliscore output file does not exist. Check results for BUSCO: 285488at7742
## Thu Jul 14 02:33:36 CEST 2022 - The aliscore output appears to be empty. Check results for BUSCO: 285488at7742
## Thu Jul 14 02:33:39 CEST 2022 - The aliscore output file does not exist. Check results for BUSCO: 353461at7742
## Thu Jul 14 02:33:39 CEST 2022 - The aliscore output appears to be empty. Check results for BUSCO: 353461at7742
## Thu Jul 14 02:33:39 CEST 2022 - The aliscore output file does not exist. Check results for BUSCO: 405236at7742
## Thu Jul 14 02:33:39 CEST 2022 - The aliscore output appears to be empty. Check results for BUSCO: 405236at7742
## Thu Jul 14 02:33:40 CEST 2022 - The aliscore output file does not exist. Check results for BUSCO: 298518at7742
## Thu Jul 14 02:33:40 CEST 2022 - The aliscore output appears to be empty. Check results for BUSCO: 298518at7742
## Thu Jul 14 02:33:41 CEST 2022 - The aliscore output file does not exist. Check results for BUSCO: 321832at7742
## Thu Jul 14 02:33:41 CEST 2022 - The aliscore output appears to be empty. Check results for BUSCO: 321832at7742
## Thu Jul 14 02:33:41 CEST 2022 - The aliscore output file does not exist. Check results for BUSCO: 283060at7742
## Thu Jul 14 02:33:41 CEST 2022 - The aliscore output appears to be empty. Check results for BUSCO: 283060at7742
## Thu Jul 14 02:33:42 CEST 2022 - The aliscore output file does not exist. Check results for BUSCO: 235347at7742
## Thu Jul 14 02:33:42 CEST 2022 - The aliscore output appears to be empty. Check results for BUSCO: 235347at7742
## Thu Jul 14 02:33:43 CEST 2022 - The aliscore output file does not exist. Check results for BUSCO: 285488at7742
## Thu Jul 14 02:33:43 CEST 2022 - The aliscore output appears to be empty. Check results for BUSCO: 285488at7742
## Thu Jul 14 02:33:44 CEST 2022 - The aliscore output file does not exist. Check results for BUSCO: 366454at7742
## Thu Jul 14 02:33:44 CEST 2022 - The aliscore output appears to be empty. Check results for BUSCO: 366454at7742
## Thu Jul 14 02:33:45 CEST 2022 - The aliscore output file does not exist. Check results for BUSCO: 407088at7742
## Thu Jul 14 02:33:45 CEST 2022 - The aliscore output appears to be empty. Check results for BUSCO: 407088at7742
## Thu Jul 14 02:33:46 CEST 2022 - The aliscore output file does not exist. Check results for BUSCO: 417968at7742
## Thu Jul 14 02:33:46 CEST 2022 - The aliscore output appears to be empty. Check results for BUSCO: 417968at7742
## Thu Jul 14 02:33:47 CEST 2022 - The aliscore output file does not exist. Check results for BUSCO: 312646at7742
## Thu Jul 14 02:33:47 CEST 2022 - The aliscore output appears to be empty. Check results for BUSCO: 312646at7742
## Thu Jul 14 02:33:47 CEST 2022 - The aliscore output file does not exist. Check results for BUSCO: 362133at7742
## Thu Jul 14 02:33:47 CEST 2022 - The aliscore output appears to be empty. Check results for BUSCO: 362133at7742
## Thu Jul 14 02:33:49 CEST 2022 - The aliscore output file does not exist. Check results for BUSCO: 399133at7742
## Thu Jul 14 02:33:49 CEST 2022 - The aliscore output appears to be empty. Check results for BUSCO: 399133at7742
## Thu Jul 14 02:33:50 CEST 2022 - The aliscore output file does not exist. Check results for BUSCO: 361767at7742
## Thu Jul 14 02:33:50 CEST 2022 - The aliscore output appears to be empty. Check results for BUSCO: 361767at7742
## Thu Jul 14 02:33:51 CEST 2022 - The aliscore output file does not exist. Check results for BUSCO: 409625at7742
## Thu Jul 14 02:33:51 CEST 2022 - The aliscore output appears to be empty. Check results for BUSCO: 409625at7742
## Thu Jul 14 02:33:52 CEST 2022 - The aliscore output file does not exist. Check results for BUSCO: 293276at7742
## Thu Jul 14 02:33:52 CEST 2022 - The aliscore output appears to be empty. Check results for BUSCO: 293276at7742
## Thu Jul 14 02:33:52 CEST 2022 - The aliscore output file does not exist. Check results for BUSCO: 235347at7742
## Thu Jul 14 02:33:52 CEST 2022 - The aliscore output appears to be empty. Check results for BUSCO: 235347at7742
## Thu Jul 14 02:33:54 CEST 2022 - The aliscore output file does not exist. Check results for BUSCO: 359273at7742
## Thu Jul 14 02:33:54 CEST 2022 - The aliscore output appears to be empty. Check results for BUSCO: 359273at7742
## Thu Jul 14 02:33:54 CEST 2022 - The aliscore output file does not exist. Check results for BUSCO: 342286at7742
## Thu Jul 14 02:33:54 CEST 2022 - The aliscore output file does not exist. Check results for BUSCO: 342286at7742
## Thu Jul 14 02:33:54 CEST 2022 - The aliscore output appears to be empty. Check results for BUSCO: 342286at7742
## Thu Jul 14 02:33:54 CEST 2022 - The aliscore output appears to be empty. Check results for BUSCO: 342286at7742
## Thu Jul 14 02:33:57 CEST 2022 - The aliscore output file does not exist. Check results for BUSCO: 412561at7742
## Thu Jul 14 02:33:57 CEST 2022 - The aliscore output appears to be empty. Check results for BUSCO: 412561at7742
## Thu Jul 14 02:33:59 CEST 2022 - The aliscore output file does not exist. Check results for BUSCO: 385398at7742
## Thu Jul 14 02:33:59 CEST 2022 - The aliscore output appears to be empty. Check results for BUSCO: 385398at7742
## Thu Jul 14 02:33:59 CEST 2022 - The aliscore output file does not exist. Check results for BUSCO: 372023at7742
## Thu Jul 14 02:33:59 CEST 2022 - The aliscore output appears to be empty. Check results for BUSCO: 372023at7742
## Thu Jul 14 02:33:59 CEST 2022 - The aliscore output file does not exist. Check results for BUSCO: 372023at7742
## Thu Jul 14 02:33:59 CEST 2022 - The aliscore output appears to be empty. Check results for BUSCO: 372023at7742
## Thu Jul 14 02:33:59 CEST 2022 - The aliscore output file does not exist. Check results for BUSCO: 388786at7742
## Thu Jul 14 02:33:59 CEST 2022 - The aliscore output appears to be empty. Check results for BUSCO: 388786at7742
## Thu Jul 14 02:34:01 CEST 2022 - The aliscore output file does not exist. Check results for BUSCO: 397721at7742
## Thu Jul 14 02:34:01 CEST 2022 - The aliscore output appears to be empty. Check results for BUSCO: 397721at7742
## Thu Jul 14 02:34:02 CEST 2022 - The aliscore output file does not exist. Check results for BUSCO: 346436at7742
## Thu Jul 14 02:34:02 CEST 2022 - The aliscore output appears to be empty. Check results for BUSCO: 346436at7742
## Thu Jul 14 02:34:04 CEST 2022 - The aliscore output file does not exist. Check results for BUSCO: 398403at7742
## Thu Jul 14 02:34:04 CEST 2022 - The aliscore output appears to be empty. Check results for BUSCO: 398403at7742
## Thu Jul 14 02:34:04 CEST 2022 - The aliscore output file does not exist. Check results for BUSCO: 342286at7742
## Thu Jul 14 02:34:05 CEST 2022 - The aliscore output appears to be empty. Check results for BUSCO: 342286at7742
## Thu Jul 14 02:34:06 CEST 2022 - The aliscore output file does not exist. Check results for BUSCO: 407321at7742
## Thu Jul 14 02:34:06 CEST 2022 - The aliscore output appears to be empty. Check results for BUSCO: 407321at7742
## Thu Jul 14 02:34:06 CEST 2022 - The aliscore output file does not exist. Check results for BUSCO: 303403at7742
## Thu Jul 14 02:34:06 CEST 2022 - The aliscore output appears to be empty. Check results for BUSCO: 303403at7742
## Thu Jul 14 02:34:08 CEST 2022 - The aliscore output file does not exist. Check results for BUSCO: 266654at7742
## Thu Jul 14 02:34:08 CEST 2022 - The aliscore output appears to be empty. Check results for BUSCO: 266654at7742
## Thu Jul 14 02:34:08 CEST 2022 - The aliscore output file does not exist. Check results for BUSCO: 266654at7742
## Thu Jul 14 02:34:08 CEST 2022 - The aliscore output appears to be empty. Check results for BUSCO: 266654at7742
## Thu Jul 14 02:34:08 CEST 2022 - The aliscore output file does not exist. Check results for BUSCO: 347954at7742
## Thu Jul 14 02:34:08 CEST 2022 - The aliscore output appears to be empty. Check results for BUSCO: 347954at7742
## Thu Jul 14 02:34:10 CEST 2022 - The aliscore output file does not exist. Check results for BUSCO: 397721at7742
## Thu Jul 14 02:34:10 CEST 2022 - The aliscore output appears to be empty. Check results for BUSCO: 397721at7742
## Thu Jul 14 02:34:10 CEST 2022 - The aliscore output file does not exist. Check results for BUSCO: 398727at7742
## Thu Jul 14 02:34:10 CEST 2022 - The aliscore output appears to be empty. Check results for BUSCO: 398727at7742
## Thu Jul 14 02:34:12 CEST 2022 - The aliscore output file does not exist. Check results for BUSCO: 356030at7742
## Thu Jul 14 02:34:12 CEST 2022 - The aliscore output appears to be empty. Check results for BUSCO: 356030at7742
## Thu Jul 14 02:34:12 CEST 2022 - The aliscore output file does not exist. Check results for BUSCO: 317760at7742
## Thu Jul 14 02:34:12 CEST 2022 - The aliscore output appears to be empty. Check results for BUSCO: 317760at7742
## Thu Jul 14 02:34:14 CEST 2022 - The aliscore output file does not exist. Check results for BUSCO: 407064at7742
## Thu Jul 14 02:34:14 CEST 2022 - The aliscore output appears to be empty. Check results for BUSCO: 407064at7742
## Thu Jul 14 02:34:14 CEST 2022 - The aliscore output file does not exist. Check results for BUSCO: 287059at7742
## Thu Jul 14 02:34:14 CEST 2022 - The aliscore output appears to be empty. Check results for BUSCO: 287059at7742
## Thu Jul 14 02:34:15 CEST 2022 - The aliscore output file does not exist. Check results for BUSCO: 355312at7742
## Thu Jul 14 02:34:15 CEST 2022 - The aliscore output appears to be empty. Check results for BUSCO: 355312at7742
## Thu Jul 14 02:34:16 CEST 2022 - The aliscore output file does not exist. Check results for BUSCO: 329799at7742
## Thu Jul 14 02:34:16 CEST 2022 - The aliscore output appears to be empty. Check results for BUSCO: 329799at7742
## Thu Jul 14 02:34:17 CEST 2022 - The aliscore output file does not exist. Check results for BUSCO: 398379at7742
## Thu Jul 14 02:34:17 CEST 2022 - The aliscore output appears to be empty. Check results for BUSCO: 398379at7742
## Thu Jul 14 02:34:18 CEST 2022 - The aliscore output file does not exist. Check results for BUSCO: 344798at7742
## Thu Jul 14 02:34:19 CEST 2022 - The aliscore output appears to be empty. Check results for BUSCO: 344798at7742
## Thu Jul 14 02:34:19 CEST 2022 - The aliscore output file does not exist. Check results for BUSCO: 285301at7742
## Thu Jul 14 02:34:19 CEST 2022 - The aliscore output appears to be empty. Check results for BUSCO: 285301at7742
## Thu Jul 14 02:34:20 CEST 2022 - The aliscore output file does not exist. Check results for BUSCO: 293276at7742
## Thu Jul 14 02:34:20 CEST 2022 - The aliscore output appears to be empty. Check results for BUSCO: 293276at7742
## Thu Jul 14 02:34:20 CEST 2022 - The aliscore output file does not exist. Check results for BUSCO: 353461at7742
## Thu Jul 14 02:34:20 CEST 2022 - The aliscore output appears to be empty. Check results for BUSCO: 353461at7742
## Thu Jul 14 02:34:23 CEST 2022 - The aliscore output file does not exist. Check results for BUSCO: 278384at7742
## Thu Jul 14 02:34:23 CEST 2022 - The aliscore output appears to be empty. Check results for BUSCO: 278384at7742
## Thu Jul 14 02:34:23 CEST 2022 - The aliscore output file does not exist. Check results for BUSCO: 312646at7742
## Thu Jul 14 02:34:23 CEST 2022 - The aliscore output appears to be empty. Check results for BUSCO: 312646at7742
## Thu Jul 14 02:34:25 CEST 2022 - The aliscore output file does not exist. Check results for BUSCO: 348744at7742
## Thu Jul 14 02:34:25 CEST 2022 - The aliscore output appears to be empty. Check results for BUSCO: 348744at7742
## Thu Jul 14 02:34:25 CEST 2022 - The aliscore output file does not exist. Check results for BUSCO: 420085at7742
## Thu Jul 14 02:34:25 CEST 2022 - The aliscore output file does not exist. Check results for BUSCO: 327515at7742
## Thu Jul 14 02:34:25 CEST 2022 - The aliscore output appears to be empty. Check results for BUSCO: 420085at7742
## Thu Jul 14 02:34:25 CEST 2022 - The aliscore output appears to be empty. Check results for BUSCO: 327515at7742
## Thu Jul 14 02:34:26 CEST 2022 - The aliscore output file does not exist. Check results for BUSCO: 366454at7742
## Thu Jul 14 02:34:26 CEST 2022 - The aliscore output appears to be empty. Check results for BUSCO: 366454at7742
## Thu Jul 14 02:34:27 CEST 2022 - The aliscore output file does not exist. Check results for BUSCO: 406535at7742
## Thu Jul 14 02:34:27 CEST 2022 - The aliscore output appears to be empty. Check results for BUSCO: 406535at7742
## Thu Jul 14 02:34:30 CEST 2022 - The aliscore output file does not exist. Check results for BUSCO: 278384at7742
## Thu Jul 14 02:34:30 CEST 2022 - The aliscore output appears to be empty. Check results for BUSCO: 278384at7742
## Thu Jul 14 02:34:30 CEST 2022 - The aliscore output file does not exist. Check results for BUSCO: 355312at7742
## Thu Jul 14 02:34:30 CEST 2022 - The aliscore output appears to be empty. Check results for BUSCO: 355312at7742
## Thu Jul 14 02:34:31 CEST 2022 - The aliscore output file does not exist. Check results for BUSCO: 392383at7742
## Thu Jul 14 02:34:31 CEST 2022 - The aliscore output appears to be empty. Check results for BUSCO: 392383at7742
## Thu Jul 14 02:34:31 CEST 2022 - The aliscore output file does not exist. Check results for BUSCO: 293643at7742
## Thu Jul 14 02:34:31 CEST 2022 - The aliscore output appears to be empty. Check results for BUSCO: 293643at7742
## Thu Jul 14 02:34:32 CEST 2022 - The aliscore output file does not exist. Check results for BUSCO: 349482at7742
## Thu Jul 14 02:34:32 CEST 2022 - The aliscore output appears to be empty. Check results for BUSCO: 349482at7742
## Thu Jul 14 02:34:32 CEST 2022 - The aliscore output file does not exist. Check results for BUSCO: 393341at7742
## Thu Jul 14 02:34:33 CEST 2022 - The aliscore output appears to be empty. Check results for BUSCO: 393341at7742
## Thu Jul 14 02:34:33 CEST 2022 - The aliscore output file does not exist. Check results for BUSCO: 405886at7742
## Thu Jul 14 02:34:33 CEST 2022 - The aliscore output appears to be empty. Check results for BUSCO: 405886at7742
## Thu Jul 14 02:34:34 CEST 2022 - The aliscore output file does not exist. Check results for BUSCO: 355312at7742
## Thu Jul 14 02:34:34 CEST 2022 - The aliscore output appears to be empty. Check results for BUSCO: 355312at7742
## Thu Jul 14 02:34:36 CEST 2022 - The aliscore output file does not exist. Check results for BUSCO: 316607at7742
## Thu Jul 14 02:34:36 CEST 2022 - The aliscore output appears to be empty. Check results for BUSCO: 316607at7742
## Thu Jul 14 02:34:38 CEST 2022 - The aliscore output file does not exist. Check results for BUSCO: 352153at7742
## Thu Jul 14 02:34:38 CEST 2022 - The aliscore output appears to be empty. Check results for BUSCO: 352153at7742
## Thu Jul 14 02:34:38 CEST 2022 - The aliscore output file does not exist. Check results for BUSCO: 412708at7742
## Thu Jul 14 02:34:38 CEST 2022 - The aliscore output appears to be empty. Check results for BUSCO: 412708at7742
## Thu Jul 14 02:34:39 CEST 2022 - The aliscore output file does not exist. Check results for BUSCO: 371207at7742
## Thu Jul 14 02:34:39 CEST 2022 - The aliscore output appears to be empty. Check results for BUSCO: 371207at7742
## Thu Jul 14 02:34:40 CEST 2022 - The aliscore output file does not exist. Check results for BUSCO: 328506at7742
## Thu Jul 14 02:34:40 CEST 2022 - The aliscore output appears to be empty. Check results for BUSCO: 328506at7742
## Thu Jul 14 02:34:40 CEST 2022 - The aliscore output file does not exist. Check results for BUSCO: 390104at7742
## Thu Jul 14 02:34:40 CEST 2022 - The aliscore output appears to be empty. Check results for BUSCO: 390104at7742
## Thu Jul 14 02:34:41 CEST 2022 - The aliscore output file does not exist. Check results for BUSCO: 377323at7742
## Thu Jul 14 02:34:41 CEST 2022 - The aliscore output appears to be empty. Check results for BUSCO: 377323at7742
## Thu Jul 14 02:34:42 CEST 2022 - The aliscore output file does not exist. Check results for BUSCO: 331632at7742
## Thu Jul 14 02:34:42 CEST 2022 - The aliscore output appears to be empty. Check results for BUSCO: 331632at7742
## Thu Jul 14 02:34:44 CEST 2022 - The aliscore output file does not exist. Check results for BUSCO: 331632at7742
## Thu Jul 14 02:34:44 CEST 2022 - The aliscore output appears to be empty. Check results for BUSCO: 331632at7742
## Thu Jul 14 02:34:44 CEST 2022 - The aliscore output file does not exist. Check results for BUSCO: 413027at7742
## Thu Jul 14 02:34:44 CEST 2022 - The aliscore output appears to be empty. Check results for BUSCO: 413027at7742
## Thu Jul 14 02:34:45 CEST 2022 - The aliscore output file does not exist. Check results for BUSCO: 323567at7742
## Thu Jul 14 02:34:45 CEST 2022 - The aliscore output appears to be empty. Check results for BUSCO: 323567at7742
## Thu Jul 14 02:34:46 CEST 2022 - The aliscore output file does not exist. Check results for BUSCO: 388786at7742
## Thu Jul 14 02:34:46 CEST 2022 - The aliscore output appears to be empty. Check results for BUSCO: 388786at7742
## Thu Jul 14 02:34:46 CEST 2022 - The aliscore output file does not exist. Check results for BUSCO: 332775at7742
## Thu Jul 14 02:34:47 CEST 2022 - The aliscore output appears to be empty. Check results for BUSCO: 332775at7742
## Thu Jul 14 02:34:48 CEST 2022 - The aliscore output file does not exist. Check results for BUSCO: 349482at7742
## Thu Jul 14 02:34:48 CEST 2022 - The aliscore output appears to be empty. Check results for BUSCO: 349482at7742
## Thu Jul 14 02:34:50 CEST 2022 - The aliscore output file does not exist. Check results for BUSCO: 322712at7742
## Thu Jul 14 02:34:50 CEST 2022 - The aliscore output appears to be empty. Check results for BUSCO: 322712at7742
## Thu Jul 14 02:34:51 CEST 2022 - The aliscore output file does not exist. Check results for BUSCO: 398727at7742
## Thu Jul 14 02:34:51 CEST 2022 - The aliscore output appears to be empty. Check results for BUSCO: 398727at7742
## Thu Jul 14 02:34:51 CEST 2022 - The aliscore output file does not exist. Check results for BUSCO: 398727at7742
## Thu Jul 14 02:34:51 CEST 2022 - The aliscore output appears to be empty. Check results for BUSCO: 398727at7742
## Thu Jul 14 02:34:52 CEST 2022 - The aliscore output file does not exist. Check results for BUSCO: 371714at7742
## Thu Jul 14 02:34:52 CEST 2022 - The aliscore output appears to be empty. Check results for BUSCO: 371714at7742
## Thu Jul 14 02:34:53 CEST 2022 - The aliscore output file does not exist. Check results for BUSCO: 385758at7742
## Thu Jul 14 02:34:53 CEST 2022 - The aliscore output appears to be empty. Check results for BUSCO: 385758at7742
## Thu Jul 14 02:34:54 CEST 2022 - The aliscore output file does not exist. Check results for BUSCO: 331632at7742
## Thu Jul 14 02:34:54 CEST 2022 - The aliscore output appears to be empty. Check results for BUSCO: 331632at7742
## Thu Jul 14 02:34:55 CEST 2022 - The aliscore output file does not exist. Check results for BUSCO: 415130at7742
## Thu Jul 14 02:34:55 CEST 2022 - The aliscore output appears to be empty. Check results for BUSCO: 415130at7742
## Thu Jul 14 02:34:55 CEST 2022 - The aliscore output file does not exist. Check results for BUSCO: 363317at7742
## Thu Jul 14 02:34:55 CEST 2022 - The aliscore output appears to be empty. Check results for BUSCO: 363317at7742
## Thu Jul 14 02:34:56 CEST 2022 - The aliscore output file does not exist. Check results for BUSCO: 377176at7742
## Thu Jul 14 02:34:56 CEST 2022 - The aliscore output appears to be empty. Check results for BUSCO: 377176at7742
## Thu Jul 14 02:34:58 CEST 2022 - The aliscore output file does not exist. Check results for BUSCO: 395764at7742
## Thu Jul 14 02:34:58 CEST 2022 - The aliscore output appears to be empty. Check results for BUSCO: 395764at7742
## Thu Jul 14 02:34:58 CEST 2022 - The aliscore output file does not exist. Check results for BUSCO: 371207at7742
## Thu Jul 14 02:34:58 CEST 2022 - The aliscore output appears to be empty. Check results for BUSCO: 371207at7742
## Thu Jul 14 02:34:59 CEST 2022 - The aliscore output file does not exist. Check results for BUSCO: 351113at7742
## Thu Jul 14 02:34:59 CEST 2022 - The aliscore output appears to be empty. Check results for BUSCO: 351113at7742
## Thu Jul 14 02:35:00 CEST 2022 - The aliscore output file does not exist. Check results for BUSCO: 354349at7742
## Thu Jul 14 02:35:00 CEST 2022 - The aliscore output appears to be empty. Check results for BUSCO: 354349at7742
## Thu Jul 14 02:35:01 CEST 2022 - The aliscore output file does not exist. Check results for BUSCO: 414818at7742
## Thu Jul 14 02:35:02 CEST 2022 - The aliscore output appears to be empty. Check results for BUSCO: 414818at7742
## Thu Jul 14 02:35:02 CEST 2022 - The aliscore output file does not exist. Check results for BUSCO: 398379at7742
## Thu Jul 14 02:35:02 CEST 2022 - The aliscore output appears to be empty. Check results for BUSCO: 398379at7742
## Thu Jul 14 02:35:03 CEST 2022 - The aliscore output file does not exist. Check results for BUSCO: 285301at7742
## Thu Jul 14 02:35:03 CEST 2022 - The aliscore output appears to be empty. Check results for BUSCO: 285301at7742
## Thu Jul 14 02:35:04 CEST 2022 - The aliscore output file does not exist. Check results for BUSCO: 407064at7742
## Thu Jul 14 02:35:04 CEST 2022 - The aliscore output appears to be empty. Check results for BUSCO: 407064at7742
## Thu Jul 14 02:35:04 CEST 2022 - The aliscore output file does not exist. Check results for BUSCO: 409736at7742
## Thu Jul 14 02:35:04 CEST 2022 - The aliscore output appears to be empty. Check results for BUSCO: 409736at7742
## Thu Jul 14 02:35:05 CEST 2022 - The aliscore output file does not exist. Check results for BUSCO: 341187at7742
## Thu Jul 14 02:35:05 CEST 2022 - The aliscore output appears to be empty. Check results for BUSCO: 341187at7742
## Thu Jul 14 02:35:07 CEST 2022 - The aliscore output file does not exist. Check results for BUSCO: 437625at7742
## Thu Jul 14 02:35:07 CEST 2022 - The aliscore output appears to be empty. Check results for BUSCO: 437625at7742
## Thu Jul 14 02:35:09 CEST 2022 - The aliscore output file does not exist. Check results for BUSCO: 283060at7742
## Thu Jul 14 02:35:09 CEST 2022 - The aliscore output appears to be empty. Check results for BUSCO: 283060at7742
## Thu Jul 14 02:35:09 CEST 2022 - The aliscore output file does not exist. Check results for BUSCO: 415928at7742
## Thu Jul 14 02:35:09 CEST 2022 - The aliscore output appears to be empty. Check results for BUSCO: 415928at7742
## Thu Jul 14 02:35:11 CEST 2022 - The aliscore output file does not exist. Check results for BUSCO: 273166at7742
## Thu Jul 14 02:35:11 CEST 2022 - The aliscore output appears to be empty. Check results for BUSCO: 273166at7742
## Thu Jul 14 02:35:11 CEST 2022 - The aliscore output file does not exist. Check results for BUSCO: 393599at7742
## Thu Jul 14 02:35:11 CEST 2022 - The aliscore output appears to be empty. Check results for BUSCO: 393599at7742
## Thu Jul 14 02:35:12 CEST 2022 - The aliscore output file does not exist. Check results for BUSCO: 422259at7742
## Thu Jul 14 02:35:12 CEST 2022 - The aliscore output appears to be empty. Check results for BUSCO: 422259at7742
## Thu Jul 14 02:35:12 CEST 2022 - The aliscore output file does not exist. Check results for BUSCO: 391345at7742
## Thu Jul 14 02:35:12 CEST 2022 - The aliscore output appears to be empty. Check results for BUSCO: 391345at7742
## Thu Jul 14 02:35:14 CEST 2022 - The aliscore output file does not exist. Check results for BUSCO: 381431at7742
## Thu Jul 14 02:35:14 CEST 2022 - The aliscore output appears to be empty. Check results for BUSCO: 381431at7742
## Thu Jul 14 02:35:14 CEST 2022 - The aliscore output file does not exist. Check results for BUSCO: 385022at7742
## Thu Jul 14 02:35:14 CEST 2022 - The aliscore output appears to be empty. Check results for BUSCO: 385022at7742
## Thu Jul 14 02:35:15 CEST 2022 - The aliscore output file does not exist. Check results for BUSCO: 304456at7742
## Thu Jul 14 02:35:15 CEST 2022 - The aliscore output appears to be empty. Check results for BUSCO: 304456at7742
## Thu Jul 14 02:35:16 CEST 2022 - The aliscore output file does not exist. Check results for BUSCO: 412561at7742
## Thu Jul 14 02:35:16 CEST 2022 - The aliscore output appears to be empty. Check results for BUSCO: 412561at7742
## Thu Jul 14 02:35:17 CEST 2022 - The aliscore output file does not exist. Check results for BUSCO: 332775at7742
## Thu Jul 14 02:35:17 CEST 2022 - The aliscore output appears to be empty. Check results for BUSCO: 332775at7742
## Thu Jul 14 02:35:18 CEST 2022 - The aliscore output file does not exist. Check results for BUSCO: 361078at7742
## Thu Jul 14 02:35:18 CEST 2022 - The aliscore output appears to be empty. Check results for BUSCO: 361078at7742
## Thu Jul 14 02:35:18 CEST 2022 - The aliscore output file does not exist. Check results for BUSCO: 392048at7742
## Thu Jul 14 02:35:18 CEST 2022 - The aliscore output appears to be empty. Check results for BUSCO: 392048at7742
## Thu Jul 14 02:35:19 CEST 2022 - The aliscore output file does not exist. Check results for BUSCO: 406306at7742
## Thu Jul 14 02:35:19 CEST 2022 - The aliscore output appears to be empty. Check results for BUSCO: 406306at7742
## Thu Jul 14 02:35:21 CEST 2022 - The aliscore output file does not exist. Check results for BUSCO: 287059at7742
## Thu Jul 14 02:35:21 CEST 2022 - The aliscore output appears to be empty. Check results for BUSCO: 287059at7742
## Thu Jul 14 02:35:22 CEST 2022 - The aliscore output file does not exist. Check results for BUSCO: 437625at7742
## Thu Jul 14 02:35:22 CEST 2022 - The aliscore output appears to be empty. Check results for BUSCO: 437625at7742
## Thu Jul 14 02:35:22 CEST 2022 - The aliscore output file does not exist. Check results for BUSCO: 251459at7742
## Thu Jul 14 02:35:22 CEST 2022 - The aliscore output appears to be empty. Check results for BUSCO: 251459at7742
## Thu Jul 14 02:35:23 CEST 2022 - The aliscore output file does not exist. Check results for BUSCO: 304456at7742
## Thu Jul 14 02:35:23 CEST 2022 - The aliscore output appears to be empty. Check results for BUSCO: 304456at7742
## Thu Jul 14 02:35:23 CEST 2022 - The aliscore output file does not exist. Check results for BUSCO: 397721at7742
## Thu Jul 14 02:35:23 CEST 2022 - The aliscore output appears to be empty. Check results for BUSCO: 397721at7742
## Thu Jul 14 02:35:24 CEST 2022 - The aliscore output file does not exist. Check results for BUSCO: 364898at7742
## Thu Jul 14 02:35:24 CEST 2022 - The aliscore output appears to be empty. Check results for BUSCO: 364898at7742
## Thu Jul 14 02:35:25 CEST 2022 - The aliscore output file does not exist. Check results for BUSCO: 406935at7742
## Thu Jul 14 02:35:26 CEST 2022 - The aliscore output appears to be empty. Check results for BUSCO: 406935at7742
## Thu Jul 14 02:35:27 CEST 2022 - The aliscore output file does not exist. Check results for BUSCO: 317760at7742
## Thu Jul 14 02:35:27 CEST 2022 - The aliscore output appears to be empty. Check results for BUSCO: 317760at7742
## Thu Jul 14 02:35:28 CEST 2022 - The aliscore output file does not exist. Check results for BUSCO: 347416at7742
## Thu Jul 14 02:35:28 CEST 2022 - The aliscore output appears to be empty. Check results for BUSCO: 347416at7742
## Thu Jul 14 02:35:28 CEST 2022 - The aliscore output file does not exist. Check results for BUSCO: 368773at7742
## Thu Jul 14 02:35:28 CEST 2022 - The aliscore output appears to be empty. Check results for BUSCO: 368773at7742
## Thu Jul 14 02:35:29 CEST 2022 - The aliscore output file does not exist. Check results for BUSCO: 322470at7742
## Thu Jul 14 02:35:29 CEST 2022 - The aliscore output appears to be empty. Check results for BUSCO: 322470at7742
## Thu Jul 14 02:35:31 CEST 2022 - The aliscore output file does not exist. Check results for BUSCO: 386508at7742
## Thu Jul 14 02:35:31 CEST 2022 - The aliscore output appears to be empty. Check results for BUSCO: 386508at7742
## Thu Jul 14 02:35:31 CEST 2022 - The aliscore output file does not exist. Check results for BUSCO: 251459at7742
## Thu Jul 14 02:35:31 CEST 2022 - The ALICUT output appears to be empty. Will touch an empty file so the pipeline will continue. Check results for BUSCO: 251459at7742
## Thu Jul 14 02:35:32 CEST 2022 - The aliscore output file does not exist. Check results for BUSCO: 304456at7742
## Thu Jul 14 02:35:32 CEST 2022 - The aliscore output appears to be empty. Check results for BUSCO: 304456at7742
## Thu Jul 14 02:35:33 CEST 2022 - The aliscore output file does not exist. Check results for BUSCO: 399667at7742
## Thu Jul 14 02:35:33 CEST 2022 - The aliscore output appears to be empty. Check results for BUSCO: 399667at7742
## Thu Jul 14 02:35:33 CEST 2022 - The aliscore output file does not exist. Check results for BUSCO: 393341at7742
## Thu Jul 14 02:35:33 CEST 2022 - The aliscore output appears to be empty. Check results for BUSCO: 393341at7742
## Thu Jul 14 02:35:34 CEST 2022 - The aliscore output file does not exist. Check results for BUSCO: 322712at7742
## Thu Jul 14 02:35:34 CEST 2022 - The aliscore output appears to be empty. Check results for BUSCO: 322712at7742
## Thu Jul 14 02:35:35 CEST 2022 - The aliscore output file does not exist. Check results for BUSCO: 363317at7742
## Thu Jul 14 02:35:35 CEST 2022 - The aliscore output appears to be empty. Check results for BUSCO: 363317at7742
## Thu Jul 14 02:35:37 CEST 2022 - The aliscore output file does not exist. Check results for BUSCO: 341385at7742
## Thu Jul 14 02:35:37 CEST 2022 - The aliscore output appears to be empty. Check results for BUSCO: 341385at7742
## Thu Jul 14 02:35:38 CEST 2022 - The aliscore output file does not exist. Check results for BUSCO: 350839at7742
## Thu Jul 14 02:35:38 CEST 2022 - The aliscore output appears to be empty. Check results for BUSCO: 350839at7742
## Thu Jul 14 02:35:38 CEST 2022 - The aliscore output file does not exist. Check results for BUSCO: 358276at7742
## Thu Jul 14 02:35:38 CEST 2022 - The aliscore output appears to be empty. Check results for BUSCO: 358276at7742
## Thu Jul 14 02:35:39 CEST 2022 - The aliscore output file does not exist. Check results for BUSCO: 251459at7742
## Thu Jul 14 02:35:39 CEST 2022 - The aliscore output appears to be empty. Check results for BUSCO: 251459at7742
## Thu Jul 14 02:35:41 CEST 2022 - The aliscore output file does not exist. Check results for BUSCO: 393852at7742
## Thu Jul 14 02:35:41 CEST 2022 - The aliscore output appears to be empty. Check results for BUSCO: 393852at7742
## Thu Jul 14 02:35:41 CEST 2022 - The aliscore output file does not exist. Check results for BUSCO: 367430at7742
## Thu Jul 14 02:35:41 CEST 2022 - The aliscore output appears to be empty. Check results for BUSCO: 367430at7742
## Thu Jul 14 02:35:41 CEST 2022 - The aliscore output file does not exist. Check results for BUSCO: 398584at7742
## Thu Jul 14 02:35:42 CEST 2022 - The aliscore output appears to be empty. Check results for BUSCO: 398584at7742
## Thu Jul 14 02:35:43 CEST 2022 - The aliscore output file does not exist. Check results for BUSCO: 388786at7742
## Thu Jul 14 02:35:43 CEST 2022 - The aliscore output appears to be empty. Check results for BUSCO: 388786at7742
## Thu Jul 14 02:35:44 CEST 2022 - The aliscore output file does not exist. Check results for BUSCO: 358562at7742
## Thu Jul 14 02:35:44 CEST 2022 - The aliscore output appears to be empty. Check results for BUSCO: 358562at7742
## Thu Jul 14 02:35:45 CEST 2022 - The aliscore output file does not exist. Check results for BUSCO: 358562at7742
## Thu Jul 14 02:35:45 CEST 2022 - The aliscore output appears to be empty. Check results for BUSCO: 358562at7742
## Thu Jul 14 02:35:46 CEST 2022 - The aliscore output file does not exist. Check results for BUSCO: 363627at7742
## Thu Jul 14 02:35:46 CEST 2022 - The aliscore output appears to be empty. Check results for BUSCO: 363627at7742
## Thu Jul 14 02:35:46 CEST 2022 - The aliscore output file does not exist. Check results for BUSCO: 322712at7742
## Thu Jul 14 02:35:46 CEST 2022 - The aliscore output appears to be empty. Check results for BUSCO: 322712at7742
## Thu Jul 14 02:35:48 CEST 2022 - The aliscore output file does not exist. Check results for BUSCO: 259882at7742
## Thu Jul 14 02:35:48 CEST 2022 - The aliscore output appears to be empty. Check results for BUSCO: 259882at7742
## Thu Jul 14 02:35:48 CEST 2022 - The aliscore output file does not exist. Check results for BUSCO: 368773at7742
## Thu Jul 14 02:35:48 CEST 2022 - The aliscore output appears to be empty. Check results for BUSCO: 368773at7742
## Thu Jul 14 02:35:49 CEST 2022 - The aliscore output file does not exist. Check results for BUSCO: 345605at7742
## Thu Jul 14 02:35:49 CEST 2022 - The aliscore output appears to be empty. Check results for BUSCO: 345605at7742
## Thu Jul 14 02:35:50 CEST 2022 - The aliscore output file does not exist. Check results for BUSCO: 388019at7742
## Thu Jul 14 02:35:50 CEST 2022 - The aliscore output appears to be empty. Check results for BUSCO: 388019at7742
## Thu Jul 14 02:35:50 CEST 2022 - The aliscore output file does not exist. Check results for BUSCO: 345605at7742
## Thu Jul 14 02:35:50 CEST 2022 - The aliscore output appears to be empty. Check results for BUSCO: 345605at7742
## Thu Jul 14 02:35:52 CEST 2022 - The aliscore output file does not exist. Check results for BUSCO: 332775at7742
## Thu Jul 14 02:35:52 CEST 2022 - The aliscore output appears to be empty. Check results for BUSCO: 332775at7742
## Thu Jul 14 02:35:53 CEST 2022 - The aliscore output file does not exist. Check results for BUSCO: 336175at7742
## Thu Jul 14 02:35:53 CEST 2022 - The aliscore output appears to be empty. Check results for BUSCO: 336175at7742
## Thu Jul 14 02:35:53 CEST 2022 - The aliscore output file does not exist. Check results for BUSCO: 273166at7742
## Thu Jul 14 02:35:54 CEST 2022 - The aliscore output appears to be empty. Check results for BUSCO: 273166at7742
## Thu Jul 14 02:35:55 CEST 2022 - The aliscore output file does not exist. Check results for BUSCO: 285352at7742
## Thu Jul 14 02:35:55 CEST 2022 - The aliscore output appears to be empty. Check results for BUSCO: 285352at7742
## Thu Jul 14 02:35:56 CEST 2022 - The aliscore output file does not exist. Check results for BUSCO: 380785at7742
## Thu Jul 14 02:35:56 CEST 2022 - The aliscore output appears to be empty. Check results for BUSCO: 380785at7742
## Thu Jul 14 02:35:58 CEST 2022 - The aliscore output file does not exist. Check results for BUSCO: 417968at7742
## Thu Jul 14 02:35:58 CEST 2022 - The aliscore output appears to be empty. Check results for BUSCO: 417968at7742
## Thu Jul 14 02:35:58 CEST 2022 - The aliscore output file does not exist. Check results for BUSCO: 350839at7742
## Thu Jul 14 02:35:58 CEST 2022 - The aliscore output appears to be empty. Check results for BUSCO: 350839at7742
## Thu Jul 14 02:35:59 CEST 2022 - The aliscore output file does not exist. Check results for BUSCO: 417968at7742
## Thu Jul 14 02:35:59 CEST 2022 - The aliscore output appears to be empty. Check results for BUSCO: 417968at7742
## Thu Jul 14 02:36:00 CEST 2022 - The aliscore output file does not exist. Check results for BUSCO: 301105at7742
## Thu Jul 14 02:36:00 CEST 2022 - The aliscore output appears to be empty. Check results for BUSCO: 301105at7742
## Thu Jul 14 02:36:00 CEST 2022 - The aliscore output file does not exist. Check results for BUSCO: 387095at7742
## Thu Jul 14 02:36:00 CEST 2022 - The aliscore output appears to be empty. Check results for BUSCO: 387095at7742
## Thu Jul 14 02:36:01 CEST 2022 - The aliscore output file does not exist. Check results for BUSCO: 279246at7742
## Thu Jul 14 02:36:01 CEST 2022 - The aliscore output appears to be empty. Check results for BUSCO: 279246at7742
## Thu Jul 14 02:36:02 CEST 2022 - The aliscore output file does not exist. Check results for BUSCO: 303403at7742
## Thu Jul 14 02:36:02 CEST 2022 - The aliscore output appears to be empty. Check results for BUSCO: 303403at7742
## Thu Jul 14 02:36:04 CEST 2022 - The aliscore output file does not exist. Check results for BUSCO: 414818at7742
## Thu Jul 14 02:36:04 CEST 2022 - The aliscore output appears to be empty. Check results for BUSCO: 414818at7742
## Thu Jul 14 02:36:04 CEST 2022 - The aliscore output file does not exist. Check results for BUSCO: 330116at7742
## Thu Jul 14 02:36:04 CEST 2022 - The aliscore output appears to be empty. Check results for BUSCO: 330116at7742
## Thu Jul 14 02:36:05 CEST 2022 - The aliscore output file does not exist. Check results for BUSCO: 361358at7742
## Thu Jul 14 02:36:05 CEST 2022 - The aliscore output appears to be empty. Check results for BUSCO: 361358at7742
## Thu Jul 14 02:36:07 CEST 2022 - The aliscore output file does not exist. Check results for BUSCO: 285352at7742
## Thu Jul 14 02:36:07 CEST 2022 - The aliscore output appears to be empty. Check results for BUSCO: 285352at7742
## Thu Jul 14 02:36:07 CEST 2022 - The aliscore output file does not exist. Check results for BUSCO: 341385at7742
## Thu Jul 14 02:36:07 CEST 2022 - The aliscore output appears to be empty. Check results for BUSCO: 341385at7742
## Thu Jul 14 02:36:07 CEST 2022 - The aliscore output file does not exist. Check results for BUSCO: 259882at7742
## Thu Jul 14 02:36:07 CEST 2022 - The aliscore output appears to be empty. Check results for BUSCO: 259882at7742
## Thu Jul 14 02:36:08 CEST 2022 - The aliscore output file does not exist. Check results for BUSCO: 406535at7742
## Thu Jul 14 02:36:08 CEST 2022 - The aliscore output appears to be empty. Check results for BUSCO: 406535at7742
## Thu Jul 14 02:36:09 CEST 2022 - The aliscore output file does not exist. Check results for BUSCO: 381528at7742
## Thu Jul 14 02:36:09 CEST 2022 - The aliscore output appears to be empty. Check results for BUSCO: 381528at7742
## Thu Jul 14 02:36:12 CEST 2022 - The aliscore output file does not exist. Check results for BUSCO: 345605at7742
## Thu Jul 14 02:36:12 CEST 2022 - The aliscore output appears to be empty. Check results for BUSCO: 345605at7742
## Thu Jul 14 02:36:12 CEST 2022 - The aliscore output file does not exist. Check results for BUSCO: 414818at7742
## Thu Jul 14 02:36:12 CEST 2022 - The aliscore output appears to be empty. Check results for BUSCO: 414818at7742
## Thu Jul 14 02:36:13 CEST 2022 - The aliscore output file does not exist. Check results for BUSCO: 285352at7742
## Thu Jul 14 02:36:13 CEST 2022 - The aliscore output appears to be empty. Check results for BUSCO: 285352at7742
## Thu Jul 14 02:36:13 CEST 2022 - The aliscore output file does not exist. Check results for BUSCO: 381830at7742
## Thu Jul 14 02:36:14 CEST 2022 - The aliscore output appears to be empty. Check results for BUSCO: 381830at7742
## Thu Jul 14 02:36:14 CEST 2022 - The aliscore output file does not exist. Check results for BUSCO: 371714at7742
## Thu Jul 14 02:36:14 CEST 2022 - The aliscore output appears to be empty. Check results for BUSCO: 371714at7742
## Thu Jul 14 02:36:15 CEST 2022 - The aliscore output file does not exist. Check results for BUSCO: 397031at7742
## Thu Jul 14 02:36:15 CEST 2022 - The aliscore output appears to be empty. Check results for BUSCO: 397031at7742
## Thu Jul 14 02:36:16 CEST 2022 - The aliscore output file does not exist. Check results for BUSCO: 326283at7742
## Thu Jul 14 02:36:16 CEST 2022 - The aliscore output appears to be empty. Check results for BUSCO: 326283at7742
## Thu Jul 14 02:36:17 CEST 2022 - The aliscore output file does not exist. Check results for BUSCO: 346605at7742
## Thu Jul 14 02:36:17 CEST 2022 - The aliscore output appears to be empty. Check results for BUSCO: 346605at7742
## Thu Jul 14 02:36:18 CEST 2022 - The aliscore output file does not exist. Check results for BUSCO: 336175at7742
## Thu Jul 14 02:36:18 CEST 2022 - The aliscore output appears to be empty. Check results for BUSCO: 336175at7742
## Thu Jul 14 02:36:19 CEST 2022 - The aliscore output file does not exist. Check results for BUSCO: 279246at7742
## Thu Jul 14 02:36:19 CEST 2022 - The aliscore output appears to be empty. Check results for BUSCO: 279246at7742
## Thu Jul 14 02:36:19 CEST 2022 - The aliscore output file does not exist. Check results for BUSCO: 279246at7742
## Thu Jul 14 02:36:20 CEST 2022 - The aliscore output appears to be empty. Check results for BUSCO: 279246at7742
## Thu Jul 14 02:36:20 CEST 2022 - The aliscore output file does not exist. Check results for BUSCO: 319099at7742
## Thu Jul 14 02:36:20 CEST 2022 - The aliscore output appears to be empty. Check results for BUSCO: 319099at7742
## Thu Jul 14 02:36:21 CEST 2022 - The aliscore output file does not exist. Check results for BUSCO: 367430at7742
## Thu Jul 14 02:36:21 CEST 2022 - The aliscore output appears to be empty. Check results for BUSCO: 367430at7742
## Thu Jul 14 02:36:22 CEST 2022 - The aliscore output file does not exist. Check results for BUSCO: 367430at7742
## Thu Jul 14 02:36:22 CEST 2022 - The aliscore output appears to be empty. Check results for BUSCO: 367430at7742
## Thu Jul 14 02:36:23 CEST 2022 - The aliscore output file does not exist. Check results for BUSCO: 377176at7742
## Thu Jul 14 02:36:23 CEST 2022 - The aliscore output appears to be empty. Check results for BUSCO: 377176at7742
## Thu Jul 14 02:36:24 CEST 2022 - The aliscore output file does not exist. Check results for BUSCO: 377176at7742
## Thu Jul 14 02:36:24 CEST 2022 - The aliscore output appears to be empty. Check results for BUSCO: 377176at7742
## Thu Jul 14 02:36:25 CEST 2022 - The aliscore output file does not exist. Check results for BUSCO: 259882at7742
## Thu Jul 14 02:36:25 CEST 2022 - The aliscore output appears to be empty. Check results for BUSCO: 259882at7742
## Thu Jul 14 02:36:25 CEST 2022 - The aliscore output file does not exist. Check results for BUSCO: 381106at7742
## Thu Jul 14 02:36:25 CEST 2022 - The aliscore output appears to be empty. Check results for BUSCO: 381106at7742
## Thu Jul 14 02:36:27 CEST 2022 - The aliscore output file does not exist. Check results for BUSCO: 391345at7742
## Thu Jul 14 02:36:27 CEST 2022 - The aliscore output file does not exist. Check results for BUSCO: 425988at7742
## Thu Jul 14 02:36:27 CEST 2022 - The aliscore output appears to be empty. Check results for BUSCO: 391345at7742
## Thu Jul 14 02:36:27 CEST 2022 - The aliscore output appears to be empty. Check results for BUSCO: 425988at7742
## Thu Jul 14 02:36:28 CEST 2022 - The aliscore output file does not exist. Check results for BUSCO: 323438at7742
## Thu Jul 14 02:36:28 CEST 2022 - The aliscore output appears to be empty. Check results for BUSCO: 323438at7742
## Thu Jul 14 02:36:30 CEST 2022 - The aliscore output file does not exist. Check results for BUSCO: 317760at7742
## Thu Jul 14 02:36:30 CEST 2022 - The aliscore output appears to be empty. Check results for BUSCO: 317760at7742
## Thu Jul 14 02:36:31 CEST 2022 - The aliscore output file does not exist. Check results for BUSCO: 368773at7742
## Thu Jul 14 02:36:31 CEST 2022 - The aliscore output file does not exist. Check results for BUSCO: 401723at7742
## Thu Jul 14 02:36:31 CEST 2022 - The aliscore output appears to be empty. Check results for BUSCO: 368773at7742
## Thu Jul 14 02:36:31 CEST 2022 - The aliscore output appears to be empty. Check results for BUSCO: 401723at7742
## Thu Jul 14 02:36:33 CEST 2022 - The aliscore output file does not exist. Check results for BUSCO: 395170at7742
## Thu Jul 14 02:36:33 CEST 2022 - The aliscore output appears to be empty. Check results for BUSCO: 395170at7742
## Thu Jul 14 02:36:33 CEST 2022 - The aliscore output file does not exist. Check results for BUSCO: 416829at7742
## Thu Jul 14 02:36:33 CEST 2022 - The aliscore output appears to be empty. Check results for BUSCO: 416829at7742
## Thu Jul 14 02:36:34 CEST 2022 - The aliscore output file does not exist. Check results for BUSCO: 349482at7742
## Thu Jul 14 02:36:35 CEST 2022 - The aliscore output appears to be empty. Check results for BUSCO: 349482at7742
## Thu Jul 14 02:36:35 CEST 2022 - The aliscore output file does not exist. Check results for BUSCO: 381106at7742
## Thu Jul 14 02:36:35 CEST 2022 - The aliscore output appears to be empty. Check results for BUSCO: 381106at7742
## Thu Jul 14 02:36:36 CEST 2022 - The aliscore output file does not exist. Check results for BUSCO: 421992at7742
## Thu Jul 14 02:36:36 CEST 2022 - The aliscore output appears to be empty. Check results for BUSCO: 421992at7742
## Thu Jul 14 02:36:37 CEST 2022 - The aliscore output file does not exist. Check results for BUSCO: 398268at7742
## Thu Jul 14 02:36:37 CEST 2022 - The aliscore output appears to be empty. Check results for BUSCO: 398268at7742
## Thu Jul 14 02:36:38 CEST 2022 - The aliscore output file does not exist. Check results for BUSCO: 288056at7742
## Thu Jul 14 02:36:38 CEST 2022 - The aliscore output appears to be empty. Check results for BUSCO: 288056at7742
## Thu Jul 14 02:36:39 CEST 2022 - The aliscore output file does not exist. Check results for BUSCO: 422508at7742
## Thu Jul 14 02:36:39 CEST 2022 - The aliscore output appears to be empty. Check results for BUSCO: 422508at7742
## Thu Jul 14 02:36:40 CEST 2022 - The aliscore output file does not exist. Check results for BUSCO: 398268at7742
## Thu Jul 14 02:36:40 CEST 2022 - The aliscore output appears to be empty. Check results for BUSCO: 398268at7742
## Thu Jul 14 02:36:41 CEST 2022 - The aliscore output file does not exist. Check results for BUSCO: 301105at7742
## Thu Jul 14 02:36:41 CEST 2022 - The aliscore output appears to be empty. Check results for BUSCO: 301105at7742
## Thu Jul 14 02:36:41 CEST 2022 - The aliscore output file does not exist. Check results for BUSCO: 369368at7742
## Thu Jul 14 02:36:41 CEST 2022 - The aliscore output appears to be empty. Check results for BUSCO: 369368at7742
## Thu Jul 14 02:36:41 CEST 2022 - The aliscore output file does not exist. Check results for BUSCO: 415928at7742
## Thu Jul 14 02:36:41 CEST 2022 - The aliscore output appears to be empty. Check results for BUSCO: 415928at7742
## Thu Jul 14 02:36:42 CEST 2022 - The aliscore output file does not exist. Check results for BUSCO: 385758at7742
## Thu Jul 14 02:36:42 CEST 2022 - The aliscore output appears to be empty. Check results for BUSCO: 385758at7742
## Thu Jul 14 02:36:42 CEST 2022 - The aliscore output file does not exist. Check results for BUSCO: 390104at7742
## Thu Jul 14 02:36:42 CEST 2022 - The aliscore output appears to be empty. Check results for BUSCO: 390104at7742
## Thu Jul 14 02:36:43 CEST 2022 - The aliscore output file does not exist. Check results for BUSCO: 390104at7742
## Thu Jul 14 02:36:43 CEST 2022 - The aliscore output appears to be empty. Check results for BUSCO: 390104at7742
## Thu Jul 14 02:36:46 CEST 2022 - The aliscore output file does not exist. Check results for BUSCO: 347416at7742
## Thu Jul 14 02:36:46 CEST 2022 - The aliscore output appears to be empty. Check results for BUSCO: 347416at7742
## Thu Jul 14 02:36:47 CEST 2022 - The aliscore output file does not exist. Check results for BUSCO: 347954at7742
## Thu Jul 14 02:36:47 CEST 2022 - The aliscore output appears to be empty. Check results for BUSCO: 347954at7742
## Thu Jul 14 02:36:47 CEST 2022 - The aliscore output file does not exist. Check results for BUSCO: 304775at7742
## Thu Jul 14 02:36:47 CEST 2022 - The aliscore output appears to be empty. Check results for BUSCO: 304775at7742
## Thu Jul 14 02:36:48 CEST 2022 - The aliscore output file does not exist. Check results for BUSCO: 381371at7742
## Thu Jul 14 02:36:48 CEST 2022 - The aliscore output appears to be empty. Check results for BUSCO: 381371at7742
## Thu Jul 14 02:36:48 CEST 2022 - The aliscore output file does not exist. Check results for BUSCO: 400282at7742
## Thu Jul 14 02:36:48 CEST 2022 - The aliscore output appears to be empty. Check results for BUSCO: 400282at7742
## Thu Jul 14 02:36:49 CEST 2022 - The aliscore output file does not exist. Check results for BUSCO: 361767at7742
## Thu Jul 14 02:36:49 CEST 2022 - The aliscore output appears to be empty. Check results for BUSCO: 361767at7742
## Thu Jul 14 02:36:52 CEST 2022 - The aliscore output file does not exist. Check results for BUSCO: 400325at7742
## Thu Jul 14 02:36:52 CEST 2022 - The aliscore output appears to be empty. Check results for BUSCO: 400325at7742
## Thu Jul 14 02:36:52 CEST 2022 - The aliscore output file does not exist. Check results for BUSCO: 400874at7742
## Thu Jul 14 02:36:52 CEST 2022 - The aliscore output appears to be empty. Check results for BUSCO: 400874at7742
## Thu Jul 14 02:36:53 CEST 2022 - The aliscore output file does not exist. Check results for BUSCO: 371714at7742
## Thu Jul 14 02:36:53 CEST 2022 - The aliscore output appears to be empty. Check results for BUSCO: 371714at7742
## Thu Jul 14 02:36:54 CEST 2022 - The aliscore output file does not exist. Check results for BUSCO: 301105at7742
## Thu Jul 14 02:36:54 CEST 2022 - The aliscore output appears to be empty. Check results for BUSCO: 301105at7742
## Thu Jul 14 02:36:55 CEST 2022 - The aliscore output file does not exist. Check results for BUSCO: 405980at7742
## Thu Jul 14 02:36:55 CEST 2022 - The aliscore output appears to be empty. Check results for BUSCO: 405980at7742
## Thu Jul 14 02:36:55 CEST 2022 - The aliscore output file does not exist. Check results for BUSCO: 264273at7742
## Thu Jul 14 02:36:55 CEST 2022 - The aliscore output appears to be empty. Check results for BUSCO: 264273at7742
## Thu Jul 14 02:36:56 CEST 2022 - The aliscore output file does not exist. Check results for BUSCO: 407064at7742
## Thu Jul 14 02:36:56 CEST 2022 - The aliscore output appears to be empty. Check results for BUSCO: 407064at7742
## Thu Jul 14 02:36:57 CEST 2022 - The aliscore output file does not exist. Check results for BUSCO: 264273at7742
## Thu Jul 14 02:36:58 CEST 2022 - The aliscore output appears to be empty. Check results for BUSCO: 264273at7742
## Thu Jul 14 02:36:59 CEST 2022 - The aliscore output file does not exist. Check results for BUSCO: 371710at7742
## Thu Jul 14 02:36:59 CEST 2022 - The aliscore output appears to be empty. Check results for BUSCO: 371710at7742
## Thu Jul 14 02:37:00 CEST 2022 - The aliscore output file does not exist. Check results for BUSCO: 418345at7742
## Thu Jul 14 02:37:00 CEST 2022 - The aliscore output appears to be empty. Check results for BUSCO: 418345at7742
## Thu Jul 14 02:37:01 CEST 2022 - The aliscore output file does not exist. Check results for BUSCO: 368156at7742
## Thu Jul 14 02:37:01 CEST 2022 - The aliscore output appears to be empty. Check results for BUSCO: 368156at7742
## Thu Jul 14 02:37:01 CEST 2022 - The aliscore output file does not exist. Check results for BUSCO: 398268at7742
## Thu Jul 14 02:37:01 CEST 2022 - The aliscore output file does not exist. Check results for BUSCO: 419163at7742
## Thu Jul 14 02:37:01 CEST 2022 - The aliscore output appears to be empty. Check results for BUSCO: 398268at7742
## Thu Jul 14 02:37:01 CEST 2022 - The aliscore output appears to be empty. Check results for BUSCO: 419163at7742
## Thu Jul 14 02:37:03 CEST 2022 - The aliscore output file does not exist. Check results for BUSCO: 305009at7742
## Thu Jul 14 02:37:03 CEST 2022 - The aliscore output appears to be empty. Check results for BUSCO: 305009at7742
## Thu Jul 14 02:37:03 CEST 2022 - The aliscore output file does not exist. Check results for BUSCO: 264273at7742
## Thu Jul 14 02:37:03 CEST 2022 - The aliscore output appears to be empty. Check results for BUSCO: 264273at7742
## Thu Jul 14 02:37:05 CEST 2022 - The aliscore output file does not exist. Check results for BUSCO: 422508at7742
## Thu Jul 14 02:37:05 CEST 2022 - The aliscore output appears to be empty. Check results for BUSCO: 422508at7742
## Thu Jul 14 02:37:06 CEST 2022 - The aliscore output file does not exist. Check results for BUSCO: 363317at7742
## Thu Jul 14 02:37:06 CEST 2022 - The aliscore output appears to be empty. Check results for BUSCO: 363317at7742
## Thu Jul 14 02:37:06 CEST 2022 - The aliscore output file does not exist. Check results for BUSCO: 373225at7742
## Thu Jul 14 02:37:06 CEST 2022 - The aliscore output appears to be empty. Check results for BUSCO: 373225at7742
## Thu Jul 14 02:37:07 CEST 2022 - The aliscore output file does not exist. Check results for BUSCO: 378341at7742
## Thu Jul 14 02:37:07 CEST 2022 - The aliscore output appears to be empty. Check results for BUSCO: 378341at7742
## Thu Jul 14 02:37:08 CEST 2022 - The aliscore output file does not exist. Check results for BUSCO: 371710at7742
## Thu Jul 14 02:37:08 CEST 2022 - The aliscore output appears to be empty. Check results for BUSCO: 371710at7742
## Thu Jul 14 02:37:09 CEST 2022 - The aliscore output file does not exist. Check results for BUSCO: 398584at7742
## Thu Jul 14 02:37:09 CEST 2022 - The aliscore output appears to be empty. Check results for BUSCO: 398584at7742
## Thu Jul 14 02:37:09 CEST 2022 - The aliscore output file does not exist. Check results for BUSCO: 381106at7742
## Thu Jul 14 02:37:09 CEST 2022 - The aliscore output appears to be empty. Check results for BUSCO: 381106at7742
## Thu Jul 14 02:37:10 CEST 2022 - The aliscore output file does not exist. Check results for BUSCO: 421992at7742
## Thu Jul 14 02:37:10 CEST 2022 - The aliscore output appears to be empty. Check results for BUSCO: 421992at7742
## Thu Jul 14 02:37:12 CEST 2022 - The aliscore output file does not exist. Check results for BUSCO: 394099at7742
## Thu Jul 14 02:37:12 CEST 2022 - The aliscore output appears to be empty. Check results for BUSCO: 394099at7742
## Thu Jul 14 02:37:14 CEST 2022 - The aliscore output file does not exist. Check results for BUSCO: 385469at7742
## Thu Jul 14 02:37:14 CEST 2022 - The aliscore output file does not exist. Check results for BUSCO: 384470at7742
## Thu Jul 14 02:37:14 CEST 2022 - The aliscore output appears to be empty. Check results for BUSCO: 385469at7742
## Thu Jul 14 02:37:14 CEST 2022 - The aliscore output appears to be empty. Check results for BUSCO: 384470at7742
## Thu Jul 14 02:37:14 CEST 2022 - The aliscore output file does not exist. Check results for BUSCO: 401745at7742
## Thu Jul 14 02:37:14 CEST 2022 - The aliscore output appears to be empty. Check results for BUSCO: 401745at7742
## Thu Jul 14 02:37:16 CEST 2022 - The aliscore output file does not exist. Check results for BUSCO: 382432at7742
## Thu Jul 14 02:37:16 CEST 2022 - The aliscore output appears to be empty. Check results for BUSCO: 382432at7742
## Thu Jul 14 02:37:17 CEST 2022 - The aliscore output file does not exist. Check results for BUSCO: 406935at7742
## Thu Jul 14 02:37:17 CEST 2022 - The aliscore output appears to be empty. Check results for BUSCO: 406935at7742
## Thu Jul 14 02:37:18 CEST 2022 - The aliscore output file does not exist. Check results for BUSCO: 420868at7742
## Thu Jul 14 02:37:18 CEST 2022 - The aliscore output appears to be empty. Check results for BUSCO: 420868at7742
## Thu Jul 14 02:37:18 CEST 2022 - The aliscore output file does not exist. Check results for BUSCO: 401723at7742
## Thu Jul 14 02:37:18 CEST 2022 - The aliscore output appears to be empty. Check results for BUSCO: 401723at7742
## Thu Jul 14 02:37:19 CEST 2022 - The aliscore output file does not exist. Check results for BUSCO: 360409at7742
## Thu Jul 14 02:37:20 CEST 2022 - The aliscore output appears to be empty. Check results for BUSCO: 360409at7742
## Thu Jul 14 02:37:20 CEST 2022 - The aliscore output file does not exist. Check results for BUSCO: 319099at7742
## Thu Jul 14 02:37:20 CEST 2022 - The aliscore output appears to be empty. Check results for BUSCO: 319099at7742
## Thu Jul 14 02:37:20 CEST 2022 - The aliscore output file does not exist. Check results for BUSCO: 371581at7742
## Thu Jul 14 02:37:20 CEST 2022 - The aliscore output appears to be empty. Check results for BUSCO: 371581at7742
## Thu Jul 14 02:37:21 CEST 2022 - The aliscore output file does not exist. Check results for BUSCO: 337697at7742
## Thu Jul 14 02:37:21 CEST 2022 - The aliscore output appears to be empty. Check results for BUSCO: 337697at7742
## Thu Jul 14 02:37:22 CEST 2022 - The aliscore output file does not exist. Check results for BUSCO: 394099at7742
## Thu Jul 14 02:37:22 CEST 2022 - The aliscore output appears to be empty. Check results for BUSCO: 394099at7742
## Thu Jul 14 02:37:23 CEST 2022 - The aliscore output file does not exist. Check results for BUSCO: 330116at7742
## Thu Jul 14 02:37:23 CEST 2022 - The aliscore output appears to be empty. Check results for BUSCO: 330116at7742
## Thu Jul 14 02:37:23 CEST 2022 - The aliscore output file does not exist. Check results for BUSCO: 357862at7742
## Thu Jul 14 02:37:24 CEST 2022 - The aliscore output appears to be empty. Check results for BUSCO: 357862at7742
## Thu Jul 14 02:37:25 CEST 2022 - The aliscore output file does not exist. Check results for BUSCO: 304775at7742
## Thu Jul 14 02:37:25 CEST 2022 - The aliscore output appears to be empty. Check results for BUSCO: 304775at7742
## Thu Jul 14 02:37:26 CEST 2022 - The aliscore output file does not exist. Check results for BUSCO: 351113at7742
## Thu Jul 14 02:37:26 CEST 2022 - The aliscore output appears to be empty. Check results for BUSCO: 351113at7742
## Thu Jul 14 02:37:26 CEST 2022 - The aliscore output file does not exist. Check results for BUSCO: 336175at7742
## Thu Jul 14 02:37:26 CEST 2022 - The aliscore output appears to be empty. Check results for BUSCO: 336175at7742
## Thu Jul 14 02:37:28 CEST 2022 - The aliscore output file does not exist. Check results for BUSCO: 402833at7742
## Thu Jul 14 02:37:28 CEST 2022 - The aliscore output appears to be empty. Check results for BUSCO: 402833at7742
## Thu Jul 14 02:37:30 CEST 2022 - The aliscore output file does not exist. Check results for BUSCO: 345592at7742
## Thu Jul 14 02:37:30 CEST 2022 - The aliscore output appears to be empty. Check results for BUSCO: 345592at7742
## Thu Jul 14 02:37:30 CEST 2022 - The aliscore output file does not exist. Check results for BUSCO: 328338at7742
## Thu Jul 14 02:37:30 CEST 2022 - The aliscore output appears to be empty. Check results for BUSCO: 328338at7742
## Thu Jul 14 02:37:30 CEST 2022 - The aliscore output file does not exist. Check results for BUSCO: 289723at7742
## Thu Jul 14 02:37:30 CEST 2022 - The aliscore output appears to be empty. Check results for BUSCO: 289723at7742
## Thu Jul 14 02:37:32 CEST 2022 - The aliscore output file does not exist. Check results for BUSCO: 308505at7742
## Thu Jul 14 02:37:32 CEST 2022 - The aliscore output appears to be empty. Check results for BUSCO: 308505at7742
## Thu Jul 14 02:37:33 CEST 2022 - The aliscore output file does not exist. Check results for BUSCO: 346605at7742
## Thu Jul 14 02:37:33 CEST 2022 - The aliscore output appears to be empty. Check results for BUSCO: 346605at7742
## Thu Jul 14 02:37:33 CEST 2022 - The aliscore output file does not exist. Check results for BUSCO: 400282at7742
## Thu Jul 14 02:37:33 CEST 2022 - The aliscore output appears to be empty. Check results for BUSCO: 400282at7742
## Thu Jul 14 02:37:34 CEST 2022 - The aliscore output file does not exist. Check results for BUSCO: 361358at7742
## Thu Jul 14 02:37:34 CEST 2022 - The aliscore output appears to be empty. Check results for BUSCO: 361358at7742
## Thu Jul 14 02:37:35 CEST 2022 - The aliscore output file does not exist. Check results for BUSCO: 342980at7742
## Thu Jul 14 02:37:35 CEST 2022 - The aliscore output appears to be empty. Check results for BUSCO: 342980at7742
## Thu Jul 14 02:37:36 CEST 2022 - The aliscore output file does not exist. Check results for BUSCO: 368156at7742
## Thu Jul 14 02:37:36 CEST 2022 - The aliscore output appears to be empty. Check results for BUSCO: 368156at7742
## Thu Jul 14 02:37:37 CEST 2022 - The aliscore output file does not exist. Check results for BUSCO: 395764at7742
## Thu Jul 14 02:37:37 CEST 2022 - The aliscore output appears to be empty. Check results for BUSCO: 395764at7742
## Thu Jul 14 02:37:38 CEST 2022 - The aliscore output file does not exist. Check results for BUSCO: 356826at7742
## Thu Jul 14 02:37:38 CEST 2022 - The aliscore output appears to be empty. Check results for BUSCO: 356826at7742
## Thu Jul 14 02:37:38 CEST 2022 - The aliscore output file does not exist. Check results for BUSCO: 342980at7742
## Thu Jul 14 02:37:38 CEST 2022 - The aliscore output appears to be empty. Check results for BUSCO: 342980at7742
## Thu Jul 14 02:37:40 CEST 2022 - The aliscore output file does not exist. Check results for BUSCO: 394099at7742
## Thu Jul 14 02:37:40 CEST 2022 - The aliscore output appears to be empty. Check results for BUSCO: 394099at7742
## Thu Jul 14 02:37:42 CEST 2022 - The aliscore output file does not exist. Check results for BUSCO: 329907at7742
## Thu Jul 14 02:37:42 CEST 2022 - The aliscore output appears to be empty. Check results for BUSCO: 329907at7742
## Thu Jul 14 02:37:42 CEST 2022 - The aliscore output file does not exist. Check results for BUSCO: 395170at7742
## Thu Jul 14 02:37:42 CEST 2022 - The aliscore output appears to be empty. Check results for BUSCO: 395170at7742
## Thu Jul 14 02:37:42 CEST 2022 - The aliscore output file does not exist. Check results for BUSCO: 393852at7742
## Thu Jul 14 02:37:42 CEST 2022 - The aliscore output appears to be empty. Check results for BUSCO: 393852at7742
## Thu Jul 14 02:37:43 CEST 2022 - The aliscore output file does not exist. Check results for BUSCO: 351113at7742
## Thu Jul 14 02:37:43 CEST 2022 - The aliscore output appears to be empty. Check results for BUSCO: 351113at7742
## Thu Jul 14 02:37:44 CEST 2022 - The aliscore output file does not exist. Check results for BUSCO: 402833at7742
## Thu Jul 14 02:37:44 CEST 2022 - The aliscore output appears to be empty. Check results for BUSCO: 402833at7742
## Thu Jul 14 02:37:44 CEST 2022 - The aliscore output file does not exist. Check results for BUSCO: 381371at7742
## Thu Jul 14 02:37:44 CEST 2022 - The aliscore output appears to be empty. Check results for BUSCO: 381371at7742
## Thu Jul 14 02:37:45 CEST 2022 - The aliscore output file does not exist. Check results for BUSCO: 337603at7742
## Thu Jul 14 02:37:45 CEST 2022 - The aliscore output appears to be empty. Check results for BUSCO: 337603at7742
## Thu Jul 14 02:37:47 CEST 2022 - The aliscore output file does not exist. Check results for BUSCO: 360409at7742
## Thu Jul 14 02:37:47 CEST 2022 - The aliscore output appears to be empty. Check results for BUSCO: 360409at7742
## Thu Jul 14 02:37:47 CEST 2022 - The aliscore output file does not exist. Check results for BUSCO: 328338at7742
## Thu Jul 14 02:37:47 CEST 2022 - The aliscore output appears to be empty. Check results for BUSCO: 328338at7742
## Thu Jul 14 02:37:49 CEST 2022 - The aliscore output file does not exist. Check results for BUSCO: 422508at7742
## Thu Jul 14 02:37:49 CEST 2022 - The aliscore output appears to be empty. Check results for BUSCO: 422508at7742
## Thu Jul 14 02:37:49 CEST 2022 - The aliscore output file does not exist. Check results for BUSCO: 370331at7742
## Thu Jul 14 02:37:49 CEST 2022 - The aliscore output appears to be empty. Check results for BUSCO: 370331at7742
## Thu Jul 14 02:37:49 CEST 2022 - The aliscore output file does not exist. Check results for BUSCO: 365811at7742
## Thu Jul 14 02:37:49 CEST 2022 - The aliscore output appears to be empty. Check results for BUSCO: 365811at7742
## Thu Jul 14 02:37:51 CEST 2022 - The aliscore output file does not exist. Check results for BUSCO: 405236at7742
## Thu Jul 14 02:37:52 CEST 2022 - The aliscore output appears to be empty. Check results for BUSCO: 405236at7742
## Thu Jul 14 02:37:52 CEST 2022 - The aliscore output file does not exist. Check results for BUSCO: 412561at7742
## Thu Jul 14 02:37:52 CEST 2022 - The aliscore output appears to be empty. Check results for BUSCO: 412561at7742
## Thu Jul 14 02:37:53 CEST 2022 - The aliscore output file does not exist. Check results for BUSCO: 304775at7742
## Thu Jul 14 02:37:53 CEST 2022 - The aliscore output appears to be empty. Check results for BUSCO: 304775at7742
## Thu Jul 14 02:37:53 CEST 2022 - The aliscore output file does not exist. Check results for BUSCO: 418345at7742
## Thu Jul 14 02:37:53 CEST 2022 - The aliscore output appears to be empty. Check results for BUSCO: 418345at7742
## Thu Jul 14 02:37:55 CEST 2022 - The aliscore output file does not exist. Check results for BUSCO: 330116at7742
## Thu Jul 14 02:37:55 CEST 2022 - The aliscore output appears to be empty. Check results for BUSCO: 330116at7742
## Thu Jul 14 02:37:55 CEST 2022 - The aliscore output file does not exist. Check results for BUSCO: 356826at7742
## Thu Jul 14 02:37:55 CEST 2022 - The aliscore output appears to be empty. Check results for BUSCO: 356826at7742
## Thu Jul 14 02:37:55 CEST 2022 - The aliscore output file does not exist. Check results for BUSCO: 398584at7742
## Thu Jul 14 02:37:55 CEST 2022 - The aliscore output appears to be empty. Check results for BUSCO: 398584at7742
## Thu Jul 14 02:37:57 CEST 2022 - The aliscore output file does not exist. Check results for BUSCO: 398761at7742
## Thu Jul 14 02:37:57 CEST 2022 - The aliscore output appears to be empty. Check results for BUSCO: 398761at7742
## Thu Jul 14 02:37:58 CEST 2022 - The aliscore output file does not exist. Check results for BUSCO: 378341at7742
## Thu Jul 14 02:37:58 CEST 2022 - The aliscore output appears to be empty. Check results for BUSCO: 378341at7742
## Thu Jul 14 02:37:58 CEST 2022 - The aliscore output file does not exist. Check results for BUSCO: 398761at7742
## Thu Jul 14 02:37:58 CEST 2022 - The aliscore output appears to be empty. Check results for BUSCO: 398761at7742
## Thu Jul 14 02:37:58 CEST 2022 - The aliscore output file does not exist. Check results for BUSCO: 323438at7742
## Thu Jul 14 02:37:58 CEST 2022 - The aliscore output appears to be empty. Check results for BUSCO: 323438at7742
## Thu Jul 14 02:38:00 CEST 2022 - The aliscore output file does not exist. Check results for BUSCO: 361767at7742
## Thu Jul 14 02:38:00 CEST 2022 - The aliscore output appears to be empty. Check results for BUSCO: 361767at7742
## Thu Jul 14 02:38:00 CEST 2022 - The aliscore output file does not exist. Check results for BUSCO: 385398at7742
## Thu Jul 14 02:38:00 CEST 2022 - The aliscore output appears to be empty. Check results for BUSCO: 385398at7742
## Thu Jul 14 02:38:02 CEST 2022 - The aliscore output file does not exist. Check results for BUSCO: 271346at7742
## Thu Jul 14 02:38:02 CEST 2022 - The aliscore output appears to be empty. Check results for BUSCO: 271346at7742
## Thu Jul 14 02:38:02 CEST 2022 - The aliscore output file does not exist. Check results for BUSCO: 404437at7742
## Thu Jul 14 02:38:02 CEST 2022 - The aliscore output appears to be empty. Check results for BUSCO: 404437at7742
## Thu Jul 14 02:38:03 CEST 2022 - The aliscore output file does not exist. Check results for BUSCO: 361802at7742
## Thu Jul 14 02:38:03 CEST 2022 - The aliscore output appears to be empty. Check results for BUSCO: 361802at7742
## Thu Jul 14 02:38:06 CEST 2022 - The aliscore output file does not exist. Check results for BUSCO: 412708at7742
## Thu Jul 14 02:38:06 CEST 2022 - The aliscore output appears to be empty. Check results for BUSCO: 412708at7742
## Thu Jul 14 02:38:06 CEST 2022 - The aliscore output file does not exist. Check results for BUSCO: 397807at7742
## Thu Jul 14 02:38:07 CEST 2022 - The aliscore output appears to be empty. Check results for BUSCO: 397807at7742
## Thu Jul 14 02:38:07 CEST 2022 - The aliscore output file does not exist. Check results for BUSCO: 308505at7742
## Thu Jul 14 02:38:07 CEST 2022 - The aliscore output appears to be empty. Check results for BUSCO: 308505at7742
## Thu Jul 14 02:38:07 CEST 2022 - The aliscore output file does not exist. Check results for BUSCO: 352153at7742
## Thu Jul 14 02:38:07 CEST 2022 - The aliscore output appears to be empty. Check results for BUSCO: 352153at7742
## Thu Jul 14 02:38:07 CEST 2022 - The aliscore output file does not exist. Check results for BUSCO: 397807at7742
## Thu Jul 14 02:38:07 CEST 2022 - The aliscore output appears to be empty. Check results for BUSCO: 397807at7742
## Thu Jul 14 02:38:09 CEST 2022 - The aliscore output file does not exist. Check results for BUSCO: 381431at7742
## Thu Jul 14 02:38:09 CEST 2022 - The aliscore output appears to be empty. Check results for BUSCO: 381431at7742
## Thu Jul 14 02:38:10 CEST 2022 - The aliscore output file does not exist. Check results for BUSCO: 395170at7742
## Thu Jul 14 02:38:10 CEST 2022 - The aliscore output appears to be empty. Check results for BUSCO: 395170at7742
## Thu Jul 14 02:38:11 CEST 2022 - The aliscore output file does not exist. Check results for BUSCO: 337697at7742
## Thu Jul 14 02:38:11 CEST 2022 - The aliscore output file does not exist. Check results for BUSCO: 345592at7742
## Thu Jul 14 02:38:11 CEST 2022 - The aliscore output appears to be empty. Check results for BUSCO: 337697at7742
## Thu Jul 14 02:38:11 CEST 2022 - The aliscore output appears to be empty. Check results for BUSCO: 345592at7742
## Thu Jul 14 02:38:12 CEST 2022 - The aliscore output file does not exist. Check results for BUSCO: 384470at7742
## Thu Jul 14 02:38:12 CEST 2022 - The aliscore output appears to be empty. Check results for BUSCO: 384470at7742
## Thu Jul 14 02:38:13 CEST 2022 - The aliscore output file does not exist. Check results for BUSCO: 398761at7742
## Thu Jul 14 02:38:13 CEST 2022 - The aliscore output appears to be empty. Check results for BUSCO: 398761at7742
## Thu Jul 14 02:38:13 CEST 2022 - The aliscore output file does not exist. Check results for BUSCO: 407321at7742
## Thu Jul 14 02:38:13 CEST 2022 - The aliscore output appears to be empty. Check results for BUSCO: 407321at7742
## Thu Jul 14 02:38:15 CEST 2022 - The aliscore output file does not exist. Check results for BUSCO: 351522at7742
## Thu Jul 14 02:38:15 CEST 2022 - The aliscore output appears to be empty. Check results for BUSCO: 351522at7742
## Thu Jul 14 02:38:15 CEST 2022 - The aliscore output file does not exist. Check results for BUSCO: 399667at7742
## Thu Jul 14 02:38:15 CEST 2022 - The aliscore output appears to be empty. Check results for BUSCO: 399667at7742
## Thu Jul 14 02:38:16 CEST 2022 - The aliscore output file does not exist. Check results for BUSCO: 357862at7742
## Thu Jul 14 02:38:16 CEST 2022 - The aliscore output appears to be empty. Check results for BUSCO: 357862at7742
## Thu Jul 14 02:38:18 CEST 2022 - The aliscore output file does not exist. Check results for BUSCO: 327681at7742
## Thu Jul 14 02:38:18 CEST 2022 - The aliscore output appears to be empty. Check results for BUSCO: 327681at7742
## Thu Jul 14 02:38:19 CEST 2022 - The aliscore output file does not exist. Check results for BUSCO: 323438at7742
## Thu Jul 14 02:38:19 CEST 2022 - The aliscore output file does not exist. Check results for BUSCO: 390017at7742
## Thu Jul 14 02:38:19 CEST 2022 - The aliscore output appears to be empty. Check results for BUSCO: 323438at7742
## Thu Jul 14 02:38:19 CEST 2022 - The aliscore output appears to be empty. Check results for BUSCO: 390017at7742
## Thu Jul 14 02:38:21 CEST 2022 - The aliscore output file does not exist. Check results for BUSCO: 331131at7742
## Thu Jul 14 02:38:21 CEST 2022 - The aliscore output appears to be empty. Check results for BUSCO: 331131at7742
## Thu Jul 14 02:38:21 CEST 2022 - The aliscore output file does not exist. Check results for BUSCO: 332363at7742
## Thu Jul 14 02:38:21 CEST 2022 - The aliscore output appears to be empty. Check results for BUSCO: 332363at7742
## Thu Jul 14 02:38:22 CEST 2022 - The aliscore output file does not exist. Check results for BUSCO: 406935at7742
## Thu Jul 14 02:38:22 CEST 2022 - The aliscore output appears to be empty. Check results for BUSCO: 406935at7742
## Thu Jul 14 02:38:23 CEST 2022 - The aliscore output file does not exist. Check results for BUSCO: 339259at7742
## Thu Jul 14 02:38:23 CEST 2022 - The aliscore output appears to be empty. Check results for BUSCO: 339259at7742
## Thu Jul 14 02:38:23 CEST 2022 - The aliscore output file does not exist. Check results for BUSCO: 293761at7742
## Thu Jul 14 02:38:23 CEST 2022 - The aliscore output appears to be empty. Check results for BUSCO: 293761at7742
## Thu Jul 14 02:38:24 CEST 2022 - The aliscore output file does not exist. Check results for BUSCO: 288056at7742
## Thu Jul 14 02:38:24 CEST 2022 - The aliscore output file does not exist. Check results for BUSCO: 293761at7742
## Thu Jul 14 02:38:24 CEST 2022 - The aliscore output appears to be empty. Check results for BUSCO: 288056at7742
## Thu Jul 14 02:38:24 CEST 2022 - The aliscore output appears to be empty. Check results for BUSCO: 293761at7742
## Thu Jul 14 02:38:24 CEST 2022 - The aliscore output file does not exist. Check results for BUSCO: 293761at7742
## Thu Jul 14 02:38:25 CEST 2022 - The aliscore output appears to be empty. Check results for BUSCO: 293761at7742
## Thu Jul 14 02:38:27 CEST 2022 - The aliscore output file does not exist. Check results for BUSCO: 399133at7742
## Thu Jul 14 02:38:27 CEST 2022 - The aliscore output appears to be empty. Check results for BUSCO: 399133at7742
## Thu Jul 14 02:38:27 CEST 2022 - The aliscore output file does not exist. Check results for BUSCO: 324813at7742
## Thu Jul 14 02:38:27 CEST 2022 - The aliscore output appears to be empty. Check results for BUSCO: 324813at7742
## Thu Jul 14 02:38:29 CEST 2022 - The aliscore output file does not exist. Check results for BUSCO: 381371at7742
## Thu Jul 14 02:38:29 CEST 2022 - The aliscore output appears to be empty. Check results for BUSCO: 381371at7742
## Thu Jul 14 02:38:29 CEST 2022 - The aliscore output file does not exist. Check results for BUSCO: 437625at7742
## Thu Jul 14 02:38:29 CEST 2022 - The aliscore output appears to be empty. Check results for BUSCO: 437625at7742
## Thu Jul 14 02:38:30 CEST 2022 - The aliscore output file does not exist. Check results for BUSCO: 337603at7742
## Thu Jul 14 02:38:30 CEST 2022 - The aliscore output appears to be empty. Check results for BUSCO: 337603at7742
## Thu Jul 14 02:38:31 CEST 2022 - The aliscore output file does not exist. Check results for BUSCO: 365924at7742
## Thu Jul 14 02:38:31 CEST 2022 - The aliscore output appears to be empty. Check results for BUSCO: 365924at7742
## Thu Jul 14 02:38:33 CEST 2022 - The aliscore output file does not exist. Check results for BUSCO: 379007at7742
## Thu Jul 14 02:38:33 CEST 2022 - The aliscore output appears to be empty. Check results for BUSCO: 379007at7742
## Thu Jul 14 02:38:33 CEST 2022 - The aliscore output file does not exist. Check results for BUSCO: 402520at7742
## Thu Jul 14 02:38:33 CEST 2022 - The aliscore output appears to be empty. Check results for BUSCO: 402520at7742
## Thu Jul 14 02:38:35 CEST 2022 - The aliscore output file does not exist. Check results for BUSCO: 407871at7742
## Thu Jul 14 02:38:35 CEST 2022 - The aliscore output appears to be empty. Check results for BUSCO: 407871at7742
## Thu Jul 14 02:38:36 CEST 2022 - The aliscore output file does not exist. Check results for BUSCO: 388019at7742
## Thu Jul 14 02:38:36 CEST 2022 - The aliscore output appears to be empty. Check results for BUSCO: 388019at7742
## Thu Jul 14 02:38:36 CEST 2022 - The aliscore output file does not exist. Check results for BUSCO: 416829at7742
## Thu Jul 14 02:38:36 CEST 2022 - The aliscore output appears to be empty. Check results for BUSCO: 416829at7742
## Thu Jul 14 02:38:37 CEST 2022 - The aliscore output file does not exist. Check results for BUSCO: 329907at7742
## Thu Jul 14 02:38:37 CEST 2022 - The aliscore output appears to be empty. Check results for BUSCO: 329907at7742
## Thu Jul 14 02:38:37 CEST 2022 - The aliscore output file does not exist. Check results for BUSCO: 288056at7742
## Thu Jul 14 02:38:37 CEST 2022 - The aliscore output appears to be empty. Check results for BUSCO: 288056at7742
## Thu Jul 14 02:38:39 CEST 2022 - The aliscore output file does not exist. Check results for BUSCO: 405980at7742
## Thu Jul 14 02:38:39 CEST 2022 - The aliscore output appears to be empty. Check results for BUSCO: 405980at7742
## Thu Jul 14 02:38:40 CEST 2022 - The aliscore output file does not exist. Check results for BUSCO: 379007at7742
## Thu Jul 14 02:38:40 CEST 2022 - The aliscore output appears to be empty. Check results for BUSCO: 379007at7742
## Thu Jul 14 02:38:40 CEST 2022 - The aliscore output file does not exist. Check results for BUSCO: 379661at7742
## Thu Jul 14 02:38:40 CEST 2022 - The aliscore output appears to be empty. Check results for BUSCO: 379661at7742
## Thu Jul 14 02:38:42 CEST 2022 - The aliscore output file does not exist. Check results for BUSCO: 364898at7742
## Thu Jul 14 02:38:42 CEST 2022 - The aliscore output appears to be empty. Check results for BUSCO: 364898at7742
## Thu Jul 14 02:38:43 CEST 2022 - The aliscore output file does not exist. Check results for BUSCO: 407871at7742
## Thu Jul 14 02:38:43 CEST 2022 - The aliscore output appears to be empty. Check results for BUSCO: 407871at7742
## Thu Jul 14 02:38:44 CEST 2022 - The aliscore output file does not exist. Check results for BUSCO: 427300at7742
## Thu Jul 14 02:38:44 CEST 2022 - The aliscore output appears to be empty. Check results for BUSCO: 427300at7742
## Thu Jul 14 02:38:44 CEST 2022 - The aliscore output file does not exist. Check results for BUSCO: 323567at7742
## Thu Jul 14 02:38:44 CEST 2022 - The aliscore output appears to be empty. Check results for BUSCO: 323567at7742
## Thu Jul 14 02:38:45 CEST 2022 - The aliscore output file does not exist. Check results for BUSCO: 418711at7742
## Thu Jul 14 02:38:45 CEST 2022 - The aliscore output appears to be empty. Check results for BUSCO: 418711at7742
## Thu Jul 14 02:38:46 CEST 2022 - The aliscore output file does not exist. Check results for BUSCO: 325447at7742
## Thu Jul 14 02:38:46 CEST 2022 - The aliscore output appears to be empty. Check results for BUSCO: 325447at7742
## Thu Jul 14 02:38:46 CEST 2022 - The aliscore output file does not exist. Check results for BUSCO: 357862at7742
## Thu Jul 14 02:38:46 CEST 2022 - The aliscore output appears to be empty. Check results for BUSCO: 357862at7742
## Thu Jul 14 02:38:46 CEST 2022 - The aliscore output file does not exist. Check results for BUSCO: 342980at7742
## Thu Jul 14 02:38:46 CEST 2022 - The aliscore output appears to be empty. Check results for BUSCO: 342980at7742
## Thu Jul 14 02:38:48 CEST 2022 - The aliscore output file does not exist. Check results for BUSCO: 329907at7742
## Thu Jul 14 02:38:48 CEST 2022 - The aliscore output appears to be empty. Check results for BUSCO: 329907at7742
## Thu Jul 14 02:38:49 CEST 2022 - The aliscore output file does not exist. Check results for BUSCO: 356826at7742
## Thu Jul 14 02:38:49 CEST 2022 - The aliscore output appears to be empty. Check results for BUSCO: 356826at7742
## Thu Jul 14 02:38:50 CEST 2022 - The aliscore output file does not exist. Check results for BUSCO: 289723at7742
## Thu Jul 14 02:38:50 CEST 2022 - The aliscore output appears to be empty. Check results for BUSCO: 289723at7742
## Thu Jul 14 02:38:50 CEST 2022 - The aliscore output file does not exist. Check results for BUSCO: 350289at7742
## Thu Jul 14 02:38:50 CEST 2022 - The aliscore output appears to be empty. Check results for BUSCO: 350289at7742
## Thu Jul 14 02:38:51 CEST 2022 - The aliscore output file does not exist. Check results for BUSCO: 271346at7742
## Thu Jul 14 02:38:51 CEST 2022 - The aliscore output appears to be empty. Check results for BUSCO: 271346at7742
## Thu Jul 14 02:38:52 CEST 2022 - The aliscore output file does not exist. Check results for BUSCO: 386476at7742
## Thu Jul 14 02:38:52 CEST 2022 - The aliscore output appears to be empty. Check results for BUSCO: 386476at7742
## Thu Jul 14 02:38:52 CEST 2022 - The aliscore output file does not exist. Check results for BUSCO: 370331at7742
## Thu Jul 14 02:38:52 CEST 2022 - The aliscore output appears to be empty. Check results for BUSCO: 370331at7742
## Thu Jul 14 02:38:53 CEST 2022 - The aliscore output file does not exist. Check results for BUSCO: 393599at7742
## Thu Jul 14 02:38:53 CEST 2022 - The aliscore output appears to be empty. Check results for BUSCO: 393599at7742
## Thu Jul 14 02:38:55 CEST 2022 - The aliscore output file does not exist. Check results for BUSCO: 407871at7742
## Thu Jul 14 02:38:55 CEST 2022 - The aliscore output appears to be empty. Check results for BUSCO: 407871at7742
## Thu Jul 14 02:38:55 CEST 2022 - The aliscore output file does not exist. Check results for BUSCO: 400282at7742
## Thu Jul 14 02:38:55 CEST 2022 - The aliscore output appears to be empty. Check results for BUSCO: 400282at7742
## Thu Jul 14 02:38:55 CEST 2022 - The aliscore output file does not exist. Check results for BUSCO: 377681at7742
## Thu Jul 14 02:38:55 CEST 2022 - The aliscore output appears to be empty. Check results for BUSCO: 377681at7742
## Thu Jul 14 02:38:58 CEST 2022 - The aliscore output file does not exist. Check results for BUSCO: 381830at7742
## Thu Jul 14 02:38:58 CEST 2022 - The aliscore output appears to be empty. Check results for BUSCO: 381830at7742
## Thu Jul 14 02:38:58 CEST 2022 - The aliscore output file does not exist. Check results for BUSCO: 386476at7742
## Thu Jul 14 02:38:58 CEST 2022 - The aliscore output appears to be empty. Check results for BUSCO: 386476at7742
## Thu Jul 14 02:38:59 CEST 2022 - The aliscore output file does not exist. Check results for BUSCO: 339259at7742
## Thu Jul 14 02:38:59 CEST 2022 - The aliscore output appears to be empty. Check results for BUSCO: 339259at7742
## Thu Jul 14 02:39:01 CEST 2022 - The aliscore output file does not exist. Check results for BUSCO: 430342at7742
## Thu Jul 14 02:39:01 CEST 2022 - The aliscore output appears to be empty. Check results for BUSCO: 430342at7742
## Thu Jul 14 02:39:02 CEST 2022 - The aliscore output file does not exist. Check results for BUSCO: 371710at7742
## Thu Jul 14 02:39:02 CEST 2022 - The aliscore output appears to be empty. Check results for BUSCO: 371710at7742
## Thu Jul 14 02:39:02 CEST 2022 - The aliscore output file does not exist. Check results for BUSCO: 325703at7742
## Thu Jul 14 02:39:02 CEST 2022 - The aliscore output appears to be empty. Check results for BUSCO: 325703at7742
## Thu Jul 14 02:39:02 CEST 2022 - The aliscore output file does not exist. Check results for BUSCO: 379007at7742
## Thu Jul 14 02:39:02 CEST 2022 - The aliscore output file does not exist. Check results for BUSCO: 402833at7742
## Thu Jul 14 02:39:02 CEST 2022 - The aliscore output appears to be empty. Check results for BUSCO: 379007at7742
## Thu Jul 14 02:39:02 CEST 2022 - The aliscore output appears to be empty. Check results for BUSCO: 402833at7742
## Thu Jul 14 02:39:03 CEST 2022 - The aliscore output file does not exist. Check results for BUSCO: 339259at7742
## Thu Jul 14 02:39:03 CEST 2022 - The aliscore output appears to be empty. Check results for BUSCO: 339259at7742
## Thu Jul 14 02:39:04 CEST 2022 - The aliscore output file does not exist. Check results for BUSCO: 419468at7742
## Thu Jul 14 02:39:05 CEST 2022 - The aliscore output appears to be empty. Check results for BUSCO: 419468at7742
## Thu Jul 14 02:39:05 CEST 2022 - The aliscore output file does not exist. Check results for BUSCO: 430342at7742
## Thu Jul 14 02:39:05 CEST 2022 - The aliscore output appears to be empty. Check results for BUSCO: 430342at7742
## Thu Jul 14 02:39:07 CEST 2022 - The aliscore output file does not exist. Check results for BUSCO: 386476at7742
## Thu Jul 14 02:39:07 CEST 2022 - The aliscore output appears to be empty. Check results for BUSCO: 386476at7742
## Thu Jul 14 02:39:07 CEST 2022 - The aliscore output file does not exist. Check results for BUSCO: 388097at7742
## Thu Jul 14 02:39:08 CEST 2022 - The aliscore output appears to be empty. Check results for BUSCO: 388097at7742
## Thu Jul 14 02:39:08 CEST 2022 - The aliscore output file does not exist. Check results for BUSCO: 418711at7742
## Thu Jul 14 02:39:08 CEST 2022 - The aliscore output appears to be empty. Check results for BUSCO: 418711at7742
## Thu Jul 14 02:39:09 CEST 2022 - The aliscore output file does not exist. Check results for BUSCO: 415130at7742
## Thu Jul 14 02:39:09 CEST 2022 - The aliscore output appears to be empty. Check results for BUSCO: 415130at7742
## Thu Jul 14 02:39:09 CEST 2022 - The aliscore output file does not exist. Check results for BUSCO: 390810at7742
## Thu Jul 14 02:39:09 CEST 2022 - The aliscore output appears to be empty. Check results for BUSCO: 390810at7742
## Thu Jul 14 02:39:09 CEST 2022 - The aliscore output file does not exist. Check results for BUSCO: 369390at7742
## Thu Jul 14 02:39:09 CEST 2022 - The aliscore output appears to be empty. Check results for BUSCO: 369390at7742
## Thu Jul 14 02:39:10 CEST 2022 - The aliscore output file does not exist. Check results for BUSCO: 377681at7742
## Thu Jul 14 02:39:10 CEST 2022 - The aliscore output appears to be empty. Check results for BUSCO: 377681at7742
## Thu Jul 14 02:39:12 CEST 2022 - The aliscore output file does not exist. Check results for BUSCO: 337917at7742
## Thu Jul 14 02:39:12 CEST 2022 - The aliscore output appears to be empty. Check results for BUSCO: 337917at7742
## Thu Jul 14 02:39:14 CEST 2022 - The aliscore output file does not exist. Check results for BUSCO: 289723at7742
## Thu Jul 14 02:39:14 CEST 2022 - The aliscore output appears to be empty. Check results for BUSCO: 289723at7742
## Thu Jul 14 02:39:14 CEST 2022 - The aliscore output file does not exist. Check results for BUSCO: 399531at7742
## Thu Jul 14 02:39:14 CEST 2022 - The aliscore output appears to be empty. Check results for BUSCO: 399531at7742
## Thu Jul 14 02:39:14 CEST 2022 - The aliscore output file does not exist. Check results for BUSCO: 390017at7742
## Thu Jul 14 02:39:14 CEST 2022 - The aliscore output appears to be empty. Check results for BUSCO: 390017at7742
## Thu Jul 14 02:39:14 CEST 2022 - The aliscore output file does not exist. Check results for BUSCO: 372023at7742
## Thu Jul 14 02:39:14 CEST 2022 - The aliscore output appears to be empty. Check results for BUSCO: 372023at7742
## Thu Jul 14 02:39:16 CEST 2022 - The aliscore output file does not exist. Check results for BUSCO: 270182at7742
## Thu Jul 14 02:39:16 CEST 2022 - The aliscore output appears to be empty. Check results for BUSCO: 270182at7742
## Thu Jul 14 02:39:17 CEST 2022 - The aliscore output file does not exist. Check results for BUSCO: 324813at7742
## Thu Jul 14 02:39:17 CEST 2022 - The aliscore output appears to be empty. Check results for BUSCO: 324813at7742
## Thu Jul 14 02:39:18 CEST 2022 - The aliscore output file does not exist. Check results for BUSCO: 421992at7742
## Thu Jul 14 02:39:18 CEST 2022 - The aliscore output appears to be empty. Check results for BUSCO: 421992at7742
## Thu Jul 14 02:39:19 CEST 2022 - The aliscore output file does not exist. Check results for BUSCO: 308505at7742
## Thu Jul 14 02:39:19 CEST 2022 - The aliscore output appears to be empty. Check results for BUSCO: 308505at7742
## Thu Jul 14 02:39:20 CEST 2022 - The aliscore output file does not exist. Check results for BUSCO: 370331at7742
## Thu Jul 14 02:39:20 CEST 2022 - The aliscore output appears to be empty. Check results for BUSCO: 370331at7742
## Thu Jul 14 02:39:20 CEST 2022 - The aliscore output file does not exist. Check results for BUSCO: 271346at7742
## Thu Jul 14 02:39:20 CEST 2022 - The aliscore output appears to be empty. Check results for BUSCO: 271346at7742
## Thu Jul 14 02:39:21 CEST 2022 - The aliscore output file does not exist. Check results for BUSCO: 378120at7742
## Thu Jul 14 02:39:21 CEST 2022 - The aliscore output appears to be empty. Check results for BUSCO: 378120at7742
## Thu Jul 14 02:39:22 CEST 2022 - The aliscore output file does not exist. Check results for BUSCO: 368156at7742
## Thu Jul 14 02:39:22 CEST 2022 - The aliscore output appears to be empty. Check results for BUSCO: 368156at7742
## Thu Jul 14 02:39:23 CEST 2022 - The aliscore output file does not exist. Check results for BUSCO: 390810at7742
## Thu Jul 14 02:39:23 CEST 2022 - The aliscore output appears to be empty. Check results for BUSCO: 390810at7742
## Thu Jul 14 02:39:24 CEST 2022 - The aliscore output file does not exist. Check results for BUSCO: 420868at7742
## Thu Jul 14 02:39:24 CEST 2022 - The aliscore output appears to be empty. Check results for BUSCO: 420868at7742
## Thu Jul 14 02:39:26 CEST 2022 - The aliscore output file does not exist. Check results for BUSCO: 361078at7742
## Thu Jul 14 02:39:26 CEST 2022 - The aliscore output appears to be empty. Check results for BUSCO: 361078at7742
## Thu Jul 14 02:39:26 CEST 2022 - The aliscore output file does not exist. Check results for BUSCO: 325542at7742
## Thu Jul 14 02:39:26 CEST 2022 - The aliscore output appears to be empty. Check results for BUSCO: 325542at7742
## Thu Jul 14 02:39:26 CEST 2022 - The aliscore output file does not exist. Check results for BUSCO: 377681at7742
## Thu Jul 14 02:39:26 CEST 2022 - The aliscore output appears to be empty. Check results for BUSCO: 377681at7742
## Thu Jul 14 02:39:28 CEST 2022 - The aliscore output file does not exist. Check results for BUSCO: 327681at7742
## Thu Jul 14 02:39:28 CEST 2022 - The aliscore output appears to be empty. Check results for BUSCO: 327681at7742
## Thu Jul 14 02:39:29 CEST 2022 - The aliscore output file does not exist. Check results for BUSCO: 327681at7742
## Thu Jul 14 02:39:29 CEST 2022 - The aliscore output file does not exist. Check results for BUSCO: 337917at7742
## Thu Jul 14 02:39:29 CEST 2022 - The aliscore output appears to be empty. Check results for BUSCO: 327681at7742
## Thu Jul 14 02:39:29 CEST 2022 - The aliscore output appears to be empty. Check results for BUSCO: 337917at7742
## Thu Jul 14 02:39:30 CEST 2022 - The aliscore output file does not exist. Check results for BUSCO: 404316at7742
## Thu Jul 14 02:39:30 CEST 2022 - The aliscore output appears to be empty. Check results for BUSCO: 404316at7742
## Thu Jul 14 02:39:31 CEST 2022 - The aliscore output file does not exist. Check results for BUSCO: 270182at7742
## Thu Jul 14 02:39:31 CEST 2022 - The aliscore output appears to be empty. Check results for BUSCO: 270182at7742
## Thu Jul 14 02:39:32 CEST 2022 - The aliscore output file does not exist. Check results for BUSCO: 270182at7742
## Thu Jul 14 02:39:32 CEST 2022 - The aliscore output appears to be empty. Check results for BUSCO: 270182at7742
## Thu Jul 14 02:39:33 CEST 2022 - The aliscore output file does not exist. Check results for BUSCO: 361078at7742
## Thu Jul 14 02:39:33 CEST 2022 - The aliscore output appears to be empty. Check results for BUSCO: 361078at7742
## Thu Jul 14 02:39:33 CEST 2022 - The aliscore output file does not exist. Check results for BUSCO: 427300at7742
## Thu Jul 14 02:39:33 CEST 2022 - The aliscore output appears to be empty. Check results for BUSCO: 427300at7742
## Thu Jul 14 02:39:35 CEST 2022 - The aliscore output file does not exist. Check results for BUSCO: 392048at7742
## Thu Jul 14 02:39:35 CEST 2022 - The aliscore output appears to be empty. Check results for BUSCO: 392048at7742
## Thu Jul 14 02:39:35 CEST 2022 - The aliscore output file does not exist. Check results for BUSCO: 325542at7742
## Thu Jul 14 02:39:35 CEST 2022 - The aliscore output appears to be empty. Check results for BUSCO: 325542at7742
## Thu Jul 14 02:39:35 CEST 2022 - The aliscore output file does not exist. Check results for BUSCO: 332363at7742
## Thu Jul 14 02:39:35 CEST 2022 - The aliscore output appears to be empty. Check results for BUSCO: 332363at7742
## Thu Jul 14 02:39:37 CEST 2022 - The aliscore output file does not exist. Check results for BUSCO: 406535at7742
## Thu Jul 14 02:39:37 CEST 2022 - The aliscore output appears to be empty. Check results for BUSCO: 406535at7742
## Thu Jul 14 02:39:38 CEST 2022 - The aliscore output file does not exist. Check results for BUSCO: 407088at7742
## Thu Jul 14 02:39:38 CEST 2022 - The aliscore output appears to be empty. Check results for BUSCO: 407088at7742
## Thu Jul 14 02:39:40 CEST 2022 - The aliscore output file does not exist. Check results for BUSCO: 408438at7742
## Thu Jul 14 02:39:40 CEST 2022 - The aliscore output file does not exist. Check results for BUSCO: 348185at7742
## Thu Jul 14 02:39:40 CEST 2022 - The aliscore output appears to be empty. Check results for BUSCO: 408438at7742
## Thu Jul 14 02:39:40 CEST 2022 - The aliscore output appears to be empty. Check results for BUSCO: 348185at7742
## Thu Jul 14 02:39:40 CEST 2022 - The aliscore output file does not exist. Check results for BUSCO: 381830at7742
## Thu Jul 14 02:39:40 CEST 2022 - The aliscore output appears to be empty. Check results for BUSCO: 381830at7742
## Thu Jul 14 02:39:43 CEST 2022 - The aliscore output file does not exist. Check results for BUSCO: 356030at7742
## Thu Jul 14 02:39:43 CEST 2022 - The aliscore output appears to be empty. Check results for BUSCO: 356030at7742
## Thu Jul 14 02:39:43 CEST 2022 - The aliscore output file does not exist. Check results for BUSCO: 332363at7742
## Thu Jul 14 02:39:43 CEST 2022 - The aliscore output appears to be empty. Check results for BUSCO: 332363at7742
## Thu Jul 14 02:39:43 CEST 2022 - The aliscore output file does not exist. Check results for BUSCO: 405980at7742
## Thu Jul 14 02:39:43 CEST 2022 - The aliscore output appears to be empty. Check results for BUSCO: 405980at7742
## Thu Jul 14 02:39:44 CEST 2022 - The aliscore output file does not exist. Check results for BUSCO: 388430at7742
## Thu Jul 14 02:39:44 CEST 2022 - The aliscore output file does not exist. Check results for BUSCO: 344785at7742
## Thu Jul 14 02:39:44 CEST 2022 - The aliscore output appears to be empty. Check results for BUSCO: 388430at7742
## Thu Jul 14 02:39:44 CEST 2022 - The aliscore output appears to be empty. Check results for BUSCO: 344785at7742
## Thu Jul 14 02:39:46 CEST 2022 - The aliscore output file does not exist. Check results for BUSCO: 315750at7742
## Thu Jul 14 02:39:46 CEST 2022 - The aliscore output appears to be empty. Check results for BUSCO: 315750at7742
## Thu Jul 14 02:39:46 CEST 2022 - The aliscore output file does not exist. Check results for BUSCO: 351522at7742
## Thu Jul 14 02:39:46 CEST 2022 - The aliscore output appears to be empty. Check results for BUSCO: 351522at7742
## Thu Jul 14 02:39:47 CEST 2022 - The aliscore output file does not exist. Check results for BUSCO: 311774at7742
## Thu Jul 14 02:39:47 CEST 2022 - The aliscore output appears to be empty. Check results for BUSCO: 311774at7742
## Thu Jul 14 02:39:47 CEST 2022 - The aliscore output file does not exist. Check results for BUSCO: 399133at7742
## Thu Jul 14 02:39:47 CEST 2022 - The aliscore output appears to be empty. Check results for BUSCO: 399133at7742
## Thu Jul 14 02:39:48 CEST 2022 - The aliscore output file does not exist. Check results for BUSCO: 325542at7742
## Thu Jul 14 02:39:49 CEST 2022 - The aliscore output appears to be empty. Check results for BUSCO: 325542at7742
## Thu Jul 14 02:39:50 CEST 2022 - The aliscore output file does not exist. Check results for BUSCO: 341385at7742
## Thu Jul 14 02:39:50 CEST 2022 - The aliscore output appears to be empty. Check results for BUSCO: 341385at7742
## Thu Jul 14 02:39:51 CEST 2022 - The aliscore output file does not exist. Check results for BUSCO: 401745at7742
## Thu Jul 14 02:39:51 CEST 2022 - The aliscore output appears to be empty. Check results for BUSCO: 401745at7742
## Thu Jul 14 02:39:52 CEST 2022 - The aliscore output file does not exist. Check results for BUSCO: 388097at7742
## Thu Jul 14 02:39:52 CEST 2022 - The aliscore output appears to be empty. Check results for BUSCO: 388097at7742
## Thu Jul 14 02:39:52 CEST 2022 - The aliscore output file does not exist. Check results for BUSCO: 344785at7742
## Thu Jul 14 02:39:52 CEST 2022 - The aliscore output appears to be empty. Check results for BUSCO: 344785at7742
## Thu Jul 14 02:39:54 CEST 2022 - The aliscore output file does not exist. Check results for BUSCO: 372079at7742
## Thu Jul 14 02:39:54 CEST 2022 - The aliscore output appears to be empty. Check results for BUSCO: 372079at7742
## Thu Jul 14 02:39:54 CEST 2022 - The aliscore output file does not exist. Check results for BUSCO: 402520at7742
## Thu Jul 14 02:39:54 CEST 2022 - The aliscore output appears to be empty. Check results for BUSCO: 402520at7742
## Thu Jul 14 02:39:55 CEST 2022 - The aliscore output file does not exist. Check results for BUSCO: 372079at7742
## Thu Jul 14 02:39:55 CEST 2022 - The aliscore output appears to be empty. Check results for BUSCO: 372079at7742
## Thu Jul 14 02:39:56 CEST 2022 - The aliscore output file does not exist. Check results for BUSCO: 295790at7742
## Thu Jul 14 02:39:56 CEST 2022 - The aliscore output appears to be empty. Check results for BUSCO: 295790at7742
## Thu Jul 14 02:39:57 CEST 2022 - The aliscore output file does not exist. Check results for BUSCO: 295790at7742
## Thu Jul 14 02:39:57 CEST 2022 - The aliscore output file does not exist. Check results for BUSCO: 392048at7742
## Thu Jul 14 02:39:57 CEST 2022 - The aliscore output appears to be empty. Check results for BUSCO: 295790at7742
## Thu Jul 14 02:39:57 CEST 2022 - The aliscore output appears to be empty. Check results for BUSCO: 392048at7742
## Thu Jul 14 02:39:58 CEST 2022 - The aliscore output file does not exist. Check results for BUSCO: 369390at7742
## Thu Jul 14 02:39:58 CEST 2022 - The aliscore output file does not exist. Check results for BUSCO: 407323at7742
## Thu Jul 14 02:39:58 CEST 2022 - The aliscore output appears to be empty. Check results for BUSCO: 369390at7742
## Thu Jul 14 02:39:58 CEST 2022 - The aliscore output appears to be empty. Check results for BUSCO: 407323at7742
## Thu Jul 14 02:40:00 CEST 2022 - The aliscore output file does not exist. Check results for BUSCO: 355988at7742
## Thu Jul 14 02:40:00 CEST 2022 - The aliscore output file does not exist. Check results for BUSCO: 336050at7742
## Thu Jul 14 02:40:00 CEST 2022 - The aliscore output appears to be empty. Check results for BUSCO: 355988at7742
## Thu Jul 14 02:40:00 CEST 2022 - The aliscore output appears to be empty. Check results for BUSCO: 336050at7742
## Thu Jul 14 02:40:01 CEST 2022 - The aliscore output file does not exist. Check results for BUSCO: 350289at7742
## Thu Jul 14 02:40:01 CEST 2022 - The aliscore output appears to be empty. Check results for BUSCO: 350289at7742
## Thu Jul 14 02:40:03 CEST 2022 - The aliscore output file does not exist. Check results for BUSCO: 343091at7742
## Thu Jul 14 02:40:03 CEST 2022 - The aliscore output appears to be empty. Check results for BUSCO: 343091at7742
## Thu Jul 14 02:40:04 CEST 2022 - The aliscore output file does not exist. Check results for BUSCO: 343091at7742
## Thu Jul 14 02:40:04 CEST 2022 - The aliscore output appears to be empty. Check results for BUSCO: 343091at7742
## Thu Jul 14 02:40:04 CEST 2022 - The aliscore output file does not exist. Check results for BUSCO: 398403at7742
## Thu Jul 14 02:40:04 CEST 2022 - The aliscore output appears to be empty. Check results for BUSCO: 398403at7742
## Thu Jul 14 02:40:05 CEST 2022 - The aliscore output file does not exist. Check results for BUSCO: 337917at7742
## Thu Jul 14 02:40:05 CEST 2022 - The aliscore output appears to be empty. Check results for BUSCO: 337917at7742
## Thu Jul 14 02:40:05 CEST 2022 - The aliscore output file does not exist. Check results for BUSCO: 324813at7742
## Thu Jul 14 02:40:05 CEST 2022 - The aliscore output appears to be empty. Check results for BUSCO: 324813at7742
## Thu Jul 14 02:40:07 CEST 2022 - The aliscore output file does not exist. Check results for BUSCO: 407323at7742
## Thu Jul 14 02:40:07 CEST 2022 - The aliscore output file does not exist. Check results for BUSCO: 362666at7742
## Thu Jul 14 02:40:07 CEST 2022 - The aliscore output appears to be empty. Check results for BUSCO: 407323at7742
## Thu Jul 14 02:40:07 CEST 2022 - The aliscore output appears to be empty. Check results for BUSCO: 362666at7742
## Thu Jul 14 02:40:09 CEST 2022 - The aliscore output file does not exist. Check results for BUSCO: 389339at7742
## Thu Jul 14 02:40:09 CEST 2022 - The aliscore output appears to be empty. Check results for BUSCO: 389339at7742
## Thu Jul 14 02:40:10 CEST 2022 - The aliscore output file does not exist. Check results for BUSCO: 397455at7742
## Thu Jul 14 02:40:10 CEST 2022 - The aliscore output appears to be empty. Check results for BUSCO: 397455at7742
## Thu Jul 14 02:40:10 CEST 2022 - The aliscore output file does not exist. Check results for BUSCO: 372079at7742
## Thu Jul 14 02:40:10 CEST 2022 - The aliscore output appears to be empty. Check results for BUSCO: 372079at7742
## Thu Jul 14 02:40:12 CEST 2022 - The aliscore output file does not exist. Check results for BUSCO: 298413at7742
## Thu Jul 14 02:40:12 CEST 2022 - The aliscore output appears to be empty. Check results for BUSCO: 298413at7742
## Thu Jul 14 02:40:12 CEST 2022 - The aliscore output file does not exist. Check results for BUSCO: 400845at7742
## Thu Jul 14 02:40:12 CEST 2022 - The aliscore output appears to be empty. Check results for BUSCO: 400845at7742
## Thu Jul 14 02:40:13 CEST 2022 - The aliscore output file does not exist. Check results for BUSCO: 350289at7742
## Thu Jul 14 02:40:13 CEST 2022 - The aliscore output appears to be empty. Check results for BUSCO: 350289at7742
## Thu Jul 14 02:40:13 CEST 2022 - The aliscore output file does not exist. Check results for BUSCO: 331131at7742
## Thu Jul 14 02:40:13 CEST 2022 - The aliscore output appears to be empty. Check results for BUSCO: 331131at7742
## Thu Jul 14 02:40:14 CEST 2022 - The aliscore output file does not exist. Check results for BUSCO: 359566at7742
## Thu Jul 14 02:40:14 CEST 2022 - The aliscore output appears to be empty. Check results for BUSCO: 359566at7742
## Thu Jul 14 02:40:15 CEST 2022 - The aliscore output file does not exist. Check results for BUSCO: 295790at7742
## Thu Jul 14 02:40:15 CEST 2022 - The aliscore output appears to be empty. Check results for BUSCO: 295790at7742
## Thu Jul 14 02:40:15 CEST 2022 - The aliscore output file does not exist. Check results for BUSCO: 343091at7742
## Thu Jul 14 02:40:15 CEST 2022 - The aliscore output appears to be empty. Check results for BUSCO: 343091at7742
## Thu Jul 14 02:40:17 CEST 2022 - The aliscore output file does not exist. Check results for BUSCO: 420868at7742
## Thu Jul 14 02:40:17 CEST 2022 - The aliscore output appears to be empty. Check results for BUSCO: 420868at7742
## Thu Jul 14 02:40:17 CEST 2022 - The aliscore output file does not exist. Check results for BUSCO: 393595at7742
## Thu Jul 14 02:40:17 CEST 2022 - The aliscore output appears to be empty. Check results for BUSCO: 393595at7742
## Thu Jul 14 02:40:18 CEST 2022 - The aliscore output file does not exist. Check results for BUSCO: 382095at7742
## Thu Jul 14 02:40:18 CEST 2022 - The aliscore output appears to be empty. Check results for BUSCO: 382095at7742
## Thu Jul 14 02:40:20 CEST 2022 - The aliscore output file does not exist. Check results for BUSCO: 399531at7742
## Thu Jul 14 02:40:20 CEST 2022 - The aliscore output appears to be empty. Check results for BUSCO: 399531at7742
## Thu Jul 14 02:40:21 CEST 2022 - The aliscore output file does not exist. Check results for BUSCO: 411659at7742
## Thu Jul 14 02:40:21 CEST 2022 - The aliscore output file does not exist. Check results for BUSCO: 379661at7742
## Thu Jul 14 02:40:21 CEST 2022 - The aliscore output appears to be empty. Check results for BUSCO: 411659at7742
## Thu Jul 14 02:40:21 CEST 2022 - The aliscore output file does not exist. Check results for BUSCO: 344060at7742
## Thu Jul 14 02:40:21 CEST 2022 - The aliscore output appears to be empty. Check results for BUSCO: 379661at7742
## Thu Jul 14 02:40:21 CEST 2022 - The aliscore output appears to be empty. Check results for BUSCO: 344060at7742
## Thu Jul 14 02:40:23 CEST 2022 - The aliscore output file does not exist. Check results for BUSCO: 359566at7742
## Thu Jul 14 02:40:23 CEST 2022 - The aliscore output appears to be empty. Check results for BUSCO: 359566at7742
## Thu Jul 14 02:40:23 CEST 2022 - The aliscore output file does not exist. Check results for BUSCO: 322958at7742
## Thu Jul 14 02:40:23 CEST 2022 - The aliscore output appears to be empty. Check results for BUSCO: 322958at7742
## Thu Jul 14 02:40:24 CEST 2022 - The aliscore output file does not exist. Check results for BUSCO: 328338at7742
## Thu Jul 14 02:40:24 CEST 2022 - The aliscore output appears to be empty. Check results for BUSCO: 328338at7742
## Thu Jul 14 02:40:24 CEST 2022 - The aliscore output file does not exist. Check results for BUSCO: 355988at7742
## Thu Jul 14 02:40:24 CEST 2022 - The aliscore output appears to be empty. Check results for BUSCO: 355988at7742
## Thu Jul 14 02:40:25 CEST 2022 - The aliscore output file does not exist. Check results for BUSCO: 377323at7742
## Thu Jul 14 02:40:26 CEST 2022 - The aliscore output appears to be empty. Check results for BUSCO: 377323at7742
## Thu Jul 14 02:40:27 CEST 2022 - The aliscore output file does not exist. Check results for BUSCO: 286190at7742
## Thu Jul 14 02:40:27 CEST 2022 - The aliscore output appears to be empty. Check results for BUSCO: 286190at7742
## Thu Jul 14 02:40:27 CEST 2022 - The aliscore output file does not exist. Check results for BUSCO: 298413at7742
## Thu Jul 14 02:40:27 CEST 2022 - The aliscore output appears to be empty. Check results for BUSCO: 298413at7742
## Thu Jul 14 02:40:29 CEST 2022 - The aliscore output file does not exist. Check results for BUSCO: 344060at7742
## Thu Jul 14 02:40:29 CEST 2022 - The aliscore output appears to be empty. Check results for BUSCO: 344060at7742
## Thu Jul 14 02:40:29 CEST 2022 - The aliscore output file does not exist. Check results for BUSCO: 311774at7742
## Thu Jul 14 02:40:29 CEST 2022 - The aliscore output appears to be empty. Check results for BUSCO: 311774at7742
## Thu Jul 14 02:40:31 CEST 2022 - The aliscore output file does not exist. Check results for BUSCO: 351522at7742
## Thu Jul 14 02:40:31 CEST 2022 - The aliscore output appears to be empty. Check results for BUSCO: 351522at7742
## Thu Jul 14 02:40:32 CEST 2022 - The aliscore output file does not exist. Check results for BUSCO: 337603at7742
## Thu Jul 14 02:40:32 CEST 2022 - The aliscore output appears to be empty. Check results for BUSCO: 337603at7742
## Thu Jul 14 02:40:32 CEST 2022 - The aliscore output file does not exist. Check results for BUSCO: 298413at7742
## Thu Jul 14 02:40:32 CEST 2022 - The aliscore output appears to be empty. Check results for BUSCO: 298413at7742
## Thu Jul 14 02:40:33 CEST 2022 - The aliscore output file does not exist. Check results for BUSCO: 413027at7742
## Thu Jul 14 02:40:33 CEST 2022 - The aliscore output appears to be empty. Check results for BUSCO: 413027at7742
## Thu Jul 14 02:40:34 CEST 2022 - The aliscore output file does not exist. Check results for BUSCO: 315750at7742
## Thu Jul 14 02:40:34 CEST 2022 - The aliscore output appears to be empty. Check results for BUSCO: 315750at7742
## Thu Jul 14 02:40:34 CEST 2022 - The aliscore output file does not exist. Check results for BUSCO: 346606at7742
## Thu Jul 14 02:40:34 CEST 2022 - The aliscore output appears to be empty. Check results for BUSCO: 346606at7742
## Thu Jul 14 02:40:35 CEST 2022 - The aliscore output file does not exist. Check results for BUSCO: 383748at7742
## Thu Jul 14 02:40:35 CEST 2022 - The aliscore output appears to be empty. Check results for BUSCO: 383748at7742
## Thu Jul 14 02:40:35 CEST 2022 - The aliscore output file does not exist. Check results for BUSCO: 359566at7742
## Thu Jul 14 02:40:35 CEST 2022 - The aliscore output appears to be empty. Check results for BUSCO: 359566at7742
## Thu Jul 14 02:40:37 CEST 2022 - The aliscore output file does not exist. Check results for BUSCO: 389339at7742
## Thu Jul 14 02:40:37 CEST 2022 - The aliscore output appears to be empty. Check results for BUSCO: 389339at7742
## Thu Jul 14 02:40:38 CEST 2022 - The aliscore output file does not exist. Check results for BUSCO: 286190at7742
## Thu Jul 14 02:40:38 CEST 2022 - The aliscore output appears to be empty. Check results for BUSCO: 286190at7742
## Thu Jul 14 02:40:39 CEST 2022 - The aliscore output file does not exist. Check results for BUSCO: 385791at7742
## Thu Jul 14 02:40:39 CEST 2022 - The aliscore output appears to be empty. Check results for BUSCO: 385791at7742
## Thu Jul 14 02:40:40 CEST 2022 - The aliscore output file does not exist. Check results for BUSCO: 379661at7742
## Thu Jul 14 02:40:40 CEST 2022 - The aliscore output appears to be empty. Check results for BUSCO: 379661at7742
## Thu Jul 14 02:40:40 CEST 2022 - The aliscore output file does not exist. Check results for BUSCO: 416829at7742
## Thu Jul 14 02:40:40 CEST 2022 - The aliscore output appears to be empty. Check results for BUSCO: 416829at7742
## Thu Jul 14 02:40:41 CEST 2022 - The aliscore output file does not exist. Check results for BUSCO: 419468at7742
## Thu Jul 14 02:40:41 CEST 2022 - The aliscore output appears to be empty. Check results for BUSCO: 419468at7742
## Thu Jul 14 02:40:42 CEST 2022 - The aliscore output file does not exist. Check results for BUSCO: 361802at7742
## Thu Jul 14 02:40:42 CEST 2022 - The aliscore output appears to be empty. Check results for BUSCO: 361802at7742
## Thu Jul 14 02:40:43 CEST 2022 - The aliscore output file does not exist. Check results for BUSCO: 339111at7742
## Thu Jul 14 02:40:43 CEST 2022 - The aliscore output appears to be empty. Check results for BUSCO: 339111at7742
## Thu Jul 14 02:40:43 CEST 2022 - The aliscore output file does not exist. Check results for BUSCO: 409625at7742
## Thu Jul 14 02:40:43 CEST 2022 - The aliscore output appears to be empty. Check results for BUSCO: 409625at7742
## Thu Jul 14 02:40:44 CEST 2022 - The aliscore output file does not exist. Check results for BUSCO: 393595at7742
## Thu Jul 14 02:40:44 CEST 2022 - The aliscore output appears to be empty. Check results for BUSCO: 393595at7742
## Thu Jul 14 02:40:44 CEST 2022 - The aliscore output file does not exist. Check results for BUSCO: 369390at7742
## Thu Jul 14 02:40:44 CEST 2022 - The aliscore output appears to be empty. Check results for BUSCO: 369390at7742
## Thu Jul 14 02:40:45 CEST 2022 - The aliscore output file does not exist. Check results for BUSCO: 402520at7742
## Thu Jul 14 02:40:45 CEST 2022 - The aliscore output appears to be empty. Check results for BUSCO: 402520at7742
## Thu Jul 14 02:40:46 CEST 2022 - The aliscore output file does not exist. Check results for BUSCO: 365924at7742
## Thu Jul 14 02:40:46 CEST 2022 - The aliscore output appears to be empty. Check results for BUSCO: 365924at7742
## Thu Jul 14 02:40:47 CEST 2022 - The aliscore output file does not exist. Check results for BUSCO: 344060at7742
## Thu Jul 14 02:40:47 CEST 2022 - The aliscore output appears to be empty. Check results for BUSCO: 344060at7742
## Thu Jul 14 02:40:47 CEST 2022 - The aliscore output file does not exist. Check results for BUSCO: 385469at7742
## Thu Jul 14 02:40:47 CEST 2022 - The aliscore output appears to be empty. Check results for BUSCO: 385469at7742
## Thu Jul 14 02:40:50 CEST 2022 - The aliscore output file does not exist. Check results for BUSCO: 348185at7742
## Thu Jul 14 02:40:50 CEST 2022 - The aliscore output appears to be empty. Check results for BUSCO: 348185at7742
## Thu Jul 14 02:40:50 CEST 2022 - The aliscore output file does not exist. Check results for BUSCO: 384962at7742
## Thu Jul 14 02:40:50 CEST 2022 - The aliscore output appears to be empty. Check results for BUSCO: 384962at7742
## Thu Jul 14 02:40:51 CEST 2022 - The aliscore output file does not exist. Check results for BUSCO: 382095at7742
## Thu Jul 14 02:40:51 CEST 2022 - The aliscore output appears to be empty. Check results for BUSCO: 382095at7742
## Thu Jul 14 02:40:53 CEST 2022 - The aliscore output file does not exist. Check results for BUSCO: 416838at7742
## Thu Jul 14 02:40:53 CEST 2022 - The aliscore output appears to be empty. Check results for BUSCO: 416838at7742
## Thu Jul 14 02:40:53 CEST 2022 - The aliscore output file does not exist. Check results for BUSCO: 404316at7742
## Thu Jul 14 02:40:53 CEST 2022 - The aliscore output appears to be empty. Check results for BUSCO: 404316at7742
## Thu Jul 14 02:40:53 CEST 2022 - The aliscore output file does not exist. Check results for BUSCO: 345592at7742
## Thu Jul 14 02:40:53 CEST 2022 - The aliscore output appears to be empty. Check results for BUSCO: 345592at7742
## Thu Jul 14 02:40:54 CEST 2022 - The aliscore output file does not exist. Check results for BUSCO: 291391at7742
## Thu Jul 14 02:40:54 CEST 2022 - The aliscore output appears to be empty. Check results for BUSCO: 291391at7742
## Thu Jul 14 02:40:55 CEST 2022 - The aliscore output file does not exist. Check results for BUSCO: 362666at7742
## Thu Jul 14 02:40:55 CEST 2022 - The aliscore output appears to be empty. Check results for BUSCO: 362666at7742
## Thu Jul 14 02:40:55 CEST 2022 - The aliscore output file does not exist. Check results for BUSCO: 409526at7742
## Thu Jul 14 02:40:55 CEST 2022 - The aliscore output appears to be empty. Check results for BUSCO: 409526at7742
## Thu Jul 14 02:40:56 CEST 2022 - The aliscore output file does not exist. Check results for BUSCO: 311774at7742
## Thu Jul 14 02:40:56 CEST 2022 - The aliscore output appears to be empty. Check results for BUSCO: 311774at7742
## Thu Jul 14 02:40:56 CEST 2022 - The aliscore output file does not exist. Check results for BUSCO: 392330at7742
## Thu Jul 14 02:40:56 CEST 2022 - The aliscore output appears to be empty. Check results for BUSCO: 392330at7742
## Thu Jul 14 02:40:57 CEST 2022 - The aliscore output file does not exist. Check results for BUSCO: 407321at7742
## Thu Jul 14 02:40:57 CEST 2022 - The aliscore output appears to be empty. Check results for BUSCO: 407321at7742
## Thu Jul 14 02:40:59 CEST 2022 - The aliscore output file does not exist. Check results for BUSCO: 366658at7742
## Thu Jul 14 02:40:59 CEST 2022 - The aliscore output appears to be empty. Check results for BUSCO: 366658at7742
## Thu Jul 14 02:41:00 CEST 2022 - The aliscore output file does not exist. Check results for BUSCO: 378120at7742
## Thu Jul 14 02:41:00 CEST 2022 - The aliscore output appears to be empty. Check results for BUSCO: 378120at7742
## Thu Jul 14 02:41:02 CEST 2022 - The aliscore output file does not exist. Check results for BUSCO: 346606at7742
## Thu Jul 14 02:41:02 CEST 2022 - The aliscore output appears to be empty. Check results for BUSCO: 346606at7742
## Thu Jul 14 02:41:02 CEST 2022 - The aliscore output file does not exist. Check results for BUSCO: 346606at7742
## Thu Jul 14 02:41:02 CEST 2022 - The aliscore output appears to be empty. Check results for BUSCO: 346606at7742
## Thu Jul 14 02:41:02 CEST 2022 - The aliscore output file does not exist. Check results for BUSCO: 382432at7742
## Thu Jul 14 02:41:02 CEST 2022 - The aliscore output appears to be empty. Check results for BUSCO: 382432at7742
## Thu Jul 14 02:41:03 CEST 2022 - The aliscore output file does not exist. Check results for BUSCO: 419468at7742
## Thu Jul 14 02:41:03 CEST 2022 - The aliscore output appears to be empty. Check results for BUSCO: 419468at7742
## Thu Jul 14 02:41:05 CEST 2022 - The aliscore output file does not exist. Check results for BUSCO: 365811at7742
## Thu Jul 14 02:41:05 CEST 2022 - The aliscore output appears to be empty. Check results for BUSCO: 365811at7742
## Thu Jul 14 02:41:05 CEST 2022 - The aliscore output file does not exist. Check results for BUSCO: 325447at7742
## Thu Jul 14 02:41:05 CEST 2022 - The aliscore output appears to be empty. Check results for BUSCO: 325447at7742
## Thu Jul 14 02:41:06 CEST 2022 - The aliscore output file does not exist. Check results for BUSCO: 385469at7742
## Thu Jul 14 02:41:06 CEST 2022 - The aliscore output appears to be empty. Check results for BUSCO: 385469at7742
## Thu Jul 14 02:41:06 CEST 2022 - The aliscore output file does not exist. Check results for BUSCO: 388430at7742
## Thu Jul 14 02:41:06 CEST 2022 - The aliscore output appears to be empty. Check results for BUSCO: 388430at7742
## Thu Jul 14 02:41:07 CEST 2022 - The aliscore output file does not exist. Check results for BUSCO: 389183at7742
## Thu Jul 14 02:41:07 CEST 2022 - The aliscore output appears to be empty. Check results for BUSCO: 389183at7742
## Thu Jul 14 02:41:08 CEST 2022 - The aliscore output file does not exist. Check results for BUSCO: 399109at7742
## Thu Jul 14 02:41:08 CEST 2022 - The aliscore output appears to be empty. Check results for BUSCO: 399109at7742
## Thu Jul 14 02:41:09 CEST 2022 - The aliscore output file does not exist. Check results for BUSCO: 400874at7742
## Thu Jul 14 02:41:09 CEST 2022 - The aliscore output appears to be empty. Check results for BUSCO: 400874at7742
## Thu Jul 14 02:41:09 CEST 2022 - The aliscore output file does not exist. Check results for BUSCO: 355988at7742
## Thu Jul 14 02:41:09 CEST 2022 - The aliscore output appears to be empty. Check results for BUSCO: 355988at7742
## Thu Jul 14 02:41:10 CEST 2022 - The aliscore output file does not exist. Check results for BUSCO: 286190at7742
## Thu Jul 14 02:41:10 CEST 2022 - The aliscore output appears to be empty. Check results for BUSCO: 286190at7742
## Thu Jul 14 02:41:11 CEST 2022 - The aliscore output file does not exist. Check results for BUSCO: 382432at7742
## Thu Jul 14 02:41:11 CEST 2022 - The aliscore output appears to be empty. Check results for BUSCO: 382432at7742
## Thu Jul 14 02:41:12 CEST 2022 - The aliscore output file does not exist. Check results for BUSCO: 342096at7742
## Thu Jul 14 02:41:12 CEST 2022 - The aliscore output appears to be empty. Check results for BUSCO: 342096at7742
## Thu Jul 14 02:41:13 CEST 2022 - The aliscore output file does not exist. Check results for BUSCO: 371106at7742
## Thu Jul 14 02:41:13 CEST 2022 - The aliscore output file does not exist. Check results for BUSCO: 404333at7742
## Thu Jul 14 02:41:13 CEST 2022 - The aliscore output appears to be empty. Check results for BUSCO: 371106at7742
## Thu Jul 14 02:41:13 CEST 2022 - The aliscore output appears to be empty. Check results for BUSCO: 404333at7742
## Thu Jul 14 02:41:15 CEST 2022 - The aliscore output file does not exist. Check results for BUSCO: 389183at7742
## Thu Jul 14 02:41:15 CEST 2022 - The aliscore output appears to be empty. Check results for BUSCO: 389183at7742
## Thu Jul 14 02:41:16 CEST 2022 - The aliscore output file does not exist. Check results for BUSCO: 404316at7742
## Thu Jul 14 02:41:16 CEST 2022 - The aliscore output appears to be empty. Check results for BUSCO: 404316at7742
## Thu Jul 14 02:41:16 CEST 2022 - The aliscore output file does not exist. Check results for BUSCO: 408072at7742
## Thu Jul 14 02:41:16 CEST 2022 - The aliscore output appears to be empty. Check results for BUSCO: 408072at7742
## Thu Jul 14 02:41:18 CEST 2022 - The aliscore output file does not exist. Check results for BUSCO: 421281at7742
## Thu Jul 14 02:41:18 CEST 2022 - The aliscore output appears to be empty. Check results for BUSCO: 421281at7742
## Thu Jul 14 02:41:18 CEST 2022 - The aliscore output file does not exist. Check results for BUSCO: 291391at7742
## Thu Jul 14 02:41:18 CEST 2022 - The aliscore output appears to be empty. Check results for BUSCO: 291391at7742
## Thu Jul 14 02:41:18 CEST 2022 - The aliscore output file does not exist. Check results for BUSCO: 380785at7742
## Thu Jul 14 02:41:18 CEST 2022 - The aliscore output appears to be empty. Check results for BUSCO: 380785at7742
## Thu Jul 14 02:41:20 CEST 2022 - The aliscore output file does not exist. Check results for BUSCO: 369736at7742
## Thu Jul 14 02:41:20 CEST 2022 - The aliscore output appears to be empty. Check results for BUSCO: 369736at7742
## Thu Jul 14 02:41:20 CEST 2022 - The aliscore output file does not exist. Check results for BUSCO: 357298at7742
## Thu Jul 14 02:41:20 CEST 2022 - The aliscore output appears to be empty. Check results for BUSCO: 357298at7742
## Thu Jul 14 02:41:21 CEST 2022 - The aliscore output file does not exist. Check results for BUSCO: 399667at7742
## Thu Jul 14 02:41:21 CEST 2022 - The aliscore output appears to be empty. Check results for BUSCO: 399667at7742
## Thu Jul 14 02:41:22 CEST 2022 - The aliscore output file does not exist. Check results for BUSCO: 427300at7742
## Thu Jul 14 02:41:22 CEST 2022 - The aliscore output appears to be empty. Check results for BUSCO: 427300at7742
## Thu Jul 14 02:41:24 CEST 2022 - The aliscore output file does not exist. Check results for BUSCO: 342096at7742
## Thu Jul 14 02:41:24 CEST 2022 - The aliscore output appears to be empty. Check results for BUSCO: 342096at7742
## Thu Jul 14 02:41:24 CEST 2022 - The aliscore output file does not exist. Check results for BUSCO: 362666at7742
## Thu Jul 14 02:41:24 CEST 2022 - The aliscore output appears to be empty. Check results for BUSCO: 362666at7742
## Thu Jul 14 02:41:25 CEST 2022 - The aliscore output file does not exist. Check results for BUSCO: 411659at7742
## Thu Jul 14 02:41:25 CEST 2022 - The aliscore output appears to be empty. Check results for BUSCO: 411659at7742
## Thu Jul 14 02:41:26 CEST 2022 - The aliscore output file does not exist. Check results for BUSCO: 424089at7742
## Thu Jul 14 02:41:26 CEST 2022 - The aliscore output appears to be empty. Check results for BUSCO: 424089at7742
## Thu Jul 14 02:41:27 CEST 2022 - The aliscore output file does not exist. Check results for BUSCO: 291391at7742
## Thu Jul 14 02:41:27 CEST 2022 - The aliscore output appears to be empty. Check results for BUSCO: 291391at7742
## Thu Jul 14 02:41:28 CEST 2022 - The aliscore output file does not exist. Check results for BUSCO: 399109at7742
## Thu Jul 14 02:41:28 CEST 2022 - The aliscore output file does not exist. Check results for BUSCO: 414796at7742
## Thu Jul 14 02:41:28 CEST 2022 - The aliscore output appears to be empty. Check results for BUSCO: 399109at7742
## Thu Jul 14 02:41:28 CEST 2022 - The aliscore output appears to be empty. Check results for BUSCO: 414796at7742
## Thu Jul 14 02:41:29 CEST 2022 - The aliscore output file does not exist. Check results for BUSCO: 409625at7742
## Thu Jul 14 02:41:29 CEST 2022 - The aliscore output appears to be empty. Check results for BUSCO: 409625at7742
## Thu Jul 14 02:41:30 CEST 2022 - The aliscore output file does not exist. Check results for BUSCO: 378341at7742
## Thu Jul 14 02:41:30 CEST 2022 - The aliscore output appears to be empty. Check results for BUSCO: 378341at7742
## Thu Jul 14 02:41:31 CEST 2022 - The aliscore output file does not exist. Check results for BUSCO: 425988at7742
## Thu Jul 14 02:41:31 CEST 2022 - The aliscore output appears to be empty. Check results for BUSCO: 425988at7742
## Thu Jul 14 02:41:32 CEST 2022 - The aliscore output file does not exist. Check results for BUSCO: 365924at7742
## Thu Jul 14 02:41:32 CEST 2022 - The aliscore output appears to be empty. Check results for BUSCO: 365924at7742
## Thu Jul 14 02:41:32 CEST 2022 - The aliscore output file does not exist. Check results for BUSCO: 304251at7742
## Thu Jul 14 02:41:32 CEST 2022 - The aliscore output appears to be empty. Check results for BUSCO: 304251at7742
## Thu Jul 14 02:41:34 CEST 2022 - The aliscore output file does not exist. Check results for BUSCO: 392901at7742
## Thu Jul 14 02:41:34 CEST 2022 - The aliscore output appears to be empty. Check results for BUSCO: 392901at7742
## Thu Jul 14 02:41:35 CEST 2022 - The aliscore output file does not exist. Check results for BUSCO: 388430at7742
## Thu Jul 14 02:41:35 CEST 2022 - The aliscore output appears to be empty. Check results for BUSCO: 388430at7742
## Thu Jul 14 02:41:36 CEST 2022 - The aliscore output file does not exist. Check results for BUSCO: 400196at7742
## Thu Jul 14 02:41:36 CEST 2022 - The aliscore output appears to be empty. Check results for BUSCO: 400196at7742
## Thu Jul 14 02:41:37 CEST 2022 - The aliscore output file does not exist. Check results for BUSCO: 322958at7742
## Thu Jul 14 02:41:37 CEST 2022 - The aliscore output appears to be empty. Check results for BUSCO: 322958at7742
## Thu Jul 14 02:41:38 CEST 2022 - The aliscore output file does not exist. Check results for BUSCO: 291326at7742
## Thu Jul 14 02:41:38 CEST 2022 - The aliscore output appears to be empty. Check results for BUSCO: 291326at7742
## Thu Jul 14 02:41:38 CEST 2022 - The aliscore output file does not exist. Check results for BUSCO: 322958at7742
## Thu Jul 14 02:41:38 CEST 2022 - The aliscore output appears to be empty. Check results for BUSCO: 322958at7742
## Thu Jul 14 02:41:39 CEST 2022 - The aliscore output file does not exist. Check results for BUSCO: 363461at7742
## Thu Jul 14 02:41:39 CEST 2022 - The aliscore output appears to be empty. Check results for BUSCO: 363461at7742
## Thu Jul 14 02:41:40 CEST 2022 - The aliscore output file does not exist. Check results for BUSCO: 339111at7742
## Thu Jul 14 02:41:40 CEST 2022 - The aliscore output appears to be empty. Check results for BUSCO: 339111at7742
## Thu Jul 14 02:41:40 CEST 2022 - The aliscore output file does not exist. Check results for BUSCO: 412708at7742
## Thu Jul 14 02:41:41 CEST 2022 - The aliscore output appears to be empty. Check results for BUSCO: 412708at7742
## Thu Jul 14 02:41:41 CEST 2022 - The aliscore output file does not exist. Check results for BUSCO: 304251at7742
## Thu Jul 14 02:41:41 CEST 2022 - The aliscore output appears to be empty. Check results for BUSCO: 304251at7742
## Thu Jul 14 02:41:42 CEST 2022 - The aliscore output file does not exist. Check results for BUSCO: 368039at7742
## Thu Jul 14 02:41:42 CEST 2022 - The aliscore output appears to be empty. Check results for BUSCO: 368039at7742
## Thu Jul 14 02:41:43 CEST 2022 - The aliscore output file does not exist. Check results for BUSCO: 418675at7742
## Thu Jul 14 02:41:43 CEST 2022 - The aliscore output appears to be empty. Check results for BUSCO: 418675at7742
## Thu Jul 14 02:41:44 CEST 2022 - The aliscore output file does not exist. Check results for BUSCO: 325638at7742
## Thu Jul 14 02:41:44 CEST 2022 - The aliscore output appears to be empty. Check results for BUSCO: 325638at7742
## Thu Jul 14 02:41:44 CEST 2022 - The aliscore output file does not exist. Check results for BUSCO: 325638at7742
## Thu Jul 14 02:41:44 CEST 2022 - The aliscore output appears to be empty. Check results for BUSCO: 325638at7742
## Thu Jul 14 02:41:45 CEST 2022 - The aliscore output file does not exist. Check results for BUSCO: 425988at7742
## Thu Jul 14 02:41:45 CEST 2022 - The aliscore output appears to be empty. Check results for BUSCO: 425988at7742
## Thu Jul 14 02:41:46 CEST 2022 - The aliscore output file does not exist. Check results for BUSCO: 365811at7742
## Thu Jul 14 02:41:46 CEST 2022 - The aliscore output appears to be empty. Check results for BUSCO: 365811at7742
## Thu Jul 14 02:41:47 CEST 2022 - The aliscore output file does not exist. Check results for BUSCO: 389339at7742
## Thu Jul 14 02:41:47 CEST 2022 - The aliscore output appears to be empty. Check results for BUSCO: 389339at7742
## Thu Jul 14 02:41:47 CEST 2022 - The aliscore output file does not exist. Check results for BUSCO: 369735at7742
## Thu Jul 14 02:41:47 CEST 2022 - The aliscore output appears to be empty. Check results for BUSCO: 369735at7742
## Thu Jul 14 02:41:49 CEST 2022 - The aliscore output file does not exist. Check results for BUSCO: 364336at7742
## Thu Jul 14 02:41:49 CEST 2022 - The aliscore output appears to be empty. Check results for BUSCO: 364336at7742
## Thu Jul 14 02:41:49 CEST 2022 - The aliscore output file does not exist. Check results for BUSCO: 432038at7742
## Thu Jul 14 02:41:49 CEST 2022 - The aliscore output appears to be empty. Check results for BUSCO: 432038at7742
## Thu Jul 14 02:41:51 CEST 2022 - The aliscore output file does not exist. Check results for BUSCO: 367717at7742
## Thu Jul 14 02:41:51 CEST 2022 - The aliscore output appears to be empty. Check results for BUSCO: 367717at7742
## Thu Jul 14 02:41:51 CEST 2022 - The aliscore output file does not exist. Check results for BUSCO: 361503at7742
## Thu Jul 14 02:41:51 CEST 2022 - The aliscore output appears to be empty. Check results for BUSCO: 361503at7742
## Thu Jul 14 02:41:51 CEST 2022 - The aliscore output file does not exist. Check results for BUSCO: 396678at7742
## Thu Jul 14 02:41:52 CEST 2022 - The aliscore output appears to be empty. Check results for BUSCO: 396678at7742
## Thu Jul 14 02:41:52 CEST 2022 - The aliscore output file does not exist. Check results for BUSCO: 325638at7742
## Thu Jul 14 02:41:53 CEST 2022 - The aliscore output appears to be empty. Check results for BUSCO: 325638at7742
## Thu Jul 14 02:41:53 CEST 2022 - The aliscore output file does not exist. Check results for BUSCO: 392330at7742
## Thu Jul 14 02:41:53 CEST 2022 - The aliscore output appears to be empty. Check results for BUSCO: 392330at7742
## Thu Jul 14 02:41:55 CEST 2022 - The aliscore output file does not exist. Check results for BUSCO: 405886at7742
## Thu Jul 14 02:41:55 CEST 2022 - The aliscore output file does not exist. Check results for BUSCO: 405886at7742
## Thu Jul 14 02:41:55 CEST 2022 - The aliscore output appears to be empty. Check results for BUSCO: 405886at7742
## Thu Jul 14 02:41:55 CEST 2022 - The aliscore output appears to be empty. Check results for BUSCO: 405886at7742
## Thu Jul 14 02:41:56 CEST 2022 - The aliscore output file does not exist. Check results for BUSCO: 421575at7742
## Thu Jul 14 02:41:56 CEST 2022 - The aliscore output appears to be empty. Check results for BUSCO: 421575at7742
## Thu Jul 14 02:41:57 CEST 2022 - The aliscore output file does not exist. Check results for BUSCO: 382946at7742
## Thu Jul 14 02:41:57 CEST 2022 - The aliscore output appears to be empty. Check results for BUSCO: 382946at7742
## Thu Jul 14 02:41:57 CEST 2022 - The aliscore output file does not exist. Check results for BUSCO: 422259at7742
## Thu Jul 14 02:41:57 CEST 2022 - The aliscore output appears to be empty. Check results for BUSCO: 422259at7742
## Thu Jul 14 02:41:59 CEST 2022 - The aliscore output file does not exist. Check results for BUSCO: 424089at7742
## Thu Jul 14 02:41:59 CEST 2022 - The aliscore output appears to be empty. Check results for BUSCO: 424089at7742
## Thu Jul 14 02:42:01 CEST 2022 - The aliscore output file does not exist. Check results for BUSCO: 304251at7742
## Thu Jul 14 02:42:01 CEST 2022 - The aliscore output appears to be empty. Check results for BUSCO: 304251at7742
## Thu Jul 14 02:42:01 CEST 2022 - The aliscore output file does not exist. Check results for BUSCO: 400196at7742
## Thu Jul 14 02:42:01 CEST 2022 - The aliscore output appears to be empty. Check results for BUSCO: 400196at7742
## Thu Jul 14 02:42:01 CEST 2022 - The aliscore output file does not exist. Check results for BUSCO: 369736at7742
## Thu Jul 14 02:42:01 CEST 2022 - The aliscore output appears to be empty. Check results for BUSCO: 369736at7742
## Thu Jul 14 02:42:02 CEST 2022 - The aliscore output file does not exist. Check results for BUSCO: 388097at7742
## Thu Jul 14 02:42:02 CEST 2022 - The aliscore output appears to be empty. Check results for BUSCO: 388097at7742
